# Single-nucleotide Differences and Cell Type Decide the Subcellular Localization of miRNA Isoforms (isomiRs), tRNA-derived Fragments (tRFs) and rRNA-derived Fragments (rRFs)

**DOI:** 10.1101/2022.08.22.503746

**Authors:** Tess Cherlin, Yi Jing, Venetia Pliatsika, Haley Wilson, Lily Thompson, Panagiotis I. Vlantis, Phillipe Loher, Benjamin Leiby, Isidore Rigoutsos

**Affiliations:** Computational Medicine Center, Thomas Jefferson University, Philadelphia, PA 19017; Division of Biostatistics, Thomas Jefferson University, Philadelphia, PA 19017

## Abstract

**Background:** MicroRNAs (miRNAs) and their isoforms (isomiRs), tRNA-derived fragments (tRFs), and rRNA-derived fragments (rRFs) represent ~95% of all short RNAs found in cells. All three types modulate mRNA and protein abundance and are dysregulated in diseases. Experimental studies to date assumed that the subcellular localization of these molecules is well understood and constant across cell types.

**Results:** We investigated the localization of isomiRs, tRFs, and rRFs in biological replicates from three frequently-used model cell lines. In each case, we analyzed the contents of the nucleus, cytoplasm, whole mitochondrion, mitoplast, and the whole cell. We used a rigorous mathematical model to account for cross-fraction contamination and technical errors and adjusted abundances accordingly. We found that isomiRs, tRFs, and rRFs exhibit complex and unexpected patterns of subcellular localization. These patterns depend on the type of the RNA molecule, its exact sequence, and the cell type. Even for “sibling” RNAs from the same parental RNA whose sequences differ by only a few nucleotides, their subcellular localization depends on each sibling’s exact sequence and the cell type.

**Conclusions:** Previous studies of isomiRs, tRFs, and rRFs that used ectopic expression without accounting for isoforms may need to be re-evaluated. Future experiments with these molecules will need to distinguish among the multiple isoforms and account for the fact that each isoform’s abundance and destination depend on its exact sequence and cell type. The findings additionally suggest the existence of an intracellular trafficking program for isomiRs, tRFs, and rRFs and, by extension, expanded roles for these molecules – both dimensions await characterization. To help design future experiments, we compiled a first-of-its-kind Atlas to catalogue the subcellular localization and abundance of 5,898 isomiRs, tRFs, and rRFs across three model cell lines.

**Results Summary:**

- We analyzed the distribution of microRNA isoforms (isomiRs), tRNA-derived fragments (tRFs), and rRNA-derived fragments (rRFs) in the

◦ nucleus
◦ cytoplasm
◦ mitochondrion, and
◦ mitoplast

of biological replicates from three cell lines from the same tissue.
- We corrected the measured abundances by accounting for cross-fraction contamination and technical errors through a rigorous mathematical model.
- Our analysis revealed complex localization patterns involving numerous isomiRs, tRFs, and rRFs.
- The subcellular localization of these RNAs depends on their exact sequence and differs even for molecules that arise from the same parental miRNA, tRNA, or rRNA.
- For a given RNA, its subcellular localization additionally depends on cell type.
- The findings have implications for previous and future molecular studies of the function of isomiRs, tRFs, and rRFs.
- The findings suggest the existence of a complex subcellular trafficking program, and hint at expanded functions for these RNA molecules that differ by compartment.
- To assist with the design of future experiments, we created a first-of-its-kind Atlas that catalogues the subcellular distribution and abundance of 5,898 isomiRs, tRFs, and rRFs across three cell lines.

## INTRODUCTION

Most experimental studies to date assumed that the destination of isomiRs, tRFs, and rRFs within the cell is well understood and unchanged across cell types, and that the isoforms of these molecules merit no dedicated study. However, as we showed in previous work, personal attributes (e.g., sex, genetic ancestry, and age)^1–6^ and context (e.g., cell type, tissue type and disease)^4–10^ modulate the profiles of all three molecular types. The differences are consequential because they include differing 5’ or 3’ termini and changes in abundance, which affect the identity of the targeted mRNAs and the level of suppression^1,3–5^.

Considering these dependencies, it is important to know which isomiRs, tRFs, and rRFs are produced in a given cellular context. Also important is to establish where in the cell these short RNAs localize, as this can provide important clues as to their roles. The localization question need not have an obvious answer considering that these molecules arise from both the nuclear and the mitochondrial (MT) genomes.

The nuclear genome hosts more than five thousand miRNA precursors^11–18^, 610 nuclear tRNA genes, and multiple instances of four nuclear rRNA genes (5S, 5.8S, 18S and 28S)^19–21^. These precursor RNAs produce isomiRs, tRFs and rRFs, respectively. The nucleus-encoded miRNAs, tRNAs and rRNAs carry out their canonical functions in the cytoplasm where they are shuttled *via* dedicated mechanisms. Mature tRNAs are charged with amino acids, interact with the ribosome, and contribute their amino acids to the nascent polypeptides during protein translation^22^. As for rRNAs, they serve as components of the ribosome’s scaffold and specific rRNA motifs are essential for the translation process^23,24^. MiRNA precursors produce mature miRNAs and multiple isomiRs that enter the RNA interference pathway (RNAi)^25^. Nuclear tRNAs and rRNAs produce tRFs and rRFs, respectively, through mechanisms that are not well understood^26–28^. At least some tRFs and rRFs are known to also enter the RNAi pathway{Kumar, 2016 #8156;Lambert, 2019 #1572}, just like isomiRs.

The MT genome encodes 22 tRNA and two rRNA genes (12S and 16S)^29^ all of which produce tRFs and rRFs. For more details, we refer the reader to recent reviews that summarize the current knowledge on these RNA types^27,30–32^. The MT-encoded tRNAs and rRNAs are initially transcribed as polycistrons from the MT’s circular genome in the inner membrane of the mitochondrion and remain inside the inner membrane of the MT where they carry out their canonical functions during protein translation^29,33^. In analogy to their nuclear counterparts, the mechanism by which MT tRNAs and rRNA produce tRFs and rRFs is not understood.

Based on previous studies, we know that *long* non-coding RNAs (lncRNAs) localize preferentially to subcellular compartments^34–41^, sometimes in response to internal or external signaling^8,34,35,38,39,42–45^. However, the question of *short* RNA localization has received comparatively less attention. The localization of miRNAs is the one notable exception^36,46–56^.

Conventionally, <u>miRNAs</u> are expected in the cytoplasm. But previous studies showed that miRNAs can also localize and function in the nucleus^48,52,55–61^, and *at* or *in* the mitochondrion^36,62,63^. One important limitation of these studies is that they neither distinguished among isomiRs nor examined the dependence of isomiR localization on cellular context. In regards tRFs, our analytical pan-cancer studies suggest that tRFs must localize to the nucleus, cytoplasm, and MT in various combinations, where they potentially play various roles^27^. This is in agreement with a previous report of a full-length MT tRNA^Met^ in the cytoplasm, where it plays an unknown role^64^. Even less is known about the localization patterns of <u>rRFs</u>, the most recent addition to the pantheon of short RNA classes{Cherlin, 2020 #894;Lambert, 2019 #1572}. One relevant piece of evidence is that the nuclearly-encoded 5S rRNA localizes to both the cytoplasm and the MT^65,66^, raising the possibility that rRFs have multiple destinations. Of note, some rRFs are loaded on Argonaute^28^ indicating that at least some of them are in the cytoplasm.

We are not aware of a systematic study that addressed the question of how isomiRs, tRFs, and rRFs are distributed within the cell or how cellular context affects that distribution. In what follows, we present our findings from such a systematic exploration. We used as a testbed three commercially available cell lines (BT-20, MDA-MB-231, MDA-MB-468) that model triple-negative breast cancer (TNBC). For each cell line, we generated and analyzed data from several biological replicates and four subcellular compartments. Being mindful of the possibility of cross-fraction contamination and technical errors, we used a rigorous mixed-effects model to estimate both and reconstruct the true abundance of each molecule separately in each fraction. We also compared our findings to those from three earlier reports on the subcellular localization of short RNAs^51,62,63^. Lastly, for a select set of dozens of short isomiRs, tRFs, and rRFs we generated their profiles across 6,203 public datasets from different tissue sources.

## METHODS

For the purposes of this project, we developed a new method, “Fraction-seq.” Fraction-seq combines experimental and analytical steps. We describe each component of Fraction-seq below, followed by information on other methods that we used in this project.

### Cell Culture

We grow BT-20 (ATCC #HTB-19), MDA-MB231 (ATCC #HTB-26), and MDA-MB468 (ATCC #HTB-132) cells according to standard cell growth conditions using Complete DMEM (Gibco Dulbecco’s Modified Eagle Medium). We test all cell lines frequently to ensure that they are mycoplasma-free.

### Fraction-seq: cellular fractionation step

Our subcellular localization method leverages deep-sequencing and profiles short RNAs in the nuclear, cytoplasmic, mitochondrial, and mitoplast fractions of a cell starting from the same material. Specifically, we grow cells in 20-40 10 cm dishes depending on cell type in Complete DMEM Media. When cells reach 90% confluence, we wash the plates 2 times with ice cold 1X PBS. We add 2 ml of 1X ice cold PBS to each plate and then mechanically scrape cells using a cell scraper. We pool cells into two 50 ml ice cold conical tubes that we spin at 500 g for 10 minutes. We remove the supernatant and resuspend the cells in 2x packed cell volume (PCV) of 0.9% NaCl followed by incubation on ice for 10 minutes. We then aliquot cells into 10 ice cold 1.5 ml microcentrifuge tubes and spin them at 500 g for 10 minutes. We remove the supernatant by aspirating. Using the Qiagen Mitochondria Isolation Kit (Qiagen #37612), we resuspend cell pellets in ice cold Lysis Buffer + Protease Inhibitor and 1 mM EGTA and incubate for 10 minutes rotating (end-over-end shaker) at 4°C. We save 0.4% of lysate from each tube for **total cell sample**. We centrifuge the lysate at 1,000 g for 10 minutes at 4°C then carefully remove the supernatant and transfer it to fresh 1.5 ml microcentrifuge tubes – this is the *crude* cytoplasmic fraction. We spin the cytoplasmic fraction at 16,000 g for 10 minutes at 4°C to remove additional cell debris and transfer the supernatant to ultracentrifuge tubes and centrifuge them at 100,000 g for 30 minutes at 4°C. The supernatant now contains the ***pure* cytoplasmic fraction**. We resuspend the pellet from the cell lysate in 1.5 ml ice-cold Disruption Buffer (+ protease inhibitor). We pass the pellet slowly through a blunt-ended 26-gauge needle syringe, 20x on ice (making sure to avoid bubbles). We centrifuge the lysate at 1,000 g for 10 minutes at 4°C then transfer the supernatant to a clean 1.5 ml microcentrifuge tube. We save the pellet as *crude* nuclear fraction and centrifuge the supernatant at 6,000 g for 10 minutes at 4°C. The pellet that remains after we remove and discard the supernatant is the *crude* mitochondrial fraction. We combine the two crude mitochondrial pellets into one and carefully resuspend them in 750 μl Mitochondria Purification Buffer (+ protease inhibitor) using a 1 ml pipette. 750 μl Mitochondria Purification Buffer is also added to a 2 ml microcentrifuge tube and 500 μl Disruption Buffer is carefully added underneath (see image on page 13 of QProteome Mitochondria Isolation Handbook). Finally, we add the mitochondrial suspension carefully to the top. We centrifuge this multilayer solution at 14,000 g for 15 minutes at 4°C which produces a delicate, soft, white band at the bottom of the tube. We carefully remove and discard 1.5 ml of supernatant then resuspend the remaining 500 μl (mitochondrial pellet and buffer) and transfer it to clean 1.5 ml tubes. We resuspend 1 ml of Mitochondria Storage Buffer into the solution and centrifuge at 8,000 g for 10 minutes at 4°C. We carefully remove and discard 1.5 ml of the supernatant and resuspend it in 1.5 ml Mitochondria Storage Buffer, then centrifuge it at 8,000 g for 10 minutes at 4°C. We repeat this process until a pellet forms at the bottom of the tube. We combine these ***pure* mitochondrial fraction** pellets in 100 μl Mitochondrial Storage Buffer (+ protease inhibitor). We measure the crude nuclear fraction pellets and resuspend them in the appropriate volume of CERI buffer (+ protease inhibitors) from the NE-PER Nuclear Cytoplasmic Extraction Kit (Thermo Fisher Scientific #78833). We vortex the tubes at the highest setting for 15 seconds and incubate on ice for 10 minutes. We add ice-cold CER II buffer to the tubes. We vortex the pellet for 5 seconds at the highest setting and then incubate on ice for 1 additional minute. Again, we vortex the pellet for 5 seconds on the highest setting then centrifuge it at maximum speed (~16,000 g). We remove and discard the supernatant. We then resuspend the *insoluble* pellet in NER buffer (+ protease inhibitors), vortex the pellet at the highest setting for 15 seconds, incubate on ice for 10 minutes (one time only), and, finally, centrifuge it at max speed (~16,000 g) for 10 minutes at 4°C. The supernatant contains the ***pure* nuclear fraction**. Following validation of the mitochondrial fraction’s purity using protein methods (described below), we divide the pure mitochondrial fraction into thirds: 1/3^rd^ represents the pure mitochondrial fraction; the remaining 2/3^rds^ are transferred to a new tube to be purified into the mitoplast fraction. We centrifuge the pre-mitoplast pellet at 8,000 g for 10 minutes, then discard the supernatant. We resuspend the pellet in 10x pellet volume 100 mM Swelling Buffer (NaPO4, pH 7.4) (+ protease inhibitor) and incubate on ice for 20 minutes. The hypotonic reaction is stopped by adding 3.75 x pellet volume (60%) to solution. We centrifuge the solution at 12,000 g for 15 minutes, followed by removal of the supernatant. We wash the pellet twice in 1 ml Mitochondrial Storage Buffer (+ protease inhibitor). Lastly, we resuspend the ***pure* mitoplast fraction** in 50 μl Mitochondrial Storage Buffer (+ protease inhibitor). We store all fractions in −20°C.

### Fraction-seq: WES “blotting” step

We subject all fraction lysates (total, nuclear, cytoplasmic, mitochondrial, and mitoplast) to Bradford Protein Assay (BioRad #5000006) to calculate protein concentration. We use the Simple Western size-based Assay from Protein Simple (WES) to validate purity of the cell fractions based on protein profiling. We carry out WES validation according to the company’s specifications (https://proteinsimple.com) by loading 0.2 mg/ml of each sample to a 12-230 kDa Jess/Wes Separation Module, 8 x 25 Capillary Cartridge (Protein Simple #SM-W004). Primary antibodies, Cytochrome C -- Mitoplast (BD Pharmagen #556433), GAPDH -- Cytoplasm (NEB #2118), LAMIN A/C – Nucleus (NEB #4777), SDHA – Mitochondria/Mitoplast (NEB #11998), and TFAM – Mitochondria/Mitoplast (NEB #8076) are used as cell fraction markers. Secondary Anti-Rabbit (Protein Simple #DM-001) and Anti-Mouse (Protein Simple #DM-002) antibodies are used. To analyze the WES results, we follow the company’s protocol and calculate the area under the curve for each antibody.

### Fraction-seq: RNA isolation step

After we validate the protein purity of the cell fractions, we subject the mitochondrial fraction to incubation with 2 mg/ml RNase A for 15 minutes on ice. Immediately, 900 μl of Trizol (Invitrogen) is added to Total, Nuclear, Cytoplasmic, Mitochondrial, and Mitoplast fractions. A 1 ml pipette is used to resuspend samples in Trizol, and samples are incubated at room temperature for 5 minutes to equilibrate. We add 180 μl chloroform to the samples. The Trizol/chloroform solution is shaken vigorously for 30 seconds. We incubate the samples at room temperature for 3 minutes and then centrifuge them at 12,000 g for 15 minutes. We transfer the clear supernatant from each sample to clean tubes and added 500 μl 100% isopropanol to each tube. We incubate the samples at 20°C for 2 hrs allowing the RNA to precipitate out of solution. We centrifuge the samples at 12,000 g for 10 minutes at 4°C and remove the supernatant. We wash the samples twice with 1 ml 70% ethanol and centrifuged them at 7,600 g for 5 minutes. We remove all ethanol and allow the pellets to dry for 15 minutes. We then resuspend the pellets in molecular grade water (Fisher #BP2819-1). We detect the RNA concentrations using a spectrophotometer.

### Fraction-seq: next generation sequencing step

We assess the integrity and purity of the RNA of the validated preparations (above) using the Agilent 2100 Bioanalyzer. We prepare the validated RNAs for sequencing using the NEBNext Library preparation method (#E7330) with the kit’s standard protocol. The NEBNext 3’-adapter is AGATCGGAAGAGCACACGTCT. All samples are sequenced using the Illumina NextSeq 500 platform at 75 cycles and an average depth of 30 million reads.

### Fraction-seq: profiling isomiRs, tRFs and rRFs step

We profile isomiRs using isoMiRmap^67^, tRFs using MINTmap^68,69^, and rRFs using the brute-force approach that we described previously^1^. Prior to mapping, we used *cutadapt* to remove the 3’ adapter^70^. For all three RNA classes, the profiling methods ensure exhaustive and deterministic identification and quantification of the respective molecules. We threshold the mapped short RNAs using the adaptive, sample-specific Threshold-seq tool^71^ and default parameter settings. Since the mitoplast fraction is not treated with RNase A because of its limiting yield, we apply additional computational filters: specifically, for a given biological replicate, we remove any short RNA that is present in the Mitoplast fraction but absent from the mitochondrial fraction. We then maintain isomiRs, tRFs, and rRFs whose sequence lengths are ≥ 18 nts. This is a more stringent length threshold than what is used by MINTmap and isoMiRmap. Finally, we normalize the abundances of all isomiRs, tRFs and rRFs and express them in reads-per-million (RPM).

### Differential Abundance of short RNAs

We use the DESeq2 package^72^ in R to determine short RNAs that are differentially abundant between two cell lines, or two cell fractions. For the values input into the DESeq2 package, we use a mean raw read threshold of 300 for either of the two groups being compared. For normalization, we used DESeq2’s default parameters which calculates size factors based on the median of ratios. Differential Abundance (DA) is measured in |log_2_| ≥ 1 change. We determine statistical significance using a false discovery rate (FDR) of 0.05.

### Northern blotting

We run 5 or 10 μg of RNA from each cell line on a 15% acrylamide/8M urea gel at 300V until the lower dye front reaches the bottom of the gel. We transfer the gel to Hybond^TM^-N^+^ membrane (Amersham Biosciences, catalog number: RPN303B) at 400 mA for 30 minutes. We dry the membrane and then cross-link twice at 120,000 μJ/cm^2^. We pre-hybridize the membranes in hybridization buffer (PerfectHyb™ Plus Hybridization Buffer: H7033-1L) for 30 minutes rotating at 37°C. We prepare northern probes using the DIG labeling kit (DIG Oligonucleotide 3’-End Labeling Kit, 2nd Gen: 3353575910). Detection of membranes is done using the DIG detection kit (DIG Wash and Block Buffer Set: 11585762001, Anti-Digoxigenin-AP, Fab fragments: 11093274910, CDP-Star Chemiluminescent Substrate: C0712-100ML) following the manufacturer’s instructions. **Supplemental Table S1** lists the northern probes we used in this study.

### Mixed-effects modeling

We model the fractionation process by distinguishing among the following seven groups of short RNAs: isomiRs; tRFs derived from nuclear tRNAs; tRFs derived from mitochondrial tRNAs; ambiguous tRFs whose sequences are present in both nuclear and mitochondrial tRNAs; rRFs derived from the nuclear 5S rRNA; rRFs derived from the nuclear 45S rRNA cassette; and rRFs derived from the two mitochondrial rRNAs. Each short RNA, including those that are adjacent or overlapping on the parental RNA from which they arise, is treated as an independent molecule. Furthermore, we assume the possibility of contamination during the fractionation process, and of errors that differed from replicate to replicate.

For the estimation process, and separately for each cell line:

- We assume that the cytoplasmic fraction (“cyto”) is uncontaminated and the measured abundance of the *i^th^* molecule in the *j^th^* replicate is linked to the *i^th^* molecule’s true value as follows:

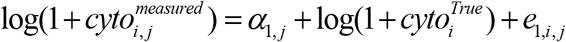

where 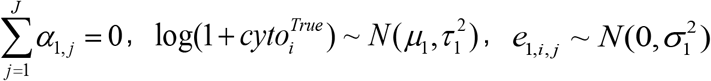, and *J* is the number of replicates. This is a mixed effects model where 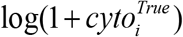 is the random effect. The mean of the log-transformed true value (*μ*_1*i*_) is a function of group membership. We fit this model using restricted maximum likelihood estimation in SAS Proc Mixed. From the results of fitting this model, we can estimate the expected true value for the *i^th^* molecule from 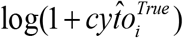, and the *j^th^* replicate of the *i^th^* molecule from 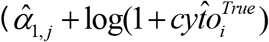).
- We assume that the nuclear fraction (“nuc”), mitochondrial fraction (“mito”) and mitoplast fraction (“MP”) are contaminating one another and the measured abundance of the *i^th^* molecule in the *j^th^* replicate is linked to the *i^th^* molecule’s true value as follows:

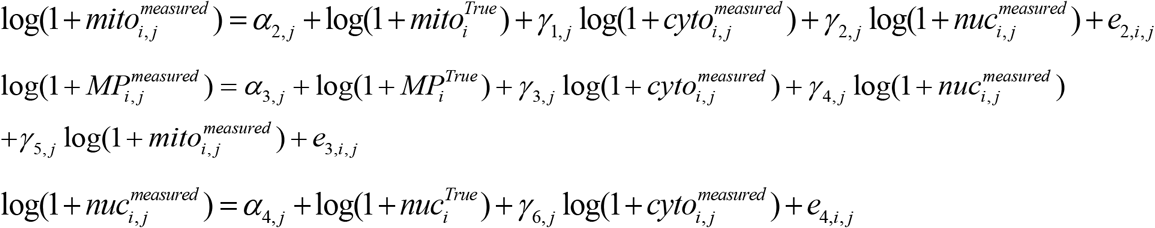

where 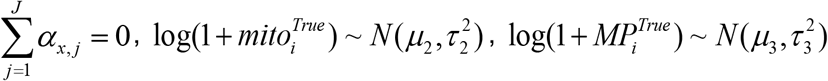, 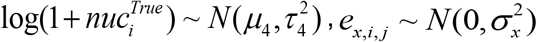, and *J* is the number of replicates. Fitting these models directly would result in high estimates for the various *γ*’s as they would capture contamination and true correlation. We tackle this complication using a three-step process:

1. We fit a model with estimated *True* values as predictors. E.g., for the nuclear fraction

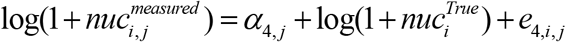

where 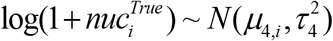 and 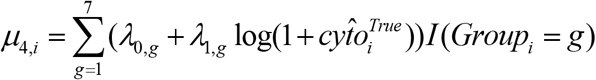. That is, we allow the mean of the *i^th^* molecule’s true value in the nuclear fraction to depend on the group type of the *i^th^* molecule and the molecule’s estimated true cytoplasmic value which captures the correlation.
2. Using the estimated residuals from step 1, we fit the model

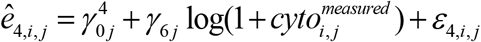 The idea here is that if all association between measured the *i^th^* molecule’s measured value in the nuclear and cytoplasmic fractions is due to real correlation, then there should be no association between the residuals of step 1 and the measured cytoplasmic value. Any association may be present indicates contamination.
3. We subtract the contamination from the measured value to estimate the uncontaminated value of the *i^th^* molecule in the *j^th^* replicate as:

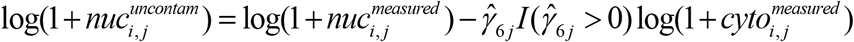 Any non-positive estimates for *γ* are ignored. We then refit the mixed effects model using the uncontaminated estimates as the dependent variable:

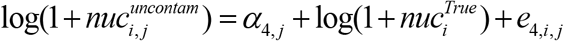 Using the results from the model we estimate the true abundance of the *i^th^* molecule in the *j^th^* replicate in the nuclear fraction 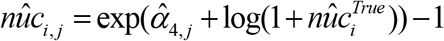. We repeat the last three steps for the mitochondrial and mitoplast fractions. In these two models, the estimated mean true abundance of the *i^th^* molecule in the nuclear and cytoplasmic fractions are included in step 1 (including the mitochondrial fraction in the mitoplast results in nonconvergence). To estimate the contamination, all factors in the contamination model are included.
- Finally, for each replicate, we fit the following equation for rescaling purposes:

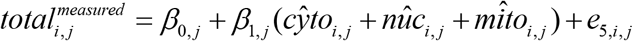

where 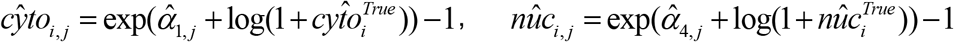, and 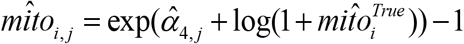. This calculation excluded the top 1% and the bottom 1% of the molecules (outliers).

## RESULTS

In these experiments we used three cell lines that model TNBC: BT-20 (derived from a pleural effusion of the mammary gland/breast epithelial tissue from a 74-year-old White/European American female), MDA-MB-231 (derived from a pleural effusion from the metastatic site from a 51-year-old White/European American female), and MDA-MB-468 (derived from a pleural effusion from the metastatic site from a 51-year-old Black/African American female). From each cell line, we isolated the nuclear, cytoplasmic, mitochondrial and mitoplast fractions, and confirmed their purity using fraction-specific protein markers (see Methods and **Supplemental Figure S1**). We generated fractions from three biological replicates for BT-20 and MDA-MB-231 cells and four replicates for MDA-MB-468 cells. For each replicate, we deep-sequenced five RNA preparations that included the nuclear, cytoplasmic, mitochondrial and mitoplast fractions and the corresponding total RNA. We profiled the isomiRs, tRFs and rRFs separately for each preparation. **Figure 1** captures the workflow. Throughout this paper we refer to the physical material that we isolated as cell “fractions” and the pre-defined organelles (nucleus, cytoplasm, mitochondrion, and mitoplast) as cell “compartments”.

**Figure 1.**
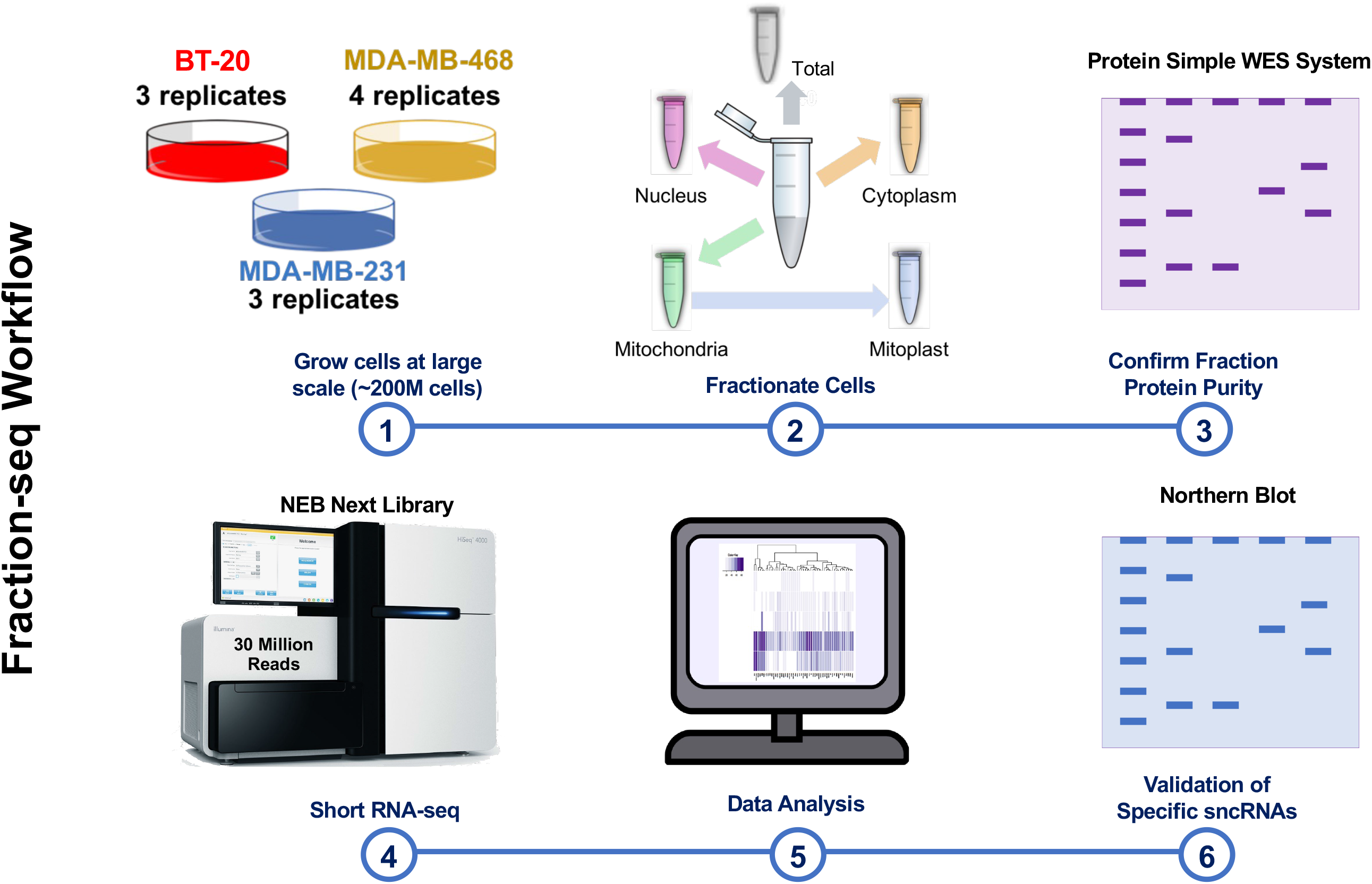
Overview of the Fraction-seq workflow. **1.** Three Triple Negative Breast Cancer (TNBC) cell lines BT-20, MDA-MB-231, and MDA-MB-468 were grown to ~200M cells and harvested. **2.** From the same starting material, cells were separated into total, nuclear, cytoplasmic, mitochondrial, and mitoplast subcellular compartments. **3.** The WES protein detection assay was used to analyze cell compartment markers in each cell fraction. **4.** Short RNA-seq library preparation was carried out using NEBNext with 100 ng RNA from each sample followed by RNA sequencing using the Illumina NextSeq 500 platform at 75 cycles and an average depth of 30 million reads. **5.** High quality short RNAs reads were mapped to isomiRs, tRFs, and rRFs and short RNAs between 18 and 50 nts and normalized to sequencing depth. Normalized reads with abundances greater than 10 RPM were further analyzed for the purpose of this study. **6.** Northern blot analysis was used to confirm short RNAs identified by this study. Steps 1-5 were done using biological replicates (BT-20 – three replicates, MDA-MB-231 – three replicates, MDA-MB-468 4 replicates).

### TNBC model cell lines exhibit cell-type specific features

We reasoned that if the cell lines *truly* differ from one another, their difference should be evident if we considered *only* their abundant molecules. Therefore, we focused *only* on short RNAs whose abundance in total RNA was ≥ 10 RPM when averaged across each cell line’s replicates. **Supplemental Table S2** catalogues the average abundance and standard deviations for the short RNAs that satisfy this criterion.

**Figure 2A** shows a Principal Component Analysis (PCA) of the three cell lines using the 7,165 short RNAs that satisfy the 10 RPM threshold. We see that the replicates of each cell line – BT-20 (blue), MDA-MB-231 (yellow), MDA-MB-468 (red) – cluster with one another, as expected. In **Figure 2B**, we show a hierarchical clustering of the cell line replicates after running a Pearson correlation on the short RNAs with an abundance > 10 RPM in each sample (10,070 short RNAs). **Figures 2C-E** shows that the clustering of each cell line’s replicates persists when we apply the PCA method to each RNA class in turn (1,421 isomiRs, 567 tRFs, and 5,177 rRFs, respectively).

**Figure 2.**
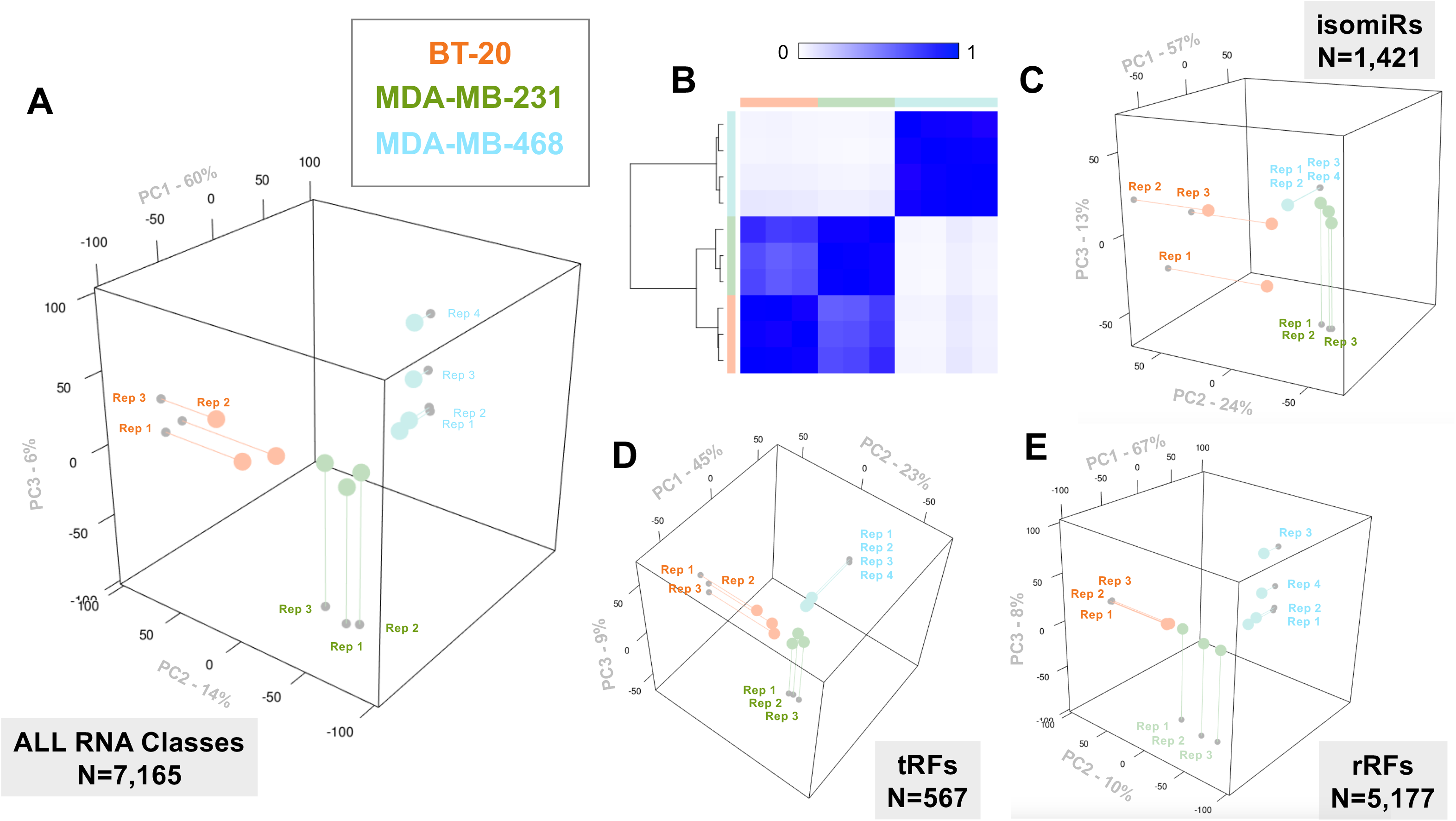
Cell replicates are reproducible and characterized by RNA classes. **A.** Principal Component Analysis (PCA) visualization of the short RNAs in each sample with an average abundance of 10 RPM or greater across replicates (n=7,165). **B.** Hierarchical clustering of the Pearson correlations for replicates from each cell line with short RNA abundances greater than 10 RPM (n=10,070). Color bar represents the Pearson correlation values which range from 0 (white, not correlated) to 1 (blue, exactly correlated). **C.** PCA visualization of isomiRs in each sample with an average abundance of 10 RPM or greater across replicates (n=1,421). **D.** PCA visualization of the tRFs in each sample with an average abundance of 10 RPM or greater across replicates (n=567). **E.** PCA of the rRFs in each sample with an average abundance of 10 RPM or greater across replicates (n=5,177). **A,C-E.** All PCA used components 1,2, and 3. **A-E.** BT-20 samples are represented by orange, MDA-MB-231 samples are represented by green, and MDA-MB-468 samples are represented by blue.

### IsomiRs, tRFs and rRFs can be unique to a cell line or shared among cell lines

After confirming the reproducibility of the cell line replicates for this study, we sought to determine the intercellular similarities and differences of the short RNA populations. We did this by comparing the top 10% most abundant short RNAs (isomiRs, tRFs, or rRFs) from each cell line’s total RNA based on average abundance (**Supplemental Table S3**): the average abundances of these RNAs range between ~189 and ~64,137 RPM. **Figure 3A** shows the overlap among the top 10% most abundant short RNAs found in BT-20, MBDA-MB-231, and MDA-MB-468. Clearly, each cell line expresses its own unique profile of abundant short RNAs with only 17 short RNAs being common to all three cell lines (**Supplemental Figure S2**). BT-20 and MDA-MB-231 cells share 140 of their top 10% most abundant RNAs. On the other hand, MDA-MB-468 cells are unlike BT-20 and MDA-MB-231 with 416 of their top 10% most abundant RNAs being unique to the MDA-MB-468 cell line.

**Figure 3.**
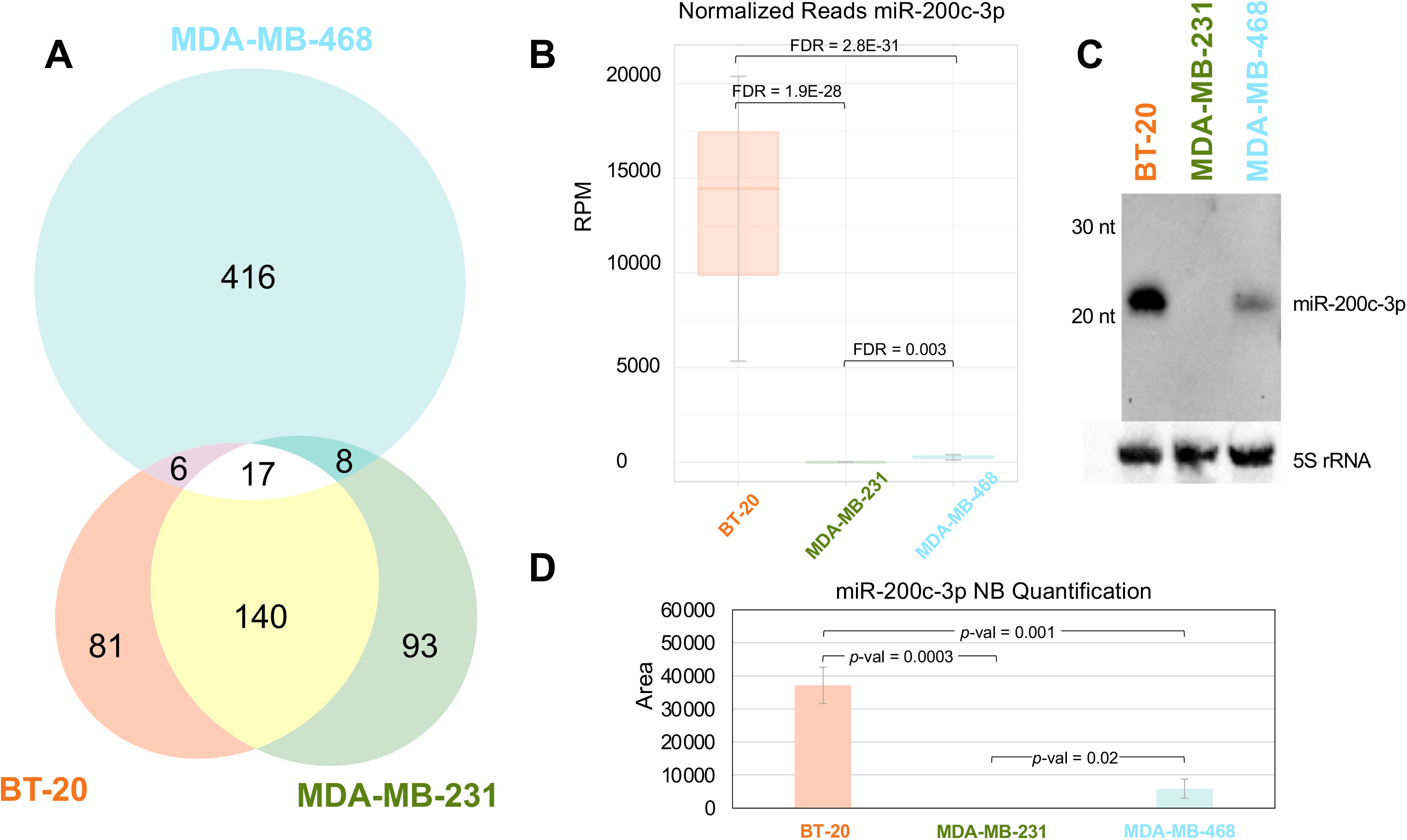
Short RNAs show cell line specific features. **A.** Venn Diagram of the top 10% most abundant short RNAs in each the three TNBC cell lines (BT-20, MDA-MB-231, and MDA-MB468) based on average abundance. **B.** Boxplots represent the normalized reads for miR-200c-3p in BT-20, MDA-MB-231, and MDA-MB-468 cells. **C.** Northern blot to detect miR-200c-3p in BT-20, MDA-MB-231, and MDA-MB-468 cell line RNA using 5μg RNA from each cell line. Northern detection of 5S rRNA at the bottom is the loading control. **D.** Quantification of northern blot analysis for miR-200c-3p done in triplicate. Colors of cell lines: BT-20 (orange), MDA-MB-231 (green), and MDA-MB-468 (blue).

For those short RNAs that passed Threshold-seq and 18-nt length cutoffs (Methods), we calculated pairwise differential abundances using DESeq2 (see Methods), and determined the log_2_ fold-change and associated adjusted *p*-values. We found 6,593 short RNAs that are differentially abundant (DA) in various pairwise combinations: they comprised 2,983 isomiRs, 532 tRFs, and 4,765 rRFs (**Supplemental Table S4**). When we compared BT-20 and MDA-MB-231 cells, which share many of their most abundant short RNAs, we identified 901 molecules that are differentially abundant at statistically-significant levels (FDR ≤ 0.05) with a | log_2_ *(fold change)* | of ≥ 1. Not surprisingly, when we compared MDA-MB-468 with BT-20 and MDA-MB-231, we identified 5,327 and 5,253 short RNAs, respectively (FDR ≤ 0.05, | log_2_ | fold change ≥ 1).

#### A specific example

One of the most abundant and differentially abundant short RNA from our analysis is the 0|0 isoform of miR-200c-3p. This isomiR’s abundance in BT-20 cells is more than 1000x higher than in MDA-MB-231 cells (adjusted *p-val=1.31E-62)* and more than 200x higher than in MDA-MB-231 cells (adjusted *p-val*=1.17E-43) (**Supplemental Table S4** and **Figure 3B**). Probing for this isomiR on a northern blot (**Figure 3C**) revealed a much more intense band in BT-20 cells compared to MDA-MB-231 and MDA-MB-468 cells. The quantification for these differences in technical replicates of the northern is shown in **Figure 3D**.

### IsomiRs, tRFs and rRFs can be enriched in one subcellular fraction or multiple fractions

In analogy to our total RNA analyses, we also compared fractions within a cell line and across cell lines.

We first identified the top 10% most abundant among the thresholded short RNAs for each fraction based on average abundance and compared them across cell-lines (**Supplemental Table S5)**. As the Venn diagrams of **Supplemental Figures S3A-D** show, each fraction’s contents largely mirror the similarities and differences that we observed in the total RNA comparisons shown in **Figure 3**. We also carried out a differential abundance analysis, separately for each fraction. **Supplemental Table S6** lists molecules whose abundance in a given fraction differs by at least 2-fold between cell lines. Only molecules with average abundance ≥ 10 RPM in total RNA were considered in this analysis.

### IsomiRs, tRFs and rRFs can be enriched in a specific cell line and fraction combination

The differential abundance comparisons listed in **Supplemental Table S6** reveal complex patterns of expression and preferential localization for isomiRs, tRFs and rRFs. We highlight this point with the help of several 5’ tRNA-halves (5’-tRHs) from the nuclear tRNA^LysCTT^. **Figure 4A** shows 20 5’-tRHs from two different isodecoders of tRNA^LysCTT^: the Lys^CTT^ isodecoder chr1.trna119-LysCTT located between positions 145395522 and 145395594 on the reverse strand of chromosome 1 (human genome assembly GRCh37); and the Lys^CTT^ isodecoder chr16.trna10-LysCTT located between positions 3241501 and 3241573 on the forward strand of chromosome 16. As shown in **Figure 4A**, the two isodecoders differ at exactly one position, position 29 (indicated in red/blue. Because of the extensive sequence similarity, two of the 20 5’-tRFs, those with lengths 27 and 28 nts, cannot be assigned unambiguously to one of the two isodecoders.

**Figure 4.**
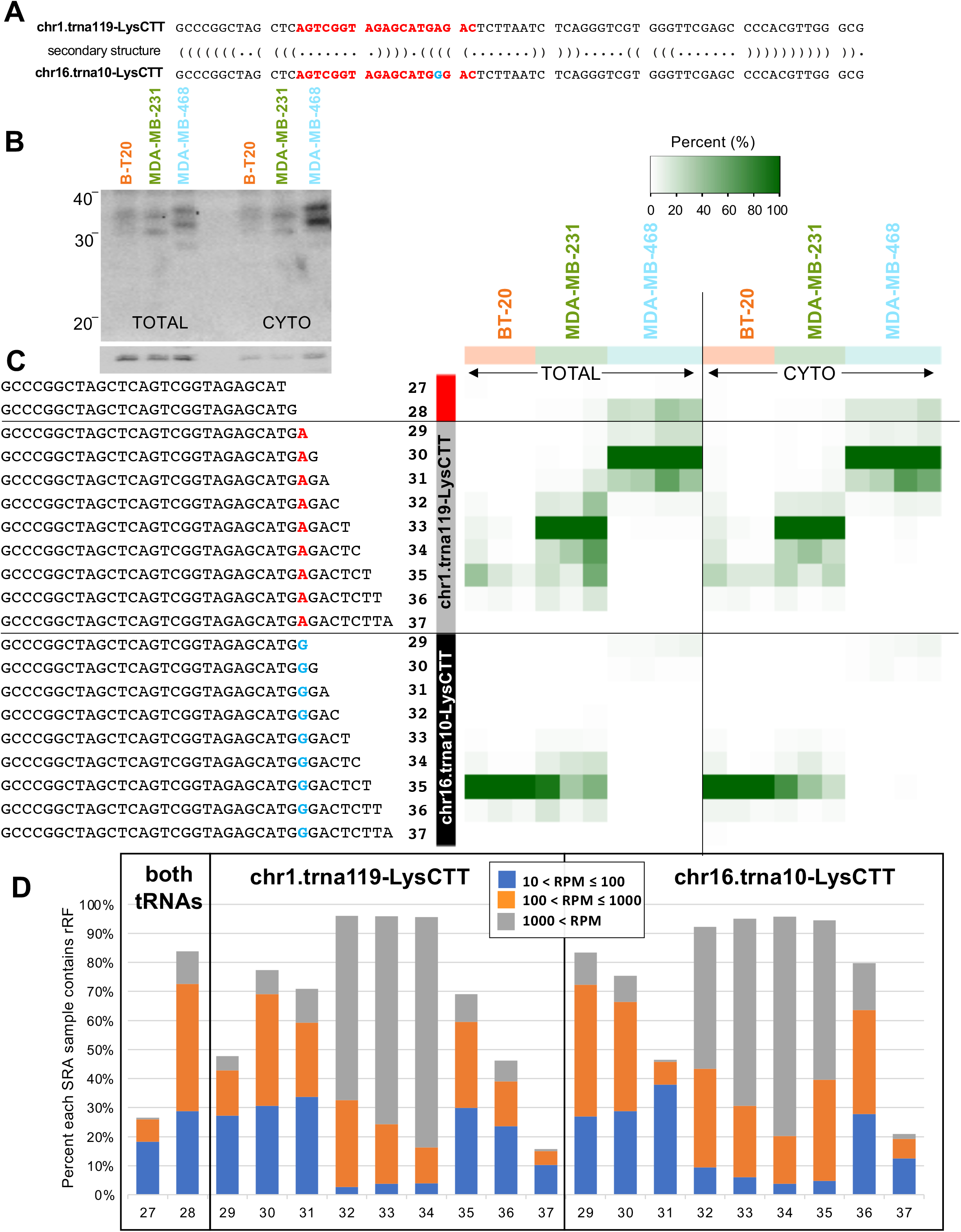
Lys^CTT^ tRNA-halves present differently when it comes to cell line and cell compartment. **A.** The primary and secondary structures of two Lys^CTT^ tRNA isodecoders (tRNA-Lys-CTT-2-1 and tRNA-LysCTT-4-1) that differ by one nucleotide (A – red /G – blue at position 29). The red text in the sequence shows the region targeted by the northern blot probe. **B.** The northern blot shows the short RNAs (range: 20-40 nts) detected using 5 μg of Total and Cytoplasmic (CYTO) RNA and Lys^CTT^ probe. Full-length Lys^CTT^ was used as a loading control. **C.** The heatmap shows normalized abundances of each short RNA (row) where the denominator is the most abundant RNA from the 20 short RNAs (originating from the two Lys^CTT^ isodecoders) in Total and cytoplasmic (CYTO) replicates for BT-20 (orange, n=3), MDA-MB-231 (green, n=3), and MDA-MB-468 (blue, n=4) short RNA-seq. The red/blue A/G denotes the single nucleotide difference between the two isodecoders at position 29. **D.** The percent of the 20 tRFs from the two Lys^CTT^ isodecoders (Figure 4C) in 6203 publicly available datasets/samples (SRA). The barplot shows the fraction of the public samples containing the corresponding molecule at an abundance between 10 and 100 RPM (blue), between 100 and 1000 RPM (orange), or above 1,000 RPM (grey).

The 5’-tRFs and 5’-tRHs from these two tRNA^LysCTT^ isodecoders are examples of RNAs with cell-line-dependent enrichment profiles. Deep sequencing can discern these differences. For independent validation, we also investigated these length-dependent differences using a northern blot. In **Figure 4B**, we observe tRNA^LysCTT^ fragments of differing lengths in total RNA and the cytoplasmic fractions, in the three cell lines. Moreover, the northern blots reveal the presence of molecules that are not captured by standard deep sequencing; we will return to this point below.

**Figure 4C** shows a heatmap of each tRF’s relative abundance to the 35-mer from chr16.trna10-LysCTT, which is the most abundant among these 20 tRFs (BT-20 cells). Note how this 35-mer is much less abundant in MDA-MB-231 and absent from MDA-MB-468 cells. On the other hand, the 33-mer from chr1.trna119-LysCTT is most abundant in MDA-MB-231 cells, much less so in BT-20 cells, and absent from MDA-MB-468 cells. In MDA-MB-468 cells, the most abundant tRF is a 30-mer 5’-tRF from chr1.trna119-LysCTT; the fragment is absent from BT-20 and MDA-MB-231 cells. These differences persist in the cytoplasmic fraction (**Figure 4C**). The complete collection of differentially abundant tRNA^LysCTT^ molecules and associated adjusted *p*-values are listed in **Supplemental Table S7**. These short RNAs are among numerous molecules whose abundance in a given fraction differs statistically significantly among cell lines (data not shown).

We also searched for these same 20 tRFs in public datasets. Given the lengths of the Lys^CTT^ tRFs discussed here, we restricted the search to 6,203 datasets that were sequenced at a minimum of 45 cycles. All 20 tRFs from the chr1.trna119-LysCTT and chr16.trna10-LysCTT isodecoders are highly abundant (**Supplemental Table S8** and **Figure 4D**). From **Figure 4D** we can also see that the 32-, 33-, and 34-mer tRFs from chr1.trna119-LysCTT and the 32-, 33-, 34-, and 35-mer tRFs from chr16.trna10-LysCTT are the most abundant: their abundance exceeds 10 RPM in 90% of the 6,203 examined datasets. In **Supplemental Table S8**, note how the 35-mer tRFs from chr16.trna10-LysCTT is more abundant in more datasets than its counterpart 35-mer from chr1.trna119-LysCTT. On the other hand, the 28-mer 5’-tRFs, which can arise from either isodecoder, is present in 84% of the samples.

### The enrichment patterns of isomiRs, tRFs, and rRFs can be cell-type specific

In addition to studying the *intercellular* profiles of the short RNAs across cell lines, we also investigated their *intracellular* profiles in each cell line. We first compared the top 10% most abundant short RNAs in each cell compartment (nucleus, cytoplasm, mitochondrion, and mitoplast) from a given cell line based on average abundance of the replicates (**Figure 5A-C**). By using the top 10% most abundant RNAs, we focus on short RNAs that are more likely to be consequential in the respective compartment while minimizing potential crosstalk from contamination. The Venn diagrams in **Figures 5A-C** show that many of the most abundant RNAs in each cell compartment are actually shared. In BT-20 cells, the four compartments share 106 short RNAs, whereas in MDA-MB-231 and MDA-MB-468 cells they share 94 and 145, respectively. **Figures 5A-C** also show that there are many RNAs that are unique to each cell compartment.

**Figure 5.**
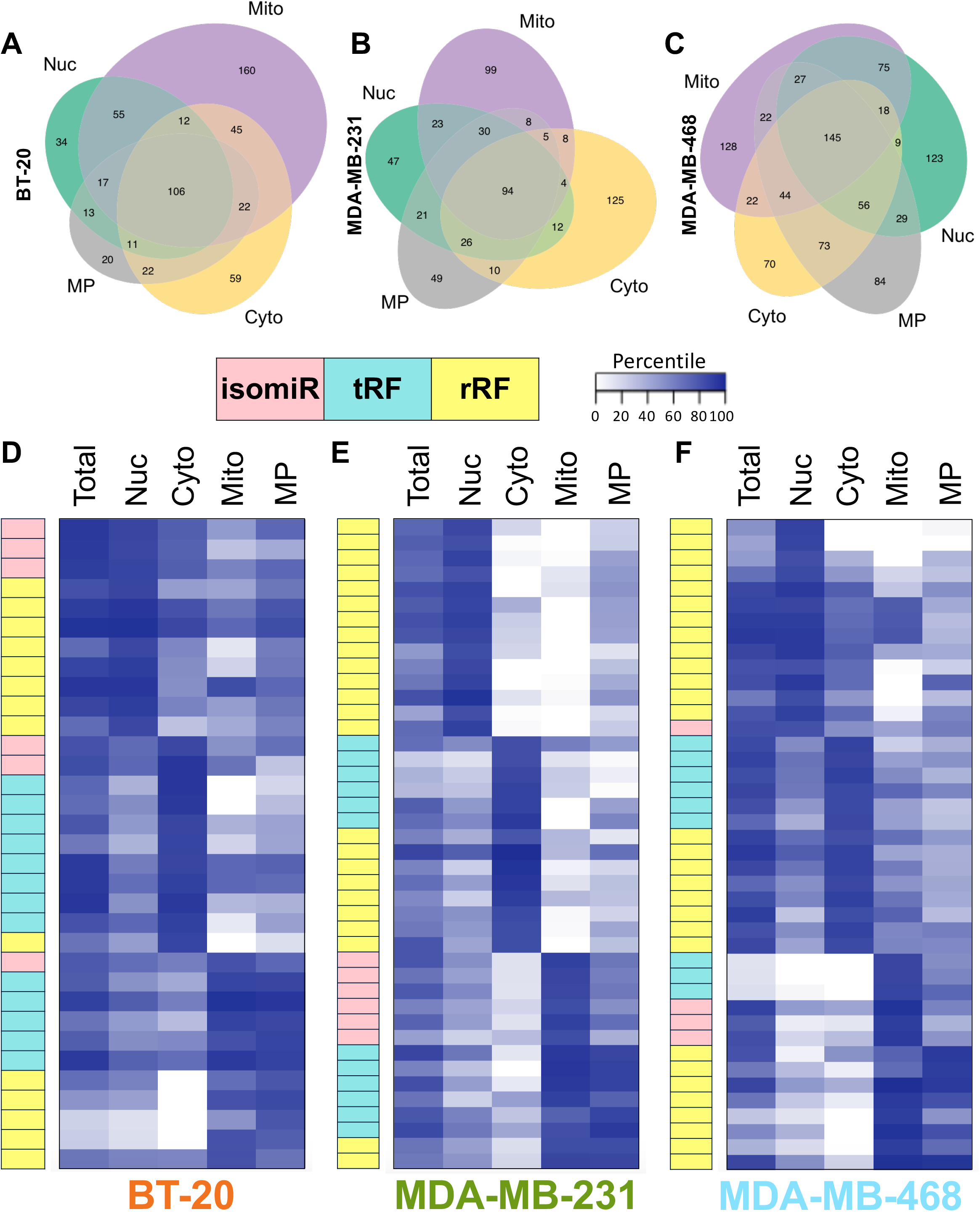
Short RNA are enriched in specific cell compartments. **A-C.** The Venn diagrams show the top 10% most abundant short RNAs in each cell compartment of BT-20 (A), MDA-MB-231 (B), and MDA-MB-468 (C) cells based on average abundance. Colors for A-C correspond to the different cell compartments: nucleus (Nuc) – green, cytoplasm (Cyto) – yellow, mitochondrion (Mito) – purple, and mitoplast (MP) grey. **D-F.** Heatmaps of the of the percentiles of the average normalized abundance (RPM) of short RNAs in total, nucleus (NUC), cytoplasm (CYTO), mitochondrion (MITO), and mitoplast (MP) samples for BT-20 (D), MDA-MB-231 (E), and MDA-MB-468 (F) cells. The heatmap colors range from white (lowest percentile) to dark-purple (highest percentile). The rows of the heatmap correspond to specific short RNA, where the colored bars to the left of each heatmap represent the short RNA class (pink – isomiRs, teal – tRFs, and yellow – rRFs). The sequences of the corresponding short RNAs and their type are listed in **Appendix C Figure 4 A-C**.

We next wanted to know more granularly, where a given short RNA goes in a cell and whether the localization is cell-type specific. Unlike the comparisons of total RNA or of individual fractions across cell lines, we cannot directly compare the abundances of a given short RNA in two different fractions from the same cell line^73^. By sequencing a given fraction (e.g., nucleus, cytoplasm, mitochondrion, or mitoplast) at the same depth as total RNA, we effectively *amplify* the abundance of each molecule in that fraction by an *unknown* factor that will differ for each fraction. Despite this limitation, and even without the benefit of modeling, we can reach important conclusions. To do so, we rank the RNAs in each sample from most abundant to least abundant, assign the ranks to percentiles (100=most abundant RNA, 0=least abundant RNA), and compare the percentile of specific collections of short RNAs in different fractions. We will return to this below when we describe a mathematical model that aims to solve this problem by estimating each molecule’s abundance while accounting for technical errors and possible contamination.

This qualitative method helped demonstrate that some short RNAs are enriched differently in different compartments. We discuss several such examples below. We also found that even short RNAs that arise from the same precursor (i.e., the same miRNA arm, tRNA isodecoder, or rRNA) can exhibit different preference in their subcellular localization. It is worth reiterating that we treated the mitochondrial fraction with RNase A to remove RNAs from the outer mitochondrial membrane. This allows to focus on short RNAs that are protected by, e.g., Argonaute proteins bound to the mitochondrial membrane, and short RNAs that are *inside* the mitochondrion. We did not treat the mitoplast fraction with RNAse A due to the very small yields we obtained. Consequently, in our analyses of the mitoplast fraction we considered only molecules that are present in both the mitoplast *<u>and</u>* mitochondrial fractions.

The heatmaps in **Figures 5D-F** show examples of short RNAs that exhibit compartment-specific enrichments. The heatmaps use average percentile values. The values range from the 100^th^ percentile (high enrichment / darker shade) to the 0^th^ percentile (no enrichment / white). Color coding indicates the short RNA class (isomiRs, tRFs or rRFs). The identities of the molecules shown in **Figures 5D-F** here are listed in **Supplemental Figures S4A-C**. As expected, we can observe many RNAs whose fraction enrichment is the one expected. For example, tRFs arising from nuclear tRNAs are enriched in the cytoplasmic fraction and tRFs arising from MT tRNAs are enriched in the mitochondrial fraction. However, the heatmaps also show some RNAs being enriched in fractions where they would not be expected. For example: we found that some isomiRs are enriched in the nuclear and mitochondrial fractions; also, some rRFs from the same rRNA are enriched in different compartments independently. We discuss some of these observations below.

**Figure 5D** and accompanying **Supplemental Figure S4A** show examples of 33 short RNAs from BT-20 cells. Seven of the shown rRFs are from the 5’ ETS region of the 45S rRNA. They have overlapping sequences but different enrichments in the various fractions. All eight tRFs from nuclear tRNAs are enriched in the cytoplasmic fraction, as expected. Of these eight, the two tRFs from the nuclear tRNA^GlyGCC^ are also enriched (62^nd^-74^th^ percentiles) in the nuclear, mitochondrial, and mitoplast fractions. Conversely, tRFs from the mitochondrial tRNA^ValTAC^ are enriched in the nuclear and cytoplasmic fractions, but not the mitochondrial/mitoplast fractions. The two short RNAs that arise from 5.8S and 18S rRNA, respectively, offer another interesting example: the first is enriched in the nuclear and the second in the cytoplasmic fraction even though both parental rRNAs are part of the 45S rRNA cassette.

**Figure 5E** and **Supplemental Figure S4B** show analogous examples from MDA-MB-231 cells. Again, we see multiple rRFs from the 5’ ETS and 5.8S regions of the 45S cassette that are highly enriched in the nuclear fraction. On the other hand, multiple rRFs from the 18S and 28S regions of 45S are highly enriched in the cytoplasmic fraction. Perhaps the most notable observation here involves six isomiRs from different miRNAs that are much more enriched in the mitochondrial than the cytoplasmic fraction.

Lastly, **Figure 5F** and **Supplemental Figure S4C** show some examples from MDA-MB-468 cells. Two groups of molecules are worth noting here. The first comprises four isomiRs from miR-30a, miR-30c, miR-30e, and miR-200c. Of these, miR-30c is comparatively more enriched in the nuclear fraction whereas the other three are enriched in the mitochondrial fraction. The second group comprises rRFs from several regions of 28S rRNA. We observe that the rRFs form subgroups each of which is enriched in a different fraction. Notably, rRFs that are immediately adjacent to one another on the 28S rRNA show different enrichments in each compartment.

### Previously unreported short RNAs show preferential subcellular enrichments

The rRFs that arise from the 45S precursor rRNA and have been discussed in the literature are fragments of the precursor’s three component rRNA (18S, 5.8S, and 28S)^1^. In addition, evidence supports short RNAs, including miRNAs, that are encoded in both the rRNA-coding regions and the spacer regions of the 45S rRNA^74–76^. These reports are interesting because the spacer regions are believed to be removed quickly by a variety of nucleases and then degraded in order to release the mature rRNAs and maintain cell homeostasis^30,77,78^.

Based on our analysis, the 5’ ETS region of the 45S rRNA is unlike the remaining three spacers, i.e., ITS1, ITS2, and 3’ ETS: 5’ ETS is the only spacer that produces short RNAs in high abundance. In **Figures 5D-F** and **Supplemental Figures S4A-C** we highlighted several rRFs from this region: while they are consistently enriched in the nuclear fraction, their abundances in total RNA differ widely among the three cell lines. One of them, the 31-mer CTTCGTGATCGATGTGGTGACGTCGTGCTCT begins 11 nucleotides after the A’ position and is consistently the most abundant of all fragments in total RNA, in all three cell lines. Its absolute abundance varies across the three cell lines: it is more abundant in BT-20 (median: ~950 RPM) and MDA-MB-231 (median: ~1,430 RPM) than in MDA-MB-468 (median ~35.5 RPM) cells. These differences are statistically significant: the adjusted *p*-values are < 1.32E-28 for BT-20 vs. MDA-MB-468, and – 8.21E-28 for MDA-MB-231 vs. MDA-MB-468 (**Figure 6A**). Beyond the three TNBC cell lines, we find the 31-mer in 4,420 of the 6,203 public datasets we also analyzed (**Supplemental Table S8**) at an abundance ≥ 10 RPM.

**Figure 6.**
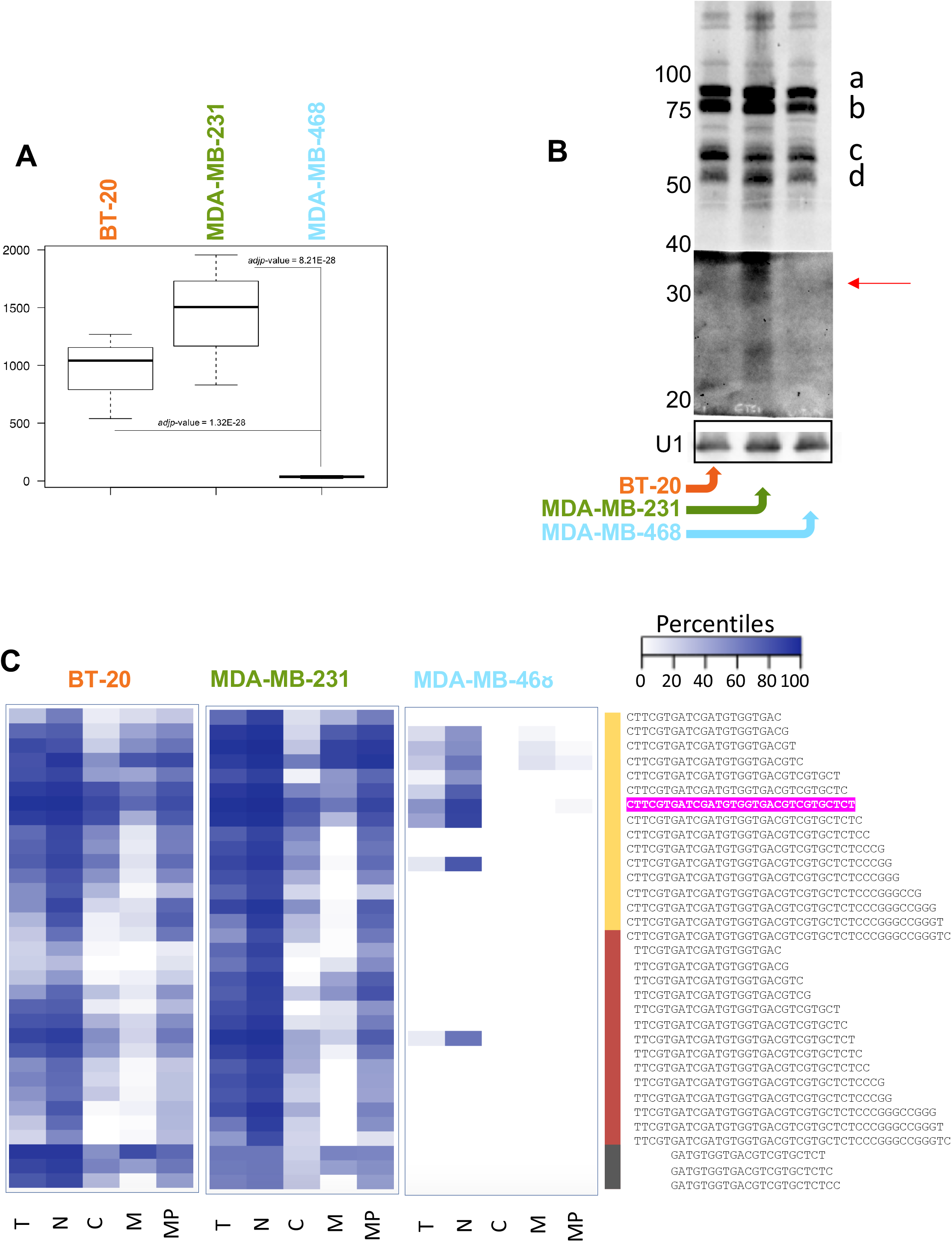
Short RNA from 5’ ETS rRNA are present, abundant, and have cell-line and cellcompartment features. **A.** Boxplot of 31-mer 5’ ETS rRF abundance (RPM) in BT-20 (n=3), MDA-MB-231 (n=3), and MDA-MB-468 (n=4) cell line RNA. Significance is shown by FDR (adjusted *p*-value) from differential expression analysis (DESeq2). BT-20/MDA-MB-468 FDR = 1.32E-28, MDA-MB-231/MDA-MB468 FDR = 8.21 E-28. **B.** Northern blot of 31-mer 5’ ETS rRF using two oligo probes overlapping the full-length of the short RNAs in 20 μg BT-20, MDA-MB-231, and MDA-MB-468 cell line RNA. U1 is used as loading control. The red arrow points to the RNA length region of interest on the northern blot where the 31-mer 5’ ETS rRF is expected. Additional bands above 50 nts are indicated by a, b, c, and d and are discussed in the results. **C.** Heatmaps show percentiles the 5’ ETS rRFs beginning at position +425 of the 45S rRNA based on average abundance for the cell fractions (T (total), N (nucleus), C (cytoplasm), M (mitochondrion), MP (mitoplast) in each cell line (BT-20, MDA-MB-231, MDA-MB-468). The heatmap colors range from white (lowest percentile) to dark-purple (highest percentile). The 34 5’ ETS rRFs are located to the right of the heatmaps and are delineated by three groups based on start position (yellow – +425, red – +426, dark-grey – +432). The 31-mer rRF is highlighted in hot pink. Colors of cell lines: BT-20 (orange), MDA-MB-231 (green), and MDA-Mb-468 (blue).

Intrigued by the absence of previous reports on 5’ ETS-derived fragments, we sought to validate the above 31-mer further using northern blots (**Figure 6B**). Qualitatively, the blots show that an rRF in the length range of 31 nts is present in MDA-MB-231 cells (indicated by the red arrow). The blots also reveal many additional molecules, both shorter and longer. MDA-MB-231 produces short (22-50 nts) rRFs from the 5’ ETS locus in higher abundance than BT-20 or MDA-MB-468. The blots also reveal four major and several minor rRFs from the 5’ ETS, all longer than 50 nts. The molecules corresponding to bands *a*-*d* are highly abundant and present in all three TNBC cell lines. The two molecules in the 50-70 nts range appear to have cell-line-specific length preferences: the molecules of band *c* are more abundant in MDA-MB-231 cells whereas those of band *d* are more abundant in BT-20 cells.

Considering the distinct 5’ ETS bands detecting in the northern blot, we computationally investigated fragments from this region further. We find that 5’ ETS produces 161 short RNAs with lengths between 18 and 50 nts and abundances between ~10 and ~1,955 RPM. Most of these short RNAs arise from the interval between positions +424 and +484 of 45S rRNA (**Supplemental Figure S5A-B**). This interval is an unstructured region slightly downstream from the A’ position (+414) of 5’ ETS^79^.

Of these 161 5’ ETS-derived rRFs, 33 are present in the top 15^th^ percentile of at least one cellular fraction, in at least one of the three cell lines (BT-20, MDA-MB-231, and MDA-MB-468). The heatmap in **Figure 6C** shows the average percentiles of these 33 rRFs across the cell compartments in each cell line. The 31-mer is highlighted. 16 of the 33 rRFs begin at position +425 of the 45S rRNA (yellow), 14 start at position +426 (red), and three start at position +435 (grey). Many of the 33 molecules are highly enriched in the nuclear fraction in all three cell lines. It is clear that these 5’ ETS-derived rRFs are enriched in the various fractions in a cell-line- and length-dependent manner. For example, the cytoplasmic fraction of BT-20 cells contains primarily medium length rRFs from this locus, whereas in MDA-MB-231 cells it contains shorter rRFs as well. MDA-MB-468 stands apart in that 5’ ETS-derived rRFs appear to be generally at lower abundance than the other two cell lines but present primarily in the nucleus.

### “Standard” RNA-seq offers only a partial picture of the subcellular enrichment preferences of isomiRs, tRFs, and rRFs

We already presented example northern blots that revealed abundant tRFs or rRFs that are not “visible” to the standard RNA-seq to which we subjected our preparations (see **Figures 4B** and **6B**). Such observations have been documented in the literature and sometimes result from the fact that these molecules have unexpected terminal phosphate modifications^32,80^. The inability of standard RNA-seq to reveal such molecules becomes a consideration given that the specific localization preferences of isomiRs, tRFs, and rRFs are expected to translate to functional differences in different cell lines and tissues. We revisit the point of cell-line-specific localization preferences by focusing on rRFs from the 5’ end of 28S rRNA.

Others and we previously showed that the 5’ region of the 28S rRNA is a hotspot for the production of highly abundant rRFs in multiple cell types and tissues^1,81,82^. One of the most abundant short RNAs in our data is the rRF CGCGACCTCAGA<u>TCAGACGT GGCGACCCGCTGA</u>ATTT that corresponds to the first 37 nts of the 28S rRNA. The top boxplot in **Figure 7A** shows this rRF’s abundances in total RNA from BT-20, MDA-MB-231, and MDA-MB468 cells. The bottom boxplot shows the same rRF’s abundance in the nuclear fractions from the three cell lines.

**Figure 7.**
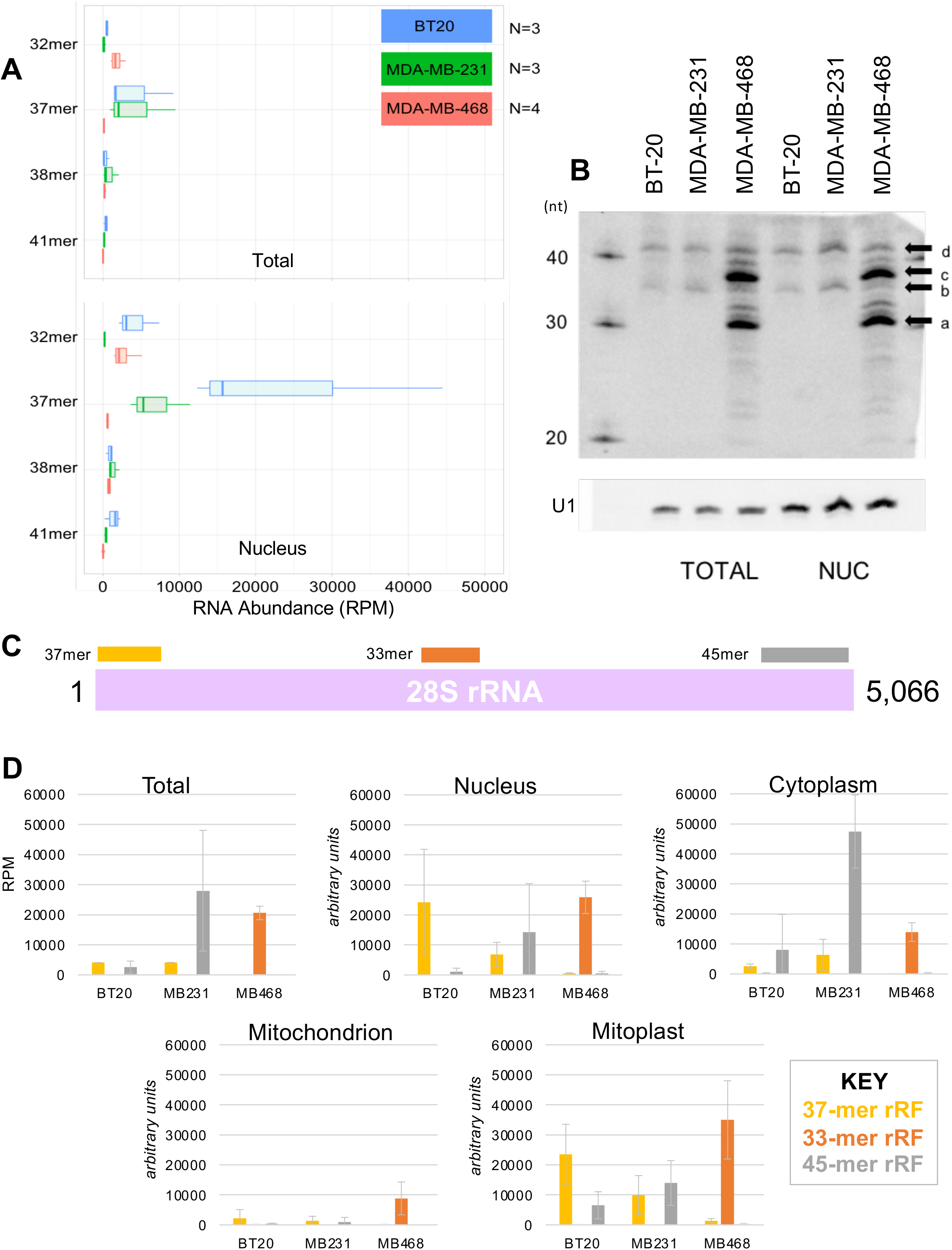
28S rRFs have cell type- and cell compartment-specific features that are not fully accounted for by traditional RNA-seq. **A.** Boxplots showing the relative abundance of four 28S rRFs from the 5’ end of the 28S rRNA in BT-20 (blue, n=3), MDA-MB-231 (green, n=3), and MDA-MB-468 (red, n=4) RNA-seq data in total cell RNA and nuclear RNA. **B.** Northern blot probing for 37-mer 28S rRF in total cell RNA and nuclear RNA from the three cell lines (BT-20, MDA-MB-231, MDA-MB-468). U1 RNA is used as a loading control. Specific bands of interest are indicated by a, b, c, d (see text). **C.** Schematic of full length 5,066 nt 28S rRNA (magenta, not to scale) with annotations of the most abundant rRF from each cell line BT-20 (37-mer, yellow), MDA-MB-231 (45-mer, grey), and MDA-MB-468 (33-mer, orange). **D.** Barplots show the average abundance of the three 28S rRF from Figure 7C in total cell RNA (Total) and the cell fractions (Nucleus, Cytoplasm, Mitochondrion, and Mitoplast). Colors represent the different 28S rRF: yellow (37-mer), orange (33-mer), and grey (45-mer).

In total RNA, this 37-mer rRF is abundant in both BT-20 and MDA-MB-231 cells, but not in MDA-MB-468. At the same time, the 37-mer is characteristically more enriched in the nuclear fraction of BT-20, compared to the nuclear fraction of MDA-MB-231 or MDA-MB-468 cells. Considering the nearly 35-fold difference in abundance between BT-20 cells and MDA-MB-231, we sought to validate it with a northern blot. Our probe targeted the underlined region near the center of the 37mer rRF (see above).

The northern blot revealed several cell-line specific rRFs that are not present in the RNA-seq data. As can be seen in **Figure 7B**, the 37-mer (arrow *b*) is enriched in the nuclear fraction relative to total RNA and slightly more abundant in MDA-MB-468 compared to BT-20 and MDA-MB-231. This difference and the richer rRF profile observed in the northern is a clear departure from what from the RNA-seq allows us to deduce. Indeed, the 37-mer rRF is overshadowed by two other rRFs in MDA-MB-468 cells, a 32-mer (arrow *a*) and a 38-mer (arrow c). These fragments are clearly very abundant in the nucleus of MDA-MB-468. Additionally, there is a 41-mer rRF (arrow *d*) whose band intensity matches that of the 37-mer rRF, but is not recapitulated by the RNA-seq data (**Figure 7A**).

### The most abundant rRFs from a parental rRNA can differ by cell line and can have different enrichment preferences

A comparison between RNA-seq and northern blotting reveals differences in the profiles of the short RNAs that are present in a given cell-line/cell-fraction combination. Nonetheless, examination of RNA-seq data can provide useful insights and serve a hypothesis generation role. In the previous section, we focused on the 5’ region of 28S rRNA, which is a hotspot of rRF production, according to the literature^1,31,81,82^. Given our findings on the dependence of rRFs on context, we asked whether this region of 28S remains a hotspot independent of cell type.

We searched our RNA-seq data from total RNA to determine the 28S regions that produce the most abundant rRFs in each of the three cell lines we studied. **Figure 7C** summarizes our findings and shows that the location and length of the most abundant rRFs differ characteristically among BT-20, MDA-MB-231, and MDA-MB-468. **Figure 7D** shows the relative average abundance for these fragments in total RNA and the four cell fractions. In BT-20 cells, the most abundant rRF is a 37-mer from the 5’ end of 28S rRNA (blue) that is enriched in the nuclear and mitoplast fractions, and whose average abundance is 4,105 RPM. Its sequence is CGCGACCTCAGATCAGACGTGGCGACCCGCTGAATTT. In MDA-MB-231 cells, the most abundant rRF is a 45-mer from the 3’ end of 28S (gray) that is enriched in the cytoplasm, and whose average abundance is 27,983 RPM. Its sequence is CTCGCTGCGATCTATTGAAAGTCAGCCCTCGACACAAGGGTTTGT. In MDA-MB-468 cells, the most abundant rRF is a 33-mer from the middle of the 28S rRNA (orange) that is enriched in the nucleus and mitoplast and has an average abundance of 20,587 RPM. Its sequence is CGGTTCCGGCGGCGTCCGGTGAGCTCTCGCTGG.

### Public datasets independently support the preferential subcellular enrichment of short RNAs

Several previous studies examined the subcellular localization of proteins and messenger RNAs (mRNAs) in mouse and human cells^83–87^. These studies confirmed known localization preferences for these molecules but also unveiled previously unsuspected ones. Additionally, lncRNAs have become a topic of subcellular exploration. Scientists have shown that understanding the localization profile of a lncRNA can aid in uncovering its function in the cell^73,88,89^. Recent work extended these analyses to short RNAs that localize to the nucleus and the mitochondrion^36,48,52,55,56,62,63^ but did not attempt to account for cross-fraction contamination. To the best of our knowledge, only three studies attempted to study short RNAs in different compartments in an unbiased manner. The first study looked at mitochondrial short RNAs in HEK293 cells and HeLa cells^63^. The second study examined the short RNA contents of the mitochondrion and the mitoplast of 143B cells^62^. The third study examined short RNAs in the nucleus and the cytoplasm of 5-8F cells^51^. All three studies used deep sequencing to profile short RNAs between 18 and 30 nts in various cell fractions^51,62,63^. Notably, all these experiments used a single replicate for each cell line and cell fraction, did not account for cross-fraction contamination and were carried out at a time that isomiRs, tRFs, and rRFs were considered cell oddities.

We reanalyzed the data of these three studies using the same normalization and analysis protocol that we used to analyze the BT-20, MDA-MB-231 and MDA-MB-468 datasets (Methods). For each dataset, we identified and analyzed only isomiRs, tRFs, and rRFs between 18 and 30 nts with abundances greater than 10 RPM.

**Supplemental Figures S6A-B** summarize our findings. First, we see that all previous studies consistently support the presence of isomiRs, tRFs, and rRFs in the nucleus, cytoplasm and mitochondrion by observing the percentage of unique RNAs for each RNA type in **Supplemental Figure S6A**. Proportionally, tRFs (yellow) and rRFs (blue) account for nearly 85-90% of the unique short RNA identities in each of the cell fractions. IsomiRs (green) are more abundant in HEK293 mitochondrion than HeLa or 143B. IsomiRs comprise nearly 10% of the distinct molecules found in nuclear and cytoplasmic fractions of 5-8F cells. tRFs and rRFs derived from mitochondrial tRNAs and rRNAs are more abundant in the mitochondrial fraction of 143B cells, compared to HEK293 and HeLa. Interestingly, a considerable portion of the nuclear and cytoplasmic fractions of 5-8F cells corresponds to mitochondrial tRFs and rRFs (light yellow and light blue respectively). When we look at the number of unique RNAs for each RNA type in the five fractions samples in **Supplemental Figure SF6B**, we see that the 5-8F nuclear and cytoplasmic samples contain approximately 5,000 more reads than the HEK293 and HeLa mitochondrial samples, while near-similar reads to the 143B mitochondrial samples.

Next, we computed the same profiles for the short RNAs from our fractions from BT-20, MDA-MB-231, and MDA-MB-468 cells. To allow for consistent comparisons, we counted the unique short RNAs with lengths 18-30 nts whose abundance ≥ 10 RPM in at least two thirds of the replicates, and separately for each cell fraction (**Supplemental Figure S6C-D**). When we consider the percent of unique RNAs for each RNA type in each fraction of each cell line (**Supplemental Figure S6C**) we see that the nuclear, cytoplasmic, and mitochondrial fractions of BT-20 cells contain mostly nuclear-tRNA-derived tRFs (~40-50%), many isomiRs (~25%), and fewer nuclear-rRNA-derived rRFs (~15%). In MDA-MB-231 cells, we see more nuclear-rRNA-derived rRFs in the nucleus and cytoplasm (~25-30%). Lastly, in MDA-MB-468 cells, the nuclear, cytoplasmic, and mitochondrial fractions contain primarily rRFs (~50%), and tRFs (~50%) and very few isomiRs. We note that the mitochondrion of all three TNBC cell lines contain the most diverse populations of short RNA, which is similar to the mitochondrion of 143B cells. When we look at the number of unique RNAs for each RNA type (**Supplemental Figure S6D**) we see that there are 2X as many short RNAs in the fractions of MDA-MB-468 cells compared with BT-20 and MDA-MB-231, a side-effect of the consistency thresholds we employed (above). However, we are also able to see that overall, the nuclear fractions of BT-20 and MDA-MB-231 cells contain the most unique short RNA molecules, while the cytoplasm of MDA-MB-468 cells contains the most unique short RNA molecules.

### Mathematical modeling adjusts for technical errors and cross-fraction contamination

By applying qualitative arguments to our RNA-seq data (**Figure 5** and **Supplemental Figure 4**), we showed multiple isomiRs, tRFs, and rRFs are enriched preferentially in different fractions in different cell lines. However, this approach can be used only with short RNAs whose abundances and rankings show large changes among the various fractions. These molecules represent only a small fraction of the many molecules that are present at statistically-significant levels. To determine in a uniform manner all molecules that exhibit preferential localization, independent of their abundance levels, we created a mixed-effects model (see Methods). The model distinguishes among seven molecular groups (a combination of RNA class and whether the RNA is encoded in the nuclear or the mitochondrial genome – see Methods), and accounts for cross-fraction contamination and possible technical errors in the various replicates. Using the model, we reconstructed the *true* value of each molecule in each fraction from the values we measured in each replicate while correcting for contamination and errors. Finally, we fitted each molecule’s reconstructed values in the nuclear, cytoplasmic, and mitochondrial fractions to their measured values in total RNA. By doing so, we were able to determine each fraction’s relative contribution to total RNA. We find the following nuclear:cytoplasm:mitochondrion ratios: BT-20 = 21%: 52%: 27%, MDA-MB-231 = 25%: 35%: 40%, and MDA-MB-468 = 39%: 44%: 17%.

The scaling also allows us to directly compare a molecule’s abundance across fractions. **Supplemental Table S9** lists the reconstructed average values for each molecule in the nuclear, cytoplasmic, mitochondrial and mitoplast fractions. Each cell line is shown on a separate sheet. All listed molecules exceed a threshold of 10 RPM (reconstructed abundance) in at least one of the fractions. In these tables, we also mark “outlier” molecules, which we define as molecules that are at the top 1% or bottom 1% in at least one of the fractions. In BT-20, we found 2,081 molecules ≥ 10 RPM (72 outliers) of which 881 are isomiRs, 292 tRFs, and 908 rRFs. In MDA-MB-231, we found 2,417 molecules ≥ 10 RPM (69 outliers) of which 896 are isomiRs, 325 tRFs, and 1,196 rRFs. Lastly, in MDA-MB-468, we found 3,534 molecules ≥ 10 RPM (160 outliers) of which 140 isomiRs, 165 tRFs, and 3,229 rRFs.

Using the reconstructed abundances for the short RNAs in each fraction and excluding the outlier RNAs, we updated **Supplemental Figure S6C-D**. In **Figure 8A-B**, we show the new version. As can be seen, the stringent thresholding did not alter the relative proportions of the various RNA types in any of the three cell lines. However, the number of unique RNAs that exceed the threshold in each cell type has changed. For example, in BT-20 cells, there are fewer short RNAs in the nucleus and mitochondrion that survive thresholding. The number of RNAs in the cytoplasm of BT-20 cells remains virtually unchanged (~2,100 molecules). In MDA-MB-231 cells, the number of short RNAs in the nucleus and cytoplasm decreases by ~50% but remains unchanged in the mitochondrion (~2000). Lastly, in MDA-MB-468 cells, there are ~36% fewer short RNAs in the nucleus and cytoplasm, and ~12% fewer in the mitochondrion.

**Figure 8.**
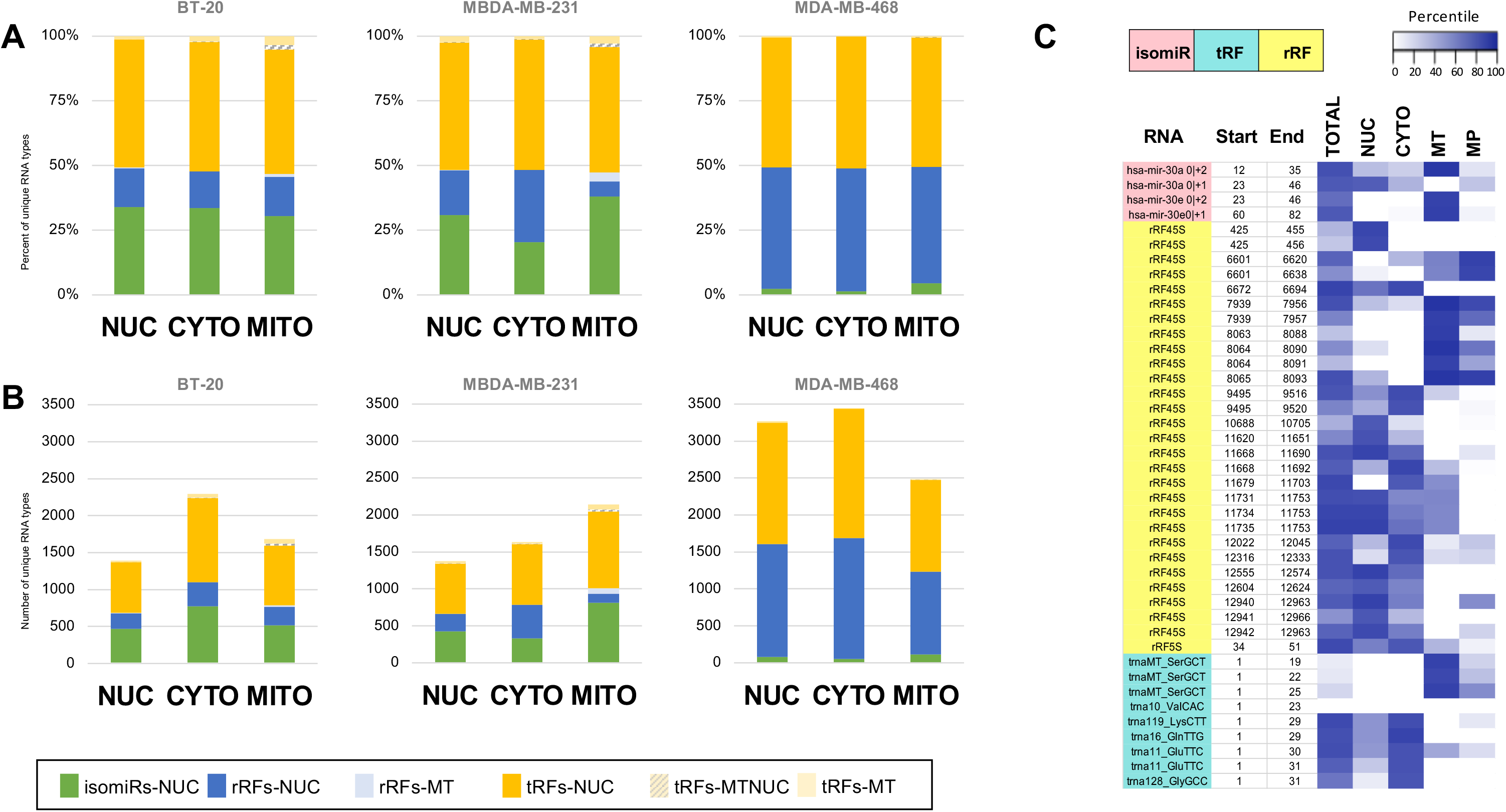
Nuclear and mitochondrial genomes produce short RNAs that can be found in nuclear, cytoplasmic, and mitochondrial fractions. **A-B.** Stacked barplots represent the percent of unique short RNA **(A)** and the number of unique short RNA based on average abundance **(B)** that are found in cell fractions from three cell lines (BT20, MDA-MB-231, and MDA-MB-468) Colors denote the different RNA classes: nuclear isomiRs (green), nuclear rRFs (dark blue), nuclear tRFs (dark yellow), mitochondrial rRFs (light blue), mitochondrial tRFs (light yellow), and tRFs that map to both nuclear and mitochondrial genomes (diagonal yellow lines). **C.** Heatmaps of the of the percentiles of the average normalized abundance (RPM) of short RNA in total, nucleus (NUC), cytoplasm (CYTO), mitochondrion (MITO), and mitoplast (MP) samples for MDA-MB-468 cells after reconstructing short RNA abundances. Outliers are removed. The heatmap colors range from white (lowest percentile) to dark-purple (highest percentile). The rows of the heatmap correspond to specific short RNA, where the colored bars to the left of each heatmap represent the short RNA class (pink – isomiRs, teal – tRFs, and yellow – rRFs).

We also recalculated the percentile values for the 42 short RNAs in MDA-MB-468 cell fractions (**Figure 5F)** using their reconstructed abundances (**Figure 8C**). Additionally, we reordered the rows so that molecules that are progressively further from the 5’ terminus of the parental RNA appear on consecutive rows. This reorganization allows several observations. For example, we see more pronounced differences in each molecule’s relative presence across the fractions. Additionally, we can see very distinct enrichment profiles for RNAs with near-similar sequences. For example, the shorter of the two isomiRs of miR-30a (miR-30a|0|+1|) is enriched in the nucleus whereas miR-30a|0|+2|, which is longer by one nucleotide, is enriched in the mitochondrion. The short RNAs from 45S provide even more striking examples of enrichment profile biases. As one moves away from the 5’ terminus of 45S, the corresponding rRFs show enrichments that favor a single fraction or specific combinations of fractions.

Given the above observation of the miR-30a isomiRs, we sought to examine the distribution of isomiRs arising from the miR-17/92 cluster^90–93^. The cluster, also known as “oncomiR-1,” spans several hundred nucleotides and harbors six miRNA precursors (miR-17, miR-18, miR-19a, miR-19b, miR-20a and miR-92a). **Figure 9** shows the reconstructed abundances (in RPM values, as well as color-coded) of the cluster’s isomiR abundances in BT-20 and MDA-MB-231 cells. IsomiRs with reconstructed abundance below threshold are not listed. The isomiRs highlighted in green are the ones found in miRBase and are also known as “archetype” or “reference” isoforms^3,32^. The archetype isoforms of a miRNA are those for which off-the-shelf commercial assays are available; these are also the isoforms that have been primarily studied in the last twenty years. As can be seen from **Figure 9**, the abundance and enrichment profiles of the various isomiRs, including the archetype miRNAs, differ greatly even though they all arise from a single polycistronic transcript^92^. Moreover, a given isomiR’s enrichment in a subcellular compartment can differ by cell line. In several instances, even the archetype isoform can be absent in a given cell line: e.g., miR-92a|0|0| is absent from MDA-MB-231, yet is very abundant in BT-20 cells. Another unexpected finding is the prevalence of many isomiRs with non-templated additions (nucleotides in red, boldface).

**Figure 9.**
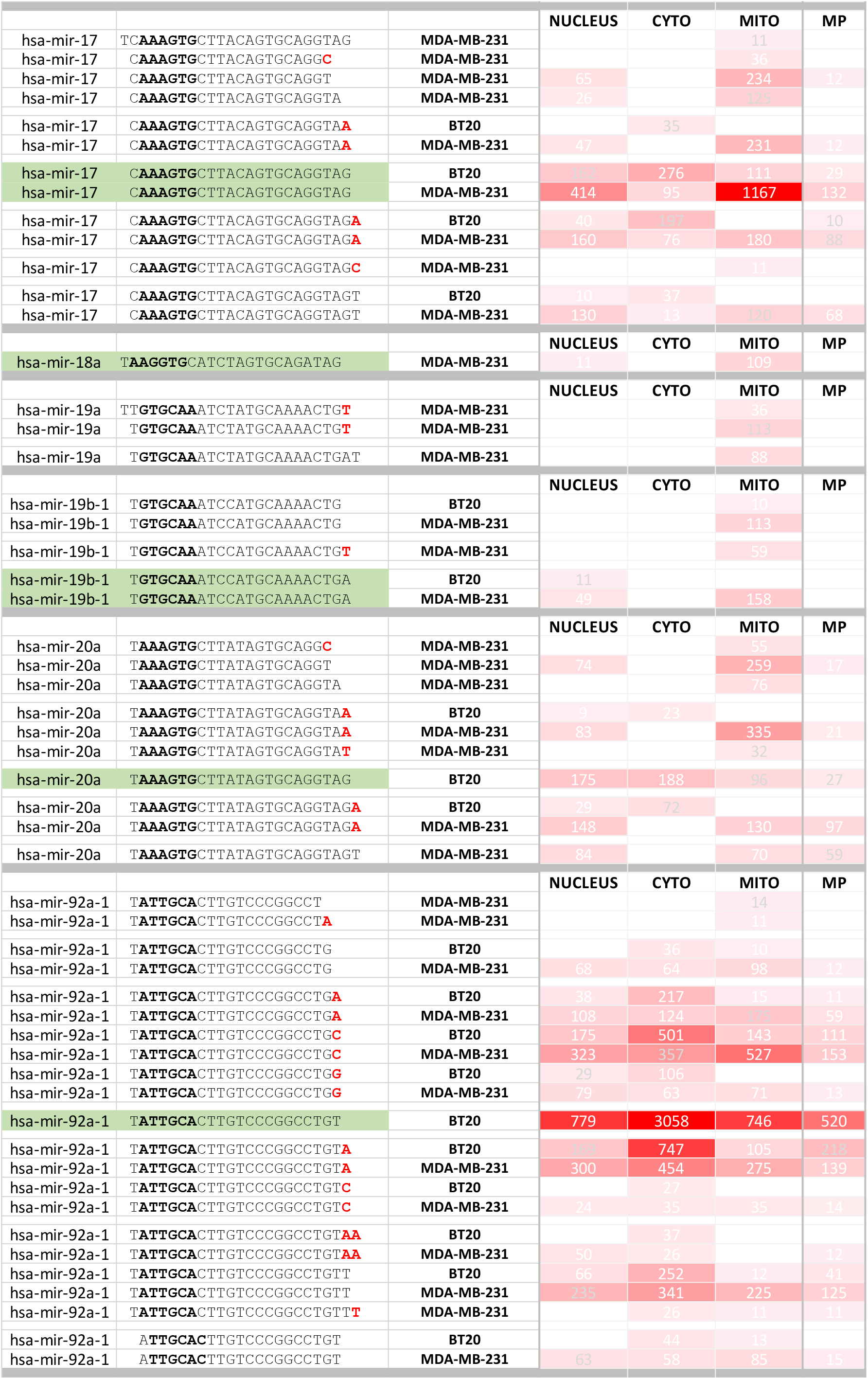
Specific isomiRs from miR-19/92 cluster localize to unique cell compartments. **A.** Heatmap of average reconstructed abundances of miR-17/92 cluster isomiRs in BT-20 and MDA-MB-231 cell fractions. Heatmaps colors range from 0 RPM (white) to 1000 RPM (red). Green highlighting indicates reference or miRBase miRNA sequences (0|0).

## DISCUSSION

We developed a cell fractionation and sequencing method to experimentally explore the localization of short RNAs in any cell line. We also developed a computational pipeline to systematically analyze the fractionation data and applied the pipeline to model cell lines in a specific disease.

The study reported our findings on the subcellular enrichment preferences of short RNAs (isomiRs, tRFs, and rRFs) in three TNBC cell lines. We uncovered complex abundance profiles that are RNA class- and cell-type specific. Knowledge of the enrichment patterns of these molecules can prove instrumental in designing better-targeted experiments to uncover the function of these molecules and prioritizing molecules that are easier to manipulate experimentally. Moreover, given the regulatory nature of isomiRs, tRFs, and rRFs and their context-specific (sex, ancestry, age, tissue type, and disease) enrichment patterns, accounting for short RNAs can improve personalized medicine. We believe that short RNA localization plays an important role in this ongoing inquiry.

In recent years, thorough and systematic studies examined the localization of <u>long</u> non-coding RNAs using data from the ENCODE project^73,88,89,94^. These studies assumed a continuum of enrichments across compartments, which led to computational methods that report the *relative* and *absolute presence* of a given molecule in different compartments^73,88,89,95–97^. Interestingly, some studies considered lncRNA localization separately for each cell line – a cell-specific feature, while others did not make this distinction.

By comparison, <u>short</u> RNA localization studies have received less attention. Arguably, the problem of subcellular localization is more difficult for short RNAs. First, there are many more short RNAs than lncRNAs: in recent studies, we described a combined total of more than 50,000 isomiRs^6,67^, tRFs^9,69^, and rRFs^1^. The second complicating factor is that short RNAs are produced by the nuclear and the mitochondrial genomes^1,6,9,67,69^. Of the handful of localization studies to date, the experimental designs did not account for cross-fraction contamination, did not include biological replicates^62,63,82^, or examined one or two cell fractions^62,63,82,94,98^. For example, the ENCODE project generated short RNA-seq datasets using replicates from 11 different cell lines but focused on the nuclear and cytoplasmic fractions only. We are not aware of any studies on short RNAs that used replicates from two or more model cell lines <u>from the same tissue</u> and/or disease to study subcellular localization in nuclear, cytoplasmic, mitochondrion, and mitoplast fractions while also controlling for cross-fraction contamination.

### Fraction-Seq: a new method for profiling RNAs in subcellular fractions

In this project, we analyzed multiple biological replicates from three cell lines that model TNBC (BT-20, MDA-MB-231, MDA-MB-468). For each replicate, we profiled the short RNA populations in total RNA, and the nuclear, cytoplasmic, mitochondrial, and mitoplast fractions, for a total of five preparations. These preparations were derived from the same starting cell material using “Fraction-seq” (see Methods). Fraction-seq is a hybrid approach that combines a novel cell fractionation method, which we developed for this project, with deep sequencing. After confirming the purity of each fraction using standard compartment-specific protein markers, Fraction-seq calls for deep-sequencing (short RNA-seq) of total RNA and RNA from the four fractions, for a total of five datasets per biological replicate. **Figure 1** summarizes the approach. We worked with three replicates for BT-20 and MDA-MB-231, and four for MDA-MB-468, and analyzed a total of 50 datasets. In each dataset, we identified the isomiRs, tRFs and rRFs that are present using established methods. The methods include candidate ranking, differential expression analysis, and mixed-effects modeling.

### Different abundant isomiRs, tRFs, and rRFs in different cell lines

Through our analysis of the short RNA profiles in total RNA from BT-20, MDA-MB-231, and MDA-MB-468 cell lines, we confirmed the congruence of the biological replicates (**Figure 2**) and the unique character of each cell line’s short RNA profiles (**Figure 3A**). The three cell lines shared only 17 short RNAs of the top-10% (244 for BT-20 cells, 258 for MDA-MB-231 cells, and 447 for MDA-MB-468 cells) most abundant molecules in total RNA. This was unexpected given that the cell lines are from the same tissue type and disease (**Supplemental Figure S2**). However, two of the cell lines, BT-20 and MDA-MB-231, share more than half of their top-10% most abundant molecules (**Figure 3A**). Indeed, a comprehensive differential abundance analysis of total RNA shows that only 2-12% of the top 10% most abundant RNAs from each fraction are shared across cell lines. We experimentally validated a few short RNAs that exhibited cell-type-specific or fraction-specific enrichment. We also computed differential expression across cell lines for a given cell compartment and quantified the differences statistically **(Supplemental Table 4** and **Supplemental Table 6)**. One of the differentially abundant RNAs is the miRNA miR-200c-3p 0|0. This miRNA is a characteristic example of a cell-type dependent molecule. Indeed, RNA-seq indicates that the miRNA is significantly more abundant in BT-20 cells than in MDA-MB231 or MDA-MB-468 cells. Northern blot analysis independently confirmed its differential abundance in these cell lines (**Figure 12B-C**). This result is important because it points out that different cell lines modeling the same disease cannot be used interchangeably.

In addition to miRNAs, tRNA halves have been shown to be important in disease contexts^27,99^. In our analysis, we explored tRFs produced from two isodecoders of tRNA^LysCTT^, chr1.trna119-LysCTT and chr16.trna10-LysCTT, which provide several more notable examples of cell-type-specific and fraction-specific enrichment. The sequences of the two isodecoders differ by a single nucleotide at position 29 (**Figure 4**). Both isodecoders produce multiple 5’-tRFs and 5’-tRHs, each with different abundance and fraction-enrichment profiles. For example, a 30-mer 5’ tRF chr1.trna119-LysCTT is exclusively present in MDA-MB-468 cells and enriched in the cytoplasm. On the other hand, the slightly longer 33-mer 5’ tRFs from tRNA-Lys-CTT-2-1 is also enriched in the cytoplasm but present exclusively in MDA-MB-231 cells (**Figure 4C**). Analogously, the even longer 35-mer 5’-tRFs from chr16.trna10-LysCTT is also enriched in the cytoplasm but present predominantly in BT-20 cells. These findings additionally indicate that different isodecoders of the same isoacceptor produce different fragments (tRFs or tRHs) in different cell lines, with the produced fragments having specific subcellular enrichments. This observation is potentially important for a number of contexts such as translational efficiency^100^ or targeted therapeutics^5,9,10^. We stress that these tRFs arise from distinct isodecoders of the same isoacceptor on two chromosomes, differ by a single nucleotide, and have distinct expression patterns. Further studies are needed to determine whether these near-identical tRFs also have different functional roles. We also note that the abundance differences among the tRFs produced by these two tRNA^LysCTT^ isodecoders are not unique to the three TNBC cell lines we analyzed. Examination of 6,203 public datasets from NIH’s Sequence Read Archive revealed that these two isodecoders produce highly abundant tRFs across different tissues, cell lines, disease states, and other contexts **(Figure 4D and Supplemental Table 8**). There appear to be context-specific circumstances contributing to the preferential production of these tRFs.

### Percentiles reveal cell fraction-specific enrichment profiles

One key attribute of the Fraction-seq method is its use of equal amounts of RNA from each cell fraction. This is intentional and is meant to avoid the documented problems of spike-ins, which are technically challenging to accurately reproduce^101^, or cell equivalents, which are operating on estimates. Such limitations become particularly acute given the multiple biological replicates and the fact that for each replicate, Fraction-seq generates five RNA preparations from the same starting material. The limitation of this strategy is that it requires an additional analytical step (discussed below) to determine how much each fraction contributes to a cell’s total RNA. Without this additional step, we cannot *quantitatively* compare fractions from the same cell line to each other^73^. Of course, by replacing a molecule’s abundance in a fraction by a percentile, we can still observe different short RNAs being enriched in different cell compartments (**Figure 5**). Ideally however, we would like to be able to directly compare within a cell line.

### Mixed-effect modeling corrects for technical and contamination errors

One of the main reasons to isolate each cell compartment is to compare the localization preferences and abundances of molecules across the fractions of a cell. This means determining how many of the isomiRs, tRFs, and rRFs that exceed a reasonable RPM threshold exhibit unexpected enrichment patterns. However, experimental errors introduced by biological replicates and cross-fraction contamination, which may occur during the fractionation stage, can affect the results. **Supplemental Figure S1** shows that based on fraction-specific protein markers, our fractions are free of contamination. However, there remains the possibility that contamination is present but below the sensitivity capabilities of these assays. We therefore built a mathematical model of the process. For a specific combination of fraction and biological replicate, the model assumes that the measured abundance of a molecule is the sum of its true abundance, a contribution from a replicate-specific technical error, and a contribution from contamination of the fraction at hand by other fractions. We applied the model to each cell line separately, using all available biological replicates simultaneously and constraining its calculations by the measured abundances of the total RNA preparation. Through a mixed-effect modeling approach, the model can recover each molecule’s true abundance in each fraction of a cell line while correcting for errors and contamination. Our mathematical model can be applied after fractionation and enable any kind of analysis that requires knowledge of the true abundance of each molecule. Future work will be necessary to validate this model experimentally.

### Reconstructed profiles reveal the molecules’ true abundance in each cell fraction

Having recovered the true abundances of short RNA molecules, we carried out exhaustive within-a-cell-line comparisons to determine each molecule’s enrichment across the various subcellular compartments. We found a diverse canvas of enrichments that preferentially place various combinations of isomiRs, tRFs, and rRFs in different compartments (**Figure 8A**). Depending on the samples being evaluated, we see changes in the number of reconstructed short RNAs (**Figure 8B**) compared to the short RNAs that we started with (**Supplemental Figure S6D**) for all short RNA types. These enrichment patterns can shed light on various cellular processes and deepen our understanding of short RNA biology. For example, in **Figure 9A**, we can see that miR-17, a miRNA involved in cancer progression and inflammation^90–93^, is most enriched in the mitochondrion of MDA-MB-231 cells and the cytoplasm of BT-20 cells. Results from this work can deepen our understanding of RNA biology and help improve experimental design. Eventually, it is possible that the findings will contribute to developing more targeted disease treatments.

### Intracellular trafficking of short RNAs occurs from the nucleus to the mitochondrion

It is well established that the majority of mitochondrial proteins are encoded by the nuclear genome, and the mitochondrial-bound mRNAs must be localized to the mitochondrion^102^. Evidence suggests translation occurs either before localization or in a process of co-localization allowing for the negatively-charged amino acids to traverse the positively charged mitochondrial membrane^102^. The full-length 5S rRNA has been shown to enter the mitochondrion encapsulated by the proteins Rhodenase and MRP-L18^46,65,66^. However, a functional role for 5S rRNA in the mitochondrion has yet to be identified. Nevertheless, it is still not clear if nuclear-encoded short RNAs enter the mitochondrion like their longer RNA counterparts.

Our contribution to these established localization processes is that short RNAs encoded by the nuclear genome also localize to the mitochondrion. For example, our data shows strong enrichment in the mitochondrial <u>and</u> mitoplast fractions for several rRFs derived from nuclear rRNAs (**Figure 8C**). This suggests that at least some of these short RNAs are *inside* the mitochondrion. However, it remains unclear whether the fragments are processed locally from copies of imported nuclear rRNAs, or imported as rRFs from the cytoplasm. A mechanism for the translocation of small RNAs into the mitochondrion from the cytoplasm has yet to be described^46^. Exceptions like 5S rRNA-derived rRFs notwithstanding, it remains formally possible that the majority of the short RNAs we see enriched in the mitochondrial fraction are simply “attached” to the outer membrane of the mitochondrion and never enter it. For example, miRNA-loaded Argonaute complexes attached to the mitochondrial membrane would be unaffected by our RNAse A treatment due to their size^103,104^. Nevertheless, our data uncovers many isomiRs that are enriched in the mitochondrial fraction but depleted from the cytoplasmic fraction (**Figure 8C**). This disparity suggests that the process is not accidental and further studies are needed to independently validate these findings.

### The 5’ ETS rRNA spacer is a rich source of short RNAs

Previous work has shown miRNA genes to be embedded in both the coding regions and rRNA spacer regions of the 45S rRNA; mainly from the 5’ ETS and 28S^74–76^. While many of the rRNA-derived miRNAs are predicted through bioinformatic methods, others have been experimentally validated in humans and other species. The findings that miRNAs are embedded within the spacer regions of the 45S rRNA suggests a regulatory role for some rRFs, which also may be important for rRNA biogenesis and the regulation of protein translation^11,30^. Murine miR-712 for example (and its human counterpart miR-205 on chr1: 209432133-209432242), is a miRNA derived from the ITS2 spacer region of the mouse genome and is involved in regulating cardiovascular disease pathways^76^. Human miR-663 (chr20: 26208186-26208278), which plays an important role as an inappropriately silenced tumor suppressor in cancer progression, is an example of a functional short RNA encoded in the ETS1 spacer region^105–108^. Because of these results, we decided to explore the short RNAs of the spacer regions of 45S.

Our analysis identified multiple rRFs from the A’ region of the 5’ ETS of 45S rRNA, which has no previously identified miRNA annotations. This region was previously thought to be degraded^19,77^. One such identified rRF, a 31-mer, is more abundant in MDA-MB-231 cells than in BT-20 or MDA-MB-468 cells (**Figure 6A**), a finding that is supported by northern blotting (**Figure 6B**). It is important to point out that while we do see a 31-nt band based on the RNA marker, northern blotting is still a qualitative method. Due to binding affinities of the northern probe, it remains possible that the band we see is the 31-mer of interest or a secondary 31-mer rRF with slightly different endpoints. Further validation experiments will need to be done to quantitatively confirm the presence of this fragment in the nuclear fraction of cells and its cell-line specific abundances. Unfortunately, there are currently no sufficiently accurate or sufficiently sensitive methods for detecting a short RNA with specific endpoints^109^. This technology gap is currently a limiting feature of the fast-evolving short RNA field.

Of note, the northern blot also revealed longer abundant rRFs from the same locus as the 31-mer in all three cell lines (**Figure 6B**). In various combinations, subsets of the 33-mer 5’ ETS-derived rRF with abundances ≥ 10 RPM exhibit very distinct cell-type-dependent patterns (**Figure 6C**). This cell-/tissue-specificity extends beyond the three cell lines, as evidenced by the abundance profile of the 31-mer rRF across 6,203 publicly available RNA-seq datasets (**Supplemental Table 8**).

The biogenesis and function of these 5’ ETS-derived rRFs is currently unknown. The region of the 5’ ETS where we find these short RNAs is slightly downstream of the A’ position, an essential cleavage location for early internal processing of the 45S rRNA^79^. Studies have reported that functional defects in nuclear exosome-associated nucleases contribute to incomplete degradation of 5’ ETS RNA^77,78^, leading to the production of long rRNA fragments (~1,000 nts) beginning at the A’ position of the 5’ ETS with varying 3’ endpoints^77^. However, we are not aware of previous studies that reported short RNAs that arise from this region.

Interestingly, this locus, which lies directly between the A’ position and is a two-branch stem-loop, contains the vertebrate box motif, a single-stranded stretch of the 5’ ETS that is evolutionarily conserved between mammals and amphibians^79,110^. In addition, this region contains a binding motif for both the protein nucleolin (TCGA), which resides within the ~11 nt evolutionary conserved region (ECM), and the U3 snoRNA binding motif (GTGCTCT)^111–113^. Nucleolin is required to recruit snoRNA U3 for rRNA processing and has been linked to promoting carcinogenesis, metastasis, and aging^112–116^. The relevance of this region of the 45S being a hotspot for rRNA processing is interesting in light of the observation that this locus can produce short RNAs (**Supplemental Figure S5**). Nucleolin primarily localizes to the nucleus^117^ while our analyses show that rRFs from 5’ ETS RNA are enriched in the nuclear fraction and have abundances that place them well above the 50^th^ percentile (**Figure 6C**). Their co-localization makes it conceivable that these short RNAs regulate rRNA synthesis by displacing nucleolin from its rRNA binding site in a fashion analogous to YBX1’s displacement by tRFs^118^.

### The most abundant short RNAs in a cell are cell-line-specific

Another unexpected result pertains to the fact that the most abundant rRFs from a given precursor rRNA is a feature of cell type. This extends to rRFs an observation we reported previously for isomiRs^5,6,10^ and tRFs^7,9^. Our RNA-seq data shows a 37-mer rRF from the 5’-end of the 28S rRNA to be significantly more abundant in the nuclear fraction of BT-20 cells as compared to both MDA-MB-231 and MDA-MB-468 cells (**Figure 7B** and **Supplemental Table 6**). However, when we probed for this sequence in both total and nuclear RNA from the three cell lines, we saw the 37-mer being more abundant in MDA-MB-231, not BT-20 (**Figure 7B**). Interestingly, MDA-MB-468 has two extremely strong bands at approximately 32 and 38 nts. The presence of these RNAs was surprising as they are not supported by our RNA-seq data. The disparity between northern blot and RNA-seq is not unexpected, especially for longer rRNA fragments, and may be the result of a 5’ or 3’-terminal modification that interferes with adapter ligation during the library preparation step of RNA-sequencing^80^.

When we checked to see which were the most abundant rRFs within the three cell lines, we found that all were 28S rRFs. However, each had originated from a different location on the 28S rRNA (**Figure 7C**). The most abundant rRF in BT-20 cells is the 37-mer 5’ 28S rRF (yellow, **Figure 7CD**), the most abundant rRF in MDA-MB-231 cells comes from the 3’ end of the 28S rRNA (grey, **Figure 7C-D**), and the most abundant rRF in MDA-MB-468 cells comes from the center region of the 28S rRNA (orange, **Figure 7C-D**). Moreover, the most abundant rRF of each cell line is enriched in a different compartment.

Interestingly, it has been suggested that both short and long rRFs from the 3’-end of the 28S have the potential to function in biological pathways in humans and zebrafish^119,120^. Taken together, the rRFs referenced above could underlie cell-line specific processes and future work could investigate what is regulating the differential localization profiles.

To summarize, this research has broken new ground in terms of our understanding of the enrichment patterns of short RNAs. Our analysis of the short RNAs found in our Fraction-seq results both agreed with and extended the results from earlier work (**Supplemental Figure 3**). Prompted by our RNA-seq-based analyses, we pursued a few select isomiRs, tRFs, and rRFs, and we experimentally corroborated previously uncharacterized cell type- and cell-compartment-dependent behavior. We also expanded our profiling of a select group of isomiRs, tRFs, and rRFs to 6,203 publicly available short RNA-seq datasets and showed that these molecules’ contextdependent expression extends to most other datasets (**Supplemental Table 8**).

Most importantly, we devised a multi-faceted experimental and analytical approach to analyze the short RNA profiles of subcellular compartments. We found that each compartment contains its own distinct profile of isomiRs, tRFs, and rRFs. The short RNA profiles are rich in short RNA molecule diversity that depends on cell type. Unexpectedly, we found that the enrichment patterns of a molecule do *not* depend on the precursor RNA from which it arises or the RNA class to which the molecule belongs (**Figure 6C**). These findings are important because isomiRs, tRFs, and rRFs modulate protein abundance.

While our findings are derived from the study of three cancer cell lines modeling TNBC, the findings extend to cell contexts beyond cancer. Moreover, the experimental and analytical protocol we devised can be applied to any cell line. It is our hope that Fraction-seq will facilitate the creation of an “Atlas” of subcellular localization for short RNAs in multiple cells and tissues. Such a cellular cartograph for isomiRs, tRFs, and rRFs would highlight useful constraints on their possible functions, help prioritize among molecules arising from the same precursor RNA, and could be leveraged to design better-targeted diagnostics, prognostics, and therapeutics.

## Authors’ contributions

IR conceived and supervised the study. TC and IR designed the experiments. TC, YJ, HW, and LT performed the experiments. VP and TC managed the submission and maintenance of original data deposited to NCBI. TC, IR, and BL designed the analysis methodology. TC, IR, BL, VP, PV, and PL contributed the analytical tools. TC and IR analyzed the data and designed the figures. TC and IR wrote the manuscript. All authors read and approved the final manuscript.

## Acknowledgments

The authors thank the TNBC patients who donated their samples for the three cell lines (BT-20, MDA-MB-231, and MDA-MB-468) used in this study. The authors would like to also thank the thousands of anonymous donors without whose contributions, this study would not be possible and the NCBI for supporting the SRA infrastructure allowing us to access thousands of publicly available datasets. The authors thank the members of the CMC for their helpful discussions, assistance and suggestions. The authors especially would like to thank Drs. Yohei Kirino, Megumi Hamasaki, Erin Seifert, Gyorgy Csordas, and Sergio De La Fuente Perez for their helpful conversations and guidance regarding the cell fractionation method. This work was supported partially by a William M. Keck Foundation grant (IR), and by Institutional Funds.

## Availability of data

All data generated or analyzed during this study are included in this published article, its supplementary information files, and publicly available repositories. The 50 RNA-seq datasets that we generated can be found in the NCBI SRA repository, https://www.ncbi.nlm.nih.gov/bioproject/PRJNA816866, and will be available upon publication of this article.

## Supplemental Figure Legends

**Supplemental Figure 1 Protein marker validation of cell fractionation via WES**

**A-C.** WES blots showing detection of cell fraction protein markers Lamin A/C (nucleus), GAPDH (cytoplasm), SDHA (mitochondrion), Cytochrome C (mitochondrion) TFAM (mitoplast). 2mg/ml protein lysate from each sample was loaded into a given lane. Absence of Cytochrome C in the mitoplast fraction indicates removal of intermembrane material from the mitochondrion. **A.** BT-20 – three replicates. **B.** MDA-MB-231 – three replicates **C.** MDA-MB-468 – four replicates.

**Supplemental Figure 2 Abundances of 17 short RNAs shared between TNBC cell lines**

The identities of the 17 short RNAs that are in common between BT-20, MDA-MB-231, and MDA-MB-468 cell lines when comparing the top 10% most abundant RNAs from each cell line based on average abundance. **A-Q.** Boxplot of the abundance (RPM) of the 17 short RNAs shared by the top 10% most abundant short RNAs across TNBC cell lines: BT-20 (n=3), MDA-MB-231 (n=3), and MDA-MB-468 (n=4).

**Supplemental Figure 3 Comparisons of short RNAs across cell lines reveal cell-line specific features**

**A-D.** Venn Diagrams of the top 10% most abundant short RNA in BT20 (red), MDA-MB-231 (green), and MDA-MB-468 (blue) cells based on average abundance in each cell fraction (nucleus, cytoplasm, mitochondrion, and mitoplast). **A.** Nucleus. **B.** Cytoplasm. **C.** Mitochondrion. **D.** Mitoplast.

**Supplemental Figure 4 Heatmaps of short RNAs in TNBC cell line fractions with labels**

**A-C.** Heatmaps with corresponding short RNA labels of the of the percentiles of the average normalized abundance (RPM) of short RNAs in total (Total), nucleus (Nuc), cytoplasm (Cyto), mitochondrion (Mito), and mitoplast (MP) samples for BT-20 (A), MDA-MB-231 (B), and MDA-MB-468 (C) cells. The heatmap colors range from white (lowest percentile) to dark-purple (highest percentile). Each row of the heatmap corresponds to specific the rRNA class, RNA precursor molecule, RNA start position, RNA end position, and RNA sequence of the short RNA, where the colors represent the short RNA class (pink – isomiRs, teal – tRF, and yellow – rRF).

**Supplemental Figure 5 Hotspot 5’ ETS region produces short RNAs**

**A.** Schematic of the full 13,351 45S rRNA with annotations of the three rRNA (18S, 5.8S, 28S) and four spacer regions (5’ ETS, ITS1, ITS2, 3’ ETS). The 5’ ETS is annotated with a pileup of the short RNAs mapping to this region for BT-20 (blue), MDA-MB-231 (orange), and MDA-MB-468 (grey) based on average. **B.** Zoomed-in image of the 5’ ETS region of the 45S rRNA for BT-20 (blue), MDA-MB-231 (orange), and MDA-MB-468 (grey) based on average. Pileup shows the position along the 3,654 nt 5’ ETS where short RNAs are being produced. A-B) The A’ position (+414) and the start (+425) and end (+484) positions of the main pileups are indicated by black arrows. Error for the pileups are shown as shaded area above and below pileup lines.

**Supplemental Figure 6 Nuclear and mitochondrial genomes produce short RNA that can be found in nuclear, cytoplasmic, and mitochondrial fractions**

**A.** Stacked barplots representing the percent of unique short RNA types found in cell fractions from three independent studies: nucleus and cytoplasm (5-8F cells, doi: 10.1371/journal.pone.0010563), mitochondrion (HEK293 cells, HeLa cells doi: 10.1371/journal.pone.0044873, 143B cells doi: 10.1016/j.cell.2011.06.051). **B.** Stacked barplots representing the number of unique RNA types found in the same samples as A. **C.** Stacked barplots representing the percent of unique short RNA that is found in cell fractions from three cell lines (BT20, MDA-MB-231, and MDA-MB-468) based on average abundance. **D.** Stacked barplots representing the number of unique short RNA that is found in cell fractions from three cell lines (BT20, MDA-MB-231, and MDA-MB-468) based on average abundance. Colors denote the different RNA classes: nuclear isomiRs (green), nuclear rRFs (dark blue), nuclear tRFs (dark yellow), mitochondrial rRFs (light blue), mitochondrial tRFs (light yellow), and tRFs that map to both nuclear and mitochondrial genomes (diagonal yellow lines).

