## Supplementary figures and images for "Single-nucleotide Differences and Cell Type Decide the Subcellular Localization of miRNA Isoforms (isomiRs), tRNA-derived Fragments (tRFs) and rRNA-derived Fragments (rRFs)"

### Supp. Figure S1

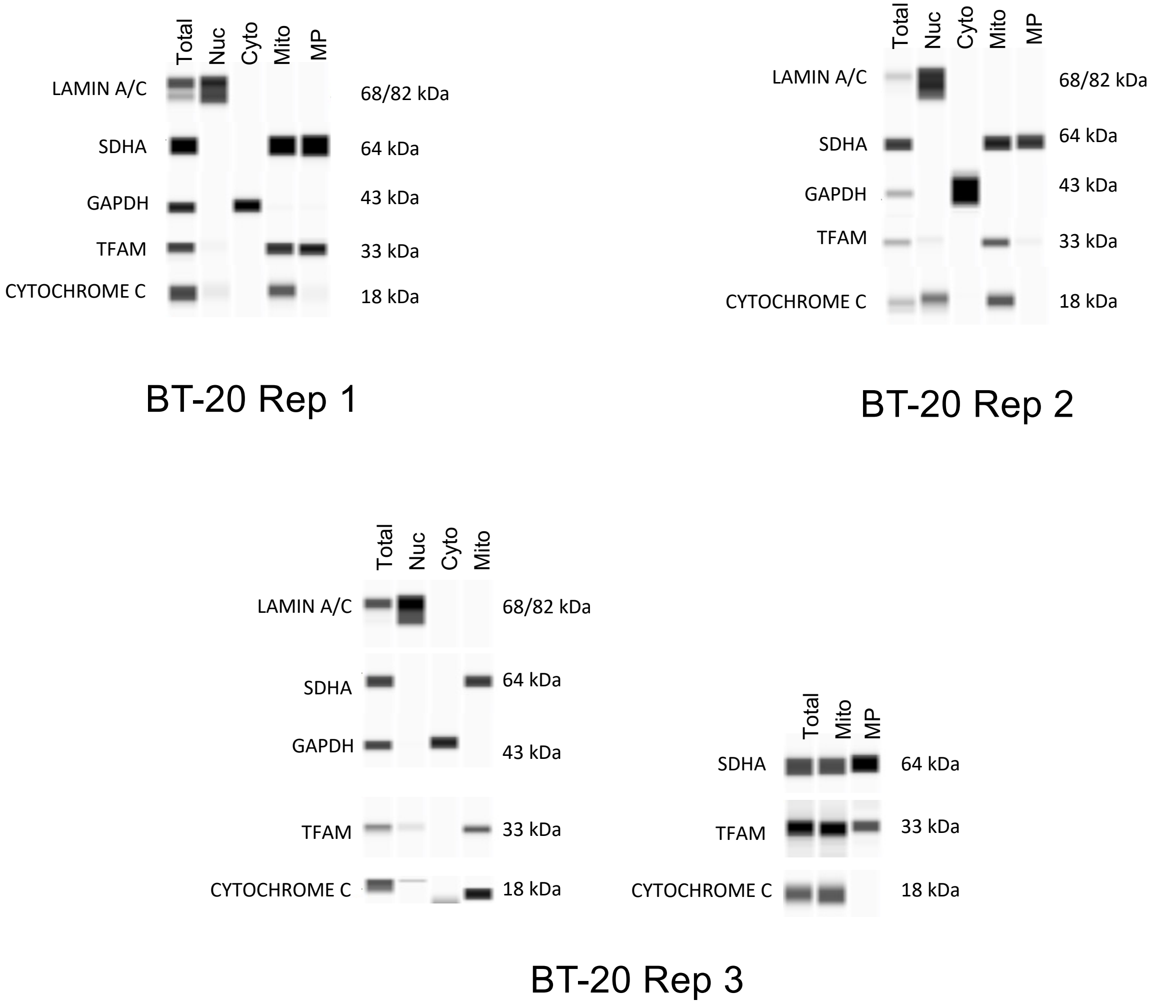

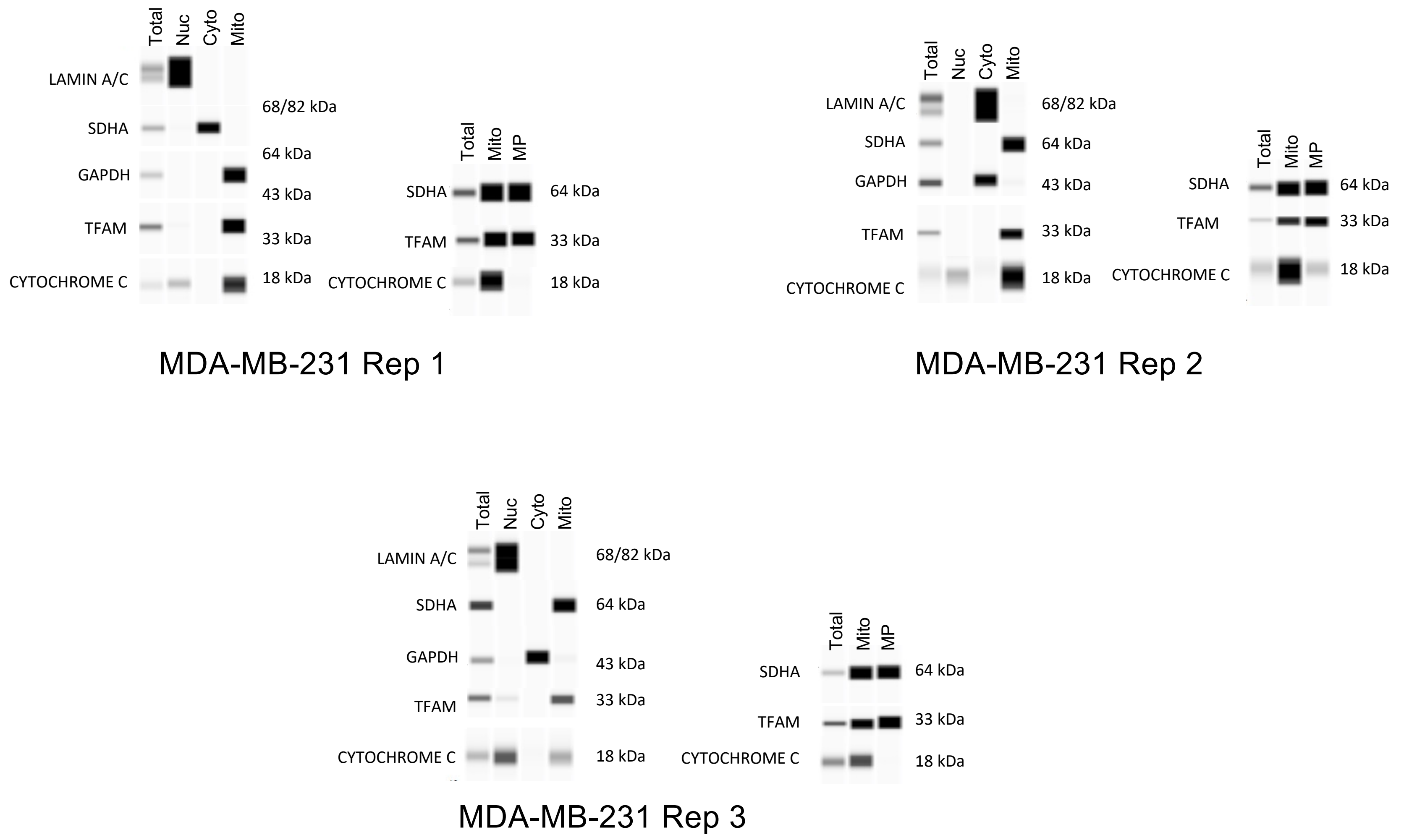

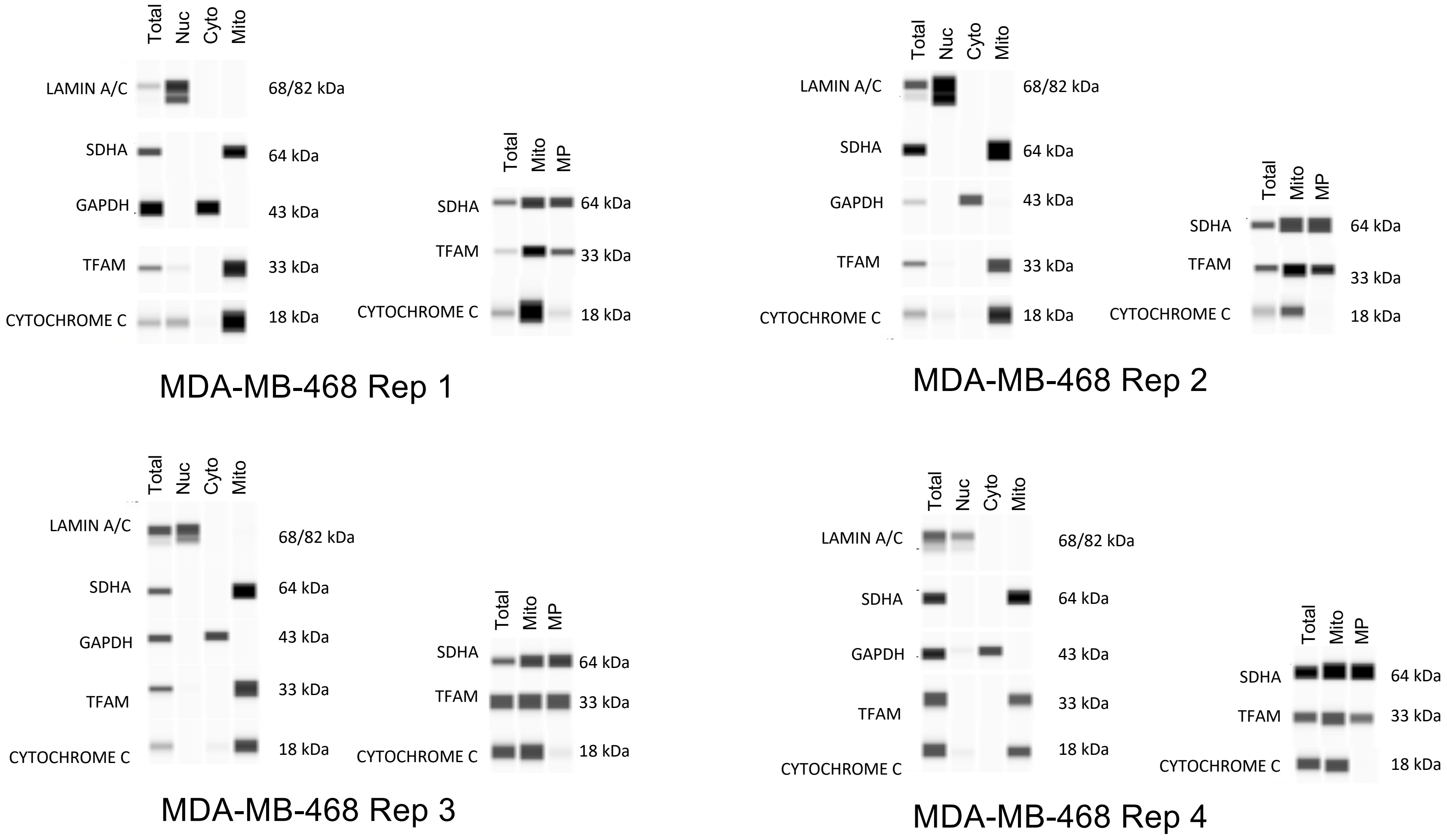

### Supp. Figure S3

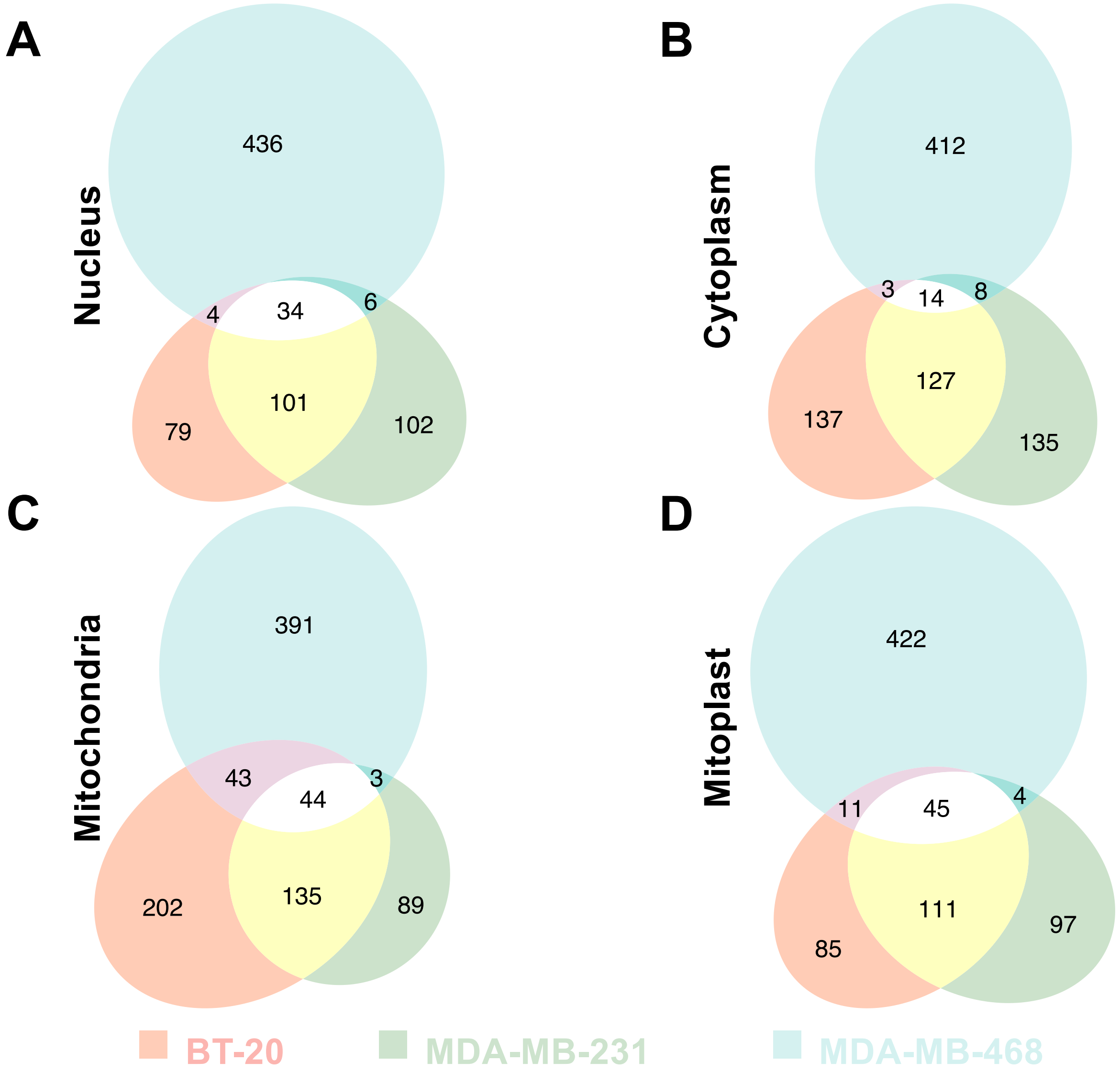

### Supp. Figure S5

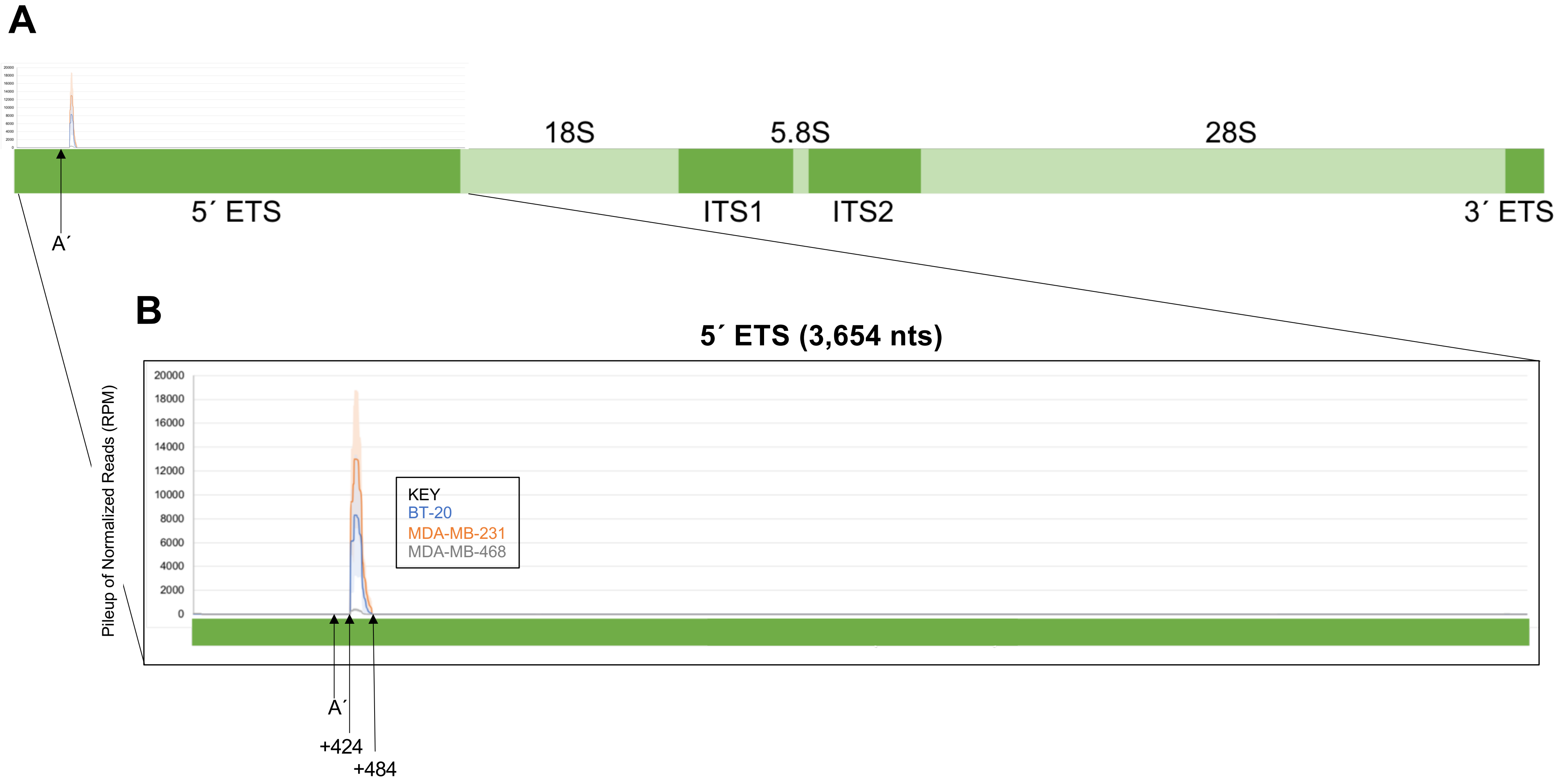

### Supp. Figure S6

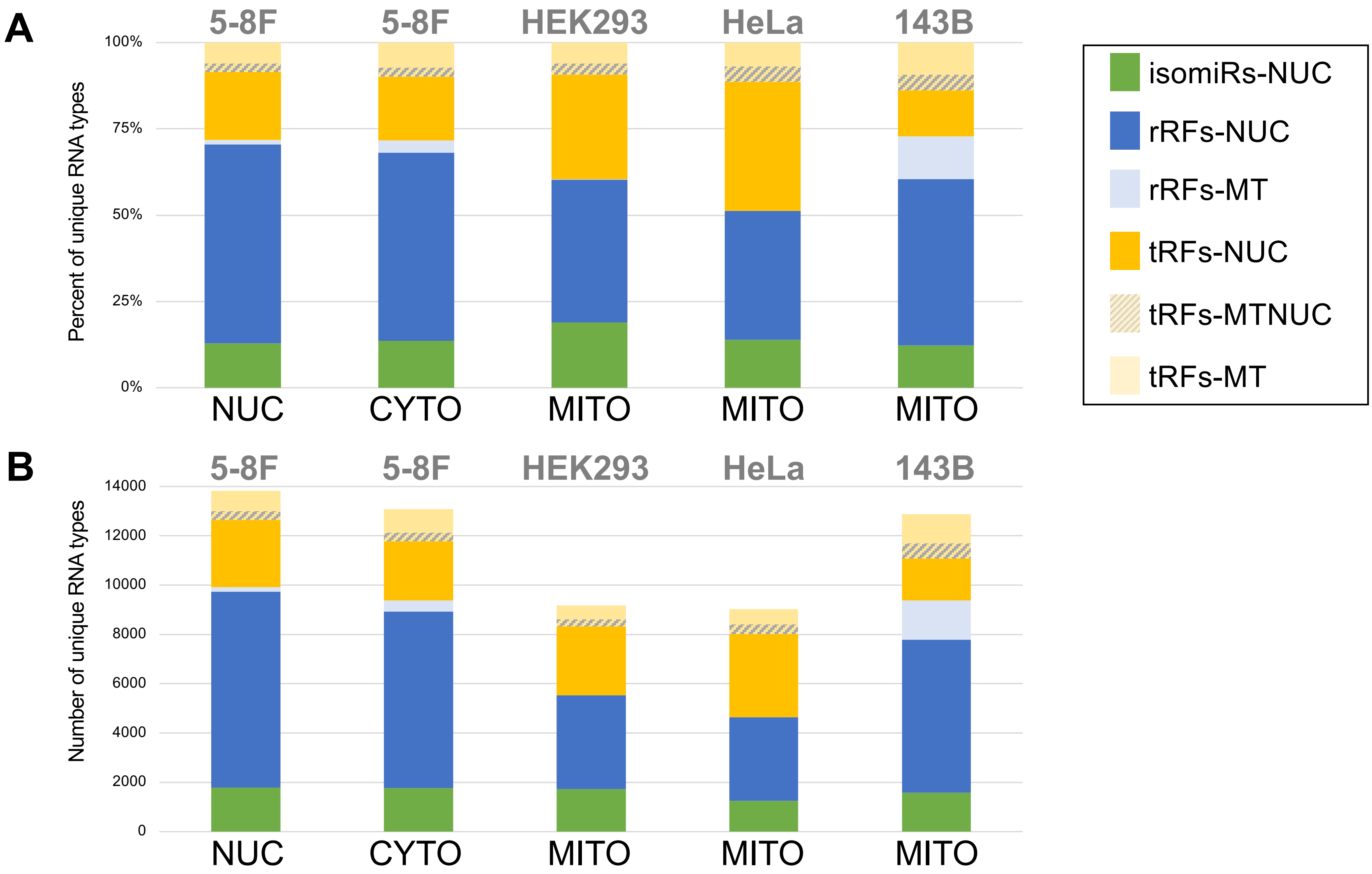

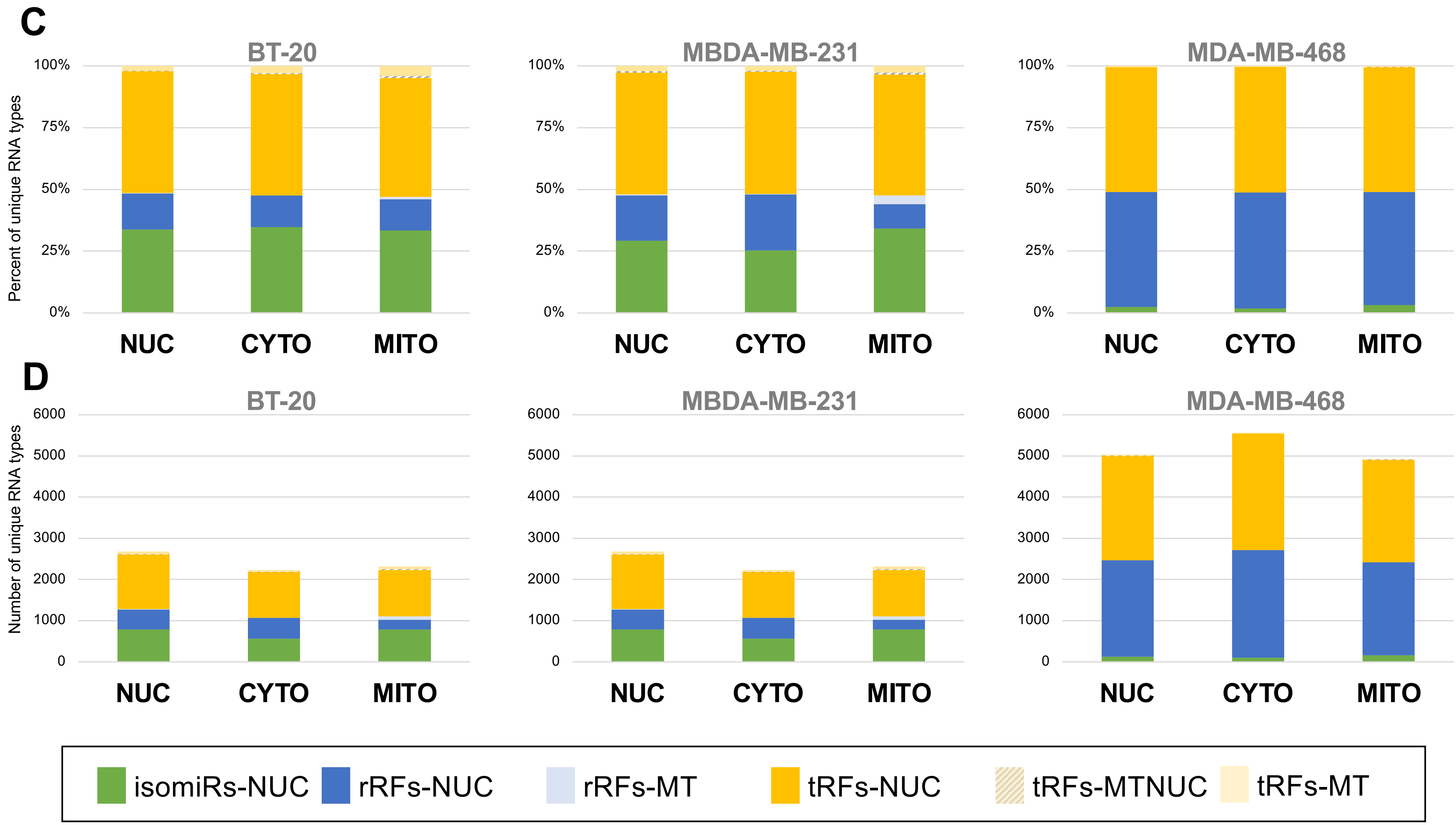
