## Supplementary material for "Single-nucleotide Differences and Cell Type Decide the Subcellular Localization of miRNA Isoforms (isomiRs), tRNA-derived Fragments (tRFs) and rRNA-derived Fragments (rRFs)": Supp. Figure S2

**Supp. Fig. S2**

### 17 Shared short RNAs from Venn Diagram

| | Short RNAs | Sequence |
|---|---|---|
| 1 | 5S 82.150.39 | GATGGGAGACCGCCTGGGAATACCGGGTGCTGTAGGCTT |
| 2 | 5.8S 6596.6619.24 | TCGTACGACTCTTAGCGGTGGATC |
| 3 | 28S 7925.7944.20 | CGCGACCTCAGATCAGACGT |
| 4 | 28S 7925.7958.34 | CGCGACCTCAGATCAGACGTGGCGACCCGCTGAA |
| 5 | 28S 7925.7962.38 | CGCGACCTCAGATCAGACGTGGCGACCCGCTGAATTTA |
| 6 | let-7a-5p 0\|0 | TGAGGTAGTAGGTTGTATAGTT |
| 7 | let-7f-5p 0\|0 | TGAGGTAGTAGATTGTATAGTT |
| 8 | miR-21-5p 0\|0 | TAGCTTATCAGACTGATGTTGA |
| 9 | miR-21-5p 0\|+1(+1A) | TAGCTTATCAGACTGATGTTGACA |
| 10 | miR-21-5p 0\|+1 | TAGCTTATCAGACTGATGTTGAC |
| 11 | miR-21-5p 0\|+2 | TAGCTTATCAGACTGATGTTGACT |
| 12 | miR-30a-5p 0\|+2 | TGTAAACATCCTCGACTGGAAGCT |
| 13 | miR-148a-3p 0\|0 | TCAGTGCACTACAGAACTTTGT |
| 14 | miR-7-5p 0\|0 | TGGAAGACTAGTGATTTTGTTGTT |
| 15 | let-7g-5p 0\|0 | TGAGGTAGTAGTTTGTACAGTT |
| 16 | let-7i-5p 0\|0 | TGAGGTAGTAGTTTGTGCTGTT |
| 17 | miR-26a-5p 0\|0 | TTCAAGTAATCCAGGATAGGCT |

**A**

1. 5S 82.120.39 GATGGGAGACCGCCTGGGAATACCGGGTGCTGTAGGCTT

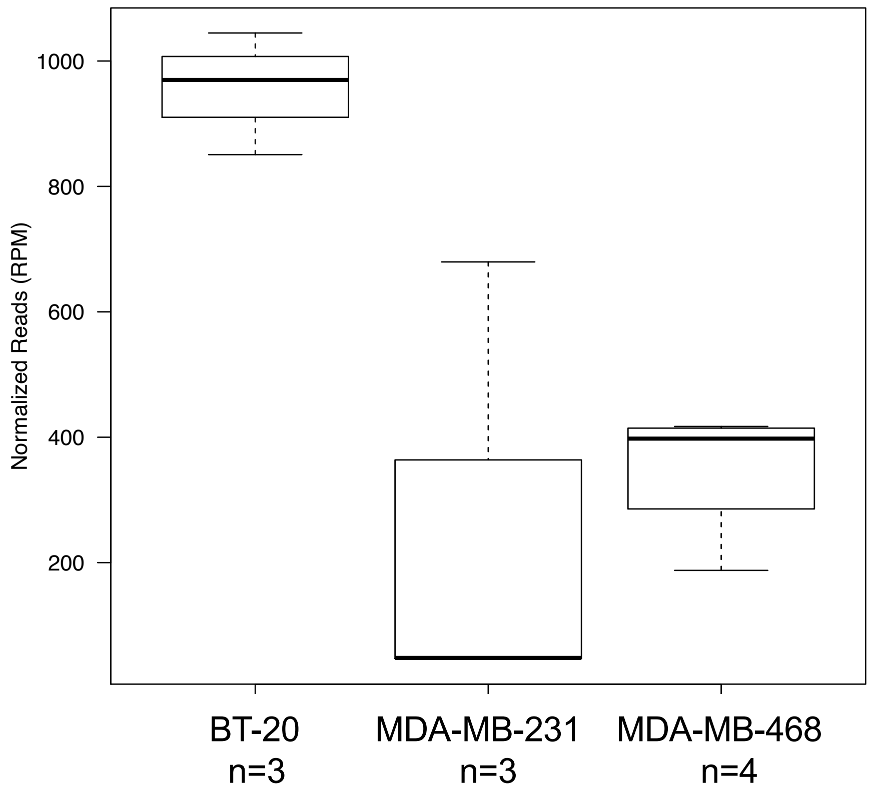

**B**

2. 5.8S 6596.6619.24 TCGTACGACTCTTAGCGGTGGATC

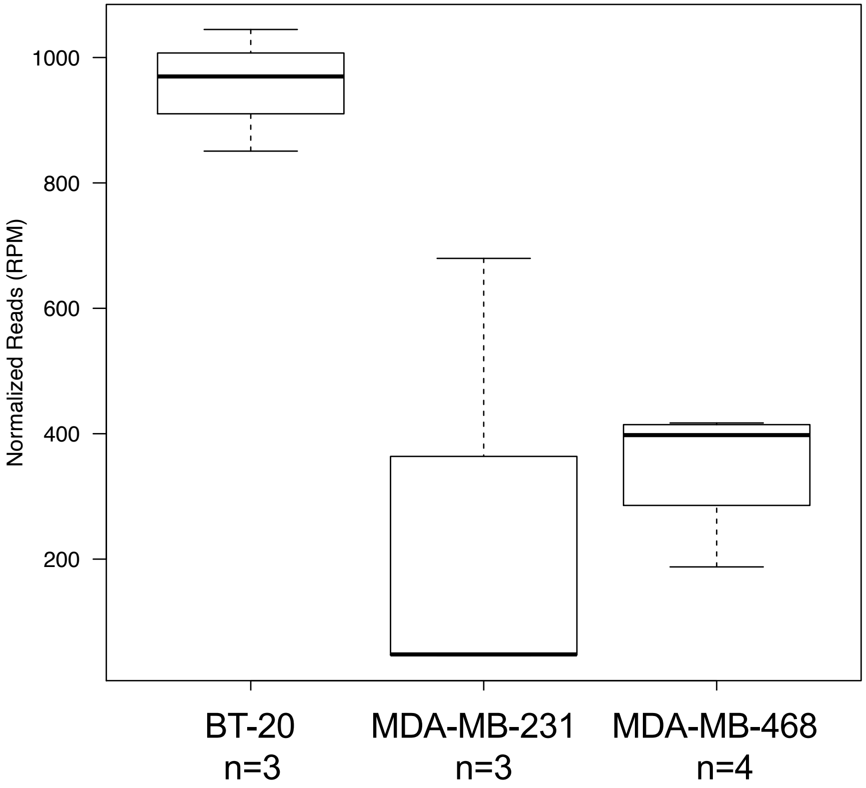

**C**

3. 28S 7925.7944.20 CGCGACCTCAGATCAGACGT

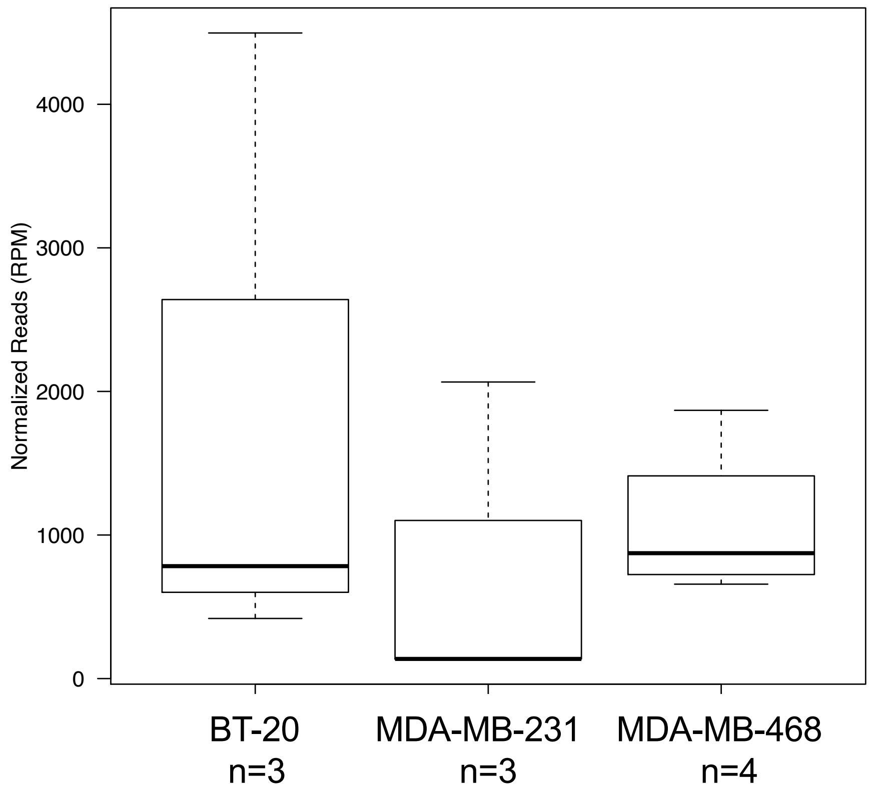

**D**

4. 28S 7925.7958.34 CGCGACCTCAGATCAGACGTGGCGACCCGCTGAA

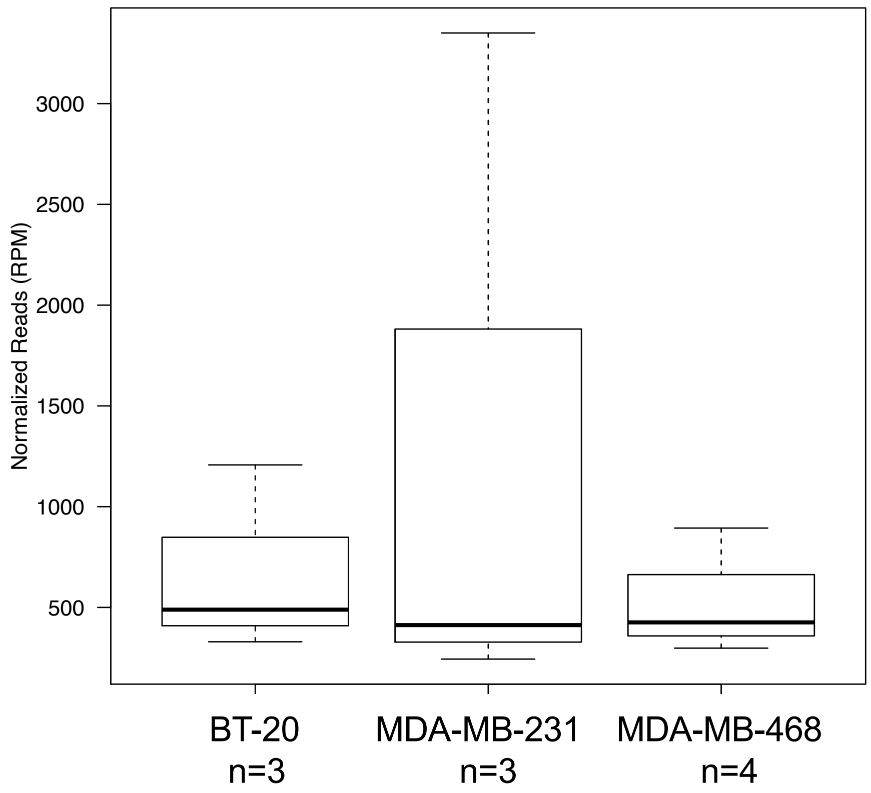

**E**

5. 28S 7925.7962.38 CGCGACCTCAGATCAGACGTGGCGACCCGCTGAATTTA

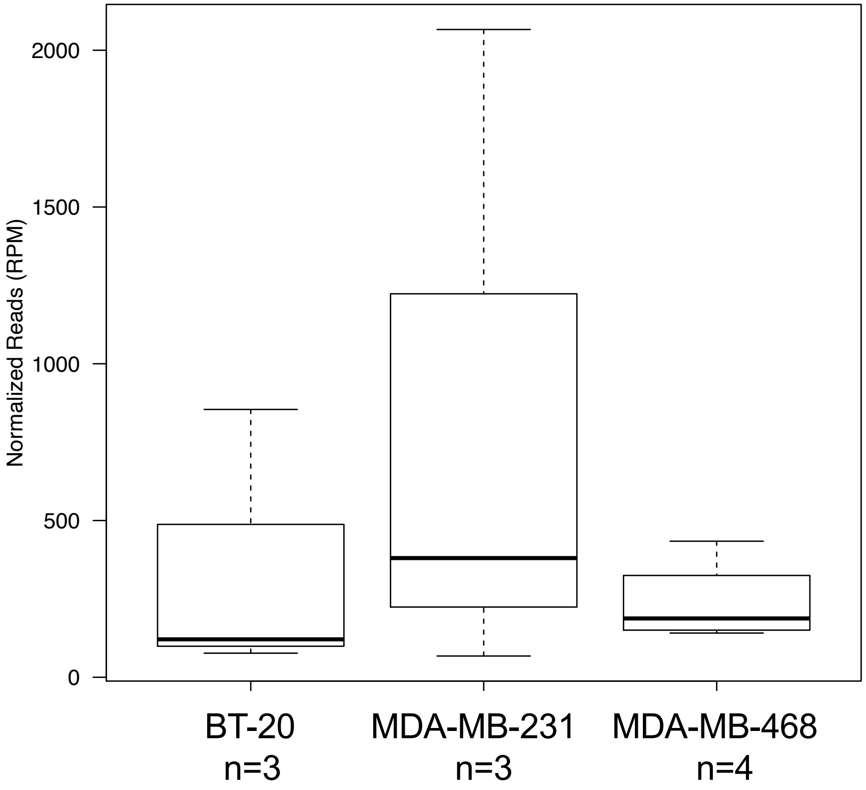

**F**

6. let-7a-5p 0|0 TGAGGTAGTAGGTTGTATAGTT

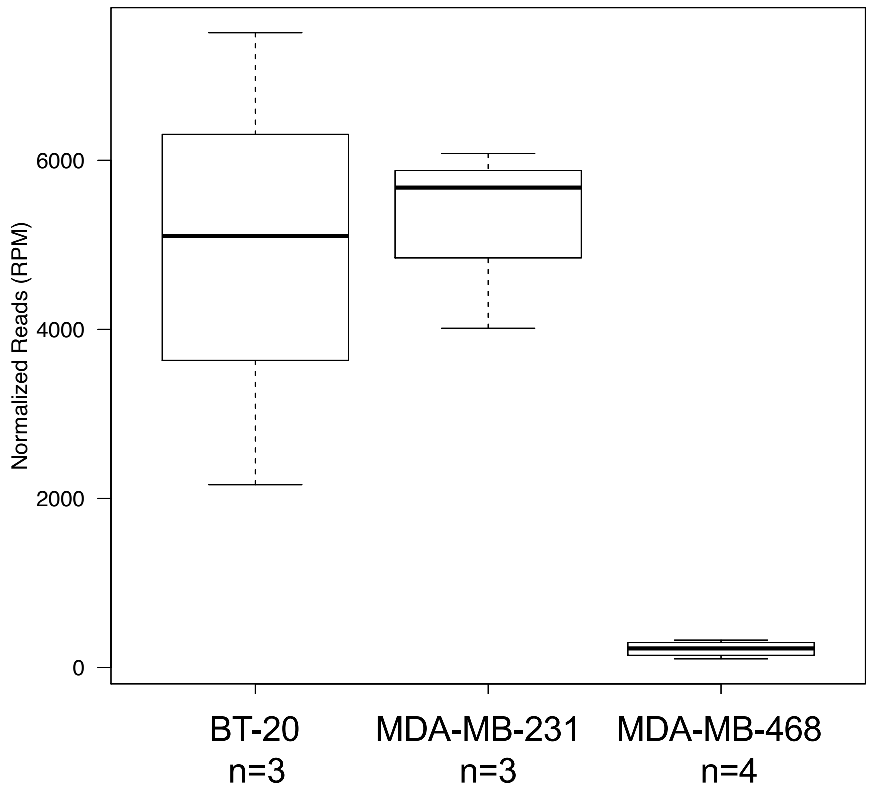

G

7. let-7f-5p 0|0 TGAGGTAGTAGATTGTATAGTT

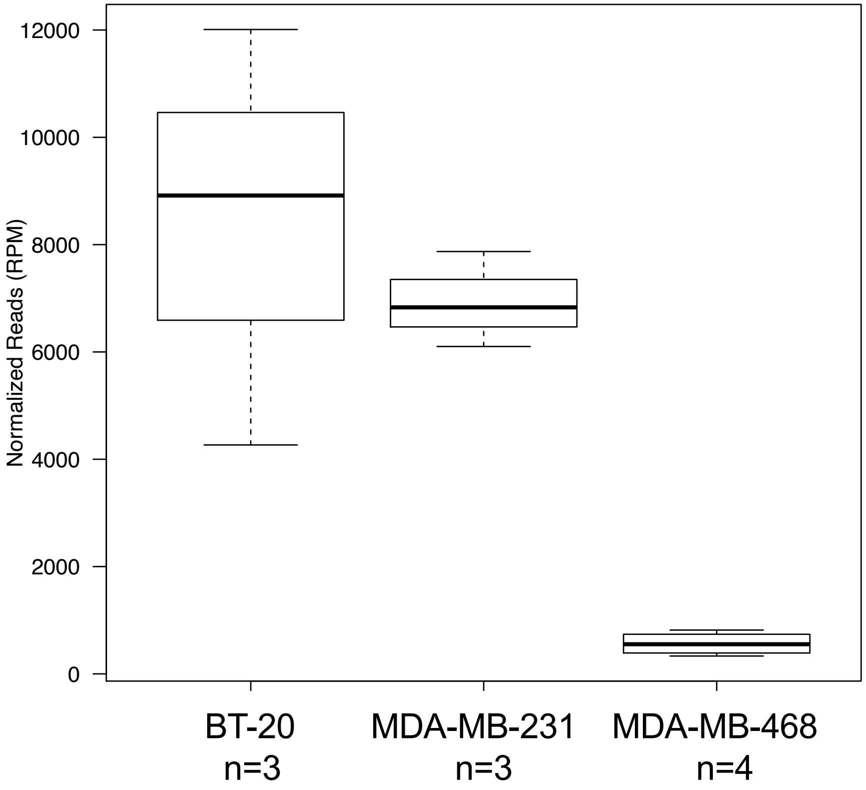

H

8. miR-21-5p 0|0 TAGCTTATCAGACTGATGTTGA

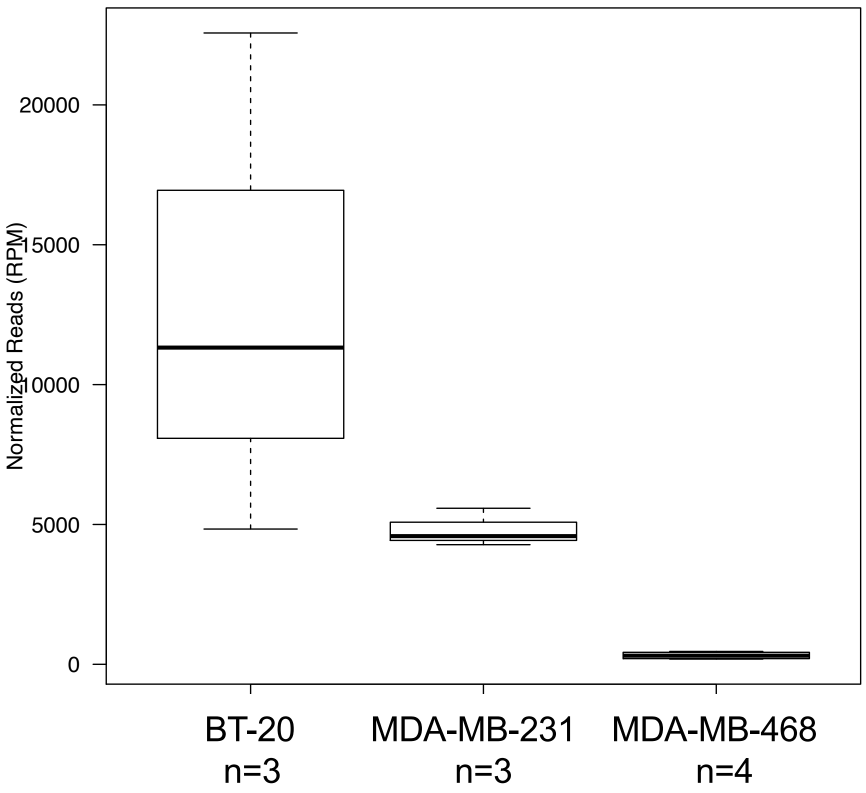

I

9. miR-21-5p 0|+1(+1A) TAGCTTATCAGACTGATGTTGACA

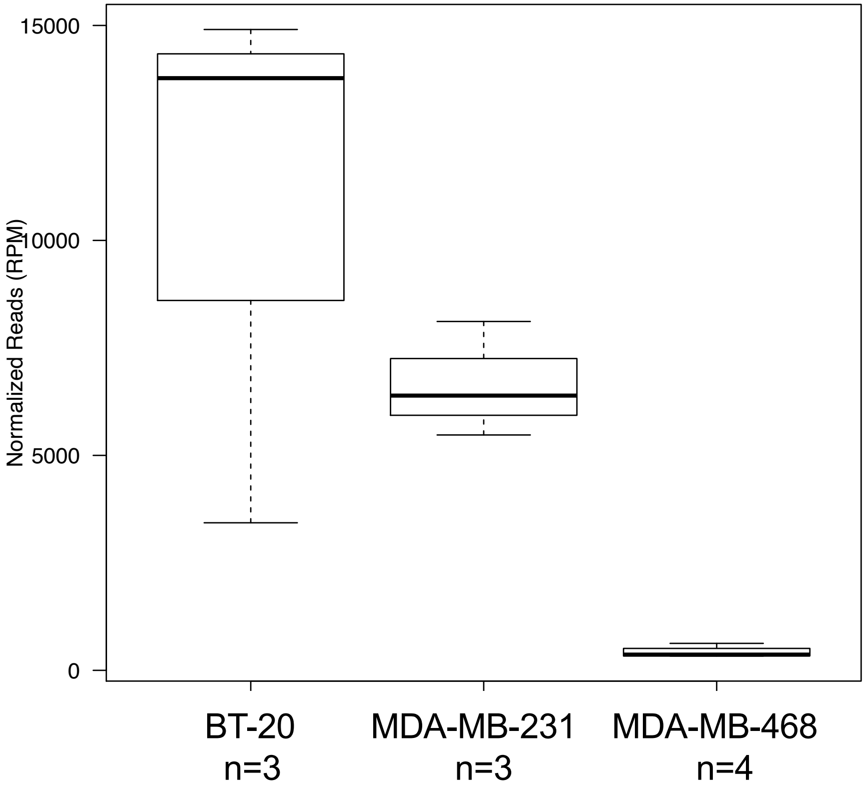

**J**

10. miR-21-5p 0|+1 TAGCTTATCAGACTGATGTTGAC

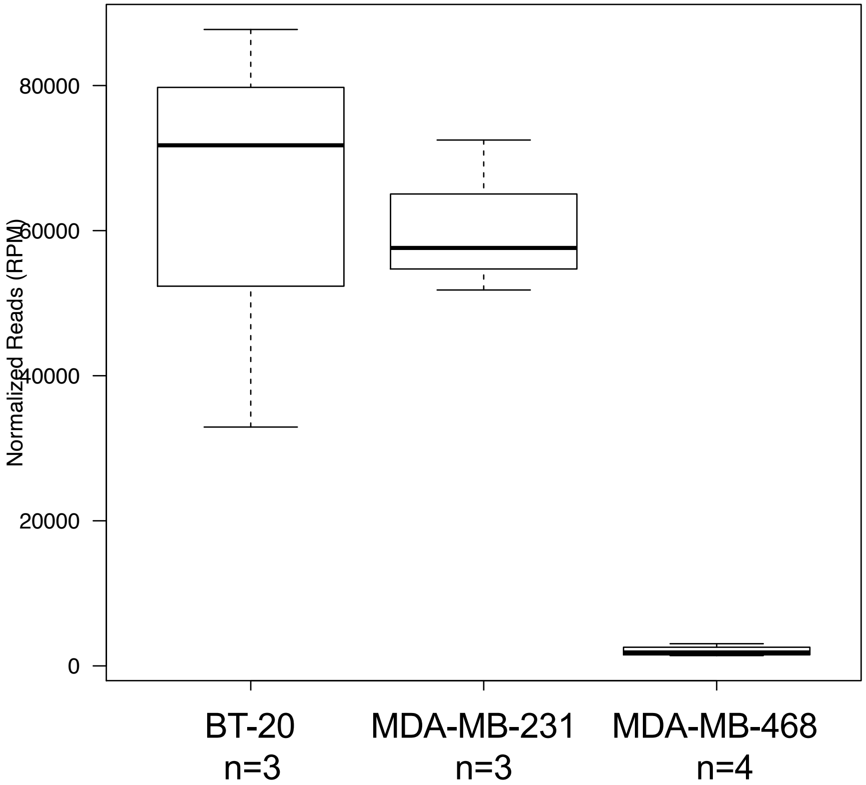

**K**

11. miR-21-5p 0|+2 TAGCTTATCAGACTGATGTTGACT

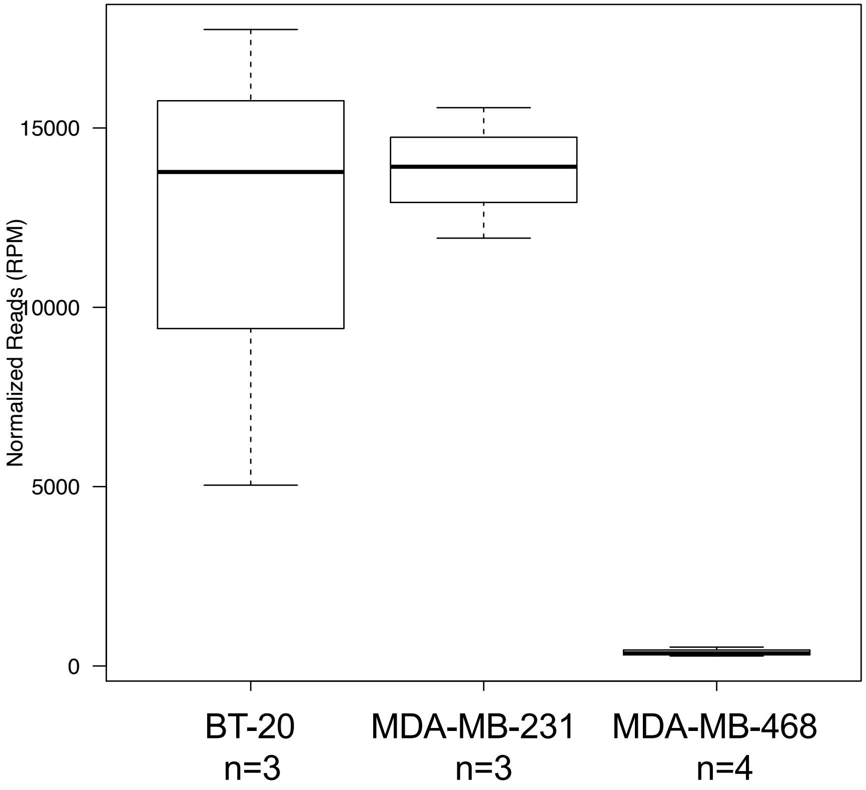

**L**

12. miR-30a-5p 0|+2 TGTAAACATCCTCGACTGGAAGCT

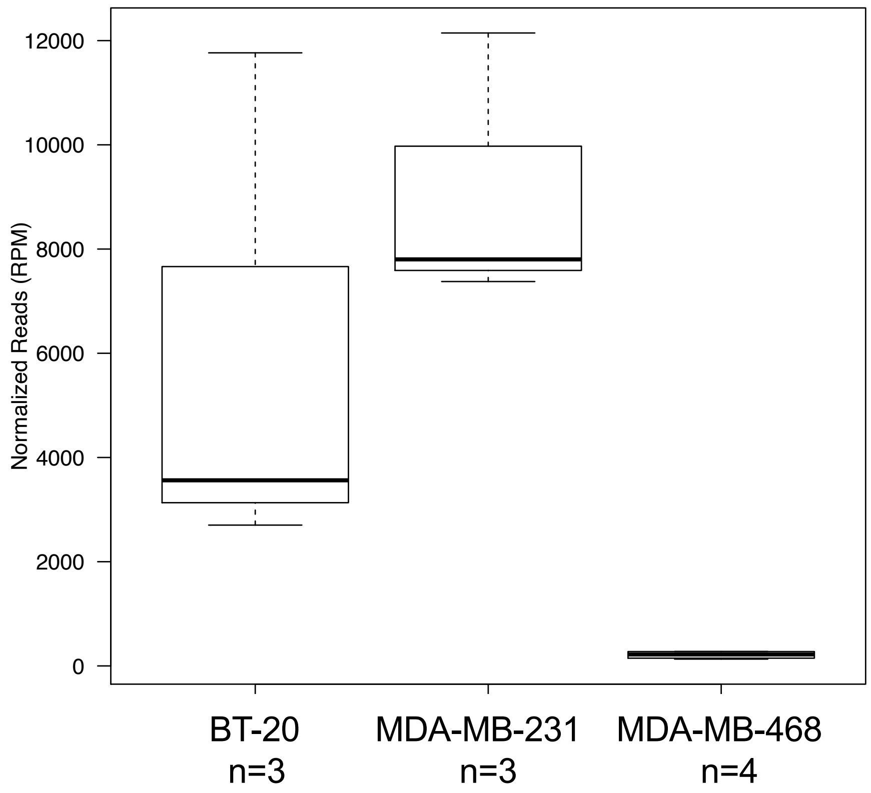

M

13. miR-148a-3p 0|0 TCAGTGCACTACAGAACTTTGT

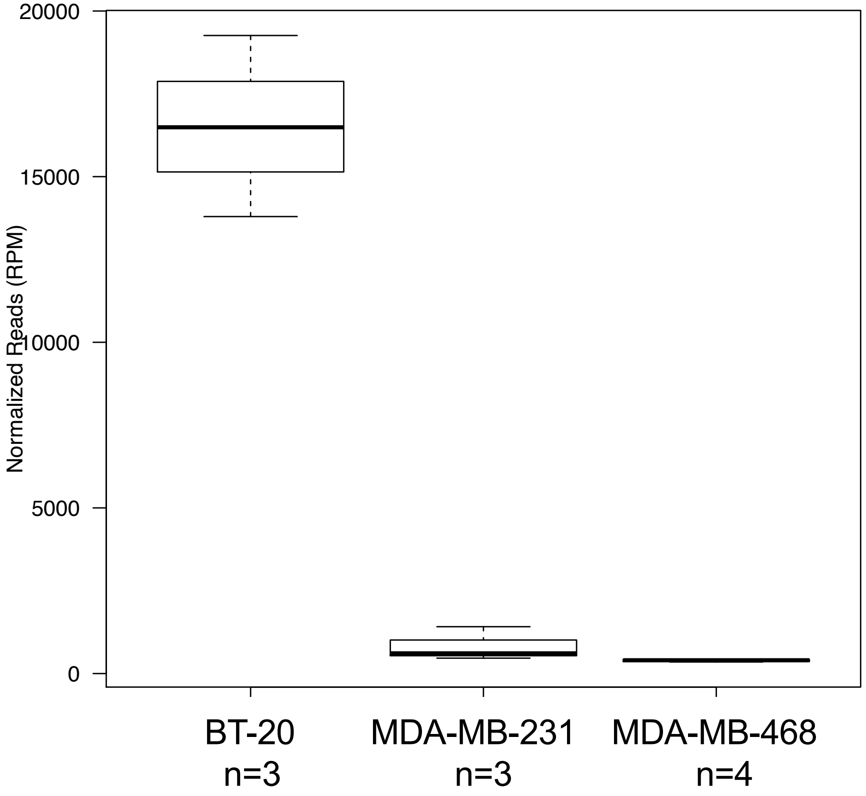

**N**

14. miR-7 -5p 0|0 TGGAAGACTAGTGATTTTGTTGTT

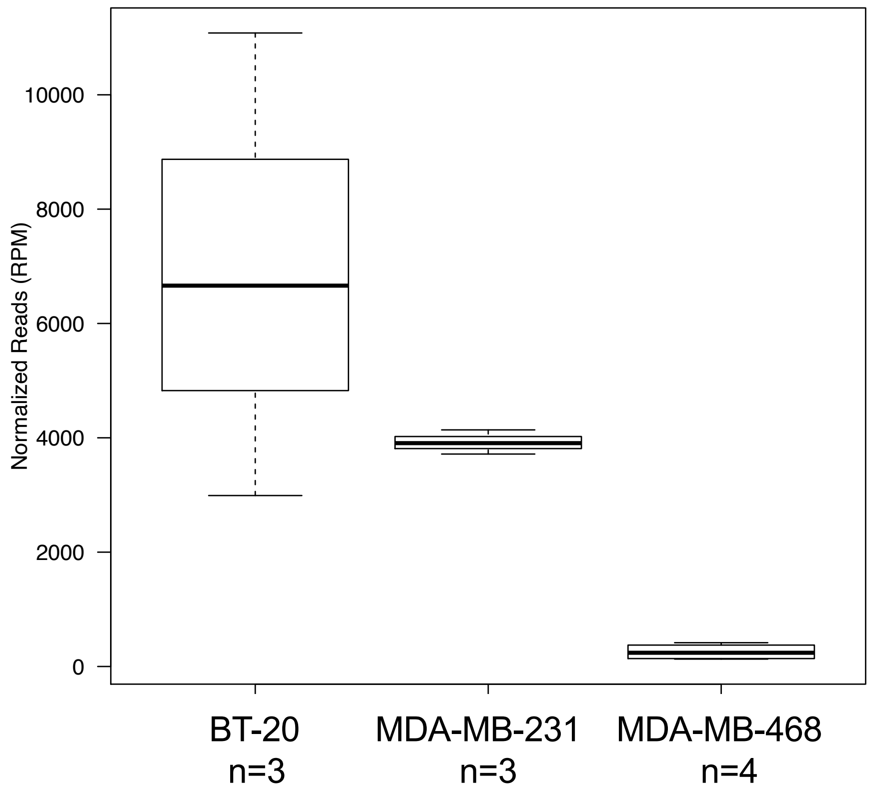

**O**

15. let-7g-5p 0|0 TGAGGTAGTAGTTTGTACAGTT

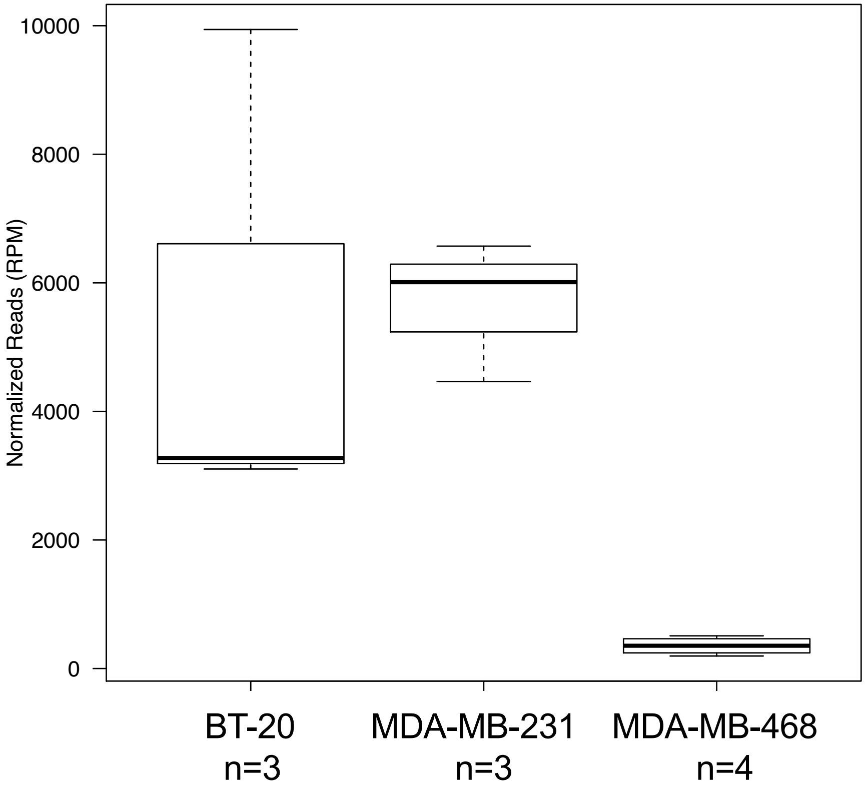

P

16. let-7i-5p 0|0 TGAGGTAGTAGTTTGTGCTGTT

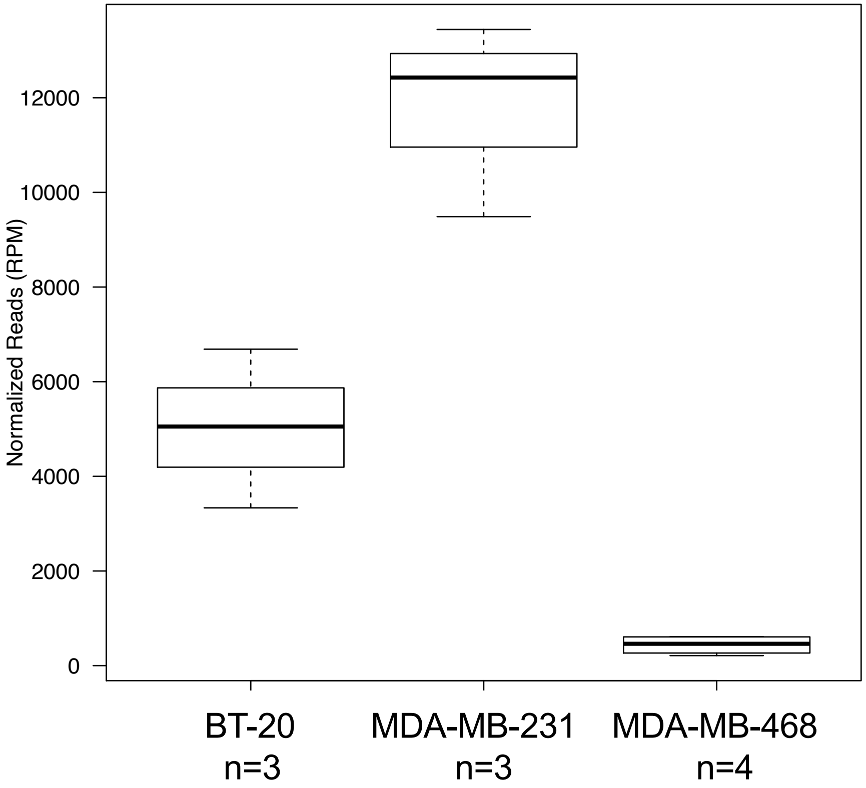

**Q**

17. miR-26a-5p 0|0 TTCAAGTAATCCAGGATAGGCT

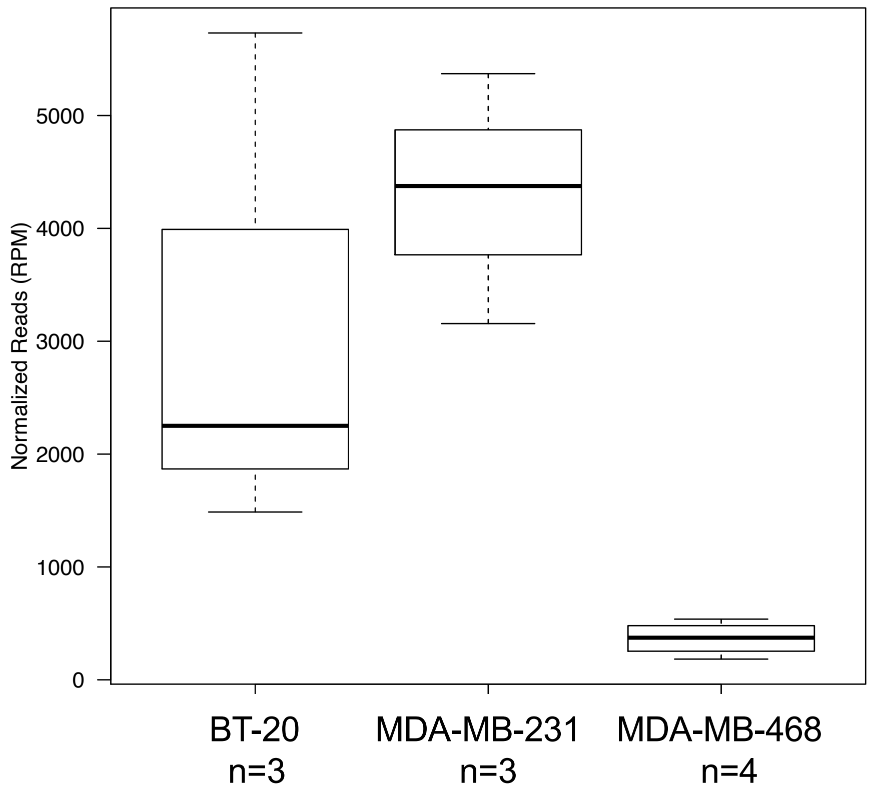
