## Supplementary material for "Single-nucleotide Differences and Cell Type Decide the Subcellular Localization of miRNA Isoforms (isomiRs), tRNA-derived Fragments (tRFs) and rRNA-derived Fragments (rRFs)": Supp. Figure S4

**Supp. Fig. S4**

## BT-20

| isomiR | tRF | rRF |
|---|---|---|

| Class | RNA | Start | End | Sequence |
|---|---|---|---|---|
| isomiR | mir-96 | 15 | 37 | TTTGGCACTAGCACATTTTTGCT |
| isomiR | mir-30b | 23 | 44 | TGTAAACATCCTACACTCAGCT |
| isomiR | mir-24 | 56 | 77 | TGGCTCAGTTCAGCAGGAACAGT |
| rRF | 5' ETS | 425 | 453 | CTTCGTGATCGATGTGGTGACGTCGTGCT |
| rRF | 5' ETS | 425 | 454 | CTTCGTGATCGATGTGGTGACGTCGTGCTC |
| rRF | 5' ETS | 425 | 455 | CTTCGTGATCGATGTGGTGACGTCGTGCTCT |
| rRF | 5' ETS | 425 | 457 | CTTCGTGATCGATGTGGTGACGTCGTGCTCTCC |
| rRF | 5' ETS | 426 | 455 | TTCGTGATCGATGTGGTGACGTCGTGCTCT |
| rRF | 5' ETS | 435 | 455 | GATGTGGTGACGTCGTGCTCT |
| rRF | 5' ETS | 435 | 456 | GATGTGGTGACGTCGTGCTCTC |
| rRF | 5.8S | 6601 | 6633 | CGACTCTTAGCGGTGGATCACTCGGCTCGTGCG |
| isomiR | mir-221 | 71 | 93 | AGCTACATTGTCTGCTGGGTTTCT |
| isomiR | mir-320a | 48 | 69 | AAAAGCTGGGTTGAGAGGGCGATT |
| tRF | trna10_AlaCGC | 58 | 75 | TCCCCGGCATCTCCACCA |
| tRF | trna101_AlaCGC | 58 | 75 | TCCCCGGCACCTCCACCA |
| tRF | trna11_GluTTC | 1 | 33 | TCCCACATGGTCTAGCGGTTAGGATTCCTGGTT |
| tRF | trna134_GluTTC | 1 | 33 | TCCCTGGTGGTCTAGTGGCTAGGATTCGGCGCT |
| tRF | trna128_GlyGCC | 1 | 32 | GCATTGGTGGTTCAGTGGTAGAATTCTCGCCT |
| tRF | trna2_GlyGCC | 1 | 32 | GCATGGGTGGTTCAGTGGTAGAATTCTCGCCT |
| tRF | trna119_LysCTT | 1 | 35 | GCCCGGCTAGCTCAGTCGGTAGAGCATGAGACTCT |
| tRF | trnaMT_ValTAC | 48 | 67 | CAACTTAACTTGACCGCTCT |
| rRF | 18S | 3655 | 3678 | TACCTGGTTGATCCTGCCAGTAGC |
| isomiR | mir-584 | 22 | 42 | TTATGGTTTGCCTGGGACTGA |
| tRF | trnaMT_GluTTC | 38 | 72 | CATTGGTCGTGGTTGTAGTCCGTGCGAGAATACCA |
| tRF | trnaMT_GluTTC | 40 | 72 | TTGGTCGTGGTTGTAGTCCGTGCGAGAATACCA |
| tRF | trnaMT_GluTTC | 41 | 72 | TGGTCGTGGTTGTAGTCCGTGCGAGAATACCA |
| tRF | trnaMT_PheGAA | 39 | 56 | ATGTTTAGACGGGCTCAC |
| tRF | trnaMT_SerGCT | 1 | 25 | GAGAAAGCTCACAAGAACTGCTAAC |
| rRF | 16S | 51 | 74 | GCTAAACCTAGCCCCAAACCCACT |
| rRF | 16S | 51 | 84 | GCTAAACCTAGCCCCAAACCCACTCCACCTTACT |
| rRF | 16S | 1 | 45 | GCTAAACCTAGCCCCAAACCCACTCCACCTTACTACCAGACAACC |
| rRF | 12S | 2 | 27 | ATAGGTTTGGTCCTAGCCTTTCTATT |
| rRF | 5S | 86 | 120 | GGAGACCGCCTGGGAATACCGGGTGCTGTAGGCTT |

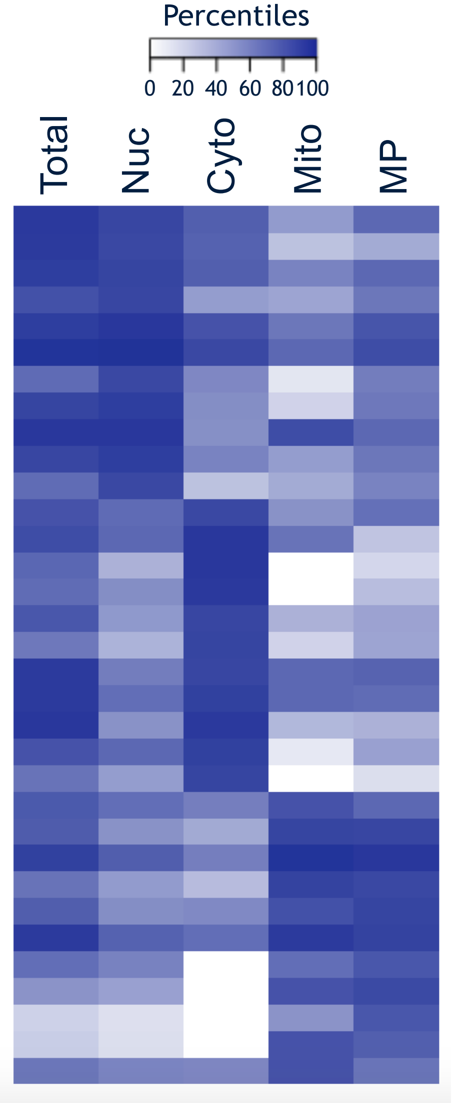

### MDA-MB-231

| isomiR | tRF | rRF |
|---|---|---|

| Class | RNA | Start | End | Sequence |
|---|---|---|---|---|
| rRF | 5' ETS | 425 | 464 | CTTCGTGATCGATGTGGTGACGTCGTGCTCTCCCGGGCCG |
| rRF | 5' ETS | 425 | 465 | CTTCGTGATCGATGTGGTGACGTCGTGCTCTCCCGGGCCGG |
| rRF | 5' ETS | 426 | 445 | TTCGTGATCGATGTGGTGAC |
| rRF | 5' ETS | 426 | 451 | TTCGTGATCGATGTGGTGACGTCGTG |
| rRF | 5' ETS | 426 | 453 | TTCGTGATCGATGTGGTGACGTCGTGCT |
| rRF | 5' ETS | 426 | 457 | TTCGTGATCGATGTGGTGACGTCGTGCTCTCC |
| rRF | 5' ETS | 426 | 459 | TTCGTGATCGATGTGGTGACGTCGTGCTCTCCCG |
| rRF | 5' ETS | 426 | 460 | TTCGTGATCGATGTGGTGACGTCGTGCTCTCCCGG |
| rRF | 5' ETS | 426 | 468 | TTCGTGATCGATGTGGTGACGTCGTGCTCTCCCGGGCCGGGTC |
| rRF | 5.8S | 6599 | 6633 | TACGACTCTTAGCGGTGGATCACTCGGCTCGTGCG |
| rRF | 5.8S | 6600 | 6633 | ACGACTCTTAGCGGTGGATCACTCGGCTCGTGCG |
| rRF | 5.8S | 6602 | 6633 | GACTCTTAGCGGTGGATCACTCGGCTCGTGCG |
| rRF | 5.8S | 6603 | 6633 | ACTCTTAGCGGTGGATCACTCGGCTCGTGCG |
| rRF | 5.8S | 6611 | 6633 | CGGTGGATCACTCGGCTCGTGCG |
| tRF | trna111_HisGTG | 1 | 33 | GCCGTGATCGTATAGTGGTTAGTACTCTGCGTT |
| tRF | trna10_AspGTC | 36 | 74 | CACGCGGGAGACCGGGGTTCGATTCCCCGACGGGGAGCC |
| tRF | trna10_AspGTC | 36 | 75 | CACGCGGGAGACCGGGGTTCGATTCCCCGACGGGGAGCCA |
| tRF | trna10_AspGTC | 37 | 73 | ACGCGGGAGACCGGGGTTCGATTCCCCGACGGGGAGC |
| tRF | trna10_AspGTC | 58 | 75 | TCCCCGGCATCTCCACCA |
| tRF | trna101_AlaAGC | 58 | 75 | TCCCCGGCACCTCCACCA |
| rRF | 5S | 82 | 119 | GATGGGAGACCGCCTGGGAATACCGGGTGCTGTAGGCT |
| rRF | 18S | 5422 | 5466 | ACGGCCCTGGCGGAGCGCTGAGAAGACGGTCGAACTTGACTATCT |
| rRF | 18S | 5423 | 5466 | CGGCCCTGGCGGAGCGCTGAGAAGACGGTCGAACTTGACTATCT |
| rRF | 18S | 5439 | 5466 | CTGAGAAGACGGTCGAACTTGACTATCT |
| rRF | 18S | 5441 | 5466 | GAGAAGACGGTCGAACTTGACTATCT |
| rRF | 28S | 12366 | 12383 | TTCGATGTCGGCTCTTCC |
| rRF | 28S | 12369 | 12398 | GATGTCGGCTCTTCCTATCATTGTGAAGCA |
| rRF | 28S | 12943 | 12989 | CCCTCGCTGCGATCTATTGAAAGTCAGCCCTCGACACAAGGGTTTGT |
| isomiR | miR-21 | 14 | 34 | TAGCTTATCAGACTGATGTTGT |
| isomiR | mir-29a | 48 | 67 | TAGCACCATCTGAAATCGGTC |
| isomiR | mir-30a | 12 | 33 | TGTAAACATCCTCGACTGGAAGA |
| isomiR | mir-93 | 17 | 38 | CAAAGTGCTGTTCGTGCAGGTAA |
| isomiR | mir-181a | 30 | 53 | AACATTCAACGCTGTCGGTGAGTT |
| isomiR | mir-629 | 28 | 49 | TGGGTTTACGTTGGGAGAACTT |
| tRF | trna10_LysCTT | 1 | 34 | GCCCGGCTAGCTCAGTCGGTAGAGCATGGGACTC |
| tRF | trnaMT_SerGCT | 1 | 19 | GAGAAAGCTCACAAGAACT |
| tRF | trnaMT_SerGCT | 1 | 21 | GAGAAAGCTCACAAGAACTGC |
| tRF | trnaMT_SerGCT | 1 | 22 | GAGAAAGCTCACAAGAACTGCT |
| tRF | trnaMT_HisGTG | 32 | 70 | TGAATCTGACAACAGAGGCTTACGACCCCTTATTTACCC |
| tRF | trnaMT_HisGTG | 32 | 71 | TGAATCTGACAACAGAGGCTTACGACCCCTTATTTACCCC |
| rRF | 16S | 51 | 87 | GCTAAACCTAGCCCCAAACCCACTCCACCTTACTACC |
| rRF | 16S | 51 | 88 | GCTAAACCTAGCCCCAAACCCACTCCACCTTACTACCA |

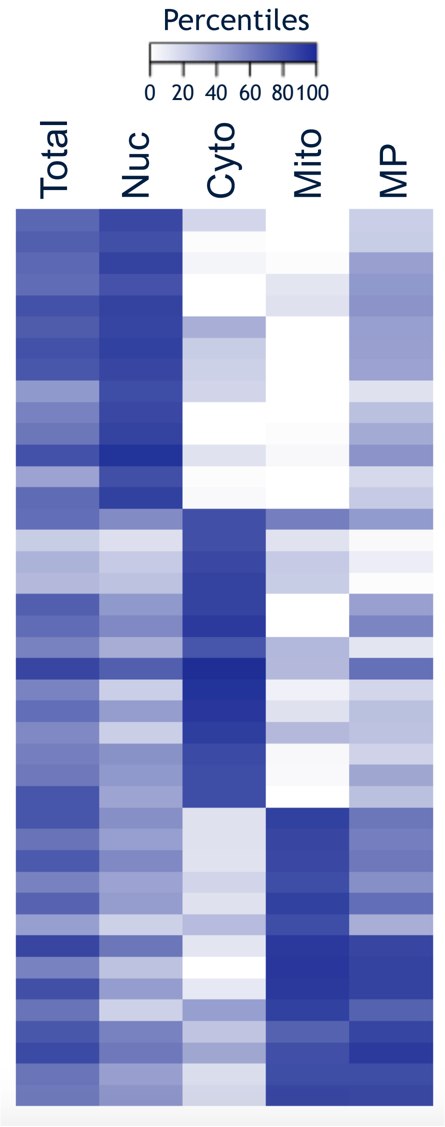

### MDA-MB-468

isomiR | tRF | rRF

| Class | RNA | Start | End | Sequence |
|---|---|---|---|---|
| rRF | 5' ETS | 425 | 455 | CTTCGTGATCGATGTGGTGACGTCGTGCTCT |
| rRF | 5' ETS | 425 | 456 | CTTCGTGATCGATGTGGTGACGTCGTGCTCTC |
| rRF | 28S | 10688 | 10705 | AAATCTCGCGCCGGGCCG |
| rRF | 28S | 11620 | 11651 | CCCAGTGCTCTGAATGTCAAAGTGAAGAAATT |
| rRF | 28S | 11668 | 11690 | AACGGCGGGAGTAACTATGACTC |
| rRF | 28S | 11731 | 11753 | CGCATGAATGGATGAACGAGATT |
| rRF | 28S | 11734 | 11753 | ATGAATGGATGAACGAGATT |
| rRF | 28S | 11735 | 11753 | TGAATGGATGAACGAGATT |
| rRF | 28S | 12555 | 12574 | TATGTGCTTGGCTGAGGAGC |
| rRF | 28S | 12604 | 12624 | TATGACTGAACGCCTCTAAGT |
| rRF | 28S | 12940 | 12963 | GCTCCCTCGCTGCGATCTATTGAA |
| rRF | 28S | 12941 | 12966 | CTCCCTCGCTGCGATCTATTGAAAGT |
| rRF | 28S | 12942 | 12963 | TCCCTCGCTGCGATCTATTGAA |
| isomiR | mir-30c | 23 | 46 | TGTAAACATCCTACACTCTCAGCT |
| tRF | trna10_ValCAC | 1 | 23 | GTTTCCGTAGTGTAGTGGTTATC |
| tRF | trna119_LysCTT | 1 | 29 | GCCCGGCTAGCTCAGTCGGTAGAGCATGA |
| tRF | trna16_GlnTTG | 1 | 29 | GGTCCCATGGTGTAATGGTTAGCACTCTG |
| tRF | trna11_GluTTC | 1 | 30 | TCCCACATGGTCTAGCGGTTAGGATTCCTG |
| tRF | trna11_GluTTC | 1 | 31 | TCCCACATGGTCTAGCGGTTAGGATTCCTGG |
| tRF | trna128_GlyGCC | 1 | 31 | GCATTGGTGGTTCAGTGGTAGAATTCTCGCC |
| rRF | 5S | 34 | 51 | CTCGTCTGATCTCGGAAG |
| rRF | 5.8S | 6672 | 6694 | ATTGCAGGACACATTGATCATCG |
| rRF | 28S | 9495 | 9516 | GTGGAGGTCCGTAGCGGTCCTG |
| rRF | 28S | 9495 | 9520 | GTGGAGGTCCGTAGCGGTCCTGACGT |
| rRF | 28S | 11668 | 11692 | AACGGCGGGAGTAACTATGACTCTC |
| rRF | 28S | 11679 | 11703 | TAACTATGACTCTCTTAAGGTAGCC |
| rRF | 28S | 12022 | 12045 | CCCGAGGGGCTCTCGCTTCTGGCG |
| rRF | 28S | 12316 | 12333 | TAACTGGCTTGTGGCGGC |
| tRF | trnaMT_SerGCT | 1 | 19 | GAGAAAGCTCACAAGAACT |
| tRF | trnaMT_SerGCT | 1 | 22 | GAGAAAGCTCACAAGAACTGCT |
| tRF | trnaMT_SerGCT | 1 | 25 | GAGAAAGCTCACAAGAACTGCTAAC |
| isomiR | mir-30a | 12 | 35 | TGTAAACATCCTCGACTGGAAGCT |
| isomiR | mir-30e | 23 | 46 | TGTAAACATCCTTGACTGGAAGCT |
| isomiR | mir-200a | 60 | 82 | TAACACTGTCTGGTAACGATGTT |
| rRF | 5.8S | 6601 | 6620 | CGACTCTTAGCGGTGGATCA |
| rRF | 5.8S | 6601 | 6638 | CGACTCTTAGCGGTGGATCACTCGGCTCGTGCGTCGAT |
| rRF | 28S | 7939 | 7956 | AGACGTGGCGACCCGCTG |
| rRF | 28S | 7939 | 7957 | AGACGTGGCGACCCGCTGA |
| rRF | 28S | 8063 | 8088 | GGCGCGGGACATGTGGCGTACGGAAG |
| rRF | 28S | 8064 | 8090 | GCGCGGGACATGTGGCGTACGGAAGAC |
| rRF | 28S | 8064 | 8091 | GCGCGGGACATGTGGCGTACGGAAGACC |
| rRF | 28S | 8065 | 8093 | CGCGGGACATGTGGCGTACGGAAGACCCG |

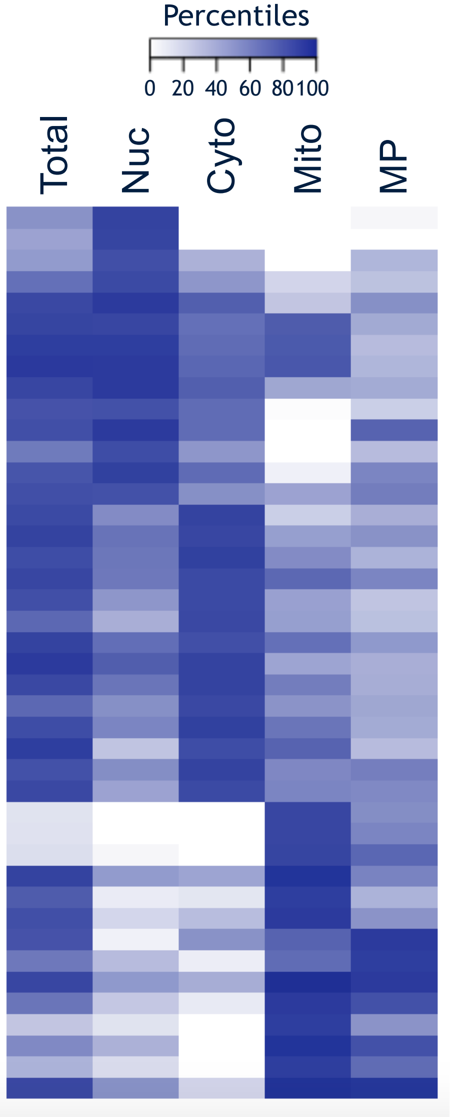
