## Supplementary material for "Single-nucleotide Differences and Cell Type Decide the Subcellular Localization of miRNA Isoforms (isomiRs), tRNA-derived Fragments (tRFs) and rRNA-derived Fragments (rRFs)": Supp. Table S1

**Supplemental Table S1:** DNA oligo sequences ordered from Fisher/Invitrogen used for the DIG northern blot probes n this study. The separation below indicates the oligos that were used to target specfic short RNA vs. RNA markers .

| **Probe Name** | **Sequence** |
|---|---|
| **Short RNA targets** | |
| miR_200c_5p | TCCATCATTACCCGGCAGTATTA |
| LysCTT tRF | GTCTCATGCTCTACCGACT |
| ETS1_1 rRF | AGAGCACGACGTCACCACAT |
| ETS1_2 rRF | GTCACCACATCGATCACGAAG |
| 28S rRF | TCGCCACGTCTGATCTGA |
| **RNA markers** | |
| 5S rRNA | AGCCTACAGCACCCGGTATTC |
| Nuclear_RNA_marker_U1 | GTGATCATGGTATCTCCCCTGCCAG |
| Mito_RNA_marker_ tRNAValUAC | GTGTTAAGCTACACTCTG |
| Cyto_RNA_marker_tRNALysCUU | GTCTCATGCTCTACCGACT |
