## Supplementary material for "Single-nucleotide Differences and Cell Type Decide the Subcellular Localization of miRNA Isoforms (isomiRs), tRNA-derived Fragments (tRFs) and rRNA-derived Fragments (rRFs)": Supp. Table S2

**Supplemental Table S2:** Data table of the average abundances (RPM) for each short RNA found in RNA-seq greater than 10 RPM in a given cell line and cell fraction. The annotation "am" is used to indicate abiguous short RNA, those that can be mapped to a noncoding RNA gene as well as somewhere else on the geneome. The annotation "multi" referrs to short RNA that map to locations in both the mitochondrial and nuclear genomes. The rRF are flanked by 50 nucleotdies upstream and downstream and their coordiantes reflect this. The coordiantes for the isomiR and rRF are from the hg38 human genome assembly while the coordintes for the tRF are from the hg19 human genome assembly.

<table>
<tr><th>Full Name</th><th>Shortened Name</th><th>Start</th><th>End</th><th>Length</th><th>Sequence</th><th>Cell Lines</th><th>Total Ave</th><th>Nuc Ave</th><th>Cyto Ave</th><th>Mito Ave</th><th>MP Ave</th><th>Total SD</th><th>Nuc SD</th><th>Cyto SD</th><th>Mito SD</th><th>MP SD</th></tr>
<tr><td>>am_MI0000748|hsa-mir-130b&WithFlank&22|+|21653298|21653391@19.39.21&[MIMAT0004680&hsa-miR-130b-5p&offsets|0|0;m-234&22|+|21653316|21653337&offsets|0|-1]|ACTCTTTCCCTGTTGCACTAC</td><td>>am_MI0000748|hsa-mir-130b&WithFlank&22|+|21653298|21653391</td><td>19</td><td>39</td><td>21</td><td>ACTCTTTCCCTGTTGCACTAC</td><td>BT20</td><td>12</td><td>6</td><td>24</td><td>0</td><td>3</td><td>16.7</td><td>2.4</td><td>13.0</td><td>0.4</td><td>4.7</td></tr>
<tr><td>>am_MI0000748|hsa-mir-130b&WithFlank&22|+|21653298|21653391@19.39.21&[MIMAT0004680&hsa-miR-130b-5p&offsets|0|0;m-234&22|+|21653316|21653337&offsets|0|-1]|ACTCTTTCCCTGTTGCACTAC</td><td>>am_MI0000748|hsa-mir-130b&WithFlank&22|+|21653298|21653391</td><td>19</td><td>39</td><td>21</td><td>ACTCTTTCCCTGTTGCACTAC</td><td>MB231</td><td>25</td><td>35</td><td>36</td><td>1</td><td>14</td><td>21.5</td><td>12.7</td><td>10.4</td><td>0.8</td><td>12.7</td></tr>
<tr><td>>am_MI0000786|hsa-mir-378a&WithFlank&5|+|149732819|149732896@49.69.21(+1A)&[MIMAT0000732&hsa-miR-378a-3p&offsets|0|-1(+1A);m-44&5|+|149732867|149732888&offsets|0|-1(+1A)]|ACTGGACTTGGAGTCAGAAGGA</td><td>>am_MI0000786|hsa-mir-378a&WithFlank&5|+|149732819|149732896</td><td>49</td><td>69</td><td>21(+1A)</td><td>ACTGGACTTGGAGTCAGAAGGA</td><td>BT20</td><td>77</td><td>53</td><td>122</td><td>257</td><td>33</td><td>27.6</td><td>23.2</td><td>25.1</td><td>214.3</td><td>6.6</td></tr>
<tr><td>>am_MI0000786|hsa-mir-378a&WithFlank&5|+|149732819|149732896@49.69.21(+1A)&[MIMAT0000732&hsa-miR-378a-3p&offsets|0|-1(+1A);m-44&5|+|149732867|149732888&offsets|0|-1(+1A)]|ACTGGACTTGGAGTCAGAAGGA</td><td>>am_MI0000786|hsa-mir-378a&WithFlank&5|+|149732819|149732896</td><td>49</td><td>69</td><td>21(+1A)</td><td>ACTGGACTTGGAGTCAGAAGGA</td><td>MB231</td><td>32</td><td>33</td><td>18</td><td>265</td><td>36</td><td>11.7</td><td>3.0</td><td>15.7</td><td>25.7</td><td>11.9</td></tr>
<tr><td>>am_MI0000786|hsa-mir-378a&WithFlank&5|+|149732819|149732896@49.69.21(+1U)&[MIMAT0000732&hsa-miR-378a-3p&offsets|0|-1(+1U);m-44&5|+|149732867|149732888&offsets|0|-1(+1U)]|ACTGGACTTGGAGTCAGAAGGT</td><td>>am_MI0000786|hsa-mir-378a&WithFlank&5|+|149732819|149732896</td><td>49</td><td>69</td><td>21(+1U)</td><td>ACTGGACTTGGAGTCAGAAGGT</td><td>BT20</td><td>230</td><td>166</td><td>318</td><td>945</td><td>106</td><td>69.2</td><td>73.8</td><td>41.2</td><td>807.1</td><td>28.7</td></tr>
<tr><td>>am_MI0000786|hsa-mir-378a&WithFlank&5|+|149732819|149732896@49.69.21(+1U)&[MIMAT0000732&hsa-miR-378a-3p&offsets|0|-1(+1U);m-44&5|+|149732867|149732888&offsets|0|-1(+1U)]|ACTGGACTTGGAGTCAGAAGGT</td><td>>am_MI0000786|hsa-mir-378a&WithFlank&5|+|149732819|149732896</td><td>49</td><td>69</td><td>21(+1U)</td><td>ACTGGACTTGGAGTCAGAAGGT</td><td>MB231</td><td>118</td><td>109</td><td>58</td><td>995</td><td>129</td><td>44.1</td><td>9.4</td><td>37.0</td><td>135.9</td><td>52.9</td></tr>
<tr><td>>am_MI0000786|hsa-mir-378a&WithFlank&5|+|149732819|149732896@49.69.21(+1U)&[MIMAT0000732&hsa-miR-378a-3p&offsets|0|-1(+1U);m-44&5|+|149732867|149732888&offsets|0|-1(+1U)]|ACTGGACTTGGAGTCAGAAGGT</td><td>>am_MI0000786|hsa-mir-378a&WithFlank&5|+|149732819|149732896</td><td>49</td><td>69</td><td>21(+1U)</td><td>ACTGGACTTGGAGTCAGAAGGT</td><td>MB468</td><td>23</td><td>20</td><td>14</td><td>179</td><td>9</td><td>4.2</td><td>2.3</td><td>4.6</td><td>91.5</td><td>2.9</td></tr>
<tr><td>>am_MI0000786|hsa-mir-378a&WithFlank&5|+|149732819|149732896@49.69.21&[MIMAT0000732&hsa-miR-378a-3p&offsets|0|-1;m-44&5|+|149732867|149732888&offsets|0|-1]|ACTGGACTTGGAGTCAGAAGG</td><td>>am_MI0000786|hsa-mir-378a&WithFlank&5|+|149732819|149732896</td><td>49</td><td>69</td><td>21</td><td>ACTGGACTTGGAGTCAGAAGG</td><td>BT20</td><td>143</td><td>115</td><td>112</td><td>369</td><td>61</td><td>51.0</td><td>53.3</td><td>35.7</td><td>297.4</td><td>21.0</td></tr>
<tr><td>>am_MI0000786|hsa-mir-378a&WithFlank&5|+|149732819|149732896@49.69.21&[MIMAT0000732&hsa-miR-378a-3p&offsets|0|-1;m-44&5|+|149732867|149732888&offsets|0|-1]|ACTGGACTTGGAGTCAGAAGG</td><td>>am_MI0000786|hsa-mir-378a&WithFlank&5|+|149732819|149732896</td><td>49</td><td>69</td><td>21</td><td>ACTGGACTTGGAGTCAGAAGG</td><td>MB231</td><td>91</td><td>109</td><td>36</td><td>726</td><td>95</td><td>26.2</td><td>19.9</td><td>28.9</td><td>397.1</td><td>28.1</td></tr>
<tr><td>>am_MI0000786|hsa-mir-378a&WithFlank&5|+|149732819|149732896@49.69.21&[MIMAT0000732&hsa-miR-378a-3p&offsets|0|-1;m-44&5|+|149732867|149732888&offsets|0|-1]|ACTGGACTTGGAGTCAGAAGG</td><td>>am_MI0000786|hsa-mir-378a&WithFlank&5|+|149732819|149732896</td><td>49</td><td>69</td><td>21</td><td>ACTGGACTTGGAGTCAGAAGG</td><td>MB468</td><td>21</td><td>13</td><td>11</td><td>189</td><td>7</td><td>6.6</td><td>2.3</td><td>3.2</td><td>178.8</td><td>1.8</td></tr>
<tr><td>>am_MI0000786|hsa-mir-378a&WithFlank&5|+|149732819|149732896@49.70.22(+1A)&[MIMAT0000732&hsa-miR-378a-3p&offsets|0|0(+1A);m-44&5|+|149732867|149732888&offsets|0|0(+1A)]|ACTGGACTTGGAGTCAGAAGGCA</td><td>>am_MI0000786|hsa-mir-378a&WithFlank&5|+|149732819|149732896</td><td>49</td><td>70</td><td>22(+1A)</td><td>ACTGGACTTGGAGTCAGAAGGCA</td><td>BT20</td><td>74</td><td>58</td><td>170</td><td>399</td><td>41</td><td>36.3</td><td>16.5</td><td>52.9</td><td>322.5</td><td>5.7</td></tr>
<tr><td>>am_MI0000786|hsa-mir-378a&WithFlank&5|+|149732819|149732896@49.70.22(+1A)&[MIMAT0000732&hsa-miR-378a-3p&offsets|0|0(+1A);m-44&5|+|149732867|149732888&offsets|0|0(+1A)]|ACTGGACTTGGAGTCAGAAGGCA</td><td>>am_MI0000786|hsa-mir-378a&WithFlank&5|+|149732819|149732896</td><td>49</td><td>70</td><td>22(+1A)</td><td>ACTGGACTTGGAGTCAGAAGGCA</td><td>MB231</td><td>37</td><td>37</td><td>29</td><td>240</td><td>30</td><td>21.7</td><td>3.6</td><td>13.0</td><td>46.8</td><td>7.8</td></tr>
<tr><td>>am_MI0000786|hsa-mir-378a&WithFlank&5|+|149732819|149732896@49.70.22(+1A)&[MIMAT0000732&hsa-miR-378a-3p&offsets|0|0(+1A);m-44&5|+|149732867|149732888&offsets|0|0(+1A)]|ACTGGACTTGGAGTCAGAAGGCA</td><td>>am_MI0000786|hsa-mir-378a&WithFlank&5|+|149732819|149732896</td><td>49</td><td>70</td><td>22(+1A)</td><td>ACTGGACTTGGAGTCAGAAGGCA</td><td>MB468</td><td>14</td><td>17</td><td>11</td><td>99</td><td>5</td><td>4.2</td><td>3.9</td><td>4.8</td><td>46.9</td><td>2.1</td></tr>
<tr><td>>am_MI0000786|hsa-mir-378a&WithFlank&5|+|149732819|149732896@49.70.22(+1G)&[MIMAT0000732&hsa-miR-378a-3p&offsets|0|0(+1G);m-44&5|+|149732867|149732888&offsets|0|0(+1G)]|ACTGGACTTGGAGTCAGAAGGCG</td><td>>am_MI0000786|hsa-mir-378a&WithFlank&5|+|149732819|149732896</td><td>49</td><td>70</td><td>22(+1G)</td><td>ACTGGACTTGGAGTCAGAAGGCG</td><td>BT20</td><td>15</td><td>9</td><td>28</td><td>114</td><td>7</td><td>3.2</td><td>1.5</td><td>15.0</td><td>90.0</td><td>0.7</td></tr>
<tr><td>>am_MI0000786|hsa-mir-378a&WithFlank&5|+|149732819|149732896@49.70.22(+1U)&[MIMAT0000732&hsa-miR-378a-3p&offsets|0|0(+1U);m-44&5|+|149732867|149732888&offsets|0|0(+1U)]|ACTGGACTTGGAGTCAGAAGGCT</td><td>>am_MI0000786|hsa-mir-378a&WithFlank&5|+|149732819|149732896</td><td>49</td><td>70</td><td>22(+1U)</td><td>ACTGGACTTGGAGTCAGAAGGCT</td><td>BT20</td><td>50</td><td>36</td><td>107</td><td>455</td><td>43</td><td>31.3</td><td>9.3</td><td>56.8</td><td>353.0</td><td>5.4</td></tr>
<tr><td>>am_MI0000786|hsa-mir-378a&WithFlank&5|+|149732819|149732896@49.70.22(+1U)&[MIMAT0000732&hsa-miR-378a-3p&offsets|0|0(+1U);m-44&5|+|149732867|149732888&offsets|0|0(+1U)]|ACTGGACTTGGAGTCAGAAGGCT</td><td>>am_MI0000786|hsa-mir-378a&WithFlank&5|+|149732819|149732896</td><td>49</td><td>70</td><td>22(+1U)</td><td>ACTGGACTTGGAGTCAGAAGGCT</td><td>MB231</td><td>66</td><td>68</td><td>39</td><td>442</td><td>50</td><td>40.5</td><td>11.1</td><td>11.3</td><td>51.1</td><td>6.5</td></tr>
<tr><td>>am_MI0000786|hsa-mir-378a&WithFlank&5|+|149732819|149732896@49.70.22(+1U)&[MIMAT0000732&hsa-miR-378a-3p&offsets|0|0(+1U);m-44&5|+|149732867|149732888&offsets|0|0(+1U)]|ACTGGACTTGGAGTCAGAAGGCT</td><td>>am_MI0000786|hsa-mir-378a&WithFlank&5|+|149732819|149732896</td><td>49</td><td>70</td><td>22(+1U)</td><td>ACTGGACTTGGAGTCAGAAGGCT</td><td>MB468</td><td>13</td><td>21</td><td>10</td><td>93</td><td>6</td><td>2.2</td><td>9.3</td><td>5.2</td><td>44.8</td><td>5.2</td></tr>
<tr><td>>am_MI0000786|hsa-mir-378a&WithFlank&5|+|149732819|149732896@49.70.22&[MIMAT0000732&hsa-miR-378a-3p&offsets|0|0;m-44&5|+|149732867|149732888&offsets|0|0]|ACTGGACTTGGAGTCAGAAGGC</td><td>>am_MI0000786|hsa-mir-378a&WithFlank&5|+|149732819|149732896</td><td>49</td><td>70</td><td>22</td><td>ACTGGACTTGGAGTCAGAAGGC</td><td>BT20</td><td>615</td><td>499</td><td>1005</td><td>3140</td><td>400</td><td>368.4</td><td>156.9</td><td>205.6</td><td>2560.7</td><td>122.0</td></tr>
<tr><td>>am_MI0000786|hsa-mir-378a&WithFlank&5|+|149732819|149732896@49.70.22&[MIMAT0000732&hsa-miR-378a-3p&offsets|0|0;m-44&5|+|149732867|149732888&offsets|0|0]|ACTGGACTTGGAGTCAGAAGGC</td><td>>am_MI0000786|hsa-mir-378a&WithFlank&5|+|149732819|149732896</td><td>49</td><td>70</td><td>22</td><td>ACTGGACTTGGAGTCAGAAGGC</td><td>MB231</td><td>457</td><td>536</td><td>264</td><td>3064</td><td>454</td><td>203.0</td><td>92.1</td><td>165.2</td><td>1093.3</td><td>78.7</td></tr>
<tr><td>>am_MI0000786|hsa-mir-378a&WithFlank&5|+|149732819|149732896@49.70.22&[MIMAT0000732&hsa-miR-378a-3p&offsets|0|0;m-44&5|+|149732867|149732888&offsets|0|0]|ACTGGACTTGGAGTCAGAAGGC</td><td>>am_MI0000786|hsa-mir-378a&WithFlank&5|+|149732819|149732896</td><td>49</td><td>70</td><td>22</td><td>ACTGGACTTGGAGTCAGAAGGC</td><td>MB468</td><td>112</td><td>126</td><td>80</td><td>807</td><td>43</td><td>41.1</td><td>31.2</td><td>28.1</td><td>392.0</td><td>19.5</td></tr>
<tr><td>>am_MI0000786|hsa-mir-378a&WithFlank&5|+|149732819|149732896@50.69.20(+1U)&[MIMAT0000732&hsa-miR-378a-3p&offsets|+1|-1(+1U);m-44&5|+|149732867|149732888&offsets|+1|-1(+1U)]|CTGGACTTGGAGTCAGAAGGT</td><td>>am_MI0000786|hsa-mir-378a&WithFlank&5|+|149732819|149732896</td><td>50</td><td>69</td><td>20(+1U)</td><td>CTGGACTTGGAGTCAGAAGGT</td><td>BT20</td><td>22</td><td>16</td><td>41</td><td>116</td><td>12</td><td>5.3</td><td>5.2</td><td>6.4</td><td>111.9</td><td>4.3</td></tr>
<tr><td>>am_MI0000786|hsa-mir-378a&WithFlank&5|+|149732819|149732896@50.69.20&[MIMAT0000732&hsa-miR-378a-3p&offsets|+1|-1;m-44&5|+|149732867|149732888&offsets|+1|-1]|CTGGACTTGGAGTCAGAAGG</td><td>>am_MI0000786|hsa-mir-378a&WithFlank&5|+|149732819|149732896</td><td>50</td><td>69</td><td>20</td><td>CTGGACTTGGAGTCAGAAGG</td><td>BT20</td><td>10</td><td>8</td><td>11</td><td>30</td><td>5</td><td>3.7</td><td>3.0</td><td>2.9</td><td>24.4</td><td>2.2</td></tr>
<tr><td>>am_MI0000786|hsa-mir-378a&WithFlank&5|+|149732819|149732896@50.70.21(+1U)&[MIMAT0000732&hsa-miR-378a-3p&offsets|+1|0(+1U);m-44&5|+|149732867|149732888&offsets|+1|0(+1U)]|CTGGACTTGGAGTCAGAAGGCT</td><td>>am_MI0000786|hsa-mir-378a&WithFlank&5|+|149732819|149732896</td><td>50</td><td>70</td><td>21(+1U)</td><td>CTGGACTTGGAGTCAGAAGGCT</td><td>BT20</td><td>21</td><td>16</td><td>48</td><td>241</td><td>20</td><td>12.8</td><td>1.2</td><td>26.1</td><td>227.2</td><td>6.9</td></tr>
<tr><td>>am_MI0000786|hsa-mir-378a&WithFlank&5|+|149732819|149732896@50.70.21(+1U)&[MIMAT0000732&hsa-miR-378a-3p&offsets|+1|0(+1U);m-44&5|+|149732867|149732888&offsets|+1|0(+1U)]|CTGGACTTGGAGTCAGAAGGCT</td><td>>am_MI0000786|hsa-mir-378a&WithFlank&5|+|149732819|149732896</td><td>50</td><td>70</td><td>21(+1U)</td><td>CTGGACTTGGAGTCAGAAGGCT</td><td>MB231</td><td>20</td><td>19</td><td>19</td><td>147</td><td>16</td><td>8.5</td><td>2.5</td><td>5.9</td><td>23.5</td><td>2.4</td></tr>
<tr><td>>am_MI0000786|hsa-mir-378a&WithFlank&5|+|149732819|149732896@50.70.21&[MIMAT0000732&hsa-miR-378a-3p&offsets|+1|0;m-44&5|+|149732867|149732888&offsets|+1|0]|CTGGACTTGGAGTCAGAAGGC</td><td>>am_MI0000786|hsa-mir-378a&WithFlank&5|+|149732819|149732896</td><td>50</td><td>70</td><td>21</td><td>CTGGACTTGGAGTCAGAAGGC</td><td>BT20</td><td>59</td><td>50</td><td>130</td><td>369</td><td>44</td><td>32.8</td><td>9.6</td><td>19.5</td><td>295.3</td><td>14.9</td></tr>
<tr><td>>am_MI0000786|hsa-mir-378a&WithFlank&5|+|149732819|149732896@50.70.21&[MIMAT0000732&hsa-miR-378a-3p&offsets|+1|0;m-44&5|+|149732867|149732888&offsets|+1|0]|CTGGACTTGGAGTCAGAAGGC</td><td>>am_MI0000786|hsa-mir-378a&WithFlank&5|+|149732819|149732896</td><td>50</td><td>70</td><td>21</td><td>CTGGACTTGGAGTCAGAAGGC</td><td>MB231</td><td>36</td><td>38</td><td>41</td><td>254</td><td>35</td><td>14.3</td><td>6.0</td><td>24.9</td><td>85.9</td><td>7.5</td></tr>
</table>

<table>
<tr><td>>am_MI0000786|hsa-mir-378a&WithFlank&5|+|149732819|149732896@50.70.21&[MIMAT0000732&hsa-miR-378a-3p&offsets|+1|0;m-44&5|+|149732867|149732888&offsets|+1|0]|CTGGACTTGGAGTCAGAAGGC</td><td>>am_MI0000786|hsa-mir-378a&WithFlank&5|+|149732819|149732896</td><td>50</td><td>70</td><td>21</td><td>CTGGACTTGGAGTCAGAAGGC</td><td>MB468</td><td>10</td><td>15</td><td>6</td><td>97</td><td>6</td><td>3.7</td><td>3.8</td><td>2.3</td><td>51.4</td><td>2.4</td></tr>
<tr><td>>am_MI0000809|hsa-mir-151a&WithFlank&8|-|140732558|140732659@17.37.21&[MIMAT0004697&hsa-miR-151a-5p&offsets|0|0;m-75&8|-|140732623|140732643&offsets|0|0]|TCGAGGAGCTCACAGTCTAGT</td><td>>am_MI0000809|hsa-mir-151a&WithFlank&8|-|140732558|140732659</td><td>17</td><td>37</td><td>21</td><td>TCGAGGAGCTCACAGTCTAGT</td><td>BT20</td><td>190</td><td>179</td><td>159</td><td>46</td><td>28</td><td>154.5</td><td>117.1</td><td>53.0</td><td>39.9</td><td>21.8</td></tr>
<tr><td>>am_MI0000809|hsa-mir-151a&WithFlank&8|-|140732558|140732659@17.37.21&[MIMAT0004697&hsa-miR-151a-5p&offsets|0|0;m-75&8|-|140732623|140732643&offsets|0|0]|TCGAGGAGCTCACAGTCTAGT</td><td>>am_MI0000809|hsa-mir-151a&WithFlank&8|-|140732558|140732659</td><td>17</td><td>37</td><td>21</td><td>TCGAGGAGCTCACAGTCTAGT</td><td>MB231</td><td>31</td><td>62</td><td>25</td><td>54</td><td>52</td><td>13.2</td><td>34.5</td><td>27.3</td><td>16.1</td><td>37.5</td></tr>
<tr><td>>am_MI0000809|hsa-mir-151a&WithFlank&8|-|140732558|140732659@17.38.22&[MIMAT0004697&hsa-miR-151a-5p&offsets|0|+1;m-75&8|-|140732623|140732643&offsets|0|+1]|TCGAGGAGCTCACAGTCTAGTA</td><td>>am_MI0000809|hsa-mir-151a&WithFlank&8|-|140732558|140732659</td><td>17</td><td>38</td><td>22</td><td>TCGAGGAGCTCACAGTCTAGTA</td><td>BT20</td><td>87</td><td>66</td><td>54</td><td>6</td><td>10</td><td>81.8</td><td>43.5</td><td>15.8</td><td>3.1</td><td>5.5</td></tr>
<tr><td>>am_MI0000809|hsa-mir-151a&WithFlank&8|-|140732558|140732659@17.38.22&[MIMAT0004697&hsa-miR-151a-5p&offsets|0|+1;m-75&8|-|140732623|140732643&offsets|0|+1]|TCGAGGAGCTCACAGTCTAGTA</td><td>>am_MI0000809|hsa-mir-151a&WithFlank&8|-|140732558|140732659</td><td>17</td><td>38</td><td>22</td><td>TCGAGGAGCTCACAGTCTAGTA</td><td>MB231</td><td>13</td><td>25</td><td>8</td><td>11</td><td>18</td><td>4.1</td><td>12.3</td><td>6.6</td><td>2.7</td><td>8.3</td></tr>
<tr><td>>am_MI0000809|hsa-mir-151a&WithFlank&8|-|140732558|140732659@52.72.21(+1A)&[MIMAT0000757&hsa-miR-151a-3p&offsets|-1|-1(+1A);m-30&8|-|140732587|140732607&offsets|-1|-1(+1A)]|ACTAGACTGAAGCTCCTTGAGA</td><td>>am_MI0000809|hsa-mir-151a&WithFlank&8|-|140732558|140732659</td><td>52</td><td>72</td><td>21(+1A)</td><td>ACTAGACTGAAGCTCCTTGAGA</td><td>BT20</td><td>14</td><td>9</td><td>25</td><td>8</td><td>7</td><td>4.3</td><td>5.2</td><td>6.1</td><td>3.6</td><td>1.5</td></tr>
<tr><td>>am_MI0000809|hsa-mir-151a&WithFlank&8|-|140732558|140732659@52.73.22&[MIMAT0000757&hsa-miR-151a-3p&offsets|-1|0;m-30&8|-|140732587|140732607&offsets|-1|0]|ACTAGACTGAAGCTCCTTGAGG</td><td>>am_MI0000809|hsa-mir-151a&WithFlank&8|-|140732558|140732659</td><td>52</td><td>73</td><td>22</td><td>ACTAGACTGAAGCTCCTTGAGG</td><td>BT20</td><td>85</td><td>64</td><td>139</td><td>69</td><td>61</td><td>51.8</td><td>33.7</td><td>33.2</td><td>19.6</td><td>8.2</td></tr>
<tr><td>>am_MI0000809|hsa-mir-151a&WithFlank&8|-|140732558|140732659@52.73.22&[MIMAT0000757&hsa-miR-151a-3p&offsets|-1|0;m-30&8|-|140732587|140732607&offsets|-1|0]|ACTAGACTGAAGCTCCTTGAGG</td><td>>am_MI0000809|hsa-mir-151a&WithFlank&8|-|140732558|140732659</td><td>52</td><td>73</td><td>22</td><td>ACTAGACTGAAGCTCCTTGAGG</td><td>MB231</td><td>54</td><td>65</td><td>36</td><td>85</td><td>50</td><td>20.6</td><td>8.4</td><td>18.1</td><td>43.2</td><td>8.8</td></tr>
<tr><td>>am_MI0000809|hsa-mir-151a&WithFlank&8|-|140732558|140732659@53.70.18(+3G)&[MIMAT0000757&hsa-miR-151a-3p&offsets|0|-3(+3G);m-30&8|-|140732587|140732607&offsets|0|-3(+3G)]|CTAGACTGAAGCTCCTTGGGG</td><td>>am_MI0000809|hsa-mir-151a&WithFlank&8|-|140732558|140732659</td><td>53</td><td>70</td><td>18(+3G)</td><td>CTAGACTGAAGCTCCTTGGGG</td><td>BT20</td><td>21</td><td>15</td><td>29</td><td>35</td><td>16</td><td>2.2</td><td>6.3</td><td>11.7</td><td>24.6</td><td>4.1</td></tr>
<tr><td>>am_MI0000809|hsa-mir-151a&WithFlank&8|-|140732558|140732659@53.70.18(+3G)&[MIMAT0000757&hsa-miR-151a-3p&offsets|0|-3(+3G);m-30&8|-|140732587|140732607&offsets|0|-3(+3G)]|CTAGACTGAAGCTCCTTGGGG</td><td>>am_MI0000809|hsa-mir-151a&WithFlank&8|-|140732558|140732659</td><td>53</td><td>70</td><td>18(+3G)</td><td>CTAGACTGAAGCTCCTTGGGG</td><td>MB231</td><td>11</td><td>10</td><td>7</td><td>16</td><td>8</td><td>2.4</td><td>2.3</td><td>3.5</td><td>2.8</td><td>1.4</td></tr>
<tr><td>>am_MI0000809|hsa-mir-151a&WithFlank&8|-|140732558|140732659@53.70.18&[MIMAT0000757&hsa-miR-151a-3p&offsets|0|-3;m-30&8|-|140732587|140732607&offsets|0|-3]|CTAGACTGAAGCTCCTTG</td><td>>am_MI0000809|hsa-mir-151a&WithFlank&8|-|140732558|140732659</td><td>53</td><td>70</td><td>18</td><td>CTAGACTGAAGCTCCTTG</td><td>BT20</td><td>87</td><td>67</td><td>94</td><td>212</td><td>78</td><td>42.2</td><td>29.4</td><td>42.8</td><td>147.7</td><td>9.1</td></tr>
<tr><td>>am_MI0000809|hsa-mir-151a&WithFlank&8|-|140732558|140732659@53.70.18&[MIMAT0000757&hsa-miR-151a-3p&offsets|0|-3;m-30&8|-|140732587|140732607&offsets|0|-3]|CTAGACTGAAGCTCCTTG</td><td>>am_MI0000809|hsa-mir-151a&WithFlank&8|-|140732558|140732659</td><td>53</td><td>70</td><td>18</td><td>CTAGACTGAAGCTCCTTG</td><td>MB231</td><td>34</td><td>40</td><td>21</td><td>112</td><td>37</td><td>10.5</td><td>17.5</td><td>7.6</td><td>20.1</td><td>11.2</td></tr>
<tr><td>>am_MI0000809|hsa-mir-151a&WithFlank&8|-|140732558|140732659@53.71.19(+1A)&[MIMAT0000757&hsa-miR-151a-3p&offsets|0|-2(+1A);m-30&8|-|140732587|140732607&offsets|0|-2(+1A)]|CTAGACTGAAGCTCCTTGAA</td><td>>am_MI0000809|hsa-mir-151a&WithFlank&8|-|140732558|140732659</td><td>53</td><td>71</td><td>19(+1A)</td><td>CTAGACTGAAGCTCCTTGAA</td><td>BT20</td><td>42</td><td>17</td><td>30</td><td>45</td><td>13</td><td>21.8</td><td>6.6</td><td>2.8</td><td>32.9</td><td>1.3</td></tr>
<tr><td>>am_MI0000809|hsa-mir-151a&WithFlank&8|-|140732558|140732659@53.71.19&[MIMAT0000757&hsa-miR-151a-3p&offsets|0|-2;m-30&8|-|140732587|140732607&offsets|0|-2]|CTAGACTGAAGCTCCTTGA</td><td>>am_MI0000809|hsa-mir-151a&WithFlank&8|-|140732558|140732659</td><td>53</td><td>71</td><td>19</td><td>CTAGACTGAAGCTCCTTGA</td><td>BT20</td><td>72</td><td>53</td><td>64</td><td>117</td><td>59</td><td>33.1</td><td>18.7</td><td>28.9</td><td>69.0</td><td>12.0</td></tr>
<tr><td>>am_MI0000809|hsa-mir-151a&WithFlank&8|-|140732558|140732659@53.71.19&[MIMAT0000757&hsa-miR-151a-3p&offsets|0|-2;m-30&8|-|140732587|140732607&offsets|0|-2]|CTAGACTGAAGCTCCTTGA</td><td>>am_MI0000809|hsa-mir-151a&WithFlank&8|-|140732558|140732659</td><td>53</td><td>71</td><td>19</td><td>CTAGACTGAAGCTCCTTGA</td><td>MB231</td><td>23</td><td>27</td><td>9</td><td>45</td><td>23</td><td>5.8</td><td>12.3</td><td>3.8</td><td>7.4</td><td>7.6</td></tr>
<tr><td>>am_MI0000809|hsa-mir-151a&WithFlank&8|-|140732558|140732659@53.72.20(+1A)&[MIMAT0000757&hsa-miR-151a-3p&offsets|0|-1(+1A);m-30&8|-|140732587|140732607&offsets|0|-1(+1A)]|CTAGACTGAAGCTCCTTGAGA</td><td>>am_MI0000809|hsa-mir-151a&WithFlank&8|-|140732558|140732659</td><td>53</td><td>72</td><td>20(+1A)</td><td>CTAGACTGAAGCTCCTTGAGA</td><td>BT20</td><td>801</td><td>495</td><td>1191</td><td>951</td><td>475</td><td>130.6</td><td>188.1</td><td>429.6</td><td>677.2</td><td>134.4</td></tr>
<tr><td>>am_MI0000809|hsa-mir-151a&WithFlank&8|-|140732558|140732659@53.72.20(+1A)&[MIMAT0000757&hsa-miR-151a-3p&offsets|0|-1(+1A);m-30&8|-|140732587|140732607&offsets|0|-1(+1A)]|CTAGACTGAAGCTCCTTGAGA</td><td>>am_MI0000809|hsa-mir-151a&WithFlank&8|-|140732558|140732659</td><td>53</td><td>72</td><td>20(+1A)</td><td>CTAGACTGAAGCTCCTTGAGA</td><td>MB231</td><td>276</td><td>227</td><td>186</td><td>557</td><td>268</td><td>77.2</td><td>7.4</td><td>86.5</td><td>41.8</td><td>64.9</td></tr>
<tr><td>>am_MI0000809|hsa-mir-151a&WithFlank&8|-|140732558|140732659@53.72.20(+1C)&[MIMAT0000757&hsa-miR-151a-3p&offsets|0|-1(+1C);m-30&8|-|140732587|140732607&offsets|0|-1(+1C)]|CTAGACTGAAGCTCCTTGAGC</td><td>>am_MI0000809|hsa-mir-151a&WithFlank&8|-|140732558|140732659</td><td>53</td><td>72</td><td>20(+1C)</td><td>CTAGACTGAAGCTCCTTGAGC</td><td>BT20</td><td>63</td><td>38</td><td>58</td><td>69</td><td>31</td><td>9.5</td><td>10.8</td><td>8.0</td><td>56.7</td><td>8.0</td></tr>
<tr><td>>am_MI0000809|hsa-mir-151a&WithFlank&8|-|140732558|140732659@53.72.20(+1C)&[MIMAT0000757&hsa-miR-151a-3p&offsets|0|-1(+1C);m-30&8|-|140732587|140732607&offsets|0|-1(+1C)]|CTAGACTGAAGCTCCTTGAGC</td><td>>am_MI0000809|hsa-mir-151a&WithFlank&8|-|140732558|140732659</td><td>53</td><td>72</td><td>20(+1C)</td><td>CTAGACTGAAGCTCCTTGAGC</td><td>MB231</td><td>30</td><td>26</td><td>11</td><td>59</td><td>30</td><td>6.9</td><td>10.7</td><td>2.9</td><td>18.7</td><td>13.2</td></tr>
<tr><td>>am_MI0000809|hsa-mir-151a&WithFlank&8|-|140732558|140732659@53.72.20(+1U)&[MIMAT0000757&hsa-miR-151a-3p&offsets|0|-1(+1U);m-30&8|-|140732587|140732607&offsets|0|-1(+1U)]|CTAGACTGAAGCTCCTTGAGT</td><td>>am_MI0000809|hsa-mir-151a&WithFlank&8|-|140732558|140732659</td><td>53</td><td>72</td><td>20(+1U)</td><td>CTAGACTGAAGCTCCTTGAGT</td><td>BT20</td><td>374</td><td>196</td><td>298</td><td>332</td><td>183</td><td>76.2</td><td>50.5</td><td>54.0</td><td>210.6</td><td>43.3</td></tr>
<tr><td>>am_MI0000809|hsa-mir-151a&WithFlank&8|-|140732558|140732659@53.72.20(+1U)&[MIMAT0000757&hsa-miR-151a-3p&offsets|0|-1(+1U);m-30&8|-|140732587|140732607&offsets|0|-1(+1U)]|CTAGACTGAAGCTCCTTGAGT</td><td>>am_MI0000809|hsa-mir-151a&WithFlank&8|-|140732558|140732659</td><td>53</td><td>72</td><td>20(+1U)</td><td>CTAGACTGAAGCTCCTTGAGT</td><td>MB231</td><td>197</td><td>178</td><td>68</td><td>302</td><td>181</td><td>51.5</td><td>69.8</td><td>25.4</td><td>79.9</td><td>75.3</td></tr>
<tr><td>>am_MI0000809|hsa-mir-151a&WithFlank&8|-|140732558|140732659@53.72.20(+2U)&[MIMAT0000757&hsa-miR-151a-3p&offsets|0|-1(+2U);m-30&8|-|140732587|140732607&offsets|0|-1(+2U)]|CTAGACTGAAGCTCCTTGAGTT</td><td>>am_MI0000809|hsa-mir-151a&WithFlank&8|-|140732558|140732659</td><td>53</td><td>72</td><td>20(+2U)</td><td>CTAGACTGAAGCTCCTTGAGTT</td><td>BT20</td><td>15</td><td>7</td><td>10</td><td>8</td><td>5</td><td>4.6</td><td>3.0</td><td>1.4</td><td>5.3</td><td>0.5</td></tr>
<tr><td>>am_MI0000809|hsa-mir-151a&WithFlank&8|-|140732558|140732659@53.72.20(+2U)&[MIMAT0000757&hsa-miR-151a-3p&offsets|0|-1(+2U);m-30&8|-|140732587|140732607&offsets|0|-1(+2U)]|CTAGACTGAAGCTCCTTGAGTT</td><td>>am_MI0000809|hsa-mir-151a&WithFlank&8|-|140732558|140732659</td><td>53</td><td>72</td><td>20(+2U)</td><td>CTAGACTGAAGCTCCTTGAGTT</td><td>MB231</td><td>16</td><td>11</td><td>5</td><td>18</td><td>12</td><td>6.0</td><td>3.2</td><td>1.8</td><td>7.3</td><td>6.9</td></tr>
<tr><td>>am_MI0000809|hsa-mir-151a&WithFlank&8|-|140732558|140732659@53.72.20&[MIMAT0000757&hsa-miR-151a-3p&offsets|0|-1;m-30&8|-|140732587|140732607&offsets|0|-1]|CTAGACTGAAGCTCCTTGAG</td><td>>am_MI0000809|hsa-mir-151a&WithFlank&8|-|140732558|140732659</td><td>53</td><td>72</td><td>20</td><td>CTAGACTGAAGCTCCTTGAG</td><td>BT20</td><td>254</td><td>149</td><td>205</td><td>429</td><td>123</td><td>53.1</td><td>50.5</td><td>53.3</td><td>278.6</td><td>6.6</td></tr>
<tr><td>>am_MI0000809|hsa-mir-151a&WithFlank&8|-|140732558|140732659@53.72.20&[MIMAT0000757&hsa-miR-151a-3p&offsets|0|-1;m-30&8|-|140732587|140732607&offsets|0|-1]|CTAGACTGAAGCTCCTTGAG</td><td>>am_MI0000809|hsa-mir-151a&WithFlank&8|-|140732558|140732659</td><td>53</td><td>72</td><td>20</td><td>CTAGACTGAAGCTCCTTGAG</td><td>MB231</td><td>85</td><td>93</td><td>34</td><td>182</td><td>74</td><td>12.9</td><td>39.5</td><td>12.8</td><td>11.5</td><td>21.1</td></tr>
<tr><td>>am_MI0000809|hsa-mir-151a&WithFlank&8|-|140732558|140732659@53.73.21(+1C)&[MIMAT0000757&hsa-miR-151a-3p&offsets|0|0(+1C);m-30&8|-|140732587|140732607&offsets|0|0(+1C)]|CTAGACTGAAGCTCCTTGAGGC</td><td>>am_MI0000809|hsa-mir-151a&WithFlank&8|-|140732558|140732659</td><td>53</td><td>73</td><td>21(+1C)</td><td>CTAGACTGAAGCTCCTTGAGGC</td><td>BT20</td><td>84</td><td>58</td><td>113</td><td>79</td><td>72</td><td>36.2</td><td>15.1</td><td>39.6</td><td>44.5</td><td>41.0</td></tr>
<tr><td>>am_MI0000809|hsa-mir-151a&WithFlank&8|-|140732558|140732659@53.73.21(+1C)&[MIMAT0000757&hsa-miR-151a-3p&offsets|0|0(+1C);m-30&8|-|140732587|140732607&offsets|0|0(+1C)]|CTAGACTGAAGCTCCTTGAGGC</td><td>>am_MI0000809|hsa-mir-151a&WithFlank&8|-|140732558|140732659</td><td>53</td><td>73</td><td>21(+1C)</td><td>CTAGACTGAAGCTCCTTGAGGC</td><td>MB231</td><td>55</td><td>55</td><td>28</td><td>106</td><td>63</td><td>13.6</td><td>16.7</td><td>6.0</td><td>33.9</td><td>29.2</td></tr>
<tr><td>>am_MI0000809|hsa-mir-151a&WithFlank&8|-|140732558|140732659@53.73.21(+1G)&[MIMAT0000757&hsa-miR-151a-3p&offsets|0|0(+1G);m-30&8|-|140732587|140732607&offsets|0|0(+1G)]|CTAGACTGAAGCTCCTTGAGGG</td><td>>am_MI0000809|hsa-mir-151a&WithFlank&8|-|140732558|140732659</td><td>53</td><td>73</td><td>21(+1G)</td><td>CTAGACTGAAGCTCCTTGAGGG</td><td>BT20</td><td>75</td><td>24</td><td>47</td><td>23</td><td>13</td><td>43.1</td><td>2.8</td><td>23.4</td><td>19.6</td><td>4.2</td></tr>
<tr><td>>am_MI0000809|hsa-mir-151a&WithFlank&8|-|140732558|140732659@53.73.21(+1G)&[MIMAT0000757&hsa-miR-151a-3p&offsets|0|0(+1G);m-30&8|-|140732587|140732607&offsets|0|0(+1G)]|CTAGACTGAAGCTCCTTGAGGG</td><td>>am_MI0000809|hsa-mir-151a&WithFlank&8|-|140732558|140732659</td><td>53</td><td>73</td><td>21(+1G)</td><td>CTAGACTGAAGCTCCTTGAGGG</td><td>MB231</td><td>36</td><td>16</td><td>7</td><td>20</td><td>10</td><td>18.4</td><td>6.1</td><td>2.4</td><td>6.5</td><td>3.7</td></tr>
<tr><td>>am_MI0000809|hsa-mir-151a&WithFlank&8|-|140732558|140732659@53.73.21(+1U)&[MIMAT0000757&hsa-miR-151a-3p&offsets|0|0(+1U);m-30&8|-|140732587|140732607&offsets|0|0(+1U)]|CTAGACTGAAGCTCCTTGAGGT</td><td>>am_MI0000809|hsa-mir-151a&WithFlank&8|-|140732558|140732659</td><td>53</td><td>73</td><td>21(+1U)</td><td>CTAGACTGAAGCTCCTTGAGGT</td><td>BT20</td><td>547</td><td>358</td><td>747</td><td>505</td><td>451</td><td>250.7</td><td>52.6</td><td>316.1</td><td>257.4</td><td>230.0</td></tr>
<tr><td>>am_MI0000809|hsa-mir-151a&WithFlank&8|-|140732558|140732659@53.73.21(+1U)&[MIMAT0000757&hsa-miR-151a-3p&offsets|0|0(+1U);m-30&8|-|140732587|140732607&offsets|0|0(+1U)]|CTAGACTGAAGCTCCTTGAGGT</td><td>>am_MI0000809|hsa-mir-151a&WithFlank&8|-|140732558|140732659</td><td>53</td><td>73</td><td>21(+1U)</td><td>CTAGACTGAAGCTCCTTGAGGT</td><td>MB231</td><td>415</td><td>433</td><td>221</td><td>601</td><td>416</td><td>92.6</td><td>127.4</td><td>57.6</td><td>125.4</td><td>132.8</td></tr>
</table>

| | | | | | | | | | | | | | | | | |
|---|---|---|---|---|---|---|---|---|---|---|---|---|---|---|---|---|
| >am_MI0000809\|hsa-mir-151a&WithFlank&8\|-\|140732558\|140732659@53.73.21(+2U)&[MIMAT0000757&hsa-miR-151a-3p&offsets\|0\|0(+2U);m-30&8\|-\|140732587\|140732607&offsets\|0\|0(+2U)]\|CTAGACTGAAGCTCCTTGAGGTT | >am_MI0000809\|hsa-mir-151a&WithFlank&8\|-\|140732558\|140732659 | 53 | 73 | 21(+2U) | CTAGACTGAAGCTCCTTGAGGTT | BT20 | 56 | 35 | 59 | 42 | 33 | 19.6 | 12.9 | 16.3 | 30.3 | 16.0 |
| >am_MI0000809\|hsa-mir-151a&WithFlank&8\|-\|140732558\|140732659@53.73.21(+2U)&[MIMAT0000757&hsa-miR-151a-3p&offsets\|0\|0(+2U);m-30&8\|-\|140732587\|140732607&offsets\|0\|0(+2U)]\|CTAGACTGAAGCTCCTTGAGGTT | >am_MI0000809\|hsa-mir-151a&WithFlank&8\|-\|140732558\|140732659 | 53 | 73 | 21(+2U) | CTAGACTGAAGCTCCTTGAGGTT | MB231 | 52 | 54 | 21 | 80 | 55 | 8.2 | 20.7 | 11.6 | 10.9 | 28.4 |
| >am_MI0000809\|hsa-mir-151a&WithFlank&8\|-\|140732558\|140732659@53.73.21(+3U)&[MIMAT0000757&hsa-miR-151a-3p&offsets\|0\|0(+3U);m-30&8\|-\|140732587\|140732607&offsets\|0\|0(+3U)]\|CTAGACTGAAGCTCCTTGAGGTTT | >am_MI0000809\|hsa-mir-151a&WithFlank&8\|-\|140732558\|140732659 | 53 | 73 | 21(+3U) | CTAGACTGAAGCTCCTTGAGGTTT | BT20 | 26 | 15 | 16 | 45 | 15 | 10.9 | 10.3 | 4.4 | 40.7 | 9.1 |
| >am_MI0000809\|hsa-mir-151a&WithFlank&8\|-\|140732558\|140732659@53.73.21(+3U)&[MIMAT0000757&hsa-miR-151a-3p&offsets\|0\|0(+3U);m-30&8\|-\|140732587\|140732607&offsets\|0\|0(+3U)]\|CTAGACTGAAGCTCCTTGAGGTTT | >am_MI0000809\|hsa-mir-151a&WithFlank&8\|-\|140732558\|140732659 | 53 | 73 | 21(+3U) | CTAGACTGAAGCTCCTTGAGGTTT | MB231 | 31 | 25 | 10 | 100 | 35 | 10.1 | 4.7 | 5.3 | 36.6 | 18.9 |
| >am_MI0000809\|hsa-mir-151a&WithFlank&8\|-\|140732558\|140732659@53.73.21&[MIMAT0000757&hsa-miR-151a-3p&offsets\|0\|0;m-30&8\|-\|140732587\|140732607&offsets\|0\|0]\|CTAGACTGAAGCTCCTTGAGG | >am_MI0000809\|hsa-mir-151a&WithFlank&8\|-\|140732558\|140732659 | 53 | 73 | 21 | CTAGACTGAAGCTCCTTGAGG | BT20 | 4538 | 3385 | 6711 | 7546 | 3732 | 1857.4 | 1343.6 | 2541.3 | 4341.8 | 895.0 |
| >am_MI0000809\|hsa-mir-151a&WithFlank&8\|-\|140732558\|140732659@53.73.21&[MIMAT0000757&hsa-miR-151a-3p&offsets\|0\|0;m-30&8\|-\|140732587\|140732607&offsets\|0\|0]\|CTAGACTGAAGCTCCTTGAGG | >am_MI0000809\|hsa-mir-151a&WithFlank&8\|-\|140732558\|140732659 | 53 | 73 | 21 | CTAGACTGAAGCTCCTTGAGG | MB231 | 2521 | 2475 | 1818 | 4114 | 2057 | 795.9 | 261.1 | 1143.0 | 1614.3 | 139.0 |
| >am_MI0000809\|hsa-mir-151a&WithFlank&8\|-\|140732558\|140732659@53.73.21&[MIMAT0000757&hsa-miR-151a-3p&offsets\|0\|0;m-30&8\|-\|140732587\|140732607&offsets\|0\|0]\|CTAGACTGAAGCTCCTTGAGG | >am_MI0000809\|hsa-mir-151a&WithFlank&8\|-\|140732558\|140732659 | 53 | 73 | 21 | CTAGACTGAAGCTCCTTGAGG | MB468 | 55 | 98 | 45 | 166 | 27 | 12.8 | 44.1 | 42.6 | 92.1 | 12.8 |
| >am_MI0000815\|hsa-mir-339&WithFlank&7\|-\|1022927\|1023032@21.41.21&[MIMAT0000764&hsa-miR-339-5p&offsets\|0\|-2;m-163&7\|-\|1022990\|1023012&offsets\|0\|-2]\|TCCCTGTCCTCCAGGAGCTCA | >am_MI0000815\|hsa-mir-339&WithFlank&7\|-\|1022927\|1023032 | 21 | 41 | 21 | TCCCTGTCCTCCAGGAGCTCA | BT20 | 17 | 12 | 15 | 14 | 2 | 19.0 | 12.1 | 1.5 | 11.0 | 1.9 |
| >am_MI0000815\|hsa-mir-339&WithFlank&7\|-\|1022927\|1023032@21.42.22(+1A)&[MIMAT0000764&hsa-miR-339-5p&offsets\|0\|-1(+1A);m-163&7\|-\|1022990\|1023012&offsets\|0\|-1(+1A)]\|TCCCTGTCCTCCAGGAGCTCACA | >am_MI0000815\|hsa-mir-339&WithFlank&7\|-\|1022927\|1023032 | 21 | 42 | 22(+1A) | TCCCTGTCCTCCAGGAGCTCACA | BT20 | 19 | 13 | 25 | 1 | 5 | 14.7 | 13.6 | 3.2 | 0.1 | 5.4 |
| >am_MI0000815\|hsa-mir-339&WithFlank&7\|-\|1022927\|1023032@21.42.22&[MIMAT0000764&hsa-miR-339-5p&offsets\|0\|-1;m-163&7\|-\|1022990\|1023012&offsets\|0\|-1]\|TCCCTGTCCTCCAGGAGCTCAC | >am_MI0000815\|hsa-mir-339&WithFlank&7\|-\|1022927\|1023032 | 21 | 42 | 22 | TCCCTGTCCTCCAGGAGCTCAC | BT20 | 15 | 14 | 17 | 3 | 5 | 10.9 | 13.6 | 7.6 | 2.6 | 4.0 |
| >am_MI0003581\|hsa-mir-574&WithFlank&4\|+\|38868026\|38868133@31.53.23&[MIMAT0004795&hsa-miR-574-5p&offsets\|0\|0;m-213&4\|+\|38868056\|38868078&offsets\|0\|0]\|TGAGTGTGTGTGTGTGAGTGTGT | >am_MI0003581\|hsa-mir-574&WithFlank&4\|+\|38868026\|38868133 | 31 | 53 | 23 | TGAGTGTGTGTGTGTGAGTGTGT | BT20 | 31 | 21 | 47 | 73 | 15 | 18.6 | 5.4 | 30.5 | 72.1 | 9.8 |
| >am_MI0003581\|hsa-mir-574&WithFlank&4\|+\|38868026\|38868133@31.53.23&[MIMAT0004795&hsa-miR-574-5p&offsets\|0\|0;m-213&4\|+\|38868056\|38868078&offsets\|0\|0]\|TGAGTGTGTGTGTGTGAGTGTGT | >am_MI0003581\|hsa-mir-574&WithFlank&4\|+\|38868026\|38868133 | 31 | 53 | 23 | TGAGTGTGTGTGTGTGAGTGTGT | MB231 | 34 | 30 | 25 | 165 | 48 | 9.0 | 4.6 | 16.7 | 29.2 | 26.4 |
| >am_MI0003581\|hsa-mir-574&WithFlank&4\|+\|38868026\|38868133@31.54.24&[MIMAT0004795&hsa-miR-574-5p&offsets\|0\|+1;m-213&4\|+\|38868056\|38868078&offsets\|0\|+1]\|TGAGTGTGTGTGTGTGAGTGTGTG | >am_MI0003581\|hsa-mir-574&WithFlank&4\|+\|38868026\|38868133 | 31 | 54 | 24 | TGAGTGTGTGTGTGTGAGTGTGTG | BT20 | 15 | 11 | 10 | 8 | 8 | 8.3 | 6.1 | 3.2 | 6.0 | 3.6 |
| >am_MI0003581\|hsa-mir-574&WithFlank&4\|+\|38868026\|38868133@31.54.24&[MIMAT0004795&hsa-miR-574-5p&offsets\|0\|+1;m-213&4\|+\|38868056\|38868078&offsets\|0\|+1]\|TGAGTGTGTGTGTGTGAGTGTGTG | >am_MI0003581\|hsa-mir-574&WithFlank&4\|+\|38868026\|38868133 | 31 | 54 | 24 | TGAGTGTGTGTGTGTGAGTGTGTG | MB231 | 26 | 18 | 7 | 56 | 30 | 15.9 | 6.3 | 2.8 | 17.7 | 11.4 |
| >am_MI0003581\|hsa-mir-574&WithFlank&4\|+\|38868026\|38868133@31.55.25&[MIMAT0004795&hsa-miR-574-5p&offsets\|0\|+2;m-213&4\|+\|38868056\|38868078&offsets\|0\|+2]\|TGAGTGTGTGTGTGTGAGTGTGTGT | >am_MI0003581\|hsa-mir-574&WithFlank&4\|+\|38868026\|38868133 | 31 | 55 | 25 | TGAGTGTGTGTGTGTGAGTGTGTGT | BT20 | 10 | 7 | 6 | 5 | 5 | 3.9 | 1.4 | 1.8 | 6.6 | 3.4 |
| >am_MI0003581\|hsa-mir-574&WithFlank&4\|+\|38868026\|38868133@31.55.25&[MIMAT0004795&hsa-miR-574-5p&offsets\|0\|+2;m-213&4\|+\|38868056\|38868078&offsets\|0\|+2]\|TGAGTGTGTGTGTGTGAGTGTGTGT | >am_MI0003581\|hsa-mir-574&WithFlank&4\|+\|38868026\|38868133 | 31 | 55 | 25 | TGAGTGTGTGTGTGTGAGTGTGTGT | MB231 | 33 | 21 | 9 | 76 | 48 | 12.4 | 1.3 | 4.2 | 20.2 | 33.2 |
| >am_MI0003583\|hsa-mir-576&WithFlank&4\|+\|109488692\|109488801@61.83.23&[MIMAT0004796&hsa-miR-576-3p&offsets\|0\|+1;m-524&4\|+\|109488752\|109488772&offsets\|0\|+2]\|AAGATGTGGAAAAATTGGAATCC | >am_MI0003583\|hsa-mir-576&WithFlank&4\|+\|109488692\|109488801 | 61 | 83 | 23 | AAGATGTGGAAAAATTGGAATCC | BT20 | 11 | 14 | 15 | 73 | 7 | 8.5 | 6.8 | 8.1 | 61.8 | 1.1 |
| >am_MI0003639\|hsa-mir-625&WithFlank&14\|+\|65471096\|65471192@58.78.21(+1U)&[MIMAT0004808&hsa-miR-625-3p&offsets\|0\|-1(+1U);m-137&14\|+\|65471153\|65471174&offsets\|0\|-1(+1U)]\|GACTATAGAACTTTCCCCCTCT | >am_MI0003639\|hsa-mir-625&WithFlank&14\|+\|65471096\|65471192 | 58 | 78 | 21(+1U) | GACTATAGAACTTTCCCCCTCT | MB231 | 11 | 15 | 14 | 22 | 11 | 5.6 | 1.5 | 4.2 | 6.2 | 1.3 |
| >am_MI0003639\|hsa-mir-625&WithFlank&14\|+\|65471096\|65471192@58.80.23&[MIMAT0004808&hsa-miR-625-3p&offsets\|0\|+1;m-137&14\|+\|65471153\|65471174&offsets\|0\|+1]\|GACTATAGAACTTTCCCCCTCAT | >am_MI0003639\|hsa-mir-625&WithFlank&14\|+\|65471096\|65471192 | 58 | 80 | 23 | GACTATAGAACTTTCCCCCTCAT | MB231 | 26 | 30 | 31 | 40 | 25 | 17.8 | 2.1 | 5.3 | 1.7 | 5.8 |
| >am_MI0003772\|hsa-mir-151b&WithFlank&14\|-\|100109413\|100109520@66.85.20(+1C);MI0000809\|hsa-mir-151a&WithFlank&8\|-\|140732558\|140732659@17.36.20(+1C)&[MIMAT0010214&hsa-miR-151b&offsets\|0\|+2(+1C);m-179&14\|-\|100109436\|100109455&offsets\|0\|0(+1C)];[MIMAT0004697&hsa-miR-151a-5p&offsets\|0\|-1(+1C);m-75&8\|-\|140732623\|140732643&offsets\|0\|-1(+1C)]\|TCGAGGAGCTCACAGTCTAGC | >am_MI0003772\|hsa-mir-151b&WithFlank&14\|-\|100109413\|100109520 | 66 | 85 | 20(+1C) | TCGAGGAGCTCACAGTCTAGC | BT20 | 24 | 24 | 20 | 6 | 3 | 18.4 | 17.3 | 8.5 | 5.9 | 2.9 |
| >am_MI0003772\|hsa-mir-151b&WithFlank&14\|-\|100109413\|100109520@66.85.20(+1G);MI0000809\|hsa-mir-151a&WithFlank&8\|-\|140732558\|140732659@17.36.20(+1G)&[MIMAT0010214&hsa-miR-151b&offsets\|0\|+2(+1G);m-179&14\|-\|100109436\|100109455&offsets\|0\|0(+1G)];[MIMAT0004697&hsa-miR-151a-5p&offsets\|0\|-1(+1G);m-75&8\|-\|140732623\|140732643&offsets\|0\|-1(+1G)]\|TCGAGGAGCTCACAGTCTAGG | >am_MI0003772\|hsa-mir-151b&WithFlank&14\|-\|100109413\|100109520 | 66 | 85 | 20(+1G) | TCGAGGAGCTCACAGTCTAGG | BT20 | 16 | 4 | 5 | 1 | 0 | 19.7 | 3.2 | 3.0 | 1.3 | 0.0 |
| >am_MI0003776\|hsa-mir-320b-1&WithFlank&1\|+\|116671740\|116671823@48.65.18;MI0003839\|hsa-mir-320b-2&WithFlank&1\|-\|224257034\|224257116@47.64.18;MI0008190\|hsa-mir-320d-1&WithFlank&13\|-\|40727810\|40727893@48.65.18;MI0003778\|hsa-mir-320c-1&WithFlank&18\|+\|21683512\|21683595@48.65.18;MI0008191\|hsa-mir-320c-2&WithFlank&18\|+\|24321669\|24321752@48.65.18;MI0000542\|hsa-mir-320a&WithFlank&8\|-\|22244960\|22245043@48.65.18;MI0008192\|hsa-mir-320d-2&WithFlank&X\|-\|140926154\|140926237@48.65.18&[MIMAT0005792&hsa-miR-320b&offsets\|0\|-4;m-183&1\|+\|116671787\|116671806&offsets\|0\|-2];[MIMAT0005792_1&hsa-miR-320b&offsets\|0\|-4;m-183&1\|-\|224257051\|224257070&offsets\|0\|-2];[MIMAT0006764&hsa-miR-320d&offsets\|0\|-1;m-246&13\|-\|40727829\|40727846&offsets\|0\|0];[MIMAT0005793&hsa-miR-320c&offsets\|0\|-2;m-222&18\|+\|21683559\|21683578&offsets\|0\|-2];[MIMAT0005793_1&hsa-miR-320c&offsets\|0\|-2;m-219&18\|+\|24321716\|24321735&offsets\|0\|-2];[MIMAT0000510&hsa-miR-320a-3p&offsets\|0\|-4;m-60&8\|-\|22244975\|22244996&offsets\|0\|-4];[MIMAT0006764_1&hsa-miR-320d&offsets\|0\|-1;m-253&X\|-\|140926173\|140926190&offsets\|0\|0]\|AAAAGCTGGGTTGAGAGG | >am_MI0003776\|hsa-mir-320b-1&WithFlank&1\|+\|116671740\|116671823 | 48 | 65 | 18 | AAAAGCTGGGTTGAGAGG | BT20 | 19 | 18 | 56 | 173 | 9 | 6.2 | 2.8 | 28.4 | 131.0 | 1.4 |

<table>
<tr><td>>am_MI0005763|hsa-mir-941-1&WithFlank&20|+|63919443|63919526@53.74.22;MI0005764|hsa-mir-941-2&WithFlank&20|+|63919499|63919582@53.74.22;MI0005765|hsa-mir-941-3&WithFlank&20|+|63919555|63919638@53.74.22;MI0005766|hsa-mir-941-4&WithFlank&20|+|63919750|63919833@53.74.22;MI0031520|hsa-mir-941-5&WithFlank&20|+|63919862|63919945@53.74.22&[MIMAT0004984&hsa-miR-941&offsets|0|-1;m-263&20|+|63919495|63919517&offsets|0|-1];[MIMAT0004984_1&hsa-miR-941&offsets|0|-1;m-263&20|+|63919551|63919573&offsets|0|-1];[MIMAT0004984_2&hsa-miR-941&offsets|0|-1;m-263&20|+|63919607|63919629&offsets|0|-1];[MIMAT0004984_3&hsa-miR-941&offsets|0|-1;m-263&20|+|63919802|63919824&offsets|0|-1];[MIMAT0004984_4&hsa-miR-941&offsets|0|-1;m-263&20|+|63919914|63919936&offsets|0|-1]|CACCCGGCTGTGTGCACATGTG</td><td>>am_MI0005763|hsa-mir-941-1&WithFlank&20|+|63919443|63919526</td><td>53</td><td>74</td><td>22</td><td>CACCCGGCTGTGTGCACATGTG</td><td>BT20</td><td>45</td><td>23</td><td>63</td><td>30</td><td>16</td><td>19.9</td><td>10.2</td><td>5.2</td><td>13.3</td><td>3.6</td></tr>
<tr><td>>am_MI0005763|hsa-mir-941-1&WithFlank&20|+|63919443|63919526@53.74.22;MI0005764|hsa-mir-941-2&WithFlank&20|+|63919499|63919582@53.74.22;MI0005765|hsa-mir-941-3&WithFlank&20|+|63919555|63919638@53.74.22;MI0005766|hsa-mir-941-4&WithFlank&20|+|63919750|63919833@53.74.22;MI0031520|hsa-mir-941-5&WithFlank&20|+|63919862|63919945@53.74.22&[MIMAT0004984&hsa-miR-941&offsets|0|-1;m-263&20|+|63919495|63919517&offsets|0|-1];[MIMAT0004984_1&hsa-miR-941&offsets|0|-1;m-263&20|+|63919551|63919573&offsets|0|-1];[MIMAT0004984_2&hsa-miR-941&offsets|0|-1;m-263&20|+|63919607|63919629&offsets|0|-1];[MIMAT0004984_3&hsa-miR-941&offsets|0|-1;m-263&20|+|63919802|63919824&offsets|0|-1];[MIMAT0004984_4&hsa-miR-941&offsets|0|-1;m-263&20|+|63919914|63919936&offsets|0|-1]|CACCCGGCTGTGTGCACATGTG</td><td>>am_MI0005763|hsa-mir-941-1&WithFlank&20|+|63919443|63919526</td><td>53</td><td>74</td><td>22</td><td>CACCCGGCTGTGTGCACATGTG</td><td>MB231</td><td>35</td><td>26</td><td>25</td><td>57</td><td>27</td><td>5.4</td><td>8.0</td><td>22.4</td><td>8.5</td><td>12.3</td></tr>
<tr><td>>am_MI0005763|hsa-mir-941-1&WithFlank&20|+|63919443|63919526@53.74.22(+1A);MI0005764|hsa-mir-941-2&WithFlank&20|+|63919499|63919582@53.74.22(+1A);MI0005765|hsa-mir-941-3&WithFlank&20|+|63919555|63919638@53.74.22(+1A);MI0005766|hsa-mir-941-4&WithFlank&20|+|63919750|63919833@53.74.22(+1A);MI0031520|hsa-mir-941-5&WithFlank&20|+|63919862|63919945@53.74.22(+1A)&[MIMAT0004984&hsa-miR-941&offsets|0|-1(+1A);m-263&20|+|63919495|63919517&offsets|0|-1(+1A)];[MIMAT0004984_1&hsa-miR-941&offsets|0|-1(+1A);m-263&20|+|63919551|63919573&offsets|0|-1(+1A)];[MIMAT0004984_2&hsa-miR-941&offsets|0|-1(+1A);m-263&20|+|63919607|63919629&offsets|0|-1(+1A)];[MIMAT0004984_3&hsa-miR-941&offsets|0|-1(+1A);m-263&20|+|63919802|63919824&offsets|0|-1(+1A)];[MIMAT0004984_4&hsa-miR-941&offsets|0|-1(+1A);m-263&20|+|63919914|63919936&offsets|0|-1(+1A)]|CACCCGGCTGTGTGCACATGTGA</td><td>>am_MI0005763|hsa-mir-941-1&WithFlank&20|+|63919443|63919526</td><td>53</td><td>74</td><td>22(+1A)</td><td>CACCCGGCTGTGTGCACATGTGA</td><td>BT20</td><td>57</td><td>24</td><td>96</td><td>27</td><td>32</td><td>15.6</td><td>11.7</td><td>25.6</td><td>10.3</td><td>12.8</td></tr>
<tr><td>>am_MI0005763|hsa-mir-941-1&WithFlank&20|+|63919443|63919526@53.74.22(+1A);MI0005764|hsa-mir-941-2&WithFlank&20|+|63919499|63919582@53.74.22(+1A);MI0005765|hsa-mir-941-3&WithFlank&20|+|63919555|63919638@53.74.22(+1A);MI0005766|hsa-mir-941-4&WithFlank&20|+|63919750|63919833@53.74.22(+1A);MI0031520|hsa-mir-941-5&WithFlank&20|+|63919862|63919945@53.74.22(+1A)&[MIMAT0004984&hsa-miR-941&offsets|0|-1(+1A);m-263&20|+|63919495|63919517&offsets|0|-1(+1A)];[MIMAT0004984_1&hsa-miR-941&offsets|0|-1(+1A);m-263&20|+|63919551|63919573&offsets|0|-1(+1A)];[MIMAT0004984_2&hsa-miR-941&offsets|0|-1(+1A);m-263&20|+|63919607|63919629&offsets|0|-1(+1A)];[MIMAT0004984_3&hsa-miR-941&offsets|0|-1(+1A);m-263&20|+|63919802|63919824&offsets|0|-1(+1A)];[MIMAT0004984_4&hsa-miR-941&offsets|0|-1(+1A);m-263&20|+|63919914|63919936&offsets|0|-1(+1A)]|CACCCGGCTGTGTGCACATGTGA</td><td>>am_MI0005763|hsa-mir-941-1&WithFlank&20|+|63919443|63919526</td><td>53</td><td>74</td><td>22(+1A)</td><td>CACCCGGCTGTGTGCACATGTGA</td><td>MB231</td><td>37</td><td>29</td><td>39</td><td>30</td><td>30</td><td>15.1</td><td>11.4</td><td>20.0</td><td>7.5</td><td>11.4</td></tr>
<tr><td>>am_MI0005763|hsa-mir-941-1&WithFlank&20|+|63919443|63919526@53.74.22(+1U);MI0005764|hsa-mir-941-2&WithFlank&20|+|63919499|63919582@53.74.22(+1U);MI0005765|hsa-mir-941-3&WithFlank&20|+|63919555|63919638@53.74.22(+1U);MI0005766|hsa-mir-941-4&WithFlank&20|+|63919750|63919833@53.74.22(+1U);MI0031520|hsa-mir-941-5&WithFlank&20|+|63919862|63919945@53.74.22(+1U)&[MIMAT0004984&hsa-miR-941&offsets|0|-1(+1U);m-263&20|+|63919495|63919517&offsets|0|-1(+1U)];[MIMAT0004984_1&hsa-miR-941&offsets|0|-1(+1U);m-263&20|+|63919551|63919573&offsets|0|-1(+1U)];[MIMAT0004984_2&hsa-miR-941&offsets|0|-1(+1U);m-263&20|+|63919607|63919629&offsets|0|-1(+1U)];[MIMAT0004984_3&hsa-miR-941&offsets|0|-1(+1U);m-263&20|+|63919802|63919824&offsets|0|-1(+1U)];[MIMAT0004984_4&hsa-miR-941&offsets|0|-1(+1U);m-263&20|+|63919914|63919936&offsets|0|-1(+1U)]|CACCCGGCTGTGTGCACATGTGT</td><td>>am_MI0005763|hsa-mir-941-1&WithFlank&20|+|63919443|63919526</td><td>53</td><td>74</td><td>22(+1U)</td><td>CACCCGGCTGTGTGCACATGTGT</td><td>BT20</td><td>211</td><td>107</td><td>300</td><td>111</td><td>141</td><td>65.7</td><td>47.0</td><td>68.4</td><td>49.6</td><td>50.3</td></tr>
<tr><td>>am_MI0005763|hsa-mir-941-1&WithFlank&20|+|63919443|63919526@53.74.22(+1U);MI0005764|hsa-mir-941-2&WithFlank&20|+|63919499|63919582@53.74.22(+1U);MI0005765|hsa-mir-941-3&WithFlank&20|+|63919555|63919638@53.74.22(+1U);MI0005766|hsa-mir-941-4&WithFlank&20|+|63919750|63919833@53.74.22(+1U);MI0031520|hsa-mir-941-5&WithFlank&20|+|63919862|63919945@53.74.22(+1U)&[MIMAT0004984&hsa-miR-941&offsets|0|-1(+1U);m-263&20|+|63919495|63919517&offsets|0|-1(+1U)];[MIMAT0004984_1&hsa-miR-941&offsets|0|-1(+1U);m-263&20|+|63919551|63919573&offsets|0|-1(+1U)];[MIMAT0004984_2&hsa-miR-941&offsets|0|-1(+1U);m-263&20|+|63919607|63919629&offsets|0|-1(+1U)];[MIMAT0004984_3&hsa-miR-941&offsets|0|-1(+1U);m-263&20|+|63919802|63919824&offsets|0|-1(+1U)];[MIMAT0004984_4&hsa-miR-941&offsets|0|-1(+1U);m-263&20|+|63919914|63919936&offsets|0|-1(+1U)]|CACCCGGCTGTGTGCACATGTGT</td><td>>am_MI0005763|hsa-mir-941-1&WithFlank&20|+|63919443|63919526</td><td>53</td><td>74</td><td>22(+1U)</td><td>CACCCGGCTGTGTGCACATGTGT</td><td>MB231</td><td>167</td><td>128</td><td>148</td><td>124</td><td>137</td><td>58.9</td><td>36.6</td><td>80.5</td><td>32.3</td><td>50.6</td></tr>
<tr><td>>am_MI0005763|hsa-mir-941-1&WithFlank&20|+|63919443|63919526@53.75.23;MI0005764|hsa-mir-941-2&WithFlank&20|+|63919499|63919582@53.75.23;MI0005765|hsa-mir-941-3&WithFlank&20|+|63919555|63919638@53.75.23;MI0005766|hsa-mir-941-4&WithFlank&20|+|63919750|63919833@53.75.23;MI0031520|hsa-mir-941-5&WithFlank&20|+|63919862|63919945@53.75.23&[MIMAT0004984&hsa-miR-941&offsets|0|0;m-263&20|+|63919495|63919517&offsets|0|0];[MIMAT0004984_1&hsa-miR-941&offsets|0|0;m-263&20|+|63919551|63919573&offsets|0|0];[MIMAT0004984_2&hsa-miR-941&offsets|0|0;m-263&20|+|63919607|63919629&offsets|0|0];[MIMAT0004984_3&hsa-miR-941&offsets|0|0;m-263&20|+|63919802|63919824&offsets|0|0];[MIMAT0004984_4&hsa-miR-941&offsets|0|0;m-263&20|+|63919914|63919936&offsets|0|0]|CACCCGGCTGTGTGCACATGTGC</td><td>>am_MI0005763|hsa-mir-941-1&WithFlank&20|+|63919443|63919526</td><td>53</td><td>75</td><td>23</td><td>CACCCGGCTGTGTGCACATGTGC</td><td>BT20</td><td>564</td><td>336</td><td>949</td><td>380</td><td>555</td><td>385.3</td><td>138.0</td><td>397.4</td><td>95.6</td><td>156.4</td></tr>
<tr><td>>am_MI0005763|hsa-mir-941-1&WithFlank&20|+|63919443|63919526@53.75.23;MI0005764|hsa-mir-941-2&WithFlank&20|+|63919499|63919582@53.75.23;MI0005765|hsa-mir-941-3&WithFlank&20|+|63919555|63919638@53.75.23;MI0005766|hsa-mir-941-4&WithFlank&20|+|63919750|63919833@53.75.23;MI0031520|hsa-mir-941-5&WithFlank&20|+|63919862|63919945@53.75.23&[MIMAT0004984&hsa-miR-941&offsets|0|0;m-263&20|+|63919495|63919517&offsets|0|0];[MIMAT0004984_1&hsa-miR-941&offsets|0|0;m-263&20|+|63919551|63919573&offsets|0|0];[MIMAT0004984_2&hsa-miR-941&offsets|0|0;m-263&20|+|63919607|63919629&offsets|0|0];[MIMAT0004984_3&hsa-miR-941&offsets|0|0;m-263&20|+|63919802|63919824&offsets|0|0];[MIMAT0004984_4&hsa-miR-941&offsets|0|0;m-263&20|+|63919914|63919936&offsets|0|0]|CACCCGGCTGTGTGCACATGTGC</td><td>>am_MI0005763|hsa-mir-941-1&WithFlank&20|+|63919443|63919526</td><td>53</td><td>75</td><td>23</td><td>CACCCGGCTGTGTGCACATGTGC</td><td>MB231</td><td>608</td><td>637</td><td>671</td><td>363</td><td>456</td><td>337.2</td><td>253.4</td><td>464.5</td><td>98.0</td><td>125.5</td></tr>
</table>

<table>
<tr><th></th><th></th><th></th><th></th><th></th><th></th><th></th><th></th><th></th><th></th><th></th><th></th><th></th><th></th><th></th><th></th><th></th></tr>
<tr><td>>am_MI0005763|hsa-mir-941-1&WithFlank&20|+|63919443|63919526@53.75.23;MI0005764|hsa-mir-941-2&WithFlank&20|+|63919499|63919582@53.75.23;MI0005765|hsa-mir-941-3&WithFlank&20|+|63919555|63919638@53.75.23;MI0005766|hsa-mir-941-4&WithFlank&20|+|63919750|63919833@53.75.23;MI0031520|hsa-mir-941-5&WithFlank&20|+|63919862|63919945@53.75.23&[MIMAT0004984&hsa-miR-941&offsets|0|0;m-263&20|+|63919495|63919517&offsets|0|0];[MIMAT0004984_1&hsa-miR-941&offsets|0|0;m-263&20|+|63919551|63919573&offsets|0|0];[MIMAT0004984_2&hsa-miR-941&offsets|0|0;m-263&20|+|63919607|63919629&offsets|0|0];[MIMAT0004984_3&hsa-miR-941&offsets|0|0;m-263&20|+|63919802|63919824&offsets|0|0];[MIMAT0004984_4&hsa-miR-941&offsets|0|0;m-263&20|+|63919914|63919936&offsets|0|0]|CACCCGGCTGTGTGCACATGTGC</td><td>>am_MI0005763|hsa-mir-941-1&WithFlank&20|+|63919443|63919526</td><td>53</td><td>75</td><td>23</td><td>CACCCGGCTGTGTGCACATGTGC</td><td>MB468</td><td>16</td><td>33</td><td>18</td><td>17</td><td>8</td><td>4.4</td><td>15.6</td><td>12.0</td><td>6.3</td><td>2.8</td></tr>
<tr><td>>am_MI0005763|hsa-mir-941-1&WithFlank&20|+|63919443|63919526@53.75.23(+1U);MI0005764|hsa-mir-941-2&WithFlank&20|+|63919499|63919582@53.75.23(+1U);MI0005765|hsa-mir-941-3&WithFlank&20|+|63919555|63919638@53.75.23(+1U);MI0005766|hsa-mir-941-4&WithFlank&20|+|63919750|63919833@53.75.23(+1U);MI0031520|hsa-mir-941-5&WithFlank&20|+|63919862|63919945@53.75.23(+1U)&[MIMAT0004984&hsa-miR-941&offsets|0|0(+1U);m-263&20|+|63919495|63919517&offsets|0|0(+1U)];[MIMAT0004984_1&hsa-miR-941&offsets|0|0(+1U);m-263&20|+|63919551|63919573&offsets|0|0(+1U)];[MIMAT0004984_2&hsa-miR-941&offsets|0|0(+1U);m-263&20|+|63919607|63919629&offsets|0|0(+1U)];[MIMAT0004984_3&hsa-miR-941&offsets|0|0(+1U);m-263&20|+|63919802|63919824&offsets|0|0(+1U)];[MIMAT0004984_4&hsa-miR-941&offsets|0|0(+1U);m-263&20|+|63919914|63919936&offsets|0|0(+1U)]|CACCCGGCTGTGTGCACATGTGCT</td><td>>am_MI0005763|hsa-mir-941-1&WithFlank&20|+|63919443|63919526</td><td>53</td><td>75</td><td>23(+1U)</td><td>CACCCGGCTGTGTGCACATGTGCT</td><td>BT20</td><td>24</td><td>10</td><td>26</td><td>3</td><td>17</td><td>12.4</td><td>4.5</td><td>11.4</td><td>0.3</td><td>10.3</td></tr>
<tr><td>>am_MI0005763|hsa-mir-941-1&WithFlank&20|+|63919443|63919526@53.75.23(+1U);MI0005764|hsa-mir-941-2&WithFlank&20|+|63919499|63919582@53.75.23(+1U);MI0005765|hsa-mir-941-3&WithFlank&20|+|63919555|63919638@53.75.23(+1U);MI0005766|hsa-mir-941-4&WithFlank&20|+|63919750|63919833@53.75.23(+1U);MI0031520|hsa-mir-941-5&WithFlank&20|+|63919862|63919945@53.75.23(+1U)&[MIMAT0004984&hsa-miR-941&offsets|0|0(+1U);m-263&20|+|63919495|63919517&offsets|0|0(+1U)];[MIMAT0004984_1&hsa-miR-941&offsets|0|0(+1U);m-263&20|+|63919551|63919573&offsets|0|0(+1U)];[MIMAT0004984_2&hsa-miR-941&offsets|0|0(+1U);m-263&20|+|63919607|63919629&offsets|0|0(+1U)];[MIMAT0004984_3&hsa-miR-941&offsets|0|0(+1U);m-263&20|+|63919802|63919824&offsets|0|0(+1U)];[MIMAT0004984_4&hsa-miR-941&offsets|0|0(+1U);m-263&20|+|63919914|63919936&offsets|0|0(+1U)]|CACCCGGCTGTGTGCACATGTGCT</td><td>>am_MI0005763|hsa-mir-941-1&WithFlank&20|+|63919443|63919526</td><td>53</td><td>75</td><td>23(+1U)</td><td>CACCCGGCTGTGTGCACATGTGCT</td><td>MB231</td><td>34</td><td>20</td><td>17</td><td>9</td><td>26</td><td>12.5</td><td>5.5</td><td>5.9</td><td>2.0</td><td>7.8</td></tr>
<tr><td>>am_MI0006370|hsa-mir-1303&WithFlank&5|+|154685770|154685867@57.79.23&[MIMAT0005891&hsa-miR-1303&offsets|-1|0;m-516&5|+|154685826|154685848&offsets|0|0]|TTTTAGAGACGGGGTCTTGCTCT</td><td>>am_MI0006370|hsa-mir-1303&WithFlank&5|+|154685770|154685867</td><td>57</td><td>79</td><td>23</td><td>TTTTAGAGACGGGGTCTTGCTCT</td><td>BT20</td><td>11</td><td>7</td><td>10</td><td>12</td><td>10</td><td>4.0</td><td>4.8</td><td>3.0</td><td>5.5</td><td>0.7</td></tr>
<tr><td>>am_MI0014244|hsa-mir-548x&WithFlank&21|-|18686084|18686170@52.73.22&[MIMAT0015081_1&hsa-miR-548x-3p&offsets|0|+2;m-1121&21|-|18686099|18686119&offsets|0|+1]|TAAAAACTGCAATTACTTTCAC</td><td>>am_MI0014244|hsa-mir-548x&WithFlank&21|-|18686084|18686170</td><td>52</td><td>73</td><td>22</td><td>TAAAAACTGCAATTACTTTCAC</td><td>BT20</td><td>16</td><td>7</td><td>11</td><td>2</td><td>3</td><td>10.4</td><td>5.1</td><td>11.5</td><td>1.3</td><td>2.3</td></tr>
<tr><td>>am_MI0015825|hsa-mir-378c&WithFlank&10|-|130962582|130962674@17.34.18;MI0016749|hsa-mir-378d-1&WithFlank&4|-|5923269|5923334@37.54.18;MI0000786|hsa-mir-378a&WithFlank&5|+|149732819|149732896@49.66.18;MI0003840|hsa-mir-378d-2&WithFlank&8|-|93916016|93916125@28.45.18&[MIMAT0016847&hsa-miR-378c&offsets|0|-7;m-130&10|-|130962639|130962658&offsets|0|-2];[MIMAT0018926&hsa-miR-378d&offsets|0|-2;m-228&4|-|5923281|5923298&offsets|0|0];[MIMAT0000732&hsa-miR-378a-3p&offsets|0|-4;m-44&5|+|149732867|149732888&offsets|0|-4];[MIMAT0018926_1&hsa-miR-378d&offsets|0|-2;m-227&8|-|93916081|93916098&offsets|0|0]|ACTGGACTTGGAGTCAGA</td><td>>am_MI0015825|hsa-mir-378c&WithFlank&10|-|130962582|130962674</td><td>17</td><td>34</td><td>18</td><td>ACTGGACTTGGAGTCAGA</td><td>BT20</td><td>16</td><td>12</td><td>19</td><td>82</td><td>9</td><td>12.8</td><td>8.2</td><td>9.6</td><td>79.3</td><td>2.7</td></tr>
<tr><td>>am_MI0015825|hsa-mir-378c&WithFlank&10|-|130962582|130962674@17.35.19;MI0016749|hsa-mir-378d-1&WithFlank&4|-|5923269|5923334@37.55.19;MI0000786|hsa-mir-378a&WithFlank&5|+|149732819|149732896@49.67.19;MI0003840|hsa-mir-378d-2&WithFlank&8|-|93916016|93916125@28.46.19&[MIMAT0016847&hsa-miR-378c&offsets|0|-6;m-130&10|-|130962639|130962658&offsets|0|-1];[MIMAT0018926&hsa-miR-378d&offsets|0|-1;m-228&4|-|5923281|5923298&offsets|0|+1];[MIMAT0000732&hsa-miR-378a-3p&offsets|0|-3;m-44&5|+|149732867|149732888&offsets|0|-3];[MIMAT0018926_1&hsa-miR-378d&offsets|0|-1;m-227&8|-|93916081|93916098&offsets|0|+1]|ACTGGACTTGGAGTCAGAA</td><td>>am_MI0015825|hsa-mir-378c&WithFlank&10|-|130962582|130962674</td><td>17</td><td>35</td><td>19</td><td>ACTGGACTTGGAGTCAGAA</td><td>BT20</td><td>12</td><td>12</td><td>16</td><td>85</td><td>8</td><td>8.4</td><td>6.0</td><td>8.6</td><td>70.6</td><td>1.8</td></tr>
<tr><td>>am_MI0015825|hsa-mir-378c&WithFlank&10|-|130962582|130962674@17.36.20;MI0000786|hsa-mir-378a&WithFlank&5|+|149732819|149732896@49.68.20&[MIMAT0016847&hsa-miR-378c&offsets|0|-5;m-130&10|-|130962639|130962658&offsets|0|0];[MIMAT0000732&hsa-miR-378a-3p&offsets|0|-2;m-44&5|+|149732867|149732888&offsets|0|-2]|ACTGGACTTGGAGTCAGAAG</td><td>>am_MI0015825|hsa-mir-378c&WithFlank&10|-|130962582|130962674</td><td>17</td><td>36</td><td>20</td><td>ACTGGACTTGGAGTCAGAAG</td><td>BT20</td><td>27</td><td>14</td><td>9</td><td>25</td><td>5</td><td>16.1</td><td>11.7</td><td>2.7</td><td>21.6</td><td>1.8</td></tr>
<tr><td>>am_MI0015825|hsa-mir-378c&WithFlank&10|-|130962582|130962674@17.36.20;MI0000786|hsa-mir-378a&WithFlank&5|+|149732819|149732896@49.68.20&[MIMAT0016847&hsa-miR-378c&offsets|0|-5;m-130&10|-|130962639|130962658&offsets|0|0];[MIMAT0000732&hsa-miR-378a-3p&offsets|0|-2;m-44&5|+|149732867|149732888&offsets|0|-2]|ACTGGACTTGGAGTCAGAAG</td><td>>am_MI0015825|hsa-mir-378c&WithFlank&10|-|130962582|130962674</td><td>17</td><td>36</td><td>20</td><td>ACTGGACTTGGAGTCAGAAG</td><td>MB231</td><td>13</td><td>12</td><td>3</td><td>88</td><td>16</td><td>1.9</td><td>3.4</td><td>2.1</td><td>73.5</td><td>10.0</td></tr>
<tr><td>>am_MI0015825|hsa-mir-378c&WithFlank&10|-|130962582|130962674@17.37.21&[MIMAT0016847&hsa-miR-378c&offsets|0|-4;m-130&10|-|130962639|130962658&offsets|0|+1]|ACTGGACTTGGAGTCAGAAGA</td><td>>am_MI0015825|hsa-mir-378c&WithFlank&10|-|130962582|130962674</td><td>17</td><td>37</td><td>21</td><td>ACTGGACTTGGAGTCAGAAGA</td><td>BT20</td><td>27</td><td>17</td><td>20</td><td>47</td><td>7</td><td>12.4</td><td>8.8</td><td>3.6</td><td>40.3</td><td>3.0</td></tr>
<tr><td>>am_MI0015825|hsa-mir-378c&WithFlank&10|-|130962582|130962674@17.37.21&[MIMAT0016847&hsa-miR-378c&offsets|0|-4;m-130&10|-|130962639|130962658&offsets|0|+1]|ACTGGACTTGGAGTCAGAAGA</td><td>>am_MI0015825|hsa-mir-378c&WithFlank&10|-|130962582|130962674</td><td>17</td><td>37</td><td>21</td><td>ACTGGACTTGGAGTCAGAAGA</td><td>MB231</td><td>10</td><td>10</td><td>4</td><td>100</td><td>13</td><td>2.6</td><td>1.1</td><td>4.2</td><td>20.4</td><td>6.0</td></tr>
<tr><td>>am_MI0016746|hsa-mir-548o-2&WithFlank&20|+|38516557|38516638@51.71.21(+1U);MI0006402|hsa-mir-548o&WithFlank&7|-|102405736|102405861@93.113.21(+1U)&[MIMAT0005919&hsa-miR-548o-3p&offsets|0|-1(+1U);m-431&20|+|38516607|38516628&offsets|0|-1(+1U)];[MIMAT0005919_1&hsa-miR-548o-3p&offsets|0|-1(+1U);m-430&7|-|102405748|102405769&offsets|0|-1(+1U)]|CCAAAACTGCAGTTACTTTTGT</td><td>>am_MI0016746|hsa-mir-548o-2&WithFlank&20|+|38516557|38516638</td><td>51</td><td>71</td><td>21(+1U)</td><td>CCAAAACTGCAGTTACTTTTGT</td><td>BT20</td><td>12</td><td>6</td><td>22</td><td>11</td><td>3</td><td>4.1</td><td>2.9</td><td>4.5</td><td>11.2</td><td>1.8</td></tr>
<tr><td>>am_MI0016846|hsa-mir-4485&WithFlank&11|-|10508264|10508332@34.52.19&[MIMAT0019019&hsa-miR-4485-3p&offsets|-3|-4;m-604&11|-|10508277|10508296&offsets|-3|-4]|GTTTAACGGCCGCGGTACC</td><td>>am_MI0016846|hsa-mir-4485&WithFlank&11|-|10508264|10508332</td><td>34</td><td>52</td><td>19</td><td>GTTTAACGGCCGCGGTACC</td><td>MB231</td><td>14</td><td>5</td><td>3</td><td>10</td><td>16</td><td>15.9</td><td>4.1</td><td>2.3</td><td>10.5</td><td>9.1</td></tr>
<tr><td>>am_MI0016846|hsa-mir-4485&WithFlank&11|-|10508264|10508332@35.52.18&[MIMAT0019019&hsa-miR-4485-3p&offsets|-2|-4;m-604&11|-|10508277|10508296&offsets|-2|-4]|TTTAACGGCCGCGGTACC</td><td>>am_MI0016846|hsa-mir-4485&WithFlank&11|-|10508264|10508332</td><td>35</td><td>52</td><td>18</td><td>TTTAACGGCCGCGGTACC</td><td>MB231</td><td>17</td><td>4</td><td>4</td><td>6</td><td>12</td><td>21.4</td><td>3.3</td><td>3.1</td><td>7.1</td><td>7.9</td></tr>
<tr><td>>am_MI0016846|hsa-mir-4485&WithFlank&11|-|10508264|10508332@35.53.19&[MIMAT0019019&hsa-miR-4485-3p&offsets|-2|-3;m-604&11|-|10508277|10508296&offsets|-2|-3]|TTTAACGGCCGCGGTACCC</td><td>>am_MI0016846|hsa-mir-4485&WithFlank&11|-|10508264|10508332</td><td>35</td><td>53</td><td>19</td><td>TTTAACGGCCGCGGTACCC</td><td>MB231</td><td>10</td><td>2</td><td>2</td><td>2</td><td>6</td><td>11.9</td><td>1.7</td><td>1.5</td><td>2.9</td><td>6.1</td></tr>
</table>

| | | | | | | | | | | | | | | | | |
|---|---|---|---|---|---|---|---|---|---|---|---|---|---|---|---|---|
| >am_MI0039740\|hsa-mir-12136&WithFlank&1\|-\|632609\|632691@7.32.26&[MIMAT0049032&hsa-miR-12136&offsets\|0\|+8]\|GAAAAAGTCATGGAGGCCATGGGGTT | >am_MI0039740\|hsa-mir-12136&WithFlank&1\|-\|632609\|632691 | 7 | 32 | 26 | GAAAAAGTCATGGAGGCCATGGGGTT | BT20 | 19 | 56 | 9 | 361 | 34 | 8.1 | 31.9 | 4.1 | 513.1 | 47.3 |
| >am_trna10_LysCTT_16_+_3241501_3241573@11.28.18,trna119_LysCTT_1_-_145395522_145395594@11.28.18,trna11_LysCTT_5_-_180648979_180649051@11.28.18,trna128_LysCTT_1_-_55423542_55423614@11.28.18,trna13_LysCTT_14_-_58706613_58706685@11.28.18,trna13_LysCTT_6_+_26556774_26556846@11.28.18,trna2_LysCTT_15_+_79152904_79152976@11.28.18,trna32_LysCTT_16_-_3207406_3207478@11.28.18,trna4_LysCTT_18_-_43669271_43669343@11.28.18,trna6_LysCTT_19_-_52425396_52425468@11.28.18,trna7_LysCTT_16_+_3225692_3225764@11.28.18,trna9_LysCTT_5_+_180634755_180634827@11.28.18\|CTCAGTCGGTAGAGCATG | >am_trna10_LysCTT_16_+_3241501_3241573 | 11 | 28 | 18 | CTCAGTCGGTAGAGCATG | MB468 | 53 | 24 | 60 | 37 | 8 | 29.0 | 15.1 | 47.5 | 13.5 | 2.3 |
| >am_trna10_LysCTT_16_+_3241501_3241573@31.51.21,trna119_LysCTT_1_-_145395522_145395594@31.51.21,trna11_LysCTT_5_-_180648979_180649051@31.51.21,trna13_LysCTT_16_+_3246153_3246225@31.51.21,trna13_LysCTT_6_+_26556774_26556846@31.51.21,trna30_LysCTT_16_-_3230555_3230627@31.51.21,trna5_LysCTT_16_+_3214939_3215011@31.51.21,trna7_LysCTT_16_+_3225692_3225764@31.51.21,trna8_LysCTT_1_+_39970195_39970267@31.51.21,trna9_LysCTT_5_+_180634755_180634827@31.51.21\|ACTCTTAATCTCAGGGTCGTG | >am_trna10_LysCTT_16_+_3241501_3241573 | 31 | 51 | 21 | ACTCTTAATCTCAGGGTCGTG | MB468 | 72 | 22 | 71 | 23 | 10 | 18.7 | 17.5 | 24.8 | 6.1 | 4.5 |
| >am_trna10_LysCTT_16_+_3241501_3241573@32.51.20,trna119_LysCTT_1_-_145395522_145395594@32.51.20,trna11_LysCTT_5_-_180648979_180649051@32.51.20,trna13_LysCTT_16_+_3246153_3246225@32.51.20,trna13_LysCTT_6_+_26556774_26556846@32.51.20,trna30_LysCTT_16_-_3230555_3230627@32.51.20,trna5_LysCTT_16_+_3214939_3215011@32.51.20,trna7_LysCTT_16_+_3225692_3225764@32.51.20,trna8_LysCTT_1_+_39970195_39970267@32.51.20,trna9_LysCTT_5_+_180634755_180634827@32.51.20\|CTCTTAATCTCAGGGTCGTG | >am_trna10_LysCTT_16_+_3241501_3241573 | 32 | 51 | 20 | CTCTTAATCTCAGGGTCGTG | MB468 | 20 | 10 | 11 | 12 | 4 | 9.7 | 8.9 | 6.6 | 6.9 | 1.4 |
| >am_trna10_LysCTT_16_+_3241501_3241573@34.51.18,trna119_LysCTT_1_-_145395522_145395594@34.51.18,trna11_LysCTT_5_-_180648979_180649051@34.51.18,trna13_LysCTT_16_+_3246153_3246225@34.51.18,trna13_LysCTT_6_+_26556774_26556846@34.51.18,trna30_LysCTT_16_-_3230555_3230627@34.51.18,trna32_LysCTT_16_-_3207406_3207478@34.51.18,trna5_LysCTT_16_+_3214939_3215011@34.51.18,trna7_LysCTT_16_+_3225692_3225764@34.51.18,trna8_LysCTT_1_+_39970195_39970267@34.51.18,trna9_LysCTT_5_+_180634755_180634827@34.51.18\|CTTAATCTCAGGGTCGTG | >am_trna10_LysCTT_16_+_3241501_3241573 | 34 | 51 | 18 | CTTAATCTCAGGGTCGTG | MB468 | 18 | 3 | 15 | 16 | 2 | 19.9 | 1.8 | 23.2 | 15.1 | 1.1 |
| >am_trna10_LysCTT_16_+_3241501_3241573@6.24.19,trna119_LysCTT_1_-_145395522_145395594@6.24.19,trna11_LysCTT_5_-_180648979_180649051@6.24.19,trna128_LysCTT_1_-_55423542_55423614@6.24.19,trna13_LysCTT_14_-_58706613_58706685@6.24.19,trna13_LysCTT_6_+_26556774_26556846@6.24.19,trna2_LysCTT_15_+_79152904_79152976@6.24.19,trna32_LysCTT_16_-_3207406_3207478@6.24.19,trna4_LysCTT_18_-_43669271_43669343@6.24.19,trna5_LysCTT_19_+_36066750_36066822@6.24.19,trna6_LysCTT_19_-_52425396_52425468@6.24.19,trna7_LysCTT_16_+_3225692_3225764@6.24.19,trna9_LysCTT_5_+_180634755_180634827@6.24.19\|GCTAGCTCAGTCGGTAGAG | >am_trna10_LysCTT_16_+_3241501_3241573 | 6 | 24 | 19 | GCTAGCTCAGTCGGTAGAG | MB468 | 11 | 1 | 7 | 3 | 1 | 11.6 | 0.8 | 9.3 | 1.7 | 0.2 |
| >am_trna10_ValCAC_5_-_180649395_180649467@1.19.19,trna12_ValAAC_5_-_180645270_180645342@1.19.19,trna132_ValAAC_6_-_27721179_27721251@1.19.19,trna133_ValCAC_6_-_27696327_27696399@1.19.19,trna136_ValAAC_6_-_27648885_27648957@1.19.19,trna139_ValAAC_6_-_27618707_27618779@1.19.19,trna15_ValAAC_5_-_180615416_180615488@1.19.19,trna18_ValCAC_5_-_180529253_180529325@1.19.19,trna2_ValAAC_3_+_169490018_169490090@1.19.19,trna2_ValCAC_5_+_180524070_180524142@1.19.19,trna37_ValAAC_6_+_27203288_27203360@1.19.19,trna4_ValAAC_5_+_180591154_180591226@1.19.19,trna5_ValAAC_5_+_180596610_180596682@1.19.19,trna6_ValCAC_5_+_180600650_180600722@1.19.19,trna85_ValCAC_1_-_161369490_161369562@1.19.19,trna90_ValCAC_1_-_149684088_149684161@1.19.19,trna98_ValCAC_1_-_149298555_149298627@1.19.19,trna9_ValCAC_6_+_26538282_26538354@1.19.19\|GTTTCCGTAGTGTAGTGGT | >am_trna10_ValCAC_5_-_180649395_180649467 | 1 | 19 | 19 | GTTTCCGTAGTGTAGTGGT | BT20 | 13 | 4 | 21 | 6 | 5 | 12.7 | 4.7 | 28.4 | 5.4 | 8.2 |
| >am_trna10_ValCAC_5_-_180649395_180649467@1.23.23,trna12_ValAAC_5_-_180645270_180645342@1.23.23,trna132_ValAAC_6_-_27721179_27721251@1.23.23,trna136_ValAAC_6_-_27648885_27648957@1.23.23,trna139_ValAAC_6_-_27618707_27618779@1.23.23,trna18_ValCAC_5_-_180529253_180529325@1.23.23,trna2_ValAAC_3_+_169490018_169490090@1.23.23,trna2_ValCAC_5_+_180524070_180524142@1.23.23,trna37_ValAAC_6_+_27203288_27203360@1.23.23,trna4_ValAAC_5_+_180591154_180591226@1.23.23,trna5_ValAAC_5_+_180596610_180596682@1.23.23,trna6_ValCAC_5_+_180600650_180600722@1.23.23,trna85_ValCAC_1_-_161369490_161369562@1.23.23,trna90_ValCAC_1_-_149684088_149684161@1.23.23,trna98_ValCAC_1_-_149298555_149298627@1.23.23,trna9_ValCAC_6_+_26538282_26538354@1.23.23\|GTTTCCGTAGTGTAGTGGTTATC | >am_trna10_ValCAC_5_-_180649395_180649467 | 1 | 23 | 23 | GTTTCCGTAGTGTAGTGGTTATC | BT20 | 102 | 30 | 141 | 301 | 139 | 149.5 | 18.2 | 171.3 | 342.0 | 198.5 |
| >am_trna10_ValCAC_5_-_180649395_180649467@1.23.23,trna12_ValAAC_5_-_180645270_180645342@1.23.23,trna132_ValAAC_6_-_27721179_27721251@1.23.23,trna136_ValAAC_6_-_27648885_27648957@1.23.23,trna139_ValAAC_6_-_27618707_27618779@1.23.23,trna18_ValCAC_5_-_180529253_180529325@1.23.23,trna2_ValAAC_3_+_169490018_169490090@1.23.23,trna2_ValCAC_5_+_180524070_180524142@1.23.23,trna37_ValAAC_6_+_27203288_27203360@1.23.23,trna4_ValAAC_5_+_180591154_180591226@1.23.23,trna5_ValAAC_5_+_180596610_180596682@1.23.23,trna6_ValCAC_5_+_180600650_180600722@1.23.23,trna85_ValCAC_1_-_161369490_161369562@1.23.23,trna90_ValCAC_1_-_149684088_149684161@1.23.23,trna98_ValCAC_1_-_149298555_149298627@1.23.23,trna9_ValCAC_6_+_26538282_26538354@1.23.23\|GTTTCCGTAGTGTAGTGGTTATC | >am_trna10_ValCAC_5_-_180649395_180649467 | 1 | 23 | 23 | GTTTCCGTAGTGTAGTGGTTATC | MB231 | 31 | 33 | 49 | 24 | 66 | 24.0 | 3.8 | 26.7 | 20.2 | 27.1 |
| >am_trna10_ValCAC_5_-_180649395_180649467@1.23.23,trna12_ValAAC_5_-_180645270_180645342@1.23.23,trna132_ValAAC_6_-_27721179_27721251@1.23.23,trna136_ValAAC_6_-_27648885_27648957@1.23.23,trna139_ValAAC_6_-_27618707_27618779@1.23.23,trna18_ValCAC_5_-_180529253_180529325@1.23.23,trna2_ValAAC_3_+_169490018_169490090@1.23.23,trna2_ValCAC_5_+_180524070_180524142@1.23.23,trna37_ValAAC_6_+_27203288_27203360@1.23.23,trna4_ValAAC_5_+_180591154_180591226@1.23.23,trna5_ValAAC_5_+_180596610_180596682@1.23.23,trna6_ValCAC_5_+_180600650_180600722@1.23.23,trna85_ValCAC_1_-_161369490_161369562@1.23.23,trna90_ValCAC_1_-_149684088_149684161@1.23.23,trna98_ValCAC_1_-_149298555_149298627@1.23.23,trna9_ValCAC_6_+_26538282_26538354@1.23.23\|GTTTCCGTAGTGTAGTGGTTATC | >am_trna10_ValCAC_5_-_180649395_180649467 | 1 | 23 | 23 | GTTTCCGTAGTGTAGTGGTTATC | MB468 | 153 | 49 | 229 | 21 | 29 | 24.1 | 19.1 | 81.4 | 25.1 | 10.2 |

| | | | | | | | | | | | | | | | | |
|---|---|---|---|---|---|---|---|---|---|---|---|---|---|---|---|---|
| >am_trna10_ValCAC_5_-_180649395_180649467@1.24.24,trna12_ValAAC_5_-_180645270_180645342@1.24.24,trna132_ValAAC_6_-_27721179_27721251@1.24.24,trna136_ValAAC_6_-_27648885_27648957@1.24.24,trna139_ValAAC_6_-_27618707_27618779@1.24.24,trna18_ValCAC_5_-_180529253_180529325@1.24.24,trna2_ValAAC_3_+_169490018_169490090@1.24.24,trna2_ValCAC_5_+_180524070_180524142@1.24.24,trna37_ValAAC_6_+_27203288_27203360@1.24.24,trna4_ValAAC_5_+_180591154_180591226@1.24.24,trna5_ValAAC_5_+_180596610_180596682@1.24.24,trna6_ValCAC_5_+_180600650_180600722@1.24.24,trna85_ValCAC_1_-_161369490_161369562@1.24.24,trna90_ValCAC_1_-_149684088_149684161@1.24.24,trna98_ValCAC_1_-_149298555_149298627@1.24.24,trna9_ValCAC_6_+_26538282_26538354@1.24.24\|GTTTCCGTAGTGTAGTGGTTATCA | >am_trna10_ValCAC_5_-_180649395_180649467 | 1 | 24 | 24 | GTTTCCGTAGTGTAGTGGTTATCA | MB468 | 15 | 5 | 16 | 5 | 3 | 5.9 | 4.4 | 14.1 | 5.8 | 2.4 |
| >am_trna10_ValCAC_5_-_180649395_180649467@1.29.29,trna12_ValAAC_5_-_180645270_180645342@1.29.29,trna132_ValAAC_6_-_27721179_27721251@1.29.29,trna136_ValAAC_6_-_27648885_27648957@1.29.29,trna139_ValAAC_6_-_27618707_27618779@1.29.29,trna18_ValCAC_5_-_180529253_180529325@1.29.29,trna2_ValAAC_3_+_169490018_169490090@1.29.29,trna2_ValCAC_5_+_180524070_180524142@1.29.29,trna4_ValAAC_5_+_180591154_180591226@1.29.29,trna5_ValAAC_5_+_180596610_180596682@1.29.29,trna6_ValCAC_5_+_180600650_180600722@1.29.29,trna85_ValCAC_1_-_161369490_161369562@1.29.29,trna90_ValCAC_1_-_149684088_149684161@1.29.29,trna98_ValCAC_1_-_149298555_149298627@1.29.29,trna9_ValCAC_6_+_26538282_26538354@1.29.29\|GTTTCCGTAGTGTAGTGGTTATCACGTTC | >am_trna10_ValCAC_5_-_180649395_180649467 | 1 | 29 | 29 | GTTTCCGTAGTGTAGTGGTTATCACGTTC | MB468 | 2181 | 403 | 1793 | 1284 | 433 | 1939.6 | 148.5 | 637.2 | 754.4 | 407.0 |
| >am_trna10_ValCAC_5_-_180649395_180649467@1.30.30,trna12_ValAAC_5_-_180645270_180645342@1.30.30,trna132_ValAAC_6_-_27721179_27721251@1.30.30,trna136_ValAAC_6_-_27648885_27648957@1.30.30,trna139_ValAAC_6_-_27618707_27618779@1.30.30,trna18_ValCAC_5_-_180529253_180529325@1.30.30,trna2_ValAAC_3_+_169490018_169490090@1.30.30,trna2_ValCAC_5_+_180524070_180524142@1.30.30,trna4_ValAAC_5_+_180591154_180591226@1.30.30,trna5_ValAAC_5_+_180596610_180596682@1.30.30,trna6_ValCAC_5_+_180600650_180600722@1.30.30,trna85_ValCAC_1_-_161369490_161369562@1.30.30,trna90_ValCAC_1_-_149684088_149684161@1.30.30,trna98_ValCAC_1_-_149298555_149298627@1.30.30,trna9_ValCAC_6_+_26538282_26538354@1.30.30\|GTTTCCGTAGTGTAGTGGTTATCACGTTCG | >am_trna10_ValCAC_5_-_180649395_180649467 | 1 | 30 | 30 | GTTTCCGTAGTGTAGTGGTTATCACGTTCG | MB468 | 24 | 5 | 21 | 5 | 5 | 26.7 | 1.6 | 9.4 | 1.9 | 5.2 |
| >am_trna10_ValCAC_5_-_180649395_180649467@1.31.31,trna12_ValAAC_5_-_180645270_180645342@1.31.31,trna132_ValAAC_6_-_27721179_27721251@1.31.31,trna136_ValAAC_6_-_27648885_27648957@1.31.31,trna139_ValAAC_6_-_27618707_27618779@1.31.31,trna18_ValCAC_5_-_180529253_180529325@1.31.31,trna2_ValAAC_3_+_169490018_169490090@1.31.31,trna2_ValCAC_5_+_180524070_180524142@1.31.31,trna4_ValAAC_5_+_180591154_180591226@1.31.31,trna5_ValAAC_5_+_180596610_180596682@1.31.31,trna6_ValCAC_5_+_180600650_180600722@1.31.31,trna85_ValCAC_1_-_161369490_161369562@1.31.31,trna90_ValCAC_1_-_149684088_149684161@1.31.31,trna98_ValCAC_1_-_149298555_149298627@1.31.31,trna9_ValCAC_6_+_26538282_26538354@1.31.31\|GTTTCCGTAGTGTAGTGGTTATCACGTTCGC | >am_trna10_ValCAC_5_-_180649395_180649467 | 1 | 31 | 31 | GTTTCCGTAGTGTAGTGGTTATCACGTTCGC | MB231 | 14 | 12 | 41 | 39 | 29 | 12.9 | 15.1 | 59.6 | 46.6 | 21.9 |
| >am_trna10_ValCAC_5_-_180649395_180649467@1.31.31,trna12_ValAAC_5_-_180645270_180645342@1.31.31,trna132_ValAAC_6_-_27721179_27721251@1.31.31,trna136_ValAAC_6_-_27648885_27648957@1.31.31,trna139_ValAAC_6_-_27618707_27618779@1.31.31,trna18_ValCAC_5_-_180529253_180529325@1.31.31,trna2_ValAAC_3_+_169490018_169490090@1.31.31,trna2_ValCAC_5_+_180524070_180524142@1.31.31,trna4_ValAAC_5_+_180591154_180591226@1.31.31,trna5_ValAAC_5_+_180596610_180596682@1.31.31,trna6_ValCAC_5_+_180600650_180600722@1.31.31,trna85_ValCAC_1_-_161369490_161369562@1.31.31,trna90_ValCAC_1_-_149684088_149684161@1.31.31,trna98_ValCAC_1_-_149298555_149298627@1.31.31,trna9_ValCAC_6_+_26538282_26538354@1.31.31\|GTTTCCGTAGTGTAGTGGTTATCACGTTCGC | >am_trna10_ValCAC_5_-_180649395_180649467 | 1 | 31 | 31 | GTTTCCGTAGTGTAGTGGTTATCACGTTCGC | MB468 | 50 | 29 | 53 | 31 | 15 | 40.9 | 13.7 | 12.7 | 8.1 | 14.2 |
| >am_trna10_ValCAC_5_-_180649395_180649467@1.32.32,trna12_ValAAC_5_-_180645270_180645342@1.32.32,trna132_ValAAC_6_-_27721179_27721251@1.32.32,trna136_ValAAC_6_-_27648885_27648957@1.32.32,trna139_ValAAC_6_-_27618707_27618779@1.32.32,trna18_ValCAC_5_-_180529253_180529325@1.32.32,trna2_ValAAC_3_+_169490018_169490090@1.32.32,trna2_ValCAC_5_+_180524070_180524142@1.32.32,trna4_ValAAC_5_+_180591154_180591226@1.32.32,trna5_ValAAC_5_+_180596610_180596682@1.32.32,trna6_ValCAC_5_+_180600650_180600722@1.32.32,trna85_ValCAC_1_-_161369490_161369562@1.32.32,trna90_ValCAC_1_-_149684088_149684161@1.32.32,trna98_ValCAC_1_-_149298555_149298627@1.32.32,trna9_ValCAC_6_+_26538282_26538354@1.32.32\|GTTTCCGTAGTGTAGTGGTTATCACGTTCGCC | >am_trna10_ValCAC_5_-_180649395_180649467 | 1 | 32 | 32 | GTTTCCGTAGTGTAGTGGTTATCACGTTCGCC | BT20 | 112 | 22 | 188 | 58 | 147 | 142.8 | 5.4 | 158.4 | 74.2 | 210.3 |
| >am_trna10_ValCAC_5_-_180649395_180649467@1.32.32,trna12_ValAAC_5_-_180645270_180645342@1.32.32,trna132_ValAAC_6_-_27721179_27721251@1.32.32,trna136_ValAAC_6_-_27648885_27648957@1.32.32,trna139_ValAAC_6_-_27618707_27618779@1.32.32,trna18_ValCAC_5_-_180529253_180529325@1.32.32,trna2_ValAAC_3_+_169490018_169490090@1.32.32,trna2_ValCAC_5_+_180524070_180524142@1.32.32,trna4_ValAAC_5_+_180591154_180591226@1.32.32,trna5_ValAAC_5_+_180596610_180596682@1.32.32,trna6_ValCAC_5_+_180600650_180600722@1.32.32,trna85_ValCAC_1_-_161369490_161369562@1.32.32,trna90_ValCAC_1_-_149684088_149684161@1.32.32,trna98_ValCAC_1_-_149298555_149298627@1.32.32,trna9_ValCAC_6_+_26538282_26538354@1.32.32\|GTTTCCGTAGTGTAGTGGTTATCACGTTCGCC | >am_trna10_ValCAC_5_-_180649395_180649467 | 1 | 32 | 32 | GTTTCCGTAGTGTAGTGGTTATCACGTTCGCC | MB231 | 255 | 206 | 842 | 141 | 323 | 169.1 | 238.8 | 1004.7 | 119.3 | 124.8 |

| | | | | | | | | | | | | | | | | |
|---|---|---|---|---|---|---|---|---|---|---|---|---|---|---|---|---|
| >am_trna10_ValCAC_5_-_180649395_180649467@1.33.33,trna12_ValAAC_5_-_180645270_180645342@1.33.33,trna132_ValAAC_6_-_27721179_27721251@1.33.33,trna136_ValAAC_6_-_27648885_27648957@1.33.33,trna139_ValAAC_6_-_27618707_27618779@1.33.33,trna18_ValCAC_5_-_180529253_180529325@1.33.33,trna2_ValAAC_3_+_169490018_169490090@1.33.33,trna2_ValCAC_5_+_180524070_180524142@1.33.33,trna4_ValAAC_5_+_180591154_180591226@1.33.33,trna5_ValAAC_5_+_180596610_180596682@1.33.33,trna6_ValCAC_5_+_180600650_180600722@1.33.33,trna85_ValCAC_1_-_161369490_161369562@1.33.33,trna90_ValCAC_1_-_149684088_149684161@1.33.33,trna98_ValCAC_1_-_149298555_149298627@1.33.33,trna9_ValCAC_6_+_26538282_26538354@1.33.33\|GTTTCCGTAGTGTAGTGGTTATCACGTTCGCCT | >am_trna10_ValCAC_5_-_180649395_180649467 | 1 | 33 | 33 | GTTTCCGTAGTGTAGTGGTTATCACGTTCGCCT | BT20 | 51 | 20 | 67 | 81 | 136 | 50.3 | 8.7 | 47.0 | 128.8 | 217.3 |
| >am_trna10_ValCAC_5_-_180649395_180649467@1.33.33,trna12_ValAAC_5_-_180645270_180645342@1.33.33,trna132_ValAAC_6_-_27721179_27721251@1.33.33,trna136_ValAAC_6_-_27648885_27648957@1.33.33,trna139_ValAAC_6_-_27618707_27618779@1.33.33,trna18_ValCAC_5_-_180529253_180529325@1.33.33,trna2_ValAAC_3_+_169490018_169490090@1.33.33,trna2_ValCAC_5_+_180524070_180524142@1.33.33,trna4_ValAAC_5_+_180591154_180591226@1.33.33,trna5_ValAAC_5_+_180596610_180596682@1.33.33,trna6_ValCAC_5_+_180600650_180600722@1.33.33,trna85_ValCAC_1_-_161369490_161369562@1.33.33,trna90_ValCAC_1_-_149684088_149684161@1.33.33,trna98_ValCAC_1_-_149298555_149298627@1.33.33,trna9_ValCAC_6_+_26538282_26538354@1.33.33\|GTTTCCGTAGTGTAGTGGTTATCACGTTCGCCT | >am_trna10_ValCAC_5_-_180649395_180649467 | 1 | 33 | 33 | GTTTCCGTAGTGTAGTGGTTATCACGTTCGCCT | MB231 | 93 | 90 | 362 | 99 | 121 | 78.3 | 96.5 | 358.0 | 95.8 | 81.6 |
| >am_trna10_ValCAC_5_-_180649395_180649467@1.34.34,trna18_ValCAC_5_-_180529253_180529325@1.34.34,trna2_ValCAC_5_+_180524070_180524142@1.34.34,trna6_ValCAC_5_+_180600650_180600722@1.34.34,trna85_ValCAC_1_-_161369490_161369562@1.34.34,trna90_ValCAC_1_-_149684088_149684161@1.34.34,trna98_ValCAC_1_-_149298555_149298627@1.34.34,trna9_ValCAC_6_+_26538282_26538354@1.34.34\|GTTTCCGTAGTGTAGTGGTTATCACGTTCGCCTC | >am_trna10_ValCAC_5_-_180649395_180649467 | 1 | 34 | 34 | GTTTCCGTAGTGTAGTGGTTATCACGTTCGCCTC | BT20 | 13 | 5 | 22 | 125 | 29 | 9.6 | 2.0 | 14.4 | 209.2 | 45.5 |
| >am_trna10_ValCAC_5_-_180649395_180649467@1.34.34,trna18_ValCAC_5_-_180529253_180529325@1.34.34,trna2_ValCAC_5_+_180524070_180524142@1.34.34,trna6_ValCAC_5_+_180600650_180600722@1.34.34,trna85_ValCAC_1_-_161369490_161369562@1.34.34,trna90_ValCAC_1_-_149684088_149684161@1.34.34,trna98_ValCAC_1_-_149298555_149298627@1.34.34,trna9_ValCAC_6_+_26538282_26538354@1.34.34\|GTTTCCGTAGTGTAGTGGTTATCACGTTCGCCTC | >am_trna10_ValCAC_5_-_180649395_180649467 | 1 | 34 | 34 | GTTTCCGTAGTGTAGTGGTTATCACGTTCGCCTC | MB231 | 51 | 50 | 280 | 49 | 35 | 55.1 | 63.9 | 325.7 | 47.6 | 30.2 |
| >am_trna10_ValCAC_5_-_180649395_180649467@1.36.36,trna18_ValCAC_5_-_180529253_180529325@1.36.36,trna2_ValCAC_5_+_180524070_180524142@1.36.36,trna6_ValCAC_5_+_180600650_180600722@1.36.36,trna85_ValCAC_1_-_161369490_161369562@1.36.36,trna90_ValCAC_1_-_149684088_149684161@1.36.36,trna98_ValCAC_1_-_149298555_149298627@1.36.36,trna9_ValCAC_6_+_26538282_26538354@1.36.36\|GTTTCCGTAGTGTAGTGGTTATCACGTTCGCCTCAC | >am_trna10_ValCAC_5_-_180649395_180649467 | 1 | 36 | 36 | GTTTCCGTAGTGTAGTGGTTATCACGTTCGCCTCAC | MB231 | 10 | 16 | 53 | 6 | 31 | 5.4 | 16.1 | 48.0 | 5.4 | 9.9 |
| >am_trna10_ValCAC_5_-_180649395_180649467@4.23.20,trna12_ValAAC_5_-_180645270_180645342@4.23.20,trna132_ValAAC_6_-_27721179_27721251@4.23.20,trna136_ValAAC_6_-_27648885_27648957@4.23.20,trna139_ValAAC_6_-_27618707_27618779@4.23.20,trna18_ValCAC_5_-_180529253_180529325@4.23.20,trna2_ValAAC_3_+_169490018_169490090@4.23.20,trna2_ValCAC_5_+_180524070_180524142@4.23.20,trna37_ValAAC_6_+_27203288_27203360@4.23.20,trna4_ValAAC_5_+_180591154_180591226@4.23.20,trna5_ValAAC_5_+_180596610_180596682@4.23.20,trna6_ValCAC_5_+_180600650_180600722@4.23.20,trna85_ValCAC_1_-_161369490_161369562@4.23.20,trna90_ValCAC_1_-_149684088_149684161@4.23.20,trna98_ValCAC_1_-_149298555_149298627@4.23.20,trna9_ValCAC_6_+_26538282_26538354@4.23.20\|TCCGTAGTGTAGTGGTTATC | >am_trna10_ValCAC_5_-_180649395_180649467 | 4 | 23 | 20 | TCCGTAGTGTAGTGGTTATC | BT20 | 10 | 1 | 13 | 5 | 6 | 14.3 | 1.1 | 19.5 | 5.0 | 7.1 |
| >am_trna10_ValCAC_5_-_180649395_180649467@5.23.19,trna12_ValAAC_5_-_180645270_180645342@5.23.19,trna132_ValAAC_6_-_27721179_27721251@5.23.19,trna136_ValAAC_6_-_27648885_27648957@5.23.19,trna139_ValAAC_6_-_27618707_27618779@5.23.19,trna18_ValCAC_5_-_180529253_180529325@5.23.19,trna2_ValAAC_3_+_169490018_169490090@5.23.19,trna2_ValCAC_5_+_180524070_180524142@5.23.19,trna37_ValAAC_6_+_27203288_27203360@5.23.19,trna4_ValAAC_5_+_180591154_180591226@5.23.19,trna5_ValAAC_5_+_180596610_180596682@5.23.19,trna6_ValCAC_5_+_180600650_180600722@5.23.19,trna85_ValCAC_1_-_161369490_161369562@5.23.19,trna90_ValCAC_1_-_149684088_149684161@5.23.19,trna98_ValCAC_1_-_149298555_149298627@5.23.19,trna9_ValCAC_6_+_26538282_26538354@5.23.19\|CCGTAGTGTAGTGGTTATC | >am_trna10_ValCAC_5_-_180649395_180649467 | 5 | 23 | 19 | CCGTAGTGTAGTGGTTATC | BT20 | 13 | 1 | 22 | 7 | 5 | 20.4 | 0.9 | 32.4 | 8.2 | 5.4 |
| >am_trna10_ValCAC_5_-_180649395_180649467@5.23.19,trna12_ValAAC_5_-_180645270_180645342@5.23.19,trna132_ValAAC_6_-_27721179_27721251@5.23.19,trna136_ValAAC_6_-_27648885_27648957@5.23.19,trna139_ValAAC_6_-_27618707_27618779@5.23.19,trna18_ValCAC_5_-_180529253_180529325@5.23.19,trna2_ValAAC_3_+_169490018_169490090@5.23.19,trna2_ValCAC_5_+_180524070_180524142@5.23.19,trna37_ValAAC_6_+_27203288_27203360@5.23.19,trna4_ValAAC_5_+_180591154_180591226@5.23.19,trna5_ValAAC_5_+_180596610_180596682@5.23.19,trna6_ValCAC_5_+_180600650_180600722@5.23.19,trna85_ValCAC_1_-_161369490_161369562@5.23.19,trna90_ValCAC_1_-_149684088_149684161@5.23.19,trna98_ValCAC_1_-_149298555_149298627@5.23.19,trna9_ValCAC_6_+_26538282_26538354@5.23.19\|CCGTAGTGTAGTGGTTATC | >am_trna10_ValCAC_5_-_180649395_180649467 | 5 | 23 | 19 | CCGTAGTGTAGTGGTTATC | MB468 | 10 | 1 | 21 | 2 | 2 | 3.7 | 1.0 | 10.1 | 1.9 | 1.6 |
| >am_trna100_LeuCAA_6_-_28864000_28864105@69.86.18,trna140_LeuCAA_6_-_27573417_27573524@70.87.18,trna141_LeuCAA_6_-_27570348_27570454@70.87.18,trna17_LeuCAG_16_+_57333863_57333945@69.86.18,trna26_LeuCAG_16_-_57334392_57334474@69.86.18,trna58_LeuCAA_1_+_249168054_249168159@70.87.18,trna74_LeuCAA_6_+_28908830_28908934@69.86.18\|ATCCCACTTCTGACACCA | >am_trna100_LeuCAA_6_-_28864000_28864105 | 69 | 86 | 18 | ATCCCACTTCTGACACCA | MB231 | 18 | 30 | 51 | 0 | 9 | 30.5 | 51.7 | 88.5 | 0.5 | 15.6 |
| >am_trna106_HisGTG_1_-_149155828_149155899@11.29.19,trna111_HisGTG_1_-_147774845_147774916@11.29.19,trna118_HisGTG_1_-_145396881_145396952@11.29.19,trna16_HisGTG_1_+_146544773_146544844@11.29.19,trna1_HisGTG_15_+_45493349_45493420@11.29.19,trna21_HisGTG_1_+_147753471_147753542@11.29.19,trna33_HisGTG_6_+_27125906_27125977@11.29.19,trna7_HisGTG_9_-_14433938_14434009@11.29.19,trna8_HisGTG_15_-_45492611_45492682@11.29.19,trna9_HisGTG_15_-_45490804_45490875@11.29.19\|TATAGTGGTTAGTACTCTG | >am_trna106_HisGTG_1_-_149155828_149155899 | 11 | 29 | 19 | TATAGTGGTTAGTACTCTG | MB468 | 10 | 1 | 6 | 7 | 3 | 2.4 | 0.6 | 4.6 | 2.1 | 2.0 |

| | | | | | | | | | | | | | | | | |
|---|---|---|---|---|---|---|---|---|---|---|---|---|---|---|---|---|
| >am_trna106_HisGTG_1_-_149155828_149155899@7.28.22,trna111_HisGTG_1_-_147774845_147774916@7.28.22,trna118_HisGTG_1_-_145396881_145396952@7.28.22,trna16_HisGTG_1_+_146544773_146544844@7.28.22,trna1_HisGTG_15_+_45493349_45493420@7.28.22,trna21_HisGTG_1_+_147753471_147753542@7.28.22,trna33_HisGTG_6_+_27125906_27125977@7.28.22,trna7_HisGTG_9_-_14433938_14434009@7.28.22,trna8_HisGTG_15_-_45492611_45492682@7.28.22,trna9_HisGTG_15_-_45490804_45490875@7.28.22\|ATCGTATAGTGGTTAGTACTCT | >am_trna106_HisGTG_1_-_149155828_149155899 | 7 | 28 | 22 | ATCGTATAGTGGTTAGTACTCT | MB468 | 60 | 7 | 65 | 19 | 9 | 30.6 | 4.4 | 52.2 | 4.4 | 2.3 |
| >am_trna106_HisGTG_1_-_149155828_149155899@7.29.23,trna111_HisGTG_1_-_147774845_147774916@7.29.23,trna118_HisGTG_1_-_145396881_145396952@7.29.23,trna16_HisGTG_1_+_146544773_146544844@7.29.23,trna1_HisGTG_15_+_45493349_45493420@7.29.23,trna21_HisGTG_1_+_147753471_147753542@7.29.23,trna33_HisGTG_6_+_27125906_27125977@7.29.23,trna7_HisGTG_9_-_14433938_14434009@7.29.23,trna8_HisGTG_15_-_45492611_45492682@7.29.23,trna9_HisGTG_15_-_45490804_45490875@7.29.23\|ATCGTATAGTGGTTAGTACTCTG | >am_trna106_HisGTG_1_-_149155828_149155899 | 7 | 29 | 23 | ATCGTATAGTGGTTAGTACTCTG | MB468 | 51 | 7 | 46 | 37 | 26 | 12.8 | 2.9 | 30.9 | 12.9 | 6.8 |
| >am_trna106_PheGAA_6_-_28775610_28775682@31.52.22,trna109_PheGAA_6_-_28758499_28758571@31.52.22,trna11_PheGAA_12_-_125412389_125412461@31.52.22,trna13_PheGAA_11_-_59333853_59333925@31.52.22,trna14_PheGAA_19_-_1383361_1383433@31.52.22,trna15_PheGAA_11_-_59324970_59325042@31.52.22,trna1_PheGAA_13_-_95201904_95201976@31.52.22,trna72_PheGAA_6_+_28732164_28732236@31.52.22,trna96_PheGAA_6_-_28949449_28949521@31.52.22\|ACTGAAGATCTAAAGGTCCCTG | >am_trna106_PheGAA_6_-_28775610_28775682 | 31 | 52 | 22 | ACTGAAGATCTAAAGGTCCCTG | MB468 | 23 | 17 | 15 | 9 | 4 | 11.6 | 15.3 | 7.1 | 1.3 | 1.5 |
| >am_trna11_ArgACG_3_-_45730491_45730563@39.56.18,trna138_ArgACG_6_-_27638344_27638416@39.56.18,trna156_ArgACG_6_-_27181623_27181695@39.56.18,trna36_ArgACG_6_+_27182952_27183024@39.56.18\|TCAGAAGATTCTAGGTTC | >am_trna11_ArgACG_3_-_45730491_45730563 | 39 | 56 | 18 | TCAGAAGATTCTAGGTTC | MB468 | 18 | 10 | 12 | 2 | 1 | 5.1 | 7.8 | 0.5 | 0.6 | 0.5 |
| >am_trna11_ArgACG_3_-_45730491_45730563@59.76.18,trna138_ArgACG_6_-_27638344_27638416@59.76.18,trna156_ArgACG_6_-_27181623_27181695@59.76.18,trna36_ArgACG_6_+_27182952_27183024@59.76.18,trna6_ArgACG_6_+_26328368_26328440@59.76.18,trna7_ArgACG_14_+_23398910_23398982@59.76.18,trna8_ArgACG_6_+_26537726_26537798@59.76.18\|CTCCTGGCTGGCTCGCCA | >am_trna11_ArgACG_3_-_45730491_45730563 | 59 | 76 | 18 | CTCCTGGCTGGCTCGCCA | MB468 | 10 | 9 | 11 | 2 | 8 | 4.8 | 5.6 | 5.6 | 0.6 | 1.5 |
| >am_trna11_ProAGG_16_+_3241989_3242060@1.27.27,trna12_ProAGG_6_+_26555498_26555569@1.27.27,trna12_ProTGG_11_-_75946869_75946940@1.27.27,trna14_ProTGG_5_-_180615854_180615925@1.27.27,trna22_ProAGG_14_-_21081560_21081631@1.27.27,trna23_ProAGG_14_-_21077495_21077566@1.27.27,trna28_ProTGG_16_-_3234133_3234204@1.27.27,trna29_ProAGG_16_-_3232635_3232706@1.27.27,trna2_ProAGG_7_+_128423504_128423575@1.27.27,trna30_ProCGG_6_+_27059521_27059592@1.27.27,trna37_ProCGG_17_-_8126151_8126222@1.27.27,trna3_ProTGG_16_+_3208923_3208994@1.27.27,trna52_ProCGG_1_+_167683962_167684033@1.27.27,trna65_ProAGG_1_-_167684725_167684796@1.27.27,trna6_ProCGG_16_+_3222049_3222120@1.27.27,trna6_ProTGG_14_+_21152175_21152246@1.27.27,trna8_ProTGG_16_+_3238094_3238165@1.27.27,trna9_ProAGG_11_+_75946557_75946628@1.27.27,trna9_ProAGG_16_+_3239634_3239705@1.27.27\|GGCTCGTTGGTCTAGGGGTATGATTCT | >am_trna11_ProAGG_16_+_3241989_3242060 | 1 | 27 | 27 | GGCTCGTTGGTCTAGGGGTATGATTCT | MB468 | 13 | 48 | 12 | 1 | 2 | 7.0 | 38.5 | 6.5 | 0.6 | 1.4 |
| >am_trna11_ProAGG_16_+_3241989_3242060@1.28.28,trna12_ProAGG_6_+_26555498_26555569@1.28.28,trna12_ProTGG_11_-_75946869_75946940@1.28.28,trna14_ProTGG_5_-_180615854_180615925@1.28.28,trna22_ProAGG_14_-_21081560_21081631@1.28.28,trna23_ProAGG_14_-_21077495_21077566@1.28.28,trna28_ProTGG_16_-_3234133_3234204@1.28.28,trna29_ProAGG_16_-_3232635_3232706@1.28.28,trna2_ProAGG_7_+_128423504_128423575@1.28.28,trna30_ProCGG_6_+_27059521_27059592@1.28.28,trna37_ProCGG_17_-_8126151_8126222@1.28.28,trna3_ProTGG_16_+_3208923_3208994@1.28.28,trna52_ProCGG_1_+_167683962_167684033@1.28.28,trna65_ProAGG_1_-_167684725_167684796@1.28.28,trna6_ProCGG_16_+_3222049_3222120@1.28.28,trna6_ProTGG_14_+_21152175_21152246@1.28.28,trna8_ProTGG_16_+_3238094_3238165@1.28.28,trna9_ProAGG_11_+_75946557_75946628@1.28.28,trna9_ProAGG_16_+_3239634_3239705@1.28.28\|GGCTCGTTGGTCTAGGGGTATGATTCTC | >am_trna11_ProAGG_16_+_3241989_3242060 | 1 | 28 | 28 | GGCTCGTTGGTCTAGGGGTATGATTCTC | MB468 | 76 | 189 | 249 | 10 | 23 | 32.1 | 176.5 | 125.6 | 9.2 | 16.2 |
| >am_trna11_ProAGG_16_+_3241989_3242060@1.29.29,trna12_ProAGG_6_+_26555498_26555569@1.29.29,trna12_ProTGG_11_-_75946869_75946940@1.29.29,trna14_ProTGG_5_-_180615854_180615925@1.29.29,trna22_ProAGG_14_-_21081560_21081631@1.29.29,trna23_ProAGG_14_-_21077495_21077566@1.29.29,trna28_ProTGG_16_-_3234133_3234204@1.29.29,trna29_ProAGG_16_-_3232635_3232706@1.29.29,trna2_ProAGG_7_+_128423504_128423575@1.29.29,trna30_ProCGG_6_+_27059521_27059592@1.29.29,trna37_ProCGG_17_-_8126151_8126222@1.29.29,trna3_ProTGG_16_+_3208923_3208994@1.29.29,trna52_ProCGG_1_+_167683962_167684033@1.29.29,trna65_ProAGG_1_-_167684725_167684796@1.29.29,trna6_ProCGG_16_+_3222049_3222120@1.29.29,trna6_ProTGG_14_+_21152175_21152246@1.29.29,trna8_ProTGG_16_+_3238094_3238165@1.29.29,trna9_ProAGG_11_+_75946557_75946628@1.29.29,trna9_ProAGG_16_+_3239634_3239705@1.29.29\|GGCTCGTTGGTCTAGGGGTATGATTCTCG | >am_trna11_ProAGG_16_+_3241989_3242060 | 1 | 29 | 29 | GGCTCGTTGGTCTAGGGGTATGATTCTCG | MB468 | 192 | 201 | 617 | 17 | 39 | 75.4 | 74.8 | 320.6 | 13.7 | 29.5 |
| >am_trna11_ProAGG_16_+_3241989_3242060@58.75.18,trna12_ProAGG_6_+_26555498_26555569@58.75.18,trna12_ProTGG_11_-_75946869_75946940@58.75.18,trna14_ProTGG_5_-_180615854_180615925@58.75.18,trna22_ProAGG_14_-_21081560_21081631@58.75.18,trna23_ProAGG_14_-_21077495_21077566@58.75.18,trna28_ProTGG_16_-_3234133_3234204@58.75.18,trna29_ProAGG_16_-_3232635_3232706@58.75.18,trna2_ProAGG_7_+_128423504_128423575@58.75.18,trna30_ProCGG_6_+_27059521_27059592@58.75.18,trna37_ProCGG_17_-_8126151_8126222@58.75.18,trna3_ProTGG_14_+_21101165_21101236@58.75.18,trna3_ProTGG_16_+_3208923_3208994@58.75.18,trna4_ProAGG_16_+_3210386_3210480@65.82.18,trna52_ProCGG_1_+_167683962_167684033@58.75.18,trna65_ProAGG_1_-_167684725_167684796@58.75.18,trna6_ProCGG_16_+_3222049_3222120@58.75.18,trna6_ProTGG_14_+_21152175_21152246@58.75.18,trna8_ProTGG_16_+_3238094_3238165@58.75.18,trna9_ProAGG_11_+_75946557_75946628@58.75.18,trna9_ProAGG_16_+_3239634_3239705@58.75.18\|ATCCCGGACGAGCCCCCA | >am_trna11_ProAGG_16_+_3241989_3242060 | 58 | 75 | 18 | ATCCCGGACGAGCCCCCA | MB231 | 26 | 49 | 95 | 3 | 13 | 45.7 | 85.0 | 163.8 | 5.4 | 23.1 |
| >am_trna111_HisGTG_1_-_147774845_147774916@-1G.23.24,trna118_HisGTG_1_-_145396881_145396952@-1G.23.24,trna16_HisGTG_1_+_146544773_146544844@-1G.23.24,trna1_HisGTG_15_+_45493349_45493420@-1G.23.24,trna21_HisGTG_1_+_147753471_147753542@-1G.23.24,trna33_HisGTG_6_+_27125906_27125977@-1G.23.24,trna7_HisGTG_9_-_14433938_14434009@-1G.23.24,trna8_HisGTG_15_-_45492611_45492682@-1G.23.24,trna9_HisGTG_15_-_45490804_45490875@-1G.23.24\|GGCCGTGATCGTATAGTGGTTAGT | >am_trna111_HisGTG_1_-_147774845_147774916 | -1G | 23 | 24 | GGCCGTGATCGTATAGTGGTTAGT | MB231 | 60 | 93 | 31 | 51 | 76 | 25.8 | 38.6 | 9.4 | 40.0 | 26.9 |

<table>
<tr><td>>am_trna111_HisGTG_1_-_147774845_147774916@-1G.26.27,trna118_HisGTG_1_-_145396881_145396952@-1G.26.27,trna16_HisGTG_1_+_146544773_146544844@-1G.26.27,trna1_HisGTG_15_+_45493349_45493420@-1G.26.27,trna21_HisGTG_1_+_147753471_147753542@-1G.26.27,trna33_HisGTG_6_+_27125906_27125977@-1G.26.27,trna7_HisGTG_9_-_14433938_14434009@-1G.26.27,trna8_HisGTG_15_-_45492611_45492682@-1G.26.27,trna9_HisGTG_15_-_45490804_45490875@-1G.26.27|GGCCGTGATCGTATAGTGGTTAGTACT</td><td>>am_trna111_HisGTG_1_-_147774845_147774916</td><td>-1G</td><td>26</td><td>27</td><td>GGCCGTGATCGTATAGTGGTTAGTACT</td><td>BT20</td><td>32</td><td>9</td><td>54</td><td>54</td><td>10</td><td>51.6</td><td>9.2</td><td>82.6</td><td>74.3</td><td>9.8</td></tr>
<tr><td>>am_trna111_HisGTG_1_-_147774845_147774916@-1G.26.27,trna118_HisGTG_1_-_145396881_145396952@-1G.26.27,trna16_HisGTG_1_+_146544773_146544844@-1G.26.27,trna1_HisGTG_15_+_45493349_45493420@-1G.26.27,trna21_HisGTG_1_+_147753471_147753542@-1G.26.27,trna33_HisGTG_6_+_27125906_27125977@-1G.26.27,trna7_HisGTG_9_-_14433938_14434009@-1G.26.27,trna8_HisGTG_15_-_45492611_45492682@-1G.26.27,trna9_HisGTG_15_-_45490804_45490875@-1G.26.27|GGCCGTGATCGTATAGTGGTTAGTACT</td><td>>am_trna111_HisGTG_1_-_147774845_147774916</td><td>-1G</td><td>26</td><td>27</td><td>GGCCGTGATCGTATAGTGGTTAGTACT</td><td>MB231</td><td>98</td><td>118</td><td>767</td><td>67</td><td>70</td><td>104.3</td><td>75.2</td><td>1262.3</td><td>57.5</td><td>6.2</td></tr>
<tr><td>>am_trna111_HisGTG_1_-_147774845_147774916@-1G.28.29,trna118_HisGTG_1_-_145396881_145396952@-1G.28.29,trna16_HisGTG_1_+_146544773_146544844@-1G.28.29,trna1_HisGTG_15_+_45493349_45493420@-1G.28.29,trna21_HisGTG_1_+_147753471_147753542@-1G.28.29,trna33_HisGTG_6_+_27125906_27125977@-1G.28.29,trna7_HisGTG_9_-_14433938_14434009@-1G.28.29,trna8_HisGTG_15_-_45492611_45492682@-1G.28.29,trna9_HisGTG_15_-_45490804_45490875@-1G.28.29|GGCCGTGATCGTATAGTGGTTAGTACTCT</td><td>>am_trna111_HisGTG_1_-_147774845_147774916</td><td>-1G</td><td>28</td><td>29</td><td>GGCCGTGATCGTATAGTGGTTAGTACTCT</td><td>BT20</td><td>14</td><td>5</td><td>27</td><td>37</td><td>6</td><td>21.0</td><td>4.3</td><td>37.6</td><td>28.3</td><td>4.5</td></tr>
<tr><td>>am_trna111_HisGTG_1_-_147774845_147774916@-1G.28.29,trna118_HisGTG_1_-_145396881_145396952@-1G.28.29,trna16_HisGTG_1_+_146544773_146544844@-1G.28.29,trna1_HisGTG_15_+_45493349_45493420@-1G.28.29,trna21_HisGTG_1_+_147753471_147753542@-1G.28.29,trna33_HisGTG_6_+_27125906_27125977@-1G.28.29,trna7_HisGTG_9_-_14433938_14434009@-1G.28.29,trna8_HisGTG_15_-_45492611_45492682@-1G.28.29,trna9_HisGTG_15_-_45490804_45490875@-1G.28.29|GGCCGTGATCGTATAGTGGTTAGTACTCT</td><td>>am_trna111_HisGTG_1_-_147774845_147774916</td><td>-1G</td><td>28</td><td>29</td><td>GGCCGTGATCGTATAGTGGTTAGTACTCT</td><td>MB231</td><td>36</td><td>39</td><td>166</td><td>40</td><td>25</td><td>41.3</td><td>18.7</td><td>227.5</td><td>24.0</td><td>6.0</td></tr>
<tr><td>>am_trna111_HisGTG_1_-_147774845_147774916@-1G.28.29,trna118_HisGTG_1_-_145396881_145396952@-1G.28.29,trna16_HisGTG_1_+_146544773_146544844@-1G.28.29,trna1_HisGTG_15_+_45493349_45493420@-1G.28.29,trna21_HisGTG_1_+_147753471_147753542@-1G.28.29,trna33_HisGTG_6_+_27125906_27125977@-1G.28.29,trna7_HisGTG_9_-_14433938_14434009@-1G.28.29,trna8_HisGTG_15_-_45492611_45492682@-1G.28.29,trna9_HisGTG_15_-_45490804_45490875@-1G.28.29|GGCCGTGATCGTATAGTGGTTAGTACTCT</td><td>>am_trna111_HisGTG_1_-_147774845_147774916</td><td>-1G</td><td>28</td><td>29</td><td>GGCCGTGATCGTATAGTGGTTAGTACTCT</td><td>MB468</td><td>79</td><td>161</td><td>126</td><td>11</td><td>9</td><td>14.4</td><td>102.5</td><td>45.9</td><td>4.0</td><td>3.0</td></tr>
<tr><td>>am_trna111_HisGTG_1_-_147774845_147774916@-1G.29.30,trna118_HisGTG_1_-_145396881_145396952@-1G.29.30,trna16_HisGTG_1_+_146544773_146544844@-1G.29.30,trna1_HisGTG_15_+_45493349_45493420@-1G.29.30,trna21_HisGTG_1_+_147753471_147753542@-1G.29.30,trna33_HisGTG_6_+_27125906_27125977@-1G.29.30,trna7_HisGTG_9_-_14433938_14434009@-1G.29.30,trna8_HisGTG_15_-_45492611_45492682@-1G.29.30,trna9_HisGTG_15_-_45490804_45490875@-1G.29.30|GGCCGTGATCGTATAGTGGTTAGTACTCTG</td><td>>am_trna111_HisGTG_1_-_147774845_147774916</td><td>-1G</td><td>29</td><td>30</td><td>GGCCGTGATCGTATAGTGGTTAGTACTCTG</td><td>MB468</td><td>495</td><td>1041</td><td>1010</td><td>271</td><td>184</td><td>160.1</td><td>571.4</td><td>641.8</td><td>153.1</td><td>88.8</td></tr>
<tr><td>>am_trna111_HisGTG_1_-_147774845_147774916@1.26.26,trna118_HisGTG_1_-_145396881_145396952@1.26.26,trna16_HisGTG_1_+_146544773_146544844@1.26.26,trna1_HisGTG_15_+_45493349_45493420@1.26.26,trna21_HisGTG_1_+_147753471_147753542@1.26.26,trna33_HisGTG_6_+_27125906_27125977@1.26.26,trna7_HisGTG_9_-_14433938_14434009@1.26.26,trna8_HisGTG_15_-_45492611_45492682@1.26.26,trna9_HisGTG_15_-_45490804_45490875@1.26.26|GCCGTGATCGTATAGTGGTTAGTACT</td><td>>am_trna111_HisGTG_1_-_147774845_147774916</td><td>1</td><td>26</td><td>26</td><td>GCCGTGATCGTATAGTGGTTAGTACT</td><td>BT20</td><td>20</td><td>7</td><td>43</td><td>25</td><td>4</td><td>29.1</td><td>5.3</td><td>44.0</td><td>29.3</td><td>2.4</td></tr>
<tr><td>>am_trna111_HisGTG_1_-_147774845_147774916@1.28.28,trna118_HisGTG_1_-_145396881_145396952@1.28.28,trna16_HisGTG_1_+_146544773_146544844@1.28.28,trna1_HisGTG_15_+_45493349_45493420@1.28.28,trna21_HisGTG_1_+_147753471_147753542@1.28.28,trna33_HisGTG_6_+_27125906_27125977@1.28.28,trna7_HisGTG_9_-_14433938_14434009@1.28.28,trna8_HisGTG_15_-_45492611_45492682@1.28.28,trna9_HisGTG_15_-_45490804_45490875@1.28.28|GCCGTGATCGTATAGTGGTTAGTACTCT</td><td>>am_trna111_HisGTG_1_-_147774845_147774916</td><td>1</td><td>28</td><td>28</td><td>GCCGTGATCGTATAGTGGTTAGTACTCT</td><td>MB468</td><td>57</td><td>64</td><td>102</td><td>7</td><td>5</td><td>10.9</td><td>40.6</td><td>22.0</td><td>1.6</td><td>2.1</td></tr>
<tr><td>>am_trna111_HisGTG_1_-_147774845_147774916@1.29.29,trna118_HisGTG_1_-_145396881_145396952@1.29.29,trna16_HisGTG_1_+_146544773_146544844@1.29.29,trna1_HisGTG_15_+_45493349_45493420@1.29.29,trna21_HisGTG_1_+_147753471_147753542@1.29.29,trna33_HisGTG_6_+_27125906_27125977@1.29.29,trna7_HisGTG_9_-_14433938_14434009@1.29.29,trna8_HisGTG_15_-_45492611_45492682@1.29.29,trna9_HisGTG_15_-_45490804_45490875@1.29.29|GCCGTGATCGTATAGTGGTTAGTACTCTG</td><td>>am_trna111_HisGTG_1_-_147774845_147774916</td><td>1</td><td>29</td><td>29</td><td>GCCGTGATCGTATAGTGGTTAGTACTCTG</td><td>MB468</td><td>548</td><td>670</td><td>1815</td><td>234</td><td>127</td><td>278.7</td><td>247.7</td><td>1463.2</td><td>109.8</td><td>80.4</td></tr>
<tr><td>>am_trna111_HisGTG_1_-_147774845_147774916@2.29.28,trna118_HisGTG_1_-_145396881_145396952@2.29.28,trna16_HisGTG_1_+_146544773_146544844@2.29.28,trna1_HisGTG_15_+_45493349_45493420@2.29.28,trna21_HisGTG_1_+_147753471_147753542@2.29.28,trna33_HisGTG_6_+_27125906_27125977@2.29.28,trna7_HisGTG_9_-_14433938_14434009@2.29.28,trna8_HisGTG_15_-_45492611_45492682@2.29.28,trna9_HisGTG_15_-_45490804_45490875@2.29.28|CCGTGATCGTATAGTGGTTAGTACTCTG</td><td>>am_trna111_HisGTG_1_-_147774845_147774916</td><td>2</td><td>29</td><td>28</td><td>CCGTGATCGTATAGTGGTTAGTACTCTG</td><td>MB468</td><td>11</td><td>11</td><td>22</td><td>30</td><td>15</td><td>3.4</td><td>4.4</td><td>4.3</td><td>10.7</td><td>9.0</td></tr>
<tr><td>>am_trna111_HisGTG_1_-_147774845_147774916@3.29.27,trna118_HisGTG_1_-_145396881_145396952@3.29.27,trna16_HisGTG_1_+_146544773_146544844@3.29.27,trna1_HisGTG_15_+_45493349_45493420@3.29.27,trna21_HisGTG_1_+_147753471_147753542@3.29.27,trna33_HisGTG_6_+_27125906_27125977@3.29.27,trna7_HisGTG_9_-_14433938_14434009@3.29.27,trna8_HisGTG_15_-_45492611_45492682@3.29.27,trna9_HisGTG_15_-_45490804_45490875@3.29.27|CGTGATCGTATAGTGGTTAGTACTCTG</td><td>>am_trna111_HisGTG_1_-_147774845_147774916</td><td>3</td><td>29</td><td>27</td><td>CGTGATCGTATAGTGGTTAGTACTCTG</td><td>MB468</td><td>11</td><td>9</td><td>22</td><td>39</td><td>18</td><td>2.9</td><td>3.5</td><td>7.4</td><td>18.6</td><td>8.7</td></tr>
<tr><td>>am_trna116_GluCTC_1_-_145399233_145399304@1.26.26,trna59_GluCTC_1_+_249168447_249168518@1.26.26,trna71_GluCTC_1_-_161439189_161439260@1.26.26,trna74_GluCTC_1_-_161431809_161431880@1.26.26,trna77_GluCTC_1_-_161424398_161424469@1.26.26,trna77_GluCTC_6_+_28949976_28950047@1.26.26,trna80_GluCTC_1_-_161417018_161417089@1.26.26,trna87_GluCTC_6_-_126101393_126101464@1.26.26|TCCCTGGTGGTCTAGTGGTTAGGATT</td><td>>am_trna116_GluCTC_1_-_145399233_145399304</td><td>1</td><td>26</td><td>26</td><td>TCCCTGGTGGTCTAGTGGTTAGGATT</td><td>BT20</td><td>54</td><td>9</td><td>252</td><td>69</td><td>24</td><td>69.3</td><td>1.6</td><td>125.1</td><td>70.5</td><td>18.5</td></tr>
<tr><td>>am_trna116_GluCTC_1_-_145399233_145399304@1.26.26,trna59_GluCTC_1_+_249168447_249168518@1.26.26,trna71_GluCTC_1_-_161439189_161439260@1.26.26,trna74_GluCTC_1_-_161431809_161431880@1.26.26,trna77_GluCTC_1_-_161424398_161424469@1.26.26,trna77_GluCTC_6_+_28949976_28950047@1.26.26,trna80_GluCTC_1_-_161417018_161417089@1.26.26,trna87_GluCTC_6_-_126101393_126101464@1.26.26|TCCCTGGTGGTCTAGTGGTTAGGATT</td><td>>am_trna116_GluCTC_1_-_145399233_145399304</td><td>1</td><td>26</td><td>26</td><td>TCCCTGGTGGTCTAGTGGTTAGGATT</td><td>MB231</td><td>14</td><td>7</td><td>74</td><td>15</td><td>13</td><td>19.5</td><td>0.9</td><td>33.2</td><td>9.1</td><td>10.1</td></tr>
</table>

| | | | | | | | | | | | | | | | | |
|---|---|---|---|---|---|---|---|---|---|---|---|---|---|---|---|---|
| >am_trna116_GluCTC_1_-_145399233_145399304@1.26.26,trna59_GluCTC_1_+_249168447_249168518@1.26.26,trna71_GluCTC_1_-_161439189_161439260@1.26.26,trna74_GluCTC_1_-_161431809_161431880@1.26.26,trna77_GluCTC_1_-_161424398_161424469@1.26.26,trna77_GluCTC_6_+_28949976_28950047@1.26.26,trna80_GluCTC_1_-_161417018_161417089@1.26.26,trna87_GluCTC_6_-_126101393_126101464@1.26.26\|TCCCTGGTGGTCTAGTGGTTAGGATT | >am_trna116_GluCTC_1_-_145399233_145399304 | 1 | 26 | 26 | TCCCTGGTGGTCTAGTGGTTAGGATT | MB468 | 108 | 50 | 64 | 13 | 8 | 71.0 | 34.5 | 24.9 | 4.7 | 1.8 |
| >am_trna116_GluCTC_1_-_145399233_145399304@1.27.27,trna59_GluCTC_1_+_249168447_249168518@1.27.27,trna71_GluCTC_1_-_161439189_161439260@1.27.27,trna74_GluCTC_1_-_161431809_161431880@1.27.27,trna77_GluCTC_1_-_161424398_161424469@1.27.27,trna77_GluCTC_6_+_28949976_28950047@1.27.27,trna80_GluCTC_1_-_161417018_161417089@1.27.27,trna87_GluCTC_6_-_126101393_126101464@1.27.27\|TCCCTGGTGGTCTAGTGGTTAGGATTC | >am_trna116_GluCTC_1_-_145399233_145399304 | 1 | 27 | 27 | TCCCTGGTGGTCTAGTGGTTAGGATTC | BT20 | 30 | 9 | 160 | 78 | 33 | 26.1 | 4.4 | 23.9 | 66.6 | 23.2 |
| >am_trna116_GluCTC_1_-_145399233_145399304@1.27.27,trna59_GluCTC_1_+_249168447_249168518@1.27.27,trna71_GluCTC_1_-_161439189_161439260@1.27.27,trna74_GluCTC_1_-_161431809_161431880@1.27.27,trna77_GluCTC_1_-_161424398_161424469@1.27.27,trna77_GluCTC_6_+_28949976_28950047@1.27.27,trna80_GluCTC_1_-_161417018_161417089@1.27.27,trna87_GluCTC_6_-_126101393_126101464@1.27.27\|TCCCTGGTGGTCTAGTGGTTAGGATTC | >am_trna116_GluCTC_1_-_145399233_145399304 | 1 | 27 | 27 | TCCCTGGTGGTCTAGTGGTTAGGATTC | MB231 | 11 | 7 | 61 | 20 | 17 | 9.1 | 2.3 | 12.0 | 11.9 | 10.1 |
| >am_trna116_GluCTC_1_-_145399233_145399304@1.27.27,trna59_GluCTC_1_+_249168447_249168518@1.27.27,trna71_GluCTC_1_-_161439189_161439260@1.27.27,trna74_GluCTC_1_-_161431809_161431880@1.27.27,trna77_GluCTC_1_-_161424398_161424469@1.27.27,trna77_GluCTC_6_+_28949976_28950047@1.27.27,trna80_GluCTC_1_-_161417018_161417089@1.27.27,trna87_GluCTC_6_-_126101393_126101464@1.27.27\|TCCCTGGTGGTCTAGTGGTTAGGATTC | >am_trna116_GluCTC_1_-_145399233_145399304 | 1 | 27 | 27 | TCCCTGGTGGTCTAGTGGTTAGGATTC | MB468 | 416 | 218 | 324 | 73 | 32 | 242.8 | 132.0 | 207.9 | 43.2 | 10.6 |
| >am_trna116_GluCTC_1_-_145399233_145399304@1.28.28,trna59_GluCTC_1_+_249168447_249168518@1.28.28,trna71_GluCTC_1_-_161439189_161439260@1.28.28,trna74_GluCTC_1_-_161431809_161431880@1.28.28,trna77_GluCTC_1_-_161424398_161424469@1.28.28,trna77_GluCTC_6_+_28949976_28950047@1.28.28,trna80_GluCTC_1_-_161417018_161417089@1.28.28,trna87_GluCTC_6_-_126101393_126101464@1.28.28\|TCCCTGGTGGTCTAGTGGTTAGGATTCG | >am_trna116_GluCTC_1_-_145399233_145399304 | 1 | 28 | 28 | TCCCTGGTGGTCTAGTGGTTAGGATTCG | BT20 | 44 | 17 | 83 | 133 | 53 | 44.3 | 18.7 | 45.5 | 180.0 | 43.9 |
| >am_trna116_GluCTC_1_-_145399233_145399304@1.28.28,trna59_GluCTC_1_+_249168447_249168518@1.28.28,trna71_GluCTC_1_-_161439189_161439260@1.28.28,trna74_GluCTC_1_-_161431809_161431880@1.28.28,trna77_GluCTC_1_-_161424398_161424469@1.28.28,trna77_GluCTC_6_+_28949976_28950047@1.28.28,trna80_GluCTC_1_-_161417018_161417089@1.28.28,trna87_GluCTC_6_-_126101393_126101464@1.28.28\|TCCCTGGTGGTCTAGTGGTTAGGATTCG | >am_trna116_GluCTC_1_-_145399233_145399304 | 1 | 28 | 28 | TCCCTGGTGGTCTAGTGGTTAGGATTCG | MB231 | 22 | 14 | 61 | 14 | 22 | 6.0 | 8.9 | 26.3 | 5.5 | 13.0 |
| >am_trna116_GluCTC_1_-_145399233_145399304@1.28.28,trna59_GluCTC_1_+_249168447_249168518@1.28.28,trna71_GluCTC_1_-_161439189_161439260@1.28.28,trna74_GluCTC_1_-_161431809_161431880@1.28.28,trna77_GluCTC_1_-_161424398_161424469@1.28.28,trna77_GluCTC_6_+_28949976_28950047@1.28.28,trna80_GluCTC_1_-_161417018_161417089@1.28.28,trna87_GluCTC_6_-_126101393_126101464@1.28.28\|TCCCTGGTGGTCTAGTGGTTAGGATTCG | >am_trna116_GluCTC_1_-_145399233_145399304 | 1 | 28 | 28 | TCCCTGGTGGTCTAGTGGTTAGGATTCG | MB468 | 2223 | 1286 | 1808 | 303 | 219 | 993.4 | 686.6 | 897.6 | 122.6 | 91.6 |
| >am_trna116_GluCTC_1_-_145399233_145399304@1.29.29,trna59_GluCTC_1_+_249168447_249168518@1.29.29,trna71_GluCTC_1_-_161439189_161439260@1.29.29,trna74_GluCTC_1_-_161431809_161431880@1.29.29,trna77_GluCTC_1_-_161424398_161424469@1.29.29,trna77_GluCTC_6_+_28949976_28950047@1.29.29,trna80_GluCTC_1_-_161417018_161417089@1.29.29,trna87_GluCTC_6_-_126101393_126101464@1.29.29\|TCCCTGGTGGTCTAGTGGTTAGGATTCGG | >am_trna116_GluCTC_1_-_145399233_145399304 | 1 | 29 | 29 | TCCCTGGTGGTCTAGTGGTTAGGATTCGG | BT20 | 62 | 35 | 128 | 128 | 80 | 80.3 | 48.6 | 68.9 | 189.9 | 103.6 |
| >am_trna116_GluCTC_1_-_145399233_145399304@1.29.29,trna59_GluCTC_1_+_249168447_249168518@1.29.29,trna71_GluCTC_1_-_161439189_161439260@1.29.29,trna74_GluCTC_1_-_161431809_161431880@1.29.29,trna77_GluCTC_1_-_161424398_161424469@1.29.29,trna77_GluCTC_6_+_28949976_28950047@1.29.29,trna80_GluCTC_1_-_161417018_161417089@1.29.29,trna87_GluCTC_6_-_126101393_126101464@1.29.29\|TCCCTGGTGGTCTAGTGGTTAGGATTCGG | >am_trna116_GluCTC_1_-_145399233_145399304 | 1 | 29 | 29 | TCCCTGGTGGTCTAGTGGTTAGGATTCGG | MB231 | 31 | 27 | 192 | 22 | 41 | 16.8 | 18.6 | 124.8 | 13.5 | 38.2 |
| >am_trna116_GluCTC_1_-_145399233_145399304@1.29.29,trna59_GluCTC_1_+_249168447_249168518@1.29.29,trna71_GluCTC_1_-_161439189_161439260@1.29.29,trna74_GluCTC_1_-_161431809_161431880@1.29.29,trna77_GluCTC_1_-_161424398_161424469@1.29.29,trna77_GluCTC_6_+_28949976_28950047@1.29.29,trna80_GluCTC_1_-_161417018_161417089@1.29.29,trna87_GluCTC_6_-_126101393_126101464@1.29.29\|TCCCTGGTGGTCTAGTGGTTAGGATTCGG | >am_trna116_GluCTC_1_-_145399233_145399304 | 1 | 29 | 29 | TCCCTGGTGGTCTAGTGGTTAGGATTCGG | MB468 | 3262 | 2213 | 5852 | 877 | 458 | 919.3 | 1420.8 | 2737.2 | 631.7 | 227.5 |
| >am_trna116_GluCTC_1_-_145399233_145399304@1.30.30,trna59_GluCTC_1_+_249168447_249168518@1.30.30,trna71_GluCTC_1_-_161439189_161439260@1.30.30,trna74_GluCTC_1_-_161431809_161431880@1.30.30,trna77_GluCTC_1_-_161424398_161424469@1.30.30,trna77_GluCTC_6_+_28949976_28950047@1.30.30,trna80_GluCTC_1_-_161417018_161417089@1.30.30,trna87_GluCTC_6_-_126101393_126101464@1.30.30\|TCCCTGGTGGTCTAGTGGTTAGGATTCGGC | >am_trna116_GluCTC_1_-_145399233_145399304 | 1 | 30 | 30 | TCCCTGGTGGTCTAGTGGTTAGGATTCGGC | BT20 | 154 | 50 | 723 | 392 | 172 | 117.2 | 44.9 | 281.5 | 534.0 | 149.1 |
| >am_trna116_GluCTC_1_-_145399233_145399304@1.30.30,trna59_GluCTC_1_+_249168447_249168518@1.30.30,trna71_GluCTC_1_-_161439189_161439260@1.30.30,trna74_GluCTC_1_-_161431809_161431880@1.30.30,trna77_GluCTC_1_-_161424398_161424469@1.30.30,trna77_GluCTC_6_+_28949976_28950047@1.30.30,trna80_GluCTC_1_-_161417018_161417089@1.30.30,trna87_GluCTC_6_-_126101393_126101464@1.30.30\|TCCCTGGTGGTCTAGTGGTTAGGATTCGGC | >am_trna116_GluCTC_1_-_145399233_145399304 | 1 | 30 | 30 | TCCCTGGTGGTCTAGTGGTTAGGATTCGGC | MB231 | 150 | 61 | 450 | 100 | 99 | 99.0 | 49.9 | 150.7 | 49.4 | 5.1 |
| >am_trna116_GluCTC_1_-_145399233_145399304@1.30.30,trna59_GluCTC_1_+_249168447_249168518@1.30.30,trna71_GluCTC_1_-_161439189_161439260@1.30.30,trna74_GluCTC_1_-_161431809_161431880@1.30.30,trna77_GluCTC_1_-_161424398_161424469@1.30.30,trna77_GluCTC_6_+_28949976_28950047@1.30.30,trna80_GluCTC_1_-_161417018_161417089@1.30.30,trna87_GluCTC_6_-_126101393_126101464@1.30.30\|TCCCTGGTGGTCTAGTGGTTAGGATTCGGC | >am_trna116_GluCTC_1_-_145399233_145399304 | 1 | 30 | 30 | TCCCTGGTGGTCTAGTGGTTAGGATTCGGC | MB468 | 1123 | 1264 | 2466 | 307 | 165 | 399.7 | 917.0 | 1173.5 | 227.2 | 98.1 |
| >am_trna116_GluCTC_1_-_145399233_145399304@13.30.18,trna25_GluCTC_2_-_71273488_71273560@13.30.18,trna59_GluCTC_1_+_249168447_249168518@13.30.18,trna71_GluCTC_1_-_161439189_161439260@13.30.18,trna74_GluCTC_1_-_161431809_161431880@13.30.18,trna77_GluCTC_1_-_161424398_161424469@13.30.18,trna77_GluCTC_6_+_28949976_28950047@13.30.18,trna80_GluCTC_1_-_161417018_161417089@13.30.18,trna87_GluCTC_6_-_126101393_126101464@13.30.18\|TAGTGGTTAGGATTCGGC | >am_trna116_GluCTC_1_-_145399233_145399304 | 13 | 30 | 18 | TAGTGGTTAGGATTCGGC | BT20 | 12 | 0 | 87 | 5 | 1 | 14.4 | 0.5 | 84.7 | 5.2 | 2.0 |

| | | | | | | | | | | | | | | | | |
|---|---|---|---|---|---|---|---|---|---|---|---|---|---|---|---|---|
| >am_trna116_GluCTC_1_-_145399233_145399304@7.28.22,trna59_GluCTC_1_+_249168447_249168518@7.28.22,trna71_GluCTC_1_-_161439189_161439260@7.28.22,trna74_GluCTC_1_-_161431809_161431880@7.28.22,trna77_GluCTC_1_-_161424398_161424469@7.28.22,trna77_GluCTC_6_+_28949976_28950047@7.28.22,trna80_GluCTC_1_-_161417018_161417089@7.28.22,trna87_GluCTC_6_-_126101393_126101464@7.28.22\|GTGGTCTAGTGGTTAGGATTCG | >am_trna116_GluCTC_1_-_145399233_145399304 | 7 | 28 | 22 | GTGGTCTAGTGGTTAGGATTCG | MB468 | 16 | 4 | 10 | 10 | 3 | 9.5 | 2.1 | 5.0 | 8.5 | 1.6 |
| >am_trna117_GlyTCC_1_-_145397864_145397935@1.28.28,trna45_GlyTCC_1_+_161500903_161500974@1.28.28,trna70_GlyTCC_1_-_161439547_161439618@1.28.28,trna73_GlyTCC_1_-_161432166_161432237@1.28.28,trna76_GlyTCC_1_-_161424756_161424827@1.28.28,trna79_GlyTCC_1_-_161417375_161417446@1.28.28\|GCGTTGGTGGTATAGTGGTGAGCATAGC | >am_trna117_GlyTCC_1_-_145397864_145397935 | 1 | 28 | 28 | GCGTTGGTGGTATAGTGGTGAGCATAGC | MB468 | 12 | 21 | 14 | 3 | 1 | 5.9 | 18.5 | 7.4 | 1.6 | 1.4 |
| >am_trna117_GlyTCC_1_-_145397864_145397935@1.29.29,trna45_GlyTCC_1_+_161500903_161500974@1.29.29,trna70_GlyTCC_1_-_161439547_161439618@1.29.29,trna73_GlyTCC_1_-_161432166_161432237@1.29.29,trna76_GlyTCC_1_-_161424756_161424827@1.29.29,trna79_GlyTCC_1_-_161417375_161417446@1.29.29\|GCGTTGGTGGTATAGTGGTGAGCATAGCT | >am_trna117_GlyTCC_1_-_145397864_145397935 | 1 | 29 | 29 | GCGTTGGTGGTATAGTGGTGAGCATAGCT | MB468 | 132 | 144 | 308 | 81 | 39 | 77.2 | 135.7 | 157.8 | 15.3 | 30.4 |
| >am_trna117_GlyTCC_1_-_145397864_145397935@1.30.30,trna45_GlyTCC_1_+_161500903_161500974@1.30.30,trna70_GlyTCC_1_-_161439547_161439618@1.30.30,trna73_GlyTCC_1_-_161432166_161432237@1.30.30,trna76_GlyTCC_1_-_161424756_161424827@1.30.30,trna79_GlyTCC_1_-_161417375_161417446@1.30.30\|GCGTTGGTGGTATAGTGGTGAGCATAGCTG | >am_trna117_GlyTCC_1_-_145397864_145397935 | 1 | 30 | 30 | GCGTTGGTGGTATAGTGGTGAGCATAGCTG | MB468 | 162 | 132 | 486 | 152 | 78 | 94.2 | 82.2 | 144.4 | 57.6 | 74.5 |
| >am_trna119_LysCTT_1_-_145395522_145395594@13.30.18,trna11_LysCTT_5_-_180648979_180649051@13.30.18,trna128_LysCTT_1_-_55423542_55423614@13.30.18,trna13_LysCTT_6_+_26556774_26556846@13.30.18,trna32_LysCTT_16_-_3207406_3207478@13.30.18,trna6_LysCTT_19_-_52425396_52425468@13.30.18,trna7_LysCTT_16_+_3225692_3225764@13.30.18,trna9_LysCTT_5_+_180634755_180634827@13.30.18\|CAGTCGGTAGAGCATGAG | >am_trna119_LysCTT_1_-_145395522_145395594 | 13 | 30 | 18 | CAGTCGGTAGAGCATGAG | MB468 | 17 | 17 | 11 | 5 | 5 | 3.2 | 9.9 | 3.3 | 2.8 | 3.5 |
| >am_trna119_LysCTT_1_-_145395522_145395594@14.31.18,trna11_LysCTT_5_-_180648979_180649051@14.31.18,trna128_LysCTT_1_-_55423542_55423614@14.31.18,trna13_LysCTT_6_+_26556774_26556846@14.31.18,trna32_LysCTT_16_-_3207406_3207478@14.31.18,trna6_LysCTT_19_-_52425396_52425468@14.31.18,trna7_LysCTT_16_+_3225692_3225764@14.31.18,trna9_LysCTT_5_+_180634755_180634827@14.31.18\|AGTCGGTAGAGCATGAGA | >am_trna119_LysCTT_1_-_145395522_145395594 | 14 | 31 | 18 | AGTCGGTAGAGCATGAGA | MB468 | 40 | 45 | 18 | 13 | 3 | 18.6 | 27.8 | 1.7 | 6.2 | 1.3 |
| >am_trna119_LysCTT_1_-_145395522_145395594@16.33.18,trna11_LysCTT_5_-_180648979_180649051@16.33.18,trna128_LysCTT_1_-_55423542_55423614@16.33.18,trna13_LysCTT_6_+_26556774_26556846@16.33.18,trna6_LysCTT_19_-_52425396_52425468@16.33.18,trna7_LysCTT_16_+_3225692_3225764@16.33.18,trna9_LysCTT_5_+_180634755_180634827@16.33.18\|TCGGTAGAGCATGAGACT | >am_trna119_LysCTT_1_-_145395522_145395594 | 16 | 33 | 18 | TCGGTAGAGCATGAGACT | MB231 | 19 | 24 | 12 | 5 | 4 | 17.0 | 24.0 | 13.1 | 1.4 | 3.0 |
| >am_trna128_GlyGCC_6_-_27870686_27870756@1.25.25,trna133_GlyCCC_1_-_16872434_16872504@1.25.25,trna18_GlyGCC_16_+_70822597_70822667@1.25.25,trna19_GlyGCC_16_+_70823410_70823480@1.25.25,trna19_GlyGCC_2_-_157257659_157257729@1.25.25,trna24_GlyGCC_16_-_70812942_70813012@1.25.25,trna25_GlyGCC_16_-_70812114_70812184@1.25.25,trna4_GlyCCC_1_+_17188416_17188486@1.25.25,trna5_GlyGCC_17_+_8029064_8029134@1.25.25,trna68_GlyGCC_1_-_161493637_161493707@1.25.25\|GCATTGGTGGTTCAGTGGTAGAATT | >am_trna128_GlyGCC_6_-_27870686_27870756 | 1 | 25 | 25 | GCATTGGTGGTTCAGTGGTAGAATT | BT20 | 12 | 3 | 46 | 29 | 8 | 12.5 | 1.3 | 18.2 | 25.5 | 4.2 |
| >am_trna128_GlyGCC_6_-_27870686_27870756@1.25.25,trna133_GlyCCC_1_-_16872434_16872504@1.25.25,trna18_GlyGCC_16_+_70822597_70822667@1.25.25,trna19_GlyGCC_16_+_70823410_70823480@1.25.25,trna19_GlyGCC_2_-_157257659_157257729@1.25.25,trna24_GlyGCC_16_-_70812942_70813012@1.25.25,trna25_GlyGCC_16_-_70812114_70812184@1.25.25,trna4_GlyCCC_1_+_17188416_17188486@1.25.25,trna5_GlyGCC_17_+_8029064_8029134@1.25.25,trna68_GlyGCC_1_-_161493637_161493707@1.25.25\|GCATTGGTGGTTCAGTGGTAGAATT | >am_trna128_GlyGCC_6_-_27870686_27870756 | 1 | 25 | 25 | GCATTGGTGGTTCAGTGGTAGAATT | MB468 | 45 | 46 | 34 | 14 | 7 | 30.9 | 40.9 | 11.5 | 6.8 | 4.0 |
| >am_trna128_GlyGCC_6_-_27870686_27870756@1.26.26,trna133_GlyCCC_1_-_16872434_16872504@1.26.26,trna18_GlyGCC_16_+_70822597_70822667@1.26.26,trna19_GlyGCC_16_+_70823410_70823480@1.26.26,trna19_GlyGCC_2_-_157257659_157257729@1.26.26,trna24_GlyGCC_16_-_70812942_70813012@1.26.26,trna25_GlyGCC_16_-_70812114_70812184@1.26.26,trna4_GlyCCC_1_+_17188416_17188486@1.26.26,trna5_GlyGCC_17_+_8029064_8029134@1.26.26,trna68_GlyGCC_1_-_161493637_161493707@1.26.26\|GCATTGGTGGTTCAGTGGTAGAATTC | >am_trna128_GlyGCC_6_-_27870686_27870756 | 1 | 26 | 26 | GCATTGGTGGTTCAGTGGTAGAATTC | MB468 | 34 | 34 | 20 | 16 | 3 | 16.9 | 19.5 | 8.8 | 12.2 | 2.3 |
| >am_trna128_GlyGCC_6_-_27870686_27870756@1.27.27,trna133_GlyCCC_1_-_16872434_16872504@1.27.27,trna18_GlyGCC_16_+_70822597_70822667@1.27.27,trna19_GlyGCC_16_+_70823410_70823480@1.27.27,trna19_GlyGCC_2_-_157257659_157257729@1.27.27,trna24_GlyGCC_16_-_70812942_70813012@1.27.27,trna25_GlyGCC_16_-_70812114_70812184@1.27.27,trna4_GlyCCC_1_+_17188416_17188486@1.27.27,trna5_GlyGCC_17_+_8029064_8029134@1.27.27,trna68_GlyGCC_1_-_161493637_161493707@1.27.27\|GCATTGGTGGTTCAGTGGTAGAATTCT | >am_trna128_GlyGCC_6_-_27870686_27870756 | 1 | 27 | 27 | GCATTGGTGGTTCAGTGGTAGAATTCT | BT20 | 21 | 10 | 34 | 31 | 21 | 13.3 | 12.9 | 11.4 | 28.4 | 17.0 |
| >am_trna128_GlyGCC_6_-_27870686_27870756@1.27.27,trna133_GlyCCC_1_-_16872434_16872504@1.27.27,trna18_GlyGCC_16_+_70822597_70822667@1.27.27,trna19_GlyGCC_16_+_70823410_70823480@1.27.27,trna19_GlyGCC_2_-_157257659_157257729@1.27.27,trna24_GlyGCC_16_-_70812942_70813012@1.27.27,trna25_GlyGCC_16_-_70812114_70812184@1.27.27,trna4_GlyCCC_1_+_17188416_17188486@1.27.27,trna5_GlyGCC_17_+_8029064_8029134@1.27.27,trna68_GlyGCC_1_-_161493637_161493707@1.27.27\|GCATTGGTGGTTCAGTGGTAGAATTCT | >am_trna128_GlyGCC_6_-_27870686_27870756 | 1 | 27 | 27 | GCATTGGTGGTTCAGTGGTAGAATTCT | MB231 | 29 | 14 | 62 | 38 | 28 | 2.0 | 4.2 | 17.4 | 6.7 | 8.3 |
| >am_trna128_GlyGCC_6_-_27870686_27870756@1.27.27,trna133_GlyCCC_1_-_16872434_16872504@1.27.27,trna18_GlyGCC_16_+_70822597_70822667@1.27.27,trna19_GlyGCC_16_+_70823410_70823480@1.27.27,trna19_GlyGCC_2_-_157257659_157257729@1.27.27,trna24_GlyGCC_16_-_70812942_70813012@1.27.27,trna25_GlyGCC_16_-_70812114_70812184@1.27.27,trna4_GlyCCC_1_+_17188416_17188486@1.27.27,trna5_GlyGCC_17_+_8029064_8029134@1.27.27,trna68_GlyGCC_1_-_161493637_161493707@1.27.27\|GCATTGGTGGTTCAGTGGTAGAATTCT | >am_trna128_GlyGCC_6_-_27870686_27870756 | 1 | 27 | 27 | GCATTGGTGGTTCAGTGGTAGAATTCT | MB468 | 148 | 147 | 123 | 26 | 12 | 49.5 | 57.4 | 59.8 | 11.1 | 7.3 |
| >am_trna128_GlyGCC_6_-_27870686_27870756@1.28.28,trna133_GlyCCC_1_-_16872434_16872504@1.28.28,trna18_GlyGCC_16_+_70822597_70822667@1.28.28,trna19_GlyGCC_16_+_70823410_70823480@1.28.28,trna19_GlyGCC_2_-_157257659_157257729@1.28.28,trna24_GlyGCC_16_-_70812942_70813012@1.28.28,trna25_GlyGCC_16_-_70812114_70812184@1.28.28,trna4_GlyCCC_1_+_17188416_17188486@1.28.28,trna5_GlyGCC_17_+_8029064_8029134@1.28.28,trna68_GlyGCC_1_-_161493637_161493707@1.28.28\|GCATTGGTGGTTCAGTGGTAGAATTCTC | >am_trna128_GlyGCC_6_-_27870686_27870756 | 1 | 28 | 28 | GCATTGGTGGTTCAGTGGTAGAATTCTC | BT20 | 38 | 14 | 105 | 171 | 45 | 38.0 | 18.1 | 47.1 | 153.9 | 54.2 |

| | | | | | | | | | | | | | | | | |
|---|---|---|---|---|---|---|---|---|---|---|---|---|---|---|---|---|
| >am_trna128_GlyGCC_6_-_27870686_27870756@1.28.28,trna133_GlyCCC_1_-_16872434_16872504@1.28.28,trna18_GlyGCC_16_+_70822597_70822667@1.28.28,trna19_GlyGCC_16_+_70823410_70823480@1.28.28,trna19_GlyGCC_2_-_157257659_157257729@1.28.28,trna24_GlyGCC_16_-_70812942_70813012@1.28.28,trna25_GlyGCC_16_-_70812114_70812184@1.28.28,trna4_GlyCCC_1_+_17188416_17188486@1.28.28,trna5_GlyGCC_17_+_8029064_8029134@1.28.28,trna68_GlyGCC_1_-_161493637_161493707@1.28.28\|GCATTGGTGGTTCAGTGGTAGAATTCTC | >am_trna128_GlyGCC_6_-_27870686_27870756 | 1 | 28 | 28 | GCATTGGTGGTTCAGTGGTAGAATTCTC | MB231 | 39 | 21 | 173 | 68 | 36 | 17.0 | 9.0 | 12.3 | 21.1 | 12.6 |
| >am_trna128_GlyGCC_6_-_27870686_27870756@1.28.28,trna133_GlyCCC_1_-_16872434_16872504@1.28.28,trna18_GlyGCC_16_+_70822597_70822667@1.28.28,trna19_GlyGCC_16_+_70823410_70823480@1.28.28,trna19_GlyGCC_2_-_157257659_157257729@1.28.28,trna24_GlyGCC_16_-_70812942_70813012@1.28.28,trna25_GlyGCC_16_-_70812114_70812184@1.28.28,trna4_GlyCCC_1_+_17188416_17188486@1.28.28,trna5_GlyGCC_17_+_8029064_8029134@1.28.28,trna68_GlyGCC_1_-_161493637_161493707@1.28.28\|GCATTGGTGGTTCAGTGGTAGAATTCTC | >am_trna128_GlyGCC_6_-_27870686_27870756 | 1 | 28 | 28 | GCATTGGTGGTTCAGTGGTAGAATTCTC | MB468 | 4150 | 3503 | 5089 | 1431 | 657 | 1524.6 | 2449.3 | 1879.1 | 475.5 | 409.9 |
| >am_trna128_GlyGCC_6_-_27870686_27870756@1.29.29,trna133_GlyCCC_1_-_16872434_16872504@1.29.29,trna18_GlyGCC_16_+_70822597_70822667@1.29.29,trna19_GlyGCC_16_+_70823410_70823480@1.29.29,trna19_GlyGCC_2_-_157257659_157257729@1.29.29,trna24_GlyGCC_16_-_70812942_70813012@1.29.29,trna25_GlyGCC_16_-_70812114_70812184@1.29.29,trna4_GlyCCC_1_+_17188416_17188486@1.29.29,trna5_GlyGCC_17_+_8029064_8029134@1.29.29,trna68_GlyGCC_1_-_161493637_161493707@1.29.29\|GCATTGGTGGTTCAGTGGTAGAATTCTCG | >am_trna128_GlyGCC_6_-_27870686_27870756 | 1 | 29 | 29 | GCATTGGTGGTTCAGTGGTAGAATTCTCG | MB231 | 27 | 13 | 122 | 39 | 28 | 14.4 | 8.4 | 49.5 | 28.6 | 7.7 |
| >am_trna128_GlyGCC_6_-_27870686_27870756@1.29.29,trna133_GlyCCC_1_-_16872434_16872504@1.29.29,trna18_GlyGCC_16_+_70822597_70822667@1.29.29,trna19_GlyGCC_16_+_70823410_70823480@1.29.29,trna19_GlyGCC_2_-_157257659_157257729@1.29.29,trna24_GlyGCC_16_-_70812942_70813012@1.29.29,trna25_GlyGCC_16_-_70812114_70812184@1.29.29,trna4_GlyCCC_1_+_17188416_17188486@1.29.29,trna5_GlyGCC_17_+_8029064_8029134@1.29.29,trna68_GlyGCC_1_-_161493637_161493707@1.29.29\|GCATTGGTGGTTCAGTGGTAGAATTCTCG | >am_trna128_GlyGCC_6_-_27870686_27870756 | 1 | 29 | 29 | GCATTGGTGGTTCAGTGGTAGAATTCTCG | MB468 | 2772 | 1392 | 3623 | 893 | 373 | 1234.6 | 495.5 | 1710.4 | 377.9 | 285.1 |
| >am_trna128_GlyGCC_6_-_27870686_27870756@1.30.30,trna133_GlyCCC_1_-_16872434_16872504@1.30.30,trna18_GlyGCC_16_+_70822597_70822667@1.30.30,trna19_GlyGCC_16_+_70823410_70823480@1.30.30,trna19_GlyGCC_2_-_157257659_157257729@1.30.30,trna24_GlyGCC_16_-_70812942_70813012@1.30.30,trna25_GlyGCC_16_-_70812114_70812184@1.30.30,trna4_GlyCCC_1_+_17188416_17188486@1.30.30,trna5_GlyGCC_17_+_8029064_8029134@1.30.30,trna68_GlyGCC_1_-_161493637_161493707@1.30.30\|GCATTGGTGGTTCAGTGGTAGAATTCTCGC | >am_trna128_GlyGCC_6_-_27870686_27870756 | 1 | 30 | 30 | GCATTGGTGGTTCAGTGGTAGAATTCTCGC | BT20 | 26 | 8 | 74 | 149 | 22 | 17.3 | 3.6 | 30.8 | 68.9 | 18.2 |
| >am_trna128_GlyGCC_6_-_27870686_27870756@1.30.30,trna133_GlyCCC_1_-_16872434_16872504@1.30.30,trna18_GlyGCC_16_+_70822597_70822667@1.30.30,trna19_GlyGCC_16_+_70823410_70823480@1.30.30,trna19_GlyGCC_2_-_157257659_157257729@1.30.30,trna24_GlyGCC_16_-_70812942_70813012@1.30.30,trna25_GlyGCC_16_-_70812114_70812184@1.30.30,trna4_GlyCCC_1_+_17188416_17188486@1.30.30,trna5_GlyGCC_17_+_8029064_8029134@1.30.30,trna68_GlyGCC_1_-_161493637_161493707@1.30.30\|GCATTGGTGGTTCAGTGGTAGAATTCTCGC | >am_trna128_GlyGCC_6_-_27870686_27870756 | 1 | 30 | 30 | GCATTGGTGGTTCAGTGGTAGAATTCTCGC | MB231 | 292 | 114 | 621 | 2113 | 349 | 210.1 | 116.5 | 496.3 | 1383.5 | 167.6 |
| >am_trna128_GlyGCC_6_-_27870686_27870756@1.30.30,trna133_GlyCCC_1_-_16872434_16872504@1.30.30,trna18_GlyGCC_16_+_70822597_70822667@1.30.30,trna19_GlyGCC_16_+_70823410_70823480@1.30.30,trna19_GlyGCC_2_-_157257659_157257729@1.30.30,trna24_GlyGCC_16_-_70812942_70813012@1.30.30,trna25_GlyGCC_16_-_70812114_70812184@1.30.30,trna4_GlyCCC_1_+_17188416_17188486@1.30.30,trna5_GlyGCC_17_+_8029064_8029134@1.30.30,trna68_GlyGCC_1_-_161493637_161493707@1.30.30\|GCATTGGTGGTTCAGTGGTAGAATTCTCGC | >am_trna128_GlyGCC_6_-_27870686_27870756 | 1 | 30 | 30 | GCATTGGTGGTTCAGTGGTAGAATTCTCGC | MB468 | 1043 | 483 | 2548 | 526 | 137 | 542.3 | 198.5 | 1461.9 | 159.0 | 120.6 |
| >am_trna128_GlyGCC_6_-_27870686_27870756@1.31.31,trna133_GlyCCC_1_-_16872434_16872504@1.31.31,trna18_GlyGCC_16_+_70822597_70822667@1.31.31,trna19_GlyGCC_16_+_70823410_70823480@1.31.31,trna19_GlyGCC_2_-_157257659_157257729@1.31.31,trna24_GlyGCC_16_-_70812942_70813012@1.31.31,trna25_GlyGCC_16_-_70812114_70812184@1.31.31,trna4_GlyCCC_1_+_17188416_17188486@1.31.31,trna5_GlyGCC_17_+_8029064_8029134@1.31.31,trna68_GlyGCC_1_-_161493637_161493707@1.31.31\|GCATTGGTGGTTCAGTGGTAGAATTCTCGCC | >am_trna128_GlyGCC_6_-_27870686_27870756 | 1 | 31 | 31 | GCATTGGTGGTTCAGTGGTAGAATTCTCGCC | BT20 | 1140 | 115 | 1085 | 1470 | 686 | 600.7 | 52.3 | 207.7 | 1616.0 | 777.4 |
| >am_trna128_GlyGCC_6_-_27870686_27870756@1.31.31,trna133_GlyCCC_1_-_16872434_16872504@1.31.31,trna18_GlyGCC_16_+_70822597_70822667@1.31.31,trna19_GlyGCC_16_+_70823410_70823480@1.31.31,trna19_GlyGCC_2_-_157257659_157257729@1.31.31,trna24_GlyGCC_16_-_70812942_70813012@1.31.31,trna25_GlyGCC_16_-_70812114_70812184@1.31.31,trna4_GlyCCC_1_+_17188416_17188486@1.31.31,trna5_GlyGCC_17_+_8029064_8029134@1.31.31,trna68_GlyGCC_1_-_161493637_161493707@1.31.31\|GCATTGGTGGTTCAGTGGTAGAATTCTCGCC | >am_trna128_GlyGCC_6_-_27870686_27870756 | 1 | 31 | 31 | GCATTGGTGGTTCAGTGGTAGAATTCTCGCC | MB231 | 2374 | 718 | 3091 | 2832 | 1827 | 1144.8 | 722.9 | 1749.0 | 1432.6 | 921.5 |
| >am_trna128_GlyGCC_6_-_27870686_27870756@1.31.31,trna133_GlyCCC_1_-_16872434_16872504@1.31.31,trna18_GlyGCC_16_+_70822597_70822667@1.31.31,trna19_GlyGCC_16_+_70823410_70823480@1.31.31,trna19_GlyGCC_2_-_157257659_157257729@1.31.31,trna24_GlyGCC_16_-_70812942_70813012@1.31.31,trna25_GlyGCC_16_-_70812114_70812184@1.31.31,trna4_GlyCCC_1_+_17188416_17188486@1.31.31,trna5_GlyGCC_17_+_8029064_8029134@1.31.31,trna68_GlyGCC_1_-_161493637_161493707@1.31.31\|GCATTGGTGGTTCAGTGGTAGAATTCTCGCC | >am_trna128_GlyGCC_6_-_27870686_27870756 | 1 | 31 | 31 | GCATTGGTGGTTCAGTGGTAGAATTCTCGCC | MB468 | 71 | 36 | 168 | 54 | 24 | 29.0 | 24.0 | 41.5 | 43.7 | 12.3 |
| >am_trna128_GlyGCC_6_-_27870686_27870756@1.32.32,trna133_GlyCCC_1_-_16872434_16872504@1.32.32,trna18_GlyGCC_16_+_70822597_70822667@1.32.32,trna19_GlyGCC_16_+_70823410_70823480@1.32.32,trna19_GlyGCC_2_-_157257659_157257729@1.32.32,trna24_GlyGCC_16_-_70812942_70813012@1.32.32,trna25_GlyGCC_16_-_70812114_70812184@1.32.32,trna4_GlyCCC_1_+_17188416_17188486@1.32.32,trna5_GlyGCC_17_+_8029064_8029134@1.32.32,trna68_GlyGCC_1_-_161493637_161493707@1.32.32\|GCATTGGTGGTTCAGTGGTAGAATTCTCGCCT | >am_trna128_GlyGCC_6_-_27870686_27870756 | 1 | 32 | 32 | GCATTGGTGGTTCAGTGGTAGAATTCTCGCCT | BT20 | 466 | 63 | 324 | 1081 | 486 | 23.9 | 18.0 | 105.2 | 1652.6 | 706.6 |
| >am_trna128_GlyGCC_6_-_27870686_27870756@1.32.32,trna133_GlyCCC_1_-_16872434_16872504@1.32.32,trna18_GlyGCC_16_+_70822597_70822667@1.32.32,trna19_GlyGCC_16_+_70823410_70823480@1.32.32,trna19_GlyGCC_2_-_157257659_157257729@1.32.32,trna24_GlyGCC_16_-_70812942_70813012@1.32.32,trna25_GlyGCC_16_-_70812114_70812184@1.32.32,trna4_GlyCCC_1_+_17188416_17188486@1.32.32,trna5_GlyGCC_17_+_8029064_8029134@1.32.32,trna68_GlyGCC_1_-_161493637_161493707@1.32.32\|GCATTGGTGGTTCAGTGGTAGAATTCTCGCCT | >am_trna128_GlyGCC_6_-_27870686_27870756 | 1 | 32 | 32 | GCATTGGTGGTTCAGTGGTAGAATTCTCGCCT | MB231 | 1644 | 394 | 1290 | 1391 | 920 | 1270.9 | 426.7 | 1262.8 | 905.2 | 510.3 |

| | | | | | | | | | | | | | | | | |
|---|---|---|---|---|---|---|---|---|---|---|---|---|---|---|---|---|
| >am_trna128_GlyGCC_6_-_27870686_27870756@1.33.33,trna18_GlyGCC_16_+_70822597_70822667@1.33.33,trna19_GlyGCC_16_+_70823410_70823480@1.33.33,trna19_GlyGCC_2_-_157257659_157257729@1.33.33,trna24_GlyGCC_16_-_70812942_70813012@1.33.33,trna25_GlyGCC_16_-_70812114_70812184@1.33.33,trna5_GlyGCC_17_+_8029064_8029134@1.33.33,trna68_GlyGCC_1_-_161493637_161493707@1.33.33\|GCATTGGTGGTTCAGTGGTAGAATTCTCGCCTG | >am_trna128_GlyGCC_6_-_27870686_27870756 | 1 | 33 | 33 | GCATTGGTGGTTCAGTGGTAGAATTCTCGCCTG | BT20 | 142 | 31 | 147 | 136 | 60 | 144.9 | 45.0 | 121.2 | 112.0 | 39.5 |
| >am_trna128_GlyGCC_6_-_27870686_27870756@1.33.33,trna18_GlyGCC_16_+_70822597_70822667@1.33.33,trna19_GlyGCC_16_+_70823410_70823480@1.33.33,trna19_GlyGCC_2_-_157257659_157257729@1.33.33,trna24_GlyGCC_16_-_70812942_70813012@1.33.33,trna25_GlyGCC_16_-_70812114_70812184@1.33.33,trna5_GlyGCC_17_+_8029064_8029134@1.33.33,trna68_GlyGCC_1_-_161493637_161493707@1.33.33\|GCATTGGTGGTTCAGTGGTAGAATTCTCGCCTG | >am_trna128_GlyGCC_6_-_27870686_27870756 | 1 | 33 | 33 | GCATTGGTGGTTCAGTGGTAGAATTCTCGCCTG | MB231 | 263 | 76 | 193 | 400 | 238 | 76.3 | 73.1 | 108.9 | 270.2 | 102.5 |
| >am_trna128_GlyGCC_6_-_27870686_27870756@1.34.34,trna18_GlyGCC_16_+_70822597_70822667@1.34.34,trna19_GlyGCC_16_+_70823410_70823480@1.34.34,trna19_GlyGCC_2_-_157257659_157257729@1.34.34,trna24_GlyGCC_16_-_70812942_70813012@1.34.34,trna25_GlyGCC_16_-_70812114_70812184@1.34.34,trna5_GlyGCC_17_+_8029064_8029134@1.34.34,trna68_GlyGCC_1_-_161493637_161493707@1.34.34\|GCATTGGTGGTTCAGTGGTAGAATTCTCGCCTGC | >am_trna128_GlyGCC_6_-_27870686_27870756 | 1 | 34 | 34 | GCATTGGTGGTTCAGTGGTAGAATTCTCGCCTGC | BT20 | 10 | 1 | 7 | 17 | 7 | 3.7 | 0.8 | 2.5 | 16.5 | 2.5 |
| >am_trna128_GlyGCC_6_-_27870686_27870756@1.34.34,trna18_GlyGCC_16_+_70822597_70822667@1.34.34,trna19_GlyGCC_16_+_70823410_70823480@1.34.34,trna19_GlyGCC_2_-_157257659_157257729@1.34.34,trna24_GlyGCC_16_-_70812942_70813012@1.34.34,trna25_GlyGCC_16_-_70812114_70812184@1.34.34,trna5_GlyGCC_17_+_8029064_8029134@1.34.34,trna68_GlyGCC_1_-_161493637_161493707@1.34.34\|GCATTGGTGGTTCAGTGGTAGAATTCTCGCCTGC | >am_trna128_GlyGCC_6_-_27870686_27870756 | 1 | 34 | 34 | GCATTGGTGGTTCAGTGGTAGAATTCTCGCCTGC | MB231 | 183 | 69 | 180 | 165 | 158 | 121.2 | 78.1 | 160.5 | 137.1 | 131.3 |
| >am_trna128_GlyGCC_6_-_27870686_27870756@1.35.35,trna18_GlyGCC_16_+_70822597_70822667@1.35.35,trna19_GlyGCC_16_+_70823410_70823480@1.35.35,trna19_GlyGCC_2_-_157257659_157257729@1.35.35,trna24_GlyGCC_16_-_70812942_70813012@1.35.35,trna25_GlyGCC_16_-_70812114_70812184@1.35.35,trna5_GlyGCC_17_+_8029064_8029134@1.35.35,trna68_GlyGCC_1_-_161493637_161493707@1.35.35\|GCATTGGTGGTTCAGTGGTAGAATTCTCGCCTGCC | >am_trna128_GlyGCC_6_-_27870686_27870756 | 1 | 35 | 35 | GCATTGGTGGTTCAGTGGTAGAATTCTCGCCTGCC | BT20 | 134 | 6 | 70 | 155 | 27 | 150.1 | 1.6 | 10.6 | 154.5 | 29.2 |
| >am_trna128_GlyGCC_6_-_27870686_27870756@1.35.35,trna18_GlyGCC_16_+_70822597_70822667@1.35.35,trna19_GlyGCC_16_+_70823410_70823480@1.35.35,trna19_GlyGCC_2_-_157257659_157257729@1.35.35,trna24_GlyGCC_16_-_70812942_70813012@1.35.35,trna25_GlyGCC_16_-_70812114_70812184@1.35.35,trna5_GlyGCC_17_+_8029064_8029134@1.35.35,trna68_GlyGCC_1_-_161493637_161493707@1.35.35\|GCATTGGTGGTTCAGTGGTAGAATTCTCGCCTGCC | >am_trna128_GlyGCC_6_-_27870686_27870756 | 1 | 35 | 35 | GCATTGGTGGTTCAGTGGTAGAATTCTCGCCTGCC | MB231 | 430 | 180 | 517 | 374 | 303 | 218.6 | 200.6 | 381.3 | 238.8 | 158.3 |
| >am_trna128_GlyGCC_6_-_27870686_27870756@1.36.36,trna18_GlyGCC_16_+_70822597_70822667@1.36.36,trna19_GlyGCC_16_+_70823410_70823480@1.36.36,trna19_GlyGCC_2_-_157257659_157257729@1.36.36,trna24_GlyGCC_16_-_70812942_70813012@1.36.36,trna25_GlyGCC_16_-_70812114_70812184@1.36.36,trna5_GlyGCC_17_+_8029064_8029134@1.36.36,trna68_GlyGCC_1_-_161493637_161493707@1.36.36\|GCATTGGTGGTTCAGTGGTAGAATTCTCGCCTGCCA | >am_trna128_GlyGCC_6_-_27870686_27870756 | 1 | 36 | 36 | GCATTGGTGGTTCAGTGGTAGAATTCTCGCCTGCCA | BT20 | 11 | 3 | 11 | 43 | 12 | 5.4 | 1.6 | 5.2 | 50.5 | 13.0 |
| >am_trna128_GlyGCC_6_-_27870686_27870756@1.36.36,trna18_GlyGCC_16_+_70822597_70822667@1.36.36,trna19_GlyGCC_16_+_70823410_70823480@1.36.36,trna19_GlyGCC_2_-_157257659_157257729@1.36.36,trna24_GlyGCC_16_-_70812942_70813012@1.36.36,trna25_GlyGCC_16_-_70812114_70812184@1.36.36,trna5_GlyGCC_17_+_8029064_8029134@1.36.36,trna68_GlyGCC_1_-_161493637_161493707@1.36.36\|GCATTGGTGGTTCAGTGGTAGAATTCTCGCCTGCCA | >am_trna128_GlyGCC_6_-_27870686_27870756 | 1 | 36 | 36 | GCATTGGTGGTTCAGTGGTAGAATTCTCGCCTGCCA | MB231 | 54 | 24 | 59 | 38 | 37 | 42.1 | 10.0 | 17.7 | 9.4 | 20.2 |
| >am_trna128_GlyGCC_6_-_27870686_27870756@3.31.29,trna133_GlyCCC_1_-_16872434_16872504@3.31.29,trna18_GlyGCC_16_+_70822597_70822667@3.31.29,trna19_GlyGCC_16_+_70823410_70823480@3.31.29,trna19_GlyGCC_2_-_157257659_157257729@3.31.29,trna24_GlyGCC_16_-_70812942_70813012@3.31.29,trna25_GlyGCC_16_-_70812114_70812184@3.31.29,trna4_GlyCCC_1_+_17188416_17188486@3.31.29,trna5_GlyGCC_17_+_8029064_8029134@3.31.29,trna68_GlyGCC_1_-_161493637_161493707@3.31.29\|ATTGGTGGTTCAGTGGTAGAATTCTCGCC | >am_trna128_GlyGCC_6_-_27870686_27870756 | 3 | 31 | 29 | ATTGGTGGTTCAGTGGTAGAATTCTCGCC | MB231 | 11 | 9 | 3 | 15 | 16 | 3.8 | 6.7 | 2.3 | 5.2 | 6.1 |
| >am_trna128_GlyGCC_6_-_27870686_27870756@3.34.32,trna18_GlyGCC_16_+_70822597_70822667@3.34.32,trna19_GlyGCC_16_+_70823410_70823480@3.34.32,trna19_GlyGCC_2_-_157257659_157257729@3.34.32,trna24_GlyGCC_16_-_70812942_70813012@3.34.32,trna25_GlyGCC_16_-_70812114_70812184@3.34.32,trna5_GlyGCC_17_+_8029064_8029134@3.34.32,trna68_GlyGCC_1_-_161493637_161493707@3.34.32\|ATTGGTGGTTCAGTGGTAGAATTCTCGCCTGC | >am_trna128_GlyGCC_6_-_27870686_27870756 | 3 | 34 | 32 | ATTGGTGGTTCAGTGGTAGAATTCTCGCCTGC | MB231 | 15 | 7 | 1 | 9 | 33 | 2.7 | 5.6 | 0.7 | 6.0 | 8.9 |
| >am_trna128_GlyGCC_6_-_27870686_27870756@3.35.33,trna18_GlyGCC_16_+_70822597_70822667@3.35.33,trna19_GlyGCC_16_+_70823410_70823480@3.35.33,trna19_GlyGCC_2_-_157257659_157257729@3.35.33,trna24_GlyGCC_16_-_70812942_70813012@3.35.33,trna25_GlyGCC_16_-_70812114_70812184@3.35.33,trna5_GlyGCC_17_+_8029064_8029134@3.35.33,trna68_GlyGCC_1_-_161493637_161493707@3.35.33\|ATTGGTGGTTCAGTGGTAGAATTCTCGCCTGCC | >am_trna128_GlyGCC_6_-_27870686_27870756 | 3 | 35 | 33 | ATTGGTGGTTCAGTGGTAGAATTCTCGCCTGCC | MB231 | 39 | 27 | 5 | 29 | 96 | 18.0 | 15.1 | 2.5 | 16.9 | 16.5 |
| >am_trna128_GlyGCC_6_-_27870686_27870756@4.28.25,trna133_GlyCCC_1_-_16872434_16872504@4.28.25,trna18_GlyGCC_16_+_70822597_70822667@4.28.25,trna19_GlyGCC_16_+_70823410_70823480@4.28.25,trna19_GlyGCC_2_-_157257659_157257729@4.28.25,trna24_GlyGCC_16_-_70812942_70813012@4.28.25,trna25_GlyGCC_16_-_70812114_70812184@4.28.25,trna4_GlyCCC_1_+_17188416_17188486@4.28.25,trna5_GlyGCC_17_+_8029064_8029134@4.28.25,trna68_GlyGCC_1_-_161493637_161493707@4.28.25\|TTGGTGGTTCAGTGGTAGAATTCTC | >am_trna128_GlyGCC_6_-_27870686_27870756 | 4 | 28 | 25 | TTGGTGGTTCAGTGGTAGAATTCTC | MB468 | 15 | 15 | 17 | 6 | 2 | 5.0 | 9.6 | 8.9 | 3.9 | 1.3 |

| | | | | | | | | | | | | | | | | |
|---|---|---|---|---|---|---|---|---|---|---|---|---|---|---|---|---|
| >am_trna128_GlyGCC_6_-_27870686_27870756@4.31.28,trna133_GlyCCC_1_-_16872434_16872504@4.31.28,trna18_GlyGCC_16_+_70822597_70822667@4.31.28,trna19_GlyGCC_16_+_70823410_70823480@4.31.28,trna19_GlyGCC_2_-_157257659_157257729@4.31.28,trna24_GlyGCC_16_-_70812942_70813012@4.31.28,trna25_GlyGCC_16_-_70812114_70812184@4.31.28,trna4_GlyCCC_1_+_17188416_17188486@4.31.28,trna5_GlyGCC_17_+_8029064_8029134@4.31.28,trna68_GlyGCC_1_-_161493637_161493707@4.31.28\|TTGGTGGTTCAGTGGTAGAATTCTCGCC | >am_trna128_GlyGCC_6_-_27870686_27870756 | 4 | 31 | 28 | TTGGTGGTTCAGTGGTAGAATTCTCGCC | MB231 | 10 | 11 | 3 | 7 | 8 | 2.3 | 9.7 | 1.0 | 2.1 | 3.9 |
| >am_trna128_GlyGCC_6_-_27870686_27870756@6.28.23,trna133_GlyCCC_1_-_16872434_16872504@6.28.23,trna17_GluTTC_2_-_204229131_204229201@6.28.23,trna18_GlyGCC_16_+_70822597_70822667@6.28.23,trna19_GlyGCC_16_+_70823410_70823480@6.28.23,trna19_GlyGCC_2_-_157257659_157257729@6.28.23,trna24_GlyGCC_16_-_70812942_70813012@6.28.23,trna25_GlyGCC_16_-_70812114_70812184@6.28.23,trna2_GlyGCC_21_-_18827107_18827177@6.28.23,trna35_GlyGCC_1_+_161413094_161413164@6.28.23,trna37_GlyGCC_1_+_161420467_161420537@6.28.23,trna39_GlyGCC_1_+_161427898_161427968@6.28.23,trna41_GlyGCC_1_+_161435258_161435328@6.28.23,trna4_GlyCCC_1_+_17188416_17188486@6.28.23,trna5_GlyGCC_17_+_8029064_8029134@6.28.23,trna68_GlyGCC_1_-_161493637_161493707@6.28.23,trna91_GlyCCC_1_-_149680210_149680280@6.28.23,trna99_ValCAC_1_-_149294666_149294736@6.28.23\|GGTGGTTCAGTGGTAGAATTCTC | >am_trna128_GlyGCC_6_-_27870686_27870756 | 6 | 28 | 23 | GGTGGTTCAGTGGTAGAATTCTC | MB468 | 12 | 11 | 10 | 9 | 6 | 4.7 | 4.1 | 5.0 | 5.3 | 3.9 |
| >am_trna128_GlyGCC_6_-_27870686_27870756@7.28.22,trna133_GlyCCC_1_-_16872434_16872504@7.28.22,trna17_GluTTC_2_-_204229131_204229201@7.28.22,trna18_GlyGCC_16_+_70822597_70822667@7.28.22,trna19_GlyGCC_16_+_70823410_70823480@7.28.22,trna19_GlyGCC_2_-_157257659_157257729@7.28.22,trna24_GlyGCC_16_-_70812942_70813012@7.28.22,trna25_GlyGCC_16_-_70812114_70812184@7.28.22,trna2_GlyGCC_21_-_18827107_18827177@7.28.22,trna35_GlyGCC_1_+_161413094_161413164@7.28.22,trna37_GlyGCC_1_+_161420467_161420537@7.28.22,trna39_GlyGCC_1_+_161427898_161427968@7.28.22,trna41_GlyGCC_1_+_161435258_161435328@7.28.22,trna4_GlyCCC_1_+_17188416_17188486@7.28.22,trna5_GlyGCC_17_+_8029064_8029134@7.28.22,trna68_GlyGCC_1_-_161493637_161493707@7.28.22,trna91_GlyCCC_1_-_149680210_149680280@7.28.22,trna99_ValCAC_1_-_149294666_149294736@7.28.22\|GTGGTTCAGTGGTAGAATTCTC | >am_trna128_GlyGCC_6_-_27870686_27870756 | 7 | 28 | 22 | GTGGTTCAGTGGTAGAATTCTC | MB468 | 21 | 11 | 16 | 15 | 7 | 11.2 | 4.5 | 12.6 | 8.6 | 3.6 |
| >am_trna129_MetCAT_6_-_27870271_27870342@34.51.18,trna142_MetCAT_6_-_27560600_27560671@34.51.18,trna150_MetCAT_6_-_27300764_27300835@34.51.18,trna169_MetCAT_6_-_26330529_26330600@34.51.18,trna171_MetCAT_6_-_26313352_26313423@34.51.18,trna20_MetCAT_17_-_80452597_80452668@34.51.18,trna2_MetCAT_6_+_26286754_26286825@34.51.18,trna32_MetCAT_1_+_153643726_153643797@34.51.18,trna61_MetCAT_6_+_27745664_27745735@34.51.18\|ATAACCCAGAGGTCGATG | >am_trna129_MetCAT_6_-_27870271_27870342 | 34 | 51 | 18 | ATAACCCAGAGGTCGATG | MB468 | 29 | 5 | 7 | 29 | 5 | 20.5 | 2.7 | 6.1 | 23.1 | 1.8 |
| >am_trna13_LysCTT_14_-_58706613_58706685@31.51.21,trna2_LysCTT_15_+_79152904_79152976@31.51.21,trna4_LysCTT_18_-_43669271_43669343@31.51.21\|ACTCTTAATCCCAGGGTCGTG | >am_trna13_LysCTT_14_-_58706613_58706685 | 31 | 51 | 21 | ACTCTTAATCCCAGGGTCGTG | MB468 | 37 | 13 | 49 | 19 | 6 | 14.4 | 7.9 | 27.6 | 7.1 | 2.2 |
| >am_trna13_LysCTT_14_-_58706613_58706685@32.51.20,trna2_LysCTT_15_+_79152904_79152976@32.51.20,trna4_LysCTT_18_-_43669271_43669343@32.51.20\|CTCTTAATCCCAGGGTCGTG | >am_trna13_LysCTT_14_-_58706613_58706685 | 32 | 51 | 20 | CTCTTAATCCCAGGGTCGTG | MB468 | 10 | 6 | 10 | 17 | 2 | 4.5 | 3.8 | 6.3 | 7.6 | 0.8 |
| >am_trna130_GlnTTG_6_-_27763640_27763711@13.30.18,trna146_GlnCTG_6_-_27515531_27515602@13.30.18,trna16_GlnTTG_17_+_47269890_47269961@13.30.18,trna173_GlnTTG_6_-_26311975_26312046@13.30.18,trna174_GlnTTG_6_-_26311424_26311495@13.30.18,trna1_GlnCTG_6_+_18836402_18836473@13.30.18,trna3_GlnCTG_17_+_8023070_8023141@13.30.18,trna42_GlnCTG_6_+_27263212_27263283@13.30.18,trna49_GlnCTG_6_+_27487308_27487379@13.30.18,trna64_GlnTTG_6_+_28557156_28557227@13.30.18,trna7_GlnCTG_15_-_66161400_66161471@13.30.18,trna84_GlnTTG_6_+_145503859_145503930@13.30.18,trna99_GlnCTG_6_-_28909378_28909449@13.30.18\|TAATGGTTAGCACTCTGG | >am_trna130_GlnTTG_6_-_27763640_27763711 | 13 | 30 | 18 | TAATGGTTAGCACTCTGG | MB468 | 22 | 18 | 14 | 2 | 1 | 6.6 | 14.0 | 3.6 | 0.8 | 0.2 |
| >am_trna134_GluTTC_1_-_16861774_16861845@1.28.28,trna5_GluTTC_1_+_17199078_17199149@1.28.28,trna84_GluTTC_1_-_161391883_161391954@1.28.28,trna94_GluTTC_1_-_149664355_149664427@1.28.28\|TCCCTGGTGGTCTAGTGGCTAGGATTCG | >am_trna134_GluTTC_1_-_16861774_16861845 | 1 | 28 | 28 | TCCCTGGTGGTCTAGTGGCTAGGATTCG | MB468 | 46 | 58 | 92 | 9 | 8 | 21.5 | 33.4 | 37.6 | 5.8 | 4.5 |
| >am_trna134_GluTTC_1_-_16861774_16861845@1.29.29,trna5_GluTTC_1_+_17199078_17199149@1.29.29,trna84_GluTTC_1_-_161391883_161391954@1.29.29,trna94_GluTTC_1_-_149664355_149664427@1.29.29\|TCCCTGGTGGTCTAGTGGCTAGGATTCGG | >am_trna134_GluTTC_1_-_16861774_16861845 | 1 | 29 | 29 | TCCCTGGTGGTCTAGTGGCTAGGATTCGG | MB468 | 119 | 195 | 354 | 32 | 24 | 38.4 | 138.2 | 166.6 | 23.6 | 13.6 |
| >am_trna134_GluTTC_1_-_16861774_16861845@1.30.30,trna5_GluTTC_1_+_17199078_17199149@1.30.30,trna84_GluTTC_1_-_161391883_161391954@1.30.30,trna94_GluTTC_1_-_149664355_149664427@1.30.30\|TCCCTGGTGGTCTAGTGGCTAGGATTCGGC | >am_trna134_GluTTC_1_-_16861774_16861845 | 1 | 30 | 30 | TCCCTGGTGGTCTAGTGGCTAGGATTCGGC | MB468 | 44 | 100 | 98 | 14 | 10 | 13.8 | 73.5 | 46.0 | 9.9 | 6.2 |
| >am_trna143_LysTTT_6_-_27559593_27559665@6.24.19,trna14_LysTTT_11_-_59327808_59327880@6.24.19,trna2_LysTTT_17_+_8022473_8022545@6.24.19,trna54_LysTTT_1_+_204475655_204475727@6.24.19,trna5_LysTTT_11_+_59323902_59323974@6.24.19,trna62_LysTTT_1_-_204476158_204476230@6.24.19,trna76_LysTTT_6_+_28918806_28918878@6.24.19\|GATAGCTCAGTCGGTAGAG | >am_trna143_LysTTT_6_-_27559593_27559665 | 6 | 24 | 19 | GATAGCTCAGTCGGTAGAG | MB468 | 43 | 4 | 36 | 10 | 5 | 18.9 | 3.6 | 37.7 | 6.4 | 1.8 |
| >am_trna146_GlnCTG_6_-_27515531_27515602@1.28.28,trna1_GlnCTG_6_+_18836402_18836473@1.28.28,trna3_GlnCTG_17_+_8023070_8023141@1.28.28,trna42_GlnCTG_6_+_27263212_27263283@1.28.28,trna49_GlnCTG_6_+_27487308_27487379@1.28.28,trna7_GlnCTG_15_-_66161400_66161471@1.28.28,trna99_GlnCTG_6_-_28909378_28909449@1.28.28\|GGTTCCATGGTGTAATGGTTAGCACTCT | >am_trna146_GlnCTG_6_-_27515531_27515602 | 1 | 28 | 28 | GGTTCCATGGTGTAATGGTTAGCACTCT | MB468 | 19 | 2 | 23 | 6 | 2 | 23.1 | 1.8 | 28.0 | 6.9 | 3.1 |
| >am_trna146_GlnCTG_6_-_27515531_27515602@1.29.29,trna1_GlnCTG_6_+_18836402_18836473@1.29.29,trna3_GlnCTG_17_+_8023070_8023141@1.29.29,trna42_GlnCTG_6_+_27263212_27263283@1.29.29,trna49_GlnCTG_6_+_27487308_27487379@1.29.29,trna7_GlnCTG_15_-_66161400_66161471@1.29.29,trna99_GlnCTG_6_-_28909378_28909449@1.29.29\|GGTTCCATGGTGTAATGGTTAGCACTCTG | >am_trna146_GlnCTG_6_-_27515531_27515602 | 1 | 29 | 29 | GGTTCCATGGTGTAATGGTTAGCACTCTG | MB468 | 1326 | 226 | 1508 | 424 | 153 | 840.0 | 82.9 | 817.0 | 169.5 | 128.3 |
| >am_trna162_MetCAT_6_-_26758550_26758622@31.51.21,trna164_MetCAT_6_-_26735574_26735646@31.51.21,trna20_MetCAT_16_+_71460396_71460468@31.51.21,trna21_MetCAT_6_+_26701712_26701784@31.51.21,trna27_MetCAT_6_+_26766444_26766516@31.51.21,trna75_MetCAT_6_+_28912352_28912424@31.51.21,trna97_MetCAT_6_-_28921042_28921114@31.51.21\|TCTCATAATCTGAAGGTCCTG | >am_trna162_MetCAT_6_-_26758550_26758622 | 31 | 51 | 21 | TCTCATAATCTGAAGGTCCTG | MB468 | 52 | 16 | 25 | 12 | 12 | 28.5 | 8.0 | 19.7 | 4.2 | 5.2 |
| >am_trna162_MetCAT_6_-_26758550_26758622@31.52.22,trna164_MetCAT_6_-_26735574_26735646@31.52.22,trna20_MetCAT_16_+_71460396_71460468@31.52.22,trna21_MetCAT_6_+_26701712_26701784@31.52.22,trna27_MetCAT_6_+_26766444_26766516@31.52.22,trna75_MetCAT_6_+_28912352_28912424@31.52.22,trna97_MetCAT_6_-_28921042_28921114@31.52.22\|TCTCATAATCTGAAGGTCCTGA | >am_trna162_MetCAT_6_-_26758550_26758622 | 31 | 52 | 22 | TCTCATAATCTGAAGGTCCTGA | MB468 | 14 | 5 | 8 | 2 | 1 | 4.1 | 2.6 | 3.1 | 2.0 | 1.2 |

| | | | | | | | | | | | | | | | |
|---|---|---|---|---|---|---|---|---|---|---|---|---|---|---|---|
| >am_trna162_MetCAT_6_-_26758550_26758622@31.53.23,trna164_MetCAT_6_-_26735574_26735646@31.53.23,trna20_MetCAT_16_+_71460396_71460468@31.53.23,trna21_MetCAT_6_+_26701712_26701784@31.53.23,trna27_MetCAT_6_+_26766444_26766516@31.53.23,trna75_MetCAT_6_+_28912352_28912424@31.53.23,trna97_MetCAT_6_-_28921042_28921114@31.53.23\|TCTCATAATCTGAAGGTCCTGAG | >am_trna162_MetCAT_6_-_26758550_26758622 | 31 | 53 | 23 | TCTCATAATCTGAAGGTCCTGAG | MB468 | 43 | 13 | 20 | 12 | 10 | 19.9 | 6.4 | 12.5 | 6.6 | 3.5 |
| >am_trna162_MetCAT_6_-_26758550_26758622@31.54.24,trna164_MetCAT_6_-_26735574_26735646@31.54.24,trna20_MetCAT_16_+_71460396_71460468@31.54.24,trna21_MetCAT_6_+_26701712_26701784@31.54.24,trna27_MetCAT_6_+_26766444_26766516@31.54.24,trna75_MetCAT_6_+_28912352_28912424@31.54.24,trna97_MetCAT_6_-_28921042_28921114@31.54.24\|TCTCATAATCTGAAGGTCCTGAGT | >am_trna162_MetCAT_6_-_26758550_26758622 | 31 | 54 | 24 | TCTCATAATCTGAAGGTCCTGAGT | MB468 | 62 | 17 | 39 | 19 | 7 | 16.7 | 7.8 | 16.5 | 7.7 | 3.1 |
| >am_trna162_MetCAT_6_-_26758550_26758622@34.54.21,trna164_MetCAT_6_-_26735574_26735646@34.54.21,trna20_MetCAT_16_+_71460396_71460468@34.54.21,trna21_MetCAT_6_+_26701712_26701784@34.54.21,trna27_MetCAT_6_+_26766444_26766516@34.54.21,trna75_MetCAT_6_+_28912352_28912424@34.54.21,trna97_MetCAT_6_-_28921042_28921114@34.54.21\|CATAATCTGAAGGTCCTGAGT | >am_trna162_MetCAT_6_-_26758550_26758622 | 34 | 54 | 21 | CATAATCTGAAGGTCCTGAGT | BT20 | 63 | 18 | 32 | 11 | 7 | 14.4 | 7.1 | 11.1 | 12.8 | 0.9 |
| >am_trna162_MetCAT_6_-_26758550_26758622@34.54.21,trna164_MetCAT_6_-_26735574_26735646@34.54.21,trna20_MetCAT_16_+_71460396_71460468@34.54.21,trna21_MetCAT_6_+_26701712_26701784@34.54.21,trna27_MetCAT_6_+_26766444_26766516@34.54.21,trna75_MetCAT_6_+_28912352_28912424@34.54.21,trna97_MetCAT_6_-_28921042_28921114@34.54.21\|CATAATCTGAAGGTCCTGAGT | >am_trna162_MetCAT_6_-_26758550_26758622 | 34 | 54 | 21 | CATAATCTGAAGGTCCTGAGT | MB231 | 51 | 27 | 11 | 19 | 22 | 19.9 | 11.1 | 6.1 | 5.8 | 20.6 |
| >am_trna162_MetCAT_6_-_26758550_26758622@34.54.21,trna164_MetCAT_6_-_26735574_26735646@34.54.21,trna20_MetCAT_16_+_71460396_71460468@34.54.21,trna21_MetCAT_6_+_26701712_26701784@34.54.21,trna27_MetCAT_6_+_26766444_26766516@34.54.21,trna75_MetCAT_6_+_28912352_28912424@34.54.21,trna97_MetCAT_6_-_28921042_28921114@34.54.21\|CATAATCTGAAGGTCCTGAGT | >am_trna162_MetCAT_6_-_26758550_26758622 | 34 | 54 | 21 | CATAATCTGAAGGTCCTGAGT | MB468 | 12 | 6 | 4 | 3 | 1 | 3.4 | 3.9 | 0.7 | 1.7 | 0.7 |
| >am_trna2_ArgCCT_16_+_3202901_3202973@14.31.18,trna3_ArgCCT_7_+_139025446_139025518@14.31.18\|AATGGATAAGGCATTGGC | >am_trna2_ArgCCT_16_+_3202901_3202973 | 14 | 31 | 18 | AATGGATAAGGCATTGGC | MB468 | 34 | 46 | 14 | 7 | 2 | 18.7 | 31.6 | 4.6 | 1.8 | 0.5 |
| >am_trna2_GlyGCC_21_-_18827107_18827177@1.25.25,trna35_GlyGCC_1_+_161413094_161413164@1.25.25,trna37_GlyGCC_1_+_161420467_161420537@1.25.25,trna39_GlyGCC_1_+_161427898_161427968@1.25.25,trna41_GlyGCC_1_+_161435258_161435328@1.25.25\|GCATGGGTGGTTCAGTGGTAGAATT | >am_trna2_GlyGCC_21_-_18827107_18827177 | 1 | 25 | 25 | GCATGGGTGGTTCAGTGGTAGAATT | BT20 | 19 | 6 | 83 | 284 | 15 | 20.2 | 4.6 | 27.3 | 344.1 | 11.3 |
| >am_trna2_GlyGCC_21_-_18827107_18827177@1.27.27,trna35_GlyGCC_1_+_161413094_161413164@1.27.27,trna37_GlyGCC_1_+_161420467_161420537@1.27.27,trna39_GlyGCC_1_+_161427898_161427968@1.27.27,trna41_GlyGCC_1_+_161435258_161435328@1.27.27\|GCATGGGTGGTTCAGTGGTAGAATTCT | >am_trna2_GlyGCC_21_-_18827107_18827177 | 1 | 27 | 27 | GCATGGGTGGTTCAGTGGTAGAATTCT | BT20 | 59 | 52 | 148 | 223 | 112 | 57.3 | 79.6 | 150.6 | 235.7 | 108.5 |
| >am_trna2_GlyGCC_21_-_18827107_18827177@1.27.27,trna35_GlyGCC_1_+_161413094_161413164@1.27.27,trna37_GlyGCC_1_+_161420467_161420537@1.27.27,trna39_GlyGCC_1_+_161427898_161427968@1.27.27,trna41_GlyGCC_1_+_161435258_161435328@1.27.27\|GCATGGGTGGTTCAGTGGTAGAATTCT | >am_trna2_GlyGCC_21_-_18827107_18827177 | 1 | 27 | 27 | GCATGGGTGGTTCAGTGGTAGAATTCT | MB468 | 46 | 77 | 75 | 23 | 6 | 15.0 | 33.2 | 40.8 | 15.5 | 4.1 |
| >am_trna2_GlyGCC_21_-_18827107_18827177@4.27.24,trna35_GlyGCC_1_+_161413094_161413164@4.27.24,trna37_GlyGCC_1_+_161420467_161420537@4.27.24,trna39_GlyGCC_1_+_161427898_161427968@4.27.24,trna41_GlyGCC_1_+_161435258_161435328@4.27.24\|TGGGTGGTTCAGTGGTAGAATTCT | >am_trna2_GlyGCC_21_-_18827107_18827177 | 4 | 27 | 24 | TGGGTGGTTCAGTGGTAGAATTCT | BT20 | 10 | 10 | 14 | 5 | 6 | 16.0 | 16.5 | 20.5 | 5.3 | 9.2 |
| >am_trna2_GlyGCC_21_-_18827107_18827177@57.74.18,trna35_GlyGCC_1_+_161413094_161413164@57.74.18,trna37_GlyGCC_1_+_161420467_161420537@57.74.18,trna39_GlyGCC_1_+_161427898_161427968@57.74.18,trna41_GlyGCC_1_+_161435258_161435328@57.74.18\|TTCCCGGCCCATGCACCA | >am_trna2_GlyGCC_21_-_18827107_18827177 | 57 | 74 | 18 | TTCCCGGCCCATGCACCA | BT20 | 28 | 12 | 72 | 2 | 11 | 14.9 | 3.0 | 3.9 | 1.2 | 6.1 |
| >am_trna2_GlyTCC_19_+_4724082_4724153@1.29.29\|GCGTTGGTGGTATAGTGGTTAGCATAGCT | >am_trna2_GlyTCC_19_+_4724082_4724153 | 1 | 29 | 29 | GCGTTGGTGGTATAGTGGTTAGCATAGCT | MB468 | 14 | 9 | 39 | 9 | 4 | 7.7 | 10.7 | 20.3 | 2.6 | 4.4 |
| >am_trna2_GlyTCC_19_+_4724082_4724153@1.30.30\|GCGTTGGTGGTATAGTGGTTAGCATAGCTG | >am_trna2_GlyTCC_19_+_4724082_4724153 | 1 | 30 | 30 | GCGTTGGTGGTATAGTGGTTAGCATAGCTG | MB468 | 28 | 14 | 99 | 26 | 13 | 17.4 | 10.6 | 31.7 | 8.6 | 11.4 |
| >am_trna34_LeuCAG_1_+_161411323_161411405@69.86.18,trna36_LeuCAG_1_+_161418741_161418823@69.86.18,trna38_LeuCAG_1_+_161426122_161426204@69.86.18,trna40_LeuCAG_1_+_161433532_161433614@69.86.18,trna42_LeuCAG_1_+_161440913_161440995@69.86.18,trna67_LeuCAG_1_-_161500132_161500214@69.86.18,trna7_LeuCAG_6_+_26521436_26521518@69.86.18\|ATCCCACTCCTGACACCA | >am_trna34_LeuCAG_1_+_161411323_161411405 | 69 | 86 | 18 | ATCCCACTCCTGACACCA | MB231 | 12 | 18 | 22 | 1 | 15 | 20.7 | 30.6 | 38.0 | 1.7 | 26.4 |
| >Antisense_hg38_21_-_8433172_8446622_RNA45SN1_45S_50ntFlanks@7674.7692.19\|CTCTCTCTCTCTCTCCCCC | >Antisense_hg38_21_-_8433172_8446622_RNA45SN1_45S_50ntFlanks | 7674 | 7692 | 19 | CTCTCTCTCTCTCTCCCCC | MB468 | 13 | 36 | 29 | 0 | 7 | 7.4 | 34.0 | 26.6 | 0.6 | 14.7 |
| >Antisense_hg38_21_-_8433172_8446622_RNA45SN1_45S_50ntFlanks@7674.7693.20\|CTCTCTCTCTCTCTCCCCCG | >Antisense_hg38_21_-_8433172_8446622_RNA45SN1_45S_50ntFlanks | 7674 | 7693 | 20 | CTCTCTCTCTCTCTCCCCCG | MB468 | 29 | 113 | 77 | 0 | 18 | 16.0 | 134.6 | 49.1 | 0.5 | 22.8 |
| >hg38_1_-_228634819_228635039_RNA5S12_5S_50ntFlanks@122.147.26\|TTAGTACTTGGATGGGAGACCGCCTG | >hg38_1_-_228634819_228635039_RNA5S12_5S_50ntFlanks | 122 | 147 | 26 | TTAGTACTTGGATGGGAGACCGCCTG | MB468 | 15 | 16 | 9 | 2 | 5 | 8.5 | 15.6 | 4.7 | 2.6 | 0.8 |
| >hg38_1_-_228634819_228635039_RNA5S12_5S_50ntFlanks@122.170.49\|TTAGTACTTGGATGGGAGACCGCCTGGGAATACCGGGTGCTGTAGGCTT | >hg38_1_-_228634819_228635039_RNA5S12_5S_50ntFlanks | 122 | 170 | 49 | TTAGTACTTGGATGGGAGACCGCCTGGGAATACCGGGTGCTGTAGGCTT | MB231 | 11 | 16 | 42 | 0 | 2 | 11.0 | 26.3 | 54.4 | 0.6 | 3.2 |
| >hg38_1_-_228634819_228635039_RNA5S12_5S_50ntFlanks@124.147.24\|AGTACTTGGATGGGAGACCGCCTG | >hg38_1_-_228634819_228635039_RNA5S12_5S_50ntFlanks | 124 | 147 | 24 | AGTACTTGGATGGGAGACCGCCTG | MB468 | 29 | 32 | 12 | 17 | 14 | 10.1 | 23.6 | 4.4 | 15.3 | 1.7 |
| >hg38_1_-_228634819_228635039_RNA5S12_5S_50ntFlanks@124.170.47\|AGTACTTGGATGGGAGACCGCCTGGGAATACCGGGTGCTGTAGGCTT | >hg38_1_-_228634819_228635039_RNA5S12_5S_50ntFlanks | 124 | 170 | 47 | AGTACTTGGATGGGAGACCGCCTGGGAATACCGGGTGCTGTAGGCTT | MB231 | 11 | 19 | 55 | 2 | 2 | 8.5 | 30.6 | 66.4 | 2.8 | 4.0 |
| >hg38_1_-_228634819_228635039_RNA5S12_5S_50ntFlanks@125.170.46\|GTACTTGGATGGGAGACCGCCTGGGAATACCGGGTGCTGTAGGCTT | >hg38_1_-_228634819_228635039_RNA5S12_5S_50ntFlanks | 125 | 170 | 46 | GTACTTGGATGGGAGACCGCCTGGGAATACCGGGTGCTGTAGGCTT | MB231 | 12 | 23 | 64 | 5 | 4 | 10.7 | 35.7 | 89.2 | 6.8 | 5.0 |
| >hg38_1_-_228634819_228635039_RNA5S12_5S_50ntFlanks@126.170.45\|TACTTGGATGGGAGACCGCCTGGGAATACCGGGTGCTGTAGGCTT | >hg38_1_-_228634819_228635039_RNA5S12_5S_50ntFlanks | 126 | 170 | 45 | TACTTGGATGGGAGACCGCCTGGGAATACCGGGTGCTGTAGGCTT | BT20 | 16 | 6 | 64 | 40 | 47 | 18.3 | 4.4 | 96.8 | 33.9 | 45.3 |
| >hg38_1_-_228634819_228635039_RNA5S12_5S_50ntFlanks@126.170.45\|TACTTGGATGGGAGACCGCCTGGGAATACCGGGTGCTGTAGGCTT | >hg38_1_-_228634819_228635039_RNA5S12_5S_50ntFlanks | 126 | 170 | 45 | TACTTGGATGGGAGACCGCCTGGGAATACCGGGTGCTGTAGGCTT | MB231 | 31 | 61 | 223 | 7 | 9 | 26.4 | 90.9 | 242.9 | 8.7 | 8.8 |
| >hg38_1_-_228634819_228635039_RNA5S12_5S_50ntFlanks@126.171.46\|TACTTGGATGGGAGACCGCCTGGGAATACCGGGTGCTGTAGGCTTT | >hg38_1_-_228634819_228635039_RNA5S12_5S_50ntFlanks | 126 | 171 | 46 | TACTTGGATGGGAGACCGCCTGGGAATACCGGGTGCTGTAGGCTTT | MB231 | 28 | 47 | 153 | 6 | 9 | 20.1 | 73.8 | 170.6 | 8.3 | 10.2 |
| >hg38_1_-_228634819_228635039_RNA5S12_5S_50ntFlanks@127.170.44\|ACTTGGATGGGAGACCGCCTGGGAATACCGGGTGCTGTAGGCTT | >hg38_1_-_228634819_228635039_RNA5S12_5S_50ntFlanks | 127 | 170 | 44 | ACTTGGATGGGAGACCGCCTGGGAATACCGGGTGCTGTAGGCTT | BT20 | 18 | 10 | 63 | 47 | 67 | 17.4 | 8.0 | 88.4 | 25.9 | 56.1 |
| >hg38_1_-_228634819_228635039_RNA5S12_5S_50ntFlanks@127.170.44\|ACTTGGATGGGAGACCGCCTGGGAATACCGGGTGCTGTAGGCTT | >hg38_1_-_228634819_228635039_RNA5S12_5S_50ntFlanks | 127 | 170 | 44 | ACTTGGATGGGAGACCGCCTGGGAATACCGGGTGCTGTAGGCTT | MB231 | 33 | 53 | 183 | 10 | 11 | 25.5 | 60.8 | 185.0 | 8.6 | 6.2 |
| >hg38_1_-_228634819_228635039_RNA5S12_5S_50ntFlanks@127.171.45\|ACTTGGATGGGAGACCGCCTGGGAATACCGGGTGCTGTAGGCTTT | >hg38_1_-_228634819_228635039_RNA5S12_5S_50ntFlanks | 127 | 171 | 45 | ACTTGGATGGGAGACCGCCTGGGAATACCGGGTGCTGTAGGCTTT | MB231 | 29 | 42 | 148 | 6 | 9 | 20.8 | 55.0 | 147.8 | 7.5 | 8.2 |

| | | | | | | | | | | | | | | | |
|---|---|---|---|---|---|---|---|---|---|---|---|---|---|---|---|
| >hg38_1_-_228634819_228635039_RNA5S12_5S_50ntFlanks@128.169.42\|CTTGGATGGGAGACCGCCTGGGAATACCGGGTGCTGTAGGCT | >hg38_1_-_228634819_228635039_RNA5S12_5S_50ntFlanks | 128 | 169 | 42 | CTTGGATGGGAGACCGCCTGGGAATACCGGGTGCTGTAGGCT | BT20 | 10 | 5 | 55 | 22 | 14 | 8.4 | 2.5 | 73.5 | 9.5 | 9.2 |
| >hg38_1_-_228634819_228635039_RNA5S12_5S_50ntFlanks@128.170.43\|CTTGGATGGGAGACCGCCTGGGAATACCGGGTGCTGTAGGCTT | >hg38_1_-_228634819_228635039_RNA5S12_5S_50ntFlanks | 128 | 170 | 43 | CTTGGATGGGAGACCGCCTGGGAATACCGGGTGCTGTAGGCTT | BT20 | 23 | 13 | 106 | 68 | 63 | 20.4 | 10.4 | 153.2 | 20.6 | 49.6 |
| >hg38_1_-_228634819_228635039_RNA5S12_5S_50ntFlanks@128.170.43\|CTTGGATGGGAGACCGCCTGGGAATACCGGGTGCTGTAGGCTT | >hg38_1_-_228634819_228635039_RNA5S12_5S_50ntFlanks | 128 | 170 | 43 | CTTGGATGGGAGACCGCCTGGGAATACCGGGTGCTGTAGGCTT | MB231 | 28 | 40 | 143 | 11 | 7 | 28.2 | 41.8 | 143.8 | 11.8 | 5.4 |
| >hg38_1_-_228634819_228635039_RNA5S12_5S_50ntFlanks@128.171.44\|CTTGGATGGGAGACCGCCTGGGAATACCGGGTGCTGTAGGCTTT | >hg38_1_-_228634819_228635039_RNA5S12_5S_50ntFlanks | 128 | 171 | 44 | CTTGGATGGGAGACCGCCTGGGAATACCGGGTGCTGTAGGCTTT | BT20 | 13 | 6 | 72 | 19 | 26 | 16.9 | 6.9 | 115.8 | 13.5 | 22.6 |
| >hg38_1_-_228634819_228635039_RNA5S12_5S_50ntFlanks@128.171.44\|CTTGGATGGGAGACCGCCTGGGAATACCGGGTGCTGTAGGCTTT | >hg38_1_-_228634819_228635039_RNA5S12_5S_50ntFlanks | 128 | 171 | 44 | CTTGGATGGGAGACCGCCTGGGAATACCGGGTGCTGTAGGCTTT | MB231 | 27 | 40 | 148 | 8 | 8 | 21.1 | 51.2 | 163.1 | 11.4 | 6.5 |
| >hg38_1_-_228634819_228635039_RNA5S12_5S_50ntFlanks@129.169.41\|TTGGATGGGAGACCGCCTGGGAATACCGGGTGCTGTAGGCT | >hg38_1_-_228634819_228635039_RNA5S12_5S_50ntFlanks | 129 | 169 | 41 | TTGGATGGGAGACCGCCTGGGAATACCGGGTGCTGTAGGCT | BT20 | 14 | 6 | 63 | 46 | 18 | 9.5 | 1.5 | 69.1 | 39.7 | 10.2 |
| >hg38_1_-_228634819_228635039_RNA5S12_5S_50ntFlanks@129.170.42\|TTGGATGGGAGACCGCCTGGGAATACCGGGTGCTGTAGGCTT | >hg38_1_-_228634819_228635039_RNA5S12_5S_50ntFlanks | 129 | 170 | 42 | TTGGATGGGAGACCGCCTGGGAATACCGGGTGCTGTAGGCTT | BT20 | 38 | 17 | 140 | 250 | 103 | 26.9 | 8.6 | 175.5 | 226.7 | 71.8 |
| >hg38_1_-_228634819_228635039_RNA5S12_5S_50ntFlanks@129.170.42\|TTGGATGGGAGACCGCCTGGGAATACCGGGTGCTGTAGGCTT | >hg38_1_-_228634819_228635039_RNA5S12_5S_50ntFlanks | 129 | 170 | 42 | TTGGATGGGAGACCGCCTGGGAATACCGGGTGCTGTAGGCTT | MB231 | 57 | 80 | 306 | 41 | 16 | 47.9 | 81.8 | 322.5 | 34.1 | 8.5 |
| >hg38_1_-_228634819_228635039_RNA5S12_5S_50ntFlanks@129.171.43\|TTGGATGGGAGACCGCCTGGGAATACCGGGTGCTGTAGGCTTT | >hg38_1_-_228634819_228635039_RNA5S12_5S_50ntFlanks | 129 | 171 | 43 | TTGGATGGGAGACCGCCTGGGAATACCGGGTGCTGTAGGCTTT | BT20 | 19 | 7 | 83 | 48 | 39 | 21.9 | 4.1 | 129.3 | 52.1 | 28.4 |
| >hg38_1_-_228634819_228635039_RNA5S12_5S_50ntFlanks@129.171.43\|TTGGATGGGAGACCGCCTGGGAATACCGGGTGCTGTAGGCTTT | >hg38_1_-_228634819_228635039_RNA5S12_5S_50ntFlanks | 129 | 171 | 43 | TTGGATGGGAGACCGCCTGGGAATACCGGGTGCTGTAGGCTTT | MB231 | 47 | 62 | 229 | 20 | 15 | 28.5 | 72.4 | 227.1 | 22.8 | 10.1 |
| >hg38_1_-_228634819_228635039_RNA5S12_5S_50ntFlanks@130.169.40\|TGGATGGGAGACCGCCTGGGAATACCGGGTGCTGTAGGCT | >hg38_1_-_228634819_228635039_RNA5S12_5S_50ntFlanks | 130 | 169 | 40 | TGGATGGGAGACCGCCTGGGAATACCGGGTGCTGTAGGCT | BT20 | 17 | 9 | 91 | 67 | 24 | 5.6 | 3.4 | 88.6 | 42.5 | 17.7 |
| >hg38_1_-_228634819_228635039_RNA5S12_5S_50ntFlanks@130.170.41\|TGGATGGGAGACCGCCTGGGAATACCGGGTGCTGTAGGCTT | >hg38_1_-_228634819_228635039_RNA5S12_5S_50ntFlanks | 130 | 170 | 41 | TGGATGGGAGACCGCCTGGGAATACCGGGTGCTGTAGGCTT | BT20 | 46 | 29 | 211 | 394 | 140 | 17.0 | 11.3 | 231.3 | 291.3 | 110.4 |
| >hg38_1_-_228634819_228635039_RNA5S12_5S_50ntFlanks@130.170.41\|TGGATGGGAGACCGCCTGGGAATACCGGGTGCTGTAGGCTT | >hg38_1_-_228634819_228635039_RNA5S12_5S_50ntFlanks | 130 | 170 | 41 | TGGATGGGAGACCGCCTGGGAATACCGGGTGCTGTAGGCTT | MB231 | 59 | 76 | 333 | 59 | 15 | 55.8 | 71.9 | 353.8 | 49.8 | 7.8 |
| >hg38_1_-_228634819_228635039_RNA5S12_5S_50ntFlanks@130.171.42\|TGGATGGGAGACCGCCTGGGAATACCGGGTGCTGTAGGCTTT | >hg38_1_-_228634819_228635039_RNA5S12_5S_50ntFlanks | 130 | 171 | 42 | TGGATGGGAGACCGCCTGGGAATACCGGGTGCTGTAGGCTTT | BT20 | 21 | 11 | 118 | 70 | 53 | 17.3 | 5.8 | 173.8 | 63.3 | 40.7 |
| >hg38_1_-_228634819_228635039_RNA5S12_5S_50ntFlanks@130.171.42\|TGGATGGGAGACCGCCTGGGAATACCGGGTGCTGTAGGCTTT | >hg38_1_-_228634819_228635039_RNA5S12_5S_50ntFlanks | 130 | 171 | 42 | TGGATGGGAGACCGCCTGGGAATACCGGGTGCTGTAGGCTTT | MB231 | 49 | 57 | 240 | 29 | 14 | 34.4 | 58.4 | 236.7 | 32.9 | 9.6 |
| >hg38_1_-_228634819_228635039_RNA5S12_5S_50ntFlanks@131.170.40\|GGATGGGAGACCGCCTGGGAATACCGGGTGCTGTAGGCTT | >hg38_1_-_228634819_228635039_RNA5S12_5S_50ntFlanks | 131 | 170 | 40 | GGATGGGAGACCGCCTGGGAATACCGGGTGCTGTAGGCTT | BT20 | 23 | 20 | 95 | 319 | 93 | 1.8 | 12.6 | 92.2 | 208.0 | 99.6 |
| >hg38_1_-_228634819_228635039_RNA5S12_5S_50ntFlanks@131.170.40\|GGATGGGAGACCGCCTGGGAATACCGGGTGCTGTAGGCTT | >hg38_1_-_228634819_228635039_RNA5S12_5S_50ntFlanks | 131 | 170 | 40 | GGATGGGAGACCGCCTGGGAATACCGGGTGCTGTAGGCTT | MB231 | 39 | 44 | 171 | 35 | 8 | 34.8 | 34.4 | 160.6 | 22.3 | 3.8 |
| >hg38_1_-_228634819_228635039_RNA5S12_5S_50ntFlanks@131.171.41\|GGATGGGAGACCGCCTGGGAATACCGGGTGCTGTAGGCTTT | >hg38_1_-_228634819_228635039_RNA5S12_5S_50ntFlanks | 131 | 171 | 41 | GGATGGGAGACCGCCTGGGAATACCGGGTGCTGTAGGCTTT | BT20 | 18 | 18 | 72 | 66 | 40 | 10.2 | 9.5 | 101.4 | 45.5 | 42.6 |
| >hg38_1_-_228634819_228635039_RNA5S12_5S_50ntFlanks@131.171.41\|GGATGGGAGACCGCCTGGGAATACCGGGTGCTGTAGGCTTT | >hg38_1_-_228634819_228635039_RNA5S12_5S_50ntFlanks | 131 | 171 | 41 | GGATGGGAGACCGCCTGGGAATACCGGGTGCTGTAGGCTTT | MB231 | 35 | 38 | 140 | 16 | 8 | 20.6 | 28.2 | 121.8 | 12.6 | 3.5 |
| >hg38_1_-_228634819_228635039_RNA5S12_5S_50ntFlanks@132.169.38\|GATGGGAGACCGCCTGGGAATACCGGGTGCTGTAGGCT | >hg38_1_-_228634819_228635039_RNA5S12_5S_50ntFlanks | 132 | 169 | 38 | GATGGGAGACCGCCTGGGAATACCGGGTGCTGTAGGCT | BT20 | 113 | 96 | 456 | 851 | 209 | 79.8 | 112.5 | 285.9 | 908.7 | 262.7 |
| >hg38_1_-_228634819_228635039_RNA5S12_5S_50ntFlanks@132.169.38\|GATGGGAGACCGCCTGGGAATACCGGGTGCTGTAGGCT | >hg38_1_-_228634819_228635039_RNA5S12_5S_50ntFlanks | 132 | 169 | 38 | GATGGGAGACCGCCTGGGAATACCGGGTGCTGTAGGCT | MB231 | 48 | 29 | 156 | 30 | 13 | 18.3 | 8.9 | 56.0 | 11.2 | 5.4 |
| >hg38_1_-_228634819_228635039_RNA5S12_5S_50ntFlanks@132.169.38\|GATGGGAGACCGCCTGGGAATACCGGGTGCTGTAGGCT | >hg38_1_-_228634819_228635039_RNA5S12_5S_50ntFlanks | 132 | 169 | 38 | GATGGGAGACCGCCTGGGAATACCGGGTGCTGTAGGCT | MB468 | 30 | 60 | 128 | 86 | 66 | 23.7 | 34.0 | 120.7 | 56.9 | 62.4 |
| >hg38_1_-_228634819_228635039_RNA5S12_5S_50ntFlanks@132.170.39\|GATGGGAGACCGCCTGGGAATACCGGGTGCTGTAGGCTT | >hg38_1_-_228634819_228635039_RNA5S12_5S_50ntFlanks | 132 | 170 | 39 | GATGGGAGACCGCCTGGGAATACCGGGTGCTGTAGGCTT | BT20 | 578 | 724 | 1598 | 9665 | 2424 | 392.9 | 725.6 | 937.7 | 10602.6 | 3159.8 |
| >hg38_1_-_228634819_228635039_RNA5S12_5S_50ntFlanks@132.170.39\|GATGGGAGACCGCCTGGGAATACCGGGTGCTGTAGGCTT | >hg38_1_-_228634819_228635039_RNA5S12_5S_50ntFlanks | 132 | 170 | 39 | GATGGGAGACCGCCTGGGAATACCGGGTGCTGTAGGCTT | MB231 | 534 | 602 | 1845 | 460 | 230 | 309.6 | 255.8 | 1625.4 | 224.0 | 145.2 |
| >hg38_1_-_228634819_228635039_RNA5S12_5S_50ntFlanks@132.170.39\|GATGGGAGACCGCCTGGGAATACCGGGTGCTGTAGGCTT | >hg38_1_-_228634819_228635039_RNA5S12_5S_50ntFlanks | 132 | 170 | 39 | GATGGGAGACCGCCTGGGAATACCGGGTGCTGTAGGCTT | MB468 | 236 | 557 | 1013 | 697 | 674 | 178.5 | 307.9 | 1060.3 | 383.5 | 624.5 |
| >hg38_1_-_228634819_228635039_RNA5S12_5S_50ntFlanks@132.171.40\|GATGGGAGACCGCCTGGGAATACCGGGTGCTGTAGGCTTT | >hg38_1_-_228634819_228635039_RNA5S12_5S_50ntFlanks | 132 | 171 | 40 | GATGGGAGACCGCCTGGGAATACCGGGTGCTGTAGGCTTT | BT20 | 170 | 149 | 442 | 1396 | 470 | 75.5 | 110.7 | 440.2 | 1679.5 | 568.4 |
| >hg38_1_-_228634819_228635039_RNA5S12_5S_50ntFlanks@132.171.40\|GATGGGAGACCGCCTGGGAATACCGGGTGCTGTAGGCTTT | >hg38_1_-_228634819_228635039_RNA5S12_5S_50ntFlanks | 132 | 171 | 40 | GATGGGAGACCGCCTGGGAATACCGGGTGCTGTAGGCTTT | MB231 | 268 | 247 | 646 | 146 | 119 | 85.4 | 112.2 | 379.8 | 95.3 | 52.6 |
| >hg38_1_-_228634819_228635039_RNA5S12_5S_50ntFlanks@132.171.40\|GATGGGAGACCGCCTGGGAATACCGGGTGCTGTAGGCTTT | >hg38_1_-_228634819_228635039_RNA5S12_5S_50ntFlanks | 132 | 171 | 40 | GATGGGAGACCGCCTGGGAATACCGGGTGCTGTAGGCTTT | MB468 | 48 | 116 | 255 | 148 | 212 | 28.0 | 78.5 | 333.6 | 96.6 | 211.2 |
| >hg38_1_-_228634819_228635039_RNA5S12_5S_50ntFlanks@133.169.37\|ATGGGAGACCGCCTGGGAATACCGGGTGCTGTAGGCT | >hg38_1_-_228634819_228635039_RNA5S12_5S_50ntFlanks | 133 | 169 | 37 | ATGGGAGACCGCCTGGGAATACCGGGTGCTGTAGGCT | BT20 | 92 | 63 | 302 | 316 | 102 | 57.6 | 65.2 | 191.2 | 330.8 | 110.2 |
| >hg38_1_-_228634819_228635039_RNA5S12_5S_50ntFlanks@133.169.37\|ATGGGAGACCGCCTGGGAATACCGGGTGCTGTAGGCT | >hg38_1_-_228634819_228635039_RNA5S12_5S_50ntFlanks | 133 | 169 | 37 | ATGGGAGACCGCCTGGGAATACCGGGTGCTGTAGGCT | MB231 | 38 | 26 | 84 | 15 | 9 | 9.9 | 4.7 | 30.3 | 7.9 | 3.9 |
| >hg38_1_-_228634819_228635039_RNA5S12_5S_50ntFlanks@133.170.38\|ATGGGAGACCGCCTGGGAATACCGGGTGCTGTAGGCTT | >hg38_1_-_228634819_228635039_RNA5S12_5S_50ntFlanks | 133 | 170 | 38 | ATGGGAGACCGCCTGGGAATACCGGGTGCTGTAGGCTT | BT20 | 317 | 307 | 637 | 2882 | 918 | 141.5 | 246.5 | 384.2 | 3000.6 | 997.5 |
| >hg38_1_-_228634819_228635039_RNA5S12_5S_50ntFlanks@133.170.38\|ATGGGAGACCGCCTGGGAATACCGGGTGCTGTAGGCTT | >hg38_1_-_228634819_228635039_RNA5S12_5S_50ntFlanks | 133 | 170 | 38 | ATGGGAGACCGCCTGGGAATACCGGGTGCTGTAGGCTT | MB231 | 292 | 366 | 480 | 165 | 122 | 145.7 | 91.6 | 354.3 | 78.2 | 74.3 |
| >hg38_1_-_228634819_228635039_RNA5S12_5S_50ntFlanks@133.170.38\|ATGGGAGACCGCCTGGGAATACCGGGTGCTGTAGGCTT | >hg38_1_-_228634819_228635039_RNA5S12_5S_50ntFlanks | 133 | 170 | 38 | ATGGGAGACCGCCTGGGAATACCGGGTGCTGTAGGCTT | MB468 | 53 | 74 | 137 | 203 | 189 | 34.8 | 37.5 | 131.3 | 119.7 | 176.1 |
| >hg38_1_-_228634819_228635039_RNA5S12_5S_50ntFlanks@133.171.39\|ATGGGAGACCGCCTGGGAATACCGGGTGCTGTAGGCTTT | >hg38_1_-_228634819_228635039_RNA5S12_5S_50ntFlanks | 133 | 171 | 39 | ATGGGAGACCGCCTGGGAATACCGGGTGCTGTAGGCTTT | BT20 | 128 | 81 | 253 | 438 | 204 | 61.7 | 45.9 | 249.4 | 509.6 | 216.2 |

| | | | | | | | | | | | | | | | | |
|---|---|---|---|---|---|---|---|---|---|---|---|---|---|---|---|---|
| >hg38_1_-_228634819_228635039_RNA5S12_5S_50ntFlanks@133.171.39\|ATGGGAGACCGCCTGGGAATACCGGGTGCTGTAGGCTTT | >hg38_1_-_228634819_228635039_RNA5S12_5S_50ntFlanks | 133 | 171 | 39 | ATGGGAGACCGCCTGGGAATACCGGGTGCTGTAGGCTTT | MB231 | 198 | 197 | 357 | 70 | 78 | 67.6 | 82.2 | 289.5 | 62.8 | 36.2 |
| >hg38_1_-_228634819_228635039_RNA5S12_5S_50ntFlanks@133.171.39\|ATGGGAGACCGCCTGGGAATACCGGGTGCTGTAGGCTTT | >hg38_1_-_228634819_228635039_RNA5S12_5S_50ntFlanks | 133 | 171 | 39 | ATGGGAGACCGCCTGGGAATACCGGGTGCTGTAGGCTTT | MB468 | 14 | 20 | 48 | 65 | 78 | 6.4 | 13.4 | 57.8 | 55.0 | 79.5 |
| >hg38_1_-_228634819_228635039_RNA5S12_5S_50ntFlanks@134.168.35\|TGGGAGACCGCCTGGGAATACCGGGTGCTGTAGGC | >hg38_1_-_228634819_228635039_RNA5S12_5S_50ntFlanks | 134 | 168 | 35 | TGGGAGACCGCCTGGGAATACCGGGTGCTGTAGGC | MB231 | 19 | 11 | 28 | 3 | 4 | 21.1 | 11.9 | 24.6 | 3.9 | 5.6 |
| >hg38_1_-_228634819_228635039_RNA5S12_5S_50ntFlanks@134.169.36\|TGGGAGACCGCCTGGGAATACCGGGTGCTGTAGGCT | >hg38_1_-_228634819_228635039_RNA5S12_5S_50ntFlanks | 134 | 169 | 36 | TGGGAGACCGCCTGGGAATACCGGGTGCTGTAGGCT | BT20 | 294 | 223 | 631 | 198 | 324 | 305.5 | 274.2 | 370.1 | 181.0 | 423.6 |
| >hg38_1_-_228634819_228635039_RNA5S12_5S_50ntFlanks@134.169.36\|TGGGAGACCGCCTGGGAATACCGGGTGCTGTAGGCT | >hg38_1_-_228634819_228635039_RNA5S12_5S_50ntFlanks | 134 | 169 | 36 | TGGGAGACCGCCTGGGAATACCGGGTGCTGTAGGCT | MB231 | 431 | 284 | 768 | 161 | 142 | 396.5 | 242.6 | 667.6 | 199.6 | 151.2 |
| >hg38_1_-_228634819_228635039_RNA5S12_5S_50ntFlanks@134.169.36\|TGGGAGACCGCCTGGGAATACCGGGTGCTGTAGGCT | >hg38_1_-_228634819_228635039_RNA5S12_5S_50ntFlanks | 134 | 169 | 36 | TGGGAGACCGCCTGGGAATACCGGGTGCTGTAGGCT | MB468 | 12 | 23 | 13 | 15 | 9 | 3.9 | 18.1 | 7.3 | 9.0 | 7.4 |
| >hg38_1_-_228634819_228635039_RNA5S12_5S_50ntFlanks@134.170.37\|TGGGAGACCGCCTGGGAATACCGGGTGCTGTAGGCTT | >hg38_1_-_228634819_228635039_RNA5S12_5S_50ntFlanks | 134 | 170 | 37 | TGGGAGACCGCCTGGGAATACCGGGTGCTGTAGGCTT | BT20 | 1370 | 1235 | 1649 | 2661 | 3372 | 1187.0 | 1315.6 | 515.8 | 2121.9 | 4623.0 |
| >hg38_1_-_228634819_228635039_RNA5S12_5S_50ntFlanks@134.170.37\|TGGGAGACCGCCTGGGAATACCGGGTGCTGTAGGCTT | >hg38_1_-_228634819_228635039_RNA5S12_5S_50ntFlanks | 134 | 170 | 37 | TGGGAGACCGCCTGGGAATACCGGGTGCTGTAGGCTT | MB231 | 5679 | 4965 | 6894 | 2899 | 2129 | 7115.6 | 4509.5 | 8512.1 | 3514.5 | 2155.0 |
| >hg38_1_-_228634819_228635039_RNA5S12_5S_50ntFlanks@134.170.37\|TGGGAGACCGCCTGGGAATACCGGGTGCTGTAGGCTT | >hg38_1_-_228634819_228635039_RNA5S12_5S_50ntFlanks | 134 | 170 | 37 | TGGGAGACCGCCTGGGAATACCGGGTGCTGTAGGCTT | MB468 | 95 | 194 | 68 | 104 | 70 | 34.1 | 181.3 | 36.9 | 68.1 | 48.7 |
| >hg38_1_-_228634819_228635039_RNA5S12_5S_50ntFlanks@134.171.38\|TGGGAGACCGCCTGGGAATACCGGGTGCTGTAGGCTTT | >hg38_1_-_228634819_228635039_RNA5S12_5S_50ntFlanks | 134 | 171 | 38 | TGGGAGACCGCCTGGGAATACCGGGTGCTGTAGGCTTT | BT20 | 427 | 255 | 468 | 331 | 575 | 306.6 | 225.5 | 206.8 | 241.3 | 742.6 |
| >hg38_1_-_228634819_228635039_RNA5S12_5S_50ntFlanks@134.171.38\|TGGGAGACCGCCTGGGAATACCGGGTGCTGTAGGCTTT | >hg38_1_-_228634819_228635039_RNA5S12_5S_50ntFlanks | 134 | 171 | 38 | TGGGAGACCGCCTGGGAATACCGGGTGCTGTAGGCTTT | MB231 | 2704 | 2116 | 2480 | 1535 | 1360 | 2667.4 | 1905.2 | 2279.2 | 2204.0 | 1560.9 |
| >hg38_1_-_228634819_228635039_RNA5S12_5S_50ntFlanks@134.171.38\|TGGGAGACCGCCTGGGAATACCGGGTGCTGTAGGCTTT | >hg38_1_-_228634819_228635039_RNA5S12_5S_50ntFlanks | 134 | 171 | 38 | TGGGAGACCGCCTGGGAATACCGGGTGCTGTAGGCTTT | MB468 | 23 | 38 | 19 | 26 | 24 | 12.3 | 30.8 | 14.2 | 19.9 | 20.4 |
| >hg38_1_-_228634819_228635039_RNA5S12_5S_50ntFlanks@134.172.39\|TGGGAGACCGCCTGGGAATACCGGGTGCTGTAGGCTTTT | >hg38_1_-_228634819_228635039_RNA5S12_5S_50ntFlanks | 134 | 172 | 39 | TGGGAGACCGCCTGGGAATACCGGGTGCTGTAGGCTTTT | MB231 | 68 | 45 | 86 | 64 | 38 | 62.2 | 46.1 | 73.1 | 104.7 | 51.2 |
| >hg38_1_-_228634819_228635039_RNA5S12_5S_50ntFlanks@134.173.40\|TGGGAGACCGCCTGGGAATACCGGGTGCTGTAGGCTTTTT | >hg38_1_-_228634819_228635039_RNA5S12_5S_50ntFlanks | 134 | 173 | 40 | TGGGAGACCGCCTGGGAATACCGGGTGCTGTAGGCTTTTT | MB231 | 12 | 7 | 20 | 6 | 4 | 8.4 | 7.3 | 17.2 | 9.5 | 7.2 |
| >hg38_1_-_228634819_228635039_RNA5S12_5S_50ntFlanks@135.169.35\|GGGAGACCGCCTGGGAATACCGGGTGCTGTAGGCT | >hg38_1_-_228634819_228635039_RNA5S12_5S_50ntFlanks | 135 | 169 | 35 | GGGAGACCGCCTGGGAATACCGGGTGCTGTAGGCT | BT20 | 59 | 62 | 167 | 77 | 74 | 39.8 | 71.8 | 99.4 | 73.3 | 90.9 |
| >hg38_1_-_228634819_228635039_RNA5S12_5S_50ntFlanks@135.169.35\|GGGAGACCGCCTGGGAATACCGGGTGCTGTAGGCT | >hg38_1_-_228634819_228635039_RNA5S12_5S_50ntFlanks | 135 | 169 | 35 | GGGAGACCGCCTGGGAATACCGGGTGCTGTAGGCT | MB231 | 55 | 36 | 65 | 32 | 19 | 43.7 | 16.2 | 30.2 | 17.3 | 14.0 |
| >hg38_1_-_228634819_228635039_RNA5S12_5S_50ntFlanks@135.170.36\|GGGAGACCGCCTGGGAATACCGGGTGCTGTAGGCTT | >hg38_1_-_228634819_228635039_RNA5S12_5S_50ntFlanks | 135 | 170 | 36 | GGGAGACCGCCTGGGAATACCGGGTGCTGTAGGCTT | BT20 | 469 | 655 | 803 | 1923 | 1451 | 306.6 | 672.8 | 260.1 | 1518.3 | 1886.4 |
| >hg38_1_-_228634819_228635039_RNA5S12_5S_50ntFlanks@135.170.36\|GGGAGACCGCCTGGGAATACCGGGTGCTGTAGGCTT | >hg38_1_-_228634819_228635039_RNA5S12_5S_50ntFlanks | 135 | 170 | 36 | GGGAGACCGCCTGGGAATACCGGGTGCTGTAGGCTT | MB231 | 1302 | 1115 | 1241 | 896 | 459 | 1636.3 | 687.0 | 1337.8 | 532.1 | 329.5 |
| >hg38_1_-_228634819_228635039_RNA5S12_5S_50ntFlanks@135.170.36\|GGGAGACCGCCTGGGAATACCGGGTGCTGTAGGCTT | >hg38_1_-_228634819_228635039_RNA5S12_5S_50ntFlanks | 135 | 170 | 36 | GGGAGACCGCCTGGGAATACCGGGTGCTGTAGGCTT | MB468 | 48 | 103 | 53 | 186 | 131 | 21.5 | 58.1 | 26.7 | 85.1 | 86.8 |
| >hg38_1_-_228634819_228635039_RNA5S12_5S_50ntFlanks@135.171.37\|GGGAGACCGCCTGGGAATACCGGGTGCTGTAGGCTTT | >hg38_1_-_228634819_228635039_RNA5S12_5S_50ntFlanks | 135 | 171 | 37 | GGGAGACCGCCTGGGAATACCGGGTGCTGTAGGCTTT | BT20 | 321 | 251 | 474 | 280 | 320 | 107.6 | 101.4 | 352.6 | 163.9 | 375.3 |
| >hg38_1_-_228634819_228635039_RNA5S12_5S_50ntFlanks@135.171.37\|GGGAGACCGCCTGGGAATACCGGGTGCTGTAGGCTTT | >hg38_1_-_228634819_228635039_RNA5S12_5S_50ntFlanks | 135 | 171 | 37 | GGGAGACCGCCTGGGAATACCGGGTGCTGTAGGCTTT | MB231 | 777 | 572 | 690 | 414 | 330 | 665.1 | 350.8 | 590.5 | 458.5 | 289.1 |
| >hg38_1_-_228634819_228635039_RNA5S12_5S_50ntFlanks@135.171.37\|GGGAGACCGCCTGGGAATACCGGGTGCTGTAGGCTTT | >hg38_1_-_228634819_228635039_RNA5S12_5S_50ntFlanks | 135 | 171 | 37 | GGGAGACCGCCTGGGAATACCGGGTGCTGTAGGCTTT | MB468 | 13 | 23 | 19 | 49 | 43 | 5.3 | 9.2 | 11.6 | 30.5 | 34.3 |
| >hg38_1_-_228634819_228635039_RNA5S12_5S_50ntFlanks@135.172.38\|GGGAGACCGCCTGGGAATACCGGGTGCTGTAGGCTTTT | >hg38_1_-_228634819_228635039_RNA5S12_5S_50ntFlanks | 135 | 172 | 38 | GGGAGACCGCCTGGGAATACCGGGTGCTGTAGGCTTTT | BT20 | 15 | 10 | 22 | 3 | 1 | 17.7 | 7.4 | 30.5 | 1.8 | 1.0 |
| >hg38_1_-_228634819_228635039_RNA5S12_5S_50ntFlanks@135.172.38\|GGGAGACCGCCTGGGAATACCGGGTGCTGTAGGCTTTT | >hg38_1_-_228634819_228635039_RNA5S12_5S_50ntFlanks | 135 | 172 | 38 | GGGAGACCGCCTGGGAATACCGGGTGCTGTAGGCTTTT | MB231 | 20 | 12 | 19 | 13 | 7 | 13.9 | 9.8 | 13.3 | 19.8 | 8.7 |
| >hg38_1_-_228634819_228635039_RNA5S12_5S_50ntFlanks@136.169.34\|GGAGACCGCCTGGGAATACCGGGTGCTGTAGGCT | >hg38_1_-_228634819_228635039_RNA5S12_5S_50ntFlanks | 136 | 169 | 34 | GGAGACCGCCTGGGAATACCGGGTGCTGTAGGCT | BT20 | 12 | 9 | 14 | 93 | 20 | 9.7 | 7.9 | 8.9 | 113.3 | 24.1 |
| >hg38_1_-_228634819_228635039_RNA5S12_5S_50ntFlanks@136.170.35\|GGAGACCGCCTGGGAATACCGGGTGCTGTAGGCTT | >hg38_1_-_228634819_228635039_RNA5S12_5S_50ntFlanks | 136 | 170 | 35 | GGAGACCGCCTGGGAATACCGGGTGCTGTAGGCTT | BT20 | 89 | 65 | 71 | 789 | 152 | 75.5 | 44.7 | 41.5 | 990.3 | 169.1 |
| >hg38_1_-_228634819_228635039_RNA5S12_5S_50ntFlanks@136.170.35\|GGAGACCGCCTGGGAATACCGGGTGCTGTAGGCTT | >hg38_1_-_228634819_228635039_RNA5S12_5S_50ntFlanks | 136 | 170 | 35 | GGAGACCGCCTGGGAATACCGGGTGCTGTAGGCTT | MB231 | 96 | 81 | 61 | 48 | 37 | 82.0 | 11.4 | 42.2 | 17.4 | 16.9 |
| >hg38_1_-_228634819_228635039_RNA5S12_5S_50ntFlanks@136.170.35\|GGAGACCGCCTGGGAATACCGGGTGCTGTAGGCTT | >hg38_1_-_228634819_228635039_RNA5S12_5S_50ntFlanks | 136 | 170 | 35 | GGAGACCGCCTGGGAATACCGGGTGCTGTAGGCTT | MB468 | 14 | 7 | 10 | 57 | 37 | 10.2 | 3.7 | 11.5 | 27.7 | 26.1 |
| >hg38_1_-_228634819_228635039_RNA5S12_5S_50ntFlanks@136.171.36\|GGAGACCGCCTGGGAATACCGGGTGCTGTAGGCTTT | >hg38_1_-_228634819_228635039_RNA5S12_5S_50ntFlanks | 136 | 171 | 36 | GGAGACCGCCTGGGAATACCGGGTGCTGTAGGCTTT | BT20 | 77 | 51 | 65 | 972 | 164 | 54.8 | 23.9 | 44.5 | 1519.4 | 184.2 |
| >hg38_1_-_228634819_228635039_RNA5S12_5S_50ntFlanks@136.171.36\|GGAGACCGCCTGGGAATACCGGGTGCTGTAGGCTTT | >hg38_1_-_228634819_228635039_RNA5S12_5S_50ntFlanks | 136 | 171 | 36 | GGAGACCGCCTGGGAATACCGGGTGCTGTAGGCTTT | MB231 | 183 | 144 | 110 | 56 | 77 | 161.8 | 31.9 | 75.6 | 36.1 | 36.1 |
| >hg38_1_-_228634819_228635039_RNA5S12_5S_50ntFlanks@136.171.36\|GGAGACCGCCTGGGAATACCGGGTGCTGTAGGCTTT | >hg38_1_-_228634819_228635039_RNA5S12_5S_50ntFlanks | 136 | 171 | 36 | GGAGACCGCCTGGGAATACCGGGTGCTGTAGGCTTT | MB468 | 17 | 7 | 15 | 55 | 45 | 11.6 | 4.3 | 19.5 | 38.2 | 34.6 |
| >hg38_1_-_228634819_228635039_RNA5S12_5S_50ntFlanks@137.169.33\|GAGACCGCCTGGGAATACCGGGTGCTGTAGGCT | >hg38_1_-_228634819_228635039_RNA5S12_5S_50ntFlanks | 137 | 169 | 33 | GAGACCGCCTGGGAATACCGGGTGCTGTAGGCT | BT20 | 12 | 5 | 21 | 45 | 10 | 6.8 | 2.6 | 12.1 | 38.8 | 5.6 |
| >hg38_1_-_228634819_228635039_RNA5S12_5S_50ntFlanks@137.169.33\|GAGACCGCCTGGGAATACCGGGTGCTGTAGGCT | >hg38_1_-_228634819_228635039_RNA5S12_5S_50ntFlanks | 137 | 169 | 33 | GAGACCGCCTGGGAATACCGGGTGCTGTAGGCT | MB231 | 11 | 9 | 8 | 6 | 4 | 7.4 | 4.4 | 4.1 | 3.5 | 1.0 |
| >hg38_1_-_228634819_228635039_RNA5S12_5S_50ntFlanks@137.170.34\|GAGACCGCCTGGGAATACCGGGTGCTGTAGGCTT | >hg38_1_-_228634819_228635039_RNA5S12_5S_50ntFlanks | 137 | 170 | 34 | GAGACCGCCTGGGAATACCGGGTGCTGTAGGCTT | BT20 | 23 | 10 | 34 | 45 | 11 | 18.6 | 2.5 | 20.2 | 18.9 | 4.9 |

| | | | | | | | | | | | | | | | | |
|---|---|---|---|---|---|---|---|---|---|---|---|---|---|---|---|---|
| >hg38_1_-_228634819_228635039_RNA5S12_5S_50ntFlanks@137.170.34\|GAGACCGCCTGGGAATACCGGGTGCTGTAGGCTT | >hg38_1_-_228634819_228635039_RNA5S12_5S_50ntFlanks | 137 | 170 | 34 | GAGACCGCCTGGGAATACCGGGTGCTGTAGGCTT | MB231 | 30 | 23 | 31 | 22 | 11 | 26.7 | 9.4 | 18.4 | 15.2 | 5.2 |
| >hg38_1_-_228634819_228635039_RNA5S12_5S_50ntFlanks@137.171.35\|GAGACCGCCTGGGAATACCGGGTGCTGTAGGCTTT | >hg38_1_-_228634819_228635039_RNA5S12_5S_50ntFlanks | 137 | 171 | 35 | GAGACCGCCTGGGAATACCGGGTGCTGTAGGCTTT | BT20 | 50 | 17 | 58 | 125 | 27 | 42.4 | 3.5 | 36.4 | 143.0 | 13.8 |
| >hg38_1_-_228634819_228635039_RNA5S12_5S_50ntFlanks@137.171.35\|GAGACCGCCTGGGAATACCGGGTGCTGTAGGCTTT | >hg38_1_-_228634819_228635039_RNA5S12_5S_50ntFlanks | 137 | 171 | 35 | GAGACCGCCTGGGAATACCGGGTGCTGTAGGCTTT | MB231 | 120 | 89 | 101 | 52 | 42 | 113.5 | 36.1 | 73.4 | 41.4 | 19.4 |
| >hg38_1_-_228634819_228635039_RNA5S12_5S_50ntFlanks@138.169.32\|AGACCGCCTGGGAATACCGGGTGCTGTAGGCT | >hg38_1_-_228634819_228635039_RNA5S12_5S_50ntFlanks | 138 | 169 | 32 | AGACCGCCTGGGAATACCGGGTGCTGTAGGCT | BT20 | 19 | 8 | 37 | 68 | 18 | 9.4 | 5.1 | 24.4 | 51.0 | 9.6 |
| >hg38_1_-_228634819_228635039_RNA5S12_5S_50ntFlanks@138.169.32\|AGACCGCCTGGGAATACCGGGTGCTGTAGGCT | >hg38_1_-_228634819_228635039_RNA5S12_5S_50ntFlanks | 138 | 169 | 32 | AGACCGCCTGGGAATACCGGGTGCTGTAGGCT | MB231 | 22 | 20 | 18 | 31 | 31 | 20.1 | 11.3 | 9.6 | 15.0 | 14.3 |
| >hg38_1_-_228634819_228635039_RNA5S12_5S_50ntFlanks@138.170.33\|AGACCGCCTGGGAATACCGGGTGCTGTAGGCTT | >hg38_1_-_228634819_228635039_RNA5S12_5S_50ntFlanks | 138 | 170 | 33 | AGACCGCCTGGGAATACCGGGTGCTGTAGGCTT | BT20 | 37 | 17 | 64 | 119 | 31 | 27.4 | 9.4 | 42.8 | 82.5 | 16.6 |
| >hg38_1_-_228634819_228635039_RNA5S12_5S_50ntFlanks@138.170.33\|AGACCGCCTGGGAATACCGGGTGCTGTAGGCTT | >hg38_1_-_228634819_228635039_RNA5S12_5S_50ntFlanks | 138 | 170 | 33 | AGACCGCCTGGGAATACCGGGTGCTGTAGGCTT | MB231 | 62 | 63 | 57 | 95 | 55 | 64.4 | 37.8 | 31.4 | 33.9 | 27.9 |
| >hg38_1_-_228634819_228635039_RNA5S12_5S_50ntFlanks@138.171.34\|AGACCGCCTGGGAATACCGGGTGCTGTAGGCTTT | >hg38_1_-_228634819_228635039_RNA5S12_5S_50ntFlanks | 138 | 171 | 34 | AGACCGCCTGGGAATACCGGGTGCTGTAGGCTTT | BT20 | 42 | 12 | 54 | 65 | 16 | 41.9 | 3.0 | 39.7 | 55.5 | 4.9 |
| >hg38_1_-_228634819_228635039_RNA5S12_5S_50ntFlanks@138.171.34\|AGACCGCCTGGGAATACCGGGTGCTGTAGGCTTT | >hg38_1_-_228634819_228635039_RNA5S12_5S_50ntFlanks | 138 | 171 | 34 | AGACCGCCTGGGAATACCGGGTGCTGTAGGCTTT | MB231 | 81 | 64 | 83 | 86 | 86 | 78.9 | 30.9 | 54.1 | 53.1 | 50.4 |
| >hg38_1_-_228634819_228635039_RNA5S12_5S_50ntFlanks@139.169.31\|GACCGCCTGGGAATACCGGGTGCTGTAGGCT | >hg38_1_-_228634819_228635039_RNA5S12_5S_50ntFlanks | 139 | 169 | 31 | GACCGCCTGGGAATACCGGGTGCTGTAGGCT | BT20 | 37 | 17 | 78 | 137 | 27 | 23.2 | 12.4 | 59.6 | 131.7 | 11.7 |
| >hg38_1_-_228634819_228635039_RNA5S12_5S_50ntFlanks@139.169.31\|GACCGCCTGGGAATACCGGGTGCTGTAGGCT | >hg38_1_-_228634819_228635039_RNA5S12_5S_50ntFlanks | 139 | 169 | 31 | GACCGCCTGGGAATACCGGGTGCTGTAGGCT | MB231 | 33 | 30 | 28 | 16 | 14 | 29.7 | 16.9 | 13.7 | 9.0 | 6.9 |
| >hg38_1_-_228634819_228635039_RNA5S12_5S_50ntFlanks@139.170.32\|GACCGCCTGGGAATACCGGGTGCTGTAGGCTT | >hg38_1_-_228634819_228635039_RNA5S12_5S_50ntFlanks | 139 | 170 | 32 | GACCGCCTGGGAATACCGGGTGCTGTAGGCTT | BT20 | 45 | 21 | 81 | 138 | 27 | 28.0 | 10.3 | 61.0 | 99.9 | 12.4 |
| >hg38_1_-_228634819_228635039_RNA5S12_5S_50ntFlanks@139.170.32\|GACCGCCTGGGAATACCGGGTGCTGTAGGCTT | >hg38_1_-_228634819_228635039_RNA5S12_5S_50ntFlanks | 139 | 170 | 32 | GACCGCCTGGGAATACCGGGTGCTGTAGGCTT | MB231 | 61 | 59 | 57 | 42 | 28 | 58.5 | 30.2 | 29.5 | 20.6 | 13.8 |
| >hg38_1_-_228634819_228635039_RNA5S12_5S_50ntFlanks@139.171.33\|GACCGCCTGGGAATACCGGGTGCTGTAGGCTTT | >hg38_1_-_228634819_228635039_RNA5S12_5S_50ntFlanks | 139 | 171 | 33 | GACCGCCTGGGAATACCGGGTGCTGTAGGCTTT | BT20 | 43 | 12 | 70 | 70 | 10 | 43.9 | 2.3 | 49.0 | 66.8 | 3.0 |
| >hg38_1_-_228634819_228635039_RNA5S12_5S_50ntFlanks@139.171.33\|GACCGCCTGGGAATACCGGGTGCTGTAGGCTTT | >hg38_1_-_228634819_228635039_RNA5S12_5S_50ntFlanks | 139 | 171 | 33 | GACCGCCTGGGAATACCGGGTGCTGTAGGCTTT | MB231 | 67 | 48 | 73 | 47 | 31 | 65.6 | 23.1 | 52.6 | 42.2 | 17.9 |
| >hg38_1_-_228634819_228635039_RNA5S12_5S_50ntFlanks@140.169.30\|ACCGCCTGGGAATACCGGGTGCTGTAGGCT | >hg38_1_-_228634819_228635039_RNA5S12_5S_50ntFlanks | 140 | 169 | 30 | ACCGCCTGGGAATACCGGGTGCTGTAGGCT | BT20 | 207 | 67 | 376 | 216 | 93 | 150.7 | 68.2 | 382.1 | 150.6 | 38.9 |
| >hg38_1_-_228634819_228635039_RNA5S12_5S_50ntFlanks@140.169.30\|ACCGCCTGGGAATACCGGGTGCTGTAGGCT | >hg38_1_-_228634819_228635039_RNA5S12_5S_50ntFlanks | 140 | 169 | 30 | ACCGCCTGGGAATACCGGGTGCTGTAGGCT | MB231 | 159 | 113 | 194 | 63 | 72 | 110.1 | 35.6 | 183.2 | 55.3 | 24.5 |
| >hg38_1_-_228634819_228635039_RNA5S12_5S_50ntFlanks@140.169.30\|ACCGCCTGGGAATACCGGGTGCTGTAGGCT | >hg38_1_-_228634819_228635039_RNA5S12_5S_50ntFlanks | 140 | 169 | 30 | ACCGCCTGGGAATACCGGGTGCTGTAGGCT | MB468 | 64 | 24 | 49 | 433 | 118 | 43.7 | 24.5 | 25.8 | 250.1 | 77.1 |
| >hg38_1_-_228634819_228635039_RNA5S12_5S_50ntFlanks@140.170.31\|ACCGCCTGGGAATACCGGGTGCTGTAGGCTT | >hg38_1_-_228634819_228635039_RNA5S12_5S_50ntFlanks | 140 | 170 | 31 | ACCGCCTGGGAATACCGGGTGCTGTAGGCTT | BT20 | 486 | 227 | 908 | 633 | 307 | 548.2 | 259.6 | 934.2 | 436.3 | 214.1 |
| >hg38_1_-_228634819_228635039_RNA5S12_5S_50ntFlanks@140.170.31\|ACCGCCTGGGAATACCGGGTGCTGTAGGCTT | >hg38_1_-_228634819_228635039_RNA5S12_5S_50ntFlanks | 140 | 170 | 31 | ACCGCCTGGGAATACCGGGTGCTGTAGGCTT | MB231 | 760 | 662 | 567 | 212 | 270 | 549.7 | 263.4 | 327.8 | 74.2 | 79.4 |
| >hg38_1_-_228634819_228635039_RNA5S12_5S_50ntFlanks@140.170.31\|ACCGCCTGGGAATACCGGGTGCTGTAGGCTT | >hg38_1_-_228634819_228635039_RNA5S12_5S_50ntFlanks | 140 | 170 | 31 | ACCGCCTGGGAATACCGGGTGCTGTAGGCTT | MB468 | 190 | 54 | 138 | 1015 | 427 | 118.0 | 28.7 | 84.2 | 513.6 | 269.0 |
| >hg38_1_-_228634819_228635039_RNA5S12_5S_50ntFlanks@140.171.32\|ACCGCCTGGGAATACCGGGTGCTGTAGGCTTT | >hg38_1_-_228634819_228635039_RNA5S12_5S_50ntFlanks | 140 | 171 | 32 | ACCGCCTGGGAATACCGGGTGCTGTAGGCTTT | BT20 | 154 | 46 | 280 | 120 | 58 | 114.1 | 43.7 | 229.2 | 83.7 | 38.3 |
| >hg38_1_-_228634819_228635039_RNA5S12_5S_50ntFlanks@140.171.32\|ACCGCCTGGGAATACCGGGTGCTGTAGGCTTT | >hg38_1_-_228634819_228635039_RNA5S12_5S_50ntFlanks | 140 | 171 | 32 | ACCGCCTGGGAATACCGGGTGCTGTAGGCTTT | MB231 | 293 | 240 | 292 | 111 | 117 | 210.6 | 87.1 | 202.8 | 86.8 | 49.6 |
| >hg38_1_-_228634819_228635039_RNA5S12_5S_50ntFlanks@140.171.32\|ACCGCCTGGGAATACCGGGTGCTGTAGGCTTT | >hg38_1_-_228634819_228635039_RNA5S12_5S_50ntFlanks | 140 | 171 | 32 | ACCGCCTGGGAATACCGGGTGCTGTAGGCTTT | MB468 | 36 | 11 | 31 | 414 | 97 | 23.8 | 7.0 | 21.8 | 400.1 | 75.5 |
| >hg38_1_-_228634819_228635039_RNA5S12_5S_50ntFlanks@141.169.29\|CCGCCTGGGAATACCGGGTGCTGTAGGCT | >hg38_1_-_228634819_228635039_RNA5S12_5S_50ntFlanks | 141 | 169 | 29 | CCGCCTGGGAATACCGGGTGCTGTAGGCT | BT20 | 54 | 23 | 97 | 64 | 23 | 29.1 | 21.4 | 75.3 | 28.3 | 7.5 |
| >hg38_1_-_228634819_228635039_RNA5S12_5S_50ntFlanks@141.169.29\|CCGCCTGGGAATACCGGGTGCTGTAGGCT | >hg38_1_-_228634819_228635039_RNA5S12_5S_50ntFlanks | 141 | 169 | 29 | CCGCCTGGGAATACCGGGTGCTGTAGGCT | MB231 | 48 | 47 | 50 | 16 | 19 | 21.0 | 10.2 | 22.8 | 7.5 | 4.0 |
| >hg38_1_-_228634819_228635039_RNA5S12_5S_50ntFlanks@141.169.29\|CCGCCTGGGAATACCGGGTGCTGTAGGCT | >hg38_1_-_228634819_228635039_RNA5S12_5S_50ntFlanks | 141 | 169 | 29 | CCGCCTGGGAATACCGGGTGCTGTAGGCT | MB468 | 12 | 6 | 10 | 56 | 31 | 8.0 | 3.2 | 4.7 | 23.6 | 22.7 |
| >hg38_1_-_228634819_228635039_RNA5S12_5S_50ntFlanks@141.170.30\|CCGCCTGGGAATACCGGGTGCTGTAGGCTT | >hg38_1_-_228634819_228635039_RNA5S12_5S_50ntFlanks | 141 | 170 | 30 | CCGCCTGGGAATACCGGGTGCTGTAGGCTT | BT20 | 103 | 38 | 159 | 99 | 31 | 85.3 | 24.7 | 127.8 | 36.5 | 5.0 |
| >hg38_1_-_228634819_228635039_RNA5S12_5S_50ntFlanks@141.170.30\|CCGCCTGGGAATACCGGGTGCTGTAGGCTT | >hg38_1_-_228634819_228635039_RNA5S12_5S_50ntFlanks | 141 | 170 | 30 | CCGCCTGGGAATACCGGGTGCTGTAGGCTT | MB231 | 117 | 100 | 147 | 58 | 53 | 65.3 | 21.5 | 107.7 | 21.8 | 17.1 |
| >hg38_1_-_228634819_228635039_RNA5S12_5S_50ntFlanks@141.170.30\|CCGCCTGGGAATACCGGGTGCTGTAGGCTT | >hg38_1_-_228634819_228635039_RNA5S12_5S_50ntFlanks | 141 | 170 | 30 | CCGCCTGGGAATACCGGGTGCTGTAGGCTT | MB468 | 11 | 8 | 10 | 148 | 42 | 6.7 | 4.6 | 5.2 | 87.4 | 36.0 |
| >hg38_1_-_228634819_228635039_RNA5S12_5S_50ntFlanks@141.171.31\|CCGCCTGGGAATACCGGGTGCTGTAGGCTTT | >hg38_1_-_228634819_228635039_RNA5S12_5S_50ntFlanks | 141 | 171 | 31 | CCGCCTGGGAATACCGGGTGCTGTAGGCTTT | BT20 | 97 | 24 | 145 | 54 | 15 | 100.2 | 9.7 | 91.9 | 20.0 | 2.1 |
| >hg38_1_-_228634819_228635039_RNA5S12_5S_50ntFlanks@141.171.31\|CCGCCTGGGAATACCGGGTGCTGTAGGCTTT | >hg38_1_-_228634819_228635039_RNA5S12_5S_50ntFlanks | 141 | 171 | 31 | CCGCCTGGGAATACCGGGTGCTGTAGGCTTT | MB231 | 164 | 108 | 237 | 77 | 72 | 77.3 | 42.2 | 227.0 | 57.0 | 29.3 |
| >hg38_1_-_228634819_228635039_RNA5S12_5S_50ntFlanks@142.169.28\|CGCCTGGGAATACCGGGTGCTGTAGGCT | >hg38_1_-_228634819_228635039_RNA5S12_5S_50ntFlanks | 142 | 169 | 28 | CGCCTGGGAATACCGGGTGCTGTAGGCT | BT20 | 11 | 6 | 37 | 8 | 5 | 6.0 | 4.8 | 39.8 | 5.3 | 2.7 |
| >hg38_1_-_228634819_228635039_RNA5S12_5S_50ntFlanks@142.170.29\|CGCCTGGGAATACCGGGTGCTGTAGGCTT | >hg38_1_-_228634819_228635039_RNA5S12_5S_50ntFlanks | 142 | 170 | 29 | CGCCTGGGAATACCGGGTGCTGTAGGCTT | BT20 | 36 | 14 | 66 | 43 | 14 | 28.9 | 4.6 | 55.2 | 19.2 | 4.8 |

| | | | | | | | | | | | | | | | | |
|---|---|---|---|---|---|---|---|---|---|---|---|---|---|---|---|---|
| >hg38_1_-_228634819_228635039_RNA5S12_5S_50ntFlanks@142.170.29\|CGCCTGGGAATACCGGGTGCTGTAGGCTT | >hg38_1_-_228634819_228635039_RNA5S12_5S_50ntFlanks | 142 | 170 | 29 | CGCCTGGGAATACCGGGTGCTGTAGGCTT | MB231 | 55 | 41 | 67 | 23 | 21 | 29.5 | 13.5 | 41.6 | 6.6 | 8.0 |
| >hg38_1_-_228634819_228635039_RNA5S12_5S_50ntFlanks@142.171.30\|CGCCTGGGAATACCGGGTGCTGTAGGCTTT | >hg38_1_-_228634819_228635039_RNA5S12_5S_50ntFlanks | 142 | 171 | 30 | CGCCTGGGAATACCGGGTGCTGTAGGCTTT | BT20 | 46 | 13 | 55 | 34 | 13 | 46.7 | 2.1 | 32.6 | 14.6 | 5.2 |
| >hg38_1_-_228634819_228635039_RNA5S12_5S_50ntFlanks@142.171.30\|CGCCTGGGAATACCGGGTGCTGTAGGCTTT | >hg38_1_-_228634819_228635039_RNA5S12_5S_50ntFlanks | 142 | 171 | 30 | CGCCTGGGAATACCGGGTGCTGTAGGCTTT | MB231 | 114 | 72 | 132 | 31 | 48 | 48.4 | 25.1 | 92.1 | 14.6 | 18.3 |
| >hg38_1_-_228634819_228635039_RNA5S12_5S_50ntFlanks@143.171.29\|GCCTGGGAATACCGGGTGCTGTAGGCTTT | >hg38_1_-_228634819_228635039_RNA5S12_5S_50ntFlanks | 143 | 171 | 29 | GCCTGGGAATACCGGGTGCTGTAGGCTTT | MB231 | 12 | 8 | 14 | 5 | 6 | 7.4 | 3.0 | 7.7 | 1.2 | 3.2 |
| >hg38_1_-_228634819_228635039_RNA5S12_5S_50ntFlanks@144.169.26\|CCTGGGAATACCGGGTGCTGTAGGCT | >hg38_1_-_228634819_228635039_RNA5S12_5S_50ntFlanks | 144 | 169 | 26 | CCTGGGAATACCGGGTGCTGTAGGCT | BT20 | 18 | 7 | 34 | 8 | 4 | 13.7 | 3.8 | 31.7 | 8.2 | 1.7 |
| >hg38_1_-_228634819_228635039_RNA5S12_5S_50ntFlanks@144.169.26\|CCTGGGAATACCGGGTGCTGTAGGCT | >hg38_1_-_228634819_228635039_RNA5S12_5S_50ntFlanks | 144 | 169 | 26 | CCTGGGAATACCGGGTGCTGTAGGCT | MB231 | 15 | 7 | 17 | 4 | 4 | 5.1 | 4.1 | 7.3 | 2.3 | 1.9 |
| >hg38_1_-_228634819_228635039_RNA5S12_5S_50ntFlanks@144.170.27\|CCTGGGAATACCGGGTGCTGTAGGCTT | >hg38_1_-_228634819_228635039_RNA5S12_5S_50ntFlanks | 144 | 170 | 27 | CCTGGGAATACCGGGTGCTGTAGGCTT | BT20 | 133 | 80 | 113 | 86 | 33 | 65.2 | 43.1 | 93.6 | 65.7 | 15.6 |
| >hg38_1_-_228634819_228635039_RNA5S12_5S_50ntFlanks@144.170.27\|CCTGGGAATACCGGGTGCTGTAGGCTT | >hg38_1_-_228634819_228635039_RNA5S12_5S_50ntFlanks | 144 | 170 | 27 | CCTGGGAATACCGGGTGCTGTAGGCTT | MB231 | 248 | 177 | 171 | 69 | 80 | 89.3 | 84.8 | 111.4 | 24.9 | 45.9 |
| >hg38_1_-_228634819_228635039_RNA5S12_5S_50ntFlanks@144.171.28\|CCTGGGAATACCGGGTGCTGTAGGCTTT | >hg38_1_-_228634819_228635039_RNA5S12_5S_50ntFlanks | 144 | 171 | 28 | CCTGGGAATACCGGGTGCTGTAGGCTTT | BT20 | 104 | 57 | 102 | 44 | 31 | 52.9 | 26.9 | 71.8 | 19.9 | 12.0 |
| >hg38_1_-_228634819_228635039_RNA5S12_5S_50ntFlanks@144.171.28\|CCTGGGAATACCGGGTGCTGTAGGCTTT | >hg38_1_-_228634819_228635039_RNA5S12_5S_50ntFlanks | 144 | 171 | 28 | CCTGGGAATACCGGGTGCTGTAGGCTTT | MB231 | 402 | 271 | 286 | 69 | 151 | 112.8 | 130.3 | 186.7 | 45.8 | 101.0 |
| >hg38_1_-_228634819_228635039_RNA5S12_5S_50ntFlanks@144.171.28\|CCTGGGAATACCGGGTGCTGTAGGCTTT | >hg38_1_-_228634819_228635039_RNA5S12_5S_50ntFlanks | 144 | 171 | 28 | CCTGGGAATACCGGGTGCTGTAGGCTTT | MB468 | 13 | 10 | 12 | 211 | 85 | 6.8 | 11.0 | 6.1 | 93.9 | 74.9 |
| >hg38_1_-_228634819_228635039_RNA5S12_5S_50ntFlanks@145.169.25\|CTGGGAATACCGGGTGCTGTAGGCT | >hg38_1_-_228634819_228635039_RNA5S12_5S_50ntFlanks | 145 | 169 | 25 | CTGGGAATACCGGGTGCTGTAGGCT | BT20 | 13 | 9 | 53 | 19 | 6 | 4.5 | 3.0 | 41.5 | 8.7 | 1.3 |
| >hg38_1_-_228634819_228635039_RNA5S12_5S_50ntFlanks@145.170.26\|CTGGGAATACCGGGTGCTGTAGGCTT | >hg38_1_-_228634819_228635039_RNA5S12_5S_50ntFlanks | 145 | 170 | 26 | CTGGGAATACCGGGTGCTGTAGGCTT | BT20 | 33 | 33 | 120 | 76 | 30 | 12.0 | 20.3 | 103.2 | 22.2 | 4.7 |
| >hg38_1_-_228634819_228635039_RNA5S12_5S_50ntFlanks@145.170.26\|CTGGGAATACCGGGTGCTGTAGGCTT | >hg38_1_-_228634819_228635039_RNA5S12_5S_50ntFlanks | 145 | 170 | 26 | CTGGGAATACCGGGTGCTGTAGGCTT | MB231 | 49 | 53 | 163 | 26 | 16 | 43.6 | 34.9 | 109.3 | 12.8 | 8.8 |
| >hg38_1_-_228634819_228635039_RNA5S12_5S_50ntFlanks@145.171.27\|CTGGGAATACCGGGTGCTGTAGGCTTT | >hg38_1_-_228634819_228635039_RNA5S12_5S_50ntFlanks | 145 | 171 | 27 | CTGGGAATACCGGGTGCTGTAGGCTTT | BT20 | 24 | 14 | 89 | 34 | 16 | 11.3 | 5.5 | 68.1 | 23.2 | 0.9 |
| >hg38_1_-_228634819_228635039_RNA5S12_5S_50ntFlanks@145.171.27\|CTGGGAATACCGGGTGCTGTAGGCTTT | >hg38_1_-_228634819_228635039_RNA5S12_5S_50ntFlanks | 145 | 171 | 27 | CTGGGAATACCGGGTGCTGTAGGCTTT | MB231 | 65 | 61 | 214 | 15 | 19 | 62.0 | 45.0 | 172.7 | 11.6 | 13.1 |
| >hg38_1_-_228634819_228635039_RNA5S12_5S_50ntFlanks@146.169.24\|TGGGAATACCGGGTGCTGTAGGCT | >hg38_1_-_228634819_228635039_RNA5S12_5S_50ntFlanks | 146 | 169 | 24 | TGGGAATACCGGGTGCTGTAGGCT | BT20 | 22 | 16 | 102 | 88 | 17 | 1.3 | 10.5 | 71.9 | 84.6 | 6.8 |
| >hg38_1_-_228634819_228635039_RNA5S12_5S_50ntFlanks@146.170.25\|TGGGAATACCGGGTGCTGTAGGCTT | >hg38_1_-_228634819_228635039_RNA5S12_5S_50ntFlanks | 146 | 170 | 25 | TGGGAATACCGGGTGCTGTAGGCTT | BT20 | 62 | 61 | 307 | 380 | 89 | 11.9 | 36.3 | 249.3 | 406.5 | 32.3 |
| >hg38_1_-_228634819_228635039_RNA5S12_5S_50ntFlanks@146.170.25\|TGGGAATACCGGGTGCTGTAGGCTT | >hg38_1_-_228634819_228635039_RNA5S12_5S_50ntFlanks | 146 | 170 | 25 | TGGGAATACCGGGTGCTGTAGGCTT | MB231 | 55 | 72 | 236 | 41 | 19 | 49.8 | 37.9 | 137.3 | 15.3 | 7.0 |
| >hg38_1_-_228634819_228635039_RNA5S12_5S_50ntFlanks@146.171.26\|TGGGAATACCGGGTGCTGTAGGCTTT | >hg38_1_-_228634819_228635039_RNA5S12_5S_50ntFlanks | 146 | 171 | 26 | TGGGAATACCGGGTGCTGTAGGCTTT | BT20 | 34 | 27 | 164 | 144 | 36 | 3.1 | 16.6 | 122.8 | 178.4 | 9.2 |
| >hg38_1_-_228634819_228635039_RNA5S12_5S_50ntFlanks@146.171.26\|TGGGAATACCGGGTGCTGTAGGCTTT | >hg38_1_-_228634819_228635039_RNA5S12_5S_50ntFlanks | 146 | 171 | 26 | TGGGAATACCGGGTGCTGTAGGCTTT | MB231 | 51 | 58 | 217 | 25 | 20 | 49.1 | 35.3 | 182.5 | 17.3 | 10.0 |
| >hg38_1_-_228634819_228635039_RNA5S12_5S_50ntFlanks@147.169.23\|GGGAATACCGGGTGCTGTAGGCT | >hg38_1_-_228634819_228635039_RNA5S12_5S_50ntFlanks | 147 | 169 | 23 | GGGAATACCGGGTGCTGTAGGCT | BT20 | 24 | 25 | 95 | 537 | 34 | 11.6 | 21.4 | 63.9 | 736.5 | 26.5 |
| >hg38_1_-_228634819_228635039_RNA5S12_5S_50ntFlanks@147.169.23\|GGGAATACCGGGTGCTGTAGGCT | >hg38_1_-_228634819_228635039_RNA5S12_5S_50ntFlanks | 147 | 169 | 23 | GGGAATACCGGGTGCTGTAGGCT | MB231 | 12 | 12 | 56 | 24 | 3 | 5.9 | 4.5 | 8.5 | 21.8 | 1.5 |
| >hg38_1_-_228634819_228635039_RNA5S12_5S_50ntFlanks@147.170.24\|GGGAATACCGGGTGCTGTAGGCTT | >hg38_1_-_228634819_228635039_RNA5S12_5S_50ntFlanks | 147 | 170 | 24 | GGGAATACCGGGTGCTGTAGGCTT | BT20 | 81 | 100 | 288 | 1873 | 140 | 49.7 | 79.3 | 224.6 | 2546.3 | 93.0 |
| >hg38_1_-_228634819_228635039_RNA5S12_5S_50ntFlanks@147.170.24\|GGGAATACCGGGTGCTGTAGGCTT | >hg38_1_-_228634819_228635039_RNA5S12_5S_50ntFlanks | 147 | 170 | 24 | GGGAATACCGGGTGCTGTAGGCTT | MB231 | 65 | 81 | 288 | 152 | 23 | 48.5 | 34.2 | 149.3 | 134.8 | 7.0 |
| >hg38_1_-_228634819_228635039_RNA5S12_5S_50ntFlanks@147.171.25\|GGGAATACCGGGTGCTGTAGGCTTT | >hg38_1_-_228634819_228635039_RNA5S12_5S_50ntFlanks | 147 | 171 | 25 | GGGAATACCGGGTGCTGTAGGCTTT | BT20 | 45 | 51 | 147 | 837 | 57 | 19.5 | 32.2 | 91.1 | 1325.1 | 27.6 |
| >hg38_1_-_228634819_228635039_RNA5S12_5S_50ntFlanks@147.171.25\|GGGAATACCGGGTGCTGTAGGCTTT | >hg38_1_-_228634819_228635039_RNA5S12_5S_50ntFlanks | 147 | 171 | 25 | GGGAATACCGGGTGCTGTAGGCTTT | MB231 | 62 | 67 | 263 | 48 | 26 | 45.9 | 34.0 | 203.0 | 16.9 | 9.9 |
| >hg38_1_-_228634819_228635039_RNA5S12_5S_50ntFlanks@148.169.22\|GGAATACCGGGTGCTGTAGGCT | >hg38_1_-_228634819_228635039_RNA5S12_5S_50ntFlanks | 148 | 169 | 22 | GGAATACCGGGTGCTGTAGGCT | BT20 | 27 | 47 | 110 | 2413 | 36 | 3.5 | 45.7 | 65.2 | 4052.9 | 15.6 |
| >hg38_1_-_228634819_228635039_RNA5S12_5S_50ntFlanks@148.169.22\|GGAATACCGGGTGCTGTAGGCT | >hg38_1_-_228634819_228635039_RNA5S12_5S_50ntFlanks | 148 | 169 | 22 | GGAATACCGGGTGCTGTAGGCT | MB231 | 12 | 13 | 42 | 9 | 4 | 9.4 | 6.6 | 13.9 | 4.3 | 1.1 |
| >hg38_1_-_228634819_228635039_RNA5S12_5S_50ntFlanks@148.169.22\|GGAATACCGGGTGCTGTAGGCT | >hg38_1_-_228634819_228635039_RNA5S12_5S_50ntFlanks | 148 | 169 | 22 | GGAATACCGGGTGCTGTAGGCT | MB468 | 16 | 25 | 28 | 287 | 68 | 8.0 | 20.8 | 3.3 | 160.6 | 41.0 |
| >hg38_1_-_228634819_228635039_RNA5S12_5S_50ntFlanks@148.170.23\|GGAATACCGGGTGCTGTAGGCTT | >hg38_1_-_228634819_228635039_RNA5S12_5S_50ntFlanks | 148 | 170 | 23 | GGAATACCGGGTGCTGTAGGCTT | BT20 | 58 | 115 | 211 | 4716 | 101 | 6.8 | 109.2 | 133.5 | 7856.3 | 33.5 |
| >hg38_1_-_228634819_228635039_RNA5S12_5S_50ntFlanks@148.170.23\|GGAATACCGGGTGCTGTAGGCTT | >hg38_1_-_228634819_228635039_RNA5S12_5S_50ntFlanks | 148 | 170 | 23 | GGAATACCGGGTGCTGTAGGCTT | MB231 | 49 | 59 | 186 | 43 | 16 | 46.1 | 33.3 | 138.5 | 21.5 | 5.5 |
| >hg38_1_-_228634819_228635039_RNA5S12_5S_50ntFlanks@148.170.23\|GGAATACCGGGTGCTGTAGGCTT | >hg38_1_-_228634819_228635039_RNA5S12_5S_50ntFlanks | 148 | 170 | 23 | GGAATACCGGGTGCTGTAGGCTT | MB468 | 35 | 58 | 55 | 859 | 191 | 13.9 | 48.0 | 7.3 | 560.3 | 98.4 |
| >hg38_1_-_228634819_228635039_RNA5S12_5S_50ntFlanks@148.171.24\|GGAATACCGGGTGCTGTAGGCTTT | >hg38_1_-_228634819_228635039_RNA5S12_5S_50ntFlanks | 148 | 171 | 24 | GGAATACCGGGTGCTGTAGGCTTT | BT20 | 44 | 99 | 187 | 3636 | 60 | 7.5 | 113.6 | 106.6 | 6197.0 | 26.3 |

| | | | | | | | | | | | | | | | | |
|---|---|---|---|---|---|---|---|---|---|---|---|---|---|---|---|---|
| >hg38_1_-_228634819_228635039_RNA5S12_5S_50ntFlanks@148.171.24\|GGAATACCGGGTGCTGTAGGCTTT | >hg38_1_-_228634819_228635039_RNA5S12_5S_50ntFlanks | 148 | 171 | 24 | GGAATACCGGGTGCTGTAGGCTTT | MB231 | 66 | 74 | 280 | 37 | 22 | 61.0 | 42.6 | 252.5 | 28.0 | 10.5 |
| >hg38_1_-_228634819_228635039_RNA5S12_5S_50ntFlanks@148.171.24\|GGAATACCGGGTGCTGTAGGCTTT | >hg38_1_-_228634819_228635039_RNA5S12_5S_50ntFlanks | 148 | 171 | 24 | GGAATACCGGGTGCTGTAGGCTTT | MB468 | 35 | 53 | 63 | 1004 | 178 | 13.9 | 45.0 | 7.1 | 672.4 | 106.8 |
| >hg38_1_-_228634819_228635039_RNA5S12_5S_50ntFlanks@149.169.21\|GAATACCGGGTGCTGTAGGCT | >hg38_1_-_228634819_228635039_RNA5S12_5S_50ntFlanks | 149 | 169 | 21 | GAATACCGGGTGCTGTAGGCT | BT20 | 20 | 11 | 77 | 107 | 16 | 3.5 | 8.7 | 58.0 | 60.4 | 5.4 |
| >hg38_1_-_228634819_228635039_RNA5S12_5S_50ntFlanks@149.170.22\|GAATACCGGGTGCTGTAGGCTT | >hg38_1_-_228634819_228635039_RNA5S12_5S_50ntFlanks | 149 | 170 | 22 | GAATACCGGGTGCTGTAGGCTT | BT20 | 52 | 38 | 203 | 345 | 58 | 8.1 | 22.3 | 164.7 | 139.7 | 17.2 |
| >hg38_1_-_228634819_228635039_RNA5S12_5S_50ntFlanks@149.170.22\|GAATACCGGGTGCTGTAGGCTT | >hg38_1_-_228634819_228635039_RNA5S12_5S_50ntFlanks | 149 | 170 | 22 | GAATACCGGGTGCTGTAGGCTT | MB231 | 50 | 68 | 212 | 41 | 17 | 47.2 | 41.5 | 160.8 | 19.5 | 7.6 |
| >hg38_1_-_228634819_228635039_RNA5S12_5S_50ntFlanks@149.171.23\|GAATACCGGGTGCTGTAGGCTTT | >hg38_1_-_228634819_228635039_RNA5S12_5S_50ntFlanks | 149 | 171 | 23 | GAATACCGGGTGCTGTAGGCTTT | BT20 | 41 | 27 | 151 | 207 | 41 | 9.3 | 14.1 | 102.8 | 271.2 | 13.1 |
| >hg38_1_-_228634819_228635039_RNA5S12_5S_50ntFlanks@149.171.23\|GAATACCGGGTGCTGTAGGCTTT | >hg38_1_-_228634819_228635039_RNA5S12_5S_50ntFlanks | 149 | 171 | 23 | GAATACCGGGTGCTGTAGGCTTT | MB231 | 71 | 88 | 296 | 33 | 23 | 66.7 | 55.6 | 268.2 | 25.3 | 10.6 |
| >hg38_1_-_228634819_228635039_RNA5S12_5S_50ntFlanks@150.169.20\|AATACCGGGTGCTGTAGGCT | >hg38_1_-_228634819_228635039_RNA5S12_5S_50ntFlanks | 150 | 169 | 20 | AATACCGGGTGCTGTAGGCT | BT20 | 22 | 12 | 78 | 114 | 13 | 7.1 | 8.0 | 60.5 | 12.8 | 4.1 |
| >hg38_1_-_228634819_228635039_RNA5S12_5S_50ntFlanks@150.169.20\|AATACCGGGTGCTGTAGGCT | >hg38_1_-_228634819_228635039_RNA5S12_5S_50ntFlanks | 150 | 169 | 20 | AATACCGGGTGCTGTAGGCT | MB231 | 11 | 11 | 31 | 11 | 3 | 5.6 | 4.5 | 13.4 | 2.8 | 1.2 |
| >hg38_1_-_228634819_228635039_RNA5S12_5S_50ntFlanks@150.170.21\|AATACCGGGTGCTGTAGGCTT | >hg38_1_-_228634819_228635039_RNA5S12_5S_50ntFlanks | 150 | 170 | 21 | AATACCGGGTGCTGTAGGCTT | BT20 | 57 | 36 | 192 | 412 | 43 | 18.1 | 19.1 | 155.9 | 119.8 | 9.4 |
| >hg38_1_-_228634819_228635039_RNA5S12_5S_50ntFlanks@150.170.21\|AATACCGGGTGCTGTAGGCTT | >hg38_1_-_228634819_228635039_RNA5S12_5S_50ntFlanks | 150 | 170 | 21 | AATACCGGGTGCTGTAGGCTT | MB231 | 58 | 71 | 204 | 67 | 22 | 49.1 | 49.8 | 147.9 | 19.7 | 9.4 |
| >hg38_1_-_228634819_228635039_RNA5S12_5S_50ntFlanks@150.170.21\|AATACCGGGTGCTGTAGGCTT | >hg38_1_-_228634819_228635039_RNA5S12_5S_50ntFlanks | 150 | 170 | 21 | AATACCGGGTGCTGTAGGCTT | MB468 | 20 | 27 | 23 | 754 | 187 | 13.4 | 34.9 | 8.1 | 514.6 | 125.0 |
| >hg38_1_-_228634819_228635039_RNA5S12_5S_50ntFlanks@150.171.22\|AATACCGGGTGCTGTAGGCTTT | >hg38_1_-_228634819_228635039_RNA5S12_5S_50ntFlanks | 150 | 171 | 22 | AATACCGGGTGCTGTAGGCTTT | BT20 | 48 | 28 | 145 | 153 | 37 | 18.1 | 12.0 | 104.9 | 174.6 | 10.1 |
| >hg38_1_-_228634819_228635039_RNA5S12_5S_50ntFlanks@150.171.22\|AATACCGGGTGCTGTAGGCTTT | >hg38_1_-_228634819_228635039_RNA5S12_5S_50ntFlanks | 150 | 171 | 22 | AATACCGGGTGCTGTAGGCTTT | MB231 | 90 | 101 | 284 | 52 | 34 | 70.8 | 66.7 | 241.2 | 29.3 | 15.2 |
| >hg38_1_-_228634819_228635039_RNA5S12_5S_50ntFlanks@150.171.22\|AATACCGGGTGCTGTAGGCTTT | >hg38_1_-_228634819_228635039_RNA5S12_5S_50ntFlanks | 150 | 171 | 22 | AATACCGGGTGCTGTAGGCTTT | MB468 | 22 | 30 | 27 | 1152 | 213 | 14.7 | 42.3 | 9.0 | 842.2 | 146.9 |
| >hg38_1_-_228634819_228635039_RNA5S12_5S_50ntFlanks@151.169.19\|ATACCGGGTGCTGTAGGCT | >hg38_1_-_228634819_228635039_RNA5S12_5S_50ntFlanks | 151 | 169 | 19 | ATACCGGGTGCTGTAGGCT | BT20 | 20 | 11 | 64 | 127 | 12 | 3.8 | 7.3 | 51.8 | 49.0 | 5.4 |
| >hg38_1_-_228634819_228635039_RNA5S12_5S_50ntFlanks@151.170.20\|ATACCGGGTGCTGTAGGCTT | >hg38_1_-_228634819_228635039_RNA5S12_5S_50ntFlanks | 151 | 170 | 20 | ATACCGGGTGCTGTAGGCTT | BT20 | 50 | 30 | 162 | 478 | 46 | 7.7 | 17.3 | 149.9 | 232.2 | 11.5 |
| >hg38_1_-_228634819_228635039_RNA5S12_5S_50ntFlanks@151.170.20\|ATACCGGGTGCTGTAGGCTT | >hg38_1_-_228634819_228635039_RNA5S12_5S_50ntFlanks | 151 | 170 | 20 | ATACCGGGTGCTGTAGGCTT | MB231 | 48 | 68 | 179 | 86 | 20 | 40.4 | 38.1 | 144.7 | 29.0 | 6.0 |
| >hg38_1_-_228634819_228635039_RNA5S12_5S_50ntFlanks@151.171.21\|ATACCGGGTGCTGTAGGCTTT | >hg38_1_-_228634819_228635039_RNA5S12_5S_50ntFlanks | 151 | 171 | 21 | ATACCGGGTGCTGTAGGCTTT | BT20 | 36 | 20 | 107 | 85 | 30 | 9.2 | 9.1 | 85.9 | 68.1 | 8.8 |
| >hg38_1_-_228634819_228635039_RNA5S12_5S_50ntFlanks@151.171.21\|ATACCGGGTGCTGTAGGCTTT | >hg38_1_-_228634819_228635039_RNA5S12_5S_50ntFlanks | 151 | 171 | 21 | ATACCGGGTGCTGTAGGCTTT | MB231 | 62 | 82 | 235 | 45 | 26 | 50.5 | 47.6 | 219.2 | 23.0 | 9.0 |
| >hg38_1_-_228634819_228635039_RNA5S12_5S_50ntFlanks@152.169.18\|TACCGGGTGCTGTAGGCT | >hg38_1_-_228634819_228635039_RNA5S12_5S_50ntFlanks | 152 | 169 | 18 | TACCGGGTGCTGTAGGCT | BT20 | 20 | 11 | 61 | 63 | 12 | 7.3 | 11.5 | 49.1 | 27.4 | 7.2 |
| >hg38_1_-_228634819_228635039_RNA5S12_5S_50ntFlanks@152.169.18\|TACCGGGTGCTGTAGGCT | >hg38_1_-_228634819_228635039_RNA5S12_5S_50ntFlanks | 152 | 169 | 18 | TACCGGGTGCTGTAGGCT | MB231 | 12 | 13 | 44 | 12 | 4 | 5.4 | 4.8 | 17.5 | 2.7 | 0.7 |
| >hg38_1_-_228634819_228635039_RNA5S12_5S_50ntFlanks@152.170.19\|TACCGGGTGCTGTAGGCTT | >hg38_1_-_228634819_228635039_RNA5S12_5S_50ntFlanks | 152 | 170 | 19 | TACCGGGTGCTGTAGGCTT | BT20 | 59 | 35 | 161 | 226 | 49 | 32.6 | 34.5 | 142.7 | 102.1 | 28.8 |
| >hg38_1_-_228634819_228635039_RNA5S12_5S_50ntFlanks@152.170.19\|TACCGGGTGCTGTAGGCTT | >hg38_1_-_228634819_228635039_RNA5S12_5S_50ntFlanks | 152 | 170 | 19 | TACCGGGTGCTGTAGGCTT | MB231 | 58 | 74 | 250 | 65 | 23 | 37.1 | 34.8 | 139.1 | 14.7 | 6.3 |
| >hg38_1_-_228634819_228635039_RNA5S12_5S_50ntFlanks@152.171.20\|TACCGGGTGCTGTAGGCTTT | >hg38_1_-_228634819_228635039_RNA5S12_5S_50ntFlanks | 152 | 171 | 20 | TACCGGGTGCTGTAGGCTTT | BT20 | 42 | 22 | 126 | 60 | 33 | 20.0 | 20.1 | 104.9 | 41.2 | 18.5 |
| >hg38_1_-_228634819_228635039_RNA5S12_5S_50ntFlanks@152.171.20\|TACCGGGTGCTGTAGGCTTT | >hg38_1_-_228634819_228635039_RNA5S12_5S_50ntFlanks | 152 | 171 | 20 | TACCGGGTGCTGTAGGCTTT | MB231 | 71 | 85 | 297 | 48 | 30 | 47.6 | 47.1 | 199.3 | 26.8 | 11.1 |
| >hg38_1_-_228634819_228635039_RNA5S12_5S_50ntFlanks@153.170.18\|ACCGGGTGCTGTAGGCTT | >hg38_1_-_228634819_228635039_RNA5S12_5S_50ntFlanks | 153 | 170 | 18 | ACCGGGTGCTGTAGGCTT | BT20 | 1367 | 883 | 1339 | 5119 | 1154 | 1902.7 | 1281.7 | 1693.7 | 6957.0 | 1487.3 |
| >hg38_1_-_228634819_228635039_RNA5S12_5S_50ntFlanks@153.170.18\|ACCGGGTGCTGTAGGCTT | >hg38_1_-_228634819_228635039_RNA5S12_5S_50ntFlanks | 153 | 170 | 18 | ACCGGGTGCTGTAGGCTT | MB231 | 338 | 409 | 1132 | 910 | 216 | 128.5 | 169.5 | 676.1 | 292.5 | 58.6 |
| >hg38_1_-_228634819_228635039_RNA5S12_5S_50ntFlanks@153.170.18\|ACCGGGTGCTGTAGGCTT | >hg38_1_-_228634819_228635039_RNA5S12_5S_50ntFlanks | 153 | 170 | 18 | ACCGGGTGCTGTAGGCTT | MB468 | 21 | 30 | 26 | 231 | 39 | 11.5 | 19.6 | 12.2 | 125.6 | 13.8 |
| >hg38_1_-_228634819_228635039_RNA5S12_5S_50ntFlanks@153.171.19\|ACCGGGTGCTGTAGGCTTT | >hg38_1_-_228634819_228635039_RNA5S12_5S_50ntFlanks | 153 | 171 | 19 | ACCGGGTGCTGTAGGCTTT | BT20 | 825 | 514 | 818 | 1612 | 666 | 1127.1 | 746.2 | 996.2 | 2357.5 | 848.4 |
| >hg38_1_-_228634819_228635039_RNA5S12_5S_50ntFlanks@153.171.19\|ACCGGGTGCTGTAGGCTTT | >hg38_1_-_228634819_228635039_RNA5S12_5S_50ntFlanks | 153 | 171 | 19 | ACCGGGTGCTGTAGGCTTT | MB231 | 349 | 381 | 1134 | 578 | 232 | 65.5 | 111.1 | 549.8 | 246.2 | 53.4 |
| >hg38_1_-_228634819_228635039_RNA5S12_5S_50ntFlanks@153.171.19\|ACCGGGTGCTGTAGGCTTT | >hg38_1_-_228634819_228635039_RNA5S12_5S_50ntFlanks | 153 | 171 | 19 | ACCGGGTGCTGTAGGCTTT | MB468 | 17 | 19 | 21 | 233 | 36 | 8.2 | 10.8 | 8.9 | 160.4 | 16.2 |
| >hg38_1_-_228634819_228635039_RNA5S12_5S_50ntFlanks@154.171.18\|CCGGGTGCTGTAGGCTTT | >hg38_1_-_228634819_228635039_RNA5S12_5S_50ntFlanks | 154 | 171 | 18 | CCGGGTGCTGTAGGCTTT | BT20 | 81 | 54 | 213 | 365 | 55 | 66.8 | 49.7 | 175.4 | 433.7 | 50.0 |
| >hg38_1_-_228634819_228635039_RNA5S12_5S_50ntFlanks@154.171.18\|CCGGGTGCTGTAGGCTTT | >hg38_1_-_228634819_228635039_RNA5S12_5S_50ntFlanks | 154 | 171 | 18 | CCGGGTGCTGTAGGCTTT | MB231 | 68 | 76 | 286 | 55 | 29 | 31.6 | 34.3 | 136.9 | 15.5 | 6.9 |
| >hg38_1_-_228634819_228635039_RNA5S12_5S_50ntFlanks@154.171.18\|CCGGGTGCTGTAGGCTTT | >hg38_1_-_228634819_228635039_RNA5S12_5S_50ntFlanks | 154 | 171 | 18 | CCGGGTGCTGTAGGCTTT | MB468 | 16 | 28 | 27 | 1102 | 160 | 9.6 | 29.0 | 10.9 | 862.7 | 103.5 |
| >hg38_1_-_228634819_228635039_RNA5S12_5S_50ntFlanks@51.70.20\|GTCTACGGCCATACCACCCT | >hg38_1_-_228634819_228635039_RNA5S12_5S_50ntFlanks | 51 | 70 | 20 | GTCTACGGCCATACCACCCT | MB468 | 12 | 19 | 17 | 8 | 8 | 8.7 | 8.9 | 13.3 | 10.4 | 6.1 |
| >hg38_1_-_228634819_228635039_RNA5S12_5S_50ntFlanks@51.71.21\|GTCTACGGCCATACCACCCTG | >hg38_1_-_228634819_228635039_RNA5S12_5S_50ntFlanks | 51 | 71 | 21 | GTCTACGGCCATACCACCCTG | MB468 | 41 | 75 | 46 | 10 | 29 | 26.9 | 45.0 | 42.7 | 7.7 | 15.4 |

| | | | | | | | | | | | | | | | | |
|---|---|---|---|---|---|---|---|---|---|---|---|---|---|---|---|---|
| >hg38_1_-_228634819_228635039_RNA5S12_5S_50ntFlanks@51.78.28\|GTCTACGGCCATACCACCCTGAACGCGC | >hg38_1_-_228634819_228635039_RNA5S12_5S_50ntFlanks | 51 | 78 | 28 | GTCTACGGCCATACCACCCTGAACGCGC | BT20 | 10 | 10 | 34 | 9 | 3 | 10.6 | 4.2 | 23.0 | 5.3 | 1.6 |
| >hg38_1_-_228634819_228635039_RNA5S12_5S_50ntFlanks@51.79.29\|GTCTACGGCCATACCACCCTGAACGCGCC | >hg38_1_-_228634819_228635039_RNA5S12_5S_50ntFlanks | 51 | 79 | 29 | GTCTACGGCCATACCACCCTGAACGCGCC | BT20 | 17 | 36 | 62 | 64 | 15 | 11.3 | 19.8 | 35.6 | 29.7 | 8.4 |
| >hg38_1_-_228634819_228635039_RNA5S12_5S_50ntFlanks@51.79.29\|GTCTACGGCCATACCACCCTGAACGCGCC | >hg38_1_-_228634819_228635039_RNA5S12_5S_50ntFlanks | 51 | 79 | 29 | GTCTACGGCCATACCACCCTGAACGCGCC | MB231 | 11 | 89 | 76 | 16 | 53 | 6.7 | 68.7 | 48.2 | 17.0 | 67.1 |
| >hg38_1_-_228634819_228635039_RNA5S12_5S_50ntFlanks@51.80.30\|GTCTACGGCCATACCACCCTGAACGCGCCC | >hg38_1_-_228634819_228635039_RNA5S12_5S_50ntFlanks | 51 | 80 | 30 | GTCTACGGCCATACCACCCTGAACGCGCCC | BT20 | 15 | 23 | 71 | 99 | 8 | 13.6 | 13.7 | 54.5 | 32.1 | 3.4 |
| >hg38_1_-_228634819_228635039_RNA5S12_5S_50ntFlanks@51.81.31\|GTCTACGGCCATACCACCCTGAACGCGCCCG | >hg38_1_-_228634819_228635039_RNA5S12_5S_50ntFlanks | 51 | 81 | 31 | GTCTACGGCCATACCACCCTGAACGCGCCCG | BT20 | 14 | 31 | 61 | 37 | 9 | 10.5 | 16.2 | 37.4 | 15.9 | 3.9 |
| >hg38_1_-_228634819_228635039_RNA5S12_5S_50ntFlanks@51.82.32\|GTCTACGGCCATACCACCCTGAACGCGCCCGA | >hg38_1_-_228634819_228635039_RNA5S12_5S_50ntFlanks | 51 | 82 | 32 | GTCTACGGCCATACCACCCTGAACGCGCCCGA | BT20 | 23 | 33 | 103 | 21 | 12 | 18.0 | 19.0 | 69.3 | 6.6 | 6.2 |
| >hg38_1_-_228634819_228635039_RNA5S12_5S_50ntFlanks@51.82.32\|GTCTACGGCCATACCACCCTGAACGCGCCCGA | >hg38_1_-_228634819_228635039_RNA5S12_5S_50ntFlanks | 51 | 82 | 32 | GTCTACGGCCATACCACCCTGAACGCGCCCGA | MB231 | 12 | 87 | 83 | 6 | 75 | 12.8 | 57.2 | 56.3 | 1.7 | 98.4 |
| >hg38_1_-_228634819_228635039_RNA5S12_5S_50ntFlanks@51.83.33\|GTCTACGGCCATACCACCCTGAACGCGCCCGAT | >hg38_1_-_228634819_228635039_RNA5S12_5S_50ntFlanks | 51 | 83 | 33 | GTCTACGGCCATACCACCCTGAACGCGCCCGAT | BT20 | 28 | 48 | 118 | 102 | 16 | 10.5 | 19.5 | 77.1 | 111.6 | 5.8 |
| >hg38_1_-_228634819_228635039_RNA5S12_5S_50ntFlanks@51.83.33\|GTCTACGGCCATACCACCCTGAACGCGCCCGAT | >hg38_1_-_228634819_228635039_RNA5S12_5S_50ntFlanks | 51 | 83 | 33 | GTCTACGGCCATACCACCCTGAACGCGCCCGAT | MB231 | 15 | 117 | 106 | 17 | 60 | 7.2 | 84.6 | 67.1 | 4.6 | 64.1 |
| >hg38_1_-_228634819_228635039_RNA5S12_5S_50ntFlanks@51.83.33\|GTCTACGGCCATACCACCCTGAACGCGCCCGAT | >hg38_1_-_228634819_228635039_RNA5S12_5S_50ntFlanks | 51 | 83 | 33 | GTCTACGGCCATACCACCCTGAACGCGCCCGAT | MB468 | 139 | 849 | 109 | 52 | 127 | 53.3 | 341.8 | 65.4 | 70.2 | 44.4 |
| >hg38_1_-_228634819_228635039_RNA5S12_5S_50ntFlanks@51.84.34\|GTCTACGGCCATACCACCCTGAACGCGCCCGATC | >hg38_1_-_228634819_228635039_RNA5S12_5S_50ntFlanks | 51 | 84 | 34 | GTCTACGGCCATACCACCCTGAACGCGCCCGATC | BT20 | 126 | 363 | 1364 | 512 | 152 | 60.1 | 375.0 | 270.6 | 105.5 | 81.8 |
| >hg38_1_-_228634819_228635039_RNA5S12_5S_50ntFlanks@51.84.34\|GTCTACGGCCATACCACCCTGAACGCGCCCGATC | >hg38_1_-_228634819_228635039_RNA5S12_5S_50ntFlanks | 51 | 84 | 34 | GTCTACGGCCATACCACCCTGAACGCGCCCGATC | MB231 | 76 | 298 | 634 | 163 | 194 | 90.8 | 76.0 | 226.8 | 242.8 | 124.0 |
| >hg38_1_-_228634819_228635039_RNA5S12_5S_50ntFlanks@51.84.34\|GTCTACGGCCATACCACCCTGAACGCGCCCGATC | >hg38_1_-_228634819_228635039_RNA5S12_5S_50ntFlanks | 51 | 84 | 34 | GTCTACGGCCATACCACCCTGAACGCGCCCGATC | MB468 | 12 | 130 | 33 | 52 | 21 | 7.0 | 77.3 | 28.1 | 69.0 | 11.4 |
| >hg38_1_-_228634819_228635039_RNA5S12_5S_50ntFlanks@51.85.35\|GTCTACGGCCATACCACCCTGAACGCGCCCGATCT | >hg38_1_-_228634819_228635039_RNA5S12_5S_50ntFlanks | 51 | 85 | 35 | GTCTACGGCCATACCACCCTGAACGCGCCCGATCT | BT20 | 20 | 37 | 83 | 32 | 18 | 11.7 | 15.7 | 36.9 | 21.1 | 13.1 |
| >hg38_1_-_228634819_228635039_RNA5S12_5S_50ntFlanks@51.85.35\|GTCTACGGCCATACCACCCTGAACGCGCCCGATCT | >hg38_1_-_228634819_228635039_RNA5S12_5S_50ntFlanks | 51 | 85 | 35 | GTCTACGGCCATACCACCCTGAACGCGCCCGATCT | MB231 | 10 | 68 | 65 | 7 | 36 | 6.3 | 44.2 | 31.8 | 1.8 | 30.2 |
| >hg38_1_-_228634819_228635039_RNA5S12_5S_50ntFlanks@51.86.36\|GTCTACGGCCATACCACCCTGAACGCGCCCGATCTC | >hg38_1_-_228634819_228635039_RNA5S12_5S_50ntFlanks | 51 | 86 | 36 | GTCTACGGCCATACCACCCTGAACGCGCCCGATCTC | BT20 | 17 | 43 | 111 | 77 | 12 | 11.1 | 21.7 | 40.7 | 31.4 | 6.0 |
| >hg38_1_-_228634819_228635039_RNA5S12_5S_50ntFlanks@51.87.37\|GTCTACGGCCATACCACCCTGAACGCGCCCGATCTCG | >hg38_1_-_228634819_228635039_RNA5S12_5S_50ntFlanks | 51 | 87 | 37 | GTCTACGGCCATACCACCCTGAACGCGCCCGATCTCG | BT20 | 42 | 158 | 173 | 17 | 25 | 31.7 | 223.8 | 156.0 | 5.6 | 21.0 |
| >hg38_1_-_228634819_228635039_RNA5S12_5S_50ntFlanks@51.87.37\|GTCTACGGCCATACCACCCTGAACGCGCCCGATCTCG | >hg38_1_-_228634819_228635039_RNA5S12_5S_50ntFlanks | 51 | 87 | 37 | GTCTACGGCCATACCACCCTGAACGCGCCCGATCTCG | MB231 | 10 | 67 | 71 | 8 | 48 | 6.4 | 41.8 | 37.9 | 3.2 | 29.2 |
| >hg38_1_-_228634819_228635039_RNA5S12_5S_50ntFlanks@51.88.38\|GTCTACGGCCATACCACCCTGAACGCGCCCGATCTCGT | >hg38_1_-_228634819_228635039_RNA5S12_5S_50ntFlanks | 51 | 88 | 38 | GTCTACGGCCATACCACCCTGAACGCGCCCGATCTCGT | BT20 | 75 | 195 | 256 | 51 | 72 | 31.3 | 179.8 | 143.8 | 24.9 | 48.8 |
| >hg38_1_-_228634819_228635039_RNA5S12_5S_50ntFlanks@51.88.38\|GTCTACGGCCATACCACCCTGAACGCGCCCGATCTCGT | >hg38_1_-_228634819_228635039_RNA5S12_5S_50ntFlanks | 51 | 88 | 38 | GTCTACGGCCATACCACCCTGAACGCGCCCGATCTCGT | MB231 | 44 | 186 | 256 | 48 | 109 | 14.4 | 181.1 | 261.2 | 22.5 | 50.5 |
| >hg38_1_-_228634819_228635039_RNA5S12_5S_50ntFlanks@51.89.39\|GTCTACGGCCATACCACCCTGAACGCGCCCGATCTCGTC | >hg38_1_-_228634819_228635039_RNA5S12_5S_50ntFlanks | 51 | 89 | 39 | GTCTACGGCCATACCACCCTGAACGCGCCCGATCTCGTC | BT20 | 926 | 5614 | 2282 | 321 | 700 | 780.2 | 9299.0 | 2167.3 | 193.1 | 426.9 |
| >hg38_1_-_228634819_228635039_RNA5S12_5S_50ntFlanks@51.89.39\|GTCTACGGCCATACCACCCTGAACGCGCCCGATCTCGTC | >hg38_1_-_228634819_228635039_RNA5S12_5S_50ntFlanks | 51 | 89 | 39 | GTCTACGGCCATACCACCCTGAACGCGCCCGATCTCGTC | MB231 | 345 | 1243 | 2621 | 341 | 1048 | 148.2 | 1409.8 | 2833.4 | 186.2 | 289.7 |
| >hg38_1_-_228634819_228635039_RNA5S12_5S_50ntFlanks@51.90.40\|GTCTACGGCCATACCACCCTGAACGCGCCCGATCTCGTCT | >hg38_1_-_228634819_228635039_RNA5S12_5S_50ntFlanks | 51 | 90 | 40 | GTCTACGGCCATACCACCCTGAACGCGCCCGATCTCGTCT | BT20 | 13 | 78 | 69 | 10 | 13 | 8.1 | 87.7 | 75.3 | 0.7 | 6.3 |
| >hg38_1_-_228634819_228635039_RNA5S12_5S_50ntFlanks@51.91.41\|GTCTACGGCCATACCACCCTGAACGCGCCCGATCTCGTCTG | >hg38_1_-_228634819_228635039_RNA5S12_5S_50ntFlanks | 51 | 91 | 41 | GTCTACGGCCATACCACCCTGAACGCGCCCGATCTCGTCTG | BT20 | 30 | 57 | 199 | 267 | 31 | 28.0 | 51.9 | 256.7 | 209.0 | 11.0 |
| >hg38_1_-_228634819_228635039_RNA5S12_5S_50ntFlanks@51.92.42\|GTCTACGGCCATACCACCCTGAACGCGCCCGATCTCGTCTGA | >hg38_1_-_228634819_228635039_RNA5S12_5S_50ntFlanks | 51 | 92 | 42 | GTCTACGGCCATACCACCCTGAACGCGCCCGATCTCGTCTGA | BT20 | 12 | 111 | 62 | 19 | 30 | 11.1 | 177.2 | 52.4 | 24.4 | 5.6 |
| >hg38_1_-_228634819_228635039_RNA5S12_5S_50ntFlanks@51.94.44\|GTCTACGGCCATACCACCCTGAACGCGCCCGATCTCGTCTGATC | >hg38_1_-_228634819_228635039_RNA5S12_5S_50ntFlanks | 51 | 94 | 44 | GTCTACGGCCATACCACCCTGAACGCGCCCGATCTCGTCTGATC | BT20 | 23 | 19 | 305 | 24 | 10 | 28.2 | 24.7 | 513.4 | 18.5 | 4.6 |
| >hg38_1_-_228634819_228635039_RNA5S12_5S_50ntFlanks@84.101.18\|CTCGTCTGATCTCGGAAG | >hg38_1_-_228634819_228635039_RNA5S12_5S_50ntFlanks | 84 | 101 | 18 | CTCGTCTGATCTCGGAAG | MB468 | 224 | 102 | 194 | 76 | 39 | 79.3 | 59.8 | 148.3 | 23.5 | 10.8 |
| >hg38_1_-_228634819_228635039_RNA5S12_5S_50ntFlanks@84.102.19\|CTCGTCTGATCTCGGAAGC | >hg38_1_-_228634819_228635039_RNA5S12_5S_50ntFlanks | 84 | 102 | 19 | CTCGTCTGATCTCGGAAGC | MB468 | 49 | 34 | 47 | 8 | 7 | 27.4 | 17.2 | 38.7 | 2.6 | 2.5 |
| >hg38_1_-_228634819_228635039_RNA5S12_5S_50ntFlanks@84.103.20\|CTCGTCTGATCTCGGAAGCT | >hg38_1_-_228634819_228635039_RNA5S12_5S_50ntFlanks | 84 | 103 | 20 | CTCGTCTGATCTCGGAAGCT | MB468 | 127 | 83 | 91 | 14 | 21 | 46.2 | 36.1 | 35.2 | 2.8 | 3.5 |
| >hg38_1_-_228634819_228635039_RNA5S12_5S_50ntFlanks@84.109.26\|CTCGTCTGATCTCGGAAGCTAAGCAG | >hg38_1_-_228634819_228635039_RNA5S12_5S_50ntFlanks | 84 | 109 | 26 | CTCGTCTGATCTCGGAAGCTAAGCAG | MB468 | 16 | 18 | 7 | 5 | 6 | 6.2 | 11.7 | 5.6 | 1.2 | 1.8 |
| >hg38_1_-_228634819_228635039_RNA5S12_5S_50ntFlanks@84.111.28\|CTCGTCTGATCTCGGAAGCTAAGCAGGG | >hg38_1_-_228634819_228635039_RNA5S12_5S_50ntFlanks | 84 | 111 | 28 | CTCGTCTGATCTCGGAAGCTAAGCAGGG | MB468 | 11 | 21 | 6 | 4 | 4 | 4.4 | 10.3 | 4.6 | 0.9 | 2.0 |
| >hg38_1_-_228634819_228635039_RNA5S12_5S_50ntFlanks@84.112.29\|CTCGTCTGATCTCGGAAGCTAAGCAGGGT | >hg38_1_-_228634819_228635039_RNA5S12_5S_50ntFlanks | 84 | 112 | 29 | CTCGTCTGATCTCGGAAGCTAAGCAGGGT | MB468 | 37 | 65 | 25 | 8 | 9 | 11.7 | 19.4 | 16.2 | 1.6 | 4.7 |
| >hg38_1_-_228634819_228635039_RNA5S12_5S_50ntFlanks@86.103.18\|CGTCTGATCTCGGAAGCT | >hg38_1_-_228634819_228635039_RNA5S12_5S_50ntFlanks | 86 | 103 | 18 | CGTCTGATCTCGGAAGCT | MB468 | 22 | 9 | 20 | 5 | 3 | 13.6 | 5.2 | 13.7 | 3.2 | 0.9 |
| >hg38_21_+_8433172_8446622_RNA45SN1_45S_50ntFlanks@10099.10117.19\|CGCGCGCGCGTGTGGTGTG | >hg38_21_+_8433172_8446622_RNA45SN1_45S_50ntFlanks | 10099 | 10117 | 19 | CGCGCGCGCGTGTGGTGTG | MB468 | 47 | 20 | 31 | 206 | 88 | 5.8 | 7.2 | 6.9 | 61.9 | 38.9 |
| >hg38_21_+_8433172_8446622_RNA45SN1_45S_50ntFlanks@10099.10127.29\|CGCGCGCGCGTGTGGTGTGCGTCGGAGGG | >hg38_21_+_8433172_8446622_RNA45SN1_45S_50ntFlanks | 10099 | 10127 | 29 | CGCGCGCGCGTGTGGTGTGCGTCGGAGGG | MB468 | 14 | 10 | 10 | 14 | 26 | 5.8 | 6.1 | 4.0 | 10.1 | 15.1 |
| >hg38_21_+_8433172_8446622_RNA45SN1_45S_50ntFlanks@10101.10125.25\|CGCGCGCGTGTGGTGTGCGTCGGAG | >hg38_21_+_8433172_8446622_RNA45SN1_45S_50ntFlanks | 10101 | 10125 | 25 | CGCGCGCGTGTGGTGTGCGTCGGAG | MB468 | 37 | 6 | 21 | 50 | 59 | 23.9 | 2.7 | 15.7 | 35.2 | 37.7 |

| | | | | | | | | | | | | | | | | |
|---|---|---|---|---|---|---|---|---|---|---|---|---|---|---|---|---|
| >hg38_21_+_8433172_8446622_RNA45SN1_45S_50ntFlanks@10101.10126.26\|CGCGCGCGTGTGGTGTGCGTCGGAGG | >hg38_21_+_8433172_8446622_RNA45SN1_45S_50ntFlanks | 10101 | 10126 | 26 | CGCGCGCGTGTGGTGTGCGTCGGAGG | MB468 | 38 | 19 | 27 | 34 | 58 | 17.4 | 13.1 | 14.3 | 23.0 | 39.6 |
| >hg38_21_+_8433172_8446622_RNA45SN1_45S_50ntFlanks@10101.10127.27\|CGCGCGCGTGTGGTGTGCGTCGGAGGG | >hg38_21_+_8433172_8446622_RNA45SN1_45S_50ntFlanks | 10101 | 10127 | 27 | CGCGCGCGTGTGGTGTGCGTCGGAGGG | MB468 | 56 | 51 | 57 | 51 | 102 | 24.4 | 41.5 | 27.3 | 31.5 | 59.1 |
| >hg38_21_+_8433172_8446622_RNA45SN1_45S_50ntFlanks@10107.10125.19\|CGTGTGGTGTGCGTCGGAG | >hg38_21_+_8433172_8446622_RNA45SN1_45S_50ntFlanks | 10107 | 10125 | 19 | CGTGTGGTGTGCGTCGGAG | MB468 | 29 | 31 | 43 | 51 | 24 | 10.0 | 17.5 | 18.7 | 23.1 | 5.8 |
| >hg38_21_+_8433172_8446622_RNA45SN1_45S_50ntFlanks@10107.10127.21\|CGTGTGGTGTGCGTCGGAGGG | >hg38_21_+_8433172_8446622_RNA45SN1_45S_50ntFlanks | 10107 | 10127 | 21 | CGTGTGGTGTGCGTCGGAGGG | MB468 | 28 | 34 | 38 | 50 | 31 | 6.5 | 15.4 | 13.6 | 14.9 | 8.0 |
| >hg38_21_+_8433172_8446622_RNA45SN1_45S_50ntFlanks@10116.10134.19\|TGCGTCGGAGGGCGGCGGC | >hg38_21_+_8433172_8446622_RNA45SN1_45S_50ntFlanks | 10116 | 10134 | 19 | TGCGTCGGAGGGCGGCGGC | MB468 | 14 | 5 | 17 | 13 | 14 | 4.0 | 2.2 | 2.4 | 11.6 | 4.7 |
| >hg38_21_+_8433172_8446622_RNA45SN1_45S_50ntFlanks@10116.10135.20\|TGCGTCGGAGGGCGGCGGCG | >hg38_21_+_8433172_8446622_RNA45SN1_45S_50ntFlanks | 10116 | 10135 | 20 | TGCGTCGGAGGGCGGCGGCG | MB468 | 19 | 14 | 32 | 18 | 28 | 2.8 | 7.5 | 8.5 | 21.0 | 7.6 |
| >hg38_21_+_8433172_8446622_RNA45SN1_45S_50ntFlanks@10116.10136.21\|TGCGTCGGAGGGCGGCGGCGG | >hg38_21_+_8433172_8446622_RNA45SN1_45S_50ntFlanks | 10116 | 10136 | 21 | TGCGTCGGAGGGCGGCGGCGG | MB468 | 22 | 25 | 38 | 32 | 47 | 2.3 | 14.9 | 14.3 | 32.8 | 16.5 |
| >hg38_21_+_8433172_8446622_RNA45SN1_45S_50ntFlanks@10186.10207.22\|TCCTCCCCTCCTCCCGCCCACG | >hg38_21_+_8433172_8446622_RNA45SN1_45S_50ntFlanks | 10186 | 10207 | 22 | TCCTCCCCTCCTCCCGCCCACG | MB468 | 20 | 77 | 34 | 8 | 19 | 5.1 | 59.3 | 14.6 | 11.8 | 15.5 |
| >hg38_21_+_8433172_8446622_RNA45SN1_45S_50ntFlanks@10186.10208.23\|TCCTCCCCTCCTCCCGCCCACGC | >hg38_21_+_8433172_8446622_RNA45SN1_45S_50ntFlanks | 10186 | 10208 | 23 | TCCTCCCCTCCTCCCGCCCACGC | MB468 | 12 | 34 | 19 | 8 | 8 | 3.1 | 18.9 | 4.6 | 12.7 | 7.5 |
| >hg38_21_+_8433172_8446622_RNA45SN1_45S_50ntFlanks@10240.10259.20\|CTACGCCGCGACGAGTAGGA | >hg38_21_+_8433172_8446622_RNA45SN1_45S_50ntFlanks | 10240 | 10259 | 20 | CTACGCCGCGACGAGTAGGA | BT20 | 11 | 23 | 15 | 286 | 54 | 3.8 | 10.4 | 1.3 | 141.7 | 17.8 |
| >hg38_21_+_8433172_8446622_RNA45SN1_45S_50ntFlanks@10320.10341.22\|CAGATCTTGGTGGTAGTAGCAA | >hg38_21_+_8433172_8446622_RNA45SN1_45S_50ntFlanks | 10320 | 10341 | 22 | CAGATCTTGGTGGTAGTAGCAA | MB468 | 11 | 6 | 6 | 15 | 13 | 5.7 | 3.4 | 2.6 | 6.5 | 5.3 |
| >hg38_21_+_8433172_8446622_RNA45SN1_45S_50ntFlanks@10341.10361.21\|AATATTCAAACGAGAACTTTG | >hg38_21_+_8433172_8446622_RNA45SN1_45S_50ntFlanks | 10341 | 10361 | 21 | AATATTCAAACGAGAACTTTG | MB468 | 11 | 3 | 9 | 12 | 7 | 2.6 | 2.2 | 5.5 | 3.0 | 3.2 |
| >hg38_21_+_8433172_8446622_RNA45SN1_45S_50ntFlanks@10342.10361.20\|ATATTCAAACGAGAACTTTG | >hg38_21_+_8433172_8446622_RNA45SN1_45S_50ntFlanks | 10342 | 10361 | 20 | ATATTCAAACGAGAACTTTG | MB468 | 85 | 17 | 56 | 85 | 39 | 14.9 | 11.9 | 30.6 | 19.8 | 18.4 |
| >hg38_21_+_8433172_8446622_RNA45SN1_45S_50ntFlanks@10342.10373.32\|ATATTCAAACGAGAACTTTGAAGGCCGAAGTG | >hg38_21_+_8433172_8446622_RNA45SN1_45S_50ntFlanks | 10342 | 10373 | 32 | ATATTCAAACGAGAACTTTGAAGGCCGAAGTG | MB231 | 10 | 10 | 17 | 3 | 4 | 8.3 | 5.0 | 13.5 | 1.4 | 2.6 |
| >hg38_21_+_8433172_8446622_RNA45SN1_45S_50ntFlanks@10343.10361.19\|TATTCAAACGAGAACTTTG | >hg38_21_+_8433172_8446622_RNA45SN1_45S_50ntFlanks | 10343 | 10361 | 19 | TATTCAAACGAGAACTTTG | MB468 | 25 | 4 | 20 | 18 | 12 | 8.4 | 2.1 | 17.7 | 5.7 | 6.7 |
| >hg38_21_+_8433172_8446622_RNA45SN1_45S_50ntFlanks@10343.10373.31\|TATTCAAACGAGAACTTTGAAGGCCGAAGTG | >hg38_21_+_8433172_8446622_RNA45SN1_45S_50ntFlanks | 10343 | 10373 | 31 | TATTCAAACGAGAACTTTGAAGGCCGAAGTG | MB231 | 10 | 7 | 24 | 3 | 3 | 7.7 | 2.7 | 18.8 | 2.4 | 2.6 |
| >hg38_21_+_8433172_8446622_RNA45SN1_45S_50ntFlanks@10344.10361.18\|ATTCAAACGAGAACTTTG | >hg38_21_+_8433172_8446622_RNA45SN1_45S_50ntFlanks | 10344 | 10361 | 18 | ATTCAAACGAGAACTTTG | MB468 | 19 | 4 | 17 | 29 | 10 | 8.4 | 2.8 | 14.6 | 9.8 | 3.7 |
| >hg38_21_+_8433172_8446622_RNA45SN1_45S_50ntFlanks@10344.10373.30\|ATTCAAACGAGAACTTTGAAGGCCGAAGTG | >hg38_21_+_8433172_8446622_RNA45SN1_45S_50ntFlanks | 10344 | 10373 | 30 | ATTCAAACGAGAACTTTGAAGGCCGAAGTG | MB231 | 15 | 12 | 29 | 4 | 4 | 11.6 | 7.2 | 24.4 | 3.5 | 3.2 |
| >hg38_21_+_8433172_8446622_RNA45SN1_45S_50ntFlanks@10345.10373.29\|TTCAAACGAGAACTTTGAAGGCCGAAGTG | >hg38_21_+_8433172_8446622_RNA45SN1_45S_50ntFlanks | 10345 | 10373 | 29 | TTCAAACGAGAACTTTGAAGGCCGAAGTG | MB231 | 18 | 14 | 43 | 3 | 5 | 13.9 | 9.4 | 34.2 | 2.4 | 3.6 |
| >hg38_21_+_8433172_8446622_RNA45SN1_45S_50ntFlanks@10346.10367.22\|TCAAACGAGAACTTTGAAGGCC | >hg38_21_+_8433172_8446622_RNA45SN1_45S_50ntFlanks | 10346 | 10367 | 22 | TCAAACGAGAACTTTGAAGGCC | MB231 | 12 | 19 | 15 | 2 | 2 | 12.4 | 18.6 | 12.6 | 1.7 | 1.4 |
| >hg38_21_+_8433172_8446622_RNA45SN1_45S_50ntFlanks@10346.10367.22\|TCAAACGAGAACTTTGAAGGCC | >hg38_21_+_8433172_8446622_RNA45SN1_45S_50ntFlanks | 10346 | 10367 | 22 | TCAAACGAGAACTTTGAAGGCC | MB468 | 14 | 11 | 16 | 19 | 15 | 7.8 | 3.1 | 5.9 | 12.4 | 7.6 |
| >hg38_21_+_8433172_8446622_RNA45SN1_45S_50ntFlanks@10346.10368.23\|TCAAACGAGAACTTTGAAGGCCG | >hg38_21_+_8433172_8446622_RNA45SN1_45S_50ntFlanks | 10346 | 10368 | 23 | TCAAACGAGAACTTTGAAGGCCG | MB231 | 13 | 30 | 14 | 1 | 3 | 11.8 | 28.9 | 9.4 | 1.0 | 3.4 |
| >hg38_21_+_8433172_8446622_RNA45SN1_45S_50ntFlanks@10346.10368.23\|TCAAACGAGAACTTTGAAGGCCG | >hg38_21_+_8433172_8446622_RNA45SN1_45S_50ntFlanks | 10346 | 10368 | 23 | TCAAACGAGAACTTTGAAGGCCG | MB468 | 239 | 176 | 237 | 224 | 258 | 79.8 | 55.2 | 78.3 | 137.1 | 82.2 |
| >hg38_21_+_8433172_8446622_RNA45SN1_45S_50ntFlanks@10346.10369.24\|TCAAACGAGAACTTTGAAGGCCGA | >hg38_21_+_8433172_8446622_RNA45SN1_45S_50ntFlanks | 10346 | 10369 | 24 | TCAAACGAGAACTTTGAAGGCCGA | MB468 | 36 | 26 | 30 | 91 | 55 | 17.7 | 17.0 | 16.7 | 49.1 | 23.7 |
| >hg38_21_+_8433172_8446622_RNA45SN1_45S_50ntFlanks@10346.10373.28\|TCAAACGAGAACTTTGAAGGCCGAAGTG | >hg38_21_+_8433172_8446622_RNA45SN1_45S_50ntFlanks | 10346 | 10373 | 28 | TCAAACGAGAACTTTGAAGGCCGAAGTG | MB231 | 13 | 9 | 28 | 4 | 4 | 10.6 | 5.5 | 24.8 | 3.6 | 3.1 |
| >hg38_21_+_8433172_8446622_RNA45SN1_45S_50ntFlanks@10346.10373.28\|TCAAACGAGAACTTTGAAGGCCGAAGTG | >hg38_21_+_8433172_8446622_RNA45SN1_45S_50ntFlanks | 10346 | 10373 | 28 | TCAAACGAGAACTTTGAAGGCCGAAGTG | MB468 | 21 | 24 | 19 | 27 | 30 | 13.0 | 5.5 | 9.9 | 9.8 | 5.3 |
| >hg38_21_+_8433172_8446622_RNA45SN1_45S_50ntFlanks@10346.10374.29\|TCAAACGAGAACTTTGAAGGCCGAAGTGG | >hg38_21_+_8433172_8446622_RNA45SN1_45S_50ntFlanks | 10346 | 10374 | 29 | TCAAACGAGAACTTTGAAGGCCGAAGTGG | MB468 | 67 | 81 | 58 | 164 | 127 | 24.9 | 24.1 | 13.4 | 82.5 | 13.4 |
| >hg38_21_+_8433172_8446622_RNA45SN1_45S_50ntFlanks@10346.10383.38\|TCAAACGAGAACTTTGAAGGCCGAAGTGGAGAAGGGTT | >hg38_21_+_8433172_8446622_RNA45SN1_45S_50ntFlanks | 10346 | 10383 | 38 | TCAAACGAGAACTTTGAAGGCCGAAGTGGAGAAGGGTT | MB468 | 10 | 35 | 16 | 18 | 49 | 3.6 | 20.3 | 10.3 | 11.6 | 30.5 |
| >hg38_21_+_8433172_8446622_RNA45SN1_45S_50ntFlanks@10347.10368.22\|CAAACGAGAACTTTGAAGGCCG | >hg38_21_+_8433172_8446622_RNA45SN1_45S_50ntFlanks | 10347 | 10368 | 22 | CAAACGAGAACTTTGAAGGCCG | MB468 | 53 | 30 | 46 | 38 | 21 | 17.2 | 13.9 | 10.2 | 30.3 | 9.0 |
| >hg38_21_+_8433172_8446622_RNA45SN1_45S_50ntFlanks@10347.10374.28\|CAAACGAGAACTTTGAAGGCCGAAGTGG | >hg38_21_+_8433172_8446622_RNA45SN1_45S_50ntFlanks | 10347 | 10374 | 28 | CAAACGAGAACTTTGAAGGCCGAAGTGG | MB468 | 17 | 21 | 16 | 62 | 16 | 5.6 | 9.7 | 3.8 | 60.4 | 4.9 |
| >hg38_21_+_8433172_8446622_RNA45SN1_45S_50ntFlanks@10348.10365.18\|AAACGAGAACTTTGAAGG | >hg38_21_+_8433172_8446622_RNA45SN1_45S_50ntFlanks | 10348 | 10365 | 18 | AAACGAGAACTTTGAAGG | MB468 | 31 | 20 | 15 | 25 | 5 | 6.7 | 16.2 | 3.7 | 17.2 | 2.3 |
| >hg38_21_+_8433172_8446622_RNA45SN1_45S_50ntFlanks@10348.10366.19\|AAACGAGAACTTTGAAGGC | >hg38_21_+_8433172_8446622_RNA45SN1_45S_50ntFlanks | 10348 | 10366 | 19 | AAACGAGAACTTTGAAGGC | MB231 | 27 | 16 | 21 | 11 | 4 | 28.6 | 12.5 | 23.7 | 8.9 | 2.0 |
| >hg38_21_+_8433172_8446622_RNA45SN1_45S_50ntFlanks@10348.10366.19\|AAACGAGAACTTTGAAGGC | >hg38_21_+_8433172_8446622_RNA45SN1_45S_50ntFlanks | 10348 | 10366 | 19 | AAACGAGAACTTTGAAGGC | MB468 | 67 | 46 | 45 | 65 | 12 | 18.0 | 35.3 | 10.8 | 48.7 | 3.0 |
| >hg38_21_+_8433172_8446622_RNA45SN1_45S_50ntFlanks@10348.10367.20\|AAACGAGAACTTTGAAGGCC | >hg38_21_+_8433172_8446622_RNA45SN1_45S_50ntFlanks | 10348 | 10367 | 20 | AAACGAGAACTTTGAAGGCC | MB231 | 26 | 15 | 30 | 15 | 4 | 27.2 | 12.0 | 26.5 | 13.0 | 2.4 |
| >hg38_21_+_8433172_8446622_RNA45SN1_45S_50ntFlanks@10348.10367.20\|AAACGAGAACTTTGAAGGCC | >hg38_21_+_8433172_8446622_RNA45SN1_45S_50ntFlanks | 10348 | 10367 | 20 | AAACGAGAACTTTGAAGGCC | MB468 | 24 | 16 | 23 | 29 | 6 | 8.7 | 11.3 | 7.5 | 25.5 | 4.5 |
| >hg38_21_+_8433172_8446622_RNA45SN1_45S_50ntFlanks@10348.10368.21\|AAACGAGAACTTTGAAGGCCG | >hg38_21_+_8433172_8446622_RNA45SN1_45S_50ntFlanks | 10348 | 10368 | 21 | AAACGAGAACTTTGAAGGCCG | MB231 | 12 | 7 | 5 | 4 | 2 | 16.7 | 6.5 | 6.0 | 3.1 | 2.3 |
| >hg38_21_+_8433172_8446622_RNA45SN1_45S_50ntFlanks@10348.10368.21\|AAACGAGAACTTTGAAGGCCG | >hg38_21_+_8433172_8446622_RNA45SN1_45S_50ntFlanks | 10348 | 10368 | 21 | AAACGAGAACTTTGAAGGCCG | MB468 | 181 | 96 | 102 | 167 | 51 | 38.1 | 63.1 | 24.1 | 144.9 | 30.3 |
| >hg38_21_+_8433172_8446622_RNA45SN1_45S_50ntFlanks@10348.10369.22\|AAACGAGAACTTTGAAGGCCGA | >hg38_21_+_8433172_8446622_RNA45SN1_45S_50ntFlanks | 10348 | 10369 | 22 | AAACGAGAACTTTGAAGGCCGA | MB231 | 24 | 31 | 26 | 23 | 28 | 17.5 | 25.9 | 20.8 | 15.1 | 32.6 |
| >hg38_21_+_8433172_8446622_RNA45SN1_45S_50ntFlanks@10348.10369.22\|AAACGAGAACTTTGAAGGCCGA | >hg38_21_+_8433172_8446622_RNA45SN1_45S_50ntFlanks | 10348 | 10369 | 22 | AAACGAGAACTTTGAAGGCCGA | MB468 | 11 | 8 | 7 | 16 | 5 | 2.3 | 3.8 | 1.9 | 9.7 | 1.7 |
| >hg38_21_+_8433172_8446622_RNA45SN1_45S_50ntFlanks@10348.10370.23\|AAACGAGAACTTTGAAGGCCGAA | >hg38_21_+_8433172_8446622_RNA45SN1_45S_50ntFlanks | 10348 | 10370 | 23 | AAACGAGAACTTTGAAGGCCGAA | MB231 | 10 | 6 | 4 | 8 | 5 | 10.7 | 5.4 | 3.9 | 6.0 | 3.2 |
| >hg38_21_+_8433172_8446622_RNA45SN1_45S_50ntFlanks@10348.10371.24\|AAACGAGAACTTTGAAGGCCGAAG | >hg38_21_+_8433172_8446622_RNA45SN1_45S_50ntFlanks | 10348 | 10371 | 24 | AAACGAGAACTTTGAAGGCCGAAG | BT20 | 11 | 9 | 12 | 51 | 19 | 10.3 | 2.3 | 10.9 | 41.8 | 9.4 |
| >hg38_21_+_8433172_8446622_RNA45SN1_45S_50ntFlanks@10348.10371.24\|AAACGAGAACTTTGAAGGCCGAAG | >hg38_21_+_8433172_8446622_RNA45SN1_45S_50ntFlanks | 10348 | 10371 | 24 | AAACGAGAACTTTGAAGGCCGAAG | MB231 | 36 | 32 | 29 | 29 | 29 | 32.4 | 32.8 | 31.7 | 23.3 | 35.2 |
| >hg38_21_+_8433172_8446622_RNA45SN1_45S_50ntFlanks@10348.10371.24\|AAACGAGAACTTTGAAGGCCGAAG | >hg38_21_+_8433172_8446622_RNA45SN1_45S_50ntFlanks | 10348 | 10371 | 24 | AAACGAGAACTTTGAAGGCCGAAG | MB468 | 27 | 25 | 13 | 37 | 19 | 11.3 | 9.7 | 6.9 | 20.4 | 7.2 |
| >hg38_21_+_8433172_8446622_RNA45SN1_45S_50ntFlanks@10348.10372.25\|AAACGAGAACTTTGAAGGCCGAAGT | >hg38_21_+_8433172_8446622_RNA45SN1_45S_50ntFlanks | 10348 | 10372 | 25 | AAACGAGAACTTTGAAGGCCGAAGT | BT20 | 15 | 13 | 12 | 38 | 19 | 6.5 | 1.3 | 1.8 | 20.4 | 7.1 |
| >hg38_21_+_8433172_8446622_RNA45SN1_45S_50ntFlanks@10348.10372.25\|AAACGAGAACTTTGAAGGCCGAAGT | >hg38_21_+_8433172_8446622_RNA45SN1_45S_50ntFlanks | 10348 | 10372 | 25 | AAACGAGAACTTTGAAGGCCGAAGT | MB231 | 61 | 50 | 35 | 43 | 38 | 55.7 | 41.8 | 35.2 | 31.5 | 33.9 |
| >hg38_21_+_8433172_8446622_RNA45SN1_45S_50ntFlanks@10348.10372.25\|AAACGAGAACTTTGAAGGCCGAAGT | >hg38_21_+_8433172_8446622_RNA45SN1_45S_50ntFlanks | 10348 | 10372 | 25 | AAACGAGAACTTTGAAGGCCGAAGT | MB468 | 14 | 16 | 10 | 27 | 11 | 8.0 | 5.4 | 4.3 | 13.6 | 2.8 |
| >hg38_21_+_8433172_8446622_RNA45SN1_45S_50ntFlanks@10348.10373.26\|AAACGAGAACTTTGAAGGCCGAAGTG | >hg38_21_+_8433172_8446622_RNA45SN1_45S_50ntFlanks | 10348 | 10373 | 26 | AAACGAGAACTTTGAAGGCCGAAGTG | MB231 | 21 | 12 | 16 | 6 | 6 | 23.6 | 12.0 | 16.2 | 6.1 | 5.5 |
| >hg38_21_+_8433172_8446622_RNA45SN1_45S_50ntFlanks@10348.10373.26\|AAACGAGAACTTTGAAGGCCGAAGTG | >hg38_21_+_8433172_8446622_RNA45SN1_45S_50ntFlanks | 10348 | 10373 | 26 | AAACGAGAACTTTGAAGGCCGAAGTG | MB468 | 11 | 12 | 10 | 26 | 6 | 5.9 | 7.5 | 4.9 | 22.3 | 2.5 |
| >hg38_21_+_8433172_8446622_RNA45SN1_45S_50ntFlanks@10348.10374.27\|AAACGAGAACTTTGAAGGCCGAAGTGG | >hg38_21_+_8433172_8446622_RNA45SN1_45S_50ntFlanks | 10348 | 10374 | 27 | AAACGAGAACTTTGAAGGCCGAAGTGG | MB468 | 58 | 76 | 54 | 130 | 43 | 19.1 | 34.3 | 14.7 | 113.5 | 11.4 |
| >hg38_21_+_8433172_8446622_RNA45SN1_45S_50ntFlanks@10348.10376.29\|AAACGAGAACTTTGAAGGCCGAAGTGGAG | >hg38_21_+_8433172_8446622_RNA45SN1_45S_50ntFlanks | 10348 | 10376 | 29 | AAACGAGAACTTTGAAGGCCGAAGTGGAG | MB468 | 15 | 20 | 13 | 25 | 9 | 7.2 | 6.9 | 4.9 | 26.7 | 5.1 |

| | | | | | | | | | | | | | | | | |
|---|---|---|---|---|---|---|---|---|---|---|---|---|---|---|---|---|
| >hg38_21_+_8433172_8446622_RNA45SN1_45S_50ntFlanks@10348.10377.30\|AAACGAGAACTTTGAAGGCCGAAGTGGAGA | >hg38_21_+_8433172_8446622_RNA45SN1_45S_50ntFlanks | 10348 | 10377 | 30 | AAACGAGAACTTTGAAGGCCGAAGTGGAGA | MB468 | 17 | 24 | 11 | 28 | 8 | 4.3 | 7.2 | 4.9 | 24.9 | 3.0 |
| >hg38_21_+_8433172_8446622_RNA45SN1_45S_50ntFlanks@10348.10382.35\|AAACGAGAACTTTGAAGGCCGAAGTGGAGAAGGGT | >hg38_21_+_8433172_8446622_RNA45SN1_45S_50ntFlanks | 10348 | 10382 | 35 | AAACGAGAACTTTGAAGGCCGAAGTGGAGAAGGGT | MB231 | 21 | 10 | 67 | 19 | 20 | 18.8 | 9.1 | 53.1 | 14.4 | 15.7 |
| >hg38_21_+_8433172_8446622_RNA45SN1_45S_50ntFlanks@10348.10383.36\|AAACGAGAACTTTGAAGGCCGAAGTGGAGAAGGGTT | >hg38_21_+_8433172_8446622_RNA45SN1_45S_50ntFlanks | 10348 | 10383 | 36 | AAACGAGAACTTTGAAGGCCGAAGTGGAGAAGGGTT | MB231 | 11 | 6 | 31 | 9 | 12 | 9.1 | 4.2 | 24.7 | 8.0 | 9.0 |
| >hg38_21_+_8433172_8446622_RNA45SN1_45S_50ntFlanks@10348.10383.36\|AAACGAGAACTTTGAAGGCCGAAGTGGAGAAGGGTT | >hg38_21_+_8433172_8446622_RNA45SN1_45S_50ntFlanks | 10348 | 10383 | 36 | AAACGAGAACTTTGAAGGCCGAAGTGGAGAAGGGTT | MB468 | 42 | 118 | 52 | 61 | 58 | 18.2 | 73.7 | 21.5 | 53.5 | 35.9 |
| >hg38_21_+_8433172_8446622_RNA45SN1_45S_50ntFlanks@10348.10388.41\|AAACGAGAACTTTGAAGGCCGAAGTGGAGAAGGGTTCCATG | >hg38_21_+_8433172_8446622_RNA45SN1_45S_50ntFlanks | 10348 | 10388 | 41 | AAACGAGAACTTTGAAGGCCGAAGTGGAGAAGGGTTCCATG | MB468 | 13 | 26 | 19 | 10 | 32 | 7.9 | 20.6 | 10.0 | 6.1 | 19.4 |
| >hg38_21_+_8433172_8446622_RNA45SN1_45S_50ntFlanks@10349.10366.18\|AACGAGAACTTTGAAGGC | >hg38_21_+_8433172_8446622_RNA45SN1_45S_50ntFlanks | 10349 | 10366 | 18 | AACGAGAACTTTGAAGGC | MB231 | 37 | 20 | 24 | 12 | 4 | 40.2 | 15.9 | 27.0 | 11.1 | 3.5 |
| >hg38_21_+_8433172_8446622_RNA45SN1_45S_50ntFlanks@10349.10366.18\|AACGAGAACTTTGAAGGC | >hg38_21_+_8433172_8446622_RNA45SN1_45S_50ntFlanks | 10349 | 10366 | 18 | AACGAGAACTTTGAAGGC | MB468 | 125 | 71 | 80 | 90 | 14 | 13.8 | 57.9 | 12.4 | 68.8 | 4.2 |
| >hg38_21_+_8433172_8446622_RNA45SN1_45S_50ntFlanks@10349.10367.19\|AACGAGAACTTTGAAGGCC | >hg38_21_+_8433172_8446622_RNA45SN1_45S_50ntFlanks | 10349 | 10367 | 19 | AACGAGAACTTTGAAGGCC | MB231 | 33 | 17 | 35 | 18 | 5 | 32.6 | 13.3 | 31.3 | 15.1 | 3.8 |
| >hg38_21_+_8433172_8446622_RNA45SN1_45S_50ntFlanks@10349.10367.19\|AACGAGAACTTTGAAGGCC | >hg38_21_+_8433172_8446622_RNA45SN1_45S_50ntFlanks | 10349 | 10367 | 19 | AACGAGAACTTTGAAGGCC | MB468 | 54 | 32 | 41 | 39 | 7 | 5.0 | 22.5 | 3.0 | 30.3 | 3.9 |
| >hg38_21_+_8433172_8446622_RNA45SN1_45S_50ntFlanks@10349.10368.20\|AACGAGAACTTTGAAGGCCG | >hg38_21_+_8433172_8446622_RNA45SN1_45S_50ntFlanks | 10349 | 10368 | 20 | AACGAGAACTTTGAAGGCCG | MB231 | 18 | 10 | 8 | 5 | 5 | 22.7 | 9.9 | 8.0 | 4.1 | 4.9 |
| >hg38_21_+_8433172_8446622_RNA45SN1_45S_50ntFlanks@10349.10368.20\|AACGAGAACTTTGAAGGCCG | >hg38_21_+_8433172_8446622_RNA45SN1_45S_50ntFlanks | 10349 | 10368 | 20 | AACGAGAACTTTGAAGGCCG | MB468 | 406 | 203 | 177 | 234 | 78 | 38.9 | 126.0 | 19.0 | 165.9 | 29.5 |
| >hg38_21_+_8433172_8446622_RNA45SN1_45S_50ntFlanks@10349.10369.21\|AACGAGAACTTTGAAGGCCGA | >hg38_21_+_8433172_8446622_RNA45SN1_45S_50ntFlanks | 10349 | 10369 | 21 | AACGAGAACTTTGAAGGCCGA | BT20 | 13 | 7 | 9 | 49 | 33 | 13.3 | 1.2 | 6.8 | 29.9 | 13.4 |
| >hg38_21_+_8433172_8446622_RNA45SN1_45S_50ntFlanks@10349.10369.21\|AACGAGAACTTTGAAGGCCGA | >hg38_21_+_8433172_8446622_RNA45SN1_45S_50ntFlanks | 10349 | 10369 | 21 | AACGAGAACTTTGAAGGCCGA | MB231 | 40 | 48 | 36 | 44 | 54 | 32.0 | 47.1 | 28.9 | 32.8 | 61.5 |
| >hg38_21_+_8433172_8446622_RNA45SN1_45S_50ntFlanks@10349.10369.21\|AACGAGAACTTTGAAGGCCGA | >hg38_21_+_8433172_8446622_RNA45SN1_45S_50ntFlanks | 10349 | 10369 | 21 | AACGAGAACTTTGAAGGCCGA | MB468 | 25 | 16 | 13 | 25 | 8 | 4.3 | 6.7 | 4.1 | 14.8 | 2.2 |
| >hg38_21_+_8433172_8446622_RNA45SN1_45S_50ntFlanks@10349.10370.22\|AACGAGAACTTTGAAGGCCGAA | >hg38_21_+_8433172_8446622_RNA45SN1_45S_50ntFlanks | 10349 | 10370 | 22 | AACGAGAACTTTGAAGGCCGAA | MB231 | 16 | 9 | 6 | 12 | 9 | 17.2 | 8.3 | 5.6 | 9.1 | 7.2 |
| >hg38_21_+_8433172_8446622_RNA45SN1_45S_50ntFlanks@10349.10370.22\|AACGAGAACTTTGAAGGCCGAA | >hg38_21_+_8433172_8446622_RNA45SN1_45S_50ntFlanks | 10349 | 10370 | 22 | AACGAGAACTTTGAAGGCCGAA | MB468 | 15 | 7 | 7 | 21 | 5 | 4.6 | 4.7 | 2.9 | 18.3 | 2.1 |
| >hg38_21_+_8433172_8446622_RNA45SN1_45S_50ntFlanks@10349.10371.23\|AACGAGAACTTTGAAGGCCGAAG | >hg38_21_+_8433172_8446622_RNA45SN1_45S_50ntFlanks | 10349 | 10371 | 23 | AACGAGAACTTTGAAGGCCGAAG | BT20 | 14 | 11 | 11 | 47 | 24 | 12.6 | 1.6 | 11.1 | 33.1 | 6.4 |
| >hg38_21_+_8433172_8446622_RNA45SN1_45S_50ntFlanks@10349.10371.23\|AACGAGAACTTTGAAGGCCGAAG | >hg38_21_+_8433172_8446622_RNA45SN1_45S_50ntFlanks | 10349 | 10371 | 23 | AACGAGAACTTTGAAGGCCGAAG | MB231 | 50 | 43 | 34 | 42 | 45 | 48.9 | 46.5 | 37.4 | 35.4 | 53.3 |
| >hg38_21_+_8433172_8446622_RNA45SN1_45S_50ntFlanks@10349.10371.23\|AACGAGAACTTTGAAGGCCGAAG | >hg38_21_+_8433172_8446622_RNA45SN1_45S_50ntFlanks | 10349 | 10371 | 23 | AACGAGAACTTTGAAGGCCGAAG | MB468 | 67 | 61 | 30 | 80 | 31 | 21.8 | 28.3 | 9.7 | 37.7 | 7.2 |
| >hg38_21_+_8433172_8446622_RNA45SN1_45S_50ntFlanks@10349.10372.24\|AACGAGAACTTTGAAGGCCGAAGT | >hg38_21_+_8433172_8446622_RNA45SN1_45S_50ntFlanks | 10349 | 10372 | 24 | AACGAGAACTTTGAAGGCCGAAGT | BT20 | 23 | 14 | 14 | 44 | 30 | 8.8 | 3.1 | 5.5 | 21.6 | 8.0 |
| >hg38_21_+_8433172_8446622_RNA45SN1_45S_50ntFlanks@10349.10372.24\|AACGAGAACTTTGAAGGCCGAAGT | >hg38_21_+_8433172_8446622_RNA45SN1_45S_50ntFlanks | 10349 | 10372 | 24 | AACGAGAACTTTGAAGGCCGAAGT | MB231 | 92 | 79 | 44 | 70 | 65 | 86.6 | 76.1 | 42.7 | 53.4 | 60.1 |
| >hg38_21_+_8433172_8446622_RNA45SN1_45S_50ntFlanks@10349.10372.24\|AACGAGAACTTTGAAGGCCGAAGT | >hg38_21_+_8433172_8446622_RNA45SN1_45S_50ntFlanks | 10349 | 10372 | 24 | AACGAGAACTTTGAAGGCCGAAGT | MB468 | 17 | 18 | 13 | 29 | 11 | 8.3 | 6.1 | 4.5 | 17.4 | 3.9 |
| >hg38_21_+_8433172_8446622_RNA45SN1_45S_50ntFlanks@10349.10373.25\|AACGAGAACTTTGAAGGCCGAAGTG | >hg38_21_+_8433172_8446622_RNA45SN1_45S_50ntFlanks | 10349 | 10373 | 25 | AACGAGAACTTTGAAGGCCGAAGTG | MB231 | 30 | 17 | 21 | 12 | 10 | 32.9 | 14.8 | 20.8 | 12.5 | 9.0 |
| >hg38_21_+_8433172_8446622_RNA45SN1_45S_50ntFlanks@10349.10373.25\|AACGAGAACTTTGAAGGCCGAAGTG | >hg38_21_+_8433172_8446622_RNA45SN1_45S_50ntFlanks | 10349 | 10373 | 25 | AACGAGAACTTTGAAGGCCGAAGTG | MB468 | 18 | 16 | 18 | 40 | 9 | 4.4 | 9.2 | 3.7 | 34.9 | 5.0 |
| >hg38_21_+_8433172_8446622_RNA45SN1_45S_50ntFlanks@10349.10374.26\|AACGAGAACTTTGAAGGCCGAAGTGG | >hg38_21_+_8433172_8446622_RNA45SN1_45S_50ntFlanks | 10349 | 10374 | 26 | AACGAGAACTTTGAAGGCCGAAGTGG | MB468 | 72 | 85 | 64 | 151 | 52 | 15.5 | 36.2 | 13.0 | 126.5 | 15.1 |
| >hg38_21_+_8433172_8446622_RNA45SN1_45S_50ntFlanks@10349.10376.28\|AACGAGAACTTTGAAGGCCGAAGTGGAG | >hg38_21_+_8433172_8446622_RNA45SN1_45S_50ntFlanks | 10349 | 10376 | 28 | AACGAGAACTTTGAAGGCCGAAGTGGAG | MB468 | 65 | 81 | 50 | 90 | 23 | 17.7 | 26.3 | 16.4 | 87.8 | 14.6 |
| >hg38_21_+_8433172_8446622_RNA45SN1_45S_50ntFlanks@10349.10377.29\|AACGAGAACTTTGAAGGCCGAAGTGGAGA | >hg38_21_+_8433172_8446622_RNA45SN1_45S_50ntFlanks | 10349 | 10377 | 29 | AACGAGAACTTTGAAGGCCGAAGTGGAGA | MB468 | 108 | 134 | 63 | 166 | 27 | 21.3 | 39.7 | 28.5 | 137.0 | 13.7 |
| >hg38_21_+_8433172_8446622_RNA45SN1_45S_50ntFlanks@10349.10378.30\|AACGAGAACTTTGAAGGCCGAAGTGGAGAA | >hg38_21_+_8433172_8446622_RNA45SN1_45S_50ntFlanks | 10349 | 10378 | 30 | AACGAGAACTTTGAAGGCCGAAGTGGAGAA | MB468 | 18 | 39 | 10 | 20 | 6 | 2.1 | 10.7 | 3.6 | 19.5 | 2.9 |
| >hg38_21_+_8433172_8446622_RNA45SN1_45S_50ntFlanks@10349.10379.31\|AACGAGAACTTTGAAGGCCGAAGTGGAGAAG | >hg38_21_+_8433172_8446622_RNA45SN1_45S_50ntFlanks | 10349 | 10379 | 31 | AACGAGAACTTTGAAGGCCGAAGTGGAGAAG | MB231 | 12 | 8 | 18 | 6 | 13 | 12.2 | 7.4 | 20.2 | 4.1 | 12.9 |
| >hg38_21_+_8433172_8446622_RNA45SN1_45S_50ntFlanks@10349.10379.31\|AACGAGAACTTTGAAGGCCGAAGTGGAGAAG | >hg38_21_+_8433172_8446622_RNA45SN1_45S_50ntFlanks | 10349 | 10379 | 31 | AACGAGAACTTTGAAGGCCGAAGTGGAGAAG | MB468 | 14 | 21 | 12 | 22 | 9 | 5.0 | 8.2 | 8.0 | 17.3 | 3.3 |
| >hg38_21_+_8433172_8446622_RNA45SN1_45S_50ntFlanks@10349.10382.34\|AACGAGAACTTTGAAGGCCGAAGTGGAGAAGGGT | >hg38_21_+_8433172_8446622_RNA45SN1_45S_50ntFlanks | 10349 | 10382 | 34 | AACGAGAACTTTGAAGGCCGAAGTGGAGAAGGGT | BT20 | 12 | 10 | 39 | 29 | 22 | 4.2 | 4.6 | 3.1 | 8.5 | 8.7 |
| >hg38_21_+_8433172_8446622_RNA45SN1_45S_50ntFlanks@10349.10382.34\|AACGAGAACTTTGAAGGCCGAAGTGGAGAAGGGT | >hg38_21_+_8433172_8446622_RNA45SN1_45S_50ntFlanks | 10349 | 10382 | 34 | AACGAGAACTTTGAAGGCCGAAGTGGAGAAGGGT | MB231 | 34 | 16 | 105 | 33 | 33 | 29.8 | 14.5 | 83.1 | 28.9 | 26.1 |
| >hg38_21_+_8433172_8446622_RNA45SN1_45S_50ntFlanks@10349.10382.34\|AACGAGAACTTTGAAGGCCGAAGTGGAGAAGGGT | >hg38_21_+_8433172_8446622_RNA45SN1_45S_50ntFlanks | 10349 | 10382 | 34 | AACGAGAACTTTGAAGGCCGAAGTGGAGAAGGGT | MB468 | 20 | 37 | 24 | 35 | 14 | 5.5 | 15.9 | 9.3 | 24.9 | 8.3 |
| >hg38_21_+_8433172_8446622_RNA45SN1_45S_50ntFlanks@10349.10383.35\|AACGAGAACTTTGAAGGCCGAAGTGGAGAAGGGTT | >hg38_21_+_8433172_8446622_RNA45SN1_45S_50ntFlanks | 10349 | 10383 | 35 | AACGAGAACTTTGAAGGCCGAAGTGGAGAAGGGTT | MB231 | 19 | 10 | 53 | 16 | 20 | 15.9 | 7.6 | 43.1 | 14.4 | 15.8 |
| >hg38_21_+_8433172_8446622_RNA45SN1_45S_50ntFlanks@10349.10383.35\|AACGAGAACTTTGAAGGCCGAAGTGGAGAAGGGTT | >hg38_21_+_8433172_8446622_RNA45SN1_45S_50ntFlanks | 10349 | 10383 | 35 | AACGAGAACTTTGAAGGCCGAAGTGGAGAAGGGTT | MB468 | 305 | 731 | 336 | 336 | 217 | 110.0 | 355.0 | 67.4 | 276.8 | 155.3 |
| >hg38_21_+_8433172_8446622_RNA45SN1_45S_50ntFlanks@10349.10384.36\|AACGAGAACTTTGAAGGCCGAAGTGGAGAAGGGTTC | >hg38_21_+_8433172_8446622_RNA45SN1_45S_50ntFlanks | 10349 | 10384 | 36 | AACGAGAACTTTGAAGGCCGAAGTGGAGAAGGGTTC | MB468 | 58 | 131 | 56 | 71 | 40 | 16.4 | 66.8 | 10.5 | 73.3 | 34.6 |
| >hg38_21_+_8433172_8446622_RNA45SN1_45S_50ntFlanks@10349.10388.40\|AACGAGAACTTTGAAGGCCGAAGTGGAGAAGGGTTCCATG | >hg38_21_+_8433172_8446622_RNA45SN1_45S_50ntFlanks | 10349 | 10388 | 40 | AACGAGAACTTTGAAGGCCGAAGTGGAGAAGGGTTCCATG | MB468 | 79 | 88 | 95 | 48 | 82 | 78.5 | 58.9 | 62.8 | 27.3 | 47.3 |
| >hg38_21_+_8433172_8446622_RNA45SN1_45S_50ntFlanks@10349.10390.42\|AACGAGAACTTTGAAGGCCGAAGTGGAGAAGGGTTCCATGTG | >hg38_21_+_8433172_8446622_RNA45SN1_45S_50ntFlanks | 10349 | 10390 | 42 | AACGAGAACTTTGAAGGCCGAAGTGGAGAAGGGTTCCATGTG | MB231 | 20 | 10 | 24 | 7 | 16 | 19.8 | 5.7 | 26.6 | 5.4 | 11.8 |
| >hg38_21_+_8433172_8446622_RNA45SN1_45S_50ntFlanks@10349.10390.42\|AACGAGAACTTTGAAGGCCGAAGTGGAGAAGGGTTCCATGTG | >hg38_21_+_8433172_8446622_RNA45SN1_45S_50ntFlanks | 10349 | 10390 | 42 | AACGAGAACTTTGAAGGCCGAAGTGGAGAAGGGTTCCATGTG | MB468 | 50 | 110 | 94 | 33 | 108 | 28.2 | 97.6 | 51.1 | 18.3 | 88.0 |
| >hg38_21_+_8433172_8446622_RNA45SN1_45S_50ntFlanks@10350.10367.18\|ACGAGAACTTTGAAGGCC | >hg38_21_+_8433172_8446622_RNA45SN1_45S_50ntFlanks | 10350 | 10367 | 18 | ACGAGAACTTTGAAGGCC | MB231 | 11 | 8 | 9 | 5 | 2 | 8.7 | 7.3 | 6.5 | 6.3 | 1.6 |
| >hg38_21_+_8433172_8446622_RNA45SN1_45S_50ntFlanks@10350.10367.18\|ACGAGAACTTTGAAGGCC | >hg38_21_+_8433172_8446622_RNA45SN1_45S_50ntFlanks | 10350 | 10367 | 18 | ACGAGAACTTTGAAGGCC | MB468 | 134 | 66 | 97 | 88 | 17 | 19.5 | 50.0 | 17.5 | 60.0 | 6.5 |
| >hg38_21_+_8433172_8446622_RNA45SN1_45S_50ntFlanks@10350.10368.19\|ACGAGAACTTTGAAGGCCG | >hg38_21_+_8433172_8446622_RNA45SN1_45S_50ntFlanks | 10350 | 10368 | 19 | ACGAGAACTTTGAAGGCCG | MB468 | 1044 | 434 | 416 | 585 | 224 | 183.8 | 252.2 | 60.3 | 309.7 | 49.2 |
| >hg38_21_+_8433172_8446622_RNA45SN1_45S_50ntFlanks@10350.10369.20\|ACGAGAACTTTGAAGGCCGA | >hg38_21_+_8433172_8446622_RNA45SN1_45S_50ntFlanks | 10350 | 10369 | 20 | ACGAGAACTTTGAAGGCCGA | MB231 | 15 | 18 | 17 | 15 | 14 | 11.4 | 11.3 | 14.5 | 11.4 | 12.2 |
| >hg38_21_+_8433172_8446622_RNA45SN1_45S_50ntFlanks@10350.10369.20\|ACGAGAACTTTGAAGGCCGA | >hg38_21_+_8433172_8446622_RNA45SN1_45S_50ntFlanks | 10350 | 10369 | 20 | ACGAGAACTTTGAAGGCCGA | MB468 | 55 | 32 | 28 | 58 | 22 | 9.6 | 11.1 | 8.0 | 30.1 | 3.9 |
| >hg38_21_+_8433172_8446622_RNA45SN1_45S_50ntFlanks@10350.10370.21\|ACGAGAACTTTGAAGGCCGAA | >hg38_21_+_8433172_8446622_RNA45SN1_45S_50ntFlanks | 10350 | 10370 | 21 | ACGAGAACTTTGAAGGCCGAA | MB468 | 28 | 12 | 15 | 46 | 11 | 7.3 | 7.4 | 7.4 | 40.6 | 3.5 |
| >hg38_21_+_8433172_8446622_RNA45SN1_45S_50ntFlanks@10350.10371.22\|ACGAGAACTTTGAAGGCCGAAG | >hg38_21_+_8433172_8446622_RNA45SN1_45S_50ntFlanks | 10350 | 10371 | 22 | ACGAGAACTTTGAAGGCCGAAG | MB231 | 18 | 17 | 11 | 17 | 13 | 13.1 | 11.4 | 6.9 | 10.1 | 13.1 |
| >hg38_21_+_8433172_8446622_RNA45SN1_45S_50ntFlanks@10350.10371.22\|ACGAGAACTTTGAAGGCCGAAG | >hg38_21_+_8433172_8446622_RNA45SN1_45S_50ntFlanks | 10350 | 10371 | 22 | ACGAGAACTTTGAAGGCCGAAG | MB468 | 127 | 110 | 58 | 159 | 73 | 28.5 | 41.7 | 14.7 | 74.1 | 30.5 |
| >hg38_21_+_8433172_8446622_RNA45SN1_45S_50ntFlanks@10350.10372.23\|ACGAGAACTTTGAAGGCCGAAGT | >hg38_21_+_8433172_8446622_RNA45SN1_45S_50ntFlanks | 10350 | 10372 | 23 | ACGAGAACTTTGAAGGCCGAAGT | MB231 | 32 | 27 | 17 | 17 | 18 | 25.0 | 20.6 | 12.1 | 14.0 | 13.9 |
| >hg38_21_+_8433172_8446622_RNA45SN1_45S_50ntFlanks@10350.10372.23\|ACGAGAACTTTGAAGGCCGAAGT | >hg38_21_+_8433172_8446622_RNA45SN1_45S_50ntFlanks | 10350 | 10372 | 23 | ACGAGAACTTTGAAGGCCGAAGT | MB468 | 32 | 30 | 25 | 56 | 24 | 6.3 | 8.3 | 6.2 | 31.8 | 3.4 |
| >hg38_21_+_8433172_8446622_RNA45SN1_45S_50ntFlanks@10350.10373.24\|ACGAGAACTTTGAAGGCCGAAGTG | >hg38_21_+_8433172_8446622_RNA45SN1_45S_50ntFlanks | 10350 | 10373 | 24 | ACGAGAACTTTGAAGGCCGAAGTG | MB231 | 11 | 9 | 7 | 5 | 4 | 9.5 | 6.6 | 5.2 | 5.9 | 3.3 |

| | | | | | | | | | | | | | | | | |
|---|---|---|---|---|---|---|---|---|---|---|---|---|---|---|---|---|
| >hg38_21_+_8433172_8446622_RNA45SN1_45S_50ntFlanks@10350.10373.24\|ACGAGAACTTTGAAGGCCGAAGTG | >hg38_21_+_8433172_8446622_RNA45SN1_45S_50ntFlanks | 10350 | 10373 | 24 | ACGAGAACTTTGAAGGCCGAAGTG | MB468 | 45 | 35 | 40 | 88 | 22 | 5.0 | 17.1 | 4.7 | 73.9 | 7.0 |
| >hg38_21_+_8433172_8446622_RNA45SN1_45S_50ntFlanks@10350.10374.25\|ACGAGAACTTTGAAGGCCGAAGTGG | >hg38_21_+_8433172_8446622_RNA45SN1_45S_50ntFlanks | 10350 | 10374 | 25 | ACGAGAACTTTGAAGGCCGAAGTGG | MB468 | 178 | 186 | 148 | 356 | 150 | 26.6 | 67.5 | 17.6 | 279.4 | 19.6 |
| >hg38_21_+_8433172_8446622_RNA45SN1_45S_50ntFlanks@10350.10376.27\|ACGAGAACTTTGAAGGCCGAAGTGGAG | >hg38_21_+_8433172_8446622_RNA45SN1_45S_50ntFlanks | 10350 | 10376 | 27 | ACGAGAACTTTGAAGGCCGAAGTGGAG | MB468 | 141 | 166 | 97 | 235 | 59 | 32.4 | 53.4 | 22.1 | 239.5 | 36.0 |
| >hg38_21_+_8433172_8446622_RNA45SN1_45S_50ntFlanks@10350.10377.28\|ACGAGAACTTTGAAGGCCGAAGTGGAGA | >hg38_21_+_8433172_8446622_RNA45SN1_45S_50ntFlanks | 10350 | 10377 | 28 | ACGAGAACTTTGAAGGCCGAAGTGGAGA | MB468 | 235 | 293 | 117 | 447 | 74 | 36.8 | 70.0 | 47.2 | 397.7 | 36.3 |
| >hg38_21_+_8433172_8446622_RNA45SN1_45S_50ntFlanks@10350.10378.29\|ACGAGAACTTTGAAGGCCGAAGTGGAGAA | >hg38_21_+_8433172_8446622_RNA45SN1_45S_50ntFlanks | 10350 | 10378 | 29 | ACGAGAACTTTGAAGGCCGAAGTGGAGAA | MB468 | 44 | 88 | 18 | 57 | 15 | 4.5 | 16.9 | 6.6 | 56.9 | 7.9 |
| >hg38_21_+_8433172_8446622_RNA45SN1_45S_50ntFlanks@10350.10379.30\|ACGAGAACTTTGAAGGCCGAAGTGGAGAAG | >hg38_21_+_8433172_8446622_RNA45SN1_45S_50ntFlanks | 10350 | 10379 | 30 | ACGAGAACTTTGAAGGCCGAAGTGGAGAAG | MB468 | 33 | 50 | 26 | 62 | 22 | 9.4 | 15.7 | 16.4 | 48.3 | 7.5 |
| >hg38_21_+_8433172_8446622_RNA45SN1_45S_50ntFlanks@10350.10380.31\|ACGAGAACTTTGAAGGCCGAAGTGGAGAAGG | >hg38_21_+_8433172_8446622_RNA45SN1_45S_50ntFlanks | 10350 | 10380 | 31 | ACGAGAACTTTGAAGGCCGAAGTGGAGAAGG | MB468 | 16 | 24 | 18 | 33 | 10 | 5.6 | 8.0 | 11.4 | 27.7 | 4.4 |
| >hg38_21_+_8433172_8446622_RNA45SN1_45S_50ntFlanks@10350.10381.32\|ACGAGAACTTTGAAGGCCGAAGTGGAGAAGGG | >hg38_21_+_8433172_8446622_RNA45SN1_45S_50ntFlanks | 10350 | 10381 | 32 | ACGAGAACTTTGAAGGCCGAAGTGGAGAAGGG | MB468 | 22 | 49 | 23 | 45 | 19 | 1.1 | 7.5 | 8.2 | 31.3 | 7.2 |
| >hg38_21_+_8433172_8446622_RNA45SN1_45S_50ntFlanks@10350.10382.33\|ACGAGAACTTTGAAGGCCGAAGTGGAGAAGGGT | >hg38_21_+_8433172_8446622_RNA45SN1_45S_50ntFlanks | 10350 | 10382 | 33 | ACGAGAACTTTGAAGGCCGAAGTGGAGAAGGGT | MB231 | 11 | 6 | 34 | 10 | 10 | 8.9 | 4.4 | 31.6 | 9.8 | 7.9 |
| >hg38_21_+_8433172_8446622_RNA45SN1_45S_50ntFlanks@10350.10382.33\|ACGAGAACTTTGAAGGCCGAAGTGGAGAAGGGT | >hg38_21_+_8433172_8446622_RNA45SN1_45S_50ntFlanks | 10350 | 10382 | 33 | ACGAGAACTTTGAAGGCCGAAGTGGAGAAGGGT | MB468 | 49 | 89 | 49 | 96 | 37 | 6.8 | 26.9 | 23.4 | 63.7 | 14.0 |
| >hg38_21_+_8433172_8446622_RNA45SN1_45S_50ntFlanks@10350.10383.34\|ACGAGAACTTTGAAGGCCGAAGTGGAGAAGGGTT | >hg38_21_+_8433172_8446622_RNA45SN1_45S_50ntFlanks | 10350 | 10383 | 34 | ACGAGAACTTTGAAGGCCGAAGTGGAGAAGGGTT | MB468 | 658 | 1604 | 562 | 862 | 478 | 223.3 | 714.5 | 79.5 | 729.3 | 278.7 |
| >hg38_21_+_8433172_8446622_RNA45SN1_45S_50ntFlanks@10350.10384.35\|ACGAGAACTTTGAAGGCCGAAGTGGAGAAGGGTTC | >hg38_21_+_8433172_8446622_RNA45SN1_45S_50ntFlanks | 10350 | 10384 | 35 | ACGAGAACTTTGAAGGCCGAAGTGGAGAAGGGTTC | MB468 | 144 | 311 | 113 | 207 | 94 | 34.3 | 128.5 | 13.5 | 215.3 | 66.2 |
| >hg38_21_+_8433172_8446622_RNA45SN1_45S_50ntFlanks@10350.10388.39\|ACGAGAACTTTGAAGGCCGAAGTGGAGAAGGGTTCCATG | >hg38_21_+_8433172_8446622_RNA45SN1_45S_50ntFlanks | 10350 | 10388 | 39 | ACGAGAACTTTGAAGGCCGAAGTGGAGAAGGGTTCCATG | MB468 | 267 | 218 | 225 | 137 | 234 | 292.5 | 161.0 | 171.3 | 74.5 | 123.8 |
| >hg38_21_+_8433172_8446622_RNA45SN1_45S_50ntFlanks@10350.10390.41\|ACGAGAACTTTGAAGGCCGAAGTGGAGAAGGGTTCCATGTG | >hg38_21_+_8433172_8446622_RNA45SN1_45S_50ntFlanks | 10350 | 10390 | 41 | ACGAGAACTTTGAAGGCCGAAGTGGAGAAGGGTTCCATGTG | MB468 | 162 | 296 | 236 | 117 | 296 | 111.6 | 202.2 | 106.7 | 68.6 | 194.0 |
| >hg38_21_+_8433172_8446622_RNA45SN1_45S_50ntFlanks@10350.10391.42\|ACGAGAACTTTGAAGGCCGAAGTGGAGAAGGGTTCCATGTGA | >hg38_21_+_8433172_8446622_RNA45SN1_45S_50ntFlanks | 10350 | 10391 | 42 | ACGAGAACTTTGAAGGCCGAAGTGGAGAAGGGTTCCATGTGA | MB468 | 14 | 35 | 12 | 11 | 31 | 15.5 | 25.8 | 8.4 | 7.0 | 21.9 |
| >hg38_21_+_8433172_8446622_RNA45SN1_45S_50ntFlanks@10351.10368.18\|CGAGAACTTTGAAGGCCG | >hg38_21_+_8433172_8446622_RNA45SN1_45S_50ntFlanks | 10351 | 10368 | 18 | CGAGAACTTTGAAGGCCG | MB468 | 3497 | 1151 | 1243 | 1423 | 744 | 910.5 | 554.9 | 102.5 | 376.6 | 261.6 |
| >hg38_21_+_8433172_8446622_RNA45SN1_45S_50ntFlanks@10351.10369.19\|CGAGAACTTTGAAGGCCGA | >hg38_21_+_8433172_8446622_RNA45SN1_45S_50ntFlanks | 10351 | 10369 | 19 | CGAGAACTTTGAAGGCCGA | MB468 | 90 | 39 | 49 | 97 | 36 | 24.0 | 15.3 | 13.9 | 56.5 | 10.4 |
| >hg38_21_+_8433172_8446622_RNA45SN1_45S_50ntFlanks@10351.10370.20\|CGAGAACTTTGAAGGCCGAA | >hg38_21_+_8433172_8446622_RNA45SN1_45S_50ntFlanks | 10351 | 10370 | 20 | CGAGAACTTTGAAGGCCGAA | MB468 | 43 | 15 | 23 | 73 | 16 | 12.0 | 10.3 | 9.8 | 57.5 | 5.8 |
| >hg38_21_+_8433172_8446622_RNA45SN1_45S_50ntFlanks@10351.10371.21\|CGAGAACTTTGAAGGCCGAAG | >hg38_21_+_8433172_8446622_RNA45SN1_45S_50ntFlanks | 10351 | 10371 | 21 | CGAGAACTTTGAAGGCCGAAG | MB468 | 103 | 70 | 54 | 132 | 76 | 16.9 | 38.8 | 8.5 | 75.0 | 23.8 |
| >hg38_21_+_8433172_8446622_RNA45SN1_45S_50ntFlanks@10351.10372.22\|CGAGAACTTTGAAGGCCGAAGT | >hg38_21_+_8433172_8446622_RNA45SN1_45S_50ntFlanks | 10351 | 10372 | 22 | CGAGAACTTTGAAGGCCGAAGT | MB468 | 36 | 27 | 29 | 67 | 24 | 9.6 | 8.6 | 7.3 | 41.9 | 7.8 |
| >hg38_21_+_8433172_8446622_RNA45SN1_45S_50ntFlanks@10351.10373.23\|CGAGAACTTTGAAGGCCGAAGTG | >hg38_21_+_8433172_8446622_RNA45SN1_45S_50ntFlanks | 10351 | 10373 | 23 | CGAGAACTTTGAAGGCCGAAGTG | MB468 | 71 | 43 | 69 | 151 | 32 | 3.2 | 22.1 | 11.8 | 118.3 | 9.6 |
| >hg38_21_+_8433172_8446622_RNA45SN1_45S_50ntFlanks@10351.10374.24\|CGAGAACTTTGAAGGCCGAAGTGG | >hg38_21_+_8433172_8446622_RNA45SN1_45S_50ntFlanks | 10351 | 10374 | 24 | CGAGAACTTTGAAGGCCGAAGTGG | MB468 | 233 | 207 | 186 | 513 | 207 | 26.8 | 88.2 | 10.7 | 416.5 | 35.8 |
| >hg38_21_+_8433172_8446622_RNA45SN1_45S_50ntFlanks@10351.10375.25\|CGAGAACTTTGAAGGCCGAAGTGGA | >hg38_21_+_8433172_8446622_RNA45SN1_45S_50ntFlanks | 10351 | 10375 | 25 | CGAGAACTTTGAAGGCCGAAGTGGA | MB468 | 20 | 16 | 9 | 48 | 6 | 6.3 | 5.9 | 4.7 | 54.4 | 4.6 |
| >hg38_21_+_8433172_8446622_RNA45SN1_45S_50ntFlanks@10351.10376.26\|CGAGAACTTTGAAGGCCGAAGTGGAG | >hg38_21_+_8433172_8446622_RNA45SN1_45S_50ntFlanks | 10351 | 10376 | 26 | CGAGAACTTTGAAGGCCGAAGTGGAG | MB468 | 175 | 143 | 111 | 536 | 88 | 40.2 | 61.7 | 21.1 | 634.3 | 71.8 |
| >hg38_21_+_8433172_8446622_RNA45SN1_45S_50ntFlanks@10351.10377.27\|CGAGAACTTTGAAGGCCGAAGTGGAGA | >hg38_21_+_8433172_8446622_RNA45SN1_45S_50ntFlanks | 10351 | 10377 | 27 | CGAGAACTTTGAAGGCCGAAGTGGAGA | MB468 | 725 | 633 | 362 | 1650 | 256 | 145.1 | 159.0 | 95.6 | 1502.1 | 130.3 |
| >hg38_21_+_8433172_8446622_RNA45SN1_45S_50ntFlanks@10351.10378.28\|CGAGAACTTTGAAGGCCGAAGTGGAGAA | >hg38_21_+_8433172_8446622_RNA45SN1_45S_50ntFlanks | 10351 | 10378 | 28 | CGAGAACTTTGAAGGCCGAAGTGGAGAA | MB468 | 109 | 173 | 50 | 174 | 43 | 13.0 | 45.7 | 14.9 | 177.8 | 24.2 |
| >hg38_21_+_8433172_8446622_RNA45SN1_45S_50ntFlanks@10351.10379.29\|CGAGAACTTTGAAGGCCGAAGTGGAGAAG | >hg38_21_+_8433172_8446622_RNA45SN1_45S_50ntFlanks | 10351 | 10379 | 29 | CGAGAACTTTGAAGGCCGAAGTGGAGAAG | MB468 | 68 | 86 | 53 | 154 | 56 | 21.8 | 29.2 | 33.8 | 138.2 | 25.5 |
| >hg38_21_+_8433172_8446622_RNA45SN1_45S_50ntFlanks@10351.10380.30\|CGAGAACTTTGAAGGCCGAAGTGGAGAAGG | >hg38_21_+_8433172_8446622_RNA45SN1_45S_50ntFlanks | 10351 | 10380 | 30 | CGAGAACTTTGAAGGCCGAAGTGGAGAAGG | MB468 | 39 | 56 | 41 | 98 | 29 | 13.1 | 17.6 | 20.4 | 84.3 | 15.2 |
| >hg38_21_+_8433172_8446622_RNA45SN1_45S_50ntFlanks@10351.10381.31\|CGAGAACTTTGAAGGCCGAAGTGGAGAAGGG | >hg38_21_+_8433172_8446622_RNA45SN1_45S_50ntFlanks | 10351 | 10381 | 31 | CGAGAACTTTGAAGGCCGAAGTGGAGAAGGG | MB468 | 50 | 100 | 52 | 111 | 43 | 2.3 | 16.3 | 15.4 | 74.5 | 15.1 |
| >hg38_21_+_8433172_8446622_RNA45SN1_45S_50ntFlanks@10351.10382.32\|CGAGAACTTTGAAGGCCGAAGTGGAGAAGGGT | >hg38_21_+_8433172_8446622_RNA45SN1_45S_50ntFlanks | 10351 | 10382 | 32 | CGAGAACTTTGAAGGCCGAAGTGGAGAAGGGT | MB468 | 97 | 167 | 97 | 212 | 83 | 6.4 | 37.0 | 34.8 | 136.1 | 26.4 |
| >hg38_21_+_8433172_8446622_RNA45SN1_45S_50ntFlanks@10351.10383.33\|CGAGAACTTTGAAGGCCGAAGTGGAGAAGGGTT | >hg38_21_+_8433172_8446622_RNA45SN1_45S_50ntFlanks | 10351 | 10383 | 33 | CGAGAACTTTGAAGGCCGAAGTGGAGAAGGGTT | MB468 | 1600 | 3450 | 1440 | 2524 | 1243 | 513.4 | 1221.1 | 206.8 | 2223.5 | 757.3 |
| >hg38_21_+_8433172_8446622_RNA45SN1_45S_50ntFlanks@10351.10384.34\|CGAGAACTTTGAAGGCCGAAGTGGAGAAGGGTTC | >hg38_21_+_8433172_8446622_RNA45SN1_45S_50ntFlanks | 10351 | 10384 | 34 | CGAGAACTTTGAAGGCCGAAGTGGAGAAGGGTTC | MB468 | 277 | 489 | 223 | 544 | 216 | 69.0 | 224.9 | 41.9 | 631.8 | 171.3 |
| >hg38_21_+_8433172_8446622_RNA45SN1_45S_50ntFlanks@10351.10385.35\|CGAGAACTTTGAAGGCCGAAGTGGAGAAGGGTTCC | >hg38_21_+_8433172_8446622_RNA45SN1_45S_50ntFlanks | 10351 | 10385 | 35 | CGAGAACTTTGAAGGCCGAAGTGGAGAAGGGTTCC | MB468 | 17 | 24 | 17 | 28 | 16 | 6.4 | 9.7 | 5.5 | 32.1 | 8.7 |
| >hg38_21_+_8433172_8446622_RNA45SN1_45S_50ntFlanks@10351.10387.37\|CGAGAACTTTGAAGGCCGAAGTGGAGAAGGGTTCCAT | >hg38_21_+_8433172_8446622_RNA45SN1_45S_50ntFlanks | 10351 | 10387 | 37 | CGAGAACTTTGAAGGCCGAAGTGGAGAAGGGTTCCAT | MB468 | 11 | 14 | 11 | 10 | 16 | 10.1 | 7.5 | 5.7 | 6.6 | 8.9 |
| >hg38_21_+_8433172_8446622_RNA45SN1_45S_50ntFlanks@10351.10388.38\|CGAGAACTTTGAAGGCCGAAGTGGAGAAGGGTTCCATG | >hg38_21_+_8433172_8446622_RNA45SN1_45S_50ntFlanks | 10351 | 10388 | 38 | CGAGAACTTTGAAGGCCGAAGTGGAGAAGGGTTCCATG | MB468 | 682 | 537 | 569 | 413 | 610 | 743.0 | 385.3 | 497.5 | 235.2 | 306.5 |
| >hg38_21_+_8433172_8446622_RNA45SN1_45S_50ntFlanks@10351.10390.40\|CGAGAACTTTGAAGGCCGAAGTGGAGAAGGGTTCCATGTG | >hg38_21_+_8433172_8446622_RNA45SN1_45S_50ntFlanks | 10351 | 10390 | 40 | CGAGAACTTTGAAGGCCGAAGTGGAGAAGGGTTCCATGTG | MB468 | 549 | 1075 | 749 | 494 | 1113 | 292.3 | 691.0 | 278.4 | 317.1 | 652.8 |
| >hg38_21_+_8433172_8446622_RNA45SN1_45S_50ntFlanks@10351.10391.41\|CGAGAACTTTGAAGGCCGAAGTGGAGAAGGGTTCCATGTGA | >hg38_21_+_8433172_8446622_RNA45SN1_45S_50ntFlanks | 10351 | 10391 | 41 | CGAGAACTTTGAAGGCCGAAGTGGAGAAGGGTTCCATGTGA | MB468 | 39 | 86 | 33 | 33 | 78 | 38.8 | 56.5 | 22.5 | 20.5 | 42.6 |
| >hg38_21_+_8433172_8446622_RNA45SN1_45S_50ntFlanks@10352.10369.18\|GAGAACTTTGAAGGCCGA | >hg38_21_+_8433172_8446622_RNA45SN1_45S_50ntFlanks | 10352 | 10369 | 18 | GAGAACTTTGAAGGCCGA | MB231 | 19 | 29 | 15 | 28 | 15 | 2.9 | 9.5 | 3.9 | 14.3 | 13.2 |
| >hg38_21_+_8433172_8446622_RNA45SN1_45S_50ntFlanks@10352.10371.20\|GAGAACTTTGAAGGCCGAAG | >hg38_21_+_8433172_8446622_RNA45SN1_45S_50ntFlanks | 10352 | 10371 | 20 | GAGAACTTTGAAGGCCGAAG | MB231 | 12 | 15 | 8 | 6 | 5 | 7.8 | 5.5 | 5.7 | 2.8 | 3.3 |
| >hg38_21_+_8433172_8446622_RNA45SN1_45S_50ntFlanks@10352.10372.21\|GAGAACTTTGAAGGCCGAAGT | >hg38_21_+_8433172_8446622_RNA45SN1_45S_50ntFlanks | 10352 | 10372 | 21 | GAGAACTTTGAAGGCCGAAGT | MB231 | 15 | 21 | 8 | 14 | 9 | 8.3 | 8.9 | 5.1 | 8.3 | 4.5 |
| >hg38_21_+_8433172_8446622_RNA45SN1_45S_50ntFlanks@10353.10377.25\|AGAACTTTGAAGGCCGAAGTGGAGA | >hg38_21_+_8433172_8446622_RNA45SN1_45S_50ntFlanks | 10353 | 10377 | 25 | AGAACTTTGAAGGCCGAAGTGGAGA | MB468 | 10 | 7 | 4 | 12 | 2 | 2.3 | 3.2 | 2.1 | 9.6 | 0.9 |
| >hg38_21_+_8433172_8446622_RNA45SN1_45S_50ntFlanks@10353.10383.31\|AGAACTTTGAAGGCCGAAGTGGAGAAGGGTT | >hg38_21_+_8433172_8446622_RNA45SN1_45S_50ntFlanks | 10353 | 10383 | 31 | AGAACTTTGAAGGCCGAAGTGGAGAAGGGTT | MB468 | 23 | 39 | 16 | 28 | 11 | 3.1 | 2.5 | 3.2 | 14.6 | 4.7 |
| >hg38_21_+_8433172_8446622_RNA45SN1_45S_50ntFlanks@10353.10388.36\|AGAACTTTGAAGGCCGAAGTGGAGAAGGGTTCCATG | >hg38_21_+_8433172_8446622_RNA45SN1_45S_50ntFlanks | 10353 | 10388 | 36 | AGAACTTTGAAGGCCGAAGTGGAGAAGGGTTCCATG | MB468 | 32 | 28 | 29 | 18 | 22 | 32.1 | 26.4 | 26.1 | 14.1 | 18.2 |
| >hg38_21_+_8433172_8446622_RNA45SN1_45S_50ntFlanks@10353.10390.38\|AGAACTTTGAAGGCCGAAGTGGAGAAGGGTTCCATGTG | >hg38_21_+_8433172_8446622_RNA45SN1_45S_50ntFlanks | 10353 | 10390 | 38 | AGAACTTTGAAGGCCGAAGTGGAGAAGGGTTCCATGTG | MB231 | 17 | 10 | 22 | 4 | 9 | 16.5 | 9.8 | 26.2 | 2.7 | 10.3 |
| >hg38_21_+_8433172_8446622_RNA45SN1_45S_50ntFlanks@10353.10390.38\|AGAACTTTGAAGGCCGAAGTGGAGAAGGGTTCCATGTG | >hg38_21_+_8433172_8446622_RNA45SN1_45S_50ntFlanks | 10353 | 10390 | 38 | AGAACTTTGAAGGCCGAAGTGGAGAAGGGTTCCATGTG | MB468 | 20 | 32 | 26 | 16 | 20 | 9.0 | 21.9 | 10.1 | 12.5 | 13.1 |
| >hg38_21_+_8433172_8446622_RNA45SN1_45S_50ntFlanks@10355.10383.29\|AACTTTGAAGGCCGAAGTGGAGAAGGGTT | >hg38_21_+_8433172_8446622_RNA45SN1_45S_50ntFlanks | 10355 | 10383 | 29 | AACTTTGAAGGCCGAAGTGGAGAAGGGTT | MB468 | 20 | 39 | 14 | 24 | 10 | 2.3 | 11.9 | 3.6 | 18.0 | 6.2 |
| >hg38_21_+_8433172_8446622_RNA45SN1_45S_50ntFlanks@10355.10388.34\|AACTTTGAAGGCCGAAGTGGAGAAGGGTTCCATG | >hg38_21_+_8433172_8446622_RNA45SN1_45S_50ntFlanks | 10355 | 10388 | 34 | AACTTTGAAGGCCGAAGTGGAGAAGGGTTCCATG | MB468 | 50 | 64 | 54 | 23 | 24 | 43.9 | 55.2 | 56.8 | 21.0 | 19.7 |
| >hg38_21_+_8433172_8446622_RNA45SN1_45S_50ntFlanks@10355.10390.36\|AACTTTGAAGGCCGAAGTGGAGAAGGGTTCCATGTG | >hg38_21_+_8433172_8446622_RNA45SN1_45S_50ntFlanks | 10355 | 10390 | 36 | AACTTTGAAGGCCGAAGTGGAGAAGGGTTCCATGTG | MB468 | 20 | 34 | 28 | 11 | 14 | 11.3 | 20.1 | 15.5 | 6.0 | 8.1 |
| >hg38_21_+_8433172_8446622_RNA45SN1_45S_50ntFlanks@10356.10388.33\|ACTTTGAAGGCCGAAGTGGAGAAGGGTTCCATG | >hg38_21_+_8433172_8446622_RNA45SN1_45S_50ntFlanks | 10356 | 10388 | 33 | ACTTTGAAGGCCGAAGTGGAGAAGGGTTCCATG | MB468 | 10 | 12 | 10 | 5 | 4 | 8.8 | 8.9 | 8.8 | 4.3 | 2.6 |
| >hg38_21_+_8433172_8446622_RNA45SN1_45S_50ntFlanks@10357.10374.18\|CTTTGAAGGCCGAAGTGG | >hg38_21_+_8433172_8446622_RNA45SN1_45S_50ntFlanks | 10357 | 10374 | 18 | CTTTGAAGGCCGAAGTGG | MB468 | 11 | 4 | 10 | 9 | 2 | 5.5 | 1.0 | 5.6 | 5.6 | 0.6 |

| | | | | | | | | | | | | | | | |
|---|---|---|---|---|---|---|---|---|---|---|---|---|---|---|---|
| >hg38_21_+_8433172_8446622_RNA45SN1_45S_50ntFlanks@10357.10376.20\|CTTTGAAGGCCGAAGTGGAG | >hg38_21_+_8433172_8446622_RNA45SN1_45S_50ntFlanks | 10357 | 10376 | 20 | CTTTGAAGGCCGAAGTGGAG | MB468 | 13 | 9 | 11 | 7 | 3 | 4.1 | 3.1 | 2.8 | 3.0 | 0.1 |
| >hg38_21_+_8433172_8446622_RNA45SN1_45S_50ntFlanks@10357.10377.21\|CTTTGAAGGCCGAAGTGGAGA | >hg38_21_+_8433172_8446622_RNA45SN1_45S_50ntFlanks | 10357 | 10377 | 21 | CTTTGAAGGCCGAAGTGGAGA | MB468 | 26 | 22 | 13 | 14 | 5 | 12.9 | 7.7 | 5.9 | 5.8 | 0.8 |
| >hg38_21_+_8433172_8446622_RNA45SN1_45S_50ntFlanks@10357.10383.27\|CTTTGAAGGCCGAAGTGGAGAAGGGTT | >hg38_21_+_8433172_8446622_RNA45SN1_45S_50ntFlanks | 10357 | 10383 | 27 | CTTTGAAGGCCGAAGTGGAGAAGGGTT | MB468 | 36 | 72 | 25 | 39 | 16 | 4.7 | 12.8 | 6.1 | 22.2 | 6.1 |
| >hg38_21_+_8433172_8446622_RNA45SN1_45S_50ntFlanks@10357.10384.28\|CTTTGAAGGCCGAAGTGGAGAAGGGTTC | >hg38_21_+_8433172_8446622_RNA45SN1_45S_50ntFlanks | 10357 | 10384 | 28 | CTTTGAAGGCCGAAGTGGAGAAGGGTTC | MB468 | 11 | 15 | 8 | 12 | 4 | 2.9 | 2.0 | 1.8 | 7.4 | 1.1 |
| >hg38_21_+_8433172_8446622_RNA45SN1_45S_50ntFlanks@10357.10388.32\|CTTTGAAGGCCGAAGTGGAGAAGGGTTCCATG | >hg38_21_+_8433172_8446622_RNA45SN1_45S_50ntFlanks | 10357 | 10388 | 32 | CTTTGAAGGCCGAAGTGGAGAAGGGTTCCATG | MB468 | 47 | 57 | 51 | 31 | 17 | 32.9 | 35.4 | 45.4 | 19.5 | 9.2 |
| >hg38_21_+_8433172_8446622_RNA45SN1_45S_50ntFlanks@10357.10390.34\|CTTTGAAGGCCGAAGTGGAGAAGGGTTCCATGTG | >hg38_21_+_8433172_8446622_RNA45SN1_45S_50ntFlanks | 10357 | 10390 | 34 | CTTTGAAGGCCGAAGTGGAGAAGGGTTCCATGTG | MB468 | 25 | 42 | 35 | 18 | 13 | 10.4 | 22.2 | 18.6 | 9.1 | 4.3 |
| >hg38_21_+_8433172_8446622_RNA45SN1_45S_50ntFlanks@10360.10388.29\|TGAAGGCCGAAGTGGAGAAGGGTTCCATG | >hg38_21_+_8433172_8446622_RNA45SN1_45S_50ntFlanks | 10360 | 10388 | 29 | TGAAGGCCGAAGTGGAGAAGGGTTCCATG | MB468 | 13 | 14 | 16 | 10 | 3 | 10.1 | 10.9 | 14.7 | 8.3 | 1.7 |
| >hg38_21_+_8433172_8446622_RNA45SN1_45S_50ntFlanks@10361.10383.23\|GAAGGCCGAAGTGGAGAAGGGTT | >hg38_21_+_8433172_8446622_RNA45SN1_45S_50ntFlanks | 10361 | 10383 | 23 | GAAGGCCGAAGTGGAGAAGGGTT | MB468 | 37 | 60 | 31 | 44 | 19 | 2.5 | 9.2 | 5.6 | 17.6 | 5.3 |
| >hg38_21_+_8433172_8446622_RNA45SN1_45S_50ntFlanks@10361.10388.28\|GAAGGCCGAAGTGGAGAAGGGTTCCATG | >hg38_21_+_8433172_8446622_RNA45SN1_45S_50ntFlanks | 10361 | 10388 | 28 | GAAGGCCGAAGTGGAGAAGGGTTCCATG | MB468 | 53 | 44 | 64 | 43 | 17 | 51.8 | 40.5 | 67.1 | 43.8 | 10.5 |
| >hg38_21_+_8433172_8446622_RNA45SN1_45S_50ntFlanks@10361.10390.30\|GAAGGCCGAAGTGGAGAAGGGTTCCATGTG | >hg38_21_+_8433172_8446622_RNA45SN1_45S_50ntFlanks | 10361 | 10390 | 30 | GAAGGCCGAAGTGGAGAAGGGTTCCATGTG | MB468 | 29 | 28 | 39 | 29 | 14 | 19.6 | 21.1 | 28.5 | 22.1 | 4.9 |
| >hg38_21_+_8433172_8446622_RNA45SN1_45S_50ntFlanks@10362.10379.18\|AAGGCCGAAGTGGAGAAG | >hg38_21_+_8433172_8446622_RNA45SN1_45S_50ntFlanks | 10362 | 10379 | 18 | AAGGCCGAAGTGGAGAAG | MB468 | 17 | 17 | 15 | 14 | 6 | 10.6 | 10.2 | 6.3 | 3.6 | 1.7 |
| >hg38_21_+_8433172_8446622_RNA45SN1_45S_50ntFlanks@10362.10382.21\|AAGGCCGAAGTGGAGAAGGGT | >hg38_21_+_8433172_8446622_RNA45SN1_45S_50ntFlanks | 10362 | 10382 | 21 | AAGGCCGAAGTGGAGAAGGGT | MB468 | 19 | 24 | 21 | 26 | 8 | 5.9 | 6.3 | 13.3 | 6.7 | 2.2 |
| >hg38_21_+_8433172_8446622_RNA45SN1_45S_50ntFlanks@10362.10383.22\|AAGGCCGAAGTGGAGAAGGGTT | >hg38_21_+_8433172_8446622_RNA45SN1_45S_50ntFlanks | 10362 | 10383 | 22 | AAGGCCGAAGTGGAGAAGGGTT | MB468 | 228 | 307 | 183 | 186 | 103 | 0.7 | 39.5 | 38.7 | 67.2 | 30.4 |
| >hg38_21_+_8433172_8446622_RNA45SN1_45S_50ntFlanks@10362.10384.23\|AAGGCCGAAGTGGAGAAGGGTTC | >hg38_21_+_8433172_8446622_RNA45SN1_45S_50ntFlanks | 10362 | 10384 | 23 | AAGGCCGAAGTGGAGAAGGGTTC | MB468 | 74 | 73 | 52 | 73 | 28 | 18.9 | 22.5 | 17.3 | 34.3 | 6.0 |
| >hg38_21_+_8433172_8446622_RNA45SN1_45S_50ntFlanks@10362.10388.27\|AAGGCCGAAGTGGAGAAGGGTTCCATG | >hg38_21_+_8433172_8446622_RNA45SN1_45S_50ntFlanks | 10362 | 10388 | 27 | AAGGCCGAAGTGGAGAAGGGTTCCATG | MB468 | 327 | 281 | 335 | 203 | 84 | 319.5 | 253.1 | 390.7 | 202.1 | 49.1 |
| >hg38_21_+_8433172_8446622_RNA45SN1_45S_50ntFlanks@10362.10390.29\|AAGGCCGAAGTGGAGAAGGGTTCCATGTG | >hg38_21_+_8433172_8446622_RNA45SN1_45S_50ntFlanks | 10362 | 10390 | 29 | AAGGCCGAAGTGGAGAAGGGTTCCATGTG | MB468 | 182 | 187 | 207 | 144 | 83 | 112.1 | 147.7 | 159.0 | 92.9 | 24.1 |
| >hg38_21_+_8433172_8446622_RNA45SN1_45S_50ntFlanks@10362.10391.30\|AAGGCCGAAGTGGAGAAGGGTTCCATGTGA | >hg38_21_+_8433172_8446622_RNA45SN1_45S_50ntFlanks | 10362 | 10391 | 30 | AAGGCCGAAGTGGAGAAGGGTTCCATGTGA | MB468 | 15 | 18 | 11 | 11 | 8 | 14.7 | 14.9 | 14.1 | 4.3 | 2.0 |
| >hg38_21_+_8433172_8446622_RNA45SN1_45S_50ntFlanks@10366.10383.18\|CCGAAGTGGAGAAGGGTT | >hg38_21_+_8433172_8446622_RNA45SN1_45S_50ntFlanks | 10366 | 10383 | 18 | CCGAAGTGGAGAAGGGTT | MB468 | 12 | 10 | 11 | 6 | 3 | 1.2 | 1.5 | 2.6 | 1.7 | 1.0 |
| >hg38_21_+_8433172_8446622_RNA45SN1_45S_50ntFlanks@10366.10388.23\|CCGAAGTGGAGAAGGGTTCCATG | >hg38_21_+_8433172_8446622_RNA45SN1_45S_50ntFlanks | 10366 | 10388 | 23 | CCGAAGTGGAGAAGGGTTCCATG | MB468 | 11 | 2 | 8 | 8 | 3 | 7.9 | 0.9 | 9.0 | 1.3 | 1.8 |
| >hg38_21_+_8433172_8446622_RNA45SN1_45S_50ntFlanks@10369.10407.39\|AAGTGGAGAAGGGTTCCATGTGAACAGCAGTTGAACATG | >hg38_21_+_8433172_8446622_RNA45SN1_45S_50ntFlanks | 10369 | 10407 | 39 | AAGTGGAGAAGGGTTCCATGTGAACAGCAGTTGAACATG | MB468 | 37 | 75 | 85 | 10 | 14 | 14.6 | 48.9 | 29.1 | 5.0 | 6.6 |
| >hg38_21_+_8433172_8446622_RNA45SN1_45S_50ntFlanks@10375.10406.32\|AGAAGGGTTCCATGTGAACAGCAGTTGAACAT | >hg38_21_+_8433172_8446622_RNA45SN1_45S_50ntFlanks | 10375 | 10406 | 32 | AGAAGGGTTCCATGTGAACAGCAGTTGAACAT | MB468 | 32 | 36 | 23 | 11 | 9 | 8.6 | 7.0 | 12.7 | 5.9 | 1.5 |
| >hg38_21_+_8433172_8446622_RNA45SN1_45S_50ntFlanks@10375.10407.33\|AGAAGGGTTCCATGTGAACAGCAGTTGAACATG | >hg38_21_+_8433172_8446622_RNA45SN1_45S_50ntFlanks | 10375 | 10407 | 33 | AGAAGGGTTCCATGTGAACAGCAGTTGAACATG | MB468 | 42 | 30 | 44 | 13 | 15 | 14.9 | 8.1 | 18.1 | 8.5 | 3.4 |
| >hg38_21_+_8433172_8446622_RNA45SN1_45S_50ntFlanks@10377.10407.31\|AAGGGTTCCATGTGAACAGCAGTTGAACATG | >hg38_21_+_8433172_8446622_RNA45SN1_45S_50ntFlanks | 10377 | 10407 | 31 | AAGGGTTCCATGTGAACAGCAGTTGAACATG | MB468 | 15 | 7 | 11 | 4 | 3 | 1.9 | 2.3 | 4.2 | 1.3 | 0.9 |
| >hg38_21_+_8433172_8446622_RNA45SN1_45S_50ntFlanks@10378.10406.29\|AGGGTTCCATGTGAACAGCAGTTGAACAT | >hg38_21_+_8433172_8446622_RNA45SN1_45S_50ntFlanks | 10378 | 10406 | 29 | AGGGTTCCATGTGAACAGCAGTTGAACAT | MB468 | 13 | 9 | 5 | 4 | 2 | 4.2 | 1.1 | 1.8 | 1.3 | 0.7 |
| >hg38_21_+_8433172_8446622_RNA45SN1_45S_50ntFlanks@10378.10407.30\|AGGGTTCCATGTGAACAGCAGTTGAACATG | >hg38_21_+_8433172_8446622_RNA45SN1_45S_50ntFlanks | 10378 | 10407 | 30 | AGGGTTCCATGTGAACAGCAGTTGAACATG | MB468 | 15 | 6 | 10 | 4 | 2 | 3.8 | 1.6 | 6.0 | 1.1 | 0.7 |
| >hg38_21_+_8433172_8446622_RNA45SN1_45S_50ntFlanks@10379.10407.29\|GGGTTCCATGTGAACAGCAGTTGAACATG | >hg38_21_+_8433172_8446622_RNA45SN1_45S_50ntFlanks | 10379 | 10407 | 29 | GGGTTCCATGTGAACAGCAGTTGAACATG | MB468 | 10 | 3 | 11 | 3 | 1 | 4.6 | 0.2 | 8.9 | 0.5 | 0.3 |
| >hg38_21_+_8433172_8446622_RNA45SN1_45S_50ntFlanks@10381.10424.44\|GTTCCATGTGAACAGCAGTTGAACATGGGTCAGTCGGTCCTGAG | >hg38_21_+_8433172_8446622_RNA45SN1_45S_50ntFlanks | 10381 | 10424 | 44 | GTTCCATGTGAACAGCAGTTGAACATGGGTCAGTCGGTCCTGAG | MB231 | 14 | 51 | 57 | 6 | 29 | 4.4 | 26.2 | 31.8 | 5.7 | 14.2 |
| >hg38_21_+_8433172_8446622_RNA45SN1_45S_50ntFlanks@10382.10424.43\|TTCCATGTGAACAGCAGTTGAACATGGGTCAGTCGGTCCTGAG | >hg38_21_+_8433172_8446622_RNA45SN1_45S_50ntFlanks | 10382 | 10424 | 43 | TTCCATGTGAACAGCAGTTGAACATGGGTCAGTCGGTCCTGAG | BT20 | 10 | 22 | 35 | 80 | 371 | 10.2 | 13.5 | 47.5 | 14.3 | 290.3 |
| >hg38_21_+_8433172_8446622_RNA45SN1_45S_50ntFlanks@10382.10424.43\|TTCCATGTGAACAGCAGTTGAACATGGGTCAGTCGGTCCTGAG | >hg38_21_+_8433172_8446622_RNA45SN1_45S_50ntFlanks | 10382 | 10424 | 43 | TTCCATGTGAACAGCAGTTGAACATGGGTCAGTCGGTCCTGAG | MB231 | 15 | 70 | 95 | 6 | 30 | 6.1 | 38.1 | 67.5 | 5.3 | 12.2 |
| >hg38_21_+_8433172_8446622_RNA45SN1_45S_50ntFlanks@10383.10424.42\|TCCATGTGAACAGCAGTTGAACATGGGTCAGTCGGTCCTGAG | >hg38_21_+_8433172_8446622_RNA45SN1_45S_50ntFlanks | 10383 | 10424 | 42 | TCCATGTGAACAGCAGTTGAACATGGGTCAGTCGGTCCTGAG | BT20 | 24 | 38 | 39 | 57 | 298 | 26.8 | 15.1 | 29.8 | 15.7 | 177.2 |
| >hg38_21_+_8433172_8446622_RNA45SN1_45S_50ntFlanks@10383.10424.42\|TCCATGTGAACAGCAGTTGAACATGGGTCAGTCGGTCCTGAG | >hg38_21_+_8433172_8446622_RNA45SN1_45S_50ntFlanks | 10383 | 10424 | 42 | TCCATGTGAACAGCAGTTGAACATGGGTCAGTCGGTCCTGAG | MB231 | 74 | 107 | 191 | 12 | 116 | 49.2 | 26.9 | 118.3 | 4.2 | 102.1 |
| >hg38_21_+_8433172_8446622_RNA45SN1_45S_50ntFlanks@10384.10401.18\|CCATGTGAACAGCAGTTG | >hg38_21_+_8433172_8446622_RNA45SN1_45S_50ntFlanks | 10384 | 10401 | 18 | CCATGTGAACAGCAGTTG | MB468 | 34 | 4 | 5 | 11 | 2 | 14.3 | 2.1 | 3.7 | 10.3 | 1.5 |
| >hg38_21_+_8433172_8446622_RNA45SN1_45S_50ntFlanks@10384.10403.20\|CCATGTGAACAGCAGTTGAA | >hg38_21_+_8433172_8446622_RNA45SN1_45S_50ntFlanks | 10384 | 10403 | 20 | CCATGTGAACAGCAGTTGAA | MB468 | 28 | 7 | 3 | 9 | 3 | 16.8 | 4.2 | 2.4 | 8.7 | 2.0 |
| >hg38_21_+_8433172_8446622_RNA45SN1_45S_50ntFlanks@10384.10404.21\|CCATGTGAACAGCAGTTGAAC | >hg38_21_+_8433172_8446622_RNA45SN1_45S_50ntFlanks | 10384 | 10404 | 21 | CCATGTGAACAGCAGTTGAAC | MB468 | 24 | 11 | 6 | 10 | 3 | 1.8 | 8.4 | 2.4 | 8.6 | 2.6 |
| >hg38_21_+_8433172_8446622_RNA45SN1_45S_50ntFlanks@10384.10405.22\|CCATGTGAACAGCAGTTGAACA | >hg38_21_+_8433172_8446622_RNA45SN1_45S_50ntFlanks | 10384 | 10405 | 22 | CCATGTGAACAGCAGTTGAACA | MB468 | 70 | 37 | 17 | 47 | 9 | 12.7 | 31.8 | 10.2 | 45.4 | 7.6 |
| >hg38_21_+_8433172_8446622_RNA45SN1_45S_50ntFlanks@10384.10406.23\|CCATGTGAACAGCAGTTGAACAT | >hg38_21_+_8433172_8446622_RNA45SN1_45S_50ntFlanks | 10384 | 10406 | 23 | CCATGTGAACAGCAGTTGAACAT | MB468 | 23 | 4 | 10 | 16 | 5 | 4.9 | 2.5 | 7.4 | 15.4 | 3.0 |
| >hg38_21_+_8433172_8446622_RNA45SN1_45S_50ntFlanks@10384.10424.41\|CCATGTGAACAGCAGTTGAACATGGGTCAGTCGGTCCTGAG | >hg38_21_+_8433172_8446622_RNA45SN1_45S_50ntFlanks | 10384 | 10424 | 41 | CCATGTGAACAGCAGTTGAACATGGGTCAGTCGGTCCTGAG | MB231 | 15 | 34 | 45 | 4 | 22 | 4.8 | 9.5 | 9.0 | 2.4 | 17.0 |
| >hg38_21_+_8433172_8446622_RNA45SN1_45S_50ntFlanks@10384.10424.41\|CCATGTGAACAGCAGTTGAACATGGGTCAGTCGGTCCTGAG | >hg38_21_+_8433172_8446622_RNA45SN1_45S_50ntFlanks | 10384 | 10424 | 41 | CCATGTGAACAGCAGTTGAACATGGGTCAGTCGGTCCTGAG | MB468 | 10 | 74 | 22 | 4 | 19 | 12.0 | 46.2 | 13.3 | 5.0 | 30.4 |
| >hg38_21_+_8433172_8446622_RNA45SN1_45S_50ntFlanks@10385.10405.21\|CATGTGAACAGCAGTTGAACA | >hg38_21_+_8433172_8446622_RNA45SN1_45S_50ntFlanks | 10385 | 10405 | 21 | CATGTGAACAGCAGTTGAACA | MB468 | 15 | 6 | 4 | 19 | 3 | 3.5 | 5.9 | 3.0 | 18.2 | 2.8 |
| >hg38_21_+_8433172_8446622_RNA45SN1_45S_50ntFlanks@10385.10406.22\|CATGTGAACAGCAGTTGAACAT | >hg38_21_+_8433172_8446622_RNA45SN1_45S_50ntFlanks | 10385 | 10406 | 22 | CATGTGAACAGCAGTTGAACAT | MB468 | 11 | 1 | 4 | 5 | 2 | 4.3 | 0.7 | 3.5 | 5.3 | 1.9 |
| >hg38_21_+_8433172_8446622_RNA45SN1_45S_50ntFlanks@10386.10410.25\|ATGTGAACAGCAGTTGAACATGGGT | >hg38_21_+_8433172_8446622_RNA45SN1_45S_50ntFlanks | 10386 | 10410 | 25 | ATGTGAACAGCAGTTGAACATGGGT | MB468 | 27 | 20 | 20 | 58 | 22 | 4.0 | 3.5 | 8.7 | 61.4 | 23.1 |
| >hg38_21_+_8433172_8446622_RNA45SN1_45S_50ntFlanks@10386.10424.39\|ATGTGAACAGCAGTTGAACATGGGTCAGTCGGTCCTGAG | >hg38_21_+_8433172_8446622_RNA45SN1_45S_50ntFlanks | 10386 | 10424 | 39 | ATGTGAACAGCAGTTGAACATGGGTCAGTCGGTCCTGAG | BT20 | 30 | 57 | 60 | 314 | 672 | 35.0 | 16.2 | 43.2 | 161.3 | 378.8 |
| >hg38_21_+_8433172_8446622_RNA45SN1_45S_50ntFlanks@10386.10424.39\|ATGTGAACAGCAGTTGAACATGGGTCAGTCGGTCCTGAG | >hg38_21_+_8433172_8446622_RNA45SN1_45S_50ntFlanks | 10386 | 10424 | 39 | ATGTGAACAGCAGTTGAACATGGGTCAGTCGGTCCTGAG | MB231 | 22 | 91 | 96 | 32 | 59 | 14.1 | 33.6 | 62.6 | 23.0 | 53.2 |
| >hg38_21_+_8433172_8446622_RNA45SN1_45S_50ntFlanks@10387.10407.21\|TGTGAACAGCAGTTGAACATG | >hg38_21_+_8433172_8446622_RNA45SN1_45S_50ntFlanks | 10387 | 10407 | 21 | TGTGAACAGCAGTTGAACATG | MB468 | 20 | 3 | 9 | 26 | 14 | 11.2 | 0.6 | 8.4 | 25.2 | 9.6 |
| >hg38_21_+_8433172_8446622_RNA45SN1_45S_50ntFlanks@10387.10410.24\|TGTGAACAGCAGTTGAACATGGGT | >hg38_21_+_8433172_8446622_RNA45SN1_45S_50ntFlanks | 10387 | 10410 | 24 | TGTGAACAGCAGTTGAACATGGGT | MB468 | 35 | 39 | 19 | 22 | 6 | 5.2 | 14.8 | 5.5 | 17.5 | 4.3 |
| >hg38_21_+_8433172_8446622_RNA45SN1_45S_50ntFlanks@10387.10424.38\|TGTGAACAGCAGTTGAACATGGGTCAGTCGGTCCTGAG | >hg38_21_+_8433172_8446622_RNA45SN1_45S_50ntFlanks | 10387 | 10424 | 38 | TGTGAACAGCAGTTGAACATGGGTCAGTCGGTCCTGAG | BT20 | 32 | 59 | 67 | 299 | 676 | 27.4 | 32.7 | 28.8 | 189.9 | 379.6 |
| >hg38_21_+_8433172_8446622_RNA45SN1_45S_50ntFlanks@10387.10424.38\|TGTGAACAGCAGTTGAACATGGGTCAGTCGGTCCTGAG | >hg38_21_+_8433172_8446622_RNA45SN1_45S_50ntFlanks | 10387 | 10424 | 38 | TGTGAACAGCAGTTGAACATGGGTCAGTCGGTCCTGAG | MB231 | 28 | 80 | 81 | 25 | 57 | 28.7 | 30.0 | 52.7 | 19.6 | 39.7 |
| >hg38_21_+_8433172_8446622_RNA45SN1_45S_50ntFlanks@10388.10407.20\|GTGAACAGCAGTTGAACATG | >hg38_21_+_8433172_8446622_RNA45SN1_45S_50ntFlanks | 10388 | 10407 | 20 | GTGAACAGCAGTTGAACATG | MB468 | 24 | 6 | 12 | 10 | 4 | 13.7 | 1.1 | 9.9 | 3.2 | 1.2 |
| >hg38_21_+_8433172_8446622_RNA45SN1_45S_50ntFlanks@10388.10409.22\|GTGAACAGCAGTTGAACATGGG | >hg38_21_+_8433172_8446622_RNA45SN1_45S_50ntFlanks | 10388 | 10409 | 22 | GTGAACAGCAGTTGAACATGGG | MB468 | 14 | 13 | 15 | 8 | 1 | 6.4 | 5.8 | 6.4 | 4.7 | 0.3 |
| >hg38_21_+_8433172_8446622_RNA45SN1_45S_50ntFlanks@10388.10410.23\|GTGAACAGCAGTTGAACATGGGT | >hg38_21_+_8433172_8446622_RNA45SN1_45S_50ntFlanks | 10388 | 10410 | 23 | GTGAACAGCAGTTGAACATGGGT | MB468 | 58 | 63 | 39 | 23 | 4 | 19.5 | 25.8 | 14.2 | 11.6 | 1.1 |

| | | | | | | | | | | | | | | | | |
|---|---|---|---|---|---|---|---|---|---|---|---|---|---|---|---|---|
| >hg38_21_+_8433172_8446622_RNA45SN1_45S_50ntFlanks@10388.10424.37\|GTGAACAGCAGTTGAACATGGGTCAGTCGGTCCTGAG | >hg38_21_+_8433172_8446622_RNA45SN1_45S_50ntFlanks | 10388 | 10424 | 37 | GTGAACAGCAGTTGAACATGGGTCAGTCGGTCCTGAG | BT20 | 56 | 94 | 89 | 516 | 1016 | 46.4 | 86.8 | 32.1 | 340.4 | 701.8 |
| >hg38_21_+_8433172_8446622_RNA45SN1_45S_50ntFlanks@10388.10424.37\|GTGAACAGCAGTTGAACATGGGTCAGTCGGTCCTGAG | >hg38_21_+_8433172_8446622_RNA45SN1_45S_50ntFlanks | 10388 | 10424 | 37 | GTGAACAGCAGTTGAACATGGGTCAGTCGGTCCTGAG | MB231 | 22 | 69 | 93 | 48 | 49 | 20.3 | 22.6 | 82.6 | 32.0 | 48.4 |
| >hg38_21_+_8433172_8446622_RNA45SN1_45S_50ntFlanks@10389.10406.18\|TGAACAGCAGTTGAACAT | >hg38_21_+_8433172_8446622_RNA45SN1_45S_50ntFlanks | 10389 | 10406 | 18 | TGAACAGCAGTTGAACAT | MB468 | 13 | 7 | 6 | 5 | 2 | 5.9 | 1.8 | 4.6 | 1.6 | 0.3 |
| >hg38_21_+_8433172_8446622_RNA45SN1_45S_50ntFlanks@10389.10407.19\|TGAACAGCAGTTGAACATG | >hg38_21_+_8433172_8446622_RNA45SN1_45S_50ntFlanks | 10389 | 10407 | 19 | TGAACAGCAGTTGAACATG | MB468 | 355 | 139 | 211 | 94 | 33 | 195.7 | 44.8 | 228.4 | 26.6 | 10.6 |
| >hg38_21_+_8433172_8446622_RNA45SN1_45S_50ntFlanks@10389.10408.20\|TGAACAGCAGTTGAACATGG | >hg38_21_+_8433172_8446622_RNA45SN1_45S_50ntFlanks | 10389 | 10408 | 20 | TGAACAGCAGTTGAACATGG | MB468 | 31 | 20 | 15 | 14 | 3 | 7.6 | 6.5 | 7.1 | 5.9 | 0.6 |
| >hg38_21_+_8433172_8446622_RNA45SN1_45S_50ntFlanks@10389.10409.21\|TGAACAGCAGTTGAACATGGG | >hg38_21_+_8433172_8446622_RNA45SN1_45S_50ntFlanks | 10389 | 10409 | 21 | TGAACAGCAGTTGAACATGGG | MB468 | 89 | 90 | 78 | 30 | 5 | 28.1 | 40.9 | 33.7 | 12.4 | 3.3 |
| >hg38_21_+_8433172_8446622_RNA45SN1_45S_50ntFlanks@10389.10410.22\|TGAACAGCAGTTGAACATGGGT | >hg38_21_+_8433172_8446622_RNA45SN1_45S_50ntFlanks | 10389 | 10410 | 22 | TGAACAGCAGTTGAACATGGGT | MB468 | 295 | 274 | 167 | 79 | 26 | 57.9 | 92.7 | 54.5 | 39.3 | 11.8 |
| >hg38_21_+_8433172_8446622_RNA45SN1_45S_50ntFlanks@10389.10411.23\|TGAACAGCAGTTGAACATGGGTC | >hg38_21_+_8433172_8446622_RNA45SN1_45S_50ntFlanks | 10389 | 10411 | 23 | TGAACAGCAGTTGAACATGGGTC | MB468 | 15 | 16 | 10 | 6 | 2 | 3.1 | 4.2 | 3.8 | 2.4 | 0.7 |
| >hg38_21_+_8433172_8446622_RNA45SN1_45S_50ntFlanks@10389.10424.36\|TGAACAGCAGTTGAACATGGGTCAGTCGGTCCTGAG | >hg38_21_+_8433172_8446622_RNA45SN1_45S_50ntFlanks | 10389 | 10424 | 36 | TGAACAGCAGTTGAACATGGGTCAGTCGGTCCTGAG | BT20 | 41 | 62 | 88 | 579 | 843 | 44.8 | 23.1 | 22.2 | 358.6 | 444.3 |
| >hg38_21_+_8433172_8446622_RNA45SN1_45S_50ntFlanks@10389.10424.36\|TGAACAGCAGTTGAACATGGGTCAGTCGGTCCTGAG | >hg38_21_+_8433172_8446622_RNA45SN1_45S_50ntFlanks | 10389 | 10424 | 36 | TGAACAGCAGTTGAACATGGGTCAGTCGGTCCTGAG | MB231 | 27 | 97 | 127 | 54 | 66 | 21.6 | 32.0 | 114.6 | 31.4 | 61.0 |
| >hg38_21_+_8433172_8446622_RNA45SN1_45S_50ntFlanks@10389.10424.36\|TGAACAGCAGTTGAACATGGGTCAGTCGGTCCTGAG | >hg38_21_+_8433172_8446622_RNA45SN1_45S_50ntFlanks | 10389 | 10424 | 36 | TGAACAGCAGTTGAACATGGGTCAGTCGGTCCTGAG | MB468 | 47 | 139 | 67 | 49 | 99 | 22.5 | 63.3 | 17.5 | 49.9 | 74.1 |
| >hg38_21_+_8433172_8446622_RNA45SN1_45S_50ntFlanks@10389.10429.41\|TGAACAGCAGTTGAACATGGGTCAGTCGGTCCTGAGAGATG | >hg38_21_+_8433172_8446622_RNA45SN1_45S_50ntFlanks | 10389 | 10429 | 41 | TGAACAGCAGTTGAACATGGGTCAGTCGGTCCTGAGAGATG | MB468 | 32 | 71 | 57 | 22 | 29 | 32.9 | 52.3 | 43.8 | 21.6 | 17.7 |
| >hg38_21_+_8433172_8446622_RNA45SN1_45S_50ntFlanks@10389.10430.42\|TGAACAGCAGTTGAACATGGGTCAGTCGGTCCTGAGAGATGG | >hg38_21_+_8433172_8446622_RNA45SN1_45S_50ntFlanks | 10389 | 10430 | 42 | TGAACAGCAGTTGAACATGGGTCAGTCGGTCCTGAGAGATGG | MB468 | 13 | 34 | 22 | 7 | 16 | 13.0 | 28.9 | 13.7 | 5.7 | 10.2 |
| >hg38_21_+_8433172_8446622_RNA45SN1_45S_50ntFlanks@10390.10424.35\|GAACAGCAGTTGAACATGGGTCAGTCGGTCCTGAG | >hg38_21_+_8433172_8446622_RNA45SN1_45S_50ntFlanks | 10390 | 10424 | 35 | GAACAGCAGTTGAACATGGGTCAGTCGGTCCTGAG | BT20 | 17 | 20 | 27 | 179 | 262 | 20.6 | 5.3 | 7.9 | 147.8 | 114.7 |
| >hg38_21_+_8433172_8446622_RNA45SN1_45S_50ntFlanks@10390.10424.35\|GAACAGCAGTTGAACATGGGTCAGTCGGTCCTGAG | >hg38_21_+_8433172_8446622_RNA45SN1_45S_50ntFlanks | 10390 | 10424 | 35 | GAACAGCAGTTGAACATGGGTCAGTCGGTCCTGAG | MB231 | 12 | 34 | 50 | 23 | 33 | 10.2 | 7.8 | 47.7 | 13.9 | 33.1 |
| >hg38_21_+_8433172_8446622_RNA45SN1_45S_50ntFlanks@10391.10408.18\|AACAGCAGTTGAACATGG | >hg38_21_+_8433172_8446622_RNA45SN1_45S_50ntFlanks | 10391 | 10408 | 18 | AACAGCAGTTGAACATGG | MB468 | 83 | 31 | 42 | 29 | 7 | 18.2 | 12.9 | 17.5 | 12.7 | 1.8 |
| >hg38_21_+_8433172_8446622_RNA45SN1_45S_50ntFlanks@10391.10409.19\|AACAGCAGTTGAACATGGG | >hg38_21_+_8433172_8446622_RNA45SN1_45S_50ntFlanks | 10391 | 10409 | 19 | AACAGCAGTTGAACATGGG | MB468 | 283 | 162 | 269 | 55 | 15 | 99.4 | 50.2 | 86.6 | 25.0 | 5.5 |
| >hg38_21_+_8433172_8446622_RNA45SN1_45S_50ntFlanks@10391.10410.20\|AACAGCAGTTGAACATGGGT | >hg38_21_+_8433172_8446622_RNA45SN1_45S_50ntFlanks | 10391 | 10410 | 20 | AACAGCAGTTGAACATGGGT | MB468 | 664 | 399 | 363 | 127 | 64 | 81.3 | 59.5 | 33.0 | 48.8 | 17.0 |
| >hg38_21_+_8433172_8446622_RNA45SN1_45S_50ntFlanks@10391.10411.21\|AACAGCAGTTGAACATGGGTC | >hg38_21_+_8433172_8446622_RNA45SN1_45S_50ntFlanks | 10391 | 10411 | 21 | AACAGCAGTTGAACATGGGTC | MB468 | 19 | 18 | 19 | 9 | 3 | 3.2 | 3.0 | 4.8 | 4.5 | 1.0 |
| >hg38_21_+_8433172_8446622_RNA45SN1_45S_50ntFlanks@10391.10419.29\|AACAGCAGTTGAACATGGGTCAGTCGGTC | >hg38_21_+_8433172_8446622_RNA45SN1_45S_50ntFlanks | 10391 | 10419 | 29 | AACAGCAGTTGAACATGGGTCAGTCGGTC | MB468 | 41 | 55 | 42 | 21 | 15 | 11.9 | 22.5 | 13.5 | 9.2 | 9.8 |
| >hg38_21_+_8433172_8446622_RNA45SN1_45S_50ntFlanks@10391.10420.30\|AACAGCAGTTGAACATGGGTCAGTCGGTCC | >hg38_21_+_8433172_8446622_RNA45SN1_45S_50ntFlanks | 10391 | 10420 | 30 | AACAGCAGTTGAACATGGGTCAGTCGGTCC | MB468 | 46 | 58 | 47 | 21 | 38 | 7.8 | 18.2 | 10.4 | 9.0 | 17.7 |
| >hg38_21_+_8433172_8446622_RNA45SN1_45S_50ntFlanks@10391.10422.32\|AACAGCAGTTGAACATGGGTCAGTCGGTCCTG | >hg38_21_+_8433172_8446622_RNA45SN1_45S_50ntFlanks | 10391 | 10422 | 32 | AACAGCAGTTGAACATGGGTCAGTCGGTCCTG | MB468 | 27 | 34 | 35 | 15 | 12 | 13.8 | 7.3 | 11.4 | 6.7 | 5.5 |
| >hg38_21_+_8433172_8446622_RNA45SN1_45S_50ntFlanks@10391.10424.34\|AACAGCAGTTGAACATGGGTCAGTCGGTCCTGAG | >hg38_21_+_8433172_8446622_RNA45SN1_45S_50ntFlanks | 10391 | 10424 | 34 | AACAGCAGTTGAACATGGGTCAGTCGGTCCTGAG | BT20 | 72 | 119 | 102 | 589 | 976 | 100.4 | 64.0 | 69.1 | 491.6 | 342.3 |
| >hg38_21_+_8433172_8446622_RNA45SN1_45S_50ntFlanks@10391.10424.34\|AACAGCAGTTGAACATGGGTCAGTCGGTCCTGAG | >hg38_21_+_8433172_8446622_RNA45SN1_45S_50ntFlanks | 10391 | 10424 | 34 | AACAGCAGTTGAACATGGGTCAGTCGGTCCTGAG | MB231 | 21 | 61 | 84 | 44 | 71 | 20.4 | 8.3 | 90.7 | 27.5 | 77.4 |
| >hg38_21_+_8433172_8446622_RNA45SN1_45S_50ntFlanks@10391.10424.34\|AACAGCAGTTGAACATGGGTCAGTCGGTCCTGAG | >hg38_21_+_8433172_8446622_RNA45SN1_45S_50ntFlanks | 10391 | 10424 | 34 | AACAGCAGTTGAACATGGGTCAGTCGGTCCTGAG | MB468 | 74 | 194 | 102 | 52 | 165 | 25.7 | 91.7 | 64.4 | 44.0 | 113.3 |
| >hg38_21_+_8433172_8446622_RNA45SN1_45S_50ntFlanks@10391.10426.36\|AACAGCAGTTGAACATGGGTCAGTCGGTCCTGAGAG | >hg38_21_+_8433172_8446622_RNA45SN1_45S_50ntFlanks | 10391 | 10426 | 36 | AACAGCAGTTGAACATGGGTCAGTCGGTCCTGAGAG | MB468 | 21 | 35 | 24 | 16 | 22 | 13.8 | 17.1 | 12.9 | 9.3 | 13.4 |
| >hg38_21_+_8433172_8446622_RNA45SN1_45S_50ntFlanks@10391.10429.39\|AACAGCAGTTGAACATGGGTCAGTCGGTCCTGAGAGATG | >hg38_21_+_8433172_8446622_RNA45SN1_45S_50ntFlanks | 10391 | 10429 | 39 | AACAGCAGTTGAACATGGGTCAGTCGGTCCTGAGAGATG | MB468 | 94 | 162 | 141 | 54 | 100 | 61.5 | 94.3 | 74.7 | 43.3 | 60.9 |
| >hg38_21_+_8433172_8446622_RNA45SN1_45S_50ntFlanks@10391.10430.40\|AACAGCAGTTGAACATGGGTCAGTCGGTCCTGAGAGATGG | >hg38_21_+_8433172_8446622_RNA45SN1_45S_50ntFlanks | 10391 | 10430 | 40 | AACAGCAGTTGAACATGGGTCAGTCGGTCCTGAGAGATGG | MB468 | 38 | 75 | 55 | 18 | 65 | 21.4 | 53.2 | 22.3 | 11.0 | 51.2 |
| >hg38_21_+_8433172_8446622_RNA45SN1_45S_50ntFlanks@10391.10433.43\|AACAGCAGTTGAACATGGGTCAGTCGGTCCTGAGAGATGGGCG | >hg38_21_+_8433172_8446622_RNA45SN1_45S_50ntFlanks | 10391 | 10433 | 43 | AACAGCAGTTGAACATGGGTCAGTCGGTCCTGAGAGATGGGCG | MB468 | 15 | 40 | 36 | 8 | 35 | 13.7 | 41.3 | 20.0 | 6.7 | 29.5 |
| >hg38_21_+_8433172_8446622_RNA45SN1_45S_50ntFlanks@10391.10435.45\|AACAGCAGTTGAACATGGGTCAGTCGGTCCTGAGAGATGGGCGAG | >hg38_21_+_8433172_8446622_RNA45SN1_45S_50ntFlanks | 10391 | 10435 | 45 | AACAGCAGTTGAACATGGGTCAGTCGGTCCTGAGAGATGGGCGAG | MB468 | 18 | 55 | 46 | 8 | 59 | 20.3 | 65.7 | 26.7 | 6.6 | 51.0 |
| >hg38_21_+_8433172_8446622_RNA45SN1_45S_50ntFlanks@10392.10409.18\|ACAGCAGTTGAACATGGG | >hg38_21_+_8433172_8446622_RNA45SN1_45S_50ntFlanks | 10392 | 10409 | 18 | ACAGCAGTTGAACATGGG | MB468 | 310 | 171 | 251 | 49 | 14 | 167.0 | 77.0 | 143.7 | 31.3 | 9.3 |
| >hg38_21_+_8433172_8446622_RNA45SN1_45S_50ntFlanks@10392.10410.19\|ACAGCAGTTGAACATGGGT | >hg38_21_+_8433172_8446622_RNA45SN1_45S_50ntFlanks | 10392 | 10410 | 19 | ACAGCAGTTGAACATGGGT | MB468 | 942 | 421 | 319 | 197 | 73 | 330.5 | 105.6 | 132.6 | 71.1 | 21.5 |
| >hg38_21_+_8433172_8446622_RNA45SN1_45S_50ntFlanks@10392.10411.20\|ACAGCAGTTGAACATGGGTC | >hg38_21_+_8433172_8446622_RNA45SN1_45S_50ntFlanks | 10392 | 10411 | 20 | ACAGCAGTTGAACATGGGTC | MB468 | 37 | 34 | 24 | 13 | 4 | 14.8 | 14.4 | 9.3 | 6.2 | 1.4 |
| >hg38_21_+_8433172_8446622_RNA45SN1_45S_50ntFlanks@10392.10418.27\|ACAGCAGTTGAACATGGGTCAGTCGGT | >hg38_21_+_8433172_8446622_RNA45SN1_45S_50ntFlanks | 10392 | 10418 | 27 | ACAGCAGTTGAACATGGGTCAGTCGGT | MB468 | 17 | 26 | 17 | 6 | 3 | 2.9 | 8.7 | 12.4 | 3.8 | 1.0 |
| >hg38_21_+_8433172_8446622_RNA45SN1_45S_50ntFlanks@10392.10419.28\|ACAGCAGTTGAACATGGGTCAGTCGGTC | >hg38_21_+_8433172_8446622_RNA45SN1_45S_50ntFlanks | 10392 | 10419 | 28 | ACAGCAGTTGAACATGGGTCAGTCGGTC | MB468 | 57 | 92 | 41 | 27 | 14 | 12.6 | 34.4 | 10.0 | 12.6 | 6.9 |
| >hg38_21_+_8433172_8446622_RNA45SN1_45S_50ntFlanks@10392.10420.29\|ACAGCAGTTGAACATGGGTCAGTCGGTCC | >hg38_21_+_8433172_8446622_RNA45SN1_45S_50ntFlanks | 10392 | 10420 | 29 | ACAGCAGTTGAACATGGGTCAGTCGGTCC | MB468 | 44 | 66 | 34 | 27 | 26 | 10.8 | 40.0 | 14.4 | 14.7 | 12.7 |
| >hg38_21_+_8433172_8446622_RNA45SN1_45S_50ntFlanks@10392.10422.31\|ACAGCAGTTGAACATGGGTCAGTCGGTCCTG | >hg38_21_+_8433172_8446622_RNA45SN1_45S_50ntFlanks | 10392 | 10422 | 31 | ACAGCAGTTGAACATGGGTCAGTCGGTCCTG | MB468 | 66 | 82 | 53 | 31 | 18 | 45.9 | 38.3 | 37.3 | 20.3 | 8.5 |
| >hg38_21_+_8433172_8446622_RNA45SN1_45S_50ntFlanks@10392.10424.33\|ACAGCAGTTGAACATGGGTCAGTCGGTCCTGAG | >hg38_21_+_8433172_8446622_RNA45SN1_45S_50ntFlanks | 10392 | 10424 | 33 | ACAGCAGTTGAACATGGGTCAGTCGGTCCTGAG | BT20 | 35 | 31 | 41 | 390 | 474 | 48.2 | 3.2 | 33.7 | 366.0 | 171.3 |
| >hg38_21_+_8433172_8446622_RNA45SN1_45S_50ntFlanks@10392.10424.33\|ACAGCAGTTGAACATGGGTCAGTCGGTCCTGAG | >hg38_21_+_8433172_8446622_RNA45SN1_45S_50ntFlanks | 10392 | 10424 | 33 | ACAGCAGTTGAACATGGGTCAGTCGGTCCTGAG | MB231 | 24 | 43 | 82 | 32 | 66 | 30.3 | 6.2 | 98.9 | 18.3 | 72.1 |
| >hg38_21_+_8433172_8446622_RNA45SN1_45S_50ntFlanks@10392.10424.33\|ACAGCAGTTGAACATGGGTCAGTCGGTCCTGAG | >hg38_21_+_8433172_8446622_RNA45SN1_45S_50ntFlanks | 10392 | 10424 | 33 | ACAGCAGTTGAACATGGGTCAGTCGGTCCTGAG | MB468 | 107 | 228 | 99 | 86 | 178 | 47.6 | 138.7 | 46.5 | 92.5 | 140.3 |
| >hg38_21_+_8433172_8446622_RNA45SN1_45S_50ntFlanks@10392.10426.35\|ACAGCAGTTGAACATGGGTCAGTCGGTCCTGAGAG | >hg38_21_+_8433172_8446622_RNA45SN1_45S_50ntFlanks | 10392 | 10426 | 35 | ACAGCAGTTGAACATGGGTCAGTCGGTCCTGAGAG | MB468 | 45 | 72 | 40 | 29 | 29 | 40.6 | 54.3 | 37.4 | 17.2 | 13.9 |
| >hg38_21_+_8433172_8446622_RNA45SN1_45S_50ntFlanks@10392.10428.37\|ACAGCAGTTGAACATGGGTCAGTCGGTCCTGAGAGAT | >hg38_21_+_8433172_8446622_RNA45SN1_45S_50ntFlanks | 10392 | 10428 | 37 | ACAGCAGTTGAACATGGGTCAGTCGGTCCTGAGAGAT | MB468 | 13 | 25 | 10 | 10 | 10 | 9.6 | 10.1 | 6.2 | 8.7 | 4.9 |
| >hg38_21_+_8433172_8446622_RNA45SN1_45S_50ntFlanks@10392.10429.38\|ACAGCAGTTGAACATGGGTCAGTCGGTCCTGAGAGATG | >hg38_21_+_8433172_8446622_RNA45SN1_45S_50ntFlanks | 10392 | 10429 | 38 | ACAGCAGTTGAACATGGGTCAGTCGGTCCTGAGAGATG | MB468 | 189 | 266 | 199 | 109 | 127 | 173.5 | 151.4 | 172.2 | 97.3 | 81.6 |
| >hg38_21_+_8433172_8446622_RNA45SN1_45S_50ntFlanks@10392.10430.39\|ACAGCAGTTGAACATGGGTCAGTCGGTCCTGAGAGATGG | >hg38_21_+_8433172_8446622_RNA45SN1_45S_50ntFlanks | 10392 | 10430 | 39 | ACAGCAGTTGAACATGGGTCAGTCGGTCCTGAGAGATGG | MB468 | 73 | 111 | 68 | 35 | 63 | 66.7 | 53.3 | 48.6 | 22.5 | 38.2 |
| >hg38_21_+_8433172_8446622_RNA45SN1_45S_50ntFlanks@10392.10431.40\|ACAGCAGTTGAACATGGGTCAGTCGGTCCTGAGAGATGGG | >hg38_21_+_8433172_8446622_RNA45SN1_45S_50ntFlanks | 10392 | 10431 | 40 | ACAGCAGTTGAACATGGGTCAGTCGGTCCTGAGAGATGGG | MB468 | 19 | 36 | 24 | 12 | 18 | 17.4 | 23.5 | 15.9 | 9.6 | 11.2 |
| >hg38_21_+_8433172_8446622_RNA45SN1_45S_50ntFlanks@10392.10432.41\|ACAGCAGTTGAACATGGGTCAGTCGGTCCTGAGAGATGGGC | >hg38_21_+_8433172_8446622_RNA45SN1_45S_50ntFlanks | 10392 | 10432 | 41 | ACAGCAGTTGAACATGGGTCAGTCGGTCCTGAGAGATGGGC | MB468 | 12 | 29 | 17 | 7 | 13 | 11.9 | 19.6 | 11.9 | 5.9 | 8.5 |
| >hg38_21_+_8433172_8446622_RNA45SN1_45S_50ntFlanks@10392.10433.42\|ACAGCAGTTGAACATGGGTCAGTCGGTCCTGAGAGATGGGCG | >hg38_21_+_8433172_8446622_RNA45SN1_45S_50ntFlanks | 10392 | 10433 | 42 | ACAGCAGTTGAACATGGGTCAGTCGGTCCTGAGAGATGGGCG | MB468 | 35 | 65 | 50 | 18 | 44 | 38.5 | 49.2 | 31.9 | 14.0 | 29.4 |
| >hg38_21_+_8433172_8446622_RNA45SN1_45S_50ntFlanks@10392.10435.44\|ACAGCAGTTGAACATGGGTCAGTCGGTCCTGAGAGATGGGCGAG | >hg38_21_+_8433172_8446622_RNA45SN1_45S_50ntFlanks | 10392 | 10435 | 44 | ACAGCAGTTGAACATGGGTCAGTCGGTCCTGAGAGATGGGCGAG | MB468 | 46 | 93 | 68 | 18 | 75 | 56.6 | 83.7 | 49.1 | 16.0 | 50.1 |
| >hg38_21_+_8433172_8446622_RNA45SN1_45S_50ntFlanks@10393.10410.18\|CAGCAGTTGAACATGGGT | >hg38_21_+_8433172_8446622_RNA45SN1_45S_50ntFlanks | 10393 | 10410 | 18 | CAGCAGTTGAACATGGGT | MB468 | 423 | 236 | 204 | 101 | 37 | 139.8 | 103.7 | 29.6 | 27.7 | 23.3 |
| >hg38_21_+_8433172_8446622_RNA45SN1_45S_50ntFlanks@10393.10411.19\|CAGCAGTTGAACATGGGTC | >hg38_21_+_8433172_8446622_RNA45SN1_45S_50ntFlanks | 10393 | 10411 | 19 | CAGCAGTTGAACATGGGTC | MB468 | 13 | 13 | 10 | 6 | 2 | 2.9 | 4.1 | 2.5 | 2.2 | 1.1 |

| | | | | | | | | | | | | | | | | |
|---|---|---|---|---|---|---|---|---|---|---|---|---|---|---|---|---|
| >hg38_21_+_8433172_8446622_RNA45SN1_45S_50ntFlanks@10393.10419.27\|CAGCAGTTGAACATGGGTCAGTCGGTC | >hg38_21_+_8433172_8446622_RNA45SN1_45S_50ntFlanks | 10393 | 10419 | 27 | CAGCAGTTGAACATGGGTCAGTCGGTC | MB468 | 31 | 46 | 24 | 19 | 12 | 6.0 | 13.0 | 6.7 | 8.6 | 5.9 |
| >hg38_21_+_8433172_8446622_RNA45SN1_45S_50ntFlanks@10393.10420.28\|CAGCAGTTGAACATGGGTCAGTCGGTCC | >hg38_21_+_8433172_8446622_RNA45SN1_45S_50ntFlanks | 10393 | 10420 | 28 | CAGCAGTTGAACATGGGTCAGTCGGTCC | MB468 | 22 | 30 | 17 | 17 | 21 | 3.4 | 11.7 | 4.7 | 7.8 | 12.3 |
| >hg38_21_+_8433172_8446622_RNA45SN1_45S_50ntFlanks@10393.10422.30\|CAGCAGTTGAACATGGGTCAGTCGGTCCTG | >hg38_21_+_8433172_8446622_RNA45SN1_45S_50ntFlanks | 10393 | 10422 | 30 | CAGCAGTTGAACATGGGTCAGTCGGTCCTG | MB468 | 43 | 46 | 32 | 24 | 17 | 28.2 | 21.6 | 19.8 | 13.2 | 7.1 |
| >hg38_21_+_8433172_8446622_RNA45SN1_45S_50ntFlanks@10393.10424.32\|CAGCAGTTGAACATGGGTCAGTCGGTCCTGAG | >hg38_21_+_8433172_8446622_RNA45SN1_45S_50ntFlanks | 10393 | 10424 | 32 | CAGCAGTTGAACATGGGTCAGTCGGTCCTGAG | BT20 | 39 | 41 | 41 | 550 | 494 | 50.5 | 5.0 | 30.3 | 645.6 | 165.9 |
| >hg38_21_+_8433172_8446622_RNA45SN1_45S_50ntFlanks@10393.10424.32\|CAGCAGTTGAACATGGGTCAGTCGGTCCTGAG | >hg38_21_+_8433172_8446622_RNA45SN1_45S_50ntFlanks | 10393 | 10424 | 32 | CAGCAGTTGAACATGGGTCAGTCGGTCCTGAG | MB231 | 25 | 59 | 85 | 42 | 81 | 23.9 | 11.6 | 87.6 | 27.3 | 96.7 |
| >hg38_21_+_8433172_8446622_RNA45SN1_45S_50ntFlanks@10393.10424.32\|CAGCAGTTGAACATGGGTCAGTCGGTCCTGAG | >hg38_21_+_8433172_8446622_RNA45SN1_45S_50ntFlanks | 10393 | 10424 | 32 | CAGCAGTTGAACATGGGTCAGTCGGTCCTGAG | MB468 | 61 | 138 | 58 | 48 | 119 | 23.4 | 61.9 | 33.1 | 50.4 | 84.7 |
| >hg38_21_+_8433172_8446622_RNA45SN1_45S_50ntFlanks@10393.10426.34\|CAGCAGTTGAACATGGGTCAGTCGGTCCTGAGAG | >hg38_21_+_8433172_8446622_RNA45SN1_45S_50ntFlanks | 10393 | 10426 | 34 | CAGCAGTTGAACATGGGTCAGTCGGTCCTGAGAG | MB468 | 24 | 39 | 22 | 18 | 19 | 19.4 | 24.8 | 16.6 | 10.9 | 8.8 |
| >hg38_21_+_8433172_8446622_RNA45SN1_45S_50ntFlanks@10393.10429.37\|CAGCAGTTGAACATGGGTCAGTCGGTCCTGAGAGATG | >hg38_21_+_8433172_8446622_RNA45SN1_45S_50ntFlanks | 10393 | 10429 | 37 | CAGCAGTTGAACATGGGTCAGTCGGTCCTGAGAGATG | MB468 | 108 | 160 | 107 | 75 | 84 | 79.1 | 70.8 | 76.4 | 61.9 | 50.6 |
| >hg38_21_+_8433172_8446622_RNA45SN1_45S_50ntFlanks@10393.10430.38\|CAGCAGTTGAACATGGGTCAGTCGGTCCTGAGAGATGG | >hg38_21_+_8433172_8446622_RNA45SN1_45S_50ntFlanks | 10393 | 10430 | 38 | CAGCAGTTGAACATGGGTCAGTCGGTCCTGAGAGATGG | MB468 | 44 | 68 | 42 | 25 | 46 | 31.3 | 31.7 | 22.5 | 14.7 | 26.5 |
| >hg38_21_+_8433172_8446622_RNA45SN1_45S_50ntFlanks@10393.10431.39\|CAGCAGTTGAACATGGGTCAGTCGGTCCTGAGAGATGGG | >hg38_21_+_8433172_8446622_RNA45SN1_45S_50ntFlanks | 10393 | 10431 | 39 | CAGCAGTTGAACATGGGTCAGTCGGTCCTGAGAGATGGG | MB468 | 12 | 23 | 14 | 8 | 14 | 9.7 | 12.8 | 8.1 | 6.0 | 8.1 |
| >hg38_21_+_8433172_8446622_RNA45SN1_45S_50ntFlanks@10393.10433.41\|CAGCAGTTGAACATGGGTCAGTCGGTCCTGAGAGATGGGCG | >hg38_21_+_8433172_8446622_RNA45SN1_45S_50ntFlanks | 10393 | 10433 | 41 | CAGCAGTTGAACATGGGTCAGTCGGTCCTGAGAGATGGGCG | MB468 | 19 | 33 | 25 | 12 | 30 | 17.6 | 26.4 | 13.8 | 7.7 | 21.4 |
| >hg38_21_+_8433172_8446622_RNA45SN1_45S_50ntFlanks@10393.10435.43\|CAGCAGTTGAACATGGGTCAGTCGGTCCTGAGAGATGGGCGAG | >hg38_21_+_8433172_8446622_RNA45SN1_45S_50ntFlanks | 10393 | 10435 | 43 | CAGCAGTTGAACATGGGTCAGTCGGTCCTGAGAGATGGGCGAG | MB468 | 25 | 47 | 35 | 11 | 50 | 28.3 | 45.8 | 22.7 | 9.0 | 35.4 |
| >hg38_21_+_8433172_8446622_RNA45SN1_45S_50ntFlanks@10394.10424.31\|AGCAGTTGAACATGGGTCAGTCGGTCCTGAG | >hg38_21_+_8433172_8446622_RNA45SN1_45S_50ntFlanks | 10394 | 10424 | 31 | AGCAGTTGAACATGGGTCAGTCGGTCCTGAG | BT20 | 29 | 35 | 32 | 550 | 445 | 36.6 | 8.6 | 22.5 | 525.0 | 202.9 |
| >hg38_21_+_8433172_8446622_RNA45SN1_45S_50ntFlanks@10394.10424.31\|AGCAGTTGAACATGGGTCAGTCGGTCCTGAG | >hg38_21_+_8433172_8446622_RNA45SN1_45S_50ntFlanks | 10394 | 10424 | 31 | AGCAGTTGAACATGGGTCAGTCGGTCCTGAG | MB231 | 19 | 48 | 57 | 69 | 59 | 20.2 | 14.3 | 54.5 | 53.8 | 78.4 |
| >hg38_21_+_8433172_8446622_RNA45SN1_45S_50ntFlanks@10394.10424.31\|AGCAGTTGAACATGGGTCAGTCGGTCCTGAG | >hg38_21_+_8433172_8446622_RNA45SN1_45S_50ntFlanks | 10394 | 10424 | 31 | AGCAGTTGAACATGGGTCAGTCGGTCCTGAG | MB468 | 10 | 25 | 9 | 12 | 16 | 3.9 | 12.6 | 5.7 | 13.7 | 11.8 |
| >hg38_21_+_8433172_8446622_RNA45SN1_45S_50ntFlanks@10394.10429.36\|AGCAGTTGAACATGGGTCAGTCGGTCCTGAGAGATG | >hg38_21_+_8433172_8446622_RNA45SN1_45S_50ntFlanks | 10394 | 10429 | 36 | AGCAGTTGAACATGGGTCAGTCGGTCCTGAGAGATG | MB468 | 20 | 24 | 16 | 16 | 12 | 14.7 | 7.8 | 11.3 | 14.2 | 8.0 |
| >hg38_21_+_8433172_8446622_RNA45SN1_45S_50ntFlanks@10395.10424.30\|GCAGTTGAACATGGGTCAGTCGGTCCTGAG | >hg38_21_+_8433172_8446622_RNA45SN1_45S_50ntFlanks | 10395 | 10424 | 30 | GCAGTTGAACATGGGTCAGTCGGTCCTGAG | BT20 | 34 | 40 | 37 | 443 | 533 | 41.4 | 11.7 | 25.3 | 323.0 | 213.8 |
| >hg38_21_+_8433172_8446622_RNA45SN1_45S_50ntFlanks@10395.10424.30\|GCAGTTGAACATGGGTCAGTCGGTCCTGAG | >hg38_21_+_8433172_8446622_RNA45SN1_45S_50ntFlanks | 10395 | 10424 | 30 | GCAGTTGAACATGGGTCAGTCGGTCCTGAG | MB231 | 31 | 62 | 89 | 54 | 67 | 35.8 | 8.8 | 102.4 | 35.8 | 74.5 |
| >hg38_21_+_8433172_8446622_RNA45SN1_45S_50ntFlanks@10395.10429.35\|GCAGTTGAACATGGGTCAGTCGGTCCTGAGAGATG | >hg38_21_+_8433172_8446622_RNA45SN1_45S_50ntFlanks | 10395 | 10429 | 35 | GCAGTTGAACATGGGTCAGTCGGTCCTGAGAGATG | MB468 | 14 | 15 | 12 | 18 | 7 | 13.7 | 6.2 | 10.4 | 16.1 | 4.3 |
| >hg38_21_+_8433172_8446622_RNA45SN1_45S_50ntFlanks@10396.10422.27\|CAGTTGAACATGGGTCAGTCGGTCCTG | >hg38_21_+_8433172_8446622_RNA45SN1_45S_50ntFlanks | 10396 | 10422 | 27 | CAGTTGAACATGGGTCAGTCGGTCCTG | MB468 | 11 | 6 | 8 | 12 | 2 | 8.2 | 2.4 | 5.0 | 9.2 | 1.1 |
| >hg38_21_+_8433172_8446622_RNA45SN1_45S_50ntFlanks@10396.10424.29\|CAGTTGAACATGGGTCAGTCGGTCCTGAG | >hg38_21_+_8433172_8446622_RNA45SN1_45S_50ntFlanks | 10396 | 10424 | 29 | CAGTTGAACATGGGTCAGTCGGTCCTGAG | BT20 | 69 | 111 | 134 | 647 | 978 | 61.0 | 40.2 | 67.6 | 315.4 | 387.1 |
| >hg38_21_+_8433172_8446622_RNA45SN1_45S_50ntFlanks@10396.10424.29\|CAGTTGAACATGGGTCAGTCGGTCCTGAG | >hg38_21_+_8433172_8446622_RNA45SN1_45S_50ntFlanks | 10396 | 10424 | 29 | CAGTTGAACATGGGTCAGTCGGTCCTGAG | MB231 | 70 | 189 | 189 | 101 | 122 | 68.7 | 29.7 | 190.2 | 64.8 | 111.9 |
| >hg38_21_+_8433172_8446622_RNA45SN1_45S_50ntFlanks@10396.10424.29\|CAGTTGAACATGGGTCAGTCGGTCCTGAG | >hg38_21_+_8433172_8446622_RNA45SN1_45S_50ntFlanks | 10396 | 10424 | 29 | CAGTTGAACATGGGTCAGTCGGTCCTGAG | MB468 | 18 | 45 | 17 | 21 | 19 | 8.3 | 23.6 | 5.6 | 22.2 | 13.5 |
| >hg38_21_+_8433172_8446622_RNA45SN1_45S_50ntFlanks@10396.10429.34\|CAGTTGAACATGGGTCAGTCGGTCCTGAGAGATG | >hg38_21_+_8433172_8446622_RNA45SN1_45S_50ntFlanks | 10396 | 10429 | 34 | CAGTTGAACATGGGTCAGTCGGTCCTGAGAGATG | MB468 | 19 | 11 | 13 | 19 | 7 | 18.8 | 4.3 | 9.0 | 7.0 | 3.1 |
| >hg38_21_+_8433172_8446622_RNA45SN1_45S_50ntFlanks@10396.10430.35\|CAGTTGAACATGGGTCAGTCGGTCCTGAGAGATGG | >hg38_21_+_8433172_8446622_RNA45SN1_45S_50ntFlanks | 10396 | 10430 | 35 | CAGTTGAACATGGGTCAGTCGGTCCTGAGAGATGG | MB468 | 11 | 10 | 8 | 8 | 4 | 9.1 | 2.4 | 4.5 | 5.1 | 2.4 |
| >hg38_21_+_8433172_8446622_RNA45SN1_45S_50ntFlanks@10398.10424.27\|GTTGAACATGGGTCAGTCGGTCCTGAG | >hg38_21_+_8433172_8446622_RNA45SN1_45S_50ntFlanks | 10398 | 10424 | 27 | GTTGAACATGGGTCAGTCGGTCCTGAG | BT20 | 38 | 66 | 66 | 463 | 660 | 30.7 | 28.2 | 9.6 | 261.9 | 299.2 |
| >hg38_21_+_8433172_8446622_RNA45SN1_45S_50ntFlanks@10398.10424.27\|GTTGAACATGGGTCAGTCGGTCCTGAG | >hg38_21_+_8433172_8446622_RNA45SN1_45S_50ntFlanks | 10398 | 10424 | 27 | GTTGAACATGGGTCAGTCGGTCCTGAG | MB231 | 23 | 97 | 68 | 63 | 50 | 15.4 | 34.2 | 52.3 | 43.0 | 52.5 |
| >hg38_21_+_8433172_8446622_RNA45SN1_45S_50ntFlanks@10400.10424.25\|TGAACATGGGTCAGTCGGTCCTGAG | >hg38_21_+_8433172_8446622_RNA45SN1_45S_50ntFlanks | 10400 | 10424 | 25 | TGAACATGGGTCAGTCGGTCCTGAG | BT20 | 31 | 52 | 56 | 406 | 571 | 30.8 | 10.3 | 6.6 | 275.4 | 250.4 |
| >hg38_21_+_8433172_8446622_RNA45SN1_45S_50ntFlanks@10400.10424.25\|TGAACATGGGTCAGTCGGTCCTGAG | >hg38_21_+_8433172_8446622_RNA45SN1_45S_50ntFlanks | 10400 | 10424 | 25 | TGAACATGGGTCAGTCGGTCCTGAG | MB231 | 16 | 86 | 57 | 44 | 48 | 12.8 | 33.1 | 44.9 | 24.9 | 37.8 |
| >hg38_21_+_8433172_8446622_RNA45SN1_45S_50ntFlanks@10401.10420.20\|GAACATGGGTCAGTCGGTCC | >hg38_21_+_8433172_8446622_RNA45SN1_45S_50ntFlanks | 10401 | 10420 | 20 | GAACATGGGTCAGTCGGTCC | MB468 | 15 | 11 | 13 | 17 | 12 | 2.4 | 7.1 | 4.9 | 12.5 | 5.2 |
| >hg38_21_+_8433172_8446622_RNA45SN1_45S_50ntFlanks@10401.10424.24\|GAACATGGGTCAGTCGGTCCTGAG | >hg38_21_+_8433172_8446622_RNA45SN1_45S_50ntFlanks | 10401 | 10424 | 24 | GAACATGGGTCAGTCGGTCCTGAG | BT20 | 54 | 72 | 66 | 582 | 698 | 48.1 | 13.6 | 18.7 | 289.0 | 259.3 |
| >hg38_21_+_8433172_8446622_RNA45SN1_45S_50ntFlanks@10401.10424.24\|GAACATGGGTCAGTCGGTCCTGAG | >hg38_21_+_8433172_8446622_RNA45SN1_45S_50ntFlanks | 10401 | 10424 | 24 | GAACATGGGTCAGTCGGTCCTGAG | MB231 | 49 | 151 | 124 | 400 | 156 | 30.5 | 79.9 | 89.3 | 400.1 | 186.5 |
| >hg38_21_+_8433172_8446622_RNA45SN1_45S_50ntFlanks@10401.10424.24\|GAACATGGGTCAGTCGGTCCTGAG | >hg38_21_+_8433172_8446622_RNA45SN1_45S_50ntFlanks | 10401 | 10424 | 24 | GAACATGGGTCAGTCGGTCCTGAG | MB468 | 12 | 20 | 10 | 29 | 20 | 6.9 | 11.0 | 5.5 | 21.3 | 13.1 |
| >hg38_21_+_8433172_8446622_RNA45SN1_45S_50ntFlanks@10401.10429.29\|GAACATGGGTCAGTCGGTCCTGAGAGATG | >hg38_21_+_8433172_8446622_RNA45SN1_45S_50ntFlanks | 10401 | 10429 | 29 | GAACATGGGTCAGTCGGTCCTGAGAGATG | MB231 | 17 | 11 | 18 | 4 | 11 | 12.1 | 7.1 | 12.6 | 2.7 | 10.3 |
| >hg38_21_+_8433172_8446622_RNA45SN1_45S_50ntFlanks@10401.10429.29\|GAACATGGGTCAGTCGGTCCTGAGAGATG | >hg38_21_+_8433172_8446622_RNA45SN1_45S_50ntFlanks | 10401 | 10429 | 29 | GAACATGGGTCAGTCGGTCCTGAGAGATG | MB468 | 46 | 65 | 40 | 74 | 28 | 34.0 | 50.2 | 35.4 | 75.5 | 26.0 |
| >hg38_21_+_8433172_8446622_RNA45SN1_45S_50ntFlanks@10401.10430.30\|GAACATGGGTCAGTCGGTCCTGAGAGATGG | >hg38_21_+_8433172_8446622_RNA45SN1_45S_50ntFlanks | 10401 | 10430 | 30 | GAACATGGGTCAGTCGGTCCTGAGAGATGG | MB468 | 18 | 26 | 15 | 17 | 9 | 12.6 | 17.3 | 11.1 | 11.8 | 6.3 |
| >hg38_21_+_8433172_8446622_RNA45SN1_45S_50ntFlanks@10401.10439.39\|GAACATGGGTCAGTCGGTCCTGAGAGATGGGCGAGCGCC | >hg38_21_+_8433172_8446622_RNA45SN1_45S_50ntFlanks | 10401 | 10439 | 39 | GAACATGGGTCAGTCGGTCCTGAGAGATGGGCGAGCGCC | MB231 | 18 | 18 | 21 | 9 | 26 | 17.4 | 12.1 | 18.6 | 4.0 | 21.0 |
| >hg38_21_+_8433172_8446622_RNA45SN1_45S_50ntFlanks@10402.10419.18\|AACATGGGTCAGTCGGTC | >hg38_21_+_8433172_8446622_RNA45SN1_45S_50ntFlanks | 10402 | 10419 | 18 | AACATGGGTCAGTCGGTC | MB468 | 79 | 101 | 94 | 72 | 22 | 28.2 | 12.2 | 33.4 | 55.2 | 10.5 |
| >hg38_21_+_8433172_8446622_RNA45SN1_45S_50ntFlanks@10402.10420.19\|AACATGGGTCAGTCGGTCC | >hg38_21_+_8433172_8446622_RNA45SN1_45S_50ntFlanks | 10402 | 10420 | 19 | AACATGGGTCAGTCGGTCC | MB468 | 167 | 61 | 148 | 124 | 107 | 43.9 | 19.9 | 31.9 | 25.6 | 10.3 |
| >hg38_21_+_8433172_8446622_RNA45SN1_45S_50ntFlanks@10402.10422.21\|AACATGGGTCAGTCGGTCCTG | >hg38_21_+_8433172_8446622_RNA45SN1_45S_50ntFlanks | 10402 | 10422 | 21 | AACATGGGTCAGTCGGTCCTG | MB468 | 34 | 4 | 25 | 66 | 11 | 14.2 | 1.0 | 8.6 | 33.4 | 3.1 |
| >hg38_21_+_8433172_8446622_RNA45SN1_45S_50ntFlanks@10402.10424.23\|AACATGGGTCAGTCGGTCCTGAG | >hg38_21_+_8433172_8446622_RNA45SN1_45S_50ntFlanks | 10402 | 10424 | 23 | AACATGGGTCAGTCGGTCCTGAG | BT20 | 49 | 65 | 65 | 503 | 651 | 44.1 | 8.0 | 23.1 | 273.9 | 298.1 |
| >hg38_21_+_8433172_8446622_RNA45SN1_45S_50ntFlanks@10402.10424.23\|AACATGGGTCAGTCGGTCCTGAG | >hg38_21_+_8433172_8446622_RNA45SN1_45S_50ntFlanks | 10402 | 10424 | 23 | AACATGGGTCAGTCGGTCCTGAG | MB231 | 24 | 101 | 82 | 130 | 86 | 15.5 | 33.8 | 67.8 | 131.2 | 93.9 |
| >hg38_21_+_8433172_8446622_RNA45SN1_45S_50ntFlanks@10402.10424.23\|AACATGGGTCAGTCGGTCCTGAG | >hg38_21_+_8433172_8446622_RNA45SN1_45S_50ntFlanks | 10402 | 10424 | 23 | AACATGGGTCAGTCGGTCCTGAG | MB468 | 101 | 203 | 107 | 110 | 168 | 34.8 | 109.7 | 71.2 | 30.3 | 67.6 |
| >hg38_21_+_8433172_8446622_RNA45SN1_45S_50ntFlanks@10402.10426.25\|AACATGGGTCAGTCGGTCCTGAGAG | >hg38_21_+_8433172_8446622_RNA45SN1_45S_50ntFlanks | 10402 | 10426 | 25 | AACATGGGTCAGTCGGTCCTGAGAG | MB468 | 51 | 29 | 37 | 45 | 27 | 27.3 | 12.3 | 22.0 | 8.8 | 6.8 |
| >hg38_21_+_8433172_8446622_RNA45SN1_45S_50ntFlanks@10402.10427.26\|AACATGGGTCAGTCGGTCCTGAGAGA | >hg38_21_+_8433172_8446622_RNA45SN1_45S_50ntFlanks | 10402 | 10427 | 26 | AACATGGGTCAGTCGGTCCTGAGAGA | MB468 | 12 | 8 | 10 | 10 | 6 | 5.6 | 3.2 | 3.7 | 1.7 | 0.7 |
| >hg38_21_+_8433172_8446622_RNA45SN1_45S_50ntFlanks@10402.10428.27\|AACATGGGTCAGTCGGTCCTGAGAGAT | >hg38_21_+_8433172_8446622_RNA45SN1_45S_50ntFlanks | 10402 | 10428 | 27 | AACATGGGTCAGTCGGTCCTGAGAGAT | MB468 | 17 | 22 | 14 | 13 | 10 | 5.0 | 5.7 | 3.1 | 3.4 | 3.6 |
| >hg38_21_+_8433172_8446622_RNA45SN1_45S_50ntFlanks@10402.10429.28\|AACATGGGTCAGTCGGTCCTGAGAGATG | >hg38_21_+_8433172_8446622_RNA45SN1_45S_50ntFlanks | 10402 | 10429 | 28 | AACATGGGTCAGTCGGTCCTGAGAGATG | MB468 | 383 | 461 | 301 | 278 | 196 | 93.9 | 155.3 | 109.7 | 148.5 | 69.2 |
| >hg38_21_+_8433172_8446622_RNA45SN1_45S_50ntFlanks@10402.10430.29\|AACATGGGTCAGTCGGTCCTGAGAGATGG | >hg38_21_+_8433172_8446622_RNA45SN1_45S_50ntFlanks | 10402 | 10430 | 29 | AACATGGGTCAGTCGGTCCTGAGAGATGG | MB468 | 187 | 246 | 160 | 109 | 109 | 42.5 | 30.2 | 23.6 | 23.7 | 51.1 |
| >hg38_21_+_8433172_8446622_RNA45SN1_45S_50ntFlanks@10402.10431.30\|AACATGGGTCAGTCGGTCCTGAGAGATGGG | >hg38_21_+_8433172_8446622_RNA45SN1_45S_50ntFlanks | 10402 | 10431 | 30 | AACATGGGTCAGTCGGTCCTGAGAGATGGG | MB468 | 76 | 174 | 106 | 52 | 41 | 1.2 | 66.4 | 20.4 | 29.1 | 21.8 |
| >hg38_21_+_8433172_8446622_RNA45SN1_45S_50ntFlanks@10402.10432.31\|AACATGGGTCAGTCGGTCCTGAGAGATGGGC | >hg38_21_+_8433172_8446622_RNA45SN1_45S_50ntFlanks | 10402 | 10432 | 31 | AACATGGGTCAGTCGGTCCTGAGAGATGGGC | MB468 | 16 | 39 | 19 | 15 | 9 | 2.2 | 13.1 | 3.9 | 5.0 | 3.5 |

| | | | | | | | | | | | | | | | | |
|---|---|---|---|---|---|---|---|---|---|---|---|---|---|---|---|---|
| >hg38_21_+_8433172_8446622_RNA45SN1_45S_50ntFlanks@10402.10433.32\|AACATGGTCAGTCGGTCCTGAGAGATGGGCG | >hg38_21_+_8433172_8446622_RNA45SN1_45S_50ntFlanks | 10402 | 10433 | 32 | AACATGGTCAGTCGGTCCTGAGAGATGGGCG | MB468 | 38 | 75 | 47 | 40 | 24 | 2.6 | 39.8 | 5.5 | 19.6 | 9.7 |
| >hg38_21_+_8433172_8446622_RNA45SN1_45S_50ntFlanks@10402.10435.34\|AACATGGTCAGTCGGTCCTGAGAGATGGGCGAG | >hg38_21_+_8433172_8446622_RNA45SN1_45S_50ntFlanks | 10402 | 10435 | 34 | AACATGGTCAGTCGGTCCTGAGAGATGGGCGAG | MB468 | 58 | 124 | 71 | 58 | 62 | 9.3 | 49.9 | 8.1 | 31.2 | 23.1 |
| >hg38_21_+_8433172_8446622_RNA45SN1_45S_50ntFlanks@10403.10420.18\|ACATGGTCAGTCGGTCC | >hg38_21_+_8433172_8446622_RNA45SN1_45S_50ntFlanks | 10403 | 10420 | 18 | ACATGGTCAGTCGGTCC | MB468 | 15 | 4 | 16 | 13 | 7 | 3.2 | 1.7 | 3.7 | 2.0 | 1.2 |
| >hg38_21_+_8433172_8446622_RNA45SN1_45S_50ntFlanks@10403.10424.22\|ACATGGTCAGTCGGTCCTGAG | >hg38_21_+_8433172_8446622_RNA45SN1_45S_50ntFlanks | 10403 | 10424 | 22 | ACATGGTCAGTCGGTCCTGAG | BT20 | 32 | 44 | 44 | 535 | 477 | 30.8 | 0.8 | 11.4 | 236.3 | 228.0 |
| >hg38_21_+_8433172_8446622_RNA45SN1_45S_50ntFlanks@10403.10424.22\|ACATGGTCAGTCGGTCCTGAG | >hg38_21_+_8433172_8446622_RNA45SN1_45S_50ntFlanks | 10403 | 10424 | 22 | ACATGGTCAGTCGGTCCTGAG | MB231 | 12 | 69 | 52 | 64 | 37 | 7.2 | 24.3 | 43.7 | 23.4 | 32.4 |
| >hg38_21_+_8433172_8446622_RNA45SN1_45S_50ntFlanks@10403.10429.27\|ACATGGTCAGTCGGTCCTGAGAGATG | >hg38_21_+_8433172_8446622_RNA45SN1_45S_50ntFlanks | 10403 | 10429 | 27 | ACATGGTCAGTCGGTCCTGAGAGATG | MB468 | 42 | 63 | 37 | 32 | 17 | 5.2 | 19.9 | 10.4 | 16.2 | 6.1 |
| >hg38_21_+_8433172_8446622_RNA45SN1_45S_50ntFlanks@10403.10430.28\|ACATGGTCAGTCGGTCCTGAGAGATGG | >hg38_21_+_8433172_8446622_RNA45SN1_45S_50ntFlanks | 10403 | 10430 | 28 | ACATGGTCAGTCGGTCCTGAGAGATGG | MB468 | 18 | 30 | 15 | 10 | 6 | 1.9 | 5.3 | 1.2 | 4.5 | 1.9 |
| >hg38_21_+_8433172_8446622_RNA45SN1_45S_50ntFlanks@10404.10422.19\|CATGGTCAGTCGGTCCTG | >hg38_21_+_8433172_8446622_RNA45SN1_45S_50ntFlanks | 10404 | 10422 | 19 | CATGGTCAGTCGGTCCTG | MB468 | 37 | 2 | 25 | 32 | 12 | 19.2 | 0.4 | 10.4 | 14.1 | 10.8 |
| >hg38_21_+_8433172_8446622_RNA45SN1_45S_50ntFlanks@10404.10424.21\|CATGGTCAGTCGGTCCTGAG | >hg38_21_+_8433172_8446622_RNA45SN1_45S_50ntFlanks | 10404 | 10424 | 21 | CATGGTCAGTCGGTCCTGAG | BT20 | 32 | 42 | 51 | 520 | 461 | 27.3 | 4.9 | 17.4 | 215.4 | 237.4 |
| >hg38_21_+_8433172_8446622_RNA45SN1_45S_50ntFlanks@10404.10424.21\|CATGGTCAGTCGGTCCTGAG | >hg38_21_+_8433172_8446622_RNA45SN1_45S_50ntFlanks | 10404 | 10424 | 21 | CATGGTCAGTCGGTCCTGAG | MB231 | 14 | 73 | 63 | 490 | 45 | 7.3 | 21.5 | 55.9 | 384.7 | 41.1 |
| >hg38_21_+_8433172_8446622_RNA45SN1_45S_50ntFlanks@10404.10424.21\|CATGGTCAGTCGGTCCTGAG | >hg38_21_+_8433172_8446622_RNA45SN1_45S_50ntFlanks | 10404 | 10424 | 21 | CATGGTCAGTCGGTCCTGAG | MB468 | 20 | 51 | 24 | 25 | 37 | 4.5 | 21.0 | 13.5 | 5.3 | 13.0 |
| >hg38_21_+_8433172_8446622_RNA45SN1_45S_50ntFlanks@10404.10429.26\|CATGGTCAGTCGGTCCTGAGAGATG | >hg38_21_+_8433172_8446622_RNA45SN1_45S_50ntFlanks | 10404 | 10429 | 26 | CATGGTCAGTCGGTCCTGAGAGATG | MB468 | 116 | 155 | 106 | 89 | 74 | 20.1 | 38.2 | 25.7 | 30.7 | 34.9 |
| >hg38_21_+_8433172_8446622_RNA45SN1_45S_50ntFlanks@10404.10430.27\|CATGGTCAGTCGGTCCTGAGAGATGG | >hg38_21_+_8433172_8446622_RNA45SN1_45S_50ntFlanks | 10404 | 10430 | 27 | CATGGTCAGTCGGTCCTGAGAGATGG | MB468 | 37 | 73 | 34 | 23 | 21 | 3.5 | 23.6 | 6.4 | 7.6 | 8.5 |
| >hg38_21_+_8433172_8446622_RNA45SN1_45S_50ntFlanks@10404.10431.28\|CATGGTCAGTCGGTCCTGAGAGATGGG | >hg38_21_+_8433172_8446622_RNA45SN1_45S_50ntFlanks | 10404 | 10431 | 28 | CATGGTCAGTCGGTCCTGAGAGATGGG | MB468 | 10 | 28 | 16 | 10 | 6 | 2.4 | 13.9 | 5.4 | 4.4 | 2.9 |
| >hg38_21_+_8433172_8446622_RNA45SN1_45S_50ntFlanks@10404.10435.32\|CATGGTCAGTCGGTCCTGAGAGATGGGCGAG | >hg38_21_+_8433172_8446622_RNA45SN1_45S_50ntFlanks | 10404 | 10435 | 32 | CATGGTCAGTCGGTCCTGAGAGATGGGCGAG | MB468 | 14 | 29 | 17 | 13 | 15 | 2.2 | 13.0 | 6.5 | 5.6 | 3.6 |
| >hg38_21_+_8433172_8446622_RNA45SN1_45S_50ntFlanks@10405.10424.20\|ATGGTCAGTCGGTCCTGAG | >hg38_21_+_8433172_8446622_RNA45SN1_45S_50ntFlanks | 10405 | 10424 | 20 | ATGGTCAGTCGGTCCTGAG | BT20 | 39 | 62 | 52 | 969 | 527 | 33.7 | 9.9 | 10.3 | 278.5 | 262.8 |
| >hg38_21_+_8433172_8446622_RNA45SN1_45S_50ntFlanks@10405.10424.20\|ATGGTCAGTCGGTCCTGAG | >hg38_21_+_8433172_8446622_RNA45SN1_45S_50ntFlanks | 10405 | 10424 | 20 | ATGGTCAGTCGGTCCTGAG | MB231 | 17 | 83 | 55 | 2000 | 60 | 8.6 | 28.8 | 43.5 | 1618.1 | 48.8 |
| >hg38_21_+_8433172_8446622_RNA45SN1_45S_50ntFlanks@10407.10429.23\|GGGTCAGTCGGTCCTGAGAGATG | >hg38_21_+_8433172_8446622_RNA45SN1_45S_50ntFlanks | 10407 | 10429 | 23 | GGGTCAGTCGGTCCTGAGAGATG | MB468 | 25 | 33 | 19 | 18 | 9 | 7.0 | 17.9 | 7.8 | 15.8 | 8.2 |
| >hg38_21_+_8433172_8446622_RNA45SN1_45S_50ntFlanks@10408.10425.18\|GGTCAGTCGGTCCTGAGA | >hg38_21_+_8433172_8446622_RNA45SN1_45S_50ntFlanks | 10408 | 10425 | 18 | GGTCAGTCGGTCCTGAGA | MB468 | 25 | 25 | 29 | 10 | 7 | 7.8 | 10.8 | 10.2 | 5.3 | 4.0 |
| >hg38_21_+_8433172_8446622_RNA45SN1_45S_50ntFlanks@10408.10426.19\|GGTCAGTCGGTCCTGAGAG | >hg38_21_+_8433172_8446622_RNA45SN1_45S_50ntFlanks | 10408 | 10426 | 19 | GGTCAGTCGGTCCTGAGAG | MB468 | 59 | 93 | 59 | 26 | 18 | 27.6 | 44.2 | 26.5 | 14.7 | 8.5 |
| >hg38_21_+_8433172_8446622_RNA45SN1_45S_50ntFlanks@10408.10427.20\|GGTCAGTCGGTCCTGAGAGA | >hg38_21_+_8433172_8446622_RNA45SN1_45S_50ntFlanks | 10408 | 10427 | 20 | GGTCAGTCGGTCCTGAGAGA | MB468 | 13 | 12 | 13 | 8 | 6 | 6.0 | 5.7 | 5.8 | 4.2 | 3.8 |
| >hg38_21_+_8433172_8446622_RNA45SN1_45S_50ntFlanks@10408.10428.21\|GGTCAGTCGGTCCTGAGAGAT | >hg38_21_+_8433172_8446622_RNA45SN1_45S_50ntFlanks | 10408 | 10428 | 21 | GGTCAGTCGGTCCTGAGAGAT | MB468 | 26 | 54 | 24 | 13 | 7 | 4.9 | 24.6 | 9.5 | 10.3 | 5.7 |
| >hg38_21_+_8433172_8446622_RNA45SN1_45S_50ntFlanks@10408.10429.22\|GGTCAGTCGGTCCTGAGAGATG | >hg38_21_+_8433172_8446622_RNA45SN1_45S_50ntFlanks | 10408 | 10429 | 22 | GGTCAGTCGGTCCTGAGAGATG | MB468 | 287 | 475 | 261 | 111 | 63 | 72.4 | 260.0 | 91.2 | 111.5 | 56.0 |
| >hg38_21_+_8433172_8446622_RNA45SN1_45S_50ntFlanks@10408.10430.23\|GGTCAGTCGGTCCTGAGAGATGG | >hg38_21_+_8433172_8446622_RNA45SN1_45S_50ntFlanks | 10408 | 10430 | 23 | GGTCAGTCGGTCCTGAGAGATGG | MB468 | 124 | 241 | 158 | 54 | 27 | 28.6 | 116.2 | 49.6 | 32.2 | 13.3 |
| >hg38_21_+_8433172_8446622_RNA45SN1_45S_50ntFlanks@10408.10431.24\|GGTCAGTCGGTCCTGAGAGATGGG | >hg38_21_+_8433172_8446622_RNA45SN1_45S_50ntFlanks | 10408 | 10431 | 24 | GGTCAGTCGGTCCTGAGAGATGGG | MB468 | 45 | 117 | 76 | 24 | 14 | 14.8 | 71.6 | 21.0 | 15.5 | 9.7 |
| >hg38_21_+_8433172_8446622_RNA45SN1_45S_50ntFlanks@10408.10432.25\|GGTCAGTCGGTCCTGAGAGATGGGC | >hg38_21_+_8433172_8446622_RNA45SN1_45S_50ntFlanks | 10408 | 10432 | 25 | GGTCAGTCGGTCCTGAGAGATGGGC | MB468 | 43 | 94 | 58 | 20 | 11 | 11.4 | 60.0 | 19.6 | 14.0 | 10.4 |
| >hg38_21_+_8433172_8446622_RNA45SN1_45S_50ntFlanks@10408.10433.26\|GGTCAGTCGGTCCTGAGAGATGGGCG | >hg38_21_+_8433172_8446622_RNA45SN1_45S_50ntFlanks | 10408 | 10433 | 26 | GGTCAGTCGGTCCTGAGAGATGGGCG | MB468 | 119 | 148 | 170 | 63 | 43 | 25.9 | 109.5 | 26.2 | 50.8 | 44.3 |
| >hg38_21_+_8433172_8446622_RNA45SN1_45S_50ntFlanks@10408.10434.27\|GGTCAGTCGGTCCTGAGAGATGGGCGA | >hg38_21_+_8433172_8446622_RNA45SN1_45S_50ntFlanks | 10408 | 10434 | 27 | GGTCAGTCGGTCCTGAGAGATGGGCGA | MB468 | 19 | 25 | 20 | 12 | 10 | 3.3 | 19.0 | 3.7 | 11.9 | 11.0 |
| >hg38_21_+_8433172_8446622_RNA45SN1_45S_50ntFlanks@10408.10435.28\|GGTCAGTCGGTCCTGAGAGATGGGCGAG | >hg38_21_+_8433172_8446622_RNA45SN1_45S_50ntFlanks | 10408 | 10435 | 28 | GGTCAGTCGGTCCTGAGAGATGGGCGAG | MB468 | 161 | 173 | 174 | 102 | 87 | 61.1 | 131.8 | 40.5 | 59.0 | 75.8 |
| >hg38_21_+_8433172_8446622_RNA45SN1_45S_50ntFlanks@10408.10437.30\|GGTCAGTCGGTCCTGAGAGATGGGCGAGCG | >hg38_21_+_8433172_8446622_RNA45SN1_45S_50ntFlanks | 10408 | 10437 | 30 | GGTCAGTCGGTCCTGAGAGATGGGCGAGCG | MB468 | 12 | 11 | 12 | 12 | 9 | 2.9 | 6.0 | 1.9 | 4.0 | 3.3 |
| >hg38_21_+_8433172_8446622_RNA45SN1_45S_50ntFlanks@10409.10426.18\|GTCAGTCGGTCCTGAGAG | >hg38_21_+_8433172_8446622_RNA45SN1_45S_50ntFlanks | 10409 | 10426 | 18 | GTCAGTCGGTCCTGAGAG | MB468 | 13 | 12 | 14 | 8 | 3 | 3.6 | 6.3 | 3.0 | 2.2 | 0.7 |
| >hg38_21_+_8433172_8446622_RNA45SN1_45S_50ntFlanks@10409.10429.21\|GTCAGTCGGTCCTGAGAGATG | >hg38_21_+_8433172_8446622_RNA45SN1_45S_50ntFlanks | 10409 | 10429 | 21 | GTCAGTCGGTCCTGAGAGATG | MB468 | 102 | 143 | 95 | 33 | 22 | 16.9 | 41.0 | 19.4 | 26.3 | 9.3 |
| >hg38_21_+_8433172_8446622_RNA45SN1_45S_50ntFlanks@10409.10430.22\|GTCAGTCGGTCCTGAGAGATGG | >hg38_21_+_8433172_8446622_RNA45SN1_45S_50ntFlanks | 10409 | 10430 | 22 | GTCAGTCGGTCCTGAGAGATGG | MB468 | 55 | 85 | 67 | 15 | 11 | 12.0 | 17.8 | 27.6 | 8.0 | 3.1 |
| >hg38_21_+_8433172_8446622_RNA45SN1_45S_50ntFlanks@10409.10431.23\|GTCAGTCGGTCCTGAGAGATGGG | >hg38_21_+_8433172_8446622_RNA45SN1_45S_50ntFlanks | 10409 | 10431 | 23 | GTCAGTCGGTCCTGAGAGATGGG | MB468 | 21 | 42 | 28 | 7 | 6 | 3.6 | 18.5 | 7.8 | 3.5 | 3.3 |
| >hg38_21_+_8433172_8446622_RNA45SN1_45S_50ntFlanks@10409.10432.24\|GTCAGTCGGTCCTGAGAGATGGGC | >hg38_21_+_8433172_8446622_RNA45SN1_45S_50ntFlanks | 10409 | 10432 | 24 | GTCAGTCGGTCCTGAGAGATGGGC | MB468 | 16 | 17 | 17 | 6 | 5 | 1.1 | 5.6 | 3.3 | 1.9 | 1.6 |
| >hg38_21_+_8433172_8446622_RNA45SN1_45S_50ntFlanks@10409.10433.25\|GTCAGTCGGTCCTGAGAGATGGGCG | >hg38_21_+_8433172_8446622_RNA45SN1_45S_50ntFlanks | 10409 | 10433 | 25 | GTCAGTCGGTCCTGAGAGATGGGCG | MB468 | 25 | 11 | 25 | 14 | 11 | 3.2 | 2.1 | 7.4 | 3.8 | 4.2 |
| >hg38_21_+_8433172_8446622_RNA45SN1_45S_50ntFlanks@10409.10435.27\|GTCAGTCGGTCCTGAGAGATGGGCGAG | >hg38_21_+_8433172_8446622_RNA45SN1_45S_50ntFlanks | 10409 | 10435 | 27 | GTCAGTCGGTCCTGAGAGATGGGCGAG | MB468 | 22 | 5 | 17 | 17 | 10 | 10.1 | 1.6 | 4.5 | 7.7 | 3.7 |
| >hg38_21_+_8433172_8446622_RNA45SN1_45S_50ntFlanks@10410.10428.19\|TCAGTCGGTCCTGAGAGAT | >hg38_21_+_8433172_8446622_RNA45SN1_45S_50ntFlanks | 10410 | 10428 | 19 | TCAGTCGGTCCTGAGAGAT | MB468 | 70 | 104 | 69 | 22 | 22 | 18.9 | 39.7 | 34.8 | 15.1 | 10.0 |
| >hg38_21_+_8433172_8446622_RNA45SN1_45S_50ntFlanks@10410.10429.20\|TCAGTCGGTCCTGAGAGATG | >hg38_21_+_8433172_8446622_RNA45SN1_45S_50ntFlanks | 10410 | 10429 | 20 | TCAGTCGGTCCTGAGAGATG | MB468 | 762 | 771 | 606 | 214 | 167 | 150.6 | 282.0 | 132.8 | 124.2 | 45.6 |
| >hg38_21_+_8433172_8446622_RNA45SN1_45S_50ntFlanks@10410.10430.21\|TCAGTCGGTCCTGAGAGATGG | >hg38_21_+_8433172_8446622_RNA45SN1_45S_50ntFlanks | 10410 | 10430 | 21 | TCAGTCGGTCCTGAGAGATGG | MB468 | 538 | 595 | 559 | 137 | 121 | 123.8 | 173.7 | 233.3 | 65.6 | 31.2 |
| >hg38_21_+_8433172_8446622_RNA45SN1_45S_50ntFlanks@10410.10431.22\|TCAGTCGGTCCTGAGAGATGGG | >hg38_21_+_8433172_8446622_RNA45SN1_45S_50ntFlanks | 10410 | 10431 | 22 | TCAGTCGGTCCTGAGAGATGGG | MB468 | 189 | 308 | 224 | 60 | 50 | 28.6 | 141.0 | 58.5 | 38.3 | 25.8 |
| >hg38_21_+_8433172_8446622_RNA45SN1_45S_50ntFlanks@10410.10432.23\|TCAGTCGGTCCTGAGAGATGGGC | >hg38_21_+_8433172_8446622_RNA45SN1_45S_50ntFlanks | 10410 | 10432 | 23 | TCAGTCGGTCCTGAGAGATGGGC | MB468 | 79 | 49 | 82 | 51 | 31 | 8.9 | 8.2 | 18.9 | 15.4 | 4.4 |
| >hg38_21_+_8433172_8446622_RNA45SN1_45S_50ntFlanks@10410.10433.24\|TCAGTCGGTCCTGAGAGATGGGCG | >hg38_21_+_8433172_8446622_RNA45SN1_45S_50ntFlanks | 10410 | 10433 | 24 | TCAGTCGGTCCTGAGAGATGGGCG | MB468 | 122 | 27 | 95 | 107 | 49 | 35.2 | 6.7 | 31.3 | 55.3 | 23.9 |
| >hg38_21_+_8433172_8446622_RNA45SN1_45S_50ntFlanks@10410.10435.26\|TCAGTCGGTCCTGAGAGATGGGCGAG | >hg38_21_+_8433172_8446622_RNA45SN1_45S_50ntFlanks | 10410 | 10435 | 26 | TCAGTCGGTCCTGAGAGATGGGCGAG | MB468 | 39 | 6 | 21 | 97 | 25 | 23.1 | 2.7 | 8.4 | 71.3 | 15.6 |
| >hg38_21_+_8433172_8446622_RNA45SN1_45S_50ntFlanks@10410.10440.31\|TCAGTCGGTCCTGAGAGATGGGCGAGCGCCG | >hg38_21_+_8433172_8446622_RNA45SN1_45S_50ntFlanks | 10410 | 10440 | 31 | TCAGTCGGTCCTGAGAGATGGGCGAGCGCCG | MB468 | 16 | 12 | 18 | 8 | 12 | 3.5 | 4.4 | 5.4 | 1.9 | 3.2 |
| >hg38_21_+_8433172_8446622_RNA45SN1_45S_50ntFlanks@10410.10442.33\|TCAGTCGGTCCTGAGAGATGGGCGAGCGCCGTT | >hg38_21_+_8433172_8446622_RNA45SN1_45S_50ntFlanks | 10410 | 10442 | 33 | TCAGTCGGTCCTGAGAGATGGGCGAGCGCCGTT | MB468 | 26 | 22 | 33 | 18 | 21 | 10.9 | 5.5 | 8.1 | 11.2 | 14.1 |
| >hg38_21_+_8433172_8446622_RNA45SN1_45S_50ntFlanks@10410.10443.34\|TCAGTCGGTCCTGAGAGATGGGCGAGCGCCGTTC | >hg38_21_+_8433172_8446622_RNA45SN1_45S_50ntFlanks | 10410 | 10443 | 34 | TCAGTCGGTCCTGAGAGATGGGCGAGCGCCGTTC | MB468 | 28 | 27 | 46 | 61 | 35 | 10.9 | 8.0 | 13.3 | 47.0 | 31.3 |
| >hg38_21_+_8433172_8446622_RNA45SN1_45S_50ntFlanks@10410.10453.44\|TCAGTCGGTCCTGAGAGATGGGCGAGCGCCGTTCCGAAGGGACG | >hg38_21_+_8433172_8446622_RNA45SN1_45S_50ntFlanks | 10410 | 10453 | 44 | TCAGTCGGTCCTGAGAGATGGGCGAGCGCCGTTCCGAAGGGACG | MB468 | 13 | 50 | 55 | 11 | 54 | 4.3 | 50.8 | 37.7 | 8.7 | 45.4 |
| >hg38_21_+_8433172_8446622_RNA45SN1_45S_50ntFlanks@10411.10428.18\|CAGTCGGTCCTGAGAGAT | >hg38_21_+_8433172_8446622_RNA45SN1_45S_50ntFlanks | 10411 | 10428 | 18 | CAGTCGGTCCTGAGAGAT | MB468 | 316 | 430 | 307 | 82 | 175 | 90.0 | 163.5 | 165.6 | 53.7 | 133.0 |
| >hg38_21_+_8433172_8446622_RNA45SN1_45S_50ntFlanks@10411.10429.19\|CAGTCGGTCCTGAGAGATG | >hg38_21_+_8433172_8446622_RNA45SN1_45S_50ntFlanks | 10411 | 10429 | 19 | CAGTCGGTCCTGAGAGATG | MB468 | 1452 | 1338 | 1075 | 426 | 467 | 235.3 | 457.8 | 274.1 | 176.1 | 203.9 |

| | | | | | | | | | | | | | | | | |
|---|---|---|---|---|---|---|---|---|---|---|---|---|---|---|---|---|
| >hg38_21_+_8433172_8446622_RNA45SN1_45S_50ntFlanks@10411.10430.20\|CAGTCGGTCCTGAGAGATGG | >hg38_21_+_8433172_8446622_RNA45SN1_45S_50ntFlanks | 10411 | 10430 | 20 | CAGTCGGTCCTGAGAGATGG | MB468 | 880 | 694 | 1033 | 329 | 270 | 281.1 | 223.8 | 504.9 | 82.7 | 67.2 |
| >hg38_21_+_8433172_8446622_RNA45SN1_45S_50ntFlanks@10411.10431.21\|CAGTCGGTCCTGAGAGATGGG | >hg38_21_+_8433172_8446622_RNA45SN1_45S_50ntFlanks | 10411 | 10431 | 21 | CAGTCGGTCCTGAGAGATGGG | MB468 | 383 | 411 | 488 | 163 | 119 | 74.7 | 124.7 | 175.7 | 51.1 | 48.7 |
| >hg38_21_+_8433172_8446622_RNA45SN1_45S_50ntFlanks@10411.10432.22\|CAGTCGGTCCTGAGAGATGGGC | >hg38_21_+_8433172_8446622_RNA45SN1_45S_50ntFlanks | 10411 | 10432 | 22 | CAGTCGGTCCTGAGAGATGGGC | MB468 | 121 | 35 | 146 | 111 | 53 | 14.6 | 4.9 | 39.1 | 23.2 | 25.3 |
| >hg38_21_+_8433172_8446622_RNA45SN1_45S_50ntFlanks@10411.10433.23\|CAGTCGGTCCTGAGAGATGGGCG | >hg38_21_+_8433172_8446622_RNA45SN1_45S_50ntFlanks | 10411 | 10433 | 23 | CAGTCGGTCCTGAGAGATGGGCG | MB468 | 151 | 18 | 104 | 183 | 88 | 45.5 | 8.8 | 52.4 | 122.2 | 81.6 |
| >hg38_21_+_8433172_8446622_RNA45SN1_45S_50ntFlanks@10411.10435.25\|CAGTCGGTCCTGAGAGATGGGCGAG | >hg38_21_+_8433172_8446622_RNA45SN1_45S_50ntFlanks | 10411 | 10435 | 25 | CAGTCGGTCCTGAGAGATGGGCGAG | MB468 | 57 | 8 | 30 | 152 | 67 | 27.6 | 4.1 | 10.4 | 124.9 | 57.4 |
| >hg38_21_+_8433172_8446622_RNA45SN1_45S_50ntFlanks@10411.10440.30\|CAGTCGGTCCTGAGAGATGGGCGAGCGCCG | >hg38_21_+_8433172_8446622_RNA45SN1_45S_50ntFlanks | 10411 | 10440 | 30 | CAGTCGGTCCTGAGAGATGGGCGAGCGCCG | MB468 | 21 | 9 | 22 | 25 | 22 | 4.9 | 2.7 | 5.1 | 19.7 | 11.5 |
| >hg38_21_+_8433172_8446622_RNA45SN1_45S_50ntFlanks@10411.10441.31\|CAGTCGGTCCTGAGAGATGGGCGAGCGCCGT | >hg38_21_+_8433172_8446622_RNA45SN1_45S_50ntFlanks | 10411 | 10441 | 31 | CAGTCGGTCCTGAGAGATGGGCGAGCGCCGT | MB468 | 14 | 9 | 14 | 22 | 13 | 3.9 | 1.7 | 1.6 | 19.0 | 5.9 |
| >hg38_21_+_8433172_8446622_RNA45SN1_45S_50ntFlanks@10411.10442.32\|CAGTCGGTCCTGAGAGATGGGCGAGCGCCGTT | >hg38_21_+_8433172_8446622_RNA45SN1_45S_50ntFlanks | 10411 | 10442 | 32 | CAGTCGGTCCTGAGAGATGGGCGAGCGCCGTT | MB468 | 55 | 30 | 60 | 79 | 60 | 27.9 | 13.9 | 21.1 | 72.3 | 45.9 |
| >hg38_21_+_8433172_8446622_RNA45SN1_45S_50ntFlanks@10411.10443.33\|CAGTCGGTCCTGAGAGATGGGCGAGCGCCGTTC | >hg38_21_+_8433172_8446622_RNA45SN1_45S_50ntFlanks | 10411 | 10443 | 33 | CAGTCGGTCCTGAGAGATGGGCGAGCGCCGTTC | MB468 | 82 | 63 | 109 | 296 | 133 | 37.5 | 32.2 | 42.1 | 275.0 | 111.8 |
| >hg38_21_+_8433172_8446622_RNA45SN1_45S_50ntFlanks@10411.10445.35\|CAGTCGGTCCTGAGAGATGGGCGAGCGCCGTTCCG | >hg38_21_+_8433172_8446622_RNA45SN1_45S_50ntFlanks | 10411 | 10445 | 35 | CAGTCGGTCCTGAGAGATGGGCGAGCGCCGTTCCG | MB468 | 11 | 20 | 15 | 9 | 38 | 2.9 | 4.9 | 1.2 | 5.1 | 10.0 |
| >hg38_21_+_8433172_8446622_RNA45SN1_45S_50ntFlanks@10411.10453.43\|CAGTCGGTCCTGAGAGATGGGCGAGCGCCGTTCCGAAGGGACG | >hg38_21_+_8433172_8446622_RNA45SN1_45S_50ntFlanks | 10411 | 10453 | 43 | CAGTCGGTCCTGAGAGATGGGCGAGCGCCGTTCCGAAGGGACG | MB468 | 37 | 85 | 109 | 45 | 169 | 4.2 | 87.7 | 79.5 | 34.0 | 133.3 |
| >hg38_21_+_8433172_8446622_RNA45SN1_45S_50ntFlanks@10412.10429.18\|AGTCGGTCCTGAGAGATG | >hg38_21_+_8433172_8446622_RNA45SN1_45S_50ntFlanks | 10412 | 10429 | 18 | AGTCGGTCCTGAGAGATG | MB468 | 22 | 18 | 16 | 14 | 6 | 2.5 | 9.2 | 2.1 | 7.5 | 2.3 |
| >hg38_21_+_8433172_8446622_RNA45SN1_45S_50ntFlanks@10412.10430.19\|AGTCGGTCCTGAGAGATGG | >hg38_21_+_8433172_8446622_RNA45SN1_45S_50ntFlanks | 10412 | 10430 | 19 | AGTCGGTCCTGAGAGATGG | MB468 | 21 | 13 | 28 | 19 | 4 | 3.3 | 4.0 | 12.1 | 10.2 | 0.3 |
| >hg38_21_+_8433172_8446622_RNA45SN1_45S_50ntFlanks@10412.10431.20\|AGTCGGTCCTGAGAGATGGG | >hg38_21_+_8433172_8446622_RNA45SN1_45S_50ntFlanks | 10412 | 10431 | 20 | AGTCGGTCCTGAGAGATGGG | MB468 | 10 | 9 | 15 | 9 | 3 | 2.9 | 4.8 | 6.0 | 2.8 | 1.2 |
| >hg38_21_+_8433172_8446622_RNA45SN1_45S_50ntFlanks@10414.10431.18\|TCGGTCCTGAGAGATGG | >hg38_21_+_8433172_8446622_RNA45SN1_45S_50ntFlanks | 10414 | 10431 | 18 | TCGGTCCTGAGAGATGG | MB468 | 44 | 8 | 84 | 27 | 5 | 21.9 | 3.9 | 68.6 | 4.4 | 2.0 |
| >hg38_21_+_8433172_8446622_RNA45SN1_45S_50ntFlanks@10420.10443.24\|CTGAGAGATGGGCGAGCGCCGTTC | >hg38_21_+_8433172_8446622_RNA45SN1_45S_50ntFlanks | 10420 | 10443 | 24 | CTGAGAGATGGGCGAGCGCCGTTC | MB468 | 41 | 36 | 65 | 54 | 12 | 20.7 | 7.0 | 38.6 | 62.3 | 6.9 |
| >hg38_21_+_8433172_8446622_RNA45SN1_45S_50ntFlanks@10420.10453.34\|CTGAGAGATGGGCGAGCGCCGTTCCGAAGGGACG | >hg38_21_+_8433172_8446622_RNA45SN1_45S_50ntFlanks | 10420 | 10453 | 34 | CTGAGAGATGGGCGAGCGCCGTTCCGAAGGGACG | MB468 | 63 | 137 | 137 | 20 | 25 | 40.3 | 91.7 | 84.5 | 12.5 | 13.8 |
| >hg38_21_+_8433172_8446622_RNA45SN1_45S_50ntFlanks@10420.10455.36\|CTGAGAGATGGGCGAGCGCCGTTCCGAAGGGACGGG | >hg38_21_+_8433172_8446622_RNA45SN1_45S_50ntFlanks | 10420 | 10455 | 36 | CTGAGAGATGGGCGAGCGCCGTTCCGAAGGGACGGG | MB468 | 13 | 29 | 38 | 3 | 3 | 8.3 | 23.1 | 24.0 | 2.3 | 1.6 |
| >hg38_21_+_8433172_8446622_RNA45SN1_45S_50ntFlanks@10420.10456.37\|CTGAGAGATGGGCGAGCGCCGTTCCGAAGGGACGGGC | >hg38_21_+_8433172_8446622_RNA45SN1_45S_50ntFlanks | 10420 | 10456 | 37 | CTGAGAGATGGGCGAGCGCCGTTCCGAAGGGACGGGC | MB468 | 104 | 300 | 344 | 26 | 32 | 66.3 | 238.7 | 234.7 | 11.4 | 15.2 |
| >hg38_21_+_8433172_8446622_RNA45SN1_45S_50ntFlanks@10420.10457.38\|CTGAGAGATGGGCGAGCGCCGTTCCGAAGGGACGGGCG | >hg38_21_+_8433172_8446622_RNA45SN1_45S_50ntFlanks | 10420 | 10457 | 38 | CTGAGAGATGGGCGAGCGCCGTTCCGAAGGGACGGGCG | MB468 | 23 | 53 | 94 | 6 | 7 | 11.2 | 39.0 | 41.2 | 4.1 | 4.1 |
| >hg38_21_+_8433172_8446622_RNA45SN1_45S_50ntFlanks@10421.10443.23\|TGAGAGATGGGCGAGCGCCGTTC | >hg38_21_+_8433172_8446622_RNA45SN1_45S_50ntFlanks | 10421 | 10443 | 23 | TGAGAGATGGGCGAGCGCCGTTC | MB468 | 12 | 10 | 19 | 17 | 3 | 4.2 | 0.9 | 9.6 | 21.5 | 1.2 |
| >hg38_21_+_8433172_8446622_RNA45SN1_45S_50ntFlanks@10421.10453.33\|TGAGAGATGGGCGAGCGCCGTTCCGAAGGGACG | >hg38_21_+_8433172_8446622_RNA45SN1_45S_50ntFlanks | 10421 | 10453 | 33 | TGAGAGATGGGCGAGCGCCGTTCCGAAGGGACG | MB468 | 25 | 56 | 62 | 5 | 7 | 7.4 | 14.4 | 39.4 | 1.8 | 3.5 |
| >hg38_21_+_8433172_8446622_RNA45SN1_45S_50ntFlanks@10421.10456.36\|TGAGAGATGGGCGAGCGCCGTTCCGAAGGGACGGGC | >hg38_21_+_8433172_8446622_RNA45SN1_45S_50ntFlanks | 10421 | 10456 | 36 | TGAGAGATGGGCGAGCGCCGTTCCGAAGGGACGGGC | MB468 | 51 | 127 | 196 | 10 | 16 | 22.9 | 51.3 | 134.1 | 5.4 | 9.9 |
| >hg38_21_+_8433172_8446622_RNA45SN1_45S_50ntFlanks@10421.10457.37\|TGAGAGATGGGCGAGCGCCGTTCCGAAGGGACGGGCG | >hg38_21_+_8433172_8446622_RNA45SN1_45S_50ntFlanks | 10421 | 10457 | 37 | TGAGAGATGGGCGAGCGCCGTTCCGAAGGGACGGGCG | MB468 | 16 | 36 | 67 | 2 | 4 | 4.5 | 16.6 | 27.1 | 1.2 | 2.3 |
| >hg38_21_+_8433172_8446622_RNA45SN1_45S_50ntFlanks@10423.10453.31\|AGAGATGGGCGAGCGCCGTTCCGAAGGGACG | >hg38_21_+_8433172_8446622_RNA45SN1_45S_50ntFlanks | 10423 | 10453 | 31 | AGAGATGGGCGAGCGCCGTTCCGAAGGGACG | MB468 | 20 | 48 | 45 | 5 | 6 | 0.8 | 10.8 | 22.0 | 1.9 | 2.4 |
| >hg38_21_+_8433172_8446622_RNA45SN1_45S_50ntFlanks@10423.10456.34\|AGAGATGGGCGAGCGCCGTTCCGAAGGGACGGGC | >hg38_21_+_8433172_8446622_RNA45SN1_45S_50ntFlanks | 10423 | 10456 | 34 | AGAGATGGGCGAGCGCCGTTCCGAAGGGACGGGC | MB468 | 29 | 70 | 81 | 6 | 10 | 5.0 | 13.3 | 43.6 | 2.2 | 6.0 |
| >hg38_21_+_8433172_8446622_RNA45SN1_45S_50ntFlanks@10423.10457.35\|AGAGATGGGCGAGCGCCGTTCCGAAGGGACGGGCG | >hg38_21_+_8433172_8446622_RNA45SN1_45S_50ntFlanks | 10423 | 10457 | 35 | AGAGATGGGCGAGCGCCGTTCCGAAGGGACGGGCG | MB468 | 18 | 48 | 61 | 3 | 5 | 4.5 | 29.7 | 23.9 | 1.1 | 4.1 |
| >hg38_21_+_8433172_8446622_RNA45SN1_45S_50ntFlanks@10425.10453.29\|AGATGGGCGAGCGCCGTTCCGAAGGGACG | >hg38_21_+_8433172_8446622_RNA45SN1_45S_50ntFlanks | 10425 | 10453 | 29 | AGATGGGCGAGCGCCGTTCCGAAGGGACG | MB468 | 21 | 36 | 52 | 6 | 5 | 1.8 | 10.2 | 25.9 | 2.4 | 1.3 |
| >hg38_21_+_8433172_8446622_RNA45SN1_45S_50ntFlanks@10425.10456.32\|AGATGGGCGAGCGCCGTTCCGAAGGGACGGGC | >hg38_21_+_8433172_8446622_RNA45SN1_45S_50ntFlanks | 10425 | 10456 | 32 | AGATGGGCGAGCGCCGTTCCGAAGGGACGGGC | MB468 | 26 | 42 | 75 | 9 | 8 | 2.3 | 15.5 | 34.1 | 6.3 | 3.4 |
| >hg38_21_+_8433172_8446622_RNA45SN1_45S_50ntFlanks@10425.10457.33\|AGATGGGCGAGCGCCGTTCCGAAGGGACGGGCG | >hg38_21_+_8433172_8446622_RNA45SN1_45S_50ntFlanks | 10425 | 10457 | 33 | AGATGGGCGAGCGCCGTTCCGAAGGGACGGGCG | MB468 | 13 | 17 | 47 | 3 | 3 | 4.8 | 11.9 | 17.8 | 1.3 | 1.5 |
| >hg38_21_+_8433172_8446622_RNA45SN1_45S_50ntFlanks@10427.10453.27\|ATGGGCGAGCGCCGTTCCGAAGGGACG | >hg38_21_+_8433172_8446622_RNA45SN1_45S_50ntFlanks | 10427 | 10453 | 27 | ATGGGCGAGCGCCGTTCCGAAGGGACG | MB468 | 46 | 67 | 86 | 19 | 12 | 15.5 | 36.0 | 28.9 | 10.5 | 4.7 |
| >hg38_21_+_8433172_8446622_RNA45SN1_45S_50ntFlanks@10427.10456.30\|ATGGGCGAGCGCCGTTCCGAAGGGACGGGC | >hg38_21_+_8433172_8446622_RNA45SN1_45S_50ntFlanks | 10427 | 10456 | 30 | ATGGGCGAGCGCCGTTCCGAAGGGACGGGC | MB468 | 70 | 66 | 134 | 42 | 24 | 13.4 | 37.2 | 51.7 | 22.7 | 13.0 |
| >hg38_21_+_8433172_8446622_RNA45SN1_45S_50ntFlanks@10427.10457.31\|ATGGGCGAGCGCCGTTCCGAAGGGACGGGCG | >hg38_21_+_8433172_8446622_RNA45SN1_45S_50ntFlanks | 10427 | 10457 | 31 | ATGGGCGAGCGCCGTTCCGAAGGGACGGGCG | MB468 | 20 | 20 | 36 | 8 | 6 | 7.5 | 17.6 | 16.0 | 2.8 | 2.5 |
| >hg38_21_+_8433172_8446622_RNA45SN1_45S_50ntFlanks@10431.10456.26\|GCGAGCGCCGTTCCGAAGGGACGGGC | >hg38_21_+_8433172_8446622_RNA45SN1_45S_50ntFlanks | 10431 | 10456 | 26 | GCGAGCGCCGTTCCGAAGGGACGGGC | MB468 | 12 | 1 | 15 | 15 | 2 | 8.1 | 0.3 | 4.1 | 10.8 | 1.7 |
| >hg38_21_+_8433172_8446622_RNA45SN1_45S_50ntFlanks@10432.10453.22\|CGAGCGCCGTTCCGAAGGGACG | >hg38_21_+_8433172_8446622_RNA45SN1_45S_50ntFlanks | 10432 | 10453 | 22 | CGAGCGCCGTTCCGAAGGGACG | MB468 | 12 | 3 | 8 | 27 | 13 | 3.6 | 1.4 | 1.7 | 15.8 | 6.7 |
| >hg38_21_+_8433172_8446622_RNA45SN1_45S_50ntFlanks@10436.10453.18\|CGCCGTTCCGAAGGGACG | >hg38_21_+_8433172_8446622_RNA45SN1_45S_50ntFlanks | 10436 | 10453 | 18 | CGCCGTTCCGAAGGGACG | MB468 | 41 | 0 | 48 | 28 | 5 | 28.9 | 0.2 | 29.3 | 12.2 | 3.9 |
| >hg38_21_+_8433172_8446622_RNA45SN1_45S_50ntFlanks@10441.10460.20\|TTCCGAAGGGACGGGCGATG | >hg38_21_+_8433172_8446622_RNA45SN1_45S_50ntFlanks | 10441 | 10460 | 20 | TTCCGAAGGGACGGGCGATG | MB468 | 26 | 39 | 20 | 15 | 4 | 11.9 | 29.4 | 11.3 | 14.9 | 2.5 |
| >hg38_21_+_8433172_8446622_RNA45SN1_45S_50ntFlanks@10443.10460.18\|CCGAAGGGACGGGCGATG | >hg38_21_+_8433172_8446622_RNA45SN1_45S_50ntFlanks | 10443 | 10460 | 18 | CCGAAGGGACGGGCGATG | MB468 | 13 | 16 | 12 | 13 | 3 | 3.5 | 8.5 | 5.0 | 6.6 | 1.3 |
| >hg38_21_+_8433172_8446622_RNA45SN1_45S_50ntFlanks@10444.10462.19\|CGAAGGGACGGGCGATGGC | >hg38_21_+_8433172_8446622_RNA45SN1_45S_50ntFlanks | 10444 | 10462 | 19 | CGAAGGGACGGGCGATGGC | MB468 | 19 | 35 | 32 | 15 | 8 | 8.1 | 6.8 | 29.9 | 7.3 | 1.3 |
| >hg38_21_+_8433172_8446622_RNA45SN1_45S_50ntFlanks@10458.10477.20\|ATGGCCTCCGTTGCCCTCGG | >hg38_21_+_8433172_8446622_RNA45SN1_45S_50ntFlanks | 10458 | 10477 | 20 | ATGGCCTCCGTTGCCCTCGG | MB468 | 261 | 107 | 413 | 122 | 95 | 124.2 | 38.6 | 358.9 | 42.8 | 13.1 |
| >hg38_21_+_8433172_8446622_RNA45SN1_45S_50ntFlanks@10458.10478.21\|ATGGCCTCCGTTGCCCTCGGC | >hg38_21_+_8433172_8446622_RNA45SN1_45S_50ntFlanks | 10458 | 10478 | 21 | ATGGCCTCCGTTGCCCTCGGC | MB468 | 298 | 310 | 264 | 20 | 119 | 80.7 | 149.1 | 173.0 | 7.4 | 10.7 |
| >hg38_21_+_8433172_8446622_RNA45SN1_45S_50ntFlanks@10458.10483.26\|ATGGCCTCCGTTGCCCTCGGCCGATC | >hg38_21_+_8433172_8446622_RNA45SN1_45S_50ntFlanks | 10458 | 10483 | 26 | ATGGCCTCCGTTGCCCTCGGCCGATC | MB468 | 10 | 3 | 16 | 10 | 5 | 5.4 | 0.8 | 11.2 | 10.8 | 3.0 |
| >hg38_21_+_8433172_8446622_RNA45SN1_45S_50ntFlanks@10458.10484.27\|ATGGCCTCCGTTGCCCTCGGCCGATCG | >hg38_21_+_8433172_8446622_RNA45SN1_45S_50ntFlanks | 10458 | 10484 | 27 | ATGGCCTCCGTTGCCCTCGGCCGATCG | MB468 | 31 | 23 | 47 | 14 | 21 | 8.1 | 5.7 | 22.2 | 11.7 | 9.3 |
| >hg38_21_+_8433172_8446622_RNA45SN1_45S_50ntFlanks@10458.10486.29\|ATGGCCTCCGTTGCCCTCGGCCGATCGAA | >hg38_21_+_8433172_8446622_RNA45SN1_45S_50ntFlanks | 10458 | 10486 | 29 | ATGGCCTCCGTTGCCCTCGGCCGATCGAA | MB468 | 18 | 24 | 14 | 3 | 22 | 9.4 | 13.7 | 9.9 | 0.9 | 9.3 |
| >hg38_21_+_8433172_8446622_RNA45SN1_45S_50ntFlanks@10458.10487.30\|ATGGCCTCCGTTGCCCTCGGCCGATCGAAA | >hg38_21_+_8433172_8446622_RNA45SN1_45S_50ntFlanks | 10458 | 10487 | 30 | ATGGCCTCCGTTGCCCTCGGCCGATCGAAA | MB468 | 41 | 48 | 38 | 8 | 34 | 26.8 | 30.3 | 29.6 | 0.8 | 8.4 |
| >hg38_21_+_8433172_8446622_RNA45SN1_45S_50ntFlanks@10458.10488.31\|ATGGCCTCCGTTGCCCTCGGCCGATCGAAAG | >hg38_21_+_8433172_8446622_RNA45SN1_45S_50ntFlanks | 10458 | 10488 | 31 | ATGGCCTCCGTTGCCCTCGGCCGATCGAAAG | MB468 | 78 | 103 | 52 | 18 | 67 | 55.2 | 82.7 | 46.0 | 3.3 | 23.9 |
| >hg38_21_+_8433172_8446622_RNA45SN1_45S_50ntFlanks@10458.10489.32\|ATGGCCTCCGTTGCCCTCGGCCGATCGAAAGG | >hg38_21_+_8433172_8446622_RNA45SN1_45S_50ntFlanks | 10458 | 10489 | 32 | ATGGCCTCCGTTGCCCTCGGCCGATCGAAAGG | MB468 | 10 | 14 | 11 | 7 | 9 | 6.2 | 7.5 | 10.6 | 2.6 | 2.9 |
| >hg38_21_+_8433172_8446622_RNA45SN1_45S_50ntFlanks@10458.10490.33\|ATGGCCTCCGTTGCCCTCGGCCGATCGAAAGGG | >hg38_21_+_8433172_8446622_RNA45SN1_45S_50ntFlanks | 10458 | 10490 | 33 | ATGGCCTCCGTTGCCCTCGGCCGATCGAAAGGG | MB468 | 13 | 5 | 7 | 12 | 6 | 6.9 | 2.6 | 3.8 | 4.5 | 2.4 |
| >hg38_21_+_8433172_8446622_RNA45SN1_45S_50ntFlanks@10458.10492.35\|ATGGCCTCCGTTGCCCTCGGCCGATCGAAAGGGAG | >hg38_21_+_8433172_8446622_RNA45SN1_45S_50ntFlanks | 10458 | 10492 | 35 | ATGGCCTCCGTTGCCCTCGGCCGATCGAAAGGGAG | MB468 | 10 | 3 | 3 | 9 | 4 | 5.9 | 1.2 | 1.3 | 7.3 | 1.5 |
| >hg38_21_+_8433172_8446622_RNA45SN1_45S_50ntFlanks@10458.10495.38\|ATGGCCTCCGTTGCCCTCGGCCGATCGAAAGGGAGTCG | >hg38_21_+_8433172_8446622_RNA45SN1_45S_50ntFlanks | 10458 | 10495 | 38 | ATGGCCTCCGTTGCCCTCGGCCGATCGAAAGGGAGTCG | MB468 | 28 | 8 | 16 | 28 | 25 | 18.1 | 3.4 | 8.5 | 15.1 | 8.5 |

| | | | | | | | | | | | | | | | |
|---|---|---|---|---|---|---|---|---|---|---|---|---|---|---|---|
| >hg38_21_+_8433172_8446622_RNA45SN1_45S_50ntFlanks@10458.10497.40\|ATGGCCTCCGTTGCCCTCGGCCGATCGAAAGGGAGTCGGG | >hg38_21_+_8433172_8446622_RNA45SN1_45S_50ntFlanks | 10458 | 10497 | 40 | ATGGCCTCCGTTGCCCTCGGCCGATCGAAAGGGAGTCGGG | MB468 | 28 | 29 | 42 | 55 | 100 | 4.7 | 22.3 | 24.4 | 38.2 | 79.7 |
| >hg38_21_+_8433172_8446622_RNA45SN1_45S_50ntFlanks@10461.10480.20\|GCCTCCGTTGCCCTCGGCCG | >hg38_21_+_8433172_8446622_RNA45SN1_45S_50ntFlanks | 10461 | 10480 | 20 | GCCTCCGTTGCCCTCGGCCG | MB468 | 10 | 12 | 8 | 2 | 10 | 2.3 | 5.2 | 2.1 | 1.1 | 1.8 |
| >hg38_21_+_8433172_8446622_RNA45SN1_45S_50ntFlanks@10461.10497.37\|GCCTCCGTTGCCCTCGGCCGATCGAAAGGGAGTCGGG | >hg38_21_+_8433172_8446622_RNA45SN1_45S_50ntFlanks | 10461 | 10497 | 37 | GCCTCCGTTGCCCTCGGCCGATCGAAAGGGAGTCGGG | MB468 | 15 | 9 | 14 | 7 | 25 | 2.9 | 5.2 | 5.8 | 3.9 | 10.4 |
| >hg38_21_+_8433172_8446622_RNA45SN1_45S_50ntFlanks@10463.10483.21\|CTCCGTTGCCCTCGGCCGATC | >hg38_21_+_8433172_8446622_RNA45SN1_45S_50ntFlanks | 10463 | 10483 | 21 | CTCCGTTGCCCTCGGCCGATC | MB468 | 17 | 9 | 20 | 6 | 9 | 5.8 | 3.2 | 11.2 | 3.6 | 3.7 |
| >hg38_21_+_8433172_8446622_RNA45SN1_45S_50ntFlanks@10463.10484.22\|CTCCGTTGCCCTCGGCCGATCG | >hg38_21_+_8433172_8446622_RNA45SN1_45S_50ntFlanks | 10463 | 10484 | 22 | CTCCGTTGCCCTCGGCCGATCG | MB468 | 11 | 10 | 14 | 4 | 9 | 2.8 | 3.2 | 5.0 | 2.1 | 4.0 |
| >hg38_21_+_8433172_8446622_RNA45SN1_45S_50ntFlanks@10463.10487.25\|CTCCGTTGCCCTCGGCCGATCGAAA | >hg38_21_+_8433172_8446622_RNA45SN1_45S_50ntFlanks | 10463 | 10487 | 25 | CTCCGTTGCCCTCGGCCGATCGAAA | MB468 | 10 | 12 | 10 | 3 | 10 | 6.0 | 7.1 | 5.9 | 1.1 | 4.6 |
| >hg38_21_+_8433172_8446622_RNA45SN1_45S_50ntFlanks@10463.10497.35\|CTCCGTTGCCCTCGGCCGATCGAAAGGGAGTCGGG | >hg38_21_+_8433172_8446622_RNA45SN1_45S_50ntFlanks | 10463 | 10497 | 35 | CTCCGTTGCCCTCGGCCGATCGAAAGGGAGTCGGG | MB468 | 16 | 10 | 24 | 29 | 24 | 4.0 | 2.6 | 9.7 | 9.1 | 12.2 |
| >hg38_21_+_8433172_8446622_RNA45SN1_45S_50ntFlanks@10466.10487.22\|CGTTGCCCTCGGCCGATCGAAA | >hg38_21_+_8433172_8446622_RNA45SN1_45S_50ntFlanks | 10466 | 10487 | 22 | CGTTGCCCTCGGCCGATCGAAA | MB468 | 14 | 12 | 17 | 4 | 16 | 10.1 | 7.6 | 11.2 | 1.3 | 10.0 |
| >hg38_21_+_8433172_8446622_RNA45SN1_45S_50ntFlanks@10466.10488.23\|CGTTGCCCTCGGCCGATCGAAAG | >hg38_21_+_8433172_8446622_RNA45SN1_45S_50ntFlanks | 10466 | 10488 | 23 | CGTTGCCCTCGGCCGATCGAAAG | MB468 | 23 | 25 | 23 | 6 | 29 | 18.2 | 18.9 | 17.5 | 3.1 | 20.8 |
| >hg38_21_+_8433172_8446622_RNA45SN1_45S_50ntFlanks@10466.10497.32\|CGTTGCCCTCGGCCGATCGAAAGGGAGTCGGG | >hg38_21_+_8433172_8446622_RNA45SN1_45S_50ntFlanks | 10466 | 10497 | 32 | CGTTGCCCTCGGCCGATCGAAAGGGAGTCGGG | MB468 | 67 | 65 | 74 | 45 | 75 | 6.9 | 12.2 | 22.6 | 11.1 | 10.7 |
| >hg38_21_+_8433172_8446622_RNA45SN1_45S_50ntFlanks@10468.10487.20\|TTGCCCTCGGCCGATCGAAA | >hg38_21_+_8433172_8446622_RNA45SN1_45S_50ntFlanks | 10468 | 10487 | 20 | TTGCCCTCGGCCGATCGAAA | MB468 | 14 | 32 | 14 | 1 | 8 | 7.3 | 21.0 | 9.1 | 0.5 | 3.3 |
| >hg38_21_+_8433172_8446622_RNA45SN1_45S_50ntFlanks@10468.10488.21\|TTGCCCTCGGCCGATCGAAAG | >hg38_21_+_8433172_8446622_RNA45SN1_45S_50ntFlanks | 10468 | 10488 | 21 | TTGCCCTCGGCCGATCGAAAG | MB468 | 21 | 54 | 18 | 2 | 12 | 12.4 | 43.4 | 12.7 | 0.5 | 6.2 |
| >hg38_21_+_8433172_8446622_RNA45SN1_45S_50ntFlanks@10468.10495.28\|TTGCCCTCGGCCGATCGAAAGGGAGTCG | >hg38_21_+_8433172_8446622_RNA45SN1_45S_50ntFlanks | 10468 | 10495 | 28 | TTGCCCTCGGCCGATCGAAAGGGAGTCG | MB468 | 18 | 9 | 17 | 5 | 8 | 14.7 | 6.4 | 13.7 | 2.1 | 1.9 |
| >hg38_21_+_8433172_8446622_RNA45SN1_45S_50ntFlanks@10468.10497.30\|TTGCCCTCGGCCGATCGAAAGGGAGTCGGG | >hg38_21_+_8433172_8446622_RNA45SN1_45S_50ntFlanks | 10468 | 10497 | 30 | TTGCCCTCGGCCGATCGAAAGGGAGTCGGG | MB468 | 34 | 18 | 41 | 18 | 15 | 10.2 | 4.3 | 13.2 | 5.2 | 3.0 |
| >hg38_21_+_8433172_8446622_RNA45SN1_45S_50ntFlanks@10471.10488.18\|CCCTCGGCCGATCGAAAG | >hg38_21_+_8433172_8446622_RNA45SN1_45S_50ntFlanks | 10471 | 10488 | 18 | CCCTCGGCCGATCGAAAG | MB468 | 58 | 119 | 50 | 8 | 62 | 23.5 | 77.9 | 28.5 | 7.4 | 48.8 |
| >hg38_21_+_8433172_8446622_RNA45SN1_45S_50ntFlanks@10471.10495.25\|CCCTCGGCCGATCGAAAGGGAGTCG | >hg38_21_+_8433172_8446622_RNA45SN1_45S_50ntFlanks | 10471 | 10495 | 25 | CCCTCGGCCGATCGAAAGGGAGTCG | MB468 | 14 | 7 | 12 | 3 | 14 | 9.6 | 4.6 | 9.6 | 1.8 | 4.6 |
| >hg38_21_+_8433172_8446622_RNA45SN1_45S_50ntFlanks@10473.10490.18\|CTCGGCCGATCGAAAGGG | >hg38_21_+_8433172_8446622_RNA45SN1_45S_50ntFlanks | 10473 | 10490 | 18 | CTCGGCCGATCGAAAGGG | MB468 | 14 | 13 | 11 | 6 | 4 | 5.6 | 6.6 | 5.9 | 1.4 | 0.6 |
| >hg38_21_+_8433172_8446622_RNA45SN1_45S_50ntFlanks@10473.10494.22\|CTCGGCCGATCGAAAGGGAGTC | >hg38_21_+_8433172_8446622_RNA45SN1_45S_50ntFlanks | 10473 | 10494 | 22 | CTCGGCCGATCGAAAGGGAGTC | MB468 | 13 | 12 | 6 | 6 | 4 | 2.1 | 2.6 | 1.4 | 2.8 | 0.8 |
| >hg38_21_+_8433172_8446622_RNA45SN1_45S_50ntFlanks@10477.10495.19\|GCCGATCGAAAGGGAGTCG | >hg38_21_+_8433172_8446622_RNA45SN1_45S_50ntFlanks | 10477 | 10495 | 19 | GCCGATCGAAAGGGAGTCG | MB468 | 11 | 5 | 9 | 16 | 4 | 3.2 | 2.1 | 5.1 | 7.4 | 0.4 |
| >hg38_21_+_8433172_8446622_RNA45SN1_45S_50ntFlanks@10478.10512.35\|CCGATCGAAAGGGAGTCGGGTTCAGATCCCCGAAT | >hg38_21_+_8433172_8446622_RNA45SN1_45S_50ntFlanks | 10478 | 10512 | 35 | CCGATCGAAAGGGAGTCGGGTTCAGATCCCCGAAT | MB468 | 12 | 2 | 24 | 6 | 10 | 2.7 | 0.7 | 6.3 | 3.0 | 5.6 |
| >hg38_21_+_8433172_8446622_RNA45SN1_45S_50ntFlanks@10479.10497.19\|CGATCGAAAGGGAGTCGGG | >hg38_21_+_8433172_8446622_RNA45SN1_45S_50ntFlanks | 10479 | 10497 | 19 | CGATCGAAAGGGAGTCGGG | MB468 | 51 | 15 | 66 | 91 | 8 | 9.9 | 6.4 | 15.2 | 40.9 | 5.0 |
| >hg38_21_+_8433172_8446622_RNA45SN1_45S_50ntFlanks@10479.10512.34\|CGATCGAAAGGGAGTCGGGTTCAGATCCCCGAAT | >hg38_21_+_8433172_8446622_RNA45SN1_45S_50ntFlanks | 10479 | 10512 | 34 | CGATCGAAAGGGAGTCGGGTTCAGATCCCCGAAT | MB468 | 17 | 4 | 67 | 7 | 3 | 9.8 | 1.9 | 18.9 | 3.4 | 1.5 |
| >hg38_21_+_8433172_8446622_RNA45SN1_45S_50ntFlanks@10481.10498.18\|ATCGAAAGGGAGTCGGGT | >hg38_21_+_8433172_8446622_RNA45SN1_45S_50ntFlanks | 10481 | 10498 | 18 | ATCGAAAGGGAGTCGGGT | MB468 | 14 | 8 | 27 | 12 | 2 | 3.8 | 4.5 | 9.1 | 6.6 | 0.6 |
| >hg38_21_+_8433172_8446622_RNA45SN1_45S_50ntFlanks@10482.10511.30\|TCGAAAGGGAGTCGGGTTCAGATCCCCGAA | >hg38_21_+_8433172_8446622_RNA45SN1_45S_50ntFlanks | 10482 | 10511 | 30 | TCGAAAGGGAGTCGGGTTCAGATCCCCGAA | MB468 | 11 | 1 | 36 | 3 | 1 | 5.2 | 0.4 | 22.4 | 1.7 | 0.4 |
| >hg38_21_+_8433172_8446622_RNA45SN1_45S_50ntFlanks@10482.10512.31\|TCGAAAGGGAGTCGGGTTCAGATCCCCGAAT | >hg38_21_+_8433172_8446622_RNA45SN1_45S_50ntFlanks | 10482 | 10512 | 31 | TCGAAAGGGAGTCGGGTTCAGATCCCCGAAT | MB468 | 19 | 2 | 61 | 5 | 3 | 8.6 | 0.5 | 33.9 | 2.9 | 1.2 |
| >hg38_21_+_8433172_8446622_RNA45SN1_45S_50ntFlanks@10498.10516.19\|TTCAGATCCCCGAATCCGG | >hg38_21_+_8433172_8446622_RNA45SN1_45S_50ntFlanks | 10498 | 10516 | 19 | TTCAGATCCCCGAATCCGG | MB468 | 25 | 7 | 38 | 13 | 4 | 4.0 | 1.7 | 14.4 | 7.4 | 1.7 |
| >hg38_21_+_8433172_8446622_RNA45SN1_45S_50ntFlanks@10498.10521.24\|TTCAGATCCCCGAATCCGGAGTGG | >hg38_21_+_8433172_8446622_RNA45SN1_45S_50ntFlanks | 10498 | 10521 | 24 | TTCAGATCCCCGAATCCGGAGTGG | MB468 | 28 | 8 | 25 | 13 | 4 | 27.8 | 7.1 | 34.0 | 13.4 | 3.7 |
| >hg38_21_+_8433172_8446622_RNA45SN1_45S_50ntFlanks@10498.10524.27\|TTCAGATCCCCGAATCCGGAGTGGCGG | >hg38_21_+_8433172_8446622_RNA45SN1_45S_50ntFlanks | 10498 | 10524 | 27 | TTCAGATCCCCGAATCCGGAGTGGCGG | MB468 | 11 | 10 | 13 | 6 | 3 | 5.9 | 5.0 | 10.6 | 2.7 | 0.3 |
| >hg38_21_+_8433172_8446622_RNA45SN1_45S_50ntFlanks@10508.10533.26\|CGAATCCGGAGTGGCGGAGATGGGCG | >hg38_21_+_8433172_8446622_RNA45SN1_45S_50ntFlanks | 10508 | 10533 | 26 | CGAATCCGGAGTGGCGGAGATGGGCG | MB468 | 17 | 21 | 12 | 84 | 28 | 6.4 | 14.7 | 5.4 | 42.4 | 15.3 |
| >hg38_21_+_8433172_8446622_RNA45SN1_45S_50ntFlanks@10509.10533.25\|GAATCCGGAGTGGCGGAGATGGGCG | >hg38_21_+_8433172_8446622_RNA45SN1_45S_50ntFlanks | 10509 | 10533 | 25 | GAATCCGGAGTGGCGGAGATGGGCG | MB468 | 13 | 19 | 11 | 80 | 15 | 6.6 | 18.5 | 5.3 | 62.0 | 10.8 |
| >hg38_21_+_8433172_8446622_RNA45SN1_45S_50ntFlanks@10510.10533.24\|AATCCGGAGTGGCGGAGATGGGCG | >hg38_21_+_8433172_8446622_RNA45SN1_45S_50ntFlanks | 10510 | 10533 | 24 | AATCCGGAGTGGCGGAGATGGGCG | MB468 | 81 | 138 | 62 | 601 | 145 | 35.1 | 113.2 | 28.0 | 393.2 | 90.9 |
| >hg38_21_+_8433172_8446622_RNA45SN1_45S_50ntFlanks@10511.10533.23\|ATCCGGAGTGGCGGAGATGGGCG | >hg38_21_+_8433172_8446622_RNA45SN1_45S_50ntFlanks | 10511 | 10533 | 23 | ATCCGGAGTGGCGGAGATGGGCG | MB468 | 16 | 28 | 11 | 126 | 28 | 7.0 | 25.8 | 6.0 | 83.3 | 19.9 |
| >hg38_21_+_8433172_8446622_RNA45SN1_45S_50ntFlanks@10512.10533.22\|TCCGGAGTGGCGGAGATGGGCG | >hg38_21_+_8433172_8446622_RNA45SN1_45S_50ntFlanks | 10512 | 10533 | 22 | TCCGGAGTGGCGGAGATGGGCG | MB468 | 114 | 202 | 88 | 814 | 174 | 69.4 | 198.4 | 55.7 | 512.8 | 137.5 |
| >hg38_21_+_8433172_8446622_RNA45SN1_45S_50ntFlanks@10512.10534.23\|TCCGGAGTGGCGGAGATGGGCGC | >hg38_21_+_8433172_8446622_RNA45SN1_45S_50ntFlanks | 10512 | 10534 | 23 | TCCGGAGTGGCGGAGATGGGCGC | MB468 | 18 | 21 | 11 | 106 | 25 | 8.6 | 14.9 | 5.1 | 41.1 | 12.4 |
| >hg38_21_+_8433172_8446622_RNA45SN1_45S_50ntFlanks@10513.10533.21\|CCGGAGTGGCGGAGATGGGCG | >hg38_21_+_8433172_8446622_RNA45SN1_45S_50ntFlanks | 10513 | 10533 | 21 | CCGGAGTGGCGGAGATGGGCG | MB468 | 110 | 217 | 78 | 990 | 263 | 53.7 | 162.3 | 37.6 | 703.3 | 190.3 |
| >hg38_21_+_8433172_8446622_RNA45SN1_45S_50ntFlanks@10513.10534.22\|CCGGAGTGGCGGAGATGGGCGC | >hg38_21_+_8433172_8446622_RNA45SN1_45S_50ntFlanks | 10513 | 10534 | 22 | CCGGAGTGGCGGAGATGGGCGC | MB468 | 18 | 25 | 10 | 157 | 36 | 7.7 | 16.7 | 4.5 | 86.3 | 22.7 |
| >hg38_21_+_8433172_8446622_RNA45SN1_45S_50ntFlanks@10514.10533.20\|CGGAGTGGCGGAGATGGGCG | >hg38_21_+_8433172_8446622_RNA45SN1_45S_50ntFlanks | 10514 | 10533 | 20 | CGGAGTGGCGGAGATGGGCG | MB468 | 24 | 34 | 16 | 164 | 49 | 11.7 | 28.5 | 7.1 | 94.3 | 32.0 |
| >hg38_21_+_8433172_8446622_RNA45SN1_45S_50ntFlanks@10516.10533.18\|GAGTGGCGGAGATGGGCG | >hg38_21_+_8433172_8446622_RNA45SN1_45S_50ntFlanks | 10516 | 10533 | 18 | GAGTGGCGGAGATGGGCG | MB468 | 37 | 35 | 24 | 284 | 61 | 18.0 | 23.8 | 10.2 | 189.8 | 26.1 |
| >hg38_21_+_8433172_8446622_RNA45SN1_45S_50ntFlanks@10534.10570.37\|CCGCGAGGCGTCCAGTGCGGTAACGCGACCGATCCCG | >hg38_21_+_8433172_8446622_RNA45SN1_45S_50ntFlanks | 10534 | 10570 | 37 | CCGCGAGGCGTCCAGTGCGGTAACGCGACCGATCCCG | MB468 | 27 | 11 | 19 | 47 | 85 | 22.1 | 3.8 | 13.3 | 28.7 | 62.9 |
| >hg38_21_+_8433172_8446622_RNA45SN1_45S_50ntFlanks@10571.10608.38\|GAGAAGCCGGCGGGAGCCCCGGGGAGAGTTCTCTTTTC | >hg38_21_+_8433172_8446622_RNA45SN1_45S_50ntFlanks | 10571 | 10608 | 38 | GAGAAGCCGGCGGGAGCCCCGGGGAGAGTTCTCTTTTC | MB468 | 17 | 25 | 30 | 7 | 17 | 6.7 | 12.3 | 9.8 | 2.0 | 16.8 |
| >hg38_21_+_8433172_8446622_RNA45SN1_45S_50ntFlanks@10572.10607.36\|AGAAGCCGGCGGGAGCCCCGGGGAGAGTTCTCTTTT | >hg38_21_+_8433172_8446622_RNA45SN1_45S_50ntFlanks | 10572 | 10607 | 36 | AGAAGCCGGCGGGAGCCCCGGGGAGAGTTCTCTTTT | MB468 | 28 | 52 | 36 | 5 | 18 | 7.0 | 21.2 | 9.1 | 2.3 | 9.2 |
| >hg38_21_+_8433172_8446622_RNA45SN1_45S_50ntFlanks@10572.10608.37\|AGAAGCCGGCGGGAGCCCCGGGGAGAGTTCTCTTTTC | >hg38_21_+_8433172_8446622_RNA45SN1_45S_50ntFlanks | 10572 | 10608 | 37 | AGAAGCCGGCGGGAGCCCCGGGGAGAGTTCTCTTTTC | MB468 | 54 | 89 | 107 | 11 | 31 | 22.2 | 37.4 | 34.8 | 7.3 | 19.3 |
| >hg38_21_+_8433172_8446622_RNA45SN1_45S_50ntFlanks@10574.10607.34\|AAGCCGGCGGGAGCCCCGGGGAGAGTTCTCTTTT | >hg38_21_+_8433172_8446622_RNA45SN1_45S_50ntFlanks | 10574 | 10607 | 34 | AAGCCGGCGGGAGCCCCGGGGAGAGTTCTCTTTT | MB468 | 25 | 24 | 34 | 3 | 12 | 8.7 | 6.0 | 20.1 | 0.7 | 6.8 |
| >hg38_21_+_8433172_8446622_RNA45SN1_45S_50ntFlanks@10574.10608.35\|AAGCCGGCGGGAGCCCCGGGGAGAGTTCTCTTTTC | >hg38_21_+_8433172_8446622_RNA45SN1_45S_50ntFlanks | 10574 | 10608 | 35 | AAGCCGGCGGGAGCCCCGGGGAGAGTTCTCTTTTC | MB468 | 71 | 84 | 132 | 13 | 32 | 31.8 | 31.3 | 55.5 | 7.8 | 12.6 |
| >hg38_21_+_8433172_8446622_RNA45SN1_45S_50ntFlanks@10587.10607.21\|CCCCGGGGAGAGTTCTCTTTT | >hg38_21_+_8433172_8446622_RNA45SN1_45S_50ntFlanks | 10587 | 10607 | 21 | CCCCGGGGAGAGTTCTCTTTT | MB468 | 41 | 20 | 23 | 1 | 13 | 16.5 | 4.9 | 10.2 | 0.7 | 3.5 |
| >hg38_21_+_8433172_8446622_RNA45SN1_45S_50ntFlanks@10587.10608.22\|CCCCGGGGAGAGTTCTCTTTTC | >hg38_21_+_8433172_8446622_RNA45SN1_45S_50ntFlanks | 10587 | 10608 | 22 | CCCCGGGGAGAGTTCTCTTTTC | MB468 | 17 | 13 | 22 | 2 | 8 | 6.0 | 4.5 | 10.0 | 0.9 | 4.7 |
| >hg38_21_+_8433172_8446622_RNA45SN1_45S_50ntFlanks@10588.10607.20\|CCCGGGGAGAGTTCTCTTTT | >hg38_21_+_8433172_8446622_RNA45SN1_45S_50ntFlanks | 10588 | 10607 | 20 | CCCGGGGAGAGTTCTCTTTT | MB468 | 13 | 3 | 10 | 2 | 4 | 2.4 | 1.5 | 2.4 | 0.7 | 1.4 |
| >hg38_21_+_8433172_8446622_RNA45SN1_45S_50ntFlanks@10597.10617.21\|AGTTCTCTTTTCTTTGTGAAG | >hg38_21_+_8433172_8446622_RNA45SN1_45S_50ntFlanks | 10597 | 10617 | 21 | AGTTCTCTTTTCTTTGTGAAG | MB468 | 17 | 9 | 4 | 8 | 1 | 5.1 | 7.3 | 1.2 | 2.5 | 0.4 |
| >hg38_21_+_8433172_8446622_RNA45SN1_45S_50ntFlanks@10664.10706.43\|CTTGGAAAGCGTCGCGGTTCCGGCGGCGTCCGGTGAGCTCTCG | >hg38_21_+_8433172_8446622_RNA45SN1_45S_50ntFlanks | 10664 | 10706 | 43 | CTTGGAAAGCGTCGCGGTTCCGGCGGCGTCCGGTGAGCTCTCG | MB468 | 26 | 108 | 54 | 16 | 71 | 18.1 | 67.3 | 43.4 | 15.8 | 45.8 |
| >hg38_21_+_8433172_8446622_RNA45SN1_45S_50ntFlanks@10668.10706.39\|GAAAGCGTCGCGGTTCCGGCGGCGTCCGGTGAGCTCTCG | >hg38_21_+_8433172_8446622_RNA45SN1_45S_50ntFlanks | 10668 | 10706 | 39 | GAAAGCGTCGCGGTTCCGGCGGCGTCCGGTGAGCTCTCG | MB468 | 28 | 86 | 49 | 12 | 33 | 29.3 | 73.9 | 44.8 | 13.8 | 32.1 |

| | | | | | | | | | | | | | | | | |
|---|---|---|---|---|---|---|---|---|---|---|---|---|---|---|---|---|
| >hg38_21_+_8433172_8446622_RNA45SN1_45S_50ntFlanks@10669.10706.38\|AAAGCGTCGC GGTTCCGGCGGCGTCCGGTGAGCTCTCG | >hg38_21_+_8433172_8446622_RNA45SN1_45S_50ntFlanks | 10669 | 10706 | 38 | AAAGCGTCGCGGTTCCGGCGGCGTCCGGTGAGCTCTCG | MB468 | 24 | 90 | 45 | 10 | 31 | 27.6 | 102.6 | 43.2 | 13.7 | 43.2 |
| >hg38_21_+_8433172_8446622_RNA45SN1_45S_50ntFlanks@10677.10706.30\|GCGGTTCCGG CGGCGTCCGGTGAGCTCTCG | >hg38_21_+_8433172_8446622_RNA45SN1_45S_50ntFlanks | 10677 | 10706 | 30 | GCGGTTCCGGCGGCGTCCGGTGAGCTCTCG | BT20 | 29 | 22 | 6 | 18 | 27 | 47.4 | 36.9 | 9.0 | 20.9 | 40.2 |
| >hg38_21_+_8433172_8446622_RNA45SN1_45S_50ntFlanks@10677.10706.30\|GCGGTTCCGG CGGCGTCCGGTGAGCTCTCG | >hg38_21_+_8433172_8446622_RNA45SN1_45S_50ntFlanks | 10677 | 10706 | 30 | GCGGTTCCGGCGGCGTCCGGTGAGCTCTCG | MB468 | 20 | 13 | 22 | 11 | 13 | 5.4 | 2.4 | 8.4 | 8.0 | 6.6 |
| >hg38_21_+_8433172_8446622_RNA45SN1_45S_50ntFlanks@10677.10707.31\|GCGGTTCCGG CGGCGTCCGGTGAGCTCTCGC | >hg38_21_+_8433172_8446622_RNA45SN1_45S_50ntFlanks | 10677 | 10707 | 31 | GCGGTTCCGGCGGCGTCCGGTGAGCTCTCGC | BT20 | 15 | 21 | 4 | 32 | 42 | 22.6 | 34.8 | 5.7 | 38.3 | 64.2 |
| >hg38_21_+_8433172_8446622_RNA45SN1_45S_50ntFlanks@10677.10708.32\|GCGGTTCCGG CGGCGTCCGGTGAGCTCTCGCT | >hg38_21_+_8433172_8446622_RNA45SN1_45S_50ntFlanks | 10677 | 10708 | 32 | GCGGTTCCGGCGGCGTCCGGTGAGCTCTCGCT | BT20 | 46 | 41 | 11 | 24 | 64 | 72.1 | 65.3 | 15.9 | 25.5 | 92.5 |
| >hg38_21_+_8433172_8446622_RNA45SN1_45S_50ntFlanks@10677.10708.32\|GCGGTTCCGG CGGCGTCCGGTGAGCTCTCGCT | >hg38_21_+_8433172_8446622_RNA45SN1_45S_50ntFlanks | 10677 | 10708 | 32 | GCGGTTCCGGCGGCGTCCGGTGAGCTCTCGCT | MB231 | 11 | 7 | 4 | 5 | 13 | 5.8 | 2.6 | 3.2 | 2.3 | 2.6 |
| >hg38_21_+_8433172_8446622_RNA45SN1_45S_50ntFlanks@10677.10708.32\|GCGGTTCCGG CGGCGTCCGGTGAGCTCTCGCT | >hg38_21_+_8433172_8446622_RNA45SN1_45S_50ntFlanks | 10677 | 10708 | 32 | GCGGTTCCGGCGGCGTCCGGTGAGCTCTCGCT | MB468 | 11 | 31 | 13 | 3 | 15 | 2.1 | 9.2 | 5.8 | 2.4 | 5.4 |
| >hg38_21_+_8433172_8446622_RNA45SN1_45S_50ntFlanks@10677.10709.33\|GCGGTTCCGG CGGCGTCCGGTGAGCTCTCGCTG | >hg38_21_+_8433172_8446622_RNA45SN1_45S_50ntFlanks | 10677 | 10709 | 33 | GCGGTTCCGGCGGCGTCCGGTGAGCTCTCGCTG | BT20 | 48 | 53 | 13 | 8 | 89 | 75.9 | 86.6 | 21.0 | 5.9 | 132.9 |
| >hg38_21_+_8433172_8446622_RNA45SN1_45S_50ntFlanks@10677.10709.33\|GCGGTTCCGG CGGCGTCCGGTGAGCTCTCGCTG | >hg38_21_+_8433172_8446622_RNA45SN1_45S_50ntFlanks | 10677 | 10709 | 33 | GCGGTTCCGGCGGCGTCCGGTGAGCTCTCGCTG | MB468 | 26 | 50 | 26 | 7 | 33 | 8.6 | 22.6 | 7.9 | 2.8 | 19.8 |
| >hg38_21_+_8433172_8446622_RNA45SN1_45S_50ntFlanks@10677.10710.34\|GCGGTTCCGG CGGCGTCCGGTGAGCTCTCGCTGG | >hg38_21_+_8433172_8446622_RNA45SN1_45S_50ntFlanks | 10677 | 10710 | 34 | GCGGTTCCGGCGGCGTCCGGTGAGCTCTCGCTGG | BT20 | 33 | 61 | 15 | 13 | 150 | 49.4 | 99.7 | 20.6 | 8.1 | 232.2 |
| >hg38_21_+_8433172_8446622_RNA45SN1_45S_50ntFlanks@10677.10710.34\|GCGGTTCCGG CGGCGTCCGGTGAGCTCTCGCTGG | >hg38_21_+_8433172_8446622_RNA45SN1_45S_50ntFlanks | 10677 | 10710 | 34 | GCGGTTCCGGCGGCGTCCGGTGAGCTCTCGCTGG | MB468 | 116 | 347 | 146 | 31 | 160 | 8.6 | 139.3 | 81.0 | 26.0 | 63.1 |
| >hg38_21_+_8433172_8446622_RNA45SN1_45S_50ntFlanks@10677.10711.35\|GCGGTTCCGG CGGCGTCCGGTGAGCTCTCGCTGGC | >hg38_21_+_8433172_8446622_RNA45SN1_45S_50ntFlanks | 10677 | 10711 | 35 | GCGGTTCCGGCGGCGTCCGGTGAGCTCTCGCTGGC | MB468 | 31 | 93 | 38 | 7 | 62 | 5.4 | 66.7 | 16.4 | 3.9 | 28.6 |
| >hg38_21_+_8433172_8446622_RNA45SN1_45S_50ntFlanks@10677.10712.36\|GCGGTTCCGG CGGCGTCCGGTGAGCTCTCGCTGGCC | >hg38_21_+_8433172_8446622_RNA45SN1_45S_50ntFlanks | 10677 | 10712 | 36 | GCGGTTCCGGCGGCGTCCGGTGAGCTCTCGCTGGCC | BT20 | 12 | 25 | 7 | 11 | 57 | 13.7 | 35.9 | 6.9 | 8.0 | 74.1 |
| >hg38_21_+_8433172_8446622_RNA45SN1_45S_50ntFlanks@10677.10712.36\|GCGGTTCCGG CGGCGTCCGGTGAGCTCTCGCTGGCC | >hg38_21_+_8433172_8446622_RNA45SN1_45S_50ntFlanks | 10677 | 10712 | 36 | GCGGTTCCGGCGGCGTCCGGTGAGCTCTCGCTGGCC | MB231 | 16 | 17 | 16 | 6 | 33 | 17.1 | 8.1 | 13.2 | 1.8 | 22.1 |
| >hg38_21_+_8433172_8446622_RNA45SN1_45S_50ntFlanks@10677.10712.36\|GCGGTTCCGG CGGCGTCCGGTGAGCTCTCGCTGGCC | >hg38_21_+_8433172_8446622_RNA45SN1_45S_50ntFlanks | 10677 | 10712 | 36 | GCGGTTCCGGCGGCGTCCGGTGAGCTCTCGCTGGCC | MB468 | 22 | 67 | 36 | 10 | 51 | 4.5 | 55.0 | 21.1 | 5.7 | 31.9 |
| >hg38_21_+_8433172_8446622_RNA45SN1_45S_50ntFlanks@10678.10695.18\|CGGTTCCGGC GGCGTCCG | >hg38_21_+_8433172_8446622_RNA45SN1_45S_50ntFlanks | 10678 | 10695 | 18 | CGGTTCCGGCGGCGTCCG | MB468 | 13 | 10 | 12 | 12 | 9 | 3.4 | 4.5 | 2.5 | 4.0 | 4.2 |
| >hg38_21_+_8433172_8446622_RNA45SN1_45S_50ntFlanks@10678.10696.19\|CGGTTCCGGC GGCGTCCGG | >hg38_21_+_8433172_8446622_RNA45SN1_45S_50ntFlanks | 10678 | 10696 | 19 | CGGTTCCGGCGGCGTCCGG | MB468 | 10 | 1 | 19 | 16 | 5 | 1.0 | 0.3 | 5.5 | 9.2 | 1.2 |
| >hg38_21_+_8433172_8446622_RNA45SN1_45S_50ntFlanks@10678.10705.28\|CGGTTCCGGC GGCGTCCGGTGAGCTCTC | >hg38_21_+_8433172_8446622_RNA45SN1_45S_50ntFlanks | 10678 | 10705 | 28 | CGGTTCCGGCGGCGTCCGGTGAGCTCTC | MB468 | 63 | 47 | 92 | 74 | 53 | 22.2 | 14.2 | 47.7 | 18.5 | 34.8 |
| >hg38_21_+_8433172_8446622_RNA45SN1_45S_50ntFlanks@10678.10706.29\|CGGTTCCGGC GGCGTCCGGTGAGCTCTCG | >hg38_21_+_8433172_8446622_RNA45SN1_45S_50ntFlanks | 10678 | 10706 | 29 | CGGTTCCGGCGGCGTCCGGTGAGCTCTCG | BT20 | 15 | 12 | 14 | 14 | 16 | 24.1 | 20.9 | 15.3 | 10.8 | 22.7 |
| >hg38_21_+_8433172_8446622_RNA45SN1_45S_50ntFlanks@10678.10706.29\|CGGTTCCGGC GGCGTCCGGTGAGCTCTCG | >hg38_21_+_8433172_8446622_RNA45SN1_45S_50ntFlanks | 10678 | 10706 | 29 | CGGTTCCGGCGGCGTCCGGTGAGCTCTCG | MB468 | 2241 | 1184 | 1691 | 1979 | 1784 | 754.4 | 195.4 | 733.6 | 714.7 | 476.9 |
| >hg38_21_+_8433172_8446622_RNA45SN1_45S_50ntFlanks@10678.10707.30\|CGGTTCCGGC GGCGTCCGGTGAGCTCTCGC | >hg38_21_+_8433172_8446622_RNA45SN1_45S_50ntFlanks | 10678 | 10707 | 30 | CGGTTCCGGCGGCGTCCGGTGAGCTCTCGC | MB468 | 36 | 49 | 30 | 26 | 75 | 9.4 | 8.0 | 15.2 | 18.9 | 33.7 |
| >hg38_21_+_8433172_8446622_RNA45SN1_45S_50ntFlanks@10678.10708.31\|CGGTTCCGGC GGCGTCCGGTGAGCTCTCGCT | >hg38_21_+_8433172_8446622_RNA45SN1_45S_50ntFlanks | 10678 | 10708 | 31 | CGGTTCCGGCGGCGTCCGGTGAGCTCTCGCT | BT20 | 44 | 31 | 27 | 26 | 48 | 68.8 | 50.7 | 26.1 | 23.6 | 69.6 |
| >hg38_21_+_8433172_8446622_RNA45SN1_45S_50ntFlanks@10678.10708.31\|CGGTTCCGGC GGCGTCCGGTGAGCTCTCGCT | >hg38_21_+_8433172_8446622_RNA45SN1_45S_50ntFlanks | 10678 | 10708 | 31 | CGGTTCCGGCGGCGTCCGGTGAGCTCTCGCT | MB468 | 1289 | 2547 | 898 | 512 | 2302 | 366.9 | 1121.0 | 333.4 | 402.7 | 790.8 |
| >hg38_21_+_8433172_8446622_RNA45SN1_45S_50ntFlanks@10678.10709.32\|CGGTTCCGGC GGCGTCCGGTGAGCTCTCGCTG | >hg38_21_+_8433172_8446622_RNA45SN1_45S_50ntFlanks | 10678 | 10709 | 32 | CGGTTCCGGCGGCGTCCGGTGAGCTCTCGCTG | BT20 | 23 | 20 | 20 | 15 | 39 | 36.2 | 33.4 | 20.1 | 20.6 | 58.8 |
| >hg38_21_+_8433172_8446622_RNA45SN1_45S_50ntFlanks@10678.10709.32\|CGGTTCCGGC GGCGTCCGGTGAGCTCTCGCTG | >hg38_21_+_8433172_8446622_RNA45SN1_45S_50ntFlanks | 10678 | 10709 | 32 | CGGTTCCGGCGGCGTCCGGTGAGCTCTCGCTG | MB468 | 2619 | 4437 | 1570 | 916 | 4527 | 859.9 | 3824.4 | 692.9 | 339.8 | 1961.2 |
| >hg38_21_+_8433172_8446622_RNA45SN1_45S_50ntFlanks@10678.10710.33\|CGGTTCCGGC GGCGTCCGGTGAGCTCTCGCTGG | >hg38_21_+_8433172_8446622_RNA45SN1_45S_50ntFlanks | 10678 | 10710 | 33 | CGGTTCCGGCGGCGTCCGGTGAGCTCTCGCTGG | BT20 | 40 | 64 | 158 | 138 | 115 | 62.1 | 103.4 | 216.1 | 211.9 | 167.4 |
| >hg38_21_+_8433172_8446622_RNA45SN1_45S_50ntFlanks@10678.10710.33\|CGGTTCCGGC GGCGTCCGGTGAGCTCTCGCTGG | >hg38_21_+_8433172_8446622_RNA45SN1_45S_50ntFlanks | 10678 | 10710 | 33 | CGGTTCCGGCGGCGTCCGGTGAGCTCTCGCTGG | MB468 | 20587 | 25857 | 13922 | 8833 | 34998 | 2244.8 | 5392.1 | 3114.5 | 5482.7 | 13027.4 |
| >hg38_21_+_8433172_8446622_RNA45SN1_45S_50ntFlanks@10678.10711.34\|CGGTTCCGGC GGCGTCCGGTGAGCTCTCGCTGGC | >hg38_21_+_8433172_8446622_RNA45SN1_45S_50ntFlanks | 10678 | 10711 | 34 | CGGTTCCGGCGGCGTCCGGTGAGCTCTCGCTGGC | MB468 | 4006 | 6323 | 3043 | 1073 | 8692 | 336.3 | 3814.1 | 1107.1 | 550.1 | 3247.5 |
| >hg38_21_+_8433172_8446622_RNA45SN1_45S_50ntFlanks@10678.10712.35\|CGGTTCCGGC GGCGTCCGGTGAGCTCTCGCTGGCC | >hg38_21_+_8433172_8446622_RNA45SN1_45S_50ntFlanks | 10678 | 10712 | 35 | CGGTTCCGGCGGCGTCCGGTGAGCTCTCGCTGGCC | MB468 | 849 | 1354 | 723 | 976 | 2223 | 205.3 | 821.9 | 292.8 | 336.3 | 1263.9 |
| >hg38_21_+_8433172_8446622_RNA45SN1_45S_50ntFlanks@10679.10705.27\|GGTTCCGGCG GCGTCCGGTGAGCTCTC | >hg38_21_+_8433172_8446622_RNA45SN1_45S_50ntFlanks | 10679 | 10705 | 27 | GGTTCCGGCGGCGTCCGGTGAGCTCTC | MB468 | 33 | 18 | 45 | 43 | 25 | 12.0 | 6.1 | 26.3 | 12.9 | 18.8 |
| >hg38_21_+_8433172_8446622_RNA45SN1_45S_50ntFlanks@10679.10706.28\|GGTTCCGGCG GCGTCCGGTGAGCTCTCG | >hg38_21_+_8433172_8446622_RNA45SN1_45S_50ntFlanks | 10679 | 10706 | 28 | GGTTCCGGCGGCGTCCGGTGAGCTCTCG | MB468 | 567 | 238 | 378 | 592 | 369 | 170.0 | 31.4 | 114.1 | 159.3 | 220.1 |
| >hg38_21_+_8433172_8446622_RNA45SN1_45S_50ntFlanks@10679.10707.29\|GGTTCCGGCG GCGTCCGGTGAGCTCTCGC | >hg38_21_+_8433172_8446622_RNA45SN1_45S_50ntFlanks | 10679 | 10707 | 29 | GGTTCCGGCGGCGTCCGGTGAGCTCTCGC | MB468 | 19 | 10 | 10 | 9 | 17 | 5.7 | 4.0 | 2.5 | 3.6 | 9.0 |
| >hg38_21_+_8433172_8446622_RNA45SN1_45S_50ntFlanks@10679.10708.30\|GGTTCCGGCG GCGTCCGGTGAGCTCTCGCT | >hg38_21_+_8433172_8446622_RNA45SN1_45S_50ntFlanks | 10679 | 10708 | 30 | GGTTCCGGCGGCGTCCGGTGAGCTCTCGCT | MB468 | 651 | 638 | 323 | 155 | 769 | 63.2 | 181.1 | 79.0 | 115.6 | 219.2 |
| >hg38_21_+_8433172_8446622_RNA45SN1_45S_50ntFlanks@10679.10709.31\|GGTTCCGGCG GCGTCCGGTGAGCTCTCGCTG | >hg38_21_+_8433172_8446622_RNA45SN1_45S_50ntFlanks | 10679 | 10709 | 31 | GGTTCCGGCGGCGTCCGGTGAGCTCTCGCTG | MB468 | 1044 | 746 | 420 | 268 | 1212 | 267.2 | 473.2 | 176.2 | 78.9 | 619.3 |
| >hg38_21_+_8433172_8446622_RNA45SN1_45S_50ntFlanks@10679.10710.32\|GGTTCCGGCG GCGTCCGGTGAGCTCTCGCTGG | >hg38_21_+_8433172_8446622_RNA45SN1_45S_50ntFlanks | 10679 | 10710 | 32 | GGTTCCGGCGGCGTCCGGTGAGCTCTCGCTGG | MB468 | 4099 | 2843 | 2264 | 1460 | 4434 | 382.4 | 627.0 | 272.5 | 634.8 | 1657.1 |
| >hg38_21_+_8433172_8446622_RNA45SN1_45S_50ntFlanks@10679.10711.33\|GGTTCCGGCG GCGTCCGGTGAGCTCTCGCTGGC | >hg38_21_+_8433172_8446622_RNA45SN1_45S_50ntFlanks | 10679 | 10711 | 33 | GGTTCCGGCGGCGTCCGGTGAGCTCTCGCTGGC | MB468 | 735 | 598 | 424 | 193 | 1076 | 152.7 | 457.5 | 100.2 | 60.0 | 595.3 |
| >hg38_21_+_8433172_8446622_RNA45SN1_45S_50ntFlanks@10679.10712.34\|GGTTCCGGCG GCGTCCGGTGAGCTCTCGCTGGCC | >hg38_21_+_8433172_8446622_RNA45SN1_45S_50ntFlanks | 10679 | 10712 | 34 | GGTTCCGGCGGCGTCCGGTGAGCTCTCGCTGGCC | MB468 | 36 | 38 | 28 | 153 | 62 | 19.4 | 28.1 | 7.2 | 101.1 | 53.7 |
| >hg38_21_+_8433172_8446622_RNA45SN1_45S_50ntFlanks@10680.10705.26\|GTTCCGGCGG CGTCCGGTGAGCTCTC | >hg38_21_+_8433172_8446622_RNA45SN1_45S_50ntFlanks | 10680 | 10705 | 26 | GTTCCGGCGGCGTCCGGTGAGCTCTC | MB468 | 39 | 49 | 75 | 34 | 20 | 9.1 | 3.6 | 31.4 | 7.6 | 9.6 |
| >hg38_21_+_8433172_8446622_RNA45SN1_45S_50ntFlanks@10680.10706.27\|GTTCCGGCGG CGTCCGGTGAGCTCTCG | >hg38_21_+_8433172_8446622_RNA45SN1_45S_50ntFlanks | 10680 | 10706 | 27 | GTTCCGGCGGCGTCCGGTGAGCTCTCG | MB468 | 985 | 1070 | 1039 | 506 | 504 | 357.7 | 444.6 | 618.1 | 234.6 | 103.9 |
| >hg38_21_+_8433172_8446622_RNA45SN1_45S_50ntFlanks@10680.10707.28\|GTTCCGGCGG CGTCCGGTGAGCTCTCGC | >hg38_21_+_8433172_8446622_RNA45SN1_45S_50ntFlanks | 10680 | 10707 | 28 | GTTCCGGCGGCGTCCGGTGAGCTCTCGC | MB468 | 12 | 9 | 11 | 7 | 10 | 0.9 | 0.9 | 2.5 | 3.7 | 1.5 |
| >hg38_21_+_8433172_8446622_RNA45SN1_45S_50ntFlanks@10680.10708.29\|GTTCCGGCGG CGTCCGGTGAGCTCTCGCT | >hg38_21_+_8433172_8446622_RNA45SN1_45S_50ntFlanks | 10680 | 10708 | 29 | GTTCCGGCGGCGTCCGGTGAGCTCTCGCT | BT20 | 11 | 10 | 15 | 27 | 21 | 15.3 | 13.8 | 18.3 | 38.3 | 25.0 |
| >hg38_21_+_8433172_8446622_RNA45SN1_45S_50ntFlanks@10680.10708.29\|GTTCCGGCGG CGTCCGGTGAGCTCTCGCT | >hg38_21_+_8433172_8446622_RNA45SN1_45S_50ntFlanks | 10680 | 10708 | 29 | GTTCCGGCGGCGTCCGGTGAGCTCTCGCT | MB468 | 1802 | 1541 | 1268 | 482 | 1684 | 496.5 | 409.6 | 446.1 | 352.3 | 389.1 |
| >hg38_21_+_8433172_8446622_RNA45SN1_45S_50ntFlanks@10680.10709.30\|GTTCCGGCGG CGTCCGGTGAGCTCTCGCTG | >hg38_21_+_8433172_8446622_RNA45SN1_45S_50ntFlanks | 10680 | 10709 | 30 | GTTCCGGCGGCGTCCGGTGAGCTCTCGCTG | MB468 | 582 | 208 | 287 | 160 | 447 | 85.6 | 82.7 | 69.1 | 35.4 | 88.5 |
| >hg38_21_+_8433172_8446622_RNA45SN1_45S_50ntFlanks@10680.10710.31\|GTTCCGGCGG CGTCCGGTGAGCTCTCGCTGG | >hg38_21_+_8433172_8446622_RNA45SN1_45S_50ntFlanks | 10680 | 10710 | 31 | GTTCCGGCGGCGTCCGGTGAGCTCTCGCTGG | MB468 | 2203 | 1108 | 1704 | 822 | 1712 | 696.4 | 425.2 | 896.5 | 146.1 | 394.4 |
| >hg38_21_+_8433172_8446622_RNA45SN1_45S_50ntFlanks@10680.10711.32\|GTTCCGGCGG CGTCCGGTGAGCTCTCGCTGGC | >hg38_21_+_8433172_8446622_RNA45SN1_45S_50ntFlanks | 10680 | 10711 | 32 | GTTCCGGCGGCGTCCGGTGAGCTCTCGCTGGC | MB468 | 944 | 695 | 926 | 185 | 882 | 353.0 | 442.4 | 645.8 | 73.5 | 182.4 |
| >hg38_21_+_8433172_8446622_RNA45SN1_45S_50ntFlanks@10680.10712.33\|GTTCCGGCGG CGTCCGGTGAGCTCTCGCTGGCC | >hg38_21_+_8433172_8446622_RNA45SN1_45S_50ntFlanks | 10680 | 10712 | 33 | GTTCCGGCGGCGTCCGGTGAGCTCTCGCTGGCC | MB468 | 365 | 472 | 431 | 147 | 445 | 67.3 | 208.0 | 203.7 | 43.7 | 163.7 |
| >hg38_21_+_8433172_8446622_RNA45SN1_45S_50ntFlanks@10681.10706.26\|TTCCGGCGGC GTCCGGTGAGCTCTCG | >hg38_21_+_8433172_8446622_RNA45SN1_45S_50ntFlanks | 10681 | 10706 | 26 | TTCCGGCGGCGTCCGGTGAGCTCTCG | MB468 | 219 | 279 | 234 | 71 | 85 | 71.5 | 117.5 | 158.5 | 30.3 | 21.9 |
| >hg38_21_+_8433172_8446622_RNA45SN1_45S_50ntFlanks@10681.10708.28\|TTCCGGCGGC GTCCGGTGAGCTCTCGCT | >hg38_21_+_8433172_8446622_RNA45SN1_45S_50ntFlanks | 10681 | 10708 | 28 | TTCCGGCGGCGTCCGGTGAGCTCTCGCT | MB468 | 47 | 22 | 28 | 9 | 30 | 17.8 | 11.4 | 15.9 | 2.4 | 4.7 |
| >hg38_21_+_8433172_8446622_RNA45SN1_45S_50ntFlanks@10681.10709.29\|TTCCGGCGGC GTCCGGTGAGCTCTCGCTG | >hg38_21_+_8433172_8446622_RNA45SN1_45S_50ntFlanks | 10681 | 10709 | 29 | TTCCGGCGGCGTCCGGTGAGCTCTCGCTG | MB468 | 57 | 9 | 18 | 13 | 31 | 13.8 | 3.6 | 6.9 | 2.5 | 9.0 |
| >hg38_21_+_8433172_8446622_RNA45SN1_45S_50ntFlanks@10681.10710.30\|TTCCGGCGGC GTCCGGTGAGCTCTCGCTGG | >hg38_21_+_8433172_8446622_RNA45SN1_45S_50ntFlanks | 10681 | 10710 | 30 | TTCCGGCGGCGTCCGGTGAGCTCTCGCTGG | MB468 | 265 | 84 | 169 | 85 | 142 | 121.3 | 47.6 | 139.0 | 28.3 | 31.6 |

| | | | | | | | | | | | | | | | | |
|---|---|---|---|---|---|---|---|---|---|---|---|---|---|---|---|---|
| >hg38_21_+_8433172_8446622_RNA45SN1_45S_50ntFlanks@10681.10711.31\|TTCCGGCGGCGTCCGGTGAGCTCTCGCTGGC | >hg38_21_+_8433172_8446622_RNA45SN1_45S_50ntFlanks | 10681 | 10711 | 31 | TTCCGGCGGCGTCCGGTGAGCTCTCGCTGGC | MB468 | 112 | 75 | 116 | 16 | 72 | 54.5 | 50.2 | 98.0 | 2.2 | 23.2 |
| >hg38_21_+_8433172_8446622_RNA45SN1_45S_50ntFlanks@10681.10712.32\|TTCCGGCGGCGTCCGGTGAGCTCTCGCTGGCC | >hg38_21_+_8433172_8446622_RNA45SN1_45S_50ntFlanks | 10681 | 10712 | 32 | TTCCGGCGGCGTCCGGTGAGCTCTCGCTGGCC | MB468 | 159 | 219 | 185 | 23 | 133 | 35.7 | 102.5 | 97.5 | 4.7 | 52.0 |
| >hg38_21_+_8433172_8446622_RNA45SN1_45S_50ntFlanks@10682.10706.25\|TCCGGCGGCGTCCGGTGAGCTCTCG | >hg38_21_+_8433172_8446622_RNA45SN1_45S_50ntFlanks | 10682 | 10706 | 25 | TCCGGCGGCGTCCGGTGAGCTCTCG | MB468 | 148 | 201 | 168 | 58 | 99 | 53.1 | 104.1 | 114.4 | 33.1 | 36.2 |
| >hg38_21_+_8433172_8446622_RNA45SN1_45S_50ntFlanks@10682.10709.28\|TCCGGCGGCGTCCGGTGAGCTCTCGCTG | >hg38_21_+_8433172_8446622_RNA45SN1_45S_50ntFlanks | 10682 | 10709 | 28 | TCCGGCGGCGTCCGGTGAGCTCTCGCTG | MB468 | 13 | 1 | 3 | 4 | 8 | 5.2 | 0.2 | 1.5 | 0.7 | 3.6 |
| >hg38_21_+_8433172_8446622_RNA45SN1_45S_50ntFlanks@10682.10710.29\|TCCGGCGGCGTCCGGTGAGCTCTCGCTGG | >hg38_21_+_8433172_8446622_RNA45SN1_45S_50ntFlanks | 10682 | 10710 | 29 | TCCGGCGGCGTCCGGTGAGCTCTCGCTGG | MB468 | 20 | 5 | 12 | 9 | 13 | 11.1 | 2.6 | 9.7 | 3.0 | 4.1 |
| >hg38_21_+_8433172_8446622_RNA45SN1_45S_50ntFlanks@10682.10711.30\|TCCGGCGGCGTCCGGTGAGCTCTCGCTGGC | >hg38_21_+_8433172_8446622_RNA45SN1_45S_50ntFlanks | 10682 | 10711 | 30 | TCCGGCGGCGTCCGGTGAGCTCTCGCTGGC | MB468 | 17 | 9 | 17 | 5 | 14 | 7.2 | 4.7 | 14.1 | 1.2 | 4.0 |
| >hg38_21_+_8433172_8446622_RNA45SN1_45S_50ntFlanks@10682.10712.31\|TCCGGCGGCGTCCGGTGAGCTCTCGCTGGCC | >hg38_21_+_8433172_8446622_RNA45SN1_45S_50ntFlanks | 10682 | 10712 | 31 | TCCGGCGGCGTCCGGTGAGCTCTCGCTGGCC | MB468 | 25 | 27 | 30 | 8 | 29 | 6.7 | 9.0 | 17.7 | 2.9 | 8.4 |
| >hg38_21_+_8433172_8446622_RNA45SN1_45S_50ntFlanks@10683.10705.23\|CCGGCGGCGTCCGGTGAGCTCTC | >hg38_21_+_8433172_8446622_RNA45SN1_45S_50ntFlanks | 10683 | 10705 | 23 | CCGGCGGCGTCCGGTGAGCTCTC | MB468 | 46 | 72 | 61 | 32 | 53 | 12.0 | 7.7 | 21.9 | 6.2 | 16.1 |
| >hg38_21_+_8433172_8446622_RNA45SN1_45S_50ntFlanks@10683.10706.24\|CCGGCGGCGTCCGGTGAGCTCTCG | >hg38_21_+_8433172_8446622_RNA45SN1_45S_50ntFlanks | 10683 | 10706 | 24 | CCGGCGGCGTCCGGTGAGCTCTCG | MB468 | 687 | 979 | 693 | 424 | 742 | 186.2 | 320.7 | 393.7 | 204.6 | 139.6 |
| >hg38_21_+_8433172_8446622_RNA45SN1_45S_50ntFlanks@10683.10708.26\|CCGGCGGCGTCCGGTGAGCTCTCGCT | >hg38_21_+_8433172_8446622_RNA45SN1_45S_50ntFlanks | 10683 | 10708 | 26 | CCGGCGGCGTCCGGTGAGCTCTCGCT | MB468 | 25 | 8 | 15 | 13 | 23 | 8.9 | 3.9 | 7.1 | 4.0 | 6.2 |
| >hg38_21_+_8433172_8446622_RNA45SN1_45S_50ntFlanks@10683.10709.27\|CCGGCGGCGTCCGGTGAGCTCTCGCTG | >hg38_21_+_8433172_8446622_RNA45SN1_45S_50ntFlanks | 10683 | 10709 | 27 | CCGGCGGCGTCCGGTGAGCTCTCGCTG | MB468 | 174 | 21 | 48 | 56 | 155 | 63.0 | 6.8 | 14.9 | 15.6 | 60.8 |
| >hg38_21_+_8433172_8446622_RNA45SN1_45S_50ntFlanks@10683.10710.28\|CCGGCGGCGTCCGGTGAGCTCTCGCTGG | >hg38_21_+_8433172_8446622_RNA45SN1_45S_50ntFlanks | 10683 | 10710 | 28 | CCGGCGGCGTCCGGTGAGCTCTCGCTGG | MB468 | 40 | 10 | 26 | 23 | 31 | 14.4 | 5.5 | 20.9 | 7.8 | 9.9 |
| >hg38_21_+_8433172_8446622_RNA45SN1_45S_50ntFlanks@10683.10711.29\|CCGGCGGCGTCCGGTGAGCTCTCGCTGGC | >hg38_21_+_8433172_8446622_RNA45SN1_45S_50ntFlanks | 10683 | 10711 | 29 | CCGGCGGCGTCCGGTGAGCTCTCGCTGGC | MB468 | 137 | 72 | 139 | 66 | 150 | 53.8 | 34.3 | 92.6 | 9.8 | 47.8 |
| >hg38_21_+_8433172_8446622_RNA45SN1_45S_50ntFlanks@10683.10712.30\|CCGGCGGCGTCCGGTGAGCTCTCGCTGGCC | >hg38_21_+_8433172_8446622_RNA45SN1_45S_50ntFlanks | 10683 | 10712 | 30 | CCGGCGGCGTCCGGTGAGCTCTCGCTGGCC | MB468 | 158 | 157 | 185 | 60 | 180 | 30.6 | 76.8 | 91.6 | 17.2 | 52.9 |
| >hg38_21_+_8433172_8446622_RNA45SN1_45S_50ntFlanks@10684.10702.19\|CGGCGGCGTCCGGTGAGCT | >hg38_21_+_8433172_8446622_RNA45SN1_45S_50ntFlanks | 10684 | 10702 | 19 | CGGCGGCGTCCGGTGAGCT | MB468 | 21 | 10 | 9 | 11 | 10 | 13.2 | 6.7 | 5.3 | 5.9 | 3.3 |
| >hg38_21_+_8433172_8446622_RNA45SN1_45S_50ntFlanks@10684.10705.22\|CGGCGGCGTCCGGTGAGCTCTC | >hg38_21_+_8433172_8446622_RNA45SN1_45S_50ntFlanks | 10684 | 10705 | 22 | CGGCGGCGTCCGGTGAGCTCTC | MB468 | 23 | 36 | 21 | 13 | 18 | 7.7 | 10.8 | 7.7 | 3.6 | 4.6 |
| >hg38_21_+_8433172_8446622_RNA45SN1_45S_50ntFlanks@10684.10706.23\|CGGCGGCGTCCGGTGAGCTCTCG | >hg38_21_+_8433172_8446622_RNA45SN1_45S_50ntFlanks | 10684 | 10706 | 23 | CGGCGGCGTCCGGTGAGCTCTCG | MB468 | 580 | 777 | 468 | 262 | 509 | 212.0 | 415.6 | 205.4 | 120.2 | 164.2 |
| >hg38_21_+_8433172_8446622_RNA45SN1_45S_50ntFlanks@10684.10708.25\|CGGCGGCGTCCGGTGAGCTCTCGCT | >hg38_21_+_8433172_8446622_RNA45SN1_45S_50ntFlanks | 10684 | 10708 | 25 | CGGCGGCGTCCGGTGAGCTCTCGCT | MB468 | 71 | 66 | 52 | 26 | 76 | 13.2 | 9.5 | 14.6 | 12.2 | 18.3 |
| >hg38_21_+_8433172_8446622_RNA45SN1_45S_50ntFlanks@10684.10709.26\|CGGCGGCGTCCGGTGAGCTCTCGCTG | >hg38_21_+_8433172_8446622_RNA45SN1_45S_50ntFlanks | 10684 | 10709 | 26 | CGGCGGCGTCCGGTGAGCTCTCGCTG | MB468 | 269 | 79 | 133 | 88 | 224 | 65.4 | 18.9 | 11.1 | 6.5 | 63.5 |
| >hg38_21_+_8433172_8446622_RNA45SN1_45S_50ntFlanks@10684.10710.27\|CGGCGGCGTCCGGTGAGCTCTCGCTGG | >hg38_21_+_8433172_8446622_RNA45SN1_45S_50ntFlanks | 10684 | 10710 | 27 | CGGCGGCGTCCGGTGAGCTCTCGCTGG | MB468 | 883 | 861 | 747 | 195 | 785 | 135.7 | 189.3 | 319.7 | 62.0 | 181.9 |
| >hg38_21_+_8433172_8446622_RNA45SN1_45S_50ntFlanks@10684.10711.28\|CGGCGGCGTCCGGTGAGCTCTCGCTGGC | >hg38_21_+_8433172_8446622_RNA45SN1_45S_50ntFlanks | 10684 | 10711 | 28 | CGGCGGCGTCCGGTGAGCTCTCGCTGGC | MB468 | 326 | 439 | 416 | 89 | 350 | 59.4 | 135.0 | 225.4 | 19.9 | 37.9 |
| >hg38_21_+_8433172_8446622_RNA45SN1_45S_50ntFlanks@10684.10712.29\|CGGCGGCGTCCGGTGAGCTCTCGCTGGCC | >hg38_21_+_8433172_8446622_RNA45SN1_45S_50ntFlanks | 10684 | 10712 | 29 | CGGCGGCGTCCGGTGAGCTCTCGCTGGCC | MB468 | 37 | 41 | 50 | 25 | 38 | 7.0 | 23.2 | 24.8 | 9.5 | 10.5 |
| >hg38_21_+_8433172_8446622_RNA45SN1_45S_50ntFlanks@10685.10706.22\|GGCGGCGTCCGGTGAGCTCTCG | >hg38_21_+_8433172_8446622_RNA45SN1_45S_50ntFlanks | 10685 | 10706 | 22 | GGCGGCGTCCGGTGAGCTCTCG | MB468 | 20 | 35 | 17 | 12 | 15 | 7.8 | 21.9 | 7.6 | 6.4 | 5.8 |
| >hg38_21_+_8433172_8446622_RNA45SN1_45S_50ntFlanks@10685.10708.24\|GGCGGCGTCCGGTGAGCTCTCGCT | >hg38_21_+_8433172_8446622_RNA45SN1_45S_50ntFlanks | 10685 | 10708 | 24 | GGCGGCGTCCGGTGAGCTCTCGCT | BT20 | 10 | 10 | 3 | 14 | 32 | 12.7 | 14.4 | 2.9 | 5.2 | 37.4 |
| >hg38_21_+_8433172_8446622_RNA45SN1_45S_50ntFlanks@10685.10709.25\|GGCGGCGTCCGGTGAGCTCTCGCTG | >hg38_21_+_8433172_8446622_RNA45SN1_45S_50ntFlanks | 10685 | 10709 | 25 | GGCGGCGTCCGGTGAGCTCTCGCTG | BT20 | 11 | 10 | 3 | 5 | 27 | 15.6 | 16.2 | 4.5 | 4.4 | 34.6 |
| >hg38_21_+_8433172_8446622_RNA45SN1_45S_50ntFlanks@10685.10709.25\|GGCGGCGTCCGGTGAGCTCTCGCTG | >hg38_21_+_8433172_8446622_RNA45SN1_45S_50ntFlanks | 10685 | 10709 | 25 | GGCGGCGTCCGGTGAGCTCTCGCTG | MB468 | 17 | 10 | 11 | 3 | 13 | 3.5 | 1.2 | 2.5 | 0.7 | 3.8 |
| >hg38_21_+_8433172_8446622_RNA45SN1_45S_50ntFlanks@10685.10710.26\|GGCGGCGTCCGGTGAGCTCTCGCTGG | >hg38_21_+_8433172_8446622_RNA45SN1_45S_50ntFlanks | 10685 | 10710 | 26 | GGCGGCGTCCGGTGAGCTCTCGCTGG | MB468 | 48 | 57 | 33 | 13 | 39 | 17.3 | 19.3 | 10.6 | 4.4 | 10.5 |
| >hg38_21_+_8433172_8446622_RNA45SN1_45S_50ntFlanks@10685.10711.27\|GGCGGCGTCCGGTGAGCTCTCGCTGGC | >hg38_21_+_8433172_8446622_RNA45SN1_45S_50ntFlanks | 10685 | 10711 | 27 | GGCGGCGTCCGGTGAGCTCTCGCTGGC | MB468 | 15 | 18 | 16 | 6 | 15 | 4.7 | 4.5 | 5.9 | 1.4 | 3.6 |
| >hg38_21_+_8433172_8446622_RNA45SN1_45S_50ntFlanks@10686.10706.21\|GCGGCGTCCGGTGAGCTCTCG | >hg38_21_+_8433172_8446622_RNA45SN1_45S_50ntFlanks | 10686 | 10706 | 21 | GCGGCGTCCGGTGAGCTCTCG | MB468 | 83 | 112 | 69 | 58 | 59 | 32.6 | 60.9 | 36.8 | 34.8 | 14.0 |
| >hg38_21_+_8433172_8446622_RNA45SN1_45S_50ntFlanks@10686.10707.22\|GCGGCGTCCGGTGAGCTCTCGC | >hg38_21_+_8433172_8446622_RNA45SN1_45S_50ntFlanks | 10686 | 10707 | 22 | GCGGCGTCCGGTGAGCTCTCGC | BT20 | 12 | 13 | 7 | 33 | 46 | 16.0 | 18.7 | 6.1 | 40.1 | 62.7 |
| >hg38_21_+_8433172_8446622_RNA45SN1_45S_50ntFlanks@10686.10708.23\|GCGGCGTCCGGTGAGCTCTCGCT | >hg38_21_+_8433172_8446622_RNA45SN1_45S_50ntFlanks | 10686 | 10708 | 23 | GCGGCGTCCGGTGAGCTCTCGCT | BT20 | 44 | 43 | 17 | 16 | 152 | 59.6 | 67.2 | 19.6 | 6.8 | 203.4 |
| >hg38_21_+_8433172_8446622_RNA45SN1_45S_50ntFlanks@10686.10708.23\|GCGGCGTCCGGTGAGCTCTCGCT | >hg38_21_+_8433172_8446622_RNA45SN1_45S_50ntFlanks | 10686 | 10708 | 23 | GCGGCGTCCGGTGAGCTCTCGCT | MB231 | 19 | 25 | 10 | 3 | 24 | 4.8 | 7.6 | 4.3 | 4.1 | 33.9 |
| >hg38_21_+_8433172_8446622_RNA45SN1_45S_50ntFlanks@10686.10708.23\|GCGGCGTCCGGTGAGCTCTCGCT | >hg38_21_+_8433172_8446622_RNA45SN1_45S_50ntFlanks | 10686 | 10708 | 23 | GCGGCGTCCGGTGAGCTCTCGCT | MB468 | 44 | 70 | 42 | 12 | 40 | 8.3 | 12.5 | 17.5 | 10.1 | 10.6 |
| >hg38_21_+_8433172_8446622_RNA45SN1_45S_50ntFlanks@10686.10709.24\|GCGGCGTCCGGTGAGCTCTCGCTG | >hg38_21_+_8433172_8446622_RNA45SN1_45S_50ntFlanks | 10686 | 10709 | 24 | GCGGCGTCCGGTGAGCTCTCGCTG | BT20 | 23 | 21 | 8 | 11 | 67 | 33.6 | 33.7 | 10.5 | 11.1 | 93.6 |
| >hg38_21_+_8433172_8446622_RNA45SN1_45S_50ntFlanks@10686.10709.24\|GCGGCGTCCGGTGAGCTCTCGCTG | >hg38_21_+_8433172_8446622_RNA45SN1_45S_50ntFlanks | 10686 | 10709 | 24 | GCGGCGTCCGGTGAGCTCTCGCTG | MB468 | 94 | 63 | 67 | 21 | 70 | 23.5 | 11.3 | 27.2 | 9.2 | 13.0 |
| >hg38_21_+_8433172_8446622_RNA45SN1_45S_50ntFlanks@10686.10710.25\|GCGGCGTCCGGTGAGCTCTCGCTGG | >hg38_21_+_8433172_8446622_RNA45SN1_45S_50ntFlanks | 10686 | 10710 | 25 | GCGGCGTCCGGTGAGCTCTCGCTGG | MB468 | 266 | 245 | 214 | 64 | 207 | 81.0 | 69.6 | 84.7 | 30.3 | 46.6 |
| >hg38_21_+_8433172_8446622_RNA45SN1_45S_50ntFlanks@10686.10711.26\|GCGGCGTCCGGTGAGCTCTCGCTGGC | >hg38_21_+_8433172_8446622_RNA45SN1_45S_50ntFlanks | 10686 | 10711 | 26 | GCGGCGTCCGGTGAGCTCTCGCTGGC | MB468 | 50 | 45 | 54 | 25 | 47 | 13.2 | 16.7 | 21.9 | 3.5 | 9.5 |
| >hg38_21_+_8433172_8446622_RNA45SN1_45S_50ntFlanks@10686.10712.27\|GCGGCGTCCGGTGAGCTCTCGCTGGCC | >hg38_21_+_8433172_8446622_RNA45SN1_45S_50ntFlanks | 10686 | 10712 | 27 | GCGGCGTCCGGTGAGCTCTCGCTGGCC | MB468 | 25 | 29 | 32 | 11 | 26 | 8.6 | 12.2 | 14.3 | 1.7 | 8.2 |
| >hg38_21_+_8433172_8446622_RNA45SN1_45S_50ntFlanks@10687.10705.19\|CGGCGTCCGGTGAGCTCTC | >hg38_21_+_8433172_8446622_RNA45SN1_45S_50ntFlanks | 10687 | 10705 | 19 | CGGCGTCCGGTGAGCTCTC | MB468 | 18 | 30 | 23 | 12 | 15 | 5.9 | 10.7 | 9.4 | 4.7 | 3.1 |
| >hg38_21_+_8433172_8446622_RNA45SN1_45S_50ntFlanks@10687.10706.20\|CGGCGTCCGGTGAGCTCTCG | >hg38_21_+_8433172_8446622_RNA45SN1_45S_50ntFlanks | 10687 | 10706 | 20 | CGGCGTCCGGTGAGCTCTCG | MB468 | 395 | 384 | 398 | 200 | 264 | 86.7 | 99.1 | 201.3 | 56.0 | 36.8 |
| >hg38_21_+_8433172_8446622_RNA45SN1_45S_50ntFlanks@10687.10708.22\|CGGCGTCCGGTGAGCTCTCGCT | >hg38_21_+_8433172_8446622_RNA45SN1_45S_50ntFlanks | 10687 | 10708 | 22 | CGGCGTCCGGTGAGCTCTCGCT | MB468 | 167 | 178 | 196 | 47 | 118 | 54.3 | 59.5 | 99.5 | 21.2 | 13.1 |
| >hg38_21_+_8433172_8446622_RNA45SN1_45S_50ntFlanks@10687.10709.23\|CGGCGTCCGGTGAGCTCTCGCTG | >hg38_21_+_8433172_8446622_RNA45SN1_45S_50ntFlanks | 10687 | 10709 | 23 | CGGCGTCCGGTGAGCTCTCGCTG | MB468 | 359 | 166 | 281 | 99 | 221 | 111.7 | 71.9 | 120.5 | 30.8 | 57.0 |
| >hg38_21_+_8433172_8446622_RNA45SN1_45S_50ntFlanks@10687.10710.24\|CGGCGTCCGGTGAGCTCTCGCTGG | >hg38_21_+_8433172_8446622_RNA45SN1_45S_50ntFlanks | 10687 | 10710 | 24 | CGGCGTCCGGTGAGCTCTCGCTGG | MB468 | 575 | 364 | 612 | 252 | 370 | 161.6 | 125.6 | 242.4 | 51.7 | 73.7 |
| >hg38_21_+_8433172_8446622_RNA45SN1_45S_50ntFlanks@10687.10711.25\|CGGCGTCCGGTGAGCTCTCGCTGGC | >hg38_21_+_8433172_8446622_RNA45SN1_45S_50ntFlanks | 10687 | 10711 | 25 | CGGCGTCCGGTGAGCTCTCGCTGGC | MB468 | 338 | 365 | 524 | 136 | 291 | 104.0 | 137.0 | 274.8 | 35.5 | 99.7 |
| >hg38_21_+_8433172_8446622_RNA45SN1_45S_50ntFlanks@10687.10712.26\|CGGCGTCCGGTGAGCTCTCGCTGGCC | >hg38_21_+_8433172_8446622_RNA45SN1_45S_50ntFlanks | 10687 | 10712 | 26 | CGGCGTCCGGTGAGCTCTCGCTGGCC | MB468 | 93 | 69 | 163 | 51 | 60 | 28.5 | 37.6 | 86.1 | 20.3 | 20.6 |
| >hg38_21_+_8433172_8446622_RNA45SN1_45S_50ntFlanks@10689.10706.18\|GCGTCCGGTGAGCTCTCG | >hg38_21_+_8433172_8446622_RNA45SN1_45S_50ntFlanks | 10689 | 10706 | 18 | GCGTCCGGTGAGCTCTCG | MB468 | 153 | 191 | 155 | 104 | 85 | 67.6 | 143.4 | 100.3 | 56.4 | 21.2 |
| >hg38_21_+_8433172_8446622_RNA45SN1_45S_50ntFlanks@10689.10708.20\|GCGTCCGGTGAGCTCTCGCT | >hg38_21_+_8433172_8446622_RNA45SN1_45S_50ntFlanks | 10689 | 10708 | 20 | GCGTCCGGTGAGCTCTCGCT | MB468 | 89 | 94 | 113 | 37 | 54 | 15.2 | 15.6 | 32.9 | 18.4 | 9.3 |
| >hg38_21_+_8433172_8446622_RNA45SN1_45S_50ntFlanks@10689.10709.21\|GCGTCCGGTGAGCTCTCGCTG | >hg38_21_+_8433172_8446622_RNA45SN1_45S_50ntFlanks | 10689 | 10709 | 21 | GCGTCCGGTGAGCTCTCGCTG | MB468 | 59 | 62 | 69 | 13 | 40 | 15.6 | 23.9 | 35.0 | 3.4 | 9.2 |
| >hg38_21_+_8433172_8446622_RNA45SN1_45S_50ntFlanks@10689.10710.22\|GCGTCCGGTGAGCTCTCGCTGG | >hg38_21_+_8433172_8446622_RNA45SN1_45S_50ntFlanks | 10689 | 10710 | 22 | GCGTCCGGTGAGCTCTCGCTGG | MB468 | 15 | 8 | 19 | 14 | 6 | 2.9 | 1.6 | 9.2 | 8.9 | 1.9 |
| >hg38_21_+_8433172_8446622_RNA45SN1_45S_50ntFlanks@10689.10711.23\|GCGTCCGGTGAGCTCTCGCTGGC | >hg38_21_+_8433172_8446622_RNA45SN1_45S_50ntFlanks | 10689 | 10711 | 23 | GCGTCCGGTGAGCTCTCGCTGGC | MB468 | 12 | 5 | 20 | 21 | 8 | 4.1 | 2.6 | 11.7 | 17.5 | 3.5 |

| | | | | | | | | | | | | | | | | |
|---|---|---|---|---|---|---|---|---|---|---|---|---|---|---|---|---|
| >hg38_21_+_8433172_8446622_RNA45SN1_45S_50ntFlanks@10690.10708.19\|CGTCCGGTGAGCTCTCGCT | >hg38_21_+_8433172_8446622_RNA45SN1_45S_50ntFlanks | 10690 | 10708 | 19 | CGTCCGGTGAGCTCTCGCT | MB468 | 589 | 443 | 793 | 357 | 350 | 112.6 | 119.4 | 171.3 | 69.4 | 120.9 |
| >hg38_21_+_8433172_8446622_RNA45SN1_45S_50ntFlanks@10690.10709.20\|CGTCCGGTGAGCTCTCGCTG | >hg38_21_+_8433172_8446622_RNA45SN1_45S_50ntFlanks | 10690 | 10709 | 20 | CGTCCGGTGAGCTCTCGCTG | MB468 | 155 | 130 | 203 | 59 | 106 | 31.1 | 62.9 | 82.5 | 4.9 | 16.5 |
| >hg38_21_+_8433172_8446622_RNA45SN1_45S_50ntFlanks@10690.10710.21\|CGTCCGGTGAGCTCTCGCTGG | >hg38_21_+_8433172_8446622_RNA45SN1_45S_50ntFlanks | 10690 | 10710 | 21 | CGTCCGGTGAGCTCTCGCTGG | MB468 | 79 | 27 | 105 | 87 | 36 | 17.8 | 12.9 | 47.5 | 59.4 | 15.6 |
| >hg38_21_+_8433172_8446622_RNA45SN1_45S_50ntFlanks@10690.10711.22\|CGTCCGGTGAGCTCTCGCTGGC | >hg38_21_+_8433172_8446622_RNA45SN1_45S_50ntFlanks | 10690 | 10711 | 22 | CGTCCGGTGAGCTCTCGCTGGC | MB468 | 13 | 4 | 20 | 25 | 9 | 5.0 | 2.6 | 11.0 | 22.5 | 6.6 |
| >hg38_21_+_8433172_8446622_RNA45SN1_45S_50ntFlanks@10692.10709.18\|TCCGGTGAGCTCTCGCTG | >hg38_21_+_8433172_8446622_RNA45SN1_45S_50ntFlanks | 10692 | 10709 | 18 | TCCGGTGAGCTCTCGCTG | MB468 | 17 | 3 | 13 | 19 | 9 | 4.6 | 0.4 | 4.5 | 9.1 | 2.6 |
| >hg38_21_+_8433172_8446622_RNA45SN1_45S_50ntFlanks@10694.10711.18\|CGGTGAGCTCTCGCTGGC | >hg38_21_+_8433172_8446622_RNA45SN1_45S_50ntFlanks | 10694 | 10711 | 18 | CGGTGAGCTCTCGCTGGC | MB468 | 50 | 12 | 58 | 65 | 48 | 13.3 | 2.7 | 17.9 | 16.2 | 18.7 |
| >hg38_21_+_8433172_8446622_RNA45SN1_45S_50ntFlanks@10713.10736.24\|CTTGAAAATCCGGGGGAGAGGGTG | >hg38_21_+_8433172_8446622_RNA45SN1_45S_50ntFlanks | 10713 | 10736 | 24 | CTTGAAAATCCGGGGGAGAGGGTG | MB468 | 16 | 18 | 27 | 42 | 12 | 4.0 | 5.8 | 10.1 | 15.9 | 3.4 |
| >hg38_21_+_8433172_8446622_RNA45SN1_45S_50ntFlanks@10717.10745.29\|AAAATCCGGGGGAGAGGGTGTAAATCTCG | >hg38_21_+_8433172_8446622_RNA45SN1_45S_50ntFlanks | 10717 | 10745 | 29 | AAAATCCGGGGGAGAGGGTGTAAATCTCG | MB231 | 13 | 7 | 23 | 8 | 8 | 12.7 | 6.2 | 32.7 | 8.0 | 7.2 |
| >hg38_21_+_8433172_8446622_RNA45SN1_45S_50ntFlanks@10727.10745.19\|GGAGAGGGTGTAAATCTCG | >hg38_21_+_8433172_8446622_RNA45SN1_45S_50ntFlanks | 10727 | 10745 | 19 | GGAGAGGGTGTAAATCTCG | MB468 | 18 | 2 | 11 | 11 | 5 | 6.1 | 1.0 | 3.4 | 8.4 | 2.3 |
| >hg38_21_+_8433172_8446622_RNA45SN1_45S_50ntFlanks@10727.10747.21\|GGAGAGGGTGTAAATCTCGCG | >hg38_21_+_8433172_8446622_RNA45SN1_45S_50ntFlanks | 10727 | 10747 | 21 | GGAGAGGGTGTAAATCTCGCG | MB468 | 11 | 2 | 7 | 9 | 4 | 4.6 | 0.7 | 1.4 | 5.8 | 2.5 |
| >hg38_21_+_8433172_8446622_RNA45SN1_45S_50ntFlanks@10727.10752.26\|GGAGAGGGTGTAAATCTCGCGCCGGG | >hg38_21_+_8433172_8446622_RNA45SN1_45S_50ntFlanks | 10727 | 10752 | 26 | GGAGAGGGTGTAAATCTCGCGCCGGG | MB468 | 16 | 5 | 15 | 50 | 4 | 6.5 | 1.8 | 5.4 | 22.2 | 1.8 |
| >hg38_21_+_8433172_8446622_RNA45SN1_45S_50ntFlanks@10727.10755.29\|GGAGAGGGTGTAAATCTCGCGCCGGGCCG | >hg38_21_+_8433172_8446622_RNA45SN1_45S_50ntFlanks | 10727 | 10755 | 29 | GGAGAGGGTGTAAATCTCGCGCCGGGCCG | MB468 | 70 | 130 | 58 | 86 | 38 | 36.8 | 84.8 | 28.1 | 29.1 | 16.3 |
| >hg38_21_+_8433172_8446622_RNA45SN1_45S_50ntFlanks@10727.10757.31\|GGAGAGGGTGTAAATCTCGCGCCGGGCCGTA | >hg38_21_+_8433172_8446622_RNA45SN1_45S_50ntFlanks | 10727 | 10757 | 31 | GGAGAGGGTGTAAATCTCGCGCCGGGCCGTA | MB468 | 27 | 62 | 29 | 18 | 20 | 8.7 | 24.2 | 16.1 | 5.1 | 9.1 |
| >hg38_21_+_8433172_8446622_RNA45SN1_45S_50ntFlanks@10728.10755.28\|GAGAGGGTGTAAATCTCGCGCCGGGCCG | >hg38_21_+_8433172_8446622_RNA45SN1_45S_50ntFlanks | 10728 | 10755 | 28 | GAGAGGGTGTAAATCTCGCGCCGGGCCG | MB468 | 16 | 47 | 15 | 7 | 8 | 6.9 | 26.9 | 5.6 | 2.2 | 3.8 |
| >hg38_21_+_8433172_8446622_RNA45SN1_45S_50ntFlanks@10729.10750.22\|AGAGGGTGTAAATCTCGCGCCG | >hg38_21_+_8433172_8446622_RNA45SN1_45S_50ntFlanks | 10729 | 10750 | 22 | AGAGGGTGTAAATCTCGCGCCG | MB468 | 11 | 1 | 8 | 12 | 4 | 5.8 | 0.5 | 2.3 | 7.3 | 1.9 |
| >hg38_21_+_8433172_8446622_RNA45SN1_45S_50ntFlanks@10729.10752.24\|AGAGGGTGTAAATCTCGCGCCGGG | >hg38_21_+_8433172_8446622_RNA45SN1_45S_50ntFlanks | 10729 | 10752 | 24 | AGAGGGTGTAAATCTCGCGCCGGG | MB468 | 15 | 10 | 17 | 16 | 5 | 6.7 | 4.1 | 6.1 | 3.0 | 1.5 |
| >hg38_21_+_8433172_8446622_RNA45SN1_45S_50ntFlanks@10729.10755.27\|AGAGGGTGTAAATCTCGCGCCGGGCCG | >hg38_21_+_8433172_8446622_RNA45SN1_45S_50ntFlanks | 10729 | 10755 | 27 | AGAGGGTGTAAATCTCGCGCCGGGCCG | MB468 | 27 | 59 | 26 | 22 | 15 | 8.8 | 30.2 | 10.4 | 5.7 | 4.2 |
| >hg38_21_+_8433172_8446622_RNA45SN1_45S_50ntFlanks@10731.10752.22\|AGGGTGTAAATCTCGCGCCGGG | >hg38_21_+_8433172_8446622_RNA45SN1_45S_50ntFlanks | 10731 | 10752 | 22 | AGGGTGTAAATCTCGCGCCGGG | MB468 | 25 | 7 | 29 | 35 | 11 | 9.4 | 2.0 | 6.4 | 3.8 | 3.6 |
| >hg38_21_+_8433172_8446622_RNA45SN1_45S_50ntFlanks@10731.10755.25\|AGGGTGTAAATCTCGCGCCGGGCCG | >hg38_21_+_8433172_8446622_RNA45SN1_45S_50ntFlanks | 10731 | 10755 | 25 | AGGGTGTAAATCTCGCGCCGGGCCG | MB468 | 31 | 68 | 32 | 35 | 24 | 12.2 | 26.7 | 9.0 | 7.7 | 7.9 |
| >hg38_21_+_8433172_8446622_RNA45SN1_45S_50ntFlanks@10734.10755.22\|GTGTAAATCTCGCGCCGGGCCG | >hg38_21_+_8433172_8446622_RNA45SN1_45S_50ntFlanks | 10734 | 10755 | 22 | GTGTAAATCTCGCGCCGGGCCG | MB468 | 12 | 36 | 14 | 5 | 10 | 6.3 | 16.4 | 5.3 | 2.0 | 4.4 |
| >hg38_21_+_8433172_8446622_RNA45SN1_45S_50ntFlanks@10735.10752.18\|TGTAAATCTCGCGCCGGG | >hg38_21_+_8433172_8446622_RNA45SN1_45S_50ntFlanks | 10735 | 10752 | 18 | TGTAAATCTCGCGCCGGG | MB468 | 20 | 16 | 25 | 10 | 13 | 8.5 | 6.0 | 9.6 | 2.4 | 5.0 |
| >hg38_21_+_8433172_8446622_RNA45SN1_45S_50ntFlanks@10735.10755.21\|TGTAAATCTCGCGCCGGGCCG | >hg38_21_+_8433172_8446622_RNA45SN1_45S_50ntFlanks | 10735 | 10755 | 21 | TGTAAATCTCGCGCCGGGCCG | MB468 | 75 | 245 | 85 | 17 | 55 | 39.7 | 107.9 | 41.1 | 6.6 | 25.7 |
| >hg38_21_+_8433172_8446622_RNA45SN1_45S_50ntFlanks@10735.10757.23\|TGTAAATCTCGCGCCGGGCCGTA | >hg38_21_+_8433172_8446622_RNA45SN1_45S_50ntFlanks | 10735 | 10757 | 23 | TGTAAATCTCGCGCCGGGCCGTA | MB468 | 15 | 47 | 19 | 5 | 13 | 3.5 | 25.0 | 6.4 | 0.6 | 4.0 |
| >hg38_21_+_8433172_8446622_RNA45SN1_45S_50ntFlanks@10736.10755.20\|GTAAATCTCGCGCCGGGCCG | >hg38_21_+_8433172_8446622_RNA45SN1_45S_50ntFlanks | 10736 | 10755 | 20 | GTAAATCTCGCGCCGGGCCG | MB468 | 12 | 46 | 13 | 2 | 12 | 6.2 | 26.0 | 6.6 | 0.8 | 6.7 |
| >hg38_21_+_8433172_8446622_RNA45SN1_45S_50ntFlanks@10737.10755.19\|TAAATCTCGCGCCGGGCCG | >hg38_21_+_8433172_8446622_RNA45SN1_45S_50ntFlanks | 10737 | 10755 | 19 | TAAATCTCGCGCCGGGCCG | MB468 | 77 | 297 | 73 | 10 | 74 | 39.3 | 126.1 | 40.6 | 2.2 | 32.5 |
| >hg38_21_+_8433172_8446622_RNA45SN1_45S_50ntFlanks@10737.10757.21\|TAAATCTCGCGCCGGGCCGTA | >hg38_21_+_8433172_8446622_RNA45SN1_45S_50ntFlanks | 10737 | 10757 | 21 | TAAATCTCGCGCCGGGCCGTA | MB468 | 16 | 58 | 17 | 3 | 18 | 3.1 | 30.0 | 6.8 | 0.5 | 5.2 |
| >hg38_21_+_8433172_8446622_RNA45SN1_45S_50ntFlanks@10737.10767.31\|TAAATCTCGCGCCGGGCCGTACCCATATCCG | >hg38_21_+_8433172_8446622_RNA45SN1_45S_50ntFlanks | 10737 | 10767 | 31 | TAAATCTCGCGCCGGGCCGTACCCATATCCG | MB468 | 10 | 38 | 15 | 1 | 9 | 2.9 | 18.6 | 6.3 | 1.6 | 10.5 |
| >hg38_21_+_8433172_8446622_RNA45SN1_45S_50ntFlanks@10738.10755.18\|AAATCTCGCGCCGGGCCG | >hg38_21_+_8433172_8446622_RNA45SN1_45S_50ntFlanks | 10738 | 10755 | 18 | AAATCTCGCGCCGGGCCG | MB468 | 34 | 171 | 29 | 3 | 27 | 17.6 | 71.1 | 18.9 | 1.3 | 11.5 |
| >hg38_21_+_8433172_8446622_RNA45SN1_45S_50ntFlanks@10738.10756.19\|AAATCTCGCGCCGGGCCGT | >hg38_21_+_8433172_8446622_RNA45SN1_45S_50ntFlanks | 10738 | 10756 | 19 | AAATCTCGCGCCGGGCCGT | MB231 | 14 | 39 | 37 | 1 | 6 | 9.1 | 35.8 | 34.6 | 1.1 | 5.2 |
| >hg38_21_+_8433172_8446622_RNA45SN1_45S_50ntFlanks@10738.10757.20\|AAATCTCGCGCCGGGCCGTA | >hg38_21_+_8433172_8446622_RNA45SN1_45S_50ntFlanks | 10738 | 10757 | 20 | AAATCTCGCGCCGGGCCGTA | MB468 | 11 | 43 | 9 | 1 | 6 | 4.2 | 17.4 | 4.9 | 0.8 | 4.1 |
| >hg38_21_+_8433172_8446622_RNA45SN1_45S_50ntFlanks@10756.10785.30\|TACCCATATCCGCAGCAGGTCTCCAAGGTG | >hg38_21_+_8433172_8446622_RNA45SN1_45S_50ntFlanks | 10756 | 10785 | 30 | TACCCATATCCGCAGCAGGTCTCCAAGGTG | MB468 | 11 | 8 | 8 | 6 | 11 | 4.6 | 3.1 | 2.0 | 2.6 | 8.8 |
| >hg38_21_+_8433172_8446622_RNA45SN1_45S_50ntFlanks@10760.10785.26\|CATATCCGCAGCAGGTCTCCAAGGTG | >hg38_21_+_8433172_8446622_RNA45SN1_45S_50ntFlanks | 10760 | 10785 | 26 | CATATCCGCAGCAGGTCTCCAAGGTG | MB468 | 11 | 11 | 7 | 14 | 14 | 1.8 | 2.3 | 0.7 | 8.9 | 7.0 |
| >hg38_21_+_8433172_8446622_RNA45SN1_45S_50ntFlanks@10761.10780.20\|ATATCCGCAGCAGGTCTCCA | >hg38_21_+_8433172_8446622_RNA45SN1_45S_50ntFlanks | 10761 | 10780 | 20 | ATATCCGCAGCAGGTCTCCA | MB468 | 13 | 7 | 6 | 1 | 2 | 7.1 | 2.4 | 5.7 | 1.0 | 1.5 |
| >hg38_21_+_8433172_8446622_RNA45SN1_45S_50ntFlanks@10761.10785.25\|ATATCCGCAGCAGGTCTCCAAGGTG | >hg38_21_+_8433172_8446622_RNA45SN1_45S_50ntFlanks | 10761 | 10785 | 25 | ATATCCGCAGCAGGTCTCCAAGGTG | MB468 | 14 | 15 | 9 | 9 | 14 | 1.8 | 3.8 | 3.2 | 3.5 | 4.6 |
| >hg38_21_+_8433172_8446622_RNA45SN1_45S_50ntFlanks@10764.10785.22\|TCCGCAGCAGGTCTCCAAGGTG | >hg38_21_+_8433172_8446622_RNA45SN1_45S_50ntFlanks | 10764 | 10785 | 22 | TCCGCAGCAGGTCTCCAAGGTG | MB468 | 10 | 13 | 9 | 5 | 12 | 2.7 | 3.7 | 3.0 | 1.7 | 1.1 |
| >hg38_21_+_8433172_8446622_RNA45SN1_45S_50ntFlanks@10766.10785.20\|CGCAGCAGGTCTCCAAGGTG | >hg38_21_+_8433172_8446622_RNA45SN1_45S_50ntFlanks | 10766 | 10785 | 20 | CGCAGCAGGTCTCCAAGGTG | MB468 | 18 | 19 | 15 | 16 | 14 | 4.1 | 6.9 | 3.1 | 8.7 | 4.0 |
| >hg38_21_+_8433172_8446622_RNA45SN1_45S_50ntFlanks@10766.10797.32\|CGCAGCAGGTCTCCAAGGTGAACAGCCTCTGG | >hg38_21_+_8433172_8446622_RNA45SN1_45S_50ntFlanks | 10766 | 10797 | 32 | CGCAGCAGGTCTCCAAGGTGAACAGCCTCTGG | MB468 | 14 | 11 | 12 | 12 | 9 | 3.8 | 5.8 | 5.0 | 9.7 | 7.4 |
| >hg38_21_+_8433172_8446622_RNA45SN1_45S_50ntFlanks@10767.10785.19\|GCAGCAGGTCTCCAAGGTG | >hg38_21_+_8433172_8446622_RNA45SN1_45S_50ntFlanks | 10767 | 10785 | 19 | GCAGCAGGTCTCCAAGGTG | MB468 | 24 | 17 | 18 | 15 | 13 | 4.2 | 8.3 | 10.3 | 12.7 | 5.2 |
| >hg38_21_+_8433172_8446622_RNA45SN1_45S_50ntFlanks@10768.10785.18\|CAGCAGGTCTCCAAGGTG | >hg38_21_+_8433172_8446622_RNA45SN1_45S_50ntFlanks | 10768 | 10785 | 18 | CAGCAGGTCTCCAAGGTG | MB468 | 86 | 36 | 47 | 55 | 47 | 14.2 | 19.9 | 23.4 | 30.7 | 11.8 |
| >hg38_21_+_8433172_8446622_RNA45SN1_45S_50ntFlanks@10768.10793.26\|CAGCAGGTCTCCAAGGTGAACAGCCT | >hg38_21_+_8433172_8446622_RNA45SN1_45S_50ntFlanks | 10768 | 10793 | 26 | CAGCAGGTCTCCAAGGTGAACAGCCT | MB468 | 14 | 7 | 4 | 4 | 5 | 6.6 | 3.6 | 2.3 | 1.9 | 1.8 |
| >hg38_21_+_8433172_8446622_RNA45SN1_45S_50ntFlanks@10768.10797.30\|CAGCAGGTCTCCAAGGTGAACAGCCTCTGG | >hg38_21_+_8433172_8446622_RNA45SN1_45S_50ntFlanks | 10768 | 10797 | 30 | CAGCAGGTCTCCAAGGTGAACAGCCTCTGG | MB468 | 20 | 5 | 16 | 18 | 6 | 3.3 | 1.9 | 4.0 | 6.1 | 2.5 |
| >hg38_21_+_8433172_8446622_RNA45SN1_45S_50ntFlanks@10769.10796.28\|AGCAGGTCTCCAAGGTGAACAGCCTCTG | >hg38_21_+_8433172_8446622_RNA45SN1_45S_50ntFlanks | 10769 | 10796 | 28 | AGCAGGTCTCCAAGGTGAACAGCCTCTG | MB231 | 26 | 15 | 8 | 4 | 10 | 27.4 | 14.8 | 6.6 | 4.9 | 12.1 |
| >hg38_21_+_8433172_8446622_RNA45SN1_45S_50ntFlanks@10769.10796.28\|AGCAGGTCTCCAAGGTGAACAGCCTCTG | >hg38_21_+_8433172_8446622_RNA45SN1_45S_50ntFlanks | 10769 | 10796 | 28 | AGCAGGTCTCCAAGGTGAACAGCCTCTG | MB468 | 36 | 15 | 14 | 12 | 24 | 7.2 | 4.2 | 1.7 | 4.8 | 4.3 |
| >hg38_21_+_8433172_8446622_RNA45SN1_45S_50ntFlanks@10770.10796.27\|GCAGGTCTCCAAGGTGAACAGCCTCTG | >hg38_21_+_8433172_8446622_RNA45SN1_45S_50ntFlanks | 10770 | 10796 | 27 | GCAGGTCTCCAAGGTGAACAGCCTCTG | MB231 | 18 | 13 | 13 | 4 | 8 | 17.0 | 10.7 | 9.8 | 4.5 | 9.9 |
| >hg38_21_+_8433172_8446622_RNA45SN1_45S_50ntFlanks@10770.10796.27\|GCAGGTCTCCAAGGTGAACAGCCTCTG | >hg38_21_+_8433172_8446622_RNA45SN1_45S_50ntFlanks | 10770 | 10796 | 27 | GCAGGTCTCCAAGGTGAACAGCCTCTG | MB468 | 62 | 40 | 44 | 19 | 31 | 13.4 | 16.9 | 14.1 | 8.2 | 5.1 |
| >hg38_21_+_8433172_8446622_RNA45SN1_45S_50ntFlanks@10770.10797.28\|GCAGGTCTCCAAGGTGAACAGCCTCTGG | >hg38_21_+_8433172_8446622_RNA45SN1_45S_50ntFlanks | 10770 | 10797 | 28 | GCAGGTCTCCAAGGTGAACAGCCTCTGG | MB468 | 13 | 9 | 10 | 4 | 3 | 2.0 | 5.1 | 4.1 | 1.3 | 1.6 |
| >hg38_21_+_8433172_8446622_RNA45SN1_45S_50ntFlanks@10770.10805.36\|GCAGGTCTCCAAGGTGAACAGCCTCTGGCATGTTGG | >hg38_21_+_8433172_8446622_RNA45SN1_45S_50ntFlanks | 10770 | 10805 | 36 | GCAGGTCTCCAAGGTGAACAGCCTCTGGCATGTTGG | MB468 | 16 | 13 | 9 | 5 | 16 | 5.2 | 2.6 | 0.6 | 2.1 | 5.8 |
| >hg38_21_+_8433172_8446622_RNA45SN1_45S_50ntFlanks@10771.10788.18\|CAGGTCTCCAAGGTGAAC | >hg38_21_+_8433172_8446622_RNA45SN1_45S_50ntFlanks | 10771 | 10788 | 18 | CAGGTCTCCAAGGTGAAC | MB468 | 13 | 7 | 4 | 4 | 3 | 6.0 | 3.7 | 1.1 | 1.7 | 0.9 |
| >hg38_21_+_8433172_8446622_RNA45SN1_45S_50ntFlanks@10771.10790.20\|CAGGTCTCCAAGGTGAACAG | >hg38_21_+_8433172_8446622_RNA45SN1_45S_50ntFlanks | 10771 | 10790 | 20 | CAGGTCTCCAAGGTGAACAG | MB468 | 27 | 8 | 5 | 7 | 6 | 19.3 | 5.3 | 1.4 | 1.8 | 3.0 |
| >hg38_21_+_8433172_8446622_RNA45SN1_45S_50ntFlanks@10771.10791.21\|CAGGTCTCCAAGGTGAACAGC | >hg38_21_+_8433172_8446622_RNA45SN1_45S_50ntFlanks | 10771 | 10791 | 21 | CAGGTCTCCAAGGTGAACAGC | MB468 | 46 | 27 | 10 | 11 | 8 | 24.2 | 13.4 | 3.7 | 0.6 | 1.6 |

| | | | | | | | | | | | | | | | | |
|---|---|---|---|---|---|---|---|---|---|---|---|---|---|---|---|---|
| >hg38_21_+_8433172_8446622_RNA45SN1_45S_50ntFlanks@10771.10792.22\|CAGGTCTCCAAGGTGAACAGCC | >hg38_21_+_8433172_8446622_RNA45SN1_45S_50ntFlanks | 10771 | 10792 | 22 | CAGGTCTCCAAGGTGAACAGCC | MB468 | 20 | 13 | 4 | 4 | 5 | 9.3 | 7.8 | 1.0 | 0.8 | 1.3 |
| >hg38_21_+_8433172_8446622_RNA45SN1_45S_50ntFlanks@10771.10793.23\|CAGGTCTCCAAGGTGAACAGCCT | >hg38_21_+_8433172_8446622_RNA45SN1_45S_50ntFlanks | 10771 | 10793 | 23 | CAGGTCTCCAAGGTGAACAGCCT | MB468 | 13 | 2 | 3 | 4 | 3 | 8.8 | 1.4 | 1.9 | 1.0 | 0.9 |
| >hg38_21_+_8433172_8446622_RNA45SN1_45S_50ntFlanks@10771.10796.26\|CAGGTCTCCAAGGTGAACAGCCTCTG | >hg38_21_+_8433172_8446622_RNA45SN1_45S_50ntFlanks | 10771 | 10796 | 26 | CAGGTCTCCAAGGTGAACAGCCTCTG | MB468 | 76 | 39 | 43 | 38 | 27 | 29.7 | 20.9 | 19.8 | 4.8 | 4.3 |
| >hg38_21_+_8433172_8446622_RNA45SN1_45S_50ntFlanks@10771.10797.27\|CAGGTCTCCAAGGTGAACAGCCTCTGG | >hg38_21_+_8433172_8446622_RNA45SN1_45S_50ntFlanks | 10771 | 10797 | 27 | CAGGTCTCCAAGGTGAACAGCCTCTGG | MB468 | 64 | 23 | 55 | 47 | 13 | 17.9 | 13.6 | 24.8 | 17.8 | 3.6 |
| >hg38_21_+_8433172_8446622_RNA45SN1_45S_50ntFlanks@10771.10798.28\|CAGGTCTCCAAGGTGAACAGCCTCTGGC | >hg38_21_+_8433172_8446622_RNA45SN1_45S_50ntFlanks | 10771 | 10798 | 28 | CAGGTCTCCAAGGTGAACAGCCTCTGGC | MB468 | 10 | 3 | 6 | 5 | 4 | 4.3 | 1.3 | 3.4 | 0.9 | 1.7 |
| >hg38_21_+_8433172_8446622_RNA45SN1_45S_50ntFlanks@10771.10805.35\|CAGGTCTCCAAGGTGAACAGCCTCTGGCATGTTGG | >hg38_21_+_8433172_8446622_RNA45SN1_45S_50ntFlanks | 10771 | 10805 | 35 | CAGGTCTCCAAGGTGAACAGCCTCTGGCATGTTGG | MB468 | 99 | 89 | 64 | 44 | 73 | 18.1 | 14.7 | 8.6 | 20.5 | 17.1 |
| >hg38_21_+_8433172_8446622_RNA45SN1_45S_50ntFlanks@10771.10806.36\|CAGGTCTCCAAGGTGAACAGCCTCTGGCATGTTGGA | >hg38_21_+_8433172_8446622_RNA45SN1_45S_50ntFlanks | 10771 | 10806 | 36 | CAGGTCTCCAAGGTGAACAGCCTCTGGCATGTTGGA | MB468 | 64 | 65 | 53 | 40 | 69 | 6.3 | 31.3 | 27.7 | 25.7 | 14.6 |
| >hg38_21_+_8433172_8446622_RNA45SN1_45S_50ntFlanks@10771.10807.37\|CAGGTCTCCAAGGTGAACAGCCTCTGGCATGTTGGAA | >hg38_21_+_8433172_8446622_RNA45SN1_45S_50ntFlanks | 10771 | 10807 | 37 | CAGGTCTCCAAGGTGAACAGCCTCTGGCATGTTGGAA | MB468 | 10 | 12 | 8 | 6 | 11 | 4.8 | 4.7 | 5.3 | 7.2 | 6.5 |
| >hg38_21_+_8433172_8446622_RNA45SN1_45S_50ntFlanks@10771.10818.48\|CAGGTCTCCAAGGTGAACAGCCTCTGGCATGTTGGAACAATGTAGGTA | >hg38_21_+_8433172_8446622_RNA45SN1_45S_50ntFlanks | 10771 | 10818 | 48 | CAGGTCTCCAAGGTGAACAGCCTCTGGCATGTTGGAACAATGTAGGTA | MB468 | 23 | 49 | 77 | 9 | 35 | 27.3 | 55.5 | 69.7 | 12.3 | 39.2 |
| >hg38_21_+_8433172_8446622_RNA45SN1_45S_50ntFlanks@10772.10796.25\|AGGTCTCCAAGGTGAACAGCCTCTG | >hg38_21_+_8433172_8446622_RNA45SN1_45S_50ntFlanks | 10772 | 10796 | 25 | AGGTCTCCAAGGTGAACAGCCTCTG | MB468 | 47 | 47 | 26 | 20 | 21 | 19.0 | 26.0 | 10.1 | 12.0 | 6.4 |
| >hg38_21_+_8433172_8446622_RNA45SN1_45S_50ntFlanks@10772.10805.34\|AGGTCTCCAAGGTGAACAGCCTCTGGCATGTTGG | >hg38_21_+_8433172_8446622_RNA45SN1_45S_50ntFlanks | 10772 | 10805 | 34 | AGGTCTCCAAGGTGAACAGCCTCTGGCATGTTGG | MB468 | 12 | 11 | 8 | 5 | 12 | 2.5 | 2.8 | 0.7 | 2.0 | 2.5 |
| >hg38_21_+_8433172_8446622_RNA45SN1_45S_50ntFlanks@10774.10796.23\|GTCTCCAAGGTGAACAGCCTCTG | >hg38_21_+_8433172_8446622_RNA45SN1_45S_50ntFlanks | 10774 | 10796 | 23 | GTCTCCAAGGTGAACAGCCTCTG | MB468 | 14 | 4 | 8 | 4 | 3 | 7.2 | 2.4 | 4.8 | 2.9 | 1.2 |
| >hg38_21_+_8433172_8446622_RNA45SN1_45S_50ntFlanks@10775.10796.22\|TCTCCAAGGTGAACAGCCTCTG | >hg38_21_+_8433172_8446622_RNA45SN1_45S_50ntFlanks | 10775 | 10796 | 22 | TCTCCAAGGTGAACAGCCTCTG | MB468 | 12 | 1 | 6 | 5 | 2 | 7.3 | 0.1 | 3.7 | 0.7 | 1.3 |
| >hg38_21_+_8433172_8446622_RNA45SN1_45S_50ntFlanks@10777.10818.42\|TCCAAGGTGAACAGCCTCTGGCATGTTGGAACAATGTAGGTA | >hg38_21_+_8433172_8446622_RNA45SN1_45S_50ntFlanks | 10777 | 10818 | 42 | TCCAAGGTGAACAGCCTCTGGCATGTTGGAACAATGTAGGTA | MB468 | 18 | 10 | 35 | 7 | 21 | 11.0 | 4.8 | 24.1 | 7.6 | 12.2 |
| >hg38_21_+_8433172_8446622_RNA45SN1_45S_50ntFlanks@10778.10818.41\|CCAAGGTGAACAGCCTCTGGCATGTTGGAACAATGTAGGTA | >hg38_21_+_8433172_8446622_RNA45SN1_45S_50ntFlanks | 10778 | 10818 | 41 | CCAAGGTGAACAGCCTCTGGCATGTTGGAACAATGTAGGTA | MB468 | 19 | 8 | 34 | 7 | 24 | 11.2 | 3.5 | 22.5 | 7.1 | 12.9 |
| >hg38_21_+_8433172_8446622_RNA45SN1_45S_50ntFlanks@10780.10801.22\|AAGGTGAACAGCCTCTGGCATG | >hg38_21_+_8433172_8446622_RNA45SN1_45S_50ntFlanks | 10780 | 10801 | 22 | AAGGTGAACAGCCTCTGGCATG | MB468 | 25 | 10 | 16 | 11 | 9 | 4.4 | 3.4 | 1.7 | 4.8 | 3.0 |
| >hg38_21_+_8433172_8446622_RNA45SN1_45S_50ntFlanks@10780.10818.39\|AAGGTGAACAGCCTCTGGCATGTTGGAACAATGTAGGTA | >hg38_21_+_8433172_8446622_RNA45SN1_45S_50ntFlanks | 10780 | 10818 | 39 | AAGGTGAACAGCCTCTGGCATGTTGGAACAATGTAGGTA | MB468 | 10 | 4 | 12 | 4 | 9 | 5.4 | 1.2 | 5.2 | 2.6 | 5.5 |
| >hg38_21_+_8433172_8446622_RNA45SN1_45S_50ntFlanks@10781.10799.19\|AGGTGAACAGCCTCTGGCA | >hg38_21_+_8433172_8446622_RNA45SN1_45S_50ntFlanks | 10781 | 10799 | 19 | AGGTGAACAGCCTCTGGCA | MB468 | 16 | 2 | 17 | 19 | 4 | 6.2 | 1.4 | 10.6 | 7.7 | 1.8 |
| >hg38_21_+_8433172_8446622_RNA45SN1_45S_50ntFlanks@10781.10800.20\|AGGTGAACAGCCTCTGGCAT | >hg38_21_+_8433172_8446622_RNA45SN1_45S_50ntFlanks | 10781 | 10800 | 20 | AGGTGAACAGCCTCTGGCAT | MB468 | 13 | 3 | 9 | 10 | 2 | 2.7 | 1.8 | 3.2 | 3.7 | 0.7 |
| >hg38_21_+_8433172_8446622_RNA45SN1_45S_50ntFlanks@10781.10801.21\|AGGTGAACAGCCTCTGGCATG | >hg38_21_+_8433172_8446622_RNA45SN1_45S_50ntFlanks | 10781 | 10801 | 21 | AGGTGAACAGCCTCTGGCATG | MB468 | 81 | 20 | 64 | 68 | 27 | 18.8 | 7.4 | 21.3 | 14.3 | 8.2 |
| >hg38_21_+_8433172_8446622_RNA45SN1_45S_50ntFlanks@10781.10802.22\|AGGTGAACAGCCTCTGGCATGT | >hg38_21_+_8433172_8446622_RNA45SN1_45S_50ntFlanks | 10781 | 10802 | 22 | AGGTGAACAGCCTCTGGCATGT | MB468 | 27 | 9 | 31 | 11 | 6 | 1.8 | 3.6 | 14.9 | 4.3 | 1.4 |
| >hg38_21_+_8433172_8446622_RNA45SN1_45S_50ntFlanks@10781.10803.23\|AGGTGAACAGCCTCTGGCATGTT | >hg38_21_+_8433172_8446622_RNA45SN1_45S_50ntFlanks | 10781 | 10803 | 23 | AGGTGAACAGCCTCTGGCATGTT | MB468 | 32 | 6 | 38 | 13 | 8 | 13.2 | 1.1 | 23.1 | 2.4 | 2.8 |
| >hg38_21_+_8433172_8446622_RNA45SN1_45S_50ntFlanks@10781.10804.24\|AGGTGAACAGCCTCTGGCATGTTG | >hg38_21_+_8433172_8446622_RNA45SN1_45S_50ntFlanks | 10781 | 10804 | 24 | AGGTGAACAGCCTCTGGCATGTTG | MB468 | 32 | 7 | 33 | 26 | 11 | 19.9 | 0.7 | 30.9 | 11.4 | 3.4 |
| >hg38_21_+_8433172_8446622_RNA45SN1_45S_50ntFlanks@10781.10805.25\|AGGTGAACAGCCTCTGGCATGTTGG | >hg38_21_+_8433172_8446622_RNA45SN1_45S_50ntFlanks | 10781 | 10805 | 25 | AGGTGAACAGCCTCTGGCATGTTGG | MB468 | 36 | 11 | 38 | 19 | 10 | 11.3 | 1.5 | 24.9 | 6.6 | 4.4 |
| >hg38_21_+_8433172_8446622_RNA45SN1_45S_50ntFlanks@10781.10806.26\|AGGTGAACAGCCTCTGGCATGTTGGA | >hg38_21_+_8433172_8446622_RNA45SN1_45S_50ntFlanks | 10781 | 10806 | 26 | AGGTGAACAGCCTCTGGCATGTTGGA | MB468 | 28 | 15 | 29 | 17 | 9 | 12.0 | 4.7 | 16.2 | 9.1 | 3.6 |
| >hg38_21_+_8433172_8446622_RNA45SN1_45S_50ntFlanks@10781.10807.27\|AGGTGAACAGCCTCTGGCATGTTGGAA | >hg38_21_+_8433172_8446622_RNA45SN1_45S_50ntFlanks | 10781 | 10807 | 27 | AGGTGAACAGCCTCTGGCATGTTGGAA | MB468 | 39 | 28 | 33 | 15 | 12 | 20.5 | 6.1 | 25.6 | 10.9 | 4.4 |
| >hg38_21_+_8433172_8446622_RNA45SN1_45S_50ntFlanks@10781.10808.28\|AGGTGAACAGCCTCTGGCATGTTGGAAC | >hg38_21_+_8433172_8446622_RNA45SN1_45S_50ntFlanks | 10781 | 10808 | 28 | AGGTGAACAGCCTCTGGCATGTTGGAAC | MB468 | 14 | 11 | 14 | 6 | 4 | 5.2 | 4.3 | 11.7 | 1.3 | 1.1 |
| >hg38_21_+_8433172_8446622_RNA45SN1_45S_50ntFlanks@10781.10809.29\|AGGTGAACAGCCTCTGGCATGTTGGAACA | >hg38_21_+_8433172_8446622_RNA45SN1_45S_50ntFlanks | 10781 | 10809 | 29 | AGGTGAACAGCCTCTGGCATGTTGGAACA | MB468 | 13 | 13 | 16 | 11 | 14 | 3.1 | 5.0 | 10.3 | 2.7 | 3.5 |
| >hg38_21_+_8433172_8446622_RNA45SN1_45S_50ntFlanks@10781.10814.34\|AGGTGAACAGCCTCTGGCATGTTGGAACAATGTA | >hg38_21_+_8433172_8446622_RNA45SN1_45S_50ntFlanks | 10781 | 10814 | 34 | AGGTGAACAGCCTCTGGCATGTTGGAACAATGTA | MB468 | 14 | 11 | 14 | 7 | 5 | 2.8 | 1.6 | 9.1 | 3.3 | 2.4 |
| >hg38_21_+_8433172_8446622_RNA45SN1_45S_50ntFlanks@10781.10817.37\|AGGTGAACAGCCTCTGGCATGTTGGAACAATGTAGGT | >hg38_21_+_8433172_8446622_RNA45SN1_45S_50ntFlanks | 10781 | 10817 | 37 | AGGTGAACAGCCTCTGGCATGTTGGAACAATGTAGGT | MB468 | 37 | 19 | 24 | 23 | 23 | 14.4 | 4.1 | 11.4 | 14.9 | 10.7 |
| >hg38_21_+_8433172_8446622_RNA45SN1_45S_50ntFlanks@10781.10818.38\|AGGTGAACAGCCTCTGGCATGTTGGAACAATGTAGGTA | >hg38_21_+_8433172_8446622_RNA45SN1_45S_50ntFlanks | 10781 | 10818 | 38 | AGGTGAACAGCCTCTGGCATGTTGGAACAATGTAGGTA | MB468 | 194 | 68 | 200 | 70 | 138 | 110.4 | 9.2 | 95.7 | 54.4 | 68.4 |
| >hg38_21_+_8433172_8446622_RNA45SN1_45S_50ntFlanks@10782.10801.20\|GGTGAACAGCCTCTGGCATG | >hg38_21_+_8433172_8446622_RNA45SN1_45S_50ntFlanks | 10782 | 10801 | 20 | GGTGAACAGCCTCTGGCATG | MB468 | 63 | 38 | 55 | 29 | 26 | 8.1 | 11.9 | 13.8 | 13.7 | 2.7 |
| >hg38_21_+_8433172_8446622_RNA45SN1_45S_50ntFlanks@10782.10818.37\|GGTGAACAGCCTCTGGCATGTTGGAACAATGTAGGTA | >hg38_21_+_8433172_8446622_RNA45SN1_45S_50ntFlanks | 10782 | 10818 | 37 | GGTGAACAGCCTCTGGCATGTTGGAACAATGTAGGTA | MB468 | 40 | 10 | 34 | 15 | 29 | 19.6 | 3.5 | 16.2 | 10.6 | 12.5 |
| >hg38_21_+_8433172_8446622_RNA45SN1_45S_50ntFlanks@10783.10801.19\|GTGAACAGCCTCTGGCATG | >hg38_21_+_8433172_8446622_RNA45SN1_45S_50ntFlanks | 10783 | 10801 | 19 | GTGAACAGCCTCTGGCATG | MB468 | 28 | 6 | 31 | 37 | 7 | 9.8 | 1.4 | 20.8 | 16.3 | 2.3 |
| >hg38_21_+_8433172_8446622_RNA45SN1_45S_50ntFlanks@10783.10816.34\|GTGAACAGCCTCTGGCATGTTGGAACAATGTAGG | >hg38_21_+_8433172_8446622_RNA45SN1_45S_50ntFlanks | 10783 | 10816 | 34 | GTGAACAGCCTCTGGCATGTTGGAACAATGTAGG | MB468 | 11 | 14 | 27 | 5 | 4 | 4.1 | 8.4 | 22.2 | 2.1 | 1.9 |
| >hg38_21_+_8433172_8446622_RNA45SN1_45S_50ntFlanks@10783.10817.35\|GTGAACAGCCTCTGGCATGTTGGAACAATGTAGGT | >hg38_21_+_8433172_8446622_RNA45SN1_45S_50ntFlanks | 10783 | 10817 | 35 | GTGAACAGCCTCTGGCATGTTGGAACAATGTAGGT | MB468 | 30 | 22 | 33 | 21 | 13 | 9.6 | 7.2 | 19.4 | 11.4 | 4.9 |
| >hg38_21_+_8433172_8446622_RNA45SN1_45S_50ntFlanks@10783.10818.36\|GTGAACAGCCTCTGGCATGTTGGAACAATGTAGGTA | >hg38_21_+_8433172_8446622_RNA45SN1_45S_50ntFlanks | 10783 | 10818 | 36 | GTGAACAGCCTCTGGCATGTTGGAACAATGTAGGTA | MB468 | 138 | 53 | 163 | 70 | 74 | 65.6 | 20.8 | 99.3 | 47.6 | 35.0 |
| >hg38_21_+_8433172_8446622_RNA45SN1_45S_50ntFlanks@10784.10817.34\|TGAACAGCCTCTGGCATGTTGGAACAATGTAGGT | >hg38_21_+_8433172_8446622_RNA45SN1_45S_50ntFlanks | 10784 | 10817 | 34 | TGAACAGCCTCTGGCATGTTGGAACAATGTAGGT | MB468 | 11 | 4 | 9 | 8 | 7 | 2.1 | 1.2 | 3.5 | 3.8 | 3.3 |
| >hg38_21_+_8433172_8446622_RNA45SN1_45S_50ntFlanks@10784.10818.35\|TGAACAGCCTCTGGCATGTTGGAACAATGTAGGTA | >hg38_21_+_8433172_8446622_RNA45SN1_45S_50ntFlanks | 10784 | 10818 | 35 | TGAACAGCCTCTGGCATGTTGGAACAATGTAGGTA | MB468 | 50 | 11 | 42 | 24 | 35 | 6.6 | 0.9 | 17.5 | 14.5 | 11.7 |
| >hg38_21_+_8433172_8446622_RNA45SN1_45S_50ntFlanks@10786.10812.27\|AACAGCCTCTGGCATGTTGGAACAATG | >hg38_21_+_8433172_8446622_RNA45SN1_45S_50ntFlanks | 10786 | 10812 | 27 | AACAGCCTCTGGCATGTTGGAACAATG | MB468 | 11 | 6 | 27 | 16 | 8 | 2.9 | 0.8 | 17.2 | 10.2 | 1.5 |
| >hg38_21_+_8433172_8446622_RNA45SN1_45S_50ntFlanks@10786.10813.28\|AACAGCCTCTGGCATGTTGGAACAATGT | >hg38_21_+_8433172_8446622_RNA45SN1_45S_50ntFlanks | 10786 | 10813 | 28 | AACAGCCTCTGGCATGTTGGAACAATGT | MB468 | 10 | 4 | 6 | 15 | 12 | 6.0 | 1.8 | 2.5 | 11.3 | 8.9 |
| >hg38_21_+_8433172_8446622_RNA45SN1_45S_50ntFlanks@10786.10815.30\|AACAGCCTCTGGCATGTTGGAACAATGTAG | >hg38_21_+_8433172_8446622_RNA45SN1_45S_50ntFlanks | 10786 | 10815 | 30 | AACAGCCTCTGGCATGTTGGAACAATGTAG | MB468 | 25 | 26 | 38 | 18 | 14 | 3.2 | 10.0 | 20.0 | 5.9 | 1.2 |
| >hg38_21_+_8433172_8446622_RNA45SN1_45S_50ntFlanks@10786.10816.31\|AACAGCCTCTGGCATGTTGGAACAATGTAGG | >hg38_21_+_8433172_8446622_RNA45SN1_45S_50ntFlanks | 10786 | 10816 | 31 | AACAGCCTCTGGCATGTTGGAACAATGTAGG | MB468 | 117 | 88 | 212 | 50 | 41 | 17.6 | 35.6 | 92.2 | 17.5 | 19.6 |
| >hg38_21_+_8433172_8446622_RNA45SN1_45S_50ntFlanks@10786.10817.32\|AACAGCCTCTGGCATGTTGGAACAATGTAGGT | >hg38_21_+_8433172_8446622_RNA45SN1_45S_50ntFlanks | 10786 | 10817 | 32 | AACAGCCTCTGGCATGTTGGAACAATGTAGGT | MB468 | 145 | 33 | 72 | 94 | 61 | 26.7 | 14.1 | 28.0 | 33.4 | 6.8 |
| >hg38_21_+_8433172_8446622_RNA45SN1_45S_50ntFlanks@10786.10818.33\|AACAGCCTCTGGCATGTTGGAACAATGTAGGTA | >hg38_21_+_8433172_8446622_RNA45SN1_45S_50ntFlanks | 10786 | 10818 | 33 | AACAGCCTCTGGCATGTTGGAACAATGTAGGTA | MB468 | 799 | 98 | 399 | 378 | 387 | 254.8 | 19.4 | 137.1 | 163.1 | 149.1 |
| >hg38_21_+_8433172_8446622_RNA45SN1_45S_50ntFlanks@10786.10819.34\|AACAGCCTCTGGCATGTTGGAACAATGTAGGTAA | >hg38_21_+_8433172_8446622_RNA45SN1_45S_50ntFlanks | 10786 | 10819 | 34 | AACAGCCTCTGGCATGTTGGAACAATGTAGGTAA | MB468 | 11 | 3 | 5 | 10 | 13 | 0.9 | 0.3 | 2.6 | 5.3 | 3.6 |
| >hg38_21_+_8433172_8446622_RNA45SN1_45S_50ntFlanks@10786.10820.35\|AACAGCCTCTGGCATGTTGGAACAATGTAGGTAAG | >hg38_21_+_8433172_8446622_RNA45SN1_45S_50ntFlanks | 10786 | 10820 | 35 | AACAGCCTCTGGCATGTTGGAACAATGTAGGTAAG | MB468 | 34 | 9 | 26 | 25 | 47 | 1.4 | 2.7 | 10.8 | 10.4 | 17.4 |
| >hg38_21_+_8433172_8446622_RNA45SN1_45S_50ntFlanks@10786.10821.36\|AACAGCCTCTGGCATGTTGGAACAATGTAGGTAAGG | >hg38_21_+_8433172_8446622_RNA45SN1_45S_50ntFlanks | 10786 | 10821 | 36 | AACAGCCTCTGGCATGTTGGAACAATGTAGGTAAGG | MB468 | 36 | 14 | 26 | 27 | 61 | 4.7 | 1.8 | 11.9 | 12.0 | 27.4 |
| >hg38_21_+_8433172_8446622_RNA45SN1_45S_50ntFlanks@10786.10822.37\|AACAGCCTCTGGCATGTTGGAACAATGTAGGTAAGGG | >hg38_21_+_8433172_8446622_RNA45SN1_45S_50ntFlanks | 10786 | 10822 | 37 | AACAGCCTCTGGCATGTTGGAACAATGTAGGTAAGGG | MB468 | 64 | 30 | 45 | 59 | 121 | 10.5 | 11.9 | 22.2 | 33.5 | 58.4 |
| >hg38_21_+_8433172_8446622_RNA45SN1_45S_50ntFlanks@10786.10823.38\|AACAGCCTCTGGCATGTTGGAACAATGTAGGTAAGGGA | >hg38_21_+_8433172_8446622_RNA45SN1_45S_50ntFlanks | 10786 | 10823 | 38 | AACAGCCTCTGGCATGTTGGAACAATGTAGGTAAGGGA | MB468 | 43 | 27 | 30 | 47 | 123 | 5.8 | 7.7 | 17.3 | 29.7 | 63.8 |

| | | | | | | | | | | | | | | | |
|---|---|---|---|---|---|---|---|---|---|---|---|---|---|---|---|
| >hg38_21_+_8433172_8446622_RNA45SN1_45S_50ntFlanks@10786.10825.40\|AACAGCCTCTGGCATGTTGGAACAATGTAGGTAAGGGAAG | >hg38_21_+_8433172_8446622_RNA45SN1_45S_50ntFlanks | 10786 | 10825 | 40 | AACAGCCTCTGGCATGTTGGAACAATGTAGGTAAGGGAAG | MB468 | 44 | 36 | 37 | 30 | 126 | 14.9 | 17.0 | 22.4 | 21.8 | 69.9 |
| >hg38_21_+_8433172_8446622_RNA45SN1_45S_50ntFlanks@10786.10826.41\|AACAGCCTCTGGCATGTTGGAACAATGTAGGTAAGGGAAGT | >hg38_21_+_8433172_8446622_RNA45SN1_45S_50ntFlanks | 10786 | 10826 | 41 | AACAGCCTCTGGCATGTTGGAACAATGTAGGTAAGGGAAGT | MB468 | 49 | 54 | 51 | 40 | 209 | 8.3 | 37.3 | 39.2 | 29.9 | 153.6 |
| >hg38_21_+_8433172_8446622_RNA45SN1_45S_50ntFlanks@10786.10828.43\|AACAGCCTCTGGCATGTTGGAACAATGTAGGTAAGGGAAGTCG | >hg38_21_+_8433172_8446622_RNA45SN1_45S_50ntFlanks | 10786 | 10828 | 43 | AACAGCCTCTGGCATGTTGGAACAATGTAGGTAAGGGAAGTCG | MB468 | 12 | 17 | 19 | 10 | 90 | 4.1 | 15.8 | 14.5 | 7.1 | 66.1 |
| >hg38_21_+_8433172_8446622_RNA45SN1_45S_50ntFlanks@10787.10815.29\|ACAGCCTCTGGCATGTTGGAACAATGTAG | >hg38_21_+_8433172_8446622_RNA45SN1_45S_50ntFlanks | 10787 | 10815 | 29 | ACAGCCTCTGGCATGTTGGAACAATGTAG | MB468 | 19 | 23 | 18 | 6 | 5 | 3.2 | 11.1 | 10.4 | 3.1 | 1.6 |
| >hg38_21_+_8433172_8446622_RNA45SN1_45S_50ntFlanks@10787.10816.30\|ACAGCCTCTGGCATGTTGGAACAATGTAGG | >hg38_21_+_8433172_8446622_RNA45SN1_45S_50ntFlanks | 10787 | 10816 | 30 | ACAGCCTCTGGCATGTTGGAACAATGTAGG | MB468 | 37 | 38 | 45 | 18 | 12 | 5.1 | 14.2 | 23.3 | 7.7 | 6.9 |
| >hg38_21_+_8433172_8446622_RNA45SN1_45S_50ntFlanks@10787.10817.31\|ACAGCCTCTGGCATGTTGGAACAATGTAGGT | >hg38_21_+_8433172_8446622_RNA45SN1_45S_50ntFlanks | 10787 | 10817 | 31 | ACAGCCTCTGGCATGTTGGAACAATGTAGGT | MB468 | 31 | 6 | 11 | 23 | 14 | 9.3 | 3.5 | 3.5 | 7.1 | 1.4 |
| >hg38_21_+_8433172_8446622_RNA45SN1_45S_50ntFlanks@10787.10818.32\|ACAGCCTCTGGCATGTTGGAACAATGTAGGTA | >hg38_21_+_8433172_8446622_RNA45SN1_45S_50ntFlanks | 10787 | 10818 | 32 | ACAGCCTCTGGCATGTTGGAACAATGTAGGTA | MB468 | 210 | 25 | 86 | 109 | 111 | 93.3 | 7.7 | 26.8 | 39.1 | 56.4 |
| >hg38_21_+_8433172_8446622_RNA45SN1_45S_50ntFlanks@10787.10822.36\|ACAGCCTCTGGCATGTTGGAACAATGTAGGTAAGGG | >hg38_21_+_8433172_8446622_RNA45SN1_45S_50ntFlanks | 10787 | 10822 | 36 | ACAGCCTCTGGCATGTTGGAACAATGTAGGTAAGGG | MB468 | 13 | 7 | 9 | 13 | 20 | 2.0 | 3.0 | 3.4 | 5.3 | 10.6 |
| >hg38_21_+_8433172_8446622_RNA45SN1_45S_50ntFlanks@10787.10826.40\|ACAGCCTCTGGCATGTTGGAACAATGTAGGTAAGGGAAGT | >hg38_21_+_8433172_8446622_RNA45SN1_45S_50ntFlanks | 10787 | 10826 | 40 | ACAGCCTCTGGCATGTTGGAACAATGTAGGTAAGGGAAGT | MB468 | 10 | 11 | 9 | 11 | 35 | 2.2 | 6.0 | 6.1 | 7.6 | 20.0 |
| >hg38_21_+_8433172_8446622_RNA45SN1_45S_50ntFlanks@10788.10806.19\|CAGCCTCTGGCATGTTGGA | >hg38_21_+_8433172_8446622_RNA45SN1_45S_50ntFlanks | 10788 | 10806 | 19 | CAGCCTCTGGCATGTTGGA | MB468 | 12 | 2 | 24 | 8 | 4 | 5.2 | 0.6 | 13.8 | 0.8 | 2.2 |
| >hg38_21_+_8433172_8446622_RNA45SN1_45S_50ntFlanks@10788.10807.20\|CAGCCTCTGGCATGTTGGAA | >hg38_21_+_8433172_8446622_RNA45SN1_45S_50ntFlanks | 10788 | 10807 | 20 | CAGCCTCTGGCATGTTGGAA | MB468 | 16 | 8 | 22 | 6 | 5 | 8.7 | 1.6 | 16.2 | 2.1 | 1.9 |
| >hg38_21_+_8433172_8446622_RNA45SN1_45S_50ntFlanks@10788.10815.28\|CAGCCTCTGGCATGTTGGAACAATGTAG | >hg38_21_+_8433172_8446622_RNA45SN1_45S_50ntFlanks | 10788 | 10815 | 28 | CAGCCTCTGGCATGTTGGAACAATGTAG | MB468 | 33 | 53 | 40 | 9 | 11 | 6.6 | 31.5 | 22.8 | 5.4 | 7.9 |
| >hg38_21_+_8433172_8446622_RNA45SN1_45S_50ntFlanks@10788.10816.29\|CAGCCTCTGGCATGTTGGAACAATGTAGG | >hg38_21_+_8433172_8446622_RNA45SN1_45S_50ntFlanks | 10788 | 10816 | 29 | CAGCCTCTGGCATGTTGGAACAATGTAGG | MB468 | 55 | 44 | 79 | 27 | 22 | 4.3 | 18.1 | 38.8 | 12.1 | 11.5 |
| >hg38_21_+_8433172_8446622_RNA45SN1_45S_50ntFlanks@10788.10817.30\|CAGCCTCTGGCATGTTGGAACAATGTAGGT | >hg38_21_+_8433172_8446622_RNA45SN1_45S_50ntFlanks | 10788 | 10817 | 30 | CAGCCTCTGGCATGTTGGAACAATGTAGGT | MB468 | 30 | 2 | 10 | 27 | 16 | 15.4 | 0.8 | 3.8 | 10.4 | 4.0 |
| >hg38_21_+_8433172_8446622_RNA45SN1_45S_50ntFlanks@10788.10818.31\|CAGCCTCTGGCATGTTGGAACAATGTAGGTA | >hg38_21_+_8433172_8446622_RNA45SN1_45S_50ntFlanks | 10788 | 10818 | 31 | CAGCCTCTGGCATGTTGGAACAATGTAGGTA | MB468 | 298 | 21 | 105 | 177 | 184 | 173.9 | 7.0 | 40.9 | 80.7 | 110.8 |
| >hg38_21_+_8433172_8446622_RNA45SN1_45S_50ntFlanks@10788.10822.35\|CAGCCTCTGGCATGTTGGAACAATGTAGGTAAGGG | >hg38_21_+_8433172_8446622_RNA45SN1_45S_50ntFlanks | 10788 | 10822 | 35 | CAGCCTCTGGCATGTTGGAACAATGTAGGTAAGGG | MB468 | 15 | 5 | 9 | 14 | 20 | 5.0 | 1.5 | 2.6 | 6.4 | 9.4 |
| >hg38_21_+_8433172_8446622_RNA45SN1_45S_50ntFlanks@10788.10825.38\|CAGCCTCTGGCATGTTGGAACAATGTAGGTAAGGGAAG | >hg38_21_+_8433172_8446622_RNA45SN1_45S_50ntFlanks | 10788 | 10825 | 38 | CAGCCTCTGGCATGTTGGAACAATGTAGGTAAGGGAAG | MB468 | 10 | 7 | 7 | 8 | 20 | 5.6 | 4.2 | 2.9 | 5.4 | 8.5 |
| >hg38_21_+_8433172_8446622_RNA45SN1_45S_50ntFlanks@10789.10818.30\|AGCCTCTGGCATGTTGGAACAATGTAGGTA | >hg38_21_+_8433172_8446622_RNA45SN1_45S_50ntFlanks | 10789 | 10818 | 30 | AGCCTCTGGCATGTTGGAACAATGTAGGTA | MB468 | 54 | 7 | 12 | 39 | 35 | 30.2 | 2.3 | 5.6 | 14.3 | 23.7 |
| >hg38_21_+_8433172_8446622_RNA45SN1_45S_50ntFlanks@10791.10816.26\|CCTCTGGCATGTTGGAACAATGTAGG | >hg38_21_+_8433172_8446622_RNA45SN1_45S_50ntFlanks | 10791 | 10816 | 26 | CCTCTGGCATGTTGGAACAATGTAGG | MB468 | 13 | 10 | 7 | 13 | 6 | 1.8 | 5.1 | 3.2 | 4.5 | 2.0 |
| >hg38_21_+_8433172_8446622_RNA45SN1_45S_50ntFlanks@10791.10818.28\|CCTCTGGCATGTTGGAACAATGTAGGTA | >hg38_21_+_8433172_8446622_RNA45SN1_45S_50ntFlanks | 10791 | 10818 | 28 | CCTCTGGCATGTTGGAACAATGTAGGTA | MB468 | 32 | 5 | 16 | 15 | 14 | 17.7 | 2.2 | 2.5 | 4.5 | 0.2 |
| >hg38_21_+_8433172_8446622_RNA45SN1_45S_50ntFlanks@10791.10823.33\|CCTCTGGCATGTTGGAACAATGTAGGTAAGGGA | >hg38_21_+_8433172_8446622_RNA45SN1_45S_50ntFlanks | 10791 | 10823 | 33 | CCTCTGGCATGTTGGAACAATGTAGGTAAGGGA | MB468 | 12 | 4 | 4 | 15 | 19 | 1.8 | 0.7 | 0.7 | 4.9 | 7.7 |
| >hg38_21_+_8433172_8446622_RNA45SN1_45S_50ntFlanks@10792.10815.24\|CTCTGGCATGTTGGAACAATGTAG | >hg38_21_+_8433172_8446622_RNA45SN1_45S_50ntFlanks | 10792 | 10815 | 24 | CTCTGGCATGTTGGAACAATGTAG | MB468 | 21 | 16 | 15 | 10 | 3 | 7.7 | 9.9 | 9.7 | 3.2 | 1.4 |
| >hg38_21_+_8433172_8446622_RNA45SN1_45S_50ntFlanks@10792.10816.25\|CTCTGGCATGTTGGAACAATGTAGG | >hg38_21_+_8433172_8446622_RNA45SN1_45S_50ntFlanks | 10792 | 10816 | 25 | CTCTGGCATGTTGGAACAATGTAGG | MB468 | 24 | 15 | 9 | 22 | 7 | 8.1 | 10.2 | 5.0 | 9.2 | 2.2 |
| >hg38_21_+_8433172_8446622_RNA45SN1_45S_50ntFlanks@10792.10818.27\|CTCTGGCATGTTGGAACAATGTAGGTA | >hg38_21_+_8433172_8446622_RNA45SN1_45S_50ntFlanks | 10792 | 10818 | 27 | CTCTGGCATGTTGGAACAATGTAGGTA | MB468 | 53 | 8 | 18 | 25 | 21 | 32.4 | 3.4 | 5.3 | 11.4 | 6.8 |
| >hg38_21_+_8433172_8446622_RNA45SN1_45S_50ntFlanks@10792.10822.31\|CTCTGGCATGTTGGAACAATGTAGGTAAGGG | >hg38_21_+_8433172_8446622_RNA45SN1_45S_50ntFlanks | 10792 | 10822 | 31 | CTCTGGCATGTTGGAACAATGTAGGTAAGGG | MB468 | 17 | 11 | 8 | 19 | 16 | 6.4 | 6.2 | 3.6 | 6.9 | 7.5 |
| >hg38_21_+_8433172_8446622_RNA45SN1_45S_50ntFlanks@10793.10816.24\|TCTGGCATGTTGGAACAATGTAGG | >hg38_21_+_8433172_8446622_RNA45SN1_45S_50ntFlanks | 10793 | 10816 | 24 | TCTGGCATGTTGGAACAATGTAGG | MB468 | 19 | 6 | 5 | 10 | 4 | 7.4 | 3.4 | 2.1 | 2.6 | 1.7 |
| >hg38_21_+_8433172_8446622_RNA45SN1_45S_50ntFlanks@10793.10818.26\|TCTGGCATGTTGGAACAATGTAGGTA | >hg38_21_+_8433172_8446622_RNA45SN1_45S_50ntFlanks | 10793 | 10818 | 26 | TCTGGCATGTTGGAACAATGTAGGTA | MB468 | 51 | 6 | 11 | 25 | 20 | 37.1 | 2.8 | 2.6 | 10.6 | 8.2 |
| >hg38_21_+_8433172_8446622_RNA45SN1_45S_50ntFlanks@10794.10812.19\|CTGGCATGTTGGAACAATG | >hg38_21_+_8433172_8446622_RNA45SN1_45S_50ntFlanks | 10794 | 10812 | 19 | CTGGCATGTTGGAACAATG | MB468 | 13 | 9 | 14 | 9 | 2 | 4.1 | 5.6 | 8.7 | 1.9 | 0.7 |
| >hg38_21_+_8433172_8446622_RNA45SN1_45S_50ntFlanks@10794.10813.20\|CTGGCATGTTGGAACAATGT | >hg38_21_+_8433172_8446622_RNA45SN1_45S_50ntFlanks | 10794 | 10813 | 20 | CTGGCATGTTGGAACAATGT | MB468 | 14 | 3 | 5 | 10 | 6 | 9.2 | 1.8 | 4.9 | 3.3 | 4.3 |
| >hg38_21_+_8433172_8446622_RNA45SN1_45S_50ntFlanks@10794.10814.21\|CTGGCATGTTGGAACAATGTA | >hg38_21_+_8433172_8446622_RNA45SN1_45S_50ntFlanks | 10794 | 10814 | 21 | CTGGCATGTTGGAACAATGTA | MB468 | 28 | 5 | 16 | 9 | 3 | 18.5 | 2.5 | 15.2 | 1.4 | 1.5 |
| >hg38_21_+_8433172_8446622_RNA45SN1_45S_50ntFlanks@10794.10815.22\|CTGGCATGTTGGAACAATGTAG | >hg38_21_+_8433172_8446622_RNA45SN1_45S_50ntFlanks | 10794 | 10815 | 22 | CTGGCATGTTGGAACAATGTAG | MB468 | 53 | 10 | 22 | 16 | 6 | 35.6 | 4.7 | 24.4 | 3.6 | 4.4 |
| >hg38_21_+_8433172_8446622_RNA45SN1_45S_50ntFlanks@10794.10816.23\|CTGGCATGTTGGAACAATGTAGG | >hg38_21_+_8433172_8446622_RNA45SN1_45S_50ntFlanks | 10794 | 10816 | 23 | CTGGCATGTTGGAACAATGTAGG | MB468 | 200 | 49 | 54 | 111 | 43 | 84.8 | 24.8 | 16.6 | 25.7 | 21.8 |
| >hg38_21_+_8433172_8446622_RNA45SN1_45S_50ntFlanks@10794.10817.24\|CTGGCATGTTGGAACAATGTAGGT | >hg38_21_+_8433172_8446622_RNA45SN1_45S_50ntFlanks | 10794 | 10817 | 24 | CTGGCATGTTGGAACAATGTAGGT | MB468 | 78 | 16 | 22 | 52 | 33 | 36.4 | 9.7 | 6.4 | 23.3 | 12.3 |
| >hg38_21_+_8433172_8446622_RNA45SN1_45S_50ntFlanks@10794.10818.25\|CTGGCATGTTGGAACAATGTAGGTA | >hg38_21_+_8433172_8446622_RNA45SN1_45S_50ntFlanks | 10794 | 10818 | 25 | CTGGCATGTTGGAACAATGTAGGTA | MB468 | 402 | 65 | 167 | 154 | 148 | 247.7 | 37.4 | 60.8 | 72.4 | 47.6 |
| >hg38_21_+_8433172_8446622_RNA45SN1_45S_50ntFlanks@10794.10820.27\|CTGGCATGTTGGAACAATGTAGGTAAG | >hg38_21_+_8433172_8446622_RNA45SN1_45S_50ntFlanks | 10794 | 10820 | 27 | CTGGCATGTTGGAACAATGTAGGTAAG | MB468 | 15 | 5 | 8 | 12 | 11 | 5.2 | 2.1 | 1.0 | 4.6 | 0.9 |
| >hg38_21_+_8433172_8446622_RNA45SN1_45S_50ntFlanks@10794.10821.28\|CTGGCATGTTGGAACAATGTAGGTAAGG | >hg38_21_+_8433172_8446622_RNA45SN1_45S_50ntFlanks | 10794 | 10821 | 28 | CTGGCATGTTGGAACAATGTAGGTAAGG | MB468 | 17 | 6 | 10 | 13 | 11 | 4.7 | 1.8 | 0.5 | 5.2 | 2.2 |
| >hg38_21_+_8433172_8446622_RNA45SN1_45S_50ntFlanks@10794.10822.29\|CTGGCATGTTGGAACAATGTAGGTAAGGG | >hg38_21_+_8433172_8446622_RNA45SN1_45S_50ntFlanks | 10794 | 10822 | 29 | CTGGCATGTTGGAACAATGTAGGTAAGGG | MB468 | 38 | 21 | 26 | 37 | 29 | 13.2 | 9.5 | 8.5 | 7.1 | 8.9 |
| >hg38_21_+_8433172_8446622_RNA45SN1_45S_50ntFlanks@10794.10823.30\|CTGGCATGTTGGAACAATGTAGGTAAGGGA | >hg38_21_+_8433172_8446622_RNA45SN1_45S_50ntFlanks | 10794 | 10823 | 30 | CTGGCATGTTGGAACAATGTAGGTAAGGGA | MB468 | 26 | 17 | 14 | 32 | 26 | 7.1 | 7.4 | 4.6 | 6.4 | 6.3 |
| >hg38_21_+_8433172_8446622_RNA45SN1_45S_50ntFlanks@10794.10825.32\|CTGGCATGTTGGAACAATGTAGGTAAGGGAAG | >hg38_21_+_8433172_8446622_RNA45SN1_45S_50ntFlanks | 10794 | 10825 | 32 | CTGGCATGTTGGAACAATGTAGGTAAGGGAAG | MB468 | 37 | 28 | 24 | 31 | 39 | 18.7 | 15.9 | 17.0 | 9.0 | 16.4 |
| >hg38_21_+_8433172_8446622_RNA45SN1_45S_50ntFlanks@10794.10826.33\|CTGGCATGTTGGAACAATGTAGGTAAGGGAAGT | >hg38_21_+_8433172_8446622_RNA45SN1_45S_50ntFlanks | 10794 | 10826 | 33 | CTGGCATGTTGGAACAATGTAGGTAAGGGAAGT | MB468 | 30 | 21 | 16 | 39 | 38 | 10.8 | 7.2 | 7.4 | 15.1 | 12.1 |
| >hg38_21_+_8433172_8446622_RNA45SN1_45S_50ntFlanks@10794.10828.35\|CTGGCATGTTGGAACAATGTAGGTAAGGGAAGTCG | >hg38_21_+_8433172_8446622_RNA45SN1_45S_50ntFlanks | 10794 | 10828 | 35 | CTGGCATGTTGGAACAATGTAGGTAAGGGAAGTCG | MB468 | 11 | 7 | 7 | 12 | 20 | 5.0 | 3.7 | 3.8 | 4.1 | 9.8 |
| >hg38_21_+_8433172_8446622_RNA45SN1_45S_50ntFlanks@10795.10816.22\|TGGCATGTTGGAACAATGTAGG | >hg38_21_+_8433172_8446622_RNA45SN1_45S_50ntFlanks | 10795 | 10816 | 22 | TGGCATGTTGGAACAATGTAGG | MB468 | 15 | 3 | 9 | 4 | 2 | 6.1 | 2.2 | 4.6 | 1.3 | 0.6 |
| >hg38_21_+_8433172_8446622_RNA45SN1_45S_50ntFlanks@10795.10817.23\|TGGCATGTTGGAACAATGTAGGT | >hg38_21_+_8433172_8446622_RNA45SN1_45S_50ntFlanks | 10795 | 10817 | 23 | TGGCATGTTGGAACAATGTAGGT | MB468 | 25 | 6 | 12 | 13 | 8 | 8.4 | 3.6 | 3.9 | 6.2 | 2.0 |
| >hg38_21_+_8433172_8446622_RNA45SN1_45S_50ntFlanks@10795.10818.24\|TGGCATGTTGGAACAATGTAGGTA | >hg38_21_+_8433172_8446622_RNA45SN1_45S_50ntFlanks | 10795 | 10818 | 24 | TGGCATGTTGGAACAATGTAGGTA | MB468 | 119 | 31 | 90 | 37 | 39 | 50.6 | 14.0 | 35.7 | 10.6 | 11.8 |
| >hg38_21_+_8433172_8446622_RNA45SN1_45S_50ntFlanks@10796.10816.21\|GGCATGTTGGAACAATGTAGG | >hg38_21_+_8433172_8446622_RNA45SN1_45S_50ntFlanks | 10796 | 10816 | 21 | GGCATGTTGGAACAATGTAGG | MB468 | 10 | 2 | 6 | 4 | 2 | 5.2 | 1.6 | 2.4 | 1.6 | 0.9 |
| >hg38_21_+_8433172_8446622_RNA45SN1_45S_50ntFlanks@10796.10817.22\|GGCATGTTGGAACAATGTAGGT | >hg38_21_+_8433172_8446622_RNA45SN1_45S_50ntFlanks | 10796 | 10817 | 22 | GGCATGTTGGAACAATGTAGGT | MB468 | 14 | 4 | 7 | 12 | 6 | 6.1 | 2.1 | 2.6 | 5.7 | 2.1 |
| >hg38_21_+_8433172_8446622_RNA45SN1_45S_50ntFlanks@10796.10818.23\|GGCATGTTGGAACAATGTAGGTA | >hg38_21_+_8433172_8446622_RNA45SN1_45S_50ntFlanks | 10796 | 10818 | 23 | GGCATGTTGGAACAATGTAGGTA | MB468 | 75 | 19 | 49 | 41 | 26 | 38.8 | 9.8 | 19.5 | 12.1 | 11.4 |
| >hg38_21_+_8433172_8446622_RNA45SN1_45S_50ntFlanks@10797.10816.20\|GCATGTTGGAACAATGTAGG | >hg38_21_+_8433172_8446622_RNA45SN1_45S_50ntFlanks | 10797 | 10816 | 20 | GCATGTTGGAACAATGTAGG | MB468 | 28 | 3 | 17 | 11 | 2 | 19.4 | 2.3 | 13.4 | 3.9 | 1.0 |
| >hg38_21_+_8433172_8446622_RNA45SN1_45S_50ntFlanks@10797.10817.21\|GCATGTTGGAACAATGTAGGT | >hg38_21_+_8433172_8446622_RNA45SN1_45S_50ntFlanks | 10797 | 10817 | 21 | GCATGTTGGAACAATGTAGGT | MB468 | 69 | 10 | 33 | 49 | 15 | 38.9 | 8.9 | 15.9 | 16.5 | 5.8 |
| >hg38_21_+_8433172_8446622_RNA45SN1_45S_50ntFlanks@10797.10818.22\|GCATGTTGGAACAATGTAGGTA | >hg38_21_+_8433172_8446622_RNA45SN1_45S_50ntFlanks | 10797 | 10818 | 22 | GCATGTTGGAACAATGTAGGTA | MB468 | 388 | 48 | 255 | 187 | 89 | 225.1 | 31.3 | 141.1 | 43.6 | 40.1 |

| | | | | | | | | | | | | | | | | |
|---|---|---|---|---|---|---|---|---|---|---|---|---|---|---|---|---|
| >hg38_21_+_8433172_8446622_RNA45SN1_45S_50ntFlanks@10797.10820.24\|GCATGTTGGAACAATGTAGGTAAG | >hg38_21_+_8433172_8446622_RNA45SN1_45S_50ntFlanks | 10797 | 10820 | 24 | GCATGTTGGAACAATGTAGGTAAG | MB468 | 11 | 3 | 10 | 13 | 6 | 3.4 | 1.3 | 2.5 | 7.8 | 1.1 |
| >hg38_21_+_8433172_8446622_RNA45SN1_45S_50ntFlanks@10797.10822.26\|GCATGTTGGAACAATGTAGGTAAGGG | >hg38_21_+_8433172_8446622_RNA45SN1_45S_50ntFlanks | 10797 | 10822 | 26 | GCATGTTGGAACAATGTAGGTAAGGG | MB468 | 21 | 11 | 20 | 49 | 16 | 7.1 | 4.8 | 5.8 | 16.9 | 1.9 |
| >hg38_21_+_8433172_8446622_RNA45SN1_45S_50ntFlanks@10797.10823.27\|GCATGTTGGAACAATGTAGGTAAGGGA | >hg38_21_+_8433172_8446622_RNA45SN1_45S_50ntFlanks | 10797 | 10823 | 27 | GCATGTTGGAACAATGTAGGTAAGGGA | MB468 | 14 | 8 | 11 | 38 | 14 | 3.4 | 2.8 | 3.3 | 15.0 | 1.9 |
| >hg38_21_+_8433172_8446622_RNA45SN1_45S_50ntFlanks@10797.10825.29\|GCATGTTGGAACAATGTAGGTAAGGGAAG | >hg38_21_+_8433172_8446622_RNA45SN1_45S_50ntFlanks | 10797 | 10825 | 29 | GCATGTTGGAACAATGTAGGTAAGGGAAG | MB468 | 20 | 13 | 15 | 29 | 21 | 10.2 | 8.0 | 11.2 | 7.9 | 4.3 |
| >hg38_21_+_8433172_8446622_RNA45SN1_45S_50ntFlanks@10797.10826.30\|GCATGTTGGAACAATGTAGGTAAGGGAAGT | >hg38_21_+_8433172_8446622_RNA45SN1_45S_50ntFlanks | 10797 | 10826 | 30 | GCATGTTGGAACAATGTAGGTAAGGGAAGT | MB468 | 18 | 11 | 11 | 48 | 22 | 6.2 | 3.3 | 5.3 | 12.7 | 5.2 |
| >hg38_21_+_8433172_8446622_RNA45SN1_45S_50ntFlanks@10798.10815.18\|CATGTTGGAACAATGTAG | >hg38_21_+_8433172_8446622_RNA45SN1_45S_50ntFlanks | 10798 | 10815 | 18 | CATGTTGGAACAATGTAG | MB468 | 14 | 1 | 3 | 7 | 1 | 9.7 | 1.2 | 2.1 | 2.0 | 0.8 |
| >hg38_21_+_8433172_8446622_RNA45SN1_45S_50ntFlanks@10798.10816.19\|CATGTTGGAACAATGTAGG | >hg38_21_+_8433172_8446622_RNA45SN1_45S_50ntFlanks | 10798 | 10816 | 19 | CATGTTGGAACAATGTAGG | MB468 | 61 | 7 | 39 | 17 | 5 | 31.3 | 4.6 | 16.9 | 5.7 | 2.1 |
| >hg38_21_+_8433172_8446622_RNA45SN1_45S_50ntFlanks@10798.10817.20\|CATGTTGGAACAATGTAGGT | >hg38_21_+_8433172_8446622_RNA45SN1_45S_50ntFlanks | 10798 | 10817 | 20 | CATGTTGGAACAATGTAGGT | MB468 | 78 | 9 | 54 | 44 | 19 | 37.4 | 5.3 | 17.8 | 22.1 | 7.1 |
| >hg38_21_+_8433172_8446622_RNA45SN1_45S_50ntFlanks@10798.10818.21\|CATGTTGGAACAATGTAGGTA | >hg38_21_+_8433172_8446622_RNA45SN1_45S_50ntFlanks | 10798 | 10818 | 21 | CATGTTGGAACAATGTAGGTA | MB468 | 321 | 39 | 310 | 109 | 81 | 182.1 | 22.7 | 163.7 | 34.6 | 36.8 |
| >hg38_21_+_8433172_8446622_RNA45SN1_45S_50ntFlanks@10798.10822.25\|CATGTTGGAACAATGTAGGTAAGGG | >hg38_21_+_8433172_8446622_RNA45SN1_45S_50ntFlanks | 10798 | 10822 | 25 | CATGTTGGAACAATGTAGGTAAGGG | MB468 | 21 | 14 | 19 | 38 | 21 | 7.0 | 4.5 | 6.2 | 11.8 | 4.3 |
| >hg38_21_+_8433172_8446622_RNA45SN1_45S_50ntFlanks@10798.10823.26\|CATGTTGGAACAATGTAGGTAAGGGA | >hg38_21_+_8433172_8446622_RNA45SN1_45S_50ntFlanks | 10798 | 10823 | 26 | CATGTTGGAACAATGTAGGTAAGGGA | MB468 | 14 | 11 | 11 | 29 | 18 | 4.0 | 3.6 | 2.7 | 11.6 | 3.7 |
| >hg38_21_+_8433172_8446622_RNA45SN1_45S_50ntFlanks@10798.10825.28\|CATGTTGGAACAATGTAGGTAAGGGAAG | >hg38_21_+_8433172_8446622_RNA45SN1_45S_50ntFlanks | 10798 | 10825 | 28 | CATGTTGGAACAATGTAGGTAAGGGAAG | MB468 | 22 | 18 | 17 | 24 | 23 | 9.5 | 7.9 | 10.1 | 6.9 | 3.3 |
| >hg38_21_+_8433172_8446622_RNA45SN1_45S_50ntFlanks@10798.10826.29\|CATGTTGGAACAATGTAGGTAAGGGAAGT | >hg38_21_+_8433172_8446622_RNA45SN1_45S_50ntFlanks | 10798 | 10826 | 29 | CATGTTGGAACAATGTAGGTAAGGGAAGT | MB468 | 21 | 16 | 13 | 36 | 26 | 5.5 | 4.2 | 5.1 | 11.0 | 6.4 |
| >hg38_21_+_8433172_8446622_RNA45SN1_45S_50ntFlanks@10800.10818.19\|TGTTGGAACAATGTAGGTA | >hg38_21_+_8433172_8446622_RNA45SN1_45S_50ntFlanks | 10800 | 10818 | 19 | TGTTGGAACAATGTAGGTA | MB468 | 26 | 5 | 31 | 6 | 5 | 17.1 | 3.0 | 26.1 | 3.7 | 3.1 |
| >hg38_21_+_8433172_8446622_RNA45SN1_45S_50ntFlanks@10801.10818.18\|GTTGGAACAATGTAGGTA | >hg38_21_+_8433172_8446622_RNA45SN1_45S_50ntFlanks | 10801 | 10818 | 18 | GTTGGAACAATGTAGGTA | MB468 | 41 | 9 | 56 | 9 | 7 | 29.7 | 6.1 | 47.3 | 5.0 | 5.6 |
| >hg38_21_+_8433172_8446622_RNA45SN1_45S_50ntFlanks@10802.10822.21\|TTGGAACAATGTAGGTAAGGG | >hg38_21_+_8433172_8446622_RNA45SN1_45S_50ntFlanks | 10802 | 10822 | 21 | TTGGAACAATGTAGGTAAGGG | MB468 | 15 | 8 | 18 | 27 | 8 | 2.7 | 2.0 | 2.3 | 6.7 | 0.6 |
| >hg38_21_+_8433172_8446622_RNA45SN1_45S_50ntFlanks@10802.10825.24\|TTGGAACAATGTAGGTAAGGGAAG | >hg38_21_+_8433172_8446622_RNA45SN1_45S_50ntFlanks | 10802 | 10825 | 24 | TTGGAACAATGTAGGTAAGGGAAG | MB468 | 11 | 11 | 12 | 16 | 10 | 4.4 | 4.7 | 5.9 | 4.9 | 1.4 |
| >hg38_21_+_8433172_8446622_RNA45SN1_45S_50ntFlanks@10802.10826.25\|TTGGAACAATGTAGGTAAGGGAAGT | >hg38_21_+_8433172_8446622_RNA45SN1_45S_50ntFlanks | 10802 | 10826 | 25 | TTGGAACAATGTAGGTAAGGGAAGT | MB468 | 11 | 10 | 10 | 27 | 11 | 1.9 | 1.9 | 2.2 | 7.4 | 3.6 |
| >hg38_21_+_8433172_8446622_RNA45SN1_45S_50ntFlanks@10805.10822.18\|GAACAATGTAGGTAAGGG | >hg38_21_+_8433172_8446622_RNA45SN1_45S_50ntFlanks | 10805 | 10822 | 18 | GAACAATGTAGGTAAGGG | MB468 | 29 | 15 | 26 | 12 | 17 | 8.1 | 7.6 | 3.6 | 3.2 | 3.5 |
| >hg38_21_+_8433172_8446622_RNA45SN1_45S_50ntFlanks@10805.10823.19\|GAACAATGTAGGTAAGGGA | >hg38_21_+_8433172_8446622_RNA45SN1_45S_50ntFlanks | 10805 | 10823 | 19 | GAACAATGTAGGTAAGGGA | MB468 | 26 | 16 | 22 | 16 | 23 | 7.1 | 7.7 | 5.1 | 4.6 | 1.9 |
| >hg38_21_+_8433172_8446622_RNA45SN1_45S_50ntFlanks@10805.10825.21\|GAACAATGTAGGTAAGGGAAG | >hg38_21_+_8433172_8446622_RNA45SN1_45S_50ntFlanks | 10805 | 10825 | 21 | GAACAATGTAGGTAAGGGAAG | MB468 | 18 | 18 | 16 | 8 | 17 | 7.9 | 12.5 | 5.7 | 1.1 | 4.3 |
| >hg38_21_+_8433172_8446622_RNA45SN1_45S_50ntFlanks@10805.10826.22\|GAACAATGTAGGTAAGGGAAGT | >hg38_21_+_8433172_8446622_RNA45SN1_45S_50ntFlanks | 10805 | 10826 | 22 | GAACAATGTAGGTAAGGGAAGT | MB468 | 20 | 18 | 15 | 10 | 23 | 3.8 | 8.2 | 1.1 | 2.1 | 2.7 |
| >hg38_21_+_8433172_8446622_RNA45SN1_45S_50ntFlanks@10805.10828.24\|GAACAATGTAGGTAAGGGAAGTCG | >hg38_21_+_8433172_8446622_RNA45SN1_45S_50ntFlanks | 10805 | 10828 | 24 | GAACAATGTAGGTAAGGGAAGTCG | MB468 | 14 | 12 | 14 | 7 | 15 | 3.3 | 4.9 | 3.2 | 2.0 | 2.8 |
| >hg38_21_+_8433172_8446622_RNA45SN1_45S_50ntFlanks@10805.10840.36\|GAACAATGTAGGTAAGGGAAGTCGGCAAGCCGGATC | >hg38_21_+_8433172_8446622_RNA45SN1_45S_50ntFlanks | 10805 | 10840 | 36 | GAACAATGTAGGTAAGGGAAGTCGGCAAGCCGGATC | MB468 | 14 | 16 | 19 | 12 | 32 | 2.6 | 4.5 | 2.9 | 4.4 | 15.2 |
| >hg38_21_+_8433172_8446622_RNA45SN1_45S_50ntFlanks@10806.10823.18\|AACAATGTAGGTAAGGGA | >hg38_21_+_8433172_8446622_RNA45SN1_45S_50ntFlanks | 10806 | 10823 | 18 | AACAATGTAGGTAAGGGA | MB468 | 53 | 39 | 44 | 33 | 50 | 12.0 | 22.0 | 15.5 | 13.7 | 9.2 |
| >hg38_21_+_8433172_8446622_RNA45SN1_45S_50ntFlanks@10806.10825.20\|AACAATGTAGGTAAGGGAAG | >hg38_21_+_8433172_8446622_RNA45SN1_45S_50ntFlanks | 10806 | 10825 | 20 | AACAATGTAGGTAAGGGAAG | MB468 | 36 | 42 | 32 | 18 | 40 | 14.8 | 34.2 | 15.7 | 5.7 | 8.5 |
| >hg38_21_+_8433172_8446622_RNA45SN1_45S_50ntFlanks@10806.10826.21\|AACAATGTAGGTAAGGGAAGT | >hg38_21_+_8433172_8446622_RNA45SN1_45S_50ntFlanks | 10806 | 10826 | 21 | AACAATGTAGGTAAGGGAAGT | MB468 | 40 | 38 | 29 | 20 | 48 | 5.4 | 21.4 | 5.9 | 6.7 | 8.1 |
| >hg38_21_+_8433172_8446622_RNA45SN1_45S_50ntFlanks@10806.10827.22\|AACAATGTAGGTAAGGGAAGTC | >hg38_21_+_8433172_8446622_RNA45SN1_45S_50ntFlanks | 10806 | 10827 | 22 | AACAATGTAGGTAAGGGAAGTC | MB468 | 14 | 16 | 13 | 11 | 16 | 3.4 | 12.8 | 5.6 | 3.7 | 3.6 |
| >hg38_21_+_8433172_8446622_RNA45SN1_45S_50ntFlanks@10806.10828.23\|AACAATGTAGGTAAGGGAAGTCG | >hg38_21_+_8433172_8446622_RNA45SN1_45S_50ntFlanks | 10806 | 10828 | 23 | AACAATGTAGGTAAGGGAAGTCG | MB468 | 27 | 28 | 28 | 15 | 36 | 4.0 | 13.6 | 6.5 | 4.4 | 5.8 |
| >hg38_21_+_8433172_8446622_RNA45SN1_45S_50ntFlanks@10806.10829.24\|AACAATGTAGGTAAGGGAAGTCGG | >hg38_21_+_8433172_8446622_RNA45SN1_45S_50ntFlanks | 10806 | 10829 | 24 | AACAATGTAGGTAAGGGAAGTCGG | MB468 | 12 | 22 | 15 | 12 | 20 | 3.2 | 9.3 | 2.8 | 5.8 | 5.8 |
| >hg38_21_+_8433172_8446622_RNA45SN1_45S_50ntFlanks@10806.10830.25\|AACAATGTAGGTAAGGGAAGTCGGC | >hg38_21_+_8433172_8446622_RNA45SN1_45S_50ntFlanks | 10806 | 10830 | 25 | AACAATGTAGGTAAGGGAAGTCGGC | MB468 | 11 | 14 | 14 | 10 | 13 | 3.8 | 3.6 | 3.9 | 3.8 | 4.5 |
| >hg38_21_+_8433172_8446622_RNA45SN1_45S_50ntFlanks@10806.10840.35\|AACAATGTAGGTAAGGGAAGTCGGCAAGCCGGATC | >hg38_21_+_8433172_8446622_RNA45SN1_45S_50ntFlanks | 10806 | 10840 | 35 | AACAATGTAGGTAAGGGAAGTCGGCAAGCCGGATC | MB468 | 30 | 37 | 37 | 32 | 76 | 4.0 | 11.8 | 5.5 | 11.5 | 32.5 |
| >hg38_21_+_8433172_8446622_RNA45SN1_45S_50ntFlanks@10807.10825.19\|ACAATGTAGGTAAGGGAAG | >hg38_21_+_8433172_8446622_RNA45SN1_45S_50ntFlanks | 10807 | 10825 | 19 | ACAATGTAGGTAAGGGAAG | MB468 | 24 | 23 | 27 | 10 | 29 | 10.1 | 20.3 | 16.3 | 3.6 | 7.8 |
| >hg38_21_+_8433172_8446622_RNA45SN1_45S_50ntFlanks@10807.10826.20\|ACAATGTAGGTAAGGGAAGT | >hg38_21_+_8433172_8446622_RNA45SN1_45S_50ntFlanks | 10807 | 10826 | 20 | ACAATGTAGGTAAGGGAAGT | MB468 | 24 | 19 | 22 | 12 | 34 | 4.2 | 12.2 | 6.8 | 3.7 | 6.3 |
| >hg38_21_+_8433172_8446622_RNA45SN1_45S_50ntFlanks@10807.10828.22\|ACAATGTAGGTAAGGGAAGTCG | >hg38_21_+_8433172_8446622_RNA45SN1_45S_50ntFlanks | 10807 | 10828 | 22 | ACAATGTAGGTAAGGGAAGTCG | MB468 | 16 | 15 | 19 | 10 | 27 | 2.2 | 9.9 | 4.6 | 2.7 | 8.1 |
| >hg38_21_+_8433172_8446622_RNA45SN1_45S_50ntFlanks@10807.10840.34\|ACAATGTAGGTAAGGGAAGTCGGCAAGCCGGATC | >hg38_21_+_8433172_8446622_RNA45SN1_45S_50ntFlanks | 10807 | 10840 | 34 | ACAATGTAGGTAAGGGAAGTCGGCAAGCCGGATC | MB468 | 18 | 21 | 21 | 24 | 52 | 2.0 | 8.3 | 2.3 | 9.8 | 23.1 |
| >hg38_21_+_8433172_8446622_RNA45SN1_45S_50ntFlanks@10808.10825.18\|CAATGTAGGTAAGGGAAG | >hg38_21_+_8433172_8446622_RNA45SN1_45S_50ntFlanks | 10808 | 10825 | 18 | CAATGTAGGTAAGGGAAG | MB468 | 99 | 105 | 128 | 37 | 137 | 49.7 | 102.5 | 77.2 | 14.6 | 70.1 |
| >hg38_21_+_8433172_8446622_RNA45SN1_45S_50ntFlanks@10808.10826.19\|CAATGTAGGTAAGGGAAGT | >hg38_21_+_8433172_8446622_RNA45SN1_45S_50ntFlanks | 10808 | 10826 | 19 | CAATGTAGGTAAGGGAAGT | MB468 | 95 | 87 | 106 | 41 | 139 | 28.8 | 64.5 | 30.0 | 14.1 | 49.5 |
| >hg38_21_+_8433172_8446622_RNA45SN1_45S_50ntFlanks@10808.10827.20\|CAATGTAGGTAAGGGAAGTC | >hg38_21_+_8433172_8446622_RNA45SN1_45S_50ntFlanks | 10808 | 10827 | 20 | CAATGTAGGTAAGGGAAGTC | MB468 | 32 | 37 | 46 | 22 | 51 | 11.0 | 32.8 | 24.6 | 6.8 | 29.2 |
| >hg38_21_+_8433172_8446622_RNA45SN1_45S_50ntFlanks@10808.10828.21\|CAATGTAGGTAAGGGAAGTCG | >hg38_21_+_8433172_8446622_RNA45SN1_45S_50ntFlanks | 10808 | 10828 | 21 | CAATGTAGGTAAGGGAAGTCG | MB468 | 59 | 68 | 88 | 30 | 119 | 11.9 | 50.6 | 18.9 | 8.9 | 67.7 |
| >hg38_21_+_8433172_8446622_RNA45SN1_45S_50ntFlanks@10808.10829.22\|CAATGTAGGTAAGGGAAGTCGG | >hg38_21_+_8433172_8446622_RNA45SN1_45S_50ntFlanks | 10808 | 10829 | 22 | CAATGTAGGTAAGGGAAGTCGG | MB468 | 23 | 40 | 38 | 21 | 48 | 6.4 | 29.6 | 7.0 | 8.3 | 23.6 |
| >hg38_21_+_8433172_8446622_RNA45SN1_45S_50ntFlanks@10808.10830.23\|CAATGTAGGTAAGGGAAGTCGGC | >hg38_21_+_8433172_8446622_RNA45SN1_45S_50ntFlanks | 10808 | 10830 | 23 | CAATGTAGGTAAGGGAAGTCGGC | MB468 | 24 | 30 | 43 | 25 | 37 | 6.4 | 14.5 | 18.9 | 7.1 | 12.1 |
| >hg38_21_+_8433172_8446622_RNA45SN1_45S_50ntFlanks@10808.10831.24\|CAATGTAGGTAAGGGAAGTCGGCA | >hg38_21_+_8433172_8446622_RNA45SN1_45S_50ntFlanks | 10808 | 10831 | 24 | CAATGTAGGTAAGGGAAGTCGGCA | MB468 | 13 | 14 | 18 | 13 | 25 | 3.9 | 7.5 | 4.2 | 3.3 | 9.2 |
| >hg38_21_+_8433172_8446622_RNA45SN1_45S_50ntFlanks@10808.10837.30\|CAATGTAGGTAAGGGAAGTCGGCAAGCCGG | >hg38_21_+_8433172_8446622_RNA45SN1_45S_50ntFlanks | 10808 | 10837 | 30 | CAATGTAGGTAAGGGAAGTCGGCAAGCCGG | MB468 | 14 | 19 | 19 | 18 | 29 | 1.6 | 11.3 | 4.6 | 7.9 | 12.8 |
| >hg38_21_+_8433172_8446622_RNA45SN1_45S_50ntFlanks@10808.10840.33\|CAATGTAGGTAAGGGAAGTCGGCAAGCCGGATC | >hg38_21_+_8433172_8446622_RNA45SN1_45S_50ntFlanks | 10808 | 10840 | 33 | CAATGTAGGTAAGGGAAGTCGGCAAGCCGGATC | MB468 | 75 | 105 | 103 | 104 | 227 | 10.6 | 44.2 | 14.8 | 41.6 | 125.6 |
| >hg38_21_+_8433172_8446622_RNA45SN1_45S_50ntFlanks@10808.10844.37\|CAATGTAGGTAAGGGAAGTCGGCAAGCCGGATCCGTA | >hg38_21_+_8433172_8446622_RNA45SN1_45S_50ntFlanks | 10808 | 10844 | 37 | CAATGTAGGTAAGGGAAGTCGGCAAGCCGGATCCGTA | MB468 | 10 | 12 | 14 | 6 | 37 | 3.2 | 5.2 | 3.2 | 3.2 | 20.1 |
| >hg38_21_+_8433172_8446622_RNA45SN1_45S_50ntFlanks@10809.10826.18\|AATGTAGGTAAGGGAAGT | >hg38_21_+_8433172_8446622_RNA45SN1_45S_50ntFlanks | 10809 | 10826 | 18 | AATGTAGGTAAGGGAAGT | MB231 | 10 | 28 | 20 | 4 | 3 | 6.8 | 21.1 | 22.6 | 2.4 | 1.6 |
| >hg38_21_+_8433172_8446622_RNA45SN1_45S_50ntFlanks@10809.10826.18\|AATGTAGGTAAGGGAAGT | >hg38_21_+_8433172_8446622_RNA45SN1_45S_50ntFlanks | 10809 | 10826 | 18 | AATGTAGGTAAGGGAAGT | MB468 | 11 | 11 | 14 | 4 | 4 | 5.5 | 7.6 | 8.8 | 1.3 | 1.3 |
| >hg38_21_+_8433172_8446622_RNA45SN1_45S_50ntFlanks@10816.10840.25\|GTAAGGGAAGTCGGCAAGCCGGATC | >hg38_21_+_8433172_8446622_RNA45SN1_45S_50ntFlanks | 10816 | 10840 | 25 | GTAAGGGAAGTCGGCAAGCCGGATC | MB468 | 19 | 23 | 21 | 16 | 11 | 2.6 | 5.7 | 7.0 | 5.7 | 4.5 |
| >hg38_21_+_8433172_8446622_RNA45SN1_45S_50ntFlanks@10817.10840.24\|TAAGGGAAGTCGGCAAGCCGGATC | >hg38_21_+_8433172_8446622_RNA45SN1_45S_50ntFlanks | 10817 | 10840 | 24 | TAAGGGAAGTCGGCAAGCCGGATC | MB468 | 21 | 27 | 24 | 15 | 15 | 2.3 | 5.3 | 7.0 | 4.7 | 5.6 |

| | | | | | | | | | | | | | | | |
|---|---|---|---|---|---|---|---|---|---|---|---|---|---|---|---|
| >hg38_21_+_8433172_8446622_RNA45SN1_45S_50ntFlanks@10818.10840.23\|AAGGGAAGTCGGCAAGCCGGATC | >hg38_21_+_8433172_8446622_RNA45SN1_45S_50ntFlanks | 10818 | 10840 | 23 | AAGGGAAGTCGGCAAGCCGGATC | MB468 | 12 | 16 | 12 | 18 | 7 | 2.9 | 5.2 | 4.8 | 7.4 | 2.4 |
| >hg38_21_+_8433172_8446622_RNA45SN1_45S_50ntFlanks@10819.10836.18\|AGGGAAGTCGGCAAGCCG | >hg38_21_+_8433172_8446622_RNA45SN1_45S_50ntFlanks | 10819 | 10836 | 18 | AGGGAAGTCGGCAAGCCG | MB468 | 11 | 6 | 16 | 26 | 7 | 6.5 | 2.6 | 14.2 | 7.3 | 2.0 |
| >hg38_21_+_8433172_8446622_RNA45SN1_45S_50ntFlanks@10819.10837.19\|AGGGAAGTCGGCAAGCCGG | >hg38_21_+_8433172_8446622_RNA45SN1_45S_50ntFlanks | 10819 | 10837 | 19 | AGGGAAGTCGGCAAGCCGG | MB468 | 32 | 21 | 43 | 63 | 11 | 13.4 | 16.6 | 20.1 | 31.0 | 3.6 |
| >hg38_21_+_8433172_8446622_RNA45SN1_45S_50ntFlanks@10819.10838.20\|AGGGAAGTCGGCAAGCCGGA | >hg38_21_+_8433172_8446622_RNA45SN1_45S_50ntFlanks | 10819 | 10838 | 20 | AGGGAAGTCGGCAAGCCGGA | MB468 | 21 | 16 | 20 | 45 | 9 | 8.5 | 9.0 | 6.8 | 22.0 | 3.7 |
| >hg38_21_+_8433172_8446622_RNA45SN1_45S_50ntFlanks@10819.10839.21\|AGGGAAGTCGGCAAGCCGGAT | >hg38_21_+_8433172_8446622_RNA45SN1_45S_50ntFlanks | 10819 | 10839 | 21 | AGGGAAGTCGGCAAGCCGGAT | MB468 | 18 | 16 | 15 | 57 | 8 | 2.8 | 6.0 | 2.4 | 39.6 | 0.4 |
| >hg38_21_+_8433172_8446622_RNA45SN1_45S_50ntFlanks@10819.10840.22\|AGGGAAGTCGGCAAGCCGGATC | >hg38_21_+_8433172_8446622_RNA45SN1_45S_50ntFlanks | 10819 | 10840 | 22 | AGGGAAGTCGGCAAGCCGGATC | MB468 | 392 | 446 | 399 | 513 | 180 | 126.0 | 166.2 | 123.9 | 236.8 | 38.9 |
| >hg38_21_+_8433172_8446622_RNA45SN1_45S_50ntFlanks@10820.10840.21\|GGGAAGTCGGCAAGCCGGATC | >hg38_21_+_8433172_8446622_RNA45SN1_45S_50ntFlanks | 10820 | 10840 | 21 | GGGAAGTCGGCAAGCCGGATC | MB468 | 118 | 116 | 120 | 123 | 47 | 43.2 | 43.5 | 49.0 | 63.2 | 6.4 |
| >hg38_21_+_8433172_8446622_RNA45SN1_45S_50ntFlanks@10821.10838.18\|GGAAGTCGGCAAGCCGGA | >hg38_21_+_8433172_8446622_RNA45SN1_45S_50ntFlanks | 10821 | 10838 | 18 | GGAAGTCGGCAAGCCGGA | MB468 | 68 | 36 | 56 | 68 | 16 | 27.2 | 15.6 | 32.8 | 36.7 | 4.2 |
| >hg38_21_+_8433172_8446622_RNA45SN1_45S_50ntFlanks@10821.10839.19\|GGAAGTCGGCAAGCCGGAT | >hg38_21_+_8433172_8446622_RNA45SN1_45S_50ntFlanks | 10821 | 10839 | 19 | GGAAGTCGGCAAGCCGGAT | MB468 | 18 | 9 | 13 | 36 | 4 | 6.7 | 4.5 | 5.5 | 29.4 | 2.7 |
| >hg38_21_+_8433172_8446622_RNA45SN1_45S_50ntFlanks@10821.10840.20\|GGAAGTCGGCAAGCCGGATC | >hg38_21_+_8433172_8446622_RNA45SN1_45S_50ntFlanks | 10821 | 10840 | 20 | GGAAGTCGGCAAGCCGGATC | MB468 | 266 | 121 | 274 | 284 | 77 | 142.5 | 18.2 | 186.7 | 219.0 | 22.9 |
| >hg38_21_+_8433172_8446622_RNA45SN1_45S_50ntFlanks@10822.10840.19\|GAAGTCGGCAAGCCGGATC | >hg38_21_+_8433172_8446622_RNA45SN1_45S_50ntFlanks | 10822 | 10840 | 19 | GAAGTCGGCAAGCCGGATC | MB468 | 32 | 5 | 37 | 29 | 8 | 17.3 | 0.9 | 37.8 | 22.4 | 5.9 |
| >hg38_21_+_8433172_8446622_RNA45SN1_45S_50ntFlanks@10822.10844.23\|GAAGTCGGCAAGCCGGATCCGTA | >hg38_21_+_8433172_8446622_RNA45SN1_45S_50ntFlanks | 10822 | 10844 | 23 | GAAGTCGGCAAGCCGGATCCGTA | MB468 | 12 | 4 | 16 | 8 | 7 | 5.0 | 0.7 | 7.3 | 4.9 | 2.2 |
| >hg38_21_+_8433172_8446622_RNA45SN1_45S_50ntFlanks@10823.10840.18\|AAGTCGGCAAGCCGGATC | >hg38_21_+_8433172_8446622_RNA45SN1_45S_50ntFlanks | 10823 | 10840 | 18 | AAGTCGGCAAGCCGGATC | MB468 | 16 | 1 | 20 | 34 | 4 | 15.7 | 0.3 | 21.0 | 35.5 | 2.7 |
| >hg38_21_+_8433172_8446622_RNA45SN1_45S_50ntFlanks@10823.10842.20\|AAGTCGGCAAGCCGGATCCG | >hg38_21_+_8433172_8446622_RNA45SN1_45S_50ntFlanks | 10823 | 10842 | 20 | AAGTCGGCAAGCCGGATCCG | MB468 | 11 | 4 | 11 | 10 | 6 | 6.4 | 1.0 | 9.1 | 2.5 | 2.1 |
| >hg38_21_+_8433172_8446622_RNA45SN1_45S_50ntFlanks@10823.10844.22\|AAGTCGGCAAGCCGGATCCGTA | >hg38_21_+_8433172_8446622_RNA45SN1_45S_50ntFlanks | 10823 | 10844 | 22 | AAGTCGGCAAGCCGGATCCGTA | MB468 | 20 | 6 | 15 | 8 | 7 | 11.0 | 0.9 | 10.6 | 4.2 | 2.6 |
| >hg38_21_+_8433172_8446622_RNA45SN1_45S_50ntFlanks@10826.10844.19\|TCGGCAAGCCGGATCCGTA | >hg38_21_+_8433172_8446622_RNA45SN1_45S_50ntFlanks | 10826 | 10844 | 19 | TCGGCAAGCCGGATCCGTA | MB468 | 49 | 8 | 50 | 8 | 9 | 35.1 | 4.0 | 55.5 | 2.2 | 4.0 |
| >hg38_21_+_8433172_8446622_RNA45SN1_45S_50ntFlanks@10826.10848.23\|TCGGCAAGCCGGATCCGTAACTT | >hg38_21_+_8433172_8446622_RNA45SN1_45S_50ntFlanks | 10826 | 10848 | 23 | TCGGCAAGCCGGATCCGTAACTT | MB468 | 11 | 3 | 20 | 5 | 1 | 12.3 | 1.3 | 18.4 | 3.6 | 0.5 |
| >hg38_21_+_8433172_8446622_RNA45SN1_45S_50ntFlanks@10826.10858.33\|TCGGCAAGCCGGATCCGTAACTTCGGGATAAGG | >hg38_21_+_8433172_8446622_RNA45SN1_45S_50ntFlanks | 10826 | 10858 | 33 | TCGGCAAGCCGGATCCGTAACTTCGGGATAAGG | MB468 | 45 | 10 | 37 | 28 | 14 | 38.5 | 4.2 | 39.9 | 21.6 | 7.0 |
| >hg38_21_+_8433172_8446622_RNA45SN1_45S_50ntFlanks@10826.10865.40\|TCGGCAAGCCGGATCCGTAACTTCGGGATAAGGATTGGCT | >hg38_21_+_8433172_8446622_RNA45SN1_45S_50ntFlanks | 10826 | 10865 | 40 | TCGGCAAGCCGGATCCGTAACTTCGGGATAAGGATTGGCT | MB468 | 16 | 8 | 82 | 5 | 11 | 13.5 | 5.2 | 58.1 | 3.7 | 5.0 |
| >hg38_21_+_8433172_8446622_RNA45SN1_45S_50ntFlanks@10827.10844.18\|CGGCAAGCCGGATCCGTA | >hg38_21_+_8433172_8446622_RNA45SN1_45S_50ntFlanks | 10827 | 10844 | 18 | CGGCAAGCCGGATCCGTA | MB468 | 16 | 5 | 14 | 4 | 5 | 6.3 | 1.5 | 9.4 | 2.3 | 2.1 |
| >hg38_21_+_8433172_8446622_RNA45SN1_45S_50ntFlanks@10827.10858.32\|CGGCAAGCCGGATCCGTAACTTCGGGATAAGG | >hg38_21_+_8433172_8446622_RNA45SN1_45S_50ntFlanks | 10827 | 10858 | 32 | CGGCAAGCCGGATCCGTAACTTCGGGATAAGG | MB468 | 14 | 3 | 9 | 18 | 9 | 8.9 | 1.3 | 6.0 | 20.7 | 5.3 |
| >hg38_21_+_8433172_8446622_RNA45SN1_45S_50ntFlanks@10829.10851.23\|GCAAGCCGGATCCGTAACTTCGG | >hg38_21_+_8433172_8446622_RNA45SN1_45S_50ntFlanks | 10829 | 10851 | 23 | GCAAGCCGGATCCGTAACTTCGG | MB468 | 18 | 10 | 14 | 9 | 7 | 3.9 | 5.3 | 4.3 | 2.5 | 2.6 |
| >hg38_21_+_8433172_8446622_RNA45SN1_45S_50ntFlanks@10829.10852.24\|GCAAGCCGGATCCGTAACTTCGGG | >hg38_21_+_8433172_8446622_RNA45SN1_45S_50ntFlanks | 10829 | 10852 | 24 | GCAAGCCGGATCCGTAACTTCGGG | MB468 | 37 | 14 | 28 | 12 | 10 | 13.7 | 5.7 | 19.5 | 7.0 | 3.8 |
| >hg38_21_+_8433172_8446622_RNA45SN1_45S_50ntFlanks@10829.10853.25\|GCAAGCCGGATCCGTAACTTCGGGA | >hg38_21_+_8433172_8446622_RNA45SN1_45S_50ntFlanks | 10829 | 10853 | 25 | GCAAGCCGGATCCGTAACTTCGGGA | MB468 | 19 | 6 | 17 | 10 | 8 | 7.9 | 1.7 | 9.4 | 4.4 | 4.3 |
| >hg38_21_+_8433172_8446622_RNA45SN1_45S_50ntFlanks@10829.10855.27\|GCAAGCCGGATCCGTAACTTCGGGATA | >hg38_21_+_8433172_8446622_RNA45SN1_45S_50ntFlanks | 10829 | 10855 | 27 | GCAAGCCGGATCCGTAACTTCGGGATA | MB468 | 19 | 10 | 18 | 6 | 6 | 9.0 | 2.6 | 13.6 | 3.4 | 5.1 |
| >hg38_21_+_8433172_8446622_RNA45SN1_45S_50ntFlanks@10829.10858.30\|GCAAGCCGGATCCGTAACTTCGGGATAAGG | >hg38_21_+_8433172_8446622_RNA45SN1_45S_50ntFlanks | 10829 | 10858 | 30 | GCAAGCCGGATCCGTAACTTCGGGATAAGG | MB468 | 24 | 20 | 34 | 27 | 13 | 7.0 | 7.0 | 20.7 | 30.1 | 10.5 |
| >hg38_21_+_8433172_8446622_RNA45SN1_45S_50ntFlanks@10830.10848.19\|CAAGCCGGATCCGTAACTT | >hg38_21_+_8433172_8446622_RNA45SN1_45S_50ntFlanks | 10830 | 10848 | 19 | CAAGCCGGATCCGTAACTT | MB468 | 18 | 9 | 15 | 7 | 6 | 9.5 | 4.0 | 5.3 | 2.7 | 2.2 |
| >hg38_21_+_8433172_8446622_RNA45SN1_45S_50ntFlanks@10830.10851.22\|CAAGCCGGATCCGTAACTTCGG | >hg38_21_+_8433172_8446622_RNA45SN1_45S_50ntFlanks | 10830 | 10851 | 22 | CAAGCCGGATCCGTAACTTCGG | MB468 | 24 | 5 | 24 | 22 | 10 | 8.5 | 3.3 | 9.1 | 7.8 | 4.3 |
| >hg38_21_+_8433172_8446622_RNA45SN1_45S_50ntFlanks@10830.10852.23\|CAAGCCGGATCCGTAACTTCGGG | >hg38_21_+_8433172_8446622_RNA45SN1_45S_50ntFlanks | 10830 | 10852 | 23 | CAAGCCGGATCCGTAACTTCGGG | MB468 | 29 | 3 | 25 | 15 | 7 | 15.6 | 0.9 | 15.1 | 3.6 | 3.5 |
| >hg38_21_+_8433172_8446622_RNA45SN1_45S_50ntFlanks@10830.10853.24\|CAAGCCGGATCCGTAACTTCGGGA | >hg38_21_+_8433172_8446622_RNA45SN1_45S_50ntFlanks | 10830 | 10853 | 24 | CAAGCCGGATCCGTAACTTCGGGA | MB468 | 21 | 2 | 16 | 18 | 9 | 11.2 | 0.4 | 8.0 | 13.8 | 5.4 |
| >hg38_21_+_8433172_8446622_RNA45SN1_45S_50ntFlanks@10830.10855.26\|CAAGCCGGATCCGTAACTTCGGGATA | >hg38_21_+_8433172_8446622_RNA45SN1_45S_50ntFlanks | 10830 | 10855 | 26 | CAAGCCGGATCCGTAACTTCGGGATA | MB468 | 16 | 3 | 12 | 13 | 10 | 10.9 | 1.0 | 8.7 | 13.3 | 6.4 |
| >hg38_21_+_8433172_8446622_RNA45SN1_45S_50ntFlanks@10830.10858.29\|CAAGCCGGATCCGTAACTTCGGGATAAGG | >hg38_21_+_8433172_8446622_RNA45SN1_45S_50ntFlanks | 10830 | 10858 | 29 | CAAGCCGGATCCGTAACTTCGGGATAAGG | MB468 | 35 | 13 | 40 | 61 | 41 | 11.1 | 5.4 | 25.5 | 63.7 | 13.5 |
| >hg38_21_+_8433172_8446622_RNA45SN1_45S_50ntFlanks@10832.10852.21\|AGCCGGATCCGTAACTTCGGG | >hg38_21_+_8433172_8446622_RNA45SN1_45S_50ntFlanks | 10832 | 10852 | 21 | AGCCGGATCCGTAACTTCGGG | MB468 | 11 | 1 | 9 | 5 | 2 | 7.8 | 0.2 | 8.1 | 2.1 | 1.6 |
| >hg38_21_+_8433172_8446622_RNA45SN1_45S_50ntFlanks@10832.10862.31\|AGCCGGATCCGTAACTTCGGGATAAGGATTG | >hg38_21_+_8433172_8446622_RNA45SN1_45S_50ntFlanks | 10832 | 10862 | 31 | AGCCGGATCCGTAACTTCGGGATAAGGATTG | MB468 | 17 | 4 | 37 | 11 | 7 | 5.9 | 1.7 | 30.9 | 8.6 | 5.4 |
| >hg38_21_+_8433172_8446622_RNA45SN1_45S_50ntFlanks@10834.10867.34\|CCGGATCCGTAACTTCGGGATAAGGATTGGCTCT | >hg38_21_+_8433172_8446622_RNA45SN1_45S_50ntFlanks | 10834 | 10867 | 34 | CCGGATCCGTAACTTCGGGATAAGGATTGGCTCT | MB231 | 19 | 6 | 72 | 6 | 5 | 18.8 | 4.8 | 93.1 | 9.0 | 5.0 |
| >hg38_21_+_8433172_8446622_RNA45SN1_45S_50ntFlanks@10835.10858.24\|CGGATCCGTAACTTCGGGATAAGG | >hg38_21_+_8433172_8446622_RNA45SN1_45S_50ntFlanks | 10835 | 10858 | 24 | CGGATCCGTAACTTCGGGATAAGG | MB468 | 25 | 13 | 27 | 24 | 7 | 6.8 | 9.8 | 4.0 | 19.6 | 4.6 |
| >hg38_21_+_8433172_8446622_RNA45SN1_45S_50ntFlanks@10835.10867.33\|CGGATCCGTAACTTCGGGATAAGGATTGGCTCT | >hg38_21_+_8433172_8446622_RNA45SN1_45S_50ntFlanks | 10835 | 10867 | 33 | CGGATCCGTAACTTCGGGATAAGGATTGGCTCT | MB231 | 10 | 3 | 31 | 2 | 2 | 11.0 | 0.9 | 33.1 | 1.3 | 0.7 |
| >hg38_21_+_8433172_8446622_RNA45SN1_45S_50ntFlanks@10838.10865.28\|ATCCGTAACTTCGGGATAAGGATTGGCT | >hg38_21_+_8433172_8446622_RNA45SN1_45S_50ntFlanks | 10838 | 10865 | 28 | ATCCGTAACTTCGGGATAAGGATTGGCT | MB468 | 21 | 11 | 49 | 17 | 8 | 7.5 | 6.3 | 36.8 | 4.3 | 2.4 |
| >hg38_21_+_8433172_8446622_RNA45SN1_45S_50ntFlanks@10838.10867.30\|ATCCGTAACTTCGGGATAAGGATTGGCTCT | >hg38_21_+_8433172_8446622_RNA45SN1_45S_50ntFlanks | 10838 | 10867 | 30 | ATCCGTAACTTCGGGATAAGGATTGGCTCT | MB231 | 12 | 13 | 37 | 2 | 2 | 20.4 | 22.3 | 63.5 | 2.9 | 4.1 |
| >hg38_21_+_8433172_8446622_RNA45SN1_45S_50ntFlanks@10838.10868.31\|ATCCGTAACTTCGGGATAAGGATTGGCTCTA | >hg38_21_+_8433172_8446622_RNA45SN1_45S_50ntFlanks | 10838 | 10868 | 31 | ATCCGTAACTTCGGGATAAGGATTGGCTCTA | MB468 | 15 | 15 | 48 | 8 | 4 | 10.0 | 3.4 | 39.9 | 4.6 | 1.1 |
| >hg38_21_+_8433172_8446622_RNA45SN1_45S_50ntFlanks@10839.10865.27\|TCCGTAACTTCGGGATAAGGATTGGCT | >hg38_21_+_8433172_8446622_RNA45SN1_45S_50ntFlanks | 10839 | 10865 | 27 | TCCGTAACTTCGGGATAAGGATTGGCT | MB468 | 68 | 31 | 115 | 25 | 17 | 22.0 | 11.4 | 67.4 | 10.8 | 6.5 |
| >hg38_21_+_8433172_8446622_RNA45SN1_45S_50ntFlanks@10839.10868.30\|TCCGTAACTTCGGGATAAGGATTGGCTCTA | >hg38_21_+_8433172_8446622_RNA45SN1_45S_50ntFlanks | 10839 | 10868 | 30 | TCCGTAACTTCGGGATAAGGATTGGCTCTA | MB468 | 23 | 25 | 31 | 7 | 5 | 11.8 | 6.5 | 29.0 | 3.5 | 1.7 |
| >hg38_21_+_8433172_8446622_RNA45SN1_45S_50ntFlanks@10840.10864.25\|CCGTAACTTCGGGATAAGGATTGGC | >hg38_21_+_8433172_8446622_RNA45SN1_45S_50ntFlanks | 10840 | 10864 | 25 | CCGTAACTTCGGGATAAGGATTGGC | MB468 | 11 | 6 | 18 | 16 | 6 | 2.7 | 0.8 | 6.0 | 11.7 | 4.6 |
| >hg38_21_+_8433172_8446622_RNA45SN1_45S_50ntFlanks@10840.10865.26\|CCGTAACTTCGGGATAAGGATTGGCT | >hg38_21_+_8433172_8446622_RNA45SN1_45S_50ntFlanks | 10840 | 10865 | 26 | CCGTAACTTCGGGATAAGGATTGGCT | MB468 | 233 | 159 | 306 | 136 | 93 | 49.1 | 39.5 | 59.8 | 61.6 | 50.6 |
| >hg38_21_+_8433172_8446622_RNA45SN1_45S_50ntFlanks@10840.10867.28\|CCGTAACTTCGGGATAAGGATTGGCTCT | >hg38_21_+_8433172_8446622_RNA45SN1_45S_50ntFlanks | 10840 | 10867 | 28 | CCGTAACTTCGGGATAAGGATTGGCTCT | MB468 | 20 | 27 | 25 | 6 | 7 | 6.7 | 5.0 | 7.7 | 5.0 | 2.0 |
| >hg38_21_+_8433172_8446622_RNA45SN1_45S_50ntFlanks@10840.10868.29\|CCGTAACTTCGGGATAAGGATTGGCTCTA | >hg38_21_+_8433172_8446622_RNA45SN1_45S_50ntFlanks | 10840 | 10868 | 29 | CCGTAACTTCGGGATAAGGATTGGCTCTA | MB468 | 63 | 80 | 48 | 31 | 26 | 16.2 | 23.5 | 29.2 | 19.4 | 10.8 |
| >hg38_21_+_8433172_8446622_RNA45SN1_45S_50ntFlanks@10841.10858.18\|CGTAACTTCGGGATAAGG | >hg38_21_+_8433172_8446622_RNA45SN1_45S_50ntFlanks | 10841 | 10858 | 18 | CGTAACTTCGGGATAAGG | MB468 | 115 | 22 | 137 | 32 | 13 | 74.2 | 14.8 | 130.7 | 7.6 | 6.2 |
| >hg38_21_+_8433172_8446622_RNA45SN1_45S_50ntFlanks@10841.10859.19\|CGTAACTTCGGGATAAGGA | >hg38_21_+_8433172_8446622_RNA45SN1_45S_50ntFlanks | 10841 | 10859 | 19 | CGTAACTTCGGGATAAGGA | MB468 | 113 | 16 | 87 | 21 | 13 | 86.9 | 11.9 | 81.1 | 5.6 | 9.6 |
| >hg38_21_+_8433172_8446622_RNA45SN1_45S_50ntFlanks@10841.10860.20\|CGTAACTTCGGGATAAGGAT | >hg38_21_+_8433172_8446622_RNA45SN1_45S_50ntFlanks | 10841 | 10860 | 20 | CGTAACTTCGGGATAAGGAT | MB468 | 27 | 6 | 38 | 6 | 6 | 14.6 | 2.6 | 19.1 | 3.4 | 3.5 |
| >hg38_21_+_8433172_8446622_RNA45SN1_45S_50ntFlanks@10841.10861.21\|CGTAACTTCGGGATAAGGATT | >hg38_21_+_8433172_8446622_RNA45SN1_45S_50ntFlanks | 10841 | 10861 | 21 | CGTAACTTCGGGATAAGGATT | MB468 | 171 | 60 | 266 | 92 | 44 | 41.5 | 25.4 | 124.1 | 45.6 | 39.0 |

| | | | | | | | | | | | | | | | | |
|---|---|---|---|---|---|---|---|---|---|---|---|---|---|---|---|---|
| >hg38_21_+_8433172_8446622_RNA45SN1_45S_50ntFlanks@10841.10862.22\|CGTAACTTCGGGATAAGGATTG | >hg38_21_+_8433172_8446622_RNA45SN1_45S_50ntFlanks | 10841 | 10862 | 22 | CGTAACTTCGGGATAAGGATTG | MB468 | 433 | 167 | 491 | 186 | 63 | 256.5 | 66.9 | 300.2 | 130.5 | 39.1 |
| >hg38_21_+_8433172_8446622_RNA45SN1_45S_50ntFlanks@10841.10863.23\|CGTAACTTCGGGATAAGGATTGG | >hg38_21_+_8433172_8446622_RNA45SN1_45S_50ntFlanks | 10841 | 10863 | 23 | CGTAACTTCGGGATAAGGATTGG | MB468 | 249 | 177 | 454 | 227 | 38 | 94.8 | 82.8 | 176.5 | 133.8 | 16.4 |
| >hg38_21_+_8433172_8446622_RNA45SN1_45S_50ntFlanks@10841.10864.24\|CGTAACTTCGGGATAAGGATTGGC | >hg38_21_+_8433172_8446622_RNA45SN1_45S_50ntFlanks | 10841 | 10864 | 24 | CGTAACTTCGGGATAAGGATTGGC | MB468 | 353 | 342 | 522 | 211 | 62 | 56.9 | 85.0 | 109.7 | 200.6 | 45.1 |
| >hg38_21_+_8433172_8446622_RNA45SN1_45S_50ntFlanks@10841.10865.25\|CGTAACTTCGGGATAAGGATTGGCT | >hg38_21_+_8433172_8446622_RNA45SN1_45S_50ntFlanks | 10841 | 10865 | 25 | CGTAACTTCGGGATAAGGATTGGCT | MB468 | 4399 | 3730 | 7064 | 1890 | 751 | 674.6 | 1161.5 | 2250.8 | 1557.9 | 439.5 |
| >hg38_21_+_8433172_8446622_RNA45SN1_45S_50ntFlanks@10841.10866.26\|CGTAACTTCGGGATAAGGATTGGCTC | >hg38_21_+_8433172_8446622_RNA45SN1_45S_50ntFlanks | 10841 | 10866 | 26 | CGTAACTTCGGGATAAGGATTGGCTC | MB468 | 116 | 131 | 123 | 35 | 16 | 49.8 | 48.5 | 58.7 | 18.4 | 7.2 |
| >hg38_21_+_8433172_8446622_RNA45SN1_45S_50ntFlanks@10841.10867.27\|CGTAACTTCGGGATAAGGATTGGCTCT | >hg38_21_+_8433172_8446622_RNA45SN1_45S_50ntFlanks | 10841 | 10867 | 27 | CGTAACTTCGGGATAAGGATTGGCTCT | BT20 | 26 | 42 | 125 | 27 | 10 | 14.0 | 33.3 | 77.6 | 35.9 | 3.7 |
| >hg38_21_+_8433172_8446622_RNA45SN1_45S_50ntFlanks@10841.10867.27\|CGTAACTTCGGGATAAGGATTGGCTCT | >hg38_21_+_8433172_8446622_RNA45SN1_45S_50ntFlanks | 10841 | 10867 | 27 | CGTAACTTCGGGATAAGGATTGGCTCT | MB231 | 41 | 33 | 183 | 6 | 8 | 58.7 | 46.8 | 241.5 | 6.2 | 8.9 |
| >hg38_21_+_8433172_8446622_RNA45SN1_45S_50ntFlanks@10841.10867.27\|CGTAACTTCGGGATAAGGATTGGCTCT | >hg38_21_+_8433172_8446622_RNA45SN1_45S_50ntFlanks | 10841 | 10867 | 27 | CGTAACTTCGGGATAAGGATTGGCTCT | MB468 | 214 | 268 | 244 | 60 | 28 | 61.3 | 57.2 | 87.5 | 55.1 | 15.6 |
| >hg38_21_+_8433172_8446622_RNA45SN1_45S_50ntFlanks@10841.10868.28\|CGTAACTTCGGGATAAGGATTGGCTCTA | >hg38_21_+_8433172_8446622_RNA45SN1_45S_50ntFlanks | 10841 | 10868 | 28 | CGTAACTTCGGGATAAGGATTGGCTCTA | MB231 | 13 | 8 | 38 | 3 | 2 | 21.1 | 10.3 | 50.0 | 3.6 | 2.6 |
| >hg38_21_+_8433172_8446622_RNA45SN1_45S_50ntFlanks@10841.10868.28\|CGTAACTTCGGGATAAGGATTGGCTCTA | >hg38_21_+_8433172_8446622_RNA45SN1_45S_50ntFlanks | 10841 | 10868 | 28 | CGTAACTTCGGGATAAGGATTGGCTCTA | MB468 | 774 | 899 | 615 | 237 | 103 | 230.5 | 255.3 | 425.4 | 200.7 | 48.9 |
| >hg38_21_+_8433172_8446622_RNA45SN1_45S_50ntFlanks@10841.10869.29\|CGTAACTTCGGGATAAGGATTGGCTCTAA | >hg38_21_+_8433172_8446622_RNA45SN1_45S_50ntFlanks | 10841 | 10869 | 29 | CGTAACTTCGGGATAAGGATTGGCTCTAA | MB468 | 12 | 21 | 13 | 2 | 1 | 3.2 | 6.1 | 6.9 | 2.1 | 0.7 |
| >hg38_21_+_8433172_8446622_RNA45SN1_45S_50ntFlanks@10841.10870.30\|CGTAACTTCGGGATAAGGATTGGCTCTAAG | >hg38_21_+_8433172_8446622_RNA45SN1_45S_50ntFlanks | 10841 | 10870 | 30 | CGTAACTTCGGGATAAGGATTGGCTCTAAG | MB468 | 13 | 21 | 15 | 2 | 2 | 3.4 | 4.9 | 6.7 | 1.0 | 0.5 |
| >hg38_21_+_8433172_8446622_RNA45SN1_45S_50ntFlanks@10841.10871.31\|CGTAACTTCGGGATAAGGATTGGCTCTAAGG | >hg38_21_+_8433172_8446622_RNA45SN1_45S_50ntFlanks | 10841 | 10871 | 31 | CGTAACTTCGGGATAAGGATTGGCTCTAAGG | MB468 | 22 | 41 | 25 | 4 | 3 | 2.3 | 12.7 | 6.5 | 2.4 | 1.4 |
| >hg38_21_+_8433172_8446622_RNA45SN1_45S_50ntFlanks@10841.10872.32\|CGTAACTTCGGGATAAGGATTGGCTCTAAGGG | >hg38_21_+_8433172_8446622_RNA45SN1_45S_50ntFlanks | 10841 | 10872 | 32 | CGTAACTTCGGGATAAGGATTGGCTCTAAGGG | MB468 | 43 | 80 | 82 | 8 | 5 | 13.9 | 27.8 | 33.7 | 5.1 | 1.6 |
| >hg38_21_+_8433172_8446622_RNA45SN1_45S_50ntFlanks@10841.10873.33\|CGTAACTTCGGGATAAGGATTGGCTCTAAGGGC | >hg38_21_+_8433172_8446622_RNA45SN1_45S_50ntFlanks | 10841 | 10873 | 33 | CGTAACTTCGGGATAAGGATTGGCTCTAAGGGC | MB468 | 31 | 79 | 42 | 9 | 5 | 8.7 | 19.3 | 16.3 | 6.6 | 1.7 |
| >hg38_21_+_8433172_8446622_RNA45SN1_45S_50ntFlanks@10841.10874.34\|CGTAACTTCGGGATAAGGATTGGCTCTAAGGGCT | >hg38_21_+_8433172_8446622_RNA45SN1_45S_50ntFlanks | 10841 | 10874 | 34 | CGTAACTTCGGGATAAGGATTGGCTCTAAGGGCT | MB468 | 46 | 89 | 71 | 14 | 8 | 17.8 | 15.8 | 45.8 | 13.1 | 5.7 |
| >hg38_21_+_8433172_8446622_RNA45SN1_45S_50ntFlanks@10841.10875.35\|CGTAACTTCGGGATAAGGATTGGCTCTAAGGGCTG | >hg38_21_+_8433172_8446622_RNA45SN1_45S_50ntFlanks | 10841 | 10875 | 35 | CGTAACTTCGGGATAAGGATTGGCTCTAAGGGCTG | MB468 | 33 | 68 | 69 | 13 | 7 | 15.5 | 26.8 | 45.0 | 9.4 | 4.2 |
| >hg38_21_+_8433172_8446622_RNA45SN1_45S_50ntFlanks@10842.10865.24\|GTAACTTCGGGATAAGGATTGGCT | >hg38_21_+_8433172_8446622_RNA45SN1_45S_50ntFlanks | 10842 | 10865 | 24 | GTAACTTCGGGATAAGGATTGGCT | MB468 | 24 | 21 | 51 | 6 | 4 | 10.4 | 10.2 | 25.6 | 3.1 | 1.6 |
| >hg38_21_+_8433172_8446622_RNA45SN1_45S_50ntFlanks@10842.10868.27\|GTAACTTCGGGATAAGGATTGGCTCTA | >hg38_21_+_8433172_8446622_RNA45SN1_45S_50ntFlanks | 10842 | 10868 | 27 | GTAACTTCGGGATAAGGATTGGCTCTA | MB468 | 11 | 11 | 13 | 3 | 1 | 7.2 | 5.0 | 15.0 | 1.4 | 0.6 |
| >hg38_21_+_8433172_8446622_RNA45SN1_45S_50ntFlanks@10843.10861.19\|TAACTTCGGGATAAGGATT | >hg38_21_+_8433172_8446622_RNA45SN1_45S_50ntFlanks | 10843 | 10861 | 19 | TAACTTCGGGATAAGGATT | MB468 | 22 | 7 | 30 | 4 | 3 | 6.0 | 2.8 | 9.0 | 1.5 | 1.6 |
| >hg38_21_+_8433172_8446622_RNA45SN1_45S_50ntFlanks@10843.10862.20\|TAACTTCGGGATAAGGATTG | >hg38_21_+_8433172_8446622_RNA45SN1_45S_50ntFlanks | 10843 | 10862 | 20 | TAACTTCGGGATAAGGATTG | MB468 | 59 | 22 | 74 | 12 | 7 | 17.6 | 7.1 | 24.8 | 6.3 | 2.2 |
| >hg38_21_+_8433172_8446622_RNA45SN1_45S_50ntFlanks@10843.10863.21\|TAACTTCGGGATAAGGATTGG | >hg38_21_+_8433172_8446622_RNA45SN1_45S_50ntFlanks | 10843 | 10863 | 21 | TAACTTCGGGATAAGGATTGG | MB468 | 22 | 13 | 55 | 9 | 3 | 7.6 | 6.2 | 19.7 | 4.5 | 0.3 |
| >hg38_21_+_8433172_8446622_RNA45SN1_45S_50ntFlanks@10843.10864.22\|TAACTTCGGGATAAGGATTGGC | >hg38_21_+_8433172_8446622_RNA45SN1_45S_50ntFlanks | 10843 | 10864 | 22 | TAACTTCGGGATAAGGATTGGC | MB468 | 14 | 11 | 27 | 3 | 1 | 6.4 | 3.7 | 13.5 | 2.1 | 0.7 |
| >hg38_21_+_8433172_8446622_RNA45SN1_45S_50ntFlanks@10843.10865.23\|TAACTTCGGGATAAGGATTGGCT | >hg38_21_+_8433172_8446622_RNA45SN1_45S_50ntFlanks | 10843 | 10865 | 23 | TAACTTCGGGATAAGGATTGGCT | MB468 | 103 | 72 | 178 | 21 | 13 | 47.1 | 31.2 | 83.8 | 10.4 | 3.8 |
| >hg38_21_+_8433172_8446622_RNA45SN1_45S_50ntFlanks@10843.10866.24\|TAACTTCGGGATAAGGATTGGCTC | >hg38_21_+_8433172_8446622_RNA45SN1_45S_50ntFlanks | 10843 | 10866 | 24 | TAACTTCGGGATAAGGATTGGCTC | MB468 | 15 | 14 | 17 | 2 | 1 | 10.1 | 7.7 | 14.8 | 1.5 | 0.7 |
| >hg38_21_+_8433172_8446622_RNA45SN1_45S_50ntFlanks@10843.10867.25\|TAACTTCGGGATAAGGATTGGCTCT | >hg38_21_+_8433172_8446622_RNA45SN1_45S_50ntFlanks | 10843 | 10867 | 25 | TAACTTCGGGATAAGGATTGGCTCT | MB468 | 25 | 24 | 31 | 5 | 3 | 11.8 | 5.0 | 18.7 | 3.8 | 0.9 |
| >hg38_21_+_8433172_8446622_RNA45SN1_45S_50ntFlanks@10843.10868.26\|TAACTTCGGGATAAGGATTGGCTCTA | >hg38_21_+_8433172_8446622_RNA45SN1_45S_50ntFlanks | 10843 | 10868 | 26 | TAACTTCGGGATAAGGATTGGCTCTA | MB468 | 93 | 84 | 78 | 20 | 10 | 46.5 | 24.2 | 79.7 | 12.3 | 3.8 |
| >hg38_21_+_8433172_8446622_RNA45SN1_45S_50ntFlanks@10844.10865.22\|AACTTCGGGATAAGGATTGGCT | >hg38_21_+_8433172_8446622_RNA45SN1_45S_50ntFlanks | 10844 | 10865 | 22 | AACTTCGGGATAAGGATTGGCT | MB468 | 40 | 34 | 53 | 12 | 6 | 17.2 | 19.8 | 21.9 | 5.2 | 1.8 |
| >hg38_21_+_8433172_8446622_RNA45SN1_45S_50ntFlanks@10844.10868.25\|AACTTCGGGATAAGGATTGGCTCTA | >hg38_21_+_8433172_8446622_RNA45SN1_45S_50ntFlanks | 10844 | 10868 | 25 | AACTTCGGGATAAGGATTGGCTCTA | MB468 | 13 | 13 | 8 | 4 | 2 | 8.8 | 7.6 | 9.3 | 2.1 | 0.4 |
| >hg38_21_+_8433172_8446622_RNA45SN1_45S_50ntFlanks@10845.10862.18\|ACTTCGGGATAAGGATTG | >hg38_21_+_8433172_8446622_RNA45SN1_45S_50ntFlanks | 10845 | 10862 | 18 | ACTTCGGGATAAGGATTG | MB468 | 89 | 39 | 100 | 17 | 10 | 46.2 | 18.1 | 47.6 | 5.6 | 4.1 |
| >hg38_21_+_8433172_8446622_RNA45SN1_45S_50ntFlanks@10845.10863.19\|ACTTCGGGATAAGGATTGG | >hg38_21_+_8433172_8446622_RNA45SN1_45S_50ntFlanks | 10845 | 10863 | 19 | ACTTCGGGATAAGGATTGG | MB468 | 46 | 33 | 102 | 23 | 6 | 15.2 | 17.1 | 42.6 | 10.2 | 1.7 |
| >hg38_21_+_8433172_8446622_RNA45SN1_45S_50ntFlanks@10845.10864.20\|ACTTCGGGATAAGGATTGGC | >hg38_21_+_8433172_8446622_RNA45SN1_45S_50ntFlanks | 10845 | 10864 | 20 | ACTTCGGGATAAGGATTGGC | MB468 | 56 | 50 | 88 | 25 | 8 | 5.6 | 12.1 | 22.3 | 18.9 | 4.7 |
| >hg38_21_+_8433172_8446622_RNA45SN1_45S_50ntFlanks@10845.10865.21\|ACTTCGGGATAAGGATTGGCT | >hg38_21_+_8433172_8446622_RNA45SN1_45S_50ntFlanks | 10845 | 10865 | 21 | ACTTCGGGATAAGGATTGGCT | MB468 | 596 | 503 | 831 | 180 | 99 | 123.0 | 219.8 | 213.5 | 112.0 | 41.2 |
| >hg38_21_+_8433172_8446622_RNA45SN1_45S_50ntFlanks@10845.10866.22\|ACTTCGGGATAAGGATTGGCTC | >hg38_21_+_8433172_8446622_RNA45SN1_45S_50ntFlanks | 10845 | 10866 | 22 | ACTTCGGGATAAGGATTGGCTC | MB468 | 32 | 32 | 30 | 7 | 5 | 13.7 | 17.4 | 16.2 | 2.3 | 1.2 |
| >hg38_21_+_8433172_8446622_RNA45SN1_45S_50ntFlanks@10845.10867.23\|ACTTCGGGATAAGGATTGGCTCT | >hg38_21_+_8433172_8446622_RNA45SN1_45S_50ntFlanks | 10845 | 10867 | 23 | ACTTCGGGATAAGGATTGGCTCT | MB468 | 42 | 39 | 40 | 11 | 6 | 19.8 | 16.8 | 21.5 | 7.6 | 2.2 |
| >hg38_21_+_8433172_8446622_RNA45SN1_45S_50ntFlanks@10845.10868.24\|ACTTCGGGATAAGGATTGGCTCTA | >hg38_21_+_8433172_8446622_RNA45SN1_45S_50ntFlanks | 10845 | 10868 | 24 | ACTTCGGGATAAGGATTGGCTCTA | MB468 | 129 | 118 | 85 | 42 | 21 | 67.0 | 54.8 | 77.5 | 25.3 | 5.2 |
| >hg38_21_+_8433172_8446622_RNA45SN1_45S_50ntFlanks@10845.10872.28\|ACTTCGGGATAAGGATTGGCTCTAAGGG | >hg38_21_+_8433172_8446622_RNA45SN1_45S_50ntFlanks | 10845 | 10872 | 28 | ACTTCGGGATAAGGATTGGCTCTAAGGG | MB468 | 12 | 17 | 18 | 2 | 2 | 4.9 | 6.1 | 7.6 | 1.2 | 0.2 |
| >hg38_21_+_8433172_8446622_RNA45SN1_45S_50ntFlanks@10846.10863.18\|CTTCGGGATAAGGATTGG | >hg38_21_+_8433172_8446622_RNA45SN1_45S_50ntFlanks | 10846 | 10863 | 18 | CTTCGGGATAAGGATTGG | MB468 | 34 | 22 | 77 | 21 | 4 | 19.2 | 12.5 | 42.3 | 6.9 | 0.9 |
| >hg38_21_+_8433172_8446622_RNA45SN1_45S_50ntFlanks@10846.10864.19\|CTTCGGGATAAGGATTGGC | >hg38_21_+_8433172_8446622_RNA45SN1_45S_50ntFlanks | 10846 | 10864 | 19 | CTTCGGGATAAGGATTGGC | MB468 | 41 | 34 | 70 | 21 | 5 | 12.8 | 11.0 | 26.9 | 16.0 | 2.5 |
| >hg38_21_+_8433172_8446622_RNA45SN1_45S_50ntFlanks@10846.10865.20\|CTTCGGGATAAGGATTGGCT | >hg38_21_+_8433172_8446622_RNA45SN1_45S_50ntFlanks | 10846 | 10865 | 20 | CTTCGGGATAAGGATTGGCT | MB468 | 318 | 224 | 490 | 135 | 54 | 103.8 | 82.3 | 199.0 | 85.6 | 22.9 |
| >hg38_21_+_8433172_8446622_RNA45SN1_45S_50ntFlanks@10846.10866.21\|CTTCGGGATAAGGATTGGCTC | >hg38_21_+_8433172_8446622_RNA45SN1_45S_50ntFlanks | 10846 | 10866 | 21 | CTTCGGGATAAGGATTGGCTC | MB468 | 19 | 18 | 22 | 5 | 3 | 10.6 | 8.1 | 13.0 | 1.5 | 0.4 |
| >hg38_21_+_8433172_8446622_RNA45SN1_45S_50ntFlanks@10846.10867.22\|CTTCGGGATAAGGATTGGCTCT | >hg38_21_+_8433172_8446622_RNA45SN1_45S_50ntFlanks | 10846 | 10867 | 22 | CTTCGGGATAAGGATTGGCTCT | MB468 | 30 | 27 | 32 | 6 | 4 | 17.4 | 13.2 | 20.6 | 3.4 | 1.4 |
| >hg38_21_+_8433172_8446622_RNA45SN1_45S_50ntFlanks@10846.10868.23\|CTTCGGGATAAGGATTGGCTCTA | >hg38_21_+_8433172_8446622_RNA45SN1_45S_50ntFlanks | 10846 | 10868 | 23 | CTTCGGGATAAGGATTGGCTCTA | MB468 | 100 | 89 | 71 | 28 | 17 | 59.9 | 47.1 | 73.3 | 12.6 | 3.0 |
| >hg38_21_+_8433172_8446622_RNA45SN1_45S_50ntFlanks@10848.10865.18\|TCGGGATAAGGATTGGCT | >hg38_21_+_8433172_8446622_RNA45SN1_45S_50ntFlanks | 10848 | 10865 | 18 | TCGGGATAAGGATTGGCT | MB468 | 13 | 9 | 29 | 4 | 1 | 4.7 | 4.7 | 16.8 | 2.3 | 0.6 |
| >hg38_21_+_8433172_8446622_RNA45SN1_45S_50ntFlanks@10849.10866.18\|CGGGATAAGGATTGGCTC | >hg38_21_+_8433172_8446622_RNA45SN1_45S_50ntFlanks | 10849 | 10866 | 18 | CGGGATAAGGATTGGCTC | MB468 | 18 | 14 | 24 | 4 | 2 | 11.9 | 9.2 | 19.9 | 1.3 | 0.3 |
| >hg38_21_+_8433172_8446622_RNA45SN1_45S_50ntFlanks@10849.10867.19\|CGGGATAAGGATTGGCTCT | >hg38_21_+_8433172_8446622_RNA45SN1_45S_50ntFlanks | 10849 | 10867 | 19 | CGGGATAAGGATTGGCTCT | MB468 | 25 | 15 | 28 | 4 | 2 | 18.9 | 7.0 | 21.2 | 2.2 | 0.6 |
| >hg38_21_+_8433172_8446622_RNA45SN1_45S_50ntFlanks@10849.10868.20\|CGGGATAAGGATTGGCTCTA | >hg38_21_+_8433172_8446622_RNA45SN1_45S_50ntFlanks | 10849 | 10868 | 20 | CGGGATAAGGATTGGCTCTA | MB468 | 73 | 50 | 52 | 19 | 7 | 59.5 | 22.9 | 64.4 | 9.0 | 1.2 |
| >hg38_21_+_8433172_8446622_RNA45SN1_45S_50ntFlanks@10850.10868.19\|GGATAAGGATTGGCTCTA | >hg38_21_+_8433172_8446622_RNA45SN1_45S_50ntFlanks | 10850 | 10868 | 19 | GGATAAGGATTGGCTCTA | MB468 | 14 | 13 | 10 | 2 | 1 | 1.5 | 4.4 | 4.2 | 2.0 | 0.9 |
| >hg38_21_+_8433172_8446622_RNA45SN1_45S_50ntFlanks@10853.10873.21\|ATAAGGATTGGCTCTAAGGGC | >hg38_21_+_8433172_8446622_RNA45SN1_45S_50ntFlanks | 10853 | 10873 | 21 | ATAAGGATTGGCTCTAAGGGC | MB468 | 17 | 15 | 27 | 11 | 4 | 5.1 | 7.1 | 11.7 | 9.5 | 1.8 |
| >hg38_21_+_8433172_8446622_RNA45SN1_45S_50ntFlanks@10853.10874.22\|ATAAGGATTGGCTCTAAGGGCT | >hg38_21_+_8433172_8446622_RNA45SN1_45S_50ntFlanks | 10853 | 10874 | 22 | ATAAGGATTGGCTCTAAGGGCT | MB468 | 11 | 4 | 11 | 7 | 2 | 6.1 | 1.8 | 7.2 | 1.9 | 0.4 |

| | | | | | | | | | | | | | | | | |
|---|---|---|---|---|---|---|---|---|---|---|---|---|---|---|---|---|
| >hg38_21_+_8433172_8446622_RNA45SN1_45S_50ntFlanks@10853.10875.23\|ATAAGGATTGGCTCTAAGGGCTG | >hg38_21_+_8433172_8446622_RNA45SN1_45S_50ntFlanks | 10853 | 10875 | 23 | ATAAGGATTGGCTCTAAGGGCTG | MB468 | 11 | 4 | 11 | 8 | 2 | 6.2 | 1.5 | 7.2 | 2.9 | 0.4 |
| >hg38_21_+_8433172_8446622_RNA45SN1_45S_50ntFlanks@10853.10879.27\|ATAAGGATTGGCTCTAAGGGCTGGGTC | >hg38_21_+_8433172_8446622_RNA45SN1_45S_50ntFlanks | 10853 | 10879 | 27 | ATAAGGATTGGCTCTAAGGGCTGGGTC | MB468 | 11 | 15 | 16 | 9 | 2 | 8.2 | 11.4 | 14.2 | 9.2 | 1.6 |
| >hg38_21_+_8433172_8446622_RNA45SN1_45S_50ntFlanks@10855.10873.19\|AAGGATTGGCTCTAAGGGC | >hg38_21_+_8433172_8446622_RNA45SN1_45S_50ntFlanks | 10855 | 10873 | 19 | AAGGATTGGCTCTAAGGGC | MB468 | 11 | 9 | 9 | 9 | 2 | 4.0 | 5.1 | 4.0 | 5.4 | 1.0 |
| >hg38_21_+_8433172_8446622_RNA45SN1_45S_50ntFlanks@10856.10873.18\|AGGATTGGCTCTAAGGGC | >hg38_21_+_8433172_8446622_RNA45SN1_45S_50ntFlanks | 10856 | 10873 | 18 | AGGATTGGCTCTAAGGGC | MB468 | 12 | 11 | 13 | 9 | 3 | 1.9 | 4.1 | 4.0 | 5.6 | 0.9 |
| >hg38_21_+_8433172_8446622_RNA45SN1_45S_50ntFlanks@10866.10883.18\|CTAAGGGCTGGGTCGGTC | >hg38_21_+_8433172_8446622_RNA45SN1_45S_50ntFlanks | 10866 | 10883 | 18 | CTAAGGGCTGGGTCGGTC | MB468 | 12 | 17 | 23 | 14 | 7 | 1.5 | 6.1 | 3.6 | 4.5 | 1.2 |
| >hg38_21_+_8433172_8446622_RNA45SN1_45S_50ntFlanks@10866.10884.19\|CTAAGGGCTGGGTCGGTCG | >hg38_21_+_8433172_8446622_RNA45SN1_45S_50ntFlanks | 10866 | 10884 | 19 | CTAAGGGCTGGGTCGGTCG | MB468 | 58 | 64 | 70 | 45 | 27 | 11.6 | 31.7 | 9.1 | 21.8 | 5.3 |
| >hg38_21_+_8433172_8446622_RNA45SN1_45S_50ntFlanks@10866.10886.21\|CTAAGGGCTGGGTCGGTCGGG | >hg38_21_+_8433172_8446622_RNA45SN1_45S_50ntFlanks | 10866 | 10886 | 21 | CTAAGGGCTGGGTCGGTCGGG | MB468 | 21 | 36 | 44 | 27 | 14 | 5.3 | 21.1 | 15.5 | 11.3 | 5.8 |
| >hg38_21_+_8433172_8446622_RNA45SN1_45S_50ntFlanks@10866.10889.24\|CTAAGGGCTGGGTCGGTCGGGCTG | >hg38_21_+_8433172_8446622_RNA45SN1_45S_50ntFlanks | 10866 | 10889 | 24 | CTAAGGGCTGGGTCGGTCGGGCTG | MB468 | 27 | 71 | 42 | 30 | 31 | 9.0 | 36.2 | 15.6 | 16.4 | 14.3 |
| >hg38_21_+_8433172_8446622_RNA45SN1_45S_50ntFlanks@10866.10890.25\|CTAAGGGCTGGGTCGGTCGGGCTGG | >hg38_21_+_8433172_8446622_RNA45SN1_45S_50ntFlanks | 10866 | 10890 | 25 | CTAAGGGCTGGGTCGGTCGGGCTGG | MB468 | 38 | 60 | 72 | 39 | 38 | 9.9 | 18.5 | 32.2 | 18.5 | 14.4 |
| >hg38_21_+_8433172_8446622_RNA45SN1_45S_50ntFlanks@10866.10891.26\|CTAAGGGCTGGGTCGGTCGGGCTGGG | >hg38_21_+_8433172_8446622_RNA45SN1_45S_50ntFlanks | 10866 | 10891 | 26 | CTAAGGGCTGGGTCGGTCGGGCTGGG | MB468 | 49 | 40 | 53 | 59 | 48 | 9.4 | 8.3 | 6.1 | 35.3 | 16.5 |
| >hg38_21_+_8433172_8446622_RNA45SN1_45S_50ntFlanks@10866.10892.27\|CTAAGGGCTGGGTCGGTCGGGCTGGGG | >hg38_21_+_8433172_8446622_RNA45SN1_45S_50ntFlanks | 10866 | 10892 | 27 | CTAAGGGCTGGGTCGGTCGGGCTGGGG | MB468 | 46 | 40 | 44 | 77 | 52 | 14.2 | 8.9 | 7.2 | 50.0 | 20.5 |
| >hg38_21_+_8433172_8446622_RNA45SN1_45S_50ntFlanks@10866.10894.29\|CTAAGGGCTGGGTCGGTCGGGCTGGGGCG | >hg38_21_+_8433172_8446622_RNA45SN1_45S_50ntFlanks | 10866 | 10894 | 29 | CTAAGGGCTGGGTCGGTCGGGCTGGGGCG | MB468 | 30 | 57 | 33 | 55 | 46 | 11.9 | 16.6 | 12.3 | 24.4 | 11.1 |
| >hg38_21_+_8433172_8446622_RNA45SN1_45S_50ntFlanks@10866.10901.36\|CTAAGGGCTGGGTCGGTCGGGCTGGGGCGCGAAGCG | >hg38_21_+_8433172_8446622_RNA45SN1_45S_50ntFlanks | 10866 | 10901 | 36 | CTAAGGGCTGGGTCGGTCGGGCTGGGGCGCGAAGCG | MB468 | 38 | 131 | 52 | 34 | 78 | 23.4 | 83.3 | 32.1 | 13.8 | 52.4 |
| >hg38_21_+_8433172_8446622_RNA45SN1_45S_50ntFlanks@10866.10902.37\|CTAAGGGCTGGGTCGGTCGGGCTGGGGCGCGAAGCGG | >hg38_21_+_8433172_8446622_RNA45SN1_45S_50ntFlanks | 10866 | 10902 | 37 | CTAAGGGCTGGGTCGGTCGGGCTGGGGCGCGAAGCGG | MB468 | 49 | 219 | 81 | 61 | 133 | 25.4 | 123.0 | 43.8 | 22.0 | 84.1 |
| >hg38_21_+_8433172_8446622_RNA45SN1_45S_50ntFlanks@10869.10886.18\|AGGGCTGGGTCGGTCGGG | >hg38_21_+_8433172_8446622_RNA45SN1_45S_50ntFlanks | 10869 | 10886 | 18 | AGGGCTGGGTCGGTCGGG | MB468 | 25 | 41 | 56 | 43 | 10 | 17.8 | 33.4 | 38.1 | 23.3 | 5.4 |
| >hg38_21_+_8433172_8446622_RNA45SN1_45S_50ntFlanks@10869.10889.21\|AGGGCTGGGTCGGTCGGGCTG | >hg38_21_+_8433172_8446622_RNA45SN1_45S_50ntFlanks | 10869 | 10889 | 21 | AGGGCTGGGTCGGTCGGGCTG | MB468 | 37 | 105 | 63 | 42 | 25 | 17.3 | 98.4 | 35.4 | 25.0 | 18.4 |
| >hg38_21_+_8433172_8446622_RNA45SN1_45S_50ntFlanks@10869.10890.22\|AGGGCTGGGTCGGTCGGGCTGG | >hg38_21_+_8433172_8446622_RNA45SN1_45S_50ntFlanks | 10869 | 10890 | 22 | AGGGCTGGGTCGGTCGGGCTGG | MB468 | 34 | 75 | 86 | 54 | 22 | 21.6 | 58.2 | 56.3 | 27.5 | 13.2 |
| >hg38_21_+_8433172_8446622_RNA45SN1_45S_50ntFlanks@10869.10891.23\|AGGGCTGGGTCGGTCGGGCTGGG | >hg38_21_+_8433172_8446622_RNA45SN1_45S_50ntFlanks | 10869 | 10891 | 23 | AGGGCTGGGTCGGTCGGGCTGGG | MB468 | 45 | 47 | 78 | 76 | 31 | 25.1 | 32.9 | 68.9 | 41.4 | 22.2 |
| >hg38_21_+_8433172_8446622_RNA45SN1_45S_50ntFlanks@10869.10892.24\|AGGGCTGGGTCGGTCGGGCTGGGG | >hg38_21_+_8433172_8446622_RNA45SN1_45S_50ntFlanks | 10869 | 10892 | 24 | AGGGCTGGGTCGGTCGGGCTGGGG | MB468 | 39 | 40 | 52 | 108 | 33 | 24.7 | 25.6 | 47.9 | 64.0 | 28.5 |
| >hg38_21_+_8433172_8446622_RNA45SN1_45S_50ntFlanks@10869.10894.26\|AGGGCTGGGTCGGTCGGGCTGGGGCG | >hg38_21_+_8433172_8446622_RNA45SN1_45S_50ntFlanks | 10869 | 10894 | 26 | AGGGCTGGGTCGGTCGGGCTGGGGCG | MB468 | 20 | 47 | 23 | 77 | 27 | 3.2 | 24.7 | 10.3 | 41.1 | 20.4 |
| >hg38_21_+_8433172_8446622_RNA45SN1_45S_50ntFlanks@10869.10901.33\|AGGGCTGGGTCGGTCGGGCTGGGGCGCGAAGCG | >hg38_21_+_8433172_8446622_RNA45SN1_45S_50ntFlanks | 10869 | 10901 | 33 | AGGGCTGGGTCGGTCGGGCTGGGGCGCGAAGCG | MB468 | 30 | 102 | 43 | 57 | 49 | 2.7 | 38.3 | 12.3 | 30.6 | 27.2 |
| >hg38_21_+_8433172_8446622_RNA45SN1_45S_50ntFlanks@10869.10902.34\|AGGGCTGGGTCGGTCGGGCTGGGGCGCGAAGCGG | >hg38_21_+_8433172_8446622_RNA45SN1_45S_50ntFlanks | 10869 | 10902 | 34 | AGGGCTGGGTCGGTCGGGCTGGGGCGCGAAGCGG | MB468 | 50 | 231 | 80 | 136 | 106 | 3.3 | 101.9 | 18.6 | 61.0 | 66.6 |
| >hg38_21_+_8433172_8446622_RNA45SN1_45S_50ntFlanks@10869.10904.36\|AGGGCTGGGTCGGTCGGGCTGGGGCGCGAAGCGGGG | >hg38_21_+_8433172_8446622_RNA45SN1_45S_50ntFlanks | 10869 | 10904 | 36 | AGGGCTGGGTCGGTCGGGCTGGGGCGCGAAGCGGGG | MB468 | 11 | 57 | 30 | 29 | 17 | 1.4 | 15.4 | 6.2 | 17.9 | 13.8 |
| >hg38_21_+_8433172_8446622_RNA45SN1_45S_50ntFlanks@10873.10890.18\|CTGGGTCGGTCGGGCTGG | >hg38_21_+_8433172_8446622_RNA45SN1_45S_50ntFlanks | 10873 | 10890 | 18 | CTGGGTCGGTCGGGCTGG | MB468 | 22 | 30 | 59 | 21 | 9 | 10.8 | 15.1 | 31.4 | 8.5 | 4.1 |
| >hg38_21_+_8433172_8446622_RNA45SN1_45S_50ntFlanks@10873.10891.19\|CTGGGTCGGTCGGGCTGGG | >hg38_21_+_8433172_8446622_RNA45SN1_45S_50ntFlanks | 10873 | 10891 | 19 | CTGGGTCGGTCGGGCTGGG | MB468 | 30 | 47 | 68 | 44 | 15 | 12.2 | 28.9 | 32.2 | 19.3 | 6.0 |
| >hg38_21_+_8433172_8446622_RNA45SN1_45S_50ntFlanks@10873.10892.20\|CTGGGTCGGTCGGGCTGGGG | >hg38_21_+_8433172_8446622_RNA45SN1_45S_50ntFlanks | 10873 | 10892 | 20 | CTGGGTCGGTCGGGCTGGGG | MB468 | 34 | 61 | 75 | 79 | 22 | 11.6 | 25.2 | 31.7 | 34.8 | 5.9 |
| >hg38_21_+_8433172_8446622_RNA45SN1_45S_50ntFlanks@10873.10894.22\|CTGGGTCGGTCGGGCTGGGGCG | >hg38_21_+_8433172_8446622_RNA45SN1_45S_50ntFlanks | 10873 | 10894 | 22 | CTGGGTCGGTCGGGCTGGGGCG | MB468 | 21 | 44 | 32 | 54 | 20 | 4.2 | 15.2 | 8.0 | 24.5 | 2.2 |
| >hg38_21_+_8433172_8446622_RNA45SN1_45S_50ntFlanks@10873.10901.29\|CTGGGTCGGTCGGGCTGGGGCGCGAAGCG | >hg38_21_+_8433172_8446622_RNA45SN1_45S_50ntFlanks | 10873 | 10901 | 29 | CTGGGTCGGTCGGGCTGGGGCGCGAAGCG | MB468 | 14 | 27 | 21 | 22 | 12 | 1.9 | 6.8 | 5.6 | 11.7 | 3.4 |
| >hg38_21_+_8433172_8446622_RNA45SN1_45S_50ntFlanks@10873.10902.30\|CTGGGTCGGTCGGGCTGGGGCGCGAAGCGG | >hg38_21_+_8433172_8446622_RNA45SN1_45S_50ntFlanks | 10873 | 10902 | 30 | CTGGGTCGGTCGGGCTGGGGCGCGAAGCGG | MB468 | 38 | 86 | 60 | 66 | 32 | 9.0 | 35.1 | 22.5 | 27.1 | 14.2 |
| >hg38_21_+_8433172_8446622_RNA45SN1_45S_50ntFlanks@10878.10901.24\|TCGGTCGGGCTGGGGCGCGAAGCG | >hg38_21_+_8433172_8446622_RNA45SN1_45S_50ntFlanks | 10878 | 10901 | 24 | TCGGTCGGGCTGGGGCGCGAAGCG | MB468 | 20 | 41 | 32 | 16 | 10 | 4.7 | 13.4 | 3.2 | 6.9 | 0.4 |
| >hg38_21_+_8433172_8446622_RNA45SN1_45S_50ntFlanks@10878.10902.25\|TCGGTCGGGCTGGGGCGCGAAGCGG | >hg38_21_+_8433172_8446622_RNA45SN1_45S_50ntFlanks | 10878 | 10902 | 25 | TCGGTCGGGCTGGGGCGCGAAGCGG | MB468 | 23 | 40 | 36 | 21 | 12 | 5.1 | 14.9 | 5.0 | 9.5 | 2.4 |
| >hg38_21_+_8433172_8446622_RNA45SN1_45S_50ntFlanks@10879.10902.24\|CGGTCGGGCTGGGGCGCGAAGCGG | >hg38_21_+_8433172_8446622_RNA45SN1_45S_50ntFlanks | 10879 | 10902 | 24 | CGGTCGGGCTGGGGCGCGAAGCGG | MB468 | 12 | 21 | 18 | 19 | 9 | 3.5 | 5.2 | 3.7 | 6.4 | 2.6 |
| >hg38_21_+_8433172_8446622_RNA45SN1_45S_50ntFlanks@10882.10902.21\|TCGGGCTGGGGCGCGAAGCGG | >hg38_21_+_8433172_8446622_RNA45SN1_45S_50ntFlanks | 10882 | 10902 | 21 | TCGGGCTGGGGCGCGAAGCGG | MB468 | 20 | 24 | 25 | 18 | 9 | 4.8 | 9.9 | 5.2 | 7.3 | 1.3 |
| >hg38_21_+_8433172_8446622_RNA45SN1_45S_50ntFlanks@10905.10933.29\|CTGGGCGCGCGCCGCGGCTGGACGAGGCG | >hg38_21_+_8433172_8446622_RNA45SN1_45S_50ntFlanks | 10905 | 10933 | 29 | CTGGGCGCGCGCCGCGGCTGGACGAGGCG | MB468 | 17 | 50 | 24 | 26 | 20 | 11.3 | 43.2 | 13.8 | 12.4 | 13.4 |
| >hg38_21_+_8433172_8446622_RNA45SN1_45S_50ntFlanks@10905.10935.31\|CTGGGCGCGCGCCGCGGCTGGACGAGGCGCC | >hg38_21_+_8433172_8446622_RNA45SN1_45S_50ntFlanks | 10905 | 10935 | 31 | CTGGGCGCGCGCCGCGGCTGGACGAGGCGCC | MB468 | 11 | 12 | 19 | 10 | 11 | 5.4 | 5.5 | 13.3 | 4.2 | 4.3 |
| >hg38_21_+_8433172_8446622_RNA45SN1_45S_50ntFlanks@10905.10936.32\|CTGGGCGCGCGCCGCGGCTGGACGAGGCGCCG | >hg38_21_+_8433172_8446622_RNA45SN1_45S_50ntFlanks | 10905 | 10936 | 32 | CTGGGCGCGCGCCGCGGCTGGACGAGGCGCCG | MB468 | 22 | 17 | 33 | 20 | 22 | 11.4 | 11.7 | 25.0 | 5.3 | 7.3 |
| >hg38_21_+_8433172_8446622_RNA45SN1_45S_50ntFlanks@10905.10939.35\|CTGGGCGCGCGCCGCGGCTGGACGAGGCGCCGCCG | >hg38_21_+_8433172_8446622_RNA45SN1_45S_50ntFlanks | 10905 | 10939 | 35 | CTGGGCGCGCGCCGCGGCTGGACGAGGCGCCGCCG | MB468 | 14 | 7 | 19 | 9 | 11 | 6.2 | 8.2 | 11.9 | 8.2 | 5.8 |
| >hg38_21_+_8433172_8446622_RNA45SN1_45S_50ntFlanks@10905.10941.37\|CTGGGCGCGCGCCGCGGCTGGACGAGGCGCCGCCGCC | >hg38_21_+_8433172_8446622_RNA45SN1_45S_50ntFlanks | 10905 | 10941 | 37 | CTGGGCGCGCGCCGCGGCTGGACGAGGCGCCGCCGCC | MB468 | 12 | 8 | 31 | 6 | 15 | 3.8 | 8.3 | 19.1 | 3.3 | 11.0 |
| >hg38_21_+_8433172_8446622_RNA45SN1_45S_50ntFlanks@10907.10930.24\|GGGCGCGCGCCGCGGCTGGACGAG | >hg38_21_+_8433172_8446622_RNA45SN1_45S_50ntFlanks | 10907 | 10930 | 24 | GGGCGCGCGCCGCGGCTGGACGAG | BT20 | 15 | 42 | 4 | 18 | 180 | 19.2 | 69.9 | 4.9 | 22.4 | 300.7 |
| >hg38_21_+_8433172_8446622_RNA45SN1_45S_50ntFlanks@10909.10930.22\|GCGCGCGCCGCGGCTGGACGAG | >hg38_21_+_8433172_8446622_RNA45SN1_45S_50ntFlanks | 10909 | 10930 | 22 | GCGCGCGCCGCGGCTGGACGAG | BT20 | 11 | 25 | 2 | 4 | 159 | 15.6 | 41.3 | 2.4 | 2.5 | 269.0 |
| >hg38_21_+_8433172_8446622_RNA45SN1_45S_50ntFlanks@10910.10928.19\|CGCGCGCCGCGGCTGGACG | >hg38_21_+_8433172_8446622_RNA45SN1_45S_50ntFlanks | 10910 | 10928 | 19 | CGCGCGCCGCGGCTGGACG | MB468 | 13 | 19 | 18 | 9 | 14 | 7.0 | 14.9 | 11.0 | 7.2 | 7.4 |
| >hg38_21_+_8433172_8446622_RNA45SN1_45S_50ntFlanks@10910.10932.23\|CGCGCGCCGCGGCTGGACGAGGC | >hg38_21_+_8433172_8446622_RNA45SN1_45S_50ntFlanks | 10910 | 10932 | 23 | CGCGCGCCGCGGCTGGACGAGGC | MB468 | 15 | 7 | 22 | 25 | 6 | 9.3 | 2.6 | 17.5 | 12.6 | 2.8 |
| >hg38_21_+_8433172_8446622_RNA45SN1_45S_50ntFlanks@10912.10931.20\|CGCGCCGCGGCTGGACGAGG | >hg38_21_+_8433172_8446622_RNA45SN1_45S_50ntFlanks | 10912 | 10931 | 20 | CGCGCCGCGGCTGGACGAGG | MB468 | 26 | 40 | 17 | 20 | 12 | 16.9 | 31.8 | 8.8 | 11.2 | 5.7 |
| >hg38_21_+_8433172_8446622_RNA45SN1_45S_50ntFlanks@10912.10932.21\|CGCGCCGCGGCTGGACGAGGC | >hg38_21_+_8433172_8446622_RNA45SN1_45S_50ntFlanks | 10912 | 10932 | 21 | CGCGCCGCGGCTGGACGAGGC | MB468 | 34 | 12 | 37 | 39 | 10 | 25.9 | 5.7 | 33.1 | 10.0 | 2.1 |
| >hg38_21_+_8433172_8446622_RNA45SN1_45S_50ntFlanks@10914.10931.18\|CGCCGCGGCTGGACGAGG | >hg38_21_+_8433172_8446622_RNA45SN1_45S_50ntFlanks | 10914 | 10931 | 18 | CGCCGCGGCTGGACGAGG | MB468 | 103 | 86 | 56 | 83 | 41 | 65.7 | 85.1 | 43.1 | 26.9 | 16.4 |
| >hg38_21_+_8433172_8446622_RNA45SN1_45S_50ntFlanks@10914.10932.19\|CGCCGCGGCTGGACGAGGC | >hg38_21_+_8433172_8446622_RNA45SN1_45S_50ntFlanks | 10914 | 10932 | 19 | CGCCGCGGCTGGACGAGGC | MB468 | 84 | 10 | 48 | 64 | 26 | 76.9 | 7.1 | 43.9 | 21.4 | 14.8 |
| >hg38_21_+_8433172_8446622_RNA45SN1_45S_50ntFlanks@10914.10935.22\|CGCCGCGGCTGGACGAGGCGCC | >hg38_21_+_8433172_8446622_RNA45SN1_45S_50ntFlanks | 10914 | 10935 | 22 | CGCCGCGGCTGGACGAGGCGCC | MB468 | 12 | 1 | 9 | 18 | 10 | 8.3 | 0.5 | 7.0 | 9.1 | 6.4 |
| >hg38_21_+_8433172_8446622_RNA45SN1_45S_50ntFlanks@10914.10936.23\|CGCCGCGGCTGGACGAGGCGCCG | >hg38_21_+_8433172_8446622_RNA45SN1_45S_50ntFlanks | 10914 | 10936 | 23 | CGCCGCGGCTGGACGAGGCGCCG | MB468 | 55 | 14 | 47 | 62 | 73 | 22.3 | 5.9 | 25.1 | 16.3 | 38.2 |
| >hg38_21_+_8433172_8446622_RNA45SN1_45S_50ntFlanks@10914.10946.33\|CGCCGCGGCTGGACGAGGCGCCGCCGCCCCCCC | >hg38_21_+_8433172_8446622_RNA45SN1_45S_50ntFlanks | 10914 | 10946 | 33 | CGCCGCGGCTGGACGAGGCGCCGCCGCCCCCCC | MB468 | 14 | 26 | 28 | 22 | 52 | 5.6 | 20.2 | 2.6 | 10.2 | 56.7 |
| >hg38_21_+_8433172_8446622_RNA45SN1_45S_50ntFlanks@10916.10933.18\|CCGCGGCTGGACGAGGCG | >hg38_21_+_8433172_8446622_RNA45SN1_45S_50ntFlanks | 10916 | 10933 | 18 | CCGCGGCTGGACGAGGCG | MB468 | 30 | 47 | 46 | 49 | 20 | 12.4 | 41.5 | 24.3 | 27.9 | 7.6 |

| | | | | | | | | | | | | | | | | |
|---|---|---|---|---|---|---|---|---|---|---|---|---|---|---|---|---|
| >hg38_21_+_8433172_8446622_RNA45SN1_45S_50ntFlanks@10916.10935.20\|CCGCGGCTG GACGAGGCGCC | >hg38_21_+_8433172_8446622_RNA45SN1_45S_50ntFlanks | 10916 | 10935 | 20 | CCGCGGCTGGACGAGGCGCC | MB468 | 13 | 10 | 24 | 10 | 9 | 6.5 | 5.9 | 15.0 | 4.8 | 4.2 |
| >hg38_21_+_8433172_8446622_RNA45SN1_45S_50ntFlanks@10916.10936.21\|CCGCGGCTG GACGAGGCGCCG | >hg38_21_+_8433172_8446622_RNA45SN1_45S_50ntFlanks | 10916 | 10936 | 21 | CCGCGGCTGGACGAGGCGCCG | MB468 | 27 | 16 | 34 | 21 | 23 | 15.1 | 12.6 | 18.0 | 7.3 | 13.6 |
| >hg38_21_+_8433172_8446622_RNA45SN1_45S_50ntFlanks@10921.10939.19\|GCTGGACGAG GCGCCGCCG | >hg38_21_+_8433172_8446622_RNA45SN1_45S_50ntFlanks | 10921 | 10939 | 19 | GCTGGACGAGGCGCCGCCG | MB468 | 20 | 24 | 19 | 16 | 21 | 12.9 | 21.1 | 12.8 | 7.9 | 12.1 |
| >hg38_21_+_8433172_8446622_RNA45SN1_45S_50ntFlanks@10921.10946.26\|GCTGGACGAG GCGCCGCCGCCCCCCCC | >hg38_21_+_8433172_8446622_RNA45SN1_45S_50ntFlanks | 10921 | 10946 | 26 | GCTGGACGAGGCGCCGCCGCCCCCCCC | MB468 | 12 | 20 | 30 | 18 | 13 | 2.4 | 7.0 | 14.5 | 16.4 | 5.6 |
| >hg38_21_+_8433172_8446622_RNA45SN1_45S_50ntFlanks@10922.10939.18\|CTGGACGAGG CGCCGCCG | >hg38_21_+_8433172_8446622_RNA45SN1_45S_50ntFlanks | 10922 | 10939 | 18 | CTGGACGAGGCGCCGCCG | MB468 | 112 | 142 | 93 | 110 | 129 | 75.5 | 128.4 | 57.3 | 62.8 | 75.5 |
| >hg38_21_+_8433172_8446622_RNA45SN1_45S_50ntFlanks@10922.10941.20\|CTGGACGAGG CGCCGCCGCC | >hg38_21_+_8433172_8446622_RNA45SN1_45S_50ntFlanks | 10922 | 10941 | 20 | CTGGACGAGGCGCCGCCGCC | MB468 | 38 | 64 | 59 | 14 | 37 | 18.7 | 44.0 | 42.8 | 11.7 | 19.4 |
| >hg38_21_+_8433172_8446622_RNA45SN1_45S_50ntFlanks@10922.10944.23\|CTGGACGAGG CGCCGCCGCCCCC | >hg38_21_+_8433172_8446622_RNA45SN1_45S_50ntFlanks | 10922 | 10944 | 23 | CTGGACGAGGCGCCGCCGCCCCC | MB468 | 10 | 11 | 19 | 14 | 6 | 2.4 | 2.7 | 9.3 | 8.8 | 2.8 |
| >hg38_21_+_8433172_8446622_RNA45SN1_45S_50ntFlanks@10922.10945.24\|CTGGACGAGG CGCCGCCGCCCCCC | >hg38_21_+_8433172_8446622_RNA45SN1_45S_50ntFlanks | 10922 | 10945 | 24 | CTGGACGAGGCGCCGCCGCCCCCC | MB468 | 38 | 43 | 97 | 74 | 36 | 6.6 | 3.7 | 46.4 | 52.5 | 15.8 |
| >hg38_21_+_8433172_8446622_RNA45SN1_45S_50ntFlanks@10922.10946.25\|CTGGACGAGG CGCCGCCGCCCCCCC | >hg38_21_+_8433172_8446622_RNA45SN1_45S_50ntFlanks | 10922 | 10946 | 25 | CTGGACGAGGCGCCGCCGCCCCCCC | MB468 | 55 | 69 | 158 | 105 | 59 | 8.8 | 22.8 | 78.1 | 89.2 | 24.8 |
| >hg38_21_+_8433172_8446622_RNA45SN1_45S_50ntFlanks@10922.10947.26\|CTGGACGAGG CGCCGCCGCCCCCCCC | >hg38_21_+_8433172_8446622_RNA45SN1_45S_50ntFlanks | 10922 | 10947 | 26 | CTGGACGAGGCGCCGCCGCCCCCCCC | MB468 | 11 | 16 | 25 | 17 | 12 | 2.3 | 8.3 | 14.2 | 13.1 | 7.0 |
| >hg38_21_+_8433172_8446622_RNA45SN1_45S_50ntFlanks@11027.11045.19\|CTCTCTCTCT CTCTCCCCC | >hg38_21_+_8433172_8446622_RNA45SN1_45S_50ntFlanks | 11027 | 11045 | 19 | CTCTCTCTCTCTCTCCCCC | MB468 | 13 | 36 | 29 | 0 | 7 | 7.4 | 34.0 | 26.6 | 0.6 | 14.7 |
| >hg38_21_+_8433172_8446622_RNA45SN1_45S_50ntFlanks@11027.11046.20\|CTCTCTCTCT CTCTCCCCCG | >hg38_21_+_8433172_8446622_RNA45SN1_45S_50ntFlanks | 11027 | 11046 | 20 | CTCTCTCTCTCTCTCCCCCG | MB468 | 29 | 113 | 77 | 0 | 18 | 16.0 | 134.6 | 49.1 | 0.5 | 22.8 |
| >hg38_21_+_8433172_8446622_RNA45SN1_45S_50ntFlanks@11027.11049.23\|CTCTCTCTCT CTCTCCCCCGCTC | >hg38_21_+_8433172_8446622_RNA45SN1_45S_50ntFlanks | 11027 | 11049 | 23 | CTCTCTCTCTCTCTCCCCCGCTC | MB468 | 43 | 85 | 71 | 2 | 35 | 6.5 | 59.7 | 18.6 | 1.3 | 25.1 |
| >hg38_21_+_8433172_8446622_RNA45SN1_45S_50ntFlanks@11027.11050.24\|CTCTCTCTCT CTCTCCCCCGCTCC | >hg38_21_+_8433172_8446622_RNA45SN1_45S_50ntFlanks | 11027 | 11050 | 24 | CTCTCTCTCTCTCTCCCCCGCTCC | MB468 | 64 | 126 | 122 | 5 | 55 | 16.1 | 91.8 | 45.5 | 3.3 | 35.3 |
| >hg38_21_+_8433172_8446622_RNA45SN1_45S_50ntFlanks@11027.11051.25\|CTCTCTCTCT CTCTCCCCCGCTCCC | >hg38_21_+_8433172_8446622_RNA45SN1_45S_50ntFlanks | 11027 | 11051 | 25 | CTCTCTCTCTCTCTCCCCCGCTCCC | MB468 | 67 | 111 | 144 | 5 | 63 | 11.5 | 75.1 | 23.4 | 4.2 | 37.7 |
| >hg38_21_+_8433172_8446622_RNA45SN1_45S_50ntFlanks@11027.11053.27\|CTCTCTCTCT CTCTCCCCCGCTCCCCG | >hg38_21_+_8433172_8446622_RNA45SN1_45S_50ntFlanks | 11027 | 11053 | 27 | CTCTCTCTCTCTCTCCCCCGCTCCCCG | MB468 | 47 | 148 | 104 | 2 | 33 | 17.1 | 156.5 | 40.4 | 3.7 | 38.6 |
| >hg38_21_+_8433172_8446622_RNA45SN1_45S_50ntFlanks@11027.11060.34\|CTCTCTCTCT CTCTCCCCCGCTCCCCGTCCTCCC | >hg38_21_+_8433172_8446622_RNA45SN1_45S_50ntFlanks | 11027 | 11060 | 34 | CTCTCTCTCTCTCTCCCCCGCTCCCCGTCCTCCC | MB468 | 40 | 217 | 92 | 4 | 126 | 15.8 | 223.0 | 37.9 | 2.8 | 47.5 |
| >hg38_21_+_8433172_8446622_RNA45SN1_45S_50ntFlanks@11027.11061.35\|CTCTCTCTCT CTCTCCCCCGCTCCCCGTCCTCCCC | >hg38_21_+_8433172_8446622_RNA45SN1_45S_50ntFlanks | 11027 | 11061 | 35 | CTCTCTCTCTCTCTCCCCCGCTCCCCGTCCTCCCC | MB468 | 11 | 53 | 29 | 1 | 26 | 6.0 | 49.9 | 11.9 | 0.7 | 21.1 |
| >hg38_21_+_8433172_8446622_RNA45SN1_45S_50ntFlanks@11031.11055.25\|CTCTCTCTCT CCCCCGCTCCCCGTC | >hg38_21_+_8433172_8446622_RNA45SN1_45S_50ntFlanks | 11031 | 11055 | 25 | CTCTCTCTCTCCCCCGCTCCCCGTC | BT20 | 20 | 36 | 6 | 1 | 104 | 33.5 | 58.5 | 9.6 | 1.0 | 180.0 |
| >hg38_21_+_8433172_8446622_RNA45SN1_45S_50ntFlanks@11033.11055.23\|CTCTCTCTCC CCCGCTCCCCGTC | >hg38_21_+_8433172_8446622_RNA45SN1_45S_50ntFlanks | 11033 | 11055 | 23 | CTCTCTCTCCCCCGCTCCCCGTC | BT20 | 16 | 28 | 3 | 1 | 61 | 26.2 | 47.2 | 5.6 | 0.9 | 106.3 |
| >hg38_21_+_8433172_8446622_RNA45SN1_45S_50ntFlanks@11035.11055.21\|CTCTCTCCCC CGCTCCCCGTC | >hg38_21_+_8433172_8446622_RNA45SN1_45S_50ntFlanks | 11035 | 11055 | 21 | CTCTCTCCCCCGCTCCCCGTC | BT20 | 17 | 29 | 4 | 1 | 53 | 28.0 | 48.3 | 6.3 | 1.6 | 91.7 |
| >hg38_21_+_8433172_8446622_RNA45SN1_45S_50ntFlanks@11037.11055.19\|CTCTCCCCCG CTCCCCGTC | >hg38_21_+_8433172_8446622_RNA45SN1_45S_50ntFlanks | 11037 | 11055 | 19 | CTCTCCCCCGCTCCCCGTC | BT20 | 13 | 28 | 4 | 1 | 68 | 22.1 | 46.7 | 6.6 | 2.1 | 117.9 |
| >hg38_21_+_8433172_8446622_RNA45SN1_45S_50ntFlanks@11037.11060.24\|CTCTCCCCCG CTCCCCGTCCTCCC | >hg38_21_+_8433172_8446622_RNA45SN1_45S_50ntFlanks | 11037 | 11060 | 24 | CTCTCCCCCGCTCCCCGTCCTCCC | MB468 | 17 | 115 | 17 | 7 | 38 | 4.9 | 112.5 | 9.9 | 8.8 | 8.4 |
| >hg38_21_+_8433172_8446622_RNA45SN1_45S_50ntFlanks@11038.11060.23\|TCTCCCCCGC TCCCCGTCCTCCC | >hg38_21_+_8433172_8446622_RNA45SN1_45S_50ntFlanks | 11038 | 11060 | 23 | TCTCCCCCGCTCCCCGTCCTCCC | MB468 | 13 | 83 | 26 | 5 | 10 | 6.0 | 91.9 | 16.8 | 6.8 | 7.1 |
| >hg38_21_+_8433172_8446622_RNA45SN1_45S_50ntFlanks@11051.11075.25\|CCGTCCTCCC CCCTCCCCGGGGGAG | >hg38_21_+_8433172_8446622_RNA45SN1_45S_50ntFlanks | 11051 | 11075 | 25 | CCGTCCTCCCCCCTCCCCGGGGGAG | MB468 | 19 | 2 | 39 | 8 | 3 | 12.3 | 0.4 | 12.2 | 7.3 | 1.8 |
| >hg38_21_+_8433172_8446622_RNA45SN1_45S_50ntFlanks@11076.11103.28\|CGCCGCGTG GGGGCGGCGGCGGGGGGAG | >hg38_21_+_8433172_8446622_RNA45SN1_45S_50ntFlanks | 11076 | 11103 | 28 | CGCCGCGTGGGGGCGGCGGCGGGGGGAG | MB468 | 16 | 65 | 17 | 11 | 24 | 11.2 | 38.5 | 9.5 | 12.5 | 14.7 |
| >hg38_21_+_8433172_8446622_RNA45SN1_45S_50ntFlanks@11104.11123.20\|AAGGGTCGGG GCGGCAGGGG | >hg38_21_+_8433172_8446622_RNA45SN1_45S_50ntFlanks | 11104 | 11123 | 20 | AAGGGTCGGGGCGGCAGGGG | MB468 | 19 | 76 | 42 | 49 | 16 | 5.9 | 31.3 | 17.7 | 28.1 | 6.4 |
| >hg38_21_+_8433172_8446622_RNA45SN1_45S_50ntFlanks@11104.11133.30\|AAGGGTCGGG GCGGCAGGGGCCGGCGGCGG | >hg38_21_+_8433172_8446622_RNA45SN1_45S_50ntFlanks | 11104 | 11133 | 30 | AAGGGTCGGGGCGGCAGGGGCCGGCGGCGG | MB468 | 10 | 49 | 23 | 29 | 11 | 2.2 | 11.5 | 8.7 | 24.7 | 5.8 |
| >hg38_21_+_8433172_8446622_RNA45SN1_45S_50ntFlanks@11161.11178.18\|CACGGTCCC CCGCGAGGG | >hg38_21_+_8433172_8446622_RNA45SN1_45S_50ntFlanks | 11161 | 11178 | 18 | CACGGTCCCCCGCGAGGG | MB468 | 165 | 117 | 211 | 95 | 99 | 51.1 | 16.9 | 88.4 | 61.2 | 44.2 |
| >hg38_21_+_8433172_8446622_RNA45SN1_45S_50ntFlanks@11161.11179.19\|CACGGTCCC CCGCGAGGGG | >hg38_21_+_8433172_8446622_RNA45SN1_45S_50ntFlanks | 11161 | 11179 | 19 | CACGGTCCCCCGCGAGGGG | MB468 | 91 | 66 | 128 | 72 | 47 | 44.8 | 20.2 | 52.0 | 62.4 | 28.2 |
| >hg38_21_+_8433172_8446622_RNA45SN1_45S_50ntFlanks@11161.11188.28\|CACGGTCCC CCGCGAGGGGGGGCCCGGGC | >hg38_21_+_8433172_8446622_RNA45SN1_45S_50ntFlanks | 11161 | 11188 | 28 | CACGGTCCCCCGCGAGGGGGGGCCCGGGC | MB468 | 30 | 70 | 49 | 46 | 64 | 8.9 | 52.8 | 30.5 | 43.3 | 38.0 |
| >hg38_21_+_8433172_8446622_RNA45SN1_45S_50ntFlanks@11161.11189.29\|CACGGTCCC CCGCGAGGGGGGGCCCGGGCA | >hg38_21_+_8433172_8446622_RNA45SN1_45S_50ntFlanks | 11161 | 11189 | 29 | CACGGTCCCCCGCGAGGGGGGGCCCGGGCA | MB468 | 14 | 21 | 22 | 21 | 29 | 6.3 | 18.3 | 14.2 | 23.0 | 26.8 |
| >hg38_21_+_8433172_8446622_RNA45SN1_45S_50ntFlanks@11190.11225.36\|CCCGGGGGG CCGGCGGCGGCGGCGACTCTGGACGCG | >hg38_21_+_8433172_8446622_RNA45SN1_45S_50ntFlanks | 11190 | 11225 | 36 | CCCGGGGGGCCGGCGGCGGCGGCGACTCTGGACGCG | MB468 | 10 | 84 | 45 | 42 | 45 | 5.8 | 72.7 | 33.9 | 48.9 | 61.7 |
| >hg38_21_+_8433172_8446622_RNA45SN1_45S_50ntFlanks@11199.11220.22\|CCGGCGGCG GCGGCGACTCTGG | >hg38_21_+_8433172_8446622_RNA45SN1_45S_50ntFlanks | 11199 | 11220 | 22 | CCGGCGGCGGCGGCGACTCTGG | MB468 | 28 | 50 | 43 | 14 | 25 | 6.3 | 31.0 | 6.6 | 8.0 | 6.7 |
| >hg38_21_+_8433172_8446622_RNA45SN1_45S_50ntFlanks@11199.11223.25\|CCGGCGGCG GCGGCGACTCTGGACG | >hg38_21_+_8433172_8446622_RNA45SN1_45S_50ntFlanks | 11199 | 11223 | 25 | CCGGCGGCGGCGGCGACTCTGGACG | MB468 | 22 | 57 | 50 | 26 | 27 | 2.0 | 49.2 | 13.8 | 16.0 | 11.2 |
| >hg38_21_+_8433172_8446622_RNA45SN1_45S_50ntFlanks@11199.11225.27\|CCGGCGGCG GCGGCGACTCTGGACGCG | >hg38_21_+_8433172_8446622_RNA45SN1_45S_50ntFlanks | 11199 | 11225 | 27 | CCGGCGGCGGCGGCGACTCTGGACGCG | MB468 | 127 | 195 | 278 | 83 | 104 | 21.9 | 69.3 | 66.2 | 71.2 | 7.9 |
| >hg38_21_+_8433172_8446622_RNA45SN1_45S_50ntFlanks@11199.11227.29\|CCGGCGGCG GCGGCGACTCTGGACGCGAG | >hg38_21_+_8433172_8446622_RNA45SN1_45S_50ntFlanks | 11199 | 11227 | 29 | CCGGCGGCGGCGGCGACTCTGGACGCGAG | MB468 | 38 | 62 | 88 | 16 | 29 | 12.6 | 22.6 | 33.3 | 10.5 | 4.1 |
| >hg38_21_+_8433172_8446622_RNA45SN1_45S_50ntFlanks@11200.11223.24\|CGGCGGCGG CGGCGACTCTGGACG | >hg38_21_+_8433172_8446622_RNA45SN1_45S_50ntFlanks | 11200 | 11223 | 24 | CGGCGGCGGCGGCGACTCTGGACG | MB468 | 15 | 38 | 29 | 20 | 16 | 2.0 | 33.7 | 15.5 | 10.1 | 4.9 |
| >hg38_21_+_8433172_8446622_RNA45SN1_45S_50ntFlanks@11200.11225.26\|CGGCGGCGG CGGCGACTCTGGACGCG | >hg38_21_+_8433172_8446622_RNA45SN1_45S_50ntFlanks | 11200 | 11225 | 26 | CGGCGGCGGCGGCGACTCTGGACGCG | MB468 | 69 | 54 | 123 | 63 | 51 | 5.8 | 10.2 | 28.1 | 22.7 | 22.3 |
| >hg38_21_+_8433172_8446622_RNA45SN1_45S_50ntFlanks@11200.11227.28\|CGGCGGCGG CGGCGACTCTGGACGCGAG | >hg38_21_+_8433172_8446622_RNA45SN1_45S_50ntFlanks | 11200 | 11227 | 28 | CGGCGGCGGCGGCGACTCTGGACGCGAG | MB468 | 15 | 13 | 30 | 12 | 11 | 2.4 | 3.7 | 13.3 | 4.3 | 5.7 |
| >hg38_21_+_8433172_8446622_RNA45SN1_45S_50ntFlanks@11200.11232.33\|CGGCGGCGG CGGCGACTCTGGACGCGAGCCGGG | >hg38_21_+_8433172_8446622_RNA45SN1_45S_50ntFlanks | 11200 | 11232 | 33 | CGGCGGCGGCGGCGACTCTGGACGCGAGCCGGG | MB468 | 155 | 151 | 388 | 111 | 74 | 48.0 | 26.3 | 125.4 | 49.0 | 55.4 |
| >hg38_21_+_8433172_8446622_RNA45SN1_45S_50ntFlanks@11200.11233.34\|CGGCGGCGG CGGCGACTCTGGACGCGAGCCGGGC | >hg38_21_+_8433172_8446622_RNA45SN1_45S_50ntFlanks | 11200 | 11233 | 34 | CGGCGGCGGCGGCGACTCTGGACGCGAGCCGGGC | MB468 | 11 | 11 | 29 | 9 | 6 | 5.4 | 2.1 | 9.8 | 6.5 | 5.7 |
| >hg38_21_+_8433172_8446622_RNA45SN1_45S_50ntFlanks@11200.11234.35\|CGGCGGCGG CGGCGACTCTGGACGCGAGCCGGGCC | >hg38_21_+_8433172_8446622_RNA45SN1_45S_50ntFlanks | 11200 | 11234 | 35 | CGGCGGCGGCGGCGACTCTGGACGCGAGCCGGGCC | MB468 | 10 | 6 | 33 | 8 | 5 | 4.7 | 4.5 | 3.0 | 4.5 | 5.7 |
| >hg38_21_+_8433172_8446622_RNA45SN1_45S_50ntFlanks@11203.11225.23\|CGGCGGCGG CGACTCTGGACGCG | >hg38_21_+_8433172_8446622_RNA45SN1_45S_50ntFlanks | 11203 | 11225 | 23 | CGGCGGCGGCGACTCTGGACGCG | MB468 | 17 | 12 | 25 | 18 | 11 | 5.1 | 7.3 | 6.5 | 4.4 | 1.0 |
| >hg38_21_+_8433172_8446622_RNA45SN1_45S_50ntFlanks@11203.11232.30\|CGGCGGCGG CGACTCTGGACGCGAGCCGGG | >hg38_21_+_8433172_8446622_RNA45SN1_45S_50ntFlanks | 11203 | 11232 | 30 | CGGCGGCGGCGACTCTGGACGCGAGCCGGG | MB468 | 20 | 13 | 40 | 23 | 10 | 5.9 | 1.6 | 15.4 | 17.0 | 4.7 |
| >hg38_21_+_8433172_8446622_RNA45SN1_45S_50ntFlanks@11205.11225.21\|GCGGCGGCG ACTCTGGACGCG | >hg38_21_+_8433172_8446622_RNA45SN1_45S_50ntFlanks | 11205 | 11225 | 21 | GCGGCGGCGACTCTGGACGCG | MB468 | 29 | 13 | 49 | 45 | 16 | 14.0 | 5.4 | 25.6 | 18.0 | 3.9 |
| >hg38_21_+_8433172_8446622_RNA45SN1_45S_50ntFlanks@11205.11232.28\|GCGGCGGCG ACTCTGGACGCGAGCCGGG | >hg38_21_+_8433172_8446622_RNA45SN1_45S_50ntFlanks | 11205 | 11232 | 28 | GCGGCGGCGACTCTGGACGCGAGCCGGG | MB468 | 44 | 33 | 90 | 43 | 20 | 13.2 | 9.3 | 60.4 | 26.3 | 5.3 |
| >hg38_21_+_8433172_8446622_RNA45SN1_45S_50ntFlanks@11206.11225.20\|CGGCGGCGA CTCTGGACGCG | >hg38_21_+_8433172_8446622_RNA45SN1_45S_50ntFlanks | 11206 | 11225 | 20 | CGGCGGCGACTCTGGACGCG | MB468 | 18 | 7 | 23 | 25 | 9 | 7.3 | 3.4 | 7.6 | 6.5 | 1.9 |
| >hg38_21_+_8433172_8446622_RNA45SN1_45S_50ntFlanks@11206.11232.27\|CGGCGGCGA CTCTGGACGCGAGCCGGG | >hg38_21_+_8433172_8446622_RNA45SN1_45S_50ntFlanks | 11206 | 11232 | 27 | CGGCGGCGACTCTGGACGCGAGCCGGG | MB468 | 39 | 21 | 75 | 48 | 19 | 7.8 | 3.5 | 39.5 | 33.7 | 4.2 |
| >hg38_21_+_8433172_8446622_RNA45SN1_45S_50ntFlanks@11207.11225.19\|GGCGGCGAC TCTGGACGCG | >hg38_21_+_8433172_8446622_RNA45SN1_45S_50ntFlanks | 11207 | 11225 | 19 | GGCGGCGACTCTGGACGCG | MB468 | 40 | 14 | 55 | 64 | 16 | 12.1 | 4.2 | 24.2 | 22.1 | 3.5 |

| | | | | | | | | | | | | | | | | |
|---|---|---|---|---|---|---|---|---|---|---|---|---|---|---|---|---|
| >hg38_21_+_8433172_8446622_RNA45SN1_45S_50ntFlanks@11207.11227.21\|GGCGGCGACTCTGGACGCGAG | >hg38_21_+_8433172_8446622_RNA45SN1_45S_50ntFlanks | 11207 | 11227 | 21 | GGCGGCGACTCTGGACGCGAG | MB468 | 12 | 11 | 15 | 20 | 6 | 3.1 | 4.4 | 5.6 | 7.0 | 0.7 |
| >hg38_21_+_8433172_8446622_RNA45SN1_45S_50ntFlanks@11207.11232.26\|GGCGGCGACTCTGGACGCGAGCCGGG | >hg38_21_+_8433172_8446622_RNA45SN1_45S_50ntFlanks | 11207 | 11232 | 26 | GGCGGCGACTCTGGACGCGAGCCGGG | MB468 | 39 | 41 | 79 | 35 | 16 | 9.8 | 7.5 | 42.9 | 23.6 | 6.0 |
| >hg38_21_+_8433172_8446622_RNA45SN1_45S_50ntFlanks@11207.11234.28\|GGCGGCGACTCTGGACGCGAGCCGGGCC | >hg38_21_+_8433172_8446622_RNA45SN1_45S_50ntFlanks | 11207 | 11234 | 28 | GGCGGCGACTCTGGACGCGAGCCGGGCC | MB468 | 12 | 4 | 24 | 12 | 5 | 4.6 | 2.2 | 15.6 | 8.8 | 2.6 |
| >hg38_21_+_8433172_8446622_RNA45SN1_45S_50ntFlanks@11208.11225.18\|GCGGCGACTCTGGACGCG | >hg38_21_+_8433172_8446622_RNA45SN1_45S_50ntFlanks | 11208 | 11225 | 18 | GCGGCGACTCTGGACGCG | MB468 | 369 | 58 | 579 | 874 | 123 | 185.2 | 19.9 | 250.7 | 388.5 | 30.0 |
| >hg38_21_+_8433172_8446622_RNA45SN1_45S_50ntFlanks@11208.11227.20\|GCGGCGACTCTGGACGCGAG | >hg38_21_+_8433172_8446622_RNA45SN1_45S_50ntFlanks | 11208 | 11227 | 20 | GCGGCGACTCTGGACGCGAG | MB468 | 133 | 52 | 232 | 184 | 56 | 60.3 | 17.9 | 118.6 | 73.8 | 18.6 |
| >hg38_21_+_8433172_8446622_RNA45SN1_45S_50ntFlanks@11208.11228.21\|GCGGCGACTCTGGACGCGAGC | >hg38_21_+_8433172_8446622_RNA45SN1_45S_50ntFlanks | 11208 | 11228 | 21 | GCGGCGACTCTGGACGCGAGC | BT20 | 17 | 15 | 7 | 24 | 35 | 27.6 | 23.8 | 6.6 | 28.9 | 55.0 |
| >hg38_21_+_8433172_8446622_RNA45SN1_45S_50ntFlanks@11208.11228.21\|GCGGCGACTCTGGACGCGAGC | >hg38_21_+_8433172_8446622_RNA45SN1_45S_50ntFlanks | 11208 | 11228 | 21 | GCGGCGACTCTGGACGCGAGC | MB468 | 34 | 23 | 60 | 60 | 16 | 4.5 | 4.2 | 23.7 | 42.6 | 6.8 |
| >hg38_21_+_8433172_8446622_RNA45SN1_45S_50ntFlanks@11208.11229.22\|GCGGCGACTCTGGACGCGAGCC | >hg38_21_+_8433172_8446622_RNA45SN1_45S_50ntFlanks | 11208 | 11229 | 22 | GCGGCGACTCTGGACGCGAGCC | MB468 | 47 | 56 | 88 | 38 | 22 | 12.0 | 23.0 | 36.6 | 18.3 | 4.8 |
| >hg38_21_+_8433172_8446622_RNA45SN1_45S_50ntFlanks@11208.11230.23\|GCGGCGACTCTGGACGCGAGCCG | >hg38_21_+_8433172_8446622_RNA45SN1_45S_50ntFlanks | 11208 | 11230 | 23 | GCGGCGACTCTGGACGCGAGCCG | MB468 | 73 | 106 | 129 | 72 | 42 | 19.6 | 48.0 | 68.7 | 27.3 | 8.2 |
| >hg38_21_+_8433172_8446622_RNA45SN1_45S_50ntFlanks@11208.11231.24\|GCGGCGACTCTGGACGCGAGCCGG | >hg38_21_+_8433172_8446622_RNA45SN1_45S_50ntFlanks | 11208 | 11231 | 24 | GCGGCGACTCTGGACGCGAGCCGG | MB468 | 109 | 151 | 191 | 57 | 46 | 27.1 | 56.5 | 92.4 | 25.0 | 8.7 |
| >hg38_21_+_8433172_8446622_RNA45SN1_45S_50ntFlanks@11208.11232.25\|GCGGCGACTCTGGACGCGAGCCGGG | >hg38_21_+_8433172_8446622_RNA45SN1_45S_50ntFlanks | 11208 | 11232 | 25 | GCGGCGACTCTGGACGCGAGCCGGG | MB468 | 764 | 1305 | 1442 | 632 | 372 | 236.0 | 742.2 | 760.3 | 247.5 | 100.4 |
| >hg38_21_+_8433172_8446622_RNA45SN1_45S_50ntFlanks@11208.11233.26\|GCGGCGACTCTGGACGCGAGCCGGGC | >hg38_21_+_8433172_8446622_RNA45SN1_45S_50ntFlanks | 11208 | 11233 | 26 | GCGGCGACTCTGGACGCGAGCCGGGC | MB468 | 255 | 320 | 569 | 145 | 123 | 57.3 | 104.2 | 305.5 | 42.3 | 38.4 |
| >hg38_21_+_8433172_8446622_RNA45SN1_45S_50ntFlanks@11208.11234.27\|GCGGCGACTCTGGACGCGAGCCGGGCC | >hg38_21_+_8433172_8446622_RNA45SN1_45S_50ntFlanks | 11208 | 11234 | 27 | GCGGCGACTCTGGACGCGAGCCGGGCC | MB468 | 275 | 199 | 642 | 135 | 118 | 122.3 | 87.5 | 413.2 | 60.0 | 39.7 |
| >hg38_21_+_8433172_8446622_RNA45SN1_45S_50ntFlanks@11208.11235.28\|GCGGCGACTCTGGACGCGAGCCGGGCCC | >hg38_21_+_8433172_8446622_RNA45SN1_45S_50ntFlanks | 11208 | 11235 | 28 | GCGGCGACTCTGGACGCGAGCCGGGCCC | MB468 | 11 | 6 | 22 | 2 | 4 | 3.6 | 1.5 | 9.0 | 1.2 | 1.4 |
| >hg38_21_+_8433172_8446622_RNA45SN1_45S_50ntFlanks@11208.11238.31\|GCGGCGACTCTGGACGCGAGCCGGGCCCTTC | >hg38_21_+_8433172_8446622_RNA45SN1_45S_50ntFlanks | 11208 | 11238 | 31 | GCGGCGACTCTGGACGCGAGCCGGGCCCTTC | MB468 | 12 | 9 | 35 | 7 | 7 | 4.3 | 4.1 | 15.1 | 3.2 | 5.7 |
| >hg38_21_+_8433172_8446622_RNA45SN1_45S_50ntFlanks@11208.11239.32\|GCGGCGACTCTGGACGCGAGCCGGGCCCTTCC | >hg38_21_+_8433172_8446622_RNA45SN1_45S_50ntFlanks | 11208 | 11239 | 32 | GCGGCGACTCTGGACGCGAGCCGGGCCCTTCC | MB468 | 19 | 21 | 55 | 3 | 12 | 6.9 | 5.7 | 27.5 | 1.9 | 5.4 |
| >hg38_21_+_8433172_8446622_RNA45SN1_45S_50ntFlanks@11208.11252.45\|GCGGCGACTCTGGACGCGAGCCGGGCCCTTCCCGTGGATCGCCCC | >hg38_21_+_8433172_8446622_RNA45SN1_45S_50ntFlanks | 11208 | 11252 | 45 | GCGGCGACTCTGGACGCGAGCCGGGCCCTTCCCGTGGATCGCCCC | MB468 | 12 | 138 | 74 | 3 | 50 | 6.5 | 121.4 | 59.6 | 2.9 | 41.9 |
| >hg38_21_+_8433172_8446622_RNA45SN1_45S_50ntFlanks@11208.11254.47\|GCGGCGACTCTGGACGCGAGCCGGGCCCTTCCCGTGGATCGCCCCAG | >hg38_21_+_8433172_8446622_RNA45SN1_45S_50ntFlanks | 11208 | 11254 | 47 | GCGGCGACTCTGGACGCGAGCCGGGCCCTTCCCGTGGATCGCCCCAG | MB468 | 128 | 2075 | 872 | 46 | 630 | 73.6 | 2291.2 | 831.8 | 55.5 | 462.5 |
| >hg38_21_+_8433172_8446622_RNA45SN1_45S_50ntFlanks@11209.11227.19\|CGGCGACTCTGGACGCGAG | >hg38_21_+_8433172_8446622_RNA45SN1_45S_50ntFlanks | 11209 | 11227 | 19 | CGGCGACTCTGGACGCGAG | MB468 | 97 | 31 | 161 | 141 | 38 | 42.5 | 10.8 | 92.4 | 74.6 | 7.2 |
| >hg38_21_+_8433172_8446622_RNA45SN1_45S_50ntFlanks@11209.11229.21\|CGGCGACTCTGGACGCGAGCC | >hg38_21_+_8433172_8446622_RNA45SN1_45S_50ntFlanks | 11209 | 11229 | 21 | CGGCGACTCTGGACGCGAGCC | BT20 | 10 | 12 | 7 | 20 | 27 | 16.1 | 17.8 | 3.2 | 27.6 | 33.8 |
| >hg38_21_+_8433172_8446622_RNA45SN1_45S_50ntFlanks@11209.11229.21\|CGGCGACTCTGGACGCGAGCC | >hg38_21_+_8433172_8446622_RNA45SN1_45S_50ntFlanks | 11209 | 11229 | 21 | CGGCGACTCTGGACGCGAGCC | MB468 | 43 | 38 | 58 | 37 | 23 | 12.6 | 21.1 | 24.0 | 13.4 | 4.7 |
| >hg38_21_+_8433172_8446622_RNA45SN1_45S_50ntFlanks@11209.11230.22\|CGGCGACTCTGGACGCGAGCCG | >hg38_21_+_8433172_8446622_RNA45SN1_45S_50ntFlanks | 11209 | 11230 | 22 | CGGCGACTCTGGACGCGAGCCG | MB468 | 77 | 50 | 105 | 77 | 44 | 24.2 | 19.3 | 57.6 | 20.8 | 14.1 |
| >hg38_21_+_8433172_8446622_RNA45SN1_45S_50ntFlanks@11209.11231.23\|CGGCGACTCTGGACGCGAGCCGG | >hg38_21_+_8433172_8446622_RNA45SN1_45S_50ntFlanks | 11209 | 11231 | 23 | CGGCGACTCTGGACGCGAGCCGG | MB468 | 99 | 84 | 149 | 57 | 48 | 18.8 | 28.2 | 63.2 | 27.3 | 8.7 |
| >hg38_21_+_8433172_8446622_RNA45SN1_45S_50ntFlanks@11209.11232.24\|CGGCGACTCTGGACGCGAGCCGGG | >hg38_21_+_8433172_8446622_RNA45SN1_45S_50ntFlanks | 11209 | 11232 | 24 | CGGCGACTCTGGACGCGAGCCGGG | BT20 | 11 | 10 | 14 | 15 | 27 | 17.3 | 16.4 | 15.3 | 17.4 | 31.2 |
| >hg38_21_+_8433172_8446622_RNA45SN1_45S_50ntFlanks@11209.11232.24\|CGGCGACTCTGGACGCGAGCCGGG | >hg38_21_+_8433172_8446622_RNA45SN1_45S_50ntFlanks | 11209 | 11232 | 24 | CGGCGACTCTGGACGCGAGCCGGG | MB468 | 1251 | 1230 | 1973 | 993 | 675 | 313.8 | 460.9 | 1051.9 | 270.1 | 132.4 |
| >hg38_21_+_8433172_8446622_RNA45SN1_45S_50ntFlanks@11209.11233.25\|CGGCGACTCTGGACGCGAGCCGGGC | >hg38_21_+_8433172_8446622_RNA45SN1_45S_50ntFlanks | 11209 | 11233 | 25 | CGGCGACTCTGGACGCGAGCCGGGC | MB468 | 115 | 96 | 231 | 74 | 59 | 27.8 | 21.5 | 123.3 | 33.2 | 22.6 |
| >hg38_21_+_8433172_8446622_RNA45SN1_45S_50ntFlanks@11209.11234.26\|CGGCGACTCTGGACGCGAGCCGGGCC | >hg38_21_+_8433172_8446622_RNA45SN1_45S_50ntFlanks | 11209 | 11234 | 26 | CGGCGACTCTGGACGCGAGCCGGGCC | MB468 | 200 | 73 | 356 | 113 | 91 | 131.2 | 23.9 | 243.1 | 37.0 | 46.9 |
| >hg38_21_+_8433172_8446622_RNA45SN1_45S_50ntFlanks@11209.11254.46\|CGGCGACTCTGGACGCGAGCCGGGCCCTTCCCGTGGATCGCCCCAG | >hg38_21_+_8433172_8446622_RNA45SN1_45S_50ntFlanks | 11209 | 11254 | 46 | CGGCGACTCTGGACGCGAGCCGGGCCCTTCCCGTGGATCGCCCCAG | MB468 | 167 | 2630 | 1026 | 87 | 986 | 95.0 | 2890.8 | 976.0 | 103.7 | 767.3 |
| >hg38_21_+_8433172_8446622_RNA45SN1_45S_50ntFlanks@11210.11227.18\|GGCGACTCTGGACGCGAG | >hg38_21_+_8433172_8446622_RNA45SN1_45S_50ntFlanks | 11210 | 11227 | 18 | GGCGACTCTGGACGCGAG | MB468 | 19 | 7 | 27 | 21 | 7 | 12.1 | 3.1 | 22.7 | 6.6 | 1.3 |
| >hg38_21_+_8433172_8446622_RNA45SN1_45S_50ntFlanks@11210.11230.21\|GGCGACTCTGGACGCGAGCCG | >hg38_21_+_8433172_8446622_RNA45SN1_45S_50ntFlanks | 11210 | 11230 | 21 | GGCGACTCTGGACGCGAGCCG | MB468 | 11 | 8 | 11 | 9 | 6 | 6.6 | 4.9 | 6.3 | 5.0 | 4.0 |
| >hg38_21_+_8433172_8446622_RNA45SN1_45S_50ntFlanks@11210.11231.22\|GGCGACTCTGGACGCGAGCCGG | >hg38_21_+_8433172_8446622_RNA45SN1_45S_50ntFlanks | 11210 | 11231 | 22 | GGCGACTCTGGACGCGAGCCGG | MB468 | 19 | 15 | 17 | 14 | 8 | 8.6 | 6.3 | 8.2 | 5.9 | 3.7 |
| >hg38_21_+_8433172_8446622_RNA45SN1_45S_50ntFlanks@11210.11232.23\|GGCGACTCTGGACGCGAGCCGGG | >hg38_21_+_8433172_8446622_RNA45SN1_45S_50ntFlanks | 11210 | 11232 | 23 | GGCGACTCTGGACGCGAGCCGGG | MB468 | 94 | 105 | 97 | 105 | 51 | 46.6 | 71.5 | 64.1 | 49.2 | 29.2 |
| >hg38_21_+_8433172_8446622_RNA45SN1_45S_50ntFlanks@11210.11233.24\|GGCGACTCTGGACGCGAGCCGGGC | >hg38_21_+_8433172_8446622_RNA45SN1_45S_50ntFlanks | 11210 | 11233 | 24 | GGCGACTCTGGACGCGAGCCGGGC | MB468 | 30 | 24 | 41 | 14 | 13 | 12.6 | 11.7 | 24.4 | 4.8 | 2.8 |
| >hg38_21_+_8433172_8446622_RNA45SN1_45S_50ntFlanks@11210.11234.25\|GGCGACTCTGGACGCGAGCCGGGCC | >hg38_21_+_8433172_8446622_RNA45SN1_45S_50ntFlanks | 11210 | 11234 | 25 | GGCGACTCTGGACGCGAGCCGGGCC | MB468 | 12 | 2 | 14 | 10 | 4 | 10.7 | 1.0 | 12.5 | 4.9 | 3.5 |
| >hg38_21_+_8433172_8446622_RNA45SN1_45S_50ntFlanks@11210.11254.45\|GGCGACTCTGGACGCGAGCCGGGCCCTTCCCGTGGATCGCCCCAG | >hg38_21_+_8433172_8446622_RNA45SN1_45S_50ntFlanks | 11210 | 11254 | 45 | GGCGACTCTGGACGCGAGCCGGGCCCTTCCCGTGGATCGCCCCAG | MB468 | 18 | 148 | 63 | 10 | 72 | 11.3 | 152.8 | 51.4 | 11.1 | 62.0 |
| >hg38_21_+_8433172_8446622_RNA45SN1_45S_50ntFlanks@11211.11230.20\|GCGACTCTGGACGCGAGCCG | >hg38_21_+_8433172_8446622_RNA45SN1_45S_50ntFlanks | 11211 | 11230 | 20 | GCGACTCTGGACGCGAGCCG | MB468 | 11 | 3 | 11 | 7 | 3 | 4.9 | 1.1 | 4.4 | 1.3 | 1.4 |
| >hg38_21_+_8433172_8446622_RNA45SN1_45S_50ntFlanks@11211.11231.21\|GCGACTCTGGACGCGAGCCGG | >hg38_21_+_8433172_8446622_RNA45SN1_45S_50ntFlanks | 11211 | 11231 | 21 | GCGACTCTGGACGCGAGCCGG | MB468 | 11 | 3 | 12 | 7 | 3 | 4.0 | 1.2 | 3.8 | 1.8 | 1.0 |
| >hg38_21_+_8433172_8446622_RNA45SN1_45S_50ntFlanks@11211.11232.22\|GCGACTCTGGACGCGAGCCGGG | >hg38_21_+_8433172_8446622_RNA45SN1_45S_50ntFlanks | 11211 | 11232 | 22 | GCGACTCTGGACGCGAGCCGGG | MB468 | 82 | 36 | 102 | 74 | 29 | 30.9 | 13.4 | 43.6 | 28.2 | 10.3 |
| >hg38_21_+_8433172_8446622_RNA45SN1_45S_50ntFlanks@11212.11229.18\|CGACTCTGGACGCGAGCC | >hg38_21_+_8433172_8446622_RNA45SN1_45S_50ntFlanks | 11212 | 11229 | 18 | CGACTCTGGACGCGAGCC | MB468 | 30 | 7 | 36 | 14 | 7 | 13.3 | 2.9 | 7.3 | 1.8 | 2.1 |
| >hg38_21_+_8433172_8446622_RNA45SN1_45S_50ntFlanks@11212.11230.19\|CGACTCTGGACGCGAGCCG | >hg38_21_+_8433172_8446622_RNA45SN1_45S_50ntFlanks | 11212 | 11230 | 19 | CGACTCTGGACGCGAGCCG | MB468 | 18 | 2 | 21 | 17 | 4 | 6.1 | 0.6 | 8.2 | 6.0 | 2.4 |
| >hg38_21_+_8433172_8446622_RNA45SN1_45S_50ntFlanks@11212.11231.20\|CGACTCTGGACGCGAGCCGG | >hg38_21_+_8433172_8446622_RNA45SN1_45S_50ntFlanks | 11212 | 11231 | 20 | CGACTCTGGACGCGAGCCGG | MB468 | 23 | 3 | 33 | 19 | 5 | 6.5 | 1.1 | 4.6 | 6.3 | 1.5 |
| >hg38_21_+_8433172_8446622_RNA45SN1_45S_50ntFlanks@11212.11232.21\|CGACTCTGGACGCGAGCCGGG | >hg38_21_+_8433172_8446622_RNA45SN1_45S_50ntFlanks | 11212 | 11232 | 21 | CGACTCTGGACGCGAGCCGGG | MB468 | 161 | 32 | 288 | 127 | 44 | 40.5 | 8.3 | 68.2 | 18.8 | 15.5 |
| >hg38_21_+_8433172_8446622_RNA45SN1_45S_50ntFlanks@11212.11233.22\|CGACTCTGGACGCGAGCCGGGC | >hg38_21_+_8433172_8446622_RNA45SN1_45S_50ntFlanks | 11212 | 11233 | 22 | CGACTCTGGACGCGAGCCGGGC | MB468 | 14 | 2 | 20 | 5 | 3 | 5.4 | 0.7 | 4.2 | 0.3 | 1.0 |
| >hg38_21_+_8433172_8446622_RNA45SN1_45S_50ntFlanks@11212.11252.41\|CGACTCTGGACGCGAGCCGGGCCCTTCCCGTGGATCGCCCC | >hg38_21_+_8433172_8446622_RNA45SN1_45S_50ntFlanks | 11212 | 11252 | 41 | CGACTCTGGACGCGAGCCGGGCCCTTCCCGTGGATCGCCCC | MB468 | 11 | 77 | 51 | 3 | 48 | 5.6 | 51.3 | 21.2 | 2.6 | 46.7 |
| >hg38_21_+_8433172_8446622_RNA45SN1_45S_50ntFlanks@11215.11234.20\|CTCTGGACGCGAGCCGGGCC | >hg38_21_+_8433172_8446622_RNA45SN1_45S_50ntFlanks | 11215 | 11234 | 20 | CTCTGGACGCGAGCCGGGCC | MB468 | 15 | 9 | 36 | 9 | 5 | 1.4 | 1.8 | 8.2 | 6.6 | 1.9 |
| >hg38_21_+_8433172_8446622_RNA45SN1_45S_50ntFlanks@11217.11234.18\|CTGGACGCGAGCCGGGCC | >hg38_21_+_8433172_8446622_RNA45SN1_45S_50ntFlanks | 11217 | 11234 | 18 | CTGGACGCGAGCCGGGCC | MB468 | 104 | 43 | 257 | 99 | 58 | 86.3 | 29.0 | 245.5 | 68.3 | 28.8 |
| >hg38_21_+_8433172_8446622_RNA45SN1_45S_50ntFlanks@11217.11239.23\|CTGGACGCGAGCCGGGCCCTTCC | >hg38_21_+_8433172_8446622_RNA45SN1_45S_50ntFlanks | 11217 | 11239 | 23 | CTGGACGCGAGCCGGGCCCTTCC | MB468 | 10 | 10 | 18 | 18 | 26 | 4.0 | 7.0 | 9.7 | 20.1 | 20.6 |
| >hg38_21_+_8433172_8446622_RNA45SN1_45S_50ntFlanks@11217.11246.30\|CTGGACGCGAGCCGGGCCCTTCCCGTGGAT | >hg38_21_+_8433172_8446622_RNA45SN1_45S_50ntFlanks | 11217 | 11246 | 30 | CTGGACGCGAGCCGGGCCCTTCCCGTGGAT | BT20 | 11 | 14 | 5 | 3 | 22 | 12.7 | 18.3 | 3.7 | 1.6 | 22.7 |
| >hg38_21_+_8433172_8446622_RNA45SN1_45S_50ntFlanks@11217.11246.30\|CTGGACGCGAGCCGGGCCCTTCCCGTGGAT | >hg38_21_+_8433172_8446622_RNA45SN1_45S_50ntFlanks | 11217 | 11246 | 30 | CTGGACGCGAGCCGGGCCCTTCCCGTGGAT | MB231 | 12 | 15 | 13 | 4 | 17 | 8.6 | 9.6 | 9.9 | 3.2 | 13.5 |
| >hg38_21_+_8433172_8446622_RNA45SN1_45S_50ntFlanks@11233.11250.18\|CCCTTCCCGTGGATCGCC | >hg38_21_+_8433172_8446622_RNA45SN1_45S_50ntFlanks | 11233 | 11250 | 18 | CCCTTCCCGTGGATCGCC | MB468 | 70 | 39 | 113 | 12 | 35 | 46.0 | 23.5 | 88.4 | 5.7 | 12.8 |

| | | | | | | | | | | | | | | | | |
|---|---|---|---|---|---|---|---|---|---|---|---|---|---|---|---|---|
| >hg38_21_+_8433172_8446622_RNA45SN1_45S_50ntFlanks@11233.11251.19\|CCCTTCCCGTGGATCGCCC | >hg38_21_+_8433172_8446622_RNA45SN1_45S_50ntFlanks | 11233 | 11251 | 19 | CCCTTCCCGTGGATCGCCC | MB468 | 12 | 14 | 18 | 4 | 10 | 6.3 | 9.8 | 14.9 | 1.0 | 4.4 |
| >hg38_21_+_8433172_8446622_RNA45SN1_45S_50ntFlanks@11233.11252.20\|CCCTTCCCGTGGATCGCCCC | >hg38_21_+_8433172_8446622_RNA45SN1_45S_50ntFlanks | 11233 | 11252 | 20 | CCCTTCCCGTGGATCGCCCC | MB468 | 30 | 49 | 40 | 15 | 41 | 13.4 | 28.2 | 22.5 | 10.5 | 21.0 |
| >hg38_21_+_8433172_8446622_RNA45SN1_45S_50ntFlanks@11233.11253.21\|CCCTTCCCGTGGATCGCCCCA | >hg38_21_+_8433172_8446622_RNA45SN1_45S_50ntFlanks | 11233 | 11253 | 21 | CCCTTCCCGTGGATCGCCCCA | MB468 | 29 | 27 | 32 | 9 | 36 | 12.9 | 14.7 | 20.8 | 6.2 | 18.5 |
| >hg38_21_+_8433172_8446622_RNA45SN1_45S_50ntFlanks@11233.11254.22\|CCCTTCCCGTGGATCGCCCCAG | >hg38_21_+_8433172_8446622_RNA45SN1_45S_50ntFlanks | 11233 | 11254 | 22 | CCCTTCCCGTGGATCGCCCCAG | MB468 | 116 | 124 | 125 | 49 | 143 | 45.3 | 56.5 | 73.6 | 42.4 | 45.8 |
| >hg38_21_+_8433172_8446622_RNA45SN1_45S_50ntFlanks@11234.11252.19\|CCTTCCCGTGGATCGCCCC | >hg38_21_+_8433172_8446622_RNA45SN1_45S_50ntFlanks | 11234 | 11252 | 19 | CCTTCCCGTGGATCGCCCC | MB468 | 11 | 13 | 12 | 4 | 12 | 3.3 | 6.5 | 5.7 | 2.0 | 5.3 |
| >hg38_21_+_8433172_8446622_RNA45SN1_45S_50ntFlanks@11234.11253.20\|CCTTCCCGTGGATCGCCCCA | >hg38_21_+_8433172_8446622_RNA45SN1_45S_50ntFlanks | 11234 | 11253 | 20 | CCTTCCCGTGGATCGCCCCA | MB468 | 10 | 5 | 8 | 2 | 9 | 3.6 | 2.6 | 4.4 | 1.3 | 3.8 |
| >hg38_21_+_8433172_8446622_RNA45SN1_45S_50ntFlanks@11234.11254.21\|CCTTCCCGTGGATCGCCCCAG | >hg38_21_+_8433172_8446622_RNA45SN1_45S_50ntFlanks | 11234 | 11254 | 21 | CCTTCCCGTGGATCGCCCCAG | MB468 | 34 | 19 | 31 | 14 | 31 | 11.5 | 3.7 | 14.4 | 9.7 | 8.7 |
| >hg38_21_+_8433172_8446622_RNA45SN1_45S_50ntFlanks@11235.11252.18\|CTTCCCGTGGATCGCCCC | >hg38_21_+_8433172_8446622_RNA45SN1_45S_50ntFlanks | 11235 | 11252 | 18 | CTTCCCGTGGATCGCCCC | MB468 | 142 | 150 | 180 | 41 | 102 | 55.6 | 70.4 | 90.5 | 10.4 | 27.8 |
| >hg38_21_+_8433172_8446622_RNA45SN1_45S_50ntFlanks@11235.11253.19\|CTTCCCGTGGATCGCCCCA | >hg38_21_+_8433172_8446622_RNA45SN1_45S_50ntFlanks | 11235 | 11253 | 19 | CTTCCCGTGGATCGCCCCA | MB468 | 127 | 68 | 123 | 30 | 89 | 46.3 | 30.8 | 72.9 | 12.7 | 29.5 |
| >hg38_21_+_8433172_8446622_RNA45SN1_45S_50ntFlanks@11235.11254.20\|CTTCCCGTGGATCGCCCCAG | >hg38_21_+_8433172_8446622_RNA45SN1_45S_50ntFlanks | 11235 | 11254 | 20 | CTTCCCGTGGATCGCCCCAG | MB468 | 322 | 124 | 281 | 113 | 186 | 164.5 | 37.2 | 181.5 | 66.9 | 49.2 |
| >hg38_21_+_8433172_8446622_RNA45SN1_45S_50ntFlanks@11235.11255.21\|CTTCCCGTGGATCGCCCCAGC | >hg38_21_+_8433172_8446622_RNA45SN1_45S_50ntFlanks | 11235 | 11255 | 21 | CTTCCCGTGGATCGCCCCAGC | MB468 | 30 | 28 | 28 | 11 | 20 | 6.2 | 9.3 | 7.4 | 7.4 | 6.3 |
| >hg38_21_+_8433172_8446622_RNA45SN1_45S_50ntFlanks@11235.11268.34\|CTTCCCGTGGATCGCCCCAGCTGCGGCGGGCGTC | >hg38_21_+_8433172_8446622_RNA45SN1_45S_50ntFlanks | 11235 | 11268 | 34 | CTTCCCGTGGATCGCCCCAGCTGCGGCGGGCGTC | MB468 | 16 | 31 | 14 | 5 | 29 | 4.4 | 24.0 | 1.1 | 0.5 | 15.1 |
| >hg38_21_+_8433172_8446622_RNA45SN1_45S_50ntFlanks@11235.11269.35\|CTTCCCGTGGATCGCCCCAGCTGCGGCGGGCGTCG | >hg38_21_+_8433172_8446622_RNA45SN1_45S_50ntFlanks | 11235 | 11269 | 35 | CTTCCCGTGGATCGCCCCAGCTGCGGCGGGCGTCG | MB468 | 53 | 98 | 46 | 15 | 107 | 10.4 | 74.2 | 16.7 | 7.3 | 35.7 |
| >hg38_21_+_8433172_8446622_RNA45SN1_45S_50ntFlanks@11236.11254.19\|TTCCCGTGGATCGCCCCAG | >hg38_21_+_8433172_8446622_RNA45SN1_45S_50ntFlanks | 11236 | 11254 | 19 | TTCCCGTGGATCGCCCCAG | MB468 | 15 | 3 | 10 | 4 | 5 | 11.6 | 0.9 | 7.9 | 2.4 | 1.6 |
| >hg38_21_+_8433172_8446622_RNA45SN1_45S_50ntFlanks@11255.11281.27\|CTGCGGCGGGCGTCGCGGCCGCCCCCG | >hg38_21_+_8433172_8446622_RNA45SN1_45S_50ntFlanks | 11255 | 11281 | 27 | CTGCGGCGGGCGTCGCGGCCGCCCCCG | MB468 | 13 | 9 | 15 | 6 | 12 | 4.8 | 4.5 | 8.7 | 4.0 | 4.1 |
| >hg38_21_+_8433172_8446622_RNA45SN1_45S_50ntFlanks@11314.11344.31\|CCCACCCCACGTCTCGTCGCGCGCGCGTCCG | >hg38_21_+_8433172_8446622_RNA45SN1_45S_50ntFlanks | 11314 | 11344 | 31 | CCCACCCCACGTCTCGTCGCGCGCGCGTCCG | MB468 | 32 | 167 | 55 | 5 | 76 | 10.5 | 37.5 | 28.2 | 2.0 | 30.1 |
| >hg38_21_+_8433172_8446622_RNA45SN1_45S_50ntFlanks@11315.11332.18\|CCACCCCACGTCTCGTCG | >hg38_21_+_8433172_8446622_RNA45SN1_45S_50ntFlanks | 11315 | 11332 | 18 | CCACCCCACGTCTCGTCG | MB468 | 40 | 82 | 45 | 6 | 26 | 31.8 | 103.2 | 47.7 | 2.1 | 3.1 |
| >hg38_21_+_8433172_8446622_RNA45SN1_45S_50ntFlanks@11315.11334.20\|CCACCCCACGTCTCGTCGCG | >hg38_21_+_8433172_8446622_RNA45SN1_45S_50ntFlanks | 11315 | 11334 | 20 | CCACCCCACGTCTCGTCGCG | MB468 | 26 | 34 | 36 | 4 | 22 | 15.9 | 33.9 | 30.0 | 1.9 | 11.3 |
| >hg38_21_+_8433172_8446622_RNA45SN1_45S_50ntFlanks@11315.11336.22\|CCACCCCACGTCTCGTCGCGCG | >hg38_21_+_8433172_8446622_RNA45SN1_45S_50ntFlanks | 11315 | 11336 | 22 | CCACCCCACGTCTCGTCGCGCG | MB468 | 30 | 84 | 48 | 8 | 53 | 16.4 | 76.0 | 30.6 | 4.5 | 38.5 |
| >hg38_21_+_8433172_8446622_RNA45SN1_45S_50ntFlanks@11315.11338.24\|CCACCCCACGTCTCGTCGCGCGCG | >hg38_21_+_8433172_8446622_RNA45SN1_45S_50ntFlanks | 11315 | 11338 | 24 | CCACCCCACGTCTCGTCGCGCGCG | MB468 | 34 | 93 | 51 | 9 | 58 | 17.0 | 62.2 | 24.5 | 3.4 | 37.4 |
| >hg38_21_+_8433172_8446622_RNA45SN1_45S_50ntFlanks@11315.11342.28\|CCACCCCACGTCTCGTCGCGCGCGCGTC | >hg38_21_+_8433172_8446622_RNA45SN1_45S_50ntFlanks | 11315 | 11342 | 28 | CCACCCCACGTCTCGTCGCGCGCGCGTC | MB468 | 33 | 108 | 51 | 15 | 60 | 7.5 | 32.1 | 19.8 | 11.8 | 28.4 |
| >hg38_21_+_8433172_8446622_RNA45SN1_45S_50ntFlanks@11315.11343.29\|CCACCCCACGTCTCGTCGCGCGCGCGTCC | >hg38_21_+_8433172_8446622_RNA45SN1_45S_50ntFlanks | 11315 | 11343 | 29 | CCACCCCACGTCTCGTCGCGCGCGCGTCC | MB468 | 12 | 67 | 15 | 2 | 20 | 5.4 | 42.7 | 5.0 | 1.4 | 7.5 |
| >hg38_21_+_8433172_8446622_RNA45SN1_45S_50ntFlanks@11315.11344.30\|CCACCCCACGTCTCGTCGCGCGCGCGTCCG | >hg38_21_+_8433172_8446622_RNA45SN1_45S_50ntFlanks | 11315 | 11344 | 30 | CCACCCCACGTCTCGTCGCGCGCGCGTCCG | MB468 | 139 | 641 | 213 | 21 | 330 | 51.9 | 230.4 | 93.0 | 6.7 | 142.6 |
| >hg38_21_+_8433172_8446622_RNA45SN1_45S_50ntFlanks@11316.11334.19\|CACCCCACGTCTCGTCGCG | >hg38_21_+_8433172_8446622_RNA45SN1_45S_50ntFlanks | 11316 | 11334 | 19 | CACCCCACGTCTCGTCGCG | MB468 | 109 | 66 | 131 | 13 | 78 | 55.3 | 57.0 | 86.9 | 5.6 | 39.5 |
| >hg38_21_+_8433172_8446622_RNA45SN1_45S_50ntFlanks@11316.11335.20\|CACCCCACGTCTCGTCGCGC | >hg38_21_+_8433172_8446622_RNA45SN1_45S_50ntFlanks | 11316 | 11335 | 20 | CACCCCACGTCTCGTCGCGC | MB468 | 11 | 12 | 12 | 2 | 14 | 4.0 | 6.4 | 5.0 | 1.2 | 6.1 |
| >hg38_21_+_8433172_8446622_RNA45SN1_45S_50ntFlanks@11316.11336.21\|CACCCCACGTCTCGTCGCGCG | >hg38_21_+_8433172_8446622_RNA45SN1_45S_50ntFlanks | 11316 | 11336 | 21 | CACCCCACGTCTCGTCGCGCG | MB468 | 132 | 220 | 189 | 22 | 244 | 53.1 | 151.1 | 75.5 | 11.6 | 172.3 |
| >hg38_21_+_8433172_8446622_RNA45SN1_45S_50ntFlanks@11316.11338.23\|CACCCCACGTCTCGTCGCGCGCG | >hg38_21_+_8433172_8446622_RNA45SN1_45S_50ntFlanks | 11316 | 11338 | 23 | CACCCCACGTCTCGTCGCGCGCG | MB468 | 152 | 325 | 238 | 25 | 308 | 54.8 | 129.6 | 68.9 | 12.3 | 184.3 |
| >hg38_21_+_8433172_8446622_RNA45SN1_45S_50ntFlanks@11316.11340.25\|CACCCCACGTCTCGTCGCGCGCGCG | >hg38_21_+_8433172_8446622_RNA45SN1_45S_50ntFlanks | 11316 | 11340 | 25 | CACCCCACGTCTCGTCGCGCGCGCG | MB468 | 29 | 77 | 38 | 10 | 133 | 13.2 | 22.9 | 13.1 | 6.2 | 109.2 |
| >hg38_21_+_8433172_8446622_RNA45SN1_45S_50ntFlanks@11316.11342.27\|CACCCCACGTCTCGTCGCGCGCGCGTC | >hg38_21_+_8433172_8446622_RNA45SN1_45S_50ntFlanks | 11316 | 11342 | 27 | CACCCCACGTCTCGTCGCGCGCGCGTC | MB468 | 224 | 605 | 345 | 72 | 532 | 122.2 | 222.8 | 228.6 | 66.3 | 313.8 |
| >hg38_21_+_8433172_8446622_RNA45SN1_45S_50ntFlanks@11316.11343.28\|CACCCCACGTCTCGTCGCGCGCGCGTCC | >hg38_21_+_8433172_8446622_RNA45SN1_45S_50ntFlanks | 11316 | 11343 | 28 | CACCCCACGTCTCGTCGCGCGCGCGTCC | MB468 | 50 | 248 | 56 | 7 | 109 | 18.1 | 139.5 | 14.0 | 4.9 | 38.3 |
| >hg38_21_+_8433172_8446622_RNA45SN1_45S_50ntFlanks@11316.11344.29\|CACCCCACGTCTCGTCGCGCGCGCGTCCG | >hg38_21_+_8433172_8446622_RNA45SN1_45S_50ntFlanks | 11316 | 11344 | 29 | CACCCCACGTCTCGTCGCGCGCGCGTCCG | MB468 | 653 | 2675 | 980 | 81 | 1968 | 180.8 | 279.3 | 492.1 | 29.2 | 856.3 |
| >hg38_21_+_8433172_8446622_RNA45SN1_45S_50ntFlanks@11317.11334.18\|ACCCCACGTCTCGTCGCG | >hg38_21_+_8433172_8446622_RNA45SN1_45S_50ntFlanks | 11317 | 11334 | 18 | ACCCCACGTCTCGTCGCG | MB468 | 52 | 21 | 51 | 2 | 36 | 33.3 | 14.2 | 41.0 | 0.5 | 30.9 |
| >hg38_21_+_8433172_8446622_RNA45SN1_45S_50ntFlanks@11317.11336.20\|ACCCCACGTCTCGTCGCGCG | >hg38_21_+_8433172_8446622_RNA45SN1_45S_50ntFlanks | 11317 | 11336 | 20 | ACCCCACGTCTCGTCGCGCG | MB468 | 64 | 96 | 82 | 3 | 112 | 34.3 | 58.8 | 45.6 | 1.6 | 97.8 |
| >hg38_21_+_8433172_8446622_RNA45SN1_45S_50ntFlanks@11317.11338.22\|ACCCCACGTCTCGTCGCGCGCG | >hg38_21_+_8433172_8446622_RNA45SN1_45S_50ntFlanks | 11317 | 11338 | 22 | ACCCCACGTCTCGTCGCGCGCG | MB468 | 74 | 160 | 101 | 4 | 147 | 37.2 | 59.9 | 42.0 | 1.7 | 105.3 |
| >hg38_21_+_8433172_8446622_RNA45SN1_45S_50ntFlanks@11317.11340.24\|ACCCCACGTCTCGTCGCGCGCGCG | >hg38_21_+_8433172_8446622_RNA45SN1_45S_50ntFlanks | 11317 | 11340 | 24 | ACCCCACGTCTCGTCGCGCGCGCG | MB468 | 14 | 40 | 17 | 2 | 58 | 7.5 | 14.5 | 6.7 | 0.9 | 48.8 |
| >hg38_21_+_8433172_8446622_RNA45SN1_45S_50ntFlanks@11317.11342.26\|ACCCCACGTCTCGTCGCGCGCGCGTC | >hg38_21_+_8433172_8446622_RNA45SN1_45S_50ntFlanks | 11317 | 11342 | 26 | ACCCCACGTCTCGTCGCGCGCGCGTC | MB468 | 103 | 266 | 139 | 13 | 201 | 39.8 | 62.3 | 55.7 | 8.3 | 116.1 |
| >hg38_21_+_8433172_8446622_RNA45SN1_45S_50ntFlanks@11317.11343.27\|ACCCCACGTCTCGTCGCGCGCGCGTCC | >hg38_21_+_8433172_8446622_RNA45SN1_45S_50ntFlanks | 11317 | 11343 | 27 | ACCCCACGTCTCGTCGCGCGCGCGTCC | MB468 | 39 | 169 | 41 | 2 | 65 | 17.1 | 96.4 | 10.8 | 1.3 | 25.0 |
| >hg38_21_+_8433172_8446622_RNA45SN1_45S_50ntFlanks@11317.11344.28\|ACCCCACGTCTCGTCGCGCGCGCGTCCG | >hg38_21_+_8433172_8446622_RNA45SN1_45S_50ntFlanks | 11317 | 11344 | 28 | ACCCCACGTCTCGTCGCGCGCGCGTCCG | MB468 | 423 | 1674 | 558 | 24 | 1000 | 156.3 | 337.5 | 233.7 | 13.6 | 479.5 |
| >hg38_21_+_8433172_8446622_RNA45SN1_45S_50ntFlanks@11318.11336.19\|CCCCACGTCTCGTCGCGCG | >hg38_21_+_8433172_8446622_RNA45SN1_45S_50ntFlanks | 11318 | 11336 | 19 | CCCCACGTCTCGTCGCGCG | MB468 | 46 | 44 | 65 | 4 | 64 | 26.5 | 25.7 | 41.3 | 2.3 | 63.3 |
| >hg38_21_+_8433172_8446622_RNA45SN1_45S_50ntFlanks@11318.11338.21\|CCCCACGTCTCGTCGCGCGCG | >hg38_21_+_8433172_8446622_RNA45SN1_45S_50ntFlanks | 11318 | 11338 | 21 | CCCCACGTCTCGTCGCGCGCG | MB468 | 45 | 62 | 71 | 4 | 68 | 19.8 | 29.4 | 41.0 | 2.5 | 46.1 |
| >hg38_21_+_8433172_8446622_RNA45SN1_45S_50ntFlanks@11318.11342.25\|CCCCACGTCTCGTCGCGCGCGCGTC | >hg38_21_+_8433172_8446622_RNA45SN1_45S_50ntFlanks | 11318 | 11342 | 25 | CCCCACGTCTCGTCGCGCGCGCGTC | MB468 | 80 | 199 | 116 | 9 | 140 | 28.3 | 50.1 | 39.2 | 4.7 | 70.3 |
| >hg38_21_+_8433172_8446622_RNA45SN1_45S_50ntFlanks@11318.11343.26\|CCCCACGTCTCGTCGCGCGCGCGTCC | >hg38_21_+_8433172_8446622_RNA45SN1_45S_50ntFlanks | 11318 | 11343 | 26 | CCCCACGTCTCGTCGCGCGCGCGTCC | MB468 | 37 | 156 | 41 | 2 | 49 | 16.7 | 85.5 | 10.7 | 1.1 | 18.1 |
| >hg38_21_+_8433172_8446622_RNA45SN1_45S_50ntFlanks@11318.11344.27\|CCCCACGTCTCGTCGCGCGCGCGTCCG | >hg38_21_+_8433172_8446622_RNA45SN1_45S_50ntFlanks | 11318 | 11344 | 27 | CCCCACGTCTCGTCGCGCGCGCGTCCG | MB468 | 342 | 1147 | 500 | 16 | 747 | 113.1 | 131.9 | 183.6 | 9.8 | 351.8 |
| >hg38_21_+_8433172_8446622_RNA45SN1_45S_50ntFlanks@11319.11336.18\|CCCACGTCTCGTCGCGCG | >hg38_21_+_8433172_8446622_RNA45SN1_45S_50ntFlanks | 11319 | 11336 | 18 | CCCACGTCTCGTCGCGCG | MB468 | 12 | 7 | 22 | 3 | 10 | 7.3 | 3.7 | 12.2 | 1.1 | 8.8 |
| >hg38_21_+_8433172_8446622_RNA45SN1_45S_50ntFlanks@11319.11338.20\|CCCACGTCTCGTCGCGCGCG | >hg38_21_+_8433172_8446622_RNA45SN1_45S_50ntFlanks | 11319 | 11338 | 20 | CCCACGTCTCGTCGCGCGCG | MB468 | 11 | 10 | 22 | 3 | 11 | 3.5 | 4.7 | 10.2 | 1.0 | 5.7 |
| >hg38_21_+_8433172_8446622_RNA45SN1_45S_50ntFlanks@11319.11342.24\|CCCACGTCTCGTCGCGCGCGCGTC | >hg38_21_+_8433172_8446622_RNA45SN1_45S_50ntFlanks | 11319 | 11342 | 24 | CCCACGTCTCGTCGCGCGCGCGTC | MB468 | 37 | 68 | 72 | 7 | 42 | 11.2 | 12.1 | 24.6 | 3.8 | 20.9 |
| >hg38_21_+_8433172_8446622_RNA45SN1_45S_50ntFlanks@11319.11343.25\|CCCACGTCTCGTCGCGCGCGCGTCC | >hg38_21_+_8433172_8446622_RNA45SN1_45S_50ntFlanks | 11319 | 11343 | 25 | CCCACGTCTCGTCGCGCGCGCGTCC | MB468 | 13 | 47 | 20 | 1 | 13 | 4.3 | 25.9 | 6.2 | 0.5 | 4.0 |
| >hg38_21_+_8433172_8446622_RNA45SN1_45S_50ntFlanks@11319.11344.26\|CCCACGTCTCGTCGCGCGCGCGTCCG | >hg38_21_+_8433172_8446622_RNA45SN1_45S_50ntFlanks | 11319 | 11344 | 26 | CCCACGTCTCGTCGCGCGCGCGTCCG | MB468 | 128 | 365 | 238 | 9 | 217 | 29.3 | 61.5 | 77.8 | 4.4 | 93.1 |
| >hg38_21_+_8433172_8446622_RNA45SN1_45S_50ntFlanks@11320.11338.19\|CCACGTCTCGTCGCGCGCG | >hg38_21_+_8433172_8446622_RNA45SN1_45S_50ntFlanks | 11320 | 11338 | 19 | CCACGTCTCGTCGCGCGCG | MB468 | 12 | 11 | 22 | 10 | 14 | 3.2 | 4.5 | 7.7 | 5.1 | 6.4 |
| >hg38_21_+_8433172_8446622_RNA45SN1_45S_50ntFlanks@11320.11342.23\|CCACGTCTCGTCGCGCGCGCGTC | >hg38_21_+_8433172_8446622_RNA45SN1_45S_50ntFlanks | 11320 | 11342 | 23 | CCACGTCTCGTCGCGCGCGCGTC | MB468 | 75 | 122 | 141 | 22 | 107 | 35.5 | 46.5 | 65.2 | 15.4 | 67.3 |

| | | | | | | | | | | | | | | | | |
|---|---|---|---|---|---|---|---|---|---|---|---|---|---|---|---|---|
| >hg38_21_+_8433172_8446622_RNA45SN1_45S_50ntFlanks@11320.11343.24\|CCACGTCTCGTCGCGCGCGTCC | >hg38_21_+_8433172_8446622_RNA45SN1_45S_50ntFlanks | 11320 | 11343 | 24 | CCACGTCTCGTCGCGCGCGTCC | MB468 | 21 | 65 | 30 | 2 | 24 | 6.3 | 26.8 | 4.2 | 1.5 | 4.4 |
| >hg38_21_+_8433172_8446622_RNA45SN1_45S_50ntFlanks@11320.11344.25\|CCACGTCTCGTCGCGCGCGTCCG | >hg38_21_+_8433172_8446622_RNA45SN1_45S_50ntFlanks | 11320 | 11344 | 25 | CCACGTCTCGTCGCGCGCGTCCG | MB468 | 233 | 503 | 379 | 29 | 408 | 20.4 | 18.0 | 88.0 | 12.1 | 105.4 |
| >hg38_21_+_8433172_8446622_RNA45SN1_45S_50ntFlanks@11321.11338.18\|CACGTCTCGTCGCGCGCG | >hg38_21_+_8433172_8446622_RNA45SN1_45S_50ntFlanks | 11321 | 11338 | 18 | CACGTCTCGTCGCGCGCG | MB468 | 58 | 47 | 104 | 16 | 43 | 18.7 | 18.3 | 55.4 | 3.0 | 18.5 |
| >hg38_21_+_8433172_8446622_RNA45SN1_45S_50ntFlanks@11321.11340.20\|CACGTCTCGTCGCGCGCGCG | >hg38_21_+_8433172_8446622_RNA45SN1_45S_50ntFlanks | 11321 | 11340 | 20 | CACGTCTCGTCGCGCGCGCG | MB468 | 21 | 28 | 30 | 7 | 31 | 8.0 | 8.6 | 15.1 | 1.6 | 18.3 |
| >hg38_21_+_8433172_8446622_RNA45SN1_45S_50ntFlanks@11321.11342.22\|CACGTCTCGTCGCGCGCGCGTC | >hg38_21_+_8433172_8446622_RNA45SN1_45S_50ntFlanks | 11321 | 11342 | 22 | CACGTCTCGTCGCGCGCGCGTC | MB468 | 107 | 159 | 186 | 30 | 131 | 37.2 | 55.2 | 59.3 | 10.1 | 78.3 |
| >hg38_21_+_8433172_8446622_RNA45SN1_45S_50ntFlanks@11321.11343.23\|CACGTCTCGTCGCGCGCGCGTCC | >hg38_21_+_8433172_8446622_RNA45SN1_45S_50ntFlanks | 11321 | 11343 | 23 | CACGTCTCGTCGCGCGCGCGTCC | MB468 | 48 | 141 | 60 | 5 | 59 | 13.0 | 43.2 | 17.2 | 1.9 | 19.0 |
| >hg38_21_+_8433172_8446622_RNA45SN1_45S_50ntFlanks@11321.11344.24\|CACGTCTCGTCGCGCGCGCGTCCG | >hg38_21_+_8433172_8446622_RNA45SN1_45S_50ntFlanks | 11321 | 11344 | 24 | CACGTCTCGTCGCGCGCGCGTCCG | MB468 | 420 | 809 | 601 | 92 | 671 | 67.0 | 96.6 | 166.5 | 51.8 | 251.9 |
| >hg38_21_+_8433172_8446622_RNA45SN1_45S_50ntFlanks@11322.11340.19\|ACGTCTCGTCGCGCGCGCG | >hg38_21_+_8433172_8446622_RNA45SN1_45S_50ntFlanks | 11322 | 11340 | 19 | ACGTCTCGTCGCGCGCGCG | MB468 | 28 | 34 | 37 | 11 | 32 | 13.5 | 6.0 | 18.0 | 2.9 | 20.5 |
| >hg38_21_+_8433172_8446622_RNA45SN1_45S_50ntFlanks@11322.11342.21\|ACGTCTCGTCGCGCGCGCGTC | >hg38_21_+_8433172_8446622_RNA45SN1_45S_50ntFlanks | 11322 | 11342 | 21 | ACGTCTCGTCGCGCGCGCGTC | BT20 | 13 | 26 | 9 | 3 | 34 | 21.9 | 42.0 | 11.6 | 2.2 | 51.5 |
| >hg38_21_+_8433172_8446622_RNA45SN1_45S_50ntFlanks@11322.11342.21\|ACGTCTCGTCGCGCGCGCGTC | >hg38_21_+_8433172_8446622_RNA45SN1_45S_50ntFlanks | 11322 | 11342 | 21 | ACGTCTCGTCGCGCGCGCGTC | MB468 | 112 | 183 | 181 | 47 | 114 | 19.4 | 53.5 | 38.1 | 9.6 | 63.3 |
| >hg38_21_+_8433172_8446622_RNA45SN1_45S_50ntFlanks@11322.11343.22\|ACGTCTCGTCGCGCGCGCGTCC | >hg38_21_+_8433172_8446622_RNA45SN1_45S_50ntFlanks | 11322 | 11343 | 22 | ACGTCTCGTCGCGCGCGCGTCC | BT20 | 14 | 36 | 12 | 1 | 58 | 20.0 | 55.1 | 11.7 | 1.2 | 96.3 |
| >hg38_21_+_8433172_8446622_RNA45SN1_45S_50ntFlanks@11322.11343.22\|ACGTCTCGTCGCGCGCGCGTCC | >hg38_21_+_8433172_8446622_RNA45SN1_45S_50ntFlanks | 11322 | 11343 | 22 | ACGTCTCGTCGCGCGCGCGTCC | MB468 | 83 | 227 | 100 | 12 | 83 | 27.1 | 68.0 | 32.5 | 4.9 | 39.1 |
| >hg38_21_+_8433172_8446622_RNA45SN1_45S_50ntFlanks@11322.11344.23\|ACGTCTCGTCGCGCGCGCGTCCG | >hg38_21_+_8433172_8446622_RNA45SN1_45S_50ntFlanks | 11322 | 11344 | 23 | ACGTCTCGTCGCGCGCGCGTCCG | MB468 | 550 | 989 | 742 | 171 | 728 | 145.4 | 242.4 | 262.4 | 96.5 | 344.1 |
| >hg38_21_+_8433172_8446622_RNA45SN1_45S_50ntFlanks@11322.11346.25\|ACGTCTCGTCGCGCGCGCGTCCGCT | >hg38_21_+_8433172_8446622_RNA45SN1_45S_50ntFlanks | 11322 | 11346 | 25 | ACGTCTCGTCGCGCGCGCGTCCGCT | BT20 | 11 | 30 | 7 | 1 | 66 | 17.1 | 50.0 | 10.6 | 1.5 | 106.8 |
| >hg38_21_+_8433172_8446622_RNA45SN1_45S_50ntFlanks@11322.11347.26\|ACGTCTCGTCGCGCGCGCGTCCGCTG | >hg38_21_+_8433172_8446622_RNA45SN1_45S_50ntFlanks | 11322 | 11347 | 26 | ACGTCTCGTCGCGCGCGCGTCCGCTG | BT20 | 15 | 46 | 12 | 8 | 4 | 22.6 | 75.7 | 19.2 | 13.4 | 6.1 |
| >hg38_21_+_8433172_8446622_RNA45SN1_45S_50ntFlanks@11323.11340.18\|CGTCTCGTCGCGCGCGCG | >hg38_21_+_8433172_8446622_RNA45SN1_45S_50ntFlanks | 11323 | 11340 | 18 | CGTCTCGTCGCGCGCGCG | MB468 | 49 | 69 | 81 | 39 | 95 | 19.2 | 14.0 | 31.2 | 10.2 | 62.8 |
| >hg38_21_+_8433172_8446622_RNA45SN1_45S_50ntFlanks@11323.11341.19\|CGTCTCGTCGCGCGCGCGT | >hg38_21_+_8433172_8446622_RNA45SN1_45S_50ntFlanks | 11323 | 11341 | 19 | CGTCTCGTCGCGCGCGCGT | MB468 | 11 | 16 | 15 | 4 | 10 | 2.4 | 4.1 | 2.7 | 2.1 | 3.7 |
| >hg38_21_+_8433172_8446622_RNA45SN1_45S_50ntFlanks@11323.11342.20\|CGTCTCGTCGCGCGCGCGTC | >hg38_21_+_8433172_8446622_RNA45SN1_45S_50ntFlanks | 11323 | 11342 | 20 | CGTCTCGTCGCGCGCGCGTC | BT20 | 17 | 38 | 12 | 4 | 50 | 27.9 | 63.0 | 14.2 | 3.7 | 84.4 |
| >hg38_21_+_8433172_8446622_RNA45SN1_45S_50ntFlanks@11323.11342.20\|CGTCTCGTCGCGCGCGCGTC | >hg38_21_+_8433172_8446622_RNA45SN1_45S_50ntFlanks | 11323 | 11342 | 20 | CGTCTCGTCGCGCGCGCGTC | MB468 | 450 | 650 | 864 | 259 | 555 | 135.4 | 257.4 | 270.5 | 105.5 | 326.0 |
| >hg38_21_+_8433172_8446622_RNA45SN1_45S_50ntFlanks@11323.11343.21\|CGTCTCGTCGCGCGCGCGTCC | >hg38_21_+_8433172_8446622_RNA45SN1_45S_50ntFlanks | 11323 | 11343 | 21 | CGTCTCGTCGCGCGCGCGTCC | BT20 | 22 | 66 | 17 | 3 | 127 | 36.5 | 111.2 | 17.5 | 3.8 | 215.5 |
| >hg38_21_+_8433172_8446622_RNA45SN1_45S_50ntFlanks@11323.11343.21\|CGTCTCGTCGCGCGCGCGTCC | >hg38_21_+_8433172_8446622_RNA45SN1_45S_50ntFlanks | 11323 | 11343 | 21 | CGTCTCGTCGCGCGCGCGTCC | MB468 | 407 | 996 | 534 | 62 | 520 | 117.4 | 230.2 | 158.4 | 36.6 | 193.0 |
| >hg38_21_+_8433172_8446622_RNA45SN1_45S_50ntFlanks@11323.11344.22\|CGTCTCGTCGCGCGCGCGTCCG | >hg38_21_+_8433172_8446622_RNA45SN1_45S_50ntFlanks | 11323 | 11344 | 22 | CGTCTCGTCGCGCGCGCGTCCG | MB468 | 830 | 1412 | 1416 | 582 | 1526 | 137.2 | 570.1 | 406.4 | 263.0 | 899.8 |
| >hg38_21_+_8433172_8446622_RNA45SN1_45S_50ntFlanks@11323.11346.24\|CGTCTCGTCGCGCGCGCGTCCGCT | >hg38_21_+_8433172_8446622_RNA45SN1_45S_50ntFlanks | 11323 | 11346 | 24 | CGTCTCGTCGCGCGCGCGTCCGCT | BT20 | 21 | 53 | 10 | 3 | 113 | 33.8 | 87.4 | 15.0 | 3.2 | 179.3 |
| >hg38_21_+_8433172_8446622_RNA45SN1_45S_50ntFlanks@11323.11347.25\|CGTCTCGTCGCGCGCGCGTCCGCTG | >hg38_21_+_8433172_8446622_RNA45SN1_45S_50ntFlanks | 11323 | 11347 | 25 | CGTCTCGTCGCGCGCGCGTCCGCTG | BT20 | 20 | 62 | 10 | 13 | 215 | 29.5 | 101.9 | 15.8 | 21.2 | 362.9 |
| >hg38_21_+_8433172_8446622_RNA45SN1_45S_50ntFlanks@11324.11344.21\|GTCTCGTCGCGCGCGCGTCCG | >hg38_21_+_8433172_8446622_RNA45SN1_45S_50ntFlanks | 11324 | 11344 | 21 | GTCTCGTCGCGCGCGCGTCCG | MB468 | 15 | 28 | 27 | 8 | 21 | 3.8 | 17.7 | 9.5 | 4.1 | 14.9 |
| >hg38_21_+_8433172_8446622_RNA45SN1_45S_50ntFlanks@11325.11344.20\|TCTCGTCGCGCGCGCGTCCG | >hg38_21_+_8433172_8446622_RNA45SN1_45S_50ntFlanks | 11325 | 11344 | 20 | TCTCGTCGCGCGCGCGTCCG | MB468 | 37 | 61 | 65 | 23 | 47 | 6.2 | 30.6 | 18.7 | 17.1 | 22.8 |
| >hg38_21_+_8433172_8446622_RNA45SN1_45S_50ntFlanks@11326.11345.20\|CTCGTCGCGCGCGCGTCCGC | >hg38_21_+_8433172_8446622_RNA45SN1_45S_50ntFlanks | 11326 | 11345 | 20 | CTCGTCGCGCGCGCGTCCGC | MB231 | 21 | 93 | 65 | 4 | 61 | 16.6 | 95.1 | 83.8 | 3.5 | 67.4 |
| >hg38_21_+_8433172_8446622_RNA45SN1_45S_50ntFlanks@11452.11471.20\|CTCCTCGGCGCGCGGCGGCG | >hg38_21_+_8433172_8446622_RNA45SN1_45S_50ntFlanks | 11452 | 11471 | 20 | CTCCTCGGCGCGCGGCGGCG | MB468 | 11 | 41 | 25 | 8 | 13 | 2.8 | 32.9 | 4.6 | 4.5 | 5.4 |
| >hg38_21_+_8433172_8446622_RNA45SN1_45S_50ntFlanks@11452.11472.21\|CTCCTCGGCGCGCGGCGGCGG | >hg38_21_+_8433172_8446622_RNA45SN1_45S_50ntFlanks | 11452 | 11472 | 21 | CTCCTCGGCGCGCGGCGGCGG | MB468 | 25 | 81 | 52 | 11 | 22 | 7.5 | 75.6 | 8.9 | 5.2 | 8.4 |
| >hg38_21_+_8433172_8446622_RNA45SN1_45S_50ntFlanks@11452.11478.27\|CTCCTCGGCGCGCGGCGGCGGCGGCGG | >hg38_21_+_8433172_8446622_RNA45SN1_45S_50ntFlanks | 11452 | 11478 | 27 | CTCCTCGGCGCGCGGCGGCGGCGGCGG | MB468 | 23 | 67 | 81 | 57 | 26 | 8.4 | 29.3 | 26.1 | 23.1 | 6.3 |
| >hg38_21_+_8433172_8446622_RNA45SN1_45S_50ntFlanks@11517.11539.23\|CCGCCCCCGGGGCCGCGGTTCCG | >hg38_21_+_8433172_8446622_RNA45SN1_45S_50ntFlanks | 11517 | 11539 | 23 | CCGCCCCCGGGGCCGCGGTTCCG | MB468 | 12 | 40 | 23 | 1 | 15 | 2.0 | 20.6 | 9.1 | 1.6 | 13.3 |
| >hg38_21_+_8433172_8446622_RNA45SN1_45S_50ntFlanks@11534.11551.18\|GTTCCGCGCGGCGCCTCG | >hg38_21_+_8433172_8446622_RNA45SN1_45S_50ntFlanks | 11534 | 11551 | 18 | GTTCCGCGCGGCGCCTCG | MB468 | 119 | 310 | 121 | 7 | 103 | 79.6 | 247.9 | 60.9 | 3.8 | 48.0 |
| >hg38_21_+_8433172_8446622_RNA45SN1_45S_50ntFlanks@11534.11552.19\|GTTCCGCGCGGCGCCTCGC | >hg38_21_+_8433172_8446622_RNA45SN1_45S_50ntFlanks | 11534 | 11552 | 19 | GTTCCGCGCGGCGCCTCGC | MB468 | 72 | 138 | 58 | 5 | 60 | 39.3 | 112.6 | 29.6 | 2.1 | 37.5 |
| >hg38_21_+_8433172_8446622_RNA45SN1_45S_50ntFlanks@11534.11553.20\|GTTCCGCGCGGCGCCTCGCC | >hg38_21_+_8433172_8446622_RNA45SN1_45S_50ntFlanks | 11534 | 11553 | 20 | GTTCCGCGCGGCGCCTCGCC | MB468 | 78 | 65 | 100 | 16 | 37 | 40.8 | 41.9 | 50.3 | 8.0 | 13.3 |
| >hg38_21_+_8433172_8446622_RNA45SN1_45S_50ntFlanks@11534.11555.22\|GTTCCGCGCGGCGCCTCGCCTC | >hg38_21_+_8433172_8446622_RNA45SN1_45S_50ntFlanks | 11534 | 11555 | 22 | GTTCCGCGCGGCGCCTCGCCTC | MB468 | 17 | 24 | 34 | 2 | 7 | 5.2 | 14.1 | 14.9 | 1.2 | 1.7 |
| >hg38_21_+_8433172_8446622_RNA45SN1_45S_50ntFlanks@11534.11557.24\|GTTCCGCGCGGCGCCTCGCCTCGG | >hg38_21_+_8433172_8446622_RNA45SN1_45S_50ntFlanks | 11534 | 11557 | 24 | GTTCCGCGCGGCGCCTCGCCTCGG | MB468 | 12 | 10 | 27 | 4 | 3 | 6.5 | 5.0 | 21.9 | 1.3 | 0.6 |
| >hg38_21_+_8433172_8446622_RNA45SN1_45S_50ntFlanks@11534.11560.27\|GTTCCGCGCGGCGCCTCGCCTCGGCCG | >hg38_21_+_8433172_8446622_RNA45SN1_45S_50ntFlanks | 11534 | 11560 | 27 | GTTCCGCGCGGCGCCTCGCCTCGGCCG | MB468 | 35 | 38 | 54 | 13 | 54 | 8.4 | 15.6 | 28.8 | 6.2 | 20.6 |
| >hg38_21_+_8433172_8446622_RNA45SN1_45S_50ntFlanks@11535.11552.18\|TTCCGCGCGGCGCCTCGC | >hg38_21_+_8433172_8446622_RNA45SN1_45S_50ntFlanks | 11535 | 11552 | 18 | TTCCGCGCGGCGCCTCGC | MB468 | 18 | 39 | 14 | 1 | 9 | 5.6 | 24.5 | 5.5 | 0.6 | 6.7 |
| >hg38_21_+_8433172_8446622_RNA45SN1_45S_50ntFlanks@11535.11553.19\|TTCCGCGCGGCGCCTCGCC | >hg38_21_+_8433172_8446622_RNA45SN1_45S_50ntFlanks | 11535 | 11553 | 19 | TTCCGCGCGGCGCCTCGCC | MB468 | 15 | 17 | 18 | 3 | 8 | 4.0 | 7.1 | 5.7 | 1.4 | 2.6 |
| >hg38_21_+_8433172_8446622_RNA45SN1_45S_50ntFlanks@11536.11553.18\|TCCGCGCGGCGCCTCGCC | >hg38_21_+_8433172_8446622_RNA45SN1_45S_50ntFlanks | 11536 | 11553 | 18 | TCCGCGCGGCGCCTCGCC | MB468 | 11 | 14 | 15 | 1 | 6 | 5.8 | 6.8 | 7.5 | 0.7 | 1.6 |
| >hg38_21_+_8433172_8446622_RNA45SN1_45S_50ntFlanks@11537.11560.24\|CCGCGCGGCGCCTCGCCTCGGCCG | >hg38_21_+_8433172_8446622_RNA45SN1_45S_50ntFlanks | 11537 | 11560 | 24 | CCGCGCGGCGCCTCGCCTCGGCCG | MB468 | 12 | 23 | 20 | 8 | 24 | 5.1 | 19.6 | 5.3 | 4.9 | 19.6 |
| >hg38_21_+_8433172_8446622_RNA45SN1_45S_50ntFlanks@11538.11584.47\|CGCGCGGCGCCTCGCCTCGGCCGGCGCCTAGCAGCCGACTTAGAACT | >hg38_21_+_8433172_8446622_RNA45SN1_45S_50ntFlanks | 11538 | 11584 | 47 | CGCGCGGCGCCTCGCCTCGGCCGGCGCCTAGCAGCCGACTTAGAACT | MB231 | 11 | 11 | 22 | 0 | 0 | 9.5 | 7.3 | 16.3 | 0.0 | 0.0 |
| >hg38_21_+_8433172_8446622_RNA45SN1_45S_50ntFlanks@11539.11583.45\|GCGCGGCGCCTCGCCTCGGCCGGCGCCTAGCAGCCGACTTAGAAC | >hg38_21_+_8433172_8446622_RNA45SN1_45S_50ntFlanks | 11539 | 11583 | 45 | GCGCGGCGCCTCGCCTCGGCCGGCGCCTAGCAGCCGACTTAGAAC | MB231 | 14 | 15 | 27 | 0 | 0 | 11.3 | 9.1 | 21.5 | 0.0 | 0.0 |
| >hg38_21_+_8433172_8446622_RNA45SN1_45S_50ntFlanks@11539.11584.46\|GCGCGGCGCCTCGCCTCGGCCGGCGCCTAGCAGCCGACTTAGAACT | >hg38_21_+_8433172_8446622_RNA45SN1_45S_50ntFlanks | 11539 | 11584 | 46 | GCGCGGCGCCTCGCCTCGGCCGGCGCCTAGCAGCCGACTTAGAACT | MB231 | 28 | 28 | 48 | 0 | 22 | 21.0 | 22.4 | 47.0 | 0.7 | 37.5 |
| >hg38_21_+_8433172_8446622_RNA45SN1_45S_50ntFlanks@11540.11580.41\|CGCGGCGCCTCGCCTCGGCCGGCGCCTAGCAGCCGACTTAG | >hg38_21_+_8433172_8446622_RNA45SN1_45S_50ntFlanks | 11540 | 11580 | 41 | CGCGGCGCCTCGCCTCGGCCGGCGCCTAGCAGCCGACTTAG | MB231 | 17 | 23 | 22 | 0 | 16 | 14.2 | 23.9 | 27.1 | 0.2 | 28.6 |
| >hg38_21_+_8433172_8446622_RNA45SN1_45S_50ntFlanks@11540.11580.41\|CGCGGCGCCTCGCCTCGGCCGGCGCCTAGCAGCCGACTTAG | >hg38_21_+_8433172_8446622_RNA45SN1_45S_50ntFlanks | 11540 | 11580 | 41 | CGCGGCGCCTCGCCTCGGCCGGCGCCTAGCAGCCGACTTAG | MB468 | 10 | 82 | 16 | 1 | 35 | 4.9 | 93.5 | 10.1 | 1.3 | 48.4 |
| >hg38_21_+_8433172_8446622_RNA45SN1_45S_50ntFlanks@11540.11581.42\|CGCGGCGCCTCGCCTCGGCCGGCGCCTAGCAGCCGACTTAGA | >hg38_21_+_8433172_8446622_RNA45SN1_45S_50ntFlanks | 11540 | 11581 | 42 | CGCGGCGCCTCGCCTCGGCCGGCGCCTAGCAGCCGACTTAGA | MB231 | 14 | 19 | 20 | 0 | 11 | 10.7 | 12.1 | 16.7 | 0.3 | 18.8 |
| >hg38_21_+_8433172_8446622_RNA45SN1_45S_50ntFlanks@11540.11582.43\|CGCGGCGCCTCGCCTCGGCCGGCGCCTAGCAGCCGACTTAGAA | >hg38_21_+_8433172_8446622_RNA45SN1_45S_50ntFlanks | 11540 | 11582 | 43 | CGCGGCGCCTCGCCTCGGCCGGCGCCTAGCAGCCGACTTAGAA | MB231 | 31 | 43 | 52 | 0 | 22 | 26.2 | 29.0 | 46.7 | 0.3 | 38.6 |
| >hg38_21_+_8433172_8446622_RNA45SN1_45S_50ntFlanks@11540.11583.44\|CGCGGCGCCTCGCCTCGGCCGGCGCCTAGCAGCCGACTTAGAAC | >hg38_21_+_8433172_8446622_RNA45SN1_45S_50ntFlanks | 11540 | 11583 | 44 | CGCGGCGCCTCGCCTCGGCCGGCGCCTAGCAGCCGACTTAGAAC | MB231 | 46 | 53 | 107 | 1 | 71 | 39.1 | 34.6 | 81.6 | 1.0 | 61.7 |
| >hg38_21_+_8433172_8446622_RNA45SN1_45S_50ntFlanks@11540.11584.45\|CGCGGCGCCTCGCCTCGGCCGGCGCCTAGCAGCCGACTTAGAACT | >hg38_21_+_8433172_8446622_RNA45SN1_45S_50ntFlanks | 11540 | 11584 | 45 | CGCGGCGCCTCGCCTCGGCCGGCGCCTAGCAGCCGACTTAGAACT | BT20 | 12 | 12 | 10 | 3 | 39 | 6.7 | 2.3 | 1.8 | 0.8 | 21.4 |

| | | | | | | | | | | | | | | | | |
|---|---|---|---|---|---|---|---|---|---|---|---|---|---|---|---|---|
| >hg38_21_+_8433172_8446622_RNA45SN1_45S_50ntFlanks@11540.11584.45\|CGCGGCGCCTCGCCTCGGCCGGCGCCTAGCAGCCGACTTAGAACT | >hg38_21_+_8433172_8446622_RNA45SN1_45S_50ntFlanks | 11540 | 11584 | 45 | CGCGGCGCCTCGCCTCGGCCGGCGCCTAGCAGCCGACTTAGAACT | MB231 | 110 | 123 | 255 | 2 | 141 | 92.1 | 101.1 | 191.3 | 2.3 | 128.9 |
| >hg38_21_+_8433172_8446622_RNA45SN1_45S_50ntFlanks@11540.11586.47\|CGCGGCGCCTCGCCTCGGCCGGCGCCTAGCAGCCGACTTAGAACTGG | >hg38_21_+_8433172_8446622_RNA45SN1_45S_50ntFlanks | 11540 | 11586 | 47 | CGCGGCGCCTCGCCTCGGCCGGCGCCTAGCAGCCGACTTAGAACTGG | MB231 | 20 | 16 | 28 | 1 | 30 | 18.8 | 14.5 | 25.3 | 1.2 | 27.9 |
| >hg38_21_+_8433172_8446622_RNA45SN1_45S_50ntFlanks@11541.11584.44\|GCGGCGCCTCGCCTCGGCCGGCGCCTAGCAGCCGACTTAGAACT | >hg38_21_+_8433172_8446622_RNA45SN1_45S_50ntFlanks | 11541 | 11584 | 44 | GCGGCGCCTCGCCTCGGCCGGCGCCTAGCAGCCGACTTAGAACT | MB231 | 13 | 15 | 23 | 0 | 9 | 10.0 | 6.6 | 15.7 | 0.4 | 14.8 |
| >hg38_21_+_8433172_8446622_RNA45SN1_45S_50ntFlanks@11542.11584.43\|CGGCGCCTCGCCTCGGCCGGCGCCTAGCAGCCGACTTAGAACT | >hg38_21_+_8433172_8446622_RNA45SN1_45S_50ntFlanks | 11542 | 11584 | 43 | CGGCGCCTCGCCTCGGCCGGCGCCTAGCAGCCGACTTAGAACT | MB231 | 13 | 16 | 19 | 0 | 0 | 10.9 | 6.4 | 11.3 | 0.0 | 0.0 |
| >hg38_21_+_8433172_8446622_RNA45SN1_45S_50ntFlanks@11552.11569.18\|CCTCGGCCGGCGCCTAGC | >hg38_21_+_8433172_8446622_RNA45SN1_45S_50ntFlanks | 11552 | 11569 | 18 | CCTCGGCCGGCGCCTAGC | MB468 | 28 | 46 | 30 | 2 | 49 | 11.8 | 15.3 | 17.2 | 1.1 | 20.2 |
| >hg38_21_+_8433172_8446622_RNA45SN1_45S_50ntFlanks@11552.11574.23\|CCTCGGCCGGCGCCTAGCAGCCG | >hg38_21_+_8433172_8446622_RNA45SN1_45S_50ntFlanks | 11552 | 11574 | 23 | CCTCGGCCGGCGCCTAGCAGCCG | MB468 | 12 | 12 | 19 | 8 | 19 | 2.5 | 3.6 | 6.5 | 3.9 | 12.3 |
| >hg38_21_+_8433172_8446622_RNA45SN1_45S_50ntFlanks@11552.11588.37\|CCTCGGCCGGCGCCTAGCAGCCGACTTAGAACTGGTG | >hg38_21_+_8433172_8446622_RNA45SN1_45S_50ntFlanks | 11552 | 11588 | 37 | CCTCGGCCGGCGCCTAGCAGCCGACTTAGAACTGGTG | MB468 | 14 | 9 | 15 | 6 | 30 | 2.5 | 5.7 | 5.8 | 3.8 | 18.1 |
| >hg38_21_+_8433172_8446622_RNA45SN1_45S_50ntFlanks@11552.11590.39\|CCTCGGCCGGCGCCTAGCAGCCGACTTAGAACTGGTGCG | >hg38_21_+_8433172_8446622_RNA45SN1_45S_50ntFlanks | 11552 | 11590 | 39 | CCTCGGCCGGCGCCTAGCAGCCGACTTAGAACTGGTGCG | MB231 | 11 | 10 | 4 | 0 | 5 | 10.3 | 8.0 | 3.9 | 0.4 | 9.4 |
| >hg38_21_+_8433172_8446622_RNA45SN1_45S_50ntFlanks@11552.11591.40\|CCTCGGCCGGCGCCTAGCAGCCGACTTAGAACTGGTGCGG | >hg38_21_+_8433172_8446622_RNA45SN1_45S_50ntFlanks | 11552 | 11591 | 40 | CCTCGGCCGGCGCCTAGCAGCCGACTTAGAACTGGTGCGG | MB468 | 17 | 63 | 34 | 4 | 33 | 4.6 | 64.4 | 19.2 | 3.9 | 28.5 |
| >hg38_21_+_8433172_8446622_RNA45SN1_45S_50ntFlanks@11552.11592.41\|CCTCGGCCGGCGCCTAGCAGCCGACTTAGAACTGGTGCGGA | >hg38_21_+_8433172_8446622_RNA45SN1_45S_50ntFlanks | 11552 | 11592 | 41 | CCTCGGCCGGCGCCTAGCAGCCGACTTAGAACTGGTGCGGA | MB468 | 11 | 39 | 27 | 2 | 19 | 3.1 | 45.8 | 21.1 | 2.4 | 21.1 |
| >hg38_21_+_8433172_8446622_RNA45SN1_45S_50ntFlanks@11561.11580.20\|GCGCCTAGCAGCCGACTTAG | >hg38_21_+_8433172_8446622_RNA45SN1_45S_50ntFlanks | 11561 | 11580 | 20 | GCGCCTAGCAGCCGACTTAG | MB468 | 13 | 2 | 10 | 8 | 15 | 7.1 | 0.6 | 8.6 | 6.7 | 6.4 |
| >hg38_21_+_8433172_8446622_RNA45SN1_45S_50ntFlanks@11561.11582.22\|GCGCCTAGCAGCCGACTTAGAA | >hg38_21_+_8433172_8446622_RNA45SN1_45S_50ntFlanks | 11561 | 11582 | 22 | GCGCCTAGCAGCCGACTTAGAA | MB468 | 28 | 12 | 13 | 8 | 41 | 16.2 | 6.9 | 13.9 | 2.9 | 26.7 |
| >hg38_21_+_8433172_8446622_RNA45SN1_45S_50ntFlanks@11561.11584.24\|GCGCCTAGCAGCCGACTTAGAACT | >hg38_21_+_8433172_8446622_RNA45SN1_45S_50ntFlanks | 11561 | 11584 | 24 | GCGCCTAGCAGCCGACTTAGAACT | MB468 | 12 | 17 | 11 | 14 | 27 | 3.0 | 7.8 | 2.6 | 5.4 | 9.7 |
| >hg38_21_+_8433172_8446622_RNA45SN1_45S_50ntFlanks@11561.11585.25\|GCGCCTAGCAGCCGACTTAGAACTG | >hg38_21_+_8433172_8446622_RNA45SN1_45S_50ntFlanks | 11561 | 11585 | 25 | GCGCCTAGCAGCCGACTTAGAACTG | MB468 | 169 | 119 | 100 | 116 | 198 | 42.3 | 41.2 | 24.8 | 22.6 | 59.8 |
| >hg38_21_+_8433172_8446622_RNA45SN1_45S_50ntFlanks@11561.11586.26\|GCGCCTAGCAGCCGACTTAGAACTGG | >hg38_21_+_8433172_8446622_RNA45SN1_45S_50ntFlanks | 11561 | 11586 | 26 | GCGCCTAGCAGCCGACTTAGAACTGG | MB468 | 63 | 52 | 27 | 73 | 107 | 11.1 | 19.8 | 5.0 | 10.3 | 35.8 |
| >hg38_21_+_8433172_8446622_RNA45SN1_45S_50ntFlanks@11561.11587.27\|GCGCCTAGCAGCCGACTTAGAACTGGT | >hg38_21_+_8433172_8446622_RNA45SN1_45S_50ntFlanks | 11561 | 11587 | 27 | GCGCCTAGCAGCCGACTTAGAACTGGT | MB468 | 14 | 11 | 13 | 20 | 17 | 2.6 | 5.1 | 6.4 | 8.5 | 9.0 |
| >hg38_21_+_8433172_8446622_RNA45SN1_45S_50ntFlanks@11561.11588.28\|GCGCCTAGCAGCCGACTTAGAACTGGTG | >hg38_21_+_8433172_8446622_RNA45SN1_45S_50ntFlanks | 11561 | 11588 | 28 | GCGCCTAGCAGCCGACTTAGAACTGGTG | MB468 | 64 | 28 | 89 | 63 | 49 | 11.9 | 11.3 | 26.9 | 20.2 | 22.4 |
| >hg38_21_+_8433172_8446622_RNA45SN1_45S_50ntFlanks@11561.11589.29\|GCGCCTAGCAGCCGACTTAGAACTGGTGC | >hg38_21_+_8433172_8446622_RNA45SN1_45S_50ntFlanks | 11561 | 11589 | 29 | GCGCCTAGCAGCCGACTTAGAACTGGTGC | MB468 | 20 | 6 | 11 | 15 | 19 | 6.3 | 2.8 | 0.8 | 5.4 | 7.8 |
| >hg38_21_+_8433172_8446622_RNA45SN1_45S_50ntFlanks@11561.11590.30\|GCGCCTAGCAGCCGACTTAGAACTGGTGCG | >hg38_21_+_8433172_8446622_RNA45SN1_45S_50ntFlanks | 11561 | 11590 | 30 | GCGCCTAGCAGCCGACTTAGAACTGGTGCG | MB468 | 41 | 5 | 21 | 23 | 23 | 21.3 | 2.0 | 3.6 | 9.1 | 3.1 |
| >hg38_21_+_8433172_8446622_RNA45SN1_45S_50ntFlanks@11561.11591.31\|GCGCCTAGCAGCCGACTTAGAACTGGTGCGG | >hg38_21_+_8433172_8446622_RNA45SN1_45S_50ntFlanks | 11561 | 11591 | 31 | GCGCCTAGCAGCCGACTTAGAACTGGTGCGG | MB468 | 135 | 20 | 73 | 71 | 93 | 73.1 | 10.2 | 22.7 | 40.4 | 16.5 |
| >hg38_21_+_8433172_8446622_RNA45SN1_45S_50ntFlanks@11561.11592.32\|GCGCCTAGCAGCCGACTTAGAACTGGTGCGGA | >hg38_21_+_8433172_8446622_RNA45SN1_45S_50ntFlanks | 11561 | 11592 | 32 | GCGCCTAGCAGCCGACTTAGAACTGGTGCGGA | MB468 | 155 | 32 | 105 | 130 | 131 | 56.6 | 12.0 | 34.9 | 118.8 | 47.0 |
| >hg38_21_+_8433172_8446622_RNA45SN1_45S_50ntFlanks@11561.11593.33\|GCGCCTAGCAGCCGACTTAGAACTGGTGCGGAC | >hg38_21_+_8433172_8446622_RNA45SN1_45S_50ntFlanks | 11561 | 11593 | 33 | GCGCCTAGCAGCCGACTTAGAACTGGTGCGGAC | MB468 | 35 | 9 | 39 | 28 | 41 | 13.2 | 3.1 | 16.3 | 22.3 | 17.8 |
| >hg38_21_+_8433172_8446622_RNA45SN1_45S_50ntFlanks@11561.11594.34\|GCGCCTAGCAGCCGACTTAGAACTGGTGCGGACC | >hg38_21_+_8433172_8446622_RNA45SN1_45S_50ntFlanks | 11561 | 11594 | 34 | GCGCCTAGCAGCCGACTTAGAACTGGTGCGGACC | MB468 | 25 | 18 | 37 | 15 | 34 | 9.2 | 8.9 | 15.1 | 6.5 | 11.4 |
| >hg38_21_+_8433172_8446622_RNA45SN1_45S_50ntFlanks@11561.11600.40\|GCGCCTAGCAGCCGACTTAGAACTGGTGCGGACCAGGGGA | >hg38_21_+_8433172_8446622_RNA45SN1_45S_50ntFlanks | 11561 | 11600 | 40 | GCGCCTAGCAGCCGACTTAGAACTGGTGCGGACCAGGGGA | MB468 | 23 | 116 | 30 | 38 | 77 | 10.5 | 48.9 | 18.0 | 32.9 | 46.9 |
| >hg38_21_+_8433172_8446622_RNA45SN1_45S_50ntFlanks@11561.11601.41\|GCGCCTAGCAGCCGACTTAGAACTGGTGCGGACCAGGGGAA | >hg38_21_+_8433172_8446622_RNA45SN1_45S_50ntFlanks | 11561 | 11601 | 41 | GCGCCTAGCAGCCGACTTAGAACTGGTGCGGACCAGGGGAA | MB468 | 48 | 248 | 62 | 92 | 282 | 30.4 | 155.4 | 38.8 | 93.3 | 194.6 |
| >hg38_21_+_8433172_8446622_RNA45SN1_45S_50ntFlanks@11561.11602.42\|GCGCCTAGCAGCCGACTTAGAACTGGTGCGGACCAGGGGAAT | >hg38_21_+_8433172_8446622_RNA45SN1_45S_50ntFlanks | 11561 | 11602 | 42 | GCGCCTAGCAGCCGACTTAGAACTGGTGCGGACCAGGGGAAT | MB468 | 30 | 150 | 39 | 48 | 190 | 17.8 | 74.1 | 26.8 | 49.4 | 113.7 |
| >hg38_21_+_8433172_8446622_RNA45SN1_45S_50ntFlanks@11561.11603.43\|GCGCCTAGCAGCCGACTTAGAACTGGTGCGGACCAGGGGAATC | >hg38_21_+_8433172_8446622_RNA45SN1_45S_50ntFlanks | 11561 | 11603 | 43 | GCGCCTAGCAGCCGACTTAGAACTGGTGCGGACCAGGGGAATC | MB468 | 40 | 229 | 42 | 59 | 468 | 21.4 | 142.1 | 22.4 | 61.3 | 260.3 |
| >hg38_21_+_8433172_8446622_RNA45SN1_45S_50ntFlanks@11562.11585.24\|CGCCTAGCAGCCGACTTAGAACTG | >hg38_21_+_8433172_8446622_RNA45SN1_45S_50ntFlanks | 11562 | 11585 | 24 | CGCCTAGCAGCCGACTTAGAACTG | MB468 | 12 | 6 | 5 | 14 | 15 | 4.8 | 2.1 | 0.9 | 2.8 | 8.4 |
| >hg38_21_+_8433172_8446622_RNA45SN1_45S_50ntFlanks@11562.11588.27\|CGCCTAGCAGCCGACTTAGAACTGGTG | >hg38_21_+_8433172_8446622_RNA45SN1_45S_50ntFlanks | 11562 | 11588 | 27 | CGCCTAGCAGCCGACTTAGAACTGGTG | MB468 | 46 | 29 | 29 | 43 | 22 | 8.7 | 10.4 | 8.4 | 21.5 | 7.4 |
| >hg38_21_+_8433172_8446622_RNA45SN1_45S_50ntFlanks@11562.11591.30\|CGCCTAGCAGCCGACTTAGAACTGGTGCGG | >hg38_21_+_8433172_8446622_RNA45SN1_45S_50ntFlanks | 11562 | 11591 | 30 | CGCCTAGCAGCCGACTTAGAACTGGTGCGG | MB468 | 19 | 2 | 10 | 29 | 11 | 5.8 | 1.5 | 1.6 | 29.9 | 5.4 |
| >hg38_21_+_8433172_8446622_RNA45SN1_45S_50ntFlanks@11562.11592.31\|CGCCTAGCAGCCGACTTAGAACTGGTGCGGA | >hg38_21_+_8433172_8446622_RNA45SN1_45S_50ntFlanks | 11562 | 11592 | 31 | CGCCTAGCAGCCGACTTAGAACTGGTGCGGA | MB468 | 108 | 20 | 70 | 199 | 66 | 31.3 | 10.0 | 25.0 | 211.7 | 34.6 |
| >hg38_21_+_8433172_8446622_RNA45SN1_45S_50ntFlanks@11563.11584.22\|GCCTAGCAGCCGACTTAGAACT | >hg38_21_+_8433172_8446622_RNA45SN1_45S_50ntFlanks | 11563 | 11584 | 22 | GCCTAGCAGCCGACTTAGAACT | MB231 | 19 | 18 | 10 | 6 | 12 | 14.5 | 17.1 | 9.4 | 4.3 | 11.4 |
| >hg38_21_+_8433172_8446622_RNA45SN1_45S_50ntFlanks@11564.11581.18\|CCTAGCAGCCGACTTAGA | >hg38_21_+_8433172_8446622_RNA45SN1_45S_50ntFlanks | 11564 | 11581 | 18 | CCTAGCAGCCGACTTAGA | MB468 | 113 | 90 | 71 | 23 | 114 | 14.7 | 39.8 | 20.0 | 5.4 | 23.2 |
| >hg38_21_+_8433172_8446622_RNA45SN1_45S_50ntFlanks@11564.11582.19\|CCTAGCAGCCGACTTAGAA | >hg38_21_+_8433172_8446622_RNA45SN1_45S_50ntFlanks | 11564 | 11582 | 19 | CCTAGCAGCCGACTTAGAA | MB468 | 152 | 206 | 84 | 33 | 162 | 21.7 | 95.9 | 41.4 | 7.6 | 55.0 |
| >hg38_21_+_8433172_8446622_RNA45SN1_45S_50ntFlanks@11564.11584.21\|CCTAGCAGCCGACTTAGAACT | >hg38_21_+_8433172_8446622_RNA45SN1_45S_50ntFlanks | 11564 | 11584 | 21 | CCTAGCAGCCGACTTAGAACT | MB468 | 15 | 18 | 17 | 11 | 18 | 9.0 | 4.0 | 8.3 | 1.7 | 1.5 |
| >hg38_21_+_8433172_8446622_RNA45SN1_45S_50ntFlanks@11564.11585.22\|CCTAGCAGCCGACTTAGAACTG | >hg38_21_+_8433172_8446622_RNA45SN1_45S_50ntFlanks | 11564 | 11585 | 22 | CCTAGCAGCCGACTTAGAACTG | MB468 | 189 | 188 | 159 | 125 | 156 | 13.6 | 42.2 | 17.2 | 23.0 | 53.2 |
| >hg38_21_+_8433172_8446622_RNA45SN1_45S_50ntFlanks@11564.11586.23\|CCTAGCAGCCGACTTAGAACTGG | >hg38_21_+_8433172_8446622_RNA45SN1_45S_50ntFlanks | 11564 | 11586 | 23 | CCTAGCAGCCGACTTAGAACTGG | MB468 | 67 | 45 | 39 | 74 | 66 | 7.5 | 8.6 | 2.1 | 22.2 | 25.9 |
| >hg38_21_+_8433172_8446622_RNA45SN1_45S_50ntFlanks@11564.11587.24\|CCTAGCAGCCGACTTAGAACTGGT | >hg38_21_+_8433172_8446622_RNA45SN1_45S_50ntFlanks | 11564 | 11587 | 24 | CCTAGCAGCCGACTTAGAACTGGT | MB468 | 22 | 13 | 16 | 20 | 13 | 5.8 | 4.3 | 3.9 | 6.3 | 3.6 |
| >hg38_21_+_8433172_8446622_RNA45SN1_45S_50ntFlanks@11564.11588.25\|CCTAGCAGCCGACTTAGAACTGGTG | >hg38_21_+_8433172_8446622_RNA45SN1_45S_50ntFlanks | 11564 | 11588 | 25 | CCTAGCAGCCGACTTAGAACTGGTG | MB468 | 117 | 61 | 85 | 105 | 65 | 19.8 | 12.1 | 3.0 | 29.1 | 8.3 |
| >hg38_21_+_8433172_8446622_RNA45SN1_45S_50ntFlanks@11564.11590.27\|CCTAGCAGCCGACTTAGAACTGGTGCG | >hg38_21_+_8433172_8446622_RNA45SN1_45S_50ntFlanks | 11564 | 11590 | 27 | CCTAGCAGCCGACTTAGAACTGGTGCG | MB468 | 30 | 9 | 16 | 48 | 23 | 7.2 | 2.4 | 6.3 | 36.2 | 5.2 |
| >hg38_21_+_8433172_8446622_RNA45SN1_45S_50ntFlanks@11564.11591.28\|CCTAGCAGCCGACTTAGAACTGGTGCGG | >hg38_21_+_8433172_8446622_RNA45SN1_45S_50ntFlanks | 11564 | 11591 | 28 | CCTAGCAGCCGACTTAGAACTGGTGCGG | MB468 | 182 | 53 | 85 | 164 | 146 | 66.6 | 4.7 | 55.2 | 109.8 | 39.4 |
| >hg38_21_+_8433172_8446622_RNA45SN1_45S_50ntFlanks@11564.11592.29\|CCTAGCAGCCGACTTAGAACTGGTGCGGA | >hg38_21_+_8433172_8446622_RNA45SN1_45S_50ntFlanks | 11564 | 11592 | 29 | CCTAGCAGCCGACTTAGAACTGGTGCGGA | MB468 | 47 | 11 | 33 | 132 | 33 | 22.7 | 5.0 | 14.5 | 154.9 | 20.2 |
| >hg38_21_+_8433172_8446622_RNA45SN1_45S_50ntFlanks@11564.11593.30\|CCTAGCAGCCGACTTAGAACTGGTGCGGAC | >hg38_21_+_8433172_8446622_RNA45SN1_45S_50ntFlanks | 11564 | 11593 | 30 | CCTAGCAGCCGACTTAGAACTGGTGCGGAC | MB468 | 11 | 4 | 12 | 19 | 13 | 4.2 | 0.9 | 6.0 | 17.5 | 2.9 |
| >hg38_21_+_8433172_8446622_RNA45SN1_45S_50ntFlanks@11564.11600.37\|CCTAGCAGCCGACTTAGAACTGGTGCGGACCAGGGGA | >hg38_21_+_8433172_8446622_RNA45SN1_45S_50ntFlanks | 11564 | 11600 | 37 | CCTAGCAGCCGACTTAGAACTGGTGCGGACCAGGGGA | MB468 | 55 | 103 | 50 | 94 | 103 | 14.5 | 31.3 | 9.4 | 55.3 | 47.1 |
| >hg38_21_+_8433172_8446622_RNA45SN1_45S_50ntFlanks@11564.11601.38\|CCTAGCAGCCGACTTAGAACTGGTGCGGACCAGGGGAA | >hg38_21_+_8433172_8446622_RNA45SN1_45S_50ntFlanks | 11564 | 11601 | 38 | CCTAGCAGCCGACTTAGAACTGGTGCGGACCAGGGGAA | MB468 | 135 | 321 | 125 | 230 | 490 | 60.3 | 162.1 | 38.3 | 163.3 | 247.0 |
| >hg38_21_+_8433172_8446622_RNA45SN1_45S_50ntFlanks@11564.11602.39\|CCTAGCAGCCGACTTAGAACTGGTGCGGACCAGGGGAAT | >hg38_21_+_8433172_8446622_RNA45SN1_45S_50ntFlanks | 11564 | 11602 | 39 | CCTAGCAGCCGACTTAGAACTGGTGCGGACCAGGGGAAT | MB468 | 74 | 171 | 72 | 112 | 343 | 31.3 | 58.4 | 22.4 | 86.3 | 183.5 |
| >hg38_21_+_8433172_8446622_RNA45SN1_45S_50ntFlanks@11564.11603.40\|CCTAGCAGCCGACTTAGAACTGGTGCGGACCAGGGGAATC | >hg38_21_+_8433172_8446622_RNA45SN1_45S_50ntFlanks | 11564 | 11603 | 40 | CCTAGCAGCCGACTTAGAACTGGTGCGGACCAGGGGAATC | MB468 | 85 | 212 | 67 | 122 | 778 | 26.0 | 106.7 | 33.0 | 95.7 | 447.7 |
| >hg38_21_+_8433172_8446622_RNA45SN1_45S_50ntFlanks@11564.11611.48\|CCTAGCAGCCGACTTAGAACTGGTGCGGACCAGGGGAATCCGACTGTT | >hg38_21_+_8433172_8446622_RNA45SN1_45S_50ntFlanks | 11564 | 11611 | 48 | CCTAGCAGCCGACTTAGAACTGGTGCGGACCAGGGGAATCCGACTGTT | MB468 | 24 | 92 | 43 | 24 | 366 | 21.6 | 124.2 | 28.5 | 29.3 | 350.4 |
| >hg38_21_+_8433172_8446622_RNA45SN1_45S_50ntFlanks@11565.11582.18\|CTAGCAGCCGACTTAGAA | >hg38_21_+_8433172_8446622_RNA45SN1_45S_50ntFlanks | 11565 | 11582 | 18 | CTAGCAGCCGACTTAGAA | MB468 | 15 | 15 | 9 | 3 | 9 | 3.1 | 5.8 | 2.9 | 1.0 | 2.0 |
| >hg38_21_+_8433172_8446622_RNA45SN1_45S_50ntFlanks@11565.11585.21\|CTAGCAGCCGACTTAGAACTG | >hg38_21_+_8433172_8446622_RNA45SN1_45S_50ntFlanks | 11565 | 11585 | 21 | CTAGCAGCCGACTTAGAACTG | MB468 | 24 | 14 | 18 | 10 | 13 | 3.4 | 3.1 | 1.3 | 1.0 | 2.6 |

| | | | | | | | | | | | | | | | | |
|---|---|---|---|---|---|---|---|---|---|---|---|---|---|---|---|---|
| >hg38_21_+_8433172_8446622_RNA45SN1_45S_50ntFlanks@11565.11586.22\|CTAGCAGCCGACTTAGAACTGG | >hg38_21_+_8433172_8446622_RNA45SN1_45S_50ntFlanks | 11565 | 11586 | 22 | CTAGCAGCCGACTTAGAACTGG | MB468 | 11 | 5 | 5 | 7 | 6 | 2.4 | 1.5 | 1.1 | 1.3 | 1.8 |
| >hg38_21_+_8433172_8446622_RNA45SN1_45S_50ntFlanks@11565.11588.24\|CTAGCAGCCGACTTAGAACTGGTG | >hg38_21_+_8433172_8446622_RNA45SN1_45S_50ntFlanks | 11565 | 11588 | 24 | CTAGCAGCCGACTTAGAACTGGTG | MB468 | 16 | 4 | 11 | 9 | 5 | 3.4 | 1.4 | 1.3 | 2.5 | 1.4 |
| >hg38_21_+_8433172_8446622_RNA45SN1_45S_50ntFlanks@11566.11583.18\|TAGCAGCCGACTTAGAAC | >hg38_21_+_8433172_8446622_RNA45SN1_45S_50ntFlanks | 11566 | 11583 | 18 | TAGCAGCCGACTTAGAAC | MB231 | 19 | 15 | 36 | 3 | 6 | 14.9 | 10.3 | 22.9 | 3.0 | 4.9 |
| >hg38_21_+_8433172_8446622_RNA45SN1_45S_50ntFlanks@11566.11584.19\|TAGCAGCCGACTTAGAACT | >hg38_21_+_8433172_8446622_RNA45SN1_45S_50ntFlanks | 11566 | 11584 | 19 | TAGCAGCCGACTTAGAACT | BT20 | 27 | 9 | 36 | 11 | 13 | 9.6 | 4.3 | 9.3 | 7.3 | 3.9 |
| >hg38_21_+_8433172_8446622_RNA45SN1_45S_50ntFlanks@11566.11584.19\|TAGCAGCCGACTTAGAACT | >hg38_21_+_8433172_8446622_RNA45SN1_45S_50ntFlanks | 11566 | 11584 | 19 | TAGCAGCCGACTTAGAACT | MB231 | 97 | 66 | 121 | 10 | 24 | 81.0 | 53.0 | 89.1 | 9.0 | 19.2 |
| >hg38_21_+_8433172_8446622_RNA45SN1_45S_50ntFlanks@11566.11585.20\|TAGCAGCCGACTTAGAACTG | >hg38_21_+_8433172_8446622_RNA45SN1_45S_50ntFlanks | 11566 | 11585 | 20 | TAGCAGCCGACTTAGAACTG | MB231 | 32 | 22 | 17 | 6 | 10 | 27.1 | 21.9 | 13.1 | 6.0 | 10.4 |
| >hg38_21_+_8433172_8446622_RNA45SN1_45S_50ntFlanks@11566.11585.20\|TAGCAGCCGACTTAGAACTG | >hg38_21_+_8433172_8446622_RNA45SN1_45S_50ntFlanks | 11566 | 11585 | 20 | TAGCAGCCGACTTAGAACTG | MB468 | 12 | 6 | 8 | 4 | 3 | 2.3 | 1.8 | 1.6 | 0.3 | 0.9 |
| >hg38_21_+_8433172_8446622_RNA45SN1_45S_50ntFlanks@11566.11586.21\|TAGCAGCCGACTTAGAACTGG | >hg38_21_+_8433172_8446622_RNA45SN1_45S_50ntFlanks | 11566 | 11586 | 21 | TAGCAGCCGACTTAGAACTGG | MB231 | 32 | 21 | 14 | 9 | 13 | 25.5 | 20.8 | 11.0 | 8.3 | 13.6 |
| >hg38_21_+_8433172_8446622_RNA45SN1_45S_50ntFlanks@11566.11611.46\|TAGCAGCCGACTTAGAACTGGTGCGGACCAGGGGAATCCGACTGTT | >hg38_21_+_8433172_8446622_RNA45SN1_45S_50ntFlanks | 11566 | 11611 | 46 | TAGCAGCCGACTTAGAACTGGTGCGGACCAGGGGAATCCGACTGTT | BT20 | 12 | 17 | 19 | 37 | 187 | 12.4 | 18.1 | 29.2 | 48.0 | 151.8 |
| >hg38_21_+_8433172_8446622_RNA45SN1_45S_50ntFlanks@11566.11611.46\|TAGCAGCCGACTTAGAACTGGTGCGGACCAGGGGAATCCGACTGTT | >hg38_21_+_8433172_8446622_RNA45SN1_45S_50ntFlanks | 11566 | 11611 | 46 | TAGCAGCCGACTTAGAACTGGTGCGGACCAGGGGAATCCGACTGTT | MB231 | 24 | 70 | 79 | 9 | 84 | 31.8 | 120.5 | 132.0 | 14.9 | 145.9 |
| >hg38_21_+_8433172_8446622_RNA45SN1_45S_50ntFlanks@11567.11584.18\|AGCAGCCGACTTAGAACT | >hg38_21_+_8433172_8446622_RNA45SN1_45S_50ntFlanks | 11567 | 11584 | 18 | AGCAGCCGACTTAGAACT | BT20 | 120 | 58 | 91 | 39 | 82 | 85.2 | 32.6 | 35.1 | 23.1 | 62.6 |
| >hg38_21_+_8433172_8446622_RNA45SN1_45S_50ntFlanks@11567.11584.18\|AGCAGCCGACTTAGAACT | >hg38_21_+_8433172_8446622_RNA45SN1_45S_50ntFlanks | 11567 | 11584 | 18 | AGCAGCCGACTTAGAACT | MB231 | 264 | 225 | 223 | 44 | 129 | 230.7 | 177.8 | 164.5 | 35.3 | 110.8 |
| >hg38_21_+_8433172_8446622_RNA45SN1_45S_50ntFlanks@11567.11585.19\|AGCAGCCGACTTAGAACTG | >hg38_21_+_8433172_8446622_RNA45SN1_45S_50ntFlanks | 11567 | 11585 | 19 | AGCAGCCGACTTAGAACTG | BT20 | 29 | 12 | 14 | 18 | 29 | 13.7 | 1.4 | 3.1 | 14.8 | 12.2 |
| >hg38_21_+_8433172_8446622_RNA45SN1_45S_50ntFlanks@11567.11585.19\|AGCAGCCGACTTAGAACTG | >hg38_21_+_8433172_8446622_RNA45SN1_45S_50ntFlanks | 11567 | 11585 | 19 | AGCAGCCGACTTAGAACTG | MB231 | 103 | 87 | 43 | 25 | 68 | 98.2 | 96.3 | 36.7 | 23.8 | 78.2 |
| >hg38_21_+_8433172_8446622_RNA45SN1_45S_50ntFlanks@11567.11585.19\|AGCAGCCGACTTAGAACTG | >hg38_21_+_8433172_8446622_RNA45SN1_45S_50ntFlanks | 11567 | 11585 | 19 | AGCAGCCGACTTAGAACTG | MB468 | 109 | 56 | 48 | 60 | 35 | 15.2 | 27.1 | 6.0 | 15.8 | 7.8 |
| >hg38_21_+_8433172_8446622_RNA45SN1_45S_50ntFlanks@11567.11586.20\|AGCAGCCGACTTAGAACTGG | >hg38_21_+_8433172_8446622_RNA45SN1_45S_50ntFlanks | 11567 | 11586 | 20 | AGCAGCCGACTTAGAACTGG | BT20 | 21 | 8 | 10 | 17 | 21 | 13.9 | 2.9 | 5.3 | 9.8 | 1.8 |
| >hg38_21_+_8433172_8446622_RNA45SN1_45S_50ntFlanks@11567.11586.20\|AGCAGCCGACTTAGAACTGG | >hg38_21_+_8433172_8446622_RNA45SN1_45S_50ntFlanks | 11567 | 11586 | 20 | AGCAGCCGACTTAGAACTGG | MB231 | 96 | 82 | 41 | 42 | 72 | 76.8 | 87.4 | 30.0 | 38.3 | 80.5 |
| >hg38_21_+_8433172_8446622_RNA45SN1_45S_50ntFlanks@11567.11586.20\|AGCAGCCGACTTAGAACTGG | >hg38_21_+_8433172_8446622_RNA45SN1_45S_50ntFlanks | 11567 | 11586 | 20 | AGCAGCCGACTTAGAACTGG | MB468 | 47 | 20 | 19 | 37 | 18 | 7.3 | 9.9 | 3.3 | 9.3 | 5.1 |
| >hg38_21_+_8433172_8446622_RNA45SN1_45S_50ntFlanks@11567.11587.21\|AGCAGCCGACTTAGAACTGGT | >hg38_21_+_8433172_8446622_RNA45SN1_45S_50ntFlanks | 11567 | 11587 | 21 | AGCAGCCGACTTAGAACTGGT | MB231 | 20 | 21 | 8 | 13 | 17 | 11.6 | 18.6 | 3.4 | 10.6 | 17.7 |
| >hg38_21_+_8433172_8446622_RNA45SN1_45S_50ntFlanks@11567.11587.21\|AGCAGCCGACTTAGAACTGGT | >hg38_21_+_8433172_8446622_RNA45SN1_45S_50ntFlanks | 11567 | 11587 | 21 | AGCAGCCGACTTAGAACTGGT | MB468 | 13 | 4 | 7 | 9 | 3 | 3.6 | 2.2 | 1.7 | 3.3 | 0.5 |
| >hg38_21_+_8433172_8446622_RNA45SN1_45S_50ntFlanks@11567.11588.22\|AGCAGCCGACTTAGAACTGGTG | >hg38_21_+_8433172_8446622_RNA45SN1_45S_50ntFlanks | 11567 | 11588 | 22 | AGCAGCCGACTTAGAACTGGTG | MB231 | 16 | 12 | 5 | 10 | 17 | 14.1 | 11.6 | 2.7 | 6.8 | 19.9 |
| >hg38_21_+_8433172_8446622_RNA45SN1_45S_50ntFlanks@11567.11588.22\|AGCAGCCGACTTAGAACTGGTG | >hg38_21_+_8433172_8446622_RNA45SN1_45S_50ntFlanks | 11567 | 11588 | 22 | AGCAGCCGACTTAGAACTGGTG | MB468 | 53 | 17 | 26 | 45 | 13 | 13.5 | 7.9 | 2.5 | 20.9 | 2.1 |
| >hg38_21_+_8433172_8446622_RNA45SN1_45S_50ntFlanks@11567.11590.24\|AGCAGCCGACTTAGAACTGGTGCG | >hg38_21_+_8433172_8446622_RNA45SN1_45S_50ntFlanks | 11567 | 11590 | 24 | AGCAGCCGACTTAGAACTGGTGCG | MB231 | 10 | 5 | 2 | 5 | 7 | 9.7 | 5.1 | 1.6 | 4.3 | 7.6 |
| >hg38_21_+_8433172_8446622_RNA45SN1_45S_50ntFlanks@11567.11592.26\|AGCAGCCGACTTAGAACTGGTGCGGA | >hg38_21_+_8433172_8446622_RNA45SN1_45S_50ntFlanks | 11567 | 11592 | 26 | AGCAGCCGACTTAGAACTGGTGCGGA | MB468 | 17 | 4 | 15 | 55 | 8 | 4.7 | 2.7 | 6.6 | 54.3 | 5.0 |
| >hg38_21_+_8433172_8446622_RNA45SN1_45S_50ntFlanks@11567.11594.28\|AGCAGCCGACTTAGAACTGGTGCGGACC | >hg38_21_+_8433172_8446622_RNA45SN1_45S_50ntFlanks | 11567 | 11594 | 28 | AGCAGCCGACTTAGAACTGGTGCGGACC | MB231 | 14 | 10 | 7 | 6 | 11 | 10.3 | 5.4 | 3.8 | 4.5 | 8.4 |
| >hg38_21_+_8433172_8446622_RNA45SN1_45S_50ntFlanks@11567.11600.34\|AGCAGCCGACTTAGAACTGGTGCGGACCAGGGGA | >hg38_21_+_8433172_8446622_RNA45SN1_45S_50ntFlanks | 11567 | 11600 | 34 | AGCAGCCGACTTAGAACTGGTGCGGACCAGGGGA | MB468 | 11 | 47 | 12 | 17 | 11 | 1.2 | 9.4 | 5.5 | 5.8 | 5.0 |
| >hg38_21_+_8433172_8446622_RNA45SN1_45S_50ntFlanks@11567.11601.35\|AGCAGCCGACTTAGAACTGGTGCGGACCAGGGGAA | >hg38_21_+_8433172_8446622_RNA45SN1_45S_50ntFlanks | 11567 | 11601 | 35 | AGCAGCCGACTTAGAACTGGTGCGGACCAGGGGAA | MB468 | 20 | 92 | 20 | 33 | 33 | 1.3 | 19.3 | 8.4 | 16.3 | 16.5 |
| >hg38_21_+_8433172_8446622_RNA45SN1_45S_50ntFlanks@11567.11602.36\|AGCAGCCGACTTAGAACTGGTGCGGACCAGGGGAAT | >hg38_21_+_8433172_8446622_RNA45SN1_45S_50ntFlanks | 11567 | 11602 | 36 | AGCAGCCGACTTAGAACTGGTGCGGACCAGGGGAAT | MB468 | 10 | 40 | 9 | 16 | 22 | 1.5 | 9.5 | 2.8 | 8.7 | 11.1 |
| >hg38_21_+_8433172_8446622_RNA45SN1_45S_50ntFlanks@11567.11603.37\|AGCAGCCGACTTAGAACTGGTGCGGACCAGGGGAATC | >hg38_21_+_8433172_8446622_RNA45SN1_45S_50ntFlanks | 11567 | 11603 | 37 | AGCAGCCGACTTAGAACTGGTGCGGACCAGGGGAATC | MB468 | 17 | 64 | 15 | 31 | 81 | 1.7 | 24.7 | 3.1 | 15.5 | 43.4 |
| >hg38_21_+_8433172_8446622_RNA45SN1_45S_50ntFlanks@11567.11611.45\|AGCAGCCGACTTAGAACTGGTGCGGACCAGGGGAATCCGACTGTT | >hg38_21_+_8433172_8446622_RNA45SN1_45S_50ntFlanks | 11567 | 11611 | 45 | AGCAGCCGACTTAGAACTGGTGCGGACCAGGGGAATCCGACTGTT | BT20 | 10 | 21 | 15 | 38 | 184 | 5.1 | 12.2 | 19.1 | 48.1 | 160.3 |
| >hg38_21_+_8433172_8446622_RNA45SN1_45S_50ntFlanks@11567.11611.45\|AGCAGCCGACTTAGAACTGGTGCGGACCAGGGGAATCCGACTGTT | >hg38_21_+_8433172_8446622_RNA45SN1_45S_50ntFlanks | 11567 | 11611 | 45 | AGCAGCCGACTTAGAACTGGTGCGGACCAGGGGAATCCGACTGTT | MB231 | 16 | 35 | 39 | 7 | 57 | 18.4 | 58.6 | 61.0 | 11.4 | 93.8 |
| >hg38_21_+_8433172_8446622_RNA45SN1_45S_50ntFlanks@11568.11585.18\|GCAGCCGACTTAGAACTG | >hg38_21_+_8433172_8446622_RNA45SN1_45S_50ntFlanks | 11568 | 11585 | 18 | GCAGCCGACTTAGAACTG | BT20 | 16 | 8 | 9 | 19 | 17 | 11.1 | 5.1 | 1.3 | 16.4 | 8.2 |
| >hg38_21_+_8433172_8446622_RNA45SN1_45S_50ntFlanks@11568.11585.18\|GCAGCCGACTTAGAACTG | >hg38_21_+_8433172_8446622_RNA45SN1_45S_50ntFlanks | 11568 | 11585 | 18 | GCAGCCGACTTAGAACTG | MB231 | 42 | 40 | 25 | 14 | 18 | 33.1 | 31.9 | 18.3 | 11.4 | 18.0 |
| >hg38_21_+_8433172_8446622_RNA45SN1_45S_50ntFlanks@11568.11585.18\|GCAGCCGACTTAGAACTG | >hg38_21_+_8433172_8446622_RNA45SN1_45S_50ntFlanks | 11568 | 11585 | 18 | GCAGCCGACTTAGAACTG | MB468 | 195 | 105 | 101 | 122 | 61 | 45.4 | 62.4 | 35.1 | 47.3 | 8.3 |
| >hg38_21_+_8433172_8446622_RNA45SN1_45S_50ntFlanks@11568.11586.19\|GCAGCCGACTTAGAACTGG | >hg38_21_+_8433172_8446622_RNA45SN1_45S_50ntFlanks | 11568 | 11586 | 19 | GCAGCCGACTTAGAACTGG | BT20 | 12 | 6 | 6 | 17 | 12 | 6.4 | 2.3 | 2.4 | 8.7 | 7.1 |
| >hg38_21_+_8433172_8446622_RNA45SN1_45S_50ntFlanks@11568.11586.19\|GCAGCCGACTTAGAACTGG | >hg38_21_+_8433172_8446622_RNA45SN1_45S_50ntFlanks | 11568 | 11586 | 19 | GCAGCCGACTTAGAACTGG | MB231 | 49 | 42 | 27 | 23 | 22 | 38.4 | 33.4 | 27.2 | 18.1 | 22.5 |
| >hg38_21_+_8433172_8446622_RNA45SN1_45S_50ntFlanks@11568.11586.19\|GCAGCCGACTTAGAACTGG | >hg38_21_+_8433172_8446622_RNA45SN1_45S_50ntFlanks | 11568 | 11586 | 19 | GCAGCCGACTTAGAACTGG | MB468 | 77 | 29 | 30 | 68 | 27 | 18.3 | 17.6 | 13.1 | 25.1 | 3.6 |
| >hg38_21_+_8433172_8446622_RNA45SN1_45S_50ntFlanks@11568.11587.20\|GCAGCCGACTTAGAACTGGT | >hg38_21_+_8433172_8446622_RNA45SN1_45S_50ntFlanks | 11568 | 11587 | 20 | GCAGCCGACTTAGAACTGGT | MB231 | 10 | 9 | 5 | 7 | 9 | 5.9 | 6.4 | 3.1 | 4.4 | 8.5 |
| >hg38_21_+_8433172_8446622_RNA45SN1_45S_50ntFlanks@11568.11587.20\|GCAGCCGACTTAGAACTGGT | >hg38_21_+_8433172_8446622_RNA45SN1_45S_50ntFlanks | 11568 | 11587 | 20 | GCAGCCGACTTAGAACTGGT | MB468 | 17 | 5 | 11 | 15 | 4 | 5.8 | 3.2 | 3.2 | 6.3 | 1.1 |
| >hg38_21_+_8433172_8446622_RNA45SN1_45S_50ntFlanks@11568.11588.21\|GCAGCCGACTTAGAACTGGTG | >hg38_21_+_8433172_8446622_RNA45SN1_45S_50ntFlanks | 11568 | 11588 | 21 | GCAGCCGACTTAGAACTGGTG | MB468 | 68 | 23 | 39 | 71 | 18 | 13.4 | 13.4 | 4.6 | 43.1 | 4.1 |
| >hg38_21_+_8433172_8446622_RNA45SN1_45S_50ntFlanks@11568.11591.24\|GCAGCCGACTTAGAACTGGTGCGG | >hg38_21_+_8433172_8446622_RNA45SN1_45S_50ntFlanks | 11568 | 11591 | 24 | GCAGCCGACTTAGAACTGGTGCGG | MB468 | 12 | 3 | 12 | 37 | 6 | 2.9 | 2.1 | 3.1 | 34.5 | 3.6 |
| >hg38_21_+_8433172_8446622_RNA45SN1_45S_50ntFlanks@11568.11592.25\|GCAGCCGACTTAGAACTGGTGCGGA | >hg38_21_+_8433172_8446622_RNA45SN1_45S_50ntFlanks | 11568 | 11592 | 25 | GCAGCCGACTTAGAACTGGTGCGGA | MB468 | 29 | 8 | 27 | 111 | 18 | 3.6 | 5.1 | 8.0 | 112.3 | 12.4 |
| >hg38_21_+_8433172_8446622_RNA45SN1_45S_50ntFlanks@11568.11600.33\|GCAGCCGACTTAGAACTGGTGCGGACCAGGGGA | >hg38_21_+_8433172_8446622_RNA45SN1_45S_50ntFlanks | 11568 | 11600 | 33 | GCAGCCGACTTAGAACTGGTGCGGACCAGGGGA | MB468 | 17 | 66 | 19 | 35 | 16 | 3.9 | 19.0 | 11.9 | 13.9 | 8.0 |
| >hg38_21_+_8433172_8446622_RNA45SN1_45S_50ntFlanks@11568.11601.34\|GCAGCCGACTTAGAACTGGTGCGGACCAGGGGAA | >hg38_21_+_8433172_8446622_RNA45SN1_45S_50ntFlanks | 11568 | 11601 | 34 | GCAGCCGACTTAGAACTGGTGCGGACCAGGGGAA | MB468 | 32 | 139 | 33 | 71 | 51 | 4.6 | 32.8 | 18.3 | 30.3 | 24.3 |
| >hg38_21_+_8433172_8446622_RNA45SN1_45S_50ntFlanks@11568.11602.35\|GCAGCCGACTTAGAACTGGTGCGGACCAGGGGAAT | >hg38_21_+_8433172_8446622_RNA45SN1_45S_50ntFlanks | 11568 | 11602 | 35 | GCAGCCGACTTAGAACTGGTGCGGACCAGGGGAAT | MB468 | 18 | 69 | 17 | 34 | 37 | 4.6 | 23.7 | 7.0 | 16.6 | 18.8 |
| >hg38_21_+_8433172_8446622_RNA45SN1_45S_50ntFlanks@11568.11603.36\|GCAGCCGACTTAGAACTGGTGCGGACCAGGGGAATC | >hg38_21_+_8433172_8446622_RNA45SN1_45S_50ntFlanks | 11568 | 11603 | 36 | GCAGCCGACTTAGAACTGGTGCGGACCAGGGGAATC | MB468 | 34 | 126 | 32 | 59 | 144 | 11.1 | 64.0 | 14.0 | 26.2 | 89.5 |
| >hg38_21_+_8433172_8446622_RNA45SN1_45S_50ntFlanks@11568.11611.44\|GCAGCCGACTTAGAACTGGTGCGGACCAGGGGAATCCGACTGTT | >hg38_21_+_8433172_8446622_RNA45SN1_45S_50ntFlanks | 11568 | 11611 | 44 | GCAGCCGACTTAGAACTGGTGCGGACCAGGGGAATCCGACTGTT | BT20 | 71 | 88 | 82 | 67 | 639 | 41.3 | 50.7 | 108.9 | 90.2 | 705.5 |
| >hg38_21_+_8433172_8446622_RNA45SN1_45S_50ntFlanks@11568.11611.44\|GCAGCCGACTTAGAACTGGTGCGGACCAGGGGAATCCGACTGTT | >hg38_21_+_8433172_8446622_RNA45SN1_45S_50ntFlanks | 11568 | 11611 | 44 | GCAGCCGACTTAGAACTGGTGCGGACCAGGGGAATCCGACTGTT | MB231 | 23 | 28 | 49 | 5 | 82 | 21.3 | 46.1 | 72.6 | 7.3 | 108.3 |
| >hg38_21_+_8433172_8446622_RNA45SN1_45S_50ntFlanks@11569.11586.18\|CAGCCGACTTAGAACTGG | >hg38_21_+_8433172_8446622_RNA45SN1_45S_50ntFlanks | 11569 | 11586 | 18 | CAGCCGACTTAGAACTGG | MB231 | 16 | 9 | 10 | 7 | 6 | 12.0 | 9.4 | 7.5 | 5.4 | 5.5 |
| >hg38_21_+_8433172_8446622_RNA45SN1_45S_50ntFlanks@11569.11586.18\|CAGCCGACTTAGAACTGG | >hg38_21_+_8433172_8446622_RNA45SN1_45S_50ntFlanks | 11569 | 11586 | 18 | CAGCCGACTTAGAACTGG | MB468 | 138 | 25 | 87 | 111 | 56 | 46.6 | 9.1 | 29.6 | 21.4 | 34.8 |

| | | | | | | | | | | | | | | | | |
|---|---|---|---|---|---|---|---|---|---|---|---|---|---|---|---|---|
| >hg38_21_+_8433172_8446622_RNA45SN1_45S_50ntFlanks@11569.11587.19\|CAGCCGACTTAGAACTGGT | >hg38_21_+_8433172_8446622_RNA45SN1_45S_50ntFlanks | 11569 | 11587 | 19 | CAGCCGACTTAGAACTGGT | MB468 | 10 | 1 | 10 | 13 | 3 | 4.1 | 0.9 | 4.1 | 4.6 | 1.6 |
| >hg38_21_+_8433172_8446622_RNA45SN1_45S_50ntFlanks@11569.11588.20\|CAGCCGACTTAGAACTGGTG | >hg38_21_+_8433172_8446622_RNA45SN1_45S_50ntFlanks | 11569 | 11588 | 20 | CAGCCGACTTAGAACTGGTG | MB468 | 17 | 1 | 13 | 28 | 4 | 9.8 | 0.7 | 7.5 | 22.4 | 1.8 |
| >hg38_21_+_8433172_8446622_RNA45SN1_45S_50ntFlanks@11569.11590.22\|CAGCCGACTTAGAACTGGTGCG | >hg38_21_+_8433172_8446622_RNA45SN1_45S_50ntFlanks | 11569 | 11590 | 22 | CAGCCGACTTAGAACTGGTGCG | MB468 | 19 | 4 | 12 | 56 | 7 | 7.0 | 2.7 | 4.7 | 53.1 | 1.3 |
| >hg38_21_+_8433172_8446622_RNA45SN1_45S_50ntFlanks@11569.11591.23\|CAGCCGACTTAGAACTGGTGCGG | >hg38_21_+_8433172_8446622_RNA45SN1_45S_50ntFlanks | 11569 | 11591 | 23 | CAGCCGACTTAGAACTGGTGCGG | MB468 | 25 | 5 | 22 | 65 | 11 | 13.6 | 1.7 | 17.3 | 62.4 | 3.4 |
| >hg38_21_+_8433172_8446622_RNA45SN1_45S_50ntFlanks@11569.11592.24\|CAGCCGACTTAGAACTGGTGCGGA | >hg38_21_+_8433172_8446622_RNA45SN1_45S_50ntFlanks | 11569 | 11592 | 24 | CAGCCGACTTAGAACTGGTGCGGA | MB468 | 61 | 9 | 31 | 268 | 28 | 31.9 | 6.0 | 18.0 | 297.3 | 19.5 |
| >hg38_21_+_8433172_8446622_RNA45SN1_45S_50ntFlanks@11569.11593.25\|CAGCCGACTTAGAACTGGTGCGGAC | >hg38_21_+_8433172_8446622_RNA45SN1_45S_50ntFlanks | 11569 | 11593 | 25 | CAGCCGACTTAGAACTGGTGCGGAC | MB468 | 23 | 4 | 14 | 51 | 14 | 10.8 | 1.8 | 9.3 | 49.0 | 4.2 |
| >hg38_21_+_8433172_8446622_RNA45SN1_45S_50ntFlanks@11569.11594.26\|CAGCCGACTTAGAACTGGTGCGGACC | >hg38_21_+_8433172_8446622_RNA45SN1_45S_50ntFlanks | 11569 | 11594 | 26 | CAGCCGACTTAGAACTGGTGCGGACC | MB468 | 35 | 13 | 36 | 49 | 24 | 20.2 | 5.5 | 24.1 | 29.3 | 3.8 |
| >hg38_21_+_8433172_8446622_RNA45SN1_45S_50ntFlanks@11569.11596.28\|CAGCCGACTTAGAACTGGTGCGGACCAG | >hg38_21_+_8433172_8446622_RNA45SN1_45S_50ntFlanks | 11569 | 11596 | 28 | CAGCCGACTTAGAACTGGTGCGGACCAG | MB468 | 11 | 8 | 12 | 6 | 11 | 5.8 | 5.8 | 6.4 | 2.1 | 8.0 |
| >hg38_21_+_8433172_8446622_RNA45SN1_45S_50ntFlanks@11569.11597.29\|CAGCCGACTTAGAACTGGTGCGGACCAGG | >hg38_21_+_8433172_8446622_RNA45SN1_45S_50ntFlanks | 11569 | 11597 | 29 | CAGCCGACTTAGAACTGGTGCGGACCAGG | MB468 | 20 | 20 | 36 | 17 | 18 | 17.4 | 15.0 | 42.7 | 7.7 | 2.4 |
| >hg38_21_+_8433172_8446622_RNA45SN1_45S_50ntFlanks@11569.11599.31\|CAGCCGACTTAGAACTGGTGCGGACCAGGGG | >hg38_21_+_8433172_8446622_RNA45SN1_45S_50ntFlanks | 11569 | 11599 | 31 | CAGCCGACTTAGAACTGGTGCGGACCAGGGG | MB468 | 23 | 45 | 38 | 27 | 13 | 7.3 | 12.7 | 22.7 | 10.6 | 3.5 |
| >hg38_21_+_8433172_8446622_RNA45SN1_45S_50ntFlanks@11569.11600.32\|CAGCCGACTTAGAACTGGTGCGGACCAGGGGA | >hg38_21_+_8433172_8446622_RNA45SN1_45S_50ntFlanks | 11569 | 11600 | 32 | CAGCCGACTTAGAACTGGTGCGGACCAGGGGA | MB468 | 130 | 470 | 143 | 202 | 98 | 17.6 | 93.0 | 66.6 | 85.7 | 43.4 |
| >hg38_21_+_8433172_8446622_RNA45SN1_45S_50ntFlanks@11569.11601.33\|CAGCCGACTTAGAACTGGTGCGGACCAGGGGAA | >hg38_21_+_8433172_8446622_RNA45SN1_45S_50ntFlanks | 11569 | 11601 | 33 | CAGCCGACTTAGAACTGGTGCGGACCAGGGGAA | MB468 | 281 | 1097 | 276 | 469 | 369 | 79.9 | 296.7 | 139.7 | 229.3 | 269.3 |
| >hg38_21_+_8433172_8446622_RNA45SN1_45S_50ntFlanks@11569.11602.34\|CAGCCGACTTAGAACTGGTGCGGACCAGGGGAAT | >hg38_21_+_8433172_8446622_RNA45SN1_45S_50ntFlanks | 11569 | 11602 | 34 | CAGCCGACTTAGAACTGGTGCGGACCAGGGGAAT | MB468 | 168 | 558 | 147 | 231 | 288 | 45.1 | 48.5 | 54.2 | 116.2 | 191.5 |
| >hg38_21_+_8433172_8446622_RNA45SN1_45S_50ntFlanks@11569.11603.35\|CAGCCGACTTAGAACTGGTGCGGACCAGGGGAATC | >hg38_21_+_8433172_8446622_RNA45SN1_45S_50ntFlanks | 11569 | 11603 | 35 | CAGCCGACTTAGAACTGGTGCGGACCAGGGGAATC | MB468 | 226 | 653 | 172 | 315 | 818 | 28.0 | 96.4 | 33.3 | 161.7 | 474.4 |
| >hg38_21_+_8433172_8446622_RNA45SN1_45S_50ntFlanks@11569.11605.37\|CAGCCGACTTAGAACTGGTGCGGACCAGGGGAATCCG | >hg38_21_+_8433172_8446622_RNA45SN1_45S_50ntFlanks | 11569 | 11605 | 37 | CAGCCGACTTAGAACTGGTGCGGACCAGGGGAATCCG | MB468 | 35 | 99 | 32 | 64 | 173 | 8.3 | 31.3 | 8.7 | 49.0 | 146.7 |
| >hg38_21_+_8433172_8446622_RNA45SN1_45S_50ntFlanks@11569.11609.41\|CAGCCGACTTAGAACTGGTGCGGACCAGGGGAATCCGACTG | >hg38_21_+_8433172_8446622_RNA45SN1_45S_50ntFlanks | 11569 | 11609 | 41 | CAGCCGACTTAGAACTGGTGCGGACCAGGGGAATCCGACTG | MB468 | 15 | 16 | 26 | 16 | 89 | 14.4 | 14.7 | 14.7 | 11.0 | 56.2 |
| >hg38_21_+_8433172_8446622_RNA45SN1_45S_50ntFlanks@11569.11611.43\|CAGCCGACTTAGAACTGGTGCGGACCAGGGGAATCCGACTGTT | >hg38_21_+_8433172_8446622_RNA45SN1_45S_50ntFlanks | 11569 | 11611 | 43 | CAGCCGACTTAGAACTGGTGCGGACCAGGGGAATCCGACTGTT | MB468 | 16 | 38 | 31 | 14 | 180 | 17.2 | 50.3 | 21.5 | 15.0 | 160.2 |
| >hg38_21_+_8433172_8446622_RNA45SN1_45S_50ntFlanks@11569.11612.44\|CAGCCGACTTAGAACTGGTGCGGACCAGGGGAATCCGACTGTTT | >hg38_21_+_8433172_8446622_RNA45SN1_45S_50ntFlanks | 11569 | 11612 | 44 | CAGCCGACTTAGAACTGGTGCGGACCAGGGGAATCCGACTGTTT | MB468 | 25 | 132 | 61 | 46 | 324 | 26.3 | 147.2 | 53.2 | 56.3 | 221.5 |
| >hg38_21_+_8433172_8446622_RNA45SN1_45S_50ntFlanks@11570.11588.19\|AGCCGACTTAGAACTGGTG | >hg38_21_+_8433172_8446622_RNA45SN1_45S_50ntFlanks | 11570 | 11588 | 19 | AGCCGACTTAGAACTGGTG | MB468 | 19 | 2 | 13 | 23 | 4 | 7.9 | 0.9 | 4.9 | 14.9 | 1.1 |
| >hg38_21_+_8433172_8446622_RNA45SN1_45S_50ntFlanks@11570.11591.22\|AGCCGACTTAGAACTGGTGCGG | >hg38_21_+_8433172_8446622_RNA45SN1_45S_50ntFlanks | 11570 | 11591 | 22 | AGCCGACTTAGAACTGGTGCGG | MB468 | 62 | 21 | 30 | 65 | 25 | 20.4 | 5.3 | 12.8 | 32.5 | 4.5 |
| >hg38_21_+_8433172_8446622_RNA45SN1_45S_50ntFlanks@11570.11592.23\|AGCCGACTTAGAACTGGTGCGGA | >hg38_21_+_8433172_8446622_RNA45SN1_45S_50ntFlanks | 11570 | 11592 | 23 | AGCCGACTTAGAACTGGTGCGGA | MB468 | 41 | 9 | 18 | 71 | 17 | 14.9 | 3.0 | 8.3 | 50.9 | 4.3 |
| >hg38_21_+_8433172_8446622_RNA45SN1_45S_50ntFlanks@11570.11600.31\|AGCCGACTTAGAACTGGTGCGGACCAGGGGA | >hg38_21_+_8433172_8446622_RNA45SN1_45S_50ntFlanks | 11570 | 11600 | 31 | AGCCGACTTAGAACTGGTGCGGACCAGGGGA | MB468 | 24 | 80 | 23 | 35 | 16 | 2.8 | 14.0 | 9.7 | 10.8 | 7.1 |
| >hg38_21_+_8433172_8446622_RNA45SN1_45S_50ntFlanks@11570.11601.32\|AGCCGACTTAGAACTGGTGCGGACCAGGGGAA | >hg38_21_+_8433172_8446622_RNA45SN1_45S_50ntFlanks | 11570 | 11601 | 32 | AGCCGACTTAGAACTGGTGCGGACCAGGGGAA | MB468 | 65 | 220 | 56 | 103 | 84 | 13.4 | 60.0 | 27.7 | 42.1 | 57.9 |
| >hg38_21_+_8433172_8446622_RNA45SN1_45S_50ntFlanks@11570.11602.33\|AGCCGACTTAGAACTGGTGCGGACCAGGGGAAT | >hg38_21_+_8433172_8446622_RNA45SN1_45S_50ntFlanks | 11570 | 11602 | 33 | AGCCGACTTAGAACTGGTGCGGACCAGGGGAAT | MB468 | 36 | 104 | 28 | 51 | 60 | 9.2 | 8.1 | 9.4 | 21.2 | 37.1 |
| >hg38_21_+_8433172_8446622_RNA45SN1_45S_50ntFlanks@11570.11603.34\|AGCCGACTTAGAACTGGTGCGGACCAGGGGAATC | >hg38_21_+_8433172_8446622_RNA45SN1_45S_50ntFlanks | 11570 | 11603 | 34 | AGCCGACTTAGAACTGGTGCGGACCAGGGGAATC | MB468 | 50 | 139 | 32 | 79 | 187 | 9.8 | 18.6 | 6.6 | 31.4 | 99.8 |
| >hg38_21_+_8433172_8446622_RNA45SN1_45S_50ntFlanks@11571.11588.18\|GCCGACTTAGAACTGGTG | >hg38_21_+_8433172_8446622_RNA45SN1_45S_50ntFlanks | 11571 | 11588 | 18 | GCCGACTTAGAACTGGTG | MB468 | 11 | 1 | 13 | 11 | 3 | 6.0 | 0.8 | 8.0 | 2.9 | 1.1 |
| >hg38_21_+_8433172_8446622_RNA45SN1_45S_50ntFlanks@11571.11590.20\|GCCGACTTAGAACTGGTGCG | >hg38_21_+_8433172_8446622_RNA45SN1_45S_50ntFlanks | 11571 | 11590 | 20 | GCCGACTTAGAACTGGTGCG | MB231 | 16 | 9 | 9 | 5 | 9 | 13.0 | 7.4 | 5.2 | 4.7 | 9.1 |
| >hg38_21_+_8433172_8446622_RNA45SN1_45S_50ntFlanks@11571.11591.21\|GCCGACTTAGAACTGGTGCGG | >hg38_21_+_8433172_8446622_RNA45SN1_45S_50ntFlanks | 11571 | 11591 | 21 | GCCGACTTAGAACTGGTGCGG | MB468 | 29 | 9 | 17 | 31 | 10 | 13.6 | 2.2 | 9.5 | 12.4 | 2.0 |
| >hg38_21_+_8433172_8446622_RNA45SN1_45S_50ntFlanks@11571.11592.22\|GCCGACTTAGAACTGGTGCGGA | >hg38_21_+_8433172_8446622_RNA45SN1_45S_50ntFlanks | 11571 | 11592 | 22 | GCCGACTTAGAACTGGTGCGGA | MB468 | 35 | 7 | 18 | 48 | 12 | 16.3 | 1.7 | 12.1 | 23.8 | 2.9 |
| >hg38_21_+_8433172_8446622_RNA45SN1_45S_50ntFlanks@11571.11594.24\|GCCGACTTAGAACTGGTGCGGACC | >hg38_21_+_8433172_8446622_RNA45SN1_45S_50ntFlanks | 11571 | 11594 | 24 | GCCGACTTAGAACTGGTGCGGACC | MB231 | 25 | 18 | 15 | 8 | 15 | 21.6 | 9.4 | 9.2 | 5.5 | 11.6 |
| >hg38_21_+_8433172_8446622_RNA45SN1_45S_50ntFlanks@11571.11600.30\|GCCGACTTAGAACTGGTGCGGACCAGGGGA | >hg38_21_+_8433172_8446622_RNA45SN1_45S_50ntFlanks | 11571 | 11600 | 30 | GCCGACTTAGAACTGGTGCGGACCAGGGGA | MB468 | 14 | 45 | 14 | 22 | 9 | 4.1 | 14.1 | 6.9 | 6.6 | 4.0 |
| >hg38_21_+_8433172_8446622_RNA45SN1_45S_50ntFlanks@11571.11601.31\|GCCGACTTAGAACTGGTGCGGACCAGGGGAA | >hg38_21_+_8433172_8446622_RNA45SN1_45S_50ntFlanks | 11571 | 11601 | 31 | GCCGACTTAGAACTGGTGCGGACCAGGGGAA | MB468 | 26 | 88 | 24 | 46 | 29 | 11.7 | 36.7 | 14.8 | 22.2 | 24.1 |
| >hg38_21_+_8433172_8446622_RNA45SN1_45S_50ntFlanks@11571.11602.32\|GCCGACTTAGAACTGGTGCGGACCAGGGGAAT | >hg38_21_+_8433172_8446622_RNA45SN1_45S_50ntFlanks | 11571 | 11602 | 32 | GCCGACTTAGAACTGGTGCGGACCAGGGGAAT | MB468 | 14 | 40 | 11 | 23 | 22 | 5.6 | 8.9 | 6.1 | 12.2 | 17.4 |
| >hg38_21_+_8433172_8446622_RNA45SN1_45S_50ntFlanks@11571.11603.33\|GCCGACTTAGAACTGGTGCGGACCAGGGGAATC | >hg38_21_+_8433172_8446622_RNA45SN1_45S_50ntFlanks | 11571 | 11603 | 33 | GCCGACTTAGAACTGGTGCGGACCAGGGGAATC | MB468 | 19 | 48 | 12 | 33 | 62 | 5.3 | 7.5 | 4.9 | 16.9 | 45.8 |
| >hg38_21_+_8433172_8446622_RNA45SN1_45S_50ntFlanks@11572.11589.18\|CCGACTTAGAACTGGTGC | >hg38_21_+_8433172_8446622_RNA45SN1_45S_50ntFlanks | 11572 | 11589 | 18 | CCGACTTAGAACTGGTGC | MB468 | 34 | 18 | 25 | 35 | 7 | 8.0 | 7.3 | 5.4 | 12.8 | 1.7 |
| >hg38_21_+_8433172_8446622_RNA45SN1_45S_50ntFlanks@11572.11590.19\|CCGACTTAGAACTGGTGCG | >hg38_21_+_8433172_8446622_RNA45SN1_45S_50ntFlanks | 11572 | 11590 | 19 | CCGACTTAGAACTGGTGCG | MB468 | 263 | 206 | 194 | 249 | 47 | 41.5 | 96.2 | 24.9 | 110.2 | 8.3 |
| >hg38_21_+_8433172_8446622_RNA45SN1_45S_50ntFlanks@11572.11591.20\|CCGACTTAGAACTGGTGCGG | >hg38_21_+_8433172_8446622_RNA45SN1_45S_50ntFlanks | 11572 | 11591 | 20 | CCGACTTAGAACTGGTGCGG | MB468 | 907 | 126 | 433 | 702 | 182 | 539.8 | 19.0 | 355.4 | 202.4 | 62.7 |
| >hg38_21_+_8433172_8446622_RNA45SN1_45S_50ntFlanks@11572.11592.21\|CCGACTTAGAACTGGTGCGGA | >hg38_21_+_8433172_8446622_RNA45SN1_45S_50ntFlanks | 11572 | 11592 | 21 | CCGACTTAGAACTGGTGCGGA | MB468 | 108 | 10 | 66 | 308 | 26 | 61.7 | 6.0 | 32.8 | 263.8 | 9.4 |
| >hg38_21_+_8433172_8446622_RNA45SN1_45S_50ntFlanks@11572.11593.22\|CCGACTTAGAACTGGTGCGGAC | >hg38_21_+_8433172_8446622_RNA45SN1_45S_50ntFlanks | 11572 | 11593 | 22 | CCGACTTAGAACTGGTGCGGAC | MB468 | 61 | 7 | 62 | 75 | 34 | 31.6 | 1.8 | 55.0 | 27.7 | 16.0 |
| >hg38_21_+_8433172_8446622_RNA45SN1_45S_50ntFlanks@11572.11594.23\|CCGACTTAGAACTGGTGCGGACC | >hg38_21_+_8433172_8446622_RNA45SN1_45S_50ntFlanks | 11572 | 11594 | 23 | CCGACTTAGAACTGGTGCGGACC | MB468 | 28 | 7 | 49 | 29 | 13 | 20.4 | 1.8 | 46.3 | 8.6 | 2.4 |
| >hg38_21_+_8433172_8446622_RNA45SN1_45S_50ntFlanks@11572.11597.26\|CCGACTTAGAACTGGTGCGGACCAGG | >hg38_21_+_8433172_8446622_RNA45SN1_45S_50ntFlanks | 11572 | 11597 | 26 | CCGACTTAGAACTGGTGCGGACCAGG | MB468 | 35 | 10 | 60 | 24 | 15 | 24.8 | 5.1 | 69.8 | 8.4 | 3.5 |
| >hg38_21_+_8433172_8446622_RNA45SN1_45S_50ntFlanks@11572.11599.28\|CCGACTTAGAACTGGTGCGGACCAGGGG | >hg38_21_+_8433172_8446622_RNA45SN1_45S_50ntFlanks | 11572 | 11599 | 28 | CCGACTTAGAACTGGTGCGGACCAGGGG | MB468 | 38 | 27 | 62 | 44 | 15 | 7.3 | 5.3 | 24.2 | 21.8 | 4.2 |
| >hg38_21_+_8433172_8446622_RNA45SN1_45S_50ntFlanks@11572.11600.29\|CCGACTTAGAACTGGTGCGGACCAGGGGA | >hg38_21_+_8433172_8446622_RNA45SN1_45S_50ntFlanks | 11572 | 11600 | 29 | CCGACTTAGAACTGGTGCGGACCAGGGGA | MB468 | 279 | 467 | 373 | 476 | 126 | 84.1 | 146.3 | 161.4 | 204.7 | 46.7 |
| >hg38_21_+_8433172_8446622_RNA45SN1_45S_50ntFlanks@11572.11601.30\|CCGACTTAGAACTGGTGCGGACCAGGGGAA | >hg38_21_+_8433172_8446622_RNA45SN1_45S_50ntFlanks | 11572 | 11601 | 30 | CCGACTTAGAACTGGTGCGGACCAGGGGAA | MB468 | 796 | 1391 | 996 | 1128 | 444 | 383.6 | 507.7 | 530.1 | 470.3 | 182.0 |
| >hg38_21_+_8433172_8446622_RNA45SN1_45S_50ntFlanks@11572.11602.31\|CCGACTTAGAACTGGTGCGGACCAGGGGAAT | >hg38_21_+_8433172_8446622_RNA45SN1_45S_50ntFlanks | 11572 | 11602 | 31 | CCGACTTAGAACTGGTGCGGACCAGGGGAAT | MB468 | 424 | 635 | 471 | 506 | 305 | 181.6 | 138.9 | 193.1 | 199.4 | 139.8 |
| >hg38_21_+_8433172_8446622_RNA45SN1_45S_50ntFlanks@11572.11603.32\|CCGACTTAGAACTGGTGCGGACCAGGGGAATC | >hg38_21_+_8433172_8446622_RNA45SN1_45S_50ntFlanks | 11572 | 11603 | 32 | CCGACTTAGAACTGGTGCGGACCAGGGGAATC | MB468 | 528 | 771 | 465 | 660 | 910 | 147.6 | 67.4 | 131.6 | 229.1 | 439.0 |
| >hg38_21_+_8433172_8446622_RNA45SN1_45S_50ntFlanks@11572.11604.33\|CCGACTTAGAACTGGTGCGGACCAGGGGAATCC | >hg38_21_+_8433172_8446622_RNA45SN1_45S_50ntFlanks | 11572 | 11604 | 33 | CCGACTTAGAACTGGTGCGGACCAGGGGAATCC | MB468 | 12 | 25 | 13 | 14 | 24 | 3.5 | 5.2 | 3.2 | 7.5 | 10.3 |
| >hg38_21_+_8433172_8446622_RNA45SN1_45S_50ntFlanks@11572.11605.34\|CCGACTTAGAACTGGTGCGGACCAGGGGAATCCG | >hg38_21_+_8433172_8446622_RNA45SN1_45S_50ntFlanks | 11572 | 11605 | 34 | CCGACTTAGAACTGGTGCGGACCAGGGGAATCCG | MB468 | 175 | 377 | 220 | 174 | 452 | 32.2 | 52.0 | 55.9 | 93.3 | 319.3 |
| >hg38_21_+_8433172_8446622_RNA45SN1_45S_50ntFlanks@11572.11606.35\|CCGACTTAGAACTGGTGCGGACCAGGGGAATCCGA | >hg38_21_+_8433172_8446622_RNA45SN1_45S_50ntFlanks | 11572 | 11606 | 35 | CCGACTTAGAACTGGTGCGGACCAGGGGAATCCGA | MB468 | 16 | 31 | 22 | 20 | 39 | 4.6 | 11.4 | 4.2 | 11.9 | 20.7 |
| >hg38_21_+_8433172_8446622_RNA45SN1_45S_50ntFlanks@11572.11607.36\|CCGACTTAGAACTGGTGCGGACCAGGGGAATCCGAC | >hg38_21_+_8433172_8446622_RNA45SN1_45S_50ntFlanks | 11572 | 11607 | 36 | CCGACTTAGAACTGGTGCGGACCAGGGGAATCCGAC | MB468 | 13 | 18 | 20 | 12 | 28 | 7.7 | 8.7 | 7.0 | 9.5 | 15.4 |

| | | | | | | | | | | | | | | | | |
|---|---|---|---|---|---|---|---|---|---|---|---|---|---|---|---|---|
| >hg38_21_+_8433172_8446622_RNA45SN1_45S_50ntFlanks@11572.11609.38\|CCGACTTAGAACTGGTGCGGACCAGGGGAATCCGACTG | >hg38_21_+_8433172_8446622_RNA45SN1_45S_50ntFlanks | 11572 | 11609 | 38 | CCGACTTAGAACTGGTGCGGACCAGGGGAATCCGACTG | MB468 | 22 | 34 | 25 | 32 | 81 | 12.0 | 14.0 | 9.6 | 19.2 | 47.6 |
| >hg38_21_+_8433172_8446622_RNA45SN1_45S_50ntFlanks@11572.11611.40\|CCGACTTAGAACTGGTGCGGACCAGGGGAATCCGACTGTT | >hg38_21_+_8433172_8446622_RNA45SN1_45S_50ntFlanks | 11572 | 11611 | 40 | CCGACTTAGAACTGGTGCGGACCAGGGGAATCCGACTGTT | MB468 | 42 | 86 | 47 | 55 | 292 | 35.7 | 54.2 | 24.6 | 41.9 | 174.7 |
| >hg38_21_+_8433172_8446622_RNA45SN1_45S_50ntFlanks@11572.11612.41\|CCGACTTAGAACTGGTGCGGACCAGGGGAATCCGACTGTTT | >hg38_21_+_8433172_8446622_RNA45SN1_45S_50ntFlanks | 11572 | 11612 | 41 | CCGACTTAGAACTGGTGCGGACCAGGGGAATCCGACTGTTT | MB468 | 57 | 230 | 92 | 94 | 472 | 58.5 | 192.0 | 54.7 | 94.8 | 302.0 |
| >hg38_21_+_8433172_8446622_RNA45SN1_45S_50ntFlanks@11573.11590.18\|CGACTTAGAACTGGTGCG | >hg38_21_+_8433172_8446622_RNA45SN1_45S_50ntFlanks | 11573 | 11590 | 18 | CGACTTAGAACTGGTGCG | MB231 | 11 | 7 | 6 | 6 | 3 | 8.5 | 5.2 | 4.2 | 4.8 | 2.2 |
| >hg38_21_+_8433172_8446622_RNA45SN1_45S_50ntFlanks@11573.11590.18\|CGACTTAGAACTGGTGCG | >hg38_21_+_8433172_8446622_RNA45SN1_45S_50ntFlanks | 11573 | 11590 | 18 | CGACTTAGAACTGGTGCG | MB468 | 33 | 13 | 17 | 33 | 5 | 8.0 | 5.1 | 4.7 | 16.0 | 1.3 |
| >hg38_21_+_8433172_8446622_RNA45SN1_45S_50ntFlanks@11573.11591.19\|CGACTTAGAACTGGTGCGG | >hg38_21_+_8433172_8446622_RNA45SN1_45S_50ntFlanks | 11573 | 11591 | 19 | CGACTTAGAACTGGTGCGG | MB468 | 45 | 20 | 27 | 50 | 8 | 14.6 | 8.4 | 10.0 | 39.8 | 3.6 |
| >hg38_21_+_8433172_8446622_RNA45SN1_45S_50ntFlanks@11573.11592.20\|CGACTTAGAACTGGTGCGGA | >hg38_21_+_8433172_8446622_RNA45SN1_45S_50ntFlanks | 11573 | 11592 | 20 | CGACTTAGAACTGGTGCGGA | MB468 | 60 | 28 | 34 | 86 | 16 | 13.8 | 8.5 | 10.8 | 73.7 | 7.6 |
| >hg38_21_+_8433172_8446622_RNA45SN1_45S_50ntFlanks@11573.11593.21\|CGACTTAGAACTGGTGCGGAC | >hg38_21_+_8433172_8446622_RNA45SN1_45S_50ntFlanks | 11573 | 11593 | 21 | CGACTTAGAACTGGTGCGGAC | MB468 | 32 | 28 | 24 | 28 | 11 | 4.9 | 11.1 | 7.8 | 16.3 | 2.6 |
| >hg38_21_+_8433172_8446622_RNA45SN1_45S_50ntFlanks@11573.11594.22\|CGACTTAGAACTGGTGCGGACC | >hg38_21_+_8433172_8446622_RNA45SN1_45S_50ntFlanks | 11573 | 11594 | 22 | CGACTTAGAACTGGTGCGGACC | MB468 | 11 | 21 | 11 | 10 | 4 | 2.3 | 7.9 | 4.4 | 4.0 | 1.4 |
| >hg38_21_+_8433172_8446622_RNA45SN1_45S_50ntFlanks@11573.11600.28\|CGACTTAGAACTGGTGCGGACCAGGGGA | >hg38_21_+_8433172_8446622_RNA45SN1_45S_50ntFlanks | 11573 | 11600 | 28 | CGACTTAGAACTGGTGCGGACCAGGGGA | MB468 | 31 | 52 | 33 | 56 | 13 | 5.8 | 15.1 | 10.9 | 32.1 | 5.1 |
| >hg38_21_+_8433172_8446622_RNA45SN1_45S_50ntFlanks@11573.11601.29\|CGACTTAGAACTGGTGCGGACCAGGGGAA | >hg38_21_+_8433172_8446622_RNA45SN1_45S_50ntFlanks | 11573 | 11601 | 29 | CGACTTAGAACTGGTGCGGACCAGGGGAA | MB468 | 77 | 151 | 78 | 114 | 31 | 25.1 | 59.2 | 35.8 | 71.0 | 11.1 |
| >hg38_21_+_8433172_8446622_RNA45SN1_45S_50ntFlanks@11573.11602.30\|CGACTTAGAACTGGTGCGGACCAGGGGAAT | >hg38_21_+_8433172_8446622_RNA45SN1_45S_50ntFlanks | 11573 | 11602 | 30 | CGACTTAGAACTGGTGCGGACCAGGGGAAT | MB468 | 43 | 80 | 44 | 53 | 22 | 13.5 | 23.7 | 14.4 | 33.3 | 9.7 |
| >hg38_21_+_8433172_8446622_RNA45SN1_45S_50ntFlanks@11573.11603.31\|CGACTTAGAACTGGTGCGGACCAGGGGAATC | >hg38_21_+_8433172_8446622_RNA45SN1_45S_50ntFlanks | 11573 | 11603 | 31 | CGACTTAGAACTGGTGCGGACCAGGGGAATC | MB468 | 54 | 80 | 53 | 61 | 42 | 15.1 | 11.6 | 11.2 | 44.6 | 18.3 |
| >hg38_21_+_8433172_8446622_RNA45SN1_45S_50ntFlanks@11573.11605.33\|CGACTTAGAACTGGTGCGGACCAGGGGAATCCG | >hg38_21_+_8433172_8446622_RNA45SN1_45S_50ntFlanks | 11573 | 11605 | 33 | CGACTTAGAACTGGTGCGGACCAGGGGAATCCG | MB468 | 25 | 46 | 37 | 27 | 27 | 5.0 | 14.3 | 7.3 | 17.6 | 13.4 |
| >hg38_21_+_8433172_8446622_RNA45SN1_45S_50ntFlanks@11573.11613.41\|CGACTTAGAACTGGTGCGGACCAGGGGAATCCGACTGTTTA | >hg38_21_+_8433172_8446622_RNA45SN1_45S_50ntFlanks | 11573 | 11613 | 41 | CGACTTAGAACTGGTGCGGACCAGGGGAATCCGACTGTTTA | MB231 | 12 | 17 | 40 | 4 | 16 | 8.9 | 23.3 | 46.9 | 5.0 | 16.1 |
| >hg38_21_+_8433172_8446622_RNA45SN1_45S_50ntFlanks@11574.11603.30\|GACTTAGAACTGGTGCGGACCAGGGGAATC | >hg38_21_+_8433172_8446622_RNA45SN1_45S_50ntFlanks | 11574 | 11603 | 30 | GACTTAGAACTGGTGCGGACCAGGGGAATC | MB468 | 14 | 18 | 17 | 9 | 6 | 5.0 | 5.8 | 7.5 | 1.5 | 3.4 |
| >hg38_21_+_8433172_8446622_RNA45SN1_45S_50ntFlanks@11575.11592.18\|ACTTAGAACTGGTGCGGA | >hg38_21_+_8433172_8446622_RNA45SN1_45S_50ntFlanks | 11575 | 11592 | 18 | ACTTAGAACTGGTGCGGA | MB468 | 413 | 193 | 322 | 239 | 81 | 110.8 | 65.6 | 110.6 | 142.0 | 23.6 |
| >hg38_21_+_8433172_8446622_RNA45SN1_45S_50ntFlanks@11575.11593.19\|ACTTAGAACTGGTGCGGAC | >hg38_21_+_8433172_8446622_RNA45SN1_45S_50ntFlanks | 11575 | 11593 | 19 | ACTTAGAACTGGTGCGGAC | MB468 | 164 | 117 | 196 | 71 | 40 | 25.1 | 38.9 | 44.7 | 33.8 | 5.2 |
| >hg38_21_+_8433172_8446622_RNA45SN1_45S_50ntFlanks@11575.11594.20\|ACTTAGAACTGGTGCGGACC | >hg38_21_+_8433172_8446622_RNA45SN1_45S_50ntFlanks | 11575 | 11594 | 20 | ACTTAGAACTGGTGCGGACC | MB468 | 119 | 200 | 136 | 53 | 26 | 15.4 | 75.9 | 50.2 | 23.1 | 7.1 |
| >hg38_21_+_8433172_8446622_RNA45SN1_45S_50ntFlanks@11575.11599.25\|ACTTAGAACTGGTGCGGACCAGGGG | >hg38_21_+_8433172_8446622_RNA45SN1_45S_50ntFlanks | 11575 | 11599 | 25 | ACTTAGAACTGGTGCGGACCAGGGG | MB468 | 21 | 6 | 30 | 11 | 4 | 6.9 | 2.0 | 21.2 | 8.8 | 1.8 |
| >hg38_21_+_8433172_8446622_RNA45SN1_45S_50ntFlanks@11575.11600.26\|ACTTAGAACTGGTGCGGACCAGGGGA | >hg38_21_+_8433172_8446622_RNA45SN1_45S_50ntFlanks | 11575 | 11600 | 26 | ACTTAGAACTGGTGCGGACCAGGGGA | MB468 | 75 | 50 | 72 | 45 | 16 | 25.8 | 5.8 | 33.3 | 29.3 | 7.3 |
| >hg38_21_+_8433172_8446622_RNA45SN1_45S_50ntFlanks@11575.11601.27\|ACTTAGAACTGGTGCGGACCAGGGGAA | >hg38_21_+_8433172_8446622_RNA45SN1_45S_50ntFlanks | 11575 | 11601 | 27 | ACTTAGAACTGGTGCGGACCAGGGGAA | MB468 | 136 | 150 | 119 | 72 | 35 | 62.1 | 19.7 | 65.9 | 41.3 | 17.7 |
| >hg38_21_+_8433172_8446622_RNA45SN1_45S_50ntFlanks@11575.11602.28\|ACTTAGAACTGGTGCGGACCAGGGGAAT | >hg38_21_+_8433172_8446622_RNA45SN1_45S_50ntFlanks | 11575 | 11602 | 28 | ACTTAGAACTGGTGCGGACCAGGGGAAT | MB468 | 111 | 84 | 138 | 45 | 34 | 48.7 | 11.5 | 39.3 | 27.5 | 18.1 |
| >hg38_21_+_8433172_8446622_RNA45SN1_45S_50ntFlanks@11575.11603.29\|ACTTAGAACTGGTGCGGACCAGGGGAATC | >hg38_21_+_8433172_8446622_RNA45SN1_45S_50ntFlanks | 11575 | 11603 | 29 | ACTTAGAACTGGTGCGGACCAGGGGAATC | MB468 | 507 | 443 | 693 | 313 | 221 | 141.9 | 166.2 | 138.2 | 230.5 | 169.9 |
| >hg38_21_+_8433172_8446622_RNA45SN1_45S_50ntFlanks@11575.11604.30\|ACTTAGAACTGGTGCGGACCAGGGGAATCC | >hg38_21_+_8433172_8446622_RNA45SN1_45S_50ntFlanks | 11575 | 11604 | 30 | ACTTAGAACTGGTGCGGACCAGGGGAATCC | MB468 | 13 | 18 | 13 | 5 | 6 | 5.9 | 9.8 | 4.4 | 3.8 | 5.2 |
| >hg38_21_+_8433172_8446622_RNA45SN1_45S_50ntFlanks@11575.11605.31\|ACTTAGAACTGGTGCGGACCAGGGGAATCCG | >hg38_21_+_8433172_8446622_RNA45SN1_45S_50ntFlanks | 11575 | 11605 | 31 | ACTTAGAACTGGTGCGGACCAGGGGAATCCG | MB468 | 140 | 137 | 207 | 75 | 90 | 25.7 | 28.6 | 11.8 | 52.0 | 61.4 |
| >hg38_21_+_8433172_8446622_RNA45SN1_45S_50ntFlanks@11575.11606.32\|ACTTAGAACTGGTGCGGACCAGGGGAATCCGA | >hg38_21_+_8433172_8446622_RNA45SN1_45S_50ntFlanks | 11575 | 11606 | 32 | ACTTAGAACTGGTGCGGACCAGGGGAATCCGA | MB468 | 20 | 14 | 28 | 11 | 12 | 5.1 | 4.5 | 6.6 | 8.6 | 7.7 |
| >hg38_21_+_8433172_8446622_RNA45SN1_45S_50ntFlanks@11575.11607.33\|ACTTAGAACTGGTGCGGACCAGGGGAATCCGAC | >hg38_21_+_8433172_8446622_RNA45SN1_45S_50ntFlanks | 11575 | 11607 | 33 | ACTTAGAACTGGTGCGGACCAGGGGAATCCGAC | MB468 | 35 | 28 | 50 | 19 | 27 | 11.7 | 8.6 | 11.2 | 18.1 | 19.9 |
| >hg38_21_+_8433172_8446622_RNA45SN1_45S_50ntFlanks@11575.11609.35\|ACTTAGAACTGGTGCGGACCAGGGGAATCCGACTG | >hg38_21_+_8433172_8446622_RNA45SN1_45S_50ntFlanks | 11575 | 11609 | 35 | ACTTAGAACTGGTGCGGACCAGGGGAATCCGACTG | MB468 | 48 | 26 | 38 | 46 | 46 | 19.7 | 8.5 | 15.5 | 28.6 | 22.5 |
| >hg38_21_+_8433172_8446622_RNA45SN1_45S_50ntFlanks@11575.11611.37\|ACTTAGAACTGGTGCGGACCAGGGGAATCCGACTGTT | >hg38_21_+_8433172_8446622_RNA45SN1_45S_50ntFlanks | 11575 | 11611 | 37 | ACTTAGAACTGGTGCGGACCAGGGGAATCCGACTGTT | MB468 | 31 | 31 | 25 | 52 | 128 | 17.5 | 9.0 | 13.3 | 30.9 | 92.0 |
| >hg38_21_+_8433172_8446622_RNA45SN1_45S_50ntFlanks@11575.11612.38\|ACTTAGAACTGGTGCGGACCAGGGGAATCCGACTGTTT | >hg38_21_+_8433172_8446622_RNA45SN1_45S_50ntFlanks | 11575 | 11612 | 38 | ACTTAGAACTGGTGCGGACCAGGGGAATCCGACTGTTT | MB468 | 18 | 14 | 18 | 10 | 60 | 18.2 | 10.4 | 10.7 | 6.6 | 54.7 |
| >hg38_21_+_8433172_8446622_RNA45SN1_45S_50ntFlanks@11575.11613.39\|ACTTAGAACTGGTGCGGACCAGGGGAATCCGACTGTTTA | >hg38_21_+_8433172_8446622_RNA45SN1_45S_50ntFlanks | 11575 | 11613 | 39 | ACTTAGAACTGGTGCGGACCAGGGGAATCCGACTGTTTA | MB468 | 18 | 31 | 23 | 15 | 93 | 15.9 | 23.9 | 13.3 | 11.4 | 84.7 |
| >hg38_21_+_8433172_8446622_RNA45SN1_45S_50ntFlanks@11576.11593.18\|CTTAGAACTGGTGCGGAC | >hg38_21_+_8433172_8446622_RNA45SN1_45S_50ntFlanks | 11576 | 11593 | 18 | CTTAGAACTGGTGCGGAC | MB468 | 120 | 88 | 149 | 39 | 27 | 35.3 | 47.9 | 53.1 | 11.5 | 5.8 |
| >hg38_21_+_8433172_8446622_RNA45SN1_45S_50ntFlanks@11576.11594.19\|CTTAGAACTGGTGCGGACC | >hg38_21_+_8433172_8446622_RNA45SN1_45S_50ntFlanks | 11576 | 11594 | 19 | CTTAGAACTGGTGCGGACC | MB468 | 92 | 129 | 113 | 34 | 20 | 21.9 | 50.8 | 41.8 | 11.8 | 3.1 |
| >hg38_21_+_8433172_8446622_RNA45SN1_45S_50ntFlanks@11576.11599.24\|CTTAGAACTGGTGCGGACCAGGGG | >hg38_21_+_8433172_8446622_RNA45SN1_45S_50ntFlanks | 11576 | 11599 | 24 | CTTAGAACTGGTGCGGACCAGGGG | MB468 | 11 | 2 | 14 | 6 | 2 | 3.7 | 0.6 | 7.8 | 4.5 | 0.8 |
| >hg38_21_+_8433172_8446622_RNA45SN1_45S_50ntFlanks@11576.11600.25\|CTTAGAACTGGTGCGGACCAGGGGA | >hg38_21_+_8433172_8446622_RNA45SN1_45S_50ntFlanks | 11576 | 11600 | 25 | CTTAGAACTGGTGCGGACCAGGGGA | MB468 | 69 | 34 | 66 | 30 | 11 | 32.3 | 2.2 | 46.6 | 18.3 | 3.2 |
| >hg38_21_+_8433172_8446622_RNA45SN1_45S_50ntFlanks@11576.11601.26\|CTTAGAACTGGTGCGGACCAGGGGAA | >hg38_21_+_8433172_8446622_RNA45SN1_45S_50ntFlanks | 11576 | 11601 | 26 | CTTAGAACTGGTGCGGACCAGGGGAA | MB468 | 141 | 127 | 127 | 51 | 25 | 79.4 | 13.7 | 108.0 | 28.2 | 10.0 |
| >hg38_21_+_8433172_8446622_RNA45SN1_45S_50ntFlanks@11576.11602.27\|CTTAGAACTGGTGCGGACCAGGGGAAT | >hg38_21_+_8433172_8446622_RNA45SN1_45S_50ntFlanks | 11576 | 11602 | 27 | CTTAGAACTGGTGCGGACCAGGGGAAT | MB468 | 102 | 64 | 121 | 31 | 23 | 63.9 | 11.1 | 82.0 | 14.5 | 9.2 |
| >hg38_21_+_8433172_8446622_RNA45SN1_45S_50ntFlanks@11576.11603.28\|CTTAGAACTGGTGCGGACCAGGGGAATC | >hg38_21_+_8433172_8446622_RNA45SN1_45S_50ntFlanks | 11576 | 11603 | 28 | CTTAGAACTGGTGCGGACCAGGGGAATC | MB468 | 353 | 301 | 480 | 187 | 107 | 153.0 | 68.6 | 201.6 | 103.3 | 47.2 |
| >hg38_21_+_8433172_8446622_RNA45SN1_45S_50ntFlanks@11576.11605.30\|CTTAGAACTGGTGCGGACCAGGGGAATCCG | >hg38_21_+_8433172_8446622_RNA45SN1_45S_50ntFlanks | 11576 | 11605 | 30 | CTTAGAACTGGTGCGGACCAGGGGAATCCG | MB468 | 105 | 107 | 138 | 48 | 50 | 39.2 | 18.2 | 40.8 | 29.3 | 21.9 |
| >hg38_21_+_8433172_8446622_RNA45SN1_45S_50ntFlanks@11576.11606.31\|CTTAGAACTGGTGCGGACCAGGGGAATCCGA | >hg38_21_+_8433172_8446622_RNA45SN1_45S_50ntFlanks | 11576 | 11606 | 31 | CTTAGAACTGGTGCGGACCAGGGGAATCCGA | MB468 | 12 | 10 | 15 | 6 | 5 | 4.7 | 4.0 | 7.5 | 5.2 | 2.4 |
| >hg38_21_+_8433172_8446622_RNA45SN1_45S_50ntFlanks@11576.11607.32\|CTTAGAACTGGTGCGGACCAGGGGAATCCGAC | >hg38_21_+_8433172_8446622_RNA45SN1_45S_50ntFlanks | 11576 | 11607 | 32 | CTTAGAACTGGTGCGGACCAGGGGAATCCGAC | MB468 | 32 | 25 | 38 | 14 | 16 | 16.6 | 7.7 | 16.5 | 12.7 | 7.1 |
| >hg38_21_+_8433172_8446622_RNA45SN1_45S_50ntFlanks@11576.11609.34\|CTTAGAACTGGTGCGGACCAGGGGAATCCGACTG | >hg38_21_+_8433172_8446622_RNA45SN1_45S_50ntFlanks | 11576 | 11609 | 34 | CTTAGAACTGGTGCGGACCAGGGGAATCCGACTG | MB468 | 70 | 36 | 52 | 55 | 43 | 32.4 | 7.3 | 13.5 | 40.0 | 24.6 |
| >hg38_21_+_8433172_8446622_RNA45SN1_45S_50ntFlanks@11576.11611.36\|CTTAGAACTGGTGCGGACCAGGGGAATCCGACTGTT | >hg38_21_+_8433172_8446622_RNA45SN1_45S_50ntFlanks | 11576 | 11611 | 36 | CTTAGAACTGGTGCGGACCAGGGGAATCCGACTGTT | MB468 | 39 | 22 | 30 | 36 | 71 | 31.6 | 5.7 | 13.9 | 22.5 | 31.0 |
| >hg38_21_+_8433172_8446622_RNA45SN1_45S_50ntFlanks@11576.11612.37\|CTTAGAACTGGTGCGGACCAGGGGAATCCGACTGTTT | >hg38_21_+_8433172_8446622_RNA45SN1_45S_50ntFlanks | 11576 | 11612 | 37 | CTTAGAACTGGTGCGGACCAGGGGAATCCGACTGTTT | MB468 | 21 | 9 | 22 | 6 | 27 | 26.8 | 5.9 | 14.2 | 4.1 | 19.3 |
| >hg38_21_+_8433172_8446622_RNA45SN1_45S_50ntFlanks@11576.11613.38\|CTTAGAACTGGTGCGGACCAGGGGAATCCGACTGTTTA | >hg38_21_+_8433172_8446622_RNA45SN1_45S_50ntFlanks | 11576 | 11613 | 38 | CTTAGAACTGGTGCGGACCAGGGGAATCCGACTGTTTA | MB468 | 17 | 18 | 25 | 10 | 42 | 17.9 | 9.1 | 12.6 | 7.3 | 26.2 |
| >hg38_21_+_8433172_8446622_RNA45SN1_45S_50ntFlanks@11577.11594.18\|TTAGAACTGGTGCGGACC | >hg38_21_+_8433172_8446622_RNA45SN1_45S_50ntFlanks | 11577 | 11594 | 18 | TTAGAACTGGTGCGGACC | MB468 | 23 | 24 | 32 | 8 | 5 | 2.3 | 9.4 | 13.5 | 2.2 | 1.7 |
| >hg38_21_+_8433172_8446622_RNA45SN1_45S_50ntFlanks@11577.11600.24\|TTAGAACTGGTGCGGACCAGGGGA | >hg38_21_+_8433172_8446622_RNA45SN1_45S_50ntFlanks | 11577 | 11600 | 24 | TTAGAACTGGTGCGGACCAGGGGA | MB468 | 14 | 6 | 15 | 4 | 3 | 6.0 | 1.8 | 8.4 | 4.0 | 1.4 |
| >hg38_21_+_8433172_8446622_RNA45SN1_45S_50ntFlanks@11577.11601.25\|TTAGAACTGGTGCGGACCAGGGGAA | >hg38_21_+_8433172_8446622_RNA45SN1_45S_50ntFlanks | 11577 | 11601 | 25 | TTAGAACTGGTGCGGACCAGGGGAA | MB468 | 33 | 26 | 31 | 10 | 7 | 14.0 | 3.9 | 16.6 | 6.9 | 3.7 |
| >hg38_21_+_8433172_8446622_RNA45SN1_45S_50ntFlanks@11577.11602.26\|TTAGAACTGGTGCGGACCAGGGGAAT | >hg38_21_+_8433172_8446622_RNA45SN1_45S_50ntFlanks | 11577 | 11602 | 26 | TTAGAACTGGTGCGGACCAGGGGAAT | MB468 | 22 | 13 | 24 | 7 | 7 | 10.0 | 4.5 | 7.5 | 4.6 | 3.8 |

| | | | | | | | | | | | | | | | | |
|---|---|---|---|---|---|---|---|---|---|---|---|---|---|---|---|---|
| >hg38_21_+_8433172_8446622_RNA45SN1_45S_50ntFlanks@11577.11603.27\|TTAGAACTGG TGCGGACCAGGGGAATC | >hg38_21_+_8433172_8446622_RNA45SN1_45S_50ntFlanks | 11577 | 11603 | 27 | TTAGAACTGGTGCGGACCAGGGGAATC | MB468 | 115 | 80 | 165 | 70 | 44 | 34.5 | 33.8 | 28.8 | 49.4 | 32.0 |
| >hg38_21_+_8433172_8446622_RNA45SN1_45S_50ntFlanks@11577.11605.29\|TTAGAACTGG TGCGGACCAGGGGAATCCG | >hg38_21_+_8433172_8446622_RNA45SN1_45S_50ntFlanks | 11577 | 11605 | 29 | TTAGAACTGGTGCGGACCAGGGGAATCCG | MB468 | 50 | 45 | 64 | 17 | 24 | 10.0 | 7.5 | 6.3 | 9.2 | 13.8 |
| >hg38_21_+_8433172_8446622_RNA45SN1_45S_50ntFlanks@11577.11609.33\|TTAGAACTGG TGCGGACCAGGGGAATCCGACTG | >hg38_21_+_8433172_8446622_RNA45SN1_45S_50ntFlanks | 11577 | 11609 | 33 | TTAGAACTGGTGCGGACCAGGGGAATCCGACTG | MB468 | 20 | 8 | 13 | 14 | 10 | 9.2 | 2.4 | 3.4 | 7.3 | 5.7 |
| >hg38_21_+_8433172_8446622_RNA45SN1_45S_50ntFlanks@11577.11611.35\|TTAGAACTGG TGCGGACCAGGGGAATCCGACTGTT | >hg38_21_+_8433172_8446622_RNA45SN1_45S_50ntFlanks | 11577 | 11611 | 35 | TTAGAACTGGTGCGGACCAGGGGAATCCGACTGTT | MB468 | 28 | 9 | 19 | 19 | 34 | 12.8 | 3.6 | 5.8 | 13.7 | 17.1 |
| >hg38_21_+_8433172_8446622_RNA45SN1_45S_50ntFlanks@11578.11600.23\|TAGAACTGGT GCGGACCAGGGGA | >hg38_21_+_8433172_8446622_RNA45SN1_45S_50ntFlanks | 11578 | 11600 | 23 | TAGAACTGGTGCGGACCAGGGGA | MB468 | 10 | 6 | 10 | 3 | 2 | 3.1 | 1.1 | 6.3 | 2.1 | 0.3 |
| >hg38_21_+_8433172_8446622_RNA45SN1_45S_50ntFlanks@11578.11601.24\|TAGAACTGGT GCGGACCAGGGGAA | >hg38_21_+_8433172_8446622_RNA45SN1_45S_50ntFlanks | 11578 | 11601 | 24 | TAGAACTGGTGCGGACCAGGGGAA | MB468 | 24 | 22 | 22 | 8 | 10 | 8.7 | 4.3 | 9.7 | 3.5 | 3.6 |
| >hg38_21_+_8433172_8446622_RNA45SN1_45S_50ntFlanks@11578.11602.25\|TAGAACTGGT GCGGACCAGGGGAAT | >hg38_21_+_8433172_8446622_RNA45SN1_45S_50ntFlanks | 11578 | 11602 | 25 | TAGAACTGGTGCGGACCAGGGGAAT | MB468 | 16 | 10 | 17 | 5 | 7 | 7.0 | 2.4 | 5.6 | 1.8 | 2.0 |
| >hg38_21_+_8433172_8446622_RNA45SN1_45S_50ntFlanks@11578.11603.26\|TAGAACTGGT GCGGACCAGGGGAATC | >hg38_21_+_8433172_8446622_RNA45SN1_45S_50ntFlanks | 11578 | 11603 | 26 | TAGAACTGGTGCGGACCAGGGGAATC | MB468 | 78 | 54 | 117 | 56 | 33 | 23.2 | 24.4 | 16.9 | 40.9 | 16.1 |
| >hg38_21_+_8433172_8446622_RNA45SN1_45S_50ntFlanks@11578.11605.28\|TAGAACTGGT GCGGACCAGGGGAATCCG | >hg38_21_+_8433172_8446622_RNA45SN1_45S_50ntFlanks | 11578 | 11605 | 28 | TAGAACTGGTGCGGACCAGGGGAATCCG | MB468 | 41 | 39 | 51 | 15 | 21 | 6.9 | 3.4 | 5.4 | 7.2 | 8.8 |
| >hg38_21_+_8433172_8446622_RNA45SN1_45S_50ntFlanks@11578.11609.32\|TAGAACTGGT GCGGACCAGGGGAATCCGACTG | >hg38_21_+_8433172_8446622_RNA45SN1_45S_50ntFlanks | 11578 | 11609 | 32 | TAGAACTGGTGCGGACCAGGGGAATCCGACTG | MB468 | 14 | 6 | 9 | 11 | 9 | 6.0 | 1.7 | 2.1 | 5.9 | 5.1 |
| >hg38_21_+_8433172_8446622_RNA45SN1_45S_50ntFlanks@11578.11611.34\|TAGAACTGGT GCGGACCAGGGGAATCCGACTGTT | >hg38_21_+_8433172_8446622_RNA45SN1_45S_50ntFlanks | 11578 | 11611 | 34 | TAGAACTGGTGCGGACCAGGGGAATCCGACTGTT | MB468 | 22 | 9 | 15 | 14 | 28 | 9.7 | 2.6 | 3.8 | 9.6 | 9.8 |
| >hg38_21_+_8433172_8446622_RNA45SN1_45S_50ntFlanks@11578.11613.36\|TAGAACTGGT GCGGACCAGGGGAATCCGACTGTTTA | >hg38_21_+_8433172_8446622_RNA45SN1_45S_50ntFlanks | 11578 | 11613 | 36 | TAGAACTGGTGCGGACCAGGGGAATCCGACTGTTTA | BT20 | 17 | 51 | 8 | 7 | 68 | 20.5 | 25.6 | 4.6 | 2.6 | 12.6 |
| >hg38_21_+_8433172_8446622_RNA45SN1_45S_50ntFlanks@11579.11600.22\|AGAACTGGTG CGGACCAGGGGA | >hg38_21_+_8433172_8446622_RNA45SN1_45S_50ntFlanks | 11579 | 11600 | 22 | AGAACTGGTGCGGACCAGGGGA | MB468 | 21 | 4 | 21 | 14 | 5 | 11.8 | 0.9 | 18.0 | 12.3 | 1.6 |
| >hg38_21_+_8433172_8446622_RNA45SN1_45S_50ntFlanks@11579.11601.23\|AGAACTGGTG CGGACCAGGGGAA | >hg38_21_+_8433172_8446622_RNA45SN1_45S_50ntFlanks | 11579 | 11601 | 23 | AGAACTGGTGCGGACCAGGGGAA | MB468 | 77 | 40 | 79 | 51 | 24 | 45.3 | 6.8 | 52.6 | 42.5 | 11.7 |
| >hg38_21_+_8433172_8446622_RNA45SN1_45S_50ntFlanks@11579.11602.24\|AGAACTGGTG CGGACCAGGGGAAT | >hg38_21_+_8433172_8446622_RNA45SN1_45S_50ntFlanks | 11579 | 11602 | 24 | AGAACTGGTGCGGACCAGGGGAAT | MB468 | 55 | 22 | 62 | 45 | 24 | 34.9 | 7.6 | 34.8 | 31.9 | 12.0 |
| >hg38_21_+_8433172_8446622_RNA45SN1_45S_50ntFlanks@11579.11603.25\|AGAACTGGTG CGGACCAGGGGAATC | >hg38_21_+_8433172_8446622_RNA45SN1_45S_50ntFlanks | 11579 | 11603 | 25 | AGAACTGGTGCGGACCAGGGGAATC | MB468 | 315 | 173 | 438 | 704 | 184 | 107.2 | 82.3 | 81.7 | 706.8 | 88.7 |
| >hg38_21_+_8433172_8446622_RNA45SN1_45S_50ntFlanks@11579.11604.26\|AGAACTGGTG CGGACCAGGGGAATCC | >hg38_21_+_8433172_8446622_RNA45SN1_45S_50ntFlanks | 11579 | 11604 | 26 | AGAACTGGTGCGGACCAGGGGAATCC | MB468 | 13 | 14 | 10 | 7 | 7 | 7.4 | 7.2 | 3.0 | 4.7 | 4.7 |
| >hg38_21_+_8433172_8446622_RNA45SN1_45S_50ntFlanks@11579.11605.27\|AGAACTGGTG CGGACCAGGGGAATCCG | >hg38_21_+_8433172_8446622_RNA45SN1_45S_50ntFlanks | 11579 | 11605 | 27 | AGAACTGGTGCGGACCAGGGGAATCCG | MB468 | 138 | 117 | 153 | 116 | 104 | 29.6 | 17.2 | 16.2 | 62.7 | 40.4 |
| >hg38_21_+_8433172_8446622_RNA45SN1_45S_50ntFlanks@11579.11606.28\|AGAACTGGTG CGGACCAGGGGAATCCGA | >hg38_21_+_8433172_8446622_RNA45SN1_45S_50ntFlanks | 11579 | 11606 | 28 | AGAACTGGTGCGGACCAGGGGAATCCGA | MB468 | 17 | 11 | 15 | 17 | 13 | 3.0 | 2.8 | 3.9 | 9.7 | 4.1 |
| >hg38_21_+_8433172_8446622_RNA45SN1_45S_50ntFlanks@11579.11607.29\|AGAACTGGTG CGGACCAGGGGAATCCGAC | >hg38_21_+_8433172_8446622_RNA45SN1_45S_50ntFlanks | 11579 | 11607 | 29 | AGAACTGGTGCGGACCAGGGGAATCCGAC | MB468 | 38 | 21 | 32 | 36 | 35 | 14.9 | 5.9 | 6.1 | 25.8 | 11.4 |
| >hg38_21_+_8433172_8446622_RNA45SN1_45S_50ntFlanks@11579.11609.31\|AGAACTGGTG CGGACCAGGGGAATCCGACTG | >hg38_21_+_8433172_8446622_RNA45SN1_45S_50ntFlanks | 11579 | 11609 | 31 | AGAACTGGTGCGGACCAGGGGAATCCGACTG | MB468 | 59 | 21 | 23 | 71 | 46 | 30.5 | 8.1 | 2.6 | 37.0 | 30.6 |
| >hg38_21_+_8433172_8446622_RNA45SN1_45S_50ntFlanks@11579.11611.33\|AGAACTGGTG CGGACCAGGGGAATCCGACTGTT | >hg38_21_+_8433172_8446622_RNA45SN1_45S_50ntFlanks | 11579 | 11611 | 33 | AGAACTGGTGCGGACCAGGGGAATCCGACTGTT | MB468 | 93 | 27 | 43 | 86 | 126 | 56.0 | 8.8 | 14.4 | 69.0 | 26.2 |
| >hg38_21_+_8433172_8446622_RNA45SN1_45S_50ntFlanks@11579.11612.34\|AGAACTGGTG CGGACCAGGGGAATCCGACTGTTT | >hg38_21_+_8433172_8446622_RNA45SN1_45S_50ntFlanks | 11579 | 11612 | 34 | AGAACTGGTGCGGACCAGGGGAATCCGACTGTTT | MB468 | 17 | 3 | 13 | 6 | 19 | 14.0 | 1.5 | 8.0 | 5.0 | 9.3 |
| >hg38_21_+_8433172_8446622_RNA45SN1_45S_50ntFlanks@11580.11600.21\|GAACTGGTGC GGACCAGGGGA | >hg38_21_+_8433172_8446622_RNA45SN1_45S_50ntFlanks | 11580 | 11600 | 21 | GAACTGGTGCGGACCAGGGGA | MB468 | 38 | 10 | 53 | 33 | 9 | 16.1 | 2.1 | 30.3 | 29.2 | 3.8 |
| >hg38_21_+_8433172_8446622_RNA45SN1_45S_50ntFlanks@11580.11601.22\|GAACTGGTGC GGACCAGGGGAA | >hg38_21_+_8433172_8446622_RNA45SN1_45S_50ntFlanks | 11580 | 11601 | 22 | GAACTGGTGCGGACCAGGGGAA | MB468 | 90 | 47 | 114 | 78 | 30 | 46.3 | 11.1 | 71.6 | 59.4 | 13.8 |
| >hg38_21_+_8433172_8446622_RNA45SN1_45S_50ntFlanks@11580.11602.23\|GAACTGGTGC GGACCAGGGGAAT | >hg38_21_+_8433172_8446622_RNA45SN1_45S_50ntFlanks | 11580 | 11602 | 23 | GAACTGGTGCGGACCAGGGGAAT | MB468 | 70 | 27 | 92 | 59 | 28 | 42.7 | 11.1 | 52.8 | 45.7 | 14.1 |
| >hg38_21_+_8433172_8446622_RNA45SN1_45S_50ntFlanks@11580.11603.24\|GAACTGGTGC GGACCAGGGGAATC | >hg38_21_+_8433172_8446622_RNA45SN1_45S_50ntFlanks | 11580 | 11603 | 24 | GAACTGGTGCGGACCAGGGGAATC | MB468 | 350 | 187 | 605 | 748 | 175 | 116.6 | 101.6 | 111.1 | 787.7 | 97.3 |
| >hg38_21_+_8433172_8446622_RNA45SN1_45S_50ntFlanks@11580.11604.25\|GAACTGGTGC GGACCAGGGGAATCC | >hg38_21_+_8433172_8446622_RNA45SN1_45S_50ntFlanks | 11580 | 11604 | 25 | GAACTGGTGCGGACCAGGGGAATCC | MB468 | 11 | 11 | 12 | 9 | 7 | 4.9 | 4.8 | 1.8 | 6.7 | 4.3 |
| >hg38_21_+_8433172_8446622_RNA45SN1_45S_50ntFlanks@11580.11605.26\|GAACTGGTGC GGACCAGGGGAATCCG | >hg38_21_+_8433172_8446622_RNA45SN1_45S_50ntFlanks | 11580 | 11605 | 26 | GAACTGGTGCGGACCAGGGGAATCCG | MB468 | 131 | 95 | 161 | 114 | 94 | 24.3 | 8.9 | 28.7 | 69.3 | 31.8 |
| >hg38_21_+_8433172_8446622_RNA45SN1_45S_50ntFlanks@11580.11606.27\|GAACTGGTGC GGACCAGGGGAATCCGA | >hg38_21_+_8433172_8446622_RNA45SN1_45S_50ntFlanks | 11580 | 11606 | 27 | GAACTGGTGCGGACCAGGGGAATCCGA | MB468 | 17 | 8 | 15 | 18 | 11 | 2.8 | 2.3 | 6.2 | 11.0 | 2.8 |
| >hg38_21_+_8433172_8446622_RNA45SN1_45S_50ntFlanks@11580.11607.28\|GAACTGGTGC GGACCAGGGGAATCCGAC | >hg38_21_+_8433172_8446622_RNA45SN1_45S_50ntFlanks | 11580 | 11607 | 28 | GAACTGGTGCGGACCAGGGGAATCCGAC | MB468 | 33 | 14 | 27 | 31 | 28 | 11.4 | 5.6 | 9.0 | 25.4 | 12.6 |
| >hg38_21_+_8433172_8446622_RNA45SN1_45S_50ntFlanks@11580.11609.30\|GAACTGGTGC GGACCAGGGGAATCCGACTG | >hg38_21_+_8433172_8446622_RNA45SN1_45S_50ntFlanks | 11580 | 11609 | 30 | GAACTGGTGCGGACCAGGGGAATCCGACTG | MB468 | 88 | 35 | 51 | 102 | 63 | 35.3 | 12.7 | 7.0 | 46.0 | 25.0 |
| >hg38_21_+_8433172_8446622_RNA45SN1_45S_50ntFlanks@11580.11611.32\|GAACTGGTGC GGACCAGGGGAATCCGACTGTT | >hg38_21_+_8433172_8446622_RNA45SN1_45S_50ntFlanks | 11580 | 11611 | 32 | GAACTGGTGCGGACCAGGGGAATCCGACTGTT | MB231 | 13 | 8 | 12 | 5 | 10 | 7.9 | 2.4 | 3.6 | 2.9 | 6.4 |
| >hg38_21_+_8433172_8446622_RNA45SN1_45S_50ntFlanks@11580.11611.32\|GAACTGGTGC GGACCAGGGGAATCCGACTGTT | >hg38_21_+_8433172_8446622_RNA45SN1_45S_50ntFlanks | 11580 | 11611 | 32 | GAACTGGTGCGGACCAGGGGAATCCGACTGTT | MB468 | 206 | 58 | 97 | 168 | 234 | 89.5 | 22.0 | 34.8 | 134.4 | 69.2 |
| >hg38_21_+_8433172_8446622_RNA45SN1_45S_50ntFlanks@11580.11612.33\|GAACTGGTGC GGACCAGGGGAATCCGACTGTTT | >hg38_21_+_8433172_8446622_RNA45SN1_45S_50ntFlanks | 11580 | 11612 | 33 | GAACTGGTGCGGACCAGGGGAATCCGACTGTTT | MB468 | 13 | 2 | 8 | 5 | 12 | 8.0 | 1.0 | 3.0 | 4.3 | 6.3 |
| >hg38_21_+_8433172_8446622_RNA45SN1_45S_50ntFlanks@11580.11613.34\|GAACTGGTGC GGACCAGGGGAATCCGACTGTTTA | >hg38_21_+_8433172_8446622_RNA45SN1_45S_50ntFlanks | 11580 | 11613 | 34 | GAACTGGTGCGGACCAGGGGAATCCGACTGTTTA | BT20 | 38 | 58 | 45 | 22 | 143 | 23.6 | 26.4 | 16.9 | 5.8 | 25.7 |
| >hg38_21_+_8433172_8446622_RNA45SN1_45S_50ntFlanks@11580.11613.34\|GAACTGGTGC GGACCAGGGGAATCCGACTGTTTA | >hg38_21_+_8433172_8446622_RNA45SN1_45S_50ntFlanks | 11580 | 11613 | 34 | GAACTGGTGCGGACCAGGGGAATCCGACTGTTTA | MB231 | 33 | 18 | 44 | 8 | 27 | 39.9 | 21.1 | 43.8 | 5.2 | 29.0 |
| >hg38_21_+_8433172_8446622_RNA45SN1_45S_50ntFlanks@11580.11613.34\|GAACTGGTGC GGACCAGGGGAATCCGACTGTTTA | >hg38_21_+_8433172_8446622_RNA45SN1_45S_50ntFlanks | 11580 | 11613 | 34 | GAACTGGTGCGGACCAGGGGAATCCGACTGTTTA | MB468 | 53 | 32 | 45 | 26 | 101 | 21.5 | 25.3 | 8.4 | 21.6 | 75.5 |
| >hg38_21_+_8433172_8446622_RNA45SN1_45S_50ntFlanks@11581.11599.19\|AACTGGTGCG GACCAGGGG | >hg38_21_+_8433172_8446622_RNA45SN1_45S_50ntFlanks | 11581 | 11599 | 19 | AACTGGTGCGGACCAGGGG | MB468 | 12 | 1 | 18 | 9 | 2 | 3.4 | 0.2 | 10.3 | 7.9 | 1.0 |
| >hg38_21_+_8433172_8446622_RNA45SN1_45S_50ntFlanks@11581.11600.20\|AACTGGTGCG GACCAGGGGA | >hg38_21_+_8433172_8446622_RNA45SN1_45S_50ntFlanks | 11581 | 11600 | 20 | AACTGGTGCGGACCAGGGGA | MB468 | 57 | 10 | 71 | 45 | 12 | 22.5 | 2.3 | 35.5 | 37.8 | 4.0 |
| >hg38_21_+_8433172_8446622_RNA45SN1_45S_50ntFlanks@11581.11601.21\|AACTGGTGCG GACCAGGGGAA | >hg38_21_+_8433172_8446622_RNA45SN1_45S_50ntFlanks | 11581 | 11601 | 21 | AACTGGTGCGGACCAGGGGAA | MB468 | 133 | 48 | 146 | 89 | 37 | 74.0 | 11.8 | 97.1 | 74.3 | 16.5 |
| >hg38_21_+_8433172_8446622_RNA45SN1_45S_50ntFlanks@11581.11602.22\|AACTGGTGCG GACCAGGGGAAT | >hg38_21_+_8433172_8446622_RNA45SN1_45S_50ntFlanks | 11581 | 11602 | 22 | AACTGGTGCGGACCAGGGGAAT | MB468 | 103 | 30 | 123 | 73 | 39 | 64.1 | 11.1 | 68.3 | 54.1 | 15.8 |
| >hg38_21_+_8433172_8446622_RNA45SN1_45S_50ntFlanks@11581.11603.23\|AACTGGTGCG GACCAGGGGAATC | >hg38_21_+_8433172_8446622_RNA45SN1_45S_50ntFlanks | 11581 | 11603 | 23 | AACTGGTGCGGACCAGGGGAATC | MB468 | 473 | 195 | 683 | 942 | 208 | 152.8 | 102.7 | 128.9 | 939.7 | 100.0 |
| >hg38_21_+_8433172_8446622_RNA45SN1_45S_50ntFlanks@11581.11604.24\|AACTGGTGCG GACCAGGGGAATCC | >hg38_21_+_8433172_8446622_RNA45SN1_45S_50ntFlanks | 11581 | 11604 | 24 | AACTGGTGCGGACCAGGGGAATCC | MB468 | 20 | 15 | 17 | 13 | 10 | 9.7 | 8.1 | 3.3 | 9.8 | 5.8 |
| >hg38_21_+_8433172_8446622_RNA45SN1_45S_50ntFlanks@11581.11605.25\|AACTGGTGCG GACCAGGGGAATCCG | >hg38_21_+_8433172_8446622_RNA45SN1_45S_50ntFlanks | 11581 | 11605 | 25 | AACTGGTGCGGACCAGGGGAATCCG | MB468 | 214 | 116 | 226 | 176 | 139 | 31.0 | 14.9 | 29.5 | 97.5 | 40.0 |
| >hg38_21_+_8433172_8446622_RNA45SN1_45S_50ntFlanks@11581.11606.26\|AACTGGTGCG GACCAGGGGAATCCGA | >hg38_21_+_8433172_8446622_RNA45SN1_45S_50ntFlanks | 11581 | 11606 | 26 | AACTGGTGCGGACCAGGGGAATCCGA | MB468 | 29 | 11 | 22 | 27 | 18 | 5.0 | 1.3 | 8.6 | 12.5 | 2.5 |
| >hg38_21_+_8433172_8446622_RNA45SN1_45S_50ntFlanks@11581.11607.27\|AACTGGTGCG GACCAGGGGAATCCGAC | >hg38_21_+_8433172_8446622_RNA45SN1_45S_50ntFlanks | 11581 | 11607 | 27 | AACTGGTGCGGACCAGGGGAATCCGAC | MB468 | 35 | 11 | 29 | 31 | 27 | 14.4 | 3.7 | 10.5 | 21.3 | 10.5 |
| >hg38_21_+_8433172_8446622_RNA45SN1_45S_50ntFlanks@11581.11608.28\|AACTGGTGCG GACCAGGGGAATCCGACT | >hg38_21_+_8433172_8446622_RNA45SN1_45S_50ntFlanks | 11581 | 11608 | 28 | AACTGGTGCGGACCAGGGGAATCCGACT | MB468 | 11 | 5 | 7 | 9 | 8 | 3.0 | 1.7 | 1.5 | 3.5 | 1.4 |
| >hg38_21_+_8433172_8446622_RNA45SN1_45S_50ntFlanks@11581.11609.29\|AACTGGTGCG GACCAGGGGAATCCGACTG | >hg38_21_+_8433172_8446622_RNA45SN1_45S_50ntFlanks | 11581 | 11609 | 29 | AACTGGTGCGGACCAGGGGAATCCGACTG | MB468 | 179 | 74 | 131 | 166 | 115 | 51.8 | 19.1 | 14.0 | 51.1 | 32.0 |
| >hg38_21_+_8433172_8446622_RNA45SN1_45S_50ntFlanks@11581.11610.30\|AACTGGTGCG GACCAGGGGAATCCGACTGT | >hg38_21_+_8433172_8446622_RNA45SN1_45S_50ntFlanks | 11581 | 11610 | 30 | AACTGGTGCGGACCAGGGGAATCCGACTGT | MB468 | 11 | 7 | 9 | 5 | 4 | 4.0 | 3.5 | 2.2 | 6.1 | 2.9 |

| | | | | | | | | | | | | | | | | |
|---|---|---|---|---|---|---|---|---|---|---|---|---|---|---|---|---|
| >hg38_21_+_8433172_8446622_RNA45SN1_45S_50ntFlanks@11581.11611.31\|AACTGGTGCGGACCAGGGGAATCCGACTGTT | >hg38_21_+_8433172_8446622_RNA45SN1_45S_50ntFlanks | 11581 | 11611 | 31 | AACTGGTGCGGACCAGGGGAATCCGACTGTT | MB468 | 203 | 41 | 87 | 138 | 180 | 82.4 | 11.4 | 41.9 | 97.9 | 33.2 |
| >hg38_21_+_8433172_8446622_RNA45SN1_45S_50ntFlanks@11581.11612.32\|AACTGGTGCGGACCAGGGGAATCCGACTGTTT | >hg38_21_+_8433172_8446622_RNA45SN1_45S_50ntFlanks | 11581 | 11612 | 32 | AACTGGTGCGGACCAGGGGAATCCGACTGTTT | MB231 | 11 | 6 | 14 | 3 | 8 | 6.5 | 1.6 | 5.6 | 0.8 | 1.4 |
| >hg38_21_+_8433172_8446622_RNA45SN1_45S_50ntFlanks@11581.11612.32\|AACTGGTGCGGACCAGGGGAATCCGACTGTTT | >hg38_21_+_8433172_8446622_RNA45SN1_45S_50ntFlanks | 11581 | 11612 | 32 | AACTGGTGCGGACCAGGGGAATCCGACTGTTT | MB468 | 217 | 65 | 130 | 134 | 235 | 121.3 | 23.6 | 37.1 | 107.0 | 95.2 |
| >hg38_21_+_8433172_8446622_RNA45SN1_45S_50ntFlanks@11581.11613.33\|AACTGGTGCGGACCAGGGGAATCCGACTGTTTA | >hg38_21_+_8433172_8446622_RNA45SN1_45S_50ntFlanks | 11581 | 11613 | 33 | AACTGGTGCGGACCAGGGGAATCCGACTGTTTA | MB468 | 27 | 16 | 18 | 28 | 63 | 7.7 | 6.5 | 3.9 | 16.7 | 30.1 |
| >hg38_21_+_8433172_8446622_RNA45SN1_45S_50ntFlanks@11581.11616.36\|AACTGGTGCGGACCAGGGGAATCCGACTGTTTAATT | >hg38_21_+_8433172_8446622_RNA45SN1_45S_50ntFlanks | 11581 | 11616 | 36 | AACTGGTGCGGACCAGGGGAATCCGACTGTTTAATT | MB231 | 15 | 8 | 15 | 5 | 11 | 12.4 | 3.9 | 10.5 | 0.9 | 4.4 |
| >hg38_21_+_8433172_8446622_RNA45SN1_45S_50ntFlanks@11582.11600.19\|ACTGGTGCGGACCAGGGGA | >hg38_21_+_8433172_8446622_RNA45SN1_45S_50ntFlanks | 11582 | 11600 | 19 | ACTGGTGCGGACCAGGGGA | MB468 | 16 | 5 | 23 | 14 | 3 | 6.9 | 1.5 | 11.8 | 10.2 | 0.4 |
| >hg38_21_+_8433172_8446622_RNA45SN1_45S_50ntFlanks@11582.11601.20\|ACTGGTGCGGACCAGGGGAA | >hg38_21_+_8433172_8446622_RNA45SN1_45S_50ntFlanks | 11582 | 11601 | 20 | ACTGGTGCGGACCAGGGGAA | MB468 | 29 | 18 | 35 | 23 | 8 | 12.6 | 5.4 | 22.3 | 18.0 | 3.0 |
| >hg38_21_+_8433172_8446622_RNA45SN1_45S_50ntFlanks@11582.11602.21\|ACTGGTGCGGACCAGGGGAAT | >hg38_21_+_8433172_8446622_RNA45SN1_45S_50ntFlanks | 11582 | 11602 | 21 | ACTGGTGCGGACCAGGGGAAT | BT20 | 16 | 9 | 19 | 4 | 6 | 6.0 | 3.6 | 3.0 | 2.7 | 2.0 |
| >hg38_21_+_8433172_8446622_RNA45SN1_45S_50ntFlanks@11582.11602.21\|ACTGGTGCGGACCAGGGGAAT | >hg38_21_+_8433172_8446622_RNA45SN1_45S_50ntFlanks | 11582 | 11602 | 21 | ACTGGTGCGGACCAGGGGAAT | MB468 | 27 | 11 | 35 | 29 | 17 | 16.4 | 3.3 | 17.4 | 15.2 | 7.7 |
| >hg38_21_+_8433172_8446622_RNA45SN1_45S_50ntFlanks@11582.11603.22\|ACTGGTGCGGACCAGGGGAATC | >hg38_21_+_8433172_8446622_RNA45SN1_45S_50ntFlanks | 11582 | 11603 | 22 | ACTGGTGCGGACCAGGGGAATC | MB468 | 76 | 48 | 107 | 159 | 32 | 24.5 | 21.2 | 35.0 | 130.9 | 8.0 |
| >hg38_21_+_8433172_8446622_RNA45SN1_45S_50ntFlanks@11582.11605.24\|ACTGGTGCGGACCAGGGGAATCCG | >hg38_21_+_8433172_8446622_RNA45SN1_45S_50ntFlanks | 11582 | 11605 | 24 | ACTGGTGCGGACCAGGGGAATCCG | MB468 | 29 | 15 | 29 | 22 | 16 | 6.9 | 2.6 | 8.9 | 11.7 | 2.6 |
| >hg38_21_+_8433172_8446622_RNA45SN1_45S_50ntFlanks@11582.11609.28\|ACTGGTGCGGACCAGGGGAATCCGACTG | >hg38_21_+_8433172_8446622_RNA45SN1_45S_50ntFlanks | 11582 | 11609 | 28 | ACTGGTGCGGACCAGGGGAATCCGACTG | MB468 | 25 | 9 | 17 | 21 | 14 | 12.4 | 3.1 | 5.9 | 6.9 | 5.8 |
| >hg38_21_+_8433172_8446622_RNA45SN1_45S_50ntFlanks@11582.11611.30\|ACTGGTGCGGACCAGGGGAATCCGACTGTT | >hg38_21_+_8433172_8446622_RNA45SN1_45S_50ntFlanks | 11582 | 11611 | 30 | ACTGGTGCGGACCAGGGGAATCCGACTGTT | MB231 | 14 | 9 | 17 | 3 | 8 | 5.4 | 2.6 | 1.3 | 1.7 | 1.2 |
| >hg38_21_+_8433172_8446622_RNA45SN1_45S_50ntFlanks@11582.11611.30\|ACTGGTGCGGACCAGGGGAATCCGACTGTT | >hg38_21_+_8433172_8446622_RNA45SN1_45S_50ntFlanks | 11582 | 11611 | 30 | ACTGGTGCGGACCAGGGGAATCCGACTGTT | MB468 | 94 | 35 | 52 | 76 | 94 | 34.7 | 19.3 | 16.1 | 35.2 | 24.0 |
| >hg38_21_+_8433172_8446622_RNA45SN1_45S_50ntFlanks@11582.11612.31\|ACTGGTGCGGACCAGGGGAATCCGACTGTTT | >hg38_21_+_8433172_8446622_RNA45SN1_45S_50ntFlanks | 11582 | 11612 | 31 | ACTGGTGCGGACCAGGGGAATCCGACTGTTT | MB468 | 36 | 16 | 21 | 25 | 36 | 18.5 | 8.3 | 5.7 | 14.2 | 15.2 |
| >hg38_21_+_8433172_8446622_RNA45SN1_45S_50ntFlanks@11583.11600.18\|CTGGTGCGGACCAGGGGA | >hg38_21_+_8433172_8446622_RNA45SN1_45S_50ntFlanks | 11583 | 11600 | 18 | CTGGTGCGGACCAGGGGA | MB468 | 41 | 22 | 69 | 28 | 8 | 17.7 | 4.5 | 40.3 | 16.8 | 1.1 |
| >hg38_21_+_8433172_8446622_RNA45SN1_45S_50ntFlanks@11583.11601.19\|CTGGTGCGGACCAGGGGAA | >hg38_21_+_8433172_8446622_RNA45SN1_45S_50ntFlanks | 11583 | 11601 | 19 | CTGGTGCGGACCAGGGGAA | MB468 | 85 | 89 | 112 | 58 | 32 | 48.5 | 23.0 | 71.0 | 29.9 | 10.2 |
| >hg38_21_+_8433172_8446622_RNA45SN1_45S_50ntFlanks@11583.11602.20\|CTGGTGCGGACCAGGGGAAT | >hg38_21_+_8433172_8446622_RNA45SN1_45S_50ntFlanks | 11583 | 11602 | 20 | CTGGTGCGGACCAGGGGAAT | MB468 | 66 | 52 | 80 | 65 | 42 | 45.0 | 11.0 | 52.4 | 17.5 | 29.0 |
| >hg38_21_+_8433172_8446622_RNA45SN1_45S_50ntFlanks@11583.11603.21\|CTGGTGCGGACCAGGGGAATC | >hg38_21_+_8433172_8446622_RNA45SN1_45S_50ntFlanks | 11583 | 11603 | 21 | CTGGTGCGGACCAGGGGAATC | MB468 | 166 | 135 | 207 | 330 | 104 | 67.1 | 34.5 | 89.3 | 173.1 | 29.5 |
| >hg38_21_+_8433172_8446622_RNA45SN1_45S_50ntFlanks@11583.11605.23\|CTGGTGCGGACCAGGGGAATCCG | >hg38_21_+_8433172_8446622_RNA45SN1_45S_50ntFlanks | 11583 | 11605 | 23 | CTGGTGCGGACCAGGGGAATCCG | MB468 | 79 | 50 | 82 | 62 | 81 | 30.4 | 10.4 | 30.9 | 15.4 | 38.5 |
| >hg38_21_+_8433172_8446622_RNA45SN1_45S_50ntFlanks@11583.11606.24\|CTGGTGCGGACCAGGGGAATCCGA | >hg38_21_+_8433172_8446622_RNA45SN1_45S_50ntFlanks | 11583 | 11606 | 24 | CTGGTGCGGACCAGGGGAATCCGA | MB468 | 10 | 4 | 8 | 7 | 8 | 4.0 | 0.5 | 4.7 | 2.0 | 3.1 |
| >hg38_21_+_8433172_8446622_RNA45SN1_45S_50ntFlanks@11583.11607.25\|CTGGTGCGGACCAGGGGAATCCGAC | >hg38_21_+_8433172_8446622_RNA45SN1_45S_50ntFlanks | 11583 | 11607 | 25 | CTGGTGCGGACCAGGGGAATCCGAC | MB468 | 24 | 7 | 21 | 15 | 18 | 14.8 | 2.2 | 12.4 | 7.0 | 2.9 |
| >hg38_21_+_8433172_8446622_RNA45SN1_45S_50ntFlanks@11583.11609.27\|CTGGTGCGGACCAGGGGAATCCGACTG | >hg38_21_+_8433172_8446622_RNA45SN1_45S_50ntFlanks | 11583 | 11609 | 27 | CTGGTGCGGACCAGGGGAATCCGACTG | MB468 | 66 | 29 | 51 | 54 | 62 | 36.1 | 7.4 | 24.1 | 8.0 | 30.8 |
| >hg38_21_+_8433172_8446622_RNA45SN1_45S_50ntFlanks@11583.11611.29\|CTGGTGCGGACCAGGGGAATCCGACTGTT | >hg38_21_+_8433172_8446622_RNA45SN1_45S_50ntFlanks | 11583 | 11611 | 29 | CTGGTGCGGACCAGGGGAATCCGACTGTT | BT20 | 22 | 59 | 17 | 22 | 48 | 26.1 | 54.3 | 8.6 | 17.1 | 32.7 |
| >hg38_21_+_8433172_8446622_RNA45SN1_45S_50ntFlanks@11583.11611.29\|CTGGTGCGGACCAGGGGAATCCGACTGTT | >hg38_21_+_8433172_8446622_RNA45SN1_45S_50ntFlanks | 11583 | 11611 | 29 | CTGGTGCGGACCAGGGGAATCCGACTGTT | MB231 | 17 | 10 | 22 | 3 | 10 | 15.3 | 3.6 | 10.9 | 2.1 | 5.5 |
| >hg38_21_+_8433172_8446622_RNA45SN1_45S_50ntFlanks@11583.11611.29\|CTGGTGCGGACCAGGGGAATCCGACTGTT | >hg38_21_+_8433172_8446622_RNA45SN1_45S_50ntFlanks | 11583 | 11611 | 29 | CTGGTGCGGACCAGGGGAATCCGACTGTT | MB468 | 236 | 133 | 159 | 205 | 298 | 100.4 | 43.7 | 58.3 | 34.5 | 48.1 |
| >hg38_21_+_8433172_8446622_RNA45SN1_45S_50ntFlanks@11583.11612.30\|CTGGTGCGGACCAGGGGAATCCGACTGTTT | >hg38_21_+_8433172_8446622_RNA45SN1_45S_50ntFlanks | 11583 | 11612 | 30 | CTGGTGCGGACCAGGGGAATCCGACTGTTT | BT20 | 10 | 51 | 9 | 8 | 20 | 8.2 | 65.9 | 3.4 | 5.5 | 10.1 |
| >hg38_21_+_8433172_8446622_RNA45SN1_45S_50ntFlanks@11583.11612.30\|CTGGTGCGGACCAGGGGAATCCGACTGTTT | >hg38_21_+_8433172_8446622_RNA45SN1_45S_50ntFlanks | 11583 | 11612 | 30 | CTGGTGCGGACCAGGGGAATCCGACTGTTT | MB468 | 100 | 64 | 72 | 74 | 128 | 61.7 | 17.8 | 32.2 | 15.5 | 40.4 |
| >hg38_21_+_8433172_8446622_RNA45SN1_45S_50ntFlanks@11583.11613.31\|CTGGTGCGGACCAGGGGAATCCGACTGTTTA | >hg38_21_+_8433172_8446622_RNA45SN1_45S_50ntFlanks | 11583 | 11613 | 31 | CTGGTGCGGACCAGGGGAATCCGACTGTTTA | MB468 | 17 | 17 | 12 | 18 | 37 | 8.8 | 7.3 | 6.0 | 10.7 | 24.0 |
| >hg38_21_+_8433172_8446622_RNA45SN1_45S_50ntFlanks@11584.11611.28\|TGGTGCGGACCAGGGGAATCCGACTGTT | >hg38_21_+_8433172_8446622_RNA45SN1_45S_50ntFlanks | 11584 | 11611 | 28 | TGGTGCGGACCAGGGGAATCCGACTGTT | BT20 | 16 | 46 | 9 | 16 | 47 | 14.3 | 25.0 | 3.8 | 12.1 | 24.7 |
| >hg38_21_+_8433172_8446622_RNA45SN1_45S_50ntFlanks@11585.11616.32\|GGTGCGGACCAGGGGAATCCGACTGTTTAATT | >hg38_21_+_8433172_8446622_RNA45SN1_45S_50ntFlanks | 11585 | 11616 | 32 | GGTGCGGACCAGGGGAATCCGACTGTTTAATT | MB231 | 11 | 8 | 16 | 2 | 7 | 9.4 | 5.0 | 13.9 | 3.0 | 6.3 |
| >hg38_21_+_8433172_8446622_RNA45SN1_45S_50ntFlanks@11586.11603.18\|GTGCGGACCAGGGGAATC | >hg38_21_+_8433172_8446622_RNA45SN1_45S_50ntFlanks | 11586 | 11603 | 18 | GTGCGGACCAGGGGAATC | MB468 | 145 | 221 | 133 | 154 | 54 | 19.3 | 35.1 | 33.1 | 111.4 | 12.3 |
| >hg38_21_+_8433172_8446622_RNA45SN1_45S_50ntFlanks@11586.11605.20\|GTGCGGACCAGGGGAATCCG | >hg38_21_+_8433172_8446622_RNA45SN1_45S_50ntFlanks | 11586 | 11605 | 20 | GTGCGGACCAGGGGAATCCG | MB468 | 30 | 19 | 25 | 43 | 10 | 10.4 | 6.8 | 8.2 | 35.4 | 1.9 |
| >hg38_21_+_8433172_8446622_RNA45SN1_45S_50ntFlanks@11586.11609.24\|GTGCGGACCAGGGGAATCCGACTG | >hg38_21_+_8433172_8446622_RNA45SN1_45S_50ntFlanks | 11586 | 11609 | 24 | GTGCGGACCAGGGGAATCCGACTG | MB468 | 55 | 22 | 47 | 46 | 23 | 32.0 | 8.3 | 24.4 | 23.4 | 2.8 |
| >hg38_21_+_8433172_8446622_RNA45SN1_45S_50ntFlanks@11586.11611.26\|GTGCGGACCAGGGGAATCCGACTGTT | >hg38_21_+_8433172_8446622_RNA45SN1_45S_50ntFlanks | 11586 | 11611 | 26 | GTGCGGACCAGGGGAATCCGACTGTT | MB468 | 232 | 137 | 202 | 156 | 148 | 102.5 | 35.4 | 98.4 | 60.1 | 32.3 |
| >hg38_21_+_8433172_8446622_RNA45SN1_45S_50ntFlanks@11586.11612.27\|GTGCGGACCAGGGGAATCCGACTGTTT | >hg38_21_+_8433172_8446622_RNA45SN1_45S_50ntFlanks | 11586 | 11612 | 27 | GTGCGGACCAGGGGAATCCGACTGTTT | MB468 | 111 | 73 | 89 | 90 | 85 | 75.8 | 27.7 | 48.9 | 38.4 | 21.1 |
| >hg38_21_+_8433172_8446622_RNA45SN1_45S_50ntFlanks@11586.11613.28\|GTGCGGACCAGGGGAATCCGACTGTTTA | >hg38_21_+_8433172_8446622_RNA45SN1_45S_50ntFlanks | 11586 | 11613 | 28 | GTGCGGACCAGGGGAATCCGACTGTTTA | MB468 | 17 | 14 | 13 | 17 | 19 | 7.6 | 6.2 | 7.0 | 6.9 | 8.1 |
| >hg38_21_+_8433172_8446622_RNA45SN1_45S_50ntFlanks@11587.11605.19\|TGCGGACCAGGGGAATCCG | >hg38_21_+_8433172_8446622_RNA45SN1_45S_50ntFlanks | 11587 | 11605 | 19 | TGCGGACCAGGGGAATCCG | MB468 | 17 | 5 | 10 | 17 | 6 | 5.4 | 0.9 | 2.0 | 15.2 | 1.9 |
| >hg38_21_+_8433172_8446622_RNA45SN1_45S_50ntFlanks@11587.11609.23\|TGCGGACCAGGGGAATCCGACTG | >hg38_21_+_8433172_8446622_RNA45SN1_45S_50ntFlanks | 11587 | 11609 | 23 | TGCGGACCAGGGGAATCCGACTG | MB468 | 11 | 5 | 8 | 15 | 6 | 4.3 | 2.5 | 2.4 | 9.6 | 0.7 |
| >hg38_21_+_8433172_8446622_RNA45SN1_45S_50ntFlanks@11587.11611.25\|TGCGGACCAGGGGAATCCGACTGTT | >hg38_21_+_8433172_8446622_RNA45SN1_45S_50ntFlanks | 11587 | 11611 | 25 | TGCGGACCAGGGGAATCCGACTGTT | MB468 | 54 | 28 | 39 | 44 | 41 | 14.6 | 11.0 | 11.0 | 27.5 | 11.2 |
| >hg38_21_+_8433172_8446622_RNA45SN1_45S_50ntFlanks@11587.11612.26\|TGCGGACCAGGGGAATCCGACTGTTT | >hg38_21_+_8433172_8446622_RNA45SN1_45S_50ntFlanks | 11587 | 11612 | 26 | TGCGGACCAGGGGAATCCGACTGTTT | MB468 | 19 | 12 | 15 | 27 | 16 | 8.0 | 4.1 | 5.3 | 19.3 | 5.5 |
| >hg38_21_+_8433172_8446622_RNA45SN1_45S_50ntFlanks@11587.11616.30\|TGCGGACCAGGGGAATCCGACTGTTTAATT | >hg38_21_+_8433172_8446622_RNA45SN1_45S_50ntFlanks | 11587 | 11616 | 30 | TGCGGACCAGGGGAATCCGACTGTTTAATT | MB231 | 12 | 8 | 10 | 2 | 11 | 11.2 | 6.0 | 8.2 | 2.1 | 10.4 |
| >hg38_21_+_8433172_8446622_RNA45SN1_45S_50ntFlanks@11588.11611.24\|GCGGACCAGGGGAATCCGACTGTT | >hg38_21_+_8433172_8446622_RNA45SN1_45S_50ntFlanks | 11588 | 11611 | 24 | GCGGACCAGGGGAATCCGACTGTT | MB231 | 15 | 16 | 19 | 5 | 11 | 8.3 | 1.1 | 5.4 | 3.0 | 7.1 |
| >hg38_21_+_8433172_8446622_RNA45SN1_45S_50ntFlanks@11589.11609.21\|CGGACCAGGGGAATCCGACTG | >hg38_21_+_8433172_8446622_RNA45SN1_45S_50ntFlanks | 11589 | 11609 | 21 | CGGACCAGGGGAATCCGACTG | MB468 | 21 | 11 | 24 | 28 | 21 | 7.2 | 3.9 | 5.6 | 8.1 | 4.8 |
| >hg38_21_+_8433172_8446622_RNA45SN1_45S_50ntFlanks@11589.11611.23\|CGGACCAGGGGAATCCGACTGTT | >hg38_21_+_8433172_8446622_RNA45SN1_45S_50ntFlanks | 11589 | 11611 | 23 | CGGACCAGGGGAATCCGACTGTT | MB468 | 44 | 38 | 48 | 58 | 41 | 10.1 | 10.3 | 15.5 | 28.7 | 4.3 |
| >hg38_21_+_8433172_8446622_RNA45SN1_45S_50ntFlanks@11589.11612.24\|CGGACCAGGGGAATCCGACTGTTT | >hg38_21_+_8433172_8446622_RNA45SN1_45S_50ntFlanks | 11589 | 11612 | 24 | CGGACCAGGGGAATCCGACTGTTT | MB468 | 20 | 21 | 22 | 29 | 21 | 7.9 | 8.9 | 8.4 | 15.7 | 6.4 |
| >hg38_21_+_8433172_8446622_RNA45SN1_45S_50ntFlanks@11591.11611.21\|GACCAGGGGAATCCGACTGTT | >hg38_21_+_8433172_8446622_RNA45SN1_45S_50ntFlanks | 11591 | 11611 | 21 | GACCAGGGGAATCCGACTGTT | MB468 | 16 | 8 | 8 | 15 | 9 | 4.6 | 2.1 | 5.9 | 12.6 | 3.2 |
| >hg38_21_+_8433172_8446622_RNA45SN1_45S_50ntFlanks@11592.11609.18\|ACCAGGGGAATCCGACTG | >hg38_21_+_8433172_8446622_RNA45SN1_45S_50ntFlanks | 11592 | 11609 | 18 | ACCAGGGGAATCCGACTG | MB468 | 12 | 5 | 8 | 13 | 5 | 6.1 | 1.8 | 6.0 | 7.2 | 1.4 |
| >hg38_21_+_8433172_8446622_RNA45SN1_45S_50ntFlanks@11592.11611.20\|ACCAGGGGAATCCGACTGTT | >hg38_21_+_8433172_8446622_RNA45SN1_45S_50ntFlanks | 11592 | 11611 | 20 | ACCAGGGGAATCCGACTGTT | MB468 | 57 | 19 | 41 | 69 | 41 | 29.1 | 4.4 | 39.0 | 54.4 | 16.7 |
| >hg38_21_+_8433172_8446622_RNA45SN1_45S_50ntFlanks@11592.11612.21\|ACCAGGGGAATCCGACTGTTT | >hg38_21_+_8433172_8446622_RNA45SN1_45S_50ntFlanks | 11592 | 11612 | 21 | ACCAGGGGAATCCGACTGTTT | MB468 | 52 | 15 | 37 | 40 | 38 | 41.8 | 2.3 | 41.2 | 29.7 | 16.3 |

| | | | | | | | | | | | | | | | | |
|---|---|---|---|---|---|---|---|---|---|---|---|---|---|---|---|---|
| >hg38_21_+_8433172_8446622_RNA45SN1_45S_50ntFlanks@11592.11613.22\|ACCAGGGGAATCCGACTGTTTA | >hg38_21_+_8433172_8446622_RNA45SN1_45S_50ntFlanks | 11592 | 11613 | 22 | ACCAGGGGAATCCGACTGTTTA | MB468 | 23 | 13 | 15 | 13 | 29 | 13.1 | 8.9 | 17.5 | 5.0 | 6.2 |
| >hg38_21_+_8433172_8446622_RNA45SN1_45S_50ntFlanks@11592.11616.25\|ACCAGGGGAATCCGACTGTTTAATT | >hg38_21_+_8433172_8446622_RNA45SN1_45S_50ntFlanks | 11592 | 11616 | 25 | ACCAGGGGAATCCGACTGTTTAATT | MB468 | 11 | 11 | 6 | 4 | 13 | 7.5 | 5.3 | 6.3 | 1.6 | 2.9 |
| >hg38_21_+_8433172_8446622_RNA45SN1_45S_50ntFlanks@11593.11611.19\|CCAGGGGAATCCGACTGTT | >hg38_21_+_8433172_8446622_RNA45SN1_45S_50ntFlanks | 11593 | 11611 | 19 | CCAGGGGAATCCGACTGTT | MB468 | 312 | 167 | 262 | 174 | 183 | 67.4 | 14.4 | 105.6 | 70.4 | 46.9 |
| >hg38_21_+_8433172_8446622_RNA45SN1_45S_50ntFlanks@11593.11612.20\|CCAGGGGAATCCGACTGTTT | >hg38_21_+_8433172_8446622_RNA45SN1_45S_50ntFlanks | 11593 | 11612 | 20 | CCAGGGGAATCCGACTGTTT | MB468 | 126 | 68 | 103 | 78 | 82 | 51.1 | 6.6 | 57.9 | 40.8 | 16.1 |
| >hg38_21_+_8433172_8446622_RNA45SN1_45S_50ntFlanks@11593.11613.21\|CCAGGGGAATCCGACTGTTTA | >hg38_21_+_8433172_8446622_RNA45SN1_45S_50ntFlanks | 11593 | 11613 | 21 | CCAGGGGAATCCGACTGTTTA | MB468 | 40 | 36 | 31 | 25 | 51 | 7.7 | 17.7 | 17.1 | 8.4 | 16.6 |
| >hg38_21_+_8433172_8446622_RNA45SN1_45S_50ntFlanks@11594.11611.18\|CAGGGGAATCCGACTGTT | >hg38_21_+_8433172_8446622_RNA45SN1_45S_50ntFlanks | 11594 | 11611 | 18 | CAGGGGAATCCGACTGTT | MB468 | 205 | 125 | 172 | 99 | 166 | 35.3 | 21.9 | 54.3 | 25.3 | 78.6 |
| >hg38_21_+_8433172_8446622_RNA45SN1_45S_50ntFlanks@11594.11612.19\|CAGGGGAATCCGACTGTTT | >hg38_21_+_8433172_8446622_RNA45SN1_45S_50ntFlanks | 11594 | 11612 | 19 | CAGGGGAATCCGACTGTTT | MB468 | 84 | 53 | 68 | 41 | 75 | 27.8 | 7.8 | 29.8 | 13.8 | 34.9 |
| >hg38_21_+_8433172_8446622_RNA45SN1_45S_50ntFlanks@11594.11613.20\|CAGGGGAATCCGACTGTTTA | >hg38_21_+_8433172_8446622_RNA45SN1_45S_50ntFlanks | 11594 | 11613 | 20 | CAGGGGAATCCGACTGTTTA | MB468 | 26 | 25 | 20 | 16 | 41 | 2.9 | 9.8 | 5.5 | 8.6 | 22.0 |
| >hg38_21_+_8433172_8446622_RNA45SN1_45S_50ntFlanks@11595.11612.18\|AGGGGAATCCGACTGTTT | >hg38_21_+_8433172_8446622_RNA45SN1_45S_50ntFlanks | 11595 | 11612 | 18 | AGGGGAATCCGACTGTTT | MB468 | 10 | 11 | 8 | 5 | 4 | 3.6 | 5.5 | 4.5 | 1.3 | 1.4 |
| >hg38_21_+_8433172_8446622_RNA45SN1_45S_50ntFlanks@11601.11619.19\|ATCCGACTGTTTAATTAAA | >hg38_21_+_8433172_8446622_RNA45SN1_45S_50ntFlanks | 11601 | 11619 | 19 | ATCCGACTGTTTAATTAAA | MB468 | 15 | 22 | 19 | 6 | 7 | 2.0 | 13.1 | 9.5 | 1.3 | 2.4 |
| >hg38_21_+_8433172_8446622_RNA45SN1_45S_50ntFlanks@11601.11620.20\|ATCCGACTGTTTAATTAAAA | >hg38_21_+_8433172_8446622_RNA45SN1_45S_50ntFlanks | 11601 | 11620 | 20 | ATCCGACTGTTTAATTAAAA | MB468 | 14 | 24 | 13 | 5 | 6 | 1.3 | 14.8 | 7.4 | 1.0 | 2.1 |
| >hg38_21_+_8433172_8446622_RNA45SN1_45S_50ntFlanks@11602.11619.18\|TCCGACTGTTTAATTAAA | >hg38_21_+_8433172_8446622_RNA45SN1_45S_50ntFlanks | 11602 | 11619 | 18 | TCCGACTGTTTAATTAAA | MB468 | 82 | 94 | 112 | 31 | 39 | 27.3 | 50.6 | 74.1 | 2.0 | 12.1 |
| >hg38_21_+_8433172_8446622_RNA45SN1_45S_50ntFlanks@11602.11620.19\|TCCGACTGTTTAATTAAAA | >hg38_21_+_8433172_8446622_RNA45SN1_45S_50ntFlanks | 11602 | 11620 | 19 | TCCGACTGTTTAATTAAAA | MB468 | 68 | 101 | 87 | 25 | 32 | 19.2 | 54.8 | 61.9 | 1.3 | 8.2 |
| >hg38_21_+_8433172_8446622_RNA45SN1_45S_50ntFlanks@11602.11621.20\|TCCGACTGTTTAATTAAAAC | >hg38_21_+_8433172_8446622_RNA45SN1_45S_50ntFlanks | 11602 | 11621 | 20 | TCCGACTGTTTAATTAAAAC | MB468 | 17 | 20 | 32 | 14 | 9 | 4.4 | 12.3 | 13.1 | 2.3 | 1.0 |
| >hg38_21_+_8433172_8446622_RNA45SN1_45S_50ntFlanks@11603.11620.18\|CCGACTGTTTAATTAAAA | >hg38_21_+_8433172_8446622_RNA45SN1_45S_50ntFlanks | 11603 | 11620 | 18 | CCGACTGTTTAATTAAAA | MB468 | 39 | 54 | 50 | 16 | 19 | 8.5 | 34.4 | 34.7 | 1.8 | 5.4 |
| >hg38_21_+_8433172_8446622_RNA45SN1_45S_50ntFlanks@11604.11621.18\|CGACTGTTTAATTAAAAC | >hg38_21_+_8433172_8446622_RNA45SN1_45S_50ntFlanks | 11604 | 11621 | 18 | CGACTGTTTAATTAAAAC | MB468 | 39 | 30 | 62 | 38 | 27 | 9.6 | 23.6 | 18.3 | 4.0 | 2.7 |
| >hg38_21_+_8433172_8446622_RNA45SN1_45S_50ntFlanks@11612.11640.29\|TAATTAAAACAAAGCATCGCGAAGGCCCG | >hg38_21_+_8433172_8446622_RNA45SN1_45S_50ntFlanks | 11612 | 11640 | 29 | TAATTAAAACAAAGCATCGCGAAGGCCCG | MB468 | 56 | 24 | 11 | 66 | 165 | 21.8 | 8.8 | 5.0 | 25.3 | 143.0 |
| >hg38_21_+_8433172_8446622_RNA45SN1_45S_50ntFlanks@11613.11640.28\|AATTAAAACAAAGCATCGCGAAGGCCCG | >hg38_21_+_8433172_8446622_RNA45SN1_45S_50ntFlanks | 11613 | 11640 | 28 | AATTAAAACAAAGCATCGCGAAGGCCCG | MB468 | 70 | 29 | 11 | 171 | 213 | 25.3 | 12.1 | 6.3 | 48.1 | 160.1 |
| >hg38_21_+_8433172_8446622_RNA45SN1_45S_50ntFlanks@11614.11640.27\|ATTAAAACAAAGCATCGCGAAGGCCCG | >hg38_21_+_8433172_8446622_RNA45SN1_45S_50ntFlanks | 11614 | 11640 | 27 | ATTAAAACAAAGCATCGCGAAGGCCCG | MB468 | 21 | 9 | 3 | 56 | 65 | 8.4 | 3.1 | 2.6 | 16.0 | 42.0 |
| >hg38_21_+_8433172_8446622_RNA45SN1_45S_50ntFlanks@11617.11640.24\|AAAACAAAGCATCGCGAAGGCCCG | >hg38_21_+_8433172_8446622_RNA45SN1_45S_50ntFlanks | 11617 | 11640 | 24 | AAAACAAAGCATCGCGAAGGCCCG | MB468 | 12 | 5 | 2 | 61 | 42 | 4.2 | 3.4 | 1.2 | 29.6 | 25.0 |
| >hg38_21_+_8433172_8446622_RNA45SN1_45S_50ntFlanks@11618.11640.23\|AAACAAAGCATCGCGAAGGCCCG | >hg38_21_+_8433172_8446622_RNA45SN1_45S_50ntFlanks | 11618 | 11640 | 23 | AAACAAAGCATCGCGAAGGCCCG | MB468 | 18 | 8 | 3 | 107 | 73 | 7.1 | 6.3 | 1.4 | 52.0 | 44.3 |
| >hg38_21_+_8433172_8446622_RNA45SN1_45S_50ntFlanks@11619.11640.22\|AACAAAGCATCGCGAAGGCCCG | >hg38_21_+_8433172_8446622_RNA45SN1_45S_50ntFlanks | 11619 | 11640 | 22 | AACAAAGCATCGCGAAGGCCCG | MB468 | 23 | 9 | 3 | 131 | 92 | 9.1 | 6.7 | 1.8 | 64.8 | 56.5 |
| >hg38_21_+_8433172_8446622_RNA45SN1_45S_50ntFlanks@11620.11640.21\|ACAAAGCATCGCGAAGGCCCG | >hg38_21_+_8433172_8446622_RNA45SN1_45S_50ntFlanks | 11620 | 11640 | 21 | ACAAAGCATCGCGAAGGCCCG | MB468 | 39 | 14 | 6 | 213 | 169 | 17.8 | 11.0 | 2.5 | 110.8 | 111.8 |
| >hg38_21_+_8433172_8446622_RNA45SN1_45S_50ntFlanks@11621.11640.20\|CAAAGCATCGCGAAGGCCCG | >hg38_21_+_8433172_8446622_RNA45SN1_45S_50ntFlanks | 11621 | 11640 | 20 | CAAAGCATCGCGAAGGCCCG | MB468 | 43 | 13 | 6 | 182 | 182 | 19.5 | 10.4 | 3.0 | 107.6 | 119.7 |
| >hg38_21_+_8433172_8446622_RNA45SN1_45S_50ntFlanks@11641.11659.19\|CGGCGGGTGTTGACGCGAT | >hg38_21_+_8433172_8446622_RNA45SN1_45S_50ntFlanks | 11641 | 11659 | 19 | CGGCGGGTGTTGACGCGAT | MB468 | 31 | 6 | 3 | 186 | 98 | 12.8 | 4.6 | 0.9 | 111.1 | 58.8 |
| >hg38_21_+_8433172_8446622_RNA45SN1_45S_50ntFlanks@11641.11660.20\|CGGCGGGTGTTGACGCGATG | >hg38_21_+_8433172_8446622_RNA45SN1_45S_50ntFlanks | 11641 | 11660 | 20 | CGGCGGGTGTTGACGCGATG | MB468 | 1300 | 206 | 137 | 11819 | 3483 | 592.4 | 135.0 | 51.2 | 9350.3 | 2172.6 |
| >hg38_21_+_8433172_8446622_RNA45SN1_45S_50ntFlanks@11641.11661.21\|CGGCGGGTGTTGACGCGATGT | >hg38_21_+_8433172_8446622_RNA45SN1_45S_50ntFlanks | 11641 | 11661 | 21 | CGGCGGGTGTTGACGCGATGT | MB468 | 17 | 7 | 3 | 52 | 44 | 6.8 | 5.2 | 1.2 | 18.7 | 21.0 |
| >hg38_21_+_8433172_8446622_RNA45SN1_45S_50ntFlanks@11641.11662.22\|CGGCGGGTGTTGACGCGATGTG | >hg38_21_+_8433172_8446622_RNA45SN1_45S_50ntFlanks | 11641 | 11662 | 22 | CGGCGGGTGTTGACGCGATGTG | MB468 | 209 | 103 | 41 | 644 | 399 | 112.2 | 70.3 | 17.2 | 227.0 | 179.0 |
| >hg38_21_+_8433172_8446622_RNA45SN1_45S_50ntFlanks@11641.11663.23\|CGGCGGGTGTTGACGCGATGTGA | >hg38_21_+_8433172_8446622_RNA45SN1_45S_50ntFlanks | 11641 | 11663 | 23 | CGGCGGGTGTTGACGCGATGTGA | MB468 | 50 | 30 | 9 | 104 | 74 | 36.3 | 19.2 | 3.6 | 43.3 | 37.4 |
| >hg38_21_+_8433172_8446622_RNA45SN1_45S_50ntFlanks@11641.11664.24\|CGGCGGGTGTTGACGCGATGTGAT | >hg38_21_+_8433172_8446622_RNA45SN1_45S_50ntFlanks | 11641 | 11664 | 24 | CGGCGGGTGTTGACGCGATGTGAT | MB468 | 99 | 70 | 25 | 207 | 145 | 64.0 | 42.6 | 11.0 | 190.3 | 99.4 |
| >hg38_21_+_8433172_8446622_RNA45SN1_45S_50ntFlanks@11641.11665.25\|CGGCGGGTGTTGACGCGATGTGATT | >hg38_21_+_8433172_8446622_RNA45SN1_45S_50ntFlanks | 11641 | 11665 | 25 | CGGCGGGTGTTGACGCGATGTGATT | MB468 | 447 | 244 | 101 | 680 | 697 | 210.3 | 109.7 | 58.1 | 374.3 | 314.2 |
| >hg38_21_+_8433172_8446622_RNA45SN1_45S_50ntFlanks@11641.11666.26\|CGGCGGGTGTTGACGCGATGTGATTT | >hg38_21_+_8433172_8446622_RNA45SN1_45S_50ntFlanks | 11641 | 11666 | 26 | CGGCGGGTGTTGACGCGATGTGATTT | MB468 | 15 | 8 | 3 | 20 | 29 | 8.6 | 4.9 | 2.2 | 8.3 | 15.1 |
| >hg38_21_+_8433172_8446622_RNA45SN1_45S_50ntFlanks@11641.11667.27\|CGGCGGGTGTTGACGCGATGTGATTTC | >hg38_21_+_8433172_8446622_RNA45SN1_45S_50ntFlanks | 11641 | 11667 | 27 | CGGCGGGTGTTGACGCGATGTGATTTC | MB468 | 55 | 48 | 17 | 121 | 100 | 30.8 | 32.3 | 10.5 | 116.9 | 96.7 |
| >hg38_21_+_8433172_8446622_RNA45SN1_45S_50ntFlanks@11641.11668.28\|CGGCGGGTGTTGACGCGATGTGATTTCT | >hg38_21_+_8433172_8446622_RNA45SN1_45S_50ntFlanks | 11641 | 11668 | 28 | CGGCGGGTGTTGACGCGATGTGATTTCT | MB468 | 40 | 37 | 13 | 97 | 77 | 14.8 | 17.5 | 5.7 | 88.9 | 62.4 |
| >hg38_21_+_8433172_8446622_RNA45SN1_45S_50ntFlanks@11641.11669.29\|CGGCGGGTGTTGACGCGATGTGATTTCTG | >hg38_21_+_8433172_8446622_RNA45SN1_45S_50ntFlanks | 11641 | 11669 | 29 | CGGCGGGTGTTGACGCGATGTGATTTCTG | MB468 | 205 | 149 | 72 | 436 | 314 | 64.3 | 54.3 | 27.1 | 294.0 | 211.3 |
| >hg38_21_+_8433172_8446622_RNA45SN1_45S_50ntFlanks@11641.11670.30\|CGGCGGGTGTTGACGCGATGTGATTTCTGC | >hg38_21_+_8433172_8446622_RNA45SN1_45S_50ntFlanks | 11641 | 11670 | 30 | CGGCGGGTGTTGACGCGATGTGATTTCTGC | MB468 | 317 | 298 | 84 | 479 | 808 | 164.5 | 218.7 | 63.2 | 600.6 | 1028.1 |
| >hg38_21_+_8433172_8446622_RNA45SN1_45S_50ntFlanks@11641.11679.39\|CGGCGGGTGTTGACGCGATGTGATTTCTGCCCAGTGCTC | >hg38_21_+_8433172_8446622_RNA45SN1_45S_50ntFlanks | 11641 | 11679 | 39 | CGGCGGGTGTTGACGCGATGTGATTTCTGCCCAGTGCTC | MB468 | 18 | 9 | 7 | 24 | 87 | 11.5 | 5.6 | 5.9 | 19.5 | 49.9 |
| >hg38_21_+_8433172_8446622_RNA45SN1_45S_50ntFlanks@11641.11680.40\|CGGCGGGTGTTGACGCGATGTGATTTCTGCCCAGTGCTCT | >hg38_21_+_8433172_8446622_RNA45SN1_45S_50ntFlanks | 11641 | 11680 | 40 | CGGCGGGTGTTGACGCGATGTGATTTCTGCCCAGTGCTCT | MB468 | 22 | 15 | 9 | 16 | 138 | 22.0 | 13.9 | 9.1 | 18.7 | 147.6 |
| >hg38_21_+_8433172_8446622_RNA45SN1_45S_50ntFlanks@11641.11681.41\|CGGCGGGTGTTGACGCGATGTGATTTCTGCCCAGTGCTCTG | >hg38_21_+_8433172_8446622_RNA45SN1_45S_50ntFlanks | 11641 | 11681 | 41 | CGGCGGGTGTTGACGCGATGTGATTTCTGCCCAGTGCTCTG | MB468 | 16 | 7 | 8 | 21 | 91 | 12.3 | 4.0 | 5.5 | 16.6 | 58.8 |
| >hg38_21_+_8433172_8446622_RNA45SN1_45S_50ntFlanks@11641.11685.45\|CGGCGGGTGTTGACGCGATGTGATTTCTGCCCAGTGCTCTGAATG | >hg38_21_+_8433172_8446622_RNA45SN1_45S_50ntFlanks | 11641 | 11685 | 45 | CGGCGGGTGTTGACGCGATGTGATTTCTGCCCAGTGCTCTGAATG | MB468 | 33 | 112 | 39 | 44 | 522 | 20.4 | 99.2 | 27.7 | 48.1 | 365.5 |
| >hg38_21_+_8433172_8446622_RNA45SN1_45S_50ntFlanks@11643.11660.18\|GCGGGTGTTGACGCGATG | >hg38_21_+_8433172_8446622_RNA45SN1_45S_50ntFlanks | 11643 | 11660 | 18 | GCGGGTGTTGACGCGATG | MB468 | 26 | 4 | 3 | 314 | 59 | 15.3 | 3.4 | 1.5 | 323.9 | 43.6 |
| >hg38_21_+_8433172_8446622_RNA45SN1_45S_50ntFlanks@11660.11685.26\|GTGATTTCTGCCCAGTGCTCTGAATG | >hg38_21_+_8433172_8446622_RNA45SN1_45S_50ntFlanks | 11660 | 11685 | 26 | GTGATTTCTGCCCAGTGCTCTGAATG | MB468 | 30 | 57 | 32 | 43 | 195 | 7.2 | 11.6 | 6.5 | 14.7 | 149.8 |
| >hg38_21_+_8433172_8446622_RNA45SN1_45S_50ntFlanks@11661.11678.18\|TGATTTCTGCCCAGTGCT | >hg38_21_+_8433172_8446622_RNA45SN1_45S_50ntFlanks | 11661 | 11678 | 18 | TGATTTCTGCCCAGTGCT | MB468 | 34 | 9 | 31 | 6 | 5 | 13.8 | 4.0 | 22.9 | 3.4 | 3.4 |
| >hg38_21_+_8433172_8446622_RNA45SN1_45S_50ntFlanks@11661.11679.19\|TGATTTCTGCCCAGTGCTC | >hg38_21_+_8433172_8446622_RNA45SN1_45S_50ntFlanks | 11661 | 11679 | 19 | TGATTTCTGCCCAGTGCTC | MB468 | 227 | 158 | 255 | 14 | 43 | 126.5 | 92.6 | 267.4 | 6.7 | 8.5 |
| >hg38_21_+_8433172_8446622_RNA45SN1_45S_50ntFlanks@11661.11680.20\|TGATTTCTGCCCAGTGCTCT | >hg38_21_+_8433172_8446622_RNA45SN1_45S_50ntFlanks | 11661 | 11680 | 20 | TGATTTCTGCCCAGTGCTCT | MB468 | 62 | 24 | 49 | 5 | 13 | 53.0 | 8.4 | 54.1 | 4.1 | 4.1 |
| >hg38_21_+_8433172_8446622_RNA45SN1_45S_50ntFlanks@11661.11681.21\|TGATTTCTGCCCAGTGCTCTG | >hg38_21_+_8433172_8446622_RNA45SN1_45S_50ntFlanks | 11661 | 11681 | 21 | TGATTTCTGCCCAGTGCTCTG | MB468 | 444 | 535 | 453 | 71 | 159 | 122.4 | 267.2 | 217.6 | 58.5 | 91.5 |
| >hg38_21_+_8433172_8446622_RNA45SN1_45S_50ntFlanks@11661.11682.22\|TGATTTCTGCCCAGTGCTCTGA | >hg38_21_+_8433172_8446622_RNA45SN1_45S_50ntFlanks | 11661 | 11682 | 22 | TGATTTCTGCCCAGTGCTCTGA | MB468 | 67 | 59 | 58 | 15 | 13 | 17.7 | 10.0 | 16.8 | 5.2 | 2.0 |
| >hg38_21_+_8433172_8446622_RNA45SN1_45S_50ntFlanks@11661.11684.24\|TGATTTCTGCCCAGTGCTCTGAAT | >hg38_21_+_8433172_8446622_RNA45SN1_45S_50ntFlanks | 11661 | 11684 | 24 | TGATTTCTGCCCAGTGCTCTGAAT | MB468 | 16 | 8 | 15 | 8 | 6 | 7.9 | 2.1 | 7.3 | 3.0 | 0.7 |
| >hg38_21_+_8433172_8446622_RNA45SN1_45S_50ntFlanks@11661.11685.25\|TGATTTCTGCCCAGTGCTCTGAATG | >hg38_21_+_8433172_8446622_RNA45SN1_45S_50ntFlanks | 11661 | 11685 | 25 | TGATTTCTGCCCAGTGCTCTGAATG | MB468 | 582 | 501 | 569 | 318 | 367 | 145.2 | 103.0 | 195.7 | 93.7 | 99.3 |
| >hg38_21_+_8433172_8446622_RNA45SN1_45S_50ntFlanks@11661.11687.27\|TGATTTCTGCCCAGTGCTCTGAATGTC | >hg38_21_+_8433172_8446622_RNA45SN1_45S_50ntFlanks | 11661 | 11687 | 27 | TGATTTCTGCCCAGTGCTCTGAATGTC | MB468 | 13 | 13 | 13 | 8 | 6 | 5.6 | 4.7 | 10.1 | 3.2 | 2.8 |

| | | | | | | | | | | | | | | | | |
|---|---|---|---|---|---|---|---|---|---|---|---|---|---|---|---|---|
| >hg38_21_+_8433172_8446622_RNA45SN1_45S_50ntFlanks@11662.11681.20\|GATTTCTGCCCAGTGCTCTG | >hg38_21_+_8433172_8446622_RNA45SN1_45S_50ntFlanks | 11662 | 11681 | 20 | GATTTCTGCCCAGTGCTCTG | MB468 | 13 | 18 | 13 | 4 | 9 | 2.7 | 8.8 | 3.0 | 3.3 | 10.2 |
| >hg38_21_+_8433172_8446622_RNA45SN1_45S_50ntFlanks@11662.11685.24\|GATTTCTGCCCAGTGCTCTGAATG | >hg38_21_+_8433172_8446622_RNA45SN1_45S_50ntFlanks | 11662 | 11685 | 24 | GATTTCTGCCCAGTGCTCTGAATG | MB468 | 11 | 11 | 9 | 12 | 9 | 2.7 | 1.4 | 3.9 | 5.8 | 2.2 |
| >hg38_21_+_8433172_8446622_RNA45SN1_45S_50ntFlanks@11663.11681.19\|ATTTCTGCCCAGTGCTCTG | >hg38_21_+_8433172_8446622_RNA45SN1_45S_50ntFlanks | 11663 | 11681 | 19 | ATTTCTGCCCAGTGCTCTG | MB468 | 17 | 6 | 21 | 7 | 4 | 5.3 | 2.0 | 14.4 | 5.3 | 1.3 |
| >hg38_21_+_8433172_8446622_RNA45SN1_45S_50ntFlanks@11663.11685.23\|ATTTCTGCCCAGTGCTCTGAATG | >hg38_21_+_8433172_8446622_RNA45SN1_45S_50ntFlanks | 11663 | 11685 | 23 | ATTTCTGCCCAGTGCTCTGAATG | MB468 | 57 | 27 | 55 | 76 | 32 | 21.3 | 15.3 | 17.9 | 84.6 | 22.7 |
| >hg38_21_+_8433172_8446622_RNA45SN1_45S_50ntFlanks@11664.11685.22\|TTTCTGCCCAGTGCTCTGAATG | >hg38_21_+_8433172_8446622_RNA45SN1_45S_50ntFlanks | 11664 | 11685 | 22 | TTTCTGCCCAGTGCTCTGAATG | MB468 | 19 | 13 | 26 | 21 | 12 | 9.9 | 7.1 | 14.7 | 16.0 | 7.1 |
| >hg38_21_+_8433172_8446622_RNA45SN1_45S_50ntFlanks@11665.11685.21\|TTCTGCCCAGTGCTCTGAATG | >hg38_21_+_8433172_8446622_RNA45SN1_45S_50ntFlanks | 11665 | 11685 | 21 | TTCTGCCCAGTGCTCTGAATG | MB468 | 30 | 18 | 37 | 25 | 18 | 13.0 | 2.4 | 14.3 | 8.6 | 4.0 |
| >hg38_21_+_8433172_8446622_RNA45SN1_45S_50ntFlanks@11665.11687.23\|TTCTGCCCAGTGCTCTGAATGTC | >hg38_21_+_8433172_8446622_RNA45SN1_45S_50ntFlanks | 11665 | 11687 | 23 | TTCTGCCCAGTGCTCTGAATGTC | MB231 | 22 | 18 | 42 | 5 | 6 | 17.5 | 11.7 | 30.6 | 2.7 | 3.2 |
| >hg38_21_+_8433172_8446622_RNA45SN1_45S_50ntFlanks@11665.11702.38\|TTCTGCCCAGTGCTCTGAATGTCAAAGTGAAGAAATTC | >hg38_21_+_8433172_8446622_RNA45SN1_45S_50ntFlanks | 11665 | 11702 | 38 | TTCTGCCCAGTGCTCTGAATGTCAAAGTGAAGAAATTC | MB231 | 19 | 23 | 91 | 2 | 6 | 16.5 | 20.9 | 84.1 | 3.0 | 5.1 |
| >hg38_21_+_8433172_8446622_RNA45SN1_45S_50ntFlanks@11666.11684.19\|TCTGCCCAGTGCTCTGAAT | >hg38_21_+_8433172_8446622_RNA45SN1_45S_50ntFlanks | 11666 | 11684 | 19 | TCTGCCCAGTGCTCTGAAT | BT20 | 14 | 26 | 6 | 13 | 4 | 17.3 | 42.3 | 5.8 | 13.7 | 2.0 |
| >hg38_21_+_8433172_8446622_RNA45SN1_45S_50ntFlanks@11666.11685.20\|TCTGCCCAGTGCTCTGAATG | >hg38_21_+_8433172_8446622_RNA45SN1_45S_50ntFlanks | 11666 | 11685 | 20 | TCTGCCCAGTGCTCTGAATG | BT20 | 24 | 41 | 11 | 23 | 6 | 33.8 | 65.7 | 12.9 | 14.1 | 1.6 |
| >hg38_21_+_8433172_8446622_RNA45SN1_45S_50ntFlanks@11666.11685.20\|TCTGCCCAGTGCTCTGAATG | >hg38_21_+_8433172_8446622_RNA45SN1_45S_50ntFlanks | 11666 | 11685 | 20 | TCTGCCCAGTGCTCTGAATG | MB231 | 13 | 12 | 14 | 4 | 4 | 10.8 | 9.1 | 11.3 | 0.7 | 0.7 |
| >hg38_21_+_8433172_8446622_RNA45SN1_45S_50ntFlanks@11666.11685.20\|TCTGCCCAGTGCTCTGAATG | >hg38_21_+_8433172_8446622_RNA45SN1_45S_50ntFlanks | 11666 | 11685 | 20 | TCTGCCCAGTGCTCTGAATG | MB468 | 175 | 101 | 254 | 177 | 176 | 97.8 | 21.2 | 173.8 | 108.5 | 116.6 |
| >hg38_21_+_8433172_8446622_RNA45SN1_45S_50ntFlanks@11666.11686.21\|TCTGCCCAGTGCTCTGAATGT | >hg38_21_+_8433172_8446622_RNA45SN1_45S_50ntFlanks | 11666 | 11686 | 21 | TCTGCCCAGTGCTCTGAATGT | BT20 | 14 | 18 | 4 | 6 | 1 | 22.6 | 30.7 | 5.7 | 7.4 | 1.2 |
| >hg38_21_+_8433172_8446622_RNA45SN1_45S_50ntFlanks@11666.11687.22\|TCTGCCCAGTGCTCTGAATGTC | >hg38_21_+_8433172_8446622_RNA45SN1_45S_50ntFlanks | 11666 | 11687 | 22 | TCTGCCCAGTGCTCTGAATGTC | BT20 | 12 | 8 | 28 | 35 | 9 | 6.2 | 3.0 | 14.2 | 46.5 | 2.1 |
| >hg38_21_+_8433172_8446622_RNA45SN1_45S_50ntFlanks@11666.11687.22\|TCTGCCCAGTGCTCTGAATGTC | >hg38_21_+_8433172_8446622_RNA45SN1_45S_50ntFlanks | 11666 | 11687 | 22 | TCTGCCCAGTGCTCTGAATGTC | MB231 | 38 | 38 | 94 | 10 | 9 | 28.3 | 23.2 | 61.2 | 6.3 | 5.3 |
| >hg38_21_+_8433172_8446622_RNA45SN1_45S_50ntFlanks@11666.11688.23\|TCTGCCCAGTGCTCTGAATGTCA | >hg38_21_+_8433172_8446622_RNA45SN1_45S_50ntFlanks | 11666 | 11688 | 23 | TCTGCCCAGTGCTCTGAATGTCA | BT20 | 12 | 10 | 10 | 3 | 4 | 17.0 | 16.8 | 9.4 | 1.2 | 1.1 |
| >hg38_21_+_8433172_8446622_RNA45SN1_45S_50ntFlanks@11666.11688.23\|TCTGCCCAGTGCTCTGAATGTCA | >hg38_21_+_8433172_8446622_RNA45SN1_45S_50ntFlanks | 11666 | 11688 | 23 | TCTGCCCAGTGCTCTGAATGTCA | MB231 | 11 | 10 | 20 | 2 | 3 | 8.4 | 6.9 | 15.5 | 1.5 | 2.8 |
| >hg38_21_+_8433172_8446622_RNA45SN1_45S_50ntFlanks@11666.11690.25\|TCTGCCCAGTGCTCTGAATGTCAAA | >hg38_21_+_8433172_8446622_RNA45SN1_45S_50ntFlanks | 11666 | 11690 | 25 | TCTGCCCAGTGCTCTGAATGTCAAA | BT20 | 18 | 12 | 13 | 4 | 6 | 28.9 | 19.0 | 17.6 | 2.7 | 5.0 |
| >hg38_21_+_8433172_8446622_RNA45SN1_45S_50ntFlanks@11666.11691.26\|TCTGCCCAGTGCTCTGAATGTCAAAG | >hg38_21_+_8433172_8446622_RNA45SN1_45S_50ntFlanks | 11666 | 11691 | 26 | TCTGCCCAGTGCTCTGAATGTCAAAG | BT20 | 16 | 14 | 18 | 10 | 15 | 19.6 | 15.1 | 19.0 | 8.6 | 8.2 |
| >hg38_21_+_8433172_8446622_RNA45SN1_45S_50ntFlanks@11666.11692.27\|TCTGCCCAGTGCTCTGAATGTCAAAGT | >hg38_21_+_8433172_8446622_RNA45SN1_45S_50ntFlanks | 11666 | 11692 | 27 | TCTGCCCAGTGCTCTGAATGTCAAAGT | BT20 | 13 | 12 | 29 | 13 | 14 | 13.0 | 9.7 | 42.6 | 9.3 | 9.3 |
| >hg38_21_+_8433172_8446622_RNA45SN1_45S_50ntFlanks@11666.11693.28\|TCTGCCCAGTGCTCTGAATGTCAAAGTG | >hg38_21_+_8433172_8446622_RNA45SN1_45S_50ntFlanks | 11666 | 11693 | 28 | TCTGCCCAGTGCTCTGAATGTCAAAGTG | MB231 | 11 | 10 | 22 | 3 | 5 | 8.8 | 7.8 | 19.0 | 1.0 | 3.2 |
| >hg38_21_+_8433172_8446622_RNA45SN1_45S_50ntFlanks@11666.11697.32\|TCTGCCCAGTGCTCTGAATGTCAAAGTGAAGA | >hg38_21_+_8433172_8446622_RNA45SN1_45S_50ntFlanks | 11666 | 11697 | 32 | TCTGCCCAGTGCTCTGAATGTCAAAGTGAAGA | MB231 | 10 | 11 | 11 | 3 | 4 | 6.9 | 5.2 | 8.7 | 1.3 | 2.5 |
| >hg38_21_+_8433172_8446622_RNA45SN1_45S_50ntFlanks@11666.11702.37\|TCTGCCCAGTGCTCTGAATGTCAAAGTGAAGAAATTC | >hg38_21_+_8433172_8446622_RNA45SN1_45S_50ntFlanks | 11666 | 11702 | 37 | TCTGCCCAGTGCTCTGAATGTCAAAGTGAAGAAATTC | MB231 | 29 | 26 | 109 | 5 | 7 | 26.0 | 19.9 | 74.7 | 5.5 | 5.6 |
| >hg38_21_+_8433172_8446622_RNA45SN1_45S_50ntFlanks@11666.11706.41\|TCTGCCCAGTGCTCTGAATGTCAAAGTGAAGAAATTCAATG | >hg38_21_+_8433172_8446622_RNA45SN1_45S_50ntFlanks | 11666 | 11706 | 41 | TCTGCCCAGTGCTCTGAATGTCAAAGTGAAGAAATTCAATG | MB231 | 12 | 14 | 80 | 2 | 2 | 8.2 | 17.9 | 80.4 | 2.6 | 3.0 |
| >hg38_21_+_8433172_8446622_RNA45SN1_45S_50ntFlanks@11667.11685.19\|CTGCCCAGTGCTCTGAATG | >hg38_21_+_8433172_8446622_RNA45SN1_45S_50ntFlanks | 11667 | 11685 | 19 | CTGCCCAGTGCTCTGAATG | MB468 | 41 | 36 | 51 | 32 | 38 | 19.3 | 12.9 | 39.9 | 23.6 | 17.3 |
| >hg38_21_+_8433172_8446622_RNA45SN1_45S_50ntFlanks@11667.11687.21\|CTGCCCAGTGCTCTGAATGTC | >hg38_21_+_8433172_8446622_RNA45SN1_45S_50ntFlanks | 11667 | 11687 | 21 | CTGCCCAGTGCTCTGAATGTC | MB231 | 26 | 26 | 64 | 6 | 8 | 21.0 | 15.5 | 45.0 | 3.8 | 5.5 |
| >hg38_21_+_8433172_8446622_RNA45SN1_45S_50ntFlanks@11667.11702.36\|CTGCCCAGTGCTCTGAATGTCAAAGTGAAGAAATTC | >hg38_21_+_8433172_8446622_RNA45SN1_45S_50ntFlanks | 11667 | 11702 | 36 | CTGCCCAGTGCTCTGAATGTCAAAGTGAAGAAATTC | MB231 | 20 | 20 | 123 | 3 | 7 | 16.7 | 15.8 | 131.3 | 2.9 | 5.3 |
| >hg38_21_+_8433172_8446622_RNA45SN1_45S_50ntFlanks@11668.11685.18\|TGCCCAGTGCTCTGAATG | >hg38_21_+_8433172_8446622_RNA45SN1_45S_50ntFlanks | 11668 | 11685 | 18 | TGCCCAGTGCTCTGAATG | MB231 | 31 | 33 | 24 | 6 | 8 | 29.0 | 32.1 | 24.7 | 3.3 | 6.9 |
| >hg38_21_+_8433172_8446622_RNA45SN1_45S_50ntFlanks@11668.11685.18\|TGCCCAGTGCTCTGAATG | >hg38_21_+_8433172_8446622_RNA45SN1_45S_50ntFlanks | 11668 | 11685 | 18 | TGCCCAGTGCTCTGAATG | MB468 | 46 | 33 | 37 | 27 | 25 | 17.0 | 10.7 | 11.8 | 6.0 | 6.4 |
| >hg38_21_+_8433172_8446622_RNA45SN1_45S_50ntFlanks@11668.11686.19\|TGCCCAGTGCTCTGAATGT | >hg38_21_+_8433172_8446622_RNA45SN1_45S_50ntFlanks | 11668 | 11686 | 19 | TGCCCAGTGCTCTGAATGT | MB231 | 15 | 12 | 13 | 1 | 3 | 15.4 | 12.7 | 15.1 | 1.7 | 4.2 |
| >hg38_21_+_8433172_8446622_RNA45SN1_45S_50ntFlanks@11668.11687.20\|TGCCCAGTGCTCTGAATGTC | >hg38_21_+_8433172_8446622_RNA45SN1_45S_50ntFlanks | 11668 | 11687 | 20 | TGCCCAGTGCTCTGAATGTC | BT20 | 23 | 11 | 34 | 29 | 10 | 12.2 | 2.7 | 15.0 | 38.3 | 2.8 |
| >hg38_21_+_8433172_8446622_RNA45SN1_45S_50ntFlanks@11668.11687.20\|TGCCCAGTGCTCTGAATGTC | >hg38_21_+_8433172_8446622_RNA45SN1_45S_50ntFlanks | 11668 | 11687 | 20 | TGCCCAGTGCTCTGAATGTC | MB231 | 90 | 67 | 139 | 19 | 23 | 72.7 | 50.7 | 106.8 | 13.2 | 18.0 |
| >hg38_21_+_8433172_8446622_RNA45SN1_45S_50ntFlanks@11668.11688.21\|TGCCCAGTGCTCTGAATGTCA | >hg38_21_+_8433172_8446622_RNA45SN1_45S_50ntFlanks | 11668 | 11688 | 21 | TGCCCAGTGCTCTGAATGTCA | MB231 | 38 | 25 | 44 | 4 | 13 | 34.2 | 18.7 | 38.3 | 2.9 | 9.2 |
| >hg38_21_+_8433172_8446622_RNA45SN1_45S_50ntFlanks@11668.11691.24\|TGCCCAGTGCTCTGAATGTCAAAG | >hg38_21_+_8433172_8446622_RNA45SN1_45S_50ntFlanks | 11668 | 11691 | 24 | TGCCCAGTGCTCTGAATGTCAAAG | MB231 | 20 | 21 | 32 | 5 | 17 | 16.3 | 13.5 | 28.0 | 4.6 | 11.5 |
| >hg38_21_+_8433172_8446622_RNA45SN1_45S_50ntFlanks@11668.11692.25\|TGCCCAGTGCTCTGAATGTCAAAGT | >hg38_21_+_8433172_8446622_RNA45SN1_45S_50ntFlanks | 11668 | 11692 | 25 | TGCCCAGTGCTCTGAATGTCAAAGT | MB231 | 12 | 14 | 23 | 3 | 9 | 8.2 | 8.2 | 15.0 | 2.8 | 9.0 |
| >hg38_21_+_8433172_8446622_RNA45SN1_45S_50ntFlanks@11668.11693.26\|TGCCCAGTGCTCTGAATGTCAAAGTG | >hg38_21_+_8433172_8446622_RNA45SN1_45S_50ntFlanks | 11668 | 11693 | 26 | TGCCCAGTGCTCTGAATGTCAAAGTG | MB231 | 30 | 25 | 43 | 6 | 14 | 24.2 | 22.6 | 41.2 | 3.6 | 12.4 |
| >hg38_21_+_8433172_8446622_RNA45SN1_45S_50ntFlanks@11668.11701.34\|TGCCCAGTGCTCTGAATGTCAAAGTGAAGAAATT | >hg38_21_+_8433172_8446622_RNA45SN1_45S_50ntFlanks | 11668 | 11701 | 34 | TGCCCAGTGCTCTGAATGTCAAAGTGAAGAAATT | MB231 | 11 | 10 | 29 | 2 | 5 | 8.2 | 9.4 | 26.7 | 1.3 | 6.4 |
| >hg38_21_+_8433172_8446622_RNA45SN1_45S_50ntFlanks@11668.11701.34\|TGCCCAGTGCTCTGAATGTCAAAGTGAAGAAATT | >hg38_21_+_8433172_8446622_RNA45SN1_45S_50ntFlanks | 11668 | 11701 | 34 | TGCCCAGTGCTCTGAATGTCAAAGTGAAGAAATT | MB468 | 14 | 58 | 10 | 2 | 6 | 6.9 | 21.4 | 3.7 | 1.5 | 1.9 |
| >hg38_21_+_8433172_8446622_RNA45SN1_45S_50ntFlanks@11668.11702.35\|TGCCCAGTGCTCTGAATGTCAAAGTGAAGAAATTC | >hg38_21_+_8433172_8446622_RNA45SN1_45S_50ntFlanks | 11668 | 11702 | 35 | TGCCCAGTGCTCTGAATGTCAAAGTGAAGAAATTC | BT20 | 18 | 11 | 54 | 15 | 15 | 15.9 | 7.1 | 19.1 | 16.4 | 5.1 |
| >hg38_21_+_8433172_8446622_RNA45SN1_45S_50ntFlanks@11668.11702.35\|TGCCCAGTGCTCTGAATGTCAAAGTGAAGAAATTC | >hg38_21_+_8433172_8446622_RNA45SN1_45S_50ntFlanks | 11668 | 11702 | 35 | TGCCCAGTGCTCTGAATGTCAAAGTGAAGAAATTC | MB231 | 85 | 84 | 386 | 12 | 27 | 75.5 | 74.3 | 397.2 | 12.3 | 20.7 |
| >hg38_21_+_8433172_8446622_RNA45SN1_45S_50ntFlanks@11668.11702.35\|TGCCCAGTGCTCTGAATGTCAAAGTGAAGAAATTC | >hg38_21_+_8433172_8446622_RNA45SN1_45S_50ntFlanks | 11668 | 11702 | 35 | TGCCCAGTGCTCTGAATGTCAAAGTGAAGAAATTC | MB468 | 30 | 210 | 44 | 15 | 57 | 19.2 | 129.1 | 21.2 | 10.4 | 21.0 |
| >hg38_21_+_8433172_8446622_RNA45SN1_45S_50ntFlanks@11668.11703.36\|TGCCCAGTGCTCTGAATGTCAAAGTGAAGAAATTCA | >hg38_21_+_8433172_8446622_RNA45SN1_45S_50ntFlanks | 11668 | 11703 | 36 | TGCCCAGTGCTCTGAATGTCAAAGTGAAGAAATTCA | MB231 | 35 | 30 | 121 | 4 | 12 | 28.9 | 20.9 | 89.5 | 2.4 | 9.1 |
| >hg38_21_+_8433172_8446622_RNA45SN1_45S_50ntFlanks@11668.11704.37\|TGCCCAGTGCTCTGAATGTCAAAGTGAAGAAATTCAA | >hg38_21_+_8433172_8446622_RNA45SN1_45S_50ntFlanks | 11668 | 11704 | 37 | TGCCCAGTGCTCTGAATGTCAAAGTGAAGAAATTCAA | MB231 | 10 | 10 | 34 | 1 | 4 | 9.0 | 7.3 | 23.3 | 1.3 | 3.7 |
| >hg38_21_+_8433172_8446622_RNA45SN1_45S_50ntFlanks@11668.11706.39\|TGCCCAGTGCTCTGAATGTCAAAGTGAAGAAATTCAATG | >hg38_21_+_8433172_8446622_RNA45SN1_45S_50ntFlanks | 11668 | 11706 | 39 | TGCCCAGTGCTCTGAATGTCAAAGTGAAGAAATTCAATG | MB231 | 38 | 42 | 182 | 6 | 9 | 28.1 | 43.6 | 136.1 | 7.2 | 6.2 |
| >hg38_21_+_8433172_8446622_RNA45SN1_45S_50ntFlanks@11668.11707.40\|TGCCCAGTGCTCTGAATGTCAAAGTGAAGAAATTCAATGA | >hg38_21_+_8433172_8446622_RNA45SN1_45S_50ntFlanks | 11668 | 11707 | 40 | TGCCCAGTGCTCTGAATGTCAAAGTGAAGAAATTCAATGA | MB231 | 24 | 32 | 109 | 3 | 6 | 17.5 | 37.7 | 72.3 | 3.5 | 6.1 |
| >hg38_21_+_8433172_8446622_RNA45SN1_45S_50ntFlanks@11668.11709.42\|TGCCCAGTGCTCTGAATGTCAAAGTGAAGAAATTCAATGAAG | >hg38_21_+_8433172_8446622_RNA45SN1_45S_50ntFlanks | 11668 | 11709 | 42 | TGCCCAGTGCTCTGAATGTCAAAGTGAAGAAATTCAATGAAG | MB231 | 10 | 25 | 53 | 1 | 2 | 7.2 | 35.4 | 51.5 | 1.3 | 3.7 |
| >hg38_21_+_8433172_8446622_RNA45SN1_45S_50ntFlanks@11669.11686.18\|GCCCAGTGCTCTGAATGT | >hg38_21_+_8433172_8446622_RNA45SN1_45S_50ntFlanks | 11669 | 11686 | 18 | GCCCAGTGCTCTGAATGT | MB231 | 15 | 12 | 15 | 2 | 3 | 17.0 | 12.6 | 19.3 | 2.6 | 3.5 |
| >hg38_21_+_8433172_8446622_RNA45SN1_45S_50ntFlanks@11669.11687.19\|GCCCAGTGCTCTGAATGTC | >hg38_21_+_8433172_8446622_RNA45SN1_45S_50ntFlanks | 11669 | 11687 | 19 | GCCCAGTGCTCTGAATGTC | BT20 | 21 | 12 | 35 | 28 | 12 | 12.6 | 6.3 | 17.1 | 37.3 | 4.1 |
| >hg38_21_+_8433172_8446622_RNA45SN1_45S_50ntFlanks@11669.11687.19\|GCCCAGTGCTCTGAATGTC | >hg38_21_+_8433172_8446622_RNA45SN1_45S_50ntFlanks | 11669 | 11687 | 19 | GCCCAGTGCTCTGAATGTC | MB231 | 84 | 64 | 163 | 14 | 21 | 72.9 | 47.5 | 134.0 | 9.8 | 16.3 |
| >hg38_21_+_8433172_8446622_RNA45SN1_45S_50ntFlanks@11669.11688.20\|GCCCAGTGCTCTGAATGTCA | >hg38_21_+_8433172_8446622_RNA45SN1_45S_50ntFlanks | 11669 | 11688 | 20 | GCCCAGTGCTCTGAATGTCA | MB231 | 24 | 20 | 40 | 4 | 9 | 22.3 | 14.6 | 34.9 | 3.3 | 7.5 |

| | | | | | | | | | | | | | | | |
|---|---|---|---|---|---|---|---|---|---|---|---|---|---|---|---|
| >hg38_21_+_8433172_8446622_RNA45SN1_45S_50ntFlanks@11669.11691.23\|GCCCAGTGCTCTGAATGTCAAAG | >hg38_21_+_8433172_8446622_RNA45SN1_45S_50ntFlanks | 11669 | 11691 | 23 | GCCCAGTGCTCTGAATGTCAAAG | MB231 | 13 | 18 | 25 | 4 | 13 | 10.0 | 10.2 | 17.6 | 2.2 | 9.1 |
| >hg38_21_+_8433172_8446622_RNA45SN1_45S_50ntFlanks@11669.11692.24\|GCCCAGTGCTCTGAATGTCAAAGT | >hg38_21_+_8433172_8446622_RNA45SN1_45S_50ntFlanks | 11669 | 11692 | 24 | GCCCAGTGCTCTGAATGTCAAAGT | MB231 | 11 | 14 | 19 | 2 | 8 | 5.7 | 3.5 | 10.8 | 2.2 | 6.5 |
| >hg38_21_+_8433172_8446622_RNA45SN1_45S_50ntFlanks@11669.11693.25\|GCCCAGTGCTCTGAATGTCAAAGTG | >hg38_21_+_8433172_8446622_RNA45SN1_45S_50ntFlanks | 11669 | 11693 | 25 | GCCCAGTGCTCTGAATGTCAAAGTG | MB231 | 22 | 20 | 39 | 4 | 10 | 18.1 | 18.1 | 39.1 | 4.7 | 9.6 |
| >hg38_21_+_8433172_8446622_RNA45SN1_45S_50ntFlanks@11669.11702.34\|GCCCAGTGCTCTGAATGTCAAAGTGAAGAAATTC | >hg38_21_+_8433172_8446622_RNA45SN1_45S_50ntFlanks | 11669 | 11702 | 34 | GCCCAGTGCTCTGAATGTCAAAGTGAAGAAATTC | BT20 | 16 | 10 | 44 | 11 | 16 | 14.4 | 8.4 | 21.8 | 14.7 | 7.0 |
| >hg38_21_+_8433172_8446622_RNA45SN1_45S_50ntFlanks@11669.11702.34\|GCCCAGTGCTCTGAATGTCAAAGTGAAGAAATTC | >hg38_21_+_8433172_8446622_RNA45SN1_45S_50ntFlanks | 11669 | 11702 | 34 | GCCCAGTGCTCTGAATGTCAAAGTGAAGAAATTC | MB231 | 69 | 74 | 307 | 7 | 22 | 59.8 | 60.4 | 303.8 | 5.4 | 17.7 |
| >hg38_21_+_8433172_8446622_RNA45SN1_45S_50ntFlanks@11669.11703.35\|GCCCAGTGCTCTGAATGTCAAAGTGAAGAAATTCA | >hg38_21_+_8433172_8446622_RNA45SN1_45S_50ntFlanks | 11669 | 11703 | 35 | GCCCAGTGCTCTGAATGTCAAAGTGAAGAAATTCA | MB231 | 23 | 19 | 86 | 2 | 8 | 18.4 | 12.3 | 64.1 | 1.7 | 7.5 |
| >hg38_21_+_8433172_8446622_RNA45SN1_45S_50ntFlanks@11669.11706.38\|GCCCAGTGCTCTGAATGTCAAAGTGAAGAAATTCAATG | >hg38_21_+_8433172_8446622_RNA45SN1_45S_50ntFlanks | 11669 | 11706 | 38 | GCCCAGTGCTCTGAATGTCAAAGTGAAGAAATTCAATG | MB231 | 31 | 31 | 176 | 3 | 8 | 22.4 | 26.0 | 136.0 | 3.6 | 6.7 |
| >hg38_21_+_8433172_8446622_RNA45SN1_45S_50ntFlanks@11669.11707.39\|GCCCAGTGCTCTGAATGTCAAAGTGAAGAAATTCAATGA | >hg38_21_+_8433172_8446622_RNA45SN1_45S_50ntFlanks | 11669 | 11707 | 39 | GCCCAGTGCTCTGAATGTCAAAGTGAAGAAATTCAATGA | MB231 | 18 | 16 | 97 | 2 | 4 | 13.7 | 13.7 | 73.8 | 2.4 | 4.0 |
| >hg38_21_+_8433172_8446622_RNA45SN1_45S_50ntFlanks@11669.11709.41\|GCCCAGTGCTCTGAATGTCAAAGTGAAGAAATTCAATGAAG | >hg38_21_+_8433172_8446622_RNA45SN1_45S_50ntFlanks | 11669 | 11709 | 41 | GCCCAGTGCTCTGAATGTCAAAGTGAAGAAATTCAATGAAG | MB231 | 11 | 20 | 60 | 0 | 3 | 6.8 | 25.1 | 57.5 | 0.5 | 2.4 |
| >hg38_21_+_8433172_8446622_RNA45SN1_45S_50ntFlanks@11670.11687.18\|CCCAGTGCTCTGAATGTC | >hg38_21_+_8433172_8446622_RNA45SN1_45S_50ntFlanks | 11670 | 11687 | 18 | CCCAGTGCTCTGAATGTC | MB468 | 77 | 73 | 93 | 38 | 22 | 21.0 | 20.5 | 60.5 | 13.7 | 8.3 |
| >hg38_21_+_8433172_8446622_RNA45SN1_45S_50ntFlanks@11670.11688.19\|CCCAGTGCTCTGAATGTCA | >hg38_21_+_8433172_8446622_RNA45SN1_45S_50ntFlanks | 11670 | 11688 | 19 | CCCAGTGCTCTGAATGTCA | MB468 | 63 | 61 | 51 | 15 | 12 | 36.9 | 23.5 | 56.2 | 5.3 | 4.2 |
| >hg38_21_+_8433172_8446622_RNA45SN1_45S_50ntFlanks@11670.11690.21\|CCCAGTGCTCTGAATGTCAAA | >hg38_21_+_8433172_8446622_RNA45SN1_45S_50ntFlanks | 11670 | 11690 | 21 | CCCAGTGCTCTGAATGTCAAA | MB468 | 11 | 14 | 11 | 4 | 6 | 2.2 | 5.5 | 2.1 | 0.9 | 1.7 |
| >hg38_21_+_8433172_8446622_RNA45SN1_45S_50ntFlanks@11670.11691.22\|CCCAGTGCTCTGAATGTCAAAG | >hg38_21_+_8433172_8446622_RNA45SN1_45S_50ntFlanks | 11670 | 11691 | 22 | CCCAGTGCTCTGAATGTCAAAG | MB468 | 44 | 91 | 32 | 34 | 60 | 8.4 | 49.6 | 12.8 | 15.2 | 68.0 |
| >hg38_21_+_8433172_8446622_RNA45SN1_45S_50ntFlanks@11670.11692.23\|CCCAGTGCTCTGAATGTCAAAGT | >hg38_21_+_8433172_8446622_RNA45SN1_45S_50ntFlanks | 11670 | 11692 | 23 | CCCAGTGCTCTGAATGTCAAAGT | MB468 | 21 | 39 | 15 | 8 | 5 | 2.3 | 14.3 | 3.5 | 2.5 | 1.4 |
| >hg38_21_+_8433172_8446622_RNA45SN1_45S_50ntFlanks@11670.11693.24\|CCCAGTGCTCTGAATGTCAAAGTG | >hg38_21_+_8433172_8446622_RNA45SN1_45S_50ntFlanks | 11670 | 11693 | 24 | CCCAGTGCTCTGAATGTCAAAGTG | MB468 | 53 | 72 | 48 | 24 | 15 | 17.0 | 20.8 | 26.6 | 14.4 | 8.6 |
| >hg38_21_+_8433172_8446622_RNA45SN1_45S_50ntFlanks@11670.11697.28\|CCCAGTGCTCTGAATGTCAAAGTGAAGA | >hg38_21_+_8433172_8446622_RNA45SN1_45S_50ntFlanks | 11670 | 11697 | 28 | CCCAGTGCTCTGAATGTCAAAGTGAAGA | MB468 | 13 | 21 | 10 | 7 | 4 | 4.8 | 8.3 | 4.8 | 2.2 | 0.7 |
| >hg38_21_+_8433172_8446622_RNA45SN1_45S_50ntFlanks@11670.11698.29\|CCCAGTGCTCTGAATGTCAAAGTGAAGAA | >hg38_21_+_8433172_8446622_RNA45SN1_45S_50ntFlanks | 11670 | 11698 | 29 | CCCAGTGCTCTGAATGTCAAAGTGAAGAA | MB468 | 28 | 45 | 19 | 9 | 9 | 7.8 | 11.5 | 9.0 | 2.9 | 2.0 |
| >hg38_21_+_8433172_8446622_RNA45SN1_45S_50ntFlanks@11670.11699.30\|CCCAGTGCTCTGAATGTCAAAGTGAAGAAA | >hg38_21_+_8433172_8446622_RNA45SN1_45S_50ntFlanks | 11670 | 11699 | 30 | CCCAGTGCTCTGAATGTCAAAGTGAAGAAA | MB468 | 26 | 48 | 20 | 8 | 11 | 7.1 | 10.9 | 10.5 | 2.9 | 3.0 |
| >hg38_21_+_8433172_8446622_RNA45SN1_45S_50ntFlanks@11670.11700.31\|CCCAGTGCTCTGAATGTCAAAGTGAAGAAAT | >hg38_21_+_8433172_8446622_RNA45SN1_45S_50ntFlanks | 11670 | 11700 | 31 | CCCAGTGCTCTGAATGTCAAAGTGAAGAAAT | MB468 | 33 | 69 | 13 | 8 | 13 | 15.5 | 18.3 | 4.1 | 3.1 | 2.6 |
| >hg38_21_+_8433172_8446622_RNA45SN1_45S_50ntFlanks@11670.11701.32\|CCCAGTGCTCTGAATGTCAAAGTGAAGAAATT | >hg38_21_+_8433172_8446622_RNA45SN1_45S_50ntFlanks | 11670 | 11701 | 32 | CCCAGTGCTCTGAATGTCAAAGTGAAGAAATT | MB468 | 71 | 205 | 39 | 16 | 21 | 37.8 | 69.7 | 21.2 | 7.7 | 4.7 |
| >hg38_21_+_8433172_8446622_RNA45SN1_45S_50ntFlanks@11670.11702.33\|CCCAGTGCTCTGAATGTCAAAGTGAAGAAATTC | >hg38_21_+_8433172_8446622_RNA45SN1_45S_50ntFlanks | 11670 | 11702 | 33 | CCCAGTGCTCTGAATGTCAAAGTGAAGAAATTC | MB468 | 131 | 506 | 155 | 60 | 117 | 78.4 | 247.7 | 66.4 | 26.1 | 14.7 |
| >hg38_21_+_8433172_8446622_RNA45SN1_45S_50ntFlanks@11670.11703.34\|CCCAGTGCTCTGAATGTCAAAGTGAAGAAATTCA | >hg38_21_+_8433172_8446622_RNA45SN1_45S_50ntFlanks | 11670 | 11703 | 34 | CCCAGTGCTCTGAATGTCAAAGTGAAGAAATTCA | MB468 | 35 | 115 | 45 | 15 | 41 | 17.7 | 49.4 | 15.6 | 7.6 | 13.3 |
| >hg38_21_+_8433172_8446622_RNA45SN1_45S_50ntFlanks@11670.11704.35\|CCCAGTGCTCTGAATGTCAAAGTGAAGAAATTCAA | >hg38_21_+_8433172_8446622_RNA45SN1_45S_50ntFlanks | 11670 | 11704 | 35 | CCCAGTGCTCTGAATGTCAAAGTGAAGAAATTCAA | MB468 | 17 | 51 | 19 | 5 | 13 | 4.4 | 8.2 | 3.3 | 3.0 | 4.1 |
| >hg38_21_+_8433172_8446622_RNA45SN1_45S_50ntFlanks@11671.11688.18\|CCAGTGCTCTGAATGTCA | >hg38_21_+_8433172_8446622_RNA45SN1_45S_50ntFlanks | 11671 | 11688 | 18 | CCAGTGCTCTGAATGTCA | MB468 | 72 | 31 | 59 | 13 | 7 | 78.6 | 18.4 | 94.6 | 7.6 | 5.6 |
| >hg38_21_+_8433172_8446622_RNA45SN1_45S_50ntFlanks@11671.11690.20\|CCAGTGCTCTGAATGTCAAA | >hg38_21_+_8433172_8446622_RNA45SN1_45S_50ntFlanks | 11671 | 11690 | 20 | CCAGTGCTCTGAATGTCAAA | MB468 | 10 | 8 | 9 | 4 | 2 | 4.5 | 4.2 | 7.1 | 2.4 | 1.0 |
| >hg38_21_+_8433172_8446622_RNA45SN1_45S_50ntFlanks@11671.11691.21\|CCAGTGCTCTGAATGTCAAAG | >hg38_21_+_8433172_8446622_RNA45SN1_45S_50ntFlanks | 11671 | 11691 | 21 | CCAGTGCTCTGAATGTCAAAG | MB468 | 39 | 52 | 22 | 22 | 13 | 17.0 | 33.7 | 18.8 | 14.4 | 12.5 |
| >hg38_21_+_8433172_8446622_RNA45SN1_45S_50ntFlanks@11671.11692.22\|CCAGTGCTCTGAATGTCAAAGT | >hg38_21_+_8433172_8446622_RNA45SN1_45S_50ntFlanks | 11671 | 11692 | 22 | CCAGTGCTCTGAATGTCAAAGT | MB468 | 29 | 28 | 17 | 9 | 3 | 18.0 | 16.1 | 16.2 | 6.4 | 1.3 |
| >hg38_21_+_8433172_8446622_RNA45SN1_45S_50ntFlanks@11671.11693.23\|CCAGTGCTCTGAATGTCAAAGTG | >hg38_21_+_8433172_8446622_RNA45SN1_45S_50ntFlanks | 11671 | 11693 | 23 | CCAGTGCTCTGAATGTCAAAGTG | MB468 | 106 | 66 | 88 | 34 | 11 | 109.2 | 42.0 | 128.6 | 35.7 | 11.6 |
| >hg38_21_+_8433172_8446622_RNA45SN1_45S_50ntFlanks@11671.11698.28\|CCAGTGCTCTGAATGTCAAAGTGAAGAA | >hg38_21_+_8433172_8446622_RNA45SN1_45S_50ntFlanks | 11671 | 11698 | 28 | CCAGTGCTCTGAATGTCAAAGTGAAGAA | MB468 | 14 | 15 | 7 | 6 | 3 | 2.6 | 5.8 | 1.3 | 1.7 | 0.7 |
| >hg38_21_+_8433172_8446622_RNA45SN1_45S_50ntFlanks@11671.11699.29\|CCAGTGCTCTGAATGTCAAAGTGAAGAAA | >hg38_21_+_8433172_8446622_RNA45SN1_45S_50ntFlanks | 11671 | 11699 | 29 | CCAGTGCTCTGAATGTCAAAGTGAAGAAA | MB468 | 20 | 25 | 11 | 7 | 5 | 5.0 | 9.6 | 4.0 | 4.8 | 2.7 |
| >hg38_21_+_8433172_8446622_RNA45SN1_45S_50ntFlanks@11671.11700.30\|CCAGTGCTCTGAATGTCAAAGTGAAGAAAT | >hg38_21_+_8433172_8446622_RNA45SN1_45S_50ntFlanks | 11671 | 11700 | 30 | CCAGTGCTCTGAATGTCAAAGTGAAGAAAT | MB468 | 25 | 37 | 14 | 8 | 6 | 5.2 | 9.5 | 3.8 | 6.0 | 2.7 |
| >hg38_21_+_8433172_8446622_RNA45SN1_45S_50ntFlanks@11671.11701.31\|CCAGTGCTCTGAATGTCAAAGTGAAGAAATT | >hg38_21_+_8433172_8446622_RNA45SN1_45S_50ntFlanks | 11671 | 11701 | 31 | CCAGTGCTCTGAATGTCAAAGTGAAGAAATT | MB468 | 41 | 84 | 27 | 12 | 9 | 10.5 | 24.8 | 4.6 | 9.4 | 2.9 |
| >hg38_21_+_8433172_8446622_RNA45SN1_45S_50ntFlanks@11671.11702.32\|CCAGTGCTCTGAATGTCAAAGTGAAGAAATTC | >hg38_21_+_8433172_8446622_RNA45SN1_45S_50ntFlanks | 11671 | 11702 | 32 | CCAGTGCTCTGAATGTCAAAGTGAAGAAATTC | MB468 | 101 | 252 | 93 | 52 | 47 | 40.1 | 118.0 | 24.9 | 36.1 | 15.4 |
| >hg38_21_+_8433172_8446622_RNA45SN1_45S_50ntFlanks@11671.11703.33\|CCAGTGCTCTGAATGTCAAAGTGAAGAAATTCA | >hg38_21_+_8433172_8446622_RNA45SN1_45S_50ntFlanks | 11671 | 11703 | 33 | CCAGTGCTCTGAATGTCAAAGTGAAGAAATTCA | MB468 | 54 | 96 | 46 | 23 | 18 | 33.8 | 38.5 | 24.0 | 19.4 | 8.1 |
| >hg38_21_+_8433172_8446622_RNA45SN1_45S_50ntFlanks@11671.11704.34\|CCAGTGCTCTGAATGTCAAAGTGAAGAAATTCAA | >hg38_21_+_8433172_8446622_RNA45SN1_45S_50ntFlanks | 11671 | 11704 | 34 | CCAGTGCTCTGAATGTCAAAGTGAAGAAATTCAA | MB468 | 33 | 58 | 26 | 10 | 8 | 29.1 | 22.7 | 22.0 | 9.6 | 5.2 |
| >hg38_21_+_8433172_8446622_RNA45SN1_45S_50ntFlanks@11671.11706.36\|CCAGTGCTCTGAATGTCAAAGTGAAGAAATTCAATG | >hg38_21_+_8433172_8446622_RNA45SN1_45S_50ntFlanks | 11671 | 11706 | 36 | CCAGTGCTCTGAATGTCAAAGTGAAGAAATTCAATG | MB468 | 28 | 68 | 34 | 7 | 5 | 31.9 | 22.5 | 37.0 | 8.2 | 2.8 |
| >hg38_21_+_8433172_8446622_RNA45SN1_45S_50ntFlanks@11671.11707.37\|CCAGTGCTCTGAATGTCAAAGTGAAGAAATTCAATGA | >hg38_21_+_8433172_8446622_RNA45SN1_45S_50ntFlanks | 11671 | 11707 | 37 | CCAGTGCTCTGAATGTCAAAGTGAAGAAATTCAATGA | MB468 | 12 | 37 | 20 | 3 | 1 | 15.7 | 15.3 | 25.6 | 3.8 | 1.7 |
| >hg38_21_+_8433172_8446622_RNA45SN1_45S_50ntFlanks@11672.11691.20\|CAGTGCTCTGAATGTCAAAG | >hg38_21_+_8433172_8446622_RNA45SN1_45S_50ntFlanks | 11672 | 11691 | 20 | CAGTGCTCTGAATGTCAAAG | MB231 | 10 | 10 | 9 | 4 | 7 | 4.7 | 4.5 | 7.1 | 3.6 | 2.9 |
| >hg38_21_+_8433172_8446622_RNA45SN1_45S_50ntFlanks@11672.11691.20\|CAGTGCTCTGAATGTCAAAG | >hg38_21_+_8433172_8446622_RNA45SN1_45S_50ntFlanks | 11672 | 11691 | 20 | CAGTGCTCTGAATGTCAAAG | MB468 | 24 | 22 | 10 | 12 | 5 | 8.3 | 18.9 | 6.3 | 3.4 | 3.5 |
| >hg38_21_+_8433172_8446622_RNA45SN1_45S_50ntFlanks@11672.11692.21\|CAGTGCTCTGAATGTCAAAGT | >hg38_21_+_8433172_8446622_RNA45SN1_45S_50ntFlanks | 11672 | 11692 | 21 | CAGTGCTCTGAATGTCAAAGT | MB231 | 12 | 13 | 10 | 3 | 6 | 3.3 | 4.3 | 6.0 | 1.3 | 2.9 |
| >hg38_21_+_8433172_8446622_RNA45SN1_45S_50ntFlanks@11672.11692.21\|CAGTGCTCTGAATGTCAAAGT | >hg38_21_+_8433172_8446622_RNA45SN1_45S_50ntFlanks | 11672 | 11692 | 21 | CAGTGCTCTGAATGTCAAAGT | MB468 | 18 | 11 | 7 | 4 | 1 | 7.8 | 8.0 | 3.4 | 1.0 | 0.1 |
| >hg38_21_+_8433172_8446622_RNA45SN1_45S_50ntFlanks@11672.11693.22\|CAGTGCTCTGAATGTCAAAGTG | >hg38_21_+_8433172_8446622_RNA45SN1_45S_50ntFlanks | 11672 | 11693 | 22 | CAGTGCTCTGAATGTCAAAGTG | MB231 | 12 | 10 | 8 | 2 | 2 | 10.4 | 10.7 | 6.9 | 1.6 | 1.7 |
| >hg38_21_+_8433172_8446622_RNA45SN1_45S_50ntFlanks@11672.11693.22\|CAGTGCTCTGAATGTCAAAGTG | >hg38_21_+_8433172_8446622_RNA45SN1_45S_50ntFlanks | 11672 | 11693 | 22 | CAGTGCTCTGAATGTCAAAGTG | MB468 | 44 | 16 | 23 | 12 | 3 | 31.7 | 8.1 | 25.5 | 7.5 | 2.5 |
| >hg38_21_+_8433172_8446622_RNA45SN1_45S_50ntFlanks@11672.11699.28\|CAGTGCTCTGAATGTCAAAGTGAAGAAA | >hg38_21_+_8433172_8446622_RNA45SN1_45S_50ntFlanks | 11672 | 11699 | 28 | CAGTGCTCTGAATGTCAAAGTGAAGAAA | MB468 | 15 | 13 | 6 | 4 | 2 | 3.5 | 5.2 | 2.9 | 1.9 | 1.1 |
| >hg38_21_+_8433172_8446622_RNA45SN1_45S_50ntFlanks@11672.11700.29\|CAGTGCTCTGAATGTCAAAGTGAAGAAAT | >hg38_21_+_8433172_8446622_RNA45SN1_45S_50ntFlanks | 11672 | 11700 | 29 | CAGTGCTCTGAATGTCAAAGTGAAGAAAT | MB468 | 17 | 18 | 8 | 4 | 2 | 3.6 | 6.2 | 2.4 | 1.4 | 0.8 |
| >hg38_21_+_8433172_8446622_RNA45SN1_45S_50ntFlanks@11672.11701.30\|CAGTGCTCTGAATGTCAAAGTGAAGAAATT | >hg38_21_+_8433172_8446622_RNA45SN1_45S_50ntFlanks | 11672 | 11701 | 30 | CAGTGCTCTGAATGTCAAAGTGAAGAAATT | MB231 | 12 | 17 | 9 | 2 | 4 | 9.4 | 13.8 | 9.0 | 1.2 | 3.2 |
| >hg38_21_+_8433172_8446622_RNA45SN1_45S_50ntFlanks@11672.11701.30\|CAGTGCTCTGAATGTCAAAGTGAAGAAATT | >hg38_21_+_8433172_8446622_RNA45SN1_45S_50ntFlanks | 11672 | 11701 | 30 | CAGTGCTCTGAATGTCAAAGTGAAGAAATT | MB468 | 26 | 42 | 15 | 5 | 3 | 8.7 | 20.2 | 2.1 | 2.6 | 1.3 |
| >hg38_21_+_8433172_8446622_RNA45SN1_45S_50ntFlanks@11672.11702.31\|CAGTGCTCTGAATGTCAAAGTGAAGAAATTC | >hg38_21_+_8433172_8446622_RNA45SN1_45S_50ntFlanks | 11672 | 11702 | 31 | CAGTGCTCTGAATGTCAAAGTGAAGAAATTC | MB231 | 28 | 37 | 29 | 5 | 6 | 20.8 | 22.4 | 23.7 | 4.6 | 4.4 |
| >hg38_21_+_8433172_8446622_RNA45SN1_45S_50ntFlanks@11672.11702.31\|CAGTGCTCTGAATGTCAAAGTGAAGAAATTC | >hg38_21_+_8433172_8446622_RNA45SN1_45S_50ntFlanks | 11672 | 11702 | 31 | CAGTGCTCTGAATGTCAAAGTGAAGAAATTC | MB468 | 72 | 131 | 42 | 22 | 12 | 20.7 | 91.0 | 7.1 | 13.0 | 3.9 |
| >hg38_21_+_8433172_8446622_RNA45SN1_45S_50ntFlanks@11672.11703.32\|CAGTGCTCTGAATGTCAAAGTGAAGAAATTCA | >hg38_21_+_8433172_8446622_RNA45SN1_45S_50ntFlanks | 11672 | 11703 | 32 | CAGTGCTCTGAATGTCAAAGTGAAGAAATTCA | MB231 | 16 | 19 | 15 | 3 | 4 | 10.5 | 6.1 | 13.4 | 3.0 | 3.5 |

| | | | | | | | | | | | | | | | |
|---|---|---|---|---|---|---|---|---|---|---|---|---|---|---|---|
| >hg38_21_+_8433172_8446622_RNA45SN1_45S_50ntFlanks@11672.11703.32\|CAGTGCTCTGAATGTCAAAGTGAAGAAATTCA | >hg38_21_+_8433172_8446622_RNA45SN1_45S_50ntFlanks | 11672 | 11703 | 32 | CAGTGCTCTGAATGTCAAAGTGAAGAAATTCA | MB468 | 29 | 39 | 16 | 10 | 5 | 12.1 | 23.7 | 3.3 | 6.3 | 2.2 |
| >hg38_21_+_8433172_8446622_RNA45SN1_45S_50ntFlanks@11672.11704.33\|CAGTGCTCTGAATGTCAAAGTGAAGAAATTCAA | >hg38_21_+_8433172_8446622_RNA45SN1_45S_50ntFlanks | 11672 | 11704 | 33 | CAGTGCTCTGAATGTCAAAGTGAAGAAATTCAA | MB468 | 27 | 36 | 14 | 7 | 4 | 14.6 | 22.4 | 5.1 | 5.0 | 2.5 |
| >hg38_21_+_8433172_8446622_RNA45SN1_45S_50ntFlanks@11672.11706.35\|CAGTGCTCTGAATGTCAAAGTGAAGAAATTCAATG | >hg38_21_+_8433172_8446622_RNA45SN1_45S_50ntFlanks | 11672 | 11706 | 35 | CAGTGCTCTGAATGTCAAAGTGAAGAAATTCAATG | MB231 | 21 | 17 | 33 | 3 | 3 | 20.5 | 8.3 | 24.3 | 4.5 | 3.0 |
| >hg38_21_+_8433172_8446622_RNA45SN1_45S_50ntFlanks@11672.11706.35\|CAGTGCTCTGAATGTCAAAGTGAAGAAATTCAATG | >hg38_21_+_8433172_8446622_RNA45SN1_45S_50ntFlanks | 11672 | 11706 | 35 | CAGTGCTCTGAATGTCAAAGTGAAGAAATTCAATG | MB468 | 24 | 51 | 23 | 6 | 3 | 19.2 | 24.6 | 13.9 | 5.5 | 2.1 |
| >hg38_21_+_8433172_8446622_RNA45SN1_45S_50ntFlanks@11672.11707.36\|CAGTGCTCTGAATGTCAAAGTGAAGAAATTCAATGA | >hg38_21_+_8433172_8446622_RNA45SN1_45S_50ntFlanks | 11672 | 11707 | 36 | CAGTGCTCTGAATGTCAAAGTGAAGAAATTCAATGA | MB231 | 17 | 16 | 35 | 3 | 4 | 15.1 | 8.6 | 25.1 | 1.9 | 2.8 |
| >hg38_21_+_8433172_8446622_RNA45SN1_45S_50ntFlanks@11672.11709.38\|CAGTGCTCTGAATGTCAAAGTGAAGAAATTCAATGAAG | >hg38_21_+_8433172_8446622_RNA45SN1_45S_50ntFlanks | 11672 | 11709 | 38 | CAGTGCTCTGAATGTCAAAGTGAAGAAATTCAATGAAG | MB231 | 19 | 23 | 33 | 2 | 3 | 16.3 | 16.0 | 22.4 | 2.8 | 2.7 |
| >hg38_21_+_8433172_8446622_RNA45SN1_45S_50ntFlanks@11675.11702.28\|TGCTCTGAATGTCAAAGTGAAGAAATTC | >hg38_21_+_8433172_8446622_RNA45SN1_45S_50ntFlanks | 11675 | 11702 | 28 | TGCTCTGAATGTCAAAGTGAAGAAATTC | BT20 | 12 | 6 | 21 | 5 | 8 | 6.9 | 2.4 | 3.8 | 3.4 | 2.9 |
| >hg38_21_+_8433172_8446622_RNA45SN1_45S_50ntFlanks@11675.11702.28\|TGCTCTGAATGTCAAAGTGAAGAAATTC | >hg38_21_+_8433172_8446622_RNA45SN1_45S_50ntFlanks | 11675 | 11702 | 28 | TGCTCTGAATGTCAAAGTGAAGAAATTC | MB231 | 67 | 67 | 58 | 16 | 12 | 51.9 | 52.7 | 41.6 | 15.7 | 8.2 |
| >hg38_21_+_8433172_8446622_RNA45SN1_45S_50ntFlanks@11675.11702.28\|TGCTCTGAATGTCAAAGTGAAGAAATTC | >hg38_21_+_8433172_8446622_RNA45SN1_45S_50ntFlanks | 11675 | 11702 | 28 | TGCTCTGAATGTCAAAGTGAAGAAATTC | MB468 | 14 | 8 | 14 | 15 | 22 | 3.8 | 4.5 | 1.9 | 7.3 | 13.7 |
| >hg38_21_+_8433172_8446622_RNA45SN1_45S_50ntFlanks@11675.11706.32\|TGCTCTGAATGTCAAAGTGAAGAAATTCAATG | >hg38_21_+_8433172_8446622_RNA45SN1_45S_50ntFlanks | 11675 | 11706 | 32 | TGCTCTGAATGTCAAAGTGAAGAAATTCAATG | MB231 | 34 | 29 | 39 | 8 | 7 | 25.5 | 20.9 | 27.3 | 7.3 | 4.3 |
| >hg38_21_+_8433172_8446622_RNA45SN1_45S_50ntFlanks@11675.11707.33\|TGCTCTGAATGTCAAAGTGAAGAAATTCAATGA | >hg38_21_+_8433172_8446622_RNA45SN1_45S_50ntFlanks | 11675 | 11707 | 33 | TGCTCTGAATGTCAAAGTGAAGAAATTCAATGA | MB231 | 23 | 28 | 30 | 6 | 5 | 17.4 | 20.5 | 23.7 | 4.7 | 4.1 |
| >hg38_21_+_8433172_8446622_RNA45SN1_45S_50ntFlanks@11675.11709.35\|TGCTCTGAATGTCAAAGTGAAGAAATTCAATGAAG | >hg38_21_+_8433172_8446622_RNA45SN1_45S_50ntFlanks | 11675 | 11709 | 35 | TGCTCTGAATGTCAAAGTGAAGAAATTCAATGAAG | MB231 | 20 | 27 | 25 | 3 | 5 | 15.9 | 18.5 | 20.4 | 3.3 | 4.0 |
| >hg38_21_+_8433172_8446622_RNA45SN1_45S_50ntFlanks@11677.11702.26\|CTCTGAATGTCAAAGTGAAGAAATTC | >hg38_21_+_8433172_8446622_RNA45SN1_45S_50ntFlanks | 11677 | 11702 | 26 | CTCTGAATGTCAAAGTGAAGAAATTC | MB468 | 57 | 26 | 39 | 38 | 24 | 14.3 | 14.7 | 12.4 | 3.9 | 21.9 |
| >hg38_21_+_8433172_8446622_RNA45SN1_45S_50ntFlanks@11679.11702.24\|CTGAATGTCAAAGTGAAGAAATTC | >hg38_21_+_8433172_8446622_RNA45SN1_45S_50ntFlanks | 11679 | 11702 | 24 | CTGAATGTCAAAGTGAAGAAATTC | MB468 | 12 | 7 | 15 | 13 | 7 | 2.6 | 1.9 | 9.8 | 3.9 | 6.2 |
| >hg38_21_+_8433172_8446622_RNA45SN1_45S_50ntFlanks@11680.11700.21\|TGAATGTCAAAGTGAAGAAAT | >hg38_21_+_8433172_8446622_RNA45SN1_45S_50ntFlanks | 11680 | 11700 | 21 | TGAATGTCAAAGTGAAGAAAT | MB468 | 11 | 4 | 5 | 1 | 0 | 7.0 | 1.3 | 7.8 | 0.7 | 0.5 |
| >hg38_21_+_8433172_8446622_RNA45SN1_45S_50ntFlanks@11680.11701.22\|TGAATGTCAAAGTGAAGAAATT | >hg38_21_+_8433172_8446622_RNA45SN1_45S_50ntFlanks | 11680 | 11701 | 22 | TGAATGTCAAAGTGAAGAAATT | MB468 | 14 | 7 | 9 | 2 | 1 | 9.4 | 2.3 | 14.8 | 1.0 | 0.4 |
| >hg38_21_+_8433172_8446622_RNA45SN1_45S_50ntFlanks@11682.11700.19\|AATGTCAAAGTGAAGAAAT | >hg38_21_+_8433172_8446622_RNA45SN1_45S_50ntFlanks | 11682 | 11700 | 19 | AATGTCAAAGTGAAGAAAT | MB468 | 11 | 3 | 5 | 5 | 1 | 6.9 | 1.0 | 7.4 | 3.8 | 0.5 |
| >hg38_21_+_8433172_8446622_RNA45SN1_45S_50ntFlanks@11682.11702.21\|AATGTCAAAGTGAAGAAATTC | >hg38_21_+_8433172_8446622_RNA45SN1_45S_50ntFlanks | 11682 | 11702 | 21 | AATGTCAAAGTGAAGAAATTC | MB468 | 44 | 11 | 41 | 52 | 7 | 42.6 | 3.4 | 65.0 | 42.7 | 4.2 |
| >hg38_21_+_8433172_8446622_RNA45SN1_45S_50ntFlanks@11682.11703.22\|AATGTCAAAGTGAAGAAATTCA | >hg38_21_+_8433172_8446622_RNA45SN1_45S_50ntFlanks | 11682 | 11703 | 22 | AATGTCAAAGTGAAGAAATTCA | MB468 | 11 | 1 | 9 | 5 | 1 | 11.4 | 0.4 | 14.6 | 2.5 | 0.8 |
| >hg38_21_+_8433172_8446622_RNA45SN1_45S_50ntFlanks@11682.11704.23\|AATGTCAAAGTGAAGAAATTCAA | >hg38_21_+_8433172_8446622_RNA45SN1_45S_50ntFlanks | 11682 | 11704 | 23 | AATGTCAAAGTGAAGAAATTCAA | MB468 | 11 | 2 | 8 | 3 | 1 | 13.1 | 1.1 | 13.2 | 1.8 | 0.4 |
| >hg38_21_+_8433172_8446622_RNA45SN1_45S_50ntFlanks@11682.11706.25\|AATGTCAAAGTGAAGAAATTCAATG | >hg38_21_+_8433172_8446622_RNA45SN1_45S_50ntFlanks | 11682 | 11706 | 25 | AATGTCAAAGTGAAGAAATTCAATG | MB468 | 65 | 7 | 50 | 14 | 4 | 90.5 | 2.6 | 86.1 | 7.1 | 1.7 |
| >hg38_21_+_8433172_8446622_RNA45SN1_45S_50ntFlanks@11683.11701.19\|ATGTCAAAGTGAAGAAATT | >hg38_21_+_8433172_8446622_RNA45SN1_45S_50ntFlanks | 11683 | 11701 | 19 | ATGTCAAAGTGAAGAAATT | MB468 | 13 | 4 | 6 | 3 | 1 | 7.7 | 1.5 | 8.6 | 2.5 | 0.5 |
| >hg38_21_+_8433172_8446622_RNA45SN1_45S_50ntFlanks@11683.11702.20\|ATGTCAAAGTGAAGAAATTC | >hg38_21_+_8433172_8446622_RNA45SN1_45S_50ntFlanks | 11683 | 11702 | 20 | ATGTCAAAGTGAAGAAATTC | MB468 | 29 | 6 | 23 | 52 | 2 | 17.7 | 3.1 | 31.1 | 52.1 | 1.0 |
| >hg38_21_+_8433172_8446622_RNA45SN1_45S_50ntFlanks@11683.11706.24\|ATGTCAAAGTGAAGAAATTCAATG | >hg38_21_+_8433172_8446622_RNA45SN1_45S_50ntFlanks | 11683 | 11706 | 24 | ATGTCAAAGTGAAGAAATTCAATG | MB468 | 15 | 2 | 17 | 7 | 2 | 11.5 | 0.6 | 23.0 | 3.3 | 0.4 |
| >hg38_21_+_8433172_8446622_RNA45SN1_45S_50ntFlanks@11686.11717.32\|TCAAAGTGAAGAAATTCAATGAAGCGCGGGTA | >hg38_21_+_8433172_8446622_RNA45SN1_45S_50ntFlanks | 11686 | 11717 | 32 | TCAAAGTGAAGAAATTCAATGAAGCGCGGGTA | MB468 | 10 | 13 | 19 | 1 | 9 | 3.3 | 7.8 | 12.7 | 0.4 | 5.3 |
| >hg38_21_+_8433172_8446622_RNA45SN1_45S_50ntFlanks@11694.11711.18\|AAGAAATTCAATGAAGCG | >hg38_21_+_8433172_8446622_RNA45SN1_45S_50ntFlanks | 11694 | 11711 | 18 | AAGAAATTCAATGAAGCG | MB468 | 27 | 7 | 16 | 2 | 1 | 22.1 | 4.3 | 23.8 | 1.1 | 0.5 |
| >hg38_21_+_8433172_8446622_RNA45SN1_45S_50ntFlanks@11694.11717.24\|AAGAAATTCAATGAAGCGCGGGTA | >hg38_21_+_8433172_8446622_RNA45SN1_45S_50ntFlanks | 11694 | 11717 | 24 | AAGAAATTCAATGAAGCGCGGGTA | MB468 | 30 | 22 | 27 | 27 | 9 | 12.9 | 12.6 | 25.3 | 21.1 | 4.6 |
| >hg38_21_+_8433172_8446622_RNA45SN1_45S_50ntFlanks@11694.11728.35\|AAGAAATTCAATGAAGCGCGGGTAAACGGCGGGAG | >hg38_21_+_8433172_8446622_RNA45SN1_45S_50ntFlanks | 11694 | 11728 | 35 | AAGAAATTCAATGAAGCGCGGGTAAACGGCGGGAG | MB468 | 10 | 27 | 14 | 9 | 25 | 2.0 | 20.4 | 4.0 | 5.5 | 22.7 |
| >hg38_21_+_8433172_8446622_RNA45SN1_45S_50ntFlanks@11702.11728.27\|CAATGAAGCGCGGGTAAACGGCGGGAG | >hg38_21_+_8433172_8446622_RNA45SN1_45S_50ntFlanks | 11702 | 11728 | 27 | CAATGAAGCGCGGGTAAACGGCGGGAG | MB468 | 13 | 19 | 10 | 12 | 11 | 2.4 | 5.4 | 2.7 | 8.0 | 4.2 |
| >hg38_21_+_8433172_8446622_RNA45SN1_45S_50ntFlanks@11703.11725.23\|AATGAAGCGCGGGTAAACGGCGG | >hg38_21_+_8433172_8446622_RNA45SN1_45S_50ntFlanks | 11703 | 11725 | 23 | AATGAAGCGCGGGTAAACGGCGG | MB468 | 14 | 7 | 7 | 11 | 4 | 4.6 | 2.0 | 2.2 | 10.1 | 2.4 |
| >hg38_21_+_8433172_8446622_RNA45SN1_45S_50ntFlanks@11703.11726.24\|AATGAAGCGCGGGTAAACGGCGGG | >hg38_21_+_8433172_8446622_RNA45SN1_45S_50ntFlanks | 11703 | 11726 | 24 | AATGAAGCGCGGGTAAACGGCGGG | MB468 | 26 | 28 | 18 | 25 | 15 | 6.5 | 6.5 | 3.9 | 16.5 | 3.8 |
| >hg38_21_+_8433172_8446622_RNA45SN1_45S_50ntFlanks@11703.11727.25\|AATGAAGCGCGGGTAAACGGCGGGA | >hg38_21_+_8433172_8446622_RNA45SN1_45S_50ntFlanks | 11703 | 11727 | 25 | AATGAAGCGCGGGTAAACGGCGGGA | MB468 | 19 | 16 | 12 | 13 | 11 | 2.9 | 3.7 | 3.5 | 11.1 | 4.5 |
| >hg38_21_+_8433172_8446622_RNA45SN1_45S_50ntFlanks@11703.11728.26\|AATGAAGCGCGGGTAAACGGCGGGAG | >hg38_21_+_8433172_8446622_RNA45SN1_45S_50ntFlanks | 11703 | 11728 | 26 | AATGAAGCGCGGGTAAACGGCGGGAG | MB468 | 210 | 272 | 145 | 190 | 177 | 33.0 | 73.2 | 45.4 | 127.0 | 80.9 |
| >hg38_21_+_8433172_8446622_RNA45SN1_45S_50ntFlanks@11703.11736.34\|AATGAAGCGCGGGTAAACGGCGGGAGTAACTATG | >hg38_21_+_8433172_8446622_RNA45SN1_45S_50ntFlanks | 11703 | 11736 | 34 | AATGAAGCGCGGGTAAACGGCGGGAGTAACTATG | MB468 | 19 | 24 | 16 | 12 | 13 | 10.9 | 10.4 | 8.1 | 6.8 | 9.4 |
| >hg38_21_+_8433172_8446622_RNA45SN1_45S_50ntFlanks@11703.11740.38\|AATGAAGCGCGGGTAAACGGCGGGAGTAACTATGACTC | >hg38_21_+_8433172_8446622_RNA45SN1_45S_50ntFlanks | 11703 | 11740 | 38 | AATGAAGCGCGGGTAAACGGCGGGAGTAACTATGACTC | MB468 | 15 | 32 | 17 | 5 | 17 | 8.9 | 12.5 | 6.4 | 4.0 | 10.7 |
| >hg38_21_+_8433172_8446622_RNA45SN1_45S_50ntFlanks@11704.11726.23\|ATGAAGCGCGGGTAAACGGCGGG | >hg38_21_+_8433172_8446622_RNA45SN1_45S_50ntFlanks | 11704 | 11726 | 23 | ATGAAGCGCGGGTAAACGGCGGG | MB468 | 12 | 12 | 7 | 11 | 7 | 2.1 | 1.5 | 1.3 | 6.5 | 2.3 |
| >hg38_21_+_8433172_8446622_RNA45SN1_45S_50ntFlanks@11704.11728.25\|ATGAAGCGCGGGTAAACGGCGGGAG | >hg38_21_+_8433172_8446622_RNA45SN1_45S_50ntFlanks | 11704 | 11728 | 25 | ATGAAGCGCGGGTAAACGGCGGGAG | MB468 | 97 | 136 | 59 | 86 | 88 | 14.8 | 24.5 | 17.6 | 49.6 | 26.8 |
| >hg38_21_+_8433172_8446622_RNA45SN1_45S_50ntFlanks@11704.11736.33\|ATGAAGCGCGGGTAAACGGCGGGAGTAACTATG | >hg38_21_+_8433172_8446622_RNA45SN1_45S_50ntFlanks | 11704 | 11736 | 33 | ATGAAGCGCGGGTAAACGGCGGGAGTAACTATG | MB468 | 15 | 17 | 12 | 9 | 11 | 10.4 | 9.3 | 7.5 | 6.6 | 11.3 |
| >hg38_21_+_8433172_8446622_RNA45SN1_45S_50ntFlanks@11704.11740.37\|ATGAAGCGCGGGTAAACGGCGGGAGTAACTATGACTC | >hg38_21_+_8433172_8446622_RNA45SN1_45S_50ntFlanks | 11704 | 11740 | 37 | ATGAAGCGCGGGTAAACGGCGGGAGTAACTATGACTC | MB468 | 13 | 23 | 12 | 4 | 14 | 9.8 | 11.4 | 6.6 | 3.9 | 11.7 |
| >hg38_21_+_8433172_8446622_RNA45SN1_45S_50ntFlanks@11705.11728.24\|TGAAGCGCGGGTAAACGGCGGGAG | >hg38_21_+_8433172_8446622_RNA45SN1_45S_50ntFlanks | 11705 | 11728 | 24 | TGAAGCGCGGGTAAACGGCGGGAG | MB468 | 65 | 102 | 49 | 49 | 47 | 3.1 | 18.5 | 12.8 | 25.8 | 13.8 |
| >hg38_21_+_8433172_8446622_RNA45SN1_45S_50ntFlanks@11705.11736.32\|TGAAGCGCGGGTAAACGGCGGGAGTAACTATG | >hg38_21_+_8433172_8446622_RNA45SN1_45S_50ntFlanks | 11705 | 11736 | 32 | TGAAGCGCGGGTAAACGGCGGGAGTAACTATG | MB468 | 11 | 12 | 7 | 6 | 6 | 9.0 | 7.0 | 7.2 | 5.1 | 6.3 |
| >hg38_21_+_8433172_8446622_RNA45SN1_45S_50ntFlanks@11705.11740.36\|TGAAGCGCGGGTAAACGGCGGGAGTAACTATGACTC | >hg38_21_+_8433172_8446622_RNA45SN1_45S_50ntFlanks | 11705 | 11740 | 36 | TGAAGCGCGGGTAAACGGCGGGAGTAACTATGACTC | MB468 | 12 | 22 | 11 | 4 | 12 | 10.2 | 11.1 | 6.6 | 3.1 | 11.6 |
| >hg38_21_+_8433172_8446622_RNA45SN1_45S_50ntFlanks@11707.11728.22\|AAGCGCGGGTAAACGGCGGGAG | >hg38_21_+_8433172_8446622_RNA45SN1_45S_50ntFlanks | 11707 | 11728 | 22 | AAGCGCGGGTAAACGGCGGGAG | MB468 | 27 | 40 | 24 | 26 | 18 | 2.3 | 8.2 | 3.8 | 10.9 | 4.9 |
| >hg38_21_+_8433172_8446622_RNA45SN1_45S_50ntFlanks@11707.11736.30\|AAGCGCGGGTAAACGGCGGGAGTAACTATG | >hg38_21_+_8433172_8446622_RNA45SN1_45S_50ntFlanks | 11707 | 11736 | 30 | AAGCGCGGGTAAACGGCGGGAGTAACTATG | MB468 | 12 | 15 | 8 | 8 | 8 | 11.6 | 11.2 | 9.4 | 7.5 | 8.5 |
| >hg38_21_+_8433172_8446622_RNA45SN1_45S_50ntFlanks@11707.11740.34\|AAGCGCGGGTAAACGGCGGGAGTAACTATGACTC | >hg38_21_+_8433172_8446622_RNA45SN1_45S_50ntFlanks | 11707 | 11740 | 34 | AAGCGCGGGTAAACGGCGGGAGTAACTATGACTC | MB468 | 17 | 25 | 13 | 4 | 11 | 15.1 | 16.9 | 14.3 | 4.2 | 11.0 |
| >hg38_21_+_8433172_8446622_RNA45SN1_45S_50ntFlanks@11710.11728.19\|CGCGGGTAAACGGCGGGAG | >hg38_21_+_8433172_8446622_RNA45SN1_45S_50ntFlanks | 11710 | 11728 | 19 | CGCGGGTAAACGGCGGGAG | MB468 | 38 | 47 | 45 | 47 | 20 | 7.4 | 14.4 | 11.8 | 20.2 | 8.7 |
| >hg38_21_+_8433172_8446622_RNA45SN1_45S_50ntFlanks@11710.11736.27\|CGCGGGTAAACGGCGGGAGTAACTATG | >hg38_21_+_8433172_8446622_RNA45SN1_45S_50ntFlanks | 11710 | 11736 | 27 | CGCGGGTAAACGGCGGGAGTAACTATG | MB468 | 23 | 21 | 17 | 16 | 13 | 26.7 | 15.5 | 19.7 | 13.1 | 15.5 |
| >hg38_21_+_8433172_8446622_RNA45SN1_45S_50ntFlanks@11710.11738.29\|CGCGGGTAAACGGCGGGAGTAACTATGAC | >hg38_21_+_8433172_8446622_RNA45SN1_45S_50ntFlanks | 11710 | 11738 | 29 | CGCGGGTAAACGGCGGGAGTAACTATGAC | MB468 | 13 | 9 | 10 | 4 | 6 | 15.4 | 4.7 | 10.1 | 1.6 | 4.2 |
| >hg38_21_+_8433172_8446622_RNA45SN1_45S_50ntFlanks@11710.11740.31\|CGCGGGTAAACGGCGGGAGTAACTATGACTC | >hg38_21_+_8433172_8446622_RNA45SN1_45S_50ntFlanks | 11710 | 11740 | 31 | CGCGGGTAAACGGCGGGAGTAACTATGACTC | MB468 | 29 | 27 | 27 | 10 | 16 | 32.9 | 18.9 | 29.4 | 11.2 | 21.3 |
| >hg38_21_+_8433172_8446622_RNA45SN1_45S_50ntFlanks@11710.11742.33\|CGCGGGTAAACGGCGGGAGTAACTATGACTCTC | >hg38_21_+_8433172_8446622_RNA45SN1_45S_50ntFlanks | 11710 | 11742 | 33 | CGCGGGTAAACGGCGGGAGTAACTATGACTCTC | MB468 | 14 | 9 | 34 | 5 | 5 | 13.5 | 3.4 | 40.5 | 5.3 | 3.6 |

| | | | | | | | | | | | | | | | | |
|---|---|---|---|---|---|---|---|---|---|---|---|---|---|---|---|---|
| >hg38_21_+_8433172_8446622_RNA45SN1_45S_50ntFlanks@11710.11744.35\|CGCGGGTAAACGGCGGGAGTAACTATGACTCTCTT | >hg38_21_+_8433172_8446622_RNA45SN1_45S_50ntFlanks | 11710 | 11744 | 35 | CGCGGGTAAACGGCGGGAGTAACTATGACTCTCTT | MB468 | 13 | 7 | 18 | 4 | 6 | 6.9 | 1.4 | 11.6 | 1.7 | 2.6 |
| >hg38_21_+_8433172_8446622_RNA45SN1_45S_50ntFlanks@11712.11736.25\|CGGGTAAACGGCGGGAGTAACTATG | >hg38_21_+_8433172_8446622_RNA45SN1_45S_50ntFlanks | 11712 | 11736 | 25 | CGGGTAAACGGCGGGAGTAACTATG | MB468 | 13 | 10 | 11 | 8 | 4 | 14.2 | 4.9 | 11.9 | 5.1 | 4.1 |
| >hg38_21_+_8433172_8446622_RNA45SN1_45S_50ntFlanks@11712.11740.29\|CGGGTAAACGGCGGGAGTAACTATGACTC | >hg38_21_+_8433172_8446622_RNA45SN1_45S_50ntFlanks | 11712 | 11740 | 29 | CGGGTAAACGGCGGGAGTAACTATGACTC | MB468 | 15 | 22 | 13 | 4 | 6 | 14.1 | 14.8 | 14.5 | 3.2 | 7.9 |
| >hg38_21_+_8433172_8446622_RNA45SN1_45S_50ntFlanks@11712.11743.32\|CGGGTAAACGGCGGGAGTAACTATGACTCTCT | >hg38_21_+_8433172_8446622_RNA45SN1_45S_50ntFlanks | 11712 | 11743 | 32 | CGGGTAAACGGCGGGAGTAACTATGACTCTCT | BT20 | 12 | 4 | 4 | 0 | 0 | 20.1 | 7.7 | 2.9 | 0.8 | 0.4 |
| >hg38_21_+_8433172_8446622_RNA45SN1_45S_50ntFlanks@11714.11744.31\|GGTAAACGGCGGGAGTAACTATGACTCTCTT | >hg38_21_+_8433172_8446622_RNA45SN1_45S_50ntFlanks | 11714 | 11744 | 31 | GGTAAACGGCGGGAGTAACTATGACTCTCTT | MB468 | 12 | 2 | 19 | 4 | 3 | 5.5 | 0.9 | 14.5 | 1.7 | 0.7 |
| >hg38_21_+_8433172_8446622_RNA45SN1_45S_50ntFlanks@11717.11743.27\|AAACGGCGGGAGTAACTATGACTCTCT | >hg38_21_+_8433172_8446622_RNA45SN1_45S_50ntFlanks | 11717 | 11743 | 27 | AAACGGCGGGAGTAACTATGACTCTCT | BT20 | 17 | 13 | 2 | 1 | 1 | 29.5 | 22.7 | 3.6 | 2.2 | 1.0 |
| >hg38_21_+_8433172_8446622_RNA45SN1_45S_50ntFlanks@11717.11744.28\|AAACGGCGGGAGTAACTATGACTCTCTT | >hg38_21_+_8433172_8446622_RNA45SN1_45S_50ntFlanks | 11717 | 11744 | 28 | AAACGGCGGGAGTAACTATGACTCTCTT | BT20 | 15 | 9 | 2 | 2 | 1 | 23.5 | 11.9 | 1.6 | 1.3 | 0.6 |
| >hg38_21_+_8433172_8446622_RNA45SN1_45S_50ntFlanks@11718.11736.19\|AACGGCGGGAGTAACTATG | >hg38_21_+_8433172_8446622_RNA45SN1_45S_50ntFlanks | 11718 | 11736 | 19 | AACGGCGGGAGTAACTATG | MB468 | 125 | 121 | 62 | 41 | 34 | 121.0 | 83.0 | 70.3 | 38.7 | 30.7 |
| >hg38_21_+_8433172_8446622_RNA45SN1_45S_50ntFlanks@11718.11737.20\|AACGGCGGGAGTAACTATGA | >hg38_21_+_8433172_8446622_RNA45SN1_45S_50ntFlanks | 11718 | 11737 | 20 | AACGGCGGGAGTAACTATGA | MB468 | 21 | 25 | 9 | 10 | 6 | 15.7 | 11.4 | 9.1 | 4.0 | 2.2 |
| >hg38_21_+_8433172_8446622_RNA45SN1_45S_50ntFlanks@11718.11738.21\|AACGGCGGGAGTAACTATGAC | >hg38_21_+_8433172_8446622_RNA45SN1_45S_50ntFlanks | 11718 | 11738 | 21 | AACGGCGGGAGTAACTATGAC | MB468 | 82 | 98 | 52 | 25 | 27 | 76.5 | 60.6 | 55.6 | 7.2 | 10.9 |
| >hg38_21_+_8433172_8446622_RNA45SN1_45S_50ntFlanks@11718.11739.22\|AACGGCGGGAGTAACTATGACT | >hg38_21_+_8433172_8446622_RNA45SN1_45S_50ntFlanks | 11718 | 11739 | 22 | AACGGCGGGAGTAACTATGACT | MB468 | 28 | 37 | 14 | 4 | 8 | 24.2 | 24.3 | 17.6 | 2.3 | 5.0 |
| >hg38_21_+_8433172_8446622_RNA45SN1_45S_50ntFlanks@11718.11740.23\|AACGGCGGGAGTAACTATGACTC | >hg38_21_+_8433172_8446622_RNA45SN1_45S_50ntFlanks | 11718 | 11740 | 23 | AACGGCGGGAGTAACTATGACTC | MB468 | 190 | 471 | 156 | 26 | 53 | 132.8 | 344.4 | 173.8 | 22.6 | 41.4 |
| >hg38_21_+_8433172_8446622_RNA45SN1_45S_50ntFlanks@11718.11742.25\|AACGGCGGGAGTAACTATGACTCTC | >hg38_21_+_8433172_8446622_RNA45SN1_45S_50ntFlanks | 11718 | 11742 | 25 | AACGGCGGGAGTAACTATGACTCTC | MB468 | 151 | 50 | 319 | 68 | 30 | 79.6 | 10.6 | 242.1 | 21.3 | 7.8 |
| >hg38_21_+_8433172_8446622_RNA45SN1_45S_50ntFlanks@11718.11743.26\|AACGGCGGGAGTAACTATGACTCTCT | >hg38_21_+_8433172_8446622_RNA45SN1_45S_50ntFlanks | 11718 | 11743 | 26 | AACGGCGGGAGTAACTATGACTCTCT | BT20 | 26 | 17 | 3 | 3 | 1 | 44.9 | 30.1 | 4.3 | 5.0 | 1.3 |
| >hg38_21_+_8433172_8446622_RNA45SN1_45S_50ntFlanks@11718.11743.26\|AACGGCGGGAGTAACTATGACTCTCT | >hg38_21_+_8433172_8446622_RNA45SN1_45S_50ntFlanks | 11718 | 11743 | 26 | AACGGCGGGAGTAACTATGACTCTCT | MB468 | 36 | 4 | 59 | 19 | 8 | 30.3 | 0.6 | 69.7 | 6.1 | 4.0 |
| >hg38_21_+_8433172_8446622_RNA45SN1_45S_50ntFlanks@11718.11744.27\|AACGGCGGGAGTAACTATGACTCTCTT | >hg38_21_+_8433172_8446622_RNA45SN1_45S_50ntFlanks | 11718 | 11744 | 27 | AACGGCGGGAGTAACTATGACTCTCTT | BT20 | 18 | 7 | 2 | 2 | 0 | 29.1 | 12.4 | 1.2 | 2.3 | 0.6 |
| >hg38_21_+_8433172_8446622_RNA45SN1_45S_50ntFlanks@11718.11744.27\|AACGGCGGGAGTAACTATGACTCTCTT | >hg38_21_+_8433172_8446622_RNA45SN1_45S_50ntFlanks | 11718 | 11744 | 27 | AACGGCGGGAGTAACTATGACTCTCTT | MB468 | 62 | 3 | 50 | 45 | 16 | 53.5 | 0.3 | 57.3 | 23.3 | 6.5 |
| >hg38_21_+_8433172_8446622_RNA45SN1_45S_50ntFlanks@11718.11745.28\|AACGGCGGGAGTAACTATGACTCTCTTA | >hg38_21_+_8433172_8446622_RNA45SN1_45S_50ntFlanks | 11718 | 11745 | 28 | AACGGCGGGAGTAACTATGACTCTCTTA | MB468 | 11 | 1 | 9 | 6 | 3 | 11.1 | 0.2 | 11.1 | 3.9 | 0.8 |
| >hg38_21_+_8433172_8446622_RNA45SN1_45S_50ntFlanks@11718.11751.34\|AACGGCGGGAGTAACTATGACTCTCTTAAGGTAG | >hg38_21_+_8433172_8446622_RNA45SN1_45S_50ntFlanks | 11718 | 11751 | 34 | AACGGCGGGAGTAACTATGACTCTCTTAAGGTAG | MB468 | 79 | 37 | 59 | 24 | 20 | 49.0 | 11.0 | 44.5 | 15.3 | 6.4 |
| >hg38_21_+_8433172_8446622_RNA45SN1_45S_50ntFlanks@11719.11740.22\|ACGGCGGGAGTAACTATGACTC | >hg38_21_+_8433172_8446622_RNA45SN1_45S_50ntFlanks | 11719 | 11740 | 22 | ACGGCGGGAGTAACTATGACTC | MB468 | 11 | 28 | 9 | 2 | 5 | 5.0 | 18.8 | 7.3 | 1.6 | 3.4 |
| >hg38_21_+_8433172_8446622_RNA45SN1_45S_50ntFlanks@11719.11742.24\|ACGGCGGGAGTAACTATGACTCTC | >hg38_21_+_8433172_8446622_RNA45SN1_45S_50ntFlanks | 11719 | 11742 | 24 | ACGGCGGGAGTAACTATGACTCTC | MB468 | 12 | 5 | 19 | 7 | 3 | 3.5 | 1.1 | 10.6 | 1.9 | 1.0 |
| >hg38_21_+_8433172_8446622_RNA45SN1_45S_50ntFlanks@11719.11743.25\|ACGGCGGGAGTAACTATGACTCTCT | >hg38_21_+_8433172_8446622_RNA45SN1_45S_50ntFlanks | 11719 | 11743 | 25 | ACGGCGGGAGTAACTATGACTCTCT | BT20 | 20 | 15 | 2 | 3 | 1 | 34.5 | 26.6 | 2.8 | 3.8 | 1.0 |
| >hg38_21_+_8433172_8446622_RNA45SN1_45S_50ntFlanks@11719.11751.33\|ACGGCGGGAGTAACTATGACTCTCTTAAGGTAG | >hg38_21_+_8433172_8446622_RNA45SN1_45S_50ntFlanks | 11719 | 11751 | 33 | ACGGCGGGAGTAACTATGACTCTCTTAAGGTAG | MB468 | 12 | 6 | 7 | 4 | 3 | 4.6 | 1.7 | 3.3 | 1.8 | 1.0 |
| >hg38_21_+_8433172_8446622_RNA45SN1_45S_50ntFlanks@11726.11751.26\|GAGTAACTATGACTCTCTTAAGGTAG | >hg38_21_+_8433172_8446622_RNA45SN1_45S_50ntFlanks | 11726 | 11751 | 26 | GAGTAACTATGACTCTCTTAAGGTAG | MB468 | 15 | 5 | 5 | 2 | 1 | 9.1 | 2.6 | 4.9 | 1.4 | 0.7 |
| >hg38_21_+_8433172_8446622_RNA45SN1_45S_50ntFlanks@11727.11744.18\|AGTAACTATGACTCTCTT | >hg38_21_+_8433172_8446622_RNA45SN1_45S_50ntFlanks | 11727 | 11744 | 18 | AGTAACTATGACTCTCTT | MB231 | 21 | 10 | 36 | 0 | 0 | 25.3 | 7.3 | 41.9 | 0.0 | 0.0 |
| >hg38_21_+_8433172_8446622_RNA45SN1_45S_50ntFlanks@11727.11744.18\|AGTAACTATGACTCTCTT | >hg38_21_+_8433172_8446622_RNA45SN1_45S_50ntFlanks | 11727 | 11744 | 18 | AGTAACTATGACTCTCTT | MB468 | 50 | 54 | 22 | 8 | 9 | 10.2 | 29.7 | 4.5 | 1.9 | 4.7 |
| >hg38_21_+_8433172_8446622_RNA45SN1_45S_50ntFlanks@11727.11745.19\|AGTAACTATGACTCTCTTA | >hg38_21_+_8433172_8446622_RNA45SN1_45S_50ntFlanks | 11727 | 11745 | 19 | AGTAACTATGACTCTCTTA | MB231 | 16 | 11 | 23 | 0 | 1 | 18.3 | 7.7 | 24.0 | 0.3 | 2.0 |
| >hg38_21_+_8433172_8446622_RNA45SN1_45S_50ntFlanks@11727.11748.22\|AGTAACTATGACTCTCTTAAGG | >hg38_21_+_8433172_8446622_RNA45SN1_45S_50ntFlanks | 11727 | 11748 | 22 | AGTAACTATGACTCTCTTAAGG | MB231 | 12 | 13 | 22 | 0 | 2 | 10.1 | 8.3 | 19.5 | 0.5 | 3.4 |
| >hg38_21_+_8433172_8446622_RNA45SN1_45S_50ntFlanks@11727.11749.23\|AGTAACTATGACTCTCTTAAGGT | >hg38_21_+_8433172_8446622_RNA45SN1_45S_50ntFlanks | 11727 | 11749 | 23 | AGTAACTATGACTCTCTTAAGGT | MB468 | 12 | 12 | 3 | 2 | 2 | 2.3 | 3.6 | 0.4 | 0.7 | 0.4 |
| >hg38_21_+_8433172_8446622_RNA45SN1_45S_50ntFlanks@11727.11751.25\|AGTAACTATGACTCTCTTAAGGTAG | >hg38_21_+_8433172_8446622_RNA45SN1_45S_50ntFlanks | 11727 | 11751 | 25 | AGTAACTATGACTCTCTTAAGGTAG | MB468 | 91 | 91 | 12 | 8 | 10 | 28.3 | 37.9 | 3.3 | 4.2 | 2.8 |
| >hg38_21_+_8433172_8446622_RNA45SN1_45S_50ntFlanks@11727.11752.26\|AGTAACTATGACTCTCTTAAGGTAGC | >hg38_21_+_8433172_8446622_RNA45SN1_45S_50ntFlanks | 11727 | 11752 | 26 | AGTAACTATGACTCTCTTAAGGTAGC | MB231 | 13 | 5 | 9 | 1 | 1 | 10.7 | 3.0 | 6.7 | 0.8 | 1.3 |
| >hg38_21_+_8433172_8446622_RNA45SN1_45S_50ntFlanks@11727.11753.27\|AGTAACTATGACTCTCTTAAGGTAGCC | >hg38_21_+_8433172_8446622_RNA45SN1_45S_50ntFlanks | 11727 | 11753 | 27 | AGTAACTATGACTCTCTTAAGGTAGCC | MB468 | 34 | 6 | 5 | 8 | 2 | 24.5 | 3.3 | 2.2 | 3.6 | 0.7 |
| >hg38_21_+_8433172_8446622_RNA45SN1_45S_50ntFlanks@11728.11745.18\|GTAACTATGACTCTCTTA | >hg38_21_+_8433172_8446622_RNA45SN1_45S_50ntFlanks | 11728 | 11745 | 18 | GTAACTATGACTCTCTTA | MB468 | 14 | 26 | 4 | 1 | 2 | 3.8 | 9.6 | 0.9 | 0.9 | 1.6 |
| >hg38_21_+_8433172_8446622_RNA45SN1_45S_50ntFlanks@11728.11749.22\|GTAACTATGACTCTCTTAAGGT | >hg38_21_+_8433172_8446622_RNA45SN1_45S_50ntFlanks | 11728 | 11749 | 22 | GTAACTATGACTCTCTTAAGGT | MB468 | 15 | 18 | 3 | 1 | 1 | 5.6 | 11.3 | 0.6 | 1.2 | 0.7 |
| >hg38_21_+_8433172_8446622_RNA45SN1_45S_50ntFlanks@11728.11751.24\|GTAACTATGACTCTCTTAAGGTAG | >hg38_21_+_8433172_8446622_RNA45SN1_45S_50ntFlanks | 11728 | 11751 | 24 | GTAACTATGACTCTCTTAAGGTAG | MB468 | 140 | 177 | 17 | 5 | 10 | 73.3 | 131.4 | 3.0 | 4.2 | 3.8 |
| >hg38_21_+_8433172_8446622_RNA45SN1_45S_50ntFlanks@11728.11753.26\|GTAACTATGACTCTCTTAAGGTAGCC | >hg38_21_+_8433172_8446622_RNA45SN1_45S_50ntFlanks | 11728 | 11753 | 26 | GTAACTATGACTCTCTTAAGGTAGCC | MB468 | 22 | 4 | 3 | 4 | 1 | 16.2 | 1.4 | 1.1 | 2.0 | 0.3 |
| >hg38_21_+_8433172_8446622_RNA45SN1_45S_50ntFlanks@11729.11746.18\|TAACTATGACTCTCTTAA | >hg38_21_+_8433172_8446622_RNA45SN1_45S_50ntFlanks | 11729 | 11746 | 18 | TAACTATGACTCTCTTAA | MB468 | 21 | 14 | 22 | 3 | 7 | 7.7 | 5.9 | 10.7 | 2.1 | 4.6 |
| >hg38_21_+_8433172_8446622_RNA45SN1_45S_50ntFlanks@11729.11747.19\|TAACTATGACTCTCTTAAG | >hg38_21_+_8433172_8446622_RNA45SN1_45S_50ntFlanks | 11729 | 11747 | 19 | TAACTATGACTCTCTTAAG | MB468 | 418 | 579 | 549 | 24 | 127 | 167.5 | 352.4 | 487.8 | 15.5 | 75.1 |
| >hg38_21_+_8433172_8446622_RNA45SN1_45S_50ntFlanks@11729.11748.20\|TAACTATGACTCTCTTAAGG | >hg38_21_+_8433172_8446622_RNA45SN1_45S_50ntFlanks | 11729 | 11748 | 20 | TAACTATGACTCTCTTAAGG | MB231 | 13 | 15 | 32 | 0 | 2 | 10.1 | 9.4 | 25.8 | 0.3 | 4.3 |
| >hg38_21_+_8433172_8446622_RNA45SN1_45S_50ntFlanks@11729.11748.20\|TAACTATGACTCTCTTAAGG | >hg38_21_+_8433172_8446622_RNA45SN1_45S_50ntFlanks | 11729 | 11748 | 20 | TAACTATGACTCTCTTAAGG | MB468 | 351 | 526 | 382 | 43 | 121 | 59.0 | 256.4 | 208.1 | 20.0 | 11.3 |
| >hg38_21_+_8433172_8446622_RNA45SN1_45S_50ntFlanks@11729.11749.21\|TAACTATGACTCTCTTAAGGT | >hg38_21_+_8433172_8446622_RNA45SN1_45S_50ntFlanks | 11729 | 11749 | 21 | TAACTATGACTCTCTTAAGGT | MB468 | 762 | 517 | 460 | 75 | 107 | 130.8 | 130.3 | 76.9 | 32.1 | 19.6 |
| >hg38_21_+_8433172_8446622_RNA45SN1_45S_50ntFlanks@11729.11750.22\|TAACTATGACTCTCTTAAGGTA | >hg38_21_+_8433172_8446622_RNA45SN1_45S_50ntFlanks | 11729 | 11750 | 22 | TAACTATGACTCTCTTAAGGTA | MB468 | 157 | 111 | 150 | 26 | 26 | 21.2 | 41.2 | 37.2 | 4.6 | 11.6 |
| >hg38_21_+_8433172_8446622_RNA45SN1_45S_50ntFlanks@11729.11751.23\|TAACTATGACTCTCTTAAGGTAG | >hg38_21_+_8433172_8446622_RNA45SN1_45S_50ntFlanks | 11729 | 11751 | 23 | TAACTATGACTCTCTTAAGGTAG | MB468 | 7977 | 6038 | 3564 | 405 | 898 | 2085.4 | 1991.0 | 803.1 | 250.6 | 84.9 |
| >hg38_21_+_8433172_8446622_RNA45SN1_45S_50ntFlanks@11729.11752.24\|TAACTATGACTCTCTTAAGGTAGC | >hg38_21_+_8433172_8446622_RNA45SN1_45S_50ntFlanks | 11729 | 11752 | 24 | TAACTATGACTCTCTTAAGGTAGC | MB468 | 236 | 38 | 75 | 28 | 27 | 98.8 | 6.3 | 9.2 | 10.2 | 11.2 |
| >hg38_21_+_8433172_8446622_RNA45SN1_45S_50ntFlanks@11729.11753.25\|TAACTATGACTCTCTTAAGGTAGCC | >hg38_21_+_8433172_8446622_RNA45SN1_45S_50ntFlanks | 11729 | 11753 | 25 | TAACTATGACTCTCTTAAGGTAGCC | MB468 | 291 | 22 | 171 | 105 | 27 | 165.3 | 7.9 | 87.2 | 44.9 | 16.1 |
| >hg38_21_+_8433172_8446622_RNA45SN1_45S_50ntFlanks@11729.11754.26\|TAACTATGACTCTCTTAAGGTAGCCA | >hg38_21_+_8433172_8446622_RNA45SN1_45S_50ntFlanks | 11729 | 11754 | 26 | TAACTATGACTCTCTTAAGGTAGCCA | MB468 | 29 | 0 | 17 | 20 | 2 | 28.9 | 0.5 | 12.6 | 15.7 | 1.5 |
| >hg38_21_+_8433172_8446622_RNA45SN1_45S_50ntFlanks@11729.11761.33\|TAACTATGACTCTCTTAAGGTAGCCAAATGCCT | >hg38_21_+_8433172_8446622_RNA45SN1_45S_50ntFlanks | 11729 | 11761 | 33 | TAACTATGACTCTCTTAAGGTAGCCAAATGCCT | MB468 | 39 | 10 | 37 | 8 | 13 | 19.4 | 4.7 | 13.4 | 5.5 | 7.0 |
| >hg38_21_+_8433172_8446622_RNA45SN1_45S_50ntFlanks@11729.11764.36\|TAACTATGACTCTCTTAAGGTAGCCAAATGCCTCGT | >hg38_21_+_8433172_8446622_RNA45SN1_45S_50ntFlanks | 11729 | 11764 | 36 | TAACTATGACTCTCTTAAGGTAGCCAAATGCCTCGT | MB468 | 47 | 34 | 158 | 4 | 16 | 18.2 | 22.7 | 142.5 | 2.4 | 12.7 |
| >hg38_21_+_8433172_8446622_RNA45SN1_45S_50ntFlanks@11729.11765.37\|TAACTATGACTCTCTTAAGGTAGCCAAATGCCTCGTC | >hg38_21_+_8433172_8446622_RNA45SN1_45S_50ntFlanks | 11729 | 11765 | 37 | TAACTATGACTCTCTTAAGGTAGCCAAATGCCTCGTC | MB468 | 357 | 560 | 808 | 45 | 158 | 106.9 | 351.7 | 434.8 | 28.7 | 111.5 |
| >hg38_21_+_8433172_8446622_RNA45SN1_45S_50ntFlanks@11729.11766.38\|TAACTATGACTCTCTTAAGGTAGCCAAATGCCTCGTCA | >hg38_21_+_8433172_8446622_RNA45SN1_45S_50ntFlanks | 11729 | 11766 | 38 | TAACTATGACTCTCTTAAGGTAGCCAAATGCCTCGTCA | MB468 | 18 | 27 | 51 | 3 | 16 | 7.8 | 27.7 | 30.8 | 1.7 | 13.2 |

| | | | | | | | | | | | | | | | | |
|---|---|---|---|---|---|---|---|---|---|---|---|---|---|---|---|---|
| >hg38_21_+_8433172_8446622_RNA45SN1_45S_50ntFlanks@11729.11767.39\|TAACTATGACTCTCTTAAGGTAGCCAAATGCCTCGTCAT | >hg38_21_+_8433172_8446622_RNA45SN1_45S_50ntFlanks | 11729 | 11767 | 39 | TAACTATGACTCTCTTAAGGTAGCCAAATGCCTCGTCAT | MB468 | 108 | 128 | 369 | 80 | 92 | 31.1 | 97.9 | 237.4 | 48.2 | 58.9 |
| >hg38_21_+_8433172_8446622_RNA45SN1_45S_50ntFlanks@11730.11751.22\|AACTATGACTCTCTTAAGGTAG | >hg38_21_+_8433172_8446622_RNA45SN1_45S_50ntFlanks | 11730 | 11751 | 22 | AACTATGACTCTCTTAAGGTAG | MB468 | 51 | 21 | 13 | 4 | 5 | 24.8 | 8.4 | 4.8 | 2.1 | 1.1 |
| >hg38_21_+_8433172_8446622_RNA45SN1_45S_50ntFlanks@11731.11751.21\|ACTATGACTCTCTTAAGGTAG | >hg38_21_+_8433172_8446622_RNA45SN1_45S_50ntFlanks | 11731 | 11751 | 21 | ACTATGACTCTCTTAAGGTAG | MB468 | 90 | 31 | 19 | 8 | 9 | 25.7 | 7.3 | 3.8 | 2.2 | 2.1 |
| >hg38_21_+_8433172_8446622_RNA45SN1_45S_50ntFlanks@11733.11751.19\|TATGACTCTCTTAAGGTAG | >hg38_21_+_8433172_8446622_RNA45SN1_45S_50ntFlanks | 11733 | 11751 | 19 | TATGACTCTCTTAAGGTAG | MB468 | 38 | 24 | 19 | 6 | 6 | 8.3 | 11.4 | 1.8 | 1.3 | 0.9 |
| >hg38_21_+_8433172_8446622_RNA45SN1_45S_50ntFlanks@11734.11751.18\|ATGACTCTCTTAAGGTAG | >hg38_21_+_8433172_8446622_RNA45SN1_45S_50ntFlanks | 11734 | 11751 | 18 | ATGACTCTCTTAAGGTAG | MB468 | 14 | 6 | 5 | 3 | 2 | 6.2 | 3.5 | 3.7 | 1.3 | 0.6 |
| >hg38_21_+_8433172_8446622_RNA45SN1_45S_50ntFlanks@11737.11768.32\|ACTCTCTTAAGGTAGCCAAATGCCTCGTCATC | >hg38_21_+_8433172_8446622_RNA45SN1_45S_50ntFlanks | 11737 | 11768 | 32 | ACTCTCTTAAGGTAGCCAAATGCCTCGTCATC | MB468 | 12 | 10 | 3 | 1 | 3 | 2.4 | 2.0 | 0.9 | 0.9 | 1.8 |
| >hg38_21_+_8433172_8446622_RNA45SN1_45S_50ntFlanks@11739.11758.20\|TCTCTTAAGGTAGCCAAATG | >hg38_21_+_8433172_8446622_RNA45SN1_45S_50ntFlanks | 11739 | 11758 | 20 | TCTCTTAAGGTAGCCAAATG | MB468 | 31 | 15 | 33 | 3 | 3 | 15.5 | 10.8 | 24.6 | 1.6 | 1.0 |
| >hg38_21_+_8433172_8446622_RNA45SN1_45S_50ntFlanks@11739.11767.29\|TCTCTTAAGGTAGCCAAATGCCTCGTCAT | >hg38_21_+_8433172_8446622_RNA45SN1_45S_50ntFlanks | 11739 | 11767 | 29 | TCTCTTAAGGTAGCCAAATGCCTCGTCAT | MB468 | 18 | 7 | 4 | 4 | 3 | 6.7 | 3.2 | 1.3 | 1.7 | 0.8 |
| >hg38_21_+_8433172_8446622_RNA45SN1_45S_50ntFlanks@11739.11768.30\|TCTCTTAAGGTAGCCAAATGCCTCGTCATC | >hg38_21_+_8433172_8446622_RNA45SN1_45S_50ntFlanks | 11739 | 11768 | 30 | TCTCTTAAGGTAGCCAAATGCCTCGTCATC | MB468 | 32 | 18 | 12 | 7 | 6 | 4.9 | 4.7 | 3.6 | 2.9 | 3.4 |
| >hg38_21_+_8433172_8446622_RNA45SN1_45S_50ntFlanks@11740.11758.19\|CTCTTAAGGTAGCCAAATG | >hg38_21_+_8433172_8446622_RNA45SN1_45S_50ntFlanks | 11740 | 11758 | 19 | CTCTTAAGGTAGCCAAATG | MB468 | 10 | 3 | 10 | 1 | 0 | 7.0 | 2.2 | 10.2 | 0.9 | 0.5 |
| >hg38_21_+_8433172_8446622_RNA45SN1_45S_50ntFlanks@11741.11758.18\|TCTTAAGGTAGCCAAATG | >hg38_21_+_8433172_8446622_RNA45SN1_45S_50ntFlanks | 11741 | 11758 | 18 | TCTTAAGGTAGCCAAATG | MB468 | 32 | 6 | 38 | 2 | 1 | 23.9 | 4.5 | 48.7 | 1.8 | 1.4 |
| >hg38_21_+_8433172_8446622_RNA45SN1_45S_50ntFlanks@11741.11759.19\|TCTTAAGGTAGCCAAATGC | >hg38_21_+_8433172_8446622_RNA45SN1_45S_50ntFlanks | 11741 | 11759 | 19 | TCTTAAGGTAGCCAAATGC | MB468 | 20 | 5 | 23 | 1 | 1 | 13.8 | 2.8 | 23.8 | 0.6 | 0.5 |
| >hg38_21_+_8433172_8446622_RNA45SN1_45S_50ntFlanks@11752.11777.26\|CCAAATGCCTCGTCATCTAATTAGTG | >hg38_21_+_8433172_8446622_RNA45SN1_45S_50ntFlanks | 11752 | 11777 | 26 | CCAAATGCCTCGTCATCTAATTAGTG | MB468 | 11 | 10 | 9 | 2 | 5 | 0.9 | 4.8 | 2.0 | 0.5 | 1.7 |
| >hg38_21_+_8433172_8446622_RNA45SN1_45S_50ntFlanks@11752.11780.29\|CCAAATGCCTCGTCATCTAATTAGTGACG | >hg38_21_+_8433172_8446622_RNA45SN1_45S_50ntFlanks | 11752 | 11780 | 29 | CCAAATGCCTCGTCATCTAATTAGTGACG | MB468 | 21 | 6 | 36 | 4 | 3 | 6.6 | 2.1 | 12.2 | 1.7 | 1.5 |
| >hg38_21_+_8433172_8446622_RNA45SN1_45S_50ntFlanks@11755.11777.23\|AATGCCTCGTCATCTAATTAGTG | >hg38_21_+_8433172_8446622_RNA45SN1_45S_50ntFlanks | 11755 | 11777 | 23 | AATGCCTCGTCATCTAATTAGTG | MB468 | 31 | 18 | 19 | 4 | 15 | 2.8 | 7.0 | 8.2 | 0.9 | 4.9 |
| >hg38_21_+_8433172_8446622_RNA45SN1_45S_50ntFlanks@11755.11780.26\|AATGCCTCGTCATCTAATTAGTGACG | >hg38_21_+_8433172_8446622_RNA45SN1_45S_50ntFlanks | 11755 | 11780 | 26 | AATGCCTCGTCATCTAATTAGTGACG | MB468 | 11 | 1 | 12 | 4 | 1 | 7.4 | 0.4 | 10.2 | 2.5 | 0.1 |
| >hg38_21_+_8433172_8446622_RNA45SN1_45S_50ntFlanks@11759.11777.19\|CCTCGTCATCTAATTAGTG | >hg38_21_+_8433172_8446622_RNA45SN1_45S_50ntFlanks | 11759 | 11777 | 19 | CCTCGTCATCTAATTAGTG | MB468 | 44 | 12 | 37 | 8 | 16 | 19.3 | 4.8 | 25.3 | 2.1 | 3.9 |
| >hg38_21_+_8433172_8446622_RNA45SN1_45S_50ntFlanks@11759.11780.22\|CCTCGTCATCTAATTAGTGACG | >hg38_21_+_8433172_8446622_RNA45SN1_45S_50ntFlanks | 11759 | 11780 | 22 | CCTCGTCATCTAATTAGTGACG | MB468 | 23 | 0 | 20 | 8 | 1 | 20.9 | 0.2 | 12.2 | 7.3 | 0.7 |
| >hg38_21_+_8433172_8446622_RNA45SN1_45S_50ntFlanks@11760.11777.18\|CTCGTCATCTAATTAGTG | >hg38_21_+_8433172_8446622_RNA45SN1_45S_50ntFlanks | 11760 | 11777 | 18 | CTCGTCATCTAATTAGTG | MB468 | 17 | 2 | 10 | 7 | 9 | 2.9 | 0.6 | 4.5 | 2.8 | 3.9 |
| >hg38_21_+_8433172_8446622_RNA45SN1_45S_50ntFlanks@11766.11784.19\|ATCTAATTAGTGACGCGCA | >hg38_21_+_8433172_8446622_RNA45SN1_45S_50ntFlanks | 11766 | 11784 | 19 | ATCTAATTAGTGACGCGCA | MB468 | 22 | 7 | 17 | 2 | 5 | 21.2 | 4.6 | 16.0 | 1.8 | 3.0 |
| >hg38_21_+_8433172_8446622_RNA45SN1_45S_50ntFlanks@11766.11786.21\|ATCTAATTAGTGACGCGCATG | >hg38_21_+_8433172_8446622_RNA45SN1_45S_50ntFlanks | 11766 | 11786 | 21 | ATCTAATTAGTGACGCGCATG | MB468 | 13 | 13 | 14 | 3 | 6 | 4.3 | 8.2 | 6.9 | 1.4 | 2.0 |
| >hg38_21_+_8433172_8446622_RNA45SN1_45S_50ntFlanks@11766.11789.24\|ATCTAATTAGTGACGCGCATGAAT | >hg38_21_+_8433172_8446622_RNA45SN1_45S_50ntFlanks | 11766 | 11789 | 24 | ATCTAATTAGTGACGCGCATGAAT | MB468 | 10 | 20 | 7 | 3 | 3 | 3.6 | 9.7 | 2.4 | 1.9 | 0.7 |
| >hg38_21_+_8433172_8446622_RNA45SN1_45S_50ntFlanks@11766.11790.25\|ATCTAATTAGTGACGCGCATGAATG | >hg38_21_+_8433172_8446622_RNA45SN1_45S_50ntFlanks | 11766 | 11790 | 25 | ATCTAATTAGTGACGCGCATGAATG | MB468 | 27 | 34 | 35 | 14 | 9 | 6.3 | 15.0 | 14.5 | 13.3 | 6.4 |
| >hg38_21_+_8433172_8446622_RNA45SN1_45S_50ntFlanks@11766.11791.26\|ATCTAATTAGTGACGCGCATGAATGG | >hg38_21_+_8433172_8446622_RNA45SN1_45S_50ntFlanks | 11766 | 11791 | 26 | ATCTAATTAGTGACGCGCATGAATGG | MB468 | 35 | 44 | 23 | 9 | 9 | 4.5 | 23.2 | 7.2 | 1.7 | 3.5 |
| >hg38_21_+_8433172_8446622_RNA45SN1_45S_50ntFlanks@11766.11794.29\|ATCTAATTAGTGACGCGCATGAATGGATG | >hg38_21_+_8433172_8446622_RNA45SN1_45S_50ntFlanks | 11766 | 11794 | 29 | ATCTAATTAGTGACGCGCATGAATGGATG | MB468 | 15 | 11 | 19 | 7 | 7 | 3.8 | 5.0 | 8.5 | 2.2 | 5.0 |
| >hg38_21_+_8433172_8446622_RNA45SN1_45S_50ntFlanks@11766.11803.38\|ATCTAATTAGTGACGCGCATGAATGGATGAACGAGATT | >hg38_21_+_8433172_8446622_RNA45SN1_45S_50ntFlanks | 11766 | 11803 | 38 | ATCTAATTAGTGACGCGCATGAATGGATGAACGAGATT | MB468 | 26 | 31 | 24 | 9 | 9 | 9.6 | 5.3 | 13.0 | 6.6 | 3.8 |
| >hg38_21_+_8433172_8446622_RNA45SN1_45S_50ntFlanks@11767.11784.18\|TCTAATTAGTGACGCGCA | >hg38_21_+_8433172_8446622_RNA45SN1_45S_50ntFlanks | 11767 | 11784 | 18 | TCTAATTAGTGACGCGCA | MB468 | 13 | 4 | 12 | 1 | 4 | 9.9 | 1.7 | 10.0 | 0.5 | 2.2 |
| >hg38_21_+_8433172_8446622_RNA45SN1_45S_50ntFlanks@11767.11790.24\|TCTAATTAGTGACGCGCATGAATG | >hg38_21_+_8433172_8446622_RNA45SN1_45S_50ntFlanks | 11767 | 11790 | 24 | TCTAATTAGTGACGCGCATGAATG | MB468 | 21 | 24 | 28 | 6 | 6 | 6.7 | 8.8 | 10.7 | 3.1 | 0.9 |
| >hg38_21_+_8433172_8446622_RNA45SN1_45S_50ntFlanks@11767.11791.25\|TCTAATTAGTGACGCGCATGAATGG | >hg38_21_+_8433172_8446622_RNA45SN1_45S_50ntFlanks | 11767 | 11791 | 25 | TCTAATTAGTGACGCGCATGAATGG | MB468 | 23 | 29 | 17 | 5 | 7 | 2.3 | 16.4 | 5.5 | 1.3 | 3.0 |
| >hg38_21_+_8433172_8446622_RNA45SN1_45S_50ntFlanks@11767.11794.28\|TCTAATTAGTGACGCGCATGAATGGATG | >hg38_21_+_8433172_8446622_RNA45SN1_45S_50ntFlanks | 11767 | 11794 | 28 | TCTAATTAGTGACGCGCATGAATGGATG | MB468 | 142 | 201 | 91 | 39 | 25 | 16.9 | 67.2 | 13.8 | 8.6 | 8.2 |
| >hg38_21_+_8433172_8446622_RNA45SN1_45S_50ntFlanks@11767.11803.37\|TCTAATTAGTGACGCGCATGAATGGATGAACGAGATT | >hg38_21_+_8433172_8446622_RNA45SN1_45S_50ntFlanks | 11767 | 11803 | 37 | TCTAATTAGTGACGCGCATGAATGGATGAACGAGATT | MB468 | 52 | 51 | 36 | 15 | 11 | 7.2 | 8.8 | 18.4 | 10.7 | 2.4 |
| >hg38_21_+_8433172_8446622_RNA45SN1_45S_50ntFlanks@11767.11804.38\|TCTAATTAGTGACGCGCATGAATGGATGAACGAGATTC | >hg38_21_+_8433172_8446622_RNA45SN1_45S_50ntFlanks | 11767 | 11804 | 38 | TCTAATTAGTGACGCGCATGAATGGATGAACGAGATTC | MB468 | 12 | 15 | 11 | 4 | 6 | 1.6 | 3.5 | 5.0 | 3.1 | 1.8 |
| >hg38_21_+_8433172_8446622_RNA45SN1_45S_50ntFlanks@11768.11785.18\|CTAATTAGTGACGCGCAT | >hg38_21_+_8433172_8446622_RNA45SN1_45S_50ntFlanks | 11768 | 11785 | 18 | CTAATTAGTGACGCGCAT | MB468 | 14 | 6 | 14 | 1 | 2 | 9.0 | 2.6 | 11.9 | 0.5 | 1.9 |
| >hg38_21_+_8433172_8446622_RNA45SN1_45S_50ntFlanks@11768.11786.19\|CTAATTAGTGACGCGCATG | >hg38_21_+_8433172_8446622_RNA45SN1_45S_50ntFlanks | 11768 | 11786 | 19 | CTAATTAGTGACGCGCATG | MB468 | 88 | 59 | 97 | 18 | 30 | 28.9 | 35.1 | 63.0 | 6.9 | 8.2 |
| >hg38_21_+_8433172_8446622_RNA45SN1_45S_50ntFlanks@11768.11790.23\|CTAATTAGTGACGCGCATGAATG | >hg38_21_+_8433172_8446622_RNA45SN1_45S_50ntFlanks | 11768 | 11790 | 23 | CTAATTAGTGACGCGCATGAATG | MB468 | 144 | 141 | 173 | 37 | 28 | 31.1 | 52.5 | 36.8 | 18.8 | 7.6 |
| >hg38_21_+_8433172_8446622_RNA45SN1_45S_50ntFlanks@11768.11791.24\|CTAATTAGTGACGCGCATGAATGG | >hg38_21_+_8433172_8446622_RNA45SN1_45S_50ntFlanks | 11768 | 11791 | 24 | CTAATTAGTGACGCGCATGAATGG | MB468 | 75 | 92 | 57 | 19 | 25 | 9.0 | 58.2 | 20.1 | 4.8 | 10.0 |
| >hg38_21_+_8433172_8446622_RNA45SN1_45S_50ntFlanks@11768.11792.25\|CTAATTAGTGACGCGCATGAATGGA | >hg38_21_+_8433172_8446622_RNA45SN1_45S_50ntFlanks | 11768 | 11792 | 25 | CTAATTAGTGACGCGCATGAATGGA | MB468 | 225 | 263 | 191 | 64 | 43 | 41.4 | 93.3 | 67.1 | 41.9 | 21.8 |
| >hg38_21_+_8433172_8446622_RNA45SN1_45S_50ntFlanks@11768.11794.27\|CTAATTAGTGACGCGCATGAATGGATG | >hg38_21_+_8433172_8446622_RNA45SN1_45S_50ntFlanks | 11768 | 11794 | 27 | CTAATTAGTGACGCGCATGAATGGATG | MB468 | 310 | 492 | 234 | 145 | 62 | 14.1 | 114.2 | 93.1 | 41.3 | 10.8 |
| >hg38_21_+_8433172_8446622_RNA45SN1_45S_50ntFlanks@11768.11798.31\|CTAATTAGTGACGCGCATGAATGGATGAACG | >hg38_21_+_8433172_8446622_RNA45SN1_45S_50ntFlanks | 11768 | 11798 | 31 | CTAATTAGTGACGCGCATGAATGGATGAACG | MB468 | 11 | 8 | 7 | 5 | 4 | 3.8 | 2.5 | 3.1 | 1.7 | 1.5 |
| >hg38_21_+_8433172_8446622_RNA45SN1_45S_50ntFlanks@11768.11803.36\|CTAATTAGTGACGCGCATGAATGGATGAACGAGATT | >hg38_21_+_8433172_8446622_RNA45SN1_45S_50ntFlanks | 11768 | 11803 | 36 | CTAATTAGTGACGCGCATGAATGGATGAACGAGATT | MB468 | 213 | 152 | 111 | 76 | 46 | 47.4 | 27.9 | 46.7 | 50.3 | 20.7 |
| >hg38_21_+_8433172_8446622_RNA45SN1_45S_50ntFlanks@11768.11804.37\|CTAATTAGTGACGCGCATGAATGGATGAACGAGATTC | >hg38_21_+_8433172_8446622_RNA45SN1_45S_50ntFlanks | 11768 | 11804 | 37 | CTAATTAGTGACGCGCATGAATGGATGAACGAGATTC | MB468 | 63 | 60 | 37 | 33 | 31 | 19.2 | 22.5 | 12.5 | 20.8 | 18.5 |
| >hg38_21_+_8433172_8446622_RNA45SN1_45S_50ntFlanks@11768.11806.39\|CTAATTAGTGACGCGCATGAATGGATGAACGAGATTCCC | >hg38_21_+_8433172_8446622_RNA45SN1_45S_50ntFlanks | 11768 | 11806 | 39 | CTAATTAGTGACGCGCATGAATGGATGAACGAGATTCCC | MB468 | 13 | 28 | 18 | 8 | 22 | 4.0 | 14.8 | 6.3 | 5.5 | 10.9 |
| >hg38_21_+_8433172_8446622_RNA45SN1_45S_50ntFlanks@11769.11786.18\|TAATTAGTGACGCGCATG | >hg38_21_+_8433172_8446622_RNA45SN1_45S_50ntFlanks | 11769 | 11786 | 18 | TAATTAGTGACGCGCATG | MB468 | 42 | 22 | 47 | 3 | 8 | 11.2 | 12.5 | 37.3 | 2.2 | 1.8 |
| >hg38_21_+_8433172_8446622_RNA45SN1_45S_50ntFlanks@11769.11790.22\|TAATTAGTGACGCGCATGAATG | >hg38_21_+_8433172_8446622_RNA45SN1_45S_50ntFlanks | 11769 | 11790 | 22 | TAATTAGTGACGCGCATGAATG | MB468 | 63 | 49 | 75 | 11 | 9 | 15.9 | 10.7 | 23.3 | 7.4 | 3.8 |
| >hg38_21_+_8433172_8446622_RNA45SN1_45S_50ntFlanks@11769.11791.23\|TAATTAGTGACGCGCATGAATGG | >hg38_21_+_8433172_8446622_RNA45SN1_45S_50ntFlanks | 11769 | 11791 | 23 | TAATTAGTGACGCGCATGAATGG | MB468 | 111 | 137 | 85 | 16 | 13 | 18.3 | 44.1 | 22.2 | 7.3 | 2.9 |
| >hg38_21_+_8433172_8446622_RNA45SN1_45S_50ntFlanks@11769.11792.24\|TAATTAGTGACGCGCATGAATGGA | >hg38_21_+_8433172_8446622_RNA45SN1_45S_50ntFlanks | 11769 | 11792 | 24 | TAATTAGTGACGCGCATGAATGGA | MB468 | 76 | 104 | 62 | 13 | 10 | 11.4 | 21.3 | 20.0 | 7.5 | 3.5 |
| >hg38_21_+_8433172_8446622_RNA45SN1_45S_50ntFlanks@11769.11794.26\|TAATTAGTGACGCGCATGAATGGATG | >hg38_21_+_8433172_8446622_RNA45SN1_45S_50ntFlanks | 11769 | 11794 | 26 | TAATTAGTGACGCGCATGAATGGATG | MB468 | 114 | 190 | 92 | 31 | 17 | 17.4 | 42.2 | 25.0 | 17.3 | 5.1 |
| >hg38_21_+_8433172_8446622_RNA45SN1_45S_50ntFlanks@11769.11803.35\|TAATTAGTGACGCGCATGAATGGATGAACGAGATT | >hg38_21_+_8433172_8446622_RNA45SN1_45S_50ntFlanks | 11769 | 11803 | 35 | TAATTAGTGACGCGCATGAATGGATGAACGAGATT | MB468 | 69 | 62 | 59 | 10 | 10 | 11.9 | 9.1 | 18.0 | 5.8 | 4.6 |
| >hg38_21_+_8433172_8446622_RNA45SN1_45S_50ntFlanks@11769.11804.36\|TAATTAGTGACGCGCATGAATGGATGAACGAGATTC | >hg38_21_+_8433172_8446622_RNA45SN1_45S_50ntFlanks | 11769 | 11804 | 36 | TAATTAGTGACGCGCATGAATGGATGAACGAGATTC | MB468 | 48 | 70 | 53 | 13 | 16 | 7.1 | 18.4 | 8.7 | 7.9 | 8.2 |
| >hg38_21_+_8433172_8446622_RNA45SN1_45S_50ntFlanks@11770.11794.25\|AATTAGTGACGCGCATGAATGGATG | >hg38_21_+_8433172_8446622_RNA45SN1_45S_50ntFlanks | 11770 | 11794 | 25 | AATTAGTGACGCGCATGAATGGATG | MB231 | 22 | 10 | 80 | 12 | 4 | 18.0 | 7.4 | 59.0 | 9.3 | 4.1 |

| | | | | | | | | | | | | | | | | |
|---|---|---|---|---|---|---|---|---|---|---|---|---|---|---|---|---|
| >hg38_21_+_8433172_8446622_RNA45SN1_45S_50ntFlanks@11770.11794.25\|AATTAGTGACGCGCATGAATGGATG | >hg38_21_+_8433172_8446622_RNA45SN1_45S_50ntFlanks | 11770 | 11794 | 25 | AATTAGTGACGCGCATGAATGGATG | MB468 | 17 | 34 | 11 | 12 | 4 | 3.0 | 5.8 | 4.0 | 5.6 | 0.7 |
| >hg38_21_+_8433172_8446622_RNA45SN1_45S_50ntFlanks@11770.11803.34\|AATTAGTGACGCGCATGAATGGATGAACGAGATT | >hg38_21_+_8433172_8446622_RNA45SN1_45S_50ntFlanks | 11770 | 11803 | 34 | AATTAGTGACGCGCATGAATGGATGAACGAGATT | MB231 | 10 | 4 | 8 | 4 | 3 | 9.1 | 4.0 | 11.4 | 2.2 | 3.2 |
| >hg38_21_+_8433172_8446622_RNA45SN1_45S_50ntFlanks@11771.11790.20\|ATTAGTGACGCGCATGAATG | >hg38_21_+_8433172_8446622_RNA45SN1_45S_50ntFlanks | 11771 | 11790 | 20 | ATTAGTGACGCGCATGAATG | MB468 | 11 | 13 | 12 | 6 | 3 | 2.0 | 5.3 | 3.1 | 5.5 | 1.8 |
| >hg38_21_+_8433172_8446622_RNA45SN1_45S_50ntFlanks@11771.11791.21\|ATTAGTGACGCGCATGAATGG | >hg38_21_+_8433172_8446622_RNA45SN1_45S_50ntFlanks | 11771 | 11791 | 21 | ATTAGTGACGCGCATGAATGG | MB468 | 17 | 26 | 11 | 7 | 3 | 4.2 | 13.2 | 5.5 | 1.9 | 0.9 |
| >hg38_21_+_8433172_8446622_RNA45SN1_45S_50ntFlanks@11771.11792.22\|ATTAGTGACGCGCATGAATGGA | >hg38_21_+_8433172_8446622_RNA45SN1_45S_50ntFlanks | 11771 | 11792 | 22 | ATTAGTGACGCGCATGAATGGA | MB468 | 10 | 19 | 7 | 6 | 3 | 1.7 | 6.4 | 3.5 | 1.7 | 0.6 |
| >hg38_21_+_8433172_8446622_RNA45SN1_45S_50ntFlanks@11771.11794.24\|ATTAGTGACGCGCATGAATGGATG | >hg38_21_+_8433172_8446622_RNA45SN1_45S_50ntFlanks | 11771 | 11794 | 24 | ATTAGTGACGCGCATGAATGGATG | MB231 | 11 | 5 | 41 | 4 | 2 | 8.0 | 3.6 | 29.1 | 2.9 | 1.1 |
| >hg38_21_+_8433172_8446622_RNA45SN1_45S_50ntFlanks@11771.11794.24\|ATTAGTGACGCGCATGAATGGATG | >hg38_21_+_8433172_8446622_RNA45SN1_45S_50ntFlanks | 11771 | 11794 | 24 | ATTAGTGACGCGCATGAATGGATG | MB468 | 28 | 66 | 19 | 22 | 8 | 3.6 | 19.0 | 8.7 | 9.9 | 2.3 |
| >hg38_21_+_8433172_8446622_RNA45SN1_45S_50ntFlanks@11771.11803.33\|ATTAGTGACGCGCATGAATGGATGAACGAGATT | >hg38_21_+_8433172_8446622_RNA45SN1_45S_50ntFlanks | 11771 | 11803 | 33 | ATTAGTGACGCGCATGAATGGATGAACGAGATT | MB468 | 10 | 8 | 6 | 4 | 2 | 3.2 | 3.6 | 2.3 | 1.5 | 1.6 |
| >hg38_21_+_8433172_8446622_RNA45SN1_45S_50ntFlanks@11772.11794.23\|TTAGTGACGCGCATGAATGGATG | >hg38_21_+_8433172_8446622_RNA45SN1_45S_50ntFlanks | 11772 | 11794 | 23 | TTAGTGACGCGCATGAATGGATG | MB468 | 23 | 50 | 18 | 14 | 4 | 2.7 | 14.5 | 6.9 | 6.7 | 1.0 |
| >hg38_21_+_8433172_8446622_RNA45SN1_45S_50ntFlanks@11773.11793.21\|TAGTGACGCGCATGAATGGAT | >hg38_21_+_8433172_8446622_RNA45SN1_45S_50ntFlanks | 11773 | 11793 | 21 | TAGTGACGCGCATGAATGGAT | BT20 | 16 | 8 | 4 | 0 | 0 | 28.1 | 14.3 | 6.6 | 0.4 | 0.0 |
| >hg38_21_+_8433172_8446622_RNA45SN1_45S_50ntFlanks@11774.11793.20\|AGTGACGCGCATGAATGGAT | >hg38_21_+_8433172_8446622_RNA45SN1_45S_50ntFlanks | 11774 | 11793 | 20 | AGTGACGCGCATGAATGGAT | BT20 | 19 | 12 | 8 | 3 | 1 | 27.9 | 19.1 | 8.9 | 3.4 | 0.9 |
| >hg38_21_+_8433172_8446622_RNA45SN1_45S_50ntFlanks@11774.11794.21\|AGTGACGCGCATGAATGGATG | >hg38_21_+_8433172_8446622_RNA45SN1_45S_50ntFlanks | 11774 | 11794 | 21 | AGTGACGCGCATGAATGGATG | MB231 | 13 | 8 | 40 | 8 | 3 | 11.9 | 6.7 | 31.9 | 7.5 | 3.5 |
| >hg38_21_+_8433172_8446622_RNA45SN1_45S_50ntFlanks@11775.11793.19\|GTGACGCGCATGAATGGAT | >hg38_21_+_8433172_8446622_RNA45SN1_45S_50ntFlanks | 11775 | 11793 | 19 | GTGACGCGCATGAATGGAT | BT20 | 22 | 15 | 6 | 2 | 0 | 37.4 | 25.5 | 8.2 | 2.6 | 0.8 |
| >hg38_21_+_8433172_8446622_RNA45SN1_45S_50ntFlanks@11778.11798.21\|ACGCGCATGAATGGATGAACG | >hg38_21_+_8433172_8446622_RNA45SN1_45S_50ntFlanks | 11778 | 11798 | 21 | ACGCGCATGAATGGATGAACG | MB468 | 22 | 5 | 9 | 17 | 5 | 6.1 | 2.1 | 2.1 | 8.9 | 3.4 |
| >hg38_21_+_8433172_8446622_RNA45SN1_45S_50ntFlanks@11778.11803.26\|ACGCGCATGAATGGATGAACGAGATT | >hg38_21_+_8433172_8446622_RNA45SN1_45S_50ntFlanks | 11778 | 11803 | 26 | ACGCGCATGAATGGATGAACGAGATT | MB468 | 78 | 21 | 26 | 59 | 12 | 26.2 | 7.8 | 8.1 | 25.3 | 5.3 |
| >hg38_21_+_8433172_8446622_RNA45SN1_45S_50ntFlanks@11778.11804.27\|ACGCGCATGAATGGATGAACGAGATTC | >hg38_21_+_8433172_8446622_RNA45SN1_45S_50ntFlanks | 11778 | 11804 | 27 | ACGCGCATGAATGGATGAACGAGATTC | MB468 | 48 | 21 | 18 | 73 | 17 | 17.7 | 8.9 | 7.0 | 38.8 | 5.1 |
| >hg38_21_+_8433172_8446622_RNA45SN1_45S_50ntFlanks@11778.11805.28\|ACGCGCATGAATGGATGAACGAGATTCC | >hg38_21_+_8433172_8446622_RNA45SN1_45S_50ntFlanks | 11778 | 11805 | 28 | ACGCGCATGAATGGATGAACGAGATTCC | MB468 | 12 | 7 | 9 | 6 | 5 | 1.7 | 2.4 | 2.0 | 0.9 | 1.4 |
| >hg38_21_+_8433172_8446622_RNA45SN1_45S_50ntFlanks@11778.11806.29\|ACGCGCATGAATGGATGAACGAGATTCCC | >hg38_21_+_8433172_8446622_RNA45SN1_45S_50ntFlanks | 11778 | 11806 | 29 | ACGCGCATGAATGGATGAACGAGATTCCC | MB468 | 16 | 14 | 15 | 13 | 11 | 1.7 | 4.9 | 2.7 | 6.1 | 1.3 |
| >hg38_21_+_8433172_8446622_RNA45SN1_45S_50ntFlanks@11781.11798.18\|CGCATGAATGGATGAACG | >hg38_21_+_8433172_8446622_RNA45SN1_45S_50ntFlanks | 11781 | 11798 | 18 | CGCATGAATGGATGAACG | MB468 | 89 | 59 | 66 | 47 | 17 | 25.1 | 23.1 | 43.1 | 17.1 | 2.5 |
| >hg38_21_+_8433172_8446622_RNA45SN1_45S_50ntFlanks@11781.11803.23\|CGCATGAATGGATGAACGAGATT | >hg38_21_+_8433172_8446622_RNA45SN1_45S_50ntFlanks | 11781 | 11803 | 23 | CGCATGAATGGATGAACGAGATT | MB468 | 196 | 217 | 68 | 127 | 32 | 16.3 | 66.4 | 32.3 | 62.4 | 13.4 |
| >hg38_21_+_8433172_8446622_RNA45SN1_45S_50ntFlanks@11781.11804.24\|CGCATGAATGGATGAACGAGATTC | >hg38_21_+_8433172_8446622_RNA45SN1_45S_50ntFlanks | 11781 | 11804 | 24 | CGCATGAATGGATGAACGAGATTC | MB468 | 355 | 491 | 151 | 339 | 102 | 38.1 | 175.6 | 52.5 | 174.0 | 25.0 |
| >hg38_21_+_8433172_8446622_RNA45SN1_45S_50ntFlanks@11781.11805.25\|CGCATGAATGGATGAACGAGATTCC | >hg38_21_+_8433172_8446622_RNA45SN1_45S_50ntFlanks | 11781 | 11805 | 25 | CGCATGAATGGATGAACGAGATTCC | MB468 | 58 | 61 | 36 | 31 | 18 | 2.1 | 16.4 | 12.7 | 8.5 | 2.9 |
| >hg38_21_+_8433172_8446622_RNA45SN1_45S_50ntFlanks@11781.11806.26\|CGCATGAATGGATGAACGAGATTCCC | >hg38_21_+_8433172_8446622_RNA45SN1_45S_50ntFlanks | 11781 | 11806 | 26 | CGCATGAATGGATGAACGAGATTCCC | MB468 | 83 | 60 | 46 | 57 | 49 | 3.9 | 10.3 | 14.0 | 26.2 | 4.1 |
| >hg38_21_+_8433172_8446622_RNA45SN1_45S_50ntFlanks@11781.11807.27\|CGCATGAATGGATGAACGAGATTCCCA | >hg38_21_+_8433172_8446622_RNA45SN1_45S_50ntFlanks | 11781 | 11807 | 27 | CGCATGAATGGATGAACGAGATTCCCA | MB468 | 12 | 5 | 5 | 12 | 7 | 1.7 | 1.9 | 2.8 | 11.2 | 4.3 |
| >hg38_21_+_8433172_8446622_RNA45SN1_45S_50ntFlanks@11782.11804.23\|GCATGAATGGATGAACGAGATTC | >hg38_21_+_8433172_8446622_RNA45SN1_45S_50ntFlanks | 11782 | 11804 | 23 | GCATGAATGGATGAACGAGATTC | MB468 | 20 | 32 | 10 | 17 | 5 | 4.2 | 14.5 | 3.0 | 7.4 | 0.3 |
| >hg38_21_+_8433172_8446622_RNA45SN1_45S_50ntFlanks@11783.11800.18\|CATGAATGGATGAACGAG | >hg38_21_+_8433172_8446622_RNA45SN1_45S_50ntFlanks | 11783 | 11800 | 18 | CATGAATGGATGAACGAG | MB468 | 48 | 70 | 30 | 52 | 17 | 17.1 | 39.0 | 15.5 | 17.3 | 4.6 |
| >hg38_21_+_8433172_8446622_RNA45SN1_45S_50ntFlanks@11783.11802.20\|CATGAATGGATGAACGAGAT | >hg38_21_+_8433172_8446622_RNA45SN1_45S_50ntFlanks | 11783 | 11802 | 20 | CATGAATGGATGAACGAGAT | MB468 | 33 | 30 | 16 | 52 | 13 | 6.7 | 8.7 | 11.2 | 38.6 | 7.6 |
| >hg38_21_+_8433172_8446622_RNA45SN1_45S_50ntFlanks@11783.11803.21\|CATGAATGGATGAACGAGATT | >hg38_21_+_8433172_8446622_RNA45SN1_45S_50ntFlanks | 11783 | 11803 | 21 | CATGAATGGATGAACGAGATT | MB468 | 1005 | 1213 | 283 | 433 | 105 | 325.5 | 385.9 | 225.2 | 162.1 | 50.8 |
| >hg38_21_+_8433172_8446622_RNA45SN1_45S_50ntFlanks@11783.11804.22\|CATGAATGGATGAACGAGATTC | >hg38_21_+_8433172_8446622_RNA45SN1_45S_50ntFlanks | 11783 | 11804 | 22 | CATGAATGGATGAACGAGATTC | MB468 | 2949 | 2843 | 1463 | 1537 | 564 | 637.1 | 1194.5 | 1013.2 | 362.3 | 83.8 |
| >hg38_21_+_8433172_8446622_RNA45SN1_45S_50ntFlanks@11783.11805.23\|CATGAATGGATGAACGAGATTCC | >hg38_21_+_8433172_8446622_RNA45SN1_45S_50ntFlanks | 11783 | 11805 | 23 | CATGAATGGATGAACGAGATTCC | MB468 | 275 | 141 | 160 | 115 | 63 | 105.3 | 52.2 | 153.4 | 24.5 | 12.1 |
| >hg38_21_+_8433172_8446622_RNA45SN1_45S_50ntFlanks@11783.11806.24\|CATGAATGGATGAACGAGATTCCC | >hg38_21_+_8433172_8446622_RNA45SN1_45S_50ntFlanks | 11783 | 11806 | 24 | CATGAATGGATGAACGAGATTCCC | MB468 | 363 | 108 | 150 | 211 | 156 | 145.9 | 16.6 | 138.4 | 78.5 | 46.5 |
| >hg38_21_+_8433172_8446622_RNA45SN1_45S_50ntFlanks@11783.11807.25\|CATGAATGGATGAACGAGATTCCCA | >hg38_21_+_8433172_8446622_RNA45SN1_45S_50ntFlanks | 11783 | 11807 | 25 | CATGAATGGATGAACGAGATTCCCA | MB468 | 32 | 6 | 9 | 28 | 11 | 16.9 | 1.8 | 7.4 | 28.4 | 6.2 |
| >hg38_21_+_8433172_8446622_RNA45SN1_45S_50ntFlanks@11783.11810.28\|CATGAATGGATGAACGAGATTCCCACTG | >hg38_21_+_8433172_8446622_RNA45SN1_45S_50ntFlanks | 11783 | 11810 | 28 | CATGAATGGATGAACGAGATTCCCACTG | MB468 | 14 | 2 | 3 | 15 | 7 | 7.3 | 0.9 | 2.8 | 14.0 | 6.1 |
| >hg38_21_+_8433172_8446622_RNA45SN1_45S_50ntFlanks@11783.11816.34\|CATGAATGGATGAACGAGATTCCCACTGTCCCTA | >hg38_21_+_8433172_8446622_RNA45SN1_45S_50ntFlanks | 11783 | 11816 | 34 | CATGAATGGATGAACGAGATTCCCACTGTCCCTA | MB468 | 11 | 6 | 7 | 18 | 11 | 2.4 | 1.5 | 2.9 | 16.8 | 9.3 |
| >hg38_21_+_8433172_8446622_RNA45SN1_45S_50ntFlanks@11784.11803.20\|ATGAATGGATGAACGAGATT | >hg38_21_+_8433172_8446622_RNA45SN1_45S_50ntFlanks | 11784 | 11803 | 20 | ATGAATGGATGAACGAGATT | MB468 | 295 | 353 | 96 | 119 | 24 | 120.4 | 145.2 | 89.0 | 31.2 | 5.7 |
| >hg38_21_+_8433172_8446622_RNA45SN1_45S_50ntFlanks@11784.11804.21\|ATGAATGGATGAACGAGATTC | >hg38_21_+_8433172_8446622_RNA45SN1_45S_50ntFlanks | 11784 | 11804 | 21 | ATGAATGGATGAACGAGATTC | MB468 | 702 | 751 | 408 | 449 | 97 | 197.7 | 296.7 | 360.1 | 151.4 | 21.7 |
| >hg38_21_+_8433172_8446622_RNA45SN1_45S_50ntFlanks@11784.11805.22\|ATGAATGGATGAACGAGATTCC | >hg38_21_+_8433172_8446622_RNA45SN1_45S_50ntFlanks | 11784 | 11805 | 22 | ATGAATGGATGAACGAGATTCC | MB468 | 82 | 36 | 47 | 43 | 12 | 38.4 | 11.2 | 54.8 | 14.7 | 3.0 |
| >hg38_21_+_8433172_8446622_RNA45SN1_45S_50ntFlanks@11784.11806.23\|ATGAATGGATGAACGAGATTCCC | >hg38_21_+_8433172_8446622_RNA45SN1_45S_50ntFlanks | 11784 | 11806 | 23 | ATGAATGGATGAACGAGATTCCC | MB468 | 117 | 28 | 45 | 81 | 34 | 58.0 | 5.7 | 47.7 | 29.3 | 14.3 |
| >hg38_21_+_8433172_8446622_RNA45SN1_45S_50ntFlanks@11785.11802.18\|TGAATGGATGAACGAGAT | >hg38_21_+_8433172_8446622_RNA45SN1_45S_50ntFlanks | 11785 | 11802 | 18 | TGAATGGATGAACGAGAT | MB468 | 14 | 14 | 5 | 24 | 5 | 1.7 | 6.0 | 2.7 | 31.3 | 4.7 |
| >hg38_21_+_8433172_8446622_RNA45SN1_45S_50ntFlanks@11785.11803.19\|TGAATGGATGAACGAGATT | >hg38_21_+_8433172_8446622_RNA45SN1_45S_50ntFlanks | 11785 | 11803 | 19 | TGAATGGATGAACGAGATT | MB468 | 397 | 479 | 131 | 124 | 26 | 213.6 | 239.3 | 145.6 | 39.4 | 6.1 |
| >hg38_21_+_8433172_8446622_RNA45SN1_45S_50ntFlanks@11785.11804.20\|TGAATGGATGAACGAGATTC | >hg38_21_+_8433172_8446622_RNA45SN1_45S_50ntFlanks | 11785 | 11804 | 20 | TGAATGGATGAACGAGATTC | MB468 | 1158 | 1029 | 610 | 481 | 116 | 512.7 | 557.0 | 635.0 | 175.8 | 31.1 |
| >hg38_21_+_8433172_8446622_RNA45SN1_45S_50ntFlanks@11785.11805.21\|TGAATGGATGAACGAGATTCC | >hg38_21_+_8433172_8446622_RNA45SN1_45S_50ntFlanks | 11785 | 11805 | 21 | TGAATGGATGAACGAGATTCC | MB468 | 115 | 35 | 73 | 46 | 13 | 81.9 | 13.0 | 98.9 | 16.7 | 4.7 |
| >hg38_21_+_8433172_8446622_RNA45SN1_45S_50ntFlanks@11785.11806.22\|TGAATGGATGAACGAGATTCCC | >hg38_21_+_8433172_8446622_RNA45SN1_45S_50ntFlanks | 11785 | 11806 | 22 | TGAATGGATGAACGAGATTCCC | MB468 | 120 | 19 | 51 | 63 | 24 | 93.5 | 6.6 | 67.3 | 23.3 | 15.7 |
| >hg38_21_+_8433172_8446622_RNA45SN1_45S_50ntFlanks@11786.11803.18\|GAATGGATGAACGAGATT | >hg38_21_+_8433172_8446622_RNA45SN1_45S_50ntFlanks | 11786 | 11803 | 18 | GAATGGATGAACGAGATT | MB468 | 134 | 138 | 36 | 65 | 11 | 63.5 | 70.9 | 37.4 | 13.2 | 2.9 |
| >hg38_21_+_8433172_8446622_RNA45SN1_45S_50ntFlanks@11786.11804.19\|GAATGGATGAACGAGATTC | >hg38_21_+_8433172_8446622_RNA45SN1_45S_50ntFlanks | 11786 | 11804 | 19 | GAATGGATGAACGAGATTC | MB468 | 172 | 88 | 76 | 111 | 22 | 86.2 | 53.3 | 96.0 | 47.6 | 3.1 |
| >hg38_21_+_8433172_8446622_RNA45SN1_45S_50ntFlanks@11786.11805.20\|GAATGGATGAACGAGATTCC | >hg38_21_+_8433172_8446622_RNA45SN1_45S_50ntFlanks | 11786 | 11805 | 20 | GAATGGATGAACGAGATTCC | MB468 | 11 | 3 | 7 | 7 | 2 | 7.1 | 1.1 | 9.1 | 2.7 | 0.6 |
| >hg38_21_+_8433172_8446622_RNA45SN1_45S_50ntFlanks@11787.11804.18\|AATGGATGAACGAGATTC | >hg38_21_+_8433172_8446622_RNA45SN1_45S_50ntFlanks | 11787 | 11804 | 18 | AATGGATGAACGAGATTC | MB468 | 927 | 594 | 286 | 701 | 113 | 321.3 | 309.9 | 254.6 | 239.3 | 23.9 |
| >hg38_21_+_8433172_8446622_RNA45SN1_45S_50ntFlanks@11787.11805.19\|AATGGATGAACGAGATTCC | >hg38_21_+_8433172_8446622_RNA45SN1_45S_50ntFlanks | 11787 | 11805 | 19 | AATGGATGAACGAGATTCC | MB468 | 16 | 1 | 7 | 9 | 2 | 12.5 | 0.7 | 10.1 | 3.1 | 1.2 |
| >hg38_21_+_8433172_8446622_RNA45SN1_45S_50ntFlanks@11787.11806.20\|AATGGATGAACGAGATTCCC | >hg38_21_+_8433172_8446622_RNA45SN1_45S_50ntFlanks | 11787 | 11806 | 20 | AATGGATGAACGAGATTCCC | MB468 | 22 | 1 | 6 | 24 | 4 | 18.1 | 0.4 | 8.3 | 21.2 | 3.2 |
| >hg38_21_+_8433172_8446622_RNA45SN1_45S_50ntFlanks@11792.11810.19\|ATGAACGAGATTCCCACTG | >hg38_21_+_8433172_8446622_RNA45SN1_45S_50ntFlanks | 11792 | 11810 | 19 | ATGAACGAGATTCCCACTG | MB468 | 25 | 11 | 23 | 30 | 10 | 9.6 | 2.9 | 21.8 | 22.6 | 2.7 |

| | | | | | | | | | | | | | | | | |
|---|---|---|---|---|---|---|---|---|---|---|---|---|---|---|---|---|
| >hg38_21_+_8433172_8446622_RNA45SN1_45S_50ntFlanks@11792.11820.29\|ATGAACGAGATTCCCACTGTCCCTACCTA | >hg38_21_+_8433172_8446622_RNA45SN1_45S_50ntFlanks | 11792 | 11820 | 29 | ATGAACGAGATTCCCACTGTCCCTACCTA | MB468 | 10 | 16 | 11 | 15 | 21 | 2.0 | 3.8 | 3.7 | 8.6 | 10.3 |
| >hg38_21_+_8433172_8446622_RNA45SN1_45S_50ntFlanks@11795.11820.26\|AACGAGATTCCCACTGTCCCTACCTA | >hg38_21_+_8433172_8446622_RNA45SN1_45S_50ntFlanks | 11795 | 11820 | 26 | AACGAGATTCCCACTGTCCCTACCTA | MB468 | 12 | 21 | 8 | 5 | 19 | 2.2 | 5.9 | 3.0 | 4.3 | 7.3 |
| >hg38_21_+_8433172_8446622_RNA45SN1_45S_50ntFlanks@11804.11821.18\|CCCACTGTCCCTACCTAC | >hg38_21_+_8433172_8446622_RNA45SN1_45S_50ntFlanks | 11804 | 11821 | 18 | CCCACTGTCCCTACCTAC | MB468 | 14 | 36 | 18 | 1 | 16 | 5.8 | 27.5 | 5.5 | 0.6 | 13.5 |
| >hg38_21_+_8433172_8446622_RNA45SN1_45S_50ntFlanks@11805.11823.19\|CCACTGTCCCTACCTACTA | >hg38_21_+_8433172_8446622_RNA45SN1_45S_50ntFlanks | 11805 | 11823 | 19 | CCACTGTCCCTACCTACTA | MB468 | 11 | 20 | 11 | 3 | 17 | 4.8 | 8.1 | 5.0 | 2.1 | 6.0 |
| >hg38_21_+_8433172_8446622_RNA45SN1_45S_50ntFlanks@11821.11842.22\|CTATCCAGCGAAACCACAGCCA | >hg38_21_+_8433172_8446622_RNA45SN1_45S_50ntFlanks | 11821 | 11842 | 22 | CTATCCAGCGAAACCACAGCCA | MB468 | 19 | 17 | 14 | 21 | 17 | 2.8 | 6.0 | 4.9 | 16.5 | 9.5 |
| >hg38_21_+_8433172_8446622_RNA45SN1_45S_50ntFlanks@11821.11844.24\|CTATCCAGCGAAACCACAGCCAAG | >hg38_21_+_8433172_8446622_RNA45SN1_45S_50ntFlanks | 11821 | 11844 | 24 | CTATCCAGCGAAACCACAGCCAAG | MB468 | 28 | 23 | 15 | 45 | 29 | 2.9 | 8.0 | 2.1 | 37.3 | 12.0 |
| >hg38_21_+_8433172_8446622_RNA45SN1_45S_50ntFlanks@11843.11872.30\|AGGGAACGGGCTTGGCGGAATCAGCGGGGA | >hg38_21_+_8433172_8446622_RNA45SN1_45S_50ntFlanks | 11843 | 11872 | 30 | AGGGAACGGGCTTGGCGGAATCAGCGGGGA | MB468 | 17 | 19 | 19 | 79 | 23 | 5.0 | 4.4 | 9.1 | 19.3 | 7.5 |
| >hg38_21_+_8433172_8446622_RNA45SN1_45S_50ntFlanks@11844.11872.29\|GGGAACGGGCTTGGCGGAATCAGCGGGGA | >hg38_21_+_8433172_8446622_RNA45SN1_45S_50ntFlanks | 11844 | 11872 | 29 | GGGAACGGGCTTGGCGGAATCAGCGGGGA | MB468 | 11 | 11 | 11 | 63 | 12 | 4.3 | 2.6 | 4.3 | 14.2 | 6.4 |
| >hg38_21_+_8433172_8446622_RNA45SN1_45S_50ntFlanks@11845.11872.28\|GGAACGGGCTTGGCGGAATCAGCGGGGA | >hg38_21_+_8433172_8446622_RNA45SN1_45S_50ntFlanks | 11845 | 11872 | 28 | GGAACGGGCTTGGCGGAATCAGCGGGGA | MB468 | 30 | 31 | 19 | 168 | 50 | 9.0 | 2.7 | 6.7 | 30.5 | 14.9 |
| >hg38_21_+_8433172_8446622_RNA45SN1_45S_50ntFlanks@11847.11872.26\|AACGGGCTTGGCGGAATCAGCGGGGA | >hg38_21_+_8433172_8446622_RNA45SN1_45S_50ntFlanks | 11847 | 11872 | 26 | AACGGGCTTGGCGGAATCAGCGGGGA | MB468 | 29 | 35 | 19 | 117 | 47 | 6.7 | 1.7 | 5.6 | 40.4 | 16.0 |
| >hg38_21_+_8433172_8446622_RNA45SN1_45S_50ntFlanks@11848.11872.25\|ACGGGCTTGGCGGAATCAGCGGGGA | >hg38_21_+_8433172_8446622_RNA45SN1_45S_50ntFlanks | 11848 | 11872 | 25 | ACGGGCTTGGCGGAATCAGCGGGGA | MB468 | 16 | 21 | 11 | 68 | 27 | 4.4 | 0.4 | 2.8 | 20.4 | 10.6 |
| >hg38_21_+_8433172_8446622_RNA45SN1_45S_50ntFlanks@11849.11872.24\|CGGGCTTGGCGGAATCAGCGGGGA | >hg38_21_+_8433172_8446622_RNA45SN1_45S_50ntFlanks | 11849 | 11872 | 24 | CGGGCTTGGCGGAATCAGCGGGGA | MB468 | 31 | 45 | 23 | 113 | 58 | 6.9 | 5.1 | 5.1 | 41.7 | 21.7 |
| >hg38_21_+_8433172_8446622_RNA45SN1_45S_50ntFlanks@11853.11872.20\|CTTGGCGGAATCAGCGGGGA | >hg38_21_+_8433172_8446622_RNA45SN1_45S_50ntFlanks | 11853 | 11872 | 20 | CTTGGCGGAATCAGCGGGGA | MB468 | 15 | 15 | 9 | 22 | 14 | 9.5 | 6.9 | 3.5 | 4.1 | 6.8 |
| >hg38_21_+_8433172_8446622_RNA45SN1_45S_50ntFlanks@11872.11893.22\|AAAGAAGACCCTGTTGAGCTTG | >hg38_21_+_8433172_8446622_RNA45SN1_45S_50ntFlanks | 11872 | 11893 | 22 | AAAGAAGACCCTGTTGAGCTTG | MB468 | 28 | 18 | 22 | 25 | 11 | 5.6 | 8.3 | 8.8 | 8.9 | 3.9 |
| >hg38_21_+_8433172_8446622_RNA45SN1_45S_50ntFlanks@11873.11890.18\|AAGAAGACCCTGTTGAGC | >hg38_21_+_8433172_8446622_RNA45SN1_45S_50ntFlanks | 11873 | 11890 | 18 | AAGAAGACCCTGTTGAGC | MB468 | 119 | 113 | 67 | 96 | 35 | 24.5 | 58.0 | 25.0 | 46.6 | 10.5 |
| >hg38_21_+_8433172_8446622_RNA45SN1_45S_50ntFlanks@11873.11892.20\|AAGAAGACCCTGTTGAGCTT | >hg38_21_+_8433172_8446622_RNA45SN1_45S_50ntFlanks | 11873 | 11892 | 20 | AAGAAGACCCTGTTGAGCTT | MB468 | 99 | 51 | 43 | 54 | 29 | 6.0 | 23.3 | 6.9 | 16.0 | 8.9 |
| >hg38_21_+_8433172_8446622_RNA45SN1_45S_50ntFlanks@11873.11893.21\|AAGAAGACCCTGTTGAGCTTG | >hg38_21_+_8433172_8446622_RNA45SN1_45S_50ntFlanks | 11873 | 11893 | 21 | AAGAAGACCCTGTTGAGCTTG | MB468 | 541 | 282 | 371 | 433 | 133 | 16.1 | 171.4 | 104.9 | 153.0 | 39.3 |
| >hg38_21_+_8433172_8446622_RNA45SN1_45S_50ntFlanks@11873.11894.22\|AAGAAGACCCTGTTGAGCTTGA | >hg38_21_+_8433172_8446622_RNA45SN1_45S_50ntFlanks | 11873 | 11894 | 22 | AAGAAGACCCTGTTGAGCTTGA | MB468 | 17 | 7 | 5 | 15 | 7 | 3.5 | 4.2 | 0.7 | 4.5 | 3.3 |
| >hg38_21_+_8433172_8446622_RNA45SN1_45S_50ntFlanks@11873.11897.25\|AAGAAGACCCTGTTGAGCTTGACTC | >hg38_21_+_8433172_8446622_RNA45SN1_45S_50ntFlanks | 11873 | 11897 | 25 | AAGAAGACCCTGTTGAGCTTGACTC | MB468 | 75 | 51 | 54 | 33 | 30 | 13.8 | 8.4 | 14.6 | 9.6 | 6.3 |
| >hg38_21_+_8433172_8446622_RNA45SN1_45S_50ntFlanks@11873.11898.26\|AAGAAGACCCTGTTGAGCTTGACTCT | >hg38_21_+_8433172_8446622_RNA45SN1_45S_50ntFlanks | 11873 | 11898 | 26 | AAGAAGACCCTGTTGAGCTTGACTCT | MB468 | 132 | 118 | 98 | 53 | 63 | 15.9 | 15.6 | 17.7 | 14.1 | 9.2 |
| >hg38_21_+_8433172_8446622_RNA45SN1_45S_50ntFlanks@11873.11899.27\|AAGAAGACCCTGTTGAGCTTGACTCTA | >hg38_21_+_8433172_8446622_RNA45SN1_45S_50ntFlanks | 11873 | 11899 | 27 | AAGAAGACCCTGTTGAGCTTGACTCTA | MB468 | 35 | 34 | 29 | 12 | 24 | 8.2 | 3.3 | 7.3 | 3.1 | 3.3 |
| >hg38_21_+_8433172_8446622_RNA45SN1_45S_50ntFlanks@11873.11900.28\|AAGAAGACCCTGTTGAGCTTGACTCTAG | >hg38_21_+_8433172_8446622_RNA45SN1_45S_50ntFlanks | 11873 | 11900 | 28 | AAGAAGACCCTGTTGAGCTTGACTCTAG | MB468 | 92 | 83 | 82 | 45 | 42 | 24.3 | 23.4 | 42.2 | 15.2 | 11.8 |
| >hg38_21_+_8433172_8446622_RNA45SN1_45S_50ntFlanks@11873.11901.29\|AAGAAGACCCTGTTGAGCTTGACTCTAGT | >hg38_21_+_8433172_8446622_RNA45SN1_45S_50ntFlanks | 11873 | 11901 | 29 | AAGAAGACCCTGTTGAGCTTGACTCTAGT | MB468 | 14 | 13 | 13 | 5 | 6 | 2.0 | 3.6 | 4.1 | 0.7 | 3.0 |
| >hg38_21_+_8433172_8446622_RNA45SN1_45S_50ntFlanks@11873.11902.30\|AAGAAGACCCTGTTGAGCTTGACTCTAGTC | >hg38_21_+_8433172_8446622_RNA45SN1_45S_50ntFlanks | 11873 | 11902 | 30 | AAGAAGACCCTGTTGAGCTTGACTCTAGTC | MB468 | 27 | 20 | 26 | 13 | 9 | 7.6 | 6.1 | 9.2 | 1.4 | 3.9 |
| >hg38_21_+_8433172_8446622_RNA45SN1_45S_50ntFlanks@11873.11903.31\|AAGAAGACCCTGTTGAGCTTGACTCTAGTCT | >hg38_21_+_8433172_8446622_RNA45SN1_45S_50ntFlanks | 11873 | 11903 | 31 | AAGAAGACCCTGTTGAGCTTGACTCTAGTCT | MB468 | 34 | 12 | 19 | 32 | 33 | 10.4 | 4.1 | 5.3 | 9.0 | 7.6 |
| >hg38_21_+_8433172_8446622_RNA45SN1_45S_50ntFlanks@11873.11904.32\|AAGAAGACCCTGTTGAGCTTGACTCTAGTCTG | >hg38_21_+_8433172_8446622_RNA45SN1_45S_50ntFlanks | 11873 | 11904 | 32 | AAGAAGACCCTGTTGAGCTTGACTCTAGTCTG | MB468 | 75 | 10 | 27 | 162 | 55 | 36.2 | 2.9 | 7.8 | 87.7 | 16.0 |
| >hg38_21_+_8433172_8446622_RNA45SN1_45S_50ntFlanks@11873.11905.33\|AAGAAGACCCTGTTGAGCTTGACTCTAGTCTGG | >hg38_21_+_8433172_8446622_RNA45SN1_45S_50ntFlanks | 11873 | 11905 | 33 | AAGAAGACCCTGTTGAGCTTGACTCTAGTCTGG | MB468 | 71 | 15 | 24 | 80 | 71 | 7.3 | 3.3 | 4.4 | 42.2 | 16.5 |
| >hg38_21_+_8433172_8446622_RNA45SN1_45S_50ntFlanks@11873.11906.34\|AAGAAGACCCTGTTGAGCTTGACTCTAGTCTGGC | >hg38_21_+_8433172_8446622_RNA45SN1_45S_50ntFlanks | 11873 | 11906 | 34 | AAGAAGACCCTGTTGAGCTTGACTCTAGTCTGGC | MB468 | 131 | 46 | 96 | 72 | 143 | 28.3 | 12.0 | 37.7 | 29.8 | 34.6 |
| >hg38_21_+_8433172_8446622_RNA45SN1_45S_50ntFlanks@11873.11907.35\|AAGAAGACCCTGTTGAGCTTGACTCTAGTCTGGCA | >hg38_21_+_8433172_8446622_RNA45SN1_45S_50ntFlanks | 11873 | 11907 | 35 | AAGAAGACCCTGTTGAGCTTGACTCTAGTCTGGCA | MB468 | 32 | 9 | 26 | 18 | 43 | 7.8 | 3.4 | 5.5 | 9.2 | 22.8 |
| >hg38_21_+_8433172_8446622_RNA45SN1_45S_50ntFlanks@11873.11908.36\|AAGAAGACCCTGTTGAGCTTGACTCTAGTCTGGCAC | >hg38_21_+_8433172_8446622_RNA45SN1_45S_50ntFlanks | 11873 | 11908 | 36 | AAGAAGACCCTGTTGAGCTTGACTCTAGTCTGGCAC | MB468 | 29 | 20 | 27 | 9 | 51 | 9.5 | 8.0 | 14.3 | 7.1 | 21.3 |
| >hg38_21_+_8433172_8446622_RNA45SN1_45S_50ntFlanks@11873.11909.37\|AAGAAGACCCTGTTGAGCTTGACTCTAGTCTGGCACG | >hg38_21_+_8433172_8446622_RNA45SN1_45S_50ntFlanks | 11873 | 11909 | 37 | AAGAAGACCCTGTTGAGCTTGACTCTAGTCTGGCACG | MB468 | 17 | 11 | 18 | 9 | 37 | 4.5 | 5.3 | 5.2 | 5.7 | 25.4 |
| >hg38_21_+_8433172_8446622_RNA45SN1_45S_50ntFlanks@11873.11910.38\|AAGAAGACCCTGTTGAGCTTGACTCTAGTCTGGCACGG | >hg38_21_+_8433172_8446622_RNA45SN1_45S_50ntFlanks | 11873 | 11910 | 38 | AAGAAGACCCTGTTGAGCTTGACTCTAGTCTGGCACGG | MB468 | 14 | 10 | 12 | 11 | 51 | 5.0 | 6.4 | 4.4 | 6.9 | 21.4 |
| >hg38_21_+_8433172_8446622_RNA45SN1_45S_50ntFlanks@11873.11912.40\|AAGAAGACCCTGTTGAGCTTGACTCTAGTCTGGCACGGTG | >hg38_21_+_8433172_8446622_RNA45SN1_45S_50ntFlanks | 11873 | 11912 | 40 | AAGAAGACCCTGTTGAGCTTGACTCTAGTCTGGCACGGTG | MB468 | 14 | 6 | 15 | 25 | 30 | 4.5 | 3.5 | 7.4 | 20.5 | 27.7 |
| >hg38_21_+_8433172_8446622_RNA45SN1_45S_50ntFlanks@11874.11892.19\|AGAAGACCCTGTTGAGCTT | >hg38_21_+_8433172_8446622_RNA45SN1_45S_50ntFlanks | 11874 | 11892 | 19 | AGAAGACCCTGTTGAGCTT | MB468 | 79 | 34 | 39 | 44 | 24 | 9.5 | 21.2 | 12.7 | 15.0 | 7.3 |
| >hg38_21_+_8433172_8446622_RNA45SN1_45S_50ntFlanks@11874.11893.20\|AGAAGACCCTGTTGAGCTTG | >hg38_21_+_8433172_8446622_RNA45SN1_45S_50ntFlanks | 11874 | 11893 | 20 | AGAAGACCCTGTTGAGCTTG | MB468 | 578 | 307 | 461 | 405 | 161 | 57.9 | 186.6 | 164.5 | 140.9 | 39.9 |
| >hg38_21_+_8433172_8446622_RNA45SN1_45S_50ntFlanks@11874.11894.21\|AGAAGACCCTGTTGAGCTTGA | >hg38_21_+_8433172_8446622_RNA45SN1_45S_50ntFlanks | 11874 | 11894 | 21 | AGAAGACCCTGTTGAGCTTGA | MB468 | 17 | 10 | 8 | 14 | 9 | 2.2 | 7.4 | 2.2 | 4.8 | 4.0 |
| >hg38_21_+_8433172_8446622_RNA45SN1_45S_50ntFlanks@11874.11897.24\|AGAAGACCCTGTTGAGCTTGACTC | >hg38_21_+_8433172_8446622_RNA45SN1_45S_50ntFlanks | 11874 | 11897 | 24 | AGAAGACCCTGTTGAGCTTGACTC | MB468 | 86 | 65 | 74 | 33 | 39 | 29.1 | 11.3 | 29.5 | 5.9 | 9.1 |
| >hg38_21_+_8433172_8446622_RNA45SN1_45S_50ntFlanks@11874.11898.25\|AGAAGACCCTGTTGAGCTTGACTCT | >hg38_21_+_8433172_8446622_RNA45SN1_45S_50ntFlanks | 11874 | 11898 | 25 | AGAAGACCCTGTTGAGCTTGACTCT | MB468 | 207 | 169 | 169 | 81 | 117 | 29.0 | 21.0 | 24.0 | 25.4 | 28.4 |
| >hg38_21_+_8433172_8446622_RNA45SN1_45S_50ntFlanks@11874.11899.26\|AGAAGACCCTGTTGAGCTTGACTCTA | >hg38_21_+_8433172_8446622_RNA45SN1_45S_50ntFlanks | 11874 | 11899 | 26 | AGAAGACCCTGTTGAGCTTGACTCTA | MB468 | 43 | 40 | 43 | 13 | 35 | 10.0 | 5.6 | 11.9 | 2.7 | 4.7 |
| >hg38_21_+_8433172_8446622_RNA45SN1_45S_50ntFlanks@11874.11900.27\|AGAAGACCCTGTTGAGCTTGACTCTAG | >hg38_21_+_8433172_8446622_RNA45SN1_45S_50ntFlanks | 11874 | 11900 | 27 | AGAAGACCCTGTTGAGCTTGACTCTAG | MB468 | 83 | 67 | 84 | 40 | 48 | 18.7 | 21.8 | 30.7 | 9.4 | 8.0 |
| >hg38_21_+_8433172_8446622_RNA45SN1_45S_50ntFlanks@11874.11901.28\|AGAAGACCCTGTTGAGCTTGACTCTAGT | >hg38_21_+_8433172_8446622_RNA45SN1_45S_50ntFlanks | 11874 | 11901 | 28 | AGAAGACCCTGTTGAGCTTGACTCTAGT | MB468 | 13 | 11 | 13 | 4 | 7 | 2.9 | 3.7 | 3.9 | 1.1 | 3.3 |
| >hg38_21_+_8433172_8446622_RNA45SN1_45S_50ntFlanks@11874.11902.29\|AGAAGACCCTGTTGAGCTTGACTCTAGTC | >hg38_21_+_8433172_8446622_RNA45SN1_45S_50ntFlanks | 11874 | 11902 | 29 | AGAAGACCCTGTTGAGCTTGACTCTAGTC | MB468 | 33 | 26 | 29 | 15 | 13 | 6.1 | 10.9 | 5.9 | 2.0 | 5.0 |
| >hg38_21_+_8433172_8446622_RNA45SN1_45S_50ntFlanks@11874.11903.30\|AGAAGACCCTGTTGAGCTTGACTCTAGTCT | >hg38_21_+_8433172_8446622_RNA45SN1_45S_50ntFlanks | 11874 | 11903 | 30 | AGAAGACCCTGTTGAGCTTGACTCTAGTCT | MB468 | 31 | 8 | 16 | 33 | 37 | 5.1 | 3.1 | 3.6 | 12.1 | 13.2 |
| >hg38_21_+_8433172_8446622_RNA45SN1_45S_50ntFlanks@11874.11904.31\|AGAAGACCCTGTTGAGCTTGACTCTAGTCTG | >hg38_21_+_8433172_8446622_RNA45SN1_45S_50ntFlanks | 11874 | 11904 | 31 | AGAAGACCCTGTTGAGCTTGACTCTAGTCTG | MB468 | 62 | 8 | 20 | 159 | 58 | 19.9 | 2.3 | 6.0 | 98.4 | 18.9 |
| >hg38_21_+_8433172_8446622_RNA45SN1_45S_50ntFlanks@11874.11905.32\|AGAAGACCCTGTTGAGCTTGACTCTAGTCTGG | >hg38_21_+_8433172_8446622_RNA45SN1_45S_50ntFlanks | 11874 | 11905 | 32 | AGAAGACCCTGTTGAGCTTGACTCTAGTCTGG | MB468 | 63 | 12 | 19 | 83 | 74 | 3.7 | 3.6 | 6.7 | 48.8 | 20.7 |
| >hg38_21_+_8433172_8446622_RNA45SN1_45S_50ntFlanks@11874.11906.33\|AGAAGACCCTGTTGAGCTTGACTCTAGTCTGGC | >hg38_21_+_8433172_8446622_RNA45SN1_45S_50ntFlanks | 11874 | 11906 | 33 | AGAAGACCCTGTTGAGCTTGACTCTAGTCTGGC | MB468 | 157 | 47 | 103 | 75 | 198 | 27.7 | 10.0 | 20.6 | 27.1 | 15.1 |
| >hg38_21_+_8433172_8446622_RNA45SN1_45S_50ntFlanks@11874.11907.34\|AGAAGACCCTGTTGAGCTTGACTCTAGTCTGGCA | >hg38_21_+_8433172_8446622_RNA45SN1_45S_50ntFlanks | 11874 | 11907 | 34 | AGAAGACCCTGTTGAGCTTGACTCTAGTCTGGCA | MB468 | 35 | 11 | 26 | 19 | 57 | 6.8 | 2.9 | 6.2 | 10.4 | 19.6 |
| >hg38_21_+_8433172_8446622_RNA45SN1_45S_50ntFlanks@11874.11908.35\|AGAAGACCCTGTTGAGCTTGACTCTAGTCTGGCAC | >hg38_21_+_8433172_8446622_RNA45SN1_45S_50ntFlanks | 11874 | 11908 | 35 | AGAAGACCCTGTTGAGCTTGACTCTAGTCTGGCAC | MB468 | 44 | 25 | 37 | 11 | 79 | 15.3 | 7.1 | 14.5 | 8.4 | 23.1 |
| >hg38_21_+_8433172_8446622_RNA45SN1_45S_50ntFlanks@11874.11909.36\|AGAAGACCCTGTTGAGCTTGACTCTAGTCTGGCACG | >hg38_21_+_8433172_8446622_RNA45SN1_45S_50ntFlanks | 11874 | 11909 | 36 | AGAAGACCCTGTTGAGCTTGACTCTAGTCTGGCACG | MB468 | 26 | 17 | 23 | 11 | 59 | 8.5 | 5.4 | 5.7 | 7.4 | 28.5 |
| >hg38_21_+_8433172_8446622_RNA45SN1_45S_50ntFlanks@11874.11910.37\|AGAAGACCCTGTTGAGCTTGACTCTAGTCTGGCACGG | >hg38_21_+_8433172_8446622_RNA45SN1_45S_50ntFlanks | 11874 | 11910 | 37 | AGAAGACCCTGTTGAGCTTGACTCTAGTCTGGCACGG | MB468 | 23 | 16 | 18 | 15 | 89 | 9.5 | 8.0 | 8.5 | 8.0 | 28.1 |

| | | | | | | | | | | | | | | |
|---|---|---|---|---|---|---|---|---|---|---|---|---|---|---|
| >hg38_21_+_8433172_8446622_RNA45SN1_45S_50ntFlanks@11874.11912.39\|AGAAGACCCTGTTGAGCTTGACTCTAGTCTGGCACGGTG | >hg38_21_+_8433172_8446622_RNA45SN1_45S_50ntFlanks | 11874 | 11912 | 39 | AGAAGACCCTGTTGAGCTTGACTCTAGTCTGGCACGGTG | MB468 | 16 | 7 | 17 | 30 | 40 | 3.2 | 3.5 | 8.9 | 26.2 | 30.2 |
| >hg38_21_+_8433172_8446622_RNA45SN1_45S_50ntFlanks@11875.11892.18\|GAAGACCCTGTTGAGCTT | >hg38_21_+_8433172_8446622_RNA45SN1_45S_50ntFlanks | 11875 | 11892 | 18 | GAAGACCCTGTTGAGCTT | MB468 | 13 | 6 | 8 | 7 | 4 | 1.8 | 3.3 | 1.3 | 3.3 | 1.0 |
| >hg38_21_+_8433172_8446622_RNA45SN1_45S_50ntFlanks@11875.11893.19\|GAAGACCCTGTTGAGCTTG | >hg38_21_+_8433172_8446622_RNA45SN1_45S_50ntFlanks | 11875 | 11893 | 19 | GAAGACCCTGTTGAGCTTG | MB468 | 88 | 52 | 84 | 60 | 22 | 5.0 | 32.0 | 22.9 | 25.3 | 2.5 |
| >hg38_21_+_8433172_8446622_RNA45SN1_45S_50ntFlanks@11875.11897.23\|GAAGACCCTGTTGAGCTTGACTC | >hg38_21_+_8433172_8446622_RNA45SN1_45S_50ntFlanks | 11875 | 11897 | 23 | GAAGACCCTGTTGAGCTTGACTC | MB468 | 17 | 16 | 20 | 6 | 8 | 5.0 | 3.4 | 6.6 | 1.5 | 2.1 |
| >hg38_21_+_8433172_8446622_RNA45SN1_45S_50ntFlanks@11875.11898.24\|GAAGACCCTGTTGAGCTTGACTCT | >hg38_21_+_8433172_8446622_RNA45SN1_45S_50ntFlanks | 11875 | 11898 | 24 | GAAGACCCTGTTGAGCTTGACTCT | MB468 | 23 | 23 | 23 | 6 | 14 | 2.0 | 7.1 | 6.4 | 2.3 | 3.3 |
| >hg38_21_+_8433172_8446622_RNA45SN1_45S_50ntFlanks@11875.11900.26\|GAAGACCCTGTTGAGCTTGACTCTAG | >hg38_21_+_8433172_8446622_RNA45SN1_45S_50ntFlanks | 11875 | 11900 | 26 | GAAGACCCTGTTGAGCTTGACTCTAG | MB468 | 11 | 8 | 11 | 5 | 6 | 3.9 | 2.0 | 7.4 | 1.2 | 0.8 |
| >hg38_21_+_8433172_8446622_RNA45SN1_45S_50ntFlanks@11875.11906.32\|GAAGACCCTGTTGAGCTTGACTCTAGTCTGGC | >hg38_21_+_8433172_8446622_RNA45SN1_45S_50ntFlanks | 11875 | 11906 | 32 | GAAGACCCTGTTGAGCTTGACTCTAGTCTGGC | MB468 | 14 | 2 | 8 | 7 | 15 | 4.7 | 1.1 | 4.3 | 1.7 | 4.7 |
| >hg38_21_+_8433172_8446622_RNA45SN1_45S_50ntFlanks@11876.11893.18\|AAGACCCTGTTGAGCTTG | >hg38_21_+_8433172_8446622_RNA45SN1_45S_50ntFlanks | 11876 | 11893 | 18 | AAGACCCTGTTGAGCTTG | MB468 | 266 | 112 | 324 | 220 | 70 | 43.4 | 78.1 | 77.0 | 86.2 | 11.6 |
| >hg38_21_+_8433172_8446622_RNA45SN1_45S_50ntFlanks@11876.11897.22\|AAGACCCTGTTGAGCTTGACTC | >hg38_21_+_8433172_8446622_RNA45SN1_45S_50ntFlanks | 11876 | 11897 | 22 | AAGACCCTGTTGAGCTTGACTC | MB468 | 44 | 38 | 55 | 20 | 23 | 5.5 | 10.3 | 12.2 | 5.6 | 9.3 |
| >hg38_21_+_8433172_8446622_RNA45SN1_45S_50ntFlanks@11876.11898.23\|AAGACCCTGTTGAGCTTGACTCT | >hg38_21_+_8433172_8446622_RNA45SN1_45S_50ntFlanks | 11876 | 11898 | 23 | AAGACCCTGTTGAGCTTGACTCT | MB468 | 74 | 89 | 77 | 24 | 53 | 12.2 | 30.2 | 29.9 | 12.7 | 17.6 |
| >hg38_21_+_8433172_8446622_RNA45SN1_45S_50ntFlanks@11876.11899.24\|AAGACCCTGTTGAGCTTGACTCTA | >hg38_21_+_8433172_8446622_RNA45SN1_45S_50ntFlanks | 11876 | 11899 | 24 | AAGACCCTGTTGAGCTTGACTCTA | MB468 | 29 | 36 | 32 | 8 | 25 | 5.4 | 11.5 | 11.0 | 3.7 | 5.5 |
| >hg38_21_+_8433172_8446622_RNA45SN1_45S_50ntFlanks@11876.11900.25\|AAGACCCTGTTGAGCTTGACTCTAG | >hg38_21_+_8433172_8446622_RNA45SN1_45S_50ntFlanks | 11876 | 11900 | 25 | AAGACCCTGTTGAGCTTGACTCTAG | MB468 | 38 | 35 | 41 | 17 | 23 | 13.4 | 10.8 | 30.1 | 5.7 | 3.1 |
| >hg38_21_+_8433172_8446622_RNA45SN1_45S_50ntFlanks@11876.11904.29\|AAGACCCTGTTGAGCTTGACTCTAGTCTG | >hg38_21_+_8433172_8446622_RNA45SN1_45S_50ntFlanks | 11876 | 11904 | 29 | AAGACCCTGTTGAGCTTGACTCTAGTCTG | MB468 | 18 | 2 | 5 | 48 | 18 | 10.1 | 0.6 | 3.7 | 31.2 | 14.2 |
| >hg38_21_+_8433172_8446622_RNA45SN1_45S_50ntFlanks@11876.11905.30\|AAGACCCTGTTGAGCTTGACTCTAGTCTGG | >hg38_21_+_8433172_8446622_RNA45SN1_45S_50ntFlanks | 11876 | 11905 | 30 | AAGACCCTGTTGAGCTTGACTCTAGTCTGG | MB468 | 32 | 5 | 10 | 39 | 42 | 10.7 | 0.8 | 3.6 | 11.0 | 25.3 |
| >hg38_21_+_8433172_8446622_RNA45SN1_45S_50ntFlanks@11876.11906.31\|AAGACCCTGTTGAGCTTGACTCTAGTCTGGC | >hg38_21_+_8433172_8446622_RNA45SN1_45S_50ntFlanks | 11876 | 11906 | 31 | AAGACCCTGTTGAGCTTGACTCTAGTCTGGC | MB468 | 46 | 9 | 29 | 25 | 58 | 14.3 | 2.5 | 18.9 | 7.9 | 19.3 |
| >hg38_21_+_8433172_8446622_RNA45SN1_45S_50ntFlanks@11876.11908.33\|AAGACCCTGTTGAGCTTGACTCTAGTCTGGCAC | >hg38_21_+_8433172_8446622_RNA45SN1_45S_50ntFlanks | 11876 | 11908 | 33 | AAGACCCTGTTGAGCTTGACTCTAGTCTGGCAC | MB468 | 19 | 9 | 13 | 6 | 34 | 6.4 | 3.6 | 6.0 | 4.6 | 4.4 |
| >hg38_21_+_8433172_8446622_RNA45SN1_45S_50ntFlanks@11876.11909.34\|AAGACCCTGTTGAGCTTGACTCTAGTCTGGCACG | >hg38_21_+_8433172_8446622_RNA45SN1_45S_50ntFlanks | 11876 | 11909 | 34 | AAGACCCTGTTGAGCTTGACTCTAGTCTGGCACG | MB468 | 13 | 8 | 9 | 6 | 24 | 2.8 | 2.4 | 1.7 | 3.4 | 5.2 |
| >hg38_21_+_8433172_8446622_RNA45SN1_45S_50ntFlanks@11877.11898.22\|AGACCCTGTTGAGCTTGACTCT | >hg38_21_+_8433172_8446622_RNA45SN1_45S_50ntFlanks | 11877 | 11898 | 22 | AGACCCTGTTGAGCTTGACTCT | BT20 | 11 | 10 | 4 | 11 | 10 | 16.5 | 15.2 | 3.2 | 16.5 | 7.7 |
| >hg38_21_+_8433172_8446622_RNA45SN1_45S_50ntFlanks@11877.11898.22\|AGACCCTGTTGAGCTTGACTCT | >hg38_21_+_8433172_8446622_RNA45SN1_45S_50ntFlanks | 11877 | 11898 | 22 | AGACCCTGTTGAGCTTGACTCT | MB468 | 15 | 14 | 16 | 4 | 7 | 2.0 | 3.6 | 3.1 | 2.1 | 1.7 |
| >hg38_21_+_8433172_8446622_RNA45SN1_45S_50ntFlanks@11877.11906.30\|AGACCCTGTTGAGCTTGACTCTAGTCTGGC | >hg38_21_+_8433172_8446622_RNA45SN1_45S_50ntFlanks | 11877 | 11906 | 30 | AGACCCTGTTGAGCTTGACTCTAGTCTGGC | MB468 | 12 | 3 | 7 | 4 | 13 | 3.2 | 0.6 | 2.8 | 1.2 | 3.0 |
| >hg38_21_+_8433172_8446622_RNA45SN1_45S_50ntFlanks@11878.11898.21\|GACCCTGTTGAGCTTGACTCT | >hg38_21_+_8433172_8446622_RNA45SN1_45S_50ntFlanks | 11878 | 11898 | 21 | GACCCTGTTGAGCTTGACTCT | BT20 | 13 | 14 | 4 | 4 | 5 | 19.5 | 22.5 | 3.2 | 6.5 | 5.3 |
| >hg38_21_+_8433172_8446622_RNA45SN1_45S_50ntFlanks@11879.11897.19\|ACCCTGTTGAGCTTGACTC | >hg38_21_+_8433172_8446622_RNA45SN1_45S_50ntFlanks | 11879 | 11897 | 19 | ACCCTGTTGAGCTTGACTC | MB468 | 11 | 7 | 10 | 3 | 2 | 0.8 | 3.5 | 1.5 | 1.4 | 0.4 |
| >hg38_21_+_8433172_8446622_RNA45SN1_45S_50ntFlanks@11879.11898.20\|ACCCTGTTGAGCTTGACTCT | >hg38_21_+_8433172_8446622_RNA45SN1_45S_50ntFlanks | 11879 | 11898 | 20 | ACCCTGTTGAGCTTGACTCT | BT20 | 39 | 34 | 7 | 5 | 11 | 64.1 | 56.9 | 7.9 | 6.2 | 6.9 |
| >hg38_21_+_8433172_8446622_RNA45SN1_45S_50ntFlanks@11879.11898.20\|ACCCTGTTGAGCTTGACTCT | >hg38_21_+_8433172_8446622_RNA45SN1_45S_50ntFlanks | 11879 | 11898 | 20 | ACCCTGTTGAGCTTGACTCT | MB468 | 11 | 8 | 13 | 3 | 4 | 1.1 | 2.0 | 3.0 | 1.5 | 0.8 |
| >hg38_21_+_8433172_8446622_RNA45SN1_45S_50ntFlanks@11879.11904.26\|ACCCTGTTGAGCTTGACTCTAGTCTG | >hg38_21_+_8433172_8446622_RNA45SN1_45S_50ntFlanks | 11879 | 11904 | 26 | ACCCTGTTGAGCTTGACTCTAGTCTG | MB468 | 17 | 14 | 11 | 8 | 9 | 5.5 | 5.0 | 2.1 | 5.6 | 0.6 |
| >hg38_21_+_8433172_8446622_RNA45SN1_45S_50ntFlanks@11880.11898.19\|CCCTGTTGAGCTTGACTCT | >hg38_21_+_8433172_8446622_RNA45SN1_45S_50ntFlanks | 11880 | 11898 | 19 | CCCTGTTGAGCTTGACTCT | BT20 | 76 | 28 | 16 | 4 | 5 | 128.9 | 48.0 | 25.5 | 5.7 | 4.6 |
| >hg38_21_+_8433172_8446622_RNA45SN1_45S_50ntFlanks@11880.11899.20\|CCCTGTTGAGCTTGACTCTA | >hg38_21_+_8433172_8446622_RNA45SN1_45S_50ntFlanks | 11880 | 11899 | 20 | CCCTGTTGAGCTTGACTCTA | BT20 | 13 | 6 | 3 | 2 | 2 | 21.2 | 10.9 | 3.6 | 2.9 | 2.2 |
| >hg38_21_+_8433172_8446622_RNA45SN1_45S_50ntFlanks@11880.11904.25\|CCCTGTTGAGCTTGACTCTAGTCTG | >hg38_21_+_8433172_8446622_RNA45SN1_45S_50ntFlanks | 11880 | 11904 | 25 | CCCTGTTGAGCTTGACTCTAGTCTG | MB468 | 11 | 7 | 9 | 8 | 7 | 1.8 | 2.3 | 0.9 | 5.3 | 1.1 |
| >hg38_21_+_8433172_8446622_RNA45SN1_45S_50ntFlanks@11881.11898.18\|CCTGTTGAGCTTGACTCT | >hg38_21_+_8433172_8446622_RNA45SN1_45S_50ntFlanks | 11881 | 11898 | 18 | CCTGTTGAGCTTGACTCT | MB468 | 12 | 2 | 13 | 10 | 4 | 2.8 | 0.9 | 3.4 | 3.1 | 2.4 |
| >hg38_21_+_8433172_8446622_RNA45SN1_45S_50ntFlanks@11881.11899.19\|CCTGTTGAGCTTGACTCTA | >hg38_21_+_8433172_8446622_RNA45SN1_45S_50ntFlanks | 11881 | 11899 | 19 | CCTGTTGAGCTTGACTCTA | MB468 | 14 | 1 | 15 | 6 | 4 | 8.7 | 0.5 | 8.9 | 3.2 | 2.5 |
| >hg38_21_+_8433172_8446622_RNA45SN1_45S_50ntFlanks@11881.11900.20\|CCTGTTGAGCTTGACTCTAG | >hg38_21_+_8433172_8446622_RNA45SN1_45S_50ntFlanks | 11881 | 11900 | 20 | CCTGTTGAGCTTGACTCTAG | MB468 | 19 | 1 | 17 | 18 | 7 | 7.8 | 0.7 | 8.5 | 12.8 | 4.3 |
| >hg38_21_+_8433172_8446622_RNA45SN1_45S_50ntFlanks@11881.11903.23\|CCTGTTGAGCTTGACTCTAGTCT | >hg38_21_+_8433172_8446622_RNA45SN1_45S_50ntFlanks | 11881 | 11903 | 23 | CCTGTTGAGCTTGACTCTAGTCT | MB468 | 40 | 15 | 38 | 19 | 17 | 7.2 | 4.7 | 18.5 | 10.3 | 7.6 |
| >hg38_21_+_8433172_8446622_RNA45SN1_45S_50ntFlanks@11881.11904.24\|CCTGTTGAGCTTGACTCTAGTCTG | >hg38_21_+_8433172_8446622_RNA45SN1_45S_50ntFlanks | 11881 | 11904 | 24 | CCTGTTGAGCTTGACTCTAGTCTG | MB468 | 190 | 74 | 129 | 136 | 77 | 20.7 | 21.8 | 49.0 | 61.7 | 29.6 |
| >hg38_21_+_8433172_8446622_RNA45SN1_45S_50ntFlanks@11881.11905.25\|CCTGTTGAGCTTGACTCTAGTCTGG | >hg38_21_+_8433172_8446622_RNA45SN1_45S_50ntFlanks | 11881 | 11905 | 25 | CCTGTTGAGCTTGACTCTAGTCTGG | MB468 | 21 | 7 | 19 | 15 | 7 | 5.1 | 2.5 | 10.1 | 5.0 | 5.3 |
| >hg38_21_+_8433172_8446622_RNA45SN1_45S_50ntFlanks@11881.11906.26\|CCTGTTGAGCTTGACTCTAGTCTGGC | >hg38_21_+_8433172_8446622_RNA45SN1_45S_50ntFlanks | 11881 | 11906 | 26 | CCTGTTGAGCTTGACTCTAGTCTGGC | MB468 | 21 | 5 | 41 | 8 | 6 | 7.5 | 1.1 | 20.8 | 1.5 | 3.3 |
| >hg38_21_+_8433172_8446622_RNA45SN1_45S_50ntFlanks@11882.11900.19\|CTGTTGAGCTTGACTCTAG | >hg38_21_+_8433172_8446622_RNA45SN1_45S_50ntFlanks | 11882 | 11900 | 19 | CTGTTGAGCTTGACTCTAG | MB468 | 30 | 1 | 11 | 13 | 7 | 20.9 | 0.5 | 5.9 | 9.2 | 7.5 |
| >hg38_21_+_8433172_8446622_RNA45SN1_45S_50ntFlanks@11882.11903.22\|CTGTTGAGCTTGACTCTAGTCT | >hg38_21_+_8433172_8446622_RNA45SN1_45S_50ntFlanks | 11882 | 11903 | 22 | CTGTTGAGCTTGACTCTAGTCT | MB468 | 13 | 3 | 12 | 8 | 6 | 3.1 | 1.3 | 4.6 | 2.6 | 2.6 |
| >hg38_21_+_8433172_8446622_RNA45SN1_45S_50ntFlanks@11882.11904.23\|CTGTTGAGCTTGACTCTAGTCTG | >hg38_21_+_8433172_8446622_RNA45SN1_45S_50ntFlanks | 11882 | 11904 | 23 | CTGTTGAGCTTGACTCTAGTCTG | MB468 | 125 | 35 | 82 | 73 | 56 | 40.7 | 6.4 | 29.2 | 25.9 | 32.2 |
| >hg38_21_+_8433172_8446622_RNA45SN1_45S_50ntFlanks@11883.11904.22\|TGTTGAGCTTGACTCTAGTCTG | >hg38_21_+_8433172_8446622_RNA45SN1_45S_50ntFlanks | 11883 | 11904 | 22 | TGTTGAGCTTGACTCTAGTCTG | MB468 | 13 | 4 | 10 | 9 | 4 | 5.8 | 0.9 | 5.2 | 2.4 | 1.3 |
| >hg38_21_+_8433172_8446622_RNA45SN1_45S_50ntFlanks@11883.11905.23\|TGTTGAGCTTGACTCTAGTCTGG | >hg38_21_+_8433172_8446622_RNA45SN1_45S_50ntFlanks | 11883 | 11905 | 23 | TGTTGAGCTTGACTCTAGTCTGG | MB468 | 12 | 8 | 10 | 5 | 4 | 4.3 | 3.3 | 2.3 | 2.2 | 1.6 |
| >hg38_21_+_8433172_8446622_RNA45SN1_45S_50ntFlanks@11884.11904.21\|GTTGAGCTTGACTCTAGTCTG | >hg38_21_+_8433172_8446622_RNA45SN1_45S_50ntFlanks | 11884 | 11904 | 21 | GTTGAGCTTGACTCTAGTCTG | MB468 | 20 | 6 | 16 | 13 | 9 | 8.0 | 2.0 | 10.8 | 6.9 | 3.4 |
| >hg38_21_+_8433172_8446622_RNA45SN1_45S_50ntFlanks@11884.11905.22\|GTTGAGCTTGACTCTAGTCTGG | >hg38_21_+_8433172_8446622_RNA45SN1_45S_50ntFlanks | 11884 | 11905 | 22 | GTTGAGCTTGACTCTAGTCTGG | MB468 | 10 | 6 | 8 | 4 | 3 | 2.3 | 1.7 | 2.5 | 1.8 | 1.0 |
| >hg38_21_+_8433172_8446622_RNA45SN1_45S_50ntFlanks@11885.11903.19\|TTGAGCTTGACTCTAGTCT | >hg38_21_+_8433172_8446622_RNA45SN1_45S_50ntFlanks | 11885 | 11903 | 19 | TTGAGCTTGACTCTAGTCT | MB468 | 52 | 11 | 53 | 25 | 20 | 23.9 | 2.2 | 37.7 | 5.4 | 10.1 |
| >hg38_21_+_8433172_8446622_RNA45SN1_45S_50ntFlanks@11885.11904.20\|TTGAGCTTGACTCTAGTCTG | >hg38_21_+_8433172_8446622_RNA45SN1_45S_50ntFlanks | 11885 | 11904 | 20 | TTGAGCTTGACTCTAGTCTG | MB468 | 181 | 28 | 205 | 132 | 76 | 98.3 | 4.9 | 151.8 | 29.2 | 29.1 |
| >hg38_21_+_8433172_8446622_RNA45SN1_45S_50ntFlanks@11885.11905.21\|TTGAGCTTGACTCTAGTCTGG | >hg38_21_+_8433172_8446622_RNA45SN1_45S_50ntFlanks | 11885 | 11905 | 21 | TTGAGCTTGACTCTAGTCTGG | MB468 | 106 | 32 | 85 | 43 | 30 | 29.5 | 7.3 | 47.0 | 12.3 | 20.3 |
| >hg38_21_+_8433172_8446622_RNA45SN1_45S_50ntFlanks@11885.11912.28\|TTGAGCTTGACTCTAGTCTGGCACGGTG | >hg38_21_+_8433172_8446622_RNA45SN1_45S_50ntFlanks | 11885 | 11912 | 28 | TTGAGCTTGACTCTAGTCTGGCACGGTG | MB468 | 20 | 3 | 11 | 34 | 15 | 1.2 | 0.7 | 4.6 | 21.8 | 4.6 |
| >hg38_21_+_8433172_8446622_RNA45SN1_45S_50ntFlanks@11886.11903.18\|TGAGCTTGACTCTAGTCT | >hg38_21_+_8433172_8446622_RNA45SN1_45S_50ntFlanks | 11886 | 11903 | 18 | TGAGCTTGACTCTAGTCT | MB468 | 25 | 4 | 25 | 12 | 11 | 7.6 | 0.9 | 16.3 | 2.1 | 6.1 |
| >hg38_21_+_8433172_8446622_RNA45SN1_45S_50ntFlanks@11886.11904.19\|TGAGCTTGACTCTAGTCTG | >hg38_21_+_8433172_8446622_RNA45SN1_45S_50ntFlanks | 11886 | 11904 | 19 | TGAGCTTGACTCTAGTCTG | MB468 | 94 | 9 | 111 | 68 | 42 | 38.9 | 1.4 | 72.8 | 21.8 | 16.2 |
| >hg38_21_+_8433172_8446622_RNA45SN1_45S_50ntFlanks@11886.11905.20\|TGAGCTTGACTCTAGTCTGG | >hg38_21_+_8433172_8446622_RNA45SN1_45S_50ntFlanks | 11886 | 11905 | 20 | TGAGCTTGACTCTAGTCTGG | MB468 | 54 | 13 | 43 | 22 | 16 | 11.6 | 3.1 | 22.7 | 10.2 | 11.4 |
| >hg38_21_+_8433172_8446622_RNA45SN1_45S_50ntFlanks@11887.11904.18\|GAGCTTGACTCTAGTCTG | >hg38_21_+_8433172_8446622_RNA45SN1_45S_50ntFlanks | 11887 | 11904 | 18 | GAGCTTGACTCTAGTCTG | MB468 | 12 | 2 | 10 | 6 | 4 | 4.6 | 0.8 | 5.2 | 1.2 | 2.3 |

| | | | | | | | | | | | | | | | | |
|---|---|---|---|---|---|---|---|---|---|---|---|---|---|---|---|---|
| >hg38_21_+_8433172_8446622_RNA45SN1_45S_50ntFlanks@11888.11905.18\|AGCTTGACTCTAGTCTGG | >hg38_21_+_8433172_8446622_RNA45SN1_45S_50ntFlanks | 11888 | 11905 | 18 | AGCTTGACTCTAGTCTGG | MB468 | 22 | 3 | 16 | 11 | 11 | 7.1 | 1.1 | 8.9 | 3.2 | 14.8 |
| >hg38_21_+_8433172_8446622_RNA45SN1_45S_50ntFlanks@11890.11907.18\|CTTGACTCTAGTCTGGCA | >hg38_21_+_8433172_8446622_RNA45SN1_45S_50ntFlanks | 11890 | 11907 | 18 | CTTGACTCTAGTCTGGCA | MB468 | 80 | 6 | 95 | 87 | 42 | 16.7 | 3.6 | 17.0 | 42.7 | 30.7 |
| >hg38_21_+_8433172_8446622_RNA45SN1_45S_50ntFlanks@11890.11912.23\|CTTGACTCTAGTCTGGCACGGTG | >hg38_21_+_8433172_8446622_RNA45SN1_45S_50ntFlanks | 11890 | 11912 | 23 | CTTGACTCTAGTCTGGCACGGTG | MB468 | 75 | 24 | 132 | 74 | 45 | 5.3 | 4.6 | 29.0 | 29.3 | 12.2 |
| >hg38_21_+_8433172_8446622_RNA45SN1_45S_50ntFlanks@11891.11909.19\|TTGACTCTAGTCTGGCACG | >hg38_21_+_8433172_8446622_RNA45SN1_45S_50ntFlanks | 11891 | 11909 | 19 | TTGACTCTAGTCTGGCACG | MB468 | 17 | 5 | 28 | 12 | 8 | 4.7 | 1.4 | 10.2 | 4.4 | 2.7 |
| >hg38_21_+_8433172_8446622_RNA45SN1_45S_50ntFlanks@11891.11910.20\|TTGACTCTAGTCTGGCACGG | >hg38_21_+_8433172_8446622_RNA45SN1_45S_50ntFlanks | 11891 | 11910 | 20 | TTGACTCTAGTCTGGCACGG | MB468 | 21 | 18 | 40 | 9 | 24 | 9.2 | 4.7 | 22.2 | 0.9 | 12.4 |
| >hg38_21_+_8433172_8446622_RNA45SN1_45S_50ntFlanks@11891.11912.22\|TTGACTCTAGTCTGGCACGGTG | >hg38_21_+_8433172_8446622_RNA45SN1_45S_50ntFlanks | 11891 | 11912 | 22 | TTGACTCTAGTCTGGCACGGTG | MB468 | 99 | 92 | 185 | 51 | 48 | 29.0 | 34.9 | 60.2 | 13.3 | 8.7 |
| >hg38_21_+_8433172_8446622_RNA45SN1_45S_50ntFlanks@11894.11911.18\|ACTCTAGTCTGGCACGGT | >hg38_21_+_8433172_8446622_RNA45SN1_45S_50ntFlanks | 11894 | 11911 | 18 | ACTCTAGTCTGGCACGGT | MB468 | 12 | 10 | 9 | 2 | 3 | 3.3 | 4.2 | 2.8 | 1.0 | 1.0 |
| >hg38_21_+_8433172_8446622_RNA45SN1_45S_50ntFlanks@11894.11912.19\|ACTCTAGTCTGGCACGGTG | >hg38_21_+_8433172_8446622_RNA45SN1_45S_50ntFlanks | 11894 | 11912 | 19 | ACTCTAGTCTGGCACGGTG | MB468 | 156 | 93 | 137 | 32 | 25 | 46.1 | 37.2 | 31.2 | 7.7 | 5.6 |
| >hg38_21_+_8433172_8446622_RNA45SN1_45S_50ntFlanks@11894.11916.23\|ACTCTAGTCTGGCACGGTGAAGA | >hg38_21_+_8433172_8446622_RNA45SN1_45S_50ntFlanks | 11894 | 11916 | 23 | ACTCTAGTCTGGCACGGTGAAGA | MB468 | 40 | 49 | 19 | 3 | 4 | 18.7 | 27.3 | 9.5 | 1.1 | 2.1 |
| >hg38_21_+_8433172_8446622_RNA45SN1_45S_50ntFlanks@11894.11920.27\|ACTCTAGTCTGGCACGGTGAAGAGACA | >hg38_21_+_8433172_8446622_RNA45SN1_45S_50ntFlanks | 11894 | 11920 | 27 | ACTCTAGTCTGGCACGGTGAAGAGACA | MB468 | 17 | 4 | 13 | 4 | 4 | 7.3 | 0.6 | 6.1 | 1.6 | 1.5 |
| >hg38_21_+_8433172_8446622_RNA45SN1_45S_50ntFlanks@11895.11912.18\|CTCTAGTCTGGCACGGTG | >hg38_21_+_8433172_8446622_RNA45SN1_45S_50ntFlanks | 11895 | 11912 | 18 | CTCTAGTCTGGCACGGTG | MB468 | 12 | 5 | 9 | 5 | 2 | 3.5 | 3.0 | 3.3 | 2.3 | 0.7 |
| >hg38_21_+_8433172_8446622_RNA45SN1_45S_50ntFlanks@11901.11920.20\|TCTGGCACGGTGAAGAGACA | >hg38_21_+_8433172_8446622_RNA45SN1_45S_50ntFlanks | 11901 | 11920 | 20 | TCTGGCACGGTGAAGAGACA | MB468 | 11 | 14 | 7 | 1 | 1 | 5.0 | 6.9 | 4.3 | 0.6 | 0.7 |
| >hg38_21_+_8433172_8446622_RNA45SN1_45S_50ntFlanks@11906.11929.24\|CACGGTGAAGAGACATGAGAGGTG | >hg38_21_+_8433172_8446622_RNA45SN1_45S_50ntFlanks | 11906 | 11929 | 24 | CACGGTGAAGAGACATGAGAGGTG | MB468 | 12 | 14 | 11 | 16 | 7 | 5.1 | 2.6 | 7.9 | 9.0 | 4.8 |
| >hg38_21_+_8433172_8446622_RNA45SN1_45S_50ntFlanks@11918.11948.31\|ACATGAGAGGTGTAGAATAAGTGGGAGGCCC | >hg38_21_+_8433172_8446622_RNA45SN1_45S_50ntFlanks | 11918 | 11948 | 31 | ACATGAGAGGTGTAGAATAAGTGGGAGGCCC | MB468 | 19 | 12 | 77 | 96 | 14 | 12.9 | 6.8 | 64.8 | 70.4 | 7.5 |
| >hg38_21_+_8433172_8446622_RNA45SN1_45S_50ntFlanks@11918.11949.32\|ACATGAGAGGTGTAGAATAAGTGGGAGGCCCC | >hg38_21_+_8433172_8446622_RNA45SN1_45S_50ntFlanks | 11918 | 11949 | 32 | ACATGAGAGGTGTAGAATAAGTGGGAGGCCCC | MB468 | 11 | 7 | 49 | 27 | 6 | 5.9 | 4.0 | 40.3 | 19.1 | 3.5 |
| >hg38_21_+_8433172_8446622_RNA45SN1_45S_50ntFlanks@11918.11950.33\|ACATGAGAGGTGTAGAATAAGTGGGAGGCCCCC | >hg38_21_+_8433172_8446622_RNA45SN1_45S_50ntFlanks | 11918 | 11950 | 33 | ACATGAGAGGTGTAGAATAAGTGGGAGGCCCCC | MB468 | 36 | 25 | 159 | 47 | 25 | 12.5 | 15.0 | 108.3 | 38.8 | 15.6 |
| >hg38_21_+_8433172_8446622_RNA45SN1_45S_50ntFlanks@11918.11951.34\|ACATGAGAGGTGTAGAATAAGTGGGAGGCCCCCG | >hg38_21_+_8433172_8446622_RNA45SN1_45S_50ntFlanks | 11918 | 11951 | 34 | ACATGAGAGGTGTAGAATAAGTGGGAGGCCCCCG | MB468 | 34 | 22 | 138 | 50 | 23 | 11.8 | 6.2 | 78.0 | 33.6 | 15.9 |
| >hg38_21_+_8433172_8446622_RNA45SN1_45S_50ntFlanks@11918.11954.37\|ACATGAGAGGTGTAGAATAAGTGGGAGGCCCCCGGCG | >hg38_21_+_8433172_8446622_RNA45SN1_45S_50ntFlanks | 11918 | 11954 | 37 | ACATGAGAGGTGTAGAATAAGTGGGAGGCCCCCGGCG | MB468 | 10 | 16 | 29 | 12 | 6 | 5.3 | 6.5 | 28.6 | 7.0 | 4.0 |
| >hg38_21_+_8433172_8446622_RNA45SN1_45S_50ntFlanks@11920.11950.31\|ATGAGAGGTGTAGAATAAGTGGGAGGCCCCC | >hg38_21_+_8433172_8446622_RNA45SN1_45S_50ntFlanks | 11920 | 11950 | 31 | ATGAGAGGTGTAGAATAAGTGGGAGGCCCCC | MB468 | 11 | 14 | 40 | 41 | 10 | 4.1 | 9.0 | 21.0 | 34.4 | 6.3 |
| >hg38_21_+_8433172_8446622_RNA45SN1_45S_50ntFlanks@11920.11951.32\|ATGAGAGGTGTAGAATAAGTGGGAGGCCCCCG | >hg38_21_+_8433172_8446622_RNA45SN1_45S_50ntFlanks | 11920 | 11951 | 32 | ATGAGAGGTGTAGAATAAGTGGGAGGCCCCCG | MB468 | 12 | 10 | 41 | 41 | 9 | 2.7 | 3.8 | 18.0 | 17.9 | 5.2 |
| >hg38_21_+_8433172_8446622_RNA45SN1_45S_50ntFlanks@11921.11950.30\|TGAGAGGTGTAGAATAAGTGGGAGGCCCCC | >hg38_21_+_8433172_8446622_RNA45SN1_45S_50ntFlanks | 11921 | 11950 | 30 | TGAGAGGTGTAGAATAAGTGGGAGGCCCCC | MB468 | 11 | 12 | 37 | 32 | 8 | 6.2 | 9.6 | 24.6 | 29.2 | 5.8 |
| >hg38_21_+_8433172_8446622_RNA45SN1_45S_50ntFlanks@11921.11951.31\|TGAGAGGTGTAGAATAAGTGGGAGGCCCCCG | >hg38_21_+_8433172_8446622_RNA45SN1_45S_50ntFlanks | 11921 | 11951 | 31 | TGAGAGGTGTAGAATAAGTGGGAGGCCCCCG | MB468 | 11 | 9 | 35 | 29 | 6 | 5.5 | 5.1 | 16.1 | 12.9 | 3.7 |
| >hg38_21_+_8433172_8446622_RNA45SN1_45S_50ntFlanks@11998.12032.35\|GCCCTGCGGGCCGCCGGTGAAATACCACTACTCTG | >hg38_21_+_8433172_8446622_RNA45SN1_45S_50ntFlanks | 11998 | 12032 | 35 | GCCCTGCGGGCCGCCGGTGAAATACCACTACTCTG | MB468 | 12 | 31 | 18 | 4 | 29 | 3.2 | 15.8 | 20.0 | 3.3 | 21.3 |
| >hg38_21_+_8433172_8446622_RNA45SN1_45S_50ntFlanks@11999.12029.31\|CCCTGCGGGCCGCCGGTGAAATACCACTACT | >hg38_21_+_8433172_8446622_RNA45SN1_45S_50ntFlanks | 11999 | 12029 | 31 | CCCTGCGGGCCGCCGGTGAAATACCACTACT | MB468 | 16 | 46 | 16 | 2 | 10 | 4.8 | 16.6 | 4.9 | 0.9 | 7.6 |
| >hg38_21_+_8433172_8446622_RNA45SN1_45S_50ntFlanks@11999.12030.32\|CCCTGCGGGCCGCCGGTGAAATACCACTACTC | >hg38_21_+_8433172_8446622_RNA45SN1_45S_50ntFlanks | 11999 | 12030 | 32 | CCCTGCGGGCCGCCGGTGAAATACCACTACTC | MB468 | 29 | 118 | 35 | 4 | 31 | 8.7 | 43.0 | 16.3 | 1.6 | 14.2 |
| >hg38_21_+_8433172_8446622_RNA45SN1_45S_50ntFlanks@11999.12031.33\|CCCTGCGGGCCGCCGGTGAAATACCACTACTCT | >hg38_21_+_8433172_8446622_RNA45SN1_45S_50ntFlanks | 11999 | 12031 | 33 | CCCTGCGGGCCGCCGGTGAAATACCACTACTCT | MB468 | 82 | 241 | 121 | 13 | 125 | 17.7 | 57.3 | 58.9 | 6.2 | 57.8 |
| >hg38_21_+_8433172_8446622_RNA45SN1_45S_50ntFlanks@11999.12032.34\|CCCTGCGGGCCGCCGGTGAAATACCACTACTCTG | >hg38_21_+_8433172_8446622_RNA45SN1_45S_50ntFlanks | 11999 | 12032 | 34 | CCCTGCGGGCCGCCGGTGAAATACCACTACTCTG | MB468 | 606 | 1753 | 1147 | 183 | 1070 | 159.3 | 399.9 | 844.3 | 96.4 | 714.1 |
| >hg38_21_+_8433172_8446622_RNA45SN1_45S_50ntFlanks@11999.12033.35\|CCCTGCGGGCCGCCGGTGAAATACCACTACTCTGA | >hg38_21_+_8433172_8446622_RNA45SN1_45S_50ntFlanks | 11999 | 12033 | 35 | CCCTGCGGGCCGCCGGTGAAATACCACTACTCTGA | MB468 | 17 | 54 | 33 | 8 | 54 | 6.3 | 12.8 | 20.3 | 4.1 | 27.2 |
| >hg38_21_+_8433172_8446622_RNA45SN1_45S_50ntFlanks@11999.12035.37\|CCCTGCGGGCCGCCGGTGAAATACCACTACTCTGATC | >hg38_21_+_8433172_8446622_RNA45SN1_45S_50ntFlanks | 11999 | 12035 | 37 | CCCTGCGGGCCGCCGGTGAAATACCACTACTCTGATC | MB468 | 13 | 32 | 18 | 9 | 67 | 9.5 | 17.6 | 7.6 | 6.6 | 68.4 |
| >hg38_21_+_8433172_8446622_RNA45SN1_45S_50ntFlanks@11999.12043.45\|CCCTGCGGGCCGCCGGTGAAATACCACTACTCTGATCGTTTTTTC | >hg38_21_+_8433172_8446622_RNA45SN1_45S_50ntFlanks | 11999 | 12043 | 45 | CCCTGCGGGCCGCCGGTGAAATACCACTACTCTGATCGTTTTTTC | MB468 | 18 | 158 | 55 | 18 | 195 | 12.4 | 144.7 | 70.5 | 26.8 | 137.1 |
| >hg38_21_+_8433172_8446622_RNA45SN1_45S_50ntFlanks@12000.12032.33\|CCTGCGGGCCGCCGGTGAAATACCACTACTCTG | >hg38_21_+_8433172_8446622_RNA45SN1_45S_50ntFlanks | 12000 | 12032 | 33 | CCTGCGGGCCGCCGGTGAAATACCACTACTCTG | MB468 | 30 | 94 | 50 | 14 | 44 | 8.7 | 47.7 | 25.7 | 4.5 | 20.1 |
| >hg38_21_+_8433172_8446622_RNA45SN1_45S_50ntFlanks@12001.12030.30\|CTGCGGGCCGCCGGTGAAATACCACTACTC | >hg38_21_+_8433172_8446622_RNA45SN1_45S_50ntFlanks | 12001 | 12030 | 30 | CTGCGGGCCGCCGGTGAAATACCACTACTC | MB468 | 13 | 45 | 19 | 8 | 14 | 4.9 | 17.0 | 9.4 | 1.9 | 2.5 |
| >hg38_21_+_8433172_8446622_RNA45SN1_45S_50ntFlanks@12001.12031.31\|CTGCGGGCCGCCGGTGAAATACCACTACTCT | >hg38_21_+_8433172_8446622_RNA45SN1_45S_50ntFlanks | 12001 | 12031 | 31 | CTGCGGGCCGCCGGTGAAATACCACTACTCT | MB468 | 29 | 48 | 43 | 16 | 34 | 4.6 | 15.7 | 21.0 | 5.6 | 8.0 |
| >hg38_21_+_8433172_8446622_RNA45SN1_45S_50ntFlanks@12001.12032.32\|CTGCGGGCCGCCGGTGAAATACCACTACTCTG | >hg38_21_+_8433172_8446622_RNA45SN1_45S_50ntFlanks | 12001 | 12032 | 32 | CTGCGGGCCGCCGGTGAAATACCACTACTCTG | MB468 | 196 | 311 | 326 | 135 | 246 | 27.2 | 104.2 | 195.9 | 49.1 | 76.0 |
| >hg38_21_+_8433172_8446622_RNA45SN1_45S_50ntFlanks@12002.12029.28\|TGCGGGCCGCCGGTGAAATACCACTACT | >hg38_21_+_8433172_8446622_RNA45SN1_45S_50ntFlanks | 12002 | 12029 | 28 | TGCGGGCCGCCGGTGAAATACCACTACT | MB468 | 11 | 24 | 13 | 11 | 10 | 2.4 | 8.7 | 6.3 | 4.3 | 3.9 |
| >hg38_21_+_8433172_8446622_RNA45SN1_45S_50ntFlanks@12002.12030.29\|TGCGGGCCGCCGGTGAAATACCACTACTC | >hg38_21_+_8433172_8446622_RNA45SN1_45S_50ntFlanks | 12002 | 12030 | 29 | TGCGGGCCGCCGGTGAAATACCACTACTC | MB468 | 32 | 66 | 40 | 18 | 29 | 12.7 | 28.4 | 20.9 | 6.5 | 4.9 |
| >hg38_21_+_8433172_8446622_RNA45SN1_45S_50ntFlanks@12002.12031.30\|TGCGGGCCGCCGGTGAAATACCACTACTCT | >hg38_21_+_8433172_8446622_RNA45SN1_45S_50ntFlanks | 12002 | 12031 | 30 | TGCGGGCCGCCGGTGAAATACCACTACTCT | MB468 | 71 | 68 | 88 | 33 | 62 | 22.2 | 23.5 | 49.6 | 8.8 | 8.8 |
| >hg38_21_+_8433172_8446622_RNA45SN1_45S_50ntFlanks@12002.12032.31\|TGCGGGCCGCCGGTGAAATACCACTACTCTG | >hg38_21_+_8433172_8446622_RNA45SN1_45S_50ntFlanks | 12002 | 12032 | 31 | TGCGGGCCGCCGGTGAAATACCACTACTCTG | MB468 | 325 | 412 | 510 | 236 | 361 | 49.4 | 166.8 | 350.7 | 59.2 | 95.7 |
| >hg38_21_+_8433172_8446622_RNA45SN1_45S_50ntFlanks@12002.12043.42\|TGCGGGCCGCCGGTGAAATACCACTACTCTGATCGTTTTTTC | >hg38_21_+_8433172_8446622_RNA45SN1_45S_50ntFlanks | 12002 | 12043 | 42 | TGCGGGCCGCCGGTGAAATACCACTACTCTGATCGTTTTTTC | MB468 | 18 | 145 | 53 | 21 | 144 | 14.9 | 113.0 | 60.4 | 25.7 | 106.2 |
| >hg38_21_+_8433172_8446622_RNA45SN1_45S_50ntFlanks@12003.12029.27\|GCGGGCCGCCGGTGAAATACCACTACT | >hg38_21_+_8433172_8446622_RNA45SN1_45S_50ntFlanks | 12003 | 12029 | 27 | GCGGGCCGCCGGTGAAATACCACTACT | MB231 | 18 | 32 | 61 | 1 | 4 | 15.1 | 20.5 | 41.7 | 0.7 | 1.8 |
| >hg38_21_+_8433172_8446622_RNA45SN1_45S_50ntFlanks@12003.12030.28\|GCGGGCCGCCGGTGAAATACCACTACTC | >hg38_21_+_8433172_8446622_RNA45SN1_45S_50ntFlanks | 12003 | 12030 | 28 | GCGGGCCGCCGGTGAAATACCACTACTC | MB468 | 20 | 45 | 19 | 15 | 30 | 9.5 | 29.0 | 8.1 | 3.6 | 5.7 |
| >hg38_21_+_8433172_8446622_RNA45SN1_45S_50ntFlanks@12003.12031.29\|GCGGGCCGCCGGTGAAATACCACTACTCT | >hg38_21_+_8433172_8446622_RNA45SN1_45S_50ntFlanks | 12003 | 12031 | 29 | GCGGGCCGCCGGTGAAATACCACTACTCT | MB468 | 37 | 36 | 42 | 34 | 57 | 13.5 | 13.8 | 18.4 | 8.4 | 8.9 |
| >hg38_21_+_8433172_8446622_RNA45SN1_45S_50ntFlanks@12003.12032.30\|GCGGGCCGCCGGTGAAATACCACTACTCTG | >hg38_21_+_8433172_8446622_RNA45SN1_45S_50ntFlanks | 12003 | 12032 | 30 | GCGGGCCGCCGGTGAAATACCACTACTCTG | MB468 | 162 | 199 | 273 | 342 | 306 | 29.9 | 67.9 | 151.8 | 70.4 | 38.9 |
| >hg38_21_+_8433172_8446622_RNA45SN1_45S_50ntFlanks@12003.12033.31\|GCGGGCCGCCGGTGAAATACCACTACTCTGA | >hg38_21_+_8433172_8446622_RNA45SN1_45S_50ntFlanks | 12003 | 12033 | 31 | GCGGGCCGCCGGTGAAATACCACTACTCTGA | BT20 | 12 | 11 | 15 | 12 | 37 | 7.8 | 12.9 | 10.4 | 19.5 | 15.9 |
| >hg38_21_+_8433172_8446622_RNA45SN1_45S_50ntFlanks@12003.12033.31\|GCGGGCCGCCGGTGAAATACCACTACTCTGA | >hg38_21_+_8433172_8446622_RNA45SN1_45S_50ntFlanks | 12003 | 12033 | 31 | GCGGGCCGCCGGTGAAATACCACTACTCTGA | MB231 | 21 | 40 | 64 | 3 | 13 | 12.1 | 40.6 | 82.9 | 1.1 | 8.8 |
| >hg38_21_+_8433172_8446622_RNA45SN1_45S_50ntFlanks@12003.12034.32\|GCGGGCCGCCGGTGAAATACCACTACTCTGAT | >hg38_21_+_8433172_8446622_RNA45SN1_45S_50ntFlanks | 12003 | 12034 | 32 | GCGGGCCGCCGGTGAAATACCACTACTCTGAT | MB231 | 28 | 42 | 70 | 3 | 9 | 16.6 | 36.7 | 71.3 | 1.5 | 4.2 |
| >hg38_21_+_8433172_8446622_RNA45SN1_45S_50ntFlanks@12004.12032.29\|CGGGCCGCCGGTGAAATACCACTACTCTG | >hg38_21_+_8433172_8446622_RNA45SN1_45S_50ntFlanks | 12004 | 12032 | 29 | CGGGCCGCCGGTGAAATACCACTACTCTG | MB468 | 24 | 8 | 22 | 79 | 39 | 6.5 | 2.0 | 7.7 | 9.9 | 15.0 |
| >hg38_21_+_8433172_8446622_RNA45SN1_45S_50ntFlanks@12004.12033.30\|CGGGCCGCCGGTGAAATACCACTACTCTGA | >hg38_21_+_8433172_8446622_RNA45SN1_45S_50ntFlanks | 12004 | 12033 | 30 | CGGGCCGCCGGTGAAATACCACTACTCTGA | BT20 | 67 | 27 | 35 | 35 | 79 | 75.0 | 33.5 | 27.3 | 54.7 | 52.0 |
| >hg38_21_+_8433172_8446622_RNA45SN1_45S_50ntFlanks@12004.12034.31\|CGGGCCGCCGGTGAAATACCACTACTCTGAT | >hg38_21_+_8433172_8446622_RNA45SN1_45S_50ntFlanks | 12004 | 12034 | 31 | CGGGCCGCCGGTGAAATACCACTACTCTGAT | BT20 | 15 | 9 | 8 | 4 | 24 | 12.8 | 10.1 | 5.1 | 4.5 | 15.1 |
| >hg38_21_+_8433172_8446622_RNA45SN1_45S_50ntFlanks@12004.12035.32\|CGGGCCGCCGGTGAAATACCACTACTCTGATC | >hg38_21_+_8433172_8446622_RNA45SN1_45S_50ntFlanks | 12004 | 12035 | 32 | CGGGCCGCCGGTGAAATACCACTACTCTGATC | BT20 | 32 | 41 | 25 | 80 | 123 | 25.0 | 49.2 | 15.3 | 97.0 | 87.3 |

| | | | | | | | | | | | | | | | | |
|---|---|---|---|---|---|---|---|---|---|---|---|---|---|---|---|---|
| >hg38_21_+_8433172_8446622_RNA45SN1_45S_50ntFlanks@12006.12032.27\|GGCCGCCGGTGAAATACCACTACTCTG | >hg38_21_+_8433172_8446622_RNA45SN1_45S_50ntFlanks | 12006 | 12032 | 27 | GGCCGCCGGTGAAATACCACTACTCTG | MB468 | 24 | 53 | 35 | 44 | 59 | 7.5 | 42.4 | 16.9 | 28.7 | 29.3 |
| >hg38_21_+_8433172_8446622_RNA45SN1_45S_50ntFlanks@12006.12033.28\|GGCCGCCGGTGAAATACCACTACTCTGA | >hg38_21_+_8433172_8446622_RNA45SN1_45S_50ntFlanks | 12006 | 12033 | 28 | GGCCGCCGGTGAAATACCACTACTCTGA | BT20 | 12 | 11 | 14 | 3 | 18 | 5.3 | 8.9 | 10.1 | 4.6 | 18.4 |
| >hg38_21_+_8433172_8446622_RNA45SN1_45S_50ntFlanks@12007.12029.23\|GCCGCCGGTGAAATACCACTACT | >hg38_21_+_8433172_8446622_RNA45SN1_45S_50ntFlanks | 12007 | 12029 | 23 | GCCGCCGGTGAAATACCACTACT | BT20 | 28 | 27 | 9 | 3 | 45 | 44.2 | 43.1 | 12.3 | 5.3 | 74.9 |
| >hg38_21_+_8433172_8446622_RNA45SN1_45S_50ntFlanks@12007.12030.24\|GCCGCCGGTGAAATACCACTACTC | >hg38_21_+_8433172_8446622_RNA45SN1_45S_50ntFlanks | 12007 | 12030 | 24 | GCCGCCGGTGAAATACCACTACTC | MB468 | 14 | 32 | 8 | 32 | 174 | 5.1 | 18.5 | 3.1 | 14.1 | 47.0 |
| >hg38_21_+_8433172_8446622_RNA45SN1_45S_50ntFlanks@12007.12031.25\|GCCGCCGGTGAAATACCACTACTCT | >hg38_21_+_8433172_8446622_RNA45SN1_45S_50ntFlanks | 12007 | 12031 | 25 | GCCGCCGGTGAAATACCACTACTCT | MB468 | 50 | 91 | 45 | 73 | 329 | 15.2 | 54.4 | 14.8 | 45.9 | 102.6 |
| >hg38_21_+_8433172_8446622_RNA45SN1_45S_50ntFlanks@12007.12032.26\|GCCGCCGGTGAAATACCACTACTCTG | >hg38_21_+_8433172_8446622_RNA45SN1_45S_50ntFlanks | 12007 | 12032 | 26 | GCCGCCGGTGAAATACCACTACTCTG | MB468 | 208 | 464 | 271 | 776 | 1026 | 46.3 | 286.4 | 133.5 | 346.4 | 405.6 |
| >hg38_21_+_8433172_8446622_RNA45SN1_45S_50ntFlanks@12007.12035.29\|GCCGCCGGTGAAATACCACTACTCTGATC | >hg38_21_+_8433172_8446622_RNA45SN1_45S_50ntFlanks | 12007 | 12035 | 29 | GCCGCCGGTGAAATACCACTACTCTGATC | BT20 | 10 | 21 | 6 | 21 | 47 | 10.6 | 25.4 | 4.3 | 17.5 | 49.9 |
| >hg38_21_+_8433172_8446622_RNA45SN1_45S_50ntFlanks@12007.12043.37\|GCCGCCGGTGAAATACCACTACTCTGATCGTTTTTTC | >hg38_21_+_8433172_8446622_RNA45SN1_45S_50ntFlanks | 12007 | 12043 | 37 | GCCGCCGGTGAAATACCACTACTCTGATCGTTTTTTC | MB468 | 26 | 220 | 75 | 77 | 489 | 15.7 | 171.5 | 61.4 | 89.5 | 274.2 |
| >hg38_21_+_8433172_8446622_RNA45SN1_45S_50ntFlanks@12008.12029.22\|CCGCCGGTGAAATACCACTACT | >hg38_21_+_8433172_8446622_RNA45SN1_45S_50ntFlanks | 12008 | 12029 | 22 | CCGCCGGTGAAATACCACTACT | MB468 | 20 | 5 | 11 | 114 | 95 | 6.5 | 3.8 | 4.6 | 30.6 | 31.3 |
| >hg38_21_+_8433172_8446622_RNA45SN1_45S_50ntFlanks@12008.12030.23\|CCGCCGGTGAAATACCACTACTC | >hg38_21_+_8433172_8446622_RNA45SN1_45S_50ntFlanks | 12008 | 12030 | 23 | CCGCCGGTGAAATACCACTACTC | MB468 | 100 | 42 | 86 | 343 | 369 | 40.5 | 20.9 | 42.0 | 143.6 | 110.0 |
| >hg38_21_+_8433172_8446622_RNA45SN1_45S_50ntFlanks@12008.12031.24\|CCGCCGGTGAAATACCACTACTCT | >hg38_21_+_8433172_8446622_RNA45SN1_45S_50ntFlanks | 12008 | 12031 | 24 | CCGCCGGTGAAATACCACTACTCT | MB468 | 169 | 85 | 211 | 500 | 476 | 42.5 | 43.4 | 133.4 | 267.3 | 60.2 |
| >hg38_21_+_8433172_8446622_RNA45SN1_45S_50ntFlanks@12008.12032.25\|CCGCCGGTGAAATACCACTACTCTG | >hg38_21_+_8433172_8446622_RNA45SN1_45S_50ntFlanks | 12008 | 12032 | 25 | CCGCCGGTGAAATACCACTACTCTG | MB468 | 873 | 839 | 1508 | 3473 | 2563 | 129.2 | 504.0 | 1159.1 | 1372.2 | 549.9 |
| >hg38_21_+_8433172_8446622_RNA45SN1_45S_50ntFlanks@12008.12033.26\|CCGCCGGTGAAATACCACTACTCTGA | >hg38_21_+_8433172_8446622_RNA45SN1_45S_50ntFlanks | 12008 | 12033 | 26 | CCGCCGGTGAAATACCACTACTCTGA | MB468 | 36 | 32 | 59 | 49 | 58 | 6.1 | 17.8 | 42.5 | 21.3 | 13.7 |
| >hg38_21_+_8433172_8446622_RNA45SN1_45S_50ntFlanks@12008.12034.27\|CCGCCGGTGAAATACCACTACTCTGAT | >hg38_21_+_8433172_8446622_RNA45SN1_45S_50ntFlanks | 12008 | 12034 | 27 | CCGCCGGTGAAATACCACTACTCTGAT | MB468 | 25 | 18 | 49 | 23 | 30 | 3.2 | 5.4 | 20.4 | 8.0 | 7.6 |
| >hg38_21_+_8433172_8446622_RNA45SN1_45S_50ntFlanks@12008.12035.28\|CCGCCGGTGAAATACCACTACTCTGATC | >hg38_21_+_8433172_8446622_RNA45SN1_45S_50ntFlanks | 12008 | 12035 | 28 | CCGCCGGTGAAATACCACTACTCTGATC | MB468 | 12 | 17 | 20 | 45 | 32 | 3.9 | 5.3 | 9.2 | 13.7 | 9.7 |
| >hg38_21_+_8433172_8446622_RNA45SN1_45S_50ntFlanks@12008.12039.32\|CCGCCGGTGAAATACCACTACTCTGATCGTTT | >hg38_21_+_8433172_8446622_RNA45SN1_45S_50ntFlanks | 12008 | 12039 | 32 | CCGCCGGTGAAATACCACTACTCTGATCGTTT | MB468 | 14 | 37 | 36 | 26 | 40 | 1.3 | 19.4 | 25.5 | 28.3 | 12.0 |
| >hg38_21_+_8433172_8446622_RNA45SN1_45S_50ntFlanks@12008.12042.35\|CCGCCGGTGAAATACCACTACTCTGATCGTTTTTT | >hg38_21_+_8433172_8446622_RNA45SN1_45S_50ntFlanks | 12008 | 12042 | 35 | CCGCCGGTGAAATACCACTACTCTGATCGTTTTTT | MB468 | 14 | 68 | 54 | 20 | 119 | 11.8 | 52.7 | 36.6 | 19.3 | 82.8 |
| >hg38_21_+_8433172_8446622_RNA45SN1_45S_50ntFlanks@12008.12043.36\|CCGCCGGTGAAATACCACTACTCTGATCGTTTTTTC | >hg38_21_+_8433172_8446622_RNA45SN1_45S_50ntFlanks | 12008 | 12043 | 36 | CCGCCGGTGAAATACCACTACTCTGATCGTTTTTTC | MB468 | 121 | 751 | 378 | 284 | 1232 | 90.0 | 496.5 | 384.4 | 348.5 | 720.7 |
| >hg38_21_+_8433172_8446622_RNA45SN1_45S_50ntFlanks@12025.12043.19\|CTACTCTGATCGTTTTTTC | >hg38_21_+_8433172_8446622_RNA45SN1_45S_50ntFlanks | 12025 | 12043 | 19 | CTACTCTGATCGTTTTTTC | MB468 | 10 | 9 | 12 | 3 | 5 | 5.5 | 4.9 | 9.8 | 0.3 | 1.9 |
| >hg38_21_+_8433172_8446622_RNA45SN1_45S_50ntFlanks@12033.12052.20\|ATCGTTTTTTCACTGACCCG | >hg38_21_+_8433172_8446622_RNA45SN1_45S_50ntFlanks | 12033 | 12052 | 20 | ATCGTTTTTTCACTGACCCG | MB468 | 22 | 3 | 8 | 7 | 13 | 9.1 | 0.7 | 4.8 | 3.2 | 5.4 |
| >hg38_21_+_8433172_8446622_RNA45SN1_45S_50ntFlanks@12033.12053.21\|ATCGTTTTTTCACTGACCCGG | >hg38_21_+_8433172_8446622_RNA45SN1_45S_50ntFlanks | 12033 | 12053 | 21 | ATCGTTTTTTCACTGACCCGG | MB468 | 69 | 17 | 35 | 36 | 81 | 25.7 | 5.1 | 18.3 | 17.2 | 13.4 |
| >hg38_21_+_8433172_8446622_RNA45SN1_45S_50ntFlanks@12033.12055.23\|ATCGTTTTTTCACTGACCCGGTG | >hg38_21_+_8433172_8446622_RNA45SN1_45S_50ntFlanks | 12033 | 12055 | 23 | ATCGTTTTTTCACTGACCCGGTG | MB468 | 13 | 2 | 7 | 25 | 14 | 3.0 | 0.8 | 5.1 | 12.3 | 3.5 |
| >hg38_21_+_8433172_8446622_RNA45SN1_45S_50ntFlanks@12035.12053.19\|CGTTTTTTCACTGACCCGG | >hg38_21_+_8433172_8446622_RNA45SN1_45S_50ntFlanks | 12035 | 12053 | 19 | CGTTTTTTCACTGACCCGG | MB468 | 15 | 4 | 12 | 4 | 7 | 5.8 | 1.3 | 5.6 | 2.6 | 2.2 |
| >hg38_21_+_8433172_8446622_RNA45SN1_45S_50ntFlanks@12044.12068.25\|ACTGACCCGGTGAGGCGGGGGGGCG | >hg38_21_+_8433172_8446622_RNA45SN1_45S_50ntFlanks | 12044 | 12068 | 25 | ACTGACCCGGTGAGGCGGGGGGGCG | MB468 | 32 | 54 | 20 | 64 | 120 | 3.0 | 34.2 | 5.6 | 40.2 | 63.5 |
| >hg38_21_+_8433172_8446622_RNA45SN1_45S_50ntFlanks@12044.12070.27\|ACTGACCCGGTGAGGCGGGGGGGCGAG | >hg38_21_+_8433172_8446622_RNA45SN1_45S_50ntFlanks | 12044 | 12070 | 27 | ACTGACCCGGTGAGGCGGGGGGGCGAG | MB468 | 44 | 100 | 25 | 72 | 164 | 5.3 | 44.5 | 7.6 | 46.2 | 95.4 |
| >hg38_21_+_8433172_8446622_RNA45SN1_45S_50ntFlanks@12044.12071.28\|ACTGACCCGGTGAGGCGGGGGGGCGAGC | >hg38_21_+_8433172_8446622_RNA45SN1_45S_50ntFlanks | 12044 | 12071 | 28 | ACTGACCCGGTGAGGCGGGGGGGCGAGC | MB468 | 21 | 43 | 12 | 51 | 104 | 5.9 | 11.3 | 5.9 | 39.2 | 56.5 |
| >hg38_21_+_8433172_8446622_RNA45SN1_45S_50ntFlanks@12044.12072.29\|ACTGACCCGGTGAGGCGGGGGGGCGAGCC | >hg38_21_+_8433172_8446622_RNA45SN1_45S_50ntFlanks | 12044 | 12072 | 29 | ACTGACCCGGTGAGGCGGGGGGGCGAGCC | MB468 | 24 | 61 | 19 | 43 | 141 | 2.7 | 29.6 | 10.4 | 28.3 | 73.1 |
| >hg38_21_+_8433172_8446622_RNA45SN1_45S_50ntFlanks@12045.12070.26\|CTGACCCGGTGAGGCGGGGGGGCGAG | >hg38_21_+_8433172_8446622_RNA45SN1_45S_50ntFlanks | 12045 | 12070 | 26 | CTGACCCGGTGAGGCGGGGGGGCGAG | MB468 | 13 | 34 | 10 | 21 | 45 | 2.4 | 19.5 | 3.3 | 13.1 | 30.8 |
| >hg38_21_+_8433172_8446622_RNA45SN1_45S_50ntFlanks@12071.12090.20\|CCCCGAGGGGCTCTCGCTTC | >hg38_21_+_8433172_8446622_RNA45SN1_45S_50ntFlanks | 12071 | 12090 | 20 | CCCCGAGGGGCTCTCGCTTC | MB468 | 10 | 5 | 11 | 2 | 7 | 3.7 | 1.7 | 4.8 | 1.2 | 2.0 |
| >hg38_21_+_8433172_8446622_RNA45SN1_45S_50ntFlanks@12071.12092.22\|CCCCGAGGGGCTCTCGCTTCTG | >hg38_21_+_8433172_8446622_RNA45SN1_45S_50ntFlanks | 12071 | 12092 | 22 | CCCCGAGGGGCTCTCGCTTCTG | MB468 | 15 | 9 | 30 | 2 | 12 | 4.1 | 2.2 | 14.4 | 1.3 | 3.1 |
| >hg38_21_+_8433172_8446622_RNA45SN1_45S_50ntFlanks@12071.12093.23\|CCCCGAGGGGCTCTCGCTTCTGG | >hg38_21_+_8433172_8446622_RNA45SN1_45S_50ntFlanks | 12071 | 12093 | 23 | CCCCGAGGGGCTCTCGCTTCTGG | MB468 | 14 | 3 | 20 | 3 | 4 | 3.1 | 1.0 | 11.2 | 1.4 | 0.5 |
| >hg38_21_+_8433172_8446622_RNA45SN1_45S_50ntFlanks@12071.12094.24\|CCCCGAGGGGCTCTCGCTTCTGGC | >hg38_21_+_8433172_8446622_RNA45SN1_45S_50ntFlanks | 12071 | 12094 | 24 | CCCCGAGGGGCTCTCGCTTCTGGC | MB468 | 25 | 7 | 45 | 3 | 6 | 7.2 | 4.1 | 15.9 | 2.8 | 3.1 |
| >hg38_21_+_8433172_8446622_RNA45SN1_45S_50ntFlanks@12071.12095.25\|CCCCGAGGGGCTCTCGCTTCTGGCG | >hg38_21_+_8433172_8446622_RNA45SN1_45S_50ntFlanks | 12071 | 12095 | 25 | CCCCGAGGGGCTCTCGCTTCTGGCG | MB468 | 156 | 60 | 250 | 56 | 91 | 56.1 | 12.1 | 150.8 | 62.8 | 36.5 |
| >hg38_21_+_8433172_8446622_RNA45SN1_45S_50ntFlanks@12071.12096.26\|CCCCGAGGGGCTCTCGCTTCTGGCGC | >hg38_21_+_8433172_8446622_RNA45SN1_45S_50ntFlanks | 12071 | 12096 | 26 | CCCCGAGGGGCTCTCGCTTCTGGCGC | MB468 | 48 | 17 | 59 | 8 | 25 | 25.8 | 8.9 | 56.7 | 10.4 | 8.2 |
| >hg38_21_+_8433172_8446622_RNA45SN1_45S_50ntFlanks@12071.12097.27\|CCCCGAGGGGCTCTCGCTTCTGGCGCC | >hg38_21_+_8433172_8446622_RNA45SN1_45S_50ntFlanks | 12071 | 12097 | 27 | CCCCGAGGGGCTCTCGCTTCTGGCGCC | MB468 | 27 | 24 | 33 | 2 | 23 | 12.3 | 14.5 | 22.2 | 2.9 | 21.9 |
| >hg38_21_+_8433172_8446622_RNA45SN1_45S_50ntFlanks@12071.12106.36\|CCCCGAGGGGCTCTCGCTTCTGGCGCCAAGCGCCCG | >hg38_21_+_8433172_8446622_RNA45SN1_45S_50ntFlanks | 12071 | 12106 | 36 | CCCCGAGGGGCTCTCGCTTCTGGCGCCAAGCGCCCG | MB468 | 24 | 32 | 28 | 2 | 43 | 10.2 | 20.9 | 19.0 | 2.4 | 35.9 |
| >hg38_21_+_8433172_8446622_RNA45SN1_45S_50ntFlanks@12072.12089.18\|CCCGAGGGGCTCTCGCTT | >hg38_21_+_8433172_8446622_RNA45SN1_45S_50ntFlanks | 12072 | 12089 | 18 | CCCGAGGGGCTCTCGCTT | MB468 | 14 | 4 | 12 | 3 | 12 | 2.6 | 1.7 | 2.3 | 1.4 | 3.8 |
| >hg38_21_+_8433172_8446622_RNA45SN1_45S_50ntFlanks@12072.12090.19\|CCCGAGGGGCTCTCGCTTC | >hg38_21_+_8433172_8446622_RNA45SN1_45S_50ntFlanks | 12072 | 12090 | 19 | CCCGAGGGGCTCTCGCTTC | MB468 | 22 | 8 | 26 | 6 | 12 | 4.8 | 2.1 | 6.2 | 2.2 | 4.3 |
| >hg38_21_+_8433172_8446622_RNA45SN1_45S_50ntFlanks@12072.12092.21\|CCCGAGGGGCTCTCGCTTCTG | >hg38_21_+_8433172_8446622_RNA45SN1_45S_50ntFlanks | 12072 | 12092 | 21 | CCCGAGGGGCTCTCGCTTCTG | MB468 | 19 | 4 | 50 | 5 | 8 | 2.6 | 1.4 | 12.4 | 2.3 | 2.9 |
| >hg38_21_+_8433172_8446622_RNA45SN1_45S_50ntFlanks@12072.12093.22\|CCCGAGGGGCTCTCGCTTCTGG | >hg38_21_+_8433172_8446622_RNA45SN1_45S_50ntFlanks | 12072 | 12093 | 22 | CCCGAGGGGCTCTCGCTTCTGG | MB468 | 12 | 2 | 22 | 6 | 4 | 3.7 | 0.8 | 6.0 | 1.8 | 0.9 |
| >hg38_21_+_8433172_8446622_RNA45SN1_45S_50ntFlanks@12072.12094.23\|CCCGAGGGGCTCTCGCTTCTGGC | >hg38_21_+_8433172_8446622_RNA45SN1_45S_50ntFlanks | 12072 | 12094 | 23 | CCCGAGGGGCTCTCGCTTCTGGC | MB468 | 16 | 4 | 31 | 5 | 5 | 5.0 | 2.6 | 8.6 | 1.5 | 3.4 |
| >hg38_21_+_8433172_8446622_RNA45SN1_45S_50ntFlanks@12072.12095.24\|CCCGAGGGGCTCTCGCTTCTGGCG | >hg38_21_+_8433172_8446622_RNA45SN1_45S_50ntFlanks | 12072 | 12095 | 24 | CCCGAGGGGCTCTCGCTTCTGGCG | MB468 | 118 | 46 | 229 | 53 | 60 | 28.1 | 11.2 | 76.1 | 26.4 | 21.5 |
| >hg38_21_+_8433172_8446622_RNA45SN1_45S_50ntFlanks@12072.12096.25\|CCCGAGGGGCTCTCGCTTCTGGCGC | >hg38_21_+_8433172_8446622_RNA45SN1_45S_50ntFlanks | 12072 | 12096 | 25 | CCCGAGGGGCTCTCGCTTCTGGCGC | MB468 | 34 | 11 | 59 | 9 | 17 | 13.3 | 5.8 | 42.1 | 10.2 | 5.0 |
| >hg38_21_+_8433172_8446622_RNA45SN1_45S_50ntFlanks@12072.12097.26\|CCCGAGGGGCTCTCGCTTCTGGCGCC | >hg38_21_+_8433172_8446622_RNA45SN1_45S_50ntFlanks | 12072 | 12097 | 26 | CCCGAGGGGCTCTCGCTTCTGGCGCC | MB468 | 40 | 42 | 57 | 6 | 39 | 12.9 | 15.2 | 26.7 | 5.5 | 13.0 |
| >hg38_21_+_8433172_8446622_RNA45SN1_45S_50ntFlanks@12072.12106.35\|CCCGAGGGGCTCTCGCTTCTGGCGCCAAGCGCCCG | >hg38_21_+_8433172_8446622_RNA45SN1_45S_50ntFlanks | 12072 | 12106 | 35 | CCCGAGGGGCTCTCGCTTCTGGCGCCAAGCGCCCG | MB468 | 20 | 9 | 15 | 5 | 33 | 7.5 | 4.2 | 6.6 | 3.9 | 8.9 |
| >hg38_21_+_8433172_8446622_RNA45SN1_45S_50ntFlanks@12073.12090.18\|CCGAGGGGCTCTCGCTTC | >hg38_21_+_8433172_8446622_RNA45SN1_45S_50ntFlanks | 12073 | 12090 | 18 | CCGAGGGGCTCTCGCTTC | MB468 | 52 | 11 | 50 | 34 | 29 | 10.1 | 3.2 | 13.8 | 14.1 | 12.1 |
| >hg38_21_+_8433172_8446622_RNA45SN1_45S_50ntFlanks@12073.12091.19\|CCGAGGGGCTCTCGCTTCT | >hg38_21_+_8433172_8446622_RNA45SN1_45S_50ntFlanks | 12073 | 12091 | 19 | CCGAGGGGCTCTCGCTTCT | MB468 | 14 | 2 | 19 | 7 | 9 | 6.3 | 0.6 | 12.5 | 1.7 | 5.6 |
| >hg38_21_+_8433172_8446622_RNA45SN1_45S_50ntFlanks@12073.12092.20\|CCGAGGGGCTCTCGCTTCTG | >hg38_21_+_8433172_8446622_RNA45SN1_45S_50ntFlanks | 12073 | 12092 | 20 | CCGAGGGGCTCTCGCTTCTG | MB468 | 36 | 3 | 72 | 21 | 16 | 4.6 | 0.8 | 24.9 | 3.5 | 6.6 |
| >hg38_21_+_8433172_8446622_RNA45SN1_45S_50ntFlanks@12073.12093.21\|CCGAGGGGCTCTCGCTTCTGG | >hg38_21_+_8433172_8446622_RNA45SN1_45S_50ntFlanks | 12073 | 12093 | 21 | CCGAGGGGCTCTCGCTTCTGG | MB468 | 14 | 2 | 19 | 16 | 5 | 3.3 | 0.7 | 6.7 | 12.5 | 1.4 |
| >hg38_21_+_8433172_8446622_RNA45SN1_45S_50ntFlanks@12073.12094.22\|CCGAGGGGCTCTCGCTTCTGGC | >hg38_21_+_8433172_8446622_RNA45SN1_45S_50ntFlanks | 12073 | 12094 | 22 | CCGAGGGGCTCTCGCTTCTGGC | MB468 | 34 | 7 | 61 | 25 | 16 | 9.3 | 4.4 | 20.0 | 15.5 | 10.1 |

| | | | | | | | | | | | | | | | | |
|---|---|---|---|---|---|---|---|---|---|---|---|---|---|---|---|---|
| >hg38_21_+_8433172_8446622_RNA45SN1_45S_50ntFlanks@12073.12095.23\|CCGAGGGGCTCTCGCTTCTGGCG | >hg38_21_+_8433172_8446622_RNA45SN1_45S_50ntFlanks | 12073 | 12095 | 23 | CCGAGGGGCTCTCGCTTCTGGCG | MB468 | 222 | 78 | 430 | 186 | 121 | 61.7 | 20.5 | 197.4 | 85.9 | 56.4 |
| >hg38_21_+_8433172_8446622_RNA45SN1_45S_50ntFlanks@12073.12096.24\|CCGAGGGGCTCTCGCTTCTGGCGC | >hg38_21_+_8433172_8446622_RNA45SN1_45S_50ntFlanks | 12073 | 12096 | 24 | CCGAGGGGCTCTCGCTTCTGGCGC | MB468 | 98 | 39 | 154 | 38 | 54 | 37.4 | 18.3 | 94.6 | 23.6 | 18.1 |
| >hg38_21_+_8433172_8446622_RNA45SN1_45S_50ntFlanks@12073.12097.25\|CCGAGGGGCTCTCGCTTCTGGCGCC | >hg38_21_+_8433172_8446622_RNA45SN1_45S_50ntFlanks | 12073 | 12097 | 25 | CCGAGGGGCTCTCGCTTCTGGCGCC | MB468 | 156 | 143 | 189 | 37 | 159 | 44.9 | 40.6 | 77.6 | 22.8 | 42.7 |
| >hg38_21_+_8433172_8446622_RNA45SN1_45S_50ntFlanks@12073.12098.26\|CCGAGGGGCTCTCGCTTCTGGCGCCA | >hg38_21_+_8433172_8446622_RNA45SN1_45S_50ntFlanks | 12073 | 12098 | 26 | CCGAGGGGCTCTCGCTTCTGGCGCCA | MB468 | 25 | 15 | 27 | 4 | 20 | 7.5 | 5.5 | 10.4 | 1.4 | 7.2 |
| >hg38_21_+_8433172_8446622_RNA45SN1_45S_50ntFlanks@12073.12106.34\|CCGAGGGGCTCTCGCTTCTGGCGCCAAGCGCCCG | >hg38_21_+_8433172_8446622_RNA45SN1_45S_50ntFlanks | 12073 | 12106 | 34 | CCGAGGGGCTCTCGCTTCTGGCGCCAAGCGCCCG | MB468 | 32 | 9 | 20 | 14 | 49 | 15.3 | 3.8 | 11.0 | 6.5 | 14.9 |
| >hg38_21_+_8433172_8446622_RNA45SN1_45S_50ntFlanks@12073.12108.36\|CCGAGGGGCTCTCGCTTCTGGCGCCAAGCGCCCGGC | >hg38_21_+_8433172_8446622_RNA45SN1_45S_50ntFlanks | 12073 | 12108 | 36 | CCGAGGGGCTCTCGCTTCTGGCGCCAAGCGCCCGGC | MB468 | 10 | 4 | 6 | 18 | 28 | 3.6 | 1.8 | 3.5 | 6.3 | 11.1 |
| >hg38_21_+_8433172_8446622_RNA45SN1_45S_50ntFlanks@12073.12110.38\|CCGAGGGGCTCTCGCTTCTGGCGCCAAGCGCCCGGCCG | >hg38_21_+_8433172_8446622_RNA45SN1_45S_50ntFlanks | 12073 | 12110 | 38 | CCGAGGGGCTCTCGCTTCTGGCGCCAAGCGCCCGGCCG | MB468 | 27 | 19 | 17 | 22 | 83 | 4.9 | 17.0 | 8.3 | 14.4 | 41.0 |
| >hg38_21_+_8433172_8446622_RNA45SN1_45S_50ntFlanks@12074.12095.22\|CGAGGGGCTCTCGCTTCTGGCG | >hg38_21_+_8433172_8446622_RNA45SN1_45S_50ntFlanks | 12074 | 12095 | 22 | CGAGGGGCTCTCGCTTCTGGCG | MB468 | 69 | 41 | 96 | 45 | 68 | 13.5 | 10.2 | 31.0 | 15.2 | 30.4 |
| >hg38_21_+_8433172_8446622_RNA45SN1_45S_50ntFlanks@12075.12095.21\|GAGGGGCTCTCGCTTCTGGCG | >hg38_21_+_8433172_8446622_RNA45SN1_45S_50ntFlanks | 12075 | 12095 | 21 | GAGGGGCTCTCGCTTCTGGCG | MB468 | 18 | 21 | 30 | 8 | 12 | 4.2 | 6.3 | 15.1 | 6.0 | 5.3 |
| >hg38_21_+_8433172_8446622_RNA45SN1_45S_50ntFlanks@12076.12095.20\|AGGGGCTCTCGCTTCTGGCG | >hg38_21_+_8433172_8446622_RNA45SN1_45S_50ntFlanks | 12076 | 12095 | 20 | AGGGGCTCTCGCTTCTGGCG | MB468 | 333 | 286 | 728 | 104 | 122 | 74.7 | 99.1 | 278.8 | 100.8 | 52.7 |
| >hg38_21_+_8433172_8446622_RNA45SN1_45S_50ntFlanks@12076.12096.21\|AGGGGCTCTCGCTTCTGGCGC | >hg38_21_+_8433172_8446622_RNA45SN1_45S_50ntFlanks | 12076 | 12096 | 21 | AGGGGCTCTCGCTTCTGGCGC | MB468 | 24 | 27 | 42 | 6 | 10 | 7.4 | 9.7 | 20.3 | 5.7 | 2.7 |
| >hg38_21_+_8433172_8446622_RNA45SN1_45S_50ntFlanks@12076.12097.22\|AGGGGCTCTCGCTTCTGGCGCC | >hg38_21_+_8433172_8446622_RNA45SN1_45S_50ntFlanks | 12076 | 12097 | 22 | AGGGGCTCTCGCTTCTGGCGCC | MB468 | 24 | 20 | 32 | 5 | 12 | 9.4 | 2.8 | 12.4 | 3.8 | 3.2 |
| >hg38_21_+_8433172_8446622_RNA45SN1_45S_50ntFlanks@12077.12095.19\|GGGGCTCTCGCTTCTGGCG | >hg38_21_+_8433172_8446622_RNA45SN1_45S_50ntFlanks | 12077 | 12095 | 19 | GGGGCTCTCGCTTCTGGCG | MB468 | 25 | 49 | 31 | 4 | 9 | 6.3 | 17.3 | 15.3 | 3.8 | 3.3 |
| >hg38_21_+_8433172_8446622_RNA45SN1_45S_50ntFlanks@12078.12095.18\|GGGCTCTCGCTTCTGGCG | >hg38_21_+_8433172_8446622_RNA45SN1_45S_50ntFlanks | 12078 | 12095 | 18 | GGGCTCTCGCTTCTGGCG | MB468 | 42 | 58 | 46 | 6 | 12 | 15.4 | 25.0 | 27.5 | 4.5 | 4.3 |
| >hg38_21_+_8433172_8446622_RNA45SN1_45S_50ntFlanks@12081.12106.26\|CTCTCGCTTCTGGCGCCAAGCGCCCG | >hg38_21_+_8433172_8446622_RNA45SN1_45S_50ntFlanks | 12081 | 12106 | 26 | CTCTCGCTTCTGGCGCCAAGCGCCCG | MB468 | 51 | 17 | 50 | 6 | 53 | 22.0 | 7.1 | 29.2 | 3.2 | 17.3 |
| >hg38_21_+_8433172_8446622_RNA45SN1_45S_50ntFlanks@12081.12107.27\|CTCTCGCTTCTGGCGCCAAGCGCCCGG | >hg38_21_+_8433172_8446622_RNA45SN1_45S_50ntFlanks | 12081 | 12107 | 27 | CTCTCGCTTCTGGCGCCAAGCGCCCGG | MB468 | 50 | 17 | 48 | 15 | 50 | 14.8 | 3.8 | 18.1 | 4.4 | 11.8 |
| >hg38_21_+_8433172_8446622_RNA45SN1_45S_50ntFlanks@12081.12108.28\|CTCTCGCTTCTGGCGCCAAGCGCCCGGC | >hg38_21_+_8433172_8446622_RNA45SN1_45S_50ntFlanks | 12081 | 12108 | 28 | CTCTCGCTTCTGGCGCCAAGCGCCCGGC | MB468 | 54 | 15 | 49 | 17 | 58 | 23.1 | 6.5 | 27.9 | 6.5 | 13.9 |
| >hg38_21_+_8433172_8446622_RNA45SN1_45S_50ntFlanks@12081.12109.29\|CTCTCGCTTCTGGCGCCAAGCGCCCGGCC | >hg38_21_+_8433172_8446622_RNA45SN1_45S_50ntFlanks | 12081 | 12109 | 29 | CTCTCGCTTCTGGCGCCAAGCGCCCGGCC | MB468 | 12 | 2 | 9 | 6 | 13 | 3.9 | 0.8 | 4.0 | 2.6 | 2.8 |
| >hg38_21_+_8433172_8446622_RNA45SN1_45S_50ntFlanks@12081.12110.30\|CTCTCGCTTCTGGCGCCAAGCGCCCGGCCG | >hg38_21_+_8433172_8446622_RNA45SN1_45S_50ntFlanks | 12081 | 12110 | 30 | CTCTCGCTTCTGGCGCCAAGCGCCCGGCCG | MB468 | 92 | 27 | 76 | 43 | 107 | 12.8 | 10.9 | 17.6 | 21.3 | 37.8 |
| >hg38_21_+_8433172_8446622_RNA45SN1_45S_50ntFlanks@12083.12106.24\|CTCGCTTCTGGCGCCAAGCGCCCG | >hg38_21_+_8433172_8446622_RNA45SN1_45S_50ntFlanks | 12083 | 12106 | 24 | CTCGCTTCTGGCGCCAAGCGCCCG | MB468 | 44 | 4 | 21 | 14 | 46 | 26.0 | 1.6 | 12.7 | 6.5 | 26.0 |
| >hg38_21_+_8433172_8446622_RNA45SN1_45S_50ntFlanks@12083.12107.25\|CTCGCTTCTGGCGCCAAGCGCCCGG | >hg38_21_+_8433172_8446622_RNA45SN1_45S_50ntFlanks | 12083 | 12107 | 25 | CTCGCTTCTGGCGCCAAGCGCCCGG | MB468 | 35 | 5 | 19 | 24 | 37 | 14.4 | 2.1 | 9.1 | 7.3 | 10.1 |
| >hg38_21_+_8433172_8446622_RNA45SN1_45S_50ntFlanks@12083.12108.26\|CTCGCTTCTGGCGCCAAGCGCCCGGC | >hg38_21_+_8433172_8446622_RNA45SN1_45S_50ntFlanks | 12083 | 12108 | 26 | CTCGCTTCTGGCGCCAAGCGCCCGGC | MB468 | 27 | 3 | 14 | 22 | 32 | 15.0 | 0.8 | 10.0 | 7.3 | 13.2 |
| >hg38_21_+_8433172_8446622_RNA45SN1_45S_50ntFlanks@12083.12110.28\|CTCGCTTCTGGCGCCAAGCGCCCGGCCG | >hg38_21_+_8433172_8446622_RNA45SN1_45S_50ntFlanks | 12083 | 12110 | 28 | CTCGCTTCTGGCGCCAAGCGCCCGGCCG | MB468 | 13 | 4 | 8 | 14 | 17 | 3.8 | 2.6 | 2.0 | 3.4 | 3.3 |
| >hg38_21_+_8433172_8446622_RNA45SN1_45S_50ntFlanks@12126.12146.21\|CGACCCGCTCCGGGGACAGTG | >hg38_21_+_8433172_8446622_RNA45SN1_45S_50ntFlanks | 12126 | 12146 | 21 | CGACCCGCTCCGGGGACAGTG | MB468 | 43 | 11 | 43 | 171 | 70 | 8.3 | 3.1 | 16.7 | 86.5 | 63.3 |
| >hg38_21_+_8433172_8446622_RNA45SN1_45S_50ntFlanks@12126.12148.23\|CGACCCGCTCCGGGGACAGTGCC | >hg38_21_+_8433172_8446622_RNA45SN1_45S_50ntFlanks | 12126 | 12148 | 23 | CGACCCGCTCCGGGGACAGTGCC | MB468 | 20 | 10 | 15 | 29 | 44 | 4.7 | 4.6 | 5.9 | 20.9 | 24.2 |
| >hg38_21_+_8433172_8446622_RNA45SN1_45S_50ntFlanks@12126.12150.25\|CGACCCGCTCCGGGGACAGTGCCAG | >hg38_21_+_8433172_8446622_RNA45SN1_45S_50ntFlanks | 12126 | 12150 | 25 | CGACCCGCTCCGGGGACAGTGCCAG | MB468 | 51 | 36 | 51 | 41 | 131 | 13.5 | 14.3 | 18.0 | 23.6 | 56.7 |
| >hg38_21_+_8433172_8446622_RNA45SN1_45S_50ntFlanks@12126.12151.26\|CGACCCGCTCCGGGGACAGTGCCAGG | >hg38_21_+_8433172_8446622_RNA45SN1_45S_50ntFlanks | 12126 | 12151 | 26 | CGACCCGCTCCGGGGACAGTGCCAGG | MB468 | 14 | 17 | 10 | 13 | 52 | 3.8 | 11.1 | 3.8 | 4.7 | 32.7 |
| >hg38_21_+_8433172_8446622_RNA45SN1_45S_50ntFlanks@12128.12146.19\|ACCCGCTCCGGGGACAGTG | >hg38_21_+_8433172_8446622_RNA45SN1_45S_50ntFlanks | 12128 | 12146 | 19 | ACCCGCTCCGGGGACAGTG | MB468 | 11 | 8 | 20 | 7 | 14 | 1.6 | 3.5 | 6.0 | 2.5 | 7.8 |
| >hg38_21_+_8433172_8446622_RNA45SN1_45S_50ntFlanks@12133.12150.18\|CTCCGGGGACAGTGCCAG | >hg38_21_+_8433172_8446622_RNA45SN1_45S_50ntFlanks | 12133 | 12150 | 18 | CTCCGGGGACAGTGCCAG | MB468 | 135 | 138 | 138 | 61 | 210 | 80.7 | 129.6 | 70.1 | 47.3 | 146.1 |
| >hg38_21_+_8433172_8446622_RNA45SN1_45S_50ntFlanks@12151.12179.29\|GTGGGGAGTTTGACTGGGGCGGTACACCT | >hg38_21_+_8433172_8446622_RNA45SN1_45S_50ntFlanks | 12151 | 12179 | 29 | GTGGGGAGTTTGACTGGGGCGGTACACCT | MB468 | 29 | 18 | 16 | 39 | 16 | 13.1 | 7.6 | 7.9 | 12.3 | 6.4 |
| >hg38_21_+_8433172_8446622_RNA45SN1_45S_50ntFlanks@12151.12180.30\|GTGGGGAGTTTGACTGGGGCGGTACACCTG | >hg38_21_+_8433172_8446622_RNA45SN1_45S_50ntFlanks | 12151 | 12180 | 30 | GTGGGGAGTTTGACTGGGGCGGTACACCTG | MB468 | 51 | 30 | 21 | 56 | 28 | 21.4 | 13.5 | 8.2 | 10.5 | 9.0 |
| >hg38_21_+_8433172_8446622_RNA45SN1_45S_50ntFlanks@12151.12181.31\|GTGGGGAGTTTGACTGGGGCGGTACACCTGT | >hg38_21_+_8433172_8446622_RNA45SN1_45S_50ntFlanks | 12151 | 12181 | 31 | GTGGGGAGTTTGACTGGGGCGGTACACCTGT | MB468 | 23 | 16 | 9 | 26 | 17 | 5.8 | 6.0 | 1.7 | 4.8 | 4.7 |
| >hg38_21_+_8433172_8446622_RNA45SN1_45S_50ntFlanks@12151.12182.32\|GTGGGGAGTTTGACTGGGGCGGTACACCTGTC | >hg38_21_+_8433172_8446622_RNA45SN1_45S_50ntFlanks | 12151 | 12182 | 32 | GTGGGGAGTTTGACTGGGGCGGTACACCTGTC | MB468 | 46 | 26 | 20 | 56 | 32 | 14.2 | 13.1 | 9.7 | 18.0 | 16.7 |
| >hg38_21_+_8433172_8446622_RNA45SN1_45S_50ntFlanks@12151.12183.33\|GTGGGGAGTTTGACTGGGGCGGTACACCTGTCA | >hg38_21_+_8433172_8446622_RNA45SN1_45S_50ntFlanks | 12151 | 12183 | 33 | GTGGGGAGTTTGACTGGGGCGGTACACCTGTCA | MB468 | 19 | 14 | 8 | 20 | 16 | 4.0 | 10.8 | 2.8 | 10.8 | 11.4 |
| >hg38_21_+_8433172_8446622_RNA45SN1_45S_50ntFlanks@12152.12180.29\|TGGGGAGTTTGACTGGGGCGGTACACCTG | >hg38_21_+_8433172_8446622_RNA45SN1_45S_50ntFlanks | 12152 | 12180 | 29 | TGGGGAGTTTGACTGGGGCGGTACACCTG | MB468 | 13 | 8 | 5 | 22 | 7 | 4.7 | 4.3 | 2.8 | 3.8 | 2.0 |
| >hg38_21_+_8433172_8446622_RNA45SN1_45S_50ntFlanks@12152.12182.31\|TGGGGAGTTTGACTGGGGCGGTACACCTGTC | >hg38_21_+_8433172_8446622_RNA45SN1_45S_50ntFlanks | 12152 | 12182 | 31 | TGGGGAGTTTGACTGGGGCGGTACACCTGTC | MB468 | 11 | 6 | 5 | 28 | 7 | 3.8 | 2.7 | 2.2 | 7.3 | 2.4 |
| >hg38_21_+_8433172_8446622_RNA45SN1_45S_50ntFlanks@12154.12180.27\|GGGAGTTTGACTGGGGCGGTACACCTG | >hg38_21_+_8433172_8446622_RNA45SN1_45S_50ntFlanks | 12154 | 12180 | 27 | GGGAGTTTGACTGGGGCGGTACACCTG | MB468 | 13 | 11 | 7 | 47 | 12 | 5.0 | 5.3 | 3.7 | 13.4 | 3.3 |
| >hg38_21_+_8433172_8446622_RNA45SN1_45S_50ntFlanks@12154.12182.29\|GGGAGTTTGACTGGGGCGGTACACCTGTC | >hg38_21_+_8433172_8446622_RNA45SN1_45S_50ntFlanks | 12154 | 12182 | 29 | GGGAGTTTGACTGGGGCGGTACACCTGTC | MB468 | 13 | 7 | 5 | 53 | 12 | 5.1 | 2.2 | 2.7 | 30.0 | 4.1 |
| >hg38_21_+_8433172_8446622_RNA45SN1_45S_50ntFlanks@12156.12180.25\|GAGTTTGACTGGGGCGGTACACCTG | >hg38_21_+_8433172_8446622_RNA45SN1_45S_50ntFlanks | 12156 | 12180 | 25 | GAGTTTGACTGGGGCGGTACACCTG | MB468 | 13 | 6 | 4 | 23 | 11 | 6.4 | 4.1 | 1.4 | 8.0 | 2.1 |
| >hg38_21_+_8433172_8446622_RNA45SN1_45S_50ntFlanks@12157.12178.22\|AGTTTGACTGGGGCGGTACACC | >hg38_21_+_8433172_8446622_RNA45SN1_45S_50ntFlanks | 12157 | 12178 | 22 | AGTTTGACTGGGGCGGTACACC | MB468 | 11 | 7 | 6 | 25 | 9 | 8.4 | 2.4 | 6.0 | 13.1 | 3.4 |
| >hg38_21_+_8433172_8446622_RNA45SN1_45S_50ntFlanks@12157.12179.23\|AGTTTGACTGGGGCGGTACACCT | >hg38_21_+_8433172_8446622_RNA45SN1_45S_50ntFlanks | 12157 | 12179 | 23 | AGTTTGACTGGGGCGGTACACCT | MB468 | 76 | 19 | 25 | 139 | 57 | 38.4 | 8.6 | 7.4 | 49.0 | 19.9 |
| >hg38_21_+_8433172_8446622_RNA45SN1_45S_50ntFlanks@12157.12180.24\|AGTTTGACTGGGGCGGTACACCTG | >hg38_21_+_8433172_8446622_RNA45SN1_45S_50ntFlanks | 12157 | 12180 | 24 | AGTTTGACTGGGGCGGTACACCTG | MB468 | 129 | 41 | 38 | 234 | 115 | 54.9 | 20.1 | 11.3 | 60.9 | 31.2 |
| >hg38_21_+_8433172_8446622_RNA45SN1_45S_50ntFlanks@12157.12181.25\|AGTTTGACTGGGGCGGTACACCTGT | >hg38_21_+_8433172_8446622_RNA45SN1_45S_50ntFlanks | 12157 | 12181 | 25 | AGTTTGACTGGGGCGGTACACCTGT | MB468 | 73 | 25 | 20 | 153 | 83 | 26.2 | 10.9 | 7.3 | 21.4 | 18.6 |
| >hg38_21_+_8433172_8446622_RNA45SN1_45S_50ntFlanks@12157.12182.26\|AGTTTGACTGGGGCGGTACACCTGTC | >hg38_21_+_8433172_8446622_RNA45SN1_45S_50ntFlanks | 12157 | 12182 | 26 | AGTTTGACTGGGGCGGTACACCTGTC | MB468 | 71 | 7 | 11 | 244 | 81 | 19.6 | 1.6 | 6.2 | 198.5 | 33.2 |
| >hg38_21_+_8433172_8446622_RNA45SN1_45S_50ntFlanks@12157.12183.27\|AGTTTGACTGGGGCGGTACACCTGTCA | >hg38_21_+_8433172_8446622_RNA45SN1_45S_50ntFlanks | 12157 | 12183 | 27 | AGTTTGACTGGGGCGGTACACCTGTCA | MB468 | 13 | 1 | 2 | 41 | 15 | 3.0 | 0.3 | 1.5 | 23.3 | 5.1 |
| >hg38_21_+_8433172_8446622_RNA45SN1_45S_50ntFlanks@12158.12180.23\|GTTTGACTGGGGCGGTACACCTG | >hg38_21_+_8433172_8446622_RNA45SN1_45S_50ntFlanks | 12158 | 12180 | 23 | GTTTGACTGGGGCGGTACACCTG | MB468 | 13 | 4 | 4 | 30 | 13 | 7.7 | 2.1 | 0.9 | 8.6 | 5.0 |
| >hg38_21_+_8433172_8446622_RNA45SN1_45S_50ntFlanks@12159.12178.20\|TTTGACTGGGGCGGTACACC | >hg38_21_+_8433172_8446622_RNA45SN1_45S_50ntFlanks | 12159 | 12178 | 20 | TTTGACTGGGGCGGTACACC | MB468 | 52 | 36 | 33 | 78 | 29 | 26.9 | 23.5 | 41.2 | 23.5 | 5.7 |
| >hg38_21_+_8433172_8446622_RNA45SN1_45S_50ntFlanks@12159.12179.21\|TTTGACTGGGGCGGTACACCT | >hg38_21_+_8433172_8446622_RNA45SN1_45S_50ntFlanks | 12159 | 12179 | 21 | TTTGACTGGGGCGGTACACCT | MB468 | 358 | 58 | 84 | 433 | 191 | 169.2 | 23.0 | 22.9 | 155.9 | 81.1 |
| >hg38_21_+_8433172_8446622_RNA45SN1_45S_50ntFlanks@12159.12180.22\|TTTGACTGGGGCGGTACACCTG | >hg38_21_+_8433172_8446622_RNA45SN1_45S_50ntFlanks | 12159 | 12180 | 22 | TTTGACTGGGGCGGTACACCTG | MB468 | 300 | 49 | 57 | 440 | 165 | 137.5 | 23.0 | 19.5 | 80.2 | 62.5 |
| >hg38_21_+_8433172_8446622_RNA45SN1_45S_50ntFlanks@12159.12181.23\|TTTGACTGGGGCGGTACACCTGT | >hg38_21_+_8433172_8446622_RNA45SN1_45S_50ntFlanks | 12159 | 12181 | 23 | TTTGACTGGGGCGGTACACCTGT | MB468 | 172 | 29 | 29 | 293 | 127 | 65.5 | 12.2 | 12.4 | 85.1 | 46.6 |

| | | | | | | | | | | | | | | | | |
|---|---|---|---|---|---|---|---|---|---|---|---|---|---|---|---|---|
| >hg38_21_+_8433172_8446622_RNA45SN1_45S_50ntFlanks@12159.12182.24\|TTTGACTGGGGCGGTACACCTGTC | >hg38_21_+_8433172_8446622_RNA45SN1_45S_50ntFlanks | 12159 | 12182 | 24 | TTTGACTGGGGCGGTACACCTGTC | MB468 | 161 | 10 | 20 | 358 | 128 | 56.2 | 1.4 | 11.6 | 249.0 | 62.9 |
| >hg38_21_+_8433172_8446622_RNA45SN1_45S_50ntFlanks@12159.12183.25\|TTTGACTGGGGCGGTACACCTGTCA | >hg38_21_+_8433172_8446622_RNA45SN1_45S_50ntFlanks | 12159 | 12183 | 25 | TTTGACTGGGGCGGTACACCTGTCA | MB468 | 27 | 1 | 3 | 46 | 19 | 12.5 | 0.3 | 1.8 | 27.8 | 11.8 |
| >hg38_21_+_8433172_8446622_RNA45SN1_45S_50ntFlanks@12160.12179.20\|TTGACTGGGGCGGTACACCT | >hg38_21_+_8433172_8446622_RNA45SN1_45S_50ntFlanks | 12160 | 12179 | 20 | TTGACTGGGGCGGTACACCT | MB468 | 18 | 4 | 4 | 30 | 14 | 9.5 | 1.6 | 1.2 | 10.9 | 9.4 |
| >hg38_21_+_8433172_8446622_RNA45SN1_45S_50ntFlanks@12160.12180.21\|TTGACTGGGGCGGTACACCTG | >hg38_21_+_8433172_8446622_RNA45SN1_45S_50ntFlanks | 12160 | 12180 | 21 | TTGACTGGGGCGGTACACCTG | MB468 | 15 | 3 | 3 | 29 | 12 | 6.7 | 1.4 | 1.2 | 3.8 | 7.2 |
| >hg38_21_+_8433172_8446622_RNA45SN1_45S_50ntFlanks@12161.12179.19\|TGACTGGGGCGGTACACCT | >hg38_21_+_8433172_8446622_RNA45SN1_45S_50ntFlanks | 12161 | 12179 | 19 | TGACTGGGGCGGTACACCT | MB468 | 50 | 7 | 9 | 84 | 43 | 31.8 | 3.1 | 3.1 | 40.4 | 31.8 |
| >hg38_21_+_8433172_8446622_RNA45SN1_45S_50ntFlanks@12161.12180.20\|TGACTGGGGCGGTACACCTG | >hg38_21_+_8433172_8446622_RNA45SN1_45S_50ntFlanks | 12161 | 12180 | 20 | TGACTGGGGCGGTACACCTG | MB468 | 45 | 7 | 8 | 90 | 36 | 23.5 | 3.4 | 3.6 | 13.4 | 22.3 |
| >hg38_21_+_8433172_8446622_RNA45SN1_45S_50ntFlanks@12161.12181.21\|TGACTGGGGCGGTACACCTGT | >hg38_21_+_8433172_8446622_RNA45SN1_45S_50ntFlanks | 12161 | 12181 | 21 | TGACTGGGGCGGTACACCTGT | MB468 | 22 | 3 | 4 | 45 | 21 | 9.7 | 1.7 | 1.6 | 7.9 | 11.2 |
| >hg38_21_+_8433172_8446622_RNA45SN1_45S_50ntFlanks@12162.12179.18\|GACTGGGGCGGTACACCT | >hg38_21_+_8433172_8446622_RNA45SN1_45S_50ntFlanks | 12162 | 12179 | 18 | GACTGGGGCGGTACACCT | MB468 | 25 | 3 | 4 | 46 | 26 | 20.5 | 1.9 | 1.1 | 32.1 | 24.8 |
| >hg38_21_+_8433172_8446622_RNA45SN1_45S_50ntFlanks@12162.12180.19\|GACTGGGGCGGTACACCTG | >hg38_21_+_8433172_8446622_RNA45SN1_45S_50ntFlanks | 12162 | 12180 | 19 | GACTGGGGCGGTACACCTG | MB468 | 46 | 8 | 8 | 112 | 45 | 24.2 | 3.9 | 3.3 | 31.4 | 31.3 |
| >hg38_21_+_8433172_8446622_RNA45SN1_45S_50ntFlanks@12162.12181.20\|GACTGGGGCGGTACACCTGT | >hg38_21_+_8433172_8446622_RNA45SN1_45S_50ntFlanks | 12162 | 12181 | 20 | GACTGGGGCGGTACACCTGT | MB468 | 20 | 3 | 3 | 44 | 21 | 10.1 | 1.3 | 0.8 | 14.0 | 14.1 |
| >hg38_21_+_8433172_8446622_RNA45SN1_45S_50ntFlanks@12163.12180.18\|ACTGGGGCGGTACACCTG | >hg38_21_+_8433172_8446622_RNA45SN1_45S_50ntFlanks | 12163 | 12180 | 18 | ACTGGGGCGGTACACCTG | MB468 | 293 | 36 | 46 | 521 | 212 | 162.5 | 19.4 | 28.3 | 139.2 | 176.5 |
| >hg38_21_+_8433172_8446622_RNA45SN1_45S_50ntFlanks@12163.12181.19\|ACTGGGGCGGTACACCTGT | >hg38_21_+_8433172_8446622_RNA45SN1_45S_50ntFlanks | 12163 | 12181 | 19 | ACTGGGGCGGTACACCTGT | MB468 | 204 | 19 | 23 | 318 | 194 | 124.9 | 6.6 | 11.8 | 100.7 | 189.1 |
| >hg38_21_+_8433172_8446622_RNA45SN1_45S_50ntFlanks@12163.12182.20\|ACTGGGGCGGTACACCTGTC | >hg38_21_+_8433172_8446622_RNA45SN1_45S_50ntFlanks | 12163 | 12182 | 20 | ACTGGGGCGGTACACCTGTC | MB468 | 402 | 49 | 46 | 616 | 430 | 223.5 | 21.5 | 26.5 | 148.8 | 275.5 |
| >hg38_21_+_8433172_8446622_RNA45SN1_45S_50ntFlanks@12163.12183.21\|ACTGGGGCGGTACACCTGTCA | >hg38_21_+_8433172_8446622_RNA45SN1_45S_50ntFlanks | 12163 | 12183 | 21 | ACTGGGGCGGTACACCTGTCA | MB468 | 56 | 6 | 8 | 59 | 54 | 27.2 | 3.1 | 3.9 | 23.9 | 23.8 |
| >hg38_21_+_8433172_8446622_RNA45SN1_45S_50ntFlanks@12164.12182.19\|CTGGGGCGGTACACCTGTC | >hg38_21_+_8433172_8446622_RNA45SN1_45S_50ntFlanks | 12164 | 12182 | 19 | CTGGGGCGGTACACCTGTC | MB468 | 17 | 2 | 2 | 29 | 19 | 10.0 | 1.0 | 1.2 | 12.5 | 16.0 |
| >hg38_21_+_8433172_8446622_RNA45SN1_45S_50ntFlanks@12165.12182.18\|TGGGGCGGTACACCTGTC | >hg38_21_+_8433172_8446622_RNA45SN1_45S_50ntFlanks | 12165 | 12182 | 18 | TGGGGCGGTACACCTGTC | MB468 | 13 | 1 | 2 | 24 | 13 | 8.7 | 0.6 | 0.8 | 10.6 | 11.9 |
| >hg38_21_+_8433172_8446622_RNA45SN1_45S_50ntFlanks@12202.12220.19\|CTAAGGCGAGCTCAGGGAG | >hg38_21_+_8433172_8446622_RNA45SN1_45S_50ntFlanks | 12202 | 12220 | 19 | CTAAGGCGAGCTCAGGGAG | MB468 | 14 | 36 | 13 | 47 | 59 | 8.6 | 12.4 | 13.4 | 28.9 | 58.6 |
| >hg38_21_+_8433172_8446622_RNA45SN1_45S_50ntFlanks@12202.12221.20\|CTAAGGCGAGCTCAGGGAGG | >hg38_21_+_8433172_8446622_RNA45SN1_45S_50ntFlanks | 12202 | 12221 | 20 | CTAAGGCGAGCTCAGGGAGG | MB468 | 11 | 27 | 18 | 39 | 45 | 8.1 | 7.9 | 17.9 | 18.1 | 40.2 |
| >hg38_21_+_8433172_8446622_RNA45SN1_45S_50ntFlanks@12241.12271.31\|CAGAAGGGCAAAAGCTCGCTTGATCTTGATT | >hg38_21_+_8433172_8446622_RNA45SN1_45S_50ntFlanks | 12241 | 12271 | 31 | CAGAAGGGCAAAAGCTCGCTTGATCTTGATT | MB231 | 12 | 6 | 11 | 1 | 4 | 16.0 | 6.7 | 13.0 | 1.9 | 4.2 |
| >hg38_21_+_8433172_8446622_RNA45SN1_45S_50ntFlanks@12241.12272.32\|CAGAAGGGCAAAAGCTCGCTTGATCTTGATTT | >hg38_21_+_8433172_8446622_RNA45SN1_45S_50ntFlanks | 12241 | 12272 | 32 | CAGAAGGGCAAAAGCTCGCTTGATCTTGATTT | MB468 | 10 | 21 | 12 | 25 | 100 | 4.3 | 6.4 | 9.8 | 14.9 | 40.7 |
| >hg38_21_+_8433172_8446622_RNA45SN1_45S_50ntFlanks@12249.12272.24\|CAAAAGCTCGCTTGATCTTGATTT | >hg38_21_+_8433172_8446622_RNA45SN1_45S_50ntFlanks | 12249 | 12272 | 24 | CAAAAGCTCGCTTGATCTTGATTT | MB468 | 12 | 21 | 17 | 21 | 122 | 4.8 | 12.4 | 3.7 | 12.4 | 138.5 |
| >hg38_21_+_8433172_8446622_RNA45SN1_45S_50ntFlanks@12258.12284.27\|GCTTGATCTTGATTTTCAGTACGAATA | >hg38_21_+_8433172_8446622_RNA45SN1_45S_50ntFlanks | 12258 | 12284 | 27 | GCTTGATCTTGATTTTCAGTACGAATA | MB468 | 19 | 50 | 11 | 31 | 46 | 11.3 | 20.1 | 8.1 | 31.4 | 40.6 |
| >hg38_21_+_8433172_8446622_RNA45SN1_45S_50ntFlanks@12259.12284.26\|CTTGATCTTGATTTTCAGTACGAATA | >hg38_21_+_8433172_8446622_RNA45SN1_45S_50ntFlanks | 12259 | 12284 | 26 | CTTGATCTTGATTTTCAGTACGAATA | MB468 | 35 | 59 | 23 | 123 | 135 | 21.1 | 47.7 | 19.7 | 93.6 | 133.2 |
| >hg38_21_+_8433172_8446622_RNA45SN1_45S_50ntFlanks@12274.12293.20\|CAGTACGAATACAGACCGTG | >hg38_21_+_8433172_8446622_RNA45SN1_45S_50ntFlanks | 12274 | 12293 | 20 | CAGTACGAATACAGACCGTG | MB468 | 13 | 4 | 7 | 57 | 18 | 7.6 | 3.2 | 6.2 | 47.6 | 11.9 |
| >hg38_21_+_8433172_8446622_RNA45SN1_45S_50ntFlanks@12324.12342.19\|TTGGGTTTTAAGCAGGAGG | >hg38_21_+_8433172_8446622_RNA45SN1_45S_50ntFlanks | 12324 | 12342 | 19 | TTGGGTTTTAAGCAGGAGG | MB468 | 14 | 4 | 39 | 17 | 4 | 6.1 | 0.6 | 20.4 | 8.7 | 2.2 |
| >hg38_21_+_8433172_8446622_RNA45SN1_45S_50ntFlanks@12324.12343.20\|TTGGGTTTTAAGCAGGAGGT | >hg38_21_+_8433172_8446622_RNA45SN1_45S_50ntFlanks | 12324 | 12343 | 20 | TTGGGTTTTAAGCAGGAGGT | MB468 | 13 | 6 | 23 | 11 | 3 | 3.1 | 0.9 | 4.6 | 2.3 | 1.2 |
| >hg38_21_+_8433172_8446622_RNA45SN1_45S_50ntFlanks@12324.12344.21\|TTGGGTTTTAAGCAGGAGGTG | >hg38_21_+_8433172_8446622_RNA45SN1_45S_50ntFlanks | 12324 | 12344 | 21 | TTGGGTTTTAAGCAGGAGGTG | MB468 | 134 | 106 | 276 | 130 | 51 | 38.1 | 32.0 | 81.5 | 47.3 | 24.5 |
| >hg38_21_+_8433172_8446622_RNA45SN1_45S_50ntFlanks@12324.12345.22\|TTGGGTTTTAAGCAGGAGGTGT | >hg38_21_+_8433172_8446622_RNA45SN1_45S_50ntFlanks | 12324 | 12345 | 22 | TTGGGTTTTAAGCAGGAGGTGT | MB468 | 36 | 23 | 67 | 17 | 8 | 22.9 | 12.8 | 42.6 | 3.9 | 2.3 |
| >hg38_21_+_8433172_8446622_RNA45SN1_45S_50ntFlanks@12324.12346.23\|TTGGGTTTTAAGCAGGAGGTGTC | >hg38_21_+_8433172_8446622_RNA45SN1_45S_50ntFlanks | 12324 | 12346 | 23 | TTGGGTTTTAAGCAGGAGGTGTC | MB468 | 232 | 154 | 207 | 236 | 88 | 39.3 | 46.1 | 35.4 | 50.9 | 41.3 |
| >hg38_21_+_8433172_8446622_RNA45SN1_45S_50ntFlanks@12324.12347.24\|TTGGGTTTTAAGCAGGAGGTGTCA | >hg38_21_+_8433172_8446622_RNA45SN1_45S_50ntFlanks | 12324 | 12347 | 24 | TTGGGTTTTAAGCAGGAGGTGTCA | MB468 | 12 | 8 | 13 | 11 | 4 | 3.5 | 2.2 | 3.1 | 2.5 | 1.5 |
| >hg38_21_+_8433172_8446622_RNA45SN1_45S_50ntFlanks@12324.12348.25\|TTGGGTTTTAAGCAGGAGGTGTCAG | >hg38_21_+_8433172_8446622_RNA45SN1_45S_50ntFlanks | 12324 | 12348 | 25 | TTGGGTTTTAAGCAGGAGGTGTCAG | MB468 | 61 | 43 | 73 | 58 | 23 | 11.4 | 7.9 | 9.9 | 5.1 | 7.7 |
| >hg38_21_+_8433172_8446622_RNA45SN1_45S_50ntFlanks@12324.12349.26\|TTGGGTTTTAAGCAGGAGGTGTCAGA | >hg38_21_+_8433172_8446622_RNA45SN1_45S_50ntFlanks | 12324 | 12349 | 26 | TTGGGTTTTAAGCAGGAGGTGTCAGA | MB468 | 18 | 10 | 17 | 25 | 9 | 5.6 | 2.8 | 2.8 | 3.9 | 3.5 |
| >hg38_21_+_8433172_8446622_RNA45SN1_45S_50ntFlanks@12324.12355.32\|TTGGGTTTTAAGCAGGAGGTGTCAGAAAAGTT | >hg38_21_+_8433172_8446622_RNA45SN1_45S_50ntFlanks | 12324 | 12355 | 32 | TTGGGTTTTAAGCAGGAGGTGTCAGAAAAGTT | MB468 | 15 | 15 | 16 | 18 | 11 | 4.1 | 9.2 | 4.8 | 10.3 | 7.8 |
| >hg38_21_+_8433172_8446622_RNA45SN1_45S_50ntFlanks@12324.12356.33\|TTGGGTTTTAAGCAGGAGGTGTCAGAAAAGTTA | >hg38_21_+_8433172_8446622_RNA45SN1_45S_50ntFlanks | 12324 | 12356 | 33 | TTGGGTTTTAAGCAGGAGGTGTCAGAAAAGTTA | MB468 | 13 | 8 | 10 | 23 | 12 | 1.9 | 5.2 | 3.3 | 16.0 | 9.1 |
| >hg38_21_+_8433172_8446622_RNA45SN1_45S_50ntFlanks@12328.12346.19\|GTTTTAAGCAGGAGGTGTC | >hg38_21_+_8433172_8446622_RNA45SN1_45S_50ntFlanks | 12328 | 12346 | 19 | GTTTTAAGCAGGAGGTGTC | MB468 | 18 | 19 | 14 | 9 | 4 | 7.0 | 5.5 | 4.1 | 6.5 | 2.2 |
| >hg38_21_+_8433172_8446622_RNA45SN1_45S_50ntFlanks@12329.12346.18\|TTTTAAGCAGGAGGTGTC | >hg38_21_+_8433172_8446622_RNA45SN1_45S_50ntFlanks | 12329 | 12346 | 18 | TTTTAAGCAGGAGGTGTC | MB468 | 31 | 33 | 19 | 10 | 4 | 11.0 | 12.2 | 8.3 | 5.1 | 1.4 |
| >hg38_21_+_8433172_8446622_RNA45SN1_45S_50ntFlanks@12332.12349.18\|TAAGCAGGAGGTGTCAGA | >hg38_21_+_8433172_8446622_RNA45SN1_45S_50ntFlanks | 12332 | 12349 | 18 | TAAGCAGGAGGTGTCAGA | MB468 | 11 | 6 | 6 | 5 | 3 | 5.4 | 2.6 | 1.7 | 1.4 | 0.6 |
| >hg38_21_+_8433172_8446622_RNA45SN1_45S_50ntFlanks@12347.12365.19\|AGAAAAGTTACCACAGGGA | >hg38_21_+_8433172_8446622_RNA45SN1_45S_50ntFlanks | 12347 | 12365 | 19 | AGAAAAGTTACCACAGGGA | MB468 | 43 | 20 | 62 | 26 | 7 | 6.6 | 7.6 | 14.7 | 18.9 | 4.1 |
| >hg38_21_+_8433172_8446622_RNA45SN1_45S_50ntFlanks@12347.12366.20\|AGAAAAGTTACCACAGGGAT | >hg38_21_+_8433172_8446622_RNA45SN1_45S_50ntFlanks | 12347 | 12366 | 20 | AGAAAAGTTACCACAGGGAT | MB468 | 12 | 5 | 14 | 6 | 3 | 2.2 | 0.9 | 4.5 | 4.2 | 2.4 |
| >hg38_21_+_8433172_8446622_RNA45SN1_45S_50ntFlanks@12353.12376.24\|GTTACCACAGGGATAACTGGCTTG | >hg38_21_+_8433172_8446622_RNA45SN1_45S_50ntFlanks | 12353 | 12376 | 24 | GTTACCACAGGGATAACTGGCTTG | MB468 | 11 | 6 | 5 | 5 | 5 | 6.2 | 4.0 | 4.3 | 1.8 | 2.9 |
| >hg38_21_+_8433172_8446622_RNA45SN1_45S_50ntFlanks@12354.12376.23\|TTACCACAGGGATAACTGGCTTG | >hg38_21_+_8433172_8446622_RNA45SN1_45S_50ntFlanks | 12354 | 12376 | 23 | TTACCACAGGGATAACTGGCTTG | MB468 | 15 | 2 | 4 | 5 | 4 | 7.8 | 0.4 | 1.4 | 1.0 | 1.0 |
| >hg38_21_+_8433172_8446622_RNA45SN1_45S_50ntFlanks@12356.12375.20\|ACCACAGGGATAACTGGCTT | >hg38_21_+_8433172_8446622_RNA45SN1_45S_50ntFlanks | 12356 | 12375 | 20 | ACCACAGGGATAACTGGCTT | MB468 | 15 | 3 | 4 | 6 | 4 | 5.0 | 1.4 | 0.7 | 1.7 | 2.2 |
| >hg38_21_+_8433172_8446622_RNA45SN1_45S_50ntFlanks@12356.12376.21\|ACCACAGGGATAACTGGCTTG | >hg38_21_+_8433172_8446622_RNA45SN1_45S_50ntFlanks | 12356 | 12376 | 21 | ACCACAGGGATAACTGGCTTG | MB468 | 38 | 8 | 11 | 16 | 11 | 15.9 | 2.8 | 3.2 | 4.7 | 5.1 |
| >hg38_21_+_8433172_8446622_RNA45SN1_45S_50ntFlanks@12357.12375.19\|CCACAGGGATAACTGGCTT | >hg38_21_+_8433172_8446622_RNA45SN1_45S_50ntFlanks | 12357 | 12375 | 19 | CCACAGGGATAACTGGCTT | MB468 | 61 | 19 | 26 | 24 | 22 | 14.0 | 6.1 | 6.3 | 8.4 | 10.7 |
| >hg38_21_+_8433172_8446622_RNA45SN1_45S_50ntFlanks@12357.12376.20\|CCACAGGGATAACTGGCTTG | >hg38_21_+_8433172_8446622_RNA45SN1_45S_50ntFlanks | 12357 | 12376 | 20 | CCACAGGGATAACTGGCTTG | MB468 | 121 | 24 | 47 | 66 | 42 | 53.4 | 6.0 | 16.1 | 18.5 | 29.4 |
| >hg38_21_+_8433172_8446622_RNA45SN1_45S_50ntFlanks@12358.12375.18\|CACAGGGATAACTGGCTT | >hg38_21_+_8433172_8446622_RNA45SN1_45S_50ntFlanks | 12358 | 12375 | 18 | CACAGGGATAACTGGCTT | MB468 | 11 | 3 | 6 | 6 | 4 | 1.1 | 1.4 | 2.4 | 1.2 | 2.2 |
| >hg38_21_+_8433172_8446622_RNA45SN1_45S_50ntFlanks@12358.12376.19\|CACAGGGATAACTGGCTTG | >hg38_21_+_8433172_8446622_RNA45SN1_45S_50ntFlanks | 12358 | 12376 | 19 | CACAGGGATAACTGGCTTG | MB468 | 17 | 4 | 10 | 12 | 6 | 7.1 | 0.8 | 5.5 | 6.1 | 3.7 |
| >hg38_21_+_8433172_8446622_RNA45SN1_45S_50ntFlanks@12360.12382.23\|CAGGGATAACTGGCTTGTGGCGG | >hg38_21_+_8433172_8446622_RNA45SN1_45S_50ntFlanks | 12360 | 12382 | 23 | CAGGGATAACTGGCTTGTGGCGG | MB468 | 24 | 36 | 19 | 17 | 17 | 10.2 | 12.1 | 9.0 | 4.5 | 3.0 |
| >hg38_21_+_8433172_8446622_RNA45SN1_45S_50ntFlanks@12360.12384.25\|CAGGGATAACTGGCTTGTGGCGGCC | >hg38_21_+_8433172_8446622_RNA45SN1_45S_50ntFlanks | 12360 | 12384 | 25 | CAGGGATAACTGGCTTGTGGCGGCC | MB468 | 17 | 13 | 25 | 13 | 22 | 6.9 | 2.4 | 7.8 | 6.4 | 10.5 |
| >hg38_21_+_8433172_8446622_RNA45SN1_45S_50ntFlanks@12363.12382.20\|GGATAACTGGCTTGTGGCGG | >hg38_21_+_8433172_8446622_RNA45SN1_45S_50ntFlanks | 12363 | 12382 | 20 | GGATAACTGGCTTGTGGCGG | MB468 | 43 | 37 | 36 | 7 | 10 | 24.7 | 21.7 | 23.0 | 5.8 | 5.2 |

| | | | | | | | | | | | | | | | | |
|---|---|---|---|---|---|---|---|---|---|---|---|---|---|---|---|---|
| >hg38_21_+_8433172_8446622_RNA45SN1_45S_50ntFlanks@12363.12383.21\|GGATAACTGGCTTGTGGCGGC | >hg38_21_+_8433172_8446622_RNA45SN1_45S_50ntFlanks | 12363 | 12383 | 21 | GGATAACTGGCTTGTGGCGGC | MB468 | 12 | 7 | 11 | 2 | 3 | 6.9 | 4.4 | 7.9 | 1.3 | 2.1 |
| >hg38_21_+_8433172_8446622_RNA45SN1_45S_50ntFlanks@12365.12382.18\|ATAACTGGCTTGTGGCGG | >hg38_21_+_8433172_8446622_RNA45SN1_45S_50ntFlanks | 12365 | 12382 | 18 | ATAACTGGCTTGTGGCGG | MB468 | 98 | 32 | 99 | 40 | 38 | 22.4 | 4.1 | 28.6 | 15.3 | 14.2 |
| >hg38_21_+_8433172_8446622_RNA45SN1_45S_50ntFlanks@12365.12383.19\|ATAACTGGCTTGTGGCGGC | >hg38_21_+_8433172_8446622_RNA45SN1_45S_50ntFlanks | 12365 | 12383 | 19 | ATAACTGGCTTGTGGCGGC | MB468 | 26 | 6 | 28 | 10 | 12 | 5.2 | 1.0 | 6.4 | 3.7 | 3.8 |
| >hg38_21_+_8433172_8446622_RNA45SN1_45S_50ntFlanks@12366.12383.18\|TAACTGGCTTGTGGCGGC | >hg38_21_+_8433172_8446622_RNA45SN1_45S_50ntFlanks | 12366 | 12383 | 18 | TAACTGGCTTGTGGCGGC | MB468 | 172 | 32 | 188 | 50 | 52 | 65.0 | 4.8 | 104.6 | 8.3 | 19.7 |
| >hg38_21_+_8433172_8446622_RNA45SN1_45S_50ntFlanks@12366.12384.19\|TAACTGGCTTGTGGCGGCC | >hg38_21_+_8433172_8446622_RNA45SN1_45S_50ntFlanks | 12366 | 12384 | 19 | TAACTGGCTTGTGGCGGCC | MB468 | 52 | 8 | 68 | 45 | 41 | 9.6 | 2.7 | 37.0 | 21.9 | 27.1 |
| >hg38_21_+_8433172_8446622_RNA45SN1_45S_50ntFlanks@12366.12404.39\|TAACTGGCTTGTGGCGGCCAAGCGTTCATAGCGACGTCG | >hg38_21_+_8433172_8446622_RNA45SN1_45S_50ntFlanks | 12366 | 12404 | 39 | TAACTGGCTTGTGGCGGCCAAGCGTTCATAGCGACGTCG | MB468 | 16 | 4 | 24 | 3 | 16 | 16.2 | 2.4 | 26.9 | 2.1 | 8.4 |
| >hg38_21_+_8433172_8446622_RNA45SN1_45S_50ntFlanks@12367.12384.18\|AACTGGCTTGTGGCGGCC | >hg38_21_+_8433172_8446622_RNA45SN1_45S_50ntFlanks | 12367 | 12384 | 18 | AACTGGCTTGTGGCGGCC | MB468 | 19 | 4 | 20 | 42 | 13 | 1.4 | 1.5 | 6.0 | 37.4 | 6.8 |
| >hg38_21_+_8433172_8446622_RNA45SN1_45S_50ntFlanks@12386.12414.29\|AGCGTTCATAGCGACGTCGCTTTTTGATC | >hg38_21_+_8433172_8446622_RNA45SN1_45S_50ntFlanks | 12386 | 12414 | 29 | AGCGTTCATAGCGACGTCGCTTTTTGATC | MB468 | 19 | 7 | 33 | 31 | 99 | 4.1 | 1.0 | 16.5 | 7.5 | 91.9 |
| >hg38_21_+_8433172_8446622_RNA45SN1_45S_50ntFlanks@12388.12433.46\|CGTTCATAGCGACGTCGCTTTTTGATCCTTCGATGTCGGCTCTTCC | >hg38_21_+_8433172_8446622_RNA45SN1_45S_50ntFlanks | 12388 | 12433 | 46 | CGTTCATAGCGACGTCGCTTTTTGATCCTTCGATGTCGGCTCTTCC | MB231 | 17 | 16 | 115 | 0 | 0 | 12.2 | 24.6 | 78.7 | 0.0 | 0.0 |
| >hg38_21_+_8433172_8446622_RNA45SN1_45S_50ntFlanks@12393.12414.22\|ATAGCGACGTCGCTTTTTGATC | >hg38_21_+_8433172_8446622_RNA45SN1_45S_50ntFlanks | 12393 | 12414 | 22 | ATAGCGACGTCGCTTTTTGATC | MB231 | 12 | 15 | 15 | 2 | 7 | 9.5 | 8.7 | 11.2 | 0.9 | 7.7 |
| >hg38_21_+_8433172_8446622_RNA45SN1_45S_50ntFlanks@12393.12414.22\|ATAGCGACGTCGCTTTTTGATC | >hg38_21_+_8433172_8446622_RNA45SN1_45S_50ntFlanks | 12393 | 12414 | 22 | ATAGCGACGTCGCTTTTTGATC | MB468 | 15 | 15 | 19 | 15 | 37 | 6.4 | 3.5 | 7.3 | 8.8 | 39.0 |
| >hg38_21_+_8433172_8446622_RNA45SN1_45S_50ntFlanks@12394.12414.21\|TAGCGACGTCGCTTTTTGATC | >hg38_21_+_8433172_8446622_RNA45SN1_45S_50ntFlanks | 12394 | 12414 | 21 | TAGCGACGTCGCTTTTTGATC | MB231 | 30 | 35 | 46 | 2 | 8 | 29.9 | 34.5 | 43.7 | 2.7 | 10.5 |
| >hg38_21_+_8433172_8446622_RNA45SN1_45S_50ntFlanks@12394.12414.21\|TAGCGACGTCGCTTTTTGATC | >hg38_21_+_8433172_8446622_RNA45SN1_45S_50ntFlanks | 12394 | 12414 | 21 | TAGCGACGTCGCTTTTTGATC | MB468 | 86 | 80 | 106 | 37 | 140 | 38.6 | 28.1 | 40.7 | 28.6 | 121.7 |
| >hg38_21_+_8433172_8446622_RNA45SN1_45S_50ntFlanks@12394.12417.24\|TAGCGACGTCGCTTTTTGATCCTT | >hg38_21_+_8433172_8446622_RNA45SN1_45S_50ntFlanks | 12394 | 12417 | 24 | TAGCGACGTCGCTTTTTGATCCTT | MB468 | 14 | 17 | 9 | 13 | 72 | 5.6 | 6.2 | 3.2 | 11.7 | 63.3 |
| >hg38_21_+_8433172_8446622_RNA45SN1_45S_50ntFlanks@12394.12418.25\|TAGCGACGTCGCTTTTTGATCCTTC | >hg38_21_+_8433172_8446622_RNA45SN1_45S_50ntFlanks | 12394 | 12418 | 25 | TAGCGACGTCGCTTTTTGATCCTTC | MB231 | 14 | 20 | 18 | 0 | 4 | 14.6 | 16.8 | 14.7 | 0.7 | 6.2 |
| >hg38_21_+_8433172_8446622_RNA45SN1_45S_50ntFlanks@12394.12419.26\|TAGCGACGTCGCTTTTTGATCCTTCG | >hg38_21_+_8433172_8446622_RNA45SN1_45S_50ntFlanks | 12394 | 12419 | 26 | TAGCGACGTCGCTTTTTGATCCTTCG | MB468 | 49 | 58 | 51 | 11 | 98 | 25.1 | 26.6 | 19.5 | 7.7 | 44.7 |
| >hg38_21_+_8433172_8446622_RNA45SN1_45S_50ntFlanks@12395.12414.20\|AGCGACGTCGCTTTTTGATC | >hg38_21_+_8433172_8446622_RNA45SN1_45S_50ntFlanks | 12395 | 12414 | 20 | AGCGACGTCGCTTTTTGATC | BT20 | 11 | 10 | 12 | 12 | 17 | 6.5 | 6.4 | 6.7 | 11.0 | 8.6 |
| >hg38_21_+_8433172_8446622_RNA45SN1_45S_50ntFlanks@12395.12414.20\|AGCGACGTCGCTTTTTGATC | >hg38_21_+_8433172_8446622_RNA45SN1_45S_50ntFlanks | 12395 | 12414 | 20 | AGCGACGTCGCTTTTTGATC | MB231 | 44 | 44 | 50 | 7 | 14 | 26.7 | 23.2 | 26.4 | 4.3 | 12.6 |
| >hg38_21_+_8433172_8446622_RNA45SN1_45S_50ntFlanks@12395.12415.21\|AGCGACGTCGCTTTTTGATCC | >hg38_21_+_8433172_8446622_RNA45SN1_45S_50ntFlanks | 12395 | 12415 | 21 | AGCGACGTCGCTTTTTGATCC | MB231 | 10 | 7 | 6 | 1 | 4 | 8.3 | 5.7 | 5.4 | 1.6 | 5.2 |
| >hg38_21_+_8433172_8446622_RNA45SN1_45S_50ntFlanks@12395.12418.24\|AGCGACGTCGCTTTTTGATCCTTC | >hg38_21_+_8433172_8446622_RNA45SN1_45S_50ntFlanks | 12395 | 12418 | 24 | AGCGACGTCGCTTTTTGATCCTTC | MB231 | 27 | 27 | 29 | 1 | 8 | 32.0 | 11.1 | 27.2 | 0.6 | 7.6 |
| >hg38_21_+_8433172_8446622_RNA45SN1_45S_50ntFlanks@12396.12414.19\|GCGACGTCGCTTTTTGATC | >hg38_21_+_8433172_8446622_RNA45SN1_45S_50ntFlanks | 12396 | 12414 | 19 | GCGACGTCGCTTTTTGATC | BT20 | 13 | 12 | 19 | 8 | 18 | 8.3 | 11.5 | 8.2 | 10.7 | 6.2 |
| >hg38_21_+_8433172_8446622_RNA45SN1_45S_50ntFlanks@12396.12414.19\|GCGACGTCGCTTTTTGATC | >hg38_21_+_8433172_8446622_RNA45SN1_45S_50ntFlanks | 12396 | 12414 | 19 | GCGACGTCGCTTTTTGATC | MB231 | 47 | 35 | 105 | 3 | 16 | 31.4 | 19.0 | 84.1 | 2.7 | 13.7 |
| >hg38_21_+_8433172_8446622_RNA45SN1_45S_50ntFlanks@12396.12414.19\|GCGACGTCGCTTTTTGATC | >hg38_21_+_8433172_8446622_RNA45SN1_45S_50ntFlanks | 12396 | 12414 | 19 | GCGACGTCGCTTTTTGATC | MB468 | 120 | 114 | 140 | 43 | 73 | 29.8 | 33.3 | 36.6 | 47.1 | 30.3 |
| >hg38_21_+_8433172_8446622_RNA45SN1_45S_50ntFlanks@12396.12416.21\|GCGACGTCGCTTTTTGATCCT | >hg38_21_+_8433172_8446622_RNA45SN1_45S_50ntFlanks | 12396 | 12416 | 21 | GCGACGTCGCTTTTTGATCCT | MB468 | 22 | 35 | 11 | 4 | 27 | 5.7 | 15.4 | 1.6 | 4.4 | 19.9 |
| >hg38_21_+_8433172_8446622_RNA45SN1_45S_50ntFlanks@12396.12417.22\|GCGACGTCGCTTTTTGATCCTT | >hg38_21_+_8433172_8446622_RNA45SN1_45S_50ntFlanks | 12396 | 12417 | 22 | GCGACGTCGCTTTTTGATCCTT | MB468 | 55 | 102 | 28 | 6 | 87 | 16.1 | 44.3 | 2.4 | 6.8 | 85.8 |
| >hg38_21_+_8433172_8446622_RNA45SN1_45S_50ntFlanks@12396.12418.23\|GCGACGTCGCTTTTTGATCCTTC | >hg38_21_+_8433172_8446622_RNA45SN1_45S_50ntFlanks | 12396 | 12418 | 23 | GCGACGTCGCTTTTTGATCCTTC | MB231 | 23 | 17 | 40 | 0 | 3 | 31.0 | 14.3 | 42.1 | 0.3 | 4.5 |
| >hg38_21_+_8433172_8446622_RNA45SN1_45S_50ntFlanks@12397.12414.18\|CGACGTCGCTTTTTGATC | >hg38_21_+_8433172_8446622_RNA45SN1_45S_50ntFlanks | 12397 | 12414 | 18 | CGACGTCGCTTTTTGATC | MB231 | 16 | 13 | 36 | 2 | 8 | 9.5 | 5.4 | 22.2 | 1.4 | 4.1 |
| >hg38_21_+_8433172_8446622_RNA45SN1_45S_50ntFlanks@12397.12414.18\|CGACGTCGCTTTTTGATC | >hg38_21_+_8433172_8446622_RNA45SN1_45S_50ntFlanks | 12397 | 12414 | 18 | CGACGTCGCTTTTTGATC | MB468 | 115 | 53 | 166 | 81 | 123 | 53.2 | 10.6 | 46.8 | 64.4 | 60.3 |
| >hg38_21_+_8433172_8446622_RNA45SN1_45S_50ntFlanks@12397.12416.20\|CGACGTCGCTTTTTGATCCT | >hg38_21_+_8433172_8446622_RNA45SN1_45S_50ntFlanks | 12397 | 12416 | 20 | CGACGTCGCTTTTTGATCCT | MB468 | 16 | 13 | 17 | 5 | 25 | 9.1 | 3.1 | 6.2 | 2.9 | 14.6 |
| >hg38_21_+_8433172_8446622_RNA45SN1_45S_50ntFlanks@12397.12417.21\|CGACGTCGCTTTTTGATCCTT | >hg38_21_+_8433172_8446622_RNA45SN1_45S_50ntFlanks | 12397 | 12417 | 21 | CGACGTCGCTTTTTGATCCTT | MB468 | 69 | 68 | 72 | 17 | 114 | 46.0 | 16.9 | 27.1 | 10.5 | 82.5 |
| >hg38_21_+_8433172_8446622_RNA45SN1_45S_50ntFlanks@12397.12418.22\|CGACGTCGCTTTTTGATCCTTC | >hg38_21_+_8433172_8446622_RNA45SN1_45S_50ntFlanks | 12397 | 12418 | 22 | CGACGTCGCTTTTTGATCCTTC | MB231 | 36 | 25 | 133 | 1 | 9 | 33.0 | 18.1 | 122.4 | 0.5 | 7.6 |
| >hg38_21_+_8433172_8446622_RNA45SN1_45S_50ntFlanks@12397.12418.22\|CGACGTCGCTTTTTGATCCTTC | >hg38_21_+_8433172_8446622_RNA45SN1_45S_50ntFlanks | 12397 | 12418 | 22 | CGACGTCGCTTTTTGATCCTTC | MB468 | 22 | 23 | 32 | 3 | 11 | 14.9 | 6.2 | 15.2 | 0.9 | 3.8 |
| >hg38_21_+_8433172_8446622_RNA45SN1_45S_50ntFlanks@12397.12419.23\|CGACGTCGCTTTTTGATCCTTCG | >hg38_21_+_8433172_8446622_RNA45SN1_45S_50ntFlanks | 12397 | 12419 | 23 | CGACGTCGCTTTTTGATCCTTCG | MB468 | 11 | 5 | 11 | 7 | 16 | 2.5 | 2.1 | 4.2 | 1.4 | 7.5 |
| >hg38_21_+_8433172_8446622_RNA45SN1_45S_50ntFlanks@12399.12419.21\|ACGTCGCTTTTTGATCCTTCG | >hg38_21_+_8433172_8446622_RNA45SN1_45S_50ntFlanks | 12399 | 12419 | 21 | ACGTCGCTTTTTGATCCTTCG | MB468 | 21 | 8 | 26 | 5 | 22 | 8.3 | 2.7 | 12.8 | 1.5 | 11.8 |
| >hg38_21_+_8433172_8446622_RNA45SN1_45S_50ntFlanks@12399.12433.35\|ACGTCGCTTTTTGATCCTTCGATGTCGGCTCTTCC | >hg38_21_+_8433172_8446622_RNA45SN1_45S_50ntFlanks | 12399 | 12433 | 35 | ACGTCGCTTTTTGATCCTTCGATGTCGGCTCTTCC | MB231 | 18 | 19 | 20 | 1 | 10 | 20.7 | 10.6 | 18.3 | 0.6 | 8.3 |
| >hg38_21_+_8433172_8446622_RNA45SN1_45S_50ntFlanks@12404.12431.28\|GCTTTTTGATCCTTCGATGTCGGCTCTT | >hg38_21_+_8433172_8446622_RNA45SN1_45S_50ntFlanks | 12404 | 12431 | 28 | GCTTTTTGATCCTTCGATGTCGGCTCTT | MB468 | 21 | 17 | 37 | 5 | 5 | 6.9 | 8.1 | 5.0 | 2.3 | 2.3 |
| >hg38_21_+_8433172_8446622_RNA45SN1_45S_50ntFlanks@12404.12432.29\|GCTTTTTGATCCTTCGATGTCGGCTCTTC | >hg38_21_+_8433172_8446622_RNA45SN1_45S_50ntFlanks | 12404 | 12432 | 29 | GCTTTTTGATCCTTCGATGTCGGCTCTTC | MB468 | 18 | 20 | 31 | 7 | 6 | 4.5 | 7.3 | 9.9 | 2.5 | 3.9 |
| >hg38_21_+_8433172_8446622_RNA45SN1_45S_50ntFlanks@12404.12433.30\|GCTTTTTGATCCTTCGATGTCGGCTCTTCC | >hg38_21_+_8433172_8446622_RNA45SN1_45S_50ntFlanks | 12404 | 12433 | 30 | GCTTTTTGATCCTTCGATGTCGGCTCTTCC | MB231 | 11 | 15 | 26 | 0 | 3 | 13.6 | 9.7 | 35.3 | 0.3 | 4.9 |
| >hg38_21_+_8433172_8446622_RNA45SN1_45S_50ntFlanks@12404.12434.31\|GCTTTTTGATCCTTCGATGTCGGCTCTTCCT | >hg38_21_+_8433172_8446622_RNA45SN1_45S_50ntFlanks | 12404 | 12434 | 31 | GCTTTTTGATCCTTCGATGTCGGCTCTTCCT | MB468 | 16 | 71 | 19 | 2 | 12 | 7.2 | 47.8 | 6.6 | 0.9 | 6.5 |
| >hg38_21_+_8433172_8446622_RNA45SN1_45S_50ntFlanks@12404.12436.33\|GCTTTTTGATCCTTCGATGTCGGCTCTTCCTAT | >hg38_21_+_8433172_8446622_RNA45SN1_45S_50ntFlanks | 12404 | 12436 | 33 | GCTTTTTGATCCTTCGATGTCGGCTCTTCCTAT | MB468 | 13 | 70 | 10 | 1 | 12 | 3.6 | 64.8 | 2.8 | 0.5 | 8.4 |
| >hg38_21_+_8433172_8446622_RNA45SN1_45S_50ntFlanks@12405.12422.18\|CTTTTTGATCCTTCGATG | >hg38_21_+_8433172_8446622_RNA45SN1_45S_50ntFlanks | 12405 | 12422 | 18 | CTTTTTGATCCTTCGATG | MB468 | 103 | 14 | 151 | 39 | 28 | 42.5 | 6.3 | 104.5 | 14.7 | 9.8 |
| >hg38_21_+_8433172_8446622_RNA45SN1_45S_50ntFlanks@12405.12423.19\|CTTTTTGATCCTTCGATGT | >hg38_21_+_8433172_8446622_RNA45SN1_45S_50ntFlanks | 12405 | 12423 | 19 | CTTTTTGATCCTTCGATGT | MB468 | 34 | 6 | 42 | 11 | 5 | 13.0 | 2.5 | 17.3 | 3.2 | 2.2 |
| >hg38_21_+_8433172_8446622_RNA45SN1_45S_50ntFlanks@12405.12424.20\|CTTTTTGATCCTTCGATGTC | >hg38_21_+_8433172_8446622_RNA45SN1_45S_50ntFlanks | 12405 | 12424 | 20 | CTTTTTGATCCTTCGATGTC | MB468 | 164 | 44 | 209 | 72 | 42 | 79.2 | 12.6 | 138.4 | 26.7 | 32.2 |
| >hg38_21_+_8433172_8446622_RNA45SN1_45S_50ntFlanks@12405.12425.21\|CTTTTTGATCCTTCGATGTCG | >hg38_21_+_8433172_8446622_RNA45SN1_45S_50ntFlanks | 12405 | 12425 | 21 | CTTTTTGATCCTTCGATGTCG | MB468 | 135 | 34 | 127 | 42 | 36 | 45.1 | 5.4 | 47.0 | 9.0 | 15.4 |
| >hg38_21_+_8433172_8446622_RNA45SN1_45S_50ntFlanks@12405.12426.22\|CTTTTTGATCCTTCGATGTCGG | >hg38_21_+_8433172_8446622_RNA45SN1_45S_50ntFlanks | 12405 | 12426 | 22 | CTTTTTGATCCTTCGATGTCGG | MB468 | 413 | 213 | 548 | 183 | 116 | 84.3 | 62.8 | 185.4 | 88.4 | 27.9 |
| >hg38_21_+_8433172_8446622_RNA45SN1_45S_50ntFlanks@12405.12427.23\|CTTTTTGATCCTTCGATGTCGGC | >hg38_21_+_8433172_8446622_RNA45SN1_45S_50ntFlanks | 12405 | 12427 | 23 | CTTTTTGATCCTTCGATGTCGGC | MB468 | 17 | 17 | 22 | 8 | 4 | 3.0 | 3.4 | 7.6 | 2.9 | 0.3 |
| >hg38_21_+_8433172_8446622_RNA45SN1_45S_50ntFlanks@12405.12428.24\|CTTTTTGATCCTTCGATGTCGGCT | >hg38_21_+_8433172_8446622_RNA45SN1_45S_50ntFlanks | 12405 | 12428 | 24 | CTTTTTGATCCTTCGATGTCGGCT | MB468 | 16 | 14 | 25 | 13 | 9 | 3.0 | 1.7 | 8.3 | 7.5 | 3.5 |
| >hg38_21_+_8433172_8446622_RNA45SN1_45S_50ntFlanks@12405.12429.25\|CTTTTTGATCCTTCGATGTCGGCTC | >hg38_21_+_8433172_8446622_RNA45SN1_45S_50ntFlanks | 12405 | 12429 | 25 | CTTTTTGATCCTTCGATGTCGGCTC | MB468 | 353 | 506 | 543 | 77 | 75 | 37.1 | 93.5 | 148.9 | 37.6 | 19.5 |
| >hg38_21_+_8433172_8446622_RNA45SN1_45S_50ntFlanks@12405.12430.26\|CTTTTTGATCCTTCGATGTCGGCTCT | >hg38_21_+_8433172_8446622_RNA45SN1_45S_50ntFlanks | 12405 | 12430 | 26 | CTTTTTGATCCTTCGATGTCGGCTCT | MB468 | 90 | 76 | 170 | 36 | 22 | 17.7 | 22.6 | 53.4 | 9.7 | 7.6 |
| >hg38_21_+_8433172_8446622_RNA45SN1_45S_50ntFlanks@12405.12431.27\|CTTTTTGATCCTTCGATGTCGGCTCTT | >hg38_21_+_8433172_8446622_RNA45SN1_45S_50ntFlanks | 12405 | 12431 | 27 | CTTTTTGATCCTTCGATGTCGGCTCTT | MB468 | 924 | 851 | 1693 | 235 | 204 | 284.1 | 192.1 | 811.0 | 113.8 | 38.0 |

| | | | | | | | | | | | | | | | | |
|---|---|---|---|---|---|---|---|---|---|---|---|---|---|---|---|---|
| >hg38_21_+_8433172_8446622_RNA45SN1_45S_50ntFlanks@12405.12432.28\|CTTTTTGATCCTTCGATGTCGGCTCTTC | >hg38_21_+_8433172_8446622_RNA45SN1_45S_50ntFlanks | 12405 | 12432 | 28 | CTTTTTGATCCTTCGATGTCGGCTCTTC | MB468 | 696 | 850 | 1126 | 320 | 246 | 323.6 | 226.7 | 818.6 | 121.5 | 66.2 |
| >hg38_21_+_8433172_8446622_RNA45SN1_45S_50ntFlanks@12405.12433.29\|CTTTTTGATCCTTCGATGTCGGCTCTTCC | >hg38_21_+_8433172_8446622_RNA45SN1_45S_50ntFlanks | 12405 | 12433 | 29 | CTTTTTGATCCTTCGATGTCGGCTCTTCC | MB468 | 232 | 910 | 354 | 64 | 151 | 54.3 | 189.4 | 177.1 | 23.7 | 14.7 |
| >hg38_21_+_8433172_8446622_RNA45SN1_45S_50ntFlanks@12405.12434.30\|CTTTTTGATCCTTCGATGTCGGCTCTTCCT | >hg38_21_+_8433172_8446622_RNA45SN1_45S_50ntFlanks | 12405 | 12434 | 30 | CTTTTTGATCCTTCGATGTCGGCTCTTCCT | MB468 | 795 | 3223 | 1167 | 136 | 691 | 214.7 | 1314.0 | 455.9 | 51.2 | 84.7 |
| >hg38_21_+_8433172_8446622_RNA45SN1_45S_50ntFlanks@12405.12435.31\|CTTTTTGATCCTTCGATGTCGGCTCTTCCTA | >hg38_21_+_8433172_8446622_RNA45SN1_45S_50ntFlanks | 12405 | 12435 | 31 | CTTTTTGATCCTTCGATGTCGGCTCTTCCTA | MB468 | 362 | 1425 | 368 | 94 | 369 | 67.3 | 867.7 | 161.0 | 40.8 | 64.2 |
| >hg38_21_+_8433172_8446622_RNA45SN1_45S_50ntFlanks@12405.12436.32\|CTTTTTGATCCTTCGATGTCGGCTCTTCCTAT | >hg38_21_+_8433172_8446622_RNA45SN1_45S_50ntFlanks | 12405 | 12436 | 32 | CTTTTTGATCCTTCGATGTCGGCTCTTCCTAT | MB468 | 665 | 3424 | 630 | 89 | 1048 | 129.7 | 2128.7 | 424.6 | 39.5 | 236.9 |
| >hg38_21_+_8433172_8446622_RNA45SN1_45S_50ntFlanks@12405.12437.33\|CTTTTTGATCCTTCGATGTCGGCTCTTCCTATC | >hg38_21_+_8433172_8446622_RNA45SN1_45S_50ntFlanks | 12405 | 12437 | 33 | CTTTTTGATCCTTCGATGTCGGCTCTTCCTATC | MB468 | 216 | 1175 | 258 | 49 | 350 | 48.9 | 787.7 | 143.1 | 20.2 | 54.1 |
| >hg38_21_+_8433172_8446622_RNA45SN1_45S_50ntFlanks@12405.12438.34\|CTTTTTGATCCTTCGATGTCGGCTCTTCCTATCA | >hg38_21_+_8433172_8446622_RNA45SN1_45S_50ntFlanks | 12405 | 12438 | 34 | CTTTTTGATCCTTCGATGTCGGCTCTTCCTATCA | MB468 | 12 | 109 | 14 | 5 | 81 | 3.9 | 76.3 | 3.1 | 4.2 | 50.9 |
| >hg38_21_+_8433172_8446622_RNA45SN1_45S_50ntFlanks@12405.12439.35\|CTTTTTGATCCTTCGATGTCGGCTCTTCCTATCAT | >hg38_21_+_8433172_8446622_RNA45SN1_45S_50ntFlanks | 12405 | 12439 | 35 | CTTTTTGATCCTTCGATGTCGGCTCTTCCTATCAT | MB468 | 11 | 79 | 17 | 3 | 20 | 1.2 | 41.3 | 10.0 | 2.4 | 13.5 |
| >hg38_21_+_8433172_8446622_RNA45SN1_45S_50ntFlanks@12405.12440.36\|CTTTTTGATCCTTCGATGTCGGCTCTTCCTATCATT | >hg38_21_+_8433172_8446622_RNA45SN1_45S_50ntFlanks | 12405 | 12440 | 36 | CTTTTTGATCCTTCGATGTCGGCTCTTCCTATCATT | MB468 | 17 | 144 | 35 | 4 | 27 | 6.3 | 83.5 | 4.7 | 2.5 | 9.7 |
| >hg38_21_+_8433172_8446622_RNA45SN1_45S_50ntFlanks@12405.12441.37\|CTTTTTGATCCTTCGATGTCGGCTCTTCCTATCATTG | >hg38_21_+_8433172_8446622_RNA45SN1_45S_50ntFlanks | 12405 | 12441 | 37 | CTTTTTGATCCTTCGATGTCGGCTCTTCCTATCATTG | MB468 | 136 | 1294 | 207 | 25 | 231 | 41.9 | 872.7 | 37.8 | 22.3 | 91.7 |
| >hg38_21_+_8433172_8446622_RNA45SN1_45S_50ntFlanks@12405.12443.39\|CTTTTTGATCCTTCGATGTCGGCTCTTCCTATCATTGTG | >hg38_21_+_8433172_8446622_RNA45SN1_45S_50ntFlanks | 12405 | 12443 | 39 | CTTTTTGATCCTTCGATGTCGGCTCTTCCTATCATTGTG | MB468 | 248 | 3160 | 434 | 70 | 569 | 80.6 | 2448.9 | 120.9 | 63.1 | 321.5 |
| >hg38_21_+_8433172_8446622_RNA45SN1_45S_50ntFlanks@12405.12444.40\|CTTTTTGATCCTTCGATGTCGGCTCTTCCTATCATTGTGA | >hg38_21_+_8433172_8446622_RNA45SN1_45S_50ntFlanks | 12405 | 12444 | 40 | CTTTTTGATCCTTCGATGTCGGCTCTTCCTATCATTGTGA | MB468 | 30 | 459 | 54 | 7 | 72 | 9.2 | 454.1 | 20.1 | 7.2 | 43.8 |
| >hg38_21_+_8433172_8446622_RNA45SN1_45S_50ntFlanks@12405.12445.41\|CTTTTTGATCCTTCGATGTCGGCTCTTCCTATCATTGTGAA | >hg38_21_+_8433172_8446622_RNA45SN1_45S_50ntFlanks | 12405 | 12445 | 41 | CTTTTTGATCCTTCGATGTCGGCTCTTCCTATCATTGTGAA | MB468 | 10 | 53 | 13 | 2 | 21 | 3.0 | 40.3 | 7.2 | 2.8 | 24.7 |
| >hg38_21_+_8433172_8446622_RNA45SN1_45S_50ntFlanks@12405.12446.42\|CTTTTTGATCCTTCGATGTCGGCTCTTCCTATCATTGTGAAG | >hg38_21_+_8433172_8446622_RNA45SN1_45S_50ntFlanks | 12405 | 12446 | 42 | CTTTTTGATCCTTCGATGTCGGCTCTTCCTATCATTGTGAAG | MB468 | 48 | 528 | 128 | 12 | 135 | 13.5 | 578.1 | 85.9 | 15.8 | 99.6 |
| >hg38_21_+_8433172_8446622_RNA45SN1_45S_50ntFlanks@12406.12429.24\|TTTTTGATCCTTCGATGTCGGCTC | >hg38_21_+_8433172_8446622_RNA45SN1_45S_50ntFlanks | 12406 | 12429 | 24 | TTTTTGATCCTTCGATGTCGGCTC | MB468 | 10 | 10 | 15 | 2 | 1 | 2.3 | 1.1 | 1.6 | 1.1 | 0.1 |
| >hg38_21_+_8433172_8446622_RNA45SN1_45S_50ntFlanks@12406.12431.26\|TTTTTGATCCTTCGATGTCGGCTCTT | >hg38_21_+_8433172_8446622_RNA45SN1_45S_50ntFlanks | 12406 | 12431 | 26 | TTTTTGATCCTTCGATGTCGGCTCTT | MB468 | 31 | 22 | 59 | 8 | 8 | 12.8 | 6.0 | 31.0 | 4.8 | 3.0 |
| >hg38_21_+_8433172_8446622_RNA45SN1_45S_50ntFlanks@12406.12432.27\|TTTTTGATCCTTCGATGTCGGCTCTTC | >hg38_21_+_8433172_8446622_RNA45SN1_45S_50ntFlanks | 12406 | 12432 | 27 | TTTTTGATCCTTCGATGTCGGCTCTTC | MB468 | 17 | 18 | 32 | 8 | 6 | 7.4 | 3.3 | 21.8 | 4.7 | 0.9 |
| >hg38_21_+_8433172_8446622_RNA45SN1_45S_50ntFlanks@12406.12434.29\|TTTTTGATCCTTCGATGTCGGCTCTTCCT | >hg38_21_+_8433172_8446622_RNA45SN1_45S_50ntFlanks | 12406 | 12434 | 29 | TTTTTGATCCTTCGATGTCGGCTCTTCCT | MB468 | 31 | 144 | 49 | 4 | 19 | 10.7 | 52.1 | 21.1 | 2.2 | 5.4 |
| >hg38_21_+_8433172_8446622_RNA45SN1_45S_50ntFlanks@12406.12435.30\|TTTTTGATCCTTCGATGTCGGCTCTTCCTA | >hg38_21_+_8433172_8446622_RNA45SN1_45S_50ntFlanks | 12406 | 12435 | 30 | TTTTTGATCCTTCGATGTCGGCTCTTCCTA | MB468 | 13 | 52 | 14 | 3 | 8 | 3.4 | 33.0 | 4.8 | 1.2 | 1.9 |
| >hg38_21_+_8433172_8446622_RNA45SN1_45S_50ntFlanks@12406.12436.31\|TTTTTGATCCTTCGATGTCGGCTCTTCCTAT | >hg38_21_+_8433172_8446622_RNA45SN1_45S_50ntFlanks | 12406 | 12436 | 31 | TTTTTGATCCTTCGATGTCGGCTCTTCCTAT | MB468 | 26 | 134 | 25 | 2 | 20 | 4.8 | 82.4 | 15.4 | 1.0 | 5.6 |
| >hg38_21_+_8433172_8446622_RNA45SN1_45S_50ntFlanks@12407.12424.18\|TTTTGATCCTTCGATGTC | >hg38_21_+_8433172_8446622_RNA45SN1_45S_50ntFlanks | 12407 | 12424 | 18 | TTTTGATCCTTCGATGTC | MB468 | 12 | 2 | 13 | 7 | 3 | 6.5 | 0.4 | 11.7 | 2.3 | 1.7 |
| >hg38_21_+_8433172_8446622_RNA45SN1_45S_50ntFlanks@12407.12425.19\|TTTTGATCCTTCGATGTCG | >hg38_21_+_8433172_8446622_RNA45SN1_45S_50ntFlanks | 12407 | 12425 | 19 | TTTTGATCCTTCGATGTCG | MB468 | 14 | 2 | 11 | 5 | 2 | 7.1 | 0.7 | 7.3 | 2.0 | 0.9 |
| >hg38_21_+_8433172_8446622_RNA45SN1_45S_50ntFlanks@12407.12426.20\|TTTTGATCCTTCGATGTCGG | >hg38_21_+_8433172_8446622_RNA45SN1_45S_50ntFlanks | 12407 | 12426 | 20 | TTTTGATCCTTCGATGTCGG | MB468 | 43 | 12 | 46 | 25 | 9 | 14.5 | 1.9 | 9.4 | 18.7 | 2.1 |
| >hg38_21_+_8433172_8446622_RNA45SN1_45S_50ntFlanks@12407.12429.23\|TTTTGATCCTTCGATGTCGGCTC | >hg38_21_+_8433172_8446622_RNA45SN1_45S_50ntFlanks | 12407 | 12429 | 23 | TTTTGATCCTTCGATGTCGGCTC | MB468 | 19 | 14 | 35 | 8 | 6 | 5.3 | 2.1 | 4.5 | 4.8 | 1.1 |
| >hg38_21_+_8433172_8446622_RNA45SN1_45S_50ntFlanks@12407.12431.25\|TTTTGATCCTTCGATGTCGGCTCTT | >hg38_21_+_8433172_8446622_RNA45SN1_45S_50ntFlanks | 12407 | 12431 | 25 | TTTTGATCCTTCGATGTCGGCTCTT | MB468 | 89 | 46 | 198 | 35 | 28 | 36.6 | 13.6 | 84.8 | 21.6 | 12.4 |
| >hg38_21_+_8433172_8446622_RNA45SN1_45S_50ntFlanks@12407.12432.26\|TTTTGATCCTTCGATGTCGGCTCTTC | >hg38_21_+_8433172_8446622_RNA45SN1_45S_50ntFlanks | 12407 | 12432 | 26 | TTTTGATCCTTCGATGTCGGCTCTTC | MB468 | 35 | 28 | 75 | 26 | 19 | 16.9 | 5.5 | 41.4 | 10.7 | 10.1 |
| >hg38_21_+_8433172_8446622_RNA45SN1_45S_50ntFlanks@12407.12433.27\|TTTTGATCCTTCGATGTCGGCTCTTCC | >hg38_21_+_8433172_8446622_RNA45SN1_45S_50ntFlanks | 12407 | 12433 | 27 | TTTTGATCCTTCGATGTCGGCTCTTCC | MB231 | 27 | 28 | 109 | 2 | 8 | 26.0 | 13.9 | 137.8 | 1.6 | 7.8 |
| >hg38_21_+_8433172_8446622_RNA45SN1_45S_50ntFlanks@12407.12433.27\|TTTTGATCCTTCGATGTCGGCTCTTCC | >hg38_21_+_8433172_8446622_RNA45SN1_45S_50ntFlanks | 12407 | 12433 | 27 | TTTTGATCCTTCGATGTCGGCTCTTCC | MB468 | 26 | 89 | 49 | 8 | 19 | 8.8 | 17.3 | 24.8 | 3.3 | 3.7 |
| >hg38_21_+_8433172_8446622_RNA45SN1_45S_50ntFlanks@12407.12434.28\|TTTTGATCCTTCGATGTCGGCTCTTCCT | >hg38_21_+_8433172_8446622_RNA45SN1_45S_50ntFlanks | 12407 | 12434 | 28 | TTTTGATCCTTCGATGTCGGCTCTTCCT | MB231 | 18 | 22 | 54 | 1 | 8 | 19.6 | 7.0 | 67.7 | 0.9 | 7.1 |
| >hg38_21_+_8433172_8446622_RNA45SN1_45S_50ntFlanks@12407.12434.28\|TTTTGATCCTTCGATGTCGGCTCTTCCT | >hg38_21_+_8433172_8446622_RNA45SN1_45S_50ntFlanks | 12407 | 12434 | 28 | TTTTGATCCTTCGATGTCGGCTCTTCCT | MB468 | 74 | 283 | 124 | 14 | 87 | 28.7 | 108.9 | 59.2 | 4.9 | 21.4 |
| >hg38_21_+_8433172_8446622_RNA45SN1_45S_50ntFlanks@12407.12435.29\|TTTTGATCCTTCGATGTCGGCTCTTCCTA | >hg38_21_+_8433172_8446622_RNA45SN1_45S_50ntFlanks | 12407 | 12435 | 29 | TTTTGATCCTTCGATGTCGGCTCTTCCTA | MB468 | 38 | 117 | 45 | 10 | 46 | 8.6 | 67.3 | 17.6 | 1.8 | 16.4 |
| >hg38_21_+_8433172_8446622_RNA45SN1_45S_50ntFlanks@12407.12436.30\|TTTTGATCCTTCGATGTCGGCTCTTCCTAT | >hg38_21_+_8433172_8446622_RNA45SN1_45S_50ntFlanks | 12407 | 12436 | 30 | TTTTGATCCTTCGATGTCGGCTCTTCCTAT | MB468 | 109 | 404 | 94 | 12 | 166 | 19.7 | 214.0 | 42.2 | 3.7 | 56.4 |
| >hg38_21_+_8433172_8446622_RNA45SN1_45S_50ntFlanks@12407.12437.31\|TTTTGATCCTTCGATGTCGGCTCTTCCTATC | >hg38_21_+_8433172_8446622_RNA45SN1_45S_50ntFlanks | 12407 | 12437 | 31 | TTTTGATCCTTCGATGTCGGCTCTTCCTATC | MB231 | 13 | 19 | 29 | 1 | 7 | 13.8 | 10.2 | 31.6 | 0.6 | 5.8 |
| >hg38_21_+_8433172_8446622_RNA45SN1_45S_50ntFlanks@12407.12437.31\|TTTTGATCCTTCGATGTCGGCTCTTCCTATC | >hg38_21_+_8433172_8446622_RNA45SN1_45S_50ntFlanks | 12407 | 12437 | 31 | TTTTGATCCTTCGATGTCGGCTCTTCCTATC | MB468 | 39 | 168 | 40 | 9 | 62 | 13.7 | 84.0 | 17.7 | 1.5 | 16.1 |
| >hg38_21_+_8433172_8446622_RNA45SN1_45S_50ntFlanks@12407.12438.32\|TTTTGATCCTTCGATGTCGGCTCTTCCTATCA | >hg38_21_+_8433172_8446622_RNA45SN1_45S_50ntFlanks | 12407 | 12438 | 32 | TTTTGATCCTTCGATGTCGGCTCTTCCTATCA | MB231 | 13 | 18 | 16 | 0 | 4 | 14.5 | 10.3 | 15.7 | 0.4 | 6.1 |
| >hg38_21_+_8433172_8446622_RNA45SN1_45S_50ntFlanks@12407.12441.35\|TTTTGATCCTTCGATGTCGGCTCTTCCTATCATTG | >hg38_21_+_8433172_8446622_RNA45SN1_45S_50ntFlanks | 12407 | 12441 | 35 | TTTTGATCCTTCGATGTCGGCTCTTCCTATCATTG | MB468 | 22 | 144 | 24 | 4 | 35 | 13.5 | 57.5 | 11.7 | 2.8 | 14.8 |
| >hg38_21_+_8433172_8446622_RNA45SN1_45S_50ntFlanks@12407.12443.37\|TTTTGATCCTTCGATGTCGGCTCTTCCTATCATTGTG | >hg38_21_+_8433172_8446622_RNA45SN1_45S_50ntFlanks | 12407 | 12443 | 37 | TTTTGATCCTTCGATGTCGGCTCTTCCTATCATTGTG | MB231 | 14 | 16 | 34 | 1 | 11 | 15.0 | 10.3 | 43.5 | 0.5 | 11.4 |
| >hg38_21_+_8433172_8446622_RNA45SN1_45S_50ntFlanks@12407.12443.37\|TTTTGATCCTTCGATGTCGGCTCTTCCTATCATTGTG | >hg38_21_+_8433172_8446622_RNA45SN1_45S_50ntFlanks | 12407 | 12443 | 37 | TTTTGATCCTTCGATGTCGGCTCTTCCTATCATTGTG | MB468 | 49 | 364 | 48 | 16 | 114 | 26.4 | 185.5 | 16.5 | 12.8 | 67.8 |
| >hg38_21_+_8433172_8446622_RNA45SN1_45S_50ntFlanks@12407.12446.40\|TTTTGATCCTTCGATGTCGGCTCTTCCTATCATTGTGAAG | >hg38_21_+_8433172_8446622_RNA45SN1_45S_50ntFlanks | 12407 | 12446 | 40 | TTTTGATCCTTCGATGTCGGCTCTTCCTATCATTGTGAAG | MB468 | 14 | 83 | 27 | 3 | 17 | 6.7 | 55.5 | 13.8 | 3.8 | 19.2 |
| >hg38_21_+_8433172_8446622_RNA45SN1_45S_50ntFlanks@12407.12448.42\|TTTTGATCCTTCGATGTCGGCTCTTCCTATCATTGTGAAGCA | >hg38_21_+_8433172_8446622_RNA45SN1_45S_50ntFlanks | 12407 | 12448 | 42 | TTTTGATCCTTCGATGTCGGCTCTTCCTATCATTGTGAAGCA | MB231 | 17 | 19 | 39 | 0 | 6 | 14.4 | 20.6 | 45.4 | 0.5 | 10.0 |
| >hg38_21_+_8433172_8446622_RNA45SN1_45S_50ntFlanks@12408.12431.24\|TTTGATCCTTCGATGTCGGCTCTT | >hg38_21_+_8433172_8446622_RNA45SN1_45S_50ntFlanks | 12408 | 12431 | 24 | TTTGATCCTTCGATGTCGGCTCTT | MB468 | 34 | 8 | 76 | 12 | 6 | 21.6 | 1.9 | 58.5 | 7.2 | 0.8 |
| >hg38_21_+_8433172_8446622_RNA45SN1_45S_50ntFlanks@12408.12432.25\|TTTGATCCTTCGATGTCGGCTCTTC | >hg38_21_+_8433172_8446622_RNA45SN1_45S_50ntFlanks | 12408 | 12432 | 25 | TTTGATCCTTCGATGTCGGCTCTTC | MB468 | 14 | 6 | 29 | 9 | 4 | 12.3 | 1.7 | 27.3 | 5.1 | 1.0 |
| >hg38_21_+_8433172_8446622_RNA45SN1_45S_50ntFlanks@12408.12433.26\|TTTGATCCTTCGATGTCGGCTCTTCC | >hg38_21_+_8433172_8446622_RNA45SN1_45S_50ntFlanks | 12408 | 12433 | 26 | TTTGATCCTTCGATGTCGGCTCTTCC | MB231 | 13 | 15 | 61 | 1 | 5 | 12.6 | 3.2 | 78.9 | 0.7 | 4.4 |
| >hg38_21_+_8433172_8446622_RNA45SN1_45S_50ntFlanks@12408.12434.27\|TTTGATCCTTCGATGTCGGCTCTTCCT | >hg38_21_+_8433172_8446622_RNA45SN1_45S_50ntFlanks | 12408 | 12434 | 27 | TTTGATCCTTCGATGTCGGCTCTTCCT | BT20 | 14 | 27 | 10 | 6 | 3 | 17.8 | 28.8 | 9.3 | 10.6 | 5.7 |
| >hg38_21_+_8433172_8446622_RNA45SN1_45S_50ntFlanks@12408.12434.27\|TTTGATCCTTCGATGTCGGCTCTTCCT | >hg38_21_+_8433172_8446622_RNA45SN1_45S_50ntFlanks | 12408 | 12434 | 27 | TTTGATCCTTCGATGTCGGCTCTTCCT | MB468 | 14 | 41 | 22 | 2 | 13 | 5.1 | 12.8 | 12.3 | 0.8 | 1.6 |
| >hg38_21_+_8433172_8446622_RNA45SN1_45S_50ntFlanks@12408.12436.29\|TTTGATCCTTCGATGTCGGCTCTTCCTAT | >hg38_21_+_8433172_8446622_RNA45SN1_45S_50ntFlanks | 12408 | 12436 | 29 | TTTGATCCTTCGATGTCGGCTCTTCCTAT | MB468 | 13 | 41 | 15 | 1 | 13 | 4.7 | 17.2 | 8.9 | 0.4 | 1.2 |
| >hg38_21_+_8433172_8446622_RNA45SN1_45S_50ntFlanks@12409.12431.23\|TTGATCCTTCGATGTCGGCTCTT | >hg38_21_+_8433172_8446622_RNA45SN1_45S_50ntFlanks | 12409 | 12431 | 23 | TTGATCCTTCGATGTCGGCTCTT | MB468 | 10 | 1 | 22 | 5 | 2 | 9.2 | 0.3 | 23.1 | 2.4 | 0.5 |
| >hg38_21_+_8433172_8446622_RNA45SN1_45S_50ntFlanks@12409.12433.25\|TTGATCCTTCGATGTCGGCTCTTCC | >hg38_21_+_8433172_8446622_RNA45SN1_45S_50ntFlanks | 12409 | 12433 | 25 | TTGATCCTTCGATGTCGGCTCTTCC | MB231 | 17 | 14 | 43 | 1 | 7 | 15.4 | 4.5 | 38.7 | 1.0 | 6.3 |
| >hg38_21_+_8433172_8446622_RNA45SN1_45S_50ntFlanks@12409.12434.26\|TTGATCCTTCGATGTCGGCTCTTCCT | >hg38_21_+_8433172_8446622_RNA45SN1_45S_50ntFlanks | 12409 | 12434 | 26 | TTGATCCTTCGATGTCGGCTCTTCCT | BT20 | 22 | 48 | 15 | 21 | 2 | 32.5 | 54.1 | 13.6 | 35.8 | 4.3 |
| >hg38_21_+_8433172_8446622_RNA45SN1_45S_50ntFlanks@12409.12434.26\|TTGATCCTTCGATGTCGGCTCTTCCT | >hg38_21_+_8433172_8446622_RNA45SN1_45S_50ntFlanks | 12409 | 12434 | 26 | TTGATCCTTCGATGTCGGCTCTTCCT | MB231 | 13 | 15 | 23 | 1 | 6 | 13.0 | 1.2 | 22.3 | 0.4 | 6.1 |

| | | | | | | | | | | | | | | | |
|---|---|---|---|---|---|---|---|---|---|---|---|---|---|---|---|
| >hg38_21_+_8433172_8446622_RNA45SN1_45S_50ntFlanks@12409.12437.29\|TTGATCCTTCGATGTCGGCTCTTCCTATC | >hg38_21_+_8433172_8446622_RNA45SN1_45S_50ntFlanks | 12409 | 12437 | 29 | TTGATCCTTCGATGTCGGCTCTTCCTATC | MB231 | 10 | 14 | 15 | 0 | 0 | 11.6 | 7.5 | 13.3 | 0.0 | 0.0 |
| >hg38_21_+_8433172_8446622_RNA45SN1_45S_50ntFlanks@12409.12448.40\|TTGATCCTTCGATGTCGGCTCTTCCTATCATTGTGAAGCA | >hg38_21_+_8433172_8446622_RNA45SN1_45S_50ntFlanks | 12409 | 12448 | 40 | TTGATCCTTCGATGTCGGCTCTTCCTATCATTGTGAAGCA | MB231 | 15 | 15 | 17 | 0 | 8 | 16.0 | 12.4 | 15.9 | 0.3 | 7.2 |
| >hg38_21_+_8433172_8446622_RNA45SN1_45S_50ntFlanks@12410.12433.24\|TGATCCTTCGATGTCGGCTCTTCC | >hg38_21_+_8433172_8446622_RNA45SN1_45S_50ntFlanks | 12410 | 12433 | 24 | TGATCCTTCGATGTCGGCTCTTCC | MB231 | 11 | 8 | 29 | 0 | 3 | 8.4 | 3.9 | 22.4 | 0.8 | 5.0 |
| >hg38_21_+_8433172_8446622_RNA45SN1_45S_50ntFlanks@12410.12434.25\|TGATCCTTCGATGTCGGCTCTTCCT | >hg38_21_+_8433172_8446622_RNA45SN1_45S_50ntFlanks | 12410 | 12434 | 25 | TGATCCTTCGATGTCGGCTCTTCCT | BT20 | 41 | 78 | 21 | 2 | 1 | 65.7 | 122.9 | 24.9 | 3.4 | 1.9 |
| >hg38_21_+_8433172_8446622_RNA45SN1_45S_50ntFlanks@12411.12432.22\|GATCCTTCGATGTCGGCTCTTC | >hg38_21_+_8433172_8446622_RNA45SN1_45S_50ntFlanks | 12411 | 12432 | 22 | GATCCTTCGATGTCGGCTCTTC | MB468 | 11 | 8 | 23 | 4 | 3 | 8.7 | 3.0 | 23.9 | 2.6 | 0.5 |
| >hg38_21_+_8433172_8446622_RNA45SN1_45S_50ntFlanks@12411.12433.23\|GATCCTTCGATGTCGGCTCTTCC | >hg38_21_+_8433172_8446622_RNA45SN1_45S_50ntFlanks | 12411 | 12433 | 23 | GATCCTTCGATGTCGGCTCTTCC | MB231 | 11 | 10 | 23 | 0 | 3 | 19.4 | 16.9 | 40.7 | 0.5 | 4.4 |
| >hg38_21_+_8433172_8446622_RNA45SN1_45S_50ntFlanks@12411.12434.24\|GATCCTTCGATGTCGGCTCTTCCT | >hg38_21_+_8433172_8446622_RNA45SN1_45S_50ntFlanks | 12411 | 12434 | 24 | GATCCTTCGATGTCGGCTCTTCCT | MB231 | 12 | 12 | 17 | 0 | 4 | 20.5 | 21.0 | 29.6 | 0.3 | 7.0 |
| >hg38_21_+_8433172_8446622_RNA45SN1_45S_50ntFlanks@12411.12448.38\|GATCCTTCGATGTCGGCTCTTCCTATCATTGTGAAGCA | >hg38_21_+_8433172_8446622_RNA45SN1_45S_50ntFlanks | 12411 | 12448 | 38 | GATCCTTCGATGTCGGCTCTTCCTATCATTGTGAAGCA | MB231 | 17 | 9 | 18 | 0 | 0 | 29.7 | 15.6 | 30.8 | 0.0 | 0.0 |
| >hg38_21_+_8433172_8446622_RNA45SN1_45S_50ntFlanks@12412.12429.18\|ATCCTTCGATGTCGGCTC | >hg38_21_+_8433172_8446622_RNA45SN1_45S_50ntFlanks | 12412 | 12429 | 18 | ATCCTTCGATGTCGGCTC | MB468 | 13 | 6 | 24 | 2 | 2 | 6.0 | 1.0 | 11.6 | 0.8 | 0.6 |
| >hg38_21_+_8433172_8446622_RNA45SN1_45S_50ntFlanks@12412.12430.19\|ATCCTTCGATGTCGGCTCT | >hg38_21_+_8433172_8446622_RNA45SN1_45S_50ntFlanks | 12412 | 12430 | 19 | ATCCTTCGATGTCGGCTCT | MB468 | 11 | 9 | 25 | 2 | 3 | 3.8 | 1.4 | 10.9 | 1.0 | 0.7 |
| >hg38_21_+_8433172_8446622_RNA45SN1_45S_50ntFlanks@12412.12431.20\|ATCCTTCGATGTCGGCTCTT | >hg38_21_+_8433172_8446622_RNA45SN1_45S_50ntFlanks | 12412 | 12431 | 20 | ATCCTTCGATGTCGGCTCTT | MB468 | 34 | 26 | 47 | 6 | 11 | 15.7 | 6.3 | 23.3 | 1.5 | 2.4 |
| >hg38_21_+_8433172_8446622_RNA45SN1_45S_50ntFlanks@12412.12432.21\|ATCCTTCGATGTCGGCTCTTC | >hg38_21_+_8433172_8446622_RNA45SN1_45S_50ntFlanks | 12412 | 12432 | 21 | ATCCTTCGATGTCGGCTCTTC | MB468 | 71 | 130 | 107 | 14 | 36 | 21.2 | 67.1 | 54.5 | 5.9 | 8.7 |
| >hg38_21_+_8433172_8446622_RNA45SN1_45S_50ntFlanks@12412.12433.22\|ATCCTTCGATGTCGGCTCTTCC | >hg38_21_+_8433172_8446622_RNA45SN1_45S_50ntFlanks | 12412 | 12433 | 22 | ATCCTTCGATGTCGGCTCTTCC | MB468 | 15 | 37 | 22 | 2 | 7 | 5.9 | 10.4 | 12.3 | 0.6 | 1.0 |
| >hg38_21_+_8433172_8446622_RNA45SN1_45S_50ntFlanks@12412.12434.23\|ATCCTTCGATGTCGGCTCTTCCT | >hg38_21_+_8433172_8446622_RNA45SN1_45S_50ntFlanks | 12412 | 12434 | 23 | ATCCTTCGATGTCGGCTCTTCCT | MB468 | 41 | 105 | 50 | 4 | 28 | 16.6 | 34.4 | 31.3 | 1.6 | 6.1 |
| >hg38_21_+_8433172_8446622_RNA45SN1_45S_50ntFlanks@12412.12435.24\|ATCCTTCGATGTCGGCTCTTCCTA | >hg38_21_+_8433172_8446622_RNA45SN1_45S_50ntFlanks | 12412 | 12435 | 24 | ATCCTTCGATGTCGGCTCTTCCTA | MB468 | 23 | 70 | 20 | 3 | 21 | 9.1 | 47.6 | 15.6 | 1.0 | 11.1 |
| >hg38_21_+_8433172_8446622_RNA45SN1_45S_50ntFlanks@12412.12436.25\|ATCCTTCGATGTCGGCTCTTCCTAT | >hg38_21_+_8433172_8446622_RNA45SN1_45S_50ntFlanks | 12412 | 12436 | 25 | ATCCTTCGATGTCGGCTCTTCCTAT | MB468 | 40 | 146 | 35 | 2 | 55 | 15.6 | 84.6 | 23.7 | 0.4 | 39.1 |
| >hg38_21_+_8433172_8446622_RNA45SN1_45S_50ntFlanks@12412.12437.26\|ATCCTTCGATGTCGGCTCTTCCTATC | >hg38_21_+_8433172_8446622_RNA45SN1_45S_50ntFlanks | 12412 | 12437 | 26 | ATCCTTCGATGTCGGCTCTTCCTATC | MB468 | 16 | 41 | 16 | 2 | 20 | 6.6 | 13.1 | 8.6 | 0.2 | 10.9 |
| >hg38_21_+_8433172_8446622_RNA45SN1_45S_50ntFlanks@12412.12441.30\|ATCCTTCGATGTCGGCTCTTCCTATCATTG | >hg38_21_+_8433172_8446622_RNA45SN1_45S_50ntFlanks | 12412 | 12441 | 30 | ATCCTTCGATGTCGGCTCTTCCTATCATTG | MB468 | 18 | 56 | 21 | 1 | 20 | 5.4 | 16.4 | 4.9 | 0.5 | 6.7 |
| >hg38_21_+_8433172_8446622_RNA45SN1_45S_50ntFlanks@12412.12443.32\|ATCCTTCGATGTCGGCTCTTCCTATCATTGTG | >hg38_21_+_8433172_8446622_RNA45SN1_45S_50ntFlanks | 12412 | 12443 | 32 | ATCCTTCGATGTCGGCTCTTCCTATCATTGTG | MB468 | 42 | 133 | 71 | 4 | 49 | 14.8 | 44.3 | 15.6 | 2.2 | 13.2 |
| >hg38_21_+_8433172_8446622_RNA45SN1_45S_50ntFlanks@12412.12446.35\|ATCCTTCGATGTCGGCTCTTCCTATCATTGTGAAG | >hg38_21_+_8433172_8446622_RNA45SN1_45S_50ntFlanks | 12412 | 12446 | 35 | ATCCTTCGATGTCGGCTCTTCCTATCATTGTGAAG | MB468 | 11 | 16 | 33 | 1 | 8 | 2.8 | 10.2 | 15.0 | 1.5 | 5.4 |
| >hg38_21_+_8433172_8446622_RNA45SN1_45S_50ntFlanks@12413.12430.18\|TCCTTCGATGTCGGCTCT | >hg38_21_+_8433172_8446622_RNA45SN1_45S_50ntFlanks | 12413 | 12430 | 18 | TCCTTCGATGTCGGCTCT | BT20 | 12 | 24 | 8 | 0 | 0 | 19.8 | 40.9 | 11.4 | 0.2 | 0.7 |
| >hg38_21_+_8433172_8446622_RNA45SN1_45S_50ntFlanks@12413.12431.19\|TCCTTCGATGTCGGCTCTT | >hg38_21_+_8433172_8446622_RNA45SN1_45S_50ntFlanks | 12413 | 12431 | 19 | TCCTTCGATGTCGGCTCTT | MB468 | 20 | 17 | 24 | 4 | 6 | 6.5 | 3.1 | 6.3 | 1.5 | 1.8 |
| >hg38_21_+_8433172_8446622_RNA45SN1_45S_50ntFlanks@12413.12432.20\|TCCTTCGATGTCGGCTCTTC | >hg38_21_+_8433172_8446622_RNA45SN1_45S_50ntFlanks | 12413 | 12432 | 20 | TCCTTCGATGTCGGCTCTTC | MB468 | 42 | 71 | 57 | 9 | 17 | 11.6 | 32.5 | 20.2 | 4.0 | 3.9 |
| >hg38_21_+_8433172_8446622_RNA45SN1_45S_50ntFlanks@12413.12434.22\|TCCTTCGATGTCGGCTCTTCCT | >hg38_21_+_8433172_8446622_RNA45SN1_45S_50ntFlanks | 12413 | 12434 | 22 | TCCTTCGATGTCGGCTCTTCCT | BT20 | 51 | 44 | 34 | 2 | 4 | 84.8 | 73.5 | 54.8 | 2.2 | 4.3 |
| >hg38_21_+_8433172_8446622_RNA45SN1_45S_50ntFlanks@12413.12434.22\|TCCTTCGATGTCGGCTCTTCCT | >hg38_21_+_8433172_8446622_RNA45SN1_45S_50ntFlanks | 12413 | 12434 | 22 | TCCTTCGATGTCGGCTCTTCCT | MB468 | 21 | 50 | 24 | 3 | 11 | 8.7 | 16.0 | 14.7 | 0.8 | 0.7 |
| >hg38_21_+_8433172_8446622_RNA45SN1_45S_50ntFlanks@12413.12435.23\|TCCTTCGATGTCGGCTCTTCCTA | >hg38_21_+_8433172_8446622_RNA45SN1_45S_50ntFlanks | 12413 | 12435 | 23 | TCCTTCGATGTCGGCTCTTCCTA | MB468 | 11 | 35 | 9 | 2 | 6 | 4.0 | 22.4 | 6.0 | 1.7 | 4.1 |
| >hg38_21_+_8433172_8446622_RNA45SN1_45S_50ntFlanks@12413.12436.24\|TCCTTCGATGTCGGCTCTTCCTAT | >hg38_21_+_8433172_8446622_RNA45SN1_45S_50ntFlanks | 12413 | 12436 | 24 | TCCTTCGATGTCGGCTCTTCCTAT | MB468 | 21 | 68 | 16 | 1 | 19 | 8.5 | 40.2 | 10.3 | 0.6 | 8.0 |
| >hg38_21_+_8433172_8446622_RNA45SN1_45S_50ntFlanks@12413.12443.31\|TCCTTCGATGTCGGCTCTTCCTATCATTGTG | >hg38_21_+_8433172_8446622_RNA45SN1_45S_50ntFlanks | 12413 | 12443 | 31 | TCCTTCGATGTCGGCTCTTCCTATCATTGTG | MB468 | 18 | 55 | 35 | 2 | 13 | 7.7 | 20.8 | 9.4 | 0.8 | 3.0 |
| >hg38_21_+_8433172_8446622_RNA45SN1_45S_50ntFlanks@12414.12434.21\|CCTTCGATGTCGGCTCTTCCT | >hg38_21_+_8433172_8446622_RNA45SN1_45S_50ntFlanks | 12414 | 12434 | 21 | CCTTCGATGTCGGCTCTTCCT | BT20 | 10 | 16 | 18 | 4 | 4 | 12.1 | 19.8 | 23.7 | 6.3 | 3.7 |
| >hg38_21_+_8433172_8446622_RNA45SN1_45S_50ntFlanks@12414.12448.35\|CCTTCGATGTCGGCTCTTCCTATCATTGTGAAGCA | >hg38_21_+_8433172_8446622_RNA45SN1_45S_50ntFlanks | 12414 | 12448 | 35 | CCTTCGATGTCGGCTCTTCCTATCATTGTGAAGCA | MB231 | 20 | 14 | 50 | 0 | 8 | 14.3 | 6.0 | 38.2 | 0.4 | 13.6 |
| >hg38_21_+_8433172_8446622_RNA45SN1_45S_50ntFlanks@12415.12432.18\|CTTCGATGTCGGCTCTTC | >hg38_21_+_8433172_8446622_RNA45SN1_45S_50ntFlanks | 12415 | 12432 | 18 | CTTCGATGTCGGCTCTTC | MB231 | 16 | 16 | 89 | 1 | 4 | 15.3 | 10.7 | 65.2 | 1.0 | 6.6 |
| >hg38_21_+_8433172_8446622_RNA45SN1_45S_50ntFlanks@12415.12432.18\|CTTCGATGTCGGCTCTTC | >hg38_21_+_8433172_8446622_RNA45SN1_45S_50ntFlanks | 12415 | 12432 | 18 | CTTCGATGTCGGCTCTTC | MB468 | 339 | 405 | 540 | 93 | 127 | 79.0 | 119.2 | 73.5 | 39.5 | 19.1 |
| >hg38_21_+_8433172_8446622_RNA45SN1_45S_50ntFlanks@12415.12433.19\|CTTCGATGTCGGCTCTTCC | >hg38_21_+_8433172_8446622_RNA45SN1_45S_50ntFlanks | 12415 | 12433 | 19 | CTTCGATGTCGGCTCTTCC | BT20 | 23 | 21 | 61 | 19 | 18 | 19.0 | 14.0 | 66.6 | 28.3 | 3.2 |
| >hg38_21_+_8433172_8446622_RNA45SN1_45S_50ntFlanks@12415.12433.19\|CTTCGATGTCGGCTCTTCC | >hg38_21_+_8433172_8446622_RNA45SN1_45S_50ntFlanks | 12415 | 12433 | 19 | CTTCGATGTCGGCTCTTCC | MB231 | 30 | 42 | 159 | 2 | 14 | 28.1 | 26.7 | 115.7 | 2.0 | 12.9 |
| >hg38_21_+_8433172_8446622_RNA45SN1_45S_50ntFlanks@12415.12433.19\|CTTCGATGTCGGCTCTTCC | >hg38_21_+_8433172_8446622_RNA45SN1_45S_50ntFlanks | 12415 | 12433 | 19 | CTTCGATGTCGGCTCTTCC | MB468 | 53 | 91 | 88 | 19 | 28 | 16.3 | 23.4 | 31.8 | 6.6 | 4.8 |
| >hg38_21_+_8433172_8446622_RNA45SN1_45S_50ntFlanks@12415.12434.20\|CTTCGATGTCGGCTCTTCCT | >hg38_21_+_8433172_8446622_RNA45SN1_45S_50ntFlanks | 12415 | 12434 | 20 | CTTCGATGTCGGCTCTTCCT | BT20 | 198 | 233 | 120 | 17 | 33 | 306.0 | 321.5 | 160.7 | 25.5 | 13.3 |
| >hg38_21_+_8433172_8446622_RNA45SN1_45S_50ntFlanks@12415.12434.20\|CTTCGATGTCGGCTCTTCCT | >hg38_21_+_8433172_8446622_RNA45SN1_45S_50ntFlanks | 12415 | 12434 | 20 | CTTCGATGTCGGCTCTTCCT | MB231 | 25 | 69 | 113 | 2 | 19 | 14.9 | 54.2 | 72.9 | 1.3 | 15.5 |
| >hg38_21_+_8433172_8446622_RNA45SN1_45S_50ntFlanks@12415.12434.20\|CTTCGATGTCGGCTCTTCCT | >hg38_21_+_8433172_8446622_RNA45SN1_45S_50ntFlanks | 12415 | 12434 | 20 | CTTCGATGTCGGCTCTTCCT | MB468 | 206 | 368 | 309 | 48 | 166 | 67.6 | 55.0 | 116.0 | 14.6 | 54.1 |
| >hg38_21_+_8433172_8446622_RNA45SN1_45S_50ntFlanks@12415.12435.21\|CTTCGATGTCGGCTCTTCCTA | >hg38_21_+_8433172_8446622_RNA45SN1_45S_50ntFlanks | 12415 | 12435 | 21 | CTTCGATGTCGGCTCTTCCTA | BT20 | 11 | 13 | 6 | 2 | 1 | 15.7 | 19.4 | 4.9 | 2.7 | 2.1 |
| >hg38_21_+_8433172_8446622_RNA45SN1_45S_50ntFlanks@12415.12435.21\|CTTCGATGTCGGCTCTTCCTA | >hg38_21_+_8433172_8446622_RNA45SN1_45S_50ntFlanks | 12415 | 12435 | 21 | CTTCGATGTCGGCTCTTCCTA | MB468 | 177 | 387 | 192 | 34 | 201 | 28.4 | 169.5 | 54.0 | 5.9 | 102.1 |
| >hg38_21_+_8433172_8446622_RNA45SN1_45S_50ntFlanks@12415.12436.22\|CTTCGATGTCGGCTCTTCCTAT | >hg38_21_+_8433172_8446622_RNA45SN1_45S_50ntFlanks | 12415 | 12436 | 22 | CTTCGATGTCGGCTCTTCCTAT | BT20 | 19 | 16 | 10 | 2 | 1 | 27.5 | 24.7 | 9.3 | 3.6 | 1.3 |
| >hg38_21_+_8433172_8446622_RNA45SN1_45S_50ntFlanks@12415.12436.22\|CTTCGATGTCGGCTCTTCCTAT | >hg38_21_+_8433172_8446622_RNA45SN1_45S_50ntFlanks | 12415 | 12436 | 22 | CTTCGATGTCGGCTCTTCCTAT | MB468 | 314 | 786 | 344 | 35 | 602 | 68.8 | 274.5 | 85.9 | 13.3 | 301.6 |
| >hg38_21_+_8433172_8446622_RNA45SN1_45S_50ntFlanks@12415.12437.23\|CTTCGATGTCGGCTCTTCCTATC | >hg38_21_+_8433172_8446622_RNA45SN1_45S_50ntFlanks | 12415 | 12437 | 23 | CTTCGATGTCGGCTCTTCCTATC | BT20 | 25 | 20 | 38 | 5 | 27 | 17.3 | 13.4 | 31.7 | 5.0 | 2.9 |
| >hg38_21_+_8433172_8446622_RNA45SN1_45S_50ntFlanks@12415.12437.23\|CTTCGATGTCGGCTCTTCCTATC | >hg38_21_+_8433172_8446622_RNA45SN1_45S_50ntFlanks | 12415 | 12437 | 23 | CTTCGATGTCGGCTCTTCCTATC | MB231 | 29 | 36 | 86 | 1 | 14 | 24.0 | 29.5 | 70.1 | 1.9 | 25.1 |
| >hg38_21_+_8433172_8446622_RNA45SN1_45S_50ntFlanks@12415.12437.23\|CTTCGATGTCGGCTCTTCCTATC | >hg38_21_+_8433172_8446622_RNA45SN1_45S_50ntFlanks | 12415 | 12437 | 23 | CTTCGATGTCGGCTCTTCCTATC | MB468 | 101 | 221 | 152 | 18 | 217 | 22.5 | 31.1 | 43.4 | 5.6 | 84.3 |
| >hg38_21_+_8433172_8446622_RNA45SN1_45S_50ntFlanks@12415.12438.24\|CTTCGATGTCGGCTCTTCCTATCA | >hg38_21_+_8433172_8446622_RNA45SN1_45S_50ntFlanks | 12415 | 12438 | 24 | CTTCGATGTCGGCTCTTCCTATCA | BT20 | 30 | 34 | 22 | 1 | 9 | 32.9 | 28.6 | 17.4 | 2.5 | 14.9 |
| >hg38_21_+_8433172_8446622_RNA45SN1_45S_50ntFlanks@12415.12438.24\|CTTCGATGTCGGCTCTTCCTATCA | >hg38_21_+_8433172_8446622_RNA45SN1_45S_50ntFlanks | 12415 | 12438 | 24 | CTTCGATGTCGGCTCTTCCTATCA | MB231 | 17 | 26 | 46 | 0 | 12 | 12.3 | 24.1 | 48.2 | 0.7 | 20.2 |
| >hg38_21_+_8433172_8446622_RNA45SN1_45S_50ntFlanks@12415.12439.25\|CTTCGATGTCGGCTCTTCCTATCAT | >hg38_21_+_8433172_8446622_RNA45SN1_45S_50ntFlanks | 12415 | 12439 | 25 | CTTCGATGTCGGCTCTTCCTATCAT | BT20 | 30 | 26 | 24 | 2 | 3 | 34.7 | 27.6 | 22.7 | 3.0 | 4.8 |
| >hg38_21_+_8433172_8446622_RNA45SN1_45S_50ntFlanks@12415.12439.25\|CTTCGATGTCGGCTCTTCCTATCAT | >hg38_21_+_8433172_8446622_RNA45SN1_45S_50ntFlanks | 12415 | 12439 | 25 | CTTCGATGTCGGCTCTTCCTATCAT | MB231 | 11 | 12 | 35 | 0 | 6 | 9.1 | 11.4 | 28.4 | 0.5 | 10.4 |
| >hg38_21_+_8433172_8446622_RNA45SN1_45S_50ntFlanks@12415.12440.26\|CTTCGATGTCGGCTCTTCCTATCATT | >hg38_21_+_8433172_8446622_RNA45SN1_45S_50ntFlanks | 12415 | 12440 | 26 | CTTCGATGTCGGCTCTTCCTATCATT | BT20 | 13 | 10 | 22 | 2 | 7 | 9.0 | 5.6 | 18.9 | 3.1 | 6.5 |

| | | | | | | | | | | | | | | | | |
|---|---|---|---|---|---|---|---|---|---|---|---|---|---|---|---|---|
| >hg38_21_+_8433172_8446622_RNA45SN1_45S_50ntFlanks@12415.12440.26\|CTTCGATGTCGGCTCTTCCTATCATT | >hg38_21_+_8433172_8446622_RNA45SN1_45S_50ntFlanks | 12415 | 12440 | 26 | CTTCGATGTCGGCTCTTCCTATCATT | MB231 | 18 | 17 | 56 | 0 | 10 | 17.8 | 19.3 | 58.5 | 0.6 | 17.7 |
| >hg38_21_+_8433172_8446622_RNA45SN1_45S_50ntFlanks@12415.12441.27\|CTTCGATGTCGGCTCTTCCTATCATTG | >hg38_21_+_8433172_8446622_RNA45SN1_45S_50ntFlanks | 12415 | 12441 | 27 | CTTCGATGTCGGCTCTTCCTATCATTG | MB231 | 15 | 19 | 51 | 0 | 11 | 16.8 | 22.9 | 62.0 | 0.7 | 19.5 |
| >hg38_21_+_8433172_8446622_RNA45SN1_45S_50ntFlanks@12415.12441.27\|CTTCGATGTCGGCTCTTCCTATCATTG | >hg38_21_+_8433172_8446622_RNA45SN1_45S_50ntFlanks | 12415 | 12441 | 27 | CTTCGATGTCGGCTCTTCCTATCATTG | MB468 | 84 | 196 | 140 | 11 | 206 | 30.0 | 28.4 | 46.2 | 7.0 | 85.2 |
| >hg38_21_+_8433172_8446622_RNA45SN1_45S_50ntFlanks@12415.12442.28\|CTTCGATGTCGGCTCTTCCTATCATTGT | >hg38_21_+_8433172_8446622_RNA45SN1_45S_50ntFlanks | 12415 | 12442 | 28 | CTTCGATGTCGGCTCTTCCTATCATTGT | MB231 | 16 | 19 | 66 | 0 | 10 | 15.4 | 18.9 | 69.9 | 0.7 | 16.9 |
| >hg38_21_+_8433172_8446622_RNA45SN1_45S_50ntFlanks@12415.12443.29\|CTTCGATGTCGGCTCTTCCTATCATTGTG | >hg38_21_+_8433172_8446622_RNA45SN1_45S_50ntFlanks | 12415 | 12443 | 29 | CTTCGATGTCGGCTCTTCCTATCATTGTG | BT20 | 12 | 10 | 21 | 8 | 10 | 8.2 | 5.9 | 10.7 | 13.1 | 9.0 |
| >hg38_21_+_8433172_8446622_RNA45SN1_45S_50ntFlanks@12415.12443.29\|CTTCGATGTCGGCTCTTCCTATCATTGTG | >hg38_21_+_8433172_8446622_RNA45SN1_45S_50ntFlanks | 12415 | 12443 | 29 | CTTCGATGTCGGCTCTTCCTATCATTGTG | MB231 | 27 | 32 | 81 | 1 | 22 | 32.1 | 39.0 | 102.5 | 1.6 | 37.9 |
| >hg38_21_+_8433172_8446622_RNA45SN1_45S_50ntFlanks@12415.12443.29\|CTTCGATGTCGGCTCTTCCTATCATTGTG | >hg38_21_+_8433172_8446622_RNA45SN1_45S_50ntFlanks | 12415 | 12443 | 29 | CTTCGATGTCGGCTCTTCCTATCATTGTG | MB468 | 145 | 235 | 324 | 42 | 508 | 47.2 | 43.4 | 61.8 | 29.2 | 214.6 |
| >hg38_21_+_8433172_8446622_RNA45SN1_45S_50ntFlanks@12415.12444.30\|CTTCGATGTCGGCTCTTCCTATCATTGTGA | >hg38_21_+_8433172_8446622_RNA45SN1_45S_50ntFlanks | 12415 | 12444 | 30 | CTTCGATGTCGGCTCTTCCTATCATTGTGA | BT20 | 13 | 10 | 26 | 17 | 15 | 9.7 | 4.8 | 18.3 | 28.1 | 13.1 |
| >hg38_21_+_8433172_8446622_RNA45SN1_45S_50ntFlanks@12415.12444.30\|CTTCGATGTCGGCTCTTCCTATCATTGTGA | >hg38_21_+_8433172_8446622_RNA45SN1_45S_50ntFlanks | 12415 | 12444 | 30 | CTTCGATGTCGGCTCTTCCTATCATTGTGA | MB231 | 24 | 31 | 67 | 1 | 22 | 22.9 | 32.4 | 68.5 | 0.9 | 37.2 |
| >hg38_21_+_8433172_8446622_RNA45SN1_45S_50ntFlanks@12415.12444.30\|CTTCGATGTCGGCTCTTCCTATCATTGTGA | >hg38_21_+_8433172_8446622_RNA45SN1_45S_50ntFlanks | 12415 | 12444 | 30 | CTTCGATGTCGGCTCTTCCTATCATTGTGA | MB468 | 36 | 74 | 80 | 9 | 90 | 14.7 | 21.1 | 27.1 | 6.4 | 32.8 |
| >hg38_21_+_8433172_8446622_RNA45SN1_45S_50ntFlanks@12415.12445.31\|CTTCGATGTCGGCTCTTCCTATCATTGTGAA | >hg38_21_+_8433172_8446622_RNA45SN1_45S_50ntFlanks | 12415 | 12445 | 31 | CTTCGATGTCGGCTCTTCCTATCATTGTGAA | MB468 | 19 | 28 | 33 | 3 | 38 | 7.6 | 6.9 | 9.2 | 2.1 | 10.4 |
| >hg38_21_+_8433172_8446622_RNA45SN1_45S_50ntFlanks@12415.12446.32\|CTTCGATGTCGGCTCTTCCTATCATTGTGAAG | >hg38_21_+_8433172_8446622_RNA45SN1_45S_50ntFlanks | 12415 | 12446 | 32 | CTTCGATGTCGGCTCTTCCTATCATTGTGAAG | MB231 | 21 | 16 | 41 | 0 | 11 | 27.0 | 14.5 | 47.6 | 0.7 | 18.4 |
| >hg38_21_+_8433172_8446622_RNA45SN1_45S_50ntFlanks@12415.12446.32\|CTTCGATGTCGGCTCTTCCTATCATTGTGAAG | >hg38_21_+_8433172_8446622_RNA45SN1_45S_50ntFlanks | 12415 | 12446 | 32 | CTTCGATGTCGGCTCTTCCTATCATTGTGAAG | MB468 | 42 | 41 | 75 | 9 | 99 | 8.5 | 16.5 | 26.9 | 8.2 | 32.9 |
| >hg38_21_+_8433172_8446622_RNA45SN1_45S_50ntFlanks@12415.12447.33\|CTTCGATGTCGGCTCTTCCTATCATTGTGAAGC | >hg38_21_+_8433172_8446622_RNA45SN1_45S_50ntFlanks | 12415 | 12447 | 33 | CTTCGATGTCGGCTCTTCCTATCATTGTGAAGC | MB231 | 26 | 21 | 53 | 1 | 16 | 25.8 | 12.7 | 40.5 | 0.5 | 18.6 |
| >hg38_21_+_8433172_8446622_RNA45SN1_45S_50ntFlanks@12415.12448.34\|CTTCGATGTCGGCTCTTCCTATCATTGTGAAGCA | >hg38_21_+_8433172_8446622_RNA45SN1_45S_50ntFlanks | 12415 | 12448 | 34 | CTTCGATGTCGGCTCTTCCTATCATTGTGAAGCA | BT20 | 20 | 21 | 41 | 7 | 47 | 9.6 | 11.9 | 29.3 | 8.4 | 19.9 |
| >hg38_21_+_8433172_8446622_RNA45SN1_45S_50ntFlanks@12415.12448.34\|CTTCGATGTCGGCTCTTCCTATCATTGTGAAGCA | >hg38_21_+_8433172_8446622_RNA45SN1_45S_50ntFlanks | 12415 | 12448 | 34 | CTTCGATGTCGGCTCTTCCTATCATTGTGAAGCA | MB231 | 67 | 38 | 139 | 2 | 41 | 65.7 | 32.3 | 102.1 | 1.1 | 27.8 |
| >hg38_21_+_8433172_8446622_RNA45SN1_45S_50ntFlanks@12415.12449.35\|CTTCGATGTCGGCTCTTCCTATCATTGTGAAGCAG | >hg38_21_+_8433172_8446622_RNA45SN1_45S_50ntFlanks | 12415 | 12449 | 35 | CTTCGATGTCGGCTCTTCCTATCATTGTGAAGCAG | MB231 | 31 | 18 | 52 | 2 | 18 | 28.8 | 15.6 | 43.6 | 1.8 | 17.2 |
| >hg38_21_+_8433172_8446622_RNA45SN1_45S_50ntFlanks@12415.12450.36\|CTTCGATGTCGGCTCTTCCTATCATTGTGAAGCAGA | >hg38_21_+_8433172_8446622_RNA45SN1_45S_50ntFlanks | 12415 | 12450 | 36 | CTTCGATGTCGGCTCTTCCTATCATTGTGAAGCAGA | MB231 | 11 | 7 | 10 | 0 | 5 | 7.8 | 4.8 | 6.1 | 0.6 | 9.0 |
| >hg38_21_+_8433172_8446622_RNA45SN1_45S_50ntFlanks@12415.12452.38\|CTTCGATGTCGGCTCTTCCTATCATTGTGAAGCAGAAT | >hg38_21_+_8433172_8446622_RNA45SN1_45S_50ntFlanks | 12415 | 12452 | 38 | CTTCGATGTCGGCTCTTCCTATCATTGTGAAGCAGAAT | MB231 | 12 | 7 | 11 | 1 | 7 | 8.1 | 3.9 | 8.3 | 0.5 | 7.0 |
| >hg38_21_+_8433172_8446622_RNA45SN1_45S_50ntFlanks@12415.12453.39\|CTTCGATGTCGGCTCTTCCTATCATTGTGAAGCAGAATT | >hg38_21_+_8433172_8446622_RNA45SN1_45S_50ntFlanks | 12415 | 12453 | 39 | CTTCGATGTCGGCTCTTCCTATCATTGTGAAGCAGAATT | MB231 | 10 | 6 | 10 | 0 | 4 | 8.5 | 2.3 | 7.7 | 0.3 | 6.7 |
| >hg38_21_+_8433172_8446622_RNA45SN1_45S_50ntFlanks@12415.12455.41\|CTTCGATGTCGGCTCTTCCTATCATTGTGAAGCAGAATTCA | >hg38_21_+_8433172_8446622_RNA45SN1_45S_50ntFlanks | 12415 | 12455 | 41 | CTTCGATGTCGGCTCTTCCTATCATTGTGAAGCAGAATTCA | MB468 | 19 | 24 | 39 | 10 | 154 | 3.7 | 22.1 | 47.5 | 12.8 | 123.7 |
| >hg38_21_+_8433172_8446622_RNA45SN1_45S_50ntFlanks@12415.12457.43\|CTTCGATGTCGGCTCTTCCTATCATTGTGAAGCAGAATTCACC | >hg38_21_+_8433172_8446622_RNA45SN1_45S_50ntFlanks | 12415 | 12457 | 43 | CTTCGATGTCGGCTCTTCCTATCATTGTGAAGCAGAATTCACC | MB231 | 11 | 7 | 18 | 0 | 0 | 6.6 | 5.0 | 15.6 | 0.0 | 0.0 |
| >hg38_21_+_8433172_8446622_RNA45SN1_45S_50ntFlanks@12416.12433.18\|TTCGATGTCGGCTCTTCC | >hg38_21_+_8433172_8446622_RNA45SN1_45S_50ntFlanks | 12416 | 12433 | 18 | TTCGATGTCGGCTCTTCC | BT20 | 26 | 12 | 111 | 14 | 12 | 16.8 | 8.9 | 123.8 | 12.0 | 3.6 |
| >hg38_21_+_8433172_8446622_RNA45SN1_45S_50ntFlanks@12416.12433.18\|TTCGATGTCGGCTCTTCC | >hg38_21_+_8433172_8446622_RNA45SN1_45S_50ntFlanks | 12416 | 12433 | 18 | TTCGATGTCGGCTCTTCC | MB231 | 103 | 65 | 569 | 8 | 18 | 93.3 | 46.8 | 474.4 | 4.8 | 12.5 |
| >hg38_21_+_8433172_8446622_RNA45SN1_45S_50ntFlanks@12416.12433.18\|TTCGATGTCGGCTCTTCC | >hg38_21_+_8433172_8446622_RNA45SN1_45S_50ntFlanks | 12416 | 12433 | 18 | TTCGATGTCGGCTCTTCC | MB468 | 10 | 12 | 21 | 4 | 3 | 4.3 | 7.0 | 9.4 | 1.8 | 0.5 |
| >hg38_21_+_8433172_8446622_RNA45SN1_45S_50ntFlanks@12416.12434.19\|TTCGATGTCGGCTCTTCCT | >hg38_21_+_8433172_8446622_RNA45SN1_45S_50ntFlanks | 12416 | 12434 | 19 | TTCGATGTCGGCTCTTCCT | BT20 | 28 | 22 | 93 | 11 | 14 | 16.7 | 9.9 | 97.4 | 11.6 | 1.2 |
| >hg38_21_+_8433172_8446622_RNA45SN1_45S_50ntFlanks@12416.12434.19\|TTCGATGTCGGCTCTTCCT | >hg38_21_+_8433172_8446622_RNA45SN1_45S_50ntFlanks | 12416 | 12434 | 19 | TTCGATGTCGGCTCTTCCT | MB231 | 85 | 59 | 400 | 7 | 24 | 66.9 | 37.2 | 366.1 | 4.5 | 9.9 |
| >hg38_21_+_8433172_8446622_RNA45SN1_45S_50ntFlanks@12416.12434.19\|TTCGATGTCGGCTCTTCCT | >hg38_21_+_8433172_8446622_RNA45SN1_45S_50ntFlanks | 12416 | 12434 | 19 | TTCGATGTCGGCTCTTCCT | MB468 | 39 | 48 | 76 | 10 | 15 | 16.0 | 19.9 | 31.5 | 3.0 | 1.2 |
| >hg38_21_+_8433172_8446622_RNA45SN1_45S_50ntFlanks@12416.12435.20\|TTCGATGTCGGCTCTTCCTA | >hg38_21_+_8433172_8446622_RNA45SN1_45S_50ntFlanks | 12416 | 12435 | 20 | TTCGATGTCGGCTCTTCCTA | MB468 | 58 | 71 | 81 | 9 | 24 | 15.9 | 15.3 | 16.8 | 2.5 | 1.7 |
| >hg38_21_+_8433172_8446622_RNA45SN1_45S_50ntFlanks@12416.12436.21\|TTCGATGTCGGCTCTTCCTAT | >hg38_21_+_8433172_8446622_RNA45SN1_45S_50ntFlanks | 12416 | 12436 | 21 | TTCGATGTCGGCTCTTCCTAT | MB231 | 20 | 21 | 51 | 1 | 7 | 15.5 | 20.6 | 46.4 | 0.9 | 11.7 |
| >hg38_21_+_8433172_8446622_RNA45SN1_45S_50ntFlanks@12416.12436.21\|TTCGATGTCGGCTCTTCCTAT | >hg38_21_+_8433172_8446622_RNA45SN1_45S_50ntFlanks | 12416 | 12436 | 21 | TTCGATGTCGGCTCTTCCTAT | MB468 | 80 | 113 | 95 | 8 | 57 | 31.0 | 18.1 | 13.8 | 1.9 | 14.1 |
| >hg38_21_+_8433172_8446622_RNA45SN1_45S_50ntFlanks@12416.12437.22\|TTCGATGTCGGCTCTTCCTATC | >hg38_21_+_8433172_8446622_RNA45SN1_45S_50ntFlanks | 12416 | 12437 | 22 | TTCGATGTCGGCTCTTCCTATC | BT20 | 30 | 14 | 75 | 6 | 20 | 15.5 | 7.3 | 67.5 | 4.6 | 6.3 |
| >hg38_21_+_8433172_8446622_RNA45SN1_45S_50ntFlanks@12416.12437.22\|TTCGATGTCGGCTCTTCCTATC | >hg38_21_+_8433172_8446622_RNA45SN1_45S_50ntFlanks | 12416 | 12437 | 22 | TTCGATGTCGGCTCTTCCTATC | MB231 | 130 | 100 | 375 | 4 | 51 | 113.7 | 89.8 | 328.5 | 4.4 | 49.3 |
| >hg38_21_+_8433172_8446622_RNA45SN1_45S_50ntFlanks@12416.12437.22\|TTCGATGTCGGCTCTTCCTATC | >hg38_21_+_8433172_8446622_RNA45SN1_45S_50ntFlanks | 12416 | 12437 | 22 | TTCGATGTCGGCTCTTCCTATC | MB468 | 25 | 36 | 38 | 3 | 20 | 10.7 | 5.9 | 10.5 | 1.4 | 3.8 |
| >hg38_21_+_8433172_8446622_RNA45SN1_45S_50ntFlanks@12416.12438.23\|TTCGATGTCGGCTCTTCCTATCA | >hg38_21_+_8433172_8446622_RNA45SN1_45S_50ntFlanks | 12416 | 12438 | 23 | TTCGATGTCGGCTCTTCCTATCA | BT20 | 22 | 17 | 42 | 3 | 22 | 13.2 | 7.5 | 23.3 | 2.6 | 5.1 |
| >hg38_21_+_8433172_8446622_RNA45SN1_45S_50ntFlanks@12416.12438.23\|TTCGATGTCGGCTCTTCCTATCA | >hg38_21_+_8433172_8446622_RNA45SN1_45S_50ntFlanks | 12416 | 12438 | 23 | TTCGATGTCGGCTCTTCCTATCA | MB231 | 83 | 85 | 231 | 2 | 52 | 70.8 | 97.4 | 268.4 | 3.2 | 64.9 |
| >hg38_21_+_8433172_8446622_RNA45SN1_45S_50ntFlanks@12416.12439.24\|TTCGATGTCGGCTCTTCCTATCAT | >hg38_21_+_8433172_8446622_RNA45SN1_45S_50ntFlanks | 12416 | 12439 | 24 | TTCGATGTCGGCTCTTCCTATCAT | BT20 | 18 | 10 | 36 | 2 | 6 | 7.6 | 4.1 | 23.1 | 2.7 | 5.8 |
| >hg38_21_+_8433172_8446622_RNA45SN1_45S_50ntFlanks@12416.12439.24\|TTCGATGTCGGCTCTTCCTATCAT | >hg38_21_+_8433172_8446622_RNA45SN1_45S_50ntFlanks | 12416 | 12439 | 24 | TTCGATGTCGGCTCTTCCTATCAT | MB231 | 54 | 40 | 158 | 1 | 20 | 53.4 | 40.4 | 141.5 | 1.4 | 34.9 |
| >hg38_21_+_8433172_8446622_RNA45SN1_45S_50ntFlanks@12416.12440.25\|TTCGATGTCGGCTCTTCCTATCATT | >hg38_21_+_8433172_8446622_RNA45SN1_45S_50ntFlanks | 12416 | 12440 | 25 | TTCGATGTCGGCTCTTCCTATCATT | BT20 | 23 | 10 | 45 | 3 | 16 | 13.3 | 4.3 | 28.5 | 1.8 | 8.6 |
| >hg38_21_+_8433172_8446622_RNA45SN1_45S_50ntFlanks@12416.12440.25\|TTCGATGTCGGCTCTTCCTATCATT | >hg38_21_+_8433172_8446622_RNA45SN1_45S_50ntFlanks | 12416 | 12440 | 25 | TTCGATGTCGGCTCTTCCTATCATT | MB231 | 84 | 58 | 229 | 1 | 39 | 94.8 | 65.6 | 256.3 | 2.1 | 52.3 |
| >hg38_21_+_8433172_8446622_RNA45SN1_45S_50ntFlanks@12416.12441.26\|TTCGATGTCGGCTCTTCCTATCATTG | >hg38_21_+_8433172_8446622_RNA45SN1_45S_50ntFlanks | 12416 | 12441 | 26 | TTCGATGTCGGCTCTTCCTATCATTG | BT20 | 11 | 7 | 22 | 2 | 6 | 6.2 | 2.2 | 8.2 | 2.7 | 4.9 |
| >hg38_21_+_8433172_8446622_RNA45SN1_45S_50ntFlanks@12416.12441.26\|TTCGATGTCGGCTCTTCCTATCATTG | >hg38_21_+_8433172_8446622_RNA45SN1_45S_50ntFlanks | 12416 | 12441 | 26 | TTCGATGTCGGCTCTTCCTATCATTG | MB231 | 53 | 47 | 138 | 1 | 25 | 61.0 | 57.2 | 172.1 | 2.2 | 42.7 |
| >hg38_21_+_8433172_8446622_RNA45SN1_45S_50ntFlanks@12416.12441.26\|TTCGATGTCGGCTCTTCCTATCATTG | >hg38_21_+_8433172_8446622_RNA45SN1_45S_50ntFlanks | 12416 | 12441 | 26 | TTCGATGTCGGCTCTTCCTATCATTG | MB468 | 20 | 26 | 38 | 3 | 17 | 8.7 | 8.7 | 14.4 | 1.6 | 3.4 |
| >hg38_21_+_8433172_8446622_RNA45SN1_45S_50ntFlanks@12416.12442.27\|TTCGATGTCGGCTCTTCCTATCATTGT | >hg38_21_+_8433172_8446622_RNA45SN1_45S_50ntFlanks | 12416 | 12442 | 27 | TTCGATGTCGGCTCTTCCTATCATTGT | BT20 | 14 | 7 | 37 | 2 | 7 | 8.5 | 2.2 | 20.6 | 3.3 | 6.9 |
| >hg38_21_+_8433172_8446622_RNA45SN1_45S_50ntFlanks@12416.12442.27\|TTCGATGTCGGCTCTTCCTATCATTGT | >hg38_21_+_8433172_8446622_RNA45SN1_45S_50ntFlanks | 12416 | 12442 | 27 | TTCGATGTCGGCTCTTCCTATCATTGT | MB231 | 58 | 40 | 164 | 1 | 20 | 53.9 | 43.1 | 183.5 | 2.0 | 35.4 |
| >hg38_21_+_8433172_8446622_RNA45SN1_45S_50ntFlanks@12416.12443.28\|TTCGATGTCGGCTCTTCCTATCATTGTG | >hg38_21_+_8433172_8446622_RNA45SN1_45S_50ntFlanks | 12416 | 12443 | 28 | TTCGATGTCGGCTCTTCCTATCATTGTG | BT20 | 14 | 9 | 28 | 3 | 14 | 8.7 | 4.0 | 14.8 | 1.7 | 6.3 |
| >hg38_21_+_8433172_8446622_RNA45SN1_45S_50ntFlanks@12416.12443.28\|TTCGATGTCGGCTCTTCCTATCATTGTG | >hg38_21_+_8433172_8446622_RNA45SN1_45S_50ntFlanks | 12416 | 12443 | 28 | TTCGATGTCGGCTCTTCCTATCATTGTG | MB231 | 73 | 47 | 171 | 2 | 42 | 78.0 | 59.0 | 226.6 | 3.5 | 54.3 |
| >hg38_21_+_8433172_8446622_RNA45SN1_45S_50ntFlanks@12416.12443.28\|TTCGATGTCGGCTCTTCCTATCATTGTG | >hg38_21_+_8433172_8446622_RNA45SN1_45S_50ntFlanks | 12416 | 12443 | 28 | TTCGATGTCGGCTCTTCCTATCATTGTG | MB468 | 33 | 30 | 75 | 8 | 41 | 13.0 | 7.8 | 19.6 | 4.5 | 16.5 |
| >hg38_21_+_8433172_8446622_RNA45SN1_45S_50ntFlanks@12416.12444.29\|TTCGATGTCGGCTCTTCCTATCATTGTGA | >hg38_21_+_8433172_8446622_RNA45SN1_45S_50ntFlanks | 12416 | 12444 | 29 | TTCGATGTCGGCTCTTCCTATCATTGTGA | BT20 | 18 | 11 | 46 | 5 | 11 | 11.3 | 4.5 | 34.7 | 6.6 | 10.4 |
| >hg38_21_+_8433172_8446622_RNA45SN1_45S_50ntFlanks@12416.12444.29\|TTCGATGTCGGCTCTTCCTATCATTGTGA | >hg38_21_+_8433172_8446622_RNA45SN1_45S_50ntFlanks | 12416 | 12444 | 29 | TTCGATGTCGGCTCTTCCTATCATTGTGA | MB231 | 72 | 54 | 190 | 2 | 52 | 63.2 | 57.7 | 193.0 | 2.9 | 58.8 |

| | | | | | | | | | | | | | | | | |
|---|---|---|---|---|---|---|---|---|---|---|---|---|---|---|---|---|
| >hg38_21_+_8433172_8446622_RNA45SN1_45S_50ntFlanks@12416.12444.29\|TTCGATGTCGGCTCTTCCTATCATTGTGA | >hg38_21_+_8433172_8446622_RNA45SN1_45S_50ntFlanks | 12416 | 12444 | 29 | TTCGATGTCGGCTCTTCCTATCATTGTGA | MB468 | 11 | 15 | 24 | 2 | 11 | 5.8 | 8.4 | 10.6 | 1.2 | 4.2 |
| >hg38_21_+_8433172_8446622_RNA45SN1_45S_50ntFlanks@12416.12445.30\|TTCGATGTCGGCTCTTCCTATCATTGTGAA | >hg38_21_+_8433172_8446622_RNA45SN1_45S_50ntFlanks | 12416 | 12445 | 30 | TTCGATGTCGGCTCTTCCTATCATTGTGAA | MB231 | 28 | 22 | 65 | 1 | 15 | 22.9 | 18.4 | 63.2 | 1.1 | 25.2 |
| >hg38_21_+_8433172_8446622_RNA45SN1_45S_50ntFlanks@12416.12446.31\|TTCGATGTCGGCTCTTCCTATCATTGTGAAG | >hg38_21_+_8433172_8446622_RNA45SN1_45S_50ntFlanks | 12416 | 12446 | 31 | TTCGATGTCGGCTCTTCCTATCATTGTGAAG | MB231 | 73 | 28 | 163 | 3 | 30 | 85.7 | 25.1 | 127.0 | 2.9 | 29.1 |
| >hg38_21_+_8433172_8446622_RNA45SN1_45S_50ntFlanks@12416.12446.31\|TTCGATGTCGGCTCTTCCTATCATTGTGAAG | >hg38_21_+_8433172_8446622_RNA45SN1_45S_50ntFlanks | 12416 | 12446 | 31 | TTCGATGTCGGCTCTTCCTATCATTGTGAAG | MB468 | 13 | 21 | 32 | 4 | 10 | 7.7 | 14.4 | 17.6 | 1.2 | 3.1 |
| >hg38_21_+_8433172_8446622_RNA45SN1_45S_50ntFlanks@12416.12447.32\|TTCGATGTCGGCTCTTCCTATCATTGTGAAGC | >hg38_21_+_8433172_8446622_RNA45SN1_45S_50ntFlanks | 12416 | 12447 | 32 | TTCGATGTCGGCTCTTCCTATCATTGTGAAGC | MB231 | 73 | 33 | 134 | 2 | 32 | 70.6 | 25.5 | 105.9 | 2.1 | 29.4 |
| >hg38_21_+_8433172_8446622_RNA45SN1_45S_50ntFlanks@12416.12448.33\|TTCGATGTCGGCTCTTCCTATCATTGTGAAGCA | >hg38_21_+_8433172_8446622_RNA45SN1_45S_50ntFlanks | 12416 | 12448 | 33 | TTCGATGTCGGCTCTTCCTATCATTGTGAAGCA | BT20 | 33 | 20 | 84 | 6 | 19 | 13.9 | 8.2 | 84.8 | 2.6 | 5.7 |
| >hg38_21_+_8433172_8446622_RNA45SN1_45S_50ntFlanks@12416.12448.33\|TTCGATGTCGGCTCTTCCTATCATTGTGAAGCA | >hg38_21_+_8433172_8446622_RNA45SN1_45S_50ntFlanks | 12416 | 12448 | 33 | TTCGATGTCGGCTCTTCCTATCATTGTGAAGCA | MB231 | 195 | 79 | 335 | 6 | 69 | 187.1 | 63.1 | 254.7 | 3.8 | 49.9 |
| >hg38_21_+_8433172_8446622_RNA45SN1_45S_50ntFlanks@12416.12449.34\|TTCGATGTCGGCTCTTCCTATCATTGTGAAGCAG | >hg38_21_+_8433172_8446622_RNA45SN1_45S_50ntFlanks | 12416 | 12449 | 34 | TTCGATGTCGGCTCTTCCTATCATTGTGAAGCAG | BT20 | 12 | 4 | 18 | 2 | 5 | 7.3 | 2.8 | 11.8 | 1.7 | 2.7 |
| >hg38_21_+_8433172_8446622_RNA45SN1_45S_50ntFlanks@12416.12449.34\|TTCGATGTCGGCTCTTCCTATCATTGTGAAGCAG | >hg38_21_+_8433172_8446622_RNA45SN1_45S_50ntFlanks | 12416 | 12449 | 34 | TTCGATGTCGGCTCTTCCTATCATTGTGAAGCAG | MB231 | 85 | 33 | 68 | 7 | 42 | 75.6 | 26.6 | 60.2 | 4.8 | 29.6 |
| >hg38_21_+_8433172_8446622_RNA45SN1_45S_50ntFlanks@12416.12450.35\|TTCGATGTCGGCTCTTCCTATCATTGTGAAGCAGA | >hg38_21_+_8433172_8446622_RNA45SN1_45S_50ntFlanks | 12416 | 12450 | 35 | TTCGATGTCGGCTCTTCCTATCATTGTGAAGCAGA | MB231 | 28 | 13 | 21 | 2 | 18 | 22.4 | 10.1 | 14.8 | 2.0 | 15.5 |
| >hg38_21_+_8433172_8446622_RNA45SN1_45S_50ntFlanks@12416.12452.37\|TTCGATGTCGGCTCTTCCTATCATTGTGAAGCAGAAT | >hg38_21_+_8433172_8446622_RNA45SN1_45S_50ntFlanks | 12416 | 12452 | 37 | TTCGATGTCGGCTCTTCCTATCATTGTGAAGCAGAAT | MB231 | 39 | 18 | 29 | 3 | 24 | 27.6 | 10.8 | 22.0 | 2.7 | 21.2 |
| >hg38_21_+_8433172_8446622_RNA45SN1_45S_50ntFlanks@12416.12453.38\|TTCGATGTCGGCTCTTCCTATCATTGTGAAGCAGAATT | >hg38_21_+_8433172_8446622_RNA45SN1_45S_50ntFlanks | 12416 | 12453 | 38 | TTCGATGTCGGCTCTTCCTATCATTGTGAAGCAGAATT | MB231 | 32 | 16 | 28 | 2 | 19 | 26.5 | 10.2 | 21.1 | 1.4 | 13.0 |
| >hg38_21_+_8433172_8446622_RNA45SN1_45S_50ntFlanks@12416.12454.39\|TTCGATGTCGGCTCTTCCTATCATTGTGAAGCAGAATTC | >hg38_21_+_8433172_8446622_RNA45SN1_45S_50ntFlanks | 12416 | 12454 | 39 | TTCGATGTCGGCTCTTCCTATCATTGTGAAGCAGAATTC | MB231 | 26 | 16 | 27 | 2 | 16 | 19.2 | 9.3 | 19.7 | 2.7 | 13.6 |
| >hg38_21_+_8433172_8446622_RNA45SN1_45S_50ntFlanks@12416.12455.40\|TTCGATGTCGGCTCTTCCTATCATTGTGAAGCAGAATTCA | >hg38_21_+_8433172_8446622_RNA45SN1_45S_50ntFlanks | 12416 | 12455 | 40 | TTCGATGTCGGCTCTTCCTATCATTGTGAAGCAGAATTCA | MB231 | 16 | 10 | 15 | 1 | 15 | 13.1 | 6.0 | 11.2 | 0.7 | 13.5 |
| >hg38_21_+_8433172_8446622_RNA45SN1_45S_50ntFlanks@12416.12456.41\|TTCGATGTCGGCTCTTCCTATCATTGTGAAGCAGAATTCAC | >hg38_21_+_8433172_8446622_RNA45SN1_45S_50ntFlanks | 12416 | 12456 | 41 | TTCGATGTCGGCTCTTCCTATCATTGTGAAGCAGAATTCAC | MB231 | 15 | 9 | 21 | 0 | 13 | 9.6 | 6.6 | 16.2 | 0.6 | 22.1 |
| >hg38_21_+_8433172_8446622_RNA45SN1_45S_50ntFlanks@12416.12457.42\|TTCGATGTCGGCTCTTCCTATCATTGTGAAGCAGAATTCACC | >hg38_21_+_8433172_8446622_RNA45SN1_45S_50ntFlanks | 12416 | 12457 | 42 | TTCGATGTCGGCTCTTCCTATCATTGTGAAGCAGAATTCACC | MB231 | 53 | 33 | 135 | 1 | 22 | 40.3 | 35.0 | 95.5 | 1.2 | 23.1 |
| >hg38_21_+_8433172_8446622_RNA45SN1_45S_50ntFlanks@12416.12458.43\|TTCGATGTCGGCTCTTCCTATCATTGTGAAGCAGAATTCACCA | >hg38_21_+_8433172_8446622_RNA45SN1_45S_50ntFlanks | 12416 | 12458 | 43 | TTCGATGTCGGCTCTTCCTATCATTGTGAAGCAGAATTCACCA | MB231 | 16 | 13 | 44 | 0 | 0 | 9.9 | 15.9 | 32.7 | 0.0 | 0.0 |
| >hg38_21_+_8433172_8446622_RNA45SN1_45S_50ntFlanks@12417.12434.18\|TCGATGTCGGCTCTTCCT | >hg38_21_+_8433172_8446622_RNA45SN1_45S_50ntFlanks | 12417 | 12434 | 18 | TCGATGTCGGCTCTTCCT | BT20 | 60 | 93 | 73 | 11 | 12 | 91.6 | 147.8 | 103.4 | 14.9 | 3.9 |
| >hg38_21_+_8433172_8446622_RNA45SN1_45S_50ntFlanks@12417.12434.18\|TCGATGTCGGCTCTTCCT | >hg38_21_+_8433172_8446622_RNA45SN1_45S_50ntFlanks | 12417 | 12434 | 18 | TCGATGTCGGCTCTTCCT | MB231 | 32 | 21 | 147 | 2 | 10 | 22.6 | 10.0 | 150.0 | 1.0 | 5.5 |
| >hg38_21_+_8433172_8446622_RNA45SN1_45S_50ntFlanks@12417.12434.18\|TCGATGTCGGCTCTTCCT | >hg38_21_+_8433172_8446622_RNA45SN1_45S_50ntFlanks | 12417 | 12434 | 18 | TCGATGTCGGCTCTTCCT | MB468 | 334 | 350 | 638 | 65 | 126 | 137.1 | 140.4 | 318.8 | 17.9 | 12.0 |
| >hg38_21_+_8433172_8446622_RNA45SN1_45S_50ntFlanks@12417.12435.19\|TCGATGTCGGCTCTTCCTA | >hg38_21_+_8433172_8446622_RNA45SN1_45S_50ntFlanks | 12417 | 12435 | 19 | TCGATGTCGGCTCTTCCTA | MB468 | 368 | 449 | 489 | 57 | 159 | 89.0 | 117.9 | 114.1 | 11.8 | 22.9 |
| >hg38_21_+_8433172_8446622_RNA45SN1_45S_50ntFlanks@12417.12436.20\|TCGATGTCGGCTCTTCCTAT | >hg38_21_+_8433172_8446622_RNA45SN1_45S_50ntFlanks | 12417 | 12436 | 20 | TCGATGTCGGCTCTTCCTAT | MB468 | 585 | 817 | 707 | 53 | 428 | 173.8 | 152.6 | 164.9 | 20.0 | 132.7 |
| >hg38_21_+_8433172_8446622_RNA45SN1_45S_50ntFlanks@12417.12437.21\|TCGATGTCGGCTCTTCCTATC | >hg38_21_+_8433172_8446622_RNA45SN1_45S_50ntFlanks | 12417 | 12437 | 21 | TCGATGTCGGCTCTTCCTATC | BT20 | 12 | 7 | 29 | 4 | 9 | 7.9 | 4.4 | 24.2 | 5.9 | 7.8 |
| >hg38_21_+_8433172_8446622_RNA45SN1_45S_50ntFlanks@12417.12437.21\|TCGATGTCGGCTCTTCCTATC | >hg38_21_+_8433172_8446622_RNA45SN1_45S_50ntFlanks | 12417 | 12437 | 21 | TCGATGTCGGCTCTTCCTATC | MB231 | 45 | 34 | 122 | 1 | 18 | 38.3 | 24.8 | 86.8 | 1.3 | 18.4 |
| >hg38_21_+_8433172_8446622_RNA45SN1_45S_50ntFlanks@12417.12437.21\|TCGATGTCGGCTCTTCCTATC | >hg38_21_+_8433172_8446622_RNA45SN1_45S_50ntFlanks | 12417 | 12437 | 21 | TCGATGTCGGCTCTTCCTATC | MB468 | 171 | 244 | 273 | 23 | 146 | 53.1 | 35.4 | 87.2 | 5.9 | 30.7 |
| >hg38_21_+_8433172_8446622_RNA45SN1_45S_50ntFlanks@12417.12438.22\|TCGATGTCGGCTCTTCCTATCA | >hg38_21_+_8433172_8446622_RNA45SN1_45S_50ntFlanks | 12417 | 12438 | 22 | TCGATGTCGGCTCTTCCTATCA | BT20 | 19 | 18 | 23 | 2 | 6 | 20.2 | 21.3 | 16.4 | 3.9 | 10.2 |
| >hg38_21_+_8433172_8446622_RNA45SN1_45S_50ntFlanks@12417.12438.22\|TCGATGTCGGCTCTTCCTATCA | >hg38_21_+_8433172_8446622_RNA45SN1_45S_50ntFlanks | 12417 | 12438 | 22 | TCGATGTCGGCTCTTCCTATCA | MB231 | 32 | 31 | 78 | 1 | 13 | 24.3 | 28.1 | 73.9 | 1.1 | 22.6 |
| >hg38_21_+_8433172_8446622_RNA45SN1_45S_50ntFlanks@12417.12438.22\|TCGATGTCGGCTCTTCCTATCA | >hg38_21_+_8433172_8446622_RNA45SN1_45S_50ntFlanks | 12417 | 12438 | 22 | TCGATGTCGGCTCTTCCTATCA | MB468 | 14 | 33 | 20 | 5 | 81 | 5.2 | 13.3 | 4.2 | 4.5 | 65.2 |
| >hg38_21_+_8433172_8446622_RNA45SN1_45S_50ntFlanks@12417.12439.23\|TCGATGTCGGCTCTTCCTATCAT | >hg38_21_+_8433172_8446622_RNA45SN1_45S_50ntFlanks | 12417 | 12439 | 23 | TCGATGTCGGCTCTTCCTATCAT | BT20 | 14 | 12 | 20 | 1 | 3 | 16.9 | 16.0 | 17.4 | 1.5 | 4.4 |
| >hg38_21_+_8433172_8446622_RNA45SN1_45S_50ntFlanks@12417.12439.23\|TCGATGTCGGCTCTTCCTATCAT | >hg38_21_+_8433172_8446622_RNA45SN1_45S_50ntFlanks | 12417 | 12439 | 23 | TCGATGTCGGCTCTTCCTATCAT | MB231 | 19 | 14 | 50 | 0 | 7 | 17.0 | 12.3 | 39.0 | 0.5 | 12.4 |
| >hg38_21_+_8433172_8446622_RNA45SN1_45S_50ntFlanks@12417.12439.23\|TCGATGTCGGCTCTTCCTATCAT | >hg38_21_+_8433172_8446622_RNA45SN1_45S_50ntFlanks | 12417 | 12439 | 23 | TCGATGTCGGCTCTTCCTATCAT | MB468 | 14 | 22 | 25 | 2 | 12 | 3.6 | 7.2 | 13.5 | 0.6 | 4.8 |
| >hg38_21_+_8433172_8446622_RNA45SN1_45S_50ntFlanks@12417.12440.24\|TCGATGTCGGCTCTTCCTATCATT | >hg38_21_+_8433172_8446622_RNA45SN1_45S_50ntFlanks | 12417 | 12440 | 24 | TCGATGTCGGCTCTTCCTATCATT | MB231 | 25 | 18 | 66 | 0 | 8 | 27.8 | 18.0 | 65.4 | 0.8 | 14.7 |
| >hg38_21_+_8433172_8446622_RNA45SN1_45S_50ntFlanks@12417.12440.24\|TCGATGTCGGCTCTTCCTATCATT | >hg38_21_+_8433172_8446622_RNA45SN1_45S_50ntFlanks | 12417 | 12440 | 24 | TCGATGTCGGCTCTTCCTATCATT | MB468 | 17 | 27 | 31 | 2 | 12 | 5.9 | 8.0 | 8.0 | 1.0 | 2.3 |
| >hg38_21_+_8433172_8446622_RNA45SN1_45S_50ntFlanks@12417.12441.25\|TCGATGTCGGCTCTTCCTATCATTG | >hg38_21_+_8433172_8446622_RNA45SN1_45S_50ntFlanks | 12417 | 12441 | 25 | TCGATGTCGGCTCTTCCTATCATTG | MB468 | 94 | 100 | 145 | 17 | 87 | 22.1 | 11.9 | 17.8 | 10.4 | 35.7 |
| >hg38_21_+_8433172_8446622_RNA45SN1_45S_50ntFlanks@12417.12442.26\|TCGATGTCGGCTCTTCCTATCATTGT | >hg38_21_+_8433172_8446622_RNA45SN1_45S_50ntFlanks | 12417 | 12442 | 26 | TCGATGTCGGCTCTTCCTATCATTGT | MB231 | 17 | 13 | 46 | 0 | 7 | 14.4 | 11.5 | 44.3 | 0.5 | 12.7 |
| >hg38_21_+_8433172_8446622_RNA45SN1_45S_50ntFlanks@12417.12443.27\|TCGATGTCGGCTCTTCCTATCATTGTG | >hg38_21_+_8433172_8446622_RNA45SN1_45S_50ntFlanks | 12417 | 12443 | 27 | TCGATGTCGGCTCTTCCTATCATTGTG | MB231 | 26 | 16 | 70 | 1 | 11 | 24.4 | 16.2 | 77.8 | 1.0 | 19.8 |
| >hg38_21_+_8433172_8446622_RNA45SN1_45S_50ntFlanks@12417.12443.27\|TCGATGTCGGCTCTTCCTATCATTGTG | >hg38_21_+_8433172_8446622_RNA45SN1_45S_50ntFlanks | 12417 | 12443 | 27 | TCGATGTCGGCTCTTCCTATCATTGTG | MB468 | 239 | 226 | 647 | 52 | 304 | 75.9 | 46.6 | 185.3 | 35.2 | 135.9 |
| >hg38_21_+_8433172_8446622_RNA45SN1_45S_50ntFlanks@12417.12444.28\|TCGATGTCGGCTCTTCCTATCATTGTGA | >hg38_21_+_8433172_8446622_RNA45SN1_45S_50ntFlanks | 12417 | 12444 | 28 | TCGATGTCGGCTCTTCCTATCATTGTGA | MB231 | 20 | 15 | 63 | 1 | 11 | 17.2 | 11.3 | 53.4 | 1.1 | 19.4 |
| >hg38_21_+_8433172_8446622_RNA45SN1_45S_50ntFlanks@12417.12444.28\|TCGATGTCGGCTCTTCCTATCATTGTGA | >hg38_21_+_8433172_8446622_RNA45SN1_45S_50ntFlanks | 12417 | 12444 | 28 | TCGATGTCGGCTCTTCCTATCATTGTGA | MB468 | 61 | 76 | 171 | 13 | 61 | 22.5 | 29.5 | 63.6 | 9.1 | 20.8 |
| >hg38_21_+_8433172_8446622_RNA45SN1_45S_50ntFlanks@12417.12445.29\|TCGATGTCGGCTCTTCCTATCATTGTGAA | >hg38_21_+_8433172_8446622_RNA45SN1_45S_50ntFlanks | 12417 | 12445 | 29 | TCGATGTCGGCTCTTCCTATCATTGTGAA | MB468 | 28 | 30 | 71 | 5 | 23 | 9.3 | 8.7 | 25.9 | 3.3 | 4.8 |
| >hg38_21_+_8433172_8446622_RNA45SN1_45S_50ntFlanks@12417.12446.30\|TCGATGTCGGCTCTTCCTATCATTGTGAAG | >hg38_21_+_8433172_8446622_RNA45SN1_45S_50ntFlanks | 12417 | 12446 | 30 | TCGATGTCGGCTCTTCCTATCATTGTGAAG | MB231 | 18 | 8 | 46 | 1 | 13 | 19.3 | 4.9 | 33.1 | 1.0 | 12.1 |
| >hg38_21_+_8433172_8446622_RNA45SN1_45S_50ntFlanks@12417.12446.30\|TCGATGTCGGCTCTTCCTATCATTGTGAAG | >hg38_21_+_8433172_8446622_RNA45SN1_45S_50ntFlanks | 12417 | 12446 | 30 | TCGATGTCGGCTCTTCCTATCATTGTGAAG | MB468 | 69 | 106 | 195 | 26 | 51 | 31.4 | 51.2 | 113.5 | 9.9 | 8.4 |
| >hg38_21_+_8433172_8446622_RNA45SN1_45S_50ntFlanks@12417.12447.31\|TCGATGTCGGCTCTTCCTATCATTGTGAAGC | >hg38_21_+_8433172_8446622_RNA45SN1_45S_50ntFlanks | 12417 | 12447 | 31 | TCGATGTCGGCTCTTCCTATCATTGTGAAGC | MB231 | 18 | 10 | 50 | 1 | 17 | 16.0 | 5.1 | 33.1 | 1.0 | 15.5 |
| >hg38_21_+_8433172_8446622_RNA45SN1_45S_50ntFlanks@12417.12448.32\|TCGATGTCGGCTCTTCCTATCATTGTGAAGCA | >hg38_21_+_8433172_8446622_RNA45SN1_45S_50ntFlanks | 12417 | 12448 | 32 | TCGATGTCGGCTCTTCCTATCATTGTGAAGCA | MB231 | 47 | 21 | 147 | 1 | 25 | 45.0 | 12.9 | 106.9 | 1.2 | 21.6 |
| >hg38_21_+_8433172_8446622_RNA45SN1_45S_50ntFlanks@12417.12449.33\|TCGATGTCGGCTCTTCCTATCATTGTGAAGCAG | >hg38_21_+_8433172_8446622_RNA45SN1_45S_50ntFlanks | 12417 | 12449 | 33 | TCGATGTCGGCTCTTCCTATCATTGTGAAGCAG | MB231 | 12 | 4 | 14 | 2 | 10 | 11.1 | 2.8 | 12.2 | 1.8 | 9.0 |
| >hg38_21_+_8433172_8446622_RNA45SN1_45S_50ntFlanks@12417.12452.36\|TCGATGTCGGCTCTTCCTATCATTGTGAAGCAGAAT | >hg38_21_+_8433172_8446622_RNA45SN1_45S_50ntFlanks | 12417 | 12452 | 36 | TCGATGTCGGCTCTTCCTATCATTGTGAAGCAGAAT | MB231 | 15 | 10 | 20 | 2 | 19 | 10.0 | 4.1 | 15.1 | 1.8 | 16.5 |
| >hg38_21_+_8433172_8446622_RNA45SN1_45S_50ntFlanks@12417.12453.37\|TCGATGTCGGCTCTTCCTATCATTGTGAAGCAGAATT | >hg38_21_+_8433172_8446622_RNA45SN1_45S_50ntFlanks | 12417 | 12453 | 37 | TCGATGTCGGCTCTTCCTATCATTGTGAAGCAGAATT | MB231 | 12 | 9 | 16 | 1 | 11 | 10.0 | 3.5 | 11.5 | 1.1 | 9.8 |
| >hg38_21_+_8433172_8446622_RNA45SN1_45S_50ntFlanks@12417.12455.39\|TCGATGTCGGCTCTTCCTATCATTGTGAAGCAGAATTCA | >hg38_21_+_8433172_8446622_RNA45SN1_45S_50ntFlanks | 12417 | 12455 | 39 | TCGATGTCGGCTCTTCCTATCATTGTGAAGCAGAATTCA | MB468 | 50 | 124 | 131 | 20 | 195 | 28.2 | 104.8 | 107.3 | 17.6 | 168.1 |
| >hg38_21_+_8433172_8446622_RNA45SN1_45S_50ntFlanks@12417.12457.41\|TCGATGTCGGCTCTTCCTATCATTGTGAAGCAGAATTCACC | >hg38_21_+_8433172_8446622_RNA45SN1_45S_50ntFlanks | 12417 | 12457 | 41 | TCGATGTCGGCTCTTCCTATCATTGTGAAGCAGAATTCACC | MB231 | 17 | 13 | 35 | 1 | 13 | 13.0 | 10.3 | 24.2 | 0.6 | 12.6 |
| >hg38_21_+_8433172_8446622_RNA45SN1_45S_50ntFlanks@12418.12435.18\|CGATGTCGGCTCTTCCTA | >hg38_21_+_8433172_8446622_RNA45SN1_45S_50ntFlanks | 12418 | 12435 | 18 | CGATGTCGGCTCTTCCTA | BT20 | 24 | 60 | 17 | 3 | 3 | 40.4 | 103.1 | 21.6 | 4.5 | 2.5 |

| | | | | | | | | | | | | | | | | |
|---|---|---|---|---|---|---|---|---|---|---|---|---|---|---|---|---|
| >hg38_21_+_8433172_8446622_RNA45SN1_45S_50ntFlanks@12418.12435.18\|CGATGTCGGCTCTTCCTA | >hg38_21_+_8433172_8446622_RNA45SN1_45S_50ntFlanks | 12418 | 12435 | 18 | CGATGTCGGCTCTTCCTA | MB468 | 1000 | 1344 | 1084 | 143 | 529 | 331.3 | 283.3 | 266.1 | 30.4 | 126.2 |
| >hg38_21_+_8433172_8446622_RNA45SN1_45S_50ntFlanks@12418.12436.19\|CGATGTCGGCTCTTCCTAT | >hg38_21_+_8433172_8446622_RNA45SN1_45S_50ntFlanks | 12418 | 12436 | 19 | CGATGTCGGCTCTTCCTAT | BT20 | 28 | 50 | 17 | 4 | 1 | 46.7 | 84.8 | 19.8 | 6.8 | 1.9 |
| >hg38_21_+_8433172_8446622_RNA45SN1_45S_50ntFlanks@12418.12436.19\|CGATGTCGGCTCTTCCTAT | >hg38_21_+_8433172_8446622_RNA45SN1_45S_50ntFlanks | 12418 | 12436 | 19 | CGATGTCGGCTCTTCCTAT | MB468 | 1535 | 2345 | 1548 | 140 | 1280 | 546.3 | 477.3 | 324.5 | 48.4 | 487.4 |
| >hg38_21_+_8433172_8446622_RNA45SN1_45S_50ntFlanks@12418.12437.20\|CGATGTCGGCTCTTCCTATC | >hg38_21_+_8433172_8446622_RNA45SN1_45S_50ntFlanks | 12418 | 12437 | 20 | CGATGTCGGCTCTTCCTATC | BT20 | 16 | 10 | 23 | 11 | 12 | 14.2 | 7.8 | 15.8 | 16.0 | 5.5 |
| >hg38_21_+_8433172_8446622_RNA45SN1_45S_50ntFlanks@12418.12437.20\|CGATGTCGGCTCTTCCTATC | >hg38_21_+_8433172_8446622_RNA45SN1_45S_50ntFlanks | 12418 | 12437 | 20 | CGATGTCGGCTCTTCCTATC | MB231 | 38 | 31 | 79 | 1 | 17 | 28.9 | 24.9 | 55.5 | 1.5 | 18.4 |
| >hg38_21_+_8433172_8446622_RNA45SN1_45S_50ntFlanks@12418.12437.20\|CGATGTCGGCTCTTCCTATC | >hg38_21_+_8433172_8446622_RNA45SN1_45S_50ntFlanks | 12418 | 12437 | 20 | CGATGTCGGCTCTTCCTATC | MB468 | 429 | 628 | 577 | 56 | 424 | 176.7 | 98.2 | 171.4 | 22.1 | 114.1 |
| >hg38_21_+_8433172_8446622_RNA45SN1_45S_50ntFlanks@12418.12438.21\|CGATGTCGGCTCTTCCTATCA | >hg38_21_+_8433172_8446622_RNA45SN1_45S_50ntFlanks | 12418 | 12438 | 21 | CGATGTCGGCTCTTCCTATCA | BT20 | 30 | 37 | 21 | 2 | 4 | 43.6 | 56.7 | 19.9 | 4.3 | 6.8 |
| >hg38_21_+_8433172_8446622_RNA45SN1_45S_50ntFlanks@12418.12438.21\|CGATGTCGGCTCTTCCTATCA | >hg38_21_+_8433172_8446622_RNA45SN1_45S_50ntFlanks | 12418 | 12438 | 21 | CGATGTCGGCTCTTCCTATCA | MB231 | 19 | 24 | 37 | 1 | 12 | 15.0 | 23.5 | 38.7 | 0.9 | 20.9 |
| >hg38_21_+_8433172_8446622_RNA45SN1_45S_50ntFlanks@12418.12438.21\|CGATGTCGGCTCTTCCTATCA | >hg38_21_+_8433172_8446622_RNA45SN1_45S_50ntFlanks | 12418 | 12438 | 21 | CGATGTCGGCTCTTCCTATCA | MB468 | 25 | 54 | 29 | 7 | 176 | 13.7 | 16.0 | 10.2 | 6.5 | 145.1 |
| >hg38_21_+_8433172_8446622_RNA45SN1_45S_50ntFlanks@12418.12439.22\|CGATGTCGGCTCTTCCTATCAT | >hg38_21_+_8433172_8446622_RNA45SN1_45S_50ntFlanks | 12418 | 12439 | 22 | CGATGTCGGCTCTTCCTATCAT | BT20 | 31 | 37 | 24 | 2 | 4 | 45.6 | 58.0 | 26.0 | 2.8 | 3.6 |
| >hg38_21_+_8433172_8446622_RNA45SN1_45S_50ntFlanks@12418.12439.22\|CGATGTCGGCTCTTCCTATCAT | >hg38_21_+_8433172_8446622_RNA45SN1_45S_50ntFlanks | 12418 | 12439 | 22 | CGATGTCGGCTCTTCCTATCAT | MB231 | 14 | 13 | 30 | 0 | 7 | 12.6 | 10.7 | 23.2 | 0.5 | 12.4 |
| >hg38_21_+_8433172_8446622_RNA45SN1_45S_50ntFlanks@12418.12439.22\|CGATGTCGGCTCTTCCTATCAT | >hg38_21_+_8433172_8446622_RNA45SN1_45S_50ntFlanks | 12418 | 12439 | 22 | CGATGTCGGCTCTTCCTATCAT | MB468 | 27 | 49 | 45 | 4 | 33 | 8.0 | 17.7 | 24.0 | 1.1 | 13.6 |
| >hg38_21_+_8433172_8446622_RNA45SN1_45S_50ntFlanks@12418.12440.23\|CGATGTCGGCTCTTCCTATCATT | >hg38_21_+_8433172_8446622_RNA45SN1_45S_50ntFlanks | 12418 | 12440 | 23 | CGATGTCGGCTCTTCCTATCATT | MB231 | 16 | 16 | 34 | 0 | 7 | 17.6 | 18.0 | 35.9 | 0.8 | 12.8 |
| >hg38_21_+_8433172_8446622_RNA45SN1_45S_50ntFlanks@12418.12440.23\|CGATGTCGGCTCTTCCTATCATT | >hg38_21_+_8433172_8446622_RNA45SN1_45S_50ntFlanks | 12418 | 12440 | 23 | CGATGTCGGCTCTTCCTATCATT | MB468 | 26 | 39 | 41 | 4 | 23 | 9.6 | 13.4 | 14.1 | 1.2 | 5.4 |
| >hg38_21_+_8433172_8446622_RNA45SN1_45S_50ntFlanks@12418.12441.24\|CGATGTCGGCTCTTCCTATCATTG | >hg38_21_+_8433172_8446622_RNA45SN1_45S_50ntFlanks | 12418 | 12441 | 24 | CGATGTCGGCTCTTCCTATCATTG | MB468 | 106 | 103 | 140 | 21 | 123 | 42.1 | 9.1 | 24.3 | 9.1 | 69.6 |
| >hg38_21_+_8433172_8446622_RNA45SN1_45S_50ntFlanks@12418.12442.25\|CGATGTCGGCTCTTCCTATCATTGT | >hg38_21_+_8433172_8446622_RNA45SN1_45S_50ntFlanks | 12418 | 12442 | 25 | CGATGTCGGCTCTTCCTATCATTGT | MB231 | 12 | 12 | 34 | 0 | 7 | 9.3 | 10.8 | 29.9 | 0.5 | 12.2 |
| >hg38_21_+_8433172_8446622_RNA45SN1_45S_50ntFlanks@12418.12442.25\|CGATGTCGGCTCTTCCTATCATTGT | >hg38_21_+_8433172_8446622_RNA45SN1_45S_50ntFlanks | 12418 | 12442 | 25 | CGATGTCGGCTCTTCCTATCATTGT | MB468 | 16 | 28 | 31 | 3 | 14 | 11.9 | 16.5 | 19.5 | 0.9 | 4.5 |
| >hg38_21_+_8433172_8446622_RNA45SN1_45S_50ntFlanks@12418.12443.26\|CGATGTCGGCTCTTCCTATCATTGTG | >hg38_21_+_8433172_8446622_RNA45SN1_45S_50ntFlanks | 12418 | 12443 | 26 | CGATGTCGGCTCTTCCTATCATTGTG | MB231 | 15 | 17 | 48 | 1 | 12 | 13.9 | 18.6 | 38.1 | 1.0 | 19.9 |
| >hg38_21_+_8433172_8446622_RNA45SN1_45S_50ntFlanks@12418.12443.26\|CGATGTCGGCTCTTCCTATCATTGTG | >hg38_21_+_8433172_8446622_RNA45SN1_45S_50ntFlanks | 12418 | 12443 | 26 | CGATGTCGGCTCTTCCTATCATTGTG | MB468 | 463 | 1079 | 869 | 97 | 618 | 272.1 | 380.8 | 530.8 | 47.0 | 250.5 |
| >hg38_21_+_8433172_8446622_RNA45SN1_45S_50ntFlanks@12418.12444.27\|CGATGTCGGCTCTTCCTATCATTGTGA | >hg38_21_+_8433172_8446622_RNA45SN1_45S_50ntFlanks | 12418 | 12444 | 27 | CGATGTCGGCTCTTCCTATCATTGTGA | MB231 | 11 | 12 | 37 | 1 | 12 | 8.5 | 10.1 | 25.0 | 0.7 | 13.9 |
| >hg38_21_+_8433172_8446622_RNA45SN1_45S_50ntFlanks@12418.12444.27\|CGATGTCGGCTCTTCCTATCATTGTGA | >hg38_21_+_8433172_8446622_RNA45SN1_45S_50ntFlanks | 12418 | 12444 | 27 | CGATGTCGGCTCTTCCTATCATTGTGA | MB468 | 85 | 246 | 160 | 20 | 106 | 52.6 | 97.5 | 101.4 | 5.3 | 39.8 |
| >hg38_21_+_8433172_8446622_RNA45SN1_45S_50ntFlanks@12418.12445.28\|CGATGTCGGCTCTTCCTATCATTGTGAA | >hg38_21_+_8433172_8446622_RNA45SN1_45S_50ntFlanks | 12418 | 12445 | 28 | CGATGTCGGCTCTTCCTATCATTGTGAA | MB468 | 46 | 121 | 80 | 9 | 51 | 24.8 | 37.9 | 42.8 | 2.1 | 11.1 |
| >hg38_21_+_8433172_8446622_RNA45SN1_45S_50ntFlanks@12418.12446.29\|CGATGTCGGCTCTTCCTATCATTGTGAAG | >hg38_21_+_8433172_8446622_RNA45SN1_45S_50ntFlanks | 12418 | 12446 | 29 | CGATGTCGGCTCTTCCTATCATTGTGAAG | MB231 | 12 | 6 | 29 | 2 | 10 | 11.6 | 3.5 | 19.1 | 1.5 | 9.1 |
| >hg38_21_+_8433172_8446622_RNA45SN1_45S_50ntFlanks@12418.12446.29\|CGATGTCGGCTCTTCCTATCATTGTGAAG | >hg38_21_+_8433172_8446622_RNA45SN1_45S_50ntFlanks | 12418 | 12446 | 29 | CGATGTCGGCTCTTCCTATCATTGTGAAG | MB468 | 145 | 278 | 305 | 56 | 135 | 80.0 | 142.8 | 177.4 | 7.9 | 28.1 |
| >hg38_21_+_8433172_8446622_RNA45SN1_45S_50ntFlanks@12418.12447.30\|CGATGTCGGCTCTTCCTATCATTGTGAAGC | >hg38_21_+_8433172_8446622_RNA45SN1_45S_50ntFlanks | 12418 | 12447 | 30 | CGATGTCGGCTCTTCCTATCATTGTGAAGC | MB231 | 13 | 8 | 26 | 1 | 16 | 9.8 | 3.6 | 15.1 | 0.9 | 14.7 |
| >hg38_21_+_8433172_8446622_RNA45SN1_45S_50ntFlanks@12418.12448.31\|CGATGTCGGCTCTTCCTATCATTGTGAAGCA | >hg38_21_+_8433172_8446622_RNA45SN1_45S_50ntFlanks | 12418 | 12448 | 31 | CGATGTCGGCTCTTCCTATCATTGTGAAGCA | MB231 | 34 | 18 | 91 | 2 | 29 | 27.5 | 10.7 | 63.3 | 1.0 | 19.9 |
| >hg38_21_+_8433172_8446622_RNA45SN1_45S_50ntFlanks@12418.12449.32\|CGATGTCGGCTCTTCCTATCATTGTGAAGCAG | >hg38_21_+_8433172_8446622_RNA45SN1_45S_50ntFlanks | 12418 | 12449 | 32 | CGATGTCGGCTCTTCCTATCATTGTGAAGCAG | MB231 | 19 | 10 | 29 | 3 | 25 | 15.2 | 5.2 | 20.5 | 2.1 | 19.3 |
| >hg38_21_+_8433172_8446622_RNA45SN1_45S_50ntFlanks@12418.12449.32\|CGATGTCGGCTCTTCCTATCATTGTGAAGCAG | >hg38_21_+_8433172_8446622_RNA45SN1_45S_50ntFlanks | 12418 | 12449 | 32 | CGATGTCGGCTCTTCCTATCATTGTGAAGCAG | MB468 | 22 | 33 | 43 | 11 | 26 | 10.0 | 17.0 | 20.1 | 4.0 | 6.6 |
| >hg38_21_+_8433172_8446622_RNA45SN1_45S_50ntFlanks@12418.12451.34\|CGATGTCGGCTCTTCCTATCATTGTGAAGCAGAA | >hg38_21_+_8433172_8446622_RNA45SN1_45S_50ntFlanks | 12418 | 12451 | 34 | CGATGTCGGCTCTTCCTATCATTGTGAAGCAGAA | MB468 | 10 | 14 | 19 | 6 | 22 | 4.5 | 7.3 | 11.5 | 3.6 | 5.4 |
| >hg38_21_+_8433172_8446622_RNA45SN1_45S_50ntFlanks@12418.12452.35\|CGATGTCGGCTCTTCCTATCATTGTGAAGCAGAAT | >hg38_21_+_8433172_8446622_RNA45SN1_45S_50ntFlanks | 12418 | 12452 | 35 | CGATGTCGGCTCTTCCTATCATTGTGAAGCAGAAT | MB231 | 14 | 11 | 18 | 3 | 22 | 8.0 | 4.6 | 13.1 | 1.8 | 18.3 |
| >hg38_21_+_8433172_8446622_RNA45SN1_45S_50ntFlanks@12418.12453.36\|CGATGTCGGCTCTTCCTATCATTGTGAAGCAGAATT | >hg38_21_+_8433172_8446622_RNA45SN1_45S_50ntFlanks | 12418 | 12453 | 36 | CGATGTCGGCTCTTCCTATCATTGTGAAGCAGAATT | MB231 | 11 | 10 | 13 | 1 | 12 | 7.8 | 4.8 | 8.4 | 1.1 | 10.8 |
| >hg38_21_+_8433172_8446622_RNA45SN1_45S_50ntFlanks@12418.12454.37\|CGATGTCGGCTCTTCCTATCATTGTGAAGCAGAATTC | >hg38_21_+_8433172_8446622_RNA45SN1_45S_50ntFlanks | 12418 | 12454 | 37 | CGATGTCGGCTCTTCCTATCATTGTGAAGCAGAATTC | MB468 | 12 | 29 | 25 | 8 | 31 | 6.8 | 18.7 | 15.2 | 5.9 | 23.8 |
| >hg38_21_+_8433172_8446622_RNA45SN1_45S_50ntFlanks@12418.12455.38\|CGATGTCGGCTCTTCCTATCATTGTGAAGCAGAATTCA | >hg38_21_+_8433172_8446622_RNA45SN1_45S_50ntFlanks | 12418 | 12455 | 38 | CGATGTCGGCTCTTCCTATCATTGTGAAGCAGAATTCA | MB468 | 123 | 277 | 321 | 44 | 541 | 79.9 | 249.9 | 224.1 | 36.3 | 420.7 |
| >hg38_21_+_8433172_8446622_RNA45SN1_45S_50ntFlanks@12418.12456.39\|CGATGTCGGCTCTTCCTATCATTGTGAAGCAGAATTCAC | >hg38_21_+_8433172_8446622_RNA45SN1_45S_50ntFlanks | 12418 | 12456 | 39 | CGATGTCGGCTCTTCCTATCATTGTGAAGCAGAATTCAC | MB468 | 28 | 156 | 99 | 5 | 86 | 14.7 | 127.4 | 61.2 | 4.3 | 70.2 |
| >hg38_21_+_8433172_8446622_RNA45SN1_45S_50ntFlanks@12418.12457.40\|CGATGTCGGCTCTTCCTATCATTGTGAAGCAGAATTCACC | >hg38_21_+_8433172_8446622_RNA45SN1_45S_50ntFlanks | 12418 | 12457 | 40 | CGATGTCGGCTCTTCCTATCATTGTGAAGCAGAATTCACC | MB231 | 17 | 16 | 49 | 1 | 15 | 11.2 | 13.6 | 30.0 | 0.6 | 14.3 |
| >hg38_21_+_8433172_8446622_RNA45SN1_45S_50ntFlanks@12419.12436.18\|GATGTCGGCTCTTCCTAT | >hg38_21_+_8433172_8446622_RNA45SN1_45S_50ntFlanks | 12419 | 12436 | 18 | GATGTCGGCTCTTCCTAT | MB231 | 26 | 26 | 39 | 1 | 6 | 17.2 | 19.8 | 28.8 | 1.4 | 10.8 |
| >hg38_21_+_8433172_8446622_RNA45SN1_45S_50ntFlanks@12419.12436.18\|GATGTCGGCTCTTCCTAT | >hg38_21_+_8433172_8446622_RNA45SN1_45S_50ntFlanks | 12419 | 12436 | 18 | GATGTCGGCTCTTCCTAT | MB468 | 212 | 332 | 176 | 17 | 67 | 85.0 | 50.3 | 47.8 | 4.7 | 16.7 |
| >hg38_21_+_8433172_8446622_RNA45SN1_45S_50ntFlanks@12419.12437.19\|GATGTCGGCTCTTCCTATC | >hg38_21_+_8433172_8446622_RNA45SN1_45S_50ntFlanks | 12419 | 12437 | 19 | GATGTCGGCTCTTCCTATC | BT20 | 43 | 32 | 100 | 41 | 33 | 19.5 | 9.6 | 48.1 | 65.7 | 10.5 |
| >hg38_21_+_8433172_8446622_RNA45SN1_45S_50ntFlanks@12419.12437.19\|GATGTCGGCTCTTCCTATC | >hg38_21_+_8433172_8446622_RNA45SN1_45S_50ntFlanks | 12419 | 12437 | 19 | GATGTCGGCTCTTCCTATC | MB231 | 221 | 181 | 429 | 5 | 61 | 181.8 | 147.5 | 333.1 | 4.6 | 47.4 |
| >hg38_21_+_8433172_8446622_RNA45SN1_45S_50ntFlanks@12419.12437.19\|GATGTCGGCTCTTCCTATC | >hg38_21_+_8433172_8446622_RNA45SN1_45S_50ntFlanks | 12419 | 12437 | 19 | GATGTCGGCTCTTCCTATC | MB468 | 77 | 111 | 84 | 6 | 27 | 36.0 | 27.1 | 18.6 | 2.0 | 7.3 |
| >hg38_21_+_8433172_8446622_RNA45SN1_45S_50ntFlanks@12419.12438.20\|GATGTCGGCTCTTCCTATCA | >hg38_21_+_8433172_8446622_RNA45SN1_45S_50ntFlanks | 12419 | 12438 | 20 | GATGTCGGCTCTTCCTATCA | BT20 | 25 | 22 | 43 | 8 | 30 | 17.2 | 12.7 | 15.4 | 10.7 | 3.2 |
| >hg38_21_+_8433172_8446622_RNA45SN1_45S_50ntFlanks@12419.12438.20\|GATGTCGGCTCTTCCTATCA | >hg38_21_+_8433172_8446622_RNA45SN1_45S_50ntFlanks | 12419 | 12438 | 20 | GATGTCGGCTCTTCCTATCA | MB231 | 113 | 125 | 221 | 3 | 50 | 89.4 | 134.3 | 240.4 | 4.6 | 56.6 |
| >hg38_21_+_8433172_8446622_RNA45SN1_45S_50ntFlanks@12419.12439.21\|GATGTCGGCTCTTCCTATCAT | >hg38_21_+_8433172_8446622_RNA45SN1_45S_50ntFlanks | 12419 | 12439 | 21 | GATGTCGGCTCTTCCTATCAT | BT20 | 25 | 16 | 45 | 7 | 5 | 5.2 | 7.8 | 14.0 | 12.5 | 8.0 |
| >hg38_21_+_8433172_8446622_RNA45SN1_45S_50ntFlanks@12419.12439.21\|GATGTCGGCTCTTCCTATCAT | >hg38_21_+_8433172_8446622_RNA45SN1_45S_50ntFlanks | 12419 | 12439 | 21 | GATGTCGGCTCTTCCTATCAT | MB231 | 84 | 67 | 183 | 1 | 19 | 71.8 | 61.2 | 152.9 | 2.1 | 32.6 |
| >hg38_21_+_8433172_8446622_RNA45SN1_45S_50ntFlanks@12419.12440.22\|GATGTCGGCTCTTCCTATCATT | >hg38_21_+_8433172_8446622_RNA45SN1_45S_50ntFlanks | 12419 | 12440 | 22 | GATGTCGGCTCTTCCTATCATT | BT20 | 11 | 7 | 26 | 4 | 6 | 5.8 | 2.5 | 4.5 | 6.2 | 5.5 |
| >hg38_21_+_8433172_8446622_RNA45SN1_45S_50ntFlanks@12419.12440.22\|GATGTCGGCTCTTCCTATCATT | >hg38_21_+_8433172_8446622_RNA45SN1_45S_50ntFlanks | 12419 | 12440 | 22 | GATGTCGGCTCTTCCTATCATT | MB231 | 44 | 39 | 99 | 2 | 12 | 40.8 | 43.2 | 111.0 | 2.7 | 20.3 |
| >hg38_21_+_8433172_8446622_RNA45SN1_45S_50ntFlanks@12419.12441.23\|GATGTCGGCTCTTCCTATCATTG | >hg38_21_+_8433172_8446622_RNA45SN1_45S_50ntFlanks | 12419 | 12441 | 23 | GATGTCGGCTCTTCCTATCATTG | MB231 | 45 | 49 | 136 | 1 | 16 | 44.9 | 51.1 | 128.6 | 1.7 | 27.3 |
| >hg38_21_+_8433172_8446622_RNA45SN1_45S_50ntFlanks@12419.12441.23\|GATGTCGGCTCTTCCTATCATTG | >hg38_21_+_8433172_8446622_RNA45SN1_45S_50ntFlanks | 12419 | 12441 | 23 | GATGTCGGCTCTTCCTATCATTG | MB468 | 25 | 56 | 43 | 2 | 11 | 14.2 | 22.0 | 24.7 | 1.2 | 2.9 |
| >hg38_21_+_8433172_8446622_RNA45SN1_45S_50ntFlanks@12419.12442.24\|GATGTCGGCTCTTCCTATCATTGT | >hg38_21_+_8433172_8446622_RNA45SN1_45S_50ntFlanks | 12419 | 12442 | 24 | GATGTCGGCTCTTCCTATCATTGT | BT20 | 13 | 10 | 43 | 32 | 6 | 7.6 | 1.4 | 20.9 | 55.0 | 5.2 |
| >hg38_21_+_8433172_8446622_RNA45SN1_45S_50ntFlanks@12419.12442.24\|GATGTCGGCTCTTCCTATCATTGT | >hg38_21_+_8433172_8446622_RNA45SN1_45S_50ntFlanks | 12419 | 12442 | 24 | GATGTCGGCTCTTCCTATCATTGT | MB231 | 53 | 54 | 225 | 1 | 16 | 42.6 | 50.7 | 166.3 | 2.0 | 26.9 |

| | | | | | | | | | | | | | | | | |
|---|---|---|---|---|---|---|---|---|---|---|---|---|---|---|---|---|
| >hg38_21_+_8433172_8446622_RNA45SN1_45S_50ntFlanks@12419.12443.25\|GATGTCGGCTCTTCCTATCATTGTG | >hg38_21_+_8433172_8446622_RNA45SN1_45S_50ntFlanks | 12419 | 12443 | 25 | GATGTCGGCTCTTCCTATCATTGTG | BT20 | 14 | 10 | 27 | 36 | 11 | 9.9 | 4.2 | 13.4 | 56.0 | 4.3 |
| >hg38_21_+_8433172_8446622_RNA45SN1_45S_50ntFlanks@12419.12443.25\|GATGTCGGCTCTTCCTATCATTGTG | >hg38_21_+_8433172_8446622_RNA45SN1_45S_50ntFlanks | 12419 | 12443 | 25 | GATGTCGGCTCTTCCTATCATTGTG | MB231 | 50 | 50 | 165 | 3 | 28 | 43.5 | 53.1 | 136.5 | 3.5 | 35.8 |
| >hg38_21_+_8433172_8446622_RNA45SN1_45S_50ntFlanks@12419.12443.25\|GATGTCGGCTCTTCCTATCATTGTG | >hg38_21_+_8433172_8446622_RNA45SN1_45S_50ntFlanks | 12419 | 12443 | 25 | GATGTCGGCTCTTCCTATCATTGTG | MB468 | 36 | 114 | 63 | 8 | 16 | 22.7 | 36.9 | 39.8 | 4.2 | 6.4 |
| >hg38_21_+_8433172_8446622_RNA45SN1_45S_50ntFlanks@12419.12444.26\|GATGTCGGCTCTTCCTATCATTGTGA | >hg38_21_+_8433172_8446622_RNA45SN1_45S_50ntFlanks | 12419 | 12444 | 26 | GATGTCGGCTCTTCCTATCATTGTGA | BT20 | 12 | 10 | 30 | 77 | 11 | 7.7 | 1.8 | 16.9 | 129.3 | 1.8 |
| >hg38_21_+_8433172_8446622_RNA45SN1_45S_50ntFlanks@12419.12444.26\|GATGTCGGCTCTTCCTATCATTGTGA | >hg38_21_+_8433172_8446622_RNA45SN1_45S_50ntFlanks | 12419 | 12444 | 26 | GATGTCGGCTCTTCCTATCATTGTGA | MB231 | 38 | 40 | 134 | 2 | 25 | 28.6 | 36.9 | 93.1 | 2.9 | 28.5 |
| >hg38_21_+_8433172_8446622_RNA45SN1_45S_50ntFlanks@12419.12444.26\|GATGTCGGCTCTTCCTATCATTGTGA | >hg38_21_+_8433172_8446622_RNA45SN1_45S_50ntFlanks | 12419 | 12444 | 26 | GATGTCGGCTCTTCCTATCATTGTGA | MB468 | 10 | 37 | 16 | 2 | 4 | 6.1 | 12.5 | 9.7 | 0.8 | 2.0 |
| >hg38_21_+_8433172_8446622_RNA45SN1_45S_50ntFlanks@12419.12445.27\|GATGTCGGCTCTTCCTATCATTGTGAA | >hg38_21_+_8433172_8446622_RNA45SN1_45S_50ntFlanks | 12419 | 12445 | 27 | GATGTCGGCTCTTCCTATCATTGTGAA | MB231 | 13 | 14 | 45 | 1 | 9 | 9.6 | 7.7 | 30.2 | 0.9 | 9.5 |
| >hg38_21_+_8433172_8446622_RNA45SN1_45S_50ntFlanks@12419.12446.28\|GATGTCGGCTCTTCCTATCATTGTGAAG | >hg38_21_+_8433172_8446622_RNA45SN1_45S_50ntFlanks | 12419 | 12446 | 28 | GATGTCGGCTCTTCCTATCATTGTGAAG | MB231 | 38 | 17 | 104 | 4 | 15 | 36.9 | 9.4 | 68.9 | 2.5 | 11.2 |
| >hg38_21_+_8433172_8446622_RNA45SN1_45S_50ntFlanks@12419.12446.28\|GATGTCGGCTCTTCCTATCATTGTGAAG | >hg38_21_+_8433172_8446622_RNA45SN1_45S_50ntFlanks | 12419 | 12446 | 28 | GATGTCGGCTCTTCCTATCATTGTGAAG | MB468 | 27 | 53 | 45 | 7 | 8 | 17.3 | 30.7 | 26.1 | 1.6 | 3.6 |
| >hg38_21_+_8433172_8446622_RNA45SN1_45S_50ntFlanks@12419.12447.29\|GATGTCGGCTCTTCCTATCATTGTGAAGC | >hg38_21_+_8433172_8446622_RNA45SN1_45S_50ntFlanks | 12419 | 12447 | 29 | GATGTCGGCTCTTCCTATCATTGTGAAGC | MB231 | 52 | 29 | 187 | 4 | 21 | 37.3 | 10.0 | 120.3 | 2.5 | 15.4 |
| >hg38_21_+_8433172_8446622_RNA45SN1_45S_50ntFlanks@12419.12448.30\|GATGTCGGCTCTTCCTATCATTGTGAAGCA | >hg38_21_+_8433172_8446622_RNA45SN1_45S_50ntFlanks | 12419 | 12448 | 30 | GATGTCGGCTCTTCCTATCATTGTGAAGCA | BT20 | 22 | 12 | 74 | 13 | 20 | 7.3 | 5.9 | 53.7 | 12.5 | 1.6 |
| >hg38_21_+_8433172_8446622_RNA45SN1_45S_50ntFlanks@12419.12448.30\|GATGTCGGCTCTTCCTATCATTGTGAAGCA | >hg38_21_+_8433172_8446622_RNA45SN1_45S_50ntFlanks | 12419 | 12448 | 30 | GATGTCGGCTCTTCCTATCATTGTGAAGCA | MB231 | 141 | 57 | 436 | 8 | 45 | 115.8 | 38.8 | 338.9 | 5.3 | 32.0 |
| >hg38_21_+_8433172_8446622_RNA45SN1_45S_50ntFlanks@12419.12449.31\|GATGTCGGCTCTTCCTATCATTGTGAAGCAG | >hg38_21_+_8433172_8446622_RNA45SN1_45S_50ntFlanks | 12419 | 12449 | 31 | GATGTCGGCTCTTCCTATCATTGTGAAGCAG | MB231 | 76 | 29 | 154 | 7 | 30 | 62.0 | 20.2 | 107.9 | 4.3 | 23.1 |
| >hg38_21_+_8433172_8446622_RNA45SN1_45S_50ntFlanks@12419.12450.32\|GATGTCGGCTCTTCCTATCATTGTGAAGCAGA | >hg38_21_+_8433172_8446622_RNA45SN1_45S_50ntFlanks | 12419 | 12450 | 32 | GATGTCGGCTCTTCCTATCATTGTGAAGCAGA | MB231 | 22 | 11 | 38 | 3 | 11 | 15.7 | 5.3 | 23.9 | 2.5 | 9.8 |
| >hg38_21_+_8433172_8446622_RNA45SN1_45S_50ntFlanks@12419.12451.33\|GATGTCGGCTCTTCCTATCATTGTGAAGCAGAA | >hg38_21_+_8433172_8446622_RNA45SN1_45S_50ntFlanks | 12419 | 12451 | 33 | GATGTCGGCTCTTCCTATCATTGTGAAGCAGAA | MB231 | 11 | 7 | 13 | 1 | 10 | 7.8 | 3.4 | 8.8 | 0.8 | 8.8 |
| >hg38_21_+_8433172_8446622_RNA45SN1_45S_50ntFlanks@12419.12452.34\|GATGTCGGCTCTTCCTATCATTGTGAAGCAGAAT | >hg38_21_+_8433172_8446622_RNA45SN1_45S_50ntFlanks | 12419 | 12452 | 34 | GATGTCGGCTCTTCCTATCATTGTGAAGCAGAAT | MB231 | 63 | 32 | 80 | 8 | 30 | 47.4 | 20.7 | 68.9 | 5.9 | 23.9 |
| >hg38_21_+_8433172_8446622_RNA45SN1_45S_50ntFlanks@12419.12453.35\|GATGTCGGCTCTTCCTATCATTGTGAAGCAGAATT | >hg38_21_+_8433172_8446622_RNA45SN1_45S_50ntFlanks | 12419 | 12453 | 35 | GATGTCGGCTCTTCCTATCATTGTGAAGCAGAATT | MB231 | 41 | 25 | 45 | 6 | 19 | 29.0 | 13.9 | 32.5 | 4.6 | 14.1 |
| >hg38_21_+_8433172_8446622_RNA45SN1_45S_50ntFlanks@12419.12454.36\|GATGTCGGCTCTTCCTATCATTGTGAAGCAGAATTC | >hg38_21_+_8433172_8446622_RNA45SN1_45S_50ntFlanks | 12419 | 12454 | 36 | GATGTCGGCTCTTCCTATCATTGTGAAGCAGAATTC | MB231 | 37 | 28 | 47 | 8 | 19 | 27.0 | 16.6 | 31.3 | 7.8 | 15.0 |
| >hg38_21_+_8433172_8446622_RNA45SN1_45S_50ntFlanks@12419.12455.37\|GATGTCGGCTCTTCCTATCATTGTGAAGCAGAATTCA | >hg38_21_+_8433172_8446622_RNA45SN1_45S_50ntFlanks | 12419 | 12455 | 37 | GATGTCGGCTCTTCCTATCATTGTGAAGCAGAATTCA | MB231 | 17 | 14 | 23 | 2 | 10 | 10.8 | 7.9 | 15.9 | 2.0 | 8.7 |
| >hg38_21_+_8433172_8446622_RNA45SN1_45S_50ntFlanks@12419.12455.37\|GATGTCGGCTCTTCCTATCATTGTGAAGCAGAATTCA | >hg38_21_+_8433172_8446622_RNA45SN1_45S_50ntFlanks | 12419 | 12455 | 37 | GATGTCGGCTCTTCCTATCATTGTGAAGCAGAATTCA | MB468 | 18 | 51 | 33 | 7 | 35 | 13.4 | 56.2 | 26.8 | 5.0 | 35.1 |
| >hg38_21_+_8433172_8446622_RNA45SN1_45S_50ntFlanks@12419.12456.38\|GATGTCGGCTCTTCCTATCATTGTGAAGCAGAATTCAC | >hg38_21_+_8433172_8446622_RNA45SN1_45S_50ntFlanks | 12419 | 12456 | 38 | GATGTCGGCTCTTCCTATCATTGTGAAGCAGAATTCAC | MB231 | 23 | 13 | 64 | 1 | 13 | 15.3 | 8.4 | 50.7 | 0.5 | 12.7 |
| >hg38_21_+_8433172_8446622_RNA45SN1_45S_50ntFlanks@12419.12457.39\|GATGTCGGCTCTTCCTATCATTGTGAAGCAGAATTCACC | >hg38_21_+_8433172_8446622_RNA45SN1_45S_50ntFlanks | 12419 | 12457 | 39 | GATGTCGGCTCTTCCTATCATTGTGAAGCAGAATTCACC | MB231 | 57 | 47 | 141 | 4 | 21 | 41.6 | 37.7 | 89.9 | 3.0 | 14.9 |
| >hg38_21_+_8433172_8446622_RNA45SN1_45S_50ntFlanks@12419.12458.40\|GATGTCGGCTCTTCCTATCATTGTGAAGCAGAATTCACCA | >hg38_21_+_8433172_8446622_RNA45SN1_45S_50ntFlanks | 12419 | 12458 | 40 | GATGTCGGCTCTTCCTATCATTGTGAAGCAGAATTCACCA | MB231 | 23 | 19 | 71 | 1 | 14 | 15.4 | 15.3 | 44.5 | 0.6 | 13.0 |
| >hg38_21_+_8433172_8446622_RNA45SN1_45S_50ntFlanks@12420.12437.18\|ATGTCGGCTCTTCCTATC | >hg38_21_+_8433172_8446622_RNA45SN1_45S_50ntFlanks | 12420 | 12437 | 18 | ATGTCGGCTCTTCCTATC | BT20 | 18 | 12 | 41 | 5 | 9 | 11.3 | 8.3 | 37.5 | 4.1 | 1.4 |
| >hg38_21_+_8433172_8446622_RNA45SN1_45S_50ntFlanks@12420.12437.18\|ATGTCGGCTCTTCCTATC | >hg38_21_+_8433172_8446622_RNA45SN1_45S_50ntFlanks | 12420 | 12437 | 18 | ATGTCGGCTCTTCCTATC | MB231 | 58 | 60 | 111 | 2 | 40 | 45.3 | 50.8 | 90.7 | 2.7 | 48.8 |
| >hg38_21_+_8433172_8446622_RNA45SN1_45S_50ntFlanks@12420.12437.18\|ATGTCGGCTCTTCCTATC | >hg38_21_+_8433172_8446622_RNA45SN1_45S_50ntFlanks | 12420 | 12437 | 18 | ATGTCGGCTCTTCCTATC | MB468 | 548 | 808 | 568 | 68 | 206 | 207.9 | 65.1 | 209.8 | 33.1 | 52.3 |
| >hg38_21_+_8433172_8446622_RNA45SN1_45S_50ntFlanks@12420.12438.19\|ATGTCGGCTCTTCCTATCA | >hg38_21_+_8433172_8446622_RNA45SN1_45S_50ntFlanks | 12420 | 12438 | 19 | ATGTCGGCTCTTCCTATCA | BT20 | 13 | 20 | 17 | 1 | 5 | 8.6 | 22.1 | 11.3 | 2.5 | 8.7 |
| >hg38_21_+_8433172_8446622_RNA45SN1_45S_50ntFlanks@12420.12438.19\|ATGTCGGCTCTTCCTATCA | >hg38_21_+_8433172_8446622_RNA45SN1_45S_50ntFlanks | 12420 | 12438 | 19 | ATGTCGGCTCTTCCTATCA | MB231 | 31 | 44 | 56 | 1 | 33 | 24.7 | 47.2 | 63.4 | 1.8 | 56.3 |
| >hg38_21_+_8433172_8446622_RNA45SN1_45S_50ntFlanks@12420.12438.19\|ATGTCGGCTCTTCCTATCA | >hg38_21_+_8433172_8446622_RNA45SN1_45S_50ntFlanks | 12420 | 12438 | 19 | ATGTCGGCTCTTCCTATCA | MB468 | 32 | 68 | 29 | 4 | 29 | 17.0 | 18.2 | 9.4 | 2.1 | 9.8 |
| >hg38_21_+_8433172_8446622_RNA45SN1_45S_50ntFlanks@12420.12439.20\|ATGTCGGCTCTTCCTATCAT | >hg38_21_+_8433172_8446622_RNA45SN1_45S_50ntFlanks | 12420 | 12439 | 20 | ATGTCGGCTCTTCCTATCAT | BT20 | 13 | 24 | 21 | 3 | 6 | 7.8 | 28.8 | 4.0 | 4.0 | 5.4 |
| >hg38_21_+_8433172_8446622_RNA45SN1_45S_50ntFlanks@12420.12439.20\|ATGTCGGCTCTTCCTATCAT | >hg38_21_+_8433172_8446622_RNA45SN1_45S_50ntFlanks | 12420 | 12439 | 20 | ATGTCGGCTCTTCCTATCAT | MB231 | 34 | 37 | 70 | 1 | 26 | 30.4 | 37.5 | 75.5 | 2.1 | 45.1 |
| >hg38_21_+_8433172_8446622_RNA45SN1_45S_50ntFlanks@12420.12439.20\|ATGTCGGCTCTTCCTATCAT | >hg38_21_+_8433172_8446622_RNA45SN1_45S_50ntFlanks | 12420 | 12439 | 20 | ATGTCGGCTCTTCCTATCAT | MB468 | 69 | 83 | 92 | 7 | 37 | 18.9 | 20.1 | 36.6 | 3.9 | 4.9 |
| >hg38_21_+_8433172_8446622_RNA45SN1_45S_50ntFlanks@12420.12440.21\|ATGTCGGCTCTTCCTATCATT | >hg38_21_+_8433172_8446622_RNA45SN1_45S_50ntFlanks | 12420 | 12440 | 21 | ATGTCGGCTCTTCCTATCATT | MB231 | 31 | 30 | 74 | 1 | 20 | 32.7 | 33.2 | 78.5 | 1.8 | 34.7 |
| >hg38_21_+_8433172_8446622_RNA45SN1_45S_50ntFlanks@12420.12440.21\|ATGTCGGCTCTTCCTATCATT | >hg38_21_+_8433172_8446622_RNA45SN1_45S_50ntFlanks | 12420 | 12440 | 21 | ATGTCGGCTCTTCCTATCATT | MB468 | 38 | 68 | 55 | 5 | 20 | 15.5 | 18.9 | 20.5 | 2.4 | 6.9 |
| >hg38_21_+_8433172_8446622_RNA45SN1_45S_50ntFlanks@12420.12441.22\|ATGTCGGCTCTTCCTATCATTG | >hg38_21_+_8433172_8446622_RNA45SN1_45S_50ntFlanks | 12420 | 12441 | 22 | ATGTCGGCTCTTCCTATCATTG | MB231 | 12 | 17 | 37 | 0 | 13 | 11.4 | 19.4 | 33.6 | 0.6 | 22.1 |
| >hg38_21_+_8433172_8446622_RNA45SN1_45S_50ntFlanks@12420.12441.22\|ATGTCGGCTCTTCCTATCATTG | >hg38_21_+_8433172_8446622_RNA45SN1_45S_50ntFlanks | 12420 | 12441 | 22 | ATGTCGGCTCTTCCTATCATTG | MB468 | 195 | 504 | 262 | 23 | 96 | 107.6 | 110.9 | 142.4 | 10.6 | 34.5 |
| >hg38_21_+_8433172_8446622_RNA45SN1_45S_50ntFlanks@12420.12442.23\|ATGTCGGCTCTTCCTATCATTGT | >hg38_21_+_8433172_8446622_RNA45SN1_45S_50ntFlanks | 12420 | 12442 | 23 | ATGTCGGCTCTTCCTATCATTGT | MB231 | 13 | 19 | 56 | 1 | 14 | 11.0 | 19.5 | 45.1 | 1.4 | 24.0 |
| >hg38_21_+_8433172_8446622_RNA45SN1_45S_50ntFlanks@12420.12442.23\|ATGTCGGCTCTTCCTATCATTGT | >hg38_21_+_8433172_8446622_RNA45SN1_45S_50ntFlanks | 12420 | 12442 | 23 | ATGTCGGCTCTTCCTATCATTGT | MB468 | 16 | 39 | 26 | 3 | 6 | 11.2 | 20.5 | 15.2 | 1.4 | 2.7 |
| >hg38_21_+_8433172_8446622_RNA45SN1_45S_50ntFlanks@12420.12443.24\|ATGTCGGCTCTTCCTATCATTGTG | >hg38_21_+_8433172_8446622_RNA45SN1_45S_50ntFlanks | 12420 | 12443 | 24 | ATGTCGGCTCTTCCTATCATTGTG | MB231 | 18 | 22 | 50 | 1 | 18 | 18.1 | 23.5 | 48.0 | 1.8 | 26.3 |
| >hg38_21_+_8433172_8446622_RNA45SN1_45S_50ntFlanks@12420.12443.24\|ATGTCGGCTCTTCCTATCATTGTG | >hg38_21_+_8433172_8446622_RNA45SN1_45S_50ntFlanks | 12420 | 12443 | 24 | ATGTCGGCTCTTCCTATCATTGTG | MB468 | 332 | 816 | 452 | 89 | 174 | 188.8 | 227.4 | 241.0 | 59.3 | 69.7 |
| >hg38_21_+_8433172_8446622_RNA45SN1_45S_50ntFlanks@12420.12444.25\|ATGTCGGCTCTTCCTATCATTGTGA | >hg38_21_+_8433172_8446622_RNA45SN1_45S_50ntFlanks | 12420 | 12444 | 25 | ATGTCGGCTCTTCCTATCATTGTGA | MB231 | 17 | 19 | 49 | 1 | 22 | 13.6 | 18.9 | 40.9 | 2.4 | 37.6 |
| >hg38_21_+_8433172_8446622_RNA45SN1_45S_50ntFlanks@12420.12444.25\|ATGTCGGCTCTTCCTATCATTGTGA | >hg38_21_+_8433172_8446622_RNA45SN1_45S_50ntFlanks | 12420 | 12444 | 25 | ATGTCGGCTCTTCCTATCATTGTGA | MB468 | 107 | 316 | 148 | 22 | 48 | 52.9 | 98.1 | 71.3 | 9.2 | 17.5 |
| >hg38_21_+_8433172_8446622_RNA45SN1_45S_50ntFlanks@12420.12445.26\|ATGTCGGCTCTTCCTATCATTGTGAA | >hg38_21_+_8433172_8446622_RNA45SN1_45S_50ntFlanks | 12420 | 12445 | 26 | ATGTCGGCTCTTCCTATCATTGTGAA | MB468 | 82 | 208 | 94 | 12 | 35 | 35.6 | 30.2 | 31.6 | 3.6 | 11.7 |
| >hg38_21_+_8433172_8446622_RNA45SN1_45S_50ntFlanks@12420.12446.27\|ATGTCGGCTCTTCCTATCATTGTGAAG | >hg38_21_+_8433172_8446622_RNA45SN1_45S_50ntFlanks | 12420 | 12446 | 27 | ATGTCGGCTCTTCCTATCATTGTGAAG | MB231 | 25 | 13 | 58 | 3 | 31 | 26.5 | 9.6 | 38.8 | 2.3 | 28.1 |
| >hg38_21_+_8433172_8446622_RNA45SN1_45S_50ntFlanks@12420.12446.27\|ATGTCGGCTCTTCCTATCATTGTGAAG | >hg38_21_+_8433172_8446622_RNA45SN1_45S_50ntFlanks | 12420 | 12446 | 27 | ATGTCGGCTCTTCCTATCATTGTGAAG | MB468 | 490 | 828 | 810 | 127 | 154 | 238.6 | 396.9 | 346.7 | 30.4 | 62.8 |
| >hg38_21_+_8433172_8446622_RNA45SN1_45S_50ntFlanks@12420.12447.28\|ATGTCGGCTCTTCCTATCATTGTGAAGC | >hg38_21_+_8433172_8446622_RNA45SN1_45S_50ntFlanks | 12420 | 12447 | 28 | ATGTCGGCTCTTCCTATCATTGTGAAGC | MB231 | 26 | 18 | 56 | 3 | 62 | 18.5 | 9.3 | 37.6 | 2.8 | 66.1 |
| >hg38_21_+_8433172_8446622_RNA45SN1_45S_50ntFlanks@12420.12447.28\|ATGTCGGCTCTTCCTATCATTGTGAAGC | >hg38_21_+_8433172_8446622_RNA45SN1_45S_50ntFlanks | 12420 | 12447 | 28 | ATGTCGGCTCTTCCTATCATTGTGAAGC | MB468 | 15 | 24 | 21 | 4 | 10 | 6.8 | 13.6 | 9.4 | 0.8 | 4.9 |
| >hg38_21_+_8433172_8446622_RNA45SN1_45S_50ntFlanks@12420.12448.29\|ATGTCGGCTCTTCCTATCATTGTGAAGCA | >hg38_21_+_8433172_8446622_RNA45SN1_45S_50ntFlanks | 12420 | 12448 | 29 | ATGTCGGCTCTTCCTATCATTGTGAAGCA | MB231 | 68 | 37 | 137 | 4 | 122 | 54.4 | 26.2 | 99.2 | 3.4 | 94.2 |
| >hg38_21_+_8433172_8446622_RNA45SN1_45S_50ntFlanks@12420.12448.29\|ATGTCGGCTCTTCCTATCATTGTGAAGCA | >hg38_21_+_8433172_8446622_RNA45SN1_45S_50ntFlanks | 12420 | 12448 | 29 | ATGTCGGCTCTTCCTATCATTGTGAAGCA | MB468 | 19 | 36 | 27 | 4 | 24 | 10.3 | 32.6 | 16.9 | 1.5 | 11.1 |
| >hg38_21_+_8433172_8446622_RNA45SN1_45S_50ntFlanks@12420.12449.30\|ATGTCGGCTCTTCCTATCATTGTGAAGCAG | >hg38_21_+_8433172_8446622_RNA45SN1_45S_50ntFlanks | 12420 | 12449 | 30 | ATGTCGGCTCTTCCTATCATTGTGAAGCAG | MB231 | 30 | 16 | 40 | 5 | 69 | 23.8 | 12.6 | 32.7 | 3.8 | 55.4 |

| | | | | | | | | | | | | | | | | |
|---|---|---|---|---|---|---|---|---|---|---|---|---|---|---|---|---|
| >hg38_21_+_8433172_8446622_RNA45SN1_45S_50ntFlanks@12420.12449.30\|ATGTCGGCTCTTCCTATCATTGTGAAGCAG | >hg38_21_+_8433172_8446622_RNA45SN1_45S_50ntFlanks | 12420 | 12449 | 30 | ATGTCGGCTCTTCCTATCATTGTGAAGCAG | MB468 | 61 | 72 | 79 | 17 | 32 | 21.4 | 31.0 | 19.9 | 5.5 | 13.6 |
| >hg38_21_+_8433172_8446622_RNA45SN1_45S_50ntFlanks@12420.12450.31\|ATGTCGGCTCTTCCTATCATTGTGAAGCAGA | >hg38_21_+_8433172_8446622_RNA45SN1_45S_50ntFlanks | 12420 | 12450 | 31 | ATGTCGGCTCTTCCTATCATTGTGAAGCAGA | MB231 | 12 | 7 | 9 | 2 | 37 | 8.4 | 4.5 | 5.6 | 2.3 | 33.6 |
| >hg38_21_+_8433172_8446622_RNA45SN1_45S_50ntFlanks@12420.12450.31\|ATGTCGGCTCTTCCTATCATTGTGAAGCAGA | >hg38_21_+_8433172_8446622_RNA45SN1_45S_50ntFlanks | 12420 | 12450 | 31 | ATGTCGGCTCTTCCTATCATTGTGAAGCAGA | MB468 | 23 | 20 | 28 | 8 | 12 | 9.3 | 12.9 | 11.3 | 3.0 | 5.2 |
| >hg38_21_+_8433172_8446622_RNA45SN1_45S_50ntFlanks@12420.12451.32\|ATGTCGGCTCTTCCTATCATTGTGAAGCAGAA | >hg38_21_+_8433172_8446622_RNA45SN1_45S_50ntFlanks | 12420 | 12451 | 32 | ATGTCGGCTCTTCCTATCATTGTGAAGCAGAA | MB468 | 17 | 13 | 12 | 4 | 11 | 3.6 | 6.0 | 3.9 | 1.8 | 6.0 |
| >hg38_21_+_8433172_8446622_RNA45SN1_45S_50ntFlanks@12420.12452.33\|ATGTCGGCTCTTCCTATCATTGTGAAGCAGAAT | >hg38_21_+_8433172_8446622_RNA45SN1_45S_50ntFlanks | 12420 | 12452 | 33 | ATGTCGGCTCTTCCTATCATTGTGAAGCAGAAT | MB231 | 20 | 12 | 12 | 6 | 51 | 16.2 | 6.3 | 7.3 | 4.0 | 36.7 |
| >hg38_21_+_8433172_8446622_RNA45SN1_45S_50ntFlanks@12420.12453.34\|ATGTCGGCTCTTCCTATCATTGTGAAGCAGAATT | >hg38_21_+_8433172_8446622_RNA45SN1_45S_50ntFlanks | 12420 | 12453 | 34 | ATGTCGGCTCTTCCTATCATTGTGAAGCAGAATT | MB231 | 17 | 11 | 13 | 2 | 30 | 12.0 | 6.0 | 8.0 | 1.1 | 21.8 |
| >hg38_21_+_8433172_8446622_RNA45SN1_45S_50ntFlanks@12420.12454.35\|ATGTCGGCTCTTCCTATCATTGTGAAGCAGAATTC | >hg38_21_+_8433172_8446622_RNA45SN1_45S_50ntFlanks | 12420 | 12454 | 35 | ATGTCGGCTCTTCCTATCATTGTGAAGCAGAATTC | MB231 | 13 | 10 | 9 | 3 | 22 | 8.4 | 5.8 | 7.0 | 3.0 | 18.9 |
| >hg38_21_+_8433172_8446622_RNA45SN1_45S_50ntFlanks@12420.12454.35\|ATGTCGGCTCTTCCTATCATTGTGAAGCAGAATTC | >hg38_21_+_8433172_8446622_RNA45SN1_45S_50ntFlanks | 12420 | 12454 | 35 | ATGTCGGCTCTTCCTATCATTGTGAAGCAGAATTC | MB468 | 19 | 21 | 19 | 7 | 21 | 7.7 | 13.9 | 8.6 | 3.5 | 17.8 |
| >hg38_21_+_8433172_8446622_RNA45SN1_45S_50ntFlanks@12420.12455.36\|ATGTCGGCTCTTCCTATCATTGTGAAGCAGAATTCA | >hg38_21_+_8433172_8446622_RNA45SN1_45S_50ntFlanks | 12420 | 12455 | 36 | ATGTCGGCTCTTCCTATCATTGTGAAGCAGAATTCA | MB468 | 230 | 325 | 272 | 59 | 425 | 117.2 | 325.4 | 193.0 | 39.1 | 403.9 |
| >hg38_21_+_8433172_8446622_RNA45SN1_45S_50ntFlanks@12420.12456.37\|ATGTCGGCTCTTCCTATCATTGTGAAGCAGAATTCAC | >hg38_21_+_8433172_8446622_RNA45SN1_45S_50ntFlanks | 12420 | 12456 | 37 | ATGTCGGCTCTTCCTATCATTGTGAAGCAGAATTCAC | MB231 | 18 | 15 | 20 | 1 | 94 | 11.8 | 7.7 | 14.2 | 1.2 | 82.6 |
| >hg38_21_+_8433172_8446622_RNA45SN1_45S_50ntFlanks@12420.12456.37\|ATGTCGGCTCTTCCTATCATTGTGAAGCAGAATTCAC | >hg38_21_+_8433172_8446622_RNA45SN1_45S_50ntFlanks | 12420 | 12456 | 37 | ATGTCGGCTCTTCCTATCATTGTGAAGCAGAATTCAC | MB468 | 50 | 143 | 69 | 6 | 58 | 24.7 | 100.9 | 16.0 | 4.8 | 41.6 |
| >hg38_21_+_8433172_8446622_RNA45SN1_45S_50ntFlanks@12420.12457.38\|ATGTCGGCTCTTCCTATCATTGTGAAGCAGAATTCACC | >hg38_21_+_8433172_8446622_RNA45SN1_45S_50ntFlanks | 12420 | 12457 | 38 | ATGTCGGCTCTTCCTATCATTGTGAAGCAGAATTCACC | MB231 | 50 | 43 | 150 | 2 | 78 | 35.9 | 29.6 | 103.5 | 1.7 | 67.6 |
| >hg38_21_+_8433172_8446622_RNA45SN1_45S_50ntFlanks@12420.12458.39\|ATGTCGGCTCTTCCTATCATTGTGAAGCAGAATTCACCA | >hg38_21_+_8433172_8446622_RNA45SN1_45S_50ntFlanks | 12420 | 12458 | 39 | ATGTCGGCTCTTCCTATCATTGTGAAGCAGAATTCACCA | MB231 | 22 | 20 | 63 | 1 | 87 | 14.7 | 14.7 | 45.4 | 0.9 | 75.9 |
| >hg38_21_+_8433172_8446622_RNA45SN1_45S_50ntFlanks@12420.12459.40\|ATGTCGGCTCTTCCTATCATTGTGAAGCAGAATTCACCAA | >hg38_21_+_8433172_8446622_RNA45SN1_45S_50ntFlanks | 12420 | 12459 | 40 | ATGTCGGCTCTTCCTATCATTGTGAAGCAGAATTCACCAA | MB231 | 16 | 15 | 16 | 1 | 168 | 11.6 | 13.1 | 11.5 | 0.7 | 176.2 |
| >hg38_21_+_8433172_8446622_RNA45SN1_45S_50ntFlanks@12420.12460.41\|ATGTCGGCTCTTCCTATCATTGTGAAGCAGAATTCACCAAG | >hg38_21_+_8433172_8446622_RNA45SN1_45S_50ntFlanks | 12420 | 12460 | 41 | ATGTCGGCTCTTCCTATCATTGTGAAGCAGAATTCACCAAG | MB231 | 15 | 14 | 15 | 1 | 149 | 10.3 | 13.4 | 11.3 | 1.1 | 159.6 |
| >hg38_21_+_8433172_8446622_RNA45SN1_45S_50ntFlanks@12423.12441.19\|TCGGCTCTTCCTATCATTG | >hg38_21_+_8433172_8446622_RNA45SN1_45S_50ntFlanks | 12423 | 12441 | 19 | TCGGCTCTTCCTATCATTG | MB468 | 15 | 27 | 15 | 0 | 4 | 5.0 | 6.8 | 5.3 | 0.6 | 5.0 |
| >hg38_21_+_8433172_8446622_RNA45SN1_45S_50ntFlanks@12423.12443.21\|TCGGCTCTTCCTATCATTGTG | >hg38_21_+_8433172_8446622_RNA45SN1_45S_50ntFlanks | 12423 | 12443 | 21 | TCGGCTCTTCCTATCATTGTG | MB468 | 36 | 43 | 36 | 3 | 17 | 9.1 | 7.8 | 13.0 | 1.1 | 7.2 |
| >hg38_21_+_8433172_8446622_RNA45SN1_45S_50ntFlanks@12423.12445.23\|TCGGCTCTTCCTATCATTGTGAA | >hg38_21_+_8433172_8446622_RNA45SN1_45S_50ntFlanks | 12423 | 12445 | 23 | TCGGCTCTTCCTATCATTGTGAA | MB468 | 12 | 17 | 12 | 0 | 2 | 5.0 | 3.4 | 3.3 | 0.3 | 2.3 |
| >hg38_21_+_8433172_8446622_RNA45SN1_45S_50ntFlanks@12423.12455.33\|TCGGCTCTTCCTATCATTGTGAAGCAGAATTCA | >hg38_21_+_8433172_8446622_RNA45SN1_45S_50ntFlanks | 12423 | 12455 | 33 | TCGGCTCTTCCTATCATTGTGAAGCAGAATTCA | MB468 | 13 | 9 | 14 | 5 | 15 | 5.0 | 6.4 | 8.9 | 2.5 | 9.4 |
| >hg38_21_+_8433172_8446622_RNA45SN1_45S_50ntFlanks@12423.12457.35\|TCGGCTCTTCCTATCATTGTGAAGCAGAATTCACC | >hg38_21_+_8433172_8446622_RNA45SN1_45S_50ntFlanks | 12423 | 12457 | 35 | TCGGCTCTTCCTATCATTGTGAAGCAGAATTCACC | MB231 | 16 | 16 | 40 | 0 | 3 | 12.2 | 9.6 | 28.0 | 0.5 | 5.9 |
| >hg38_21_+_8433172_8446622_RNA45SN1_45S_50ntFlanks@12431.12455.25\|TCCTATCATTGTGAAGCAGAATTCA | >hg38_21_+_8433172_8446622_RNA45SN1_45S_50ntFlanks | 12431 | 12455 | 25 | TCCTATCATTGTGAAGCAGAATTCA | MB468 | 10 | 21 | 6 | 5 | 22 | 3.7 | 4.5 | 2.8 | 2.1 | 8.6 |
| >hg38_21_+_8433172_8446622_RNA45SN1_45S_50ntFlanks@12432.12455.24\|CCTATCATTGTGAAGCAGAATTCA | >hg38_21_+_8433172_8446622_RNA45SN1_45S_50ntFlanks | 12432 | 12455 | 24 | CCTATCATTGTGAAGCAGAATTCA | MB468 | 32 | 32 | 19 | 35 | 24 | 5.1 | 12.8 | 6.7 | 25.7 | 8.9 |
| >hg38_21_+_8433172_8446622_RNA45SN1_45S_50ntFlanks@12433.12452.20\|CTATCATTGTGAAGCAGAAT | >hg38_21_+_8433172_8446622_RNA45SN1_45S_50ntFlanks | 12433 | 12452 | 20 | CTATCATTGTGAAGCAGAAT | MB468 | 12 | 7 | 8 | 5 | 3 | 3.0 | 3.6 | 2.8 | 1.3 | 1.4 |
| >hg38_21_+_8433172_8446622_RNA45SN1_45S_50ntFlanks@12433.12454.22\|CTATCATTGTGAAGCAGAATTC | >hg38_21_+_8433172_8446622_RNA45SN1_45S_50ntFlanks | 12433 | 12454 | 22 | CTATCATTGTGAAGCAGAATTC | MB468 | 19 | 20 | 12 | 34 | 4 | 9.4 | 10.8 | 8.6 | 20.5 | 1.4 |
| >hg38_21_+_8433172_8446622_RNA45SN1_45S_50ntFlanks@12433.12455.23\|CTATCATTGTGAAGCAGAATTCA | >hg38_21_+_8433172_8446622_RNA45SN1_45S_50ntFlanks | 12433 | 12455 | 23 | CTATCATTGTGAAGCAGAATTCA | MB468 | 23 | 10 | 12 | 34 | 16 | 4.1 | 5.7 | 6.0 | 21.8 | 8.1 |
| >hg38_21_+_8433172_8446622_RNA45SN1_45S_50ntFlanks@12434.12455.22\|TATCATTGTGAAGCAGAATTCA | >hg38_21_+_8433172_8446622_RNA45SN1_45S_50ntFlanks | 12434 | 12455 | 22 | TATCATTGTGAAGCAGAATTCA | MB468 | 31 | 23 | 18 | 11 | 16 | 5.9 | 4.0 | 14.6 | 2.2 | 5.5 |
| >hg38_21_+_8433172_8446622_RNA45SN1_45S_50ntFlanks@12435.12454.20\|ATCATTGTGAAGCAGAATTC | >hg38_21_+_8433172_8446622_RNA45SN1_45S_50ntFlanks | 12435 | 12454 | 20 | ATCATTGTGAAGCAGAATTC | MB468 | 48 | 34 | 29 | 52 | 5 | 37.3 | 13.6 | 37.9 | 31.5 | 1.4 |
| >hg38_21_+_8433172_8446622_RNA45SN1_45S_50ntFlanks@12435.12455.21\|ATCATTGTGAAGCAGAATTCA | >hg38_21_+_8433172_8446622_RNA45SN1_45S_50ntFlanks | 12435 | 12455 | 21 | ATCATTGTGAAGCAGAATTCA | MB468 | 235 | 81 | 116 | 147 | 51 | 67.9 | 24.4 | 119.0 | 79.6 | 16.0 |
| >hg38_21_+_8433172_8446622_RNA45SN1_45S_50ntFlanks@12436.12454.19\|TCATTGTGAAGCAGAATTC | >hg38_21_+_8433172_8446622_RNA45SN1_45S_50ntFlanks | 12436 | 12454 | 19 | TCATTGTGAAGCAGAATTC | MB468 | 136 | 80 | 91 | 137 | 17 | 88.8 | 37.2 | 104.9 | 79.9 | 6.1 |
| >hg38_21_+_8433172_8446622_RNA45SN1_45S_50ntFlanks@12436.12455.20\|TCATTGTGAAGCAGAATTCA | >hg38_21_+_8433172_8446622_RNA45SN1_45S_50ntFlanks | 12436 | 12455 | 20 | TCATTGTGAAGCAGAATTCA | MB468 | 545 | 121 | 274 | 284 | 137 | 211.4 | 40.3 | 279.9 | 138.0 | 42.0 |
| >hg38_21_+_8433172_8446622_RNA45SN1_45S_50ntFlanks@12436.12473.38\|TCATTGTGAAGCAGAATTCACCAAGCGTTGGATTGTTC | >hg38_21_+_8433172_8446622_RNA45SN1_45S_50ntFlanks | 12436 | 12473 | 38 | TCATTGTGAAGCAGAATTCACCAAGCGTTGGATTGTTC | MB468 | 29 | 32 | 88 | 17 | 11 | 11.7 | 20.6 | 44.4 | 10.5 | 9.3 |
| >hg38_21_+_8433172_8446622_RNA45SN1_45S_50ntFlanks@12437.12454.18\|CATTGTGAAGCAGAATTC | >hg38_21_+_8433172_8446622_RNA45SN1_45S_50ntFlanks | 12437 | 12454 | 18 | CATTGTGAAGCAGAATTC | MB468 | 307 | 191 | 182 | 321 | 45 | 70.4 | 129.8 | 147.8 | 234.0 | 18.2 |
| >hg38_21_+_8433172_8446622_RNA45SN1_45S_50ntFlanks@12437.12455.19\|CATTGTGAAGCAGAATTCA | >hg38_21_+_8433172_8446622_RNA45SN1_45S_50ntFlanks | 12437 | 12455 | 19 | CATTGTGAAGCAGAATTCA | MB468 | 2567 | 497 | 1369 | 1337 | 617 | 937.4 | 173.0 | 1185.4 | 622.6 | 356.0 |
| >hg38_21_+_8433172_8446622_RNA45SN1_45S_50ntFlanks@12437.12456.20\|CATTGTGAAGCAGAATTCAC | >hg38_21_+_8433172_8446622_RNA45SN1_45S_50ntFlanks | 12437 | 12456 | 20 | CATTGTGAAGCAGAATTCAC | MB468 | 11 | 0 | 3 | 3 | 3 | 11.2 | 0.1 | 2.8 | 1.2 | 3.5 |
| >hg38_21_+_8433172_8446622_RNA45SN1_45S_50ntFlanks@12437.12472.36\|CATTGTGAAGCAGAATTCACCAAGCGTTGGATTGTT | >hg38_21_+_8433172_8446622_RNA45SN1_45S_50ntFlanks | 12437 | 12472 | 36 | CATTGTGAAGCAGAATTCACCAAGCGTTGGATTGTT | MB468 | 15 | 12 | 30 | 7 | 4 | 7.9 | 7.4 | 19.9 | 7.6 | 3.6 |
| >hg38_21_+_8433172_8446622_RNA45SN1_45S_50ntFlanks@12437.12473.37\|CATTGTGAAGCAGAATTCACCAAGCGTTGGATTGTTC | >hg38_21_+_8433172_8446622_RNA45SN1_45S_50ntFlanks | 12437 | 12473 | 37 | CATTGTGAAGCAGAATTCACCAAGCGTTGGATTGTTC | MB468 | 170 | 141 | 480 | 141 | 60 | 59.9 | 86.6 | 174.0 | 137.9 | 50.3 |
| >hg38_21_+_8433172_8446622_RNA45SN1_45S_50ntFlanks@12437.12474.38\|CATTGTGAAGCAGAATTCACCAAGCGTTGGATTGTTCA | >hg38_21_+_8433172_8446622_RNA45SN1_45S_50ntFlanks | 12437 | 12474 | 38 | CATTGTGAAGCAGAATTCACCAAGCGTTGGATTGTTCA | MB468 | 29 | 25 | 47 | 16 | 15 | 11.7 | 24.8 | 39.2 | 20.5 | 14.9 |
| >hg38_21_+_8433172_8446622_RNA45SN1_45S_50ntFlanks@12438.12455.18\|ATTGTGAAGCAGAATTCA | >hg38_21_+_8433172_8446622_RNA45SN1_45S_50ntFlanks | 12438 | 12455 | 18 | ATTGTGAAGCAGAATTCA | MB468 | 628 | 80 | 228 | 379 | 106 | 353.7 | 30.1 | 239.0 | 262.4 | 65.1 |
| >hg38_21_+_8433172_8446622_RNA45SN1_45S_50ntFlanks@12438.12472.35\|ATTGTGAAGCAGAATTCACCAAGCGTTGGATTGTT | >hg38_21_+_8433172_8446622_RNA45SN1_45S_50ntFlanks | 12438 | 12472 | 35 | ATTGTGAAGCAGAATTCACCAAGCGTTGGATTGTT | MB231 | 13 | 7 | 12 | 2 | 2 | 12.5 | 6.5 | 14.5 | 0.9 | 1.9 |
| >hg38_21_+_8433172_8446622_RNA45SN1_45S_50ntFlanks@12438.12473.36\|ATTGTGAAGCAGAATTCACCAAGCGTTGGATTGTTC | >hg38_21_+_8433172_8446622_RNA45SN1_45S_50ntFlanks | 12438 | 12473 | 36 | ATTGTGAAGCAGAATTCACCAAGCGTTGGATTGTTC | MB231 | 23 | 12 | 32 | 10 | 4 | 21.7 | 12.8 | 38.3 | 5.6 | 3.0 |
| >hg38_21_+_8433172_8446622_RNA45SN1_45S_50ntFlanks@12438.12473.36\|ATTGTGAAGCAGAATTCACCAAGCGTTGGATTGTTC | >hg38_21_+_8433172_8446622_RNA45SN1_45S_50ntFlanks | 12438 | 12473 | 36 | ATTGTGAAGCAGAATTCACCAAGCGTTGGATTGTTC | MB468 | 39 | 29 | 86 | 15 | 7 | 12.9 | 15.8 | 28.3 | 7.0 | 4.0 |
| >hg38_21_+_8433172_8446622_RNA45SN1_45S_50ntFlanks@12438.12474.37\|ATTGTGAAGCAGAATTCACCAAGCGTTGGATTGTTCA | >hg38_21_+_8433172_8446622_RNA45SN1_45S_50ntFlanks | 12438 | 12474 | 37 | ATTGTGAAGCAGAATTCACCAAGCGTTGGATTGTTCA | MB468 | 11 | 7 | 13 | 2 | 2 | 3.5 | 6.5 | 9.4 | 1.8 | 1.6 |
| >hg38_21_+_8433172_8446622_RNA45SN1_45S_50ntFlanks@12439.12472.34\|TTGTGAAGCAGAATTCACCAAGCGTTGGATTGTT | >hg38_21_+_8433172_8446622_RNA45SN1_45S_50ntFlanks | 12439 | 12472 | 34 | TTGTGAAGCAGAATTCACCAAGCGTTGGATTGTT | MB231 | 13 | 6 | 8 | 2 | 3 | 12.0 | 5.9 | 7.3 | 1.0 | 1.5 |
| >hg38_21_+_8433172_8446622_RNA45SN1_45S_50ntFlanks@12439.12473.35\|TTGTGAAGCAGAATTCACCAAGCGTTGGATTGTTC | >hg38_21_+_8433172_8446622_RNA45SN1_45S_50ntFlanks | 12439 | 12473 | 35 | TTGTGAAGCAGAATTCACCAAGCGTTGGATTGTTC | MB231 | 29 | 15 | 26 | 16 | 6 | 24.9 | 16.6 | 27.8 | 8.5 | 3.5 |
| >hg38_21_+_8433172_8446622_RNA45SN1_45S_50ntFlanks@12440.12473.34\|TGTGAAGCAGAATTCACCAAGCGTTGGATTGTTC | >hg38_21_+_8433172_8446622_RNA45SN1_45S_50ntFlanks | 12440 | 12473 | 34 | TGTGAAGCAGAATTCACCAAGCGTTGGATTGTTC | MB231 | 13 | 7 | 10 | 7 | 3 | 10.8 | 7.1 | 9.7 | 4.2 | 1.8 |
| >hg38_21_+_8433172_8446622_RNA45SN1_45S_50ntFlanks@12442.12473.32\|TGAAGCAGAATTCACCAAGCGTTGGATTGTTC | >hg38_21_+_8433172_8446622_RNA45SN1_45S_50ntFlanks | 12442 | 12473 | 32 | TGAAGCAGAATTCACCAAGCGTTGGATTGTTC | MB468 | 21 | 3 | 9 | 12 | 3 | 10.4 | 1.7 | 5.9 | 8.3 | 2.4 |
| >hg38_21_+_8433172_8446622_RNA45SN1_45S_50ntFlanks@12444.12466.23\|AAGCAGAATTCACCAAGCGTTGG | >hg38_21_+_8433172_8446622_RNA45SN1_45S_50ntFlanks | 12444 | 12466 | 23 | AAGCAGAATTCACCAAGCGTTGG | MB468 | 10 | 5 | 8 | 3 | 2 | 4.9 | 1.6 | 4.1 | 0.9 | 0.5 |
| >hg38_21_+_8433172_8446622_RNA45SN1_45S_50ntFlanks@12444.12467.24\|AAGCAGAATTCACCAAGCGTTGGA | >hg38_21_+_8433172_8446622_RNA45SN1_45S_50ntFlanks | 12444 | 12467 | 24 | AAGCAGAATTCACCAAGCGTTGGA | MB468 | 18 | 9 | 8 | 8 | 3 | 14.1 | 3.9 | 9.5 | 4.5 | 0.3 |
| >hg38_21_+_8433172_8446622_RNA45SN1_45S_50ntFlanks@12444.12473.30\|AAGCAGAATTCACCAAGCGTTGGATTGTTC | >hg38_21_+_8433172_8446622_RNA45SN1_45S_50ntFlanks | 12444 | 12473 | 30 | AAGCAGAATTCACCAAGCGTTGGATTGTTC | MB468 | 10 | 2 | 4 | 28 | 2 | 8.3 | 1.2 | 2.4 | 16.5 | 1.3 |
| >hg38_21_+_8433172_8446622_RNA45SN1_45S_50ntFlanks@12444.12475.32\|AAGCAGAATTCACCAAGCGTTGGATTGTTCAC | >hg38_21_+_8433172_8446622_RNA45SN1_45S_50ntFlanks | 12444 | 12475 | 32 | AAGCAGAATTCACCAAGCGTTGGATTGTTCAC | MB468 | 15 | 3 | 4 | 10 | 7 | 10.3 | 2.6 | 3.8 | 9.6 | 3.5 |

| | | | | | | | | | | | | | | | | |
|---|---|---|---|---|---|---|---|---|---|---|---|---|---|---|---|---|
| >hg38_21_+_8433172_8446622_RNA45SN1_45S_50ntFlanks@12444.12476.33\|AAGCAGAATTCACCAAGCGTTGGATTGTTCACC | >hg38_21_+_8433172_8446622_RNA45SN1_45S_50ntFlanks | 12444 | 12476 | 33 | AAGCAGAATTCACCAAGCGTTGGATTGTTCACC | MB231 | 26 | 17 | 16 | 11 | 4 | 21.5 | 14.8 | 12.2 | 6.9 | 1.9 |
| >hg38_21_+_8433172_8446622_RNA45SN1_45S_50ntFlanks@12444.12476.33\|AAGCAGAATTCACCAAGCGTTGGATTGTTCACC | >hg38_21_+_8433172_8446622_RNA45SN1_45S_50ntFlanks | 12444 | 12476 | 33 | AAGCAGAATTCACCAAGCGTTGGATTGTTCACC | MB468 | 37 | 12 | 23 | 43 | 26 | 16.7 | 10.1 | 10.2 | 39.0 | 18.4 |
| >hg38_21_+_8433172_8446622_RNA45SN1_45S_50ntFlanks@12444.12477.34\|AAGCAGAATTCACCAAGCGTTGGATTGTTCACCC | >hg38_21_+_8433172_8446622_RNA45SN1_45S_50ntFlanks | 12444 | 12477 | 34 | AAGCAGAATTCACCAAGCGTTGGATTGTTCACCC | MB231 | 18 | 13 | 17 | 6 | 2 | 16.2 | 11.1 | 19.7 | 3.6 | 0.9 |
| >hg38_21_+_8433172_8446622_RNA45SN1_45S_50ntFlanks@12444.12478.35\|AAGCAGAATTCACCAAGCGTTGGATTGTTCACCCA | >hg38_21_+_8433172_8446622_RNA45SN1_45S_50ntFlanks | 12444 | 12478 | 35 | AAGCAGAATTCACCAAGCGTTGGATTGTTCACCCA | MB468 | 17 | 16 | 15 | 12 | 14 | 9.1 | 9.0 | 7.0 | 15.8 | 11.4 |
| >hg38_21_+_8433172_8446622_RNA45SN1_45S_50ntFlanks@12444.12480.37\|AAGCAGAATTCACCAAGCGTTGGATTGTTCACCCACT | >hg38_21_+_8433172_8446622_RNA45SN1_45S_50ntFlanks | 12444 | 12480 | 37 | AAGCAGAATTCACCAAGCGTTGGATTGTTCACCCACT | MB231 | 16 | 20 | 24 | 5 | 6 | 14.1 | 15.4 | 22.7 | 4.0 | 4.5 |
| >hg38_21_+_8433172_8446622_RNA45SN1_45S_50ntFlanks@12444.12480.37\|AAGCAGAATTCACCAAGCGTTGGATTGTTCACCCACT | >hg38_21_+_8433172_8446622_RNA45SN1_45S_50ntFlanks | 12444 | 12480 | 37 | AAGCAGAATTCACCAAGCGTTGGATTGTTCACCCACT | MB468 | 14 | 23 | 19 | 14 | 20 | 6.6 | 7.5 | 10.1 | 13.0 | 15.2 |
| >hg38_21_+_8433172_8446622_RNA45SN1_45S_50ntFlanks@12444.12484.41\|AAGCAGAATTCACCAAGCGTTGGATTGTTCACCCACTAATA | >hg38_21_+_8433172_8446622_RNA45SN1_45S_50ntFlanks | 12444 | 12484 | 41 | AAGCAGAATTCACCAAGCGTTGGATTGTTCACCCACTAATA | MB468 | 14 | 69 | 20 | 6 | 28 | 10.3 | 52.5 | 12.9 | 4.4 | 27.8 |
| >hg38_21_+_8433172_8446622_RNA45SN1_45S_50ntFlanks@12444.12486.43\|AAGCAGAATTCACCAAGCGTTGGATTGTTCACCCACTAATAGG | >hg38_21_+_8433172_8446622_RNA45SN1_45S_50ntFlanks | 12444 | 12486 | 43 | AAGCAGAATTCACCAAGCGTTGGATTGTTCACCCACTAATAGG | MB468 | 11 | 58 | 20 | 3 | 11 | 6.0 | 70.2 | 15.4 | 3.2 | 19.0 |
| >hg38_21_+_8433172_8446622_RNA45SN1_45S_50ntFlanks@12445.12472.28\|AGCAGAATTCACCAAGCGTTGGATTGTT | >hg38_21_+_8433172_8446622_RNA45SN1_45S_50ntFlanks | 12445 | 12472 | 28 | AGCAGAATTCACCAAGCGTTGGATTGTT | MB468 | 13 | 1 | 5 | 4 | 2 | 6.6 | 1.1 | 3.2 | 1.7 | 2.8 |
| >hg38_21_+_8433172_8446622_RNA45SN1_45S_50ntFlanks@12445.12476.32\|AGCAGAATTCACCAAGCGTTGGATTGTTCACC | >hg38_21_+_8433172_8446622_RNA45SN1_45S_50ntFlanks | 12445 | 12476 | 32 | AGCAGAATTCACCAAGCGTTGGATTGTTCACC | MB231 | 15 | 10 | 8 | 10 | 3 | 13.7 | 8.9 | 6.4 | 6.5 | 1.4 |
| >hg38_21_+_8433172_8446622_RNA45SN1_45S_50ntFlanks@12445.12476.32\|AGCAGAATTCACCAAGCGTTGGATTGTTCACC | >hg38_21_+_8433172_8446622_RNA45SN1_45S_50ntFlanks | 12445 | 12476 | 32 | AGCAGAATTCACCAAGCGTTGGATTGTTCACC | MB468 | 11 | 3 | 7 | 14 | 7 | 4.7 | 2.5 | 3.4 | 12.8 | 4.2 |
| >hg38_21_+_8433172_8446622_RNA45SN1_45S_50ntFlanks@12445.12480.36\|AGCAGAATTCACCAAGCGTTGGATTGTTCACCCACT | >hg38_21_+_8433172_8446622_RNA45SN1_45S_50ntFlanks | 12445 | 12480 | 36 | AGCAGAATTCACCAAGCGTTGGATTGTTCACCCACT | MB231 | 11 | 13 | 14 | 6 | 4 | 8.8 | 10.6 | 12.2 | 4.9 | 3.6 |
| >hg38_21_+_8433172_8446622_RNA45SN1_45S_50ntFlanks@12446.12476.31\|GCAGAATTCACCAAGCGTTGGATTGTTCACC | >hg38_21_+_8433172_8446622_RNA45SN1_45S_50ntFlanks | 12446 | 12476 | 31 | GCAGAATTCACCAAGCGTTGGATTGTTCACC | MB231 | 26 | 11 | 19 | 11 | 2 | 23.6 | 8.3 | 15.2 | 8.3 | 1.0 |
| >hg38_21_+_8433172_8446622_RNA45SN1_45S_50ntFlanks@12446.12477.32\|GCAGAATTCACCAAGCGTTGGATTGTTCACCC | >hg38_21_+_8433172_8446622_RNA45SN1_45S_50ntFlanks | 12446 | 12477 | 32 | GCAGAATTCACCAAGCGTTGGATTGTTCACCC | MB231 | 11 | 5 | 16 | 4 | 1 | 11.3 | 3.7 | 18.0 | 3.3 | 1.0 |
| >hg38_21_+_8433172_8446622_RNA45SN1_45S_50ntFlanks@12446.12480.35\|GCAGAATTCACCAAGCGTTGGATTGTTCACCCACT | >hg38_21_+_8433172_8446622_RNA45SN1_45S_50ntFlanks | 12446 | 12480 | 35 | GCAGAATTCACCAAGCGTTGGATTGTTCACCCACT | MB231 | 12 | 14 | 26 | 7 | 4 | 11.0 | 10.0 | 19.9 | 9.5 | 4.4 |
| >hg38_21_+_8433172_8446622_RNA45SN1_45S_50ntFlanks@12447.12475.29\|CAGAATTCACCAAGCGTTGGATTGTTCAC | >hg38_21_+_8433172_8446622_RNA45SN1_45S_50ntFlanks | 12447 | 12475 | 29 | CAGAATTCACCAAGCGTTGGATTGTTCAC | MB468 | 17 | 1 | 6 | 14 | 5 | 15.0 | 1.1 | 6.6 | 11.7 | 2.8 |
| >hg38_21_+_8433172_8446622_RNA45SN1_45S_50ntFlanks@12447.12476.30\|CAGAATTCACCAAGCGTTGGATTGTTCACC | >hg38_21_+_8433172_8446622_RNA45SN1_45S_50ntFlanks | 12447 | 12476 | 30 | CAGAATTCACCAAGCGTTGGATTGTTCACC | MB468 | 21 | 3 | 15 | 33 | 10 | 12.9 | 2.3 | 9.9 | 26.8 | 5.1 |
| >hg38_21_+_8433172_8446622_RNA45SN1_45S_50ntFlanks@12448.12476.29\|AGAATTCACCAAGCGTTGGATTGTTCACC | >hg38_21_+_8433172_8446622_RNA45SN1_45S_50ntFlanks | 12448 | 12476 | 29 | AGAATTCACCAAGCGTTGGATTGTTCACC | MB231 | 12 | 6 | 7 | 6 | 2 | 13.6 | 4.9 | 5.2 | 3.8 | 1.0 |
| >hg38_21_+_8433172_8446622_RNA45SN1_45S_50ntFlanks@12448.12480.33\|AGAATTCACCAAGCGTTGGATTGTTCACCCACT | >hg38_21_+_8433172_8446622_RNA45SN1_45S_50ntFlanks | 12448 | 12480 | 33 | AGAATTCACCAAGCGTTGGATTGTTCACCCACT | MB231 | 14 | 16 | 28 | 3 | 4 | 13.4 | 10.7 | 23.3 | 2.1 | 2.1 |
| >hg38_21_+_8433172_8446622_RNA45SN1_45S_50ntFlanks@12449.12476.28\|GAATTCACCAAGCGTTGGATTGTTCACC | >hg38_21_+_8433172_8446622_RNA45SN1_45S_50ntFlanks | 12449 | 12476 | 28 | GAATTCACCAAGCGTTGGATTGTTCACC | MB231 | 36 | 5 | 24 | 16 | 3 | 39.7 | 4.1 | 24.0 | 11.5 | 1.3 |
| >hg38_21_+_8433172_8446622_RNA45SN1_45S_50ntFlanks@12449.12477.29\|GAATTCACCAAGCGTTGGATTGTTCACCC | >hg38_21_+_8433172_8446622_RNA45SN1_45S_50ntFlanks | 12449 | 12477 | 29 | GAATTCACCAAGCGTTGGATTGTTCACCC | MB231 | 15 | 2 | 20 | 6 | 1 | 19.0 | 2.0 | 26.3 | 4.0 | 0.4 |
| >hg38_21_+_8433172_8446622_RNA45SN1_45S_50ntFlanks@12449.12480.32\|GAATTCACCAAGCGTTGGATTGTTCACCCACT | >hg38_21_+_8433172_8446622_RNA45SN1_45S_50ntFlanks | 12449 | 12480 | 32 | GAATTCACCAAGCGTTGGATTGTTCACCCACT | BT20 | 16 | 19 | 45 | 5 | 2 | 9.9 | 11.9 | 42.4 | 4.4 | 0.9 |
| >hg38_21_+_8433172_8446622_RNA45SN1_45S_50ntFlanks@12449.12480.32\|GAATTCACCAAGCGTTGGATTGTTCACCCACT | >hg38_21_+_8433172_8446622_RNA45SN1_45S_50ntFlanks | 12449 | 12480 | 32 | GAATTCACCAAGCGTTGGATTGTTCACCCACT | MB231 | 22 | 12 | 48 | 8 | 3 | 23.8 | 8.0 | 43.2 | 4.4 | 1.6 |
| >hg38_21_+_8433172_8446622_RNA45SN1_45S_50ntFlanks@12449.12486.38\|GAATTCACCAAGCGTTGGATTGTTCACCCACTAATAGG | >hg38_21_+_8433172_8446622_RNA45SN1_45S_50ntFlanks | 12449 | 12486 | 38 | GAATTCACCAAGCGTTGGATTGTTCACCCACTAATAGG | MB231 | 11 | 10 | 20 | 2 | 3 | 8.6 | 7.7 | 22.4 | 2.4 | 3.5 |
| >hg38_21_+_8433172_8446622_RNA45SN1_45S_50ntFlanks@12450.12476.27\|AATTCACCAAGCGTTGGATTGTTCACC | >hg38_21_+_8433172_8446622_RNA45SN1_45S_50ntFlanks | 12450 | 12476 | 27 | AATTCACCAAGCGTTGGATTGTTCACC | MB231 | 15 | 2 | 12 | 8 | 2 | 15.6 | 2.4 | 9.4 | 6.1 | 0.7 |
| >hg38_21_+_8433172_8446622_RNA45SN1_45S_50ntFlanks@12450.12480.31\|AATTCACCAAGCGTTGGATTGTTCACCCACT | >hg38_21_+_8433172_8446622_RNA45SN1_45S_50ntFlanks | 12450 | 12480 | 31 | AATTCACCAAGCGTTGGATTGTTCACCCACT | BT20 | 11 | 14 | 38 | 3 | 1 | 5.8 | 8.5 | 31.9 | 1.8 | 0.2 |
| >hg38_21_+_8433172_8446622_RNA45SN1_45S_50ntFlanks@12450.12480.31\|AATTCACCAAGCGTTGGATTGTTCACCCACT | >hg38_21_+_8433172_8446622_RNA45SN1_45S_50ntFlanks | 12450 | 12480 | 31 | AATTCACCAAGCGTTGGATTGTTCACCCACT | MB231 | 14 | 8 | 38 | 4 | 3 | 13.2 | 6.0 | 31.5 | 2.4 | 1.7 |
| >hg38_21_+_8433172_8446622_RNA45SN1_45S_50ntFlanks@12451.12476.26\|ATTCACCAAGCGTTGGATTGTTCACC | >hg38_21_+_8433172_8446622_RNA45SN1_45S_50ntFlanks | 12451 | 12476 | 26 | ATTCACCAAGCGTTGGATTGTTCACC | MB231 | 22 | 15 | 20 | 5 | 6 | 16.0 | 10.4 | 14.7 | 3.5 | 4.6 |
| >hg38_21_+_8433172_8446622_RNA45SN1_45S_50ntFlanks@12451.12477.27\|ATTCACCAAGCGTTGGATTGTTCACCC | >hg38_21_+_8433172_8446622_RNA45SN1_45S_50ntFlanks | 12451 | 12477 | 27 | ATTCACCAAGCGTTGGATTGTTCACCC | MB231 | 14 | 8 | 20 | 2 | 3 | 10.2 | 6.1 | 20.8 | 1.6 | 3.1 |
| >hg38_21_+_8433172_8446622_RNA45SN1_45S_50ntFlanks@12453.12476.24\|TCACCAAGCGTTGGATTGTTCACC | >hg38_21_+_8433172_8446622_RNA45SN1_45S_50ntFlanks | 12453 | 12476 | 24 | TCACCAAGCGTTGGATTGTTCACC | MB231 | 23 | 13 | 34 | 5 | 5 | 16.9 | 9.1 | 22.9 | 3.8 | 3.7 |
| >hg38_21_+_8433172_8446622_RNA45SN1_45S_50ntFlanks@12453.12477.25\|TCACCAAGCGTTGGATTGTTCACCC | >hg38_21_+_8433172_8446622_RNA45SN1_45S_50ntFlanks | 12453 | 12477 | 25 | TCACCAAGCGTTGGATTGTTCACCC | MB231 | 14 | 7 | 25 | 1 | 2 | 10.8 | 5.2 | 22.5 | 1.2 | 2.1 |
| >hg38_21_+_8433172_8446622_RNA45SN1_45S_50ntFlanks@12454.12476.23\|CACCAAGCGTTGGATTGTTCACC | >hg38_21_+_8433172_8446622_RNA45SN1_45S_50ntFlanks | 12454 | 12476 | 23 | CACCAAGCGTTGGATTGTTCACC | MB231 | 17 | 10 | 36 | 4 | 6 | 12.1 | 6.5 | 25.2 | 2.7 | 3.8 |
| >hg38_21_+_8433172_8446622_RNA45SN1_45S_50ntFlanks@12454.12476.23\|CACCAAGCGTTGGATTGTTCACC | >hg38_21_+_8433172_8446622_RNA45SN1_45S_50ntFlanks | 12454 | 12476 | 23 | CACCAAGCGTTGGATTGTTCACC | MB468 | 21 | 8 | 18 | 19 | 10 | 7.4 | 4.5 | 6.1 | 15.4 | 0.5 |
| >hg38_21_+_8433172_8446622_RNA45SN1_45S_50ntFlanks@12455.12473.19\|ACCAAGCGTTGGATTGTTC | >hg38_21_+_8433172_8446622_RNA45SN1_45S_50ntFlanks | 12455 | 12473 | 19 | ACCAAGCGTTGGATTGTTC | MB468 | 18 | 2 | 20 | 57 | 2 | 16.6 | 1.8 | 22.5 | 36.4 | 0.8 |
| >hg38_21_+_8433172_8446622_RNA45SN1_45S_50ntFlanks@12455.12474.20\|ACCAAGCGTTGGATTGTTCA | >hg38_21_+_8433172_8446622_RNA45SN1_45S_50ntFlanks | 12455 | 12474 | 20 | ACCAAGCGTTGGATTGTTCA | MB468 | 11 | 1 | 17 | 10 | 1 | 10.5 | 0.8 | 21.3 | 7.5 | 0.4 |
| >hg38_21_+_8433172_8446622_RNA45SN1_45S_50ntFlanks@12455.12475.21\|ACCAAGCGTTGGATTGTTCAC | >hg38_21_+_8433172_8446622_RNA45SN1_45S_50ntFlanks | 12455 | 12475 | 21 | ACCAAGCGTTGGATTGTTCAC | MB468 | 18 | 4 | 14 | 5 | 5 | 12.5 | 2.7 | 14.0 | 3.4 | 2.5 |
| >hg38_21_+_8433172_8446622_RNA45SN1_45S_50ntFlanks@12455.12476.22\|ACCAAGCGTTGGATTGTTCACC | >hg38_21_+_8433172_8446622_RNA45SN1_45S_50ntFlanks | 12455 | 12476 | 22 | ACCAAGCGTTGGATTGTTCACC | MB231 | 23 | 15 | 25 | 6 | 7 | 15.5 | 9.1 | 16.3 | 3.7 | 5.2 |
| >hg38_21_+_8433172_8446622_RNA45SN1_45S_50ntFlanks@12455.12476.22\|ACCAAGCGTTGGATTGTTCACC | >hg38_21_+_8433172_8446622_RNA45SN1_45S_50ntFlanks | 12455 | 12476 | 22 | ACCAAGCGTTGGATTGTTCACC | MB468 | 74 | 19 | 54 | 40 | 22 | 66.0 | 7.3 | 56.9 | 24.0 | 7.4 |
| >hg38_21_+_8433172_8446622_RNA45SN1_45S_50ntFlanks@12455.12477.23\|ACCAAGCGTTGGATTGTTCACCC | >hg38_21_+_8433172_8446622_RNA45SN1_45S_50ntFlanks | 12455 | 12477 | 23 | ACCAAGCGTTGGATTGTTCACCC | MB231 | 11 | 7 | 14 | 2 | 3 | 8.1 | 5.4 | 12.5 | 1.6 | 3.3 |
| >hg38_21_+_8433172_8446622_RNA45SN1_45S_50ntFlanks@12455.12477.23\|ACCAAGCGTTGGATTGTTCACCC | >hg38_21_+_8433172_8446622_RNA45SN1_45S_50ntFlanks | 12455 | 12477 | 23 | ACCAAGCGTTGGATTGTTCACCC | MB468 | 21 | 10 | 17 | 8 | 8 | 18.8 | 3.3 | 16.2 | 3.9 | 2.5 |
| >hg38_21_+_8433172_8446622_RNA45SN1_45S_50ntFlanks@12455.12478.24\|ACCAAGCGTTGGATTGTTCACCCA | >hg38_21_+_8433172_8446622_RNA45SN1_45S_50ntFlanks | 12455 | 12478 | 24 | ACCAAGCGTTGGATTGTTCACCCA | MB468 | 20 | 5 | 10 | 7 | 5 | 20.0 | 1.3 | 10.6 | 2.3 | 0.4 |
| >hg38_21_+_8433172_8446622_RNA45SN1_45S_50ntFlanks@12455.12484.30\|ACCAAGCGTTGGATTGTTCACCCACTAATA | >hg38_21_+_8433172_8446622_RNA45SN1_45S_50ntFlanks | 12455 | 12484 | 30 | ACCAAGCGTTGGATTGTTCACCCACTAATA | MB468 | 17 | 11 | 10 | 3 | 7 | 16.9 | 8.2 | 14.4 | 1.7 | 4.3 |
| >hg38_21_+_8433172_8446622_RNA45SN1_45S_50ntFlanks@12455.12486.32\|ACCAAGCGTTGGATTGTTCACCCACTAATAGG | >hg38_21_+_8433172_8446622_RNA45SN1_45S_50ntFlanks | 12455 | 12486 | 32 | ACCAAGCGTTGGATTGTTCACCCACTAATAGG | MB468 | 10 | 5 | 4 | 1 | 2 | 8.0 | 1.8 | 3.9 | 1.1 | 2.2 |
| >hg38_21_+_8433172_8446622_RNA45SN1_45S_50ntFlanks@12455.12487.33\|ACCAAGCGTTGGATTGTTCACCCACTAATAGGG | >hg38_21_+_8433172_8446622_RNA45SN1_45S_50ntFlanks | 12455 | 12487 | 33 | ACCAAGCGTTGGATTGTTCACCCACTAATAGGG | MB468 | 34 | 22 | 17 | 7 | 9 | 23.3 | 7.8 | 10.7 | 5.8 | 7.1 |
| >hg38_21_+_8433172_8446622_RNA45SN1_45S_50ntFlanks@12456.12473.18\|CCAAGCGTTGGATTGTTC | >hg38_21_+_8433172_8446622_RNA45SN1_45S_50ntFlanks | 12456 | 12473 | 18 | CCAAGCGTTGGATTGTTC | MB468 | 778 | 131 | 864 | 2287 | 104 | 409.9 | 135.2 | 462.4 | 1681.3 | 63.2 |
| >hg38_21_+_8433172_8446622_RNA45SN1_45S_50ntFlanks@12456.12474.19\|CCAAGCGTTGGATTGTTCA | >hg38_21_+_8433172_8446622_RNA45SN1_45S_50ntFlanks | 12456 | 12474 | 19 | CCAAGCGTTGGATTGTTCA | MB468 | 649 | 116 | 917 | 542 | 87 | 205.0 | 111.1 | 565.5 | 420.5 | 37.8 |
| >hg38_21_+_8433172_8446622_RNA45SN1_45S_50ntFlanks@12456.12475.20\|CCAAGCGTTGGATTGTTCAC | >hg38_21_+_8433172_8446622_RNA45SN1_45S_50ntFlanks | 12456 | 12475 | 20 | CCAAGCGTTGGATTGTTCAC | MB468 | 2648 | 600 | 1933 | 687 | 406 | 760.8 | 534.3 | 1234.7 | 457.0 | 197.7 |
| >hg38_21_+_8433172_8446622_RNA45SN1_45S_50ntFlanks@12456.12476.21\|CCAAGCGTTGGATTGTTCACC | >hg38_21_+_8433172_8446622_RNA45SN1_45S_50ntFlanks | 12456 | 12476 | 21 | CCAAGCGTTGGATTGTTCACC | MB231 | 12 | 7 | 35 | 4 | 4 | 8.5 | 6.7 | 40.3 | 2.3 | 3.1 |
| >hg38_21_+_8433172_8446622_RNA45SN1_45S_50ntFlanks@12456.12476.21\|CCAAGCGTTGGATTGTTCACC | >hg38_21_+_8433172_8446622_RNA45SN1_45S_50ntFlanks | 12456 | 12476 | 21 | CCAAGCGTTGGATTGTTCACC | MB468 | 4718 | 1256 | 2776 | 2520 | 983 | 1147.3 | 886.2 | 839.8 | 1747.0 | 286.7 |
| >hg38_21_+_8433172_8446622_RNA45SN1_45S_50ntFlanks@12456.12477.22\|CCAAGCGTTGGATTGTTCACCC | >hg38_21_+_8433172_8446622_RNA45SN1_45S_50ntFlanks | 12456 | 12477 | 22 | CCAAGCGTTGGATTGTTCACCC | MB468 | 894 | 343 | 635 | 473 | 266 | 168.4 | 125.5 | 115.6 | 309.6 | 58.6 |

| | | | | | | | | | | | | | | | |
|---|---|---|---|---|---|---|---|---|---|---|---|---|---|---|---|
| >hg38_21_+_8433172_8446622_RNA45SN1_45S_50ntFlanks@12456.12478.23\|CCAAGCGTTGGATTGTTCACCCA | >hg38_21_+_8433172_8446622_RNA45SN1_45S_50ntFlanks | 12456 | 12478 | 23 | CCAAGCGTTGGATTGTTCACCCA | MB468 | 1356 | 766 | 894 | 431 | 500 | 231.8 | 158.1 | 261.5 | 165.4 | 115.0 |
| >hg38_21_+_8433172_8446622_RNA45SN1_45S_50ntFlanks@12456.12479.24\|CCAAGCGTTGGATTGTTCACCCAC | >hg38_21_+_8433172_8446622_RNA45SN1_45S_50ntFlanks | 12456 | 12479 | 24 | CCAAGCGTTGGATTGTTCACCCAC | MB468 | 77 | 30 | 42 | 81 | 39 | 27.0 | 16.4 | 21.2 | 41.1 | 5.5 |
| >hg38_21_+_8433172_8446622_RNA45SN1_45S_50ntFlanks@12456.12480.25\|CCAAGCGTTGGATTGTTCACCCACT | >hg38_21_+_8433172_8446622_RNA45SN1_45S_50ntFlanks | 12456 | 12480 | 25 | CCAAGCGTTGGATTGTTCACCCACT | MB468 | 257 | 90 | 202 | 244 | 75 | 141.0 | 100.6 | 105.8 | 139.9 | 20.0 |
| >hg38_21_+_8433172_8446622_RNA45SN1_45S_50ntFlanks@12456.12481.26\|CCAAGCGTTGGATTGTTCACCCACTA | >hg38_21_+_8433172_8446622_RNA45SN1_45S_50ntFlanks | 12456 | 12481 | 26 | CCAAGCGTTGGATTGTTCACCCACTA | MB468 | 139 | 55 | 91 | 66 | 41 | 61.7 | 60.1 | 52.6 | 36.5 | 6.8 |
| >hg38_21_+_8433172_8446622_RNA45SN1_45S_50ntFlanks@12456.12482.27\|CCAAGCGTTGGATTGTTCACCCACTAA | >hg38_21_+_8433172_8446622_RNA45SN1_45S_50ntFlanks | 12456 | 12482 | 27 | CCAAGCGTTGGATTGTTCACCCACTAA | MB468 | 22 | 8 | 12 | 15 | 12 | 3.3 | 9.6 | 4.0 | 11.2 | 4.0 |
| >hg38_21_+_8433172_8446622_RNA45SN1_45S_50ntFlanks@12456.12483.28\|CCAAGCGTTGGATTGTTCACCCACTAAT | >hg38_21_+_8433172_8446622_RNA45SN1_45S_50ntFlanks | 12456 | 12483 | 28 | CCAAGCGTTGGATTGTTCACCCACTAAT | MB468 | 207 | 196 | 103 | 65 | 84 | 24.4 | 122.5 | 24.1 | 12.2 | 5.9 |
| >hg38_21_+_8433172_8446622_RNA45SN1_45S_50ntFlanks@12456.12484.29\|CCAAGCGTTGGATTGTTCACCCACTAATA | >hg38_21_+_8433172_8446622_RNA45SN1_45S_50ntFlanks | 12456 | 12484 | 29 | CCAAGCGTTGGATTGTTCACCCACTAATA | MB468 | 1224 | 953 | 575 | 281 | 531 | 170.1 | 543.0 | 337.5 | 43.2 | 184.5 |
| >hg38_21_+_8433172_8446622_RNA45SN1_45S_50ntFlanks@12456.12485.30\|CCAAGCGTTGGATTGTTCACCCACTAATAG | >hg38_21_+_8433172_8446622_RNA45SN1_45S_50ntFlanks | 12456 | 12485 | 30 | CCAAGCGTTGGATTGTTCACCCACTAATAG | MB468 | 483 | 410 | 185 | 110 | 129 | 83.5 | 304.8 | 40.3 | 40.3 | 38.8 |
| >hg38_21_+_8433172_8446622_RNA45SN1_45S_50ntFlanks@12456.12486.31\|CCAAGCGTTGGATTGTTCACCCACTAATAGG | >hg38_21_+_8433172_8446622_RNA45SN1_45S_50ntFlanks | 12456 | 12486 | 31 | CCAAGCGTTGGATTGTTCACCCACTAATAGG | MB468 | 1518 | 1052 | 495 | 356 | 327 | 418.4 | 698.7 | 181.6 | 263.6 | 106.6 |
| >hg38_21_+_8433172_8446622_RNA45SN1_45S_50ntFlanks@12456.12487.32\|CCAAGCGTTGGATTGTTCACCCACTAATAGGG | >hg38_21_+_8433172_8446622_RNA45SN1_45S_50ntFlanks | 12456 | 12487 | 32 | CCAAGCGTTGGATTGTTCACCCACTAATAGGG | MB468 | 5678 | 5408 | 3310 | 1179 | 1070 | 2326.1 | 3134.3 | 505.1 | 264.7 | 437.9 |
| >hg38_21_+_8433172_8446622_RNA45SN1_45S_50ntFlanks@12456.12488.33\|CCAAGCGTTGGATTGTTCACCCACTAATAGGGA | >hg38_21_+_8433172_8446622_RNA45SN1_45S_50ntFlanks | 12456 | 12488 | 33 | CCAAGCGTTGGATTGTTCACCCACTAATAGGGA | MB468 | 780 | 724 | 519 | 384 | 267 | 277.4 | 368.3 | 71.1 | 339.7 | 172.2 |
| >hg38_21_+_8433172_8446622_RNA45SN1_45S_50ntFlanks@12456.12489.34\|CCAAGCGTTGGATTGTTCACCCACTAATAGGGAA | >hg38_21_+_8433172_8446622_RNA45SN1_45S_50ntFlanks | 12456 | 12489 | 34 | CCAAGCGTTGGATTGTTCACCCACTAATAGGGAA | MB468 | 131 | 171 | 91 | 89 | 76 | 48.5 | 117.6 | 16.9 | 107.8 | 59.9 |
| >hg38_21_+_8433172_8446622_RNA45SN1_45S_50ntFlanks@12456.12490.35\|CCAAGCGTTGGATTGTTCACCCACTAATAGGGAAC | >hg38_21_+_8433172_8446622_RNA45SN1_45S_50ntFlanks | 12456 | 12490 | 35 | CCAAGCGTTGGATTGTTCACCCACTAATAGGGAAC | MB468 | 67 | 161 | 88 | 73 | 80 | 38.9 | 162.0 | 34.0 | 42.6 | 52.4 |
| >hg38_21_+_8433172_8446622_RNA45SN1_45S_50ntFlanks@12456.12491.36\|CCAAGCGTTGGATTGTTCACCCACTAATAGGGAACG | >hg38_21_+_8433172_8446622_RNA45SN1_45S_50ntFlanks | 12456 | 12491 | 36 | CCAAGCGTTGGATTGTTCACCCACTAATAGGGAACG | MB468 | 12 | 19 | 18 | 17 | 19 | 8.6 | 16.3 | 15.6 | 18.5 | 6.9 |
| >hg38_21_+_8433172_8446622_RNA45SN1_45S_50ntFlanks@12456.12493.38\|CCAAGCGTTGGATTGTTCACCCACTAATAGGGAACGTG | >hg38_21_+_8433172_8446622_RNA45SN1_45S_50ntFlanks | 12456 | 12493 | 38 | CCAAGCGTTGGATTGTTCACCCACTAATAGGGAACGTG | MB468 | 28 | 57 | 41 | 32 | 44 | 13.3 | 44.1 | 20.3 | 21.3 | 34.0 |
| >hg38_21_+_8433172_8446622_RNA45SN1_45S_50ntFlanks@12456.12496.41\|CCAAGCGTTGGATTGTTCACCCACTAATAGGGAACGTGAGC | >hg38_21_+_8433172_8446622_RNA45SN1_45S_50ntFlanks | 12456 | 12496 | 41 | CCAAGCGTTGGATTGTTCACCCACTAATAGGGAACGTGAGC | MB468 | 14 | 63 | 56 | 16 | 41 | 9.3 | 51.7 | 38.5 | 11.3 | 34.6 |
| >hg38_21_+_8433172_8446622_RNA45SN1_45S_50ntFlanks@12456.12499.44\|CCAAGCGTTGGATTGTTCACCCACTAATAGGGAACGTGAGCTGG | >hg38_21_+_8433172_8446622_RNA45SN1_45S_50ntFlanks | 12456 | 12499 | 44 | CCAAGCGTTGGATTGTTCACCCACTAATAGGGAACGTGAGCTGG | MB468 | 12 | 27 | 59 | 9 | 35 | 5.8 | 31.8 | 45.5 | 9.1 | 37.6 |
| >hg38_21_+_8433172_8446622_RNA45SN1_45S_50ntFlanks@12457.12474.18\|CAAGCGTTGGATTGTTCA | >hg38_21_+_8433172_8446622_RNA45SN1_45S_50ntFlanks | 12457 | 12474 | 18 | CAAGCGTTGGATTGTTCA | MB468 | 87 | 15 | 143 | 28 | 11 | 55.1 | 6.5 | 147.7 | 12.3 | 0.7 |
| >hg38_21_+_8433172_8446622_RNA45SN1_45S_50ntFlanks@12457.12475.19\|CAAGCGTTGGATTGTTCAC | >hg38_21_+_8433172_8446622_RNA45SN1_45S_50ntFlanks | 12457 | 12475 | 19 | CAAGCGTTGGATTGTTCAC | MB468 | 131 | 28 | 122 | 23 | 41 | 63.0 | 13.5 | 101.8 | 9.1 | 11.4 |
| >hg38_21_+_8433172_8446622_RNA45SN1_45S_50ntFlanks@12457.12476.20\|CAAGCGTTGGATTGTTCACC | >hg38_21_+_8433172_8446622_RNA45SN1_45S_50ntFlanks | 12457 | 12476 | 20 | CAAGCGTTGGATTGTTCACC | MB231 | 23 | 20 | 41 | 4 | 10 | 20.6 | 18.9 | 31.9 | 3.1 | 9.7 |
| >hg38_21_+_8433172_8446622_RNA45SN1_45S_50ntFlanks@12457.12476.20\|CAAGCGTTGGATTGTTCACC | >hg38_21_+_8433172_8446622_RNA45SN1_45S_50ntFlanks | 12457 | 12476 | 20 | CAAGCGTTGGATTGTTCACC | MB468 | 623 | 283 | 764 | 186 | 201 | 377.0 | 95.5 | 689.4 | 70.8 | 73.6 |
| >hg38_21_+_8433172_8446622_RNA45SN1_45S_50ntFlanks@12457.12477.21\|CAAGCGTTGGATTGTTCACCC | >hg38_21_+_8433172_8446622_RNA45SN1_45S_50ntFlanks | 12457 | 12477 | 21 | CAAGCGTTGGATTGTTCACCC | MB468 | 342 | 217 | 376 | 63 | 94 | 201.8 | 80.3 | 331.7 | 25.6 | 48.9 |
| >hg38_21_+_8433172_8446622_RNA45SN1_45S_50ntFlanks@12457.12478.22\|CAAGCGTTGGATTGTTCACCCA | >hg38_21_+_8433172_8446622_RNA45SN1_45S_50ntFlanks | 12457 | 12478 | 22 | CAAGCGTTGGATTGTTCACCCA | MB468 | 418 | 228 | 330 | 68 | 142 | 153.8 | 59.6 | 261.0 | 33.1 | 63.3 |
| >hg38_21_+_8433172_8446622_RNA45SN1_45S_50ntFlanks@12457.12479.23\|CAAGCGTTGGATTGTTCACCCAC | >hg38_21_+_8433172_8446622_RNA45SN1_45S_50ntFlanks | 12457 | 12479 | 23 | CAAGCGTTGGATTGTTCACCCAC | MB468 | 15 | 3 | 8 | 6 | 5 | 16.5 | 1.2 | 9.4 | 2.2 | 1.4 |
| >hg38_21_+_8433172_8446622_RNA45SN1_45S_50ntFlanks@12457.12480.24\|CAAGCGTTGGATTGTTCACCCACT | >hg38_21_+_8433172_8446622_RNA45SN1_45S_50ntFlanks | 12457 | 12480 | 24 | CAAGCGTTGGATTGTTCACCCACT | MB468 | 45 | 5 | 14 | 13 | 11 | 52.0 | 3.9 | 14.2 | 5.1 | 4.2 |
| >hg38_21_+_8433172_8446622_RNA45SN1_45S_50ntFlanks@12457.12481.25\|CAAGCGTTGGATTGTTCACCCACTA | >hg38_21_+_8433172_8446622_RNA45SN1_45S_50ntFlanks | 12457 | 12481 | 25 | CAAGCGTTGGATTGTTCACCCACTA | MB468 | 20 | 2 | 7 | 3 | 4 | 25.7 | 1.5 | 9.2 | 1.3 | 1.0 |
| >hg38_21_+_8433172_8446622_RNA45SN1_45S_50ntFlanks@12457.12483.27\|CAAGCGTTGGATTGTTCACCCACTAAT | >hg38_21_+_8433172_8446622_RNA45SN1_45S_50ntFlanks | 12457 | 12483 | 27 | CAAGCGTTGGATTGTTCACCCACTAAT | MB468 | 16 | 10 | 6 | 3 | 8 | 11.7 | 6.1 | 4.7 | 2.5 | 5.7 |
| >hg38_21_+_8433172_8446622_RNA45SN1_45S_50ntFlanks@12457.12484.28\|CAAGCGTTGGATTGTTCACCCACTAATA | >hg38_21_+_8433172_8446622_RNA45SN1_45S_50ntFlanks | 12457 | 12484 | 28 | CAAGCGTTGGATTGTTCACCCACTAATA | MB468 | 119 | 37 | 54 | 13 | 57 | 107.1 | 17.2 | 75.3 | 6.7 | 42.4 |
| >hg38_21_+_8433172_8446622_RNA45SN1_45S_50ntFlanks@12457.12485.29\|CAAGCGTTGGATTGTTCACCCACTAATAG | >hg38_21_+_8433172_8446622_RNA45SN1_45S_50ntFlanks | 12457 | 12485 | 29 | CAAGCGTTGGATTGTTCACCCACTAATAG | MB468 | 32 | 18 | 12 | 5 | 10 | 27.5 | 7.9 | 10.1 | 2.4 | 5.3 |
| >hg38_21_+_8433172_8446622_RNA45SN1_45S_50ntFlanks@12457.12486.30\|CAAGCGTTGGATTGTTCACCCACTAATAGG | >hg38_21_+_8433172_8446622_RNA45SN1_45S_50ntFlanks | 12457 | 12486 | 30 | CAAGCGTTGGATTGTTCACCCACTAATAGG | MB468 | 55 | 36 | 19 | 8 | 17 | 28.4 | 18.4 | 9.5 | 3.3 | 13.5 |
| >hg38_21_+_8433172_8446622_RNA45SN1_45S_50ntFlanks@12457.12487.31\|CAAGCGTTGGATTGTTCACCCACTAATAGGG | >hg38_21_+_8433172_8446622_RNA45SN1_45S_50ntFlanks | 12457 | 12487 | 31 | CAAGCGTTGGATTGTTCACCCACTAATAGGG | MB468 | 230 | 214 | 141 | 40 | 80 | 103.4 | 75.0 | 66.7 | 31.1 | 76.0 |
| >hg38_21_+_8433172_8446622_RNA45SN1_45S_50ntFlanks@12457.12488.32\|CAAGCGTTGGATTGTTCACCCACTAATAGGGA | >hg38_21_+_8433172_8446622_RNA45SN1_45S_50ntFlanks | 12457 | 12488 | 32 | CAAGCGTTGGATTGTTCACCCACTAATAGGGA | MB468 | 74 | 70 | 47 | 17 | 18 | 43.7 | 12.0 | 36.2 | 10.1 | 9.5 |
| >hg38_21_+_8433172_8446622_RNA45SN1_45S_50ntFlanks@12457.12489.33\|CAAGCGTTGGATTGTTCACCCACTAATAGGGAA | >hg38_21_+_8433172_8446622_RNA45SN1_45S_50ntFlanks | 12457 | 12489 | 33 | CAAGCGTTGGATTGTTCACCCACTAATAGGGAA | MB468 | 10 | 12 | 7 | 4 | 5 | 5.2 | 6.1 | 5.2 | 4.5 | 4.0 |
| >hg38_21_+_8433172_8446622_RNA45SN1_45S_50ntFlanks@12458.12475.18\|AAGCGTTGGATTGTTCAC | >hg38_21_+_8433172_8446622_RNA45SN1_45S_50ntFlanks | 12458 | 12475 | 18 | AAGCGTTGGATTGTTCAC | MB468 | 53 | 17 | 55 | 11 | 9 | 17.8 | 10.6 | 36.3 | 7.1 | 3.0 |
| >hg38_21_+_8433172_8446622_RNA45SN1_45S_50ntFlanks@12458.12476.19\|AAGCGTTGGATTGTTCACC | >hg38_21_+_8433172_8446622_RNA45SN1_45S_50ntFlanks | 12458 | 12476 | 19 | AAGCGTTGGATTGTTCACC | BT20 | 15 | 17 | 26 | 77 | 9 | 5.8 | 3.8 | 1.1 | 124.4 | 4.2 |
| >hg38_21_+_8433172_8446622_RNA45SN1_45S_50ntFlanks@12458.12476.19\|AAGCGTTGGATTGTTCACC | >hg38_21_+_8433172_8446622_RNA45SN1_45S_50ntFlanks | 12458 | 12476 | 19 | AAGCGTTGGATTGTTCACC | MB231 | 63 | 62 | 85 | 13 | 19 | 47.2 | 60.6 | 64.2 | 8.8 | 19.6 |
| >hg38_21_+_8433172_8446622_RNA45SN1_45S_50ntFlanks@12458.12476.19\|AAGCGTTGGATTGTTCACC | >hg38_21_+_8433172_8446622_RNA45SN1_45S_50ntFlanks | 12458 | 12476 | 19 | AAGCGTTGGATTGTTCACC | MB468 | 248 | 136 | 306 | 93 | 57 | 68.0 | 69.0 | 169.0 | 45.3 | 10.3 |
| >hg38_21_+_8433172_8446622_RNA45SN1_45S_50ntFlanks@12458.12477.20\|AAGCGTTGGATTGTTCACCC | >hg38_21_+_8433172_8446622_RNA45SN1_45S_50ntFlanks | 12458 | 12477 | 20 | AAGCGTTGGATTGTTCACCC | MB231 | 29 | 29 | 46 | 5 | 11 | 22.0 | 31.4 | 51.4 | 3.7 | 11.6 |
| >hg38_21_+_8433172_8446622_RNA45SN1_45S_50ntFlanks@12458.12477.20\|AAGCGTTGGATTGTTCACCC | >hg38_21_+_8433172_8446622_RNA45SN1_45S_50ntFlanks | 12458 | 12477 | 20 | AAGCGTTGGATTGTTCACCC | MB468 | 89 | 60 | 102 | 23 | 21 | 30.4 | 20.1 | 57.5 | 11.2 | 8.3 |
| >hg38_21_+_8433172_8446622_RNA45SN1_45S_50ntFlanks@12458.12478.21\|AAGCGTTGGATTGTTCACCCA | >hg38_21_+_8433172_8446622_RNA45SN1_45S_50ntFlanks | 12458 | 12478 | 21 | AAGCGTTGGATTGTTCACCCA | MB231 | 11 | 11 | 11 | 1 | 4 | 8.3 | 12.0 | 10.9 | 1.6 | 4.9 |
| >hg38_21_+_8433172_8446622_RNA45SN1_45S_50ntFlanks@12458.12478.21\|AAGCGTTGGATTGTTCACCCA | >hg38_21_+_8433172_8446622_RNA45SN1_45S_50ntFlanks | 12458 | 12478 | 21 | AAGCGTTGGATTGTTCACCCA | MB468 | 110 | 66 | 92 | 25 | 32 | 22.3 | 19.6 | 53.2 | 14.8 | 11.7 |
| >hg38_21_+_8433172_8446622_RNA45SN1_45S_50ntFlanks@12458.12480.23\|AAGCGTTGGATTGTTCACCCACT | >hg38_21_+_8433172_8446622_RNA45SN1_45S_50ntFlanks | 12458 | 12480 | 23 | AAGCGTTGGATTGTTCACCCACT | MB468 | 21 | 3 | 6 | 6 | 4 | 19.7 | 1.3 | 4.2 | 2.3 | 1.8 |
| >hg38_21_+_8433172_8446622_RNA45SN1_45S_50ntFlanks@12458.12484.27\|AAGCGTTGGATTGTTCACCCACTAATA | >hg38_21_+_8433172_8446622_RNA45SN1_45S_50ntFlanks | 12458 | 12484 | 27 | AAGCGTTGGATTGTTCACCCACTAATA | MB468 | 31 | 9 | 9 | 4 | 7 | 16.9 | 4.9 | 8.6 | 2.2 | 3.9 |
| >hg38_21_+_8433172_8446622_RNA45SN1_45S_50ntFlanks@12458.12485.28\|AAGCGTTGGATTGTTCACCCACTAATAG | >hg38_21_+_8433172_8446622_RNA45SN1_45S_50ntFlanks | 12458 | 12485 | 28 | AAGCGTTGGATTGTTCACCCACTAATAG | MB468 | 10 | 7 | 4 | 1 | 1 | 4.9 | 3.5 | 2.0 | 0.7 | 0.9 |
| >hg38_21_+_8433172_8446622_RNA45SN1_45S_50ntFlanks@12458.12486.29\|AAGCGTTGGATTGTTCACCCACTAATAGG | >hg38_21_+_8433172_8446622_RNA45SN1_45S_50ntFlanks | 12458 | 12486 | 29 | AAGCGTTGGATTGTTCACCCACTAATAGG | MB468 | 18 | 13 | 7 | 2 | 3 | 4.9 | 8.8 | 1.9 | 0.7 | 1.2 |
| >hg38_21_+_8433172_8446622_RNA45SN1_45S_50ntFlanks@12458.12487.30\|AAGCGTTGGATTGTTCACCCACTAATAGGG | >hg38_21_+_8433172_8446622_RNA45SN1_45S_50ntFlanks | 12458 | 12487 | 30 | AAGCGTTGGATTGTTCACCCACTAATAGGG | MB468 | 76 | 88 | 55 | 11 | 13 | 27.3 | 32.0 | 16.5 | 8.5 | 8.9 |
| >hg38_21_+_8433172_8446622_RNA45SN1_45S_50ntFlanks@12458.12488.31\|AAGCGTTGGATTGTTCACCCACTAATAGGGA | >hg38_21_+_8433172_8446622_RNA45SN1_45S_50ntFlanks | 12458 | 12488 | 31 | AAGCGTTGGATTGTTCACCCACTAATAGGGA | MB468 | 15 | 12 | 9 | 3 | 2 | 5.3 | 2.3 | 4.0 | 1.0 | 1.6 |
| >hg38_21_+_8433172_8446622_RNA45SN1_45S_50ntFlanks@12458.12496.39\|AAGCGTTGGATTGTTCACCCACTAATAGGGAACGTGAGC | >hg38_21_+_8433172_8446622_RNA45SN1_45S_50ntFlanks | 12458 | 12496 | 39 | AAGCGTTGGATTGTTCACCCACTAATAGGGAACGTGAGC | MB231 | 17 | 14 | 29 | 3 | 5 | 11.4 | 7.3 | 22.2 | 1.6 | 3.6 |
| >hg38_21_+_8433172_8446622_RNA45SN1_45S_50ntFlanks@12458.12497.40\|AAGCGTTGGATTGTTCACCCACTAATAGGGAACGTGAGCT | >hg38_21_+_8433172_8446622_RNA45SN1_45S_50ntFlanks | 12458 | 12497 | 40 | AAGCGTTGGATTGTTCACCCACTAATAGGGAACGTGAGCT | MB231 | 11 | 5 | 8 | 3 | 2 | 9.0 | 3.4 | 7.3 | 1.4 | 1.2 |
| >hg38_21_+_8433172_8446622_RNA45SN1_45S_50ntFlanks@12459.12476.18\|AGCGTTGGATTGTTCACC | >hg38_21_+_8433172_8446622_RNA45SN1_45S_50ntFlanks | 12459 | 12476 | 18 | AGCGTTGGATTGTTCACC | MB468 | 290 | 154 | 264 | 118 | 62 | 68.1 | 91.6 | 114.9 | 68.6 | 13.5 |

| | | | | | | | | | | | | | | | |
|---|---|---|---|---|---|---|---|---|---|---|---|---|---|---|---|
| >hg38_21_+_8433172_8446622_RNA45SN1_45S_50ntFlanks@12459.12477.19\|AGCGTTGGATTGTTCACCC | >hg38_21_+_8433172_8446622_RNA45SN1_45S_50ntFlanks | 12459 | 12477 | 19 | AGCGTTGGATTGTTCACCC | MB468 | 79 | 42 | 69 | 23 | 19 | 25.7 | 20.1 | 31.9 | 11.1 | 8.4 |
| >hg38_21_+_8433172_8446622_RNA45SN1_45S_50ntFlanks@12459.12478.20\|AGCGTTGGATTGTTCACCCA | >hg38_21_+_8433172_8446622_RNA45SN1_45S_50ntFlanks | 12459 | 12478 | 20 | AGCGTTGGATTGTTCACCCA | MB468 | 84 | 35 | 62 | 22 | 24 | 21.3 | 15.3 | 35.5 | 12.4 | 11.5 |
| >hg38_21_+_8433172_8446622_RNA45SN1_45S_50ntFlanks@12459.12480.22\|AGCGTTGGATTGTTCACCCACT | >hg38_21_+_8433172_8446622_RNA45SN1_45S_50ntFlanks | 12459 | 12480 | 22 | AGCGTTGGATTGTTCACCCACT | MB468 | 29 | 3 | 7 | 6 | 5 | 28.6 | 1.7 | 5.4 | 2.5 | 3.2 |
| >hg38_21_+_8433172_8446622_RNA45SN1_45S_50ntFlanks@12459.12484.26\|AGCGTTGGATTGTTCACCCACTAATA | >hg38_21_+_8433172_8446622_RNA45SN1_45S_50ntFlanks | 12459 | 12484 | 26 | AGCGTTGGATTGTTCACCCACTAATA | MB468 | 21 | 4 | 6 | 3 | 5 | 11.5 | 2.2 | 6.0 | 1.7 | 3.9 |
| >hg38_21_+_8433172_8446622_RNA45SN1_45S_50ntFlanks@12459.12486.28\|AGCGTTGGATTGTTCACCCACTAATAGG | >hg38_21_+_8433172_8446622_RNA45SN1_45S_50ntFlanks | 12459 | 12486 | 28 | AGCGTTGGATTGTTCACCCACTAATAGG | MB468 | 17 | 9 | 4 | 2 | 2 | 6.2 | 6.0 | 1.2 | 0.9 | 1.5 |
| >hg38_21_+_8433172_8446622_RNA45SN1_45S_50ntFlanks@12459.12487.29\|AGCGTTGGATTGTTCACCCACTAATAGGG | >hg38_21_+_8433172_8446622_RNA45SN1_45S_50ntFlanks | 12459 | 12487 | 29 | AGCGTTGGATTGTTCACCCACTAATAGGG | MB468 | 89 | 93 | 41 | 13 | 15 | 39.3 | 45.3 | 11.6 | 11.0 | 14.0 |
| >hg38_21_+_8433172_8446622_RNA45SN1_45S_50ntFlanks@12459.12488.30\|AGCGTTGGATTGTTCACCCACTAATAGGGA | >hg38_21_+_8433172_8446622_RNA45SN1_45S_50ntFlanks | 12459 | 12488 | 30 | AGCGTTGGATTGTTCACCCACTAATAGGGA | MB468 | 18 | 19 | 10 | 3 | 3 | 6.9 | 5.8 | 4.0 | 1.4 | 2.3 |
| >hg38_21_+_8433172_8446622_RNA45SN1_45S_50ntFlanks@12461.12478.18\|CGTTGGATTGTTCACCCA | >hg38_21_+_8433172_8446622_RNA45SN1_45S_50ntFlanks | 12461 | 12478 | 18 | CGTTGGATTGTTCACCCA | MB468 | 11 | 1 | 3 | 5 | 2 | 9.9 | 0.3 | 2.8 | 2.4 | 0.9 |
| >hg38_21_+_8433172_8446622_RNA45SN1_45S_50ntFlanks@12461.12487.27\|CGTTGGATTGTTCACCCACTAATAGGG | >hg38_21_+_8433172_8446622_RNA45SN1_45S_50ntFlanks | 12461 | 12487 | 27 | CGTTGGATTGTTCACCCACTAATAGGG | MB468 | 26 | 12 | 10 | 7 | 5 | 10.6 | 5.9 | 4.5 | 3.8 | 2.8 |
| >hg38_21_+_8433172_8446622_RNA45SN1_45S_50ntFlanks@12461.12488.28\|CGTTGGATTGTTCACCCACTAATAGGGA | >hg38_21_+_8433172_8446622_RNA45SN1_45S_50ntFlanks | 12461 | 12488 | 28 | CGTTGGATTGTTCACCCACTAATAGGGA | MB468 | 16 | 7 | 6 | 4 | 2 | 11.1 | 3.4 | 4.1 | 0.9 | 1.0 |
| >hg38_21_+_8433172_8446622_RNA45SN1_45S_50ntFlanks@12463.12480.18\|TTGGATTGTTCACCCACT | >hg38_21_+_8433172_8446622_RNA45SN1_45S_50ntFlanks | 12463 | 12480 | 18 | TTGGATTGTTCACCCACT | BT20 | 20 | 7 | 3 | 4 | 1 | 29.2 | 9.7 | 3.1 | 7.0 | 1.1 |
| >hg38_21_+_8433172_8446622_RNA45SN1_45S_50ntFlanks@12463.12486.24\|TTGGATTGTTCACCCACTAATAGG | >hg38_21_+_8433172_8446622_RNA45SN1_45S_50ntFlanks | 12463 | 12486 | 24 | TTGGATTGTTCACCCACTAATAGG | MB231 | 11 | 4 | 3 | 1 | 2 | 7.5 | 3.6 | 2.1 | 1.2 | 1.8 |
| >hg38_21_+_8433172_8446622_RNA45SN1_45S_50ntFlanks@12463.12487.25\|TTGGATTGTTCACCCACTAATAGGG | >hg38_21_+_8433172_8446622_RNA45SN1_45S_50ntFlanks | 12463 | 12487 | 25 | TTGGATTGTTCACCCACTAATAGGG | MB468 | 19 | 8 | 15 | 5 | 1 | 9.5 | 6.8 | 9.0 | 2.0 | 0.4 |
| >hg38_21_+_8433172_8446622_RNA45SN1_45S_50ntFlanks@12463.12488.26\|TTGGATTGTTCACCCACTAATAGGGA | >hg38_21_+_8433172_8446622_RNA45SN1_45S_50ntFlanks | 12463 | 12488 | 26 | TTGGATTGTTCACCCACTAATAGGGA | MB468 | 11 | 4 | 4 | 2 | 1 | 5.7 | 1.4 | 3.1 | 1.3 | 0.4 |
| >hg38_21_+_8433172_8446622_RNA45SN1_45S_50ntFlanks@12463.12496.34\|TTGGATTGTTCACCCACTAATAGGGAACGTGAGC | >hg38_21_+_8433172_8446622_RNA45SN1_45S_50ntFlanks | 12463 | 12496 | 34 | TTGGATTGTTCACCCACTAATAGGGAACGTGAGC | MB231 | 23 | 14 | 26 | 5 | 8 | 13.3 | 5.2 | 6.3 | 4.9 | 4.7 |
| >hg38_21_+_8433172_8446622_RNA45SN1_45S_50ntFlanks@12464.12488.25\|TGGATTGTTCACCCACTAATAGGGA | >hg38_21_+_8433172_8446622_RNA45SN1_45S_50ntFlanks | 12464 | 12488 | 25 | TGGATTGTTCACCCACTAATAGGGA | BT20 | 12 | 1 | 1 | 1 | 0 | 20.4 | 1.5 | 0.9 | 0.9 | 0.6 |
| >hg38_21_+_8433172_8446622_RNA45SN1_45S_50ntFlanks@12473.12496.24\|CACCCACTAATAGGGAACGTGAGC | >hg38_21_+_8433172_8446622_RNA45SN1_45S_50ntFlanks | 12473 | 12496 | 24 | CACCCACTAATAGGGAACGTGAGC | MB231 | 26 | 13 | 26 | 2 | 3 | 19.1 | 8.1 | 22.3 | 2.2 | 2.9 |
| >hg38_21_+_8433172_8446622_RNA45SN1_45S_50ntFlanks@12473.12496.24\|CACCCACTAATAGGGAACGTGAGC | >hg38_21_+_8433172_8446622_RNA45SN1_45S_50ntFlanks | 12473 | 12496 | 24 | CACCCACTAATAGGGAACGTGAGC | MB468 | 11 | 5 | 8 | 3 | 2 | 4.4 | 4.4 | 5.0 | 0.2 | 0.6 |
| >hg38_21_+_8433172_8446622_RNA45SN1_45S_50ntFlanks@12473.12497.25\|CACCCACTAATAGGGAACGTGAGCT | >hg38_21_+_8433172_8446622_RNA45SN1_45S_50ntFlanks | 12473 | 12497 | 25 | CACCCACTAATAGGGAACGTGAGCT | MB231 | 23 | 7 | 10 | 1 | 3 | 22.9 | 4.7 | 7.6 | 0.9 | 2.6 |
| >hg38_21_+_8433172_8446622_RNA45SN1_45S_50ntFlanks@12474.12496.23\|ACCCACTAATAGGGAACGTGAGC | >hg38_21_+_8433172_8446622_RNA45SN1_45S_50ntFlanks | 12474 | 12496 | 23 | ACCCACTAATAGGGAACGTGAGC | BT20 | 13 | 4 | 19 | 5 | 7 | 5.0 | 0.5 | 7.1 | 4.0 | 0.8 |
| >hg38_21_+_8433172_8446622_RNA45SN1_45S_50ntFlanks@12474.12496.23\|ACCCACTAATAGGGAACGTGAGC | >hg38_21_+_8433172_8446622_RNA45SN1_45S_50ntFlanks | 12474 | 12496 | 23 | ACCCACTAATAGGGAACGTGAGC | MB231 | 81 | 46 | 74 | 5 | 10 | 63.1 | 37.2 | 68.5 | 5.0 | 10.8 |
| >hg38_21_+_8433172_8446622_RNA45SN1_45S_50ntFlanks@12474.12496.23\|ACCCACTAATAGGGAACGTGAGC | >hg38_21_+_8433172_8446622_RNA45SN1_45S_50ntFlanks | 12474 | 12496 | 23 | ACCCACTAATAGGGAACGTGAGC | MB468 | 33 | 18 | 26 | 11 | 9 | 11.3 | 17.4 | 15.0 | 1.6 | 4.4 |
| >hg38_21_+_8433172_8446622_RNA45SN1_45S_50ntFlanks@12474.12497.24\|ACCCACTAATAGGGAACGTGAGCT | >hg38_21_+_8433172_8446622_RNA45SN1_45S_50ntFlanks | 12474 | 12497 | 24 | ACCCACTAATAGGGAACGTGAGCT | MB231 | 57 | 21 | 26 | 2 | 8 | 55.0 | 15.9 | 21.2 | 1.8 | 7.5 |
| >hg38_21_+_8433172_8446622_RNA45SN1_45S_50ntFlanks@12474.12497.24\|ACCCACTAATAGGGAACGTGAGCT | >hg38_21_+_8433172_8446622_RNA45SN1_45S_50ntFlanks | 12474 | 12497 | 24 | ACCCACTAATAGGGAACGTGAGCT | MB468 | 20 | 7 | 9 | 3 | 7 | 6.4 | 4.8 | 3.4 | 0.9 | 2.5 |
| >hg38_21_+_8433172_8446622_RNA45SN1_45S_50ntFlanks@12474.12498.25\|ACCCACTAATAGGGAACGTGAGCTG | >hg38_21_+_8433172_8446622_RNA45SN1_45S_50ntFlanks | 12474 | 12498 | 25 | ACCCACTAATAGGGAACGTGAGCTG | MB468 | 71 | 21 | 28 | 8 | 14 | 20.6 | 4.7 | 4.3 | 1.8 | 2.9 |
| >hg38_21_+_8433172_8446622_RNA45SN1_45S_50ntFlanks@12474.12499.26\|ACCCACTAATAGGGAACGTGAGCTGG | >hg38_21_+_8433172_8446622_RNA45SN1_45S_50ntFlanks | 12474 | 12499 | 26 | ACCCACTAATAGGGAACGTGAGCTGG | MB468 | 16 | 6 | 11 | 3 | 2 | 5.5 | 2.2 | 3.8 | 1.6 | 0.8 |
| >hg38_21_+_8433172_8446622_RNA45SN1_45S_50ntFlanks@12475.12496.22\|CCCACTAATAGGGAACGTGAGC | >hg38_21_+_8433172_8446622_RNA45SN1_45S_50ntFlanks | 12475 | 12496 | 22 | CCCACTAATAGGGAACGTGAGC | MB231 | 36 | 32 | 42 | 3 | 6 | 28.3 | 21.9 | 32.4 | 2.6 | 5.9 |
| >hg38_21_+_8433172_8446622_RNA45SN1_45S_50ntFlanks@12475.12496.22\|CCCACTAATAGGGAACGTGAGC | >hg38_21_+_8433172_8446622_RNA45SN1_45S_50ntFlanks | 12475 | 12496 | 22 | CCCACTAATAGGGAACGTGAGC | MB468 | 27 | 23 | 27 | 13 | 9 | 12.1 | 14.9 | 12.1 | 2.9 | 2.5 |
| >hg38_21_+_8433172_8446622_RNA45SN1_45S_50ntFlanks@12475.12497.23\|CCCACTAATAGGGAACGTGAGCT | >hg38_21_+_8433172_8446622_RNA45SN1_45S_50ntFlanks | 12475 | 12497 | 23 | CCCACTAATAGGGAACGTGAGCT | MB231 | 28 | 14 | 27 | 2 | 5 | 22.7 | 9.1 | 19.3 | 2.1 | 4.3 |
| >hg38_21_+_8433172_8446622_RNA45SN1_45S_50ntFlanks@12475.12497.23\|CCCACTAATAGGGAACGTGAGCT | >hg38_21_+_8433172_8446622_RNA45SN1_45S_50ntFlanks | 12475 | 12497 | 23 | CCCACTAATAGGGAACGTGAGCT | MB468 | 19 | 13 | 18 | 6 | 9 | 7.9 | 6.8 | 6.4 | 1.5 | 1.9 |
| >hg38_21_+_8433172_8446622_RNA45SN1_45S_50ntFlanks@12475.12498.24\|CCCACTAATAGGGAACGTGAGCTG | >hg38_21_+_8433172_8446622_RNA45SN1_45S_50ntFlanks | 12475 | 12498 | 24 | CCCACTAATAGGGAACGTGAGCTG | MB468 | 74 | 50 | 43 | 14 | 28 | 27.1 | 16.9 | 8.6 | 5.5 | 8.9 |
| >hg38_21_+_8433172_8446622_RNA45SN1_45S_50ntFlanks@12475.12499.25\|CCCACTAATAGGGAACGTGAGCTGG | >hg38_21_+_8433172_8446622_RNA45SN1_45S_50ntFlanks | 12475 | 12499 | 25 | CCCACTAATAGGGAACGTGAGCTGG | MB468 | 13 | 6 | 15 | 3 | 3 | 3.7 | 1.8 | 7.5 | 1.6 | 1.2 |
| >hg38_21_+_8433172_8446622_RNA45SN1_45S_50ntFlanks@12476.12493.18\|CCACTAATAGGGAACGTG | >hg38_21_+_8433172_8446622_RNA45SN1_45S_50ntFlanks | 12476 | 12493 | 18 | CCACTAATAGGGAACGTG | MB468 | 44 | 19 | 36 | 26 | 15 | 14.4 | 8.4 | 11.2 | 5.5 | 6.5 |
| >hg38_21_+_8433172_8446622_RNA45SN1_45S_50ntFlanks@12476.12496.21\|CCACTAATAGGGAACGTGAGC | >hg38_21_+_8433172_8446622_RNA45SN1_45S_50ntFlanks | 12476 | 12496 | 21 | CCACTAATAGGGAACGTGAGC | MB468 | 220 | 162 | 166 | 137 | 96 | 107.6 | 92.0 | 107.5 | 41.6 | 21.8 |
| >hg38_21_+_8433172_8446622_RNA45SN1_45S_50ntFlanks@12476.12497.22\|CCACTAATAGGGAACGTGAGCT | >hg38_21_+_8433172_8446622_RNA45SN1_45S_50ntFlanks | 12476 | 12497 | 22 | CCACTAATAGGGAACGTGAGCT | MB231 | 11 | 5 | 16 | 2 | 3 | 7.2 | 2.3 | 11.2 | 0.9 | 2.5 |
| >hg38_21_+_8433172_8446622_RNA45SN1_45S_50ntFlanks@12476.12497.22\|CCACTAATAGGGAACGTGAGCT | >hg38_21_+_8433172_8446622_RNA45SN1_45S_50ntFlanks | 12476 | 12497 | 22 | CCACTAATAGGGAACGTGAGCT | MB468 | 186 | 120 | 137 | 87 | 57 | 87.0 | 47.5 | 69.8 | 26.3 | 22.9 |
| >hg38_21_+_8433172_8446622_RNA45SN1_45S_50ntFlanks@12476.12498.23\|CCACTAATAGGGAACGTGAGCTG | >hg38_21_+_8433172_8446622_RNA45SN1_45S_50ntFlanks | 12476 | 12498 | 23 | CCACTAATAGGGAACGTGAGCTG | MB468 | 257 | 168 | 189 | 131 | 69 | 100.7 | 39.3 | 75.0 | 54.5 | 22.9 |
| >hg38_21_+_8433172_8446622_RNA45SN1_45S_50ntFlanks@12476.12499.24\|CCACTAATAGGGAACGTGAGCTGG | >hg38_21_+_8433172_8446622_RNA45SN1_45S_50ntFlanks | 12476 | 12499 | 24 | CCACTAATAGGGAACGTGAGCTGG | MB468 | 46 | 27 | 34 | 15 | 10 | 12.1 | 8.1 | 13.4 | 6.8 | 4.3 |
| >hg38_21_+_8433172_8446622_RNA45SN1_45S_50ntFlanks@12476.12500.25\|CCACTAATAGGGAACGTGAGCTGGG | >hg38_21_+_8433172_8446622_RNA45SN1_45S_50ntFlanks | 12476 | 12500 | 25 | CCACTAATAGGGAACGTGAGCTGGG | MB468 | 73 | 72 | 64 | 29 | 14 | 28.4 | 16.6 | 38.6 | 13.2 | 3.7 |
| >hg38_21_+_8433172_8446622_RNA45SN1_45S_50ntFlanks@12476.12509.34\|CCACTAATAGGGAACGTGAGCTGGGTTTAGACCG | >hg38_21_+_8433172_8446622_RNA45SN1_45S_50ntFlanks | 12476 | 12509 | 34 | CCACTAATAGGGAACGTGAGCTGGGTTTAGACCG | MB468 | 14 | 15 | 10 | 2 | 3 | 10.3 | 10.4 | 5.2 | 0.4 | 2.4 |
| >hg38_21_+_8433172_8446622_RNA45SN1_45S_50ntFlanks@12477.12496.20\|CACTAATAGGGAACGTGAGC | >hg38_21_+_8433172_8446622_RNA45SN1_45S_50ntFlanks | 12477 | 12496 | 20 | CACTAATAGGGAACGTGAGC | BT20 | 23 | 5 | 83 | 9 | 6 | 10.7 | 0.7 | 60.3 | 5.9 | 0.4 |
| >hg38_21_+_8433172_8446622_RNA45SN1_45S_50ntFlanks@12477.12496.20\|CACTAATAGGGAACGTGAGC | >hg38_21_+_8433172_8446622_RNA45SN1_45S_50ntFlanks | 12477 | 12496 | 20 | CACTAATAGGGAACGTGAGC | MB231 | 61 | 25 | 141 | 15 | 10 | 50.6 | 16.8 | 116.8 | 9.9 | 8.8 |
| >hg38_21_+_8433172_8446622_RNA45SN1_45S_50ntFlanks@12477.12496.20\|CACTAATAGGGAACGTGAGC | >hg38_21_+_8433172_8446622_RNA45SN1_45S_50ntFlanks | 12477 | 12496 | 20 | CACTAATAGGGAACGTGAGC | MB468 | 43 | 26 | 46 | 19 | 9 | 13.0 | 9.0 | 17.7 | 7.9 | 3.6 |
| >hg38_21_+_8433172_8446622_RNA45SN1_45S_50ntFlanks@12477.12497.21\|CACTAATAGGGAACGTGAGCT | >hg38_21_+_8433172_8446622_RNA45SN1_45S_50ntFlanks | 12477 | 12497 | 21 | CACTAATAGGGAACGTGAGCT | BT20 | 36 | 8 | 84 | 15 | 12 | 17.7 | 2.6 | 59.0 | 7.2 | 1.7 |
| >hg38_21_+_8433172_8446622_RNA45SN1_45S_50ntFlanks@12477.12497.21\|CACTAATAGGGAACGTGAGCT | >hg38_21_+_8433172_8446622_RNA45SN1_45S_50ntFlanks | 12477 | 12497 | 21 | CACTAATAGGGAACGTGAGCT | MB231 | 70 | 19 | 152 | 18 | 14 | 55.5 | 15.5 | 129.3 | 12.3 | 11.3 |
| >hg38_21_+_8433172_8446622_RNA45SN1_45S_50ntFlanks@12477.12497.21\|CACTAATAGGGAACGTGAGCT | >hg38_21_+_8433172_8446622_RNA45SN1_45S_50ntFlanks | 12477 | 12497 | 21 | CACTAATAGGGAACGTGAGCT | MB468 | 37 | 21 | 38 | 12 | 8 | 9.6 | 4.6 | 13.5 | 4.5 | 2.5 |
| >hg38_21_+_8433172_8446622_RNA45SN1_45S_50ntFlanks@12477.12498.22\|CACTAATAGGGAACGTGAGCTG | >hg38_21_+_8433172_8446622_RNA45SN1_45S_50ntFlanks | 12477 | 12498 | 22 | CACTAATAGGGAACGTGAGCTG | MB468 | 126 | 90 | 132 | 36 | 19 | 35.5 | 25.4 | 42.4 | 12.1 | 4.6 |
| >hg38_21_+_8433172_8446622_RNA45SN1_45S_50ntFlanks@12477.12499.23\|CACTAATAGGGAACGTGAGCTGG | >hg38_21_+_8433172_8446622_RNA45SN1_45S_50ntFlanks | 12477 | 12499 | 23 | CACTAATAGGGAACGTGAGCTGG | MB468 | 25 | 21 | 39 | 8 | 4 | 5.7 | 4.9 | 20.1 | 2.3 | 1.2 |
| >hg38_21_+_8433172_8446622_RNA45SN1_45S_50ntFlanks@12477.12500.24\|CACTAATAGGGAACGTGAGCTGGG | >hg38_21_+_8433172_8446622_RNA45SN1_45S_50ntFlanks | 12477 | 12500 | 24 | CACTAATAGGGAACGTGAGCTGGG | MB468 | 11 | 13 | 14 | 5 | 2 | 3.6 | 2.2 | 6.2 | 2.6 | 0.8 |
| >hg38_21_+_8433172_8446622_RNA45SN1_45S_50ntFlanks@12478.12496.19\|ACTAATAGGGAACGTGAGC | >hg38_21_+_8433172_8446622_RNA45SN1_45S_50ntFlanks | 12478 | 12496 | 19 | ACTAATAGGGAACGTGAGC | BT20 | 18 | 8 | 77 | 20 | 8 | 8.9 | 3.3 | 77.5 | 15.1 | 1.7 |

| | | | | | | | | | | | | | | | |
|---|---|---|---|---|---|---|---|---|---|---|---|---|---|---|---|
| >hg38_21_+_8433172_8446622_RNA45SN1_45S_50ntFlanks@12478.12496.19\|ACTAATAGGGAACGTGAGC | >hg38_21_+_8433172_8446622_RNA45SN1_45S_50ntFlanks | 12478 | 12496 | 19 | ACTAATAGGGAACGTGAGC | MB231 | 30 | 21 | 67 | 18 | 9 | 23.5 | 15.2 | 51.8 | 11.0 | 8.6 |
| >hg38_21_+_8433172_8446622_RNA45SN1_45S_50ntFlanks@12478.12496.19\|ACTAATAGGGAACGTGAGC | >hg38_21_+_8433172_8446622_RNA45SN1_45S_50ntFlanks | 12478 | 12496 | 19 | ACTAATAGGGAACGTGAGC | MB468 | 29 | 31 | 25 | 11 | 4 | 9.8 | 9.1 | 8.4 | 5.1 | 1.0 |
| >hg38_21_+_8433172_8446622_RNA45SN1_45S_50ntFlanks@12478.12497.20\|ACTAATAGGGAACGTGAGCT | >hg38_21_+_8433172_8446622_RNA45SN1_45S_50ntFlanks | 12478 | 12497 | 20 | ACTAATAGGGAACGTGAGCT | BT20 | 37 | 12 | 107 | 35 | 19 | 19.5 | 7.4 | 110.3 | 33.3 | 4.5 |
| >hg38_21_+_8433172_8446622_RNA45SN1_45S_50ntFlanks@12478.12497.20\|ACTAATAGGGAACGTGAGCT | >hg38_21_+_8433172_8446622_RNA45SN1_45S_50ntFlanks | 12478 | 12497 | 20 | ACTAATAGGGAACGTGAGCT | MB231 | 55 | 25 | 117 | 16 | 14 | 40.9 | 18.6 | 85.7 | 10.8 | 12.8 |
| >hg38_21_+_8433172_8446622_RNA45SN1_45S_50ntFlanks@12478.12497.20\|ACTAATAGGGAACGTGAGCT | >hg38_21_+_8433172_8446622_RNA45SN1_45S_50ntFlanks | 12478 | 12497 | 20 | ACTAATAGGGAACGTGAGCT | MB468 | 30 | 27 | 28 | 10 | 4 | 8.9 | 6.3 | 9.0 | 4.8 | 0.8 |
| >hg38_21_+_8433172_8446622_RNA45SN1_45S_50ntFlanks@12478.12498.21\|ACTAATAGGGAACGTGAGCTG | >hg38_21_+_8433172_8446622_RNA45SN1_45S_50ntFlanks | 12478 | 12498 | 21 | ACTAATAGGGAACGTGAGCTG | MB468 | 60 | 59 | 52 | 19 | 6 | 19.0 | 13.7 | 14.2 | 10.8 | 3.2 |
| >hg38_21_+_8433172_8446622_RNA45SN1_45S_50ntFlanks@12478.12499.22\|ACTAATAGGGAACGTGAGCTGG | >hg38_21_+_8433172_8446622_RNA45SN1_45S_50ntFlanks | 12478 | 12499 | 22 | ACTAATAGGGAACGTGAGCTGG | MB468 | 24 | 29 | 27 | 8 | 2 | 8.5 | 8.6 | 14.4 | 3.4 | 1.1 |
| >hg38_21_+_8433172_8446622_RNA45SN1_45S_50ntFlanks@12478.12500.23\|ACTAATAGGGAACGTGAGCTGGG | >hg38_21_+_8433172_8446622_RNA45SN1_45S_50ntFlanks | 12478 | 12500 | 23 | ACTAATAGGGAACGTGAGCTGGG | MB468 | 15 | 20 | 15 | 6 | 2 | 5.1 | 3.6 | 5.3 | 3.8 | 0.8 |
| >hg38_21_+_8433172_8446622_RNA45SN1_45S_50ntFlanks@12479.12496.18\|CTAATAGGGAACGTGAGC | >hg38_21_+_8433172_8446622_RNA45SN1_45S_50ntFlanks | 12479 | 12496 | 18 | CTAATAGGGAACGTGAGC | BT20 | 15 | 5 | 86 | 17 | 4 | 6.6 | 2.5 | 97.7 | 17.9 | 1.2 |
| >hg38_21_+_8433172_8446622_RNA45SN1_45S_50ntFlanks@12479.12496.18\|CTAATAGGGAACGTGAGC | >hg38_21_+_8433172_8446622_RNA45SN1_45S_50ntFlanks | 12479 | 12496 | 18 | CTAATAGGGAACGTGAGC | MB231 | 33 | 16 | 81 | 13 | 7 | 30.2 | 12.2 | 60.6 | 8.7 | 6.0 |
| >hg38_21_+_8433172_8446622_RNA45SN1_45S_50ntFlanks@12479.12496.18\|CTAATAGGGAACGTGAGC | >hg38_21_+_8433172_8446622_RNA45SN1_45S_50ntFlanks | 12479 | 12496 | 18 | CTAATAGGGAACGTGAGC | MB468 | 56 | 48 | 61 | 23 | 10 | 18.5 | 11.1 | 25.1 | 8.3 | 3.1 |
| >hg38_21_+_8433172_8446622_RNA45SN1_45S_50ntFlanks@12479.12497.19\|CTAATAGGGAACGTGAGCT | >hg38_21_+_8433172_8446622_RNA45SN1_45S_50ntFlanks | 12479 | 12497 | 19 | CTAATAGGGAACGTGAGCT | BT20 | 36 | 10 | 139 | 28 | 12 | 20.1 | 9.7 | 156.9 | 29.0 | 3.5 |
| >hg38_21_+_8433172_8446622_RNA45SN1_45S_50ntFlanks@12479.12497.19\|CTAATAGGGAACGTGAGCT | >hg38_21_+_8433172_8446622_RNA45SN1_45S_50ntFlanks | 12479 | 12497 | 19 | CTAATAGGGAACGTGAGCT | MB231 | 59 | 18 | 157 | 12 | 10 | 49.1 | 13.8 | 121.1 | 8.6 | 10.0 |
| >hg38_21_+_8433172_8446622_RNA45SN1_45S_50ntFlanks@12479.12497.19\|CTAATAGGGAACGTGAGCT | >hg38_21_+_8433172_8446622_RNA45SN1_45S_50ntFlanks | 12479 | 12497 | 19 | CTAATAGGGAACGTGAGCT | MB468 | 63 | 48 | 70 | 21 | 11 | 18.1 | 9.3 | 20.5 | 9.2 | 3.2 |
| >hg38_21_+_8433172_8446622_RNA45SN1_45S_50ntFlanks@12479.12498.20\|CTAATAGGGAACGTGAGCTG | >hg38_21_+_8433172_8446622_RNA45SN1_45S_50ntFlanks | 12479 | 12498 | 20 | CTAATAGGGAACGTGAGCTG | MB468 | 101 | 77 | 102 | 32 | 14 | 30.9 | 6.7 | 27.2 | 15.2 | 5.8 |
| >hg38_21_+_8433172_8446622_RNA45SN1_45S_50ntFlanks@12479.12499.21\|CTAATAGGGAACGTGAGCTGG | >hg38_21_+_8433172_8446622_RNA45SN1_45S_50ntFlanks | 12479 | 12499 | 21 | CTAATAGGGAACGTGAGCTGG | MB468 | 36 | 37 | 43 | 11 | 4 | 9.9 | 2.6 | 16.8 | 4.6 | 1.2 |
| >hg38_21_+_8433172_8446622_RNA45SN1_45S_50ntFlanks@12479.12500.22\|CTAATAGGGAACGTGAGCTGGG | >hg38_21_+_8433172_8446622_RNA45SN1_45S_50ntFlanks | 12479 | 12500 | 22 | CTAATAGGGAACGTGAGCTGGG | MB468 | 35 | 38 | 37 | 15 | 5 | 11.0 | 15.8 | 12.0 | 9.6 | 2.2 |
| >hg38_21_+_8433172_8446622_RNA45SN1_45S_50ntFlanks@12479.12509.31\|CTAATAGGGAACGTGAGCTGGGTTTAGACCG | >hg38_21_+_8433172_8446622_RNA45SN1_45S_50ntFlanks | 12479 | 12509 | 31 | CTAATAGGGAACGTGAGCTGGGTTTAGACCG | MB468 | 19 | 18 | 17 | 2 | 2 | 13.1 | 17.3 | 7.5 | 1.0 | 1.7 |
| >hg38_21_+_8433172_8446622_RNA45SN1_45S_50ntFlanks@12480.12497.18\|TAATAGGGAACGTGAGCT | >hg38_21_+_8433172_8446622_RNA45SN1_45S_50ntFlanks | 12480 | 12497 | 18 | TAATAGGGAACGTGAGCT | BT20 | 24 | 6 | 86 | 18 | 7 | 11.9 | 4.3 | 101.8 | 16.1 | 1.1 |
| >hg38_21_+_8433172_8446622_RNA45SN1_45S_50ntFlanks@12480.12497.18\|TAATAGGGAACGTGAGCT | >hg38_21_+_8433172_8446622_RNA45SN1_45S_50ntFlanks | 12480 | 12497 | 18 | TAATAGGGAACGTGAGCT | MB231 | 45 | 16 | 103 | 10 | 9 | 36.8 | 12.0 | 79.1 | 7.3 | 7.1 |
| >hg38_21_+_8433172_8446622_RNA45SN1_45S_50ntFlanks@12480.12498.19\|TAATAGGGAACGTGAGCTG | >hg38_21_+_8433172_8446622_RNA45SN1_45S_50ntFlanks | 12480 | 12498 | 19 | TAATAGGGAACGTGAGCTG | MB468 | 19 | 15 | 17 | 5 | 2 | 5.0 | 3.1 | 3.3 | 1.8 | 0.7 |
| >hg38_21_+_8433172_8446622_RNA45SN1_45S_50ntFlanks@12481.12498.18\|AATAGGGAACGTGAGCTG | >hg38_21_+_8433172_8446622_RNA45SN1_45S_50ntFlanks | 12481 | 12498 | 18 | AATAGGGAACGTGAGCTG | MB468 | 31 | 28 | 26 | 15 | 4 | 8.6 | 8.6 | 4.7 | 8.5 | 2.0 |
| >hg38_21_+_8433172_8446622_RNA45SN1_45S_50ntFlanks@12481.12499.19\|AATAGGGAACGTGAGCTGG | >hg38_21_+_8433172_8446622_RNA45SN1_45S_50ntFlanks | 12481 | 12499 | 19 | AATAGGGAACGTGAGCTGG | BT20 | 19 | 21 | 21 | 16 | 4 | 30.4 | 34.4 | 26.2 | 18.9 | 3.0 |
| >hg38_21_+_8433172_8446622_RNA45SN1_45S_50ntFlanks@12481.12499.19\|AATAGGGAACGTGAGCTGG | >hg38_21_+_8433172_8446622_RNA45SN1_45S_50ntFlanks | 12481 | 12499 | 19 | AATAGGGAACGTGAGCTGG | MB468 | 13 | 16 | 13 | 7 | 2 | 3.3 | 4.5 | 4.9 | 3.0 | 0.4 |
| >hg38_21_+_8433172_8446622_RNA45SN1_45S_50ntFlanks@12481.12500.20\|AATAGGGAACGTGAGCTGGG | >hg38_21_+_8433172_8446622_RNA45SN1_45S_50ntFlanks | 12481 | 12500 | 20 | AATAGGGAACGTGAGCTGGG | BT20 | 18 | 16 | 9 | 10 | 2 | 27.3 | 26.2 | 9.4 | 9.5 | 2.5 |
| >hg38_21_+_8433172_8446622_RNA45SN1_45S_50ntFlanks@12481.12500.20\|AATAGGGAACGTGAGCTGGG | >hg38_21_+_8433172_8446622_RNA45SN1_45S_50ntFlanks | 12481 | 12500 | 20 | AATAGGGAACGTGAGCTGGG | MB468 | 11 | 14 | 10 | 8 | 2 | 3.9 | 2.9 | 2.7 | 5.6 | 0.8 |
| >hg38_21_+_8433172_8446622_RNA45SN1_45S_50ntFlanks@12481.12501.21\|AATAGGGAACGTGAGCTGGGT | >hg38_21_+_8433172_8446622_RNA45SN1_45S_50ntFlanks | 12481 | 12501 | 21 | AATAGGGAACGTGAGCTGGGT | BT20 | 57 | 27 | 10 | 16 | 3 | 98.6 | 46.1 | 16.6 | 27.6 | 4.6 |
| >hg38_21_+_8433172_8446622_RNA45SN1_45S_50ntFlanks@12481.12502.22\|AATAGGGAACGTGAGCTGGGTT | >hg38_21_+_8433172_8446622_RNA45SN1_45S_50ntFlanks | 12481 | 12502 | 22 | AATAGGGAACGTGAGCTGGGTT | BT20 | 33 | 17 | 6 | 8 | 2 | 56.2 | 30.0 | 9.3 | 13.6 | 2.7 |
| >hg38_21_+_8433172_8446622_RNA45SN1_45S_50ntFlanks@12482.12499.18\|ATAGGGAACGTGAGCTGG | >hg38_21_+_8433172_8446622_RNA45SN1_45S_50ntFlanks | 12482 | 12499 | 18 | ATAGGGAACGTGAGCTGG | BT20 | 28 | 48 | 19 | 6 | 2 | 48.2 | 83.1 | 30.9 | 4.8 | 1.7 |
| >hg38_21_+_8433172_8446622_RNA45SN1_45S_50ntFlanks@12482.12499.18\|ATAGGGAACGTGAGCTGG | >hg38_21_+_8433172_8446622_RNA45SN1_45S_50ntFlanks | 12482 | 12499 | 18 | ATAGGGAACGTGAGCTGG | MB468 | 12 | 13 | 14 | 7 | 2 | 3.4 | 4.5 | 6.0 | 3.8 | 0.5 |
| >hg38_21_+_8433172_8446622_RNA45SN1_45S_50ntFlanks@12482.12500.19\|ATAGGGAACGTGAGCTGGG | >hg38_21_+_8433172_8446622_RNA45SN1_45S_50ntFlanks | 12482 | 12500 | 19 | ATAGGGAACGTGAGCTGGG | BT20 | 29 | 37 | 8 | 11 | 2 | 49.0 | 64.3 | 12.6 | 15.5 | 2.5 |
| >hg38_21_+_8433172_8446622_RNA45SN1_45S_50ntFlanks@12482.12500.19\|ATAGGGAACGTGAGCTGGG | >hg38_21_+_8433172_8446622_RNA45SN1_45S_50ntFlanks | 12482 | 12500 | 19 | ATAGGGAACGTGAGCTGGG | MB468 | 15 | 19 | 14 | 13 | 2 | 3.7 | 1.4 | 4.8 | 8.8 | 0.9 |
| >hg38_21_+_8433172_8446622_RNA45SN1_45S_50ntFlanks@12482.12501.20\|ATAGGGAACGTGAGCTGGGT | >hg38_21_+_8433172_8446622_RNA45SN1_45S_50ntFlanks | 12482 | 12501 | 20 | ATAGGGAACGTGAGCTGGGT | BT20 | 107 | 74 | 16 | 32 | 4 | 184.5 | 128.0 | 28.1 | 55.3 | 6.7 |
| >hg38_21_+_8433172_8446622_RNA45SN1_45S_50ntFlanks@12482.12502.21\|ATAGGGAACGTGAGCTGGGTT | >hg38_21_+_8433172_8446622_RNA45SN1_45S_50ntFlanks | 12482 | 12502 | 21 | ATAGGGAACGTGAGCTGGGTT | BT20 | 55 | 42 | 8 | 16 | 3 | 95.2 | 73.3 | 13.6 | 28.4 | 4.3 |
| >hg38_21_+_8433172_8446622_RNA45SN1_45S_50ntFlanks@12482.12503.22\|ATAGGGAACGTGAGCTGGGTTT | >hg38_21_+_8433172_8446622_RNA45SN1_45S_50ntFlanks | 12482 | 12503 | 22 | ATAGGGAACGTGAGCTGGGTTT | BT20 | 14 | 9 | 1 | 3 | 1 | 23.5 | 15.8 | 2.4 | 4.7 | 1.2 |
| >hg38_21_+_8433172_8446622_RNA45SN1_45S_50ntFlanks@12487.12509.23\|GAACGTGAGCTGGGTTTAGACCG | >hg38_21_+_8433172_8446622_RNA45SN1_45S_50ntFlanks | 12487 | 12509 | 23 | GAACGTGAGCTGGGTTTAGACCG | MB468 | 27 | 20 | 18 | 7 | 3 | 5.5 | 5.4 | 5.3 | 4.7 | 0.7 |
| >hg38_21_+_8433172_8446622_RNA45SN1_45S_50ntFlanks@12488.12505.18\|AACGTGAGCTGGGTTTAG | >hg38_21_+_8433172_8446622_RNA45SN1_45S_50ntFlanks | 12488 | 12505 | 18 | AACGTGAGCTGGGTTTAG | MB468 | 24 | 27 | 12 | 8 | 2 | 6.4 | 20.5 | 7.2 | 5.7 | 1.4 |
| >hg38_21_+_8433172_8446622_RNA45SN1_45S_50ntFlanks@12488.12508.21\|AACGTGAGCTGGGTTTAGACC | >hg38_21_+_8433172_8446622_RNA45SN1_45S_50ntFlanks | 12488 | 12508 | 21 | AACGTGAGCTGGGTTTAGACC | MB468 | 18 | 10 | 8 | 7 | 4 | 16.3 | 2.8 | 4.7 | 8.3 | 3.9 |
| >hg38_21_+_8433172_8446622_RNA45SN1_45S_50ntFlanks@12488.12509.22\|AACGTGAGCTGGGTTTAGACCG | >hg38_21_+_8433172_8446622_RNA45SN1_45S_50ntFlanks | 12488 | 12509 | 22 | AACGTGAGCTGGGTTTAGACCG | MB468 | 652 | 397 | 309 | 155 | 50 | 182.7 | 137.2 | 39.1 | 94.7 | 17.3 |
| >hg38_21_+_8433172_8446622_RNA45SN1_45S_50ntFlanks@12488.12512.25\|AACGTGAGCTGGGTTTAGACCGTCG | >hg38_21_+_8433172_8446622_RNA45SN1_45S_50ntFlanks | 12488 | 12512 | 25 | AACGTGAGCTGGGTTTAGACCGTCG | MB468 | 27 | 3 | 19 | 7 | 3 | 6.2 | 1.6 | 8.2 | 6.0 | 1.3 |
| >hg38_21_+_8433172_8446622_RNA45SN1_45S_50ntFlanks@12488.12518.31\|AACGTGAGCTGGGTTTAGACCGTCGTGAGAC | >hg38_21_+_8433172_8446622_RNA45SN1_45S_50ntFlanks | 12488 | 12518 | 31 | AACGTGAGCTGGGTTTAGACCGTCGTGAGAC | MB468 | 43 | 15 | 20 | 21 | 16 | 11.9 | 2.0 | 8.0 | 7.9 | 6.7 |
| >hg38_21_+_8433172_8446622_RNA45SN1_45S_50ntFlanks@12488.12519.32\|AACGTGAGCTGGGTTTAGACCGTCGTGAGACA | >hg38_21_+_8433172_8446622_RNA45SN1_45S_50ntFlanks | 12488 | 12519 | 32 | AACGTGAGCTGGGTTTAGACCGTCGTGAGACA | MB468 | 23 | 9 | 11 | 9 | 13 | 9.8 | 1.6 | 4.2 | 2.4 | 8.2 |
| >hg38_21_+_8433172_8446622_RNA45SN1_45S_50ntFlanks@12488.12520.33\|AACGTGAGCTGGGTTTAGACCGTCGTGAGACAG | >hg38_21_+_8433172_8446622_RNA45SN1_45S_50ntFlanks | 12488 | 12520 | 33 | AACGTGAGCTGGGTTTAGACCGTCGTGAGACAG | MB468 | 47 | 16 | 19 | 14 | 23 | 21.2 | 6.2 | 5.5 | 4.4 | 11.9 |
| >hg38_21_+_8433172_8446622_RNA45SN1_45S_50ntFlanks@12488.12521.34\|AACGTGAGCTGGGTTTAGACCGTCGTGAGACAGG | >hg38_21_+_8433172_8446622_RNA45SN1_45S_50ntFlanks | 12488 | 12521 | 34 | AACGTGAGCTGGGTTTAGACCGTCGTGAGACAGG | MB468 | 78 | 37 | 36 | 28 | 49 | 46.5 | 12.0 | 15.1 | 7.2 | 38.0 |
| >hg38_21_+_8433172_8446622_RNA45SN1_45S_50ntFlanks@12488.12522.35\|AACGTGAGCTGGGTTTAGACCGTCGTGAGACAGGT | >hg38_21_+_8433172_8446622_RNA45SN1_45S_50ntFlanks | 12488 | 12522 | 35 | AACGTGAGCTGGGTTTAGACCGTCGTGAGACAGGT | MB468 | 25 | 29 | 18 | 7 | 14 | 18.4 | 16.1 | 7.1 | 3.6 | 9.2 |
| >hg38_21_+_8433172_8446622_RNA45SN1_45S_50ntFlanks@12488.12524.37\|AACGTGAGCTGGGTTTAGACCGTCGTGAGACAGGTTA | >hg38_21_+_8433172_8446622_RNA45SN1_45S_50ntFlanks | 12488 | 12524 | 37 | AACGTGAGCTGGGTTTAGACCGTCGTGAGACAGGTTA | MB468 | 18 | 30 | 16 | 4 | 12 | 15.4 | 25.4 | 7.0 | 1.5 | 13.0 |
| >hg38_21_+_8433172_8446622_RNA45SN1_45S_50ntFlanks@12488.12525.38\|AACGTGAGCTGGGTTTAGACCGTCGTGAGACAGGTTAG | >hg38_21_+_8433172_8446622_RNA45SN1_45S_50ntFlanks | 12488 | 12525 | 38 | AACGTGAGCTGGGTTTAGACCGTCGTGAGACAGGTTAG | MB468 | 17 | 36 | 16 | 4 | 11 | 10.3 | 36.8 | 8.3 | 1.7 | 8.9 |
| >hg38_21_+_8433172_8446622_RNA45SN1_45S_50ntFlanks@12489.12509.21\|ACGTGAGCTGGGTTTAGACCG | >hg38_21_+_8433172_8446622_RNA45SN1_45S_50ntFlanks | 12489 | 12509 | 21 | ACGTGAGCTGGGTTTAGACCG | MB468 | 110 | 46 | 57 | 27 | 9 | 24.9 | 16.3 | 14.1 | 16.8 | 3.0 |
| >hg38_21_+_8433172_8446622_RNA45SN1_45S_50ntFlanks@12489.12518.30\|ACGTGAGCTGGGTTTAGACCGTCGTGAGAC | >hg38_21_+_8433172_8446622_RNA45SN1_45S_50ntFlanks | 12489 | 12518 | 30 | ACGTGAGCTGGGTTTAGACCGTCGTGAGAC | MB468 | 11 | 4 | 6 | 5 | 4 | 3.5 | 0.5 | 1.8 | 2.3 | 1.8 |
| >hg38_21_+_8433172_8446622_RNA45SN1_45S_50ntFlanks@12489.12521.33\|ACGTGAGCTGGGTTTAGACCGTCGTGAGACAGG | >hg38_21_+_8433172_8446622_RNA45SN1_45S_50ntFlanks | 12489 | 12521 | 33 | ACGTGAGCTGGGTTTAGACCGTCGTGAGACAGG | MB468 | 17 | 7 | 8 | 6 | 10 | 9.0 | 2.2 | 2.9 | 1.8 | 6.5 |

| | | | | | | | | | | | | | | | | |
|---|---|---|---|---|---|---|---|---|---|---|---|---|---|---|---|---|
| >hg38_21_+_8433172_8446622_RNA45SN1_45S_50ntFlanks@12490.12509.20\|CGTGAGCTGGGTTTAGACCG | >hg38_21_+_8433172_8446622_RNA45SN1_45S_50ntFlanks | 12490 | 12509 | 20 | CGTGAGCTGGGTTTAGACCG | MB468 | 15 | 4 | 8 | 6 | 2 | 2.8 | 1.6 | 3.4 | 4.4 | 0.6 |
| >hg38_21_+_8433172_8446622_RNA45SN1_45S_50ntFlanks@12490.12520.31\|CGTGAGCTGGGTTTAGACCGTCGTGAGACAG | >hg38_21_+_8433172_8446622_RNA45SN1_45S_50ntFlanks | 12490 | 12520 | 31 | CGTGAGCTGGGTTTAGACCGTCGTGAGACAG | MB468 | 17 | 6 | 9 | 8 | 12 | 5.8 | 2.2 | 3.3 | 3.0 | 7.4 |
| >hg38_21_+_8433172_8446622_RNA45SN1_45S_50ntFlanks@12490.12521.32\|CGTGAGCTGGGTTTAGACCGTCGTGAGACAGG | >hg38_21_+_8433172_8446622_RNA45SN1_45S_50ntFlanks | 12490 | 12521 | 32 | CGTGAGCTGGGTTTAGACCGTCGTGAGACAGG | MB468 | 19 | 8 | 10 | 8 | 16 | 9.4 | 2.4 | 3.8 | 3.9 | 11.2 |
| >hg38_21_+_8433172_8446622_RNA45SN1_45S_50ntFlanks@12499.12516.18\|GGTTTAGACCGTCGTGAG | >hg38_21_+_8433172_8446622_RNA45SN1_45S_50ntFlanks | 12499 | 12516 | 18 | GGTTTAGACCGTCGTGAG | MB468 | 11 | 1 | 14 | 4 | 1 | 6.7 | 0.4 | 3.2 | 4.4 | 1.1 |
| >hg38_21_+_8433172_8446622_RNA45SN1_45S_50ntFlanks@12499.12517.19\|GGTTTAGACCGTCGTGAGA | >hg38_21_+_8433172_8446622_RNA45SN1_45S_50ntFlanks | 12499 | 12517 | 19 | GGTTTAGACCGTCGTGAGA | MB468 | 11 | 2 | 15 | 6 | 1 | 3.1 | 0.4 | 6.0 | 6.0 | 0.7 |
| >hg38_21_+_8433172_8446622_RNA45SN1_45S_50ntFlanks@12499.12518.20\|GGTTTAGACCGTCGTGAGAC | >hg38_21_+_8433172_8446622_RNA45SN1_45S_50ntFlanks | 12499 | 12518 | 20 | GGTTTAGACCGTCGTGAGAC | MB468 | 33 | 1 | 16 | 14 | 13 | 14.5 | 0.8 | 9.8 | 7.7 | 8.2 |
| >hg38_21_+_8433172_8446622_RNA45SN1_45S_50ntFlanks@12499.12521.23\|GGTTTAGACCGTCGTGAGACAGG | >hg38_21_+_8433172_8446622_RNA45SN1_45S_50ntFlanks | 12499 | 12521 | 23 | GGTTTAGACCGTCGTGAGACAGG | MB468 | 13 | 0 | 7 | 10 | 4 | 2.1 | 0.3 | 3.3 | 10.9 | 2.0 |
| >hg38_21_+_8433172_8446622_RNA45SN1_45S_50ntFlanks@12500.12518.19\|GTTTAGACCGTCGTGAGAC | >hg38_21_+_8433172_8446622_RNA45SN1_45S_50ntFlanks | 12500 | 12518 | 19 | GTTTAGACCGTCGTGAGAC | MB468 | 14 | 0 | 5 | 7 | 4 | 6.6 | 0.2 | 3.6 | 3.6 | 2.9 |
| >hg38_21_+_8433172_8446622_RNA45SN1_45S_50ntFlanks@12510.12527.18\|TCGTGAGACAGGTTAGTT | >hg38_21_+_8433172_8446622_RNA45SN1_45S_50ntFlanks | 12510 | 12527 | 18 | TCGTGAGACAGGTTAGTT | MB468 | 46 | 8 | 107 | 26 | 14 | 25.6 | 5.9 | 83.8 | 8.4 | 4.1 |
| >hg38_21_+_8433172_8446622_RNA45SN1_45S_50ntFlanks@12510.12538.29\|TCGTGAGACAGGTTAGTTTTACCCTACTG | >hg38_21_+_8433172_8446622_RNA45SN1_45S_50ntFlanks | 12510 | 12538 | 29 | TCGTGAGACAGGTTAGTTTTACCCTACTG | MB468 | 13 | 2 | 5 | 12 | 9 | 5.7 | 0.7 | 2.3 | 8.1 | 4.8 |
| >hg38_21_+_8433172_8446622_RNA45SN1_45S_50ntFlanks@12511.12538.28\|CGTGAGACAGGTTAGTTTTACCCTACTG | >hg38_21_+_8433172_8446622_RNA45SN1_45S_50ntFlanks | 12511 | 12538 | 28 | CGTGAGACAGGTTAGTTTTACCCTACTG | MB468 | 13 | 3 | 4 | 17 | 20 | 2.6 | 0.7 | 2.1 | 4.2 | 7.0 |
| >hg38_21_+_8433172_8446622_RNA45SN1_45S_50ntFlanks@12513.12538.26\|TGAGACAGGTTAGTTTTACCCTACTG | >hg38_21_+_8433172_8446622_RNA45SN1_45S_50ntFlanks | 12513 | 12538 | 26 | TGAGACAGGTTAGTTTTACCCTACTG | MB468 | 14 | 1 | 5 | 15 | 8 | 8.7 | 0.3 | 3.3 | 10.1 | 4.0 |
| >hg38_21_+_8433172_8446622_RNA45SN1_45S_50ntFlanks@12539.12557.19\|ATGATGTGTTGTTGCCATG | >hg38_21_+_8433172_8446622_RNA45SN1_45S_50ntFlanks | 12539 | 12557 | 19 | ATGATGTGTTGTTGCCATG | MB468 | 23 | 8 | 11 | 39 | 31 | 9.5 | 6.7 | 6.4 | 30.2 | 15.3 |
| >hg38_21_+_8433172_8446622_RNA45SN1_45S_50ntFlanks@12539.12558.20\|ATGATGTGTTGTTGCCATGG | >hg38_21_+_8433172_8446622_RNA45SN1_45S_50ntFlanks | 12539 | 12558 | 20 | ATGATGTGTTGTTGCCATGG | MB468 | 12 | 7 | 5 | 8 | 9 | 4.5 | 4.6 | 1.4 | 5.0 | 3.4 |
| >hg38_21_+_8433172_8446622_RNA45SN1_45S_50ntFlanks@12539.12560.22\|ATGATGTGTTGTTGCCATGGTA | >hg38_21_+_8433172_8446622_RNA45SN1_45S_50ntFlanks | 12539 | 12560 | 22 | ATGATGTGTTGTTGCCATGGTA | MB468 | 17 | 19 | 11 | 18 | 20 | 6.5 | 7.2 | 3.3 | 12.8 | 7.3 |
| >hg38_21_+_8433172_8446622_RNA45SN1_45S_50ntFlanks@12539.12563.25\|ATGATGTGTTGTTGCCATGGTAATC | >hg38_21_+_8433172_8446622_RNA45SN1_45S_50ntFlanks | 12539 | 12563 | 25 | ATGATGTGTTGTTGCCATGGTAATC | MB468 | 56 | 54 | 36 | 43 | 34 | 17.9 | 25.9 | 11.0 | 19.8 | 17.8 |
| >hg38_21_+_8433172_8446622_RNA45SN1_45S_50ntFlanks@12539.12565.27\|ATGATGTGTTGTTGCCATGGTAATCCT | >hg38_21_+_8433172_8446622_RNA45SN1_45S_50ntFlanks | 12539 | 12565 | 27 | ATGATGTGTTGTTGCCATGGTAATCCT | MB468 | 27 | 31 | 18 | 22 | 23 | 13.4 | 20.8 | 6.7 | 7.5 | 8.8 |
| >hg38_21_+_8433172_8446622_RNA45SN1_45S_50ntFlanks@12539.12566.28\|ATGATGTGTTGTTGCCATGGTAATCCTG | >hg38_21_+_8433172_8446622_RNA45SN1_45S_50ntFlanks | 12539 | 12566 | 28 | ATGATGTGTTGTTGCCATGGTAATCCTG | MB468 | 127 | 104 | 87 | 108 | 71 | 62.3 | 30.2 | 36.9 | 33.5 | 18.5 |
| >hg38_21_+_8433172_8446622_RNA45SN1_45S_50ntFlanks@12539.12567.29\|ATGATGTGTTGTTGCCATGGTAATCCTGC | >hg38_21_+_8433172_8446622_RNA45SN1_45S_50ntFlanks | 12539 | 12567 | 29 | ATGATGTGTTGTTGCCATGGTAATCCTGC | MB468 | 64 | 77 | 54 | 58 | 49 | 38.1 | 16.5 | 27.3 | 16.0 | 22.5 |
| >hg38_21_+_8433172_8446622_RNA45SN1_45S_50ntFlanks@12539.12568.30\|ATGATGTGTTGTTGCCATGGTAATCCTGCT | >hg38_21_+_8433172_8446622_RNA45SN1_45S_50ntFlanks | 12539 | 12568 | 30 | ATGATGTGTTGTTGCCATGGTAATCCTGCT | MB468 | 13 | 11 | 9 | 41 | 19 | 5.6 | 3.7 | 2.9 | 14.8 | 1.5 |
| >hg38_21_+_8433172_8446622_RNA45SN1_45S_50ntFlanks@12539.12569.31\|ATGATGTGTTGTTGCCATGGTAATCCTGCTC | >hg38_21_+_8433172_8446622_RNA45SN1_45S_50ntFlanks | 12539 | 12569 | 31 | ATGATGTGTTGTTGCCATGGTAATCCTGCTC | MB468 | 169 | 272 | 162 | 118 | 174 | 83.7 | 91.2 | 49.4 | 42.5 | 73.5 |
| >hg38_21_+_8433172_8446622_RNA45SN1_45S_50ntFlanks@12539.12573.35\|ATGATGTGTTGTTGCCATGGTAATCCTGCTCAGTA | >hg38_21_+_8433172_8446622_RNA45SN1_45S_50ntFlanks | 12539 | 12573 | 35 | ATGATGTGTTGTTGCCATGGTAATCCTGCTCAGTA | MB468 | 21 | 24 | 17 | 45 | 51 | 5.7 | 13.5 | 5.9 | 18.8 | 28.2 |
| >hg38_21_+_8433172_8446622_RNA45SN1_45S_50ntFlanks@12540.12569.30\|TGATGTGTTGTTGCCATGGTAATCCTGCTC | >hg38_21_+_8433172_8446622_RNA45SN1_45S_50ntFlanks | 12540 | 12569 | 30 | TGATGTGTTGTTGCCATGGTAATCCTGCTC | MB468 | 12 | 19 | 13 | 8 | 14 | 5.5 | 7.0 | 4.6 | 1.6 | 5.6 |
| >hg38_21_+_8433172_8446622_RNA45SN1_45S_50ntFlanks@12567.12587.21\|CTCAGTACGAGAGGAACCGCA | >hg38_21_+_8433172_8446622_RNA45SN1_45S_50ntFlanks | 12567 | 12587 | 21 | CTCAGTACGAGAGGAACCGCA | MB468 | 59 | 29 | 31 | 26 | 61 | 13.1 | 9.8 | 6.3 | 9.5 | 5.0 |
| >hg38_21_+_8433172_8446622_RNA45SN1_45S_50ntFlanks@12568.12587.20\|TCAGTACGAGAGGAACCGCA | >hg38_21_+_8433172_8446622_RNA45SN1_45S_50ntFlanks | 12568 | 12587 | 20 | TCAGTACGAGAGGAACCGCA | MB468 | 17 | 9 | 9 | 10 | 9 | 4.8 | 4.4 | 3.8 | 3.8 | 4.0 |
| >hg38_21_+_8433172_8446622_RNA45SN1_45S_50ntFlanks@12569.12587.19\|CAGTACGAGAGGAACCGCA | >hg38_21_+_8433172_8446622_RNA45SN1_45S_50ntFlanks | 12569 | 12587 | 19 | CAGTACGAGAGGAACCGCA | MB468 | 17 | 8 | 9 | 16 | 15 | 5.0 | 4.2 | 5.0 | 8.9 | 3.0 |
| >hg38_21_+_8433172_8446622_RNA45SN1_45S_50ntFlanks@12570.12587.18\|AGTACGAGAGGAACCGCA | >hg38_21_+_8433172_8446622_RNA45SN1_45S_50ntFlanks | 12570 | 12587 | 18 | AGTACGAGAGGAACCGCA | MB468 | 122 | 44 | 50 | 101 | 58 | 44.3 | 22.2 | 21.8 | 57.7 | 14.4 |
| >hg38_21_+_8433172_8446622_RNA45SN1_45S_50ntFlanks@12574.12604.31\|CGAGAGGAACCGCAGGTTCAGACATTTGGTG | >hg38_21_+_8433172_8446622_RNA45SN1_45S_50ntFlanks | 12574 | 12604 | 31 | CGAGAGGAACCGCAGGTTCAGACATTTGGTG | MB468 | 13 | 8 | 7 | 48 | 20 | 4.5 | 3.0 | 2.7 | 6.8 | 3.8 |
| >hg38_21_+_8433172_8446622_RNA45SN1_45S_50ntFlanks@12588.12610.23\|GGTTCAGACATTTGGTGTATGTG | >hg38_21_+_8433172_8446622_RNA45SN1_45S_50ntFlanks | 12588 | 12610 | 23 | GGTTCAGACATTTGGTGTATGTG | MB468 | 15 | 4 | 8 | 30 | 6 | 4.3 | 0.9 | 2.6 | 17.6 | 2.0 |
| >hg38_21_+_8433172_8446622_RNA45SN1_45S_50ntFlanks@12588.12614.27\|GGTTCAGACATTTGGTGTATGTGCTTG | >hg38_21_+_8433172_8446622_RNA45SN1_45S_50ntFlanks | 12588 | 12614 | 27 | GGTTCAGACATTTGGTGTATGTGCTTG | MB468 | 16 | 2 | 6 | 27 | 10 | 8.2 | 0.7 | 4.4 | 21.0 | 3.5 |
| >hg38_21_+_8433172_8446622_RNA45SN1_45S_50ntFlanks@12588.12616.29\|GGTTCAGACATTTGGTGTATGTGCTTGGC | >hg38_21_+_8433172_8446622_RNA45SN1_45S_50ntFlanks | 12588 | 12616 | 29 | GGTTCAGACATTTGGTGTATGTGCTTGGC | MB468 | 13 | 4 | 10 | 9 | 6 | 3.5 | 1.5 | 4.4 | 4.1 | 1.4 |
| >hg38_21_+_8433172_8446622_RNA45SN1_45S_50ntFlanks@12589.12614.26\|GTTCAGACATTTGGTGTATGTGCTTG | >hg38_21_+_8433172_8446622_RNA45SN1_45S_50ntFlanks | 12589 | 12614 | 26 | GTTCAGACATTTGGTGTATGTGCTTG | MB468 | 59 | 9 | 24 | 47 | 22 | 29.1 | 2.0 | 17.3 | 16.7 | 7.2 |
| >hg38_21_+_8433172_8446622_RNA45SN1_45S_50ntFlanks@12597.12614.18\|ATTTGGTGTATGTGCTTG | >hg38_21_+_8433172_8446622_RNA45SN1_45S_50ntFlanks | 12597 | 12614 | 18 | ATTTGGTGTATGTGCTTG | MB468 | 12 | 8 | 20 | 14 | 19 | 8.5 | 3.7 | 18.8 | 12.2 | 15.2 |
| >hg38_21_+_8433172_8446622_RNA45SN1_45S_50ntFlanks@12605.12622.18\|TATGTGCTTGGCTGAGGA | >hg38_21_+_8433172_8446622_RNA45SN1_45S_50ntFlanks | 12605 | 12622 | 18 | TATGTGCTTGGCTGAGGA | MB468 | 25 | 35 | 13 | 4 | 4 | 10.0 | 12.2 | 5.6 | 1.3 | 1.1 |
| >hg38_21_+_8433172_8446622_RNA45SN1_45S_50ntFlanks@12605.12623.19\|TATGTGCTTGGCTGAGGAG | >hg38_21_+_8433172_8446622_RNA45SN1_45S_50ntFlanks | 12605 | 12623 | 19 | TATGTGCTTGGCTGAGGAG | MB468 | 37 | 76 | 39 | 8 | 12 | 14.9 | 19.6 | 19.6 | 2.0 | 2.5 |
| >hg38_21_+_8433172_8446622_RNA45SN1_45S_50ntFlanks@12605.12624.20\|TATGTGCTTGGCTGAGGAGC | >hg38_21_+_8433172_8446622_RNA45SN1_45S_50ntFlanks | 12605 | 12624 | 20 | TATGTGCTTGGCTGAGGAGC | MB231 | 12 | 8 | 35 | 2 | 3 | 10.8 | 4.7 | 27.5 | 1.0 | 2.3 |
| >hg38_21_+_8433172_8446622_RNA45SN1_45S_50ntFlanks@12605.12624.20\|TATGTGCTTGGCTGAGGAGC | >hg38_21_+_8433172_8446622_RNA45SN1_45S_50ntFlanks | 12605 | 12624 | 20 | TATGTGCTTGGCTGAGGAGC | MB468 | 206 | 438 | 111 | 32 | 29 | 92.0 | 160.8 | 57.8 | 11.8 | 6.2 |
| >hg38_21_+_8433172_8446622_RNA45SN1_45S_50ntFlanks@12605.12625.21\|TATGTGCTTGGCTGAGGAGCC | >hg38_21_+_8433172_8446622_RNA45SN1_45S_50ntFlanks | 12605 | 12625 | 21 | TATGTGCTTGGCTGAGGAGCC | MB468 | 861 | 705 | 588 | 185 | 115 | 236.2 | 351.9 | 231.8 | 82.5 | 40.8 |
| >hg38_21_+_8433172_8446622_RNA45SN1_45S_50ntFlanks@12605.12626.22\|TATGTGCTTGGCTGAGGAGCCA | >hg38_21_+_8433172_8446622_RNA45SN1_45S_50ntFlanks | 12605 | 12626 | 22 | TATGTGCTTGGCTGAGGAGCCA | MB468 | 568 | 313 | 427 | 139 | 82 | 178.6 | 189.2 | 87.0 | 21.5 | 52.4 |
| >hg38_21_+_8433172_8446622_RNA45SN1_45S_50ntFlanks@12605.12634.30\|TATGTGCTTGGCTGAGGAGCCAATGGGGCG | >hg38_21_+_8433172_8446622_RNA45SN1_45S_50ntFlanks | 12605 | 12634 | 30 | TATGTGCTTGGCTGAGGAGCCAATGGGGCG | MB468 | 11 | 21 | 17 | 7 | 5 | 7.8 | 12.1 | 10.8 | 3.5 | 2.0 |
| >hg38_21_+_8433172_8446622_RNA45SN1_45S_50ntFlanks@12606.12624.19\|ATGTGCTTGGCTGAGGAGC | >hg38_21_+_8433172_8446622_RNA45SN1_45S_50ntFlanks | 12606 | 12624 | 19 | ATGTGCTTGGCTGAGGAGC | MB468 | 19 | 49 | 9 | 5 | 3 | 8.1 | 21.2 | 3.8 | 1.0 | 0.4 |
| >hg38_21_+_8433172_8446622_RNA45SN1_45S_50ntFlanks@12606.12625.20\|ATGTGCTTGGCTGAGGAGCC | >hg38_21_+_8433172_8446622_RNA45SN1_45S_50ntFlanks | 12606 | 12625 | 20 | ATGTGCTTGGCTGAGGAGCC | MB468 | 73 | 58 | 46 | 24 | 11 | 26.5 | 31.8 | 23.2 | 4.1 | 5.3 |
| >hg38_21_+_8433172_8446622_RNA45SN1_45S_50ntFlanks@12606.12626.21\|ATGTGCTTGGCTGAGGAGCCA | >hg38_21_+_8433172_8446622_RNA45SN1_45S_50ntFlanks | 12606 | 12626 | 21 | ATGTGCTTGGCTGAGGAGCCA | MB468 | 44 | 25 | 28 | 21 | 8 | 20.0 | 15.8 | 8.9 | 8.4 | 7.5 |
| >hg38_21_+_8433172_8446622_RNA45SN1_45S_50ntFlanks@12607.12624.18\|TGTGCTTGGCTGAGGAGC | >hg38_21_+_8433172_8446622_RNA45SN1_45S_50ntFlanks | 12607 | 12624 | 18 | TGTGCTTGGCTGAGGAGC | MB231 | 23 | 20 | 31 | 4 | 4 | 17.6 | 14.2 | 20.1 | 2.2 | 3.1 |
| >hg38_21_+_8433172_8446622_RNA45SN1_45S_50ntFlanks@12607.12625.19\|TGTGCTTGGCTGAGGAGCC | >hg38_21_+_8433172_8446622_RNA45SN1_45S_50ntFlanks | 12607 | 12625 | 19 | TGTGCTTGGCTGAGGAGCC | MB468 | 34 | 25 | 23 | 15 | 11 | 4.2 | 9.4 | 5.1 | 4.9 | 7.8 |
| >hg38_21_+_8433172_8446622_RNA45SN1_45S_50ntFlanks@12607.12626.20\|TGTGCTTGGCTGAGGAGCCA | >hg38_21_+_8433172_8446622_RNA45SN1_45S_50ntFlanks | 12607 | 12626 | 20 | TGTGCTTGGCTGAGGAGCCA | MB468 | 25 | 11 | 18 | 12 | 10 | 6.5 | 6.8 | 1.1 | 4.1 | 11.7 |
| >hg38_21_+_8433172_8446622_RNA45SN1_45S_50ntFlanks@12615.12653.39\|GCTGAGGAGCCAATGGGGCGAAGCTACCATCTGTGGGAT | >hg38_21_+_8433172_8446622_RNA45SN1_45S_50ntFlanks | 12615 | 12653 | 39 | GCTGAGGAGCCAATGGGGCGAAGCTACCATCTGTGGGAT | MB468 | 102 | 131 | 92 | 77 | 245 | 22.9 | 104.1 | 55.4 | 51.9 | 147.1 |
| >hg38_21_+_8433172_8446622_RNA45SN1_45S_50ntFlanks@12615.12654.40\|GCTGAGGAGCCAATGGGGCGAAGCTACCATCTGTGGGATT | >hg38_21_+_8433172_8446622_RNA45SN1_45S_50ntFlanks | 12615 | 12654 | 40 | GCTGAGGAGCCAATGGGGCGAAGCTACCATCTGTGGGATT | MB468 | 294 | 707 | 305 | 138 | 824 | 78.5 | 712.3 | 192.0 | 84.6 | 546.1 |
| >hg38_21_+_8433172_8446622_RNA45SN1_45S_50ntFlanks@12615.12661.47\|GCTGAGGAGCCAATGGGGCGAAGCTACCATCTGTGGGATTATGACTG | >hg38_21_+_8433172_8446622_RNA45SN1_45S_50ntFlanks | 12615 | 12661 | 47 | GCTGAGGAGCCAATGGGGCGAAGCTACCATCTGTGGGATTATGACTG | MB468 | 11 | 60 | 11 | 15 | 132 | 5.1 | 66.9 | 7.1 | 17.9 | 94.5 |
| >hg38_21_+_8433172_8446622_RNA45SN1_45S_50ntFlanks@12616.12653.38\|CTGAGGAGCCAATGGGGCGAAGCTACCATCTGTGGGAT | >hg38_21_+_8433172_8446622_RNA45SN1_45S_50ntFlanks | 12616 | 12653 | 38 | CTGAGGAGCCAATGGGGCGAAGCTACCATCTGTGGGAT | MB468 | 28 | 37 | 25 | 31 | 82 | 7.5 | 19.2 | 13.3 | 16.9 | 46.5 |

| | | | | | | | | | | | | | | | | |
|---|---|---|---|---|---|---|---|---|---|---|---|---|---|---|---|---|
| >hg38_21_+_8433172_8446622_RNA45SN1_45S_50ntFlanks@12616.12654.39\|CTGAGGAGCCAATGGGGCGAAGCTACCATCTGTGGGATT | >hg38_21_+_8433172_8446622_RNA45SN1_45S_50ntFlanks | 12616 | 12654 | 39 | CTGAGGAGCCAATGGGGCGAAGCTACCATCTGTGGGATT | MB468 | 61 | 113 | 61 | 45 | 223 | 19.7 | 88.3 | 37.0 | 23.9 | 132.3 |
| >hg38_21_+_8433172_8446622_RNA45SN1_45S_50ntFlanks@12617.12653.37\|TGAGGAGCCAATGGGGCGAAGCTACCATCTGTGGGAT | >hg38_21_+_8433172_8446622_RNA45SN1_45S_50ntFlanks | 12617 | 12653 | 37 | TGAGGAGCCAATGGGGCGAAGCTACCATCTGTGGGAT | MB468 | 37 | 40 | 32 | 32 | 60 | 7.4 | 11.6 | 15.5 | 15.2 | 36.3 |
| >hg38_21_+_8433172_8446622_RNA45SN1_45S_50ntFlanks@12617.12654.38\|TGAGGAGCCAATGGGGCGAAGCTACCATCTGTGGGATT | >hg38_21_+_8433172_8446622_RNA45SN1_45S_50ntFlanks | 12617 | 12654 | 38 | TGAGGAGCCAATGGGGCGAAGCTACCATCTGTGGGATT | MB468 | 77 | 112 | 76 | 48 | 167 | 20.7 | 67.4 | 39.9 | 24.6 | 103.4 |
| >hg38_21_+_8433172_8446622_RNA45SN1_45S_50ntFlanks@12618.12654.37\|GAGGAGCCAATGGGGCGAAGCTACCATCTGTGGGATT | >hg38_21_+_8433172_8446622_RNA45SN1_45S_50ntFlanks | 12618 | 12654 | 37 | GAGGAGCCAATGGGGCGAAGCTACCATCTGTGGGATT | MB468 | 12 | 16 | 12 | 9 | 17 | 3.7 | 6.3 | 7.5 | 4.0 | 11.2 |
| >hg38_21_+_8433172_8446622_RNA45SN1_45S_50ntFlanks@12619.12653.35\|AGGAGCCAATGGGGCGAAGCTACCATCTGTGGGAT | >hg38_21_+_8433172_8446622_RNA45SN1_45S_50ntFlanks | 12619 | 12653 | 35 | AGGAGCCAATGGGGCGAAGCTACCATCTGTGGGAT | MB468 | 60 | 55 | 49 | 53 | 72 | 12.2 | 9.5 | 27.0 | 21.2 | 36.2 |
| >hg38_21_+_8433172_8446622_RNA45SN1_45S_50ntFlanks@12619.12654.36\|AGGAGCCAATGGGGCGAAGCTACCATCTGTGGGATT | >hg38_21_+_8433172_8446622_RNA45SN1_45S_50ntFlanks | 12619 | 12654 | 36 | AGGAGCCAATGGGGCGAAGCTACCATCTGTGGGATT | MB468 | 125 | 148 | 113 | 76 | 203 | 45.8 | 72.0 | 57.1 | 31.8 | 108.7 |
| >hg38_21_+_8433172_8446622_RNA45SN1_45S_50ntFlanks@12621.12654.34\|GAGCCAATGGGGCGAAGCTACCATCTGTGGGATT | >hg38_21_+_8433172_8446622_RNA45SN1_45S_50ntFlanks | 12621 | 12654 | 34 | GAGCCAATGGGGCGAAGCTACCATCTGTGGGATT | MB468 | 19 | 10 | 10 | 13 | 28 | 2.3 | 3.6 | 4.3 | 4.5 | 11.1 |
| >hg38_21_+_8433172_8446622_RNA45SN1_45S_50ntFlanks@12622.12654.33\|AGCCAATGGGGCGAAGCTACCATCTGTGGGATT | >hg38_21_+_8433172_8446622_RNA45SN1_45S_50ntFlanks | 12622 | 12654 | 33 | AGCCAATGGGGCGAAGCTACCATCTGTGGGATT | MB468 | 36 | 14 | 17 | 22 | 55 | 7.2 | 6.5 | 8.1 | 6.7 | 27.2 |
| >hg38_21_+_8433172_8446622_RNA45SN1_45S_50ntFlanks@12623.12654.32\|GCCAATGGGGCGAAGCTACCATCTGTGGGATT | >hg38_21_+_8433172_8446622_RNA45SN1_45S_50ntFlanks | 12623 | 12654 | 32 | GCCAATGGGGCGAAGCTACCATCTGTGGGATT | MB468 | 26 | 8 | 12 | 22 | 43 | 5.2 | 2.8 | 4.9 | 3.3 | 15.0 |
| >hg38_21_+_8433172_8446622_RNA45SN1_45S_50ntFlanks@12624.12652.29\|CCAATGGGGCGAAGCTACCATCTGTGGGA | >hg38_21_+_8433172_8446622_RNA45SN1_45S_50ntFlanks | 12624 | 12652 | 29 | CCAATGGGGCGAAGCTACCATCTGTGGGA | MB468 | 14 | 2 | 6 | 38 | 23 | 6.6 | 0.9 | 2.8 | 29.5 | 9.3 |
| >hg38_21_+_8433172_8446622_RNA45SN1_45S_50ntFlanks@12624.12653.30\|CCAATGGGGCGAAGCTACCATCTGTGGGAT | >hg38_21_+_8433172_8446622_RNA45SN1_45S_50ntFlanks | 12624 | 12653 | 30 | CCAATGGGGCGAAGCTACCATCTGTGGGAT | MB468 | 56 | 7 | 23 | 97 | 98 | 12.9 | 2.0 | 9.2 | 59.8 | 42.5 |
| >hg38_21_+_8433172_8446622_RNA45SN1_45S_50ntFlanks@12624.12654.31\|CCAATGGGGCGAAGCTACCATCTGTGGGATT | >hg38_21_+_8433172_8446622_RNA45SN1_45S_50ntFlanks | 12624 | 12654 | 31 | CCAATGGGGCGAAGCTACCATCTGTGGGATT | MB468 | 464 | 108 | 169 | 511 | 951 | 123.4 | 46.0 | 94.8 | 247.2 | 398.3 |
| >hg38_21_+_8433172_8446622_RNA45SN1_45S_50ntFlanks@12624.12657.34\|CCAATGGGGCGAAGCTACCATCTGTGGGATTATG | >hg38_21_+_8433172_8446622_RNA45SN1_45S_50ntFlanks | 12624 | 12657 | 34 | CCAATGGGGCGAAGCTACCATCTGTGGGATTATG | MB468 | 65 | 40 | 26 | 79 | 231 | 39.2 | 16.7 | 20.9 | 35.6 | 98.4 |
| >hg38_21_+_8433172_8446622_RNA45SN1_45S_50ntFlanks@12624.12659.36\|CCAATGGGGCGAAGCTACCATCTGTGGGATTATGAC | >hg38_21_+_8433172_8446622_RNA45SN1_45S_50ntFlanks | 12624 | 12659 | 36 | CCAATGGGGCGAAGCTACCATCTGTGGGATTATGAC | MB468 | 24 | 22 | 11 | 27 | 119 | 7.4 | 11.6 | 4.1 | 13.5 | 53.4 |
| >hg38_21_+_8433172_8446622_RNA45SN1_45S_50ntFlanks@12624.12661.38\|CCAATGGGGCGAAGCTACCATCTGTGGGATTATGACTG | >hg38_21_+_8433172_8446622_RNA45SN1_45S_50ntFlanks | 12624 | 12661 | 38 | CCAATGGGGCGAAGCTACCATCTGTGGGATTATGACTG | MB468 | 93 | 136 | 35 | 122 | 602 | 26.6 | 26.0 | 10.6 | 109.0 | 492.6 |
| >hg38_21_+_8433172_8446622_RNA45SN1_45S_50ntFlanks@12625.12652.28\|CAATGGGGCGAAGCTACCATCTGTGGGA | >hg38_21_+_8433172_8446622_RNA45SN1_45S_50ntFlanks | 12625 | 12652 | 28 | CAATGGGGCGAAGCTACCATCTGTGGGA | MB468 | 11 | 4 | 7 | 22 | 13 | 4.4 | 0.6 | 1.7 | 15.0 | 6.1 |
| >hg38_21_+_8433172_8446622_RNA45SN1_45S_50ntFlanks@12625.12653.29\|CAATGGGGCGAAGCTACCATCTGTGGGAT | >hg38_21_+_8433172_8446622_RNA45SN1_45S_50ntFlanks | 12625 | 12653 | 29 | CAATGGGGCGAAGCTACCATCTGTGGGAT | MB468 | 176 | 73 | 92 | 126 | 209 | 26.1 | 18.1 | 22.9 | 19.0 | 78.3 |
| >hg38_21_+_8433172_8446622_RNA45SN1_45S_50ntFlanks@12625.12654.30\|CAATGGGGCGAAGCTACCATCTGTGGGATT | >hg38_21_+_8433172_8446622_RNA45SN1_45S_50ntFlanks | 12625 | 12654 | 30 | CAATGGGGCGAAGCTACCATCTGTGGGATT | MB468 | 490 | 342 | 245 | 270 | 647 | 132.4 | 28.2 | 84.9 | 40.6 | 383.1 |
| >hg38_21_+_8433172_8446622_RNA45SN1_45S_50ntFlanks@12625.12657.33\|CAATGGGGCGAAGCTACCATCTGTGGGATTATG | >hg38_21_+_8433172_8446622_RNA45SN1_45S_50ntFlanks | 12625 | 12657 | 33 | CAATGGGGCGAAGCTACCATCTGTGGGATTATG | MB468 | 18 | 18 | 10 | 21 | 57 | 5.7 | 5.7 | 5.9 | 10.4 | 44.9 |
| >hg38_21_+_8433172_8446622_RNA45SN1_45S_50ntFlanks@12625.12659.35\|CAATGGGGCGAAGCTACCATCTGTGGGATTATGAC | >hg38_21_+_8433172_8446622_RNA45SN1_45S_50ntFlanks | 12625 | 12659 | 35 | CAATGGGGCGAAGCTACCATCTGTGGGATTATGAC | MB468 | 15 | 20 | 7 | 18 | 76 | 2.2 | 4.0 | 3.7 | 17.3 | 89.6 |
| >hg38_21_+_8433172_8446622_RNA45SN1_45S_50ntFlanks@12625.12661.37\|CAATGGGGCGAAGCTACCATCTGTGGGATTATGACTG | >hg38_21_+_8433172_8446622_RNA45SN1_45S_50ntFlanks | 12625 | 12661 | 37 | CAATGGGGCGAAGCTACCATCTGTGGGATTATGACTG | MB468 | 26 | 23 | 9 | 39 | 111 | 8.8 | 4.7 | 2.3 | 29.1 | 81.9 |
| >hg38_21_+_8433172_8446622_RNA45SN1_45S_50ntFlanks@12625.12667.43\|CAATGGGGCGAAGCTACCATCTGTGGGATTATGACTGAACGCC | >hg38_21_+_8433172_8446622_RNA45SN1_45S_50ntFlanks | 12625 | 12667 | 43 | CAATGGGGCGAAGCTACCATCTGTGGGATTATGACTGAACGCC | MB231 | 25 | 10 | 31 | 3 | 20 | 24.3 | 9.9 | 28.4 | 3.7 | 18.9 |
| >hg38_21_+_8433172_8446622_RNA45SN1_45S_50ntFlanks@12626.12652.27\|AATGGGGCGAAGCTACCATCTGTGGGA | >hg38_21_+_8433172_8446622_RNA45SN1_45S_50ntFlanks | 12626 | 12652 | 27 | AATGGGGCGAAGCTACCATCTGTGGGA | MB468 | 30 | 13 | 22 | 88 | 30 | 10.1 | 1.8 | 6.0 | 83.2 | 11.6 |
| >hg38_21_+_8433172_8446622_RNA45SN1_45S_50ntFlanks@12626.12653.28\|AATGGGGCGAAGCTACCATCTGTGGGAT | >hg38_21_+_8433172_8446622_RNA45SN1_45S_50ntFlanks | 12626 | 12653 | 28 | AATGGGGCGAAGCTACCATCTGTGGGAT | MB468 | 491 | 211 | 269 | 565 | 480 | 163.7 | 88.3 | 107.9 | 159.8 | 204.2 |
| >hg38_21_+_8433172_8446622_RNA45SN1_45S_50ntFlanks@12626.12654.29\|AATGGGGCGAAGCTACCATCTGTGGGATT | >hg38_21_+_8433172_8446622_RNA45SN1_45S_50ntFlanks | 12626 | 12654 | 29 | AATGGGGCGAAGCTACCATCTGTGGGATT | MB468 | 966 | 658 | 510 | 790 | 1105 | 126.1 | 145.0 | 161.0 | 296.8 | 690.1 |
| >hg38_21_+_8433172_8446622_RNA45SN1_45S_50ntFlanks@12626.12657.32\|AATGGGGCGAAGCTACCATCTGTGGGATTATG | >hg38_21_+_8433172_8446622_RNA45SN1_45S_50ntFlanks | 12626 | 12657 | 32 | AATGGGGCGAAGCTACCATCTGTGGGATTATG | MB468 | 42 | 39 | 23 | 55 | 119 | 5.2 | 15.1 | 14.2 | 23.1 | 99.5 |
| >hg38_21_+_8433172_8446622_RNA45SN1_45S_50ntFlanks@12626.12659.34\|AATGGGGCGAAGCTACCATCTGTGGGATTATGAC | >hg38_21_+_8433172_8446622_RNA45SN1_45S_50ntFlanks | 12626 | 12659 | 34 | AATGGGGCGAAGCTACCATCTGTGGGATTATGAC | MB468 | 57 | 74 | 29 | 90 | 277 | 12.3 | 30.7 | 15.7 | 97.6 | 365.4 |
| >hg38_21_+_8433172_8446622_RNA45SN1_45S_50ntFlanks@12626.12661.36\|AATGGGGCGAAGCTACCATCTGTGGGATTATGACTG | >hg38_21_+_8433172_8446622_RNA45SN1_45S_50ntFlanks | 12626 | 12661 | 36 | AATGGGGCGAAGCTACCATCTGTGGGATTATGACTG | MB468 | 123 | 94 | 40 | 193 | 416 | 35.9 | 23.5 | 9.5 | 139.3 | 344.0 |
| >hg38_21_+_8433172_8446622_RNA45SN1_45S_50ntFlanks@12626.12667.42\|AATGGGGCGAAGCTACCATCTGTGGGATTATGACTGAACGCC | >hg38_21_+_8433172_8446622_RNA45SN1_45S_50ntFlanks | 12626 | 12667 | 42 | AATGGGGCGAAGCTACCATCTGTGGGATTATGACTGAACGCC | MB231 | 56 | 32 | 68 | 13 | 52 | 50.0 | 26.1 | 54.2 | 12.4 | 41.0 |
| >hg38_21_+_8433172_8446622_RNA45SN1_45S_50ntFlanks@12626.12668.43\|AATGGGGCGAAGCTACCATCTGTGGGATTATGACTGAACGCCT | >hg38_21_+_8433172_8446622_RNA45SN1_45S_50ntFlanks | 12626 | 12668 | 43 | AATGGGGCGAAGCTACCATCTGTGGGATTATGACTGAACGCCT | MB231 | 13 | 4 | 6 | 2 | 9 | 14.8 | 3.0 | 7.9 | 2.8 | 9.5 |
| >hg38_21_+_8433172_8446622_RNA45SN1_45S_50ntFlanks@12626.12669.44\|AATGGGGCGAAGCTACCATCTGTGGGATTATGACTGAACGCCTC | >hg38_21_+_8433172_8446622_RNA45SN1_45S_50ntFlanks | 12626 | 12669 | 44 | AATGGGGCGAAGCTACCATCTGTGGGATTATGACTGAACGCCTC | MB231 | 11 | 7 | 3 | 13 | 16 | 8.9 | 8.8 | 3.1 | 20.1 | 18.1 |
| >hg38_21_+_8433172_8446622_RNA45SN1_45S_50ntFlanks@12627.12652.26\|ATGGGGCGAAGCTACCATCTGTGGGA | >hg38_21_+_8433172_8446622_RNA45SN1_45S_50ntFlanks | 12627 | 12652 | 26 | ATGGGGCGAAGCTACCATCTGTGGGA | MB468 | 40 | 14 | 28 | 76 | 40 | 16.9 | 4.9 | 9.0 | 69.8 | 19.3 |
| >hg38_21_+_8433172_8446622_RNA45SN1_45S_50ntFlanks@12627.12653.27\|ATGGGGCGAAGCTACCATCTGTGGGAT | >hg38_21_+_8433172_8446622_RNA45SN1_45S_50ntFlanks | 12627 | 12653 | 27 | ATGGGGCGAAGCTACCATCTGTGGGAT | MB468 | 434 | 164 | 208 | 539 | 507 | 114.7 | 54.4 | 71.7 | 136.4 | 165.1 |
| >hg38_21_+_8433172_8446622_RNA45SN1_45S_50ntFlanks@12627.12654.28\|ATGGGGCGAAGCTACCATCTGTGGGATT | >hg38_21_+_8433172_8446622_RNA45SN1_45S_50ntFlanks | 12627 | 12654 | 28 | ATGGGGCGAAGCTACCATCTGTGGGATT | MB468 | 847 | 528 | 395 | 742 | 1138 | 64.2 | 70.3 | 111.0 | 308.8 | 541.8 |
| >hg38_21_+_8433172_8446622_RNA45SN1_45S_50ntFlanks@12627.12657.31\|ATGGGGCGAAGCTACCATCTGTGGGATTATG | >hg38_21_+_8433172_8446622_RNA45SN1_45S_50ntFlanks | 12627 | 12657 | 31 | ATGGGGCGAAGCTACCATCTGTGGGATTATG | MB468 | 62 | 54 | 27 | 68 | 212 | 13.1 | 17.5 | 14.2 | 28.2 | 175.5 |
| >hg38_21_+_8433172_8446622_RNA45SN1_45S_50ntFlanks@12627.12659.33\|ATGGGGCGAAGCTACCATCTGTGGGATTATGAC | >hg38_21_+_8433172_8446622_RNA45SN1_45S_50ntFlanks | 12627 | 12659 | 33 | ATGGGGCGAAGCTACCATCTGTGGGATTATGAC | MB468 | 109 | 139 | 45 | 170 | 649 | 16.3 | 62.8 | 23.2 | 196.5 | 872.5 |
| >hg38_21_+_8433172_8446622_RNA45SN1_45S_50ntFlanks@12627.12661.35\|ATGGGGCGAAGCTACCATCTGTGGGATTATGACTG | >hg38_21_+_8433172_8446622_RNA45SN1_45S_50ntFlanks | 12627 | 12661 | 35 | ATGGGGCGAAGCTACCATCTGTGGGATTATGACTG | MB468 | 105 | 57 | 30 | 167 | 376 | 43.4 | 8.9 | 4.6 | 65.0 | 230.0 |
| >hg38_21_+_8433172_8446622_RNA45SN1_45S_50ntFlanks@12627.12667.41\|ATGGGGCGAAGCTACCATCTGTGGGATTATGACTGAACGCC | >hg38_21_+_8433172_8446622_RNA45SN1_45S_50ntFlanks | 12627 | 12667 | 41 | ATGGGGCGAAGCTACCATCTGTGGGATTATGACTGAACGCC | MB231 | 23 | 13 | 20 | 5 | 21 | 19.6 | 10.3 | 16.9 | 4.9 | 16.8 |
| >hg38_21_+_8433172_8446622_RNA45SN1_45S_50ntFlanks@12628.12652.25\|TGGGGCGAAGCTACCATCTGTGGGA | >hg38_21_+_8433172_8446622_RNA45SN1_45S_50ntFlanks | 12628 | 12652 | 25 | TGGGGCGAAGCTACCATCTGTGGGA | MB468 | 11 | 5 | 10 | 15 | 10 | 2.5 | 1.1 | 4.3 | 11.1 | 3.8 |
| >hg38_21_+_8433172_8446622_RNA45SN1_45S_50ntFlanks@12628.12653.26\|TGGGGCGAAGCTACCATCTGTGGGAT | >hg38_21_+_8433172_8446622_RNA45SN1_45S_50ntFlanks | 12628 | 12653 | 26 | TGGGGCGAAGCTACCATCTGTGGGAT | MB468 | 145 | 55 | 78 | 120 | 148 | 50.5 | 18.2 | 25.2 | 30.7 | 62.2 |
| >hg38_21_+_8433172_8446622_RNA45SN1_45S_50ntFlanks@12628.12654.27\|TGGGGCGAAGCTACCATCTGTGGGATT | >hg38_21_+_8433172_8446622_RNA45SN1_45S_50ntFlanks | 12628 | 12654 | 27 | TGGGGCGAAGCTACCATCTGTGGGATT | MB468 | 307 | 238 | 159 | 189 | 400 | 33.7 | 37.6 | 58.6 | 70.9 | 241.8 |
| >hg38_21_+_8433172_8446622_RNA45SN1_45S_50ntFlanks@12628.12657.30\|TGGGGCGAAGCTACCATCTGTGGGATTATG | >hg38_21_+_8433172_8446622_RNA45SN1_45S_50ntFlanks | 12628 | 12657 | 30 | TGGGGCGAAGCTACCATCTGTGGGATTATG | MB468 | 18 | 17 | 8 | 18 | 58 | 3.7 | 7.8 | 5.3 | 13.3 | 57.6 |
| >hg38_21_+_8433172_8446622_RNA45SN1_45S_50ntFlanks@12628.12659.32\|TGGGGCGAAGCTACCATCTGTGGGATTATGAC | >hg38_21_+_8433172_8446622_RNA45SN1_45S_50ntFlanks | 12628 | 12659 | 32 | TGGGGCGAAGCTACCATCTGTGGGATTATGAC | MB468 | 27 | 37 | 11 | 43 | 146 | 7.5 | 18.6 | 8.6 | 50.6 | 203.5 |
| >hg38_21_+_8433172_8446622_RNA45SN1_45S_50ntFlanks@12628.12661.34\|TGGGGCGAAGCTACCATCTGTGGGATTATGACTG | >hg38_21_+_8433172_8446622_RNA45SN1_45S_50ntFlanks | 12628 | 12661 | 34 | TGGGGCGAAGCTACCATCTGTGGGATTATGACTG | MB468 | 60 | 77 | 17 | 114 | 335 | 13.7 | 43.2 | 5.1 | 127.2 | 458.6 |
| >hg38_21_+_8433172_8446622_RNA45SN1_45S_50ntFlanks@12630.12654.25\|GGGCGAAGCTACCATCTGTGGGATT | >hg38_21_+_8433172_8446622_RNA45SN1_45S_50ntFlanks | 12630 | 12654 | 25 | GGGCGAAGCTACCATCTGTGGGATT | MB468 | 24 | 15 | 12 | 19 | 31 | 6.8 | 5.9 | 2.3 | 9.9 | 24.0 |
| >hg38_21_+_8433172_8446622_RNA45SN1_45S_50ntFlanks@12631.12653.23\|GGCGAAGCTACCATCTGTGGGAT | >hg38_21_+_8433172_8446622_RNA45SN1_45S_50ntFlanks | 12631 | 12653 | 23 | GGCGAAGCTACCATCTGTGGGAT | MB468 | 16 | 2 | 3 | 12 | 12 | 9.2 | 0.8 | 1.1 | 2.9 | 5.2 |
| >hg38_21_+_8433172_8446622_RNA45SN1_45S_50ntFlanks@12631.12654.24\|GGCGAAGCTACCATCTGTGGGATT | >hg38_21_+_8433172_8446622_RNA45SN1_45S_50ntFlanks | 12631 | 12654 | 24 | GGCGAAGCTACCATCTGTGGGATT | MB468 | 56 | 18 | 19 | 31 | 54 | 15.3 | 5.3 | 2.9 | 15.0 | 24.5 |
| >hg38_21_+_8433172_8446622_RNA45SN1_45S_50ntFlanks@12631.12661.31\|GGCGAAGCTACCATCTGTGGGATTATGACTG | >hg38_21_+_8433172_8446622_RNA45SN1_45S_50ntFlanks | 12631 | 12661 | 31 | GGCGAAGCTACCATCTGTGGGATTATGACTG | MB468 | 12 | 12 | 3 | 35 | 56 | 2.9 | 5.6 | 1.8 | 45.6 | 63.9 |
| >hg38_21_+_8433172_8446622_RNA45SN1_45S_50ntFlanks@12632.12653.22\|GCGAAGCTACCATCTGTGGGAT | >hg38_21_+_8433172_8446622_RNA45SN1_45S_50ntFlanks | 12632 | 12653 | 22 | GCGAAGCTACCATCTGTGGGAT | MB468 | 117 | 20 | 38 | 140 | 108 | 67.3 | 11.2 | 16.0 | 37.9 | 41.2 |
| >hg38_21_+_8433172_8446622_RNA45SN1_45S_50ntFlanks@12632.12654.23\|GCGAAGCTACCATCTGTGGGATT | >hg38_21_+_8433172_8446622_RNA45SN1_45S_50ntFlanks | 12632 | 12654 | 23 | GCGAAGCTACCATCTGTGGGATT | MB468 | 241 | 86 | 72 | 228 | 271 | 95.7 | 40.9 | 17.3 | 62.2 | 82.8 |

| | | | | | | | | | | | | | | | | |
|---|---|---|---|---|---|---|---|---|---|---|---|---|---|---|---|---|
| >hg38_21_+_8433172_8446622_RNA45SN1_45S_50ntFlanks@12632.12657.26\|GCGAAGCTACCATCTGTGGGATTATG | >hg38_21_+_8433172_8446622_RNA45SN1_45S_50ntFlanks | 12632 | 12657 | 26 | GCGAAGCTACCATCTGTGGGATTATG | MB468 | 14 | 6 | 3 | 27 | 36 | 2.1 | 3.6 | 1.7 | 15.6 | 20.7 |
| >hg38_21_+_8433172_8446622_RNA45SN1_45S_50ntFlanks@12632.12659.28\|GCGAAGCTACCATCTGTGGGATTATGAC | >hg38_21_+_8433172_8446622_RNA45SN1_45S_50ntFlanks | 12632 | 12659 | 28 | GCGAAGCTACCATCTGTGGGATTATGAC | MB468 | 29 | 29 | 9 | 89 | 133 | 10.0 | 17.8 | 7.1 | 88.8 | 112.0 |
| >hg38_21_+_8433172_8446622_RNA45SN1_45S_50ntFlanks@12632.12661.30\|GCGAAGCTACCATCTGTGGGATTATGACTG | >hg38_21_+_8433172_8446622_RNA45SN1_45S_50ntFlanks | 12632 | 12661 | 30 | GCGAAGCTACCATCTGTGGGATTATGACTG | MB468 | 34 | 30 | 9 | 126 | 143 | 5.9 | 13.8 | 3.9 | 138.6 | 121.7 |
| >hg38_21_+_8433172_8446622_RNA45SN1_45S_50ntFlanks@12632.12667.36\|GCGAAGCTACCATCTGTGGGATTATGACTGAACGCC | >hg38_21_+_8433172_8446622_RNA45SN1_45S_50ntFlanks | 12632 | 12667 | 36 | GCGAAGCTACCATCTGTGGGATTATGACTGAACGCC | MB231 | 14 | 6 | 8 | 2 | 12 | 15.6 | 4.9 | 10.5 | 2.1 | 11.3 |
| >hg38_21_+_8433172_8446622_RNA45SN1_45S_50ntFlanks@12633.12651.19\|CGAAGCTACCATCTGTGGG | >hg38_21_+_8433172_8446622_RNA45SN1_45S_50ntFlanks | 12633 | 12651 | 19 | CGAAGCTACCATCTGTGGG | MB468 | 88 | 26 | 50 | 107 | 45 | 34.3 | 10.8 | 23.5 | 13.5 | 13.5 |
| >hg38_21_+_8433172_8446622_RNA45SN1_45S_50ntFlanks@12633.12652.20\|CGAAGCTACCATCTGTGGGA | >hg38_21_+_8433172_8446622_RNA45SN1_45S_50ntFlanks | 12633 | 12652 | 20 | CGAAGCTACCATCTGTGGGA | MB468 | 99 | 14 | 35 | 155 | 56 | 33.4 | 4.9 | 11.1 | 99.5 | 8.6 |
| >hg38_21_+_8433172_8446622_RNA45SN1_45S_50ntFlanks@12633.12653.21\|CGAAGCTACCATCTGTGGGAT | >hg38_21_+_8433172_8446622_RNA45SN1_45S_50ntFlanks | 12633 | 12653 | 21 | CGAAGCTACCATCTGTGGGAT | MB468 | 959 | 110 | 196 | 1024 | 665 | 538.9 | 40.7 | 60.4 | 286.2 | 367.8 |
| >hg38_21_+_8433172_8446622_RNA45SN1_45S_50ntFlanks@12633.12654.22\|CGAAGCTACCATCTGTGGGATT | >hg38_21_+_8433172_8446622_RNA45SN1_45S_50ntFlanks | 12633 | 12654 | 22 | CGAAGCTACCATCTGTGGGATT | MB468 | 1415 | 201 | 279 | 1333 | 1230 | 592.2 | 77.9 | 129.8 | 319.8 | 523.5 |
| >hg38_21_+_8433172_8446622_RNA45SN1_45S_50ntFlanks@12633.12656.24\|CGAAGCTACCATCTGTGGGATTAT | >hg38_21_+_8433172_8446622_RNA45SN1_45S_50ntFlanks | 12633 | 12656 | 24 | CGAAGCTACCATCTGTGGGATTAT | MB468 | 16 | 3 | 5 | 17 | 28 | 5.2 | 1.3 | 4.0 | 5.5 | 19.4 |
| >hg38_21_+_8433172_8446622_RNA45SN1_45S_50ntFlanks@12633.12657.25\|CGAAGCTACCATCTGTGGGATTATG | >hg38_21_+_8433172_8446622_RNA45SN1_45S_50ntFlanks | 12633 | 12657 | 25 | CGAAGCTACCATCTGTGGGATTATG | MB468 | 71 | 17 | 14 | 126 | 121 | 16.9 | 12.4 | 11.1 | 33.2 | 54.5 |
| >hg38_21_+_8433172_8446622_RNA45SN1_45S_50ntFlanks@12633.12659.27\|CGAAGCTACCATCTGTGGGATTATGAC | >hg38_21_+_8433172_8446622_RNA45SN1_45S_50ntFlanks | 12633 | 12659 | 27 | CGAAGCTACCATCTGTGGGATTATGAC | MB468 | 118 | 70 | 31 | 174 | 392 | 30.8 | 54.1 | 23.5 | 95.4 | 283.4 |
| >hg38_21_+_8433172_8446622_RNA45SN1_45S_50ntFlanks@12633.12661.29\|CGAAGCTACCATCTGTGGGATTATGACTG | >hg38_21_+_8433172_8446622_RNA45SN1_45S_50ntFlanks | 12633 | 12661 | 29 | CGAAGCTACCATCTGTGGGATTATGACTG | MB468 | 200 | 103 | 44 | 429 | 589 | 27.6 | 45.8 | 17.7 | 226.1 | 329.6 |
| >hg38_21_+_8433172_8446622_RNA45SN1_45S_50ntFlanks@12634.12653.20\|GAAGCTACCATCTGTGGGAT | >hg38_21_+_8433172_8446622_RNA45SN1_45S_50ntFlanks | 12634 | 12653 | 20 | GAAGCTACCATCTGTGGGAT | MB468 | 25 | 1 | 4 | 35 | 13 | 16.8 | 0.5 | 1.3 | 9.6 | 10.1 |
| >hg38_21_+_8433172_8446622_RNA45SN1_45S_50ntFlanks@12634.12654.21\|GAAGCTACCATCTGTGGGATT | >hg38_21_+_8433172_8446622_RNA45SN1_45S_50ntFlanks | 12634 | 12654 | 21 | GAAGCTACCATCTGTGGGATT | MB468 | 48 | 3 | 6 | 61 | 36 | 30.8 | 1.3 | 2.6 | 13.5 | 29.2 |
| >hg38_21_+_8433172_8446622_RNA45SN1_45S_50ntFlanks@12634.12659.26\|GAAGCTACCATCTGTGGGATTATGAC | >hg38_21_+_8433172_8446622_RNA45SN1_45S_50ntFlanks | 12634 | 12659 | 26 | GAAGCTACCATCTGTGGGATTATGAC | MB468 | 11 | 5 | 2 | 25 | 35 | 3.7 | 4.0 | 1.2 | 19.5 | 27.7 |
| >hg38_21_+_8433172_8446622_RNA45SN1_45S_50ntFlanks@12634.12661.28\|GAAGCTACCATCTGTGGGATTATGACTG | >hg38_21_+_8433172_8446622_RNA45SN1_45S_50ntFlanks | 12634 | 12661 | 28 | GAAGCTACCATCTGTGGGATTATGACTG | MB468 | 19 | 9 | 4 | 63 | 57 | 3.3 | 4.1 | 2.0 | 52.9 | 35.0 |
| >hg38_21_+_8433172_8446622_RNA45SN1_45S_50ntFlanks@12635.12652.18\|AAGCTACCATCTGTGGGA | >hg38_21_+_8433172_8446622_RNA45SN1_45S_50ntFlanks | 12635 | 12652 | 18 | AAGCTACCATCTGTGGGA | MB468 | 46 | 3 | 12 | 86 | 17 | 20.0 | 1.7 | 3.5 | 57.8 | 8.3 |
| >hg38_21_+_8433172_8446622_RNA45SN1_45S_50ntFlanks@12635.12653.19\|AAGCTACCATCTGTGGGAT | >hg38_21_+_8433172_8446622_RNA45SN1_45S_50ntFlanks | 12635 | 12653 | 19 | AAGCTACCATCTGTGGGAT | MB468 | 443 | 16 | 58 | 534 | 188 | 332.4 | 6.3 | 15.4 | 198.5 | 164.1 |
| >hg38_21_+_8433172_8446622_RNA45SN1_45S_50ntFlanks@12635.12654.20\|AAGCTACCATCTGTGGGATT | >hg38_21_+_8433172_8446622_RNA45SN1_45S_50ntFlanks | 12635 | 12654 | 20 | AAGCTACCATCTGTGGGATT | MB468 | 958 | 54 | 118 | 1249 | 780 | 657.6 | 15.6 | 60.2 | 325.0 | 787.2 |
| >hg38_21_+_8433172_8446622_RNA45SN1_45S_50ntFlanks@12635.12657.23\|AAGCTACCATCTGTGGGATTATG | >hg38_21_+_8433172_8446622_RNA45SN1_45S_50ntFlanks | 12635 | 12657 | 23 | AAGCTACCATCTGTGGGATTATG | MB468 | 83 | 17 | 17 | 147 | 114 | 17.3 | 15.3 | 14.1 | 14.7 | 67.0 |
| >hg38_21_+_8433172_8446622_RNA45SN1_45S_50ntFlanks@12635.12659.25\|AAGCTACCATCTGTGGGATTATGAC | >hg38_21_+_8433172_8446622_RNA45SN1_45S_50ntFlanks | 12635 | 12659 | 25 | AAGCTACCATCTGTGGGATTATGAC | MB468 | 188 | 80 | 40 | 324 | 553 | 51.8 | 59.5 | 26.5 | 227.5 | 432.4 |
| >hg38_21_+_8433172_8446622_RNA45SN1_45S_50ntFlanks@12635.12660.26\|AAGCTACCATCTGTGGGATTATGACT | >hg38_21_+_8433172_8446622_RNA45SN1_45S_50ntFlanks | 12635 | 12660 | 26 | AAGCTACCATCTGTGGGATTATGACT | MB468 | 15 | 7 | 3 | 41 | 40 | 6.2 | 3.6 | 1.9 | 22.0 | 14.1 |
| >hg38_21_+_8433172_8446622_RNA45SN1_45S_50ntFlanks@12635.12661.27\|AAGCTACCATCTGTGGGATTATGACTG | >hg38_21_+_8433172_8446622_RNA45SN1_45S_50ntFlanks | 12635 | 12661 | 27 | AAGCTACCATCTGTGGGATTATGACTG | MB468 | 366 | 153 | 69 | 1010 | 1011 | 57.1 | 68.8 | 23.4 | 685.8 | 603.1 |
| >hg38_21_+_8433172_8446622_RNA45SN1_45S_50ntFlanks@12635.12667.33\|AAGCTACCATCTGTGGGATTATGACTGAACGCC | >hg38_21_+_8433172_8446622_RNA45SN1_45S_50ntFlanks | 12635 | 12667 | 33 | AAGCTACCATCTGTGGGATTATGACTGAACGCC | MB231 | 17 | 11 | 30 | 3 | 19 | 14.9 | 9.9 | 26.6 | 2.7 | 23.0 |
| >hg38_21_+_8433172_8446622_RNA45SN1_45S_50ntFlanks@12636.12653.18\|AGCTACCATCTGTGGGAT | >hg38_21_+_8433172_8446622_RNA45SN1_45S_50ntFlanks | 12636 | 12653 | 18 | AGCTACCATCTGTGGGAT | MB468 | 142 | 4 | 20 | 131 | 64 | 109.3 | 1.7 | 5.1 | 55.7 | 62.5 |
| >hg38_21_+_8433172_8446622_RNA45SN1_45S_50ntFlanks@12636.12654.19\|AGCTACCATCTGTGGGATT | >hg38_21_+_8433172_8446622_RNA45SN1_45S_50ntFlanks | 12636 | 12654 | 19 | AGCTACCATCTGTGGGATT | MB468 | 308 | 18 | 45 | 329 | 247 | 188.8 | 7.3 | 21.8 | 86.4 | 244.8 |
| >hg38_21_+_8433172_8446622_RNA45SN1_45S_50ntFlanks@12636.12657.22\|AGCTACCATCTGTGGGATTATG | >hg38_21_+_8433172_8446622_RNA45SN1_45S_50ntFlanks | 12636 | 12657 | 22 | AGCTACCATCTGTGGGATTATG | MB468 | 32 | 6 | 7 | 54 | 48 | 5.4 | 5.1 | 5.3 | 6.0 | 34.5 |
| >hg38_21_+_8433172_8446622_RNA45SN1_45S_50ntFlanks@12636.12659.24\|AGCTACCATCTGTGGGATTATGAC | >hg38_21_+_8433172_8446622_RNA45SN1_45S_50ntFlanks | 12636 | 12659 | 24 | AGCTACCATCTGTGGGATTATGAC | MB468 | 81 | 37 | 19 | 136 | 285 | 21.5 | 28.9 | 13.1 | 90.0 | 246.7 |
| >hg38_21_+_8433172_8446622_RNA45SN1_45S_50ntFlanks@12636.12661.26\|AGCTACCATCTGTGGGATTATGACTG | >hg38_21_+_8433172_8446622_RNA45SN1_45S_50ntFlanks | 12636 | 12661 | 26 | AGCTACCATCTGTGGGATTATGACTG | MB468 | 111 | 52 | 22 | 218 | 351 | 16.4 | 27.2 | 10.4 | 167.7 | 276.2 |
| >hg38_21_+_8433172_8446622_RNA45SN1_45S_50ntFlanks@12637.12654.18\|GCTACCATCTGTGGGATT | >hg38_21_+_8433172_8446622_RNA45SN1_45S_50ntFlanks | 12637 | 12654 | 18 | GCTACCATCTGTGGGATT | MB468 | 128 | 6 | 20 | 199 | 91 | 81.3 | 2.8 | 6.8 | 30.6 | 74.3 |
| >hg38_21_+_8433172_8446622_RNA45SN1_45S_50ntFlanks@12637.12657.21\|GCTACCATCTGTGGGATTATG | >hg38_21_+_8433172_8446622_RNA45SN1_45S_50ntFlanks | 12637 | 12657 | 21 | GCTACCATCTGTGGGATTATG | MB468 | 14 | 3 | 3 | 38 | 25 | 3.3 | 2.3 | 2.3 | 7.6 | 17.8 |
| >hg38_21_+_8433172_8446622_RNA45SN1_45S_50ntFlanks@12637.12659.23\|GCTACCATCTGTGGGATTATGAC | >hg38_21_+_8433172_8446622_RNA45SN1_45S_50ntFlanks | 12637 | 12659 | 23 | GCTACCATCTGTGGGATTATGAC | MB468 | 39 | 19 | 10 | 115 | 156 | 11.0 | 15.9 | 6.5 | 91.9 | 142.6 |
| >hg38_21_+_8433172_8446622_RNA45SN1_45S_50ntFlanks@12637.12661.25\|GCTACCATCTGTGGGATTATGACTG | >hg38_21_+_8433172_8446622_RNA45SN1_45S_50ntFlanks | 12637 | 12661 | 25 | GCTACCATCTGTGGGATTATGACTG | MB468 | 50 | 24 | 9 | 162 | 181 | 6.5 | 11.5 | 3.7 | 152.0 | 156.9 |
| >hg38_21_+_8433172_8446622_RNA45SN1_45S_50ntFlanks@12638.12657.20\|CTACCATCTGTGGGATTATG | >hg38_21_+_8433172_8446622_RNA45SN1_45S_50ntFlanks | 12638 | 12657 | 20 | CTACCATCTGTGGGATTATG | MB468 | 37 | 3 | 8 | 78 | 59 | 17.5 | 2.6 | 6.7 | 10.2 | 70.0 |
| >hg38_21_+_8433172_8446622_RNA45SN1_45S_50ntFlanks@12638.12659.22\|CTACCATCTGTGGGATTATGAC | >hg38_21_+_8433172_8446622_RNA45SN1_45S_50ntFlanks | 12638 | 12659 | 22 | CTACCATCTGTGGGATTATGAC | MB468 | 95 | 15 | 20 | 136 | 290 | 43.1 | 13.0 | 12.1 | 79.2 | 257.8 |
| >hg38_21_+_8433172_8446622_RNA45SN1_45S_50ntFlanks@12638.12661.24\|CTACCATCTGTGGGATTATGACTG | >hg38_21_+_8433172_8446622_RNA45SN1_45S_50ntFlanks | 12638 | 12661 | 24 | CTACCATCTGTGGGATTATGACTG | MB468 | 121 | 36 | 23 | 316 | 376 | 17.6 | 17.9 | 9.7 | 217.7 | 304.2 |
| >hg38_21_+_8433172_8446622_RNA45SN1_45S_50ntFlanks@12639.12659.21\|TACCATCTGTGGGATTATGAC | >hg38_21_+_8433172_8446622_RNA45SN1_45S_50ntFlanks | 12639 | 12659 | 21 | TACCATCTGTGGGATTATGAC | MB468 | 12 | 2 | 4 | 14 | 23 | 3.7 | 1.4 | 1.9 | 10.5 | 16.2 |
| >hg38_21_+_8433172_8446622_RNA45SN1_45S_50ntFlanks@12639.12661.23\|TACCATCTGTGGGATTATGACTG | >hg38_21_+_8433172_8446622_RNA45SN1_45S_50ntFlanks | 12639 | 12661 | 23 | TACCATCTGTGGGATTATGACTG | MB468 | 22 | 6 | 4 | 47 | 46 | 2.5 | 3.8 | 2.1 | 32.3 | 26.1 |
| >hg38_21_+_8433172_8446622_RNA45SN1_45S_50ntFlanks@12640.12659.20\|ACCATCTGTGGGATTATGAC | >hg38_21_+_8433172_8446622_RNA45SN1_45S_50ntFlanks | 12640 | 12659 | 20 | ACCATCTGTGGGATTATGAC | MB468 | 36 | 15 | 14 | 87 | 97 | 4.7 | 16.0 | 9.3 | 77.4 | 63.6 |
| >hg38_21_+_8433172_8446622_RNA45SN1_45S_50ntFlanks@12640.12661.22\|ACCATCTGTGGGATTATGACTG | >hg38_21_+_8433172_8446622_RNA45SN1_45S_50ntFlanks | 12640 | 12661 | 22 | ACCATCTGTGGGATTATGACTG | MB468 | 38 | 9 | 4 | 181 | 100 | 10.0 | 7.0 | 3.2 | 151.5 | 72.2 |
| >hg38_21_+_8433172_8446622_RNA45SN1_45S_50ntFlanks@12640.12667.28\|ACCATCTGTGGGATTATGACTGAACGCC | >hg38_21_+_8433172_8446622_RNA45SN1_45S_50ntFlanks | 12640 | 12667 | 28 | ACCATCTGTGGGATTATGACTGAACGCC | MB231 | 18 | 15 | 33 | 4 | 18 | 14.6 | 12.3 | 26.2 | 3.6 | 21.6 |
| >hg38_21_+_8433172_8446622_RNA45SN1_45S_50ntFlanks@12640.12668.29\|ACCATCTGTGGGATTATGACTGAACGCCT | >hg38_21_+_8433172_8446622_RNA45SN1_45S_50ntFlanks | 12640 | 12668 | 29 | ACCATCTGTGGGATTATGACTGAACGCCT | MB231 | 13 | 11 | 19 | 3 | 14 | 12.1 | 9.3 | 16.0 | 3.4 | 16.8 |
| >hg38_21_+_8433172_8446622_RNA45SN1_45S_50ntFlanks@12640.12669.30\|ACCATCTGTGGGATTATGACTGAACGCCTC | >hg38_21_+_8433172_8446622_RNA45SN1_45S_50ntFlanks | 12640 | 12669 | 30 | ACCATCTGTGGGATTATGACTGAACGCCTC | MB231 | 19 | 15 | 65 | 4 | 21 | 17.7 | 10.6 | 74.5 | 3.3 | 19.4 |
| >hg38_21_+_8433172_8446622_RNA45SN1_45S_50ntFlanks@12641.12659.19\|CCATCTGTGGGATTATGAC | >hg38_21_+_8433172_8446622_RNA45SN1_45S_50ntFlanks | 12641 | 12659 | 19 | CCATCTGTGGGATTATGAC | MB468 | 86 | 37 | 64 | 95 | 353 | 36.4 | 37.9 | 39.8 | 69.4 | 300.4 |
| >hg38_21_+_8433172_8446622_RNA45SN1_45S_50ntFlanks@12641.12661.21\|CCATCTGTGGGATTATGACTG | >hg38_21_+_8433172_8446622_RNA45SN1_45S_50ntFlanks | 12641 | 12661 | 21 | CCATCTGTGGGATTATGACTG | MB468 | 48 | 27 | 27 | 142 | 207 | 25.3 | 16.8 | 14.9 | 90.5 | 127.2 |
| >hg38_21_+_8433172_8446622_RNA45SN1_45S_50ntFlanks@12642.12659.18\|CATCTGTGGGATTATGAC | >hg38_21_+_8433172_8446622_RNA45SN1_45S_50ntFlanks | 12642 | 12659 | 18 | CATCTGTGGGATTATGAC | MB468 | 25 | 15 | 29 | 20 | 84 | 13.1 | 9.0 | 16.4 | 14.1 | 77.4 |
| >hg38_21_+_8433172_8446622_RNA45SN1_45S_50ntFlanks@12642.12661.20\|CATCTGTGGGATTATGACTG | >hg38_21_+_8433172_8446622_RNA45SN1_45S_50ntFlanks | 12642 | 12661 | 20 | CATCTGTGGGATTATGACTG | MB468 | 18 | 11 | 12 | 46 | 71 | 9.8 | 5.6 | 6.1 | 15.3 | 53.1 |
| >hg38_21_+_8433172_8446622_RNA45SN1_45S_50ntFlanks@12643.12661.19\|ATCTGTGGGATTATGACTG | >hg38_21_+_8433172_8446622_RNA45SN1_45S_50ntFlanks | 12643 | 12661 | 19 | ATCTGTGGGATTATGACTG | MB468 | 19 | 11 | 13 | 49 | 68 | 10.1 | 5.7 | 6.3 | 26.9 | 50.9 |
| >hg38_21_+_8433172_8446622_RNA45SN1_45S_50ntFlanks@12644.12661.18\|TCTGTGGGATTATGACTG | >hg38_21_+_8433172_8446622_RNA45SN1_45S_50ntFlanks | 12644 | 12661 | 18 | TCTGTGGGATTATGACTG | MB468 | 40 | 19 | 29 | 121 | 121 | 25.1 | 12.7 | 15.6 | 68.7 | 80.0 |
| >hg38_21_+_8433172_8446622_RNA45SN1_45S_50ntFlanks@12652.12675.24\|ATTATGACTGAACGCCTCTAAGTC | >hg38_21_+_8433172_8446622_RNA45SN1_45S_50ntFlanks | 12652 | 12675 | 24 | ATTATGACTGAACGCCTCTAAGTC | MB468 | 14 | 10 | 18 | 2 | 5 | 6.9 | 4.1 | 12.2 | 1.0 | 4.7 |

| | | | | | | | | | | | | | | |
|---|---|---|---|---|---|---|---|---|---|---|---|---|---|---|
| >hg38_21_+_8433172_8446622_RNA45SN1_45S_50ntFlanks@12654.12671.18\|TATGACTGAACGCCTCTA | >hg38_21_+_8433172_8446622_RNA45SN1_45S_50ntFlanks | 12654 | 12671 | 18 | TATGACTGAACGCCTCTA | MB468 | 19 | 20 | 12 | 6 | 6 | 8.1 | 6.6 | 4.7 | 1.7 | 1.1 |
| >hg38_21_+_8433172_8446622_RNA45SN1_45S_50ntFlanks@12654.12673.20\|TATGACTGAACGCCTCTAAG | >hg38_21_+_8433172_8446622_RNA45SN1_45S_50ntFlanks | 12654 | 12673 | 20 | TATGACTGAACGCCTCTAAG | MB468 | 14 | 22 | 7 | 3 | 4 | 5.8 | 10.8 | 2.1 | 0.5 | 0.8 |
| >hg38_21_+_8433172_8446622_RNA45SN1_45S_50ntFlanks@12654.12674.21\|TATGACTGAACGCCTCTAAGT | >hg38_21_+_8433172_8446622_RNA45SN1_45S_50ntFlanks | 12654 | 12674 | 21 | TATGACTGAACGCCTCTAAGT | MB468 | 133 | 172 | 71 | 8 | 20 | 56.2 | 80.9 | 23.7 | 2.1 | 11.1 |
| >hg38_21_+_8433172_8446622_RNA45SN1_45S_50ntFlanks@12654.12675.22\|TATGACTGAACGCCTCTAAGTC | >hg38_21_+_8433172_8446622_RNA45SN1_45S_50ntFlanks | 12654 | 12675 | 22 | TATGACTGAACGCCTCTAAGTC | MB468 | 410 | 372 | 535 | 42 | 148 | 145.7 | 82.5 | 293.4 | 15.0 | 66.8 |
| >hg38_21_+_8433172_8446622_RNA45SN1_45S_50ntFlanks@12655.12673.19\|ATGACTGAACGCCTCTAAG | >hg38_21_+_8433172_8446622_RNA45SN1_45S_50ntFlanks | 12655 | 12673 | 19 | ATGACTGAACGCCTCTAAG | MB468 | 52 | 90 | 23 | 25 | 19 | 23.7 | 54.6 | 10.5 | 11.7 | 3.2 |
| >hg38_21_+_8433172_8446622_RNA45SN1_45S_50ntFlanks@12655.12674.20\|ATGACTGAACGCCTCTAAGT | >hg38_21_+_8433172_8446622_RNA45SN1_45S_50ntFlanks | 12655 | 12674 | 20 | ATGACTGAACGCCTCTAAGT | MB468 | 530 | 707 | 255 | 58 | 98 | 245.9 | 335.5 | 80.5 | 21.0 | 51.0 |
| >hg38_21_+_8433172_8446622_RNA45SN1_45S_50ntFlanks@12655.12675.21\|ATGACTGAACGCCTCTAAGTC | >hg38_21_+_8433172_8446622_RNA45SN1_45S_50ntFlanks | 12655 | 12675 | 21 | ATGACTGAACGCCTCTAAGTC | MB468 | 1032 | 673 | 1328 | 78 | 371 | 242.9 | 128.6 | 389.0 | 6.3 | 210.4 |
| >hg38_21_+_8433172_8446622_RNA45SN1_45S_50ntFlanks@12656.12674.19\|TGACTGAACGCCTCTAAGT | >hg38_21_+_8433172_8446622_RNA45SN1_45S_50ntFlanks | 12656 | 12674 | 19 | TGACTGAACGCCTCTAAGT | MB468 | 26 | 35 | 16 | 3 | 5 | 6.6 | 16.6 | 4.5 | 0.7 | 2.4 |
| >hg38_21_+_8433172_8446622_RNA45SN1_45S_50ntFlanks@12656.12675.20\|TGACTGAACGCCTCTAAGTC | >hg38_21_+_8433172_8446622_RNA45SN1_45S_50ntFlanks | 12656 | 12675 | 20 | TGACTGAACGCCTCTAAGTC | MB468 | 32 | 19 | 37 | 2 | 13 | 17.5 | 9.9 | 23.7 | 0.4 | 8.0 |
| >hg38_21_+_8433172_8446622_RNA45SN1_45S_50ntFlanks@12657.12674.18\|GACTGAACGCCTCTAAGT | >hg38_21_+_8433172_8446622_RNA45SN1_45S_50ntFlanks | 12657 | 12674 | 18 | GACTGAACGCCTCTAAGT | MB468 | 61 | 72 | 31 | 12 | 14 | 10.2 | 25.9 | 4.5 | 7.1 | 5.2 |
| >hg38_21_+_8433172_8446622_RNA45SN1_45S_50ntFlanks@12658.12675.18\|ACTGAACGCCTCTAAGTC | >hg38_21_+_8433172_8446622_RNA45SN1_45S_50ntFlanks | 12658 | 12675 | 18 | ACTGAACGCCTCTAAGTC | MB468 | 86 | 104 | 59 | 31 | 55 | 9.0 | 40.3 | 16.6 | 13.4 | 20.4 |
| >hg38_21_+_8433172_8446622_RNA45SN1_45S_50ntFlanks@12676.12703.28\|AGAATCCCGCCCAGGCGGAACGATACGG | >hg38_21_+_8433172_8446622_RNA45SN1_45S_50ntFlanks | 12676 | 12703 | 28 | AGAATCCCGCCCAGGCGGAACGATACGG | MB468 | 52 | 6 | 34 | 181 | 37 | 19.1 | 2.9 | 15.8 | 145.7 | 14.8 |
| >hg38_21_+_8433172_8446622_RNA45SN1_45S_50ntFlanks@12676.12708.33\|AGAATCCCGCCCAGGCGGAACGATACGGCAGCG | >hg38_21_+_8433172_8446622_RNA45SN1_45S_50ntFlanks | 12676 | 12708 | 33 | AGAATCCCGCCCAGGCGGAACGATACGGCAGCG | MB468 | 21 | 18 | 22 | 58 | 27 | 5.2 | 3.5 | 9.6 | 6.4 | 12.4 |
| >hg38_21_+_8433172_8446622_RNA45SN1_45S_50ntFlanks@12678.12703.26\|AATCCCGCCCAGGCGGAACGATACGG | >hg38_21_+_8433172_8446622_RNA45SN1_45S_50ntFlanks | 12678 | 12703 | 26 | AATCCCGCCCAGGCGGAACGATACGG | MB468 | 37 | 5 | 27 | 132 | 31 | 12.0 | 1.5 | 16.7 | 96.4 | 12.7 |
| >hg38_21_+_8433172_8446622_RNA45SN1_45S_50ntFlanks@12685.12703.19\|CCCAGGCGGAACGATACGG | >hg38_21_+_8433172_8446622_RNA45SN1_45S_50ntFlanks | 12685 | 12703 | 19 | CCCAGGCGGAACGATACGG | MB468 | 14 | 11 | 9 | 12 | 15 | 6.5 | 3.7 | 1.6 | 8.2 | 8.2 |
| >hg38_21_+_8433172_8446622_RNA45SN1_45S_50ntFlanks@12685.12711.27\|CCCAGGCGGAACGATACGGCAGCGCCG | >hg38_21_+_8433172_8446622_RNA45SN1_45S_50ntFlanks | 12685 | 12711 | 27 | CCCAGGCGGAACGATACGGCAGCGCCG | MB468 | 14 | 5 | 6 | 12 | 43 | 10.9 | 2.1 | 1.3 | 10.0 | 28.0 |
| >hg38_21_+_8433172_8446622_RNA45SN1_45S_50ntFlanks@12686.12703.18\|CCAGGCGGAACGATACGG | >hg38_21_+_8433172_8446622_RNA45SN1_45S_50ntFlanks | 12686 | 12703 | 18 | CCAGGCGGAACGATACGG | MB468 | 16 | 10 | 10 | 14 | 12 | 8.0 | 2.5 | 6.1 | 7.3 | 7.9 |
| >hg38_21_+_8433172_8446622_RNA45SN1_45S_50ntFlanks@12709.12742.34\|CCGCGGAGCCTCGGTTGGCCTCGGATAGCCGGTC | >hg38_21_+_8433172_8446622_RNA45SN1_45S_50ntFlanks | 12709 | 12742 | 34 | CCGCGGAGCCTCGGTTGGCCTCGGATAGCCGGTC | MB468 | 12 | 7 | 3 | 43 | 68 | 6.9 | 2.5 | 1.5 | 15.0 | 42.2 |
| >hg38_21_+_8433172_8446622_RNA45SN1_45S_50ntFlanks@12712.12742.31\|CGGAGCCTCGGTTGGCCTCGGATAGCCGGTC | >hg38_21_+_8433172_8446622_RNA45SN1_45S_50ntFlanks | 12712 | 12742 | 31 | CGGAGCCTCGGTTGGCCTCGGATAGCCGGTC | MB468 | 12 | 4 | 5 | 266 | 39 | 6.7 | 2.6 | 4.9 | 134.3 | 30.6 |
| >hg38_21_+_8433172_8446622_RNA45SN1_45S_50ntFlanks@12712.12751.40\|CGGAGCCTCGGTTGGCCTCGGATAGCCGGTCCCCCGCCTG | >hg38_21_+_8433172_8446622_RNA45SN1_45S_50ntFlanks | 12712 | 12751 | 40 | CGGAGCCTCGGTTGGCCTCGGATAGCCGGTCCCCCGCCTG | MB468 | 13 | 8 | 4 | 62 | 117 | 6.1 | 2.2 | 1.4 | 70.9 | 98.2 |
| >hg38_21_+_8433172_8446622_RNA45SN1_45S_50ntFlanks@12714.12733.20\|GAGCCTCGGTTGGCCTCGGA | >hg38_21_+_8433172_8446622_RNA45SN1_45S_50ntFlanks | 12714 | 12733 | 20 | GAGCCTCGGTTGGCCTCGGA | BT20 | 12 | 1 | 1 | 2 | 0 | 20.0 | 1.0 | 0.8 | 1.6 | 0.8 |
| >hg38_21_+_8433172_8446622_RNA45SN1_45S_50ntFlanks@12715.12733.19\|AGCCTCGGTTGGCCTCGGA | >hg38_21_+_8433172_8446622_RNA45SN1_45S_50ntFlanks | 12715 | 12733 | 19 | AGCCTCGGTTGGCCTCGGA | BT20 | 16 | 3 | 4 | 7 | 3 | 26.1 | 4.0 | 3.3 | 8.1 | 0.9 |
| >hg38_21_+_8433172_8446622_RNA45SN1_45S_50ntFlanks@12804.12821.18\|CGCGTGCCCCGCCGCGCG | >hg38_21_+_8433172_8446622_RNA45SN1_45S_50ntFlanks | 12804 | 12821 | 18 | CGCGTGCCCCGCCGCGCG | MB468 | 12 | 1 | 1 | 16 | 20 | 2.8 | 0.4 | 0.2 | 17.5 | 7.7 |
| >hg38_21_+_8433172_8446622_RNA45SN1_45S_50ntFlanks@12874.12891.18\|GCTGGAAAGGCGGCCGCC | >hg38_21_+_8433172_8446622_RNA45SN1_45S_50ntFlanks | 12874 | 12891 | 18 | GCTGGAAAGGCGGCCGCC | MB468 | 22 | 13 | 31 | 13 | 4 | 10.4 | 8.6 | 13.4 | 5.9 | 1.1 |
| >hg38_21_+_8433172_8446622_RNA45SN1_45S_50ntFlanks@12874.12892.19\|GCTGGAAAGGCGGCCGCCC | >hg38_21_+_8433172_8446622_RNA45SN1_45S_50ntFlanks | 12874 | 12892 | 19 | GCTGGAAAGGCGGCCGCCC | MB468 | 11 | 8 | 14 | 5 | 4 | 3.8 | 5.4 | 6.2 | 3.8 | 0.6 |
| >hg38_21_+_8433172_8446622_RNA45SN1_45S_50ntFlanks@12875.12892.18\|CTGGAAAGGCGGCCGCCC | >hg38_21_+_8433172_8446622_RNA45SN1_45S_50ntFlanks | 12875 | 12892 | 18 | CTGGAAAGGCGGCCGCCC | MB468 | 12 | 5 | 11 | 24 | 6 | 4.9 | 3.2 | 3.8 | 17.4 | 2.2 |
| >hg38_21_+_8433172_8446622_RNA45SN1_45S_50ntFlanks@12892.12930.39\|CCCTCGCCCGTCACGCACCGCACGTTCGTGGGGAACCTG | >hg38_21_+_8433172_8446622_RNA45SN1_45S_50ntFlanks | 12892 | 12930 | 39 | CCCTCGCCCGTCACGCACCGCACGTTCGTGGGGAACCTG | MB468 | 11 | 5 | 3 | 21 | 62 | 7.0 | 3.7 | 2.3 | 25.1 | 28.7 |
| >hg38_21_+_8433172_8446622_RNA45SN1_45S_50ntFlanks@12892.12932.41\|CCCTCGCCCGTCACGCACCGCACGTTCGTGGGGAACCTGGC | >hg38_21_+_8433172_8446622_RNA45SN1_45S_50ntFlanks | 12892 | 12932 | 41 | CCCTCGCCCGTCACGCACCGCACGTTCGTGGGGAACCTGGC | MB468 | 48 | 46 | 14 | 27 | 391 | 27.4 | 21.4 | 8.4 | 35.5 | 180.8 |
| >hg38_21_+_8433172_8446622_RNA45SN1_45S_50ntFlanks@12892.12933.42\|CCCTCGCCCGTCACGCACCGCACGTTCGTGGGGAACCTGGCG | >hg38_21_+_8433172_8446622_RNA45SN1_45S_50ntFlanks | 12892 | 12933 | 42 | CCCTCGCCCGTCACGCACCGCACGTTCGTGGGGAACCTGGCG | MB468 | 18 | 16 | 6 | 29 | 158 | 8.0 | 7.7 | 3.3 | 38.4 | 86.4 |
| >hg38_21_+_8433172_8446622_RNA45SN1_45S_50ntFlanks@12893.12930.38\|CCTCGCCCGTCACGCACCGCACGTTCGTGGGGAACCTG | >hg38_21_+_8433172_8446622_RNA45SN1_45S_50ntFlanks | 12893 | 12930 | 38 | CCTCGCCCGTCACGCACCGCACGTTCGTGGGGAACCTG | MB468 | 34 | 13 | 7 | 88 | 183 | 20.4 | 10.0 | 5.4 | 101.2 | 62.9 |
| >hg38_21_+_8433172_8446622_RNA45SN1_45S_50ntFlanks@12893.12932.40\|CCTCGCCCGTCACGCACCGCACGTTCGTGGGGAACCTGGC | >hg38_21_+_8433172_8446622_RNA45SN1_45S_50ntFlanks | 12893 | 12932 | 40 | CCTCGCCCGTCACGCACCGCACGTTCGTGGGGAACCTGGC | MB468 | 46 | 33 | 14 | 81 | 307 | 21.0 | 8.7 | 7.1 | 101.8 | 167.4 |
| >hg38_21_+_8433172_8446622_RNA45SN1_45S_50ntFlanks@12893.12933.41\|CCTCGCCCGTCACGCACCGCACGTTCGTGGGGAACCTGGCG | >hg38_21_+_8433172_8446622_RNA45SN1_45S_50ntFlanks | 12893 | 12933 | 41 | CCTCGCCCGTCACGCACCGCACGTTCGTGGGGAACCTGGCG | MB468 | 40 | 37 | 15 | 115 | 328 | 15.6 | 16.6 | 6.2 | 127.8 | 192.3 |
| >hg38_21_+_8433172_8446622_RNA45SN1_45S_50ntFlanks@12907.12931.25\|CACCGCACGTTCGTGGGGAACCTGG | >hg38_21_+_8433172_8446622_RNA45SN1_45S_50ntFlanks | 12907 | 12931 | 25 | CACCGCACGTTCGTGGGGAACCTGG | MB468 | 10 | 8 | 15 | 13 | 15 | 8.5 | 9.9 | 18.6 | 9.7 | 10.6 |
| >hg38_21_+_8433172_8446622_RNA45SN1_45S_50ntFlanks@12907.12932.26\|CACCGCACGTTCGTGGGGAACCTGGC | >hg38_21_+_8433172_8446622_RNA45SN1_45S_50ntFlanks | 12907 | 12932 | 26 | CACCGCACGTTCGTGGGGAACCTGGC | MB468 | 15 | 17 | 23 | 15 | 25 | 14.8 | 18.9 | 33.0 | 13.5 | 17.6 |
| >hg38_21_+_8433172_8446622_RNA45SN1_45S_50ntFlanks@12907.12933.27\|CACCGCACGTTCGTGGGGAACCTGGCG | >hg38_21_+_8433172_8446622_RNA45SN1_45S_50ntFlanks | 12907 | 12933 | 27 | CACCGCACGTTCGTGGGGAACCTGGCG | MB468 | 74 | 81 | 87 | 60 | 128 | 75.2 | 86.3 | 113.4 | 33.8 | 51.7 |
| >hg38_21_+_8433172_8446622_RNA45SN1_45S_50ntFlanks@12907.12935.29\|CACCGCACGTTCGTGGGGAACCTGGCGCT | >hg38_21_+_8433172_8446622_RNA45SN1_45S_50ntFlanks | 12907 | 12935 | 29 | CACCGCACGTTCGTGGGGAACCTGGCGCT | MB468 | 19 | 20 | 19 | 12 | 47 | 18.7 | 26.1 | 23.5 | 6.1 | 23.2 |
| >hg38_21_+_8433172_8446622_RNA45SN1_45S_50ntFlanks@12909.12933.25\|CCGCACGTTCGTGGGGAACCTGGCG | >hg38_21_+_8433172_8446622_RNA45SN1_45S_50ntFlanks | 12909 | 12933 | 25 | CCGCACGTTCGTGGGGAACCTGGCG | MB468 | 16 | 11 | 19 | 23 | 25 | 14.4 | 8.0 | 21.1 | 9.8 | 5.0 |
| >hg38_21_+_8433172_8446622_RNA45SN1_45S_50ntFlanks@12912.12932.21\|CACGTTCGTGGGGAACCTGGC | >hg38_21_+_8433172_8446622_RNA45SN1_45S_50ntFlanks | 12912 | 12932 | 21 | CACGTTCGTGGGGAACCTGGC | MB468 | 14 | 14 | 33 | 15 | 12 | 14.8 | 13.5 | 42.6 | 9.7 | 5.7 |
| >hg38_21_+_8433172_8446622_RNA45SN1_45S_50ntFlanks@12912.12933.22\|CACGTTCGTGGGGAACCTGGCG | >hg38_21_+_8433172_8446622_RNA45SN1_45S_50ntFlanks | 12912 | 12933 | 22 | CACGTTCGTGGGGAACCTGGCG | MB468 | 61 | 41 | 103 | 76 | 63 | 60.6 | 48.0 | 121.2 | 55.3 | 40.2 |
| >hg38_21_+_8433172_8446622_RNA45SN1_45S_50ntFlanks@12912.12935.24\|CACGTTCGTGGGGAACCTGGCGCT | >hg38_21_+_8433172_8446622_RNA45SN1_45S_50ntFlanks | 12912 | 12935 | 24 | CACGTTCGTGGGGAACCTGGCGCT | MB468 | 17 | 8 | 19 | 27 | 29 | 17.2 | 11.3 | 20.6 | 20.7 | 19.3 |
| >hg38_21_+_8433172_8446622_RNA45SN1_45S_50ntFlanks@12913.12933.21\|ACGTTCGTGGGGAACCTGGCG | >hg38_21_+_8433172_8446622_RNA45SN1_45S_50ntFlanks | 12913 | 12933 | 21 | ACGTTCGTGGGGAACCTGGCG | MB468 | 20 | 7 | 29 | 34 | 16 | 19.5 | 6.1 | 30.9 | 26.8 | 9.6 |
| >hg38_21_+_8433172_8446622_RNA45SN1_45S_50ntFlanks@12914.12933.20\|CGTTCGTGGGGAACCTGGCG | >hg38_21_+_8433172_8446622_RNA45SN1_45S_50ntFlanks | 12914 | 12933 | 20 | CGTTCGTGGGGAACCTGGCG | MB468 | 27 | 11 | 33 | 77 | 31 | 24.2 | 5.8 | 34.6 | 50.8 | 11.0 |
| >hg38_21_+_8433172_8446622_RNA45SN1_45S_50ntFlanks@12916.12933.18\|TTCGTGGGGAACCTGGCG | >hg38_21_+_8433172_8446622_RNA45SN1_45S_50ntFlanks | 12916 | 12933 | 18 | TTCGTGGGGAACCTGGCG | MB468 | 13 | 13 | 26 | 10 | 8 | 13.0 | 7.8 | 32.2 | 8.6 | 1.0 |
| >hg38_21_+_8433172_8446622_RNA45SN1_45S_50ntFlanks@12926.12956.31\|ACCTGGCGCTAAACCATTCGTAGACGACCTG | >hg38_21_+_8433172_8446622_RNA45SN1_45S_50ntFlanks | 12926 | 12956 | 31 | ACCTGGCGCTAAACCATTCGTAGACGACCTG | MB468 | 14 | 7 | 10 | 12 | 47 | 10.2 | 5.2 | 6.3 | 2.9 | 27.0 |
| >hg38_21_+_8433172_8446622_RNA45SN1_45S_50ntFlanks@12927.12953.27\|CCTGGCGCTAAACCATTCGTAGACGAC | >hg38_21_+_8433172_8446622_RNA45SN1_45S_50ntFlanks | 12927 | 12953 | 27 | CCTGGCGCTAAACCATTCGTAGACGAC | MB468 | 14 | 8 | 9 | 14 | 35 | 6.4 | 3.1 | 5.1 | 7.3 | 18.1 |
| >hg38_21_+_8433172_8446622_RNA45SN1_45S_50ntFlanks@12927.12956.30\|CCTGGCGCTAAACCATTCGTAGACGACCTG | >hg38_21_+_8433172_8446622_RNA45SN1_45S_50ntFlanks | 12927 | 12956 | 30 | CCTGGCGCTAAACCATTCGTAGACGACCTG | MB468 | 24 | 4 | 10 | 26 | 58 | 5.1 | 0.7 | 3.5 | 6.3 | 10.2 |
| >hg38_21_+_8433172_8446622_RNA45SN1_45S_50ntFlanks@12928.12956.29\|CTGGCGCTAAACCATTCGTAGACGACCTG | >hg38_21_+_8433172_8446622_RNA45SN1_45S_50ntFlanks | 12928 | 12956 | 29 | CTGGCGCTAAACCATTCGTAGACGACCTG | MB468 | 13 | 2 | 5 | 14 | 30 | 3.2 | 0.6 | 2.1 | 4.2 | 6.4 |
| >hg38_21_+_8433172_8446622_RNA45SN1_45S_50ntFlanks@12931.12956.26\|GCGCTAAACCATTCGTAGACGACCTG | >hg38_21_+_8433172_8446622_RNA45SN1_45S_50ntFlanks | 12931 | 12956 | 26 | GCGCTAAACCATTCGTAGACGACCTG | MB468 | 51 | 80 | 45 | 48 | 77 | 11.0 | 13.1 | 22.4 | 21.1 | 13.0 |
| >hg38_21_+_8433172_8446622_RNA45SN1_45S_50ntFlanks@12932.12953.22\|CGCTAAACCATTCGTAGACGAC | >hg38_21_+_8433172_8446622_RNA45SN1_45S_50ntFlanks | 12932 | 12953 | 22 | CGCTAAACCATTCGTAGACGAC | MB468 | 12 | 4 | 10 | 11 | 6 | 3.9 | 1.1 | 2.8 | 3.8 | 0.6 |
| >hg38_21_+_8433172_8446622_RNA45SN1_45S_50ntFlanks@12932.12956.25\|CGCTAAACCATTCGTAGACGACCTG | >hg38_21_+_8433172_8446622_RNA45SN1_45S_50ntFlanks | 12932 | 12956 | 25 | CGCTAAACCATTCGTAGACGACCTG | MB468 | 59 | 75 | 53 | 49 | 61 | 11.8 | 6.6 | 22.5 | 5.5 | 14.3 |

| | | | | | | | | | | | | | | | | |
|---|---|---|---|---|---|---|---|---|---|---|---|---|---|---|---|---|
| >hg38_21_+_8433172_8446622_RNA45SN1_45S_50ntFlanks@12932.12972.41\|CGCTAAACCATTCGTAGACGACCTGCTTCTGGGTCGGGGTT | >hg38_21_+_8433172_8446622_RNA45SN1_45S_50ntFlanks | 12932 | 12972 | 41 | CGCTAAACCATTCGTAGACGACCTGCTTCTGGGTCGGGGTT | MB468 | 26 | 35 | 89 | 21 | 48 | 16.2 | 21.7 | 55.4 | 21.0 | 29.5 |
| >hg38_21_+_8433172_8446622_RNA45SN1_45S_50ntFlanks@12933.12951.19\|GCTAAACCATTCGTAGACG | >hg38_21_+_8433172_8446622_RNA45SN1_45S_50ntFlanks | 12933 | 12951 | 19 | GCTAAACCATTCGTAGACG | MB468 | 15 | 11 | 14 | 4 | 5 | 5.7 | 6.5 | 9.0 | 1.4 | 0.9 |
| >hg38_21_+_8433172_8446622_RNA45SN1_45S_50ntFlanks@12933.12952.20\|GCTAAACCATTCGTAGACGA | >hg38_21_+_8433172_8446622_RNA45SN1_45S_50ntFlanks | 12933 | 12952 | 20 | GCTAAACCATTCGTAGACGA | MB468 | 13 | 10 | 10 | 3 | 3 | 5.6 | 6.1 | 5.7 | 0.6 | 0.8 |
| >hg38_21_+_8433172_8446622_RNA45SN1_45S_50ntFlanks@12933.12953.21\|GCTAAACCATTCGTAGACGAC | >hg38_21_+_8433172_8446622_RNA45SN1_45S_50ntFlanks | 12933 | 12953 | 21 | GCTAAACCATTCGTAGACGAC | MB468 | 21 | 24 | 23 | 7 | 6 | 8.6 | 14.5 | 14.0 | 4.4 | 1.1 |
| >hg38_21_+_8433172_8446622_RNA45SN1_45S_50ntFlanks@12933.12956.24\|GCTAAACCATTCGTAGACGACCTG | >hg38_21_+_8433172_8446622_RNA45SN1_45S_50ntFlanks | 12933 | 12956 | 24 | GCTAAACCATTCGTAGACGACCTG | MB468 | 92 | 222 | 94 | 25 | 37 | 9.4 | 102.5 | 33.6 | 11.0 | 6.7 |
| >hg38_21_+_8433172_8446622_RNA45SN1_45S_50ntFlanks@12933.12965.33\|GCTAAACCATTCGTAGACGACCTGCTTCTGGGT | >hg38_21_+_8433172_8446622_RNA45SN1_45S_50ntFlanks | 12933 | 12965 | 33 | GCTAAACCATTCGTAGACGACCTGCTTCTGGGT | MB468 | 12 | 23 | 21 | 7 | 9 | 5.5 | 9.5 | 18.1 | 6.8 | 3.0 |
| >hg38_21_+_8433172_8446622_RNA45SN1_45S_50ntFlanks@12934.12951.18\|CTAAACCATTCGTAGACG | >hg38_21_+_8433172_8446622_RNA45SN1_45S_50ntFlanks | 12934 | 12951 | 18 | CTAAACCATTCGTAGACG | MB468 | 96 | 37 | 124 | 58 | 40 | 24.1 | 22.4 | 48.5 | 9.3 | 6.9 |
| >hg38_21_+_8433172_8446622_RNA45SN1_45S_50ntFlanks@12934.12952.19\|CTAAACCATTCGTAGACGA | >hg38_21_+_8433172_8446622_RNA45SN1_45S_50ntFlanks | 12934 | 12952 | 19 | CTAAACCATTCGTAGACGA | MB468 | 62 | 22 | 72 | 34 | 19 | 18.3 | 9.8 | 24.4 | 9.2 | 7.4 |
| >hg38_21_+_8433172_8446622_RNA45SN1_45S_50ntFlanks@12934.12953.20\|CTAAACCATTCGTAGACGAC | >hg38_21_+_8433172_8446622_RNA45SN1_45S_50ntFlanks | 12934 | 12953 | 20 | CTAAACCATTCGTAGACGAC | MB468 | 91 | 60 | 130 | 48 | 30 | 30.5 | 27.3 | 59.0 | 18.3 | 4.1 |
| >hg38_21_+_8433172_8446622_RNA45SN1_45S_50ntFlanks@12934.12954.21\|CTAAACCATTCGTAGACGACC | >hg38_21_+_8433172_8446622_RNA45SN1_45S_50ntFlanks | 12934 | 12954 | 21 | CTAAACCATTCGTAGACGACC | MB468 | 24 | 24 | 40 | 6 | 10 | 5.7 | 6.4 | 15.0 | 2.7 | 0.7 |
| >hg38_21_+_8433172_8446622_RNA45SN1_45S_50ntFlanks@12934.12955.22\|CTAAACCATTCGTAGACGACCT | >hg38_21_+_8433172_8446622_RNA45SN1_45S_50ntFlanks | 12934 | 12955 | 22 | CTAAACCATTCGTAGACGACCT | MB468 | 42 | 64 | 53 | 11 | 19 | 7.1 | 5.7 | 14.6 | 1.8 | 4.2 |
| >hg38_21_+_8433172_8446622_RNA45SN1_45S_50ntFlanks@12934.12956.23\|CTAAACCATTCGTAGACGACCTG | >hg38_21_+_8433172_8446622_RNA45SN1_45S_50ntFlanks | 12934 | 12956 | 23 | CTAAACCATTCGTAGACGACCTG | MB468 | 701 | 1169 | 842 | 226 | 280 | 82.2 | 310.3 | 243.1 | 44.4 | 40.8 |
| >hg38_21_+_8433172_8446622_RNA45SN1_45S_50ntFlanks@12934.12957.24\|CTAAACCATTCGTAGACGACCTGC | >hg38_21_+_8433172_8446622_RNA45SN1_45S_50ntFlanks | 12934 | 12957 | 24 | CTAAACCATTCGTAGACGACCTGC | MB468 | 27 | 43 | 31 | 5 | 10 | 5.8 | 11.0 | 15.7 | 0.8 | 2.6 |
| >hg38_21_+_8433172_8446622_RNA45SN1_45S_50ntFlanks@12934.12958.25\|CTAAACCATTCGTAGACGACCTGCT | >hg38_21_+_8433172_8446622_RNA45SN1_45S_50ntFlanks | 12934 | 12958 | 25 | CTAAACCATTCGTAGACGACCTGCT | MB468 | 34 | 49 | 22 | 9 | 23 | 9.3 | 2.9 | 6.5 | 2.3 | 3.6 |
| >hg38_21_+_8433172_8446622_RNA45SN1_45S_50ntFlanks@12934.12959.26\|CTAAACCATTCGTAGACGACCTGCTT | >hg38_21_+_8433172_8446622_RNA45SN1_45S_50ntFlanks | 12934 | 12959 | 26 | CTAAACCATTCGTAGACGACCTGCTT | MB468 | 67 | 134 | 42 | 13 | 36 | 14.8 | 19.7 | 13.5 | 1.4 | 5.1 |
| >hg38_21_+_8433172_8446622_RNA45SN1_45S_50ntFlanks@12934.12960.27\|CTAAACCATTCGTAGACGACCTGCTTC | >hg38_21_+_8433172_8446622_RNA45SN1_45S_50ntFlanks | 12934 | 12960 | 27 | CTAAACCATTCGTAGACGACCTGCTTC | MB468 | 42 | 73 | 25 | 17 | 24 | 17.5 | 18.2 | 11.6 | 1.9 | 2.7 |
| >hg38_21_+_8433172_8446622_RNA45SN1_45S_50ntFlanks@12934.12961.28\|CTAAACCATTCGTAGACGACCTGCTTCT | >hg38_21_+_8433172_8446622_RNA45SN1_45S_50ntFlanks | 12934 | 12961 | 28 | CTAAACCATTCGTAGACGACCTGCTTCT | MB468 | 50 | 117 | 35 | 9 | 18 | 18.0 | 26.5 | 18.8 | 2.1 | 4.3 |
| >hg38_21_+_8433172_8446622_RNA45SN1_45S_50ntFlanks@12934.12962.29\|CTAAACCATTCGTAGACGACCTGCTTCTG | >hg38_21_+_8433172_8446622_RNA45SN1_45S_50ntFlanks | 12934 | 12962 | 29 | CTAAACCATTCGTAGACGACCTGCTTCTG | MB468 | 28 | 63 | 30 | 11 | 14 | 12.4 | 18.0 | 16.3 | 2.7 | 1.5 |
| >hg38_21_+_8433172_8446622_RNA45SN1_45S_50ntFlanks@12934.12965.32\|CTAAACCATTCGTAGACGACCTGCTTCTGGGT | >hg38_21_+_8433172_8446622_RNA45SN1_45S_50ntFlanks | 12934 | 12965 | 32 | CTAAACCATTCGTAGACGACCTGCTTCTGGGT | MB468 | 50 | 47 | 93 | 35 | 27 | 15.5 | 19.3 | 69.5 | 23.7 | 8.2 |
| >hg38_21_+_8433172_8446622_RNA45SN1_45S_50ntFlanks@12934.12966.33\|CTAAACCATTCGTAGACGACCTGCTTCTGGGTC | >hg38_21_+_8433172_8446622_RNA45SN1_45S_50ntFlanks | 12934 | 12966 | 33 | CTAAACCATTCGTAGACGACCTGCTTCTGGGTC | MB468 | 38 | 23 | 107 | 16 | 17 | 9.1 | 12.6 | 57.7 | 13.5 | 5.9 |
| >hg38_21_+_8433172_8446622_RNA45SN1_45S_50ntFlanks@12934.12969.36\|CTAAACCATTCGTAGACGACCTGCTTCTGGGTCGGG | >hg38_21_+_8433172_8446622_RNA45SN1_45S_50ntFlanks | 12934 | 12969 | 36 | CTAAACCATTCGTAGACGACCTGCTTCTGGGTCGGG | MB468 | 23 | 10 | 56 | 16 | 18 | 11.7 | 4.5 | 43.7 | 14.5 | 6.2 |
| >hg38_21_+_8433172_8446622_RNA45SN1_45S_50ntFlanks@12934.12970.37\|CTAAACCATTCGTAGACGACCTGCTTCTGGGTCGGGG | >hg38_21_+_8433172_8446622_RNA45SN1_45S_50ntFlanks | 12934 | 12970 | 37 | CTAAACCATTCGTAGACGACCTGCTTCTGGGTCGGGG | MB468 | 27 | 6 | 57 | 12 | 12 | 14.3 | 2.0 | 34.7 | 10.8 | 4.5 |
| >hg38_21_+_8433172_8446622_RNA45SN1_45S_50ntFlanks@12934.12972.39\|CTAAACCATTCGTAGACGACCTGCTTCTGGGTCGGGGTT | >hg38_21_+_8433172_8446622_RNA45SN1_45S_50ntFlanks | 12934 | 12972 | 39 | CTAAACCATTCGTAGACGACCTGCTTCTGGGTCGGGGTT | MB468 | 143 | 87 | 647 | 86 | 126 | 67.1 | 48.2 | 463.3 | 76.2 | 64.3 |
| >hg38_21_+_8433172_8446622_RNA45SN1_45S_50ntFlanks@12935.12952.18\|TAAACCATTCGTAGACGA | >hg38_21_+_8433172_8446622_RNA45SN1_45S_50ntFlanks | 12935 | 12952 | 18 | TAAACCATTCGTAGACGA | MB468 | 19 | 5 | 20 | 5 | 4 | 4.7 | 2.2 | 9.3 | 1.8 | 1.8 |
| >hg38_21_+_8433172_8446622_RNA45SN1_45S_50ntFlanks@12935.12953.19\|TAAACCATTCGTAGACGAC | >hg38_21_+_8433172_8446622_RNA45SN1_45S_50ntFlanks | 12935 | 12953 | 19 | TAAACCATTCGTAGACGAC | MB468 | 27 | 13 | 37 | 8 | 6 | 6.1 | 5.6 | 18.7 | 3.8 | 1.0 |
| >hg38_21_+_8433172_8446622_RNA45SN1_45S_50ntFlanks@12935.12955.21\|TAAACCATTCGTAGACGACCT | >hg38_21_+_8433172_8446622_RNA45SN1_45S_50ntFlanks | 12935 | 12955 | 21 | TAAACCATTCGTAGACGACCT | MB468 | 13 | 17 | 16 | 2 | 4 | 3.2 | 1.8 | 2.9 | 0.3 | 1.4 |
| >hg38_21_+_8433172_8446622_RNA45SN1_45S_50ntFlanks@12935.12956.22\|TAAACCATTCGTAGACGACCTG | >hg38_21_+_8433172_8446622_RNA45SN1_45S_50ntFlanks | 12935 | 12956 | 22 | TAAACCATTCGTAGACGACCTG | MB468 | 213 | 299 | 270 | 44 | 64 | 39.3 | 70.6 | 52.8 | 10.1 | 12.3 |
| >hg38_21_+_8433172_8446622_RNA45SN1_45S_50ntFlanks@12935.12959.25\|TAAACCATTCGTAGACGACCTGCTT | >hg38_21_+_8433172_8446622_RNA45SN1_45S_50ntFlanks | 12935 | 12959 | 25 | TAAACCATTCGTAGACGACCTGCTT | MB468 | 14 | 25 | 10 | 2 | 5 | 2.8 | 4.1 | 2.9 | 0.2 | 1.9 |
| >hg38_21_+_8433172_8446622_RNA45SN1_45S_50ntFlanks@12935.12961.27\|TAAACCATTCGTAGACGACCTGCTTCT | >hg38_21_+_8433172_8446622_RNA45SN1_45S_50ntFlanks | 12935 | 12961 | 27 | TAAACCATTCGTAGACGACCTGCTTCT | MB468 | 13 | 29 | 9 | 1 | 3 | 2.7 | 4.8 | 5.1 | 0.8 | 2.5 |
| >hg38_21_+_8433172_8446622_RNA45SN1_45S_50ntFlanks@12935.12966.32\|TAAACCATTCGTAGACGACCTGCTTCTGGGTC | >hg38_21_+_8433172_8446622_RNA45SN1_45S_50ntFlanks | 12935 | 12966 | 32 | TAAACCATTCGTAGACGACCTGCTTCTGGGTC | MB468 | 11 | 4 | 33 | 3 | 3 | 2.1 | 2.0 | 18.9 | 2.3 | 0.6 |
| >hg38_21_+_8433172_8446622_RNA45SN1_45S_50ntFlanks@12936.12953.18\|AAACCATTCGTAGACGAC | >hg38_21_+_8433172_8446622_RNA45SN1_45S_50ntFlanks | 12936 | 12953 | 18 | AAACCATTCGTAGACGAC | MB468 | 76 | 43 | 60 | 32 | 21 | 16.8 | 20.8 | 25.6 | 9.5 | 6.0 |
| >hg38_21_+_8433172_8446622_RNA45SN1_45S_50ntFlanks@12936.12954.19\|AAACCATTCGTAGACGACC | >hg38_21_+_8433172_8446622_RNA45SN1_45S_50ntFlanks | 12936 | 12954 | 19 | AAACCATTCGTAGACGACC | MB468 | 19 | 18 | 19 | 3 | 7 | 4.9 | 5.8 | 5.1 | 1.4 | 0.6 |
| >hg38_21_+_8433172_8446622_RNA45SN1_45S_50ntFlanks@12936.12955.20\|AAACCATTCGTAGACGACCT | >hg38_21_+_8433172_8446622_RNA45SN1_45S_50ntFlanks | 12936 | 12955 | 20 | AAACCATTCGTAGACGACCT | MB468 | 38 | 51 | 30 | 8 | 15 | 9.2 | 4.8 | 5.7 | 1.6 | 4.1 |
| >hg38_21_+_8433172_8446622_RNA45SN1_45S_50ntFlanks@12936.12956.21\|AAACCATTCGTAGACGACCTG | >hg38_21_+_8433172_8446622_RNA45SN1_45S_50ntFlanks | 12936 | 12956 | 21 | AAACCATTCGTAGACGACCTG | MB468 | 623 | 987 | 481 | 215 | 237 | 143.7 | 259.6 | 93.5 | 45.5 | 56.8 |
| >hg38_21_+_8433172_8446622_RNA45SN1_45S_50ntFlanks@12936.12957.22\|AAACCATTCGTAGACGACCTGC | >hg38_21_+_8433172_8446622_RNA45SN1_45S_50ntFlanks | 12936 | 12957 | 22 | AAACCATTCGTAGACGACCTGC | MB468 | 22 | 36 | 18 | 3 | 8 | 6.0 | 11.4 | 8.6 | 0.5 | 1.3 |
| >hg38_21_+_8433172_8446622_RNA45SN1_45S_50ntFlanks@12936.12958.23\|AAACCATTCGTAGACGACCTGCT | >hg38_21_+_8433172_8446622_RNA45SN1_45S_50ntFlanks | 12936 | 12958 | 23 | AAACCATTCGTAGACGACCTGCT | MB468 | 22 | 36 | 12 | 4 | 13 | 6.3 | 3.8 | 3.9 | 1.4 | 2.9 |
| >hg38_21_+_8433172_8446622_RNA45SN1_45S_50ntFlanks@12936.12959.24\|AAACCATTCGTAGACGACCTGCTT | >hg38_21_+_8433172_8446622_RNA45SN1_45S_50ntFlanks | 12936 | 12959 | 24 | AAACCATTCGTAGACGACCTGCTT | MB468 | 39 | 74 | 20 | 5 | 15 | 7.5 | 11.3 | 5.0 | 1.1 | 5.7 |
| >hg38_21_+_8433172_8446622_RNA45SN1_45S_50ntFlanks@12936.12960.25\|AAACCATTCGTAGACGACCTGCTTC | >hg38_21_+_8433172_8446622_RNA45SN1_45S_50ntFlanks | 12936 | 12960 | 25 | AAACCATTCGTAGACGACCTGCTTC | MB468 | 30 | 59 | 13 | 8 | 15 | 11.9 | 14.4 | 4.8 | 2.2 | 3.2 |
| >hg38_21_+_8433172_8446622_RNA45SN1_45S_50ntFlanks@12936.12961.26\|AAACCATTCGTAGACGACCTGCTTCT | >hg38_21_+_8433172_8446622_RNA45SN1_45S_50ntFlanks | 12936 | 12961 | 26 | AAACCATTCGTAGACGACCTGCTTCT | MB468 | 31 | 77 | 14 | 3 | 11 | 7.9 | 11.6 | 6.7 | 0.8 | 2.7 |
| >hg38_21_+_8433172_8446622_RNA45SN1_45S_50ntFlanks@12936.12962.27\|AAACCATTCGTAGACGACCTGCTTCTG | >hg38_21_+_8433172_8446622_RNA45SN1_45S_50ntFlanks | 12936 | 12962 | 27 | AAACCATTCGTAGACGACCTGCTTCTG | MB468 | 17 | 40 | 12 | 5 | 8 | 7.5 | 11.5 | 6.9 | 1.2 | 1.7 |
| >hg38_21_+_8433172_8446622_RNA45SN1_45S_50ntFlanks@12936.12966.31\|AAACCATTCGTAGACGACCTGCTTCTGGGTC | >hg38_21_+_8433172_8446622_RNA45SN1_45S_50ntFlanks | 12936 | 12966 | 31 | AAACCATTCGTAGACGACCTGCTTCTGGGTC | MB468 | 29 | 13 | 50 | 9 | 9 | 10.9 | 7.4 | 30.2 | 6.2 | 3.8 |
| >hg38_21_+_8433172_8446622_RNA45SN1_45S_50ntFlanks@12936.12969.34\|AAACCATTCGTAGACGACCTGCTTCTGGGTCGGG | >hg38_21_+_8433172_8446622_RNA45SN1_45S_50ntFlanks | 12936 | 12969 | 34 | AAACCATTCGTAGACGACCTGCTTCTGGGTCGGG | MB468 | 10 | 3 | 18 | 7 | 6 | 5.8 | 1.9 | 14.3 | 5.3 | 1.7 |
| >hg38_21_+_8433172_8446622_RNA45SN1_45S_50ntFlanks@12937.12954.18\|AACCATTCGTAGACGACC | >hg38_21_+_8433172_8446622_RNA45SN1_45S_50ntFlanks | 12937 | 12954 | 18 | AACCATTCGTAGACGACC | MB468 | 15 | 12 | 17 | 4 | 5 | 2.8 | 4.2 | 5.5 | 1.9 | 0.4 |
| >hg38_21_+_8433172_8446622_RNA45SN1_45S_50ntFlanks@12937.12955.19\|AACCATTCGTAGACGACCT | >hg38_21_+_8433172_8446622_RNA45SN1_45S_50ntFlanks | 12937 | 12955 | 19 | AACCATTCGTAGACGACCT | MB468 | 30 | 37 | 27 | 7 | 11 | 8.0 | 7.5 | 5.9 | 1.3 | 3.3 |
| >hg38_21_+_8433172_8446622_RNA45SN1_45S_50ntFlanks@12937.12956.20\|AACCATTCGTAGACGACCTG | >hg38_21_+_8433172_8446622_RNA45SN1_45S_50ntFlanks | 12937 | 12956 | 20 | AACCATTCGTAGACGACCTG | MB468 | 469 | 715 | 417 | 167 | 169 | 103.8 | 215.5 | 100.3 | 32.5 | 41.8 |
| >hg38_21_+_8433172_8446622_RNA45SN1_45S_50ntFlanks@12937.12957.21\|AACCATTCGTAGACGACCTGC | >hg38_21_+_8433172_8446622_RNA45SN1_45S_50ntFlanks | 12937 | 12957 | 21 | AACCATTCGTAGACGACCTGC | MB468 | 17 | 28 | 15 | 3 | 6 | 4.8 | 12.0 | 9.3 | 0.2 | 1.2 |
| >hg38_21_+_8433172_8446622_RNA45SN1_45S_50ntFlanks@12937.12958.22\|AACCATTCGTAGACGACCTGCT | >hg38_21_+_8433172_8446622_RNA45SN1_45S_50ntFlanks | 12937 | 12958 | 22 | AACCATTCGTAGACGACCTGCT | MB468 | 16 | 25 | 9 | 3 | 8 | 4.5 | 6.1 | 3.6 | 0.7 | 2.2 |
| >hg38_21_+_8433172_8446622_RNA45SN1_45S_50ntFlanks@12937.12959.23\|AACCATTCGTAGACGACCTGCTT | >hg38_21_+_8433172_8446622_RNA45SN1_45S_50ntFlanks | 12937 | 12959 | 23 | AACCATTCGTAGACGACCTGCTT | MB468 | 34 | 59 | 19 | 4 | 11 | 6.9 | 16.9 | 5.3 | 0.5 | 3.6 |
| >hg38_21_+_8433172_8446622_RNA45SN1_45S_50ntFlanks@12937.12960.24\|AACCATTCGTAGACGACCTGCTTC | >hg38_21_+_8433172_8446622_RNA45SN1_45S_50ntFlanks | 12937 | 12960 | 24 | AACCATTCGTAGACGACCTGCTTC | MB468 | 19 | 36 | 9 | 6 | 9 | 8.0 | 8.8 | 3.4 | 1.5 | 2.0 |
| >hg38_21_+_8433172_8446622_RNA45SN1_45S_50ntFlanks@12937.12961.25\|AACCATTCGTAGACGACCTGCTTCT | >hg38_21_+_8433172_8446622_RNA45SN1_45S_50ntFlanks | 12937 | 12961 | 25 | AACCATTCGTAGACGACCTGCTTCT | MB468 | 25 | 59 | 12 | 3 | 9 | 6.1 | 6.7 | 4.3 | 0.6 | 2.7 |

| | | | | | | | | | | | | | | | |
|---|---|---|---|---|---|---|---|---|---|---|---|---|---|---|---|
| >hg38_21_+_8433172_8446622_RNA45SN1_45S_50ntFlanks@12937.12962.26\|AACCATTCGTAGACGACCTGCTTCTG | >hg38_21_+_8433172_8446622_RNA45SN1_45S_50ntFlanks | 12937 | 12962 | 26 | AACCATTCGTAGACGACCTGCTTCTG | MB468 | 13 | 29 | 10 | 5 | 6 | 6.3 | 10.4 | 6.2 | 0.8 | 1.3 |
| >hg38_21_+_8433172_8446622_RNA45SN1_45S_50ntFlanks@12937.12966.30\|AACCATTCGTAGACGACCTGCTTCTGGGTC | >hg38_21_+_8433172_8446622_RNA45SN1_45S_50ntFlanks | 12937 | 12966 | 30 | AACCATTCGTAGACGACCTGCTTCTGGGTC | MB468 | 27 | 11 | 46 | 10 | 8 | 6.4 | 5.2 | 22.3 | 5.8 | 1.5 |
| >hg38_21_+_8433172_8446622_RNA45SN1_45S_50ntFlanks@12938.12955.18\|ACCATTCGTAGACGACCT | >hg38_21_+_8433172_8446622_RNA45SN1_45S_50ntFlanks | 12938 | 12955 | 18 | ACCATTCGTAGACGACCT | MB468 | 12 | 11 | 12 | 3 | 4 | 2.8 | 3.4 | 3.3 | 1.6 | 1.6 |
| >hg38_21_+_8433172_8446622_RNA45SN1_45S_50ntFlanks@12938.12956.19\|ACCATTCGTAGACGACCTG | >hg38_21_+_8433172_8446622_RNA45SN1_45S_50ntFlanks | 12938 | 12956 | 19 | ACCATTCGTAGACGACCTG | MB468 | 184 | 218 | 190 | 64 | 61 | 29.8 | 82.8 | 55.5 | 15.9 | 14.3 |
| >hg38_21_+_8433172_8446622_RNA45SN1_45S_50ntFlanks@12938.12959.22\|ACCATTCGTAGACGACCTGCTT | >hg38_21_+_8433172_8446622_RNA45SN1_45S_50ntFlanks | 12938 | 12959 | 22 | ACCATTCGTAGACGACCTGCTT | MB468 | 11 | 18 | 6 | 1 | 3 | 2.5 | 5.3 | 2.4 | 1.0 | 2.0 |
| >hg38_21_+_8433172_8446622_RNA45SN1_45S_50ntFlanks@12938.12966.29\|ACCATTCGTAGACGACCTGCTTCTGGGTC | >hg38_21_+_8433172_8446622_RNA45SN1_45S_50ntFlanks | 12938 | 12966 | 29 | ACCATTCGTAGACGACCTGCTTCTGGGTC | MB468 | 14 | 5 | 23 | 7 | 5 | 5.9 | 3.5 | 12.5 | 4.8 | 1.9 |
| >hg38_21_+_8433172_8446622_RNA45SN1_45S_50ntFlanks@12939.12956.18\|CCATTCGTAGACGACCTG | >hg38_21_+_8433172_8446622_RNA45SN1_45S_50ntFlanks | 12939 | 12956 | 18 | CCATTCGTAGACGACCTG | MB468 | 121 | 73 | 137 | 55 | 46 | 16.9 | 26.5 | 36.1 | 10.8 | 12.5 |
| >hg38_21_+_8433172_8446622_RNA45SN1_45S_50ntFlanks@12939.12966.28\|CCATTCGTAGACGACCTGCTTCTGGGTC | >hg38_21_+_8433172_8446622_RNA45SN1_45S_50ntFlanks | 12939 | 12966 | 28 | CCATTCGTAGACGACCTGCTTCTGGGTC | MB468 | 15 | 3 | 24 | 10 | 6 | 3.8 | 1.9 | 11.5 | 7.0 | 2.9 |
| >hg38_21_+_8433172_8446622_RNA45SN1_45S_50ntFlanks@12939.12972.34\|CCATTCGTAGACGACCTGCTTCTGGGTCGGGGTT | >hg38_21_+_8433172_8446622_RNA45SN1_45S_50ntFlanks | 12939 | 12972 | 34 | CCATTCGTAGACGACCTGCTTCTGGGTCGGGGTT | MB468 | 10 | 3 | 17 | 10 | 5 | 3.0 | 0.3 | 8.0 | 5.7 | 2.5 |
| >hg38_21_+_8433172_8446622_RNA45SN1_45S_50ntFlanks@12940.12959.20\|CATTCGTAGACGACCTGCTT | >hg38_21_+_8433172_8446622_RNA45SN1_45S_50ntFlanks | 12940 | 12959 | 20 | CATTCGTAGACGACCTGCTT | MB468 | 13 | 7 | 12 | 3 | 4 | 3.9 | 2.8 | 4.2 | 1.2 | 1.7 |
| >hg38_21_+_8433172_8446622_RNA45SN1_45S_50ntFlanks@12940.12965.26\|CATTCGTAGACGACCTGCTTCTGGGT | >hg38_21_+_8433172_8446622_RNA45SN1_45S_50ntFlanks | 12940 | 12965 | 26 | CATTCGTAGACGACCTGCTTCTGGGT | MB468 | 17 | 12 | 19 | 10 | 5 | 2.1 | 3.5 | 7.1 | 1.5 | 1.4 |
| >hg38_21_+_8433172_8446622_RNA45SN1_45S_50ntFlanks@12940.12966.27\|CATTCGTAGACGACCTGCTTCTGGGTC | >hg38_21_+_8433172_8446622_RNA45SN1_45S_50ntFlanks | 12940 | 12966 | 27 | CATTCGTAGACGACCTGCTTCTGGGTC | MB468 | 69 | 6 | 123 | 33 | 17 | 19.6 | 2.1 | 56.7 | 20.7 | 9.7 |
| >hg38_21_+_8433172_8446622_RNA45SN1_45S_50ntFlanks@12940.12972.33\|CATTCGTAGACGACCTGCTTCTGGGTCGGGGTT | >hg38_21_+_8433172_8446622_RNA45SN1_45S_50ntFlanks | 12940 | 12972 | 33 | CATTCGTAGACGACCTGCTTCTGGGTCGGGGTT | MB468 | 31 | 7 | 51 | 30 | 10 | 9.8 | 1.6 | 26.1 | 15.1 | 4.0 |
| >hg38_21_+_8433172_8446622_RNA45SN1_45S_50ntFlanks@12941.12958.18\|ATTCGTAGACGACCTGCT | >hg38_21_+_8433172_8446622_RNA45SN1_45S_50ntFlanks | 12941 | 12958 | 18 | ATTCGTAGACGACCTGCT | MB231 | 15 | 9 | 21 | 1 | 3 | 12.8 | 2.2 | 14.4 | 1.2 | 2.7 |
| >hg38_21_+_8433172_8446622_RNA45SN1_45S_50ntFlanks@12941.12958.18\|ATTCGTAGACGACCTGCT | >hg38_21_+_8433172_8446622_RNA45SN1_45S_50ntFlanks | 12941 | 12958 | 18 | ATTCGTAGACGACCTGCT | MB468 | 12 | 5 | 11 | 4 | 5 | 2.6 | 1.7 | 4.0 | 0.5 | 1.3 |
| >hg38_21_+_8433172_8446622_RNA45SN1_45S_50ntFlanks@12941.12959.19\|ATTCGTAGACGACCTGCTT | >hg38_21_+_8433172_8446622_RNA45SN1_45S_50ntFlanks | 12941 | 12959 | 19 | ATTCGTAGACGACCTGCTT | MB231 | 14 | 9 | 20 | 1 | 1 | 13.5 | 3.5 | 14.0 | 1.2 | 2.4 |
| >hg38_21_+_8433172_8446622_RNA45SN1_45S_50ntFlanks@12941.12959.19\|ATTCGTAGACGACCTGCTT | >hg38_21_+_8433172_8446622_RNA45SN1_45S_50ntFlanks | 12941 | 12959 | 19 | ATTCGTAGACGACCTGCTT | MB468 | 14 | 7 | 14 | 4 | 4 | 3.7 | 4.1 | 2.8 | 0.5 | 2.2 |
| >hg38_21_+_8433172_8446622_RNA45SN1_45S_50ntFlanks@12941.12960.20\|ATTCGTAGACGACCTGCTTC | >hg38_21_+_8433172_8446622_RNA45SN1_45S_50ntFlanks | 12941 | 12960 | 20 | ATTCGTAGACGACCTGCTTC | BT20 | 11 | 7 | 28 | 6 | 6 | 3.3 | 2.0 | 15.8 | 3.9 | 0.6 |
| >hg38_21_+_8433172_8446622_RNA45SN1_45S_50ntFlanks@12941.12960.20\|ATTCGTAGACGACCTGCTTC | >hg38_21_+_8433172_8446622_RNA45SN1_45S_50ntFlanks | 12941 | 12960 | 20 | ATTCGTAGACGACCTGCTTC | MB231 | 19 | 16 | 32 | 4 | 5 | 16.5 | 7.8 | 23.1 | 1.6 | 2.3 |
| >hg38_21_+_8433172_8446622_RNA45SN1_45S_50ntFlanks@12941.12961.21\|ATTCGTAGACGACCTGCTTCT | >hg38_21_+_8433172_8446622_RNA45SN1_45S_50ntFlanks | 12941 | 12961 | 21 | ATTCGTAGACGACCTGCTTCT | MB231 | 17 | 16 | 23 | 3 | 5 | 14.3 | 10.5 | 21.8 | 3.2 | 2.9 |
| >hg38_21_+_8433172_8446622_RNA45SN1_45S_50ntFlanks@12941.12964.24\|ATTCGTAGACGACCTGCTTCTGGG | >hg38_21_+_8433172_8446622_RNA45SN1_45S_50ntFlanks | 12941 | 12964 | 24 | ATTCGTAGACGACCTGCTTCTGGG | MB468 | 14 | 13 | 11 | 6 | 4 | 2.7 | 5.1 | 3.4 | 2.7 | 1.5 |
| >hg38_21_+_8433172_8446622_RNA45SN1_45S_50ntFlanks@12941.12965.25\|ATTCGTAGACGACCTGCTTCTGGGT | >hg38_21_+_8433172_8446622_RNA45SN1_45S_50ntFlanks | 12941 | 12965 | 25 | ATTCGTAGACGACCTGCTTCTGGGT | MB468 | 28 | 14 | 28 | 13 | 6 | 2.1 | 6.0 | 11.2 | 7.4 | 1.7 |
| >hg38_21_+_8433172_8446622_RNA45SN1_45S_50ntFlanks@12941.12966.26\|ATTCGTAGACGACCTGCTTCTGGGTC | >hg38_21_+_8433172_8446622_RNA45SN1_45S_50ntFlanks | 12941 | 12966 | 26 | ATTCGTAGACGACCTGCTTCTGGGTC | MB468 | 78 | 4 | 112 | 40 | 13 | 40.2 | 1.4 | 54.6 | 26.6 | 9.8 |
| >hg38_21_+_8433172_8446622_RNA45SN1_45S_50ntFlanks@12941.12972.32\|ATTCGTAGACGACCTGCTTCTGGGTCGGGGTT | >hg38_21_+_8433172_8446622_RNA45SN1_45S_50ntFlanks | 12941 | 12972 | 32 | ATTCGTAGACGACCTGCTTCTGGGTCGGGGTT | MB468 | 14 | 2 | 22 | 28 | 5 | 3.8 | 0.3 | 11.7 | 16.8 | 2.2 |
| >hg38_21_+_8433172_8446622_RNA45SN1_45S_50ntFlanks@12942.12959.18\|TTCGTAGACGACCTGCTT | >hg38_21_+_8433172_8446622_RNA45SN1_45S_50ntFlanks | 12942 | 12959 | 18 | TTCGTAGACGACCTGCTT | MB468 | 65 | 23 | 70 | 14 | 13 | 20.6 | 11.4 | 18.0 | 2.4 | 6.7 |
| >hg38_21_+_8433172_8446622_RNA45SN1_45S_50ntFlanks@12942.12960.19\|TTCGTAGACGACCTGCTTC | >hg38_21_+_8433172_8446622_RNA45SN1_45S_50ntFlanks | 12942 | 12960 | 19 | TTCGTAGACGACCTGCTTC | MB231 | 12 | 12 | 18 | 2 | 2 | 10.4 | 5.9 | 10.8 | 1.6 | 2.1 |
| >hg38_21_+_8433172_8446622_RNA45SN1_45S_50ntFlanks@12942.12960.19\|TTCGTAGACGACCTGCTTC | >hg38_21_+_8433172_8446622_RNA45SN1_45S_50ntFlanks | 12942 | 12960 | 19 | TTCGTAGACGACCTGCTTC | MB468 | 35 | 19 | 34 | 20 | 9 | 13.0 | 7.6 | 13.8 | 5.5 | 2.5 |
| >hg38_21_+_8433172_8446622_RNA45SN1_45S_50ntFlanks@12942.12961.20\|TTCGTAGACGACCTGCTTCT | >hg38_21_+_8433172_8446622_RNA45SN1_45S_50ntFlanks | 12942 | 12961 | 20 | TTCGTAGACGACCTGCTTCT | MB231 | 11 | 12 | 13 | 1 | 2 | 9.3 | 7.7 | 11.0 | 1.5 | 2.1 |
| >hg38_21_+_8433172_8446622_RNA45SN1_45S_50ntFlanks@12942.12961.20\|TTCGTAGACGACCTGCTTCT | >hg38_21_+_8433172_8446622_RNA45SN1_45S_50ntFlanks | 12942 | 12961 | 20 | TTCGTAGACGACCTGCTTCT | MB468 | 30 | 22 | 28 | 10 | 8 | 12.8 | 5.6 | 14.0 | 3.5 | 1.3 |
| >hg38_21_+_8433172_8446622_RNA45SN1_45S_50ntFlanks@12942.12962.21\|TTCGTAGACGACCTGCTTCTG | >hg38_21_+_8433172_8446622_RNA45SN1_45S_50ntFlanks | 12942 | 12962 | 21 | TTCGTAGACGACCTGCTTCTG | MB468 | 18 | 20 | 24 | 12 | 7 | 5.6 | 6.5 | 10.8 | 5.4 | 1.4 |
| >hg38_21_+_8433172_8446622_RNA45SN1_45S_50ntFlanks@12942.12964.23\|TTCGTAGACGACCTGCTTCTGGG | >hg38_21_+_8433172_8446622_RNA45SN1_45S_50ntFlanks | 12942 | 12964 | 23 | TTCGTAGACGACCTGCTTCTGGG | MB468 | 23 | 10 | 16 | 10 | 6 | 5.4 | 3.2 | 4.1 | 5.2 | 2.7 |
| >hg38_21_+_8433172_8446622_RNA45SN1_45S_50ntFlanks@12942.12965.24\|TTCGTAGACGACCTGCTTCTGGGT | >hg38_21_+_8433172_8446622_RNA45SN1_45S_50ntFlanks | 12942 | 12965 | 24 | TTCGTAGACGACCTGCTTCTGGGT | MB468 | 47 | 21 | 34 | 18 | 11 | 5.4 | 8.3 | 11.3 | 10.6 | 4.2 |
| >hg38_21_+_8433172_8446622_RNA45SN1_45S_50ntFlanks@12942.12966.25\|TTCGTAGACGACCTGCTTCTGGGTC | >hg38_21_+_8433172_8446622_RNA45SN1_45S_50ntFlanks | 12942 | 12966 | 25 | TTCGTAGACGACCTGCTTCTGGGTC | MB468 | 73 | 3 | 88 | 28 | 12 | 42.0 | 1.1 | 45.3 | 21.3 | 10.6 |
| >hg38_21_+_8433172_8446622_RNA45SN1_45S_50ntFlanks@12942.12972.31\|TTCGTAGACGACCTGCTTCTGGGTCGGGGTT | >hg38_21_+_8433172_8446622_RNA45SN1_45S_50ntFlanks | 12942 | 12972 | 31 | TTCGTAGACGACCTGCTTCTGGGTCGGGGTT | MB468 | 50 | 8 | 81 | 75 | 19 | 10.7 | 2.1 | 39.8 | 40.4 | 7.5 |
| >hg38_21_+_8433172_8446622_RNA45SN1_45S_50ntFlanks@12942.12973.32\|TTCGTAGACGACCTGCTTCTGGGTCGGGGTTT | >hg38_21_+_8433172_8446622_RNA45SN1_45S_50ntFlanks | 12942 | 12973 | 32 | TTCGTAGACGACCTGCTTCTGGGTCGGGGTTT | MB468 | 24 | 5 | 27 | 19 | 11 | 8.8 | 0.6 | 14.5 | 7.7 | 3.6 |
| >hg38_21_+_8433172_8446622_RNA45SN1_45S_50ntFlanks@12944.12961.18\|CGTAGACGACCTGCTTCT | >hg38_21_+_8433172_8446622_RNA45SN1_45S_50ntFlanks | 12944 | 12961 | 18 | CGTAGACGACCTGCTTCT | MB468 | 15 | 10 | 14 | 7 | 7 | 7.8 | 4.3 | 6.0 | 4.6 | 2.4 |
| >hg38_21_+_8433172_8446622_RNA45SN1_45S_50ntFlanks@12944.12962.19\|CGTAGACGACCTGCTTCTG | >hg38_21_+_8433172_8446622_RNA45SN1_45S_50ntFlanks | 12944 | 12962 | 19 | CGTAGACGACCTGCTTCTG | MB468 | 15 | 15 | 20 | 15 | 7 | 5.5 | 6.7 | 9.1 | 6.9 | 1.1 |
| >hg38_21_+_8433172_8446622_RNA45SN1_45S_50ntFlanks@12944.12964.21\|CGTAGACGACCTGCTTCTGGG | >hg38_21_+_8433172_8446622_RNA45SN1_45S_50ntFlanks | 12944 | 12964 | 21 | CGTAGACGACCTGCTTCTGGG | MB468 | 37 | 38 | 19 | 15 | 13 | 16.6 | 20.8 | 5.9 | 11.4 | 5.7 |
| >hg38_21_+_8433172_8446622_RNA45SN1_45S_50ntFlanks@12944.12965.22\|CGTAGACGACCTGCTTCTGGGT | >hg38_21_+_8433172_8446622_RNA45SN1_45S_50ntFlanks | 12944 | 12965 | 22 | CGTAGACGACCTGCTTCTGGGT | MB468 | 16 | 3 | 11 | 9 | 5 | 6.6 | 0.7 | 4.1 | 4.5 | 2.3 |
| >hg38_21_+_8433172_8446622_RNA45SN1_45S_50ntFlanks@12944.12966.23\|CGTAGACGACCTGCTTCTGGGTC | >hg38_21_+_8433172_8446622_RNA45SN1_45S_50ntFlanks | 12944 | 12966 | 23 | CGTAGACGACCTGCTTCTGGGTC | MB468 | 15 | 0 | 9 | 6 | 5 | 13.8 | 0.1 | 4.9 | 4.2 | 5.9 |
| >hg38_21_+_8433172_8446622_RNA45SN1_45S_50ntFlanks@12944.12972.29\|CGTAGACGACCTGCTTCTGGGTCGGGGTT | >hg38_21_+_8433172_8446622_RNA45SN1_45S_50ntFlanks | 12944 | 12972 | 29 | CGTAGACGACCTGCTTCTGGGTCGGGGTT | MB468 | 21 | 3 | 27 | 39 | 12 | 6.9 | 1.2 | 13.7 | 18.7 | 5.2 |
| >hg38_21_+_8433172_8446622_RNA45SN1_45S_50ntFlanks@12946.12964.19\|TAGACGACCTGCTTCTGGG | >hg38_21_+_8433172_8446622_RNA45SN1_45S_50ntFlanks | 12946 | 12964 | 19 | TAGACGACCTGCTTCTGGG | MB468 | 23 | 11 | 9 | 8 | 11 | 8.5 | 5.0 | 2.6 | 5.2 | 7.5 |
| >hg38_21_+_8433172_8446622_RNA45SN1_45S_50ntFlanks@12949.12982.34\|ACGACCTGCTTCTGGGTCGGGGTTTCGTACGTAG | >hg38_21_+_8433172_8446622_RNA45SN1_45S_50ntFlanks | 12949 | 12982 | 34 | ACGACCTGCTTCTGGGTCGGGGTTTCGTACGTAG | MB468 | 13 | 14 | 24 | 4 | 10 | 5.9 | 5.6 | 12.0 | 2.9 | 10.4 |
| >hg38_21_+_8433172_8446622_RNA45SN1_45S_50ntFlanks@12954.12972.19\|CTGCTTCTGGGTCGGGGTT | >hg38_21_+_8433172_8446622_RNA45SN1_45S_50ntFlanks | 12954 | 12972 | 19 | CTGCTTCTGGGTCGGGGTT | MB468 | 67 | 58 | 42 | 27 | 21 | 27.9 | 30.7 | 12.8 | 8.4 | 7.5 |
| >hg38_21_+_8433172_8446622_RNA45SN1_45S_50ntFlanks@12954.12973.20\|CTGCTTCTGGGTCGGGGTTT | >hg38_21_+_8433172_8446622_RNA45SN1_45S_50ntFlanks | 12954 | 12973 | 20 | CTGCTTCTGGGTCGGGGTTT | MB468 | 19 | 16 | 11 | 11 | 7 | 9.4 | 10.8 | 4.7 | 4.1 | 2.4 |
| >hg38_21_+_8433172_8446622_RNA45SN1_45S_50ntFlanks@12954.12982.29\|CTGCTTCTGGGTCGGGGTTTCGTACGTAG | >hg38_21_+_8433172_8446622_RNA45SN1_45S_50ntFlanks | 12954 | 12982 | 29 | CTGCTTCTGGGTCGGGGTTTCGTACGTAG | MB468 | 15 | 32 | 14 | 8 | 9 | 4.1 | 14.6 | 2.5 | 1.6 | 2.4 |
| >hg38_21_+_8433172_8446622_RNA45SN1_45S_50ntFlanks@12955.12972.18\|TGCTTCTGGGTCGGGGTT | >hg38_21_+_8433172_8446622_RNA45SN1_45S_50ntFlanks | 12955 | 12972 | 18 | TGCTTCTGGGTCGGGGTT | MB468 | 12 | 9 | 8 | 2 | 3 | 4.5 | 4.0 | 1.9 | 0.8 | 0.7 |
| >hg38_21_+_8433172_8446622_RNA45SN1_45S_50ntFlanks@12957.12975.19\|CTTCTGGGTCGGGGTTTCG | >hg38_21_+_8433172_8446622_RNA45SN1_45S_50ntFlanks | 12957 | 12975 | 19 | CTTCTGGGTCGGGGTTTCG | MB468 | 13 | 23 | 15 | 6 | 13 | 2.9 | 7.9 | 5.9 | 1.8 | 5.3 |
| >hg38_21_+_8433172_8446622_RNA45SN1_45S_50ntFlanks@12957.12979.23\|CTTCTGGGTCGGGGTTTCGTACG | >hg38_21_+_8433172_8446622_RNA45SN1_45S_50ntFlanks | 12957 | 12979 | 23 | CTTCTGGGTCGGGGTTTCGTACG | MB468 | 50 | 168 | 57 | 22 | 56 | 12.3 | 70.0 | 20.9 | 12.9 | 34.7 |
| >hg38_21_+_8433172_8446622_RNA45SN1_45S_50ntFlanks@12957.12980.24\|CTTCTGGGTCGGGGTTTCGTACGT | >hg38_21_+_8433172_8446622_RNA45SN1_45S_50ntFlanks | 12957 | 12980 | 24 | CTTCTGGGTCGGGGTTTCGTACGT | MB468 | 14 | 44 | 14 | 7 | 15 | 4.3 | 13.3 | 4.5 | 2.6 | 6.1 |

| | | | | | | | | | | | | | | | |
|---|---|---|---|---|---|---|---|---|---|---|---|---|---|---|---|
| >hg38_21_+_8433172_8446622_RNA45SN1_45S_50ntFlanks@12957.12981.25\|CTTCTGGGTCGGGGTTTCGTACGTA | >hg38_21_+_8433172_8446622_RNA45SN1_45S_50ntFlanks | 12957 | 12981 | 25 | CTTCTGGGTCGGGGTTTCGTACGTA | MB468 | 16 | 31 | 17 | 10 | 17 | 6.3 | 9.6 | 7.9 | 3.8 | 5.2 |
| >hg38_21_+_8433172_8446622_RNA45SN1_45S_50ntFlanks@12957.12982.26\|CTTCTGGGTCGGGGTTTCGTACGTAG | >hg38_21_+_8433172_8446622_RNA45SN1_45S_50ntFlanks | 12957 | 12982 | 26 | CTTCTGGGTCGGGGTTTCGTACGTAG | MB468 | 151 | 417 | 149 | 63 | 135 | 28.8 | 127.7 | 46.0 | 16.6 | 32.4 |
| >hg38_21_+_8433172_8446622_RNA45SN1_45S_50ntFlanks@12957.12985.29\|CTTCTGGGTCGGGGTTTCGTACGTAGCAG | >hg38_21_+_8433172_8446622_RNA45SN1_45S_50ntFlanks | 12957 | 12985 | 29 | CTTCTGGGTCGGGGTTTCGTACGTAGCAG | MB468 | 12 | 26 | 9 | 3 | 8 | 4.2 | 10.8 | 2.0 | 0.8 | 1.2 |
| >hg38_21_+_8433172_8446622_RNA45SN1_45S_50ntFlanks@12957.12986.30\|CTTCTGGGTCGGGGTTTCGTACGTAGCAGA | >hg38_21_+_8433172_8446622_RNA45SN1_45S_50ntFlanks | 12957 | 12986 | 30 | CTTCTGGGTCGGGGTTTCGTACGTAGCAGA | MB468 | 19 | 53 | 25 | 7 | 16 | 7.9 | 22.1 | 15.1 | 5.7 | 4.3 |
| >hg38_21_+_8433172_8446622_RNA45SN1_45S_50ntFlanks@12957.12987.31\|CTTCTGGGTCGGGGTTTCGTACGTAGCAGAG | >hg38_21_+_8433172_8446622_RNA45SN1_45S_50ntFlanks | 12957 | 12987 | 31 | CTTCTGGGTCGGGGTTTCGTACGTAGCAGAG | MB468 | 32 | 70 | 24 | 8 | 17 | 11.0 | 33.9 | 5.5 | 3.1 | 4.3 |
| >hg38_21_+_8433172_8446622_RNA45SN1_45S_50ntFlanks@12957.12989.33\|CTTCTGGGTCGGGGTTTCGTACGTAGCAGAGCA | >hg38_21_+_8433172_8446622_RNA45SN1_45S_50ntFlanks | 12957 | 12989 | 33 | CTTCTGGGTCGGGGTTTCGTACGTAGCAGAGCA | MB468 | 14 | 44 | 15 | 3 | 8 | 5.6 | 13.8 | 5.1 | 1.3 | 3.5 |
| >hg38_21_+_8433172_8446622_RNA45SN1_45S_50ntFlanks@12957.12990.34\|CTTCTGGGTCGGGGTTTCGTACGTAGCAGAGCAG | >hg38_21_+_8433172_8446622_RNA45SN1_45S_50ntFlanks | 12957 | 12990 | 34 | CTTCTGGGTCGGGGTTTCGTACGTAGCAGAGCAG | MB468 | 38 | 150 | 38 | 10 | 31 | 11.3 | 30.9 | 10.6 | 5.9 | 11.1 |
| >hg38_21_+_8433172_8446622_RNA45SN1_45S_50ntFlanks@12959.12982.24\|TCTGGGTCGGGGTTTCGTACGTAG | >hg38_21_+_8433172_8446622_RNA45SN1_45S_50ntFlanks | 12959 | 12982 | 24 | TCTGGGTCGGGGTTTCGTACGTAG | MB468 | 16 | 45 | 17 | 4 | 6 | 2.8 | 12.2 | 1.4 | 2.4 | 2.4 |
| >hg38_21_+_8433172_8446622_RNA45SN1_45S_50ntFlanks@12960.12982.23\|CTGGGTCGGGGTTTCGTACGTAG | >hg38_21_+_8433172_8446622_RNA45SN1_45S_50ntFlanks | 12960 | 12982 | 23 | CTGGGTCGGGGTTTCGTACGTAG | MB468 | 17 | 43 | 19 | 10 | 15 | 3.7 | 11.2 | 3.1 | 5.9 | 9.3 |
| >hg38_21_+_8433172_8446622_RNA45SN1_45S_50ntFlanks@12970.12990.21\|GTTTCGTACGTAGCAGAGCAG | >hg38_21_+_8433172_8446622_RNA45SN1_45S_50ntFlanks | 12970 | 12990 | 21 | GTTTCGTACGTAGCAGAGCAG | MB468 | 18 | 19 | 13 | 3 | 5 | 8.1 | 7.8 | 5.7 | 2.2 | 1.9 |
| >hg38_21_+_8433172_8446622_RNA45SN1_45S_50ntFlanks@12970.12991.22\|GTTTCGTACGTAGCAGAGCAGC | >hg38_21_+_8433172_8446622_RNA45SN1_45S_50ntFlanks | 12970 | 12991 | 22 | GTTTCGTACGTAGCAGAGCAGC | MB468 | 11 | 8 | 13 | 1 | 2 | 4.9 | 2.2 | 7.5 | 0.9 | 0.2 |
| >hg38_21_+_8433172_8446622_RNA45SN1_45S_50ntFlanks@12971.12988.18\|TTTCGTACGTAGCAGAGC | >hg38_21_+_8433172_8446622_RNA45SN1_45S_50ntFlanks | 12971 | 12988 | 18 | TTTCGTACGTAGCAGAGC | MB468 | 13 | 26 | 11 | 1 | 2 | 2.2 | 7.3 | 3.4 | 0.5 | 1.4 |
| >hg38_21_+_8433172_8446622_RNA45SN1_45S_50ntFlanks@12971.12989.19\|TTTCGTACGTAGCAGAGCA | >hg38_21_+_8433172_8446622_RNA45SN1_45S_50ntFlanks | 12971 | 12989 | 19 | TTTCGTACGTAGCAGAGCA | MB468 | 31 | 37 | 28 | 1 | 4 | 4.6 | 12.5 | 6.8 | 0.7 | 2.8 |
| >hg38_21_+_8433172_8446622_RNA45SN1_45S_50ntFlanks@12971.12990.20\|TTTCGTACGTAGCAGAGCAG | >hg38_21_+_8433172_8446622_RNA45SN1_45S_50ntFlanks | 12971 | 12990 | 20 | TTTCGTACGTAGCAGAGCAG | MB468 | 63 | 131 | 47 | 4 | 15 | 11.9 | 37.0 | 10.8 | 0.8 | 3.0 |
| >hg38_21_+_8433172_8446622_RNA45SN1_45S_50ntFlanks@12971.12991.21\|TTTCGTACGTAGCAGAGCAGC | >hg38_21_+_8433172_8446622_RNA45SN1_45S_50ntFlanks | 12971 | 12991 | 21 | TTTCGTACGTAGCAGAGCAGC | MB468 | 50 | 72 | 47 | 2 | 8 | 10.1 | 12.2 | 15.5 | 0.5 | 0.9 |
| >hg38_21_+_8433172_8446622_RNA45SN1_45S_50ntFlanks@12971.12992.22\|TTTCGTACGTAGCAGAGCAGCT | >hg38_21_+_8433172_8446622_RNA45SN1_45S_50ntFlanks | 12971 | 12992 | 22 | TTTCGTACGTAGCAGAGCAGCT | MB468 | 13 | 19 | 16 | 0 | 1 | 3.3 | 4.6 | 5.5 | 0.2 | 1.2 |
| >hg38_21_+_8433172_8446622_RNA45SN1_45S_50ntFlanks@12973.12990.18\|TCGTACGTAGCAGAGCAG | >hg38_21_+_8433172_8446622_RNA45SN1_45S_50ntFlanks | 12973 | 12990 | 18 | TCGTACGTAGCAGAGCAG | MB468 | 1026 | 1807 | 501 | 189 | 201 | 174.3 | 715.4 | 112.6 | 42.5 | 49.6 |
| >hg38_21_+_8433172_8446622_RNA45SN1_45S_50ntFlanks@12973.12991.19\|TCGTACGTAGCAGAGCAGC | >hg38_21_+_8433172_8446622_RNA45SN1_45S_50ntFlanks | 12973 | 12991 | 19 | TCGTACGTAGCAGAGCAGC | MB468 | 864 | 974 | 507 | 112 | 152 | 214.4 | 246.0 | 170.8 | 11.9 | 25.9 |
| >hg38_21_+_8433172_8446622_RNA45SN1_45S_50ntFlanks@12973.12992.20\|TCGTACGTAGCAGAGCAGCT | >hg38_21_+_8433172_8446622_RNA45SN1_45S_50ntFlanks | 12973 | 12992 | 20 | TCGTACGTAGCAGAGCAGCT | MB468 | 150 | 141 | 105 | 19 | 32 | 27.4 | 35.6 | 19.7 | 4.1 | 6.1 |
| >hg38_21_+_8433172_8446622_RNA45SN1_45S_50ntFlanks@12973.12994.22\|TCGTACGTAGCAGAGCAGCTCC | >hg38_21_+_8433172_8446622_RNA45SN1_45S_50ntFlanks | 12973 | 12994 | 22 | TCGTACGTAGCAGAGCAGCTCC | MB468 | 11 | 9 | 7 | 3 | 5 | 3.3 | 2.0 | 2.2 | 0.5 | 1.8 |
| >hg38_21_+_8433172_8446622_RNA45SN1_45S_50ntFlanks@12973.13000.28\|TCGTACGTAGCAGAGCAGCTCCCTCGCT | >hg38_21_+_8433172_8446622_RNA45SN1_45S_50ntFlanks | 12973 | 13000 | 28 | TCGTACGTAGCAGAGCAGCTCCCTCGCT | MB468 | 12 | 18 | 14 | 5 | 10 | 2.3 | 4.4 | 4.0 | 0.6 | 2.1 |
| >hg38_21_+_8433172_8446622_RNA45SN1_45S_50ntFlanks@12973.13001.29\|TCGTACGTAGCAGAGCAGCTCCCTCGCTG | >hg38_21_+_8433172_8446622_RNA45SN1_45S_50ntFlanks | 12973 | 13001 | 29 | TCGTACGTAGCAGAGCAGCTCCCTCGCTG | MB468 | 29 | 46 | 34 | 14 | 21 | 10.5 | 6.5 | 14.0 | 1.2 | 7.9 |
| >hg38_21_+_8433172_8446622_RNA45SN1_45S_50ntFlanks@12974.12991.18\|CGTACGTAGCAGAGCAGC | >hg38_21_+_8433172_8446622_RNA45SN1_45S_50ntFlanks | 12974 | 12991 | 18 | CGTACGTAGCAGAGCAGC | MB468 | 629 | 768 | 275 | 85 | 124 | 163.1 | 270.0 | 89.6 | 25.9 | 11.0 |
| >hg38_21_+_8433172_8446622_RNA45SN1_45S_50ntFlanks@12974.12992.19\|CGTACGTAGCAGAGCAGCT | >hg38_21_+_8433172_8446622_RNA45SN1_45S_50ntFlanks | 12974 | 12992 | 19 | CGTACGTAGCAGAGCAGCT | MB468 | 54 | 44 | 26 | 9 | 11 | 8.5 | 9.6 | 6.8 | 1.5 | 2.4 |
| >hg38_21_+_8433172_8446622_RNA45SN1_45S_50ntFlanks@12974.13001.28\|CGTACGTAGCAGAGCAGCTCCCTCGCTG | >hg38_21_+_8433172_8446622_RNA45SN1_45S_50ntFlanks | 12974 | 13001 | 28 | CGTACGTAGCAGAGCAGCTCCCTCGCTG | MB468 | 16 | 19 | 11 | 13 | 14 | 2.8 | 4.9 | 1.6 | 2.8 | 2.5 |
| >hg38_21_+_8433172_8446622_RNA45SN1_45S_50ntFlanks@12988.13013.26\|CAGCTCCCTCGCTGCGATCTATTGAA | >hg38_21_+_8433172_8446622_RNA45SN1_45S_50ntFlanks | 12988 | 13013 | 26 | CAGCTCCCTCGCTGCGATCTATTGAA | MB231 | 18 | 25 | 45 | 1 | 7 | 15.1 | 15.9 | 26.2 | 1.3 | 6.0 |
| >hg38_21_+_8433172_8446622_RNA45SN1_45S_50ntFlanks@12988.13013.26\|CAGCTCCCTCGCTGCGATCTATTGAA | >hg38_21_+_8433172_8446622_RNA45SN1_45S_50ntFlanks | 12988 | 13013 | 26 | CAGCTCCCTCGCTGCGATCTATTGAA | MB468 | 17 | 29 | 13 | 7 | 100 | 4.8 | 4.6 | 3.1 | 2.0 | 79.2 |
| >hg38_21_+_8433172_8446622_RNA45SN1_45S_50ntFlanks@12990.13012.23\|GCTCCCTCGCTGCGATCTATTGA | >hg38_21_+_8433172_8446622_RNA45SN1_45S_50ntFlanks | 12990 | 13012 | 23 | GCTCCCTCGCTGCGATCTATTGA | MB468 | 28 | 99 | 19 | 0 | 5 | 8.9 | 85.1 | 13.0 | 0.3 | 9.0 |
| >hg38_21_+_8433172_8446622_RNA45SN1_45S_50ntFlanks@12990.13013.24\|GCTCCCTCGCTGCGATCTATTGAA | >hg38_21_+_8433172_8446622_RNA45SN1_45S_50ntFlanks | 12990 | 13013 | 24 | GCTCCCTCGCTGCGATCTATTGAA | MB231 | 20 | 20 | 49 | 0 | 0 | 21.3 | 18.3 | 36.8 | 0.0 | 0.0 |
| >hg38_21_+_8433172_8446622_RNA45SN1_45S_50ntFlanks@12990.13013.24\|GCTCCCTCGCTGCGATCTATTGAA | >hg38_21_+_8433172_8446622_RNA45SN1_45S_50ntFlanks | 12990 | 13013 | 24 | GCTCCCTCGCTGCGATCTATTGAA | MB468 | 128 | 497 | 88 | 4 | 100 | 37.4 | 361.4 | 58.0 | 1.9 | 29.6 |
| >hg38_21_+_8433172_8446622_RNA45SN1_45S_50ntFlanks@12990.13014.25\|GCTCCCTCGCTGCGATCTATTGAAA | >hg38_21_+_8433172_8446622_RNA45SN1_45S_50ntFlanks | 12990 | 13014 | 25 | GCTCCCTCGCTGCGATCTATTGAAA | MB231 | 11 | 10 | 25 | 0 | 0 | 10.4 | 7.5 | 18.0 | 0.0 | 0.0 |
| >hg38_21_+_8433172_8446622_RNA45SN1_45S_50ntFlanks@12990.13014.25\|GCTCCCTCGCTGCGATCTATTGAAA | >hg38_21_+_8433172_8446622_RNA45SN1_45S_50ntFlanks | 12990 | 13014 | 25 | GCTCCCTCGCTGCGATCTATTGAAA | MB468 | 32 | 134 | 21 | 1 | 18 | 12.0 | 88.0 | 13.6 | 0.5 | 13.0 |
| >hg38_21_+_8433172_8446622_RNA45SN1_45S_50ntFlanks@12990.13015.26\|GCTCCCTCGCTGCGATCTATTGAAAG | >hg38_21_+_8433172_8446622_RNA45SN1_45S_50ntFlanks | 12990 | 13015 | 26 | GCTCCCTCGCTGCGATCTATTGAAAG | MB468 | 22 | 68 | 13 | 0 | 8 | 9.8 | 42.4 | 10.1 | 0.2 | 15.9 |
| >hg38_21_+_8433172_8446622_RNA45SN1_45S_50ntFlanks@12990.13017.28\|GCTCCCTCGCTGCGATCTATTGAAAGTC | >hg38_21_+_8433172_8446622_RNA45SN1_45S_50ntFlanks | 12990 | 13017 | 28 | GCTCCCTCGCTGCGATCTATTGAAAGTC | MB231 | 14 | 8 | 30 | 0 | 0 | 18.1 | 7.1 | 22.3 | 0.0 | 0.0 |
| >hg38_21_+_8433172_8446622_RNA45SN1_45S_50ntFlanks@12990.13039.50\|GCTCCCTCGCTGCGATCTATTGAAAGTCAGCCCTCGACACAAGGGTTTGT | >hg38_21_+_8433172_8446622_RNA45SN1_45S_50ntFlanks | 12990 | 13039 | 50 | GCTCCCTCGCTGCGATCTATTGAAAGTCAGCCCTCGACACAAGGGTTTGT | MB231 | 12 | 27 | 6 | 0 | 0 | 6.9 | 46.2 | 3.8 | 0.0 | 0.0 |
| >hg38_21_+_8433172_8446622_RNA45SN1_45S_50ntFlanks@12991.13008.18\|CTCCCTCGCTGCGATCTA | >hg38_21_+_8433172_8446622_RNA45SN1_45S_50ntFlanks | 12991 | 13008 | 18 | CTCCCTCGCTGCGATCTA | MB468 | 36 | 44 | 28 | 2 | 17 | 19.6 | 32.2 | 27.7 | 1.0 | 4.4 |
| >hg38_21_+_8433172_8446622_RNA45SN1_45S_50ntFlanks@12991.13009.19\|CTCCCTCGCTGCGATCTAT | >hg38_21_+_8433172_8446622_RNA45SN1_45S_50ntFlanks | 12991 | 13009 | 19 | CTCCCTCGCTGCGATCTAT | MB468 | 31 | 48 | 20 | 1 | 14 | 15.3 | 46.9 | 20.9 | 0.7 | 9.8 |
| >hg38_21_+_8433172_8446622_RNA45SN1_45S_50ntFlanks@12991.13010.20\|CTCCCTCGCTGCGATCTATT | >hg38_21_+_8433172_8446622_RNA45SN1_45S_50ntFlanks | 12991 | 13010 | 20 | CTCCCTCGCTGCGATCTATT | MB468 | 49 | 61 | 44 | 2 | 33 | 16.6 | 37.1 | 29.1 | 0.8 | 5.2 |
| >hg38_21_+_8433172_8446622_RNA45SN1_45S_50ntFlanks@12991.13011.21\|CTCCCTCGCTGCGATCTATTG | >hg38_21_+_8433172_8446622_RNA45SN1_45S_50ntFlanks | 12991 | 13011 | 21 | CTCCCTCGCTGCGATCTATTG | MB468 | 152 | 368 | 122 | 5 | 90 | 55.9 | 253.3 | 64.0 | 1.0 | 18.1 |
| >hg38_21_+_8433172_8446622_RNA45SN1_45S_50ntFlanks@12991.13012.22\|CTCCCTCGCTGCGATCTATTGA | >hg38_21_+_8433172_8446622_RNA45SN1_45S_50ntFlanks | 12991 | 13012 | 22 | CTCCCTCGCTGCGATCTATTGA | MB468 | 230 | 817 | 188 | 11 | 111 | 36.8 | 448.5 | 74.1 | 3.3 | 35.8 |
| >hg38_21_+_8433172_8446622_RNA45SN1_45S_50ntFlanks@12991.13013.23\|CTCCCTCGCTGCGATCTATTGAA | >hg38_21_+_8433172_8446622_RNA45SN1_45S_50ntFlanks | 12991 | 13013 | 23 | CTCCCTCGCTGCGATCTATTGAA | MB468 | 846 | 3038 | 652 | 86 | 548 | 194.9 | 1566.3 | 402.7 | 27.8 | 225.2 |
| >hg38_21_+_8433172_8446622_RNA45SN1_45S_50ntFlanks@12991.13014.24\|CTCCCTCGCTGCGATCTATTGAAA | >hg38_21_+_8433172_8446622_RNA45SN1_45S_50ntFlanks | 12991 | 13014 | 24 | CTCCCTCGCTGCGATCTATTGAAA | MB468 | 224 | 1001 | 158 | 19 | 146 | 55.0 | 482.8 | 96.8 | 7.2 | 60.2 |
| >hg38_21_+_8433172_8446622_RNA45SN1_45S_50ntFlanks@12991.13015.25\|CTCCCTCGCTGCGATCTATTGAAAG | >hg38_21_+_8433172_8446622_RNA45SN1_45S_50ntFlanks | 12991 | 13015 | 25 | CTCCCTCGCTGCGATCTATTGAAAG | MB468 | 343 | 1268 | 208 | 14 | 371 | 108.6 | 579.5 | 110.1 | 6.9 | 226.5 |
| >hg38_21_+_8433172_8446622_RNA45SN1_45S_50ntFlanks@12991.13016.26\|CTCCCTCGCTGCGATCTATTGAAAGT | >hg38_21_+_8433172_8446622_RNA45SN1_45S_50ntFlanks | 12991 | 13016 | 26 | CTCCCTCGCTGCGATCTATTGAAAGT | MB468 | 47 | 173 | 39 | 2 | 24 | 7.5 | 49.7 | 15.7 | 0.4 | 6.6 |
| >hg38_21_+_8433172_8446622_RNA45SN1_45S_50ntFlanks@12991.13017.27\|CTCCCTCGCTGCGATCTATTGAAAGTC | >hg38_21_+_8433172_8446622_RNA45SN1_45S_50ntFlanks | 12991 | 13017 | 27 | CTCCCTCGCTGCGATCTATTGAAAGTC | MB468 | 33 | 111 | 20 | 1 | 20 | 13.7 | 16.9 | 6.8 | 0.4 | 6.0 |
| >hg38_21_+_8433172_8446622_RNA45SN1_45S_50ntFlanks@12991.13019.29\|CTCCCTCGCTGCGATCTATTGAAAGTCAG | >hg38_21_+_8433172_8446622_RNA45SN1_45S_50ntFlanks | 12991 | 13019 | 29 | CTCCCTCGCTGCGATCTATTGAAAGTCAG | MB468 | 58 | 139 | 30 | 1 | 25 | 29.2 | 57.1 | 5.7 | 0.6 | 8.8 |
| >hg38_21_+_8433172_8446622_RNA45SN1_45S_50ntFlanks@12991.13027.37\|CTCCCTCGCTGCGATCTATTGAAAGTCAGCCCTCGAC | >hg38_21_+_8433172_8446622_RNA45SN1_45S_50ntFlanks | 12991 | 13027 | 37 | CTCCCTCGCTGCGATCTATTGAAAGTCAGCCCTCGAC | MB468 | 35 | 65 | 32 | 4 | 73 | 14.2 | 39.9 | 18.3 | 3.8 | 44.6 |
| >hg38_21_+_8433172_8446622_RNA45SN1_45S_50ntFlanks@12991.13028.38\|CTCCCTCGCTGCGATCTATTGAAAGTCAGCCCTCGACA | >hg38_21_+_8433172_8446622_RNA45SN1_45S_50ntFlanks | 12991 | 13028 | 38 | CTCCCTCGCTGCGATCTATTGAAAGTCAGCCCTCGACA | MB468 | 24 | 41 | 24 | 4 | 57 | 8.3 | 42.0 | 10.8 | 3.0 | 47.2 |
| >hg38_21_+_8433172_8446622_RNA45SN1_45S_50ntFlanks@12991.13039.49\|CTCCCTCGCTGCGATCTATTGAAAGTCAGCCCTCGACACAAGGGTTTGT | >hg38_21_+_8433172_8446622_RNA45SN1_45S_50ntFlanks | 12991 | 13039 | 49 | CTCCCTCGCTGCGATCTATTGAAAGTCAGCCCTCGACACAAGGGTTTGT | MB231 | 24 | 58 | 20 | 0 | 0 | 13.4 | 97.4 | 13.7 | 0.0 | 0.0 |
| >hg38_21_+_8433172_8446622_RNA45SN1_45S_50ntFlanks@12991.13039.49\|CTCCCTCGCTGCGATCTATTGAAAGTCAGCCCTCGACACAAGGGTTTGT | >hg38_21_+_8433172_8446622_RNA45SN1_45S_50ntFlanks | 12991 | 13039 | 49 | CTCCCTCGCTGCGATCTATTGAAAGTCAGCCCTCGACACAAGGGTTTGT | MB468 | 117 | 707 | 181 | 20 | 707 | 100.2 | 1130.3 | 128.7 | 35.6 | 816.3 |

| | | | | | | | | | | | | | | | | |
|---|---|---|---|---|---|---|---|---|---|---|---|---|---|---|---|---|
| >hg38_21_+_8433172_8446622_RNA45SN1_45S_50ntFlanks@12992.13011.20\|TCCCTCGCTGCGATCTATTG | >hg38_21_+_8433172_8446622_RNA45SN1_45S_50ntFlanks | 12992 | 13011 | 20 | TCCCTCGCTGCGATCTATTG | MB468 | 26 | 43 | 21 | 0 | 5 | 10.1 | 23.9 | 11.4 | 0.4 | 5.5 |
| >hg38_21_+_8433172_8446622_RNA45SN1_45S_50ntFlanks@12992.13012.21\|TCCCTCGCTGCGATCTATTGA | >hg38_21_+_8433172_8446622_RNA45SN1_45S_50ntFlanks | 12992 | 13012 | 21 | TCCCTCGCTGCGATCTATTGA | MB468 | 32 | 86 | 31 | 2 | 12 | 6.6 | 40.1 | 12.2 | 0.7 | 1.5 |
| >hg38_21_+_8433172_8446622_RNA45SN1_45S_50ntFlanks@12992.13013.22\|TCCCTCGCTGCGATCTATTGAA | >hg38_21_+_8433172_8446622_RNA45SN1_45S_50ntFlanks | 12992 | 13013 | 22 | TCCCTCGCTGCGATCTATTGAA | MB468 | 104 | 309 | 83 | 12 | 51 | 19.8 | 141.9 | 44.7 | 1.9 | 10.8 |
| >hg38_21_+_8433172_8446622_RNA45SN1_45S_50ntFlanks@12992.13014.23\|TCCCTCGCTGCGATCTATTGAAA | >hg38_21_+_8433172_8446622_RNA45SN1_45S_50ntFlanks | 12992 | 13014 | 23 | TCCCTCGCTGCGATCTATTGAAA | MB468 | 28 | 103 | 22 | 3 | 14 | 6.2 | 40.7 | 11.4 | 1.0 | 2.7 |
| >hg38_21_+_8433172_8446622_RNA45SN1_45S_50ntFlanks@12992.13015.24\|TCCCTCGCTGCGATCTATTGAAAG | >hg38_21_+_8433172_8446622_RNA45SN1_45S_50ntFlanks | 12992 | 13015 | 24 | TCCCTCGCTGCGATCTATTGAAAG | MB468 | 41 | 132 | 32 | 3 | 35 | 11.2 | 48.3 | 15.4 | 1.5 | 15.4 |
| >hg38_21_+_8433172_8446622_RNA45SN1_45S_50ntFlanks@12992.13039.48\|TCCCTCGCTGCGATCTATTGAAAGTCAGCCCTCGACACAAGGGTTTGT | >hg38_21_+_8433172_8446622_RNA45SN1_45S_50ntFlanks | 12992 | 13039 | 48 | TCCCTCGCTGCGATCTATTGAAAGTCAGCCCTCGACACAAGGGTTTGT | MB231 | 34 | 78 | 37 | 0 | 0 | 19.1 | 131.0 | 17.0 | 0.0 | 0.0 |
| >hg38_21_+_8433172_8446622_RNA45SN1_45S_50ntFlanks@12992.13039.48\|TCCCTCGCTGCGATCTATTGAAAGTCAGCCCTCGACACAAGGGTTTGT | >hg38_21_+_8433172_8446622_RNA45SN1_45S_50ntFlanks | 12992 | 13039 | 48 | TCCCTCGCTGCGATCTATTGAAAGTCAGCCCTCGACACAAGGGTTTGT | MB468 | 25 | 181 | 40 | 4 | 11 | 19.0 | 290.5 | 32.0 | 7.0 | 17.2 |
| >hg38_21_+_8433172_8446622_RNA45SN1_45S_50ntFlanks@12993.13012.20\|CCCTCGCTGCGATCTATTGA | >hg38_21_+_8433172_8446622_RNA45SN1_45S_50ntFlanks | 12993 | 13012 | 20 | CCCTCGCTGCGATCTATTGA | MB468 | 13 | 35 | 14 | 1 | 6 | 2.5 | 10.9 | 6.8 | 0.6 | 1.3 |
| >hg38_21_+_8433172_8446622_RNA45SN1_45S_50ntFlanks@12993.13013.21\|CCCTCGCTGCGATCTATTGAA | >hg38_21_+_8433172_8446622_RNA45SN1_45S_50ntFlanks | 12993 | 13013 | 21 | CCCTCGCTGCGATCTATTGAA | MB468 | 38 | 123 | 34 | 6 | 23 | 9.6 | 50.1 | 21.5 | 2.4 | 9.8 |
| >hg38_21_+_8433172_8446622_RNA45SN1_45S_50ntFlanks@12993.13015.23\|CCCTCGCTGCGATCTATTGAAAG | >hg38_21_+_8433172_8446622_RNA45SN1_45S_50ntFlanks | 12993 | 13015 | 23 | CCCTCGCTGCGATCTATTGAAAG | MB468 | 13 | 47 | 11 | 2 | 14 | 3.2 | 16.2 | 4.9 | 1.3 | 6.4 |
| >hg38_21_+_8433172_8446622_RNA45SN1_45S_50ntFlanks@12993.13039.47\|CCCTCGCTGCGATCTATTGAAAGTCAGCCCTCGACACAAGGGTTTGT | >hg38_21_+_8433172_8446622_RNA45SN1_45S_50ntFlanks | 12993 | 13039 | 47 | CCCTCGCTGCGATCTATTGAAAGTCAGCCCTCGACACAAGGGTTTGT | BT20 | 30 | 12 | 51 | 3 | 331 | 23.2 | 9.8 | 66.5 | 0.1 | 333.2 |
| >hg38_21_+_8433172_8446622_RNA45SN1_45S_50ntFlanks@12993.13039.47\|CCCTCGCTGCGATCTATTGAAAGTCAGCCCTCGACACAAGGGTTTGT | >hg38_21_+_8433172_8446622_RNA45SN1_45S_50ntFlanks | 12993 | 13039 | 47 | CCCTCGCTGCGATCTATTGAAAGTCAGCCCTCGACACAAGGGTTTGT | MB231 | 174 | 251 | 215 | 1 | 149 | 111.1 | 405.6 | 82.4 | 1.0 | 258.5 |
| >hg38_21_+_8433172_8446622_RNA45SN1_45S_50ntFlanks@12993.13039.47\|CCCTCGCTGCGATCTATTGAAAGTCAGCCCTCGACACAAGGGTTTGT | >hg38_21_+_8433172_8446622_RNA45SN1_45S_50ntFlanks | 12993 | 13039 | 47 | CCCTCGCTGCGATCTATTGAAAGTCAGCCCTCGACACAAGGGTTTGT | MB468 | 15 | 146 | 23 | 2 | 12 | 10.2 | 228.6 | 19.0 | 4.2 | 19.3 |
| >hg38_21_+_8433172_8446622_RNA45SN1_45S_50ntFlanks@12994.13012.19\|CCTCGCTGCGATCTATTGA | >hg38_21_+_8433172_8446622_RNA45SN1_45S_50ntFlanks | 12994 | 13012 | 19 | CCTCGCTGCGATCTATTGA | MB468 | 23 | 45 | 28 | 3 | 10 | 3.1 | 11.4 | 11.2 | 1.1 | 4.5 |
| >hg38_21_+_8433172_8446622_RNA45SN1_45S_50ntFlanks@12994.13013.20\|CCTCGCTGCGATCTATTGAA | >hg38_21_+_8433172_8446622_RNA45SN1_45S_50ntFlanks | 12994 | 13013 | 20 | CCTCGCTGCGATCTATTGAA | MB468 | 82 | 197 | 90 | 16 | 57 | 25.1 | 61.9 | 60.4 | 7.8 | 35.5 |
| >hg38_21_+_8433172_8446622_RNA45SN1_45S_50ntFlanks@12994.13014.21\|CCTCGCTGCGATCTATTGAAA | >hg38_21_+_8433172_8446622_RNA45SN1_45S_50ntFlanks | 12994 | 13014 | 21 | CCTCGCTGCGATCTATTGAAA | MB468 | 21 | 68 | 21 | 4 | 12 | 5.8 | 24.3 | 13.0 | 1.8 | 6.3 |
| >hg38_21_+_8433172_8446622_RNA45SN1_45S_50ntFlanks@12994.13015.22\|CCTCGCTGCGATCTATTGAAAG | >hg38_21_+_8433172_8446622_RNA45SN1_45S_50ntFlanks | 12994 | 13015 | 22 | CCTCGCTGCGATCTATTGAAAG | MB468 | 29 | 77 | 28 | 5 | 20 | 9.2 | 34.1 | 16.5 | 2.1 | 10.6 |
| >hg38_21_+_8433172_8446622_RNA45SN1_45S_50ntFlanks@12994.13036.43\|CCTCGCTGCGATCTATTGAAAGTCAGCCCTCGACACAAGGGTT | >hg38_21_+_8433172_8446622_RNA45SN1_45S_50ntFlanks | 12994 | 13036 | 43 | CCTCGCTGCGATCTATTGAAAGTCAGCCCTCGACACAAGGGTT | MB231 | 10 | 8 | 19 | 0 | 0 | 1.3 | 8.1 | 6.4 | 0.0 | 0.0 |
| >hg38_21_+_8433172_8446622_RNA45SN1_45S_50ntFlanks@12994.13038.45\|CCTCGCTGCGATCTATTGAAAGTCAGCCCTCGACACAAGGGTTTG | >hg38_21_+_8433172_8446622_RNA45SN1_45S_50ntFlanks | 12994 | 13038 | 45 | CCTCGCTGCGATCTATTGAAAGTCAGCCCTCGACACAAGGGTTTG | MB231 | 21 | 6 | 27 | 0 | 0 | 14.9 | 3.6 | 19.6 | 0.0 | 0.0 |
| >hg38_21_+_8433172_8446622_RNA45SN1_45S_50ntFlanks@12994.13039.46\|CCTCGCTGCGATCTATTGAAAGTCAGCCCTCGACACAAGGGTTTGT | >hg38_21_+_8433172_8446622_RNA45SN1_45S_50ntFlanks | 12994 | 13039 | 46 | CCTCGCTGCGATCTATTGAAAGTCAGCCCTCGACACAAGGGTTTGT | BT20 | 86 | 25 | 210 | 19 | 676 | 72.5 | 20.2 | 290.0 | 0.8 | 712.3 |
| >hg38_21_+_8433172_8446622_RNA45SN1_45S_50ntFlanks@12994.13039.46\|CCTCGCTGCGATCTATTGAAAGTCAGCCCTCGACACAAGGGTTTGT | >hg38_21_+_8433172_8446622_RNA45SN1_45S_50ntFlanks | 12994 | 13039 | 46 | CCTCGCTGCGATCTATTGAAAGTCAGCCCTCGACACAAGGGTTTGT | MB231 | 1653 | 755 | 1712 | 8 | 747 | 1763.9 | 790.7 | 458.4 | 5.0 | 269.8 |
| >hg38_21_+_8433172_8446622_RNA45SN1_45S_50ntFlanks@12994.13039.46\|CCTCGCTGCGATCTATTGAAAGTCAGCCCTCGACACAAGGGTTTGT | >hg38_21_+_8433172_8446622_RNA45SN1_45S_50ntFlanks | 12994 | 13039 | 46 | CCTCGCTGCGATCTATTGAAAGTCAGCCCTCGACACAAGGGTTTGT | MB468 | 12 | 130 | 28 | 3 | 18 | 8.6 | 194.5 | 26.6 | 5.2 | 31.9 |
| >hg38_21_+_8433172_8446622_RNA45SN1_45S_50ntFlanks@12994.13040.47\|CCTCGCTGCGATCTATTGAAAGTCAGCCCTCGACACAAGGGTTTGTC | >hg38_21_+_8433172_8446622_RNA45SN1_45S_50ntFlanks | 12994 | 13040 | 47 | CCTCGCTGCGATCTATTGAAAGTCAGCCCTCGACACAAGGGTTTGTC | MB231 | 24 | 6 | 3 | 0 | 0 | 33.5 | 4.3 | 1.8 | 0.0 | 0.0 |
| >hg38_21_+_8433172_8446622_RNA45SN1_45S_50ntFlanks@12995.13012.18\|CTCGCTGCGATCTATTGA | >hg38_21_+_8433172_8446622_RNA45SN1_45S_50ntFlanks | 12995 | 13012 | 18 | CTCGCTGCGATCTATTGA | MB231 | 19 | 31 | 41 | 1 | 10 | 11.0 | 13.6 | 21.4 | 1.4 | 17.3 |
| >hg38_21_+_8433172_8446622_RNA45SN1_45S_50ntFlanks@12995.13012.18\|CTCGCTGCGATCTATTGA | >hg38_21_+_8433172_8446622_RNA45SN1_45S_50ntFlanks | 12995 | 13012 | 18 | CTCGCTGCGATCTATTGA | MB468 | 15 | 22 | 18 | 2 | 5 | 2.3 | 9.9 | 5.9 | 0.5 | 1.3 |
| >hg38_21_+_8433172_8446622_RNA45SN1_45S_50ntFlanks@12995.13013.19\|CTCGCTGCGATCTATTGAA | >hg38_21_+_8433172_8446622_RNA45SN1_45S_50ntFlanks | 12995 | 13013 | 19 | CTCGCTGCGATCTATTGAA | BT20 | 11 | 4 | 23 | 2 | 4 | 9.9 | 1.1 | 3.6 | 3.5 | 4.2 |
| >hg38_21_+_8433172_8446622_RNA45SN1_45S_50ntFlanks@12995.13013.19\|CTCGCTGCGATCTATTGAA | >hg38_21_+_8433172_8446622_RNA45SN1_45S_50ntFlanks | 12995 | 13013 | 19 | CTCGCTGCGATCTATTGAA | MB231 | 22 | 33 | 51 | 2 | 15 | 10.9 | 12.8 | 24.6 | 1.8 | 14.1 |
| >hg38_21_+_8433172_8446622_RNA45SN1_45S_50ntFlanks@12995.13013.19\|CTCGCTGCGATCTATTGAA | >hg38_21_+_8433172_8446622_RNA45SN1_45S_50ntFlanks | 12995 | 13013 | 19 | CTCGCTGCGATCTATTGAA | MB468 | 40 | 79 | 42 | 6 | 21 | 15.1 | 39.3 | 33.4 | 2.9 | 8.9 |
| >hg38_21_+_8433172_8446622_RNA45SN1_45S_50ntFlanks@12995.13014.20\|CTCGCTGCGATCTATTGAAA | >hg38_21_+_8433172_8446622_RNA45SN1_45S_50ntFlanks | 12995 | 13014 | 20 | CTCGCTGCGATCTATTGAAA | MB231 | 17 | 24 | 34 | 1 | 16 | 9.1 | 11.0 | 11.7 | 1.2 | 22.2 |
| >hg38_21_+_8433172_8446622_RNA45SN1_45S_50ntFlanks@12995.13014.20\|CTCGCTGCGATCTATTGAAA | >hg38_21_+_8433172_8446622_RNA45SN1_45S_50ntFlanks | 12995 | 13014 | 20 | CTCGCTGCGATCTATTGAAA | MB468 | 11 | 30 | 10 | 2 | 5 | 3.5 | 14.6 | 7.2 | 1.2 | 1.7 |
| >hg38_21_+_8433172_8446622_RNA45SN1_45S_50ntFlanks@12995.13015.21\|CTCGCTGCGATCTATTGAAAG | >hg38_21_+_8433172_8446622_RNA45SN1_45S_50ntFlanks | 12995 | 13015 | 21 | CTCGCTGCGATCTATTGAAAG | BT20 | 22 | 22 | 21 | 4 | 7 | 22.9 | 28.7 | 17.6 | 4.6 | 2.2 |
| >hg38_21_+_8433172_8446622_RNA45SN1_45S_50ntFlanks@12995.13015.21\|CTCGCTGCGATCTATTGAAAG | >hg38_21_+_8433172_8446622_RNA45SN1_45S_50ntFlanks | 12995 | 13015 | 21 | CTCGCTGCGATCTATTGAAAG | MB231 | 22 | 33 | 40 | 3 | 24 | 9.5 | 14.4 | 10.7 | 1.2 | 20.1 |
| >hg38_21_+_8433172_8446622_RNA45SN1_45S_50ntFlanks@12995.13015.21\|CTCGCTGCGATCTATTGAAAG | >hg38_21_+_8433172_8446622_RNA45SN1_45S_50ntFlanks | 12995 | 13015 | 21 | CTCGCTGCGATCTATTGAAAG | MB468 | 14 | 30 | 13 | 4 | 12 | 4.3 | 18.1 | 6.9 | 1.8 | 3.6 |
| >hg38_21_+_8433172_8446622_RNA45SN1_45S_50ntFlanks@12995.13016.22\|CTCGCTGCGATCTATTGAAAGT | >hg38_21_+_8433172_8446622_RNA45SN1_45S_50ntFlanks | 12995 | 13016 | 22 | CTCGCTGCGATCTATTGAAAGT | BT20 | 22 | 14 | 39 | 4 | 12 | 2.7 | 6.1 | 12.2 | 3.8 | 5.3 |
| >hg38_21_+_8433172_8446622_RNA45SN1_45S_50ntFlanks@12995.13016.22\|CTCGCTGCGATCTATTGAAAGT | >hg38_21_+_8433172_8446622_RNA45SN1_45S_50ntFlanks | 12995 | 13016 | 22 | CTCGCTGCGATCTATTGAAAGT | MB231 | 42 | 49 | 52 | 7 | 25 | 33.9 | 30.7 | 21.1 | 8.0 | 25.2 |
| >hg38_21_+_8433172_8446622_RNA45SN1_45S_50ntFlanks@12995.13017.23\|CTCGCTGCGATCTATTGAAAGTC | >hg38_21_+_8433172_8446622_RNA45SN1_45S_50ntFlanks | 12995 | 13017 | 23 | CTCGCTGCGATCTATTGAAAGTC | BT20 | 12 | 4 | 28 | 3 | 6 | 12.9 | 0.3 | 13.3 | 2.6 | 6.7 |
| >hg38_21_+_8433172_8446622_RNA45SN1_45S_50ntFlanks@12995.13017.23\|CTCGCTGCGATCTATTGAAAGTC | >hg38_21_+_8433172_8446622_RNA45SN1_45S_50ntFlanks | 12995 | 13017 | 23 | CTCGCTGCGATCTATTGAAAGTC | MB231 | 33 | 35 | 75 | 3 | 13 | 20.7 | 17.9 | 44.2 | 2.4 | 11.7 |
| >hg38_21_+_8433172_8446622_RNA45SN1_45S_50ntFlanks@12995.13035.41\|CTCGCTGCGATCTATTGAAAGTCAGCCCTCGACACAAGGGT | >hg38_21_+_8433172_8446622_RNA45SN1_45S_50ntFlanks | 12995 | 13035 | 41 | CTCGCTGCGATCTATTGAAAGTCAGCCCTCGACACAAGGGT | MB231 | 13 | 9 | 37 | 1 | 15 | 7.1 | 3.2 | 25.0 | 0.4 | 18.3 |
| >hg38_21_+_8433172_8446622_RNA45SN1_45S_50ntFlanks@12995.13036.42\|CTCGCTGCGATCTATTGAAAGTCAGCCCTCGACACAAGGGTT | >hg38_21_+_8433172_8446622_RNA45SN1_45S_50ntFlanks | 12995 | 13036 | 42 | CTCGCTGCGATCTATTGAAAGTCAGCCCTCGACACAAGGGTT | BT20 | 81 | 41 | 233 | 7 | 83 | 57.8 | 31.0 | 289.3 | 3.0 | 13.6 |
| >hg38_21_+_8433172_8446622_RNA45SN1_45S_50ntFlanks@12995.13036.42\|CTCGCTGCGATCTATTGAAAGTCAGCCCTCGACACAAGGGTT | >hg38_21_+_8433172_8446622_RNA45SN1_45S_50ntFlanks | 12995 | 13036 | 42 | CTCGCTGCGATCTATTGAAAGTCAGCCCTCGACACAAGGGTT | MB231 | 132 | 85 | 318 | 3 | 96 | 56.7 | 80.6 | 157.4 | 2.2 | 68.5 |
| >hg38_21_+_8433172_8446622_RNA45SN1_45S_50ntFlanks@12995.13037.43\|CTCGCTGCGATCTATTGAAAGTCAGCCCTCGACACAAGGGTTT | >hg38_21_+_8433172_8446622_RNA45SN1_45S_50ntFlanks | 12995 | 13037 | 43 | CTCGCTGCGATCTATTGAAAGTCAGCCCTCGACACAAGGGTTT | MB231 | 50 | 21 | 171 | 1 | 34 | 26.8 | 6.4 | 130.3 | 0.2 | 21.6 |
| >hg38_21_+_8433172_8446622_RNA45SN1_45S_50ntFlanks@12995.13038.44\|CTCGCTGCGATCTATTGAAAGTCAGCCCTCGACACAAGGGTTTG | >hg38_21_+_8433172_8446622_RNA45SN1_45S_50ntFlanks | 12995 | 13038 | 44 | CTCGCTGCGATCTATTGAAAGTCAGCCCTCGACACAAGGGTTTG | BT20 | 30 | 10 | 53 | 6 | 36 | 30.2 | 10.2 | 73.4 | 5.3 | 16.9 |
| >hg38_21_+_8433172_8446622_RNA45SN1_45S_50ntFlanks@12995.13038.44\|CTCGCTGCGATCTATTGAAAGTCAGCCCTCGACACAAGGGTTTG | >hg38_21_+_8433172_8446622_RNA45SN1_45S_50ntFlanks | 12995 | 13038 | 44 | CTCGCTGCGATCTATTGAAAGTCAGCCCTCGACACAAGGGTTTG | MB231 | 263 | 60 | 409 | 3 | 104 | 176.5 | 33.2 | 307.9 | 0.9 | 32.5 |
| >hg38_21_+_8433172_8446622_RNA45SN1_45S_50ntFlanks@12995.13039.45\|CTCGCTGCGATCTATTGAAAGTCAGCCCTCGACACAAGGGTTTGT | >hg38_21_+_8433172_8446622_RNA45SN1_45S_50ntFlanks | 12995 | 13039 | 45 | CTCGCTGCGATCTATTGAAAGTCAGCCCTCGACACAAGGGTTTGT | BT20 | 2574 | 1049 | 8053 | 357 | 6526 | 2020.9 | 1204.5 | 11794.7 | 204.3 | 4520.4 |
| >hg38_21_+_8433172_8446622_RNA45SN1_45S_50ntFlanks@12995.13039.45\|CTCGCTGCGATCTATTGAAAGTCAGCCCTCGACACAAGGGTTTGT | >hg38_21_+_8433172_8446622_RNA45SN1_45S_50ntFlanks | 12995 | 13039 | 45 | CTCGCTGCGATCTATTGAAAGTCAGCCCTCGACACAAGGGTTTGT | MB231 | 27983 | 14212 | 47443 | 254 | 13965 | 20072.8 | 16211.9 | 12201.0 | 163.1 | 7455.0 |
| >hg38_21_+_8433172_8446622_RNA45SN1_45S_50ntFlanks@12995.13039.45\|CTCGCTGCGATCTATTGAAAGTCAGCCCTCGACACAAGGGTTTGT | >hg38_21_+_8433172_8446622_RNA45SN1_45S_50ntFlanks | 12995 | 13039 | 45 | CTCGCTGCGATCTATTGAAAGTCAGCCCTCGACACAAGGGTTTGT | MB468 | 55 | 486 | 187 | 12 | 232 | 32.2 | 717.7 | 233.7 | 11.0 | 258.9 |
| >hg38_21_+_8433172_8446622_RNA45SN1_45S_50ntFlanks@12995.13040.46\|CTCGCTGCGATCTATTGAAAGTCAGCCCTCGACACAAGGGTTTGTC | >hg38_21_+_8433172_8446622_RNA45SN1_45S_50ntFlanks | 12995 | 13040 | 46 | CTCGCTGCGATCTATTGAAAGTCAGCCCTCGACACAAGGGTTTGTC | MB231 | 137 | 39 | 20 | 1 | 60 | 131.0 | 25.4 | 6.5 | 1.2 | 58.2 |
| >hg38_21_+_8433172_8446622_RNA45SN1_45S_50ntFlanks@12996.13013.18\|TCGCTGCGATCTATTGAA | >hg38_21_+_8433172_8446622_RNA45SN1_45S_50ntFlanks | 12996 | 13013 | 18 | TCGCTGCGATCTATTGAA | BT20 | 43 | 17 | 102 | 6 | 9 | 26.5 | 1.5 | 8.5 | 4.0 | 1.1 |
| >hg38_21_+_8433172_8446622_RNA45SN1_45S_50ntFlanks@12996.13013.18\|TCGCTGCGATCTATTGAA | >hg38_21_+_8433172_8446622_RNA45SN1_45S_50ntFlanks | 12996 | 13013 | 18 | TCGCTGCGATCTATTGAA | MB231 | 147 | 171 | 241 | 14 | 48 | 94.1 | 107.4 | 134.4 | 11.6 | 36.9 |
| >hg38_21_+_8433172_8446622_RNA45SN1_45S_50ntFlanks@12996.13013.18\|TCGCTGCGATCTATTGAA | >hg38_21_+_8433172_8446622_RNA45SN1_45S_50ntFlanks | 12996 | 13013 | 18 | TCGCTGCGATCTATTGAA | MB468 | 35 | 52 | 40 | 6 | 16 | 9.9 | 25.3 | 27.5 | 2.6 | 7.9 |

| | | | | | | | | | | | | | | | | |
|---|---|---|---|---|---|---|---|---|---|---|---|---|---|---|---|---|
| >hg38_21_+_8433172_8446622_RNA45SN1_45S_50ntFlanks@12996.13014.19\|TCGCTGCGATCTATTGAAA | >hg38_21_+_8433172_8446622_RNA45SN1_45S_50ntFlanks | 12996 | 13014 | 19 | TCGCTGCGATCTATTGAAA | BT20 | 25 | 11 | 53 | 3 | 7 | 14.6 | 2.3 | 7.3 | 1.4 | 0.8 |
| >hg38_21_+_8433172_8446622_RNA45SN1_45S_50ntFlanks@12996.13014.19\|TCGCTGCGATCTATTGAAA | >hg38_21_+_8433172_8446622_RNA45SN1_45S_50ntFlanks | 12996 | 13014 | 19 | TCGCTGCGATCTATTGAAA | MB231 | 95 | 97 | 133 | 9 | 34 | 75.4 | 59.8 | 76.3 | 8.1 | 32.5 |
| >hg38_21_+_8433172_8446622_RNA45SN1_45S_50ntFlanks@12996.13014.19\|TCGCTGCGATCTATTGAAA | >hg38_21_+_8433172_8446622_RNA45SN1_45S_50ntFlanks | 12996 | 13014 | 19 | TCGCTGCGATCTATTGAAA | MB468 | 10 | 22 | 10 | 2 | 4 | 1.4 | 11.2 | 5.1 | 0.7 | 1.2 |
| >hg38_21_+_8433172_8446622_RNA45SN1_45S_50ntFlanks@12996.13015.20\|TCGCTGCGATCTATTGAAAG | >hg38_21_+_8433172_8446622_RNA45SN1_45S_50ntFlanks | 12996 | 13015 | 20 | TCGCTGCGATCTATTGAAAG | BT20 | 20 | 16 | 19 | 5 | 7 | 16.7 | 15.7 | 10.8 | 2.1 | 1.0 |
| >hg38_21_+_8433172_8446622_RNA45SN1_45S_50ntFlanks@12996.13015.20\|TCGCTGCGATCTATTGAAAG | >hg38_21_+_8433172_8446622_RNA45SN1_45S_50ntFlanks | 12996 | 13015 | 20 | TCGCTGCGATCTATTGAAAG | MB231 | 33 | 37 | 42 | 8 | 22 | 23.8 | 23.6 | 29.7 | 3.4 | 10.4 |
| >hg38_21_+_8433172_8446622_RNA45SN1_45S_50ntFlanks@12996.13016.21\|TCGCTGCGATCTATTGAAAGT | >hg38_21_+_8433172_8446622_RNA45SN1_45S_50ntFlanks | 12996 | 13016 | 21 | TCGCTGCGATCTATTGAAAGT | BT20 | 13 | 8 | 15 | 2 | 5 | 1.6 | 2.8 | 3.0 | 0.7 | 1.6 |
| >hg38_21_+_8433172_8446622_RNA45SN1_45S_50ntFlanks@12996.13016.21\|TCGCTGCGATCTATTGAAAGT | >hg38_21_+_8433172_8446622_RNA45SN1_45S_50ntFlanks | 12996 | 13016 | 21 | TCGCTGCGATCTATTGAAAGT | MB231 | 31 | 25 | 29 | 7 | 11 | 29.4 | 19.7 | 25.7 | 8.0 | 8.0 |
| >hg38_21_+_8433172_8446622_RNA45SN1_45S_50ntFlanks@12996.13017.22\|TCGCTGCGATCTATTGAAAGTC | >hg38_21_+_8433172_8446622_RNA45SN1_45S_50ntFlanks | 12996 | 13017 | 22 | TCGCTGCGATCTATTGAAAGTC | BT20 | 28 | 11 | 62 | 3 | 8 | 30.3 | 5.4 | 52.5 | 1.1 | 4.1 |
| >hg38_21_+_8433172_8446622_RNA45SN1_45S_50ntFlanks@12996.13017.22\|TCGCTGCGATCTATTGAAAGTC | >hg38_21_+_8433172_8446622_RNA45SN1_45S_50ntFlanks | 12996 | 13017 | 22 | TCGCTGCGATCTATTGAAAGTC | MB231 | 84 | 72 | 127 | 10 | 22 | 63.9 | 47.5 | 99.2 | 8.6 | 15.3 |
| >hg38_21_+_8433172_8446622_RNA45SN1_45S_50ntFlanks@12996.13036.41\|TCGCTGCGATCTATTGAAAGTCAGCCCTCGACACAAGGGTT | >hg38_21_+_8433172_8446622_RNA45SN1_45S_50ntFlanks | 12996 | 13036 | 41 | TCGCTGCGATCTATTGAAAGTCAGCCCTCGACACAAGGGTT | BT20 | 82 | 56 | 171 | 7 | 90 | 51.9 | 22.9 | 187.4 | 3.2 | 37.8 |
| >hg38_21_+_8433172_8446622_RNA45SN1_45S_50ntFlanks@12996.13036.41\|TCGCTGCGATCTATTGAAAGTCAGCCCTCGACACAAGGGTT | >hg38_21_+_8433172_8446622_RNA45SN1_45S_50ntFlanks | 12996 | 13036 | 41 | TCGCTGCGATCTATTGAAAGTCAGCCCTCGACACAAGGGTT | MB231 | 118 | 96 | 182 | 5 | 88 | 108.0 | 130.1 | 80.4 | 4.5 | 99.2 |
| >hg38_21_+_8433172_8446622_RNA45SN1_45S_50ntFlanks@12996.13037.42\|TCGCTGCGATCTATTGAAAGTCAGCCCTCGACACAAGGGTTT | >hg38_21_+_8433172_8446622_RNA45SN1_45S_50ntFlanks | 12996 | 13037 | 42 | TCGCTGCGATCTATTGAAAGTCAGCCCTCGACACAAGGGTTT | MB231 | 19 | 11 | 60 | 1 | 16 | 6.4 | 8.2 | 49.7 | 0.0 | 9.1 |
| >hg38_21_+_8433172_8446622_RNA45SN1_45S_50ntFlanks@12996.13038.43\|TCGCTGCGATCTATTGAAAGTCAGCCCTCGACACAAGGGTTTG | >hg38_21_+_8433172_8446622_RNA45SN1_45S_50ntFlanks | 12996 | 13038 | 43 | TCGCTGCGATCTATTGAAAGTCAGCCCTCGACACAAGGGTTTG | BT20 | 25 | 10 | 41 | 6 | 19 | 25.3 | 7.4 | 56.2 | 5.6 | 23.4 |
| >hg38_21_+_8433172_8446622_RNA45SN1_45S_50ntFlanks@12996.13038.43\|TCGCTGCGATCTATTGAAAGTCAGCCCTCGACACAAGGGTTTG | >hg38_21_+_8433172_8446622_RNA45SN1_45S_50ntFlanks | 12996 | 13038 | 43 | TCGCTGCGATCTATTGAAAGTCAGCCCTCGACACAAGGGTTTG | MB231 | 135 | 48 | 193 | 5 | 80 | 62.9 | 45.3 | 142.5 | 3.5 | 66.7 |
| >hg38_21_+_8433172_8446622_RNA45SN1_45S_50ntFlanks@12996.13039.44\|TCGCTGCGATCTATTGAAAGTCAGCCCTCGACACAAGGGTTTGT | >hg38_21_+_8433172_8446622_RNA45SN1_45S_50ntFlanks | 12996 | 13039 | 44 | TCGCTGCGATCTATTGAAAGTCAGCCCTCGACACAAGGGTTTGT | BT20 | 1742 | 998 | 4233 | 370 | 4573 | 1261.7 | 836.0 | 5891.9 | 226.4 | 2741.9 |
| >hg38_21_+_8433172_8446622_RNA45SN1_45S_50ntFlanks@12996.13039.44\|TCGCTGCGATCTATTGAAAGTCAGCCCTCGACACAAGGGTTTGT | >hg38_21_+_8433172_8446622_RNA45SN1_45S_50ntFlanks | 12996 | 13039 | 44 | TCGCTGCGATCTATTGAAAGTCAGCCCTCGACACAAGGGTTTGT | MB231 | 12367 | 9985 | 15016 | 371 | 9919 | 5105.5 | 13741.6 | 2367.1 | 281.0 | 9661.3 |
| >hg38_21_+_8433172_8446622_RNA45SN1_45S_50ntFlanks@12996.13039.44\|TCGCTGCGATCTATTGAAAGTCAGCCCTCGACACAAGGGTTTGT | >hg38_21_+_8433172_8446622_RNA45SN1_45S_50ntFlanks | 12996 | 13039 | 44 | TCGCTGCGATCTATTGAAAGTCAGCCCTCGACACAAGGGTTTGT | MB468 | 63 | 349 | 158 | 12 | 242 | 36.5 | 506.2 | 171.4 | 14.1 | 265.1 |
| >hg38_21_+_8433172_8446622_RNA45SN1_45S_50ntFlanks@12996.13040.45\|TCGCTGCGATCTATTGAAAGTCAGCCCTCGACACAAGGGTTTGTC | >hg38_21_+_8433172_8446622_RNA45SN1_45S_50ntFlanks | 12996 | 13040 | 45 | TCGCTGCGATCTATTGAAAGTCAGCCCTCGACACAAGGGTTTGTC | MB231 | 42 | 20 | 6 | 2 | 30 | 20.1 | 21.2 | 2.1 | 2.0 | 25.9 |
| >hg38_21_+_8433172_8446622_RNA45SN1_45S_50ntFlanks@12997.13014.18\|CGCTGCGATCTATTGAAA | >hg38_21_+_8433172_8446622_RNA45SN1_45S_50ntFlanks | 12997 | 13014 | 18 | CGCTGCGATCTATTGAAA | BT20 | 58 | 18 | 102 | 6 | 10 | 32.6 | 7.9 | 10.3 | 3.8 | 2.8 |
| >hg38_21_+_8433172_8446622_RNA45SN1_45S_50ntFlanks@12997.13014.18\|CGCTGCGATCTATTGAAA | >hg38_21_+_8433172_8446622_RNA45SN1_45S_50ntFlanks | 12997 | 13014 | 18 | CGCTGCGATCTATTGAAA | MB231 | 240 | 205 | 256 | 18 | 74 | 203.7 | 119.4 | 143.5 | 14.7 | 69.6 |
| >hg38_21_+_8433172_8446622_RNA45SN1_45S_50ntFlanks@12997.13014.18\|CGCTGCGATCTATTGAAA | >hg38_21_+_8433172_8446622_RNA45SN1_45S_50ntFlanks | 12997 | 13014 | 18 | CGCTGCGATCTATTGAAA | MB468 | 28 | 53 | 25 | 6 | 17 | 8.1 | 29.6 | 13.3 | 2.9 | 11.0 |
| >hg38_21_+_8433172_8446622_RNA45SN1_45S_50ntFlanks@12997.13015.19\|CGCTGCGATCTATTGAAAG | >hg38_21_+_8433172_8446622_RNA45SN1_45S_50ntFlanks | 12997 | 13015 | 19 | CGCTGCGATCTATTGAAAG | BT20 | 21 | 18 | 21 | 5 | 6 | 12.0 | 17.6 | 11.4 | 2.9 | 1.2 |
| >hg38_21_+_8433172_8446622_RNA45SN1_45S_50ntFlanks@12997.13015.19\|CGCTGCGATCTATTGAAAG | >hg38_21_+_8433172_8446622_RNA45SN1_45S_50ntFlanks | 12997 | 13015 | 19 | CGCTGCGATCTATTGAAAG | MB231 | 49 | 49 | 46 | 14 | 26 | 35.8 | 28.5 | 34.9 | 11.1 | 17.4 |
| >hg38_21_+_8433172_8446622_RNA45SN1_45S_50ntFlanks@12997.13015.19\|CGCTGCGATCTATTGAAAG | >hg38_21_+_8433172_8446622_RNA45SN1_45S_50ntFlanks | 12997 | 13015 | 19 | CGCTGCGATCTATTGAAAG | MB468 | 14 | 14 | 11 | 6 | 12 | 3.3 | 6.6 | 5.8 | 2.4 | 4.9 |
| >hg38_21_+_8433172_8446622_RNA45SN1_45S_50ntFlanks@12997.13016.20\|CGCTGCGATCTATTGAAAGT | >hg38_21_+_8433172_8446622_RNA45SN1_45S_50ntFlanks | 12997 | 13016 | 20 | CGCTGCGATCTATTGAAAGT | BT20 | 10 | 6 | 11 | 2 | 2 | 1.3 | 2.0 | 3.2 | 2.0 | 2.0 |
| >hg38_21_+_8433172_8446622_RNA45SN1_45S_50ntFlanks@12997.13016.20\|CGCTGCGATCTATTGAAAGT | >hg38_21_+_8433172_8446622_RNA45SN1_45S_50ntFlanks | 12997 | 13016 | 20 | CGCTGCGATCTATTGAAAGT | MB231 | 36 | 32 | 24 | 15 | 12 | 35.0 | 23.6 | 21.8 | 15.5 | 8.7 |
| >hg38_21_+_8433172_8446622_RNA45SN1_45S_50ntFlanks@12997.13017.21\|CGCTGCGATCTATTGAAAGTC | >hg38_21_+_8433172_8446622_RNA45SN1_45S_50ntFlanks | 12997 | 13017 | 21 | CGCTGCGATCTATTGAAAGTC | BT20 | 46 | 17 | 85 | 5 | 11 | 52.2 | 10.7 | 84.8 | 1.7 | 7.0 |
| >hg38_21_+_8433172_8446622_RNA45SN1_45S_50ntFlanks@12997.13017.21\|CGCTGCGATCTATTGAAAGTC | >hg38_21_+_8433172_8446622_RNA45SN1_45S_50ntFlanks | 12997 | 13017 | 21 | CGCTGCGATCTATTGAAAGTC | MB231 | 122 | 102 | 142 | 20 | 31 | 89.0 | 67.0 | 115.4 | 19.8 | 22.7 |
| >hg38_21_+_8433172_8446622_RNA45SN1_45S_50ntFlanks@12997.13035.39\|CGCTGCGATCTATTGAAAGTCAGCCCTCGACACAAGGGT | >hg38_21_+_8433172_8446622_RNA45SN1_45S_50ntFlanks | 12997 | 13035 | 39 | CGCTGCGATCTATTGAAAGTCAGCCCTCGACACAAGGGT | MB231 | 17 | 13 | 40 | 4 | 21 | 8.7 | 8.2 | 37.0 | 3.4 | 23.9 |
| >hg38_21_+_8433172_8446622_RNA45SN1_45S_50ntFlanks@12997.13036.40\|CGCTGCGATCTATTGAAAGTCAGCCCTCGACACAAGGGTT | >hg38_21_+_8433172_8446622_RNA45SN1_45S_50ntFlanks | 12997 | 13036 | 40 | CGCTGCGATCTATTGAAAGTCAGCCCTCGACACAAGGGTT | BT20 | 79 | 51 | 150 | 7 | 78 | 62.4 | 7.1 | 142.3 | 4.7 | 35.3 |
| >hg38_21_+_8433172_8446622_RNA45SN1_45S_50ntFlanks@12997.13036.40\|CGCTGCGATCTATTGAAAGTCAGCCCTCGACACAAGGGTT | >hg38_21_+_8433172_8446622_RNA45SN1_45S_50ntFlanks | 12997 | 13036 | 40 | CGCTGCGATCTATTGAAAGTCAGCCCTCGACACAAGGGTT | MB231 | 146 | 93 | 173 | 8 | 123 | 116.6 | 96.5 | 81.8 | 5.0 | 114.4 |
| >hg38_21_+_8433172_8446622_RNA45SN1_45S_50ntFlanks@12997.13037.41\|CGCTGCGATCTATTGAAAGTCAGCCCTCGACACAAGGGTTT | >hg38_21_+_8433172_8446622_RNA45SN1_45S_50ntFlanks | 12997 | 13037 | 41 | CGCTGCGATCTATTGAAAGTCAGCCCTCGACACAAGGGTTT | MB231 | 30 | 16 | 77 | 3 | 29 | 14.5 | 5.8 | 76.4 | 1.0 | 18.0 |
| >hg38_21_+_8433172_8446622_RNA45SN1_45S_50ntFlanks@12997.13038.42\|CGCTGCGATCTATTGAAAGTCAGCCCTCGACACAAGGGTTTG | >hg38_21_+_8433172_8446622_RNA45SN1_45S_50ntFlanks | 12997 | 13038 | 42 | CGCTGCGATCTATTGAAAGTCAGCCCTCGACACAAGGGTTTG | BT20 | 24 | 11 | 37 | 8 | 25 | 16.8 | 5.3 | 45.7 | 6.8 | 12.8 |
| >hg38_21_+_8433172_8446622_RNA45SN1_45S_50ntFlanks@12997.13038.42\|CGCTGCGATCTATTGAAAGTCAGCCCTCGACACAAGGGTTTG | >hg38_21_+_8433172_8446622_RNA45SN1_45S_50ntFlanks | 12997 | 13038 | 42 | CGCTGCGATCTATTGAAAGTCAGCCCTCGACACAAGGGTTTG | MB231 | 220 | 67 | 253 | 10 | 130 | 150.0 | 37.6 | 235.0 | 1.6 | 79.4 |
| >hg38_21_+_8433172_8446622_RNA45SN1_45S_50ntFlanks@12997.13039.43\|CGCTGCGATCTATTGAAAGTCAGCCCTCGACACAAGGGTTTGT | >hg38_21_+_8433172_8446622_RNA45SN1_45S_50ntFlanks | 12997 | 13039 | 43 | CGCTGCGATCTATTGAAAGTCAGCCCTCGACACAAGGGTTTGT | BT20 | 1747 | 1021 | 4134 | 441 | 3690 | 1371.4 | 648.1 | 5368.1 | 324.4 | 1803.9 |
| >hg38_21_+_8433172_8446622_RNA45SN1_45S_50ntFlanks@12997.13039.43\|CGCTGCGATCTATTGAAAGTCAGCCCTCGACACAAGGGTTTGT | >hg38_21_+_8433172_8446622_RNA45SN1_45S_50ntFlanks | 12997 | 13039 | 43 | CGCTGCGATCTATTGAAAGTCAGCCCTCGACACAAGGGTTTGT | MB231 | 15787 | 10416 | 16542 | 846 | 14771 | 4251.0 | 10701.8 | 4816.0 | 493.5 | 11883.9 |
| >hg38_21_+_8433172_8446622_RNA45SN1_45S_50ntFlanks@12997.13039.43\|CGCTGCGATCTATTGAAAGTCAGCCCTCGACACAAGGGTTTGT | >hg38_21_+_8433172_8446622_RNA45SN1_45S_50ntFlanks | 12997 | 13039 | 43 | CGCTGCGATCTATTGAAAGTCAGCCCTCGACACAAGGGTTTGT | MB468 | 47 | 192 | 86 | 11 | 208 | 28.4 | 278.8 | 63.5 | 9.4 | 215.8 |
| >hg38_21_+_8433172_8446622_RNA45SN1_45S_50ntFlanks@12997.13040.44\|CGCTGCGATCTATTGAAAGTCAGCCCTCGACACAAGGGTTTGTC | >hg38_21_+_8433172_8446622_RNA45SN1_45S_50ntFlanks | 12997 | 13040 | 44 | CGCTGCGATCTATTGAAAGTCAGCCCTCGACACAAGGGTTTGTC | MB231 | 29 | 12 | 4 | 3 | 26 | 10.9 | 6.6 | 1.2 | 2.0 | 14.2 |
| >hg38_21_+_8433172_8446622_RNA45SN1_45S_50ntFlanks@12998.13017.20\|GCTGCGATCTATTGAAAGTC | >hg38_21_+_8433172_8446622_RNA45SN1_45S_50ntFlanks | 12998 | 13017 | 20 | GCTGCGATCTATTGAAAGTC | MB231 | 16 | 16 | 13 | 4 | 4 | 11.6 | 10.8 | 9.6 | 3.4 | 3.4 |
| >hg38_21_+_8433172_8446622_RNA45SN1_45S_50ntFlanks@12998.13036.39\|GCTGCGATCTATTGAAAGTCAGCCCTCGACACAAGGGTT | >hg38_21_+_8433172_8446622_RNA45SN1_45S_50ntFlanks | 12998 | 13036 | 39 | GCTGCGATCTATTGAAAGTCAGCCCTCGACACAAGGGTT | BT20 | 10 | 6 | 9 | 3 | 6 | 7.1 | 2.5 | 7.4 | 2.5 | 4.4 |
| >hg38_21_+_8433172_8446622_RNA45SN1_45S_50ntFlanks@12998.13039.42\|GCTGCGATCTATTGAAAGTCAGCCCTCGACACAAGGGTTTGT | >hg38_21_+_8433172_8446622_RNA45SN1_45S_50ntFlanks | 12998 | 13039 | 42 | GCTGCGATCTATTGAAAGTCAGCCCTCGACACAAGGGTTTGT | BT20 | 283 | 159 | 210 | 123 | 274 | 224.8 | 114.6 | 224.3 | 106.3 | 69.2 |
| >hg38_21_+_8433172_8446622_RNA45SN1_45S_50ntFlanks@12998.13039.42\|GCTGCGATCTATTGAAAGTCAGCCCTCGACACAAGGGTTTGT | >hg38_21_+_8433172_8446622_RNA45SN1_45S_50ntFlanks | 12998 | 13039 | 42 | GCTGCGATCTATTGAAAGTCAGCCCTCGACACAAGGGTTTGT | MB231 | 471 | 321 | 287 | 104 | 493 | 141.9 | 255.1 | 168.2 | 69.3 | 398.5 |
| >hg38_21_+_8433172_8446622_RNA45SN1_45S_50ntFlanks@12999.13027.29\|CTGCGATCTATTGAAAGTCAGCCCTCGAC | >hg38_21_+_8433172_8446622_RNA45SN1_45S_50ntFlanks | 12999 | 13027 | 29 | CTGCGATCTATTGAAAGTCAGCCCTCGAC | MB231 | 15 | 25 | 29 | 1 | 6 | 15.0 | 19.3 | 26.0 | 0.5 | 4.9 |
| >hg38_21_+_8433172_8446622_RNA45SN1_45S_50ntFlanks@12999.13027.29\|CTGCGATCTATTGAAAGTCAGCCCTCGAC | >hg38_21_+_8433172_8446622_RNA45SN1_45S_50ntFlanks | 12999 | 13027 | 29 | CTGCGATCTATTGAAAGTCAGCCCTCGAC | MB468 | 48 | 89 | 52 | 11 | 37 | 15.9 | 20.4 | 28.7 | 4.8 | 6.2 |
| >hg38_21_+_8433172_8446622_RNA45SN1_45S_50ntFlanks@12999.13028.30\|CTGCGATCTATTGAAAGTCAGCCCTCGACA | >hg38_21_+_8433172_8446622_RNA45SN1_45S_50ntFlanks | 12999 | 13028 | 30 | CTGCGATCTATTGAAAGTCAGCCCTCGACA | MB468 | 30 | 40 | 29 | 5 | 21 | 10.5 | 16.3 | 11.8 | 1.5 | 3.1 |
| >hg38_21_+_8433172_8446622_RNA45SN1_45S_50ntFlanks@12999.13039.41\|CTGCGATCTATTGAAAGTCAGCCCTCGACACAAGGGTTTGT | >hg38_21_+_8433172_8446622_RNA45SN1_45S_50ntFlanks | 12999 | 13039 | 41 | CTGCGATCTATTGAAAGTCAGCCCTCGACACAAGGGTTTGT | BT20 | 83 | 66 | 144 | 53 | 221 | 43.7 | 6.1 | 130.7 | 27.4 | 113.7 |
| >hg38_21_+_8433172_8446622_RNA45SN1_45S_50ntFlanks@12999.13039.41\|CTGCGATCTATTGAAAGTCAGCCCTCGACACAAGGGTTTGT | >hg38_21_+_8433172_8446622_RNA45SN1_45S_50ntFlanks | 12999 | 13039 | 41 | CTGCGATCTATTGAAAGTCAGCCCTCGACACAAGGGTTTGT | MB231 | 507 | 458 | 484 | 59 | 475 | 227.6 | 444.5 | 111.5 | 51.8 | 434.7 |
| >hg38_21_+_8433172_8446622_RNA45SN1_45S_50ntFlanks@12999.13039.41\|CTGCGATCTATTGAAAGTCAGCCCTCGACACAAGGGTTTGT | >hg38_21_+_8433172_8446622_RNA45SN1_45S_50ntFlanks | 12999 | 13039 | 41 | CTGCGATCTATTGAAAGTCAGCCCTCGACACAAGGGTTTGT | MB468 | 14 | 34 | 27 | 6 | 55 | 7.7 | 46.9 | 19.1 | 6.3 | 51.8 |
| >hg38_21_+_8433172_8446622_RNA45SN1_45S_50ntFlanks@13000.13027.28\|TGCGATCTATTGAAAGTCAGCCCTCGAC | >hg38_21_+_8433172_8446622_RNA45SN1_45S_50ntFlanks | 13000 | 13027 | 28 | TGCGATCTATTGAAAGTCAGCCCTCGAC | MB231 | 20 | 39 | 16 | 0 | 9 | 19.7 | 31.7 | 12.8 | 0.3 | 14.9 |
| >hg38_21_+_8433172_8446622_RNA45SN1_45S_50ntFlanks@13000.13027.28\|TGCGATCTATTGAAAGTCAGCCCTCGAC | >hg38_21_+_8433172_8446622_RNA45SN1_45S_50ntFlanks | 13000 | 13027 | 28 | TGCGATCTATTGAAAGTCAGCCCTCGAC | MB468 | 32 | 58 | 41 | 3 | 18 | 13.6 | 22.1 | 25.9 | 1.7 | 3.3 |

| | | | | | | | | | | | | | | | | |
|---|---|---|---|---|---|---|---|---|---|---|---|---|---|---|---|---|
| >hg38_21_+_8433172_8446622_RNA45SN1_45S_50ntFlanks@13000.13028.29\|TGCGATCTATTGAAAGTCAGCCCTCGACA | >hg38_21_+_8433172_8446622_RNA45SN1_45S_50ntFlanks | 13000 | 13028 | 29 | TGCGATCTATTGAAAGTCAGCCCTCGACA | MB468 | 14 | 12 | 14 | 1 | 7 | 6.5 | 4.8 | 8.5 | 0.2 | 0.8 |
| >hg38_21_+_8433172_8446622_RNA45SN1_45S_50ntFlanks@13000.13039.40\|TGCGATCTATTGAAAGTCAGCCCTCGACACAAGGGTTTGT | >hg38_21_+_8433172_8446622_RNA45SN1_45S_50ntFlanks | 13000 | 13039 | 40 | TGCGATCTATTGAAAGTCAGCCCTCGACACAAGGGTTTGT | BT20 | 11 | 7 | 12 | 8 | 23 | 8.8 | 4.7 | 12.0 | 5.4 | 5.7 |
| >hg38_21_+_8433172_8446622_RNA45SN1_45S_50ntFlanks@13000.13039.40\|TGCGATCTATTGAAAGTCAGCCCTCGACACAAGGGTTTGT | >hg38_21_+_8433172_8446622_RNA45SN1_45S_50ntFlanks | 13000 | 13039 | 40 | TGCGATCTATTGAAAGTCAGCCCTCGACACAAGGGTTTGT | MB231 | 42 | 38 | 21 | 15 | 148 | 19.4 | 14.3 | 6.1 | 2.0 | 92.8 |
| >hg38_21_+_8433172_8446622_RNA45SN1_45S_50ntFlanks@13001.13027.27\|GCGATCTATTGAAAGTCAGCCCTCGAC | >hg38_21_+_8433172_8446622_RNA45SN1_45S_50ntFlanks | 13001 | 13027 | 27 | GCGATCTATTGAAAGTCAGCCCTCGAC | MB468 | 31 | 29 | 27 | 4 | 17 | 13.6 | 6.6 | 16.8 | 0.9 | 4.8 |
| >hg38_21_+_8433172_8446622_RNA45SN1_45S_50ntFlanks@13001.13028.28\|GCGATCTATTGAAAGTCAGCCCTCGACA | >hg38_21_+_8433172_8446622_RNA45SN1_45S_50ntFlanks | 13001 | 13028 | 28 | GCGATCTATTGAAAGTCAGCCCTCGACA | MB468 | 12 | 11 | 9 | 3 | 7 | 4.7 | 3.6 | 4.9 | 1.0 | 1.3 |
| >hg38_21_+_8433172_8446622_RNA45SN1_45S_50ntFlanks@13002.13020.19\|CGATCTATTGAAAGTCAGC | >hg38_21_+_8433172_8446622_RNA45SN1_45S_50ntFlanks | 13002 | 13020 | 19 | CGATCTATTGAAAGTCAGC | MB468 | 30 | 7 | 29 | 18 | 8 | 4.3 | 4.9 | 7.5 | 7.3 | 3.3 |
| >hg38_21_+_8433172_8446622_RNA45SN1_45S_50ntFlanks@13002.13023.22\|CGATCTATTGAAAGTCAGCCCT | >hg38_21_+_8433172_8446622_RNA45SN1_45S_50ntFlanks | 13002 | 13023 | 22 | CGATCTATTGAAAGTCAGCCCT | MB468 | 64 | 60 | 70 | 53 | 25 | 33.8 | 27.8 | 36.7 | 24.0 | 18.2 |
| >hg38_21_+_8433172_8446622_RNA45SN1_45S_50ntFlanks@13002.13025.24\|CGATCTATTGAAAGTCAGCCCTCG | >hg38_21_+_8433172_8446622_RNA45SN1_45S_50ntFlanks | 13002 | 13025 | 24 | CGATCTATTGAAAGTCAGCCCTCG | MB468 | 36 | 21 | 51 | 18 | 9 | 7.8 | 3.5 | 9.7 | 9.7 | 4.4 |
| >hg38_21_+_8433172_8446622_RNA45SN1_45S_50ntFlanks@13002.13026.25\|CGATCTATTGAAAGTCAGCCCTCGA | >hg38_21_+_8433172_8446622_RNA45SN1_45S_50ntFlanks | 13002 | 13026 | 25 | CGATCTATTGAAAGTCAGCCCTCGA | MB468 | 19 | 11 | 29 | 3 | 7 | 2.8 | 3.2 | 5.8 | 1.4 | 2.4 |
| >hg38_21_+_8433172_8446622_RNA45SN1_45S_50ntFlanks@13002.13027.26\|CGATCTATTGAAAGTCAGCCCTCGAC | >hg38_21_+_8433172_8446622_RNA45SN1_45S_50ntFlanks | 13002 | 13027 | 26 | CGATCTATTGAAAGTCAGCCCTCGAC | MB468 | 757 | 653 | 1049 | 101 | 353 | 155.4 | 67.9 | 309.0 | 15.9 | 108.4 |
| >hg38_21_+_8433172_8446622_RNA45SN1_45S_50ntFlanks@13002.13028.27\|CGATCTATTGAAAGTCAGCCCTCGACA | >hg38_21_+_8433172_8446622_RNA45SN1_45S_50ntFlanks | 13002 | 13028 | 27 | CGATCTATTGAAAGTCAGCCCTCGACA | MB468 | 388 | 302 | 467 | 75 | 209 | 45.5 | 54.7 | 97.6 | 26.2 | 70.8 |
| >hg38_21_+_8433172_8446622_RNA45SN1_45S_50ntFlanks@13002.13029.28\|CGATCTATTGAAAGTCAGCCCTCGACAC | >hg38_21_+_8433172_8446622_RNA45SN1_45S_50ntFlanks | 13002 | 13029 | 28 | CGATCTATTGAAAGTCAGCCCTCGACAC | MB468 | 220 | 201 | 264 | 20 | 164 | 22.3 | 45.8 | 45.7 | 2.9 | 25.5 |
| >hg38_21_+_8433172_8446622_RNA45SN1_45S_50ntFlanks@13002.13030.29\|CGATCTATTGAAAGTCAGCCCTCGACACA | >hg38_21_+_8433172_8446622_RNA45SN1_45S_50ntFlanks | 13002 | 13030 | 29 | CGATCTATTGAAAGTCAGCCCTCGACACA | MB468 | 16 | 17 | 20 | 4 | 17 | 7.7 | 6.1 | 9.0 | 2.0 | 7.6 |
| >hg38_21_+_8433172_8446622_RNA45SN1_45S_50ntFlanks@13002.13039.38\|CGATCTATTGAAAGTCAGCCCTCGACACAAGGGTTTGT | >hg38_21_+_8433172_8446622_RNA45SN1_45S_50ntFlanks | 13002 | 13039 | 38 | CGATCTATTGAAAGTCAGCCCTCGACACAAGGGTTTGT | MB231 | 13 | 19 | 8 | 3 | 31 | 22.9 | 33.0 | 14.0 | 5.5 | 53.3 |
| >hg38_21_+_8433172_8446622_RNA45SN1_45S_50ntFlanks@13002.13039.38\|CGATCTATTGAAAGTCAGCCCTCGACACAAGGGTTTGT | >hg38_21_+_8433172_8446622_RNA45SN1_45S_50ntFlanks | 13002 | 13039 | 38 | CGATCTATTGAAAGTCAGCCCTCGACACAAGGGTTTGT | MB468 | 141 | 97 | 218 | 59 | 421 | 47.0 | 92.8 | 200.1 | 42.1 | 366.2 |
| >hg38_21_+_8433172_8446622_RNA45SN1_45S_50ntFlanks@13003.13027.25\|GATCTATTGAAAGTCAGCCCTCGAC | >hg38_21_+_8433172_8446622_RNA45SN1_45S_50ntFlanks | 13003 | 13027 | 25 | GATCTATTGAAAGTCAGCCCTCGAC | MB468 | 14 | 17 | 16 | 0 | 3 | 6.1 | 7.4 | 7.5 | 0.2 | 3.9 |
| >hg38_21_+_8433172_8446622_RNA45SN1_45S_50ntFlanks@13004.13023.20\|ATCTATTGAAAGTCAGCCCT | >hg38_21_+_8433172_8446622_RNA45SN1_45S_50ntFlanks | 13004 | 13023 | 20 | ATCTATTGAAAGTCAGCCCT | MB468 | 20 | 14 | 24 | 4 | 5 | 3.7 | 3.2 | 7.3 | 2.3 | 2.4 |
| >hg38_21_+_8433172_8446622_RNA45SN1_45S_50ntFlanks@13004.13025.22\|ATCTATTGAAAGTCAGCCCTCG | >hg38_21_+_8433172_8446622_RNA45SN1_45S_50ntFlanks | 13004 | 13025 | 22 | ATCTATTGAAAGTCAGCCCTCG | MB468 | 83 | 77 | 111 | 8 | 22 | 33.7 | 49.5 | 46.0 | 2.6 | 7.5 |
| >hg38_21_+_8433172_8446622_RNA45SN1_45S_50ntFlanks@13004.13027.24\|ATCTATTGAAAGTCAGCCCTCGAC | >hg38_21_+_8433172_8446622_RNA45SN1_45S_50ntFlanks | 13004 | 13027 | 24 | ATCTATTGAAAGTCAGCCCTCGAC | MB468 | 215 | 165 | 227 | 12 | 61 | 96.6 | 79.7 | 133.6 | 1.6 | 16.5 |
| >hg38_21_+_8433172_8446622_RNA45SN1_45S_50ntFlanks@13004.13028.25\|ATCTATTGAAAGTCAGCCCTCGACA | >hg38_21_+_8433172_8446622_RNA45SN1_45S_50ntFlanks | 13004 | 13028 | 25 | ATCTATTGAAAGTCAGCCCTCGACA | MB468 | 97 | 41 | 120 | 9 | 26 | 43.7 | 12.7 | 69.8 | 3.1 | 15.1 |
| >hg38_21_+_8433172_8446622_RNA45SN1_45S_50ntFlanks@13004.13029.26\|ATCTATTGAAAGTCAGCCCTCGACAC | >hg38_21_+_8433172_8446622_RNA45SN1_45S_50ntFlanks | 13004 | 13029 | 26 | ATCTATTGAAAGTCAGCCCTCGACAC | MB468 | 18 | 12 | 22 | 1 | 6 | 6.7 | 3.3 | 7.4 | 0.3 | 3.3 |
| >hg38_21_+_8433172_8446622_RNA45SN1_45S_50ntFlanks@13004.13039.36\|ATCTATTGAAAGTCAGCCCTCGACACAAGGGTTTGT | >hg38_21_+_8433172_8446622_RNA45SN1_45S_50ntFlanks | 13004 | 13039 | 36 | ATCTATTGAAAGTCAGCCCTCGACACAAGGGTTTGT | MB468 | 15 | 4 | 9 | 7 | 23 | 6.0 | 2.8 | 8.6 | 5.2 | 21.7 |
| >hg38_21_+_8433172_8446622_RNA45SN1_45S_50ntFlanks@13005.13025.21\|TCTATTGAAAGTCAGCCCTCG | >hg38_21_+_8433172_8446622_RNA45SN1_45S_50ntFlanks | 13005 | 13025 | 21 | TCTATTGAAAGTCAGCCCTCG | MB468 | 19 | 14 | 27 | 2 | 4 | 9.8 | 8.6 | 14.3 | 0.4 | 0.9 |
| >hg38_21_+_8433172_8446622_RNA45SN1_45S_50ntFlanks@13005.13027.23\|TCTATTGAAAGTCAGCCCTCGAC | >hg38_21_+_8433172_8446622_RNA45SN1_45S_50ntFlanks | 13005 | 13027 | 23 | TCTATTGAAAGTCAGCCCTCGAC | MB468 | 31 | 13 | 38 | 2 | 6 | 18.8 | 4.6 | 27.3 | 0.6 | 1.5 |
| >hg38_21_+_8433172_8446622_RNA45SN1_45S_50ntFlanks@13005.13028.24\|TCTATTGAAAGTCAGCCCTCGACA | >hg38_21_+_8433172_8446622_RNA45SN1_45S_50ntFlanks | 13005 | 13028 | 24 | TCTATTGAAAGTCAGCCCTCGACA | MB468 | 20 | 6 | 29 | 2 | 5 | 10.8 | 2.3 | 18.1 | 0.6 | 1.9 |
| >hg38_21_+_8433172_8446622_RNA45SN1_45S_50ntFlanks@13005.13039.35\|TCTATTGAAAGTCAGCCCTCGACACAAGGGTTTGT | >hg38_21_+_8433172_8446622_RNA45SN1_45S_50ntFlanks | 13005 | 13039 | 35 | TCTATTGAAAGTCAGCCCTCGACACAAGGGTTTGT | MB231 | 12 | 11 | 8 | 3 | 14 | 14.2 | 9.0 | 9.0 | 1.4 | 14.5 |
| >hg38_21_+_8433172_8446622_RNA45SN1_45S_50ntFlanks@13006.13023.18\|CTATTGAAAGTCAGCCCT | >hg38_21_+_8433172_8446622_RNA45SN1_45S_50ntFlanks | 13006 | 13023 | 18 | CTATTGAAAGTCAGCCCT | MB468 | 12 | 6 | 18 | 4 | 3 | 3.0 | 1.6 | 5.9 | 2.8 | 1.3 |
| >hg38_21_+_8433172_8446622_RNA45SN1_45S_50ntFlanks@13006.13025.20\|CTATTGAAAGTCAGCCCTCG | >hg38_21_+_8433172_8446622_RNA45SN1_45S_50ntFlanks | 13006 | 13025 | 20 | CTATTGAAAGTCAGCCCTCG | MB468 | 87 | 60 | 89 | 11 | 17 | 56.4 | 34.3 | 61.6 | 3.3 | 4.7 |
| >hg38_21_+_8433172_8446622_RNA45SN1_45S_50ntFlanks@13006.13027.22\|CTATTGAAAGTCAGCCCTCGAC | >hg38_21_+_8433172_8446622_RNA45SN1_45S_50ntFlanks | 13006 | 13027 | 22 | CTATTGAAAGTCAGCCCTCGAC | MB468 | 99 | 32 | 117 | 9 | 17 | 77.0 | 10.5 | 102.2 | 1.5 | 7.3 |
| >hg38_21_+_8433172_8446622_RNA45SN1_45S_50ntFlanks@13006.13028.23\|CTATTGAAAGTCAGCCCTCGACA | >hg38_21_+_8433172_8446622_RNA45SN1_45S_50ntFlanks | 13006 | 13028 | 23 | CTATTGAAAGTCAGCCCTCGACA | MB468 | 70 | 16 | 99 | 9 | 14 | 47.2 | 5.8 | 72.9 | 2.6 | 6.1 |
| >hg38_21_+_8433172_8446622_RNA45SN1_45S_50ntFlanks@13006.13039.34\|CTATTGAAAGTCAGCCCTCGACACAAGGGTTTGT | >hg38_21_+_8433172_8446622_RNA45SN1_45S_50ntFlanks | 13006 | 13039 | 34 | CTATTGAAAGTCAGCCCTCGACACAAGGGTTTGT | BT20 | 33 | 40 | 46 | 40 | 105 | 22.3 | 9.4 | 43.3 | 24.5 | 92.2 |
| >hg38_21_+_8433172_8446622_RNA45SN1_45S_50ntFlanks@13006.13039.34\|CTATTGAAAGTCAGCCCTCGACACAAGGGTTTGT | >hg38_21_+_8433172_8446622_RNA45SN1_45S_50ntFlanks | 13006 | 13039 | 34 | CTATTGAAAGTCAGCCCTCGACACAAGGGTTTGT | MB231 | 38 | 38 | 24 | 16 | 50 | 43.8 | 31.3 | 24.8 | 5.9 | 37.6 |
| >hg38_21_+_8433172_8446622_RNA45SN1_45S_50ntFlanks@13006.13039.34\|CTATTGAAAGTCAGCCCTCGACACAAGGGTTTGT | >hg38_21_+_8433172_8446622_RNA45SN1_45S_50ntFlanks | 13006 | 13039 | 34 | CTATTGAAAGTCAGCCCTCGACACAAGGGTTTGT | MB468 | 12 | 9 | 6 | 9 | 13 | 6.1 | 3.8 | 3.4 | 6.1 | 9.1 |
| >hg38_21_+_8433172_8446622_RNA45SN1_45S_50ntFlanks@13007.13025.19\|TATTGAAAGTCAGCCCTCG | >hg38_21_+_8433172_8446622_RNA45SN1_45S_50ntFlanks | 13007 | 13025 | 19 | TATTGAAAGTCAGCCCTCG | MB468 | 84 | 56 | 89 | 7 | 14 | 42.0 | 23.6 | 52.0 | 3.1 | 7.7 |
| >hg38_21_+_8433172_8446622_RNA45SN1_45S_50ntFlanks@13007.13027.21\|TATTGAAAGTCAGCCCTCGAC | >hg38_21_+_8433172_8446622_RNA45SN1_45S_50ntFlanks | 13007 | 13027 | 21 | TATTGAAAGTCAGCCCTCGAC | MB468 | 57 | 18 | 62 | 4 | 8 | 40.7 | 3.3 | 54.5 | 1.7 | 5.9 |
| >hg38_21_+_8433172_8446622_RNA45SN1_45S_50ntFlanks@13007.13028.22\|TATTGAAAGTCAGCCCTCGACA | >hg38_21_+_8433172_8446622_RNA45SN1_45S_50ntFlanks | 13007 | 13028 | 22 | TATTGAAAGTCAGCCCTCGACA | MB468 | 66 | 22 | 72 | 5 | 12 | 39.7 | 6.5 | 48.0 | 2.7 | 7.8 |
| >hg38_21_+_8433172_8446622_RNA45SN1_45S_50ntFlanks@13007.13039.33\|TATTGAAAGTCAGCCCTCGACACAAGGGTTTGT | >hg38_21_+_8433172_8446622_RNA45SN1_45S_50ntFlanks | 13007 | 13039 | 33 | TATTGAAAGTCAGCCCTCGACACAAGGGTTTGT | BT20 | 68 | 54 | 85 | 44 | 190 | 27.7 | 12.4 | 55.4 | 7.9 | 112.3 |
| >hg38_21_+_8433172_8446622_RNA45SN1_45S_50ntFlanks@13007.13039.33\|TATTGAAAGTCAGCCCTCGACACAAGGGTTTGT | >hg38_21_+_8433172_8446622_RNA45SN1_45S_50ntFlanks | 13007 | 13039 | 33 | TATTGAAAGTCAGCCCTCGACACAAGGGTTTGT | MB231 | 67 | 64 | 49 | 23 | 87 | 24.0 | 23.8 | 5.2 | 12.0 | 12.2 |
| >hg38_21_+_8433172_8446622_RNA45SN1_45S_50ntFlanks@13007.13039.33\|TATTGAAAGTCAGCCCTCGACACAAGGGTTTGT | >hg38_21_+_8433172_8446622_RNA45SN1_45S_50ntFlanks | 13007 | 13039 | 33 | TATTGAAAGTCAGCCCTCGACACAAGGGTTTGT | MB468 | 11 | 6 | 6 | 10 | 9 | 4.4 | 3.5 | 2.9 | 9.7 | 6.4 |
| >hg38_21_+_8433172_8446622_RNA45SN1_45S_50ntFlanks@13008.13025.18\|ATTGAAAGTCAGCCCTCG | >hg38_21_+_8433172_8446622_RNA45SN1_45S_50ntFlanks | 13008 | 13025 | 18 | ATTGAAAGTCAGCCCTCG | MB468 | 147 | 82 | 123 | 21 | 24 | 97.3 | 49.3 | 100.2 | 11.8 | 10.4 |
| >hg38_21_+_8433172_8446622_RNA45SN1_45S_50ntFlanks@13008.13026.19\|ATTGAAAGTCAGCCCTCGA | >hg38_21_+_8433172_8446622_RNA45SN1_45S_50ntFlanks | 13008 | 13026 | 19 | ATTGAAAGTCAGCCCTCGA | MB468 | 14 | 8 | 13 | 2 | 3 | 5.9 | 3.2 | 7.8 | 0.9 | 1.3 |
| >hg38_21_+_8433172_8446622_RNA45SN1_45S_50ntFlanks@13008.13027.20\|ATTGAAAGTCAGCCCTCGAC | >hg38_21_+_8433172_8446622_RNA45SN1_45S_50ntFlanks | 13008 | 13027 | 20 | ATTGAAAGTCAGCCCTCGAC | MB468 | 77 | 21 | 69 | 11 | 10 | 67.0 | 4.9 | 72.9 | 4.1 | 5.9 |
| >hg38_21_+_8433172_8446622_RNA45SN1_45S_50ntFlanks@13008.13028.21\|ATTGAAAGTCAGCCCTCGACA | >hg38_21_+_8433172_8446622_RNA45SN1_45S_50ntFlanks | 13008 | 13028 | 21 | ATTGAAAGTCAGCCCTCGACA | MB468 | 71 | 18 | 63 | 14 | 11 | 54.5 | 3.3 | 58.1 | 7.7 | 6.0 |
| >hg38_21_+_8433172_8446622_RNA45SN1_45S_50ntFlanks@13008.13029.22\|ATTGAAAGTCAGCCCTCGACAC | >hg38_21_+_8433172_8446622_RNA45SN1_45S_50ntFlanks | 13008 | 13029 | 22 | ATTGAAAGTCAGCCCTCGACAC | MB468 | 11 | 5 | 9 | 1 | 2 | 8.5 | 1.1 | 7.3 | 1.1 | 1.4 |
| >hg38_21_+_8433172_8446622_RNA45SN1_45S_50ntFlanks@13008.13039.32\|ATTGAAAGTCAGCCCTCGACACAAGGGTTTGT | >hg38_21_+_8433172_8446622_RNA45SN1_45S_50ntFlanks | 13008 | 13039 | 32 | ATTGAAAGTCAGCCCTCGACACAAGGGTTTGT | BT20 | 113 | 170 | 165 | 111 | 427 | 73.7 | 26.7 | 133.0 | 12.5 | 311.0 |
| >hg38_21_+_8433172_8446622_RNA45SN1_45S_50ntFlanks@13008.13039.32\|ATTGAAAGTCAGCCCTCGACACAAGGGTTTGT | >hg38_21_+_8433172_8446622_RNA45SN1_45S_50ntFlanks | 13008 | 13039 | 32 | ATTGAAAGTCAGCCCTCGACACAAGGGTTTGT | MB231 | 76 | 73 | 41 | 26 | 73 | 91.2 | 63.3 | 45.2 | 9.5 | 70.9 |
| >hg38_21_+_8433172_8446622_RNA45SN1_45S_50ntFlanks@13008.13039.32\|ATTGAAAGTCAGCCCTCGACACAAGGGTTTGT | >hg38_21_+_8433172_8446622_RNA45SN1_45S_50ntFlanks | 13008 | 13039 | 32 | ATTGAAAGTCAGCCCTCGACACAAGGGTTTGT | MB468 | 20 | 26 | 9 | 27 | 15 | 10.7 | 23.4 | 4.1 | 23.1 | 5.3 |
| >hg38_21_+_8433172_8446622_RNA45SN1_45S_50ntFlanks@13009.13027.19\|TTGAAAGTCAGCCCTCGAC | >hg38_21_+_8433172_8446622_RNA45SN1_45S_50ntFlanks | 13009 | 13027 | 19 | TTGAAAGTCAGCCCTCGAC | MB468 | 46 | 12 | 48 | 5 | 6 | 39.2 | 3.2 | 57.0 | 2.4 | 2.9 |
| >hg38_21_+_8433172_8446622_RNA45SN1_45S_50ntFlanks@13009.13028.20\|TTGAAAGTCAGCCCTCGACA | >hg38_21_+_8433172_8446622_RNA45SN1_45S_50ntFlanks | 13009 | 13028 | 20 | TTGAAAGTCAGCCCTCGACA | MB468 | 21 | 5 | 24 | 5 | 3 | 17.1 | 0.9 | 25.7 | 2.5 | 1.8 |
| >hg38_21_+_8433172_8446622_RNA45SN1_45S_50ntFlanks@13010.13027.18\|TGAAAGTCAGCCCTCGAC | >hg38_21_+_8433172_8446622_RNA45SN1_45S_50ntFlanks | 13010 | 13027 | 18 | TGAAAGTCAGCCCTCGAC | MB468 | 18 | 5 | 15 | 1 | 2 | 14.2 | 1.4 | 15.9 | 1.2 | 1.7 |

| | | | | | | | | | | | | | | | | |
|---|---|---|---|---|---|---|---|---|---|---|---|---|---|---|---|---|
| >hg38_21_+_8433172_8446622_RNA45SN1_45S_50ntFlanks@13011.13039.29\|GAAAGTCAGCCCTCGACACAAGGGTTTGT | >hg38_21_+_8433172_8446622_RNA45SN1_45S_50ntFlanks | 13011 | 13039 | 29 | GAAAGTCAGCCCTCGACACAAGGGTTTGT | BT20 | 11 | 8 | 6 | 6 | 18 | 10.4 | 4.2 | 1.9 | 1.6 | 16.4 |
| >hg38_21_+_8433172_8446622_RNA45SN1_45S_50ntFlanks@13022.13039.18\|CTCGACACAAGGGTTTGT | >hg38_21_+_8433172_8446622_RNA45SN1_45S_50ntFlanks | 13022 | 13039 | 18 | CTCGACACAAGGGTTTGT | BT20 | 33 | 18 | 31 | 17 | 39 | 7.7 | 8.0 | 10.4 | 10.2 | 6.6 |
| >hg38_21_+_8433172_8446622_RNA45SN1_45S_50ntFlanks@13022.13039.18\|CTCGACACAAGGGTTTGT | >hg38_21_+_8433172_8446622_RNA45SN1_45S_50ntFlanks | 13022 | 13039 | 18 | CTCGACACAAGGGTTTGT | MB231 | 50 | 38 | 36 | 24 | 72 | 32.6 | 17.2 | 17.8 | 13.5 | 24.2 |
| >hg38_21_+_8433172_8446622_RNA45SN1_45S_50ntFlanks@13022.13039.18\|CTCGACACAAGGGTTTGT | >hg38_21_+_8433172_8446622_RNA45SN1_45S_50ntFlanks | 13022 | 13039 | 18 | CTCGACACAAGGGTTTGT | MB468 | 18 | 4 | 8 | 6 | 10 | 6.9 | 1.3 | 2.5 | 1.4 | 5.2 |
| >hg38_21_+_8433172_8446622_RNA45SN1_45S_50ntFlanks@2959.2979.21\|CGACGTGTGGCGTGGGTCGAC | >hg38_21_+_8433172_8446622_RNA45SN1_45S_50ntFlanks | 2959 | 2979 | 21 | CGACGTGTGGCGTGGGTCGAC | MB468 | 17 | 29 | 9 | 5 | 2 | 8.6 | 11.0 | 6.0 | 4.0 | 0.6 |
| >hg38_21_+_8433172_8446622_RNA45SN1_45S_50ntFlanks@3593.3610.18\|CCCGTCCGTCCGTCCGTC | >hg38_21_+_8433172_8446622_RNA45SN1_45S_50ntFlanks | 3593 | 3610 | 18 | CCCGTCCGTCCGTCCGTC | MB468 | 16 | 16 | 33 | 19 | 22 | 5.4 | 13.0 | 7.6 | 17.8 | 6.8 |
| >hg38_21_+_8433172_8446622_RNA45SN1_45S_50ntFlanks@3705.3722.18\|TACCTGGTTGATCCTGCC | >hg38_21_+_8433172_8446622_RNA45SN1_45S_50ntFlanks | 3705 | 3722 | 18 | TACCTGGTTGATCCTGCC | BT20 | 322 | 386 | 1263 | 161 | 238 | 192.4 | 199.0 | 145.6 | 185.5 | 127.7 |
| >hg38_21_+_8433172_8446622_RNA45SN1_45S_50ntFlanks@3705.3722.18\|TACCTGGTTGATCCTGCC | >hg38_21_+_8433172_8446622_RNA45SN1_45S_50ntFlanks | 3705 | 3722 | 18 | TACCTGGTTGATCCTGCC | MB231 | 137 | 209 | 905 | 30 | 337 | 172.5 | 78.1 | 75.7 | 23.3 | 356.3 |
| >hg38_21_+_8433172_8446622_RNA45SN1_45S_50ntFlanks@3705.3722.18\|TACCTGGTTGATCCTGCC | >hg38_21_+_8433172_8446622_RNA45SN1_45S_50ntFlanks | 3705 | 3722 | 18 | TACCTGGTTGATCCTGCC | MB468 | 64 | 50 | 90 | 55 | 151 | 15.2 | 28.8 | 39.0 | 39.4 | 163.1 |
| >hg38_21_+_8433172_8446622_RNA45SN1_45S_50ntFlanks@3705.3723.19\|TACCTGGTTGATCCTGCCA | >hg38_21_+_8433172_8446622_RNA45SN1_45S_50ntFlanks | 3705 | 3723 | 19 | TACCTGGTTGATCCTGCCA | BT20 | 19 | 29 | 63 | 7 | 17 | 7.5 | 13.7 | 13.3 | 0.7 | 12.4 |
| >hg38_21_+_8433172_8446622_RNA45SN1_45S_50ntFlanks@3705.3725.21\|TACCTGGTTGATCCTGCCAGT | >hg38_21_+_8433172_8446622_RNA45SN1_45S_50ntFlanks | 3705 | 3725 | 21 | TACCTGGTTGATCCTGCCAGT | BT20 | 458 | 252 | 1217 | 44 | 104 | 445.5 | 83.2 | 827.7 | 38.4 | 77.2 |
| >hg38_21_+_8433172_8446622_RNA45SN1_45S_50ntFlanks@3705.3725.21\|TACCTGGTTGATCCTGCCAGT | >hg38_21_+_8433172_8446622_RNA45SN1_45S_50ntFlanks | 3705 | 3725 | 21 | TACCTGGTTGATCCTGCCAGT | MB231 | 28 | 11 | 35 | 12 | 106 | 25.6 | 10.4 | 16.9 | 17.5 | 93.6 |
| >hg38_21_+_8433172_8446622_RNA45SN1_45S_50ntFlanks@3705.3726.22\|TACCTGGTTGATCCTGCCAGTA | >hg38_21_+_8433172_8446622_RNA45SN1_45S_50ntFlanks | 3705 | 3726 | 22 | TACCTGGTTGATCCTGCCAGTA | BT20 | 10 | 6 | 46 | 1 | 3 | 6.2 | 1.1 | 30.4 | 0.7 | 3.0 |
| >hg38_21_+_8433172_8446622_RNA45SN1_45S_50ntFlanks@3705.3728.24\|TACCTGGTTGATCCTGCCAGTAGC | >hg38_21_+_8433172_8446622_RNA45SN1_45S_50ntFlanks | 3705 | 3728 | 24 | TACCTGGTTGATCCTGCCAGTAGC | BT20 | 106 | 37 | 406 | 6 | 20 | 107.3 | 5.8 | 221.8 | 4.2 | 18.2 |
| >hg38_21_+_8433172_8446622_RNA45SN1_45S_50ntFlanks@3705.3728.24\|TACCTGGTTGATCCTGCCAGTAGC | >hg38_21_+_8433172_8446622_RNA45SN1_45S_50ntFlanks | 3705 | 3728 | 24 | TACCTGGTTGATCCTGCCAGTAGC | MB231 | 17 | 9 | 53 | 3 | 31 | 12.8 | 4.3 | 5.2 | 5.2 | 32.2 |
| >hg38_21_+_8433172_8446622_RNA45SN1_45S_50ntFlanks@3705.3729.25\|TACCTGGTTGATCCTGCCAGTAGCA | >hg38_21_+_8433172_8446622_RNA45SN1_45S_50ntFlanks | 3705 | 3729 | 25 | TACCTGGTTGATCCTGCCAGTAGCA | BT20 | 40 | 8 | 19 | 1 | 1 | 65.4 | 8.2 | 12.4 | 1.5 | 1.4 |
| >hg38_21_+_8433172_8446622_RNA45SN1_45S_50ntFlanks@3705.3730.26\|TACCTGGTTGATCCTGCCAGTAGCAT | >hg38_21_+_8433172_8446622_RNA45SN1_45S_50ntFlanks | 3705 | 3730 | 26 | TACCTGGTTGATCCTGCCAGTAGCAT | BT20 | 57 | 12 | 13 | 0 | 0 | 92.1 | 8.0 | 7.0 | 0.0 | 0.0 |
| >hg38_21_+_8433172_8446622_RNA45SN1_45S_50ntFlanks@3705.3732.28\|TACCTGGTTGATCCTGCCAGTAGCATAT | >hg38_21_+_8433172_8446622_RNA45SN1_45S_50ntFlanks | 3705 | 3732 | 28 | TACCTGGTTGATCCTGCCAGTAGCATAT | BT20 | 54 | 4 | 19 | 0 | 0 | 89.3 | 2.0 | 23.1 | 0.0 | 0.0 |
| >hg38_21_+_8433172_8446622_RNA45SN1_45S_50ntFlanks@3705.3735.31\|TACCTGGTTGATCCTGCCAGTAGCATATGCT | >hg38_21_+_8433172_8446622_RNA45SN1_45S_50ntFlanks | 3705 | 3735 | 31 | TACCTGGTTGATCCTGCCAGTAGCATATGCT | BT20 | 31 | 19 | 103 | 0 | 15 | 28.2 | 5.5 | 63.5 | 0.6 | 25.3 |
| >hg38_21_+_8433172_8446622_RNA45SN1_45S_50ntFlanks@3705.3736.32\|TACCTGGTTGATCCTGCCAGTAGCATATGCTT | >hg38_21_+_8433172_8446622_RNA45SN1_45S_50ntFlanks | 3705 | 3736 | 32 | TACCTGGTTGATCCTGCCAGTAGCATATGCTT | BT20 | 82 | 48 | 106 | 3 | 60 | 64.5 | 17.5 | 32.5 | 1.3 | 68.4 |
| >hg38_21_+_8433172_8446622_RNA45SN1_45S_50ntFlanks@3705.3736.32\|TACCTGGTTGATCCTGCCAGTAGCATATGCTT | >hg38_21_+_8433172_8446622_RNA45SN1_45S_50ntFlanks | 3705 | 3736 | 32 | TACCTGGTTGATCCTGCCAGTAGCATATGCTT | MB231 | 14 | 8 | 32 | 1 | 88 | 12.9 | 2.3 | 7.4 | 1.0 | 23.5 |
| >hg38_21_+_8433172_8446622_RNA45SN1_45S_50ntFlanks@3705.3737.33\|TACCTGGTTGATCCTGCCAGTAGCATATGCTTG | >hg38_21_+_8433172_8446622_RNA45SN1_45S_50ntFlanks | 3705 | 3737 | 33 | TACCTGGTTGATCCTGCCAGTAGCATATGCTTG | MB468 | 12 | 4 | 12 | 7 | 24 | 5.5 | 1.4 | 6.1 | 3.3 | 17.5 |
| >hg38_21_+_8433172_8446622_RNA45SN1_45S_50ntFlanks@3705.3739.35\|TACCTGGTTGATCCTGCCAGTAGCATATGCTTGTC | >hg38_21_+_8433172_8446622_RNA45SN1_45S_50ntFlanks | 3705 | 3739 | 35 | TACCTGGTTGATCCTGCCAGTAGCATATGCTTGTC | BT20 | 64 | 85 | 164 | 19 | 34 | 33.0 | 8.7 | 91.9 | 13.4 | 29.6 |
| >hg38_21_+_8433172_8446622_RNA45SN1_45S_50ntFlanks@3705.3739.35\|TACCTGGTTGATCCTGCCAGTAGCATATGCTTGTC | >hg38_21_+_8433172_8446622_RNA45SN1_45S_50ntFlanks | 3705 | 3739 | 35 | TACCTGGTTGATCCTGCCAGTAGCATATGCTTGTC | MB231 | 12 | 12 | 45 | 12 | 54 | 8.2 | 2.7 | 15.2 | 17.2 | 16.4 |
| >hg38_21_+_8433172_8446622_RNA45SN1_45S_50ntFlanks@3705.3740.36\|TACCTGGTTGATCCTGCCAGTAGCATATGCTTGTCT | >hg38_21_+_8433172_8446622_RNA45SN1_45S_50ntFlanks | 3705 | 3740 | 36 | TACCTGGTTGATCCTGCCAGTAGCATATGCTTGTCT | BT20 | 12 | 10 | 38 | 6 | 6 | 6.2 | 2.7 | 15.2 | 5.2 | 5.2 |
| >hg38_21_+_8433172_8446622_RNA45SN1_45S_50ntFlanks@3705.3741.37\|TACCTGGTTGATCCTGCCAGTAGCATATGCTTGTCTC | >hg38_21_+_8433172_8446622_RNA45SN1_45S_50ntFlanks | 3705 | 3741 | 37 | TACCTGGTTGATCCTGCCAGTAGCATATGCTTGTCTC | BT20 | 115 | 97 | 288 | 18 | 43 | 117.4 | 21.9 | 206.1 | 8.3 | 24.7 |
| >hg38_21_+_8433172_8446622_RNA45SN1_45S_50ntFlanks@3705.3741.37\|TACCTGGTTGATCCTGCCAGTAGCATATGCTTGTCTC | >hg38_21_+_8433172_8446622_RNA45SN1_45S_50ntFlanks | 3705 | 3741 | 37 | TACCTGGTTGATCCTGCCAGTAGCATATGCTTGTCTC | MB231 | 26 | 16 | 94 | 8 | 54 | 13.8 | 4.2 | 19.9 | 8.2 | 28.7 |
| >hg38_21_+_8433172_8446622_RNA45SN1_45S_50ntFlanks@3705.3742.38\|TACCTGGTTGATCCTGCCAGTAGCATATGCTTGTCTCA | >hg38_21_+_8433172_8446622_RNA45SN1_45S_50ntFlanks | 3705 | 3742 | 38 | TACCTGGTTGATCCTGCCAGTAGCATATGCTTGTCTCA | BT20 | 98 | 95 | 362 | 8 | 35 | 65.1 | 9.7 | 243.5 | 4.2 | 16.5 |
| >hg38_21_+_8433172_8446622_RNA45SN1_45S_50ntFlanks@3705.3742.38\|TACCTGGTTGATCCTGCCAGTAGCATATGCTTGTCTCA | >hg38_21_+_8433172_8446622_RNA45SN1_45S_50ntFlanks | 3705 | 3742 | 38 | TACCTGGTTGATCCTGCCAGTAGCATATGCTTGTCTCA | MB231 | 25 | 15 | 65 | 2 | 42 | 19.9 | 13.9 | 13.4 | 1.9 | 33.1 |
| >hg38_21_+_8433172_8446622_RNA45SN1_45S_50ntFlanks@3705.3743.39\|TACCTGGTTGATCCTGCCAGTAGCATATGCTTGTCTCAA | >hg38_21_+_8433172_8446622_RNA45SN1_45S_50ntFlanks | 3705 | 3743 | 39 | TACCTGGTTGATCCTGCCAGTAGCATATGCTTGTCTCAA | BT20 | 56 | 88 | 214 | 23 | 38 | 30.2 | 50.4 | 107.8 | 21.8 | 12.8 |
| >hg38_21_+_8433172_8446622_RNA45SN1_45S_50ntFlanks@3705.3743.39\|TACCTGGTTGATCCTGCCAGTAGCATATGCTTGTCTCAA | >hg38_21_+_8433172_8446622_RNA45SN1_45S_50ntFlanks | 3705 | 3743 | 39 | TACCTGGTTGATCCTGCCAGTAGCATATGCTTGTCTCAA | MB231 | 32 | 26 | 154 | 11 | 49 | 16.3 | 8.7 | 82.0 | 8.4 | 14.7 |
| >hg38_21_+_8433172_8446622_RNA45SN1_45S_50ntFlanks@3705.3744.40\|TACCTGGTTGATCCTGCCAGTAGCATATGCTTGTCTCAAA | >hg38_21_+_8433172_8446622_RNA45SN1_45S_50ntFlanks | 3705 | 3744 | 40 | TACCTGGTTGATCCTGCCAGTAGCATATGCTTGTCTCAAA | BT20 | 43 | 74 | 80 | 5 | 14 | 42.9 | 34.5 | 74.4 | 3.3 | 4.0 |
| >hg38_21_+_8433172_8446622_RNA45SN1_45S_50ntFlanks@3705.3744.40\|TACCTGGTTGATCCTGCCAGTAGCATATGCTTGTCTCAAA | >hg38_21_+_8433172_8446622_RNA45SN1_45S_50ntFlanks | 3705 | 3744 | 40 | TACCTGGTTGATCCTGCCAGTAGCATATGCTTGTCTCAAA | MB231 | 10 | 10 | 26 | 2 | 19 | 8.3 | 7.3 | 5.8 | 2.1 | 21.8 |
| >hg38_21_+_8433172_8446622_RNA45SN1_45S_50ntFlanks@3705.3745.41\|TACCTGGTTGATCCTGCCAGTAGCATATGCTTGTCTCAAAG | >hg38_21_+_8433172_8446622_RNA45SN1_45S_50ntFlanks | 3705 | 3745 | 41 | TACCTGGTTGATCCTGCCAGTAGCATATGCTTGTCTCAAAG | BT20 | 30 | 41 | 78 | 3 | 8 | 25.8 | 23.5 | 82.2 | 1.4 | 3.0 |
| >hg38_21_+_8433172_8446622_RNA45SN1_45S_50ntFlanks@3705.3745.41\|TACCTGGTTGATCCTGCCAGTAGCATATGCTTGTCTCAAAG | >hg38_21_+_8433172_8446622_RNA45SN1_45S_50ntFlanks | 3705 | 3745 | 41 | TACCTGGTTGATCCTGCCAGTAGCATATGCTTGTCTCAAAG | MB231 | 13 | 10 | 42 | 2 | 28 | 6.8 | 4.9 | 11.4 | 2.2 | 10.0 |
| >hg38_21_+_8433172_8446622_RNA45SN1_45S_50ntFlanks@3705.3746.42\|TACCTGGTTGATCCTGCCAGTAGCATATGCTTGTCTCAAAGA | >hg38_21_+_8433172_8446622_RNA45SN1_45S_50ntFlanks | 3705 | 3746 | 42 | TACCTGGTTGATCCTGCCAGTAGCATATGCTTGTCTCAAAGA | BT20 | 227 | 376 | 410 | 9 | 39 | 361.8 | 568.1 | 651.2 | 4.9 | 6.0 |
| >hg38_21_+_8433172_8446622_RNA45SN1_45S_50ntFlanks@3705.3746.42\|TACCTGGTTGATCCTGCCAGTAGCATATGCTTGTCTCAAAGA | >hg38_21_+_8433172_8446622_RNA45SN1_45S_50ntFlanks | 3705 | 3746 | 42 | TACCTGGTTGATCCTGCCAGTAGCATATGCTTGTCTCAAAGA | MB231 | 15 | 12 | 67 | 7 | 119 | 8.3 | 9.9 | 25.3 | 6.6 | 70.4 |
| >hg38_21_+_8433172_8446622_RNA45SN1_45S_50ntFlanks@3705.3747.43\|TACCTGGTTGATCCTGCCAGTAGCATATGCTTGTCTCAAAGAT | >hg38_21_+_8433172_8446622_RNA45SN1_45S_50ntFlanks | 3705 | 3747 | 43 | TACCTGGTTGATCCTGCCAGTAGCATATGCTTGTCTCAAAGAT | BT20 | 75 | 73 | 85 | 2 | 2 | 123.7 | 108.7 | 130.5 | 2.6 | 2.2 |
| >hg38_21_+_8433172_8446622_RNA45SN1_45S_50ntFlanks@3705.3748.44\|TACCTGGTTGATCCTGCCAGTAGCATATGCTTGTCTCAAAGATT | >hg38_21_+_8433172_8446622_RNA45SN1_45S_50ntFlanks | 3705 | 3748 | 44 | TACCTGGTTGATCCTGCCAGTAGCATATGCTTGTCTCAAAGATT | BT20 | 319 | 342 | 365 | 21 | 26 | 512.2 | 487.9 | 574.7 | 22.9 | 9.4 |
| >hg38_21_+_8433172_8446622_RNA45SN1_45S_50ntFlanks@3705.3748.44\|TACCTGGTTGATCCTGCCAGTAGCATATGCTTGTCTCAAAGATT | >hg38_21_+_8433172_8446622_RNA45SN1_45S_50ntFlanks | 3705 | 3748 | 44 | TACCTGGTTGATCCTGCCAGTAGCATATGCTTGTCTCAAAGATT | MB231 | 18 | 19 | 86 | 11 | 46 | 4.9 | 21.4 | 32.2 | 15.7 | 32.6 |
| >hg38_21_+_8433172_8446622_RNA45SN1_45S_50ntFlanks@3711.3729.19\|GTTGATCCTGCCAGTAGCA | >hg38_21_+_8433172_8446622_RNA45SN1_45S_50ntFlanks | 3711 | 3729 | 19 | GTTGATCCTGCCAGTAGCA | MB468 | 19 | 21 | 10 | 10 | 24 | 8.5 | 8.6 | 4.6 | 7.7 | 9.6 |
| >hg38_21_+_8433172_8446622_RNA45SN1_45S_50ntFlanks@3711.3737.27\|GTTGATCCTGCCAGTAGCATATGCTTG | >hg38_21_+_8433172_8446622_RNA45SN1_45S_50ntFlanks | 3711 | 3737 | 27 | GTTGATCCTGCCAGTAGCATATGCTTG | MB468 | 63 | 107 | 46 | 21 | 50 | 29.7 | 76.4 | 13.0 | 20.1 | 17.8 |
| >hg38_21_+_8433172_8446622_RNA45SN1_45S_50ntFlanks@3711.3739.29\|GTTGATCCTGCCAGTAGCATATGCTTGTC | >hg38_21_+_8433172_8446622_RNA45SN1_45S_50ntFlanks | 3711 | 3739 | 29 | GTTGATCCTGCCAGTAGCATATGCTTGTC | MB468 | 11 | 20 | 13 | 4 | 8 | 5.5 | 5.6 | 4.6 | 3.1 | 2.3 |
| >hg38_21_+_8433172_8446622_RNA45SN1_45S_50ntFlanks@3728.3748.21\|CATATGCTTGTCTCAAAGATT | >hg38_21_+_8433172_8446622_RNA45SN1_45S_50ntFlanks | 3728 | 3748 | 21 | CATATGCTTGTCTCAAAGATT | MB468 | 12 | 3 | 5 | 5 | 5 | 4.7 | 1.9 | 1.9 | 1.0 | 2.9 |
| >hg38_21_+_8433172_8446622_RNA45SN1_45S_50ntFlanks@3731.3748.18\|ATGCTTGTCTCAAAGATT | >hg38_21_+_8433172_8446622_RNA45SN1_45S_50ntFlanks | 3731 | 3748 | 18 | ATGCTTGTCTCAAAGATT | BT20 | 11 | 7 | 10 | 0 | 1 | 7.9 | 2.9 | 3.9 | 0.4 | 1.5 |
| >hg38_21_+_8433172_8446622_RNA45SN1_45S_50ntFlanks@3738.3756.19\|TCTCAAAGATTAAGCCATG | >hg38_21_+_8433172_8446622_RNA45SN1_45S_50ntFlanks | 3738 | 3756 | 19 | TCTCAAAGATTAAGCCATG | MB468 | 12 | 6 | 14 | 79 | 19 | 4.9 | 2.9 | 4.9 | 30.7 | 10.8 |
| >hg38_21_+_8433172_8446622_RNA45SN1_45S_50ntFlanks@3769.3786.18\|CGCACGGCCGGTACAGTG | >hg38_21_+_8433172_8446622_RNA45SN1_45S_50ntFlanks | 3769 | 3786 | 18 | CGCACGGCCGGTACAGTG | MB468 | 18 | 14 | 10 | 13 | 20 | 5.6 | 7.2 | 3.0 | 4.7 | 17.6 |
| >hg38_21_+_8433172_8446622_RNA45SN1_45S_50ntFlanks@3769.3792.24\|CGCACGGCCGGTACAGTGAAACTG | >hg38_21_+_8433172_8446622_RNA45SN1_45S_50ntFlanks | 3769 | 3792 | 24 | CGCACGGCCGGTACAGTGAAACTG | MB468 | 24 | 57 | 14 | 8 | 42 | 8.8 | 24.0 | 2.4 | 5.6 | 37.5 |
| >hg38_21_+_8433172_8446622_RNA45SN1_45S_50ntFlanks@3769.3794.26\|CGCACGGCCGGTACAGTGAAACTGCG | >hg38_21_+_8433172_8446622_RNA45SN1_45S_50ntFlanks | 3769 | 3794 | 26 | CGCACGGCCGGTACAGTGAAACTGCG | MB468 | 22 | 27 | 16 | 11 | 27 | 6.2 | 15.5 | 4.3 | 7.1 | 21.7 |
| >hg38_21_+_8433172_8446622_RNA45SN1_45S_50ntFlanks@3771.3791.21\|CACGGCCGGTACAGTGAAACT | >hg38_21_+_8433172_8446622_RNA45SN1_45S_50ntFlanks | 3771 | 3791 | 21 | CACGGCCGGTACAGTGAAACT | MB468 | 15 | 24 | 10 | 6 | 17 | 9.2 | 5.4 | 4.1 | 7.0 | 13.2 |

| | | | | | | | | | | | | | | | | |
|---|---|---|---|---|---|---|---|---|---|---|---|---|---|---|---|---|
| >hg38_21_+_8433172_8446622_RNA45SN1_45S_50ntFlanks@3771.3792.22\|CACGGCCGGTACAGTGAAACTG | >hg38_21_+_8433172_8446622_RNA45SN1_45S_50ntFlanks | 3771 | 3792 | 22 | CACGGCCGGTACAGTGAAACTG | MB468 | 42 | 76 | 30 | 23 | 69 | 16.1 | 29.0 | 6.0 | 19.9 | 54.4 |
| >hg38_21_+_8433172_8446622_RNA45SN1_45S_50ntFlanks@3771.3794.24\|CACGGCCGGTACAGTGAAACTGCG | >hg38_21_+_8433172_8446622_RNA45SN1_45S_50ntFlanks | 3771 | 3794 | 24 | CACGGCCGGTACAGTGAAACTGCG | MB468 | 18 | 33 | 17 | 13 | 28 | 7.2 | 14.2 | 3.5 | 15.1 | 17.7 |
| >hg38_21_+_8433172_8446622_RNA45SN1_45S_50ntFlanks@3771.3799.29\|CACGGCCGGTACAGTGAAACTGCGAATGG | >hg38_21_+_8433172_8446622_RNA45SN1_45S_50ntFlanks | 3771 | 3799 | 29 | CACGGCCGGTACAGTGAAACTGCGAATGG | MB468 | 11 | 25 | 10 | 13 | 11 | 3.3 | 4.2 | 2.3 | 5.9 | 3.0 |
| >hg38_21_+_8433172_8446622_RNA45SN1_45S_50ntFlanks@3773.3792.20\|CGGCCGGTACAGTGAAACTG | >hg38_21_+_8433172_8446622_RNA45SN1_45S_50ntFlanks | 3773 | 3792 | 20 | CGGCCGGTACAGTGAAACTG | MB468 | 28 | 34 | 17 | 20 | 76 | 9.4 | 14.6 | 3.1 | 14.9 | 89.6 |
| >hg38_21_+_8433172_8446622_RNA45SN1_45S_50ntFlanks@3775.3792.18\|GCCGGTACAGTGAAACTG | >hg38_21_+_8433172_8446622_RNA45SN1_45S_50ntFlanks | 3775 | 3792 | 18 | GCCGGTACAGTGAAACTG | MB468 | 265 | 203 | 193 | 163 | 182 | 128.1 | 109.8 | 43.2 | 142.4 | 139.3 |
| >hg38_21_+_8433172_8446622_RNA45SN1_45S_50ntFlanks@3775.3794.20\|GCCGGTACAGTGAAACTGCG | >hg38_21_+_8433172_8446622_RNA45SN1_45S_50ntFlanks | 3775 | 3794 | 20 | GCCGGTACAGTGAAACTGCG | MB468 | 22 | 9 | 20 | 19 | 18 | 5.1 | 2.7 | 4.1 | 16.6 | 16.8 |
| >hg38_21_+_8433172_8446622_RNA45SN1_45S_50ntFlanks@3775.3798.24\|GCCGGTACAGTGAAACTGCGAATG | >hg38_21_+_8433172_8446622_RNA45SN1_45S_50ntFlanks | 3775 | 3798 | 24 | GCCGGTACAGTGAAACTGCGAATG | MB468 | 17 | 19 | 24 | 17 | 12 | 6.0 | 5.6 | 13.3 | 17.1 | 7.7 |
| >hg38_21_+_8433172_8446622_RNA45SN1_45S_50ntFlanks@3775.3799.25\|GCCGGTACAGTGAAACTGCGAATGG | >hg38_21_+_8433172_8446622_RNA45SN1_45S_50ntFlanks | 3775 | 3799 | 25 | GCCGGTACAGTGAAACTGCGAATGG | MB468 | 12 | 8 | 7 | 19 | 14 | 4.9 | 1.4 | 3.5 | 5.2 | 11.1 |
| >hg38_21_+_8433172_8446622_RNA45SN1_45S_50ntFlanks@3776.3801.26\|CCGGTACAGTGAAACTGCGAATGGCT | >hg38_21_+_8433172_8446622_RNA45SN1_45S_50ntFlanks | 3776 | 3801 | 26 | CCGGTACAGTGAAACTGCGAATGGCT | MB468 | 40 | 28 | 45 | 57 | 37 | 8.8 | 8.4 | 12.0 | 24.8 | 16.3 |
| >hg38_21_+_8433172_8446622_RNA45SN1_45S_50ntFlanks@3776.3805.30\|CCGGTACAGTGAAACTGCGAATGGCTCATT | >hg38_21_+_8433172_8446622_RNA45SN1_45S_50ntFlanks | 3776 | 3805 | 30 | CCGGTACAGTGAAACTGCGAATGGCTCATT | MB468 | 21 | 47 | 22 | 29 | 25 | 10.0 | 18.5 | 4.7 | 32.4 | 8.4 |
| >hg38_21_+_8433172_8446622_RNA45SN1_45S_50ntFlanks@3777.3794.18\|CGGTACAGTGAAACTGCG | >hg38_21_+_8433172_8446622_RNA45SN1_45S_50ntFlanks | 3777 | 3794 | 18 | CGGTACAGTGAAACTGCG | MB468 | 16 | 1 | 10 | 15 | 11 | 8.1 | 0.7 | 3.5 | 5.3 | 13.8 |
| >hg38_21_+_8433172_8446622_RNA45SN1_45S_50ntFlanks@3777.3801.25\|CGGTACAGTGAAACTGCGAATGGCT | >hg38_21_+_8433172_8446622_RNA45SN1_45S_50ntFlanks | 3777 | 3801 | 25 | CGGTACAGTGAAACTGCGAATGGCT | MB468 | 56 | 77 | 50 | 58 | 64 | 21.2 | 34.1 | 20.3 | 35.9 | 34.3 |
| >hg38_21_+_8433172_8446622_RNA45SN1_45S_50ntFlanks@3777.3805.29\|CGGTACAGTGAAACTGCGAATGGCTCATT | >hg38_21_+_8433172_8446622_RNA45SN1_45S_50ntFlanks | 3777 | 3805 | 29 | CGGTACAGTGAAACTGCGAATGGCTCATT | MB468 | 15 | 25 | 13 | 14 | 16 | 8.5 | 10.4 | 5.8 | 14.0 | 10.2 |
| >hg38_21_+_8433172_8446622_RNA45SN1_45S_50ntFlanks@3780.3801.22\|TACAGTGAAACTGCGAATGGCT | >hg38_21_+_8433172_8446622_RNA45SN1_45S_50ntFlanks | 3780 | 3801 | 22 | TACAGTGAAACTGCGAATGGCT | MB468 | 24 | 26 | 16 | 18 | 32 | 6.9 | 23.0 | 4.3 | 15.7 | 19.3 |
| >hg38_21_+_8433172_8446622_RNA45SN1_45S_50ntFlanks@3781.3801.21\|ACAGTGAAACTGCGAATGGCT | >hg38_21_+_8433172_8446622_RNA45SN1_45S_50ntFlanks | 3781 | 3801 | 21 | ACAGTGAAACTGCGAATGGCT | MB468 | 14 | 14 | 10 | 16 | 24 | 4.4 | 12.6 | 1.6 | 15.0 | 14.1 |
| >hg38_21_+_8433172_8446622_RNA45SN1_45S_50ntFlanks@3782.3800.19\|CAGTGAAACTGCGAATGGC | >hg38_21_+_8433172_8446622_RNA45SN1_45S_50ntFlanks | 3782 | 3800 | 19 | CAGTGAAACTGCGAATGGC | MB468 | 39 | 52 | 37 | 65 | 53 | 17.4 | 35.7 | 16.7 | 76.7 | 36.5 |
| >hg38_21_+_8433172_8446622_RNA45SN1_45S_50ntFlanks@3782.3801.20\|CAGTGAAACTGCGAATGGCT | >hg38_21_+_8433172_8446622_RNA45SN1_45S_50ntFlanks | 3782 | 3801 | 20 | CAGTGAAACTGCGAATGGCT | MB468 | 148 | 138 | 93 | 153 | 212 | 32.6 | 86.6 | 29.0 | 127.6 | 161.5 |
| >hg38_21_+_8433172_8446622_RNA45SN1_45S_50ntFlanks@3782.3802.21\|CAGTGAAACTGCGAATGGCTC | >hg38_21_+_8433172_8446622_RNA45SN1_45S_50ntFlanks | 3782 | 3802 | 21 | CAGTGAAACTGCGAATGGCTC | MB468 | 19 | 28 | 14 | 28 | 27 | 4.2 | 16.3 | 5.5 | 24.3 | 17.8 |
| >hg38_21_+_8433172_8446622_RNA45SN1_45S_50ntFlanks@3782.3805.24\|CAGTGAAACTGCGAATGGCTCATT | >hg38_21_+_8433172_8446622_RNA45SN1_45S_50ntFlanks | 3782 | 3805 | 24 | CAGTGAAACTGCGAATGGCTCATT | MB468 | 37 | 29 | 24 | 36 | 51 | 9.1 | 14.6 | 8.2 | 30.5 | 48.8 |
| >hg38_21_+_8433172_8446622_RNA45SN1_45S_50ntFlanks@3782.3810.29\|CAGTGAAACTGCGAATGGCTCATTAAATC | >hg38_21_+_8433172_8446622_RNA45SN1_45S_50ntFlanks | 3782 | 3810 | 29 | CAGTGAAACTGCGAATGGCTCATTAAATC | MB468 | 16 | 29 | 20 | 15 | 22 | 5.9 | 9.8 | 6.2 | 12.4 | 15.2 |
| >hg38_21_+_8433172_8446622_RNA45SN1_45S_50ntFlanks@3782.3812.31\|CAGTGAAACTGCGAATGGCTCATTAAATCAG | >hg38_21_+_8433172_8446622_RNA45SN1_45S_50ntFlanks | 3782 | 3812 | 31 | CAGTGAAACTGCGAATGGCTCATTAAATCAG | MB468 | 17 | 38 | 14 | 18 | 38 | 5.6 | 29.8 | 4.0 | 20.2 | 31.4 |
| >hg38_21_+_8433172_8446622_RNA45SN1_45S_50ntFlanks@3782.3814.33\|CAGTGAAACTGCGAATGGCTCATTAAATCAGTT | >hg38_21_+_8433172_8446622_RNA45SN1_45S_50ntFlanks | 3782 | 3814 | 33 | CAGTGAAACTGCGAATGGCTCATTAAATCAGTT | MB468 | 11 | 31 | 12 | 10 | 16 | 1.4 | 14.8 | 4.4 | 9.4 | 9.9 |
| >hg38_21_+_8433172_8446622_RNA45SN1_45S_50ntFlanks@3783.3800.18\|AGTGAAACTGCGAATGGC | >hg38_21_+_8433172_8446622_RNA45SN1_45S_50ntFlanks | 3783 | 3800 | 18 | AGTGAAACTGCGAATGGC | MB468 | 12 | 17 | 12 | 23 | 15 | 5.7 | 11.2 | 6.3 | 25.3 | 11.0 |
| >hg38_21_+_8433172_8446622_RNA45SN1_45S_50ntFlanks@3783.3801.19\|AGTGAAACTGCGAATGGCT | >hg38_21_+_8433172_8446622_RNA45SN1_45S_50ntFlanks | 3783 | 3801 | 19 | AGTGAAACTGCGAATGGCT | MB468 | 34 | 30 | 17 | 40 | 54 | 15.6 | 22.9 | 6.4 | 34.9 | 52.9 |
| >hg38_21_+_8433172_8446622_RNA45SN1_45S_50ntFlanks@3784.3801.18\|GTGAAACTGCGAATGGCT | >hg38_21_+_8433172_8446622_RNA45SN1_45S_50ntFlanks | 3784 | 3801 | 18 | GTGAAACTGCGAATGGCT | MB468 | 17 | 13 | 13 | 19 | 19 | 4.4 | 8.9 | 3.1 | 18.1 | 7.7 |
| >hg38_21_+_8433172_8446622_RNA45SN1_45S_50ntFlanks@3785.3802.18\|TGAAACTGCGAATGGCTC | >hg38_21_+_8433172_8446622_RNA45SN1_45S_50ntFlanks | 3785 | 3802 | 18 | TGAAACTGCGAATGGCTC | MB468 | 13 | 12 | 10 | 16 | 11 | 2.2 | 5.2 | 2.1 | 17.0 | 2.2 |
| >hg38_21_+_8433172_8446622_RNA45SN1_45S_50ntFlanks@3785.3810.26\|TGAAACTGCGAATGGCTCATTAAATC | >hg38_21_+_8433172_8446622_RNA45SN1_45S_50ntFlanks | 3785 | 3810 | 26 | TGAAACTGCGAATGGCTCATTAAATC | MB468 | 20 | 38 | 19 | 11 | 18 | 4.9 | 14.9 | 7.9 | 13.3 | 8.3 |
| >hg38_21_+_8433172_8446622_RNA45SN1_45S_50ntFlanks@3785.3812.28\|TGAAACTGCGAATGGCTCATTAAATCAG | >hg38_21_+_8433172_8446622_RNA45SN1_45S_50ntFlanks | 3785 | 3812 | 28 | TGAAACTGCGAATGGCTCATTAAATCAG | MB468 | 27 | 56 | 15 | 17 | 38 | 11.1 | 44.6 | 3.4 | 20.9 | 34.1 |
| >hg38_21_+_8433172_8446622_RNA45SN1_45S_50ntFlanks@3787.3805.19\|AAACTGCGAATGGCTCATT | >hg38_21_+_8433172_8446622_RNA45SN1_45S_50ntFlanks | 3787 | 3805 | 19 | AAACTGCGAATGGCTCATT | MB468 | 18 | 11 | 10 | 10 | 23 | 5.4 | 5.7 | 4.4 | 6.0 | 15.4 |
| >hg38_21_+_8433172_8446622_RNA45SN1_45S_50ntFlanks@3787.3810.24\|AAACTGCGAATGGCTCATTAAATC | >hg38_21_+_8433172_8446622_RNA45SN1_45S_50ntFlanks | 3787 | 3810 | 24 | AAACTGCGAATGGCTCATTAAATC | MB468 | 51 | 81 | 43 | 22 | 50 | 24.4 | 18.9 | 20.1 | 16.4 | 27.3 |
| >hg38_21_+_8433172_8446622_RNA45SN1_45S_50ntFlanks@3787.3812.26\|AAACTGCGAATGGCTCATTAAATCAG | >hg38_21_+_8433172_8446622_RNA45SN1_45S_50ntFlanks | 3787 | 3812 | 26 | AAACTGCGAATGGCTCATTAAATCAG | MB468 | 27 | 39 | 11 | 15 | 62 | 14.1 | 28.5 | 3.3 | 16.5 | 77.9 |
| >hg38_21_+_8433172_8446622_RNA45SN1_45S_50ntFlanks@3788.3810.23\|AACTGCGAATGGCTCATTAAATC | >hg38_21_+_8433172_8446622_RNA45SN1_45S_50ntFlanks | 3788 | 3810 | 23 | AACTGCGAATGGCTCATTAAATC | MB468 | 16 | 26 | 13 | 8 | 15 | 7.8 | 9.4 | 5.7 | 7.0 | 7.2 |
| >hg38_21_+_8433172_8446622_RNA45SN1_45S_50ntFlanks@3789.3810.22\|ACTGCGAATGGCTCATTAAATC | >hg38_21_+_8433172_8446622_RNA45SN1_45S_50ntFlanks | 3789 | 3810 | 22 | ACTGCGAATGGCTCATTAAATC | MB468 | 31 | 44 | 24 | 22 | 35 | 11.2 | 25.9 | 4.3 | 17.1 | 19.5 |
| >hg38_21_+_8433172_8446622_RNA45SN1_45S_50ntFlanks@3793.3810.18\|CGAATGGCTCATTAAATC | >hg38_21_+_8433172_8446622_RNA45SN1_45S_50ntFlanks | 3793 | 3810 | 18 | CGAATGGCTCATTAAATC | MB468 | 19 | 10 | 13 | 11 | 24 | 4.8 | 2.8 | 5.8 | 6.1 | 22.4 |
| >hg38_21_+_8433172_8446622_RNA45SN1_45S_50ntFlanks@3793.3814.22\|CGAATGGCTCATTAAATCAGTT | >hg38_21_+_8433172_8446622_RNA45SN1_45S_50ntFlanks | 3793 | 3814 | 22 | CGAATGGCTCATTAAATCAGTT | MB468 | 11 | 7 | 5 | 14 | 6 | 2.7 | 3.8 | 0.7 | 2.4 | 6.1 |
| >hg38_21_+_8433172_8446622_RNA45SN1_45S_50ntFlanks@3802.3829.28\|CATTAAATCAGTTATGGTTCCTTTGGTC | >hg38_21_+_8433172_8446622_RNA45SN1_45S_50ntFlanks | 3802 | 3829 | 28 | CATTAAATCAGTTATGGTTCCTTTGGTC | MB468 | 21 | 17 | 23 | 7 | 7 | 2.3 | 5.3 | 11.6 | 4.9 | 2.9 |
| >hg38_21_+_8433172_8446622_RNA45SN1_45S_50ntFlanks@3802.3834.33\|CATTAAATCAGTTATGGTTCCTTTGGTCGCTCG | >hg38_21_+_8433172_8446622_RNA45SN1_45S_50ntFlanks | 3802 | 3834 | 33 | CATTAAATCAGTTATGGTTCCTTTGGTCGCTCG | MB468 | 28 | 97 | 50 | 5 | 36 | 8.8 | 62.2 | 18.7 | 2.4 | 22.8 |
| >hg38_21_+_8433172_8446622_RNA45SN1_45S_50ntFlanks@3803.3829.27\|ATTAAATCAGTTATGGTTCCTTTGGTC | >hg38_21_+_8433172_8446622_RNA45SN1_45S_50ntFlanks | 3803 | 3829 | 27 | ATTAAATCAGTTATGGTTCCTTTGGTC | MB468 | 11 | 12 | 7 | 2 | 2 | 3.3 | 3.8 | 2.2 | 1.4 | 0.7 |
| >hg38_21_+_8433172_8446622_RNA45SN1_45S_50ntFlanks@3811.3834.24\|AGTTATGGTTCCTTTGGTCGCTCG | >hg38_21_+_8433172_8446622_RNA45SN1_45S_50ntFlanks | 3811 | 3834 | 24 | AGTTATGGTTCCTTTGGTCGCTCG | MB468 | 12 | 32 | 12 | 3 | 12 | 4.6 | 11.9 | 6.2 | 0.9 | 7.2 |
| >hg38_21_+_8433172_8446622_RNA45SN1_45S_50ntFlanks@3813.3830.18\|TTATGGTTCCTTTGGTCG | >hg38_21_+_8433172_8446622_RNA45SN1_45S_50ntFlanks | 3813 | 3830 | 18 | TTATGGTTCCTTTGGTCG | MB468 | 11 | 11 | 7 | 4 | 8 | 4.2 | 5.1 | 2.7 | 0.8 | 1.4 |
| >hg38_21_+_8433172_8446622_RNA45SN1_45S_50ntFlanks@3813.3834.22\|TTATGGTTCCTTTGGTCGCTCG | >hg38_21_+_8433172_8446622_RNA45SN1_45S_50ntFlanks | 3813 | 3834 | 22 | TTATGGTTCCTTTGGTCGCTCG | MB468 | 14 | 36 | 28 | 8 | 29 | 2.5 | 17.6 | 9.0 | 1.4 | 24.3 |
| >hg38_21_+_8433172_8446622_RNA45SN1_45S_50ntFlanks@3813.3837.25\|TTATGGTTCCTTTGGTCGCTCGCTC | >hg38_21_+_8433172_8446622_RNA45SN1_45S_50ntFlanks | 3813 | 3837 | 25 | TTATGGTTCCTTTGGTCGCTCGCTC | MB468 | 10 | 46 | 10 | 9 | 14 | 1.4 | 23.0 | 1.6 | 2.8 | 3.3 |
| >hg38_21_+_8433172_8446622_RNA45SN1_45S_50ntFlanks@3831.3860.30\|CTCGCTCCTCTCCTACTTGGATAACTGTGG | >hg38_21_+_8433172_8446622_RNA45SN1_45S_50ntFlanks | 3831 | 3860 | 30 | CTCGCTCCTCTCCTACTTGGATAACTGTGG | MB468 | 23 | 132 | 36 | 6 | 30 | 10.0 | 31.0 | 11.4 | 4.9 | 12.8 |
| >hg38_21_+_8433172_8446622_RNA45SN1_45S_50ntFlanks@3835.3859.25\|CTCCTCTCCTACTTGGATAACTGTG | >hg38_21_+_8433172_8446622_RNA45SN1_45S_50ntFlanks | 3835 | 3859 | 25 | CTCCTCTCCTACTTGGATAACTGTG | MB468 | 11 | 12 | 17 | 5 | 22 | 5.0 | 3.2 | 9.0 | 3.1 | 26.4 |
| >hg38_21_+_8433172_8446622_RNA45SN1_45S_50ntFlanks@3835.3860.26\|CTCCTCTCCTACTTGGATAACTGTGG | >hg38_21_+_8433172_8446622_RNA45SN1_45S_50ntFlanks | 3835 | 3860 | 26 | CTCCTCTCCTACTTGGATAACTGTGG | MB468 | 145 | 261 | 186 | 36 | 195 | 120.4 | 76.4 | 97.0 | 30.7 | 158.2 |
| >hg38_21_+_8433172_8446622_RNA45SN1_45S_50ntFlanks@3835.3861.27\|CTCCTCTCCTACTTGGATAACTGTGGT | >hg38_21_+_8433172_8446622_RNA45SN1_45S_50ntFlanks | 3835 | 3861 | 27 | CTCCTCTCCTACTTGGATAACTGTGGT | MB468 | 22 | 40 | 25 | 3 | 59 | 10.3 | 14.2 | 8.2 | 2.1 | 75.8 |
| >hg38_21_+_8433172_8446622_RNA45SN1_45S_50ntFlanks@3835.3862.28\|CTCCTCTCCTACTTGGATAACTGTGGTA | >hg38_21_+_8433172_8446622_RNA45SN1_45S_50ntFlanks | 3835 | 3862 | 28 | CTCCTCTCCTACTTGGATAACTGTGGTA | MB468 | 13 | 17 | 16 | 3 | 58 | 5.7 | 1.8 | 7.7 | 1.5 | 76.4 |
| >hg38_21_+_8433172_8446622_RNA45SN1_45S_50ntFlanks@3835.3868.34\|CTCCTCTCCTACTTGGATAACTGTGGTAATTCTA | >hg38_21_+_8433172_8446622_RNA45SN1_45S_50ntFlanks | 3835 | 3868 | 34 | CTCCTCTCCTACTTGGATAACTGTGGTAATTCTA | MB468 | 10 | 38 | 28 | 4 | 19 | 6.1 | 11.0 | 12.3 | 3.4 | 10.2 |
| >hg38_21_+_8433172_8446622_RNA45SN1_45S_50ntFlanks@3835.3871.37\|CTCCTCTCCTACTTGGATAACTGTGGTAATTCTAGAG | >hg38_21_+_8433172_8446622_RNA45SN1_45S_50ntFlanks | 3835 | 3871 | 37 | CTCCTCTCCTACTTGGATAACTGTGGTAATTCTAGAG | MB468 | 18 | 70 | 66 | 3 | 29 | 16.6 | 48.0 | 69.0 | 2.0 | 20.0 |
| >hg38_21_+_8433172_8446622_RNA45SN1_45S_50ntFlanks@3837.3860.24\|CCTCTCCTACTTGGATAACTGTGG | >hg38_21_+_8433172_8446622_RNA45SN1_45S_50ntFlanks | 3837 | 3860 | 24 | CCTCTCCTACTTGGATAACTGTGG | MB468 | 21 | 25 | 19 | 12 | 32 | 7.6 | 8.6 | 2.5 | 10.5 | 31.9 |

| | | | | | | | | | | | | | | | | |
|---|---|---|---|---|---|---|---|---|---|---|---|---|---|---|---|---|
| >hg38_21_+_8433172_8446622_RNA45SN1_45S_50ntFlanks@3837.3861.25\|CCTCTCCTACTTGGATAACTGTGGT | >hg38_21_+_8433172_8446622_RNA45SN1_45S_50ntFlanks | 3837 | 3861 | 25 | CCTCTCCTACTTGGATAACTGTGGT | MB468 | 19 | 27 | 16 | 3 | 19 | 7.6 | 11.4 | 2.9 | 2.7 | 21.4 |
| >hg38_21_+_8433172_8446622_RNA45SN1_45S_50ntFlanks@3837.3868.32\|CCTCTCCTACTTGGATAACTGTGGTAATTCTA | >hg38_21_+_8433172_8446622_RNA45SN1_45S_50ntFlanks | 3837 | 3868 | 32 | CCTCTCCTACTTGGATAACTGTGGTAATTCTA | MB468 | 12 | 33 | 19 | 2 | 16 | 5.5 | 9.4 | 7.5 | 2.0 | 8.8 |
| >hg38_21_+_8433172_8446622_RNA45SN1_45S_50ntFlanks@3837.3871.35\|CCTCTCCTACTTGGATAACTGTGGTAATTCTAGAG | >hg38_21_+_8433172_8446622_RNA45SN1_45S_50ntFlanks | 3837 | 3871 | 35 | CCTCTCCTACTTGGATAACTGTGGTAATTCTAGAG | MB468 | 12 | 32 | 26 | 2 | 18 | 9.0 | 8.2 | 21.1 | 2.0 | 12.9 |
| >hg38_21_+_8433172_8446622_RNA45SN1_45S_50ntFlanks@3838.3859.22\|CTCTCCTACTTGGATAACTGTG | >hg38_21_+_8433172_8446622_RNA45SN1_45S_50ntFlanks | 3838 | 3859 | 22 | CTCTCCTACTTGGATAACTGTG | MB468 | 10 | 6 | 12 | 8 | 10 | 1.1 | 2.4 | 3.9 | 8.3 | 7.0 |
| >hg38_21_+_8433172_8446622_RNA45SN1_45S_50ntFlanks@3838.3860.23\|CTCTCCTACTTGGATAACTGTGG | >hg38_21_+_8433172_8446622_RNA45SN1_45S_50ntFlanks | 3838 | 3860 | 23 | CTCTCCTACTTGGATAACTGTGG | MB468 | 72 | 125 | 98 | 41 | 46 | 27.8 | 53.2 | 36.7 | 53.4 | 26.2 |
| >hg38_21_+_8433172_8446622_RNA45SN1_45S_50ntFlanks@3838.3861.24\|CTCTCCTACTTGGATAACTGTGGT | >hg38_21_+_8433172_8446622_RNA45SN1_45S_50ntFlanks | 3838 | 3861 | 24 | CTCTCCTACTTGGATAACTGTGGT | MB468 | 14 | 24 | 16 | 4 | 9 | 4.2 | 12.4 | 4.0 | 4.2 | 6.0 |
| >hg38_21_+_8433172_8446622_RNA45SN1_45S_50ntFlanks@3838.3862.25\|CTCTCCTACTTGGATAACTGTGGTA | >hg38_21_+_8433172_8446622_RNA45SN1_45S_50ntFlanks | 3838 | 3862 | 25 | CTCTCCTACTTGGATAACTGTGGTA | MB468 | 13 | 12 | 11 | 4 | 8 | 2.6 | 2.8 | 2.3 | 3.0 | 5.5 |
| >hg38_21_+_8433172_8446622_RNA45SN1_45S_50ntFlanks@3838.3868.31\|CTCTCCTACTTGGATAACTGTGGTAATTCTA | >hg38_21_+_8433172_8446622_RNA45SN1_45S_50ntFlanks | 3838 | 3868 | 31 | CTCTCCTACTTGGATAACTGTGGTAATTCTA | MB468 | 15 | 39 | 30 | 5 | 14 | 6.4 | 17.0 | 13.5 | 5.5 | 8.0 |
| >hg38_21_+_8433172_8446622_RNA45SN1_45S_50ntFlanks@3838.3871.34\|CTCTCCTACTTGGATAACTGTGGTAATTCTAGAG | >hg38_21_+_8433172_8446622_RNA45SN1_45S_50ntFlanks | 3838 | 3871 | 34 | CTCTCCTACTTGGATAACTGTGGTAATTCTAGAG | MB468 | 20 | 36 | 38 | 9 | 18 | 15.7 | 9.6 | 26.4 | 6.5 | 2.3 |
| >hg38_21_+_8433172_8446622_RNA45SN1_45S_50ntFlanks@3839.3868.30\|TCTCCTACTTGGATAACTGTGGTAATTCTA | >hg38_21_+_8433172_8446622_RNA45SN1_45S_50ntFlanks | 3839 | 3868 | 30 | TCTCCTACTTGGATAACTGTGGTAATTCTA | BT20 | 13 | 17 | 44 | 2 | 1 | 15.6 | 22.7 | 61.4 | 3.5 | 1.1 |
| >hg38_21_+_8433172_8446622_RNA45SN1_45S_50ntFlanks@3840.3860.21\|CTCCTACTTGGATAACTGTGG | >hg38_21_+_8433172_8446622_RNA45SN1_45S_50ntFlanks | 3840 | 3860 | 21 | CTCCTACTTGGATAACTGTGG | BT20 | 13 | 23 | 59 | 4 | 5 | 5.0 | 12.3 | 19.6 | 6.1 | 4.5 |
| >hg38_21_+_8433172_8446622_RNA45SN1_45S_50ntFlanks@3840.3860.21\|CTCCTACTTGGATAACTGTGG | >hg38_21_+_8433172_8446622_RNA45SN1_45S_50ntFlanks | 3840 | 3860 | 21 | CTCCTACTTGGATAACTGTGG | MB468 | 27 | 25 | 38 | 44 | 27 | 5.2 | 15.5 | 8.4 | 58.6 | 16.8 |
| >hg38_21_+_8433172_8446622_RNA45SN1_45S_50ntFlanks@3840.3861.22\|CTCCTACTTGGATAACTGTGGT | >hg38_21_+_8433172_8446622_RNA45SN1_45S_50ntFlanks | 3840 | 3861 | 22 | CTCCTACTTGGATAACTGTGGT | BT20 | 14 | 23 | 53 | 2 | 1 | 4.7 | 10.2 | 15.5 | 3.4 | 1.6 |
| >hg38_21_+_8433172_8446622_RNA45SN1_45S_50ntFlanks@3840.3862.23\|CTCCTACTTGGATAACTGTGGTA | >hg38_21_+_8433172_8446622_RNA45SN1_45S_50ntFlanks | 3840 | 3862 | 23 | CTCCTACTTGGATAACTGTGGTA | BT20 | 16 | 23 | 114 | 3 | 5 | 6.5 | 11.1 | 61.3 | 3.3 | 4.5 |
| >hg38_21_+_8433172_8446622_RNA45SN1_45S_50ntFlanks@3840.3865.26\|CTCCTACTTGGATAACTGTGGTAATT | >hg38_21_+_8433172_8446622_RNA45SN1_45S_50ntFlanks | 3840 | 3865 | 26 | CTCCTACTTGGATAACTGTGGTAATT | BT20 | 11 | 15 | 38 | 0 | 1 | 4.8 | 7.4 | 19.9 | 0.9 | 1.5 |
| >hg38_21_+_8433172_8446622_RNA45SN1_45S_50ntFlanks@3840.3868.29\|CTCCTACTTGGATAACTGTGGTAATTCTA | >hg38_21_+_8433172_8446622_RNA45SN1_45S_50ntFlanks | 3840 | 3868 | 29 | CTCCTACTTGGATAACTGTGGTAATTCTA | BT20 | 20 | 30 | 68 | 9 | 12 | 12.8 | 6.2 | 34.2 | 15.4 | 11.3 |
| >hg38_21_+_8433172_8446622_RNA45SN1_45S_50ntFlanks@3840.3873.34\|CTCCTACTTGGATAACTGTGGTAATTCTAGAGCT | >hg38_21_+_8433172_8446622_RNA45SN1_45S_50ntFlanks | 3840 | 3873 | 34 | CTCCTACTTGGATAACTGTGGTAATTCTAGAGCT | BT20 | 15 | 27 | 37 | 1 | 1 | 7.6 | 18.8 | 21.5 | 1.6 | 2.3 |
| >hg38_21_+_8433172_8446622_RNA45SN1_45S_50ntFlanks@3840.3879.40\|CTCCTACTTGGATAACTGTGGTAATTCTAGAGCTAATACA | >hg38_21_+_8433172_8446622_RNA45SN1_45S_50ntFlanks | 3840 | 3879 | 40 | CTCCTACTTGGATAACTGTGGTAATTCTAGAGCTAATACA | BT20 | 14 | 26 | 46 | 1 | 6 | 7.0 | 29.8 | 42.2 | 1.8 | 10.2 |
| >hg38_21_+_8433172_8446622_RNA45SN1_45S_50ntFlanks@3841.3879.39\|TCCTACTTGGATAACTGTGGTAATTCTAGAGCTAATACA | >hg38_21_+_8433172_8446622_RNA45SN1_45S_50ntFlanks | 3841 | 3879 | 39 | TCCTACTTGGATAACTGTGGTAATTCTAGAGCTAATACA | BT20 | 10 | 17 | 29 | 4 | 10 | 6.3 | 23.1 | 30.9 | 2.2 | 6.6 |
| >hg38_21_+_8433172_8446622_RNA45SN1_45S_50ntFlanks@3842.3859.18\|CCTACTTGGATAACTGTG | >hg38_21_+_8433172_8446622_RNA45SN1_45S_50ntFlanks | 3842 | 3859 | 18 | CCTACTTGGATAACTGTG | MB468 | 22 | 10 | 38 | 19 | 16 | 3.5 | 6.0 | 20.7 | 20.0 | 6.0 |
| >hg38_21_+_8433172_8446622_RNA45SN1_45S_50ntFlanks@3842.3860.19\|CCTACTTGGATAACTGTGG | >hg38_21_+_8433172_8446622_RNA45SN1_45S_50ntFlanks | 3842 | 3860 | 19 | CCTACTTGGATAACTGTGG | MB468 | 15 | 7 | 14 | 13 | 8 | 3.2 | 4.0 | 6.5 | 11.6 | 5.4 |
| >hg38_21_+_8433172_8446622_RNA45SN1_45S_50ntFlanks@3842.3861.20\|CCTACTTGGATAACTGTGGT | >hg38_21_+_8433172_8446622_RNA45SN1_45S_50ntFlanks | 3842 | 3861 | 20 | CCTACTTGGATAACTGTGGT | MB468 | 13 | 6 | 16 | 9 | 7 | 2.4 | 2.4 | 4.6 | 10.2 | 4.5 |
| >hg38_21_+_8433172_8446622_RNA45SN1_45S_50ntFlanks@3842.3868.27\|CCTACTTGGATAACTGTGGTAATTCTA | >hg38_21_+_8433172_8446622_RNA45SN1_45S_50ntFlanks | 3842 | 3868 | 27 | CCTACTTGGATAACTGTGGTAATTCTA | MB468 | 25 | 22 | 37 | 14 | 19 | 9.1 | 11.1 | 11.7 | 16.8 | 11.7 |
| >hg38_21_+_8433172_8446622_RNA45SN1_45S_50ntFlanks@3843.3860.18\|CTACTTGGATAACTGTGG | >hg38_21_+_8433172_8446622_RNA45SN1_45S_50ntFlanks | 3843 | 3860 | 18 | CTACTTGGATAACTGTGG | MB468 | 30 | 11 | 28 | 24 | 19 | 9.9 | 6.7 | 12.7 | 22.3 | 17.3 |
| >hg38_21_+_8433172_8446622_RNA45SN1_45S_50ntFlanks@3843.3861.19\|CTACTTGGATAACTGTGGT | >hg38_21_+_8433172_8446622_RNA45SN1_45S_50ntFlanks | 3843 | 3861 | 19 | CTACTTGGATAACTGTGGT | MB468 | 83 | 41 | 96 | 47 | 48 | 31.5 | 21.6 | 36.1 | 57.2 | 40.5 |
| >hg38_21_+_8433172_8446622_RNA45SN1_45S_50ntFlanks@3843.3862.20\|CTACTTGGATAACTGTGGTA | >hg38_21_+_8433172_8446622_RNA45SN1_45S_50ntFlanks | 3843 | 3862 | 20 | CTACTTGGATAACTGTGGTA | MB468 | 18 | 4 | 9 | 10 | 10 | 6.5 | 2.4 | 3.9 | 7.4 | 11.4 |
| >hg38_21_+_8433172_8446622_RNA45SN1_45S_50ntFlanks@3843.3868.26\|CTACTTGGATAACTGTGGTAATTCTA | >hg38_21_+_8433172_8446622_RNA45SN1_45S_50ntFlanks | 3843 | 3868 | 26 | CTACTTGGATAACTGTGGTAATTCTA | MB468 | 57 | 21 | 57 | 37 | 53 | 22.0 | 11.0 | 19.8 | 43.3 | 42.2 |
| >hg38_21_+_8433172_8446622_RNA45SN1_45S_50ntFlanks@3843.3869.27\|CTACTTGGATAACTGTGGTAATTCTAG | >hg38_21_+_8433172_8446622_RNA45SN1_45S_50ntFlanks | 3843 | 3869 | 27 | CTACTTGGATAACTGTGGTAATTCTAG | MB468 | 16 | 7 | 19 | 14 | 18 | 3.0 | 2.4 | 7.5 | 9.3 | 15.4 |
| >hg38_21_+_8433172_8446622_RNA45SN1_45S_50ntFlanks@3843.3871.29\|CTACTTGGATAACTGTGGTAATTCTAGAG | >hg38_21_+_8433172_8446622_RNA45SN1_45S_50ntFlanks | 3843 | 3871 | 29 | CTACTTGGATAACTGTGGTAATTCTAGAG | MB468 | 40 | 13 | 45 | 30 | 33 | 33.4 | 4.8 | 44.4 | 12.3 | 27.9 |
| >hg38_21_+_8433172_8446622_RNA45SN1_45S_50ntFlanks@3843.3872.30\|CTACTTGGATAACTGTGGTAATTCTAGAGC | >hg38_21_+_8433172_8446622_RNA45SN1_45S_50ntFlanks | 3843 | 3872 | 30 | CTACTTGGATAACTGTGGTAATTCTAGAGC | MB468 | 12 | 5 | 14 | 15 | 17 | 3.0 | 0.7 | 6.4 | 7.2 | 13.0 |
| >hg38_21_+_8433172_8446622_RNA45SN1_45S_50ntFlanks@3844.3866.23\|TACTTGGATAACTGTGGTAATTC | >hg38_21_+_8433172_8446622_RNA45SN1_45S_50ntFlanks | 3844 | 3866 | 23 | TACTTGGATAACTGTGGTAATTC | MB231 | 10 | 7 | 16 | 2 | 2 | 8.2 | 6.4 | 11.7 | 2.0 | 1.9 |
| >hg38_21_+_8433172_8446622_RNA45SN1_45S_50ntFlanks@3844.3868.25\|TACTTGGATAACTGTGGTAATTCTA | >hg38_21_+_8433172_8446622_RNA45SN1_45S_50ntFlanks | 3844 | 3868 | 25 | TACTTGGATAACTGTGGTAATTCTA | MB468 | 14 | 14 | 25 | 10 | 14 | 7.2 | 5.8 | 7.3 | 11.3 | 9.9 |
| >hg38_21_+_8433172_8446622_RNA45SN1_45S_50ntFlanks@3845.3868.24\|ACTTGGATAACTGTGGTAATTCTA | >hg38_21_+_8433172_8446622_RNA45SN1_45S_50ntFlanks | 3845 | 3868 | 24 | ACTTGGATAACTGTGGTAATTCTA | MB231 | 12 | 11 | 36 | 1 | 2 | 9.4 | 8.2 | 35.1 | 1.3 | 1.9 |
| >hg38_21_+_8433172_8446622_RNA45SN1_45S_50ntFlanks@3845.3868.24\|ACTTGGATAACTGTGGTAATTCTA | >hg38_21_+_8433172_8446622_RNA45SN1_45S_50ntFlanks | 3845 | 3868 | 24 | ACTTGGATAACTGTGGTAATTCTA | MB468 | 15 | 19 | 29 | 18 | 18 | 6.6 | 8.7 | 9.8 | 20.3 | 10.7 |
| >hg38_21_+_8433172_8446622_RNA45SN1_45S_50ntFlanks@3846.3868.23\|CTTGGATAACTGTGGTAATTCTA | >hg38_21_+_8433172_8446622_RNA45SN1_45S_50ntFlanks | 3846 | 3868 | 23 | CTTGGATAACTGTGGTAATTCTA | MB468 | 37 | 32 | 79 | 45 | 40 | 14.7 | 13.2 | 30.1 | 39.5 | 26.2 |
| >hg38_21_+_8433172_8446622_RNA45SN1_45S_50ntFlanks@3846.3869.24\|CTTGGATAACTGTGGTAATTCTAG | >hg38_21_+_8433172_8446622_RNA45SN1_45S_50ntFlanks | 3846 | 3869 | 24 | CTTGGATAACTGTGGTAATTCTAG | MB468 | 36 | 24 | 56 | 38 | 40 | 17.5 | 6.6 | 19.2 | 19.7 | 35.7 |
| >hg38_21_+_8433172_8446622_RNA45SN1_45S_50ntFlanks@3846.3881.36\|CTTGGATAACTGTGGTAATTCTAGAGCTAATACATG | >hg38_21_+_8433172_8446622_RNA45SN1_45S_50ntFlanks | 3846 | 3881 | 36 | CTTGGATAACTGTGGTAATTCTAGAGCTAATACATG | MB468 | 12 | 10 | 12 | 18 | 34 | 3.3 | 3.1 | 13.1 | 14.0 | 16.2 |
| >hg38_21_+_8433172_8446622_RNA45SN1_45S_50ntFlanks@3848.3866.19\|TGGATAACTGTGGTAATTC | >hg38_21_+_8433172_8446622_RNA45SN1_45S_50ntFlanks | 3848 | 3866 | 19 | TGGATAACTGTGGTAATTC | MB231 | 12 | 10 | 21 | 4 | 2 | 8.9 | 8.5 | 16.3 | 3.7 | 1.9 |
| >hg38_21_+_8433172_8446622_RNA45SN1_45S_50ntFlanks@3849.3866.18\|GGATAACTGTGGTAATTC | >hg38_21_+_8433172_8446622_RNA45SN1_45S_50ntFlanks | 3849 | 3866 | 18 | GGATAACTGTGGTAATTC | MB231 | 12 | 10 | 18 | 3 | 1 | 10.1 | 8.6 | 14.8 | 3.1 | 1.3 |
| >hg38_21_+_8433172_8446622_RNA45SN1_45S_50ntFlanks@3850.3869.20\|GATAACTGTGGTAATTCTAG | >hg38_21_+_8433172_8446622_RNA45SN1_45S_50ntFlanks | 3850 | 3869 | 20 | GATAACTGTGGTAATTCTAG | MB468 | 11 | 5 | 17 | 11 | 7 | 3.4 | 3.8 | 5.5 | 2.6 | 4.0 |
| >hg38_21_+_8433172_8446622_RNA45SN1_45S_50ntFlanks@3850.3871.22\|GATAACTGTGGTAATTCTAGAG | >hg38_21_+_8433172_8446622_RNA45SN1_45S_50ntFlanks | 3850 | 3871 | 22 | GATAACTGTGGTAATTCTAGAG | MB468 | 17 | 6 | 17 | 13 | 10 | 14.2 | 3.3 | 14.2 | 4.7 | 7.1 |
| >hg38_21_+_8433172_8446622_RNA45SN1_45S_50ntFlanks@3850.3872.23\|GATAACTGTGGTAATTCTAGAGC | >hg38_21_+_8433172_8446622_RNA45SN1_45S_50ntFlanks | 3850 | 3872 | 23 | GATAACTGTGGTAATTCTAGAGC | MB468 | 19 | 9 | 21 | 20 | 15 | 8.0 | 4.1 | 9.4 | 9.5 | 13.9 |
| >hg38_21_+_8433172_8446622_RNA45SN1_45S_50ntFlanks@3850.3873.24\|GATAACTGTGGTAATTCTAGAGCT | >hg38_21_+_8433172_8446622_RNA45SN1_45S_50ntFlanks | 3850 | 3873 | 24 | GATAACTGTGGTAATTCTAGAGCT | MB231 | 13 | 6 | 21 | 2 | 2 | 11.9 | 5.9 | 15.1 | 1.0 | 1.3 |
| >hg38_21_+_8433172_8446622_RNA45SN1_45S_50ntFlanks@3850.3873.24\|GATAACTGTGGTAATTCTAGAGCT | >hg38_21_+_8433172_8446622_RNA45SN1_45S_50ntFlanks | 3850 | 3873 | 24 | GATAACTGTGGTAATTCTAGAGCT | MB468 | 24 | 13 | 21 | 30 | 16 | 9.9 | 7.3 | 8.0 | 16.3 | 9.5 |
| >hg38_21_+_8433172_8446622_RNA45SN1_45S_50ntFlanks@3850.3874.25\|GATAACTGTGGTAATTCTAGAGCTA | >hg38_21_+_8433172_8446622_RNA45SN1_45S_50ntFlanks | 3850 | 3874 | 25 | GATAACTGTGGTAATTCTAGAGCTA | MB468 | 11 | 4 | 9 | 17 | 11 | 4.4 | 2.8 | 6.3 | 4.4 | 6.1 |
| >hg38_21_+_8433172_8446622_RNA45SN1_45S_50ntFlanks@3850.3881.32\|GATAACTGTGGTAATTCTAGAGCTAATACATG | >hg38_21_+_8433172_8446622_RNA45SN1_45S_50ntFlanks | 3850 | 3881 | 32 | GATAACTGTGGTAATTCTAGAGCTAATACATG | MB468 | 11 | 9 | 13 | 12 | 20 | 3.5 | 2.9 | 9.2 | 7.5 | 7.1 |
| >hg38_21_+_8433172_8446622_RNA45SN1_45S_50ntFlanks@3851.3872.22\|ATAACTGTGGTAATTCTAGAGC | >hg38_21_+_8433172_8446622_RNA45SN1_45S_50ntFlanks | 3851 | 3872 | 22 | ATAACTGTGGTAATTCTAGAGC | MB468 | 11 | 3 | 10 | 8 | 9 | 4.3 | 1.4 | 5.3 | 1.7 | 10.5 |
| >hg38_21_+_8433172_8446622_RNA45SN1_45S_50ntFlanks@3851.3873.23\|ATAACTGTGGTAATTCTAGAGCT | >hg38_21_+_8433172_8446622_RNA45SN1_45S_50ntFlanks | 3851 | 3873 | 23 | ATAACTGTGGTAATTCTAGAGCT | MB468 | 16 | 4 | 12 | 17 | 11 | 4.2 | 3.0 | 7.8 | 4.8 | 10.0 |
| >hg38_21_+_8433172_8446622_RNA45SN1_45S_50ntFlanks@3851.3874.24\|ATAACTGTGGTAATTCTAGAGCTA | >hg38_21_+_8433172_8446622_RNA45SN1_45S_50ntFlanks | 3851 | 3874 | 24 | ATAACTGTGGTAATTCTAGAGCTA | MB468 | 12 | 3 | 9 | 15 | 11 | 4.0 | 0.6 | 7.6 | 5.1 | 9.4 |
| >hg38_21_+_8433172_8446622_RNA45SN1_45S_50ntFlanks@3851.3881.31\|ATAACTGTGGTAATTCTAGAGCTAATACATG | >hg38_21_+_8433172_8446622_RNA45SN1_45S_50ntFlanks | 3851 | 3881 | 31 | ATAACTGTGGTAATTCTAGAGCTAATACATG | MB468 | 12 | 6 | 12 | 8 | 16 | 3.7 | 1.6 | 8.2 | 3.7 | 6.1 |

| | | | | | | | | | | | | | | | | |
|---|---|---|---|---|---|---|---|---|---|---|---|---|---|---|---|---|
| >hg38_21_+_8433172_8446622_RNA45SN1_45S_50ntFlanks@3852.3872.21\|TAACTGTGGTAATTCTAGAGC | >hg38_21_+_8433172_8446622_RNA45SN1_45S_50ntFlanks | 3852 | 3872 | 21 | TAACTGTGGTAATTCTAGAGC | MB468 | 16 | 3 | 11 | 7 | 8 | 5.6 | 1.5 | 5.7 | 2.2 | 9.8 |
| >hg38_21_+_8433172_8446622_RNA45SN1_45S_50ntFlanks@3852.3874.23\|TAACTGTGGTAATTCTAGAGCTA | >hg38_21_+_8433172_8446622_RNA45SN1_45S_50ntFlanks | 3852 | 3874 | 23 | TAACTGTGGTAATTCTAGAGCTA | MB468 | 19 | 3 | 12 | 14 | 13 | 8.7 | 2.1 | 10.4 | 3.8 | 13.4 |
| >hg38_21_+_8433172_8446622_RNA45SN1_45S_50ntFlanks@3852.3881.30\|TAACTGTGGTAATTCTAGAGCTAATACATG | >hg38_21_+_8433172_8446622_RNA45SN1_45S_50ntFlanks | 3852 | 3881 | 30 | TAACTGTGGTAATTCTAGAGCTAATACATG | MB468 | 16 | 9 | 16 | 8 | 19 | 5.1 | 2.8 | 10.7 | 4.2 | 7.1 |
| >hg38_21_+_8433172_8446622_RNA45SN1_45S_50ntFlanks@3853.3871.19\|AACTGTGGTAATTCTAGAG | >hg38_21_+_8433172_8446622_RNA45SN1_45S_50ntFlanks | 3853 | 3871 | 19 | AACTGTGGTAATTCTAGAG | MB468 | 29 | 4 | 13 | 27 | 12 | 14.5 | 2.5 | 9.8 | 15.5 | 14.1 |
| >hg38_21_+_8433172_8446622_RNA45SN1_45S_50ntFlanks@3853.3872.20\|AACTGTGGTAATTCTAGAGC | >hg38_21_+_8433172_8446622_RNA45SN1_45S_50ntFlanks | 3853 | 3872 | 20 | AACTGTGGTAATTCTAGAGC | MB468 | 28 | 4 | 15 | 25 | 16 | 8.4 | 2.2 | 7.3 | 0.4 | 20.7 |
| >hg38_21_+_8433172_8446622_RNA45SN1_45S_50ntFlanks@3853.3873.21\|AACTGTGGTAATTCTAGAGCT | >hg38_21_+_8433172_8446622_RNA45SN1_45S_50ntFlanks | 3853 | 3873 | 21 | AACTGTGGTAATTCTAGAGCT | MB468 | 36 | 6 | 14 | 39 | 17 | 13.6 | 3.7 | 9.8 | 13.4 | 18.1 |
| >hg38_21_+_8433172_8446622_RNA45SN1_45S_50ntFlanks@3853.3874.22\|AACTGTGGTAATTCTAGAGCTA | >hg38_21_+_8433172_8446622_RNA45SN1_45S_50ntFlanks | 3853 | 3874 | 22 | AACTGTGGTAATTCTAGAGCTA | MB468 | 16 | 3 | 9 | 23 | 13 | 3.7 | 2.7 | 7.2 | 6.0 | 10.6 |
| >hg38_21_+_8433172_8446622_RNA45SN1_45S_50ntFlanks@3853.3881.29\|AACTGTGGTAATTCTAGAGCTAATACATG | >hg38_21_+_8433172_8446622_RNA45SN1_45S_50ntFlanks | 3853 | 3881 | 29 | AACTGTGGTAATTCTAGAGCTAATACATG | MB468 | 63 | 35 | 43 | 38 | 49 | 19.5 | 14.9 | 27.3 | 11.7 | 24.6 |
| >hg38_21_+_8433172_8446622_RNA45SN1_45S_50ntFlanks@3853.3883.31\|AACTGTGGTAATTCTAGAGCTAATACATGCC | >hg38_21_+_8433172_8446622_RNA45SN1_45S_50ntFlanks | 3853 | 3883 | 31 | AACTGTGGTAATTCTAGAGCTAATACATGCC | MB468 | 11 | 12 | 14 | 7 | 6 | 4.5 | 2.8 | 5.8 | 4.9 | 2.8 |
| >hg38_21_+_8433172_8446622_RNA45SN1_45S_50ntFlanks@3854.3871.18\|ACTGTGGTAATTCTAGAG | >hg38_21_+_8433172_8446622_RNA45SN1_45S_50ntFlanks | 3854 | 3871 | 18 | ACTGTGGTAATTCTAGAG | MB468 | 47 | 6 | 20 | 37 | 12 | 18.8 | 3.9 | 13.2 | 10.1 | 12.7 |
| >hg38_21_+_8433172_8446622_RNA45SN1_45S_50ntFlanks@3854.3872.19\|ACTGTGGTAATTCTAGAGC | >hg38_21_+_8433172_8446622_RNA45SN1_45S_50ntFlanks | 3854 | 3872 | 19 | ACTGTGGTAATTCTAGAGC | MB468 | 75 | 15 | 42 | 64 | 24 | 19.9 | 8.4 | 15.7 | 41.6 | 26.5 |
| >hg38_21_+_8433172_8446622_RNA45SN1_45S_50ntFlanks@3854.3873.20\|ACTGTGGTAATTCTAGAGCT | >hg38_21_+_8433172_8446622_RNA45SN1_45S_50ntFlanks | 3854 | 3873 | 20 | ACTGTGGTAATTCTAGAGCT | MB468 | 114 | 19 | 46 | 86 | 27 | 71.2 | 12.3 | 22.3 | 47.2 | 20.6 |
| >hg38_21_+_8433172_8446622_RNA45SN1_45S_50ntFlanks@3854.3874.21\|ACTGTGGTAATTCTAGAGCTA | >hg38_21_+_8433172_8446622_RNA45SN1_45S_50ntFlanks | 3854 | 3874 | 21 | ACTGTGGTAATTCTAGAGCTA | MB468 | 50 | 12 | 24 | 36 | 20 | 10.6 | 7.5 | 8.5 | 10.1 | 12.8 |
| >hg38_21_+_8433172_8446622_RNA45SN1_45S_50ntFlanks@3854.3880.27\|ACTGTGGTAATTCTAGAGCTAATACAT | >hg38_21_+_8433172_8446622_RNA45SN1_45S_50ntFlanks | 3854 | 3880 | 27 | ACTGTGGTAATTCTAGAGCTAATACAT | MB468 | 37 | 20 | 17 | 14 | 12 | 15.2 | 15.1 | 8.7 | 15.0 | 9.6 |
| >hg38_21_+_8433172_8446622_RNA45SN1_45S_50ntFlanks@3854.3881.28\|ACTGTGGTAATTCTAGAGCTAATACATG | >hg38_21_+_8433172_8446622_RNA45SN1_45S_50ntFlanks | 3854 | 3881 | 28 | ACTGTGGTAATTCTAGAGCTAATACATG | MB468 | 203 | 118 | 130 | 77 | 80 | 86.6 | 75.2 | 104.5 | 50.7 | 50.2 |
| >hg38_21_+_8433172_8446622_RNA45SN1_45S_50ntFlanks@3854.3882.29\|ACTGTGGTAATTCTAGAGCTAATACATGC | >hg38_21_+_8433172_8446622_RNA45SN1_45S_50ntFlanks | 3854 | 3882 | 29 | ACTGTGGTAATTCTAGAGCTAATACATGC | MB468 | 21 | 16 | 20 | 8 | 7 | 2.7 | 8.0 | 10.2 | 6.9 | 3.0 |
| >hg38_21_+_8433172_8446622_RNA45SN1_45S_50ntFlanks@3854.3883.30\|ACTGTGGTAATTCTAGAGCTAATACATGCC | >hg38_21_+_8433172_8446622_RNA45SN1_45S_50ntFlanks | 3854 | 3883 | 30 | ACTGTGGTAATTCTAGAGCTAATACATGCC | MB468 | 16 | 16 | 17 | 7 | 5 | 3.7 | 6.4 | 4.6 | 5.9 | 2.2 |
| >hg38_21_+_8433172_8446622_RNA45SN1_45S_50ntFlanks@3854.3884.31\|ACTGTGGTAATTCTAGAGCTAATACATGCCG | >hg38_21_+_8433172_8446622_RNA45SN1_45S_50ntFlanks | 3854 | 3884 | 31 | ACTGTGGTAATTCTAGAGCTAATACATGCCG | MB468 | 17 | 18 | 17 | 8 | 10 | 1.8 | 7.0 | 4.8 | 6.9 | 3.5 |
| >hg38_21_+_8433172_8446622_RNA45SN1_45S_50ntFlanks@3854.3889.36\|ACTGTGGTAATTCTAGAGCTAATACATGCCGACGGG | >hg38_21_+_8433172_8446622_RNA45SN1_45S_50ntFlanks | 3854 | 3889 | 36 | ACTGTGGTAATTCTAGAGCTAATACATGCCGACGGG | MB468 | 11 | 42 | 16 | 9 | 7 | 3.9 | 17.5 | 12.3 | 7.9 | 2.6 |
| >hg38_21_+_8433172_8446622_RNA45SN1_45S_50ntFlanks@3855.3872.18\|CTGTGGTAATTCTAGAGC | >hg38_21_+_8433172_8446622_RNA45SN1_45S_50ntFlanks | 3855 | 3872 | 18 | CTGTGGTAATTCTAGAGC | MB468 | 14 | 3 | 12 | 11 | 5 | 3.8 | 1.9 | 5.6 | 4.2 | 4.9 |
| >hg38_21_+_8433172_8446622_RNA45SN1_45S_50ntFlanks@3855.3873.19\|CTGTGGTAATTCTAGAGCT | >hg38_21_+_8433172_8446622_RNA45SN1_45S_50ntFlanks | 3855 | 3873 | 19 | CTGTGGTAATTCTAGAGCT | MB468 | 12 | 3 | 7 | 10 | 5 | 4.8 | 2.3 | 3.8 | 2.4 | 3.6 |
| >hg38_21_+_8433172_8446622_RNA45SN1_45S_50ntFlanks@3855.3881.27\|CTGTGGTAATTCTAGAGCTAATACATG | >hg38_21_+_8433172_8446622_RNA45SN1_45S_50ntFlanks | 3855 | 3881 | 27 | CTGTGGTAATTCTAGAGCTAATACATG | MB468 | 26 | 15 | 15 | 12 | 17 | 9.1 | 7.6 | 10.6 | 5.5 | 13.2 |
| >hg38_21_+_8433172_8446622_RNA45SN1_45S_50ntFlanks@3858.3881.24\|TGGTAATTCTAGAGCTAATACATG | >hg38_21_+_8433172_8446622_RNA45SN1_45S_50ntFlanks | 3858 | 3881 | 24 | TGGTAATTCTAGAGCTAATACATG | MB468 | 15 | 5 | 13 | 3 | 5 | 9.6 | 3.1 | 14.8 | 2.4 | 4.4 |
| >hg38_21_+_8433172_8446622_RNA45SN1_45S_50ntFlanks@3859.3881.23\|GGTAATTCTAGAGCTAATACATG | >hg38_21_+_8433172_8446622_RNA45SN1_45S_50ntFlanks | 3859 | 3881 | 23 | GGTAATTCTAGAGCTAATACATG | MB468 | 11 | 2 | 12 | 1 | 2 | 10.7 | 1.3 | 18.6 | 1.5 | 2.1 |
| >hg38_21_+_8433172_8446622_RNA45SN1_45S_50ntFlanks@3860.3881.22\|GTAATTCTAGAGCTAATACATG | >hg38_21_+_8433172_8446622_RNA45SN1_45S_50ntFlanks | 3860 | 3881 | 22 | GTAATTCTAGAGCTAATACATG | MB468 | 11 | 2 | 9 | 2 | 4 | 8.7 | 0.9 | 11.7 | 1.8 | 3.1 |
| >hg38_21_+_8433172_8446622_RNA45SN1_45S_50ntFlanks@3861.3881.21\|TAATTCTAGAGCTAATACATG | >hg38_21_+_8433172_8446622_RNA45SN1_45S_50ntFlanks | 3861 | 3881 | 21 | TAATTCTAGAGCTAATACATG | MB468 | 22 | 4 | 31 | 2 | 6 | 15.6 | 2.3 | 42.8 | 2.0 | 4.9 |
| >hg38_21_+_8433172_8446622_RNA45SN1_45S_50ntFlanks@3862.3881.20\|AATTCTAGAGCTAATACATG | >hg38_21_+_8433172_8446622_RNA45SN1_45S_50ntFlanks | 3862 | 3881 | 20 | AATTCTAGAGCTAATACATG | MB468 | 13 | 2 | 17 | 1 | 4 | 9.1 | 1.7 | 25.3 | 0.7 | 3.3 |
| >hg38_21_+_8433172_8446622_RNA45SN1_45S_50ntFlanks@3863.3881.19\|ATTCTAGAGCTAATACATG | >hg38_21_+_8433172_8446622_RNA45SN1_45S_50ntFlanks | 3863 | 3881 | 19 | ATTCTAGAGCTAATACATG | MB468 | 12 | 1 | 13 | 2 | 5 | 10.2 | 0.7 | 18.2 | 1.1 | 5.3 |
| >hg38_21_+_8433172_8446622_RNA45SN1_45S_50ntFlanks@3866.3898.33\|CTAGAGCTAATACATGCCGACGGGCGCTGACCC | >hg38_21_+_8433172_8446622_RNA45SN1_45S_50ntFlanks | 3866 | 3898 | 33 | CTAGAGCTAATACATGCCGACGGGCGCTGACCC | MB468 | 20 | 20 | 29 | 22 | 52 | 7.9 | 18.9 | 15.0 | 18.7 | 11.1 |
| >hg38_21_+_8433172_8446622_RNA45SN1_45S_50ntFlanks@3870.3898.29\|AGCTAATACATGCCGACGGGCGCTGACCC | >hg38_21_+_8433172_8446622_RNA45SN1_45S_50ntFlanks | 3870 | 3898 | 29 | AGCTAATACATGCCGACGGGCGCTGACCC | MB468 | 26 | 25 | 30 | 17 | 52 | 8.9 | 25.0 | 17.6 | 13.0 | 18.4 |
| >hg38_21_+_8433172_8446622_RNA45SN1_45S_50ntFlanks@3872.3898.27\|CTAATACATGCCGACGGGCGCTGACCC | >hg38_21_+_8433172_8446622_RNA45SN1_45S_50ntFlanks | 3872 | 3898 | 27 | CTAATACATGCCGACGGGCGCTGACCC | MB468 | 71 | 88 | 114 | 24 | 162 | 16.1 | 52.5 | 21.2 | 12.3 | 71.5 |
| >hg38_21_+_8433172_8446622_RNA45SN1_45S_50ntFlanks@3872.3899.28\|CTAATACATGCCGACGGGCGCTGACCCC | >hg38_21_+_8433172_8446622_RNA45SN1_45S_50ntFlanks | 3872 | 3899 | 28 | CTAATACATGCCGACGGGCGCTGACCCC | MB468 | 18 | 15 | 24 | 2 | 22 | 5.3 | 5.5 | 4.3 | 1.6 | 5.6 |
| >hg38_21_+_8433172_8446622_RNA45SN1_45S_50ntFlanks@3873.3898.26\|TAATACATGCCGACGGGCGCTGACCC | >hg38_21_+_8433172_8446622_RNA45SN1_45S_50ntFlanks | 3873 | 3898 | 26 | TAATACATGCCGACGGGCGCTGACCC | MB468 | 145 | 155 | 203 | 28 | 194 | 33.6 | 121.9 | 85.3 | 19.3 | 55.1 |
| >hg38_21_+_8433172_8446622_RNA45SN1_45S_50ntFlanks@3873.3899.27\|TAATACATGCCGACGGGCGCTGACCCC | >hg38_21_+_8433172_8446622_RNA45SN1_45S_50ntFlanks | 3873 | 3899 | 27 | TAATACATGCCGACGGGCGCTGACCCC | MB468 | 38 | 27 | 39 | 2 | 27 | 11.2 | 16.0 | 13.9 | 1.7 | 4.1 |
| >hg38_21_+_8433172_8446622_RNA45SN1_45S_50ntFlanks@3874.3898.25\|AATACATGCCGACGGGCGCTGACCC | >hg38_21_+_8433172_8446622_RNA45SN1_45S_50ntFlanks | 3874 | 3898 | 25 | AATACATGCCGACGGGCGCTGACCC | MB468 | 241 | 284 | 325 | 109 | 271 | 106.7 | 250.2 | 239.1 | 59.3 | 47.6 |
| >hg38_21_+_8433172_8446622_RNA45SN1_45S_50ntFlanks@3874.3899.26\|AATACATGCCGACGGGCGCTGACCCC | >hg38_21_+_8433172_8446622_RNA45SN1_45S_50ntFlanks | 3874 | 3899 | 26 | AATACATGCCGACGGGCGCTGACCCC | MB468 | 60 | 47 | 56 | 9 | 43 | 20.3 | 30.4 | 28.7 | 6.4 | 2.9 |
| >hg38_21_+_8433172_8446622_RNA45SN1_45S_50ntFlanks@3875.3898.24\|ATACATGCCGACGGGCGCTGACCC | >hg38_21_+_8433172_8446622_RNA45SN1_45S_50ntFlanks | 3875 | 3898 | 24 | ATACATGCCGACGGGCGCTGACCC | MB468 | 237 | 306 | 321 | 163 | 564 | 27.8 | 257.6 | 71.7 | 79.3 | 127.2 |
| >hg38_21_+_8433172_8446622_RNA45SN1_45S_50ntFlanks@3875.3899.25\|ATACATGCCGACGGGCGCTGACCCC | >hg38_21_+_8433172_8446622_RNA45SN1_45S_50ntFlanks | 3875 | 3899 | 25 | ATACATGCCGACGGGCGCTGACCCC | MB468 | 70 | 53 | 69 | 12 | 69 | 22.7 | 35.1 | 12.9 | 8.9 | 15.9 |
| >hg38_21_+_8433172_8446622_RNA45SN1_45S_50ntFlanks@3876.3898.23\|TACATGCCGACGGGCGCTGACCC | >hg38_21_+_8433172_8446622_RNA45SN1_45S_50ntFlanks | 3876 | 3898 | 23 | TACATGCCGACGGGCGCTGACCC | MB468 | 86 | 73 | 120 | 30 | 89 | 28.5 | 50.5 | 60.6 | 17.6 | 16.3 |
| >hg38_21_+_8433172_8446622_RNA45SN1_45S_50ntFlanks@3876.3899.24\|TACATGCCGACGGGCGCTGACCCC | >hg38_21_+_8433172_8446622_RNA45SN1_45S_50ntFlanks | 3876 | 3899 | 24 | TACATGCCGACGGGCGCTGACCCC | MB468 | 22 | 13 | 23 | 2 | 11 | 2.0 | 5.3 | 5.6 | 1.2 | 1.3 |
| >hg38_21_+_8433172_8446622_RNA45SN1_45S_50ntFlanks@3878.3898.21\|CATGCCGACGGGCGCTGACCC | >hg38_21_+_8433172_8446622_RNA45SN1_45S_50ntFlanks | 3878 | 3898 | 21 | CATGCCGACGGGCGCTGACCC | MB468 | 20 | 24 | 32 | 6 | 18 | 3.7 | 12.1 | 8.0 | 3.1 | 4.9 |
| >hg38_21_+_8433172_8446622_RNA45SN1_45S_50ntFlanks@3880.3898.19\|TGCCGACGGGCGCTGACCC | >hg38_21_+_8433172_8446622_RNA45SN1_45S_50ntFlanks | 3880 | 3898 | 19 | TGCCGACGGGCGCTGACCC | MB468 | 10 | 8 | 16 | 4 | 9 | 6.0 | 5.3 | 8.5 | 1.6 | 4.5 |
| >hg38_21_+_8433172_8446622_RNA45SN1_45S_50ntFlanks@3881.3898.18\|GCCGACGGGCGCTGACCC | >hg38_21_+_8433172_8446622_RNA45SN1_45S_50ntFlanks | 3881 | 3898 | 18 | GCCGACGGGCGCTGACCC | MB468 | 24 | 15 | 32 | 7 | 28 | 14.8 | 12.7 | 24.0 | 4.5 | 15.7 |
| >hg38_21_+_8433172_8446622_RNA45SN1_45S_50ntFlanks@3881.3899.19\|GCCGACGGGCGCTGACCCC | >hg38_21_+_8433172_8446622_RNA45SN1_45S_50ntFlanks | 3881 | 3899 | 19 | GCCGACGGGCGCTGACCCC | MB468 | 12 | 4 | 12 | 1 | 11 | 7.5 | 2.7 | 7.3 | 0.9 | 9.2 |
| >hg38_21_+_8433172_8446622_RNA45SN1_45S_50ntFlanks@3882.3899.18\|CCGACGGGCGCTGACCCC | >hg38_21_+_8433172_8446622_RNA45SN1_45S_50ntFlanks | 3882 | 3899 | 18 | CCGACGGGCGCTGACCCC | MB468 | 212 | 32 | 166 | 246 | 213 | 117.3 | 18.3 | 89.1 | 91.1 | 169.2 |
| >hg38_21_+_8433172_8446622_RNA45SN1_45S_50ntFlanks@3882.3915.34\|CCGACGGGCGCTGACCCCCTTCGCGGGGGGGATG | >hg38_21_+_8433172_8446622_RNA45SN1_45S_50ntFlanks | 3882 | 3915 | 34 | CCGACGGGCGCTGACCCCCTTCGCGGGGGGGATG | MB468 | 11 | 15 | 29 | 6 | 10 | 6.3 | 5.9 | 14.9 | 3.2 | 3.9 |
| >hg38_21_+_8433172_8446622_RNA45SN1_45S_50ntFlanks@3885.3915.31\|ACGGGCGCTGACCCCCTTCGCGGGGGGGATG | >hg38_21_+_8433172_8446622_RNA45SN1_45S_50ntFlanks | 3885 | 3915 | 31 | ACGGGCGCTGACCCCCTTCGCGGGGGGGATG | MB468 | 19 | 22 | 52 | 12 | 13 | 12.9 | 4.8 | 28.1 | 2.3 | 2.0 |
| >hg38_21_+_8433172_8446622_RNA45SN1_45S_50ntFlanks@3899.3919.21\|CCTTCGCGGGGGGGATGCGTG | >hg38_21_+_8433172_8446622_RNA45SN1_45S_50ntFlanks | 3899 | 3919 | 21 | CCTTCGCGGGGGGGATGCGTG | MB468 | 64 | 149 | 128 | 357 | 340 | 11.0 | 129.6 | 64.9 | 256.0 | 212.9 |
| >hg38_21_+_8433172_8446622_RNA45SN1_45S_50ntFlanks@3899.3920.22\|CCTTCGCGGGGGGGATGCGTGC | >hg38_21_+_8433172_8446622_RNA45SN1_45S_50ntFlanks | 3899 | 3920 | 22 | CCTTCGCGGGGGGGATGCGTGC | MB468 | 23 | 63 | 65 | 77 | 59 | 1.0 | 67.3 | 24.5 | 54.3 | 35.6 |
| >hg38_21_+_8433172_8446622_RNA45SN1_45S_50ntFlanks@3899.3927.29\|CCTTCGCGGGGGGGATGCGTGCATTTATC | >hg38_21_+_8433172_8446622_RNA45SN1_45S_50ntFlanks | 3899 | 3927 | 29 | CCTTCGCGGGGGGGATGCGTGCATTTATC | MB468 | 26 | 25 | 35 | 34 | 66 | 15.7 | 18.8 | 22.2 | 18.6 | 29.2 |

| | | | | | | | | | | | | | | | |
|---|---|---|---|---|---|---|---|---|---|---|---|---|---|---|---|
| >hg38_21_+_8433172_8446622_RNA45SN1_45S_50ntFlanks@3899.3937.39\|CCTTCGCGGGGGGGATGCGTGCATTTATCAGATCAAAAC | >hg38_21_+_8433172_8446622_RNA45SN1_45S_50ntFlanks | 3899 | 3937 | 39 | CCTTCGCGGGGGGGATGCGTGCATTTATCAGATCAAAAC | MB468 | 13 | 31 | 40 | 53 | 65 | 12.7 | 28.6 | 40.1 | 58.1 | 37.2 |
| >hg38_21_+_8433172_8446622_RNA45SN1_45S_50ntFlanks@3899.3938.40\|CCTTCGCGGGGGGGATGCGTGCATTTATCAGATCAAAACC | >hg38_21_+_8433172_8446622_RNA45SN1_45S_50ntFlanks | 3899 | 3938 | 40 | CCTTCGCGGGGGGGATGCGTGCATTTATCAGATCAAAACC | MB468 | 13 | 15 | 45 | 32 | 69 | 15.6 | 15.0 | 40.4 | 38.2 | 73.6 |
| >hg38_21_+_8433172_8446622_RNA45SN1_45S_50ntFlanks@3900.3919.20\|CTTCGCGGGGGGGATGCGTG | >hg38_21_+_8433172_8446622_RNA45SN1_45S_50ntFlanks | 3900 | 3919 | 20 | CTTCGCGGGGGGGATGCGTG | MB468 | 18 | 36 | 27 | 145 | 45 | 5.5 | 19.9 | 13.7 | 78.7 | 14.4 |
| >hg38_21_+_8433172_8446622_RNA45SN1_45S_50ntFlanks@3930.3959.30\|ATCAAAACCAACCCGGTCAGCCCCTCTCCG | >hg38_21_+_8433172_8446622_RNA45SN1_45S_50ntFlanks | 3930 | 3959 | 30 | ATCAAAACCAACCCGGTCAGCCCCTCTCCG | MB468 | 23 | 57 | 61 | 7 | 48 | 1.9 | 17.9 | 16.1 | 1.7 | 8.0 |
| >hg38_21_+_8433172_8446622_RNA45SN1_45S_50ntFlanks@3931.3959.29\|TCAAAACCAACCCGGTCAGCCCCTCTCCG | >hg38_21_+_8433172_8446622_RNA45SN1_45S_50ntFlanks | 3931 | 3959 | 29 | TCAAAACCAACCCGGTCAGCCCCTCTCCG | MB468 | 15 | 36 | 41 | 2 | 37 | 3.5 | 6.9 | 14.9 | 0.5 | 9.9 |
| >hg38_21_+_8433172_8446622_RNA45SN1_45S_50ntFlanks@3939.3958.20\|AACCCGGTCAGCCCCTCTCC | >hg38_21_+_8433172_8446622_RNA45SN1_45S_50ntFlanks | 3939 | 3958 | 20 | AACCCGGTCAGCCCCTCTCC | MB468 | 15 | 25 | 17 | 4 | 39 | 8.2 | 24.5 | 11.6 | 2.4 | 8.3 |
| >hg38_21_+_8433172_8446622_RNA45SN1_45S_50ntFlanks@3939.3959.21\|AACCCGGTCAGCCCCTCTCCG | >hg38_21_+_8433172_8446622_RNA45SN1_45S_50ntFlanks | 3939 | 3959 | 21 | AACCCGGTCAGCCCCTCTCCG | MB468 | 60 | 60 | 86 | 20 | 148 | 36.2 | 48.6 | 66.4 | 12.9 | 49.5 |
| >hg38_21_+_8433172_8446622_RNA45SN1_45S_50ntFlanks@3940.3959.20\|ACCCGGTCAGCCCCTCTCCG | >hg38_21_+_8433172_8446622_RNA45SN1_45S_50ntFlanks | 3940 | 3959 | 20 | ACCCGGTCAGCCCCTCTCCG | MB468 | 13 | 16 | 27 | 2 | 22 | 3.9 | 8.2 | 10.5 | 1.2 | 7.0 |
| >hg38_21_+_8433172_8446622_RNA45SN1_45S_50ntFlanks@3984.4001.18\|GCGGCTTTGGTGACTCTA | >hg38_21_+_8433172_8446622_RNA45SN1_45S_50ntFlanks | 3984 | 4001 | 18 | GCGGCTTTGGTGACTCTA | MB468 | 101 | 317 | 69 | 121 | 146 | 34.2 | 145.5 | 35.4 | 124.0 | 90.0 |
| >hg38_21_+_8433172_8446622_RNA45SN1_45S_50ntFlanks@3984.4002.19\|GCGGCTTTGGTGACTCTAG | >hg38_21_+_8433172_8446622_RNA45SN1_45S_50ntFlanks | 3984 | 4002 | 19 | GCGGCTTTGGTGACTCTAG | MB468 | 78 | 299 | 45 | 140 | 208 | 23.1 | 188.4 | 18.1 | 143.8 | 170.3 |
| >hg38_21_+_8433172_8446622_RNA45SN1_45S_50ntFlanks@3984.4005.22\|GCGGCTTTGGTGACTCTAGATA | >hg38_21_+_8433172_8446622_RNA45SN1_45S_50ntFlanks | 3984 | 4005 | 22 | GCGGCTTTGGTGACTCTAGATA | MB468 | 14 | 49 | 4 | 31 | 30 | 5.0 | 29.1 | 2.0 | 22.3 | 22.5 |
| >hg38_21_+_8433172_8446622_RNA45SN1_45S_50ntFlanks@3984.4024.41\|GCGGCTTTGGTGACTCTAGATAACCTCGGGCCGATCGCACG | >hg38_21_+_8433172_8446622_RNA45SN1_45S_50ntFlanks | 3984 | 4024 | 41 | GCGGCTTTGGTGACTCTAGATAACCTCGGGCCGATCGCACG | MB468 | 34 | 17 | 74 | 229 | 154 | 35.4 | 10.2 | 76.5 | 195.7 | 135.5 |
| >hg38_21_+_8433172_8446622_RNA45SN1_45S_50ntFlanks@3984.4026.43\|GCGGCTTTGGTGACTCTAGATAACCTCGGGCCGATCGCACGCC | >hg38_21_+_8433172_8446622_RNA45SN1_45S_50ntFlanks | 3984 | 4026 | 43 | GCGGCTTTGGTGACTCTAGATAACCTCGGGCCGATCGCACGCC | MB468 | 38 | 13 | 56 | 69 | 161 | 38.8 | 5.3 | 54.9 | 58.1 | 166.2 |
| >hg38_21_+_8433172_8446622_RNA45SN1_45S_50ntFlanks@3985.4002.18\|CGGCTTTGGTGACTCTAG | >hg38_21_+_8433172_8446622_RNA45SN1_45S_50ntFlanks | 3985 | 4002 | 18 | CGGCTTTGGTGACTCTAG | MB468 | 337 | 478 | 202 | 360 | 355 | 119.7 | 216.0 | 60.9 | 299.9 | 219.8 |
| >hg38_21_+_8433172_8446622_RNA45SN1_45S_50ntFlanks@3985.4003.19\|CGGCTTTGGTGACTCTAGA | >hg38_21_+_8433172_8446622_RNA45SN1_45S_50ntFlanks | 3985 | 4003 | 19 | CGGCTTTGGTGACTCTAGA | MB468 | 10 | 14 | 6 | 12 | 17 | 4.0 | 5.6 | 1.5 | 10.2 | 12.8 |
| >hg38_21_+_8433172_8446622_RNA45SN1_45S_50ntFlanks@3985.4004.20\|CGGCTTTGGTGACTCTAGAT | >hg38_21_+_8433172_8446622_RNA45SN1_45S_50ntFlanks | 3985 | 4004 | 20 | CGGCTTTGGTGACTCTAGAT | MB468 | 18 | 31 | 11 | 20 | 26 | 5.3 | 17.9 | 1.4 | 11.6 | 21.8 |
| >hg38_21_+_8433172_8446622_RNA45SN1_45S_50ntFlanks@3985.4005.21\|CGGCTTTGGTGACTCTAGATA | >hg38_21_+_8433172_8446622_RNA45SN1_45S_50ntFlanks | 3985 | 4005 | 21 | CGGCTTTGGTGACTCTAGATA | MB468 | 54 | 89 | 20 | 68 | 55 | 14.9 | 33.2 | 4.0 | 52.0 | 36.0 |
| >hg38_21_+_8433172_8446622_RNA45SN1_45S_50ntFlanks@3985.4010.26\|CGGCTTTGGTGACTCTAGATAACCTC | >hg38_21_+_8433172_8446622_RNA45SN1_45S_50ntFlanks | 3985 | 4010 | 26 | CGGCTTTGGTGACTCTAGATAACCTC | MB468 | 11 | 37 | 8 | 8 | 16 | 4.3 | 16.9 | 5.3 | 6.8 | 4.8 |
| >hg38_21_+_8433172_8446622_RNA45SN1_45S_50ntFlanks@3985.4024.40\|CGGCTTTGGTGACTCTAGATAACCTCGGGCCGATCGCACG | >hg38_21_+_8433172_8446622_RNA45SN1_45S_50ntFlanks | 3985 | 4024 | 40 | CGGCTTTGGTGACTCTAGATAACCTCGGGCCGATCGCACG | MB468 | 38 | 18 | 95 | 259 | 146 | 41.1 | 5.6 | 96.3 | 221.0 | 126.0 |
| >hg38_21_+_8433172_8446622_RNA45SN1_45S_50ntFlanks@3985.4026.42\|CGGCTTTGGTGACTCTAGATAACCTCGGGCCGATCGCACGCC | >hg38_21_+_8433172_8446622_RNA45SN1_45S_50ntFlanks | 3985 | 4026 | 42 | CGGCTTTGGTGACTCTAGATAACCTCGGGCCGATCGCACGCC | MB468 | 71 | 61 | 138 | 111 | 264 | 67.0 | 25.1 | 130.2 | 100.1 | 243.4 |
| >hg38_21_+_8433172_8446622_RNA45SN1_45S_50ntFlanks@3986.4023.38\|GGCTTTGGTGACTCTAGATAACCTCGGGCCGATCGCAC | >hg38_21_+_8433172_8446622_RNA45SN1_45S_50ntFlanks | 3986 | 4023 | 38 | GGCTTTGGTGACTCTAGATAACCTCGGGCCGATCGCAC | BT20 | 17 | 18 | 59 | 4 | 8 | 4.0 | 12.6 | 16.7 | 2.7 | 5.9 |
| >hg38_21_+_8433172_8446622_RNA45SN1_45S_50ntFlanks@3986.4023.38\|GGCTTTGGTGACTCTAGATAACCTCGGGCCGATCGCAC | >hg38_21_+_8433172_8446622_RNA45SN1_45S_50ntFlanks | 3986 | 4023 | 38 | GGCTTTGGTGACTCTAGATAACCTCGGGCCGATCGCAC | MB231 | 13 | 12 | 43 | 2 | 7 | 3.9 | 8.6 | 43.0 | 1.3 | 2.6 |
| >hg38_21_+_8433172_8446622_RNA45SN1_45S_50ntFlanks@3987.4023.37\|GCTTTGGTGACTCTAGATAACCTCGGGCCGATCGCAC | >hg38_21_+_8433172_8446622_RNA45SN1_45S_50ntFlanks | 3987 | 4023 | 37 | GCTTTGGTGACTCTAGATAACCTCGGGCCGATCGCAC | BT20 | 22 | 18 | 48 | 34 | 14 | 11.3 | 14.8 | 11.3 | 27.4 | 15.6 |
| >hg38_21_+_8433172_8446622_RNA45SN1_45S_50ntFlanks@3988.4023.36\|CTTTGGTGACTCTAGATAACCTCGGGCCGATCGCAC | >hg38_21_+_8433172_8446622_RNA45SN1_45S_50ntFlanks | 3988 | 4023 | 36 | CTTTGGTGACTCTAGATAACCTCGGGCCGATCGCAC | BT20 | 46 | 37 | 101 | 149 | 35 | 26.0 | 22.0 | 24.5 | 232.6 | 29.4 |
| >hg38_21_+_8433172_8446622_RNA45SN1_45S_50ntFlanks@3988.4023.36\|CTTTGGTGACTCTAGATAACCTCGGGCCGATCGCAC | >hg38_21_+_8433172_8446622_RNA45SN1_45S_50ntFlanks | 3988 | 4023 | 36 | CTTTGGTGACTCTAGATAACCTCGGGCCGATCGCAC | MB231 | 20 | 17 | 47 | 2 | 8 | 8.2 | 9.7 | 19.6 | 0.4 | 1.9 |
| >hg38_21_+_8433172_8446622_RNA45SN1_45S_50ntFlanks@3990.4023.34\|TTGGTGACTCTAGATAACCTCGGGCCGATCGCAC | >hg38_21_+_8433172_8446622_RNA45SN1_45S_50ntFlanks | 3990 | 4023 | 34 | TTGGTGACTCTAGATAACCTCGGGCCGATCGCAC | BT20 | 24 | 17 | 49 | 11 | 16 | 16.0 | 13.7 | 32.4 | 1.9 | 12.4 |
| >hg38_21_+_8433172_8446622_RNA45SN1_45S_50ntFlanks@3990.4023.34\|TTGGTGACTCTAGATAACCTCGGGCCGATCGCAC | >hg38_21_+_8433172_8446622_RNA45SN1_45S_50ntFlanks | 3990 | 4023 | 34 | TTGGTGACTCTAGATAACCTCGGGCCGATCGCAC | MB231 | 34 | 32 | 43 | 6 | 25 | 26.5 | 18.8 | 29.0 | 3.3 | 19.9 |
| >hg38_21_+_8433172_8446622_RNA45SN1_45S_50ntFlanks@3990.4024.35\|TTGGTGACTCTAGATAACCTCGGGCCGATCGCACG | >hg38_21_+_8433172_8446622_RNA45SN1_45S_50ntFlanks | 3990 | 4024 | 35 | TTGGTGACTCTAGATAACCTCGGGCCGATCGCACG | MB231 | 10 | 10 | 23 | 2 | 7 | 8.4 | 5.2 | 24.3 | 1.3 | 5.5 |
| >hg38_21_+_8433172_8446622_RNA45SN1_45S_50ntFlanks@3991.4023.33\|TGGTGACTCTAGATAACCTCGGGCCGATCGCAC | >hg38_21_+_8433172_8446622_RNA45SN1_45S_50ntFlanks | 3991 | 4023 | 33 | TGGTGACTCTAGATAACCTCGGGCCGATCGCAC | BT20 | 13 | 9 | 27 | 9 | 9 | 9.2 | 6.9 | 20.8 | 6.6 | 5.5 |
| >hg38_21_+_8433172_8446622_RNA45SN1_45S_50ntFlanks@3991.4023.33\|TGGTGACTCTAGATAACCTCGGGCCGATCGCAC | >hg38_21_+_8433172_8446622_RNA45SN1_45S_50ntFlanks | 3991 | 4023 | 33 | TGGTGACTCTAGATAACCTCGGGCCGATCGCAC | MB231 | 14 | 13 | 19 | 3 | 10 | 10.9 | 7.2 | 11.2 | 1.5 | 8.0 |
| >hg38_21_+_8433172_8446622_RNA45SN1_45S_50ntFlanks@3991.4024.34\|TGGTGACTCTAGATAACCTCGGGCCGATCGCACG | >hg38_21_+_8433172_8446622_RNA45SN1_45S_50ntFlanks | 3991 | 4024 | 34 | TGGTGACTCTAGATAACCTCGGGCCGATCGCACG | MB468 | 13 | 43 | 21 | 20 | 33 | 5.8 | 18.0 | 11.4 | 9.8 | 4.1 |
| >hg38_21_+_8433172_8446622_RNA45SN1_45S_50ntFlanks@3992.4023.32\|GGTGACTCTAGATAACCTCGGGCCGATCGCAC | >hg38_21_+_8433172_8446622_RNA45SN1_45S_50ntFlanks | 3992 | 4023 | 32 | GGTGACTCTAGATAACCTCGGGCCGATCGCAC | BT20 | 15 | 9 | 28 | 11 | 9 | 10.2 | 5.9 | 16.7 | 6.1 | 5.7 |
| >hg38_21_+_8433172_8446622_RNA45SN1_45S_50ntFlanks@3992.4023.32\|GGTGACTCTAGATAACCTCGGGCCGATCGCAC | >hg38_21_+_8433172_8446622_RNA45SN1_45S_50ntFlanks | 3992 | 4023 | 32 | GGTGACTCTAGATAACCTCGGGCCGATCGCAC | MB231 | 34 | 28 | 34 | 7 | 25 | 28.5 | 24.1 | 37.5 | 4.6 | 21.7 |
| >hg38_21_+_8433172_8446622_RNA45SN1_45S_50ntFlanks@3992.4024.33\|GGTGACTCTAGATAACCTCGGGCCGATCGCACG | >hg38_21_+_8433172_8446622_RNA45SN1_45S_50ntFlanks | 3992 | 4024 | 33 | GGTGACTCTAGATAACCTCGGGCCGATCGCACG | MB231 | 16 | 13 | 24 | 3 | 10 | 14.7 | 8.0 | 19.9 | 2.2 | 9.0 |
| >hg38_21_+_8433172_8446622_RNA45SN1_45S_50ntFlanks@3993.4013.21\|GTGACTCTAGATAACCTCGGG | >hg38_21_+_8433172_8446622_RNA45SN1_45S_50ntFlanks | 3993 | 4013 | 21 | GTGACTCTAGATAACCTCGGG | MB468 | 39 | 84 | 52 | 38 | 36 | 13.0 | 48.4 | 21.8 | 23.6 | 23.0 |
| >hg38_21_+_8433172_8446622_RNA45SN1_45S_50ntFlanks@3993.4022.30\|GTGACTCTAGATAACCTCGGGCCGATCGCA | >hg38_21_+_8433172_8446622_RNA45SN1_45S_50ntFlanks | 3993 | 4022 | 30 | GTGACTCTAGATAACCTCGGGCCGATCGCA | MB231 | 10 | 5 | 8 | 2 | 4 | 9.6 | 5.3 | 8.6 | 1.8 | 3.9 |
| >hg38_21_+_8433172_8446622_RNA45SN1_45S_50ntFlanks@3993.4023.31\|GTGACTCTAGATAACCTCGGGCCGATCGCAC | >hg38_21_+_8433172_8446622_RNA45SN1_45S_50ntFlanks | 3993 | 4023 | 31 | GTGACTCTAGATAACCTCGGGCCGATCGCAC | BT20 | 26 | 16 | 49 | 16 | 21 | 18.0 | 11.9 | 41.0 | 10.1 | 15.7 |
| >hg38_21_+_8433172_8446622_RNA45SN1_45S_50ntFlanks@3993.4023.31\|GTGACTCTAGATAACCTCGGGCCGATCGCAC | >hg38_21_+_8433172_8446622_RNA45SN1_45S_50ntFlanks | 3993 | 4023 | 31 | GTGACTCTAGATAACCTCGGGCCGATCGCAC | MB231 | 59 | 45 | 60 | 10 | 38 | 48.8 | 40.8 | 49.2 | 7.0 | 32.0 |
| >hg38_21_+_8433172_8446622_RNA45SN1_45S_50ntFlanks@3993.4024.32\|GTGACTCTAGATAACCTCGGGCCGATCGCACG | >hg38_21_+_8433172_8446622_RNA45SN1_45S_50ntFlanks | 3993 | 4024 | 32 | GTGACTCTAGATAACCTCGGGCCGATCGCACG | MB231 | 18 | 16 | 43 | 2 | 12 | 14.4 | 10.7 | 46.1 | 1.7 | 11.2 |
| >hg38_21_+_8433172_8446622_RNA45SN1_45S_50ntFlanks@3993.4024.32\|GTGACTCTAGATAACCTCGGGCCGATCGCACG | >hg38_21_+_8433172_8446622_RNA45SN1_45S_50ntFlanks | 3993 | 4024 | 32 | GTGACTCTAGATAACCTCGGGCCGATCGCACG | MB468 | 21 | 62 | 16 | 40 | 119 | 6.5 | 32.3 | 7.5 | 20.0 | 71.4 |
| >hg38_21_+_8433172_8446622_RNA45SN1_45S_50ntFlanks@3994.4013.20\|TGACTCTAGATAACCTCGGG | >hg38_21_+_8433172_8446622_RNA45SN1_45S_50ntFlanks | 3994 | 4013 | 20 | TGACTCTAGATAACCTCGGG | MB468 | 15 | 18 | 17 | 7 | 4 | 6.9 | 12.6 | 7.0 | 1.6 | 1.6 |
| >hg38_21_+_8433172_8446622_RNA45SN1_45S_50ntFlanks@3994.4023.30\|TGACTCTAGATAACCTCGGGCCGATCGCAC | >hg38_21_+_8433172_8446622_RNA45SN1_45S_50ntFlanks | 3994 | 4023 | 30 | TGACTCTAGATAACCTCGGGCCGATCGCAC | MB231 | 12 | 11 | 19 | 1 | 8 | 10.1 | 7.4 | 14.4 | 1.0 | 7.8 |
| >hg38_21_+_8433172_8446622_RNA45SN1_45S_50ntFlanks@3996.4023.28\|ACTCTAGATAACCTCGGGCCGATCGCAC | >hg38_21_+_8433172_8446622_RNA45SN1_45S_50ntFlanks | 3996 | 4023 | 28 | ACTCTAGATAACCTCGGGCCGATCGCAC | BT20 | 10 | 11 | 13 | 1 | 0 | 10.9 | 16.8 | 18.3 | 2.5 | 0.7 |
| >hg38_21_+_8433172_8446622_RNA45SN1_45S_50ntFlanks@3996.4024.29\|ACTCTAGATAACCTCGGGCCGATCGCACG | >hg38_21_+_8433172_8446622_RNA45SN1_45S_50ntFlanks | 3996 | 4024 | 29 | ACTCTAGATAACCTCGGGCCGATCGCACG | MB468 | 15 | 69 | 10 | 10 | 30 | 5.7 | 45.9 | 3.9 | 6.2 | 11.4 |
| >hg38_21_+_8433172_8446622_RNA45SN1_45S_50ntFlanks@3997.4023.27\|CTCTAGATAACCTCGGGCCGATCGCAC | >hg38_21_+_8433172_8446622_RNA45SN1_45S_50ntFlanks | 3997 | 4023 | 27 | CTCTAGATAACCTCGGGCCGATCGCAC | BT20 | 22 | 23 | 43 | 1 | 10 | 32.6 | 37.0 | 71.1 | 1.5 | 15.7 |
| >hg38_21_+_8433172_8446622_RNA45SN1_45S_50ntFlanks@3999.4024.26\|CTAGATAACCTCGGGCCGATCGCACG | >hg38_21_+_8433172_8446622_RNA45SN1_45S_50ntFlanks | 3999 | 4024 | 26 | CTAGATAACCTCGGGCCGATCGCACG | MB468 | 12 | 25 | 18 | 7 | 13 | 3.7 | 10.2 | 8.4 | 3.4 | 3.3 |
| >hg38_21_+_8433172_8446622_RNA45SN1_45S_50ntFlanks@4000.4024.25\|TAGATAACCTCGGGCCGATCGCACG | >hg38_21_+_8433172_8446622_RNA45SN1_45S_50ntFlanks | 4000 | 4024 | 25 | TAGATAACCTCGGGCCGATCGCACG | MB468 | 33 | 49 | 48 | 16 | 30 | 4.3 | 13.7 | 9.0 | 9.3 | 8.3 |
| >hg38_21_+_8433172_8446622_RNA45SN1_45S_50ntFlanks@4002.4020.19\|GATAACCTCGGGCCGATCG | >hg38_21_+_8433172_8446622_RNA45SN1_45S_50ntFlanks | 4002 | 4020 | 19 | GATAACCTCGGGCCGATCG | MB468 | 27 | 105 | 35 | 8 | 32 | 10.0 | 70.0 | 16.9 | 3.9 | 15.6 |
| >hg38_21_+_8433172_8446622_RNA45SN1_45S_50ntFlanks@4002.4022.21\|GATAACCTCGGGCCGATCGCA | >hg38_21_+_8433172_8446622_RNA45SN1_45S_50ntFlanks | 4002 | 4022 | 21 | GATAACCTCGGGCCGATCGCA | MB468 | 27 | 83 | 39 | 7 | 29 | 8.9 | 55.6 | 16.7 | 1.9 | 12.1 |
| >hg38_21_+_8433172_8446622_RNA45SN1_45S_50ntFlanks@4002.4023.22\|GATAACCTCGGGCCGATCGCAC | >hg38_21_+_8433172_8446622_RNA45SN1_45S_50ntFlanks | 4002 | 4023 | 22 | GATAACCTCGGGCCGATCGCAC | MB468 | 24 | 116 | 44 | 6 | 28 | 8.9 | 57.3 | 23.4 | 3.0 | 9.1 |

| | | | | | | | | | | | | | | | | |
|---|---|---|---|---|---|---|---|---|---|---|---|---|---|---|---|---|
| >hg38_21_+_8433172_8446622_RNA45SN1_45S_50ntFlanks@4002.4024.23\|GATAACCTCGGGCCGATCGCACG | >hg38_21_+_8433172_8446622_RNA45SN1_45S_50ntFlanks | 4002 | 4024 | 23 | GATAACCTCGGGCCGATCGCACG | MB468 | 113 | 182 | 232 | 58 | 101 | 28.9 | 76.0 | 62.6 | 32.1 | 25.5 |
| >hg38_21_+_8433172_8446622_RNA45SN1_45S_50ntFlanks@4002.4025.24\|GATAACCTCGGGCCGATCGCACGC | >hg38_21_+_8433172_8446622_RNA45SN1_45S_50ntFlanks | 4002 | 4025 | 24 | GATAACCTCGGGCCGATCGCACGC | MB468 | 24 | 76 | 38 | 5 | 31 | 9.8 | 60.0 | 13.8 | 1.9 | 14.2 |
| >hg38_21_+_8433172_8446622_RNA45SN1_45S_50ntFlanks@4002.4026.25\|GATAACCTCGGGCCGATCGCACGCC | >hg38_21_+_8433172_8446622_RNA45SN1_45S_50ntFlanks | 4002 | 4026 | 25 | GATAACCTCGGGCCGATCGCACGCC | MB468 | 16 | 27 | 30 | 17 | 24 | 2.4 | 17.7 | 7.1 | 7.1 | 10.5 |
| >hg38_21_+_8433172_8446622_RNA45SN1_45S_50ntFlanks@4002.4042.41\|GATAACCTCGGGCCGATCGCACGCCCCCCGTGGCGGCGACG | >hg38_21_+_8433172_8446622_RNA45SN1_45S_50ntFlanks | 4002 | 4042 | 41 | GATAACCTCGGGCCGATCGCACGCCCCCCGTGGCGGCGACG | MB468 | 18 | 51 | 106 | 2 | 15 | 9.4 | 29.7 | 73.2 | 2.0 | 12.3 |
| >hg38_21_+_8433172_8446622_RNA45SN1_45S_50ntFlanks@4003.4020.18\|ATAACCTCGGGCCGATCG | >hg38_21_+_8433172_8446622_RNA45SN1_45S_50ntFlanks | 4003 | 4020 | 18 | ATAACCTCGGGCCGATCG | MB468 | 63 | 211 | 55 | 12 | 73 | 27.5 | 113.9 | 21.8 | 3.8 | 32.9 |
| >hg38_21_+_8433172_8446622_RNA45SN1_45S_50ntFlanks@4003.4022.20\|ATAACCTCGGGCCGATCGCA | >hg38_21_+_8433172_8446622_RNA45SN1_45S_50ntFlanks | 4003 | 4022 | 20 | ATAACCTCGGGCCGATCGCA | MB468 | 53 | 137 | 51 | 10 | 57 | 19.4 | 73.0 | 13.3 | 2.3 | 25.1 |
| >hg38_21_+_8433172_8446622_RNA45SN1_45S_50ntFlanks@4003.4023.21\|ATAACCTCGGGCCGATCGCAC | >hg38_21_+_8433172_8446622_RNA45SN1_45S_50ntFlanks | 4003 | 4023 | 21 | ATAACCTCGGGCCGATCGCAC | MB468 | 35 | 154 | 44 | 7 | 44 | 13.6 | 56.9 | 22.8 | 1.2 | 19.1 |
| >hg38_21_+_8433172_8446622_RNA45SN1_45S_50ntFlanks@4003.4024.22\|ATAACCTCGGGCCGATCGCACG | >hg38_21_+_8433172_8446622_RNA45SN1_45S_50ntFlanks | 4003 | 4024 | 22 | ATAACCTCGGGCCGATCGCACG | MB468 | 216 | 228 | 335 | 93 | 199 | 77.6 | 68.6 | 80.4 | 30.3 | 73.6 |
| >hg38_21_+_8433172_8446622_RNA45SN1_45S_50ntFlanks@4003.4025.23\|ATAACCTCGGGCCGATCGCACGC | >hg38_21_+_8433172_8446622_RNA45SN1_45S_50ntFlanks | 4003 | 4025 | 23 | ATAACCTCGGGCCGATCGCACGC | MB468 | 63 | 113 | 76 | 9 | 78 | 21.7 | 61.5 | 27.1 | 2.6 | 42.3 |
| >hg38_21_+_8433172_8446622_RNA45SN1_45S_50ntFlanks@4003.4026.24\|ATAACCTCGGGCCGATCGCACGCC | >hg38_21_+_8433172_8446622_RNA45SN1_45S_50ntFlanks | 4003 | 4026 | 24 | ATAACCTCGGGCCGATCGCACGCC | MB468 | 48 | 70 | 68 | 28 | 84 | 7.6 | 45.4 | 12.4 | 9.1 | 52.6 |
| >hg38_21_+_8433172_8446622_RNA45SN1_45S_50ntFlanks@4003.4042.40\|ATAACCTCGGGCCGATCGCACGCCCCCCGTGGCGGCGACG | >hg38_21_+_8433172_8446622_RNA45SN1_45S_50ntFlanks | 4003 | 4042 | 40 | ATAACCTCGGGCCGATCGCACGCCCCCCGTGGCGGCGACG | MB468 | 56 | 151 | 230 | 6 | 79 | 31.2 | 27.3 | 146.9 | 4.9 | 36.3 |
| >hg38_21_+_8433172_8446622_RNA45SN1_45S_50ntFlanks@4004.4024.21\|TAACCTCGGGCCGATCGCACG | >hg38_21_+_8433172_8446622_RNA45SN1_45S_50ntFlanks | 4004 | 4024 | 21 | TAACCTCGGGCCGATCGCACG | MB468 | 10 | 9 | 17 | 3 | 6 | 3.1 | 3.8 | 2.8 | 1.0 | 1.7 |
| >hg38_21_+_8433172_8446622_RNA45SN1_45S_50ntFlanks@4005.4022.18\|AACCTCGGGCCGATCGCA | >hg38_21_+_8433172_8446622_RNA45SN1_45S_50ntFlanks | 4005 | 4022 | 18 | AACCTCGGGCCGATCGCA | MB468 | 20 | 55 | 19 | 5 | 23 | 4.4 | 33.7 | 6.1 | 1.8 | 9.9 |
| >hg38_21_+_8433172_8446622_RNA45SN1_45S_50ntFlanks@4005.4023.19\|AACCTCGGGCCGATCGCAC | >hg38_21_+_8433172_8446622_RNA45SN1_45S_50ntFlanks | 4005 | 4023 | 19 | AACCTCGGGCCGATCGCAC | BT20 | 11 | 11 | 44 | 7 | 2 | 8.7 | 11.5 | 58.5 | 11.6 | 1.9 |
| >hg38_21_+_8433172_8446622_RNA45SN1_45S_50ntFlanks@4005.4023.19\|AACCTCGGGCCGATCGCAC | >hg38_21_+_8433172_8446622_RNA45SN1_45S_50ntFlanks | 4005 | 4023 | 19 | AACCTCGGGCCGATCGCAC | MB231 | 12 | 15 | 69 | 1 | 8 | 10.6 | 8.3 | 70.1 | 0.5 | 6.6 |
| >hg38_21_+_8433172_8446622_RNA45SN1_45S_50ntFlanks@4005.4023.19\|AACCTCGGGCCGATCGCAC | >hg38_21_+_8433172_8446622_RNA45SN1_45S_50ntFlanks | 4005 | 4023 | 19 | AACCTCGGGCCGATCGCAC | MB468 | 11 | 49 | 14 | 2 | 13 | 2.8 | 22.8 | 5.4 | 0.6 | 4.2 |
| >hg38_21_+_8433172_8446622_RNA45SN1_45S_50ntFlanks@4005.4024.20\|AACCTCGGGCCGATCGCACG | >hg38_21_+_8433172_8446622_RNA45SN1_45S_50ntFlanks | 4005 | 4024 | 20 | AACCTCGGGCCGATCGCACG | MB468 | 63 | 67 | 102 | 30 | 48 | 14.9 | 27.5 | 26.1 | 9.1 | 18.4 |
| >hg38_21_+_8433172_8446622_RNA45SN1_45S_50ntFlanks@4005.4025.21\|AACCTCGGGCCGATCGCACGC | >hg38_21_+_8433172_8446622_RNA45SN1_45S_50ntFlanks | 4005 | 4025 | 21 | AACCTCGGGCCGATCGCACGC | MB468 | 21 | 42 | 25 | 3 | 27 | 5.8 | 30.6 | 9.8 | 0.9 | 17.7 |
| >hg38_21_+_8433172_8446622_RNA45SN1_45S_50ntFlanks@4005.4026.22\|AACCTCGGGCCGATCGCACGCC | >hg38_21_+_8433172_8446622_RNA45SN1_45S_50ntFlanks | 4005 | 4026 | 22 | AACCTCGGGCCGATCGCACGCC | MB468 | 16 | 26 | 24 | 8 | 29 | 4.7 | 20.4 | 7.0 | 3.7 | 22.0 |
| >hg38_21_+_8433172_8446622_RNA45SN1_45S_50ntFlanks@4005.4042.38\|AACCTCGGGCCGATCGCACGCCCCCCGTGGCGGCGACG | >hg38_21_+_8433172_8446622_RNA45SN1_45S_50ntFlanks | 4005 | 4042 | 38 | AACCTCGGGCCGATCGCACGCCCCCCGTGGCGGCGACG | MB468 | 21 | 49 | 57 | 2 | 26 | 7.8 | 10.5 | 39.6 | 1.3 | 13.3 |
| >hg38_21_+_8433172_8446622_RNA45SN1_45S_50ntFlanks@4006.4023.18\|ACCTCGGGCCGATCGCAC | >hg38_21_+_8433172_8446622_RNA45SN1_45S_50ntFlanks | 4006 | 4023 | 18 | ACCTCGGGCCGATCGCAC | BT20 | 14 | 11 | 41 | 4 | 2 | 16.5 | 15.1 | 57.4 | 5.2 | 1.7 |
| >hg38_21_+_8433172_8446622_RNA45SN1_45S_50ntFlanks@4006.4023.18\|ACCTCGGGCCGATCGCAC | >hg38_21_+_8433172_8446622_RNA45SN1_45S_50ntFlanks | 4006 | 4023 | 18 | ACCTCGGGCCGATCGCAC | MB231 | 11 | 11 | 54 | 0 | 5 | 9.7 | 8.2 | 56.0 | 0.4 | 4.1 |
| >hg38_21_+_8433172_8446622_RNA45SN1_45S_50ntFlanks@4006.4023.18\|ACCTCGGGCCGATCGCAC | >hg38_21_+_8433172_8446622_RNA45SN1_45S_50ntFlanks | 4006 | 4023 | 18 | ACCTCGGGCCGATCGCAC | MB468 | 18 | 74 | 25 | 4 | 18 | 4.4 | 32.3 | 11.2 | 1.2 | 6.5 |
| >hg38_21_+_8433172_8446622_RNA45SN1_45S_50ntFlanks@4006.4024.19\|ACCTCGGGCCGATCGCACG | >hg38_21_+_8433172_8446622_RNA45SN1_45S_50ntFlanks | 4006 | 4024 | 19 | ACCTCGGGCCGATCGCACG | MB468 | 72 | 61 | 113 | 38 | 46 | 8.5 | 25.3 | 26.3 | 9.9 | 14.4 |
| >hg38_21_+_8433172_8446622_RNA45SN1_45S_50ntFlanks@4006.4025.20\|ACCTCGGGCCGATCGCACGC | >hg38_21_+_8433172_8446622_RNA45SN1_45S_50ntFlanks | 4006 | 4025 | 20 | ACCTCGGGCCGATCGCACGC | MB468 | 33 | 45 | 36 | 5 | 33 | 5.9 | 30.1 | 8.1 | 1.9 | 17.5 |
| >hg38_21_+_8433172_8446622_RNA45SN1_45S_50ntFlanks@4006.4026.21\|ACCTCGGGCCGATCGCACGCC | >hg38_21_+_8433172_8446622_RNA45SN1_45S_50ntFlanks | 4006 | 4026 | 21 | ACCTCGGGCCGATCGCACGCC | MB468 | 18 | 23 | 27 | 11 | 23 | 4.6 | 16.4 | 6.5 | 4.5 | 14.8 |
| >hg38_21_+_8433172_8446622_RNA45SN1_45S_50ntFlanks@4006.4042.37\|ACCTCGGGCCGATCGCACGCCCCCCGTGGCGGCGACG | >hg38_21_+_8433172_8446622_RNA45SN1_45S_50ntFlanks | 4006 | 4042 | 37 | ACCTCGGGCCGATCGCACGCCCCCCGTGGCGGCGACG | MB468 | 26 | 48 | 70 | 3 | 19 | 10.4 | 13.6 | 36.0 | 1.8 | 5.0 |
| >hg38_21_+_8433172_8446622_RNA45SN1_45S_50ntFlanks@4007.4024.18\|CCTCGGGCCGATCGCACG | >hg38_21_+_8433172_8446622_RNA45SN1_45S_50ntFlanks | 4007 | 4024 | 18 | CCTCGGGCCGATCGCACG | MB468 | 45 | 24 | 67 | 46 | 43 | 10.1 | 9.8 | 16.4 | 13.5 | 23.2 |
| >hg38_21_+_8433172_8446622_RNA45SN1_45S_50ntFlanks@4007.4025.19\|CCTCGGGCCGATCGCACGC | >hg38_21_+_8433172_8446622_RNA45SN1_45S_50ntFlanks | 4007 | 4025 | 19 | CCTCGGGCCGATCGCACGC | MB468 | 15 | 12 | 18 | 6 | 21 | 2.1 | 6.6 | 3.3 | 3.9 | 13.7 |
| >hg38_21_+_8433172_8446622_RNA45SN1_45S_50ntFlanks@4007.4026.20\|CCTCGGGCCGATCGCACGCC | >hg38_21_+_8433172_8446622_RNA45SN1_45S_50ntFlanks | 4007 | 4026 | 20 | CCTCGGGCCGATCGCACGCC | MB468 | 12 | 10 | 17 | 15 | 22 | 5.3 | 6.3 | 6.4 | 6.9 | 16.4 |
| >hg38_21_+_8433172_8446622_RNA45SN1_45S_50ntFlanks@4007.4042.36\|CCTCGGGCCGATCGCACGCCCCCCGTGGCGGCGACG | >hg38_21_+_8433172_8446622_RNA45SN1_45S_50ntFlanks | 4007 | 4042 | 36 | CCTCGGGCCGATCGCACGCCCCCCGTGGCGGCGACG | MB468 | 17 | 19 | 34 | 3 | 15 | 3.8 | 4.5 | 19.7 | 1.7 | 2.7 |
| >hg38_21_+_8433172_8446622_RNA45SN1_45S_50ntFlanks@4008.4025.18\|CTCGGGCCGATCGCACGC | >hg38_21_+_8433172_8446622_RNA45SN1_45S_50ntFlanks | 4008 | 4025 | 18 | CTCGGGCCGATCGCACGC | MB468 | 18 | 11 | 22 | 10 | 16 | 1.7 | 4.1 | 5.7 | 5.3 | 9.3 |
| >hg38_21_+_8433172_8446622_RNA45SN1_45S_50ntFlanks@4008.4042.35\|CTCGGGCCGATCGCACGCCCCCCGTGGCGGCGACG | >hg38_21_+_8433172_8446622_RNA45SN1_45S_50ntFlanks | 4008 | 4042 | 35 | CTCGGGCCGATCGCACGCCCCCCGTGGCGGCGACG | MB468 | 21 | 13 | 33 | 6 | 14 | 5.9 | 4.8 | 15.9 | 2.7 | 6.1 |
| >hg38_21_+_8433172_8446622_RNA45SN1_45S_50ntFlanks@4027.4045.19\|CCCCGTGGCGGCGACGACC | >hg38_21_+_8433172_8446622_RNA45SN1_45S_50ntFlanks | 4027 | 4045 | 19 | CCCCGTGGCGGCGACGACC | MB468 | 10 | 15 | 5 | 8 | 26 | 4.5 | 13.7 | 1.0 | 3.9 | 20.6 |
| >hg38_21_+_8433172_8446622_RNA45SN1_45S_50ntFlanks@4043.4063.21\|ACCCATTCGAACGTCTGCCCT | >hg38_21_+_8433172_8446622_RNA45SN1_45S_50ntFlanks | 4043 | 4063 | 21 | ACCCATTCGAACGTCTGCCCT | MB468 | 13 | 38 | 19 | 2 | 24 | 1.0 | 27.1 | 3.5 | 1.6 | 21.9 |
| >hg38_21_+_8433172_8446622_RNA45SN1_45S_50ntFlanks@4044.4063.20\|CCCATTCGAACGTCTGCCCT | >hg38_21_+_8433172_8446622_RNA45SN1_45S_50ntFlanks | 4044 | 4063 | 20 | CCCATTCGAACGTCTGCCCT | MB468 | 12 | 21 | 19 | 2 | 14 | 2.9 | 6.8 | 5.1 | 0.9 | 9.7 |
| >hg38_21_+_8433172_8446622_RNA45SN1_45S_50ntFlanks@4045.4062.18\|CCATTCGAACGTCTGCCC | >hg38_21_+_8433172_8446622_RNA45SN1_45S_50ntFlanks | 4045 | 4062 | 18 | CCATTCGAACGTCTGCCC | MB468 | 35 | 44 | 59 | 14 | 26 | 5.2 | 16.1 | 16.0 | 1.6 | 14.2 |
| >hg38_21_+_8433172_8446622_RNA45SN1_45S_50ntFlanks@4045.4063.19\|CCATTCGAACGTCTGCCCT | >hg38_21_+_8433172_8446622_RNA45SN1_45S_50ntFlanks | 4045 | 4063 | 19 | CCATTCGAACGTCTGCCCT | MB468 | 117 | 149 | 191 | 40 | 107 | 17.1 | 52.4 | 26.8 | 12.9 | 60.1 |
| >hg38_21_+_8433172_8446622_RNA45SN1_45S_50ntFlanks@4045.4066.22\|CCATTCGAACGTCTGCCCTATC | >hg38_21_+_8433172_8446622_RNA45SN1_45S_50ntFlanks | 4045 | 4066 | 22 | CCATTCGAACGTCTGCCCTATC | MB468 | 31 | 92 | 38 | 10 | 94 | 8.8 | 42.2 | 10.7 | 0.9 | 100.0 |
| >hg38_21_+_8433172_8446622_RNA45SN1_45S_50ntFlanks@4046.4063.18\|CATTCGAACGTCTGCCCT | >hg38_21_+_8433172_8446622_RNA45SN1_45S_50ntFlanks | 4046 | 4063 | 18 | CATTCGAACGTCTGCCCT | MB468 | 229 | 210 | 317 | 84 | 167 | 34.3 | 69.5 | 26.8 | 28.3 | 115.3 |
| >hg38_21_+_8433172_8446622_RNA45SN1_45S_50ntFlanks@4046.4064.19\|CATTCGAACGTCTGCCCTA | >hg38_21_+_8433172_8446622_RNA45SN1_45S_50ntFlanks | 4046 | 4064 | 19 | CATTCGAACGTCTGCCCTA | MB468 | 12 | 11 | 15 | 10 | 19 | 1.3 | 3.2 | 3.1 | 3.5 | 16.9 |
| >hg38_21_+_8433172_8446622_RNA45SN1_45S_50ntFlanks@4046.4065.20\|CATTCGAACGTCTGCCCTAT | >hg38_21_+_8433172_8446622_RNA45SN1_45S_50ntFlanks | 4046 | 4065 | 20 | CATTCGAACGTCTGCCCTAT | MB468 | 15 | 15 | 19 | 3 | 12 | 1.4 | 4.4 | 1.9 | 1.4 | 9.0 |
| >hg38_21_+_8433172_8446622_RNA45SN1_45S_50ntFlanks@4046.4066.21\|CATTCGAACGTCTGCCCTATC | >hg38_21_+_8433172_8446622_RNA45SN1_45S_50ntFlanks | 4046 | 4066 | 21 | CATTCGAACGTCTGCCCTATC | MB231 | 12 | 6 | 45 | 1 | 3 | 9.7 | 4.3 | 35.6 | 0.7 | 2.5 |
| >hg38_21_+_8433172_8446622_RNA45SN1_45S_50ntFlanks@4046.4066.21\|CATTCGAACGTCTGCCCTATC | >hg38_21_+_8433172_8446622_RNA45SN1_45S_50ntFlanks | 4046 | 4066 | 21 | CATTCGAACGTCTGCCCTATC | MB468 | 66 | 154 | 76 | 31 | 165 | 7.0 | 51.5 | 22.8 | 7.9 | 166.5 |
| >hg38_21_+_8433172_8446622_RNA45SN1_45S_50ntFlanks@4046.4067.22\|CATTCGAACGTCTGCCCTATCA | >hg38_21_+_8433172_8446622_RNA45SN1_45S_50ntFlanks | 4046 | 4067 | 22 | CATTCGAACGTCTGCCCTATCA | MB468 | 14 | 29 | 16 | 5 | 25 | 2.0 | 5.0 | 3.4 | 1.2 | 21.0 |
| >hg38_21_+_8433172_8446622_RNA45SN1_45S_50ntFlanks@4047.4064.18\|ATTCGAACGTCTGCCCTA | >hg38_21_+_8433172_8446622_RNA45SN1_45S_50ntFlanks | 4047 | 4064 | 18 | ATTCGAACGTCTGCCCTA | MB468 | 23 | 21 | 31 | 26 | 40 | 7.0 | 7.3 | 5.6 | 8.1 | 38.2 |
| >hg38_21_+_8433172_8446622_RNA45SN1_45S_50ntFlanks@4047.4065.19\|ATTCGAACGTCTGCCCTAT | >hg38_21_+_8433172_8446622_RNA45SN1_45S_50ntFlanks | 4047 | 4065 | 19 | ATTCGAACGTCTGCCCTAT | MB468 | 31 | 33 | 39 | 7 | 28 | 6.6 | 11.7 | 6.7 | 3.2 | 21.8 |
| >hg38_21_+_8433172_8446622_RNA45SN1_45S_50ntFlanks@4047.4066.20\|ATTCGAACGTCTGCCCTATC | >hg38_21_+_8433172_8446622_RNA45SN1_45S_50ntFlanks | 4047 | 4066 | 20 | ATTCGAACGTCTGCCCTATC | MB231 | 11 | 8 | 41 | 1 | 3 | 8.2 | 5.3 | 34.0 | 0.7 | 2.5 |
| >hg38_21_+_8433172_8446622_RNA45SN1_45S_50ntFlanks@4047.4066.20\|ATTCGAACGTCTGCCCTATC | >hg38_21_+_8433172_8446622_RNA45SN1_45S_50ntFlanks | 4047 | 4066 | 20 | ATTCGAACGTCTGCCCTATC | MB468 | 109 | 262 | 143 | 79 | 352 | 42.1 | 92.3 | 46.0 | 26.5 | 360.9 |
| >hg38_21_+_8433172_8446622_RNA45SN1_45S_50ntFlanks@4047.4067.21\|ATTCGAACGTCTGCCCTATCA | >hg38_21_+_8433172_8446622_RNA45SN1_45S_50ntFlanks | 4047 | 4067 | 21 | ATTCGAACGTCTGCCCTATCA | MB468 | 23 | 50 | 27 | 11 | 46 | 6.1 | 18.5 | 4.1 | 2.1 | 41.4 |

| | | | | | | | | | | | | | | | | |
|---|---|---|---|---|---|---|---|---|---|---|---|---|---|---|---|---|
| >hg38_21_+_8433172_8446622_RNA45SN1_45S_50ntFlanks@4047.4073.27\|ATTCGAACGTCT GCCCTATCAACTTTC | >hg38_21_+_8433172_8446622_RNA45SN1_45S_50ntFlanks | 4047 | 4073 | 27 | ATTCGAACGTCTGCCCTATCAACTTTC | MB468 | 11 | 15 | 22 | 9 | 17 | 4.2 | 4.9 | 7.9 | 2.5 | 16.0 |
| >hg38_21_+_8433172_8446622_RNA45SN1_45S_50ntFlanks@4047.4081.35\|ATTCGAACGTCT GCCCTATCAACTTTCGATGGTAG | >hg38_21_+_8433172_8446622_RNA45SN1_45S_50ntFlanks | 4047 | 4081 | 35 | ATTCGAACGTCTGCCCTATCAACTTTCGATGGTAG | MB468 | 12 | 21 | 24 | 13 | 54 | 2.1 | 11.2 | 12.7 | 12.7 | 25.9 |
| >hg38_21_+_8433172_8446622_RNA45SN1_45S_50ntFlanks@4048.4066.19\|TTCGAACGTCTG CCCTATC | >hg38_21_+_8433172_8446622_RNA45SN1_45S_50ntFlanks | 4048 | 4066 | 19 | TTCGAACGTCTGCCCTATC | MB231 | 11 | 7 | 40 | 0 | 1 | 8.6 | 5.0 | 33.6 | 0.6 | 1.6 |
| >hg38_21_+_8433172_8446622_RNA45SN1_45S_50ntFlanks@4048.4066.19\|TTCGAACGTCTG CCCTATC | >hg38_21_+_8433172_8446622_RNA45SN1_45S_50ntFlanks | 4048 | 4066 | 19 | TTCGAACGTCTGCCCTATC | MB468 | 17 | 34 | 28 | 13 | 40 | 4.0 | 14.8 | 7.7 | 9.6 | 35.9 |
| >hg38_21_+_8433172_8446622_RNA45SN1_45S_50ntFlanks@4049.4066.18\|TCGAACGTCTGC CCTATC | >hg38_21_+_8433172_8446622_RNA45SN1_45S_50ntFlanks | 4049 | 4066 | 18 | TCGAACGTCTGCCCTATC | MB231 | 12 | 6 | 42 | 1 | 3 | 9.9 | 4.6 | 38.2 | 0.6 | 2.3 |
| >hg38_21_+_8433172_8446622_RNA45SN1_45S_50ntFlanks@4064.4081.18\|ATCAACTTTCGA TGGTAG | >hg38_21_+_8433172_8446622_RNA45SN1_45S_50ntFlanks | 4064 | 4081 | 18 | ATCAACTTTCGATGGTAG | MB468 | 67 | 3 | 47 | 35 | 32 | 45.3 | 1.4 | 39.0 | 8.2 | 42.0 |
| >hg38_21_+_8433172_8446622_RNA45SN1_45S_50ntFlanks@4064.4101.38\|ATCAACTTTCGA TGGTAGTCGCCGTGCCTACCATGGTG | >hg38_21_+_8433172_8446622_RNA45SN1_45S_50ntFlanks | 4064 | 4101 | 38 | ATCAACTTTCGATGGTAGTCGCCGTGCCTACCATGGTG | MB468 | 12 | 19 | 53 | 9 | 12 | 5.6 | 3.4 | 30.1 | 5.0 | 6.7 |
| >hg38_21_+_8433172_8446622_RNA45SN1_45S_50ntFlanks@4068.4089.22\|ACTTTCGATGGT AGTCGCCGTG | >hg38_21_+_8433172_8446622_RNA45SN1_45S_50ntFlanks | 4068 | 4089 | 22 | ACTTTCGATGGTAGTCGCCGTG | MB468 | 11 | 6 | 14 | 14 | 41 | 4.9 | 2.0 | 6.9 | 3.2 | 62.0 |
| >hg38_21_+_8433172_8446622_RNA45SN1_45S_50ntFlanks@4069.4089.21\|CTTTCGATGGTA GTCGCCGTG | >hg38_21_+_8433172_8446622_RNA45SN1_45S_50ntFlanks | 4069 | 4089 | 21 | CTTTCGATGGTAGTCGCCGTG | MB468 | 21 | 6 | 23 | 31 | 98 | 9.9 | 2.2 | 12.0 | 5.9 | 144.1 |
| >hg38_21_+_8433172_8446622_RNA45SN1_45S_50ntFlanks@4069.4094.26\|CTTTCGATGGTA GTCGCCGTGCCTAC | >hg38_21_+_8433172_8446622_RNA45SN1_45S_50ntFlanks | 4069 | 4094 | 26 | CTTTCGATGGTAGTCGCCGTGCCTAC | MB468 | 13 | 13 | 19 | 24 | 10 | 4.2 | 4.9 | 3.3 | 7.0 | 2.0 |
| >hg38_21_+_8433172_8446622_RNA45SN1_45S_50ntFlanks@4069.4095.27\|CTTTCGATGGTA GTCGCCGTGCCTACC | >hg38_21_+_8433172_8446622_RNA45SN1_45S_50ntFlanks | 4069 | 4095 | 27 | CTTTCGATGGTAGTCGCCGTGCCTACC | MB468 | 22 | 4 | 22 | 46 | 11 | 5.0 | 1.3 | 4.5 | 23.3 | 4.4 |
| >hg38_21_+_8433172_8446622_RNA45SN1_45S_50ntFlanks@4069.4098.30\|CTTTCGATGGTA GTCGCCGTGCCTACCATG | >hg38_21_+_8433172_8446622_RNA45SN1_45S_50ntFlanks | 4069 | 4098 | 30 | CTTTCGATGGTAGTCGCCGTGCCTACCATG | MB468 | 17 | 4 | 36 | 9 | 7 | 8.0 | 2.2 | 16.5 | 1.6 | 5.3 |
| >hg38_21_+_8433172_8446622_RNA45SN1_45S_50ntFlanks@4069.4101.33\|CTTTCGATGGTA GTCGCCGTGCCTACCATGGTG | >hg38_21_+_8433172_8446622_RNA45SN1_45S_50ntFlanks | 4069 | 4101 | 33 | CTTTCGATGGTAGTCGCCGTGCCTACCATGGTG | MB468 | 21 | 15 | 48 | 22 | 10 | 9.8 | 7.6 | 10.3 | 4.1 | 4.5 |
| >hg38_21_+_8433172_8446622_RNA45SN1_45S_50ntFlanks@4099.4121.23\|GTGACCACGGG TGACGGGGAATC | >hg38_21_+_8433172_8446622_RNA45SN1_45S_50ntFlanks | 4099 | 4121 | 23 | GTGACCACGGGTGACGGGGAATC | MB468 | 12 | 28 | 15 | 25 | 12 | 2.7 | 18.6 | 4.1 | 9.5 | 9.2 |
| >hg38_21_+_8433172_8446622_RNA45SN1_45S_50ntFlanks@4102.4120.19\|ACCACGGGTGA CGGGGAAT | >hg38_21_+_8433172_8446622_RNA45SN1_45S_50ntFlanks | 4102 | 4120 | 19 | ACCACGGGTGACGGGGAAT | MB468 | 15 | 19 | 24 | 14 | 11 | 4.7 | 16.9 | 9.4 | 6.3 | 11.1 |
| >hg38_21_+_8433172_8446622_RNA45SN1_45S_50ntFlanks@4102.4121.20\|ACCACGGGTGA CGGGGAATC | >hg38_21_+_8433172_8446622_RNA45SN1_45S_50ntFlanks | 4102 | 4121 | 20 | ACCACGGGTGACGGGGAATC | MB468 | 65 | 134 | 116 | 135 | 59 | 21.8 | 121.3 | 47.1 | 89.7 | 59.4 |
| >hg38_21_+_8433172_8446622_RNA45SN1_45S_50ntFlanks@4103.4121.19\|CCACGGGTGAC GGGGAATC | >hg38_21_+_8433172_8446622_RNA45SN1_45S_50ntFlanks | 4103 | 4121 | 19 | CCACGGGTGACGGGGAATC | MB468 | 27 | 67 | 56 | 76 | 32 | 5.2 | 55.6 | 9.7 | 56.1 | 28.4 |
| >hg38_21_+_8433172_8446622_RNA45SN1_45S_50ntFlanks@4122.4145.24\|AGGGTTCGATTC CGGAGAGGGAGC | >hg38_21_+_8433172_8446622_RNA45SN1_45S_50ntFlanks | 4122 | 4145 | 24 | AGGGTTCGATTCCGGAGAGGGAGC | MB468 | 12 | 22 | 9 | 38 | 11 | 4.5 | 7.8 | 5.0 | 18.0 | 9.9 |
| >hg38_21_+_8433172_8446622_RNA45SN1_45S_50ntFlanks@4125.4147.23\|GTTCGATTCCGG AGAGGGAGCCT | >hg38_21_+_8433172_8446622_RNA45SN1_45S_50ntFlanks | 4125 | 4147 | 23 | GTTCGATTCCGGAGAGGGAGCCT | BT20 | 12 | 7 | 2 | 7 | 2 | 20.0 | 12.1 | 3.2 | 8.4 | 1.8 |
| >hg38_21_+_8433172_8446622_RNA45SN1_45S_50ntFlanks@4128.4149.22\|CGATTCCGGAG AGGGAGCCTGA | >hg38_21_+_8433172_8446622_RNA45SN1_45S_50ntFlanks | 4128 | 4149 | 22 | CGATTCCGGAGAGGGAGCCTGA | BT20 | 12 | 20 | 3 | 1 | 0 | 20.4 | 34.0 | 4.3 | 1.4 | 0.0 |
| >hg38_21_+_8433172_8446622_RNA45SN1_45S_50ntFlanks@4128.4158.31\|CGATTCCGGAG AGGGAGCCTGAGAAACGGCT | >hg38_21_+_8433172_8446622_RNA45SN1_45S_50ntFlanks | 4128 | 4158 | 31 | CGATTCCGGAGAGGGAGCCTGAGAAACGGCT | MB468 | 11 | 10 | 7 | 10 | 8 | 2.5 | 3.9 | 2.3 | 5.2 | 2.1 |
| >hg38_21_+_8433172_8446622_RNA45SN1_45S_50ntFlanks@4128.4162.35\|CGATTCCGGAG AGGGAGCCTGAGAAACGGCTACCA | >hg38_21_+_8433172_8446622_RNA45SN1_45S_50ntFlanks | 4128 | 4162 | 35 | CGATTCCGGAGAGGGAGCCTGAGAAACGGCTACCA | MB468 | 19 | 44 | 42 | 10 | 34 | 5.2 | 20.6 | 14.4 | 7.5 | 15.6 |
| >hg38_21_+_8433172_8446622_RNA45SN1_45S_50ntFlanks@4129.4162.34\|GATTCCGGAGA GGGAGCCTGAGAAACGGCTACCA | >hg38_21_+_8433172_8446622_RNA45SN1_45S_50ntFlanks | 4129 | 4162 | 34 | GATTCCGGAGAGGGAGCCTGAGAAACGGCTACCA | MB468 | 19 | 32 | 39 | 7 | 15 | 3.4 | 9.1 | 5.2 | 6.3 | 6.2 |
| >hg38_21_+_8433172_8446622_RNA45SN1_45S_50ntFlanks@4130.4162.33\|ATTCCGGAGAG GGAGCCTGAGAAACGGCTACCA | >hg38_21_+_8433172_8446622_RNA45SN1_45S_50ntFlanks | 4130 | 4162 | 33 | ATTCCGGAGAGGGAGCCTGAGAAACGGCTACCA | MB468 | 51 | 54 | 61 | 24 | 42 | 19.0 | 14.0 | 25.8 | 14.2 | 18.7 |
| >hg38_21_+_8433172_8446622_RNA45SN1_45S_50ntFlanks@4160.4178.19\|CCACATCCAAG GAAGGCAG | >hg38_21_+_8433172_8446622_RNA45SN1_45S_50ntFlanks | 4160 | 4178 | 19 | CCACATCCAAGGAAGGCAG | MB468 | 13 | 8 | 7 | 14 | 6 | 4.1 | 4.2 | 2.8 | 6.7 | 3.2 |
| >hg38_21_+_8433172_8446622_RNA45SN1_45S_50ntFlanks@4160.4182.23\|CCACATCCAAG GAAGGCAGCAGG | >hg38_21_+_8433172_8446622_RNA45SN1_45S_50ntFlanks | 4160 | 4182 | 23 | CCACATCCAAGGAAGGCAGCAGG | MB468 | 21 | 29 | 12 | 25 | 7 | 10.4 | 12.6 | 4.2 | 24.3 | 1.9 |
| >hg38_21_+_8433172_8446622_RNA45SN1_45S_50ntFlanks@4160.4184.25\|CCACATCCAAG GAAGGCAGCAGGCG | >hg38_21_+_8433172_8446622_RNA45SN1_45S_50ntFlanks | 4160 | 4184 | 25 | CCACATCCAAGGAAGGCAGCAGGCG | MB468 | 15 | 26 | 11 | 28 | 9 | 6.9 | 8.5 | 2.4 | 22.0 | 2.3 |
| >hg38_21_+_8433172_8446622_RNA45SN1_45S_50ntFlanks@4161.4182.22\|CACATCCAAGG AAGGCAGCAGG | >hg38_21_+_8433172_8446622_RNA45SN1_45S_50ntFlanks | 4161 | 4182 | 22 | CACATCCAAGGAAGGCAGCAGG | MB468 | 11 | 14 | 7 | 10 | 4 | 4.9 | 5.3 | 2.1 | 7.6 | 1.1 |
| >hg38_21_+_8433172_8446622_RNA45SN1_45S_50ntFlanks@4163.4182.20\|CATCCAAGGAA GGCAGCAGG | >hg38_21_+_8433172_8446622_RNA45SN1_45S_50ntFlanks | 4163 | 4182 | 20 | CATCCAAGGAAGGCAGCAGG | MB468 | 85 | 110 | 48 | 51 | 22 | 39.2 | 48.2 | 13.7 | 32.1 | 6.2 |
| >hg38_21_+_8433172_8446622_RNA45SN1_45S_50ntFlanks@4163.4184.22\|CATCCAAGGAA GGCAGCAGGCG | >hg38_21_+_8433172_8446622_RNA45SN1_45S_50ntFlanks | 4163 | 4184 | 22 | CATCCAAGGAAGGCAGCAGGCG | MB468 | 57 | 109 | 49 | 63 | 28 | 29.1 | 40.0 | 14.1 | 32.2 | 7.1 |
| >hg38_21_+_8433172_8446622_RNA45SN1_45S_50ntFlanks@4163.4186.24\|CATCCAAGGAA GGCAGCAGGCGCG | >hg38_21_+_8433172_8446622_RNA45SN1_45S_50ntFlanks | 4163 | 4186 | 24 | CATCCAAGGAAGGCAGCAGGCGCG | MB468 | 22 | 51 | 34 | 25 | 25 | 11.7 | 11.5 | 16.0 | 9.3 | 9.6 |
| >hg38_21_+_8433172_8446622_RNA45SN1_45S_50ntFlanks@4220.4242.23\|ACGAAAAATAAC AATACAGGACT | >hg38_21_+_8433172_8446622_RNA45SN1_45S_50ntFlanks | 4220 | 4242 | 23 | ACGAAAAATAACAATACAGGACT | MB468 | 52 | 31 | 74 | 116 | 75 | 15.0 | 17.0 | 44.3 | 58.5 | 24.3 |
| >hg38_21_+_8433172_8446622_RNA45SN1_45S_50ntFlanks@4220.4243.24\|ACGAAAAATAAC AATACAGGACTC | >hg38_21_+_8433172_8446622_RNA45SN1_45S_50ntFlanks | 4220 | 4243 | 24 | ACGAAAAATAACAATACAGGACTC | MB468 | 12 | 9 | 18 | 24 | 12 | 1.7 | 5.9 | 6.2 | 7.4 | 8.0 |
| >hg38_21_+_8433172_8446622_RNA45SN1_45S_50ntFlanks@4223.4242.20\|AAAAATAACAAT ACAGGACT | >hg38_21_+_8433172_8446622_RNA45SN1_45S_50ntFlanks | 4223 | 4242 | 20 | AAAAATAACAATACAGGACT | MB468 | 21 | 12 | 23 | 38 | 19 | 14.7 | 5.8 | 15.5 | 20.1 | 6.9 |
| >hg38_21_+_8433172_8446622_RNA45SN1_45S_50ntFlanks@4224.4242.19\|AAAATAACAATA CAGGACT | >hg38_21_+_8433172_8446622_RNA45SN1_45S_50ntFlanks | 4224 | 4242 | 19 | AAAATAACAATACAGGACT | MB468 | 62 | 28 | 63 | 100 | 44 | 17.6 | 18.9 | 49.4 | 56.3 | 13.8 |
| >hg38_21_+_8433172_8446622_RNA45SN1_45S_50ntFlanks@4225.4242.18\|AAATAACAATAC AGGACT | >hg38_21_+_8433172_8446622_RNA45SN1_45S_50ntFlanks | 4225 | 4242 | 18 | AAATAACAATACAGGACT | MB468 | 42 | 16 | 43 | 80 | 32 | 14.3 | 11.9 | 43.2 | 41.5 | 9.7 |
| >hg38_21_+_8433172_8446622_RNA45SN1_45S_50ntFlanks@4240.4258.19\|ACTCTTTCGAGG CCCTGTA | >hg38_21_+_8433172_8446622_RNA45SN1_45S_50ntFlanks | 4240 | 4258 | 19 | ACTCTTTCGAGGCCCTGTA | MB468 | 28 | 12 | 48 | 12 | 32 | 5.0 | 2.8 | 18.6 | 5.2 | 21.6 |
| >hg38_21_+_8433172_8446622_RNA45SN1_45S_50ntFlanks@4241.4258.18\|CTCTTTCGAGGC CCTGTA | >hg38_21_+_8433172_8446622_RNA45SN1_45S_50ntFlanks | 4241 | 4258 | 18 | CTCTTTCGAGGCCCTGTA | MB468 | 56 | 26 | 125 | 13 | 28 | 9.3 | 8.3 | 47.0 | 4.8 | 12.4 |
| >hg38_21_+_8433172_8446622_RNA45SN1_45S_50ntFlanks@4243.4263.21\|CTTTCGAGGCC CTGTAATTGG | >hg38_21_+_8433172_8446622_RNA45SN1_45S_50ntFlanks | 4243 | 4263 | 21 | CTTTCGAGGCCCTGTAATTGG | MB468 | 25 | 35 | 27 | 12 | 28 | 7.2 | 17.8 | 7.1 | 10.8 | 12.5 |
| >hg38_21_+_8433172_8446622_RNA45SN1_45S_50ntFlanks@4243.4264.22\|CTTTCGAGGCC CTGTAATTGGA | >hg38_21_+_8433172_8446622_RNA45SN1_45S_50ntFlanks | 4243 | 4264 | 22 | CTTTCGAGGCCCTGTAATTGGA | MB468 | 13 | 20 | 20 | 9 | 15 | 4.6 | 8.1 | 6.6 | 6.8 | 6.9 |
| >hg38_21_+_8433172_8446622_RNA45SN1_45S_50ntFlanks@4257.4275.19\|TAATTGGAATGA GTCCACT | >hg38_21_+_8433172_8446622_RNA45SN1_45S_50ntFlanks | 4257 | 4275 | 19 | TAATTGGAATGAGTCCACT | MB468 | 11 | 1 | 7 | 17 | 18 | 6.4 | 0.3 | 4.9 | 5.7 | 27.0 |
| >hg38_21_+_8433172_8446622_RNA45SN1_45S_50ntFlanks@4264.4282.19\|AATGAGTCCACT TTAAATC | >hg38_21_+_8433172_8446622_RNA45SN1_45S_50ntFlanks | 4264 | 4282 | 19 | AATGAGTCCACTTTAAATC | MB468 | 18 | 9 | 11 | 9 | 15 | 6.7 | 4.2 | 4.6 | 1.6 | 12.7 |
| >hg38_21_+_8433172_8446622_RNA45SN1_45S_50ntFlanks@4279.4314.36\|AATCCTTTAACG AGGATCCATTGGAGGGCAAGTCTG | >hg38_21_+_8433172_8446622_RNA45SN1_45S_50ntFlanks | 4279 | 4314 | 36 | AATCCTTTAACGAGGATCCATTGGAGGGCAAGTCTG | MB468 | 11 | 29 | 13 | 5 | 4 | 2.2 | 14.9 | 3.6 | 3.4 | 1.4 |
| >hg38_21_+_8433172_8446622_RNA45SN1_45S_50ntFlanks@4283.4307.25\|CTTTAACGAGGA TCCATTGGAGGGC | >hg38_21_+_8433172_8446622_RNA45SN1_45S_50ntFlanks | 4283 | 4307 | 25 | CTTTAACGAGGATCCATTGGAGGGC | MB468 | 19 | 32 | 14 | 32 | 10 | 7.7 | 10.7 | 7.0 | 11.9 | 1.9 |
| >hg38_21_+_8433172_8446622_RNA45SN1_45S_50ntFlanks@4283.4314.32\|CTTTAACGAGGA TCCATTGGAGGGCAAGTCTG | >hg38_21_+_8433172_8446622_RNA45SN1_45S_50ntFlanks | 4283 | 4314 | 32 | CTTTAACGAGGATCCATTGGAGGGCAAGTCTG | MB468 | 21 | 62 | 27 | 38 | 18 | 6.5 | 35.7 | 13.0 | 33.8 | 11.2 |
| >hg38_21_+_8433172_8446622_RNA45SN1_45S_50ntFlanks@4283.4317.35\|CTTTAACGAGGA TCCATTGGAGGGCAAGTCTGGTG | >hg38_21_+_8433172_8446622_RNA45SN1_45S_50ntFlanks | 4283 | 4317 | 35 | CTTTAACGAGGATCCATTGGAGGGCAAGTCTGGTG | MB468 | 96 | 390 | 131 | 227 | 88 | 44.2 | 176.1 | 55.1 | 224.5 | 45.9 |
| >hg38_21_+_8433172_8446622_RNA45SN1_45S_50ntFlanks@4297.4314.18\|CATTGGAGGGC AAGTCTG | >hg38_21_+_8433172_8446622_RNA45SN1_45S_50ntFlanks | 4297 | 4314 | 18 | CATTGGAGGGCAAGTCTG | MB468 | 51 | 39 | 32 | 26 | 11 | 31.6 | 24.2 | 22.2 | 8.8 | 4.4 |
| >hg38_21_+_8433172_8446622_RNA45SN1_45S_50ntFlanks@4297.4316.20\|CATTGGAGGGC AAGTCTGGT | >hg38_21_+_8433172_8446622_RNA45SN1_45S_50ntFlanks | 4297 | 4316 | 20 | CATTGGAGGGCAAGTCTGGT | MB468 | 22 | 21 | 20 | 16 | 7 | 12.5 | 12.2 | 14.7 | 6.5 | 3.3 |
| >hg38_21_+_8433172_8446622_RNA45SN1_45S_50ntFlanks@4297.4317.21\|CATTGGAGGGC AAGTCTGGTG | >hg38_21_+_8433172_8446622_RNA45SN1_45S_50ntFlanks | 4297 | 4317 | 21 | CATTGGAGGGCAAGTCTGGTG | MB468 | 52 | 28 | 55 | 91 | 38 | 25.3 | 9.1 | 38.6 | 36.0 | 29.8 |
| >hg38_21_+_8433172_8446622_RNA45SN1_45S_50ntFlanks@4297.4318.22\|CATTGGAGGGC AAGTCTGGTGC | >hg38_21_+_8433172_8446622_RNA45SN1_45S_50ntFlanks | 4297 | 4318 | 22 | CATTGGAGGGCAAGTCTGGTGC | MB468 | 12 | 14 | 14 | 8 | 7 | 5.2 | 4.8 | 6.6 | 2.5 | 6.7 |

| | | | | | | | | | | | | | | | |
|---|---|---|---|---|---|---|---|---|---|---|---|---|---|---|---|
| >hg38_21_+_8433172_8446622_RNA45SN1_45S_50ntFlanks@4297.4319.23\|CATTGGAGGGCAAGTCTGGTGCC | >hg38_21_+_8433172_8446622_RNA45SN1_45S_50ntFlanks | 4297 | 4319 | 23 | CATTGGAGGGCAAGTCTGGTGCC | MB468 | 18 | 17 | 23 | 13 | 15 | 8.2 | 4.6 | 8.8 | 3.8 | 10.9 |
| >hg38_21_+_8433172_8446622_RNA45SN1_45S_50ntFlanks@4297.4320.24\|CATTGGAGGGCAAGTCTGGTGCCA | >hg38_21_+_8433172_8446622_RNA45SN1_45S_50ntFlanks | 4297 | 4320 | 24 | CATTGGAGGGCAAGTCTGGTGCCA | MB468 | 20 | 16 | 16 | 8 | 10 | 8.3 | 2.8 | 6.4 | 3.4 | 2.9 |
| >hg38_21_+_8433172_8446622_RNA45SN1_45S_50ntFlanks@4297.4321.25\|CATTGGAGGGCAAGTCTGGTGCCAG | >hg38_21_+_8433172_8446622_RNA45SN1_45S_50ntFlanks | 4297 | 4321 | 25 | CATTGGAGGGCAAGTCTGGTGCCAG | MB468 | 28 | 24 | 36 | 17 | 21 | 7.8 | 4.4 | 13.4 | 6.6 | 14.3 |
| >hg38_21_+_8433172_8446622_RNA45SN1_45S_50ntFlanks@4312.4332.21\|CTGGTGCCAGCAGCCGCGGTA | >hg38_21_+_8433172_8446622_RNA45SN1_45S_50ntFlanks | 4312 | 4332 | 21 | CTGGTGCCAGCAGCCGCGGTA | BT20 | 16 | 10 | 4 | 2 | 1 | 26.9 | 17.0 | 6.2 | 2.9 | 1.3 |
| >hg38_21_+_8433172_8446622_RNA45SN1_45S_50ntFlanks@4322.4339.18\|CAGCCGCGGTAATTCCAG | >hg38_21_+_8433172_8446622_RNA45SN1_45S_50ntFlanks | 4322 | 4339 | 18 | CAGCCGCGGTAATTCCAG | MB468 | 15 | 9 | 7 | 3 | 13 | 7.0 | 2.8 | 2.8 | 1.6 | 4.9 |
| >hg38_21_+_8433172_8446622_RNA45SN1_45S_50ntFlanks@4322.4346.25\|CAGCCGCGGTAATTCCAGCTCCAAT | >hg38_21_+_8433172_8446622_RNA45SN1_45S_50ntFlanks | 4322 | 4346 | 25 | CAGCCGCGGTAATTCCAGCTCCAAT | MB468 | 22 | 2 | 7 | 4 | 9 | 12.9 | 0.2 | 3.4 | 1.2 | 4.5 |
| >hg38_21_+_8433172_8446622_RNA45SN1_45S_50ntFlanks@4322.4352.31\|CAGCCGCGGTAATTCCAGCTCCAATAGCGTA | >hg38_21_+_8433172_8446622_RNA45SN1_45S_50ntFlanks | 4322 | 4352 | 31 | CAGCCGCGGTAATTCCAGCTCCAATAGCGTA | MB468 | 10 | 5 | 6 | 3 | 8 | 1.2 | 1.4 | 3.6 | 1.8 | 5.0 |
| >hg38_21_+_8433172_8446622_RNA45SN1_45S_50ntFlanks@4325.4346.22\|CCGCGGTAATTCCAGCTCCAAT | >hg38_21_+_8433172_8446622_RNA45SN1_45S_50ntFlanks | 4325 | 4346 | 22 | CCGCGGTAATTCCAGCTCCAAT | MB468 | 14 | 5 | 8 | 2 | 11 | 6.8 | 1.8 | 5.5 | 0.4 | 8.0 |
| >hg38_21_+_8433172_8446622_RNA45SN1_45S_50ntFlanks@4328.4346.19\|CGGTAATTCCAGCTCCAAT | >hg38_21_+_8433172_8446622_RNA45SN1_45S_50ntFlanks | 4328 | 4346 | 19 | CGGTAATTCCAGCTCCAAT | MB468 | 16 | 5 | 12 | 3 | 8 | 5.2 | 1.6 | 9.8 | 0.7 | 5.0 |
| >hg38_21_+_8433172_8446622_RNA45SN1_45S_50ntFlanks@4353.4371.19\|TATTAAAGTTGCTGCAGTT | >hg38_21_+_8433172_8446622_RNA45SN1_45S_50ntFlanks | 4353 | 4371 | 19 | TATTAAAGTTGCTGCAGTT | MB468 | 18 | 8 | 26 | 8 | 9 | 6.7 | 2.7 | 8.7 | 3.1 | 3.1 |
| >hg38_21_+_8433172_8446622_RNA45SN1_45S_50ntFlanks@4364.4391.28\|CTGCAGTTAAAAAGCTCGTAGTTGGATC | >hg38_21_+_8433172_8446622_RNA45SN1_45S_50ntFlanks | 4364 | 4391 | 28 | CTGCAGTTAAAAAGCTCGTAGTTGGATC | MB468 | 13 | 6 | 7 | 9 | 5 | 3.8 | 4.5 | 2.7 | 3.2 | 1.6 |
| >hg38_21_+_8433172_8446622_RNA45SN1_45S_50ntFlanks@4367.4388.22\|CAGTTAAAAAGCTCGTAGTTGG | >hg38_21_+_8433172_8446622_RNA45SN1_45S_50ntFlanks | 4367 | 4388 | 22 | CAGTTAAAAAGCTCGTAGTTGG | MB468 | 29 | 5 | 22 | 8 | 8 | 11.9 | 3.2 | 20.6 | 3.3 | 4.3 |
| >hg38_21_+_8433172_8446622_RNA45SN1_45S_50ntFlanks@4367.4389.23\|CAGTTAAAAAGCTCGTAGTTGGA | >hg38_21_+_8433172_8446622_RNA45SN1_45S_50ntFlanks | 4367 | 4389 | 23 | CAGTTAAAAAGCTCGTAGTTGGA | MB468 | 194 | 65 | 131 | 61 | 34 | 71.6 | 52.1 | 77.0 | 33.4 | 12.5 |
| >hg38_21_+_8433172_8446622_RNA45SN1_45S_50ntFlanks@4367.4391.25\|CAGTTAAAAAGCTCGTAGTTGGATC | >hg38_21_+_8433172_8446622_RNA45SN1_45S_50ntFlanks | 4367 | 4391 | 25 | CAGTTAAAAAGCTCGTAGTTGGATC | MB468 | 156 | 87 | 133 | 62 | 36 | 44.1 | 64.0 | 42.3 | 20.4 | 9.7 |
| >hg38_21_+_8433172_8446622_RNA45SN1_45S_50ntFlanks@4367.4393.27\|CAGTTAAAAAGCTCGTAGTTGGATCTT | >hg38_21_+_8433172_8446622_RNA45SN1_45S_50ntFlanks | 4367 | 4393 | 27 | CAGTTAAAAAGCTCGTAGTTGGATCTT | MB468 | 17 | 9 | 10 | 7 | 5 | 4.4 | 6.0 | 3.2 | 3.4 | 1.7 |
| >hg38_21_+_8433172_8446622_RNA45SN1_45S_50ntFlanks@4367.4394.28\|CAGTTAAAAAGCTCGTAGTTGGATCTTG | >hg38_21_+_8433172_8446622_RNA45SN1_45S_50ntFlanks | 4367 | 4394 | 28 | CAGTTAAAAAGCTCGTAGTTGGATCTTG | MB468 | 31 | 18 | 21 | 15 | 11 | 7.6 | 10.9 | 6.9 | 3.5 | 2.6 |
| >hg38_21_+_8433172_8446622_RNA45SN1_45S_50ntFlanks@4367.4396.30\|CAGTTAAAAAGCTCGTAGTTGGATCTTGGG | >hg38_21_+_8433172_8446622_RNA45SN1_45S_50ntFlanks | 4367 | 4396 | 30 | CAGTTAAAAAGCTCGTAGTTGGATCTTGGG | MB468 | 15 | 30 | 11 | 8 | 5 | 4.7 | 10.0 | 4.5 | 5.9 | 1.2 |
| >hg38_21_+_8433172_8446622_RNA45SN1_45S_50ntFlanks@4367.4398.32\|CAGTTAAAAAGCTCGTAGTTGGATCTTGGGAG | >hg38_21_+_8433172_8446622_RNA45SN1_45S_50ntFlanks | 4367 | 4398 | 32 | CAGTTAAAAAGCTCGTAGTTGGATCTTGGGAG | MB468 | 42 | 90 | 32 | 24 | 35 | 16.0 | 44.6 | 14.9 | 17.3 | 10.7 |
| >hg38_21_+_8433172_8446622_RNA45SN1_45S_50ntFlanks@4367.4400.34\|CAGTTAAAAAGCTCGTAGTTGGATCTTGGGAGCG | >hg38_21_+_8433172_8446622_RNA45SN1_45S_50ntFlanks | 4367 | 4400 | 34 | CAGTTAAAAAGCTCGTAGTTGGATCTTGGGAGCG | MB468 | 10 | 9 | 12 | 8 | 8 | 1.5 | 2.2 | 7.4 | 5.0 | 2.8 |
| >hg38_21_+_8433172_8446622_RNA45SN1_45S_50ntFlanks@4368.4389.22\|AGTTAAAAAGCTCGTAGTTGGA | >hg38_21_+_8433172_8446622_RNA45SN1_45S_50ntFlanks | 4368 | 4389 | 22 | AGTTAAAAAGCTCGTAGTTGGA | MB468 | 11 | 4 | 6 | 5 | 3 | 3.6 | 4.4 | 3.6 | 3.4 | 1.5 |
| >hg38_21_+_8433172_8446622_RNA45SN1_45S_50ntFlanks@4368.4391.24\|AGTTAAAAAGCTCGTAGTTGGATC | >hg38_21_+_8433172_8446622_RNA45SN1_45S_50ntFlanks | 4368 | 4391 | 24 | AGTTAAAAAGCTCGTAGTTGGATC | MB468 | 17 | 8 | 12 | 8 | 5 | 5.4 | 6.7 | 5.3 | 3.1 | 1.6 |
| >hg38_21_+_8433172_8446622_RNA45SN1_45S_50ntFlanks@4369.4389.21\|GTTAAAAAGCTCGTAGTTGGA | >hg38_21_+_8433172_8446622_RNA45SN1_45S_50ntFlanks | 4369 | 4389 | 21 | GTTAAAAAGCTCGTAGTTGGA | MB468 | 11 | 3 | 7 | 6 | 3 | 3.2 | 3.2 | 3.4 | 5.2 | 1.3 |
| >hg38_21_+_8433172_8446622_RNA45SN1_45S_50ntFlanks@4369.4391.23\|GTTAAAAAGCTCGTAGTTGGATC | >hg38_21_+_8433172_8446622_RNA45SN1_45S_50ntFlanks | 4369 | 4391 | 23 | GTTAAAAAGCTCGTAGTTGGATC | MB468 | 22 | 11 | 19 | 19 | 6 | 5.5 | 8.6 | 6.0 | 8.2 | 1.2 |
| >hg38_21_+_8433172_8446622_RNA45SN1_45S_50ntFlanks@4370.4389.20\|TTAAAAAGCTCGTAGTTGGA | >hg38_21_+_8433172_8446622_RNA45SN1_45S_50ntFlanks | 4370 | 4389 | 20 | TTAAAAAGCTCGTAGTTGGA | MB468 | 28 | 6 | 23 | 12 | 5 | 7.5 | 4.7 | 9.5 | 9.8 | 2.4 |
| >hg38_21_+_8433172_8446622_RNA45SN1_45S_50ntFlanks@4370.4391.22\|TTAAAAAGCTCGTAGTTGGATC | >hg38_21_+_8433172_8446622_RNA45SN1_45S_50ntFlanks | 4370 | 4391 | 22 | TTAAAAAGCTCGTAGTTGGATC | MB468 | 27 | 12 | 16 | 17 | 4 | 4.8 | 8.9 | 5.1 | 8.0 | 1.7 |
| >hg38_21_+_8433172_8446622_RNA45SN1_45S_50ntFlanks@4370.4398.29\|TTAAAAAGCTCGTAGTTGGATCTTGGGAG | >hg38_21_+_8433172_8446622_RNA45SN1_45S_50ntFlanks | 4370 | 4398 | 29 | TTAAAAAGCTCGTAGTTGGATCTTGGGAG | MB468 | 18 | 27 | 11 | 8 | 6 | 5.5 | 15.1 | 4.0 | 7.4 | 1.6 |
| >hg38_21_+_8433172_8446622_RNA45SN1_45S_50ntFlanks@4372.4389.18\|AAAAAGCTCGTAGTTGGA | >hg38_21_+_8433172_8446622_RNA45SN1_45S_50ntFlanks | 4372 | 4389 | 18 | AAAAAGCTCGTAGTTGGA | MB468 | 19 | 4 | 8 | 9 | 3 | 6.3 | 3.6 | 4.4 | 7.2 | 1.3 |
| >hg38_21_+_8433172_8446622_RNA45SN1_45S_50ntFlanks@4372.4391.20\|AAAAAGCTCGTAGTTGGATC | >hg38_21_+_8433172_8446622_RNA45SN1_45S_50ntFlanks | 4372 | 4391 | 20 | AAAAAGCTCGTAGTTGGATC | MB468 | 18 | 7 | 9 | 20 | 4 | 3.6 | 5.8 | 3.9 | 7.7 | 1.6 |
| >hg38_21_+_8433172_8446622_RNA45SN1_45S_50ntFlanks@4380.4408.29\|CGTAGTTGGATCTTGGGAGCGGGCGGGCG | >hg38_21_+_8433172_8446622_RNA45SN1_45S_50ntFlanks | 4380 | 4408 | 29 | CGTAGTTGGATCTTGGGAGCGGGCGGGCG | MB468 | 12 | 21 | 11 | 16 | 12 | 7.8 | 0.9 | 6.5 | 7.1 | 3.2 |
| >hg38_21_+_8433172_8446622_RNA45SN1_45S_50ntFlanks@4390.4408.19\|TCTTGGGAGCGGGCGGGCG | >hg38_21_+_8433172_8446622_RNA45SN1_45S_50ntFlanks | 4390 | 4408 | 19 | TCTTGGGAGCGGGCGGGCG | MB468 | 12 | 13 | 19 | 14 | 7 | 5.4 | 3.3 | 11.7 | 7.3 | 2.0 |
| >hg38_21_+_8433172_8446622_RNA45SN1_45S_50ntFlanks@4399.4423.25\|CGGGCGGGCGGTCCGCCGCGAGGCG | >hg38_21_+_8433172_8446622_RNA45SN1_45S_50ntFlanks | 4399 | 4423 | 25 | CGGGCGGGCGGTCCGCCGCGAGGCG | MB468 | 14 | 22 | 31 | 11 | 12 | 5.2 | 11.3 | 15.2 | 3.3 | 7.8 |
| >hg38_21_+_8433172_8446622_RNA45SN1_45S_50ntFlanks@4443.4470.28\|CCCTTGCCTCTCGGCGCCCCCTCGATGC | >hg38_21_+_8433172_8446622_RNA45SN1_45S_50ntFlanks | 4443 | 4470 | 28 | CCCTTGCCTCTCGGCGCCCCCTCGATGC | MB468 | 12 | 32 | 13 | 3 | 28 | 4.9 | 31.5 | 5.4 | 2.2 | 7.3 |
| >hg38_21_+_8433172_8446622_RNA45SN1_45S_50ntFlanks@4443.4471.29\|CCCTTGCCTCTCGGCGCCCCCTCGATGCT | >hg38_21_+_8433172_8446622_RNA45SN1_45S_50ntFlanks | 4443 | 4471 | 29 | CCCTTGCCTCTCGGCGCCCCCTCGATGCT | MB468 | 87 | 135 | 115 | 38 | 168 | 34.2 | 136.2 | 57.9 | 21.1 | 48.1 |
| >hg38_21_+_8433172_8446622_RNA45SN1_45S_50ntFlanks@4443.4472.30\|CCCTTGCCTCTCGGCGCCCCCTCGATGCTC | >hg38_21_+_8433172_8446622_RNA45SN1_45S_50ntFlanks | 4443 | 4472 | 30 | CCCTTGCCTCTCGGCGCCCCCTCGATGCTC | MB468 | 37 | 57 | 59 | 25 | 56 | 20.8 | 65.9 | 39.7 | 11.1 | 15.0 |
| >hg38_21_+_8433172_8446622_RNA45SN1_45S_50ntFlanks@4444.4471.28\|CCTTGCCTCTCGGCGCCCCCTCGATGCT | >hg38_21_+_8433172_8446622_RNA45SN1_45S_50ntFlanks | 4444 | 4471 | 28 | CCTTGCCTCTCGGCGCCCCCTCGATGCT | MB468 | 19 | 27 | 26 | 20 | 37 | 8.2 | 20.0 | 8.3 | 15.4 | 10.7 |
| >hg38_21_+_8433172_8446622_RNA45SN1_45S_50ntFlanks@4445.4471.27\|CTTGCCTCTCGGCGCCCCCTCGATGCT | >hg38_21_+_8433172_8446622_RNA45SN1_45S_50ntFlanks | 4445 | 4471 | 27 | CTTGCCTCTCGGCGCCCCCTCGATGCT | MB468 | 33 | 48 | 58 | 43 | 53 | 6.7 | 21.3 | 19.1 | 36.5 | 19.8 |
| >hg38_21_+_8433172_8446622_RNA45SN1_45S_50ntFlanks@4445.4472.28\|CTTGCCTCTCGGCGCCCCCTCGATGCTC | >hg38_21_+_8433172_8446622_RNA45SN1_45S_50ntFlanks | 4445 | 4472 | 28 | CTTGCCTCTCGGCGCCCCCTCGATGCTC | MB468 | 15 | 40 | 26 | 24 | 27 | 6.1 | 36.8 | 7.7 | 5.0 | 10.7 |
| >hg38_21_+_8433172_8446622_RNA45SN1_45S_50ntFlanks@4449.4471.23\|CCTCTCGGCGCCCCCTCGATGCT | >hg38_21_+_8433172_8446622_RNA45SN1_45S_50ntFlanks | 4449 | 4471 | 23 | CCTCTCGGCGCCCCCTCGATGCT | MB468 | 55 | 94 | 82 | 35 | 164 | 17.6 | 72.5 | 48.9 | 9.9 | 78.3 |
| >hg38_21_+_8433172_8446622_RNA45SN1_45S_50ntFlanks@4449.4472.24\|CCTCTCGGCGCCCCCTCGATGCTC | >hg38_21_+_8433172_8446622_RNA45SN1_45S_50ntFlanks | 4449 | 4472 | 24 | CCTCTCGGCGCCCCCTCGATGCTC | MB468 | 32 | 142 | 51 | 27 | 113 | 13.6 | 193.6 | 12.4 | 7.5 | 86.1 |
| >hg38_21_+_8433172_8446622_RNA45SN1_45S_50ntFlanks@4453.4473.21\|TCGGCGCCCCCTCGATGCTCT | >hg38_21_+_8433172_8446622_RNA45SN1_45S_50ntFlanks | 4453 | 4473 | 21 | TCGGCGCCCCCTCGATGCTCT | MB231 | 16 | 4 | 34 | 7 | 11 | 5.9 | 1.9 | 21.6 | 1.4 | 6.4 |
| >hg38_21_+_8433172_8446622_RNA45SN1_45S_50ntFlanks@4472.4492.21\|CTTAGCTGAGTGTCCCGCGGG | >hg38_21_+_8433172_8446622_RNA45SN1_45S_50ntFlanks | 4472 | 4492 | 21 | CTTAGCTGAGTGTCCCGCGGG | MB468 | 21 | 17 | 24 | 26 | 26 | 4.9 | 8.4 | 8.7 | 17.4 | 23.0 |
| >hg38_21_+_8433172_8446622_RNA45SN1_45S_50ntFlanks@4472.4493.22\|CTTAGCTGAGTGTCCCGCGGGG | >hg38_21_+_8433172_8446622_RNA45SN1_45S_50ntFlanks | 4472 | 4493 | 22 | CTTAGCTGAGTGTCCCGCGGGG | MB468 | 18 | 18 | 36 | 19 | 18 | 4.8 | 4.2 | 14.0 | 13.4 | 12.9 |
| >hg38_21_+_8433172_8446622_RNA45SN1_45S_50ntFlanks@4472.4494.23\|CTTAGCTGAGTGTCCCGCGGGGC | >hg38_21_+_8433172_8446622_RNA45SN1_45S_50ntFlanks | 4472 | 4494 | 23 | CTTAGCTGAGTGTCCCGCGGGGC | MB468 | 45 | 39 | 98 | 52 | 57 | 22.2 | 11.2 | 53.3 | 26.6 | 28.4 |
| >hg38_21_+_8433172_8446622_RNA45SN1_45S_50ntFlanks@4472.4496.25\|CTTAGCTGAGTGTCCCGCGGGGCCC | >hg38_21_+_8433172_8446622_RNA45SN1_45S_50ntFlanks | 4472 | 4496 | 25 | CTTAGCTGAGTGTCCCGCGGGGCCC | MB468 | 23 | 16 | 29 | 12 | 24 | 8.5 | 9.9 | 10.8 | 6.3 | 4.0 |
| >hg38_21_+_8433172_8446622_RNA45SN1_45S_50ntFlanks@4472.4497.26\|CTTAGCTGAGTGTCCCGCGGGGCCCG | >hg38_21_+_8433172_8446622_RNA45SN1_45S_50ntFlanks | 4472 | 4497 | 26 | CTTAGCTGAGTGTCCCGCGGGGCCCG | MB468 | 28 | 10 | 36 | 36 | 35 | 9.2 | 2.1 | 20.3 | 23.2 | 17.0 |
| >hg38_21_+_8433172_8446622_RNA45SN1_45S_50ntFlanks@4473.4494.22\|TTAGCTGAGTGTCCCGCGGGGC | >hg38_21_+_8433172_8446622_RNA45SN1_45S_50ntFlanks | 4473 | 4494 | 22 | TTAGCTGAGTGTCCCGCGGGGC | MB468 | 11 | 13 | 20 | 7 | 6 | 0.7 | 9.2 | 6.7 | 2.0 | 2.5 |
| >hg38_21_+_8433172_8446622_RNA45SN1_45S_50ntFlanks@4494.4511.18\|CCCGAAGCGTTTACTTTG | >hg38_21_+_8433172_8446622_RNA45SN1_45S_50ntFlanks | 4494 | 4511 | 18 | CCCGAAGCGTTTACTTTG | MB468 | 13 | 1 | 5 | 12 | 15 | 10.4 | 0.7 | 4.2 | 12.2 | 15.2 |
| >hg38_21_+_8433172_8446622_RNA45SN1_45S_50ntFlanks@4498.4517.20\|AAGCGTTTACTTTGAAAAAA | >hg38_21_+_8433172_8446622_RNA45SN1_45S_50ntFlanks | 4498 | 4517 | 20 | AAGCGTTTACTTTGAAAAAA | MB468 | 41 | 10 | 8 | 6 | 28 | 23.9 | 9.4 | 5.4 | 6.4 | 40.0 |
| >hg38_21_+_8433172_8446622_RNA45SN1_45S_50ntFlanks@4498.4519.22\|AAGCGTTTACTTTGAAAAAATT | >hg38_21_+_8433172_8446622_RNA45SN1_45S_50ntFlanks | 4498 | 4519 | 22 | AAGCGTTTACTTTGAAAAAATT | BT20 | 11 | 12 | 17 | 2 | 1 | 10.7 | 10.4 | 11.6 | 2.8 | 2.2 |
| >hg38_21_+_8433172_8446622_RNA45SN1_45S_50ntFlanks@4498.4519.22\|AAGCGTTTACTTTGAAAAAATT | >hg38_21_+_8433172_8446622_RNA45SN1_45S_50ntFlanks | 4498 | 4519 | 22 | AAGCGTTTACTTTGAAAAAATT | MB468 | 10 | 4 | 4 | 1 | 8 | 4.4 | 2.1 | 1.8 | 1.7 | 10.9 |

| | | | | | | | | | | | | | | | | |
|---|---|---|---|---|---|---|---|---|---|---|---|---|---|---|---|---|
| >hg38_21_+_8433172_8446622_RNA45SN1_45S_50ntFlanks@4498.4520.23\|AAGCGTTTACTTTGAAAAAATTA | >hg38_21_+_8433172_8446622_RNA45SN1_45S_50ntFlanks | 4498 | 4520 | 23 | AAGCGTTTACTTTGAAAAAATTA | MB468 | 13 | 6 | 6 | 2 | 8 | 4.8 | 3.1 | 4.0 | 2.0 | 11.4 |
| >hg38_21_+_8433172_8446622_RNA45SN1_45S_50ntFlanks@4499.4517.19\|AGCGTTTACTTTGAAAAAA | >hg38_21_+_8433172_8446622_RNA45SN1_45S_50ntFlanks | 4499 | 4517 | 19 | AGCGTTTACTTTGAAAAAA | MB468 | 16 | 9 | 7 | 1 | 10 | 8.2 | 8.8 | 4.0 | 1.0 | 14.0 |
| >hg38_21_+_8433172_8446622_RNA45SN1_45S_50ntFlanks@4547.4565.19\|CTGGATACCGCAGCTAGGA | >hg38_21_+_8433172_8446622_RNA45SN1_45S_50ntFlanks | 4547 | 4565 | 19 | CTGGATACCGCAGCTAGGA | MB468 | 23 | 10 | 7 | 42 | 12 | 12.9 | 3.0 | 2.9 | 32.9 | 5.6 |
| >hg38_21_+_8433172_8446622_RNA45SN1_45S_50ntFlanks@4554.4572.19\|CCGCAGCTAGGAATAATGG | >hg38_21_+_8433172_8446622_RNA45SN1_45S_50ntFlanks | 4554 | 4572 | 19 | CCGCAGCTAGGAATAATGG | MB468 | 14 | 13 | 9 | 10 | 6 | 6.3 | 7.1 | 3.7 | 4.0 | 3.2 |
| >hg38_21_+_8433172_8446622_RNA45SN1_45S_50ntFlanks@4554.4573.20\|CCGCAGCTAGGAATAATGGA | >hg38_21_+_8433172_8446622_RNA45SN1_45S_50ntFlanks | 4554 | 4573 | 20 | CCGCAGCTAGGAATAATGGA | MB468 | 10 | 17 | 5 | 12 | 5 | 5.0 | 10.2 | 2.1 | 6.1 | 1.3 |
| >hg38_21_+_8433172_8446622_RNA45SN1_45S_50ntFlanks@4555.4572.18\|CGCAGCTAGGAATAATGG | >hg38_21_+_8433172_8446622_RNA45SN1_45S_50ntFlanks | 4555 | 4572 | 18 | CGCAGCTAGGAATAATGG | MB468 | 50 | 79 | 37 | 44 | 20 | 23.2 | 53.6 | 20.1 | 16.8 | 10.8 |
| >hg38_21_+_8433172_8446622_RNA45SN1_45S_50ntFlanks@4555.4573.19\|CGCAGCTAGGAATAATGGA | >hg38_21_+_8433172_8446622_RNA45SN1_45S_50ntFlanks | 4555 | 4573 | 19 | CGCAGCTAGGAATAATGGA | MB468 | 24 | 38 | 9 | 31 | 10 | 12.8 | 22.7 | 4.4 | 11.8 | 5.2 |
| >hg38_21_+_8433172_8446622_RNA45SN1_45S_50ntFlanks@4555.4587.33\|CGCAGCTAGGAATAATGGAATAGGACCGCGGTT | >hg38_21_+_8433172_8446622_RNA45SN1_45S_50ntFlanks | 4555 | 4587 | 33 | CGCAGCTAGGAATAATGGAATAGGACCGCGGTT | MB468 | 17 | 24 | 16 | 8 | 13 | 2.8 | 4.7 | 3.5 | 4.4 | 1.4 |
| >hg38_21_+_8433172_8446622_RNA45SN1_45S_50ntFlanks@4557.4576.20\|CAGCTAGGAATAATGGAATA | >hg38_21_+_8433172_8446622_RNA45SN1_45S_50ntFlanks | 4557 | 4576 | 20 | CAGCTAGGAATAATGGAATA | MB468 | 11 | 10 | 5 | 5 | 3 | 4.4 | 5.7 | 1.5 | 1.5 | 1.0 |
| >hg38_21_+_8433172_8446622_RNA45SN1_45S_50ntFlanks@4557.4587.31\|CAGCTAGGAATAATGGAATAGGACCGCGGTT | >hg38_21_+_8433172_8446622_RNA45SN1_45S_50ntFlanks | 4557 | 4587 | 31 | CAGCTAGGAATAATGGAATAGGACCGCGGTT | MB468 | 28 | 39 | 26 | 12 | 9 | 9.2 | 14.1 | 3.5 | 5.9 | 2.9 |
| >hg38_21_+_8433172_8446622_RNA45SN1_45S_50ntFlanks@4558.4587.30\|AGCTAGGAATAATGGAATAGGACCGCGGTT | >hg38_21_+_8433172_8446622_RNA45SN1_45S_50ntFlanks | 4558 | 4587 | 30 | AGCTAGGAATAATGGAATAGGACCGCGGTT | MB468 | 15 | 20 | 10 | 7 | 5 | 7.0 | 8.0 | 2.4 | 6.6 | 2.9 |
| >hg38_21_+_8433172_8446622_RNA45SN1_45S_50ntFlanks@4560.4577.18\|CTAGGAATAATGGAATAG | >hg38_21_+_8433172_8446622_RNA45SN1_45S_50ntFlanks | 4560 | 4577 | 18 | CTAGGAATAATGGAATAG | MB468 | 21 | 9 | 9 | 14 | 15 | 11.1 | 3.8 | 7.0 | 5.5 | 19.6 |
| >hg38_21_+_8433172_8446622_RNA45SN1_45S_50ntFlanks@4560.4578.19\|CTAGGAATAATGGAATAGG | >hg38_21_+_8433172_8446622_RNA45SN1_45S_50ntFlanks | 4560 | 4578 | 19 | CTAGGAATAATGGAATAGG | MB468 | 17 | 8 | 17 | 8 | 5 | 7.8 | 5.6 | 4.1 | 3.0 | 3.1 |
| >hg38_21_+_8433172_8446622_RNA45SN1_45S_50ntFlanks@4560.4579.20\|CTAGGAATAATGGAATAGGA | >hg38_21_+_8433172_8446622_RNA45SN1_45S_50ntFlanks | 4560 | 4579 | 20 | CTAGGAATAATGGAATAGGA | MB468 | 30 | 17 | 33 | 17 | 7 | 16.5 | 12.3 | 4.7 | 7.4 | 3.1 |
| >hg38_21_+_8433172_8446622_RNA45SN1_45S_50ntFlanks@4560.4587.28\|CTAGGAATAATGGAATAGGACCGCGGTT | >hg38_21_+_8433172_8446622_RNA45SN1_45S_50ntFlanks | 4560 | 4587 | 28 | CTAGGAATAATGGAATAGGACCGCGGTT | MB468 | 35 | 51 | 54 | 36 | 15 | 10.8 | 15.5 | 14.6 | 14.5 | 5.6 |
| >hg38_21_+_8433172_8446622_RNA45SN1_45S_50ntFlanks@4560.4589.30\|CTAGGAATAATGGAATAGGACCGCGGTTCT | >hg38_21_+_8433172_8446622_RNA45SN1_45S_50ntFlanks | 4560 | 4589 | 30 | CTAGGAATAATGGAATAGGACCGCGGTTCT | MB468 | 15 | 10 | 24 | 13 | 17 | 7.7 | 2.9 | 15.5 | 9.1 | 11.9 |
| >hg38_21_+_8433172_8446622_RNA45SN1_45S_50ntFlanks@4564.4587.24\|GAATAATGGAATAGGACCGCGGTT | >hg38_21_+_8433172_8446622_RNA45SN1_45S_50ntFlanks | 4564 | 4587 | 24 | GAATAATGGAATAGGACCGCGGTT | MB468 | 18 | 28 | 15 | 6 | 7 | 7.9 | 16.2 | 3.5 | 7.9 | 3.6 |
| >hg38_21_+_8433172_8446622_RNA45SN1_45S_50ntFlanks@4565.4587.23\|AATAATGGAATAGGACCGCGGTT | >hg38_21_+_8433172_8446622_RNA45SN1_45S_50ntFlanks | 4565 | 4587 | 23 | AATAATGGAATAGGACCGCGGTT | MB468 | 55 | 78 | 44 | 18 | 21 | 21.5 | 33.4 | 13.2 | 18.4 | 6.2 |
| >hg38_21_+_8433172_8446622_RNA45SN1_45S_50ntFlanks@4566.4587.22\|ATAATGGAATAGGACCGCGGTT | >hg38_21_+_8433172_8446622_RNA45SN1_45S_50ntFlanks | 4566 | 4587 | 22 | ATAATGGAATAGGACCGCGGTT | MB468 | 30 | 39 | 27 | 8 | 9 | 16.5 | 16.6 | 7.7 | 7.3 | 3.5 |
| >hg38_21_+_8433172_8446622_RNA45SN1_45S_50ntFlanks@4567.4587.21\|TAATGGAATAGGACCGCGGTT | >hg38_21_+_8433172_8446622_RNA45SN1_45S_50ntFlanks | 4567 | 4587 | 21 | TAATGGAATAGGACCGCGGTT | MB468 | 50 | 51 | 47 | 13 | 18 | 17.3 | 20.2 | 10.4 | 9.5 | 7.5 |
| >hg38_21_+_8433172_8446622_RNA45SN1_45S_50ntFlanks@4567.4589.23\|TAATGGAATAGGACCGCGGTTCT | >hg38_21_+_8433172_8446622_RNA45SN1_45S_50ntFlanks | 4567 | 4589 | 23 | TAATGGAATAGGACCGCGGTTCT | MB468 | 23 | 7 | 25 | 14 | 25 | 9.3 | 1.8 | 9.1 | 6.6 | 27.1 |
| >hg38_21_+_8433172_8446622_RNA45SN1_45S_50ntFlanks@4568.4589.22\|AATGGAATAGGACCGCGGTTCT | >hg38_21_+_8433172_8446622_RNA45SN1_45S_50ntFlanks | 4568 | 4589 | 22 | AATGGAATAGGACCGCGGTTCT | MB231 | 14 | 27 | 13 | 9 | 7 | 7.3 | 14.2 | 4.7 | 1.9 | 4.1 |
| >hg38_21_+_8433172_8446622_RNA45SN1_45S_50ntFlanks@4569.4587.19\|ATGGAATAGGACCGCGGTT | >hg38_21_+_8433172_8446622_RNA45SN1_45S_50ntFlanks | 4569 | 4587 | 19 | ATGGAATAGGACCGCGGTT | MB468 | 31 | 40 | 22 | 14 | 11 | 12.5 | 17.9 | 5.9 | 8.1 | 1.9 |
| >hg38_21_+_8433172_8446622_RNA45SN1_45S_50ntFlanks@4569.4589.21\|ATGGAATAGGACCGCGGTTCT | >hg38_21_+_8433172_8446622_RNA45SN1_45S_50ntFlanks | 4569 | 4589 | 21 | ATGGAATAGGACCGCGGTTCT | MB468 | 11 | 2 | 9 | 11 | 11 | 5.1 | 0.7 | 3.7 | 5.3 | 11.8 |
| >hg38_21_+_8433172_8446622_RNA45SN1_45S_50ntFlanks@4570.4587.18\|TGGAATAGGACCGCGGTT | >hg38_21_+_8433172_8446622_RNA45SN1_45S_50ntFlanks | 4570 | 4587 | 18 | TGGAATAGGACCGCGGTT | MB468 | 36 | 50 | 30 | 12 | 12 | 14.4 | 21.2 | 9.2 | 5.6 | 3.5 |
| >hg38_21_+_8433172_8446622_RNA45SN1_45S_50ntFlanks@4570.4589.20\|TGGAATAGGACCGCGGTTCT | >hg38_21_+_8433172_8446622_RNA45SN1_45S_50ntFlanks | 4570 | 4589 | 20 | TGGAATAGGACCGCGGTTCT | MB468 | 12 | 3 | 12 | 9 | 19 | 5.2 | 0.4 | 4.1 | 3.7 | 29.1 |
| >hg38_21_+_8433172_8446622_RNA45SN1_45S_50ntFlanks@4588.4608.21\|CTATTTTGTTGGTTTTCGGAA | >hg38_21_+_8433172_8446622_RNA45SN1_45S_50ntFlanks | 4588 | 4608 | 21 | CTATTTTGTTGGTTTTCGGAA | MB468 | 21 | 6 | 38 | 9 | 9 | 13.5 | 4.0 | 28.0 | 4.3 | 4.7 |
| >hg38_21_+_8433172_8446622_RNA45SN1_45S_50ntFlanks@4588.4611.24\|CTATTTTGTTGGTTTTCGGAACTG | >hg38_21_+_8433172_8446622_RNA45SN1_45S_50ntFlanks | 4588 | 4611 | 24 | CTATTTTGTTGGTTTTCGGAACTG | MB468 | 13 | 4 | 19 | 30 | 9 | 3.0 | 1.8 | 7.1 | 14.8 | 2.9 |
| >hg38_21_+_8433172_8446622_RNA45SN1_45S_50ntFlanks@4588.4613.26\|CTATTTTGTTGGTTTTCGGAACTGAG | >hg38_21_+_8433172_8446622_RNA45SN1_45S_50ntFlanks | 4588 | 4613 | 26 | CTATTTTGTTGGTTTTCGGAACTGAG | MB468 | 21 | 9 | 17 | 47 | 14 | 7.6 | 3.4 | 4.2 | 17.2 | 3.7 |
| >hg38_21_+_8433172_8446622_RNA45SN1_45S_50ntFlanks@4588.4614.27\|CTATTTTGTTGGTTTTCGGAACTGAGG | >hg38_21_+_8433172_8446622_RNA45SN1_45S_50ntFlanks | 4588 | 4614 | 27 | CTATTTTGTTGGTTTTCGGAACTGAGG | MB468 | 56 | 48 | 52 | 48 | 22 | 13.4 | 25.0 | 24.0 | 18.5 | 8.0 |
| >hg38_21_+_8433172_8446622_RNA45SN1_45S_50ntFlanks@4588.4615.28\|CTATTTTGTTGGTTTTCGGAACTGAGGC | >hg38_21_+_8433172_8446622_RNA45SN1_45S_50ntFlanks | 4588 | 4615 | 28 | CTATTTTGTTGGTTTTCGGAACTGAGGC | MB468 | 19 | 26 | 25 | 16 | 6 | 8.6 | 15.5 | 18.9 | 7.1 | 3.7 |
| >hg38_21_+_8433172_8446622_RNA45SN1_45S_50ntFlanks@4588.4616.29\|CTATTTTGTTGGTTTTCGGAACTGAGGCC | >hg38_21_+_8433172_8446622_RNA45SN1_45S_50ntFlanks | 4588 | 4616 | 29 | CTATTTTGTTGGTTTTCGGAACTGAGGCC | MB468 | 121 | 147 | 212 | 115 | 72 | 46.4 | 61.6 | 122.7 | 69.4 | 16.8 |
| >hg38_21_+_8433172_8446622_RNA45SN1_45S_50ntFlanks@4588.4617.30\|CTATTTTGTTGGTTTTCGGAACTGAGGCCA | >hg38_21_+_8433172_8446622_RNA45SN1_45S_50ntFlanks | 4588 | 4617 | 30 | CTATTTTGTTGGTTTTCGGAACTGAGGCCA | MB468 | 11 | 4 | 13 | 11 | 7 | 2.5 | 1.0 | 8.6 | 7.5 | 2.6 |
| >hg38_21_+_8433172_8446622_RNA45SN1_45S_50ntFlanks@4588.4619.32\|CTATTTTGTTGGTTTTCGGAACTGAGGCCATG | >hg38_21_+_8433172_8446622_RNA45SN1_45S_50ntFlanks | 4588 | 4619 | 32 | CTATTTTGTTGGTTTTCGGAACTGAGGCCATG | MB468 | 20 | 8 | 27 | 30 | 24 | 5.3 | 2.7 | 17.1 | 19.7 | 9.8 |
| >hg38_21_+_8433172_8446622_RNA45SN1_45S_50ntFlanks@4588.4621.34\|CTATTTTGTTGGTTTTCGGAACTGAGGCCATGAT | >hg38_21_+_8433172_8446622_RNA45SN1_45S_50ntFlanks | 4588 | 4621 | 34 | CTATTTTGTTGGTTTTCGGAACTGAGGCCATGAT | MB468 | 13 | 6 | 22 | 15 | 26 | 4.4 | 2.7 | 24.4 | 8.7 | 12.7 |
| >hg38_21_+_8433172_8446622_RNA45SN1_45S_50ntFlanks@4590.4614.25\|ATTTTGTTGGTTTTCGGAACTGAGG | >hg38_21_+_8433172_8446622_RNA45SN1_45S_50ntFlanks | 4590 | 4614 | 25 | ATTTTGTTGGTTTTCGGAACTGAGG | MB468 | 16 | 13 | 11 | 14 | 5 | 1.9 | 5.6 | 4.4 | 4.4 | 1.3 |
| >hg38_21_+_8433172_8446622_RNA45SN1_45S_50ntFlanks@4590.4616.27\|ATTTTGTTGGTTTTCGGAACTGAGGCC | >hg38_21_+_8433172_8446622_RNA45SN1_45S_50ntFlanks | 4590 | 4616 | 27 | ATTTTGTTGGTTTTCGGAACTGAGGCC | MB468 | 31 | 30 | 38 | 42 | 20 | 16.7 | 8.6 | 21.4 | 15.7 | 6.6 |
| >hg38_21_+_8433172_8446622_RNA45SN1_45S_50ntFlanks@4591.4616.26\|TTTTGTTGGTTTTCGGAACTGAGGCC | >hg38_21_+_8433172_8446622_RNA45SN1_45S_50ntFlanks | 4591 | 4616 | 26 | TTTTGTTGGTTTTCGGAACTGAGGCC | MB468 | 18 | 17 | 15 | 22 | 7 | 9.5 | 6.9 | 8.3 | 6.8 | 1.0 |
| >hg38_21_+_8433172_8446622_RNA45SN1_45S_50ntFlanks@4592.4616.25\|TTTGTTGGTTTTCGGAACTGAGGCC | >hg38_21_+_8433172_8446622_RNA45SN1_45S_50ntFlanks | 4592 | 4616 | 25 | TTTGTTGGTTTTCGGAACTGAGGCC | MB468 | 15 | 12 | 16 | 18 | 7 | 6.7 | 4.4 | 5.6 | 5.8 | 1.2 |
| >hg38_21_+_8433172_8446622_RNA45SN1_45S_50ntFlanks@4594.4616.23\|TGTTGGTTTTCGGAACTGAGGCC | >hg38_21_+_8433172_8446622_RNA45SN1_45S_50ntFlanks | 4594 | 4616 | 23 | TGTTGGTTTTCGGAACTGAGGCC | MB468 | 11 | 6 | 13 | 24 | 7 | 6.7 | 3.7 | 6.0 | 11.6 | 1.8 |
| >hg38_21_+_8433172_8446622_RNA45SN1_45S_50ntFlanks@4602.4619.18\|TTCGGAACTGAGGCCATG | >hg38_21_+_8433172_8446622_RNA45SN1_45S_50ntFlanks | 4602 | 4619 | 18 | TTCGGAACTGAGGCCATG | MB468 | 12 | 5 | 4 | 14 | 5 | 4.6 | 3.1 | 0.9 | 5.2 | 1.8 |
| >hg38_21_+_8433172_8446622_RNA45SN1_45S_50ntFlanks@4657.4680.24\|CTAGAGGTGAAATTCTTGGACCGG | >hg38_21_+_8433172_8446622_RNA45SN1_45S_50ntFlanks | 4657 | 4680 | 24 | CTAGAGGTGAAATTCTTGGACCGG | MB468 | 15 | 11 | 20 | 7 | 12 | 10.9 | 2.4 | 12.3 | 4.1 | 7.9 |
| >hg38_21_+_8433172_8446622_RNA45SN1_45S_50ntFlanks@4657.4682.26\|CTAGAGGTGAAATTCTTGGACCGGCG | >hg38_21_+_8433172_8446622_RNA45SN1_45S_50ntFlanks | 4657 | 4682 | 26 | CTAGAGGTGAAATTCTTGGACCGGCG | MB468 | 53 | 103 | 58 | 14 | 56 | 37.6 | 25.1 | 42.8 | 6.8 | 33.5 |
| >hg38_21_+_8433172_8446622_RNA45SN1_45S_50ntFlanks@4657.4684.28\|CTAGAGGTGAAATTCTTGGACCGGCGCA | >hg38_21_+_8433172_8446622_RNA45SN1_45S_50ntFlanks | 4657 | 4684 | 28 | CTAGAGGTGAAATTCTTGGACCGGCGCA | MB468 | 13 | 21 | 21 | 17 | 55 | 8.2 | 2.4 | 15.3 | 10.3 | 65.4 |
| >hg38_21_+_8433172_8446622_RNA45SN1_45S_50ntFlanks@4657.4686.30\|CTAGAGGTGAAATTCTTGGACCGGCGCAAG | >hg38_21_+_8433172_8446622_RNA45SN1_45S_50ntFlanks | 4657 | 4686 | 30 | CTAGAGGTGAAATTCTTGGACCGGCGCAAG | MB468 | 17 | 16 | 23 | 23 | 36 | 11.4 | 4.3 | 13.8 | 14.9 | 28.7 |
| >hg38_21_+_8433172_8446622_RNA45SN1_45S_50ntFlanks@4657.4688.32\|CTAGAGGTGAAATTCTTGGACCGGCGCAAGAC | >hg38_21_+_8433172_8446622_RNA45SN1_45S_50ntFlanks | 4657 | 4688 | 32 | CTAGAGGTGAAATTCTTGGACCGGCGCAAGAC | MB468 | 25 | 37 | 37 | 13 | 101 | 20.1 | 29.5 | 38.8 | 9.0 | 84.5 |
| >hg38_21_+_8433172_8446622_RNA45SN1_45S_50ntFlanks@4657.4689.33\|CTAGAGGTGAAATTCTTGGACCGGCGCAAGACG | >hg38_21_+_8433172_8446622_RNA45SN1_45S_50ntFlanks | 4657 | 4689 | 33 | CTAGAGGTGAAATTCTTGGACCGGCGCAAGACG | MB468 | 298 | 301 | 386 | 267 | 1039 | 203.9 | 164.0 | 294.7 | 146.3 | 788.8 |
| >hg38_21_+_8433172_8446622_RNA45SN1_45S_50ntFlanks@4659.4678.20\|AGAGGTGAAATTCTTGGACC | >hg38_21_+_8433172_8446622_RNA45SN1_45S_50ntFlanks | 4659 | 4678 | 20 | AGAGGTGAAATTCTTGGACC | MB231 | 10 | 13 | 10 | 10 | 3 | 9.4 | 13.7 | 12.7 | 9.1 | 2.1 |
| >hg38_21_+_8433172_8446622_RNA45SN1_45S_50ntFlanks@4661.4689.29\|AGGTGAAATTCTTGGACCGGCGCAAGACG | >hg38_21_+_8433172_8446622_RNA45SN1_45S_50ntFlanks | 4661 | 4689 | 29 | AGGTGAAATTCTTGGACCGGCGCAAGACG | MB468 | 13 | 21 | 19 | 29 | 22 | 10.2 | 14.2 | 16.8 | 17.8 | 7.1 |
| >hg38_21_+_8433172_8446622_RNA45SN1_45S_50ntFlanks@4683.4707.25\|CAAGACGGACCAGAGCGAAAGCATT | >hg38_21_+_8433172_8446622_RNA45SN1_45S_50ntFlanks | 4683 | 4707 | 25 | CAAGACGGACCAGAGCGAAAGCATT | MB468 | 27 | 36 | 45 | 41 | 53 | 23.0 | 33.1 | 49.9 | 15.3 | 19.5 |

| | | | | | | | | | | | | | | | | |
|---|---|---|---|---|---|---|---|---|---|---|---|---|---|---|---|---|
| >hg38_21_+_8433172_8446622_RNA45SN1_45S_50ntFlanks@4685.4707.23\|AGACGGACCAGAGCGAAAGCATT | >hg38_21_+_8433172_8446622_RNA45SN1_45S_50ntFlanks | 4685 | 4707 | 23 | AGACGGACCAGAGCGAAAGCATT | MB468 | 17 | 29 | 22 | 20 | 30 | 13.0 | 28.7 | 23.3 | 8.5 | 22.7 |
| >hg38_21_+_8433172_8446622_RNA45SN1_45S_50ntFlanks@4687.4707.21\|ACGGACCAGAGCGAAAGCATT | >hg38_21_+_8433172_8446622_RNA45SN1_45S_50ntFlanks | 4687 | 4707 | 21 | ACGGACCAGAGCGAAAGCATT | MB468 | 15 | 22 | 18 | 15 | 38 | 11.3 | 17.9 | 19.4 | 5.2 | 32.8 |
| >hg38_21_+_8433172_8446622_RNA45SN1_45S_50ntFlanks@4722.4748.27\|TCATTAATCAAGAACGAAAGTCGGAGG | >hg38_21_+_8433172_8446622_RNA45SN1_45S_50ntFlanks | 4722 | 4748 | 27 | TCATTAATCAAGAACGAAAGTCGGAGG | MB468 | 47 | 28 | 49 | 28 | 41 | 33.3 | 9.8 | 43.2 | 9.9 | 17.6 |
| >hg38_21_+_8433172_8446622_RNA45SN1_45S_50ntFlanks@4722.4751.30\|TCATTAATCAAGAACGAAAGTCGGAGGTTC | >hg38_21_+_8433172_8446622_RNA45SN1_45S_50ntFlanks | 4722 | 4751 | 30 | TCATTAATCAAGAACGAAAGTCGGAGGTTC | MB468 | 55 | 44 | 52 | 55 | 40 | 32.6 | 10.3 | 43.8 | 14.4 | 14.9 |
| >hg38_21_+_8433172_8446622_RNA45SN1_45S_50ntFlanks@4722.4752.31\|TCATTAATCAAGAACGAAAGTCGGAGGTTCG | >hg38_21_+_8433172_8446622_RNA45SN1_45S_50ntFlanks | 4722 | 4752 | 31 | TCATTAATCAAGAACGAAAGTCGGAGGTTCG | MB468 | 69 | 51 | 92 | 46 | 86 | 43.8 | 10.9 | 85.8 | 18.3 | 10.4 |
| >hg38_21_+_8433172_8446622_RNA45SN1_45S_50ntFlanks@4722.4758.37\|TCATTAATCAAGAACGAAAGTCGGAGGTTCGAAGACG | >hg38_21_+_8433172_8446622_RNA45SN1_45S_50ntFlanks | 4722 | 4758 | 37 | TCATTAATCAAGAACGAAAGTCGGAGGTTCGAAGACG | MB468 | 360 | 814 | 523 | 432 | 723 | 242.0 | 135.4 | 420.3 | 314.3 | 291.7 |
| >hg38_21_+_8433172_8446622_RNA45SN1_45S_50ntFlanks@4722.4759.38\|TCATTAATCAAGAACGAAAGTCGGAGGTTCGAAGACGA | >hg38_21_+_8433172_8446622_RNA45SN1_45S_50ntFlanks | 4722 | 4759 | 38 | TCATTAATCAAGAACGAAAGTCGGAGGTTCGAAGACGA | MB468 | 133 | 339 | 290 | 117 | 246 | 98.3 | 208.5 | 237.2 | 69.7 | 72.6 |
| >hg38_21_+_8433172_8446622_RNA45SN1_45S_50ntFlanks@4723.4748.26\|CATTAATCAAGAACGAAAGTCGGAGG | >hg38_21_+_8433172_8446622_RNA45SN1_45S_50ntFlanks | 4723 | 4748 | 26 | CATTAATCAAGAACGAAAGTCGGAGG | MB468 | 40 | 21 | 45 | 36 | 38 | 25.5 | 7.5 | 34.0 | 16.3 | 16.9 |
| >hg38_21_+_8433172_8446622_RNA45SN1_45S_50ntFlanks@4723.4751.29\|CATTAATCAAGAACGAAAGTCGGAGGTTC | >hg38_21_+_8433172_8446622_RNA45SN1_45S_50ntFlanks | 4723 | 4751 | 29 | CATTAATCAAGAACGAAAGTCGGAGGTTC | MB468 | 56 | 37 | 47 | 67 | 38 | 22.5 | 8.4 | 35.2 | 16.4 | 14.2 |
| >hg38_21_+_8433172_8446622_RNA45SN1_45S_50ntFlanks@4723.4752.30\|CATTAATCAAGAACGAAAGTCGGAGGTTCG | >hg38_21_+_8433172_8446622_RNA45SN1_45S_50ntFlanks | 4723 | 4752 | 30 | CATTAATCAAGAACGAAAGTCGGAGGTTCG | MB468 | 70 | 43 | 77 | 65 | 100 | 31.1 | 7.8 | 62.7 | 25.3 | 54.0 |
| >hg38_21_+_8433172_8446622_RNA45SN1_45S_50ntFlanks@4723.4758.36\|CATTAATCAAGAACGAAAGTCGGAGGTTCGAAGACG | >hg38_21_+_8433172_8446622_RNA45SN1_45S_50ntFlanks | 4723 | 4758 | 36 | CATTAATCAAGAACGAAAGTCGGAGGTTCGAAGACG | MB468 | 352 | 613 | 328 | 502 | 951 | 144.0 | 32.6 | 217.9 | 341.4 | 552.9 |
| >hg38_21_+_8433172_8446622_RNA45SN1_45S_50ntFlanks@4723.4759.37\|CATTAATCAAGAACGAAAGTCGGAGGTTCGAAGACGA | >hg38_21_+_8433172_8446622_RNA45SN1_45S_50ntFlanks | 4723 | 4759 | 37 | CATTAATCAAGAACGAAAGTCGGAGGTTCGAAGACGA | MB468 | 86 | 156 | 127 | 91 | 210 | 48.2 | 71.3 | 89.3 | 53.4 | 70.4 |
| >hg38_21_+_8433172_8446622_RNA45SN1_45S_50ntFlanks@4724.4748.25\|ATTAATCAAGAACGAAAGTCGGAGG | >hg38_21_+_8433172_8446622_RNA45SN1_45S_50ntFlanks | 4724 | 4748 | 25 | ATTAATCAAGAACGAAAGTCGGAGG | MB468 | 42 | 27 | 45 | 44 | 43 | 24.8 | 10.9 | 39.5 | 12.1 | 30.6 |
| >hg38_21_+_8433172_8446622_RNA45SN1_45S_50ntFlanks@4724.4751.28\|ATTAATCAAGAACGAAAGTCGGAGGTTC | >hg38_21_+_8433172_8446622_RNA45SN1_45S_50ntFlanks | 4724 | 4751 | 28 | ATTAATCAAGAACGAAAGTCGGAGGTTC | MB468 | 62 | 48 | 52 | 93 | 45 | 23.1 | 13.3 | 37.5 | 12.9 | 31.1 |
| >hg38_21_+_8433172_8446622_RNA45SN1_45S_50ntFlanks@4724.4752.29\|ATTAATCAAGAACGAAAGTCGGAGGTTCG | >hg38_21_+_8433172_8446622_RNA45SN1_45S_50ntFlanks | 4724 | 4752 | 29 | ATTAATCAAGAACGAAAGTCGGAGGTTCG | MB468 | 66 | 51 | 75 | 68 | 91 | 27.5 | 11.5 | 63.3 | 13.0 | 56.5 |
| >hg38_21_+_8433172_8446622_RNA45SN1_45S_50ntFlanks@4724.4757.34\|ATTAATCAAGAACGAAAGTCGGAGGTTCGAAGAC | >hg38_21_+_8433172_8446622_RNA45SN1_45S_50ntFlanks | 4724 | 4757 | 34 | ATTAATCAAGAACGAAAGTCGGAGGTTCGAAGAC | MB468 | 10 | 20 | 9 | 14 | 26 | 5.6 | 0.9 | 5.8 | 11.3 | 16.7 |
| >hg38_21_+_8433172_8446622_RNA45SN1_45S_50ntFlanks@4724.4758.35\|ATTAATCAAGAACGAAAGTCGGAGGTTCGAAGACG | >hg38_21_+_8433172_8446622_RNA45SN1_45S_50ntFlanks | 4724 | 4758 | 35 | ATTAATCAAGAACGAAAGTCGGAGGTTCGAAGACG | MB468 | 576 | 1078 | 505 | 846 | 1499 | 273.9 | 87.5 | 315.7 | 625.9 | 886.8 |
| >hg38_21_+_8433172_8446622_RNA45SN1_45S_50ntFlanks@4724.4759.36\|ATTAATCAAGAACGAAAGTCGGAGGTTCGAAGACGA | >hg38_21_+_8433172_8446622_RNA45SN1_45S_50ntFlanks | 4724 | 4759 | 36 | ATTAATCAAGAACGAAAGTCGGAGGTTCGAAGACGA | MB468 | 159 | 331 | 209 | 175 | 395 | 102.6 | 96.8 | 139.7 | 109.7 | 194.2 |
| >hg38_21_+_8433172_8446622_RNA45SN1_45S_50ntFlanks@4725.4758.34\|TTAATCAAGAACGAAAGTCGGAGGTTCGAAGACG | >hg38_21_+_8433172_8446622_RNA45SN1_45S_50ntFlanks | 4725 | 4758 | 34 | TTAATCAAGAACGAAAGTCGGAGGTTCGAAGACG | MB468 | 58 | 129 | 60 | 91 | 92 | 35.7 | 14.5 | 32.2 | 63.7 | 46.6 |
| >hg38_21_+_8433172_8446622_RNA45SN1_45S_50ntFlanks@4725.4759.35\|TTAATCAAGAACGAAAGTCGGAGGTTCGAAGACGA | >hg38_21_+_8433172_8446622_RNA45SN1_45S_50ntFlanks | 4725 | 4759 | 35 | TTAATCAAGAACGAAAGTCGGAGGTTCGAAGACGA | MB468 | 18 | 39 | 24 | 20 | 29 | 13.2 | 9.0 | 13.7 | 10.4 | 11.8 |
| >hg38_21_+_8433172_8446622_RNA45SN1_45S_50ntFlanks@4726.4748.23\|TAATCAAGAACGAAAGTCGGAGG | >hg38_21_+_8433172_8446622_RNA45SN1_45S_50ntFlanks | 4726 | 4748 | 23 | TAATCAAGAACGAAAGTCGGAGG | MB468 | 18 | 9 | 15 | 12 | 9 | 8.0 | 3.8 | 11.2 | 5.7 | 5.2 |
| >hg38_21_+_8433172_8446622_RNA45SN1_45S_50ntFlanks@4726.4751.26\|TAATCAAGAACGAAAGTCGGAGGTTC | >hg38_21_+_8433172_8446622_RNA45SN1_45S_50ntFlanks | 4726 | 4751 | 26 | TAATCAAGAACGAAAGTCGGAGGTTC | MB468 | 20 | 14 | 18 | 28 | 9 | 9.2 | 4.1 | 9.5 | 9.2 | 6.7 |
| >hg38_21_+_8433172_8446622_RNA45SN1_45S_50ntFlanks@4726.4752.27\|TAATCAAGAACGAAAGTCGGAGGTTCG | >hg38_21_+_8433172_8446622_RNA45SN1_45S_50ntFlanks | 4726 | 4752 | 27 | TAATCAAGAACGAAAGTCGGAGGTTCG | MB468 | 29 | 22 | 31 | 21 | 20 | 13.2 | 5.4 | 19.5 | 5.8 | 12.0 |
| >hg38_21_+_8433172_8446622_RNA45SN1_45S_50ntFlanks@4726.4758.33\|TAATCAAGAACGAAAGTCGGAGGTTCGAAGACG | >hg38_21_+_8433172_8446622_RNA45SN1_45S_50ntFlanks | 4726 | 4758 | 33 | TAATCAAGAACGAAAGTCGGAGGTTCGAAGACG | MB468 | 202 | 344 | 177 | 212 | 243 | 100.4 | 39.3 | 85.2 | 131.9 | 108.2 |
| >hg38_21_+_8433172_8446622_RNA45SN1_45S_50ntFlanks@4726.4759.34\|TAATCAAGAACGAAAGTCGGAGGTTCGAAGACGA | >hg38_21_+_8433172_8446622_RNA45SN1_45S_50ntFlanks | 4726 | 4759 | 34 | TAATCAAGAACGAAAGTCGGAGGTTCGAAGACGA | MB468 | 80 | 132 | 91 | 68 | 91 | 53.8 | 28.4 | 49.2 | 36.1 | 31.0 |
| >hg38_21_+_8433172_8446622_RNA45SN1_45S_50ntFlanks@4727.4752.26\|AATCAAGAACGAAAGTCGGAGGTTCG | >hg38_21_+_8433172_8446622_RNA45SN1_45S_50ntFlanks | 4727 | 4752 | 26 | AATCAAGAACGAAAGTCGGAGGTTCG | MB468 | 13 | 10 | 12 | 21 | 9 | 6.2 | 4.0 | 7.7 | 6.8 | 1.4 |
| >hg38_21_+_8433172_8446622_RNA45SN1_45S_50ntFlanks@4727.4758.32\|AATCAAGAACGAAAGTCGGAGGTTCGAAGACG | >hg38_21_+_8433172_8446622_RNA45SN1_45S_50ntFlanks | 4727 | 4758 | 32 | AATCAAGAACGAAAGTCGGAGGTTCGAAGACG | MB231 | 20 | 7 | 35 | 48 | 10 | 12.3 | 1.4 | 26.5 | 38.0 | 2.6 |
| >hg38_21_+_8433172_8446622_RNA45SN1_45S_50ntFlanks@4727.4758.32\|AATCAAGAACGAAAGTCGGAGGTTCGAAGACG | >hg38_21_+_8433172_8446622_RNA45SN1_45S_50ntFlanks | 4727 | 4758 | 32 | AATCAAGAACGAAAGTCGGAGGTTCGAAGACG | MB468 | 26 | 42 | 28 | 62 | 49 | 15.0 | 11.6 | 15.5 | 15.4 | 16.7 |
| >hg38_21_+_8433172_8446622_RNA45SN1_45S_50ntFlanks@4727.4759.33\|AATCAAGAACGAAAGTCGGAGGTTCGAAGACGA | >hg38_21_+_8433172_8446622_RNA45SN1_45S_50ntFlanks | 4727 | 4759 | 33 | AATCAAGAACGAAAGTCGGAGGTTCGAAGACGA | MB468 | 12 | 17 | 15 | 24 | 21 | 8.3 | 5.0 | 8.3 | 11.0 | 5.5 |
| >hg38_21_+_8433172_8446622_RNA45SN1_45S_50ntFlanks@4727.4761.35\|AATCAAGAACGAAAGTCGGAGGTTCGAAGACGATC | >hg38_21_+_8433172_8446622_RNA45SN1_45S_50ntFlanks | 4727 | 4761 | 35 | AATCAAGAACGAAAGTCGGAGGTTCGAAGACGATC | MB231 | 18 | 12 | 58 | 7 | 5 | 8.3 | 4.0 | 27.9 | 5.8 | 1.2 |
| >hg38_21_+_8433172_8446622_RNA45SN1_45S_50ntFlanks@4727.4765.39\|AATCAAGAACGAAAGTCGGAGGTTCGAAGACGATCAGAT | >hg38_21_+_8433172_8446622_RNA45SN1_45S_50ntFlanks | 4727 | 4765 | 39 | AATCAAGAACGAAAGTCGGAGGTTCGAAGACGATCAGAT | BT20 | 15 | 27 | 93 | 61 | 14 | 8.9 | 6.4 | 59.8 | 41.7 | 2.6 |
| >hg38_21_+_8433172_8446622_RNA45SN1_45S_50ntFlanks@4727.4765.39\|AATCAAGAACGAAAGTCGGAGGTTCGAAGACGATCAGAT | >hg38_21_+_8433172_8446622_RNA45SN1_45S_50ntFlanks | 4727 | 4765 | 39 | AATCAAGAACGAAAGTCGGAGGTTCGAAGACGATCAGAT | MB231 | 24 | 18 | 125 | 19 | 10 | 11.3 | 4.3 | 70.5 | 18.4 | 4.2 |
| >hg38_21_+_8433172_8446622_RNA45SN1_45S_50ntFlanks@4727.4767.41\|AATCAAGAACGAAAGTCGGAGGTTCGAAGACGATCAGATAC | >hg38_21_+_8433172_8446622_RNA45SN1_45S_50ntFlanks | 4727 | 4767 | 41 | AATCAAGAACGAAAGTCGGAGGTTCGAAGACGATCAGATAC | MB231 | 37 | 14 | 268 | 10 | 9 | 31.6 | 13.4 | 215.4 | 12.5 | 6.7 |
| >hg38_21_+_8433172_8446622_RNA45SN1_45S_50ntFlanks@4727.4768.42\|AATCAAGAACGAAAGTCGGAGGTTCGAAGACGATCAGATACC | >hg38_21_+_8433172_8446622_RNA45SN1_45S_50ntFlanks | 4727 | 4768 | 42 | AATCAAGAACGAAAGTCGGAGGTTCGAAGACGATCAGATACC | MB231 | 21 | 10 | 174 | 5 | 5 | 17.5 | 9.3 | 149.7 | 7.1 | 4.2 |
| >hg38_21_+_8433172_8446622_RNA45SN1_45S_50ntFlanks@4728.4748.21\|ATCAAGAACGAAAGTCGGAGG | >hg38_21_+_8433172_8446622_RNA45SN1_45S_50ntFlanks | 4728 | 4748 | 21 | ATCAAGAACGAAAGTCGGAGG | MB468 | 21 | 9 | 19 | 31 | 8 | 9.5 | 5.0 | 10.9 | 17.7 | 2.6 |
| >hg38_21_+_8433172_8446622_RNA45SN1_45S_50ntFlanks@4728.4751.24\|ATCAAGAACGAAAGTCGGAGGTTC | >hg38_21_+_8433172_8446622_RNA45SN1_45S_50ntFlanks | 4728 | 4751 | 24 | ATCAAGAACGAAAGTCGGAGGTTC | MB468 | 53 | 50 | 46 | 91 | 14 | 17.3 | 25.3 | 14.1 | 36.9 | 3.9 |
| >hg38_21_+_8433172_8446622_RNA45SN1_45S_50ntFlanks@4728.4752.25\|ATCAAGAACGAAAGTCGGAGGTTCG | >hg38_21_+_8433172_8446622_RNA45SN1_45S_50ntFlanks | 4728 | 4752 | 25 | ATCAAGAACGAAAGTCGGAGGTTCG | MB468 | 46 | 33 | 38 | 45 | 17 | 12.6 | 15.2 | 16.1 | 13.0 | 2.1 |
| >hg38_21_+_8433172_8446622_RNA45SN1_45S_50ntFlanks@4728.4758.31\|ATCAAGAACGAAAGTCGGAGGTTCGAAGACG | >hg38_21_+_8433172_8446622_RNA45SN1_45S_50ntFlanks | 4728 | 4758 | 31 | ATCAAGAACGAAAGTCGGAGGTTCGAAGACG | MB468 | 153 | 263 | 129 | 329 | 134 | 75.6 | 59.7 | 72.1 | 83.4 | 29.0 |
| >hg38_21_+_8433172_8446622_RNA45SN1_45S_50ntFlanks@4728.4759.32\|ATCAAGAACGAAAGTCGGAGGTTCGAAGACGA | >hg38_21_+_8433172_8446622_RNA45SN1_45S_50ntFlanks | 4728 | 4759 | 32 | ATCAAGAACGAAAGTCGGAGGTTCGAAGACGA | MB468 | 29 | 31 | 31 | 59 | 20 | 14.1 | 7.0 | 14.3 | 30.6 | 5.0 |
| >hg38_21_+_8433172_8446622_RNA45SN1_45S_50ntFlanks@4728.4767.40\|ATCAAGAACGAAAGTCGGAGGTTCGAAGACGATCAGATAC | >hg38_21_+_8433172_8446622_RNA45SN1_45S_50ntFlanks | 4728 | 4767 | 40 | ATCAAGAACGAAAGTCGGAGGTTCGAAGACGATCAGATAC | MB231 | 11 | 5 | 28 | 5 | 2 | 18.4 | 9.3 | 48.6 | 8.7 | 4.1 |
| >hg38_21_+_8433172_8446622_RNA45SN1_45S_50ntFlanks@4729.4748.20\|TCAAGAACGAAAGTCGGAGG | >hg38_21_+_8433172_8446622_RNA45SN1_45S_50ntFlanks | 4729 | 4748 | 20 | TCAAGAACGAAAGTCGGAGG | MB468 | 13 | 4 | 10 | 17 | 6 | 4.3 | 2.0 | 5.4 | 6.3 | 1.2 |
| >hg38_21_+_8433172_8446622_RNA45SN1_45S_50ntFlanks@4729.4751.23\|TCAAGAACGAAAGTCGGAGGTTC | >hg38_21_+_8433172_8446622_RNA45SN1_45S_50ntFlanks | 4729 | 4751 | 23 | TCAAGAACGAAAGTCGGAGGTTC | MB468 | 27 | 26 | 22 | 44 | 10 | 8.2 | 9.8 | 5.8 | 11.9 | 2.7 |
| >hg38_21_+_8433172_8446622_RNA45SN1_45S_50ntFlanks@4729.4752.24\|TCAAGAACGAAAGTCGGAGGTTCG | >hg38_21_+_8433172_8446622_RNA45SN1_45S_50ntFlanks | 4729 | 4752 | 24 | TCAAGAACGAAAGTCGGAGGTTCG | MB468 | 17 | 10 | 15 | 16 | 12 | 2.8 | 4.1 | 4.4 | 1.6 | 2.4 |
| >hg38_21_+_8433172_8446622_RNA45SN1_45S_50ntFlanks@4729.4758.30\|TCAAGAACGAAAGTCGGAGGTTCGAAGACG | >hg38_21_+_8433172_8446622_RNA45SN1_45S_50ntFlanks | 4729 | 4758 | 30 | TCAAGAACGAAAGTCGGAGGTTCGAAGACG | MB468 | 82 | 159 | 68 | 218 | 136 | 35.3 | 60.3 | 32.7 | 100.7 | 82.5 |
| >hg38_21_+_8433172_8446622_RNA45SN1_45S_50ntFlanks@4729.4759.31\|TCAAGAACGAAAGTCGGAGGTTCGAAGACGA | >hg38_21_+_8433172_8446622_RNA45SN1_45S_50ntFlanks | 4729 | 4759 | 31 | TCAAGAACGAAAGTCGGAGGTTCGAAGACGA | MB468 | 28 | 40 | 30 | 55 | 32 | 14.8 | 5.0 | 11.4 | 19.1 | 6.2 |
| >hg38_21_+_8433172_8446622_RNA45SN1_45S_50ntFlanks@4729.4767.39\|TCAAGAACGAAAGTCGGAGGTTCGAAGACGATCAGATAC | >hg38_21_+_8433172_8446622_RNA45SN1_45S_50ntFlanks | 4729 | 4767 | 39 | TCAAGAACGAAAGTCGGAGGTTCGAAGACGATCAGATAC | MB231 | 10 | 4 | 36 | 3 | 2 | 10.7 | 4.0 | 27.8 | 3.4 | 1.6 |
| >hg38_21_+_8433172_8446622_RNA45SN1_45S_50ntFlanks@4730.4748.19\|CAAGAACGAAAGTCGGAGG | >hg38_21_+_8433172_8446622_RNA45SN1_45S_50ntFlanks | 4730 | 4748 | 19 | CAAGAACGAAAGTCGGAGG | MB468 | 19 | 6 | 12 | 28 | 10 | 7.4 | 3.1 | 6.6 | 14.0 | 2.8 |
| >hg38_21_+_8433172_8446622_RNA45SN1_45S_50ntFlanks@4730.4751.22\|CAAGAACGAAAGTCGGAGGTTC | >hg38_21_+_8433172_8446622_RNA45SN1_45S_50ntFlanks | 4730 | 4751 | 22 | CAAGAACGAAAGTCGGAGGTTC | MB468 | 42 | 32 | 29 | 64 | 17 | 8.5 | 12.2 | 9.4 | 22.4 | 2.5 |
| >hg38_21_+_8433172_8446622_RNA45SN1_45S_50ntFlanks@4730.4752.23\|CAAGAACGAAAGTCGGAGGTTCG | >hg38_21_+_8433172_8446622_RNA45SN1_45S_50ntFlanks | 4730 | 4752 | 23 | CAAGAACGAAAGTCGGAGGTTCG | MB468 | 27 | 13 | 22 | 22 | 16 | 2.8 | 5.1 | 6.7 | 6.4 | 1.9 |
| >hg38_21_+_8433172_8446622_RNA45SN1_45S_50ntFlanks@4730.4758.29\|CAAGAACGAAAGTCGGAGGTTCGAAGACG | >hg38_21_+_8433172_8446622_RNA45SN1_45S_50ntFlanks | 4730 | 4758 | 29 | CAAGAACGAAAGTCGGAGGTTCGAAGACG | MB231 | 13 | 5 | 33 | 31 | 7 | 9.7 | 2.4 | 18.4 | 27.4 | 2.5 |

| | | | | | | | | | | | | | | | | |
|---|---|---|---|---|---|---|---|---|---|---|---|---|---|---|---|---|
| >hg38_21_+_8433172_8446622_RNA45SN1_45S_50ntFlanks@4730.4758.29\|CAAGAACGAAAGTCGGAGGTTCGAAGACG | >hg38_21_+_8433172_8446622_RNA45SN1_45S_50ntFlanks | 4730 | 4758 | 29 | CAAGAACGAAAGTCGGAGGTTCGAAGACG | MB468 | 119 | 210 | 77 | 308 | 229 | 57.5 | 66.1 | 40.5 | 135.4 | 159.9 |
| >hg38_21_+_8433172_8446622_RNA45SN1_45S_50ntFlanks@4730.4759.30\|CAAGAACGAAAGTCGGAGGTTCGAAGACGA | >hg38_21_+_8433172_8446622_RNA45SN1_45S_50ntFlanks | 4730 | 4759 | 30 | CAAGAACGAAAGTCGGAGGTTCGAAGACGA | MB468 | 36 | 45 | 24 | 83 | 49 | 18.9 | 5.3 | 11.2 | 30.0 | 23.3 |
| >hg38_21_+_8433172_8446622_RNA45SN1_45S_50ntFlanks@4730.4765.36\|CAAGAACGAAAGTCGGAGGTTCGAAGACGATCAGAT | >hg38_21_+_8433172_8446622_RNA45SN1_45S_50ntFlanks | 4730 | 4765 | 36 | CAAGAACGAAAGTCGGAGGTTCGAAGACGATCAGAT | MB231 | 10 | 6 | 34 | 7 | 4 | 7.8 | 1.7 | 25.6 | 3.2 | 3.0 |
| >hg38_21_+_8433172_8446622_RNA45SN1_45S_50ntFlanks@4730.4767.38\|CAAGAACGAAAGTCGGAGGTTCGAAGACGATCAGATAC | >hg38_21_+_8433172_8446622_RNA45SN1_45S_50ntFlanks | 4730 | 4767 | 38 | CAAGAACGAAAGTCGGAGGTTCGAAGACGATCAGATAC | MB231 | 26 | 10 | 103 | 8 | 6 | 22.2 | 7.5 | 92.5 | 5.5 | 5.6 |
| >hg38_21_+_8433172_8446622_RNA45SN1_45S_50ntFlanks@4730.4768.39\|CAAGAACGAAAGTCGGAGGTTCGAAGACGATCAGATACC | >hg38_21_+_8433172_8446622_RNA45SN1_45S_50ntFlanks | 4730 | 4768 | 39 | CAAGAACGAAAGTCGGAGGTTCGAAGACGATCAGATACC | MB231 | 14 | 5 | 55 | 3 | 3 | 13.0 | 4.0 | 55.9 | 3.1 | 3.3 |
| >hg38_21_+_8433172_8446622_RNA45SN1_45S_50ntFlanks@4731.4748.18\|AAGAACGAAAGTCGGAGG | >hg38_21_+_8433172_8446622_RNA45SN1_45S_50ntFlanks | 4731 | 4748 | 18 | AAGAACGAAAGTCGGAGG | MB468 | 30 | 13 | 26 | 47 | 12 | 11.4 | 7.7 | 12.8 | 19.3 | 4.6 |
| >hg38_21_+_8433172_8446622_RNA45SN1_45S_50ntFlanks@4731.4750.20\|AAGAACGAAAGTCGGAGGTT | >hg38_21_+_8433172_8446622_RNA45SN1_45S_50ntFlanks | 4731 | 4750 | 20 | AAGAACGAAAGTCGGAGGTT | MB468 | 12 | 14 | 5 | 6 | 2 | 5.7 | 7.9 | 2.9 | 2.7 | 0.4 |
| >hg38_21_+_8433172_8446622_RNA45SN1_45S_50ntFlanks@4731.4751.21\|AAGAACGAAAGTCGGAGGTTC | >hg38_21_+_8433172_8446622_RNA45SN1_45S_50ntFlanks | 4731 | 4751 | 21 | AAGAACGAAAGTCGGAGGTTC | MB231 | 17 | 8 | 19 | 12 | 5 | 22.7 | 8.1 | 23.6 | 6.3 | 3.4 |
| >hg38_21_+_8433172_8446622_RNA45SN1_45S_50ntFlanks@4731.4751.21\|AAGAACGAAAGTCGGAGGTTC | >hg38_21_+_8433172_8446622_RNA45SN1_45S_50ntFlanks | 4731 | 4751 | 21 | AAGAACGAAAGTCGGAGGTTC | MB468 | 84 | 83 | 58 | 158 | 30 | 14.5 | 35.3 | 6.2 | 57.9 | 6.9 |
| >hg38_21_+_8433172_8446622_RNA45SN1_45S_50ntFlanks@4731.4752.22\|AAGAACGAAAGTCGGAGGTTCG | >hg38_21_+_8433172_8446622_RNA45SN1_45S_50ntFlanks | 4731 | 4752 | 22 | AAGAACGAAAGTCGGAGGTTCG | MB468 | 18 | 8 | 18 | 22 | 10 | 2.0 | 3.1 | 6.3 | 4.6 | 5.1 |
| >hg38_21_+_8433172_8446622_RNA45SN1_45S_50ntFlanks@4731.4757.27\|AAGAACGAAAGTCGGAGGTTCGAAGAC | >hg38_21_+_8433172_8446622_RNA45SN1_45S_50ntFlanks | 4731 | 4757 | 27 | AAGAACGAAAGTCGGAGGTTCGAAGAC | MB231 | 12 | 7 | 25 | 4 | 2 | 13.2 | 3.8 | 27.7 | 2.5 | 1.3 |
| >hg38_21_+_8433172_8446622_RNA45SN1_45S_50ntFlanks@4731.4758.28\|AAGAACGAAAGTCGGAGGTTCGAAGACG | >hg38_21_+_8433172_8446622_RNA45SN1_45S_50ntFlanks | 4731 | 4758 | 28 | AAGAACGAAAGTCGGAGGTTCGAAGACG | BT20 | 19 | 12 | 150 | 192 | 13 | 10.2 | 7.7 | 126.8 | 239.0 | 2.5 |
| >hg38_21_+_8433172_8446622_RNA45SN1_45S_50ntFlanks@4731.4758.28\|AAGAACGAAAGTCGGAGGTTCGAAGACG | >hg38_21_+_8433172_8446622_RNA45SN1_45S_50ntFlanks | 4731 | 4758 | 28 | AAGAACGAAAGTCGGAGGTTCGAAGACG | MB231 | 48 | 24 | 123 | 45 | 13 | 54.3 | 16.6 | 88.1 | 27.6 | 10.0 |
| >hg38_21_+_8433172_8446622_RNA45SN1_45S_50ntFlanks@4731.4758.28\|AAGAACGAAAGTCGGAGGTTCGAAGACG | >hg38_21_+_8433172_8446622_RNA45SN1_45S_50ntFlanks | 4731 | 4758 | 28 | AAGAACGAAAGTCGGAGGTTCGAAGACG | MB468 | 273 | 575 | 246 | 918 | 399 | 106.2 | 208.4 | 139.3 | 420.9 | 267.1 |
| >hg38_21_+_8433172_8446622_RNA45SN1_45S_50ntFlanks@4731.4759.29\|AAGAACGAAAGTCGGAGGTTCGAAGACGA | >hg38_21_+_8433172_8446622_RNA45SN1_45S_50ntFlanks | 4731 | 4759 | 29 | AAGAACGAAAGTCGGAGGTTCGAAGACGA | MB468 | 96 | 157 | 88 | 295 | 102 | 46.2 | 27.8 | 42.9 | 89.9 | 43.4 |
| >hg38_21_+_8433172_8446622_RNA45SN1_45S_50ntFlanks@4731.4761.31\|AAGAACGAAAGTCGGAGGTTCGAAGACGATC | >hg38_21_+_8433172_8446622_RNA45SN1_45S_50ntFlanks | 4731 | 4761 | 31 | AAGAACGAAAGTCGGAGGTTCGAAGACGATC | BT20 | 21 | 31 | 131 | 160 | 16 | 6.9 | 5.2 | 80.8 | 227.8 | 0.7 |
| >hg38_21_+_8433172_8446622_RNA45SN1_45S_50ntFlanks@4731.4761.31\|AAGAACGAAAGTCGGAGGTTCGAAGACGATC | >hg38_21_+_8433172_8446622_RNA45SN1_45S_50ntFlanks | 4731 | 4761 | 31 | AAGAACGAAAGTCGGAGGTTCGAAGACGATC | MB231 | 47 | 47 | 141 | 27 | 18 | 32.1 | 25.6 | 71.8 | 18.0 | 8.3 |
| >hg38_21_+_8433172_8446622_RNA45SN1_45S_50ntFlanks@4731.4765.35\|AAGAACGAAAGTCGGAGGTTCGAAGACGATCAGAT | >hg38_21_+_8433172_8446622_RNA45SN1_45S_50ntFlanks | 4731 | 4765 | 35 | AAGAACGAAAGTCGGAGGTTCGAAGACGATCAGAT | BT20 | 23 | 34 | 158 | 69 | 14 | 9.1 | 10.4 | 92.0 | 22.1 | 3.8 |
| >hg38_21_+_8433172_8446622_RNA45SN1_45S_50ntFlanks@4731.4765.35\|AAGAACGAAAGTCGGAGGTTCGAAGACGATCAGAT | >hg38_21_+_8433172_8446622_RNA45SN1_45S_50ntFlanks | 4731 | 4765 | 35 | AAGAACGAAAGTCGGAGGTTCGAAGACGATCAGAT | MB231 | 38 | 28 | 132 | 23 | 13 | 36.9 | 15.9 | 143.9 | 9.3 | 10.1 |
| >hg38_21_+_8433172_8446622_RNA45SN1_45S_50ntFlanks@4731.4766.36\|AAGAACGAAAGTCGGAGGTTCGAAGACGATCAGATA | >hg38_21_+_8433172_8446622_RNA45SN1_45S_50ntFlanks | 4731 | 4766 | 36 | AAGAACGAAAGTCGGAGGTTCGAAGACGATCAGATA | BT20 | 10 | 6 | 142 | 15 | 5 | 2.7 | 3.7 | 109.2 | 4.1 | 2.5 |
| >hg38_21_+_8433172_8446622_RNA45SN1_45S_50ntFlanks@4731.4766.36\|AAGAACGAAAGTCGGAGGTTCGAAGACGATCAGATA | >hg38_21_+_8433172_8446622_RNA45SN1_45S_50ntFlanks | 4731 | 4766 | 36 | AAGAACGAAAGTCGGAGGTTCGAAGACGATCAGATA | MB231 | 22 | 9 | 88 | 8 | 5 | 27.2 | 8.8 | 107.4 | 4.2 | 5.1 |
| >hg38_21_+_8433172_8446622_RNA45SN1_45S_50ntFlanks@4731.4767.37\|AAGAACGAAAGTCGGAGGTTCGAAGACGATCAGATAC | >hg38_21_+_8433172_8446622_RNA45SN1_45S_50ntFlanks | 4731 | 4767 | 37 | AAGAACGAAAGTCGGAGGTTCGAAGACGATCAGATAC | BT20 | 20 | 12 | 191 | 22 | 8 | 12.3 | 2.5 | 78.8 | 10.6 | 6.6 |
| >hg38_21_+_8433172_8446622_RNA45SN1_45S_50ntFlanks@4731.4767.37\|AAGAACGAAAGTCGGAGGTTCGAAGACGATCAGATAC | >hg38_21_+_8433172_8446622_RNA45SN1_45S_50ntFlanks | 4731 | 4767 | 37 | AAGAACGAAAGTCGGAGGTTCGAAGACGATCAGATAC | MB231 | 52 | 24 | 251 | 19 | 13 | 48.7 | 23.1 | 309.5 | 11.7 | 13.8 |
| >hg38_21_+_8433172_8446622_RNA45SN1_45S_50ntFlanks@4731.4768.38\|AAGAACGAAAGTCGGAGGTTCGAAGACGATCAGATACC | >hg38_21_+_8433172_8446622_RNA45SN1_45S_50ntFlanks | 4731 | 4768 | 38 | AAGAACGAAAGTCGGAGGTTCGAAGACGATCAGATACC | BT20 | 15 | 10 | 109 | 15 | 8 | 5.2 | 3.3 | 55.3 | 8.2 | 4.7 |
| >hg38_21_+_8433172_8446622_RNA45SN1_45S_50ntFlanks@4731.4768.38\|AAGAACGAAAGTCGGAGGTTCGAAGACGATCAGATACC | >hg38_21_+_8433172_8446622_RNA45SN1_45S_50ntFlanks | 4731 | 4768 | 38 | AAGAACGAAAGTCGGAGGTTCGAAGACGATCAGATACC | MB231 | 41 | 18 | 193 | 11 | 10 | 46.5 | 17.7 | 262.1 | 7.3 | 10.5 |
| >hg38_21_+_8433172_8446622_RNA45SN1_45S_50ntFlanks@4732.4751.20\|AGAACGAAAGTCGGAGGTTC | >hg38_21_+_8433172_8446622_RNA45SN1_45S_50ntFlanks | 4732 | 4751 | 20 | AGAACGAAAGTCGGAGGTTC | MB231 | 10 | 5 | 9 | 8 | 3 | 12.4 | 5.5 | 10.0 | 5.0 | 2.1 |
| >hg38_21_+_8433172_8446622_RNA45SN1_45S_50ntFlanks@4732.4751.20\|AGAACGAAAGTCGGAGGTTC | >hg38_21_+_8433172_8446622_RNA45SN1_45S_50ntFlanks | 4732 | 4751 | 20 | AGAACGAAAGTCGGAGGTTC | MB468 | 25 | 23 | 17 | 43 | 9 | 6.1 | 11.4 | 4.3 | 17.8 | 1.5 |
| >hg38_21_+_8433172_8446622_RNA45SN1_45S_50ntFlanks@4732.4752.21\|AGAACGAAAGTCGGAGGTTCG | >hg38_21_+_8433172_8446622_RNA45SN1_45S_50ntFlanks | 4732 | 4752 | 21 | AGAACGAAAGTCGGAGGTTCG | MB468 | 20 | 10 | 16 | 25 | 10 | 3.6 | 4.3 | 5.4 | 6.7 | 2.3 |
| >hg38_21_+_8433172_8446622_RNA45SN1_45S_50ntFlanks@4732.4758.27\|AGAACGAAAGTCGGAGGTTCGAAGACG | >hg38_21_+_8433172_8446622_RNA45SN1_45S_50ntFlanks | 4732 | 4758 | 27 | AGAACGAAAGTCGGAGGTTCGAAGACG | MB231 | 21 | 11 | 65 | 16 | 5 | 22.1 | 8.1 | 44.4 | 13.4 | 3.6 |
| >hg38_21_+_8433172_8446622_RNA45SN1_45S_50ntFlanks@4732.4758.27\|AGAACGAAAGTCGGAGGTTCGAAGACG | >hg38_21_+_8433172_8446622_RNA45SN1_45S_50ntFlanks | 4732 | 4758 | 27 | AGAACGAAAGTCGGAGGTTCGAAGACG | MB468 | 59 | 131 | 60 | 222 | 91 | 28.7 | 47.0 | 34.6 | 107.5 | 66.4 |
| >hg38_21_+_8433172_8446622_RNA45SN1_45S_50ntFlanks@4732.4759.28\|AGAACGAAAGTCGGAGGTTCGAAGACGA | >hg38_21_+_8433172_8446622_RNA45SN1_45S_50ntFlanks | 4732 | 4759 | 28 | AGAACGAAAGTCGGAGGTTCGAAGACGA | MB468 | 24 | 37 | 22 | 79 | 25 | 13.1 | 7.8 | 11.9 | 30.0 | 11.5 |
| >hg38_21_+_8433172_8446622_RNA45SN1_45S_50ntFlanks@4732.4761.30\|AGAACGAAAGTCGGAGGTTCGAAGACGATC | >hg38_21_+_8433172_8446622_RNA45SN1_45S_50ntFlanks | 4732 | 4761 | 30 | AGAACGAAAGTCGGAGGTTCGAAGACGATC | BT20 | 11 | 12 | 73 | 147 | 11 | 5.2 | 5.6 | 57.4 | 218.3 | 2.5 |
| >hg38_21_+_8433172_8446622_RNA45SN1_45S_50ntFlanks@4732.4761.30\|AGAACGAAAGTCGGAGGTTCGAAGACGATC | >hg38_21_+_8433172_8446622_RNA45SN1_45S_50ntFlanks | 4732 | 4761 | 30 | AGAACGAAAGTCGGAGGTTCGAAGACGATC | MB231 | 26 | 16 | 70 | 23 | 11 | 20.9 | 6.6 | 38.1 | 21.8 | 5.3 |
| >hg38_21_+_8433172_8446622_RNA45SN1_45S_50ntFlanks@4732.4765.34\|AGAACGAAAGTCGGAGGTTCGAAGACGATCAGAT | >hg38_21_+_8433172_8446622_RNA45SN1_45S_50ntFlanks | 4732 | 4765 | 34 | AGAACGAAAGTCGGAGGTTCGAAGACGATCAGAT | BT20 | 11 | 11 | 71 | 26 | 6 | 5.6 | 6.1 | 43.8 | 11.5 | 1.3 |
| >hg38_21_+_8433172_8446622_RNA45SN1_45S_50ntFlanks@4732.4765.34\|AGAACGAAAGTCGGAGGTTCGAAGACGATCAGAT | >hg38_21_+_8433172_8446622_RNA45SN1_45S_50ntFlanks | 4732 | 4765 | 34 | AGAACGAAAGTCGGAGGTTCGAAGACGATCAGAT | MB231 | 18 | 11 | 52 | 14 | 6 | 19.4 | 8.0 | 51.7 | 8.6 | 4.6 |
| >hg38_21_+_8433172_8446622_RNA45SN1_45S_50ntFlanks@4732.4766.35\|AGAACGAAAGTCGGAGGTTCGAAGACGATCAGATA | >hg38_21_+_8433172_8446622_RNA45SN1_45S_50ntFlanks | 4732 | 4766 | 35 | AGAACGAAAGTCGGAGGTTCGAAGACGATCAGATA | MB231 | 15 | 6 | 45 | 6 | 4 | 18.3 | 5.4 | 44.7 | 4.9 | 3.1 |
| >hg38_21_+_8433172_8446622_RNA45SN1_45S_50ntFlanks@4732.4767.36\|AGAACGAAAGTCGGAGGTTCGAAGACGATCAGATAC | >hg38_21_+_8433172_8446622_RNA45SN1_45S_50ntFlanks | 4732 | 4767 | 36 | AGAACGAAAGTCGGAGGTTCGAAGACGATCAGATAC | BT20 | 14 | 7 | 107 | 13 | 5 | 8.8 | 2.3 | 42.3 | 5.5 | 3.1 |
| >hg38_21_+_8433172_8446622_RNA45SN1_45S_50ntFlanks@4732.4767.36\|AGAACGAAAGTCGGAGGTTCGAAGACGATCAGATAC | >hg38_21_+_8433172_8446622_RNA45SN1_45S_50ntFlanks | 4732 | 4767 | 36 | AGAACGAAAGTCGGAGGTTCGAAGACGATCAGATAC | MB231 | 40 | 20 | 125 | 16 | 10 | 35.6 | 19.0 | 141.9 | 11.0 | 8.9 |
| >hg38_21_+_8433172_8446622_RNA45SN1_45S_50ntFlanks@4732.4768.37\|AGAACGAAAGTCGGAGGTTCGAAGACGATCAGATACC | >hg38_21_+_8433172_8446622_RNA45SN1_45S_50ntFlanks | 4732 | 4768 | 37 | AGAACGAAAGTCGGAGGTTCGAAGACGATCAGATACC | MB231 | 28 | 16 | 76 | 9 | 7 | 28.7 | 15.1 | 91.3 | 6.8 | 6.8 |
| >hg38_21_+_8433172_8446622_RNA45SN1_45S_50ntFlanks@4733.4751.19\|GAACGAAAGTCGGAGGTTC | >hg38_21_+_8433172_8446622_RNA45SN1_45S_50ntFlanks | 4733 | 4751 | 19 | GAACGAAAGTCGGAGGTTC | MB231 | 11 | 6 | 8 | 7 | 2 | 10.1 | 4.7 | 7.2 | 5.0 | 1.4 |
| >hg38_21_+_8433172_8446622_RNA45SN1_45S_50ntFlanks@4733.4751.19\|GAACGAAAGTCGGAGGTTC | >hg38_21_+_8433172_8446622_RNA45SN1_45S_50ntFlanks | 4733 | 4751 | 19 | GAACGAAAGTCGGAGGTTC | MB468 | 20 | 16 | 14 | 31 | 7 | 3.7 | 7.8 | 4.4 | 11.2 | 1.1 |
| >hg38_21_+_8433172_8446622_RNA45SN1_45S_50ntFlanks@4733.4752.20\|GAACGAAAGTCGGAGGTTCG | >hg38_21_+_8433172_8446622_RNA45SN1_45S_50ntFlanks | 4733 | 4752 | 20 | GAACGAAAGTCGGAGGTTCG | MB468 | 12 | 6 | 14 | 11 | 5 | 2.8 | 2.6 | 3.4 | 2.9 | 1.8 |
| >hg38_21_+_8433172_8446622_RNA45SN1_45S_50ntFlanks@4733.4758.26\|GAACGAAAGTCGGAGGTTCGAAGACG | >hg38_21_+_8433172_8446622_RNA45SN1_45S_50ntFlanks | 4733 | 4758 | 26 | GAACGAAAGTCGGAGGTTCGAAGACG | MB231 | 18 | 10 | 55 | 25 | 5 | 12.6 | 5.4 | 46.7 | 26.1 | 2.6 |
| >hg38_21_+_8433172_8446622_RNA45SN1_45S_50ntFlanks@4733.4758.26\|GAACGAAAGTCGGAGGTTCGAAGACG | >hg38_21_+_8433172_8446622_RNA45SN1_45S_50ntFlanks | 4733 | 4758 | 26 | GAACGAAAGTCGGAGGTTCGAAGACG | MB468 | 32 | 67 | 35 | 115 | 53 | 11.2 | 23.8 | 18.1 | 55.8 | 39.5 |
| >hg38_21_+_8433172_8446622_RNA45SN1_45S_50ntFlanks@4733.4759.27\|GAACGAAAGTCGGAGGTTCGAAGACGA | >hg38_21_+_8433172_8446622_RNA45SN1_45S_50ntFlanks | 4733 | 4759 | 27 | GAACGAAAGTCGGAGGTTCGAAGACGA | MB468 | 14 | 20 | 13 | 43 | 15 | 6.2 | 3.4 | 7.1 | 15.8 | 8.3 |
| >hg38_21_+_8433172_8446622_RNA45SN1_45S_50ntFlanks@4733.4761.29\|GAACGAAAGTCGGAGGTTCGAAGACGATC | >hg38_21_+_8433172_8446622_RNA45SN1_45S_50ntFlanks | 4733 | 4761 | 29 | GAACGAAAGTCGGAGGTTCGAAGACGATC | MB231 | 25 | 19 | 60 | 14 | 6 | 12.3 | 3.4 | 29.4 | 14.7 | 3.2 |
| >hg38_21_+_8433172_8446622_RNA45SN1_45S_50ntFlanks@4733.4765.33\|GAACGAAAGTCGGAGGTTCGAAGACGATCAGAT | >hg38_21_+_8433172_8446622_RNA45SN1_45S_50ntFlanks | 4733 | 4765 | 33 | GAACGAAAGTCGGAGGTTCGAAGACGATCAGAT | MB231 | 18 | 15 | 41 | 14 | 6 | 10.7 | 3.8 | 23.2 | 8.8 | 3.1 |
| >hg38_21_+_8433172_8446622_RNA45SN1_45S_50ntFlanks@4733.4766.34\|GAACGAAAGTCGGAGGTTCGAAGACGATCAGATA | >hg38_21_+_8433172_8446622_RNA45SN1_45S_50ntFlanks | 4733 | 4766 | 34 | GAACGAAAGTCGGAGGTTCGAAGACGATCAGATA | MB231 | 13 | 6 | 32 | 5 | 2 | 11.1 | 4.3 | 21.8 | 4.5 | 2.3 |
| >hg38_21_+_8433172_8446622_RNA45SN1_45S_50ntFlanks@4733.4767.35\|GAACGAAAGTCGGAGGTTCGAAGACGATCAGATAC | >hg38_21_+_8433172_8446622_RNA45SN1_45S_50ntFlanks | 4733 | 4767 | 35 | GAACGAAAGTCGGAGGTTCGAAGACGATCAGATAC | BT20 | 10 | 4 | 60 | 15 | 4 | 6.9 | 1.2 | 21.1 | 10.6 | 1.6 |
| >hg38_21_+_8433172_8446622_RNA45SN1_45S_50ntFlanks@4733.4767.35\|GAACGAAAGTCGGAGGTTCGAAGACGATCAGATAC | >hg38_21_+_8433172_8446622_RNA45SN1_45S_50ntFlanks | 4733 | 4767 | 35 | GAACGAAAGTCGGAGGTTCGAAGACGATCAGATAC | MB231 | 47 | 23 | 108 | 17 | 10 | 45.1 | 18.7 | 87.2 | 16.0 | 7.8 |

| | | | | | | | | | | | | | | | | |
|---|---|---|---|---|---|---|---|---|---|---|---|---|---|---|---|---|
| >hg38_21_+_8433172_8446622_RNA45SN1_45S_50ntFlanks@4733.4768.36\|GAACGAAAGTCGGAGGTTCGAAGACGATCAGATACC | >hg38_21_+_8433172_8446622_RNA45SN1_45S_50ntFlanks | 4733 | 4768 | 36 | GAACGAAAGTCGGAGGTTCGAAGACGATCAGATACC | MB231 | 23 | 13 | 37 | 7 | 5 | 19.4 | 10.8 | 34.4 | 6.6 | 4.0 |
| >hg38_21_+_8433172_8446622_RNA45SN1_45S_50ntFlanks@4734.4751.18\|AACGAAAGTCGGAGGTTC | >hg38_21_+_8433172_8446622_RNA45SN1_45S_50ntFlanks | 4734 | 4751 | 18 | AACGAAAGTCGGAGGTTC | MB468 | 51 | 62 | 33 | 63 | 18 | 16.1 | 33.3 | 8.7 | 25.7 | 1.8 |
| >hg38_21_+_8433172_8446622_RNA45SN1_45S_50ntFlanks@4734.4752.19\|AACGAAAGTCGGAGGTTCG | >hg38_21_+_8433172_8446622_RNA45SN1_45S_50ntFlanks | 4734 | 4752 | 19 | AACGAAAGTCGGAGGTTCG | MB468 | 52 | 24 | 31 | 36 | 20 | 15.8 | 16.0 | 10.4 | 11.0 | 4.4 |
| >hg38_21_+_8433172_8446622_RNA45SN1_45S_50ntFlanks@4734.4758.25\|AACGAAAGTCGGAGGTTCGAAGACG | >hg38_21_+_8433172_8446622_RNA45SN1_45S_50ntFlanks | 4734 | 4758 | 25 | AACGAAAGTCGGAGGTTCGAAGACG | BT20 | 18 | 4 | 36 | 195 | 13 | 11.7 | 1.3 | 31.2 | 10.4 | 7.4 |
| >hg38_21_+_8433172_8446622_RNA45SN1_45S_50ntFlanks@4734.4758.25\|AACGAAAGTCGGAGGTTCGAAGACG | >hg38_21_+_8433172_8446622_RNA45SN1_45S_50ntFlanks | 4734 | 4758 | 25 | AACGAAAGTCGGAGGTTCGAAGACG | MB231 | 23 | 10 | 27 | 172 | 23 | 8.2 | 4.6 | 16.0 | 177.9 | 4.7 |
| >hg38_21_+_8433172_8446622_RNA45SN1_45S_50ntFlanks@4734.4758.25\|AACGAAAGTCGGAGGTTCGAAGACG | >hg38_21_+_8433172_8446622_RNA45SN1_45S_50ntFlanks | 4734 | 4758 | 25 | AACGAAAGTCGGAGGTTCGAAGACG | MB468 | 75 | 132 | 48 | 191 | 86 | 26.5 | 51.3 | 24.6 | 55.5 | 54.6 |
| >hg38_21_+_8433172_8446622_RNA45SN1_45S_50ntFlanks@4734.4759.26\|AACGAAAGTCGGAGGTTCGAAGACGA | >hg38_21_+_8433172_8446622_RNA45SN1_45S_50ntFlanks | 4734 | 4759 | 26 | AACGAAAGTCGGAGGTTCGAAGACGA | MB468 | 41 | 64 | 28 | 100 | 32 | 20.3 | 18.2 | 14.7 | 36.4 | 14.7 |
| >hg38_21_+_8433172_8446622_RNA45SN1_45S_50ntFlanks@4734.4761.28\|AACGAAAGTCGGAGGTTCGAAGACGATC | >hg38_21_+_8433172_8446622_RNA45SN1_45S_50ntFlanks | 4734 | 4761 | 28 | AACGAAAGTCGGAGGTTCGAAGACGATC | MB231 | 16 | 12 | 29 | 15 | 6 | 10.7 | 2.4 | 10.8 | 13.9 | 0.8 |
| >hg38_21_+_8433172_8446622_RNA45SN1_45S_50ntFlanks@4734.4765.32\|AACGAAAGTCGGAGGTTCGAAGACGATCAGAT | >hg38_21_+_8433172_8446622_RNA45SN1_45S_50ntFlanks | 4734 | 4765 | 32 | AACGAAAGTCGGAGGTTCGAAGACGATCAGAT | BT20 | 11 | 10 | 29 | 61 | 9 | 3.8 | 6.0 | 18.7 | 42.5 | 4.0 |
| >hg38_21_+_8433172_8446622_RNA45SN1_45S_50ntFlanks@4734.4767.34\|AACGAAAGTCGGAGGTTCGAAGACGATCAGATAC | >hg38_21_+_8433172_8446622_RNA45SN1_45S_50ntFlanks | 4734 | 4767 | 34 | AACGAAAGTCGGAGGTTCGAAGACGATCAGATAC | MB231 | 20 | 11 | 32 | 8 | 4 | 16.5 | 9.0 | 34.7 | 7.9 | 3.0 |
| >hg38_21_+_8433172_8446622_RNA45SN1_45S_50ntFlanks@4734.4768.35\|AACGAAAGTCGGAGGTTCGAAGACGATCAGATACC | >hg38_21_+_8433172_8446622_RNA45SN1_45S_50ntFlanks | 4734 | 4768 | 35 | AACGAAAGTCGGAGGTTCGAAGACGATCAGATACC | MB231 | 14 | 7 | 21 | 5 | 3 | 12.6 | 5.9 | 22.6 | 3.8 | 1.9 |
| >hg38_21_+_8433172_8446622_RNA45SN1_45S_50ntFlanks@4735.4752.18\|ACGAAAGTCGGAGGTTCG | >hg38_21_+_8433172_8446622_RNA45SN1_45S_50ntFlanks | 4735 | 4752 | 18 | ACGAAAGTCGGAGGTTCG | MB468 | 18 | 7 | 10 | 6 | 4 | 6.8 | 5.4 | 4.8 | 1.8 | 0.8 |
| >hg38_21_+_8433172_8446622_RNA45SN1_45S_50ntFlanks@4735.4758.24\|ACGAAAGTCGGAGGTTCGAAGACG | >hg38_21_+_8433172_8446622_RNA45SN1_45S_50ntFlanks | 4735 | 4758 | 24 | ACGAAAGTCGGAGGTTCGAAGACG | MB231 | 32 | 23 | 58 | 18 | 6 | 24.7 | 14.0 | 38.0 | 17.9 | 4.1 |
| >hg38_21_+_8433172_8446622_RNA45SN1_45S_50ntFlanks@4735.4758.24\|ACGAAAGTCGGAGGTTCGAAGACG | >hg38_21_+_8433172_8446622_RNA45SN1_45S_50ntFlanks | 4735 | 4758 | 24 | ACGAAAGTCGGAGGTTCGAAGACG | MB468 | 44 | 101 | 27 | 64 | 21 | 35.0 | 69.8 | 18.9 | 22.4 | 3.2 |
| >hg38_21_+_8433172_8446622_RNA45SN1_45S_50ntFlanks@4735.4759.25\|ACGAAAGTCGGAGGTTCGAAGACGA | >hg38_21_+_8433172_8446622_RNA45SN1_45S_50ntFlanks | 4735 | 4759 | 25 | ACGAAAGTCGGAGGTTCGAAGACGA | MB468 | 22 | 37 | 15 | 31 | 8 | 17.5 | 24.4 | 10.2 | 12.6 | 1.7 |
| >hg38_21_+_8433172_8446622_RNA45SN1_45S_50ntFlanks@4735.4761.27\|ACGAAAGTCGGAGGTTCGAAGACGATC | >hg38_21_+_8433172_8446622_RNA45SN1_45S_50ntFlanks | 4735 | 4761 | 27 | ACGAAAGTCGGAGGTTCGAAGACGATC | MB231 | 31 | 25 | 43 | 23 | 9 | 13.8 | 4.5 | 13.9 | 22.4 | 3.2 |
| >hg38_21_+_8433172_8446622_RNA45SN1_45S_50ntFlanks@4735.4765.31\|ACGAAAGTCGGAGGTTCGAAGACGATCAGAT | >hg38_21_+_8433172_8446622_RNA45SN1_45S_50ntFlanks | 4735 | 4765 | 31 | ACGAAAGTCGGAGGTTCGAAGACGATCAGAT | MB231 | 14 | 9 | 20 | 7 | 4 | 11.2 | 4.8 | 18.8 | 3.7 | 1.8 |
| >hg38_21_+_8433172_8446622_RNA45SN1_45S_50ntFlanks@4735.4766.32\|ACGAAAGTCGGAGGTTCGAAGACGATCAGATA | >hg38_21_+_8433172_8446622_RNA45SN1_45S_50ntFlanks | 4735 | 4766 | 32 | ACGAAAGTCGGAGGTTCGAAGACGATCAGATA | MB231 | 13 | 6 | 22 | 7 | 3 | 11.1 | 5.0 | 21.0 | 6.9 | 1.8 |
| >hg38_21_+_8433172_8446622_RNA45SN1_45S_50ntFlanks@4735.4767.33\|ACGAAAGTCGGAGGTTCGAAGACGATCAGATAC | >hg38_21_+_8433172_8446622_RNA45SN1_45S_50ntFlanks | 4735 | 4767 | 33 | ACGAAAGTCGGAGGTTCGAAGACGATCAGATAC | MB231 | 33 | 16 | 42 | 9 | 4 | 28.8 | 13.7 | 50.0 | 8.6 | 3.8 |
| >hg38_21_+_8433172_8446622_RNA45SN1_45S_50ntFlanks@4735.4768.34\|ACGAAAGTCGGAGGTTCGAAGACGATCAGATACC | >hg38_21_+_8433172_8446622_RNA45SN1_45S_50ntFlanks | 4735 | 4768 | 34 | ACGAAAGTCGGAGGTTCGAAGACGATCAGATACC | MB231 | 25 | 15 | 40 | 6 | 4 | 21.4 | 13.4 | 45.6 | 6.5 | 3.3 |
| >hg38_21_+_8433172_8446622_RNA45SN1_45S_50ntFlanks@4736.4758.23\|CGAAAGTCGGAGGTTCGAAGACG | >hg38_21_+_8433172_8446622_RNA45SN1_45S_50ntFlanks | 4736 | 4758 | 23 | CGAAAGTCGGAGGTTCGAAGACG | MB468 | 20 | 50 | 13 | 48 | 23 | 6.4 | 27.2 | 5.2 | 31.7 | 17.2 |
| >hg38_21_+_8433172_8446622_RNA45SN1_45S_50ntFlanks@4738.4757.20\|AAAGTCGGAGGTTCGAAGAC | >hg38_21_+_8433172_8446622_RNA45SN1_45S_50ntFlanks | 4738 | 4757 | 20 | AAAGTCGGAGGTTCGAAGAC | MB231 | 14 | 8 | 28 | 11 | 5 | 12.4 | 4.8 | 27.2 | 2.9 | 1.3 |
| >hg38_21_+_8433172_8446622_RNA45SN1_45S_50ntFlanks@4738.4758.21\|AAAGTCGGAGGTTCGAAGACG | >hg38_21_+_8433172_8446622_RNA45SN1_45S_50ntFlanks | 4738 | 4758 | 21 | AAAGTCGGAGGTTCGAAGACG | BT20 | 18 | 8 | 49 | 125 | 13 | 5.9 | 0.5 | 29.3 | 13.9 | 7.1 |
| >hg38_21_+_8433172_8446622_RNA45SN1_45S_50ntFlanks@4738.4758.21\|AAAGTCGGAGGTTCGAAGACG | >hg38_21_+_8433172_8446622_RNA45SN1_45S_50ntFlanks | 4738 | 4758 | 21 | AAAGTCGGAGGTTCGAAGACG | MB231 | 46 | 28 | 61 | 279 | 45 | 30.7 | 12.3 | 47.4 | 157.3 | 7.3 |
| >hg38_21_+_8433172_8446622_RNA45SN1_45S_50ntFlanks@4738.4758.21\|AAAGTCGGAGGTTCGAAGACG | >hg38_21_+_8433172_8446622_RNA45SN1_45S_50ntFlanks | 4738 | 4758 | 21 | AAAGTCGGAGGTTCGAAGACG | MB468 | 165 | 229 | 143 | 581 | 172 | 86.2 | 69.9 | 77.5 | 132.7 | 90.2 |
| >hg38_21_+_8433172_8446622_RNA45SN1_45S_50ntFlanks@4738.4759.22\|AAAGTCGGAGGTTCGAAGACGA | >hg38_21_+_8433172_8446622_RNA45SN1_45S_50ntFlanks | 4738 | 4759 | 22 | AAAGTCGGAGGTTCGAAGACGA | MB468 | 29 | 21 | 30 | 132 | 27 | 16.1 | 11.5 | 14.1 | 64.0 | 12.7 |
| >hg38_21_+_8433172_8446622_RNA45SN1_45S_50ntFlanks@4738.4761.24\|AAAGTCGGAGGTTCGAAGACGATC | >hg38_21_+_8433172_8446622_RNA45SN1_45S_50ntFlanks | 4738 | 4761 | 24 | AAAGTCGGAGGTTCGAAGACGATC | MB231 | 34 | 21 | 67 | 24 | 10 | 29.2 | 12.4 | 36.5 | 16.5 | 6.5 |
| >hg38_21_+_8433172_8446622_RNA45SN1_45S_50ntFlanks@4738.4765.28\|AAAGTCGGAGGTTCGAAGACGATCAGAT | >hg38_21_+_8433172_8446622_RNA45SN1_45S_50ntFlanks | 4738 | 4765 | 28 | AAAGTCGGAGGTTCGAAGACGATCAGAT | MB231 | 20 | 10 | 25 | 25 | 6 | 20.2 | 9.3 | 25.7 | 15.2 | 3.8 |
| >hg38_21_+_8433172_8446622_RNA45SN1_45S_50ntFlanks@4738.4766.29\|AAAGTCGGAGGTTCGAAGACGATCAGATA | >hg38_21_+_8433172_8446622_RNA45SN1_45S_50ntFlanks | 4738 | 4766 | 29 | AAAGTCGGAGGTTCGAAGACGATCAGATA | MB231 | 20 | 9 | 27 | 11 | 4 | 22.5 | 9.9 | 25.7 | 8.9 | 3.2 |
| >hg38_21_+_8433172_8446622_RNA45SN1_45S_50ntFlanks@4738.4767.30\|AAAGTCGGAGGTTCGAAGACGATCAGATAC | >hg38_21_+_8433172_8446622_RNA45SN1_45S_50ntFlanks | 4738 | 4767 | 30 | AAAGTCGGAGGTTCGAAGACGATCAGATAC | BT20 | 15 | 6 | 65 | 12 | 4 | 7.7 | 1.3 | 34.7 | 1.9 | 1.8 |
| >hg38_21_+_8433172_8446622_RNA45SN1_45S_50ntFlanks@4738.4767.30\|AAAGTCGGAGGTTCGAAGACGATCAGATAC | >hg38_21_+_8433172_8446622_RNA45SN1_45S_50ntFlanks | 4738 | 4767 | 30 | AAAGTCGGAGGTTCGAAGACGATCAGATAC | MB231 | 46 | 23 | 56 | 25 | 10 | 38.7 | 20.8 | 60.0 | 18.6 | 8.3 |
| >hg38_21_+_8433172_8446622_RNA45SN1_45S_50ntFlanks@4738.4768.31\|AAAGTCGGAGGTTCGAAGACGATCAGATACC | >hg38_21_+_8433172_8446622_RNA45SN1_45S_50ntFlanks | 4738 | 4768 | 31 | AAAGTCGGAGGTTCGAAGACGATCAGATACC | BT20 | 15 | 9 | 72 | 18 | 5 | 7.0 | 3.1 | 43.8 | 5.3 | 0.9 |
| >hg38_21_+_8433172_8446622_RNA45SN1_45S_50ntFlanks@4738.4768.31\|AAAGTCGGAGGTTCGAAGACGATCAGATACC | >hg38_21_+_8433172_8446622_RNA45SN1_45S_50ntFlanks | 4738 | 4768 | 31 | AAAGTCGGAGGTTCGAAGACGATCAGATACC | MB231 | 45 | 27 | 71 | 24 | 10 | 42.0 | 26.6 | 74.0 | 19.1 | 9.0 |
| >hg38_21_+_8433172_8446622_RNA45SN1_45S_50ntFlanks@4739.4758.20\|AAGTCGGAGGTTCGAAGACG | >hg38_21_+_8433172_8446622_RNA45SN1_45S_50ntFlanks | 4739 | 4758 | 20 | AAGTCGGAGGTTCGAAGACG | MB468 | 14 | 11 | 13 | 55 | 17 | 4.6 | 4.6 | 5.3 | 21.0 | 10.1 |
| >hg38_21_+_8433172_8446622_RNA45SN1_45S_50ntFlanks@474.497.24\|TCTTCGTGATCGATGTGGTGACGT | >hg38_21_+_8433172_8446622_RNA45SN1_45S_50ntFlanks | 474 | 497 | 24 | TCTTCGTGATCGATGTGGTGACGT | BT20 | 11 | 24 | 1 | 8 | 6 | 12.8 | 32.6 | 1.0 | 11.3 | 5.8 |
| >hg38_21_+_8433172_8446622_RNA45SN1_45S_50ntFlanks@474.497.24\|TCTTCGTGATCGATGTGGTGACGT | >hg38_21_+_8433172_8446622_RNA45SN1_45S_50ntFlanks | 474 | 497 | 24 | TCTTCGTGATCGATGTGGTGACGT | MB231 | 50 | 159 | 2 | 27 | 39 | 23.0 | 83.5 | 1.1 | 8.9 | 16.3 |
| >hg38_21_+_8433172_8446622_RNA45SN1_45S_50ntFlanks@474.498.25\|TCTTCGTGATCGATGTGGTGACGTC | >hg38_21_+_8433172_8446622_RNA45SN1_45S_50ntFlanks | 474 | 498 | 25 | TCTTCGTGATCGATGTGGTGACGTC | MB231 | 12 | 43 | 3 | 16 | 41 | 11.1 | 26.0 | 1.6 | 7.3 | 8.4 |
| >hg38_21_+_8433172_8446622_RNA45SN1_45S_50ntFlanks@474.504.31\|TCTTCGTGATCGATGTGGTGACGTCGTGCTC | >hg38_21_+_8433172_8446622_RNA45SN1_45S_50ntFlanks | 474 | 504 | 31 | TCTTCGTGATCGATGTGGTGACGTCGTGCTC | BT20 | 14 | 29 | 6 | 4 | 8 | 6.2 | 20.3 | 2.7 | 1.7 | 4.5 |
| >hg38_21_+_8433172_8446622_RNA45SN1_45S_50ntFlanks@474.504.31\|TCTTCGTGATCGATGTGGTGACGTCGTGCTC | >hg38_21_+_8433172_8446622_RNA45SN1_45S_50ntFlanks | 474 | 504 | 31 | TCTTCGTGATCGATGTGGTGACGTCGTGCTC | MB231 | 19 | 57 | 3 | 2 | 8 | 12.0 | 23.9 | 1.6 | 1.5 | 2.4 |
| >hg38_21_+_8433172_8446622_RNA45SN1_45S_50ntFlanks@474.505.32\|TCTTCGTGATCGATGTGGTGACGTCGTGCTCT | >hg38_21_+_8433172_8446622_RNA45SN1_45S_50ntFlanks | 474 | 505 | 32 | TCTTCGTGATCGATGTGGTGACGTCGTGCTCT | BT20 | 26 | 46 | 12 | 5 | 8 | 10.9 | 26.0 | 6.3 | 2.2 | 4.1 |
| >hg38_21_+_8433172_8446622_RNA45SN1_45S_50ntFlanks@474.505.32\|TCTTCGTGATCGATGTGGTGACGTCGTGCTCT | >hg38_21_+_8433172_8446622_RNA45SN1_45S_50ntFlanks | 474 | 505 | 32 | TCTTCGTGATCGATGTGGTGACGTCGTGCTCT | MB231 | 50 | 123 | 8 | 4 | 13 | 20.0 | 62.2 | 4.2 | 1.6 | 3.2 |
| >hg38_21_+_8433172_8446622_RNA45SN1_45S_50ntFlanks@474.506.33\|TCTTCGTGATCGATGTGGTGACGTCGTGCTCTC | >hg38_21_+_8433172_8446622_RNA45SN1_45S_50ntFlanks | 474 | 506 | 33 | TCTTCGTGATCGATGTGGTGACGTCGTGCTCTC | BT20 | 10 | 23 | 9 | 2 | 5 | 4.7 | 11.3 | 4.6 | 2.1 | 4.1 |
| >hg38_21_+_8433172_8446622_RNA45SN1_45S_50ntFlanks@474.506.33\|TCTTCGTGATCGATGTGGTGACGTCGTGCTCTC | >hg38_21_+_8433172_8446622_RNA45SN1_45S_50ntFlanks | 474 | 506 | 33 | TCTTCGTGATCGATGTGGTGACGTCGTGCTCTC | MB231 | 21 | 57 | 6 | 1 | 7 | 9.2 | 26.3 | 3.3 | 0.4 | 1.0 |
| >hg38_21_+_8433172_8446622_RNA45SN1_45S_50ntFlanks@4742.4759.18\|TCGGAGGTTCGAAGACGA | >hg38_21_+_8433172_8446622_RNA45SN1_45S_50ntFlanks | 4742 | 4759 | 18 | TCGGAGGTTCGAAGACGA | MB468 | 21 | 28 | 19 | 27 | 16 | 11.3 | 10.0 | 10.6 | 12.6 | 10.1 |
| >hg38_21_+_8433172_8446622_RNA45SN1_45S_50ntFlanks@475.492.18\|CTTCGTGATCGATGTGGT | >hg38_21_+_8433172_8446622_RNA45SN1_45S_50ntFlanks | 475 | 492 | 18 | CTTCGTGATCGATGTGGT | BT20 | 122 | 116 | 10 | 12 | 19 | 40.2 | 22.9 | 9.0 | 11.4 | 18.9 |
| >hg38_21_+_8433172_8446622_RNA45SN1_45S_50ntFlanks@475.492.18\|CTTCGTGATCGATGTGGT | >hg38_21_+_8433172_8446622_RNA45SN1_45S_50ntFlanks | 475 | 492 | 18 | CTTCGTGATCGATGTGGT | MB231 | 28 | 36 | 1 | 2 | 5 | 38.6 | 28.0 | 0.4 | 1.4 | 4.2 |
| >hg38_21_+_8433172_8446622_RNA45SN1_45S_50ntFlanks@475.492.18\|CTTCGTGATCGATGTGGT | >hg38_21_+_8433172_8446622_RNA45SN1_45S_50ntFlanks | 475 | 492 | 18 | CTTCGTGATCGATGTGGT | MB468 | 17 | 12 | 0 | 1 | 1 | 20.8 | 3.8 | 0.1 | 1.0 | 0.7 |
| >hg38_21_+_8433172_8446622_RNA45SN1_45S_50ntFlanks@475.493.19\|CTTCGTGATCGATGTGGTG | >hg38_21_+_8433172_8446622_RNA45SN1_45S_50ntFlanks | 475 | 493 | 19 | CTTCGTGATCGATGTGGTG | BT20 | 13 | 21 | 2 | 7 | 5 | 11.3 | 6.9 | 0.5 | 6.3 | 2.7 |
| >hg38_21_+_8433172_8446622_RNA45SN1_45S_50ntFlanks@475.494.20\|CTTCGTGATCGATGTGGTGA | >hg38_21_+_8433172_8446622_RNA45SN1_45S_50ntFlanks | 475 | 494 | 20 | CTTCGTGATCGATGTGGTGA | BT20 | 14 | 22 | 2 | 12 | 8 | 20.8 | 19.5 | 1.9 | 12.0 | 5.9 |
| >hg38_21_+_8433172_8446622_RNA45SN1_45S_50ntFlanks@475.495.21\|CTTCGTGATCGATGTGGTGAC | >hg38_21_+_8433172_8446622_RNA45SN1_45S_50ntFlanks | 475 | 495 | 21 | CTTCGTGATCGATGTGGTGAC | BT20 | 30 | 77 | 13 | 17 | 22 | 25.9 | 41.1 | 3.2 | 12.1 | 12.3 |

| | | | | | | | | | | | | | | | | |
|---|---|---|---|---|---|---|---|---|---|---|---|---|---|---|---|---|
| >hg38_21_+_8433172_8446622_RNA45SN1_45S_50ntFlanks@475.495.21\|CTTCGTGATCGATGTGGTGAC | >hg38_21_+_8433172_8446622_RNA45SN1_45S_50ntFlanks | 475 | 495 | 21 | CTTCGTGATCGATGTGGTGAC | MB231 | 74 | 347 | 16 | 32 | 64 | 31.6 | 75.8 | 7.8 | 7.4 | 2.7 |
| >hg38_21_+_8433172_8446622_RNA45SN1_45S_50ntFlanks@475.496.22\|CTTCGTGATCGATGTGGTGACG | >hg38_21_+_8433172_8446622_RNA45SN1_45S_50ntFlanks | 475 | 496 | 22 | CTTCGTGATCGATGTGGTGACG | BT20 | 174 | 342 | 20 | 76 | 81 | 187.7 | 393.9 | 17.0 | 95.5 | 63.5 |
| >hg38_21_+_8433172_8446622_RNA45SN1_45S_50ntFlanks@475.496.22\|CTTCGTGATCGATGTGGTGACG | >hg38_21_+_8433172_8446622_RNA45SN1_45S_50ntFlanks | 475 | 496 | 22 | CTTCGTGATCGATGTGGTGACG | MB231 | 370 | 1257 | 18 | 166 | 282 | 143.8 | 499.6 | 8.3 | 23.6 | 68.4 |
| >hg38_21_+_8433172_8446622_RNA45SN1_45S_50ntFlanks@475.496.22\|CTTCGTGATCGATGTGGTGACG | >hg38_21_+_8433172_8446622_RNA45SN1_45S_50ntFlanks | 475 | 496 | 22 | CTTCGTGATCGATGTGGTGACG | MB468 | 15 | 48 | 0 | 7 | 3 | 4.0 | 31.0 | 0.4 | 7.7 | 3.5 |
| >hg38_21_+_8433172_8446622_RNA45SN1_45S_50ntFlanks@475.497.23\|CTTCGTGATCGATGTGGTGACGT | >hg38_21_+_8433172_8446622_RNA45SN1_45S_50ntFlanks | 475 | 497 | 23 | CTTCGTGATCGATGTGGTGACGT | BT20 | 387 | 914 | 28 | 153 | 178 | 420.1 | 1349.8 | 27.3 | 200.7 | 202.8 |
| >hg38_21_+_8433172_8446622_RNA45SN1_45S_50ntFlanks@475.497.23\|CTTCGTGATCGATGTGGTGACGT | >hg38_21_+_8433172_8446622_RNA45SN1_45S_50ntFlanks | 475 | 497 | 23 | CTTCGTGATCGATGTGGTGACGT | MB231 | 938 | 2814 | 26 | 493 | 700 | 416.4 | 1193.3 | 17.1 | 119.4 | 436.9 |
| >hg38_21_+_8433172_8446622_RNA45SN1_45S_50ntFlanks@475.497.23\|CTTCGTGATCGATGTGGTGACGT | >hg38_21_+_8433172_8446622_RNA45SN1_45S_50ntFlanks | 475 | 497 | 23 | CTTCGTGATCGATGTGGTGACGT | MB468 | 22 | 55 | 0 | 11 | 4 | 8.7 | 41.8 | 0.4 | 13.4 | 5.5 |
| >hg38_21_+_8433172_8446622_RNA45SN1_45S_50ntFlanks@475.498.24\|CTTCGTGATCGATGTGGTGACGTC | >hg38_21_+_8433172_8446622_RNA45SN1_45S_50ntFlanks | 475 | 498 | 24 | CTTCGTGATCGATGTGGTGACGTC | BT20 | 496 | 1175 | 74 | 471 | 542 | 691.7 | 913.7 | 69.1 | 568.6 | 557.5 |
| >hg38_21_+_8433172_8446622_RNA45SN1_45S_50ntFlanks@475.498.24\|CTTCGTGATCGATGTGGTGACGTC | >hg38_21_+_8433172_8446622_RNA45SN1_45S_50ntFlanks | 475 | 498 | 24 | CTTCGTGATCGATGTGGTGACGTC | MB231 | 481 | 1640 | 81 | 531 | 1135 | 447.5 | 912.6 | 40.7 | 203.2 | 403.8 |
| >hg38_21_+_8433172_8446622_RNA45SN1_45S_50ntFlanks@475.498.24\|CTTCGTGATCGATGTGGTGACGTC | >hg38_21_+_8433172_8446622_RNA45SN1_45S_50ntFlanks | 475 | 498 | 24 | CTTCGTGATCGATGTGGTGACGTC | MB468 | 20 | 80 | 1 | 12 | 9 | 9.9 | 41.5 | 0.6 | 10.1 | 8.3 |
| >hg38_21_+_8433172_8446622_RNA45SN1_45S_50ntFlanks@475.499.25\|CTTCGTGATCGATGTGGTGACGTCG | >hg38_21_+_8433172_8446622_RNA45SN1_45S_50ntFlanks | 475 | 499 | 25 | CTTCGTGATCGATGTGGTGACGTCG | BT20 | 20 | 50 | 5 | 8 | 13 | 20.6 | 64.8 | 2.1 | 9.1 | 13.0 |
| >hg38_21_+_8433172_8446622_RNA45SN1_45S_50ntFlanks@475.499.25\|CTTCGTGATCGATGTGGTGACGTCG | >hg38_21_+_8433172_8446622_RNA45SN1_45S_50ntFlanks | 475 | 499 | 25 | CTTCGTGATCGATGTGGTGACGTCG | MB231 | 22 | 87 | 2 | 16 | 31 | 9.7 | 70.9 | 1.6 | 7.2 | 16.6 |
| >hg38_21_+_8433172_8446622_RNA45SN1_45S_50ntFlanks@475.501.27\|CTTCGTGATCGATGTGGTGACGTCGTG | >hg38_21_+_8433172_8446622_RNA45SN1_45S_50ntFlanks | 475 | 501 | 27 | CTTCGTGATCGATGTGGTGACGTCGTG | BT20 | 44 | 84 | 8 | 12 | 34 | 30.4 | 54.0 | 3.4 | 5.7 | 23.3 |
| >hg38_21_+_8433172_8446622_RNA45SN1_45S_50ntFlanks@475.501.27\|CTTCGTGATCGATGTGGTGACGTCGTG | >hg38_21_+_8433172_8446622_RNA45SN1_45S_50ntFlanks | 475 | 501 | 27 | CTTCGTGATCGATGTGGTGACGTCGTG | MB231 | 28 | 62 | 3 | 25 | 54 | 14.8 | 26.4 | 1.6 | 8.3 | 24.5 |
| >hg38_21_+_8433172_8446622_RNA45SN1_45S_50ntFlanks@475.502.28\|CTTCGTGATCGATGTGGTGACGTCGTGC | >hg38_21_+_8433172_8446622_RNA45SN1_45S_50ntFlanks | 475 | 502 | 28 | CTTCGTGATCGATGTGGTGACGTCGTGC | BT20 | 55 | 90 | 13 | 15 | 32 | 57.4 | 67.1 | 6.1 | 13.0 | 27.6 |
| >hg38_21_+_8433172_8446622_RNA45SN1_45S_50ntFlanks@475.502.28\|CTTCGTGATCGATGTGGTGACGTCGTGC | >hg38_21_+_8433172_8446622_RNA45SN1_45S_50ntFlanks | 475 | 502 | 28 | CTTCGTGATCGATGTGGTGACGTCGTGC | MB231 | 34 | 88 | 2 | 25 | 37 | 18.5 | 28.4 | 1.2 | 3.9 | 10.3 |
| >hg38_21_+_8433172_8446622_RNA45SN1_45S_50ntFlanks@475.503.29\|CTTCGTGATCGATGTGGTGACGTCGTGCT | >hg38_21_+_8433172_8446622_RNA45SN1_45S_50ntFlanks | 475 | 503 | 29 | CTTCGTGATCGATGTGGTGACGTCGTGCT | BT20 | 225 | 378 | 43 | 44 | 108 | 169.2 | 230.3 | 17.2 | 21.0 | 62.4 |
| >hg38_21_+_8433172_8446622_RNA45SN1_45S_50ntFlanks@475.503.29\|CTTCGTGATCGATGTGGTGACGTCGTGCT | >hg38_21_+_8433172_8446622_RNA45SN1_45S_50ntFlanks | 475 | 503 | 29 | CTTCGTGATCGATGTGGTGACGTCGTGCT | MB231 | 142 | 358 | 10 | 50 | 109 | 94.1 | 100.1 | 4.0 | 7.6 | 27.1 |
| >hg38_21_+_8433172_8446622_RNA45SN1_45S_50ntFlanks@475.503.29\|CTTCGTGATCGATGTGGTGACGTCGTGCT | >hg38_21_+_8433172_8446622_RNA45SN1_45S_50ntFlanks | 475 | 503 | 29 | CTTCGTGATCGATGTGGTGACGTCGTGCT | MB468 | 12 | 49 | 0 | 2 | 2 | 3.9 | 27.2 | 0.3 | 2.4 | 2.6 |
| >hg38_21_+_8433172_8446622_RNA45SN1_45S_50ntFlanks@475.504.30\|CTTCGTGATCGATGTGGTGACGTCGTGCTC | >hg38_21_+_8433172_8446622_RNA45SN1_45S_50ntFlanks | 475 | 504 | 30 | CTTCGTGATCGATGTGGTGACGTCGTGCTC | BT20 | 442 | 867 | 201 | 126 | 274 | 210.8 | 590.3 | 79.0 | 93.2 | 187.6 |
| >hg38_21_+_8433172_8446622_RNA45SN1_45S_50ntFlanks@475.504.30\|CTTCGTGATCGATGTGGTGACGTCGTGCTC | >hg38_21_+_8433172_8446622_RNA45SN1_45S_50ntFlanks | 475 | 504 | 30 | CTTCGTGATCGATGTGGTGACGTCGTGCTC | MB231 | 490 | 1397 | 67 | 58 | 190 | 276.9 | 551.8 | 29.6 | 32.3 | 48.8 |
| >hg38_21_+_8433172_8446622_RNA45SN1_45S_50ntFlanks@475.504.30\|CTTCGTGATCGATGTGGTGACGTCGTGCTC | >hg38_21_+_8433172_8446622_RNA45SN1_45S_50ntFlanks | 475 | 504 | 30 | CTTCGTGATCGATGTGGTGACGTCGTGCTC | MB468 | 17 | 131 | 1 | 2 | 3 | 5.3 | 94.3 | 0.7 | 1.6 | 3.5 |
| >hg38_21_+_8433172_8446622_RNA45SN1_45S_50ntFlanks@475.505.31\|CTTCGTGATCGATGTGGTGACGTCGTGCTCT | >hg38_21_+_8433172_8446622_RNA45SN1_45S_50ntFlanks | 475 | 505 | 31 | CTTCGTGATCGATGTGGTGACGTCGTGCTCT | BT20 | 950 | 1576 | 360 | 157 | 323 | 373.2 | 833.6 | 188.6 | 102.3 | 149.6 |
| >hg38_21_+_8433172_8446622_RNA45SN1_45S_50ntFlanks@475.505.31\|CTTCGTGATCGATGTGGTGACGTCGTGCTCT | >hg38_21_+_8433172_8446622_RNA45SN1_45S_50ntFlanks | 475 | 505 | 31 | CTTCGTGATCGATGTGGTGACGTCGTGCTCT | MB231 | 1430 | 3435 | 183 | 93 | 380 | 566.4 | 1537.0 | 81.9 | 28.9 | 54.9 |
| >hg38_21_+_8433172_8446622_RNA45SN1_45S_50ntFlanks@475.505.31\|CTTCGTGATCGATGTGGTGACGTCGTGCTCT | >hg38_21_+_8433172_8446622_RNA45SN1_45S_50ntFlanks | 475 | 505 | 31 | CTTCGTGATCGATGTGGTGACGTCGTGCTCT | MB468 | 36 | 327 | 2 | 4 | 6 | 11.2 | 200.4 | 0.8 | 3.6 | 6.1 |
| >hg38_21_+_8433172_8446622_RNA45SN1_45S_50ntFlanks@475.506.32\|CTTCGTGATCGATGTGGTGACGTCGTGCTCTC | >hg38_21_+_8433172_8446622_RNA45SN1_45S_50ntFlanks | 475 | 506 | 32 | CTTCGTGATCGATGTGGTGACGTCGTGCTCTC | BT20 | 395 | 780 | 258 | 65 | 204 | 189.9 | 331.3 | 139.9 | 70.9 | 78.8 |
| >hg38_21_+_8433172_8446622_RNA45SN1_45S_50ntFlanks@475.506.32\|CTTCGTGATCGATGTGGTGACGTCGTGCTCTC | >hg38_21_+_8433172_8446622_RNA45SN1_45S_50ntFlanks | 475 | 506 | 32 | CTTCGTGATCGATGTGGTGACGTCGTGCTCTC | MB231 | 625 | 1645 | 149 | 30 | 189 | 229.4 | 744.8 | 76.3 | 10.0 | 30.7 |
| >hg38_21_+_8433172_8446622_RNA45SN1_45S_50ntFlanks@475.506.32\|CTTCGTGATCGATGTGGTGACGTCGTGCTCTC | >hg38_21_+_8433172_8446622_RNA45SN1_45S_50ntFlanks | 475 | 506 | 32 | CTTCGTGATCGATGTGGTGACGTCGTGCTCTC | MB468 | 29 | 276 | 2 | 2 | 4 | 9.3 | 166.7 | 1.3 | 1.3 | 4.4 |
| >hg38_21_+_8433172_8446622_RNA45SN1_45S_50ntFlanks@475.507.33\|CTTCGTGATCGATGTGGTGACGTCGTGCTCTCC | >hg38_21_+_8433172_8446622_RNA45SN1_45S_50ntFlanks | 475 | 507 | 33 | CTTCGTGATCGATGTGGTGACGTCGTGCTCTCC | BT20 | 125 | 246 | 62 | 13 | 83 | 118.6 | 102.9 | 25.8 | 13.6 | 38.4 |
| >hg38_21_+_8433172_8446622_RNA45SN1_45S_50ntFlanks@475.507.33\|CTTCGTGATCGATGTGGTGACGTCGTGCTCTCC | >hg38_21_+_8433172_8446622_RNA45SN1_45S_50ntFlanks | 475 | 507 | 33 | CTTCGTGATCGATGTGGTGACGTCGTGCTCTCC | MB231 | 107 | 299 | 87 | 7 | 93 | 70.1 | 123.5 | 80.9 | 2.1 | 18.7 |
| >hg38_21_+_8433172_8446622_RNA45SN1_45S_50ntFlanks@475.508.34\|CTTCGTGATCGATGTGGTGACGTCGTGCTCTCCC | >hg38_21_+_8433172_8446622_RNA45SN1_45S_50ntFlanks | 475 | 508 | 34 | CTTCGTGATCGATGTGGTGACGTCGTGCTCTCCC | BT20 | 43 | 83 | 21 | 6 | 31 | 40.6 | 37.2 | 10.9 | 6.0 | 27.3 |
| >hg38_21_+_8433172_8446622_RNA45SN1_45S_50ntFlanks@475.508.34\|CTTCGTGATCGATGTGGTGACGTCGTGCTCTCCC | >hg38_21_+_8433172_8446622_RNA45SN1_45S_50ntFlanks | 475 | 508 | 34 | CTTCGTGATCGATGTGGTGACGTCGTGCTCTCCC | MB231 | 34 | 98 | 25 | 2 | 35 | 22.9 | 37.6 | 25.4 | 1.2 | 7.8 |
| >hg38_21_+_8433172_8446622_RNA45SN1_45S_50ntFlanks@475.509.35\|CTTCGTGATCGATGTGGTGACGTCGTGCTCTCCCG | >hg38_21_+_8433172_8446622_RNA45SN1_45S_50ntFlanks | 475 | 509 | 35 | CTTCGTGATCGATGTGGTGACGTCGTGCTCTCCCG | BT20 | 127 | 242 | 55 | 9 | 75 | 103.3 | 116.2 | 25.5 | 11.4 | 28.8 |
| >hg38_21_+_8433172_8446622_RNA45SN1_45S_50ntFlanks@475.509.35\|CTTCGTGATCGATGTGGTGACGTCGTGCTCTCCCG | >hg38_21_+_8433172_8446622_RNA45SN1_45S_50ntFlanks | 475 | 509 | 35 | CTTCGTGATCGATGTGGTGACGTCGTGCTCTCCCG | MB231 | 161 | 444 | 55 | 5 | 75 | 93.3 | 198.9 | 44.4 | 2.5 | 16.8 |
| >hg38_21_+_8433172_8446622_RNA45SN1_45S_50ntFlanks@475.510.36\|CTTCGTGATCGATGTGGTGACGTCGTGCTCTCCCGG | >hg38_21_+_8433172_8446622_RNA45SN1_45S_50ntFlanks | 475 | 510 | 36 | CTTCGTGATCGATGTGGTGACGTCGTGCTCTCCCGG | BT20 | 151 | 297 | 90 | 15 | 108 | 132.6 | 125.4 | 36.4 | 16.4 | 25.4 |
| >hg38_21_+_8433172_8446622_RNA45SN1_45S_50ntFlanks@475.510.36\|CTTCGTGATCGATGTGGTGACGTCGTGCTCTCCCGG | >hg38_21_+_8433172_8446622_RNA45SN1_45S_50ntFlanks | 475 | 510 | 36 | CTTCGTGATCGATGTGGTGACGTCGTGCTCTCCCGG | MB231 | 240 | 597 | 73 | 8 | 99 | 147.5 | 284.3 | 59.3 | 3.6 | 25.9 |
| >hg38_21_+_8433172_8446622_RNA45SN1_45S_50ntFlanks@475.510.36\|CTTCGTGATCGATGTGGTGACGTCGTGCTCTCCCGG | >hg38_21_+_8433172_8446622_RNA45SN1_45S_50ntFlanks | 475 | 510 | 36 | CTTCGTGATCGATGTGGTGACGTCGTGCTCTCCCGG | MB468 | 13 | 140 | 1 | 0 | 2 | 4.0 | 52.6 | 0.3 | 0.7 | 4.2 |
| >hg38_21_+_8433172_8446622_RNA45SN1_45S_50ntFlanks@475.511.37\|CTTCGTGATCGATGTGGTGACGTCGTGCTCTCCCGGG | >hg38_21_+_8433172_8446622_RNA45SN1_45S_50ntFlanks | 475 | 511 | 37 | CTTCGTGATCGATGTGGTGACGTCGTGCTCTCCCGGG | BT20 | 100 | 189 | 51 | 10 | 64 | 89.9 | 86.2 | 13.3 | 11.2 | 20.2 |
| >hg38_21_+_8433172_8446622_RNA45SN1_45S_50ntFlanks@475.511.37\|CTTCGTGATCGATGTGGTGACGTCGTGCTCTCCCGGG | >hg38_21_+_8433172_8446622_RNA45SN1_45S_50ntFlanks | 475 | 511 | 37 | CTTCGTGATCGATGTGGTGACGTCGTGCTCTCCCGGG | MB231 | 142 | 311 | 36 | 6 | 53 | 78.1 | 135.7 | 24.5 | 2.7 | 12.3 |
| >hg38_21_+_8433172_8446622_RNA45SN1_45S_50ntFlanks@475.512.38\|CTTCGTGATCGATGTGGTGACGTCGTGCTCTCCCGGGC | >hg38_21_+_8433172_8446622_RNA45SN1_45S_50ntFlanks | 475 | 512 | 38 | CTTCGTGATCGATGTGGTGACGTCGTGCTCTCCCGGGC | BT20 | 25 | 56 | 11 | 3 | 10 | 24.6 | 33.1 | 1.7 | 3.0 | 9.8 |
| >hg38_21_+_8433172_8446622_RNA45SN1_45S_50ntFlanks@475.512.38\|CTTCGTGATCGATGTGGTGACGTCGTGCTCTCCCGGGC | >hg38_21_+_8433172_8446622_RNA45SN1_45S_50ntFlanks | 475 | 512 | 38 | CTTCGTGATCGATGTGGTGACGTCGTGCTCTCCCGGGC | MB231 | 25 | 73 | 6 | 2 | 13 | 19.0 | 26.4 | 2.2 | 1.3 | 5.8 |
| >hg38_21_+_8433172_8446622_RNA45SN1_45S_50ntFlanks@475.513.39\|CTTCGTGATCGATGTGGTGACGTCGTGCTCTCCCGGGCC | >hg38_21_+_8433172_8446622_RNA45SN1_45S_50ntFlanks | 475 | 513 | 39 | CTTCGTGATCGATGTGGTGACGTCGTGCTCTCCCGGGCC | BT20 | 25 | 57 | 11 | 2 | 15 | 26.1 | 31.4 | 4.6 | 2.2 | 12.9 |
| >hg38_21_+_8433172_8446622_RNA45SN1_45S_50ntFlanks@475.513.39\|CTTCGTGATCGATGTGGTGACGTCGTGCTCTCCCGGGCC | >hg38_21_+_8433172_8446622_RNA45SN1_45S_50ntFlanks | 475 | 513 | 39 | CTTCGTGATCGATGTGGTGACGTCGTGCTCTCCCGGGCC | MB231 | 24 | 73 | 9 | 0 | 0 | 20.7 | 35.2 | 7.1 | 0.0 | 0.0 |
| >hg38_21_+_8433172_8446622_RNA45SN1_45S_50ntFlanks@475.514.40\|CTTCGTGATCGATGTGGTGACGTCGTGCTCTCCCGGGCCG | >hg38_21_+_8433172_8446622_RNA45SN1_45S_50ntFlanks | 475 | 514 | 40 | CTTCGTGATCGATGTGGTGACGTCGTGCTCTCCCGGGCCG | BT20 | 74 | 175 | 32 | 5 | 28 | 69.0 | 102.7 | 15.8 | 5.8 | 24.7 |
| >hg38_21_+_8433172_8446622_RNA45SN1_45S_50ntFlanks@475.514.40\|CTTCGTGATCGATGTGGTGACGTCGTGCTCTCCCGGGCCG | >hg38_21_+_8433172_8446622_RNA45SN1_45S_50ntFlanks | 475 | 514 | 40 | CTTCGTGATCGATGTGGTGACGTCGTGCTCTCCCGGGCCG | MB231 | 86 | 311 | 18 | 1 | 20 | 47.1 | 182.7 | 8.8 | 1.1 | 19.9 |
| >hg38_21_+_8433172_8446622_RNA45SN1_45S_50ntFlanks@475.515.41\|CTTCGTGATCGATGTGGTGACGTCGTGCTCTCCCGGGCCGG | >hg38_21_+_8433172_8446622_RNA45SN1_45S_50ntFlanks | 475 | 515 | 41 | CTTCGTGATCGATGTGGTGACGTCGTGCTCTCCCGGGCCGG | BT20 | 50 | 96 | 6 | 3 | 11 | 48.9 | 58.8 | 1.4 | 2.5 | 11.3 |
| >hg38_21_+_8433172_8446622_RNA45SN1_45S_50ntFlanks@475.515.41\|CTTCGTGATCGATGTGGTGACGTCGTGCTCTCCCGGGCCGG | >hg38_21_+_8433172_8446622_RNA45SN1_45S_50ntFlanks | 475 | 515 | 41 | CTTCGTGATCGATGTGGTGACGTCGTGCTCTCCCGGGCCGG | MB231 | 96 | 219 | 8 | 3 | 21 | 43.4 | 107.2 | 2.8 | 2.4 | 22.5 |
| >hg38_21_+_8433172_8446622_RNA45SN1_45S_50ntFlanks@475.516.42\|CTTCGTGATCGATGTGGTGACGTCGTGCTCTCCCGGGCCGGG | >hg38_21_+_8433172_8446622_RNA45SN1_45S_50ntFlanks | 475 | 516 | 42 | CTTCGTGATCGATGTGGTGACGTCGTGCTCTCCCGGGCCGGG | BT20 | 156 | 381 | 20 | 14 | 120 | 227.4 | 245.4 | 10.1 | 14.6 | 12.8 |
| >hg38_21_+_8433172_8446622_RNA45SN1_45S_50ntFlanks@475.516.42\|CTTCGTGATCGATGTGGTGACGTCGTGCTCTCCCGGGCCGGG | >hg38_21_+_8433172_8446622_RNA45SN1_45S_50ntFlanks | 475 | 516 | 42 | CTTCGTGATCGATGTGGTGACGTCGTGCTCTCCCGGGCCGGG | MB231 | 249 | 1066 | 61 | 8 | 131 | 174.6 | 585.8 | 46.2 | 5.0 | 21.9 |
| >hg38_21_+_8433172_8446622_RNA45SN1_45S_50ntFlanks@475.517.43\|CTTCGTGATCGATGTGGTGACGTCGTGCTCTCCCGGGCCGGGT | >hg38_21_+_8433172_8446622_RNA45SN1_45S_50ntFlanks | 475 | 517 | 43 | CTTCGTGATCGATGTGGTGACGTCGTGCTCTCCCGGGCCGGGT | BT20 | 84 | 275 | 12 | 15 | 155 | 123.6 | 202.2 | 7.7 | 12.5 | 45.5 |

| | | | | | | | | | | | | | | | | |
|---|---|---|---|---|---|---|---|---|---|---|---|---|---|---|---|---|
| >hg38_21_+_8433172_8446622_RNA45SN1_45S_50ntFlanks@475.517.43\|CTTCGTGATCGATGTGGTGACGTCGTGCTCTCCCGGGCCGGGT | >hg38_21_+_8433172_8446622_RNA45SN1_45S_50ntFlanks | 475 | 517 | 43 | CTTCGTGATCGATGTGGTGACGTCGTGCTCTCCCGGGCCGGGT | MB231 | 100 | 517 | 57 | 7 | 113 | 108.1 | 306.6 | 77.4 | 5.7 | 38.6 |
| >hg38_21_+_8433172_8446622_RNA45SN1_45S_50ntFlanks@475.518.44\|CTTCGTGATCGATGTGGTGACGTCGTGCTCTCCCGGGCCGGGTC | >hg38_21_+_8433172_8446622_RNA45SN1_45S_50ntFlanks | 475 | 518 | 44 | CTTCGTGATCGATGTGGTGACGTCGTGCTCTCCCGGGCCGGGTC | BT20 | 39 | 169 | 9 | 12 | 89 | 53.7 | 142.6 | 5.9 | 8.4 | 33.9 |
| >hg38_21_+_8433172_8446622_RNA45SN1_45S_50ntFlanks@475.518.44\|CTTCGTGATCGATGTGGTGACGTCGTGCTCTCCCGGGCCGGGTC | >hg38_21_+_8433172_8446622_RNA45SN1_45S_50ntFlanks | 475 | 518 | 44 | CTTCGTGATCGATGTGGTGACGTCGTGCTCTCCCGGGCCGGGTC | MB231 | 68 | 303 | 33 | 6 | 82 | 71.1 | 193.3 | 34.4 | 4.9 | 48.0 |
| >hg38_21_+_8433172_8446622_RNA45SN1_45S_50ntFlanks@475.519.45\|CTTCGTGATCGATGTGGTGACGTCGTGCTCTCCCGGGCCGGGTCC | >hg38_21_+_8433172_8446622_RNA45SN1_45S_50ntFlanks | 475 | 519 | 45 | CTTCGTGATCGATGTGGTGACGTCGTGCTCTCCCGGGCCGGGTCC | MB231 | 18 | 66 | 3 | 1 | 6 | 10.5 | 36.3 | 0.9 | 1.0 | 11.2 |
| >hg38_21_+_8433172_8446622_RNA45SN1_45S_50ntFlanks@475.520.46\|CTTCGTGATCGATGTGGTGACGTCGTGCTCTCCCGGGCCGGGTCCG | >hg38_21_+_8433172_8446622_RNA45SN1_45S_50ntFlanks | 475 | 520 | 46 | CTTCGTGATCGATGTGGTGACGTCGTGCTCTCCCGGGCCGGGTCCG | BT20 | 20 | 67 | 2 | 2 | 11 | 17.6 | 62.0 | 2.0 | 2.0 | 11.3 |
| >hg38_21_+_8433172_8446622_RNA45SN1_45S_50ntFlanks@475.520.46\|CTTCGTGATCGATGTGGTGACGTCGTGCTCTCCCGGGCCGGGTCCG | >hg38_21_+_8433172_8446622_RNA45SN1_45S_50ntFlanks | 475 | 520 | 46 | CTTCGTGATCGATGTGGTGACGTCGTGCTCTCCCGGGCCGGGTCCG | MB231 | 48 | 153 | 6 | 1 | 36 | 15.9 | 78.4 | 2.3 | 1.4 | 19.8 |
| >hg38_21_+_8433172_8446622_RNA45SN1_45S_50ntFlanks@475.521.47\|CTTCGTGATCGATGTGGTGACGTCGTGCTCTCCCGGGCCGGGTCCGA | >hg38_21_+_8433172_8446622_RNA45SN1_45S_50ntFlanks | 475 | 521 | 47 | CTTCGTGATCGATGTGGTGACGTCGTGCTCTCCCGGGCCGGGTCCGA | BT20 | 31 | 151 | 10 | 6 | 39 | 28.0 | 179.7 | 13.8 | 4.4 | 27.8 |
| >hg38_21_+_8433172_8446622_RNA45SN1_45S_50ntFlanks@475.521.47\|CTTCGTGATCGATGTGGTGACGTCGTGCTCTCCCGGGCCGGGTCCGA | >hg38_21_+_8433172_8446622_RNA45SN1_45S_50ntFlanks | 475 | 521 | 47 | CTTCGTGATCGATGTGGTGACGTCGTGCTCTCCCGGGCCGGGTCCGA | MB231 | 63 | 171 | 8 | 4 | 77 | 22.6 | 96.5 | 3.7 | 2.9 | 57.3 |
| >hg38_21_+_8433172_8446622_RNA45SN1_45S_50ntFlanks@475.522.48\|CTTCGTGATCGATGTGGTGACGTCGTGCTCTCCCGGGCCGGGTCCGAG | >hg38_21_+_8433172_8446622_RNA45SN1_45S_50ntFlanks | 475 | 522 | 48 | CTTCGTGATCGATGTGGTGACGTCGTGCTCTCCCGGGCCGGGTCCGAG | BT20 | 13 | 68 | 2 | 1 | 9 | 11.1 | 75.9 | 2.4 | 0.9 | 12.1 |
| >hg38_21_+_8433172_8446622_RNA45SN1_45S_50ntFlanks@475.522.48\|CTTCGTGATCGATGTGGTGACGTCGTGCTCTCCCGGGCCGGGTCCGAG | >hg38_21_+_8433172_8446622_RNA45SN1_45S_50ntFlanks | 475 | 522 | 48 | CTTCGTGATCGATGTGGTGACGTCGTGCTCTCCCGGGCCGGGTCCGAG | MB231 | 30 | 167 | 3 | 1 | 17 | 12.5 | 207.2 | 2.0 | 0.8 | 15.4 |
| >hg38_21_+_8433172_8446622_RNA45SN1_45S_50ntFlanks@475.523.49\|CTTCGTGATCGATGTGGTGACGTCGTGCTCTCCCGGGCCGGGTCCGAGC | >hg38_21_+_8433172_8446622_RNA45SN1_45S_50ntFlanks | 475 | 523 | 49 | CTTCGTGATCGATGTGGTGACGTCGTGCTCTCCCGGGCCGGGTCCGAGC | BT20 | 11 | 53 | 1 | 2 | 13 | 9.1 | 57.6 | 1.2 | 1.6 | 11.7 |
| >hg38_21_+_8433172_8446622_RNA45SN1_45S_50ntFlanks@475.523.49\|CTTCGTGATCGATGTGGTGACGTCGTGCTCTCCCGGGCCGGGTCCGAGC | >hg38_21_+_8433172_8446622_RNA45SN1_45S_50ntFlanks | 475 | 523 | 49 | CTTCGTGATCGATGTGGTGACGTCGTGCTCTCCCGGGCCGGGTCCGAGC | MB231 | 34 | 269 | 3 | 1 | 28 | 21.6 | 390.2 | 1.0 | 1.3 | 26.3 |
| >hg38_21_+_8433172_8446622_RNA45SN1_45S_50ntFlanks@475.524.50\|CTTCGTGATCGATGTGGTGACGTCGTGCTCTCCCGGGCCGGGTCCGAGCC | >hg38_21_+_8433172_8446622_RNA45SN1_45S_50ntFlanks | 475 | 524 | 50 | CTTCGTGATCGATGTGGTGACGTCGTGCTCTCCCGGGCCGGGTCCGAGCC | MB231 | 37 | 417 | 6 | 1 | 17 | 26.8 | 652.6 | 5.4 | 0.9 | 28.8 |
| >hg38_21_+_8433172_8446622_RNA45SN1_45S_50ntFlanks@4752.4782.31\|GAAGACGATCAGATACCGTCGTAGTTCCGAC | >hg38_21_+_8433172_8446622_RNA45SN1_45S_50ntFlanks | 4752 | 4782 | 31 | GAAGACGATCAGATACCGTCGTAGTTCCGAC | MB468 | 13 | 11 | 16 | 14 | 44 | 1.9 | 5.6 | 5.9 | 9.7 | 53.3 |
| >hg38_21_+_8433172_8446622_RNA45SN1_45S_50ntFlanks@4753.4782.30\|AAGACGATCAGATACCGTCGTAGTTCCGAC | >hg38_21_+_8433172_8446622_RNA45SN1_45S_50ntFlanks | 4753 | 4782 | 30 | AAGACGATCAGATACCGTCGTAGTTCCGAC | MB468 | 30 | 32 | 40 | 27 | 72 | 4.9 | 8.7 | 20.1 | 20.4 | 83.8 |
| >hg38_21_+_8433172_8446622_RNA45SN1_45S_50ntFlanks@4753.4783.31\|AAGACGATCAGATACCGTCGTAGTTCCGACC | >hg38_21_+_8433172_8446622_RNA45SN1_45S_50ntFlanks | 4753 | 4783 | 31 | AAGACGATCAGATACCGTCGTAGTTCCGACC | MB468 | 22 | 45 | 29 | 8 | 65 | 6.3 | 26.7 | 15.9 | 7.4 | 59.7 |
| >hg38_21_+_8433172_8446622_RNA45SN1_45S_50ntFlanks@4758.4780.23\|GATCAGATACCGTCGTAGTTCCG | >hg38_21_+_8433172_8446622_RNA45SN1_45S_50ntFlanks | 4758 | 4780 | 23 | GATCAGATACCGTCGTAGTTCCG | MB468 | 11 | 12 | 10 | 7 | 9 | 3.9 | 7.4 | 3.2 | 6.3 | 6.8 |
| >hg38_21_+_8433172_8446622_RNA45SN1_45S_50ntFlanks@4759.4776.18\|ATCAGATACCGTCGTAGT | >hg38_21_+_8433172_8446622_RNA45SN1_45S_50ntFlanks | 4759 | 4776 | 18 | ATCAGATACCGTCGTAGT | MB468 | 22 | 6 | 22 | 8 | 7 | 11.7 | 2.8 | 17.3 | 2.3 | 3.7 |
| >hg38_21_+_8433172_8446622_RNA45SN1_45S_50ntFlanks@4759.4777.19\|ATCAGATACCGTCGTAGTT | >hg38_21_+_8433172_8446622_RNA45SN1_45S_50ntFlanks | 4759 | 4777 | 19 | ATCAGATACCGTCGTAGTT | MB468 | 21 | 5 | 14 | 23 | 10 | 5.0 | 2.3 | 9.9 | 9.0 | 5.9 |
| >hg38_21_+_8433172_8446622_RNA45SN1_45S_50ntFlanks@4759.4778.20\|ATCAGATACCGTCGTAGTTC | >hg38_21_+_8433172_8446622_RNA45SN1_45S_50ntFlanks | 4759 | 4778 | 20 | ATCAGATACCGTCGTAGTTC | MB468 | 145 | 109 | 110 | 89 | 76 | 34.9 | 51.2 | 28.7 | 46.3 | 46.9 |
| >hg38_21_+_8433172_8446622_RNA45SN1_45S_50ntFlanks@4759.4779.21\|ATCAGATACCGTCGTAGTTCC | >hg38_21_+_8433172_8446622_RNA45SN1_45S_50ntFlanks | 4759 | 4779 | 21 | ATCAGATACCGTCGTAGTTCC | MB468 | 73 | 73 | 64 | 24 | 39 | 9.0 | 18.8 | 11.3 | 23.7 | 24.1 |
| >hg38_21_+_8433172_8446622_RNA45SN1_45S_50ntFlanks@4759.4780.22\|ATCAGATACCGTCGTAGTTCCG | >hg38_21_+_8433172_8446622_RNA45SN1_45S_50ntFlanks | 4759 | 4780 | 22 | ATCAGATACCGTCGTAGTTCCG | MB468 | 914 | 686 | 779 | 387 | 511 | 94.6 | 294.1 | 200.1 | 362.3 | 321.6 |
| >hg38_21_+_8433172_8446622_RNA45SN1_45S_50ntFlanks@4759.4781.23\|ATCAGATACCGTCGTAGTTCCGA | >hg38_21_+_8433172_8446622_RNA45SN1_45S_50ntFlanks | 4759 | 4781 | 23 | ATCAGATACCGTCGTAGTTCCGA | MB468 | 32 | 7 | 41 | 19 | 15 | 7.3 | 0.8 | 30.6 | 12.2 | 8.9 |
| >hg38_21_+_8433172_8446622_RNA45SN1_45S_50ntFlanks@4759.4782.24\|ATCAGATACCGTCGTAGTTCCGAC | >hg38_21_+_8433172_8446622_RNA45SN1_45S_50ntFlanks | 4759 | 4782 | 24 | ATCAGATACCGTCGTAGTTCCGAC | MB468 | 211 | 38 | 159 | 250 | 251 | 15.5 | 7.6 | 85.0 | 80.3 | 338.6 |
| >hg38_21_+_8433172_8446622_RNA45SN1_45S_50ntFlanks@4759.4783.25\|ATCAGATACCGTCGTAGTTCCGACC | >hg38_21_+_8433172_8446622_RNA45SN1_45S_50ntFlanks | 4759 | 4783 | 25 | ATCAGATACCGTCGTAGTTCCGACC | MB468 | 118 | 51 | 98 | 115 | 159 | 15.5 | 4.6 | 27.5 | 93.2 | 182.0 |
| >hg38_21_+_8433172_8446622_RNA45SN1_45S_50ntFlanks@4759.4786.28\|ATCAGATACCGTCGTAGTTCCGACCATA | >hg38_21_+_8433172_8446622_RNA45SN1_45S_50ntFlanks | 4759 | 4786 | 28 | ATCAGATACCGTCGTAGTTCCGACCATA | MB468 | 73 | 16 | 46 | 65 | 83 | 23.6 | 2.3 | 21.7 | 22.6 | 68.2 |
| >hg38_21_+_8433172_8446622_RNA45SN1_45S_50ntFlanks@4759.4793.35\|ATCAGATACCGTCGTAGTTCCGACCATAAACGATG | >hg38_21_+_8433172_8446622_RNA45SN1_45S_50ntFlanks | 4759 | 4793 | 35 | ATCAGATACCGTCGTAGTTCCGACCATAAACGATG | MB468 | 22 | 41 | 77 | 33 | 38 | 4.8 | 8.0 | 37.8 | 23.4 | 12.8 |
| >hg38_21_+_8433172_8446622_RNA45SN1_45S_50ntFlanks@476.494.19\|TTCGTGATCGATGTGGTGA | >hg38_21_+_8433172_8446622_RNA45SN1_45S_50ntFlanks | 476 | 494 | 19 | TTCGTGATCGATGTGGTGA | BT20 | 10 | 19 | 1 | 1 | 4 | 15.8 | 26.5 | 1.3 | 1.2 | 5.1 |
| >hg38_21_+_8433172_8446622_RNA45SN1_45S_50ntFlanks@476.495.20\|TTCGTGATCGATGTGGTGAC | >hg38_21_+_8433172_8446622_RNA45SN1_45S_50ntFlanks | 476 | 495 | 20 | TTCGTGATCGATGTGGTGAC | BT20 | 25 | 64 | 7 | 2 | 10 | 28.0 | 69.9 | 5.3 | 1.8 | 11.2 |
| >hg38_21_+_8433172_8446622_RNA45SN1_45S_50ntFlanks@476.495.20\|TTCGTGATCGATGTGGTGAC | >hg38_21_+_8433172_8446622_RNA45SN1_45S_50ntFlanks | 476 | 495 | 20 | TTCGTGATCGATGTGGTGAC | MB231 | 73 | 337 | 8 | 9 | 37 | 16.8 | 102.9 | 5.7 | 2.2 | 5.2 |
| >hg38_21_+_8433172_8446622_RNA45SN1_45S_50ntFlanks@476.496.21\|TTCGTGATCGATGTGGTGACG | >hg38_21_+_8433172_8446622_RNA45SN1_45S_50ntFlanks | 476 | 496 | 21 | TTCGTGATCGATGTGGTGACG | BT20 | 27 | 74 | 2 | 2 | 14 | 34.0 | 113.5 | 2.5 | 3.2 | 18.4 |
| >hg38_21_+_8433172_8446622_RNA45SN1_45S_50ntFlanks@476.496.21\|TTCGTGATCGATGTGGTGACG | >hg38_21_+_8433172_8446622_RNA45SN1_45S_50ntFlanks | 476 | 496 | 21 | TTCGTGATCGATGTGGTGACG | MB231 | 63 | 255 | 2 | 16 | 52 | 28.3 | 166.4 | 1.7 | 4.9 | 22.3 |
| >hg38_21_+_8433172_8446622_RNA45SN1_45S_50ntFlanks@476.497.22\|TTCGTGATCGATGTGGTGACGT | >hg38_21_+_8433172_8446622_RNA45SN1_45S_50ntFlanks | 476 | 497 | 22 | TTCGTGATCGATGTGGTGACGT | BT20 | 31 | 90 | 3 | 2 | 19 | 28.2 | 138.7 | 3.3 | 2.7 | 28.7 |
| >hg38_21_+_8433172_8446622_RNA45SN1_45S_50ntFlanks@476.497.22\|TTCGTGATCGATGTGGTGACGT | >hg38_21_+_8433172_8446622_RNA45SN1_45S_50ntFlanks | 476 | 497 | 22 | TTCGTGATCGATGTGGTGACGT | MB231 | 66 | 183 | 3 | 25 | 79 | 22.2 | 104.0 | 2.5 | 9.2 | 66.5 |
| >hg38_21_+_8433172_8446622_RNA45SN1_45S_50ntFlanks@476.498.23\|TTCGTGATCGATGTGGTGACGTC | >hg38_21_+_8433172_8446622_RNA45SN1_45S_50ntFlanks | 476 | 498 | 23 | TTCGTGATCGATGTGGTGACGTC | BT20 | 57 | 160 | 5 | 9 | 38 | 82.2 | 227.8 | 6.9 | 9.4 | 52.7 |
| >hg38_21_+_8433172_8446622_RNA45SN1_45S_50ntFlanks@476.498.23\|TTCGTGATCGATGTGGTGACGTC | >hg38_21_+_8433172_8446622_RNA45SN1_45S_50ntFlanks | 476 | 498 | 23 | TTCGTGATCGATGTGGTGACGTC | MB231 | 136 | 501 | 6 | 45 | 144 | 76.4 | 339.0 | 4.9 | 23.5 | 84.0 |
| >hg38_21_+_8433172_8446622_RNA45SN1_45S_50ntFlanks@476.499.24\|TTCGTGATCGATGTGGTGACGTCG | >hg38_21_+_8433172_8446622_RNA45SN1_45S_50ntFlanks | 476 | 499 | 24 | TTCGTGATCGATGTGGTGACGTCG | BT20 | 98 | 204 | 30 | 16 | 74 | 132.5 | 225.8 | 24.0 | 12.0 | 75.4 |
| >hg38_21_+_8433172_8446622_RNA45SN1_45S_50ntFlanks@476.499.24\|TTCGTGATCGATGTGGTGACGTCG | >hg38_21_+_8433172_8446622_RNA45SN1_45S_50ntFlanks | 476 | 499 | 24 | TTCGTGATCGATGTGGTGACGTCG | MB231 | 126 | 416 | 23 | 57 | 185 | 93.1 | 202.9 | 13.9 | 26.9 | 65.4 |
| >hg38_21_+_8433172_8446622_RNA45SN1_45S_50ntFlanks@476.501.26\|TTCGTGATCGATGTGGTGACGTCGTG | >hg38_21_+_8433172_8446622_RNA45SN1_45S_50ntFlanks | 476 | 501 | 26 | TTCGTGATCGATGTGGTGACGTCGTG | BT20 | 62 | 110 | 7 | 7 | 24 | 44.3 | 71.3 | 3.8 | 1.2 | 10.7 |
| >hg38_21_+_8433172_8446622_RNA45SN1_45S_50ntFlanks@476.501.26\|TTCGTGATCGATGTGGTGACGTCGTG | >hg38_21_+_8433172_8446622_RNA45SN1_45S_50ntFlanks | 476 | 501 | 26 | TTCGTGATCGATGTGGTGACGTCGTG | MB231 | 75 | 171 | 3 | 15 | 42 | 35.4 | 35.0 | 1.1 | 5.0 | 12.0 |
| >hg38_21_+_8433172_8446622_RNA45SN1_45S_50ntFlanks@476.502.27\|TTCGTGATCGATGTGGTGACGTCGTGC | >hg38_21_+_8433172_8446622_RNA45SN1_45S_50ntFlanks | 476 | 502 | 27 | TTCGTGATCGATGTGGTGACGTCGTGC | BT20 | 43 | 68 | 5 | 3 | 12 | 38.2 | 59.9 | 2.5 | 1.7 | 8.5 |
| >hg38_21_+_8433172_8446622_RNA45SN1_45S_50ntFlanks@476.502.27\|TTCGTGATCGATGTGGTGACGTCGTGC | >hg38_21_+_8433172_8446622_RNA45SN1_45S_50ntFlanks | 476 | 502 | 27 | TTCGTGATCGATGTGGTGACGTCGTGC | MB231 | 55 | 140 | 1 | 10 | 19 | 23.1 | 23.6 | 0.6 | 4.5 | 4.6 |
| >hg38_21_+_8433172_8446622_RNA45SN1_45S_50ntFlanks@476.503.28\|TTCGTGATCGATGTGGTGACGTCGTGCT | >hg38_21_+_8433172_8446622_RNA45SN1_45S_50ntFlanks | 476 | 503 | 28 | TTCGTGATCGATGTGGTGACGTCGTGCT | BT20 | 117 | 168 | 13 | 8 | 31 | 66.1 | 110.4 | 6.0 | 4.8 | 17.4 |
| >hg38_21_+_8433172_8446622_RNA45SN1_45S_50ntFlanks@476.503.28\|TTCGTGATCGATGTGGTGACGTCGTGCT | >hg38_21_+_8433172_8446622_RNA45SN1_45S_50ntFlanks | 476 | 503 | 28 | TTCGTGATCGATGTGGTGACGTCGTGCT | MB231 | 147 | 332 | 4 | 16 | 44 | 66.0 | 58.0 | 1.8 | 6.0 | 9.1 |
| >hg38_21_+_8433172_8446622_RNA45SN1_45S_50ntFlanks@476.504.29\|TTCGTGATCGATGTGGTGACGTCGTGCTC | >hg38_21_+_8433172_8446622_RNA45SN1_45S_50ntFlanks | 476 | 504 | 29 | TTCGTGATCGATGTGGTGACGTCGTGCTC | BT20 | 148 | 248 | 34 | 23 | 63 | 66.3 | 171.3 | 16.9 | 23.9 | 40.0 |
| >hg38_21_+_8433172_8446622_RNA45SN1_45S_50ntFlanks@476.504.29\|TTCGTGATCGATGTGGTGACGTCGTGCTC | >hg38_21_+_8433172_8446622_RNA45SN1_45S_50ntFlanks | 476 | 504 | 29 | TTCGTGATCGATGTGGTGACGTCGTGCTC | MB231 | 210 | 557 | 13 | 17 | 67 | 111.4 | 176.7 | 6.3 | 12.1 | 18.1 |
| >hg38_21_+_8433172_8446622_RNA45SN1_45S_50ntFlanks@476.505.30\|TTCGTGATCGATGTGGTGACGTCGTGCTCT | >hg38_21_+_8433172_8446622_RNA45SN1_45S_50ntFlanks | 476 | 505 | 30 | TTCGTGATCGATGTGGTGACGTCGTGCTCT | BT20 | 311 | 449 | 63 | 29 | 88 | 133.0 | 245.8 | 35.4 | 32.3 | 38.9 |
| >hg38_21_+_8433172_8446622_RNA45SN1_45S_50ntFlanks@476.505.30\|TTCGTGATCGATGTGGTGACGTCGTGCTCT | >hg38_21_+_8433172_8446622_RNA45SN1_45S_50ntFlanks | 476 | 505 | 30 | TTCGTGATCGATGTGGTGACGTCGTGCTCT | MB231 | 568 | 1333 | 37 | 23 | 139 | 225.7 | 486.9 | 17.5 | 9.9 | 23.6 |
| >hg38_21_+_8433172_8446622_RNA45SN1_45S_50ntFlanks@476.505.30\|TTCGTGATCGATGTGGTGACGTCGTGCTCT | >hg38_21_+_8433172_8446622_RNA45SN1_45S_50ntFlanks | 476 | 505 | 30 | TTCGTGATCGATGTGGTGACGTCGTGCTCT | MB468 | 13 | 75 | 1 | 1 | 2 | 3.7 | 44.2 | 0.4 | 0.8 | 2.0 |
| >hg38_21_+_8433172_8446622_RNA45SN1_45S_50ntFlanks@476.506.31\|TTCGTGATCGATGTGGTGACGTCGTGCTCTC | >hg38_21_+_8433172_8446622_RNA45SN1_45S_50ntFlanks | 476 | 506 | 31 | TTCGTGATCGATGTGGTGACGTCGTGCTCTC | BT20 | 132 | 249 | 50 | 15 | 53 | 47.2 | 125.9 | 21.6 | 17.8 | 18.3 |

| | | | | | | | | | | | | | | | | |
|---|---|---|---|---|---|---|---|---|---|---|---|---|---|---|---|---|
| >hg38_21_+_8433172_8446622_RNA45SN1_45S_50ntFlanks@476.506.31\|TTCGTGATCGATGTGGTGACGTCGTGCTCTC | >hg38_21_+_8433172_8446622_RNA45SN1_45S_50ntFlanks | 476 | 506 | 31 | TTCGTGATCGATGTGGTGACGTCGTGCTCTC | MB231 | 272 | 725 | 36 | 11 | 76 | 100.3 | 283.0 | 17.0 | 5.8 | 17.1 |
| >hg38_21_+_8433172_8446622_RNA45SN1_45S_50ntFlanks@476.507.32\|TTCGTGATCGATGTGGTGACGTCGTGCTCTCC | >hg38_21_+_8433172_8446622_RNA45SN1_45S_50ntFlanks | 476 | 507 | 32 | TTCGTGATCGATGTGGTGACGTCGTGCTCTCC | BT20 | 58 | 111 | 21 | 5 | 21 | 47.8 | 54.1 | 7.8 | 6.3 | 19.0 |
| >hg38_21_+_8433172_8446622_RNA45SN1_45S_50ntFlanks@476.507.32\|TTCGTGATCGATGTGGTGACGTCGTGCTCTCC | >hg38_21_+_8433172_8446622_RNA45SN1_45S_50ntFlanks | 476 | 507 | 32 | TTCGTGATCGATGTGGTGACGTCGTGCTCTCC | MB231 | 111 | 336 | 41 | 4 | 39 | 60.4 | 145.2 | 35.2 | 2.2 | 10.2 |
| >hg38_21_+_8433172_8446622_RNA45SN1_45S_50ntFlanks@476.508.33\|TTCGTGATCGATGTGGTGACGTCGTGCTCTCCC | >hg38_21_+_8433172_8446622_RNA45SN1_45S_50ntFlanks | 476 | 508 | 33 | TTCGTGATCGATGTGGTGACGTCGTGCTCTCCC | BT20 | 18 | 35 | 6 | 2 | 9 | 16.6 | 17.2 | 3.0 | 2.8 | 7.9 |
| >hg38_21_+_8433172_8446622_RNA45SN1_45S_50ntFlanks@476.508.33\|TTCGTGATCGATGTGGTGACGTCGTGCTCTCCC | >hg38_21_+_8433172_8446622_RNA45SN1_45S_50ntFlanks | 476 | 508 | 33 | TTCGTGATCGATGTGGTGACGTCGTGCTCTCCC | MB231 | 26 | 80 | 12 | 1 | 15 | 18.6 | 31.2 | 11.5 | 0.2 | 4.3 |
| >hg38_21_+_8433172_8446622_RNA45SN1_45S_50ntFlanks@476.509.34\|TTCGTGATCGATGTGGTGACGTCGTGCTCTCCCG | >hg38_21_+_8433172_8446622_RNA45SN1_45S_50ntFlanks | 476 | 509 | 34 | TTCGTGATCGATGTGGTGACGTCGTGCTCTCCCG | BT20 | 62 | 110 | 18 | 4 | 24 | 41.9 | 55.3 | 7.1 | 6.5 | 20.6 |
| >hg38_21_+_8433172_8446622_RNA45SN1_45S_50ntFlanks@476.509.34\|TTCGTGATCGATGTGGTGACGTCGTGCTCTCCCG | >hg38_21_+_8433172_8446622_RNA45SN1_45S_50ntFlanks | 476 | 509 | 34 | TTCGTGATCGATGTGGTGACGTCGTGCTCTCCCG | MB231 | 142 | 394 | 24 | 2 | 38 | 64.5 | 158.9 | 20.6 | 0.9 | 10.9 |
| >hg38_21_+_8433172_8446622_RNA45SN1_45S_50ntFlanks@476.510.35\|TTCGTGATCGATGTGGTGACGTCGTGCTCTCCCGG | >hg38_21_+_8433172_8446622_RNA45SN1_45S_50ntFlanks | 476 | 510 | 35 | TTCGTGATCGATGTGGTGACGTCGTGCTCTCCCGG | BT20 | 52 | 93 | 23 | 5 | 23 | 40.7 | 42.6 | 9.8 | 7.3 | 20.3 |
| >hg38_21_+_8433172_8446622_RNA45SN1_45S_50ntFlanks@476.510.35\|TTCGTGATCGATGTGGTGACGTCGTGCTCTCCCGG | >hg38_21_+_8433172_8446622_RNA45SN1_45S_50ntFlanks | 476 | 510 | 35 | TTCGTGATCGATGTGGTGACGTCGTGCTCTCCCGG | MB231 | 119 | 313 | 22 | 3 | 38 | 65.6 | 141.7 | 18.7 | 1.5 | 10.7 |
| >hg38_21_+_8433172_8446622_RNA45SN1_45S_50ntFlanks@476.511.36\|TTCGTGATCGATGTGGTGACGTCGTGCTCTCCCGGG | >hg38_21_+_8433172_8446622_RNA45SN1_45S_50ntFlanks | 476 | 511 | 36 | TTCGTGATCGATGTGGTGACGTCGTGCTCTCCCGGG | BT20 | 39 | 75 | 18 | 4 | 18 | 33.2 | 29.6 | 7.2 | 4.5 | 15.2 |
| >hg38_21_+_8433172_8446622_RNA45SN1_45S_50ntFlanks@476.511.36\|TTCGTGATCGATGTGGTGACGTCGTGCTCTCCCGGG | >hg38_21_+_8433172_8446622_RNA45SN1_45S_50ntFlanks | 476 | 511 | 36 | TTCGTGATCGATGTGGTGACGTCGTGCTCTCCCGGG | MB231 | 81 | 195 | 20 | 2 | 23 | 44.8 | 89.6 | 14.5 | 0.8 | 6.4 |
| >hg38_21_+_8433172_8446622_RNA45SN1_45S_50ntFlanks@476.512.37\|TTCGTGATCGATGTGGTGACGTCGTGCTCTCCCGGGC | >hg38_21_+_8433172_8446622_RNA45SN1_45S_50ntFlanks | 476 | 512 | 37 | TTCGTGATCGATGTGGTGACGTCGTGCTCTCCCGGGC | BT20 | 19 | 40 | 9 | 2 | 9 | 15.4 | 18.9 | 3.2 | 2.4 | 7.8 |
| >hg38_21_+_8433172_8446622_RNA45SN1_45S_50ntFlanks@476.512.37\|TTCGTGATCGATGTGGTGACGTCGTGCTCTCCCGGGC | >hg38_21_+_8433172_8446622_RNA45SN1_45S_50ntFlanks | 476 | 512 | 37 | TTCGTGATCGATGTGGTGACGTCGTGCTCTCCCGGGC | MB231 | 37 | 106 | 11 | 1 | 6 | 25.1 | 39.1 | 9.5 | 0.8 | 5.5 |
| >hg38_21_+_8433172_8446622_RNA45SN1_45S_50ntFlanks@476.513.38\|TTCGTGATCGATGTGGTGACGTCGTGCTCTCCCGGGCC | >hg38_21_+_8433172_8446622_RNA45SN1_45S_50ntFlanks | 476 | 513 | 38 | TTCGTGATCGATGTGGTGACGTCGTGCTCTCCCGGGCC | BT20 | 13 | 29 | 5 | 1 | 3 | 12.0 | 14.7 | 1.6 | 1.4 | 5.1 |
| >hg38_21_+_8433172_8446622_RNA45SN1_45S_50ntFlanks@476.513.38\|TTCGTGATCGATGTGGTGACGTCGTGCTCTCCCGGGCC | >hg38_21_+_8433172_8446622_RNA45SN1_45S_50ntFlanks | 476 | 513 | 38 | TTCGTGATCGATGTGGTGACGTCGTGCTCTCCCGGGCC | MB231 | 24 | 77 | 6 | 0 | 0 | 18.1 | 37.8 | 5.5 | 0.0 | 0.0 |
| >hg38_21_+_8433172_8446622_RNA45SN1_45S_50ntFlanks@476.514.39\|TTCGTGATCGATGTGGTGACGTCGTGCTCTCCCGGGCCG | >hg38_21_+_8433172_8446622_RNA45SN1_45S_50ntFlanks | 476 | 514 | 39 | TTCGTGATCGATGTGGTGACGTCGTGCTCTCCCGGGCCG | BT20 | 11 | 24 | 4 | 1 | 2 | 6.2 | 14.2 | 2.2 | 0.7 | 2.2 |
| >hg38_21_+_8433172_8446622_RNA45SN1_45S_50ntFlanks@476.514.39\|TTCGTGATCGATGTGGTGACGTCGTGCTCTCCCGGGCCG | >hg38_21_+_8433172_8446622_RNA45SN1_45S_50ntFlanks | 476 | 514 | 39 | TTCGTGATCGATGTGGTGACGTCGTGCTCTCCCGGGCCG | MB231 | 17 | 46 | 2 | 0 | 3 | 5.5 | 28.1 | 1.5 | 0.6 | 5.2 |
| >hg38_21_+_8433172_8446622_RNA45SN1_45S_50ntFlanks@476.515.40\|TTCGTGATCGATGTGGTGACGTCGTGCTCTCCCGGGCCGG | >hg38_21_+_8433172_8446622_RNA45SN1_45S_50ntFlanks | 476 | 515 | 40 | TTCGTGATCGATGTGGTGACGTCGTGCTCTCCCGGGCCGG | BT20 | 11 | 20 | 1 | 1 | 2 | 4.4 | 12.3 | 0.6 | 0.5 | 1.8 |
| >hg38_21_+_8433172_8446622_RNA45SN1_45S_50ntFlanks@476.515.40\|TTCGTGATCGATGTGGTGACGTCGTGCTCTCCCGGGCCGG | >hg38_21_+_8433172_8446622_RNA45SN1_45S_50ntFlanks | 476 | 515 | 40 | TTCGTGATCGATGTGGTGACGTCGTGCTCTCCCGGGCCGG | MB231 | 28 | 52 | 2 | 1 | 5 | 8.5 | 28.3 | 1.1 | 1.4 | 6.2 |
| >hg38_21_+_8433172_8446622_RNA45SN1_45S_50ntFlanks@476.516.41\|TTCGTGATCGATGTGGTGACGTCGTGCTCTCCCGGGCCGGG | >hg38_21_+_8433172_8446622_RNA45SN1_45S_50ntFlanks | 476 | 516 | 41 | TTCGTGATCGATGTGGTGACGTCGTGCTCTCCCGGGCCGGG | BT20 | 72 | 188 | 10 | 6 | 32 | 93.4 | 106.5 | 2.8 | 8.3 | 27.9 |
| >hg38_21_+_8433172_8446622_RNA45SN1_45S_50ntFlanks@476.516.41\|TTCGTGATCGATGTGGTGACGTCGTGCTCTCCCGGGCCGGG | >hg38_21_+_8433172_8446622_RNA45SN1_45S_50ntFlanks | 476 | 516 | 41 | TTCGTGATCGATGTGGTGACGTCGTGCTCTCCCGGGCCGGG | MB231 | 179 | 786 | 37 | 3 | 62 | 104.5 | 431.9 | 28.5 | 2.1 | 12.4 |
| >hg38_21_+_8433172_8446622_RNA45SN1_45S_50ntFlanks@476.517.42\|TTCGTGATCGATGTGGTGACGTCGTGCTCTCCCGGGCCGGGT | >hg38_21_+_8433172_8446622_RNA45SN1_45S_50ntFlanks | 476 | 517 | 42 | TTCGTGATCGATGTGGTGACGTCGTGCTCTCCCGGGCCGGGT | BT20 | 45 | 142 | 6 | 5 | 31 | 62.1 | 94.2 | 2.8 | 5.6 | 27.2 |
| >hg38_21_+_8433172_8446622_RNA45SN1_45S_50ntFlanks@476.517.42\|TTCGTGATCGATGTGGTGACGTCGTGCTCTCCCGGGCCGGGT | >hg38_21_+_8433172_8446622_RNA45SN1_45S_50ntFlanks | 476 | 517 | 42 | TTCGTGATCGATGTGGTGACGTCGTGCTCTCCCGGGCCGGGT | MB231 | 96 | 553 | 30 | 4 | 49 | 94.2 | 358.1 | 38.8 | 3.2 | 18.9 |
| >hg38_21_+_8433172_8446622_RNA45SN1_45S_50ntFlanks@476.518.43\|TTCGTGATCGATGTGGTGACGTCGTGCTCTCCCGGGCCGGGTC | >hg38_21_+_8433172_8446622_RNA45SN1_45S_50ntFlanks | 476 | 518 | 43 | TTCGTGATCGATGTGGTGACGTCGTGCTCTCCCGGGCCGGGTC | BT20 | 22 | 80 | 4 | 6 | 25 | 28.8 | 55.4 | 2.5 | 5.4 | 7.0 |
| >hg38_21_+_8433172_8446622_RNA45SN1_45S_50ntFlanks@476.518.43\|TTCGTGATCGATGTGGTGACGTCGTGCTCTCCCGGGCCGGGTC | >hg38_21_+_8433172_8446622_RNA45SN1_45S_50ntFlanks | 476 | 518 | 43 | TTCGTGATCGATGTGGTGACGTCGTGCTCTCCCGGGCCGGGTC | MB231 | 63 | 343 | 21 | 3 | 13 | 64.9 | 235.8 | 22.2 | 3.8 | 10.9 |
| >hg38_21_+_8433172_8446622_RNA45SN1_45S_50ntFlanks@476.519.44\|TTCGTGATCGATGTGGTGACGTCGTGCTCTCCCGGGCCGGGTCC | >hg38_21_+_8433172_8446622_RNA45SN1_45S_50ntFlanks | 476 | 519 | 44 | TTCGTGATCGATGTGGTGACGTCGTGCTCTCCCGGGCCGGGTCC | MB231 | 20 | 93 | 3 | 0 | 2 | 13.5 | 52.7 | 1.3 | 0.6 | 4.2 |
| >hg38_21_+_8433172_8446622_RNA45SN1_45S_50ntFlanks@476.520.45\|TTCGTGATCGATGTGGTGACGTCGTGCTCTCCCGGGCCGGGTCCG | >hg38_21_+_8433172_8446622_RNA45SN1_45S_50ntFlanks | 476 | 520 | 45 | TTCGTGATCGATGTGGTGACGTCGTGCTCTCCCGGGCCGGGTCCG | BT20 | 17 | 47 | 2 | 1 | 7 | 18.8 | 38.2 | 1.1 | 0.6 | 6.7 |
| >hg38_21_+_8433172_8446622_RNA45SN1_45S_50ntFlanks@476.520.45\|TTCGTGATCGATGTGGTGACGTCGTGCTCTCCCGGGCCGGGTCCG | >hg38_21_+_8433172_8446622_RNA45SN1_45S_50ntFlanks | 476 | 520 | 45 | TTCGTGATCGATGTGGTGACGTCGTGCTCTCCCGGGCCGGGTCCG | MB231 | 49 | 207 | 5 | 1 | 15 | 21.6 | 112.6 | 2.4 | 1.0 | 14.1 |
| >hg38_21_+_8433172_8446622_RNA45SN1_45S_50ntFlanks@476.521.46\|TTCGTGATCGATGTGGTGACGTCGTGCTCTCCCGGGCCGGGTCCGA | >hg38_21_+_8433172_8446622_RNA45SN1_45S_50ntFlanks | 476 | 521 | 46 | TTCGTGATCGATGTGGTGACGTCGTGCTCTCCCGGGCCGGGTCCGA | BT20 | 17 | 62 | 4 | 1 | 15 | 13.1 | 63.0 | 4.7 | 0.6 | 8.1 |
| >hg38_21_+_8433172_8446622_RNA45SN1_45S_50ntFlanks@476.521.46\|TTCGTGATCGATGTGGTGACGTCGTGCTCTCCCGGGCCGGGTCCGA | >hg38_21_+_8433172_8446622_RNA45SN1_45S_50ntFlanks | 476 | 521 | 46 | TTCGTGATCGATGTGGTGACGTCGTGCTCTCCCGGGCCGGGTCCGA | MB231 | 40 | 166 | 4 | 1 | 25 | 10.8 | 99.4 | 1.8 | 1.0 | 23.2 |
| >hg38_21_+_8433172_8446622_RNA45SN1_45S_50ntFlanks@476.522.47\|TTCGTGATCGATGTGGTGACGTCGTGCTCTCCCGGGCCGGGTCCGAG | >hg38_21_+_8433172_8446622_RNA45SN1_45S_50ntFlanks | 476 | 522 | 47 | TTCGTGATCGATGTGGTGACGTCGTGCTCTCCCGGGCCGGGTCCGAG | MB231 | 23 | 169 | 3 | 0 | 0 | 12.7 | 209.5 | 2.7 | 0.0 | 0.0 |
| >hg38_21_+_8433172_8446622_RNA45SN1_45S_50ntFlanks@476.523.48\|TTCGTGATCGATGTGGTGACGTCGTGCTCTCCCGGGCCGGGTCCGAGC | >hg38_21_+_8433172_8446622_RNA45SN1_45S_50ntFlanks | 476 | 523 | 48 | TTCGTGATCGATGTGGTGACGTCGTGCTCTCCCGGGCCGGGTCCGAGC | BT20 | 10 | 59 | 1 | 1 | 17 | 8.3 | 69.4 | 1.4 | 0.2 | 14.7 |
| >hg38_21_+_8433172_8446622_RNA45SN1_45S_50ntFlanks@476.523.48\|TTCGTGATCGATGTGGTGACGTCGTGCTCTCCCGGGCCGGGTCCGAGC | >hg38_21_+_8433172_8446622_RNA45SN1_45S_50ntFlanks | 476 | 523 | 48 | TTCGTGATCGATGTGGTGACGTCGTGCTCTCCCGGGCCGGGTCCGAGC | MB231 | 62 | 635 | 6 | 1 | 28 | 46.8 | 901.0 | 4.4 | 0.9 | 31.0 |
| >hg38_21_+_8433172_8446622_RNA45SN1_45S_50ntFlanks@476.524.49\|TTCGTGATCGATGTGGTGACGTCGTGCTCTCCCGGGCCGGGTCCGAGCC | >hg38_21_+_8433172_8446622_RNA45SN1_45S_50ntFlanks | 476 | 524 | 49 | TTCGTGATCGATGTGGTGACGTCGTGCTCTCCCGGGCCGGGTCCGAGCC | MB231 | 38 | 497 | 6 | 0 | 14 | 28.1 | 763.3 | 6.4 | 0.4 | 15.3 |
| >hg38_21_+_8433172_8446622_RNA45SN1_45S_50ntFlanks@476.525.50\|TTCGTGATCGATGTGGTGACGTCGTGCTCTCCCGGGCCGGGTCCGAGCCG | >hg38_21_+_8433172_8446622_RNA45SN1_45S_50ntFlanks | 476 | 525 | 50 | TTCGTGATCGATGTGGTGACGTCGTGCTCTCCCGGGCCGGGTCCGAGCCG | MB231 | 38 | 118 | 1 | 0 | 0 | 27.9 | 156.9 | 0.1 | 0.0 | 0.0 |
| >hg38_21_+_8433172_8446622_RNA45SN1_45S_50ntFlanks@4760.4777.18\|TCAGATACCGTCGTAGTT | >hg38_21_+_8433172_8446622_RNA45SN1_45S_50ntFlanks | 4760 | 4777 | 18 | TCAGATACCGTCGTAGTT | MB468 | 16 | 4 | 13 | 12 | 6 | 5.0 | 2.6 | 8.2 | 5.0 | 2.9 |
| >hg38_21_+_8433172_8446622_RNA45SN1_45S_50ntFlanks@4760.4778.19\|TCAGATACCGTCGTAGTTC | >hg38_21_+_8433172_8446622_RNA45SN1_45S_50ntFlanks | 4760 | 4778 | 19 | TCAGATACCGTCGTAGTTC | MB468 | 64 | 40 | 53 | 34 | 30 | 19.9 | 15.0 | 17.3 | 11.4 | 13.6 |
| >hg38_21_+_8433172_8446622_RNA45SN1_45S_50ntFlanks@4760.4779.20\|TCAGATACCGTCGTAGTTCC | >hg38_21_+_8433172_8446622_RNA45SN1_45S_50ntFlanks | 4760 | 4779 | 20 | TCAGATACCGTCGTAGTTCC | MB468 | 38 | 33 | 38 | 7 | 16 | 10.4 | 12.7 | 12.2 | 5.5 | 7.9 |
| >hg38_21_+_8433172_8446622_RNA45SN1_45S_50ntFlanks@4760.4780.21\|TCAGATACCGTCGTAGTTCCG | >hg38_21_+_8433172_8446622_RNA45SN1_45S_50ntFlanks | 4760 | 4780 | 21 | TCAGATACCGTCGTAGTTCCG | MB468 | 443 | 281 | 418 | 129 | 196 | 110.0 | 105.4 | 166.0 | 86.4 | 103.6 |
| >hg38_21_+_8433172_8446622_RNA45SN1_45S_50ntFlanks@4760.4782.23\|TCAGATACCGTCGTAGTTCCGAC | >hg38_21_+_8433172_8446622_RNA45SN1_45S_50ntFlanks | 4760 | 4782 | 23 | TCAGATACCGTCGTAGTTCCGAC | MB468 | 115 | 33 | 107 | 84 | 116 | 27.1 | 14.8 | 47.1 | 20.3 | 156.0 |
| >hg38_21_+_8433172_8446622_RNA45SN1_45S_50ntFlanks@4760.4783.24\|TCAGATACCGTCGTAGTTCCGACC | >hg38_21_+_8433172_8446622_RNA45SN1_45S_50ntFlanks | 4760 | 4783 | 24 | TCAGATACCGTCGTAGTTCCGACC | MB468 | 67 | 26 | 53 | 45 | 63 | 21.8 | 10.2 | 18.0 | 24.4 | 67.7 |
| >hg38_21_+_8433172_8446622_RNA45SN1_45S_50ntFlanks@4760.4786.27\|TCAGATACCGTCGTAGTTCCGACCATA | >hg38_21_+_8433172_8446622_RNA45SN1_45S_50ntFlanks | 4760 | 4786 | 27 | TCAGATACCGTCGTAGTTCCGACCATA | MB468 | 42 | 7 | 25 | 36 | 39 | 9.2 | 3.8 | 10.5 | 10.8 | 31.1 |
| >hg38_21_+_8433172_8446622_RNA45SN1_45S_50ntFlanks@4760.4793.34\|TCAGATACCGTCGTAGTTCCGACCATAAACGATG | >hg38_21_+_8433172_8446622_RNA45SN1_45S_50ntFlanks | 4760 | 4793 | 34 | TCAGATACCGTCGTAGTTCCGACCATAAACGATG | MB468 | 16 | 31 | 37 | 12 | 14 | 6.0 | 17.0 | 13.7 | 6.0 | 4.6 |
| >hg38_21_+_8433172_8446622_RNA45SN1_45S_50ntFlanks@4761.4780.20\|CAGATACCGTCGTAGTTCCG | >hg38_21_+_8433172_8446622_RNA45SN1_45S_50ntFlanks | 4761 | 4780 | 20 | CAGATACCGTCGTAGTTCCG | MB468 | 27 | 7 | 22 | 13 | 11 | 9.5 | 3.6 | 8.7 | 7.1 | 3.1 |
| >hg38_21_+_8433172_8446622_RNA45SN1_45S_50ntFlanks@4762.4780.19\|AGATACCGTCGTAGTTCCG | >hg38_21_+_8433172_8446622_RNA45SN1_45S_50ntFlanks | 4762 | 4780 | 19 | AGATACCGTCGTAGTTCCG | MB468 | 69 | 31 | 55 | 18 | 30 | 15.7 | 15.7 | 13.2 | 10.8 | 7.2 |
| >hg38_21_+_8433172_8446622_RNA45SN1_45S_50ntFlanks@4763.4780.18\|GATACCGTCGTAGTTCCG | >hg38_21_+_8433172_8446622_RNA45SN1_45S_50ntFlanks | 4763 | 4780 | 18 | GATACCGTCGTAGTTCCG | MB468 | 89 | 36 | 102 | 19 | 35 | 26.9 | 20.3 | 46.4 | 12.3 | 4.9 |
| >hg38_21_+_8433172_8446622_RNA45SN1_45S_50ntFlanks@4764.4781.18\|ATACCGTCGTAGTTCCGA | >hg38_21_+_8433172_8446622_RNA45SN1_45S_50ntFlanks | 4764 | 4781 | 18 | ATACCGTCGTAGTTCCGA | MB468 | 13 | 3 | 18 | 4 | 4 | 5.0 | 0.8 | 8.5 | 1.2 | 1.3 |
| >hg38_21_+_8433172_8446622_RNA45SN1_45S_50ntFlanks@4770.4793.24\|TCGTAGTTCCGACCATAAACGATG | >hg38_21_+_8433172_8446622_RNA45SN1_45S_50ntFlanks | 4770 | 4793 | 24 | TCGTAGTTCCGACCATAAACGATG | MB468 | 24 | 19 | 42 | 16 | 14 | 11.8 | 11.5 | 13.8 | 8.9 | 2.8 |
| >hg38_21_+_8433172_8446622_RNA45SN1_45S_50ntFlanks@4772.4793.22\|GTAGTTCCGACCATAAACGATG | >hg38_21_+_8433172_8446622_RNA45SN1_45S_50ntFlanks | 4772 | 4793 | 22 | GTAGTTCCGACCATAAACGATG | MB468 | 23 | 26 | 32 | 11 | 12 | 9.6 | 16.0 | 13.0 | 7.7 | 2.7 |
| >hg38_21_+_8433172_8446622_RNA45SN1_45S_50ntFlanks@4773.4793.21\|TAGTTCCGACCATAAACGATG | >hg38_21_+_8433172_8446622_RNA45SN1_45S_50ntFlanks | 4773 | 4793 | 21 | TAGTTCCGACCATAAACGATG | MB468 | 199 | 173 | 174 | 58 | 56 | 109.3 | 153.0 | 66.7 | 29.6 | 18.6 |

| | | | | | | | | | | | | | | | | |
|---|---|---|---|---|---|---|---|---|---|---|---|---|---|---|---|---|
| >hg38_21_+_8433172_8446622_RNA45SN1_45S_50ntFlanks@4783.4800.18\|CATAAACGATGCCGACCG | >hg38_21_+_8433172_8446622_RNA45SN1_45S_50ntFlanks | 4783 | 4800 | 18 | CATAAACGATGCCGACCG | MB468 | 10 | 9 | 10 | 4 | 13 | 4.2 | 3.4 | 3.6 | 2.1 | 10.8 |
| >hg38_21_+_8433172_8446622_RNA45SN1_45S_50ntFlanks@481.505.25\|GATCGATGTGGTGACGTCGTGCTCT | >hg38_21_+_8433172_8446622_RNA45SN1_45S_50ntFlanks | 481 | 505 | 25 | GATCGATGTGGTGACGTCGTGCTCT | MB231 | 21 | 23 | 0 | 16 | 6 | 36.4 | 40.3 | 0.7 | 28.4 | 10.4 |
| >hg38_21_+_8433172_8446622_RNA45SN1_45S_50ntFlanks@482.504.23\|ATCGATGTGGTGACGTCGTGCTC | >hg38_21_+_8433172_8446622_RNA45SN1_45S_50ntFlanks | 482 | 504 | 23 | ATCGATGTGGTGACGTCGTGCTC | MB231 | 25 | 17 | 0 | 23 | 7 | 44.1 | 28.9 | 0.3 | 39.3 | 12.4 |
| >hg38_21_+_8433172_8446622_RNA45SN1_45S_50ntFlanks@482.505.24\|ATCGATGTGGTGACGTCGTGCTCT | >hg38_21_+_8433172_8446622_RNA45SN1_45S_50ntFlanks | 482 | 505 | 24 | ATCGATGTGGTGACGTCGTGCTCT | MB231 | 304 | 355 | 6 | 201 | 70 | 525.6 | 614.4 | 10.0 | 348.0 | 121.9 |
| >hg38_21_+_8433172_8446622_RNA45SN1_45S_50ntFlanks@482.505.24\|ATCGATGTGGTGACGTCGTGCTCT | >hg38_21_+_8433172_8446622_RNA45SN1_45S_50ntFlanks | 482 | 505 | 24 | ATCGATGTGGTGACGTCGTGCTCT | MB468 | 25 | 52 | 1 | 29 | 4 | 8.8 | 28.2 | 0.5 | 20.8 | 4.1 |
| >hg38_21_+_8433172_8446622_RNA45SN1_45S_50ntFlanks@482.506.25\|ATCGATGTGGTGACGTCGTGCTCTC | >hg38_21_+_8433172_8446622_RNA45SN1_45S_50ntFlanks | 482 | 506 | 25 | ATCGATGTGGTGACGTCGTGCTCTC | MB231 | 133 | 230 | 7 | 25 | 43 | 229.6 | 397.6 | 12.0 | 43.9 | 74.7 |
| >hg38_21_+_8433172_8446622_RNA45SN1_45S_50ntFlanks@482.506.25\|ATCGATGTGGTGACGTCGTGCTCTC | >hg38_21_+_8433172_8446622_RNA45SN1_45S_50ntFlanks | 482 | 506 | 25 | ATCGATGTGGTGACGTCGTGCTCTC | MB468 | 18 | 60 | 1 | 7 | 4 | 6.1 | 52.1 | 1.1 | 4.0 | 3.6 |
| >hg38_21_+_8433172_8446622_RNA45SN1_45S_50ntFlanks@482.507.26\|ATCGATGTGGTGACGTCGTGCTCTCC | >hg38_21_+_8433172_8446622_RNA45SN1_45S_50ntFlanks | 482 | 507 | 26 | ATCGATGTGGTGACGTCGTGCTCTCC | MB231 | 86 | 206 | 12 | 1 | 24 | 148.1 | 356.9 | 21.3 | 1.8 | 41.1 |
| >hg38_21_+_8433172_8446622_RNA45SN1_45S_50ntFlanks@482.508.27\|ATCGATGTGGTGACGTCGTGCTCTCCC | >hg38_21_+_8433172_8446622_RNA45SN1_45S_50ntFlanks | 482 | 508 | 27 | ATCGATGTGGTGACGTCGTGCTCTCCC | MB231 | 34 | 46 | 2 | 0 | 6 | 59.7 | 80.2 | 4.3 | 0.6 | 10.7 |
| >hg38_21_+_8433172_8446622_RNA45SN1_45S_50ntFlanks@482.509.28\|ATCGATGTGGTGACGTCGTGCTCTCCCG | >hg38_21_+_8433172_8446622_RNA45SN1_45S_50ntFlanks | 482 | 509 | 28 | ATCGATGTGGTGACGTCGTGCTCTCCCG | MB231 | 53 | 47 | 2 | 0 | 6 | 91.5 | 80.7 | 3.6 | 0.8 | 11.1 |
| >hg38_21_+_8433172_8446622_RNA45SN1_45S_50ntFlanks@482.510.29\|ATCGATGTGGTGACGTCGTGCTCTCCCGG | >hg38_21_+_8433172_8446622_RNA45SN1_45S_50ntFlanks | 482 | 510 | 29 | ATCGATGTGGTGACGTCGTGCTCTCCCGG | MB231 | 46 | 43 | 2 | 0 | 6 | 80.0 | 74.6 | 2.8 | 0.6 | 10.8 |
| >hg38_21_+_8433172_8446622_RNA45SN1_45S_50ntFlanks@482.511.30\|ATCGATGTGGTGACGTCGTGCTCTCCCGGG | >hg38_21_+_8433172_8446622_RNA45SN1_45S_50ntFlanks | 482 | 511 | 30 | ATCGATGTGGTGACGTCGTGCTCTCCCGGG | MB231 | 19 | 24 | 1 | 1 | 4 | 33.5 | 42.2 | 1.6 | 1.8 | 6.9 |
| >hg38_21_+_8433172_8446622_RNA45SN1_45S_50ntFlanks@482.514.33\|ATCGATGTGGTGACGTCGTGCTCTCCCGGGCCG | >hg38_21_+_8433172_8446622_RNA45SN1_45S_50ntFlanks | 482 | 514 | 33 | ATCGATGTGGTGACGTCGTGCTCTCCCGGGCCG | MB231 | 24 | 59 | 4 | 0 | 0 | 41.2 | 101.6 | 6.1 | 0.0 | 0.0 |
| >hg38_21_+_8433172_8446622_RNA45SN1_45S_50ntFlanks@482.515.34\|ATCGATGTGGTGACGTCGTGCTCTCCCGGGCCGG | >hg38_21_+_8433172_8446622_RNA45SN1_45S_50ntFlanks | 482 | 515 | 34 | ATCGATGTGGTGACGTCGTGCTCTCCCGGGCCGG | MB231 | 38 | 70 | 2 | 0 | 6 | 66.3 | 121.8 | 2.9 | 0.5 | 10.2 |
| >hg38_21_+_8433172_8446622_RNA45SN1_45S_50ntFlanks@482.516.35\|ATCGATGTGGTGACGTCGTGCTCTCCCGGGCCGGG | >hg38_21_+_8433172_8446622_RNA45SN1_45S_50ntFlanks | 482 | 516 | 35 | ATCGATGTGGTGACGTCGTGCTCTCCCGGGCCGGG | MB231 | 35 | 56 | 1 | 1 | 9 | 60.5 | 97.8 | 2.4 | 2.4 | 15.0 |
| >hg38_21_+_8433172_8446622_RNA45SN1_45S_50ntFlanks@482.517.36\|ATCGATGTGGTGACGTCGTGCTCTCCCGGGCCGGGT | >hg38_21_+_8433172_8446622_RNA45SN1_45S_50ntFlanks | 482 | 517 | 36 | ATCGATGTGGTGACGTCGTGCTCTCCCGGGCCGGGT | MB231 | 22 | 62 | 2 | 1 | 10 | 38.6 | 107.2 | 4.1 | 2.6 | 17.9 |
| >hg38_21_+_8433172_8446622_RNA45SN1_45S_50ntFlanks@482.518.37\|ATCGATGTGGTGACGTCGTGCTCTCCCGGGCCGGGTC | >hg38_21_+_8433172_8446622_RNA45SN1_45S_50ntFlanks | 482 | 518 | 37 | ATCGATGTGGTGACGTCGTGCTCTCCCGGGCCGGGTC | MB231 | 15 | 34 | 1 | 2 | 7 | 25.4 | 59.3 | 1.7 | 3.5 | 12.6 |
| >hg38_21_+_8433172_8446622_RNA45SN1_45S_50ntFlanks@482.519.38\|ATCGATGTGGTGACGTCGTGCTCTCCCGGGCCGGGTCC | >hg38_21_+_8433172_8446622_RNA45SN1_45S_50ntFlanks | 482 | 519 | 38 | ATCGATGTGGTGACGTCGTGCTCTCCCGGGCCGGGTCC | MB231 | 18 | 40 | 1 | 1 | 3 | 30.3 | 69.2 | 1.2 | 1.1 | 5.8 |
| >hg38_21_+_8433172_8446622_RNA45SN1_45S_50ntFlanks@482.520.39\|ATCGATGTGGTGACGTCGTGCTCTCCCGGGCCGGGTCCG | >hg38_21_+_8433172_8446622_RNA45SN1_45S_50ntFlanks | 482 | 520 | 39 | ATCGATGTGGTGACGTCGTGCTCTCCCGGGCCGGGTCCG | MB231 | 47 | 132 | 3 | 1 | 11 | 80.9 | 228.0 | 4.6 | 2.4 | 19.6 |
| >hg38_21_+_8433172_8446622_RNA45SN1_45S_50ntFlanks@482.521.40\|ATCGATGTGGTGACGTCGTGCTCTCCCGGGCCGGGTCCGA | >hg38_21_+_8433172_8446622_RNA45SN1_45S_50ntFlanks | 482 | 521 | 40 | ATCGATGTGGTGACGTCGTGCTCTCCCGGGCCGGGTCCGA | MB231 | 37 | 125 | 3 | 2 | 17 | 64.0 | 216.5 | 5.0 | 2.9 | 30.2 |
| >hg38_21_+_8433172_8446622_RNA45SN1_45S_50ntFlanks@482.522.41\|ATCGATGTGGTGACGTCGTGCTCTCCCGGGCCGGGTCCGAG | >hg38_21_+_8433172_8446622_RNA45SN1_45S_50ntFlanks | 482 | 522 | 41 | ATCGATGTGGTGACGTCGTGCTCTCCCGGGCCGGGTCCGAG | MB231 | 38 | 199 | 4 | 2 | 12 | 66.5 | 344.4 | 7.2 | 2.9 | 20.6 |
| >hg38_21_+_8433172_8446622_RNA45SN1_45S_50ntFlanks@482.524.43\|ATCGATGTGGTGACGTCGTGCTCTCCCGGGCCGGGTCCGAGCC | >hg38_21_+_8433172_8446622_RNA45SN1_45S_50ntFlanks | 482 | 524 | 43 | ATCGATGTGGTGACGTCGTGCTCTCCCGGGCCGGGTCCGAGCC | MB231 | 31 | 239 | 2 | 5 | 10 | 54.4 | 414.6 | 2.7 | 8.8 | 17.2 |
| >hg38_21_+_8433172_8446622_RNA45SN1_45S_50ntFlanks@482.525.44\|ATCGATGTGGTGACGTCGTGCTCTCCCGGGCCGGGTCCGAGCCG | >hg38_21_+_8433172_8446622_RNA45SN1_45S_50ntFlanks | 482 | 525 | 44 | ATCGATGTGGTGACGTCGTGCTCTCCCGGGCCGGGTCCGAGCCG | MB231 | 29 | 164 | 1 | 1 | 10 | 50.6 | 284.2 | 1.8 | 2.5 | 16.8 |
| >hg38_21_+_8433172_8446622_RNA45SN1_45S_50ntFlanks@482.526.45\|ATCGATGTGGTGACGTCGTGCTCTCCCGGGCCGGGTCCGAGCCGC | >hg38_21_+_8433172_8446622_RNA45SN1_45S_50ntFlanks | 482 | 526 | 45 | ATCGATGTGGTGACGTCGTGCTCTCCCGGGCCGGGTCCGAGCCGC | MB231 | 10 | 99 | 0 | 1 | 6 | 18.1 | 171.1 | 0.7 | 2.0 | 9.6 |
| >hg38_21_+_8433172_8446622_RNA45SN1_45S_50ntFlanks@482.527.46\|ATCGATGTGGTGACGTCGTGCTCTCCCGGGCCGGGTCCGAGCCGCG | >hg38_21_+_8433172_8446622_RNA45SN1_45S_50ntFlanks | 482 | 527 | 46 | ATCGATGTGGTGACGTCGTGCTCTCCCGGGCCGGGTCCGAGCCGCG | MB231 | 19 | 140 | 1 | 2 | 10 | 32.5 | 242.5 | 1.3 | 4.1 | 18.1 |
| >hg38_21_+_8433172_8446622_RNA45SN1_45S_50ntFlanks@482.529.48\|ATCGATGTGGTGACGTCGTGCTCTCCCGGGCCGGGTCCGAGCCGCGAC | >hg38_21_+_8433172_8446622_RNA45SN1_45S_50ntFlanks | 482 | 529 | 48 | ATCGATGTGGTGACGTCGTGCTCTCCCGGGCCGGGTCCGAGCCGCGAC | MB231 | 21 | 177 | 0 | 1 | 7 | 35.6 | 306.8 | 0.6 | 2.2 | 12.4 |
| >hg38_21_+_8433172_8446622_RNA45SN1_45S_50ntFlanks@482.530.49\|ATCGATGTGGTGACGTCGTGCTCTCCCGGGCCGGGTCCGAGCCGCGACG | >hg38_21_+_8433172_8446622_RNA45SN1_45S_50ntFlanks | 482 | 530 | 49 | ATCGATGTGGTGACGTCGTGCTCTCCCGGGCCGGGTCCGAGCCGCGACG | MB231 | 23 | 220 | 0 | 2 | 5 | 39.8 | 381.5 | 0.3 | 2.7 | 8.8 |
| >hg38_21_+_8433172_8446622_RNA45SN1_45S_50ntFlanks@482.531.50\|ATCGATGTGGTGACGTCGTGCTCTCCCGGGCCGGGTCCGAGCCGCGACGG | >hg38_21_+_8433172_8446622_RNA45SN1_45S_50ntFlanks | 482 | 531 | 50 | ATCGATGTGGTGACGTCGTGCTCTCCCGGGCCGGGTCCGAGCCGCGACGG | MB231 | 27 | 299 | 0 | 1 | 6 | 46.7 | 517.7 | 0.6 | 2.4 | 9.8 |
| >hg38_21_+_8433172_8446622_RNA45SN1_45S_50ntFlanks@4823.4851.29\|ATGACCCGCCGGGCAGCTTCCGGGAAACC | >hg38_21_+_8433172_8446622_RNA45SN1_45S_50ntFlanks | 4823 | 4851 | 29 | ATGACCCGCCGGGCAGCTTCCGGGAAACC | MB231 | 20 | 9 | 20 | 0 | 2 | 16.1 | 7.7 | 17.7 | 0.4 | 4.2 |
| >hg38_21_+_8433172_8446622_RNA45SN1_45S_50ntFlanks@4823.4857.35\|ATGACCCGCCGGGCAGCTTCCGGGAAACCAAAGTC | >hg38_21_+_8433172_8446622_RNA45SN1_45S_50ntFlanks | 4823 | 4857 | 35 | ATGACCCGCCGGGCAGCTTCCGGGAAACCAAAGTC | MB231 | 10 | 3 | 16 | 0 | 1 | 10.2 | 2.2 | 17.9 | 0.3 | 0.9 |
| >hg38_21_+_8433172_8446622_RNA45SN1_45S_50ntFlanks@4823.4859.37\|ATGACCCGCCGGGCAGCTTCCGGGAAACCAAAGTCTT | >hg38_21_+_8433172_8446622_RNA45SN1_45S_50ntFlanks | 4823 | 4859 | 37 | ATGACCCGCCGGGCAGCTTCCGGGAAACCAAAGTCTT | MB231 | 12 | 3 | 12 | 0 | 0 | 12.9 | 2.9 | 15.6 | 0.0 | 0.0 |
| >hg38_21_+_8433172_8446622_RNA45SN1_45S_50ntFlanks@483.505.23\|TCGATGTGGTGACGTCGTGCTCT | >hg38_21_+_8433172_8446622_RNA45SN1_45S_50ntFlanks | 483 | 505 | 23 | TCGATGTGGTGACGTCGTGCTCT | BT20 | 45 | 91 | 5 | 11 | 7 | 22.7 | 98.8 | 2.4 | 8.0 | 4.0 |
| >hg38_21_+_8433172_8446622_RNA45SN1_45S_50ntFlanks@483.505.23\|TCGATGTGGTGACGTCGTGCTCT | >hg38_21_+_8433172_8446622_RNA45SN1_45S_50ntFlanks | 483 | 505 | 23 | TCGATGTGGTGACGTCGTGCTCT | MB231 | 41 | 66 | 2 | 12 | 11 | 14.6 | 18.6 | 0.9 | 1.3 | 6.2 |
| >hg38_21_+_8433172_8446622_RNA45SN1_45S_50ntFlanks@483.506.24\|TCGATGTGGTGACGTCGTGCTCTC | >hg38_21_+_8433172_8446622_RNA45SN1_45S_50ntFlanks | 483 | 506 | 24 | TCGATGTGGTGACGTCGTGCTCTC | BT20 | 17 | 34 | 6 | 2 | 4 | 8.3 | 34.1 | 3.2 | 2.1 | 3.6 |
| >hg38_21_+_8433172_8446622_RNA45SN1_45S_50ntFlanks@483.506.24\|TCGATGTGGTGACGTCGTGCTCTC | >hg38_21_+_8433172_8446622_RNA45SN1_45S_50ntFlanks | 483 | 506 | 24 | TCGATGTGGTGACGTCGTGCTCTC | MB231 | 20 | 47 | 3 | 2 | 5 | 7.8 | 20.3 | 1.9 | 0.6 | 1.8 |
| >hg38_21_+_8433172_8446622_RNA45SN1_45S_50ntFlanks@483.507.25\|TCGATGTGGTGACGTCGTGCTCTCC | >hg38_21_+_8433172_8446622_RNA45SN1_45S_50ntFlanks | 483 | 507 | 25 | TCGATGTGGTGACGTCGTGCTCTCC | BT20 | 11 | 19 | 3 | 0 | 1 | 7.1 | 10.7 | 0.5 | 0.4 | 1.6 |
| >hg38_21_+_8433172_8446622_RNA45SN1_45S_50ntFlanks@483.507.25\|TCGATGTGGTGACGTCGTGCTCTCC | >hg38_21_+_8433172_8446622_RNA45SN1_45S_50ntFlanks | 483 | 507 | 25 | TCGATGTGGTGACGTCGTGCTCTCC | MB231 | 12 | 34 | 2 | 0 | 0 | 4.1 | 10.9 | 0.8 | 0.0 | 0.0 |
| >hg38_21_+_8433172_8446622_RNA45SN1_45S_50ntFlanks@483.529.47\|TCGATGTGGTGACGTCGTGCTCTCCCGGGCCGGGTCCGAGCCGCGAC | >hg38_21_+_8433172_8446622_RNA45SN1_45S_50ntFlanks | 483 | 529 | 47 | TCGATGTGGTGACGTCGTGCTCTCCCGGGCCGGGTCCGAGCCGCGAC | MB231 | 11 | 61 | 0 | 1 | 1 | 6.6 | 55.1 | 0.2 | 0.9 | 2.4 |
| >hg38_21_+_8433172_8446622_RNA45SN1_45S_50ntFlanks@483.532.50\|TCGATGTGGTGACGTCGTGCTCTCCCGGGCCGGGTCCGAGCCGCGACGGG | >hg38_21_+_8433172_8446622_RNA45SN1_45S_50ntFlanks | 483 | 532 | 50 | TCGATGTGGTGACGTCGTGCTCTCCCGGGCCGGGTCCGAGCCGCGACGGG | BT20 | 20 | 61 | 2 | 2 | 1 | 27.0 | 75.8 | 3.8 | 1.8 | 0.8 |
| >hg38_21_+_8433172_8446622_RNA45SN1_45S_50ntFlanks@483.532.50\|TCGATGTGGTGACGTCGTGCTCTCCCGGGCCGGGTCCGAGCCGCGACGGG | >hg38_21_+_8433172_8446622_RNA45SN1_45S_50ntFlanks | 483 | 532 | 50 | TCGATGTGGTGACGTCGTGCTCTCCCGGGCCGGGTCCGAGCCGCGACGGG | MB231 | 78 | 254 | 2 | 1 | 9 | 68.0 | 360.3 | 0.3 | 1.0 | 7.7 |
| >hg38_21_+_8433172_8446622_RNA45SN1_45S_50ntFlanks@485.503.19\|GATGTGGTGACGTCGTGCT | >hg38_21_+_8433172_8446622_RNA45SN1_45S_50ntFlanks | 485 | 503 | 19 | GATGTGGTGACGTCGTGCT | BT20 | 10 | 11 | 0 | 2 | 2 | 7.3 | 7.5 | 0.4 | 1.5 | 1.1 |
| >hg38_21_+_8433172_8446622_RNA45SN1_45S_50ntFlanks@485.504.20\|GATGTGGTGACGTCGTGCTC | >hg38_21_+_8433172_8446622_RNA45SN1_45S_50ntFlanks | 485 | 504 | 20 | GATGTGGTGACGTCGTGCTC | BT20 | 53 | 65 | 5 | 43 | 16 | 41.1 | 49.4 | 2.4 | 33.7 | 11.1 |
| >hg38_21_+_8433172_8446622_RNA45SN1_45S_50ntFlanks@485.504.20\|GATGTGGTGACGTCGTGCTC | >hg38_21_+_8433172_8446622_RNA45SN1_45S_50ntFlanks | 485 | 504 | 20 | GATGTGGTGACGTCGTGCTC | MB231 | 25 | 101 | 2 | 48 | 20 | 25.0 | 87.4 | 1.4 | 42.9 | 20.5 |
| >hg38_21_+_8433172_8446622_RNA45SN1_45S_50ntFlanks@485.505.21\|GATGTGGTGACGTCGTGCTCT | >hg38_21_+_8433172_8446622_RNA45SN1_45S_50ntFlanks | 485 | 505 | 21 | GATGTGGTGACGTCGTGCTCT | BT20 | 804 | 951 | 63 | 369 | 146 | 706.2 | 669.4 | 42.9 | 196.4 | 82.5 |
| >hg38_21_+_8433172_8446622_RNA45SN1_45S_50ntFlanks@485.505.21\|GATGTGGTGACGTCGTGCTCT | >hg38_21_+_8433172_8446622_RNA45SN1_45S_50ntFlanks | 485 | 505 | 21 | GATGTGGTGACGTCGTGCTCT | MB231 | 510 | 1223 | 18 | 285 | 200 | 521.3 | 1089.4 | 18.1 | 246.9 | 204.3 |
| >hg38_21_+_8433172_8446622_RNA45SN1_45S_50ntFlanks@485.506.22\|GATGTGGTGACGTCGTGCTCTC | >hg38_21_+_8433172_8446622_RNA45SN1_45S_50ntFlanks | 485 | 506 | 22 | GATGTGGTGACGTCGTGCTCTC | BT20 | 292 | 488 | 73 | 59 | 96 | 174.9 | 322.0 | 42.3 | 35.8 | 49.3 |
| >hg38_21_+_8433172_8446622_RNA45SN1_45S_50ntFlanks@485.506.22\|GATGTGGTGACGTCGTGCTCTC | >hg38_21_+_8433172_8446622_RNA45SN1_45S_50ntFlanks | 485 | 506 | 22 | GATGTGGTGACGTCGTGCTCTC | MB231 | 262 | 790 | 23 | 76 | 100 | 263.9 | 746.9 | 25.5 | 66.3 | 93.2 |
| >hg38_21_+_8433172_8446622_RNA45SN1_45S_50ntFlanks@485.507.23\|GATGTGGTGACGTCGTGCTCTCC | >hg38_21_+_8433172_8446622_RNA45SN1_45S_50ntFlanks | 485 | 507 | 23 | GATGTGGTGACGTCGTGCTCTCC | BT20 | 151 | 265 | 28 | 6 | 45 | 84.7 | 135.6 | 6.9 | 4.7 | 25.1 |
| >hg38_21_+_8433172_8446622_RNA45SN1_45S_50ntFlanks@485.507.23\|GATGTGGTGACGTCGTGCTCTCC | >hg38_21_+_8433172_8446622_RNA45SN1_45S_50ntFlanks | 485 | 507 | 23 | GATGTGGTGACGTCGTGCTCTCC | MB231 | 177 | 543 | 22 | 3 | 63 | 174.0 | 559.2 | 22.9 | 2.8 | 58.0 |
| >hg38_21_+_8433172_8446622_RNA45SN1_45S_50ntFlanks@485.508.24\|GATGTGGTGACGTCGTGCTCTCCC | >hg38_21_+_8433172_8446622_RNA45SN1_45S_50ntFlanks | 485 | 508 | 24 | GATGTGGTGACGTCGTGCTCTCCC | BT20 | 50 | 75 | 7 | 14 | 13 | 26.7 | 33.6 | 1.8 | 17.4 | 11.8 |

| | | | | | | | | | | | | | | | | |
|---|---|---|---|---|---|---|---|---|---|---|---|---|---|---|---|---|
| >hg38_21_+_8433172_8446622_RNA45SN1_45S_50ntFlanks@485.508.24\|GATGTGGTGACGTCGTGCTCTCCC | >hg38_21_+_8433172_8446622_RNA45SN1_45S_50ntFlanks | 485 | 508 | 24 | GATGTGGTGACGTCGTGCTCTCCC | MB231 | 47 | 143 | 4 | 4 | 19 | 46.1 | 136.7 | 4.0 | 4.0 | 17.3 |
| >hg38_21_+_8433172_8446622_RNA45SN1_45S_50ntFlanks@485.509.25\|GATGTGGTGACGTCGTGCTCTCCCG | >hg38_21_+_8433172_8446622_RNA45SN1_45S_50ntFlanks | 485 | 509 | 25 | GATGTGGTGACGTCGTGCTCTCCCG | BT20 | 67 | 85 | 6 | 2 | 18 | 37.6 | 49.6 | 2.5 | 2.2 | 16.8 |
| >hg38_21_+_8433172_8446622_RNA45SN1_45S_50ntFlanks@485.509.25\|GATGTGGTGACGTCGTGCTCTCCCG | >hg38_21_+_8433172_8446622_RNA45SN1_45S_50ntFlanks | 485 | 509 | 25 | GATGTGGTGACGTCGTGCTCTCCCG | MB231 | 61 | 170 | 3 | 3 | 23 | 56.2 | 162.4 | 2.8 | 2.5 | 20.3 |
| >hg38_21_+_8433172_8446622_RNA45SN1_45S_50ntFlanks@485.510.26\|GATGTGGTGACGTCGTGCTCTCCCGG | >hg38_21_+_8433172_8446622_RNA45SN1_45S_50ntFlanks | 485 | 510 | 26 | GATGTGGTGACGTCGTGCTCTCCCGG | BT20 | 62 | 90 | 9 | 5 | 23 | 40.3 | 52.6 | 2.7 | 4.3 | 21.2 |
| >hg38_21_+_8433172_8446622_RNA45SN1_45S_50ntFlanks@485.510.26\|GATGTGGTGACGTCGTGCTCTCCCGG | >hg38_21_+_8433172_8446622_RNA45SN1_45S_50ntFlanks | 485 | 510 | 26 | GATGTGGTGACGTCGTGCTCTCCCGG | MB231 | 51 | 157 | 3 | 5 | 22 | 48.4 | 157.3 | 3.4 | 4.1 | 18.9 |
| >hg38_21_+_8433172_8446622_RNA45SN1_45S_50ntFlanks@485.511.27\|GATGTGGTGACGTCGTGCTCTCCCGGG | >hg38_21_+_8433172_8446622_RNA45SN1_45S_50ntFlanks | 485 | 511 | 27 | GATGTGGTGACGTCGTGCTCTCCCGGG | BT20 | 36 | 67 | 9 | 6 | 16 | 21.9 | 32.5 | 4.2 | 3.9 | 6.2 |
| >hg38_21_+_8433172_8446622_RNA45SN1_45S_50ntFlanks@485.511.27\|GATGTGGTGACGTCGTGCTCTCCCGGG | >hg38_21_+_8433172_8446622_RNA45SN1_45S_50ntFlanks | 485 | 511 | 27 | GATGTGGTGACGTCGTGCTCTCCCGGG | MB231 | 30 | 92 | 3 | 6 | 11 | 28.5 | 93.5 | 2.5 | 5.3 | 9.2 |
| >hg38_21_+_8433172_8446622_RNA45SN1_45S_50ntFlanks@485.514.30\|GATGTGGTGACGTCGTGCTCTCCCGGGCCG | >hg38_21_+_8433172_8446622_RNA45SN1_45S_50ntFlanks | 485 | 514 | 30 | GATGTGGTGACGTCGTGCTCTCCCGGGCCG | BT20 | 42 | 103 | 16 | 4 | 19 | 40.9 | 56.1 | 3.4 | 3.9 | 16.3 |
| >hg38_21_+_8433172_8446622_RNA45SN1_45S_50ntFlanks@485.514.30\|GATGTGGTGACGTCGTGCTCTCCCGGGCCG | >hg38_21_+_8433172_8446622_RNA45SN1_45S_50ntFlanks | 485 | 514 | 30 | GATGTGGTGACGTCGTGCTCTCCCGGGCCG | MB231 | 24 | 161 | 6 | 1 | 10 | 22.3 | 184.3 | 5.9 | 1.1 | 9.0 |
| >hg38_21_+_8433172_8446622_RNA45SN1_45S_50ntFlanks@485.515.31\|GATGTGGTGACGTCGTGCTCTCCCGGGCCGG | >hg38_21_+_8433172_8446622_RNA45SN1_45S_50ntFlanks | 485 | 515 | 31 | GATGTGGTGACGTCGTGCTCTCCCGGGCCGG | BT20 | 58 | 105 | 6 | 4 | 23 | 66.0 | 47.5 | 2.8 | 2.6 | 9.9 |
| >hg38_21_+_8433172_8446622_RNA45SN1_45S_50ntFlanks@485.515.31\|GATGTGGTGACGTCGTGCTCTCCCGGGCCGG | >hg38_21_+_8433172_8446622_RNA45SN1_45S_50ntFlanks | 485 | 515 | 31 | GATGTGGTGACGTCGTGCTCTCCCGGGCCGG | MB231 | 47 | 240 | 5 | 5 | 17 | 48.2 | 258.2 | 4.2 | 4.6 | 14.4 |
| >hg38_21_+_8433172_8446622_RNA45SN1_45S_50ntFlanks@485.516.32\|GATGTGGTGACGTCGTGCTCTCCCGGGCCGGG | >hg38_21_+_8433172_8446622_RNA45SN1_45S_50ntFlanks | 485 | 516 | 32 | GATGTGGTGACGTCGTGCTCTCCCGGGCCGGG | BT20 | 42 | 118 | 5 | 14 | 23 | 54.8 | 70.7 | 2.3 | 10.7 | 13.1 |
| >hg38_21_+_8433172_8446622_RNA45SN1_45S_50ntFlanks@485.516.32\|GATGTGGTGACGTCGTGCTCTCCCGGGCCGGG | >hg38_21_+_8433172_8446622_RNA45SN1_45S_50ntFlanks | 485 | 516 | 32 | GATGTGGTGACGTCGTGCTCTCCCGGGCCGGG | MB231 | 39 | 324 | 5 | 13 | 24 | 37.7 | 331.5 | 4.1 | 11.6 | 21.1 |
| >hg38_21_+_8433172_8446622_RNA45SN1_45S_50ntFlanks@485.517.33\|GATGTGGTGACGTCGTGCTCTCCCGGGCCGGGT | >hg38_21_+_8433172_8446622_RNA45SN1_45S_50ntFlanks | 485 | 517 | 33 | GATGTGGTGACGTCGTGCTCTCCCGGGCCGGGT | BT20 | 39 | 160 | 6 | 24 | 42 | 51.9 | 96.7 | 2.6 | 22.5 | 14.5 |
| >hg38_21_+_8433172_8446622_RNA45SN1_45S_50ntFlanks@485.517.33\|GATGTGGTGACGTCGTGCTCTCCCGGGCCGGGT | >hg38_21_+_8433172_8446622_RNA45SN1_45S_50ntFlanks | 485 | 517 | 33 | GATGTGGTGACGTCGTGCTCTCCCGGGCCGGGT | MB231 | 20 | 258 | 2 | 20 | 25 | 21.0 | 260.8 | 2.2 | 17.0 | 21.7 |
| >hg38_21_+_8433172_8446622_RNA45SN1_45S_50ntFlanks@485.518.34\|GATGTGGTGACGTCGTGCTCTCCCGGGCCGGGTC | >hg38_21_+_8433172_8446622_RNA45SN1_45S_50ntFlanks | 485 | 518 | 34 | GATGTGGTGACGTCGTGCTCTCCCGGGCCGGGTC | BT20 | 20 | 86 | 3 | 14 | 14 | 23.4 | 67.5 | 1.6 | 13.6 | 8.2 |
| >hg38_21_+_8433172_8446622_RNA45SN1_45S_50ntFlanks@485.518.34\|GATGTGGTGACGTCGTGCTCTCCCGGGCCGGGTC | >hg38_21_+_8433172_8446622_RNA45SN1_45S_50ntFlanks | 485 | 518 | 34 | GATGTGGTGACGTCGTGCTCTCCCGGGCCGGGTC | MB231 | 13 | 125 | 2 | 11 | 14 | 14.1 | 133.5 | 1.9 | 9.8 | 13.1 |
| >hg38_21_+_8433172_8446622_RNA45SN1_45S_50ntFlanks@485.519.35\|GATGTGGTGACGTCGTGCTCTCCCGGGCCGGGTCC | >hg38_21_+_8433172_8446622_RNA45SN1_45S_50ntFlanks | 485 | 519 | 35 | GATGTGGTGACGTCGTGCTCTCCCGGGCCGGGTCC | BT20 | 12 | 44 | 1 | 5 | 5 | 15.2 | 25.7 | 0.6 | 5.8 | 5.0 |
| >hg38_21_+_8433172_8446622_RNA45SN1_45S_50ntFlanks@485.519.35\|GATGTGGTGACGTCGTGCTCTCCCGGGCCGGGTCC | >hg38_21_+_8433172_8446622_RNA45SN1_45S_50ntFlanks | 485 | 519 | 35 | GATGTGGTGACGTCGTGCTCTCCCGGGCCGGGTCC | MB231 | 16 | 154 | 1 | 5 | 10 | 16.7 | 159.2 | 0.9 | 4.4 | 9.1 |
| >hg38_21_+_8433172_8446622_RNA45SN1_45S_50ntFlanks@485.520.36\|GATGTGGTGACGTCGTGCTCTCCCGGGCCGGGTCCG | >hg38_21_+_8433172_8446622_RNA45SN1_45S_50ntFlanks | 485 | 520 | 36 | GATGTGGTGACGTCGTGCTCTCCCGGGCCGGGTCCG | BT20 | 44 | 116 | 4 | 7 | 18 | 53.2 | 73.5 | 0.6 | 7.7 | 17.0 |
| >hg38_21_+_8433172_8446622_RNA45SN1_45S_50ntFlanks@485.520.36\|GATGTGGTGACGTCGTGCTCTCCCGGGCCGGGTCCG | >hg38_21_+_8433172_8446622_RNA45SN1_45S_50ntFlanks | 485 | 520 | 36 | GATGTGGTGACGTCGTGCTCTCCCGGGCCGGGTCCG | MB231 | 43 | 441 | 3 | 8 | 24 | 38.7 | 387.0 | 2.2 | 7.5 | 20.6 |
| >hg38_21_+_8433172_8446622_RNA45SN1_45S_50ntFlanks@485.521.37\|GATGTGGTGACGTCGTGCTCTCCCGGGCCGGGTCCGA | >hg38_21_+_8433172_8446622_RNA45SN1_45S_50ntFlanks | 485 | 521 | 37 | GATGTGGTGACGTCGTGCTCTCCCGGGCCGGGTCCGA | BT20 | 56 | 195 | 12 | 10 | 30 | 47.2 | 131.1 | 6.0 | 6.8 | 9.2 |
| >hg38_21_+_8433172_8446622_RNA45SN1_45S_50ntFlanks@485.521.37\|GATGTGGTGACGTCGTGCTCTCCCGGGCCGGGTCCGA | >hg38_21_+_8433172_8446622_RNA45SN1_45S_50ntFlanks | 485 | 521 | 37 | GATGTGGTGACGTCGTGCTCTCCCGGGCCGGGTCCGA | MB231 | 44 | 403 | 5 | 9 | 24 | 38.1 | 350.4 | 4.4 | 8.1 | 21.9 |
| >hg38_21_+_8433172_8446622_RNA45SN1_45S_50ntFlanks@485.522.38\|GATGTGGTGACGTCGTGCTCTCCCGGGCCGGGTCCGAG | >hg38_21_+_8433172_8446622_RNA45SN1_45S_50ntFlanks | 485 | 522 | 38 | GATGTGGTGACGTCGTGCTCTCCCGGGCCGGGTCCGAG | BT20 | 41 | 208 | 5 | 10 | 23 | 44.5 | 161.3 | 2.5 | 7.4 | 10.8 |
| >hg38_21_+_8433172_8446622_RNA45SN1_45S_50ntFlanks@485.522.38\|GATGTGGTGACGTCGTGCTCTCCCGGGCCGGGTCCGAG | >hg38_21_+_8433172_8446622_RNA45SN1_45S_50ntFlanks | 485 | 522 | 38 | GATGTGGTGACGTCGTGCTCTCCCGGGCCGGGTCCGAG | MB231 | 29 | 346 | 2 | 9 | 16 | 25.4 | 299.9 | 2.2 | 8.4 | 14.1 |
| >hg38_21_+_8433172_8446622_RNA45SN1_45S_50ntFlanks@485.524.40\|GATGTGGTGACGTCGTGCTCTCCCGGGCCGGGTCCGAGCC | >hg38_21_+_8433172_8446622_RNA45SN1_45S_50ntFlanks | 485 | 524 | 40 | GATGTGGTGACGTCGTGCTCTCCCGGGCCGGGTCCGAGCC | BT20 | 19 | 85 | 2 | 21 | 11 | 22.4 | 60.1 | 0.8 | 23.8 | 3.3 |
| >hg38_21_+_8433172_8446622_RNA45SN1_45S_50ntFlanks@485.524.40\|GATGTGGTGACGTCGTGCTCTCCCGGGCCGGGTCCGAGCC | >hg38_21_+_8433172_8446622_RNA45SN1_45S_50ntFlanks | 485 | 524 | 40 | GATGTGGTGACGTCGTGCTCTCCCGGGCCGGGTCCGAGCC | MB231 | 20 | 304 | 2 | 4 | 14 | 20.0 | 306.9 | 1.5 | 3.8 | 12.8 |
| >hg38_21_+_8433172_8446622_RNA45SN1_45S_50ntFlanks@485.525.41\|GATGTGGTGACGTCGTGCTCTCCCGGGCCGGGTCCGAGCCG | >hg38_21_+_8433172_8446622_RNA45SN1_45S_50ntFlanks | 485 | 525 | 41 | GATGTGGTGACGTCGTGCTCTCCCGGGCCGGGTCCGAGCCG | BT20 | 32 | 142 | 2 | 7 | 15 | 26.5 | 119.3 | 1.7 | 5.9 | 6.4 |
| >hg38_21_+_8433172_8446622_RNA45SN1_45S_50ntFlanks@485.525.41\|GATGTGGTGACGTCGTGCTCTCCCGGGCCGGGTCCGAGCCG | >hg38_21_+_8433172_8446622_RNA45SN1_45S_50ntFlanks | 485 | 525 | 41 | GATGTGGTGACGTCGTGCTCTCCCGGGCCGGGTCCGAGCCG | MB231 | 51 | 353 | 3 | 6 | 22 | 44.7 | 313.6 | 2.8 | 5.7 | 20.8 |
| >hg38_21_+_8433172_8446622_RNA45SN1_45S_50ntFlanks@485.526.42\|GATGTGGTGACGTCGTGCTCTCCCGGGCCGGGTCCGAGCCGC | >hg38_21_+_8433172_8446622_RNA45SN1_45S_50ntFlanks | 485 | 526 | 42 | GATGTGGTGACGTCGTGCTCTCCCGGGCCGGGTCCGAGCCGC | BT20 | 10 | 75 | 1 | 5 | 6 | 8.0 | 68.3 | 1.2 | 6.1 | 2.7 |
| >hg38_21_+_8433172_8446622_RNA45SN1_45S_50ntFlanks@485.526.42\|GATGTGGTGACGTCGTGCTCTCCCGGGCCGGGTCCGAGCCGC | >hg38_21_+_8433172_8446622_RNA45SN1_45S_50ntFlanks | 485 | 526 | 42 | GATGTGGTGACGTCGTGCTCTCCCGGGCCGGGTCCGAGCCGC | MB231 | 19 | 141 | 1 | 4 | 11 | 21.1 | 157.5 | 1.2 | 3.9 | 13.3 |
| >hg38_21_+_8433172_8446622_RNA45SN1_45S_50ntFlanks@485.527.43\|GATGTGGTGACGTCGTGCTCTCCCGGGCCGGGTCCGAGCCGCG | >hg38_21_+_8433172_8446622_RNA45SN1_45S_50ntFlanks | 485 | 527 | 43 | GATGTGGTGACGTCGTGCTCTCCCGGGCCGGGTCCGAGCCGCG | BT20 | 20 | 131 | 1 | 10 | 14 | 15.9 | 129.8 | 1.5 | 11.9 | 8.0 |
| >hg38_21_+_8433172_8446622_RNA45SN1_45S_50ntFlanks@485.527.43\|GATGTGGTGACGTCGTGCTCTCCCGGGCCGGGTCCGAGCCGCG | >hg38_21_+_8433172_8446622_RNA45SN1_45S_50ntFlanks | 485 | 527 | 43 | GATGTGGTGACGTCGTGCTCTCCCGGGCCGGGTCCGAGCCGCG | MB231 | 33 | 145 | 2 | 6 | 17 | 36.5 | 162.3 | 2.5 | 7.5 | 20.6 |
| >hg38_21_+_8433172_8446622_RNA45SN1_45S_50ntFlanks@485.528.44\|GATGTGGTGACGTCGTGCTCTCCCGGGCCGGGTCCGAGCCGCGA | >hg38_21_+_8433172_8446622_RNA45SN1_45S_50ntFlanks | 485 | 528 | 44 | GATGTGGTGACGTCGTGCTCTCCCGGGCCGGGTCCGAGCCGCGA | BT20 | 11 | 69 | 1 | 5 | 2 | 10.6 | 70.5 | 1.8 | 6.2 | 1.9 |
| >hg38_21_+_8433172_8446622_RNA45SN1_45S_50ntFlanks@485.528.44\|GATGTGGTGACGTCGTGCTCTCCCGGGCCGGGTCCGAGCCGCGA | >hg38_21_+_8433172_8446622_RNA45SN1_45S_50ntFlanks | 485 | 528 | 44 | GATGTGGTGACGTCGTGCTCTCCCGGGCCGGGTCCGAGCCGCGA | MB231 | 21 | 57 | 1 | 3 | 9 | 23.9 | 56.7 | 1.6 | 3.6 | 13.6 |
| >hg38_21_+_8433172_8446622_RNA45SN1_45S_50ntFlanks@485.529.45\|GATGTGGTGACGTCGTGCTCTCCCGGGCCGGGTCCGAGCCGCGAC | >hg38_21_+_8433172_8446622_RNA45SN1_45S_50ntFlanks | 485 | 529 | 45 | GATGTGGTGACGTCGTGCTCTCCCGGGCCGGGTCCGAGCCGCGAC | BT20 | 11 | 79 | 1 | 7 | 9 | 9.0 | 81.9 | 0.9 | 7.7 | 7.5 |
| >hg38_21_+_8433172_8446622_RNA45SN1_45S_50ntFlanks@485.529.45\|GATGTGGTGACGTCGTGCTCTCCCGGGCCGGGTCCGAGCCGCGAC | >hg38_21_+_8433172_8446622_RNA45SN1_45S_50ntFlanks | 485 | 529 | 45 | GATGTGGTGACGTCGTGCTCTCCCGGGCCGGGTCCGAGCCGCGAC | MB231 | 36 | 94 | 1 | 4 | 13 | 46.0 | 105.8 | 1.0 | 5.9 | 18.1 |
| >hg38_21_+_8433172_8446622_RNA45SN1_45S_50ntFlanks@485.530.46\|GATGTGGTGACGTCGTGCTCTCCCGGGCCGGGTCCGAGCCGCGACG | >hg38_21_+_8433172_8446622_RNA45SN1_45S_50ntFlanks | 485 | 530 | 46 | GATGTGGTGACGTCGTGCTCTCCCGGGCCGGGTCCGAGCCGCGACG | MB231 | 31 | 42 | 1 | 3 | 7 | 38.8 | 45.8 | 1.0 | 4.0 | 9.7 |
| >hg38_21_+_8433172_8446622_RNA45SN1_45S_50ntFlanks@485.531.47\|GATGTGGTGACGTCGTGCTCTCCCGGGCCGGGTCCGAGCCGCGACGG | >hg38_21_+_8433172_8446622_RNA45SN1_45S_50ntFlanks | 485 | 531 | 47 | GATGTGGTGACGTCGTGCTCTCCCGGGCCGGGTCCGAGCCGCGACGG | BT20 | 10 | 57 | 0 | 6 | 2 | 11.6 | 73.9 | 0.6 | 6.0 | 2.1 |
| >hg38_21_+_8433172_8446622_RNA45SN1_45S_50ntFlanks@485.531.47\|GATGTGGTGACGTCGTGCTCTCCCGGGCCGGGTCCGAGCCGCGACGG | >hg38_21_+_8433172_8446622_RNA45SN1_45S_50ntFlanks | 485 | 531 | 47 | GATGTGGTGACGTCGTGCTCTCCCGGGCCGGGTCCGAGCCGCGACGG | MB231 | 37 | 41 | 1 | 4 | 7 | 52.6 | 45.1 | 0.7 | 5.2 | 10.0 |
| >hg38_21_+_8433172_8446622_RNA45SN1_45S_50ntFlanks@485.532.48\|GATGTGGTGACGTCGTGCTCTCCCGGGCCGGGTCCGAGCCGCGACGGG | >hg38_21_+_8433172_8446622_RNA45SN1_45S_50ntFlanks | 485 | 532 | 48 | GATGTGGTGACGTCGTGCTCTCCCGGGCCGGGTCCGAGCCGCGACGGG | BT20 | 62 | 279 | 3 | 12 | 32 | 83.4 | 359.1 | 5.5 | 14.6 | 43.9 |
| >hg38_21_+_8433172_8446622_RNA45SN1_45S_50ntFlanks@485.532.48\|GATGTGGTGACGTCGTGCTCTCCCGGGCCGGGTCCGAGCCGCGACGGG | >hg38_21_+_8433172_8446622_RNA45SN1_45S_50ntFlanks | 485 | 532 | 48 | GATGTGGTGACGTCGTGCTCTCCCGGGCCGGGTCCGAGCCGCGACGGG | MB231 | 324 | 135 | 4 | 4 | 33 | 490.2 | 170.8 | 3.8 | 5.5 | 45.6 |
| >hg38_21_+_8433172_8446622_RNA45SN1_45S_50ntFlanks@485.533.49\|GATGTGGTGACGTCGTGCTCTCCCGGGCCGGGTCCGAGCCGCGACGGGC | >hg38_21_+_8433172_8446622_RNA45SN1_45S_50ntFlanks | 485 | 533 | 49 | GATGTGGTGACGTCGTGCTCTCCCGGGCCGGGTCCGAGCCGCGACGGGC | MB231 | 27 | 14 | 0 | 0 | 6 | 44.4 | 18.8 | 0.1 | 0.7 | 11.2 |
| >hg38_21_+_8433172_8446622_RNA45SN1_45S_50ntFlanks@485.534.50\|GATGTGGTGACGTCGTGCTCTCCCGGGCCGGGTCCGAGCCGCGACGGGCG | >hg38_21_+_8433172_8446622_RNA45SN1_45S_50ntFlanks | 485 | 534 | 50 | GATGTGGTGACGTCGTGCTCTCCCGGGCCGGGTCCGAGCCGCGACGGGCG | MB231 | 30 | 21 | 0 | 0 | 8 | 49.6 | 30.5 | 0.0 | 0.8 | 13.2 |
| >hg38_21_+_8433172_8446622_RNA45SN1_45S_50ntFlanks@4853.4883.31\|AAGTCTTTGGGTTCCGGGGGGAGTATGGTTG | >hg38_21_+_8433172_8446622_RNA45SN1_45S_50ntFlanks | 4853 | 4883 | 31 | AAGTCTTTGGGTTCCGGGGGGAGTATGGTTG | MB468 | 16 | 44 | 18 | 13 | 17 | 5.7 | 5.5 | 7.4 | 4.8 | 8.7 |
| >hg38_21_+_8433172_8446622_RNA45SN1_45S_50ntFlanks@4884.4906.23\|CAAAGCTGAAACTTAAAGGAATT | >hg38_21_+_8433172_8446622_RNA45SN1_45S_50ntFlanks | 4884 | 4906 | 23 | CAAAGCTGAAACTTAAAGGAATT | MB468 | 14 | 8 | 13 | 15 | 10 | 1.5 | 5.0 | 4.0 | 2.6 | 6.2 |
| >hg38_21_+_8433172_8446622_RNA45SN1_45S_50ntFlanks@4971.4999.29\|CCGGCCCGGACACGGACAGGATTGACAGA | >hg38_21_+_8433172_8446622_RNA45SN1_45S_50ntFlanks | 4971 | 4999 | 29 | CCGGCCCGGACACGGACAGGATTGACAGA | MB468 | 12 | 1 | 11 | 56 | 43 | 6.8 | 0.6 | 8.4 | 45.4 | 33.0 |
| >hg38_21_+_8433172_8446622_RNA45SN1_45S_50ntFlanks@4976.4993.18\|CCGGACACGGACAGGATT | >hg38_21_+_8433172_8446622_RNA45SN1_45S_50ntFlanks | 4976 | 4993 | 18 | CCGGACACGGACAGGATT | BT20 | 10 | 16 | 23 | 9 | 4 | 2.8 | 5.6 | 6.2 | 6.5 | 0.5 |
| >hg38_21_+_8433172_8446622_RNA45SN1_45S_50ntFlanks@4995.5013.19\|ACAGATTGATAGCTCTTTC | >hg38_21_+_8433172_8446622_RNA45SN1_45S_50ntFlanks | 4995 | 5013 | 19 | ACAGATTGATAGCTCTTTC | MB468 | 17 | 10 | 14 | 117 | 53 | 9.6 | 3.6 | 12.1 | 125.7 | 64.9 |
| >hg38_21_+_8433172_8446622_RNA45SN1_45S_50ntFlanks@4995.5022.28\|ACAGATTGATAGCTCTTTCTCGATTCCG | >hg38_21_+_8433172_8446622_RNA45SN1_45S_50ntFlanks | 4995 | 5022 | 28 | ACAGATTGATAGCTCTTTCTCGATTCCG | MB468 | 14 | 66 | 8 | 34 | 66 | 3.9 | 69.7 | 5.1 | 41.6 | 69.2 |

| | | | | | | | | | | | | | | | | |
|---|---|---|---|---|---|---|---|---|---|---|---|---|---|---|---|---|
| >hg38_21_+_8433172_8446622_RNA45SN1_45S_50ntFlanks@4995.5024.30\|ACAGATTGATAG<br>CTCTTTCTCGATTCCGTG | >hg38_21_+_8433172_8446622_RNA45SN1_45S_50ntFlanks | 4995 | 5024 | 30 | ACAGATTGATAGCTCTTTCTCGATTCCGTG | MB468 | 35 | 200 | 20 | 118 | 85 | 14.3 | 217.3 | 14.7 | 127.7 | 70.4 |
| >hg38_21_+_8433172_8446622_RNA45SN1_45S_50ntFlanks@5056.5082.27\|GAGCGATTTGTC<br>TGGTTAATTCCGATA | >hg38_21_+_8433172_8446622_RNA45SN1_45S_50ntFlanks | 5056 | 5082 | 27 | GAGCGATTTGTCTGGTTAATTCCGATA | MB468 | 10 | 20 | 5 | 90 | 22 | 1.9 | 14.2 | 3.5 | 123.9 | 20.9 |
| >hg38_21_+_8433172_8446622_RNA45SN1_45S_50ntFlanks@5057.5081.25\|AGCGATTTGTCT<br>GGTTAATTCCGAT | >hg38_21_+_8433172_8446622_RNA45SN1_45S_50ntFlanks | 5057 | 5081 | 25 | AGCGATTTGTCTGGTTAATTCCGAT | BT20 | 17 | 4 | 12 | 55 | 10 | 6.4 | 1.9 | 10.0 | 59.0 | 3.5 |
| >hg38_21_+_8433172_8446622_RNA45SN1_45S_50ntFlanks@5071.5098.28\|TTAATTCCGATA<br>ACGAACGAGACTCTGG | >hg38_21_+_8433172_8446622_RNA45SN1_45S_50ntFlanks | 5071 | 5098 | 28 | TTAATTCCGATAACGAACGAGACTCTGG | MB468 | 21 | 101 | 14 | 114 | 68 | 4.7 | 97.4 | 8.1 | 147.6 | 69.2 |
| >hg38_21_+_8433172_8446622_RNA45SN1_45S_50ntFlanks@5073.5097.25\|AATTCCGATAAC<br>GAACGAGACTCTG | >hg38_21_+_8433172_8446622_RNA45SN1_45S_50ntFlanks | 5073 | 5097 | 25 | AATTCCGATAACGAACGAGACTCTG | MB468 | 12 | 55 | 10 | 78 | 51 | 5.0 | 50.0 | 5.1 | 116.9 | 50.2 |
| >hg38_21_+_8433172_8446622_RNA45SN1_45S_50ntFlanks@5073.5098.26\|AATTCCGATAAC<br>GAACGAGACTCTGG | >hg38_21_+_8433172_8446622_RNA45SN1_45S_50ntFlanks | 5073 | 5098 | 26 | AATTCCGATAACGAACGAGACTCTGG | MB468 | 38 | 178 | 26 | 239 | 182 | 8.6 | 195.5 | 13.1 | 344.8 | 202.9 |
| >hg38_21_+_8433172_8446622_RNA45SN1_45S_50ntFlanks@5078.5097.20\|CGATAACGAAC<br>GAGACTCTG | >hg38_21_+_8433172_8446622_RNA45SN1_45S_50ntFlanks | 5078 | 5097 | 20 | CGATAACGAACGAGACTCTG | MB468 | 10 | 21 | 9 | 80 | 28 | 6.6 | 17.8 | 6.2 | 98.3 | 17.3 |
| >hg38_21_+_8433172_8446622_RNA45SN1_45S_50ntFlanks@5078.5102.25\|CGATAACGAAC<br>GAGACTCTGGCATG | >hg38_21_+_8433172_8446622_RNA45SN1_45S_50ntFlanks | 5078 | 5102 | 25 | CGATAACGAACGAGACTCTGGCATG | MB468 | 11 | 49 | 12 | 73 | 44 | 4.6 | 54.6 | 8.8 | 79.3 | 40.0 |
| >hg38_21_+_8433172_8446622_RNA45SN1_45S_50ntFlanks@5084.5110.27\|CGAACGAGACT<br>CTGGCATGCTAACTAG | >hg38_21_+_8433172_8446622_RNA45SN1_45S_50ntFlanks | 5084 | 5110 | 27 | CGAACGAGACTCTGGCATGCTAACTAG | MB468 | 16 | 86 | 9 | 98 | 222 | 4.2 | 62.9 | 7.1 | 114.8 | 232.8 |
| >hg38_21_+_8433172_8446622_RNA45SN1_45S_50ntFlanks@5084.5115.32\|CGAACGAGACT<br>CTGGCATGCTAACTAGTTACG | >hg38_21_+_8433172_8446622_RNA45SN1_45S_50ntFlanks | 5084 | 5115 | 32 | CGAACGAGACTCTGGCATGCTAACTAGTTACG | MB468 | 11 | 28 | 4 | 28 | 41 | 7.2 | 23.1 | 1.9 | 25.7 | 37.0 |
| >hg38_21_+_8433172_8446622_RNA45SN1_45S_50ntFlanks@5086.5112.27\|AACGAGACTCTG<br>GCATGCTAACTAGTT | >hg38_21_+_8433172_8446622_RNA45SN1_45S_50ntFlanks | 5086 | 5112 | 27 | AACGAGACTCTGGCATGCTAACTAGTT | MB231 | 12 | 10 | 15 | 5 | 4 | 13.3 | 7.6 | 11.1 | 1.5 | 2.1 |
| >hg38_21_+_8433172_8446622_RNA45SN1_45S_50ntFlanks@51.71.21\|GCTGACACGCTGTCC<br>TCTGGC | >hg38_21_+_8433172_8446622_RNA45SN1_45S_50ntFlanks | 51 | 71 | 21 | GCTGACACGCTGTCCTCTGGC | MB231 | 15 | 31 | 4 | 0 | 1 | 9.3 | 25.3 | 2.2 | 0.4 | 1.1 |
| >hg38_21_+_8433172_8446622_RNA45SN1_45S_50ntFlanks@51.72.22\|GCTGACACGCTGTCC<br>TCTGGCG | >hg38_21_+_8433172_8446622_RNA45SN1_45S_50ntFlanks | 51 | 72 | 22 | GCTGACACGCTGTCCTCTGGCG | MB468 | 44 | 66 | 12 | 7 | 4 | 11.5 | 26.3 | 1.7 | 1.7 | 3.4 |
| >hg38_21_+_8433172_8446622_RNA45SN1_45S_50ntFlanks@51.74.24\|GCTGACACGCTGTCC<br>TCTGGCGAC | >hg38_21_+_8433172_8446622_RNA45SN1_45S_50ntFlanks | 51 | 74 | 24 | GCTGACACGCTGTCCTCTGGCGAC | MB231 | 10 | 62 | 10 | 1 | 8 | 3.6 | 48.7 | 9.1 | 0.5 | 13.3 |
| >hg38_21_+_8433172_8446622_RNA45SN1_45S_50ntFlanks@5125.5145.21\|AGCGGTCGGCG<br>TCCCCCAACT | >hg38_21_+_8433172_8446622_RNA45SN1_45S_50ntFlanks | 5125 | 5145 | 21 | AGCGGTCGGCGTCCCCCAACT | BT20 | 10 | 39 | 29 | 1 | 5 | 8.2 | 49.1 | 12.9 | 1.0 | 5.1 |
| >hg38_21_+_8433172_8446622_RNA45SN1_45S_50ntFlanks@5125.5158.34\|AGCGGTCGGCG<br>TCCCCCAACTTCTTAGAGGGACA | >hg38_21_+_8433172_8446622_RNA45SN1_45S_50ntFlanks | 5125 | 5158 | 34 | AGCGGTCGGCGTCCCCCAACTTCTTAGAGGGACA | BT20 | 14 | 14 | 107 | 5 | 10 | 6.8 | 8.6 | 77.8 | 4.3 | 6.3 |
| >hg38_21_+_8433172_8446622_RNA45SN1_45S_50ntFlanks@5152.5185.34\|AGGGACAAGTG<br>GCGTTCAGCCACCCGAGATTGAG | >hg38_21_+_8433172_8446622_RNA45SN1_45S_50ntFlanks | 5152 | 5185 | 34 | AGGGACAAGTGGCGTTCAGCCACCCGAGATTGAG | MB468 | 72 | 33 | 4 | 42 | 84 | 77.6 | 38.2 | 3.1 | 45.5 | 89.7 |
| >hg38_21_+_8433172_8446622_RNA45SN1_45S_50ntFlanks@5186.5205.20\|CAATAACAGGTC<br>TGTGATGC | >hg38_21_+_8433172_8446622_RNA45SN1_45S_50ntFlanks | 5186 | 5205 | 20 | CAATAACAGGTCTGTGATGC | MB468 | 13 | 29 | 13 | 38 | 30 | 0.9 | 20.8 | 5.5 | 53.6 | 27.2 |
| >hg38_21_+_8433172_8446622_RNA45SN1_45S_50ntFlanks@5190.5214.25\|AACAGGTCTGTG<br>ATGCCCTTAGATG | >hg38_21_+_8433172_8446622_RNA45SN1_45S_50ntFlanks | 5190 | 5214 | 25 | AACAGGTCTGTGATGCCCTTAGATG | BT20 | 10 | 8 | 2 | 6 | 1 | 16.9 | 13.7 | 1.4 | 5.7 | 0.5 |
| >hg38_21_+_8433172_8446622_RNA45SN1_45S_50ntFlanks@5190.5215.26\|AACAGGTCTGTG<br>ATGCCCTTAGATGT | >hg38_21_+_8433172_8446622_RNA45SN1_45S_50ntFlanks | 5190 | 5215 | 26 | AACAGGTCTGTGATGCCCTTAGATGT | BT20 | 12 | 5 | 1 | 3 | 1 | 19.5 | 9.0 | 1.0 | 1.5 | 0.2 |
| >hg38_21_+_8433172_8446622_RNA45SN1_45S_50ntFlanks@5191.5214.24\|ACAGGTCTGTGA<br>TGCCCTTAGATG | >hg38_21_+_8433172_8446622_RNA45SN1_45S_50ntFlanks | 5191 | 5214 | 24 | ACAGGTCTGTGATGCCCTTAGATG | BT20 | 11 | 9 | 2 | 3 | 1 | 17.6 | 15.3 | 2.1 | 1.9 | 0.5 |
| >hg38_21_+_8433172_8446622_RNA45SN1_45S_50ntFlanks@5191.5215.25\|ACAGGTCTGTGA<br>TGCCCTTAGATGT | >hg38_21_+_8433172_8446622_RNA45SN1_45S_50ntFlanks | 5191 | 5215 | 25 | ACAGGTCTGTGATGCCCTTAGATGT | BT20 | 19 | 10 | 2 | 2 | 1 | 32.4 | 17.0 | 2.5 | 2.1 | 1.0 |
| >hg38_21_+_8433172_8446622_RNA45SN1_45S_50ntFlanks@5192.5214.23\|CAGGTCTGTGAT<br>GCCCTTAGATG | >hg38_21_+_8433172_8446622_RNA45SN1_45S_50ntFlanks | 5192 | 5214 | 23 | CAGGTCTGTGATGCCCTTAGATG | BT20 | 21 | 11 | 3 | 3 | 1 | 34.7 | 18.7 | 3.6 | 1.9 | 1.0 |
| >hg38_21_+_8433172_8446622_RNA45SN1_45S_50ntFlanks@5192.5215.24\|CAGGTCTGTGAT<br>GCCCTTAGATGT | >hg38_21_+_8433172_8446622_RNA45SN1_45S_50ntFlanks | 5192 | 5215 | 24 | CAGGTCTGTGATGCCCTTAGATGT | BT20 | 28 | 6 | 3 | 2 | 1 | 48.7 | 10.8 | 3.7 | 1.8 | 0.8 |
| >hg38_21_+_8433172_8446622_RNA45SN1_45S_50ntFlanks@5237.5254.18\|ACTGACTGGCTC<br>AGCGTG | >hg38_21_+_8433172_8446622_RNA45SN1_45S_50ntFlanks | 5237 | 5254 | 18 | ACTGACTGGCTCAGCGTG | MB468 | 11 | 1 | 2 | 50 | 51 | 8.7 | 0.6 | 1.2 | 30.0 | 82.9 |
| >hg38_21_+_8433172_8446622_RNA45SN1_45S_50ntFlanks@5237.5255.19\|ACTGACTGGCTC<br>AGCGTGT | >hg38_21_+_8433172_8446622_RNA45SN1_45S_50ntFlanks | 5237 | 5255 | 19 | ACTGACTGGCTCAGCGTGT | BT20 | 12 | 1 | 4 | 76 | 8 | 6.9 | 0.3 | 2.4 | 97.8 | 2.1 |
| >hg38_21_+_8433172_8446622_RNA45SN1_45S_50ntFlanks@5417.5443.27\|CTACCGATTGGA<br>TGGTTTAGTGAGGCC | >hg38_21_+_8433172_8446622_RNA45SN1_45S_50ntFlanks | 5417 | 5443 | 27 | CTACCGATTGGATGGTTTAGTGAGGCC | MB468 | 16 | 10 | 22 | 17 | 9 | 7.0 | 3.4 | 11.5 | 5.4 | 3.9 |
| >hg38_21_+_8433172_8446622_RNA45SN1_45S_50ntFlanks@5417.5445.29\|CTACCGATTGGA<br>TGGTTTAGTGAGGCCCT | >hg38_21_+_8433172_8446622_RNA45SN1_45S_50ntFlanks | 5417 | 5445 | 29 | CTACCGATTGGATGGTTTAGTGAGGCCCT | MB468 | 23 | 39 | 46 | 16 | 26 | 10.8 | 20.2 | 19.1 | 10.9 | 17.4 |
| >hg38_21_+_8433172_8446622_RNA45SN1_45S_50ntFlanks@5419.5445.27\|ACCGATTGGATG<br>GTTTAGTGAGGCCCT | >hg38_21_+_8433172_8446622_RNA45SN1_45S_50ntFlanks | 5419 | 5445 | 27 | ACCGATTGGATGGTTTAGTGAGGCCCT | MB468 | 12 | 19 | 20 | 6 | 9 | 5.0 | 12.9 | 6.1 | 4.7 | 5.3 |
| >hg38_21_+_8433172_8446622_RNA45SN1_45S_50ntFlanks@5420.5438.19\|CCGATTGGATGG<br>TTTAGTG | >hg38_21_+_8433172_8446622_RNA45SN1_45S_50ntFlanks | 5420 | 5438 | 19 | CCGATTGGATGGTTTAGTG | MB468 | 15 | 8 | 21 | 29 | 7 | 8.9 | 6.8 | 10.6 | 15.3 | 2.5 |
| >hg38_21_+_8433172_8446622_RNA45SN1_45S_50ntFlanks@5420.5443.24\|CCGATTGGATGG<br>TTTAGTGAGGCC | >hg38_21_+_8433172_8446622_RNA45SN1_45S_50ntFlanks | 5420 | 5443 | 24 | CCGATTGGATGGTTTAGTGAGGCC | MB468 | 119 | 116 | 372 | 122 | 66 | 57.9 | 49.6 | 151.8 | 74.2 | 20.5 |
| >hg38_21_+_8433172_8446622_RNA45SN1_45S_50ntFlanks@5420.5444.25\|CCGATTGGATGG<br>TTTAGTGAGGCCC | >hg38_21_+_8433172_8446622_RNA45SN1_45S_50ntFlanks | 5420 | 5444 | 25 | CCGATTGGATGGTTTAGTGAGGCCC | MB468 | 24 | 38 | 70 | 23 | 16 | 8.8 | 39.0 | 25.3 | 19.9 | 9.6 |
| >hg38_21_+_8433172_8446622_RNA45SN1_45S_50ntFlanks@5420.5445.26\|CCGATTGGATGG<br>TTTAGTGAGGCCCT | >hg38_21_+_8433172_8446622_RNA45SN1_45S_50ntFlanks | 5420 | 5445 | 26 | CCGATTGGATGGTTTAGTGAGGCCCT | MB468 | 69 | 93 | 177 | 52 | 62 | 24.3 | 83.0 | 48.6 | 41.6 | 40.0 |
| >hg38_21_+_8433172_8446622_RNA45SN1_45S_50ntFlanks@5421.5443.23\|CGATTGGATGGT<br>TTAGTGAGGCC | >hg38_21_+_8433172_8446622_RNA45SN1_45S_50ntFlanks | 5421 | 5443 | 23 | CGATTGGATGGTTTAGTGAGGCC | MB468 | 50 | 59 | 114 | 98 | 37 | 13.5 | 34.1 | 57.0 | 80.0 | 18.3 |
| >hg38_21_+_8433172_8446622_RNA45SN1_45S_50ntFlanks@5421.5445.25\|CGATTGGATGGT<br>TTAGTGAGGCCCT | >hg38_21_+_8433172_8446622_RNA45SN1_45S_50ntFlanks | 5421 | 5445 | 25 | CGATTGGATGGTTTAGTGAGGCCCT | MB468 | 34 | 44 | 54 | 42 | 50 | 7.7 | 37.3 | 15.3 | 33.5 | 39.1 |
| >hg38_21_+_8433172_8446622_RNA45SN1_45S_50ntFlanks@5423.5443.21\|ATTGGATGGTTT<br>AGTGAGGCC | >hg38_21_+_8433172_8446622_RNA45SN1_45S_50ntFlanks | 5423 | 5443 | 21 | ATTGGATGGTTTAGTGAGGCC | MB468 | 14 | 9 | 16 | 26 | 7 | 4.0 | 6.4 | 4.2 | 21.8 | 3.6 |
| >hg38_21_+_8433172_8446622_RNA45SN1_45S_50ntFlanks@5423.5445.23\|ATTGGATGGTTT<br>AGTGAGGCCCT | >hg38_21_+_8433172_8446622_RNA45SN1_45S_50ntFlanks | 5423 | 5445 | 23 | ATTGGATGGTTTAGTGAGGCCCT | MB468 | 17 | 7 | 14 | 16 | 10 | 6.5 | 4.2 | 4.4 | 14.5 | 4.0 |
| >hg38_21_+_8433172_8446622_RNA45SN1_45S_50ntFlanks@5425.5450.26\|TGGATGGTTTAG<br>TGAGGCCCTCGGAT | >hg38_21_+_8433172_8446622_RNA45SN1_45S_50ntFlanks | 5425 | 5450 | 26 | TGGATGGTTTAGTGAGGCCCTCGGAT | MB231 | 12 | 12 | 20 | 1 | 1 | 10.9 | 10.9 | 23.3 | 1.0 | 1.4 |
| >hg38_21_+_8433172_8446622_RNA45SN1_45S_50ntFlanks@5436.5466.31\|GTGAGGCCCTC<br>GGATCGGCCCCGCCGGGGTC | >hg38_21_+_8433172_8446622_RNA45SN1_45S_50ntFlanks | 5436 | 5466 | 31 | GTGAGGCCCTCGGATCGGCCCCGCCGGGGTC | MB468 | 14 | 33 | 27 | 16 | 20 | 6.4 | 9.6 | 9.7 | 5.0 | 6.2 |
| >hg38_21_+_8433172_8446622_RNA45SN1_45S_50ntFlanks@5442.5466.25\|CCCTCGGATCG<br>GCCCCGCCGGGGTC | >hg38_21_+_8433172_8446622_RNA45SN1_45S_50ntFlanks | 5442 | 5466 | 25 | CCCTCGGATCGGCCCCGCCGGGGTC | MB468 | 21 | 35 | 34 | 6 | 31 | 10.1 | 11.4 | 24.3 | 4.1 | 7.0 |
| >hg38_21_+_8433172_8446622_RNA45SN1_45S_50ntFlanks@5443.5466.24\|CCTCGGATCGG<br>CCCCGCCGGGGTC | >hg38_21_+_8433172_8446622_RNA45SN1_45S_50ntFlanks | 5443 | 5466 | 24 | CCTCGGATCGGCCCCGCCGGGGTC | MB468 | 44 | 75 | 76 | 18 | 72 | 14.9 | 21.0 | 45.2 | 11.6 | 13.1 |
| >hg38_21_+_8433172_8446622_RNA45SN1_45S_50ntFlanks@5444.5465.22\|CTCGGATCGGC<br>CCCGCCGGGGT | >hg38_21_+_8433172_8446622_RNA45SN1_45S_50ntFlanks | 5444 | 5465 | 22 | CTCGGATCGGCCCCGCCGGGGT | MB468 | 17 | 25 | 20 | 11 | 21 | 7.0 | 5.9 | 9.1 | 3.9 | 8.4 |
| >hg38_21_+_8433172_8446622_RNA45SN1_45S_50ntFlanks@5444.5466.23\|CTCGGATCGGC<br>CCCGCCGGGGTC | >hg38_21_+_8433172_8446622_RNA45SN1_45S_50ntFlanks | 5444 | 5466 | 23 | CTCGGATCGGCCCCGCCGGGGTC | BT20 | 31 | 33 | 52 | 14 | 35 | 51.4 | 55.0 | 65.3 | 13.0 | 58.6 |
| >hg38_21_+_8433172_8446622_RNA45SN1_45S_50ntFlanks@5444.5466.23\|CTCGGATCGGC<br>CCCGCCGGGGTC | >hg38_21_+_8433172_8446622_RNA45SN1_45S_50ntFlanks | 5444 | 5466 | 23 | CTCGGATCGGCCCCGCCGGGGTC | MB468 | 1041 | 1342 | 2030 | 2049 | 1340 | 698.9 | 709.9 | 1120.0 | 803.5 | 305.2 |
| >hg38_21_+_8433172_8446622_RNA45SN1_45S_50ntFlanks@5444.5467.24\|CTCGGATCGGC<br>CCCGCCGGGGTCG | >hg38_21_+_8433172_8446622_RNA45SN1_45S_50ntFlanks | 5444 | 5467 | 24 | CTCGGATCGGCCCCGCCGGGGTCG | MB468 | 49 | 51 | 96 | 229 | 69 | 24.7 | 13.5 | 49.4 | 83.1 | 34.4 |
| >hg38_21_+_8433172_8446622_RNA45SN1_45S_50ntFlanks@5445.5466.22\|TCGGATCGGCC<br>CCGCCGGGGTC | >hg38_21_+_8433172_8446622_RNA45SN1_45S_50ntFlanks | 5445 | 5466 | 22 | TCGGATCGGCCCCGCCGGGGTC | MB468 | 267 | 262 | 482 | 929 | 327 | 72.8 | 30.5 | 187.9 | 552.1 | 95.7 |
| >hg38_21_+_8433172_8446622_RNA45SN1_45S_50ntFlanks@5445.5467.23\|TCGGATCGGCC<br>CCGCCGGGGTCG | >hg38_21_+_8433172_8446622_RNA45SN1_45S_50ntFlanks | 5445 | 5467 | 23 | TCGGATCGGCCCCGCCGGGGTCG | MB468 | 12 | 11 | 22 | 53 | 15 | 2.0 | 1.6 | 7.5 | 30.5 | 6.8 |
| >hg38_21_+_8433172_8446622_RNA45SN1_45S_50ntFlanks@5446.5466.21\|CGGATCGGCCC<br>CGCCGGGGTC | >hg38_21_+_8433172_8446622_RNA45SN1_45S_50ntFlanks | 5446 | 5466 | 21 | CGGATCGGCCCCGCCGGGGTC | MB468 | 465 | 409 | 741 | 2674 | 1050 | 247.1 | 75.7 | 332.1 | 2134.5 | 500.4 |
| >hg38_21_+_8433172_8446622_RNA45SN1_45S_50ntFlanks@5446.5467.22\|CGGATCGGCCC<br>CGCCGGGGTCG | >hg38_21_+_8433172_8446622_RNA45SN1_45S_50ntFlanks | 5446 | 5467 | 22 | CGGATCGGCCCCGCCGGGGTCG | MB468 | 17 | 14 | 30 | 96 | 33 | 9.3 | 3.5 | 15.5 | 74.4 | 15.6 |
| >hg38_21_+_8433172_8446622_RNA45SN1_45S_50ntFlanks@5449.5466.18\|ATCGGCCCCGC<br>CGGGGTC | >hg38_21_+_8433172_8446622_RNA45SN1_45S_50ntFlanks | 5449 | 5466 | 18 | ATCGGCCCCGCCGGGGTC | MB468 | 17 | 5 | 18 | 184 | 21 | 1.8 | 1.3 | 8.5 | 113.6 | 5.5 |

| | | | | | | | | | | | | | | | |
|---|---|---|---|---|---|---|---|---|---|---|---|---|---|---|---|
| >hg38_21_+_8433172_8446622_RNA45SN1_45S_50ntFlanks@5467.5484.18\|GGCCCACGGCCCTGGCGG | >hg38_21_+_8433172_8446622_RNA45SN1_45S_50ntFlanks | 5467 | 5484 | 18 | GGCCCACGGCCCTGGCGG | MB468 | 25 | 22 | 20 | 14 | 32 | 6.4 | 10.2 | 12.0 | 4.8 | 12.0 |
| >hg38_21_+_8433172_8446622_RNA45SN1_45S_50ntFlanks@5467.5491.25\|GGCCCACGGCCCTGGCGGAGCGCTG | >hg38_21_+_8433172_8446622_RNA45SN1_45S_50ntFlanks | 5467 | 5491 | 25 | GGCCCACGGCCCTGGCGGAGCGCTG | MB468 | 18 | 18 | 23 | 8 | 20 | 9.7 | 6.0 | 13.4 | 6.4 | 3.7 |
| >hg38_21_+_8433172_8446622_RNA45SN1_45S_50ntFlanks@5467.5499.33\|GGCCCACGGCCCTGGCGGAGCGCTGAGAAGACG | >hg38_21_+_8433172_8446622_RNA45SN1_45S_50ntFlanks | 5467 | 5499 | 33 | GGCCCACGGCCCTGGCGGAGCGCTGAGAAGACG | MB468 | 128 | 509 | 167 | 145 | 175 | 58.8 | 324.3 | 107.2 | 182.7 | 169.8 |
| >hg38_21_+_8433172_8446622_RNA45SN1_45S_50ntFlanks@5468.5499.32\|GCCCACGGCCCTGGCGGAGCGCTGAGAAGACG | >hg38_21_+_8433172_8446622_RNA45SN1_45S_50ntFlanks | 5468 | 5499 | 32 | GCCCACGGCCCTGGCGGAGCGCTGAGAAGACG | MB468 | 13 | 49 | 19 | 10 | 14 | 5.4 | 36.0 | 10.4 | 10.1 | 10.2 |
| >hg38_21_+_8433172_8446622_RNA45SN1_45S_50ntFlanks@5469.5499.31\|CCCACGGCCCTGGCGGAGCGCTGAGAAGACG | >hg38_21_+_8433172_8446622_RNA45SN1_45S_50ntFlanks | 5469 | 5499 | 31 | CCCACGGCCCTGGCGGAGCGCTGAGAAGACG | MB468 | 21 | 49 | 34 | 20 | 18 | 9.7 | 16.9 | 19.9 | 14.2 | 6.6 |
| >hg38_21_+_8433172_8446622_RNA45SN1_45S_50ntFlanks@5470.5499.30\|CCACGGCCCTGGCGGAGCGCTGAGAAGACG | >hg38_21_+_8433172_8446622_RNA45SN1_45S_50ntFlanks | 5470 | 5499 | 30 | CCACGGCCCTGGCGGAGCGCTGAGAAGACG | MB468 | 37 | 70 | 83 | 55 | 36 | 22.4 | 30.6 | 65.6 | 39.5 | 11.3 |
| >hg38_21_+_8433172_8446622_RNA45SN1_45S_50ntFlanks@5471.5499.29\|CACGGCCCTGGCGGAGCGCTGAGAAGACG | >hg38_21_+_8433172_8446622_RNA45SN1_45S_50ntFlanks | 5471 | 5499 | 29 | CACGGCCCTGGCGGAGCGCTGAGAAGACG | MB468 | 96 | 192 | 266 | 203 | 88 | 41.9 | 88.2 | 182.2 | 169.7 | 42.1 |
| >hg38_21_+_8433172_8446622_RNA45SN1_45S_50ntFlanks@5471.5514.44\|CACGGCCCTGGCGGAGCGCTGAGAAGACGGTCGAACTTGACTAT | >hg38_21_+_8433172_8446622_RNA45SN1_45S_50ntFlanks | 5471 | 5514 | 44 | CACGGCCCTGGCGGAGCGCTGAGAAGACGGTCGAACTTGACTAT | MB468 | 16 | 105 | 231 | 39 | 74 | 8.1 | 78.1 | 220.1 | 47.4 | 55.3 |
| >hg38_21_+_8433172_8446622_RNA45SN1_45S_50ntFlanks@5471.5516.46\|CACGGCCCTGGCGGAGCGCTGAGAAGACGGTCGAACTTGACTATCT | >hg38_21_+_8433172_8446622_RNA45SN1_45S_50ntFlanks | 5471 | 5516 | 46 | CACGGCCCTGGCGGAGCGCTGAGAAGACGGTCGAACTTGACTATCT | MB231 | 49 | 7 | 454 | 7 | 14 | 66.5 | 1.4 | 285.1 | 10.4 | 13.4 |
| >hg38_21_+_8433172_8446622_RNA45SN1_45S_50ntFlanks@5472.5489.18\|ACGGCCCTGGCGGAGCGC | >hg38_21_+_8433172_8446622_RNA45SN1_45S_50ntFlanks | 5472 | 5489 | 18 | ACGGCCCTGGCGGAGCGC | BT20 | 14 | 27 | 117 | 13 | 11 | 10.0 | 20.2 | 110.8 | 6.9 | 7.3 |
| >hg38_21_+_8433172_8446622_RNA45SN1_45S_50ntFlanks@5472.5489.18\|ACGGCCCTGGCGGAGCGC | >hg38_21_+_8433172_8446622_RNA45SN1_45S_50ntFlanks | 5472 | 5489 | 18 | ACGGCCCTGGCGGAGCGC | MB231 | 38 | 37 | 100 | 32 | 37 | 18.5 | 11.4 | 46.4 | 27.0 | 12.2 |
| >hg38_21_+_8433172_8446622_RNA45SN1_45S_50ntFlanks@5472.5490.19\|ACGGCCCTGGCGGAGCGCT | >hg38_21_+_8433172_8446622_RNA45SN1_45S_50ntFlanks | 5472 | 5490 | 19 | ACGGCCCTGGCGGAGCGCT | BT20 | 18 | 19 | 153 | 5 | 16 | 17.0 | 16.1 | 217.7 | 3.6 | 16.0 |
| >hg38_21_+_8433172_8446622_RNA45SN1_45S_50ntFlanks@5472.5490.19\|ACGGCCCTGGCGGAGCGCT | >hg38_21_+_8433172_8446622_RNA45SN1_45S_50ntFlanks | 5472 | 5490 | 19 | ACGGCCCTGGCGGAGCGCT | MB231 | 17 | 16 | 72 | 3 | 12 | 9.7 | 3.6 | 36.3 | 1.7 | 8.5 |
| >hg38_21_+_8433172_8446622_RNA45SN1_45S_50ntFlanks@5472.5491.20\|ACGGCCCTGGCGGAGCGCTG | >hg38_21_+_8433172_8446622_RNA45SN1_45S_50ntFlanks | 5472 | 5491 | 20 | ACGGCCCTGGCGGAGCGCTG | BT20 | 10 | 9 | 134 | 7 | 11 | 13.3 | 12.3 | 208.9 | 4.3 | 14.1 |
| >hg38_21_+_8433172_8446622_RNA45SN1_45S_50ntFlanks@5472.5494.23\|ACGGCCCTGGCGGAGCGCTGAGA | >hg38_21_+_8433172_8446622_RNA45SN1_45S_50ntFlanks | 5472 | 5494 | 23 | ACGGCCCTGGCGGAGCGCTGAGA | BT20 | 15 | 20 | 375 | 21 | 29 | 17.0 | 25.7 | 583.0 | 14.7 | 40.7 |
| >hg38_21_+_8433172_8446622_RNA45SN1_45S_50ntFlanks@5472.5499.28\|ACGGCCCTGGCGGAGCGCTGAGAAGACG | >hg38_21_+_8433172_8446622_RNA45SN1_45S_50ntFlanks | 5472 | 5499 | 28 | ACGGCCCTGGCGGAGCGCTGAGAAGACG | BT20 | 32 | 32 | 432 | 42 | 50 | 37.9 | 44.4 | 648.8 | 7.2 | 58.8 |
| >hg38_21_+_8433172_8446622_RNA45SN1_45S_50ntFlanks@5472.5499.28\|ACGGCCCTGGCGGAGCGCTGAGAAGACG | >hg38_21_+_8433172_8446622_RNA45SN1_45S_50ntFlanks | 5472 | 5499 | 28 | ACGGCCCTGGCGGAGCGCTGAGAAGACG | MB231 | 49 | 49 | 322 | 16 | 36 | 35.4 | 31.9 | 245.1 | 10.7 | 42.0 |
| >hg38_21_+_8433172_8446622_RNA45SN1_45S_50ntFlanks@5472.5499.28\|ACGGCCCTGGCGGAGCGCTGAGAAGACG | >hg38_21_+_8433172_8446622_RNA45SN1_45S_50ntFlanks | 5472 | 5499 | 28 | ACGGCCCTGGCGGAGCGCTGAGAAGACG | MB468 | 60 | 115 | 139 | 102 | 83 | 17.5 | 35.3 | 79.4 | 102.7 | 74.1 |
| >hg38_21_+_8433172_8446622_RNA45SN1_45S_50ntFlanks@5472.5500.29\|ACGGCCCTGGCGGAGCGCTGAGAAGACGG | >hg38_21_+_8433172_8446622_RNA45SN1_45S_50ntFlanks | 5472 | 5500 | 29 | ACGGCCCTGGCGGAGCGCTGAGAAGACGG | MB231 | 10 | 22 | 133 | 6 | 22 | 5.4 | 9.1 | 79.2 | 2.6 | 33.5 |
| >hg38_21_+_8433172_8446622_RNA45SN1_45S_50ntFlanks@5472.5501.30\|ACGGCCCTGGCGGAGCGCTGAGAAGACGGT | >hg38_21_+_8433172_8446622_RNA45SN1_45S_50ntFlanks | 5472 | 5501 | 30 | ACGGCCCTGGCGGAGCGCTGAGAAGACGGT | BT20 | 76 | 70 | 1054 | 46 | 87 | 105.3 | 102.2 | 1617.9 | 50.6 | 122.7 |
| >hg38_21_+_8433172_8446622_RNA45SN1_45S_50ntFlanks@5472.5501.30\|ACGGCCCTGGCGGAGCGCTGAGAAGACGGT | >hg38_21_+_8433172_8446622_RNA45SN1_45S_50ntFlanks | 5472 | 5501 | 30 | ACGGCCCTGGCGGAGCGCTGAGAAGACGGT | MB231 | 25 | 38 | 375 | 9 | 29 | 15.8 | 9.8 | 358.8 | 3.0 | 41.7 |
| >hg38_21_+_8433172_8446622_RNA45SN1_45S_50ntFlanks@5472.5502.31\|ACGGCCCTGGCGGAGCGCTGAGAAGACGGTC | >hg38_21_+_8433172_8446622_RNA45SN1_45S_50ntFlanks | 5472 | 5502 | 31 | ACGGCCCTGGCGGAGCGCTGAGAAGACGGTC | BT20 | 17 | 22 | 295 | 33 | 29 | 20.2 | 28.1 | 446.4 | 35.4 | 37.8 |
| >hg38_21_+_8433172_8446622_RNA45SN1_45S_50ntFlanks@5472.5506.35\|ACGGCCCTGGCGGAGCGCTGAGAAGACGGTCGAAC | >hg38_21_+_8433172_8446622_RNA45SN1_45S_50ntFlanks | 5472 | 5506 | 35 | ACGGCCCTGGCGGAGCGCTGAGAAGACGGTCGAAC | MB231 | 14 | 28 | 98 | 25 | 13 | 5.9 | 28.7 | 65.6 | 13.6 | 8.6 |
| >hg38_21_+_8433172_8446622_RNA45SN1_45S_50ntFlanks@5472.5507.36\|ACGGCCCTGGCGGAGCGCTGAGAAGACGGTCGAACT | >hg38_21_+_8433172_8446622_RNA45SN1_45S_50ntFlanks | 5472 | 5507 | 36 | ACGGCCCTGGCGGAGCGCTGAGAAGACGGTCGAACT | BT20 | 14 | 21 | 194 | 5 | 13 | 17.6 | 21.0 | 270.2 | 5.7 | 16.7 |
| >hg38_21_+_8433172_8446622_RNA45SN1_45S_50ntFlanks@5472.5509.38\|ACGGCCCTGGCGGAGCGCTGAGAAGACGGTCGAACTTG | >hg38_21_+_8433172_8446622_RNA45SN1_45S_50ntFlanks | 5472 | 5509 | 38 | ACGGCCCTGGCGGAGCGCTGAGAAGACGGTCGAACTTG | BT20 | 28 | 40 | 199 | 20 | 46 | 25.3 | 32.2 | 190.0 | 11.2 | 51.4 |
| >hg38_21_+_8433172_8446622_RNA45SN1_45S_50ntFlanks@5472.5509.38\|ACGGCCCTGGCGGAGCGCTGAGAAGACGGTCGAACTTG | >hg38_21_+_8433172_8446622_RNA45SN1_45S_50ntFlanks | 5472 | 5509 | 38 | ACGGCCCTGGCGGAGCGCTGAGAAGACGGTCGAACTTG | MB231 | 27 | 39 | 154 | 12 | 25 | 2.9 | 27.5 | 74.6 | 2.1 | 11.3 |
| >hg38_21_+_8433172_8446622_RNA45SN1_45S_50ntFlanks@5472.5511.40\|ACGGCCCTGGCGGAGCGCTGAGAAGACGGTCGAACTTGAC | >hg38_21_+_8433172_8446622_RNA45SN1_45S_50ntFlanks | 5472 | 5511 | 40 | ACGGCCCTGGCGGAGCGCTGAGAAGACGGTCGAACTTGAC | MB231 | 12 | 24 | 65 | 4 | 10 | 3.2 | 17.9 | 34.2 | 1.1 | 6.9 |
| >hg38_21_+_8433172_8446622_RNA45SN1_45S_50ntFlanks@5472.5512.41\|ACGGCCCTGGCGGAGCGCTGAGAAGACGGTCGAACTTGACT | >hg38_21_+_8433172_8446622_RNA45SN1_45S_50ntFlanks | 5472 | 5512 | 41 | ACGGCCCTGGCGGAGCGCTGAGAAGACGGTCGAACTTGACT | BT20 | 26 | 42 | 188 | 37 | 71 | 17.7 | 22.1 | 150.2 | 33.7 | 97.9 |
| >hg38_21_+_8433172_8446622_RNA45SN1_45S_50ntFlanks@5472.5512.41\|ACGGCCCTGGCGGAGCGCTGAGAAGACGGTCGAACTTGACT | >hg38_21_+_8433172_8446622_RNA45SN1_45S_50ntFlanks | 5472 | 5512 | 41 | ACGGCCCTGGCGGAGCGCTGAGAAGACGGTCGAACTTGACT | MB231 | 73 | 150 | 410 | 20 | 36 | 13.6 | 82.4 | 24.0 | 8.1 | 19.2 |
| >hg38_21_+_8433172_8446622_RNA45SN1_45S_50ntFlanks@5472.5513.42\|ACGGCCCTGGCGGAGCGCTGAGAAGACGGTCGAACTTGACTA | >hg38_21_+_8433172_8446622_RNA45SN1_45S_50ntFlanks | 5472 | 5513 | 42 | ACGGCCCTGGCGGAGCGCTGAGAAGACGGTCGAACTTGACTA | MB231 | 11 | 12 | 66 | 3 | 10 | 4.7 | 7.6 | 26.6 | 1.5 | 5.0 |
| >hg38_21_+_8433172_8446622_RNA45SN1_45S_50ntFlanks@5472.5514.43\|ACGGCCCTGGCGGAGCGCTGAGAAGACGGTCGAACTTGACTAT | >hg38_21_+_8433172_8446622_RNA45SN1_45S_50ntFlanks | 5472 | 5514 | 43 | ACGGCCCTGGCGGAGCGCTGAGAAGACGGTCGAACTTGACTAT | MB231 | 37 | 50 | 220 | 4 | 20 | 22.1 | 41.9 | 31.5 | 1.4 | 11.0 |
| >hg38_21_+_8433172_8446622_RNA45SN1_45S_50ntFlanks@5472.5514.43\|ACGGCCCTGGCGGAGCGCTGAGAAGACGGTCGAACTTGACTAT | >hg38_21_+_8433172_8446622_RNA45SN1_45S_50ntFlanks | 5472 | 5514 | 43 | ACGGCCCTGGCGGAGCGCTGAGAAGACGGTCGAACTTGACTAT | MB468 | 11 | 70 | 121 | 21 | 42 | 4.0 | 52.6 | 102.7 | 27.3 | 26.1 |
| >hg38_21_+_8433172_8446622_RNA45SN1_45S_50ntFlanks@5472.5515.44\|ACGGCCCTGGCGGAGCGCTGAGAAGACGGTCGAACTTGACTATC | >hg38_21_+_8433172_8446622_RNA45SN1_45S_50ntFlanks | 5472 | 5515 | 44 | ACGGCCCTGGCGGAGCGCTGAGAAGACGGTCGAACTTGACTATC | MB231 | 12 | 17 | 134 | 2 | 7 | 0.8 | 13.9 | 70.0 | 1.7 | 2.9 |
| >hg38_21_+_8433172_8446622_RNA45SN1_45S_50ntFlanks@5472.5516.45\|ACGGCCCTGGCGGAGCGCTGAGAAGACGGTCGAACTTGACTATCT | >hg38_21_+_8433172_8446622_RNA45SN1_45S_50ntFlanks | 5472 | 5516 | 45 | ACGGCCCTGGCGGAGCGCTGAGAAGACGGTCGAACTTGACTATCT | BT20 | 40 | 36 | 188 | 113 | 835 | 33.9 | 26.4 | 94.4 | 136.0 | 1229.3 |
| >hg38_21_+_8433172_8446622_RNA45SN1_45S_50ntFlanks@5472.5516.45\|ACGGCCCTGGCGGAGCGCTGAGAAGACGGTCGAACTTGACTATCT | >hg38_21_+_8433172_8446622_RNA45SN1_45S_50ntFlanks | 5472 | 5516 | 45 | ACGGCCCTGGCGGAGCGCTGAGAAGACGGTCGAACTTGACTATCT | MB231 | 292 | 128 | 4066 | 65 | 129 | 271.7 | 59.9 | 649.9 | 88.2 | 116.9 |
| >hg38_21_+_8433172_8446622_RNA45SN1_45S_50ntFlanks@5473.5499.27\|CGGCCCTGGCGGAGCGCTGAGAAGACG | >hg38_21_+_8433172_8446622_RNA45SN1_45S_50ntFlanks | 5473 | 5499 | 27 | CGGCCCTGGCGGAGCGCTGAGAAGACG | BT20 | 20 | 23 | 216 | 26 | 30 | 24.3 | 32.9 | 302.4 | 9.4 | 32.8 |
| >hg38_21_+_8433172_8446622_RNA45SN1_45S_50ntFlanks@5473.5499.27\|CGGCCCTGGCGGAGCGCTGAGAAGACG | >hg38_21_+_8433172_8446622_RNA45SN1_45S_50ntFlanks | 5473 | 5499 | 27 | CGGCCCTGGCGGAGCGCTGAGAAGACG | MB231 | 36 | 31 | 138 | 14 | 18 | 28.2 | 23.7 | 113.7 | 8.3 | 15.0 |
| >hg38_21_+_8433172_8446622_RNA45SN1_45S_50ntFlanks@5473.5499.27\|CGGCCCTGGCGGAGCGCTGAGAAGACG | >hg38_21_+_8433172_8446622_RNA45SN1_45S_50ntFlanks | 5473 | 5499 | 27 | CGGCCCTGGCGGAGCGCTGAGAAGACG | MB468 | 37 | 55 | 86 | 160 | 93 | 15.7 | 12.9 | 56.3 | 104.2 | 53.5 |
| >hg38_21_+_8433172_8446622_RNA45SN1_45S_50ntFlanks@5473.5501.29\|CGGCCCTGGCGGAGCGCTGAGAAGACGGT | >hg38_21_+_8433172_8446622_RNA45SN1_45S_50ntFlanks | 5473 | 5501 | 29 | CGGCCCTGGCGGAGCGCTGAGAAGACGGT | BT20 | 28 | 29 | 339 | 18 | 36 | 40.5 | 45.5 | 513.5 | 16.5 | 55.3 |
| >hg38_21_+_8433172_8446622_RNA45SN1_45S_50ntFlanks@5473.5509.37\|CGGCCCTGGCGGAGCGCTGAGAAGACGGTCGAACTTG | >hg38_21_+_8433172_8446622_RNA45SN1_45S_50ntFlanks | 5473 | 5509 | 37 | CGGCCCTGGCGGAGCGCTGAGAAGACGGTCGAACTTG | BT20 | 13 | 16 | 81 | 7 | 26 | 18.4 | 24.7 | 96.9 | 5.2 | 37.6 |
| >hg38_21_+_8433172_8446622_RNA45SN1_45S_50ntFlanks@5473.5512.40\|CGGCCCTGGCGGAGCGCTGAGAAGACGGTCGAACTTGACT | >hg38_21_+_8433172_8446622_RNA45SN1_45S_50ntFlanks | 5473 | 5512 | 40 | CGGCCCTGGCGGAGCGCTGAGAAGACGGTCGAACTTGACT | BT20 | 11 | 17 | 82 | 17 | 37 | 11.3 | 20.2 | 78.8 | 16.4 | 51.8 |
| >hg38_21_+_8433172_8446622_RNA45SN1_45S_50ntFlanks@5473.5512.40\|CGGCCCTGGCGGAGCGCTGAGAAGACGGTCGAACTTGACT | >hg38_21_+_8433172_8446622_RNA45SN1_45S_50ntFlanks | 5473 | 5512 | 40 | CGGCCCTGGCGGAGCGCTGAGAAGACGGTCGAACTTGACT | MB231 | 16 | 12 | 98 | 6 | 10 | 12.3 | 7.2 | 71.3 | 2.9 | 6.0 |
| >hg38_21_+_8433172_8446622_RNA45SN1_45S_50ntFlanks@5473.5516.44\|CGGCCCTGGCGGAGCGCTGAGAAGACGGTCGAACTTGACTATCT | >hg38_21_+_8433172_8446622_RNA45SN1_45S_50ntFlanks | 5473 | 5516 | 44 | CGGCCCTGGCGGAGCGCTGAGAAGACGGTCGAACTTGACTATCT | BT20 | 23 | 28 | 109 | 98 | 731 | 19.5 | 29.4 | 56.8 | 132.6 | 1138.8 |
| >hg38_21_+_8433172_8446622_RNA45SN1_45S_50ntFlanks@5473.5516.44\|CGGCCCTGGCGGAGCGCTGAGAAGACGGTCGAACTTGACTATCT | >hg38_21_+_8433172_8446622_RNA45SN1_45S_50ntFlanks | 5473 | 5516 | 44 | CGGCCCTGGCGGAGCGCTGAGAAGACGGTCGAACTTGACTATCT | MB231 | 62 | 26 | 1651 | 8 | 17 | 37.3 | 30.5 | 1563.1 | 9.2 | 9.8 |
| >hg38_21_+_8433172_8446622_RNA45SN1_45S_50ntFlanks@5476.5516.41\|CCCTGGCGGAGCGCTGAGAAGACGGTCGAACTTGACTATCT | >hg38_21_+_8433172_8446622_RNA45SN1_45S_50ntFlanks | 5476 | 5516 | 41 | CCCTGGCGGAGCGCTGAGAAGACGGTCGAACTTGACTATCT | BT20 | 10 | 12 | 61 | 15 | 116 | 8.8 | 15.3 | 50.4 | 10.4 | 136.2 |
| >hg38_21_+_8433172_8446622_RNA45SN1_45S_50ntFlanks@5482.5499.18\|CGGAGCGCTGAGAAGACG | >hg38_21_+_8433172_8446622_RNA45SN1_45S_50ntFlanks | 5482 | 5499 | 18 | CGGAGCGCTGAGAAGACG | MB468 | 14 | 16 | 18 | 154 | 18 | 2.6 | 8.1 | 8.1 | 74.3 | 10.2 |
| >hg38_21_+_8433172_8446622_RNA45SN1_45S_50ntFlanks@5482.5514.33\|CGGAGCGCTGAGAAGACGGTCGAACTTGACTAT | >hg38_21_+_8433172_8446622_RNA45SN1_45S_50ntFlanks | 5482 | 5514 | 33 | CGGAGCGCTGAGAAGACGGTCGAACTTGACTAT | MB468 | 11 | 26 | 24 | 51 | 27 | 2.1 | 16.9 | 8.4 | 34.2 | 21.6 |
| >hg38_21_+_8433172_8446622_RNA45SN1_45S_50ntFlanks@5484.5509.26\|GAGCGCTGAGAAGACGGTCGAACTTG | >hg38_21_+_8433172_8446622_RNA45SN1_45S_50ntFlanks | 5484 | 5509 | 26 | GAGCGCTGAGAAGACGGTCGAACTTG | MB468 | 11 | 6 | 13 | 72 | 16 | 4.0 | 2.6 | 5.6 | 25.7 | 3.0 |
| >hg38_21_+_8433172_8446622_RNA45SN1_45S_50ntFlanks@5484.5514.31\|GAGCGCTGAGAAGACGGTCGAACTTGACTAT | >hg38_21_+_8433172_8446622_RNA45SN1_45S_50ntFlanks | 5484 | 5514 | 31 | GAGCGCTGAGAAGACGGTCGAACTTGACTAT | MB468 | 214 | 330 | 386 | 600 | 391 | 78.8 | 216.4 | 176.7 | 347.1 | 231.8 |
| >hg38_21_+_8433172_8446622_RNA45SN1_45S_50ntFlanks@5484.5518.35\|GAGCGCTGAGAAGACGGTCGAACTTGACTATCTAG | >hg38_21_+_8433172_8446622_RNA45SN1_45S_50ntFlanks | 5484 | 5518 | 35 | GAGCGCTGAGAAGACGGTCGAACTTGACTATCTAG | MB468 | 10 | 19 | 32 | 28 | 21 | 6.4 | 12.3 | 23.4 | 14.3 | 10.2 |

| | | | | | | | | | | | | | | | | |
|---|---|---|---|---|---|---|---|---|---|---|---|---|---|---|---|---|
| >hg38_21_+_8433172_8446622_RNA45SN1_45S_50ntFlanks@5485.5509.25\|AGCGCTGAGAAGACGGTCGAACTTG | >hg38_21_+_8433172_8446622_RNA45SN1_45S_50ntFlanks | 5485 | 5509 | 25 | AGCGCTGAGAAGACGGTCGAACTTG | MB468 | 22 | 15 | 24 | 188 | 39 | 5.9 | 6.5 | 6.6 | 60.3 | 4.5 |
| >hg38_21_+_8433172_8446622_RNA45SN1_45S_50ntFlanks@5485.5510.26\|AGCGCTGAGAAGACGGTCGAACTTGA | >hg38_21_+_8433172_8446622_RNA45SN1_45S_50ntFlanks | 5485 | 5510 | 26 | AGCGCTGAGAAGACGGTCGAACTTGA | MB468 | 14 | 11 | 12 | 96 | 26 | 4.1 | 5.8 | 6.2 | 18.3 | 5.8 |
| >hg38_21_+_8433172_8446622_RNA45SN1_45S_50ntFlanks@5485.5512.28\|AGCGCTGAGAAGACGGTCGAACTTGACT | >hg38_21_+_8433172_8446622_RNA45SN1_45S_50ntFlanks | 5485 | 5512 | 28 | AGCGCTGAGAAGACGGTCGAACTTGACT | BT20 | 20 | 24 | 150 | 18 | 18 | 31.3 | 38.4 | 244.9 | 25.2 | 26.5 |
| >hg38_21_+_8433172_8446622_RNA45SN1_45S_50ntFlanks@5485.5513.29\|AGCGCTGAGAAGACGGTCGAACTTGACTA | >hg38_21_+_8433172_8446622_RNA45SN1_45S_50ntFlanks | 5485 | 5513 | 29 | AGCGCTGAGAAGACGGTCGAACTTGACTA | MB468 | 11 | 10 | 17 | 89 | 33 | 4.2 | 3.0 | 3.6 | 32.6 | 5.8 |
| >hg38_21_+_8433172_8446622_RNA45SN1_45S_50ntFlanks@5485.5514.30\|AGCGCTGAGAAGACGGTCGAACTTGACTAT | >hg38_21_+_8433172_8446622_RNA45SN1_45S_50ntFlanks | 5485 | 5514 | 30 | AGCGCTGAGAAGACGGTCGAACTTGACTAT | MB468 | 220 | 187 | 426 | 875 | 436 | 62.9 | 71.2 | 140.0 | 251.5 | 130.9 |
| >hg38_21_+_8433172_8446622_RNA45SN1_45S_50ntFlanks@5485.5515.31\|AGCGCTGAGAAGACGGTCGAACTTGACTATC | >hg38_21_+_8433172_8446622_RNA45SN1_45S_50ntFlanks | 5485 | 5515 | 31 | AGCGCTGAGAAGACGGTCGAACTTGACTATC | MB468 | 14 | 24 | 28 | 72 | 42 | 2.8 | 12.7 | 11.6 | 26.7 | 20.3 |
| >hg38_21_+_8433172_8446622_RNA45SN1_45S_50ntFlanks@5485.5516.32\|AGCGCTGAGAAGACGGTCGAACTTGACTATCT | >hg38_21_+_8433172_8446622_RNA45SN1_45S_50ntFlanks | 5485 | 5516 | 32 | AGCGCTGAGAAGACGGTCGAACTTGACTATCT | BT20 | 167 | 314 | 401 | 178 | 661 | 223.1 | 511.3 | 454.2 | 178.2 | 1038.5 |
| >hg38_21_+_8433172_8446622_RNA45SN1_45S_50ntFlanks@5485.5516.32\|AGCGCTGAGAAGACGGTCGAACTTGACTATCT | >hg38_21_+_8433172_8446622_RNA45SN1_45S_50ntFlanks | 5485 | 5516 | 32 | AGCGCTGAGAAGACGGTCGAACTTGACTATCT | MB468 | 43 | 39 | 77 | 70 | 66 | 16.9 | 24.7 | 41.2 | 49.6 | 35.8 |
| >hg38_21_+_8433172_8446622_RNA45SN1_45S_50ntFlanks@5485.5518.34\|AGCGCTGAGAAGACGGTCGAACTTGACTATCTAG | >hg38_21_+_8433172_8446622_RNA45SN1_45S_50ntFlanks | 5485 | 5518 | 34 | AGCGCTGAGAAGACGGTCGAACTTGACTATCTAG | MB468 | 45 | 42 | 121 | 102 | 73 | 39.6 | 24.8 | 127.9 | 55.4 | 34.6 |
| >hg38_21_+_8433172_8446622_RNA45SN1_45S_50ntFlanks@5485.5526.42\|AGCGCTGAGAAGACGGTCGAACTTGACTATCTAGAGGAAGTA | >hg38_21_+_8433172_8446622_RNA45SN1_45S_50ntFlanks | 5485 | 5526 | 42 | AGCGCTGAGAAGACGGTCGAACTTGACTATCTAGAGGAAGTA | MB468 | 19 | 46 | 36 | 35 | 128 | 15.7 | 27.7 | 22.0 | 27.4 | 55.3 |
| >hg38_21_+_8433172_8446622_RNA45SN1_45S_50ntFlanks@5486.5514.29\|GCGCTGAGAAGACGGTCGAACTTGACTAT | >hg38_21_+_8433172_8446622_RNA45SN1_45S_50ntFlanks | 5486 | 5514 | 29 | GCGCTGAGAAGACGGTCGAACTTGACTAT | MB468 | 43 | 37 | 85 | 220 | 88 | 17.7 | 17.3 | 44.9 | 82.6 | 25.9 |
| >hg38_21_+_8433172_8446622_RNA45SN1_45S_50ntFlanks@5486.5516.31\|GCGCTGAGAAGACGGTCGAACTTGACTATCT | >hg38_21_+_8433172_8446622_RNA45SN1_45S_50ntFlanks | 5486 | 5516 | 31 | GCGCTGAGAAGACGGTCGAACTTGACTATCT | BT20 | 22 | 27 | 101 | 16 | 42 | 18.7 | 36.0 | 59.9 | 4.0 | 47.1 |
| >hg38_21_+_8433172_8446622_RNA45SN1_45S_50ntFlanks@5487.5509.23\|CGCTGAGAAGACGGTCGAACTTG | >hg38_21_+_8433172_8446622_RNA45SN1_45S_50ntFlanks | 5487 | 5509 | 23 | CGCTGAGAAGACGGTCGAACTTG | MB468 | 11 | 7 | 15 | 98 | 17 | 4.7 | 4.5 | 5.3 | 45.0 | 4.8 |
| >hg38_21_+_8433172_8446622_RNA45SN1_45S_50ntFlanks@5487.5514.28\|CGCTGAGAAGACGGTCGAACTTGACTAT | >hg38_21_+_8433172_8446622_RNA45SN1_45S_50ntFlanks | 5487 | 5514 | 28 | CGCTGAGAAGACGGTCGAACTTGACTAT | MB468 | 190 | 81 | 322 | 704 | 355 | 58.1 | 42.0 | 62.8 | 173.4 | 104.5 |
| >hg38_21_+_8433172_8446622_RNA45SN1_45S_50ntFlanks@5487.5515.29\|CGCTGAGAAGACGGTCGAACTTGACTATC | >hg38_21_+_8433172_8446622_RNA45SN1_45S_50ntFlanks | 5487 | 5515 | 29 | CGCTGAGAAGACGGTCGAACTTGACTATC | MB468 | 12 | 10 | 18 | 79 | 42 | 3.6 | 4.9 | 4.1 | 18.5 | 21.7 |
| >hg38_21_+_8433172_8446622_RNA45SN1_45S_50ntFlanks@5487.5516.30\|CGCTGAGAAGACGGTCGAACTTGACTATCT | >hg38_21_+_8433172_8446622_RNA45SN1_45S_50ntFlanks | 5487 | 5516 | 30 | CGCTGAGAAGACGGTCGAACTTGACTATCT | BT20 | 57 | 50 | 309 | 46 | 76 | 50.1 | 66.3 | 166.9 | 16.4 | 34.2 |
| >hg38_21_+_8433172_8446622_RNA45SN1_45S_50ntFlanks@5487.5516.30\|CGCTGAGAAGACGGTCGAACTTGACTATCT | >hg38_21_+_8433172_8446622_RNA45SN1_45S_50ntFlanks | 5487 | 5516 | 30 | CGCTGAGAAGACGGTCGAACTTGACTATCT | MB231 | 24 | 9 | 195 | 10 | 8 | 15.4 | 0.5 | 49.5 | 9.2 | 3.4 |
| >hg38_21_+_8433172_8446622_RNA45SN1_45S_50ntFlanks@5487.5516.30\|CGCTGAGAAGACGGTCGAACTTGACTATCT | >hg38_21_+_8433172_8446622_RNA45SN1_45S_50ntFlanks | 5487 | 5516 | 30 | CGCTGAGAAGACGGTCGAACTTGACTATCT | MB468 | 39 | 25 | 77 | 60 | 48 | 21.5 | 14.6 | 52.7 | 39.9 | 28.3 |
| >hg38_21_+_8433172_8446622_RNA45SN1_45S_50ntFlanks@5487.5526.40\|CGCTGAGAAGACGGTCGAACTTGACTATCTAGAGGAAGTA | >hg38_21_+_8433172_8446622_RNA45SN1_45S_50ntFlanks | 5487 | 5526 | 40 | CGCTGAGAAGACGGTCGAACTTGACTATCTAGAGGAAGTA | MB468 | 29 | 35 | 52 | 49 | 127 | 20.6 | 16.6 | 35.3 | 30.1 | 39.4 |
| >hg38_21_+_8433172_8446622_RNA45SN1_45S_50ntFlanks@5488.5516.29\|GCTGAGAAGACGGTCGAACTTGACTATCT | >hg38_21_+_8433172_8446622_RNA45SN1_45S_50ntFlanks | 5488 | 5516 | 29 | GCTGAGAAGACGGTCGAACTTGACTATCT | BT20 | 42 | 18 | 311 | 32 | 37 | 56.7 | 6.5 | 309.1 | 17.6 | 12.4 |
| >hg38_21_+_8433172_8446622_RNA45SN1_45S_50ntFlanks@5488.5516.29\|GCTGAGAAGACGGTCGAACTTGACTATCT | >hg38_21_+_8433172_8446622_RNA45SN1_45S_50ntFlanks | 5488 | 5516 | 29 | GCTGAGAAGACGGTCGAACTTGACTATCT | MB231 | 14 | 5 | 75 | 5 | 6 | 16.9 | 1.0 | 50.3 | 3.1 | 2.5 |
| >hg38_21_+_8433172_8446622_RNA45SN1_45S_50ntFlanks@5489.5512.24\|CTGAGAAGACGGTCGAACTTGACT | >hg38_21_+_8433172_8446622_RNA45SN1_45S_50ntFlanks | 5489 | 5512 | 24 | CTGAGAAGACGGTCGAACTTGACT | BT20 | 12 | 5 | 49 | 11 | 4 | 6.2 | 3.0 | 47.1 | 5.3 | 1.6 |
| >hg38_21_+_8433172_8446622_RNA45SN1_45S_50ntFlanks@5489.5512.24\|CTGAGAAGACGGTCGAACTTGACT | >hg38_21_+_8433172_8446622_RNA45SN1_45S_50ntFlanks | 5489 | 5512 | 24 | CTGAGAAGACGGTCGAACTTGACT | MB231 | 24 | 6 | 20 | 8 | 3 | 28.7 | 4.9 | 13.9 | 5.9 | 1.1 |
| >hg38_21_+_8433172_8446622_RNA45SN1_45S_50ntFlanks@5489.5514.26\|CTGAGAAGACGGTCGAACTTGACTAT | >hg38_21_+_8433172_8446622_RNA45SN1_45S_50ntFlanks | 5489 | 5514 | 26 | CTGAGAAGACGGTCGAACTTGACTAT | MB468 | 47 | 8 | 65 | 373 | 74 | 21.7 | 5.7 | 28.9 | 96.9 | 5.9 |
| >hg38_21_+_8433172_8446622_RNA45SN1_45S_50ntFlanks@5489.5516.28\|CTGAGAAGACGGTCGAACTTGACTATCT | >hg38_21_+_8433172_8446622_RNA45SN1_45S_50ntFlanks | 5489 | 5516 | 28 | CTGAGAAGACGGTCGAACTTGACTATCT | BT20 | 162 | 126 | 1164 | 382 | 178 | 199.4 | 176.6 | 1426.7 | 538.6 | 105.4 |
| >hg38_21_+_8433172_8446622_RNA45SN1_45S_50ntFlanks@5489.5516.28\|CTGAGAAGACGGTCGAACTTGACTATCT | >hg38_21_+_8433172_8446622_RNA45SN1_45S_50ntFlanks | 5489 | 5516 | 28 | CTGAGAAGACGGTCGAACTTGACTATCT | MB231 | 70 | 37 | 647 | 15 | 28 | 38.1 | 13.8 | 269.1 | 7.0 | 23.6 |
| >hg38_21_+_8433172_8446622_RNA45SN1_45S_50ntFlanks@5489.5516.28\|CTGAGAAGACGGTCGAACTTGACTATCT | >hg38_21_+_8433172_8446622_RNA45SN1_45S_50ntFlanks | 5489 | 5516 | 28 | CTGAGAAGACGGTCGAACTTGACTATCT | MB468 | 20 | 16 | 49 | 61 | 42 | 12.3 | 8.6 | 35.3 | 56.9 | 26.7 |
| >hg38_21_+_8433172_8446622_RNA45SN1_45S_50ntFlanks@5490.5512.23\|TGAGAAGACGGTCGAACTTGACT | >hg38_21_+_8433172_8446622_RNA45SN1_45S_50ntFlanks | 5490 | 5512 | 23 | TGAGAAGACGGTCGAACTTGACT | BT20 | 15 | 7 | 68 | 11 | 5 | 9.2 | 6.8 | 77.3 | 6.9 | 2.6 |
| >hg38_21_+_8433172_8446622_RNA45SN1_45S_50ntFlanks@5490.5512.23\|TGAGAAGACGGTCGAACTTGACT | >hg38_21_+_8433172_8446622_RNA45SN1_45S_50ntFlanks | 5490 | 5512 | 23 | TGAGAAGACGGTCGAACTTGACT | MB231 | 35 | 8 | 26 | 8 | 4 | 36.4 | 5.8 | 18.1 | 6.3 | 1.3 |
| >hg38_21_+_8433172_8446622_RNA45SN1_45S_50ntFlanks@5490.5516.27\|TGAGAAGACGGTCGAACTTGACTATCT | >hg38_21_+_8433172_8446622_RNA45SN1_45S_50ntFlanks | 5490 | 5516 | 27 | TGAGAAGACGGTCGAACTTGACTATCT | BT20 | 108 | 109 | 706 | 103 | 187 | 141.6 | 165.6 | 866.8 | 69.8 | 208.5 |
| >hg38_21_+_8433172_8446622_RNA45SN1_45S_50ntFlanks@5490.5516.27\|TGAGAAGACGGTCGAACTTGACTATCT | >hg38_21_+_8433172_8446622_RNA45SN1_45S_50ntFlanks | 5490 | 5516 | 27 | TGAGAAGACGGTCGAACTTGACTATCT | MB231 | 52 | 20 | 329 | 10 | 20 | 63.0 | 9.2 | 269.5 | 1.8 | 17.9 |
| >hg38_21_+_8433172_8446622_RNA45SN1_45S_50ntFlanks@5491.5512.22\|GAGAAGACGGTCGAACTTGACT | >hg38_21_+_8433172_8446622_RNA45SN1_45S_50ntFlanks | 5491 | 5512 | 22 | GAGAAGACGGTCGAACTTGACT | BT20 | 27 | 28 | 256 | 25 | 19 | 36.6 | 40.7 | 408.5 | 30.8 | 24.7 |
| >hg38_21_+_8433172_8446622_RNA45SN1_45S_50ntFlanks@5491.5512.22\|GAGAAGACGGTCGAACTTGACT | >hg38_21_+_8433172_8446622_RNA45SN1_45S_50ntFlanks | 5491 | 5512 | 22 | GAGAAGACGGTCGAACTTGACT | MB231 | 20 | 7 | 16 | 10 | 4 | 14.9 | 2.0 | 7.1 | 6.2 | 1.2 |
| >hg38_21_+_8433172_8446622_RNA45SN1_45S_50ntFlanks@5491.5516.26\|GAGAAGACGGTCGAACTTGACTATCT | >hg38_21_+_8433172_8446622_RNA45SN1_45S_50ntFlanks | 5491 | 5516 | 26 | GAGAAGACGGTCGAACTTGACTATCT | BT20 | 540 | 712 | 3185 | 376 | 1045 | 784.9 | 1160.9 | 4653.0 | 418.4 | 1557.7 |
| >hg38_21_+_8433172_8446622_RNA45SN1_45S_50ntFlanks@5491.5516.26\|GAGAAGACGGTCGAACTTGACTATCT | >hg38_21_+_8433172_8446622_RNA45SN1_45S_50ntFlanks | 5491 | 5516 | 26 | GAGAAGACGGTCGAACTTGACTATCT | MB231 | 43 | 20 | 378 | 37 | 30 | 13.4 | 10.2 | 37.5 | 24.8 | 27.4 |
| >hg38_21_+_8433172_8446622_RNA45SN1_45S_50ntFlanks@5491.5525.35\|GAGAAGACGGTCGAACTTGACTATCTAGAGGAAGT | >hg38_21_+_8433172_8446622_RNA45SN1_45S_50ntFlanks | 5491 | 5525 | 35 | GAGAAGACGGTCGAACTTGACTATCTAGAGGAAGT | BT20 | 11 | 26 | 78 | 22 | 11 | 12.8 | 35.1 | 114.2 | 10.7 | 13.5 |
| >hg38_21_+_8433172_8446622_RNA45SN1_45S_50ntFlanks@5492.5509.18\|AGAAGACGGTCGAACTTG | >hg38_21_+_8433172_8446622_RNA45SN1_45S_50ntFlanks | 5492 | 5509 | 18 | AGAAGACGGTCGAACTTG | MB468 | 42 | 17 | 37 | 80 | 23 | 31.9 | 7.0 | 28.2 | 39.6 | 6.6 |
| >hg38_21_+_8433172_8446622_RNA45SN1_45S_50ntFlanks@5492.5510.19\|AGAAGACGGTCGAACTTGA | >hg38_21_+_8433172_8446622_RNA45SN1_45S_50ntFlanks | 5492 | 5510 | 19 | AGAAGACGGTCGAACTTGA | MB468 | 31 | 13 | 18 | 41 | 13 | 14.4 | 4.5 | 9.5 | 15.6 | 4.1 |
| >hg38_21_+_8433172_8446622_RNA45SN1_45S_50ntFlanks@5492.5512.21\|AGAAGACGGTCGAACTTGACT | >hg38_21_+_8433172_8446622_RNA45SN1_45S_50ntFlanks | 5492 | 5512 | 21 | AGAAGACGGTCGAACTTGACT | BT20 | 27 | 31 | 137 | 14 | 17 | 37.1 | 45.9 | 213.7 | 15.8 | 17.7 |
| >hg38_21_+_8433172_8446622_RNA45SN1_45S_50ntFlanks@5492.5513.22\|AGAAGACGGTCGAACTTGACTA | >hg38_21_+_8433172_8446622_RNA45SN1_45S_50ntFlanks | 5492 | 5513 | 22 | AGAAGACGGTCGAACTTGACTA | MB468 | 18 | 2 | 15 | 69 | 17 | 14.5 | 1.8 | 17.0 | 41.4 | 11.3 |
| >hg38_21_+_8433172_8446622_RNA45SN1_45S_50ntFlanks@5492.5514.23\|AGAAGACGGTCGAACTTGACTAT | >hg38_21_+_8433172_8446622_RNA45SN1_45S_50ntFlanks | 5492 | 5514 | 23 | AGAAGACGGTCGAACTTGACTAT | MB468 | 144 | 13 | 138 | 565 | 100 | 106.5 | 12.8 | 149.2 | 381.6 | 51.4 |
| >hg38_21_+_8433172_8446622_RNA45SN1_45S_50ntFlanks@5492.5516.25\|AGAAGACGGTCGAACTTGACTATCT | >hg38_21_+_8433172_8446622_RNA45SN1_45S_50ntFlanks | 5492 | 5516 | 25 | AGAAGACGGTCGAACTTGACTATCT | BT20 | 641 | 887 | 2200 | 261 | 832 | 902.2 | 1415.4 | 2880.5 | 208.6 | 1116.2 |
| >hg38_21_+_8433172_8446622_RNA45SN1_45S_50ntFlanks@5492.5516.25\|AGAAGACGGTCGAACTTGACTATCT | >hg38_21_+_8433172_8446622_RNA45SN1_45S_50ntFlanks | 5492 | 5516 | 25 | AGAAGACGGTCGAACTTGACTATCT | MB231 | 53 | 36 | 465 | 33 | 41 | 5.2 | 16.3 | 139.3 | 23.2 | 27.1 |
| >hg38_21_+_8433172_8446622_RNA45SN1_45S_50ntFlanks@5492.5525.34\|AGAAGACGGTCGAACTTGACTATCTAGAGGAAGT | >hg38_21_+_8433172_8446622_RNA45SN1_45S_50ntFlanks | 5492 | 5525 | 34 | AGAAGACGGTCGAACTTGACTATCTAGAGGAAGT | BT20 | 13 | 29 | 56 | 25 | 12 | 16.7 | 35.7 | 71.9 | 23.9 | 13.9 |
| >hg38_21_+_8433172_8446622_RNA45SN1_45S_50ntFlanks@5492.5525.34\|AGAAGACGGTCGAACTTGACTATCTAGAGGAAGT | >hg38_21_+_8433172_8446622_RNA45SN1_45S_50ntFlanks | 5492 | 5525 | 34 | AGAAGACGGTCGAACTTGACTATCTAGAGGAAGT | MB468 | 12 | 8 | 19 | 24 | 27 | 11.0 | 3.3 | 18.6 | 13.7 | 5.0 |
| >hg38_21_+_8433172_8446622_RNA45SN1_45S_50ntFlanks@5492.5526.35\|AGAAGACGGTCGAACTTGACTATCTAGAGGAAGTA | >hg38_21_+_8433172_8446622_RNA45SN1_45S_50ntFlanks | 5492 | 5526 | 35 | AGAAGACGGTCGAACTTGACTATCTAGAGGAAGTA | MB468 | 40 | 26 | 65 | 82 | 129 | 28.2 | 3.3 | 76.8 | 43.0 | 21.9 |
| >hg38_21_+_8433172_8446622_RNA45SN1_45S_50ntFlanks@5493.5516.24\|GAAGACGGTCGAACTTGACTATCT | >hg38_21_+_8433172_8446622_RNA45SN1_45S_50ntFlanks | 5493 | 5516 | 24 | GAAGACGGTCGAACTTGACTATCT | BT20 | 24 | 29 | 121 | 41 | 55 | 20.7 | 35.7 | 75.5 | 37.5 | 28.7 |
| >hg38_21_+_8433172_8446622_RNA45SN1_45S_50ntFlanks@5493.5516.24\|GAAGACGGTCGAACTTGACTATCT | >hg38_21_+_8433172_8446622_RNA45SN1_45S_50ntFlanks | 5493 | 5516 | 24 | GAAGACGGTCGAACTTGACTATCT | MB231 | 13 | 12 | 113 | 5 | 7 | 9.9 | 2.4 | 51.9 | 2.8 | 4.0 |
| >hg38_21_+_8433172_8446622_RNA45SN1_45S_50ntFlanks@5494.5514.21\|AAGACGGTCGAACTTGACTAT | >hg38_21_+_8433172_8446622_RNA45SN1_45S_50ntFlanks | 5494 | 5514 | 21 | AAGACGGTCGAACTTGACTAT | MB468 | 24 | 1 | 20 | 155 | 15 | 10.6 | 1.2 | 15.5 | 88.9 | 9.9 |
| >hg38_21_+_8433172_8446622_RNA45SN1_45S_50ntFlanks@5494.5516.23\|AAGACGGTCGAACTTGACTATCT | >hg38_21_+_8433172_8446622_RNA45SN1_45S_50ntFlanks | 5494 | 5516 | 23 | AAGACGGTCGAACTTGACTATCT | BT20 | 46 | 49 | 247 | 77 | 104 | 44.2 | 63.8 | 215.6 | 75.4 | 64.5 |

| | | | | | | | | | | | | | | | | |
|---|---|---|---|---|---|---|---|---|---|---|---|---|---|---|---|---|
| >hg38_21_+_8433172_8446622_RNA45SN1_45S_50ntFlanks@5494.5516.23\|AAGACGGTCGAACTTGACTATCT | >hg38_21_+_8433172_8446622_RNA45SN1_45S_50ntFlanks | 5494 | 5516 | 23 | AAGACGGTCGAACTTGACTATCT | MB231 | 21 | 27 | 209 | 7 | 12 | 16.1 | 3.2 | 72.3 | 5.2 | 5.9 |
| >hg38_21_+_8433172_8446622_RNA45SN1_45S_50ntFlanks@5494.5526.33\|AAGACGGTCGAACTTGACTATCTAGAGGAAGTA | >hg38_21_+_8433172_8446622_RNA45SN1_45S_50ntFlanks | 5494 | 5526 | 33 | AAGACGGTCGAACTTGACTATCTAGAGGAAGTA | MB468 | 16 | 9 | 18 | 50 | 51 | 11.1 | 3.1 | 17.1 | 22.6 | 19.4 |
| >hg38_21_+_8433172_8446622_RNA45SN1_45S_50ntFlanks@5495.5516.22\|AGACGGTCGAACTTGACTATCT | >hg38_21_+_8433172_8446622_RNA45SN1_45S_50ntFlanks | 5495 | 5516 | 22 | AGACGGTCGAACTTGACTATCT | BT20 | 29 | 44 | 103 | 29 | 54 | 37.5 | 64.0 | 102.3 | 13.7 | 51.1 |
| >hg38_21_+_8433172_8446622_RNA45SN1_45S_50ntFlanks@5496.5516.21\|GACGGTCGAACTTGACTATCT | >hg38_21_+_8433172_8446622_RNA45SN1_45S_50ntFlanks | 5496 | 5516 | 21 | GACGGTCGAACTTGACTATCT | BT20 | 16 | 31 | 41 | 17 | 32 | 20.5 | 44.9 | 44.3 | 6.9 | 41.0 |
| >hg38_21_+_8433172_8446622_RNA45SN1_45S_50ntFlanks@5497.5526.30\|ACGGTCGAACTTGACTATCTAGAGGAAGTA | >hg38_21_+_8433172_8446622_RNA45SN1_45S_50ntFlanks | 5497 | 5526 | 30 | ACGGTCGAACTTGACTATCTAGAGGAAGTA | MB468 | 12 | 8 | 11 | 24 | 19 | 9.4 | 6.0 | 7.4 | 13.7 | 7.2 |
| >hg38_21_+_8433172_8446622_RNA45SN1_45S_50ntFlanks@5515.5533.19\|CTAGAGGAAGTAAAAGTCG | >hg38_21_+_8433172_8446622_RNA45SN1_45S_50ntFlanks | 5515 | 5533 | 19 | CTAGAGGAAGTAAAAGTCG | MB468 | 12 | 5 | 17 | 22 | 11 | 4.0 | 3.1 | 9.0 | 5.2 | 8.4 |
| >hg38_21_+_8433172_8446622_RNA45SN1_45S_50ntFlanks@5515.5534.20\|CTAGAGGAAGTAAAAGTCGT | >hg38_21_+_8433172_8446622_RNA45SN1_45S_50ntFlanks | 5515 | 5534 | 20 | CTAGAGGAAGTAAAAGTCGT | MB468 | 12 | 4 | 15 | 20 | 10 | 4.7 | 1.7 | 7.5 | 6.9 | 7.8 |
| >hg38_21_+_8433172_8446622_RNA45SN1_45S_50ntFlanks@5515.5535.21\|CTAGAGGAAGTAAAAGTCGTA | >hg38_21_+_8433172_8446622_RNA45SN1_45S_50ntFlanks | 5515 | 5535 | 21 | CTAGAGGAAGTAAAAGTCGTA | MB468 | 34 | 12 | 37 | 46 | 27 | 14.2 | 4.9 | 18.6 | 6.0 | 18.1 |
| >hg38_21_+_8433172_8446622_RNA45SN1_45S_50ntFlanks@5523.5550.28\|AGTAAAAGTCGTAACAAGGTTTCCGTAG | >hg38_21_+_8433172_8446622_RNA45SN1_45S_50ntFlanks | 5523 | 5550 | 28 | AGTAAAAGTCGTAACAAGGTTTCCGTAG | MB468 | 12 | 77 | 10 | 5 | 2 | 2.8 | 47.3 | 4.6 | 2.7 | 0.8 |
| >hg38_21_+_8433172_8446622_RNA45SN1_45S_50ntFlanks@5524.5550.27\|GTAAAAGTCGTAACAAGGTTTCCGTAG | >hg38_21_+_8433172_8446622_RNA45SN1_45S_50ntFlanks | 5524 | 5550 | 27 | GTAAAAGTCGTAACAAGGTTTCCGTAG | MB468 | 20 | 45 | 23 | 9 | 4 | 2.4 | 29.0 | 8.9 | 4.8 | 0.3 |
| >hg38_21_+_8433172_8446622_RNA45SN1_45S_50ntFlanks@5525.5549.25\|TAAAAGTCGTAACAAGGTTTCCGTA | >hg38_21_+_8433172_8446622_RNA45SN1_45S_50ntFlanks | 5525 | 5549 | 25 | TAAAAGTCGTAACAAGGTTTCCGTA | MB468 | 10 | 5 | 8 | 8 | 4 | 1.2 | 0.9 | 4.2 | 3.4 | 1.1 |
| >hg38_21_+_8433172_8446622_RNA45SN1_45S_50ntFlanks@5525.5550.26\|TAAAAGTCGTAACAAGGTTTCCGTAG | >hg38_21_+_8433172_8446622_RNA45SN1_45S_50ntFlanks | 5525 | 5550 | 26 | TAAAAGTCGTAACAAGGTTTCCGTAG | MB468 | 25 | 51 | 23 | 14 | 7 | 2.0 | 31.9 | 7.0 | 7.3 | 1.3 |
| >hg38_21_+_8433172_8446622_RNA45SN1_45S_50ntFlanks@5526.5550.25\|AAAAGTCGTAACAAGGTTTCCGTAG | >hg38_21_+_8433172_8446622_RNA45SN1_45S_50ntFlanks | 5526 | 5550 | 25 | AAAAGTCGTAACAAGGTTTCCGTAG | MB468 | 19 | 37 | 13 | 11 | 5 | 2.9 | 23.4 | 6.3 | 5.5 | 1.0 |
| >hg38_21_+_8433172_8446622_RNA45SN1_45S_50ntFlanks@5527.5548.22\|AAAGTCGTAACAAGGTTTCCGT | >hg38_21_+_8433172_8446622_RNA45SN1_45S_50ntFlanks | 5527 | 5548 | 22 | AAAGTCGTAACAAGGTTTCCGT | MB468 | 12 | 14 | 8 | 20 | 4 | 2.4 | 10.5 | 3.4 | 7.0 | 2.0 |
| >hg38_21_+_8433172_8446622_RNA45SN1_45S_50ntFlanks@5527.5549.23\|AAAGTCGTAACAAGGTTTCCGTA | >hg38_21_+_8433172_8446622_RNA45SN1_45S_50ntFlanks | 5527 | 5549 | 23 | AAAGTCGTAACAAGGTTTCCGTA | MB468 | 35 | 14 | 18 | 37 | 11 | 9.4 | 5.0 | 10.0 | 11.8 | 4.8 |
| >hg38_21_+_8433172_8446622_RNA45SN1_45S_50ntFlanks@5527.5550.24\|AAAGTCGTAACAAGGTTTCCGTAG | >hg38_21_+_8433172_8446622_RNA45SN1_45S_50ntFlanks | 5527 | 5550 | 24 | AAAGTCGTAACAAGGTTTCCGTAG | MB468 | 102 | 151 | 61 | 74 | 30 | 23.0 | 115.8 | 25.2 | 30.0 | 11.8 |
| >hg38_21_+_8433172_8446622_RNA45SN1_45S_50ntFlanks@5527.5553.27\|AAAGTCGTAACAAGGTTTCCGTAGGTG | >hg38_21_+_8433172_8446622_RNA45SN1_45S_50ntFlanks | 5527 | 5553 | 27 | AAAGTCGTAACAAGGTTTCCGTAGGTG | MB468 | 10 | 10 | 10 | 5 | 5 | 2.8 | 4.8 | 3.8 | 0.7 | 3.6 |
| >hg38_21_+_8433172_8446622_RNA45SN1_45S_50ntFlanks@6643.6683.41\|ACCTCGTACGACTCTTAGCGGTGGATCACTCGGCTCGTGCG | >hg38_21_+_8433172_8446622_RNA45SN1_45S_50ntFlanks | 6643 | 6683 | 41 | ACCTCGTACGACTCTTAGCGGTGGATCACTCGGCTCGTGCG | MB231 | 14 | 42 | 0 | 2 | 2 | 3.1 | 27.3 | 0.2 | 1.9 | 0.6 |
| >hg38_21_+_8433172_8446622_RNA45SN1_45S_50ntFlanks@6644.6665.22\|CCTCGTACGACTCTTAGCGGTG | >hg38_21_+_8433172_8446622_RNA45SN1_45S_50ntFlanks | 6644 | 6665 | 22 | CCTCGTACGACTCTTAGCGGTG | BT20 | 18 | 34 | 14 | 45 | 61 | 9.2 | 21.1 | 2.3 | 17.1 | 12.3 |
| >hg38_21_+_8433172_8446622_RNA45SN1_45S_50ntFlanks@6644.6665.22\|CCTCGTACGACTCTTAGCGGTG | >hg38_21_+_8433172_8446622_RNA45SN1_45S_50ntFlanks | 6644 | 6665 | 22 | CCTCGTACGACTCTTAGCGGTG | MB231 | 14 | 39 | 14 | 12 | 119 | 19.0 | 22.9 | 14.2 | 16.6 | 154.9 |
| >hg38_21_+_8433172_8446622_RNA45SN1_45S_50ntFlanks@6644.6669.26\|CCTCGTACGACTCTTAGCGGTGGATC | >hg38_21_+_8433172_8446622_RNA45SN1_45S_50ntFlanks | 6644 | 6669 | 26 | CCTCGTACGACTCTTAGCGGTGGATC | MB231 | 11 | 11 | 9 | 5 | 35 | 16.6 | 9.6 | 11.3 | 7.2 | 42.2 |
| >hg38_21_+_8433172_8446622_RNA45SN1_45S_50ntFlanks@6644.6683.40\|CCTCGTACGACTCTTAGCGGTGGATCACTCGGCTCGTGCG | >hg38_21_+_8433172_8446622_RNA45SN1_45S_50ntFlanks | 6644 | 6683 | 40 | CCTCGTACGACTCTTAGCGGTGGATCACTCGGCTCGTGCG | MB231 | 22 | 58 | 1 | 4 | 7 | 2.6 | 34.6 | 0.6 | 4.6 | 0.8 |
| >hg38_21_+_8433172_8446622_RNA45SN1_45S_50ntFlanks@6645.6662.18\|CTCGTACGACTCTTAGCG | >hg38_21_+_8433172_8446622_RNA45SN1_45S_50ntFlanks | 6645 | 6662 | 18 | CTCGTACGACTCTTAGCG | BT20 | 132 | 178 | 205 | 532 | 390 | 61.7 | 76.1 | 101.2 | 115.1 | 18.9 |
| >hg38_21_+_8433172_8446622_RNA45SN1_45S_50ntFlanks@6645.6662.18\|CTCGTACGACTCTTAGCG | >hg38_21_+_8433172_8446622_RNA45SN1_45S_50ntFlanks | 6645 | 6662 | 18 | CTCGTACGACTCTTAGCG | MB231 | 45 | 336 | 240 | 113 | 1034 | 54.1 | 276.9 | 156.8 | 139.0 | 1520.2 |
| >hg38_21_+_8433172_8446622_RNA45SN1_45S_50ntFlanks@6645.6662.18\|CTCGTACGACTCTTAGCG | >hg38_21_+_8433172_8446622_RNA45SN1_45S_50ntFlanks | 6645 | 6662 | 18 | CTCGTACGACTCTTAGCG | MB468 | 63 | 55 | 84 | 69 | 48 | 29.2 | 51.1 | 50.5 | 22.0 | 21.6 |
| >hg38_21_+_8433172_8446622_RNA45SN1_45S_50ntFlanks@6645.6663.19\|CTCGTACGACTCTTAGCGG | >hg38_21_+_8433172_8446622_RNA45SN1_45S_50ntFlanks | 6645 | 6663 | 19 | CTCGTACGACTCTTAGCGG | BT20 | 78 | 105 | 105 | 253 | 263 | 45.8 | 39.6 | 42.2 | 16.6 | 38.9 |
| >hg38_21_+_8433172_8446622_RNA45SN1_45S_50ntFlanks@6645.6663.19\|CTCGTACGACTCTTAGCGG | >hg38_21_+_8433172_8446622_RNA45SN1_45S_50ntFlanks | 6645 | 6663 | 19 | CTCGTACGACTCTTAGCGG | MB231 | 25 | 266 | 158 | 51 | 1057 | 29.6 | 249.9 | 52.2 | 43.0 | 1658.6 |
| >hg38_21_+_8433172_8446622_RNA45SN1_45S_50ntFlanks@6645.6663.19\|CTCGTACGACTCTTAGCGG | >hg38_21_+_8433172_8446622_RNA45SN1_45S_50ntFlanks | 6645 | 6663 | 19 | CTCGTACGACTCTTAGCGG | MB468 | 44 | 49 | 43 | 58 | 66 | 19.3 | 49.9 | 31.3 | 26.2 | 22.6 |
| >hg38_21_+_8433172_8446622_RNA45SN1_45S_50ntFlanks@6645.6664.20\|CTCGTACGACTCTTAGCGGT | >hg38_21_+_8433172_8446622_RNA45SN1_45S_50ntFlanks | 6645 | 6664 | 20 | CTCGTACGACTCTTAGCGGT | BT20 | 117 | 143 | 162 | 410 | 412 | 38.5 | 57.9 | 59.2 | 71.0 | 85.2 |
| >hg38_21_+_8433172_8446622_RNA45SN1_45S_50ntFlanks@6645.6664.20\|CTCGTACGACTCTTAGCGGT | >hg38_21_+_8433172_8446622_RNA45SN1_45S_50ntFlanks | 6645 | 6664 | 20 | CTCGTACGACTCTTAGCGGT | MB231 | 56 | 119 | 123 | 115 | 423 | 69.2 | 48.8 | 110.5 | 118.0 | 421.0 |
| >hg38_21_+_8433172_8446622_RNA45SN1_45S_50ntFlanks@6645.6664.20\|CTCGTACGACTCTTAGCGGT | >hg38_21_+_8433172_8446622_RNA45SN1_45S_50ntFlanks | 6645 | 6664 | 20 | CTCGTACGACTCTTAGCGGT | MB468 | 12 | 18 | 12 | 21 | 35 | 5.4 | 12.0 | 10.3 | 7.9 | 12.5 |
| >hg38_21_+_8433172_8446622_RNA45SN1_45S_50ntFlanks@6645.6665.21\|CTCGTACGACTCTTAGCGGTG | >hg38_21_+_8433172_8446622_RNA45SN1_45S_50ntFlanks | 6645 | 6665 | 21 | CTCGTACGACTCTTAGCGGTG | BT20 | 405 | 473 | 511 | 1530 | 1158 | 73.9 | 274.3 | 119.9 | 718.2 | 209.9 |
| >hg38_21_+_8433172_8446622_RNA45SN1_45S_50ntFlanks@6645.6665.21\|CTCGTACGACTCTTAGCGGTG | >hg38_21_+_8433172_8446622_RNA45SN1_45S_50ntFlanks | 6645 | 6665 | 21 | CTCGTACGACTCTTAGCGGTG | MB231 | 103 | 272 | 184 | 180 | 907 | 131.9 | 166.0 | 142.5 | 263.1 | 1133.1 |
| >hg38_21_+_8433172_8446622_RNA45SN1_45S_50ntFlanks@6645.6665.21\|CTCGTACGACTCTTAGCGGTG | >hg38_21_+_8433172_8446622_RNA45SN1_45S_50ntFlanks | 6645 | 6665 | 21 | CTCGTACGACTCTTAGCGGTG | MB468 | 34 | 44 | 39 | 47 | 62 | 15.0 | 37.5 | 25.0 | 17.6 | 27.5 |
| >hg38_21_+_8433172_8446622_RNA45SN1_45S_50ntFlanks@6645.6666.22\|CTCGTACGACTCTTAGCGGTGG | >hg38_21_+_8433172_8446622_RNA45SN1_45S_50ntFlanks | 6645 | 6666 | 22 | CTCGTACGACTCTTAGCGGTGG | BT20 | 92 | 130 | 139 | 229 | 431 | 36.3 | 71.6 | 57.9 | 17.0 | 48.7 |
| >hg38_21_+_8433172_8446622_RNA45SN1_45S_50ntFlanks@6645.6666.22\|CTCGTACGACTCTTAGCGGTGG | >hg38_21_+_8433172_8446622_RNA45SN1_45S_50ntFlanks | 6645 | 6666 | 22 | CTCGTACGACTCTTAGCGGTGG | MB231 | 44 | 135 | 84 | 27 | 483 | 55.0 | 88.4 | 65.5 | 30.9 | 601.6 |
| >hg38_21_+_8433172_8446622_RNA45SN1_45S_50ntFlanks@6645.6666.22\|CTCGTACGACTCTTAGCGGTGG | >hg38_21_+_8433172_8446622_RNA45SN1_45S_50ntFlanks | 6645 | 6666 | 22 | CTCGTACGACTCTTAGCGGTGG | MB468 | 53 | 61 | 69 | 30 | 87 | 39.2 | 60.2 | 77.0 | 17.3 | 55.8 |
| >hg38_21_+_8433172_8446622_RNA45SN1_45S_50ntFlanks@6645.6667.23\|CTCGTACGACTCTTAGCGGTGGA | >hg38_21_+_8433172_8446622_RNA45SN1_45S_50ntFlanks | 6645 | 6667 | 23 | CTCGTACGACTCTTAGCGGTGGA | BT20 | 84 | 108 | 169 | 134 | 388 | 24.9 | 63.0 | 94.7 | 0.1 | 90.5 |
| >hg38_21_+_8433172_8446622_RNA45SN1_45S_50ntFlanks@6645.6667.23\|CTCGTACGACTCTTAGCGGTGGA | >hg38_21_+_8433172_8446622_RNA45SN1_45S_50ntFlanks | 6645 | 6667 | 23 | CTCGTACGACTCTTAGCGGTGGA | MB231 | 47 | 63 | 55 | 26 | 285 | 55.4 | 26.0 | 46.3 | 29.5 | 244.3 |
| >hg38_21_+_8433172_8446622_RNA45SN1_45S_50ntFlanks@6645.6668.24\|CTCGTACGACTCTTAGCGGTGGAT | >hg38_21_+_8433172_8446622_RNA45SN1_45S_50ntFlanks | 6645 | 6668 | 24 | CTCGTACGACTCTTAGCGGTGGAT | BT20 | 71 | 89 | 103 | 84 | 303 | 14.1 | 39.9 | 55.0 | 6.0 | 108.6 |
| >hg38_21_+_8433172_8446622_RNA45SN1_45S_50ntFlanks@6645.6668.24\|CTCGTACGACTCTTAGCGGTGGAT | >hg38_21_+_8433172_8446622_RNA45SN1_45S_50ntFlanks | 6645 | 6668 | 24 | CTCGTACGACTCTTAGCGGTGGAT | MB231 | 39 | 54 | 36 | 31 | 175 | 34.8 | 32.7 | 21.9 | 8.1 | 142.4 |
| >hg38_21_+_8433172_8446622_RNA45SN1_45S_50ntFlanks@6645.6668.24\|CTCGTACGACTCTTAGCGGTGGAT | >hg38_21_+_8433172_8446622_RNA45SN1_45S_50ntFlanks | 6645 | 6668 | 24 | CTCGTACGACTCTTAGCGGTGGAT | MB468 | 10 | 9 | 7 | 13 | 20 | 4.7 | 5.4 | 3.8 | 2.4 | 8.3 |
| >hg38_21_+_8433172_8446622_RNA45SN1_45S_50ntFlanks@6645.6669.25\|CTCGTACGACTCTTAGCGGTGGATC | >hg38_21_+_8433172_8446622_RNA45SN1_45S_50ntFlanks | 6645 | 6669 | 25 | CTCGTACGACTCTTAGCGGTGGATC | BT20 | 147 | 279 | 241 | 174 | 1006 | 41.2 | 162.8 | 75.2 | 94.9 | 230.6 |
| >hg38_21_+_8433172_8446622_RNA45SN1_45S_50ntFlanks@6645.6669.25\|CTCGTACGACTCTTAGCGGTGGATC | >hg38_21_+_8433172_8446622_RNA45SN1_45S_50ntFlanks | 6645 | 6669 | 25 | CTCGTACGACTCTTAGCGGTGGATC | MB231 | 68 | 74 | 105 | 37 | 255 | 98.8 | 49.9 | 142.6 | 54.1 | 183.3 |
| >hg38_21_+_8433172_8446622_RNA45SN1_45S_50ntFlanks@6645.6669.25\|CTCGTACGACTCTTAGCGGTGGATC | >hg38_21_+_8433172_8446622_RNA45SN1_45S_50ntFlanks | 6645 | 6669 | 25 | CTCGTACGACTCTTAGCGGTGGATC | MB468 | 57 | 53 | 33 | 60 | 222 | 17.7 | 18.7 | 17.1 | 13.5 | 46.3 |
| >hg38_21_+_8433172_8446622_RNA45SN1_45S_50ntFlanks@6645.6670.26\|CTCGTACGACTCTTAGCGGTGGATCA | >hg38_21_+_8433172_8446622_RNA45SN1_45S_50ntFlanks | 6645 | 6670 | 26 | CTCGTACGACTCTTAGCGGTGGATCA | MB468 | 51 | 29 | 20 | 37 | 199 | 12.9 | 5.4 | 3.9 | 5.3 | 16.9 |
| >hg38_21_+_8433172_8446622_RNA45SN1_45S_50ntFlanks@6645.6672.28\|CTCGTACGACTCTTAGCGGTGGATCACT | >hg38_21_+_8433172_8446622_RNA45SN1_45S_50ntFlanks | 6645 | 6672 | 28 | CTCGTACGACTCTTAGCGGTGGATCACT | BT20 | 42 | 67 | 59 | 29 | 186 | 10.8 | 10.8 | 21.0 | 22.9 | 93.9 |
| >hg38_21_+_8433172_8446622_RNA45SN1_45S_50ntFlanks@6645.6672.28\|CTCGTACGACTCTTAGCGGTGGATCACT | >hg38_21_+_8433172_8446622_RNA45SN1_45S_50ntFlanks | 6645 | 6672 | 28 | CTCGTACGACTCTTAGCGGTGGATCACT | MB231 | 16 | 43 | 38 | 7 | 175 | 21.6 | 27.3 | 28.4 | 6.7 | 211.6 |
| >hg38_21_+_8433172_8446622_RNA45SN1_45S_50ntFlanks@6645.6674.30\|CTCGTACGACTCTTAGCGGTGGATCACTCG | >hg38_21_+_8433172_8446622_RNA45SN1_45S_50ntFlanks | 6645 | 6674 | 30 | CTCGTACGACTCTTAGCGGTGGATCACTCG | BT20 | 13 | 30 | 19 | 74 | 160 | 5.8 | 10.2 | 7.1 | 81.6 | 29.4 |
| >hg38_21_+_8433172_8446622_RNA45SN1_45S_50ntFlanks@6645.6676.32\|CTCGTACGACTCTTAGCGGTGGATCACTCGGC | >hg38_21_+_8433172_8446622_RNA45SN1_45S_50ntFlanks | 6645 | 6676 | 32 | CTCGTACGACTCTTAGCGGTGGATCACTCGGC | BT20 | 11 | 32 | 42 | 62 | 120 | 4.1 | 8.7 | 44.6 | 33.7 | 22.1 |

| | | | | | | | | | | | | | | | | |
|---|---|---|---|---|---|---|---|---|---|---|---|---|---|---|---|---|
| >hg38_21_+_8433172_8446622_RNA45SN1_45S_50ntFlanks@6645.6677.33\|CTCGTACGACTCTTAGCGGTGGATCACTCGGCT | >hg38_21_+_8433172_8446622_RNA45SN1_45S_50ntFlanks | 6645 | 6677 | 33 | CTCGTACGACTCTTAGCGGTGGATCACTCGGCT | BT20 | 19 | 38 | 21 | 55 | 103 | 5.8 | 3.9 | 4.3 | 24.3 | 21.6 |
| >hg38_21_+_8433172_8446622_RNA45SN1_45S_50ntFlanks@6645.6678.34\|CTCGTACGACTCTTAGCGGTGGATCACTCGGCTC | >hg38_21_+_8433172_8446622_RNA45SN1_45S_50ntFlanks | 6645 | 6678 | 34 | CTCGTACGACTCTTAGCGGTGGATCACTCGGCTC | BT20 | 15 | 34 | 15 | 48 | 93 | 5.7 | 4.3 | 1.6 | 11.5 | 19.2 |
| >hg38_21_+_8433172_8446622_RNA45SN1_45S_50ntFlanks@6645.6679.35\|CTCGTACGACTCTTAGCGGTGGATCACTCGGCTCG | >hg38_21_+_8433172_8446622_RNA45SN1_45S_50ntFlanks | 6645 | 6679 | 35 | CTCGTACGACTCTTAGCGGTGGATCACTCGGCTCG | MB231 | 16 | 52 | 22 | 5 | 201 | 20.0 | 33.4 | 13.0 | 0.6 | 265.7 |
| >hg38_21_+_8433172_8446622_RNA45SN1_45S_50ntFlanks@6645.6679.35\|CTCGTACGACTCTTAGCGGTGGATCACTCGGCTCG | >hg38_21_+_8433172_8446622_RNA45SN1_45S_50ntFlanks | 6645 | 6679 | 35 | CTCGTACGACTCTTAGCGGTGGATCACTCGGCTCG | MB468 | 28 | 68 | 16 | 35 | 279 | 4.9 | 24.8 | 7.3 | 30.8 | 220.3 |
| >hg38_21_+_8433172_8446622_RNA45SN1_45S_50ntFlanks@6645.6680.36\|CTCGTACGACTCTTAGCGGTGGATCACTCGGCTCGT | >hg38_21_+_8433172_8446622_RNA45SN1_45S_50ntFlanks | 6645 | 6680 | 36 | CTCGTACGACTCTTAGCGGTGGATCACTCGGCTCGT | BT20 | 18 | 37 | 28 | 56 | 102 | 2.7 | 15.0 | 14.8 | 31.1 | 5.7 |
| >hg38_21_+_8433172_8446622_RNA45SN1_45S_50ntFlanks@6645.6680.36\|CTCGTACGACTCTTAGCGGTGGATCACTCGGCTCGT | >hg38_21_+_8433172_8446622_RNA45SN1_45S_50ntFlanks | 6645 | 6680 | 36 | CTCGTACGACTCTTAGCGGTGGATCACTCGGCTCGT | MB231 | 13 | 42 | 20 | 9 | 132 | 17.7 | 28.0 | 10.3 | 9.3 | 167.5 |
| >hg38_21_+_8433172_8446622_RNA45SN1_45S_50ntFlanks@6645.6680.36\|CTCGTACGACTCTTAGCGGTGGATCACTCGGCTCGT | >hg38_21_+_8433172_8446622_RNA45SN1_45S_50ntFlanks | 6645 | 6680 | 36 | CTCGTACGACTCTTAGCGGTGGATCACTCGGCTCGT | MB468 | 16 | 27 | 8 | 23 | 97 | 3.8 | 14.2 | 4.9 | 15.2 | 57.8 |
| >hg38_21_+_8433172_8446622_RNA45SN1_45S_50ntFlanks@6645.6681.37\|CTCGTACGACTCTTAGCGGTGGATCACTCGGCTCGTG | >hg38_21_+_8433172_8446622_RNA45SN1_45S_50ntFlanks | 6645 | 6681 | 37 | CTCGTACGACTCTTAGCGGTGGATCACTCGGCTCGTG | MB231 | 14 | 98 | 23 | 15 | 211 | 10.7 | 47.9 | 14.5 | 12.2 | 318.2 |
| >hg38_21_+_8433172_8446622_RNA45SN1_45S_50ntFlanks@6645.6681.37\|CTCGTACGACTCTTAGCGGTGGATCACTCGGCTCGTG | >hg38_21_+_8433172_8446622_RNA45SN1_45S_50ntFlanks | 6645 | 6681 | 37 | CTCGTACGACTCTTAGCGGTGGATCACTCGGCTCGTG | MB468 | 121 | 177 | 46 | 237 | 708 | 28.1 | 79.3 | 24.9 | 172.6 | 368.8 |
| >hg38_21_+_8433172_8446622_RNA45SN1_45S_50ntFlanks@6645.6685.41\|CTCGTACGACTCTTAGCGGTGGATCACTCGGCTCGTGCGTC | >hg38_21_+_8433172_8446622_RNA45SN1_45S_50ntFlanks | 6645 | 6685 | 41 | CTCGTACGACTCTTAGCGGTGGATCACTCGGCTCGTGCGTC | BT20 | 15 | 37 | 27 | 99 | 25 | 15.3 | 26.9 | 32.2 | 91.4 | 6.5 |
| >hg38_21_+_8433172_8446622_RNA45SN1_45S_50ntFlanks@6646.6663.18\|TCGTACGACTCTTAGCGG | >hg38_21_+_8433172_8446622_RNA45SN1_45S_50ntFlanks | 6646 | 6663 | 18 | TCGTACGACTCTTAGCGG | BT20 | 502 | 865 | 649 | 1220 | 2186 | 314.5 | 441.5 | 210.3 | 315.5 | 328.0 |
| >hg38_21_+_8433172_8446622_RNA45SN1_45S_50ntFlanks@6646.6663.18\|TCGTACGACTCTTAGCGG | >hg38_21_+_8433172_8446622_RNA45SN1_45S_50ntFlanks | 6646 | 6663 | 18 | TCGTACGACTCTTAGCGG | MB231 | 115 | 1258 | 557 | 201 | 4329 | 132.2 | 1212.8 | 246.2 | 154.9 | 6652.5 |
| >hg38_21_+_8433172_8446622_RNA45SN1_45S_50ntFlanks@6646.6663.18\|TCGTACGACTCTTAGCGG | >hg38_21_+_8433172_8446622_RNA45SN1_45S_50ntFlanks | 6646 | 6663 | 18 | TCGTACGACTCTTAGCGG | MB468 | 315 | 212 | 241 | 366 | 337 | 125.5 | 242.5 | 172.8 | 139.1 | 111.2 |
| >hg38_21_+_8433172_8446622_RNA45SN1_45S_50ntFlanks@6646.6664.19\|TCGTACGACTCTTAGCGGT | >hg38_21_+_8433172_8446622_RNA45SN1_45S_50ntFlanks | 6646 | 6664 | 19 | TCGTACGACTCTTAGCGGT | BT20 | 1097 | 1748 | 1310 | 2439 | 5882 | 361.6 | 869.0 | 360.6 | 1011.6 | 1169.1 |
| >hg38_21_+_8433172_8446622_RNA45SN1_45S_50ntFlanks@6646.6664.19\|TCGTACGACTCTTAGCGGT | >hg38_21_+_8433172_8446622_RNA45SN1_45S_50ntFlanks | 6646 | 6664 | 19 | TCGTACGACTCTTAGCGGT | MB231 | 344 | 826 | 511 | 430 | 2576 | 336.0 | 326.2 | 244.9 | 120.0 | 2317.5 |
| >hg38_21_+_8433172_8446622_RNA45SN1_45S_50ntFlanks@6646.6664.19\|TCGTACGACTCTTAGCGGT | >hg38_21_+_8433172_8446622_RNA45SN1_45S_50ntFlanks | 6646 | 6664 | 19 | TCGTACGACTCTTAGCGGT | MB468 | 103 | 160 | 80 | 127 | 322 | 39.9 | 110.6 | 60.8 | 52.0 | 115.3 |
| >hg38_21_+_8433172_8446622_RNA45SN1_45S_50ntFlanks@6646.6665.20\|TCGTACGACTCTTAGCGGTG | >hg38_21_+_8433172_8446622_RNA45SN1_45S_50ntFlanks | 6646 | 6665 | 20 | TCGTACGACTCTTAGCGGTG | BT20 | 1970 | 2421 | 2199 | 4583 | 5984 | 568.7 | 1561.1 | 128.0 | 1206.0 | 983.0 |
| >hg38_21_+_8433172_8446622_RNA45SN1_45S_50ntFlanks@6646.6665.20\|TCGTACGACTCTTAGCGGTG | >hg38_21_+_8433172_8446622_RNA45SN1_45S_50ntFlanks | 6646 | 6665 | 20 | TCGTACGACTCTTAGCGGTG | MB231 | 282 | 752 | 378 | 336 | 2326 | 357.6 | 510.1 | 211.5 | 468.0 | 2946.7 |
| >hg38_21_+_8433172_8446622_RNA45SN1_45S_50ntFlanks@6646.6665.20\|TCGTACGACTCTTAGCGGTG | >hg38_21_+_8433172_8446622_RNA45SN1_45S_50ntFlanks | 6646 | 6665 | 20 | TCGTACGACTCTTAGCGGTG | MB468 | 132 | 125 | 119 | 143 | 201 | 52.9 | 104.8 | 78.1 | 47.8 | 95.4 |
| >hg38_21_+_8433172_8446622_RNA45SN1_45S_50ntFlanks@6646.6666.21\|TCGTACGACTCTTAGCGGTGG | >hg38_21_+_8433172_8446622_RNA45SN1_45S_50ntFlanks | 6646 | 6666 | 21 | TCGTACGACTCTTAGCGGTGG | BT20 | 452 | 649 | 594 | 974 | 2216 | 165.7 | 327.9 | 125.7 | 147.7 | 582.1 |
| >hg38_21_+_8433172_8446622_RNA45SN1_45S_50ntFlanks@6646.6666.21\|TCGTACGACTCTTAGCGGTGG | >hg38_21_+_8433172_8446622_RNA45SN1_45S_50ntFlanks | 6646 | 6666 | 21 | TCGTACGACTCTTAGCGGTGG | MB231 | 110 | 378 | 146 | 113 | 1288 | 141.5 | 289.3 | 62.3 | 117.7 | 1521.9 |
| >hg38_21_+_8433172_8446622_RNA45SN1_45S_50ntFlanks@6646.6666.21\|TCGTACGACTCTTAGCGGTGG | >hg38_21_+_8433172_8446622_RNA45SN1_45S_50ntFlanks | 6646 | 6666 | 21 | TCGTACGACTCTTAGCGGTGG | MB468 | 309 | 184 | 282 | 166 | 386 | 245.7 | 188.1 | 299.9 | 92.3 | 253.3 |
| >hg38_21_+_8433172_8446622_RNA45SN1_45S_50ntFlanks@6646.6667.22\|TCGTACGACTCTTAGCGGTGGA | >hg38_21_+_8433172_8446622_RNA45SN1_45S_50ntFlanks | 6646 | 6667 | 22 | TCGTACGACTCTTAGCGGTGGA | BT20 | 419 | 520 | 680 | 604 | 1955 | 31.4 | 245.1 | 219.8 | 221.6 | 952.2 |
| >hg38_21_+_8433172_8446622_RNA45SN1_45S_50ntFlanks@6646.6667.22\|TCGTACGACTCTTAGCGGTGGA | >hg38_21_+_8433172_8446622_RNA45SN1_45S_50ntFlanks | 6646 | 6667 | 22 | TCGTACGACTCTTAGCGGTGGA | MB231 | 118 | 159 | 85 | 128 | 834 | 146.2 | 71.6 | 25.1 | 134.8 | 531.7 |
| >hg38_21_+_8433172_8446622_RNA45SN1_45S_50ntFlanks@6646.6667.22\|TCGTACGACTCTTAGCGGTGGA | >hg38_21_+_8433172_8446622_RNA45SN1_45S_50ntFlanks | 6646 | 6667 | 22 | TCGTACGACTCTTAGCGGTGGA | MB468 | 27 | 23 | 26 | 35 | 82 | 14.5 | 12.7 | 19.8 | 16.4 | 52.7 |
| >hg38_21_+_8433172_8446622_RNA45SN1_45S_50ntFlanks@6646.6668.23\|TCGTACGACTCTTAGCGGTGGAT | >hg38_21_+_8433172_8446622_RNA45SN1_45S_50ntFlanks | 6646 | 6668 | 23 | TCGTACGACTCTTAGCGGTGGAT | BT20 | 411 | 501 | 487 | 350 | 1694 | 53.9 | 217.1 | 151.2 | 46.7 | 822.3 |
| >hg38_21_+_8433172_8446622_RNA45SN1_45S_50ntFlanks@6646.6668.23\|TCGTACGACTCTTAGCGGTGGAT | >hg38_21_+_8433172_8446622_RNA45SN1_45S_50ntFlanks | 6646 | 6668 | 23 | TCGTACGACTCTTAGCGGTGGAT | MB231 | 133 | 198 | 84 | 135 | 581 | 106.0 | 133.6 | 39.3 | 38.0 | 411.4 |
| >hg38_21_+_8433172_8446622_RNA45SN1_45S_50ntFlanks@6646.6668.23\|TCGTACGACTCTTAGCGGTGGAT | >hg38_21_+_8433172_8446622_RNA45SN1_45S_50ntFlanks | 6646 | 6668 | 23 | TCGTACGACTCTTAGCGGTGGAT | MB468 | 55 | 36 | 36 | 68 | 91 | 23.7 | 20.2 | 19.0 | 9.1 | 43.9 |
| >hg38_21_+_8433172_8446622_RNA45SN1_45S_50ntFlanks@6646.6669.24\|TCGTACGACTCTTAGCGGTGGATC | >hg38_21_+_8433172_8446622_RNA45SN1_45S_50ntFlanks | 6646 | 6669 | 24 | TCGTACGACTCTTAGCGGTGGATC | BT20 | 955 | 1703 | 1340 | 945 | 5696 | 97.8 | 1013.4 | 101.1 | 390.1 | 1894.1 |
| >hg38_21_+_8433172_8446622_RNA45SN1_45S_50ntFlanks@6646.6669.24\|TCGTACGACTCTTAGCGGTGGATC | >hg38_21_+_8433172_8446622_RNA45SN1_45S_50ntFlanks | 6646 | 6669 | 24 | TCGTACGACTCTTAGCGGTGGATC | MB231 | 258 | 293 | 256 | 161 | 932 | 365.2 | 191.9 | 294.2 | 235.5 | 731.1 |
| >hg38_21_+_8433172_8446622_RNA45SN1_45S_50ntFlanks@6646.6669.24\|TCGTACGACTCTTAGCGGTGGATC | >hg38_21_+_8433172_8446622_RNA45SN1_45S_50ntFlanks | 6646 | 6669 | 24 | TCGTACGACTCTTAGCGGTGGATC | MB468 | 350 | 201 | 172 | 391 | 930 | 109.3 | 82.8 | 79.1 | 125.2 | 137.8 |
| >hg38_21_+_8433172_8446622_RNA45SN1_45S_50ntFlanks@6646.6670.25\|TCGTACGACTCTTAGCGGTGGATCA | >hg38_21_+_8433172_8446622_RNA45SN1_45S_50ntFlanks | 6646 | 6670 | 25 | TCGTACGACTCTTAGCGGTGGATCA | BT20 | 51 | 73 | 60 | 21 | 259 | 14.1 | 25.0 | 5.3 | 16.4 | 141.5 |
| >hg38_21_+_8433172_8446622_RNA45SN1_45S_50ntFlanks@6646.6670.25\|TCGTACGACTCTTAGCGGTGGATCA | >hg38_21_+_8433172_8446622_RNA45SN1_45S_50ntFlanks | 6646 | 6670 | 25 | TCGTACGACTCTTAGCGGTGGATCA | MB231 | 13 | 37 | 18 | 3 | 157 | 14.9 | 22.7 | 6.2 | 1.7 | 188.0 |
| >hg38_21_+_8433172_8446622_RNA45SN1_45S_50ntFlanks@6646.6670.25\|TCGTACGACTCTTAGCGGTGGATCA | >hg38_21_+_8433172_8446622_RNA45SN1_45S_50ntFlanks | 6646 | 6670 | 25 | TCGTACGACTCTTAGCGGTGGATCA | MB468 | 307 | 107 | 108 | 243 | 835 | 77.1 | 34.2 | 18.1 | 61.9 | 214.1 |
| >hg38_21_+_8433172_8446622_RNA45SN1_45S_50ntFlanks@6646.6671.26\|TCGTACGACTCTTAGCGGTGGATCAC | >hg38_21_+_8433172_8446622_RNA45SN1_45S_50ntFlanks | 6646 | 6671 | 26 | TCGTACGACTCTTAGCGGTGGATCAC | BT20 | 16 | 26 | 33 | 11 | 78 | 3.9 | 7.2 | 8.3 | 8.7 | 36.6 |
| >hg38_21_+_8433172_8446622_RNA45SN1_45S_50ntFlanks@6646.6672.27\|TCGTACGACTCTTAGCGGTGGATCACT | >hg38_21_+_8433172_8446622_RNA45SN1_45S_50ntFlanks | 6646 | 6672 | 27 | TCGTACGACTCTTAGCGGTGGATCACT | BT20 | 321 | 465 | 396 | 151 | 1264 | 84.2 | 139.0 | 72.0 | 119.6 | 760.5 |
| >hg38_21_+_8433172_8446622_RNA45SN1_45S_50ntFlanks@6646.6672.27\|TCGTACGACTCTTAGCGGTGGATCACT | >hg38_21_+_8433172_8446622_RNA45SN1_45S_50ntFlanks | 6646 | 6672 | 27 | TCGTACGACTCTTAGCGGTGGATCACT | MB231 | 69 | 189 | 116 | 22 | 629 | 87.2 | 123.9 | 61.2 | 20.0 | 719.8 |
| >hg38_21_+_8433172_8446622_RNA45SN1_45S_50ntFlanks@6646.6673.28\|TCGTACGACTCTTAGCGGTGGATCACTC | >hg38_21_+_8433172_8446622_RNA45SN1_45S_50ntFlanks | 6646 | 6673 | 28 | TCGTACGACTCTTAGCGGTGGATCACTC | BT20 | 45 | 88 | 58 | 93 | 245 | 15.8 | 25.3 | 13.6 | 20.2 | 121.3 |
| >hg38_21_+_8433172_8446622_RNA45SN1_45S_50ntFlanks@6646.6673.28\|TCGTACGACTCTTAGCGGTGGATCACTC | >hg38_21_+_8433172_8446622_RNA45SN1_45S_50ntFlanks | 6646 | 6673 | 28 | TCGTACGACTCTTAGCGGTGGATCACTC | MB231 | 15 | 71 | 38 | 18 | 220 | 20.0 | 57.8 | 19.0 | 20.7 | 298.9 |
| >hg38_21_+_8433172_8446622_RNA45SN1_45S_50ntFlanks@6646.6674.29\|TCGTACGACTCTTAGCGGTGGATCACTCG | >hg38_21_+_8433172_8446622_RNA45SN1_45S_50ntFlanks | 6646 | 6674 | 29 | TCGTACGACTCTTAGCGGTGGATCACTCG | BT20 | 44 | 92 | 59 | 163 | 424 | 20.1 | 36.4 | 28.1 | 161.1 | 104.6 |
| >hg38_21_+_8433172_8446622_RNA45SN1_45S_50ntFlanks@6646.6674.29\|TCGTACGACTCTTAGCGGTGGATCACTCG | >hg38_21_+_8433172_8446622_RNA45SN1_45S_50ntFlanks | 6646 | 6674 | 29 | TCGTACGACTCTTAGCGGTGGATCACTCG | MB231 | 19 | 166 | 51 | 15 | 615 | 21.0 | 162.1 | 43.1 | 6.7 | 893.3 |
| >hg38_21_+_8433172_8446622_RNA45SN1_45S_50ntFlanks@6646.6675.30\|TCGTACGACTCTTAGCGGTGGATCACTCGG | >hg38_21_+_8433172_8446622_RNA45SN1_45S_50ntFlanks | 6646 | 6675 | 30 | TCGTACGACTCTTAGCGGTGGATCACTCGG | BT20 | 43 | 97 | 54 | 221 | 347 | 28.6 | 31.4 | 32.5 | 286.2 | 145.5 |
| >hg38_21_+_8433172_8446622_RNA45SN1_45S_50ntFlanks@6646.6675.30\|TCGTACGACTCTTAGCGGTGGATCACTCGG | >hg38_21_+_8433172_8446622_RNA45SN1_45S_50ntFlanks | 6646 | 6675 | 30 | TCGTACGACTCTTAGCGGTGGATCACTCGG | MB231 | 29 | 291 | 76 | 16 | 1000 | 30.8 | 244.2 | 42.6 | 1.6 | 1633.8 |
| >hg38_21_+_8433172_8446622_RNA45SN1_45S_50ntFlanks@6646.6675.30\|TCGTACGACTCTTAGCGGTGGATCACTCGG | >hg38_21_+_8433172_8446622_RNA45SN1_45S_50ntFlanks | 6646 | 6675 | 30 | TCGTACGACTCTTAGCGGTGGATCACTCGG | MB468 | 48 | 95 | 51 | 49 | 196 | 22.5 | 80.2 | 40.5 | 37.8 | 86.2 |
| >hg38_21_+_8433172_8446622_RNA45SN1_45S_50ntFlanks@6646.6676.31\|TCGTACGACTCTTAGCGGTGGATCACTCGGC | >hg38_21_+_8433172_8446622_RNA45SN1_45S_50ntFlanks | 6646 | 6676 | 31 | TCGTACGACTCTTAGCGGTGGATCACTCGGC | BT20 | 78 | 179 | 133 | 245 | 603 | 21.4 | 63.0 | 84.1 | 146.4 | 166.4 |
| >hg38_21_+_8433172_8446622_RNA45SN1_45S_50ntFlanks@6646.6676.31\|TCGTACGACTCTTAGCGGTGGATCACTCGGC | >hg38_21_+_8433172_8446622_RNA45SN1_45S_50ntFlanks | 6646 | 6676 | 31 | TCGTACGACTCTTAGCGGTGGATCACTCGGC | MB231 | 25 | 136 | 41 | 53 | 484 | 31.2 | 120.8 | 29.7 | 25.8 | 564.9 |
| >hg38_21_+_8433172_8446622_RNA45SN1_45S_50ntFlanks@6646.6677.32\|TCGTACGACTCTTAGCGGTGGATCACTCGGCT | >hg38_21_+_8433172_8446622_RNA45SN1_45S_50ntFlanks | 6646 | 6677 | 32 | TCGTACGACTCTTAGCGGTGGATCACTCGGCT | BT20 | 133 | 237 | 124 | 281 | 651 | 44.7 | 69.3 | 12.8 | 172.8 | 217.1 |
| >hg38_21_+_8433172_8446622_RNA45SN1_45S_50ntFlanks@6646.6677.32\|TCGTACGACTCTTAGCGGTGGATCACTCGGCT | >hg38_21_+_8433172_8446622_RNA45SN1_45S_50ntFlanks | 6646 | 6677 | 32 | TCGTACGACTCTTAGCGGTGGATCACTCGGCT | MB231 | 39 | 165 | 47 | 38 | 547 | 51.8 | 123.4 | 26.7 | 40.6 | 788.8 |
| >hg38_21_+_8433172_8446622_RNA45SN1_45S_50ntFlanks@6646.6677.32\|TCGTACGACTCTTAGCGGTGGATCACTCGGCT | >hg38_21_+_8433172_8446622_RNA45SN1_45S_50ntFlanks | 6646 | 6677 | 32 | TCGTACGACTCTTAGCGGTGGATCACTCGGCT | MB468 | 16 | 41 | 13 | 31 | 91 | 4.7 | 34.6 | 9.3 | 12.9 | 46.0 |
| >hg38_21_+_8433172_8446622_RNA45SN1_45S_50ntFlanks@6646.6678.33\|TCGTACGACTCTTAGCGGTGGATCACTCGGCTC | >hg38_21_+_8433172_8446622_RNA45SN1_45S_50ntFlanks | 6646 | 6678 | 33 | TCGTACGACTCTTAGCGGTGGATCACTCGGCTC | BT20 | 89 | 177 | 77 | 219 | 473 | 36.7 | 59.4 | 27.4 | 83.3 | 159.3 |

| | | | | | | | | | | | | | | | | |
|---|---|---|---|---|---|---|---|---|---|---|---|---|---|---|---|---|
| >hg38_21_+_8433172_8446622_RNA45SN1_45S_50ntFlanks@6646.6678.33\|TCGTACGACTCTTAGCGGTGGATCACTCGGCTC | >hg38_21_+_8433172_8446622_RNA45SN1_45S_50ntFlanks | 6646 | 6678 | 33 | TCGTACGACTCTTAGCGGTGGATCACTCGGCTC | MB231 | 25 | 102 | 26 | 24 | 339 | 35.8 | 77.4 | 15.2 | 26.7 | 466.9 |
| >hg38_21_+_8433172_8446622_RNA45SN1_45S_50ntFlanks@6646.6678.33\|TCGTACGACTCTTAGCGGTGGATCACTCGGCTC | >hg38_21_+_8433172_8446622_RNA45SN1_45S_50ntFlanks | 6646 | 6678 | 33 | TCGTACGACTCTTAGCGGTGGATCACTCGGCTC | MB468 | 13 | 33 | 9 | 19 | 75 | 3.1 | 29.0 | 5.9 | 10.6 | 45.4 |
| >hg38_21_+_8433172_8446622_RNA45SN1_45S_50ntFlanks@6646.6679.34\|TCGTACGACTCTTAGCGGTGGATCACTCGGCTCG | >hg38_21_+_8433172_8446622_RNA45SN1_45S_50ntFlanks | 6646 | 6679 | 34 | TCGTACGACTCTTAGCGGTGGATCACTCGGCTCG | BT20 | 47 | 119 | 72 | 302 | 528 | 33.5 | 59.8 | 43.6 | 440.5 | 257.9 |
| >hg38_21_+_8433172_8446622_RNA45SN1_45S_50ntFlanks@6646.6679.34\|TCGTACGACTCTTAGCGGTGGATCACTCGGCTCG | >hg38_21_+_8433172_8446622_RNA45SN1_45S_50ntFlanks | 6646 | 6679 | 34 | TCGTACGACTCTTAGCGGTGGATCACTCGGCTCG | MB231 | 48 | 203 | 55 | 17 | 647 | 54.7 | 143.0 | 30.4 | 5.4 | 895.4 |
| >hg38_21_+_8433172_8446622_RNA45SN1_45S_50ntFlanks@6646.6679.34\|TCGTACGACTCTTAGCGGTGGATCACTCGGCTCG | >hg38_21_+_8433172_8446622_RNA45SN1_45S_50ntFlanks | 6646 | 6679 | 34 | TCGTACGACTCTTAGCGGTGGATCACTCGGCTCG | MB468 | 135 | 240 | 70 | 157 | 937 | 27.4 | 114.2 | 28.9 | 140.5 | 664.1 |
| >hg38_21_+_8433172_8446622_RNA45SN1_45S_50ntFlanks@6646.6680.35\|TCGTACGACTCTTAGCGGTGGATCACTCGGCTCGT | >hg38_21_+_8433172_8446622_RNA45SN1_45S_50ntFlanks | 6646 | 6680 | 35 | TCGTACGACTCTTAGCGGTGGATCACTCGGCTCGT | BT20 | 139 | 231 | 194 | 232 | 575 | 11.7 | 109.0 | 121.6 | 124.6 | 81.1 |
| >hg38_21_+_8433172_8446622_RNA45SN1_45S_50ntFlanks@6646.6680.35\|TCGTACGACTCTTAGCGGTGGATCACTCGGCTCGT | >hg38_21_+_8433172_8446622_RNA45SN1_45S_50ntFlanks | 6646 | 6680 | 35 | TCGTACGACTCTTAGCGGTGGATCACTCGGCTCGT | MB231 | 60 | 207 | 63 | 31 | 465 | 62.6 | 115.4 | 37.1 | 28.0 | 570.0 |
| >hg38_21_+_8433172_8446622_RNA45SN1_45S_50ntFlanks@6646.6680.35\|TCGTACGACTCTTAGCGGTGGATCACTCGGCTCGT | >hg38_21_+_8433172_8446622_RNA45SN1_45S_50ntFlanks | 6646 | 6680 | 35 | TCGTACGACTCTTAGCGGTGGATCACTCGGCTCGT | MB468 | 103 | 99 | 44 | 138 | 401 | 22.6 | 51.2 | 22.3 | 86.5 | 158.8 |
| >hg38_21_+_8433172_8446622_RNA45SN1_45S_50ntFlanks@6646.6681.36\|TCGTACGACTCTTAGCGGTGGATCACTCGGCTCGTG | >hg38_21_+_8433172_8446622_RNA45SN1_45S_50ntFlanks | 6646 | 6681 | 36 | TCGTACGACTCTTAGCGGTGGATCACTCGGCTCGTG | BT20 | 104 | 274 | 67 | 300 | 607 | 93.0 | 141.5 | 32.4 | 258.2 | 105.2 |
| >hg38_21_+_8433172_8446622_RNA45SN1_45S_50ntFlanks@6646.6681.36\|TCGTACGACTCTTAGCGGTGGATCACTCGGCTCGTG | >hg38_21_+_8433172_8446622_RNA45SN1_45S_50ntFlanks | 6646 | 6681 | 36 | TCGTACGACTCTTAGCGGTGGATCACTCGGCTCGTG | MB231 | 263 | 1343 | 94 | 55 | 916 | 155.1 | 454.5 | 68.6 | 42.4 | 1356.8 |
| >hg38_21_+_8433172_8446622_RNA45SN1_45S_50ntFlanks@6646.6681.36\|TCGTACGACTCTTAGCGGTGGATCACTCGGCTCGTG | >hg38_21_+_8433172_8446622_RNA45SN1_45S_50ntFlanks | 6646 | 6681 | 36 | TCGTACGACTCTTAGCGGTGGATCACTCGGCTCGTG | MB468 | 1111 | 1107 | 352 | 1630 | 4591 | 311.6 | 467.3 | 160.5 | 1198.2 | 2040.7 |
| >hg38_21_+_8433172_8446622_RNA45SN1_45S_50ntFlanks@6646.6683.38\|TCGTACGACTCTTAGCGGTGGATCACTCGGCTCGTGCG | >hg38_21_+_8433172_8446622_RNA45SN1_45S_50ntFlanks | 6646 | 6683 | 38 | TCGTACGACTCTTAGCGGTGGATCACTCGGCTCGTGCG | BT20 | 41 | 74 | 42 | 80 | 41 | 40.4 | 40.7 | 39.2 | 33.2 | 4.7 |
| >hg38_21_+_8433172_8446622_RNA45SN1_45S_50ntFlanks@6646.6683.38\|TCGTACGACTCTTAGCGGTGGATCACTCGGCTCGTGCG | >hg38_21_+_8433172_8446622_RNA45SN1_45S_50ntFlanks | 6646 | 6683 | 38 | TCGTACGACTCTTAGCGGTGGATCACTCGGCTCGTGCG | MB231 | 71 | 108 | 5 | 31 | 27 | 19.7 | 71.5 | 2.0 | 28.7 | 5.0 |
| >hg38_21_+_8433172_8446622_RNA45SN1_45S_50ntFlanks@6646.6685.40\|TCGTACGACTCTTAGCGGTGGATCACTCGGCTCGTGCGTC | >hg38_21_+_8433172_8446622_RNA45SN1_45S_50ntFlanks | 6646 | 6685 | 40 | TCGTACGACTCTTAGCGGTGGATCACTCGGCTCGTGCGTC | BT20 | 288 | 1125 | 374 | 1165 | 903 | 269.8 | 803.4 | 348.2 | 758.4 | 129.1 |
| >hg38_21_+_8433172_8446622_RNA45SN1_45S_50ntFlanks@6646.6685.40\|TCGTACGACTCTTAGCGGTGGATCACTCGGCTCGTGCGTC | >hg38_21_+_8433172_8446622_RNA45SN1_45S_50ntFlanks | 6646 | 6685 | 40 | TCGTACGACTCTTAGCGGTGGATCACTCGGCTCGTGCGTC | MB231 | 38 | 37 | 16 | 356 | 258 | 38.5 | 25.6 | 5.6 | 582.2 | 334.0 |
| >hg38_21_+_8433172_8446622_RNA45SN1_45S_50ntFlanks@6646.6688.43\|TCGTACGACTCTTAGCGGTGGATCACTCGGCTCGTGCGTCGAT | >hg38_21_+_8433172_8446622_RNA45SN1_45S_50ntFlanks | 6646 | 6688 | 43 | TCGTACGACTCTTAGCGGTGGATCACTCGGCTCGTGCGTCGAT | BT20 | 11 | 28 | 6 | 29 | 49 | 12.1 | 21.5 | 5.9 | 27.9 | 9.2 |
| >hg38_21_+_8433172_8446622_RNA45SN1_45S_50ntFlanks@6646.6689.44\|TCGTACGACTCTTAGCGGTGGATCACTCGGCTCGTGCGTCGATG | >hg38_21_+_8433172_8446622_RNA45SN1_45S_50ntFlanks | 6646 | 6689 | 44 | TCGTACGACTCTTAGCGGTGGATCACTCGGCTCGTGCGTCGATG | BT20 | 88 | 409 | 80 | 312 | 796 | 61.9 | 479.8 | 79.9 | 238.2 | 289.4 |
| >hg38_21_+_8433172_8446622_RNA45SN1_45S_50ntFlanks@6646.6689.44\|TCGTACGACTCTTAGCGGTGGATCACTCGGCTCGTGCGTCGATG | >hg38_21_+_8433172_8446622_RNA45SN1_45S_50ntFlanks | 6646 | 6689 | 44 | TCGTACGACTCTTAGCGGTGGATCACTCGGCTCGTGCGTCGATG | MB231 | 54 | 76 | 17 | 116 | 278 | 21.2 | 26.8 | 5.1 | 176.7 | 309.4 |
| >hg38_21_+_8433172_8446622_RNA45SN1_45S_50ntFlanks@6646.6689.44\|TCGTACGACTCTTAGCGGTGGATCACTCGGCTCGTGCGTCGATG | >hg38_21_+_8433172_8446622_RNA45SN1_45S_50ntFlanks | 6646 | 6689 | 44 | TCGTACGACTCTTAGCGGTGGATCACTCGGCTCGTGCGTCGATG | MB468 | 143 | 63 | 69 | 329 | 2668 | 35.6 | 46.8 | 89.5 | 241.2 | 2186.6 |
| >hg38_21_+_8433172_8446622_RNA45SN1_45S_50ntFlanks@6646.6692.47\|TCGTACGACTCTTAGCGGTGGATCACTCGGCTCGTGCGTCGATGAAG | >hg38_21_+_8433172_8446622_RNA45SN1_45S_50ntFlanks | 6646 | 6692 | 47 | TCGTACGACTCTTAGCGGTGGATCACTCGGCTCGTGCGTCGATGAAG | MB231 | 12 | 11 | 4 | 7 | 36 | 5.7 | 9.4 | 3.6 | 8.4 | 30.4 |
| >hg38_21_+_8433172_8446622_RNA45SN1_45S_50ntFlanks@6646.6693.48\|TCGTACGACTCTTAGCGGTGGATCACTCGGCTCGTGCGTCGATGAAGA | >hg38_21_+_8433172_8446622_RNA45SN1_45S_50ntFlanks | 6646 | 6693 | 48 | TCGTACGACTCTTAGCGGTGGATCACTCGGCTCGTGCGTCGATGAAGA | BT20 | 11 | 21 | 4 | 14 | 61 | 10.5 | 25.3 | 4.4 | 11.7 | 38.4 |
| >hg38_21_+_8433172_8446622_RNA45SN1_45S_50ntFlanks@6646.6695.50\|TCGTACGACTCTTAGCGGTGGATCACTCGGCTCGTGCGTCGATGAAGAAC | >hg38_21_+_8433172_8446622_RNA45SN1_45S_50ntFlanks | 6646 | 6695 | 50 | TCGTACGACTCTTAGCGGTGGATCACTCGGCTCGTGCGTCGATGAAGAAC | BT20 | 45 | 117 | 19 | 98 | 588 | 63.6 | 180.9 | 27.5 | 125.5 | 618.6 |
| >hg38_21_+_8433172_8446622_RNA45SN1_45S_50ntFlanks@6646.6695.50\|TCGTACGACTCTTAGCGGTGGATCACTCGGCTCGTGCGTCGATGAAGAAC | >hg38_21_+_8433172_8446622_RNA45SN1_45S_50ntFlanks | 6646 | 6695 | 50 | TCGTACGACTCTTAGCGGTGGATCACTCGGCTCGTGCGTCGATGAAGAAC | MB231 | 53 | 70 | 22 | 31 | 205 | 16.9 | 109.6 | 17.5 | 51.0 | 300.3 |
| >hg38_21_+_8433172_8446622_RNA45SN1_45S_50ntFlanks@6647.6664.18\|CGTACGACTCTTAGCGGT | >hg38_21_+_8433172_8446622_RNA45SN1_45S_50ntFlanks | 6647 | 6664 | 18 | CGTACGACTCTTAGCGGT | BT20 | 132 | 233 | 126 | 450 | 719 | 47.6 | 143.5 | 30.0 | 167.9 | 160.0 |
| >hg38_21_+_8433172_8446622_RNA45SN1_45S_50ntFlanks@6647.6664.18\|CGTACGACTCTTAGCGGT | >hg38_21_+_8433172_8446622_RNA45SN1_45S_50ntFlanks | 6647 | 6664 | 18 | CGTACGACTCTTAGCGGT | MB231 | 58 | 115 | 61 | 130 | 439 | 67.4 | 47.8 | 23.4 | 66.9 | 323.7 |
| >hg38_21_+_8433172_8446622_RNA45SN1_45S_50ntFlanks@6647.6664.18\|CGTACGACTCTTAGCGGT | >hg38_21_+_8433172_8446622_RNA45SN1_45S_50ntFlanks | 6647 | 6664 | 18 | CGTACGACTCTTAGCGGT | MB468 | 14 | 18 | 10 | 24 | 43 | 6.0 | 10.7 | 6.6 | 8.7 | 18.0 |
| >hg38_21_+_8433172_8446622_RNA45SN1_45S_50ntFlanks@6647.6665.19\|CGTACGACTCTTAGCGGTG | >hg38_21_+_8433172_8446622_RNA45SN1_45S_50ntFlanks | 6647 | 6665 | 19 | CGTACGACTCTTAGCGGTG | BT20 | 279 | 391 | 257 | 1127 | 1012 | 134.5 | 250.4 | 83.8 | 258.0 | 122.6 |
| >hg38_21_+_8433172_8446622_RNA45SN1_45S_50ntFlanks@6647.6665.19\|CGTACGACTCTTAGCGGTG | >hg38_21_+_8433172_8446622_RNA45SN1_45S_50ntFlanks | 6647 | 6665 | 19 | CGTACGACTCTTAGCGGTG | MB231 | 55 | 147 | 58 | 131 | 525 | 72.6 | 98.9 | 24.5 | 185.5 | 640.1 |
| >hg38_21_+_8433172_8446622_RNA45SN1_45S_50ntFlanks@6647.6665.19\|CGTACGACTCTTAGCGGTG | >hg38_21_+_8433172_8446622_RNA45SN1_45S_50ntFlanks | 6647 | 6665 | 19 | CGTACGACTCTTAGCGGTG | MB468 | 28 | 21 | 21 | 42 | 44 | 11.8 | 17.4 | 10.5 | 15.0 | 19.5 |
| >hg38_21_+_8433172_8446622_RNA45SN1_45S_50ntFlanks@6647.6666.20\|CGTACGACTCTTAGCGGTGG | >hg38_21_+_8433172_8446622_RNA45SN1_45S_50ntFlanks | 6647 | 6666 | 20 | CGTACGACTCTTAGCGGTGG | BT20 | 64 | 85 | 64 | 217 | 274 | 38.9 | 35.3 | 30.5 | 83.1 | 26.8 |
| >hg38_21_+_8433172_8446622_RNA45SN1_45S_50ntFlanks@6647.6666.20\|CGTACGACTCTTAGCGGTGG | >hg38_21_+_8433172_8446622_RNA45SN1_45S_50ntFlanks | 6647 | 6666 | 20 | CGTACGACTCTTAGCGGTGG | MB231 | 21 | 70 | 23 | 38 | 267 | 26.8 | 55.1 | 12.2 | 39.3 | 337.9 |
| >hg38_21_+_8433172_8446622_RNA45SN1_45S_50ntFlanks@6647.6666.20\|CGTACGACTCTTAGCGGTGG | >hg38_21_+_8433172_8446622_RNA45SN1_45S_50ntFlanks | 6647 | 6666 | 20 | CGTACGACTCTTAGCGGTGG | MB468 | 51 | 36 | 47 | 38 | 61 | 37.5 | 33.6 | 40.8 | 17.1 | 43.6 |
| >hg38_21_+_8433172_8446622_RNA45SN1_45S_50ntFlanks@6647.6667.21\|CGTACGACTCTTAGCGGTGGA | >hg38_21_+_8433172_8446622_RNA45SN1_45S_50ntFlanks | 6647 | 6667 | 21 | CGTACGACTCTTAGCGGTGGA | BT20 | 73 | 86 | 89 | 174 | 283 | 31.5 | 38.2 | 16.5 | 69.8 | 56.5 |
| >hg38_21_+_8433172_8446622_RNA45SN1_45S_50ntFlanks@6647.6667.21\|CGTACGACTCTTAGCGGTGGA | >hg38_21_+_8433172_8446622_RNA45SN1_45S_50ntFlanks | 6647 | 6667 | 21 | CGTACGACTCTTAGCGGTGGA | MB231 | 28 | 37 | 17 | 43 | 185 | 32.9 | 16.6 | 7.3 | 44.7 | 130.5 |
| >hg38_21_+_8433172_8446622_RNA45SN1_45S_50ntFlanks@6647.6668.22\|CGTACGACTCTTAGCGGTGGAT | >hg38_21_+_8433172_8446622_RNA45SN1_45S_50ntFlanks | 6647 | 6668 | 22 | CGTACGACTCTTAGCGGTGGAT | BT20 | 82 | 98 | 71 | 147 | 305 | 27.0 | 40.9 | 1.8 | 74.3 | 91.9 |
| >hg38_21_+_8433172_8446622_RNA45SN1_45S_50ntFlanks@6647.6668.22\|CGTACGACTCTTAGCGGTGGAT | >hg38_21_+_8433172_8446622_RNA45SN1_45S_50ntFlanks | 6647 | 6668 | 22 | CGTACGACTCTTAGCGGTGGAT | MB231 | 31 | 46 | 18 | 106 | 154 | 24.4 | 30.1 | 7.5 | 42.6 | 101.8 |
| >hg38_21_+_8433172_8446622_RNA45SN1_45S_50ntFlanks@6647.6668.22\|CGTACGACTCTTAGCGGTGGAT | >hg38_21_+_8433172_8446622_RNA45SN1_45S_50ntFlanks | 6647 | 6668 | 22 | CGTACGACTCTTAGCGGTGGAT | MB468 | 12 | 7 | 7 | 21 | 18 | 6.0 | 4.4 | 3.3 | 2.3 | 9.2 |
| >hg38_21_+_8433172_8446622_RNA45SN1_45S_50ntFlanks@6647.6669.23\|CGTACGACTCTTAGCGGTGGATC | >hg38_21_+_8433172_8446622_RNA45SN1_45S_50ntFlanks | 6647 | 6669 | 23 | CGTACGACTCTTAGCGGTGGATC | BT20 | 148 | 260 | 166 | 183 | 816 | 36.1 | 139.0 | 40.7 | 37.6 | 150.6 |
| >hg38_21_+_8433172_8446622_RNA45SN1_45S_50ntFlanks@6647.6669.23\|CGTACGACTCTTAGCGGTGGATC | >hg38_21_+_8433172_8446622_RNA45SN1_45S_50ntFlanks | 6647 | 6669 | 23 | CGTACGACTCTTAGCGGTGGATC | MB231 | 49 | 52 | 32 | 47 | 203 | 68.7 | 29.5 | 29.1 | 65.6 | 155.8 |
| >hg38_21_+_8433172_8446622_RNA45SN1_45S_50ntFlanks@6647.6669.23\|CGTACGACTCTTAGCGGTGGATC | >hg38_21_+_8433172_8446622_RNA45SN1_45S_50ntFlanks | 6647 | 6669 | 23 | CGTACGACTCTTAGCGGTGGATC | MB468 | 60 | 34 | 30 | 78 | 173 | 18.5 | 14.6 | 13.2 | 25.6 | 11.8 |
| >hg38_21_+_8433172_8446622_RNA45SN1_45S_50ntFlanks@6647.6670.24\|CGTACGACTCTTAGCGGTGGATCA | >hg38_21_+_8433172_8446622_RNA45SN1_45S_50ntFlanks | 6647 | 6670 | 24 | CGTACGACTCTTAGCGGTGGATCA | MB468 | 42 | 15 | 15 | 39 | 143 | 8.2 | 5.5 | 2.0 | 11.3 | 20.6 |
| >hg38_21_+_8433172_8446622_RNA45SN1_45S_50ntFlanks@6647.6672.26\|CGTACGACTCTTAGCGGTGGATCACT | >hg38_21_+_8433172_8446622_RNA45SN1_45S_50ntFlanks | 6647 | 6672 | 26 | CGTACGACTCTTAGCGGTGGATCACT | BT20 | 51 | 71 | 45 | 26 | 166 | 21.3 | 18.7 | 8.2 | 17.5 | 66.1 |
| >hg38_21_+_8433172_8446622_RNA45SN1_45S_50ntFlanks@6647.6672.26\|CGTACGACTCTTAGCGGTGGATCACT | >hg38_21_+_8433172_8446622_RNA45SN1_45S_50ntFlanks | 6647 | 6672 | 26 | CGTACGACTCTTAGCGGTGGATCACT | MB231 | 14 | 40 | 18 | 7 | 138 | 17.4 | 27.9 | 8.6 | 5.8 | 151.3 |
| >hg38_21_+_8433172_8446622_RNA45SN1_45S_50ntFlanks@6647.6676.30\|CGTACGACTCTTAGCGGTGGATCACTCGGC | >hg38_21_+_8433172_8446622_RNA45SN1_45S_50ntFlanks | 6647 | 6676 | 30 | CGTACGACTCTTAGCGGTGGATCACTCGGC | BT20 | 19 | 38 | 32 | 61 | 129 | 10.6 | 13.1 | 34.1 | 45.0 | 27.5 |
| >hg38_21_+_8433172_8446622_RNA45SN1_45S_50ntFlanks@6647.6677.31\|CGTACGACTCTTAGCGGTGGATCACTCGGCT | >hg38_21_+_8433172_8446622_RNA45SN1_45S_50ntFlanks | 6647 | 6677 | 31 | CGTACGACTCTTAGCGGTGGATCACTCGGCT | BT20 | 24 | 39 | 15 | 54 | 98 | 13.9 | 6.9 | 5.9 | 29.2 | 16.7 |
| >hg38_21_+_8433172_8446622_RNA45SN1_45S_50ntFlanks@6647.6678.32\|CGTACGACTCTTAGCGGTGGATCACTCGGCTC | >hg38_21_+_8433172_8446622_RNA45SN1_45S_50ntFlanks | 6647 | 6678 | 32 | CGTACGACTCTTAGCGGTGGATCACTCGGCTC | BT20 | 17 | 29 | 12 | 44 | 84 | 9.4 | 7.8 | 5.6 | 12.4 | 21.5 |
| >hg38_21_+_8433172_8446622_RNA45SN1_45S_50ntFlanks@6647.6679.33\|CGTACGACTCTTAGCGGTGGATCACTCGGCTCG | >hg38_21_+_8433172_8446622_RNA45SN1_45S_50ntFlanks | 6647 | 6679 | 33 | CGTACGACTCTTAGCGGTGGATCACTCGGCTCG | MB468 | 16 | 30 | 9 | 30 | 137 | 3.2 | 15.2 | 3.7 | 25.4 | 96.6 |
| >hg38_21_+_8433172_8446622_RNA45SN1_45S_50ntFlanks@6647.6680.34\|CGTACGACTCTTAGCGGTGGATCACTCGGCTCGT | >hg38_21_+_8433172_8446622_RNA45SN1_45S_50ntFlanks | 6647 | 6680 | 34 | CGTACGACTCTTAGCGGTGGATCACTCGGCTCGT | BT20 | 23 | 34 | 25 | 41 | 75 | 5.6 | 11.4 | 13.8 | 12.2 | 3.7 |
| >hg38_21_+_8433172_8446622_RNA45SN1_45S_50ntFlanks@6647.6680.34\|CGTACGACTCTTAGCGGTGGATCACTCGGCTCGT | >hg38_21_+_8433172_8446622_RNA45SN1_45S_50ntFlanks | 6647 | 6680 | 34 | CGTACGACTCTTAGCGGTGGATCACTCGGCTCGT | MB231 | 13 | 46 | 10 | 10 | 92 | 11.1 | 24.3 | 7.6 | 9.9 | 111.7 |
| >hg38_21_+_8433172_8446622_RNA45SN1_45S_50ntFlanks@6647.6680.34\|CGTACGACTCTTAGCGGTGGATCACTCGGCTCGT | >hg38_21_+_8433172_8446622_RNA45SN1_45S_50ntFlanks | 6647 | 6680 | 34 | CGTACGACTCTTAGCGGTGGATCACTCGGCTCGT | MB468 | 17 | 15 | 7 | 27 | 70 | 4.0 | 7.3 | 3.1 | 13.0 | 24.6 |

| | | | | | | | | | | | | | | | | |
|---|---|---|---|---|---|---|---|---|---|---|---|---|---|---|---|---|
| >hg38_21_+_8433172_8446622_RNA45SN1_45S_50ntFlanks@6647.6681.35\|CGTACGACTCTTAGCGGTGGATCACTCGGCTCGTG | >hg38_21_+_8433172_8446622_RNA45SN1_45S_50ntFlanks | 6647 | 6681 | 35 | CGTACGACTCTTAGCGGTGGATCACTCGGCTCGTG | BT20 | 44 | 92 | 10 | 62 | 95 | 50.1 | 36.5 | 5.6 | 49.8 | 12.0 |
| >hg38_21_+_8433172_8446622_RNA45SN1_45S_50ntFlanks@6647.6681.35\|CGTACGACTCTTAGCGGTGGATCACTCGGCTCGTG | >hg38_21_+_8433172_8446622_RNA45SN1_45S_50ntFlanks | 6647 | 6681 | 35 | CGTACGACTCTTAGCGGTGGATCACTCGGCTCGTG | MB231 | 88 | 445 | 17 | 19 | 185 | 52.6 | 217.1 | 12.4 | 14.3 | 257.1 |
| >hg38_21_+_8433172_8446622_RNA45SN1_45S_50ntFlanks@6647.6681.35\|CGTACGACTCTTAGCGGTGGATCACTCGGCTCGTG | >hg38_21_+_8433172_8446622_RNA45SN1_45S_50ntFlanks | 6647 | 6681 | 35 | CGTACGACTCTTAGCGGTGGATCACTCGGCTCGTG | MB468 | 199 | 201 | 69 | 343 | 911 | 51.9 | 75.9 | 26.2 | 234.1 | 405.9 |
| >hg38_21_+_8433172_8446622_RNA45SN1_45S_50ntFlanks@6647.6683.37\|CGTACGACTCTTAGCGGTGGATCACTCGGCTCGTGCG | >hg38_21_+_8433172_8446622_RNA45SN1_45S_50ntFlanks | 6647 | 6683 | 37 | CGTACGACTCTTAGCGGTGGATCACTCGGCTCGTGCG | MB231 | 13 | 15 | 1 | 4 | 5 | 2.7 | 11.5 | 0.6 | 3.2 | 1.6 |
| >hg38_21_+_8433172_8446622_RNA45SN1_45S_50ntFlanks@6647.6689.43\|CGTACGACTCTTAGCGGTGGATCACTCGGCTCGTGCGTCGATG | >hg38_21_+_8433172_8446622_RNA45SN1_45S_50ntFlanks | 6647 | 6689 | 43 | CGTACGACTCTTAGCGGTGGATCACTCGGCTCGTGCGTCGATG | MB468 | 14 | 10 | 7 | 41 | 263 | 3.0 | 6.7 | 7.9 | 26.4 | 207.1 |
| >hg38_21_+_8433172_8446622_RNA45SN1_45S_50ntFlanks@6647.6695.49\|CGTACGACTCTTAGCGGTGGATCACTCGGCTCGTGCGTCGATGAAGAAC | >hg38_21_+_8433172_8446622_RNA45SN1_45S_50ntFlanks | 6647 | 6695 | 49 | CGTACGACTCTTAGCGGTGGATCACTCGGCTCGTGCGTCGATGAAGAAC | MB231 | 17 | 22 | 7 | 10 | 65 | 7.1 | 31.9 | 5.6 | 16.6 | 91.9 |
| >hg38_21_+_8433172_8446622_RNA45SN1_45S_50ntFlanks@6647.6696.50\|CGTACGACTCTTAGCGGTGGATCACTCGGCTCGTGCGTCGATGAAGAACG | >hg38_21_+_8433172_8446622_RNA45SN1_45S_50ntFlanks | 6647 | 6696 | 50 | CGTACGACTCTTAGCGGTGGATCACTCGGCTCGTGCGTCGATGAAGAACG | MB231 | 22 | 37 | 6 | 5 | 30 | 8.9 | 50.9 | 5.6 | 7.9 | 52.6 |
| >hg38_21_+_8433172_8446622_RNA45SN1_45S_50ntFlanks@6648.6665.18\|GTACGACTCTTAGCGGTG | >hg38_21_+_8433172_8446622_RNA45SN1_45S_50ntFlanks | 6648 | 6665 | 18 | GTACGACTCTTAGCGGTG | BT20 | 37 | 54 | 26 | 108 | 150 | 18.0 | 35.0 | 17.0 | 19.6 | 17.6 |
| >hg38_21_+_8433172_8446622_RNA45SN1_45S_50ntFlanks@6648.6666.19\|GTACGACTCTTAGCGGTGG | >hg38_21_+_8433172_8446622_RNA45SN1_45S_50ntFlanks | 6648 | 6666 | 19 | GTACGACTCTTAGCGGTGG | BT20 | 11 | 16 | 10 | 20 | 45 | 4.0 | 7.7 | 5.3 | 5.2 | 7.4 |
| >hg38_21_+_8433172_8446622_RNA45SN1_45S_50ntFlanks@6648.6666.19\|GTACGACTCTTAGCGGTGG | >hg38_21_+_8433172_8446622_RNA45SN1_45S_50ntFlanks | 6648 | 6666 | 19 | GTACGACTCTTAGCGGTGG | MB468 | 13 | 4 | 9 | 9 | 15 | 9.7 | 3.9 | 10.3 | 5.9 | 11.1 |
| >hg38_21_+_8433172_8446622_RNA45SN1_45S_50ntFlanks@6648.6667.20\|GTACGACTCTTAGCGGTGGA | >hg38_21_+_8433172_8446622_RNA45SN1_45S_50ntFlanks | 6648 | 6667 | 20 | GTACGACTCTTAGCGGTGGA | BT20 | 14 | 16 | 14 | 32 | 57 | 4.4 | 6.9 | 6.4 | 16.1 | 18.3 |
| >hg38_21_+_8433172_8446622_RNA45SN1_45S_50ntFlanks@6648.6668.21\|GTACGACTCTTAGCGGTGGAT | >hg38_21_+_8433172_8446622_RNA45SN1_45S_50ntFlanks | 6648 | 6668 | 21 | GTACGACTCTTAGCGGTGGAT | BT20 | 51 | 64 | 36 | 57 | 241 | 26.8 | 25.8 | 21.1 | 4.3 | 116.0 |
| >hg38_21_+_8433172_8446622_RNA45SN1_45S_50ntFlanks@6648.6668.21\|GTACGACTCTTAGCGGTGGAT | >hg38_21_+_8433172_8446622_RNA45SN1_45S_50ntFlanks | 6648 | 6668 | 21 | GTACGACTCTTAGCGGTGGAT | MB231 | 24 | 27 | 9 | 29 | 114 | 23.1 | 19.2 | 6.0 | 11.5 | 70.9 |
| >hg38_21_+_8433172_8446622_RNA45SN1_45S_50ntFlanks@6648.6668.21\|GTACGACTCTTAGCGGTGGAT | >hg38_21_+_8433172_8446622_RNA45SN1_45S_50ntFlanks | 6648 | 6668 | 21 | GTACGACTCTTAGCGGTGGAT | MB468 | 10 | 4 | 5 | 9 | 16 | 4.4 | 1.8 | 1.9 | 1.4 | 10.8 |
| >hg38_21_+_8433172_8446622_RNA45SN1_45S_50ntFlanks@6648.6669.22\|GTACGACTCTTAGCGGTGGATC | >hg38_21_+_8433172_8446622_RNA45SN1_45S_50ntFlanks | 6648 | 6669 | 22 | GTACGACTCTTAGCGGTGGATC | BT20 | 47 | 78 | 48 | 60 | 256 | 10.1 | 36.6 | 27.6 | 20.0 | 110.0 |
| >hg38_21_+_8433172_8446622_RNA45SN1_45S_50ntFlanks@6648.6669.22\|GTACGACTCTTAGCGGTGGATC | >hg38_21_+_8433172_8446622_RNA45SN1_45S_50ntFlanks | 6648 | 6669 | 22 | GTACGACTCTTAGCGGTGGATC | MB231 | 17 | 16 | 7 | 13 | 77 | 23.1 | 9.7 | 4.1 | 18.3 | 60.5 |
| >hg38_21_+_8433172_8446622_RNA45SN1_45S_50ntFlanks@6648.6669.22\|GTACGACTCTTAGCGGTGGATC | >hg38_21_+_8433172_8446622_RNA45SN1_45S_50ntFlanks | 6648 | 6669 | 22 | GTACGACTCTTAGCGGTGGATC | MB468 | 27 | 8 | 10 | 31 | 70 | 8.2 | 2.5 | 1.8 | 23.1 | 22.3 |
| >hg38_21_+_8433172_8446622_RNA45SN1_45S_50ntFlanks@6648.6670.23\|GTACGACTCTTAGCGGTGGATCA | >hg38_21_+_8433172_8446622_RNA45SN1_45S_50ntFlanks | 6648 | 6670 | 23 | GTACGACTCTTAGCGGTGGATCA | MB468 | 13 | 2 | 5 | 10 | 38 | 3.7 | 0.6 | 2.4 | 6.8 | 13.0 |
| >hg38_21_+_8433172_8446622_RNA45SN1_45S_50ntFlanks@6648.6672.25\|GTACGACTCTTAGCGGTGGATCACT | >hg38_21_+_8433172_8446622_RNA45SN1_45S_50ntFlanks | 6648 | 6672 | 25 | GTACGACTCTTAGCGGTGGATCACT | BT20 | 19 | 31 | 15 | 10 | 87 | 8.7 | 9.9 | 8.2 | 7.0 | 48.8 |
| >hg38_21_+_8433172_8446622_RNA45SN1_45S_50ntFlanks@6648.6676.29\|GTACGACTCTTAGCGGTGGATCACTCGGC | >hg38_21_+_8433172_8446622_RNA45SN1_45S_50ntFlanks | 6648 | 6676 | 29 | GTACGACTCTTAGCGGTGGATCACTCGGC | BT20 | 26 | 49 | 39 | 82 | 212 | 13.9 | 25.8 | 42.2 | 94.5 | 46.5 |
| >hg38_21_+_8433172_8446622_RNA45SN1_45S_50ntFlanks@6648.6676.29\|GTACGACTCTTAGCGGTGGATCACTCGGC | >hg38_21_+_8433172_8446622_RNA45SN1_45S_50ntFlanks | 6648 | 6676 | 29 | GTACGACTCTTAGCGGTGGATCACTCGGC | MB231 | 13 | 74 | 17 | 13 | 290 | 15.7 | 66.3 | 15.0 | 5.5 | 352.4 |
| >hg38_21_+_8433172_8446622_RNA45SN1_45S_50ntFlanks@6648.6677.30\|GTACGACTCTTAGCGGTGGATCACTCGGCT | >hg38_21_+_8433172_8446622_RNA45SN1_45S_50ntFlanks | 6648 | 6677 | 30 | GTACGACTCTTAGCGGTGGATCACTCGGCT | BT20 | 18 | 32 | 9 | 40 | 117 | 12.1 | 14.9 | 6.3 | 37.1 | 14.6 |
| >hg38_21_+_8433172_8446622_RNA45SN1_45S_50ntFlanks@6648.6678.31\|GTACGACTCTTAGCGGTGGATCACTCGGCTC | >hg38_21_+_8433172_8446622_RNA45SN1_45S_50ntFlanks | 6648 | 6678 | 31 | GTACGACTCTTAGCGGTGGATCACTCGGCTC | BT20 | 30 | 39 | 16 | 43 | 168 | 25.3 | 11.5 | 12.2 | 37.8 | 41.2 |
| >hg38_21_+_8433172_8446622_RNA45SN1_45S_50ntFlanks@6648.6678.31\|GTACGACTCTTAGCGGTGGATCACTCGGCTC | >hg38_21_+_8433172_8446622_RNA45SN1_45S_50ntFlanks | 6648 | 6678 | 31 | GTACGACTCTTAGCGGTGGATCACTCGGCTC | MB231 | 13 | 51 | 9 | 3 | 218 | 19.5 | 38.1 | 6.0 | 3.2 | 326.8 |
| >hg38_21_+_8433172_8446622_RNA45SN1_45S_50ntFlanks@6648.6679.32\|GTACGACTCTTAGCGGTGGATCACTCGGCTCG | >hg38_21_+_8433172_8446622_RNA45SN1_45S_50ntFlanks | 6648 | 6679 | 32 | GTACGACTCTTAGCGGTGGATCACTCGGCTCG | BT20 | 16 | 24 | 13 | 57 | 123 | 15.6 | 9.3 | 15.2 | 86.1 | 57.0 |
| >hg38_21_+_8433172_8446622_RNA45SN1_45S_50ntFlanks@6648.6679.32\|GTACGACTCTTAGCGGTGGATCACTCGGCTCG | >hg38_21_+_8433172_8446622_RNA45SN1_45S_50ntFlanks | 6648 | 6679 | 32 | GTACGACTCTTAGCGGTGGATCACTCGGCTCG | MB231 | 27 | 69 | 15 | 3 | 274 | 36.4 | 39.2 | 9.9 | 1.1 | 391.6 |
| >hg38_21_+_8433172_8446622_RNA45SN1_45S_50ntFlanks@6648.6679.32\|GTACGACTCTTAGCGGTGGATCACTCGGCTCG | >hg38_21_+_8433172_8446622_RNA45SN1_45S_50ntFlanks | 6648 | 6679 | 32 | GTACGACTCTTAGCGGTGGATCACTCGGCTCG | MB468 | 40 | 67 | 17 | 25 | 281 | 11.1 | 31.5 | 7.9 | 22.8 | 221.0 |
| >hg38_21_+_8433172_8446622_RNA45SN1_45S_50ntFlanks@6648.6680.33\|GTACGACTCTTAGCGGTGGATCACTCGGCTCGT | >hg38_21_+_8433172_8446622_RNA45SN1_45S_50ntFlanks | 6648 | 6680 | 33 | GTACGACTCTTAGCGGTGGATCACTCGGCTCGT | BT20 | 32 | 40 | 33 | 36 | 101 | 13.8 | 20.4 | 23.9 | 25.4 | 18.1 |
| >hg38_21_+_8433172_8446622_RNA45SN1_45S_50ntFlanks@6648.6680.33\|GTACGACTCTTAGCGGTGGATCACTCGGCTCGT | >hg38_21_+_8433172_8446622_RNA45SN1_45S_50ntFlanks | 6648 | 6680 | 33 | GTACGACTCTTAGCGGTGGATCACTCGGCTCGT | MB231 | 20 | 56 | 10 | 7 | 123 | 20.7 | 34.5 | 9.0 | 6.5 | 142.2 |
| >hg38_21_+_8433172_8446622_RNA45SN1_45S_50ntFlanks@6648.6680.33\|GTACGACTCTTAGCGGTGGATCACTCGGCTCGT | >hg38_21_+_8433172_8446622_RNA45SN1_45S_50ntFlanks | 6648 | 6680 | 33 | GTACGACTCTTAGCGGTGGATCACTCGGCTCGT | MB468 | 24 | 15 | 8 | 24 | 88 | 5.3 | 8.2 | 4.8 | 14.4 | 27.0 |
| >hg38_21_+_8433172_8446622_RNA45SN1_45S_50ntFlanks@6648.6681.34\|GTACGACTCTTAGCGGTGGATCACTCGGCTCGTG | >hg38_21_+_8433172_8446622_RNA45SN1_45S_50ntFlanks | 6648 | 6681 | 34 | GTACGACTCTTAGCGGTGGATCACTCGGCTCGTG | BT20 | 20 | 42 | 7 | 33 | 81 | 20.6 | 16.9 | 5.0 | 29.1 | 17.2 |
| >hg38_21_+_8433172_8446622_RNA45SN1_45S_50ntFlanks@6648.6681.34\|GTACGACTCTTAGCGGTGGATCACTCGGCTCGTG | >hg38_21_+_8433172_8446622_RNA45SN1_45S_50ntFlanks | 6648 | 6681 | 34 | GTACGACTCTTAGCGGTGGATCACTCGGCTCGTG | MB231 | 81 | 376 | 15 | 9 | 189 | 57.1 | 189.3 | 12.5 | 5.0 | 268.4 |
| >hg38_21_+_8433172_8446622_RNA45SN1_45S_50ntFlanks@6648.6681.34\|GTACGACTCTTAGCGGTGGATCACTCGGCTCGTG | >hg38_21_+_8433172_8446622_RNA45SN1_45S_50ntFlanks | 6648 | 6681 | 34 | GTACGACTCTTAGCGGTGGATCACTCGGCTCGTG | MB468 | 258 | 148 | 68 | 264 | 939 | 77.6 | 61.8 | 30.5 | 213.1 | 357.9 |
| >hg38_21_+_8433172_8446622_RNA45SN1_45S_50ntFlanks@6648.6689.42\|GTACGACTCTTAGCGGTGGATCACTCGGCTCGTGCGTCGATG | >hg38_21_+_8433172_8446622_RNA45SN1_45S_50ntFlanks | 6648 | 6689 | 42 | GTACGACTCTTAGCGGTGGATCACTCGGCTCGTGCGTCGATG | BT20 | 16 | 70 | 13 | 45 | 188 | 10.2 | 64.9 | 7.8 | 8.8 | 40.0 |
| >hg38_21_+_8433172_8446622_RNA45SN1_45S_50ntFlanks@6648.6689.42\|GTACGACTCTTAGCGGTGGATCACTCGGCTCGTGCGTCGATG | >hg38_21_+_8433172_8446622_RNA45SN1_45S_50ntFlanks | 6648 | 6689 | 42 | GTACGACTCTTAGCGGTGGATCACTCGGCTCGTGCGTCGATG | MB231 | 32 | 59 | 8 | 19 | 99 | 22.2 | 20.5 | 4.3 | 23.7 | 97.8 |
| >hg38_21_+_8433172_8446622_RNA45SN1_45S_50ntFlanks@6648.6689.42\|GTACGACTCTTAGCGGTGGATCACTCGGCTCGTGCGTCGATG | >hg38_21_+_8433172_8446622_RNA45SN1_45S_50ntFlanks | 6648 | 6689 | 42 | GTACGACTCTTAGCGGTGGATCACTCGGCTCGTGCGTCGATG | MB468 | 76 | 77 | 26 | 112 | 978 | 13.9 | 39.4 | 18.9 | 88.5 | 544.4 |
| >hg38_21_+_8433172_8446622_RNA45SN1_45S_50ntFlanks@6648.6690.43\|GTACGACTCTTAGCGGTGGATCACTCGGCTCGTGCGTCGATGA | >hg38_21_+_8433172_8446622_RNA45SN1_45S_50ntFlanks | 6648 | 6690 | 43 | GTACGACTCTTAGCGGTGGATCACTCGGCTCGTGCGTCGATGA | MB231 | 11 | 22 | 4 | 3 | 27 | 9.3 | 8.5 | 3.1 | 0.5 | 19.9 |
| >hg38_21_+_8433172_8446622_RNA45SN1_45S_50ntFlanks@6648.6692.45\|GTACGACTCTTAGCGGTGGATCACTCGGCTCGTGCGTCGATGAAG | >hg38_21_+_8433172_8446622_RNA45SN1_45S_50ntFlanks | 6648 | 6692 | 45 | GTACGACTCTTAGCGGTGGATCACTCGGCTCGTGCGTCGATGAAG | MB231 | 25 | 38 | 10 | 2 | 51 | 19.7 | 30.6 | 8.6 | 0.9 | 40.8 |
| >hg38_21_+_8433172_8446622_RNA45SN1_45S_50ntFlanks@6648.6693.46\|GTACGACTCTTAGCGGTGGATCACTCGGCTCGTGCGTCGATGAAGA | >hg38_21_+_8433172_8446622_RNA45SN1_45S_50ntFlanks | 6648 | 6693 | 46 | GTACGACTCTTAGCGGTGGATCACTCGGCTCGTGCGTCGATGAAGA | MB231 | 14 | 22 | 6 | 1 | 36 | 11.0 | 18.7 | 4.6 | 1.5 | 35.2 |
| >hg38_21_+_8433172_8446622_RNA45SN1_45S_50ntFlanks@6648.6694.47\|GTACGACTCTTAGCGGTGGATCACTCGGCTCGTGCGTCGATGAAGAA | >hg38_21_+_8433172_8446622_RNA45SN1_45S_50ntFlanks | 6648 | 6694 | 47 | GTACGACTCTTAGCGGTGGATCACTCGGCTCGTGCGTCGATGAAGAA | MB231 | 11 | 19 | 5 | 1 | 26 | 7.9 | 24.2 | 5.4 | 1.3 | 36.0 |
| >hg38_21_+_8433172_8446622_RNA45SN1_45S_50ntFlanks@6648.6695.48\|GTACGACTCTTAGCGGTGGATCACTCGGCTCGTGCGTCGATGAAGAAC | >hg38_21_+_8433172_8446622_RNA45SN1_45S_50ntFlanks | 6648 | 6695 | 48 | GTACGACTCTTAGCGGTGGATCACTCGGCTCGTGCGTCGATGAAGAAC | MB231 | 45 | 62 | 23 | 11 | 148 | 33.4 | 86.1 | 17.6 | 16.8 | 203.6 |
| >hg38_21_+_8433172_8446622_RNA45SN1_45S_50ntFlanks@6648.6696.49\|GTACGACTCTTAGCGGTGGATCACTCGGCTCGTGCGTCGATGAAGAACG | >hg38_21_+_8433172_8446622_RNA45SN1_45S_50ntFlanks | 6648 | 6696 | 49 | GTACGACTCTTAGCGGTGGATCACTCGGCTCGTGCGTCGATGAAGAACG | MB231 | 68 | 171 | 32 | 5 | 84 | 26.1 | 243.0 | 22.7 | 7.6 | 132.5 |
| >hg38_21_+_8433172_8446622_RNA45SN1_45S_50ntFlanks@6648.6697.50\|GTACGACTCTTAGCGGTGGATCACTCGGCTCGTGCGTCGATGAAGAACGC | >hg38_21_+_8433172_8446622_RNA45SN1_45S_50ntFlanks | 6648 | 6697 | 50 | GTACGACTCTTAGCGGTGGATCACTCGGCTCGTGCGTCGATGAAGAACGC | MB231 | 14 | 25 | 6 | 1 | 29 | 13.5 | 37.6 | 4.4 | 1.5 | 50.4 |
| >hg38_21_+_8433172_8446622_RNA45SN1_45S_50ntFlanks@6649.6666.18\|TACGACTCTTAGCGGTGG | >hg38_21_+_8433172_8446622_RNA45SN1_45S_50ntFlanks | 6649 | 6666 | 18 | TACGACTCTTAGCGGTGG | MB468 | 13 | 2 | 9 | 7 | 11 | 9.4 | 1.6 | 7.6 | 3.8 | 9.4 |
| >hg38_21_+_8433172_8446622_RNA45SN1_45S_50ntFlanks@6649.6668.20\|TACGACTCTTAGCGGTGGAT | >hg38_21_+_8433172_8446622_RNA45SN1_45S_50ntFlanks | 6649 | 6668 | 20 | TACGACTCTTAGCGGTGGAT | BT20 | 22 | 35 | 16 | 20 | 145 | 3.1 | 16.4 | 4.2 | 9.7 | 72.8 |
| >hg38_21_+_8433172_8446622_RNA45SN1_45S_50ntFlanks@6649.6668.20\|TACGACTCTTAGCGGTGGAT | >hg38_21_+_8433172_8446622_RNA45SN1_45S_50ntFlanks | 6649 | 6668 | 20 | TACGACTCTTAGCGGTGGAT | MB231 | 13 | 14 | 5 | 5 | 60 | 9.1 | 10.1 | 2.7 | 1.3 | 35.0 |
| >hg38_21_+_8433172_8446622_RNA45SN1_45S_50ntFlanks@6649.6669.21\|TACGACTCTTAGCGGTGGATC | >hg38_21_+_8433172_8446622_RNA45SN1_45S_50ntFlanks | 6649 | 6669 | 21 | TACGACTCTTAGCGGTGGATC | BT20 | 21 | 47 | 22 | 37 | 200 | 0.4 | 20.6 | 9.1 | 15.7 | 109.3 |
| >hg38_21_+_8433172_8446622_RNA45SN1_45S_50ntFlanks@6649.6669.21\|TACGACTCTTAGCGGTGGATC | >hg38_21_+_8433172_8446622_RNA45SN1_45S_50ntFlanks | 6649 | 6669 | 21 | TACGACTCTTAGCGGTGGATC | MB468 | 29 | 7 | 13 | 19 | 58 | 6.1 | 2.0 | 2.1 | 8.4 | 19.0 |
| >hg38_21_+_8433172_8446622_RNA45SN1_45S_50ntFlanks@6649.6670.22\|TACGACTCTTAGCGGTGGATCA | >hg38_21_+_8433172_8446622_RNA45SN1_45S_50ntFlanks | 6649 | 6670 | 22 | TACGACTCTTAGCGGTGGATCA | MB468 | 12 | 2 | 5 | 7 | 30 | 2.9 | 0.5 | 2.5 | 3.4 | 11.5 |
| >hg38_21_+_8433172_8446622_RNA45SN1_45S_50ntFlanks@6649.6679.31\|TACGACTCTTAGCGGTGGATCACTCGGCTCG | >hg38_21_+_8433172_8446622_RNA45SN1_45S_50ntFlanks | 6649 | 6679 | 31 | TACGACTCTTAGCGGTGGATCACTCGGCTCG | MB231 | 19 | 40 | 9 | 2 | 161 | 22.1 | 21.2 | 5.9 | 0.5 | 207.8 |

| | | | | | | | | | | | | | | | |
|---|---|---|---|---|---|---|---|---|---|---|---|---|---|---|---|
| >hg38_21_+_8433172_8446622_RNA45SN1_45S_50ntFlanks@6649.6679.31\|TACGACTCTTAGCGGTGGATCACTCGGCTCG | >hg38_21_+_8433172_8446622_RNA45SN1_45S_50ntFlanks | 6649 | 6679 | 31 | TACGACTCTTAGCGGTGGATCACTCGGCTCG | MB468 | 39 | 50 | 22 | 24 | 183 | 8.9 | 18.3 | 9.3 | 20.4 | 88.3 |
| >hg38_21_+_8433172_8446622_RNA45SN1_45S_50ntFlanks@6649.6680.32\|TACGACTCTTAGCGGTGGATCACTCGGCTCGT | >hg38_21_+_8433172_8446622_RNA45SN1_45S_50ntFlanks | 6649 | 6680 | 32 | TACGACTCTTAGCGGTGGATCACTCGGCTCGT | MB468 | 15 | 6 | 6 | 17 | 39 | 3.7 | 3.0 | 2.5 | 8.1 | 4.7 |
| >hg38_21_+_8433172_8446622_RNA45SN1_45S_50ntFlanks@6649.6681.33\|TACGACTCTTAGCGGTGGATCACTCGGCTCGTG | >hg38_21_+_8433172_8446622_RNA45SN1_45S_50ntFlanks | 6649 | 6681 | 33 | TACGACTCTTAGCGGTGGATCACTCGGCTCGTG | MB468 | 67 | 23 | 21 | 129 | 186 | 12.7 | 7.8 | 5.3 | 56.8 | 58.3 |
| >hg38_21_+_8433172_8446622_RNA45SN1_45S_50ntFlanks@6649.6683.35\|TACGACTCTTAGCGGTGGATCACTCGGCTCGTGCG | >hg38_21_+_8433172_8446622_RNA45SN1_45S_50ntFlanks | 6649 | 6683 | 35 | TACGACTCTTAGCGGTGGATCACTCGGCTCGTGCG | BT20 | 18 | 37 | 3 | 4 | 13 | 20.5 | 25.4 | 3.0 | 4.5 | 3.3 |
| >hg38_21_+_8433172_8446622_RNA45SN1_45S_50ntFlanks@6649.6683.35\|TACGACTCTTAGCGGTGGATCACTCGGCTCGTGCG | >hg38_21_+_8433172_8446622_RNA45SN1_45S_50ntFlanks | 6649 | 6683 | 35 | TACGACTCTTAGCGGTGGATCACTCGGCTCGTGCG | MB231 | 57 | 295 | 3 | 4 | 36 | 35.2 | 215.8 | 2.1 | 2.1 | 38.8 |
| >hg38_21_+_8433172_8446622_RNA45SN1_45S_50ntFlanks@6649.6689.41\|TACGACTCTTAGCGGTGGATCACTCGGCTCGTGCGTCGATG | >hg38_21_+_8433172_8446622_RNA45SN1_45S_50ntFlanks | 6649 | 6689 | 41 | TACGACTCTTAGCGGTGGATCACTCGGCTCGTGCGTCGATG | BT20 | 10 | 25 | 7 | 15 | 57 | 5.5 | 12.8 | 4.7 | 4.6 | 10.6 |
| >hg38_21_+_8433172_8446622_RNA45SN1_45S_50ntFlanks@6649.6689.41\|TACGACTCTTAGCGGTGGATCACTCGGCTCGTGCGTCGATG | >hg38_21_+_8433172_8446622_RNA45SN1_45S_50ntFlanks | 6649 | 6689 | 41 | TACGACTCTTAGCGGTGGATCACTCGGCTCGTGCGTCGATG | MB231 | 13 | 24 | 3 | 7 | 42 | 8.7 | 7.4 | 1.8 | 6.5 | 32.9 |
| >hg38_21_+_8433172_8446622_RNA45SN1_45S_50ntFlanks@6649.6689.41\|TACGACTCTTAGCGGTGGATCACTCGGCTCGTGCGTCGATG | >hg38_21_+_8433172_8446622_RNA45SN1_45S_50ntFlanks | 6649 | 6689 | 41 | TACGACTCTTAGCGGTGGATCACTCGGCTCGTGCGTCGATG | MB468 | 44 | 41 | 15 | 69 | 539 | 8.0 | 30.6 | 11.9 | 46.3 | 375.1 |
| >hg38_21_+_8433172_8446622_RNA45SN1_45S_50ntFlanks@6649.6692.44\|TACGACTCTTAGCGGTGGATCACTCGGCTCGTGCGTCGATGAAG | >hg38_21_+_8433172_8446622_RNA45SN1_45S_50ntFlanks | 6649 | 6692 | 44 | TACGACTCTTAGCGGTGGATCACTCGGCTCGTGCGTCGATGAAG | MB231 | 12 | 18 | 4 | 2 | 28 | 9.5 | 15.0 | 3.2 | 1.3 | 22.5 |
| >hg38_21_+_8433172_8446622_RNA45SN1_45S_50ntFlanks@6649.6695.47\|TACGACTCTTAGCGGTGGATCACTCGGCTCGTGCGTCGATGAAGAAC | >hg38_21_+_8433172_8446622_RNA45SN1_45S_50ntFlanks | 6649 | 6695 | 47 | TACGACTCTTAGCGGTGGATCACTCGGCTCGTGCGTCGATGAAGAAC | MB231 | 18 | 19 | 12 | 3 | 55 | 12.7 | 25.1 | 10.6 | 4.6 | 74.4 |
| >hg38_21_+_8433172_8446622_RNA45SN1_45S_50ntFlanks@6649.6696.48\|TACGACTCTTAGCGGTGGATCACTCGGCTCGTGCGTCGATGAAGAACG | >hg38_21_+_8433172_8446622_RNA45SN1_45S_50ntFlanks | 6649 | 6696 | 48 | TACGACTCTTAGCGGTGGATCACTCGGCTCGTGCGTCGATGAAGAACG | MB231 | 20 | 43 | 10 | 1 | 29 | 7.5 | 58.7 | 6.8 | 1.9 | 49.6 |
| >hg38_21_+_8433172_8446622_RNA45SN1_45S_50ntFlanks@6649.6698.50\|TACGACTCTTAGCGGTGGATCACTCGGCTCGTGCGTCGATGAAGAACGCA | >hg38_21_+_8433172_8446622_RNA45SN1_45S_50ntFlanks | 6649 | 6698 | 50 | TACGACTCTTAGCGGTGGATCACTCGGCTCGTGCGTCGATGAAGAACGCA | MB231 | 17 | 66 | 13 | 1 | 37 | 16.8 | 107.5 | 16.5 | 1.0 | 63.4 |
| >hg38_21_+_8433172_8446622_RNA45SN1_45S_50ntFlanks@6650.6668.19\|ACGACTCTTAGCGGTGGAT | >hg38_21_+_8433172_8446622_RNA45SN1_45S_50ntFlanks | 6650 | 6668 | 19 | ACGACTCTTAGCGGTGGAT | BT20 | 43 | 99 | 38 | 63 | 284 | 10.2 | 42.5 | 8.8 | 17.9 | 137.2 |
| >hg38_21_+_8433172_8446622_RNA45SN1_45S_50ntFlanks@6650.6668.19\|ACGACTCTTAGCGGTGGAT | >hg38_21_+_8433172_8446622_RNA45SN1_45S_50ntFlanks | 6650 | 6668 | 19 | ACGACTCTTAGCGGTGGAT | MB231 | 18 | 24 | 7 | 29 | 97 | 13.0 | 17.1 | 4.4 | 13.1 | 56.3 |
| >hg38_21_+_8433172_8446622_RNA45SN1_45S_50ntFlanks@6650.6669.20\|ACGACTCTTAGCGGTGGATC | >hg38_21_+_8433172_8446622_RNA45SN1_45S_50ntFlanks | 6650 | 6669 | 20 | ACGACTCTTAGCGGTGGATC | BT20 | 39 | 123 | 42 | 108 | 374 | 6.8 | 54.7 | 17.8 | 54.8 | 162.6 |
| >hg38_21_+_8433172_8446622_RNA45SN1_45S_50ntFlanks@6650.6669.20\|ACGACTCTTAGCGGTGGATC | >hg38_21_+_8433172_8446622_RNA45SN1_45S_50ntFlanks | 6650 | 6669 | 20 | ACGACTCTTAGCGGTGGATC | MB231 | 11 | 17 | 6 | 14 | 74 | 14.2 | 9.3 | 2.8 | 20.3 | 49.1 |
| >hg38_21_+_8433172_8446622_RNA45SN1_45S_50ntFlanks@6650.6669.20\|ACGACTCTTAGCGGTGGATC | >hg38_21_+_8433172_8446622_RNA45SN1_45S_50ntFlanks | 6650 | 6669 | 20 | ACGACTCTTAGCGGTGGATC | MB468 | 32 | 8 | 16 | 28 | 73 | 10.7 | 2.3 | 1.7 | 18.6 | 21.3 |
| >hg38_21_+_8433172_8446622_RNA45SN1_45S_50ntFlanks@6650.6670.21\|ACGACTCTTAGCGGTGGATCA | >hg38_21_+_8433172_8446622_RNA45SN1_45S_50ntFlanks | 6650 | 6670 | 21 | ACGACTCTTAGCGGTGGATCA | MB468 | 12 | 2 | 5 | 8 | 33 | 2.8 | 0.2 | 2.1 | 4.1 | 12.9 |
| >hg38_21_+_8433172_8446622_RNA45SN1_45S_50ntFlanks@6650.6672.23\|ACGACTCTTAGCGGTGGATCACT | >hg38_21_+_8433172_8446622_RNA45SN1_45S_50ntFlanks | 6650 | 6672 | 23 | ACGACTCTTAGCGGTGGATCACT | BT20 | 19 | 62 | 15 | 23 | 146 | 7.6 | 33.6 | 6.4 | 21.8 | 78.3 |
| >hg38_21_+_8433172_8446622_RNA45SN1_45S_50ntFlanks@6650.6675.26\|ACGACTCTTAGCGGTGGATCACTCGG | >hg38_21_+_8433172_8446622_RNA45SN1_45S_50ntFlanks | 6650 | 6675 | 26 | ACGACTCTTAGCGGTGGATCACTCGG | BT20 | 11 | 29 | 14 | 62 | 76 | 9.1 | 11.2 | 16.7 | 59.8 | 22.1 |
| >hg38_21_+_8433172_8446622_RNA45SN1_45S_50ntFlanks@6650.6676.27\|ACGACTCTTAGCGGTGGATCACTCGGC | >hg38_21_+_8433172_8446622_RNA45SN1_45S_50ntFlanks | 6650 | 6676 | 27 | ACGACTCTTAGCGGTGGATCACTCGGC | BT20 | 18 | 54 | 141 | 104 | 193 | 10.5 | 25.6 | 225.1 | 116.3 | 58.2 |
| >hg38_21_+_8433172_8446622_RNA45SN1_45S_50ntFlanks@6650.6677.28\|ACGACTCTTAGCGGTGGATCACTCGGCT | >hg38_21_+_8433172_8446622_RNA45SN1_45S_50ntFlanks | 6650 | 6677 | 28 | ACGACTCTTAGCGGTGGATCACTCGGCT | BT20 | 17 | 80 | 14 | 79 | 173 | 5.7 | 43.0 | 4.9 | 81.9 | 26.3 |
| >hg38_21_+_8433172_8446622_RNA45SN1_45S_50ntFlanks@6650.6678.29\|ACGACTCTTAGCGGTGGATCACTCGGCTC | >hg38_21_+_8433172_8446622_RNA45SN1_45S_50ntFlanks | 6650 | 6678 | 29 | ACGACTCTTAGCGGTGGATCACTCGGCTC | BT20 | 24 | 67 | 19 | 81 | 197 | 15.1 | 31.2 | 9.6 | 63.4 | 41.3 |
| >hg38_21_+_8433172_8446622_RNA45SN1_45S_50ntFlanks@6650.6678.29\|ACGACTCTTAGCGGTGGATCACTCGGCTC | >hg38_21_+_8433172_8446622_RNA45SN1_45S_50ntFlanks | 6650 | 6678 | 29 | ACGACTCTTAGCGGTGGATCACTCGGCTC | MB231 | 10 | 48 | 13 | 5 | 220 | 14.9 | 34.5 | 8.8 | 5.0 | 331.8 |
| >hg38_21_+_8433172_8446622_RNA45SN1_45S_50ntFlanks@6650.6679.30\|ACGACTCTTAGCGGTGGATCACTCGGCTCG | >hg38_21_+_8433172_8446622_RNA45SN1_45S_50ntFlanks | 6650 | 6679 | 30 | ACGACTCTTAGCGGTGGATCACTCGGCTCG | BT20 | 17 | 49 | 91 | 103 | 180 | 13.8 | 26.6 | 140.8 | 140.0 | 92.9 |
| >hg38_21_+_8433172_8446622_RNA45SN1_45S_50ntFlanks@6650.6679.30\|ACGACTCTTAGCGGTGGATCACTCGGCTCG | >hg38_21_+_8433172_8446622_RNA45SN1_45S_50ntFlanks | 6650 | 6679 | 30 | ACGACTCTTAGCGGTGGATCACTCGGCTCG | MB231 | 27 | 71 | 19 | 5 | 278 | 29.9 | 42.3 | 10.7 | 1.1 | 366.7 |
| >hg38_21_+_8433172_8446622_RNA45SN1_45S_50ntFlanks@6650.6679.30\|ACGACTCTTAGCGGTGGATCACTCGGCTCG | >hg38_21_+_8433172_8446622_RNA45SN1_45S_50ntFlanks | 6650 | 6679 | 30 | ACGACTCTTAGCGGTGGATCACTCGGCTCG | MB468 | 50 | 63 | 29 | 39 | 305 | 10.1 | 25.9 | 10.1 | 29.9 | 159.0 |
| >hg38_21_+_8433172_8446622_RNA45SN1_45S_50ntFlanks@6650.6681.32\|ACGACTCTTAGCGGTGGATCACTCGGCTCGTG | >hg38_21_+_8433172_8446622_RNA45SN1_45S_50ntFlanks | 6650 | 6681 | 32 | ACGACTCTTAGCGGTGGATCACTCGGCTCGTG | BT20 | 25 | 80 | 11 | 63 | 121 | 19.9 | 40.3 | 5.2 | 50.1 | 17.7 |
| >hg38_21_+_8433172_8446622_RNA45SN1_45S_50ntFlanks@6650.6681.32\|ACGACTCTTAGCGGTGGATCACTCGGCTCGTG | >hg38_21_+_8433172_8446622_RNA45SN1_45S_50ntFlanks | 6650 | 6681 | 32 | ACGACTCTTAGCGGTGGATCACTCGGCTCGTG | MB231 | 35 | 178 | 15 | 10 | 176 | 21.1 | 79.7 | 11.3 | 7.8 | 253.1 |
| >hg38_21_+_8433172_8446622_RNA45SN1_45S_50ntFlanks@6650.6681.32\|ACGACTCTTAGCGGTGGATCACTCGGCTCGTG | >hg38_21_+_8433172_8446622_RNA45SN1_45S_50ntFlanks | 6650 | 6681 | 32 | ACGACTCTTAGCGGTGGATCACTCGGCTCGTG | MB468 | 252 | 144 | 87 | 305 | 847 | 70.4 | 59.8 | 33.5 | 233.7 | 260.1 |
| >hg38_21_+_8433172_8446622_RNA45SN1_45S_50ntFlanks@6650.6683.34\|ACGACTCTTAGCGGTGGATCACTCGGCTCGTGCG | >hg38_21_+_8433172_8446622_RNA45SN1_45S_50ntFlanks | 6650 | 6683 | 34 | ACGACTCTTAGCGGTGGATCACTCGGCTCGTGCG | BT20 | 33 | 80 | 8 | 12 | 24 | 39.1 | 55.8 | 10.4 | 11.3 | 5.8 |
| >hg38_21_+_8433172_8446622_RNA45SN1_45S_50ntFlanks@6650.6683.34\|ACGACTCTTAGCGGTGGATCACTCGGCTCGTGCG | >hg38_21_+_8433172_8446622_RNA45SN1_45S_50ntFlanks | 6650 | 6683 | 34 | ACGACTCTTAGCGGTGGATCACTCGGCTCGTGCG | MB231 | 65 | 373 | 4 | 8 | 57 | 38.6 | 271.9 | 3.1 | 3.2 | 66.2 |
| >hg38_21_+_8433172_8446622_RNA45SN1_45S_50ntFlanks@6650.6689.40\|ACGACTCTTAGCGGTGGATCACTCGGCTCGTGCGTCGATG | >hg38_21_+_8433172_8446622_RNA45SN1_45S_50ntFlanks | 6650 | 6689 | 40 | ACGACTCTTAGCGGTGGATCACTCGGCTCGTGCGTCGATG | BT20 | 21 | 71 | 14 | 50 | 114 | 11.0 | 33.3 | 7.9 | 17.2 | 25.8 |
| >hg38_21_+_8433172_8446622_RNA45SN1_45S_50ntFlanks@6650.6689.40\|ACGACTCTTAGCGGTGGATCACTCGGCTCGTGCGTCGATG | >hg38_21_+_8433172_8446622_RNA45SN1_45S_50ntFlanks | 6650 | 6689 | 40 | ACGACTCTTAGCGGTGGATCACTCGGCTCGTGCGTCGATG | MB231 | 21 | 38 | 5 | 20 | 62 | 13.9 | 6.0 | 3.2 | 25.1 | 44.5 |
| >hg38_21_+_8433172_8446622_RNA45SN1_45S_50ntFlanks@6650.6689.40\|ACGACTCTTAGCGGTGGATCACTCGGCTCGTGCGTCGATG | >hg38_21_+_8433172_8446622_RNA45SN1_45S_50ntFlanks | 6650 | 6689 | 40 | ACGACTCTTAGCGGTGGATCACTCGGCTCGTGCGTCGATG | MB468 | 64 | 47 | 18 | 103 | 719 | 13.9 | 25.5 | 13.9 | 58.4 | 461.2 |
| >hg38_21_+_8433172_8446622_RNA45SN1_45S_50ntFlanks@6650.6692.43\|ACGACTCTTAGCGGTGGATCACTCGGCTCGTGCGTCGATGAAG | >hg38_21_+_8433172_8446622_RNA45SN1_45S_50ntFlanks | 6650 | 6692 | 43 | ACGACTCTTAGCGGTGGATCACTCGGCTCGTGCGTCGATGAAG | BT20 | 13 | 27 | 11 | 10 | 39 | 11.4 | 12.4 | 13.1 | 3.7 | 17.7 |
| >hg38_21_+_8433172_8446622_RNA45SN1_45S_50ntFlanks@6650.6692.43\|ACGACTCTTAGCGGTGGATCACTCGGCTCGTGCGTCGATGAAG | >hg38_21_+_8433172_8446622_RNA45SN1_45S_50ntFlanks | 6650 | 6692 | 43 | ACGACTCTTAGCGGTGGATCACTCGGCTCGTGCGTCGATGAAG | MB231 | 27 | 47 | 10 | 8 | 67 | 23.6 | 32.2 | 7.7 | 5.2 | 49.8 |
| >hg38_21_+_8433172_8446622_RNA45SN1_45S_50ntFlanks@6650.6693.44\|ACGACTCTTAGCGGTGGATCACTCGGCTCGTGCGTCGATGAAGA | >hg38_21_+_8433172_8446622_RNA45SN1_45S_50ntFlanks | 6650 | 6693 | 44 | ACGACTCTTAGCGGTGGATCACTCGGCTCGTGCGTCGATGAAGA | MB231 | 12 | 20 | 5 | 4 | 37 | 9.2 | 10.9 | 3.7 | 1.9 | 24.8 |
| >hg38_21_+_8433172_8446622_RNA45SN1_45S_50ntFlanks@6650.6695.46\|ACGACTCTTAGCGGTGGATCACTCGGCTCGTGCGTCGATGAAGAAC | >hg38_21_+_8433172_8446622_RNA45SN1_45S_50ntFlanks | 6650 | 6695 | 46 | ACGACTCTTAGCGGTGGATCACTCGGCTCGTGCGTCGATGAAGAAC | BT20 | 15 | 47 | 10 | 20 | 121 | 8.0 | 49.0 | 10.1 | 11.3 | 66.5 |
| >hg38_21_+_8433172_8446622_RNA45SN1_45S_50ntFlanks@6650.6695.46\|ACGACTCTTAGCGGTGGATCACTCGGCTCGTGCGTCGATGAAGAAC | >hg38_21_+_8433172_8446622_RNA45SN1_45S_50ntFlanks | 6650 | 6695 | 46 | ACGACTCTTAGCGGTGGATCACTCGGCTCGTGCGTCGATGAAGAAC | MB231 | 39 | 40 | 26 | 11 | 109 | 27.3 | 37.7 | 26.6 | 14.8 | 115.4 |
| >hg38_21_+_8433172_8446622_RNA45SN1_45S_50ntFlanks@6650.6696.47\|ACGACTCTTAGCGGTGGATCACTCGGCTCGTGCGTCGATGAAGAACG | >hg38_21_+_8433172_8446622_RNA45SN1_45S_50ntFlanks | 6650 | 6696 | 47 | ACGACTCTTAGCGGTGGATCACTCGGCTCGTGCGTCGATGAAGAACG | BT20 | 11 | 33 | 3 | 9 | 50 | 7.5 | 30.7 | 2.0 | 4.2 | 34.8 |
| >hg38_21_+_8433172_8446622_RNA45SN1_45S_50ntFlanks@6650.6696.47\|ACGACTCTTAGCGGTGGATCACTCGGCTCGTGCGTCGATGAAGAACG | >hg38_21_+_8433172_8446622_RNA45SN1_45S_50ntFlanks | 6650 | 6696 | 47 | ACGACTCTTAGCGGTGGATCACTCGGCTCGTGCGTCGATGAAGAACG | MB231 | 58 | 124 | 24 | 5 | 96 | 25.5 | 140.1 | 20.5 | 6.7 | 85.3 |
| >hg38_21_+_8433172_8446622_RNA45SN1_45S_50ntFlanks@6650.6698.49\|ACGACTCTTAGCGGTGGATCACTCGGCTCGTGCGTCGATGAAGAACGCA | >hg38_21_+_8433172_8446622_RNA45SN1_45S_50ntFlanks | 6650 | 6698 | 49 | ACGACTCTTAGCGGTGGATCACTCGGCTCGTGCGTCGATGAAGAACGCA | MB231 | 48 | 158 | 33 | 4 | 85 | 48.4 | 245.1 | 32.1 | 6.8 | 146.9 |
| >hg38_21_+_8433172_8446622_RNA45SN1_45S_50ntFlanks@6650.6698.49\|ACGACTCTTAGCGGTGGATCACTCGGCTCGTGCGTCGATGAAGAACGCA | >hg38_21_+_8433172_8446622_RNA45SN1_45S_50ntFlanks | 6650 | 6698 | 49 | ACGACTCTTAGCGGTGGATCACTCGGCTCGTGCGTCGATGAAGAACGCA | MB468 | 17 | 99 | 13 | 29 | 313 | 9.2 | 87.1 | 9.6 | 35.0 | 192.8 |
| >hg38_21_+_8433172_8446622_RNA45SN1_45S_50ntFlanks@6650.6699.50\|ACGACTCTTAGCGGTGGATCACTCGGCTCGTGCGTCGATGAAGAACGCAG | >hg38_21_+_8433172_8446622_RNA45SN1_45S_50ntFlanks | 6650 | 6699 | 50 | ACGACTCTTAGCGGTGGATCACTCGGCTCGTGCGTCGATGAAGAACGCAG | MB231 | 27 | 111 | 10 | 1 | 17 | 18.2 | 173.7 | 9.4 | 2.1 | 30.0 |
| >hg38_21_+_8433172_8446622_RNA45SN1_45S_50ntFlanks@6650.6699.50\|ACGACTCTTAGCGGTGGATCACTCGGCTCGTGCGTCGATGAAGAACGCAG | >hg38_21_+_8433172_8446622_RNA45SN1_45S_50ntFlanks | 6650 | 6699 | 50 | ACGACTCTTAGCGGTGGATCACTCGGCTCGTGCGTCGATGAAGAACGCAG | MB468 | 10 | 68 | 11 | 20 | 203 | 5.7 | 81.2 | 9.7 | 29.1 | 163.4 |
| >hg38_21_+_8433172_8446622_RNA45SN1_45S_50ntFlanks@6651.6668.18\|CGACTCTTAGCGGTGGAT | >hg38_21_+_8433172_8446622_RNA45SN1_45S_50ntFlanks | 6651 | 6668 | 18 | CGACTCTTAGCGGTGGAT | BT20 | 386 | 824 | 478 | 397 | 2707 | 65.9 | 330.2 | 68.0 | 261.9 | 1096.1 |
| >hg38_21_+_8433172_8446622_RNA45SN1_45S_50ntFlanks@6651.6668.18\|CGACTCTTAGCGGTGGAT | >hg38_21_+_8433172_8446622_RNA45SN1_45S_50ntFlanks | 6651 | 6668 | 18 | CGACTCTTAGCGGTGGAT | MB231 | 168 | 213 | 94 | 75 | 915 | 142.6 | 118.8 | 38.4 | 20.2 | 531.5 |
| >hg38_21_+_8433172_8446622_RNA45SN1_45S_50ntFlanks@6651.6668.18\|CGACTCTTAGCGGTGGAT | >hg38_21_+_8433172_8446622_RNA45SN1_45S_50ntFlanks | 6651 | 6668 | 18 | CGACTCTTAGCGGTGGAT | MB468 | 87 | 26 | 69 | 56 | 155 | 27.8 | 12.7 | 21.4 | 13.2 | 91.6 |
| >hg38_21_+_8433172_8446622_RNA45SN1_45S_50ntFlanks@6651.6669.19\|CGACTCTTAGCGGTGGATC | >hg38_21_+_8433172_8446622_RNA45SN1_45S_50ntFlanks | 6651 | 6669 | 19 | CGACTCTTAGCGGTGGATC | BT20 | 276 | 826 | 404 | 612 | 2672 | 26.5 | 317.6 | 83.6 | 241.8 | 1189.6 |

| | | | | | | | | | | | | | | | |
|---|---|---|---|---|---|---|---|---|---|---|---|---|---|---|---|
| >hg38_21_+_8433172_8446622_RNA45SN1_45S_50ntFlanks@6651.6669.19\|CGACTCTTAGCGGTGGATC | >hg38_21_+_8433172_8446622_RNA45SN1_45S_50ntFlanks | 6651 | 6669 | 19 | CGACTCTTAGCGGTGGATC | MB231 | 94 | 122 | 72 | 77 | 578 | 124.7 | 57.8 | 25.7 | 103.7 | 402.1 |
| >hg38_21_+_8433172_8446622_RNA45SN1_45S_50ntFlanks@6651.6669.19\|CGACTCTTAGCGGTGGATC | >hg38_21_+_8433172_8446622_RNA45SN1_45S_50ntFlanks | 6651 | 6669 | 19 | CGACTCTTAGCGGTGGATC | MB468 | 201 | 38 | 103 | 243 | 571 | 47.7 | 12.9 | 16.9 | 169.7 | 214.2 |
| >hg38_21_+_8433172_8446622_RNA45SN1_45S_50ntFlanks@6651.6670.20\|CGACTCTTAGCGGTGGATCA | >hg38_21_+_8433172_8446622_RNA45SN1_45S_50ntFlanks | 6651 | 6670 | 20 | CGACTCTTAGCGGTGGATCA | BT20 | 15 | 33 | 15 | 5 | 66 | 8.2 | 12.9 | 5.4 | 2.9 | 36.7 |
| >hg38_21_+_8433172_8446622_RNA45SN1_45S_50ntFlanks@6651.6670.20\|CGACTCTTAGCGGTGGATCA | >hg38_21_+_8433172_8446622_RNA45SN1_45S_50ntFlanks | 6651 | 6670 | 20 | CGACTCTTAGCGGTGGATCA | MB468 | 112 | 11 | 39 | 103 | 439 | 33.3 | 2.5 | 16.7 | 55.4 | 204.9 |
| >hg38_21_+_8433172_8446622_RNA45SN1_45S_50ntFlanks@6651.6672.22\|CGACTCTTAGCGGTGGATCACT | >hg38_21_+_8433172_8446622_RNA45SN1_45S_50ntFlanks | 6651 | 6672 | 22 | CGACTCTTAGCGGTGGATCACT | BT20 | 155 | 458 | 170 | 112 | 1094 | 37.6 | 219.1 | 31.8 | 87.6 | 672.1 |
| >hg38_21_+_8433172_8446622_RNA45SN1_45S_50ntFlanks@6651.6672.22\|CGACTCTTAGCGGTGGATCACT | >hg38_21_+_8433172_8446622_RNA45SN1_45S_50ntFlanks | 6651 | 6672 | 22 | CGACTCTTAGCGGTGGATCACT | MB231 | 55 | 228 | 108 | 17 | 922 | 68.1 | 187.2 | 62.7 | 10.0 | 1019.0 |
| >hg38_21_+_8433172_8446622_RNA45SN1_45S_50ntFlanks@6651.6672.22\|CGACTCTTAGCGGTGGATCACT | >hg38_21_+_8433172_8446622_RNA45SN1_45S_50ntFlanks | 6651 | 6672 | 22 | CGACTCTTAGCGGTGGATCACT | MB468 | 11 | 19 | 11 | 11 | 64 | 6.7 | 14.5 | 8.9 | 6.1 | 33.8 |
| >hg38_21_+_8433172_8446622_RNA45SN1_45S_50ntFlanks@6651.6673.23\|CGACTCTTAGCGGTGGATCACTC | >hg38_21_+_8433172_8446622_RNA45SN1_45S_50ntFlanks | 6651 | 6673 | 23 | CGACTCTTAGCGGTGGATCACTC | BT20 | 50 | 162 | 67 | 117 | 487 | 17.6 | 72.1 | 24.9 | 70.5 | 153.5 |
| >hg38_21_+_8433172_8446622_RNA45SN1_45S_50ntFlanks@6651.6673.23\|CGACTCTTAGCGGTGGATCACTC | >hg38_21_+_8433172_8446622_RNA45SN1_45S_50ntFlanks | 6651 | 6673 | 23 | CGACTCTTAGCGGTGGATCACTC | MB231 | 27 | 172 | 91 | 15 | 644 | 35.8 | 149.4 | 54.0 | 12.4 | 849.8 |
| >hg38_21_+_8433172_8446622_RNA45SN1_45S_50ntFlanks@6651.6674.24\|CGACTCTTAGCGGTGGATCACTCG | >hg38_21_+_8433172_8446622_RNA45SN1_45S_50ntFlanks | 6651 | 6674 | 24 | CGACTCTTAGCGGTGGATCACTCG | BT20 | 40 | 124 | 85 | 171 | 578 | 28.8 | 58.1 | 97.5 | 183.1 | 163.8 |
| >hg38_21_+_8433172_8446622_RNA45SN1_45S_50ntFlanks@6651.6674.24\|CGACTCTTAGCGGTGGATCACTCG | >hg38_21_+_8433172_8446622_RNA45SN1_45S_50ntFlanks | 6651 | 6674 | 24 | CGACTCTTAGCGGTGGATCACTCG | MB231 | 23 | 260 | 94 | 17 | 1177 | 27.4 | 254.5 | 77.6 | 7.6 | 1716.1 |
| >hg38_21_+_8433172_8446622_RNA45SN1_45S_50ntFlanks@6651.6674.24\|CGACTCTTAGCGGTGGATCACTCG | >hg38_21_+_8433172_8446622_RNA45SN1_45S_50ntFlanks | 6651 | 6674 | 24 | CGACTCTTAGCGGTGGATCACTCG | MB468 | 14 | 27 | 18 | 19 | 89 | 5.4 | 34.2 | 13.8 | 9.1 | 58.9 |
| >hg38_21_+_8433172_8446622_RNA45SN1_45S_50ntFlanks@6651.6675.25\|CGACTCTTAGCGGTGGATCACTCGG | >hg38_21_+_8433172_8446622_RNA45SN1_45S_50ntFlanks | 6651 | 6675 | 25 | CGACTCTTAGCGGTGGATCACTCGG | BT20 | 47 | 150 | 157 | 220 | 394 | 37.5 | 63.9 | 222.2 | 289.3 | 82.1 |
| >hg38_21_+_8433172_8446622_RNA45SN1_45S_50ntFlanks@6651.6675.25\|CGACTCTTAGCGGTGGATCACTCGG | >hg38_21_+_8433172_8446622_RNA45SN1_45S_50ntFlanks | 6651 | 6675 | 25 | CGACTCTTAGCGGTGGATCACTCGG | MB231 | 28 | 549 | 140 | 18 | 2747 | 24.2 | 606.5 | 124.4 | 2.2 | 4591.6 |
| >hg38_21_+_8433172_8446622_RNA45SN1_45S_50ntFlanks@6651.6675.25\|CGACTCTTAGCGGTGGATCACTCGG | >hg38_21_+_8433172_8446622_RNA45SN1_45S_50ntFlanks | 6651 | 6675 | 25 | CGACTCTTAGCGGTGGATCACTCGG | MB468 | 64 | 132 | 117 | 67 | 414 | 31.5 | 114.0 | 94.9 | 47.2 | 310.6 |
| >hg38_21_+_8433172_8446622_RNA45SN1_45S_50ntFlanks@6651.6676.26\|CGACTCTTAGCGGTGGATCACTCGGC | >hg38_21_+_8433172_8446622_RNA45SN1_45S_50ntFlanks | 6651 | 6676 | 26 | CGACTCTTAGCGGTGGATCACTCGGC | BT20 | 188 | 570 | 2184 | 753 | 2262 | 73.2 | 273.5 | 3460.6 | 922.8 | 684.0 |
| >hg38_21_+_8433172_8446622_RNA45SN1_45S_50ntFlanks@6651.6676.26\|CGACTCTTAGCGGTGGATCACTCGGC | >hg38_21_+_8433172_8446622_RNA45SN1_45S_50ntFlanks | 6651 | 6676 | 26 | CGACTCTTAGCGGTGGATCACTCGGC | MB231 | 84 | 484 | 179 | 86 | 2396 | 76.4 | 449.0 | 135.4 | 36.3 | 2989.0 |
| >hg38_21_+_8433172_8446622_RNA45SN1_45S_50ntFlanks@6651.6676.26\|CGACTCTTAGCGGTGGATCACTCGGC | >hg38_21_+_8433172_8446622_RNA45SN1_45S_50ntFlanks | 6651 | 6676 | 26 | CGACTCTTAGCGGTGGATCACTCGGC | MB468 | 13 | 46 | 15 | 36 | 157 | 6.7 | 59.5 | 16.0 | 16.4 | 74.5 |
| >hg38_21_+_8433172_8446622_RNA45SN1_45S_50ntFlanks@6651.6677.27\|CGACTCTTAGCGGTGGATCACTCGGCT | >hg38_21_+_8433172_8446622_RNA45SN1_45S_50ntFlanks | 6651 | 6677 | 27 | CGACTCTTAGCGGTGGATCACTCGGCT | BT20 | 153 | 634 | 172 | 550 | 1594 | 15.9 | 396.1 | 62.6 | 598.7 | 307.0 |
| >hg38_21_+_8433172_8446622_RNA45SN1_45S_50ntFlanks@6651.6677.27\|CGACTCTTAGCGGTGGATCACTCGGCT | >hg38_21_+_8433172_8446622_RNA45SN1_45S_50ntFlanks | 6651 | 6677 | 27 | CGACTCTTAGCGGTGGATCACTCGGCT | MB231 | 78 | 497 | 145 | 45 | 2101 | 95.0 | 494.8 | 108.0 | 35.9 | 3165.3 |
| >hg38_21_+_8433172_8446622_RNA45SN1_45S_50ntFlanks@6651.6677.27\|CGACTCTTAGCGGTGGATCACTCGGCT | >hg38_21_+_8433172_8446622_RNA45SN1_45S_50ntFlanks | 6651 | 6677 | 27 | CGACTCTTAGCGGTGGATCACTCGGCT | MB468 | 65 | 86 | 45 | 90 | 436 | 20.5 | 103.1 | 31.2 | 49.3 | 301.9 |
| >hg38_21_+_8433172_8446622_RNA45SN1_45S_50ntFlanks@6651.6678.28\|CGACTCTTAGCGGTGGATCACTCGGCTC | >hg38_21_+_8433172_8446622_RNA45SN1_45S_50ntFlanks | 6651 | 6678 | 28 | CGACTCTTAGCGGTGGATCACTCGGCTC | BT20 | 224 | 619 | 220 | 1129 | 2265 | 130.7 | 313.8 | 122.2 | 1144.0 | 571.0 |
| >hg38_21_+_8433172_8446622_RNA45SN1_45S_50ntFlanks@6651.6678.28\|CGACTCTTAGCGGTGGATCACTCGGCTC | >hg38_21_+_8433172_8446622_RNA45SN1_45S_50ntFlanks | 6651 | 6678 | 28 | CGACTCTTAGCGGTGGATCACTCGGCTC | MB231 | 119 | 522 | 188 | 41 | 2369 | 175.9 | 419.0 | 111.7 | 43.4 | 3574.9 |
| >hg38_21_+_8433172_8446622_RNA45SN1_45S_50ntFlanks@6651.6678.28\|CGACTCTTAGCGGTGGATCACTCGGCTC | >hg38_21_+_8433172_8446622_RNA45SN1_45S_50ntFlanks | 6651 | 6678 | 28 | CGACTCTTAGCGGTGGATCACTCGGCTC | MB468 | 74 | 120 | 44 | 56 | 464 | 17.5 | 119.0 | 23.0 | 30.9 | 315.3 |
| >hg38_21_+_8433172_8446622_RNA45SN1_45S_50ntFlanks@6651.6679.29\|CGACTCTTAGCGGTGGATCACTCGGCTCG | >hg38_21_+_8433172_8446622_RNA45SN1_45S_50ntFlanks | 6651 | 6679 | 29 | CGACTCTTAGCGGTGGATCACTCGGCTCG | BT20 | 55 | 180 | 607 | 342 | 665 | 34.3 | 108.6 | 978.2 | 498.6 | 369.8 |
| >hg38_21_+_8433172_8446622_RNA45SN1_45S_50ntFlanks@6651.6679.29\|CGACTCTTAGCGGTGGATCACTCGGCTCG | >hg38_21_+_8433172_8446622_RNA45SN1_45S_50ntFlanks | 6651 | 6679 | 29 | CGACTCTTAGCGGTGGATCACTCGGCTCG | MB231 | 123 | 365 | 105 | 42 | 1606 | 142.7 | 290.0 | 76.4 | 10.3 | 2000.7 |
| >hg38_21_+_8433172_8446622_RNA45SN1_45S_50ntFlanks@6651.6679.29\|CGACTCTTAGCGGTGGATCACTCGGCTCG | >hg38_21_+_8433172_8446622_RNA45SN1_45S_50ntFlanks | 6651 | 6679 | 29 | CGACTCTTAGCGGTGGATCACTCGGCTCG | MB468 | 425 | 242 | 191 | 419 | 3089 | 60.0 | 128.0 | 59.7 | 321.3 | 1666.7 |
| >hg38_21_+_8433172_8446622_RNA45SN1_45S_50ntFlanks@6651.6680.30\|CGACTCTTAGCGGTGGATCACTCGGCTCGT | >hg38_21_+_8433172_8446622_RNA45SN1_45S_50ntFlanks | 6651 | 6680 | 30 | CGACTCTTAGCGGTGGATCACTCGGCTCGT | BT20 | 63 | 190 | 78 | 186 | 448 | 5.5 | 94.7 | 36.3 | 146.4 | 97.7 |
| >hg38_21_+_8433172_8446622_RNA45SN1_45S_50ntFlanks@6651.6680.30\|CGACTCTTAGCGGTGGATCACTCGGCTCGT | >hg38_21_+_8433172_8446622_RNA45SN1_45S_50ntFlanks | 6651 | 6680 | 30 | CGACTCTTAGCGGTGGATCACTCGGCTCGT | MB231 | 63 | 238 | 60 | 32 | 667 | 59.8 | 166.0 | 44.5 | 30.9 | 905.5 |
| >hg38_21_+_8433172_8446622_RNA45SN1_45S_50ntFlanks@6651.6680.30\|CGACTCTTAGCGGTGGATCACTCGGCTCGT | >hg38_21_+_8433172_8446622_RNA45SN1_45S_50ntFlanks | 6651 | 6680 | 30 | CGACTCTTAGCGGTGGATCACTCGGCTCGT | MB468 | 195 | 142 | 73 | 206 | 881 | 50.9 | 84.0 | 32.9 | 39.8 | 460.0 |
| >hg38_21_+_8433172_8446622_RNA45SN1_45S_50ntFlanks@6651.6681.31\|CGACTCTTAGCGGTGGATCACTCGGCTCGTG | >hg38_21_+_8433172_8446622_RNA45SN1_45S_50ntFlanks | 6651 | 6681 | 31 | CGACTCTTAGCGGTGGATCACTCGGCTCGTG | BT20 | 223 | 790 | 134 | 648 | 1578 | 182.5 | 461.4 | 68.0 | 763.1 | 265.7 |
| >hg38_21_+_8433172_8446622_RNA45SN1_45S_50ntFlanks@6651.6681.31\|CGACTCTTAGCGGTGGATCACTCGGCTCGTG | >hg38_21_+_8433172_8446622_RNA45SN1_45S_50ntFlanks | 6651 | 6681 | 31 | CGACTCTTAGCGGTGGATCACTCGGCTCGTG | MB231 | 455 | 2021 | 202 | 75 | 2116 | 308.5 | 661.4 | 137.2 | 53.3 | 3080.3 |
| >hg38_21_+_8433172_8446622_RNA45SN1_45S_50ntFlanks@6651.6681.31\|CGACTCTTAGCGGTGGATCACTCGGCTCGTG | >hg38_21_+_8433172_8446622_RNA45SN1_45S_50ntFlanks | 6651 | 6681 | 31 | CGACTCTTAGCGGTGGATCACTCGGCTCGTG | MB468 | 3080 | 2174 | 1129 | 3415 | 12410 | 838.0 | 1125.6 | 531.0 | 2094.7 | 4561.7 |
| >hg38_21_+_8433172_8446622_RNA45SN1_45S_50ntFlanks@6651.6682.32\|CGACTCTTAGCGGTGGATCACTCGGCTCGTGC | >hg38_21_+_8433172_8446622_RNA45SN1_45S_50ntFlanks | 6651 | 6682 | 32 | CGACTCTTAGCGGTGGATCACTCGGCTCGTGC | BT20 | 55 | 168 | 59 | 129 | 467 | 38.5 | 72.3 | 33.6 | 106.2 | 141.0 |
| >hg38_21_+_8433172_8446622_RNA45SN1_45S_50ntFlanks@6651.6682.32\|CGACTCTTAGCGGTGGATCACTCGGCTCGTGC | >hg38_21_+_8433172_8446622_RNA45SN1_45S_50ntFlanks | 6651 | 6682 | 32 | CGACTCTTAGCGGTGGATCACTCGGCTCGTGC | MB231 | 44 | 347 | 96 | 19 | 950 | 35.5 | 253.0 | 92.3 | 5.3 | 1449.6 |
| >hg38_21_+_8433172_8446622_RNA45SN1_45S_50ntFlanks@6651.6683.33\|CGACTCTTAGCGGTGGATCACTCGGCTCGTGCG | >hg38_21_+_8433172_8446622_RNA45SN1_45S_50ntFlanks | 6651 | 6683 | 33 | CGACTCTTAGCGGTGGATCACTCGGCTCGTGCG | BT20 | 108 | 307 | 43 | 40 | 69 | 92.0 | 231.4 | 55.3 | 20.1 | 21.7 |
| >hg38_21_+_8433172_8446622_RNA45SN1_45S_50ntFlanks@6651.6683.33\|CGACTCTTAGCGGTGGATCACTCGGCTCGTGCG | >hg38_21_+_8433172_8446622_RNA45SN1_45S_50ntFlanks | 6651 | 6683 | 33 | CGACTCTTAGCGGTGGATCACTCGGCTCGTGCG | MB231 | 289 | 1491 | 20 | 83 | 236 | 100.0 | 991.4 | 12.3 | 25.4 | 227.5 |
| >hg38_21_+_8433172_8446622_RNA45SN1_45S_50ntFlanks@6651.6683.33\|CGACTCTTAGCGGTGGATCACTCGGCTCGTGCG | >hg38_21_+_8433172_8446622_RNA45SN1_45S_50ntFlanks | 6651 | 6683 | 33 | CGACTCTTAGCGGTGGATCACTCGGCTCGTGCG | MB468 | 45 | 50 | 12 | 90 | 282 | 16.5 | 23.9 | 4.7 | 38.4 | 131.4 |
| >hg38_21_+_8433172_8446622_RNA45SN1_45S_50ntFlanks@6651.6684.34\|CGACTCTTAGCGGTGGATCACTCGGCTCGTGCGT | >hg38_21_+_8433172_8446622_RNA45SN1_45S_50ntFlanks | 6651 | 6684 | 34 | CGACTCTTAGCGGTGGATCACTCGGCTCGTGCGT | BT20 | 80 | 344 | 99 | 75 | 288 | 24.8 | 173.4 | 32.3 | 48.3 | 94.8 |
| >hg38_21_+_8433172_8446622_RNA45SN1_45S_50ntFlanks@6651.6684.34\|CGACTCTTAGCGGTGGATCACTCGGCTCGTGCGT | >hg38_21_+_8433172_8446622_RNA45SN1_45S_50ntFlanks | 6651 | 6684 | 34 | CGACTCTTAGCGGTGGATCACTCGGCTCGTGCGT | MB231 | 159 | 525 | 56 | 44 | 333 | 55.7 | 238.2 | 29.2 | 9.3 | 297.8 |
| >hg38_21_+_8433172_8446622_RNA45SN1_45S_50ntFlanks@6651.6684.34\|CGACTCTTAGCGGTGGATCACTCGGCTCGTGCGT | >hg38_21_+_8433172_8446622_RNA45SN1_45S_50ntFlanks | 6651 | 6684 | 34 | CGACTCTTAGCGGTGGATCACTCGGCTCGTGCGT | MB468 | 17 | 26 | 9 | 27 | 120 | 3.5 | 8.4 | 5.3 | 14.3 | 27.6 |
| >hg38_21_+_8433172_8446622_RNA45SN1_45S_50ntFlanks@6651.6685.35\|CGACTCTTAGCGGTGGATCACTCGGCTCGTGCGTC | >hg38_21_+_8433172_8446622_RNA45SN1_45S_50ntFlanks | 6651 | 6685 | 35 | CGACTCTTAGCGGTGGATCACTCGGCTCGTGCGTC | BT20 | 50 | 174 | 53 | 544 | 226 | 15.4 | 88.3 | 31.8 | 223.4 | 44.2 |
| >hg38_21_+_8433172_8446622_RNA45SN1_45S_50ntFlanks@6651.6685.35\|CGACTCTTAGCGGTGGATCACTCGGCTCGTGCGTC | >hg38_21_+_8433172_8446622_RNA45SN1_45S_50ntFlanks | 6651 | 6685 | 35 | CGACTCTTAGCGGTGGATCACTCGGCTCGTGCGTC | MB231 | 37 | 85 | 16 | 196 | 165 | 23.3 | 37.9 | 12.2 | 280.9 | 93.6 |
| >hg38_21_+_8433172_8446622_RNA45SN1_45S_50ntFlanks@6651.6685.35\|CGACTCTTAGCGGTGGATCACTCGGCTCGTGCGTC | >hg38_21_+_8433172_8446622_RNA45SN1_45S_50ntFlanks | 6651 | 6685 | 35 | CGACTCTTAGCGGTGGATCACTCGGCTCGTGCGTC | MB468 | 22 | 14 | 6 | 81 | 129 | 4.1 | 3.8 | 1.4 | 15.1 | 34.9 |
| >hg38_21_+_8433172_8446622_RNA45SN1_45S_50ntFlanks@6651.6686.36\|CGACTCTTAGCGGTGGATCACTCGGCTCGTGCGTCG | >hg38_21_+_8433172_8446622_RNA45SN1_45S_50ntFlanks | 6651 | 6686 | 36 | CGACTCTTAGCGGTGGATCACTCGGCTCGTGCGTCG | BT20 | 19 | 43 | 12 | 18 | 20 | 3.9 | 26.5 | 8.6 | 11.3 | 4.8 |
| >hg38_21_+_8433172_8446622_RNA45SN1_45S_50ntFlanks@6651.6686.36\|CGACTCTTAGCGGTGGATCACTCGGCTCGTGCGTCG | >hg38_21_+_8433172_8446622_RNA45SN1_45S_50ntFlanks | 6651 | 6686 | 36 | CGACTCTTAGCGGTGGATCACTCGGCTCGTGCGTCG | MB231 | 23 | 39 | 5 | 11 | 36 | 3.6 | 15.1 | 3.3 | 4.6 | 16.7 |
| >hg38_21_+_8433172_8446622_RNA45SN1_45S_50ntFlanks@6651.6686.36\|CGACTCTTAGCGGTGGATCACTCGGCTCGTGCGTCG | >hg38_21_+_8433172_8446622_RNA45SN1_45S_50ntFlanks | 6651 | 6686 | 36 | CGACTCTTAGCGGTGGATCACTCGGCTCGTGCGTCG | MB468 | 34 | 9 | 6 | 91 | 147 | 18.1 | 4.2 | 3.7 | 58.3 | 89.7 |
| >hg38_21_+_8433172_8446622_RNA45SN1_45S_50ntFlanks@6651.6687.37\|CGACTCTTAGCGGTGGATCACTCGGCTCGTGCGTCGA | >hg38_21_+_8433172_8446622_RNA45SN1_45S_50ntFlanks | 6651 | 6687 | 37 | CGACTCTTAGCGGTGGATCACTCGGCTCGTGCGTCGA | BT20 | 48 | 111 | 23 | 19 | 67 | 15.5 | 90.3 | 13.3 | 8.6 | 40.2 |
| >hg38_21_+_8433172_8446622_RNA45SN1_45S_50ntFlanks@6651.6687.37\|CGACTCTTAGCGGTGGATCACTCGGCTCGTGCGTCGA | >hg38_21_+_8433172_8446622_RNA45SN1_45S_50ntFlanks | 6651 | 6687 | 37 | CGACTCTTAGCGGTGGATCACTCGGCTCGTGCGTCGA | MB231 | 49 | 132 | 30 | 30 | 307 | 10.9 | 78.9 | 23.5 | 15.5 | 356.8 |
| >hg38_21_+_8433172_8446622_RNA45SN1_45S_50ntFlanks@6651.6688.38\|CGACTCTTAGCGGTGGATCACTCGGCTCGTGCGTCGAT | >hg38_21_+_8433172_8446622_RNA45SN1_45S_50ntFlanks | 6651 | 6688 | 38 | CGACTCTTAGCGGTGGATCACTCGGCTCGTGCGTCGAT | BT20 | 125 | 322 | 91 | 101 | 422 | 69.6 | 163.6 | 89.6 | 28.4 | 122.5 |

| | | | | | | | | | | | | | | | | |
|---|---|---|---|---|---|---|---|---|---|---|---|---|---|---|---|---|
| >hg38_21_+_8433172_8446622_RNA45SN1_45S_50ntFlanks@6651.6688.38\|CGACTCTTAGCGGTGGATCACTCGGCTCGTGCGTCGAT | >hg38_21_+_8433172_8446622_RNA45SN1_45S_50ntFlanks | 6651 | 6688 | 38 | CGACTCTTAGCGGTGGATCACTCGGCTCGTGCGTCGAT | MB231 | 75 | 117 | 12 | 42 | 199 | 46.2 | 56.0 | 5.2 | 30.5 | 142.5 |
| >hg38_21_+_8433172_8446622_RNA45SN1_45S_50ntFlanks@6651.6688.38\|CGACTCTTAGCGGTGGATCACTCGGCTCGTGCGTCGAT | >hg38_21_+_8433172_8446622_RNA45SN1_45S_50ntFlanks | 6651 | 6688 | 38 | CGACTCTTAGCGGTGGATCACTCGGCTCGTGCGTCGAT | MB468 | 48 | 26 | 11 | 102 | 412 | 8.9 | 14.3 | 6.5 | 60.9 | 204.2 |
| >hg38_21_+_8433172_8446622_RNA45SN1_45S_50ntFlanks@6651.6689.39\|CGACTCTTAGCGGTGGATCACTCGGCTCGTGCGTCGATG | >hg38_21_+_8433172_8446622_RNA45SN1_45S_50ntFlanks | 6651 | 6689 | 39 | CGACTCTTAGCGGTGGATCACTCGGCTCGTGCGTCGATG | BT20 | 224 | 717 | 192 | 395 | 1364 | 96.0 | 313.7 | 116.8 | 53.3 | 358.7 |
| >hg38_21_+_8433172_8446622_RNA45SN1_45S_50ntFlanks@6651.6689.39\|CGACTCTTAGCGGTGGATCACTCGGCTCGTGCGTCGATG | >hg38_21_+_8433172_8446622_RNA45SN1_45S_50ntFlanks | 6651 | 6689 | 39 | CGACTCTTAGCGGTGGATCACTCGGCTCGTGCGTCGATG | MB231 | 252 | 454 | 62 | 159 | 806 | 175.6 | 125.5 | 30.7 | 168.5 | 561.6 |
| >hg38_21_+_8433172_8446622_RNA45SN1_45S_50ntFlanks@6651.6689.39\|CGACTCTTAGCGGTGGATCACTCGGCTCGTGCGTCGATG | >hg38_21_+_8433172_8446622_RNA45SN1_45S_50ntFlanks | 6651 | 6689 | 39 | CGACTCTTAGCGGTGGATCACTCGGCTCGTGCGTCGATG | MB468 | 1220 | 854 | 301 | 1856 | 13024 | 240.7 | 391.8 | 188.7 | 1044.7 | 6134.6 |
| >hg38_21_+_8433172_8446622_RNA45SN1_45S_50ntFlanks@6651.6690.40\|CGACTCTTAGCGGTGGATCACTCGGCTCGTGCGTCGATGA | >hg38_21_+_8433172_8446622_RNA45SN1_45S_50ntFlanks | 6651 | 6690 | 40 | CGACTCTTAGCGGTGGATCACTCGGCTCGTGCGTCGATGA | BT20 | 70 | 139 | 52 | 92 | 235 | 70.1 | 69.3 | 51.4 | 50.4 | 81.6 |
| >hg38_21_+_8433172_8446622_RNA45SN1_45S_50ntFlanks@6651.6690.40\|CGACTCTTAGCGGTGGATCACTCGGCTCGTGCGTCGATGA | >hg38_21_+_8433172_8446622_RNA45SN1_45S_50ntFlanks | 6651 | 6690 | 40 | CGACTCTTAGCGGTGGATCACTCGGCTCGTGCGTCGATGA | MB231 | 76 | 129 | 29 | 33 | 245 | 66.6 | 14.7 | 21.9 | 1.2 | 191.4 |
| >hg38_21_+_8433172_8446622_RNA45SN1_45S_50ntFlanks@6651.6690.40\|CGACTCTTAGCGGTGGATCACTCGGCTCGTGCGTCGATGA | >hg38_21_+_8433172_8446622_RNA45SN1_45S_50ntFlanks | 6651 | 6690 | 40 | CGACTCTTAGCGGTGGATCACTCGGCTCGTGCGTCGATGA | MB468 | 22 | 27 | 6 | 38 | 162 | 7.2 | 10.2 | 2.5 | 34.1 | 51.1 |
| >hg38_21_+_8433172_8446622_RNA45SN1_45S_50ntFlanks@6651.6691.41\|CGACTCTTAGCGGTGGATCACTCGGCTCGTGCGTCGATGAA | >hg38_21_+_8433172_8446622_RNA45SN1_45S_50ntFlanks | 6651 | 6691 | 41 | CGACTCTTAGCGGTGGATCACTCGGCTCGTGCGTCGATGAA | BT20 | 15 | 43 | 10 | 11 | 35 | 6.9 | 28.4 | 5.6 | 2.5 | 5.3 |
| >hg38_21_+_8433172_8446622_RNA45SN1_45S_50ntFlanks@6651.6691.41\|CGACTCTTAGCGGTGGATCACTCGGCTCGTGCGTCGATGAA | >hg38_21_+_8433172_8446622_RNA45SN1_45S_50ntFlanks | 6651 | 6691 | 41 | CGACTCTTAGCGGTGGATCACTCGGCTCGTGCGTCGATGAA | MB231 | 21 | 40 | 7 | 10 | 53 | 13.3 | 13.1 | 3.7 | 4.3 | 20.5 |
| >hg38_21_+_8433172_8446622_RNA45SN1_45S_50ntFlanks@6651.6692.42\|CGACTCTTAGCGGTGGATCACTCGGCTCGTGCGTCGATGAAG | >hg38_21_+_8433172_8446622_RNA45SN1_45S_50ntFlanks | 6651 | 6692 | 42 | CGACTCTTAGCGGTGGATCACTCGGCTCGTGCGTCGATGAAG | BT20 | 105 | 208 | 147 | 80 | 373 | 85.5 | 68.5 | 185.2 | 23.4 | 150.3 |
| >hg38_21_+_8433172_8446622_RNA45SN1_45S_50ntFlanks@6651.6692.42\|CGACTCTTAGCGGTGGATCACTCGGCTCGTGCGTCGATGAAG | >hg38_21_+_8433172_8446622_RNA45SN1_45S_50ntFlanks | 6651 | 6692 | 42 | CGACTCTTAGCGGTGGATCACTCGGCTCGTGCGTCGATGAAG | MB231 | 253 | 378 | 91 | 54 | 646 | 213.0 | 141.3 | 71.9 | 10.9 | 456.6 |
| >hg38_21_+_8433172_8446622_RNA45SN1_45S_50ntFlanks@6651.6692.42\|CGACTCTTAGCGGTGGATCACTCGGCTCGTGCGTCGATGAAG | >hg38_21_+_8433172_8446622_RNA45SN1_45S_50ntFlanks | 6651 | 6692 | 42 | CGACTCTTAGCGGTGGATCACTCGGCTCGTGCGTCGATGAAG | MB468 | 98 | 100 | 27 | 139 | 766 | 37.9 | 67.3 | 12.1 | 129.4 | 346.3 |
| >hg38_21_+_8433172_8446622_RNA45SN1_45S_50ntFlanks@6651.6693.43\|CGACTCTTAGCGGTGGATCACTCGGCTCGTGCGTCGATGAAGA | >hg38_21_+_8433172_8446622_RNA45SN1_45S_50ntFlanks | 6651 | 6693 | 43 | CGACTCTTAGCGGTGGATCACTCGGCTCGTGCGTCGATGAAGA | BT20 | 159 | 257 | 85 | 57 | 555 | 178.1 | 139.6 | 67.9 | 16.8 | 92.6 |
| >hg38_21_+_8433172_8446622_RNA45SN1_45S_50ntFlanks@6651.6693.43\|CGACTCTTAGCGGTGGATCACTCGGCTCGTGCGTCGATGAAGA | >hg38_21_+_8433172_8446622_RNA45SN1_45S_50ntFlanks | 6651 | 6693 | 43 | CGACTCTTAGCGGTGGATCACTCGGCTCGTGCGTCGATGAAGA | MB231 | 154 | 210 | 65 | 32 | 500 | 130.4 | 78.5 | 39.9 | 14.8 | 334.4 |
| >hg38_21_+_8433172_8446622_RNA45SN1_45S_50ntFlanks@6651.6693.43\|CGACTCTTAGCGGTGGATCACTCGGCTCGTGCGTCGATGAAGA | >hg38_21_+_8433172_8446622_RNA45SN1_45S_50ntFlanks | 6651 | 6693 | 43 | CGACTCTTAGCGGTGGATCACTCGGCTCGTGCGTCGATGAAGA | MB468 | 10 | 19 | 5 | 18 | 102 | 5.5 | 20.5 | 3.1 | 17.2 | 84.0 |
| >hg38_21_+_8433172_8446622_RNA45SN1_45S_50ntFlanks@6651.6694.44\|CGACTCTTAGCGGTGGATCACTCGGCTCGTGCGTCGATGAAGAA | >hg38_21_+_8433172_8446622_RNA45SN1_45S_50ntFlanks | 6651 | 6694 | 44 | CGACTCTTAGCGGTGGATCACTCGGCTCGTGCGTCGATGAAGAA | BT20 | 52 | 135 | 30 | 21 | 167 | 30.6 | 83.7 | 20.1 | 5.4 | 52.7 |
| >hg38_21_+_8433172_8446622_RNA45SN1_45S_50ntFlanks@6651.6694.44\|CGACTCTTAGCGGTGGATCACTCGGCTCGTGCGTCGATGAAGAA | >hg38_21_+_8433172_8446622_RNA45SN1_45S_50ntFlanks | 6651 | 6694 | 44 | CGACTCTTAGCGGTGGATCACTCGGCTCGTGCGTCGATGAAGAA | MB231 | 111 | 149 | 50 | 17 | 279 | 79.5 | 105.4 | 51.3 | 15.6 | 269.2 |
| >hg38_21_+_8433172_8446622_RNA45SN1_45S_50ntFlanks@6651.6695.45\|CGACTCTTAGCGGTGGATCACTCGGCTCGTGCGTCGATGAAGAAC | >hg38_21_+_8433172_8446622_RNA45SN1_45S_50ntFlanks | 6651 | 6695 | 45 | CGACTCTTAGCGGTGGATCACTCGGCTCGTGCGTCGATGAAGAAC | BT20 | 132 | 387 | 105 | 137 | 868 | 74.1 | 391.7 | 101.0 | 65.7 | 392.4 |
| >hg38_21_+_8433172_8446622_RNA45SN1_45S_50ntFlanks@6651.6695.45\|CGACTCTTAGCGGTGGATCACTCGGCTCGTGCGTCGATGAAGAAC | >hg38_21_+_8433172_8446622_RNA45SN1_45S_50ntFlanks | 6651 | 6695 | 45 | CGACTCTTAGCGGTGGATCACTCGGCTCGTGCGTCGATGAAGAAC | MB231 | 309 | 311 | 220 | 90 | 910 | 194.2 | 212.6 | 180.6 | 120.0 | 942.1 |
| >hg38_21_+_8433172_8446622_RNA45SN1_45S_50ntFlanks@6651.6696.46\|CGACTCTTAGCGGTGGATCACTCGGCTCGTGCGTCGATGAAGAACG | >hg38_21_+_8433172_8446622_RNA45SN1_45S_50ntFlanks | 6651 | 6696 | 46 | CGACTCTTAGCGGTGGATCACTCGGCTCGTGCGTCGATGAAGAACG | BT20 | 138 | 381 | 48 | 89 | 740 | 103.5 | 343.8 | 32.1 | 4.3 | 464.0 |
| >hg38_21_+_8433172_8446622_RNA45SN1_45S_50ntFlanks@6651.6696.46\|CGACTCTTAGCGGTGGATCACTCGGCTCGTGCGTCGATGAAGAACG | >hg38_21_+_8433172_8446622_RNA45SN1_45S_50ntFlanks | 6651 | 6696 | 46 | CGACTCTTAGCGGTGGATCACTCGGCTCGTGCGTCGATGAAGAACG | MB231 | 717 | 1468 | 338 | 56 | 1268 | 345.1 | 1215.0 | 255.1 | 63.1 | 977.7 |
| >hg38_21_+_8433172_8446622_RNA45SN1_45S_50ntFlanks@6651.6696.46\|CGACTCTTAGCGGTGGATCACTCGGCTCGTGCGTCGATGAAGAACG | >hg38_21_+_8433172_8446622_RNA45SN1_45S_50ntFlanks | 6651 | 6696 | 46 | CGACTCTTAGCGGTGGATCACTCGGCTCGTGCGTCGATGAAGAACG | MB468 | 36 | 167 | 23 | 91 | 644 | 13.3 | 183.7 | 16.4 | 99.1 | 404.4 |
| >hg38_21_+_8433172_8446622_RNA45SN1_45S_50ntFlanks@6651.6697.47\|CGACTCTTAGCGGTGGATCACTCGGCTCGTGCGTCGATGAAGAACGC | >hg38_21_+_8433172_8446622_RNA45SN1_45S_50ntFlanks | 6651 | 6697 | 47 | CGACTCTTAGCGGTGGATCACTCGGCTCGTGCGTCGATGAAGAACGC | BT20 | 45 | 113 | 29 | 70 | 429 | 27.2 | 107.7 | 25.5 | 30.1 | 248.8 |
| >hg38_21_+_8433172_8446622_RNA45SN1_45S_50ntFlanks@6651.6697.47\|CGACTCTTAGCGGTGGATCACTCGGCTCGTGCGTCGATGAAGAACGC | >hg38_21_+_8433172_8446622_RNA45SN1_45S_50ntFlanks | 6651 | 6697 | 47 | CGACTCTTAGCGGTGGATCACTCGGCTCGTGCGTCGATGAAGAACGC | MB231 | 199 | 253 | 166 | 22 | 666 | 125.3 | 220.9 | 125.6 | 22.6 | 463.8 |
| >hg38_21_+_8433172_8446622_RNA45SN1_45S_50ntFlanks@6651.6697.47\|CGACTCTTAGCGGTGGATCACTCGGCTCGTGCGTCGATGAAGAACGC | >hg38_21_+_8433172_8446622_RNA45SN1_45S_50ntFlanks | 6651 | 6697 | 47 | CGACTCTTAGCGGTGGATCACTCGGCTCGTGCGTCGATGAAGAACGC | MB468 | 46 | 151 | 39 | 58 | 771 | 29.8 | 179.8 | 28.5 | 71.8 | 650.2 |
| >hg38_21_+_8433172_8446622_RNA45SN1_45S_50ntFlanks@6651.6698.48\|CGACTCTTAGCGGTGGATCACTCGGCTCGTGCGTCGATGAAGAACGCA | >hg38_21_+_8433172_8446622_RNA45SN1_45S_50ntFlanks | 6651 | 6698 | 48 | CGACTCTTAGCGGTGGATCACTCGGCTCGTGCGTCGATGAAGAACGCA | BT20 | 40 | 161 | 28 | 106 | 712 | 25.6 | 194.8 | 31.9 | 39.8 | 522.8 |
| >hg38_21_+_8433172_8446622_RNA45SN1_45S_50ntFlanks@6651.6698.48\|CGACTCTTAGCGGTGGATCACTCGGCTCGTGCGTCGATGAAGAACGCA | >hg38_21_+_8433172_8446622_RNA45SN1_45S_50ntFlanks | 6651 | 6698 | 48 | CGACTCTTAGCGGTGGATCACTCGGCTCGTGCGTCGATGAAGAACGCA | MB231 | 245 | 493 | 126 | 20 | 713 | 199.6 | 597.9 | 62.6 | 29.8 | 632.3 |
| >hg38_21_+_8433172_8446622_RNA45SN1_45S_50ntFlanks@6651.6698.48\|CGACTCTTAGCGGTGGATCACTCGGCTCGTGCGTCGATGAAGAACGCA | >hg38_21_+_8433172_8446622_RNA45SN1_45S_50ntFlanks | 6651 | 6698 | 48 | CGACTCTTAGCGGTGGATCACTCGGCTCGTGCGTCGATGAAGAACGCA | MB468 | 141 | 347 | 80 | 186 | 2336 | 82.8 | 279.9 | 51.7 | 226.5 | 1410.7 |
| >hg38_21_+_8433172_8446622_RNA45SN1_45S_50ntFlanks@6651.6699.49\|CGACTCTTAGCGGTGGATCACTCGGCTCGTGCGTCGATGAAGAACGCAG | >hg38_21_+_8433172_8446622_RNA45SN1_45S_50ntFlanks | 6651 | 6699 | 49 | CGACTCTTAGCGGTGGATCACTCGGCTCGTGCGTCGATGAAGAACGCAG | BT20 | 52 | 216 | 13 | 41 | 406 | 37.5 | 256.8 | 15.1 | 14.2 | 346.7 |
| >hg38_21_+_8433172_8446622_RNA45SN1_45S_50ntFlanks@6651.6699.49\|CGACTCTTAGCGGTGGATCACTCGGCTCGTGCGTCGATGAAGAACGCAG | >hg38_21_+_8433172_8446622_RNA45SN1_45S_50ntFlanks | 6651 | 6699 | 49 | CGACTCTTAGCGGTGGATCACTCGGCTCGTGCGTCGATGAAGAACGCAG | MB231 | 331 | 1175 | 148 | 16 | 456 | 210.2 | 1684.0 | 97.2 | 17.7 | 240.6 |
| >hg38_21_+_8433172_8446622_RNA45SN1_45S_50ntFlanks@6651.6699.49\|CGACTCTTAGCGGTGGATCACTCGGCTCGTGCGTCGATGAAGAACGCAG | >hg38_21_+_8433172_8446622_RNA45SN1_45S_50ntFlanks | 6651 | 6699 | 49 | CGACTCTTAGCGGTGGATCACTCGGCTCGTGCGTCGATGAAGAACGCAG | MB468 | 211 | 967 | 206 | 398 | 3976 | 109.7 | 1017.9 | 177.6 | 569.8 | 3095.4 |
| >hg38_21_+_8433172_8446622_RNA45SN1_45S_50ntFlanks@6651.6700.50\|CGACTCTTAGCGGTGGATCACTCGGCTCGTGCGTCGATGAAGAACGCAGC | >hg38_21_+_8433172_8446622_RNA45SN1_45S_50ntFlanks | 6651 | 6700 | 50 | CGACTCTTAGCGGTGGATCACTCGGCTCGTGCGTCGATGAAGAACGCAGC | BT20 | 37 | 145 | 33 | 205 | 1102 | 40.0 | 200.9 | 46.1 | 169.8 | 1155.7 |
| >hg38_21_+_8433172_8446622_RNA45SN1_45S_50ntFlanks@6651.6700.50\|CGACTCTTAGCGGTGGATCACTCGGCTCGTGCGTCGATGAAGAACGCAGC | >hg38_21_+_8433172_8446622_RNA45SN1_45S_50ntFlanks | 6651 | 6700 | 50 | CGACTCTTAGCGGTGGATCACTCGGCTCGTGCGTCGATGAAGAACGCAGC | MB231 | 135 | 288 | 131 | 42 | 593 | 97.1 | 413.6 | 54.7 | 63.6 | 442.0 |
| >hg38_21_+_8433172_8446622_RNA45SN1_45S_50ntFlanks@6652.6669.18\|GACTCTTAGCGGTGGATC | >hg38_21_+_8433172_8446622_RNA45SN1_45S_50ntFlanks | 6652 | 6669 | 18 | GACTCTTAGCGGTGGATC | BT20 | 10 | 26 | 22 | 39 | 50 | 3.7 | 11.7 | 7.6 | 28.6 | 19.8 |
| >hg38_21_+_8433172_8446622_RNA45SN1_45S_50ntFlanks@6652.6676.25\|GACTCTTAGCGGTGGATCACTCGGC | >hg38_21_+_8433172_8446622_RNA45SN1_45S_50ntFlanks | 6652 | 6676 | 25 | GACTCTTAGCGGTGGATCACTCGGC | BT20 | 18 | 62 | 461 | 223 | 234 | 10.0 | 35.9 | 763.4 | 143.4 | 70.4 |
| >hg38_21_+_8433172_8446622_RNA45SN1_45S_50ntFlanks@6652.6676.25\|GACTCTTAGCGGTGGATCACTCGGC | >hg38_21_+_8433172_8446622_RNA45SN1_45S_50ntFlanks | 6652 | 6676 | 25 | GACTCTTAGCGGTGGATCACTCGGC | MB231 | 19 | 81 | 51 | 31 | 307 | 12.2 | 56.5 | 29.0 | 12.7 | 408.9 |
| >hg38_21_+_8433172_8446622_RNA45SN1_45S_50ntFlanks@6652.6677.26\|GACTCTTAGCGGTGGATCACTCGGCT | >hg38_21_+_8433172_8446622_RNA45SN1_45S_50ntFlanks | 6652 | 6677 | 26 | GACTCTTAGCGGTGGATCACTCGGCT | BT20 | 16 | 58 | 28 | 270 | 155 | 2.4 | 34.8 | 9.5 | 158.9 | 34.1 |
| >hg38_21_+_8433172_8446622_RNA45SN1_45S_50ntFlanks@6652.6677.26\|GACTCTTAGCGGTGGATCACTCGGCT | >hg38_21_+_8433172_8446622_RNA45SN1_45S_50ntFlanks | 6652 | 6677 | 26 | GACTCTTAGCGGTGGATCACTCGGCT | MB231 | 15 | 71 | 32 | 29 | 243 | 10.8 | 57.5 | 18.0 | 24.9 | 366.9 |
| >hg38_21_+_8433172_8446622_RNA45SN1_45S_50ntFlanks@6652.6678.27\|GACTCTTAGCGGTGGATCACTCGGCTC | >hg38_21_+_8433172_8446622_RNA45SN1_45S_50ntFlanks | 6652 | 6678 | 27 | GACTCTTAGCGGTGGATCACTCGGCTC | BT20 | 39 | 103 | 54 | 363 | 404 | 29.3 | 62.8 | 37.4 | 373.6 | 90.8 |
| >hg38_21_+_8433172_8446622_RNA45SN1_45S_50ntFlanks@6652.6678.27\|GACTCTTAGCGGTGGATCACTCGGCTC | >hg38_21_+_8433172_8446622_RNA45SN1_45S_50ntFlanks | 6652 | 6678 | 27 | GACTCTTAGCGGTGGATCACTCGGCTC | MB231 | 30 | 118 | 55 | 12 | 495 | 43.8 | 84.7 | 26.9 | 12.0 | 759.3 |
| >hg38_21_+_8433172_8446622_RNA45SN1_45S_50ntFlanks@6652.6678.27\|GACTCTTAGCGGTGGATCACTCGGCTC | >hg38_21_+_8433172_8446622_RNA45SN1_45S_50ntFlanks | 6652 | 6678 | 27 | GACTCTTAGCGGTGGATCACTCGGCTC | MB468 | 16 | 37 | 16 | 11 | 93 | 4.2 | 32.7 | 9.2 | 9.2 | 66.8 |
| >hg38_21_+_8433172_8446622_RNA45SN1_45S_50ntFlanks@6652.6679.28\|GACTCTTAGCGGTGGATCACTCGGCTCG | >hg38_21_+_8433172_8446622_RNA45SN1_45S_50ntFlanks | 6652 | 6679 | 28 | GACTCTTAGCGGTGGATCACTCGGCTCG | BT20 | 14 | 37 | 202 | 122 | 187 | 13.9 | 20.9 | 332.5 | 170.3 | 133.5 |
| >hg38_21_+_8433172_8446622_RNA45SN1_45S_50ntFlanks@6652.6679.28\|GACTCTTAGCGGTGGATCACTCGGCTCG | >hg38_21_+_8433172_8446622_RNA45SN1_45S_50ntFlanks | 6652 | 6679 | 28 | GACTCTTAGCGGTGGATCACTCGGCTCG | MB231 | 27 | 78 | 26 | 5 | 332 | 34.2 | 52.2 | 19.3 | 1.0 | 493.3 |
| >hg38_21_+_8433172_8446622_RNA45SN1_45S_50ntFlanks@6652.6679.28\|GACTCTTAGCGGTGGATCACTCGGCTCG | >hg38_21_+_8433172_8446622_RNA45SN1_45S_50ntFlanks | 6652 | 6679 | 28 | GACTCTTAGCGGTGGATCACTCGGCTCG | MB468 | 55 | 97 | 40 | 51 | 488 | 17.1 | 69.2 | 21.2 | 57.8 | 403.6 |
| >hg38_21_+_8433172_8446622_RNA45SN1_45S_50ntFlanks@6652.6680.29\|GACTCTTAGCGGTGGATCACTCGGCTCGT | >hg38_21_+_8433172_8446622_RNA45SN1_45S_50ntFlanks | 6652 | 6680 | 29 | GACTCTTAGCGGTGGATCACTCGGCTCGT | MB468 | 20 | 23 | 10 | 25 | 85 | 5.3 | 14.8 | 6.2 | 12.7 | 45.1 |
| >hg38_21_+_8433172_8446622_RNA45SN1_45S_50ntFlanks@6652.6681.30\|GACTCTTAGCGGTGGATCACTCGGCTCGTG | >hg38_21_+_8433172_8446622_RNA45SN1_45S_50ntFlanks | 6652 | 6681 | 30 | GACTCTTAGCGGTGGATCACTCGGCTCGTG | BT20 | 45 | 97 | 21 | 186 | 147 | 45.9 | 23.0 | 13.5 | 95.4 | 33.4 |
| >hg38_21_+_8433172_8446622_RNA45SN1_45S_50ntFlanks@6652.6681.30\|GACTCTTAGCGGTGGATCACTCGGCTCGTG | >hg38_21_+_8433172_8446622_RNA45SN1_45S_50ntFlanks | 6652 | 6681 | 30 | GACTCTTAGCGGTGGATCACTCGGCTCGTG | MB231 | 91 | 449 | 40 | 25 | 355 | 54.9 | 184.2 | 35.3 | 21.1 | 552.8 |
| >hg38_21_+_8433172_8446622_RNA45SN1_45S_50ntFlanks@6652.6681.30\|GACTCTTAGCGGTGGATCACTCGGCTCGTG | >hg38_21_+_8433172_8446622_RNA45SN1_45S_50ntFlanks | 6652 | 6681 | 30 | GACTCTTAGCGGTGGATCACTCGGCTCGTG | MB468 | 434 | 403 | 194 | 439 | 1658 | 161.7 | 273.7 | 117.0 | 400.2 | 856.1 |
| >hg38_21_+_8433172_8446622_RNA45SN1_45S_50ntFlanks@6652.6682.31\|GACTCTTAGCGGTGGATCACTCGGCTCGTGC | >hg38_21_+_8433172_8446622_RNA45SN1_45S_50ntFlanks | 6652 | 6682 | 31 | GACTCTTAGCGGTGGATCACTCGGCTCGTGC | BT20 | 12 | 37 | 14 | 74 | 103 | 8.9 | 10.2 | 10.8 | 39.4 | 15.6 |

| | | | | | | | | | | | | | | | | |
|---|---|---|---|---|---|---|---|---|---|---|---|---|---|---|---|---|
| >hg38_21_+_8433172_8446622_RNA45SN1_45S_50ntFlanks@6652.6682.31\|GACTCTTAGCGG<br>TGGATCACTCGGCTCGTGC | >hg38_21_+_8433172_8446622_RNA45SN1_45S_50ntFlanks | 6652 | 6682 | 31 | GACTCTTAGCGGTGGATCACTCGGCTCGTGC | MB231 | 13 | 87 | 24 | 5 | 215 | 10.5 | 53.9 | 23.1 | 4.1 | 333.9 |
| >hg38_21_+_8433172_8446622_RNA45SN1_45S_50ntFlanks@6652.6683.32\|GACTCTTAGCGG<br>TGGATCACTCGGCTCGTGCG | >hg38_21_+_8433172_8446622_RNA45SN1_45S_50ntFlanks | 6652 | 6683 | 32 | GACTCTTAGCGGTGGATCACTCGGCTCGTGCG | BT20 | 87 | 143 | 29 | 32 | 68 | 121.7 | 63.8 | 42.6 | 34.9 | 16.9 |
| >hg38_21_+_8433172_8446622_RNA45SN1_45S_50ntFlanks@6652.6683.32\|GACTCTTAGCGG<br>TGGATCACTCGGCTCGTGCG | >hg38_21_+_8433172_8446622_RNA45SN1_45S_50ntFlanks | 6652 | 6683 | 32 | GACTCTTAGCGGTGGATCACTCGGCTCGTGCG | MB231 | 186 | 1105 | 15 | 10 | 107 | 155.6 | 715.9 | 7.4 | 3.2 | 137.9 |
| >hg38_21_+_8433172_8446622_RNA45SN1_45S_50ntFlanks@6652.6683.32\|GACTCTTAGCGG<br>TGGATCACTCGGCTCGTGCG | >hg38_21_+_8433172_8446622_RNA45SN1_45S_50ntFlanks | 6652 | 6683 | 32 | GACTCTTAGCGGTGGATCACTCGGCTCGTGCG | MB468 | 15 | 51 | 5 | 16 | 132 | 11.7 | 36.9 | 3.5 | 15.0 | 130.9 |
| >hg38_21_+_8433172_8446622_RNA45SN1_45S_50ntFlanks@6652.6684.33\|GACTCTTAGCGG<br>TGGATCACTCGGCTCGTGCGT | >hg38_21_+_8433172_8446622_RNA45SN1_45S_50ntFlanks | 6652 | 6684 | 33 | GACTCTTAGCGGTGGATCACTCGGCTCGTGCGT | BT20 | 16 | 67 | 24 | 41 | 67 | 11.1 | 33.2 | 7.7 | 22.8 | 12.9 |
| >hg38_21_+_8433172_8446622_RNA45SN1_45S_50ntFlanks@6652.6684.33\|GACTCTTAGCGG<br>TGGATCACTCGGCTCGTGCGT | >hg38_21_+_8433172_8446622_RNA45SN1_45S_50ntFlanks | 6652 | 6684 | 33 | GACTCTTAGCGGTGGATCACTCGGCTCGTGCGT | MB231 | 34 | 113 | 17 | 13 | 55 | 19.6 | 53.0 | 2.9 | 2.7 | 56.6 |
| >hg38_21_+_8433172_8446622_RNA45SN1_45S_50ntFlanks@6652.6686.35\|GACTCTTAGCGG<br>TGGATCACTCGGCTCGTGCGTCG | >hg38_21_+_8433172_8446622_RNA45SN1_45S_50ntFlanks | 6652 | 6686 | 35 | GACTCTTAGCGGTGGATCACTCGGCTCGTGCGTCG | BT20 | 30 | 59 | 22 | 33 | 66 | 30.1 | 25.1 | 25.9 | 11.6 | 15.0 |
| >hg38_21_+_8433172_8446622_RNA45SN1_45S_50ntFlanks@6652.6686.35\|GACTCTTAGCGG<br>TGGATCACTCGGCTCGTGCGTCG | >hg38_21_+_8433172_8446622_RNA45SN1_45S_50ntFlanks | 6652 | 6686 | 35 | GACTCTTAGCGGTGGATCACTCGGCTCGTGCGTCG | MB231 | 49 | 141 | 9 | 10 | 70 | 43.4 | 57.9 | 4.5 | 4.1 | 53.2 |
| >hg38_21_+_8433172_8446622_RNA45SN1_45S_50ntFlanks@6652.6686.35\|GACTCTTAGCGG<br>TGGATCACTCGGCTCGTGCGTCG | >hg38_21_+_8433172_8446622_RNA45SN1_45S_50ntFlanks | 6652 | 6686 | 35 | GACTCTTAGCGGTGGATCACTCGGCTCGTGCGTCG | MB468 | 64 | 46 | 12 | 51 | 339 | 16.1 | 10.7 | 1.9 | 40.3 | 79.9 |
| >hg38_21_+_8433172_8446622_RNA45SN1_45S_50ntFlanks@6652.6687.36\|GACTCTTAGCGG<br>TGGATCACTCGGCTCGTGCGTCGA | >hg38_21_+_8433172_8446622_RNA45SN1_45S_50ntFlanks | 6652 | 6687 | 36 | GACTCTTAGCGGTGGATCACTCGGCTCGTGCGTCGA | BT20 | 50 | 106 | 19 | 30 | 59 | 56.2 | 61.9 | 22.9 | 17.5 | 24.9 |
| >hg38_21_+_8433172_8446622_RNA45SN1_45S_50ntFlanks@6652.6687.36\|GACTCTTAGCGG<br>TGGATCACTCGGCTCGTGCGTCGA | >hg38_21_+_8433172_8446622_RNA45SN1_45S_50ntFlanks | 6652 | 6687 | 36 | GACTCTTAGCGGTGGATCACTCGGCTCGTGCGTCGA | MB231 | 51 | 166 | 11 | 9 | 115 | 35.1 | 103.5 | 8.5 | 0.7 | 142.0 |
| >hg38_21_+_8433172_8446622_RNA45SN1_45S_50ntFlanks@6652.6688.37\|GACTCTTAGCGG<br>TGGATCACTCGGCTCGTGCGTCGAT | >hg38_21_+_8433172_8446622_RNA45SN1_45S_50ntFlanks | 6652 | 6688 | 37 | GACTCTTAGCGGTGGATCACTCGGCTCGTGCGTCGAT | BT20 | 65 | 206 | 40 | 123 | 345 | 58.6 | 81.0 | 33.0 | 58.2 | 97.3 |
| >hg38_21_+_8433172_8446622_RNA45SN1_45S_50ntFlanks@6652.6688.37\|GACTCTTAGCGG<br>TGGATCACTCGGCTCGTGCGTCGAT | >hg38_21_+_8433172_8446622_RNA45SN1_45S_50ntFlanks | 6652 | 6688 | 37 | GACTCTTAGCGGTGGATCACTCGGCTCGTGCGTCGAT | MB231 | 62 | 133 | 14 | 25 | 110 | 52.2 | 68.4 | 4.0 | 13.3 | 79.2 |
| >hg38_21_+_8433172_8446622_RNA45SN1_45S_50ntFlanks@6652.6688.37\|GACTCTTAGCGG<br>TGGATCACTCGGCTCGTGCGTCGAT | >hg38_21_+_8433172_8446622_RNA45SN1_45S_50ntFlanks | 6652 | 6688 | 37 | GACTCTTAGCGGTGGATCACTCGGCTCGTGCGTCGAT | MB468 | 34 | 36 | 8 | 32 | 218 | 6.5 | 15.6 | 2.1 | 26.2 | 59.5 |
| >hg38_21_+_8433172_8446622_RNA45SN1_45S_50ntFlanks@6652.6689.38\|GACTCTTAGCGG<br>TGGATCACTCGGCTCGTGCGTCGATG | >hg38_21_+_8433172_8446622_RNA45SN1_45S_50ntFlanks | 6652 | 6689 | 38 | GACTCTTAGCGGTGGATCACTCGGCTCGTGCGTCGATG | BT20 | 75 | 230 | 72 | 462 | 618 | 72.0 | 118.2 | 68.0 | 202.4 | 122.8 |
| >hg38_21_+_8433172_8446622_RNA45SN1_45S_50ntFlanks@6652.6689.38\|GACTCTTAGCGG<br>TGGATCACTCGGCTCGTGCGTCGATG | >hg38_21_+_8433172_8446622_RNA45SN1_45S_50ntFlanks | 6652 | 6689 | 38 | GACTCTTAGCGGTGGATCACTCGGCTCGTGCGTCGATG | MB231 | 106 | 225 | 36 | 65 | 195 | 121.9 | 72.7 | 21.7 | 87.5 | 145.5 |
| >hg38_21_+_8433172_8446622_RNA45SN1_45S_50ntFlanks@6652.6689.38\|GACTCTTAGCGG<br>TGGATCACTCGGCTCGTGCGTCGATG | >hg38_21_+_8433172_8446622_RNA45SN1_45S_50ntFlanks | 6652 | 6689 | 38 | GACTCTTAGCGGTGGATCACTCGGCTCGTGCGTCGATG | MB468 | 400 | 648 | 107 | 350 | 3393 | 85.4 | 217.9 | 24.5 | 307.0 | 1601.7 |
| >hg38_21_+_8433172_8446622_RNA45SN1_45S_50ntFlanks@6652.6690.39\|GACTCTTAGCGG<br>TGGATCACTCGGCTCGTGCGTCGATGA | >hg38_21_+_8433172_8446622_RNA45SN1_45S_50ntFlanks | 6652 | 6690 | 39 | GACTCTTAGCGGTGGATCACTCGGCTCGTGCGTCGATGA | BT20 | 25 | 43 | 15 | 102 | 81 | 31.4 | 14.4 | 15.4 | 76.4 | 12.9 |
| >hg38_21_+_8433172_8446622_RNA45SN1_45S_50ntFlanks@6652.6690.39\|GACTCTTAGCGG<br>TGGATCACTCGGCTCGTGCGTCGATGA | >hg38_21_+_8433172_8446622_RNA45SN1_45S_50ntFlanks | 6652 | 6690 | 39 | GACTCTTAGCGGTGGATCACTCGGCTCGTGCGTCGATGA | MB231 | 23 | 48 | 8 | 12 | 51 | 25.1 | 12.8 | 5.0 | 2.5 | 50.9 |
| >hg38_21_+_8433172_8446622_RNA45SN1_45S_50ntFlanks@6652.6692.41\|GACTCTTAGCGG<br>TGGATCACTCGGCTCGTGCGTCGATGAAG | >hg38_21_+_8433172_8446622_RNA45SN1_45S_50ntFlanks | 6652 | 6692 | 41 | GACTCTTAGCGGTGGATCACTCGGCTCGTGCGTCGATGAAG | BT20 | 33 | 52 | 51 | 72 | 160 | 39.3 | 9.9 | 70.4 | 40.0 | 90.6 |
| >hg38_21_+_8433172_8446622_RNA45SN1_45S_50ntFlanks@6652.6692.41\|GACTCTTAGCGG<br>TGGATCACTCGGCTCGTGCGTCGATGAAG | >hg38_21_+_8433172_8446622_RNA45SN1_45S_50ntFlanks | 6652 | 6692 | 41 | GACTCTTAGCGGTGGATCACTCGGCTCGTGCGTCGATGAAG | MB231 | 71 | 122 | 35 | 13 | 135 | 79.6 | 50.8 | 24.2 | 6.7 | 128.8 |
| >hg38_21_+_8433172_8446622_RNA45SN1_45S_50ntFlanks@6652.6692.41\|GACTCTTAGCGG<br>TGGATCACTCGGCTCGTGCGTCGATGAAG | >hg38_21_+_8433172_8446622_RNA45SN1_45S_50ntFlanks | 6652 | 6692 | 41 | GACTCTTAGCGGTGGATCACTCGGCTCGTGCGTCGATGAAG | MB468 | 26 | 40 | 6 | 27 | 159 | 15.4 | 16.1 | 3.0 | 28.3 | 90.7 |
| >hg38_21_+_8433172_8446622_RNA45SN1_45S_50ntFlanks@6652.6693.42\|GACTCTTAGCGG<br>TGGATCACTCGGCTCGTGCGTCGATGAAGA | >hg38_21_+_8433172_8446622_RNA45SN1_45S_50ntFlanks | 6652 | 6693 | 42 | GACTCTTAGCGGTGGATCACTCGGCTCGTGCGTCGATGAAGA | BT20 | 38 | 54 | 18 | 39 | 146 | 52.7 | 29.9 | 18.0 | 19.9 | 12.2 |
| >hg38_21_+_8433172_8446622_RNA45SN1_45S_50ntFlanks@6652.6693.42\|GACTCTTAGCGG<br>TGGATCACTCGGCTCGTGCGTCGATGAAGA | >hg38_21_+_8433172_8446622_RNA45SN1_45S_50ntFlanks | 6652 | 6693 | 42 | GACTCTTAGCGGTGGATCACTCGGCTCGTGCGTCGATGAAGA | MB231 | 37 | 72 | 19 | 9 | 94 | 37.8 | 33.0 | 11.3 | 4.6 | 85.6 |
| >hg38_21_+_8433172_8446622_RNA45SN1_45S_50ntFlanks@6652.6694.43\|GACTCTTAGCGG<br>TGGATCACTCGGCTCGTGCGTCGATGAAGAA | >hg38_21_+_8433172_8446622_RNA45SN1_45S_50ntFlanks | 6652 | 6694 | 43 | GACTCTTAGCGGTGGATCACTCGGCTCGTGCGTCGATGAAGAA | BT20 | 14 | 30 | 9 | 11 | 63 | 14.3 | 16.9 | 6.5 | 2.2 | 27.3 |
| >hg38_21_+_8433172_8446622_RNA45SN1_45S_50ntFlanks@6652.6694.43\|GACTCTTAGCGG<br>TGGATCACTCGGCTCGTGCGTCGATGAAGAA | >hg38_21_+_8433172_8446622_RNA45SN1_45S_50ntFlanks | 6652 | 6694 | 43 | GACTCTTAGCGGTGGATCACTCGGCTCGTGCGTCGATGAAGAA | MB231 | 31 | 49 | 17 | 4 | 57 | 27.1 | 36.7 | 14.5 | 2.9 | 46.7 |
| >hg38_21_+_8433172_8446622_RNA45SN1_45S_50ntFlanks@6652.6695.44\|GACTCTTAGCGG<br>TGGATCACTCGGCTCGTGCGTCGATGAAGAAC | >hg38_21_+_8433172_8446622_RNA45SN1_45S_50ntFlanks | 6652 | 6695 | 44 | GACTCTTAGCGGTGGATCACTCGGCTCGTGCGTCGATGAAGAAC | BT20 | 35 | 100 | 27 | 135 | 319 | 33.9 | 100.2 | 16.1 | 30.7 | 150.7 |
| >hg38_21_+_8433172_8446622_RNA45SN1_45S_50ntFlanks@6652.6695.44\|GACTCTTAGCGG<br>TGGATCACTCGGCTCGTGCGTCGATGAAGAAC | >hg38_21_+_8433172_8446622_RNA45SN1_45S_50ntFlanks | 6652 | 6695 | 44 | GACTCTTAGCGGTGGATCACTCGGCTCGTGCGTCGATGAAGAAC | MB231 | 90 | 114 | 76 | 44 | 229 | 89.3 | 85.5 | 55.2 | 49.4 | 196.7 |
| >hg38_21_+_8433172_8446622_RNA45SN1_45S_50ntFlanks@6652.6696.45\|GACTCTTAGCGG<br>TGGATCACTCGGCTCGTGCGTCGATGAAGAACG | >hg38_21_+_8433172_8446622_RNA45SN1_45S_50ntFlanks | 6652 | 6696 | 45 | GACTCTTAGCGGTGGATCACTCGGCTCGTGCGTCGATGAAGAACG | BT20 | 29 | 74 | 11 | 59 | 155 | 29.8 | 64.4 | 6.4 | 15.1 | 100.8 |
| >hg38_21_+_8433172_8446622_RNA45SN1_45S_50ntFlanks@6652.6696.45\|GACTCTTAGCGG<br>TGGATCACTCGGCTCGTGCGTCGATGAAGAACG | >hg38_21_+_8433172_8446622_RNA45SN1_45S_50ntFlanks | 6652 | 6696 | 45 | GACTCTTAGCGGTGGATCACTCGGCTCGTGCGTCGATGAAGAACG | MB231 | 112 | 271 | 82 | 16 | 179 | 79.0 | 215.8 | 54.2 | 19.1 | 140.9 |
| >hg38_21_+_8433172_8446622_RNA45SN1_45S_50ntFlanks@6652.6697.46\|GACTCTTAGCGG<br>TGGATCACTCGGCTCGTGCGTCGATGAAGAACGC | >hg38_21_+_8433172_8446622_RNA45SN1_45S_50ntFlanks | 6652 | 6697 | 46 | GACTCTTAGCGGTGGATCACTCGGCTCGTGCGTCGATGAAGAACGC | BT20 | 11 | 32 | 7 | 79 | 139 | 9.5 | 33.1 | 5.6 | 37.8 | 83.0 |
| >hg38_21_+_8433172_8446622_RNA45SN1_45S_50ntFlanks@6652.6697.46\|GACTCTTAGCGG<br>TGGATCACTCGGCTCGTGCGTCGATGAAGAACGC | >hg38_21_+_8433172_8446622_RNA45SN1_45S_50ntFlanks | 6652 | 6697 | 46 | GACTCTTAGCGGTGGATCACTCGGCTCGTGCGTCGATGAAGAACGC | MB231 | 55 | 76 | 33 | 14 | 167 | 55.0 | 66.5 | 21.5 | 15.7 | 117.8 |
| >hg38_21_+_8433172_8446622_RNA45SN1_45S_50ntFlanks@6652.6697.46\|GACTCTTAGCGG<br>TGGATCACTCGGCTCGTGCGTCGATGAAGAACGC | >hg38_21_+_8433172_8446622_RNA45SN1_45S_50ntFlanks | 6652 | 6697 | 46 | GACTCTTAGCGGTGGATCACTCGGCTCGTGCGTCGATGAAGAACGC | MB468 | 16 | 50 | 11 | 20 | 168 | 9.1 | 40.8 | 7.2 | 24.4 | 174.4 |
| >hg38_21_+_8433172_8446622_RNA45SN1_45S_50ntFlanks@6652.6698.47\|GACTCTTAGCGG<br>TGGATCACTCGGCTCGTGCGTCGATGAAGAACGCA | >hg38_21_+_8433172_8446622_RNA45SN1_45S_50ntFlanks | 6652 | 6698 | 47 | GACTCTTAGCGGTGGATCACTCGGCTCGTGCGTCGATGAAGAACGCA | BT20 | 20 | 92 | 15 | 218 | 394 | 20.6 | 114.5 | 14.4 | 69.4 | 292.8 |
| >hg38_21_+_8433172_8446622_RNA45SN1_45S_50ntFlanks@6652.6698.47\|GACTCTTAGCGG<br>TGGATCACTCGGCTCGTGCGTCGATGAAGAACGCA | >hg38_21_+_8433172_8446622_RNA45SN1_45S_50ntFlanks | 6652 | 6698 | 47 | GACTCTTAGCGGTGGATCACTCGGCTCGTGCGTCGATGAAGAACGCA | MB231 | 144 | 339 | 100 | 21 | 343 | 158.0 | 421.8 | 68.0 | 32.3 | 299.1 |
| >hg38_21_+_8433172_8446622_RNA45SN1_45S_50ntFlanks@6652.6698.47\|GACTCTTAGCGG<br>TGGATCACTCGGCTCGTGCGTCGATGAAGAACGCA | >hg38_21_+_8433172_8446622_RNA45SN1_45S_50ntFlanks | 6652 | 6698 | 47 | GACTCTTAGCGGTGGATCACTCGGCTCGTGCGTCGATGAAGAACGCA | MB468 | 82 | 333 | 56 | 104 | 1233 | 46.9 | 226.8 | 45.6 | 121.8 | 735.3 |
| >hg38_21_+_8433172_8446622_RNA45SN1_45S_50ntFlanks@6652.6699.48\|GACTCTTAGCGG<br>TGGATCACTCGGCTCGTGCGTCGATGAAGAACGCAG | >hg38_21_+_8433172_8446622_RNA45SN1_45S_50ntFlanks | 6652 | 6699 | 48 | GACTCTTAGCGGTGGATCACTCGGCTCGTGCGTCGATGAAGAACGCAG | BT20 | 11 | 52 | 3 | 38 | 87 | 9.3 | 59.5 | 3.6 | 24.3 | 66.7 |
| >hg38_21_+_8433172_8446622_RNA45SN1_45S_50ntFlanks@6652.6699.48\|GACTCTTAGCGG<br>TGGATCACTCGGCTCGTGCGTCGATGAAGAACGCAG | >hg38_21_+_8433172_8446622_RNA45SN1_45S_50ntFlanks | 6652 | 6699 | 48 | GACTCTTAGCGGTGGATCACTCGGCTCGTGCGTCGATGAAGAACGCAG | MB231 | 62 | 218 | 28 | 11 | 91 | 39.9 | 286.7 | 9.9 | 13.2 | 37.5 |
| >hg38_21_+_8433172_8446622_RNA45SN1_45S_50ntFlanks@6652.6699.48\|GACTCTTAGCGG<br>TGGATCACTCGGCTCGTGCGTCGATGAAGAACGCAG | >hg38_21_+_8433172_8446622_RNA45SN1_45S_50ntFlanks | 6652 | 6699 | 48 | GACTCTTAGCGGTGGATCACTCGGCTCGTGCGTCGATGAAGAACGCAG | MB468 | 45 | 172 | 38 | 79 | 696 | 26.5 | 175.4 | 34.5 | 111.5 | 497.4 |
| >hg38_21_+_8433172_8446622_RNA45SN1_45S_50ntFlanks@6652.6700.49\|GACTCTTAGCGG<br>TGGATCACTCGGCTCGTGCGTCGATGAAGAACGCAGC | >hg38_21_+_8433172_8446622_RNA45SN1_45S_50ntFlanks | 6652 | 6700 | 49 | GACTCTTAGCGGTGGATCACTCGGCTCGTGCGTCGATGAAGAACGCAGC | MB231 | 34 | 67 | 29 | 40 | 136 | 24.5 | 93.9 | 8.9 | 63.1 | 100.3 |
| >hg38_21_+_8433172_8446622_RNA45SN1_45S_50ntFlanks@6652.6701.50\|GACTCTTAGCGG<br>TGGATCACTCGGCTCGTGCGTCGATGAAGAACGCAGCT | >hg38_21_+_8433172_8446622_RNA45SN1_45S_50ntFlanks | 6652 | 6701 | 50 | GACTCTTAGCGGTGGATCACTCGGCTCGTGCGTCGATGAAGAACGCAGCT | MB231 | 20 | 30 | 12 | 24 | 80 | 16.0 | 44.0 | 8.4 | 18.4 | 61.0 |
| >hg38_21_+_8433172_8446622_RNA45SN1_45S_50ntFlanks@6653.6679.27\|ACTCTTAGCGGT<br>GGATCACTCGGCTCG | >hg38_21_+_8433172_8446622_RNA45SN1_45S_50ntFlanks | 6653 | 6679 | 27 | ACTCTTAGCGGTGGATCACTCGGCTCG | MB468 | 10 | 18 | 15 | 3 | 21 | 2.3 | 9.0 | 6.2 | 3.3 | 13.4 |
| >hg38_21_+_8433172_8446622_RNA45SN1_45S_50ntFlanks@6653.6680.28\|ACTCTTAGCGGT<br>GGATCACTCGGCTCGT | >hg38_21_+_8433172_8446622_RNA45SN1_45S_50ntFlanks | 6653 | 6680 | 28 | ACTCTTAGCGGTGGATCACTCGGCTCGT | MB468 | 15 | 13 | 13 | 9 | 18 | 3.8 | 3.5 | 3.5 | 7.7 | 2.5 |
| >hg38_21_+_8433172_8446622_RNA45SN1_45S_50ntFlanks@6653.6681.29\|ACTCTTAGCGGT<br>GGATCACTCGGCTCGTG | >hg38_21_+_8433172_8446622_RNA45SN1_45S_50ntFlanks | 6653 | 6681 | 29 | ACTCTTAGCGGTGGATCACTCGGCTCGTG | BT20 | 25 | 41 | 5 | 20 | 24 | 26.0 | 21.1 | 3.2 | 11.7 | 8.6 |
| >hg38_21_+_8433172_8446622_RNA45SN1_45S_50ntFlanks@6653.6681.29\|ACTCTTAGCGGT<br>GGATCACTCGGCTCGTG | >hg38_21_+_8433172_8446622_RNA45SN1_45S_50ntFlanks | 6653 | 6681 | 29 | ACTCTTAGCGGTGGATCACTCGGCTCGTG | MB231 | 27 | 121 | 6 | 3 | 51 | 11.2 | 61.8 | 5.3 | 1.6 | 74.7 |
| >hg38_21_+_8433172_8446622_RNA45SN1_45S_50ntFlanks@6653.6681.29\|ACTCTTAGCGGT<br>GGATCACTCGGCTCGTG | >hg38_21_+_8433172_8446622_RNA45SN1_45S_50ntFlanks | 6653 | 6681 | 29 | ACTCTTAGCGGTGGATCACTCGGCTCGTG | MB468 | 192 | 133 | 140 | 164 | 258 | 58.8 | 56.0 | 58.4 | 176.1 | 44.9 |
| >hg38_21_+_8433172_8446622_RNA45SN1_45S_50ntFlanks@6653.6683.31\|ACTCTTAGCGGT<br>GGATCACTCGGCTCGTGCG | >hg38_21_+_8433172_8446622_RNA45SN1_45S_50ntFlanks | 6653 | 6683 | 31 | ACTCTTAGCGGTGGATCACTCGGCTCGTGCG | BT20 | 36 | 47 | 6 | 3 | 16 | 48.5 | 24.5 | 8.8 | 2.3 | 5.2 |
| >hg38_21_+_8433172_8446622_RNA45SN1_45S_50ntFlanks@6653.6683.31\|ACTCTTAGCGGT<br>GGATCACTCGGCTCGTGCG | >hg38_21_+_8433172_8446622_RNA45SN1_45S_50ntFlanks | 6653 | 6683 | 31 | ACTCTTAGCGGTGGATCACTCGGCTCGTGCG | MB231 | 42 | 216 | 8 | 1 | 18 | 34.4 | 117.1 | 3.7 | 0.5 | 19.2 |
| >hg38_21_+_8433172_8446622_RNA45SN1_45S_50ntFlanks@6653.6686.34\|ACTCTTAGCGGT<br>GGATCACTCGGCTCGTGCGTCG | >hg38_21_+_8433172_8446622_RNA45SN1_45S_50ntFlanks | 6653 | 6686 | 34 | ACTCTTAGCGGTGGATCACTCGGCTCGTGCGTCG | BT20 | 11 | 10 | 5 | 3 | 12 | 12.9 | 4.6 | 6.4 | 2.7 | 12.2 |
| >hg38_21_+_8433172_8446622_RNA45SN1_45S_50ntFlanks@6653.6686.34\|ACTCTTAGCGGT<br>GGATCACTCGGCTCGTGCGTCG | >hg38_21_+_8433172_8446622_RNA45SN1_45S_50ntFlanks | 6653 | 6686 | 34 | ACTCTTAGCGGTGGATCACTCGGCTCGTGCGTCG | MB231 | 12 | 33 | 4 | 0 | 10 | 15.1 | 10.0 | 2.0 | 0.3 | 17.4 |

| | | | | | | | | | | | | | | | | |
|---|---|---|---|---|---|---|---|---|---|---|---|---|---|---|---|---|
| >hg38_21_+_8433172_8446622_RNA45SN1_45S_50ntFlanks@6653.6686.34\|ACTCTTAGCGGT<br>GGATCACTCGGCTCGTGCGTCG | >hg38_21_+_8433172_8446622_RNA45SN1_45S_50ntFlanks | 6653 | 6686 | 34 | ACTCTTAGCGGTGGATCACTCGGCTCGTGCGTCG | MB468 | 16 | 22 | 6 | 7 | 48 | 6.5 | 7.7 | 1.8 | 7.0 | 29.8 |
| >hg38_21_+_8433172_8446622_RNA45SN1_45S_50ntFlanks@6653.6687.35\|ACTCTTAGCGGT<br>GGATCACTCGGCTCGTGCGTCGA | >hg38_21_+_8433172_8446622_RNA45SN1_45S_50ntFlanks | 6653 | 6687 | 35 | ACTCTTAGCGGTGGATCACTCGGCTCGTGCGTCGA | BT20 | 14 | 17 | 4 | 4 | 10 | 18.5 | 6.0 | 6.1 | 4.4 | 10.4 |
| >hg38_21_+_8433172_8446622_RNA45SN1_45S_50ntFlanks@6653.6687.35\|ACTCTTAGCGGT<br>GGATCACTCGGCTCGTGCGTCGA | >hg38_21_+_8433172_8446622_RNA45SN1_45S_50ntFlanks | 6653 | 6687 | 35 | ACTCTTAGCGGTGGATCACTCGGCTCGTGCGTCGA | MB231 | 16 | 43 | 4 | 1 | 14 | 18.1 | 9.3 | 1.0 | 0.7 | 18.4 |
| >hg38_21_+_8433172_8446622_RNA45SN1_45S_50ntFlanks@6653.6688.36\|ACTCTTAGCGGT<br>GGATCACTCGGCTCGTGCGTCGAT | >hg38_21_+_8433172_8446622_RNA45SN1_45S_50ntFlanks | 6653 | 6688 | 36 | ACTCTTAGCGGTGGATCACTCGGCTCGTGCGTCGAT | BT20 | 26 | 38 | 14 | 35 | 109 | 34.1 | 3.2 | 17.6 | 48.6 | 3.9 |
| >hg38_21_+_8433172_8446622_RNA45SN1_45S_50ntFlanks@6653.6688.36\|ACTCTTAGCGGT<br>GGATCACTCGGCTCGTGCGTCGAT | >hg38_21_+_8433172_8446622_RNA45SN1_45S_50ntFlanks | 6653 | 6688 | 36 | ACTCTTAGCGGTGGATCACTCGGCTCGTGCGTCGAT | MB231 | 25 | 49 | 9 | 2 | 22 | 28.9 | 16.4 | 4.7 | 0.4 | 13.7 |
| >hg38_21_+_8433172_8446622_RNA45SN1_45S_50ntFlanks@6653.6688.36\|ACTCTTAGCGGT<br>GGATCACTCGGCTCGTGCGTCGAT | >hg38_21_+_8433172_8446622_RNA45SN1_45S_50ntFlanks | 6653 | 6688 | 36 | ACTCTTAGCGGTGGATCACTCGGCTCGTGCGTCGAT | MB468 | 12 | 25 | 5 | 6 | 48 | 4.6 | 9.1 | 1.7 | 5.5 | 29.0 |
| >hg38_21_+_8433172_8446622_RNA45SN1_45S_50ntFlanks@6653.6689.37\|ACTCTTAGCGGT<br>GGATCACTCGGCTCGTGCGTCGATG | >hg38_21_+_8433172_8446622_RNA45SN1_45S_50ntFlanks | 6653 | 6689 | 37 | ACTCTTAGCGGTGGATCACTCGGCTCGTGCGTCGATG | BT20 | 14 | 26 | 13 | 41 | 71 | 13.8 | 1.8 | 14.6 | 21.2 | 13.1 |
| >hg38_21_+_8433172_8446622_RNA45SN1_45S_50ntFlanks@6653.6689.37\|ACTCTTAGCGGT<br>GGATCACTCGGCTCGTGCGTCGATG | >hg38_21_+_8433172_8446622_RNA45SN1_45S_50ntFlanks | 6653 | 6689 | 37 | ACTCTTAGCGGTGGATCACTCGGCTCGTGCGTCGATG | MB231 | 18 | 31 | 10 | 4 | 27 | 21.1 | 15.3 | 7.8 | 4.5 | 20.0 |
| >hg38_21_+_8433172_8446622_RNA45SN1_45S_50ntFlanks@6653.6689.37\|ACTCTTAGCGGT<br>GGATCACTCGGCTCGTGCGTCGATG | >hg38_21_+_8433172_8446622_RNA45SN1_45S_50ntFlanks | 6653 | 6689 | 37 | ACTCTTAGCGGTGGATCACTCGGCTCGTGCGTCGATG | MB468 | 69 | 100 | 32 | 50 | 304 | 17.3 | 26.9 | 6.8 | 45.7 | 108.8 |
| >hg38_21_+_8433172_8446622_RNA45SN1_45S_50ntFlanks@6653.6696.44\|ACTCTTAGCGGT<br>GGATCACTCGGCTCGTGCGTCGATGAAGAACG | >hg38_21_+_8433172_8446622_RNA45SN1_45S_50ntFlanks | 6653 | 6696 | 44 | ACTCTTAGCGGTGGATCACTCGGCTCGTGCGTCGATGAAGAACG | MB231 | 15 | 28 | 13 | 1 | 22 | 8.0 | 17.3 | 8.6 | 0.9 | 19.5 |
| >hg38_21_+_8433172_8446622_RNA45SN1_45S_50ntFlanks@6653.6698.46\|ACTCTTAGCGGT<br>GGATCACTCGGCTCGTGCGTCGATGAAGAACGCA | >hg38_21_+_8433172_8446622_RNA45SN1_45S_50ntFlanks | 6653 | 6698 | 46 | ACTCTTAGCGGTGGATCACTCGGCTCGTGCGTCGATGAAGAACGCA | MB231 | 15 | 31 | 11 | 1 | 23 | 15.0 | 31.9 | 6.6 | 1.4 | 39.2 |
| >hg38_21_+_8433172_8446622_RNA45SN1_45S_50ntFlanks@6654.6681.28\|CTCTTAGCGGTG<br>GATCACTCGGCTCGTG | >hg38_21_+_8433172_8446622_RNA45SN1_45S_50ntFlanks | 6654 | 6681 | 28 | CTCTTAGCGGTGGATCACTCGGCTCGTG | MB468 | 111 | 52 | 83 | 107 | 126 | 36.1 | 24.6 | 33.1 | 58.5 | 16.9 |
| >hg38_21_+_8433172_8446622_RNA45SN1_45S_50ntFlanks@6654.6683.30\|CTCTTAGCGGTG<br>GATCACTCGGCTCGTGCG | >hg38_21_+_8433172_8446622_RNA45SN1_45S_50ntFlanks | 6654 | 6683 | 30 | CTCTTAGCGGTGGATCACTCGGCTCGTGCG | BT20 | 11 | 17 | 3 | 2 | 2 | 14.1 | 7.9 | 4.7 | 2.8 | 3.6 |
| >hg38_21_+_8433172_8446622_RNA45SN1_45S_50ntFlanks@6654.6683.30\|CTCTTAGCGGTG<br>GATCACTCGGCTCGTGCG | >hg38_21_+_8433172_8446622_RNA45SN1_45S_50ntFlanks | 6654 | 6683 | 30 | CTCTTAGCGGTGGATCACTCGGCTCGTGCG | MB231 | 17 | 82 | 3 | 0 | 4 | 12.9 | 39.2 | 0.9 | 0.3 | 7.5 |
| >hg38_21_+_8433172_8446622_RNA45SN1_45S_50ntFlanks@6654.6689.36\|CTCTTAGCGGTG<br>GATCACTCGGCTCGTGCGTCGATG | >hg38_21_+_8433172_8446622_RNA45SN1_45S_50ntFlanks | 6654 | 6689 | 36 | CTCTTAGCGGTGGATCACTCGGCTCGTGCGTCGATG | MB468 | 33 | 52 | 16 | 23 | 115 | 8.7 | 13.7 | 6.2 | 15.0 | 52.0 |
| >hg38_21_+_8433172_8446622_RNA45SN1_45S_50ntFlanks@6655.6680.26\|TCTTAGCGGTGG<br>ATCACTCGGCTCGT | >hg38_21_+_8433172_8446622_RNA45SN1_45S_50ntFlanks | 6655 | 6680 | 26 | TCTTAGCGGTGGATCACTCGGCTCGT | MB468 | 12 | 8 | 10 | 6 | 12 | 2.6 | 2.9 | 1.8 | 4.1 | 2.6 |
| >hg38_21_+_8433172_8446622_RNA45SN1_45S_50ntFlanks@6655.6681.27\|TCTTAGCGGTGG<br>ATCACTCGGCTCGTG | >hg38_21_+_8433172_8446622_RNA45SN1_45S_50ntFlanks | 6655 | 6681 | 27 | TCTTAGCGGTGGATCACTCGGCTCGTG | MB468 | 180 | 114 | 133 | 115 | 189 | 52.3 | 49.0 | 53.2 | 99.8 | 34.5 |
| >hg38_21_+_8433172_8446622_RNA45SN1_45S_50ntFlanks@6655.6688.34\|TCTTAGCGGTGG<br>ATCACTCGGCTCGTGCGTCGAT | >hg38_21_+_8433172_8446622_RNA45SN1_45S_50ntFlanks | 6655 | 6688 | 34 | TCTTAGCGGTGGATCACTCGGCTCGTGCGTCGAT | BT20 | 14 | 17 | 4 | 10 | 17 | 15.0 | 4.6 | 5.0 | 11.1 | 1.9 |
| >hg38_21_+_8433172_8446622_RNA45SN1_45S_50ntFlanks@6655.6689.35\|TCTTAGCGGTGG<br>ATCACTCGGCTCGTGCGTCGATG | >hg38_21_+_8433172_8446622_RNA45SN1_45S_50ntFlanks | 6655 | 6689 | 35 | TCTTAGCGGTGGATCACTCGGCTCGTGCGTCGATG | MB468 | 39 | 63 | 16 | 30 | 136 | 11.2 | 10.6 | 5.3 | 21.9 | 60.8 |
| >hg38_21_+_8433172_8446622_RNA45SN1_45S_50ntFlanks@6656.6680.25\|CTTAGCGGTGGA<br>TCACTCGGCTCGT | >hg38_21_+_8433172_8446622_RNA45SN1_45S_50ntFlanks | 6656 | 6680 | 25 | CTTAGCGGTGGATCACTCGGCTCGT | MB468 | 13 | 8 | 10 | 14 | 13 | 3.4 | 2.5 | 1.9 | 7.4 | 2.8 |
| >hg38_21_+_8433172_8446622_RNA45SN1_45S_50ntFlanks@6656.6681.26\|CTTAGCGGTGGA<br>TCACTCGGCTCGTG | >hg38_21_+_8433172_8446622_RNA45SN1_45S_50ntFlanks | 6656 | 6681 | 26 | CTTAGCGGTGGATCACTCGGCTCGTG | MB468 | 226 | 120 | 166 | 278 | 262 | 70.7 | 49.2 | 57.7 | 216.6 | 36.1 |
| >hg38_21_+_8433172_8446622_RNA45SN1_45S_50ntFlanks@6656.6686.31\|CTTAGCGGTGGA<br>TCACTCGGCTCGTGCGTCG | >hg38_21_+_8433172_8446622_RNA45SN1_45S_50ntFlanks | 6656 | 6686 | 31 | CTTAGCGGTGGATCACTCGGCTCGTGCGTCG | MB468 | 11 | 11 | 5 | 12 | 23 | 5.9 | 3.2 | 1.9 | 7.1 | 6.8 |
| >hg38_21_+_8433172_8446622_RNA45SN1_45S_50ntFlanks@6656.6689.34\|CTTAGCGGTGGA<br>TCACTCGGCTCGTGCGTCGATG | >hg38_21_+_8433172_8446622_RNA45SN1_45S_50ntFlanks | 6656 | 6689 | 34 | CTTAGCGGTGGATCACTCGGCTCGTGCGTCGATG | MB468 | 34 | 44 | 16 | 45 | 79 | 9.9 | 8.0 | 5.5 | 29.7 | 30.5 |
| >hg38_21_+_8433172_8446622_RNA45SN1_45S_50ntFlanks@6657.6681.25\|TTAGCGGTGGAT<br>CACTCGGCTCGTG | >hg38_21_+_8433172_8446622_RNA45SN1_45S_50ntFlanks | 6657 | 6681 | 25 | TTAGCGGTGGATCACTCGGCTCGTG | MB468 | 58 | 31 | 39 | 88 | 64 | 20.1 | 10.7 | 8.3 | 64.8 | 16.1 |
| >hg38_21_+_8433172_8446622_RNA45SN1_45S_50ntFlanks@6658.6681.24\|TAGCGGTGGATC<br>ACTCGGCTCGTG | >hg38_21_+_8433172_8446622_RNA45SN1_45S_50ntFlanks | 6658 | 6681 | 24 | TAGCGGTGGATCACTCGGCTCGTG | MB468 | 16 | 6 | 11 | 23 | 20 | 5.1 | 1.6 | 0.8 | 5.1 | 3.9 |
| >hg38_21_+_8433172_8446622_RNA45SN1_45S_50ntFlanks@6659.6681.23\|AGCGGTGGATC<br>ACTCGGCTCGTG | >hg38_21_+_8433172_8446622_RNA45SN1_45S_50ntFlanks | 6659 | 6681 | 23 | AGCGGTGGATCACTCGGCTCGTG | MB468 | 75 | 32 | 41 | 139 | 121 | 29.2 | 14.2 | 25.1 | 93.9 | 24.8 |
| >hg38_21_+_8433172_8446622_RNA45SN1_45S_50ntFlanks@6659.6689.31\|AGCGGTGGATC<br>ACTCGGCTCGTGCGTCGATG | >hg38_21_+_8433172_8446622_RNA45SN1_45S_50ntFlanks | 6659 | 6689 | 31 | AGCGGTGGATCACTCGGCTCGTGCGTCGATG | MB468 | 18 | 17 | 5 | 39 | 45 | 7.1 | 1.8 | 1.8 | 20.3 | 19.1 |
| >hg38_21_+_8433172_8446622_RNA45SN1_45S_50ntFlanks@6660.6681.22\|GCGGTGGATCA<br>CTCGGCTCGTG | >hg38_21_+_8433172_8446622_RNA45SN1_45S_50ntFlanks | 6660 | 6681 | 22 | GCGGTGGATCACTCGGCTCGTG | MB468 | 55 | 38 | 25 | 91 | 85 | 28.6 | 20.2 | 13.5 | 75.8 | 41.0 |
| >hg38_21_+_8433172_8446622_RNA45SN1_45S_50ntFlanks@6660.6688.29\|GCGGTGGATCA<br>CTCGGCTCGTGCGTCGAT | >hg38_21_+_8433172_8446622_RNA45SN1_45S_50ntFlanks | 6660 | 6688 | 29 | GCGGTGGATCACTCGGCTCGTGCGTCGAT | BT20 | 11 | 10 | 5 | 24 | 24 | 15.4 | 4.3 | 6.2 | 17.5 | 5.2 |
| >hg38_21_+_8433172_8446622_RNA45SN1_45S_50ntFlanks@6660.6689.30\|GCGGTGGATCA<br>CTCGGCTCGTGCGTCGATG | >hg38_21_+_8433172_8446622_RNA45SN1_45S_50ntFlanks | 6660 | 6689 | 30 | GCGGTGGATCACTCGGCTCGTGCGTCGATG | MB468 | 34 | 33 | 9 | 70 | 94 | 13.1 | 6.6 | 3.2 | 40.1 | 38.7 |
| >hg38_21_+_8433172_8446622_RNA45SN1_45S_50ntFlanks@6661.6678.18\|CGGTGGATCACT<br>CGGCTC | >hg38_21_+_8433172_8446622_RNA45SN1_45S_50ntFlanks | 6661 | 6678 | 18 | CGGTGGATCACTCGGCTC | BT20 | 42 | 45 | 36 | 138 | 107 | 16.3 | 2.9 | 15.1 | 85.1 | 56.9 |
| >hg38_21_+_8433172_8446622_RNA45SN1_45S_50ntFlanks@6661.6679.19\|CGGTGGATCACT<br>CGGCTCG | >hg38_21_+_8433172_8446622_RNA45SN1_45S_50ntFlanks | 6661 | 6679 | 19 | CGGTGGATCACTCGGCTCG | BT20 | 10 | 15 | 44 | 25 | 21 | 3.0 | 6.2 | 57.3 | 9.4 | 3.6 |
| >hg38_21_+_8433172_8446622_RNA45SN1_45S_50ntFlanks@6661.6679.19\|CGGTGGATCACT<br>CGGCTCG | >hg38_21_+_8433172_8446622_RNA45SN1_45S_50ntFlanks | 6661 | 6679 | 19 | CGGTGGATCACTCGGCTCG | MB468 | 13 | 10 | 14 | 9 | 10 | 3.4 | 4.2 | 5.3 | 4.6 | 3.9 |
| >hg38_21_+_8433172_8446622_RNA45SN1_45S_50ntFlanks@6661.6680.20\|CGGTGGATCACT<br>CGGCTCGT | >hg38_21_+_8433172_8446622_RNA45SN1_45S_50ntFlanks | 6661 | 6680 | 20 | CGGTGGATCACTCGGCTCGT | BT20 | 15 | 19 | 16 | 50 | 36 | 2.6 | 1.4 | 3.5 | 23.6 | 13.0 |
| >hg38_21_+_8433172_8446622_RNA45SN1_45S_50ntFlanks@6661.6680.20\|CGGTGGATCACT<br>CGGCTCGT | >hg38_21_+_8433172_8446622_RNA45SN1_45S_50ntFlanks | 6661 | 6680 | 20 | CGGTGGATCACTCGGCTCGT | MB468 | 12 | 11 | 8 | 19 | 14 | 3.9 | 5.3 | 3.3 | 11.8 | 1.5 |
| >hg38_21_+_8433172_8446622_RNA45SN1_45S_50ntFlanks@6661.6681.21\|CGGTGGATCACT<br>CGGCTCGTG | >hg38_21_+_8433172_8446622_RNA45SN1_45S_50ntFlanks | 6661 | 6681 | 21 | CGGTGGATCACTCGGCTCGTG | BT20 | 25 | 38 | 22 | 84 | 64 | 19.3 | 6.3 | 9.3 | 14.2 | 1.7 |
| >hg38_21_+_8433172_8446622_RNA45SN1_45S_50ntFlanks@6661.6681.21\|CGGTGGATCACT<br>CGGCTCGTG | >hg38_21_+_8433172_8446622_RNA45SN1_45S_50ntFlanks | 6661 | 6681 | 21 | CGGTGGATCACTCGGCTCGTG | MB231 | 21 | 80 | 8 | 7 | 52 | 8.0 | 4.8 | 4.1 | 8.3 | 82.3 |
| >hg38_21_+_8433172_8446622_RNA45SN1_45S_50ntFlanks@6661.6681.21\|CGGTGGATCACT<br>CGGCTCGTG | >hg38_21_+_8433172_8446622_RNA45SN1_45S_50ntFlanks | 6661 | 6681 | 21 | CGGTGGATCACTCGGCTCGTG | MB468 | 158 | 128 | 71 | 198 | 191 | 59.7 | 51.7 | 32.9 | 73.4 | 22.5 |
| >hg38_21_+_8433172_8446622_RNA45SN1_45S_50ntFlanks@6661.6683.23\|CGGTGGATCACT<br>CGGCTCGTGCG | >hg38_21_+_8433172_8446622_RNA45SN1_45S_50ntFlanks | 6661 | 6683 | 23 | CGGTGGATCACTCGGCTCGTGCG | BT20 | 65 | 101 | 15 | 21 | 41 | 80.1 | 43.6 | 18.6 | 10.4 | 6.6 |
| >hg38_21_+_8433172_8446622_RNA45SN1_45S_50ntFlanks@6661.6683.23\|CGGTGGATCACT<br>CGGCTCGTGCG | >hg38_21_+_8433172_8446622_RNA45SN1_45S_50ntFlanks | 6661 | 6683 | 23 | CGGTGGATCACTCGGCTCGTGCG | MB231 | 73 | 410 | 8 | 5 | 40 | 26.8 | 137.8 | 3.9 | 1.6 | 52.9 |
| >hg38_21_+_8433172_8446622_RNA45SN1_45S_50ntFlanks@6661.6684.24\|CGGTGGATCACT<br>CGGCTCGTGCGT | >hg38_21_+_8433172_8446622_RNA45SN1_45S_50ntFlanks | 6661 | 6684 | 24 | CGGTGGATCACTCGGCTCGTGCGT | BT20 | 13 | 29 | 12 | 32 | 30 | 11.2 | 14.1 | 6.1 | 9.9 | 9.5 |
| >hg38_21_+_8433172_8446622_RNA45SN1_45S_50ntFlanks@6661.6685.25\|CGGTGGATCACT<br>CGGCTCGTGCGTC | >hg38_21_+_8433172_8446622_RNA45SN1_45S_50ntFlanks | 6661 | 6685 | 25 | CGGTGGATCACTCGGCTCGTGCGTC | BT20 | 23 | 64 | 26 | 571 | 263 | 20.1 | 34.5 | 22.9 | 219.4 | 51.9 |
| >hg38_21_+_8433172_8446622_RNA45SN1_45S_50ntFlanks@6661.6686.26\|CGGTGGATCACT<br>CGGCTCGTGCGTCG | >hg38_21_+_8433172_8446622_RNA45SN1_45S_50ntFlanks | 6661 | 6686 | 26 | CGGTGGATCACTCGGCTCGTGCGTCG | BT20 | 41 | 53 | 24 | 40 | 91 | 47.2 | 22.2 | 29.3 | 7.8 | 28.2 |
| >hg38_21_+_8433172_8446622_RNA45SN1_45S_50ntFlanks@6661.6686.26\|CGGTGGATCACT<br>CGGCTCGTGCGTCG | >hg38_21_+_8433172_8446622_RNA45SN1_45S_50ntFlanks | 6661 | 6686 | 26 | CGGTGGATCACTCGGCTCGTGCGTCG | MB231 | 18 | 55 | 5 | 4 | 30 | 15.1 | 9.7 | 1.9 | 2.1 | 32.5 |
| >hg38_21_+_8433172_8446622_RNA45SN1_45S_50ntFlanks@6661.6686.26\|CGGTGGATCACT<br>CGGCTCGTGCGTCG | >hg38_21_+_8433172_8446622_RNA45SN1_45S_50ntFlanks | 6661 | 6686 | 26 | CGGTGGATCACTCGGCTCGTGCGTCG | MB468 | 47 | 37 | 13 | 44 | 141 | 17.2 | 5.9 | 3.4 | 12.7 | 40.5 |
| >hg38_21_+_8433172_8446622_RNA45SN1_45S_50ntFlanks@6661.6687.27\|CGGTGGATCACT<br>CGGCTCGTGCGTCGA | >hg38_21_+_8433172_8446622_RNA45SN1_45S_50ntFlanks | 6661 | 6687 | 27 | CGGTGGATCACTCGGCTCGTGCGTCGA | BT20 | 60 | 82 | 25 | 51 | 85 | 78.4 | 47.8 | 32.6 | 25.8 | 35.5 |
| >hg38_21_+_8433172_8446622_RNA45SN1_45S_50ntFlanks@6661.6687.27\|CGGTGGATCACT<br>CGGCTCGTGCGTCGA | >hg38_21_+_8433172_8446622_RNA45SN1_45S_50ntFlanks | 6661 | 6687 | 27 | CGGTGGATCACTCGGCTCGTGCGTCGA | MB231 | 17 | 57 | 6 | 4 | 30 | 8.1 | 19.9 | 3.3 | 0.2 | 38.9 |
| >hg38_21_+_8433172_8446622_RNA45SN1_45S_50ntFlanks@6661.6688.28\|CGGTGGATCACT<br>CGGCTCGTGCGTCGAT | >hg38_21_+_8433172_8446622_RNA45SN1_45S_50ntFlanks | 6661 | 6688 | 28 | CGGTGGATCACTCGGCTCGTGCGTCGAT | BT20 | 91 | 145 | 40 | 241 | 440 | 117.3 | 51.3 | 46.4 | 179.1 | 134.6 |
| >hg38_21_+_8433172_8446622_RNA45SN1_45S_50ntFlanks@6661.6688.28\|CGGTGGATCACT<br>CGGCTCGTGCGTCGAT | >hg38_21_+_8433172_8446622_RNA45SN1_45S_50ntFlanks | 6661 | 6688 | 28 | CGGTGGATCACTCGGCTCGTGCGTCGAT | MB231 | 19 | 43 | 7 | 9 | 22 | 17.5 | 10.4 | 1.9 | 8.7 | 14.9 |
| >hg38_21_+_8433172_8446622_RNA45SN1_45S_50ntFlanks@6661.6688.28\|CGGTGGATCACT<br>CGGCTCGTGCGTCGAT | >hg38_21_+_8433172_8446622_RNA45SN1_45S_50ntFlanks | 6661 | 6688 | 28 | CGGTGGATCACTCGGCTCGTGCGTCGAT | MB468 | 26 | 40 | 8 | 47 | 98 | 7.3 | 13.5 | 2.2 | 13.3 | 45.3 |

| | | | | | | | | | | | | | | | | |
|---|---|---|---|---|---|---|---|---|---|---|---|---|---|---|---|---|
| >hg38_21_+_8433172_8446622_RNA45SN1_45S_50ntFlanks@6661.6689.29\|CGGTGGATCACTCGGCTCGTGCGTCGATG | >hg38_21_+_8433172_8446622_RNA45SN1_45S_50ntFlanks | 6661 | 6689 | 29 | CGGTGGATCACTCGGCTCGTGCGTCGATG | BT20 | 92 | 184 | 74 | 604 | 672 | 96.1 | 109.5 | 75.2 | 70.7 | 150.8 |
| >hg38_21_+_8433172_8446622_RNA45SN1_45S_50ntFlanks@6661.6689.29\|CGGTGGATCACTCGGCTCGTGCGTCGATG | >hg38_21_+_8433172_8446622_RNA45SN1_45S_50ntFlanks | 6661 | 6689 | 29 | CGGTGGATCACTCGGCTCGTGCGTCGATG | MB231 | 25 | 49 | 16 | 35 | 49 | 27.7 | 18.3 | 12.8 | 50.7 | 36.7 |
| >hg38_21_+_8433172_8446622_RNA45SN1_45S_50ntFlanks@6661.6689.29\|CGGTGGATCACTCGGCTCGTGCGTCGATG | >hg38_21_+_8433172_8446622_RNA45SN1_45S_50ntFlanks | 6661 | 6689 | 29 | CGGTGGATCACTCGGCTCGTGCGTCGATG | MB468 | 220 | 274 | 73 | 521 | 992 | 55.3 | 62.6 | 34.0 | 199.0 | 331.1 |
| >hg38_21_+_8433172_8446622_RNA45SN1_45S_50ntFlanks@6661.6692.32\|CGGTGGATCACTCGGCTCGTGCGTCGATGAAG | >hg38_21_+_8433172_8446622_RNA45SN1_45S_50ntFlanks | 6661 | 6692 | 32 | CGGTGGATCACTCGGCTCGTGCGTCGATGAAG | MB468 | 11 | 9 | 3 | 26 | 32 | 5.4 | 3.8 | 1.4 | 10.0 | 12.3 |
| >hg38_21_+_8433172_8446622_RNA45SN1_45S_50ntFlanks@6661.6693.33\|CGGTGGATCACTCGGCTCGTGCGTCGATGAAGA | >hg38_21_+_8433172_8446622_RNA45SN1_45S_50ntFlanks | 6661 | 6693 | 33 | CGGTGGATCACTCGGCTCGTGCGTCGATGAAGA | BT20 | 13 | 26 | 9 | 62 | 52 | 13.7 | 10.7 | 7.4 | 27.9 | 23.6 |
| >hg38_21_+_8433172_8446622_RNA45SN1_45S_50ntFlanks@6661.6694.34\|CGGTGGATCACTCGGCTCGTGCGTCGATGAAGAA | >hg38_21_+_8433172_8446622_RNA45SN1_45S_50ntFlanks | 6661 | 6694 | 34 | CGGTGGATCACTCGGCTCGTGCGTCGATGAAGAA | BT20 | 11 | 18 | 8 | 15 | 26 | 9.3 | 5.3 | 5.3 | 5.1 | 8.2 |
| >hg38_21_+_8433172_8446622_RNA45SN1_45S_50ntFlanks@6661.6695.35\|CGGTGGATCACTCGGCTCGTGCGTCGATGAAGAAC | >hg38_21_+_8433172_8446622_RNA45SN1_45S_50ntFlanks | 6661 | 6695 | 35 | CGGTGGATCACTCGGCTCGTGCGTCGATGAAGAAC | BT20 | 37 | 79 | 35 | 209 | 173 | 21.4 | 33.4 | 18.8 | 64.1 | 19.5 |
| >hg38_21_+_8433172_8446622_RNA45SN1_45S_50ntFlanks@6661.6695.35\|CGGTGGATCACTCGGCTCGTGCGTCGATGAAGAAC | >hg38_21_+_8433172_8446622_RNA45SN1_45S_50ntFlanks | 6661 | 6695 | 35 | CGGTGGATCACTCGGCTCGTGCGTCGATGAAGAAC | MB231 | 19 | 21 | 14 | 32 | 42 | 14.6 | 11.4 | 11.4 | 32.5 | 32.0 |
| >hg38_21_+_8433172_8446622_RNA45SN1_45S_50ntFlanks@6661.6696.36\|CGGTGGATCACTCGGCTCGTGCGTCGATGAAGAACG | >hg38_21_+_8433172_8446622_RNA45SN1_45S_50ntFlanks | 6661 | 6696 | 36 | CGGTGGATCACTCGGCTCGTGCGTCGATGAAGAACG | BT20 | 27 | 44 | 17 | 98 | 50 | 29.4 | 19.5 | 14.3 | 65.2 | 21.8 |
| >hg38_21_+_8433172_8446622_RNA45SN1_45S_50ntFlanks@6661.6696.36\|CGGTGGATCACTCGGCTCGTGCGTCGATGAAGAACG | >hg38_21_+_8433172_8446622_RNA45SN1_45S_50ntFlanks | 6661 | 6696 | 36 | CGGTGGATCACTCGGCTCGTGCGTCGATGAAGAACG | MB231 | 17 | 37 | 10 | 14 | 38 | 13.0 | 6.8 | 6.2 | 13.0 | 39.4 |
| >hg38_21_+_8433172_8446622_RNA45SN1_45S_50ntFlanks@6661.6696.36\|CGGTGGATCACTCGGCTCGTGCGTCGATGAAGAACG | >hg38_21_+_8433172_8446622_RNA45SN1_45S_50ntFlanks | 6661 | 6696 | 36 | CGGTGGATCACTCGGCTCGTGCGTCGATGAAGAACG | MB468 | 13 | 12 | 5 | 38 | 47 | 3.1 | 4.9 | 1.4 | 12.1 | 14.9 |
| >hg38_21_+_8433172_8446622_RNA45SN1_45S_50ntFlanks@6661.6697.37\|CGGTGGATCACTCGGCTCGTGCGTCGATGAAGAACGC | >hg38_21_+_8433172_8446622_RNA45SN1_45S_50ntFlanks | 6661 | 6697 | 37 | CGGTGGATCACTCGGCTCGTGCGTCGATGAAGAACGC | BT20 | 25 | 38 | 24 | 132 | 74 | 18.2 | 5.8 | 14.1 | 58.0 | 23.9 |
| >hg38_21_+_8433172_8446622_RNA45SN1_45S_50ntFlanks@6661.6697.37\|CGGTGGATCACTCGGCTCGTGCGTCGATGAAGAACGC | >hg38_21_+_8433172_8446622_RNA45SN1_45S_50ntFlanks | 6661 | 6697 | 37 | CGGTGGATCACTCGGCTCGTGCGTCGATGAAGAACGC | MB231 | 14 | 19 | 10 | 10 | 36 | 11.0 | 5.6 | 6.2 | 9.2 | 30.3 |
| >hg38_21_+_8433172_8446622_RNA45SN1_45S_50ntFlanks@6661.6697.37\|CGGTGGATCACTCGGCTCGTGCGTCGATGAAGAACGC | >hg38_21_+_8433172_8446622_RNA45SN1_45S_50ntFlanks | 6661 | 6697 | 37 | CGGTGGATCACTCGGCTCGTGCGTCGATGAAGAACGC | MB468 | 30 | 26 | 13 | 37 | 97 | 5.1 | 5.5 | 3.5 | 21.1 | 43.3 |
| >hg38_21_+_8433172_8446622_RNA45SN1_45S_50ntFlanks@6661.6698.38\|CGGTGGATCACTCGGCTCGTGCGTCGATGAAGAACGCA | >hg38_21_+_8433172_8446622_RNA45SN1_45S_50ntFlanks | 6661 | 6698 | 38 | CGGTGGATCACTCGGCTCGTGCGTCGATGAAGAACGCA | BT20 | 13 | 24 | 13 | 113 | 40 | 4.9 | 5.2 | 3.1 | 56.0 | 8.6 |
| >hg38_21_+_8433172_8446622_RNA45SN1_45S_50ntFlanks@6661.6698.38\|CGGTGGATCACTCGGCTCGTGCGTCGATGAAGAACGCA | >hg38_21_+_8433172_8446622_RNA45SN1_45S_50ntFlanks | 6661 | 6698 | 38 | CGGTGGATCACTCGGCTCGTGCGTCGATGAAGAACGCA | MB468 | 18 | 9 | 6 | 20 | 59 | 4.4 | 1.9 | 2.7 | 11.4 | 28.6 |
| >hg38_21_+_8433172_8446622_RNA45SN1_45S_50ntFlanks@6661.6699.39\|CGGTGGATCACTCGGCTCGTGCGTCGATGAAGAACGCAG | >hg38_21_+_8433172_8446622_RNA45SN1_45S_50ntFlanks | 6661 | 6699 | 39 | CGGTGGATCACTCGGCTCGTGCGTCGATGAAGAACGCAG | BT20 | 38 | 73 | 19 | 218 | 58 | 48.4 | 31.1 | 12.0 | 142.0 | 8.1 |
| >hg38_21_+_8433172_8446622_RNA45SN1_45S_50ntFlanks@6661.6699.39\|CGGTGGATCACTCGGCTCGTGCGTCGATGAAGAACGCAG | >hg38_21_+_8433172_8446622_RNA45SN1_45S_50ntFlanks | 6661 | 6699 | 39 | CGGTGGATCACTCGGCTCGTGCGTCGATGAAGAACGCAG | MB231 | 14 | 47 | 13 | 12 | 52 | 10.9 | 12.8 | 1.4 | 10.9 | 74.2 |
| >hg38_21_+_8433172_8446622_RNA45SN1_45S_50ntFlanks@6661.6699.39\|CGGTGGATCACTCGGCTCGTGCGTCGATGAAGAACGCAG | >hg38_21_+_8433172_8446622_RNA45SN1_45S_50ntFlanks | 6661 | 6699 | 39 | CGGTGGATCACTCGGCTCGTGCGTCGATGAAGAACGCAG | MB468 | 116 | 110 | 50 | 121 | 368 | 29.0 | 48.5 | 20.7 | 100.0 | 181.5 |
| >hg38_21_+_8433172_8446622_RNA45SN1_45S_50ntFlanks@6661.6700.40\|CGGTGGATCACTCGGCTCGTGCGTCGATGAAGAACGCAGC | >hg38_21_+_8433172_8446622_RNA45SN1_45S_50ntFlanks | 6661 | 6700 | 40 | CGGTGGATCACTCGGCTCGTGCGTCGATGAAGAACGCAGC | BT20 | 30 | 69 | 33 | 502 | 136 | 8.2 | 12.9 | 10.0 | 338.0 | 40.9 |
| >hg38_21_+_8433172_8446622_RNA45SN1_45S_50ntFlanks@6661.6701.41\|CGGTGGATCACTCGGCTCGTGCGTCGATGAAGAACGCAGCT | >hg38_21_+_8433172_8446622_RNA45SN1_45S_50ntFlanks | 6661 | 6701 | 41 | CGGTGGATCACTCGGCTCGTGCGTCGATGAAGAACGCAGCT | BT20 | 49 | 73 | 37 | 110 | 228 | 23.8 | 23.3 | 13.2 | 76.6 | 138.7 |
| >hg38_21_+_8433172_8446622_RNA45SN1_45S_50ntFlanks@6661.6701.41\|CGGTGGATCACTCGGCTCGTGCGTCGATGAAGAACGCAGCT | >hg38_21_+_8433172_8446622_RNA45SN1_45S_50ntFlanks | 6661 | 6701 | 41 | CGGTGGATCACTCGGCTCGTGCGTCGATGAAGAACGCAGCT | MB231 | 13 | 30 | 16 | 28 | 74 | 5.6 | 11.4 | 7.0 | 5.8 | 71.0 |
| >hg38_21_+_8433172_8446622_RNA45SN1_45S_50ntFlanks@6661.6703.43\|CGGTGGATCACTCGGCTCGTGCGTCGATGAAGAACGCAGCTAG | >hg38_21_+_8433172_8446622_RNA45SN1_45S_50ntFlanks | 6661 | 6703 | 43 | CGGTGGATCACTCGGCTCGTGCGTCGATGAAGAACGCAGCTAG | BT20 | 12 | 20 | 3 | 3 | 20 | 14.3 | 11.2 | 1.6 | 2.2 | 9.2 |
| >hg38_21_+_8433172_8446622_RNA45SN1_45S_50ntFlanks@6661.6706.46\|CGGTGGATCACTCGGCTCGTGCGTCGATGAAGAACGCAGCTAGCTG | >hg38_21_+_8433172_8446622_RNA45SN1_45S_50ntFlanks | 6661 | 6706 | 46 | CGGTGGATCACTCGGCTCGTGCGTCGATGAAGAACGCAGCTAGCTG | BT20 | 25 | 120 | 40 | 23 | 453 | 8.3 | 144.3 | 27.7 | 5.5 | 245.4 |
| >hg38_21_+_8433172_8446622_RNA45SN1_45S_50ntFlanks@6661.6706.46\|CGGTGGATCACTCGGCTCGTGCGTCGATGAAGAACGCAGCTAGCTG | >hg38_21_+_8433172_8446622_RNA45SN1_45S_50ntFlanks | 6661 | 6706 | 46 | CGGTGGATCACTCGGCTCGTGCGTCGATGAAGAACGCAGCTAGCTG | MB231 | 15 | 17 | 11 | 3 | 57 | 13.4 | 17.0 | 7.3 | 3.1 | 55.5 |
| >hg38_21_+_8433172_8446622_RNA45SN1_45S_50ntFlanks@6661.6707.47\|CGGTGGATCACTCGGCTCGTGCGTCGATGAAGAACGCAGCTAGCTGC | >hg38_21_+_8433172_8446622_RNA45SN1_45S_50ntFlanks | 6661 | 6707 | 47 | CGGTGGATCACTCGGCTCGTGCGTCGATGAAGAACGCAGCTAGCTGC | BT20 | 10 | 91 | 21 | 33 | 568 | 7.0 | 119.8 | 16.2 | 14.4 | 412.5 |
| >hg38_21_+_8433172_8446622_RNA45SN1_45S_50ntFlanks@6663.6680.18\|GTGGATCACTCGGCTCGT | >hg38_21_+_8433172_8446622_RNA45SN1_45S_50ntFlanks | 6663 | 6680 | 18 | GTGGATCACTCGGCTCGT | MB468 | 11 | 9 | 6 | 19 | 8 | 4.2 | 3.6 | 2.8 | 11.0 | 2.3 |
| >hg38_21_+_8433172_8446622_RNA45SN1_45S_50ntFlanks@6663.6681.19\|GTGGATCACTCGGCTCGTG | >hg38_21_+_8433172_8446622_RNA45SN1_45S_50ntFlanks | 6663 | 6681 | 19 | GTGGATCACTCGGCTCGTG | MB468 | 170 | 125 | 75 | 325 | 153 | 72.3 | 53.3 | 36.2 | 119.9 | 18.4 |
| >hg38_21_+_8433172_8446622_RNA45SN1_45S_50ntFlanks@6663.6686.24\|GTGGATCACTCGGCTCGTGCGTCG | >hg38_21_+_8433172_8446622_RNA45SN1_45S_50ntFlanks | 6663 | 6686 | 24 | GTGGATCACTCGGCTCGTGCGTCG | MB468 | 45 | 38 | 15 | 66 | 86 | 17.7 | 9.4 | 6.4 | 35.0 | 11.5 |
| >hg38_21_+_8433172_8446622_RNA45SN1_45S_50ntFlanks@6663.6688.26\|GTGGATCACTCGGCTCGTGCGTCGAT | >hg38_21_+_8433172_8446622_RNA45SN1_45S_50ntFlanks | 6663 | 6688 | 26 | GTGGATCACTCGGCTCGTGCGTCGAT | MB468 | 22 | 10 | 6 | 32 | 44 | 8.0 | 3.4 | 2.4 | 10.7 | 5.9 |
| >hg38_21_+_8433172_8446622_RNA45SN1_45S_50ntFlanks@6663.6689.27\|GTGGATCACTCGGCTCGTGCGTCGATG | >hg38_21_+_8433172_8446622_RNA45SN1_45S_50ntFlanks | 6663 | 6689 | 27 | GTGGATCACTCGGCTCGTGCGTCGATG | MB468 | 171 | 74 | 62 | 379 | 343 | 64.6 | 23.8 | 20.7 | 70.6 | 120.4 |
| >hg38_21_+_8433172_8446622_RNA45SN1_45S_50ntFlanks@6663.6699.37\|GTGGATCACTCGGCTCGTGCGTCGATGAAGAACGCAG | >hg38_21_+_8433172_8446622_RNA45SN1_45S_50ntFlanks | 6663 | 6699 | 37 | GTGGATCACTCGGCTCGTGCGTCGATGAAGAACGCAG | MB468 | 18 | 14 | 13 | 35 | 44 | 4.6 | 9.1 | 6.7 | 26.2 | 23.4 |
| >hg38_21_+_8433172_8446622_RNA45SN1_45S_50ntFlanks@6664.6681.18\|TGGATCACTCGGCTCGTG | >hg38_21_+_8433172_8446622_RNA45SN1_45S_50ntFlanks | 6664 | 6681 | 18 | TGGATCACTCGGCTCGTG | MB468 | 167 | 97 | 74 | 325 | 149 | 64.5 | 49.3 | 30.9 | 84.1 | 26.8 |
| >hg38_21_+_8433172_8446622_RNA45SN1_45S_50ntFlanks@6664.6686.23\|TGGATCACTCGGCTCGTGCGTCG | >hg38_21_+_8433172_8446622_RNA45SN1_45S_50ntFlanks | 6664 | 6686 | 23 | TGGATCACTCGGCTCGTGCGTCG | MB468 | 23 | 12 | 6 | 31 | 51 | 8.9 | 3.1 | 2.4 | 8.8 | 14.6 |
| >hg38_21_+_8433172_8446622_RNA45SN1_45S_50ntFlanks@6664.6688.25\|TGGATCACTCGGCTCGTGCGTCGAT | >hg38_21_+_8433172_8446622_RNA45SN1_45S_50ntFlanks | 6664 | 6688 | 25 | TGGATCACTCGGCTCGTGCGTCGAT | MB468 | 12 | 3 | 3 | 23 | 27 | 4.5 | 1.5 | 0.7 | 7.0 | 6.4 |
| >hg38_21_+_8433172_8446622_RNA45SN1_45S_50ntFlanks@6664.6689.26\|TGGATCACTCGGCTCGTGCGTCGATG | >hg38_21_+_8433172_8446622_RNA45SN1_45S_50ntFlanks | 6664 | 6689 | 26 | TGGATCACTCGGCTCGTGCGTCGATG | MB468 | 26 | 5 | 8 | 105 | 49 | 8.4 | 1.9 | 1.9 | 39.1 | 16.6 |
| >hg38_21_+_8433172_8446622_RNA45SN1_45S_50ntFlanks@6666.6686.21\|GATCACTCGGCTCGTGCGTCG | >hg38_21_+_8433172_8446622_RNA45SN1_45S_50ntFlanks | 6666 | 6686 | 21 | GATCACTCGGCTCGTGCGTCG | MB468 | 13 | 3 | 3 | 18 | 32 | 6.1 | 1.0 | 0.8 | 6.8 | 9.8 |
| >hg38_21_+_8433172_8446622_RNA45SN1_45S_50ntFlanks@6666.6689.24\|GATCACTCGGCTCGTGCGTCGATG | >hg38_21_+_8433172_8446622_RNA45SN1_45S_50ntFlanks | 6666 | 6689 | 24 | GATCACTCGGCTCGTGCGTCGATG | MB468 | 11 | 1 | 3 | 60 | 25 | 2.8 | 0.6 | 0.4 | 31.2 | 10.2 |
| >hg38_21_+_8433172_8446622_RNA45SN1_45S_50ntFlanks@6667.6686.20\|ATCACTCGGCTCGTGCGTCG | >hg38_21_+_8433172_8446622_RNA45SN1_45S_50ntFlanks | 6667 | 6686 | 20 | ATCACTCGGCTCGTGCGTCG | MB468 | 26 | 6 | 7 | 28 | 53 | 10.9 | 2.7 | 1.7 | 10.0 | 22.0 |
| >hg38_21_+_8433172_8446622_RNA45SN1_45S_50ntFlanks@6667.6689.23\|ATCACTCGGCTCGTGCGTCGATG | >hg38_21_+_8433172_8446622_RNA45SN1_45S_50ntFlanks | 6667 | 6689 | 23 | ATCACTCGGCTCGTGCGTCGATG | MB468 | 11 | 1 | 3 | 42 | 22 | 4.1 | 0.5 | 0.7 | 31.6 | 7.5 |
| >hg38_21_+_8433172_8446622_RNA45SN1_45S_50ntFlanks@6667.6699.33\|ATCACTCGGCTCGTGCGTCGATGAAGAACGCAG | >hg38_21_+_8433172_8446622_RNA45SN1_45S_50ntFlanks | 6667 | 6699 | 33 | ATCACTCGGCTCGTGCGTCGATGAAGAACGCAG | MB468 | 16 | 12 | 8 | 17 | 46 | 5.9 | 6.5 | 4.1 | 11.2 | 14.9 |
| >hg38_21_+_8433172_8446622_RNA45SN1_45S_50ntFlanks@6669.6686.18\|CACTCGGCTCGTGCGTCG | >hg38_21_+_8433172_8446622_RNA45SN1_45S_50ntFlanks | 6669 | 6686 | 18 | CACTCGGCTCGTGCGTCG | MB468 | 15 | 3 | 5 | 30 | 31 | 5.8 | 1.3 | 0.9 | 5.1 | 9.6 |
| >hg38_21_+_8433172_8446622_RNA45SN1_45S_50ntFlanks@6669.6699.31\|CACTCGGCTCGTGCGTCGATGAAGAACGCAG | >hg38_21_+_8433172_8446622_RNA45SN1_45S_50ntFlanks | 6669 | 6699 | 31 | CACTCGGCTCGTGCGTCGATGAAGAACGCAG | MB468 | 13 | 6 | 6 | 22 | 39 | 3.2 | 4.2 | 2.6 | 11.6 | 14.9 |
| >hg38_21_+_8433172_8446622_RNA45SN1_45S_50ntFlanks@6670.6696.27\|ACTCGGCTCGTGCGTCGATGAAGAACG | >hg38_21_+_8433172_8446622_RNA45SN1_45S_50ntFlanks | 6670 | 6696 | 27 | ACTCGGCTCGTGCGTCGATGAAGAACG | MB468 | 13 | 6 | 6 | 42 | 42 | 2.8 | 3.4 | 1.1 | 10.9 | 10.9 |
| >hg38_21_+_8433172_8446622_RNA45SN1_45S_50ntFlanks@6670.6698.29\|ACTCGGCTCGTGCGTCGATGAAGAACGCA | >hg38_21_+_8433172_8446622_RNA45SN1_45S_50ntFlanks | 6670 | 6698 | 29 | ACTCGGCTCGTGCGTCGATGAAGAACGCA | MB468 | 27 | 13 | 13 | 17 | 118 | 5.9 | 5.5 | 2.2 | 10.2 | 93.3 |
| >hg38_21_+_8433172_8446622_RNA45SN1_45S_50ntFlanks@6670.6699.30\|ACTCGGCTCGTGCGTCGATGAAGAACGCAG | >hg38_21_+_8433172_8446622_RNA45SN1_45S_50ntFlanks | 6670 | 6699 | 30 | ACTCGGCTCGTGCGTCGATGAAGAACGCAG | MB468 | 52 | 16 | 23 | 54 | 143 | 18.2 | 11.7 | 10.9 | 22.1 | 56.6 |
| >hg38_21_+_8433172_8446622_RNA45SN1_45S_50ntFlanks@6671.6696.26\|CTCGGCTCGTGCGTCGATGAAGAACG | >hg38_21_+_8433172_8446622_RNA45SN1_45S_50ntFlanks | 6671 | 6696 | 26 | CTCGGCTCGTGCGTCGATGAAGAACG | MB468 | 68 | 43 | 42 | 184 | 356 | 11.3 | 17.7 | 2.1 | 85.3 | 290.7 |
| >hg38_21_+_8433172_8446622_RNA45SN1_45S_50ntFlanks@6671.6697.27\|CTCGGCTCGTGCGTCGATGAAGAACGC | >hg38_21_+_8433172_8446622_RNA45SN1_45S_50ntFlanks | 6671 | 6697 | 27 | CTCGGCTCGTGCGTCGATGAAGAACGC | MB468 | 51 | 32 | 34 | 62 | 301 | 4.2 | 12.7 | 4.9 | 30.0 | 282.3 |
| >hg38_21_+_8433172_8446622_RNA45SN1_45S_50ntFlanks@6671.6698.28\|CTCGGCTCGTGCGTCGATGAAGAACGCA | >hg38_21_+_8433172_8446622_RNA45SN1_45S_50ntFlanks | 6671 | 6698 | 28 | CTCGGCTCGTGCGTCGATGAAGAACGCA | MB468 | 97 | 51 | 64 | 76 | 815 | 20.5 | 34.1 | 9.3 | 46.6 | 877.2 |
| >hg38_21_+_8433172_8446622_RNA45SN1_45S_50ntFlanks@6671.6699.29\|CTCGGCTCGTGCGTCGATGAAGAACGCAG | >hg38_21_+_8433172_8446622_RNA45SN1_45S_50ntFlanks | 6671 | 6699 | 29 | CTCGGCTCGTGCGTCGATGAAGAACGCAG | MB468 | 293 | 119 | 182 | 360 | 1517 | 86.1 | 71.8 | 51.7 | 129.0 | 1285.4 |

| | | | | | | | | | | | | | | | | |
|---|---|---|---|---|---|---|---|---|---|---|---|---|---|---|---|---|
| >hg38_21_+_8433172_8446622_RNA45SN1_45S_50ntFlanks@6676.6696.21\|CTCGTGCGTCGATGAAGAACG | >hg38_21_+_8433172_8446622_RNA45SN1_45S_50ntFlanks | 6676 | 6696 | 21 | CTCGTGCGTCGATGAAGAACG | MB468 | 21 | 11 | 23 | 18 | 10 | 10.7 | 9.1 | 15.9 | 4.1 | 3.9 |
| >hg38_21_+_8433172_8446622_RNA45SN1_45S_50ntFlanks@6687.6706.20\|ATGAAGAACGCAGCTAGCTG | >hg38_21_+_8433172_8446622_RNA45SN1_45S_50ntFlanks | 6687 | 6706 | 20 | ATGAAGAACGCAGCTAGCTG | MB468 | 10 | 6 | 7 | 9 | 3 | 3.8 | 2.3 | 2.3 | 3.0 | 1.2 |
| >hg38_21_+_8433172_8446622_RNA45SN1_45S_50ntFlanks@6689.6720.32\|GAAGAACGCAGCTAGCTGCGAGAATTAATGTG | >hg38_21_+_8433172_8446622_RNA45SN1_45S_50ntFlanks | 6689 | 6720 | 32 | GAAGAACGCAGCTAGCTGCGAGAATTAATGTG | MB468 | 16 | 9 | 30 | 5 | 10 | 5.1 | 1.9 | 11.0 | 1.2 | 3.8 |
| >hg38_21_+_8433172_8446622_RNA45SN1_45S_50ntFlanks@6690.6718.29\|AAGAACGCAGCTAGCTGCGAGAATTAATG | >hg38_21_+_8433172_8446622_RNA45SN1_45S_50ntFlanks | 6690 | 6718 | 29 | AAGAACGCAGCTAGCTGCGAGAATTAATG | MB468 | 14 | 3 | 13 | 3 | 18 | 6.4 | 1.1 | 6.6 | 1.5 | 8.4 |
| >hg38_21_+_8433172_8446622_RNA45SN1_45S_50ntFlanks@6690.6719.30\|AAGAACGCAGCTAGCTGCGAGAATTAATGT | >hg38_21_+_8433172_8446622_RNA45SN1_45S_50ntFlanks | 6690 | 6719 | 30 | AAGAACGCAGCTAGCTGCGAGAATTAATGT | MB468 | 11 | 5 | 13 | 2 | 11 | 6.2 | 2.2 | 3.3 | 0.9 | 4.9 |
| >hg38_21_+_8433172_8446622_RNA45SN1_45S_50ntFlanks@6690.6720.31\|AAGAACGCAGCTAGCTGCGAGAATTAATGTG | >hg38_21_+_8433172_8446622_RNA45SN1_45S_50ntFlanks | 6690 | 6720 | 31 | AAGAACGCAGCTAGCTGCGAGAATTAATGTG | MB468 | 154 | 64 | 223 | 55 | 168 | 57.5 | 4.3 | 64.9 | 9.6 | 77.1 |
| >hg38_21_+_8433172_8446622_RNA45SN1_45S_50ntFlanks@6697.6718.22\|CAGCTAGCTGCGAGAATTAATG | >hg38_21_+_8433172_8446622_RNA45SN1_45S_50ntFlanks | 6697 | 6718 | 22 | CAGCTAGCTGCGAGAATTAATG | MB468 | 39 | 39 | 34 | 90 | 210 | 16.7 | 2.0 | 9.4 | 42.8 | 33.8 |
| >hg38_21_+_8433172_8446622_RNA45SN1_45S_50ntFlanks@6697.6720.24\|CAGCTAGCTGCGAGAATTAATGTG | >hg38_21_+_8433172_8446622_RNA45SN1_45S_50ntFlanks | 6697 | 6720 | 24 | CAGCTAGCTGCGAGAATTAATGTG | MB231 | 38 | 33 | 72 | 12 | 7 | 31.3 | 25.9 | 65.9 | 8.9 | 6.8 |
| >hg38_21_+_8433172_8446622_RNA45SN1_45S_50ntFlanks@6697.6720.24\|CAGCTAGCTGCGAGAATTAATGTG | >hg38_21_+_8433172_8446622_RNA45SN1_45S_50ntFlanks | 6697 | 6720 | 24 | CAGCTAGCTGCGAGAATTAATGTG | MB468 | 74 | 91 | 80 | 79 | 61 | 6.8 | 27.4 | 23.9 | 37.4 | 27.1 |
| >hg38_21_+_8433172_8446622_RNA45SN1_45S_50ntFlanks@6698.6717.20\|AGCTAGCTGCGAGAATTAAT | >hg38_21_+_8433172_8446622_RNA45SN1_45S_50ntFlanks | 6698 | 6717 | 20 | AGCTAGCTGCGAGAATTAAT | MB468 | 20 | 30 | 15 | 29 | 16 | 3.8 | 5.7 | 2.9 | 25.1 | 3.8 |
| >hg38_21_+_8433172_8446622_RNA45SN1_45S_50ntFlanks@6698.6718.21\|AGCTAGCTGCGAGAATTAATG | >hg38_21_+_8433172_8446622_RNA45SN1_45S_50ntFlanks | 6698 | 6718 | 21 | AGCTAGCTGCGAGAATTAATG | MB468 | 610 | 903 | 503 | 861 | 354 | 149.7 | 261.8 | 151.2 | 646.3 | 94.6 |
| >hg38_21_+_8433172_8446622_RNA45SN1_45S_50ntFlanks@6698.6719.22\|AGCTAGCTGCGAGAATTAATGT | >hg38_21_+_8433172_8446622_RNA45SN1_45S_50ntFlanks | 6698 | 6719 | 22 | AGCTAGCTGCGAGAATTAATGT | MB468 | 69 | 124 | 64 | 69 | 31 | 16.0 | 33.8 | 22.4 | 52.1 | 15.5 |
| >hg38_21_+_8433172_8446622_RNA45SN1_45S_50ntFlanks@6698.6720.23\|AGCTAGCTGCGAGAATTAATGTG | >hg38_21_+_8433172_8446622_RNA45SN1_45S_50ntFlanks | 6698 | 6720 | 23 | AGCTAGCTGCGAGAATTAATGTG | MB231 | 54 | 50 | 107 | 34 | 12 | 44.3 | 39.4 | 84.8 | 32.5 | 11.6 |
| >hg38_21_+_8433172_8446622_RNA45SN1_45S_50ntFlanks@6698.6720.23\|AGCTAGCTGCGAGAATTAATGTG | >hg38_21_+_8433172_8446622_RNA45SN1_45S_50ntFlanks | 6698 | 6720 | 23 | AGCTAGCTGCGAGAATTAATGTG | MB468 | 1525 | 2349 | 1350 | 1563 | 648 | 222.4 | 969.7 | 382.8 | 991.0 | 407.5 |
| >hg38_21_+_8433172_8446622_RNA45SN1_45S_50ntFlanks@6698.6721.24\|AGCTAGCTGCGAGAATTAATGTGA | >hg38_21_+_8433172_8446622_RNA45SN1_45S_50ntFlanks | 6698 | 6721 | 24 | AGCTAGCTGCGAGAATTAATGTGA | MB468 | 12 | 24 | 7 | 14 | 6 | 2.1 | 7.7 | 2.3 | 5.8 | 1.0 |
| >hg38_21_+_8433172_8446622_RNA45SN1_45S_50ntFlanks@6699.6717.19\|GCTAGCTGCGAGAATTAAT | >hg38_21_+_8433172_8446622_RNA45SN1_45S_50ntFlanks | 6699 | 6717 | 19 | GCTAGCTGCGAGAATTAAT | MB468 | 45 | 68 | 38 | 66 | 36 | 7.1 | 16.7 | 6.9 | 63.5 | 13.9 |
| >hg38_21_+_8433172_8446622_RNA45SN1_45S_50ntFlanks@6699.6718.20\|GCTAGCTGCGAGAATTAATG | >hg38_21_+_8433172_8446622_RNA45SN1_45S_50ntFlanks | 6699 | 6718 | 20 | GCTAGCTGCGAGAATTAATG | MB468 | 1352 | 1842 | 1264 | 1995 | 870 | 354.3 | 577.4 | 526.0 | 1693.8 | 328.3 |
| >hg38_21_+_8433172_8446622_RNA45SN1_45S_50ntFlanks@6699.6719.21\|GCTAGCTGCGAGAATTAATGT | >hg38_21_+_8433172_8446622_RNA45SN1_45S_50ntFlanks | 6699 | 6719 | 21 | GCTAGCTGCGAGAATTAATGT | MB468 | 139 | 214 | 143 | 152 | 73 | 17.6 | 47.6 | 49.9 | 121.6 | 42.1 |
| >hg38_21_+_8433172_8446622_RNA45SN1_45S_50ntFlanks@6699.6720.22\|GCTAGCTGCGAGAATTAATGTG | >hg38_21_+_8433172_8446622_RNA45SN1_45S_50ntFlanks | 6699 | 6720 | 22 | GCTAGCTGCGAGAATTAATGTG | MB231 | 60 | 68 | 106 | 35 | 12 | 50.3 | 59.6 | 82.2 | 31.0 | 12.0 |
| >hg38_21_+_8433172_8446622_RNA45SN1_45S_50ntFlanks@6699.6720.22\|GCTAGCTGCGAGAATTAATGTG | >hg38_21_+_8433172_8446622_RNA45SN1_45S_50ntFlanks | 6699 | 6720 | 22 | GCTAGCTGCGAGAATTAATGTG | MB468 | 3020 | 4120 | 2989 | 3346 | 1482 | 645.8 | 1870.7 | 1081.4 | 2209.6 | 1048.6 |
| >hg38_21_+_8433172_8446622_RNA45SN1_45S_50ntFlanks@6699.6721.23\|GCTAGCTGCGAGAATTAATGTGA | >hg38_21_+_8433172_8446622_RNA45SN1_45S_50ntFlanks | 6699 | 6721 | 23 | GCTAGCTGCGAGAATTAATGTGA | MB468 | 22 | 40 | 14 | 29 | 12 | 5.1 | 12.1 | 4.6 | 10.2 | 2.5 |
| >hg38_21_+_8433172_8446622_RNA45SN1_45S_50ntFlanks@6700.6717.18\|CTAGCTGCGAGAATTAAT | >hg38_21_+_8433172_8446622_RNA45SN1_45S_50ntFlanks | 6700 | 6717 | 18 | CTAGCTGCGAGAATTAAT | MB468 | 21 | 27 | 21 | 15 | 18 | 3.9 | 9.7 | 4.3 | 14.3 | 8.6 |
| >hg38_21_+_8433172_8446622_RNA45SN1_45S_50ntFlanks@6700.6718.19\|CTAGCTGCGAGAATTAATG | >hg38_21_+_8433172_8446622_RNA45SN1_45S_50ntFlanks | 6700 | 6718 | 19 | CTAGCTGCGAGAATTAATG | MB468 | 790 | 749 | 824 | 536 | 529 | 148.9 | 196.8 | 183.6 | 521.5 | 266.8 |
| >hg38_21_+_8433172_8446622_RNA45SN1_45S_50ntFlanks@6700.6719.20\|CTAGCTGCGAGAATTAATGT | >hg38_21_+_8433172_8446622_RNA45SN1_45S_50ntFlanks | 6700 | 6719 | 20 | CTAGCTGCGAGAATTAATGT | MB468 | 51 | 29 | 48 | 38 | 18 | 12.3 | 10.9 | 8.4 | 7.4 | 4.4 |
| >hg38_21_+_8433172_8446622_RNA45SN1_45S_50ntFlanks@6700.6720.21\|CTAGCTGCGAGAATTAATGTG | >hg38_21_+_8433172_8446622_RNA45SN1_45S_50ntFlanks | 6700 | 6720 | 21 | CTAGCTGCGAGAATTAATGTG | MB231 | 51 | 33 | 58 | 16 | 8 | 57.1 | 31.8 | 59.3 | 13.9 | 8.6 |
| >hg38_21_+_8433172_8446622_RNA45SN1_45S_50ntFlanks@6700.6720.21\|CTAGCTGCGAGAATTAATGTG | >hg38_21_+_8433172_8446622_RNA45SN1_45S_50ntFlanks | 6700 | 6720 | 21 | CTAGCTGCGAGAATTAATGTG | MB468 | 1530 | 608 | 1423 | 1051 | 432 | 436.9 | 169.4 | 401.3 | 150.4 | 106.5 |
| >hg38_21_+_8433172_8446622_RNA45SN1_45S_50ntFlanks@6700.6721.22\|CTAGCTGCGAGAATTAATGTGA | >hg38_21_+_8433172_8446622_RNA45SN1_45S_50ntFlanks | 6700 | 6721 | 22 | CTAGCTGCGAGAATTAATGTGA | MB468 | 13 | 9 | 8 | 10 | 6 | 4.0 | 4.5 | 2.1 | 2.9 | 2.3 |
| >hg38_21_+_8433172_8446622_RNA45SN1_45S_50ntFlanks@6702.6720.19\|AGCTGCGAGAATTAATGTG | >hg38_21_+_8433172_8446622_RNA45SN1_45S_50ntFlanks | 6702 | 6720 | 19 | AGCTGCGAGAATTAATGTG | MB231 | 16 | 10 | 9 | 11 | 4 | 14.6 | 7.4 | 8.3 | 5.8 | 2.6 |
| >hg38_21_+_8433172_8446622_RNA45SN1_45S_50ntFlanks@6702.6720.19\|AGCTGCGAGAATTAATGTG | >hg38_21_+_8433172_8446622_RNA45SN1_45S_50ntFlanks | 6702 | 6720 | 19 | AGCTGCGAGAATTAATGTG | MB468 | 53 | 22 | 27 | 80 | 14 | 8.1 | 13.5 | 9.1 | 27.3 | 3.2 |
| >hg38_21_+_8433172_8446622_RNA45SN1_45S_50ntFlanks@6702.6724.23\|AGCTGCGAGAATTAATGTGAATT | >hg38_21_+_8433172_8446622_RNA45SN1_45S_50ntFlanks | 6702 | 6724 | 23 | AGCTGCGAGAATTAATGTGAATT | BT20 | 20 | 11 | 8 | 9 | 13 | 32.6 | 18.1 | 10.4 | 11.7 | 19.5 |
| >hg38_21_+_8433172_8446622_RNA45SN1_45S_50ntFlanks@6703.6720.18\|GCTGCGAGAATTAATGTG | >hg38_21_+_8433172_8446622_RNA45SN1_45S_50ntFlanks | 6703 | 6720 | 18 | GCTGCGAGAATTAATGTG | MB468 | 22 | 10 | 13 | 37 | 9 | 6.8 | 5.8 | 4.1 | 16.7 | 3.3 |
| >hg38_21_+_8433172_8446622_RNA45SN1_45S_50ntFlanks@6707.6736.30\|CGAGAATTAATGTGAATTGCAGGACACATT | >hg38_21_+_8433172_8446622_RNA45SN1_45S_50ntFlanks | 6707 | 6736 | 30 | CGAGAATTAATGTGAATTGCAGGACACATT | MB468 | 12 | 4 | 10 | 6 | 6 | 5.8 | 1.5 | 9.3 | 2.0 | 3.0 |
| >hg38_21_+_8433172_8446622_RNA45SN1_45S_50ntFlanks@6708.6731.24\|GAGAATTAATGTGAATTGCAGGAC | >hg38_21_+_8433172_8446622_RNA45SN1_45S_50ntFlanks | 6708 | 6731 | 24 | GAGAATTAATGTGAATTGCAGGAC | MB231 | 17 | 10 | 18 | 3 | 2 | 13.2 | 7.5 | 19.1 | 3.2 | 1.6 |
| >hg38_21_+_8433172_8446622_RNA45SN1_45S_50ntFlanks@6709.6734.26\|AGAATTAATGTGAATTGCAGGACACA | >hg38_21_+_8433172_8446622_RNA45SN1_45S_50ntFlanks | 6709 | 6734 | 26 | AGAATTAATGTGAATTGCAGGACACA | MB468 | 11 | 5 | 6 | 3 | 2 | 3.4 | 3.1 | 5.9 | 1.6 | 0.9 |
| >hg38_21_+_8433172_8446622_RNA45SN1_45S_50ntFlanks@6718.6739.22\|GTGAATTGCAGGACACATTGAT | >hg38_21_+_8433172_8446622_RNA45SN1_45S_50ntFlanks | 6718 | 6739 | 22 | GTGAATTGCAGGACACATTGAT | MB231 | 11 | 16 | 1 | 1 | 2 | 4.3 | 0.5 | 0.3 | 1.0 | 2.7 |
| >hg38_21_+_8433172_8446622_RNA45SN1_45S_50ntFlanks@6718.6743.26\|GTGAATTGCAGGACACATTGATCATC | >hg38_21_+_8433172_8446622_RNA45SN1_45S_50ntFlanks | 6718 | 6743 | 26 | GTGAATTGCAGGACACATTGATCATC | MB468 | 18 | 13 | 13 | 8 | 3 | 11.7 | 5.7 | 13.3 | 4.5 | 0.4 |
| >hg38_21_+_8433172_8446622_RNA45SN1_45S_50ntFlanks@6718.6744.27\|GTGAATTGCAGGACACATTGATCATCG | >hg38_21_+_8433172_8446622_RNA45SN1_45S_50ntFlanks | 6718 | 6744 | 27 | GTGAATTGCAGGACACATTGATCATCG | MB468 | 12 | 5 | 13 | 3 | 1 | 1.6 | 1.1 | 3.2 | 1.2 | 0.9 |
| >hg38_21_+_8433172_8446622_RNA45SN1_45S_50ntFlanks@6719.6736.18\|TGAATTGCAGGACACATT | >hg38_21_+_8433172_8446622_RNA45SN1_45S_50ntFlanks | 6719 | 6736 | 18 | TGAATTGCAGGACACATT | MB468 | 580 | 374 | 156 | 77 | 65 | 215.4 | 302.0 | 96.5 | 15.8 | 22.2 |
| >hg38_21_+_8433172_8446622_RNA45SN1_45S_50ntFlanks@6719.6737.19\|TGAATTGCAGGACACATTG | >hg38_21_+_8433172_8446622_RNA45SN1_45S_50ntFlanks | 6719 | 6737 | 19 | TGAATTGCAGGACACATTG | MB468 | 489 | 267 | 173 | 102 | 61 | 72.8 | 153.9 | 80.8 | 16.4 | 22.5 |
| >hg38_21_+_8433172_8446622_RNA45SN1_45S_50ntFlanks@6719.6738.20\|TGAATTGCAGGACACATTGA | >hg38_21_+_8433172_8446622_RNA45SN1_45S_50ntFlanks | 6719 | 6738 | 20 | TGAATTGCAGGACACATTGA | MB468 | 22 | 15 | 7 | 8 | 7 | 3.4 | 8.4 | 2.6 | 4.9 | 1.4 |
| >hg38_21_+_8433172_8446622_RNA45SN1_45S_50ntFlanks@6719.6739.21\|TGAATTGCAGGACACATTGAT | >hg38_21_+_8433172_8446622_RNA45SN1_45S_50ntFlanks | 6719 | 6739 | 21 | TGAATTGCAGGACACATTGAT | MB468 | 169 | 99 | 57 | 23 | 26 | 14.2 | 67.6 | 23.1 | 12.8 | 14.8 |
| >hg38_21_+_8433172_8446622_RNA45SN1_45S_50ntFlanks@6719.6740.22\|TGAATTGCAGGACACATTGATC | >hg38_21_+_8433172_8446622_RNA45SN1_45S_50ntFlanks | 6719 | 6740 | 22 | TGAATTGCAGGACACATTGATC | MB468 | 528 | 488 | 258 | 159 | 93 | 83.6 | 238.0 | 115.8 | 30.8 | 26.3 |
| >hg38_21_+_8433172_8446622_RNA45SN1_45S_50ntFlanks@6719.6741.23\|TGAATTGCAGGACACATTGATCA | >hg38_21_+_8433172_8446622_RNA45SN1_45S_50ntFlanks | 6719 | 6741 | 23 | TGAATTGCAGGACACATTGATCA | MB468 | 50 | 36 | 23 | 14 | 15 | 4.4 | 15.4 | 4.9 | 6.3 | 3.5 |
| >hg38_21_+_8433172_8446622_RNA45SN1_45S_50ntFlanks@6719.6742.24\|TGAATTGCAGGACACATTGATCAT | >hg38_21_+_8433172_8446622_RNA45SN1_45S_50ntFlanks | 6719 | 6742 | 24 | TGAATTGCAGGACACATTGATCAT | MB468 | 75 | 55 | 30 | 10 | 8 | 47.0 | 22.0 | 15.2 | 4.9 | 2.1 |
| >hg38_21_+_8433172_8446622_RNA45SN1_45S_50ntFlanks@6719.6743.25\|TGAATTGCAGGACACATTGATCATC | >hg38_21_+_8433172_8446622_RNA45SN1_45S_50ntFlanks | 6719 | 6743 | 25 | TGAATTGCAGGACACATTGATCATC | MB468 | 788 | 790 | 402 | 266 | 162 | 236.7 | 354.8 | 173.8 | 76.9 | 28.5 |
| >hg38_21_+_8433172_8446622_RNA45SN1_45S_50ntFlanks@6719.6744.26\|TGAATTGCAGGACACATTGATCATCG | >hg38_21_+_8433172_8446622_RNA45SN1_45S_50ntFlanks | 6719 | 6744 | 26 | TGAATTGCAGGACACATTGATCATCG | MB468 | 1147 | 506 | 1487 | 237 | 207 | 156.2 | 75.0 | 345.6 | 112.4 | 46.2 |
| >hg38_21_+_8433172_8446622_RNA45SN1_45S_50ntFlanks@6719.6745.27\|TGAATTGCAGGACACATTGATCATCGA | >hg38_21_+_8433172_8446622_RNA45SN1_45S_50ntFlanks | 6719 | 6745 | 27 | TGAATTGCAGGACACATTGATCATCGA | MB468 | 67 | 41 | 85 | 19 | 14 | 6.6 | 7.3 | 15.8 | 14.1 | 5.6 |
| >hg38_21_+_8433172_8446622_RNA45SN1_45S_50ntFlanks@6719.6746.28\|TGAATTGCAGGACACATTGATCATCGAC | >hg38_21_+_8433172_8446622_RNA45SN1_45S_50ntFlanks | 6719 | 6746 | 28 | TGAATTGCAGGACACATTGATCATCGAC | MB468 | 116 | 54 | 80 | 74 | 62 | 25.4 | 8.7 | 15.6 | 29.0 | 10.5 |
| >hg38_21_+_8433172_8446622_RNA45SN1_45S_50ntFlanks@6719.6747.29\|TGAATTGCAGGACACATTGATCATCGACA | >hg38_21_+_8433172_8446622_RNA45SN1_45S_50ntFlanks | 6719 | 6747 | 29 | TGAATTGCAGGACACATTGATCATCGACA | MB468 | 123 | 61 | 100 | 43 | 99 | 48.5 | 22.5 | 56.6 | 17.5 | 8.8 |
| >hg38_21_+_8433172_8446622_RNA45SN1_45S_50ntFlanks@6719.6749.31\|TGAATTGCAGGACACATTGATCATCGACACT | >hg38_21_+_8433172_8446622_RNA45SN1_45S_50ntFlanks | 6719 | 6749 | 31 | TGAATTGCAGGACACATTGATCATCGACACT | MB468 | 37 | 46 | 25 | 16 | 27 | 8.6 | 6.7 | 10.6 | 8.2 | 2.3 |

| | | | | | | | | | | | | | | | | |
|---|---|---|---|---|---|---|---|---|---|---|---|---|---|---|---|---|
| >hg38_21_+_8433172_8446622_RNA45SN1_45S_50ntFlanks@6719.6750.32\|TGAATTGCAGGACACATTGATCATCGACACTT | >hg38_21_+_8433172_8446622_RNA45SN1_45S_50ntFlanks | 6719 | 6750 | 32 | TGAATTGCAGGACACATTGATCATCGACACTT | MB468 | 14 | 14 | 9 | 8 | 13 | 4.1 | 4.3 | 4.4 | 4.7 | 3.4 |
| >hg38_21_+_8433172_8446622_RNA45SN1_45S_50ntFlanks@6719.6751.33\|TGAATTGCAGGACACATTGATCATCGACACTTC | >hg38_21_+_8433172_8446622_RNA45SN1_45S_50ntFlanks | 6719 | 6751 | 33 | TGAATTGCAGGACACATTGATCATCGACACTTC | MB468 | 35 | 35 | 17 | 30 | 28 | 10.5 | 16.1 | 7.1 | 22.5 | 3.6 |
| >hg38_21_+_8433172_8446622_RNA45SN1_45S_50ntFlanks@6719.6752.34\|TGAATTGCAGGACACATTGATCATCGACACTTCG | >hg38_21_+_8433172_8446622_RNA45SN1_45S_50ntFlanks | 6719 | 6752 | 34 | TGAATTGCAGGACACATTGATCATCGACACTTCG | MB468 | 28 | 22 | 17 | 28 | 26 | 8.4 | 19.8 | 13.8 | 28.7 | 5.8 |
| >hg38_21_+_8433172_8446622_RNA45SN1_45S_50ntFlanks@6719.6756.38\|TGAATTGCAGGACACATTGATCATCGACACTTCGAACG | >hg38_21_+_8433172_8446622_RNA45SN1_45S_50ntFlanks | 6719 | 6756 | 38 | TGAATTGCAGGACACATTGATCATCGACACTTCGAACG | MB468 | 12 | 14 | 14 | 11 | 24 | 5.2 | 12.1 | 7.3 | 9.2 | 17.6 |
| >hg38_21_+_8433172_8446622_RNA45SN1_45S_50ntFlanks@6719.6758.40\|TGAATTGCAGGACACATTGATCATCGACACTTCGAACGCA | >hg38_21_+_8433172_8446622_RNA45SN1_45S_50ntFlanks | 6719 | 6758 | 40 | TGAATTGCAGGACACATTGATCATCGACACTTCGAACGCA | MB468 | 13 | 24 | 20 | 8 | 35 | 5.8 | 25.6 | 9.9 | 5.3 | 26.6 |
| >hg38_21_+_8433172_8446622_RNA45SN1_45S_50ntFlanks@6721.6738.18\|AATTGCAGGACACATTGA | >hg38_21_+_8433172_8446622_RNA45SN1_45S_50ntFlanks | 6721 | 6738 | 18 | AATTGCAGGACACATTGA | MB468 | 52 | 21 | 15 | 18 | 8 | 12.8 | 12.3 | 8.3 | 11.7 | 1.9 |
| >hg38_21_+_8433172_8446622_RNA45SN1_45S_50ntFlanks@6721.6739.19\|AATTGCAGGACACATTGAT | >hg38_21_+_8433172_8446622_RNA45SN1_45S_50ntFlanks | 6721 | 6739 | 19 | AATTGCAGGACACATTGAT | MB468 | 327 | 126 | 125 | 52 | 41 | 59.6 | 94.6 | 78.8 | 33.4 | 26.5 |
| >hg38_21_+_8433172_8446622_RNA45SN1_45S_50ntFlanks@6721.6740.20\|AATTGCAGGACACATTGATC | >hg38_21_+_8433172_8446622_RNA45SN1_45S_50ntFlanks | 6721 | 6740 | 20 | AATTGCAGGACACATTGATC | MB468 | 1039 | 710 | 605 | 349 | 144 | 318.3 | 487.0 | 423.5 | 137.4 | 74.6 |
| >hg38_21_+_8433172_8446622_RNA45SN1_45S_50ntFlanks@6721.6741.21\|AATTGCAGGACACATTGATCA | >hg38_21_+_8433172_8446622_RNA45SN1_45S_50ntFlanks | 6721 | 6741 | 21 | AATTGCAGGACACATTGATCA | MB468 | 77 | 48 | 39 | 14 | 14 | 21.7 | 25.7 | 26.3 | 7.3 | 6.6 |
| >hg38_21_+_8433172_8446622_RNA45SN1_45S_50ntFlanks@6721.6742.22\|AATTGCAGGACACATTGATCAT | >hg38_21_+_8433172_8446622_RNA45SN1_45S_50ntFlanks | 6721 | 6742 | 22 | AATTGCAGGACACATTGATCAT | MB468 | 82 | 52 | 44 | 12 | 10 | 41.8 | 21.9 | 34.0 | 5.5 | 4.5 |
| >hg38_21_+_8433172_8446622_RNA45SN1_45S_50ntFlanks@6721.6743.23\|AATTGCAGGACACATTGATCATC | >hg38_21_+_8433172_8446622_RNA45SN1_45S_50ntFlanks | 6721 | 6743 | 23 | AATTGCAGGACACATTGATCATC | MB468 | 765 | 693 | 483 | 224 | 153 | 253.0 | 397.5 | 340.7 | 88.9 | 60.3 |
| >hg38_21_+_8433172_8446622_RNA45SN1_45S_50ntFlanks@6721.6744.24\|AATTGCAGGACACATTGATCATCG | >hg38_21_+_8433172_8446622_RNA45SN1_45S_50ntFlanks | 6721 | 6744 | 24 | AATTGCAGGACACATTGATCATCG | MB468 | 2435 | 922 | 2460 | 324 | 295 | 176.0 | 167.3 | 1102.3 | 139.4 | 112.9 |
| >hg38_21_+_8433172_8446622_RNA45SN1_45S_50ntFlanks@6721.6745.25\|AATTGCAGGACACATTGATCATCGA | >hg38_21_+_8433172_8446622_RNA45SN1_45S_50ntFlanks | 6721 | 6745 | 25 | AATTGCAGGACACATTGATCATCGA | MB468 | 200 | 104 | 198 | 37 | 28 | 21.7 | 24.8 | 87.8 | 25.5 | 15.9 |
| >hg38_21_+_8433172_8446622_RNA45SN1_45S_50ntFlanks@6721.6746.26\|AATTGCAGGACACATTGATCATCGAC | >hg38_21_+_8433172_8446622_RNA45SN1_45S_50ntFlanks | 6721 | 6746 | 26 | AATTGCAGGACACATTGATCATCGAC | MB468 | 271 | 123 | 162 | 169 | 115 | 50.1 | 20.6 | 44.3 | 70.4 | 53.0 |
| >hg38_21_+_8433172_8446622_RNA45SN1_45S_50ntFlanks@6721.6747.27\|AATTGCAGGACACATTGATCATCGACA | >hg38_21_+_8433172_8446622_RNA45SN1_45S_50ntFlanks | 6721 | 6747 | 27 | AATTGCAGGACACATTGATCATCGACA | MB468 | 284 | 123 | 164 | 85 | 148 | 82.2 | 56.9 | 79.9 | 34.8 | 41.7 |
| >hg38_21_+_8433172_8446622_RNA45SN1_45S_50ntFlanks@6721.6749.29\|AATTGCAGGACACATTGATCATCGACACT | >hg38_21_+_8433172_8446622_RNA45SN1_45S_50ntFlanks | 6721 | 6749 | 29 | AATTGCAGGACACATTGATCATCGACACT | MB468 | 19 | 25 | 8 | 8 | 20 | 2.8 | 10.0 | 2.3 | 3.3 | 8.7 |
| >hg38_21_+_8433172_8446622_RNA45SN1_45S_50ntFlanks@6721.6750.30\|AATTGCAGGACACATTGATCATCGACACTT | >hg38_21_+_8433172_8446622_RNA45SN1_45S_50ntFlanks | 6721 | 6750 | 30 | AATTGCAGGACACATTGATCATCGACACTT | MB468 | 37 | 35 | 19 | 19 | 29 | 5.8 | 12.6 | 5.5 | 8.8 | 9.7 |
| >hg38_21_+_8433172_8446622_RNA45SN1_45S_50ntFlanks@6721.6751.31\|AATTGCAGGACACATTGATCATCGACACTTC | >hg38_21_+_8433172_8446622_RNA45SN1_45S_50ntFlanks | 6721 | 6751 | 31 | AATTGCAGGACACATTGATCATCGACACTTC | MB468 | 11 | 11 | 6 | 11 | 11 | 1.7 | 3.7 | 0.8 | 5.1 | 6.1 |
| >hg38_21_+_8433172_8446622_RNA45SN1_45S_50ntFlanks@6721.6752.32\|AATTGCAGGACACATTGATCATCGACACTTCG | >hg38_21_+_8433172_8446622_RNA45SN1_45S_50ntFlanks | 6721 | 6752 | 32 | AATTGCAGGACACATTGATCATCGACACTTCG | MB468 | 38 | 44 | 26 | 32 | 42 | 5.2 | 13.0 | 6.4 | 19.4 | 6.8 |
| >hg38_21_+_8433172_8446622_RNA45SN1_45S_50ntFlanks@6721.6753.33\|AATTGCAGGACACATTGATCATCGACACTTCGA | >hg38_21_+_8433172_8446622_RNA45SN1_45S_50ntFlanks | 6721 | 6753 | 33 | AATTGCAGGACACATTGATCATCGACACTTCGA | MB468 | 15 | 16 | 10 | 17 | 22 | 4.0 | 4.9 | 1.1 | 10.9 | 4.8 |
| >hg38_21_+_8433172_8446622_RNA45SN1_45S_50ntFlanks@6721.6756.36\|AATTGCAGGACACATTGATCATCGACACTTCGAACG | >hg38_21_+_8433172_8446622_RNA45SN1_45S_50ntFlanks | 6721 | 6756 | 36 | AATTGCAGGACACATTGATCATCGACACTTCGAACG | MB468 | 42 | 51 | 39 | 49 | 64 | 12.7 | 19.0 | 6.4 | 29.3 | 33.3 |
| >hg38_21_+_8433172_8446622_RNA45SN1_45S_50ntFlanks@6721.6757.37\|AATTGCAGGACACATTGATCATCGACACTTCGAACGC | >hg38_21_+_8433172_8446622_RNA45SN1_45S_50ntFlanks | 6721 | 6757 | 37 | AATTGCAGGACACATTGATCATCGACACTTCGAACGC | MB468 | 20 | 29 | 21 | 14 | 29 | 7.7 | 20.7 | 5.4 | 10.8 | 22.8 |
| >hg38_21_+_8433172_8446622_RNA45SN1_45S_50ntFlanks@6721.6758.38\|AATTGCAGGACACATTGATCATCGACACTTCGAACGCA | >hg38_21_+_8433172_8446622_RNA45SN1_45S_50ntFlanks | 6721 | 6758 | 38 | AATTGCAGGACACATTGATCATCGACACTTCGAACGCA | MB468 | 140 | 276 | 158 | 67 | 171 | 61.0 | 224.9 | 55.8 | 44.7 | 122.9 |
| >hg38_21_+_8433172_8446622_RNA45SN1_45S_50ntFlanks@6721.6760.40\|AATTGCAGGACACATTGATCATCGACACTTCGAACGCACT | >hg38_21_+_8433172_8446622_RNA45SN1_45S_50ntFlanks | 6721 | 6760 | 40 | AATTGCAGGACACATTGATCATCGACACTTCGAACGCACT | MB468 | 21 | 85 | 35 | 9 | 33 | 3.4 | 87.2 | 23.5 | 6.2 | 23.7 |
| >hg38_21_+_8433172_8446622_RNA45SN1_45S_50ntFlanks@6721.6770.50\|AATTGCAGGACACATTGATCATCGACACTTCGAACGCACTTGCGGCCCCG | >hg38_21_+_8433172_8446622_RNA45SN1_45S_50ntFlanks | 6721 | 6770 | 50 | AATTGCAGGACACATTGATCATCGACACTTCGAACGCACTTGCGGCCCCG | MB468 | 23 | 61 | 29 | 26 | 146 | 16.4 | 80.3 | 24.0 | 36.2 | 142.1 |
| >hg38_21_+_8433172_8446622_RNA45SN1_45S_50ntFlanks@6722.6739.18\|ATTGCAGGACACATTGAT | >hg38_21_+_8433172_8446622_RNA45SN1_45S_50ntFlanks | 6722 | 6739 | 18 | ATTGCAGGACACATTGAT | MB468 | 41 | 14 | 12 | 5 | 5 | 11.7 | 10.0 | 9.0 | 2.8 | 3.2 |
| >hg38_21_+_8433172_8446622_RNA45SN1_45S_50ntFlanks@6722.6740.19\|ATTGCAGGACACATTGATC | >hg38_21_+_8433172_8446622_RNA45SN1_45S_50ntFlanks | 6722 | 6740 | 19 | ATTGCAGGACACATTGATC | MB468 | 136 | 85 | 64 | 37 | 16 | 46.6 | 60.0 | 49.2 | 15.9 | 8.1 |
| >hg38_21_+_8433172_8446622_RNA45SN1_45S_50ntFlanks@6722.6741.20\|ATTGCAGGACACATTGATCA | >hg38_21_+_8433172_8446622_RNA45SN1_45S_50ntFlanks | 6722 | 6741 | 20 | ATTGCAGGACACATTGATCA | MB468 | 14 | 8 | 6 | 2 | 2 | 4.8 | 4.5 | 4.0 | 1.1 | 1.2 |
| >hg38_21_+_8433172_8446622_RNA45SN1_45S_50ntFlanks@6722.6742.21\|ATTGCAGGACACATTGATCAT | >hg38_21_+_8433172_8446622_RNA45SN1_45S_50ntFlanks | 6722 | 6742 | 21 | ATTGCAGGACACATTGATCAT | MB468 | 13 | 9 | 6 | 2 | 1 | 7.6 | 4.1 | 4.7 | 0.8 | 0.6 |
| >hg38_21_+_8433172_8446622_RNA45SN1_45S_50ntFlanks@6722.6743.22\|ATTGCAGGACACATTGATCATC | >hg38_21_+_8433172_8446622_RNA45SN1_45S_50ntFlanks | 6722 | 6743 | 22 | ATTGCAGGACACATTGATCATC | MB468 | 120 | 96 | 61 | 27 | 18 | 47.4 | 54.4 | 49.9 | 11.2 | 5.9 |
| >hg38_21_+_8433172_8446622_RNA45SN1_45S_50ntFlanks@6722.6744.23\|ATTGCAGGACACATTGATCATCG | >hg38_21_+_8433172_8446622_RNA45SN1_45S_50ntFlanks | 6722 | 6744 | 23 | ATTGCAGGACACATTGATCATCG | MB468 | 299 | 108 | 264 | 33 | 29 | 55.8 | 28.4 | 149.1 | 13.2 | 11.9 |
| >hg38_21_+_8433172_8446622_RNA45SN1_45S_50ntFlanks@6722.6745.24\|ATTGCAGGACACATTGATCATCGA | >hg38_21_+_8433172_8446622_RNA45SN1_45S_50ntFlanks | 6722 | 6745 | 24 | ATTGCAGGACACATTGATCATCGA | MB468 | 23 | 13 | 22 | 3 | 3 | 3.7 | 4.0 | 10.7 | 2.0 | 1.8 |
| >hg38_21_+_8433172_8446622_RNA45SN1_45S_50ntFlanks@6722.6746.25\|ATTGCAGGACACATTGATCATCGAC | >hg38_21_+_8433172_8446622_RNA45SN1_45S_50ntFlanks | 6722 | 6746 | 25 | ATTGCAGGACACATTGATCATCGAC | MB468 | 29 | 13 | 16 | 17 | 11 | 7.3 | 2.5 | 4.8 | 5.4 | 3.8 |
| >hg38_21_+_8433172_8446622_RNA45SN1_45S_50ntFlanks@6722.6747.26\|ATTGCAGGACACATTGATCATCGACA | >hg38_21_+_8433172_8446622_RNA45SN1_45S_50ntFlanks | 6722 | 6747 | 26 | ATTGCAGGACACATTGATCATCGACA | MB468 | 27 | 11 | 15 | 7 | 11 | 6.3 | 4.2 | 6.0 | 2.1 | 1.9 |
| >hg38_21_+_8433172_8446622_RNA45SN1_45S_50ntFlanks@6722.6758.37\|ATTGCAGGACACATTGATCATCGACACTTCGAACGCA | >hg38_21_+_8433172_8446622_RNA45SN1_45S_50ntFlanks | 6722 | 6758 | 37 | ATTGCAGGACACATTGATCATCGACACTTCGAACGCA | MB468 | 12 | 19 | 15 | 5 | 9 | 5.2 | 11.0 | 6.1 | 3.7 | 6.5 |
| >hg38_21_+_8433172_8446622_RNA45SN1_45S_50ntFlanks@6723.6740.18\|TTGCAGGACACATTGATC | >hg38_21_+_8433172_8446622_RNA45SN1_45S_50ntFlanks | 6723 | 6740 | 18 | TTGCAGGACACATTGATC | MB468 | 68 | 34 | 26 | 14 | 6 | 28.3 | 28.9 | 24.4 | 5.8 | 4.0 |
| >hg38_21_+_8433172_8446622_RNA45SN1_45S_50ntFlanks@6723.6743.21\|TTGCAGGACACATTGATCATC | >hg38_21_+_8433172_8446622_RNA45SN1_45S_50ntFlanks | 6723 | 6743 | 21 | TTGCAGGACACATTGATCATC | MB468 | 65 | 45 | 29 | 11 | 8 | 32.8 | 32.9 | 28.2 | 5.2 | 3.2 |
| >hg38_21_+_8433172_8446622_RNA45SN1_45S_50ntFlanks@6723.6744.22\|TTGCAGGACACATTGATCATCG | >hg38_21_+_8433172_8446622_RNA45SN1_45S_50ntFlanks | 6723 | 6744 | 22 | TTGCAGGACACATTGATCATCG | MB468 | 144 | 47 | 121 | 14 | 12 | 43.6 | 16.7 | 92.0 | 6.4 | 4.5 |
| >hg38_21_+_8433172_8446622_RNA45SN1_45S_50ntFlanks@6723.6745.23\|TTGCAGGACACATTGATCATCGA | >hg38_21_+_8433172_8446622_RNA45SN1_45S_50ntFlanks | 6723 | 6745 | 23 | TTGCAGGACACATTGATCATCGA | MB468 | 13 | 7 | 12 | 1 | 1 | 2.8 | 3.2 | 6.2 | 1.6 | 1.0 |
| >hg38_21_+_8433172_8446622_RNA45SN1_45S_50ntFlanks@6723.6746.24\|TTGCAGGACACATTGATCATCGAC | >hg38_21_+_8433172_8446622_RNA45SN1_45S_50ntFlanks | 6723 | 6746 | 24 | TTGCAGGACACATTGATCATCGAC | MB468 | 15 | 7 | 9 | 7 | 5 | 3.5 | 2.3 | 2.7 | 2.9 | 1.6 |
| >hg38_21_+_8433172_8446622_RNA45SN1_45S_50ntFlanks@6723.6747.25\|TTGCAGGACACATTGATCATCGACA | >hg38_21_+_8433172_8446622_RNA45SN1_45S_50ntFlanks | 6723 | 6747 | 25 | TTGCAGGACACATTGATCATCGACA | MB468 | 13 | 6 | 7 | 3 | 5 | 1.8 | 3.1 | 2.7 | 1.1 | 0.7 |
| >hg38_21_+_8433172_8446622_RNA45SN1_45S_50ntFlanks@6724.6743.20\|TGCAGGACACATTGATCATC | >hg38_21_+_8433172_8446622_RNA45SN1_45S_50ntFlanks | 6724 | 6743 | 20 | TGCAGGACACATTGATCATC | MB468 | 10 | 8 | 3 | 3 | 3 | 3.1 | 6.7 | 1.8 | 1.1 | 0.9 |
| >hg38_21_+_8433172_8446622_RNA45SN1_45S_50ntFlanks@6724.6744.21\|TGCAGGACACATTGATCATCG | >hg38_21_+_8433172_8446622_RNA45SN1_45S_50ntFlanks | 6724 | 6744 | 21 | TGCAGGACACATTGATCATCG | MB468 | 18 | 6 | 12 | 3 | 3 | 3.8 | 2.8 | 6.0 | 1.9 | 0.7 |
| >hg38_21_+_8433172_8446622_RNA45SN1_45S_50ntFlanks@6725.6743.19\|GCAGGACACATTGATCATC | >hg38_21_+_8433172_8446622_RNA45SN1_45S_50ntFlanks | 6725 | 6743 | 19 | GCAGGACACATTGATCATC | MB231 | 39 | 38 | 28 | 6 | 43 | 51.3 | 55.5 | 42.7 | 7.8 | 70.1 |
| >hg38_21_+_8433172_8446622_RNA45SN1_45S_50ntFlanks@6725.6743.19\|GCAGGACACATTGATCATC | >hg38_21_+_8433172_8446622_RNA45SN1_45S_50ntFlanks | 6725 | 6743 | 19 | GCAGGACACATTGATCATC | MB468 | 32 | 42 | 8 | 7 | 5 | 5.6 | 42.9 | 2.5 | 5.2 | 3.7 |
| >hg38_21_+_8433172_8446622_RNA45SN1_45S_50ntFlanks@6725.6744.20\|GCAGGACACATTGATCATCG | >hg38_21_+_8433172_8446622_RNA45SN1_45S_50ntFlanks | 6725 | 6744 | 20 | GCAGGACACATTGATCATCG | MB231 | 10 | 4 | 6 | 1 | 3 | 8.3 | 3.8 | 6.4 | 0.5 | 3.3 |
| >hg38_21_+_8433172_8446622_RNA45SN1_45S_50ntFlanks@6725.6744.20\|GCAGGACACATTGATCATCG | >hg38_21_+_8433172_8446622_RNA45SN1_45S_50ntFlanks | 6725 | 6744 | 20 | GCAGGACACATTGATCATCG | MB468 | 45 | 31 | 23 | 8 | 7 | 14.5 | 25.2 | 13.4 | 8.1 | 6.0 |
| >hg38_21_+_8433172_8446622_RNA45SN1_45S_50ntFlanks@6725.6747.23\|GCAGGACACATTGATCATCGACA | >hg38_21_+_8433172_8446622_RNA45SN1_45S_50ntFlanks | 6725 | 6747 | 23 | GCAGGACACATTGATCATCGACA | BT20 | 11 | 3 | 23 | 2 | 10 | 9.6 | 0.5 | 18.9 | 0.2 | 8.5 |
| >hg38_21_+_8433172_8446622_RNA45SN1_45S_50ntFlanks@6725.6747.23\|GCAGGACACATTGATCATCGACA | >hg38_21_+_8433172_8446622_RNA45SN1_45S_50ntFlanks | 6725 | 6747 | 23 | GCAGGACACATTGATCATCGACA | MB231 | 38 | 15 | 40 | 2 | 25 | 57.3 | 22.2 | 65.4 | 3.8 | 37.5 |
| >hg38_21_+_8433172_8446622_RNA45SN1_45S_50ntFlanks@6726.6743.18\|CAGGACACATTGATCATC | >hg38_21_+_8433172_8446622_RNA45SN1_45S_50ntFlanks | 6726 | 6743 | 18 | CAGGACACATTGATCATC | MB468 | 28 | 11 | 16 | 9 | 2 | 28.2 | 8.4 | 21.2 | 4.5 | 0.6 |

| | | | | | | | | | | | | | | | |
|---|---|---|---|---|---|---|---|---|---|---|---|---|---|---|---|
| >hg38_21_+_8433172_8446622_RNA45SN1_45S_50ntFlanks@6726.6744.19\|CAGGACACATTGATCATCG | >hg38_21_+_8433172_8446622_RNA45SN1_45S_50ntFlanks | 6726 | 6744 | 19 | CAGGACACATTGATCATCG | MB468 | 33 | 5 | 37 | 11 | 3 | 22.1 | 2.1 | 41.8 | 11.5 | 2.1 |
| >hg38_21_+_8433172_8446622_RNA45SN1_45S_50ntFlanks@6728.6756.29\|GGACACATTGATCATCGACACTTCGAACG | >hg38_21_+_8433172_8446622_RNA45SN1_45S_50ntFlanks | 6728 | 6756 | 29 | GGACACATTGATCATCGACACTTCGAACG | MB231 | 14 | 6 | 19 | 4 | 8 | 11.7 | 4.4 | 15.6 | 4.6 | 8.6 |
| >hg38_21_+_8433172_8446622_RNA45SN1_45S_50ntFlanks@6729.6770.42\|GACACATTGATCATCGACACTTCGAACGCACTTGCGGCCCCG | >hg38_21_+_8433172_8446622_RNA45SN1_45S_50ntFlanks | 6729 | 6770 | 42 | GACACATTGATCATCGACACTTCGAACGCACTTGCGGCCCCG | MB468 | 20 | 32 | 21 | 13 | 33 | 8.5 | 21.4 | 13.7 | 16.0 | 31.4 |
| >hg38_21_+_8433172_8446622_RNA45SN1_45S_50ntFlanks@6729.6771.43\|GACACATTGATCATCGACACTTCGAACGCACTTGCGGCCCCGG | >hg38_21_+_8433172_8446622_RNA45SN1_45S_50ntFlanks | 6729 | 6771 | 43 | GACACATTGATCATCGACACTTCGAACGCACTTGCGGCCCCGG | MB468 | 13 | 30 | 8 | 7 | 39 | 4.6 | 27.8 | 4.7 | 8.2 | 41.5 |
| >hg38_21_+_8433172_8446622_RNA45SN1_45S_50ntFlanks@6730.6747.18\|ACACATTGATCATCGACA | >hg38_21_+_8433172_8446622_RNA45SN1_45S_50ntFlanks | 6730 | 6747 | 18 | ACACATTGATCATCGACA | MB468 | 11 | 2 | 14 | 4 | 3 | 4.0 | 0.8 | 6.1 | 0.7 | 0.9 |
| >hg38_21_+_8433172_8446622_RNA45SN1_45S_50ntFlanks@6730.6752.23\|ACACATTGATCATCGACACTTCG | >hg38_21_+_8433172_8446622_RNA45SN1_45S_50ntFlanks | 6730 | 6752 | 23 | ACACATTGATCATCGACACTTCG | MB468 | 15 | 12 | 17 | 5 | 8 | 3.7 | 3.6 | 3.8 | 2.1 | 2.6 |
| >hg38_21_+_8433172_8446622_RNA45SN1_45S_50ntFlanks@6730.6756.27\|ACACATTGATCATCGACACTTCGAACG | >hg38_21_+_8433172_8446622_RNA45SN1_45S_50ntFlanks | 6730 | 6756 | 27 | ACACATTGATCATCGACACTTCGAACG | MB468 | 18 | 19 | 15 | 9 | 12 | 3.6 | 6.6 | 3.8 | 2.1 | 3.0 |
| >hg38_21_+_8433172_8446622_RNA45SN1_45S_50ntFlanks@6730.6770.41\|ACACATTGATCATCGACACTTCGAACGCACTTGCGGCCCCG | >hg38_21_+_8433172_8446622_RNA45SN1_45S_50ntFlanks | 6730 | 6770 | 41 | ACACATTGATCATCGACACTTCGAACGCACTTGCGGCCCCG | MB468 | 25 | 39 | 26 | 17 | 61 | 9.1 | 25.1 | 13.7 | 17.0 | 28.9 |
| >hg38_21_+_8433172_8446622_RNA45SN1_45S_50ntFlanks@6730.6771.42\|ACACATTGATCATCGACACTTCGAACGCACTTGCGGCCCCGG | >hg38_21_+_8433172_8446622_RNA45SN1_45S_50ntFlanks | 6730 | 6771 | 42 | ACACATTGATCATCGACACTTCGAACGCACTTGCGGCCCCGG | MB468 | 17 | 39 | 11 | 9 | 75 | 5.8 | 36.4 | 5.8 | 9.6 | 44.7 |
| >hg38_21_+_8433172_8446622_RNA45SN1_45S_50ntFlanks@6731.6752.22\|CACATTGATCATCGACACTTCG | >hg38_21_+_8433172_8446622_RNA45SN1_45S_50ntFlanks | 6731 | 6752 | 22 | CACATTGATCATCGACACTTCG | MB468 | 50 | 28 | 54 | 18 | 32 | 16.7 | 8.9 | 17.5 | 5.1 | 12.0 |
| >hg38_21_+_8433172_8446622_RNA45SN1_45S_50ntFlanks@6731.6753.23\|CACATTGATCATCGACACTTCGA | >hg38_21_+_8433172_8446622_RNA45SN1_45S_50ntFlanks | 6731 | 6753 | 23 | CACATTGATCATCGACACTTCGA | MB468 | 37 | 14 | 29 | 14 | 22 | 14.9 | 4.9 | 8.8 | 5.4 | 12.4 |
| >hg38_21_+_8433172_8446622_RNA45SN1_45S_50ntFlanks@6731.6754.24\|CACATTGATCATCGACACTTCGAA | >hg38_21_+_8433172_8446622_RNA45SN1_45S_50ntFlanks | 6731 | 6754 | 24 | CACATTGATCATCGACACTTCGAA | MB468 | 18 | 8 | 11 | 5 | 10 | 5.4 | 3.6 | 2.8 | 2.9 | 6.0 |
| >hg38_21_+_8433172_8446622_RNA45SN1_45S_50ntFlanks@6731.6756.26\|CACATTGATCATCGACACTTCGAACG | >hg38_21_+_8433172_8446622_RNA45SN1_45S_50ntFlanks | 6731 | 6756 | 26 | CACATTGATCATCGACACTTCGAACG | MB468 | 76 | 50 | 59 | 40 | 49 | 23.7 | 29.0 | 17.5 | 13.8 | 27.5 |
| >hg38_21_+_8433172_8446622_RNA45SN1_45S_50ntFlanks@6731.6758.28\|CACATTGATCATCGACACTTCGAACGCA | >hg38_21_+_8433172_8446622_RNA45SN1_45S_50ntFlanks | 6731 | 6758 | 28 | CACATTGATCATCGACACTTCGAACGCA | MB468 | 24 | 15 | 22 | 10 | 20 | 7.3 | 5.4 | 10.3 | 5.6 | 6.8 |
| >hg38_21_+_8433172_8446622_RNA45SN1_45S_50ntFlanks@6731.6770.40\|CACATTGATCATCGACACTTCGAACGCACTTGCGGCCCCG | >hg38_21_+_8433172_8446622_RNA45SN1_45S_50ntFlanks | 6731 | 6770 | 40 | CACATTGATCATCGACACTTCGAACGCACTTGCGGCCCCG | MB468 | 82 | 131 | 80 | 53 | 193 | 27.6 | 78.2 | 42.9 | 56.8 | 66.3 |
| >hg38_21_+_8433172_8446622_RNA45SN1_45S_50ntFlanks@6731.6771.41\|CACATTGATCATCGACACTTCGAACGCACTTGCGGCCCCGG | >hg38_21_+_8433172_8446622_RNA45SN1_45S_50ntFlanks | 6731 | 6771 | 41 | CACATTGATCATCGACACTTCGAACGCACTTGCGGCCCCGG | MB468 | 49 | 105 | 29 | 24 | 201 | 13.6 | 85.9 | 15.8 | 27.2 | 89.9 |
| >hg38_21_+_8433172_8446622_RNA45SN1_45S_50ntFlanks@6732.6749.18\|ACATTGATCATCGACACT | >hg38_21_+_8433172_8446622_RNA45SN1_45S_50ntFlanks | 6732 | 6749 | 18 | ACATTGATCATCGACACT | BT20 | 11 | 6 | 6 | 2 | 9 | 8.5 | 2.0 | 0.9 | 0.8 | 4.1 |
| >hg38_21_+_8433172_8446622_RNA45SN1_45S_50ntFlanks@6732.6752.21\|ACATTGATCATCGACACTTCG | >hg38_21_+_8433172_8446622_RNA45SN1_45S_50ntFlanks | 6732 | 6752 | 21 | ACATTGATCATCGACACTTCG | MB231 | 14 | 12 | 6 | 1 | 14 | 11.0 | 7.8 | 5.4 | 1.2 | 14.9 |
| >hg38_21_+_8433172_8446622_RNA45SN1_45S_50ntFlanks@6732.6752.21\|ACATTGATCATCGACACTTCG | >hg38_21_+_8433172_8446622_RNA45SN1_45S_50ntFlanks | 6732 | 6752 | 21 | ACATTGATCATCGACACTTCG | MB468 | 39 | 20 | 33 | 23 | 21 | 8.5 | 7.9 | 9.2 | 3.0 | 5.5 |
| >hg38_21_+_8433172_8446622_RNA45SN1_45S_50ntFlanks@6732.6753.22\|ACATTGATCATCGACACTTCGA | >hg38_21_+_8433172_8446622_RNA45SN1_45S_50ntFlanks | 6732 | 6753 | 22 | ACATTGATCATCGACACTTCGA | MB468 | 28 | 12 | 17 | 18 | 14 | 6.5 | 6.2 | 1.8 | 5.7 | 7.0 |
| >hg38_21_+_8433172_8446622_RNA45SN1_45S_50ntFlanks@6732.6754.23\|ACATTGATCATCGACACTTCGAA | >hg38_21_+_8433172_8446622_RNA45SN1_45S_50ntFlanks | 6732 | 6754 | 23 | ACATTGATCATCGACACTTCGAA | MB468 | 12 | 6 | 6 | 4 | 5 | 1.7 | 3.1 | 3.4 | 1.5 | 2.9 |
| >hg38_21_+_8433172_8446622_RNA45SN1_45S_50ntFlanks@6732.6756.25\|ACATTGATCATCGACACTTCGAACG | >hg38_21_+_8433172_8446622_RNA45SN1_45S_50ntFlanks | 6732 | 6756 | 25 | ACATTGATCATCGACACTTCGAACG | MB231 | 13 | 14 | 7 | 3 | 14 | 9.6 | 8.8 | 5.5 | 2.4 | 16.9 |
| >hg38_21_+_8433172_8446622_RNA45SN1_45S_50ntFlanks@6732.6756.25\|ACATTGATCATCGACACTTCGAACG | >hg38_21_+_8433172_8446622_RNA45SN1_45S_50ntFlanks | 6732 | 6756 | 25 | ACATTGATCATCGACACTTCGAACG | MB468 | 43 | 32 | 26 | 37 | 31 | 8.7 | 17.6 | 4.5 | 9.1 | 14.4 |
| >hg38_21_+_8433172_8446622_RNA45SN1_45S_50ntFlanks@6732.6758.27\|ACATTGATCATCGACACTTCGAACGCA | >hg38_21_+_8433172_8446622_RNA45SN1_45S_50ntFlanks | 6732 | 6758 | 27 | ACATTGATCATCGACACTTCGAACGCA | MB468 | 19 | 13 | 12 | 14 | 15 | 5.2 | 4.7 | 3.8 | 3.1 | 5.8 |
| >hg38_21_+_8433172_8446622_RNA45SN1_45S_50ntFlanks@6732.6770.39\|ACATTGATCATCGACACTTCGAACGCACTTGCGGCCCCG | >hg38_21_+_8433172_8446622_RNA45SN1_45S_50ntFlanks | 6732 | 6770 | 39 | ACATTGATCATCGACACTTCGAACGCACTTGCGGCCCCG | MB468 | 67 | 102 | 54 | 71 | 162 | 19.3 | 54.5 | 27.9 | 69.5 | 65.0 |
| >hg38_21_+_8433172_8446622_RNA45SN1_45S_50ntFlanks@6732.6771.40\|ACATTGATCATCGACACTTCGAACGCACTTGCGGCCCCGG | >hg38_21_+_8433172_8446622_RNA45SN1_45S_50ntFlanks | 6732 | 6771 | 40 | ACATTGATCATCGACACTTCGAACGCACTTGCGGCCCCGG | MB468 | 56 | 107 | 26 | 38 | 214 | 14.5 | 83.2 | 12.4 | 38.5 | 105.1 |
| >hg38_21_+_8433172_8446622_RNA45SN1_45S_50ntFlanks@6732.6773.42\|ACATTGATCATCGACACTTCGAACGCACTTGCGGCCCCGGGT | >hg38_21_+_8433172_8446622_RNA45SN1_45S_50ntFlanks | 6732 | 6773 | 42 | ACATTGATCATCGACACTTCGAACGCACTTGCGGCCCCGGGT | BT20 | 34 | 34 | 25 | 9 | 93 | 39.3 | 28.1 | 17.0 | 5.9 | 9.5 |
| >hg38_21_+_8433172_8446622_RNA45SN1_45S_50ntFlanks@6732.6773.42\|ACATTGATCATCGACACTTCGAACGCACTTGCGGCCCCGGGT | >hg38_21_+_8433172_8446622_RNA45SN1_45S_50ntFlanks | 6732 | 6773 | 42 | ACATTGATCATCGACACTTCGAACGCACTTGCGGCCCCGGGT | MB231 | 41 | 41 | 32 | 3 | 78 | 34.8 | 23.9 | 25.7 | 3.1 | 38.7 |
| >hg38_21_+_8433172_8446622_RNA45SN1_45S_50ntFlanks@6732.6774.43\|ACATTGATCATCGACACTTCGAACGCACTTGCGGCCCCGGGTT | >hg38_21_+_8433172_8446622_RNA45SN1_45S_50ntFlanks | 6732 | 6774 | 43 | ACATTGATCATCGACACTTCGAACGCACTTGCGGCCCCGGGTT | BT20 | 19 | 23 | 13 | 7 | 61 | 14.2 | 18.0 | 8.9 | 4.4 | 5.4 |
| >hg38_21_+_8433172_8446622_RNA45SN1_45S_50ntFlanks@6732.6774.43\|ACATTGATCATCGACACTTCGAACGCACTTGCGGCCCCGGGTT | >hg38_21_+_8433172_8446622_RNA45SN1_45S_50ntFlanks | 6732 | 6774 | 43 | ACATTGATCATCGACACTTCGAACGCACTTGCGGCCCCGGGTT | MB231 | 22 | 22 | 19 | 3 | 58 | 10.4 | 13.6 | 5.0 | 3.6 | 28.0 |
| >hg38_21_+_8433172_8446622_RNA45SN1_45S_50ntFlanks@6733.6752.20\|CATTGATCATCGACACTTCG | >hg38_21_+_8433172_8446622_RNA45SN1_45S_50ntFlanks | 6733 | 6752 | 20 | CATTGATCATCGACACTTCG | MB468 | 79 | 22 | 62 | 68 | 40 | 28.0 | 12.6 | 11.0 | 30.5 | 19.1 |
| >hg38_21_+_8433172_8446622_RNA45SN1_45S_50ntFlanks@6733.6753.21\|CATTGATCATCGACACTTCGA | >hg38_21_+_8433172_8446622_RNA45SN1_45S_50ntFlanks | 6733 | 6753 | 21 | CATTGATCATCGACACTTCGA | MB468 | 46 | 12 | 26 | 47 | 22 | 16.5 | 8.5 | 7.2 | 31.8 | 12.7 |
| >hg38_21_+_8433172_8446622_RNA45SN1_45S_50ntFlanks@6733.6754.22\|CATTGATCATCGACACTTCGAA | >hg38_21_+_8433172_8446622_RNA45SN1_45S_50ntFlanks | 6733 | 6754 | 22 | CATTGATCATCGACACTTCGAA | MB468 | 18 | 5 | 8 | 12 | 7 | 3.3 | 2.7 | 1.6 | 9.1 | 3.7 |
| >hg38_21_+_8433172_8446622_RNA45SN1_45S_50ntFlanks@6733.6756.24\|CATTGATCATCGACACTTCGAACG | >hg38_21_+_8433172_8446622_RNA45SN1_45S_50ntFlanks | 6733 | 6756 | 24 | CATTGATCATCGACACTTCGAACG | MB468 | 98 | 43 | 53 | 108 | 50 | 33.9 | 30.3 | 13.7 | 74.2 | 32.5 |
| >hg38_21_+_8433172_8446622_RNA45SN1_45S_50ntFlanks@6733.6758.26\|CATTGATCATCGACACTTCGAACGCA | >hg38_21_+_8433172_8446622_RNA45SN1_45S_50ntFlanks | 6733 | 6758 | 26 | CATTGATCATCGACACTTCGAACGCA | MB468 | 39 | 21 | 22 | 36 | 35 | 12.8 | 9.9 | 9.7 | 21.6 | 19.3 |
| >hg38_21_+_8433172_8446622_RNA45SN1_45S_50ntFlanks@6733.6770.38\|CATTGATCATCGACACTTCGAACGCACTTGCGGCCCCG | >hg38_21_+_8433172_8446622_RNA45SN1_45S_50ntFlanks | 6733 | 6770 | 38 | CATTGATCATCGACACTTCGAACGCACTTGCGGCCCCG | MB468 | 114 | 193 | 89 | 108 | 281 | 27.3 | 101.5 | 44.0 | 103.6 | 109.7 |
| >hg38_21_+_8433172_8446622_RNA45SN1_45S_50ntFlanks@6733.6771.39\|CATTGATCATCGACACTTCGAACGCACTTGCGGCCCCGG | >hg38_21_+_8433172_8446622_RNA45SN1_45S_50ntFlanks | 6733 | 6771 | 39 | CATTGATCATCGACACTTCGAACGCACTTGCGGCCCCGG | MB468 | 95 | 196 | 45 | 62 | 384 | 24.1 | 152.3 | 21.4 | 63.0 | 173.8 |
| >hg38_21_+_8433172_8446622_RNA45SN1_45S_50ntFlanks@6733.6772.40\|CATTGATCATCGACACTTCGAACGCACTTGCGGCCCCGGG | >hg38_21_+_8433172_8446622_RNA45SN1_45S_50ntFlanks | 6733 | 6772 | 40 | CATTGATCATCGACACTTCGAACGCACTTGCGGCCCCGGG | MB468 | 12 | 29 | 6 | 14 | 69 | 3.3 | 18.3 | 3.7 | 16.0 | 31.2 |
| >hg38_21_+_8433172_8446622_RNA45SN1_45S_50ntFlanks@6733.6773.41\|CATTGATCATCGACACTTCGAACGCACTTGCGGCCCCGGGT | >hg38_21_+_8433172_8446622_RNA45SN1_45S_50ntFlanks | 6733 | 6773 | 41 | CATTGATCATCGACACTTCGAACGCACTTGCGGCCCCGGGT | BT20 | 18 | 16 | 12 | 5 | 45 | 14.7 | 10.0 | 7.7 | 3.7 | 4.2 |
| >hg38_21_+_8433172_8446622_RNA45SN1_45S_50ntFlanks@6733.6773.41\|CATTGATCATCGACACTTCGAACGCACTTGCGGCCCCGGGT | >hg38_21_+_8433172_8446622_RNA45SN1_45S_50ntFlanks | 6733 | 6773 | 41 | CATTGATCATCGACACTTCGAACGCACTTGCGGCCCCGGGT | MB231 | 21 | 22 | 14 | 2 | 43 | 18.5 | 13.3 | 10.8 | 2.4 | 18.7 |
| >hg38_21_+_8433172_8446622_RNA45SN1_45S_50ntFlanks@6733.6774.42\|CATTGATCATCGACACTTCGAACGCACTTGCGGCCCCGGGTT | >hg38_21_+_8433172_8446622_RNA45SN1_45S_50ntFlanks | 6733 | 6774 | 42 | CATTGATCATCGACACTTCGAACGCACTTGCGGCCCCGGGTT | BT20 | 10 | 11 | 7 | 4 | 32 | 5.8 | 7.0 | 4.2 | 2.0 | 2.9 |
| >hg38_21_+_8433172_8446622_RNA45SN1_45S_50ntFlanks@6733.6774.42\|CATTGATCATCGACACTTCGAACGCACTTGCGGCCCCGGGTT | >hg38_21_+_8433172_8446622_RNA45SN1_45S_50ntFlanks | 6733 | 6774 | 42 | CATTGATCATCGACACTTCGAACGCACTTGCGGCCCCGGGTT | MB231 | 13 | 13 | 9 | 3 | 33 | 6.9 | 7.6 | 3.7 | 2.3 | 15.7 |
| >hg38_21_+_8433172_8446622_RNA45SN1_45S_50ntFlanks@6734.6752.19\|ATTGATCATCGACACTTCG | >hg38_21_+_8433172_8446622_RNA45SN1_45S_50ntFlanks | 6734 | 6752 | 19 | ATTGATCATCGACACTTCG | MB468 | 15 | 4 | 11 | 18 | 9 | 5.2 | 2.6 | 3.4 | 4.7 | 2.9 |
| >hg38_21_+_8433172_8446622_RNA45SN1_45S_50ntFlanks@6734.6753.20\|ATTGATCATCGACACTTCGA | >hg38_21_+_8433172_8446622_RNA45SN1_45S_50ntFlanks | 6734 | 6753 | 20 | ATTGATCATCGACACTTCGA | MB468 | 11 | 4 | 7 | 11 | 5 | 3.7 | 3.3 | 0.9 | 2.8 | 1.6 |
| >hg38_21_+_8433172_8446622_RNA45SN1_45S_50ntFlanks@6734.6756.23\|ATTGATCATCGACACTTCGAACG | >hg38_21_+_8433172_8446622_RNA45SN1_45S_50ntFlanks | 6734 | 6756 | 23 | ATTGATCATCGACACTTCGAACG | MB468 | 21 | 14 | 14 | 22 | 10 | 4.0 | 9.4 | 1.6 | 5.4 | 2.5 |
| >hg38_21_+_8433172_8446622_RNA45SN1_45S_50ntFlanks@6734.6770.37\|ATTGATCATCGACACTTCGAACGCACTTGCGGCCCCG | >hg38_21_+_8433172_8446622_RNA45SN1_45S_50ntFlanks | 6734 | 6770 | 37 | ATTGATCATCGACACTTCGAACGCACTTGCGGCCCCG | MB468 | 52 | 77 | 34 | 49 | 117 | 15.7 | 25.0 | 18.6 | 37.9 | 42.8 |
| >hg38_21_+_8433172_8446622_RNA45SN1_45S_50ntFlanks@6734.6771.38\|ATTGATCATCGACACTTCGAACGCACTTGCGGCCCCGG | >hg38_21_+_8433172_8446622_RNA45SN1_45S_50ntFlanks | 6734 | 6771 | 38 | ATTGATCATCGACACTTCGAACGCACTTGCGGCCCCGG | MB468 | 32 | 57 | 11 | 26 | 124 | 8.4 | 26.0 | 5.8 | 21.5 | 57.6 |
| >hg38_21_+_8433172_8446622_RNA45SN1_45S_50ntFlanks@6734.6773.40\|ATTGATCATCGACACTTCGAACGCACTTGCGGCCCCGGGT | >hg38_21_+_8433172_8446622_RNA45SN1_45S_50ntFlanks | 6734 | 6773 | 40 | ATTGATCATCGACACTTCGAACGCACTTGCGGCCCCGGGT | BT20 | 37 | 51 | 31 | 16 | 124 | 30.2 | 47.6 | 18.9 | 14.1 | 18.8 |
| >hg38_21_+_8433172_8446622_RNA45SN1_45S_50ntFlanks@6734.6773.40\|ATTGATCATCGACACTTCGAACGCACTTGCGGCCCCGGGT | >hg38_21_+_8433172_8446622_RNA45SN1_45S_50ntFlanks | 6734 | 6773 | 40 | ATTGATCATCGACACTTCGAACGCACTTGCGGCCCCGGGT | MB231 | 44 | 55 | 20 | 8 | 105 | 25.4 | 38.7 | 9.0 | 8.4 | 10.0 |
| >hg38_21_+_8433172_8446622_RNA45SN1_45S_50ntFlanks@6734.6774.41\|ATTGATCATCGACACTTCGAACGCACTTGCGGCCCCGGGTT | >hg38_21_+_8433172_8446622_RNA45SN1_45S_50ntFlanks | 6734 | 6774 | 41 | ATTGATCATCGACACTTCGAACGCACTTGCGGCCCCGGGTT | BT20 | 23 | 34 | 18 | 11 | 88 | 12.4 | 29.5 | 11.1 | 5.5 | 11.6 |

<table>
<tr><td>>hg38_21_+_8433172_8446622_RNA45SN1_45S_50ntFlanks@6734.6774.41|ATTGATCATCGACACTTCGAACGCACTTGCGGCCCCGGGTT</td><td>>hg38_21_+_8433172_8446622_RNA45SN1_45S_50ntFlanks</td><td>6734</td><td>6774</td><td>41</td><td>ATTGATCATCGACACTTCGAACGCACTTGCGGCCCCGGGTT</td><td>MB231</td><td>31</td><td>33</td><td>18</td><td>8</td><td>78</td><td>10.9</td><td>18.6</td><td>2.1</td><td>8.2</td><td>12.2</td></tr>
<tr><td>>hg38_21_+_8433172_8446622_RNA45SN1_45S_50ntFlanks@6735.6770.36|TTGATCATCGACACTTCGAACGCACTTGCGGCCCCG</td><td>>hg38_21_+_8433172_8446622_RNA45SN1_45S_50ntFlanks</td><td>6735</td><td>6770</td><td>36</td><td>TTGATCATCGACACTTCGAACGCACTTGCGGCCCCG</td><td>MB468</td><td>28</td><td>32</td><td>18</td><td>21</td><td>45</td><td>8.5</td><td>14.2</td><td>7.2</td><td>16.1</td><td>16.2</td></tr>
<tr><td>>hg38_21_+_8433172_8446622_RNA45SN1_45S_50ntFlanks@6735.6771.37|TTGATCATCGACACTTCGAACGCACTTGCGGCCCCGG</td><td>>hg38_21_+_8433172_8446622_RNA45SN1_45S_50ntFlanks</td><td>6735</td><td>6771</td><td>37</td><td>TTGATCATCGACACTTCGAACGCACTTGCGGCCCCGG</td><td>MB468</td><td>21</td><td>31</td><td>8</td><td>12</td><td>60</td><td>6.1</td><td>14.1</td><td>3.2</td><td>9.8</td><td>24.7</td></tr>
<tr><td>>hg38_21_+_8433172_8446622_RNA45SN1_45S_50ntFlanks@6737.6756.20|GATCATCGACACTTCGAACG</td><td>>hg38_21_+_8433172_8446622_RNA45SN1_45S_50ntFlanks</td><td>6737</td><td>6756</td><td>20</td><td>GATCATCGACACTTCGAACG</td><td>MB468</td><td>15</td><td>7</td><td>9</td><td>10</td><td>4</td><td>4.5</td><td>5.3</td><td>4.3</td><td>3.1</td><td>0.6</td></tr>
<tr><td>>hg38_21_+_8433172_8446622_RNA45SN1_45S_50ntFlanks@6737.6770.34|GATCATCGACACTTCGAACGCACTTGCGGCCCCG</td><td>>hg38_21_+_8433172_8446622_RNA45SN1_45S_50ntFlanks</td><td>6737</td><td>6770</td><td>34</td><td>GATCATCGACACTTCGAACGCACTTGCGGCCCCG</td><td>MB468</td><td>55</td><td>47</td><td>27</td><td>34</td><td>81</td><td>21.4</td><td>24.5</td><td>11.8</td><td>29.3</td><td>28.0</td></tr>
<tr><td>>hg38_21_+_8433172_8446622_RNA45SN1_45S_50ntFlanks@6737.6771.35|GATCATCGACACTTCGAACGCACTTGCGGCCCCGG</td><td>>hg38_21_+_8433172_8446622_RNA45SN1_45S_50ntFlanks</td><td>6737</td><td>6771</td><td>35</td><td>GATCATCGACACTTCGAACGCACTTGCGGCCCCGG</td><td>MB468</td><td>35</td><td>38</td><td>10</td><td>19</td><td>88</td><td>11.4</td><td>16.9</td><td>4.6</td><td>18.3</td><td>38.2</td></tr>
<tr><td>>hg38_21_+_8433172_8446622_RNA45SN1_45S_50ntFlanks@6738.6756.19|ATCATCGACACTTCGAACG</td><td>>hg38_21_+_8433172_8446622_RNA45SN1_45S_50ntFlanks</td><td>6738</td><td>6756</td><td>19</td><td>ATCATCGACACTTCGAACG</td><td>MB468</td><td>72</td><td>10</td><td>53</td><td>45</td><td>18</td><td>49.6</td><td>6.8</td><td>45.8</td><td>22.0</td><td>4.6</td></tr>
<tr><td>>hg38_21_+_8433172_8446622_RNA45SN1_45S_50ntFlanks@6738.6758.21|ATCATCGACACTTCGAACGCA</td><td>>hg38_21_+_8433172_8446622_RNA45SN1_45S_50ntFlanks</td><td>6738</td><td>6758</td><td>21</td><td>ATCATCGACACTTCGAACGCA</td><td>MB468</td><td>22</td><td>3</td><td>20</td><td>9</td><td>6</td><td>10.1</td><td>2.4</td><td>8.7</td><td>3.5</td><td>3.2</td></tr>
<tr><td>>hg38_21_+_8433172_8446622_RNA45SN1_45S_50ntFlanks@6738.6760.23|ATCATCGACACTTCGAACGCACT</td><td>>hg38_21_+_8433172_8446622_RNA45SN1_45S_50ntFlanks</td><td>6738</td><td>6760</td><td>23</td><td>ATCATCGACACTTCGAACGCACT</td><td>MB468</td><td>13</td><td>7</td><td>11</td><td>5</td><td>5</td><td>6.1</td><td>3.2</td><td>5.8</td><td>0.5</td><td>1.9</td></tr>
<tr><td>>hg38_21_+_8433172_8446622_RNA45SN1_45S_50ntFlanks@6738.6769.32|ATCATCGACACTTCGAACGCACTTGCGGCCCC</td><td>>hg38_21_+_8433172_8446622_RNA45SN1_45S_50ntFlanks</td><td>6738</td><td>6769</td><td>32</td><td>ATCATCGACACTTCGAACGCACTTGCGGCCCC</td><td>MB468</td><td>14</td><td>9</td><td>10</td><td>11</td><td>18</td><td>3.7</td><td>1.5</td><td>2.3</td><td>7.0</td><td>2.9</td></tr>
<tr><td>>hg38_21_+_8433172_8446622_RNA45SN1_45S_50ntFlanks@6738.6770.33|ATCATCGACACTTCGAACGCACTTGCGGCCCCG</td><td>>hg38_21_+_8433172_8446622_RNA45SN1_45S_50ntFlanks</td><td>6738</td><td>6770</td><td>33</td><td>ATCATCGACACTTCGAACGCACTTGCGGCCCCG</td><td>MB468</td><td>340</td><td>198</td><td>165</td><td>292</td><td>498</td><td>118.3</td><td>84.6</td><td>49.2</td><td>195.1</td><td>83.2</td></tr>
<tr><td>>hg38_21_+_8433172_8446622_RNA45SN1_45S_50ntFlanks@6738.6771.34|ATCATCGACACTTCGAACGCACTTGCGGCCCCGG</td><td>>hg38_21_+_8433172_8446622_RNA45SN1_45S_50ntFlanks</td><td>6738</td><td>6771</td><td>34</td><td>ATCATCGACACTTCGAACGCACTTGCGGCCCCGG</td><td>MB468</td><td>318</td><td>220</td><td>97</td><td>206</td><td>714</td><td>57.4</td><td>68.7</td><td>15.4</td><td>177.3</td><td>155.1</td></tr>
<tr><td>>hg38_21_+_8433172_8446622_RNA45SN1_45S_50ntFlanks@6738.6772.35|ATCATCGACACTTCGAACGCACTTGCGGCCCCGGG</td><td>>hg38_21_+_8433172_8446622_RNA45SN1_45S_50ntFlanks</td><td>6738</td><td>6772</td><td>35</td><td>ATCATCGACACTTCGAACGCACTTGCGGCCCCGGG</td><td>MB468</td><td>65</td><td>59</td><td>20</td><td>51</td><td>210</td><td>23.6</td><td>25.7</td><td>7.9</td><td>50.5</td><td>103.3</td></tr>
<tr><td>>hg38_21_+_8433172_8446622_RNA45SN1_45S_50ntFlanks@6739.6770.32|TCATCGACACTTCGAACGCACTTGCGGCCCCG</td><td>>hg38_21_+_8433172_8446622_RNA45SN1_45S_50ntFlanks</td><td>6739</td><td>6770</td><td>32</td><td>TCATCGACACTTCGAACGCACTTGCGGCCCCG</td><td>MB468</td><td>47</td><td>23</td><td>22</td><td>34</td><td>65</td><td>17.8</td><td>9.9</td><td>8.3</td><td>22.8</td><td>9.8</td></tr>
<tr><td>>hg38_21_+_8433172_8446622_RNA45SN1_45S_50ntFlanks@6739.6771.33|TCATCGACACTTCGAACGCACTTGCGGCCCCGG</td><td>>hg38_21_+_8433172_8446622_RNA45SN1_45S_50ntFlanks</td><td>6739</td><td>6771</td><td>33</td><td>TCATCGACACTTCGAACGCACTTGCGGCCCCGG</td><td>MB468</td><td>43</td><td>25</td><td>12</td><td>26</td><td>85</td><td>8.9</td><td>8.9</td><td>2.8</td><td>21.7</td><td>14.2</td></tr>
<tr><td>>hg38_21_+_8433172_8446622_RNA45SN1_45S_50ntFlanks@6739.6772.34|TCATCGACACTTCGAACGCACTTGCGGCCCCGGG</td><td>>hg38_21_+_8433172_8446622_RNA45SN1_45S_50ntFlanks</td><td>6739</td><td>6772</td><td>34</td><td>TCATCGACACTTCGAACGCACTTGCGGCCCCGGG</td><td>MB468</td><td>12</td><td>10</td><td>3</td><td>7</td><td>32</td><td>4.0</td><td>4.6</td><td>1.7</td><td>6.6</td><td>16.0</td></tr>
<tr><td>>hg38_21_+_8433172_8446622_RNA45SN1_45S_50ntFlanks@6740.6770.31|CATCGACACTTCGAACGCACTTGCGGCCCCG</td><td>>hg38_21_+_8433172_8446622_RNA45SN1_45S_50ntFlanks</td><td>6740</td><td>6770</td><td>31</td><td>CATCGACACTTCGAACGCACTTGCGGCCCCG</td><td>MB468</td><td>177</td><td>86</td><td>81</td><td>144</td><td>224</td><td>50.6</td><td>30.9</td><td>18.9</td><td>67.5</td><td>59.8</td></tr>
<tr><td>>hg38_21_+_8433172_8446622_RNA45SN1_45S_50ntFlanks@6740.6771.32|CATCGACACTTCGAACGCACTTGCGGCCCCGG</td><td>>hg38_21_+_8433172_8446622_RNA45SN1_45S_50ntFlanks</td><td>6740</td><td>6771</td><td>32</td><td>CATCGACACTTCGAACGCACTTGCGGCCCCGG</td><td>MB468</td><td>173</td><td>97</td><td>45</td><td>102</td><td>350</td><td>40.5</td><td>34.9</td><td>7.4</td><td>68.7</td><td>115.5</td></tr>
<tr><td>>hg38_21_+_8433172_8446622_RNA45SN1_45S_50ntFlanks@6740.6772.33|CATCGACACTTCGAACGCACTTGCGGCCCCGGG</td><td>>hg38_21_+_8433172_8446622_RNA45SN1_45S_50ntFlanks</td><td>6740</td><td>6772</td><td>33</td><td>CATCGACACTTCGAACGCACTTGCGGCCCCGGG</td><td>MB468</td><td>41</td><td>35</td><td>10</td><td>25</td><td>122</td><td>13.3</td><td>13.5</td><td>3.6</td><td>22.3</td><td>36.6</td></tr>
<tr><td>>hg38_21_+_8433172_8446622_RNA45SN1_45S_50ntFlanks@6741.6770.30|ATCGACACTTCGAACGCACTTGCGGCCCCG</td><td>>hg38_21_+_8433172_8446622_RNA45SN1_45S_50ntFlanks</td><td>6741</td><td>6770</td><td>30</td><td>ATCGACACTTCGAACGCACTTGCGGCCCCG</td><td>MB468</td><td>194</td><td>75</td><td>134</td><td>149</td><td>160</td><td>71.3</td><td>46.4</td><td>54.8</td><td>42.6</td><td>34.8</td></tr>
<tr><td>>hg38_21_+_8433172_8446622_RNA45SN1_45S_50ntFlanks@6741.6771.31|ATCGACACTTCGAACGCACTTGCGGCCCCGG</td><td>>hg38_21_+_8433172_8446622_RNA45SN1_45S_50ntFlanks</td><td>6741</td><td>6771</td><td>31</td><td>ATCGACACTTCGAACGCACTTGCGGCCCCGG</td><td>MB468</td><td>165</td><td>70</td><td>50</td><td>99</td><td>213</td><td>35.4</td><td>27.5</td><td>12.9</td><td>49.9</td><td>42.5</td></tr>
<tr><td>>hg38_21_+_8433172_8446622_RNA45SN1_45S_50ntFlanks@6741.6772.32|ATCGACACTTCGAACGCACTTGCGGCCCCGGG</td><td>>hg38_21_+_8433172_8446622_RNA45SN1_45S_50ntFlanks</td><td>6741</td><td>6772</td><td>32</td><td>ATCGACACTTCGAACGCACTTGCGGCCCCGGG</td><td>MB468</td><td>43</td><td>28</td><td>10</td><td>27</td><td>97</td><td>14.4</td><td>15.1</td><td>4.2</td><td>22.8</td><td>43.4</td></tr>
<tr><td>>hg38_21_+_8433172_8446622_RNA45SN1_45S_50ntFlanks@6742.6770.29|TCGACACTTCGAACGCACTTGCGGCCCCG</td><td>>hg38_21_+_8433172_8446622_RNA45SN1_45S_50ntFlanks</td><td>6742</td><td>6770</td><td>29</td><td>TCGACACTTCGAACGCACTTGCGGCCCCG</td><td>MB468</td><td>41</td><td>12</td><td>44</td><td>39</td><td>31</td><td>17.2</td><td>5.6</td><td>23.5</td><td>11.2</td><td>10.6</td></tr>
<tr><td>>hg38_21_+_8433172_8446622_RNA45SN1_45S_50ntFlanks@6742.6771.30|TCGACACTTCGAACGCACTTGCGGCCCCGG</td><td>>hg38_21_+_8433172_8446622_RNA45SN1_45S_50ntFlanks</td><td>6742</td><td>6771</td><td>30</td><td>TCGACACTTCGAACGCACTTGCGGCCCCGG</td><td>MB468</td><td>38</td><td>13</td><td>15</td><td>22</td><td>38</td><td>9.0</td><td>5.6</td><td>5.6</td><td>10.6</td><td>11.3</td></tr>
<tr><td>>hg38_21_+_8433172_8446622_RNA45SN1_45S_50ntFlanks@6742.6772.31|TCGACACTTCGAACGCACTTGCGGCCCCGGG</td><td>>hg38_21_+_8433172_8446622_RNA45SN1_45S_50ntFlanks</td><td>6742</td><td>6772</td><td>31</td><td>TCGACACTTCGAACGCACTTGCGGCCCCGGG</td><td>MB468</td><td>10</td><td>5</td><td>3</td><td>6</td><td>17</td><td>4.4</td><td>3.0</td><td>1.4</td><td>5.1</td><td>5.7</td></tr>
<tr><td>>hg38_21_+_8433172_8446622_RNA45SN1_45S_50ntFlanks@6743.6770.28|CGACACTTCGAACGCACTTGCGGCCCCG</td><td>>hg38_21_+_8433172_8446622_RNA45SN1_45S_50ntFlanks</td><td>6743</td><td>6770</td><td>28</td><td>CGACACTTCGAACGCACTTGCGGCCCCG</td><td>MB468</td><td>170</td><td>78</td><td>193</td><td>167</td><td>137</td><td>87.9</td><td>58.5</td><td>145.0</td><td>67.2</td><td>9.7</td></tr>
<tr><td>>hg38_21_+_8433172_8446622_RNA45SN1_45S_50ntFlanks@6743.6771.29|CGACACTTCGAACGCACTTGCGGCCCCGG</td><td>>hg38_21_+_8433172_8446622_RNA45SN1_45S_50ntFlanks</td><td>6743</td><td>6771</td><td>29</td><td>CGACACTTCGAACGCACTTGCGGCCCCGG</td><td>MB468</td><td>185</td><td>65</td><td>82</td><td>131</td><td>185</td><td>69.5</td><td>26.8</td><td>51.7</td><td>90.8</td><td>37.3</td></tr>
<tr><td>>hg38_21_+_8433172_8446622_RNA45SN1_45S_50ntFlanks@6743.6772.30|CGACACTTCGAACGCACTTGCGGCCCCGGG</td><td>>hg38_21_+_8433172_8446622_RNA45SN1_45S_50ntFlanks</td><td>6743</td><td>6772</td><td>30</td><td>CGACACTTCGAACGCACTTGCGGCCCCGGG</td><td>MB468</td><td>36</td><td>21</td><td>11</td><td>27</td><td>74</td><td>16.6</td><td>14.2</td><td>9.5</td><td>25.7</td><td>50.7</td></tr>
<tr><td>>hg38_21_+_8433172_8446622_RNA45SN1_45S_50ntFlanks@6744.6770.27|GACACTTCGAACGCACTTGCGGCCCCG</td><td>>hg38_21_+_8433172_8446622_RNA45SN1_45S_50ntFlanks</td><td>6744</td><td>6770</td><td>27</td><td>GACACTTCGAACGCACTTGCGGCCCCG</td><td>MB468</td><td>159</td><td>82</td><td>217</td><td>123</td><td>120</td><td>67.0</td><td>56.8</td><td>129.7</td><td>61.9</td><td>13.0</td></tr>
<tr><td>>hg38_21_+_8433172_8446622_RNA45SN1_45S_50ntFlanks@6744.6771.28|GACACTTCGAACGCACTTGCGGCCCCGG</td><td>>hg38_21_+_8433172_8446622_RNA45SN1_45S_50ntFlanks</td><td>6744</td><td>6771</td><td>28</td><td>GACACTTCGAACGCACTTGCGGCCCCGG</td><td>MB468</td><td>104</td><td>36</td><td>65</td><td>100</td><td>96</td><td>37.0</td><td>12.8</td><td>28.4</td><td>76.0</td><td>25.1</td></tr>
<tr><td>>hg38_21_+_8433172_8446622_RNA45SN1_45S_50ntFlanks@6744.6772.29|GACACTTCGAACGCACTTGCGGCCCCGGG</td><td>>hg38_21_+_8433172_8446622_RNA45SN1_45S_50ntFlanks</td><td>6744</td><td>6772</td><td>29</td><td>GACACTTCGAACGCACTTGCGGCCCCGGG</td><td>MB468</td><td>15</td><td>7</td><td>8</td><td>19</td><td>24</td><td>8.0</td><td>3.6</td><td>4.5</td><td>19.6</td><td>14.6</td></tr>
<tr><td>>hg38_21_+_8433172_8446622_RNA45SN1_45S_50ntFlanks@6745.6762.18|ACACTTCGAACGCACTTG</td><td>>hg38_21_+_8433172_8446622_RNA45SN1_45S_50ntFlanks</td><td>6745</td><td>6762</td><td>18</td><td>ACACTTCGAACGCACTTG</td><td>MB468</td><td>33</td><td>7</td><td>20</td><td>10</td><td>10</td><td>13.5</td><td>3.4</td><td>5.8</td><td>2.6</td><td>4.7</td></tr>
<tr><td>>hg38_21_+_8433172_8446622_RNA45SN1_45S_50ntFlanks@6745.6769.25|ACACTTCGAACGCACTTGCGGCCCC</td><td>>hg38_21_+_8433172_8446622_RNA45SN1_45S_50ntFlanks</td><td>6745</td><td>6769</td><td>25</td><td>ACACTTCGAACGCACTTGCGGCCCC</td><td>MB468</td><td>22</td><td>12</td><td>31</td><td>18</td><td>27</td><td>6.8</td><td>3.9</td><td>11.8</td><td>7.5</td><td>11.6</td></tr>
<tr><td>>hg38_21_+_8433172_8446622_RNA45SN1_45S_50ntFlanks@6745.6770.26|ACACTTCGAACGCACTTGCGGCCCCG</td><td>>hg38_21_+_8433172_8446622_RNA45SN1_45S_50ntFlanks</td><td>6745</td><td>6770</td><td>26</td><td>ACACTTCGAACGCACTTGCGGCCCCG</td><td>MB468</td><td>424</td><td>174</td><td>486</td><td>586</td><td>575</td><td>75.4</td><td>70.8</td><td>149.7</td><td>176.6</td><td>231.0</td></tr>
<tr><td>>hg38_21_+_8433172_8446622_RNA45SN1_45S_50ntFlanks@6745.6771.27|ACACTTCGAACGCACTTGCGGCCCCGG</td><td>>hg38_21_+_8433172_8446622_RNA45SN1_45S_50ntFlanks</td><td>6745</td><td>6771</td><td>27</td><td>ACACTTCGAACGCACTTGCGGCCCCGG</td><td>MB468</td><td>386</td><td>110</td><td>208</td><td>342</td><td>539</td><td>95.0</td><td>24.3</td><td>65.6</td><td>114.4</td><td>344.1</td></tr>
<tr><td>>hg38_21_+_8433172_8446622_RNA45SN1_45S_50ntFlanks@6745.6772.28|ACACTTCGAACGCACTTGCGGCCCCGGG</td><td>>hg38_21_+_8433172_8446622_RNA45SN1_45S_50ntFlanks</td><td>6745</td><td>6772</td><td>28</td><td>ACACTTCGAACGCACTTGCGGCCCCGGG</td><td>MB468</td><td>48</td><td>18</td><td>25</td><td>67</td><td>81</td><td>12.8</td><td>5.4</td><td>5.6</td><td>41.3</td><td>18.9</td></tr>
<tr><td>>hg38_21_+_8433172_8446622_RNA45SN1_45S_50ntFlanks@6746.6770.25|CACTTCGAACGCACTTGCGGCCCCG</td><td>>hg38_21_+_8433172_8446622_RNA45SN1_45S_50ntFlanks</td><td>6746</td><td>6770</td><td>25</td><td>CACTTCGAACGCACTTGCGGCCCCG</td><td>MB468</td><td>118</td><td>48</td><td>138</td><td>153</td><td>140</td><td>35.9</td><td>20.7</td><td>51.4</td><td>40.5</td><td>46.8</td></tr>
<tr><td>>hg38_21_+_8433172_8446622_RNA45SN1_45S_50ntFlanks@6746.6771.26|CACTTCGAACGCACTTGCGGCCCCGG</td><td>>hg38_21_+_8433172_8446622_RNA45SN1_45S_50ntFlanks</td><td>6746</td><td>6771</td><td>26</td><td>CACTTCGAACGCACTTGCGGCCCCGG</td><td>MB468</td><td>86</td><td>26</td><td>53</td><td>93</td><td>108</td><td>18.4</td><td>6.7</td><td>10.2</td><td>25.0</td><td>41.0</td></tr>
<tr><td>>hg38_21_+_8433172_8446622_RNA45SN1_45S_50ntFlanks@6747.6770.24|ACTTCGAACGCACTTGCGGCCCCG</td><td>>hg38_21_+_8433172_8446622_RNA45SN1_45S_50ntFlanks</td><td>6747</td><td>6770</td><td>24</td><td>ACTTCGAACGCACTTGCGGCCCCG</td><td>MB468</td><td>57</td><td>21</td><td>66</td><td>88</td><td>61</td><td>13.7</td><td>9.3</td><td>30.8</td><td>30.1</td><td>20.3</td></tr>
<tr><td>>hg38_21_+_8433172_8446622_RNA45SN1_45S_50ntFlanks@6747.6771.25|ACTTCGAACGCACTTGCGGCCCCGG</td><td>>hg38_21_+_8433172_8446622_RNA45SN1_45S_50ntFlanks</td><td>6747</td><td>6771</td><td>25</td><td>ACTTCGAACGCACTTGCGGCCCCGG</td><td>MB468</td><td>36</td><td>10</td><td>25</td><td>60</td><td>41</td><td>7.3</td><td>3.4</td><td>6.6</td><td>15.4</td><td>18.4</td></tr>
<tr><td>>hg38_21_+_8433172_8446622_RNA45SN1_45S_50ntFlanks@6748.6770.23|CTTCGAACGCACTTGCGGCCCCG</td><td>>hg38_21_+_8433172_8446622_RNA45SN1_45S_50ntFlanks</td><td>6748</td><td>6770</td><td>23</td><td>CTTCGAACGCACTTGCGGCCCCG</td><td>MB468</td><td>34</td><td>11</td><td>31</td><td>69</td><td>45</td><td>9.6</td><td>4.3</td><td>12.0</td><td>19.9</td><td>14.2</td></tr>
<tr><td>>hg38_21_+_8433172_8446622_RNA45SN1_45S_50ntFlanks@6748.6771.24|CTTCGAACGCACTTGCGGCCCCGG</td><td>>hg38_21_+_8433172_8446622_RNA45SN1_45S_50ntFlanks</td><td>6748</td><td>6771</td><td>24</td><td>CTTCGAACGCACTTGCGGCCCCGG</td><td>MB468</td><td>16</td><td>4</td><td>11</td><td>45</td><td>21</td><td>4.6</td><td>1.1</td><td>2.3</td><td>11.1</td><td>8.3</td></tr>
<tr><td>>hg38_21_+_8433172_8446622_RNA45SN1_45S_50ntFlanks@6751.6770.20|CGAACGCACTTGCGGCCCCG</td><td>>hg38_21_+_8433172_8446622_RNA45SN1_45S_50ntFlanks</td><td>6751</td><td>6770</td><td>20</td><td>CGAACGCACTTGCGGCCCCG</td><td>MB468</td><td>11</td><td>2</td><td>11</td><td>44</td><td>12</td><td>1.4</td><td>0.8</td><td>5.0</td><td>25.9</td><td>4.9</td></tr>
<tr><td>>hg38_21_+_8433172_8446622_RNA45SN1_45S_50ntFlanks@6753.6770.18|AACGCACTTGCGGCCCCG</td><td>>hg38_21_+_8433172_8446622_RNA45SN1_45S_50ntFlanks</td><td>6753</td><td>6770</td><td>18</td><td>AACGCACTTGCGGCCCCG</td><td>MB468</td><td>11</td><td>3</td><td>13</td><td>19</td><td>10</td><td>1.8</td><td>1.9</td><td>4.0</td><td>12.3</td><td>5.6</td></tr>
<tr><td>>hg38_21_+_8433172_8446622_RNA45SN1_45S_50ntFlanks@6758.6806.49|ACTTGCGGCCCCGGGTTCCTCCCGGGGCTACGCCTGTCTGAGCGTCGCT</td><td>>hg38_21_+_8433172_8446622_RNA45SN1_45S_50ntFlanks</td><td>6758</td><td>6806</td><td>49</td><td>ACTTGCGGCCCCGGGTTCCTCCCGGGGCTACGCCTGTCTGAGCGTCGCT</td><td>MB231</td><td>13</td><td>2</td><td>14</td><td>0</td><td>10</td><td>11.6</td><td>0.8</td><td>11.4</td><td>0.7</td><td>16.6</td></tr>
<tr><td>>hg38_21_+_8433172_8446622_RNA45SN1_45S_50ntFlanks@6759.6806.48|CTTGCGGCCCCGGGTTCCTCCCGGGGCTACGCCTGTCTGAGCGTCGCT</td><td>>hg38_21_+_8433172_8446622_RNA45SN1_45S_50ntFlanks</td><td>6759</td><td>6806</td><td>48</td><td>CTTGCGGCCCCGGGTTCCTCCCGGGGCTACGCCTGTCTGAGCGTCGCT</td><td>MB231</td><td>20</td><td>5</td><td>31</td><td>1</td><td>23</td><td>11.2</td><td>1.4</td><td>25.4</td><td>0.9</td><td>20.2</td></tr>
<tr><td>>hg38_21_+_8433172_8446622_RNA45SN1_45S_50ntFlanks@6760.6806.47|TTGCGGCCCCGGGTTCCTCCCGGGGCTACGCCTGTCTGAGCGTCGCT</td><td>>hg38_21_+_8433172_8446622_RNA45SN1_45S_50ntFlanks</td><td>6760</td><td>6806</td><td>47</td><td>TTGCGGCCCCGGGTTCCTCCCGGGGCTACGCCTGTCTGAGCGTCGCT</td><td>MB231</td><td>19</td><td>5</td><td>30</td><td>1</td><td>26</td><td>11.5</td><td>1.3</td><td>23.9</td><td>1.4</td><td>22.8</td></tr>
<tr><td>>hg38_21_+_8433172_8446622_RNA45SN1_45S_50ntFlanks@6761.6806.46|TGCGGCCCCGGGTTCCTCCCGGGGCTACGCCTGTCTGAGCGTCGCT</td><td>>hg38_21_+_8433172_8446622_RNA45SN1_45S_50ntFlanks</td><td>6761</td><td>6806</td><td>46</td><td>TGCGGCCCCGGGTTCCTCCCGGGGCTACGCCTGTCTGAGCGTCGCT</td><td>MB231</td><td>24</td><td>6</td><td>27</td><td>1</td><td>30</td><td>24.4</td><td>2.7</td><td>25.2</td><td>0.5</td><td>13.6</td></tr>
<tr><td>>hg38_21_+_8433172_8446622_RNA45SN1_45S_50ntFlanks@6762.6806.45|GCGGCCCCGGGTTCCTCCCGGGGCTACGCCTGTCTGAGCGTCGCT</td><td>>hg38_21_+_8433172_8446622_RNA45SN1_45S_50ntFlanks</td><td>6762</td><td>6806</td><td>45</td><td>GCGGCCCCGGGTTCCTCCCGGGGCTACGCCTGTCTGAGCGTCGCT</td><td>MB231</td><td>18</td><td>5</td><td>20</td><td>1</td><td>13</td><td>21.1</td><td>3.3</td><td>19.9</td><td>0.7</td><td>12.2</td></tr>
<tr><td>>hg38_21_+_8433172_8446622_RNA45SN1_45S_50ntFlanks@6763.6806.44|CGGCCCCGGGTTCCTCCCGGGGCTACGCCTGTCTGAGCGTCGCT</td><td>>hg38_21_+_8433172_8446622_RNA45SN1_45S_50ntFlanks</td><td>6763</td><td>6806</td><td>44</td><td>CGGCCCCGGGTTCCTCCCGGGGCTACGCCTGTCTGAGCGTCGCT</td><td>MB231</td><td>11</td><td>4</td><td>7</td><td>1</td><td>15</td><td>11.7</td><td>2.1</td><td>5.6</td><td>0.8</td><td>1.8</td></tr>
</table>

| | | | | | | | | | | | | | | | |
|---|---|---|---|---|---|---|---|---|---|---|---|---|---|---|---|
| >hg38_21_+_8433172_8446622_RNA45SN1_45S_50ntFlanks@6771.6799.29\|GGTTCCTCCCGGGGCTACGCCTGTCTGAG | >hg38_21_+_8433172_8446622_RNA45SN1_45S_50ntFlanks | 6771 | 6799 | 29 | GGTTCCTCCCGGGGCTACGCCTGTCTGAG | MB468 | 19 | 9 | 12 | 35 | 43 | 7.3 | 1.3 | 4.2 | 22.4 | 16.6 |
| >hg38_21_+_8433172_8446622_RNA45SN1_45S_50ntFlanks@6771.6806.36\|GGTTCCTCCCGGGGCTACGCCTGTCTGAGCGTCGCT | >hg38_21_+_8433172_8446622_RNA45SN1_45S_50ntFlanks | 6771 | 6806 | 36 | GGTTCCTCCCGGGGCTACGCCTGTCTGAGCGTCGCT | MB468 | 32 | 18 | 13 | 83 | 148 | 8.7 | 9.6 | 4.6 | 33.4 | 89.4 |
| >hg38_21_+_8433172_8446622_RNA45SN1_45S_50ntFlanks@6772.6793.22\|GTTCCTCCCGGGGCTACGCCTG | >hg38_21_+_8433172_8446622_RNA45SN1_45S_50ntFlanks | 6772 | 6793 | 22 | GTTCCTCCCGGGGCTACGCCTG | MB468 | 20 | 20 | 15 | 3 | 30 | 11.4 | 10.6 | 3.1 | 2.1 | 18.1 |
| >hg38_21_+_8433172_8446622_RNA45SN1_45S_50ntFlanks@6772.6797.26\|GTTCCTCCCGGGGCTACGCCTGTCTG | >hg38_21_+_8433172_8446622_RNA45SN1_45S_50ntFlanks | 6772 | 6797 | 26 | GTTCCTCCCGGGGCTACGCCTGTCTG | MB468 | 25 | 17 | 17 | 13 | 41 | 9.3 | 4.2 | 2.9 | 9.2 | 13.3 |
| >hg38_21_+_8433172_8446622_RNA45SN1_45S_50ntFlanks@6772.6799.28\|GTTCCTCCCGGGGCTACGCCTGTCTGAG | >hg38_21_+_8433172_8446622_RNA45SN1_45S_50ntFlanks | 6772 | 6799 | 28 | GTTCCTCCCGGGGCTACGCCTGTCTGAG | MB468 | 109 | 55 | 56 | 40 | 187 | 42.4 | 7.7 | 11.9 | 16.1 | 55.0 |
| >hg38_21_+_8433172_8446622_RNA45SN1_45S_50ntFlanks@6772.6803.32\|GTTCCTCCCGGGGCTACGCCTGTCTGAGCGTC | >hg38_21_+_8433172_8446622_RNA45SN1_45S_50ntFlanks | 6772 | 6803 | 32 | GTTCCTCCCGGGGCTACGCCTGTCTGAGCGTC | MB468 | 45 | 13 | 17 | 71 | 108 | 11.4 | 2.6 | 3.0 | 9.5 | 52.9 |
| >hg38_21_+_8433172_8446622_RNA45SN1_45S_50ntFlanks@6772.6806.35\|GTTCCTCCCGGGGCTACGCCTGTCTGAGCGTCGCT | >hg38_21_+_8433172_8446622_RNA45SN1_45S_50ntFlanks | 6772 | 6806 | 35 | GTTCCTCCCGGGGCTACGCCTGTCTGAGCGTCGCT | MB231 | 17 | 11 | 22 | 9 | 38 | 14.8 | 3.7 | 4.2 | 3.3 | 25.9 |
| >hg38_21_+_8433172_8446622_RNA45SN1_45S_50ntFlanks@6772.6806.35\|GTTCCTCCCGGGGCTACGCCTGTCTGAGCGTCGCT | >hg38_21_+_8433172_8446622_RNA45SN1_45S_50ntFlanks | 6772 | 6806 | 35 | GTTCCTCCCGGGGCTACGCCTGTCTGAGCGTCGCT | MB468 | 93 | 48 | 25 | 117 | 362 | 27.3 | 23.5 | 7.5 | 59.8 | 196.6 |
| >hg38_21_+_8433172_8446622_RNA45SN1_45S_50ntFlanks@6772.6807.36\|GTTCCTCCCGGGGCTACGCCTGTCTGAGCGTCGCTT | >hg38_21_+_8433172_8446622_RNA45SN1_45S_50ntFlanks | 6772 | 6807 | 36 | GTTCCTCCCGGGGCTACGCCTGTCTGAGCGTCGCTT | MB468 | 42 | 41 | 12 | 24 | 200 | 11.7 | 20.2 | 4.6 | 9.8 | 42.0 |
| >hg38_21_+_8433172_8446622_RNA45SN1_45S_50ntFlanks@6773.6793.21\|TTCCTCCCGGGGCTACGCCTG | >hg38_21_+_8433172_8446622_RNA45SN1_45S_50ntFlanks | 6773 | 6793 | 21 | TTCCTCCCGGGGCTACGCCTG | MB468 | 44 | 44 | 30 | 2 | 42 | 22.3 | 26.4 | 9.2 | 1.3 | 17.3 |
| >hg38_21_+_8433172_8446622_RNA45SN1_45S_50ntFlanks@6773.6797.25\|TTCCTCCCGGGGCTACGCCTGTCTG | >hg38_21_+_8433172_8446622_RNA45SN1_45S_50ntFlanks | 6773 | 6797 | 25 | TTCCTCCCGGGGCTACGCCTGTCTG | MB468 | 27 | 16 | 19 | 8 | 35 | 10.7 | 4.7 | 6.5 | 4.6 | 13.0 |
| >hg38_21_+_8433172_8446622_RNA45SN1_45S_50ntFlanks@6773.6799.27\|TTCCTCCCGGGGCTACGCCTGTCTGAG | >hg38_21_+_8433172_8446622_RNA45SN1_45S_50ntFlanks | 6773 | 6799 | 27 | TTCCTCCCGGGGCTACGCCTGTCTGAG | MB468 | 53 | 19 | 28 | 20 | 69 | 16.0 | 2.4 | 8.2 | 7.4 | 23.8 |
| >hg38_21_+_8433172_8446622_RNA45SN1_45S_50ntFlanks@6773.6802.30\|TTCCTCCCGGGGCTACGCCTGTCTGAGCGT | >hg38_21_+_8433172_8446622_RNA45SN1_45S_50ntFlanks | 6773 | 6802 | 30 | TTCCTCCCGGGGCTACGCCTGTCTGAGCGT | MB468 | 20 | 16 | 11 | 5 | 29 | 9.4 | 4.4 | 4.8 | 0.8 | 13.9 |
| >hg38_21_+_8433172_8446622_RNA45SN1_45S_50ntFlanks@6773.6803.31\|TTCCTCCCGGGGCTACGCCTGTCTGAGCGTC | >hg38_21_+_8433172_8446622_RNA45SN1_45S_50ntFlanks | 6773 | 6803 | 31 | TTCCTCCCGGGGCTACGCCTGTCTGAGCGTC | MB468 | 158 | 59 | 65 | 64 | 297 | 62.3 | 18.1 | 26.8 | 12.8 | 79.6 |
| >hg38_21_+_8433172_8446622_RNA45SN1_45S_50ntFlanks@6773.6806.34\|TTCCTCCCGGGGCTACGCCTGTCTGAGCGTCGCT | >hg38_21_+_8433172_8446622_RNA45SN1_45S_50ntFlanks | 6773 | 6806 | 34 | TTCCTCCCGGGGCTACGCCTGTCTGAGCGTCGCT | BT20 | 30 | 34 | 31 | 16 | 97 | 11.2 | 4.6 | 13.0 | 9.1 | 69.8 |
| >hg38_21_+_8433172_8446622_RNA45SN1_45S_50ntFlanks@6773.6806.34\|TTCCTCCCGGGGCTACGCCTGTCTGAGCGTCGCT | >hg38_21_+_8433172_8446622_RNA45SN1_45S_50ntFlanks | 6773 | 6806 | 34 | TTCCTCCCGGGGCTACGCCTGTCTGAGCGTCGCT | MB231 | 24 | 15 | 20 | 8 | 64 | 14.3 | 2.3 | 6.5 | 4.0 | 27.6 |
| >hg38_21_+_8433172_8446622_RNA45SN1_45S_50ntFlanks@6773.6806.34\|TTCCTCCCGGGGCTACGCCTGTCTGAGCGTCGCT | >hg38_21_+_8433172_8446622_RNA45SN1_45S_50ntFlanks | 6773 | 6806 | 34 | TTCCTCCCGGGGCTACGCCTGTCTGAGCGTCGCT | MB468 | 75 | 22 | 18 | 62 | 199 | 30.5 | 10.7 | 9.0 | 32.1 | 157.1 |
| >hg38_21_+_8433172_8446622_RNA45SN1_45S_50ntFlanks@6773.6807.35\|TTCCTCCCGGGGCTACGCCTGTCTGAGCGTCGCTT | >hg38_21_+_8433172_8446622_RNA45SN1_45S_50ntFlanks | 6773 | 6807 | 35 | TTCCTCCCGGGGCTACGCCTGTCTGAGCGTCGCTT | MB468 | 45 | 24 | 10 | 14 | 132 | 15.6 | 13.0 | 5.3 | 6.5 | 38.5 |
| >hg38_21_+_8433172_8446622_RNA45SN1_45S_50ntFlanks@6774.6806.33\|TCCTCCCGGGGCTACGCCTGTCTGAGCGTCGCT | >hg38_21_+_8433172_8446622_RNA45SN1_45S_50ntFlanks | 6774 | 6806 | 33 | TCCTCCCGGGGCTACGCCTGTCTGAGCGTCGCT | BT20 | 132 | 137 | 111 | 30 | 352 | 56.5 | 53.3 | 75.0 | 11.9 | 366.3 |
| >hg38_21_+_8433172_8446622_RNA45SN1_45S_50ntFlanks@6774.6806.33\|TCCTCCCGGGGCTACGCCTGTCTGAGCGTCGCT | >hg38_21_+_8433172_8446622_RNA45SN1_45S_50ntFlanks | 6774 | 6806 | 33 | TCCTCCCGGGGCTACGCCTGTCTGAGCGTCGCT | MB231 | 51 | 32 | 33 | 14 | 159 | 10.1 | 9.7 | 9.6 | 8.0 | 80.2 |
| >hg38_21_+_8433172_8446622_RNA45SN1_45S_50ntFlanks@6774.6807.34\|TCCTCCCGGGGCTACGCCTGTCTGAGCGTCGCTT | >hg38_21_+_8433172_8446622_RNA45SN1_45S_50ntFlanks | 6774 | 6807 | 34 | TCCTCCCGGGGCTACGCCTGTCTGAGCGTCGCTT | BT20 | 11 | 13 | 10 | 3 | 38 | 7.1 | 5.1 | 7.4 | 4.0 | 26.5 |
| >hg38_21_+_8433172_8446622_RNA45SN1_45S_50ntFlanks@6775.6803.29\|CCTCCCGGGGCTACGCCTGTCTGAGCGTC | >hg38_21_+_8433172_8446622_RNA45SN1_45S_50ntFlanks | 6775 | 6803 | 29 | CCTCCCGGGGCTACGCCTGTCTGAGCGTC | MB468 | 31 | 12 | 16 | 12 | 71 | 15.1 | 1.8 | 4.6 | 4.3 | 27.6 |
| >hg38_21_+_8433172_8446622_RNA45SN1_45S_50ntFlanks@6775.6806.32\|CCTCCCGGGGCTACGCCTGTCTGAGCGTCGCT | >hg38_21_+_8433172_8446622_RNA45SN1_45S_50ntFlanks | 6775 | 6806 | 32 | CCTCCCGGGGCTACGCCTGTCTGAGCGTCGCT | BT20 | 27 | 25 | 21 | 14 | 53 | 11.7 | 9.4 | 6.4 | 3.4 | 26.7 |
| >hg38_21_+_8433172_8446622_RNA45SN1_45S_50ntFlanks@6775.6806.32\|CCTCCCGGGGCTACGCCTGTCTGAGCGTCGCT | >hg38_21_+_8433172_8446622_RNA45SN1_45S_50ntFlanks | 6775 | 6806 | 32 | CCTCCCGGGGCTACGCCTGTCTGAGCGTCGCT | MB231 | 21 | 12 | 13 | 11 | 56 | 11.9 | 2.8 | 1.1 | 2.5 | 17.7 |
| >hg38_21_+_8433172_8446622_RNA45SN1_45S_50ntFlanks@6776.6802.27\|CTCCCGGGGCTACGCCTGTCTGAGCGT | >hg38_21_+_8433172_8446622_RNA45SN1_45S_50ntFlanks | 6776 | 6802 | 27 | CTCCCGGGGCTACGCCTGTCTGAGCGT | BT20 | 12 | 15 | 11 | 41 | 7 | 9.1 | 6.5 | 3.6 | 70.6 | 12.6 |
| >hg38_21_+_8433172_8446622_RNA45SN1_45S_50ntFlanks@6776.6802.27\|CTCCCGGGGCTACGCCTGTCTGAGCGT | >hg38_21_+_8433172_8446622_RNA45SN1_45S_50ntFlanks | 6776 | 6802 | 27 | CTCCCGGGGCTACGCCTGTCTGAGCGT | MB231 | 25 | 37 | 28 | 1 | 47 | 18.2 | 20.2 | 19.0 | 0.5 | 54.8 |
| >hg38_21_+_8433172_8446622_RNA45SN1_45S_50ntFlanks@6776.6802.27\|CTCCCGGGGCTACGCCTGTCTGAGCGT | >hg38_21_+_8433172_8446622_RNA45SN1_45S_50ntFlanks | 6776 | 6802 | 27 | CTCCCGGGGCTACGCCTGTCTGAGCGT | MB468 | 11 | 15 | 9 | 1 | 10 | 5.7 | 5.6 | 6.3 | 0.9 | 7.3 |
| >hg38_21_+_8433172_8446622_RNA45SN1_45S_50ntFlanks@6776.6803.28\|CTCCCGGGGCTACGCCTGTCTGAGCGTC | >hg38_21_+_8433172_8446622_RNA45SN1_45S_50ntFlanks | 6776 | 6803 | 28 | CTCCCGGGGCTACGCCTGTCTGAGCGTC | MB468 | 75 | 45 | 44 | 33 | 185 | 35.7 | 16.2 | 10.6 | 18.6 | 90.1 |
| >hg38_21_+_8433172_8446622_RNA45SN1_45S_50ntFlanks@6776.6806.31\|CTCCCGGGGCTACGCCTGTCTGAGCGTCGCT | >hg38_21_+_8433172_8446622_RNA45SN1_45S_50ntFlanks | 6776 | 6806 | 31 | CTCCCGGGGCTACGCCTGTCTGAGCGTCGCT | BT20 | 62 | 61 | 61 | 31 | 245 | 38.3 | 19.8 | 40.0 | 15.0 | 192.7 |
| >hg38_21_+_8433172_8446622_RNA45SN1_45S_50ntFlanks@6776.6806.31\|CTCCCGGGGCTACGCCTGTCTGAGCGTCGCT | >hg38_21_+_8433172_8446622_RNA45SN1_45S_50ntFlanks | 6776 | 6806 | 31 | CTCCCGGGGCTACGCCTGTCTGAGCGTCGCT | MB231 | 98 | 80 | 59 | 23 | 207 | 88.1 | 9.2 | 13.9 | 8.0 | 138.6 |
| >hg38_21_+_8433172_8446622_RNA45SN1_45S_50ntFlanks@6776.6806.31\|CTCCCGGGGCTACGCCTGTCTGAGCGTCGCT | >hg38_21_+_8433172_8446622_RNA45SN1_45S_50ntFlanks | 6776 | 6806 | 31 | CTCCCGGGGCTACGCCTGTCTGAGCGTCGCT | MB468 | 38 | 18 | 10 | 27 | 105 | 13.2 | 7.4 | 3.4 | 15.0 | 30.2 |
| >hg38_21_+_8433172_8446622_RNA45SN1_45S_50ntFlanks@6776.6807.32\|CTCCCGGGGCTACGCCTGTCTGAGCGTCGCTT | >hg38_21_+_8433172_8446622_RNA45SN1_45S_50ntFlanks | 6776 | 6807 | 32 | CTCCCGGGGCTACGCCTGTCTGAGCGTCGCTT | MB468 | 13 | 8 | 3 | 4 | 44 | 5.8 | 4.4 | 1.1 | 1.4 | 19.4 |
| >hg38_21_+_8433172_8446622_RNA45SN1_45S_50ntFlanks@6777.6802.26\|TCCCGGGGCTACGCCTGTCTGAGCGT | >hg38_21_+_8433172_8446622_RNA45SN1_45S_50ntFlanks | 6777 | 6802 | 26 | TCCCGGGGCTACGCCTGTCTGAGCGT | MB231 | 17 | 14 | 15 | 2 | 27 | 13.4 | 11.3 | 10.8 | 1.9 | 27.1 |
| >hg38_21_+_8433172_8446622_RNA45SN1_45S_50ntFlanks@6777.6806.30\|TCCCGGGGCTACGCCTGTCTGAGCGTCGCT | >hg38_21_+_8433172_8446622_RNA45SN1_45S_50ntFlanks | 6777 | 6806 | 30 | TCCCGGGGCTACGCCTGTCTGAGCGTCGCT | BT20 | 18 | 18 | 18 | 9 | 81 | 9.4 | 1.1 | 11.2 | 2.6 | 51.9 |
| >hg38_21_+_8433172_8446622_RNA45SN1_45S_50ntFlanks@6777.6806.30\|TCCCGGGGCTACGCCTGTCTGAGCGTCGCT | >hg38_21_+_8433172_8446622_RNA45SN1_45S_50ntFlanks | 6777 | 6806 | 30 | TCCCGGGGCTACGCCTGTCTGAGCGTCGCT | MB231 | 29 | 22 | 22 | 15 | 70 | 27.1 | 6.9 | 8.8 | 7.9 | 51.1 |
| >hg38_21_+_8433172_8446622_RNA45SN1_45S_50ntFlanks@6778.6802.25\|CCCGGGGCTACGCCTGTCTGAGCGT | >hg38_21_+_8433172_8446622_RNA45SN1_45S_50ntFlanks | 6778 | 6802 | 25 | CCCGGGGCTACGCCTGTCTGAGCGT | BT20 | 45 | 48 | 39 | 18 | 104 | 43.7 | 53.2 | 37.2 | 22.6 | 87.1 |
| >hg38_21_+_8433172_8446622_RNA45SN1_45S_50ntFlanks@6778.6802.25\|CCCGGGGCTACGCCTGTCTGAGCGT | >hg38_21_+_8433172_8446622_RNA45SN1_45S_50ntFlanks | 6778 | 6802 | 25 | CCCGGGGCTACGCCTGTCTGAGCGT | MB231 | 103 | 112 | 164 | 6 | 171 | 86.6 | 95.3 | 137.1 | 6.5 | 192.8 |
| >hg38_21_+_8433172_8446622_RNA45SN1_45S_50ntFlanks@6778.6803.26\|CCCGGGGCTACGCCTGTCTGAGCGTC | >hg38_21_+_8433172_8446622_RNA45SN1_45S_50ntFlanks | 6778 | 6803 | 26 | CCCGGGGCTACGCCTGTCTGAGCGTC | MB231 | 15 | 11 | 15 | 2 | 20 | 12.6 | 9.1 | 11.9 | 1.8 | 18.5 |
| >hg38_21_+_8433172_8446622_RNA45SN1_45S_50ntFlanks@6778.6803.26\|CCCGGGGCTACGCCTGTCTGAGCGTC | >hg38_21_+_8433172_8446622_RNA45SN1_45S_50ntFlanks | 6778 | 6803 | 26 | CCCGGGGCTACGCCTGTCTGAGCGTC | MB468 | 19 | 14 | 12 | 11 | 53 | 8.2 | 4.4 | 2.2 | 7.7 | 33.0 |
| >hg38_21_+_8433172_8446622_RNA45SN1_45S_50ntFlanks@6778.6806.29\|CCCGGGGCTACGCCTGTCTGAGCGTCGCT | >hg38_21_+_8433172_8446622_RNA45SN1_45S_50ntFlanks | 6778 | 6806 | 29 | CCCGGGGCTACGCCTGTCTGAGCGTCGCT | BT20 | 48 | 47 | 45 | 30 | 202 | 24.4 | 12.8 | 26.6 | 14.9 | 117.3 |
| >hg38_21_+_8433172_8446622_RNA45SN1_45S_50ntFlanks@6778.6806.29\|CCCGGGGCTACGCCTGTCTGAGCGTCGCT | >hg38_21_+_8433172_8446622_RNA45SN1_45S_50ntFlanks | 6778 | 6806 | 29 | CCCGGGGCTACGCCTGTCTGAGCGTCGCT | MB231 | 83 | 63 | 52 | 46 | 200 | 69.6 | 36.3 | 26.1 | 35.6 | 146.8 |
| >hg38_21_+_8433172_8446622_RNA45SN1_45S_50ntFlanks@6778.6807.30\|CCCGGGGCTACGCCTGTCTGAGCGTCGCTT | >hg38_21_+_8433172_8446622_RNA45SN1_45S_50ntFlanks | 6778 | 6807 | 30 | CCCGGGGCTACGCCTGTCTGAGCGTCGCTT | MB231 | 15 | 16 | 8 | 3 | 35 | 15.1 | 12.2 | 5.7 | 3.3 | 28.7 |
| >hg38_21_+_8433172_8446622_RNA45SN1_45S_50ntFlanks@6779.6802.24\|CCGGGGCTACGCCTGTCTGAGCGT | >hg38_21_+_8433172_8446622_RNA45SN1_45S_50ntFlanks | 6779 | 6802 | 24 | CCGGGGCTACGCCTGTCTGAGCGT | BT20 | 15 | 15 | 14 | 28 | 29 | 15.7 | 15.2 | 9.9 | 36.1 | 23.4 |
| >hg38_21_+_8433172_8446622_RNA45SN1_45S_50ntFlanks@6779.6802.24\|CCGGGGCTACGCCTGTCTGAGCGT | >hg38_21_+_8433172_8446622_RNA45SN1_45S_50ntFlanks | 6779 | 6802 | 24 | CCGGGGCTACGCCTGTCTGAGCGT | MB231 | 21 | 23 | 27 | 5 | 28 | 16.2 | 15.9 | 18.6 | 4.2 | 28.1 |
| >hg38_21_+_8433172_8446622_RNA45SN1_45S_50ntFlanks@6779.6803.25\|CCGGGGCTACGCCTGTCTGAGCGTC | >hg38_21_+_8433172_8446622_RNA45SN1_45S_50ntFlanks | 6779 | 6803 | 25 | CCGGGGCTACGCCTGTCTGAGCGTC | MB468 | 28 | 18 | 23 | 57 | 87 | 7.2 | 5.7 | 5.1 | 28.3 | 49.3 |
| >hg38_21_+_8433172_8446622_RNA45SN1_45S_50ntFlanks@6779.6806.28\|CCGGGGCTACGCCTGTCTGAGCGTCGCT | >hg38_21_+_8433172_8446622_RNA45SN1_45S_50ntFlanks | 6779 | 6806 | 28 | CCGGGGCTACGCCTGTCTGAGCGTCGCT | BT20 | 21 | 18 | 24 | 33 | 82 | 10.4 | 3.6 | 14.0 | 11.2 | 51.5 |
| >hg38_21_+_8433172_8446622_RNA45SN1_45S_50ntFlanks@6779.6806.28\|CCGGGGCTACGCCTGTCTGAGCGTCGCT | >hg38_21_+_8433172_8446622_RNA45SN1_45S_50ntFlanks | 6779 | 6806 | 28 | CCGGGGCTACGCCTGTCTGAGCGTCGCT | MB231 | 32 | 23 | 25 | 44 | 79 | 27.2 | 6.6 | 8.2 | 17.0 | 49.4 |
| >hg38_21_+_8433172_8446622_RNA45SN1_45S_50ntFlanks@6780.6802.23\|CGGGGCTACGCCTGTCTGAGCGT | >hg38_21_+_8433172_8446622_RNA45SN1_45S_50ntFlanks | 6780 | 6802 | 23 | CGGGGCTACGCCTGTCTGAGCGT | BT20 | 12 | 13 | 10 | 66 | 22 | 14.1 | 9.9 | 8.3 | 85.2 | 17.6 |
| >hg38_21_+_8433172_8446622_RNA45SN1_45S_50ntFlanks@6780.6802.23\|CGGGGCTACGCCTGTCTGAGCGT | >hg38_21_+_8433172_8446622_RNA45SN1_45S_50ntFlanks | 6780 | 6802 | 23 | CGGGGCTACGCCTGTCTGAGCGT | MB231 | 19 | 26 | 30 | 14 | 22 | 8.4 | 5.7 | 6.6 | 8.2 | 19.5 |
| >hg38_21_+_8433172_8446622_RNA45SN1_45S_50ntFlanks@6780.6806.27\|CGGGGCTACGCCTGTCTGAGCGTCGCT | >hg38_21_+_8433172_8446622_RNA45SN1_45S_50ntFlanks | 6780 | 6806 | 27 | CGGGGCTACGCCTGTCTGAGCGTCGCT | BT20 | 12 | 11 | 15 | 24 | 44 | 5.3 | 3.7 | 10.8 | 10.2 | 32.0 |

| | | | | | | | | | | | | | | | | |
|---|---|---|---|---|---|---|---|---|---|---|---|---|---|---|---|---|
| >hg38_21_+_8433172_8446622_RNA45SN1_45S_50ntFlanks@6780.6806.27\|CGGGGCTACGCCTGTCTGAGCGTCGCT | >hg38_21_+_8433172_8446622_RNA45SN1_45S_50ntFlanks | 6780 | 6806 | 27 | CGGGGCTACGCCTGTCTGAGCGTCGCT | MB231 | 19 | 15 | 12 | 34 | 42 | 16.6 | 1.6 | 4.1 | 7.6 | 29.9 |
| >hg38_21_+_8433172_8446622_RNA45SN1_45S_50ntFlanks@6784.6801.18\|GCTACGCCTGTCTGAGCG | >hg38_21_+_8433172_8446622_RNA45SN1_45S_50ntFlanks | 6784 | 6801 | 18 | GCTACGCCTGTCTGAGCG | MB468 | 15 | 1 | 6 | 20 | 15 | 8.0 | 0.4 | 3.0 | 2.9 | 13.2 |
| >hg38_21_+_8433172_8446622_RNA45SN1_45S_50ntFlanks@6784.6803.20\|GCTACGCCTGTCTGAGCGTC | >hg38_21_+_8433172_8446622_RNA45SN1_45S_50ntFlanks | 6784 | 6803 | 20 | GCTACGCCTGTCTGAGCGTC | MB468 | 38 | 1 | 20 | 163 | 35 | 33.1 | 0.9 | 22.2 | 109.9 | 38.1 |
| >hg38_21_+_8433172_8446622_RNA45SN1_45S_50ntFlanks@6785.6806.22\|CTACGCCTGTCTGAGCGTCGCT | >hg38_21_+_8433172_8446622_RNA45SN1_45S_50ntFlanks | 6785 | 6806 | 22 | CTACGCCTGTCTGAGCGTCGCT | BT20 | 69 | 61 | 68 | 67 | 152 | 35.1 | 12.4 | 27.9 | 24.3 | 50.6 |
| >hg38_21_+_8433172_8446622_RNA45SN1_45S_50ntFlanks@6785.6806.22\|CTACGCCTGTCTGAGCGTCGCT | >hg38_21_+_8433172_8446622_RNA45SN1_45S_50ntFlanks | 6785 | 6806 | 22 | CTACGCCTGTCTGAGCGTCGCT | MB231 | 69 | 49 | 58 | 58 | 203 | 66.7 | 17.5 | 23.4 | 9.7 | 73.0 |
| >hg38_21_+_8433172_8446622_RNA45SN1_45S_50ntFlanks@6785.6806.22\|CTACGCCTGTCTGAGCGTCGCT | >hg38_21_+_8433172_8446622_RNA45SN1_45S_50ntFlanks | 6785 | 6806 | 22 | CTACGCCTGTCTGAGCGTCGCT | MB468 | 27 | 4 | 9 | 84 | 55 | 13.0 | 1.4 | 3.5 | 28.9 | 27.3 |
| >hg38_21_+_8433172_8446622_RNA45SN1_45S_50ntFlanks@6786.6806.21\|TACGCCTGTCTGAGCGTCGCT | >hg38_21_+_8433172_8446622_RNA45SN1_45S_50ntFlanks | 6786 | 6806 | 21 | TACGCCTGTCTGAGCGTCGCT | BT20 | 97 | 66 | 69 | 59 | 162 | 67.2 | 12.5 | 32.2 | 15.1 | 67.3 |
| >hg38_21_+_8433172_8446622_RNA45SN1_45S_50ntFlanks@6786.6806.21\|TACGCCTGTCTGAGCGTCGCT | >hg38_21_+_8433172_8446622_RNA45SN1_45S_50ntFlanks | 6786 | 6806 | 21 | TACGCCTGTCTGAGCGTCGCT | MB231 | 106 | 72 | 80 | 48 | 263 | 82.2 | 25.3 | 25.6 | 10.0 | 85.7 |
| >hg38_21_+_8433172_8446622_RNA45SN1_45S_50ntFlanks@6786.6807.22\|TACGCCTGTCTGAGCGTCGCTT | >hg38_21_+_8433172_8446622_RNA45SN1_45S_50ntFlanks | 6786 | 6807 | 22 | TACGCCTGTCTGAGCGTCGCTT | BT20 | 14 | 16 | 12 | 14 | 22 | 4.1 | 13.1 | 4.3 | 3.0 | 7.4 |
| >hg38_21_+_8433172_8446622_RNA45SN1_45S_50ntFlanks@6786.6807.22\|TACGCCTGTCTGAGCGTCGCTT | >hg38_21_+_8433172_8446622_RNA45SN1_45S_50ntFlanks | 6786 | 6807 | 22 | TACGCCTGTCTGAGCGTCGCTT | MB231 | 13 | 13 | 10 | 6 | 29 | 11.2 | 4.0 | 3.7 | 1.7 | 7.2 |
| >hg38_21_+_8433172_8446622_RNA45SN1_45S_50ntFlanks@6787.6806.20\|ACGCCTGTCTGAGCGTCGCT | >hg38_21_+_8433172_8446622_RNA45SN1_45S_50ntFlanks | 6787 | 6806 | 20 | ACGCCTGTCTGAGCGTCGCT | BT20 | 66 | 63 | 68 | 63 | 126 | 22.6 | 38.5 | 35.4 | 4.6 | 49.3 |
| >hg38_21_+_8433172_8446622_RNA45SN1_45S_50ntFlanks@6787.6806.20\|ACGCCTGTCTGAGCGTCGCT | >hg38_21_+_8433172_8446622_RNA45SN1_45S_50ntFlanks | 6787 | 6806 | 20 | ACGCCTGTCTGAGCGTCGCT | MB231 | 134 | 85 | 82 | 63 | 207 | 104.6 | 21.2 | 27.0 | 3.9 | 87.8 |
| >hg38_21_+_8433172_8446622_RNA45SN1_45S_50ntFlanks@6787.6807.21\|ACGCCTGTCTGAGCGTCGCTT | >hg38_21_+_8433172_8446622_RNA45SN1_45S_50ntFlanks | 6787 | 6807 | 21 | ACGCCTGTCTGAGCGTCGCTT | BT20 | 10 | 13 | 11 | 12 | 20 | 9.6 | 11.5 | 5.4 | 9.1 | 7.2 |
| >hg38_21_+_8433172_8446622_RNA45SN1_45S_50ntFlanks@6787.6807.21\|ACGCCTGTCTGAGCGTCGCTT | >hg38_21_+_8433172_8446622_RNA45SN1_45S_50ntFlanks | 6787 | 6807 | 21 | ACGCCTGTCTGAGCGTCGCTT | MB231 | 18 | 17 | 11 | 4 | 23 | 18.9 | 7.2 | 3.4 | 0.9 | 8.8 |
| >hg38_21_+_8433172_8446622_RNA45SN1_45S_50ntFlanks@6788.6806.19\|CGCCTGTCTGAGCGTCGCT | >hg38_21_+_8433172_8446622_RNA45SN1_45S_50ntFlanks | 6788 | 6806 | 19 | CGCCTGTCTGAGCGTCGCT | BT20 | 123 | 93 | 164 | 126 | 337 | 65.1 | 2.8 | 56.6 | 104.1 | 87.7 |
| >hg38_21_+_8433172_8446622_RNA45SN1_45S_50ntFlanks@6788.6806.19\|CGCCTGTCTGAGCGTCGCT | >hg38_21_+_8433172_8446622_RNA45SN1_45S_50ntFlanks | 6788 | 6806 | 19 | CGCCTGTCTGAGCGTCGCT | MB231 | 260 | 254 | 355 | 57 | 431 | 194.1 | 85.5 | 103.7 | 13.9 | 196.5 |
| >hg38_21_+_8433172_8446622_RNA45SN1_45S_50ntFlanks@6788.6806.19\|CGCCTGTCTGAGCGTCGCT | >hg38_21_+_8433172_8446622_RNA45SN1_45S_50ntFlanks | 6788 | 6806 | 19 | CGCCTGTCTGAGCGTCGCT | MB468 | 22 | 15 | 22 | 14 | 54 | 11.9 | 5.3 | 11.0 | 4.8 | 21.2 |
| >hg38_21_+_8433172_8446622_RNA45SN1_45S_50ntFlanks@6788.6807.20\|CGCCTGTCTGAGCGTCGCTT | >hg38_21_+_8433172_8446622_RNA45SN1_45S_50ntFlanks | 6788 | 6807 | 20 | CGCCTGTCTGAGCGTCGCTT | BT20 | 40 | 32 | 39 | 64 | 100 | 48.5 | 19.6 | 13.6 | 93.8 | 31.4 |
| >hg38_21_+_8433172_8446622_RNA45SN1_45S_50ntFlanks@6788.6807.20\|CGCCTGTCTGAGCGTCGCTT | >hg38_21_+_8433172_8446622_RNA45SN1_45S_50ntFlanks | 6788 | 6807 | 20 | CGCCTGTCTGAGCGTCGCTT | MB231 | 52 | 77 | 67 | 5 | 68 | 61.4 | 37.6 | 25.5 | 0.6 | 40.8 |
| >hg38_21_+_8433172_8446622_RNA45SN1_45S_50ntFlanks@6789.6806.18\|GCCTGTCTGAGCGTCGCT | >hg38_21_+_8433172_8446622_RNA45SN1_45S_50ntFlanks | 6789 | 6806 | 18 | GCCTGTCTGAGCGTCGCT | BT20 | 46 | 50 | 61 | 40 | 170 | 27.8 | 18.6 | 28.5 | 35.0 | 39.5 |
| >hg38_21_+_8433172_8446622_RNA45SN1_45S_50ntFlanks@6789.6806.18\|GCCTGTCTGAGCGTCGCT | >hg38_21_+_8433172_8446622_RNA45SN1_45S_50ntFlanks | 6789 | 6806 | 18 | GCCTGTCTGAGCGTCGCT | MB231 | 105 | 106 | 141 | 24 | 288 | 93.5 | 29.0 | 22.2 | 7.9 | 78.9 |
| >hg38_21_+_8433172_8446622_RNA45SN1_45S_50ntFlanks@6789.6807.19\|GCCTGTCTGAGCGTCGCTT | >hg38_21_+_8433172_8446622_RNA45SN1_45S_50ntFlanks | 6789 | 6807 | 19 | GCCTGTCTGAGCGTCGCTT | BT20 | 16 | 15 | 15 | 28 | 59 | 20.9 | 10.3 | 7.5 | 42.6 | 21.0 |
| >hg38_21_+_8433172_8446622_RNA45SN1_45S_50ntFlanks@6789.6807.19\|GCCTGTCTGAGCGTCGCTT | >hg38_21_+_8433172_8446622_RNA45SN1_45S_50ntFlanks | 6789 | 6807 | 19 | GCCTGTCTGAGCGTCGCTT | MB231 | 21 | 31 | 27 | 2 | 47 | 25.8 | 15.0 | 11.3 | 0.9 | 26.4 |
| >hg38_21_+_8433172_8446622_RNA45SN1_45S_50ntFlanks@6790.6807.18\|CCTGTCTGAGCGTCGCTT | >hg38_21_+_8433172_8446622_RNA45SN1_45S_50ntFlanks | 6790 | 6807 | 18 | CCTGTCTGAGCGTCGCTT | BT20 | 34 | 53 | 45 | 97 | 191 | 33.4 | 27.6 | 10.8 | 152.2 | 84.3 |
| >hg38_21_+_8433172_8446622_RNA45SN1_45S_50ntFlanks@6790.6807.18\|CCTGTCTGAGCGTCGCTT | >hg38_21_+_8433172_8446622_RNA45SN1_45S_50ntFlanks | 6790 | 6807 | 18 | CCTGTCTGAGCGTCGCTT | MB231 | 48 | 79 | 68 | 6 | 111 | 61.3 | 34.0 | 21.7 | 1.5 | 66.3 |
| >hg38_21_+_8433172_8446622_RNA45SN1_45S_50ntFlanks@7973.7991.19\|GACGCGACCTCAGATCAGA | >hg38_21_+_8433172_8446622_RNA45SN1_45S_50ntFlanks | 7973 | 7991 | 19 | GACGCGACCTCAGATCAGA | BT20 | 19 | 92 | 10 | 43 | 108 | 27.8 | 101.7 | 6.1 | 29.1 | 42.4 |
| >hg38_21_+_8433172_8446622_RNA45SN1_45S_50ntFlanks@7973.7992.20\|GACGCGACCTCAGATCAGAC | >hg38_21_+_8433172_8446622_RNA45SN1_45S_50ntFlanks | 7973 | 7992 | 20 | GACGCGACCTCAGATCAGAC | BT20 | 17 | 152 | 6 | 65 | 149 | 25.7 | 166.2 | 3.4 | 53.9 | 36.9 |
| >hg38_21_+_8433172_8446622_RNA45SN1_45S_50ntFlanks@7973.7993.21\|GACGCGACCTCAGATCAGACG | >hg38_21_+_8433172_8446622_RNA45SN1_45S_50ntFlanks | 7973 | 7993 | 21 | GACGCGACCTCAGATCAGACG | BT20 | 31 | 166 | 14 | 101 | 144 | 46.0 | 199.2 | 16.2 | 55.0 | 57.6 |
| >hg38_21_+_8433172_8446622_RNA45SN1_45S_50ntFlanks@7973.7993.21\|GACGCGACCTCAGATCAGACG | >hg38_21_+_8433172_8446622_RNA45SN1_45S_50ntFlanks | 7973 | 7993 | 21 | GACGCGACCTCAGATCAGACG | MB231 | 13 | 54 | 23 | 22 | 78 | 19.3 | 36.6 | 23.0 | 10.9 | 75.5 |
| >hg38_21_+_8433172_8446622_RNA45SN1_45S_50ntFlanks@7973.7994.22\|GACGCGACCTCAGATCAGACGT | >hg38_21_+_8433172_8446622_RNA45SN1_45S_50ntFlanks | 7973 | 7994 | 22 | GACGCGACCTCAGATCAGACGT | BT20 | 15 | 134 | 5 | 52 | 46 | 15.2 | 174.2 | 3.0 | 34.6 | 32.7 |
| >hg38_21_+_8433172_8446622_RNA45SN1_45S_50ntFlanks@7973.7995.23\|GACGCGACCTCAGATCAGACGTG | >hg38_21_+_8433172_8446622_RNA45SN1_45S_50ntFlanks | 7973 | 7995 | 23 | GACGCGACCTCAGATCAGACGTG | MB468 | 13 | 8 | 13 | 13 | 12 | 5.1 | 2.6 | 5.4 | 4.2 | 4.0 |
| >hg38_21_+_8433172_8446622_RNA45SN1_45S_50ntFlanks@7973.8005.33\|GACGCGACCTCAGATCAGACGTGGCGACCCGCT | >hg38_21_+_8433172_8446622_RNA45SN1_45S_50ntFlanks | 7973 | 8005 | 33 | GACGCGACCTCAGATCAGACGTGGCGACCCGCT | BT20 | 33 | 708 | 14 | 68 | 253 | 25.3 | 962.3 | 5.1 | 45.6 | 164.9 |
| >hg38_21_+_8433172_8446622_RNA45SN1_45S_50ntFlanks@7973.8006.34\|GACGCGACCTCAGATCAGACGTGGCGACCCGCTG | >hg38_21_+_8433172_8446622_RNA45SN1_45S_50ntFlanks | 7973 | 8006 | 34 | GACGCGACCTCAGATCAGACGTGGCGACCCGCTG | MB468 | 28 | 32 | 15 | 30 | 96 | 9.8 | 17.4 | 3.4 | 14.4 | 16.4 |
| >hg38_21_+_8433172_8446622_RNA45SN1_45S_50ntFlanks@7973.8008.36\|GACGCGACCTCAGATCAGACGTGGCGACCCGCTGAA | >hg38_21_+_8433172_8446622_RNA45SN1_45S_50ntFlanks | 7973 | 8008 | 36 | GACGCGACCTCAGATCAGACGTGGCGACCCGCTGAA | BT20 | 14 | 133 | 126 | 98 | 178 | 16.1 | 89.6 | 206.7 | 153.2 | 53.7 |
| >hg38_21_+_8433172_8446622_RNA45SN1_45S_50ntFlanks@7973.8008.36\|GACGCGACCTCAGATCAGACGTGGCGACCCGCTGAA | >hg38_21_+_8433172_8446622_RNA45SN1_45S_50ntFlanks | 7973 | 8008 | 36 | GACGCGACCTCAGATCAGACGTGGCGACCCGCTGAA | MB231 | 41 | 101 | 45 | 23 | 199 | 48.6 | 49.6 | 37.7 | 15.3 | 117.0 |
| >hg38_21_+_8433172_8446622_RNA45SN1_45S_50ntFlanks@7973.8008.36\|GACGCGACCTCAGATCAGACGTGGCGACCCGCTGAA | >hg38_21_+_8433172_8446622_RNA45SN1_45S_50ntFlanks | 7973 | 8008 | 36 | GACGCGACCTCAGATCAGACGTGGCGACCCGCTGAA | MB468 | 13 | 35 | 6 | 22 | 80 | 5.9 | 11.8 | 2.0 | 17.8 | 46.9 |
| >hg38_21_+_8433172_8446622_RNA45SN1_45S_50ntFlanks@7973.8009.37\|GACGCGACCTCAGATCAGACGTGGCGACCCGCTGAAT | >hg38_21_+_8433172_8446622_RNA45SN1_45S_50ntFlanks | 7973 | 8009 | 37 | GACGCGACCTCAGATCAGACGTGGCGACCCGCTGAAT | BT20 | 34 | 428 | 76 | 148 | 323 | 24.0 | 290.8 | 47.6 | 169.3 | 42.6 |
| >hg38_21_+_8433172_8446622_RNA45SN1_45S_50ntFlanks@7973.8009.37\|GACGCGACCTCAGATCAGACGTGGCGACCCGCTGAAT | >hg38_21_+_8433172_8446622_RNA45SN1_45S_50ntFlanks | 7973 | 8009 | 37 | GACGCGACCTCAGATCAGACGTGGCGACCCGCTGAAT | MB231 | 51 | 107 | 102 | 122 | 256 | 55.6 | 45.3 | 62.2 | 49.5 | 124.2 |
| >hg38_21_+_8433172_8446622_RNA45SN1_45S_50ntFlanks@7973.8010.38\|GACGCGACCTCAGATCAGACGTGGCGACCCGCTGAATT | >hg38_21_+_8433172_8446622_RNA45SN1_45S_50ntFlanks | 7973 | 8010 | 38 | GACGCGACCTCAGATCAGACGTGGCGACCCGCTGAATT | BT20 | 47 | 881 | 47 | 47 | 254 | 48.1 | 987.9 | 28.1 | 62.0 | 32.5 |
| >hg38_21_+_8433172_8446622_RNA45SN1_45S_50ntFlanks@7973.8010.38\|GACGCGACCTCAGATCAGACGTGGCGACCCGCTGAATT | >hg38_21_+_8433172_8446622_RNA45SN1_45S_50ntFlanks | 7973 | 8010 | 38 | GACGCGACCTCAGATCAGACGTGGCGACCCGCTGAATT | MB231 | 55 | 126 | 88 | 24 | 245 | 61.1 | 87.7 | 77.4 | 22.6 | 119.8 |
| >hg38_21_+_8433172_8446622_RNA45SN1_45S_50ntFlanks@7973.8011.39\|GACGCGACCTCAGATCAGACGTGGCGACCCGCTGAATTT | >hg38_21_+_8433172_8446622_RNA45SN1_45S_50ntFlanks | 7973 | 8011 | 39 | GACGCGACCTCAGATCAGACGTGGCGACCCGCTGAATTT | BT20 | 55 | 474 | 22 | 21 | 216 | 69.1 | 296.3 | 14.7 | 22.6 | 62.6 |
| >hg38_21_+_8433172_8446622_RNA45SN1_45S_50ntFlanks@7973.8011.39\|GACGCGACCTCAGATCAGACGTGGCGACCCGCTGAATTT | >hg38_21_+_8433172_8446622_RNA45SN1_45S_50ntFlanks | 7973 | 8011 | 39 | GACGCGACCTCAGATCAGACGTGGCGACCCGCTGAATTT | MB231 | 90 | 193 | 114 | 13 | 222 | 89.2 | 134.7 | 78.8 | 1.8 | 150.5 |
| >hg38_21_+_8433172_8446622_RNA45SN1_45S_50ntFlanks@7973.8012.40\|GACGCGACCTCAGATCAGACGTGGCGACCCGCTGAATTTA | >hg38_21_+_8433172_8446622_RNA45SN1_45S_50ntFlanks | 7973 | 8012 | 40 | GACGCGACCTCAGATCAGACGTGGCGACCCGCTGAATTTA | MB231 | 25 | 63 | 22 | 2 | 50 | 29.8 | 37.7 | 16.3 | 0.4 | 39.2 |
| >hg38_21_+_8433172_8446622_RNA45SN1_45S_50ntFlanks@7973.8013.41\|GACGCGACCTCAGATCAGACGTGGCGACCCGCTGAATTTAA | >hg38_21_+_8433172_8446622_RNA45SN1_45S_50ntFlanks | 7973 | 8013 | 41 | GACGCGACCTCAGATCAGACGTGGCGACCCGCTGAATTTAA | MB231 | 13 | 35 | 16 | 1 | 32 | 14.7 | 34.1 | 14.8 | 0.8 | 27.9 |
| >hg38_21_+_8433172_8446622_RNA45SN1_45S_50ntFlanks@7973.8016.44\|GACGCGACCTCAGATCAGACGTGGCGACCCGCTGAATTTAAGCA | >hg38_21_+_8433172_8446622_RNA45SN1_45S_50ntFlanks | 7973 | 8016 | 44 | GACGCGACCTCAGATCAGACGTGGCGACCCGCTGAATTTAAGCA | MB231 | 10 | 19 | 8 | 2 | 49 | 9.6 | 29.0 | 5.4 | 2.1 | 54.7 |
| >hg38_21_+_8433172_8446622_RNA45SN1_45S_50ntFlanks@7974.7991.18\|ACGCGACCTCAGATCAGA | >hg38_21_+_8433172_8446622_RNA45SN1_45S_50ntFlanks | 7974 | 7991 | 18 | ACGCGACCTCAGATCAGA | BT20 | 73 | 205 | 79 | 168 | 661 | 95.2 | 137.6 | 51.2 | 106.7 | 166.4 |
| >hg38_21_+_8433172_8446622_RNA45SN1_45S_50ntFlanks@7974.7991.18\|ACGCGACCTCAGATCAGA | >hg38_21_+_8433172_8446622_RNA45SN1_45S_50ntFlanks | 7974 | 7991 | 18 | ACGCGACCTCAGATCAGA | MB231 | 29 | 165 | 316 | 44 | 221 | 42.3 | 192.9 | 460.7 | 56.8 | 190.6 |
| >hg38_21_+_8433172_8446622_RNA45SN1_45S_50ntFlanks@7974.7991.18\|ACGCGACCTCAGATCAGA | >hg38_21_+_8433172_8446622_RNA45SN1_45S_50ntFlanks | 7974 | 7991 | 18 | ACGCGACCTCAGATCAGA | MB468 | 16 | 40 | 15 | 18 | 28 | 7.1 | 35.7 | 13.7 | 12.3 | 15.5 |
| >hg38_21_+_8433172_8446622_RNA45SN1_45S_50ntFlanks@7974.7992.19\|ACGCGACCTCAGATCAGAC | >hg38_21_+_8433172_8446622_RNA45SN1_45S_50ntFlanks | 7974 | 7992 | 19 | ACGCGACCTCAGATCAGAC | BT20 | 55 | 275 | 30 | 243 | 733 | 76.7 | 255.2 | 21.5 | 235.0 | 156.2 |
| >hg38_21_+_8433172_8446622_RNA45SN1_45S_50ntFlanks@7974.7992.19\|ACGCGACCTCAGATCAGAC | >hg38_21_+_8433172_8446622_RNA45SN1_45S_50ntFlanks | 7974 | 7992 | 19 | ACGCGACCTCAGATCAGAC | MB231 | 21 | 93 | 117 | 26 | 233 | 29.9 | 69.2 | 148.6 | 23.0 | 210.2 |

| | | | | | | | | | | | | | | | | |
|---|---|---|---|---|---|---|---|---|---|---|---|---|---|---|---|---|
| >hg38_21_+_8433172_8446622_RNA45SN1_45S_50ntFlanks@7974.7992.19\|ACGCGACCTCAGATCAGAC | >hg38_21_+_8433172_8446622_RNA45SN1_45S_50ntFlanks | 7974 | 7992 | 19 | ACGCGACCTCAGATCAGAC | MB468 | 15 | 44 | 11 | 12 | 47 | 6.5 | 17.7 | 6.3 | 3.9 | 35.5 |
| >hg38_21_+_8433172_8446622_RNA45SN1_45S_50ntFlanks@7974.7993.20\|ACGCGACCTCAGATCAGACG | >hg38_21_+_8433172_8446622_RNA45SN1_45S_50ntFlanks | 7974 | 7993 | 20 | ACGCGACCTCAGATCAGACG | BT20 | 116 | 459 | 169 | 592 | 785 | 147.7 | 438.8 | 207.4 | 288.1 | 205.4 |
| >hg38_21_+_8433172_8446622_RNA45SN1_45S_50ntFlanks@7974.7993.20\|ACGCGACCTCAGATCAGACG | >hg38_21_+_8433172_8446622_RNA45SN1_45S_50ntFlanks | 7974 | 7993 | 20 | ACGCGACCTCAGATCAGACG | MB231 | 47 | 134 | 86 | 131 | 343 | 69.8 | 77.2 | 67.1 | 104.3 | 303.1 |
| >hg38_21_+_8433172_8446622_RNA45SN1_45S_50ntFlanks@7974.7993.20\|ACGCGACCTCAGATCAGACG | >hg38_21_+_8433172_8446622_RNA45SN1_45S_50ntFlanks | 7974 | 7993 | 20 | ACGCGACCTCAGATCAGACG | MB468 | 42 | 65 | 29 | 58 | 115 | 23.7 | 53.2 | 11.2 | 12.2 | 58.7 |
| >hg38_21_+_8433172_8446622_RNA45SN1_45S_50ntFlanks@7974.7994.21\|ACGCGACCTCAGATCAGACGT | >hg38_21_+_8433172_8446622_RNA45SN1_45S_50ntFlanks | 7974 | 7994 | 21 | ACGCGACCTCAGATCAGACGT | BT20 | 65 | 371 | 35 | 203 | 274 | 52.5 | 415.4 | 28.3 | 110.1 | 184.4 |
| >hg38_21_+_8433172_8446622_RNA45SN1_45S_50ntFlanks@7974.7994.21\|ACGCGACCTCAGATCAGACGT | >hg38_21_+_8433172_8446622_RNA45SN1_45S_50ntFlanks | 7974 | 7994 | 21 | ACGCGACCTCAGATCAGACGT | MB231 | 11 | 39 | 26 | 94 | 131 | 14.2 | 15.4 | 18.5 | 104.8 | 70.9 |
| >hg38_21_+_8433172_8446622_RNA45SN1_45S_50ntFlanks@7974.7994.21\|ACGCGACCTCAGATCAGACGT | >hg38_21_+_8433172_8446622_RNA45SN1_45S_50ntFlanks | 7974 | 7994 | 21 | ACGCGACCTCAGATCAGACGT | MB468 | 32 | 16 | 25 | 58 | 38 | 12.4 | 5.6 | 10.2 | 37.6 | 16.2 |
| >hg38_21_+_8433172_8446622_RNA45SN1_45S_50ntFlanks@7974.7995.22\|ACGCGACCTCAGATCAGACGTG | >hg38_21_+_8433172_8446622_RNA45SN1_45S_50ntFlanks | 7974 | 7995 | 22 | ACGCGACCTCAGATCAGACGTG | BT20 | 45 | 223 | 253 | 148 | 311 | 41.2 | 203.9 | 394.8 | 75.3 | 145.9 |
| >hg38_21_+_8433172_8446622_RNA45SN1_45S_50ntFlanks@7974.7995.22\|ACGCGACCTCAGATCAGACGTG | >hg38_21_+_8433172_8446622_RNA45SN1_45S_50ntFlanks | 7974 | 7995 | 22 | ACGCGACCTCAGATCAGACGTG | MB231 | 18 | 95 | 91 | 56 | 280 | 24.3 | 37.3 | 70.9 | 60.2 | 282.4 |
| >hg38_21_+_8433172_8446622_RNA45SN1_45S_50ntFlanks@7974.7995.22\|ACGCGACCTCAGATCAGACGTG | >hg38_21_+_8433172_8446622_RNA45SN1_45S_50ntFlanks | 7974 | 7995 | 22 | ACGCGACCTCAGATCAGACGTG | MB468 | 107 | 44 | 93 | 87 | 96 | 59.8 | 13.9 | 42.2 | 15.2 | 44.0 |
| >hg38_21_+_8433172_8446622_RNA45SN1_45S_50ntFlanks@7974.7996.23\|ACGCGACCTCAGATCAGACGTGG | >hg38_21_+_8433172_8446622_RNA45SN1_45S_50ntFlanks | 7974 | 7996 | 23 | ACGCGACCTCAGATCAGACGTGG | MB468 | 16 | 6 | 15 | 22 | 23 | 6.3 | 1.4 | 4.7 | 8.9 | 12.8 |
| >hg38_21_+_8433172_8446622_RNA45SN1_45S_50ntFlanks@7974.7997.24\|ACGCGACCTCAGATCAGACGTGGC | >hg38_21_+_8433172_8446622_RNA45SN1_45S_50ntFlanks | 7974 | 7997 | 24 | ACGCGACCTCAGATCAGACGTGGC | BT20 | 56 | 662 | 57 | 219 | 708 | 53.8 | 616.2 | 43.4 | 116.9 | 260.7 |
| >hg38_21_+_8433172_8446622_RNA45SN1_45S_50ntFlanks@7974.7997.24\|ACGCGACCTCAGATCAGACGTGGC | >hg38_21_+_8433172_8446622_RNA45SN1_45S_50ntFlanks | 7974 | 7997 | 24 | ACGCGACCTCAGATCAGACGTGGC | MB231 | 26 | 90 | 71 | 124 | 260 | 34.1 | 48.3 | 52.3 | 122.8 | 189.6 |
| >hg38_21_+_8433172_8446622_RNA45SN1_45S_50ntFlanks@7974.8002.29\|ACGCGACCTCAGATCAGACGTGGCGACCC | >hg38_21_+_8433172_8446622_RNA45SN1_45S_50ntFlanks | 7974 | 8002 | 29 | ACGCGACCTCAGATCAGACGTGGCGACCC | BT20 | 11 | 101 | 8 | 110 | 54 | 2.8 | 100.6 | 1.7 | 63.1 | 30.3 |
| >hg38_21_+_8433172_8446622_RNA45SN1_45S_50ntFlanks@7974.8003.30\|ACGCGACCTCAGATCAGACGTGGCGACCCG | >hg38_21_+_8433172_8446622_RNA45SN1_45S_50ntFlanks | 7974 | 8003 | 30 | ACGCGACCTCAGATCAGACGTGGCGACCCG | MB468 | 15 | 12 | 16 | 14 | 30 | 6.9 | 2.1 | 8.6 | 0.8 | 11.8 |
| >hg38_21_+_8433172_8446622_RNA45SN1_45S_50ntFlanks@7974.8004.31\|ACGCGACCTCAGATCAGACGTGGCGACCCGC | >hg38_21_+_8433172_8446622_RNA45SN1_45S_50ntFlanks | 7974 | 8004 | 31 | ACGCGACCTCAGATCAGACGTGGCGACCCGC | BT20 | 26 | 323 | 180 | 114 | 270 | 18.5 | 350.3 | 283.6 | 58.5 | 130.7 |
| >hg38_21_+_8433172_8446622_RNA45SN1_45S_50ntFlanks@7974.8004.31\|ACGCGACCTCAGATCAGACGTGGCGACCCGC | >hg38_21_+_8433172_8446622_RNA45SN1_45S_50ntFlanks | 7974 | 8004 | 31 | ACGCGACCTCAGATCAGACGTGGCGACCCGC | MB231 | 11 | 31 | 24 | 22 | 101 | 10.5 | 2.5 | 5.4 | 23.3 | 69.9 |
| >hg38_21_+_8433172_8446622_RNA45SN1_45S_50ntFlanks@7974.8005.32\|ACGCGACCTCAGATCAGACGTGGCGACCCGCT | >hg38_21_+_8433172_8446622_RNA45SN1_45S_50ntFlanks | 7974 | 8005 | 32 | ACGCGACCTCAGATCAGACGTGGCGACCCGCT | BT20 | 200 | 2356 | 172 | 461 | 2315 | 95.2 | 2589.5 | 114.4 | 208.1 | 1186.8 |
| >hg38_21_+_8433172_8446622_RNA45SN1_45S_50ntFlanks@7974.8005.32\|ACGCGACCTCAGATCAGACGTGGCGACCCGCT | >hg38_21_+_8433172_8446622_RNA45SN1_45S_50ntFlanks | 7974 | 8005 | 32 | ACGCGACCTCAGATCAGACGTGGCGACCCGCT | MB231 | 51 | 93 | 49 | 77 | 499 | 62.5 | 72.9 | 19.7 | 106.7 | 569.5 |
| >hg38_21_+_8433172_8446622_RNA45SN1_45S_50ntFlanks@7974.8005.32\|ACGCGACCTCAGATCAGACGTGGCGACCCGCT | >hg38_21_+_8433172_8446622_RNA45SN1_45S_50ntFlanks | 7974 | 8005 | 32 | ACGCGACCTCAGATCAGACGTGGCGACCCGCT | MB468 | 18 | 19 | 12 | 28 | 71 | 5.8 | 4.2 | 2.0 | 14.8 | 44.8 |
| >hg38_21_+_8433172_8446622_RNA45SN1_45S_50ntFlanks@7974.8006.33\|ACGCGACCTCAGATCAGACGTGGCGACCCGCTG | >hg38_21_+_8433172_8446622_RNA45SN1_45S_50ntFlanks | 7974 | 8006 | 33 | ACGCGACCTCAGATCAGACGTGGCGACCCGCTG | BT20 | 55 | 678 | 322 | 138 | 811 | 20.8 | 699.9 | 525.0 | 91.9 | 407.8 |
| >hg38_21_+_8433172_8446622_RNA45SN1_45S_50ntFlanks@7974.8006.33\|ACGCGACCTCAGATCAGACGTGGCGACCCGCTG | >hg38_21_+_8433172_8446622_RNA45SN1_45S_50ntFlanks | 7974 | 8006 | 33 | ACGCGACCTCAGATCAGACGTGGCGACCCGCTG | MB231 | 23 | 43 | 27 | 19 | 184 | 29.6 | 21.4 | 8.0 | 20.2 | 153.5 |
| >hg38_21_+_8433172_8446622_RNA45SN1_45S_50ntFlanks@7974.8006.33\|ACGCGACCTCAGATCAGACGTGGCGACCCGCTG | >hg38_21_+_8433172_8446622_RNA45SN1_45S_50ntFlanks | 7974 | 8006 | 33 | ACGCGACCTCAGATCAGACGTGGCGACCCGCTG | MB468 | 270 | 357 | 123 | 198 | 1081 | 118.6 | 220.0 | 39.6 | 80.2 | 432.2 |
| >hg38_21_+_8433172_8446622_RNA45SN1_45S_50ntFlanks@7974.8007.34\|ACGCGACCTCAGATCAGACGTGGCGACCCGCTGA | >hg38_21_+_8433172_8446622_RNA45SN1_45S_50ntFlanks | 7974 | 8007 | 34 | ACGCGACCTCAGATCAGACGTGGCGACCCGCTGA | BT20 | 30 | 356 | 122 | 70 | 328 | 23.6 | 399.4 | 185.7 | 53.2 | 72.3 |
| >hg38_21_+_8433172_8446622_RNA45SN1_45S_50ntFlanks@7974.8007.34\|ACGCGACCTCAGATCAGACGTGGCGACCCGCTGA | >hg38_21_+_8433172_8446622_RNA45SN1_45S_50ntFlanks | 7974 | 8007 | 34 | ACGCGACCTCAGATCAGACGTGGCGACCCGCTGA | MB231 | 26 | 54 | 41 | 16 | 144 | 35.0 | 17.4 | 12.7 | 7.4 | 105.2 |
| >hg38_21_+_8433172_8446622_RNA45SN1_45S_50ntFlanks@7974.8007.34\|ACGCGACCTCAGATCAGACGTGGCGACCCGCTGA | >hg38_21_+_8433172_8446622_RNA45SN1_45S_50ntFlanks | 7974 | 8007 | 34 | ACGCGACCTCAGATCAGACGTGGCGACCCGCTGA | MB468 | 23 | 45 | 8 | 24 | 113 | 12.0 | 28.3 | 3.0 | 10.6 | 62.0 |
| >hg38_21_+_8433172_8446622_RNA45SN1_45S_50ntFlanks@7974.8008.35\|ACGCGACCTCAGATCAGACGTGGCGACCCGCTGAA | >hg38_21_+_8433172_8446622_RNA45SN1_45S_50ntFlanks | 7974 | 8008 | 35 | ACGCGACCTCAGATCAGACGTGGCGACCCGCTGAA | BT20 | 72 | 815 | 3198 | 691 | 1490 | 51.5 | 821.5 | 5416.0 | 1092.0 | 359.6 |
| >hg38_21_+_8433172_8446622_RNA45SN1_45S_50ntFlanks@7974.8008.35\|ACGCGACCTCAGATCAGACGTGGCGACCCGCTGAA | >hg38_21_+_8433172_8446622_RNA45SN1_45S_50ntFlanks | 7974 | 8008 | 35 | ACGCGACCTCAGATCAGACGTGGCGACCCGCTGAA | MB231 | 218 | 374 | 363 | 120 | 1376 | 280.2 | 166.4 | 252.5 | 69.5 | 875.2 |
| >hg38_21_+_8433172_8446622_RNA45SN1_45S_50ntFlanks@7974.8008.35\|ACGCGACCTCAGATCAGACGTGGCGACCCGCTGAA | >hg38_21_+_8433172_8446622_RNA45SN1_45S_50ntFlanks | 7974 | 8008 | 35 | ACGCGACCTCAGATCAGACGTGGCGACCCGCTGAA | MB468 | 108 | 257 | 42 | 152 | 704 | 49.4 | 96.8 | 14.8 | 109.4 | 478.0 |
| >hg38_21_+_8433172_8446622_RNA45SN1_45S_50ntFlanks@7974.8009.36\|ACGCGACCTCAGATCAGACGTGGCGACCCGCTGAAT | >hg38_21_+_8433172_8446622_RNA45SN1_45S_50ntFlanks | 7974 | 8009 | 36 | ACGCGACCTCAGATCAGACGTGGCGACCCGCTGAAT | BT20 | 238 | 2486 | 901 | 1074 | 2862 | 103.6 | 2293.5 | 568.7 | 1286.5 | 618.8 |
| >hg38_21_+_8433172_8446622_RNA45SN1_45S_50ntFlanks@7974.8009.36\|ACGCGACCTCAGATCAGACGTGGCGACCCGCTGAAT | >hg38_21_+_8433172_8446622_RNA45SN1_45S_50ntFlanks | 7974 | 8009 | 36 | ACGCGACCTCAGATCAGACGTGGCGACCCGCTGAAT | MB231 | 323 | 455 | 780 | 614 | 1710 | 401.1 | 246.3 | 430.6 | 143.6 | 919.0 |
| >hg38_21_+_8433172_8446622_RNA45SN1_45S_50ntFlanks@7974.8009.36\|ACGCGACCTCAGATCAGACGTGGCGACCCGCTGAAT | >hg38_21_+_8433172_8446622_RNA45SN1_45S_50ntFlanks | 7974 | 8009 | 36 | ACGCGACCTCAGATCAGACGTGGCGACCCGCTGAAT | MB468 | 20 | 70 | 10 | 65 | 211 | 5.6 | 17.5 | 3.3 | 42.8 | 152.6 |
| >hg38_21_+_8433172_8446622_RNA45SN1_45S_50ntFlanks@7974.8010.37\|ACGCGACCTCAGATCAGACGTGGCGACCCGCTGAATT | >hg38_21_+_8433172_8446622_RNA45SN1_45S_50ntFlanks | 7974 | 8010 | 37 | ACGCGACCTCAGATCAGACGTGGCGACCCGCTGAATT | BT20 | 321 | 3767 | 531 | 422 | 2461 | 227.2 | 2683.1 | 234.3 | 556.2 | 656.6 |
| >hg38_21_+_8433172_8446622_RNA45SN1_45S_50ntFlanks@7974.8010.37\|ACGCGACCTCAGATCAGACGTGGCGACCCGCTGAATT | >hg38_21_+_8433172_8446622_RNA45SN1_45S_50ntFlanks | 7974 | 8010 | 37 | ACGCGACCTCAGATCAGACGTGGCGACCCGCTGAATT | MB231 | 337 | 585 | 621 | 135 | 1635 | 395.1 | 393.0 | 490.9 | 106.6 | 865.6 |
| >hg38_21_+_8433172_8446622_RNA45SN1_45S_50ntFlanks@7974.8011.38\|ACGCGACCTCAGATCAGACGTGGCGACCCGCTGAATTT | >hg38_21_+_8433172_8446622_RNA45SN1_45S_50ntFlanks | 7974 | 8011 | 38 | ACGCGACCTCAGATCAGACGTGGCGACCCGCTGAATTT | BT20 | 384 | 2788 | 253 | 213 | 2190 | 411.1 | 2014.6 | 119.3 | 264.0 | 995.1 |
| >hg38_21_+_8433172_8446622_RNA45SN1_45S_50ntFlanks@7974.8011.38\|ACGCGACCTCAGATCAGACGTGGCGACCCGCTGAATTT | >hg38_21_+_8433172_8446622_RNA45SN1_45S_50ntFlanks | 7974 | 8011 | 38 | ACGCGACCTCAGATCAGACGTGGCGACCCGCTGAATTT | MB231 | 591 | 966 | 788 | 96 | 1484 | 630.5 | 629.2 | 452.4 | 12.1 | 1039.8 |
| >hg38_21_+_8433172_8446622_RNA45SN1_45S_50ntFlanks@7974.8011.38\|ACGCGACCTCAGATCAGACGTGGCGACCCGCTGAATTT | >hg38_21_+_8433172_8446622_RNA45SN1_45S_50ntFlanks | 7974 | 8011 | 38 | ACGCGACCTCAGATCAGACGTGGCGACCCGCTGAATTT | MB468 | 19 | 74 | 9 | 24 | 152 | 7.2 | 25.9 | 3.9 | 23.8 | 69.9 |
| >hg38_21_+_8433172_8446622_RNA45SN1_45S_50ntFlanks@7974.8012.39\|ACGCGACCTCAGATCAGACGTGGCGACCCGCTGAATTTA | >hg38_21_+_8433172_8446622_RNA45SN1_45S_50ntFlanks | 7974 | 8012 | 39 | ACGCGACCTCAGATCAGACGTGGCGACCCGCTGAATTTA | BT20 | 35 | 138 | 30 | 15 | 160 | 43.3 | 110.4 | 29.3 | 13.6 | 42.3 |
| >hg38_21_+_8433172_8446622_RNA45SN1_45S_50ntFlanks@7974.8012.39\|ACGCGACCTCAGATCAGACGTGGCGACCCGCTGAATTTA | >hg38_21_+_8433172_8446622_RNA45SN1_45S_50ntFlanks | 7974 | 8012 | 39 | ACGCGACCTCAGATCAGACGTGGCGACCCGCTGAATTTA | MB231 | 131 | 234 | 132 | 14 | 297 | 156.6 | 137.3 | 89.2 | 1.3 | 229.0 |
| >hg38_21_+_8433172_8446622_RNA45SN1_45S_50ntFlanks@7974.8012.39\|ACGCGACCTCAGATCAGACGTGGCGACCCGCTGAATTTA | >hg38_21_+_8433172_8446622_RNA45SN1_45S_50ntFlanks | 7974 | 8012 | 39 | ACGCGACCTCAGATCAGACGTGGCGACCCGCTGAATTTA | MB468 | 33 | 106 | 12 | 36 | 283 | 16.8 | 44.6 | 4.8 | 34.8 | 118.6 |
| >hg38_21_+_8433172_8446622_RNA45SN1_45S_50ntFlanks@7974.8013.40\|ACGCGACCTCAGATCAGACGTGGCGACCCGCTGAATTTAA | >hg38_21_+_8433172_8446622_RNA45SN1_45S_50ntFlanks | 7974 | 8013 | 40 | ACGCGACCTCAGATCAGACGTGGCGACCCGCTGAATTTAA | BT20 | 18 | 42 | 14 | 8 | 98 | 22.9 | 30.0 | 13.4 | 6.9 | 53.4 |
| >hg38_21_+_8433172_8446622_RNA45SN1_45S_50ntFlanks@7974.8013.40\|ACGCGACCTCAGATCAGACGTGGCGACCCGCTGAATTTAA | >hg38_21_+_8433172_8446622_RNA45SN1_45S_50ntFlanks | 7974 | 8013 | 40 | ACGCGACCTCAGATCAGACGTGGCGACCCGCTGAATTTAA | MB231 | 71 | 129 | 90 | 7 | 205 | 86.0 | 118.1 | 74.6 | 3.1 | 171.9 |
| >hg38_21_+_8433172_8446622_RNA45SN1_45S_50ntFlanks@7974.8014.41\|ACGCGACCTCAGATCAGACGTGGCGACCCGCTGAATTTAAG | >hg38_21_+_8433172_8446622_RNA45SN1_45S_50ntFlanks | 7974 | 8014 | 41 | ACGCGACCTCAGATCAGACGTGGCGACCCGCTGAATTTAAG | MB231 | 26 | 60 | 42 | 3 | 88 | 28.4 | 49.3 | 20.2 | 0.6 | 66.9 |
| >hg38_21_+_8433172_8446622_RNA45SN1_45S_50ntFlanks@7974.8015.42\|ACGCGACCTCAGATCAGACGTGGCGACCCGCTGAATTTAAGC | >hg38_21_+_8433172_8446622_RNA45SN1_45S_50ntFlanks | 7974 | 8015 | 42 | ACGCGACCTCAGATCAGACGTGGCGACCCGCTGAATTTAAGC | BT20 | 37 | 209 | 25 | 21 | 286 | 29.6 | 169.7 | 25.8 | 4.7 | 73.5 |
| >hg38_21_+_8433172_8446622_RNA45SN1_45S_50ntFlanks@7974.8015.42\|ACGCGACCTCAGATCAGACGTGGCGACCCGCTGAATTTAAGC | >hg38_21_+_8433172_8446622_RNA45SN1_45S_50ntFlanks | 7974 | 8015 | 42 | ACGCGACCTCAGATCAGACGTGGCGACCCGCTGAATTTAAGC | MB231 | 30 | 57 | 51 | 15 | 241 | 14.3 | 30.6 | 24.3 | 10.1 | 126.5 |
| >hg38_21_+_8433172_8446622_RNA45SN1_45S_50ntFlanks@7974.8016.43\|ACGCGACCTCAGATCAGACGTGGCGACCCGCTGAATTTAAGCA | >hg38_21_+_8433172_8446622_RNA45SN1_45S_50ntFlanks | 7974 | 8016 | 43 | ACGCGACCTCAGATCAGACGTGGCGACCCGCTGAATTTAAGCA | BT20 | 15 | 97 | 18 | 13 | 314 | 10.3 | 118.0 | 17.7 | 3.0 | 201.5 |
| >hg38_21_+_8433172_8446622_RNA45SN1_45S_50ntFlanks@7974.8016.43\|ACGCGACCTCAGATCAGACGTGGCGACCCGCTGAATTTAAGCA | >hg38_21_+_8433172_8446622_RNA45SN1_45S_50ntFlanks | 7974 | 8016 | 43 | ACGCGACCTCAGATCAGACGTGGCGACCCGCTGAATTTAAGCA | MB231 | 84 | 127 | 82 | 15 | 556 | 92.4 | 181.8 | 50.4 | 11.4 | 550.1 |
| >hg38_21_+_8433172_8446622_RNA45SN1_45S_50ntFlanks@7975.7992.18\|CGCGACCTCAGATCAGAC | >hg38_21_+_8433172_8446622_RNA45SN1_45S_50ntFlanks | 7975 | 7992 | 18 | CGCGACCTCAGATCAGAC | BT20 | 423 | 1440 | 292 | 1725 | 5901 | 559.0 | 882.7 | 179.8 | 1948.6 | 1160.3 |
| >hg38_21_+_8433172_8446622_RNA45SN1_45S_50ntFlanks@7975.7992.18\|CGCGACCTCAGATCAGAC | >hg38_21_+_8433172_8446622_RNA45SN1_45S_50ntFlanks | 7975 | 7992 | 18 | CGCGACCTCAGATCAGAC | MB231 | 134 | 479 | 812 | 105 | 1121 | 195.9 | 376.2 | 1057.1 | 83.6 | 971.4 |
| >hg38_21_+_8433172_8446622_RNA45SN1_45S_50ntFlanks@7975.7992.18\|CGCGACCTCAGATCAGAC | >hg38_21_+_8433172_8446622_RNA45SN1_45S_50ntFlanks | 7975 | 7992 | 18 | CGCGACCTCAGATCAGAC | MB468 | 90 | 218 | 81 | 77 | 392 | 43.1 | 97.9 | 50.4 | 32.5 | 405.1 |

| | | | | | | | | | | | | | | | | |
|---|---|---|---|---|---|---|---|---|---|---|---|---|---|---|---|---|
| >hg38_21_+_8433172_8446622_RNA45SN1_45S_50ntFlanks@7975.7993.19\|CGCGACCTCAGATCAGACG | >hg38_21_+_8433172_8446622_RNA45SN1_45S_50ntFlanks | 7975 | 7993 | 19 | CGCGACCTCAGATCAGACG | BT20 | 653 | 1735 | 1067 | 2283 | 4447 | 796.5 | 1194.4 | 1208.8 | 719.1 | 1062.6 |
| >hg38_21_+_8433172_8446622_RNA45SN1_45S_50ntFlanks@7975.7993.19\|CGCGACCTCAGATCAGACG | >hg38_21_+_8433172_8446622_RNA45SN1_45S_50ntFlanks | 7975 | 7993 | 19 | CGCGACCTCAGATCAGACG | MB231 | 223 | 513 | 394 | 411 | 1429 | 332.1 | 230.6 | 278.6 | 296.0 | 1233.7 |
| >hg38_21_+_8433172_8446622_RNA45SN1_45S_50ntFlanks@7975.7993.19\|CGCGACCTCAGATCAGACG | >hg38_21_+_8433172_8446622_RNA45SN1_45S_50ntFlanks | 7975 | 7993 | 19 | CGCGACCTCAGATCAGACG | MB468 | 185 | 257 | 146 | 284 | 669 | 97.1 | 223.8 | 52.1 | 104.0 | 556.8 |
| >hg38_21_+_8433172_8446622_RNA45SN1_45S_50ntFlanks@7975.7994.20\|CGCGACCTCAGATCAGACGT | >hg38_21_+_8433172_8446622_RNA45SN1_45S_50ntFlanks | 7975 | 7994 | 20 | CGCGACCTCAGATCAGACGT | BT20 | 1899 | 10134 | 1591 | 9760 | 22983 | 2256.6 | 5950.0 | 1539.8 | 7067.7 | 2829.1 |
| >hg38_21_+_8433172_8446622_RNA45SN1_45S_50ntFlanks@7975.7994.20\|CGCGACCTCAGATCAGACGT | >hg38_21_+_8433172_8446622_RNA45SN1_45S_50ntFlanks | 7975 | 7994 | 20 | CGCGACCTCAGATCAGACGT | MB231 | 779 | 2366 | 3029 | 1687 | 5268 | 1113.9 | 1550.4 | 3269.1 | 1404.6 | 3985.1 |
| >hg38_21_+_8433172_8446622_RNA45SN1_45S_50ntFlanks@7975.7994.20\|CGCGACCTCAGATCAGACGT | >hg38_21_+_8433172_8446622_RNA45SN1_45S_50ntFlanks | 7975 | 7994 | 20 | CGCGACCTCAGATCAGACGT | MB468 | 1068 | 1350 | 1247 | 1528 | 1750 | 547.1 | 575.3 | 705.2 | 725.7 | 814.6 |
| >hg38_21_+_8433172_8446622_RNA45SN1_45S_50ntFlanks@7975.7995.21\|CGCGACCTCAGATCAGACGTG | >hg38_21_+_8433172_8446622_RNA45SN1_45S_50ntFlanks | 7975 | 7995 | 21 | CGCGACCTCAGATCAGACGTG | BT20 | 281 | 859 | 996 | 625 | 1353 | 229.4 | 546.9 | 1347.3 | 363.9 | 814.5 |
| >hg38_21_+_8433172_8446622_RNA45SN1_45S_50ntFlanks@7975.7995.21\|CGCGACCTCAGATCAGACGTG | >hg38_21_+_8433172_8446622_RNA45SN1_45S_50ntFlanks | 7975 | 7995 | 21 | CGCGACCTCAGATCAGACGTG | MB231 | 52 | 232 | 334 | 116 | 725 | 61.4 | 65.0 | 282.3 | 128.8 | 744.3 |
| >hg38_21_+_8433172_8446622_RNA45SN1_45S_50ntFlanks@7975.7995.21\|CGCGACCTCAGATCAGACGTG | >hg38_21_+_8433172_8446622_RNA45SN1_45S_50ntFlanks | 7975 | 7995 | 21 | CGCGACCTCAGATCAGACGTG | MB468 | 278 | 124 | 369 | 359 | 293 | 101.2 | 53.9 | 161.7 | 47.5 | 167.1 |
| >hg38_21_+_8433172_8446622_RNA45SN1_45S_50ntFlanks@7975.7996.22\|CGCGACCTCAGATCAGACGTGG | >hg38_21_+_8433172_8446622_RNA45SN1_45S_50ntFlanks | 7975 | 7996 | 22 | CGCGACCTCAGATCAGACGTGG | BT20 | 82 | 341 | 774 | 322 | 733 | 86.9 | 249.0 | 1237.6 | 305.5 | 206.0 |
| >hg38_21_+_8433172_8446622_RNA45SN1_45S_50ntFlanks@7975.7996.22\|CGCGACCTCAGATCAGACGTGG | >hg38_21_+_8433172_8446622_RNA45SN1_45S_50ntFlanks | 7975 | 7996 | 22 | CGCGACCTCAGATCAGACGTGG | MB231 | 37 | 219 | 171 | 62 | 735 | 34.6 | 143.5 | 34.2 | 15.0 | 1018.3 |
| >hg38_21_+_8433172_8446622_RNA45SN1_45S_50ntFlanks@7975.7996.22\|CGCGACCTCAGATCAGACGTGG | >hg38_21_+_8433172_8446622_RNA45SN1_45S_50ntFlanks | 7975 | 7996 | 22 | CGCGACCTCAGATCAGACGTGG | MB468 | 132 | 90 | 164 | 203 | 303 | 53.8 | 40.9 | 52.9 | 30.8 | 249.5 |
| >hg38_21_+_8433172_8446622_RNA45SN1_45S_50ntFlanks@7975.7997.23\|CGCGACCTCAGATCAGACGTGGC | >hg38_21_+_8433172_8446622_RNA45SN1_45S_50ntFlanks | 7975 | 7997 | 23 | CGCGACCTCAGATCAGACGTGGC | BT20 | 537 | 4993 | 515 | 1644 | 7636 | 473.6 | 3365.7 | 340.0 | 1260.3 | 1893.6 |
| >hg38_21_+_8433172_8446622_RNA45SN1_45S_50ntFlanks@7975.7997.23\|CGCGACCTCAGATCAGACGTGGC | >hg38_21_+_8433172_8446622_RNA45SN1_45S_50ntFlanks | 7975 | 7997 | 23 | CGCGACCTCAGATCAGACGTGGC | MB231 | 172 | 510 | 540 | 509 | 1553 | 227.2 | 337.4 | 480.9 | 570.6 | 1365.9 |
| >hg38_21_+_8433172_8446622_RNA45SN1_45S_50ntFlanks@7975.7997.23\|CGCGACCTCAGATCAGACGTGGC | >hg38_21_+_8433172_8446622_RNA45SN1_45S_50ntFlanks | 7975 | 7997 | 23 | CGCGACCTCAGATCAGACGTGGC | MB468 | 27 | 72 | 41 | 117 | 178 | 15.5 | 37.0 | 32.2 | 91.4 | 134.4 |
| >hg38_21_+_8433172_8446622_RNA45SN1_45S_50ntFlanks@7975.7998.24\|CGCGACCTCAGATCAGACGTGGCG | >hg38_21_+_8433172_8446622_RNA45SN1_45S_50ntFlanks | 7975 | 7998 | 24 | CGCGACCTCAGATCAGACGTGGCG | BT20 | 29 | 91 | 23 | 26 | 56 | 20.2 | 80.3 | 15.1 | 4.2 | 63.0 |
| >hg38_21_+_8433172_8446622_RNA45SN1_45S_50ntFlanks@7975.8000.26\|CGCGACCTCAGATCAGACGTGGCGAC | >hg38_21_+_8433172_8446622_RNA45SN1_45S_50ntFlanks | 7975 | 8000 | 26 | CGCGACCTCAGATCAGACGTGGCGAC | BT20 | 19 | 104 | 18 | 30 | 85 | 4.6 | 71.6 | 7.0 | 4.7 | 39.2 |
| >hg38_21_+_8433172_8446622_RNA45SN1_45S_50ntFlanks@7975.8001.27\|CGCGACCTCAGATCAGACGTGGCGACC | >hg38_21_+_8433172_8446622_RNA45SN1_45S_50ntFlanks | 7975 | 8001 | 27 | CGCGACCTCAGATCAGACGTGGCGACC | BT20 | 39 | 151 | 33 | 63 | 95 | 15.2 | 71.5 | 3.0 | 15.2 | 52.4 |
| >hg38_21_+_8433172_8446622_RNA45SN1_45S_50ntFlanks@7975.8002.28\|CGCGACCTCAGATCAGACGTGGCGACCC | >hg38_21_+_8433172_8446622_RNA45SN1_45S_50ntFlanks | 7975 | 8002 | 28 | CGCGACCTCAGATCAGACGTGGCGACCC | BT20 | 86 | 474 | 61 | 687 | 428 | 11.9 | 289.0 | 8.0 | 343.0 | 206.7 |
| >hg38_21_+_8433172_8446622_RNA45SN1_45S_50ntFlanks@7975.8002.28\|CGCGACCTCAGATCAGACGTGGCGACCC | >hg38_21_+_8433172_8446622_RNA45SN1_45S_50ntFlanks | 7975 | 8002 | 28 | CGCGACCTCAGATCAGACGTGGCGACCC | MB231 | 12 | 42 | 33 | 124 | 170 | 12.6 | 20.0 | 0.6 | 164.7 | 89.9 |
| >hg38_21_+_8433172_8446622_RNA45SN1_45S_50ntFlanks@7975.8003.29\|CGCGACCTCAGATCAGACGTGGCGACCCG | >hg38_21_+_8433172_8446622_RNA45SN1_45S_50ntFlanks | 7975 | 8003 | 29 | CGCGACCTCAGATCAGACGTGGCGACCCG | BT20 | 41 | 233 | 120 | 172 | 223 | 27.4 | 173.7 | 136.7 | 152.3 | 131.9 |
| >hg38_21_+_8433172_8446622_RNA45SN1_45S_50ntFlanks@7975.8003.29\|CGCGACCTCAGATCAGACGTGGCGACCCG | >hg38_21_+_8433172_8446622_RNA45SN1_45S_50ntFlanks | 7975 | 8003 | 29 | CGCGACCTCAGATCAGACGTGGCGACCCG | MB231 | 16 | 61 | 59 | 22 | 228 | 8.0 | 44.6 | 21.8 | 11.7 | 276.6 |
| >hg38_21_+_8433172_8446622_RNA45SN1_45S_50ntFlanks@7975.8003.29\|CGCGACCTCAGATCAGACGTGGCGACCCG | >hg38_21_+_8433172_8446622_RNA45SN1_45S_50ntFlanks | 7975 | 8003 | 29 | CGCGACCTCAGATCAGACGTGGCGACCCG | MB468 | 84 | 51 | 108 | 103 | 162 | 42.4 | 12.3 | 53.9 | 25.7 | 83.4 |
| >hg38_21_+_8433172_8446622_RNA45SN1_45S_50ntFlanks@7975.8004.30\|CGCGACCTCAGATCAGACGTGGCGACCCGC | >hg38_21_+_8433172_8446622_RNA45SN1_45S_50ntFlanks | 7975 | 8004 | 30 | CGCGACCTCAGATCAGACGTGGCGACCCGC | BT20 | 222 | 1720 | 1160 | 705 | 2277 | 161.3 | 1376.3 | 1745.5 | 429.6 | 1012.0 |
| >hg38_21_+_8433172_8446622_RNA45SN1_45S_50ntFlanks@7975.8004.30\|CGCGACCTCAGATCAGACGTGGCGACCCGC | >hg38_21_+_8433172_8446622_RNA45SN1_45S_50ntFlanks | 7975 | 8004 | 30 | CGCGACCTCAGATCAGACGTGGCGACCCGC | MB231 | 76 | 202 | 179 | 79 | 552 | 73.7 | 43.7 | 75.5 | 66.9 | 381.0 |
| >hg38_21_+_8433172_8446622_RNA45SN1_45S_50ntFlanks@7975.8004.30\|CGCGACCTCAGATCAGACGTGGCGACCCGC | >hg38_21_+_8433172_8446622_RNA45SN1_45S_50ntFlanks | 7975 | 8004 | 30 | CGCGACCTCAGATCAGACGTGGCGACCCGC | MB468 | 27 | 46 | 20 | 46 | 174 | 14.0 | 20.8 | 11.0 | 30.4 | 94.8 |
| >hg38_21_+_8433172_8446622_RNA45SN1_45S_50ntFlanks@7975.8005.31\|CGCGACCTCAGATCAGACGTGGCGACCCGCT | >hg38_21_+_8433172_8446622_RNA45SN1_45S_50ntFlanks | 7975 | 8005 | 31 | CGCGACCTCAGATCAGACGTGGCGACCCGCT | BT20 | 1936 | 14093 | 1594 | 3182 | 23675 | 1035.7 | 11097.3 | 1135.7 | 1188.8 | 11046.5 |
| >hg38_21_+_8433172_8446622_RNA45SN1_45S_50ntFlanks@7975.8005.31\|CGCGACCTCAGATCAGACGTGGCGACCCGCT | >hg38_21_+_8433172_8446622_RNA45SN1_45S_50ntFlanks | 7975 | 8005 | 31 | CGCGACCTCAGATCAGACGTGGCGACCCGCT | MB231 | 458 | 762 | 528 | 274 | 3688 | 603.3 | 540.4 | 298.6 | 351.9 | 4220.4 |
| >hg38_21_+_8433172_8446622_RNA45SN1_45S_50ntFlanks@7975.8005.31\|CGCGACCTCAGATCAGACGTGGCGACCCGCT | >hg38_21_+_8433172_8446622_RNA45SN1_45S_50ntFlanks | 7975 | 8005 | 31 | CGCGACCTCAGATCAGACGTGGCGACCCGCT | MB468 | 147 | 196 | 113 | 144 | 639 | 59.1 | 57.0 | 40.3 | 99.5 | 264.2 |
| >hg38_21_+_8433172_8446622_RNA45SN1_45S_50ntFlanks@7975.8006.32\|CGCGACCTCAGATCAGACGTGGCGACCCGCTG | >hg38_21_+_8433172_8446622_RNA45SN1_45S_50ntFlanks | 7975 | 8006 | 32 | CGCGACCTCAGATCAGACGTGGCGACCCGCTG | BT20 | 521 | 4177 | 2109 | 1029 | 7322 | 197.4 | 2811.9 | 3372.5 | 942.2 | 2630.5 |
| >hg38_21_+_8433172_8446622_RNA45SN1_45S_50ntFlanks@7975.8006.32\|CGCGACCTCAGATCAGACGTGGCGACCCGCTG | >hg38_21_+_8433172_8446622_RNA45SN1_45S_50ntFlanks | 7975 | 8006 | 32 | CGCGACCTCAGATCAGACGTGGCGACCCGCTG | MB231 | 154 | 250 | 215 | 72 | 1235 | 207.7 | 98.9 | 84.1 | 68.9 | 1083.6 |
| >hg38_21_+_8433172_8446622_RNA45SN1_45S_50ntFlanks@7975.8006.32\|CGCGACCTCAGATCAGACGTGGCGACCCGCTG | >hg38_21_+_8433172_8446622_RNA45SN1_45S_50ntFlanks | 7975 | 8006 | 32 | CGCGACCTCAGATCAGACGTGGCGACCCGCTG | MB468 | 1819 | 2684 | 951 | 1215 | 10569 | 860.0 | 1650.4 | 352.1 | 498.1 | 7040.3 |
| >hg38_21_+_8433172_8446622_RNA45SN1_45S_50ntFlanks@7975.8007.33\|CGCGACCTCAGATCAGACGTGGCGACCCGCTGA | >hg38_21_+_8433172_8446622_RNA45SN1_45S_50ntFlanks | 7975 | 8007 | 33 | CGCGACCTCAGATCAGACGTGGCGACCCGCTGA | BT20 | 288 | 2118 | 861 | 526 | 3231 | 213.8 | 1732.6 | 1261.0 | 506.0 | 453.9 |
| >hg38_21_+_8433172_8446622_RNA45SN1_45S_50ntFlanks@7975.8007.33\|CGCGACCTCAGATCAGACGTGGCGACCCGCTGA | >hg38_21_+_8433172_8446622_RNA45SN1_45S_50ntFlanks | 7975 | 8007 | 33 | CGCGACCTCAGATCAGACGTGGCGACCCGCTGA | MB231 | 184 | 329 | 334 | 73 | 960 | 257.5 | 120.5 | 180.5 | 31.0 | 705.6 |
| >hg38_21_+_8433172_8446622_RNA45SN1_45S_50ntFlanks@7975.8007.33\|CGCGACCTCAGATCAGACGTGGCGACCCGCTGA | >hg38_21_+_8433172_8446622_RNA45SN1_45S_50ntFlanks | 7975 | 8007 | 33 | CGCGACCTCAGATCAGACGTGGCGACCCGCTGA | MB468 | 133 | 322 | 55 | 146 | 1043 | 71.2 | 223.4 | 27.7 | 66.1 | 874.9 |
| >hg38_21_+_8433172_8446622_RNA45SN1_45S_50ntFlanks@7975.8008.34\|CGCGACCTCAGATCAGACGTGGCGACCCGCTGAA | >hg38_21_+_8433172_8446622_RNA45SN1_45S_50ntFlanks | 7975 | 8008 | 34 | CGCGACCTCAGATCAGACGTGGCGACCCGCTGAA | BT20 | 675 | 5474 | 18863 | 5570 | 12693 | 467.6 | 5852.6 | 31795.6 | 8887.1 | 2998.2 |
| >hg38_21_+_8433172_8446622_RNA45SN1_45S_50ntFlanks@7975.8008.34\|CGCGACCTCAGATCAGACGTGGCGACCCGCTGAA | >hg38_21_+_8433172_8446622_RNA45SN1_45S_50ntFlanks | 7975 | 8008 | 34 | CGCGACCTCAGATCAGACGTGGCGACCCGCTGAA | MB231 | 1335 | 2211 | 2765 | 623 | 8150 | 1747.1 | 1228.4 | 2633.4 | 382.9 | 5927.5 |
| >hg38_21_+_8433172_8446622_RNA45SN1_45S_50ntFlanks@7975.8008.34\|CGCGACCTCAGATCAGACGTGGCGACCCGCTGAA | >hg38_21_+_8433172_8446622_RNA45SN1_45S_50ntFlanks | 7975 | 8008 | 34 | CGCGACCTCAGATCAGACGTGGCGACCCGCTGAA | MB468 | 511 | 1604 | 220 | 778 | 5822 | 262.3 | 787.4 | 102.2 | 584.4 | 5283.0 |
| >hg38_21_+_8433172_8446622_RNA45SN1_45S_50ntFlanks@7975.8009.35\|CGCGACCTCAGATCAGACGTGGCGACCCGCTGAAT | >hg38_21_+_8433172_8446622_RNA45SN1_45S_50ntFlanks | 7975 | 8009 | 35 | CGCGACCTCAGATCAGACGTGGCGACCCGCTGAAT | BT20 | 2254 | 19421 | 5955 | 8600 | 27709 | 948.0 | 20514.0 | 3198.9 | 11675.4 | 5048.6 |
| >hg38_21_+_8433172_8446622_RNA45SN1_45S_50ntFlanks@7975.8009.35\|CGCGACCTCAGATCAGACGTGGCGACCCGCTGAAT | >hg38_21_+_8433172_8446622_RNA45SN1_45S_50ntFlanks | 7975 | 8009 | 35 | CGCGACCTCAGATCAGACGTGGCGACCCGCTGAAT | MB231 | 2042 | 2638 | 5498 | 2089 | 10593 | 2620.4 | 1510.9 | 3971.7 | 980.1 | 5497.1 |
| >hg38_21_+_8433172_8446622_RNA45SN1_45S_50ntFlanks@7975.8009.35\|CGCGACCTCAGATCAGACGTGGCGACCCGCTGAAT | >hg38_21_+_8433172_8446622_RNA45SN1_45S_50ntFlanks | 7975 | 8009 | 35 | CGCGACCTCAGATCAGACGTGGCGACCCGCTGAAT | MB468 | 120 | 491 | 68 | 311 | 2053 | 33.9 | 180.9 | 26.6 | 218.5 | 1964.1 |
| >hg38_21_+_8433172_8446622_RNA45SN1_45S_50ntFlanks@7975.8010.36\|CGCGACCTCAGATCAGACGTGGCGACCCGCTGAATT | >hg38_21_+_8433172_8446622_RNA45SN1_45S_50ntFlanks | 7975 | 8010 | 36 | CGCGACCTCAGATCAGACGTGGCGACCCGCTGAATT | BT20 | 3199 | 25253 | 4387 | 4086 | 24499 | 2036.5 | 14415.5 | 1057.5 | 5450.8 | 6134.8 |
| >hg38_21_+_8433172_8446622_RNA45SN1_45S_50ntFlanks@7975.8010.36\|CGCGACCTCAGATCAGACGTGGCGACCCGCTGAATT | >hg38_21_+_8433172_8446622_RNA45SN1_45S_50ntFlanks | 7975 | 8010 | 36 | CGCGACCTCAGATCAGACGTGGCGACCCGCTGAATT | MB231 | 2392 | 3773 | 5237 | 820 | 10781 | 2916.0 | 2470.1 | 5225.3 | 730.8 | 5502.7 |
| >hg38_21_+_8433172_8446622_RNA45SN1_45S_50ntFlanks@7975.8010.36\|CGCGACCTCAGATCAGACGTGGCGACCCGCTGAATT | >hg38_21_+_8433172_8446622_RNA45SN1_45S_50ntFlanks | 7975 | 8010 | 36 | CGCGACCTCAGATCAGACGTGGCGACCCGCTGAATT | MB468 | 67 | 295 | 47 | 100 | 989 | 25.7 | 80.2 | 28.6 | 90.5 | 642.2 |
| >hg38_21_+_8433172_8446622_RNA45SN1_45S_50ntFlanks@7975.8011.37\|CGCGACCTCAGATCAGACGTGGCGACCCGCTGAATTT | >hg38_21_+_8433172_8446622_RNA45SN1_45S_50ntFlanks | 7975 | 8011 | 37 | CGCGACCTCAGATCAGACGTGGCGACCCGCTGAATTT | BT20 | 4105 | 24166 | 2505 | 2231 | 23550 | 4424.0 | 17676.9 | 808.8 | 2868.7 | 9955.0 |
| >hg38_21_+_8433172_8446622_RNA45SN1_45S_50ntFlanks@7975.8011.37\|CGCGACCTCAGATCAGACGTGGCGACCCGCTGAATTT | >hg38_21_+_8433172_8446622_RNA45SN1_45S_50ntFlanks | 7975 | 8011 | 37 | CGCGACCTCAGATCAGACGTGGCGACCCGCTGAATTT | MB231 | 4147 | 6773 | 6404 | 626 | 9955 | 4657.4 | 4095.8 | 5113.3 | 188.4 | 6471.2 |
| >hg38_21_+_8433172_8446622_RNA45SN1_45S_50ntFlanks@7975.8011.37\|CGCGACCTCAGATCAGACGTGGCGACCCGCTGAATTT | >hg38_21_+_8433172_8446622_RNA45SN1_45S_50ntFlanks | 7975 | 8011 | 37 | CGCGACCTCAGATCAGACGTGGCGACCCGCTGAATTT | MB468 | 136 | 583 | 68 | 165 | 1343 | 56.1 | 133.0 | 26.5 | 171.7 | 775.3 |
| >hg38_21_+_8433172_8446622_RNA45SN1_45S_50ntFlanks@7975.8012.38\|CGCGACCTCAGATCAGACGTGGCGACCCGCTGAATTTA | >hg38_21_+_8433172_8446622_RNA45SN1_45S_50ntFlanks | 7975 | 8012 | 38 | CGCGACCTCAGATCAGACGTGGCGACCCGCTGAATTTA | BT20 | 351 | 899 | 304 | 155 | 1685 | 436.7 | 448.7 | 283.0 | 141.7 | 480.6 |
| >hg38_21_+_8433172_8446622_RNA45SN1_45S_50ntFlanks@7975.8012.38\|CGCGACCTCAGATCAGACGTGGCGACCCGCTGAATTTA | >hg38_21_+_8433172_8446622_RNA45SN1_45S_50ntFlanks | 7975 | 8012 | 38 | CGCGACCTCAGATCAGACGTGGCGACCCGCTGAATTTA | MB231 | 838 | 1319 | 965 | 80 | 1897 | 1075.0 | 742.2 | 861.9 | 9.1 | 1414.6 |
| >hg38_21_+_8433172_8446622_RNA45SN1_45S_50ntFlanks@7975.8012.38\|CGCGACCTCAGATCAGACGTGGCGACCCGCTGAATTTA | >hg38_21_+_8433172_8446622_RNA45SN1_45S_50ntFlanks | 7975 | 8012 | 38 | CGCGACCTCAGATCAGACGTGGCGACCCGCTGAATTTA | MB468 | 238 | 801 | 81 | 261 | 2613 | 134.7 | 216.8 | 18.7 | 263.1 | 1587.8 |

| | | | | | | | | | | | | | | | | |
|---|---|---|---|---|---|---|---|---|---|---|---|---|---|---|---|---|
| >hg38_21_+_8433172_8446622_RNA45SN1_45S_50ntFlanks@7975.8013.39\|CGCGACCTCAGATCAGACGTGGCGACCCGCTGAATTTAA | >hg38_21_+_8433172_8446622_RNA45SN1_45S_50ntFlanks | 7975 | 8013 | 39 | CGCGACCTCAGATCAGACGTGGCGACCCGCTGAATTTAA | BT20 | 163 | 266 | 147 | 76 | 914 | 210.0 | 109.1 | 128.7 | 63.8 | 469.4 |
| >hg38_21_+_8433172_8446622_RNA45SN1_45S_50ntFlanks@7975.8013.39\|CGCGACCTCAGATCAGACGTGGCGACCCGCTGAATTTAA | >hg38_21_+_8433172_8446622_RNA45SN1_45S_50ntFlanks | 7975 | 8013 | 39 | CGCGACCTCAGATCAGACGTGGCGACCCGCTGAATTTAA | MB231 | 402 | 630 | 678 | 36 | 1180 | 525.8 | 576.2 | 723.5 | 10.1 | 981.8 |
| >hg38_21_+_8433172_8446622_RNA45SN1_45S_50ntFlanks@7975.8013.39\|CGCGACCTCAGATCAGACGTGGCGACCCGCTGAATTTAA | >hg38_21_+_8433172_8446622_RNA45SN1_45S_50ntFlanks | 7975 | 8013 | 39 | CGCGACCTCAGATCAGACGTGGCGACCCGCTGAATTTAA | MB468 | 11 | 42 | 5 | 20 | 98 | 7.1 | 5.2 | 1.2 | 23.7 | 47.6 |
| >hg38_21_+_8433172_8446622_RNA45SN1_45S_50ntFlanks@7975.8014.40\|CGCGACCTCAGATCAGACGTGGCGACCCGCTGAATTTAAG | >hg38_21_+_8433172_8446622_RNA45SN1_45S_50ntFlanks | 7975 | 8014 | 40 | CGCGACCTCAGATCAGACGTGGCGACCCGCTGAATTTAAG | BT20 | 102 | 426 | 140 | 71 | 1379 | 124.4 | 290.4 | 159.0 | 69.6 | 705.6 |
| >hg38_21_+_8433172_8446622_RNA45SN1_45S_50ntFlanks@7975.8014.40\|CGCGACCTCAGATCAGACGTGGCGACCCGCTGAATTTAAG | >hg38_21_+_8433172_8446622_RNA45SN1_45S_50ntFlanks | 7975 | 8014 | 40 | CGCGACCTCAGATCAGACGTGGCGACCCGCTGAATTTAAG | MB231 | 215 | 424 | 365 | 25 | 764 | 249.7 | 302.0 | 227.4 | 9.7 | 571.7 |
| >hg38_21_+_8433172_8446622_RNA45SN1_45S_50ntFlanks@7975.8014.40\|CGCGACCTCAGATCAGACGTGGCGACCCGCTGAATTTAAG | >hg38_21_+_8433172_8446622_RNA45SN1_45S_50ntFlanks | 7975 | 8014 | 40 | CGCGACCTCAGATCAGACGTGGCGACCCGCTGAATTTAAG | MB468 | 40 | 78 | 17 | 61 | 219 | 30.6 | 26.9 | 7.7 | 78.4 | 96.4 |
| >hg38_21_+_8433172_8446622_RNA45SN1_45S_50ntFlanks@7975.8015.41\|CGCGACCTCAGATCAGACGTGGCGACCCGCTGAATTTAAGC | >hg38_21_+_8433172_8446622_RNA45SN1_45S_50ntFlanks | 7975 | 8015 | 41 | CGCGACCTCAGATCAGACGTGGCGACCCGCTGAATTTAAGC | BT20 | 355 | 1359 | 203 | 190 | 2552 | 242.0 | 1045.3 | 157.3 | 48.8 | 458.0 |
| >hg38_21_+_8433172_8446622_RNA45SN1_45S_50ntFlanks@7975.8015.41\|CGCGACCTCAGATCAGACGTGGCGACCCGCTGAATTTAAGC | >hg38_21_+_8433172_8446622_RNA45SN1_45S_50ntFlanks | 7975 | 8015 | 41 | CGCGACCTCAGATCAGACGTGGCGACCCGCTGAATTTAAGC | MB231 | 189 | 380 | 319 | 104 | 1480 | 88.6 | 196.8 | 138.7 | 92.9 | 652.9 |
| >hg38_21_+_8433172_8446622_RNA45SN1_45S_50ntFlanks@7975.8016.42\|CGCGACCTCAGATCAGACGTGGCGACCCGCTGAATTTAAGCA | >hg38_21_+_8433172_8446622_RNA45SN1_45S_50ntFlanks | 7975 | 8016 | 42 | CGCGACCTCAGATCAGACGTGGCGACCCGCTGAATTTAAGCA | BT20 | 186 | 1004 | 192 | 135 | 3586 | 118.8 | 1365.0 | 152.6 | 42.6 | 2118.0 |
| >hg38_21_+_8433172_8446622_RNA45SN1_45S_50ntFlanks@7975.8016.42\|CGCGACCTCAGATCAGACGTGGCGACCCGCTGAATTTAAGCA | >hg38_21_+_8433172_8446622_RNA45SN1_45S_50ntFlanks | 7975 | 8016 | 42 | CGCGACCTCAGATCAGACGTGGCGACCCGCTGAATTTAAGCA | MB231 | 635 | 823 | 580 | 122 | 4162 | 692.5 | 1054.0 | 319.3 | 91.8 | 3867.6 |
| >hg38_21_+_8433172_8446622_RNA45SN1_45S_50ntFlanks@7975.8016.42\|CGCGACCTCAGATCAGACGTGGCGACCCGCTGAATTTAAGCA | >hg38_21_+_8433172_8446622_RNA45SN1_45S_50ntFlanks | 7975 | 8016 | 42 | CGCGACCTCAGATCAGACGTGGCGACCCGCTGAATTTAAGCA | MB468 | 10 | 36 | 11 | 24 | 378 | 5.1 | 34.5 | 5.2 | 30.8 | 231.1 |
| >hg38_21_+_8433172_8446622_RNA45SN1_45S_50ntFlanks@7975.8017.43\|CGCGACCTCAGATCAGACGTGGCGACCCGCTGAATTTAAGCAT | >hg38_21_+_8433172_8446622_RNA45SN1_45S_50ntFlanks | 7975 | 8017 | 43 | CGCGACCTCAGATCAGACGTGGCGACCCGCTGAATTTAAGCAT | BT20 | 93 | 325 | 45 | 32 | 717 | 97.8 | 427.3 | 59.2 | 29.0 | 391.7 |
| >hg38_21_+_8433172_8446622_RNA45SN1_45S_50ntFlanks@7975.8017.43\|CGCGACCTCAGATCAGACGTGGCGACCCGCTGAATTTAAGCAT | >hg38_21_+_8433172_8446622_RNA45SN1_45S_50ntFlanks | 7975 | 8017 | 43 | CGCGACCTCAGATCAGACGTGGCGACCCGCTGAATTTAAGCAT | MB231 | 83 | 102 | 91 | 20 | 686 | 28.3 | 85.7 | 38.1 | 21.1 | 370.4 |
| >hg38_21_+_8433172_8446622_RNA45SN1_45S_50ntFlanks@7975.8018.44\|CGCGACCTCAGATCAGACGTGGCGACCCGCTGAATTTAAGCATA | >hg38_21_+_8433172_8446622_RNA45SN1_45S_50ntFlanks | 7975 | 8018 | 44 | CGCGACCTCAGATCAGACGTGGCGACCCGCTGAATTTAAGCATA | BT20 | 40 | 198 | 30 | 42 | 1472 | 53.5 | 311.8 | 44.8 | 24.4 | 1765.7 |
| >hg38_21_+_8433172_8446622_RNA45SN1_45S_50ntFlanks@7975.8018.44\|CGCGACCTCAGATCAGACGTGGCGACCCGCTGAATTTAAGCATA | >hg38_21_+_8433172_8446622_RNA45SN1_45S_50ntFlanks | 7975 | 8018 | 44 | CGCGACCTCAGATCAGACGTGGCGACCCGCTGAATTTAAGCATA | MB231 | 73 | 123 | 53 | 34 | 1509 | 72.2 | 189.9 | 16.3 | 39.5 | 1149.8 |
| >hg38_21_+_8433172_8446622_RNA45SN1_45S_50ntFlanks@7975.8019.45\|CGCGACCTCAGATCAGACGTGGCGACCCGCTGAATTTAAGCATAT | >hg38_21_+_8433172_8446622_RNA45SN1_45S_50ntFlanks | 7975 | 8019 | 45 | CGCGACCTCAGATCAGACGTGGCGACCCGCTGAATTTAAGCATAT | MB231 | 14 | 21 | 17 | 2 | 68 | 9.8 | 31.1 | 9.3 | 0.9 | 59.4 |
| >hg38_21_+_8433172_8446622_RNA45SN1_45S_50ntFlanks@7975.8020.46\|CGCGACCTCAGATCAGACGTGGCGACCCGCTGAATTTAAGCATATT | >hg38_21_+_8433172_8446622_RNA45SN1_45S_50ntFlanks | 7975 | 8020 | 46 | CGCGACCTCAGATCAGACGTGGCGACCCGCTGAATTTAAGCATATT | MB231 | 14 | 18 | 15 | 1 | 39 | 12.9 | 28.6 | 12.7 | 0.5 | 34.1 |
| >hg38_21_+_8433172_8446622_RNA45SN1_45S_50ntFlanks@7976.8009.34\|GCGACCTCAGATCAGACGTGGCGACCCGCTGAAT | >hg38_21_+_8433172_8446622_RNA45SN1_45S_50ntFlanks | 7976 | 8009 | 34 | GCGACCTCAGATCAGACGTGGCGACCCGCTGAAT | MB231 | 13 | 35 | 65 | 25 | 30 | 9.7 | 29.3 | 41.8 | 11.6 | 16.7 |
| >hg38_21_+_8433172_8446622_RNA45SN1_45S_50ntFlanks@7976.8010.35\|GCGACCTCAGATCAGACGTGGCGACCCGCTGAATT | >hg38_21_+_8433172_8446622_RNA45SN1_45S_50ntFlanks | 7976 | 8010 | 35 | GCGACCTCAGATCAGACGTGGCGACCCGCTGAATT | BT20 | 12 | 80 | 14 | 107 | 89 | 12.6 | 42.3 | 5.3 | 174.7 | 19.8 |
| >hg38_21_+_8433172_8446622_RNA45SN1_45S_50ntFlanks@7976.8010.35\|GCGACCTCAGATCAGACGTGGCGACCCGCTGAATT | >hg38_21_+_8433172_8446622_RNA45SN1_45S_50ntFlanks | 7976 | 8010 | 35 | GCGACCTCAGATCAGACGTGGCGACCCGCTGAATT | MB231 | 14 | 32 | 51 | 6 | 27 | 12.6 | 15.7 | 36.5 | 4.2 | 17.0 |
| >hg38_21_+_8433172_8446622_RNA45SN1_45S_50ntFlanks@7976.8011.36\|GCGACCTCAGATCAGACGTGGCGACCCGCTGAATTT | >hg38_21_+_8433172_8446622_RNA45SN1_45S_50ntFlanks | 7976 | 8011 | 36 | GCGACCTCAGATCAGACGTGGCGACCCGCTGAATTT | BT20 | 15 | 80 | 11 | 241 | 89 | 18.7 | 52.2 | 2.8 | 410.1 | 30.6 |
| >hg38_21_+_8433172_8446622_RNA45SN1_45S_50ntFlanks@7976.8011.36\|GCGACCTCAGATCAGACGTGGCGACCCGCTGAATTT | >hg38_21_+_8433172_8446622_RNA45SN1_45S_50ntFlanks | 7976 | 8011 | 36 | GCGACCTCAGATCAGACGTGGCGACCCGCTGAATTT | MB231 | 27 | 65 | 86 | 13 | 38 | 26.5 | 33.0 | 62.2 | 10.2 | 26.8 |
| >hg38_21_+_8433172_8446622_RNA45SN1_45S_50ntFlanks@7977.7994.18\|CGACCTCAGATCAGACGT | >hg38_21_+_8433172_8446622_RNA45SN1_45S_50ntFlanks | 7977 | 7994 | 18 | CGACCTCAGATCAGACGT | MB468 | 22 | 18 | 25 | 23 | 14 | 11.6 | 9.4 | 15.7 | 4.6 | 4.2 |
| >hg38_21_+_8433172_8446622_RNA45SN1_45S_50ntFlanks@7977.7995.19\|CGACCTCAGATCAGACGTG | >hg38_21_+_8433172_8446622_RNA45SN1_45S_50ntFlanks | 7977 | 7995 | 19 | CGACCTCAGATCAGACGTG | MB468 | 22 | 17 | 27 | 22 | 11 | 11.8 | 6.9 | 16.5 | 2.8 | 4.0 |
| >hg38_21_+_8433172_8446622_RNA45SN1_45S_50ntFlanks@7977.8003.27\|CGACCTCAGATCAGACGTGGCGACCCG | >hg38_21_+_8433172_8446622_RNA45SN1_45S_50ntFlanks | 7977 | 8003 | 27 | CGACCTCAGATCAGACGTGGCGACCCG | MB468 | 12 | 14 | 13 | 7 | 15 | 6.8 | 2.5 | 7.9 | 2.4 | 5.7 |
| >hg38_21_+_8433172_8446622_RNA45SN1_45S_50ntFlanks@7977.8006.30\|CGACCTCAGATCAGACGTGGCGACCCGCTG | >hg38_21_+_8433172_8446622_RNA45SN1_45S_50ntFlanks | 7977 | 8006 | 30 | CGACCTCAGATCAGACGTGGCGACCCGCTG | MB468 | 98 | 135 | 42 | 85 | 207 | 55.1 | 23.6 | 23.7 | 25.9 | 47.1 |
| >hg38_21_+_8433172_8446622_RNA45SN1_45S_50ntFlanks@7977.8008.32\|CGACCTCAGATCAGACGTGGCGACCCGCTGAA | >hg38_21_+_8433172_8446622_RNA45SN1_45S_50ntFlanks | 7977 | 8008 | 32 | CGACCTCAGATCAGACGTGGCGACCCGCTGAA | MB468 | 20 | 37 | 5 | 35 | 58 | 10.7 | 6.0 | 3.0 | 19.2 | 11.1 |
| >hg38_21_+_8433172_8446622_RNA45SN1_45S_50ntFlanks@7977.8009.33\|CGACCTCAGATCAGACGTGGCGACCCGCTGAAT | >hg38_21_+_8433172_8446622_RNA45SN1_45S_50ntFlanks | 7977 | 8009 | 33 | CGACCTCAGATCAGACGTGGCGACCCGCTGAAT | MB231 | 12 | 15 | 38 | 10 | 33 | 10.2 | 4.7 | 35.2 | 5.8 | 33.2 |
| >hg38_21_+_8433172_8446622_RNA45SN1_45S_50ntFlanks@7977.8010.34\|CGACCTCAGATCAGACGTGGCGACCCGCTGAATT | >hg38_21_+_8433172_8446622_RNA45SN1_45S_50ntFlanks | 7977 | 8010 | 34 | CGACCTCAGATCAGACGTGGCGACCCGCTGAATT | MB231 | 24 | 26 | 74 | 5 | 50 | 22.0 | 14.6 | 83.6 | 1.6 | 45.4 |
| >hg38_21_+_8433172_8446622_RNA45SN1_45S_50ntFlanks@7977.8011.35\|CGACCTCAGATCAGACGTGGCGACCCGCTGAATTT | >hg38_21_+_8433172_8446622_RNA45SN1_45S_50ntFlanks | 7977 | 8011 | 35 | CGACCTCAGATCAGACGTGGCGACCCGCTGAATTT | BT20 | 18 | 39 | 19 | 108 | 50 | 13.7 | 19.2 | 9.8 | 176.3 | 30.1 |
| >hg38_21_+_8433172_8446622_RNA45SN1_45S_50ntFlanks@7977.8011.35\|CGACCTCAGATCAGACGTGGCGACCCGCTGAATTT | >hg38_21_+_8433172_8446622_RNA45SN1_45S_50ntFlanks | 7977 | 8011 | 35 | CGACCTCAGATCAGACGTGGCGACCCGCTGAATTT | MB231 | 76 | 77 | 224 | 11 | 111 | 80.7 | 60.7 | 272.3 | 5.3 | 90.3 |
| >hg38_21_+_8433172_8446622_RNA45SN1_45S_50ntFlanks@7977.8012.36\|CGACCTCAGATCAGACGTGGCGACCCGCTGAATTTA | >hg38_21_+_8433172_8446622_RNA45SN1_45S_50ntFlanks | 7977 | 8012 | 36 | CGACCTCAGATCAGACGTGGCGACCCGCTGAATTTA | MB231 | 23 | 22 | 51 | 2 | 33 | 28.0 | 20.6 | 62.4 | 1.8 | 29.3 |
| >hg38_21_+_8433172_8446622_RNA45SN1_45S_50ntFlanks@7977.8016.40\|CGACCTCAGATCAGACGTGGCGACCCGCTGAATTTAAGCA | >hg38_21_+_8433172_8446622_RNA45SN1_45S_50ntFlanks | 7977 | 8016 | 40 | CGACCTCAGATCAGACGTGGCGACCCGCTGAATTTAAGCA | MB231 | 69 | 66 | 146 | 4 | 117 | 77.2 | 90.5 | 175.9 | 3.6 | 132.1 |
| >hg38_21_+_8433172_8446622_RNA45SN1_45S_50ntFlanks@7977.8017.41\|CGACCTCAGATCAGACGTGGCGACCCGCTGAATTTAAGCAT | >hg38_21_+_8433172_8446622_RNA45SN1_45S_50ntFlanks | 7977 | 8017 | 41 | CGACCTCAGATCAGACGTGGCGACCCGCTGAATTTAAGCAT | MB231 | 27 | 28 | 46 | 1 | 51 | 27.8 | 39.1 | 41.0 | 1.1 | 59.1 |
| >hg38_21_+_8433172_8446622_RNA45SN1_45S_50ntFlanks@7977.8018.42\|CGACCTCAGATCAGACGTGGCGACCCGCTGAATTTAAGCATA | >hg38_21_+_8433172_8446622_RNA45SN1_45S_50ntFlanks | 7977 | 8018 | 42 | CGACCTCAGATCAGACGTGGCGACCCGCTGAATTTAAGCATA | MB231 | 13 | 18 | 30 | 0 | 23 | 13.6 | 28.3 | 32.8 | 0.7 | 39.6 |
| >hg38_21_+_8433172_8446622_RNA45SN1_45S_50ntFlanks@7977.8019.43\|CGACCTCAGATCAGACGTGGCGACCCGCTGAATTTAAGCATAT | >hg38_21_+_8433172_8446622_RNA45SN1_45S_50ntFlanks | 7977 | 8019 | 43 | CGACCTCAGATCAGACGTGGCGACCCGCTGAATTTAAGCATAT | MB231 | 21 | 33 | 44 | 0 | 46 | 21.4 | 52.5 | 40.6 | 0.3 | 65.0 |
| >hg38_21_+_8433172_8446622_RNA45SN1_45S_50ntFlanks@7978.8006.29\|GACCTCAGATCAGACGTGGCGACCCGCTG | >hg38_21_+_8433172_8446622_RNA45SN1_45S_50ntFlanks | 7978 | 8006 | 29 | GACCTCAGATCAGACGTGGCGACCCGCTG | MB468 | 19 | 29 | 9 | 16 | 32 | 11.6 | 2.8 | 5.1 | 8.1 | 8.1 |
| >hg38_21_+_8433172_8446622_RNA45SN1_45S_50ntFlanks@7978.8010.33\|GACCTCAGATCAGACGTGGCGACCCGCTGAATT | >hg38_21_+_8433172_8446622_RNA45SN1_45S_50ntFlanks | 7978 | 8010 | 33 | GACCTCAGATCAGACGTGGCGACCCGCTGAATT | MB231 | 12 | 13 | 39 | 4 | 28 | 9.0 | 6.5 | 35.0 | 3.5 | 23.6 |
| >hg38_21_+_8433172_8446622_RNA45SN1_45S_50ntFlanks@7978.8011.34\|GACCTCAGATCAGACGTGGCGACCCGCTGAATTT | >hg38_21_+_8433172_8446622_RNA45SN1_45S_50ntFlanks | 7978 | 8011 | 34 | GACCTCAGATCAGACGTGGCGACCCGCTGAATTT | BT20 | 13 | 25 | 12 | 33 | 40 | 10.8 | 10.6 | 7.6 | 46.7 | 20.9 |
| >hg38_21_+_8433172_8446622_RNA45SN1_45S_50ntFlanks@7978.8011.34\|GACCTCAGATCAGACGTGGCGACCCGCTGAATTT | >hg38_21_+_8433172_8446622_RNA45SN1_45S_50ntFlanks | 7978 | 8011 | 34 | GACCTCAGATCAGACGTGGCGACCCGCTGAATTT | MB231 | 35 | 36 | 113 | 9 | 74 | 31.5 | 22.7 | 114.3 | 5.0 | 61.0 |
| >hg38_21_+_8433172_8446622_RNA45SN1_45S_50ntFlanks@7978.8012.35\|GACCTCAGATCAGACGTGGCGACCCGCTGAATTTA | >hg38_21_+_8433172_8446622_RNA45SN1_45S_50ntFlanks | 7978 | 8012 | 35 | GACCTCAGATCAGACGTGGCGACCCGCTGAATTTA | MB231 | 11 | 10 | 27 | 2 | 30 | 12.6 | 8.5 | 28.7 | 2.1 | 25.6 |
| >hg38_21_+_8433172_8446622_RNA45SN1_45S_50ntFlanks@7978.8016.39\|GACCTCAGATCAGACGTGGCGACCCGCTGAATTTAAGCA | >hg38_21_+_8433172_8446622_RNA45SN1_45S_50ntFlanks | 7978 | 8016 | 39 | GACCTCAGATCAGACGTGGCGACCCGCTGAATTTAAGCA | MB231 | 25 | 22 | 47 | 3 | 50 | 29.2 | 27.3 | 50.8 | 3.6 | 56.9 |
| >hg38_21_+_8433172_8446622_RNA45SN1_45S_50ntFlanks@7978.8017.40\|GACCTCAGATCAGACGTGGCGACCCGCTGAATTTAAGCAT | >hg38_21_+_8433172_8446622_RNA45SN1_45S_50ntFlanks | 7978 | 8017 | 40 | GACCTCAGATCAGACGTGGCGACCCGCTGAATTTAAGCAT | MB231 | 17 | 17 | 26 | 2 | 77 | 19.3 | 20.6 | 22.5 | 1.6 | 96.8 |
| >hg38_21_+_8433172_8446622_RNA45SN1_45S_50ntFlanks@7978.8019.42\|GACCTCAGATCAGACGTGGCGACCCGCTGAATTTAAGCATAT | >hg38_21_+_8433172_8446622_RNA45SN1_45S_50ntFlanks | 7978 | 8019 | 42 | GACCTCAGATCAGACGTGGCGACCCGCTGAATTTAAGCATAT | MB231 | 11 | 16 | 29 | 0 | 24 | 11.4 | 23.6 | 26.8 | 0.4 | 29.8 |
| >hg38_21_+_8433172_8446622_RNA45SN1_45S_50ntFlanks@7979.7996.18\|ACCTCAGATCAGACGTGG | >hg38_21_+_8433172_8446622_RNA45SN1_45S_50ntFlanks | 7979 | 7996 | 18 | ACCTCAGATCAGACGTGG | MB468 | 13 | 3 | 19 | 13 | 3 | 4.1 | 2.3 | 11.6 | 6.9 | 1.5 |
| >hg38_21_+_8433172_8446622_RNA45SN1_45S_50ntFlanks@7979.7998.20\|ACCTCAGATCAGACGTGGCG | >hg38_21_+_8433172_8446622_RNA45SN1_45S_50ntFlanks | 7979 | 7998 | 20 | ACCTCAGATCAGACGTGGCG | MB468 | 13 | 2 | 12 | 7 | 4 | 9.1 | 1.4 | 10.7 | 1.5 | 1.5 |
| >hg38_21_+_8433172_8446622_RNA45SN1_45S_50ntFlanks@7979.7999.21\|ACCTCAGATCAGACGTGGCGA | >hg38_21_+_8433172_8446622_RNA45SN1_45S_50ntFlanks | 7979 | 7999 | 21 | ACCTCAGATCAGACGTGGCGA | MB468 | 15 | 4 | 20 | 10 | 4 | 11.0 | 2.6 | 16.9 | 2.1 | 1.6 |
| >hg38_21_+_8433172_8446622_RNA45SN1_45S_50ntFlanks@7979.8003.25\|ACCTCAGATCAGACGTGGCGACCCG | >hg38_21_+_8433172_8446622_RNA45SN1_45S_50ntFlanks | 7979 | 8003 | 25 | ACCTCAGATCAGACGTGGCGACCCG | MB468 | 45 | 92 | 52 | 25 | 35 | 24.7 | 37.5 | 41.9 | 4.6 | 16.6 |
| >hg38_21_+_8433172_8446622_RNA45SN1_45S_50ntFlanks@7979.8005.27\|ACCTCAGATCAGACGTGGCGACCCGCT | >hg38_21_+_8433172_8446622_RNA45SN1_45S_50ntFlanks | 7979 | 8005 | 27 | ACCTCAGATCAGACGTGGCGACCCGCT | MB468 | 19 | 22 | 12 | 16 | 33 | 8.2 | 2.9 | 6.8 | 4.7 | 14.9 |
| >hg38_21_+_8433172_8446622_RNA45SN1_45S_50ntFlanks@7979.8006.28\|ACCTCAGATCAGACGTGGCGACCCGCTG | >hg38_21_+_8433172_8446622_RNA45SN1_45S_50ntFlanks | 7979 | 8006 | 28 | ACCTCAGATCAGACGTGGCGACCCGCTG | MB468 | 555 | 563 | 209 | 372 | 925 | 334.8 | 85.4 | 113.4 | 158.0 | 273.3 |

| | | | | | | | | | | | | | | | | |
|---|---|---|---|---|---|---|---|---|---|---|---|---|---|---|---|---|
| >hg38_21_+_8433172_8446622_RNA45SN1_45S_50ntFlanks@7979.8007.29\|ACCTCAGATCAGACGTGGCGACCCGCTGA | >hg38_21_+_8433172_8446622_RNA45SN1_45S_50ntFlanks | 7979 | 8007 | 29 | ACCTCAGATCAGACGTGGCGACCCGCTGA | MB468 | 22 | 16 | 6 | 21 | 43 | 14.0 | 2.8 | 2.5 | 3.8 | 14.8 |
| >hg38_21_+_8433172_8446622_RNA45SN1_45S_50ntFlanks@7979.8008.30\|ACCTCAGATCAGACGTGGCGACCCGCTGAA | >hg38_21_+_8433172_8446622_RNA45SN1_45S_50ntFlanks | 7979 | 8008 | 30 | ACCTCAGATCAGACGTGGCGACCCGCTGAA | MB468 | 110 | 101 | 24 | 125 | 207 | 63.6 | 21.2 | 11.1 | 50.4 | 43.4 |
| >hg38_21_+_8433172_8446622_RNA45SN1_45S_50ntFlanks@7979.8009.31\|ACCTCAGATCAGACGTGGCGACCCGCTGAAT | >hg38_21_+_8433172_8446622_RNA45SN1_45S_50ntFlanks | 7979 | 8009 | 31 | ACCTCAGATCAGACGTGGCGACCCGCTGAAT | MB468 | 23 | 22 | 6 | 35 | 42 | 9.3 | 6.4 | 3.1 | 15.8 | 14.8 |
| >hg38_21_+_8433172_8446622_RNA45SN1_45S_50ntFlanks@7979.8011.33\|ACCTCAGATCAGACGTGGCGACCCGCTGAATTT | >hg38_21_+_8433172_8446622_RNA45SN1_45S_50ntFlanks | 7979 | 8011 | 33 | ACCTCAGATCAGACGTGGCGACCCGCTGAATTT | MB468 | 11 | 11 | 3 | 12 | 29 | 5.3 | 3.8 | 1.2 | 6.9 | 19.0 |
| >hg38_21_+_8433172_8446622_RNA45SN1_45S_50ntFlanks@7979.8012.34\|ACCTCAGATCAGACGTGGCGACCCGCTGAATTTA | >hg38_21_+_8433172_8446622_RNA45SN1_45S_50ntFlanks | 7979 | 8012 | 34 | ACCTCAGATCAGACGTGGCGACCCGCTGAATTTA | MB468 | 34 | 28 | 6 | 28 | 99 | 17.2 | 10.6 | 1.3 | 18.1 | 55.6 |
| >hg38_21_+_8433172_8446622_RNA45SN1_45S_50ntFlanks@7980.7998.19\|CCTCAGATCAGACGTGGCG | >hg38_21_+_8433172_8446622_RNA45SN1_45S_50ntFlanks | 7980 | 7998 | 19 | CCTCAGATCAGACGTGGCG | MB468 | 16 | 10 | 16 | 12 | 6 | 8.4 | 5.4 | 10.9 | 1.2 | 2.6 |
| >hg38_21_+_8433172_8446622_RNA45SN1_45S_50ntFlanks@7980.7999.20\|CCTCAGATCAGACGTGGCGA | >hg38_21_+_8433172_8446622_RNA45SN1_45S_50ntFlanks | 7980 | 7999 | 20 | CCTCAGATCAGACGTGGCGA | MB468 | 11 | 8 | 16 | 11 | 4 | 5.2 | 2.5 | 11.8 | 1.0 | 2.0 |
| >hg38_21_+_8433172_8446622_RNA45SN1_45S_50ntFlanks@7980.8003.24\|CCTCAGATCAGACGTGGCGACCCG | >hg38_21_+_8433172_8446622_RNA45SN1_45S_50ntFlanks | 7980 | 8003 | 24 | CCTCAGATCAGACGTGGCGACCCG | MB468 | 28 | 54 | 30 | 26 | 27 | 14.9 | 30.3 | 23.4 | 8.7 | 14.8 |
| >hg38_21_+_8433172_8446622_RNA45SN1_45S_50ntFlanks@7980.8004.25\|CCTCAGATCAGACGTGGCGACCCGC | >hg38_21_+_8433172_8446622_RNA45SN1_45S_50ntFlanks | 7980 | 8004 | 25 | CCTCAGATCAGACGTGGCGACCCGC | MB468 | 11 | 16 | 6 | 9 | 26 | 6.5 | 3.3 | 3.8 | 4.2 | 9.8 |
| >hg38_21_+_8433172_8446622_RNA45SN1_45S_50ntFlanks@7980.8005.26\|CCTCAGATCAGACGTGGCGACCCGCT | >hg38_21_+_8433172_8446622_RNA45SN1_45S_50ntFlanks | 7980 | 8005 | 26 | CCTCAGATCAGACGTGGCGACCCGCT | MB468 | 20 | 26 | 13 | 24 | 42 | 9.6 | 7.0 | 7.4 | 6.2 | 17.4 |
| >hg38_21_+_8433172_8446622_RNA45SN1_45S_50ntFlanks@7980.8006.27\|CCTCAGATCAGACGTGGCGACCCGCTG | >hg38_21_+_8433172_8446622_RNA45SN1_45S_50ntFlanks | 7980 | 8006 | 27 | CCTCAGATCAGACGTGGCGACCCGCTG | MB468 | 469 | 454 | 167 | 431 | 868 | 257.1 | 121.5 | 93.0 | 131.5 | 303.1 |
| >hg38_21_+_8433172_8446622_RNA45SN1_45S_50ntFlanks@7980.8007.28\|CCTCAGATCAGACGTGGCGACCCGCTGA | >hg38_21_+_8433172_8446622_RNA45SN1_45S_50ntFlanks | 7980 | 8007 | 28 | CCTCAGATCAGACGTGGCGACCCGCTGA | MB468 | 16 | 11 | 4 | 21 | 36 | 8.2 | 3.2 | 2.1 | 3.0 | 13.6 |
| >hg38_21_+_8433172_8446622_RNA45SN1_45S_50ntFlanks@7980.8008.29\|CCTCAGATCAGACGTGGCGACCCGCTGAA | >hg38_21_+_8433172_8446622_RNA45SN1_45S_50ntFlanks | 7980 | 8008 | 29 | CCTCAGATCAGACGTGGCGACCCGCTGAA | MB468 | 85 | 69 | 17 | 112 | 182 | 47.6 | 23.0 | 8.1 | 36.8 | 50.0 |
| >hg38_21_+_8433172_8446622_RNA45SN1_45S_50ntFlanks@7980.8009.30\|CCTCAGATCAGACGTGGCGACCCGCTGAAT | >hg38_21_+_8433172_8446622_RNA45SN1_45S_50ntFlanks | 7980 | 8009 | 30 | CCTCAGATCAGACGTGGCGACCCGCTGAAT | MB468 | 17 | 15 | 5 | 28 | 35 | 7.4 | 6.5 | 2.4 | 11.9 | 11.8 |
| >hg38_21_+_8433172_8446622_RNA45SN1_45S_50ntFlanks@7980.8012.33\|CCTCAGATCAGACGTGGCGACCCGCTGAATTTA | >hg38_21_+_8433172_8446622_RNA45SN1_45S_50ntFlanks | 7980 | 8012 | 33 | CCTCAGATCAGACGTGGCGACCCGCTGAATTTA | MB468 | 26 | 18 | 4 | 26 | 73 | 12.7 | 7.1 | 1.4 | 17.0 | 39.6 |
| >hg38_21_+_8433172_8446622_RNA45SN1_45S_50ntFlanks@7981.7998.18\|CTCAGATCAGACGTGGCG | >hg38_21_+_8433172_8446622_RNA45SN1_45S_50ntFlanks | 7981 | 7998 | 18 | CTCAGATCAGACGTGGCG | MB468 | 10 | 11 | 9 | 6 | 4 | 5.5 | 4.7 | 5.3 | 0.9 | 1.1 |
| >hg38_21_+_8433172_8446622_RNA45SN1_45S_50ntFlanks@7981.8003.23\|CTCAGATCAGACGTGGCGACCCG | >hg38_21_+_8433172_8446622_RNA45SN1_45S_50ntFlanks | 7981 | 8003 | 23 | CTCAGATCAGACGTGGCGACCCG | MB468 | 11 | 25 | 14 | 10 | 10 | 6.3 | 13.6 | 11.9 | 1.1 | 3.1 |
| >hg38_21_+_8433172_8446622_RNA45SN1_45S_50ntFlanks@7981.8004.24\|CTCAGATCAGACGTGGCGACCCGC | >hg38_21_+_8433172_8446622_RNA45SN1_45S_50ntFlanks | 7981 | 8004 | 24 | CTCAGATCAGACGTGGCGACCCGC | MB468 | 12 | 14 | 5 | 10 | 28 | 6.1 | 4.9 | 3.8 | 6.3 | 7.6 |
| >hg38_21_+_8433172_8446622_RNA45SN1_45S_50ntFlanks@7981.8005.25\|CTCAGATCAGACGTGGCGACCCGCT | >hg38_21_+_8433172_8446622_RNA45SN1_45S_50ntFlanks | 7981 | 8005 | 25 | CTCAGATCAGACGTGGCGACCCGCT | MB468 | 18 | 23 | 10 | 18 | 34 | 8.1 | 6.7 | 5.7 | 7.3 | 10.3 |
| >hg38_21_+_8433172_8446622_RNA45SN1_45S_50ntFlanks@7981.8006.26\|CTCAGATCAGACGTGGCGACCCGCTG | >hg38_21_+_8433172_8446622_RNA45SN1_45S_50ntFlanks | 7981 | 8006 | 26 | CTCAGATCAGACGTGGCGACCCGCTG | MB468 | 387 | 344 | 149 | 393 | 677 | 176.5 | 155.1 | 79.8 | 119.5 | 253.9 |
| >hg38_21_+_8433172_8446622_RNA45SN1_45S_50ntFlanks@7981.8007.27\|CTCAGATCAGACGTGGCGACCCGCTGA | >hg38_21_+_8433172_8446622_RNA45SN1_45S_50ntFlanks | 7981 | 8007 | 27 | CTCAGATCAGACGTGGCGACCCGCTGA | MB468 | 13 | 7 | 3 | 15 | 29 | 6.3 | 2.1 | 1.0 | 2.2 | 10.0 |
| >hg38_21_+_8433172_8446622_RNA45SN1_45S_50ntFlanks@7981.8008.28\|CTCAGATCAGACGTGGCGACCCGCTGAA | >hg38_21_+_8433172_8446622_RNA45SN1_45S_50ntFlanks | 7981 | 8008 | 28 | CTCAGATCAGACGTGGCGACCCGCTGAA | MB468 | 66 | 38 | 13 | 90 | 138 | 31.0 | 15.4 | 5.0 | 30.0 | 38.1 |
| >hg38_21_+_8433172_8446622_RNA45SN1_45S_50ntFlanks@7981.8009.29\|CTCAGATCAGACGTGGCGACCCGCTGAAT | >hg38_21_+_8433172_8446622_RNA45SN1_45S_50ntFlanks | 7981 | 8009 | 29 | CTCAGATCAGACGTGGCGACCCGCTGAAT | MB468 | 12 | 8 | 3 | 19 | 25 | 5.6 | 3.5 | 1.4 | 7.8 | 8.1 |
| >hg38_21_+_8433172_8446622_RNA45SN1_45S_50ntFlanks@7981.8012.32\|CTCAGATCAGACGTGGCGACCCGCTGAATTTA | >hg38_21_+_8433172_8446622_RNA45SN1_45S_50ntFlanks | 7981 | 8012 | 32 | CTCAGATCAGACGTGGCGACCCGCTGAATTTA | MB468 | 16 | 7 | 3 | 19 | 43 | 7.4 | 3.4 | 0.8 | 9.4 | 23.7 |
| >hg38_21_+_8433172_8446622_RNA45SN1_45S_50ntFlanks@7982.8004.23\|TCAGATCAGACGTGGCGACCCGC | >hg38_21_+_8433172_8446622_RNA45SN1_45S_50ntFlanks | 7982 | 8004 | 23 | TCAGATCAGACGTGGCGACCCGC | MB468 | 16 | 16 | 6 | 15 | 37 | 9.9 | 7.6 | 5.6 | 7.6 | 13.6 |
| >hg38_21_+_8433172_8446622_RNA45SN1_45S_50ntFlanks@7982.8005.24\|TCAGATCAGACGTGGCGACCCGCT | >hg38_21_+_8433172_8446622_RNA45SN1_45S_50ntFlanks | 7982 | 8005 | 24 | TCAGATCAGACGTGGCGACCCGCT | MB468 | 17 | 14 | 8 | 19 | 32 | 9.7 | 4.8 | 6.5 | 4.6 | 15.8 |
| >hg38_21_+_8433172_8446622_RNA45SN1_45S_50ntFlanks@7982.8006.25\|TCAGATCAGACGTGGCGACCCGCTG | >hg38_21_+_8433172_8446622_RNA45SN1_45S_50ntFlanks | 7982 | 8006 | 25 | TCAGATCAGACGTGGCGACCCGCTG | MB468 | 258 | 140 | 69 | 254 | 496 | 126.6 | 55.9 | 37.4 | 66.5 | 240.0 |
| >hg38_21_+_8433172_8446622_RNA45SN1_45S_50ntFlanks@7982.8007.26\|TCAGATCAGACGTGGCGACCCGCTGA | >hg38_21_+_8433172_8446622_RNA45SN1_45S_50ntFlanks | 7982 | 8007 | 26 | TCAGATCAGACGTGGCGACCCGCTGA | MB468 | 14 | 6 | 2 | 14 | 32 | 7.3 | 2.3 | 1.5 | 3.6 | 16.0 |
| >hg38_21_+_8433172_8446622_RNA45SN1_45S_50ntFlanks@7982.8008.27\|TCAGATCAGACGTGGCGACCCGCTGAA | >hg38_21_+_8433172_8446622_RNA45SN1_45S_50ntFlanks | 7982 | 8008 | 27 | TCAGATCAGACGTGGCGACCCGCTGAA | MB468 | 60 | 24 | 10 | 76 | 125 | 28.7 | 12.5 | 5.1 | 19.5 | 64.4 |
| >hg38_21_+_8433172_8446622_RNA45SN1_45S_50ntFlanks@7982.8009.28\|TCAGATCAGACGTGGCGACCCGCTGAAT | >hg38_21_+_8433172_8446622_RNA45SN1_45S_50ntFlanks | 7982 | 8009 | 28 | TCAGATCAGACGTGGCGACCCGCTGAAT | MB468 | 11 | 5 | 2 | 16 | 20 | 4.6 | 3.2 | 0.9 | 4.5 | 10.5 |
| >hg38_21_+_8433172_8446622_RNA45SN1_45S_50ntFlanks@7982.8012.31\|TCAGATCAGACGTGGCGACCCGCTGAATTTA | >hg38_21_+_8433172_8446622_RNA45SN1_45S_50ntFlanks | 7982 | 8012 | 31 | TCAGATCAGACGTGGCGACCCGCTGAATTTA | MB468 | 11 | 3 | 2 | 12 | 28 | 5.7 | 1.5 | 0.6 | 6.8 | 19.2 |
| >hg38_21_+_8433172_8446622_RNA45SN1_45S_50ntFlanks@7983.8003.21\|CAGATCAGACGTGGCGACCCG | >hg38_21_+_8433172_8446622_RNA45SN1_45S_50ntFlanks | 7983 | 8003 | 21 | CAGATCAGACGTGGCGACCCG | MB468 | 34 | 57 | 40 | 62 | 38 | 22.1 | 38.7 | 37.4 | 32.2 | 19.2 |
| >hg38_21_+_8433172_8446622_RNA45SN1_45S_50ntFlanks@7983.8004.22\|CAGATCAGACGTGGCGACCCGC | >hg38_21_+_8433172_8446622_RNA45SN1_45S_50ntFlanks | 7983 | 8004 | 22 | CAGATCAGACGTGGCGACCCGC | MB468 | 62 | 61 | 22 | 81 | 185 | 40.4 | 25.2 | 16.8 | 38.5 | 81.2 |
| >hg38_21_+_8433172_8446622_RNA45SN1_45S_50ntFlanks@7983.8005.23\|CAGATCAGACGTGGCGACCCGCT | >hg38_21_+_8433172_8446622_RNA45SN1_45S_50ntFlanks | 7983 | 8005 | 23 | CAGATCAGACGTGGCGACCCGCT | MB468 | 53 | 49 | 24 | 77 | 117 | 28.7 | 13.8 | 18.7 | 20.1 | 50.6 |
| >hg38_21_+_8433172_8446622_RNA45SN1_45S_50ntFlanks@7983.8006.24\|CAGATCAGACGTGGCGACCCGCTG | >hg38_21_+_8433172_8446622_RNA45SN1_45S_50ntFlanks | 7983 | 8006 | 24 | CAGATCAGACGTGGCGACCCGCTG | MB468 | 805 | 423 | 230 | 987 | 1689 | 381.2 | 146.1 | 111.0 | 266.0 | 826.9 |
| >hg38_21_+_8433172_8446622_RNA45SN1_45S_50ntFlanks@7983.8007.25\|CAGATCAGACGTGGCGACCCGCTGA | >hg38_21_+_8433172_8446622_RNA45SN1_45S_50ntFlanks | 7983 | 8007 | 25 | CAGATCAGACGTGGCGACCCGCTGA | MB468 | 41 | 15 | 8 | 54 | 94 | 19.2 | 6.9 | 4.4 | 22.0 | 46.7 |
| >hg38_21_+_8433172_8446622_RNA45SN1_45S_50ntFlanks@7983.8008.26\|CAGATCAGACGTGGCGACCCGCTGAA | >hg38_21_+_8433172_8446622_RNA45SN1_45S_50ntFlanks | 7983 | 8008 | 26 | CAGATCAGACGTGGCGACCCGCTGAA | MB468 | 200 | 79 | 37 | 288 | 482 | 90.2 | 39.4 | 16.9 | 78.0 | 232.5 |
| >hg38_21_+_8433172_8446622_RNA45SN1_45S_50ntFlanks@7983.8009.27\|CAGATCAGACGTGGCGACCCGCTGAAT | >hg38_21_+_8433172_8446622_RNA45SN1_45S_50ntFlanks | 7983 | 8009 | 27 | CAGATCAGACGTGGCGACCCGCTGAAT | MB468 | 31 | 13 | 6 | 53 | 67 | 13.7 | 6.9 | 1.8 | 14.2 | 33.8 |
| >hg38_21_+_8433172_8446622_RNA45SN1_45S_50ntFlanks@7983.8012.30\|CAGATCAGACGTGGCGACCCGCTGAATTTA | >hg38_21_+_8433172_8446622_RNA45SN1_45S_50ntFlanks | 7983 | 8012 | 30 | CAGATCAGACGTGGCGACCCGCTGAATTTA | MB468 | 29 | 8 | 4 | 36 | 79 | 12.4 | 4.0 | 1.4 | 17.5 | 49.5 |
| >hg38_21_+_8433172_8446622_RNA45SN1_45S_50ntFlanks@7984.8003.20\|AGATCAGACGTGGCGACCCG | >hg38_21_+_8433172_8446622_RNA45SN1_45S_50ntFlanks | 7984 | 8003 | 20 | AGATCAGACGTGGCGACCCG | MB468 | 21 | 37 | 27 | 49 | 25 | 17.5 | 23.3 | 27.3 | 26.1 | 9.4 |
| >hg38_21_+_8433172_8446622_RNA45SN1_45S_50ntFlanks@7984.8004.21\|AGATCAGACGTGGCGACCCGC | >hg38_21_+_8433172_8446622_RNA45SN1_45S_50ntFlanks | 7984 | 8004 | 21 | AGATCAGACGTGGCGACCCGC | MB468 | 28 | 23 | 10 | 60 | 76 | 20.8 | 9.1 | 8.5 | 30.0 | 43.1 |
| >hg38_21_+_8433172_8446622_RNA45SN1_45S_50ntFlanks@7984.8005.22\|AGATCAGACGTGGCGACCCGCT | >hg38_21_+_8433172_8446622_RNA45SN1_45S_50ntFlanks | 7984 | 8005 | 22 | AGATCAGACGTGGCGACCCGCT | MB468 | 27 | 16 | 11 | 52 | 53 | 17.8 | 5.8 | 7.3 | 20.1 | 31.4 |
| >hg38_21_+_8433172_8446622_RNA45SN1_45S_50ntFlanks@7984.8006.23\|AGATCAGACGTGGCGACCCGCTG | >hg38_21_+_8433172_8446622_RNA45SN1_45S_50ntFlanks | 7984 | 8006 | 23 | AGATCAGACGTGGCGACCCGCTG | MB468 | 340 | 123 | 105 | 664 | 611 | 162.0 | 47.4 | 45.1 | 362.7 | 433.4 |
| >hg38_21_+_8433172_8446622_RNA45SN1_45S_50ntFlanks@7984.8007.24\|AGATCAGACGTGGCGACCCGCTGA | >hg38_21_+_8433172_8446622_RNA45SN1_45S_50ntFlanks | 7984 | 8007 | 24 | AGATCAGACGTGGCGACCCGCTGA | MB468 | 23 | 7 | 4 | 51 | 50 | 10.4 | 2.6 | 1.4 | 31.8 | 26.2 |
| >hg38_21_+_8433172_8446622_RNA45SN1_45S_50ntFlanks@7984.8008.25\|AGATCAGACGTGGCGACCCGCTGAA | >hg38_21_+_8433172_8446622_RNA45SN1_45S_50ntFlanks | 7984 | 8008 | 25 | AGATCAGACGTGGCGACCCGCTGAA | MB468 | 122 | 39 | 20 | 294 | 282 | 51.1 | 19.3 | 9.0 | 145.1 | 139.4 |
| >hg38_21_+_8433172_8446622_RNA45SN1_45S_50ntFlanks@7984.8009.26\|AGATCAGACGTGGCGACCCGCTGAAT | >hg38_21_+_8433172_8446622_RNA45SN1_45S_50ntFlanks | 7984 | 8009 | 26 | AGATCAGACGTGGCGACCCGCTGAAT | MB468 | 18 | 6 | 4 | 61 | 39 | 7.5 | 3.7 | 1.3 | 26.7 | 19.9 |
| >hg38_21_+_8433172_8446622_RNA45SN1_45S_50ntFlanks@7984.8012.29\|AGATCAGACGTGGCGACCCGCTGAATTTA | >hg38_21_+_8433172_8446622_RNA45SN1_45S_50ntFlanks | 7984 | 8012 | 29 | AGATCAGACGTGGCGACCCGCTGAATTTA | MB468 | 17 | 4 | 3 | 26 | 45 | 7.4 | 1.8 | 1.0 | 11.8 | 26.6 |
| >hg38_21_+_8433172_8446622_RNA45SN1_45S_50ntFlanks@7985.8003.19\|GATCAGACGTGGCGACCCG | >hg38_21_+_8433172_8446622_RNA45SN1_45S_50ntFlanks | 7985 | 8003 | 19 | GATCAGACGTGGCGACCCG | MB468 | 11 | 22 | 21 | 33 | 15 | 8.5 | 14.8 | 23.3 | 13.8 | 5.2 |
| >hg38_21_+_8433172_8446622_RNA45SN1_45S_50ntFlanks@7985.8004.20\|GATCAGACGTGGCGACCCGC | >hg38_21_+_8433172_8446622_RNA45SN1_45S_50ntFlanks | 7985 | 8004 | 20 | GATCAGACGTGGCGACCCGC | MB468 | 18 | 13 | 8 | 44 | 45 | 12.7 | 5.6 | 5.4 | 22.6 | 24.6 |
| >hg38_21_+_8433172_8446622_RNA45SN1_45S_50ntFlanks@7985.8005.21\|GATCAGACGTGGCGACCCGCT | >hg38_21_+_8433172_8446622_RNA45SN1_45S_50ntFlanks | 7985 | 8005 | 21 | GATCAGACGTGGCGACCCGCT | MB468 | 18 | 11 | 8 | 40 | 36 | 10.8 | 4.6 | 5.5 | 11.4 | 18.7 |

| | | | | | | | | | | | | | | | | |
|---|---|---|---|---|---|---|---|---|---|---|---|---|---|---|---|---|
| >hg38_21_+_8433172_8446622_RNA45SN1_45S_50ntFlanks@7985.8006.22\|GATCAGACGTGGCGACCCGCTG | >hg38_21_+_8433172_8446622_RNA45SN1_45S_50ntFlanks | 7985 | 8006 | 22 | GATCAGACGTGGCGACCCGCTG | MB468 | 220 | 79 | 80 | 483 | 399 | 96.6 | 28.6 | 27.9 | 204.2 | 236.5 |
| >hg38_21_+_8433172_8446622_RNA45SN1_45S_50ntFlanks@7985.8007.23\|GATCAGACGTGGCGACCCGCTGA | >hg38_21_+_8433172_8446622_RNA45SN1_45S_50ntFlanks | 7985 | 8007 | 23 | GATCAGACGTGGCGACCCGCTGA | MB468 | 13 | 4 | 3 | 30 | 31 | 5.6 | 1.7 | 1.2 | 15.2 | 14.5 |
| >hg38_21_+_8433172_8446622_RNA45SN1_45S_50ntFlanks@7985.8008.24\|GATCAGACGTGGCGACCCGCTGAA | >hg38_21_+_8433172_8446622_RNA45SN1_45S_50ntFlanks | 7985 | 8008 | 24 | GATCAGACGTGGCGACCCGCTGAA | MB468 | 41 | 12 | 8 | 122 | 99 | 17.1 | 6.5 | 3.0 | 64.8 | 49.3 |
| >hg38_21_+_8433172_8446622_RNA45SN1_45S_50ntFlanks@7985.8012.28\|GATCAGACGTGGCGACCCGCTGAATTTA | >hg38_21_+_8433172_8446622_RNA45SN1_45S_50ntFlanks | 7985 | 8012 | 28 | GATCAGACGTGGCGACCCGCTGAATTTA | MB468 | 22 | 5 | 3 | 33 | 54 | 9.8 | 2.4 | 0.7 | 10.7 | 29.9 |
| >hg38_21_+_8433172_8446622_RNA45SN1_45S_50ntFlanks@7985.8016.32\|GATCAGACGTGGCGACCCGCTGAATTTAAGCA | >hg38_21_+_8433172_8446622_RNA45SN1_45S_50ntFlanks | 7985 | 8016 | 32 | GATCAGACGTGGCGACCCGCTGAATTTAAGCA | MB231 | 10 | 14 | 16 | 1 | 7 | 18.0 | 23.4 | 28.3 | 2.0 | 11.4 |
| >hg38_21_+_8433172_8446622_RNA45SN1_45S_50ntFlanks@7986.8003.18\|ATCAGACGTGGCGACCCG | >hg38_21_+_8433172_8446622_RNA45SN1_45S_50ntFlanks | 7986 | 8003 | 18 | ATCAGACGTGGCGACCCG | MB468 | 113 | 179 | 178 | 174 | 140 | 52.1 | 131.6 | 119.0 | 34.2 | 19.6 |
| >hg38_21_+_8433172_8446622_RNA45SN1_45S_50ntFlanks@7986.8004.19\|ATCAGACGTGGCGACCCGC | >hg38_21_+_8433172_8446622_RNA45SN1_45S_50ntFlanks | 7986 | 8004 | 19 | ATCAGACGTGGCGACCCGC | MB468 | 32 | 24 | 15 | 64 | 64 | 19.3 | 14.3 | 9.2 | 33.0 | 34.1 |
| >hg38_21_+_8433172_8446622_RNA45SN1_45S_50ntFlanks@7986.8005.20\|ATCAGACGTGGCGACCCGCT | >hg38_21_+_8433172_8446622_RNA45SN1_45S_50ntFlanks | 7986 | 8005 | 20 | ATCAGACGTGGCGACCCGCT | MB468 | 84 | 21 | 32 | 159 | 130 | 36.4 | 7.4 | 12.6 | 62.4 | 72.7 |
| >hg38_21_+_8433172_8446622_RNA45SN1_45S_50ntFlanks@7986.8006.21\|ATCAGACGTGGCGACCCGCTG | >hg38_21_+_8433172_8446622_RNA45SN1_45S_50ntFlanks | 7986 | 8006 | 21 | ATCAGACGTGGCGACCCGCTG | MB468 | 394 | 75 | 122 | 1179 | 694 | 153.2 | 26.0 | 24.3 | 787.5 | 373.5 |
| >hg38_21_+_8433172_8446622_RNA45SN1_45S_50ntFlanks@7986.8007.22\|ATCAGACGTGGCGACCCGCTGA | >hg38_21_+_8433172_8446622_RNA45SN1_45S_50ntFlanks | 7986 | 8007 | 22 | ATCAGACGTGGCGACCCGCTGA | MB468 | 14 | 2 | 2 | 49 | 31 | 5.6 | 0.6 | 0.3 | 40.5 | 14.4 |
| >hg38_21_+_8433172_8446622_RNA45SN1_45S_50ntFlanks@7986.8008.23\|ATCAGACGTGGCGACCCGCTGAA | >hg38_21_+_8433172_8446622_RNA45SN1_45S_50ntFlanks | 7986 | 8008 | 23 | ATCAGACGTGGCGACCCGCTGAA | MB468 | 56 | 8 | 8 | 235 | 128 | 21.7 | 4.9 | 2.8 | 174.3 | 61.3 |
| >hg38_21_+_8433172_8446622_RNA45SN1_45S_50ntFlanks@7986.8009.24\|ATCAGACGTGGCGACCCGCTGAAT | >hg38_21_+_8433172_8446622_RNA45SN1_45S_50ntFlanks | 7986 | 8009 | 24 | ATCAGACGTGGCGACCCGCTGAAT | MB468 | 10 | 2 | 2 | 65 | 23 | 4.3 | 1.6 | 0.5 | 45.8 | 10.1 |
| >hg38_21_+_8433172_8446622_RNA45SN1_45S_50ntFlanks@7986.8011.26\|ATCAGACGTGGCGACCCGCTGAATTT | >hg38_21_+_8433172_8446622_RNA45SN1_45S_50ntFlanks | 7986 | 8011 | 26 | ATCAGACGTGGCGACCCGCTGAATTT | MB468 | 20 | 3 | 3 | 49 | 51 | 8.9 | 2.0 | 1.2 | 28.4 | 24.7 |
| >hg38_21_+_8433172_8446622_RNA45SN1_45S_50ntFlanks@7986.8012.27\|ATCAGACGTGGCGACCCGCTGAATTTA | >hg38_21_+_8433172_8446622_RNA45SN1_45S_50ntFlanks | 7986 | 8012 | 27 | ATCAGACGTGGCGACCCGCTGAATTTA | MB468 | 68 | 10 | 7 | 125 | 172 | 28.8 | 4.4 | 2.2 | 62.0 | 87.5 |
| >hg38_21_+_8433172_8446622_RNA45SN1_45S_50ntFlanks@7986.8014.29\|ATCAGACGTGGCGACCCGCTGAATTTAAG | >hg38_21_+_8433172_8446622_RNA45SN1_45S_50ntFlanks | 7986 | 8014 | 29 | ATCAGACGTGGCGACCCGCTGAATTTAAG | MB468 | 16 | 5 | 3 | 35 | 41 | 6.4 | 4.3 | 0.9 | 8.9 | 24.6 |
| >hg38_21_+_8433172_8446622_RNA45SN1_45S_50ntFlanks@7986.8015.30\|ATCAGACGTGGCGACCCGCTGAATTTAAGC | >hg38_21_+_8433172_8446622_RNA45SN1_45S_50ntFlanks | 7986 | 8015 | 30 | ATCAGACGTGGCGACCCGCTGAATTTAAGC | MB468 | 11 | 7 | 2 | 23 | 33 | 4.8 | 4.9 | 1.1 | 6.1 | 20.6 |
| >hg38_21_+_8433172_8446622_RNA45SN1_45S_50ntFlanks@7986.8016.31\|ATCAGACGTGGCGACCCGCTGAATTTAAGCA | >hg38_21_+_8433172_8446622_RNA45SN1_45S_50ntFlanks | 7986 | 8016 | 31 | ATCAGACGTGGCGACCCGCTGAATTTAAGCA | MB468 | 24 | 15 | 5 | 48 | 77 | 9.7 | 11.5 | 2.0 | 13.2 | 51.9 |
| >hg38_21_+_8433172_8446622_RNA45SN1_45S_50ntFlanks@7986.8017.32\|ATCAGACGTGGCGACCCGCTGAATTTAAGCAT | >hg38_21_+_8433172_8446622_RNA45SN1_45S_50ntFlanks | 7986 | 8017 | 32 | ATCAGACGTGGCGACCCGCTGAATTTAAGCAT | MB468 | 28 | 17 | 5 | 51 | 97 | 11.4 | 8.9 | 0.6 | 23.7 | 71.2 |
| >hg38_21_+_8433172_8446622_RNA45SN1_45S_50ntFlanks@7986.8018.33\|ATCAGACGTGGCGACCCGCTGAATTTAAGCATA | >hg38_21_+_8433172_8446622_RNA45SN1_45S_50ntFlanks | 7986 | 8018 | 33 | ATCAGACGTGGCGACCCGCTGAATTTAAGCATA | MB468 | 15 | 11 | 3 | 31 | 72 | 5.3 | 6.7 | 0.8 | 17.3 | 49.8 |
| >hg38_21_+_8433172_8446622_RNA45SN1_45S_50ntFlanks@7987.8006.20\|TCAGACGTGGCGACCCGCTG | >hg38_21_+_8433172_8446622_RNA45SN1_45S_50ntFlanks | 7987 | 8006 | 20 | TCAGACGTGGCGACCCGCTG | MB468 | 17 | 2 | 4 | 58 | 31 | 7.4 | 0.7 | 0.5 | 42.8 | 17.5 |
| >hg38_21_+_8433172_8446622_RNA45SN1_45S_50ntFlanks@7987.8008.22\|TCAGACGTGGCGACCCGCTGAA | >hg38_21_+_8433172_8446622_RNA45SN1_45S_50ntFlanks | 7987 | 8008 | 22 | TCAGACGTGGCGACCCGCTGAA | MB468 | 10 | 2 | 2 | 31 | 25 | 4.7 | 1.9 | 0.9 | 19.6 | 13.0 |
| >hg38_21_+_8433172_8446622_RNA45SN1_45S_50ntFlanks@7988.8006.19\|CAGACGTGGCGACCCGCTG | >hg38_21_+_8433172_8446622_RNA45SN1_45S_50ntFlanks | 7988 | 8006 | 19 | CAGACGTGGCGACCCGCTG | MB468 | 19 | 2 | 3 | 127 | 40 | 11.1 | 1.4 | 0.9 | 132.9 | 22.2 |
| >hg38_21_+_8433172_8446622_RNA45SN1_45S_50ntFlanks@7988.8007.20\|CAGACGTGGCGACCCGCTGA | >hg38_21_+_8433172_8446622_RNA45SN1_45S_50ntFlanks | 7988 | 8007 | 20 | CAGACGTGGCGACCCGCTGA | MB468 | 62 | 14 | 11 | 141 | 163 | 31.0 | 8.8 | 6.0 | 61.6 | 81.1 |
| >hg38_21_+_8433172_8446622_RNA45SN1_45S_50ntFlanks@7988.8008.21\|CAGACGTGGCGACCCGCTGAA | >hg38_21_+_8433172_8446622_RNA45SN1_45S_50ntFlanks | 7988 | 8008 | 21 | CAGACGTGGCGACCCGCTGAA | MB231 | 14 | 24 | 49 | 5 | 8 | 5.8 | 12.4 | 35.4 | 2.3 | 1.4 |
| >hg38_21_+_8433172_8446622_RNA45SN1_45S_50ntFlanks@7988.8008.21\|CAGACGTGGCGACCCGCTGAA | >hg38_21_+_8433172_8446622_RNA45SN1_45S_50ntFlanks | 7988 | 8008 | 21 | CAGACGTGGCGACCCGCTGAA | MB468 | 299 | 102 | 47 | 800 | 814 | 139.3 | 96.2 | 30.7 | 305.1 | 390.5 |
| >hg38_21_+_8433172_8446622_RNA45SN1_45S_50ntFlanks@7988.8009.22\|CAGACGTGGCGACCCGCTGAAT | >hg38_21_+_8433172_8446622_RNA45SN1_45S_50ntFlanks | 7988 | 8009 | 22 | CAGACGTGGCGACCCGCTGAAT | BT20 | 10 | 37 | 17 | 29 | 9 | 7.1 | 44.9 | 2.8 | 31.8 | 4.1 |
| >hg38_21_+_8433172_8446622_RNA45SN1_45S_50ntFlanks@7988.8009.22\|CAGACGTGGCGACCCGCTGAAT | >hg38_21_+_8433172_8446622_RNA45SN1_45S_50ntFlanks | 7988 | 8009 | 22 | CAGACGTGGCGACCCGCTGAAT | MB468 | 56 | 26 | 10 | 194 | 155 | 25.6 | 23.7 | 6.8 | 62.2 | 65.8 |
| >hg38_21_+_8433172_8446622_RNA45SN1_45S_50ntFlanks@7988.8010.23\|CAGACGTGGCGACCCGCTGAATT | >hg38_21_+_8433172_8446622_RNA45SN1_45S_50ntFlanks | 7988 | 8010 | 23 | CAGACGTGGCGACCCGCTGAATT | MB468 | 20 | 8 | 3 | 53 | 60 | 10.4 | 7.7 | 2.3 | 23.1 | 31.7 |
| >hg38_21_+_8433172_8446622_RNA45SN1_45S_50ntFlanks@7988.8011.24\|CAGACGTGGCGACCCGCTGAATTT | >hg38_21_+_8433172_8446622_RNA45SN1_45S_50ntFlanks | 7988 | 8011 | 24 | CAGACGTGGCGACCCGCTGAATTT | BT20 | 14 | 56 | 15 | 29 | 16 | 7.7 | 41.3 | 4.7 | 41.0 | 7.7 |
| >hg38_21_+_8433172_8446622_RNA45SN1_45S_50ntFlanks@7988.8011.24\|CAGACGTGGCGACCCGCTGAATTT | >hg38_21_+_8433172_8446622_RNA45SN1_45S_50ntFlanks | 7988 | 8011 | 24 | CAGACGTGGCGACCCGCTGAATTT | MB231 | 17 | 14 | 14 | 4 | 13 | 12.8 | 6.5 | 7.4 | 2.1 | 3.8 |
| >hg38_21_+_8433172_8446622_RNA45SN1_45S_50ntFlanks@7988.8011.24\|CAGACGTGGCGACCCGCTGAATTT | >hg38_21_+_8433172_8446622_RNA45SN1_45S_50ntFlanks | 7988 | 8011 | 24 | CAGACGTGGCGACCCGCTGAATTT | MB468 | 41 | 15 | 7 | 95 | 126 | 20.1 | 11.4 | 5.4 | 30.2 | 64.1 |
| >hg38_21_+_8433172_8446622_RNA45SN1_45S_50ntFlanks@7988.8012.25\|CAGACGTGGCGACCCGCTGAATTTA | >hg38_21_+_8433172_8446622_RNA45SN1_45S_50ntFlanks | 7988 | 8012 | 25 | CAGACGTGGCGACCCGCTGAATTTA | MB468 | 198 | 65 | 26 | 333 | 575 | 89.1 | 42.8 | 14.2 | 72.0 | 286.3 |
| >hg38_21_+_8433172_8446622_RNA45SN1_45S_50ntFlanks@7988.8013.26\|CAGACGTGGCGACCCGCTGAATTTAA | >hg38_21_+_8433172_8446622_RNA45SN1_45S_50ntFlanks | 7988 | 8013 | 26 | CAGACGTGGCGACCCGCTGAATTTAA | MB468 | 10 | 5 | 1 | 22 | 32 | 4.3 | 3.5 | 0.9 | 4.8 | 16.4 |
| >hg38_21_+_8433172_8446622_RNA45SN1_45S_50ntFlanks@7988.8014.27\|CAGACGTGGCGACCCGCTGAATTTAAG | >hg38_21_+_8433172_8446622_RNA45SN1_45S_50ntFlanks | 7988 | 8014 | 27 | CAGACGTGGCGACCCGCTGAATTTAAG | MB468 | 34 | 18 | 5 | 83 | 102 | 14.6 | 13.9 | 3.0 | 27.1 | 55.9 |
| >hg38_21_+_8433172_8446622_RNA45SN1_45S_50ntFlanks@7988.8016.29\|CAGACGTGGCGACCCGCTGAATTTAAGCA | >hg38_21_+_8433172_8446622_RNA45SN1_45S_50ntFlanks | 7988 | 8016 | 29 | CAGACGTGGCGACCCGCTGAATTTAAGCA | MB231 | 18 | 13 | 28 | 1 | 10 | 13.1 | 7.1 | 23.7 | 1.0 | 8.4 |
| >hg38_21_+_8433172_8446622_RNA45SN1_45S_50ntFlanks@7988.8016.29\|CAGACGTGGCGACCCGCTGAATTTAAGCA | >hg38_21_+_8433172_8446622_RNA45SN1_45S_50ntFlanks | 7988 | 8016 | 29 | CAGACGTGGCGACCCGCTGAATTTAAGCA | MB468 | 20 | 17 | 4 | 55 | 69 | 8.3 | 12.7 | 2.7 | 16.7 | 39.4 |
| >hg38_21_+_8433172_8446622_RNA45SN1_45S_50ntFlanks@7988.8017.30\|CAGACGTGGCGACCCGCTGAATTTAAGCAT | >hg38_21_+_8433172_8446622_RNA45SN1_45S_50ntFlanks | 7988 | 8017 | 30 | CAGACGTGGCGACCCGCTGAATTTAAGCAT | MB468 | 19 | 19 | 3 | 50 | 75 | 8.8 | 12.6 | 1.8 | 24.7 | 39.6 |
| >hg38_21_+_8433172_8446622_RNA45SN1_45S_50ntFlanks@7988.8018.31\|CAGACGTGGCGACCCGCTGAATTTAAGCATA | >hg38_21_+_8433172_8446622_RNA45SN1_45S_50ntFlanks | 7988 | 8018 | 31 | CAGACGTGGCGACCCGCTGAATTTAAGCATA | MB468 | 11 | 11 | 2 | 31 | 51 | 4.4 | 7.0 | 1.1 | 18.6 | 30.5 |
| >hg38_21_+_8433172_8446622_RNA45SN1_45S_50ntFlanks@7988.8019.32\|CAGACGTGGCGACCCGCTGAATTTAAGCATAT | >hg38_21_+_8433172_8446622_RNA45SN1_45S_50ntFlanks | 7988 | 8019 | 32 | CAGACGTGGCGACCCGCTGAATTTAAGCATAT | MB231 | 12 | 8 | 22 | 1 | 4 | 9.7 | 5.7 | 18.1 | 0.9 | 7.2 |
| >hg38_21_+_8433172_8446622_RNA45SN1_45S_50ntFlanks@7989.8006.18\|AGACGTGGCGACCCGCTG | >hg38_21_+_8433172_8446622_RNA45SN1_45S_50ntFlanks | 7989 | 8006 | 18 | AGACGTGGCGACCCGCTG | MB468 | 233 | 42 | 29 | 1276 | 495 | 136.3 | 22.8 | 14.4 | 644.1 | 269.9 |
| >hg38_21_+_8433172_8446622_RNA45SN1_45S_50ntFlanks@7989.8007.19\|AGACGTGGCGACCCGCTGA | >hg38_21_+_8433172_8446622_RNA45SN1_45S_50ntFlanks | 7989 | 8007 | 19 | AGACGTGGCGACCCGCTGA | MB468 | 78 | 25 | 11 | 407 | 188 | 46.3 | 19.4 | 9.9 | 192.9 | 93.9 |
| >hg38_21_+_8433172_8446622_RNA45SN1_45S_50ntFlanks@7989.8008.20\|AGACGTGGCGACCCGCTGAA | >hg38_21_+_8433172_8446622_RNA45SN1_45S_50ntFlanks | 7989 | 8008 | 20 | AGACGTGGCGACCCGCTGAA | BT20 | 21 | 36 | 22 | 86 | 28 | 23.7 | 27.6 | 14.2 | 69.4 | 28.3 |
| >hg38_21_+_8433172_8446622_RNA45SN1_45S_50ntFlanks@7989.8008.20\|AGACGTGGCGACCCGCTGAA | >hg38_21_+_8433172_8446622_RNA45SN1_45S_50ntFlanks | 7989 | 8008 | 20 | AGACGTGGCGACCCGCTGAA | MB231 | 35 | 43 | 63 | 17 | 24 | 22.2 | 32.0 | 51.9 | 6.6 | 1.6 |
| >hg38_21_+_8433172_8446622_RNA45SN1_45S_50ntFlanks@7989.8008.20\|AGACGTGGCGACCCGCTGAA | >hg38_21_+_8433172_8446622_RNA45SN1_45S_50ntFlanks | 7989 | 8008 | 20 | AGACGTGGCGACCCGCTGAA | MB468 | 236 | 104 | 33 | 1722 | 626 | 136.4 | 101.9 | 29.1 | 864.4 | 319.1 |
| >hg38_21_+_8433172_8446622_RNA45SN1_45S_50ntFlanks@7989.8009.21\|AGACGTGGCGACCCGCTGAAT | >hg38_21_+_8433172_8446622_RNA45SN1_45S_50ntFlanks | 7989 | 8009 | 21 | AGACGTGGCGACCCGCTGAAT | BT20 | 42 | 103 | 43 | 126 | 43 | 31.5 | 87.0 | 30.0 | 43.0 | 24.8 |
| >hg38_21_+_8433172_8446622_RNA45SN1_45S_50ntFlanks@7989.8009.21\|AGACGTGGCGACCCGCTGAAT | >hg38_21_+_8433172_8446622_RNA45SN1_45S_50ntFlanks | 7989 | 8009 | 21 | AGACGTGGCGACCCGCTGAAT | MB231 | 119 | 107 | 133 | 103 | 82 | 101.6 | 90.0 | 147.1 | 28.1 | 22.0 |
| >hg38_21_+_8433172_8446622_RNA45SN1_45S_50ntFlanks@7989.8009.21\|AGACGTGGCGACCCGCTGAAT | >hg38_21_+_8433172_8446622_RNA45SN1_45S_50ntFlanks | 7989 | 8009 | 21 | AGACGTGGCGACCCGCTGAAT | MB468 | 31 | 18 | 4 | 277 | 82 | 17.8 | 17.0 | 3.4 | 135.8 | 41.1 |
| >hg38_21_+_8433172_8446622_RNA45SN1_45S_50ntFlanks@7989.8010.22\|AGACGTGGCGACCCGCTGAATT | >hg38_21_+_8433172_8446622_RNA45SN1_45S_50ntFlanks | 7989 | 8010 | 22 | AGACGTGGCGACCCGCTGAATT | BT20 | 49 | 78 | 37 | 49 | 47 | 50.3 | 42.1 | 32.0 | 38.8 | 43.3 |
| >hg38_21_+_8433172_8446622_RNA45SN1_45S_50ntFlanks@7989.8010.22\|AGACGTGGCGACCCGCTGAATT | >hg38_21_+_8433172_8446622_RNA45SN1_45S_50ntFlanks | 7989 | 8010 | 22 | AGACGTGGCGACCCGCTGAATT | MB231 | 110 | 96 | 93 | 25 | 79 | 54.5 | 47.3 | 70.4 | 12.2 | 13.8 |
| >hg38_21_+_8433172_8446622_RNA45SN1_45S_50ntFlanks@7989.8010.22\|AGACGTGGCGACCCGCTGAATT | >hg38_21_+_8433172_8446622_RNA45SN1_45S_50ntFlanks | 7989 | 8010 | 22 | AGACGTGGCGACCCGCTGAATT | MB468 | 15 | 9 | 2 | 58 | 42 | 9.7 | 9.3 | 2.1 | 36.7 | 24.6 |

| | | | | | | | | | | | | | | | | |
|---|---|---|---|---|---|---|---|---|---|---|---|---|---|---|---|---|
| >hg38_21_+_8433172_8446622_RNA45SN1_45S_50ntFlanks@7989.8011.23\|AGACGTGGCGACCCGCTGAATTT | >hg38_21_+_8433172_8446622_RNA45SN1_45S_50ntFlanks | 7989 | 8011 | 23 | AGACGTGGCGACCCGCTGAATTT | BT20 | 91 | 166 | 61 | 147 | 92 | 79.9 | 76.2 | 46.9 | 118.7 | 71.8 |
| >hg38_21_+_8433172_8446622_RNA45SN1_45S_50ntFlanks@7989.8011.23\|AGACGTGGCGACCCGCTGAATTT | >hg38_21_+_8433172_8446622_RNA45SN1_45S_50ntFlanks | 7989 | 8011 | 23 | AGACGTGGCGACCCGCTGAATTT | MB231 | 210 | 181 | 177 | 44 | 152 | 106.9 | 81.3 | 126.8 | 13.5 | 60.2 |
| >hg38_21_+_8433172_8446622_RNA45SN1_45S_50ntFlanks@7989.8011.23\|AGACGTGGCGACCCGCTGAATTT | >hg38_21_+_8433172_8446622_RNA45SN1_45S_50ntFlanks | 7989 | 8011 | 23 | AGACGTGGCGACCCGCTGAATTT | MB468 | 32 | 18 | 6 | 104 | 97 | 19.5 | 17.3 | 5.4 | 55.4 | 56.7 |
| >hg38_21_+_8433172_8446622_RNA45SN1_45S_50ntFlanks@7989.8012.24\|AGACGTGGCGACCCGCTGAATTTA | >hg38_21_+_8433172_8446622_RNA45SN1_45S_50ntFlanks | 7989 | 8012 | 24 | AGACGTGGCGACCCGCTGAATTTA | BT20 | 24 | 24 | 15 | 43 | 25 | 31.2 | 23.1 | 15.2 | 42.5 | 28.3 |
| >hg38_21_+_8433172_8446622_RNA45SN1_45S_50ntFlanks@7989.8012.24\|AGACGTGGCGACCCGCTGAATTTA | >hg38_21_+_8433172_8446622_RNA45SN1_45S_50ntFlanks | 7989 | 8012 | 24 | AGACGTGGCGACCCGCTGAATTTA | MB231 | 22 | 25 | 20 | 3 | 13 | 16.7 | 12.9 | 10.0 | 1.9 | 9.0 |
| >hg38_21_+_8433172_8446622_RNA45SN1_45S_50ntFlanks@7989.8012.24\|AGACGTGGCGACCCGCTGAATTTA | >hg38_21_+_8433172_8446622_RNA45SN1_45S_50ntFlanks | 7989 | 8012 | 24 | AGACGTGGCGACCCGCTGAATTTA | MB468 | 140 | 72 | 17 | 344 | 402 | 77.2 | 60.6 | 15.4 | 118.3 | 218.2 |
| >hg38_21_+_8433172_8446622_RNA45SN1_45S_50ntFlanks@7989.8013.25\|AGACGTGGCGACCCGCTGAATTTAA | >hg38_21_+_8433172_8446622_RNA45SN1_45S_50ntFlanks | 7989 | 8013 | 25 | AGACGTGGCGACCCGCTGAATTTAA | BT20 | 21 | 18 | 8 | 13 | 21 | 28.1 | 25.1 | 9.0 | 7.4 | 24.2 |
| >hg38_21_+_8433172_8446622_RNA45SN1_45S_50ntFlanks@7989.8013.25\|AGACGTGGCGACCCGCTGAATTTAA | >hg38_21_+_8433172_8446622_RNA45SN1_45S_50ntFlanks | 7989 | 8013 | 25 | AGACGTGGCGACCCGCTGAATTTAA | MB231 | 11 | 10 | 8 | 1 | 6 | 8.4 | 3.8 | 5.1 | 0.6 | 3.9 |
| >hg38_21_+_8433172_8446622_RNA45SN1_45S_50ntFlanks@7989.8014.26\|AGACGTGGCGACCCGCTGAATTTAAG | >hg38_21_+_8433172_8446622_RNA45SN1_45S_50ntFlanks | 7989 | 8014 | 26 | AGACGTGGCGACCCGCTGAATTTAAG | BT20 | 17 | 16 | 7 | 11 | 15 | 25.3 | 23.3 | 8.6 | 7.2 | 17.6 |
| >hg38_21_+_8433172_8446622_RNA45SN1_45S_50ntFlanks@7989.8014.26\|AGACGTGGCGACCCGCTGAATTTAAG | >hg38_21_+_8433172_8446622_RNA45SN1_45S_50ntFlanks | 7989 | 8014 | 26 | AGACGTGGCGACCCGCTGAATTTAAG | MB468 | 20 | 15 | 3 | 70 | 60 | 10.4 | 12.8 | 2.7 | 21.4 | 34.3 |
| >hg38_21_+_8433172_8446622_RNA45SN1_45S_50ntFlanks@7989.8015.27\|AGACGTGGCGACCCGCTGAATTTAAGC | >hg38_21_+_8433172_8446622_RNA45SN1_45S_50ntFlanks | 7989 | 8015 | 27 | AGACGTGGCGACCCGCTGAATTTAAGC | BT20 | 15 | 30 | 9 | 12 | 16 | 12.1 | 12.6 | 8.9 | 7.7 | 10.5 |
| >hg38_21_+_8433172_8446622_RNA45SN1_45S_50ntFlanks@7989.8016.28\|AGACGTGGCGACCCGCTGAATTTAAGCA | >hg38_21_+_8433172_8446622_RNA45SN1_45S_50ntFlanks | 7989 | 8016 | 28 | AGACGTGGCGACCCGCTGAATTTAAGCA | BT20 | 26 | 28 | 22 | 11 | 22 | 27.6 | 30.8 | 25.0 | 5.3 | 18.8 |
| >hg38_21_+_8433172_8446622_RNA45SN1_45S_50ntFlanks@7989.8016.28\|AGACGTGGCGACCCGCTGAATTTAAGCA | >hg38_21_+_8433172_8446622_RNA45SN1_45S_50ntFlanks | 7989 | 8016 | 28 | AGACGTGGCGACCCGCTGAATTTAAGCA | MB231 | 34 | 28 | 57 | 5 | 21 | 25.8 | 14.4 | 45.1 | 2.9 | 13.3 |
| >hg38_21_+_8433172_8446622_RNA45SN1_45S_50ntFlanks@7989.8017.29\|AGACGTGGCGACCCGCTGAATTTAAGCAT | >hg38_21_+_8433172_8446622_RNA45SN1_45S_50ntFlanks | 7989 | 8017 | 29 | AGACGTGGCGACCCGCTGAATTTAAGCAT | BT20 | 14 | 21 | 10 | 6 | 13 | 11.3 | 11.9 | 9.5 | 2.9 | 8.1 |
| >hg38_21_+_8433172_8446622_RNA45SN1_45S_50ntFlanks@7989.8017.29\|AGACGTGGCGACCCGCTGAATTTAAGCAT | >hg38_21_+_8433172_8446622_RNA45SN1_45S_50ntFlanks | 7989 | 8017 | 29 | AGACGTGGCGACCCGCTGAATTTAAGCAT | MB231 | 21 | 19 | 35 | 3 | 15 | 14.1 | 6.3 | 23.5 | 2.6 | 11.4 |
| >hg38_21_+_8433172_8446622_RNA45SN1_45S_50ntFlanks@7989.8018.30\|AGACGTGGCGACCCGCTGAATTTAAGCATA | >hg38_21_+_8433172_8446622_RNA45SN1_45S_50ntFlanks | 7989 | 8018 | 30 | AGACGTGGCGACCCGCTGAATTTAAGCATA | MB231 | 10 | 9 | 18 | 1 | 10 | 7.5 | 2.9 | 14.6 | 0.8 | 6.0 |
| >hg38_21_+_8433172_8446622_RNA45SN1_45S_50ntFlanks@7989.8019.31\|AGACGTGGCGACCCGCTGAATTTAAGCATAT | >hg38_21_+_8433172_8446622_RNA45SN1_45S_50ntFlanks | 7989 | 8019 | 31 | AGACGTGGCGACCCGCTGAATTTAAGCATAT | MB231 | 28 | 19 | 53 | 4 | 17 | 23.7 | 10.9 | 42.8 | 4.1 | 17.7 |
| >hg38_21_+_8433172_8446622_RNA45SN1_45S_50ntFlanks@7990.8008.19\|GACGTGGCGACCCGCTGAA | >hg38_21_+_8433172_8446622_RNA45SN1_45S_50ntFlanks | 7990 | 8008 | 19 | GACGTGGCGACCCGCTGAA | MB468 | 56 | 18 | 8 | 296 | 156 | 32.3 | 16.6 | 5.5 | 147.9 | 76.7 |
| >hg38_21_+_8433172_8446622_RNA45SN1_45S_50ntFlanks@7990.8009.20\|GACGTGGCGACCCGCTGAAT | >hg38_21_+_8433172_8446622_RNA45SN1_45S_50ntFlanks | 7990 | 8009 | 20 | GACGTGGCGACCCGCTGAAT | BT20 | 20 | 32 | 13 | 74 | 24 | 19.7 | 23.1 | 5.0 | 44.0 | 22.3 |
| >hg38_21_+_8433172_8446622_RNA45SN1_45S_50ntFlanks@7990.8009.20\|GACGTGGCGACCCGCTGAAT | >hg38_21_+_8433172_8446622_RNA45SN1_45S_50ntFlanks | 7990 | 8009 | 20 | GACGTGGCGACCCGCTGAAT | MB231 | 26 | 18 | 19 | 16 | 19 | 9.3 | 9.1 | 0.1 | 2.4 | 5.1 |
| >hg38_21_+_8433172_8446622_RNA45SN1_45S_50ntFlanks@7990.8010.21\|GACGTGGCGACCCGCTGAATT | >hg38_21_+_8433172_8446622_RNA45SN1_45S_50ntFlanks | 7990 | 8010 | 21 | GACGTGGCGACCCGCTGAATT | BT20 | 18 | 25 | 9 | 24 | 23 | 16.9 | 13.0 | 4.3 | 18.7 | 22.6 |
| >hg38_21_+_8433172_8446622_RNA45SN1_45S_50ntFlanks@7990.8010.21\|GACGTGGCGACCCGCTGAATT | >hg38_21_+_8433172_8446622_RNA45SN1_45S_50ntFlanks | 7990 | 8010 | 21 | GACGTGGCGACCCGCTGAATT | MB231 | 30 | 18 | 19 | 6 | 20 | 12.0 | 2.7 | 10.2 | 1.2 | 5.3 |
| >hg38_21_+_8433172_8446622_RNA45SN1_45S_50ntFlanks@7990.8011.22\|GACGTGGCGACCCGCTGAATTT | >hg38_21_+_8433172_8446622_RNA45SN1_45S_50ntFlanks | 7990 | 8011 | 22 | GACGTGGCGACCCGCTGAATTT | BT20 | 36 | 54 | 20 | 92 | 50 | 28.8 | 29.7 | 12.6 | 98.9 | 35.7 |
| >hg38_21_+_8433172_8446622_RNA45SN1_45S_50ntFlanks@7990.8011.22\|GACGTGGCGACCCGCTGAATTT | >hg38_21_+_8433172_8446622_RNA45SN1_45S_50ntFlanks | 7990 | 8011 | 22 | GACGTGGCGACCCGCTGAATTT | MB231 | 79 | 46 | 55 | 14 | 48 | 41.3 | 10.3 | 32.7 | 3.9 | 12.3 |
| >hg38_21_+_8433172_8446622_RNA45SN1_45S_50ntFlanks@7990.8012.23\|GACGTGGCGACCCGCTGAATTTA | >hg38_21_+_8433172_8446622_RNA45SN1_45S_50ntFlanks | 7990 | 8012 | 23 | GACGTGGCGACCCGCTGAATTTA | MB231 | 24 | 15 | 18 | 3 | 13 | 19.4 | 9.8 | 16.7 | 2.1 | 10.2 |
| >hg38_21_+_8433172_8446622_RNA45SN1_45S_50ntFlanks@7990.8012.23\|GACGTGGCGACCCGCTGAATTTA | >hg38_21_+_8433172_8446622_RNA45SN1_45S_50ntFlanks | 7990 | 8012 | 23 | GACGTGGCGACCCGCTGAATTTA | MB468 | 29 | 11 | 3 | 57 | 82 | 15.4 | 8.4 | 2.1 | 22.5 | 41.1 |
| >hg38_21_+_8433172_8446622_RNA45SN1_45S_50ntFlanks@7990.8013.24\|GACGTGGCGACCCGCTGAATTTAA | >hg38_21_+_8433172_8446622_RNA45SN1_45S_50ntFlanks | 7990 | 8013 | 24 | GACGTGGCGACCCGCTGAATTTAA | MB231 | 12 | 9 | 11 | 1 | 4 | 9.8 | 5.7 | 8.5 | 1.6 | 6.8 |
| >hg38_21_+_8433172_8446622_RNA45SN1_45S_50ntFlanks@7990.8016.27\|GACGTGGCGACCCGCTGAATTTAAGCA | >hg38_21_+_8433172_8446622_RNA45SN1_45S_50ntFlanks | 7990 | 8016 | 27 | GACGTGGCGACCCGCTGAATTTAAGCA | MB231 | 38 | 19 | 47 | 3 | 15 | 33.7 | 15.9 | 43.0 | 2.9 | 13.4 |
| >hg38_21_+_8433172_8446622_RNA45SN1_45S_50ntFlanks@7990.8017.28\|GACGTGGCGACCCGCTGAATTTAAGCAT | >hg38_21_+_8433172_8446622_RNA45SN1_45S_50ntFlanks | 7990 | 8017 | 28 | GACGTGGCGACCCGCTGAATTTAAGCAT | MB231 | 21 | 10 | 30 | 2 | 9 | 17.3 | 7.7 | 26.1 | 1.8 | 8.1 |
| >hg38_21_+_8433172_8446622_RNA45SN1_45S_50ntFlanks@7990.8018.29\|GACGTGGCGACCCGCTGAATTTAAGCATA | >hg38_21_+_8433172_8446622_RNA45SN1_45S_50ntFlanks | 7990 | 8018 | 29 | GACGTGGCGACCCGCTGAATTTAAGCATA | MB231 | 13 | 6 | 18 | 1 | 5 | 11.3 | 5.1 | 15.0 | 1.1 | 4.8 |
| >hg38_21_+_8433172_8446622_RNA45SN1_45S_50ntFlanks@7990.8019.30\|GACGTGGCGACCCGCTGAATTTAAGCATAT | >hg38_21_+_8433172_8446622_RNA45SN1_45S_50ntFlanks | 7990 | 8019 | 30 | GACGTGGCGACCCGCTGAATTTAAGCATAT | MB231 | 36 | 17 | 50 | 3 | 17 | 32.1 | 14.8 | 42.8 | 3.8 | 14.5 |
| >hg38_21_+_8433172_8446622_RNA45SN1_45S_50ntFlanks@7991.8008.18\|ACGTGGCGACCCGCTGAA | >hg38_21_+_8433172_8446622_RNA45SN1_45S_50ntFlanks | 7991 | 8008 | 18 | ACGTGGCGACCCGCTGAA | MB231 | 12 | 17 | 17 | 4 | 11 | 6.4 | 11.3 | 12.1 | 2.4 | 4.5 |
| >hg38_21_+_8433172_8446622_RNA45SN1_45S_50ntFlanks@7991.8008.18\|ACGTGGCGACCCGCTGAA | >hg38_21_+_8433172_8446622_RNA45SN1_45S_50ntFlanks | 7991 | 8008 | 18 | ACGTGGCGACCCGCTGAA | MB468 | 767 | 149 | 96 | 2336 | 2087 | 340.2 | 95.1 | 42.1 | 899.7 | 531.5 |
| >hg38_21_+_8433172_8446622_RNA45SN1_45S_50ntFlanks@7991.8009.19\|ACGTGGCGACCCGCTGAAT | >hg38_21_+_8433172_8446622_RNA45SN1_45S_50ntFlanks | 7991 | 8009 | 19 | ACGTGGCGACCCGCTGAAT | BT20 | 17 | 40 | 14 | 39 | 16 | 14.4 | 31.5 | 6.1 | 29.3 | 8.9 |
| >hg38_21_+_8433172_8446622_RNA45SN1_45S_50ntFlanks@7991.8009.19\|ACGTGGCGACCCGCTGAAT | >hg38_21_+_8433172_8446622_RNA45SN1_45S_50ntFlanks | 7991 | 8009 | 19 | ACGTGGCGACCCGCTGAAT | MB231 | 27 | 20 | 16 | 15 | 29 | 20.8 | 16.5 | 10.2 | 6.6 | 16.2 |
| >hg38_21_+_8433172_8446622_RNA45SN1_45S_50ntFlanks@7991.8009.19\|ACGTGGCGACCCGCTGAAT | >hg38_21_+_8433172_8446622_RNA45SN1_45S_50ntFlanks | 7991 | 8009 | 19 | ACGTGGCGACCCGCTGAAT | MB468 | 146 | 45 | 22 | 654 | 383 | 79.1 | 35.6 | 12.4 | 248.8 | 48.6 |
| >hg38_21_+_8433172_8446622_RNA45SN1_45S_50ntFlanks@7991.8010.20\|ACGTGGCGACCCGCTGAATT | >hg38_21_+_8433172_8446622_RNA45SN1_45S_50ntFlanks | 7991 | 8010 | 20 | ACGTGGCGACCCGCTGAATT | BT20 | 15 | 27 | 10 | 13 | 19 | 11.0 | 10.8 | 3.1 | 11.0 | 9.5 |
| >hg38_21_+_8433172_8446622_RNA45SN1_45S_50ntFlanks@7991.8010.20\|ACGTGGCGACCCGCTGAATT | >hg38_21_+_8433172_8446622_RNA45SN1_45S_50ntFlanks | 7991 | 8010 | 20 | ACGTGGCGACCCGCTGAATT | MB231 | 27 | 20 | 11 | 6 | 29 | 14.8 | 12.1 | 2.5 | 3.9 | 20.7 |
| >hg38_21_+_8433172_8446622_RNA45SN1_45S_50ntFlanks@7991.8010.20\|ACGTGGCGACCCGCTGAATT | >hg38_21_+_8433172_8446622_RNA45SN1_45S_50ntFlanks | 7991 | 8010 | 20 | ACGTGGCGACCCGCTGAATT | MB468 | 46 | 12 | 6 | 72 | 136 | 22.8 | 8.9 | 3.3 | 28.5 | 38.9 |
| >hg38_21_+_8433172_8446622_RNA45SN1_45S_50ntFlanks@7991.8011.21\|ACGTGGCGACCCGCTGAATTT | >hg38_21_+_8433172_8446622_RNA45SN1_45S_50ntFlanks | 7991 | 8011 | 21 | ACGTGGCGACCCGCTGAATTT | BT20 | 39 | 62 | 17 | 56 | 57 | 26.1 | 22.6 | 5.9 | 51.5 | 37.3 |
| >hg38_21_+_8433172_8446622_RNA45SN1_45S_50ntFlanks@7991.8011.21\|ACGTGGCGACCCGCTGAATTT | >hg38_21_+_8433172_8446622_RNA45SN1_45S_50ntFlanks | 7991 | 8011 | 21 | ACGTGGCGACCCGCTGAATTT | MB231 | 63 | 45 | 26 | 16 | 68 | 19.1 | 18.1 | 8.6 | 9.6 | 33.7 |
| >hg38_21_+_8433172_8446622_RNA45SN1_45S_50ntFlanks@7991.8011.21\|ACGTGGCGACCCGCTGAATTT | >hg38_21_+_8433172_8446622_RNA45SN1_45S_50ntFlanks | 7991 | 8011 | 21 | ACGTGGCGACCCGCTGAATTT | MB468 | 124 | 32 | 21 | 134 | 358 | 70.3 | 19.1 | 11.9 | 38.4 | 119.4 |
| >hg38_21_+_8433172_8446622_RNA45SN1_45S_50ntFlanks@7991.8012.22\|ACGTGGCGACCCGCTGAATTTA | >hg38_21_+_8433172_8446622_RNA45SN1_45S_50ntFlanks | 7991 | 8012 | 22 | ACGTGGCGACCCGCTGAATTTA | MB468 | 367 | 89 | 39 | 313 | 1124 | 178.6 | 42.2 | 16.9 | 99.9 | 335.9 |
| >hg38_21_+_8433172_8446622_RNA45SN1_45S_50ntFlanks@7991.8013.23\|ACGTGGCGACCCGCTGAATTTAA | >hg38_21_+_8433172_8446622_RNA45SN1_45S_50ntFlanks | 7991 | 8013 | 23 | ACGTGGCGACCCGCTGAATTTAA | MB468 | 19 | 7 | 2 | 33 | 63 | 8.9 | 5.1 | 1.2 | 17.6 | 18.9 |
| >hg38_21_+_8433172_8446622_RNA45SN1_45S_50ntFlanks@7991.8014.24\|ACGTGGCGACCCGCTGAATTTAAG | >hg38_21_+_8433172_8446622_RNA45SN1_45S_50ntFlanks | 7991 | 8014 | 24 | ACGTGGCGACCCGCTGAATTTAAG | MB468 | 59 | 29 | 8 | 125 | 176 | 23.6 | 20.1 | 3.7 | 66.1 | 59.3 |
| >hg38_21_+_8433172_8446622_RNA45SN1_45S_50ntFlanks@7991.8016.26\|ACGTGGCGACCCGCTGAATTTAAGCA | >hg38_21_+_8433172_8446622_RNA45SN1_45S_50ntFlanks | 7991 | 8016 | 26 | ACGTGGCGACCCGCTGAATTTAAGCA | MB231 | 25 | 15 | 28 | 2 | 14 | 18.6 | 10.8 | 23.1 | 0.6 | 8.1 |
| >hg38_21_+_8433172_8446622_RNA45SN1_45S_50ntFlanks@7991.8016.26\|ACGTGGCGACCCGCTGAATTTAAGCA | >hg38_21_+_8433172_8446622_RNA45SN1_45S_50ntFlanks | 7991 | 8016 | 26 | ACGTGGCGACCCGCTGAATTTAAGCA | MB468 | 23 | 18 | 4 | 39 | 88 | 10.1 | 12.1 | 2.5 | 14.6 | 22.3 |
| >hg38_21_+_8433172_8446622_RNA45SN1_45S_50ntFlanks@7991.8017.27\|ACGTGGCGACCCGCTGAATTTAAGCAT | >hg38_21_+_8433172_8446622_RNA45SN1_45S_50ntFlanks | 7991 | 8017 | 27 | ACGTGGCGACCCGCTGAATTTAAGCAT | MB231 | 10 | 6 | 10 | 0 | 4 | 6.8 | 2.9 | 7.9 | 0.7 | 6.1 |
| >hg38_21_+_8433172_8446622_RNA45SN1_45S_50ntFlanks@7991.8017.27\|ACGTGGCGACCCGCTGAATTTAAGCAT | >hg38_21_+_8433172_8446622_RNA45SN1_45S_50ntFlanks | 7991 | 8017 | 27 | ACGTGGCGACCCGCTGAATTTAAGCAT | MB468 | 22 | 13 | 3 | 21 | 75 | 8.6 | 6.9 | 0.8 | 10.2 | 12.6 |
| >hg38_21_+_8433172_8446622_RNA45SN1_45S_50ntFlanks@7991.8018.28\|ACGTGGCGACCCGCTGAATTTAAGCATA | >hg38_21_+_8433172_8446622_RNA45SN1_45S_50ntFlanks | 7991 | 8018 | 28 | ACGTGGCGACCCGCTGAATTTAAGCATA | MB468 | 15 | 9 | 2 | 19 | 66 | 7.0 | 5.1 | 0.6 | 8.5 | 17.9 |

| | | | | | | | | | | | | | | | |
|---|---|---|---|---|---|---|---|---|---|---|---|---|---|---|---|
| >hg38_21_+_8433172_8446622_RNA45SN1_45S_50ntFlanks@7991.8019.29\|ACGTGGCGACCCGCTGAATTTAAGCATAT | >hg38_21_+_8433172_8446622_RNA45SN1_45S_50ntFlanks | 7991 | 8019 | 29 | ACGTGGCGACCCGCTGAATTTAAGCATAT | MB231 | 16 | 9 | 19 | 0 | 5 | 14.8 | 6.7 | 17.1 | 0.8 | 8.7 |
| >hg38_21_+_8433172_8446622_RNA45SN1_45S_50ntFlanks@7992.8009.18\|CGTGGCGACCCGCTGAAT | >hg38_21_+_8433172_8446622_RNA45SN1_45S_50ntFlanks | 7992 | 8009 | 18 | CGTGGCGACCCGCTGAAT | MB231 | 13 | 14 | 7 | 5 | 15 | 9.8 | 11.3 | 4.0 | 3.2 | 7.5 |
| >hg38_21_+_8433172_8446622_RNA45SN1_45S_50ntFlanks@7992.8009.18\|CGTGGCGACCCGCTGAAT | >hg38_21_+_8433172_8446622_RNA45SN1_45S_50ntFlanks | 7992 | 8009 | 18 | CGTGGCGACCCGCTGAAT | MB468 | 28 | 7 | 3 | 71 | 73 | 13.8 | 4.0 | 1.9 | 37.4 | 17.4 |
| >hg38_21_+_8433172_8446622_RNA45SN1_45S_50ntFlanks@7992.8010.19\|CGTGGCGACCCGCTGAATT | >hg38_21_+_8433172_8446622_RNA45SN1_45S_50ntFlanks | 7992 | 8010 | 19 | CGTGGCGACCCGCTGAATT | MB231 | 18 | 16 | 6 | 3 | 19 | 10.8 | 10.7 | 3.1 | 1.9 | 13.0 |
| >hg38_21_+_8433172_8446622_RNA45SN1_45S_50ntFlanks@7992.8011.20\|CGTGGCGACCCGCTGAATTT | >hg38_21_+_8433172_8446622_RNA45SN1_45S_50ntFlanks | 7992 | 8011 | 20 | CGTGGCGACCCGCTGAATTT | BT20 | 17 | 42 | 6 | 31 | 20 | 11.2 | 7.7 | 1.5 | 47.2 | 15.9 |
| >hg38_21_+_8433172_8446622_RNA45SN1_45S_50ntFlanks@7992.8011.20\|CGTGGCGACCCGCTGAATTT | >hg38_21_+_8433172_8446622_RNA45SN1_45S_50ntFlanks | 7992 | 8011 | 20 | CGTGGCGACCCGCTGAATTT | MB231 | 42 | 35 | 14 | 8 | 51 | 14.5 | 17.4 | 10.4 | 4.9 | 25.1 |
| >hg38_21_+_8433172_8446622_RNA45SN1_45S_50ntFlanks@7992.8011.20\|CGTGGCGACCCGCTGAATTT | >hg38_21_+_8433172_8446622_RNA45SN1_45S_50ntFlanks | 7992 | 8011 | 20 | CGTGGCGACCCGCTGAATTT | MB468 | 19 | 4 | 3 | 15 | 54 | 10.3 | 2.1 | 1.7 | 5.9 | 19.6 |
| >hg38_21_+_8433172_8446622_RNA45SN1_45S_50ntFlanks@7992.8012.21\|CGTGGCGACCCGCTGAATTTA | >hg38_21_+_8433172_8446622_RNA45SN1_45S_50ntFlanks | 7992 | 8012 | 21 | CGTGGCGACCCGCTGAATTTA | MB468 | 46 | 9 | 4 | 32 | 142 | 24.2 | 4.3 | 2.2 | 11.1 | 58.3 |
| >hg38_21_+_8433172_8446622_RNA45SN1_45S_50ntFlanks@7992.8014.23\|CGTGGCGACCCGCTGAATTTAAG | >hg38_21_+_8433172_8446622_RNA45SN1_45S_50ntFlanks | 7992 | 8014 | 23 | CGTGGCGACCCGCTGAATTTAAG | MB468 | 15 | 6 | 2 | 21 | 42 | 7.6 | 4.1 | 0.9 | 14.7 | 12.8 |
| >hg38_21_+_8433172_8446622_RNA45SN1_45S_50ntFlanks@7992.8016.25\|CGTGGCGACCCGCTGAATTTAAGCA | >hg38_21_+_8433172_8446622_RNA45SN1_45S_50ntFlanks | 7992 | 8016 | 25 | CGTGGCGACCCGCTGAATTTAAGCA | MB231 | 37 | 24 | 43 | 1 | 19 | 29.6 | 15.6 | 34.5 | 1.6 | 17.6 |
| >hg38_21_+_8433172_8446622_RNA45SN1_45S_50ntFlanks@7992.8019.28\|CGTGGCGACCCGCTGAATTTAAGCATAT | >hg38_21_+_8433172_8446622_RNA45SN1_45S_50ntFlanks | 7992 | 8019 | 28 | CGTGGCGACCCGCTGAATTTAAGCATAT | MB231 | 14 | 10 | 17 | 0 | 6 | 11.7 | 8.6 | 18.7 | 0.8 | 11.1 |
| >hg38_21_+_8433172_8446622_RNA45SN1_45S_50ntFlanks@7993.8016.24\|GTGGCGACCCGCTGAATTTAAGCA | >hg38_21_+_8433172_8446622_RNA45SN1_45S_50ntFlanks | 7993 | 8016 | 24 | GTGGCGACCCGCTGAATTTAAGCA | MB231 | 18 | 11 | 11 | 1 | 4 | 20.0 | 9.0 | 9.0 | 0.9 | 6.9 |
| >hg38_21_+_8433172_8446622_RNA45SN1_45S_50ntFlanks@8104.8138.35\|CCGCGGCGGGGCGCGGGACATGTGGCGTACGGAAG | >hg38_21_+_8433172_8446622_RNA45SN1_45S_50ntFlanks | 8104 | 8138 | 35 | CCGCGGCGGGGCGCGGGACATGTGGCGTACGGAAG | MB468 | 26 | 22 | 4 | 223 | 83 | 17.7 | 8.8 | 2.3 | 187.5 | 50.9 |
| >hg38_21_+_8433172_8446622_RNA45SN1_45S_50ntFlanks@8104.8140.37\|CCGCGGCGGGGCGCGGGACATGTGGCGTACGGAAGAC | >hg38_21_+_8433172_8446622_RNA45SN1_45S_50ntFlanks | 8104 | 8140 | 37 | CCGCGGCGGGGCGCGGGACATGTGGCGTACGGAAGAC | MB468 | 43 | 47 | 8 | 573 | 186 | 18.3 | 13.7 | 4.7 | 586.8 | 117.7 |
| >hg38_21_+_8433172_8446622_RNA45SN1_45S_50ntFlanks@8104.8141.38\|CCGCGGCGGGGCGCGGGACATGTGGCGTACGGAAGACC | >hg38_21_+_8433172_8446622_RNA45SN1_45S_50ntFlanks | 8104 | 8141 | 38 | CCGCGGCGGGGCGCGGGACATGTGGCGTACGGAAGACC | MB468 | 18 | 24 | 4 | 167 | 93 | 9.5 | 8.3 | 2.6 | 180.1 | 72.6 |
| >hg38_21_+_8433172_8446622_RNA45SN1_45S_50ntFlanks@8104.8142.39\|CCGCGGCGGGGCGCGGGACATGTGGCGTACGGAAGACCC | >hg38_21_+_8433172_8446622_RNA45SN1_45S_50ntFlanks | 8104 | 8142 | 39 | CCGCGGCGGGGCGCGGGACATGTGGCGTACGGAAGACCC | MB468 | 10 | 15 | 4 | 48 | 52 | 5.6 | 6.5 | 3.0 | 47.6 | 35.7 |
| >hg38_21_+_8433172_8446622_RNA45SN1_45S_50ntFlanks@8104.8143.40\|CCGCGGCGGGGCGCGGGACATGTGGCGTACGGAAGACCCG | >hg38_21_+_8433172_8446622_RNA45SN1_45S_50ntFlanks | 8104 | 8143 | 40 | CCGCGGCGGGGCGCGGGACATGTGGCGTACGGAAGACCCG | MB468 | 321 | 447 | 101 | 1677 | 1716 | 165.2 | 271.1 | 85.2 | 1723.2 | 1112.4 |
| >hg38_21_+_8433172_8446622_RNA45SN1_45S_50ntFlanks@8105.8138.34\|CGCGGCGGGGCGCGGGACATGTGGCGTACGGAAG | >hg38_21_+_8433172_8446622_RNA45SN1_45S_50ntFlanks | 8105 | 8138 | 34 | CGCGGCGGGGCGCGGGACATGTGGCGTACGGAAG | MB468 | 36 | 40 | 5 | 578 | 202 | 24.9 | 18.6 | 4.2 | 389.5 | 102.6 |
| >hg38_21_+_8433172_8446622_RNA45SN1_45S_50ntFlanks@8105.8140.36\|CGCGGCGGGGCGCGGGACATGTGGCGTACGGAAGAC | >hg38_21_+_8433172_8446622_RNA45SN1_45S_50ntFlanks | 8105 | 8140 | 36 | CGCGGCGGGGCGCGGGACATGTGGCGTACGGAAGAC | MB468 | 39 | 52 | 7 | 753 | 274 | 24.0 | 17.6 | 4.4 | 624.2 | 156.4 |
| >hg38_21_+_8433172_8446622_RNA45SN1_45S_50ntFlanks@8105.8141.37\|CGCGGCGGGGCGCGGGACATGTGGCGTACGGAAGACC | >hg38_21_+_8433172_8446622_RNA45SN1_45S_50ntFlanks | 8105 | 8141 | 37 | CGCGGCGGGGCGCGGGACATGTGGCGTACGGAAGACC | MB468 | 11 | 17 | 3 | 258 | 98 | 6.8 | 4.0 | 2.0 | 218.2 | 64.7 |
| >hg38_21_+_8433172_8446622_RNA45SN1_45S_50ntFlanks@8105.8143.39\|CGCGGCGGGGCGCGGGACATGTGGCGTACGGAAGACCCG | >hg38_21_+_8433172_8446622_RNA45SN1_45S_50ntFlanks | 8105 | 8143 | 39 | CGCGGCGGGGCGCGGGACATGTGGCGTACGGAAGACCCG | MB468 | 323 | 639 | 144 | 3506 | 2031 | 202.1 | 106.8 | 92.0 | 3471.1 | 1957.1 |
| >hg38_21_+_8433172_8446622_RNA45SN1_45S_50ntFlanks@8106.8143.38\|GCGGCGGGGCGCGGGACATGTGGCGTACGGAAGACCCG | >hg38_21_+_8433172_8446622_RNA45SN1_45S_50ntFlanks | 8106 | 8143 | 38 | GCGGCGGGGCGCGGGACATGTGGCGTACGGAAGACCCG | MB468 | 14 | 13 | 3 | 107 | 63 | 9.6 | 2.5 | 1.1 | 81.5 | 33.8 |
| >hg38_21_+_8433172_8446622_RNA45SN1_45S_50ntFlanks@8107.8143.37\|CGGCGGGGCGCGGGACATGTGGCGTACGGAAGACCCG | >hg38_21_+_8433172_8446622_RNA45SN1_45S_50ntFlanks | 8107 | 8143 | 37 | CGGCGGGGCGCGGGACATGTGGCGTACGGAAGACCCG | MB468 | 22 | 16 | 4 | 110 | 58 | 15.5 | 1.3 | 1.8 | 49.6 | 36.7 |
| >hg38_21_+_8433172_8446622_RNA45SN1_45S_50ntFlanks@8113.8138.26\|GGCGCGGGACATGTGGCGTACGGAAG | >hg38_21_+_8433172_8446622_RNA45SN1_45S_50ntFlanks | 8113 | 8138 | 26 | GGCGCGGGACATGTGGCGTACGGAAG | MB468 | 21 | 13 | 2 | 446 | 49 | 15.5 | 8.5 | 1.1 | 230.1 | 24.3 |
| >hg38_21_+_8433172_8446622_RNA45SN1_45S_50ntFlanks@8113.8140.28\|GGCGCGGGACATGTGGCGTACGGAAGAC | >hg38_21_+_8433172_8446622_RNA45SN1_45S_50ntFlanks | 8113 | 8140 | 28 | GGCGCGGGACATGTGGCGTACGGAAGAC | MB468 | 15 | 8 | 1 | 274 | 39 | 10.7 | 3.5 | 0.7 | 99.7 | 14.7 |
| >hg38_21_+_8433172_8446622_RNA45SN1_45S_50ntFlanks@8113.8143.31\|GGCGCGGGACATGTGGCGTACGGAAGACCCG | >hg38_21_+_8433172_8446622_RNA45SN1_45S_50ntFlanks | 8113 | 8143 | 31 | GGCGCGGGACATGTGGCGTACGGAAGACCCG | MB468 | 18 | 6 | 1 | 230 | 49 | 10.9 | 2.3 | 0.6 | 77.6 | 29.5 |
| >hg38_21_+_8433172_8446622_RNA45SN1_45S_50ntFlanks@8114.8138.25\|GCGCGGGACATGTGGCGTACGGAAG | >hg38_21_+_8433172_8446622_RNA45SN1_45S_50ntFlanks | 8114 | 8138 | 25 | GCGCGGGACATGTGGCGTACGGAAG | MB468 | 52 | 38 | 5 | 1100 | 168 | 35.4 | 25.0 | 3.6 | 648.4 | 75.0 |
| >hg38_21_+_8433172_8446622_RNA45SN1_45S_50ntFlanks@8114.8139.26\|GCGCGGGACATGTGGCGTACGGAAGA | >hg38_21_+_8433172_8446622_RNA45SN1_45S_50ntFlanks | 8114 | 8139 | 26 | GCGCGGGACATGTGGCGTACGGAAGA | MB468 | 12 | 9 | 1 | 203 | 35 | 7.3 | 5.3 | 0.9 | 104.3 | 12.8 |
| >hg38_21_+_8433172_8446622_RNA45SN1_45S_50ntFlanks@8114.8140.27\|GCGCGGGACATGTGGCGTACGGAAGAC | >hg38_21_+_8433172_8446622_RNA45SN1_45S_50ntFlanks | 8114 | 8140 | 27 | GCGCGGGACATGTGGCGTACGGAAGAC | MB468 | 47 | 30 | 5 | 722 | 162 | 26.3 | 12.8 | 2.6 | 319.3 | 49.1 |
| >hg38_21_+_8433172_8446622_RNA45SN1_45S_50ntFlanks@8114.8141.28\|GCGCGGGACATGTGGCGTACGGAAGACC | >hg38_21_+_8433172_8446622_RNA45SN1_45S_50ntFlanks | 8114 | 8141 | 28 | GCGCGGGACATGTGGCGTACGGAAGACC | MB468 | 26 | 16 | 3 | 348 | 88 | 13.5 | 8.3 | 1.6 | 162.3 | 34.0 |
| >hg38_21_+_8433172_8446622_RNA45SN1_45S_50ntFlanks@8114.8143.30\|GCGCGGGACATGTGGCGTACGGAAGACCCG | >hg38_21_+_8433172_8446622_RNA45SN1_45S_50ntFlanks | 8114 | 8143 | 30 | GCGCGGGACATGTGGCGTACGGAAGACCCG | MB468 | 102 | 35 | 10 | 1055 | 350 | 52.9 | 15.4 | 2.9 | 447.8 | 185.1 |
| >hg38_21_+_8433172_8446622_RNA45SN1_45S_50ntFlanks@8115.8138.24\|CGCGGGACATGTGGCGTACGGAAG | >hg38_21_+_8433172_8446622_RNA45SN1_45S_50ntFlanks | 8115 | 8138 | 24 | CGCGGGACATGTGGCGTACGGAAG | MB468 | 25 | 17 | 2 | 457 | 85 | 17.1 | 10.4 | 1.8 | 336.7 | 42.8 |
| >hg38_21_+_8433172_8446622_RNA45SN1_45S_50ntFlanks@8115.8140.26\|CGCGGGACATGTGGCGTACGGAAGAC | >hg38_21_+_8433172_8446622_RNA45SN1_45S_50ntFlanks | 8115 | 8140 | 26 | CGCGGGACATGTGGCGTACGGAAGAC | MB468 | 28 | 16 | 3 | 337 | 82 | 15.3 | 8.3 | 1.9 | 193.1 | 28.9 |
| >hg38_21_+_8433172_8446622_RNA45SN1_45S_50ntFlanks@8115.8141.27\|CGCGGGACATGTGGCGTACGGAAGACC | >hg38_21_+_8433172_8446622_RNA45SN1_45S_50ntFlanks | 8115 | 8141 | 27 | CGCGGGACATGTGGCGTACGGAAGACC | BT20 | 11 | 12 | 3 | 42 | 26 | 15.2 | 18.1 | 4.2 | 44.5 | 32.7 |
| >hg38_21_+_8433172_8446622_RNA45SN1_45S_50ntFlanks@8115.8141.27\|CGCGGGACATGTGGCGTACGGAAGACC | >hg38_21_+_8433172_8446622_RNA45SN1_45S_50ntFlanks | 8115 | 8141 | 27 | CGCGGGACATGTGGCGTACGGAAGACC | MB468 | 20 | 12 | 2 | 208 | 62 | 11.0 | 5.2 | 1.4 | 105.5 | 24.9 |
| >hg38_21_+_8433172_8446622_RNA45SN1_45S_50ntFlanks@8115.8142.28\|CGCGGGACATGTGGCGTACGGAAGACCC | >hg38_21_+_8433172_8446622_RNA45SN1_45S_50ntFlanks | 8115 | 8142 | 28 | CGCGGGACATGTGGCGTACGGAAGACCC | MB468 | 12 | 6 | 2 | 76 | 38 | 6.4 | 2.6 | 0.7 | 34.8 | 16.0 |
| >hg38_21_+_8433172_8446622_RNA45SN1_45S_50ntFlanks@8115.8143.29\|CGCGGGACATGTGGCGTACGGAAGACCCG | >hg38_21_+_8433172_8446622_RNA45SN1_45S_50ntFlanks | 8115 | 8143 | 29 | CGCGGGACATGTGGCGTACGGAAGACCCG | MB468 | 185 | 44 | 17 | 1009 | 621 | 95.3 | 16.8 | 3.4 | 519.7 | 231.5 |
| >hg38_21_+_8433172_8446622_RNA45SN1_45S_50ntFlanks@8153.8181.29\|GCCGCTCGTGGGGGGCCCAAGTCCTTCTG | >hg38_21_+_8433172_8446622_RNA45SN1_45S_50ntFlanks | 8153 | 8181 | 29 | GCCGCTCGTGGGGGGCCCAAGTCCTTCTG | MB468 | 17 | 27 | 2 | 10 | 100 | 3.8 | 12.9 | 0.7 | 8.9 | 65.4 |
| >hg38_21_+_8433172_8446622_RNA45SN1_45S_50ntFlanks@8154.8178.25\|CCGCTCGTGGGGGGCCCAAGTCCTT | >hg38_21_+_8433172_8446622_RNA45SN1_45S_50ntFlanks | 8154 | 8178 | 25 | CCGCTCGTGGGGGGCCCAAGTCCTT | MB468 | 16 | 3 | 2 | 21 | 36 | 5.3 | 0.6 | 1.0 | 17.0 | 15.7 |
| >hg38_21_+_8433172_8446622_RNA45SN1_45S_50ntFlanks@8154.8181.28\|CCGCTCGTGGGGGGCCCAAGTCCTTCTG | >hg38_21_+_8433172_8446622_RNA45SN1_45S_50ntFlanks | 8154 | 8181 | 28 | CCGCTCGTGGGGGGCCCAAGTCCTTCTG | MB468 | 127 | 128 | 21 | 74 | 566 | 17.3 | 64.8 | 6.4 | 53.3 | 186.9 |
| >hg38_21_+_8433172_8446622_RNA45SN1_45S_50ntFlanks@8155.8181.27\|CGCTCGTGGGGGGCCCAAGTCCTTCTG | >hg38_21_+_8433172_8446622_RNA45SN1_45S_50ntFlanks | 8155 | 8181 | 27 | CGCTCGTGGGGGGCCCAAGTCCTTCTG | MB468 | 46 | 38 | 11 | 41 | 292 | 11.5 | 20.5 | 2.8 | 34.5 | 185.4 |
| >hg38_21_+_8433172_8446622_RNA45SN1_45S_50ntFlanks@8157.8178.22\|CTCGTGGGGGGCCCAAGTCCTT | >hg38_21_+_8433172_8446622_RNA45SN1_45S_50ntFlanks | 8157 | 8178 | 22 | CTCGTGGGGGGCCCAAGTCCTT | MB468 | 24 | 8 | 4 | 46 | 88 | 17.7 | 5.1 | 2.2 | 18.3 | 73.2 |
| >hg38_21_+_8433172_8446622_RNA45SN1_45S_50ntFlanks@8157.8181.25\|CTCGTGGGGGGCCCAAGTCCTTCTG | >hg38_21_+_8433172_8446622_RNA45SN1_45S_50ntFlanks | 8157 | 8181 | 25 | CTCGTGGGGGGCCCAAGTCCTTCTG | MB468 | 28 | 6 | 5 | 88 | 187 | 9.5 | 2.8 | 2.1 | 22.7 | 159.3 |
| >hg38_21_+_8433172_8446622_RNA45SN1_45S_50ntFlanks@8158.8188.31\|TCGTGGGGGGCCCAAGTCCTTCTGATCGAGG | >hg38_21_+_8433172_8446622_RNA45SN1_45S_50ntFlanks | 8158 | 8188 | 31 | TCGTGGGGGGCCCAAGTCCTTCTGATCGAGG | MB468 | 20 | 4 | 3 | 20 | 39 | 13.3 | 1.9 | 2.3 | 11.3 | 18.6 |
| >hg38_21_+_8433172_8446622_RNA45SN1_45S_50ntFlanks@8159.8190.32\|CGTGGGGGGCCCAAGTCCTTCTGATCGAGGCC | >hg38_21_+_8433172_8446622_RNA45SN1_45S_50ntFlanks | 8159 | 8190 | 32 | CGTGGGGGGCCCAAGTCCTTCTGATCGAGGCC | MB231 | 13 | 6 | 5 | 3 | 10 | 11.8 | 2.0 | 1.9 | 0.2 | 5.4 |
| >hg38_21_+_8433172_8446622_RNA45SN1_45S_50ntFlanks@8168.8185.18\|CCCAAGTCCTTCTGATCG | >hg38_21_+_8433172_8446622_RNA45SN1_45S_50ntFlanks | 8168 | 8185 | 18 | CCCAAGTCCTTCTGATCG | MB468 | 53 | 44 | 35 | 42 | 27 | 33.4 | 20.9 | 25.5 | 25.0 | 9.5 |
| >hg38_21_+_8433172_8446622_RNA45SN1_45S_50ntFlanks@8168.8186.19\|CCCAAGTCCTTCTGATCGA | >hg38_21_+_8433172_8446622_RNA45SN1_45S_50ntFlanks | 8168 | 8186 | 19 | CCCAAGTCCTTCTGATCGA | MB468 | 137 | 106 | 83 | 56 | 57 | 111.1 | 55.7 | 69.6 | 55.8 | 39.1 |
| >hg38_21_+_8433172_8446622_RNA45SN1_45S_50ntFlanks@8168.8187.20\|CCCAAGTCCTTCTGATCGAG | >hg38_21_+_8433172_8446622_RNA45SN1_45S_50ntFlanks | 8168 | 8187 | 20 | CCCAAGTCCTTCTGATCGAG | MB468 | 58 | 76 | 29 | 13 | 21 | 25.7 | 39.8 | 21.7 | 2.1 | 8.2 |
| >hg38_21_+_8433172_8446622_RNA45SN1_45S_50ntFlanks@8168.8188.21\|CCCAAGTCCTTCTGATCGAGG | >hg38_21_+_8433172_8446622_RNA45SN1_45S_50ntFlanks | 8168 | 8188 | 21 | CCCAAGTCCTTCTGATCGAGG | MB468 | 50 | 11 | 35 | 29 | 11 | 48.2 | 6.3 | 40.4 | 8.4 | 5.1 |
| >hg38_21_+_8433172_8446622_RNA45SN1_45S_50ntFlanks@8168.8195.28\|CCCAAGTCCTTCTGATCGAGGCCCAGCC | >hg38_21_+_8433172_8446622_RNA45SN1_45S_50ntFlanks | 8168 | 8195 | 28 | CCCAAGTCCTTCTGATCGAGGCCCAGCC | MB468 | 16 | 22 | 15 | 13 | 19 | 4.8 | 14.0 | 5.9 | 3.7 | 6.3 |

| | | | | | | | | | | | | | | | | |
|---|---|---|---|---|---|---|---|---|---|---|---|---|---|---|---|---|
| >hg38_21_+_8433172_8446622_RNA45SN1_45S_50ntFlanks@8168.8197.30\|CCCAAGTCCTTCTGATCGAGGCCCAGCCCG | >hg38_21_+_8433172_8446622_RNA45SN1_45S_50ntFlanks | 8168 | 8197 | 30 | CCCAAGTCCTTCTGATCGAGGCCCAGCCCG | MB468 | 19 | 98 | 12 | 6 | 56 | 7.6 | 105.3 | 6.3 | 2.9 | 51.2 |
| >hg38_21_+_8433172_8446622_RNA45SN1_45S_50ntFlanks@8168.8203.36\|CCCAAGTCCTTCTGATCGAGGCCCAGCCCGTGGACG | >hg38_21_+_8433172_8446622_RNA45SN1_45S_50ntFlanks | 8168 | 8203 | 36 | CCCAAGTCCTTCTGATCGAGGCCCAGCCCGTGGACG | MB468 | 11 | 15 | 24 | 43 | 18 | 7.1 | 6.2 | 20.5 | 39.0 | 16.1 |
| >hg38_21_+_8433172_8446622_RNA45SN1_45S_50ntFlanks@8169.8186.18\|CCAAGTCCTTCTGATCGA | >hg38_21_+_8433172_8446622_RNA45SN1_45S_50ntFlanks | 8169 | 8186 | 18 | CCAAGTCCTTCTGATCGA | MB468 | 51 | 23 | 27 | 29 | 18 | 41.7 | 9.2 | 17.0 | 32.2 | 11.3 |
| >hg38_21_+_8433172_8446622_RNA45SN1_45S_50ntFlanks@8169.8187.19\|CCAAGTCCTTCTGATCGAG | >hg38_21_+_8433172_8446622_RNA45SN1_45S_50ntFlanks | 8169 | 8187 | 19 | CCAAGTCCTTCTGATCGAG | MB468 | 25 | 19 | 12 | 8 | 9 | 11.7 | 10.2 | 7.9 | 2.5 | 3.3 |
| >hg38_21_+_8433172_8446622_RNA45SN1_45S_50ntFlanks@8169.8188.20\|CCAAGTCCTTCTGATCGAGG | >hg38_21_+_8433172_8446622_RNA45SN1_45S_50ntFlanks | 8169 | 8188 | 20 | CCAAGTCCTTCTGATCGAGG | MB468 | 23 | 3 | 11 | 13 | 4 | 22.8 | 1.6 | 12.2 | 4.7 | 2.3 |
| >hg38_21_+_8433172_8446622_RNA45SN1_45S_50ntFlanks@8170.8187.18\|CAAGTCCTTCTGATCGAG | >hg38_21_+_8433172_8446622_RNA45SN1_45S_50ntFlanks | 8170 | 8187 | 18 | CAAGTCCTTCTGATCGAG | MB468 | 95 | 46 | 48 | 33 | 32 | 36.5 | 22.1 | 23.8 | 11.1 | 13.9 |
| >hg38_21_+_8433172_8446622_RNA45SN1_45S_50ntFlanks@8170.8188.19\|CAAGTCCTTCTGATCGAGG | >hg38_21_+_8433172_8446622_RNA45SN1_45S_50ntFlanks | 8170 | 8188 | 19 | CAAGTCCTTCTGATCGAGG | MB468 | 92 | 6 | 46 | 46 | 16 | 82.1 | 2.9 | 42.2 | 16.6 | 13.1 |
| >hg38_21_+_8433172_8446622_RNA45SN1_45S_50ntFlanks@8170.8195.26\|CAAGTCCTTCTGATCGAGGCCCAGCC | >hg38_21_+_8433172_8446622_RNA45SN1_45S_50ntFlanks | 8170 | 8195 | 26 | CAAGTCCTTCTGATCGAGGCCCAGCC | MB468 | 98 | 46 | 91 | 138 | 86 | 39.5 | 20.5 | 52.6 | 92.1 | 33.7 |
| >hg38_21_+_8433172_8446622_RNA45SN1_45S_50ntFlanks@8170.8196.27\|CAAGTCCTTCTGATCGAGGCCCAGCCC | >hg38_21_+_8433172_8446622_RNA45SN1_45S_50ntFlanks | 8170 | 8196 | 27 | CAAGTCCTTCTGATCGAGGCCCAGCCC | MB468 | 11 | 20 | 9 | 8 | 29 | 2.0 | 15.1 | 1.6 | 6.6 | 13.7 |
| >hg38_21_+_8433172_8446622_RNA45SN1_45S_50ntFlanks@8170.8197.28\|CAAGTCCTTCTGATCGAGGCCCAGCCCG | >hg38_21_+_8433172_8446622_RNA45SN1_45S_50ntFlanks | 8170 | 8197 | 28 | CAAGTCCTTCTGATCGAGGCCCAGCCCG | MB468 | 97 | 266 | 76 | 41 | 384 | 16.6 | 222.7 | 23.3 | 26.3 | 333.7 |
| >hg38_21_+_8433172_8446622_RNA45SN1_45S_50ntFlanks@8170.8212.43\|CAAGTCCTTCTGATCGAGGCCCAGCCCGTGGACGGTGTGAGGC | >hg38_21_+_8433172_8446622_RNA45SN1_45S_50ntFlanks | 8170 | 8212 | 43 | CAAGTCCTTCTGATCGAGGCCCAGCCCGTGGACGGTGTGAGGC | MB468 | 50 | 452 | 81 | 125 | 487 | 18.2 | 272.9 | 58.3 | 113.3 | 275.6 |
| >hg38_21_+_8433172_8446622_RNA45SN1_45S_50ntFlanks@8170.8214.45\|CAAGTCCTTCTGATCGAGGCCCAGCCCGTGGACGGTGTGAGGCCG | >hg38_21_+_8433172_8446622_RNA45SN1_45S_50ntFlanks | 8170 | 8214 | 45 | CAAGTCCTTCTGATCGAGGCCCAGCCCGTGGACGGTGTGAGGCCG | MB468 | 36 | 209 | 164 | 52 | 421 | 17.3 | 115.7 | 162.1 | 36.7 | 449.9 |
| >hg38_21_+_8433172_8446622_RNA45SN1_45S_50ntFlanks@8170.8215.46\|CAAGTCCTTCTGATCGAGGCCCAGCCCGTGGACGGTGTGAGGCCGG | >hg38_21_+_8433172_8446622_RNA45SN1_45S_50ntFlanks | 8170 | 8215 | 46 | CAAGTCCTTCTGATCGAGGCCCAGCCCGTGGACGGTGTGAGGCCGG | MB468 | 19 | 146 | 79 | 26 | 179 | 11.0 | 94.2 | 48.2 | 30.8 | 116.3 |
| >hg38_21_+_8433172_8446622_RNA45SN1_45S_50ntFlanks@8171.8188.18\|AAGTCCTTCTGATCGAGG | >hg38_21_+_8433172_8446622_RNA45SN1_45S_50ntFlanks | 8171 | 8188 | 18 | AAGTCCTTCTGATCGAGG | MB468 | 28 | 1 | 11 | 15 | 3 | 28.7 | 1.1 | 11.3 | 7.0 | 2.6 |
| >hg38_21_+_8433172_8446622_RNA45SN1_45S_50ntFlanks@8171.8195.25\|AAGTCCTTCTGATCGAGGCCCAGCC | >hg38_21_+_8433172_8446622_RNA45SN1_45S_50ntFlanks | 8171 | 8195 | 25 | AAGTCCTTCTGATCGAGGCCCAGCC | MB468 | 14 | 6 | 15 | 25 | 11 | 4.7 | 4.0 | 6.3 | 23.7 | 6.7 |
| >hg38_21_+_8433172_8446622_RNA45SN1_45S_50ntFlanks@8171.8197.27\|AAGTCCTTCTGATCGAGGCCCAGCCCG | >hg38_21_+_8433172_8446622_RNA45SN1_45S_50ntFlanks | 8171 | 8197 | 27 | AAGTCCTTCTGATCGAGGCCCAGCCCG | MB468 | 22 | 71 | 21 | 8 | 63 | 5.6 | 67.7 | 9.3 | 8.7 | 60.7 |
| >hg38_21_+_8433172_8446622_RNA45SN1_45S_50ntFlanks@8171.8212.42\|AAGTCCTTCTGATCGAGGCCCAGCCCGTGGACGGTGTGAGGC | >hg38_21_+_8433172_8446622_RNA45SN1_45S_50ntFlanks | 8171 | 8212 | 42 | AAGTCCTTCTGATCGAGGCCCAGCCCGTGGACGGTGTGAGGC | MB468 | 20 | 192 | 25 | 38 | 58 | 4.4 | 158.4 | 11.8 | 36.8 | 28.4 |
| >hg38_21_+_8433172_8446622_RNA45SN1_45S_50ntFlanks@8171.8214.44\|AAGTCCTTCTGATCGAGGCCCAGCCCGTGGACGGTGTGAGGCCG | >hg38_21_+_8433172_8446622_RNA45SN1_45S_50ntFlanks | 8171 | 8214 | 44 | AAGTCCTTCTGATCGAGGCCCAGCCCGTGGACGGTGTGAGGCCG | MB468 | 16 | 76 | 70 | 21 | 60 | 4.8 | 27.9 | 50.7 | 14.3 | 55.3 |
| >hg38_21_+_8433172_8446622_RNA45SN1_45S_50ntFlanks@8172.8212.41\|AGTCCTTCTGATCGAGGCCCAGCCCGTGGACGGTGTGAGGC | >hg38_21_+_8433172_8446622_RNA45SN1_45S_50ntFlanks | 8172 | 8212 | 41 | AGTCCTTCTGATCGAGGCCCAGCCCGTGGACGGTGTGAGGC | MB468 | 11 | 73 | 14 | 14 | 14 | 2.4 | 45.4 | 7.4 | 13.7 | 6.4 |
| >hg38_21_+_8433172_8446622_RNA45SN1_45S_50ntFlanks@8173.8213.41\|GTCCTTCTGATCGAGGCCCAGCCCGTGGACGGTGTGAGGCC | >hg38_21_+_8433172_8446622_RNA45SN1_45S_50ntFlanks | 8173 | 8213 | 41 | GTCCTTCTGATCGAGGCCCAGCCCGTGGACGGTGTGAGGCC | MB231 | 10 | 8 | 19 | 1 | 4 | 11.8 | 6.3 | 27.7 | 1.4 | 3.8 |
| >hg38_21_+_8433172_8446622_RNA45SN1_45S_50ntFlanks@8174.8202.29\|TCCTTCTGATCGAGGCCCAGCCCGTGGAC | >hg38_21_+_8433172_8446622_RNA45SN1_45S_50ntFlanks | 8174 | 8202 | 29 | TCCTTCTGATCGAGGCCCAGCCCGTGGAC | MB231 | 17 | 10 | 23 | 2 | 8 | 22.2 | 8.6 | 18.7 | 2.1 | 7.0 |
| >hg38_21_+_8433172_8446622_RNA45SN1_45S_50ntFlanks@8174.8203.30\|TCCTTCTGATCGAGGCCCAGCCCGTGGACG | >hg38_21_+_8433172_8446622_RNA45SN1_45S_50ntFlanks | 8174 | 8203 | 30 | TCCTTCTGATCGAGGCCCAGCCCGTGGACG | MB231 | 15 | 9 | 12 | 2 | 13 | 17.7 | 6.8 | 9.1 | 1.4 | 12.0 |
| >hg38_21_+_8433172_8446622_RNA45SN1_45S_50ntFlanks@8175.8202.28\|CCTTCTGATCGAGGCCCAGCCCGTGGAC | >hg38_21_+_8433172_8446622_RNA45SN1_45S_50ntFlanks | 8175 | 8202 | 28 | CCTTCTGATCGAGGCCCAGCCCGTGGAC | MB231 | 18 | 9 | 30 | 3 | 10 | 22.0 | 6.9 | 25.1 | 3.7 | 8.3 |
| >hg38_21_+_8433172_8446622_RNA45SN1_45S_50ntFlanks@8175.8203.29\|CCTTCTGATCGAGGCCCAGCCCGTGGACG | >hg38_21_+_8433172_8446622_RNA45SN1_45S_50ntFlanks | 8175 | 8203 | 29 | CCTTCTGATCGAGGCCCAGCCCGTGGACG | MB231 | 11 | 7 | 10 | 3 | 11 | 12.7 | 4.7 | 8.6 | 2.8 | 10.1 |
| >hg38_21_+_8433172_8446622_RNA45SN1_45S_50ntFlanks@8175.8216.42\|CCTTCTGATCGAGGCCCAGCCCGTGGACGGTGTGAGGCCGGT | >hg38_21_+_8433172_8446622_RNA45SN1_45S_50ntFlanks | 8175 | 8216 | 42 | CCTTCTGATCGAGGCCCAGCCCGTGGACGGTGTGAGGCCGGT | BT20 | 11 | 5 | 7 | 68 | 31 | 8.6 | 1.9 | 3.0 | 54.0 | 15.1 |
| >hg38_21_+_8433172_8446622_RNA45SN1_45S_50ntFlanks@8175.8216.42\|CCTTCTGATCGAGGCCCAGCCCGTGGACGGTGTGAGGCCGGT | >hg38_21_+_8433172_8446622_RNA45SN1_45S_50ntFlanks | 8175 | 8216 | 42 | CCTTCTGATCGAGGCCCAGCCCGTGGACGGTGTGAGGCCGGT | MB231 | 13 | 10 | 11 | 28 | 31 | 5.6 | 6.8 | 4.7 | 20.4 | 20.3 |
| >hg38_21_+_8433172_8446622_RNA45SN1_45S_50ntFlanks@8176.8193.18\|CTTCTGATCGAGGCCCAG | >hg38_21_+_8433172_8446622_RNA45SN1_45S_50ntFlanks | 8176 | 8193 | 18 | CTTCTGATCGAGGCCCAG | MB468 | 15 | 4 | 29 | 33 | 7 | 5.1 | 0.9 | 12.2 | 13.1 | 2.3 |
| >hg38_21_+_8433172_8446622_RNA45SN1_45S_50ntFlanks@8176.8194.19\|CTTCTGATCGAGGCCCAGC | >hg38_21_+_8433172_8446622_RNA45SN1_45S_50ntFlanks | 8176 | 8194 | 19 | CTTCTGATCGAGGCCCAGC | MB231 | 12 | 8 | 36 | 3 | 4 | 12.5 | 6.9 | 36.1 | 2.8 | 4.6 |
| >hg38_21_+_8433172_8446622_RNA45SN1_45S_50ntFlanks@8176.8195.20\|CTTCTGATCGAGGCCCAGCC | >hg38_21_+_8433172_8446622_RNA45SN1_45S_50ntFlanks | 8176 | 8195 | 20 | CTTCTGATCGAGGCCCAGCC | BT20 | 30 | 15 | 123 | 8 | 13 | 20.7 | 8.1 | 86.2 | 5.1 | 7.9 |
| >hg38_21_+_8433172_8446622_RNA45SN1_45S_50ntFlanks@8176.8195.20\|CTTCTGATCGAGGCCCAGCC | >hg38_21_+_8433172_8446622_RNA45SN1_45S_50ntFlanks | 8176 | 8195 | 20 | CTTCTGATCGAGGCCCAGCC | MB231 | 98 | 67 | 250 | 11 | 24 | 84.8 | 50.3 | 202.6 | 8.4 | 18.3 |
| >hg38_21_+_8433172_8446622_RNA45SN1_45S_50ntFlanks@8176.8195.20\|CTTCTGATCGAGGCCCAGCC | >hg38_21_+_8433172_8446622_RNA45SN1_45S_50ntFlanks | 8176 | 8195 | 20 | CTTCTGATCGAGGCCCAGCC | MB468 | 95 | 172 | 98 | 66 | 74 | 42.6 | 39.9 | 44.4 | 29.2 | 10.4 |
| >hg38_21_+_8433172_8446622_RNA45SN1_45S_50ntFlanks@8176.8196.21\|CTTCTGATCGAGGCCCAGCCC | >hg38_21_+_8433172_8446622_RNA45SN1_45S_50ntFlanks | 8176 | 8196 | 21 | CTTCTGATCGAGGCCCAGCCC | MB231 | 10 | 7 | 39 | 1 | 3 | 8.9 | 3.4 | 28.7 | 0.7 | 2.6 |
| >hg38_21_+_8433172_8446622_RNA45SN1_45S_50ntFlanks@8176.8196.21\|CTTCTGATCGAGGCCCAGCCC | >hg38_21_+_8433172_8446622_RNA45SN1_45S_50ntFlanks | 8176 | 8196 | 21 | CTTCTGATCGAGGCCCAGCCC | MB468 | 19 | 38 | 22 | 9 | 18 | 8.2 | 7.9 | 12.7 | 4.3 | 1.0 |
| >hg38_21_+_8433172_8446622_RNA45SN1_45S_50ntFlanks@8176.8197.22\|CTTCTGATCGAGGCCCAGCCCG | >hg38_21_+_8433172_8446622_RNA45SN1_45S_50ntFlanks | 8176 | 8197 | 22 | CTTCTGATCGAGGCCCAGCCCG | MB468 | 61 | 54 | 71 | 50 | 52 | 11.0 | 9.1 | 21.9 | 15.4 | 14.3 |
| >hg38_21_+_8433172_8446622_RNA45SN1_45S_50ntFlanks@8176.8199.24\|CTTCTGATCGAGGCCCAGCCCGTG | >hg38_21_+_8433172_8446622_RNA45SN1_45S_50ntFlanks | 8176 | 8199 | 24 | CTTCTGATCGAGGCCCAGCCCGTG | MB468 | 16 | 12 | 21 | 27 | 16 | 5.2 | 0.9 | 7.8 | 12.6 | 6.9 |
| >hg38_21_+_8433172_8446622_RNA45SN1_45S_50ntFlanks@8176.8201.26\|CTTCTGATCGAGGCCCAGCCCGTGGA | >hg38_21_+_8433172_8446622_RNA45SN1_45S_50ntFlanks | 8176 | 8201 | 26 | CTTCTGATCGAGGCCCAGCCCGTGGA | MB468 | 33 | 32 | 38 | 60 | 22 | 5.0 | 4.8 | 9.0 | 38.7 | 7.9 |
| >hg38_21_+_8433172_8446622_RNA45SN1_45S_50ntFlanks@8176.8202.27\|CTTCTGATCGAGGCCCAGCCCGTGGAC | >hg38_21_+_8433172_8446622_RNA45SN1_45S_50ntFlanks | 8176 | 8202 | 27 | CTTCTGATCGAGGCCCAGCCCGTGGAC | BT20 | 33 | 18 | 80 | 21 | 29 | 26.3 | 1.5 | 41.3 | 22.9 | 11.9 |
| >hg38_21_+_8433172_8446622_RNA45SN1_45S_50ntFlanks@8176.8202.27\|CTTCTGATCGAGGCCCAGCCCGTGGAC | >hg38_21_+_8433172_8446622_RNA45SN1_45S_50ntFlanks | 8176 | 8202 | 27 | CTTCTGATCGAGGCCCAGCCCGTGGAC | MB231 | 68 | 37 | 134 | 9 | 35 | 76.6 | 28.1 | 106.0 | 7.7 | 29.8 |
| >hg38_21_+_8433172_8446622_RNA45SN1_45S_50ntFlanks@8176.8202.27\|CTTCTGATCGAGGCCCAGCCCGTGGAC | >hg38_21_+_8433172_8446622_RNA45SN1_45S_50ntFlanks | 8176 | 8202 | 27 | CTTCTGATCGAGGCCCAGCCCGTGGAC | MB468 | 30 | 49 | 40 | 31 | 26 | 12.3 | 8.6 | 21.2 | 9.5 | 9.6 |
| >hg38_21_+_8433172_8446622_RNA45SN1_45S_50ntFlanks@8176.8203.28\|CTTCTGATCGAGGCCCAGCCCGTGGACG | >hg38_21_+_8433172_8446622_RNA45SN1_45S_50ntFlanks | 8176 | 8203 | 28 | CTTCTGATCGAGGCCCAGCCCGTGGACG | MB231 | 14 | 9 | 24 | 8 | 12 | 12.8 | 6.5 | 22.3 | 6.0 | 10.0 |
| >hg38_21_+_8433172_8446622_RNA45SN1_45S_50ntFlanks@8176.8203.28\|CTTCTGATCGAGGCCCAGCCCGTGGACG | >hg38_21_+_8433172_8446622_RNA45SN1_45S_50ntFlanks | 8176 | 8203 | 28 | CTTCTGATCGAGGCCCAGCCCGTGGACG | MB468 | 235 | 275 | 230 | 562 | 197 | 39.7 | 73.3 | 90.5 | 415.1 | 147.9 |
| >hg38_21_+_8433172_8446622_RNA45SN1_45S_50ntFlanks@8176.8204.29\|CTTCTGATCGAGGCCCAGCCCGTGGACGG | >hg38_21_+_8433172_8446622_RNA45SN1_45S_50ntFlanks | 8176 | 8204 | 29 | CTTCTGATCGAGGCCCAGCCCGTGGACGG | MB468 | 93 | 112 | 80 | 172 | 77 | 6.5 | 12.7 | 11.2 | 100.2 | 40.3 |
| >hg38_21_+_8433172_8446622_RNA45SN1_45S_50ntFlanks@8176.8205.30\|CTTCTGATCGAGGCCCAGCCCGTGGACGGT | >hg38_21_+_8433172_8446622_RNA45SN1_45S_50ntFlanks | 8176 | 8205 | 30 | CTTCTGATCGAGGCCCAGCCCGTGGACGGT | MB468 | 11 | 25 | 13 | 18 | 9 | 0.8 | 4.9 | 6.2 | 6.9 | 2.7 |
| >hg38_21_+_8433172_8446622_RNA45SN1_45S_50ntFlanks@8176.8206.31\|CTTCTGATCGAGGCCCAGCCCGTGGACGGTG | >hg38_21_+_8433172_8446622_RNA45SN1_45S_50ntFlanks | 8176 | 8206 | 31 | CTTCTGATCGAGGCCCAGCCCGTGGACGGTG | MB468 | 49 | 95 | 68 | 67 | 33 | 6.7 | 28.2 | 28.1 | 25.3 | 5.6 |
| >hg38_21_+_8433172_8446622_RNA45SN1_45S_50ntFlanks@8176.8208.33\|CTTCTGATCGAGGCCCAGCCCGTGGACGGTGTG | >hg38_21_+_8433172_8446622_RNA45SN1_45S_50ntFlanks | 8176 | 8208 | 33 | CTTCTGATCGAGGCCCAGCCCGTGGACGGTGTG | MB468 | 30 | 74 | 24 | 34 | 18 | 6.1 | 21.7 | 7.5 | 13.6 | 7.0 |
| >hg38_21_+_8433172_8446622_RNA45SN1_45S_50ntFlanks@8176.8212.37\|CTTCTGATCGAGGCCCAGCCCGTGGACGGTGTGAGGC | >hg38_21_+_8433172_8446622_RNA45SN1_45S_50ntFlanks | 8176 | 8212 | 37 | CTTCTGATCGAGGCCCAGCCCGTGGACGGTGTGAGGC | MB231 | 15 | 5 | 6 | 10 | 7 | 14.4 | 4.4 | 4.9 | 11.3 | 8.2 |
| >hg38_21_+_8433172_8446622_RNA45SN1_45S_50ntFlanks@8176.8212.37\|CTTCTGATCGAGGCCCAGCCCGTGGACGGTGTGAGGC | >hg38_21_+_8433172_8446622_RNA45SN1_45S_50ntFlanks | 8176 | 8212 | 37 | CTTCTGATCGAGGCCCAGCCCGTGGACGGTGTGAGGC | MB468 | 14 | 15 | 16 | 67 | 26 | 4.1 | 10.6 | 8.1 | 48.5 | 8.8 |
| >hg38_21_+_8433172_8446622_RNA45SN1_45S_50ntFlanks@8176.8213.38\|CTTCTGATCGAGGCCCAGCCCGTGGACGGTGTGAGGCC | >hg38_21_+_8433172_8446622_RNA45SN1_45S_50ntFlanks | 8176 | 8213 | 38 | CTTCTGATCGAGGCCCAGCCCGTGGACGGTGTGAGGCC | MB231 | 12 | 3 | 9 | 3 | 2 | 15.2 | 2.1 | 12.8 | 2.8 | 2.3 |
| >hg38_21_+_8433172_8446622_RNA45SN1_45S_50ntFlanks@8176.8213.38\|CTTCTGATCGAGGCCCAGCCCGTGGACGGTGTGAGGCC | >hg38_21_+_8433172_8446622_RNA45SN1_45S_50ntFlanks | 8176 | 8213 | 38 | CTTCTGATCGAGGCCCAGCCCGTGGACGGTGTGAGGCC | MB468 | 17 | 15 | 17 | 48 | 13 | 11.0 | 16.4 | 5.5 | 30.8 | 4.0 |
| >hg38_21_+_8433172_8446622_RNA45SN1_45S_50ntFlanks@8176.8214.39\|CTTCTGATCGAGGCCCAGCCCGTGGACGGTGTGAGGCCG | >hg38_21_+_8433172_8446622_RNA45SN1_45S_50ntFlanks | 8176 | 8214 | 39 | CTTCTGATCGAGGCCCAGCCCGTGGACGGTGTGAGGCCG | MB468 | 41 | 22 | 66 | 135 | 34 | 15.9 | 17.5 | 71.2 | 141.5 | 11.0 |

| | | | | | | | | | | | | | | | | |
|---|---|---|---|---|---|---|---|---|---|---|---|---|---|---|---|---|
| >hg38_21_+_8433172_8446622_RNA45SN1_45S_50ntFlanks@8176.8215.40\|CTTCTGATCGAGGCCCAGCCCGTGGACGGTGTGAGGCCGG | >hg38_21_+_8433172_8446622_RNA45SN1_45S_50ntFlanks | 8176 | 8215 | 40 | CTTCTGATCGAGGCCCAGCCCGTGGACGGTGTGAGGCCGG | MB468 | 26 | 11 | 33 | 61 | 28 | 13.0 | 4.8 | 28.5 | 54.6 | 10.3 |
| >hg38_21_+_8433172_8446622_RNA45SN1_45S_50ntFlanks@8176.8217.42\|CTTCTGATCGAGGCCCAGCCCGTGGACGGTGTGAGGCCGGTA | >hg38_21_+_8433172_8446622_RNA45SN1_45S_50ntFlanks | 8176 | 8217 | 42 | CTTCTGATCGAGGCCCAGCCCGTGGACGGTGTGAGGCCGGTA | MB231 | 13 | 4 | 13 | 3 | 3 | 10.7 | 2.0 | 9.8 | 1.7 | 1.4 |
| >hg38_21_+_8433172_8446622_RNA45SN1_45S_50ntFlanks@8177.8195.19\|TTCTGATCGAGGCCCAGCC | >hg38_21_+_8433172_8446622_RNA45SN1_45S_50ntFlanks | 8177 | 8195 | 19 | TTCTGATCGAGGCCCAGCC | MB231 | 24 | 15 | 54 | 3 | 5 | 19.6 | 10.7 | 65.4 | 3.0 | 3.8 |
| >hg38_21_+_8433172_8446622_RNA45SN1_45S_50ntFlanks@8177.8202.26\|TTCTGATCGAGGCCCAGCCCGTGGAC | >hg38_21_+_8433172_8446622_RNA45SN1_45S_50ntFlanks | 8177 | 8202 | 26 | TTCTGATCGAGGCCCAGCCCGTGGAC | MB231 | 11 | 6 | 14 | 2 | 5 | 8.9 | 4.6 | 11.8 | 1.2 | 3.8 |
| >hg38_21_+_8433172_8446622_RNA45SN1_45S_50ntFlanks@8177.8203.27\|TTCTGATCGAGGCCCAGCCCGTGGACG | >hg38_21_+_8433172_8446622_RNA45SN1_45S_50ntFlanks | 8177 | 8203 | 27 | TTCTGATCGAGGCCCAGCCCGTGGACG | MB231 | 22 | 15 | 27 | 3 | 15 | 17.5 | 11.7 | 19.2 | 2.2 | 13.4 |
| >hg38_21_+_8433172_8446622_RNA45SN1_45S_50ntFlanks@8177.8203.27\|TTCTGATCGAGGCCCAGCCCGTGGACG | >hg38_21_+_8433172_8446622_RNA45SN1_45S_50ntFlanks | 8177 | 8203 | 27 | TTCTGATCGAGGCCCAGCCCGTGGACG | MB468 | 24 | 42 | 31 | 21 | 11 | 7.9 | 17.3 | 13.8 | 18.9 | 3.2 |
| >hg38_21_+_8433172_8446622_RNA45SN1_45S_50ntFlanks@8177.8204.28\|TTCTGATCGAGGCCCAGCCCGTGGACGG | >hg38_21_+_8433172_8446622_RNA45SN1_45S_50ntFlanks | 8177 | 8204 | 28 | TTCTGATCGAGGCCCAGCCCGTGGACGG | MB468 | 14 | 33 | 13 | 12 | 5 | 1.0 | 16.1 | 4.9 | 9.3 | 1.5 |
| >hg38_21_+_8433172_8446622_RNA45SN1_45S_50ntFlanks@8178.8203.26\|TCTGATCGAGGCCCAGCCCGTGGACG | >hg38_21_+_8433172_8446622_RNA45SN1_45S_50ntFlanks | 8178 | 8203 | 26 | TCTGATCGAGGCCCAGCCCGTGGACG | MB231 | 11 | 9 | 12 | 1 | 7 | 10.4 | 5.6 | 8.7 | 1.1 | 6.9 |
| >hg38_21_+_8433172_8446622_RNA45SN1_45S_50ntFlanks@8178.8203.26\|TCTGATCGAGGCCCAGCCCGTGGACG | >hg38_21_+_8433172_8446622_RNA45SN1_45S_50ntFlanks | 8178 | 8203 | 26 | TCTGATCGAGGCCCAGCCCGTGGACG | MB468 | 19 | 27 | 25 | 15 | 13 | 5.3 | 8.5 | 9.6 | 11.4 | 2.8 |
| >hg38_21_+_8433172_8446622_RNA45SN1_45S_50ntFlanks@8178.8207.30\|TCTGATCGAGGCCCAGCCCGTGGACGGTGT | >hg38_21_+_8433172_8446622_RNA45SN1_45S_50ntFlanks | 8178 | 8207 | 30 | TCTGATCGAGGCCCAGCCCGTGGACGGTGT | BT20 | 89 | 34 | 380 | 17 | 34 | 154.2 | 58.6 | 656.5 | 24.8 | 57.9 |
| >hg38_21_+_8433172_8446622_RNA45SN1_45S_50ntFlanks@8178.8208.31\|TCTGATCGAGGCCCAGCCCGTGGACGGTGTG | >hg38_21_+_8433172_8446622_RNA45SN1_45S_50ntFlanks | 8178 | 8208 | 31 | TCTGATCGAGGCCCAGCCCGTGGACGGTGTG | BT20 | 27 | 14 | 103 | 11 | 19 | 46.3 | 23.8 | 176.6 | 9.6 | 30.4 |
| >hg38_21_+_8433172_8446622_RNA45SN1_45S_50ntFlanks@8179.8197.19\|CTGATCGAGGCCCAGCCCG | >hg38_21_+_8433172_8446622_RNA45SN1_45S_50ntFlanks | 8179 | 8197 | 19 | CTGATCGAGGCCCAGCCCG | MB468 | 10 | 6 | 8 | 10 | 7 | 2.2 | 1.4 | 4.5 | 2.0 | 2.9 |
| >hg38_21_+_8433172_8446622_RNA45SN1_45S_50ntFlanks@8179.8203.25\|CTGATCGAGGCCCAGCCCGTGGACG | >hg38_21_+_8433172_8446622_RNA45SN1_45S_50ntFlanks | 8179 | 8203 | 25 | CTGATCGAGGCCCAGCCCGTGGACG | MB468 | 18 | 15 | 18 | 30 | 17 | 4.4 | 4.9 | 8.1 | 9.5 | 5.7 |
| >hg38_21_+_8433172_8446622_RNA45SN1_45S_50ntFlanks@8179.8206.28\|CTGATCGAGGCCCAGCCCGTGGACGGTG | >hg38_21_+_8433172_8446622_RNA45SN1_45S_50ntFlanks | 8179 | 8206 | 28 | CTGATCGAGGCCCAGCCCGTGGACGGTG | MB468 | 11 | 22 | 9 | 18 | 13 | 4.4 | 13.8 | 3.9 | 5.1 | 2.9 |
| >hg38_21_+_8433172_8446622_RNA45SN1_45S_50ntFlanks@8179.8207.29\|CTGATCGAGGCCCAGCCCGTGGACGGTGT | >hg38_21_+_8433172_8446622_RNA45SN1_45S_50ntFlanks | 8179 | 8207 | 29 | CTGATCGAGGCCCAGCCCGTGGACGGTGT | BT20 | 52 | 33 | 165 | 20 | 55 | 89.9 | 56.3 | 285.4 | 31.6 | 94.1 |
| >hg38_21_+_8433172_8446622_RNA45SN1_45S_50ntFlanks@8179.8208.30\|CTGATCGAGGCCCAGCCCGTGGACGGTGTG | >hg38_21_+_8433172_8446622_RNA45SN1_45S_50ntFlanks | 8179 | 8208 | 30 | CTGATCGAGGCCCAGCCCGTGGACGGTGTG | BT20 | 16 | 14 | 50 | 8 | 25 | 27.5 | 23.4 | 86.5 | 10.8 | 42.9 |
| >hg38_21_+_8433172_8446622_RNA45SN1_45S_50ntFlanks@8179.8208.30\|CTGATCGAGGCCCAGCCCGTGGACGGTGTG | >hg38_21_+_8433172_8446622_RNA45SN1_45S_50ntFlanks | 8179 | 8208 | 30 | CTGATCGAGGCCCAGCCCGTGGACGGTGTG | MB468 | 16 | 45 | 9 | 20 | 10 | 3.9 | 23.5 | 2.9 | 6.3 | 3.1 |
| >hg38_21_+_8433172_8446622_RNA45SN1_45S_50ntFlanks@8179.8210.32\|CTGATCGAGGCCCAGCCCGTGGACGGTGTGAG | >hg38_21_+_8433172_8446622_RNA45SN1_45S_50ntFlanks | 8179 | 8210 | 32 | CTGATCGAGGCCCAGCCCGTGGACGGTGTGAG | MB468 | 16 | 22 | 10 | 15 | 7 | 4.8 | 14.9 | 2.9 | 6.9 | 1.8 |
| >hg38_21_+_8433172_8446622_RNA45SN1_45S_50ntFlanks@8179.8211.33\|CTGATCGAGGCCCAGCCCGTGGACGGTGTGAGG | >hg38_21_+_8433172_8446622_RNA45SN1_45S_50ntFlanks | 8179 | 8211 | 33 | CTGATCGAGGCCCAGCCCGTGGACGGTGTGAGG | MB468 | 30 | 16 | 22 | 65 | 12 | 14.8 | 6.8 | 14.7 | 58.9 | 6.5 |
| >hg38_21_+_8433172_8446622_RNA45SN1_45S_50ntFlanks@8179.8212.34\|CTGATCGAGGCCCAGCCCGTGGACGGTGTGAGGC | >hg38_21_+_8433172_8446622_RNA45SN1_45S_50ntFlanks | 8179 | 8212 | 34 | CTGATCGAGGCCCAGCCCGTGGACGGTGTGAGGC | MB468 | 19 | 5 | 10 | 51 | 10 | 12.9 | 2.1 | 3.6 | 44.2 | 5.2 |
| >hg38_21_+_8433172_8446622_RNA45SN1_45S_50ntFlanks@8180.8211.32\|TGATCGAGGCCCAGCCCGTGGACGGTGTGAGG | >hg38_21_+_8433172_8446622_RNA45SN1_45S_50ntFlanks | 8180 | 8211 | 32 | TGATCGAGGCCCAGCCCGTGGACGGTGTGAGG | MB468 | 19 | 23 | 12 | 11 | 4 | 8.8 | 16.1 | 6.0 | 3.8 | 1.7 |
| >hg38_21_+_8433172_8446622_RNA45SN1_45S_50ntFlanks@8182.8203.22\|ATCGAGGCCCAGCCCGTGGACG | >hg38_21_+_8433172_8446622_RNA45SN1_45S_50ntFlanks | 8182 | 8203 | 22 | ATCGAGGCCCAGCCCGTGGACG | MB468 | 31 | 23 | 31 | 37 | 38 | 5.7 | 6.7 | 7.2 | 15.6 | 9.2 |
| >hg38_21_+_8433172_8446622_RNA45SN1_45S_50ntFlanks@8182.8204.23\|ATCGAGGCCCAGCCCGTGGACGG | >hg38_21_+_8433172_8446622_RNA45SN1_45S_50ntFlanks | 8182 | 8204 | 23 | ATCGAGGCCCAGCCCGTGGACGG | MB468 | 20 | 27 | 19 | 27 | 22 | 6.3 | 6.3 | 8.5 | 11.5 | 4.0 |
| >hg38_21_+_8433172_8446622_RNA45SN1_45S_50ntFlanks@8182.8206.25\|ATCGAGGCCCAGCCCGTGGACGGTG | >hg38_21_+_8433172_8446622_RNA45SN1_45S_50ntFlanks | 8182 | 8206 | 25 | ATCGAGGCCCAGCCCGTGGACGGTG | MB468 | 24 | 37 | 19 | 39 | 25 | 8.0 | 14.3 | 9.3 | 18.4 | 3.0 |
| >hg38_21_+_8433172_8446622_RNA45SN1_45S_50ntFlanks@8182.8208.27\|ATCGAGGCCCAGCCCGTGGACGGTGTG | >hg38_21_+_8433172_8446622_RNA45SN1_45S_50ntFlanks | 8182 | 8208 | 27 | ATCGAGGCCCAGCCCGTGGACGGTGTG | MB468 | 19 | 47 | 10 | 31 | 13 | 5.1 | 23.0 | 3.0 | 17.3 | 3.3 |
| >hg38_21_+_8433172_8446622_RNA45SN1_45S_50ntFlanks@8182.8210.29\|ATCGAGGCCCAGCCCGTGGACGGTGTGAG | >hg38_21_+_8433172_8446622_RNA45SN1_45S_50ntFlanks | 8182 | 8210 | 29 | ATCGAGGCCCAGCCCGTGGACGGTGTGAG | MB468 | 28 | 66 | 17 | 29 | 11 | 7.8 | 40.4 | 8.0 | 14.6 | 4.3 |
| >hg38_21_+_8433172_8446622_RNA45SN1_45S_50ntFlanks@8182.8211.30\|ATCGAGGCCCAGCCCGTGGACGGTGTGAGG | >hg38_21_+_8433172_8446622_RNA45SN1_45S_50ntFlanks | 8182 | 8211 | 30 | ATCGAGGCCCAGCCCGTGGACGGTGTGAGG | MB468 | 24 | 39 | 17 | 28 | 10 | 6.4 | 25.1 | 2.0 | 13.9 | 0.6 |
| >hg38_21_+_8433172_8446622_RNA45SN1_45S_50ntFlanks@8182.8223.42\|ATCGAGGCCCAGCCCGTGGACGGTGTGAGGCCGGTAGCGGCC | >hg38_21_+_8433172_8446622_RNA45SN1_45S_50ntFlanks | 8182 | 8223 | 42 | ATCGAGGCCCAGCCCGTGGACGGTGTGAGGCCGGTAGCGGCC | MB468 | 13 | 52 | 31 | 5 | 15 | 4.6 | 22.8 | 22.4 | 4.7 | 7.5 |
| >hg38_21_+_8433172_8446622_RNA45SN1_45S_50ntFlanks@8184.8210.27\|CGAGGCCCAGCCCGTGGACGGTGTGAG | >hg38_21_+_8433172_8446622_RNA45SN1_45S_50ntFlanks | 8184 | 8210 | 27 | CGAGGCCCAGCCCGTGGACGGTGTGAG | MB468 | 12 | 28 | 11 | 16 | 7 | 3.9 | 11.8 | 5.3 | 8.6 | 1.4 |
| >hg38_21_+_8433172_8446622_RNA45SN1_45S_50ntFlanks@8184.8211.28\|CGAGGCCCAGCCCGTGGACGGTGTGAGG | >hg38_21_+_8433172_8446622_RNA45SN1_45S_50ntFlanks | 8184 | 8211 | 28 | CGAGGCCCAGCCCGTGGACGGTGTGAGG | MB468 | 10 | 17 | 10 | 20 | 8 | 3.7 | 7.5 | 2.0 | 11.7 | 1.2 |
| >hg38_21_+_8433172_8446622_RNA45SN1_45S_50ntFlanks@8185.8207.23\|GAGGCCCAGCCCGTGGACGGTGT | >hg38_21_+_8433172_8446622_RNA45SN1_45S_50ntFlanks | 8185 | 8207 | 23 | GAGGCCCAGCCCGTGGACGGTGT | BT20 | 24 | 18 | 68 | 28 | 29 | 40.1 | 30.1 | 117.4 | 42.5 | 47.9 |
| >hg38_21_+_8433172_8446622_RNA45SN1_45S_50ntFlanks@8189.8208.20\|CCCAGCCCGTGGACGGTGTG | >hg38_21_+_8433172_8446622_RNA45SN1_45S_50ntFlanks | 8189 | 8208 | 20 | CCCAGCCCGTGGACGGTGTG | MB468 | 17 | 4 | 10 | 21 | 6 | 12.5 | 3.1 | 9.8 | 9.2 | 2.6 |
| >hg38_21_+_8433172_8446622_RNA45SN1_45S_50ntFlanks@8189.8212.24\|CCCAGCCCGTGGACGGTGTGAGGC | >hg38_21_+_8433172_8446622_RNA45SN1_45S_50ntFlanks | 8189 | 8212 | 24 | CCCAGCCCGTGGACGGTGTGAGGC | MB468 | 30 | 9 | 36 | 52 | 16 | 20.7 | 6.5 | 44.0 | 2.4 | 8.2 |
| >hg38_21_+_8433172_8446622_RNA45SN1_45S_50ntFlanks@8189.8213.25\|CCCAGCCCGTGGACGGTGTGAGGCC | >hg38_21_+_8433172_8446622_RNA45SN1_45S_50ntFlanks | 8189 | 8213 | 25 | CCCAGCCCGTGGACGGTGTGAGGCC | MB468 | 18 | 12 | 18 | 15 | 13 | 11.7 | 9.7 | 20.2 | 8.7 | 5.1 |
| >hg38_21_+_8433172_8446622_RNA45SN1_45S_50ntFlanks@8189.8215.27\|CCCAGCCCGTGGACGGTGTGAGGCCGG | >hg38_21_+_8433172_8446622_RNA45SN1_45S_50ntFlanks | 8189 | 8215 | 27 | CCCAGCCCGTGGACGGTGTGAGGCCGG | MB468 | 12 | 10 | 21 | 10 | 7 | 10.1 | 7.7 | 23.8 | 4.6 | 2.0 |
| >hg38_21_+_8433172_8446622_RNA45SN1_45S_50ntFlanks@8190.8212.23\|CCAGCCCGTGGACGGTGTGAGGC | >hg38_21_+_8433172_8446622_RNA45SN1_45S_50ntFlanks | 8190 | 8212 | 23 | CCAGCCCGTGGACGGTGTGAGGC | MB468 | 61 | 39 | 30 | 92 | 33 | 17.0 | 20.1 | 10.7 | 22.8 | 5.9 |
| >hg38_21_+_8433172_8446622_RNA45SN1_45S_50ntFlanks@8190.8213.24\|CCAGCCCGTGGACGGTGTGAGGCC | >hg38_21_+_8433172_8446622_RNA45SN1_45S_50ntFlanks | 8190 | 8213 | 24 | CCAGCCCGTGGACGGTGTGAGGCC | MB468 | 18 | 28 | 8 | 14 | 14 | 6.8 | 16.8 | 2.8 | 7.4 | 1.9 |
| >hg38_21_+_8433172_8446622_RNA45SN1_45S_50ntFlanks@8191.8211.21\|CAGCCCGTGGACGGTGTGAGG | >hg38_21_+_8433172_8446622_RNA45SN1_45S_50ntFlanks | 8191 | 8211 | 21 | CAGCCCGTGGACGGTGTGAGG | MB468 | 16 | 5 | 10 | 64 | 18 | 3.5 | 2.0 | 5.7 | 52.9 | 15.7 |
| >hg38_21_+_8433172_8446622_RNA45SN1_45S_50ntFlanks@8191.8212.22\|CAGCCCGTGGACGGTGTGAGGC | >hg38_21_+_8433172_8446622_RNA45SN1_45S_50ntFlanks | 8191 | 8212 | 22 | CAGCCCGTGGACGGTGTGAGGC | MB468 | 71 | 46 | 42 | 121 | 41 | 18.5 | 9.4 | 5.5 | 66.4 | 15.1 |
| >hg38_21_+_8433172_8446622_RNA45SN1_45S_50ntFlanks@8191.8213.23\|CAGCCCGTGGACGGTGTGAGGCC | >hg38_21_+_8433172_8446622_RNA45SN1_45S_50ntFlanks | 8191 | 8213 | 23 | CAGCCCGTGGACGGTGTGAGGCC | MB468 | 64 | 78 | 31 | 58 | 44 | 19.7 | 31.3 | 3.1 | 15.6 | 7.3 |
| >hg38_21_+_8433172_8446622_RNA45SN1_45S_50ntFlanks@8191.8214.24\|CAGCCCGTGGACGGTGTGAGGCCG | >hg38_21_+_8433172_8446622_RNA45SN1_45S_50ntFlanks | 8191 | 8214 | 24 | CAGCCCGTGGACGGTGTGAGGCCG | MB468 | 65 | 115 | 37 | 49 | 49 | 15.1 | 70.4 | 3.7 | 25.0 | 7.5 |
| >hg38_21_+_8433172_8446622_RNA45SN1_45S_50ntFlanks@8191.8215.25\|CAGCCCGTGGACGGTGTGAGGCCGG | >hg38_21_+_8433172_8446622_RNA45SN1_45S_50ntFlanks | 8191 | 8215 | 25 | CAGCCCGTGGACGGTGTGAGGCCGG | MB468 | 109 | 159 | 89 | 71 | 70 | 17.2 | 66.8 | 15.9 | 23.8 | 12.3 |
| >hg38_21_+_8433172_8446622_RNA45SN1_45S_50ntFlanks@8192.8211.20\|AGCCCGTGGACGGTGTGAGG | >hg38_21_+_8433172_8446622_RNA45SN1_45S_50ntFlanks | 8192 | 8211 | 20 | AGCCCGTGGACGGTGTGAGG | MB468 | 18 | 19 | 8 | 34 | 6 | 10.9 | 16.1 | 5.8 | 16.6 | 2.9 |
| >hg38_21_+_8433172_8446622_RNA45SN1_45S_50ntFlanks@8192.8212.21\|AGCCCGTGGACGGTGTGAGGC | >hg38_21_+_8433172_8446622_RNA45SN1_45S_50ntFlanks | 8192 | 8212 | 21 | AGCCCGTGGACGGTGTGAGGC | MB468 | 15 | 17 | 10 | 17 | 7 | 2.7 | 12.6 | 2.9 | 7.7 | 2.4 |
| >hg38_21_+_8433172_8446622_RNA45SN1_45S_50ntFlanks@8192.8213.22\|AGCCCGTGGACGGTGTGAGGCC | >hg38_21_+_8433172_8446622_RNA45SN1_45S_50ntFlanks | 8192 | 8213 | 22 | AGCCCGTGGACGGTGTGAGGCC | MB468 | 12 | 18 | 7 | 13 | 8 | 3.3 | 11.0 | 1.8 | 6.0 | 3.7 |
| >hg38_21_+_8433172_8446622_RNA45SN1_45S_50ntFlanks@8192.8214.23\|AGCCCGTGGACGGTGTGAGGCCG | >hg38_21_+_8433172_8446622_RNA45SN1_45S_50ntFlanks | 8192 | 8214 | 23 | AGCCCGTGGACGGTGTGAGGCCG | MB468 | 13 | 22 | 10 | 7 | 7 | 3.4 | 13.8 | 4.1 | 5.7 | 2.0 |
| >hg38_21_+_8433172_8446622_RNA45SN1_45S_50ntFlanks@8192.8215.24\|AGCCCGTGGACGGTGTGAGGCCGG | >hg38_21_+_8433172_8446622_RNA45SN1_45S_50ntFlanks | 8192 | 8215 | 24 | AGCCCGTGGACGGTGTGAGGCCGG | MB468 | 18 | 23 | 17 | 11 | 7 | 7.1 | 11.8 | 10.1 | 4.1 | 0.7 |
| >hg38_21_+_8433172_8446622_RNA45SN1_45S_50ntFlanks@8194.8211.18\|CCCGTGGACGGTGTGAGG | >hg38_21_+_8433172_8446622_RNA45SN1_45S_50ntFlanks | 8194 | 8211 | 18 | CCCGTGGACGGTGTGAGG | MB468 | 81 | 118 | 58 | 238 | 41 | 27.6 | 84.3 | 21.8 | 133.9 | 24.9 |
| >hg38_21_+_8433172_8446622_RNA45SN1_45S_50ntFlanks@8194.8212.19\|CCCGTGGACGGTGTGAGGC | >hg38_21_+_8433172_8446622_RNA45SN1_45S_50ntFlanks | 8194 | 8212 | 19 | CCCGTGGACGGTGTGAGGC | MB468 | 58 | 124 | 44 | 162 | 35 | 10.3 | 68.9 | 11.8 | 92.7 | 14.9 |
| >hg38_21_+_8433172_8446622_RNA45SN1_45S_50ntFlanks@8194.8213.20\|CCCGTGGACGGTGTGAGGCC | >hg38_21_+_8433172_8446622_RNA45SN1_45S_50ntFlanks | 8194 | 8213 | 20 | CCCGTGGACGGTGTGAGGCC | MB468 | 90 | 166 | 66 | 100 | 57 | 14.4 | 52.9 | 12.2 | 60.4 | 6.8 |

| | | | | | | | | | | | | | | | | |
|---|---|---|---|---|---|---|---|---|---|---|---|---|---|---|---|---|
| >hg38_21_+_8433172_8446622_RNA45SN1_45S_50ntFlanks@8194.8214.21\|CCCGTGGACGGTGTGAGGCCG | >hg38_21_+_8433172_8446622_RNA45SN1_45S_50ntFlanks | 8194 | 8214 | 21 | CCCGTGGACGGTGTGAGGCCG | MB468 | 76 | 114 | 83 | 73 | 41 | 16.5 | 35.0 | 13.8 | 62.3 | 8.8 |
| >hg38_21_+_8433172_8446622_RNA45SN1_45S_50ntFlanks@8194.8215.22\|CCCGTGGACGGTGTGAGGCCGG | >hg38_21_+_8433172_8446622_RNA45SN1_45S_50ntFlanks | 8194 | 8215 | 22 | CCCGTGGACGGTGTGAGGCCGG | MB468 | 169 | 205 | 206 | 130 | 72 | 61.6 | 113.1 | 126.6 | 111.0 | 32.8 |
| >hg38_21_+_8433172_8446622_RNA45SN1_45S_50ntFlanks@8194.8216.23\|CCCGTGGACGGTGTGAGGCCGGT | >hg38_21_+_8433172_8446622_RNA45SN1_45S_50ntFlanks | 8194 | 8216 | 23 | CCCGTGGACGGTGTGAGGCCGGT | MB468 | 16 | 15 | 17 | 18 | 10 | 4.8 | 6.1 | 6.1 | 5.8 | 1.9 |
| >hg38_21_+_8433172_8446622_RNA45SN1_45S_50ntFlanks@8194.8223.30\|CCCGTGGACGGTGTGAGGCCGGTAGCGGCC | >hg38_21_+_8433172_8446622_RNA45SN1_45S_50ntFlanks | 8194 | 8223 | 30 | CCCGTGGACGGTGTGAGGCCGGTAGCGGCC | MB468 | 12 | 1 | 17 | 8 | 3 | 9.4 | 0.4 | 15.3 | 1.7 | 2.1 |
| >hg38_21_+_8433172_8446622_RNA45SN1_45S_50ntFlanks@8194.8225.32\|CCCGTGGACGGTGTGAGGCCGGTAGCGGCCCC | >hg38_21_+_8433172_8446622_RNA45SN1_45S_50ntFlanks | 8194 | 8225 | 32 | CCCGTGGACGGTGTGAGGCCGGTAGCGGCCCC | MB468 | 15 | 11 | 38 | 11 | 13 | 3.4 | 3.8 | 14.4 | 7.1 | 7.4 |
| >hg38_21_+_8433172_8446622_RNA45SN1_45S_50ntFlanks@8195.8213.19\|CCGTGGACGGTGTGAGGCC | >hg38_21_+_8433172_8446622_RNA45SN1_45S_50ntFlanks | 8195 | 8213 | 19 | CCGTGGACGGTGTGAGGCC | BT20 | 13 | 5 | 88 | 63 | 12 | 10.5 | 3.3 | 76.8 | 35.7 | 4.8 |
| >hg38_21_+_8433172_8446622_RNA45SN1_45S_50ntFlanks@8195.8213.19\|CCGTGGACGGTGTGAGGCC | >hg38_21_+_8433172_8446622_RNA45SN1_45S_50ntFlanks | 8195 | 8213 | 19 | CCGTGGACGGTGTGAGGCC | MB231 | 15 | 8 | 100 | 17 | 6 | 7.4 | 4.1 | 61.9 | 9.7 | 2.9 |
| >hg38_21_+_8433172_8446622_RNA45SN1_45S_50ntFlanks@8195.8215.21\|CCGTGGACGGTGTGAGGCCGG | >hg38_21_+_8433172_8446622_RNA45SN1_45S_50ntFlanks | 8195 | 8215 | 21 | CCGTGGACGGTGTGAGGCCGG | MB468 | 14 | 11 | 15 | 16 | 4 | 4.5 | 5.6 | 7.5 | 11.6 | 2.4 |
| >hg38_21_+_8433172_8446622_RNA45SN1_45S_50ntFlanks@8196.8213.18\|CGTGGACGGTGTGAGGCC | >hg38_21_+_8433172_8446622_RNA45SN1_45S_50ntFlanks | 8196 | 8213 | 18 | CGTGGACGGTGTGAGGCC | BT20 | 22 | 14 | 109 | 236 | 18 | 15.2 | 10.9 | 89.3 | 339.3 | 13.1 |
| >hg38_21_+_8433172_8446622_RNA45SN1_45S_50ntFlanks@8196.8213.18\|CGTGGACGGTGTGAGGCC | >hg38_21_+_8433172_8446622_RNA45SN1_45S_50ntFlanks | 8196 | 8213 | 18 | CGTGGACGGTGTGAGGCC | MB231 | 63 | 36 | 446 | 30 | 13 | 51.0 | 35.9 | 587.3 | 22.5 | 11.3 |
| >hg38_21_+_8433172_8446622_RNA45SN1_45S_50ntFlanks@8196.8213.18\|CGTGGACGGTGTGAGGCC | >hg38_21_+_8433172_8446622_RNA45SN1_45S_50ntFlanks | 8196 | 8213 | 18 | CGTGGACGGTGTGAGGCC | MB468 | 73 | 129 | 89 | 146 | 41 | 30.4 | 45.3 | 38.0 | 53.2 | 9.9 |
| >hg38_21_+_8433172_8446622_RNA45SN1_45S_50ntFlanks@8196.8214.19\|CGTGGACGGTGTGAGGCCG | >hg38_21_+_8433172_8446622_RNA45SN1_45S_50ntFlanks | 8196 | 8214 | 19 | CGTGGACGGTGTGAGGCCG | MB468 | 121 | 70 | 215 | 217 | 36 | 83.5 | 19.7 | 96.3 | 92.5 | 4.7 |
| >hg38_21_+_8433172_8446622_RNA45SN1_45S_50ntFlanks@8196.8215.20\|CGTGGACGGTGTGAGGCCGG | >hg38_21_+_8433172_8446622_RNA45SN1_45S_50ntFlanks | 8196 | 8215 | 20 | CGTGGACGGTGTGAGGCCGG | MB468 | 150 | 91 | 323 | 241 | 40 | 36.0 | 35.9 | 90.9 | 106.2 | 17.7 |
| >hg38_21_+_8433172_8446622_RNA45SN1_45S_50ntFlanks@8196.8216.21\|CGTGGACGGTGTGAGGCCGGT | >hg38_21_+_8433172_8446622_RNA45SN1_45S_50ntFlanks | 8196 | 8216 | 21 | CGTGGACGGTGTGAGGCCGGT | MB468 | 22 | 9 | 38 | 47 | 8 | 9.7 | 2.7 | 14.4 | 22.6 | 2.5 |
| >hg38_21_+_8433172_8446622_RNA45SN1_45S_50ntFlanks@8196.8219.24\|CGTGGACGGTGTGAGGCCGGTAGC | >hg38_21_+_8433172_8446622_RNA45SN1_45S_50ntFlanks | 8196 | 8219 | 24 | CGTGGACGGTGTGAGGCCGGTAGC | MB231 | 12 | 7 | 26 | 9 | 3 | 12.8 | 5.5 | 30.5 | 7.3 | 2.4 |
| >hg38_21_+_8433172_8446622_RNA45SN1_45S_50ntFlanks@8196.8222.27\|CGTGGACGGTGTGAGGCCGGTAGCGGC | >hg38_21_+_8433172_8446622_RNA45SN1_45S_50ntFlanks | 8196 | 8222 | 27 | CGTGGACGGTGTGAGGCCGGTAGCGGC | MB468 | 20 | 4 | 35 | 26 | 6 | 7.1 | 0.9 | 23.0 | 6.5 | 1.7 |
| >hg38_21_+_8433172_8446622_RNA45SN1_45S_50ntFlanks@8196.8223.28\|CGTGGACGGTGTGAGGCCGGTAGCGGCC | >hg38_21_+_8433172_8446622_RNA45SN1_45S_50ntFlanks | 8196 | 8223 | 28 | CGTGGACGGTGTGAGGCCGGTAGCGGCC | MB468 | 44 | 43 | 74 | 55 | 19 | 19.0 | 16.9 | 33.8 | 41.0 | 12.7 |
| >hg38_21_+_8433172_8446622_RNA45SN1_45S_50ntFlanks@8196.8224.29\|CGTGGACGGTGTGAGGCCGGTAGCGGCCC | >hg38_21_+_8433172_8446622_RNA45SN1_45S_50ntFlanks | 8196 | 8224 | 29 | CGTGGACGGTGTGAGGCCGGTAGCGGCCC | MB468 | 56 | 91 | 119 | 43 | 24 | 14.8 | 16.1 | 55.3 | 28.0 | 12.1 |
| >hg38_21_+_8433172_8446622_RNA45SN1_45S_50ntFlanks@8196.8225.30\|CGTGGACGGTGTGAGGCCGGTAGCGGCCCC | >hg38_21_+_8433172_8446622_RNA45SN1_45S_50ntFlanks | 8196 | 8225 | 30 | CGTGGACGGTGTGAGGCCGGTAGCGGCCCC | MB468 | 36 | 70 | 92 | 16 | 17 | 13.0 | 30.6 | 51.6 | 6.9 | 8.4 |
| >hg38_21_+_8433172_8446622_RNA45SN1_45S_50ntFlanks@8196.8226.31\|CGTGGACGGTGTGAGGCCGGTAGCGGCCCCC | >hg38_21_+_8433172_8446622_RNA45SN1_45S_50ntFlanks | 8196 | 8226 | 31 | CGTGGACGGTGTGAGGCCGGTAGCGGCCCCC | MB468 | 21 | 23 | 59 | 4 | 9 | 6.6 | 2.4 | 37.4 | 1.0 | 4.7 |
| >hg38_21_+_8433172_8446622_RNA45SN1_45S_50ntFlanks@8197.8214.18\|GTGGACGGTGTGAGGCCG | >hg38_21_+_8433172_8446622_RNA45SN1_45S_50ntFlanks | 8197 | 8214 | 18 | GTGGACGGTGTGAGGCCG | MB468 | 13 | 6 | 22 | 24 | 3 | 3.7 | 2.6 | 6.8 | 14.0 | 1.1 |
| >hg38_21_+_8433172_8446622_RNA45SN1_45S_50ntFlanks@8197.8215.19\|GTGGACGGTGTGAGGCCGG | >hg38_21_+_8433172_8446622_RNA45SN1_45S_50ntFlanks | 8197 | 8215 | 19 | GTGGACGGTGTGAGGCCGG | MB468 | 17 | 8 | 32 | 26 | 3 | 7.7 | 3.7 | 17.4 | 14.6 | 1.6 |
| >hg38_21_+_8433172_8446622_RNA45SN1_45S_50ntFlanks@8198.8215.18\|TGGACGGTGTGAGGCCGG | >hg38_21_+_8433172_8446622_RNA45SN1_45S_50ntFlanks | 8198 | 8215 | 18 | TGGACGGTGTGAGGCCGG | MB468 | 253 | 41 | 311 | 169 | 31 | 178.2 | 11.3 | 365.4 | 54.1 | 18.5 |
| >hg38_21_+_8433172_8446622_RNA45SN1_45S_50ntFlanks@8198.8216.19\|TGGACGGTGTGAGGCCGGT | >hg38_21_+_8433172_8446622_RNA45SN1_45S_50ntFlanks | 8198 | 8216 | 19 | TGGACGGTGTGAGGCCGGT | MB468 | 12 | 2 | 13 | 13 | 3 | 6.8 | 0.2 | 10.3 | 4.9 | 2.1 |
| >hg38_21_+_8433172_8446622_RNA45SN1_45S_50ntFlanks@8198.8223.26\|TGGACGGTGTGAGGCCGGTAGCGGCC | >hg38_21_+_8433172_8446622_RNA45SN1_45S_50ntFlanks | 8198 | 8223 | 26 | TGGACGGTGTGAGGCCGGTAGCGGCC | MB468 | 36 | 56 | 56 | 40 | 10 | 16.2 | 13.9 | 30.6 | 24.8 | 4.8 |
| >hg38_21_+_8433172_8446622_RNA45SN1_45S_50ntFlanks@8198.8224.27\|TGGACGGTGTGAGGCCGGTAGCGGCCC | >hg38_21_+_8433172_8446622_RNA45SN1_45S_50ntFlanks | 8198 | 8224 | 27 | TGGACGGTGTGAGGCCGGTAGCGGCCC | MB468 | 22 | 28 | 44 | 19 | 7 | 5.6 | 7.1 | 20.7 | 7.5 | 2.8 |
| >hg38_21_+_8433172_8446622_RNA45SN1_45S_50ntFlanks@8198.8225.28\|TGGACGGTGTGAGGCCGGTAGCGGCCCC | >hg38_21_+_8433172_8446622_RNA45SN1_45S_50ntFlanks | 8198 | 8225 | 28 | TGGACGGTGTGAGGCCGGTAGCGGCCCC | MB468 | 28 | 28 | 64 | 20 | 9 | 8.9 | 11.4 | 22.7 | 6.2 | 2.7 |
| >hg38_21_+_8433172_8446622_RNA45SN1_45S_50ntFlanks@8317.8356.40\|CACGAGACCGATAGTCAACAAGTACCGTAAGGGAAAGTTG | >hg38_21_+_8433172_8446622_RNA45SN1_45S_50ntFlanks | 8317 | 8356 | 40 | CACGAGACCGATAGTCAACAAGTACCGTAAGGGAAAGTTG | MB468 | 13 | 45 | 41 | 27 | 57 | 7.5 | 16.3 | 29.2 | 28.3 | 18.2 |
| >hg38_21_+_8433172_8446622_RNA45SN1_45S_50ntFlanks@8319.8356.38\|CGAGACCGATAGTCAACAAGTACCGTAAGGGAAAGTTG | >hg38_21_+_8433172_8446622_RNA45SN1_45S_50ntFlanks | 8319 | 8356 | 38 | CGAGACCGATAGTCAACAAGTACCGTAAGGGAAAGTTG | MB468 | 17 | 22 | 43 | 18 | 27 | 7.4 | 8.5 | 26.1 | 13.4 | 7.6 |
| >hg38_21_+_8433172_8446622_RNA45SN1_45S_50ntFlanks@8324.8356.33\|CCGATAGTCAACAAGTACCGTAAGGGAAAGTTG | >hg38_21_+_8433172_8446622_RNA45SN1_45S_50ntFlanks | 8324 | 8356 | 33 | CCGATAGTCAACAAGTACCGTAAGGGAAAGTTG | MB468 | 30 | 3 | 65 | 21 | 8 | 1.9 | 2.4 | 19.5 | 6.8 | 2.3 |
| >hg38_21_+_8433172_8446622_RNA45SN1_45S_50ntFlanks@8325.8356.32\|CGATAGTCAACAAGTACCGTAAGGGAAAGTTG | >hg38_21_+_8433172_8446622_RNA45SN1_45S_50ntFlanks | 8325 | 8356 | 32 | CGATAGTCAACAAGTACCGTAAGGGAAAGTTG | MB468 | 12 | 1 | 19 | 10 | 4 | 3.3 | 0.7 | 7.4 | 3.5 | 1.7 |
| >hg38_21_+_8433172_8446622_RNA45SN1_45S_50ntFlanks@8327.8356.30\|ATAGTCAACAAGTACCGTAAGGGAAAGTTG | >hg38_21_+_8433172_8446622_RNA45SN1_45S_50ntFlanks | 8327 | 8356 | 30 | ATAGTCAACAAGTACCGTAAGGGAAAGTTG | MB468 | 14 | 0 | 12 | 7 | 5 | 10.1 | 0.1 | 9.9 | 2.0 | 6.9 |
| >hg38_21_+_8433172_8446622_RNA45SN1_45S_50ntFlanks@8342.8364.23\|CGTAAGGGAAAGTTGAAAAGAAC | >hg38_21_+_8433172_8446622_RNA45SN1_45S_50ntFlanks | 8342 | 8364 | 23 | CGTAAGGGAAAGTTGAAAAGAAC | MB231 | 14 | 10 | 49 | 7 | 4 | 11.8 | 8.2 | 46.2 | 5.8 | 3.5 |
| >hg38_21_+_8433172_8446622_RNA45SN1_45S_50ntFlanks@8342.8365.24\|CGTAAGGGAAAGTTGAAAAGAACT | >hg38_21_+_8433172_8446622_RNA45SN1_45S_50ntFlanks | 8342 | 8365 | 24 | CGTAAGGGAAAGTTGAAAAGAACT | BT20 | 21 | 11 | 56 | 34 | 15 | 11.1 | 3.5 | 22.9 | 22.8 | 7.3 |
| >hg38_21_+_8433172_8446622_RNA45SN1_45S_50ntFlanks@8342.8365.24\|CGTAAGGGAAAGTTGAAAAGAACT | >hg38_21_+_8433172_8446622_RNA45SN1_45S_50ntFlanks | 8342 | 8365 | 24 | CGTAAGGGAAAGTTGAAAAGAACT | MB231 | 92 | 50 | 205 | 55 | 18 | 76.5 | 41.0 | 219.3 | 44.8 | 16.5 |
| >hg38_21_+_8433172_8446622_RNA45SN1_45S_50ntFlanks@8342.8367.26\|CGTAAGGGAAAGTTGAAAAGAACTTT | >hg38_21_+_8433172_8446622_RNA45SN1_45S_50ntFlanks | 8342 | 8367 | 26 | CGTAAGGGAAAGTTGAAAAGAACTTT | MB231 | 10 | 4 | 7 | 1 | 1 | 10.8 | 3.6 | 9.0 | 1.2 | 1.1 |
| >hg38_21_+_8433172_8446622_RNA45SN1_45S_50ntFlanks@8342.8376.35\|CGTAAGGGAAAGTTGAAAAGAACTTTGAAGAGAGA | >hg38_21_+_8433172_8446622_RNA45SN1_45S_50ntFlanks | 8342 | 8376 | 35 | CGTAAGGGAAAGTTGAAAAGAACTTTGAAGAGAGA | MB231 | 13 | 3 | 36 | 5 | 1 | 10.1 | 2.4 | 45.4 | 4.3 | 1.3 |
| >hg38_21_+_8433172_8446622_RNA45SN1_45S_50ntFlanks@8342.8377.36\|CGTAAGGGAAAGTTGAAAAGAACTTTGAAGAGAGAG | >hg38_21_+_8433172_8446622_RNA45SN1_45S_50ntFlanks | 8342 | 8377 | 36 | CGTAAGGGAAAGTTGAAAAGAACTTTGAAGAGAGAG | MB231 | 11 | 3 | 32 | 6 | 2 | 10.3 | 1.7 | 30.4 | 4.4 | 1.7 |
| >hg38_21_+_8433172_8446622_RNA45SN1_45S_50ntFlanks@8342.8378.37\|CGTAAGGGAAAGTTGAAAAGAACTTTGAAGAGAGAGT | >hg38_21_+_8433172_8446622_RNA45SN1_45S_50ntFlanks | 8342 | 8378 | 37 | CGTAAGGGAAAGTTGAAAAGAACTTTGAAGAGAGAGT | MB231 | 34 | 8 | 146 | 23 | 4 | 29.7 | 6.4 | 132.8 | 15.1 | 2.8 |
| >hg38_21_+_8433172_8446622_RNA45SN1_45S_50ntFlanks@8342.8379.38\|CGTAAGGGAAAGTTGAAAAGAACTTTGAAGAGAGAGTT | >hg38_21_+_8433172_8446622_RNA45SN1_45S_50ntFlanks | 8342 | 8379 | 38 | CGTAAGGGAAAGTTGAAAAGAACTTTGAAGAGAGAGTT | MB231 | 28 | 7 | 115 | 17 | 7 | 26.6 | 5.2 | 115.4 | 9.6 | 5.0 |
| >hg38_21_+_8433172_8446622_RNA45SN1_45S_50ntFlanks@8352.8379.28\|AGTTGAAAAGAACTTTGAAGAGAGAGTT | >hg38_21_+_8433172_8446622_RNA45SN1_45S_50ntFlanks | 8352 | 8379 | 28 | AGTTGAAAAGAACTTTGAAGAGAGAGTT | MB468 | 19 | 15 | 7 | 8 | 2 | 6.4 | 12.5 | 2.3 | 4.8 | 1.0 |
| >hg38_21_+_8433172_8446622_RNA45SN1_45S_50ntFlanks@8354.8379.26\|TTGAAAAGAACTTTGAAGAGAGAGTT | >hg38_21_+_8433172_8446622_RNA45SN1_45S_50ntFlanks | 8354 | 8379 | 26 | TTGAAAAGAACTTTGAAGAGAGAGTT | MB468 | 16 | 10 | 6 | 4 | 2 | 6.2 | 8.8 | 3.7 | 3.8 | 1.1 |
| >hg38_21_+_8433172_8446622_RNA45SN1_45S_50ntFlanks@8356.8373.18\|GAAAAGAACTTTGAAGAG | >hg38_21_+_8433172_8446622_RNA45SN1_45S_50ntFlanks | 8356 | 8373 | 18 | GAAAAGAACTTTGAAGAG | MB468 | 24 | 8 | 5 | 45 | 7 | 16.9 | 5.1 | 1.8 | 28.2 | 4.8 |
| >hg38_21_+_8433172_8446622_RNA45SN1_45S_50ntFlanks@8356.8375.20\|GAAAAGAACTTTGAAGAGAG | >hg38_21_+_8433172_8446622_RNA45SN1_45S_50ntFlanks | 8356 | 8375 | 20 | GAAAAGAACTTTGAAGAGAG | MB468 | 105 | 25 | 17 | 133 | 17 | 76.3 | 15.8 | 9.8 | 58.6 | 10.3 |
| >hg38_21_+_8433172_8446622_RNA45SN1_45S_50ntFlanks@8356.8376.21\|GAAAAGAACTTTGAAGAGAGA | >hg38_21_+_8433172_8446622_RNA45SN1_45S_50ntFlanks | 8356 | 8376 | 21 | GAAAAGAACTTTGAAGAGAGA | MB468 | 23 | 6 | 2 | 24 | 2 | 14.7 | 6.5 | 1.0 | 16.9 | 1.3 |
| >hg38_21_+_8433172_8446622_RNA45SN1_45S_50ntFlanks@8356.8377.22\|GAAAAGAACTTTGAAGAGAGAG | >hg38_21_+_8433172_8446622_RNA45SN1_45S_50ntFlanks | 8356 | 8377 | 22 | GAAAAGAACTTTGAAGAGAGAG | MB468 | 126 | 46 | 19 | 126 | 20 | 69.1 | 32.9 | 4.4 | 99.1 | 7.9 |
| >hg38_21_+_8433172_8446622_RNA45SN1_45S_50ntFlanks@8356.8378.23\|GAAAAGAACTTTGAAGAGAGAGT | >hg38_21_+_8433172_8446622_RNA45SN1_45S_50ntFlanks | 8356 | 8378 | 23 | GAAAAGAACTTTGAAGAGAGAGT | MB468 | 567 | 189 | 60 | 520 | 36 | 272.9 | 159.2 | 32.5 | 432.1 | 15.6 |
| >hg38_21_+_8433172_8446622_RNA45SN1_45S_50ntFlanks@8356.8379.24\|GAAAAGAACTTTGAAGAGAGAGTT | >hg38_21_+_8433172_8446622_RNA45SN1_45S_50ntFlanks | 8356 | 8379 | 24 | GAAAAGAACTTTGAAGAGAGAGTT | MB468 | 2505 | 954 | 452 | 1036 | 297 | 1203.5 | 773.2 | 267.2 | 732.2 | 162.8 |
| >hg38_21_+_8433172_8446622_RNA45SN1_45S_50ntFlanks@8356.8380.25\|GAAAAGAACTTTGAAGAGAGAGTTC | >hg38_21_+_8433172_8446622_RNA45SN1_45S_50ntFlanks | 8356 | 8380 | 25 | GAAAAGAACTTTGAAGAGAGAGTTC | MB468 | 12 | 1 | 7 | 13 | 3 | 12.0 | 0.1 | 9.9 | 8.8 | 3.0 |
| >hg38_21_+_8433172_8446622_RNA45SN1_45S_50ntFlanks@8357.8375.19\|AAAAGAACTTTGAAGAGAG | >hg38_21_+_8433172_8446622_RNA45SN1_45S_50ntFlanks | 8357 | 8375 | 19 | AAAAGAACTTTGAAGAGAG | MB468 | 619 | 189 | 137 | 951 | 155 | 346.4 | 136.9 | 88.4 | 484.0 | 77.8 |

| | | | | | | | | | | | | | | | | |
|---|---|---|---|---|---|---|---|---|---|---|---|---|---|---|---|---|
| >hg38_21_+_8433172_8446622_RNA45SN1_45S_50ntFlanks@8357.8376.20\|AAAAGAACTTTGAAGAGAGA | >hg38_21_+_8433172_8446622_RNA45SN1_45S_50ntFlanks | 8357 | 8376 | 20 | AAAAGAACTTTGAAGAGAGA | MB468 | 106 | 39 | 20 | 134 | 14 | 48.0 | 38.8 | 9.3 | 116.2 | 6.4 |
| >hg38_21_+_8433172_8446622_RNA45SN1_45S_50ntFlanks@8357.8377.21\|AAAAGAACTTTGAAGAGAGAG | >hg38_21_+_8433172_8446622_RNA45SN1_45S_50ntFlanks | 8357 | 8377 | 21 | AAAAGAACTTTGAAGAGAGAG | MB468 | 565 | 358 | 151 | 659 | 177 | 149.3 | 276.6 | 64.0 | 595.4 | 84.9 |
| >hg38_21_+_8433172_8446622_RNA45SN1_45S_50ntFlanks@8357.8378.22\|AAAAGAACTTTGAAGAGAGAGT | >hg38_21_+_8433172_8446622_RNA45SN1_45S_50ntFlanks | 8357 | 8378 | 22 | AAAAGAACTTTGAAGAGAGAGT | MB231 | 13 | 6 | 4 | 4 | 3 | 6.5 | 4.3 | 2.7 | 3.0 | 0.0 |
| >hg38_21_+_8433172_8446622_RNA45SN1_45S_50ntFlanks@8357.8378.22\|AAAAGAACTTTGAAGAGAGAGT | >hg38_21_+_8433172_8446622_RNA45SN1_45S_50ntFlanks | 8357 | 8378 | 22 | AAAAGAACTTTGAAGAGAGAGT | MB468 | 2380 | 1080 | 397 | 2260 | 227 | 897.3 | 974.7 | 268.7 | 2018.7 | 112.2 |
| >hg38_21_+_8433172_8446622_RNA45SN1_45S_50ntFlanks@8357.8379.23\|AAAAGAACTTTGAAGAGAGAGTT | >hg38_21_+_8433172_8446622_RNA45SN1_45S_50ntFlanks | 8357 | 8379 | 23 | AAAAGAACTTTGAAGAGAGAGTT | MB468 | 7507 | 3091 | 2102 | 4634 | 2008 | 3235.1 | 2540.8 | 1588.2 | 2595.1 | 1370.5 |
| >hg38_21_+_8433172_8446622_RNA45SN1_45S_50ntFlanks@8357.8387.31\|AAAAGAACTTTGAAGAGAGAGTTCAAGAGGG | >hg38_21_+_8433172_8446622_RNA45SN1_45S_50ntFlanks | 8357 | 8387 | 31 | AAAAGAACTTTGAAGAGAGAGTTCAAGAGGG | MB468 | 19 | 23 | 27 | 45 | 17 | 3.4 | 9.8 | 9.5 | 34.2 | 3.7 |
| >hg38_21_+_8433172_8446622_RNA45SN1_45S_50ntFlanks@8357.8391.35\|AAAAGAACTTTGAAGAGAGAGTTCAAGAGGGCGTG | >hg38_21_+_8433172_8446622_RNA45SN1_45S_50ntFlanks | 8357 | 8391 | 35 | AAAAGAACTTTGAAGAGAGAGTTCAAGAGGGCGTG | MB468 | 29 | 88 | 45 | 49 | 52 | 18.3 | 25.3 | 23.2 | 25.2 | 30.9 |
| >hg38_21_+_8433172_8446622_RNA45SN1_45S_50ntFlanks@8358.8375.18\|AAAGAACTTTGAAGAGAG | >hg38_21_+_8433172_8446622_RNA45SN1_45S_50ntFlanks | 8358 | 8375 | 18 | AAAGAACTTTGAAGAGAG | MB468 | 18 | 5 | 4 | 24 | 3 | 10.7 | 4.7 | 3.0 | 16.1 | 1.0 |
| >hg38_21_+_8433172_8446622_RNA45SN1_45S_50ntFlanks@8358.8377.20\|AAAGAACTTTGAAGAGAGAG | >hg38_21_+_8433172_8446622_RNA45SN1_45S_50ntFlanks | 8358 | 8377 | 20 | AAAGAACTTTGAAGAGAGAG | MB468 | 15 | 9 | 3 | 17 | 4 | 3.9 | 7.6 | 1.3 | 16.6 | 2.0 |
| >hg38_21_+_8433172_8446622_RNA45SN1_45S_50ntFlanks@8358.8378.21\|AAAGAACTTTGAAGAGAGAGT | >hg38_21_+_8433172_8446622_RNA45SN1_45S_50ntFlanks | 8358 | 8378 | 21 | AAAGAACTTTGAAGAGAGAGT | MB468 | 61 | 28 | 9 | 57 | 5 | 24.7 | 29.2 | 5.8 | 58.8 | 2.7 |
| >hg38_21_+_8433172_8446622_RNA45SN1_45S_50ntFlanks@8358.8379.22\|AAAGAACTTTGAAGAGAGAGTT | >hg38_21_+_8433172_8446622_RNA45SN1_45S_50ntFlanks | 8358 | 8379 | 22 | AAAGAACTTTGAAGAGAGAGTT | MB468 | 149 | 64 | 36 | 75 | 32 | 64.3 | 65.5 | 24.9 | 64.9 | 20.7 |
| >hg38_21_+_8433172_8446622_RNA45SN1_45S_50ntFlanks@8359.8377.19\|AAGAACTTTGAAGAGAGAG | >hg38_21_+_8433172_8446622_RNA45SN1_45S_50ntFlanks | 8359 | 8377 | 19 | AAGAACTTTGAAGAGAGAG | MB468 | 21 | 10 | 4 | 23 | 4 | 9.4 | 9.5 | 1.7 | 19.5 | 1.9 |
| >hg38_21_+_8433172_8446622_RNA45SN1_45S_50ntFlanks@8359.8378.20\|AAGAACTTTGAAGAGAGAGT | >hg38_21_+_8433172_8446622_RNA45SN1_45S_50ntFlanks | 8359 | 8378 | 20 | AAGAACTTTGAAGAGAGAGT | MB468 | 64 | 23 | 10 | 43 | 4 | 32.6 | 24.9 | 4.6 | 36.9 | 1.5 |
| >hg38_21_+_8433172_8446622_RNA45SN1_45S_50ntFlanks@8359.8379.21\|AAGAACTTTGAAGAGAGAGTT | >hg38_21_+_8433172_8446622_RNA45SN1_45S_50ntFlanks | 8359 | 8379 | 21 | AAGAACTTTGAAGAGAGAGTT | MB468 | 130 | 37 | 36 | 52 | 21 | 79.6 | 37.3 | 25.6 | 36.2 | 12.5 |
| >hg38_21_+_8433172_8446622_RNA45SN1_45S_50ntFlanks@8360.8379.20\|AGAACTTTGAAGAGAGAGTT | >hg38_21_+_8433172_8446622_RNA45SN1_45S_50ntFlanks | 8360 | 8379 | 20 | AGAACTTTGAAGAGAGAGTT | MB468 | 12 | 3 | 3 | 5 | 2 | 6.2 | 3.1 | 0.6 | 4.0 | 1.0 |
| >hg38_21_+_8433172_8446622_RNA45SN1_45S_50ntFlanks@8378.8400.23\|TTCAAGAGGGCGTGAAACCGTTA | >hg38_21_+_8433172_8446622_RNA45SN1_45S_50ntFlanks | 8378 | 8400 | 23 | TTCAAGAGGGCGTGAAACCGTTA | MB468 | 13 | 2 | 8 | 12 | 3 | 3.9 | 1.2 | 3.0 | 7.2 | 0.9 |
| >hg38_21_+_8433172_8446622_RNA45SN1_45S_50ntFlanks@8378.8402.25\|TTCAAGAGGGCGTGAAACCGTTAAG | >hg38_21_+_8433172_8446622_RNA45SN1_45S_50ntFlanks | 8378 | 8402 | 25 | TTCAAGAGGGCGTGAAACCGTTAAG | MB468 | 19 | 4 | 10 | 32 | 8 | 8.4 | 1.2 | 2.3 | 14.1 | 1.5 |
| >hg38_21_+_8433172_8446622_RNA45SN1_45S_50ntFlanks@8379.8400.22\|TCAAGAGGGCGTGAAACCGTTA | >hg38_21_+_8433172_8446622_RNA45SN1_45S_50ntFlanks | 8379 | 8400 | 22 | TCAAGAGGGCGTGAAACCGTTA | MB468 | 24 | 3 | 12 | 20 | 11 | 10.3 | 1.6 | 2.5 | 13.7 | 4.6 |
| >hg38_21_+_8433172_8446622_RNA45SN1_45S_50ntFlanks@8379.8402.24\|TCAAGAGGGCGTGAAACCGTTAAG | >hg38_21_+_8433172_8446622_RNA45SN1_45S_50ntFlanks | 8379 | 8402 | 24 | TCAAGAGGGCGTGAAACCGTTAAG | MB468 | 24 | 4 | 10 | 43 | 19 | 16.4 | 1.3 | 4.1 | 18.8 | 7.0 |
| >hg38_21_+_8433172_8446622_RNA45SN1_45S_50ntFlanks@8380.8397.18\|CAAGAGGGCGTGAAACCG | >hg38_21_+_8433172_8446622_RNA45SN1_45S_50ntFlanks | 8380 | 8397 | 18 | CAAGAGGGCGTGAAACCG | MB468 | 148 | 40 | 79 | 112 | 32 | 78.6 | 21.3 | 24.7 | 34.0 | 6.5 |
| >hg38_21_+_8433172_8446622_RNA45SN1_45S_50ntFlanks@8380.8398.19\|CAAGAGGGCGTGAAACCGT | >hg38_21_+_8433172_8446622_RNA45SN1_45S_50ntFlanks | 8380 | 8398 | 19 | CAAGAGGGCGTGAAACCGT | MB468 | 16 | 2 | 9 | 15 | 6 | 11.3 | 1.3 | 3.3 | 6.8 | 3.7 |
| >hg38_21_+_8433172_8446622_RNA45SN1_45S_50ntFlanks@8380.8399.20\|CAAGAGGGCGTGAAACCGTT | >hg38_21_+_8433172_8446622_RNA45SN1_45S_50ntFlanks | 8380 | 8399 | 20 | CAAGAGGGCGTGAAACCGTT | MB468 | 89 | 11 | 47 | 106 | 37 | 29.7 | 4.0 | 14.3 | 73.3 | 17.1 |
| >hg38_21_+_8433172_8446622_RNA45SN1_45S_50ntFlanks@8380.8400.21\|CAAGAGGGCGTGAAACCGTTA | >hg38_21_+_8433172_8446622_RNA45SN1_45S_50ntFlanks | 8380 | 8400 | 21 | CAAGAGGGCGTGAAACCGTTA | MB468 | 329 | 41 | 204 | 296 | 155 | 112.3 | 18.2 | 59.1 | 189.9 | 55.5 |
| >hg38_21_+_8433172_8446622_RNA45SN1_45S_50ntFlanks@8380.8401.22\|CAAGAGGGCGTGAAACCGTTAA | >hg38_21_+_8433172_8446622_RNA45SN1_45S_50ntFlanks | 8380 | 8401 | 22 | CAAGAGGGCGTGAAACCGTTAA | MB468 | 60 | 7 | 44 | 40 | 33 | 25.6 | 2.5 | 18.9 | 20.6 | 15.8 |
| >hg38_21_+_8433172_8446622_RNA45SN1_45S_50ntFlanks@8380.8402.23\|CAAGAGGGCGTGAAACCGTTAAG | >hg38_21_+_8433172_8446622_RNA45SN1_45S_50ntFlanks | 8380 | 8402 | 23 | CAAGAGGGCGTGAAACCGTTAAG | MB468 | 255 | 45 | 118 | 500 | 291 | 144.5 | 18.9 | 55.7 | 226.6 | 166.8 |
| >hg38_21_+_8433172_8446622_RNA45SN1_45S_50ntFlanks@8380.8403.24\|CAAGAGGGCGTGAAACCGTTAAGA | >hg38_21_+_8433172_8446622_RNA45SN1_45S_50ntFlanks | 8380 | 8403 | 24 | CAAGAGGGCGTGAAACCGTTAAGA | MB468 | 70 | 23 | 51 | 113 | 131 | 32.6 | 7.3 | 11.2 | 19.7 | 84.1 |
| >hg38_21_+_8433172_8446622_RNA45SN1_45S_50ntFlanks@8380.8404.25\|CAAGAGGGCGTGAAACCGTTAAGAG | >hg38_21_+_8433172_8446622_RNA45SN1_45S_50ntFlanks | 8380 | 8404 | 25 | CAAGAGGGCGTGAAACCGTTAAGAG | MB468 | 47 | 22 | 46 | 47 | 67 | 30.0 | 7.9 | 16.2 | 7.4 | 42.7 |
| >hg38_21_+_8433172_8446622_RNA45SN1_45S_50ntFlanks@8380.8405.26\|CAAGAGGGCGTGAAACCGTTAAGAGG | >hg38_21_+_8433172_8446622_RNA45SN1_45S_50ntFlanks | 8380 | 8405 | 26 | CAAGAGGGCGTGAAACCGTTAAGAGG | MB468 | 103 | 85 | 136 | 303 | 223 | 58.1 | 43.8 | 55.7 | 154.7 | 197.5 |
| >hg38_21_+_8433172_8446622_RNA45SN1_45S_50ntFlanks@8380.8406.27\|CAAGAGGGCGTGAAACCGTTAAGAGGT | >hg38_21_+_8433172_8446622_RNA45SN1_45S_50ntFlanks | 8380 | 8406 | 27 | CAAGAGGGCGTGAAACCGTTAAGAGGT | MB468 | 40 | 43 | 41 | 137 | 113 | 15.3 | 20.9 | 16.4 | 54.7 | 76.6 |
| >hg38_21_+_8433172_8446622_RNA45SN1_45S_50ntFlanks@8380.8407.28\|CAAGAGGGCGTGAAACCGTTAAGAGGTA | >hg38_21_+_8433172_8446622_RNA45SN1_45S_50ntFlanks | 8380 | 8407 | 28 | CAAGAGGGCGTGAAACCGTTAAGAGGTA | MB468 | 14 | 12 | 11 | 27 | 33 | 5.7 | 4.3 | 2.5 | 16.9 | 18.6 |
| >hg38_21_+_8433172_8446622_RNA45SN1_45S_50ntFlanks@8381.8399.19\|AAGAGGGCGTGAAACCGTT | >hg38_21_+_8433172_8446622_RNA45SN1_45S_50ntFlanks | 8381 | 8399 | 19 | AAGAGGGCGTGAAACCGTT | MB468 | 35 | 5 | 15 | 63 | 11 | 12.2 | 2.2 | 3.3 | 50.8 | 6.2 |
| >hg38_21_+_8433172_8446622_RNA45SN1_45S_50ntFlanks@8381.8400.20\|AAGAGGGCGTGAAACCGTTA | >hg38_21_+_8433172_8446622_RNA45SN1_45S_50ntFlanks | 8381 | 8400 | 20 | AAGAGGGCGTGAAACCGTTA | MB468 | 130 | 17 | 67 | 161 | 42 | 46.2 | 7.8 | 17.1 | 129.8 | 20.7 |
| >hg38_21_+_8433172_8446622_RNA45SN1_45S_50ntFlanks@8381.8401.21\|AAGAGGGCGTGAAACCGTTAA | >hg38_21_+_8433172_8446622_RNA45SN1_45S_50ntFlanks | 8381 | 8401 | 21 | AAGAGGGCGTGAAACCGTTAA | MB468 | 20 | 2 | 11 | 18 | 8 | 8.0 | 0.7 | 3.8 | 12.5 | 4.7 |
| >hg38_21_+_8433172_8446622_RNA45SN1_45S_50ntFlanks@8381.8402.22\|AAGAGGGCGTGAAACCGTTAAG | >hg38_21_+_8433172_8446622_RNA45SN1_45S_50ntFlanks | 8381 | 8402 | 22 | AAGAGGGCGTGAAACCGTTAAG | MB468 | 191 | 41 | 83 | 425 | 135 | 98.9 | 16.0 | 23.9 | 224.3 | 74.0 |
| >hg38_21_+_8433172_8446622_RNA45SN1_45S_50ntFlanks@8381.8403.23\|AAGAGGGCGTGAAACCGTTAAGA | >hg38_21_+_8433172_8446622_RNA45SN1_45S_50ntFlanks | 8381 | 8403 | 23 | AAGAGGGCGTGAAACCGTTAAGA | MB468 | 30 | 11 | 21 | 67 | 35 | 13.7 | 3.1 | 5.8 | 26.5 | 21.3 |
| >hg38_21_+_8433172_8446622_RNA45SN1_45S_50ntFlanks@8381.8404.24\|AAGAGGGCGTGAAACCGTTAAGAG | >hg38_21_+_8433172_8446622_RNA45SN1_45S_50ntFlanks | 8381 | 8404 | 24 | AAGAGGGCGTGAAACCGTTAAGAG | MB468 | 21 | 12 | 20 | 28 | 18 | 10.6 | 3.8 | 6.0 | 10.4 | 13.2 |
| >hg38_21_+_8433172_8446622_RNA45SN1_45S_50ntFlanks@8381.8405.25\|AAGAGGGCGTGAAACCGTTAAGAGG | >hg38_21_+_8433172_8446622_RNA45SN1_45S_50ntFlanks | 8381 | 8405 | 25 | AAGAGGGCGTGAAACCGTTAAGAGG | MB468 | 34 | 30 | 51 | 155 | 44 | 16.5 | 13.0 | 23.8 | 98.8 | 33.4 |
| >hg38_21_+_8433172_8446622_RNA45SN1_45S_50ntFlanks@8381.8406.26\|AAGAGGGCGTGAAACCGTTAAGAGGT | >hg38_21_+_8433172_8446622_RNA45SN1_45S_50ntFlanks | 8381 | 8406 | 26 | AAGAGGGCGTGAAACCGTTAAGAGGT | MB468 | 14 | 16 | 15 | 75 | 25 | 5.6 | 6.7 | 6.1 | 36.4 | 16.0 |
| >hg38_21_+_8433172_8446622_RNA45SN1_45S_50ntFlanks@8382.8400.19\|AGAGGGCGTGAAACCGTTA | >hg38_21_+_8433172_8446622_RNA45SN1_45S_50ntFlanks | 8382 | 8400 | 19 | AGAGGGCGTGAAACCGTTA | MB468 | 10 | 1 | 5 | 14 | 4 | 4.9 | 0.6 | 1.0 | 10.7 | 2.0 |
| >hg38_21_+_8433172_8446622_RNA45SN1_45S_50ntFlanks@8382.8402.21\|AGAGGGCGTGAAACCGTTAAG | >hg38_21_+_8433172_8446622_RNA45SN1_45S_50ntFlanks | 8382 | 8402 | 21 | AGAGGGCGTGAAACCGTTAAG | MB468 | 15 | 3 | 7 | 35 | 12 | 9.7 | 0.9 | 1.9 | 18.5 | 7.9 |
| >hg38_21_+_8433172_8446622_RNA45SN1_45S_50ntFlanks@8384.8402.19\|AGGGCGTGAAACCGTTAAG | >hg38_21_+_8433172_8446622_RNA45SN1_45S_50ntFlanks | 8384 | 8402 | 19 | AGGGCGTGAAACCGTTAAG | MB468 | 17 | 1 | 6 | 28 | 12 | 14.2 | 0.8 | 2.8 | 14.5 | 11.1 |
| >hg38_21_+_8433172_8446622_RNA45SN1_45S_50ntFlanks@8388.8405.18\|CGTGAAACCGTTAAGAGG | >hg38_21_+_8433172_8446622_RNA45SN1_45S_50ntFlanks | 8388 | 8405 | 18 | CGTGAAACCGTTAAGAGG | MB468 | 19 | 2 | 39 | 15 | 11 | 9.1 | 2.2 | 32.2 | 5.7 | 4.0 |
| >hg38_21_+_8433172_8446622_RNA45SN1_45S_50ntFlanks@8423.8442.20\|CGCAGTCCGCCCGGAGGATT | >hg38_21_+_8433172_8446622_RNA45SN1_45S_50ntFlanks | 8423 | 8442 | 20 | CGCAGTCCGCCCGGAGGATT | MB468 | 24 | 10 | 5 | 124 | 430 | 11.9 | 7.2 | 3.8 | 37.5 | 364.0 |
| >hg38_21_+_8433172_8446622_RNA45SN1_45S_50ntFlanks@8472.8489.18\|CGGCCCGGCGGATCTTTC | >hg38_21_+_8433172_8446622_RNA45SN1_45S_50ntFlanks | 8472 | 8489 | 18 | CGGCCCGGCGGATCTTTC | MB468 | 12 | 21 | 9 | 5 | 10 | 6.7 | 7.3 | 5.9 | 2.1 | 0.9 |
| >hg38_21_+_8433172_8446622_RNA45SN1_45S_50ntFlanks@8472.8490.19\|CGGCCCGGCGGATCTTTCC | >hg38_21_+_8433172_8446622_RNA45SN1_45S_50ntFlanks | 8472 | 8490 | 19 | CGGCCCGGCGGATCTTTCC | MB468 | 17 | 41 | 13 | 2 | 24 | 8.2 | 31.5 | 5.8 | 0.7 | 15.9 |
| >hg38_21_+_8433172_8446622_RNA45SN1_45S_50ntFlanks@8472.8491.20\|CGGCCCGGCGGATCTTTCCC | >hg38_21_+_8433172_8446622_RNA45SN1_45S_50ntFlanks | 8472 | 8491 | 20 | CGGCCCGGCGGATCTTTCCC | MB468 | 41 | 52 | 27 | 5 | 63 | 22.7 | 31.7 | 12.0 | 1.8 | 39.4 |
| >hg38_21_+_8433172_8446622_RNA45SN1_45S_50ntFlanks@8472.8492.21\|CGGCCCGGCGGATCTTTCCCG | >hg38_21_+_8433172_8446622_RNA45SN1_45S_50ntFlanks | 8472 | 8492 | 21 | CGGCCCGGCGGATCTTTCCCG | MB468 | 71 | 39 | 32 | 9 | 134 | 37.9 | 24.9 | 8.9 | 3.2 | 87.7 |
| >hg38_21_+_8433172_8446622_RNA45SN1_45S_50ntFlanks@8472.8494.23\|CGGCCCGGCGGATCTTTCCCGCC | >hg38_21_+_8433172_8446622_RNA45SN1_45S_50ntFlanks | 8472 | 8494 | 23 | CGGCCCGGCGGATCTTTCCCGCC | MB468 | 46 | 6 | 11 | 3 | 116 | 22.4 | 4.2 | 3.3 | 1.5 | 112.6 |
| >hg38_21_+_8433172_8446622_RNA45SN1_45S_50ntFlanks@8472.8497.26\|CGGCCCGGCGGATCTTTCCCGCCCCC | >hg38_21_+_8433172_8446622_RNA45SN1_45S_50ntFlanks | 8472 | 8497 | 26 | CGGCCCGGCGGATCTTTCCCGCCCCC | MB468 | 29 | 2 | 5 | 20 | 72 | 10.4 | 1.1 | 2.7 | 16.4 | 23.0 |
| >hg38_21_+_8433172_8446622_RNA45SN1_45S_50ntFlanks@8475.8492.18\|CCCGGCGGATCTTTCCCG | >hg38_21_+_8433172_8446622_RNA45SN1_45S_50ntFlanks | 8475 | 8492 | 18 | CCCGGCGGATCTTTCCCG | MB468 | 15 | 1 | 4 | 12 | 23 | 7.6 | 0.2 | 1.2 | 4.3 | 17.2 |

| | | | | | | | | | | | | | | | |
|---|---|---|---|---|---|---|---|---|---|---|---|---|---|---|---|
| >hg38_21_+_8433172_8446622_RNA45SN1_45S_50ntFlanks@8552.8569.18\|CTCCCCGGAGGGGGCGGG | >hg38_21_+_8433172_8446622_RNA45SN1_45S_50ntFlanks | 8552 | 8569 | 18 | CTCCCCGGAGGGGGCGGG | MB468 | 66 | 18 | 60 | 138 | 60 | 18.0 | 8.1 | 18.2 | 114.4 | 44.8 |
| >hg38_21_+_8433172_8446622_RNA45SN1_45S_50ntFlanks@8552.8577.26\|CTCCCCGGAGGGGGCGGGCTCCGGCG | >hg38_21_+_8433172_8446622_RNA45SN1_45S_50ntFlanks | 8552 | 8577 | 26 | CTCCCCGGAGGGGGCGGGCTCCGGCG | MB468 | 24 | 30 | 45 | 26 | 36 | 6.3 | 12.3 | 14.7 | 28.3 | 16.5 |
| >hg38_21_+_8433172_8446622_RNA45SN1_45S_50ntFlanks@8552.8581.30\|CTCCCCGGAGGGGGCGGGCTCCGGCGGGTG | >hg38_21_+_8433172_8446622_RNA45SN1_45S_50ntFlanks | 8552 | 8581 | 30 | CTCCCCGGAGGGGGCGGGCTCCGGCGGGTG | MB468 | 13 | 18 | 19 | 10 | 25 | 7.0 | 10.8 | 3.1 | 11.2 | 16.7 |
| >hg38_21_+_8433172_8446622_RNA45SN1_45S_50ntFlanks@8686.8706.21\|CCGGCTCCGGGACGGCTGGGA | >hg38_21_+_8433172_8446622_RNA45SN1_45S_50ntFlanks | 8686 | 8706 | 21 | CCGGCTCCGGGACGGCTGGGA | MB231 | 14 | 20 | 62 | 1 | 4 | 12.0 | 14.5 | 42.3 | 0.2 | 2.4 |
| >hg38_21_+_8433172_8446622_RNA45SN1_45S_50ntFlanks@8687.8706.20\|CGGCTCCGGGACGGCTGGGA | >hg38_21_+_8433172_8446622_RNA45SN1_45S_50ntFlanks | 8687 | 8706 | 20 | CGGCTCCGGGACGGCTGGGA | MB231 | 13 | 34 | 78 | 2 | 2 | 9.8 | 23.7 | 48.7 | 0.6 | 1.3 |
| >hg38_21_+_8433172_8446622_RNA45SN1_45S_50ntFlanks@8737.8754.18\|CCCGTCCGTCCGTCCGTC | >hg38_21_+_8433172_8446622_RNA45SN1_45S_50ntFlanks | 8737 | 8754 | 18 | CCCGTCCGTCCGTCCGTC | MB468 | 16 | 16 | 33 | 19 | 22 | 5.4 | 13.0 | 7.6 | 17.8 | 6.8 |
| >hg38_21_+_8433172_8446622_RNA45SN1_45S_50ntFlanks@8737.8756.20\|CCCGTCCGTCCGTCCGTCCG | >hg38_21_+_8433172_8446622_RNA45SN1_45S_50ntFlanks | 8737 | 8756 | 20 | CCCGTCCGTCCGTCCGTCCG | MB468 | 10 | 16 | 18 | 4 | 16 | 6.2 | 16.6 | 6.8 | 1.9 | 5.5 |
| >hg38_21_+_8433172_8446622_RNA45SN1_45S_50ntFlanks@8737.8758.22\|CCCGTCCGTCCGTCCGTCCGTC | >hg38_21_+_8433172_8446622_RNA45SN1_45S_50ntFlanks | 8737 | 8758 | 22 | CCCGTCCGTCCGTCCGTCCGTC | MB468 | 17 | 25 | 27 | 19 | 19 | 7.5 | 28.0 | 8.5 | 19.7 | 5.4 |
| >hg38_21_+_8433172_8446622_RNA45SN1_45S_50ntFlanks@8799.8823.25\|CCTCGGGAGGGCGCGCGGGTCGGGG | >hg38_21_+_8433172_8446622_RNA45SN1_45S_50ntFlanks | 8799 | 8823 | 25 | CCTCGGGAGGGCGCGCGGGTCGGGG | MB468 | 13 | 27 | 14 | 31 | 19 | 2.8 | 13.9 | 8.3 | 17.6 | 8.5 |
| >hg38_21_+_8433172_8446622_RNA45SN1_45S_50ntFlanks@8912.8936.25\|CGCCGAATCCCGGGGCCGAGGGAGC | >hg38_21_+_8433172_8446622_RNA45SN1_45S_50ntFlanks | 8912 | 8936 | 25 | CGCCGAATCCCGGGGCCGAGGGAGC | MB231 | 11 | 40 | 27 | 4 | 4 | 7.4 | 45.9 | 27.8 | 2.0 | 1.6 |
| >hg38_21_+_8433172_8446622_RNA45SN1_45S_50ntFlanks@9004.9026.23\|TCTCCCCCGCGGGGGCGCGCCGG | >hg38_21_+_8433172_8446622_RNA45SN1_45S_50ntFlanks | 9004 | 9026 | 23 | TCTCCCCCGCGGGGGCGCGCCGG | MB468 | 94 | 76 | 19 | 9 | 329 | 54.4 | 27.0 | 10.8 | 7.1 | 243.6 |
| >hg38_21_+_8433172_8446622_RNA45SN1_45S_50ntFlanks@9005.9026.22\|CTCCCCCGCGGGGGCGCGCCGG | >hg38_21_+_8433172_8446622_RNA45SN1_45S_50ntFlanks | 9005 | 9026 | 22 | CTCCCCCGCGGGGGCGCGCCGG | MB468 | 14 | 12 | 3 | 4 | 65 | 8.2 | 4.4 | 1.6 | 2.5 | 40.6 |
| >hg38_21_+_8433172_8446622_RNA45SN1_45S_50ntFlanks@9021.9039.19\|CGCCGGCGTCTCCTCGTGG | >hg38_21_+_8433172_8446622_RNA45SN1_45S_50ntFlanks | 9021 | 9039 | 19 | CGCCGGCGTCTCCTCGTGG | MB468 | 11 | 13 | 10 | 1 | 6 | 1.7 | 6.2 | 1.9 | 0.4 | 4.2 |
| >hg38_21_+_8433172_8446622_RNA45SN1_45S_50ntFlanks@9027.9058.32\|CGTCTCCTCGTGGGGGGGCCGGGCCACCCCTC | >hg38_21_+_8433172_8446622_RNA45SN1_45S_50ntFlanks | 9027 | 9058 | 32 | CGTCTCCTCGTGGGGGGGCCGGGCCACCCCTC | MB468 | 11 | 4 | 1 | 16 | 67 | 5.6 | 1.8 | 0.5 | 9.8 | 35.4 |
| >hg38_21_+_8433172_8446622_RNA45SN1_45S_50ntFlanks@9027.9059.33\|CGTCTCCTCGTGGGGGGGCCGGGCCACCCCTCC | >hg38_21_+_8433172_8446622_RNA45SN1_45S_50ntFlanks | 9027 | 9059 | 33 | CGTCTCCTCGTGGGGGGGCCGGGCCACCCCTCC | MB468 | 12 | 7 | 2 | 18 | 85 | 6.5 | 3.6 | 0.6 | 8.8 | 38.4 |
| >hg38_21_+_8433172_8446622_RNA45SN1_45S_50ntFlanks@9027.9064.38\|CGTCTCCTCGTGGGGGGGCCGGGCCACCCCTCCCACGG | >hg38_21_+_8433172_8446622_RNA45SN1_45S_50ntFlanks | 9027 | 9064 | 38 | CGTCTCCTCGTGGGGGGGCCGGGCCACCCCTCCCACGG | MB468 | 20 | 16 | 5 | 20 | 127 | 9.6 | 4.9 | 2.0 | 13.7 | 92.1 |
| >hg38_21_+_8433172_8446622_RNA45SN1_45S_50ntFlanks@9027.9065.39\|CGTCTCCTCGTGGGGGGGCCGGGCCACCCCTCCCACGGC | >hg38_21_+_8433172_8446622_RNA45SN1_45S_50ntFlanks | 9027 | 9065 | 39 | CGTCTCCTCGTGGGGGGGCCGGGCCACCCCTCCCACGGC | MB468 | 14 | 12 | 3 | 19 | 110 | 7.7 | 2.3 | 2.2 | 14.8 | 81.1 |
| >hg38_21_+_8433172_8446622_RNA45SN1_45S_50ntFlanks@9033.9058.26\|CTCGTGGGGGGGCCGGGCCACCCCTC | >hg38_21_+_8433172_8446622_RNA45SN1_45S_50ntFlanks | 9033 | 9058 | 26 | CTCGTGGGGGGGCCGGGCCACCCCTC | MB468 | 12 | 3 | 2 | 157 | 69 | 6.2 | 1.8 | 1.2 | 77.0 | 34.5 |
| >hg38_21_+_8433172_8446622_RNA45SN1_45S_50ntFlanks@9033.9059.27\|CTCGTGGGGGGGCCGGGCCACCCCTCC | >hg38_21_+_8433172_8446622_RNA45SN1_45S_50ntFlanks | 9033 | 9059 | 27 | CTCGTGGGGGGGCCGGGCCACCCCTCC | MB468 | 20 | 6 | 2 | 126 | 109 | 10.2 | 4.2 | 1.3 | 58.4 | 66.1 |
| >hg38_21_+_8433172_8446622_RNA45SN1_45S_50ntFlanks@9166.9197.32\|CCGGGCGGGTCGCGCCGTCGGGCCCGGGGGAG | >hg38_21_+_8433172_8446622_RNA45SN1_45S_50ntFlanks | 9166 | 9197 | 32 | CCGGGCGGGTCGCGCCGTCGGGCCCGGGGGAG | MB468 | 34 | 39 | 3 | 99 | 222 | 17.4 | 30.5 | 1.9 | 68.8 | 218.6 |
| >hg38_21_+_8433172_8446622_RNA45SN1_45S_50ntFlanks@9178.9197.20\|CGCCGTCGGGCCCGGGGGAG | >hg38_21_+_8433172_8446622_RNA45SN1_45S_50ntFlanks | 9178 | 9197 | 20 | CGCCGTCGGGCCCGGGGGAG | MB468 | 12 | 19 | 4 | 39 | 86 | 5.4 | 16.1 | 2.2 | 22.1 | 59.7 |
| >hg38_21_+_8433172_8446622_RNA45SN1_45S_50ntFlanks@9180.9197.18\|CCGTCGGGCCCGGGGGAG | >hg38_21_+_8433172_8446622_RNA45SN1_45S_50ntFlanks | 9180 | 9197 | 18 | CCGTCGGGCCCGGGGGAG | MB468 | 23 | 33 | 7 | 98 | 205 | 9.9 | 26.4 | 3.9 | 58.6 | 124.8 |
| >hg38_21_+_8433172_8446622_RNA45SN1_45S_50ntFlanks@9198.9218.21\|GTTCTCTCGGGGCCACGCGCG | >hg38_21_+_8433172_8446622_RNA45SN1_45S_50ntFlanks | 9198 | 9218 | 21 | GTTCTCTCGGGGCCACGCGCG | MB468 | 19 | 18 | 4 | 45 | 88 | 12.1 | 17.4 | 2.2 | 23.1 | 56.9 |
| >hg38_21_+_8433172_8446622_RNA45SN1_45S_50ntFlanks@9198.9220.23\|GTTCTCTCGGGGCCACGCGCGCG | >hg38_21_+_8433172_8446622_RNA45SN1_45S_50ntFlanks | 9198 | 9220 | 23 | GTTCTCTCGGGGCCACGCGCGCG | MB468 | 16 | 17 | 3 | 28 | 86 | 10.3 | 15.6 | 1.8 | 12.4 | 67.6 |
| >hg38_21_+_8433172_8446622_RNA45SN1_45S_50ntFlanks@9198.9222.25\|GTTCTCTCGGGGCCACGCGCGCGTC | >hg38_21_+_8433172_8446622_RNA45SN1_45S_50ntFlanks | 9198 | 9222 | 25 | GTTCTCTCGGGGCCACGCGCGCGTC | MB468 | 44 | 45 | 10 | 105 | 251 | 20.8 | 33.4 | 5.2 | 53.3 | 188.4 |
| >hg38_21_+_8433172_8446622_RNA45SN1_45S_50ntFlanks@9198.9223.26\|GTTCTCTCGGGGCCACGCGCGCGTCC | >hg38_21_+_8433172_8446622_RNA45SN1_45S_50ntFlanks | 9198 | 9223 | 26 | GTTCTCTCGGGGCCACGCGCGCGTCC | MB468 | 10 | 5 | 2 | 40 | 40 | 5.6 | 3.4 | 1.1 | 22.1 | 21.5 |
| >hg38_21_+_8433172_8446622_RNA45SN1_45S_50ntFlanks@9198.9225.28\|GTTCTCTCGGGGCCACGCGCGCGTCCCC | >hg38_21_+_8433172_8446622_RNA45SN1_45S_50ntFlanks | 9198 | 9225 | 28 | GTTCTCTCGGGGCCACGCGCGCGTCCCC | MB468 | 17 | 3 | 2 | 25 | 66 | 8.7 | 1.7 | 0.9 | 16.3 | 20.0 |
| >hg38_21_+_8433172_8446622_RNA45SN1_45S_50ntFlanks@9201.9218.18\|CTCTCGGGGCCACGCGCG | >hg38_21_+_8433172_8446622_RNA45SN1_45S_50ntFlanks | 9201 | 9218 | 18 | CTCTCGGGGCCACGCGCG | MB468 | 11 | 8 | 1 | 53 | 54 | 6.5 | 6.3 | 1.3 | 22.9 | 27.2 |
| >hg38_21_+_8433172_8446622_RNA45SN1_45S_50ntFlanks@9266.9285.20\|CGACGTCGGCTACCCACCCG | >hg38_21_+_8433172_8446622_RNA45SN1_45S_50ntFlanks | 9266 | 9285 | 20 | CGACGTCGGCTACCCACCCG | MB468 | 18 | 23 | 37 | 4 | 22 | 17.6 | 25.3 | 45.9 | 2.3 | 12.1 |
| >hg38_21_+_8433172_8446622_RNA45SN1_45S_50ntFlanks@9304.9338.35\|ACCAAGGAGTCTAACACGTGCGCGAGTCGGGGGCT | >hg38_21_+_8433172_8446622_RNA45SN1_45S_50ntFlanks | 9304 | 9338 | 35 | ACCAAGGAGTCTAACACGTGCGCGAGTCGGGGGCT | MB468 | 12 | 10 | 4 | 79 | 68 | 7.3 | 3.5 | 2.6 | 46.4 | 35.3 |
| >hg38_21_+_8433172_8446622_RNA45SN1_45S_50ntFlanks@9305.9323.19\|CCAAGGAGTCTAACACGTG | >hg38_21_+_8433172_8446622_RNA45SN1_45S_50ntFlanks | 9305 | 9323 | 19 | CCAAGGAGTCTAACACGTG | MB468 | 12 | 6 | 9 | 36 | 35 | 8.8 | 3.1 | 5.5 | 32.8 | 18.7 |
| >hg38_21_+_8433172_8446622_RNA45SN1_45S_50ntFlanks@9305.9336.32\|CCAAGGAGTCTAACACGTGCGCGAGTCGGGGG | >hg38_21_+_8433172_8446622_RNA45SN1_45S_50ntFlanks | 9305 | 9336 | 32 | CCAAGGAGTCTAACACGTGCGCGAGTCGGGGG | MB468 | 13 | 7 | 3 | 98 | 44 | 8.5 | 2.5 | 2.4 | 24.7 | 25.6 |
| >hg38_21_+_8433172_8446622_RNA45SN1_45S_50ntFlanks@9305.9338.34\|CCAAGGAGTCTAACACGTGCGCGAGTCGGGGGCT | >hg38_21_+_8433172_8446622_RNA45SN1_45S_50ntFlanks | 9305 | 9338 | 34 | CCAAGGAGTCTAACACGTGCGCGAGTCGGGGGCT | MB468 | 38 | 25 | 10 | 178 | 171 | 22.6 | 8.2 | 6.0 | 66.8 | 105.7 |
| >hg38_21_+_8433172_8446622_RNA45SN1_45S_50ntFlanks@9306.9323.18\|CAAGGAGTCTAACACGTG | >hg38_21_+_8433172_8446622_RNA45SN1_45S_50ntFlanks | 9306 | 9323 | 18 | CAAGGAGTCTAACACGTG | MB468 | 18 | 5 | 12 | 67 | 26 | 12.9 | 2.5 | 5.9 | 72.2 | 6.9 |
| >hg38_21_+_8433172_8446622_RNA45SN1_45S_50ntFlanks@9306.9338.33\|CAAGGAGTCTAACACGTGCGCGAGTCGGGGGCT | >hg38_21_+_8433172_8446622_RNA45SN1_45S_50ntFlanks | 9306 | 9338 | 33 | CAAGGAGTCTAACACGTGCGCGAGTCGGGGGCT | MB468 | 17 | 12 | 6 | 113 | 84 | 9.9 | 4.4 | 3.8 | 33.2 | 53.5 |
| >hg38_21_+_8433172_8446622_RNA45SN1_45S_50ntFlanks@9307.9330.24\|AAGGAGTCTAACACGTGCGCGAGT | >hg38_21_+_8433172_8446622_RNA45SN1_45S_50ntFlanks | 9307 | 9330 | 24 | AAGGAGTCTAACACGTGCGCGAGT | MB231 | 11 | 21 | 16 | 20 | 2 | 6.5 | 15.6 | 18.5 | 8.3 | 1.1 |
| >hg38_21_+_8433172_8446622_RNA45SN1_45S_50ntFlanks@9308.9338.31\|AGGAGTCTAACACGTGCGCGAGTCGGGGGCT | >hg38_21_+_8433172_8446622_RNA45SN1_45S_50ntFlanks | 9308 | 9338 | 31 | AGGAGTCTAACACGTGCGCGAGTCGGGGGCT | MB468 | 12 | 7 | 2 | 240 | 60 | 6.7 | 2.8 | 2.0 | 109.6 | 28.9 |
| >hg38_21_+_8433172_8446622_RNA45SN1_45S_50ntFlanks@9313.9338.26\|TCTAACACGTGCGCGAGTCGGGGGCT | >hg38_21_+_8433172_8446622_RNA45SN1_45S_50ntFlanks | 9313 | 9338 | 26 | TCTAACACGTGCGCGAGTCGGGGGCT | MB468 | 12 | 12 | 3 | 48 | 38 | 9.3 | 3.8 | 2.9 | 36.5 | 14.5 |
| >hg38_21_+_8433172_8446622_RNA45SN1_45S_50ntFlanks@9339.9370.32\|CGCACGAAAGCCGCCGTGGCGCAATGAAGGTG | >hg38_21_+_8433172_8446622_RNA45SN1_45S_50ntFlanks | 9339 | 9370 | 32 | CGCACGAAAGCCGCCGTGGCGCAATGAAGGTG | MB468 | 11 | 3 | 1 | 489 | 54 | 8.5 | 1.5 | 0.7 | 411.7 | 43.2 |
| >hg38_21_+_8433172_8446622_RNA45SN1_45S_50ntFlanks@9343.9370.28\|CGAAAGCCGCCGTGGCGCAATGAAGGTG | >hg38_21_+_8433172_8446622_RNA45SN1_45S_50ntFlanks | 9343 | 9370 | 28 | CGAAAGCCGCCGTGGCGCAATGAAGGTG | MB468 | 16 | 7 | 3 | 117 | 64 | 9.9 | 3.3 | 1.8 | 67.0 | 33.5 |
| >hg38_21_+_8433172_8446622_RNA45SN1_45S_50ntFlanks@9352.9370.19\|CCGTGGCGCAATGAAGGTG | >hg38_21_+_8433172_8446622_RNA45SN1_45S_50ntFlanks | 9352 | 9370 | 19 | CCGTGGCGCAATGAAGGTG | MB468 | 130 | 44 | 24 | 302 | 232 | 106.7 | 20.6 | 24.4 | 197.8 | 122.6 |
| >hg38_21_+_8433172_8446622_RNA45SN1_45S_50ntFlanks@9352.9371.20\|CCGTGGCGCAATGAAGGTGA | >hg38_21_+_8433172_8446622_RNA45SN1_45S_50ntFlanks | 9352 | 9371 | 20 | CCGTGGCGCAATGAAGGTGA | MB468 | 13 | 4 | 2 | 20 | 23 | 10.1 | 1.5 | 1.3 | 13.9 | 10.0 |
| >hg38_21_+_8433172_8446622_RNA45SN1_45S_50ntFlanks@9352.9377.26\|CCGTGGCGCAATGAAGGTGAAGGCCG | >hg38_21_+_8433172_8446622_RNA45SN1_45S_50ntFlanks | 9352 | 9377 | 26 | CCGTGGCGCAATGAAGGTGAAGGCCG | MB468 | 12 | 3 | 2 | 21 | 27 | 6.9 | 1.0 | 0.7 | 18.9 | 17.7 |
| >hg38_21_+_8433172_8446622_RNA45SN1_45S_50ntFlanks@9353.9370.18\|CGTGGCGCAATGAAGGTG | >hg38_21_+_8433172_8446622_RNA45SN1_45S_50ntFlanks | 9353 | 9370 | 18 | CGTGGCGCAATGAAGGTG | MB468 | 10 | 3 | 4 | 74 | 18 | 7.7 | 1.7 | 4.1 | 65.2 | 9.9 |
| >hg38_21_+_8433172_8446622_RNA45SN1_45S_50ntFlanks@9381.9409.29\|CGCTCGCCGGCCGAGGTGGGATCCCGAGG | >hg38_21_+_8433172_8446622_RNA45SN1_45S_50ntFlanks | 9381 | 9409 | 29 | CGCTCGCCGGCCGAGGTGGGATCCCGAGG | MB468 | 23 | 39 | 5 | 28 | 225 | 16.5 | 20.4 | 4.3 | 25.8 | 232.9 |
| >hg38_21_+_8433172_8446622_RNA45SN1_45S_50ntFlanks@9454.9503.50\|CCGCGCCGGGGAGGTGGAGCACGAGCGCACGTGTTAGGACCCGAAAGATG | >hg38_21_+_8433172_8446622_RNA45SN1_45S_50ntFlanks | 9454 | 9503 | 50 | CCGCGCCGGGGAGGTGGAGCACGAGCGCACGTGTTAGGACCCGAAAGATG | MB468 | 24 | 206 | 27 | 135 | 436 | 21.6 | 216.3 | 27.7 | 157.7 | 402.3 |
| >hg38_21_+_8433172_8446622_RNA45SN1_45S_50ntFlanks@9465.9503.39\|AGGTGGAGCACGAGCGCACGTGTTAGGACCCGAAAGATG | >hg38_21_+_8433172_8446622_RNA45SN1_45S_50ntFlanks | 9465 | 9503 | 39 | AGGTGGAGCACGAGCGCACGTGTTAGGACCCGAAAGATG | MB468 | 10 | 7 | 2 | 33 | 83 | 5.2 | 1.9 | 1.1 | 30.1 | 63.3 |
| >hg38_21_+_8433172_8446622_RNA45SN1_45S_50ntFlanks@9477.9497.21\|AGCGCACGTGTTAGGACCCGA | >hg38_21_+_8433172_8446622_RNA45SN1_45S_50ntFlanks | 9477 | 9497 | 21 | AGCGCACGTGTTAGGACCCGA | MB231 | 23 | 44 | 83 | 0 | 1 | 24.3 | 35.7 | 56.1 | 0.3 | 1.4 |
| >hg38_21_+_8433172_8446622_RNA45SN1_45S_50ntFlanks@9478.9497.20\|GCGCACGTGTTAGGACCCGA | >hg38_21_+_8433172_8446622_RNA45SN1_45S_50ntFlanks | 9478 | 9497 | 20 | GCGCACGTGTTAGGACCCGA | MB231 | 11 | 18 | 51 | 0 | 0 | 10.5 | 12.9 | 34.5 | 0.0 | 0.0 |
| >hg38_21_+_8433172_8446622_RNA45SN1_45S_50ntFlanks@9503.9526.24\|GGTGAACTATGCCTGGGCAGGGCG | >hg38_21_+_8433172_8446622_RNA45SN1_45S_50ntFlanks | 9503 | 9526 | 24 | GGTGAACTATGCCTGGGCAGGGCG | MB468 | 12 | 7 | 10 | 48 | 6 | 4.5 | 2.8 | 3.9 | 46.3 | 1.8 |

| | | | | | | | | | | | | | | | | |
|---|---|---|---|---|---|---|---|---|---|---|---|---|---|---|---|---|
| >hg38_21_+_8433172_8446622_RNA45SN1_45S_50ntFlanks@9504.9521.18\|GTGAACTATGCCTGGGCA | >hg38_21_+_8433172_8446622_RNA45SN1_45S_50ntFlanks | 9504 | 9521 | 18 | GTGAACTATGCCTGGGCA | MB468 | 153 | 119 | 237 | 74 | 52 | 58.6 | 45.6 | 145.9 | 22.1 | 9.2 |
| >hg38_21_+_8433172_8446622_RNA45SN1_45S_50ntFlanks@9504.9522.19\|GTGAACTATGCCTGGGCAG | >hg38_21_+_8433172_8446622_RNA45SN1_45S_50ntFlanks | 9504 | 9522 | 19 | GTGAACTATGCCTGGGCAG | MB468 | 73 | 72 | 100 | 59 | 39 | 23.4 | 30.8 | 51.5 | 28.2 | 8.6 |
| >hg38_21_+_8433172_8446622_RNA45SN1_45S_50ntFlanks@9504.9523.20\|GTGAACTATGCCTGGGCAGG | >hg38_21_+_8433172_8446622_RNA45SN1_45S_50ntFlanks | 9504 | 9523 | 20 | GTGAACTATGCCTGGGCAGG | MB468 | 190 | 182 | 214 | 202 | 76 | 42.0 | 44.0 | 78.6 | 132.1 | 9.0 |
| >hg38_21_+_8433172_8446622_RNA45SN1_45S_50ntFlanks@9504.9524.21\|GTGAACTATGCCTGGGCAGGG | >hg38_21_+_8433172_8446622_RNA45SN1_45S_50ntFlanks | 9504 | 9524 | 21 | GTGAACTATGCCTGGGCAGGG | MB468 | 43 | 38 | 54 | 56 | 15 | 15.3 | 12.7 | 27.1 | 35.2 | 4.0 |
| >hg38_21_+_8433172_8446622_RNA45SN1_45S_50ntFlanks@9504.9525.22\|GTGAACTATGCCTGGGCAGGGC | >hg38_21_+_8433172_8446622_RNA45SN1_45S_50ntFlanks | 9504 | 9525 | 22 | GTGAACTATGCCTGGGCAGGGC | MB468 | 100 | 35 | 94 | 192 | 58 | 32.8 | 10.9 | 25.7 | 161.3 | 22.3 |
| >hg38_21_+_8433172_8446622_RNA45SN1_45S_50ntFlanks@9504.9526.23\|GTGAACTATGCCTGGGCAGGGCG | >hg38_21_+_8433172_8446622_RNA45SN1_45S_50ntFlanks | 9504 | 9526 | 23 | GTGAACTATGCCTGGGCAGGGCG | MB468 | 549 | 141 | 406 | 1562 | 329 | 145.4 | 33.5 | 77.7 | 1338.5 | 83.7 |
| >hg38_21_+_8433172_8446622_RNA45SN1_45S_50ntFlanks@9504.9527.24\|GTGAACTATGCCTGGGCAGGGCGA | >hg38_21_+_8433172_8446622_RNA45SN1_45S_50ntFlanks | 9504 | 9527 | 24 | GTGAACTATGCCTGGGCAGGGCGA | MB468 | 40 | 10 | 34 | 114 | 25 | 12.2 | 2.6 | 11.7 | 94.4 | 8.0 |
| >hg38_21_+_8433172_8446622_RNA45SN1_45S_50ntFlanks@9504.9529.26\|GTGAACTATGCCTGGGCAGGGCGAAG | >hg38_21_+_8433172_8446622_RNA45SN1_45S_50ntFlanks | 9504 | 9529 | 26 | GTGAACTATGCCTGGGCAGGGCGAAG | MB468 | 22 | 12 | 32 | 57 | 20 | 11.5 | 7.9 | 27.7 | 39.1 | 10.3 |
| >hg38_21_+_8433172_8446622_RNA45SN1_45S_50ntFlanks@9504.9531.28\|GTGAACTATGCCTGGGCAGGGCGAAGCC | >hg38_21_+_8433172_8446622_RNA45SN1_45S_50ntFlanks | 9504 | 9531 | 28 | GTGAACTATGCCTGGGCAGGGCGAAGCC | MB468 | 13 | 10 | 14 | 19 | 21 | 6.7 | 6.0 | 8.3 | 12.9 | 9.8 |
| >hg38_21_+_8433172_8446622_RNA45SN1_45S_50ntFlanks@9504.9532.29\|GTGAACTATGCCTGGGCAGGGCGAAGCCA | >hg38_21_+_8433172_8446622_RNA45SN1_45S_50ntFlanks | 9504 | 9532 | 29 | GTGAACTATGCCTGGGCAGGGCGAAGCCA | MB468 | 12 | 7 | 12 | 26 | 21 | 7.6 | 4.3 | 10.0 | 21.9 | 4.4 |
| >hg38_21_+_8433172_8446622_RNA45SN1_45S_50ntFlanks@9504.9533.30\|GTGAACTATGCCTGGGCAGGGCGAAGCCAG | >hg38_21_+_8433172_8446622_RNA45SN1_45S_50ntFlanks | 9504 | 9533 | 30 | GTGAACTATGCCTGGGCAGGGCGAAGCCAG | MB468 | 85 | 60 | 94 | 100 | 104 | 55.1 | 33.2 | 62.3 | 70.0 | 30.2 |
| >hg38_21_+_8433172_8446622_RNA45SN1_45S_50ntFlanks@9505.9523.19\|TGAACTATGCCTGGGCAGG | >hg38_21_+_8433172_8446622_RNA45SN1_45S_50ntFlanks | 9505 | 9523 | 19 | TGAACTATGCCTGGGCAGG | MB468 | 15 | 9 | 17 | 14 | 6 | 3.0 | 1.5 | 5.6 | 12.6 | 1.8 |
| >hg38_21_+_8433172_8446622_RNA45SN1_45S_50ntFlanks@9505.9526.22\|TGAACTATGCCTGGGCAGGGCG | >hg38_21_+_8433172_8446622_RNA45SN1_45S_50ntFlanks | 9505 | 9526 | 22 | TGAACTATGCCTGGGCAGGGCG | MB468 | 32 | 6 | 23 | 84 | 22 | 13.3 | 2.3 | 3.2 | 94.7 | 9.7 |
| >hg38_21_+_8433172_8446622_RNA45SN1_45S_50ntFlanks@9507.9526.20\|AACTATGCCTGGGCAGGGCG | >hg38_21_+_8433172_8446622_RNA45SN1_45S_50ntFlanks | 9507 | 9526 | 20 | AACTATGCCTGGGCAGGGCG | MB468 | 18 | 2 | 14 | 75 | 6 | 3.8 | 0.9 | 3.2 | 77.0 | 3.1 |
| >hg38_21_+_8433172_8446622_RNA45SN1_45S_50ntFlanks@9532.9571.40\|AGAGGAAACTCTGGTGGAGGTCCGTAGCGGTCCTGACGTG | >hg38_21_+_8433172_8446622_RNA45SN1_45S_50ntFlanks | 9532 | 9571 | 40 | AGAGGAAACTCTGGTGGAGGTCCGTAGCGGTCCTGACGTG | MB231 | 11 | 12 | 81 | 7 | 7 | 7.9 | 8.1 | 63.4 | 6.8 | 8.8 |
| >hg38_21_+_8433172_8446622_RNA45SN1_45S_50ntFlanks@9533.9550.18\|GAGGAAACTCTGGTGGAG | >hg38_21_+_8433172_8446622_RNA45SN1_45S_50ntFlanks | 9533 | 9550 | 18 | GAGGAAACTCTGGTGGAG | MB468 | 18 | 25 | 23 | 13 | 3 | 6.3 | 13.4 | 9.0 | 6.3 | 1.1 |
| >hg38_21_+_8433172_8446622_RNA45SN1_45S_50ntFlanks@9533.9551.19\|GAGGAAACTCTGGTGGAGG | >hg38_21_+_8433172_8446622_RNA45SN1_45S_50ntFlanks | 9533 | 9551 | 19 | GAGGAAACTCTGGTGGAGG | MB468 | 25 | 29 | 47 | 20 | 4 | 3.1 | 13.7 | 12.1 | 11.3 | 0.4 |
| >hg38_21_+_8433172_8446622_RNA45SN1_45S_50ntFlanks@9533.9552.20\|GAGGAAACTCTGGTGGAGGT | >hg38_21_+_8433172_8446622_RNA45SN1_45S_50ntFlanks | 9533 | 9552 | 20 | GAGGAAACTCTGGTGGAGGT | MB468 | 24 | 26 | 26 | 13 | 3 | 2.4 | 7.1 | 7.2 | 8.4 | 0.9 |
| >hg38_21_+_8433172_8446622_RNA45SN1_45S_50ntFlanks@9533.9553.21\|GAGGAAACTCTGGTGGAGGTC | >hg38_21_+_8433172_8446622_RNA45SN1_45S_50ntFlanks | 9533 | 9553 | 21 | GAGGAAACTCTGGTGGAGGTC | MB231 | 17 | 10 | 56 | 5 | 2 | 13.3 | 7.8 | 39.1 | 3.1 | 1.8 |
| >hg38_21_+_8433172_8446622_RNA45SN1_45S_50ntFlanks@9533.9553.21\|GAGGAAACTCTGGTGGAGGTC | >hg38_21_+_8433172_8446622_RNA45SN1_45S_50ntFlanks | 9533 | 9553 | 21 | GAGGAAACTCTGGTGGAGGTC | MB468 | 727 | 938 | 1415 | 273 | 73 | 301.5 | 479.9 | 949.6 | 138.3 | 17.6 |
| >hg38_21_+_8433172_8446622_RNA45SN1_45S_50ntFlanks@9533.9554.22\|GAGGAAACTCTGGTGGAGGTCC | >hg38_21_+_8433172_8446622_RNA45SN1_45S_50ntFlanks | 9533 | 9554 | 22 | GAGGAAACTCTGGTGGAGGTCC | MB468 | 25 | 14 | 38 | 15 | 3 | 15.0 | 7.1 | 41.2 | 8.4 | 0.6 |
| >hg38_21_+_8433172_8446622_RNA45SN1_45S_50ntFlanks@9533.9561.29\|GAGGAAACTCTGGTGGAGGTCCGTAGCGG | >hg38_21_+_8433172_8446622_RNA45SN1_45S_50ntFlanks | 9533 | 9561 | 29 | GAGGAAACTCTGGTGGAGGTCCGTAGCGG | MB468 | 23 | 69 | 37 | 21 | 8 | 10.5 | 60.7 | 28.8 | 8.5 | 5.0 |
| >hg38_21_+_8433172_8446622_RNA45SN1_45S_50ntFlanks@9533.9562.30\|GAGGAAACTCTGGTGGAGGTCCGTAGCGGT | >hg38_21_+_8433172_8446622_RNA45SN1_45S_50ntFlanks | 9533 | 9562 | 30 | GAGGAAACTCTGGTGGAGGTCCGTAGCGGT | MB468 | 16 | 45 | 26 | 21 | 6 | 5.2 | 27.9 | 13.5 | 8.4 | 2.8 |
| >hg38_21_+_8433172_8446622_RNA45SN1_45S_50ntFlanks@9533.9563.31\|GAGGAAACTCTGGTGGAGGTCCGTAGCGGTC | >hg38_21_+_8433172_8446622_RNA45SN1_45S_50ntFlanks | 9533 | 9563 | 31 | GAGGAAACTCTGGTGGAGGTCCGTAGCGGTC | MB468 | 122 | 322 | 244 | 99 | 30 | 54.4 | 186.9 | 185.6 | 53.7 | 12.3 |
| >hg38_21_+_8433172_8446622_RNA45SN1_45S_50ntFlanks@9533.9566.34\|GAGGAAACTCTGGTGGAGGTCCGTAGCGGTCCTG | >hg38_21_+_8433172_8446622_RNA45SN1_45S_50ntFlanks | 9533 | 9566 | 34 | GAGGAAACTCTGGTGGAGGTCCGTAGCGGTCCTG | MB468 | 13 | 40 | 26 | 9 | 6 | 6.3 | 21.0 | 19.7 | 5.3 | 2.4 |
| >hg38_21_+_8433172_8446622_RNA45SN1_45S_50ntFlanks@9533.9569.37\|GAGGAAACTCTGGTGGAGGTCCGTAGCGGTCCTGACG | >hg38_21_+_8433172_8446622_RNA45SN1_45S_50ntFlanks | 9533 | 9569 | 37 | GAGGAAACTCTGGTGGAGGTCCGTAGCGGTCCTGACG | MB468 | 19 | 56 | 48 | 15 | 11 | 17.5 | 31.4 | 48.8 | 10.2 | 6.0 |
| >hg38_21_+_8433172_8446622_RNA45SN1_45S_50ntFlanks@9533.9571.39\|GAGGAAACTCTGGTGGAGGTCCGTAGCGGTCCTGACGTG | >hg38_21_+_8433172_8446622_RNA45SN1_45S_50ntFlanks | 9533 | 9571 | 39 | GAGGAAACTCTGGTGGAGGTCCGTAGCGGTCCTGACGTG | MB231 | 11 | 17 | 138 | 6 | 8 | 8.2 | 12.1 | 175.4 | 6.7 | 9.9 |
| >hg38_21_+_8433172_8446622_RNA45SN1_45S_50ntFlanks@9533.9571.39\|GAGGAAACTCTGGTGGAGGTCCGTAGCGGTCCTGACGTG | >hg38_21_+_8433172_8446622_RNA45SN1_45S_50ntFlanks | 9533 | 9571 | 39 | GAGGAAACTCTGGTGGAGGTCCGTAGCGGTCCTGACGTG | MB468 | 74 | 734 | 186 | 96 | 94 | 20.1 | 318.8 | 92.5 | 78.1 | 68.6 |
| >hg38_21_+_8433172_8446622_RNA45SN1_45S_50ntFlanks@9534.9551.18\|AGGAAACTCTGGTGGAGG | >hg38_21_+_8433172_8446622_RNA45SN1_45S_50ntFlanks | 9534 | 9551 | 18 | AGGAAACTCTGGTGGAGG | MB468 | 528 | 330 | 501 | 261 | 50 | 112.1 | 116.0 | 143.9 | 128.9 | 8.5 |
| >hg38_21_+_8433172_8446622_RNA45SN1_45S_50ntFlanks@9534.9552.19\|AGGAAACTCTGGTGGAGGT | >hg38_21_+_8433172_8446622_RNA45SN1_45S_50ntFlanks | 9534 | 9552 | 19 | AGGAAACTCTGGTGGAGGT | MB468 | 406 | 272 | 259 | 146 | 32 | 16.7 | 74.8 | 62.2 | 78.0 | 8.3 |
| >hg38_21_+_8433172_8446622_RNA45SN1_45S_50ntFlanks@9534.9553.20\|AGGAAACTCTGGTGGAGGTC | >hg38_21_+_8433172_8446622_RNA45SN1_45S_50ntFlanks | 9534 | 9553 | 20 | AGGAAACTCTGGTGGAGGTC | MB468 | 8173 | 6841 | 9184 | 2633 | 656 | 3231.8 | 1530.9 | 6141.5 | 1008.5 | 55.3 |
| >hg38_21_+_8433172_8446622_RNA45SN1_45S_50ntFlanks@9534.9554.21\|AGGAAACTCTGGTGGAGGTCC | >hg38_21_+_8433172_8446622_RNA45SN1_45S_50ntFlanks | 9534 | 9554 | 21 | AGGAAACTCTGGTGGAGGTCC | MB468 | 39 | 10 | 41 | 35 | 5 | 22.7 | 2.5 | 42.3 | 31.0 | 1.9 |
| >hg38_21_+_8433172_8446622_RNA45SN1_45S_50ntFlanks@9534.9555.22\|AGGAAACTCTGGTGGAGGTCCG | >hg38_21_+_8433172_8446622_RNA45SN1_45S_50ntFlanks | 9534 | 9555 | 22 | AGGAAACTCTGGTGGAGGTCCG | MB468 | 29 | 3 | 19 | 44 | 8 | 13.8 | 1.2 | 12.3 | 44.7 | 4.6 |
| >hg38_21_+_8433172_8446622_RNA45SN1_45S_50ntFlanks@9534.9558.25\|AGGAAACTCTGGTGGAGGTCCGTAG | >hg38_21_+_8433172_8446622_RNA45SN1_45S_50ntFlanks | 9534 | 9558 | 25 | AGGAAACTCTGGTGGAGGTCCGTAG | MB468 | 38 | 14 | 40 | 54 | 19 | 20.6 | 3.2 | 15.5 | 35.6 | 4.7 |
| >hg38_21_+_8433172_8446622_RNA45SN1_45S_50ntFlanks@9534.9559.26\|AGGAAACTCTGGTGGAGGTCCGTAGC | >hg38_21_+_8433172_8446622_RNA45SN1_45S_50ntFlanks | 9534 | 9559 | 26 | AGGAAACTCTGGTGGAGGTCCGTAGC | MB468 | 13 | 10 | 19 | 79 | 14 | 1.6 | 3.7 | 7.3 | 40.0 | 7.0 |
| >hg38_21_+_8433172_8446622_RNA45SN1_45S_50ntFlanks@9534.9560.27\|AGGAAACTCTGGTGGAGGTCCGTAGCG | >hg38_21_+_8433172_8446622_RNA45SN1_45S_50ntFlanks | 9534 | 9560 | 27 | AGGAAACTCTGGTGGAGGTCCGTAGCG | MB468 | 46 | 46 | 58 | 75 | 22 | 13.1 | 20.2 | 26.8 | 28.0 | 3.1 |
| >hg38_21_+_8433172_8446622_RNA45SN1_45S_50ntFlanks@9534.9561.28\|AGGAAACTCTGGTGGAGGTCCGTAGCGG | >hg38_21_+_8433172_8446622_RNA45SN1_45S_50ntFlanks | 9534 | 9561 | 28 | AGGAAACTCTGGTGGAGGTCCGTAGCGG | MB468 | 197 | 423 | 236 | 179 | 92 | 92.8 | 326.4 | 141.0 | 81.4 | 76.6 |
| >hg38_21_+_8433172_8446622_RNA45SN1_45S_50ntFlanks@9534.9562.29\|AGGAAACTCTGGTGGAGGTCCGTAGCGGT | >hg38_21_+_8433172_8446622_RNA45SN1_45S_50ntFlanks | 9534 | 9562 | 29 | AGGAAACTCTGGTGGAGGTCCGTAGCGGT | MB468 | 135 | 254 | 166 | 163 | 65 | 33.8 | 116.9 | 59.7 | 65.2 | 31.0 |
| >hg38_21_+_8433172_8446622_RNA45SN1_45S_50ntFlanks@9534.9563.30\|AGGAAACTCTGGTGGAGGTCCGTAGCGGTC | >hg38_21_+_8433172_8446622_RNA45SN1_45S_50ntFlanks | 9534 | 9563 | 30 | AGGAAACTCTGGTGGAGGTCCGTAGCGGTC | MB468 | 1033 | 2163 | 1541 | 969 | 376 | 344.5 | 1060.4 | 643.3 | 494.6 | 177.8 |
| >hg38_21_+_8433172_8446622_RNA45SN1_45S_50ntFlanks@9534.9564.31\|AGGAAACTCTGGTGGAGGTCCGTAGCGGTCC | >hg38_21_+_8433172_8446622_RNA45SN1_45S_50ntFlanks | 9534 | 9564 | 31 | AGGAAACTCTGGTGGAGGTCCGTAGCGGTCC | MB468 | 134 | 376 | 132 | 81 | 43 | 21.4 | 167.4 | 34.3 | 43.4 | 21.2 |
| >hg38_21_+_8433172_8446622_RNA45SN1_45S_50ntFlanks@9534.9565.32\|AGGAAACTCTGGTGGAGGTCCGTAGCGGTCCT | >hg38_21_+_8433172_8446622_RNA45SN1_45S_50ntFlanks | 9534 | 9565 | 32 | AGGAAACTCTGGTGGAGGTCCGTAGCGGTCCT | MB468 | 119 | 198 | 98 | 33 | 38 | 16.6 | 61.4 | 15.4 | 19.0 | 12.5 |
| >hg38_21_+_8433172_8446622_RNA45SN1_45S_50ntFlanks@9534.9566.33\|AGGAAACTCTGGTGGAGGTCCGTAGCGGTCCTG | >hg38_21_+_8433172_8446622_RNA45SN1_45S_50ntFlanks | 9534 | 9566 | 33 | AGGAAACTCTGGTGGAGGTCCGTAGCGGTCCTG | MB468 | 184 | 309 | 206 | 103 | 76 | 49.0 | 99.4 | 57.4 | 42.5 | 28.1 |
| >hg38_21_+_8433172_8446622_RNA45SN1_45S_50ntFlanks@9534.9567.34\|AGGAAACTCTGGTGGAGGTCCGTAGCGGTCCTGA | >hg38_21_+_8433172_8446622_RNA45SN1_45S_50ntFlanks | 9534 | 9567 | 34 | AGGAAACTCTGGTGGAGGTCCGTAGCGGTCCTGA | MB468 | 84 | 160 | 100 | 31 | 21 | 71.0 | 121.2 | 84.5 | 18.7 | 9.1 |
| >hg38_21_+_8433172_8446622_RNA45SN1_45S_50ntFlanks@9534.9568.35\|AGGAAACTCTGGTGGAGGTCCGTAGCGGTCCTGAC | >hg38_21_+_8433172_8446622_RNA45SN1_45S_50ntFlanks | 9534 | 9568 | 35 | AGGAAACTCTGGTGGAGGTCCGTAGCGGTCCTGAC | MB468 | 26 | 58 | 42 | 9 | 8 | 25.7 | 51.5 | 38.7 | 5.0 | 5.1 |
| >hg38_21_+_8433172_8446622_RNA45SN1_45S_50ntFlanks@9534.9569.36\|AGGAAACTCTGGTGGAGGTCCGTAGCGGTCCTGACG | >hg38_21_+_8433172_8446622_RNA45SN1_45S_50ntFlanks | 9534 | 9569 | 36 | AGGAAACTCTGGTGGAGGTCCGTAGCGGTCCTGACG | MB468 | 216 | 604 | 379 | 116 | 118 | 146.9 | 468.2 | 264.9 | 71.8 | 75.8 |
| >hg38_21_+_8433172_8446622_RNA45SN1_45S_50ntFlanks@9534.9570.37\|AGGAAACTCTGGTGGAGGTCCGTAGCGGTCCTGACGT | >hg38_21_+_8433172_8446622_RNA45SN1_45S_50ntFlanks | 9534 | 9570 | 37 | AGGAAACTCTGGTGGAGGTCCGTAGCGGTCCTGACGT | MB468 | 78 | 393 | 196 | 60 | 67 | 25.9 | 125.1 | 90.6 | 44.0 | 44.8 |
| >hg38_21_+_8433172_8446622_RNA45SN1_45S_50ntFlanks@9534.9571.38\|AGGAAACTCTGGTGGAGGTCCGTAGCGGTCCTGACGTG | >hg38_21_+_8433172_8446622_RNA45SN1_45S_50ntFlanks | 9534 | 9571 | 38 | AGGAAACTCTGGTGGAGGTCCGTAGCGGTCCTGACGTG | MB468 | 882 | 5102 | 2419 | 827 | 822 | 210.8 | 1236.5 | 1141.3 | 730.8 | 587.9 |
| >hg38_21_+_8433172_8446622_RNA45SN1_45S_50ntFlanks@9534.9572.39\|AGGAAACTCTGGTGGAGGTCCGTAGCGGTCCTGACGTGC | >hg38_21_+_8433172_8446622_RNA45SN1_45S_50ntFlanks | 9534 | 9572 | 39 | AGGAAACTCTGGTGGAGGTCCGTAGCGGTCCTGACGTGC | MB468 | 118 | 643 | 275 | 76 | 112 | 34.4 | 283.6 | 120.3 | 51.4 | 63.8 |
| >hg38_21_+_8433172_8446622_RNA45SN1_45S_50ntFlanks@9534.9573.40\|AGGAAACTCTGGTGGAGGTCCGTAGCGGTCCTGACGTGCA | >hg38_21_+_8433172_8446622_RNA45SN1_45S_50ntFlanks | 9534 | 9573 | 40 | AGGAAACTCTGGTGGAGGTCCGTAGCGGTCCTGACGTGCA | MB468 | 54 | 237 | 82 | 28 | 65 | 12.0 | 140.0 | 34.8 | 16.6 | 44.9 |
| >hg38_21_+_8433172_8446622_RNA45SN1_45S_50ntFlanks@9534.9574.41\|AGGAAACTCTGGTGGAGGTCCGTAGCGGTCCTGACGTGCAA | >hg38_21_+_8433172_8446622_RNA45SN1_45S_50ntFlanks | 9534 | 9574 | 41 | AGGAAACTCTGGTGGAGGTCCGTAGCGGTCCTGACGTGCAA | MB468 | 22 | 146 | 41 | 13 | 32 | 4.6 | 105.9 | 29.2 | 8.7 | 19.2 |

| | | | | | | | | | | | | | | | |
|---|---|---|---|---|---|---|---|---|---|---|---|---|---|---|---|
| >hg38_21_+_8433172_8446622_RNA45SN1_45S_50ntFlanks@9535.9553.19\|GGAAACTCTGGTGGAGGTC | >hg38_21_+_8433172_8446622_RNA45SN1_45S_50ntFlanks | 9535 | 9553 | 19 | GGAAACTCTGGTGGAGGTC | MB468 | 21 | 10 | 31 | 12 | 2 | 5.6 | 0.7 | 16.0 | 11.4 | 0.5 |
| >hg38_21_+_8433172_8446622_RNA45SN1_45S_50ntFlanks@9535.9571.37\|GGAAACTCTGGTGGAGGTCCGTAGCGGTCCTGACGTG | >hg38_21_+_8433172_8446622_RNA45SN1_45S_50ntFlanks | 9535 | 9571 | 37 | GGAAACTCTGGTGGAGGTCCGTAGCGGTCCTGACGTG | MB468 | 18 | 97 | 36 | 8 | 13 | 6.0 | 31.7 | 17.0 | 5.8 | 7.8 |
| >hg38_21_+_8433172_8446622_RNA45SN1_45S_50ntFlanks@9536.9563.28\|GAAACTCTGGTGGAGGTCCGTAGCGGTC | >hg38_21_+_8433172_8446622_RNA45SN1_45S_50ntFlanks | 9536 | 9563 | 28 | GAAACTCTGGTGGAGGTCCGTAGCGGTC | MB468 | 11 | 20 | 19 | 8 | 2 | 4.6 | 10.7 | 11.4 | 4.3 | 0.7 |
| >hg38_21_+_8433172_8446622_RNA45SN1_45S_50ntFlanks@9536.9571.36\|GAAACTCTGGTGGAGGTCCGTAGCGGTCCTGACGTG | >hg38_21_+_8433172_8446622_RNA45SN1_45S_50ntFlanks | 9536 | 9571 | 36 | GAAACTCTGGTGGAGGTCCGTAGCGGTCCTGACGTG | MB468 | 17 | 87 | 37 | 8 | 10 | 1.6 | 9.8 | 23.1 | 4.2 | 5.5 |
| >hg38_21_+_8433172_8446622_RNA45SN1_45S_50ntFlanks@9537.9563.27\|AAACTCTGGTGGAGGTCCGTAGCGGTC | >hg38_21_+_8433172_8446622_RNA45SN1_45S_50ntFlanks | 9537 | 9563 | 27 | AAACTCTGGTGGAGGTCCGTAGCGGTC | MB468 | 13 | 18 | 16 | 5 | 2 | 4.5 | 8.3 | 6.9 | 2.3 | 0.7 |
| >hg38_21_+_8433172_8446622_RNA45SN1_45S_50ntFlanks@9537.9571.35\|AAACTCTGGTGGAGGTCCGTAGCGGTCCTGACGTG | >hg38_21_+_8433172_8446622_RNA45SN1_45S_50ntFlanks | 9537 | 9571 | 35 | AAACTCTGGTGGAGGTCCGTAGCGGTCCTGACGTG | MB468 | 18 | 73 | 26 | 6 | 9 | 2.6 | 6.5 | 11.8 | 3.1 | 5.3 |
| >hg38_21_+_8433172_8446622_RNA45SN1_45S_50ntFlanks@9538.9563.26\|AACTCTGGTGGAGGTCCGTAGCGGTC | >hg38_21_+_8433172_8446622_RNA45SN1_45S_50ntFlanks | 9538 | 9563 | 26 | AACTCTGGTGGAGGTCCGTAGCGGTC | MB468 | 12 | 19 | 14 | 5 | 2 | 5.7 | 10.0 | 8.6 | 2.4 | 1.0 |
| >hg38_21_+_8433172_8446622_RNA45SN1_45S_50ntFlanks@9538.9571.34\|AACTCTGGTGGAGGTCCGTAGCGGTCCTGACGTG | >hg38_21_+_8433172_8446622_RNA45SN1_45S_50ntFlanks | 9538 | 9571 | 34 | AACTCTGGTGGAGGTCCGTAGCGGTCCTGACGTG | MB468 | 31 | 128 | 45 | 8 | 15 | 1.6 | 12.8 | 22.9 | 4.9 | 6.2 |
| >hg38_21_+_8433172_8446622_RNA45SN1_45S_50ntFlanks@9539.9563.25\|ACTCTGGTGGAGGTCCGTAGCGGTC | >hg38_21_+_8433172_8446622_RNA45SN1_45S_50ntFlanks | 9539 | 9563 | 25 | ACTCTGGTGGAGGTCCGTAGCGGTC | MB468 | 17 | 34 | 23 | 7 | 4 | 9.0 | 19.0 | 16.7 | 2.7 | 1.8 |
| >hg38_21_+_8433172_8446622_RNA45SN1_45S_50ntFlanks@9539.9569.31\|ACTCTGGTGGAGGTCCGTAGCGGTCCTGACG | >hg38_21_+_8433172_8446622_RNA45SN1_45S_50ntFlanks | 9539 | 9569 | 31 | ACTCTGGTGGAGGTCCGTAGCGGTCCTGACG | MB468 | 14 | 40 | 19 | 4 | 6 | 8.7 | 29.4 | 17.2 | 4.3 | 5.3 |
| >hg38_21_+_8433172_8446622_RNA45SN1_45S_50ntFlanks@9539.9571.33\|ACTCTGGTGGAGGTCCGTAGCGGTCCTGACGTG | >hg38_21_+_8433172_8446622_RNA45SN1_45S_50ntFlanks | 9539 | 9571 | 33 | ACTCTGGTGGAGGTCCGTAGCGGTCCTGACGTG | MB468 | 63 | 249 | 94 | 17 | 29 | 4.0 | 11.8 | 49.1 | 13.2 | 11.6 |
| >hg38_21_+_8433172_8446622_RNA45SN1_45S_50ntFlanks@9540.9571.32\|CTCTGGTGGAGGTCCGTAGCGGTCCTGACGTG | >hg38_21_+_8433172_8446622_RNA45SN1_45S_50ntFlanks | 9540 | 9571 | 32 | CTCTGGTGGAGGTCCGTAGCGGTCCTGACGTG | MB468 | 44 | 130 | 61 | 14 | 20 | 5.9 | 37.7 | 26.1 | 6.3 | 8.3 |
| >hg38_21_+_8433172_8446622_RNA45SN1_45S_50ntFlanks@9543.9571.29\|TGGTGGAGGTCCGTAGCGGTCCTGACGTG | >hg38_21_+_8433172_8446622_RNA45SN1_45S_50ntFlanks | 9543 | 9571 | 29 | TGGTGGAGGTCCGTAGCGGTCCTGACGTG | MB468 | 14 | 35 | 22 | 7 | 6 | 4.3 | 11.1 | 14.4 | 4.4 | 1.5 |
| >hg38_21_+_8433172_8446622_RNA45SN1_45S_50ntFlanks@9544.9563.20\|GGTGGAGGTCCGTAGCGGTC | >hg38_21_+_8433172_8446622_RNA45SN1_45S_50ntFlanks | 9544 | 9563 | 20 | GGTGGAGGTCCGTAGCGGTC | MB468 | 17 | 35 | 29 | 7 | 4 | 5.5 | 17.5 | 14.4 | 5.7 | 1.9 |
| >hg38_21_+_8433172_8446622_RNA45SN1_45S_50ntFlanks@9544.9566.23\|GGTGGAGGTCCGTAGCGGTCCTG | >hg38_21_+_8433172_8446622_RNA45SN1_45S_50ntFlanks | 9544 | 9566 | 23 | GGTGGAGGTCCGTAGCGGTCCTG | MB468 | 14 | 15 | 16 | 4 | 3 | 4.8 | 2.4 | 6.1 | 3.8 | 0.7 |
| >hg38_21_+_8433172_8446622_RNA45SN1_45S_50ntFlanks@9544.9569.26\|GGTGGAGGTCCGTAGCGGTCCTGACG | >hg38_21_+_8433172_8446622_RNA45SN1_45S_50ntFlanks | 9544 | 9569 | 26 | GGTGGAGGTCCGTAGCGGTCCTGACG | MB468 | 15 | 20 | 22 | 8 | 7 | 4.5 | 1.8 | 8.1 | 5.1 | 1.4 |
| >hg38_21_+_8433172_8446622_RNA45SN1_45S_50ntFlanks@9544.9570.27\|GGTGGAGGTCCGTAGCGGTCCTGACGT | >hg38_21_+_8433172_8446622_RNA45SN1_45S_50ntFlanks | 9544 | 9570 | 27 | GGTGGAGGTCCGTAGCGGTCCTGACGT | MB468 | 14 | 34 | 29 | 6 | 6 | 4.4 | 19.3 | 15.2 | 2.6 | 2.1 |
| >hg38_21_+_8433172_8446622_RNA45SN1_45S_50ntFlanks@9544.9571.28\|GGTGGAGGTCCGTAGCGGTCCTGACGTG | >hg38_21_+_8433172_8446622_RNA45SN1_45S_50ntFlanks | 9544 | 9571 | 28 | GGTGGAGGTCCGTAGCGGTCCTGACGTG | MB231 | 13 | 12 | 100 | 6 | 4 | 11.1 | 9.5 | 121.6 | 6.1 | 5.0 |
| >hg38_21_+_8433172_8446622_RNA45SN1_45S_50ntFlanks@9544.9571.28\|GGTGGAGGTCCGTAGCGGTCCTGACGTG | >hg38_21_+_8433172_8446622_RNA45SN1_45S_50ntFlanks | 9544 | 9571 | 28 | GGTGGAGGTCCGTAGCGGTCCTGACGTG | MB468 | 138 | 423 | 189 | 63 | 48 | 28.9 | 144.3 | 127.9 | 35.7 | 16.1 |
| >hg38_21_+_8433172_8446622_RNA45SN1_45S_50ntFlanks@9544.9572.29\|GGTGGAGGTCCGTAGCGGTCCTGACGTGC | >hg38_21_+_8433172_8446622_RNA45SN1_45S_50ntFlanks | 9544 | 9572 | 29 | GGTGGAGGTCCGTAGCGGTCCTGACGTGC | MB468 | 19 | 59 | 22 | 6 | 7 | 3.4 | 9.3 | 8.4 | 4.2 | 2.1 |
| >hg38_21_+_8433172_8446622_RNA45SN1_45S_50ntFlanks@9544.9573.30\|GGTGGAGGTCCGTAGCGGTCCTGACGTGCA | >hg38_21_+_8433172_8446622_RNA45SN1_45S_50ntFlanks | 9544 | 9573 | 30 | GGTGGAGGTCCGTAGCGGTCCTGACGTGCA | MB468 | 15 | 35 | 9 | 3 | 5 | 4.7 | 6.8 | 5.0 | 1.7 | 1.9 |
| >hg38_21_+_8433172_8446622_RNA45SN1_45S_50ntFlanks@9545.9562.18\|GTGGAGGTCCGTAGCGGT | >hg38_21_+_8433172_8446622_RNA45SN1_45S_50ntFlanks | 9545 | 9562 | 18 | GTGGAGGTCCGTAGCGGT | MB468 | 20 | 36 | 25 | 16 | 6 | 2.6 | 11.2 | 6.6 | 6.1 | 0.9 |
| >hg38_21_+_8433172_8446622_RNA45SN1_45S_50ntFlanks@9545.9563.19\|GTGGAGGTCCGTAGCGGTC | >hg38_21_+_8433172_8446622_RNA45SN1_45S_50ntFlanks | 9545 | 9563 | 19 | GTGGAGGTCCGTAGCGGTC | MB468 | 326 | 660 | 397 | 143 | 63 | 81.5 | 393.3 | 136.6 | 148.0 | 33.0 |
| >hg38_21_+_8433172_8446622_RNA45SN1_45S_50ntFlanks@9545.9564.20\|GTGGAGGTCCGTAGCGGTCC | >hg38_21_+_8433172_8446622_RNA45SN1_45S_50ntFlanks | 9545 | 9564 | 20 | GTGGAGGTCCGTAGCGGTCC | MB468 | 69 | 72 | 62 | 17 | 11 | 21.4 | 28.8 | 24.0 | 15.6 | 2.2 |
| >hg38_21_+_8433172_8446622_RNA45SN1_45S_50ntFlanks@9545.9566.22\|GTGGAGGTCCGTAGCGGTCCTG | >hg38_21_+_8433172_8446622_RNA45SN1_45S_50ntFlanks | 9545 | 9566 | 22 | GTGGAGGTCCGTAGCGGTCCTG | MB468 | 160 | 72 | 210 | 61 | 32 | 26.7 | 24.6 | 28.3 | 19.2 | 13.5 |
| >hg38_21_+_8433172_8446622_RNA45SN1_45S_50ntFlanks@9545.9567.23\|GTGGAGGTCCGTAGCGGTCCTGA | >hg38_21_+_8433172_8446622_RNA45SN1_45S_50ntFlanks | 9545 | 9567 | 23 | GTGGAGGTCCGTAGCGGTCCTGA | MB468 | 32 | 19 | 39 | 12 | 8 | 11.4 | 4.7 | 17.3 | 3.0 | 2.5 |
| >hg38_21_+_8433172_8446622_RNA45SN1_45S_50ntFlanks@9545.9569.25\|GTGGAGGTCCGTAGCGGTCCTGACG | >hg38_21_+_8433172_8446622_RNA45SN1_45S_50ntFlanks | 9545 | 9569 | 25 | GTGGAGGTCCGTAGCGGTCCTGACG | MB468 | 45 | 14 | 85 | 37 | 29 | 16.4 | 4.6 | 45.9 | 10.7 | 13.3 |
| >hg38_21_+_8433172_8446622_RNA45SN1_45S_50ntFlanks@9545.9570.26\|GTGGAGGTCCGTAGCGGTCCTGACGT | >hg38_21_+_8433172_8446622_RNA45SN1_45S_50ntFlanks | 9545 | 9570 | 26 | GTGGAGGTCCGTAGCGGTCCTGACGT | MB468 | 72 | 66 | 181 | 41 | 33 | 30.6 | 61.6 | 47.0 | 14.0 | 12.2 |
| >hg38_21_+_8433172_8446622_RNA45SN1_45S_50ntFlanks@9545.9571.27\|GTGGAGGTCCGTAGCGGTCCTGACGTG | >hg38_21_+_8433172_8446622_RNA45SN1_45S_50ntFlanks | 9545 | 9571 | 27 | GTGGAGGTCCGTAGCGGTCCTGACGTG | MB231 | 18 | 10 | 100 | 5 | 4 | 14.2 | 7.3 | 105.6 | 3.9 | 3.1 |
| >hg38_21_+_8433172_8446622_RNA45SN1_45S_50ntFlanks@9545.9571.27\|GTGGAGGTCCGTAGCGGTCCTGACGTG | >hg38_21_+_8433172_8446622_RNA45SN1_45S_50ntFlanks | 9545 | 9571 | 27 | GTGGAGGTCCGTAGCGGTCCTGACGTG | MB468 | 869 | 1076 | 2162 | 472 | 377 | 359.9 | 626.7 | 925.2 | 231.9 | 109.7 |
| >hg38_21_+_8433172_8446622_RNA45SN1_45S_50ntFlanks@9545.9572.28\|GTGGAGGTCCGTAGCGGTCCTGACGTGC | >hg38_21_+_8433172_8446622_RNA45SN1_45S_50ntFlanks | 9545 | 9572 | 28 | GTGGAGGTCCGTAGCGGTCCTGACGTGC | MB468 | 104 | 153 | 225 | 42 | 52 | 27.3 | 32.8 | 54.0 | 25.0 | 9.4 |
| >hg38_21_+_8433172_8446622_RNA45SN1_45S_50ntFlanks@9545.9573.29\|GTGGAGGTCCGTAGCGGTCCTGACGTGCA | >hg38_21_+_8433172_8446622_RNA45SN1_45S_50ntFlanks | 9545 | 9573 | 29 | GTGGAGGTCCGTAGCGGTCCTGACGTGCA | MB468 | 102 | 145 | 133 | 32 | 69 | 12.3 | 54.7 | 33.5 | 13.8 | 23.1 |
| >hg38_21_+_8433172_8446622_RNA45SN1_45S_50ntFlanks@9545.9574.30\|GTGGAGGTCCGTAGCGGTCCTGACGTGCAA | >hg38_21_+_8433172_8446622_RNA45SN1_45S_50ntFlanks | 9545 | 9574 | 30 | GTGGAGGTCCGTAGCGGTCCTGACGTGCAA | MB468 | 76 | 158 | 74 | 21 | 41 | 20.5 | 36.3 | 22.2 | 12.2 | 7.3 |
| >hg38_21_+_8433172_8446622_RNA45SN1_45S_50ntFlanks@9545.9575.31\|GTGGAGGTCCGTAGCGGTCCTGACGTGCAAA | >hg38_21_+_8433172_8446622_RNA45SN1_45S_50ntFlanks | 9545 | 9575 | 31 | GTGGAGGTCCGTAGCGGTCCTGACGTGCAAA | MB468 | 12 | 30 | 11 | 7 | 18 | 3.4 | 9.3 | 7.4 | 4.7 | 3.7 |
| >hg38_21_+_8433172_8446622_RNA45SN1_45S_50ntFlanks@9546.9563.18\|TGGAGGTCCGTAGCGGTC | >hg38_21_+_8433172_8446622_RNA45SN1_45S_50ntFlanks | 9546 | 9563 | 18 | TGGAGGTCCGTAGCGGTC | MB468 | 13 | 21 | 14 | 7 | 3 | 3.2 | 3.9 | 5.8 | 7.1 | 0.7 |
| >hg38_21_+_8433172_8446622_RNA45SN1_45S_50ntFlanks@9546.9571.26\|TGGAGGTCCGTAGCGGTCCTGACGTG | >hg38_21_+_8433172_8446622_RNA45SN1_45S_50ntFlanks | 9546 | 9571 | 26 | TGGAGGTCCGTAGCGGTCCTGACGTG | MB231 | 25 | 20 | 87 | 8 | 7 | 20.7 | 13.6 | 71.8 | 6.2 | 7.3 |
| >hg38_21_+_8433172_8446622_RNA45SN1_45S_50ntFlanks@9546.9571.26\|TGGAGGTCCGTAGCGGTCCTGACGTG | >hg38_21_+_8433172_8446622_RNA45SN1_45S_50ntFlanks | 9546 | 9571 | 26 | TGGAGGTCCGTAGCGGTCCTGACGTG | MB468 | 28 | 47 | 58 | 26 | 19 | 9.9 | 22.5 | 28.8 | 19.6 | 3.5 |
| >hg38_21_+_8433172_8446622_RNA45SN1_45S_50ntFlanks@9546.9573.28\|TGGAGGTCCGTAGCGGTCCTGACGTGCA | >hg38_21_+_8433172_8446622_RNA45SN1_45S_50ntFlanks | 9546 | 9573 | 28 | TGGAGGTCCGTAGCGGTCCTGACGTGCA | MB231 | 14 | 20 | 30 | 4 | 5 | 12.4 | 16.4 | 21.2 | 5.2 | 5.0 |
| >hg38_21_+_8433172_8446622_RNA45SN1_45S_50ntFlanks@9547.9570.24\|GGAGGTCCGTAGCGGTCCTGACGT | >hg38_21_+_8433172_8446622_RNA45SN1_45S_50ntFlanks | 9547 | 9570 | 24 | GGAGGTCCGTAGCGGTCCTGACGT | MB231 | 10 | 11 | 14 | 7 | 3 | 9.6 | 12.0 | 8.5 | 8.5 | 2.8 |
| >hg38_21_+_8433172_8446622_RNA45SN1_45S_50ntFlanks@9548.9571.24\|GAGGTCCGTAGCGGTCCTGACGTG | >hg38_21_+_8433172_8446622_RNA45SN1_45S_50ntFlanks | 9548 | 9571 | 24 | GAGGTCCGTAGCGGTCCTGACGTG | MB468 | 42 | 39 | 88 | 30 | 34 | 14.4 | 22.2 | 23.0 | 14.5 | 5.2 |
| >hg38_21_+_8433172_8446622_RNA45SN1_45S_50ntFlanks@9548.9573.26\|GAGGTCCGTAGCGGTCCTGACGTGCA | >hg38_21_+_8433172_8446622_RNA45SN1_45S_50ntFlanks | 9548 | 9573 | 26 | GAGGTCCGTAGCGGTCCTGACGTGCA | MB231 | 12 | 22 | 27 | 2 | 4 | 10.0 | 16.1 | 19.5 | 2.5 | 3.9 |
| >hg38_21_+_8433172_8446622_RNA45SN1_45S_50ntFlanks@9548.9573.26\|GAGGTCCGTAGCGGTCCTGACGTGCA | >hg38_21_+_8433172_8446622_RNA45SN1_45S_50ntFlanks | 9548 | 9573 | 26 | GAGGTCCGTAGCGGTCCTGACGTGCA | MB468 | 15 | 23 | 17 | 5 | 13 | 4.5 | 7.7 | 5.3 | 2.5 | 1.1 |
| >hg38_21_+_8433172_8446622_RNA45SN1_45S_50ntFlanks@9549.9571.23\|AGGTCCGTAGCGGTCCTGACGTG | >hg38_21_+_8433172_8446622_RNA45SN1_45S_50ntFlanks | 9549 | 9571 | 23 | AGGTCCGTAGCGGTCCTGACGTG | MB468 | 15 | 10 | 40 | 8 | 8 | 5.5 | 5.6 | 10.6 | 2.4 | 3.2 |
| >hg38_21_+_8433172_8446622_RNA45SN1_45S_50ntFlanks@9551.9569.19\|GTCCGTAGCGGTCCTGACG | >hg38_21_+_8433172_8446622_RNA45SN1_45S_50ntFlanks | 9551 | 9569 | 19 | GTCCGTAGCGGTCCTGACG | MB231 | 17 | 14 | 41 | 0 | 2 | 15.7 | 9.7 | 38.8 | 0.8 | 3.9 |
| >hg38_21_+_8433172_8446622_RNA45SN1_45S_50ntFlanks@9551.9571.21\|GTCCGTAGCGGTCCTGACGTG | >hg38_21_+_8433172_8446622_RNA45SN1_45S_50ntFlanks | 9551 | 9571 | 21 | GTCCGTAGCGGTCCTGACGTG | MB231 | 15 | 13 | 31 | 1 | 3 | 12.3 | 8.7 | 22.3 | 1.2 | 2.8 |
| >hg38_21_+_8433172_8446622_RNA45SN1_45S_50ntFlanks@9551.9571.21\|GTCCGTAGCGGTCCTGACGTG | >hg38_21_+_8433172_8446622_RNA45SN1_45S_50ntFlanks | 9551 | 9571 | 21 | GTCCGTAGCGGTCCTGACGTG | MB468 | 13 | 13 | 21 | 5 | 7 | 1.8 | 4.7 | 6.3 | 3.0 | 2.4 |
| >hg38_21_+_8433172_8446622_RNA45SN1_45S_50ntFlanks@9552.9569.18\|TCCGTAGCGGTCCTGACG | >hg38_21_+_8433172_8446622_RNA45SN1_45S_50ntFlanks | 9552 | 9569 | 18 | TCCGTAGCGGTCCTGACG | MB231 | 12 | 12 | 20 | 0 | 2 | 6.9 | 3.0 | 6.1 | 0.4 | 2.7 |
| >hg38_21_+_8433172_8446622_RNA45SN1_45S_50ntFlanks@9552.9569.18\|TCCGTAGCGGTCCTGACG | >hg38_21_+_8433172_8446622_RNA45SN1_45S_50ntFlanks | 9552 | 9569 | 18 | TCCGTAGCGGTCCTGACG | MB468 | 26 | 30 | 27 | 4 | 10 | 4.7 | 9.8 | 10.6 | 4.0 | 1.4 |
| >hg38_21_+_8433172_8446622_RNA45SN1_45S_50ntFlanks@9552.9570.19\|TCCGTAGCGGTCCTGACGT | >hg38_21_+_8433172_8446622_RNA45SN1_45S_50ntFlanks | 9552 | 9570 | 19 | TCCGTAGCGGTCCTGACGT | MB468 | 11 | 10 | 14 | 2 | 4 | 2.8 | 4.2 | 5.6 | 1.4 | 1.0 |
| >hg38_21_+_8433172_8446622_RNA45SN1_45S_50ntFlanks@9552.9571.20\|TCCGTAGCGGTCCTGACGTG | >hg38_21_+_8433172_8446622_RNA45SN1_45S_50ntFlanks | 9552 | 9571 | 20 | TCCGTAGCGGTCCTGACGTG | MB468 | 70 | 40 | 70 | 14 | 23 | 12.7 | 10.4 | 14.1 | 8.0 | 8.3 |

| | | | | | | | | | | | | | | | | |
|---|---|---|---|---|---|---|---|---|---|---|---|---|---|---|---|---|
| >hg38_21_+_8433172_8446622_RNA45SN1_45S_50ntFlanks@9552.9572.21\|TCCGTAGCGGTCCTGACGTGC | >hg38_21_+_8433172_8446622_RNA45SN1_45S_50ntFlanks | 9552 | 9572 | 21 | TCCGTAGCGGTCCTGACGTGC | MB468 | 21 | 16 | 21 | 3 | 5 | 5.4 | 2.2 | 7.1 | 3.3 | 1.6 |
| >hg38_21_+_8433172_8446622_RNA45SN1_45S_50ntFlanks@9553.9570.18\|CCGTAGCGGTCCTGACGT | >hg38_21_+_8433172_8446622_RNA45SN1_45S_50ntFlanks | 9553 | 9570 | 18 | CCGTAGCGGTCCTGACGT | MB231 | 13 | 9 | 20 | 1 | 5 | 7.9 | 3.7 | 6.4 | 0.0 | 1.6 |
| >hg38_21_+_8433172_8446622_RNA45SN1_45S_50ntFlanks@9553.9571.19\|CCGTAGCGGTCCTGACGTG | >hg38_21_+_8433172_8446622_RNA45SN1_45S_50ntFlanks | 9553 | 9571 | 19 | CCGTAGCGGTCCTGACGTG | MB468 | 20 | 8 | 20 | 9 | 8 | 4.4 | 2.4 | 5.3 | 2.4 | 2.7 |
| >hg38_21_+_8433172_8446622_RNA45SN1_45S_50ntFlanks@9553.9572.20\|CCGTAGCGGTCCTGACGTGC | >hg38_21_+_8433172_8446622_RNA45SN1_45S_50ntFlanks | 9553 | 9572 | 20 | CCGTAGCGGTCCTGACGTGC | MB468 | 17 | 11 | 19 | 5 | 5 | 5.1 | 2.2 | 8.4 | 4.6 | 1.1 |
| >hg38_21_+_8433172_8446622_RNA45SN1_45S_50ntFlanks@9553.9573.21\|CCGTAGCGGTCCTGACGTGCA | >hg38_21_+_8433172_8446622_RNA45SN1_45S_50ntFlanks | 9553 | 9573 | 21 | CCGTAGCGGTCCTGACGTGCA | MB231 | 12 | 8 | 17 | 2 | 3 | 8.8 | 5.0 | 9.2 | 2.0 | 3.0 |
| >hg38_21_+_8433172_8446622_RNA45SN1_45S_50ntFlanks@9554.9571.18\|CGTAGCGGTCCTGACGTG | >hg38_21_+_8433172_8446622_RNA45SN1_45S_50ntFlanks | 9554 | 9571 | 18 | CGTAGCGGTCCTGACGTG | BT20 | 46 | 28 | 86 | 74 | 37 | 31.3 | 22.1 | 18.1 | 110.5 | 33.2 |
| >hg38_21_+_8433172_8446622_RNA45SN1_45S_50ntFlanks@9554.9571.18\|CGTAGCGGTCCTGACGTG | >hg38_21_+_8433172_8446622_RNA45SN1_45S_50ntFlanks | 9554 | 9571 | 18 | CGTAGCGGTCCTGACGTG | MB231 | 226 | 102 | 533 | 19 | 42 | 248.0 | 75.4 | 516.3 | 17.9 | 42.1 |
| >hg38_21_+_8433172_8446622_RNA45SN1_45S_50ntFlanks@9554.9571.18\|CGTAGCGGTCCTGACGTG | >hg38_21_+_8433172_8446622_RNA45SN1_45S_50ntFlanks | 9554 | 9571 | 18 | CGTAGCGGTCCTGACGTG | MB468 | 13879 | 11072 | 15793 | 4093 | 3802 | 2479.8 | 5132.7 | 4065.3 | 2940.2 | 619.3 |
| >hg38_21_+_8433172_8446622_RNA45SN1_45S_50ntFlanks@9554.9572.19\|CGTAGCGGTCCTGACGTGC | >hg38_21_+_8433172_8446622_RNA45SN1_45S_50ntFlanks | 9554 | 9572 | 19 | CGTAGCGGTCCTGACGTGC | BT20 | 13 | 11 | 33 | 48 | 18 | 7.8 | 6.5 | 11.6 | 71.1 | 9.1 |
| >hg38_21_+_8433172_8446622_RNA45SN1_45S_50ntFlanks@9554.9572.19\|CGTAGCGGTCCTGACGTGC | >hg38_21_+_8433172_8446622_RNA45SN1_45S_50ntFlanks | 9554 | 9572 | 19 | CGTAGCGGTCCTGACGTGC | MB231 | 53 | 27 | 113 | 8 | 13 | 54.7 | 22.9 | 89.3 | 10.1 | 12.6 |
| >hg38_21_+_8433172_8446622_RNA45SN1_45S_50ntFlanks@9554.9572.19\|CGTAGCGGTCCTGACGTGC | >hg38_21_+_8433172_8446622_RNA45SN1_45S_50ntFlanks | 9554 | 9572 | 19 | CGTAGCGGTCCTGACGTGC | MB468 | 1352 | 972 | 1482 | 402 | 346 | 77.3 | 149.1 | 293.5 | 339.6 | 73.3 |
| >hg38_21_+_8433172_8446622_RNA45SN1_45S_50ntFlanks@9554.9573.20\|CGTAGCGGTCCTGACGTGCA | >hg38_21_+_8433172_8446622_RNA45SN1_45S_50ntFlanks | 9554 | 9573 | 20 | CGTAGCGGTCCTGACGTGCA | BT20 | 28 | 15 | 68 | 24 | 14 | 13.8 | 5.9 | 25.8 | 27.6 | 7.2 |
| >hg38_21_+_8433172_8446622_RNA45SN1_45S_50ntFlanks@9554.9573.20\|CGTAGCGGTCCTGACGTGCA | >hg38_21_+_8433172_8446622_RNA45SN1_45S_50ntFlanks | 9554 | 9573 | 20 | CGTAGCGGTCCTGACGTGCA | MB231 | 112 | 51 | 190 | 13 | 23 | 104.5 | 35.9 | 142.1 | 15.7 | 19.9 |
| >hg38_21_+_8433172_8446622_RNA45SN1_45S_50ntFlanks@9554.9573.20\|CGTAGCGGTCCTGACGTGCA | >hg38_21_+_8433172_8446622_RNA45SN1_45S_50ntFlanks | 9554 | 9573 | 20 | CGTAGCGGTCCTGACGTGCA | MB468 | 1192 | 846 | 960 | 300 | 383 | 121.8 | 183.5 | 190.6 | 205.9 | 127.1 |
| >hg38_21_+_8433172_8446622_RNA45SN1_45S_50ntFlanks@9554.9574.21\|CGTAGCGGTCCTGACGTGCAA | >hg38_21_+_8433172_8446622_RNA45SN1_45S_50ntFlanks | 9554 | 9574 | 21 | CGTAGCGGTCCTGACGTGCAA | MB231 | 19 | 10 | 39 | 5 | 4 | 19.5 | 8.5 | 35.8 | 7.2 | 5.1 |
| >hg38_21_+_8433172_8446622_RNA45SN1_45S_50ntFlanks@9554.9574.21\|CGTAGCGGTCCTGACGTGCAA | >hg38_21_+_8433172_8446622_RNA45SN1_45S_50ntFlanks | 9554 | 9574 | 21 | CGTAGCGGTCCTGACGTGCAA | MB468 | 249 | 168 | 233 | 133 | 87 | 50.3 | 30.8 | 57.1 | 89.1 | 31.3 |
| >hg38_21_+_8433172_8446622_RNA45SN1_45S_50ntFlanks@9554.9575.22\|CGTAGCGGTCCTGACGTGCAAA | >hg38_21_+_8433172_8446622_RNA45SN1_45S_50ntFlanks | 9554 | 9575 | 22 | CGTAGCGGTCCTGACGTGCAAA | MB231 | 13 | 6 | 21 | 3 | 4 | 12.8 | 4.6 | 19.3 | 3.8 | 3.9 |
| >hg38_21_+_8433172_8446622_RNA45SN1_45S_50ntFlanks@9554.9575.22\|CGTAGCGGTCCTGACGTGCAAA | >hg38_21_+_8433172_8446622_RNA45SN1_45S_50ntFlanks | 9554 | 9575 | 22 | CGTAGCGGTCCTGACGTGCAAA | MB468 | 62 | 39 | 56 | 56 | 54 | 22.8 | 5.3 | 20.5 | 44.2 | 7.2 |
| >hg38_21_+_8433172_8446622_RNA45SN1_45S_50ntFlanks@9554.9576.23\|CGTAGCGGTCCTGACGTGCAAAT | >hg38_21_+_8433172_8446622_RNA45SN1_45S_50ntFlanks | 9554 | 9576 | 23 | CGTAGCGGTCCTGACGTGCAAAT | BT20 | 15 | 10 | 46 | 8 | 5 | 5.9 | 2.1 | 18.1 | 2.5 | 2.2 |
| >hg38_21_+_8433172_8446622_RNA45SN1_45S_50ntFlanks@9554.9576.23\|CGTAGCGGTCCTGACGTGCAAAT | >hg38_21_+_8433172_8446622_RNA45SN1_45S_50ntFlanks | 9554 | 9576 | 23 | CGTAGCGGTCCTGACGTGCAAAT | MB231 | 67 | 37 | 129 | 12 | 13 | 68.8 | 30.4 | 125.6 | 15.1 | 11.9 |
| >hg38_21_+_8433172_8446622_RNA45SN1_45S_50ntFlanks@9554.9576.23\|CGTAGCGGTCCTGACGTGCAAAT | >hg38_21_+_8433172_8446622_RNA45SN1_45S_50ntFlanks | 9554 | 9576 | 23 | CGTAGCGGTCCTGACGTGCAAAT | MB468 | 22 | 17 | 23 | 11 | 10 | 1.4 | 3.5 | 5.1 | 6.1 | 3.1 |
| >hg38_21_+_8433172_8446622_RNA45SN1_45S_50ntFlanks@9554.9577.24\|CGTAGCGGTCCTGACGTGCAAATC | >hg38_21_+_8433172_8446622_RNA45SN1_45S_50ntFlanks | 9554 | 9577 | 24 | CGTAGCGGTCCTGACGTGCAAATC | BT20 | 14 | 9 | 38 | 7 | 11 | 10.0 | 1.8 | 11.7 | 6.1 | 4.3 |
| >hg38_21_+_8433172_8446622_RNA45SN1_45S_50ntFlanks@9554.9577.24\|CGTAGCGGTCCTGACGTGCAAATC | >hg38_21_+_8433172_8446622_RNA45SN1_45S_50ntFlanks | 9554 | 9577 | 24 | CGTAGCGGTCCTGACGTGCAAATC | MB231 | 73 | 59 | 143 | 4 | 20 | 58.8 | 42.9 | 114.2 | 5.3 | 22.2 |
| >hg38_21_+_8433172_8446622_RNA45SN1_45S_50ntFlanks@9554.9577.24\|CGTAGCGGTCCTGACGTGCAAATC | >hg38_21_+_8433172_8446622_RNA45SN1_45S_50ntFlanks | 9554 | 9577 | 24 | CGTAGCGGTCCTGACGTGCAAATC | MB468 | 44 | 65 | 38 | 9 | 21 | 5.6 | 8.0 | 10.1 | 4.8 | 2.4 |
| >hg38_21_+_8433172_8446622_RNA45SN1_45S_50ntFlanks@9554.9578.25\|CGTAGCGGTCCTGACGTGCAAATCG | >hg38_21_+_8433172_8446622_RNA45SN1_45S_50ntFlanks | 9554 | 9578 | 25 | CGTAGCGGTCCTGACGTGCAAATCG | MB468 | 15 | 12 | 11 | 4 | 10 | 4.8 | 5.0 | 2.0 | 0.5 | 2.3 |
| >hg38_21_+_8433172_8446622_RNA45SN1_45S_50ntFlanks@9554.9579.26\|CGTAGCGGTCCTGACGTGCAAATCGG | >hg38_21_+_8433172_8446622_RNA45SN1_45S_50ntFlanks | 9554 | 9579 | 26 | CGTAGCGGTCCTGACGTGCAAATCGG | MB468 | 27 | 28 | 25 | 4 | 12 | 13.5 | 12.7 | 9.8 | 1.4 | 2.5 |
| >hg38_21_+_8433172_8446622_RNA45SN1_45S_50ntFlanks@9554.9580.27\|CGTAGCGGTCCTGACGTGCAAATCGGT | >hg38_21_+_8433172_8446622_RNA45SN1_45S_50ntFlanks | 9554 | 9580 | 27 | CGTAGCGGTCCTGACGTGCAAATCGGT | MB468 | 25 | 28 | 30 | 6 | 11 | 4.0 | 5.3 | 14.0 | 0.6 | 3.5 |
| >hg38_21_+_8433172_8446622_RNA45SN1_45S_50ntFlanks@9554.9581.28\|CGTAGCGGTCCTGACGTGCAAATCGGTC | >hg38_21_+_8433172_8446622_RNA45SN1_45S_50ntFlanks | 9554 | 9581 | 28 | CGTAGCGGTCCTGACGTGCAAATCGGTC | MB231 | 32 | 25 | 88 | 2 | 11 | 25.5 | 14.8 | 58.0 | 2.3 | 14.0 |
| >hg38_21_+_8433172_8446622_RNA45SN1_45S_50ntFlanks@9554.9581.28\|CGTAGCGGTCCTGACGTGCAAATCGGTC | >hg38_21_+_8433172_8446622_RNA45SN1_45S_50ntFlanks | 9554 | 9581 | 28 | CGTAGCGGTCCTGACGTGCAAATCGGTC | MB468 | 26 | 49 | 29 | 4 | 12 | 6.6 | 21.0 | 11.2 | 2.0 | 2.6 |
| >hg38_21_+_8433172_8446622_RNA45SN1_45S_50ntFlanks@9554.9582.29\|CGTAGCGGTCCTGACGTGCAAATCGGTCG | >hg38_21_+_8433172_8446622_RNA45SN1_45S_50ntFlanks | 9554 | 9582 | 29 | CGTAGCGGTCCTGACGTGCAAATCGGTCG | MB468 | 58 | 86 | 41 | 8 | 24 | 24.5 | 61.5 | 23.2 | 5.3 | 7.1 |
| >hg38_21_+_8433172_8446622_RNA45SN1_45S_50ntFlanks@9554.9584.31\|CGTAGCGGTCCTGACGTGCAAATCGGTCGTC | >hg38_21_+_8433172_8446622_RNA45SN1_45S_50ntFlanks | 9554 | 9584 | 31 | CGTAGCGGTCCTGACGTGCAAATCGGTCGTC | MB468 | 191 | 304 | 222 | 38 | 89 | 53.6 | 140.1 | 86.6 | 25.9 | 38.3 |
| >hg38_21_+_8433172_8446622_RNA45SN1_45S_50ntFlanks@9554.9585.32\|CGTAGCGGTCCTGACGTGCAAATCGGTCGTCC | >hg38_21_+_8433172_8446622_RNA45SN1_45S_50ntFlanks | 9554 | 9585 | 32 | CGTAGCGGTCCTGACGTGCAAATCGGTCGTCC | MB468 | 11 | 9 | 12 | 3 | 4 | 5.5 | 5.1 | 9.2 | 1.5 | 2.5 |
| >hg38_21_+_8433172_8446622_RNA45SN1_45S_50ntFlanks@9554.9595.42\|CGTAGCGGTCCTGACGTGCAAATCGGTCGTCCGACCTGGGTA | >hg38_21_+_8433172_8446622_RNA45SN1_45S_50ntFlanks | 9554 | 9595 | 42 | CGTAGCGGTCCTGACGTGCAAATCGGTCGTCCGACCTGGGTA | MB468 | 11 | 53 | 31 | 4 | 101 | 4.6 | 86.2 | 32.3 | 3.7 | 106.5 |
| >hg38_21_+_8433172_8446622_RNA45SN1_45S_50ntFlanks@9554.9596.43\|CGTAGCGGTCCTGACGTGCAAATCGGTCGTCCGACCTGGGTAT | >hg38_21_+_8433172_8446622_RNA45SN1_45S_50ntFlanks | 9554 | 9596 | 43 | CGTAGCGGTCCTGACGTGCAAATCGGTCGTCCGACCTGGGTAT | MB468 | 12 | 45 | 22 | 8 | 143 | 7.0 | 70.2 | 17.2 | 9.1 | 156.8 |
| >hg38_21_+_8433172_8446622_RNA45SN1_45S_50ntFlanks@9554.9597.44\|CGTAGCGGTCCTGACGTGCAAATCGGTCGTCCGACCTGGGTATA | >hg38_21_+_8433172_8446622_RNA45SN1_45S_50ntFlanks | 9554 | 9597 | 44 | CGTAGCGGTCCTGACGTGCAAATCGGTCGTCCGACCTGGGTATA | MB468 | 27 | 181 | 43 | 15 | 303 | 16.0 | 253.7 | 27.8 | 19.3 | 270.1 |
| >hg38_21_+_8433172_8446622_RNA45SN1_45S_50ntFlanks@9555.9573.19\|GTAGCGGTCCTGACGTGCA | >hg38_21_+_8433172_8446622_RNA45SN1_45S_50ntFlanks | 9555 | 9573 | 19 | GTAGCGGTCCTGACGTGCA | MB231 | 10 | 4 | 7 | 2 | 3 | 8.4 | 3.4 | 6.1 | 2.7 | 2.5 |
| >hg38_21_+_8433172_8446622_RNA45SN1_45S_50ntFlanks@9555.9573.19\|GTAGCGGTCCTGACGTGCA | >hg38_21_+_8433172_8446622_RNA45SN1_45S_50ntFlanks | 9555 | 9573 | 19 | GTAGCGGTCCTGACGTGCA | MB468 | 18 | 6 | 14 | 6 | 6 | 4.0 | 1.6 | 2.8 | 3.0 | 3.5 |
| >hg38_21_+_8433172_8446622_RNA45SN1_45S_50ntFlanks@9555.9584.30\|GTAGCGGTCCTGACGTGCAAATCGGTCGTC | >hg38_21_+_8433172_8446622_RNA45SN1_45S_50ntFlanks | 9555 | 9584 | 30 | GTAGCGGTCCTGACGTGCAAATCGGTCGTC | MB468 | 20 | 19 | 18 | 4 | 6 | 9.4 | 13.8 | 9.8 | 4.4 | 3.1 |
| >hg38_21_+_8433172_8446622_RNA45SN1_45S_50ntFlanks@9556.9573.18\|TAGCGGTCCTGACGTGCA | >hg38_21_+_8433172_8446622_RNA45SN1_45S_50ntFlanks | 9556 | 9573 | 18 | TAGCGGTCCTGACGTGCA | MB468 | 46 | 9 | 31 | 23 | 14 | 14.6 | 3.8 | 8.4 | 11.6 | 8.1 |
| >hg38_21_+_8433172_8446622_RNA45SN1_45S_50ntFlanks@9556.9581.26\|TAGCGGTCCTGACGTGCAAATCGGTC | >hg38_21_+_8433172_8446622_RNA45SN1_45S_50ntFlanks | 9556 | 9581 | 26 | TAGCGGTCCTGACGTGCAAATCGGTC | MB468 | 15 | 19 | 12 | 2 | 4 | 4.6 | 9.4 | 5.8 | 0.8 | 0.4 |
| >hg38_21_+_8433172_8446622_RNA45SN1_45S_50ntFlanks@9556.9582.27\|TAGCGGTCCTGACGTGCAAATCGGTCG | >hg38_21_+_8433172_8446622_RNA45SN1_45S_50ntFlanks | 9556 | 9582 | 27 | TAGCGGTCCTGACGTGCAAATCGGTCG | MB468 | 66 | 62 | 43 | 8 | 15 | 27.3 | 37.8 | 28.6 | 4.8 | 4.7 |
| >hg38_21_+_8433172_8446622_RNA45SN1_45S_50ntFlanks@9556.9584.29\|TAGCGGTCCTGACGTGCAAATCGGTCGTC | >hg38_21_+_8433172_8446622_RNA45SN1_45S_50ntFlanks | 9556 | 9584 | 29 | TAGCGGTCCTGACGTGCAAATCGGTCGTC | MB468 | 21 | 4 | 20 | 6 | 5 | 14.1 | 1.2 | 16.3 | 2.8 | 0.6 |
| >hg38_21_+_8433172_8446622_RNA45SN1_45S_50ntFlanks@9559.9584.26\|CGGTCCTGACGTGCAAATCGGTCGTC | >hg38_21_+_8433172_8446622_RNA45SN1_45S_50ntFlanks | 9559 | 9584 | 26 | CGGTCCTGACGTGCAAATCGGTCGTC | MB468 | 14 | 8 | 16 | 6 | 6 | 2.4 | 2.9 | 2.6 | 2.1 | 2.6 |
| >hg38_21_+_8433172_8446622_RNA45SN1_45S_50ntFlanks@9561.9596.36\|GTCCTGACGTGCAAATCGGTCGTCCGACCTGGGTAT | >hg38_21_+_8433172_8446622_RNA45SN1_45S_50ntFlanks | 9561 | 9596 | 36 | GTCCTGACGTGCAAATCGGTCGTCCGACCTGGGTAT | MB468 | 13 | 19 | 17 | 3 | 38 | 3.0 | 11.3 | 14.9 | 3.0 | 22.6 |
| >hg38_21_+_8433172_8446622_RNA45SN1_45S_50ntFlanks@9561.9597.37\|GTCCTGACGTGCAAATCGGTCGTCCGACCTGGGTATA | >hg38_21_+_8433172_8446622_RNA45SN1_45S_50ntFlanks | 9561 | 9597 | 37 | GTCCTGACGTGCAAATCGGTCGTCCGACCTGGGTATA | MB468 | 14 | 22 | 16 | 4 | 32 | 5.1 | 11.5 | 14.4 | 4.2 | 33.0 |
| >hg38_21_+_8433172_8446622_RNA45SN1_45S_50ntFlanks@9562.9584.23\|TCCTGACGTGCAAATCGGTCGTC | >hg38_21_+_8433172_8446622_RNA45SN1_45S_50ntFlanks | 9562 | 9584 | 23 | TCCTGACGTGCAAATCGGTCGTC | MB468 | 10 | 1 | 9 | 6 | 1 | 12.0 | 0.2 | 10.7 | 2.0 | 0.3 |
| >hg38_21_+_8433172_8446622_RNA45SN1_45S_50ntFlanks@9562.9592.31\|TCCTGACGTGCAAATCGGTCGTCCGACCTGG | >hg38_21_+_8433172_8446622_RNA45SN1_45S_50ntFlanks | 9562 | 9592 | 31 | TCCTGACGTGCAAATCGGTCGTCCGACCTGG | MB468 | 15 | 10 | 22 | 5 | 8 | 4.9 | 4.0 | 15.9 | 2.9 | 2.9 |
| >hg38_21_+_8433172_8446622_RNA45SN1_45S_50ntFlanks@9562.9595.34\|TCCTGACGTGCAAATCGGTCGTCCGACCTGGGTA | >hg38_21_+_8433172_8446622_RNA45SN1_45S_50ntFlanks | 9562 | 9595 | 34 | TCCTGACGTGCAAATCGGTCGTCCGACCTGGGTA | MB468 | 22 | 20 | 25 | 6 | 50 | 8.1 | 7.9 | 14.6 | 3.4 | 32.6 |
| >hg38_21_+_8433172_8446622_RNA45SN1_45S_50ntFlanks@9562.9596.35\|TCCTGACGTGCAAATCGGTCGTCCGACCTGGGTAT | >hg38_21_+_8433172_8446622_RNA45SN1_45S_50ntFlanks | 9562 | 9596 | 35 | TCCTGACGTGCAAATCGGTCGTCCGACCTGGGTAT | MB468 | 107 | 109 | 93 | 24 | 274 | 24.0 | 47.7 | 73.1 | 20.6 | 105.7 |
| >hg38_21_+_8433172_8446622_RNA45SN1_45S_50ntFlanks@9562.9597.36\|TCCTGACGTGCAAATCGGTCGTCCGACCTGGGTATA | >hg38_21_+_8433172_8446622_RNA45SN1_45S_50ntFlanks | 9562 | 9597 | 36 | TCCTGACGTGCAAATCGGTCGTCCGACCTGGGTATA | MB468 | 58 | 56 | 55 | 13 | 196 | 10.5 | 29.2 | 42.5 | 12.1 | 106.6 |

| | | | | | | | | | | | | | | | | |
|---|---|---|---|---|---|---|---|---|---|---|---|---|---|---|---|---|
| >hg38_21_+_8433172_8446622_RNA45SN1_45S_50ntFlanks@9563.9592.30\|CCTGACGTGCAAATCGGTCGTCCGACCTGG | >hg38_21_+_8433172_8446622_RNA45SN1_45S_50ntFlanks | 9563 | 9592 | 30 | CCTGACGTGCAAATCGGTCGTCCGACCTGG | MB468 | 11 | 6 | 16 | 6 | 6 | 3.4 | 2.2 | 10.0 | 2.3 | 2.5 |
| >hg38_21_+_8433172_8446622_RNA45SN1_45S_50ntFlanks@9563.9595.33\|CCTGACGTGCAAATCGGTCGTCCGACCTGGGTA | >hg38_21_+_8433172_8446622_RNA45SN1_45S_50ntFlanks | 9563 | 9595 | 33 | CCTGACGTGCAAATCGGTCGTCCGACCTGGGTA | MB468 | 13 | 11 | 13 | 5 | 31 | 6.1 | 3.8 | 7.0 | 1.8 | 15.1 |
| >hg38_21_+_8433172_8446622_RNA45SN1_45S_50ntFlanks@9563.9596.34\|CCTGACGTGCAAATCGGTCGTCCGACCTGGGTAT | >hg38_21_+_8433172_8446622_RNA45SN1_45S_50ntFlanks | 9563 | 9596 | 34 | CCTGACGTGCAAATCGGTCGTCCGACCTGGGTAT | MB468 | 122 | 128 | 100 | 33 | 354 | 21.6 | 38.9 | 73.9 | 26.6 | 146.9 |
| >hg38_21_+_8433172_8446622_RNA45SN1_45S_50ntFlanks@9563.9597.35\|CCTGACGTGCAAATCGGTCGTCCGACCTGGGTATA | >hg38_21_+_8433172_8446622_RNA45SN1_45S_50ntFlanks | 9563 | 9597 | 35 | CCTGACGTGCAAATCGGTCGTCCGACCTGGGTATA | MB468 | 46 | 44 | 44 | 14 | 166 | 8.6 | 13.3 | 37.2 | 12.0 | 61.9 |
| >hg38_21_+_8433172_8446622_RNA45SN1_45S_50ntFlanks@9564.9592.29\|CTGACGTGCAAATCGGTCGTCCGACCTGG | >hg38_21_+_8433172_8446622_RNA45SN1_45S_50ntFlanks | 9564 | 9592 | 29 | CTGACGTGCAAATCGGTCGTCCGACCTGG | MB468 | 136 | 97 | 246 | 60 | 68 | 38.3 | 29.9 | 135.3 | 16.5 | 33.5 |
| >hg38_21_+_8433172_8446622_RNA45SN1_45S_50ntFlanks@9564.9594.31\|CTGACGTGCAAATCGGTCGTCCGACCTGGGT | >hg38_21_+_8433172_8446622_RNA45SN1_45S_50ntFlanks | 9564 | 9594 | 31 | CTGACGTGCAAATCGGTCGTCCGACCTGGGT | MB468 | 45 | 70 | 46 | 20 | 84 | 21.7 | 19.6 | 20.0 | 3.5 | 32.7 |
| >hg38_21_+_8433172_8446622_RNA45SN1_45S_50ntFlanks@9564.9595.32\|CTGACGTGCAAATCGGTCGTCCGACCTGGGTA | >hg38_21_+_8433172_8446622_RNA45SN1_45S_50ntFlanks | 9564 | 9595 | 32 | CTGACGTGCAAATCGGTCGTCCGACCTGGGTA | MB468 | 161 | 219 | 177 | 52 | 312 | 96.6 | 100.2 | 88.4 | 17.7 | 169.9 |
| >hg38_21_+_8433172_8446622_RNA45SN1_45S_50ntFlanks@9564.9596.33\|CTGACGTGCAAATCGGTCGTCCGACCTGGGTAT | >hg38_21_+_8433172_8446622_RNA45SN1_45S_50ntFlanks | 9564 | 9596 | 33 | CTGACGTGCAAATCGGTCGTCCGACCTGGGTAT | MB468 | 1631 | 2400 | 1450 | 390 | 4079 | 435.6 | 785.0 | 634.0 | 239.8 | 1801.1 |
| >hg38_21_+_8433172_8446622_RNA45SN1_45S_50ntFlanks@9564.9597.34\|CTGACGTGCAAATCGGTCGTCCGACCTGGGTATA | >hg38_21_+_8433172_8446622_RNA45SN1_45S_50ntFlanks | 9564 | 9597 | 34 | CTGACGTGCAAATCGGTCGTCCGACCTGGGTATA | MB468 | 600 | 889 | 607 | 172 | 1945 | 61.8 | 257.3 | 326.3 | 109.2 | 794.3 |
| >hg38_21_+_8433172_8446622_RNA45SN1_45S_50ntFlanks@9564.9598.35\|CTGACGTGCAAATCGGTCGTCCGACCTGGGTATAG | >hg38_21_+_8433172_8446622_RNA45SN1_45S_50ntFlanks | 9564 | 9598 | 35 | CTGACGTGCAAATCGGTCGTCCGACCTGGGTATAG | MB468 | 12 | 12 | 7 | 5 | 36 | 1.2 | 7.5 | 3.9 | 3.7 | 15.5 |
| >hg38_21_+_8433172_8446622_RNA45SN1_45S_50ntFlanks@9564.9603.40\|CTGACGTGCAAATCGGTCGTCCGACCTGGGTATAGGGGCG | >hg38_21_+_8433172_8446622_RNA45SN1_45S_50ntFlanks | 9564 | 9603 | 40 | CTGACGTGCAAATCGGTCGTCCGACCTGGGTATAGGGGCG | MB231 | 13 | 8 | 4 | 2 | 16 | 11.2 | 5.6 | 4.0 | 1.7 | 14.5 |
| >hg38_21_+_8433172_8446622_RNA45SN1_45S_50ntFlanks@9564.9603.40\|CTGACGTGCAAATCGGTCGTCCGACCTGGGTATAGGGGCG | >hg38_21_+_8433172_8446622_RNA45SN1_45S_50ntFlanks | 9564 | 9603 | 40 | CTGACGTGCAAATCGGTCGTCCGACCTGGGTATAGGGGCG | MB468 | 13 | 13 | 13 | 7 | 112 | 3.4 | 10.3 | 14.8 | 4.9 | 95.7 |
| >hg38_21_+_8433172_8446622_RNA45SN1_45S_50ntFlanks@9565.9592.28\|TGACGTGCAAATCGGTCGTCCGACCTGG | >hg38_21_+_8433172_8446622_RNA45SN1_45S_50ntFlanks | 9565 | 9592 | 28 | TGACGTGCAAATCGGTCGTCCGACCTGG | MB468 | 58 | 35 | 121 | 24 | 19 | 21.0 | 15.9 | 66.7 | 4.1 | 14.1 |
| >hg38_21_+_8433172_8446622_RNA45SN1_45S_50ntFlanks@9565.9594.30\|TGACGTGCAAATCGGTCGTCCGACCTGGGT | >hg38_21_+_8433172_8446622_RNA45SN1_45S_50ntFlanks | 9565 | 9594 | 30 | TGACGTGCAAATCGGTCGTCCGACCTGGGT | MB468 | 17 | 25 | 20 | 7 | 18 | 10.8 | 11.0 | 10.9 | 2.4 | 10.7 |
| >hg38_21_+_8433172_8446622_RNA45SN1_45S_50ntFlanks@9565.9595.31\|TGACGTGCAAATCGGTCGTCCGACCTGGGTA | >hg38_21_+_8433172_8446622_RNA45SN1_45S_50ntFlanks | 9565 | 9595 | 31 | TGACGTGCAAATCGGTCGTCCGACCTGGGTA | MB468 | 71 | 93 | 90 | 23 | 82 | 52.0 | 50.1 | 53.4 | 10.7 | 60.9 |
| >hg38_21_+_8433172_8446622_RNA45SN1_45S_50ntFlanks@9565.9596.32\|TGACGTGCAAATCGGTCGTCCGACCTGGGTAT | >hg38_21_+_8433172_8446622_RNA45SN1_45S_50ntFlanks | 9565 | 9596 | 32 | TGACGTGCAAATCGGTCGTCCGACCTGGGTAT | MB468 | 611 | 741 | 577 | 164 | 914 | 318.9 | 303.1 | 241.1 | 80.7 | 520.8 |
| >hg38_21_+_8433172_8446622_RNA45SN1_45S_50ntFlanks@9565.9597.33\|TGACGTGCAAATCGGTCGTCCGACCTGGGTATA | >hg38_21_+_8433172_8446622_RNA45SN1_45S_50ntFlanks | 9565 | 9597 | 33 | TGACGTGCAAATCGGTCGTCCGACCTGGGTATA | MB468 | 219 | 270 | 227 | 73 | 445 | 65.4 | 87.2 | 118.4 | 39.2 | 248.4 |
| >hg38_21_+_8433172_8446622_RNA45SN1_45S_50ntFlanks@9566.9592.27\|GACGTGCAAATCGGTCGTCCGACCTGG | >hg38_21_+_8433172_8446622_RNA45SN1_45S_50ntFlanks | 9566 | 9592 | 27 | GACGTGCAAATCGGTCGTCCGACCTGG | MB468 | 20 | 11 | 38 | 10 | 6 | 5.5 | 4.8 | 24.5 | 2.0 | 2.5 |
| >hg38_21_+_8433172_8446622_RNA45SN1_45S_50ntFlanks@9566.9595.30\|GACGTGCAAATCGGTCGTCCGACCTGGGTA | >hg38_21_+_8433172_8446622_RNA45SN1_45S_50ntFlanks | 9566 | 9595 | 30 | GACGTGCAAATCGGTCGTCCGACCTGGGTA | MB468 | 27 | 29 | 34 | 10 | 33 | 17.4 | 11.2 | 18.7 | 0.8 | 18.9 |
| >hg38_21_+_8433172_8446622_RNA45SN1_45S_50ntFlanks@9566.9596.31\|GACGTGCAAATCGGTCGTCCGACCTGGGTAT | >hg38_21_+_8433172_8446622_RNA45SN1_45S_50ntFlanks | 9566 | 9596 | 31 | GACGTGCAAATCGGTCGTCCGACCTGGGTAT | MB468 | 238 | 241 | 235 | 78 | 372 | 87.7 | 77.5 | 130.3 | 41.6 | 134.7 |
| >hg38_21_+_8433172_8446622_RNA45SN1_45S_50ntFlanks@9566.9597.32\|GACGTGCAAATCGGTCGTCCGACCTGGGTATA | >hg38_21_+_8433172_8446622_RNA45SN1_45S_50ntFlanks | 9566 | 9597 | 32 | GACGTGCAAATCGGTCGTCCGACCTGGGTATA | MB468 | 94 | 93 | 102 | 36 | 187 | 19.0 | 22.0 | 72.8 | 20.0 | 67.9 |
| >hg38_21_+_8433172_8446622_RNA45SN1_45S_50ntFlanks@9567.9584.18\|ACGTGCAAATCGGTCGTC | >hg38_21_+_8433172_8446622_RNA45SN1_45S_50ntFlanks | 9567 | 9584 | 18 | ACGTGCAAATCGGTCGTC | MB468 | 40 | 7 | 49 | 38 | 7 | 23.4 | 3.4 | 36.8 | 19.7 | 1.0 |
| >hg38_21_+_8433172_8446622_RNA45SN1_45S_50ntFlanks@9567.9585.19\|ACGTGCAAATCGGTCGTCC | >hg38_21_+_8433172_8446622_RNA45SN1_45S_50ntFlanks | 9567 | 9585 | 19 | ACGTGCAAATCGGTCGTCC | MB468 | 19 | 10 | 29 | 4 | 5 | 10.0 | 7.5 | 22.2 | 1.7 | 1.1 |
| >hg38_21_+_8433172_8446622_RNA45SN1_45S_50ntFlanks@9567.9586.20\|ACGTGCAAATCGGTCGTCCG | >hg38_21_+_8433172_8446622_RNA45SN1_45S_50ntFlanks | 9567 | 9586 | 20 | ACGTGCAAATCGGTCGTCCG | MB468 | 49 | 15 | 52 | 15 | 13 | 19.9 | 9.9 | 27.2 | 3.9 | 2.6 |
| >hg38_21_+_8433172_8446622_RNA45SN1_45S_50ntFlanks@9567.9588.22\|ACGTGCAAATCGGTCGTCCGAC | >hg38_21_+_8433172_8446622_RNA45SN1_45S_50ntFlanks | 9567 | 9588 | 22 | ACGTGCAAATCGGTCGTCCGAC | MB468 | 16 | 2 | 18 | 13 | 4 | 5.6 | 1.0 | 9.6 | 4.0 | 0.5 |
| >hg38_21_+_8433172_8446622_RNA45SN1_45S_50ntFlanks@9567.9592.26\|ACGTGCAAATCGGTCGTCCGACCTGG | >hg38_21_+_8433172_8446622_RNA45SN1_45S_50ntFlanks | 9567 | 9592 | 26 | ACGTGCAAATCGGTCGTCCGACCTGG | MB468 | 93 | 23 | 137 | 51 | 30 | 36.2 | 8.5 | 65.1 | 7.4 | 8.2 |
| >hg38_21_+_8433172_8446622_RNA45SN1_45S_50ntFlanks@9567.9594.28\|ACGTGCAAATCGGTCGTCCGACCTGGGT | >hg38_21_+_8433172_8446622_RNA45SN1_45S_50ntFlanks | 9567 | 9594 | 28 | ACGTGCAAATCGGTCGTCCGACCTGGGT | MB468 | 39 | 24 | 52 | 39 | 55 | 15.5 | 9.1 | 16.3 | 14.5 | 30.0 |
| >hg38_21_+_8433172_8446622_RNA45SN1_45S_50ntFlanks@9567.9595.29\|ACGTGCAAATCGGTCGTCCGACCTGGGTA | >hg38_21_+_8433172_8446622_RNA45SN1_45S_50ntFlanks | 9567 | 9595 | 29 | ACGTGCAAATCGGTCGTCCGACCTGGGTA | MB468 | 208 | 120 | 315 | 111 | 249 | 111.0 | 59.6 | 136.1 | 50.5 | 169.9 |
| >hg38_21_+_8433172_8446622_RNA45SN1_45S_50ntFlanks@9567.9596.30\|ACGTGCAAATCGGTCGTCCGACCTGGGTAT | >hg38_21_+_8433172_8446622_RNA45SN1_45S_50ntFlanks | 9567 | 9596 | 30 | ACGTGCAAATCGGTCGTCCGACCTGGGTAT | MB468 | 541 | 265 | 472 | 294 | 783 | 101.0 | 70.4 | 128.5 | 131.4 | 335.5 |
| >hg38_21_+_8433172_8446622_RNA45SN1_45S_50ntFlanks@9567.9597.31\|ACGTGCAAATCGGTCGTCCGACCTGGGTATA | >hg38_21_+_8433172_8446622_RNA45SN1_45S_50ntFlanks | 9567 | 9597 | 31 | ACGTGCAAATCGGTCGTCCGACCTGGGTATA | MB468 | 424 | 193 | 355 | 219 | 759 | 96.1 | 66.3 | 137.5 | 104.1 | 381.4 |
| >hg38_21_+_8433172_8446622_RNA45SN1_45S_50ntFlanks@9567.9601.35\|ACGTGCAAATCGGTCGTCCGACCTGGGTATAGGGG | >hg38_21_+_8433172_8446622_RNA45SN1_45S_50ntFlanks | 9567 | 9601 | 35 | ACGTGCAAATCGGTCGTCCGACCTGGGTATAGGGG | MB468 | 11 | 9 | 7 | 12 | 25 | 3.8 | 2.7 | 4.2 | 8.8 | 10.9 |
| >hg38_21_+_8433172_8446622_RNA45SN1_45S_50ntFlanks@9567.9603.37\|ACGTGCAAATCGGTCGTCCGACCTGGGTATAGGGGCG | >hg38_21_+_8433172_8446622_RNA45SN1_45S_50ntFlanks | 9567 | 9603 | 37 | ACGTGCAAATCGGTCGTCCGACCTGGGTATAGGGGCG | MB468 | 27 | 28 | 25 | 23 | 108 | 7.1 | 9.7 | 14.1 | 18.8 | 75.4 |
| >hg38_21_+_8433172_8446622_RNA45SN1_45S_50ntFlanks@9568.9585.18\|CGTGCAAATCGGTCGTCC | >hg38_21_+_8433172_8446622_RNA45SN1_45S_50ntFlanks | 9568 | 9585 | 18 | CGTGCAAATCGGTCGTCC | MB468 | 11 | 6 | 20 | 3 | 3 | 5.2 | 4.1 | 16.1 | 1.3 | 0.7 |
| >hg38_21_+_8433172_8446622_RNA45SN1_45S_50ntFlanks@9568.9586.19\|CGTGCAAATCGGTCGTCCG | >hg38_21_+_8433172_8446622_RNA45SN1_45S_50ntFlanks | 9568 | 9586 | 19 | CGTGCAAATCGGTCGTCCG | MB468 | 28 | 8 | 33 | 9 | 9 | 10.4 | 5.2 | 17.0 | 3.3 | 2.4 |
| >hg38_21_+_8433172_8446622_RNA45SN1_45S_50ntFlanks@9568.9592.25\|CGTGCAAATCGGTCGTCCGACCTGG | >hg38_21_+_8433172_8446622_RNA45SN1_45S_50ntFlanks | 9568 | 9592 | 25 | CGTGCAAATCGGTCGTCCGACCTGG | MB468 | 76 | 15 | 92 | 51 | 29 | 28.5 | 4.3 | 25.3 | 10.9 | 6.4 |
| >hg38_21_+_8433172_8446622_RNA45SN1_45S_50ntFlanks@9568.9594.27\|CGTGCAAATCGGTCGTCCGACCTGGGT | >hg38_21_+_8433172_8446622_RNA45SN1_45S_50ntFlanks | 9568 | 9594 | 27 | CGTGCAAATCGGTCGTCCGACCTGGGT | MB468 | 41 | 23 | 44 | 44 | 54 | 16.2 | 6.6 | 17.9 | 13.5 | 24.3 |
| >hg38_21_+_8433172_8446622_RNA45SN1_45S_50ntFlanks@9568.9595.28\|CGTGCAAATCGGTCGTCCGACCTGGGTA | >hg38_21_+_8433172_8446622_RNA45SN1_45S_50ntFlanks | 9568 | 9595 | 28 | CGTGCAAATCGGTCGTCCGACCTGGGTA | MB468 | 206 | 104 | 284 | 113 | 221 | 97.4 | 47.7 | 124.4 | 49.2 | 138.4 |
| >hg38_21_+_8433172_8446622_RNA45SN1_45S_50ntFlanks@9568.9596.29\|CGTGCAAATCGGTCGTCCGACCTGGGTAT | >hg38_21_+_8433172_8446622_RNA45SN1_45S_50ntFlanks | 9568 | 9596 | 29 | CGTGCAAATCGGTCGTCCGACCTGGGTAT | MB468 | 468 | 203 | 357 | 271 | 674 | 58.5 | 43.9 | 86.9 | 89.4 | 228.3 |
| >hg38_21_+_8433172_8446622_RNA45SN1_45S_50ntFlanks@9568.9597.30\|CGTGCAAATCGGTCGTCCGACCTGGGTATA | >hg38_21_+_8433172_8446622_RNA45SN1_45S_50ntFlanks | 9568 | 9597 | 30 | CGTGCAAATCGGTCGTCCGACCTGGGTATA | MB468 | 437 | 184 | 291 | 226 | 803 | 94.4 | 46.8 | 97.7 | 78.0 | 238.5 |
| >hg38_21_+_8433172_8446622_RNA45SN1_45S_50ntFlanks@9568.9603.36\|CGTGCAAATCGGTCGTCCGACCTGGGTATAGGGGCG | >hg38_21_+_8433172_8446622_RNA45SN1_45S_50ntFlanks | 9568 | 9603 | 36 | CGTGCAAATCGGTCGTCCGACCTGGGTATAGGGGCG | MB468 | 38 | 45 | 28 | 40 | 143 | 8.5 | 9.2 | 9.9 | 36.6 | 81.6 |
| >hg38_21_+_8433172_8446622_RNA45SN1_45S_50ntFlanks@9569.9586.18\|GTGCAAATCGGTCGTCCG | >hg38_21_+_8433172_8446622_RNA45SN1_45S_50ntFlanks | 9569 | 9586 | 18 | GTGCAAATCGGTCGTCCG | MB468 | 12 | 5 | 16 | 4 | 2 | 5.3 | 2.6 | 10.9 | 1.4 | 0.4 |
| >hg38_21_+_8433172_8446622_RNA45SN1_45S_50ntFlanks@9569.9592.24\|GTGCAAATCGGTCGTCCGACCTGG | >hg38_21_+_8433172_8446622_RNA45SN1_45S_50ntFlanks | 9569 | 9592 | 24 | GTGCAAATCGGTCGTCCGACCTGG | MB468 | 17 | 3 | 25 | 11 | 3 | 6.9 | 1.2 | 9.4 | 2.8 | 0.8 |
| >hg38_21_+_8433172_8446622_RNA45SN1_45S_50ntFlanks@9569.9595.27\|GTGCAAATCGGTCGTCCGACCTGGGTA | >hg38_21_+_8433172_8446622_RNA45SN1_45S_50ntFlanks | 9569 | 9595 | 27 | GTGCAAATCGGTCGTCCGACCTGGGTA | MB468 | 31 | 13 | 30 | 19 | 22 | 6.4 | 3.5 | 6.6 | 9.4 | 13.4 |
| >hg38_21_+_8433172_8446622_RNA45SN1_45S_50ntFlanks@9569.9596.28\|GTGCAAATCGGTCGTCCGACCTGGGTAT | >hg38_21_+_8433172_8446622_RNA45SN1_45S_50ntFlanks | 9569 | 9596 | 28 | GTGCAAATCGGTCGTCCGACCTGGGTAT | MB468 | 53 | 15 | 29 | 38 | 49 | 15.8 | 3.1 | 14.9 | 20.2 | 15.3 |
| >hg38_21_+_8433172_8446622_RNA45SN1_45S_50ntFlanks@9569.9597.29\|GTGCAAATCGGTCGTCCGACCTGGGTATA | >hg38_21_+_8433172_8446622_RNA45SN1_45S_50ntFlanks | 9569 | 9597 | 29 | GTGCAAATCGGTCGTCCGACCTGGGTATA | MB468 | 99 | 31 | 49 | 57 | 111 | 30.4 | 9.0 | 30.3 | 16.9 | 28.8 |
| >hg38_21_+_8433172_8446622_RNA45SN1_45S_50ntFlanks@9569.9603.35\|GTGCAAATCGGTCGTCCGACCTGGGTATAGGGGCG | >hg38_21_+_8433172_8446622_RNA45SN1_45S_50ntFlanks | 9569 | 9603 | 35 | GTGCAAATCGGTCGTCCGACCTGGGTATAGGGGCG | MB231 | 23 | 15 | 23 | 6 | 34 | 21.9 | 11.9 | 20.2 | 4.3 | 31.4 |
| >hg38_21_+_8433172_8446622_RNA45SN1_45S_50ntFlanks@9570.9592.23\|TGCAAATCGGTCGTCCGACCTGG | >hg38_21_+_8433172_8446622_RNA45SN1_45S_50ntFlanks | 9570 | 9592 | 23 | TGCAAATCGGTCGTCCGACCTGG | MB468 | 15 | 1 | 17 | 23 | 2 | 3.5 | 0.4 | 5.7 | 17.4 | 1.1 |
| >hg38_21_+_8433172_8446622_RNA45SN1_45S_50ntFlanks@9570.9594.25\|TGCAAATCGGTCGTCCGACCTGGGT | >hg38_21_+_8433172_8446622_RNA45SN1_45S_50ntFlanks | 9570 | 9594 | 25 | TGCAAATCGGTCGTCCGACCTGGGT | MB468 | 39 | 9 | 32 | 59 | 39 | 13.1 | 4.6 | 13.0 | 39.1 | 32.5 |
| >hg38_21_+_8433172_8446622_RNA45SN1_45S_50ntFlanks@9570.9595.26\|TGCAAATCGGTCGTCCGACCTGGGTA | >hg38_21_+_8433172_8446622_RNA45SN1_45S_50ntFlanks | 9570 | 9595 | 26 | TGCAAATCGGTCGTCCGACCTGGGTA | MB468 | 253 | 61 | 229 | 169 | 201 | 72.8 | 26.5 | 62.4 | 126.6 | 144.1 |
| >hg38_21_+_8433172_8446622_RNA45SN1_45S_50ntFlanks@9570.9596.27\|TGCAAATCGGTCGTCCGACCTGGGTAT | >hg38_21_+_8433172_8446622_RNA45SN1_45S_50ntFlanks | 9570 | 9596 | 27 | TGCAAATCGGTCGTCCGACCTGGGTAT | MB468 | 333 | 58 | 184 | 246 | 369 | 77.6 | 21.1 | 60.3 | 162.4 | 298.4 |

| | | | | | | | | | | | | | | | | |
|---|---|---|---|---|---|---|---|---|---|---|---|---|---|---|---|---|
| >hg38_21_+_8433172_8446622_RNA45SN1_45S_50ntFlanks@9570.9597.28\|TGCAAATCGGTCGTCCGACCTGGGTATA | >hg38_21_+_8433172_8446622_RNA45SN1_45S_50ntFlanks | 9570 | 9597 | 28 | TGCAAATCGGTCGTCCGACCTGGGTATA | MB468 | 324 | 45 | 142 | 176 | 403 | 99.5 | 15.5 | 76.2 | 114.8 | 321.0 |
| >hg38_21_+_8433172_8446622_RNA45SN1_45S_50ntFlanks@9570.9603.34\|TGCAAATCGGTCGTCCGACCTGGGTATAGGGGCG | >hg38_21_+_8433172_8446622_RNA45SN1_45S_50ntFlanks | 9570 | 9603 | 34 | TGCAAATCGGTCGTCCGACCTGGGTATAGGGGCG | MB468 | 33 | 25 | 20 | 37 | 88 | 7.1 | 10.6 | 2.8 | 29.4 | 50.7 |
| >hg38_21_+_8433172_8446622_RNA45SN1_45S_50ntFlanks@9571.9595.25\|GCAAATCGGTCGTCCGACCTGGGTA | >hg38_21_+_8433172_8446622_RNA45SN1_45S_50ntFlanks | 9571 | 9595 | 25 | GCAAATCGGTCGTCCGACCTGGGTA | MB468 | 16 | 4 | 18 | 14 | 15 | 6.2 | 1.5 | 6.7 | 9.4 | 11.4 |
| >hg38_21_+_8433172_8446622_RNA45SN1_45S_50ntFlanks@9571.9596.26\|GCAAATCGGTCGTCCGACCTGGGTAT | >hg38_21_+_8433172_8446622_RNA45SN1_45S_50ntFlanks | 9571 | 9596 | 26 | GCAAATCGGTCGTCCGACCTGGGTAT | MB468 | 29 | 7 | 16 | 27 | 36 | 6.1 | 2.2 | 5.0 | 16.6 | 21.0 |
| >hg38_21_+_8433172_8446622_RNA45SN1_45S_50ntFlanks@9571.9597.27\|GCAAATCGGTCGTCCGACCTGGGTATA | >hg38_21_+_8433172_8446622_RNA45SN1_45S_50ntFlanks | 9571 | 9597 | 27 | GCAAATCGGTCGTCCGACCTGGGTATA | MB468 | 29 | 6 | 15 | 20 | 37 | 8.1 | 1.5 | 6.5 | 12.2 | 26.0 |
| >hg38_21_+_8433172_8446622_RNA45SN1_45S_50ntFlanks@9572.9592.21\|CAAATCGGTCGTCCGACCTGG | >hg38_21_+_8433172_8446622_RNA45SN1_45S_50ntFlanks | 9572 | 9592 | 21 | CAAATCGGTCGTCCGACCTGG | MB468 | 22 | 1 | 26 | 82 | 5 | 5.5 | 0.5 | 10.4 | 70.1 | 2.1 |
| >hg38_21_+_8433172_8446622_RNA45SN1_45S_50ntFlanks@9572.9594.23\|CAAATCGGTCGTCCGACCTGGGT | >hg38_21_+_8433172_8446622_RNA45SN1_45S_50ntFlanks | 9572 | 9594 | 23 | CAAATCGGTCGTCCGACCTGGGT | MB468 | 75 | 10 | 61 | 274 | 96 | 27.9 | 5.0 | 30.1 | 174.1 | 82.1 |
| >hg38_21_+_8433172_8446622_RNA45SN1_45S_50ntFlanks@9572.9595.24\|CAAATCGGTCGTCCGACCTGGGTA | >hg38_21_+_8433172_8446622_RNA45SN1_45S_50ntFlanks | 9572 | 9595 | 24 | CAAATCGGTCGTCCGACCTGGGTA | MB468 | 393 | 40 | 414 | 575 | 375 | 124.0 | 29.7 | 165.6 | 484.4 | 354.0 |
| >hg38_21_+_8433172_8446622_RNA45SN1_45S_50ntFlanks@9572.9596.25\|CAAATCGGTCGTCCGACCTGGGTAT | >hg38_21_+_8433172_8446622_RNA45SN1_45S_50ntFlanks | 9572 | 9596 | 25 | CAAATCGGTCGTCCGACCTGGGTAT | MB468 | 896 | 89 | 446 | 1265 | 1137 | 220.5 | 38.4 | 166.5 | 828.2 | 949.1 |
| >hg38_21_+_8433172_8446622_RNA45SN1_45S_50ntFlanks@9572.9597.26\|CAAATCGGTCGTCCGACCTGGGTATA | >hg38_21_+_8433172_8446622_RNA45SN1_45S_50ntFlanks | 9572 | 9597 | 26 | CAAATCGGTCGTCCGACCTGGGTATA | MB468 | 923 | 82 | 398 | 845 | 1321 | 285.1 | 29.4 | 207.4 | 512.7 | 1042.0 |
| >hg38_21_+_8433172_8446622_RNA45SN1_45S_50ntFlanks@9572.9598.27\|CAAATCGGTCGTCCGACCTGGGTATAG | >hg38_21_+_8433172_8446622_RNA45SN1_45S_50ntFlanks | 9572 | 9598 | 27 | CAAATCGGTCGTCCGACCTGGGTATAG | MB468 | 20 | 3 | 6 | 17 | 28 | 5.4 | 1.0 | 3.1 | 5.2 | 10.5 |
| >hg38_21_+_8433172_8446622_RNA45SN1_45S_50ntFlanks@9572.9601.30\|CAAATCGGTCGTCCGACCTGGGTATAGGGG | >hg38_21_+_8433172_8446622_RNA45SN1_45S_50ntFlanks | 9572 | 9601 | 30 | CAAATCGGTCGTCCGACCTGGGTATAGGGG | MB468 | 36 | 14 | 13 | 52 | 63 | 7.8 | 3.4 | 5.5 | 24.1 | 25.0 |
| >hg38_21_+_8433172_8446622_RNA45SN1_45S_50ntFlanks@9572.9602.31\|CAAATCGGTCGTCCGACCTGGGTATAGGGGC | >hg38_21_+_8433172_8446622_RNA45SN1_45S_50ntFlanks | 9572 | 9602 | 31 | CAAATCGGTCGTCCGACCTGGGTATAGGGGC | MB468 | 14 | 10 | 8 | 30 | 31 | 4.3 | 4.7 | 2.7 | 25.9 | 17.3 |
| >hg38_21_+_8433172_8446622_RNA45SN1_45S_50ntFlanks@9572.9603.32\|CAAATCGGTCGTCCGACCTGGGTATAGGGGCG | >hg38_21_+_8433172_8446622_RNA45SN1_45S_50ntFlanks | 9572 | 9603 | 32 | CAAATCGGTCGTCCGACCTGGGTATAGGGGCG | MB468 | 198 | 128 | 105 | 357 | 477 | 73.5 | 34.4 | 32.5 | 351.1 | 249.8 |
| >hg38_21_+_8433172_8446622_RNA45SN1_45S_50ntFlanks@9572.9605.34\|CAAATCGGTCGTCCGACCTGGGTATAGGGGCGAA | >hg38_21_+_8433172_8446622_RNA45SN1_45S_50ntFlanks | 9572 | 9605 | 34 | CAAATCGGTCGTCCGACCTGGGTATAGGGGCGAA | MB468 | 36 | 23 | 16 | 46 | 85 | 14.1 | 4.7 | 7.9 | 36.3 | 42.7 |
| >hg38_21_+_8433172_8446622_RNA45SN1_45S_50ntFlanks@9573.9594.22\|AAATCGGTCGTCCGACCTGGGT | >hg38_21_+_8433172_8446622_RNA45SN1_45S_50ntFlanks | 9573 | 9594 | 22 | AAATCGGTCGTCCGACCTGGGT | MB468 | 31 | 4 | 21 | 131 | 34 | 11.0 | 1.8 | 9.6 | 91.3 | 29.7 |
| >hg38_21_+_8433172_8446622_RNA45SN1_45S_50ntFlanks@9573.9595.23\|AAATCGGTCGTCCGACCTGGGTA | >hg38_21_+_8433172_8446622_RNA45SN1_45S_50ntFlanks | 9573 | 9595 | 23 | AAATCGGTCGTCCGACCTGGGTA | MB468 | 159 | 18 | 146 | 251 | 135 | 47.2 | 11.6 | 54.5 | 216.0 | 129.0 |
| >hg38_21_+_8433172_8446622_RNA45SN1_45S_50ntFlanks@9573.9596.24\|AAATCGGTCGTCCGACCTGGGTAT | >hg38_21_+_8433172_8446622_RNA45SN1_45S_50ntFlanks | 9573 | 9596 | 24 | AAATCGGTCGTCCGACCTGGGTAT | MB468 | 335 | 37 | 151 | 536 | 399 | 83.2 | 18.1 | 59.4 | 390.8 | 347.4 |
| >hg38_21_+_8433172_8446622_RNA45SN1_45S_50ntFlanks@9573.9597.25\|AAATCGGTCGTCCGACCTGGGTATA | >hg38_21_+_8433172_8446622_RNA45SN1_45S_50ntFlanks | 9573 | 9597 | 25 | AAATCGGTCGTCCGACCTGGGTATA | MB468 | 364 | 36 | 147 | 347 | 492 | 102.9 | 12.8 | 86.6 | 208.8 | 404.1 |
| >hg38_21_+_8433172_8446622_RNA45SN1_45S_50ntFlanks@9573.9601.29\|AAATCGGTCGTCCGACCTGGGTATAGGGG | >hg38_21_+_8433172_8446622_RNA45SN1_45S_50ntFlanks | 9573 | 9601 | 29 | AAATCGGTCGTCCGACCTGGGTATAGGGG | MB468 | 14 | 6 | 5 | 22 | 23 | 2.9 | 1.1 | 1.7 | 5.7 | 9.6 |
| >hg38_21_+_8433172_8446622_RNA45SN1_45S_50ntFlanks@9573.9603.31\|AAATCGGTCGTCCGACCTGGGTATAGGGGCG | >hg38_21_+_8433172_8446622_RNA45SN1_45S_50ntFlanks | 9573 | 9603 | 31 | AAATCGGTCGTCCGACCTGGGTATAGGGGCG | MB231 | 21 | 17 | 15 | 7 | 25 | 18.0 | 15.6 | 14.2 | 6.1 | 27.3 |
| >hg38_21_+_8433172_8446622_RNA45SN1_45S_50ntFlanks@9573.9603.31\|AAATCGGTCGTCCGACCTGGGTATAGGGGCG | >hg38_21_+_8433172_8446622_RNA45SN1_45S_50ntFlanks | 9573 | 9603 | 31 | AAATCGGTCGTCCGACCTGGGTATAGGGGCG | MB468 | 83 | 55 | 37 | 126 | 167 | 37.8 | 15.3 | 11.9 | 112.5 | 86.3 |
| >hg38_21_+_8433172_8446622_RNA45SN1_45S_50ntFlanks@9573.9605.33\|AAATCGGTCGTCCGACCTGGGTATAGGGGCGAA | >hg38_21_+_8433172_8446622_RNA45SN1_45S_50ntFlanks | 9573 | 9605 | 33 | AAATCGGTCGTCCGACCTGGGTATAGGGGCGAA | MB468 | 17 | 11 | 6 | 22 | 33 | 5.7 | 2.4 | 2.7 | 16.5 | 16.8 |
| >hg38_21_+_8433172_8446622_RNA45SN1_45S_50ntFlanks@9573.9610.38\|AAATCGGTCGTCCGACCTGGGTATAGGGGCGAAAGACT | >hg38_21_+_8433172_8446622_RNA45SN1_45S_50ntFlanks | 9573 | 9610 | 38 | AAATCGGTCGTCCGACCTGGGTATAGGGGCGAAAGACT | MB231 | 12 | 6 | 2 | 5 | 15 | 11.1 | 3.4 | 2.1 | 2.2 | 7.2 |
| >hg38_21_+_8433172_8446622_RNA45SN1_45S_50ntFlanks@9574.9592.19\|AATCGGTCGTCCGACCTGG | >hg38_21_+_8433172_8446622_RNA45SN1_45S_50ntFlanks | 9574 | 9592 | 19 | AATCGGTCGTCCGACCTGG | MB468 | 15 | 0 | 15 | 75 | 2 | 6.1 | 0.1 | 4.6 | 67.6 | 1.0 |
| >hg38_21_+_8433172_8446622_RNA45SN1_45S_50ntFlanks@9574.9594.21\|AATCGGTCGTCCGACCTGGGT | >hg38_21_+_8433172_8446622_RNA45SN1_45S_50ntFlanks | 9574 | 9594 | 21 | AATCGGTCGTCCGACCTGGGT | MB468 | 48 | 6 | 35 | 278 | 50 | 10.9 | 3.3 | 12.2 | 218.2 | 40.1 |
| >hg38_21_+_8433172_8446622_RNA45SN1_45S_50ntFlanks@9574.9595.22\|AATCGGTCGTCCGACCTGGGTA | >hg38_21_+_8433172_8446622_RNA45SN1_45S_50ntFlanks | 9574 | 9595 | 22 | AATCGGTCGTCCGACCTGGGTA | MB468 | 265 | 27 | 256 | 567 | 209 | 48.8 | 19.7 | 52.7 | 522.5 | 190.6 |
| >hg38_21_+_8433172_8446622_RNA45SN1_45S_50ntFlanks@9574.9596.23\|AATCGGTCGTCCGACCTGGGTAT | >hg38_21_+_8433172_8446622_RNA45SN1_45S_50ntFlanks | 9574 | 9596 | 23 | AATCGGTCGTCCGACCTGGGTAT | MB468 | 587 | 62 | 297 | 1219 | 646 | 134.7 | 22.3 | 117.8 | 944.1 | 498.3 |
| >hg38_21_+_8433172_8446622_RNA45SN1_45S_50ntFlanks@9574.9597.24\|AATCGGTCGTCCGACCTGGGTATA | >hg38_21_+_8433172_8446622_RNA45SN1_45S_50ntFlanks | 9574 | 9597 | 24 | AATCGGTCGTCCGACCTGGGTATA | MB468 | 655 | 60 | 294 | 766 | 842 | 198.7 | 18.5 | 181.2 | 505.3 | 623.7 |
| >hg38_21_+_8433172_8446622_RNA45SN1_45S_50ntFlanks@9574.9598.25\|AATCGGTCGTCCGACCTGGGTATAG | >hg38_21_+_8433172_8446622_RNA45SN1_45S_50ntFlanks | 9574 | 9598 | 25 | AATCGGTCGTCCGACCTGGGTATAG | MB468 | 16 | 3 | 5 | 17 | 20 | 3.0 | 1.0 | 2.2 | 5.6 | 6.7 |
| >hg38_21_+_8433172_8446622_RNA45SN1_45S_50ntFlanks@9574.9601.28\|AATCGGTCGTCCGACCTGGGTATAGGGG | >hg38_21_+_8433172_8446622_RNA45SN1_45S_50ntFlanks | 9574 | 9601 | 28 | AATCGGTCGTCCGACCTGGGTATAGGGG | MB468 | 27 | 12 | 9 | 51 | 44 | 8.4 | 4.1 | 4.6 | 10.8 | 10.7 |
| >hg38_21_+_8433172_8446622_RNA45SN1_45S_50ntFlanks@9574.9602.29\|AATCGGTCGTCCGACCTGGGTATAGGGGC | >hg38_21_+_8433172_8446622_RNA45SN1_45S_50ntFlanks | 9574 | 9602 | 29 | AATCGGTCGTCCGACCTGGGTATAGGGGC | BT20 | 10 | 11 | 7 | 47 | 20 | 7.2 | 8.5 | 4.7 | 43.2 | 8.4 |
| >hg38_21_+_8433172_8446622_RNA45SN1_45S_50ntFlanks@9574.9602.29\|AATCGGTCGTCCGACCTGGGTATAGGGGC | >hg38_21_+_8433172_8446622_RNA45SN1_45S_50ntFlanks | 9574 | 9602 | 29 | AATCGGTCGTCCGACCTGGGTATAGGGGC | MB231 | 36 | 33 | 24 | 25 | 48 | 31.6 | 33.5 | 21.6 | 21.6 | 47.2 |
| >hg38_21_+_8433172_8446622_RNA45SN1_45S_50ntFlanks@9574.9602.29\|AATCGGTCGTCCGACCTGGGTATAGGGGC | >hg38_21_+_8433172_8446622_RNA45SN1_45S_50ntFlanks | 9574 | 9602 | 29 | AATCGGTCGTCCGACCTGGGTATAGGGGC | MB468 | 12 | 9 | 6 | 28 | 21 | 3.3 | 3.8 | 2.1 | 18.4 | 8.8 |
| >hg38_21_+_8433172_8446622_RNA45SN1_45S_50ntFlanks@9574.9603.30\|AATCGGTCGTCCGACCTGGGTATAGGGGCG | >hg38_21_+_8433172_8446622_RNA45SN1_45S_50ntFlanks | 9574 | 9603 | 30 | AATCGGTCGTCCGACCTGGGTATAGGGGCG | BT20 | 21 | 19 | 15 | 95 | 43 | 14.2 | 17.5 | 9.3 | 67.4 | 22.4 |
| >hg38_21_+_8433172_8446622_RNA45SN1_45S_50ntFlanks@9574.9603.30\|AATCGGTCGTCCGACCTGGGTATAGGGGCG | >hg38_21_+_8433172_8446622_RNA45SN1_45S_50ntFlanks | 9574 | 9603 | 30 | AATCGGTCGTCCGACCTGGGTATAGGGGCG | MB231 | 82 | 62 | 49 | 38 | 96 | 69.7 | 61.1 | 44.6 | 21.9 | 99.5 |
| >hg38_21_+_8433172_8446622_RNA45SN1_45S_50ntFlanks@9574.9603.30\|AATCGGTCGTCCGACCTGGGTATAGGGGCG | >hg38_21_+_8433172_8446622_RNA45SN1_45S_50ntFlanks | 9574 | 9603 | 30 | AATCGGTCGTCCGACCTGGGTATAGGGGCG | MB468 | 151 | 93 | 65 | 308 | 274 | 39.1 | 26.5 | 14.5 | 266.7 | 110.9 |
| >hg38_21_+_8433172_8446622_RNA45SN1_45S_50ntFlanks@9574.9605.32\|AATCGGTCGTCCGACCTGGGTATAGGGGCGAA | >hg38_21_+_8433172_8446622_RNA45SN1_45S_50ntFlanks | 9574 | 9605 | 32 | AATCGGTCGTCCGACCTGGGTATAGGGGCGAA | MB231 | 14 | 14 | 5 | 15 | 20 | 10.5 | 11.8 | 3.6 | 6.3 | 18.8 |
| >hg38_21_+_8433172_8446622_RNA45SN1_45S_50ntFlanks@9574.9605.32\|AATCGGTCGTCCGACCTGGGTATAGGGGCGAA | >hg38_21_+_8433172_8446622_RNA45SN1_45S_50ntFlanks | 9574 | 9605 | 32 | AATCGGTCGTCCGACCTGGGTATAGGGGCGAA | MB468 | 38 | 21 | 12 | 56 | 58 | 12.3 | 8.2 | 6.6 | 37.5 | 31.9 |
| >hg38_21_+_8433172_8446622_RNA45SN1_45S_50ntFlanks@9574.9609.36\|AATCGGTCGTCCGACCTGGGTATAGGGGCGAAAGAC | >hg38_21_+_8433172_8446622_RNA45SN1_45S_50ntFlanks | 9574 | 9609 | 36 | AATCGGTCGTCCGACCTGGGTATAGGGGCGAAAGAC | MB231 | 14 | 8 | 4 | 6 | 22 | 12.7 | 8.8 | 5.0 | 4.4 | 15.7 |
| >hg38_21_+_8433172_8446622_RNA45SN1_45S_50ntFlanks@9574.9610.37\|AATCGGTCGTCCGACCTGGGTATAGGGGCGAAAGACT | >hg38_21_+_8433172_8446622_RNA45SN1_45S_50ntFlanks | 9574 | 9610 | 37 | AATCGGTCGTCCGACCTGGGTATAGGGGCGAAAGACT | BT20 | 15 | 8 | 7 | 5 | 18 | 3.5 | 3.3 | 2.0 | 2.1 | 11.8 |
| >hg38_21_+_8433172_8446622_RNA45SN1_45S_50ntFlanks@9574.9610.37\|AATCGGTCGTCCGACCTGGGTATAGGGGCGAAAGACT | >hg38_21_+_8433172_8446622_RNA45SN1_45S_50ntFlanks | 9574 | 9610 | 37 | AATCGGTCGTCCGACCTGGGTATAGGGGCGAAAGACT | MB231 | 44 | 18 | 6 | 18 | 57 | 39.9 | 19.0 | 5.1 | 12.6 | 36.3 |
| >hg38_21_+_8433172_8446622_RNA45SN1_45S_50ntFlanks@9575.9594.20\|ATCGGTCGTCCGACCTGGGT | >hg38_21_+_8433172_8446622_RNA45SN1_45S_50ntFlanks | 9575 | 9594 | 20 | ATCGGTCGTCCGACCTGGGT | MB468 | 18 | 2 | 12 | 113 | 19 | 3.9 | 1.2 | 3.9 | 88.4 | 13.3 |
| >hg38_21_+_8433172_8446622_RNA45SN1_45S_50ntFlanks@9575.9595.21\|ATCGGTCGTCCGACCTGGGTA | >hg38_21_+_8433172_8446622_RNA45SN1_45S_50ntFlanks | 9575 | 9595 | 21 | ATCGGTCGTCCGACCTGGGTA | MB468 | 90 | 7 | 85 | 240 | 66 | 9.0 | 6.4 | 16.9 | 220.6 | 54.2 |
| >hg38_21_+_8433172_8446622_RNA45SN1_45S_50ntFlanks@9575.9596.22\|ATCGGTCGTCCGACCTGGGTAT | >hg38_21_+_8433172_8446622_RNA45SN1_45S_50ntFlanks | 9575 | 9596 | 22 | ATCGGTCGTCCGACCTGGGTAT | MB468 | 230 | 19 | 114 | 546 | 239 | 62.9 | 8.3 | 44.7 | 417.6 | 163.0 |
| >hg38_21_+_8433172_8446622_RNA45SN1_45S_50ntFlanks@9575.9597.23\|ATCGGTCGTCCGACCTGGGTATA | >hg38_21_+_8433172_8446622_RNA45SN1_45S_50ntFlanks | 9575 | 9597 | 23 | ATCGGTCGTCCGACCTGGGTATA | MB468 | 266 | 20 | 120 | 317 | 322 | 98.8 | 8.4 | 74.7 | 205.1 | 223.5 |
| >hg38_21_+_8433172_8446622_RNA45SN1_45S_50ntFlanks@9575.9601.27\|ATCGGTCGTCCGACCTGGGTATAGGGG | >hg38_21_+_8433172_8446622_RNA45SN1_45S_50ntFlanks | 9575 | 9601 | 27 | ATCGGTCGTCCGACCTGGGTATAGGGG | MB468 | 11 | 5 | 4 | 23 | 18 | 4.8 | 1.7 | 2.3 | 9.2 | 1.8 |
| >hg38_21_+_8433172_8446622_RNA45SN1_45S_50ntFlanks@9575.9603.29\|ATCGGTCGTCCGACCTGGGTATAGGGGCG | >hg38_21_+_8433172_8446622_RNA45SN1_45S_50ntFlanks | 9575 | 9603 | 29 | ATCGGTCGTCCGACCTGGGTATAGGGGCG | MB231 | 13 | 6 | 7 | 3 | 7 | 22.0 | 10.5 | 12.1 | 5.1 | 12.5 |
| >hg38_21_+_8433172_8446622_RNA45SN1_45S_50ntFlanks@9575.9603.29\|ATCGGTCGTCCGACCTGGGTATAGGGGCG | >hg38_21_+_8433172_8446622_RNA45SN1_45S_50ntFlanks | 9575 | 9603 | 29 | ATCGGTCGTCCGACCTGGGTATAGGGGCG | MB468 | 63 | 38 | 29 | 159 | 113 | 6.3 | 15.1 | 5.9 | 156.5 | 46.1 |
| >hg38_21_+_8433172_8446622_RNA45SN1_45S_50ntFlanks@9575.9605.31\|ATCGGTCGTCCGACCTGGGTATAGGGGCGAA | >hg38_21_+_8433172_8446622_RNA45SN1_45S_50ntFlanks | 9575 | 9605 | 31 | ATCGGTCGTCCGACCTGGGTATAGGGGCGAA | MB468 | 19 | 9 | 6 | 29 | 27 | 7.5 | 3.3 | 3.5 | 21.8 | 14.4 |

| | | | | | | | | | | | | | | | | |
|---|---|---|---|---|---|---|---|---|---|---|---|---|---|---|---|---|
| >hg38_21_+_8433172_8446622_RNA45SN1_45S_50ntFlanks@9576.9596.21\|TCGGTCGTCCG ACCTGGGTAT | >hg38_21_+_8433172_8446622_RNA45SN1_45S_50ntFlanks | 9576 | 9596 | 21 | TCGGTCGTCCGACCTGGGTAT | MB468 | 21 | 1 | 10 | 39 | 24 | 7.4 | 0.7 | 5.8 | 23.7 | 11.8 |
| >hg38_21_+_8433172_8446622_RNA45SN1_45S_50ntFlanks@9576.9597.22\|TCGGTCGTCCG ACCTGGGTATA | >hg38_21_+_8433172_8446622_RNA45SN1_45S_50ntFlanks | 9576 | 9597 | 22 | TCGGTCGTCCGACCTGGGTATA | MB468 | 22 | 2 | 10 | 22 | 27 | 10.3 | 0.5 | 7.8 | 11.6 | 13.6 |
| >hg38_21_+_8433172_8446622_RNA45SN1_45S_50ntFlanks@9578.9603.26\|GGTCGTCCGAC CTGGGTATAGGGGCG | >hg38_21_+_8433172_8446622_RNA45SN1_45S_50ntFlanks | 9578 | 9603 | 26 | GGTCGTCCGACCTGGGTATAGGGGCG | MB231 | 13 | 14 | 9 | 6 | 18 | 21.5 | 21.3 | 14.5 | 8.6 | 29.2 |
| >hg38_21_+_8433172_8446622_RNA45SN1_45S_50ntFlanks@9602.9622.21\|CGAAAGACTAAT CGAACCATC | >hg38_21_+_8433172_8446622_RNA45SN1_45S_50ntFlanks | 9602 | 9622 | 21 | CGAAAGACTAATCGAACCATC | MB468 | 20 | 14 | 47 | 49 | 12 | 4.1 | 8.9 | 17.6 | 40.4 | 3.9 |
| >hg38_21_+_8433172_8446622_RNA45SN1_45S_50ntFlanks@9604.9621.18\|AAAGACTAATCG AACCAT | >hg38_21_+_8433172_8446622_RNA45SN1_45S_50ntFlanks | 9604 | 9621 | 18 | AAAGACTAATCGAACCAT | MB468 | 36 | 21 | 24 | 32 | 19 | 13.5 | 14.1 | 5.2 | 16.4 | 8.4 |
| >hg38_21_+_8433172_8446622_RNA45SN1_45S_50ntFlanks@9604.9622.19\|AAAGACTAATCG AACCATC | >hg38_21_+_8433172_8446622_RNA45SN1_45S_50ntFlanks | 9604 | 9622 | 19 | AAAGACTAATCGAACCATC | MB468 | 114 | 103 | 106 | 179 | 51 | 40.8 | 104.6 | 33.9 | 134.4 | 28.4 |
| >hg38_21_+_8433172_8446622_RNA45SN1_45S_50ntFlanks@9604.9623.20\|AAAGACTAATCG AACCATCT | >hg38_21_+_8433172_8446622_RNA45SN1_45S_50ntFlanks | 9604 | 9623 | 20 | AAAGACTAATCGAACCATCT | MB468 | 16 | 10 | 11 | 19 | 10 | 7.4 | 9.7 | 3.7 | 13.4 | 4.7 |
| >hg38_21_+_8433172_8446622_RNA45SN1_45S_50ntFlanks@9605.9622.18\|AAGACTAATCGA ACCATC | >hg38_21_+_8433172_8446622_RNA45SN1_45S_50ntFlanks | 9605 | 9622 | 18 | AAGACTAATCGAACCATC | MB468 | 59 | 51 | 53 | 81 | 21 | 18.6 | 53.3 | 11.7 | 60.0 | 9.1 |
| >hg38_21_+_8433172_8446622_RNA45SN1_45S_50ntFlanks@9606.9623.18\|AGACTAATCGAA CCATCT | >hg38_21_+_8433172_8446622_RNA45SN1_45S_50ntFlanks | 9606 | 9623 | 18 | AGACTAATCGAACCATCT | MB468 | 13 | 5 | 8 | 10 | 8 | 3.8 | 2.5 | 1.4 | 4.1 | 3.7 |
| >hg38_21_+_8433172_8446622_RNA45SN1_45S_50ntFlanks@9621.9648.28\|TCTAGTAGCTGG TTCCCTCCGAAGTTTC | >hg38_21_+_8433172_8446622_RNA45SN1_45S_50ntFlanks | 9621 | 9648 | 28 | TCTAGTAGCTGGTTCCCTCCGAAGTTTC | MB468 | 14 | 12 | 4 | 17 | 57 | 8.0 | 4.3 | 3.3 | 10.6 | 25.2 |
| >hg38_21_+_8433172_8446622_RNA45SN1_45S_50ntFlanks@9622.9648.27\|CTAGTAGCTGGT TCCCTCCGAAGTTTC | >hg38_21_+_8433172_8446622_RNA45SN1_45S_50ntFlanks | 9622 | 9648 | 27 | CTAGTAGCTGGTTCCCTCCGAAGTTTC | MB468 | 12 | 7 | 2 | 28 | 59 | 6.6 | 2.7 | 1.6 | 26.7 | 29.0 |
| >hg38_21_+_8433172_8446622_RNA45SN1_45S_50ntFlanks@9623.9648.26\|TAGTAGCTGGTT CCCTCCGAAGTTTC | >hg38_21_+_8433172_8446622_RNA45SN1_45S_50ntFlanks | 9623 | 9648 | 26 | TAGTAGCTGGTTCCCTCCGAAGTTTC | MB468 | 23 | 10 | 4 | 22 | 102 | 15.3 | 6.2 | 3.6 | 16.6 | 65.8 |
| >hg38_21_+_8433172_8446622_RNA45SN1_45S_50ntFlanks@9623.9649.27\|TAGTAGCTGGTT CCCTCCGAAGTTTCC | >hg38_21_+_8433172_8446622_RNA45SN1_45S_50ntFlanks | 9623 | 9649 | 27 | TAGTAGCTGGTTCCCTCCGAAGTTTCC | MB468 | 10 | 9 | 2 | 5 | 51 | 7.4 | 2.3 | 1.4 | 1.4 | 17.2 |
| >hg38_21_+_8433172_8446622_RNA45SN1_45S_50ntFlanks@9623.9650.28\|TAGTAGCTGGTT CCCTCCGAAGTTTCCC | >hg38_21_+_8433172_8446622_RNA45SN1_45S_50ntFlanks | 9623 | 9650 | 28 | TAGTAGCTGGTTCCCTCCGAAGTTTCCC | MB468 | 15 | 15 | 2 | 17 | 87 | 9.9 | 2.5 | 1.9 | 4.4 | 26.1 |
| >hg38_21_+_8433172_8446622_RNA45SN1_45S_50ntFlanks@9653.9677.25\|AGGATAGCTGG CGCTCTCGCAGACC | >hg38_21_+_8433172_8446622_RNA45SN1_45S_50ntFlanks | 9653 | 9677 | 25 | AGGATAGCTGGCGCTCTCGCAGACC | MB468 | 11 | 31 | 17 | 9 | 55 | 10.0 | 40.2 | 19.7 | 3.4 | 34.1 |
| >hg38_21_+_8433172_8446622_RNA45SN1_45S_50ntFlanks@9714.9735.22\|CGAATGATTAGA GGTCTTGGGG | >hg38_21_+_8433172_8446622_RNA45SN1_45S_50ntFlanks | 9714 | 9735 | 22 | CGAATGATTAGAGGTCTTGGGG | MB468 | 19 | 19 | 21 | 27 | 42 | 7.5 | 4.7 | 16.0 | 14.7 | 33.7 |
| >hg38_21_+_8433172_8446622_RNA45SN1_45S_50ntFlanks@9714.9737.24\|CGAATGATTAGA GGTCTTGGGGCC | >hg38_21_+_8433172_8446622_RNA45SN1_45S_50ntFlanks | 9714 | 9737 | 24 | CGAATGATTAGAGGTCTTGGGGCC | MB468 | 59 | 61 | 65 | 66 | 268 | 15.4 | 18.0 | 36.5 | 31.3 | 186.5 |
| >hg38_21_+_8433172_8446622_RNA45SN1_45S_50ntFlanks@9716.9737.22\|AATGATTAGAGG TCTTGGGGCC | >hg38_21_+_8433172_8446622_RNA45SN1_45S_50ntFlanks | 9716 | 9737 | 22 | AATGATTAGAGGTCTTGGGGCC | MB468 | 30 | 22 | 27 | 38 | 61 | 5.6 | 6.4 | 8.4 | 10.3 | 48.2 |
| >hg38_21_+_8433172_8446622_RNA45SN1_45S_50ntFlanks@9720.9737.18\|ATTAGAGGTCTT GGGGCC | >hg38_21_+_8433172_8446622_RNA45SN1_45S_50ntFlanks | 9720 | 9737 | 18 | ATTAGAGGTCTTGGGGCC | MB468 | 26 | 2 | 13 | 66 | 47 | 10.5 | 1.0 | 3.9 | 35.4 | 45.9 |
| >hg38_21_+_8433172_8446622_RNA45SN1_45S_50ntFlanks@9736.9753.18\|CCGAAACGATCT CAACCT | >hg38_21_+_8433172_8446622_RNA45SN1_45S_50ntFlanks | 9736 | 9753 | 18 | CCGAAACGATCTCAACCT | MB468 | 21 | 11 | 28 | 10 | 20 | 16.0 | 7.4 | 22.4 | 3.5 | 13.1 |
| >hg38_21_+_8433172_8446622_RNA45SN1_45S_50ntFlanks@9736.9754.19\|CCGAAACGATCT CAACCTA | >hg38_21_+_8433172_8446622_RNA45SN1_45S_50ntFlanks | 9736 | 9754 | 19 | CCGAAACGATCTCAACCTA | MB468 | 60 | 32 | 72 | 31 | 50 | 29.1 | 11.2 | 41.3 | 10.3 | 26.8 |
| >hg38_21_+_8433172_8446622_RNA45SN1_45S_50ntFlanks@9736.9755.20\|CCGAAACGATCT CAACCTAT | >hg38_21_+_8433172_8446622_RNA45SN1_45S_50ntFlanks | 9736 | 9755 | 20 | CCGAAACGATCTCAACCTAT | MB468 | 28 | 24 | 27 | 8 | 25 | 6.7 | 7.6 | 3.3 | 2.2 | 17.5 |
| >hg38_21_+_8433172_8446622_RNA45SN1_45S_50ntFlanks@9736.9757.22\|CCGAAACGATCT CAACCTATTC | >hg38_21_+_8433172_8446622_RNA45SN1_45S_50ntFlanks | 9736 | 9757 | 22 | CCGAAACGATCTCAACCTATTC | MB468 | 14 | 10 | 16 | 27 | 55 | 5.0 | 5.4 | 6.8 | 21.9 | 58.6 |
| >hg38_21_+_8433172_8446622_RNA45SN1_45S_50ntFlanks@9736.9758.23\|CCGAAACGATCT CAACCTATTCT | >hg38_21_+_8433172_8446622_RNA45SN1_45S_50ntFlanks | 9736 | 9758 | 23 | CCGAAACGATCTCAACCTATTCT | MB468 | 16 | 9 | 16 | 12 | 37 | 4.3 | 4.3 | 7.2 | 4.8 | 38.9 |
| >hg38_21_+_8433172_8446622_RNA45SN1_45S_50ntFlanks@9736.9765.30\|CCGAAACGATCT CAACCTATTCTCAAACTT | >hg38_21_+_8433172_8446622_RNA45SN1_45S_50ntFlanks | 9736 | 9765 | 30 | CCGAAACGATCTCAACCTATTCTCAAACTT | MB468 | 37 | 94 | 11 | 8 | 221 | 17.6 | 42.8 | 8.5 | 3.8 | 170.5 |
| >hg38_21_+_8433172_8446622_RNA45SN1_45S_50ntFlanks@9736.9767.32\|CCGAAACGATCT CAACCTATTCTCAAACTTTA | >hg38_21_+_8433172_8446622_RNA45SN1_45S_50ntFlanks | 9736 | 9767 | 32 | CCGAAACGATCTCAACCTATTCTCAAACTTTA | MB468 | 12 | 24 | 3 | 3 | 52 | 5.5 | 6.7 | 2.7 | 1.0 | 27.8 |
| >hg38_21_+_8433172_8446622_RNA45SN1_45S_50ntFlanks@9737.9754.18\|CGAAACGATCTC AACCTA | >hg38_21_+_8433172_8446622_RNA45SN1_45S_50ntFlanks | 9737 | 9754 | 18 | CGAAACGATCTCAACCTA | MB468 | 12 | 4 | 12 | 12 | 9 | 4.3 | 1.7 | 4.7 | 5.3 | 4.2 |
| >hg38_21_+_8433172_8446622_RNA45SN1_45S_50ntFlanks@9737.9765.29\|CGAAACGATCTC AACCTATTCTCAAACTT | >hg38_21_+_8433172_8446622_RNA45SN1_45S_50ntFlanks | 9737 | 9765 | 29 | CGAAACGATCTCAACCTATTCTCAAACTT | MB231 | 17 | 27 | 18 | 1 | 14 | 8.2 | 21.1 | 10.4 | 1.1 | 8.1 |
| >hg38_21_+_8433172_8446622_RNA45SN1_45S_50ntFlanks@9738.9755.18\|GAAACGATCTCA ACCTAT | >hg38_21_+_8433172_8446622_RNA45SN1_45S_50ntFlanks | 9738 | 9755 | 18 | GAAACGATCTCAACCTAT | MB468 | 79 | 45 | 66 | 46 | 42 | 35.3 | 29.7 | 48.3 | 9.8 | 22.1 |
| >hg38_21_+_8433172_8446622_RNA45SN1_45S_50ntFlanks@9738.9758.21\|GAAACGATCTCA ACCTATTCT | >hg38_21_+_8433172_8446622_RNA45SN1_45S_50ntFlanks | 9738 | 9758 | 21 | GAAACGATCTCAACCTATTCT | MB468 | 300 | 133 | 310 | 137 | 408 | 143.6 | 51.0 | 238.2 | 55.3 | 343.7 |
| >hg38_21_+_8433172_8446622_RNA45SN1_45S_50ntFlanks@9738.9759.22\|GAAACGATCTCA ACCTATTCTC | >hg38_21_+_8433172_8446622_RNA45SN1_45S_50ntFlanks | 9738 | 9759 | 22 | GAAACGATCTCAACCTATTCTC | MB468 | 61 | 54 | 67 | 42 | 267 | 14.2 | 14.5 | 21.9 | 15.4 | 260.5 |
| >hg38_21_+_8433172_8446622_RNA45SN1_45S_50ntFlanks@9738.9760.23\|GAAACGATCTCA ACCTATTCTCA | >hg38_21_+_8433172_8446622_RNA45SN1_45S_50ntFlanks | 9738 | 9760 | 23 | GAAACGATCTCAACCTATTCTCA | MB468 | 11 | 11 | 9 | 8 | 36 | 4.6 | 3.9 | 3.4 | 0.6 | 33.7 |
| >hg38_21_+_8433172_8446622_RNA45SN1_45S_50ntFlanks@9738.9762.25\|GAAACGATCTCA ACCTATTCTCAAA | >hg38_21_+_8433172_8446622_RNA45SN1_45S_50ntFlanks | 9738 | 9762 | 25 | GAAACGATCTCAACCTATTCTCAAA | MB468 | 14 | 16 | 6 | 5 | 42 | 4.0 | 4.8 | 1.6 | 3.9 | 47.4 |
| >hg38_21_+_8433172_8446622_RNA45SN1_45S_50ntFlanks@9738.9763.26\|GAAACGATCTCA ACCTATTCTCAAAC | >hg38_21_+_8433172_8446622_RNA45SN1_45S_50ntFlanks | 9738 | 9763 | 26 | GAAACGATCTCAACCTATTCTCAAAC | MB468 | 16 | 37 | 8 | 9 | 19 | 5.3 | 11.0 | 4.9 | 7.8 | 8.2 |
| >hg38_21_+_8433172_8446622_RNA45SN1_45S_50ntFlanks@9738.9764.27\|GAAACGATCTCA ACCTATTCTCAAACT | >hg38_21_+_8433172_8446622_RNA45SN1_45S_50ntFlanks | 9738 | 9764 | 27 | GAAACGATCTCAACCTATTCTCAAACT | MB468 | 18 | 28 | 8 | 5 | 34 | 6.9 | 1.4 | 4.9 | 2.5 | 11.2 |
| >hg38_21_+_8433172_8446622_RNA45SN1_45S_50ntFlanks@9738.9765.28\|GAAACGATCTCA ACCTATTCTCAAACTT | >hg38_21_+_8433172_8446622_RNA45SN1_45S_50ntFlanks | 9738 | 9765 | 28 | GAAACGATCTCAACCTATTCTCAAACTT | MB468 | 137 | 228 | 58 | 80 | 1818 | 41.8 | 49.9 | 32.9 | 43.9 | 887.8 |
| >hg38_21_+_8433172_8446622_RNA45SN1_45S_50ntFlanks@9738.9766.29\|GAAACGATCTCA ACCTATTCTCAAACTTT | >hg38_21_+_8433172_8446622_RNA45SN1_45S_50ntFlanks | 9738 | 9766 | 29 | GAAACGATCTCAACCTATTCTCAAACTTT | MB468 | 10 | 17 | 4 | 7 | 122 | 2.5 | 3.7 | 2.4 | 3.5 | 54.4 |
| >hg38_21_+_8433172_8446622_RNA45SN1_45S_50ntFlanks@9738.9767.30\|GAAACGATCTCA ACCTATTCTCAAACTTTA | >hg38_21_+_8433172_8446622_RNA45SN1_45S_50ntFlanks | 9738 | 9767 | 30 | GAAACGATCTCAACCTATTCTCAAACTTTA | MB468 | 55 | 79 | 16 | 47 | 467 | 18.7 | 16.7 | 13.9 | 14.7 | 139.6 |
| >hg38_21_+_8433172_8446622_RNA45SN1_45S_50ntFlanks@9739.9757.19\|AAACGATCTCAA CCTATTC | >hg38_21_+_8433172_8446622_RNA45SN1_45S_50ntFlanks | 9739 | 9757 | 19 | AAACGATCTCAACCTATTC | MB468 | 89 | 44 | 56 | 68 | 252 | 30.4 | 25.0 | 8.8 | 22.8 | 264.3 |
| >hg38_21_+_8433172_8446622_RNA45SN1_45S_50ntFlanks@9739.9758.20\|AAACGATCTCAA CCTATTCT | >hg38_21_+_8433172_8446622_RNA45SN1_45S_50ntFlanks | 9739 | 9758 | 20 | AAACGATCTCAACCTATTCT | MB468 | 234 | 145 | 186 | 101 | 439 | 25.4 | 31.4 | 44.1 | 30.1 | 300.1 |
| >hg38_21_+_8433172_8446622_RNA45SN1_45S_50ntFlanks@9739.9759.21\|AAACGATCTCAA CCTATTCTC | >hg38_21_+_8433172_8446622_RNA45SN1_45S_50ntFlanks | 9739 | 9759 | 21 | AAACGATCTCAACCTATTCTC | MB468 | 51 | 64 | 42 | 39 | 257 | 22.4 | 17.8 | 16.8 | 13.4 | 207.9 |
| >hg38_21_+_8433172_8446622_RNA45SN1_45S_50ntFlanks@9739.9760.22\|AAACGATCTCAA CCTATTCTCA | >hg38_21_+_8433172_8446622_RNA45SN1_45S_50ntFlanks | 9739 | 9760 | 22 | AAACGATCTCAACCTATTCTCA | MB468 | 11 | 12 | 7 | 9 | 46 | 5.6 | 4.3 | 3.1 | 3.5 | 50.6 |
| >hg38_21_+_8433172_8446622_RNA45SN1_45S_50ntFlanks@9739.9764.26\|AAACGATCTCAA CCTATTCTCAAACT | >hg38_21_+_8433172_8446622_RNA45SN1_45S_50ntFlanks | 9739 | 9764 | 26 | AAACGATCTCAACCTATTCTCAAACT | MB468 | 10 | 18 | 3 | 5 | 22 | 7.3 | 12.2 | 3.2 | 3.4 | 9.0 |
| >hg38_21_+_8433172_8446622_RNA45SN1_45S_50ntFlanks@9739.9765.27\|AAACGATCTCAA CCTATTCTCAAACTT | >hg38_21_+_8433172_8446622_RNA45SN1_45S_50ntFlanks | 9739 | 9765 | 27 | AAACGATCTCAACCTATTCTCAAACTT | MB468 | 90 | 170 | 31 | 80 | 1243 | 47.0 | 70.3 | 24.7 | 61.5 | 697.2 |
| >hg38_21_+_8433172_8446622_RNA45SN1_45S_50ntFlanks@9739.9767.29\|AAACGATCTCAA CCTATTCTCAAACTTTA | >hg38_21_+_8433172_8446622_RNA45SN1_45S_50ntFlanks | 9739 | 9767 | 29 | AAACGATCTCAACCTATTCTCAAACTTTA | MB468 | 28 | 59 | 8 | 28 | 235 | 15.9 | 47.5 | 9.4 | 7.9 | 94.5 |
| >hg38_21_+_8433172_8446622_RNA45SN1_45S_50ntFlanks@9740.9757.18\|AACGATCTCAAC CTATTC | >hg38_21_+_8433172_8446622_RNA45SN1_45S_50ntFlanks | 9740 | 9757 | 18 | AACGATCTCAACCTATTC | MB468 | 16 | 8 | 14 | 14 | 23 | 4.2 | 4.8 | 4.3 | 6.8 | 22.7 |
| >hg38_21_+_8433172_8446622_RNA45SN1_45S_50ntFlanks@9740.9758.19\|AACGATCTCAAC CTATTCT | >hg38_21_+_8433172_8446622_RNA45SN1_45S_50ntFlanks | 9740 | 9758 | 19 | AACGATCTCAACCTATTCT | MB468 | 33 | 17 | 27 | 13 | 42 | 4.7 | 5.1 | 11.3 | 1.4 | 32.4 |
| >hg38_21_+_8433172_8446622_RNA45SN1_45S_50ntFlanks@9740.9765.26\|AACGATCTCAAC CTATTCTCAAACTT | >hg38_21_+_8433172_8446622_RNA45SN1_45S_50ntFlanks | 9740 | 9765 | 26 | AACGATCTCAACCTATTCTCAAACTT | MB468 | 11 | 22 | 4 | 9 | 137 | 6.6 | 9.1 | 2.8 | 5.0 | 80.0 |
| >hg38_21_+_8433172_8446622_RNA45SN1_45S_50ntFlanks@9806.9830.25\|CGTGGAATGCG AGTGCCTAGTGGGC | >hg38_21_+_8433172_8446622_RNA45SN1_45S_50ntFlanks | 9806 | 9830 | 25 | CGTGGAATGCGAGTGCCTAGTGGGC | MB468 | 10 | 10 | 10 | 56 | 23 | 6.2 | 5.5 | 7.9 | 45.2 | 13.6 |
| >hg38_21_+_8433172_8446622_RNA45SN1_45S_50ntFlanks@9821.9850.30\|CCTAGTGGGCC ACTTTTGGTAAGCAGAACT | >hg38_21_+_8433172_8446622_RNA45SN1_45S_50ntFlanks | 9821 | 9850 | 30 | CCTAGTGGGCCACTTTTGGTAAGCAGAACT | MB468 | 31 | 43 | 24 | 22 | 133 | 18.8 | 24.0 | 21.0 | 13.8 | 98.9 |

| | | | | | | | | | | | | | | | |
|---|---|---|---|---|---|---|---|---|---|---|---|---|---|---|---|
| >hg38_21_+_8433172_8446622_RNA45SN1_45S_50ntFlanks@9822.9850.29\|CTAGTGGGCCACTTTTGGTAAGCAGAACT | >hg38_21_+_8433172_8446622_RNA45SN1_45S_50ntFlanks | 9822 | 9850 | 29 | CTAGTGGGCCACTTTTGGTAAGCAGAACT | MB468 | 33 | 40 | 22 | 12 | 32 | 19.5 | 27.8 | 18.8 | 3.9 | 14.8 |
| >hg38_21_+_8433172_8446622_RNA45SN1_45S_50ntFlanks@9903.9928.26\|ATCAGACCCCAGAAAAGGTGTTGGTT | >hg38_21_+_8433172_8446622_RNA45SN1_45S_50ntFlanks | 9903 | 9928 | 26 | ATCAGACCCCAGAAAAGGTGTTGGTT | MB468 | 11 | 15 | 1 | 7 | 20 | 10.3 | 9.9 | 1.3 | 4.2 | 14.4 |
| >hg38_21_+_8433172_8446622_RNA45SN1_45S_50ntFlanks@9937.9955.19\|CAGCAGGACGGTGGCCATG | >hg38_21_+_8433172_8446622_RNA45SN1_45S_50ntFlanks | 9937 | 9955 | 19 | CAGCAGGACGGTGGCCATG | BT20 | 39 | 20 | 29 | 20 | 5 | 67.4 | 34.5 | 32.1 | 7.4 | 3.3 |
| >hg38_21_+_8433172_8446622_RNA45SN1_45S_50ntFlanks@9960.9981.22\|TCGGAATCCGCTAAGGAGTGTG | >hg38_21_+_8433172_8446622_RNA45SN1_45S_50ntFlanks | 9960 | 9981 | 22 | TCGGAATCCGCTAAGGAGTGTG | BT20 | 12 | 12 | 1 | 1 | 1 | 20.3 | 20.3 | 0.7 | 1.1 | 0.9 |
| >hg38_21_+_8433172_8446622_RNA45SN1_45S_50ntFlanks@9960.9982.23\|TCGGAATCCGCTAAGGAGTGTGT | >hg38_21_+_8433172_8446622_RNA45SN1_45S_50ntFlanks | 9960 | 9982 | 23 | TCGGAATCCGCTAAGGAGTGTGT | BT20 | 22 | 11 | 1 | 1 | 0 | 38.8 | 19.6 | 1.0 | 1.4 | 0.7 |
| >hg38_21_+_8433172_8446622_RNA45SN1_45S_50ntFlanks@9961.9982.22\|CGGAATCCGCTAAGGAGTGTGT | >hg38_21_+_8433172_8446622_RNA45SN1_45S_50ntFlanks | 9961 | 9982 | 22 | CGGAATCCGCTAAGGAGTGTGT | BT20 | 12 | 6 | 0 | 1 | 0 | 20.0 | 10.6 | 0.5 | 2.6 | 0.4 |
| >hg38_MT_+_1621_3279_RNR2_16S_50ntFlanks@1000.1018.19\|GGGACCTGTATGAATGGCT | >hg38_MT_+_1621_3279_RNR2_16S_50ntFlanks | 1000 | 1018 | 19 | GGGACCTGTATGAATGGCT | MB231 | 14 | 1 | 0 | 23 | 7 | 18.2 | 1.9 | 0.5 | 21.3 | 3.8 |
| >hg38_MT_+_1621_3279_RNR2_16S_50ntFlanks@1175.1215.41\|TGCATTAAAAATTTCGGTTGGGGCGACCTCGGAGCAGAACC | >hg38_MT_+_1621_3279_RNR2_16S_50ntFlanks | 1175 | 1215 | 41 | TGCATTAAAAATTTCGGTTGGGGCGACCTCGGAGCAGAACC | MB231 | 14 | 4 | 5 | 10 | 16 | 9.8 | 1.8 | 1.8 | 11.3 | 9.4 |
| >hg38_MT_+_1621_3279_RNR2_16S_50ntFlanks@1182.1215.34\|AAAATTTCGGTTGGGGCGACCTCGGAGCAGAACC | >hg38_MT_+_1621_3279_RNR2_16S_50ntFlanks | 1182 | 1215 | 34 | AAAATTTCGGTTGGGGCGACCTCGGAGCAGAACC | MB231 | 11 | 5 | 3 | 16 | 14 | 12.4 | 3.7 | 1.7 | 11.0 | 11.3 |
| >hg38_MT_+_1621_3279_RNR2_16S_50ntFlanks@1184.1215.32\|AATTTCGGTTGGGGCGACCTCGGAGCAGAACC | >hg38_MT_+_1621_3279_RNR2_16S_50ntFlanks | 1184 | 1215 | 32 | AATTTCGGTTGGGGCGACCTCGGAGCAGAACC | MB231 | 15 | 8 | 5 | 26 | 19 | 15.0 | 5.7 | 1.7 | 18.4 | 14.5 |
| >hg38_MT_+_1621_3279_RNR2_16S_50ntFlanks@1184.1216.33\|AATTTCGGTTGGGGCGACCTCGGAGCAGAACCC | >hg38_MT_+_1621_3279_RNR2_16S_50ntFlanks | 1184 | 1216 | 33 | AATTTCGGTTGGGGCGACCTCGGAGCAGAACCC | MB231 | 11 | 8 | 3 | 17 | 16 | 9.5 | 6.3 | 1.4 | 11.6 | 13.6 |
| >hg38_MT_+_1621_3279_RNR2_16S_50ntFlanks@1216.1233.18\|CAACCTCCGAGCAGTACA | >hg38_MT_+_1621_3279_RNR2_16S_50ntFlanks | 1216 | 1233 | 18 | CAACCTCCGAGCAGTACA | MB231 | 11 | 2 | 1 | 1 | 11 | 10.4 | 1.6 | 0.9 | 1.2 | 10.3 |
| >hg38_MT_+_1621_3279_RNR2_16S_50ntFlanks@1216.1248.33\|CAACCTCCGAGCAGTACATGCTAAGACTTCACC | >hg38_MT_+_1621_3279_RNR2_16S_50ntFlanks | 1216 | 1248 | 33 | CAACCTCCGAGCAGTACATGCTAAGACTTCACC | BT20 | 14 | 3 | 1 | 15 | 33 | 2.9 | 0.5 | 0.5 | 18.9 | 40.3 |
| >hg38_MT_+_1621_3279_RNR2_16S_50ntFlanks@1216.1248.33\|CAACCTCCGAGCAGTACATGCTAAGACTTCACC | >hg38_MT_+_1621_3279_RNR2_16S_50ntFlanks | 1216 | 1248 | 33 | CAACCTCCGAGCAGTACATGCTAAGACTTCACC | MB231 | 31 | 7 | 3 | 43 | 52 | 27.0 | 3.4 | 2.5 | 30.5 | 30.9 |
| >hg38_MT_+_1621_3279_RNR2_16S_50ntFlanks@1216.1262.47\|CAACCTCCGAGCAGTACATGCTAAGACTTCACCAGTCAAAGCGAACT | >hg38_MT_+_1621_3279_RNR2_16S_50ntFlanks | 1216 | 1262 | 47 | CAACCTCCGAGCAGTACATGCTAAGACTTCACCAGTCAAAGCGAACT | BT20 | 12 | 3 | 1 | 5 | 48 | 7.7 | 2.5 | 1.0 | 6.8 | 7.0 |
| >hg38_MT_+_1621_3279_RNR2_16S_50ntFlanks@1216.1262.47\|CAACCTCCGAGCAGTACATGCTAAGACTTCACCAGTCAAAGCGAACT | >hg38_MT_+_1621_3279_RNR2_16S_50ntFlanks | 1216 | 1262 | 47 | CAACCTCCGAGCAGTACATGCTAAGACTTCACCAGTCAAAGCGAACT | MB231 | 67 | 32 | 28 | 7 | 223 | 21.0 | 46.2 | 18.6 | 10.3 | 263.1 |
| >hg38_MT_+_1621_3279_RNR2_16S_50ntFlanks@1217.1248.32\|AACCTCCGAGCAGTACATGCTAAGACTTCACC | >hg38_MT_+_1621_3279_RNR2_16S_50ntFlanks | 1217 | 1248 | 32 | AACCTCCGAGCAGTACATGCTAAGACTTCACC | MB231 | 10 | 3 | 1 | 16 | 19 | 9.9 | 1.5 | 0.9 | 13.2 | 13.0 |
| >hg38_MT_+_1621_3279_RNR2_16S_50ntFlanks@1217.1262.46\|AACCTCCGAGCAGTACATGCTAAGACTTCACCAGTCAAAGCGAACT | >hg38_MT_+_1621_3279_RNR2_16S_50ntFlanks | 1217 | 1262 | 46 | AACCTCCGAGCAGTACATGCTAAGACTTCACCAGTCAAAGCGAACT | MB231 | 11 | 6 | 5 | 2 | 28 | 4.5 | 6.5 | 1.8 | 2.3 | 40.6 |
| >hg38_MT_+_1621_3279_RNR2_16S_50ntFlanks@1249.1271.23\|AGTCAAAGCGAACTACTATACTC | >hg38_MT_+_1621_3279_RNR2_16S_50ntFlanks | 1249 | 1271 | 23 | AGTCAAAGCGAACTACTATACTC | MB231 | 12 | 2 | 0 | 41 | 4 | 15.2 | 2.7 | 0.6 | 36.1 | 1.0 |
| >hg38_MT_+_1621_3279_RNR2_16S_50ntFlanks@1347.1385.39\|CAACAATAGGGTTTACGACCTCGATGTTGGATCAGGACA | >hg38_MT_+_1621_3279_RNR2_16S_50ntFlanks | 1347 | 1385 | 39 | CAACAATAGGGTTTACGACCTCGATGTTGGATCAGGACA | MB231 | 11 | 3 | 1 | 7 | 6 | 14.0 | 2.2 | 1.1 | 4.0 | 3.2 |
| >hg38_MT_+_1621_3279_RNR2_16S_50ntFlanks@1347.1388.42\|CAACAATAGGGTTTACGACCTCGATGTTGGATCAGGACATCC | >hg38_MT_+_1621_3279_RNR2_16S_50ntFlanks | 1347 | 1388 | 42 | CAACAATAGGGTTTACGACCTCGATGTTGGATCAGGACATCC | MB231 | 16 | 4 | 2 | 6 | 10 | 19.6 | 2.1 | 2.1 | 3.3 | 3.7 |
| >hg38_MT_+_1621_3279_RNR2_16S_50ntFlanks@1347.1389.43\|CAACAATAGGGTTTACGACCTCGATGTTGGATCAGGACATCCC | >hg38_MT_+_1621_3279_RNR2_16S_50ntFlanks | 1347 | 1389 | 43 | CAACAATAGGGTTTACGACCTCGATGTTGGATCAGGACATCCC | MB231 | 23 | 7 | 3 | 6 | 20 | 24.0 | 3.0 | 3.3 | 5.6 | 11.1 |
| >hg38_MT_+_1621_3279_RNR2_16S_50ntFlanks@1348.1385.38\|AACAATAGGGTTTACGACCTCGATGTTGGATCAGGACA | >hg38_MT_+_1621_3279_RNR2_16S_50ntFlanks | 1348 | 1385 | 38 | AACAATAGGGTTTACGACCTCGATGTTGGATCAGGACA | MB231 | 38 | 9 | 4 | 38 | 21 | 49.2 | 5.5 | 3.4 | 23.0 | 11.9 |
| >hg38_MT_+_1621_3279_RNR2_16S_50ntFlanks@1348.1387.40\|AACAATAGGGTTTACGACCTCGATGTTGGATCAGGACATC | >hg38_MT_+_1621_3279_RNR2_16S_50ntFlanks | 1348 | 1387 | 40 | AACAATAGGGTTTACGACCTCGATGTTGGATCAGGACATC | MB231 | 16 | 4 | 2 | 9 | 8 | 21.6 | 2.4 | 1.8 | 5.1 | 1.5 |
| >hg38_MT_+_1621_3279_RNR2_16S_50ntFlanks@1348.1388.41\|AACAATAGGGTTTACGACCTCGATGTTGGATCAGGACATCC | >hg38_MT_+_1621_3279_RNR2_16S_50ntFlanks | 1348 | 1388 | 41 | AACAATAGGGTTTACGACCTCGATGTTGGATCAGGACATCC | MB231 | 11 | 2 | 2 | 2 | 4 | 15.6 | 1.2 | 1.7 | 1.2 | 1.6 |
| >hg38_MT_+_1621_3279_RNR2_16S_50ntFlanks@1348.1389.42\|AACAATAGGGTTTACGACCTCGATGTTGGATCAGGACATCCC | >hg38_MT_+_1621_3279_RNR2_16S_50ntFlanks | 1348 | 1389 | 42 | AACAATAGGGTTTACGACCTCGATGTTGGATCAGGACATCCC | MB231 | 16 | 3 | 2 | 8 | 9 | 21.8 | 1.3 | 1.9 | 7.5 | 1.9 |
| >hg38_MT_+_1621_3279_RNR2_16S_50ntFlanks@1349.1389.41\|ACAATAGGGTTTACGACCTCGATGTTGGATCAGGACATCCC | >hg38_MT_+_1621_3279_RNR2_16S_50ntFlanks | 1349 | 1389 | 41 | ACAATAGGGTTTACGACCTCGATGTTGGATCAGGACATCCC | MB231 | 23 | 4 | 2 | 14 | 14 | 30.5 | 2.4 | 1.9 | 13.3 | 7.2 |
| >hg38_MT_+_1621_3279_RNR2_16S_50ntFlanks@1367.1396.30\|TCGATGTTGGATCAGGACATCCCGATGGTG | >hg38_MT_+_1621_3279_RNR2_16S_50ntFlanks | 1367 | 1396 | 30 | TCGATGTTGGATCAGGACATCCCGATGGTG | MB231 | 19 | 6 | 3 | 61 | 16 | 26.3 | 7.4 | 1.5 | 48.7 | 17.4 |
| >hg38_MT_+_1621_3279_RNR2_16S_50ntFlanks@1369.1396.28\|GATGTTGGATCAGGACATCCCGATGGTG | >hg38_MT_+_1621_3279_RNR2_16S_50ntFlanks | 1369 | 1396 | 28 | GATGTTGGATCAGGACATCCCGATGGTG | MB231 | 11 | 4 | 1 | 42 | 10 | 14.6 | 4.7 | 1.2 | 31.2 | 8.0 |
| >hg38_MT_+_1621_3279_RNR2_16S_50ntFlanks@1370.1393.24\|ATGTTGGATCAGGACATCCCGATG | >hg38_MT_+_1621_3279_RNR2_16S_50ntFlanks | 1370 | 1393 | 24 | ATGTTGGATCAGGACATCCCGATG | MB231 | 15 | 3 | 1 | 50 | 7 | 18.7 | 3.7 | 0.3 | 38.9 | 1.2 |
| >hg38_MT_+_1621_3279_RNR2_16S_50ntFlanks@1370.1395.26\|ATGTTGGATCAGGACATCCCGATGGT | >hg38_MT_+_1621_3279_RNR2_16S_50ntFlanks | 1370 | 1395 | 26 | ATGTTGGATCAGGACATCCCGATGGT | MB231 | 17 | 3 | 1 | 42 | 8 | 23.5 | 4.0 | 0.8 | 33.6 | 3.5 |
| >hg38_MT_+_1621_3279_RNR2_16S_50ntFlanks@1370.1396.27\|ATGTTGGATCAGGACATCCCGATGGTG | >hg38_MT_+_1621_3279_RNR2_16S_50ntFlanks | 1370 | 1396 | 27 | ATGTTGGATCAGGACATCCCGATGGTG | BT20 | 18 | 4 | 1 | 81 | 19 | 6.2 | 2.3 | 0.1 | 63.2 | 15.4 |
| >hg38_MT_+_1621_3279_RNR2_16S_50ntFlanks@1370.1396.27\|ATGTTGGATCAGGACATCCCGATGGTG | >hg38_MT_+_1621_3279_RNR2_16S_50ntFlanks | 1370 | 1396 | 27 | ATGTTGGATCAGGACATCCCGATGGTG | MB231 | 43 | 11 | 3 | 121 | 19 | 59.5 | 14.5 | 2.4 | 107.8 | 19.7 |
| >hg38_MT_+_1621_3279_RNR2_16S_50ntFlanks@1370.1414.45\|ATGTTGGATCAGGACATCCCGATGGTGCAGCCGCTATTAAAGGTT | >hg38_MT_+_1621_3279_RNR2_16S_50ntFlanks | 1370 | 1414 | 45 | ATGTTGGATCAGGACATCCCGATGGTGCAGCCGCTATTAAAGGTT | MB231 | 14 | 6 | 7 | 33 | 23 | 9.9 | 6.3 | 4.6 | 48.6 | 19.0 |
| >hg38_MT_+_1621_3279_RNR2_16S_50ntFlanks@1389.1414.26\|CGATGGTGCAGCCGCTATTAAAGGTT | >hg38_MT_+_1621_3279_RNR2_16S_50ntFlanks | 1389 | 1414 | 26 | CGATGGTGCAGCCGCTATTAAAGGTT | MB231 | 12 | 3 | 0 | 60 | 8 | 14.4 | 3.6 | 0.5 | 60.1 | 3.5 |
| >hg38_MT_+_1621_3279_RNR2_16S_50ntFlanks@1438.1473.36\|TACGTGATCTGAGTTCAGACCGGAGTAATCCAGGTC | >hg38_MT_+_1621_3279_RNR2_16S_50ntFlanks | 1438 | 1473 | 36 | TACGTGATCTGAGTTCAGACCGGAGTAATCCAGGTC | MB231 | 11 | 4 | 3 | 5 | 8 | 7.9 | 1.3 | 0.9 | 3.2 | 4.7 |
| >hg38_MT_+_1621_3279_RNR2_16S_50ntFlanks@1438.1477.40\|TACGTGATCTGAGTTCAGACCGGAGTAATCCAGGTCGGTT | >hg38_MT_+_1621_3279_RNR2_16S_50ntFlanks | 1438 | 1477 | 40 | TACGTGATCTGAGTTCAGACCGGAGTAATCCAGGTCGGTT | BT20 | 12 | 5 | 2 | 23 | 30 | 3.2 | 2.6 | 0.5 | 16.6 | 17.3 |
| >hg38_MT_+_1621_3279_RNR2_16S_50ntFlanks@1438.1477.40\|TACGTGATCTGAGTTCAGACCGGAGTAATCCAGGTCGGTT | >hg38_MT_+_1621_3279_RNR2_16S_50ntFlanks | 1438 | 1477 | 40 | TACGTGATCTGAGTTCAGACCGGAGTAATCCAGGTCGGTT | MB231 | 29 | 10 | 11 | 14 | 21 | 33.0 | 5.6 | 5.5 | 13.1 | 11.0 |
| >hg38_MT_+_1621_3279_RNR2_16S_50ntFlanks@1438.1480.43\|TACGTGATCTGAGTTCAGACCGGAGTAATCCAGGTCGGTTTCT | >hg38_MT_+_1621_3279_RNR2_16S_50ntFlanks | 1438 | 1480 | 43 | TACGTGATCTGAGTTCAGACCGGAGTAATCCAGGTCGGTTTCT | MB231 | 12 | 4 | 5 | 3 | 10 | 10.6 | 3.3 | 0.8 | 3.5 | 6.1 |
| >hg38_MT_+_1621_3279_RNR2_16S_50ntFlanks@1439.1477.39\|ACGTGATCTGAGTTCAGACCGGAGTAATCCAGGTCGGTT | >hg38_MT_+_1621_3279_RNR2_16S_50ntFlanks | 1439 | 1477 | 39 | ACGTGATCTGAGTTCAGACCGGAGTAATCCAGGTCGGTT | MB231 | 11 | 6 | 4 | 10 | 12 | 12.1 | 2.0 | 0.6 | 7.3 | 5.4 |
| >hg38_MT_+_1621_3279_RNR2_16S_50ntFlanks@51.90.40\|GCTAAACCTAGCCCCAAACCCACTCCACCTTACTACCAGA | >hg38_MT_+_1621_3279_RNR2_16S_50ntFlanks | 51 | 90 | 40 | GCTAAACCTAGCCCCAAACCCACTCCACCTTACTACCAGA | MB231 | 29 | 43 | 9 | 218 | 327 | 14.3 | 46.3 | 6.5 | 223.4 | 322.9 |
| >hg38_MT_+_1621_3279_RNR2_16S_50ntFlanks@51.91.41\|GCTAAACCTAGCCCCAAACCCACTCCACCTTACTACCAGAC | >hg38_MT_+_1621_3279_RNR2_16S_50ntFlanks | 51 | 91 | 41 | GCTAAACCTAGCCCCAAACCCACTCCACCTTACTACCAGAC | MB231 | 13 | 12 | 6 | 71 | 122 | 5.2 | 13.2 | 3.3 | 80.2 | 123.0 |
| >hg38_MT_+_1621_3279_RNR2_16S_50ntFlanks@51.92.42\|GCTAAACCTAGCCCCAAACCCACTCCACCTTACTACCAGACA | >hg38_MT_+_1621_3279_RNR2_16S_50ntFlanks | 51 | 92 | 42 | GCTAAACCTAGCCCCAAACCCACTCCACCTTACTACCAGACA | BT20 | 14 | 8 | 4 | 82 | 56 | 11.7 | 7.2 | 5.2 | 80.2 | 4.9 |
| >hg38_MT_+_1621_3279_RNR2_16S_50ntFlanks@51.92.42\|GCTAAACCTAGCCCCAAACCCACTCCACCTTACTACCAGACA | >hg38_MT_+_1621_3279_RNR2_16S_50ntFlanks | 51 | 92 | 42 | GCTAAACCTAGCCCCAAACCCACTCCACCTTACTACCAGACA | MB231 | 48 | 50 | 25 | 271 | 472 | 19.3 | 62.4 | 20.1 | 390.1 | 511.3 |
| >hg38_MT_+_1621_3279_RNR2_16S_50ntFlanks@51.94.44\|GCTAAACCTAGCCCCAAACCCACTCCACCTTACTACCAGACAAC | >hg38_MT_+_1621_3279_RNR2_16S_50ntFlanks | 51 | 94 | 44 | GCTAAACCTAGCCCCAAACCCACTCCACCTTACTACCAGACAAC | BT20 | 12 | 7 | 1 | 67 | 123 | 12.1 | 9.0 | 1.2 | 68.5 | 43.4 |
| >hg38_MT_+_1621_3279_RNR2_16S_50ntFlanks@51.94.44\|GCTAAACCTAGCCCCAAACCCACTCCACCTTACTACCAGACAAC | >hg38_MT_+_1621_3279_RNR2_16S_50ntFlanks | 51 | 94 | 44 | GCTAAACCTAGCCCCAAACCCACTCCACCTTACTACCAGACAAC | MB231 | 30 | 46 | 7 | 158 | 465 | 15.8 | 65.9 | 4.9 | 209.0 | 424.4 |
| >hg38_MT_+_1621_3279_RNR2_16S_50ntFlanks@51.95.45\|GCTAAACCTAGCCCCAAACCCACTCCACCTTACTACCAGACAACC | >hg38_MT_+_1621_3279_RNR2_16S_50ntFlanks | 51 | 95 | 45 | GCTAAACCTAGCCCCAAACCCACTCCACCTTACTACCAGACAACC | BT20 | 24 | 14 | 1 | 117 | 257 | 30.5 | 19.7 | 2.2 | 109.9 | 152.9 |
| >hg38_MT_+_1621_3279_RNR2_16S_50ntFlanks@51.95.45\|GCTAAACCTAGCCCCAAACCCACTCCACCTTACTACCAGACAACC | >hg38_MT_+_1621_3279_RNR2_16S_50ntFlanks | 51 | 95 | 45 | GCTAAACCTAGCCCCAAACCCACTCCACCTTACTACCAGACAACC | MB231 | 65 | 104 | 14 | 277 | 941 | 45.0 | 159.1 | 7.7 | 420.0 | 885.5 |
| >hg38_MT_+_1621_3279_RNR2_16S_50ntFlanks@51.96.46\|GCTAAACCTAGCCCCAAACCCACTCCACCTTACTACCAGACAACCT | >hg38_MT_+_1621_3279_RNR2_16S_50ntFlanks | 51 | 96 | 46 | GCTAAACCTAGCCCCAAACCCACTCCACCTTACTACCAGACAACCT | MB231 | 12 | 16 | 2 | 27 | 151 | 10.7 | 25.2 | 1.3 | 39.3 | 139.1 |
| >hg38_MT_+_1621_3279_RNR2_16S_50ntFlanks@617.642.26\|AAACATATAACTGAACTCCTCACACC | >hg38_MT_+_1621_3279_RNR2_16S_50ntFlanks | 617 | 642 | 26 | AAACATATAACTGAACTCCTCACACC | BT20 | 11 | 0 | 2 | 9 | 11 | 3.8 | 0.8 | 0.5 | 8.8 | 15.1 |
| >hg38_MT_+_1621_3279_RNR2_16S_50ntFlanks@617.642.26\|AAACATATAACTGAACTCCTCACACC | >hg38_MT_+_1621_3279_RNR2_16S_50ntFlanks | 617 | 642 | 26 | AAACATATAACTGAACTCCTCACACC | MB231 | 18 | 4 | 5 | 20 | 20 | 15.2 | 2.6 | 2.6 | 13.6 | 9.0 |

<table>
<tr><th></th><th></th><th></th><th></th><th></th><th></th><th></th><th></th><th></th><th></th><th></th><th></th><th></th><th></th><th></th><th></th><th></th></tr>
<tr><td>>hg38_MT_+_1621_3279_RNR2_16S_50ntFlanks@618.642.25|AACATATAACTGAACTCCTCACACC</td><td>>hg38_MT_+_1621_3279_RNR2_16S_50ntFlanks</td><td>618</td><td>642</td><td>25</td><td>AACATATAACTGAACTCCTCACACC</td><td>MB231</td><td>12</td><td>3</td><td>3</td><td>13</td><td>14</td><td>10.6</td><td>1.4</td><td>2.2</td><td>7.7</td><td>6.1</td></tr>
<tr><td>>hg38_MT_+_1621_3279_RNR2_16S_50ntFlanks@622.642.21|TATAACTGAACTCCTCACACC</td><td>>hg38_MT_+_1621_3279_RNR2_16S_50ntFlanks</td><td>622</td><td>642</td><td>21</td><td>TATAACTGAACTCCTCACACC</td><td>MB231</td><td>13</td><td>4</td><td>4</td><td>7</td><td>10</td><td>14.3</td><td>2.9</td><td>2.5</td><td>5.8</td><td>3.4</td></tr>
<tr><td>>hg38_MT_+_1621_3279_RNR2_16S_50ntFlanks@625.642.18|AACTGAACTCCTCACACC</td><td>>hg38_MT_+_1621_3279_RNR2_16S_50ntFlanks</td><td>625</td><td>642</td><td>18</td><td>AACTGAACTCCTCACACC</td><td>MB231</td><td>11</td><td>3</td><td>3</td><td>6</td><td>8</td><td>13.9</td><td>2.5</td><td>1.9</td><td>3.9</td><td>3.3</td></tr>
<tr><td>>hg38_MT_+_1621_3279_RNR2_16S_50ntFlanks@643.663.21|CAATTGGACCAATCTATCACC</td><td>>hg38_MT_+_1621_3279_RNR2_16S_50ntFlanks</td><td>643</td><td>663</td><td>21</td><td>CAATTGGACCAATCTATCACC</td><td>MB231</td><td>12</td><td>3</td><td>1</td><td>13</td><td>8</td><td>11.3</td><td>2.9</td><td>0.9</td><td>8.0</td><td>2.6</td></tr>
<tr><td>>hg38_MT_+_1621_3279_RNR2_16S_50ntFlanks@643.676.34|CAATTGGACCAATCTATCACCCTATAGAAGAACT</td><td>>hg38_MT_+_1621_3279_RNR2_16S_50ntFlanks</td><td>643</td><td>676</td><td>34</td><td>CAATTGGACCAATCTATCACCCTATAGAAGAACT</td><td>MB231</td><td>13</td><td>2</td><td>2</td><td>12</td><td>11</td><td>15.7</td><td>2.0</td><td>1.6</td><td>6.0</td><td>3.0</td></tr>
<tr><td>>hg38_MT_+_1621_3279_RNR2_16S_50ntFlanks@644.663.20|AATTGGACCAATCTATCACC</td><td>>hg38_MT_+_1621_3279_RNR2_16S_50ntFlanks</td><td>644</td><td>663</td><td>20</td><td>AATTGGACCAATCTATCACC</td><td>BT20</td><td>10</td><td>0</td><td>1</td><td>11</td><td>6</td><td>0.8</td><td>0.4</td><td>0.6</td><td>8.4</td><td>6.7</td></tr>
<tr><td>>hg38_MT_+_1621_3279_RNR2_16S_50ntFlanks@644.663.20|AATTGGACCAATCTATCACC</td><td>>hg38_MT_+_1621_3279_RNR2_16S_50ntFlanks</td><td>644</td><td>663</td><td>20</td><td>AATTGGACCAATCTATCACC</td><td>MB231</td><td>16</td><td>4</td><td>1</td><td>16</td><td>10</td><td>16.3</td><td>4.7</td><td>0.7</td><td>10.3</td><td>3.6</td></tr>
<tr><td>>hg38_MT_+_1621_3279_RNR2_16S_50ntFlanks@644.664.21|AATTGGACCAATCTATCACCC</td><td>>hg38_MT_+_1621_3279_RNR2_16S_50ntFlanks</td><td>644</td><td>664</td><td>21</td><td>AATTGGACCAATCTATCACCC</td><td>MB231</td><td>11</td><td>4</td><td>1</td><td>11</td><td>8</td><td>10.6</td><td>3.6</td><td>0.8</td><td>7.5</td><td>2.9</td></tr>
<tr><td>>hg38_MT_+_1621_3279_RNR2_16S_50ntFlanks@644.665.22|AATTGGACCAATCTATCACCCT</td><td>>hg38_MT_+_1621_3279_RNR2_16S_50ntFlanks</td><td>644</td><td>665</td><td>22</td><td>AATTGGACCAATCTATCACCCT</td><td>MB231</td><td>10</td><td>10</td><td>3</td><td>8</td><td>14</td><td>10.4</td><td>10.0</td><td>2.2</td><td>7.6</td><td>11.6</td></tr>
<tr><td>>hg38_MT_+_1621_3279_RNR2_16S_50ntFlanks@644.676.33|AATTGGACCAATCTATCACCCTATAGAAGAACT</td><td>>hg38_MT_+_1621_3279_RNR2_16S_50ntFlanks</td><td>644</td><td>676</td><td>33</td><td>AATTGGACCAATCTATCACCCTATAGAAGAACT</td><td>BT20</td><td>11</td><td>2</td><td>1</td><td>17</td><td>10</td><td>4.8</td><td>1.2</td><td>0.5</td><td>15.5</td><td>6.0</td></tr>
<tr><td>>hg38_MT_+_1621_3279_RNR2_16S_50ntFlanks@644.676.33|AATTGGACCAATCTATCACCCTATAGAAGAACT</td><td>>hg38_MT_+_1621_3279_RNR2_16S_50ntFlanks</td><td>644</td><td>676</td><td>33</td><td>AATTGGACCAATCTATCACCCTATAGAAGAACT</td><td>MB231</td><td>16</td><td>3</td><td>3</td><td>23</td><td>13</td><td>21.0</td><td>3.0</td><td>2.5</td><td>15.7</td><td>5.3</td></tr>
<tr><td>>hg38_MT_+_1621_3279_RNR2_16S_50ntFlanks@713.740.28|GCATAAGCCTGCGTCAGATTAAAACACT</td><td>>hg38_MT_+_1621_3279_RNR2_16S_50ntFlanks</td><td>713</td><td>740</td><td>28</td><td>GCATAAGCCTGCGTCAGATTAAAACACT</td><td>MB231</td><td>12</td><td>5</td><td>2</td><td>18</td><td>22</td><td>11.9</td><td>4.3</td><td>2.5</td><td>12.4</td><td>6.3</td></tr>
<tr><td>>hg38_MT_+_1621_3279_RNR2_16S_50ntFlanks@714.740.27|CATAAGCCTGCGTCAGATTAAAACACT</td><td>>hg38_MT_+_1621_3279_RNR2_16S_50ntFlanks</td><td>714</td><td>740</td><td>27</td><td>CATAAGCCTGCGTCAGATTAAAACACT</td><td>MB231</td><td>13</td><td>4</td><td>4</td><td>10</td><td>15</td><td>10.4</td><td>2.7</td><td>2.7</td><td>7.8</td><td>8.2</td></tr>
<tr><td>>hg38_MT_+_1621_3279_RNR2_16S_50ntFlanks@716.740.25|TAAGCCTGCGTCAGATTAAAACACT</td><td>>hg38_MT_+_1621_3279_RNR2_16S_50ntFlanks</td><td>716</td><td>740</td><td>25</td><td>TAAGCCTGCGTCAGATTAAAACACT</td><td>MB231</td><td>10</td><td>3</td><td>2</td><td>8</td><td>11</td><td>8.8</td><td>1.7</td><td>1.5</td><td>5.9</td><td>5.8</td></tr>
<tr><td>>hg38_MT_+_1621_3279_RNR2_16S_50ntFlanks@718.740.23|AGCCTGCGTCAGATTAAAACACT</td><td>>hg38_MT_+_1621_3279_RNR2_16S_50ntFlanks</td><td>718</td><td>740</td><td>23</td><td>AGCCTGCGTCAGATTAAAACACT</td><td>MB231</td><td>12</td><td>4</td><td>3</td><td>13</td><td>13</td><td>9.5</td><td>2.2</td><td>1.2</td><td>9.7</td><td>7.4</td></tr>
<tr><td>>hg38_MT_+_1621_3279_RNR2_16S_50ntFlanks@722.740.19|TGCGTCAGATTAAAACACT</td><td>>hg38_MT_+_1621_3279_RNR2_16S_50ntFlanks</td><td>722</td><td>740</td><td>19</td><td>TGCGTCAGATTAAAACACT</td><td>MB231</td><td>13</td><td>4</td><td>0</td><td>8</td><td>4</td><td>15.6</td><td>3.5</td><td>0.7</td><td>6.5</td><td>2.1</td></tr>
<tr><td>>hg38_MT_+_1621_3279_RNR2_16S_50ntFlanks@732.759.28|TAAAACACTGAACTGACAATTAACAGCC</td><td>>hg38_MT_+_1621_3279_RNR2_16S_50ntFlanks</td><td>732</td><td>759</td><td>28</td><td>TAAAACACTGAACTGACAATTAACAGCC</td><td>MB231</td><td>12</td><td>3</td><td>0</td><td>29</td><td>6</td><td>12.1</td><td>3.1</td><td>0.6</td><td>26.0</td><td>2.6</td></tr>
<tr><td>>hg38_MT_+_1621_3279_RNR2_16S_50ntFlanks@929.958.30|CAGTGACACATGTTTAACGGCCGCGGTACC</td><td>>hg38_MT_+_1621_3279_RNR2_16S_50ntFlanks</td><td>929</td><td>958</td><td>30</td><td>CAGTGACACATGTTTAACGGCCGCGGTACC</td><td>MB231</td><td>11</td><td>3</td><td>2</td><td>26</td><td>10</td><td>10.2</td><td>2.3</td><td>1.8</td><td>15.2</td><td>3.9</td></tr>
<tr><td>>hg38_MT_+_1621_3279_RNR2_16S_50ntFlanks@930.958.29|AGTGACACATGTTTAACGGCCGCGGTACC</td><td>>hg38_MT_+_1621_3279_RNR2_16S_50ntFlanks</td><td>930</td><td>958</td><td>29</td><td>AGTGACACATGTTTAACGGCCGCGGTACC</td><td>MB231</td><td>16</td><td>4</td><td>3</td><td>40</td><td>12</td><td>13.1</td><td>3.7</td><td>2.6</td><td>27.8</td><td>4.2</td></tr>
<tr><td>>hg38_MT_+_1621_3279_RNR2_16S_50ntFlanks@999.1018.20|AGGGACCTGTATGAATGGCT</td><td>>hg38_MT_+_1621_3279_RNR2_16S_50ntFlanks</td><td>999</td><td>1018</td><td>20</td><td>AGGGACCTGTATGAATGGCT</td><td>MB231</td><td>14</td><td>2</td><td>1</td><td>33</td><td>8</td><td>20.0</td><td>2.7</td><td>0.4</td><td>35.9</td><td>6.6</td></tr>
<tr><td>>hg38_MT_+_598_1651_RNR1_12S_50ntFlanks@416.452.37|AAATAGACTACGAAAGTGGCTTTAACATATCTGAACA</td><td>>hg38_MT_+_598_1651_RNR1_12S_50ntFlanks</td><td>416</td><td>452</td><td>37</td><td>AAATAGACTACGAAAGTGGCTTTAACATATCTGAACA</td><td>BT20</td><td>13</td><td>2</td><td>0</td><td>4</td><td>3</td><td>12.8</td><td>0.4</td><td>0.4</td><td>4.1</td><td>3.5</td></tr>
<tr><td>>hg38_MT_+_598_1651_RNR1_12S_50ntFlanks@416.452.37|AAATAGACTACGAAAGTGGCTTTAACATATCTGAACA</td><td>>hg38_MT_+_598_1651_RNR1_12S_50ntFlanks</td><td>416</td><td>452</td><td>37</td><td>AAATAGACTACGAAAGTGGCTTTAACATATCTGAACA</td><td>MB231</td><td>23</td><td>8</td><td>2</td><td>8</td><td>8</td><td>33.8</td><td>9.6</td><td>2.8</td><td>6.8</td><td>0.2</td></tr>
<tr><td>>hg38_MT_+_598_1651_RNR1_12S_50ntFlanks@416.454.39|AAATAGACTACGAAAGTGGCTTTAACATATCTGAACACA</td><td>>hg38_MT_+_598_1651_RNR1_12S_50ntFlanks</td><td>416</td><td>454</td><td>39</td><td>AAATAGACTACGAAAGTGGCTTTAACATATCTGAACACA</td><td>MB231</td><td>11</td><td>3</td><td>1</td><td>6</td><td>6</td><td>15.6</td><td>3.2</td><td>0.7</td><td>3.8</td><td>0.6</td></tr>
<tr><td>>hg38_MT_+_598_1651_RNR1_12S_50ntFlanks@416.456.41|AAATAGACTACGAAAGTGGCTTTAACATATCTGAACACACA</td><td>>hg38_MT_+_598_1651_RNR1_12S_50ntFlanks</td><td>416</td><td>456</td><td>41</td><td>AAATAGACTACGAAAGTGGCTTTAACATATCTGAACACACA</td><td>MB231</td><td>25</td><td>4</td><td>3</td><td>6</td><td>13</td><td>32.9</td><td>1.9</td><td>2.6</td><td>6.0</td><td>5.5</td></tr>
<tr><td>>hg38_MT_+_598_1651_RNR1_12S_50ntFlanks@417.456.40|AATAGACTACGAAAGTGGCTTTAACATATCTGAACACACA</td><td>>hg38_MT_+_598_1651_RNR1_12S_50ntFlanks</td><td>417</td><td>456</td><td>40</td><td>AATAGACTACGAAAGTGGCTTTAACATATCTGAACACACA</td><td>MB231</td><td>10</td><td>2</td><td>1</td><td>4</td><td>4</td><td>14.3</td><td>1.2</td><td>0.5</td><td>2.4</td><td>1.7</td></tr>
<tr><td>>hg38_MT_+_598_1651_RNR1_12S_50ntFlanks@790.809.20|ACGATAGCCCTTATGAAACT</td><td>>hg38_MT_+_598_1651_RNR1_12S_50ntFlanks</td><td>790</td><td>809</td><td>20</td><td>ACGATAGCCCTTATGAAACT</td><td>MB231</td><td>18</td><td>5</td><td>3</td><td>9</td><td>14</td><td>19.4</td><td>5.1</td><td>2.7</td><td>8.9</td><td>7.1</td></tr>
<tr><td>>hg38_MT_+_598_1651_RNR1_12S_50ntFlanks@936.960.25|CCTACGCATTTATATAGAGGAGACA</td><td>>hg38_MT_+_598_1651_RNR1_12S_50ntFlanks</td><td>936</td><td>960</td><td>25</td><td>CCTACGCATTTATATAGAGGAGACA</td><td>MB231</td><td>14</td><td>3</td><td>0</td><td>3</td><td>4</td><td>21.3</td><td>4.1</td><td>0.4</td><td>2.8</td><td>3.4</td></tr>
<tr><td>>MI0000061|hsa-let-7a-2&WithFlank&11|-|122146516|122146599@11.28.18;MI0000064|hsa-let-7c&WithFlank&21|+|16539822|16539917@17.34.18;MI0000062|hsa-let-7a-3&WithFlank&22|+|46112743|46112828@10.27.18;MI0000060|hsa-let-7a-1&WithFlank&9|+|94175951|94176042@12.29.18&[MIMAT0000062&hsa-let-7a-5p&offsets|0|-4;m-6&11|-|122146568|122146589&offsets|0|-4];[MIMAT0000064&hsa-let-7c-5p&offsets|0|-4;m-37&21|+|16539838|16539859&offsets|0|-4];[MIMAT0000062_1&hsa-let-7a-5p&offsets|0|-4;m-5&22|+|46112752|46112773&offsets|0|-4];[MIMAT0000062_2&hsa-let-7a-5p&offsets|0|-4;m-4&9|+|94175962|94175983&offsets|0|-4]|TGAGGTAGTAGGTTGTAT</td><td>>MI0000061|hsa-let-7a-2&WithFlank&11|-|122146516|122146599</td><td>11</td><td>28</td><td>18</td><td>TGAGGTAGTAGGTTGTAT</td><td>BT20</td><td>116</td><td>157</td><td>238</td><td>58</td><td>66</td><td>124.1</td><td>106.8</td><td>222.2</td><td>12.6</td><td>9.5</td></tr>
<tr><td>>MI0000061|hsa-let-7a-2&WithFlank&11|-|122146516|122146599@11.28.18;MI0000064|hsa-let-7c&WithFlank&21|+|16539822|16539917@17.34.18;MI0000062|hsa-let-7a-3&WithFlank&22|+|46112743|46112828@10.27.18;MI0000060|hsa-let-7a-1&WithFlank&9|+|94175951|94176042@12.29.18&[MIMAT0000062&hsa-let-7a-5p&offsets|0|-4;m-6&11|-|122146568|122146589&offsets|0|-4];[MIMAT0000064&hsa-let-7c-5p&offsets|0|-4;m-37&21|+|16539838|16539859&offsets|0|-4];[MIMAT0000062_1&hsa-let-7a-5p&offsets|0|-4;m-5&22|+|46112752|46112773&offsets|0|-4];[MIMAT0000062_2&hsa-let-7a-5p&offsets|0|-4;m-4&9|+|94175962|94175983&offsets|0|-4]|TGAGGTAGTAGGTTGTAT</td><td>>MI0000061|hsa-let-7a-2&WithFlank&11|-|122146516|122146599</td><td>11</td><td>28</td><td>18</td><td>TGAGGTAGTAGGTTGTAT</td><td>MB231</td><td>117</td><td>169</td><td>112</td><td>231</td><td>123</td><td>38.2</td><td>75.7</td><td>67.9</td><td>179.6</td><td>47.2</td></tr>
<tr><td>>MI0000061|hsa-let-7a-2&WithFlank&11|-|122146516|122146599@11.29.19;MI0000062|hsa-let-7a-3&WithFlank&22|+|46112743|46112828@10.28.19;MI0000060|hsa-let-7a-1&WithFlank&9|+|94175951|94176042@12.30.19&[MIMAT0000062&hsa-let-7a-5p&offsets|0|-3;m-6&11|-|122146568|122146589&offsets|0|-3];[MIMAT0000062_1&hsa-let-7a-5p&offsets|0|-3;m-5&22|+|46112752|46112773&offsets|0|-3];[MIMAT0000062_2&hsa-let-7a-5p&offsets|0|-3;m-4&9|+|94175962|94175983&offsets|0|-3]|TGAGGTAGTAGGTTGTATA</td><td>>MI0000061|hsa-let-7a-2&WithFlank&11|-|122146516|122146599</td><td>11</td><td>29</td><td>19</td><td>TGAGGTAGTAGGTTGTATA</td><td>BT20</td><td>37</td><td>38</td><td>175</td><td>3</td><td>11</td><td>15.4</td><td>3.4</td><td>49.0</td><td>1.7</td><td>3.6</td></tr>
<tr><td>>MI0000061|hsa-let-7a-2&WithFlank&11|-|122146516|122146599@11.29.19;MI0000062|hsa-let-7a-3&WithFlank&22|+|46112743|46112828@10.28.19;MI0000060|hsa-let-7a-1&WithFlank&9|+|94175951|94176042@12.30.19&[MIMAT0000062&hsa-let-7a-5p&offsets|0|-3;m-6&11|-|122146568|122146589&offsets|0|-3];[MIMAT0000062_1&hsa-let-7a-5p&offsets|0|-3;m-5&22|+|46112752|46112773&offsets|0|-3];[MIMAT0000062_2&hsa-let-7a-5p&offsets|0|-3;m-4&9|+|94175962|94175983&offsets|0|-3]|TGAGGTAGTAGGTTGTATA</td><td>>MI0000061|hsa-let-7a-2&WithFlank&11|-|122146516|122146599</td><td>11</td><td>29</td><td>19</td><td>TGAGGTAGTAGGTTGTATA</td><td>MB231</td><td>20</td><td>16</td><td>27</td><td>5</td><td>6</td><td>24.5</td><td>23.2</td><td>36.1</td><td>1.5</td><td>6.8</td></tr>
<tr><td>>MI0000061|hsa-let-7a-2&WithFlank&11|-|122146516|122146599@11.30.20;MI0000062|hsa-let-7a-3&WithFlank&22|+|46112743|46112828@10.29.20;MI0000060|hsa-let-7a-1&WithFlank&9|+|94175951|94176042@12.31.20&[MIMAT0000062&hsa-let-7a-5p&offsets|0|-2;m-6&11|-|122146568|122146589&offsets|0|-2];[MIMAT0000062_1&hsa-let-7a-5p&offsets|0|-2;m-5&22|+|46112752|46112773&offsets|0|-2];[MIMAT0000062_2&hsa-let-7a-5p&offsets|0|-2;m-4&9|+|94175962|94175983&offsets|0|-2]|TGAGGTAGTAGGTTGTATAG</td><td>>MI0000061|hsa-let-7a-2&WithFlank&11|-|122146516|122146599</td><td>11</td><td>30</td><td>20</td><td>TGAGGTAGTAGGTTGTATAG</td><td>BT20</td><td>11</td><td>12</td><td>17</td><td>11</td><td>4</td><td>7.5</td><td>4.8</td><td>8.9</td><td>7.1</td><td>0.5</td></tr>
<tr><td>>MI0000061|hsa-let-7a-2&WithFlank&11|-|122146516|122146599@11.30.20(+1A);MI0000062|hsa-let-7a-3&WithFlank&22|+|46112743|46112828@10.29.20(+1A);MI0000060|hsa-let-7a-1&WithFlank&9|+|94175951|94176042@12.31.20(+1A)&[MIMAT0000062&hsa-let-7a-5p&offsets|0|-2(+1A);m-6&11|-|122146568|122146589&offsets|0|-2(+1A)];[MIMAT0000062_1&hsa-let-7a-5p&offsets|0|-2(+1A);m-5&22|+|46112752|46112773&offsets|0|-2(+1A)];[MIMAT0000062_2&hsa-let-7a-5p&offsets|0|-2(+1A);m-4&9|+|94175962|94175983&offsets|0|-2(+1A)]|TGAGGTAGTAGGTTGTATAGA</td><td>>MI0000061|hsa-let-7a-2&WithFlank&11|-|122146516|122146599</td><td>11</td><td>30</td><td>20(+1A)</td><td>TGAGGTAGTAGGTTGTATAGA</td><td>BT20</td><td>15</td><td>8</td><td>16</td><td>3</td><td>3</td><td>12.0</td><td>4.1</td><td>8.8</td><td>2.8</td><td>1.1</td></tr>
<tr><td>>MI0000061|hsa-let-7a-2&WithFlank&11|-|122146516|122146599@11.30.20(+1C);MI0000062|hsa-let-7a-3&WithFlank&22|+|46112743|46112828@10.29.20(+1C);MI0000060|hsa-let-7a-1&WithFlank&9|+|94175951|94176042@12.31.20(+1C)&[MIMAT0000062&hsa-let-7a-5p&offsets|0|-2(+1C);m-6&11|-|122146568|122146589&offsets|0|-2(+1C)];[MIMAT0000062_1&hsa-let-7a-5p&offsets|0|-2(+1C);m-5&22|+|46112752|46112773&offsets|0|-2(+1C)];[MIMAT0000062_2&hsa-let-7a-5p&offsets|0|-2(+1C);m-4&9|+|94175962|94175983&offsets|0|-2(+1C)]|TGAGGTAGTAGGTTGTATAGC</td><td>>MI0000061|hsa-let-7a-2&WithFlank&11|-|122146516|122146599</td><td>11</td><td>30</td><td>20(+1C)</td><td>TGAGGTAGTAGGTTGTATAGC</td><td>BT20</td><td>55</td><td>57</td><td>60</td><td>17</td><td>21</td><td>28.9</td><td>32.5</td><td>35.4</td><td>12.8</td><td>6.6</td></tr>
</table>

<table>
<tr><th></th><th></th><th></th><th></th><th></th><th></th><th></th><th></th><th></th><th></th><th></th><th></th><th></th><th></th><th></th><th></th><th></th></tr>
<tr><td>>MI0000061|hsa-let-7a-2&WithFlank&11|-|122146516|122146599@11.30.20(+1C);MI0000062|hsa-let-7a-3&WithFlank&22|+|46112743|46112828@10.29.20(+1C);MI0000060|hsa-let-7a-1&WithFlank&9|+|94175951|94176042@12.31.20(+1C)&[MIMAT0000062&hsa-let-7a-5p&offsets|0|-2(+1C);m-6&11|-|122146568|122146589&offsets|0|-2(+1C)];[MIMAT0000062_1&hsa-let-7a-5p&offsets|0|-2(+1C);m-5&22|+|46112752|46112773&offsets|0|-2(+1C)];[MIMAT0000062_2&hsa-let-7a-5p&offsets|0|-2(+1C);m-4&9|+|94175962|94175983&offsets|0|-2(+1C)]|TGAGGTAGTAGGTTGTATAGC</td><td>>MI0000061|hsa-let-7a-2&WithFlank&11|-|122146516|122146599</td><td>11</td><td>30</td><td>20(+1C)</td><td>TGAGGTAGTAGGTTGTATAGC</td><td>MB231</td><td>54</td><td>64</td><td>28</td><td>89</td><td>68</td><td>8.6</td><td>16.4</td><td>13.4</td><td>46.2</td><td>33.8</td></tr>
<tr><td>>MI0000061|hsa-let-7a-2&WithFlank&11|-|122146516|122146599@11.30.20(+1G);MI0000062|hsa-let-7a-3&WithFlank&22|+|46112743|46112828@10.29.20(+1G);MI0000060|hsa-let-7a-1&WithFlank&9|+|94175951|94176042@12.31.20(+1G)&[MIMAT0000062&hsa-let-7a-5p&offsets|0|-2(+1G);m-6&11|-|122146568|122146589&offsets|0|-2(+1G)];[MIMAT0000062_1&hsa-let-7a-5p&offsets|0|-2(+1G);m-5&22|+|46112752|46112773&offsets|0|-2(+1G)];[MIMAT0000062_2&hsa-let-7a-5p&offsets|0|-2(+1G);m-4&9|+|94175962|94175983&offsets|0|-2(+1G)]|TGAGGTAGTAGGTTGTATAGG</td><td>>MI0000061|hsa-let-7a-2&WithFlank&11|-|122146516|122146599</td><td>11</td><td>30</td><td>20(+1G)</td><td>TGAGGTAGTAGGTTGTATAGG</td><td>BT20</td><td>25</td><td>9</td><td>10</td><td>2</td><td>1</td><td>21.7</td><td>5.4</td><td>4.4</td><td>2.4</td><td>0.6</td></tr>
<tr><td>>MI0000061|hsa-let-7a-2&WithFlank&11|-|122146516|122146599@11.30.20(+1G);MI0000062|hsa-let-7a-3&WithFlank&22|+|46112743|46112828@10.29.20(+1G);MI0000060|hsa-let-7a-1&WithFlank&9|+|94175951|94176042@12.31.20(+1G)&[MIMAT0000062&hsa-let-7a-5p&offsets|0|-2(+1G);m-6&11|-|122146568|122146589&offsets|0|-2(+1G)];[MIMAT0000062_1&hsa-let-7a-5p&offsets|0|-2(+1G);m-5&22|+|46112752|46112773&offsets|0|-2(+1G)];[MIMAT0000062_2&hsa-let-7a-5p&offsets|0|-2(+1G);m-4&9|+|94175962|94175983&offsets|0|-2(+1G)]|TGAGGTAGTAGGTTGTATAGG</td><td>>MI0000061|hsa-let-7a-2&WithFlank&11|-|122146516|122146599</td><td>11</td><td>30</td><td>20(+1G)</td><td>TGAGGTAGTAGGTTGTATAGG</td><td>MB231</td><td>20</td><td>9</td><td>4</td><td>8</td><td>4</td><td>14.6</td><td>3.7</td><td>2.9</td><td>4.7</td><td>2.5</td></tr>
<tr><td>>MI0000061|hsa-let-7a-2&WithFlank&11|-|122146516|122146599@11.31.21;MI0000062|hsa-let-7a-3&WithFlank&22|+|46112743|46112828@10.30.21;MI0000060|hsa-let-7a-1&WithFlank&9|+|94175951|94176042@12.32.21&[MIMAT0000062&hsa-let-7a-5p&offsets|0|-1;m-6&11|-|122146568|122146589&offsets|0|-1];[MIMAT0000062_1&hsa-let-7a-5p&offsets|0|-1;m-5&22|+|46112752|46112773&offsets|0|-1];[MIMAT0000062_2&hsa-let-7a-5p&offsets|0|-1;m-4&9|+|94175962|94175983&offsets|0|-1]|TGAGGTAGTAGGTTGTATAGT</td><td>>MI0000061|hsa-let-7a-2&WithFlank&11|-|122146516|122146599</td><td>11</td><td>31</td><td>21</td><td>TGAGGTAGTAGGTTGTATAGT</td><td>BT20</td><td>453</td><td>418</td><td>497</td><td>126</td><td>163</td><td>244.1</td><td>210.2</td><td>304.4</td><td>69.1</td><td>37.5</td></tr>
<tr><td>>MI0000061|hsa-let-7a-2&WithFlank&11|-|122146516|122146599@11.31.21;MI0000062|hsa-let-7a-3&WithFlank&22|+|46112743|46112828@10.30.21;MI0000060|hsa-let-7a-1&WithFlank&9|+|94175951|94176042@12.32.21&[MIMAT0000062&hsa-let-7a-5p&offsets|0|-1;m-6&11|-|122146568|122146589&offsets|0|-1];[MIMAT0000062_1&hsa-let-7a-5p&offsets|0|-1;m-5&22|+|46112752|46112773&offsets|0|-1];[MIMAT0000062_2&hsa-let-7a-5p&offsets|0|-1;m-4&9|+|94175962|94175983&offsets|0|-1]|TGAGGTAGTAGGTTGTATAGT</td><td>>MI0000061|hsa-let-7a-2&WithFlank&11|-|122146516|122146599</td><td>11</td><td>31</td><td>21</td><td>TGAGGTAGTAGGTTGTATAGT</td><td>MB231</td><td>524</td><td>656</td><td>290</td><td>619</td><td>542</td><td>78.3</td><td>130.1</td><td>128.4</td><td>232.0</td><td>196.3</td></tr>
<tr><td>>MI0000061|hsa-let-7a-2&WithFlank&11|-|122146516|122146599@11.31.21;MI0000062|hsa-let-7a-3&WithFlank&22|+|46112743|46112828@10.30.21;MI0000060|hsa-let-7a-1&WithFlank&9|+|94175951|94176042@12.32.21&[MIMAT0000062&hsa-let-7a-5p&offsets|0|-1;m-6&11|-|122146568|122146589&offsets|0|-1];[MIMAT0000062_1&hsa-let-7a-5p&offsets|0|-1;m-5&22|+|46112752|46112773&offsets|0|-1];[MIMAT0000062_2&hsa-let-7a-5p&offsets|0|-1;m-4&9|+|94175962|94175983&offsets|0|-1]|TGAGGTAGTAGGTTGTATAGT</td><td>>MI0000061|hsa-let-7a-2&WithFlank&11|-|122146516|122146599</td><td>11</td><td>31</td><td>21</td><td>TGAGGTAGTAGGTTGTATAGT</td><td>MB468</td><td>18</td><td>23</td><td>9</td><td>16</td><td>5</td><td>7.6</td><td>5.6</td><td>2.6</td><td>5.4</td><td>1.6</td></tr>
<tr><td>>MI0000061|hsa-let-7a-2&WithFlank&11|-|122146516|122146599@11.31.21(+1A);MI0000062|hsa-let-7a-3&WithFlank&22|+|46112743|46112828@10.30.21(+1A);MI0000060|hsa-let-7a-1&WithFlank&9|+|94175951|94176042@12.32.21(+1A)&[MIMAT0000062&hsa-let-7a-5p&offsets|0|-1(+1A);m-6&11|-|122146568|122146589&offsets|0|-1(+1A)];[MIMAT0000062_1&hsa-let-7a-5p&offsets|0|-1(+1A);m-5&22|+|46112752|46112773&offsets|0|-1(+1A)];[MIMAT0000062_2&hsa-let-7a-5p&offsets|0|-1(+1A);m-4&9|+|94175962|94175983&offsets|0|-1(+1A)]|TGAGGTAGTAGGTTGTATAGTA</td><td>>MI0000061|hsa-let-7a-2&WithFlank&11|-|122146516|122146599</td><td>11</td><td>31</td><td>21(+1A)</td><td>TGAGGTAGTAGGTTGTATAGTA</td><td>BT20</td><td>228</td><td>112</td><td>211</td><td>24</td><td>41</td><td>174.0</td><td>67.6</td><td>113.5</td><td>16.1</td><td>8.9</td></tr>
<tr><td>>MI0000061|hsa-let-7a-2&WithFlank&11|-|122146516|122146599@11.31.21(+1A);MI0000062|hsa-let-7a-3&WithFlank&22|+|46112743|46112828@10.30.21(+1A);MI0000060|hsa-let-7a-1&WithFlank&9|+|94175951|94176042@12.32.21(+1A)&[MIMAT0000062&hsa-let-7a-5p&offsets|0|-1(+1A);m-6&11|-|122146568|122146589&offsets|0|-1(+1A)];[MIMAT0000062_1&hsa-let-7a-5p&offsets|0|-1(+1A);m-5&22|+|46112752|46112773&offsets|0|-1(+1A)];[MIMAT0000062_2&hsa-let-7a-5p&offsets|0|-1(+1A);m-4&9|+|94175962|94175983&offsets|0|-1(+1A)]|TGAGGTAGTAGGTTGTATAGTA</td><td>>MI0000061|hsa-let-7a-2&WithFlank&11|-|122146516|122146599</td><td>11</td><td>31</td><td>21(+1A)</td><td>TGAGGTAGTAGGTTGTATAGTA</td><td>MB231</td><td>114</td><td>95</td><td>71</td><td>168</td><td>122</td><td>9.4</td><td>11.6</td><td>46.3</td><td>95.1</td><td>63.7</td></tr>
<tr><td>>MI0000061|hsa-let-7a-2&WithFlank&11|-|122146516|122146599@11.31.21(+1C);MI0000062|hsa-let-7a-3&WithFlank&22|+|46112743|46112828@10.30.21(+1C);MI0000060|hsa-let-7a-1&WithFlank&9|+|94175951|94176042@12.32.21(+1C)&[MIMAT0000062&hsa-let-7a-5p&offsets|0|-1(+1C);m-6&11|-|122146568|122146589&offsets|0|-1(+1C)];[MIMAT0000062_1&hsa-let-7a-5p&offsets|0|-1(+1C);m-5&22|+|46112752|46112773&offsets|0|-1(+1C)];[MIMAT0000062_2&hsa-let-7a-5p&offsets|0|-1(+1C);m-4&9|+|94175962|94175983&offsets|0|-1(+1C)]|TGAGGTAGTAGGTTGTATAGTC</td><td>>MI0000061|hsa-let-7a-2&WithFlank&11|-|122146516|122146599</td><td>11</td><td>31</td><td>21(+1C)</td><td>TGAGGTAGTAGGTTGTATAGTC</td><td>BT20</td><td>672</td><td>669</td><td>697</td><td>135</td><td>265</td><td>327.3</td><td>415.2</td><td>364.7</td><td>82.0</td><td>73.5</td></tr>
<tr><td>>MI0000061|hsa-let-7a-2&WithFlank&11|-|122146516|122146599@11.31.21(+1C);MI0000062|hsa-let-7a-3&WithFlank&22|+|46112743|46112828@10.30.21(+1C);MI0000060|hsa-let-7a-1&WithFlank&9|+|94175951|94176042@12.32.21(+1C)&[MIMAT0000062&hsa-let-7a-5p&offsets|0|-1(+1C);m-6&11|-|122146568|122146589&offsets|0|-1(+1C)];[MIMAT0000062_1&hsa-let-7a-5p&offsets|0|-1(+1C);m-5&22|+|46112752|46112773&offsets|0|-1(+1C)];[MIMAT0000062_2&hsa-let-7a-5p&offsets|0|-1(+1C);m-4&9|+|94175962|94175983&offsets|0|-1(+1C)]|TGAGGTAGTAGGTTGTATAGTC</td><td>>MI0000061|hsa-let-7a-2&WithFlank&11|-|122146516|122146599</td><td>11</td><td>31</td><td>21(+1C)</td><td>TGAGGTAGTAGGTTGTATAGTC</td><td>MB231</td><td>575</td><td>645</td><td>308</td><td>871</td><td>741</td><td>125.6</td><td>110.1</td><td>151.1</td><td>458.9</td><td>410.9</td></tr>
<tr><td>>MI0000061|hsa-let-7a-2&WithFlank&11|-|122146516|122146599@11.31.21(+1C);MI0000062|hsa-let-7a-3&WithFlank&22|+|46112743|46112828@10.30.21(+1C);MI0000060|hsa-let-7a-1&WithFlank&9|+|94175951|94176042@12.32.21(+1C)&[MIMAT0000062&hsa-let-7a-5p&offsets|0|-1(+1C);m-6&11|-|122146568|122146589&offsets|0|-1(+1C)];[MIMAT0000062_1&hsa-let-7a-5p&offsets|0|-1(+1C);m-5&22|+|46112752|46112773&offsets|0|-1(+1C)];[MIMAT0000062_2&hsa-let-7a-5p&offsets|0|-1(+1C);m-4&9|+|94175962|94175983&offsets|0|-1(+1C)]|TGAGGTAGTAGGTTGTATAGTC</td><td>>MI0000061|hsa-let-7a-2&WithFlank&11|-|122146516|122146599</td><td>11</td><td>31</td><td>21(+1C)</td><td>TGAGGTAGTAGGTTGTATAGTC</td><td>MB468</td><td>22</td><td>29</td><td>13</td><td>25</td><td>7</td><td>8.6</td><td>3.6</td><td>4.5</td><td>19.1</td><td>2.2</td></tr>
<tr><td>>MI0000061|hsa-let-7a-2&WithFlank&11|-|122146516|122146599@11.31.21(+1G);MI0000062|hsa-let-7a-3&WithFlank&22|+|46112743|46112828@10.30.21(+1G);MI0000060|hsa-let-7a-1&WithFlank&9|+|94175951|94176042@12.32.21(+1G)&[MIMAT0000062&hsa-let-7a-5p&offsets|0|-1(+1G);m-6&11|-|122146568|122146589&offsets|0|-1(+1G)];[MIMAT0000062_1&hsa-let-7a-5p&offsets|0|-1(+1G);m-5&22|+|46112752|46112773&offsets|0|-1(+1G)];[MIMAT0000062_2&hsa-let-7a-5p&offsets|0|-1(+1G);m-4&9|+|94175962|94175983&offsets|0|-1(+1G)]|TGAGGTAGTAGGTTGTATAGTG</td><td>>MI0000061|hsa-let-7a-2&WithFlank&11|-|122146516|122146599</td><td>11</td><td>31</td><td>21(+1G)</td><td>TGAGGTAGTAGGTTGTATAGTG</td><td>BT20</td><td>187</td><td>63</td><td>77</td><td>10</td><td>10</td><td>170.4</td><td>37.8</td><td>38.0</td><td>10.1</td><td>3.1</td></tr>
<tr><td>>MI0000061|hsa-let-7a-2&WithFlank&11|-|122146516|122146599@11.31.21(+1G);MI0000062|hsa-let-7a-3&WithFlank&22|+|46112743|46112828@10.30.21(+1G);MI0000060|hsa-let-7a-1&WithFlank&9|+|94175951|94176042@12.32.21(+1G)&[MIMAT0000062&hsa-let-7a-5p&offsets|0|-1(+1G);m-6&11|-|122146568|122146589&offsets|0|-1(+1G)];[MIMAT0000062_1&hsa-let-7a-5p&offsets|0|-1(+1G);m-5&22|+|46112752|46112773&offsets|0|-1(+1G)];[MIMAT0000062_2&hsa-let-7a-5p&offsets|0|-1(+1G);m-4&9|+|94175962|94175983&offsets|0|-1(+1G)]|TGAGGTAGTAGGTTGTATAGTG</td><td>>MI0000061|hsa-let-7a-2&WithFlank&11|-|122146516|122146599</td><td>11</td><td>31</td><td>21(+1G)</td><td>TGAGGTAGTAGGTTGTATAGTG</td><td>MB231</td><td>123</td><td>56</td><td>26</td><td>45</td><td>31</td><td>81.8</td><td>15.6</td><td>17.9</td><td>24.8</td><td>16.6</td></tr>
</table>

<table>
<tr><th></th><th></th><th></th><th></th><th></th><th></th><th></th><th></th><th></th><th></th><th></th><th></th><th></th><th></th><th></th><th></th><th></th><th></th></tr>
<tr><td>>MI0000061|hsa-let-7a-2&WithFlank&11|-|122146516|122146599@11.32.22;MI0000062|hsa-let-7a-3&WithFlank&22|+|46112743|46112828@10.31.22;MI0000060|hsa-let-7a-1&WithFlank&9|+|94175951|94176042@12.33.22&[MIMAT0000062&hsa-let-7a-5p&offsets|0|0;m-6&11|-|122146568|122146589&offsets|0|0];[MIMAT0000062_1&hsa-let-7a-5p&offsets|0|0;m-5&22|+|46112752|46112773&offsets|0|0];[MIMAT0000062_2&hsa-let-7a-5p&offsets|0|0;m-4&9|+|94175962|94175983&offsets|0|0]|TGAGGTAGTAGGTTGTATAGTT</td><td>>MI0000061|hsa-let-7a-2&WithFlank&11|-|122146516|122146599</td><td>11</td><td>32</td><td>22</td><td>TGAGGTAGTAGGTTGTATAGTT</td><td>BT20</td><td>4925</td><td>4615</td><td>5324</td><td>938</td><td>1924</td><td>2678.3</td><td>2494.6</td><td>3036.9</td><td>329.5</td><td>494.5</td></tr>
<tr><td>>MI0000061|hsa-let-7a-2&WithFlank&11|-|122146516|122146599@11.32.22;MI0000062|hsa-let-7a-3&WithFlank&22|+|46112743|46112828@10.31.22;MI0000060|hsa-let-7a-1&WithFlank&9|+|94175951|94176042@12.33.22&[MIMAT0000062&hsa-let-7a-5p&offsets|0|0;m-6&11|-|122146568|122146589&offsets|0|0];[MIMAT0000062_1&hsa-let-7a-5p&offsets|0|0;m-5&22|+|46112752|46112773&offsets|0|0];[MIMAT0000062_2&hsa-let-7a-5p&offsets|0|0;m-4&9|+|94175962|94175983&offsets|0|0]|TGAGGTAGTAGGTTGTATAGTT</td><td>>MI0000061|hsa-let-7a-2&WithFlank&11|-|122146516|122146599</td><td>11</td><td>32</td><td>22</td><td>TGAGGTAGTAGGTTGTATAGTT</td><td>MB231</td><td>5257</td><td>6225</td><td>2848</td><td>5345</td><td>5531</td><td>1095.6</td><td>1036.8</td><td>1264.2</td><td>1976.7</td><td>2339.9</td></tr>
<tr><td>>MI0000061|hsa-let-7a-2&WithFlank&11|-|122146516|122146599@11.32.22;MI0000062|hsa-let-7a-3&WithFlank&22|+|46112743|46112828@10.31.22;MI0000060|hsa-let-7a-1&WithFlank&9|+|94175951|94176042@12.33.22&[MIMAT0000062&hsa-let-7a-5p&offsets|0|0;m-6&11|-|122146568|122146589&offsets|0|0];[MIMAT0000062_1&hsa-let-7a-5p&offsets|0|0;m-5&22|+|46112752|46112773&offsets|0|0];[MIMAT0000062_2&hsa-let-7a-5p&offsets|0|0;m-4&9|+|94175962|94175983&offsets|0|0]|TGAGGTAGTAGGTTGTATAGTT</td><td>>MI0000061|hsa-let-7a-2&WithFlank&11|-|122146516|122146599</td><td>11</td><td>32</td><td>22</td><td>TGAGGTAGTAGGTTGTATAGTT</td><td>MB468</td><td>219</td><td>301</td><td>120</td><td>199</td><td>62</td><td>96.5</td><td>73.3</td><td>21.6</td><td>156.0</td><td>15.5</td></tr>
<tr><td>>MI0000061|hsa-let-7a-2&WithFlank&11|-|122146516|122146599@11.32.22(+1A);MI0000062|hsa-let-7a-3&WithFlank&22|+|46112743|46112828@10.31.22(+1A);MI0000060|hsa-let-7a-1&WithFlank&9|+|94175951|94176042@12.33.22(+1A)&[MIMAT0000062&hsa-let-7a-5p&offsets|0|0(+1A);m-6&11|-|122146568|122146589&offsets|0|0(+1A)];[MIMAT0000062_1&hsa-let-7a-5p&offsets|0|0(+1A);m-5&22|+|46112752|46112773&offsets|0|0(+1A)];[MIMAT0000062_2&hsa-let-7a-5p&offsets|0|0(+1A);m-4&9|+|94175962|94175983&offsets|0|0(+1A)]|TGAGGTAGTAGGTTGTATAGTTA</td><td>>MI0000061|hsa-let-7a-2&WithFlank&11|-|122146516|122146599</td><td>11</td><td>32</td><td>22(+1A)</td><td>TGAGGTAGTAGGTTGTATAGTTA</td><td>BT20</td><td>93</td><td>77</td><td>93</td><td>86</td><td>27</td><td>54.1</td><td>53.3</td><td>53.5</td><td>61.2</td><td>6.8</td></tr>
<tr><td>>MI0000061|hsa-let-7a-2&WithFlank&11|-|122146516|122146599@11.32.22(+1A);MI0000062|hsa-let-7a-3&WithFlank&22|+|46112743|46112828@10.31.22(+1A);MI0000060|hsa-let-7a-1&WithFlank&9|+|94175951|94176042@12.33.22(+1A)&[MIMAT0000062&hsa-let-7a-5p&offsets|0|0(+1A);m-6&11|-|122146568|122146589&offsets|0|0(+1A)];[MIMAT0000062_1&hsa-let-7a-5p&offsets|0|0(+1A);m-5&22|+|46112752|46112773&offsets|0|0(+1A)];[MIMAT0000062_2&hsa-let-7a-5p&offsets|0|0(+1A);m-4&9|+|94175962|94175983&offsets|0|0(+1A)]|TGAGGTAGTAGGTTGTATAGTTA</td><td>>MI0000061|hsa-let-7a-2&WithFlank&11|-|122146516|122146599</td><td>11</td><td>32</td><td>22(+1A)</td><td>TGAGGTAGTAGGTTGTATAGTTA</td><td>MB231</td><td>59</td><td>65</td><td>35</td><td>148</td><td>57</td><td>11.2</td><td>10.1</td><td>23.1</td><td>60.3</td><td>24.2</td></tr>
<tr><td>>MI0000061|hsa-let-7a-2&WithFlank&11|-|122146516|122146599@11.32.22(+1C);MI0000062|hsa-let-7a-3&WithFlank&22|+|46112743|46112828@10.31.22(+1C);MI0000060|hsa-let-7a-1&WithFlank&9|+|94175951|94176042@12.33.22(+1C)&[MIMAT0000062&hsa-let-7a-5p&offsets|0|0(+1C);m-6&11|-|122146568|122146589&offsets|0|0(+1C)];[MIMAT0000062_1&hsa-let-7a-5p&offsets|0|0(+1C);m-5&22|+|46112752|46112773&offsets|0|0(+1C)];[MIMAT0000062_2&hsa-let-7a-5p&offsets|0|0(+1C);m-4&9|+|94175962|94175983&offsets|0|0(+1C)]|TGAGGTAGTAGGTTGTATAGTTC</td><td>>MI0000061|hsa-let-7a-2&WithFlank&11|-|122146516|122146599</td><td>11</td><td>32</td><td>22(+1C)</td><td>TGAGGTAGTAGGTTGTATAGTTC</td><td>BT20</td><td>15</td><td>14</td><td>16</td><td>15</td><td>6</td><td>7.9</td><td>8.0</td><td>8.5</td><td>13.9</td><td>0.5</td></tr>
<tr><td>>MI0000061|hsa-let-7a-2&WithFlank&11|-|122146516|122146599@11.32.22(+1C);MI0000062|hsa-let-7a-3&WithFlank&22|+|46112743|46112828@10.31.22(+1C);MI0000060|hsa-let-7a-1&WithFlank&9|+|94175951|94176042@12.33.22(+1C)&[MIMAT0000062&hsa-let-7a-5p&offsets|0|0(+1C);m-6&11|-|122146568|122146589&offsets|0|0(+1C)];[MIMAT0000062_1&hsa-let-7a-5p&offsets|0|0(+1C);m-5&22|+|46112752|46112773&offsets|0|0(+1C)];[MIMAT0000062_2&hsa-let-7a-5p&offsets|0|0(+1C);m-4&9|+|94175962|94175983&offsets|0|0(+1C)]|TGAGGTAGTAGGTTGTATAGTTC</td><td>>MI0000061|hsa-let-7a-2&WithFlank&11|-|122146516|122146599</td><td>11</td><td>32</td><td>22(+1C)</td><td>TGAGGTAGTAGGTTGTATAGTTC</td><td>MB231</td><td>17</td><td>17</td><td>11</td><td>73</td><td>20</td><td>2.5</td><td>4.2</td><td>3.9</td><td>33.2</td><td>10.7</td></tr>
<tr><td>>MI0000061|hsa-let-7a-2&WithFlank&11|-|122146516|122146599@11.32.22(+1G);MI0000062|hsa-let-7a-3&WithFlank&22|+|46112743|46112828@10.31.22(+1G);MI0000060|hsa-let-7a-1&WithFlank&9|+|94175951|94176042@12.33.22(+1G)&[MIMAT0000062&hsa-let-7a-5p&offsets|0|0(+1G);m-6&11|-|122146568|122146589&offsets|0|0(+1G)];[MIMAT0000062_1&hsa-let-7a-5p&offsets|0|0(+1G);m-5&22|+|46112752|46112773&offsets|0|0(+1G)];[MIMAT0000062_2&hsa-let-7a-5p&offsets|0|0(+1G);m-4&9|+|94175962|94175983&offsets|0|0(+1G)]|TGAGGTAGTAGGTTGTATAGTTG</td><td>>MI0000061|hsa-let-7a-2&WithFlank&11|-|122146516|122146599</td><td>11</td><td>32</td><td>22(+1G)</td><td>TGAGGTAGTAGGTTGTATAGTTG</td><td>BT20</td><td>13</td><td>11</td><td>13</td><td>12</td><td>4</td><td>6.5</td><td>1.9</td><td>7.0</td><td>9.0</td><td>0.5</td></tr>
<tr><td>>MI0000061|hsa-let-7a-2&WithFlank&11|-|122146516|122146599@11.33.23;MI0000062|hsa-let-7a-3&WithFlank&22|+|46112743|46112828@10.32.23;MI0000060|hsa-let-7a-1&WithFlank&9|+|94175951|94176042@12.34.23&[MIMAT0000062&hsa-let-7a-5p&offsets|0|+1;m-6&11|-|122146568|122146589&offsets|0|+1];[MIMAT0000062_1&hsa-let-7a-5p&offsets|0|+1;m-5&22|+|46112752|46112773&offsets|0|+1];[MIMAT0000062_2&hsa-let-7a-5p&offsets|0|+1;m-4&9|+|94175962|94175983&offsets|0|+1]|TGAGGTAGTAGGTTGTATAGTTT</td><td>>MI0000061|hsa-let-7a-2&WithFlank&11|-|122146516|122146599</td><td>11</td><td>33</td><td>23</td><td>TGAGGTAGTAGGTTGTATAGTTT</td><td>BT20</td><td>111</td><td>107</td><td>111</td><td>93</td><td>48</td><td>65.7</td><td>63.1</td><td>69.0</td><td>73.2</td><td>8.9</td></tr>
<tr><td>>MI0000061|hsa-let-7a-2&WithFlank&11|-|122146516|122146599@11.33.23;MI0000062|hsa-let-7a-3&WithFlank&22|+|46112743|46112828@10.32.23;MI0000060|hsa-let-7a-1&WithFlank&9|+|94175951|94176042@12.34.23&[MIMAT0000062&hsa-let-7a-5p&offsets|0|+1;m-6&11|-|122146568|122146589&offsets|0|+1];[MIMAT0000062_1&hsa-let-7a-5p&offsets|0|+1;m-5&22|+|46112752|46112773&offsets|0|+1];[MIMAT0000062_2&hsa-let-7a-5p&offsets|0|+1;m-4&9|+|94175962|94175983&offsets|0|+1]|TGAGGTAGTAGGTTGTATAGTTT</td><td>>MI0000061|hsa-let-7a-2&WithFlank&11|-|122146516|122146599</td><td>11</td><td>33</td><td>23</td><td>TGAGGTAGTAGGTTGTATAGTTT</td><td>MB231</td><td>168</td><td>178</td><td>97</td><td>485</td><td>179</td><td>19.5</td><td>40.6</td><td>43.0</td><td>160.3</td><td>76.2</td></tr>
<tr><td>>MI0000062|hsa-let-7a-3&WithFlank&22|+|46112743|46112828@58.78.21(+1U);MI0000060|hsa-let-7a-1&WithFlank&9|+|94175951|94176042@63.83.21(+1U)&[MIMAT0004481&hsa-let-7a-3p&offsets|0|0(+1U);m-180&22|+|46112800|46112821&offsets|0|-1(+1U)];[MIMAT0004481_1&hsa-let-7a-3p&offsets|0|0(+1U);m-181&9|+|94176013|94176034&offsets|0|-1(+1U)]|CTATACAATCTACTGTCTTTCT</td><td>>MI0000062|hsa-let-7a-3&WithFlank&22|+|46112743|46112828</td><td>58</td><td>78</td><td>21(+1U)</td><td>CTATACAATCTACTGTCTTTCT</td><td>BT20</td><td>76</td><td>47</td><td>48</td><td>35</td><td>22</td><td>68.3</td><td>48.2</td><td>65.0</td><td>32.3</td><td>23.3</td></tr>
<tr><td>>MI0000062|hsa-let-7a-3&WithFlank&22|+|46112743|46112828@58.78.21(+1U);MI0000060|hsa-let-7a-1&WithFlank&9|+|94175951|94176042@63.83.21(+1U)&[MIMAT0004481&hsa-let-7a-3p&offsets|0|0(+1U);m-180&22|+|46112800|46112821&offsets|0|-1(+1U)];[MIMAT0004481_1&hsa-let-7a-3p&offsets|0|0(+1U);m-181&9|+|94176013|94176034&offsets|0|-1(+1U)]|CTATACAATCTACTGTCTTTCT</td><td>>MI0000062|hsa-let-7a-3&WithFlank&22|+|46112743|46112828</td><td>58</td><td>78</td><td>21(+1U)</td><td>CTATACAATCTACTGTCTTTCT</td><td>MB231</td><td>67</td><td>85</td><td>19</td><td>83</td><td>120</td><td>18.4</td><td>26.0</td><td>4.2</td><td>37.7</td><td>78.6</td></tr>
<tr><td>>MI0000062|hsa-let-7a-3&WithFlank&22|+|46112743|46112828@58.79.22;MI0000060|hsa-let-7a-1&WithFlank&9|+|94175951|94176042@63.84.22&[MIMAT0004481&hsa-let-7a-3p&offsets|0|+1;m-180&22|+|46112800|46112821&offsets|0|0];[MIMAT0004481_1&hsa-let-7a-3p&offsets|0|+1;m-181&9|+|94176013|94176034&offsets|0|0]|CTATACAATCTACTGTCTTTCC</td><td>>MI0000062|hsa-let-7a-3&WithFlank&22|+|46112743|46112828</td><td>58</td><td>79</td><td>22</td><td>CTATACAATCTACTGTCTTTCC</td><td>BT20</td><td>19</td><td>14</td><td>15</td><td>13</td><td>5</td><td>14.0</td><td>15.8</td><td>17.7</td><td>10.7</td><td>5.3</td></tr>
<tr><td>>MI0000062|hsa-let-7a-3&WithFlank&22|+|46112743|46112828@58.79.22;MI0000060|hsa-let-7a-1&WithFlank&9|+|94175951|94176042@63.84.22&[MIMAT0004481&hsa-let-7a-3p&offsets|0|+1;m-180&22|+|46112800|46112821&offsets|0|0];[MIMAT0004481_1&hsa-let-7a-3p&offsets|0|+1;m-181&9|+|94176013|94176034&offsets|0|0]|CTATACAATCTACTGTCTTTCC</td><td>>MI0000062|hsa-let-7a-3&WithFlank&22|+|46112743|46112828</td><td>58</td><td>79</td><td>22</td><td>CTATACAATCTACTGTCTTTCC</td><td>MB231</td><td>11</td><td>15</td><td>4</td><td>21</td><td>23</td><td>2.8</td><td>4.9</td><td>2.1</td><td>10.2</td><td>16.3</td></tr>
<tr><td>>MI0000063|hsa-let-7b&WithFlank&22|+|46113680|46113774@12.29.18&[MIMAT0000063&hsa-let-7b-5p&offsets|0|-4;m-14&22|+|46113691|46113712&offsets|0|-4]|TGAGGTAGTAGGTTGTGT</td><td>>MI0000063|hsa-let-7b&WithFlank&22|+|46113680|46113774</td><td>12</td><td>29</td><td>18</td><td>TGAGGTAGTAGGTTGTGT</td><td>BT20</td><td>13</td><td>14</td><td>25</td><td>59</td><td>6</td><td>9.3</td><td>6.1</td><td>11.0</td><td>47.2</td><td>1.3</td></tr>
<tr><td>>MI0000063|hsa-let-7b&WithFlank&22|+|46113680|46113774@12.31.20(+1A)&[MIMAT0000063&hsa-let-7b-5p&offsets|0|-2(+1A);m-14&22|+|46113691|46113712&offsets|0|-2(+1A)]|TGAGGTAGTAGGTTGTGTGGA</td><td>>MI0000063|hsa-let-7b&WithFlank&22|+|46113680|46113774</td><td>12</td><td>31</td><td>20(+1A)</td><td>TGAGGTAGTAGGTTGTGTGGA</td><td>BT20</td><td>17</td><td>13</td><td>28</td><td>32</td><td>4</td><td>13.1</td><td>3.2</td><td>16.2</td><td>37.0</td><td>1.1</td></tr>
</table>

<table>
<tr><td>>MI0000063|hsa-let-7b&WithFlank&22|+|46113680|46113774@12.31.20(+1C)&[MIMAT0000063&hsa-let-7b-5p&offsets|0|-2(+1C);m-14&22|+|46113691|46113712&offsets|0|-2(+1C)]|TGAGGTAGTAGGTTGTGTGGC</td><td>>MI0000063|hsa-let-7b&WithFlank&22|+|46113680|46113774</td><td>12</td><td>31</td><td>20(+1C)</td><td>TGAGGTAGTAGGTTGTGTGGC</td><td>BT20</td><td>36</td><td>45</td><td>62</td><td>112</td><td>17</td><td>12.2</td><td>14.6</td><td>17.5</td><td>114.9</td><td>2.5</td></tr>
<tr><td>>MI0000063|hsa-let-7b&WithFlank&22|+|46113680|46113774@12.31.20(+1G)&[MIMAT0000063&hsa-let-7b-5p&offsets|0|-2(+1G);m-14&22|+|46113691|46113712&offsets|0|-2(+1G)]|TGAGGTAGTAGGTTGTGTGGG</td><td>>MI0000063|hsa-let-7b&WithFlank&22|+|46113680|46113774</td><td>12</td><td>31</td><td>20(+1G)</td><td>TGAGGTAGTAGGTTGTGTGGG</td><td>BT20</td><td>19</td><td>9</td><td>13</td><td>15</td><td>1</td><td>16.0</td><td>3.3</td><td>6.6</td><td>21.0</td><td>0.0</td></tr>
<tr><td>>MI0000063|hsa-let-7b&WithFlank&22|+|46113680|46113774@12.31.20&[MIMAT0000063&hsa-let-7b-5p&offsets|0|-2;m-14&22|+|46113691|46113712&offsets|0|-2]|TGAGGTAGTAGGTTGTGTGG</td><td>>MI0000063|hsa-let-7b&WithFlank&22|+|46113680|46113774</td><td>12</td><td>31</td><td>20</td><td>TGAGGTAGTAGGTTGTGTGG</td><td>BT20</td><td>32</td><td>42</td><td>62</td><td>178</td><td>15</td><td>13.3</td><td>8.1</td><td>7.4</td><td>125.3</td><td>3.9</td></tr>
<tr><td>>MI0000063|hsa-let-7b&WithFlank&22|+|46113680|46113774@12.32.21(+1A)&[MIMAT0000063&hsa-let-7b-5p&offsets|0|-1(+1A);m-14&22|+|46113691|46113712&offsets|0|-1(+1A)]|TGAGGTAGTAGGTTGTGTGGTA</td><td>>MI0000063|hsa-let-7b&WithFlank&22|+|46113680|46113774</td><td>12</td><td>32</td><td>21(+1A)</td><td>TGAGGTAGTAGGTTGTGTGGTA</td><td>BT20</td><td>175</td><td>118</td><td>271</td><td>186</td><td>45</td><td>115.2</td><td>42.3</td><td>119.9</td><td>191.9</td><td>10.4</td></tr>
<tr><td>>MI0000063|hsa-let-7b&WithFlank&22|+|46113680|46113774@12.32.21(+1A)&[MIMAT0000063&hsa-let-7b-5p&offsets|0|-1(+1A);m-14&22|+|46113691|46113712&offsets|0|-1(+1A)]|TGAGGTAGTAGGTTGTGTGGTA</td><td>>MI0000063|hsa-let-7b&WithFlank&22|+|46113680|46113774</td><td>12</td><td>32</td><td>21(+1A)</td><td>TGAGGTAGTAGGTTGTGTGGTA</td><td>MB231</td><td>18</td><td>16</td><td>17</td><td>117</td><td>18</td><td>1.0</td><td>3.9</td><td>11.7</td><td>58.9</td><td>9.1</td></tr>
<tr><td>>MI0000063|hsa-let-7b&WithFlank&22|+|46113680|46113774@12.32.21(+1C)&[MIMAT0000063&hsa-let-7b-5p&offsets|0|-1(+1C);m-14&22|+|46113691|46113712&offsets|0|-1(+1C)]|TGAGGTAGTAGGTTGTGTGGTC</td><td>>MI0000063|hsa-let-7b&WithFlank&22|+|46113680|46113774</td><td>12</td><td>32</td><td>21(+1C)</td><td>TGAGGTAGTAGGTTGTGTGGTC</td><td>BT20</td><td>338</td><td>390</td><td>531</td><td>602</td><td>158</td><td>107.1</td><td>162.7</td><td>81.6</td><td>606.0</td><td>28.4</td></tr>
<tr><td>>MI0000063|hsa-let-7b&WithFlank&22|+|46113680|46113774@12.32.21(+1C)&[MIMAT0000063&hsa-let-7b-5p&offsets|0|-1(+1C);m-14&22|+|46113691|46113712&offsets|0|-1(+1C)]|TGAGGTAGTAGGTTGTGTGGTC</td><td>>MI0000063|hsa-let-7b&WithFlank&22|+|46113680|46113774</td><td>12</td><td>32</td><td>21(+1C)</td><td>TGAGGTAGTAGGTTGTGTGGTC</td><td>MB231</td><td>54</td><td>61</td><td>46</td><td>382</td><td>65</td><td>9.3</td><td>15.5</td><td>27.7</td><td>196.4</td><td>34.4</td></tr>
<tr><td>>MI0000063|hsa-let-7b&WithFlank&22|+|46113680|46113774@12.32.21(+1G)&[MIMAT0000063&hsa-let-7b-5p&offsets|0|-1(+1G);m-14&22|+|46113691|46113712&offsets|0|-1(+1G)]|TGAGGTAGTAGGTTGTGTGGTG</td><td>>MI0000063|hsa-let-7b&WithFlank&22|+|46113680|46113774</td><td>12</td><td>32</td><td>21(+1G)</td><td>TGAGGTAGTAGGTTGTGTGGTG</td><td>BT20</td><td>130</td><td>54</td><td>85</td><td>63</td><td>9</td><td>110.1</td><td>21.3</td><td>38.8</td><td>83.2</td><td>2.5</td></tr>
<tr><td>>MI0000063|hsa-let-7b&WithFlank&22|+|46113680|46113774@12.32.21(+1G)&[MIMAT0000063&hsa-let-7b-5p&offsets|0|-1(+1G);m-14&22|+|46113691|46113712&offsets|0|-1(+1G)]|TGAGGTAGTAGGTTGTGTGGTG</td><td>>MI0000063|hsa-let-7b&WithFlank&22|+|46113680|46113774</td><td>12</td><td>32</td><td>21(+1G)</td><td>TGAGGTAGTAGGTTGTGTGGTG</td><td>MB231</td><td>17</td><td>7</td><td>5</td><td>27</td><td>4</td><td>11.9</td><td>2.8</td><td>4.4</td><td>16.5</td><td>1.9</td></tr>
<tr><td>>MI0000063|hsa-let-7b&WithFlank&22|+|46113680|46113774@12.32.21&[MIMAT0000063&hsa-let-7b-5p&offsets|0|-1;m-14&22|+|46113691|46113712&offsets|0|-1]|TGAGGTAGTAGGTTGTGTGGT</td><td>>MI0000063|hsa-let-7b&WithFlank&22|+|46113680|46113774</td><td>12</td><td>32</td><td>21</td><td>TGAGGTAGTAGGTTGTGTGGT</td><td>BT20</td><td>281</td><td>310</td><td>475</td><td>768</td><td>135</td><td>110.9</td><td>64.5</td><td>75.9</td><td>665.0</td><td>4.5</td></tr>
<tr><td>>MI0000063|hsa-let-7b&WithFlank&22|+|46113680|46113774@12.32.21&[MIMAT0000063&hsa-let-7b-5p&offsets|0|-1;m-14&22|+|46113691|46113712&offsets|0|-1]|TGAGGTAGTAGGTTGTGTGGT</td><td>>MI0000063|hsa-let-7b&WithFlank&22|+|46113680|46113774</td><td>12</td><td>32</td><td>21</td><td>TGAGGTAGTAGGTTGTGTGGT</td><td>MB231</td><td>60</td><td>76</td><td>52</td><td>393</td><td>57</td><td>9.8</td><td>14.3</td><td>31.6</td><td>138.0</td><td>17.4</td></tr>
<tr><td>>MI0000063|hsa-let-7b&WithFlank&22|+|46113680|46113774@12.33.22(+1A)&[MIMAT0000063&hsa-let-7b-5p&offsets|0|0(+1A);m-14&22|+|46113691|46113712&offsets|0|0(+1A)]|TGAGGTAGTAGGTTGTGTGGTTA</td><td>>MI0000063|hsa-let-7b&WithFlank&22|+|46113680|46113774</td><td>12</td><td>33</td><td>22(+1A)</td><td>TGAGGTAGTAGGTTGTGTGGTTA</td><td>BT20</td><td>183</td><td>129</td><td>190</td><td>453</td><td>68</td><td>110.1</td><td>61.9</td><td>28.2</td><td>337.0</td><td>18.6</td></tr>
<tr><td>>MI0000063|hsa-let-7b&WithFlank&22|+|46113680|46113774@12.33.22(+1A)&[MIMAT0000063&hsa-let-7b-5p&offsets|0|0(+1A);m-14&22|+|46113691|46113712&offsets|0|0(+1A)]|TGAGGTAGTAGGTTGTGTGGTTA</td><td>>MI0000063|hsa-let-7b&WithFlank&22|+|46113680|46113774</td><td>12</td><td>33</td><td>22(+1A)</td><td>TGAGGTAGTAGGTTGTGTGGTTA</td><td>MB231</td><td>27</td><td>28</td><td>16</td><td>162</td><td>30</td><td>5.7</td><td>5.1</td><td>10.0</td><td>32.5</td><td>12.6</td></tr>
<tr><td>>MI0000063|hsa-let-7b&WithFlank&22|+|46113680|46113774@12.33.22(+1C)&[MIMAT0000063&hsa-let-7b-5p&offsets|0|0(+1C);m-14&22|+|46113691|46113712&offsets|0|0(+1C)]|TGAGGTAGTAGGTTGTGTGGTTC</td><td>>MI0000063|hsa-let-7b&WithFlank&22|+|46113680|46113774</td><td>12</td><td>33</td><td>22(+1C)</td><td>TGAGGTAGTAGGTTGTGTGGTTC</td><td>BT20</td><td>47</td><td>43</td><td>62</td><td>142</td><td>22</td><td>19.4</td><td>17.7</td><td>5.2</td><td>119.7</td><td>4.4</td></tr>
<tr><td>>MI0000063|hsa-let-7b&WithFlank&22|+|46113680|46113774@12.33.22(+1C)&[MIMAT0000063&hsa-let-7b-5p&offsets|0|0(+1C);m-14&22|+|46113691|46113712&offsets|0|0(+1C)]|TGAGGTAGTAGGTTGTGTGGTTC</td><td>>MI0000063|hsa-let-7b&WithFlank&22|+|46113680|46113774</td><td>12</td><td>33</td><td>22(+1C)</td><td>TGAGGTAGTAGGTTGTGTGGTTC</td><td>MB231</td><td>10</td><td>10</td><td>10</td><td>87</td><td>12</td><td>1.6</td><td>3.3</td><td>5.6</td><td>43.6</td><td>7.2</td></tr>
<tr><td>>MI0000063|hsa-let-7b&WithFlank&22|+|46113680|46113774@12.33.22(+1G)&[MIMAT0000063&hsa-let-7b-5p&offsets|0|0(+1G);m-14&22|+|46113691|46113712&offsets|0|0(+1G)]|TGAGGTAGTAGGTTGTGTGGTTG</td><td>>MI0000063|hsa-let-7b&WithFlank&22|+|46113680|46113774</td><td>12</td><td>33</td><td>22(+1G)</td><td>TGAGGTAGTAGGTTGTGTGGTTG</td><td>BT20</td><td>28</td><td>17</td><td>26</td><td>62</td><td>9</td><td>10.4</td><td>4.0</td><td>4.1</td><td>48.2</td><td>1.4</td></tr>
<tr><td>>MI0000063|hsa-let-7b&WithFlank&22|+|46113680|46113774@12.33.22(+2A)&[MIMAT0000063&hsa-let-7b-5p&offsets|0|0(+2A);m-14&22|+|46113691|46113712&offsets|0|0(+2A)]|TGAGGTAGTAGGTTGTGTGGTTAA</td><td>>MI0000063|hsa-let-7b&WithFlank&22|+|46113680|46113774</td><td>12</td><td>33</td><td>22(+2A)</td><td>TGAGGTAGTAGGTTGTGTGGTTAA</td><td>BT20</td><td>16</td><td>12</td><td>18</td><td>30</td><td>6</td><td>14.1</td><td>7.6</td><td>5.0</td><td>20.9</td><td>3.3</td></tr>
<tr><td>>MI0000063|hsa-let-7b&WithFlank&22|+|46113680|46113774@12.33.22&[MIMAT0000063&hsa-let-7b-5p&offsets|0|0;m-14&22|+|46113691|46113712&offsets|0|0]|TGAGGTAGTAGGTTGTGTGGTT</td><td>>MI0000063|hsa-let-7b&WithFlank&22|+|46113680|46113774</td><td>12</td><td>33</td><td>22</td><td>TGAGGTAGTAGGTTGTGTGGTT</td><td>BT20</td><td>2399</td><td>2679</td><td>3955</td><td>4079</td><td>1174</td><td>989.3</td><td>866.5</td><td>46.0</td><td>3323.7</td><td>151.7</td></tr>
<tr><td>>MI0000063|hsa-let-7b&WithFlank&22|+|46113680|46113774@12.33.22&[MIMAT0000063&hsa-let-7b-5p&offsets|0|0;m-14&22|+|46113691|46113712&offsets|0|0]|TGAGGTAGTAGGTTGTGTGGTT</td><td>>MI0000063|hsa-let-7b&WithFlank&22|+|46113680|46113774</td><td>12</td><td>33</td><td>22</td><td>TGAGGTAGTAGGTTGTGTGGTT</td><td>MB231</td><td>463</td><td>564</td><td>416</td><td>2326</td><td>464</td><td>90.0</td><td>96.1</td><td>231.4</td><td>731.3</td><td>161.1</td></tr>
<tr><td>>MI0000063|hsa-let-7b&WithFlank&22|+|46113680|46113774@12.33.22&[MIMAT0000063&hsa-let-7b-5p&offsets|0|0;m-14&22|+|46113691|46113712&offsets|0|0]|TGAGGTAGTAGGTTGTGTGGTT</td><td>>MI0000063|hsa-let-7b&WithFlank&22|+|46113680|46113774</td><td>12</td><td>33</td><td>22</td><td>TGAGGTAGTAGGTTGTGTGGTT</td><td>MB468</td><td>51</td><td>91</td><td>32</td><td>163</td><td>16</td><td>16.5</td><td>20.8</td><td>6.6</td><td>67.7</td><td>4.1</td></tr>
<tr><td>>MI0000063|hsa-let-7b&WithFlank&22|+|46113680|46113774@12.34.23(+1U)&[MIMAT0000063&hsa-let-7b-5p&offsets|0|+1(+1U);m-14&22|+|46113691|46113712&offsets|0|+1(+1U)]|TGAGGTAGTAGGTTGTGTGGTTTT</td><td>>MI0000063|hsa-let-7b&WithFlank&22|+|46113680|46113774</td><td>12</td><td>34</td><td>23(+1U)</td><td>TGAGGTAGTAGGTTGTGTGGTTTT</td><td>BT20</td><td>33</td><td>27</td><td>52</td><td>52</td><td>14</td><td>18.7</td><td>11.8</td><td>3.5</td><td>36.0</td><td>4.8</td></tr>
<tr><td>>MI0000063|hsa-let-7b&WithFlank&22|+|46113680|46113774@12.34.23(+1U)&[MIMAT0000063&hsa-let-7b-5p&offsets|0|+1(+1U);m-14&22|+|46113691|46113712&offsets|0|+1(+1U)]|TGAGGTAGTAGGTTGTGTGGTTTT</td><td>>MI0000063|hsa-let-7b&WithFlank&22|+|46113680|46113774</td><td>12</td><td>34</td><td>23(+1U)</td><td>TGAGGTAGTAGGTTGTGTGGTTTT</td><td>MB231</td><td>13</td><td>12</td><td>13</td><td>56</td><td>12</td><td>1.3</td><td>3.4</td><td>5.5</td><td>16.2</td><td>5.0</td></tr>
<tr><td>>MI0000063|hsa-let-7b&WithFlank&22|+|46113680|46113774@12.34.23&[MIMAT0000063&hsa-let-7b-5p&offsets|0|+1;m-14&22|+|46113691|46113712&offsets|0|+1]|TGAGGTAGTAGGTTGTGTGGTTT</td><td>>MI0000063|hsa-let-7b&WithFlank&22|+|46113680|46113774</td><td>12</td><td>34</td><td>23</td><td>TGAGGTAGTAGGTTGTGTGGTTT</td><td>BT20</td><td>359</td><td>332</td><td>501</td><td>1023</td><td>179</td><td>185.9</td><td>116.1</td><td>40.3</td><td>729.7</td><td>39.8</td></tr>
<tr><td>>MI0000063|hsa-let-7b&WithFlank&22|+|46113680|46113774@12.34.23&[MIMAT0000063&hsa-let-7b-5p&offsets|0|+1;m-14&22|+|46113691|46113712&offsets|0|+1]|TGAGGTAGTAGGTTGTGTGGTTT</td><td>>MI0000063|hsa-let-7b&WithFlank&22|+|46113680|46113774</td><td>12</td><td>34</td><td>23</td><td>TGAGGTAGTAGGTTGTGTGGTTT</td><td>MB231</td><td>98</td><td>105</td><td>83</td><td>552</td><td>99</td><td>14.9</td><td>21.4</td><td>45.3</td><td>165.9</td><td>38.3</td></tr>
<tr><td>>MI0000063|hsa-let-7b&WithFlank&22|+|46113680|46113774@66.86.21(+1U)&[MIMAT0004482&hsa-let-7b-3p&offsets|0|-1(+1U);m-171&22|+|46113745|46113766&offsets|0|-1(+1U)]|CTATACAACCTACTGCCTTCCT</td><td>>MI0000063|hsa-let-7b&WithFlank&22|+|46113680|46113774</td><td>66</td><td>86</td><td>21(+1U)</td><td>CTATACAACCTACTGCCTTCCT</td><td>BT20</td><td>15</td><td>19</td><td>18</td><td>38</td><td>11</td><td>10.2</td><td>19.4</td><td>11.6</td><td>30.0</td><td>7.1</td></tr>
<tr><td>>MI0000064|hsa-let-7c&WithFlank&21|+|16539822|16539917@17.38.22&[MIMAT0000064&hsa-let-7c-5p&offsets|0|0;m-37&21|+|16539838|16539859&offsets|0|0]|TGAGGTAGTAGGTTGTATGGTT</td><td>>MI0000064|hsa-let-7c&WithFlank&21|+|16539822|16539917</td><td>17</td><td>38</td><td>22</td><td>TGAGGTAGTAGGTTGTATGGTT</td><td>BT20</td><td>14</td><td>15</td><td>31</td><td>6</td><td>5</td><td>6.0</td><td>6.5</td><td>7.1</td><td>5.2</td><td>1.1</td></tr>
<tr><td>>MI0000064|hsa-let-7c&WithFlank&21|+|16539822|16539917@17.38.22&[MIMAT0000064&hsa-let-7c-5p&offsets|0|0;m-37&21|+|16539838|16539859&offsets|0|0]|TGAGGTAGTAGGTTGTATGGTT</td><td>>MI0000064|hsa-let-7c&WithFlank&21|+|16539822|16539917</td><td>17</td><td>38</td><td>22</td><td>TGAGGTAGTAGGTTGTATGGTT</td><td>MB231</td><td>21</td><td>27</td><td>17</td><td>43</td><td>20</td><td>4.5</td><td>5.3</td><td>8.8</td><td>20.4</td><td>10.0</td></tr>
<tr><td>>MI0000064|hsa-let-7c&WithFlank&21|+|16539822|16539917@17.38.22&[MIMAT0000064&hsa-let-7c-5p&offsets|0|0;m-37&21|+|16539838|16539859&offsets|0|0]|TGAGGTAGTAGGTTGTATGGTT</td><td>>MI0000064|hsa-let-7c&WithFlank&21|+|16539822|16539917</td><td>17</td><td>38</td><td>22</td><td>TGAGGTAGTAGGTTGTATGGTT</td><td>MB468</td><td>28</td><td>46</td><td>25</td><td>57</td><td>6</td><td>8.2</td><td>12.2</td><td>7.9</td><td>26.8</td><td>1.9</td></tr>
</table>

| | | | | | | | | | | | | | | | | |
|---|---|---|---|---|---|---|---|---|---|---|---|---|---|---|---|---|
| >MI0000065\|hsa-let-7d&WithFlank&9\|+\|94178828\|94178926@14.31.18&[MIMAT0000065&hsa-let-7d-5p&offsets\|0\|-4;m-90&9\|+\|94178841\|94178862&offsets\|0\|-4]\|AGAGGTAGTAGGTTGCAT | >MI0000065\|hsa-let-7d&WithFlank&9\|+\|94178828\|94178926 | 14 | 31 | 18 | AGAGGTAGTAGGTTGCAT | BT20 | 16 | 22 | 44 | 16 | 11 | 17.1 | 12.4 | 53.1 | 7.0 | 1.3 |
| >MI0000065\|hsa-let-7d&WithFlank&9\|+\|94178828\|94178926@14.33.20(+1C)&[MIMAT0000065&hsa-let-7d-5p&offsets\|0\|-2(+1C);m-90&9\|+\|94178841\|94178862&offsets\|0\|-2(+1C)]\|AGAGGTAGTAGGTTGCATAGC | >MI0000065\|hsa-let-7d&WithFlank&9\|+\|94178828\|94178926 | 14 | 33 | 20(+1C) | AGAGGTAGTAGGTTGCATAGC | BT20 | 12 | 13 | 14 | 4 | 5 | 6.6 | 6.8 | 10.9 | 3.1 | 1.3 |
| >MI0000065\|hsa-let-7d&WithFlank&9\|+\|94178828\|94178926@14.34.21(+1A)&[MIMAT0000065&hsa-let-7d-5p&offsets\|0\|-1(+1A);m-90&9\|+\|94178841\|94178862&offsets\|0\|-1(+1A)]\|AGAGGTAGTAGGTTGCATAGTA | >MI0000065\|hsa-let-7d&WithFlank&9\|+\|94178828\|94178926 | 14 | 34 | 21(+1A) | AGAGGTAGTAGGTTGCATAGTA | BT20 | 31 | 16 | 31 | 6 | 5 | 32.2 | 7.6 | 18.5 | 5.6 | 0.5 |
| >MI0000065\|hsa-let-7d&WithFlank&9\|+\|94178828\|94178926@14.34.21(+1C)&[MIMAT0000065&hsa-let-7d-5p&offsets\|0\|-1(+1C);m-90&9\|+\|94178841\|94178862&offsets\|0\|-1(+1C)]\|AGAGGTAGTAGGTTGCATAGTC | >MI0000065\|hsa-let-7d&WithFlank&9\|+\|94178828\|94178926 | 14 | 34 | 21(+1C) | AGAGGTAGTAGGTTGCATAGTC | BT20 | 80 | 95 | 105 | 33 | 34 | 41.2 | 49.3 | 72.1 | 30.1 | 8.3 |
| >MI0000065\|hsa-let-7d&WithFlank&9\|+\|94178828\|94178926@14.34.21(+1C)&[MIMAT0000065&hsa-let-7d-5p&offsets\|0\|-1(+1C);m-90&9\|+\|94178841\|94178862&offsets\|0\|-1(+1C)]\|AGAGGTAGTAGGTTGCATAGTC | >MI0000065\|hsa-let-7d&WithFlank&9\|+\|94178828\|94178926 | 14 | 34 | 21(+1C) | AGAGGTAGTAGGTTGCATAGTC | MB231 | 29 | 40 | 20 | 86 | 43 | 10.1 | 9.1 | 11.1 | 60.6 | 26.9 |
| >MI0000065\|hsa-let-7d&WithFlank&9\|+\|94178828\|94178926@14.34.21(+1G)&[MIMAT0000065&hsa-let-7d-5p&offsets\|0\|-1(+1G);m-90&9\|+\|94178841\|94178862&offsets\|0\|-1(+1G)]\|AGAGGTAGTAGGTTGCATAGTG | >MI0000065\|hsa-let-7d&WithFlank&9\|+\|94178828\|94178926 | 14 | 34 | 21(+1G) | AGAGGTAGTAGGTTGCATAGTG | BT20 | 29 | 9 | 11 | 3 | 1 | 33.6 | 4.2 | 6.1 | 3.1 | 0.4 |
| >MI0000065\|hsa-let-7d&WithFlank&9\|+\|94178828\|94178926@14.34.21&[MIMAT0000065&hsa-let-7d-5p&offsets\|0\|-1;m-90&9\|+\|94178841\|94178862&offsets\|0\|-1]\|AGAGGTAGTAGGTTGCATAGT | >MI0000065\|hsa-let-7d&WithFlank&9\|+\|94178828\|94178926 | 14 | 34 | 21 | AGAGGTAGTAGGTTGCATAGT | BT20 | 92 | 98 | 116 | 29 | 39 | 46.1 | 35.5 | 99.2 | 15.2 | 6.5 |
| >MI0000065\|hsa-let-7d&WithFlank&9\|+\|94178828\|94178926@14.34.21&[MIMAT0000065&hsa-let-7d-5p&offsets\|0\|-1;m-90&9\|+\|94178841\|94178862&offsets\|0\|-1]\|AGAGGTAGTAGGTTGCATAGT | >MI0000065\|hsa-let-7d&WithFlank&9\|+\|94178828\|94178926 | 14 | 34 | 21 | AGAGGTAGTAGGTTGCATAGT | MB231 | 55 | 65 | 33 | 88 | 63 | 14.7 | 16.9 | 16.2 | 49.6 | 26.3 |
| >MI0000065\|hsa-let-7d&WithFlank&9\|+\|94178828\|94178926@14.35.22(+1A)&[MIMAT0000065&hsa-let-7d-5p&offsets\|0\|0(+1A);m-90&9\|+\|94178841\|94178862&offsets\|0\|0(+1A)]\|AGAGGTAGTAGGTTGCATAGTTA | >MI0000065\|hsa-let-7d&WithFlank&9\|+\|94178828\|94178926 | 14 | 35 | 22(+1A) | AGAGGTAGTAGGTTGCATAGTTA | BT20 | 15 | 13 | 17 | 15 | 5 | 10.5 | 9.2 | 13.1 | 12.3 | 1.2 |
| >MI0000065\|hsa-let-7d&WithFlank&9\|+\|94178828\|94178926@14.35.22&[MIMAT0000065&hsa-let-7d-5p&offsets\|0\|0;m-90&9\|+\|94178841\|94178862&offsets\|0\|0]\|AGAGGTAGTAGGTTGCATAGTT | >MI0000065\|hsa-let-7d&WithFlank&9\|+\|94178828\|94178926 | 14 | 35 | 22 | AGAGGTAGTAGGTTGCATAGTT | BT20 | 588 | 677 | 809 | 223 | 253 | 303.8 | 291.0 | 613.0 | 156.6 | 54.3 |
| >MI0000065\|hsa-let-7d&WithFlank&9\|+\|94178828\|94178926@14.35.22&[MIMAT0000065&hsa-let-7d-5p&offsets\|0\|0;m-90&9\|+\|94178841\|94178862&offsets\|0\|0]\|AGAGGTAGTAGGTTGCATAGTT | >MI0000065\|hsa-let-7d&WithFlank&9\|+\|94178828\|94178926 | 14 | 35 | 22 | AGAGGTAGTAGGTTGCATAGTT | MB231 | 269 | 374 | 177 | 526 | 322 | 92.5 | 89.9 | 92.2 | 310.6 | 153.9 |
| >MI0000065\|hsa-let-7d&WithFlank&9\|+\|94178828\|94178926@14.35.22&[MIMAT0000065&hsa-let-7d-5p&offsets\|0\|0;m-90&9\|+\|94178841\|94178862&offsets\|0\|0]\|AGAGGTAGTAGGTTGCATAGTT | >MI0000065\|hsa-let-7d&WithFlank&9\|+\|94178828\|94178926 | 14 | 35 | 22 | AGAGGTAGTAGGTTGCATAGTT | MB468 | 20 | 30 | 11 | 23 | 7 | 11.0 | 13.0 | 2.5 | 19.3 | 2.6 |
| >MI0000065\|hsa-let-7d&WithFlank&9\|+\|94178828\|94178926@14.36.23&[MIMAT0000065&hsa-let-7d-5p&offsets\|0\|+1;m-90&9\|+\|94178841\|94178862&offsets\|0\|+1]\|AGAGGTAGTAGGTTGCATAGTTT | >MI0000065\|hsa-let-7d&WithFlank&9\|+\|94178828\|94178926 | 14 | 36 | 23 | AGAGGTAGTAGGTTGCATAGTTT | BT20 | 19 | 24 | 28 | 30 | 10 | 10.8 | 13.8 | 23.2 | 27.4 | 2.1 |
| >MI0000065\|hsa-let-7d&WithFlank&9\|+\|94178828\|94178926@14.36.23&[MIMAT0000065&hsa-let-7d-5p&offsets\|0\|+1;m-90&9\|+\|94178841\|94178862&offsets\|0\|+1]\|AGAGGTAGTAGGTTGCATAGTTT | >MI0000065\|hsa-let-7d&WithFlank&9\|+\|94178828\|94178926 | 14 | 36 | 23 | AGAGGTAGTAGGTTGCATAGTTT | MB231 | 12 | 15 | 9 | 74 | 18 | 3.5 | 3.8 | 5.0 | 41.7 | 7.6 |
| >MI0000065\|hsa-let-7d&WithFlank&9\|+\|94178828\|94178926@68.88.21(+1A)&[MIMAT0004484&hsa-let-7d-3p&offsets\|0\|-1(+1A);m-98&9\|+\|94178895\|94178916&offsets\|0\|-1(+1A)]\|CTATACGACCTGCTGCCTTTCA | >MI0000065\|hsa-let-7d&WithFlank&9\|+\|94178828\|94178926 | 68 | 88 | 21(+1A) | CTATACGACCTGCTGCCTTTCA | BT20 | 19 | 13 | 21 | 21 | 9 | 11.6 | 7.3 | 1.5 | 17.1 | 2.4 |
| >MI0000065\|hsa-let-7d&WithFlank&9\|+\|94178828\|94178926@68.88.21(+1C)&[MIMAT0004484&hsa-let-7d-3p&offsets\|0\|-1(+1C);m-98&9\|+\|94178895\|94178916&offsets\|0\|-1(+1C)]\|CTATACGACCTGCTGCCTTTCC | >MI0000065\|hsa-let-7d&WithFlank&9\|+\|94178828\|94178926 | 68 | 88 | 21(+1C) | CTATACGACCTGCTGCCTTTCC | BT20 | 27 | 30 | 31 | 47 | 21 | 9.6 | 15.9 | 7.7 | 38.8 | 10.4 |
| >MI0000065\|hsa-let-7d&WithFlank&9\|+\|94178828\|94178926@68.88.21(+1C)&[MIMAT0004484&hsa-let-7d-3p&offsets\|0\|-1(+1C);m-98&9\|+\|94178895\|94178916&offsets\|0\|-1(+1C)]\|CTATACGACCTGCTGCCTTTCC | >MI0000065\|hsa-let-7d&WithFlank&9\|+\|94178828\|94178926 | 68 | 88 | 21(+1C) | CTATACGACCTGCTGCCTTTCC | MB231 | 15 | 19 | 10 | 46 | 29 | 6.7 | 2.6 | 2.8 | 23.5 | 21.5 |
| >MI0000065\|hsa-let-7d&WithFlank&9\|+\|94178828\|94178926@68.88.21(+1G)&[MIMAT0004484&hsa-let-7d-3p&offsets\|0\|-1(+1G);m-98&9\|+\|94178895\|94178916&offsets\|0\|-1(+1G)]\|CTATACGACCTGCTGCCTTTCG | >MI0000065\|hsa-let-7d&WithFlank&9\|+\|94178828\|94178926 | 68 | 88 | 21(+1G) | CTATACGACCTGCTGCCTTTCG | BT20 | 12 | 5 | 6 | 6 | 2 | 8.8 | 3.1 | 1.4 | 6.9 | 0.7 |
| >MI0000065\|hsa-let-7d&WithFlank&9\|+\|94178828\|94178926@68.88.21&[MIMAT0004484&hsa-let-7d-3p&offsets\|0\|-1;m-98&9\|+\|94178895\|94178916&offsets\|0\|-1]\|CTATACGACCTGCTGCCTTTC | >MI0000065\|hsa-let-7d&WithFlank&9\|+\|94178828\|94178926 | 68 | 88 | 21 | CTATACGACCTGCTGCCTTTC | BT20 | 20 | 24 | 26 | 74 | 15 | 18.6 | 11.5 | 7.9 | 61.0 | 3.0 |
| >MI0000065\|hsa-let-7d&WithFlank&9\|+\|94178828\|94178926@68.88.21&[MIMAT0004484&hsa-let-7d-3p&offsets\|0\|-1;m-98&9\|+\|94178895\|94178916&offsets\|0\|-1]\|CTATACGACCTGCTGCCTTTC | >MI0000065\|hsa-let-7d&WithFlank&9\|+\|94178828\|94178926 | 68 | 88 | 21 | CTATACGACCTGCTGCCTTTC | MB231 | 20 | 36 | 20 | 50 | 27 | 19.5 | 11.0 | 18.4 | 12.8 | 5.8 |
| >MI0000065\|hsa-let-7d&WithFlank&9\|+\|94178828\|94178926@68.89.22&[MIMAT0004484&hsa-let-7d-3p&offsets\|0\|0;m-98&9\|+\|94178895\|94178916&offsets\|0\|0]\|CTATACGACCTGCTGCCTTTCT | >MI0000065\|hsa-let-7d&WithFlank&9\|+\|94178828\|94178926 | 68 | 89 | 22 | CTATACGACCTGCTGCCTTTCT | BT20 | 173 | 184 | 213 | 295 | 130 | 68.9 | 81.6 | 67.0 | 219.6 | 57.2 |
| >MI0000065\|hsa-let-7d&WithFlank&9\|+\|94178828\|94178926@68.89.22&[MIMAT0004484&hsa-let-7d-3p&offsets\|0\|0;m-98&9\|+\|94178895\|94178916&offsets\|0\|0]\|CTATACGACCTGCTGCCTTTCT | >MI0000065\|hsa-let-7d&WithFlank&9\|+\|94178828\|94178926 | 68 | 89 | 22 | CTATACGACCTGCTGCCTTTCT | MB231 | 102 | 143 | 70 | 241 | 184 | 46.6 | 30.7 | 25.0 | 92.3 | 108.3 |
| >MI0000066\|hsa-let-7e&WithFlank&19\|+\|51692780\|51692870@14.31.18&[MIMAT0000066&hsa-let-7e-5p&offsets\|0\|-4;m-50&19\|+\|51692793\|51692814&offsets\|0\|-4]\|TGAGGTAGGAGGTTGTAT | >MI0000066\|hsa-let-7e&WithFlank&19\|+\|51692780\|51692870 | 14 | 31 | 18 | TGAGGTAGGAGGTTGTAT | BT20 | 17 | 17 | 55 | 11 | 9 | 15.1 | 9.0 | 22.7 | 4.3 | 2.0 |
| >MI0000066\|hsa-let-7e&WithFlank&19\|+\|51692780\|51692870@14.33.20(+1C)&[MIMAT0000066&hsa-let-7e-5p&offsets\|0\|-2(+1C);m-50&19\|+\|51692793\|51692814&offsets\|0\|-2(+1C)]\|TGAGGTAGGAGGTTGTATAGC | >MI0000066\|hsa-let-7e&WithFlank&19\|+\|51692780\|51692870 | 14 | 33 | 20(+1C) | TGAGGTAGGAGGTTGTATAGC | BT20 | 14 | 11 | 30 | 3 | 5 | 7.1 | 4.3 | 11.7 | 1.6 | 2.4 |
| >MI0000066\|hsa-let-7e&WithFlank&19\|+\|51692780\|51692870@14.34.21(+1A)&[MIMAT0000066&hsa-let-7e-5p&offsets\|0\|-1(+1A);m-50&19\|+\|51692793\|51692814&offsets\|0\|-1(+1A)]\|TGAGGTAGGAGGTTGTATAGTA | >MI0000066\|hsa-let-7e&WithFlank&19\|+\|51692780\|51692870 | 14 | 34 | 21(+1A) | TGAGGTAGGAGGTTGTATAGTA | BT20 | 38 | 15 | 68 | 4 | 6 | 31.8 | 6.5 | 54.1 | 0.7 | 0.9 |
| >MI0000066\|hsa-let-7e&WithFlank&19\|+\|51692780\|51692870@14.34.21(+1C)&[MIMAT0000066&hsa-let-7e-5p&offsets\|0\|-1(+1C);m-50&19\|+\|51692793\|51692814&offsets\|0\|-1(+1C)]\|TGAGGTAGGAGGTTGTATAGTC | >MI0000066\|hsa-let-7e&WithFlank&19\|+\|51692780\|51692870 | 14 | 34 | 21(+1C) | TGAGGTAGGAGGTTGTATAGTC | BT20 | 101 | 77 | 193 | 19 | 42 | 46.0 | 31.6 | 92.9 | 6.3 | 12.9 |
| >MI0000066\|hsa-let-7e&WithFlank&19\|+\|51692780\|51692870@14.34.21(+1C)&[MIMAT0000066&hsa-let-7e-5p&offsets\|0\|-1(+1C);m-50&19\|+\|51692793\|51692814&offsets\|0\|-1(+1C)]\|TGAGGTAGGAGGTTGTATAGTC | >MI0000066\|hsa-let-7e&WithFlank&19\|+\|51692780\|51692870 | 14 | 34 | 21(+1C) | TGAGGTAGGAGGTTGTATAGTC | MB231 | 39 | 33 | 46 | 32 | 45 | 8.8 | 6.3 | 33.6 | 15.9 | 22.2 |
| >MI0000066\|hsa-let-7e&WithFlank&19\|+\|51692780\|51692870@14.34.21(+1G)&[MIMAT0000066&hsa-let-7e-5p&offsets\|0\|-1(+1G);m-50&19\|+\|51692793\|51692814&offsets\|0\|-1(+1G)]\|TGAGGTAGGAGGTTGTATAGTG | >MI0000066\|hsa-let-7e&WithFlank&19\|+\|51692780\|51692870 | 14 | 34 | 21(+1G) | TGAGGTAGGAGGTTGTATAGTG | BT20 | 31 | 7 | 25 | 1 | 1 | 32.5 | 2.7 | 20.4 | 1.8 | 1.0 |
| >MI0000066\|hsa-let-7e&WithFlank&19\|+\|51692780\|51692870@14.34.21&[MIMAT0000066&hsa-let-7e-5p&offsets\|0\|-1;m-50&19\|+\|51692793\|51692814&offsets\|0\|-1]\|TGAGGTAGGAGGTTGTATAGT | >MI0000066\|hsa-let-7e&WithFlank&19\|+\|51692780\|51692870 | 14 | 34 | 21 | TGAGGTAGGAGGTTGTATAGT | BT20 | 113 | 83 | 234 | 30 | 41 | 54.0 | 31.7 | 56.8 | 9.0 | 13.8 |

<table>
<tr><td>>MI0000066|hsa-let-7e&WithFlank&19|+|51692780|51692870@14.34.21&[MIMAT0000066&hsa-let-7e-5p&offsets|0|-1;m-50&19|+|51692793|51692814&offsets|0|-1]|TGAGGTAGGAGGTTGTATAGT</td><td>>MI0000066|hsa-let-7e&WithFlank&19|+|51692780|51692870</td><td>14</td><td>34</td><td>21</td><td>TGAGGTAGGAGGTTGTATAGT</td><td>MB231</td><td>63</td><td>59</td><td>83</td><td>44</td><td>48</td><td>10.6</td><td>11.4</td><td>59.9</td><td>15.9</td><td>18.9</td></tr>
<tr><td>>MI0000066|hsa-let-7e&WithFlank&19|+|51692780|51692870@14.35.22(+1A)&[MIMAT0000066&hsa-let-7e-5p&offsets|0|0(+1A);m-50&19|+|51692793|51692814&offsets|0|0(+1A)]|TGAGGTAGGAGGTTGTATAGTTA</td><td>>MI0000066|hsa-let-7e&WithFlank&19|+|51692780|51692870</td><td>14</td><td>35</td><td>22(+1A)</td><td>TGAGGTAGGAGGTTGTATAGTTA</td><td>BT20</td><td>27</td><td>18</td><td>49</td><td>14</td><td>9</td><td>14.4</td><td>9.6</td><td>11.0</td><td>6.2</td><td>2.7</td></tr>
<tr><td>>MI0000066|hsa-let-7e&WithFlank&19|+|51692780|51692870@14.35.22(+1U)&[MIMAT0000066&hsa-let-7e-5p&offsets|0|0(+1U);m-50&19|+|51692793|51692814&offsets|0|0(+1U)]|TGAGGTAGGAGGTTGTATAGTTT</td><td>>MI0000066|hsa-let-7e&WithFlank&19|+|51692780|51692870</td><td>14</td><td>35</td><td>22(+1U)</td><td>TGAGGTAGGAGGTTGTATAGTTT</td><td>BT20</td><td>43</td><td>32</td><td>74</td><td>21</td><td>20</td><td>22.4</td><td>15.6</td><td>14.8</td><td>10.0</td><td>7.6</td></tr>
<tr><td>>MI0000066|hsa-let-7e&WithFlank&19|+|51692780|51692870@14.35.22(+1U)&[MIMAT0000066&hsa-let-7e-5p&offsets|0|0(+1U);m-50&19|+|51692793|51692814&offsets|0|0(+1U)]|TGAGGTAGGAGGTTGTATAGTTT</td><td>>MI0000066|hsa-let-7e&WithFlank&19|+|51692780|51692870</td><td>14</td><td>35</td><td>22(+1U)</td><td>TGAGGTAGGAGGTTGTATAGTTT</td><td>MB231</td><td>31</td><td>24</td><td>35</td><td>44</td><td>26</td><td>3.5</td><td>5.5</td><td>24.0</td><td>11.4</td><td>10.9</td></tr>
<tr><td>>MI0000066|hsa-let-7e&WithFlank&19|+|51692780|51692870@14.35.22&[MIMAT0000066&hsa-let-7e-5p&offsets|0|0;m-50&19|+|51692793|51692814&offsets|0|0]|TGAGGTAGGAGGTTGTATAGTT</td><td>>MI0000066|hsa-let-7e&WithFlank&19|+|51692780|51692870</td><td>14</td><td>35</td><td>22</td><td>TGAGGTAGGAGGTTGTATAGTT</td><td>BT20</td><td>759</td><td>543</td><td>1406</td><td>137</td><td>324</td><td>351.2</td><td>192.8</td><td>451.1</td><td>11.1</td><td>93.4</td></tr>
<tr><td>>MI0000066|hsa-let-7e&WithFlank&19|+|51692780|51692870@14.35.22&[MIMAT0000066&hsa-let-7e-5p&offsets|0|0;m-50&19|+|51692793|51692814&offsets|0|0]|TGAGGTAGGAGGTTGTATAGTT</td><td>>MI0000066|hsa-let-7e&WithFlank&19|+|51692780|51692870</td><td>14</td><td>35</td><td>22</td><td>TGAGGTAGGAGGTTGTATAGTT</td><td>MB231</td><td>364</td><td>326</td><td>419</td><td>194</td><td>332</td><td>51.9</td><td>43.2</td><td>299.2</td><td>57.4</td><td>126.1</td></tr>
<tr><td>>MI0000066|hsa-let-7e&WithFlank&19|+|51692780|51692870@14.35.22&[MIMAT0000066&hsa-let-7e-5p&offsets|0|0;m-50&19|+|51692793|51692814&offsets|0|0]|TGAGGTAGGAGGTTGTATAGTT</td><td>>MI0000066|hsa-let-7e&WithFlank&19|+|51692780|51692870</td><td>14</td><td>35</td><td>22</td><td>TGAGGTAGGAGGTTGTATAGTT</td><td>MB468</td><td>35</td><td>50</td><td>18</td><td>35</td><td>11</td><td>11.5</td><td>11.2</td><td>5.5</td><td>43.1</td><td>3.2</td></tr>
<tr><td>>MI0000066|hsa-let-7e&WithFlank&19|+|51692780|51692870@14.36.23&[MIMAT0000066&hsa-let-7e-5p&offsets|0|+1;m-50&19|+|51692793|51692814&offsets|0|+1]|TGAGGTAGGAGGTTGTATAGTTG</td><td>>MI0000066|hsa-let-7e&WithFlank&19|+|51692780|51692870</td><td>14</td><td>36</td><td>23</td><td>TGAGGTAGGAGGTTGTATAGTTG</td><td>BT20</td><td>28</td><td>22</td><td>57</td><td>18</td><td>15</td><td>9.9</td><td>0.7</td><td>8.9</td><td>8.9</td><td>1.2</td></tr>
<tr><td>>MI0000066|hsa-let-7e&WithFlank&19|+|51692780|51692870@14.36.23&[MIMAT0000066&hsa-let-7e-5p&offsets|0|+1;m-50&19|+|51692793|51692814&offsets|0|+1]|TGAGGTAGGAGGTTGTATAGTTG</td><td>>MI0000066|hsa-let-7e&WithFlank&19|+|51692780|51692870</td><td>14</td><td>36</td><td>23</td><td>TGAGGTAGGAGGTTGTATAGTTG</td><td>MB231</td><td>11</td><td>9</td><td>13</td><td>16</td><td>9</td><td>1.1</td><td>0.5</td><td>8.9</td><td>3.9</td><td>2.6</td></tr>
<tr><td>>MI0000067|hsa-let-7f-1&WithFlank&9|+|94176341|94176439@13.30.18;MI0000068|hsa-let-7f-2&WithFlank&X|-|53557186|53557280@14.31.18&[MIMAT0000067&hsa-let-7f-5p&offsets|0|-4;m-20&9|+|94176353|94176374&offsets|0|-4];[MIMAT0000067_1&hsa-let-7f-5p&offsets|0|-4;m-19&X|-|53557246|53557267&offsets|0|-4]|TGAGGTAGTAGATTGTAT</td><td>>MI0000067|hsa-let-7f-1&WithFlank&9|+|94176341|94176439</td><td>13</td><td>30</td><td>18</td><td>TGAGGTAGTAGATTGTAT</td><td>BT20</td><td>222</td><td>268</td><td>331</td><td>83</td><td>100</td><td>188.8</td><td>148.4</td><td>351.1</td><td>50.2</td><td>18.8</td></tr>
<tr><td>>MI0000067|hsa-let-7f-1&WithFlank&9|+|94176341|94176439@13.30.18;MI0000068|hsa-let-7f-2&WithFlank&X|-|53557186|53557280@14.31.18&[MIMAT0000067&hsa-let-7f-5p&offsets|0|-4;m-20&9|+|94176353|94176374&offsets|0|-4];[MIMAT0000067_1&hsa-let-7f-5p&offsets|0|-4;m-19&X|-|53557246|53557267&offsets|0|-4]|TGAGGTAGTAGATTGTAT</td><td>>MI0000067|hsa-let-7f-1&WithFlank&9|+|94176341|94176439</td><td>13</td><td>30</td><td>18</td><td>TGAGGTAGTAGATTGTAT</td><td>MB231</td><td>167</td><td>237</td><td>71</td><td>401</td><td>114</td><td>42.7</td><td>101.3</td><td>39.8</td><td>298.4</td><td>51.2</td></tr>
<tr><td>>MI0000067|hsa-let-7f-1&WithFlank&9|+|94176341|94176439@13.30.18;MI0000068|hsa-let-7f-2&WithFlank&X|-|53557186|53557280@14.31.18&[MIMAT0000067&hsa-let-7f-5p&offsets|0|-4;m-20&9|+|94176353|94176374&offsets|0|-4];[MIMAT0000067_1&hsa-let-7f-5p&offsets|0|-4;m-19&X|-|53557246|53557267&offsets|0|-4]|TGAGGTAGTAGATTGTAT</td><td>>MI0000067|hsa-let-7f-1&WithFlank&9|+|94176341|94176439</td><td>13</td><td>30</td><td>18</td><td>TGAGGTAGTAGATTGTAT</td><td>MB468</td><td>12</td><td>15</td><td>6</td><td>9</td><td>3</td><td>6.7</td><td>5.9</td><td>1.1</td><td>5.2</td><td>2.2</td></tr>
<tr><td>>MI0000067|hsa-let-7f-1&WithFlank&9|+|94176341|94176439@13.31.19;MI0000068|hsa-let-7f-2&WithFlank&X|-|53557186|53557280@14.32.19&[MIMAT0000067&hsa-let-7f-5p&offsets|0|-3;m-20&9|+|94176353|94176374&offsets|0|-3];[MIMAT0000067_1&hsa-let-7f-5p&offsets|0|-3;m-19&X|-|53557246|53557267&offsets|0|-3]|TGAGGTAGTAGATTGTATA</td><td>>MI0000067|hsa-let-7f-1&WithFlank&9|+|94176341|94176439</td><td>13</td><td>31</td><td>19</td><td>TGAGGTAGTAGATTGTATA</td><td>BT20</td><td>59</td><td>62</td><td>159</td><td>6</td><td>13</td><td>16.3</td><td>5.7</td><td>51.6</td><td>4.0</td><td>1.1</td></tr>
<tr><td>>MI0000067|hsa-let-7f-1&WithFlank&9|+|94176341|94176439@13.31.19;MI0000068|hsa-let-7f-2&WithFlank&X|-|53557186|53557280@14.32.19&[MIMAT0000067&hsa-let-7f-5p&offsets|0|-3;m-20&9|+|94176353|94176374&offsets|0|-3];[MIMAT0000067_1&hsa-let-7f-5p&offsets|0|-3;m-19&X|-|53557246|53557267&offsets|0|-3]|TGAGGTAGTAGATTGTATA</td><td>>MI0000067|hsa-let-7f-1&WithFlank&9|+|94176341|94176439</td><td>13</td><td>31</td><td>19</td><td>TGAGGTAGTAGATTGTATA</td><td>MB231</td><td>17</td><td>24</td><td>14</td><td>9</td><td>5</td><td>20.3</td><td>32.8</td><td>19.3</td><td>8.2</td><td>5.6</td></tr>
<tr><td>>MI0000067|hsa-let-7f-1&WithFlank&9|+|94176341|94176439@13.32.20;MI0000068|hsa-let-7f-2&WithFlank&X|-|53557186|53557280@14.33.20&[MIMAT0000067&hsa-let-7f-5p&offsets|0|-2;m-20&9|+|94176353|94176374&offsets|0|-2];[MIMAT0000067_1&hsa-let-7f-5p&offsets|0|-2;m-19&X|-|53557246|53557267&offsets|0|-2]|TGAGGTAGTAGATTGTATAG</td><td>>MI0000067|hsa-let-7f-1&WithFlank&9|+|94176341|94176439</td><td>13</td><td>32</td><td>20</td><td>TGAGGTAGTAGATTGTATAG</td><td>BT20</td><td>14</td><td>11</td><td>14</td><td>9</td><td>4</td><td>9.9</td><td>4.5</td><td>14.9</td><td>9.9</td><td>0.8</td></tr>
<tr><td>>MI0000067|hsa-let-7f-1&WithFlank&9|+|94176341|94176439@13.32.20(+1A);MI0000068|hsa-let-7f-2&WithFlank&X|-|53557186|53557280@14.33.20(+1A)&[MIMAT0000067&hsa-let-7f-5p&offsets|0|-2(+1A);m-20&9|+|94176353|94176374&offsets|0|-2(+1A)];[MIMAT0000067_1&hsa-let-7f-5p&offsets|0|-2(+1A);m-19&X|-|53557246|53557267&offsets|0|-2(+1A)]|TGAGGTAGTAGATTGTATAGA</td><td>>MI0000067|hsa-let-7f-1&WithFlank&9|+|94176341|94176439</td><td>13</td><td>32</td><td>20(+1A)</td><td>TGAGGTAGTAGATTGTATAGA</td><td>BT20</td><td>21</td><td>10</td><td>16</td><td>2</td><td>3</td><td>15.2</td><td>5.0</td><td>9.9</td><td>2.8</td><td>0.6</td></tr>
<tr><td>>MI0000067|hsa-let-7f-1&WithFlank&9|+|94176341|94176439@13.32.20(+1C);MI0000068|hsa-let-7f-2&WithFlank&X|-|53557186|53557280@14.33.20(+1C)&[MIMAT0000067&hsa-let-7f-5p&offsets|0|-2(+1C);m-20&9|+|94176353|94176374&offsets|0|-2(+1C)];[MIMAT0000067_1&hsa-let-7f-5p&offsets|0|-2(+1C);m-19&X|-|53557246|53557267&offsets|0|-2(+1C)]|TGAGGTAGTAGATTGTATAGC</td><td>>MI0000067|hsa-let-7f-1&WithFlank&9|+|94176341|94176439</td><td>13</td><td>32</td><td>20(+1C)</td><td>TGAGGTAGTAGATTGTATAGC</td><td>BT20</td><td>81</td><td>71</td><td>67</td><td>13</td><td>22</td><td>36.4</td><td>33.5</td><td>50.1</td><td>13.2</td><td>9.3</td></tr>
<tr><td>>MI0000067|hsa-let-7f-1&WithFlank&9|+|94176341|94176439@13.32.20(+1C);MI0000068|hsa-let-7f-2&WithFlank&X|-|53557186|53557280@14.33.20(+1C)&[MIMAT0000067&hsa-let-7f-5p&offsets|0|-2(+1C);m-20&9|+|94176353|94176374&offsets|0|-2(+1C)];[MIMAT0000067_1&hsa-let-7f-5p&offsets|0|-2(+1C);m-19&X|-|53557246|53557267&offsets|0|-2(+1C)]|TGAGGTAGTAGATTGTATAGC</td><td>>MI0000067|hsa-let-7f-1&WithFlank&9|+|94176341|94176439</td><td>13</td><td>32</td><td>20(+1C)</td><td>TGAGGTAGTAGATTGTATAGC</td><td>MB231</td><td>65</td><td>70</td><td>17</td><td>83</td><td>40</td><td>7.0</td><td>20.7</td><td>6.6</td><td>46.4</td><td>14.9</td></tr>
<tr><td>>MI0000067|hsa-let-7f-1&WithFlank&9|+|94176341|94176439@13.32.20(+1G);MI0000068|hsa-let-7f-2&WithFlank&X|-|53557186|53557280@14.33.20(+1G)&[MIMAT0000067&hsa-let-7f-5p&offsets|0|-2(+1G);m-20&9|+|94176353|94176374&offsets|0|-2(+1G)];[MIMAT0000067_1&hsa-let-7f-5p&offsets|0|-2(+1G);m-19&X|-|53557246|53557267&offsets|0|-2(+1G)]|TGAGGTAGTAGATTGTATAGG</td><td>>MI0000067|hsa-let-7f-1&WithFlank&9|+|94176341|94176439</td><td>13</td><td>32</td><td>20(+1G)</td><td>TGAGGTAGTAGATTGTATAGG</td><td>BT20</td><td>34</td><td>12</td><td>11</td><td>2</td><td>0</td><td>28.1</td><td>4.6</td><td>4.3</td><td>3.1</td><td>0.6</td></tr>
<tr><td>>MI0000067|hsa-let-7f-1&WithFlank&9|+|94176341|94176439@13.32.20(+1G);MI0000068|hsa-let-7f-2&WithFlank&X|-|53557186|53557280@14.33.20(+1G)&[MIMAT0000067&hsa-let-7f-5p&offsets|0|-2(+1G);m-20&9|+|94176353|94176374&offsets|0|-2(+1G)];[MIMAT0000067_1&hsa-let-7f-5p&offsets|0|-2(+1G);m-19&X|-|53557246|53557267&offsets|0|-2(+1G)]|TGAGGTAGTAGATTGTATAGG</td><td>>MI0000067|hsa-let-7f-1&WithFlank&9|+|94176341|94176439</td><td>13</td><td>32</td><td>20(+1G)</td><td>TGAGGTAGTAGATTGTATAGG</td><td>MB231</td><td>25</td><td>9</td><td>2</td><td>8</td><td>3</td><td>21.7</td><td>5.1</td><td>1.5</td><td>4.3</td><td>1.3</td></tr>
<tr><td>>MI0000067|hsa-let-7f-1&WithFlank&9|+|94176341|94176439@13.32.20(+2C);MI0000068|hsa-let-7f-2&WithFlank&X|-|53557186|53557280@14.33.20(+2C)&[MIMAT0000067&hsa-let-7f-5p&offsets|0|-2(+2C);m-20&9|+|94176353|94176374&offsets|0|-2(+2C)];[MIMAT0000067_1&hsa-let-7f-5p&offsets|0|-2(+2C);m-19&X|-|53557246|53557267&offsets|0|-2(+2C)]|TGAGGTAGTAGATTGTATAGCC</td><td>>MI0000067|hsa-let-7f-1&WithFlank&9|+|94176341|94176439</td><td>13</td><td>32</td><td>20(+2C)</td><td>TGAGGTAGTAGATTGTATAGCC</td><td>BT20</td><td>16</td><td>5</td><td>4</td><td>0</td><td>0</td><td>18.1</td><td>1.9</td><td>1.6</td><td>0.0</td><td>0.0</td></tr>
<tr><td>>MI0000067|hsa-let-7f-1&WithFlank&9|+|94176341|94176439@13.33.21;MI0000068|hsa-let-7f-2&WithFlank&X|-|53557186|53557280@14.34.21&[MIMAT0000067&hsa-let-7f-5p&offsets|0|-1;m-20&9|+|94176353|94176374&offsets|0|-1];[MIMAT0000067_1&hsa-let-7f-5p&offsets|0|-1;m-19&X|-|53557246|53557267&offsets|0|-1]|TGAGGTAGTAGATTGTATAGT</td><td>>MI0000067|hsa-let-7f-1&WithFlank&9|+|94176341|94176439</td><td>13</td><td>33</td><td>21</td><td>TGAGGTAGTAGATTGTATAGT</td><td>BT20</td><td>659</td><td>530</td><td>570</td><td>102</td><td>184</td><td>311.1</td><td>200.4</td><td>464.2</td><td>88.0</td><td>70.0</td></tr>
<tr><td>>MI0000067|hsa-let-7f-1&WithFlank&9|+|94176341|94176439@13.33.21;MI0000068|hsa-let-7f-2&WithFlank&X|-|53557186|53557280@14.34.21&[MIMAT0000067&hsa-let-7f-5p&offsets|0|-1;m-20&9|+|94176353|94176374&offsets|0|-1];[MIMAT0000067_1&hsa-let-7f-5p&offsets|0|-1;m-19&X|-|53557246|53557267&offsets|0|-1]|TGAGGTAGTAGATTGTATAGT</td><td>>MI0000067|hsa-let-7f-1&WithFlank&9|+|94176341|94176439</td><td>13</td><td>33</td><td>21</td><td>TGAGGTAGTAGATTGTATAGT</td><td>MB231</td><td>658</td><td>715</td><td>176</td><td>576</td><td>332</td><td>52.2</td><td>136.6</td><td>50.9</td><td>229.1</td><td>75.6</td></tr>
<tr><td>>MI0000067|hsa-let-7f-1&WithFlank&9|+|94176341|94176439@13.33.21;MI0000068|hsa-let-7f-2&WithFlank&X|-|53557186|53557280@14.34.21&[MIMAT0000067&hsa-let-7f-5p&offsets|0|-1;m-20&9|+|94176353|94176374&offsets|0|-1];[MIMAT0000067_1&hsa-let-7f-5p&offsets|0|-1;m-19&X|-|53557246|53557267&offsets|0|-1]|TGAGGTAGTAGATTGTATAGT</td><td>>MI0000067|hsa-let-7f-1&WithFlank&9|+|94176341|94176439</td><td>13</td><td>33</td><td>21</td><td>TGAGGTAGTAGATTGTATAGT</td><td>MB468</td><td>44</td><td>41</td><td>17</td><td>28</td><td>7</td><td>13.1</td><td>21.5</td><td>3.1</td><td>17.2</td><td>2.9</td></tr>
<tr><td>>MI0000067|hsa-let-7f-1&WithFlank&9|+|94176341|94176439@13.33.21(+1A);MI0000068|hsa-let-7f-2&WithFlank&X|-|53557186|53557280@14.34.21(+1A)&[MIMAT0000067&hsa-let-7f-5p&offsets|0|-1(+1A);m-20&9|+|94176353|94176374&offsets|0|-1(+1A)];[MIMAT0000067_1&hsa-let-7f-5p&offsets|0|-1(+1A);m-19&X|-|53557246|53557267&offsets|0|-1(+1A)]|TGAGGTAGTAGATTGTATAGTA</td><td>>MI0000067|hsa-let-7f-1&WithFlank&9|+|94176341|94176439</td><td>13</td><td>33</td><td>21(+1A)</td><td>TGAGGTAGTAGATTGTATAGTA</td><td>BT20</td><td>374</td><td>168</td><td>250</td><td>26</td><td>49</td><td>278.7</td><td>70.9</td><td>128.0</td><td>27.9</td><td>10.5</td></tr>
</table>

| | | | | | | | | | | | | | | | | |
|---|---|---|---|---|---|---|---|---|---|---|---|---|---|---|---|---|
| >MI0000067\|hsa-let-7f-1&WithFlank&9\|+\|94176341\|94176439@13.33.21(+1A);MI0000068\|hsa-let-7f-2&WithFlank&X\|-\|53557186\|53557280@14.34.21(+1A)&[MIMAT0000067&hsa-let-7f-5p&offsets\|0\|-1(+1A);m-20&9\|+\|94176353\|94176374&offsets\|0\|-1(+1A)];[MIMAT0000067_1&hsa-let-7f-5p&offsets\|0\|-1(+1A);m-19&X\|-\|53557246\|53557267&offsets\|0\|-1(+1A)]\|TGAGGTAGTAGATTGTATAGTA | >MI0000067\|hsa-let-7f-1&WithFlank&9\|+\|94176341\|94176439 | 13 | 33 | 21(+1A) | TGAGGTAGTAGATTGTATAGTA | MB231 | 145 | 117 | 45 | 188 | 76 | 11.5 | 40.1 | 18.9 | 108.2 | 28.5 |
| >MI0000067\|hsa-let-7f-1&WithFlank&9\|+\|94176341\|94176439@13.33.21(+1C);MI0000068\|hsa-let-7f-2&WithFlank&X\|-\|53557186\|53557280@14.34.21(+1C)&[MIMAT0000067&hsa-let-7f-5p&offsets\|0\|-1(+1C);m-20&9\|+\|94176353\|94176374&offsets\|0\|-1(+1C)];[MIMAT0000067_1&hsa-let-7f-5p&offsets\|0\|-1(+1C);m-19&X\|-\|53557246\|53557267&offsets\|0\|-1(+1C)]\|TGAGGTAGTAGATTGTATAGTC | >MI0000067\|hsa-let-7f-1&WithFlank&9\|+\|94176341\|94176439 | 13 | 33 | 21(+1C) | TGAGGTAGTAGATTGTATAGTC | BT20 | 1125 | 1025 | 881 | 150 | 349 | 469.9 | 453.5 | 565.4 | 149.0 | 141.8 |
| >MI0000067\|hsa-let-7f-1&WithFlank&9\|+\|94176341\|94176439@13.33.21(+1C);MI0000068\|hsa-let-7f-2&WithFlank&X\|-\|53557186\|53557280@14.34.21(+1C)&[MIMAT0000067&hsa-let-7f-5p&offsets\|0\|-1(+1C);m-20&9\|+\|94176353\|94176374&offsets\|0\|-1(+1C)];[MIMAT0000067_1&hsa-let-7f-5p&offsets\|0\|-1(+1C);m-19&X\|-\|53557246\|53557267&offsets\|0\|-1(+1C)]\|TGAGGTAGTAGATTGTATAGTC | >MI0000067\|hsa-let-7f-1&WithFlank&9\|+\|94176341\|94176439 | 13 | 33 | 21(+1C) | TGAGGTAGTAGATTGTATAGTC | MB231 | 737 | 816 | 213 | 1025 | 491 | 105.9 | 239.1 | 68.6 | 573.4 | 218.1 |
| >MI0000067\|hsa-let-7f-1&WithFlank&9\|+\|94176341\|94176439@13.33.21(+1C);MI0000068\|hsa-let-7f-2&WithFlank&X\|-\|53557186\|53557280@14.34.21(+1C)&[MIMAT0000067&hsa-let-7f-5p&offsets\|0\|-1(+1C);m-20&9\|+\|94176353\|94176374&offsets\|0\|-1(+1C)];[MIMAT0000067_1&hsa-let-7f-5p&offsets\|0\|-1(+1C);m-19&X\|-\|53557246\|53557267&offsets\|0\|-1(+1C)]\|TGAGGTAGTAGATTGTATAGTC | >MI0000067\|hsa-let-7f-1&WithFlank&9\|+\|94176341\|94176439 | 13 | 33 | 21(+1C) | TGAGGTAGTAGATTGTATAGTC | MB468 | 59 | 60 | 30 | 43 | 12 | 17.4 | 22.2 | 13.0 | 37.7 | 5.2 |
| >MI0000067\|hsa-let-7f-1&WithFlank&9\|+\|94176341\|94176439@13.33.21(+1G);MI0000068\|hsa-let-7f-2&WithFlank&X\|-\|53557186\|53557280@14.34.21(+1G)&[MIMAT0000067&hsa-let-7f-5p&offsets\|0\|-1(+1G);m-20&9\|+\|94176353\|94176374&offsets\|0\|-1(+1G)];[MIMAT0000067_1&hsa-let-7f-5p&offsets\|0\|-1(+1G);m-19&X\|-\|53557246\|53557267&offsets\|0\|-1(+1G)]\|TGAGGTAGTAGATTGTATAGTG | >MI0000067\|hsa-let-7f-1&WithFlank&9\|+\|94176341\|94176439 | 13 | 33 | 21(+1G) | TGAGGTAGTAGATTGTATAGTG | BT20 | 321 | 92 | 89 | 11 | 12 | 285.0 | 39.3 | 38.3 | 15.0 | 2.8 |
| >MI0000067\|hsa-let-7f-1&WithFlank&9\|+\|94176341\|94176439@13.33.21(+1G);MI0000068\|hsa-let-7f-2&WithFlank&X\|-\|53557186\|53557280@14.34.21(+1G)&[MIMAT0000067&hsa-let-7f-5p&offsets\|0\|-1(+1G);m-20&9\|+\|94176353\|94176374&offsets\|0\|-1(+1G)];[MIMAT0000067_1&hsa-let-7f-5p&offsets\|0\|-1(+1G);m-19&X\|-\|53557246\|53557267&offsets\|0\|-1(+1G)]\|TGAGGTAGTAGATTGTATAGTG | >MI0000067\|hsa-let-7f-1&WithFlank&9\|+\|94176341\|94176439 | 13 | 33 | 21(+1G) | TGAGGTAGTAGATTGTATAGTG | MB231 | 163 | 70 | 17 | 54 | 20 | 124.8 | 32.4 | 10.3 | 33.8 | 9.0 |
| >MI0000067\|hsa-let-7f-1&WithFlank&9\|+\|94176341\|94176439@13.33.21(+1G);MI0000068\|hsa-let-7f-2&WithFlank&X\|-\|53557186\|53557280@14.34.21(+1G)&[MIMAT0000067&hsa-let-7f-5p&offsets\|0\|-1(+1G);m-20&9\|+\|94176353\|94176374&offsets\|0\|-1(+1G)];[MIMAT0000067_1&hsa-let-7f-5p&offsets\|0\|-1(+1G);m-19&X\|-\|53557246\|53557267&offsets\|0\|-1(+1G)]\|TGAGGTAGTAGATTGTATAGTG | >MI0000067\|hsa-let-7f-1&WithFlank&9\|+\|94176341\|94176439 | 13 | 33 | 21(+1G) | TGAGGTAGTAGATTGTATAGTG | MB468 | 13 | 6 | 8 | 6 | 1 | 2.9 | 3.9 | 8.7 | 5.2 | 1.5 |
| >MI0000067\|hsa-let-7f-1&WithFlank&9\|+\|94176341\|94176439@13.34.22;MI0000068\|hsa-let-7f-2&WithFlank&X\|-\|53557186\|53557280@14.35.22&[MIMAT0000067&hsa-let-7f-5p&offsets\|0\|0;m-20&9\|+\|94176353\|94176374&offsets\|0\|0];[MIMAT0000067_1&hsa-let-7f-5p&offsets\|0\|0;m-19&X\|-\|53557246\|53557267&offsets\|0\|0]\|TGAGGTAGTAGATTGTATAGTT | >MI0000067\|hsa-let-7f-1&WithFlank&9\|+\|94176341\|94176439 | 13 | 34 | 22 | TGAGGTAGTAGATTGTATAGTT | BT20 | 8397 | 7227 | 6825 | 1004 | 2620 | 3897.5 | 2630.3 | 4837.9 | 796.5 | 1021.9 |
| >MI0000067\|hsa-let-7f-1&WithFlank&9\|+\|94176341\|94176439@13.34.22;MI0000068\|hsa-let-7f-2&WithFlank&X\|-\|53557186\|53557280@14.35.22&[MIMAT0000067&hsa-let-7f-5p&offsets\|0\|0;m-20&9\|+\|94176353\|94176374&offsets\|0\|0];[MIMAT0000067_1&hsa-let-7f-5p&offsets\|0\|0;m-19&X\|-\|53557246\|53557267&offsets\|0\|0]\|TGAGGTAGTAGATTGTATAGTT | >MI0000067\|hsa-let-7f-1&WithFlank&9\|+\|94176341\|94176439 | 13 | 34 | 22 | TGAGGTAGTAGATTGTATAGTT | MB231 | 6934 | 7832 | 1970 | 6229 | 3736 | 889.4 | 1400.3 | 459.5 | 2506.3 | 1170.3 |
| >MI0000067\|hsa-let-7f-1&WithFlank&9\|+\|94176341\|94176439@13.34.22;MI0000068\|hsa-let-7f-2&WithFlank&X\|-\|53557186\|53557280@14.35.22&[MIMAT0000067&hsa-let-7f-5p&offsets\|0\|0;m-20&9\|+\|94176353\|94176374&offsets\|0\|0];[MIMAT0000067_1&hsa-let-7f-5p&offsets\|0\|0;m-19&X\|-\|53557246\|53557267&offsets\|0\|0]\|TGAGGTAGTAGATTGTATAGTT | >MI0000067\|hsa-let-7f-1&WithFlank&9\|+\|94176341\|94176439 | 13 | 34 | 22 | TGAGGTAGTAGATTGTATAGTT | MB468 | 564 | 633 | 263 | 347 | 98 | 216.3 | 262.1 | 46.8 | 314.9 | 38.9 |
| >MI0000067\|hsa-let-7f-1&WithFlank&9\|+\|94176341\|94176439@13.34.22(+1A);MI0000068\|hsa-let-7f-2&WithFlank&X\|-\|53557186\|53557280@14.35.22(+1A)&[MIMAT0000067&hsa-let-7f-5p&offsets\|0\|0(+1A);m-20&9\|+\|94176353\|94176374&offsets\|0\|0(+1A)];[MIMAT0000067_1&hsa-let-7f-5p&offsets\|0\|0(+1A);m-19&X\|-\|53557246\|53557267&offsets\|0\|0(+1A)]\|TGAGGTAGTAGATTGTATAGTTA | >MI0000067\|hsa-let-7f-1&WithFlank&9\|+\|94176341\|94176439 | 13 | 34 | 22(+1A) | TGAGGTAGTAGATTGTATAGTTA | BT20 | 151 | 118 | 128 | 168 | 43 | 89.0 | 59.2 | 101.9 | 146.3 | 14.4 |
| >MI0000067\|hsa-let-7f-1&WithFlank&9\|+\|94176341\|94176439@13.34.22(+1A);MI0000068\|hsa-let-7f-2&WithFlank&X\|-\|53557186\|53557280@14.35.22(+1A)&[MIMAT0000067&hsa-let-7f-5p&offsets\|0\|0(+1A);m-20&9\|+\|94176353\|94176374&offsets\|0\|0(+1A)];[MIMAT0000067_1&hsa-let-7f-5p&offsets\|0\|0(+1A);m-19&X\|-\|53557246\|53557267&offsets\|0\|0(+1A)]\|TGAGGTAGTAGATTGTATAGTTA | >MI0000067\|hsa-let-7f-1&WithFlank&9\|+\|94176341\|94176439 | 13 | 34 | 22(+1A) | TGAGGTAGTAGATTGTATAGTTA | MB231 | 71 | 71 | 22 | 187 | 35 | 15.5 | 9.9 | 5.7 | 96.0 | 8.7 |
| >MI0000067\|hsa-let-7f-1&WithFlank&9\|+\|94176341\|94176439@13.34.22(+1A);MI0000068\|hsa-let-7f-2&WithFlank&X\|-\|53557186\|53557280@14.35.22(+1A)&[MIMAT0000067&hsa-let-7f-5p&offsets\|0\|0(+1A);m-20&9\|+\|94176353\|94176374&offsets\|0\|0(+1A)];[MIMAT0000067_1&hsa-let-7f-5p&offsets\|0\|0(+1A);m-19&X\|-\|53557246\|53557267&offsets\|0\|0(+1A)]\|TGAGGTAGTAGATTGTATAGTTA | >MI0000067\|hsa-let-7f-1&WithFlank&9\|+\|94176341\|94176439 | 13 | 34 | 22(+1A) | TGAGGTAGTAGATTGTATAGTTA | MB468 | 11 | 10 | 5 | 10 | 2 | 4.7 | 3.0 | 0.7 | 5.2 | 0.8 |
| >MI0000067\|hsa-let-7f-1&WithFlank&9\|+\|94176341\|94176439@13.34.22(+1C);MI0000068\|hsa-let-7f-2&WithFlank&X\|-\|53557186\|53557280@14.35.22(+1C)&[MIMAT0000067&hsa-let-7f-5p&offsets\|0\|0(+1C);m-20&9\|+\|94176353\|94176374&offsets\|0\|0(+1C)];[MIMAT0000067_1&hsa-let-7f-5p&offsets\|0\|0(+1C);m-19&X\|-\|53557246\|53557267&offsets\|0\|0(+1C)]\|TGAGGTAGTAGATTGTATAGTTC | >MI0000067\|hsa-let-7f-1&WithFlank&9\|+\|94176341\|94176439 | 13 | 34 | 22(+1C) | TGAGGTAGTAGATTGTATAGTTC | BT20 | 16 | 13 | 18 | 28 | 7 | 8.6 | 6.4 | 16.3 | 34.4 | 1.4 |
| >MI0000067\|hsa-let-7f-1&WithFlank&9\|+\|94176341\|94176439@13.34.22(+1C);MI0000068\|hsa-let-7f-2&WithFlank&X\|-\|53557186\|53557280@14.35.22(+1C)&[MIMAT0000067&hsa-let-7f-5p&offsets\|0\|0(+1C);m-20&9\|+\|94176353\|94176374&offsets\|0\|0(+1C)];[MIMAT0000067_1&hsa-let-7f-5p&offsets\|0\|0(+1C);m-19&X\|-\|53557246\|53557267&offsets\|0\|0(+1C)]\|TGAGGTAGTAGATTGTATAGTTC | >MI0000067\|hsa-let-7f-1&WithFlank&9\|+\|94176341\|94176439 | 13 | 34 | 22(+1C) | TGAGGTAGTAGATTGTATAGTTC | MB231 | 18 | 17 | 7 | 75 | 11 | 7.2 | 5.3 | 0.9 | 46.4 | 4.8 |
| >MI0000067\|hsa-let-7f-1&WithFlank&9\|+\|94176341\|94176439@13.35.23&[MIMAT0000067&hsa-let-7f-5p&offsets\|0\|+1;m-20&9\|+\|94176353\|94176374&offsets\|0\|+1]\|TGAGGTAGTAGATTGTATAGTTG | >MI0000067\|hsa-let-7f-1&WithFlank&9\|+\|94176341\|94176439 | 13 | 35 | 23 | TGAGGTAGTAGATTGTATAGTTG | BT20 | 22 | 15 | 23 | 21 | 8 | 9.1 | 2.3 | 16.6 | 22.0 | 0.5 |
| >MI0000067\|hsa-let-7f-1&WithFlank&9\|+\|94176341\|94176439@69.89.21(+1U)&[MIMAT0004486&hsa-let-7f-1-3p&offsets\|0\|-1(+1U);m-275&9\|+\|94176409\|94176430&offsets\|0\|-1(+1U)]\|CTATACAATCTATTGCCTTCCT | >MI0000067\|hsa-let-7f-1&WithFlank&9\|+\|94176341\|94176439 | 69 | 89 | 21(+1U) | CTATACAATCTATTGCCTTCCT | BT20 | 17 | 14 | 11 | 29 | 7 | 8.2 | 13.6 | 8.6 | 23.8 | 6.6 |
| >MI0000067\|hsa-let-7f-1&WithFlank&9\|+\|94176341\|94176439@69.89.21(+1U)&[MIMAT0004486&hsa-let-7f-1-3p&offsets\|0\|-1(+1U);m-275&9\|+\|94176409\|94176430&offsets\|0\|-1(+1U)]\|CTATACAATCTATTGCCTTCCT | >MI0000067\|hsa-let-7f-1&WithFlank&9\|+\|94176341\|94176439 | 69 | 89 | 21(+1U) | CTATACAATCTATTGCCTTCCT | MB231 | 12 | 20 | 7 | 45 | 19 | 4.6 | 1.4 | 1.1 | 18.0 | 10.6 |
| >MI0000068\|hsa-let-7f-2&WithFlank&X\|-\|53557186\|53557280@14.36.23&[MIMAT0000067_1&hsa-let-7f-5p&offsets\|0\|+1;m-19&X\|-\|53557246\|53557267&offsets\|0\|+1]\|TGAGGTAGTAGATTGTATAGTTT | >MI0000068\|hsa-let-7f-2&WithFlank&X\|-\|53557186\|53557280 | 14 | 36 | 23 | TGAGGTAGTAGATTGTATAGTTT | BT20 | 122 | 109 | 146 | 186 | 57 | 76.4 | 46.6 | 148.2 | 202.0 | 15.4 |
| >MI0000068\|hsa-let-7f-2&WithFlank&X\|-\|53557186\|53557280@14.36.23&[MIMAT0000067_1&hsa-let-7f-5p&offsets\|0\|+1;m-19&X\|-\|53557246\|53557267&offsets\|0\|+1]\|TGAGGTAGTAGATTGTATAGTTT | >MI0000068\|hsa-let-7f-2&WithFlank&X\|-\|53557186\|53557280 | 14 | 36 | 23 | TGAGGTAGTAGATTGTATAGTTT | MB231 | 189 | 173 | 74 | 500 | 91 | 73.2 | 47.2 | 15.8 | 261.3 | 31.0 |
| >MI0000068\|hsa-let-7f-2&WithFlank&X\|-\|53557186\|53557280@14.36.23&[MIMAT0000067_1&hsa-let-7f-5p&offsets\|0\|+1;m-19&X\|-\|53557246\|53557267&offsets\|0\|+1]\|TGAGGTAGTAGATTGTATAGTTT | >MI0000068\|hsa-let-7f-2&WithFlank&X\|-\|53557186\|53557280 | 14 | 36 | 23 | TGAGGTAGTAGATTGTATAGTTT | MB468 | 14 | 13 | 9 | 11 | 2 | 7.5 | 5.0 | 1.4 | 5.5 | 1.5 |
| >MI0000068\|hsa-let-7f-2&WithFlank&X\|-\|53557186\|53557280@64.84.21(+1U)&[MIMAT0004487&hsa-let-7f-2-3p&offsets\|0\|-1(+1U);m-453&X\|-\|53557196\|53557217&offsets\|0\|-1(+1U)]\|CTATACAGTCTACTGTCTTTCT | >MI0000068\|hsa-let-7f-2&WithFlank&X\|-\|53557186\|53557280 | 64 | 84 | 21(+1U) | CTATACAGTCTACTGTCTTTCT | BT20 | 14 | 8 | 2 | 8 | 1 | 15.0 | 9.8 | 1.4 | 6.4 | 0.9 |
| >MI0000069\|hsa-mir-15a&WithFlank&13\|-\|50049113\|50049207@20.40.21&[MIMAT0000068&hsa-miR-15a-5p&offsets\|0\|-1;m-116&13\|-\|50049168\|50049188&offsets\|0\|0]\|TAGCAGCACATAATGGTTTGT | >MI0000069\|hsa-mir-15a&WithFlank&13\|-\|50049113\|50049207 | 20 | 40 | 21 | TAGCAGCACATAATGGTTTGT | BT20 | 40 | 18 | 2 | 9 | 1 | 61.2 | 22.6 | 0.7 | 8.1 | 0.8 |
| >MI0000069\|hsa-mir-15a&WithFlank&13\|-\|50049113\|50049207@20.41.22&[MIMAT0000068&hsa-miR-15a-5p&offsets\|0\|0;m-116&13\|-\|50049168\|50049188&offsets\|0\|+1]\|TAGCAGCACATAATGGTTTGTG | >MI0000069\|hsa-mir-15a&WithFlank&13\|-\|50049113\|50049207 | 20 | 41 | 22 | TAGCAGCACATAATGGTTTGTG | BT20 | 13 | 7 | 1 | 1 | 0 | 16.4 | 8.4 | 0.8 | 1.6 | 0.5 |

<table>
<tr><td>>MI0000069|hsa-mir-15a&WithFlank&13|-|50049113|50049207@20.43.24&[MIMAT0000068&hsa-miR-15a-5p&offsets|0|+2;m-116&13|-|50049168|50049188&offsets|0|+3]|TAGCAGCACATAATGGTTTGTGGA</td><td>>MI0000069|hsa-mir-15a&WithFlank&13|-|50049113|50049207</td><td>20</td><td>43</td><td>24</td><td>TAGCAGCACATAATGGTTTGTGGA</td><td>BT20</td><td>11</td><td>6</td><td>2</td><td>1</td><td>1</td><td>12.7</td><td>6.7</td><td>1.6</td><td>1.0</td><td>0.9</td></tr>
<tr><td>>MI0000070|hsa-mir-16-1&WithFlank&13|-|50048967|50049067@20.39.20;MI0000115|hsa-mir-16-2&WithFlank&3|+|160404739|160404831@16.35.20&[MIMAT0000069&hsa-miR-16-5p&offsets|0|-2;m-62&13|-|50049027|50049048&offsets|0|-2];[MIMAT0000069_1&hsa-miR-16-5p&offsets|0|-2;m-61&3|+|160404754|160404775&offsets|0|-2]|TAGCAGCACGTAAATATTGG</td><td>>MI0000070|hsa-mir-16-1&WithFlank&13|-|50048967|50049067</td><td>20</td><td>39</td><td>20</td><td>TAGCAGCACGTAAATATTGG</td><td>BT20</td><td>22</td><td>26</td><td>26</td><td>40</td><td>5</td><td>13.5</td><td>27.8</td><td>23.9</td><td>35.5</td><td>3.6</td></tr>
<tr><td>>MI0000070|hsa-mir-16-1&WithFlank&13|-|50048967|50049067@20.39.20;MI0000115|hsa-mir-16-2&WithFlank&3|+|160404739|160404831@16.35.20&[MIMAT0000069&hsa-miR-16-5p&offsets|0|-2;m-62&13|-|50049027|50049048&offsets|0|-2];[MIMAT0000069_1&hsa-miR-16-5p&offsets|0|-2;m-61&3|+|160404754|160404775&offsets|0|-2]|TAGCAGCACGTAAATATTGG</td><td>>MI0000070|hsa-mir-16-1&WithFlank&13|-|50048967|50049067</td><td>20</td><td>39</td><td>20</td><td>TAGCAGCACGTAAATATTGG</td><td>MB231</td><td>10</td><td>14</td><td>6</td><td>58</td><td>12</td><td>2.8</td><td>8.1</td><td>4.9</td><td>41.0</td><td>8.1</td></tr>
<tr><td>>MI0000070|hsa-mir-16-1&WithFlank&13|-|50048967|50049067@20.39.20(+1A);MI0000115|hsa-mir-16-2&WithFlank&3|+|160404739|160404831@16.35.20(+1A)&[MIMAT0000069&hsa-miR-16-5p&offsets|0|-2(+1A);m-62&13|-|50049027|50049048&offsets|0|-2(+1A)];[MIMAT0000069_1&hsa-miR-16-5p&offsets|0|-2(+1A);m-61&3|+|160404754|160404775&offsets|0|-2(+1A)]|TAGCAGCACGTAAATATTGGA</td><td>>MI0000070|hsa-mir-16-1&WithFlank&13|-|50048967|50049067</td><td>20</td><td>39</td><td>20(+1A)</td><td>TAGCAGCACGTAAATATTGGA</td><td>BT20</td><td>16</td><td>7</td><td>11</td><td>9</td><td>1</td><td>20.8</td><td>6.5</td><td>7.5</td><td>7.6</td><td>0.5</td></tr>
<tr><td>>MI0000070|hsa-mir-16-1&WithFlank&13|-|50048967|50049067@20.39.20(+1U);MI0000115|hsa-mir-16-2&WithFlank&3|+|160404739|160404831@16.35.20(+1U)&[MIMAT0000069&hsa-miR-16-5p&offsets|0|-2(+1U);m-62&13|-|50049027|50049048&offsets|0|-2(+1U)];[MIMAT0000069_1&hsa-miR-16-5p&offsets|0|-2(+1U);m-61&3|+|160404754|160404775&offsets|0|-2(+1U)]|TAGCAGCACGTAAATATTGGT</td><td>>MI0000070|hsa-mir-16-1&WithFlank&13|-|50048967|50049067</td><td>20</td><td>39</td><td>20(+1U)</td><td>TAGCAGCACGTAAATATTGGT</td><td>BT20</td><td>49</td><td>30</td><td>35</td><td>35</td><td>4</td><td>51.9</td><td>30.0</td><td>21.4</td><td>31.4</td><td>2.4</td></tr>
<tr><td>>MI0000070|hsa-mir-16-1&WithFlank&13|-|50048967|50049067@20.39.20(+1U);MI0000115|hsa-mir-16-2&WithFlank&3|+|160404739|160404831@16.35.20(+1U)&[MIMAT0000069&hsa-miR-16-5p&offsets|0|-2(+1U);m-62&13|-|50049027|50049048&offsets|0|-2(+1U)];[MIMAT0000069_1&hsa-miR-16-5p&offsets|0|-2(+1U);m-61&3|+|160404754|160404775&offsets|0|-2(+1U)]|TAGCAGCACGTAAATATTGGT</td><td>>MI0000070|hsa-mir-16-1&WithFlank&13|-|50048967|50049067</td><td>20</td><td>39</td><td>20(+1U)</td><td>TAGCAGCACGTAAATATTGGT</td><td>MB231</td><td>11</td><td>10</td><td>5</td><td>87</td><td>18</td><td>1.3</td><td>3.7</td><td>3.9</td><td>54.0</td><td>16.4</td></tr>
<tr><td>>MI0000070|hsa-mir-16-1&WithFlank&13|-|50048967|50049067@20.40.21;MI0000115|hsa-mir-16-2&WithFlank&3|+|160404739|160404831@16.36.21&[MIMAT0000069&hsa-miR-16-5p&offsets|0|-1;m-62&13|-|50049027|50049048&offsets|0|-1];[MIMAT0000069_1&hsa-miR-16-5p&offsets|0|-1;m-61&3|+|160404754|160404775&offsets|0|-1]|TAGCAGCACGTAAATATTGGC</td><td>>MI0000070|hsa-mir-16-1&WithFlank&13|-|50048967|50049067</td><td>20</td><td>40</td><td>21</td><td>TAGCAGCACGTAAATATTGGC</td><td>BT20</td><td>104</td><td>86</td><td>98</td><td>111</td><td>15</td><td>88.3</td><td>75.1</td><td>61.0</td><td>90.1</td><td>10.0</td></tr>
<tr><td>>MI0000070|hsa-mir-16-1&WithFlank&13|-|50048967|50049067@20.40.21;MI0000115|hsa-mir-16-2&WithFlank&3|+|160404739|160404831@16.36.21&[MIMAT0000069&hsa-miR-16-5p&offsets|0|-1;m-62&13|-|50049027|50049048&offsets|0|-1];[MIMAT0000069_1&hsa-miR-16-5p&offsets|0|-1;m-61&3|+|160404754|160404775&offsets|0|-1]|TAGCAGCACGTAAATATTGGC</td><td>>MI0000070|hsa-mir-16-1&WithFlank&13|-|50048967|50049067</td><td>20</td><td>40</td><td>21</td><td>TAGCAGCACGTAAATATTGGC</td><td>MB231</td><td>40</td><td>52</td><td>22</td><td>223</td><td>58</td><td>3.0</td><td>17.1</td><td>17.1</td><td>96.6</td><td>38.1</td></tr>
<tr><td>>MI0000070|hsa-mir-16-1&WithFlank&13|-|50048967|50049067@20.40.21(+1A);MI0000115|hsa-mir-16-2&WithFlank&3|+|160404739|160404831@16.36.21(+1A)&[MIMAT0000069&hsa-miR-16-5p&offsets|0|-1(+1A);m-62&13|-|50049027|50049048&offsets|0|-1(+1A)];[MIMAT0000069_1&hsa-miR-16-5p&offsets|0|-1(+1A);m-61&3|+|160404754|160404775&offsets|0|-1(+1A)]|TAGCAGCACGTAAATATTGGCA</td><td>>MI0000070|hsa-mir-16-1&WithFlank&13|-|50048967|50049067</td><td>20</td><td>40</td><td>21(+1A)</td><td>TAGCAGCACGTAAATATTGGCA</td><td>BT20</td><td>252</td><td>92</td><td>82</td><td>134</td><td>6</td><td>375.0</td><td>89.7</td><td>62.4</td><td>123.8</td><td>3.5</td></tr>
<tr><td>>MI0000070|hsa-mir-16-1&WithFlank&13|-|50048967|50049067@20.40.21(+1A);MI0000115|hsa-mir-16-2&WithFlank&3|+|160404739|160404831@16.36.21(+1A)&[MIMAT0000069&hsa-miR-16-5p&offsets|0|-1(+1A);m-62&13|-|50049027|50049048&offsets|0|-1(+1A)];[MIMAT0000069_1&hsa-miR-16-5p&offsets|0|-1(+1A);m-61&3|+|160404754|160404775&offsets|0|-1(+1A)]|TAGCAGCACGTAAATATTGGCA</td><td>>MI0000070|hsa-mir-16-1&WithFlank&13|-|50048967|50049067</td><td>20</td><td>40</td><td>21(+1A)</td><td>TAGCAGCACGTAAATATTGGCA</td><td>MB231</td><td>20</td><td>21</td><td>9</td><td>186</td><td>32</td><td>7.7</td><td>11.2</td><td>8.3</td><td>67.7</td><td>37.5</td></tr>
<tr><td>>MI0000070|hsa-mir-16-1&WithFlank&13|-|50048967|50049067@20.40.21(+1C);MI0000115|hsa-mir-16-2&WithFlank&3|+|160404739|160404831@16.36.21(+1C)&[MIMAT0000069&hsa-miR-16-5p&offsets|0|-1(+1C);m-62&13|-|50049027|50049048&offsets|0|-1(+1C)];[MIMAT0000069_1&hsa-miR-16-5p&offsets|0|-1(+1C);m-61&3|+|160404754|160404775&offsets|0|-1(+1C)]|TAGCAGCACGTAAATATTGGCC</td><td>>MI0000070|hsa-mir-16-1&WithFlank&13|-|50048967|50049067</td><td>20</td><td>40</td><td>21(+1C)</td><td>TAGCAGCACGTAAATATTGGCC</td><td>BT20</td><td>12</td><td>4</td><td>3</td><td>6</td><td>0</td><td>18.0</td><td>3.6</td><td>2.3</td><td>6.7</td><td>0.0</td></tr>
<tr><td>>MI0000070|hsa-mir-16-1&WithFlank&13|-|50048967|50049067@20.40.21(+1U);MI0000115|hsa-mir-16-2&WithFlank&3|+|160404739|160404831@16.36.21(+1U)&[MIMAT0000069&hsa-miR-16-5p&offsets|0|-1(+1U);m-62&13|-|50049027|50049048&offsets|0|-1(+1U)];[MIMAT0000069_1&hsa-miR-16-5p&offsets|0|-1(+1U);m-61&3|+|160404754|160404775&offsets|0|-1(+1U)]|TAGCAGCACGTAAATATTGGCT</td><td>>MI0000070|hsa-mir-16-1&WithFlank&13|-|50048967|50049067</td><td>20</td><td>40</td><td>21(+1U)</td><td>TAGCAGCACGTAAATATTGGCT</td><td>BT20</td><td>58</td><td>14</td><td>10</td><td>22</td><td>1</td><td>89.1</td><td>11.7</td><td>8.0</td><td>21.8</td><td>0.6</td></tr>
<tr><td>>MI0000070|hsa-mir-16-1&WithFlank&13|-|50048967|50049067@20.41.22;MI0000115|hsa-mir-16-2&WithFlank&3|+|160404739|160404831@16.37.22&[MIMAT0000069&hsa-miR-16-5p&offsets|0|0;m-62&13|-|50049027|50049048&offsets|0|0];[MIMAT0000069_1&hsa-miR-16-5p&offsets|0|0;m-61&3|+|160404754|160404775&offsets|0|0]|TAGCAGCACGTAAATATTGGCG</td><td>>MI0000070|hsa-mir-16-1&WithFlank&13|-|50048967|50049067</td><td>20</td><td>41</td><td>22</td><td>TAGCAGCACGTAAATATTGGCG</td><td>BT20</td><td>606</td><td>346</td><td>243</td><td>677</td><td>25</td><td>786.5</td><td>287.2</td><td>117.8</td><td>561.8</td><td>13.9</td></tr>
<tr><td>>MI0000070|hsa-mir-16-1&WithFlank&13|-|50048967|50049067@20.41.22;MI0000115|hsa-mir-16-2&WithFlank&3|+|160404739|160404831@16.37.22&[MIMAT0000069&hsa-miR-16-5p&offsets|0|0;m-62&13|-|50049027|50049048&offsets|0|0];[MIMAT0000069_1&hsa-miR-16-5p&offsets|0|0;m-61&3|+|160404754|160404775&offsets|0|0]|TAGCAGCACGTAAATATTGGCG</td><td>>MI0000070|hsa-mir-16-1&WithFlank&13|-|50048967|50049067</td><td>20</td><td>41</td><td>22</td><td>TAGCAGCACGTAAATATTGGCG</td><td>MB231</td><td>103</td><td>130</td><td>41</td><td>755</td><td>118</td><td>42.5</td><td>65.0</td><td>37.5</td><td>116.7</td><td>109.9</td></tr>
<tr><td>>MI0000070|hsa-mir-16-1&WithFlank&13|-|50048967|50049067@20.41.22(+1A);MI0000115|hsa-mir-16-2&WithFlank&3|+|160404739|160404831@16.37.22(+1A)&[MIMAT0000069&hsa-miR-16-5p&offsets|0|0(+1A);m-62&13|-|50049027|50049048&offsets|0|0(+1A)];[MIMAT0000069_1&hsa-miR-16-5p&offsets|0|0(+1A);m-61&3|+|160404754|160404775&offsets|0|0(+1A)]|TAGCAGCACGTAAATATTGGCGA</td><td>>MI0000070|hsa-mir-16-1&WithFlank&13|-|50048967|50049067</td><td>20</td><td>41</td><td>22(+1A)</td><td>TAGCAGCACGTAAATATTGGCGA</td><td>BT20</td><td>38</td><td>12</td><td>10</td><td>12</td><td>1</td><td>56.1</td><td>13.6</td><td>7.9</td><td>10.3</td><td>0.6</td></tr>
<tr><td>>MI0000070|hsa-mir-16-1&WithFlank&13|-|50048967|50049067@20.41.22(+1C);MI0000115|hsa-mir-16-2&WithFlank&3|+|160404739|160404831@16.37.22(+1C)&[MIMAT0000069&hsa-miR-16-5p&offsets|0|0(+1C);m-62&13|-|50049027|50049048&offsets|0|0(+1C)];[MIMAT0000069_1&hsa-miR-16-5p&offsets|0|0(+1C);m-61&3|+|160404754|160404775&offsets|0|0(+1C)]|TAGCAGCACGTAAATATTGGCGC</td><td>>MI0000070|hsa-mir-16-1&WithFlank&13|-|50048967|50049067</td><td>20</td><td>41</td><td>22(+1C)</td><td>TAGCAGCACGTAAATATTGGCGC</td><td>BT20</td><td>48</td><td>32</td><td>16</td><td>31</td><td>3</td><td>60.4</td><td>39.6</td><td>9.4</td><td>27.0</td><td>2.0</td></tr>
<tr><td>>MI0000070|hsa-mir-16-1&WithFlank&13|-|50048967|50049067@20.41.22(+1G);MI0000115|hsa-mir-16-2&WithFlank&3|+|160404739|160404831@16.37.22(+1G)&[MIMAT0000069&hsa-miR-16-5p&offsets|0|0(+1G);m-62&13|-|50049027|50049048&offsets|0|0(+1G)];[MIMAT0000069_1&hsa-miR-16-5p&offsets|0|0(+1G);m-61&3|+|160404754|160404775&offsets|0|0(+1G)]|TAGCAGCACGTAAATATTGGCGG</td><td>>MI0000070|hsa-mir-16-1&WithFlank&13|-|50048967|50049067</td><td>20</td><td>41</td><td>22(+1G)</td><td>TAGCAGCACGTAAATATTGGCGG</td><td>BT20</td><td>40</td><td>7</td><td>4</td><td>5</td><td>0</td><td>60.0</td><td>7.7</td><td>2.3</td><td>5.4</td><td>0.3</td></tr>
<tr><td>>MI0000070|hsa-mir-16-1&WithFlank&13|-|50048967|50049067@20.42.23;MI0000115|hsa-mir-16-2&WithFlank&3|+|160404739|160404831@16.38.23&[MIMAT0000069&hsa-miR-16-5p&offsets|0|+1;m-62&13|-|50049027|50049048&offsets|0|+1];[MIMAT0000069_1&hsa-miR-16-5p&offsets|0|+1;m-61&3|+|160404754|160404775&offsets|0|+1]|TAGCAGCACGTAAATATTGGCGT</td><td>>MI0000070|hsa-mir-16-1&WithFlank&13|-|50048967|50049067</td><td>20</td><td>42</td><td>23</td><td>TAGCAGCACGTAAATATTGGCGT</td><td>BT20</td><td>357</td><td>202</td><td>117</td><td>226</td><td>19</td><td>454.0</td><td>235.0</td><td>57.4</td><td>189.4</td><td>15.3</td></tr>
<tr><td>>MI0000070|hsa-mir-16-1&WithFlank&13|-|50048967|50049067@20.42.23;MI0000115|hsa-mir-16-2&WithFlank&3|+|160404739|160404831@16.38.23&[MIMAT0000069&hsa-miR-16-5p&offsets|0|+1;m-62&13|-|50049027|50049048&offsets|0|+1];[MIMAT0000069_1&hsa-miR-16-5p&offsets|0|+1;m-61&3|+|160404754|160404775&offsets|0|+1]|TAGCAGCACGTAAATATTGGCGT</td><td>>MI0000070|hsa-mir-16-1&WithFlank&13|-|50048967|50049067</td><td>20</td><td>42</td><td>23</td><td>TAGCAGCACGTAAATATTGGCGT</td><td>MB231</td><td>85</td><td>84</td><td>21</td><td>354</td><td>117</td><td>27.1</td><td>50.4</td><td>17.7</td><td>119.1</td><td>130.8</td></tr>
<tr><td>>MI0000070|hsa-mir-16-1&WithFlank&13|-|50048967|50049067@20.43.24&[MIMAT0000069&hsa-miR-16-5p&offsets|0|+2;m-62&13|-|50049027|50049048&offsets|0|+2]|TAGCAGCACGTAAATATTGGCGTT</td><td>>MI0000070|hsa-mir-16-1&WithFlank&13|-|50048967|50049067</td><td>20</td><td>43</td><td>24</td><td>TAGCAGCACGTAAATATTGGCGTT</td><td>BT20</td><td>17</td><td>9</td><td>3</td><td>1</td><td>1</td><td>22.0</td><td>10.5</td><td>1.6</td><td>0.9</td><td>0.9</td></tr>
</table>

| | | | | | | | | | | | | | | | | |
|---|---|---|---|---|---|---|---|---|---|---|---|---|---|---|---|---|
| >MI0000071\|hsa-mir-17&WithFlank&13\|+\|91350599\|91350694@20.40.21&[MIMAT0000070&hsa-miR-17-5p&offsets\|0\|-2;m-88&13\|+\|91350618\|91350640&offsets\|0\|-2]\|CAAAGTGCTTACAGTGCAGGT | >MI0000071\|hsa-mir-17&WithFlank&13\|+\|91350599\|91350694 | 20 | 40 | 21 | CAAAGTGCTTACAGTGCAGGT | BT20 | 22 | 20 | 11 | 12 | 3 | 18.5 | 23.6 | 5.0 | 11.1 | 3.5 |
| >MI0000071\|hsa-mir-17&WithFlank&13\|+\|91350599\|91350694@20.40.21&[MIMAT0000070&hsa-miR-17-5p&offsets\|0\|-2;m-88&13\|+\|91350618\|91350640&offsets\|0\|-2]\|CAAAGTGCTTACAGTGCAGGT | >MI0000071\|hsa-mir-17&WithFlank&13\|+\|91350599\|91350694 | 20 | 40 | 21 | CAAAGTGCTTACAGTGCAGGT | MB231 | 47 | 48 | 11 | 191 | 69 | 5.4 | 19.6 | 9.6 | 75.3 | 61.0 |
| >MI0000071\|hsa-mir-17&WithFlank&13\|+\|91350599\|91350694@20.41.22(+1A)&[MIMAT0000070&hsa-miR-17-5p&offsets\|0\|-1(+1A);m-88&13\|+\|91350618\|91350640&offsets\|0\|-1(+1A)]\|CAAAGTGCTTACAGTGCAGGTAA | >MI0000071\|hsa-mir-17&WithFlank&13\|+\|91350599\|91350694 | 20 | 41 | 22(+1A) | CAAAGTGCTTACAGTGCAGGTAA | BT20 | 58 | 29 | 27 | 22 | 6 | 59.1 | 29.3 | 5.5 | 21.0 | 4.0 |
| >MI0000071\|hsa-mir-17&WithFlank&13\|+\|91350599\|91350694@20.41.22(+1A)&[MIMAT0000070&hsa-miR-17-5p&offsets\|0\|-1(+1A);m-88&13\|+\|91350618\|91350640&offsets\|0\|-1(+1A)]\|CAAAGTGCTTACAGTGCAGGTAA | >MI0000071\|hsa-mir-17&WithFlank&13\|+\|91350599\|91350694 | 20 | 41 | 22(+1A) | CAAAGTGCTTACAGTGCAGGTAA | MB231 | 51 | 31 | 10 | 182 | 75 | 9.4 | 8.1 | 9.3 | 75.4 | 69.3 |
| >MI0000071\|hsa-mir-17&WithFlank&13\|+\|91350599\|91350694@20.41.22&[MIMAT0000070&hsa-miR-17-5p&offsets\|0\|-1;m-88&13\|+\|91350618\|91350640&offsets\|0\|-1]\|CAAAGTGCTTACAGTGCAGGTA | >MI0000071\|hsa-mir-17&WithFlank&13\|+\|91350599\|91350694 | 20 | 41 | 22 | CAAAGTGCTTACAGTGCAGGTA | BT20 | 28 | 16 | 14 | 8 | 3 | 26.3 | 15.4 | 3.9 | 5.7 | 2.3 |
| >MI0000071\|hsa-mir-17&WithFlank&13\|+\|91350599\|91350694@20.41.22&[MIMAT0000070&hsa-miR-17-5p&offsets\|0\|-1;m-88&13\|+\|91350618\|91350640&offsets\|0\|-1]\|CAAAGTGCTTACAGTGCAGGTA | >MI0000071\|hsa-mir-17&WithFlank&13\|+\|91350599\|91350694 | 20 | 41 | 22 | CAAAGTGCTTACAGTGCAGGTA | MB231 | 32 | 31 | 11 | 91 | 39 | 5.9 | 11.7 | 10.5 | 25.8 | 31.2 |
| >MI0000071\|hsa-mir-17&WithFlank&13\|+\|91350599\|91350694@20.42.23(+1A)&[MIMAT0000070&hsa-miR-17-5p&offsets\|0\|0(+1A);m-88&13\|+\|91350618\|91350640&offsets\|0\|0(+1A)]\|CAAAGTGCTTACAGTGCAGGTAGA | >MI0000071\|hsa-mir-17&WithFlank&13\|+\|91350599\|91350694 | 20 | 42 | 23(+1A) | CAAAGTGCTTACAGTGCAGGTAGA | BT20 | 127 | 66 | 123 | 13 | 31 | 58.0 | 50.8 | 36.9 | 8.7 | 21.5 |
| >MI0000071\|hsa-mir-17&WithFlank&13\|+\|91350599\|91350694@20.42.23(+1A)&[MIMAT0000070&hsa-miR-17-5p&offsets\|0\|0(+1A);m-88&13\|+\|91350618\|91350640&offsets\|0\|0(+1A)]\|CAAAGTGCTTACAGTGCAGGTAGA | >MI0000071\|hsa-mir-17&WithFlank&13\|+\|91350599\|91350694 | 20 | 42 | 23(+1A) | CAAAGTGCTTACAGTGCAGGTAGA | MB231 | 257 | 158 | 73 | 165 | 267 | 28.2 | 53.7 | 69.1 | 66.7 | 215.8 |
| >MI0000071\|hsa-mir-17&WithFlank&13\|+\|91350599\|91350694@20.42.23(+1A)&[MIMAT0000070&hsa-miR-17-5p&offsets\|0\|0(+1A);m-88&13\|+\|91350618\|91350640&offsets\|0\|0(+1A)]\|CAAAGTGCTTACAGTGCAGGTAGA | >MI0000071\|hsa-mir-17&WithFlank&13\|+\|91350599\|91350694 | 20 | 42 | 23(+1A) | CAAAGTGCTTACAGTGCAGGTAGA | MB468 | 28 | 11 | 11 | 19 | 8 | 11.6 | 2.4 | 1.6 | 13.3 | 3.4 |
| >MI0000071\|hsa-mir-17&WithFlank&13\|+\|91350599\|91350694@20.42.23(+1C)&[MIMAT0000070&hsa-miR-17-5p&offsets\|0\|0(+1C);m-88&13\|+\|91350618\|91350640&offsets\|0\|0(+1C)]\|CAAAGTGCTTACAGTGCAGGTAGC | >MI0000071\|hsa-mir-17&WithFlank&13\|+\|91350599\|91350694 | 20 | 42 | 23(+1C) | CAAAGTGCTTACAGTGCAGGTAGC | MB231 | 18 | 13 | 3 | 16 | 28 | 5.2 | 5.8 | 2.8 | 8.2 | 31.6 |
| >MI0000071\|hsa-mir-17&WithFlank&13\|+\|91350599\|91350694@20.42.23(+1G)&[MIMAT0000070&hsa-miR-17-5p&offsets\|0\|0(+1G);m-88&13\|+\|91350618\|91350640&offsets\|0\|0(+1G)]\|CAAAGTGCTTACAGTGCAGGTAGG | >MI0000071\|hsa-mir-17&WithFlank&13\|+\|91350599\|91350694 | 20 | 42 | 23(+1G) | CAAAGTGCTTACAGTGCAGGTAGG | BT20 | 13 | 2 | 4 | 0 | 0 | 12.3 | 2.3 | 0.8 | 0.0 | 0.0 |
| >MI0000071\|hsa-mir-17&WithFlank&13\|+\|91350599\|91350694@20.42.23(+1G)&[MIMAT0000070&hsa-miR-17-5p&offsets\|0\|0(+1G);m-88&13\|+\|91350618\|91350640&offsets\|0\|0(+1G)]\|CAAAGTGCTTACAGTGCAGGTAGG | >MI0000071\|hsa-mir-17&WithFlank&13\|+\|91350599\|91350694 | 20 | 42 | 23(+1G) | CAAAGTGCTTACAGTGCAGGTAGG | MB231 | 19 | 5 | 2 | 3 | 6 | 14.8 | 2.5 | 2.0 | 1.8 | 6.5 |
| >MI0000071\|hsa-mir-17&WithFlank&13\|+\|91350599\|91350694@20.42.23&[MIMAT0000070&hsa-miR-17-5p&offsets\|0\|0;m-88&13\|+\|91350618\|91350640&offsets\|0\|0]\|CAAAGTGCTTACAGTGCAGGTAG | >MI0000071\|hsa-mir-17&WithFlank&13\|+\|91350599\|91350694 | 20 | 42 | 23 | CAAAGTGCTTACAGTGCAGGTAG | BT20 | 301 | 235 | 178 | 201 | 62 | 163.1 | 214.2 | 58.6 | 170.3 | 47.9 |
| >MI0000071\|hsa-mir-17&WithFlank&13\|+\|91350599\|91350694@20.42.23&[MIMAT0000070&hsa-miR-17-5p&offsets\|0\|0;m-88&13\|+\|91350618\|91350640&offsets\|0\|0]\|CAAAGTGCTTACAGTGCAGGTAG | >MI0000071\|hsa-mir-17&WithFlank&13\|+\|91350599\|91350694 | 20 | 42 | 23 | CAAAGTGCTTACAGTGCAGGTAG | MB231 | 563 | 461 | 95 | 1339 | 640 | 89.7 | 130.7 | 82.0 | 307.0 | 469.0 |
| >MI0000071\|hsa-mir-17&WithFlank&13\|+\|91350599\|91350694@20.42.23&[MIMAT0000070&hsa-miR-17-5p&offsets\|0\|0;m-88&13\|+\|91350618\|91350640&offsets\|0\|0]\|CAAAGTGCTTACAGTGCAGGTAG | >MI0000071\|hsa-mir-17&WithFlank&13\|+\|91350599\|91350694 | 20 | 42 | 23 | CAAAGTGCTTACAGTGCAGGTAG | MB468 | 85 | 26 | 22 | 141 | 21 | 44.0 | 6.1 | 9.1 | 87.7 | 8.7 |
| >MI0000071\|hsa-mir-17&WithFlank&13\|+\|91350599\|91350694@20.43.24&[MIMAT0000070&hsa-miR-17-5p&offsets\|0\|+1;m-88&13\|+\|91350618\|91350640&offsets\|0\|+1]\|CAAAGTGCTTACAGTGCAGGTAGT | >MI0000071\|hsa-mir-17&WithFlank&13\|+\|91350599\|91350694 | 20 | 43 | 24 | CAAAGTGCTTACAGTGCAGGTAGT | BT20 | 60 | 33 | 29 | 2 | 7 | 46.8 | 35.3 | 7.7 | 2.4 | 8.9 |
| >MI0000071\|hsa-mir-17&WithFlank&13\|+\|91350599\|91350694@20.43.24&[MIMAT0000070&hsa-miR-17-5p&offsets\|0\|+1;m-88&13\|+\|91350618\|91350640&offsets\|0\|+1]\|CAAAGTGCTTACAGTGCAGGTAGT | >MI0000071\|hsa-mir-17&WithFlank&13\|+\|91350599\|91350694 | 20 | 43 | 24 | CAAAGTGCTTACAGTGCAGGTAGT | MB231 | 159 | 116 | 29 | 100 | 200 | 35.9 | 49.0 | 26.8 | 42.8 | 200.5 |
| >MI0000071\|hsa-mir-17&WithFlank&13\|+\|91350599\|91350694@20.43.24&[MIMAT0000070&hsa-miR-17-5p&offsets\|0\|+1;m-88&13\|+\|91350618\|91350640&offsets\|0\|+1]\|CAAAGTGCTTACAGTGCAGGTAGT | >MI0000071\|hsa-mir-17&WithFlank&13\|+\|91350599\|91350694 | 20 | 43 | 24 | CAAAGTGCTTACAGTGCAGGTAGT | MB468 | 12 | 4 | 4 | 9 | 3 | 4.3 | 1.3 | 1.2 | 9.2 | 1.3 |
| >MI0000072\|hsa-mir-18a&WithFlank&13\|+\|91350745\|91350827@12.34.23&[MIMAT0000072&hsa-miR-18a-5p&offsets\|0\|0;m-262&13\|+\|91350756\|91350778&offsets\|0\|0]\|TAAGGTGCATCTAGTGCAGATAG | >MI0000072\|hsa-mir-18a&WithFlank&13\|+\|91350745\|91350827 | 12 | 34 | 23 | TAAGGTGCATCTAGTGCAGATAG | BT20 | 20 | 12 | 3 | 9 | 2 | 19.1 | 14.3 | 2.1 | 7.4 | 0.9 |
| >MI0000072\|hsa-mir-18a&WithFlank&13\|+\|91350745\|91350827@12.34.23&[MIMAT0000072&hsa-miR-18a-5p&offsets\|0\|0;m-262&13\|+\|91350756\|91350778&offsets\|0\|0]\|TAAGGTGCATCTAGTGCAGATAG | >MI0000072\|hsa-mir-18a&WithFlank&13\|+\|91350745\|91350827 | 12 | 34 | 23 | TAAGGTGCATCTAGTGCAGATAG | MB231 | 20 | 25 | 1 | 77 | 29 | 5.4 | 14.6 | 0.8 | 31.0 | 22.7 |
| >MI0000073\|hsa-mir-19a&WithFlank&13\|+\|91350885\|91350978@55.76.22(+1U)&[MIMAT0000073&hsa-miR-19a-3p&offsets\|0\|-1(+1U);m-201&13\|+\|91350939\|91350961&offsets\|0\|-1(+1U)]\|TGTGCAAATCTATGCAAAACTGT | >MI0000073\|hsa-mir-19a&WithFlank&13\|+\|91350885\|91350978 | 55 | 76 | 22(+1U) | TGTGCAAATCTATGCAAAACTGT | BT20 | 12 | 11 | 3 | 19 | 2 | 6.3 | 12.4 | 2.9 | 20.6 | 1.3 |
| >MI0000073\|hsa-mir-19a&WithFlank&13\|+\|91350885\|91350978@55.76.22(+1U)&[MIMAT0000073&hsa-miR-19a-3p&offsets\|0\|-1(+1U);m-201&13\|+\|91350939\|91350961&offsets\|0\|-1(+1U)]\|TGTGCAAATCTATGCAAAACTGT | >MI0000073\|hsa-mir-19a&WithFlank&13\|+\|91350885\|91350978 | 55 | 76 | 22(+1U) | TGTGCAAATCTATGCAAAACTGT | MB231 | 11 | 8 | 0 | 74 | 16 | 2.6 | 6.6 | 0.3 | 36.1 | 16.0 |
| >MI0000073\|hsa-mir-19a&WithFlank&13\|+\|91350885\|91350978@55.77.23&[MIMAT0000073&hsa-miR-19a-3p&offsets\|0\|0;m-201&13\|+\|91350939\|91350961&offsets\|0\|0]\|TGTGCAAATCTATGCAAAACTGA | >MI0000073\|hsa-mir-19a&WithFlank&13\|+\|91350885\|91350978 | 55 | 77 | 23 | TGTGCAAATCTATGCAAAACTGA | BT20 | 16 | 8 | 1 | 3 | 0 | 24.5 | 10.2 | 0.5 | 4.4 | 0.3 |
| >MI0000073\|hsa-mir-19a&WithFlank&13\|+\|91350885\|91350978@55.78.24&[MIMAT0000073&hsa-miR-19a-3p&offsets\|0\|+1;m-201&13\|+\|91350939\|91350961&offsets\|0\|+1]\|TGTGCAAATCTATGCAAAACTGAT | >MI0000073\|hsa-mir-19a&WithFlank&13\|+\|91350885\|91350978 | 55 | 78 | 24 | TGTGCAAATCTATGCAAAACTGAT | MB231 | 14 | 10 | 0 | 55 | 22 | 4.7 | 6.0 | 0.2 | 19.6 | 21.4 |
| >MI0000074\|hsa-mir-19b-1&WithFlank&13\|+\|91351186\|91351284@60.81.22;MI0000075\|hsa-mir-19b-2&WithFlank&X\|-\|134169665\|134169772@68.89.22&[MIMAT0000074&hsa-miR-19b-3p&offsets\|0\|-1;m-122&13\|+\|91351245\|91351267&offsets\|0\|-1];[MIMAT0000074_1&hsa-miR-19b-3p&offsets\|0\|-1;m-121&X\|-\|134169683\|134169705&offsets\|0\|-1]\|TGTGCAAATCCATGCAAAACTG | >MI0000074\|hsa-mir-19b-1&WithFlank&13\|+\|91351186\|91351284 | 60 | 81 | 22 | TGTGCAAATCCATGCAAAACTG | BT20 | 31 | 16 | 4 | 48 | 5 | 33.7 | 6.0 | 2.3 | 63.4 | 2.0 |
| >MI0000074\|hsa-mir-19b-1&WithFlank&13\|+\|91351186\|91351284@60.81.22;MI0000075\|hsa-mir-19b-2&WithFlank&X\|-\|134169665\|134169772@68.89.22&[MIMAT0000074&hsa-miR-19b-3p&offsets\|0\|-1;m-122&13\|+\|91351245\|91351267&offsets\|0\|-1];[MIMAT0000074_1&hsa-miR-19b-3p&offsets\|0\|-1;m-121&X\|-\|134169683\|134169705&offsets\|0\|-1]\|TGTGCAAATCCATGCAAAACTG | >MI0000074\|hsa-mir-19b-1&WithFlank&13\|+\|91351186\|91351284 | 60 | 81 | 22 | TGTGCAAATCCATGCAAAACTG | MB231 | 17 | 16 | 1 | 76 | 29 | 5.6 | 10.4 | 0.4 | 30.8 | 29.0 |
| >MI0000074\|hsa-mir-19b-1&WithFlank&13\|+\|91351186\|91351284@60.81.22(+1U);MI0000075\|hsa-mir-19b-2&WithFlank&X\|-\|134169665\|134169772@68.89.22(+1U)&[MIMAT0000074&hsa-miR-19b-3p&offsets\|0\|-1(+1U);m-122&13\|+\|91351245\|91351267&offsets\|0\|-1(+1U)];[MIMAT0000074_1&hsa-miR-19b-3p&offsets\|0\|-1(+1U);m-121&X\|-\|134169683\|134169705&offsets\|0\|-1(+1U)]\|TGTGCAAATCCATGCAAAACTGT | >MI0000074\|hsa-mir-19b-1&WithFlank&13\|+\|91351186\|91351284 | 60 | 81 | 22(+1U) | TGTGCAAATCCATGCAAAACTGT | BT20 | 16 | 9 | 3 | 13 | 1 | 16.8 | 9.3 | 1.8 | 15.4 | 1.0 |

<table>
<tr><td>>MI0000074|hsa-mir-19b-1&WithFlank&13|+|91351186|91351284@60.81.22(+1U);MI0000075|hsa-mir-19b-2&WithFlank&X|-|134169665|134169772@68.89.22(+1U)&[MIMAT0000074&hsa-miR-19b-3p&offsets|0|-1(+1U);m-122&13|+|91351245|91351267&offsets|0|-1(+1U)];[MIMAT0000074_1&hsa-miR-19b-3p&offsets|0|-1(+1U);m-121&X|-|134169683|134169705&offsets|0|-1(+1U)]|TGTGCAAATCCATGCAAAACTGT</td><td>>MI0000074|hsa-mir-19b-1&WithFlank&13|+|91351186|91351284</td><td>60</td><td>81</td><td>22(+1U)</td><td>TGTGCAAATCCATGCAAAACTGT</td><td>MB231</td><td>11</td><td>8</td><td>1</td><td>37</td><td>10</td><td>3.8</td><td>5.2</td><td>0.5</td><td>17.6</td><td>11.2</td></tr>
<tr><td>>MI0000074|hsa-mir-19b-1&WithFlank&13|+|91351186|91351284@60.82.23;MI0000075|hsa-mir-19b-2&WithFlank&X|-|134169665|134169772@68.90.23&[MIMAT0000074&hsa-miR-19b-3p&offsets|0|0;m-122&13|+|91351245|91351267&offsets|0|0];[MIMAT0000074_1&hsa-miR-19b-3p&offsets|0|0;m-121&X|-|134169683|134169705&offsets|0|0]|TGTGCAAATCCATGCAAAACTGA</td><td>>MI0000074|hsa-mir-19b-1&WithFlank&13|+|91351186|91351284</td><td>60</td><td>82</td><td>23</td><td>TGTGCAAATCCATGCAAAACTGA</td><td>BT20</td><td>76</td><td>33</td><td>7</td><td>34</td><td>2</td><td>109.0</td><td>34.2</td><td>3.4</td><td>47.3</td><td>1.7</td></tr>
<tr><td>>MI0000074|hsa-mir-19b-1&WithFlank&13|+|91351186|91351284@60.82.23;MI0000075|hsa-mir-19b-2&WithFlank&X|-|134169665|134169772@68.90.23&[MIMAT0000074&hsa-miR-19b-3p&offsets|0|0;m-122&13|+|91351245|91351267&offsets|0|0];[MIMAT0000074_1&hsa-miR-19b-3p&offsets|0|0;m-121&X|-|134169683|134169705&offsets|0|0]|TGTGCAAATCCATGCAAAACTGA</td><td>>MI0000074|hsa-mir-19b-1&WithFlank&13|+|91351186|91351284</td><td>60</td><td>82</td><td>23</td><td>TGTGCAAATCCATGCAAAACTGA</td><td>MB231</td><td>38</td><td>32</td><td>2</td><td>113</td><td>61</td><td>14.4</td><td>16.8</td><td>1.4</td><td>31.7</td><td>76.0</td></tr>
<tr><td>>MI0000076|hsa-mir-20a&WithFlank&13|+|91351059|91351141@14.34.21&[MIMAT0000075&hsa-miR-20a-5p&offsets|0|-2;m-96&13|+|91351072|91351094&offsets|0|-2]|TAAAGTGCTTATAGTGCAGGT</td><td>>MI0000076|hsa-mir-20a&WithFlank&13|+|91351059|91351141</td><td>14</td><td>34</td><td>21</td><td>TAAAGTGCTTATAGTGCAGGT</td><td>BT20</td><td>24</td><td>17</td><td>5</td><td>9</td><td>2</td><td>20.2</td><td>21.7</td><td>4.7</td><td>7.7</td><td>2.8</td></tr>
<tr><td>>MI0000076|hsa-mir-20a&WithFlank&13|+|91351059|91351141@14.34.21&[MIMAT0000075&hsa-miR-20a-5p&offsets|0|-2;m-96&13|+|91351072|91351094&offsets|0|-2]|TAAAGTGCTTATAGTGCAGGT</td><td>>MI0000076|hsa-mir-20a&WithFlank&13|+|91351059|91351141</td><td>14</td><td>34</td><td>21</td><td>TAAAGTGCTTATAGTGCAGGT</td><td>MB231</td><td>74</td><td>57</td><td>6</td><td>210</td><td>106</td><td>6.8</td><td>36.3</td><td>5.0</td><td>75.7</td><td>106.7</td></tr>
<tr><td>>MI0000076|hsa-mir-20a&WithFlank&13|+|91351059|91351141@14.35.22(+1A)&[MIMAT0000075&hsa-miR-20a-5p&offsets|0|-1(+1A);m-96&13|+|91351072|91351094&offsets|0|-1(+1A)]|TAAAGTGCTTATAGTGCAGGTAA</td><td>>MI0000076|hsa-mir-20a&WithFlank&13|+|91351059|91351141</td><td>14</td><td>35</td><td>22(+1A)</td><td>TAAAGTGCTTATAGTGCAGGTAA</td><td>BT20</td><td>127</td><td>38</td><td>20</td><td>21</td><td>5</td><td>146.9</td><td>48.3</td><td>11.0</td><td>20.0</td><td>5.9</td></tr>
<tr><td>>MI0000076|hsa-mir-20a&WithFlank&13|+|91351059|91351141@14.35.22(+1A)&[MIMAT0000075&hsa-miR-20a-5p&offsets|0|-1(+1A);m-96&13|+|91351072|91351094&offsets|0|-1(+1A)]|TAAAGTGCTTATAGTGCAGGTAA</td><td>>MI0000076|hsa-mir-20a&WithFlank&13|+|91351059|91351141</td><td>14</td><td>35</td><td>22(+1A)</td><td>TAAAGTGCTTATAGTGCAGGTAA</td><td>MB231</td><td>126</td><td>63</td><td>9</td><td>283</td><td>157</td><td>29.9</td><td>32.9</td><td>8.4</td><td>105.3</td><td>168.8</td></tr>
<tr><td>>MI0000076|hsa-mir-20a&WithFlank&13|+|91351059|91351141@14.35.22(+1U)&[MIMAT0000075&hsa-miR-20a-5p&offsets|0|-1(+1U);m-96&13|+|91351072|91351094&offsets|0|-1(+1U)]|TAAAGTGCTTATAGTGCAGGTAT</td><td>>MI0000076|hsa-mir-20a&WithFlank&13|+|91351059|91351141</td><td>14</td><td>35</td><td>22(+1U)</td><td>TAAAGTGCTTATAGTGCAGGTAT</td><td>MB231</td><td>10</td><td>7</td><td>0</td><td>29</td><td>14</td><td>2.4</td><td>5.5</td><td>0.5</td><td>14.3</td><td>15.3</td></tr>
<tr><td>>MI0000076|hsa-mir-20a&WithFlank&13|+|91351059|91351141@14.35.22&[MIMAT0000075&hsa-miR-20a-5p&offsets|0|-1;m-96&13|+|91351072|91351094&offsets|0|-1]|TAAAGTGCTTATAGTGCAGGTA</td><td>>MI0000076|hsa-mir-20a&WithFlank&13|+|91351059|91351141</td><td>14</td><td>35</td><td>22</td><td>TAAAGTGCTTATAGTGCAGGTA</td><td>BT20</td><td>18</td><td>6</td><td>3</td><td>1</td><td>1</td><td>24.0</td><td>8.2</td><td>2.3</td><td>1.2</td><td>1.0</td></tr>
<tr><td>>MI0000076|hsa-mir-20a&WithFlank&13|+|91351059|91351141@14.35.22&[MIMAT0000075&hsa-miR-20a-5p&offsets|0|-1;m-96&13|+|91351072|91351094&offsets|0|-1]|TAAAGTGCTTATAGTGCAGGTA</td><td>>MI0000076|hsa-mir-20a&WithFlank&13|+|91351059|91351141</td><td>14</td><td>35</td><td>22</td><td>TAAAGTGCTTATAGTGCAGGTA</td><td>MB231</td><td>23</td><td>16</td><td>3</td><td>49</td><td>29</td><td>2.4</td><td>10.9</td><td>3.0</td><td>14.4</td><td>29.5</td></tr>
<tr><td>>MI0000076|hsa-mir-20a&WithFlank&13|+|91351059|91351141@14.36.23(+1A)&[MIMAT0000075&hsa-miR-20a-5p&offsets|0|0(+1A);m-96&13|+|91351072|91351094&offsets|0|0(+1A)]|TAAAGTGCTTATAGTGCAGGTAGA</td><td>>MI0000076|hsa-mir-20a&WithFlank&13|+|91351059|91351141</td><td>14</td><td>36</td><td>23(+1A)</td><td>TAAAGTGCTTATAGTGCAGGTAGA</td><td>BT20</td><td>142</td><td>54</td><td>44</td><td>5</td><td>16</td><td>104.8</td><td>60.5</td><td>21.9</td><td>3.8</td><td>17.1</td></tr>
<tr><td>>MI0000076|hsa-mir-20a&WithFlank&13|+|91351059|91351141@14.36.23(+1A)&[MIMAT0000075&hsa-miR-20a-5p&offsets|0|0(+1A);m-96&13|+|91351072|91351094&offsets|0|0(+1A)]|TAAAGTGCTTATAGTGCAGGTAGA</td><td>>MI0000076|hsa-mir-20a&WithFlank&13|+|91351059|91351141</td><td>14</td><td>36</td><td>23(+1A)</td><td>TAAAGTGCTTATAGTGCAGGTAGA</td><td>MB231</td><td>244</td><td>134</td><td>25</td><td>105</td><td>277</td><td>41.9</td><td>62.2</td><td>21.5</td><td>12.5</td><td>266.2</td></tr>
<tr><td>>MI0000076|hsa-mir-20a&WithFlank&13|+|91351059|91351141@14.36.23(+1A)&[MIMAT0000075&hsa-miR-20a-5p&offsets|0|0(+1A);m-96&13|+|91351072|91351094&offsets|0|0(+1A)]|TAAAGTGCTTATAGTGCAGGTAGA</td><td>>MI0000076|hsa-mir-20a&WithFlank&13|+|91351059|91351141</td><td>14</td><td>36</td><td>23(+1A)</td><td>TAAAGTGCTTATAGTGCAGGTAGA</td><td>MB468</td><td>22</td><td>6</td><td>5</td><td>14</td><td>6</td><td>10.1</td><td>3.0</td><td>2.0</td><td>8.8</td><td>3.3</td></tr>
<tr><td>>MI0000076|hsa-mir-20a&WithFlank&13|+|91351059|91351141@14.36.23(+1C)&[MIMAT0000075&hsa-miR-20a-5p&offsets|0|0(+1C);m-96&13|+|91351072|91351094&offsets|0|0(+1C)]|TAAAGTGCTTATAGTGCAGGTAGC</td><td>>MI0000076|hsa-mir-20a&WithFlank&13|+|91351059|91351141</td><td>14</td><td>36</td><td>23(+1C)</td><td>TAAAGTGCTTATAGTGCAGGTAGC</td><td>MB231</td><td>12</td><td>7</td><td>1</td><td>8</td><td>19</td><td>3.1</td><td>4.4</td><td>0.5</td><td>2.9</td><td>23.3</td></tr>
<tr><td>>MI0000076|hsa-mir-20a&WithFlank&13|+|91351059|91351141@14.36.23(+1G)&[MIMAT0000075&hsa-miR-20a-5p&offsets|0|0(+1G);m-96&13|+|91351072|91351094&offsets|0|0(+1G)]|TAAAGTGCTTATAGTGCAGGTAGG</td><td>>MI0000076|hsa-mir-20a&WithFlank&13|+|91351059|91351141</td><td>14</td><td>36</td><td>23(+1G)</td><td>TAAAGTGCTTATAGTGCAGGTAGG</td><td>BT20</td><td>14</td><td>2</td><td>2</td><td>0</td><td>0</td><td>15.4</td><td>2.6</td><td>1.1</td><td>0.0</td><td>0.0</td></tr>
<tr><td>>MI0000076|hsa-mir-20a&WithFlank&13|+|91351059|91351141@14.36.23(+1G)&[MIMAT0000075&hsa-miR-20a-5p&offsets|0|0(+1G);m-96&13|+|91351072|91351094&offsets|0|0(+1G)]|TAAAGTGCTTATAGTGCAGGTAGG</td><td>>MI0000076|hsa-mir-20a&WithFlank&13|+|91351059|91351141</td><td>14</td><td>36</td><td>23(+1G)</td><td>TAAAGTGCTTATAGTGCAGGTAGG</td><td>MB231</td><td>18</td><td>4</td><td>1</td><td>3</td><td>6</td><td>15.1</td><td>2.1</td><td>0.8</td><td>0.8</td><td>5.8</td></tr>
<tr><td>>MI0000076|hsa-mir-20a&WithFlank&13|+|91351059|91351141@14.36.23&[MIMAT0000075&hsa-miR-20a-5p&offsets|0|0;m-96&13|+|91351072|91351094&offsets|0|0]|TAAAGTGCTTATAGTGCAGGTAG</td><td>>MI0000076|hsa-mir-20a&WithFlank&13|+|91351059|91351141</td><td>14</td><td>36</td><td>23</td><td>TAAAGTGCTTATAGTGCAGGTAG</td><td>BT20</td><td>631</td><td>304</td><td>133</td><td>199</td><td>63</td><td>456.1</td><td>356.2</td><td>85.9</td><td>169.0</td><td>61.4</td></tr>
<tr><td>>MI0000076|hsa-mir-20a&WithFlank&13|+|91351059|91351141@14.36.23&[MIMAT0000075&hsa-miR-20a-5p&offsets|0|0;m-96&13|+|91351072|91351094&offsets|0|0]|TAAAGTGCTTATAGTGCAGGTAG</td><td>>MI0000076|hsa-mir-20a&WithFlank&13|+|91351059|91351141</td><td>14</td><td>36</td><td>23</td><td>TAAAGTGCTTATAGTGCAGGTAG</td><td>MB231</td><td>1402</td><td>917</td><td>86</td><td>2094</td><td>1329</td><td>299.9</td><td>450.3</td><td>71.4</td><td>268.7</td><td>1151.4</td></tr>
<tr><td>>MI0000076|hsa-mir-20a&WithFlank&13|+|91351059|91351141@14.36.23&[MIMAT0000075&hsa-miR-20a-5p&offsets|0|0;m-96&13|+|91351072|91351094&offsets|0|0]|TAAAGTGCTTATAGTGCAGGTAG</td><td>>MI0000076|hsa-mir-20a&WithFlank&13|+|91351059|91351141</td><td>14</td><td>36</td><td>23</td><td>TAAAGTGCTTATAGTGCAGGTAG</td><td>MB468</td><td>135</td><td>30</td><td>22</td><td>176</td><td>32</td><td>83.8</td><td>15.1</td><td>11.6</td><td>62.0</td><td>12.9</td></tr>
<tr><td>>MI0000076|hsa-mir-20a&WithFlank&13|+|91351059|91351141@14.37.24&[MIMAT0000075&hsa-miR-20a-5p&offsets|0|+1;m-96&13|+|91351072|91351094&offsets|0|+1]|TAAAGTGCTTATAGTGCAGGTAGT</td><td>>MI0000076|hsa-mir-20a&WithFlank&13|+|91351059|91351141</td><td>14</td><td>37</td><td>24</td><td>TAAAGTGCTTATAGTGCAGGTAGT</td><td>BT20</td><td>40</td><td>16</td><td>7</td><td>1</td><td>3</td><td>40.4</td><td>20.5</td><td>3.7</td><td>0.7</td><td>3.7</td></tr>
<tr><td>>MI0000076|hsa-mir-20a&WithFlank&13|+|91351059|91351141@14.37.24&[MIMAT0000075&hsa-miR-20a-5p&offsets|0|+1;m-96&13|+|91351072|91351094&offsets|0|+1]|TAAAGTGCTTATAGTGCAGGTAGT</td><td>>MI0000076|hsa-mir-20a&WithFlank&13|+|91351059|91351141</td><td>14</td><td>37</td><td>24</td><td>TAAAGTGCTTATAGTGCAGGTAGT</td><td>MB231</td><td>104</td><td>67</td><td>8</td><td>50</td><td>147</td><td>27.5</td><td>39.1</td><td>6.3</td><td>12.3</td><td>161.1</td></tr>
<tr><td>>MI0000077|hsa-mir-21&WithFlank&17|+|59841260|59841343@13.34.22&[MIMAT0000076&hsa-miR-21-5p&offsets|-1|-1;m-1&17|+|59841273|59841294&offsets|-1|-1]|GTAGCTTATCAGACTGATGTTG</td><td>>MI0000077|hsa-mir-21&WithFlank&17|+|59841260|59841343</td><td>13</td><td>34</td><td>22</td><td>GTAGCTTATCAGACTGATGTTG</td><td>BT20</td><td>11</td><td>8</td><td>13</td><td>23</td><td>3</td><td>6.1</td><td>4.4</td><td>8.8</td><td>20.2</td><td>1.2</td></tr>
<tr><td>>MI0000077|hsa-mir-21&WithFlank&17|+|59841260|59841343@13.35.23&[MIMAT0000076&hsa-miR-21-5p&offsets|-1|0;m-1&17|+|59841273|59841294&offsets|-1|0]|GTAGCTTATCAGACTGATGTTGA</td><td>>MI0000077|hsa-mir-21&WithFlank&17|+|59841260|59841343</td><td>13</td><td>35</td><td>23</td><td>GTAGCTTATCAGACTGATGTTGA</td><td>BT20</td><td>50</td><td>32</td><td>41</td><td>158</td><td>15</td><td>26.0</td><td>17.7</td><td>25.6</td><td>145.5</td><td>6.5</td></tr>
<tr><td>>MI0000077|hsa-mir-21&WithFlank&17|+|59841260|59841343@13.35.23&[MIMAT0000076&hsa-miR-21-5p&offsets|-1|0;m-1&17|+|59841273|59841294&offsets|-1|0]|GTAGCTTATCAGACTGATGTTGA</td><td>>MI0000077|hsa-mir-21&WithFlank&17|+|59841260|59841343</td><td>13</td><td>35</td><td>23</td><td>GTAGCTTATCAGACTGATGTTGA</td><td>MB231</td><td>29</td><td>27</td><td>6</td><td>259</td><td>36</td><td>4.1</td><td>8.1</td><td>5.3</td><td>53.4</td><td>18.5</td></tr>
<tr><td>>MI0000077|hsa-mir-21&WithFlank&17|+|59841260|59841343@13.36.24&[MIMAT0000076&hsa-miR-21-5p&offsets|-1|+1;m-1&17|+|59841273|59841294&offsets|-1|+1]|GTAGCTTATCAGACTGATGTTGAC</td><td>>MI0000077|hsa-mir-21&WithFlank&17|+|59841260|59841343</td><td>13</td><td>36</td><td>24</td><td>GTAGCTTATCAGACTGATGTTGAC</td><td>BT20</td><td>14</td><td>11</td><td>16</td><td>2</td><td>5</td><td>11.2</td><td>7.0</td><td>13.6</td><td>0.7</td><td>3.1</td></tr>
<tr><td>>MI0000077|hsa-mir-21&WithFlank&17|+|59841260|59841343@13.36.24&[MIMAT0000076&hsa-miR-21-5p&offsets|-1|+1;m-1&17|+|59841273|59841294&offsets|-1|+1]|GTAGCTTATCAGACTGATGTTGAC</td><td>>MI0000077|hsa-mir-21&WithFlank&17|+|59841260|59841343</td><td>13</td><td>36</td><td>24</td><td>GTAGCTTATCAGACTGATGTTGAC</td><td>MB231</td><td>17</td><td>22</td><td>4</td><td>25</td><td>20</td><td>7.9</td><td>9.1</td><td>3.3</td><td>7.6</td><td>8.1</td></tr>
<tr><td>>MI0000077|hsa-mir-21&WithFlank&17|+|59841260|59841343@13.37.25&[MIMAT0000076&hsa-miR-21-5p&offsets|-1|+2;m-1&17|+|59841273|59841294&offsets|-1|+2]|GTAGCTTATCAGACTGATGTTGACT</td><td>>MI0000077|hsa-mir-21&WithFlank&17|+|59841260|59841343</td><td>13</td><td>37</td><td>25</td><td>GTAGCTTATCAGACTGATGTTGACT</td><td>BT20</td><td>11</td><td>7</td><td>9</td><td>1</td><td>3</td><td>6.3</td><td>3.6</td><td>5.4</td><td>0.6</td><td>1.3</td></tr>
<tr><td>>MI0000077|hsa-mir-21&WithFlank&17|+|59841260|59841343@13.37.25&[MIMAT0000076&hsa-miR-21-5p&offsets|-1|+2;m-1&17|+|59841273|59841294&offsets|-1|+2]|GTAGCTTATCAGACTGATGTTGACT</td><td>>MI0000077|hsa-mir-21&WithFlank&17|+|59841260|59841343</td><td>13</td><td>37</td><td>25</td><td>GTAGCTTATCAGACTGATGTTGACT</td><td>MB231</td><td>22</td><td>19</td><td>5</td><td>6</td><td>17</td><td>2.9</td><td>7.9</td><td>3.2</td><td>4.2</td><td>6.9</td></tr>
</table>

<table>
<tr><th></th><th></th><th></th><th></th><th></th><th></th><th></th><th></th><th></th><th></th><th></th><th></th><th></th><th></th><th></th><th></th><th></th></tr>
<tr><td>>MI0000077|hsa-mir-21&WithFlank&17|+|59841260|59841343@14.31.18(+1C)&[MIMAT0000076&hsa-miR-21-5p&offsets|0|-4(+1C);m-1&17|+|59841273|59841294&offsets|0|-4(+1C)]|TAGCTTATCAGACTGATGC</td><td>>MI0000077|hsa-mir-21&WithFlank&17|+|59841260|59841343</td><td>14</td><td>31</td><td>18(+1C)</td><td>TAGCTTATCAGACTGATGC</td><td>BT20</td><td>13</td><td>13</td><td>11</td><td>27</td><td>3</td><td>6.2</td><td>12.3</td><td>8.2</td><td>28.7</td><td>1.3</td></tr>
<tr><td>>MI0000077|hsa-mir-21&WithFlank&17|+|59841260|59841343@14.31.18&[MIMAT0000076&hsa-miR-21-5p&offsets|0|-4;m-1&17|+|59841273|59841294&offsets|0|-4]|TAGCTTATCAGACTGATG</td><td>>MI0000077|hsa-mir-21&WithFlank&17|+|59841260|59841343</td><td>14</td><td>31</td><td>18</td><td>TAGCTTATCAGACTGATG</td><td>BT20</td><td>114</td><td>69</td><td>59</td><td>67</td><td>13</td><td>49.0</td><td>37.9</td><td>31.0</td><td>86.6</td><td>3.5</td></tr>
<tr><td>>MI0000077|hsa-mir-21&WithFlank&17|+|59841260|59841343@14.31.18&[MIMAT0000076&hsa-miR-21-5p&offsets|0|-4;m-1&17|+|59841273|59841294&offsets|0|-4]|TAGCTTATCAGACTGATG</td><td>>MI0000077|hsa-mir-21&WithFlank&17|+|59841260|59841343</td><td>14</td><td>31</td><td>18</td><td>TAGCTTATCAGACTGATG</td><td>MB231</td><td>39</td><td>35</td><td>4</td><td>79</td><td>32</td><td>21.4</td><td>18.8</td><td>1.2</td><td>46.3</td><td>10.1</td></tr>
<tr><td>>MI0000077|hsa-mir-21&WithFlank&17|+|59841260|59841343@14.32.19(+1C)&[MIMAT0000076&hsa-miR-21-5p&offsets|0|-3(+1C);m-1&17|+|59841273|59841294&offsets|0|-3(+1C)]|TAGCTTATCAGACTGATGTC</td><td>>MI0000077|hsa-mir-21&WithFlank&17|+|59841260|59841343</td><td>14</td><td>32</td><td>19(+1C)</td><td>TAGCTTATCAGACTGATGTC</td><td>BT20</td><td>18</td><td>14</td><td>12</td><td>16</td><td>2</td><td>9.9</td><td>14.3</td><td>8.4</td><td>17.1</td><td>0.8</td></tr>
<tr><td>>MI0000077|hsa-mir-21&WithFlank&17|+|59841260|59841343@14.32.19&[MIMAT0000076&hsa-miR-21-5p&offsets|0|-3;m-1&17|+|59841273|59841294&offsets|0|-3]|TAGCTTATCAGACTGATGT</td><td>>MI0000077|hsa-mir-21&WithFlank&17|+|59841260|59841343</td><td>14</td><td>32</td><td>19</td><td>TAGCTTATCAGACTGATGT</td><td>BT20</td><td>102</td><td>84</td><td>88</td><td>186</td><td>22</td><td>56.5</td><td>80.2</td><td>66.5</td><td>174.9</td><td>9.2</td></tr>
<tr><td>>MI0000077|hsa-mir-21&WithFlank&17|+|59841260|59841343@14.32.19&[MIMAT0000076&hsa-miR-21-5p&offsets|0|-3;m-1&17|+|59841273|59841294&offsets|0|-3]|TAGCTTATCAGACTGATGT</td><td>>MI0000077|hsa-mir-21&WithFlank&17|+|59841260|59841343</td><td>14</td><td>32</td><td>19</td><td>TAGCTTATCAGACTGATGT</td><td>MB231</td><td>32</td><td>37</td><td>7</td><td>328</td><td>45</td><td>3.2</td><td>31.9</td><td>6.1</td><td>211.9</td><td>28.8</td></tr>
<tr><td>>MI0000077|hsa-mir-21&WithFlank&17|+|59841260|59841343@14.33.20(+1A)&[MIMAT0000076&hsa-miR-21-5p&offsets|0|-2(+1A);m-1&17|+|59841273|59841294&offsets|0|-2(+1A)]|TAGCTTATCAGACTGATGTTA</td><td>>MI0000077|hsa-mir-21&WithFlank&17|+|59841260|59841343</td><td>14</td><td>33</td><td>20(+1A)</td><td>TAGCTTATCAGACTGATGTTA</td><td>BT20</td><td>216</td><td>102</td><td>162</td><td>184</td><td>21</td><td>162.2</td><td>90.3</td><td>120.6</td><td>210.9</td><td>6.8</td></tr>
<tr><td>>MI0000077|hsa-mir-21&WithFlank&17|+|59841260|59841343@14.33.20(+1A)&[MIMAT0000076&hsa-miR-21-5p&offsets|0|-2(+1A);m-1&17|+|59841273|59841294&offsets|0|-2(+1A)]|TAGCTTATCAGACTGATGTTA</td><td>>MI0000077|hsa-mir-21&WithFlank&17|+|59841260|59841343</td><td>14</td><td>33</td><td>20(+1A)</td><td>TAGCTTATCAGACTGATGTTA</td><td>MB231</td><td>55</td><td>48</td><td>15</td><td>398</td><td>69</td><td>15.7</td><td>42.3</td><td>13.9</td><td>238.5</td><td>55.7</td></tr>
<tr><td>>MI0000077|hsa-mir-21&WithFlank&17|+|59841260|59841343@14.33.20(+1U)&[MIMAT0000076&hsa-miR-21-5p&offsets|0|-2(+1U);m-1&17|+|59841273|59841294&offsets|0|-2(+1U)]|TAGCTTATCAGACTGATGTTT</td><td>>MI0000077|hsa-mir-21&WithFlank&17|+|59841260|59841343</td><td>14</td><td>33</td><td>20(+1U)</td><td>TAGCTTATCAGACTGATGTTT</td><td>BT20</td><td>28</td><td>15</td><td>22</td><td>29</td><td>3</td><td>18.0</td><td>12.5</td><td>18.0</td><td>33.3</td><td>0.5</td></tr>
<tr><td>>MI0000077|hsa-mir-21&WithFlank&17|+|59841260|59841343@14.33.20(+2A)&[MIMAT0000076&hsa-miR-21-5p&offsets|0|-2(+2A);m-1&17|+|59841273|59841294&offsets|0|-2(+2A)]|TAGCTTATCAGACTGATGTTAA</td><td>>MI0000077|hsa-mir-21&WithFlank&17|+|59841260|59841343</td><td>14</td><td>33</td><td>20(+2A)</td><td>TAGCTTATCAGACTGATGTTAA</td><td>BT20</td><td>45</td><td>6</td><td>5</td><td>10</td><td>0</td><td>68.1</td><td>5.4</td><td>4.0</td><td>11.2</td><td>0.5</td></tr>
<tr><td>>MI0000077|hsa-mir-21&WithFlank&17|+|59841260|59841343@14.33.20&[MIMAT0000076&hsa-miR-21-5p&offsets|0|-2;m-1&17|+|59841273|59841294&offsets|0|-2]|TAGCTTATCAGACTGATGTT</td><td>>MI0000077|hsa-mir-21&WithFlank&17|+|59841260|59841343</td><td>14</td><td>33</td><td>20</td><td>TAGCTTATCAGACTGATGTT</td><td>BT20</td><td>131</td><td>101</td><td>87</td><td>112</td><td>18</td><td>79.5</td><td>96.9</td><td>65.5</td><td>101.9</td><td>6.7</td></tr>
<tr><td>>MI0000077|hsa-mir-21&WithFlank&17|+|59841260|59841343@14.33.20&[MIMAT0000076&hsa-miR-21-5p&offsets|0|-2;m-1&17|+|59841273|59841294&offsets|0|-2]|TAGCTTATCAGACTGATGTT</td><td>>MI0000077|hsa-mir-21&WithFlank&17|+|59841260|59841343</td><td>14</td><td>33</td><td>20</td><td>TAGCTTATCAGACTGATGTT</td><td>MB231</td><td>33</td><td>39</td><td>6</td><td>237</td><td>39</td><td>4.7</td><td>28.0</td><td>4.2</td><td>146.4</td><td>20.8</td></tr>
<tr><td>>MI0000077|hsa-mir-21&WithFlank&17|+|59841260|59841343@14.34.21(+1C)&[MIMAT0000076&hsa-miR-21-5p&offsets|0|-1(+1C);m-1&17|+|59841273|59841294&offsets|0|-1(+1C)]|TAGCTTATCAGACTGATGTTGC</td><td>>MI0000077|hsa-mir-21&WithFlank&17|+|59841260|59841343</td><td>14</td><td>34</td><td>21(+1C)</td><td>TAGCTTATCAGACTGATGTTGC</td><td>BT20</td><td>63</td><td>16</td><td>14</td><td>38</td><td>2</td><td>80.8</td><td>15.0</td><td>12.6</td><td>49.6</td><td>1.0</td></tr>
<tr><td>>MI0000077|hsa-mir-21&WithFlank&17|+|59841260|59841343@14.34.21(+1C)&[MIMAT0000076&hsa-miR-21-5p&offsets|0|-1(+1C);m-1&17|+|59841273|59841294&offsets|0|-1(+1C)]|TAGCTTATCAGACTGATGTTGC</td><td>>MI0000077|hsa-mir-21&WithFlank&17|+|59841260|59841343</td><td>14</td><td>34</td><td>21(+1C)</td><td>TAGCTTATCAGACTGATGTTGC</td><td>MB231</td><td>12</td><td>6</td><td>1</td><td>84</td><td>15</td><td>7.7</td><td>5.5</td><td>1.2</td><td>49.6</td><td>18.8</td></tr>
<tr><td>>MI0000077|hsa-mir-21&WithFlank&17|+|59841260|59841343@14.34.21(+1G)&[MIMAT0000076&hsa-miR-21-5p&offsets|0|-1(+1G);m-1&17|+|59841273|59841294&offsets|0|-1(+1G)]|TAGCTTATCAGACTGATGTTGG</td><td>>MI0000077|hsa-mir-21&WithFlank&17|+|59841260|59841343</td><td>14</td><td>34</td><td>21(+1G)</td><td>TAGCTTATCAGACTGATGTTGG</td><td>BT20</td><td>1048</td><td>207</td><td>186</td><td>514</td><td>18</td><td>1235.4</td><td>187.6</td><td>145.4</td><td>687.1</td><td>8.7</td></tr>
<tr><td>>MI0000077|hsa-mir-21&WithFlank&17|+|59841260|59841343@14.34.21(+1G)&[MIMAT0000076&hsa-miR-21-5p&offsets|0|-1(+1G);m-1&17|+|59841273|59841294&offsets|0|-1(+1G)]|TAGCTTATCAGACTGATGTTGG</td><td>>MI0000077|hsa-mir-21&WithFlank&17|+|59841260|59841343</td><td>14</td><td>34</td><td>21(+1G)</td><td>TAGCTTATCAGACTGATGTTGG</td><td>MB231</td><td>255</td><td>73</td><td>14</td><td>721</td><td>74</td><td>251.8</td><td>56.2</td><td>11.4</td><td>433.6</td><td>65.9</td></tr>
<tr><td>>MI0000077|hsa-mir-21&WithFlank&17|+|59841260|59841343@14.34.21(+1G)&[MIMAT0000076&hsa-miR-21-5p&offsets|0|-1(+1G);m-1&17|+|59841273|59841294&offsets|0|-1(+1G)]|TAGCTTATCAGACTGATGTTGG</td><td>>MI0000077|hsa-mir-21&WithFlank&17|+|59841260|59841343</td><td>14</td><td>34</td><td>21(+1G)</td><td>TAGCTTATCAGACTGATGTTGG</td><td>MB468</td><td>14</td><td>1</td><td>7</td><td>111</td><td>2</td><td>4.0</td><td>0.7</td><td>8.6</td><td>98.0</td><td>1.8</td></tr>
<tr><td>>MI0000077|hsa-mir-21&WithFlank&17|+|59841260|59841343@14.34.21(+1U)&[MIMAT0000076&hsa-miR-21-5p&offsets|0|-1(+1U);m-1&17|+|59841273|59841294&offsets|0|-1(+1U)]|TAGCTTATCAGACTGATGTTGT</td><td>>MI0000077|hsa-mir-21&WithFlank&17|+|59841260|59841343</td><td>14</td><td>34</td><td>21(+1U)</td><td>TAGCTTATCAGACTGATGTTGT</td><td>BT20</td><td>705</td><td>143</td><td>155</td><td>370</td><td>20</td><td>902.1</td><td>127.3</td><td>125.4</td><td>437.6</td><td>10.4</td></tr>
<tr><td>>MI0000077|hsa-mir-21&WithFlank&17|+|59841260|59841343@14.34.21(+1U)&[MIMAT0000076&hsa-miR-21-5p&offsets|0|-1(+1U);m-1&17|+|59841273|59841294&offsets|0|-1(+1U)]|TAGCTTATCAGACTGATGTTGT</td><td>>MI0000077|hsa-mir-21&WithFlank&17|+|59841260|59841343</td><td>14</td><td>34</td><td>21(+1U)</td><td>TAGCTTATCAGACTGATGTTGT</td><td>MB231</td><td>144</td><td>51</td><td>12</td><td>688</td><td>144</td><td>78.8</td><td>33.8</td><td>10.2</td><td>299.2</td><td>172.6</td></tr>
<tr><td>>MI0000077|hsa-mir-21&WithFlank&17|+|59841260|59841343@14.34.21(+2C)&[MIMAT0000076&hsa-miR-21-5p&offsets|0|-1(+2C);m-1&17|+|59841273|59841294&offsets|0|-1(+2C)]|TAGCTTATCAGACTGATGTTGCC</td><td>>MI0000077|hsa-mir-21&WithFlank&17|+|59841260|59841343</td><td>14</td><td>34</td><td>21(+2C)</td><td>TAGCTTATCAGACTGATGTTGCC</td><td>BT20</td><td>49</td><td>14</td><td>25</td><td>3</td><td>4</td><td>43.3</td><td>3.4</td><td>15.9</td><td>3.2</td><td>0.8</td></tr>
<tr><td>>MI0000077|hsa-mir-21&WithFlank&17|+|59841260|59841343@14.34.21(+2C)&[MIMAT0000076&hsa-miR-21-5p&offsets|0|-1(+2C);m-1&17|+|59841273|59841294&offsets|0|-1(+2C)]|TAGCTTATCAGACTGATGTTGCC</td><td>>MI0000077|hsa-mir-21&WithFlank&17|+|59841260|59841343</td><td>14</td><td>34</td><td>21(+2C)</td><td>TAGCTTATCAGACTGATGTTGCC</td><td>MB231</td><td>159</td><td>139</td><td>38</td><td>106</td><td>138</td><td>80.3</td><td>98.5</td><td>37.4</td><td>30.9</td><td>116.6</td></tr>
<tr><td>>MI0000077|hsa-mir-21&WithFlank&17|+|59841260|59841343@14.34.21(+2G)&[MIMAT0000076&hsa-miR-21-5p&offsets|0|-1(+2G);m-1&17|+|59841273|59841294&offsets|0|-1(+2G)]|TAGCTTATCAGACTGATGTTGGG</td><td>>MI0000077|hsa-mir-21&WithFlank&17|+|59841260|59841343</td><td>14</td><td>34</td><td>21(+2G)</td><td>TAGCTTATCAGACTGATGTTGGG</td><td>BT20</td><td>11</td><td>1</td><td>2</td><td>0</td><td>0</td><td>13.7</td><td>0.8</td><td>2.0</td><td>0.0</td><td>0.0</td></tr>
<tr><td>>MI0000077|hsa-mir-21&WithFlank&17|+|59841260|59841343@14.34.21(+2U)&[MIMAT0000076&hsa-miR-21-5p&offsets|0|-1(+2U);m-1&17|+|59841273|59841294&offsets|0|-1(+2U)]|TAGCTTATCAGACTGATGTTGTT</td><td>>MI0000077|hsa-mir-21&WithFlank&17|+|59841260|59841343</td><td>14</td><td>34</td><td>21(+2U)</td><td>TAGCTTATCAGACTGATGTTGTT</td><td>BT20</td><td>25</td><td>6</td><td>14</td><td>1</td><td>0</td><td>27.0</td><td>4.6</td><td>14.0</td><td>1.3</td><td>0.8</td></tr>
<tr><td>>MI0000077|hsa-mir-21&WithFlank&17|+|59841260|59841343@14.34.21(+3C)&[MIMAT0000076&hsa-miR-21-5p&offsets|0|-1(+3C);m-1&17|+|59841273|59841294&offsets|0|-1(+3C)]|TAGCTTATCAGACTGATGTTGCCC</td><td>>MI0000077|hsa-mir-21&WithFlank&17|+|59841260|59841343</td><td>14</td><td>34</td><td>21(+3C)</td><td>TAGCTTATCAGACTGATGTTGCCC</td><td>MB231</td><td>15</td><td>17</td><td>4</td><td>5</td><td>14</td><td>5.1</td><td>11.3</td><td>3.1</td><td>1.9</td><td>9.8</td></tr>
<tr><td>>MI0000077|hsa-mir-21&WithFlank&17|+|59841260|59841343@14.34.21&[MIMAT0000076&hsa-miR-21-5p&offsets|0|-1;m-1&17|+|59841273|59841294&offsets|0|-1]|TAGCTTATCAGACTGATGTTG</td><td>>MI0000077|hsa-mir-21&WithFlank&17|+|59841260|59841343</td><td>14</td><td>34</td><td>21</td><td>TAGCTTATCAGACTGATGTTG</td><td>BT20</td><td>1191</td><td>786</td><td>936</td><td>1420</td><td>204</td><td>652.6</td><td>637.6</td><td>674.0</td><td>1352.6</td><td>76.2</td></tr>
<tr><td>>MI0000077|hsa-mir-21&WithFlank&17|+|59841260|59841343@14.34.21&[MIMAT0000076&hsa-miR-21-5p&offsets|0|-1;m-1&17|+|59841273|59841294&offsets|0|-1]|TAGCTTATCAGACTGATGTTG</td><td>>MI0000077|hsa-mir-21&WithFlank&17|+|59841260|59841343</td><td>14</td><td>34</td><td>21</td><td>TAGCTTATCAGACTGATGTTG</td><td>MB231</td><td>542</td><td>590</td><td>128</td><td>2524</td><td>536</td><td>114.4</td><td>465.2</td><td>114.0</td><td>1057.0</td><td>297.7</td></tr>
<tr><td>>MI0000077|hsa-mir-21&WithFlank&17|+|59841260|59841343@14.34.21&[MIMAT0000076&hsa-miR-21-5p&offsets|0|-1;m-1&17|+|59841273|59841294&offsets|0|-1]|TAGCTTATCAGACTGATGTTG</td><td>>MI0000077|hsa-mir-21&WithFlank&17|+|59841260|59841343</td><td>14</td><td>34</td><td>21</td><td>TAGCTTATCAGACTGATGTTG</td><td>MB468</td><td>49</td><td>9</td><td>14</td><td>186</td><td>12</td><td>28.4</td><td>3.7</td><td>6.2</td><td>100.6</td><td>0.9</td></tr>
<tr><td>>MI0000077|hsa-mir-21&WithFlank&17|+|59841260|59841343@14.35.22(+1A)&[MIMAT0000076&hsa-miR-21-5p&offsets|0|0(+1A);m-1&17|+|59841273|59841294&offsets|0|0(+1A)]|TAGCTTATCAGACTGATGTTGAA</td><td>>MI0000077|hsa-mir-21&WithFlank&17|+|59841260|59841343</td><td>14</td><td>35</td><td>22(+1A)</td><td>TAGCTTATCAGACTGATGTTGAA</td><td>BT20</td><td>2888</td><td>875</td><td>2001</td><td>180</td><td>217</td><td>2384.8</td><td>528.5</td><td>1566.0</td><td>171.6</td><td>23.7</td></tr>
</table>

<table>
<tr><th></th><th></th><th></th><th></th><th></th><th></th><th></th><th></th><th></th><th></th><th></th><th></th><th></th><th></th><th></th><th></th><th></th></tr>
<tr><td>>MI0000077|hsa-mir-21&WithFlank&17|+|59841260|59841343@14.35.22(+1A)&[MIMAT0000076&hsa-miR-21-5p&offsets|0|0(+1A);m-1&17|+|59841273|59841294&offsets|0|0(+1A)]|TAGCTTATCAGACTGATGTTGAA</td><td>>MI0000077|hsa-mir-21&WithFlank&17|+|59841260|59841343</td><td>14</td><td>35</td><td>22(+1A)</td><td>TAGCTTATCAGACTGATGTTGAA</td><td>MB231</td><td>1120</td><td>719</td><td>257</td><td>1364</td><td>1106</td><td>83.8</td><td>295.7</td><td>232.9</td><td>299.4</td><td>761.8</td></tr>
<tr><td>>MI0000077|hsa-mir-21&WithFlank&17|+|59841260|59841343@14.35.22(+1A)&[MIMAT0000076&hsa-miR-21-5p&offsets|0|0(+1A);m-1&17|+|59841273|59841294&offsets|0|0(+1A)]|TAGCTTATCAGACTGATGTTGAA</td><td>>MI0000077|hsa-mir-21&WithFlank&17|+|59841260|59841343</td><td>14</td><td>35</td><td>22(+1A)</td><td>TAGCTTATCAGACTGATGTTGAA</td><td>MB468</td><td>37</td><td>10</td><td>19</td><td>80</td><td>13</td><td>9.1</td><td>3.6</td><td>21.1</td><td>71.5</td><td>10.2</td></tr>
<tr><td>>MI0000077|hsa-mir-21&WithFlank&17|+|59841260|59841343@14.35.22(+1G)&[MIMAT0000076&hsa-miR-21-5p&offsets|0|0(+1G);m-1&17|+|59841273|59841294&offsets|0|0(+1G)]|TAGCTTATCAGACTGATGTTGAG</td><td>>MI0000077|hsa-mir-21&WithFlank&17|+|59841260|59841343</td><td>14</td><td>35</td><td>22(+1G)</td><td>TAGCTTATCAGACTGATGTTGAG</td><td>BT20</td><td>488</td><td>71</td><td>155</td><td>13</td><td>12</td><td>556.5</td><td>39.9</td><td>144.9</td><td>15.7</td><td>0.8</td></tr>
<tr><td>>MI0000077|hsa-mir-21&WithFlank&17|+|59841260|59841343@14.35.22(+1G)&[MIMAT0000076&hsa-miR-21-5p&offsets|0|0(+1G);m-1&17|+|59841273|59841294&offsets|0|0(+1G)]|TAGCTTATCAGACTGATGTTGAG</td><td>>MI0000077|hsa-mir-21&WithFlank&17|+|59841260|59841343</td><td>14</td><td>35</td><td>22(+1G)</td><td>TAGCTTATCAGACTGATGTTGAG</td><td>MB231</td><td>196</td><td>109</td><td>30</td><td>133</td><td>125</td><td>108.7</td><td>80.8</td><td>27.7</td><td>69.3</td><td>113.3</td></tr>
<tr><td>>MI0000077|hsa-mir-21&WithFlank&17|+|59841260|59841343@14.35.22(+1U)&[MIMAT0000076&hsa-miR-21-5p&offsets|0|0(+1U);m-1&17|+|59841273|59841294&offsets|0|0(+1U)]|TAGCTTATCAGACTGATGTTGAT</td><td>>MI0000077|hsa-mir-21&WithFlank&17|+|59841260|59841343</td><td>14</td><td>35</td><td>22(+1U)</td><td>TAGCTTATCAGACTGATGTTGAT</td><td>BT20</td><td>12208</td><td>4790</td><td>8798</td><td>915</td><td>1193</td><td>6322.4</td><td>3423.4</td><td>5937.9</td><td>923.7</td><td>287.3</td></tr>
<tr><td>>MI0000077|hsa-mir-21&WithFlank&17|+|59841260|59841343@14.35.22(+1U)&[MIMAT0000076&hsa-miR-21-5p&offsets|0|0(+1U);m-1&17|+|59841273|59841294&offsets|0|0(+1U)]|TAGCTTATCAGACTGATGTTGAT</td><td>>MI0000077|hsa-mir-21&WithFlank&17|+|59841260|59841343</td><td>14</td><td>35</td><td>22(+1U)</td><td>TAGCTTATCAGACTGATGTTGAT</td><td>MB231</td><td>6308</td><td>4042</td><td>1310</td><td>7898</td><td>6589</td><td>889.3</td><td>1581.7</td><td>1069.6</td><td>2397.8</td><td>4704.6</td></tr>
<tr><td>>MI0000077|hsa-mir-21&WithFlank&17|+|59841260|59841343@14.35.22(+1U)&[MIMAT0000076&hsa-miR-21-5p&offsets|0|0(+1U);m-1&17|+|59841273|59841294&offsets|0|0(+1U)]|TAGCTTATCAGACTGATGTTGAT</td><td>>MI0000077|hsa-mir-21&WithFlank&17|+|59841260|59841343</td><td>14</td><td>35</td><td>22(+1U)</td><td>TAGCTTATCAGACTGATGTTGAT</td><td>MB468</td><td>174</td><td>48</td><td>83</td><td>278</td><td>64</td><td>52.1</td><td>25.0</td><td>98.5</td><td>248.0</td><td>57.5</td></tr>
<tr><td>>MI0000077|hsa-mir-21&WithFlank&17|+|59841260|59841343@14.35.22(+2A)&[MIMAT0000076&hsa-miR-21-5p&offsets|0|0(+2A);m-1&17|+|59841273|59841294&offsets|0|0(+2A)]|TAGCTTATCAGACTGATGTTGAAA</td><td>>MI0000077|hsa-mir-21&WithFlank&17|+|59841260|59841343</td><td>14</td><td>35</td><td>22(+2A)</td><td>TAGCTTATCAGACTGATGTTGAAA</td><td>BT20</td><td>97</td><td>48</td><td>73</td><td>9</td><td>17</td><td>63.4</td><td>27.4</td><td>49.2</td><td>1.5</td><td>6.1</td></tr>
<tr><td>>MI0000077|hsa-mir-21&WithFlank&17|+|59841260|59841343@14.35.22(+2A)&[MIMAT0000076&hsa-miR-21-5p&offsets|0|0(+2A);m-1&17|+|59841273|59841294&offsets|0|0(+2A)]|TAGCTTATCAGACTGATGTTGAAA</td><td>>MI0000077|hsa-mir-21&WithFlank&17|+|59841260|59841343</td><td>14</td><td>35</td><td>22(+2A)</td><td>TAGCTTATCAGACTGATGTTGAAA</td><td>MB231</td><td>36</td><td>35</td><td>8</td><td>17</td><td>35</td><td>3.5</td><td>16.1</td><td>5.9</td><td>4.7</td><td>15.6</td></tr>
<tr><td>>MI0000077|hsa-mir-21&WithFlank&17|+|59841260|59841343@14.35.22(+2U)&[MIMAT0000076&hsa-miR-21-5p&offsets|0|0(+2U);m-1&17|+|59841273|59841294&offsets|0|0(+2U)]|TAGCTTATCAGACTGATGTTGATT</td><td>>MI0000077|hsa-mir-21&WithFlank&17|+|59841260|59841343</td><td>14</td><td>35</td><td>22(+2U)</td><td>TAGCTTATCAGACTGATGTTGATT</td><td>BT20</td><td>101</td><td>25</td><td>29</td><td>1</td><td>3</td><td>121.6</td><td>10.7</td><td>27.5</td><td>0.6</td><td>1.3</td></tr>
<tr><td>>MI0000077|hsa-mir-21&WithFlank&17|+|59841260|59841343@14.35.22(+2U)&[MIMAT0000076&hsa-miR-21-5p&offsets|0|0(+2U);m-1&17|+|59841273|59841294&offsets|0|0(+2U)]|TAGCTTATCAGACTGATGTTGATT</td><td>>MI0000077|hsa-mir-21&WithFlank&17|+|59841260|59841343</td><td>14</td><td>35</td><td>22(+2U)</td><td>TAGCTTATCAGACTGATGTTGATT</td><td>MB231</td><td>57</td><td>38</td><td>5</td><td>6</td><td>19</td><td>30.7</td><td>27.0</td><td>4.7</td><td>1.8</td><td>14.3</td></tr>
<tr><td>>MI0000077|hsa-mir-21&WithFlank&17|+|59841260|59841343@14.35.22&[MIMAT0000076&hsa-miR-21-5p&offsets|0|0;m-1&17|+|59841273|59841294&offsets|0|0]|TAGCTTATCAGACTGATGTTGA</td><td>>MI0000077|hsa-mir-21&WithFlank&17|+|59841260|59841343</td><td>14</td><td>35</td><td>22</td><td>TAGCTTATCAGACTGATGTTGA</td><td>BT20</td><td>12910</td><td>7398</td><td>6890</td><td>21520</td><td>1506</td><td>8973.6</td><td>6125.5</td><td>4211.0</td><td>20542.4</td><td>835.6</td></tr>
<tr><td>>MI0000077|hsa-mir-21&WithFlank&17|+|59841260|59841343@14.35.22&[MIMAT0000076&hsa-miR-21-5p&offsets|0|0;m-1&17|+|59841273|59841294&offsets|0|0]|TAGCTTATCAGACTGATGTTGA</td><td>>MI0000077|hsa-mir-21&WithFlank&17|+|59841260|59841343</td><td>14</td><td>35</td><td>22</td><td>TAGCTTATCAGACTGATGTTGA</td><td>MB231</td><td>4813</td><td>3844</td><td>756</td><td>43896</td><td>5492</td><td>680.9</td><td>2054.5</td><td>634.3</td><td>13885.5</td><td>3977.8</td></tr>
<tr><td>>MI0000077|hsa-mir-21&WithFlank&17|+|59841260|59841343@14.35.22&[MIMAT0000076&hsa-miR-21-5p&offsets|0|0;m-1&17|+|59841273|59841294&offsets|0|0]|TAGCTTATCAGACTGATGTTGA</td><td>>MI0000077|hsa-mir-21&WithFlank&17|+|59841260|59841343</td><td>14</td><td>35</td><td>22</td><td>TAGCTTATCAGACTGATGTTGA</td><td>MB468</td><td>316</td><td>66</td><td>92</td><td>3587</td><td>84</td><td>133.4</td><td>23.8</td><td>58.3</td><td>2634.7</td><td>32.9</td></tr>
<tr><td>>MI0000077|hsa-mir-21&WithFlank&17|+|59841260|59841343@14.36.23(+1A)&[MIMAT0000076&hsa-miR-21-5p&offsets|0|+1(+1A);m-1&17|+|59841273|59841294&offsets|0|+1(+1A)]|TAGCTTATCAGACTGATGTTGACA</td><td>>MI0000077|hsa-mir-21&WithFlank&17|+|59841260|59841343</td><td>14</td><td>36</td><td>23(+1A)</td><td>TAGCTTATCAGACTGATGTTGACA</td><td>BT20</td><td>10704</td><td>4829</td><td>9209</td><td>804</td><td>2166</td><td>6323.0</td><td>2542.6</td><td>5127.9</td><td>128.5</td><td>851.2</td></tr>
<tr><td>>MI0000077|hsa-mir-21&WithFlank&17|+|59841260|59841343@14.36.23(+1A)&[MIMAT0000076&hsa-miR-21-5p&offsets|0|+1(+1A);m-1&17|+|59841273|59841294&offsets|0|+1(+1A)]|TAGCTTATCAGACTGATGTTGACA</td><td>>MI0000077|hsa-mir-21&WithFlank&17|+|59841260|59841343</td><td>14</td><td>36</td><td>23(+1A)</td><td>TAGCTTATCAGACTGATGTTGACA</td><td>MB231</td><td>6660</td><td>5317</td><td>1719</td><td>2383</td><td>5225</td><td>1340.1</td><td>938.7</td><td>1008.1</td><td>1093.5</td><td>2122.8</td></tr>
<tr><td>>MI0000077|hsa-mir-21&WithFlank&17|+|59841260|59841343@14.36.23(+1A)&[MIMAT0000076&hsa-miR-21-5p&offsets|0|+1(+1A);m-1&17|+|59841273|59841294&offsets|0|+1(+1A)]|TAGCTTATCAGACTGATGTTGACA</td><td>>MI0000077|hsa-mir-21&WithFlank&17|+|59841260|59841343</td><td>14</td><td>36</td><td>23(+1A)</td><td>TAGCTTATCAGACTGATGTTGACA</td><td>MB468</td><td>423</td><td>177</td><td>211</td><td>177</td><td>105</td><td>138.8</td><td>92.4</td><td>107.1</td><td>128.4</td><td>34.9</td></tr>
<tr><td>>MI0000077|hsa-mir-21&WithFlank&17|+|59841260|59841343@14.36.23(+1C)&[MIMAT0000076&hsa-miR-21-5p&offsets|0|+1(+1C);m-1&17|+|59841273|59841294&offsets|0|+1(+1C)]|TAGCTTATCAGACTGATGTTGACC</td><td>>MI0000077|hsa-mir-21&WithFlank&17|+|59841260|59841343</td><td>14</td><td>36</td><td>23(+1C)</td><td>TAGCTTATCAGACTGATGTTGACC</td><td>BT20</td><td>2428</td><td>1497</td><td>1950</td><td>229</td><td>645</td><td>1366.2</td><td>912.9</td><td>1185.7</td><td>87.5</td><td>347.6</td></tr>
<tr><td>>MI0000077|hsa-mir-21&WithFlank&17|+|59841260|59841343@14.36.23(+1C)&[MIMAT0000076&hsa-miR-21-5p&offsets|0|+1(+1C);m-1&17|+|59841273|59841294&offsets|0|+1(+1C)]|TAGCTTATCAGACTGATGTTGACC</td><td>>MI0000077|hsa-mir-21&WithFlank&17|+|59841260|59841343</td><td>14</td><td>36</td><td>23(+1C)</td><td>TAGCTTATCAGACTGATGTTGACC</td><td>MB231</td><td>2455</td><td>2439</td><td>609</td><td>947</td><td>2249</td><td>395.7</td><td>969.1</td><td>404.1</td><td>261.2</td><td>1137.1</td></tr>
<tr><td>>MI0000077|hsa-mir-21&WithFlank&17|+|59841260|59841343@14.36.23(+1C)&[MIMAT0000076&hsa-miR-21-5p&offsets|0|+1(+1C);m-1&17|+|59841273|59841294&offsets|0|+1(+1C)]|TAGCTTATCAGACTGATGTTGACC</td><td>>MI0000077|hsa-mir-21&WithFlank&17|+|59841260|59841343</td><td>14</td><td>36</td><td>23(+1C)</td><td>TAGCTTATCAGACTGATGTTGACC</td><td>MB468</td><td>60</td><td>26</td><td>28</td><td>29</td><td>16</td><td>15.6</td><td>10.0</td><td>18.5</td><td>17.6</td><td>5.6</td></tr>
<tr><td>>MI0000077|hsa-mir-21&WithFlank&17|+|59841260|59841343@14.36.23(+1G)&[MIMAT0000076&hsa-miR-21-5p&offsets|0|+1(+1G);m-1&17|+|59841273|59841294&offsets|0|+1(+1G)]|TAGCTTATCAGACTGATGTTGACG</td><td>>MI0000077|hsa-mir-21&WithFlank&17|+|59841260|59841343</td><td>14</td><td>36</td><td>23(+1G)</td><td>TAGCTTATCAGACTGATGTTGACG</td><td>BT20</td><td>1964</td><td>566</td><td>808</td><td>71</td><td>125</td><td>1610.9</td><td>301.2</td><td>490.9</td><td>54.7</td><td>44.9</td></tr>
<tr><td>>MI0000077|hsa-mir-21&WithFlank&17|+|59841260|59841343@14.36.23(+1G)&[MIMAT0000076&hsa-miR-21-5p&offsets|0|+1(+1G);m-1&17|+|59841273|59841294&offsets|0|+1(+1G)]|TAGCTTATCAGACTGATGTTGACG</td><td>>MI0000077|hsa-mir-21&WithFlank&17|+|59841260|59841343</td><td>14</td><td>36</td><td>23(+1G)</td><td>TAGCTTATCAGACTGATGTTGACG</td><td>MB231</td><td>1315</td><td>562</td><td>120</td><td>192</td><td>330</td><td>917.6</td><td>339.9</td><td>91.4</td><td>32.5</td><td>191.3</td></tr>
<tr><td>>MI0000077|hsa-mir-21&WithFlank&17|+|59841260|59841343@14.36.23(+1G)&[MIMAT0000076&hsa-miR-21-5p&offsets|0|+1(+1G);m-1&17|+|59841273|59841294&offsets|0|+1(+1G)]|TAGCTTATCAGACTGATGTTGACG</td><td>>MI0000077|hsa-mir-21&WithFlank&17|+|59841260|59841343</td><td>14</td><td>36</td><td>23(+1G)</td><td>TAGCTTATCAGACTGATGTTGACG</td><td>MB468</td><td>57</td><td>11</td><td>31</td><td>21</td><td>8</td><td>19.9</td><td>4.4</td><td>32.2</td><td>17.0</td><td>5.0</td></tr>
<tr><td>>MI0000077|hsa-mir-21&WithFlank&17|+|59841260|59841343@14.36.23&[MIMAT0000076&hsa-miR-21-5p&offsets|0|+1;m-1&17|+|59841273|59841294&offsets|0|+1]|TAGCTTATCAGACTGATGTTGAC</td><td>>MI0000077|hsa-mir-21&WithFlank&17|+|59841260|59841343</td><td>14</td><td>36</td><td>23</td><td>TAGCTTATCAGACTGATGTTGAC</td><td>BT20</td><td>64138</td><td>32047</td><td>52534</td><td>5826</td><td>10831</td><td>28185.2</td><td>16890.8</td><td>26229.6</td><td>3824.6</td><td>4560.0</td></tr>
<tr><td>>MI0000077|hsa-mir-21&WithFlank&17|+|59841260|59841343@14.36.23&[MIMAT0000076&hsa-miR-21-5p&offsets|0|+1;m-1&17|+|59841273|59841294&offsets|0|+1]|TAGCTTATCAGACTGATGTTGAC</td><td>>MI0000077|hsa-mir-21&WithFlank&17|+|59841260|59841343</td><td>14</td><td>36</td><td>23</td><td>TAGCTTATCAGACTGATGTTGAC</td><td>MB231</td><td>60639</td><td>50270</td><td>13541</td><td>47640</td><td>53771</td><td>10662.5</td><td>18139.2</td><td>11144.6</td><td>7120.3</td><td>27996.2</td></tr>
<tr><td>>MI0000077|hsa-mir-21&WithFlank&17|+|59841260|59841343@14.36.23&[MIMAT0000076&hsa-miR-21-5p&offsets|0|+1;m-1&17|+|59841273|59841294&offsets|0|+1]|TAGCTTATCAGACTGATGTTGAC</td><td>>MI0000077|hsa-mir-21&WithFlank&17|+|59841260|59841343</td><td>14</td><td>36</td><td>23</td><td>TAGCTTATCAGACTGATGTTGAC</td><td>MB468</td><td>2049</td><td>714</td><td>823</td><td>2468</td><td>538</td><td>728.2</td><td>291.7</td><td>337.4</td><td>1836.2</td><td>122.4</td></tr>
<tr><td>>MI0000077|hsa-mir-21&WithFlank&17|+|59841260|59841343@14.37.24(+1U)&[MIMAT0000076&hsa-miR-21-5p&offsets|0|+2(+1U);m-1&17|+|59841273|59841294&offsets|0|+2(+1U)]|TAGCTTATCAGACTGATGTTGACTT</td><td>>MI0000077|hsa-mir-21&WithFlank&17|+|59841260|59841343</td><td>14</td><td>37</td><td>24(+1U)</td><td>TAGCTTATCAGACTGATGTTGACTT</td><td>BT20</td><td>11</td><td>4</td><td>7</td><td>0</td><td>0</td><td>9.4</td><td>3.0</td><td>4.5</td><td>0.0</td><td>0.0</td></tr>
</table>

<table>
<tr><td>>MI0000077|hsa-mir-21&WithFlank&17|+|59841260|59841343@14.37.24(+1U)&[MIMAT0000076&hsa-miR-21-5p&offsets|0|+2(+1U);m-1&17|+|59841273|59841294&offsets|0|+2(+1U)]|TAGCTTATCAGACTGATGTTGACTT</td><td>>MI0000077|hsa-mir-21&WithFlank&17|+|59841260|59841343</td><td>14</td><td>37</td><td>24(+1U)</td><td>TAGCTTATCAGACTGATGTTGACTT</td><td>MB231</td><td>12</td><td>8</td><td>3</td><td>4</td><td>8</td><td>4.7</td><td>2.9</td><td>2.5</td><td>1.6</td><td>6.9</td></tr>
<tr><td>>MI0000077|hsa-mir-21&WithFlank&17|+|59841260|59841343@14.37.24&[MIMAT0000076&hsa-miR-21-5p&offsets|0|+2;m-1&17|+|59841273|59841294&offsets|0|+2]|TAGCTTATCAGACTGATGTTGACT</td><td>>MI0000077|hsa-mir-21&WithFlank&17|+|59841260|59841343</td><td>14</td><td>37</td><td>24</td><td>TAGCTTATCAGACTGATGTTGACT</td><td>BT20</td><td>12187</td><td>7140</td><td>8495</td><td>1063</td><td>3145</td><td>6499.2</td><td>4016.1</td><td>5407.9</td><td>406.9</td><td>1252.2</td></tr>
<tr><td>>MI0000077|hsa-mir-21&WithFlank&17|+|59841260|59841343@14.37.24&[MIMAT0000076&hsa-miR-21-5p&offsets|0|+2;m-1&17|+|59841273|59841294&offsets|0|+2]|TAGCTTATCAGACTGATGTTGACT</td><td>>MI0000077|hsa-mir-21&WithFlank&17|+|59841260|59841343</td><td>14</td><td>37</td><td>24</td><td>TAGCTTATCAGACTGATGTTGACT</td><td>MB231</td><td>13804</td><td>14033</td><td>2644</td><td>5018</td><td>11239</td><td>1821.5</td><td>6729.7</td><td>1751.2</td><td>2226.5</td><td>5255.0</td></tr>
<tr><td>>MI0000077|hsa-mir-21&WithFlank&17|+|59841260|59841343@14.37.24&[MIMAT0000076&hsa-miR-21-5p&offsets|0|+2;m-1&17|+|59841273|59841294&offsets|0|+2]|TAGCTTATCAGACTGATGTTGACT</td><td>>MI0000077|hsa-mir-21&WithFlank&17|+|59841260|59841343</td><td>14</td><td>37</td><td>24</td><td>TAGCTTATCAGACTGATGTTGACT</td><td>MB468</td><td>378</td><td>155</td><td>153</td><td>165</td><td>91</td><td>105.3</td><td>62.7</td><td>83.6</td><td>100.8</td><td>28.2</td></tr>
<tr><td>>MI0000077|hsa-mir-21&WithFlank&17|+|59841260|59841343@14.38.25&[MIMAT0000076&hsa-miR-21-5p&offsets|0|+3;m-1&17|+|59841273|59841294&offsets|0|+3]|TAGCTTATCAGACTGATGTTGACTG</td><td>>MI0000077|hsa-mir-21&WithFlank&17|+|59841260|59841343</td><td>14</td><td>38</td><td>25</td><td>TAGCTTATCAGACTGATGTTGACTG</td><td>BT20</td><td>13</td><td>7</td><td>7</td><td>1</td><td>2</td><td>6.4</td><td>0.9</td><td>1.8</td><td>1.2</td><td>1.6</td></tr>
<tr><td>>MI0000077|hsa-mir-21&WithFlank&17|+|59841260|59841343@14.39.26(+1U)&[MIMAT0000076&hsa-miR-21-5p&offsets|0|+4(+1U);m-1&17|+|59841273|59841294&offsets|0|+4(+1U)]|TAGCTTATCAGACTGATGTTGACTGTT</td><td>>MI0000077|hsa-mir-21&WithFlank&17|+|59841260|59841343</td><td>14</td><td>39</td><td>26(+1U)</td><td>TAGCTTATCAGACTGATGTTGACTGTT</td><td>BT20</td><td>26</td><td>25</td><td>15</td><td>6</td><td>13</td><td>9.0</td><td>12.9</td><td>2.0</td><td>8.0</td><td>8.4</td></tr>
<tr><td>>MI0000077|hsa-mir-21&WithFlank&17|+|59841260|59841343@14.39.26(+1U)&[MIMAT0000076&hsa-miR-21-5p&offsets|0|+4(+1U);m-1&17|+|59841273|59841294&offsets|0|+4(+1U)]|TAGCTTATCAGACTGATGTTGACTGTT</td><td>>MI0000077|hsa-mir-21&WithFlank&17|+|59841260|59841343</td><td>14</td><td>39</td><td>26(+1U)</td><td>TAGCTTATCAGACTGATGTTGACTGTT</td><td>MB231</td><td>12</td><td>13</td><td>3</td><td>5</td><td>13</td><td>2.5</td><td>6.2</td><td>2.1</td><td>1.4</td><td>6.5</td></tr>
<tr><td>>MI0000077|hsa-mir-21&WithFlank&17|+|59841260|59841343@14.39.26&[MIMAT0000076&hsa-miR-21-5p&offsets|0|+4;m-1&17|+|59841273|59841294&offsets|0|+4]|TAGCTTATCAGACTGATGTTGACTGT</td><td>>MI0000077|hsa-mir-21&WithFlank&17|+|59841260|59841343</td><td>14</td><td>39</td><td>26</td><td>TAGCTTATCAGACTGATGTTGACTGT</td><td>BT20</td><td>31</td><td>16</td><td>9</td><td>1</td><td>1</td><td>26.2</td><td>9.6</td><td>3.8</td><td>1.5</td><td>1.6</td></tr>
<tr><td>>MI0000077|hsa-mir-21&WithFlank&17|+|59841260|59841343@15.35.21(+1U)&[MIMAT0000076&hsa-miR-21-5p&offsets|+1|0(+1U);m-1&17|+|59841273|59841294&offsets|+1|0(+1U)]|AGCTTATCAGACTGATGTTGAT</td><td>>MI0000077|hsa-mir-21&WithFlank&17|+|59841260|59841343</td><td>15</td><td>35</td><td>21(+1U)</td><td>AGCTTATCAGACTGATGTTGAT</td><td>BT20</td><td>21</td><td>9</td><td>8</td><td>3</td><td>5</td><td>10.4</td><td>5.9</td><td>4.9</td><td>2.6</td><td>0.7</td></tr>
<tr><td>>MI0000077|hsa-mir-21&WithFlank&17|+|59841260|59841343@15.35.21&[MIMAT0000076&hsa-miR-21-5p&offsets|+1|0;m-1&17|+|59841273|59841294&offsets|+1|0]|AGCTTATCAGACTGATGTTGA</td><td>>MI0000077|hsa-mir-21&WithFlank&17|+|59841260|59841343</td><td>15</td><td>35</td><td>21</td><td>AGCTTATCAGACTGATGTTGA</td><td>BT20</td><td>12</td><td>8</td><td>5</td><td>27</td><td>3</td><td>6.4</td><td>6.9</td><td>2.6</td><td>25.6</td><td>0.8</td></tr>
<tr><td>>MI0000077|hsa-mir-21&WithFlank&17|+|59841260|59841343@15.36.22(+1A)&[MIMAT0000076&hsa-miR-21-5p&offsets|+1|+1(+1A);m-1&17|+|59841273|59841294&offsets|+1|+1(+1A)]|AGCTTATCAGACTGATGTTGACA</td><td>>MI0000077|hsa-mir-21&WithFlank&17|+|59841260|59841343</td><td>15</td><td>36</td><td>22(+1A)</td><td>AGCTTATCAGACTGATGTTGACA</td><td>BT20</td><td>36</td><td>18</td><td>18</td><td>5</td><td>16</td><td>18.8</td><td>8.0</td><td>8.1</td><td>3.1</td><td>1.6</td></tr>
<tr><td>>MI0000077|hsa-mir-21&WithFlank&17|+|59841260|59841343@15.36.22&[MIMAT0000076&hsa-miR-21-5p&offsets|+1|+1;m-1&17|+|59841273|59841294&offsets|+1|+1]|AGCTTATCAGACTGATGTTGAC</td><td>>MI0000077|hsa-mir-21&WithFlank&17|+|59841260|59841343</td><td>15</td><td>36</td><td>22</td><td>AGCTTATCAGACTGATGTTGAC</td><td>BT20</td><td>104</td><td>61</td><td>49</td><td>15</td><td>40</td><td>37.3</td><td>25.0</td><td>25.6</td><td>11.5</td><td>8.4</td></tr>
<tr><td>>MI0000077|hsa-mir-21&WithFlank&17|+|59841260|59841343@15.36.22&[MIMAT0000076&hsa-miR-21-5p&offsets|+1|+1;m-1&17|+|59841273|59841294&offsets|+1|+1]|AGCTTATCAGACTGATGTTGAC</td><td>>MI0000077|hsa-mir-21&WithFlank&17|+|59841260|59841343</td><td>15</td><td>36</td><td>22</td><td>AGCTTATCAGACTGATGTTGAC</td><td>MB231</td><td>64</td><td>54</td><td>10</td><td>95</td><td>83</td><td>12.6</td><td>25.9</td><td>8.4</td><td>17.3</td><td>57.2</td></tr>
<tr><td>>MI0000077|hsa-mir-21&WithFlank&17|+|59841260|59841343@15.37.23&[MIMAT0000076&hsa-miR-21-5p&offsets|+1|+2;m-1&17|+|59841273|59841294&offsets|+1|+2]|AGCTTATCAGACTGATGTTGACT</td><td>>MI0000077|hsa-mir-21&WithFlank&17|+|59841260|59841343</td><td>15</td><td>37</td><td>23</td><td>AGCTTATCAGACTGATGTTGACT</td><td>BT20</td><td>15</td><td>10</td><td>6</td><td>3</td><td>8</td><td>7.3</td><td>8.1</td><td>5.2</td><td>2.4</td><td>3.4</td></tr>
<tr><td>>MI0000077|hsa-mir-21&WithFlank&17|+|59841260|59841343@15.37.23&[MIMAT0000076&hsa-miR-21-5p&offsets|+1|+2;m-1&17|+|59841273|59841294&offsets|+1|+2]|AGCTTATCAGACTGATGTTGACT</td><td>>MI0000077|hsa-mir-21&WithFlank&17|+|59841260|59841343</td><td>15</td><td>37</td><td>23</td><td>AGCTTATCAGACTGATGTTGACT</td><td>MB231</td><td>12</td><td>12</td><td>1</td><td>18</td><td>18</td><td>1.4</td><td>6.7</td><td>0.9</td><td>2.5</td><td>12.6</td></tr>
<tr><td>>MI0000077|hsa-mir-21&WithFlank&17|+|59841260|59841343@16.36.21&[MIMAT0000076&hsa-miR-21-5p&offsets|+2|+1;m-1&17|+|59841273|59841294&offsets|+2|+1]|GCTTATCAGACTGATGTTGAC</td><td>>MI0000077|hsa-mir-21&WithFlank&17|+|59841260|59841343</td><td>16</td><td>36</td><td>21</td><td>GCTTATCAGACTGATGTTGAC</td><td>BT20</td><td>26</td><td>21</td><td>25</td><td>3</td><td>4</td><td>11.3</td><td>15.3</td><td>9.7</td><td>3.0</td><td>1.6</td></tr>
<tr><td>>MI0000077|hsa-mir-21&WithFlank&17|+|59841260|59841343@16.36.21&[MIMAT0000076&hsa-miR-21-5p&offsets|+2|+1;m-1&17|+|59841273|59841294&offsets|+2|+1]|GCTTATCAGACTGATGTTGAC</td><td>>MI0000077|hsa-mir-21&WithFlank&17|+|59841260|59841343</td><td>16</td><td>36</td><td>21</td><td>GCTTATCAGACTGATGTTGAC</td><td>MB231</td><td>11</td><td>14</td><td>5</td><td>9</td><td>8</td><td>1.6</td><td>5.2</td><td>3.9</td><td>0.3</td><td>4.1</td></tr>
<tr><td>>MI0000077|hsa-mir-21&WithFlank&17|+|59841260|59841343@52.71.20(+1C)&[MIMAT0004494&hsa-miR-21-3p&offsets|0|-1(+1C);m-52&17|+|59841311|59841332&offsets|0|-2(+1C)]|CAACACCAGTCGATGGGCTGC</td><td>>MI0000077|hsa-mir-21&WithFlank&17|+|59841260|59841343</td><td>52</td><td>71</td><td>20(+1C)</td><td>CAACACCAGTCGATGGGCTGC</td><td>BT20</td><td>12</td><td>4</td><td>3</td><td>23</td><td>2</td><td>12.4</td><td>5.0</td><td>1.6</td><td>22.7</td><td>0.7</td></tr>
<tr><td>>MI0000077|hsa-mir-21&WithFlank&17|+|59841260|59841343@52.72.21(+1A)&[MIMAT0004494&hsa-miR-21-3p&offsets|0|0(+1A);m-52&17|+|59841311|59841332&offsets|0|-1(+1A)]|CAACACCAGTCGATGGGCTGTA</td><td>>MI0000077|hsa-mir-21&WithFlank&17|+|59841260|59841343</td><td>52</td><td>72</td><td>21(+1A)</td><td>CAACACCAGTCGATGGGCTGTA</td><td>BT20</td><td>15</td><td>2</td><td>2</td><td>27</td><td>0</td><td>19.5</td><td>2.2</td><td>0.3</td><td>32.1</td><td>0.5</td></tr>
<tr><td>>MI0000077|hsa-mir-21&WithFlank&17|+|59841260|59841343@52.72.21(+1U)&[MIMAT0004494&hsa-miR-21-3p&offsets|0|0(+1U);m-52&17|+|59841311|59841332&offsets|0|-1(+1U)]|CAACACCAGTCGATGGGCTGTT</td><td>>MI0000077|hsa-mir-21&WithFlank&17|+|59841260|59841343</td><td>52</td><td>72</td><td>21(+1U)</td><td>CAACACCAGTCGATGGGCTGTT</td><td>BT20</td><td>11</td><td>1</td><td>2</td><td>23</td><td>0</td><td>12.5</td><td>2.6</td><td>0.9</td><td>22.4</td><td>0.8</td></tr>
<tr><td>>MI0000077|hsa-mir-21&WithFlank&17|+|59841260|59841343@52.72.21&[MIMAT0004494&hsa-miR-21-3p&offsets|0|0;m-52&17|+|59841311|59841332&offsets|0|-1]|CAACACCAGTCGATGGGCTGT</td><td>>MI0000077|hsa-mir-21&WithFlank&17|+|59841260|59841343</td><td>52</td><td>72</td><td>21</td><td>CAACACCAGTCGATGGGCTGT</td><td>BT20</td><td>80</td><td>27</td><td>17</td><td>162</td><td>10</td><td>78.4</td><td>29.4</td><td>7.7</td><td>146.8</td><td>4.0</td></tr>
<tr><td>>MI0000077|hsa-mir-21&WithFlank&17|+|59841260|59841343@52.72.21&[MIMAT0004494&hsa-miR-21-3p&offsets|0|0;m-52&17|+|59841311|59841332&offsets|0|-1]|CAACACCAGTCGATGGGCTGT</td><td>>MI0000077|hsa-mir-21&WithFlank&17|+|59841260|59841343</td><td>52</td><td>72</td><td>21</td><td>CAACACCAGTCGATGGGCTGT</td><td>MB231</td><td>27</td><td>12</td><td>5</td><td>136</td><td>31</td><td>9.1</td><td>9.0</td><td>4.5</td><td>40.4</td><td>37.2</td></tr>
<tr><td>>MI0000077|hsa-mir-21&WithFlank&17|+|59841260|59841343@52.73.22(+1A)&[MIMAT0004494&hsa-miR-21-3p&offsets|0|+1(+1A);m-52&17|+|59841311|59841332&offsets|0|0(+1A)]|CAACACCAGTCGATGGGCTGTCA</td><td>>MI0000077|hsa-mir-21&WithFlank&17|+|59841260|59841343</td><td>52</td><td>73</td><td>22(+1A)</td><td>CAACACCAGTCGATGGGCTGTCA</td><td>BT20</td><td>13</td><td>2</td><td>2</td><td>11</td><td>1</td><td>16.9</td><td>3.6</td><td>0.8</td><td>11.8</td><td>0.5</td></tr>
<tr><td>>MI0000077|hsa-mir-21&WithFlank&17|+|59841260|59841343@52.73.22&[MIMAT0004494&hsa-miR-21-3p&offsets|0|+1;m-52&17|+|59841311|59841332&offsets|0|0]|CAACACCAGTCGATGGGCTGTC</td><td>>MI0000077|hsa-mir-21&WithFlank&17|+|59841260|59841343</td><td>52</td><td>73</td><td>22</td><td>CAACACCAGTCGATGGGCTGTC</td><td>BT20</td><td>31</td><td>8</td><td>7</td><td>109</td><td>3</td><td>34.1</td><td>10.2</td><td>3.4</td><td>93.4</td><td>2.6</td></tr>
<tr><td>>MI0000077|hsa-mir-21&WithFlank&17|+|59841260|59841343@52.73.22&[MIMAT0004494&hsa-miR-21-3p&offsets|0|+1;m-52&17|+|59841311|59841332&offsets|0|0]|CAACACCAGTCGATGGGCTGTC</td><td>>MI0000077|hsa-mir-21&WithFlank&17|+|59841260|59841343</td><td>52</td><td>73</td><td>22</td><td>CAACACCAGTCGATGGGCTGTC</td><td>MB231</td><td>17</td><td>5</td><td>4</td><td>149</td><td>19</td><td>5.4</td><td>1.9</td><td>2.1</td><td>57.9</td><td>21.0</td></tr>
<tr><td>>MI0000077|hsa-mir-21&WithFlank&17|+|59841260|59841343@52.74.23&[MIMAT0004494&hsa-miR-21-3p&offsets|0|+2;m-52&17|+|59841311|59841332&offsets|0|+1]|CAACACCAGTCGATGGGCTGTCT</td><td>>MI0000077|hsa-mir-21&WithFlank&17|+|59841260|59841343</td><td>52</td><td>74</td><td>23</td><td>CAACACCAGTCGATGGGCTGTCT</td><td>BT20</td><td>13</td><td>3</td><td>1</td><td>28</td><td>1</td><td>14.3</td><td>4.1</td><td>0.6</td><td>28.1</td><td>0.2</td></tr>
<tr><td>>MI0000078|hsa-mir-22&WithFlank&17|-|1713897|1713993@21.41.21&[MIMAT0004495&hsa-miR-22-5p&offsets|0|-1;m-217&17|-|1713953|1713973&offsets|0|0]|AGTTCTTCAGTGGCAAGCTTT</td><td>>MI0000078|hsa-mir-22&WithFlank&17|-|1713897|1713993</td><td>21</td><td>41</td><td>21</td><td>AGTTCTTCAGTGGCAAGCTTT</td><td>BT20</td><td>30</td><td>27</td><td>28</td><td>60</td><td>19</td><td>25.6</td><td>17.6</td><td>18.5</td><td>40.8</td><td>7.4</td></tr>
<tr><td>>MI0000078|hsa-mir-22&WithFlank&17|-|1713897|1713993@21.41.21&[MIMAT0004495&hsa-miR-22-5p&offsets|0|-1;m-217&17|-|1713953|1713973&offsets|0|0]|AGTTCTTCAGTGGCAAGCTTT</td><td>>MI0000078|hsa-mir-22&WithFlank&17|-|1713897|1713993</td><td>21</td><td>41</td><td>21</td><td>AGTTCTTCAGTGGCAAGCTTT</td><td>MB231</td><td>27</td><td>38</td><td>12</td><td>298</td><td>41</td><td>15.5</td><td>14.2</td><td>4.3</td><td>129.8</td><td>15.9</td></tr>
</table>

| | | | | | | | | | | | | | | | | |
|---|---|---|---|---|---|---|---|---|---|---|---|---|---|---|---|---|
| >MI0000078\|hsa-mir-22&WithFlank&17\|-\|1713897\|1713993@21.42.22&[MIMAT0004495&hsa-miR-22-5p&offsets\|0\|0;m-217&17\|-\|1713953\|1713973&offsets\|0\|+1]\|AGTTCTTCAGTGGCAAGCTTTA | >MI0000078\|hsa-mir-22&WithFlank&17\|-\|1713897\|1713993 | 21 | 42 | 22 | AGTTCTTCAGTGGCAAGCTTTA | BT20 | 17 | 10 | 10 | 18 | 3 | 16.4 | 7.0 | 4.0 | 15.1 | 1.6 |
| >MI0000078\|hsa-mir-22&WithFlank&17\|-\|1713897\|1713993@59.78.20&[MIMAT0000077&hsa-miR-22-3p&offsets\|0\|-2;m-2&17\|-\|1713914\|1713935&offsets\|0\|-2]\|AAGCTGCCAGTTGAAGAACT | >MI0000078\|hsa-mir-22&WithFlank&17\|-\|1713897\|1713993 | 59 | 78 | 20 | AAGCTGCCAGTTGAAGAACT | BT20 | 37 | 32 | 48 | 26 | 24 | 24.2 | 17.8 | 27.1 | 11.2 | 8.4 |
| >MI0000078\|hsa-mir-22&WithFlank&17\|-\|1713897\|1713993@59.78.20&[MIMAT0000077&hsa-miR-22-3p&offsets\|0\|-2;m-2&17\|-\|1713914\|1713935&offsets\|0\|-2]\|AAGCTGCCAGTTGAAGAACT | >MI0000078\|hsa-mir-22&WithFlank&17\|-\|1713897\|1713993 | 59 | 78 | 20 | AAGCTGCCAGTTGAAGAACT | MB231 | 25 | 29 | 12 | 45 | 38 | 10.9 | 4.4 | 7.5 | 23.4 | 14.1 |
| >MI0000078\|hsa-mir-22&WithFlank&17\|-\|1713897\|1713993@59.79.21(+1A)&[MIMAT0000077&hsa-miR-22-3p&offsets\|0\|-1(+1A);m-2&17\|-\|1713914\|1713935&offsets\|0\|-1(+1A)]\|AAGCTGCCAGTTGAAGAACTGA | >MI0000078\|hsa-mir-22&WithFlank&17\|-\|1713897\|1713993 | 59 | 79 | 21(+1A) | AAGCTGCCAGTTGAAGAACTGA | BT20 | 34 | 19 | 34 | 53 | 8 | 28.9 | 12.9 | 31.7 | 41.4 | 1.5 |
| >MI0000078\|hsa-mir-22&WithFlank&17\|-\|1713897\|1713993@59.79.21(+1C)&[MIMAT0000077&hsa-miR-22-3p&offsets\|0\|-1(+1C);m-2&17\|-\|1713914\|1713935&offsets\|0\|-1(+1C)]\|AAGCTGCCAGTTGAAGAACTGC | >MI0000078\|hsa-mir-22&WithFlank&17\|-\|1713897\|1713993 | 59 | 79 | 21(+1C) | AAGCTGCCAGTTGAAGAACTGC | BT20 | 91 | 100 | 97 | 238 | 41 | 53.3 | 70.4 | 71.4 | 181.5 | 7.9 |
| >MI0000078\|hsa-mir-22&WithFlank&17\|-\|1713897\|1713993@59.79.21(+1C)&[MIMAT0000077&hsa-miR-22-3p&offsets\|0\|-1(+1C);m-2&17\|-\|1713914\|1713935&offsets\|0\|-1(+1C)]\|AAGCTGCCAGTTGAAGAACTGC | >MI0000078\|hsa-mir-22&WithFlank&17\|-\|1713897\|1713993 | 59 | 79 | 21(+1C) | AAGCTGCCAGTTGAAGAACTGC | MB231 | 42 | 66 | 13 | 626 | 109 | 14.8 | 45.4 | 10.1 | 459.7 | 77.7 |
| >MI0000078\|hsa-mir-22&WithFlank&17\|-\|1713897\|1713993@59.79.21(+1G)&[MIMAT0000077&hsa-miR-22-3p&offsets\|0\|-1(+1G);m-2&17\|-\|1713914\|1713935&offsets\|0\|-1(+1G)]\|AAGCTGCCAGTTGAAGAACTGG | >MI0000078\|hsa-mir-22&WithFlank&17\|-\|1713897\|1713993 | 59 | 79 | 21(+1G) | AAGCTGCCAGTTGAAGAACTGG | BT20 | 41 | 15 | 18 | 28 | 3 | 44.7 | 8.8 | 17.0 | 29.2 | 0.3 |
| >MI0000078\|hsa-mir-22&WithFlank&17\|-\|1713897\|1713993@59.79.21(+1G)&[MIMAT0000077&hsa-miR-22-3p&offsets\|0\|-1(+1G);m-2&17\|-\|1713914\|1713935&offsets\|0\|-1(+1G)]\|AAGCTGCCAGTTGAAGAACTGG | >MI0000078\|hsa-mir-22&WithFlank&17\|-\|1713897\|1713993 | 59 | 79 | 21(+1G) | AAGCTGCCAGTTGAAGAACTGG | MB231 | 14 | 11 | 2 | 58 | 8 | 7.6 | 8.8 | 1.8 | 47.2 | 7.1 |
| >MI0000078\|hsa-mir-22&WithFlank&17\|-\|1713897\|1713993@59.80.22(+1A)&[MIMAT0000077&hsa-miR-22-3p&offsets\|0\|0(+1A);m-2&17\|-\|1713914\|1713935&offsets\|0\|0(+1A)]\|AAGCTGCCAGTTGAAGAACTGTA | >MI0000078\|hsa-mir-22&WithFlank&17\|-\|1713897\|1713993 | 59 | 80 | 22(+1A) | AAGCTGCCAGTTGAAGAACTGTA | BT20 | 25 | 20 | 23 | 24 | 11 | 16.9 | 13.8 | 13.8 | 14.7 | 4.0 |
| >MI0000078\|hsa-mir-22&WithFlank&17\|-\|1713897\|1713993@59.80.22(+1A)&[MIMAT0000077&hsa-miR-22-3p&offsets\|0\|0(+1A);m-2&17\|-\|1713914\|1713935&offsets\|0\|0(+1A)]\|AAGCTGCCAGTTGAAGAACTGTA | >MI0000078\|hsa-mir-22&WithFlank&17\|-\|1713897\|1713993 | 59 | 80 | 22(+1A) | AAGCTGCCAGTTGAAGAACTGTA | MB231 | 11 | 11 | 3 | 62 | 17 | 5.7 | 5.0 | 1.9 | 33.4 | 9.8 |
| >MI0000078\|hsa-mir-22&WithFlank&17\|-\|1713897\|1713993@59.80.22&[MIMAT0000077&hsa-miR-22-3p&offsets\|0\|0;m-2&17\|-\|1713914\|1713935&offsets\|0\|0]\|AAGCTGCCAGTTGAAGAACTGT | >MI0000078\|hsa-mir-22&WithFlank&17\|-\|1713897\|1713993 | 59 | 80 | 22 | AAGCTGCCAGTTGAAGAACTGT | BT20 | 684 | 655 | 707 | 1687 | 306 | 410.7 | 426.2 | 459.6 | 1245.8 | 68.6 |
| >MI0000078\|hsa-mir-22&WithFlank&17\|-\|1713897\|1713993@59.80.22&[MIMAT0000077&hsa-miR-22-3p&offsets\|0\|0;m-2&17\|-\|1713914\|1713935&offsets\|0\|0]\|AAGCTGCCAGTTGAAGAACTGT | >MI0000078\|hsa-mir-22&WithFlank&17\|-\|1713897\|1713993 | 59 | 80 | 22 | AAGCTGCCAGTTGAAGAACTGT | MB231 | 357 | 563 | 117 | 3662 | 778 | 126.5 | 346.1 | 88.1 | 2481.3 | 448.0 |
| >MI0000078\|hsa-mir-22&WithFlank&17\|-\|1713897\|1713993@59.80.22&[MIMAT0000077&hsa-miR-22-3p&offsets\|0\|0;m-2&17\|-\|1713914\|1713935&offsets\|0\|0]\|AAGCTGCCAGTTGAAGAACTGT | >MI0000078\|hsa-mir-22&WithFlank&17\|-\|1713897\|1713993 | 59 | 80 | 22 | AAGCTGCCAGTTGAAGAACTGT | MB468 | 16 | 4 | 5 | 61 | 6 | 8.3 | 2.4 | 1.5 | 26.8 | 0.8 |
| >MI0000078\|hsa-mir-22&WithFlank&17\|-\|1713897\|1713993@59.81.23&[MIMAT0000077&hsa-miR-22-3p&offsets\|0\|+1;m-2&17\|-\|1713914\|1713935&offsets\|0\|+1]\|AAGCTGCCAGTTGAAGAACTGTT | >MI0000078\|hsa-mir-22&WithFlank&17\|-\|1713897\|1713993 | 59 | 81 | 23 | AAGCTGCCAGTTGAAGAACTGTT | BT20 | 16 | 16 | 12 | 14 | 7 | 9.7 | 11.0 | 6.8 | 11.7 | 2.6 |
| >MI0000078\|hsa-mir-22&WithFlank&17\|-\|1713897\|1713993@59.81.23&[MIMAT0000077&hsa-miR-22-3p&offsets\|0\|+1;m-2&17\|-\|1713914\|1713935&offsets\|0\|+1]\|AAGCTGCCAGTTGAAGAACTGTT | >MI0000078\|hsa-mir-22&WithFlank&17\|-\|1713897\|1713993 | 59 | 81 | 23 | AAGCTGCCAGTTGAAGAACTGTT | MB231 | 14 | 18 | 4 | 94 | 29 | 3.9 | 11.9 | 2.8 | 58.5 | 21.7 |
| >MI0000079\|hsa-mir-23a&WithFlank&19\|-\|13836581\|13836665@51.71.21&[MIMAT0000078&hsa-miR-23a-3p&offsets\|0\|0;m-31&19\|-\|13836595\|13836615&offsets\|0\|0]\|ATCACATTGCCAGGGATTTCC | >MI0000079\|hsa-mir-23a&WithFlank&19\|-\|13836581\|13836665 | 51 | 71 | 21 | ATCACATTGCCAGGGATTTCC | MB231 | 51 | 57 | 17 | 70 | 48 | 88.6 | 98.8 | 30.0 | 120.6 | 83.8 |
| >MI0000079\|hsa-mir-23a&WithFlank&19\|-\|13836581\|13836665@51.71.21&[MIMAT0000078&hsa-miR-23a-3p&offsets\|0\|0;m-31&19\|-\|13836595\|13836615&offsets\|0\|0]\|ATCACATTGCCAGGGATTTCC | >MI0000079\|hsa-mir-23a&WithFlank&19\|-\|13836581\|13836665 | 51 | 71 | 21 | ATCACATTGCCAGGGATTTCC | MB468 | 15 | 11 | 12 | 34 | 4 | 6.9 | 2.9 | 2.7 | 30.0 | 1.9 |
| >MI0000079\|hsa-mir-23a&WithFlank&19\|-\|13836581\|13836665@51.72.22&[MIMAT0000078&hsa-miR-23a-3p&offsets\|0\|+1;m-31&19\|-\|13836595\|13836615&offsets\|0\|+1]\|ATCACATTGCCAGGGATTTCCA | >MI0000079\|hsa-mir-23a&WithFlank&19\|-\|13836581\|13836665 | 51 | 72 | 22 | ATCACATTGCCAGGGATTTCCA | MB231 | 68 | 70 | 18 | 44 | 72 | 117.4 | 120.9 | 30.8 | 75.4 | 123.9 |
| >MI0000079\|hsa-mir-23a&WithFlank&19\|-\|13836581\|13836665@51.72.22&[MIMAT0000078&hsa-miR-23a-3p&offsets\|0\|+1;m-31&19\|-\|13836595\|13836615&offsets\|0\|+1]\|ATCACATTGCCAGGGATTTCCA | >MI0000079\|hsa-mir-23a&WithFlank&19\|-\|13836581\|13836665 | 51 | 72 | 22 | ATCACATTGCCAGGGATTTCCA | MB468 | 22 | 17 | 17 | 32 | 8 | 8.8 | 4.0 | 5.9 | 35.2 | 4.8 |
| >MI0000079\|hsa-mir-23a&WithFlank&19\|-\|13836581\|13836665@51.73.23(+1U)&[MIMAT0000078&hsa-miR-23a-3p&offsets\|0\|+2(+1U);m-31&19\|-\|13836595\|13836615&offsets\|0\|+2(+1U)]\|ATCACATTGCCAGGGATTTCCAAT | >MI0000079\|hsa-mir-23a&WithFlank&19\|-\|13836581\|13836665 | 51 | 73 | 23(+1U) | ATCACATTGCCAGGGATTTCCAAT | MB231 | 39 | 31 | 5 | 6 | 35 | 68.2 | 53.9 | 9.2 | 9.9 | 59.8 |
| >MI0000079\|hsa-mir-23a&WithFlank&19\|-\|13836581\|13836665@51.73.23&[MIMAT0000078&hsa-miR-23a-3p&offsets\|0\|+2;m-31&19\|-\|13836595\|13836615&offsets\|0\|+2]\|ATCACATTGCCAGGGATTTCCAA | >MI0000079\|hsa-mir-23a&WithFlank&19\|-\|13836581\|13836665 | 51 | 73 | 23 | ATCACATTGCCAGGGATTTCCAA | MB231 | 28 | 38 | 4 | 8 | 22 | 48.0 | 65.6 | 7.6 | 13.4 | 37.5 |
| >MI0000079\|hsa-mir-23a&WithFlank&19\|-\|13836581\|13836665@52.74.23&[MIMAT0000078&hsa-miR-23a-3p&offsets\|+1\|+3;m-31&19\|-\|13836595\|13836615&offsets\|+1\|+3]\|TCACATTGCCAGGGATTTCCAAC | >MI0000079\|hsa-mir-23a&WithFlank&19\|-\|13836581\|13836665 | 52 | 74 | 23 | TCACATTGCCAGGGATTTCCAAC | MB231 | 13 | 10 | 12 | 3 | 16 | 3.7 | 3.6 | 10.5 | 1.3 | 7.8 |
| >MI0000079\|hsa-mir-23a&WithFlank&19\|-\|13836581\|13836665@54.71.18(+1G)&[MIMAT0000078&hsa-miR-23a-3p&offsets\|+3\|0(+1G);m-31&19\|-\|13836595\|13836615&offsets\|+3\|0(+1G)]\|ACATTGCCAGGGATTTCCG | >MI0000079\|hsa-mir-23a&WithFlank&19\|-\|13836581\|13836665 | 54 | 71 | 18(+1G) | ACATTGCCAGGGATTTCCG | BT20 | 40 | 11 | 20 | 3 | 1 | 48.7 | 5.3 | 8.9 | 4.4 | 1.2 |
| >MI0000079\|hsa-mir-23a&WithFlank&19\|-\|13836581\|13836665@54.71.18(+1G)&[MIMAT0000078&hsa-miR-23a-3p&offsets\|+3\|0(+1G);m-31&19\|-\|13836595\|13836615&offsets\|+3\|0(+1G)]\|ACATTGCCAGGGATTTCCG | >MI0000079\|hsa-mir-23a&WithFlank&19\|-\|13836581\|13836665 | 54 | 71 | 18(+1G) | ACATTGCCAGGGATTTCCG | MB231 | 24 | 6 | 14 | 6 | 5 | 22.8 | 5.1 | 14.0 | 4.8 | 5.8 |
| >MI0000079\|hsa-mir-23a&WithFlank&19\|-\|13836581\|13836665@54.71.18(+1U)&[MIMAT0000078&hsa-miR-23a-3p&offsets\|+3\|0(+1U);m-31&19\|-\|13836595\|13836615&offsets\|+3\|0(+1U)]\|ACATTGCCAGGGATTTCCT | >MI0000079\|hsa-mir-23a&WithFlank&19\|-\|13836581\|13836665 | 54 | 71 | 18(+1U) | ACATTGCCAGGGATTTCCT | BT20 | 18 | 6 | 11 | 2 | 1 | 23.6 | 2.8 | 5.6 | 2.5 | 1.1 |
| >MI0000079\|hsa-mir-23a&WithFlank&19\|-\|13836581\|13836665@54.71.18&[MIMAT0000078&hsa-miR-23a-3p&offsets\|+3\|0;m-31&19\|-\|13836595\|13836615&offsets\|+3\|0]\|ACATTGCCAGGGATTTCC | >MI0000079\|hsa-mir-23a&WithFlank&19\|-\|13836581\|13836665 | 54 | 71 | 18 | ACATTGCCAGGGATTTCC | BT20 | 214 | 214 | 457 | 404 | 74 | 120.0 | 94.5 | 10.1 | 328.5 | 35.4 |
| >MI0000079\|hsa-mir-23a&WithFlank&19\|-\|13836581\|13836665@54.71.18&[MIMAT0000078&hsa-miR-23a-3p&offsets\|+3\|0;m-31&19\|-\|13836595\|13836615&offsets\|+3\|0]\|ACATTGCCAGGGATTTCC | >MI0000079\|hsa-mir-23a&WithFlank&19\|-\|13836581\|13836665 | 54 | 71 | 18 | ACATTGCCAGGGATTTCC | MB231 | 223 | 190 | 606 | 427 | 188 | 209.7 | 170.3 | 659.7 | 383.3 | 203.5 |
| >MI0000079\|hsa-mir-23a&WithFlank&19\|-\|13836581\|13836665@54.72.19(+1G)&[MIMAT0000078&hsa-miR-23a-3p&offsets\|+3\|+1(+1G);m-31&19\|-\|13836595\|13836615&offsets\|+3\|+1(+1G)]\|ACATTGCCAGGGATTTCCAG | >MI0000079\|hsa-mir-23a&WithFlank&19\|-\|13836581\|13836665 | 54 | 72 | 19(+1G) | ACATTGCCAGGGATTTCCAG | BT20 | 14 | 6 | 9 | 0 | 0 | 14.5 | 3.0 | 3.4 | 0.5 | 0.8 |
| >MI0000079\|hsa-mir-23a&WithFlank&19\|-\|13836581\|13836665@54.72.19(+1U)&[MIMAT0000078&hsa-miR-23a-3p&offsets\|+3\|+1(+1U);m-31&19\|-\|13836595\|13836615&offsets\|+3\|+1(+1U)]\|ACATTGCCAGGGATTTCCAT | >MI0000079\|hsa-mir-23a&WithFlank&19\|-\|13836581\|13836665 | 54 | 72 | 19(+1U) | ACATTGCCAGGGATTTCCAT | BT20 | 33 | 30 | 34 | 6 | 9 | 22.5 | 25.2 | 9.0 | 4.6 | 4.0 |
| >MI0000079\|hsa-mir-23a&WithFlank&19\|-\|13836581\|13836665@54.72.19(+1U)&[MIMAT0000078&hsa-miR-23a-3p&offsets\|+3\|+1(+1U);m-31&19\|-\|13836595\|13836615&offsets\|+3\|+1(+1U)]\|ACATTGCCAGGGATTTCCAT | >MI0000079\|hsa-mir-23a&WithFlank&19\|-\|13836581\|13836665 | 54 | 72 | 19(+1U) | ACATTGCCAGGGATTTCCAT | MB231 | 46 | 34 | 62 | 30 | 44 | 40.7 | 29.7 | 62.5 | 29.3 | 55.0 |
| >MI0000079\|hsa-mir-23a&WithFlank&19\|-\|13836581\|13836665@54.72.19(+2U)&[MIMAT0000078&hsa-miR-23a-3p&offsets\|+3\|+1(+2U);m-31&19\|-\|13836595\|13836615&offsets\|+3\|+1(+2U)]\|ACATTGCCAGGGATTTCCATT | >MI0000079\|hsa-mir-23a&WithFlank&19\|-\|13836581\|13836665 | 54 | 72 | 19(+2U) | ACATTGCCAGGGATTTCCATT | BT20 | 13 | 11 | 13 | 2 | 4 | 9.2 | 10.0 | 4.0 | 1.2 | 3.0 |
| >MI0000079\|hsa-mir-23a&WithFlank&19\|-\|13836581\|13836665@54.72.19(+2U)&[MIMAT0000078&hsa-miR-23a-3p&offsets\|+3\|+1(+2U);m-31&19\|-\|13836595\|13836615&offsets\|+3\|+1(+2U)]\|ACATTGCCAGGGATTTCCATT | >MI0000079\|hsa-mir-23a&WithFlank&19\|-\|13836581\|13836665 | 54 | 72 | 19(+2U) | ACATTGCCAGGGATTTCCATT | MB231 | 45 | 29 | 54 | 12 | 37 | 41.4 | 26.3 | 55.0 | 12.4 | 42.2 |
| >MI0000079\|hsa-mir-23a&WithFlank&19\|-\|13836581\|13836665@54.72.19&[MIMAT0000078&hsa-miR-23a-3p&offsets\|+3\|+1;m-31&19\|-\|13836595\|13836615&offsets\|+3\|+1]\|ACATTGCCAGGGATTTCCA | >MI0000079\|hsa-mir-23a&WithFlank&19\|-\|13836581\|13836665 | 54 | 72 | 19 | ACATTGCCAGGGATTTCCA | BT20 | 359 | 307 | 524 | 110 | 109 | 293.0 | 133.0 | 45.2 | 87.6 | 34.7 |

| | | | | | | | | | | | | | | | | |
|---|---|---|---|---|---|---|---|---|---|---|---|---|---|---|---|---|
| >MI0000079\|hsa-mir-23a&WithFlank&19\|-\|13836581\|13836665@54.72.19&[MIMAT0000078&hsa-miR-23a-3p&offsets\|+3\|+1;m-31&19\|-\|13836595\|13836615&offsets\|+3\|+1]\|ACATTGCCAGGGATTTCCA | >MI0000079\|hsa-mir-23a&WithFlank&19\|-\|13836581\|13836665 | 54 | 72 | 19 | ACATTGCCAGGGATTTCCA | MB231 | 265 | 229 | 546 | 234 | 254 | 242.2 | 208.5 | 626.1 | 218.4 | 291.2 |
| >MI0000079\|hsa-mir-23a&WithFlank&19\|-\|13836581\|13836665@54.73.20(+1A)&[MIMAT0000078&hsa-miR-23a-3p&offsets\|+3\|+2(+1A);m-31&19\|-\|13836595\|13836615&offsets\|+3\|+2(+1A)]\|ACATTGCCAGGGATTTCCAAA | >MI0000079\|hsa-mir-23a&WithFlank&19\|-\|13836581\|13836665 | 54 | 73 | 20(+1A) | ACATTGCCAGGGATTTCCAAA | MB231 | 13 | 9 | 18 | 3 | 9 | 11.3 | 8.4 | 18.6 | 3.0 | 10.7 |
| >MI0000079\|hsa-mir-23a&WithFlank&19\|-\|13836581\|13836665@54.73.20(+1U)&[MIMAT0000078&hsa-miR-23a-3p&offsets\|+3\|+2(+1U);m-31&19\|-\|13836595\|13836615&offsets\|+3\|+2(+1U)]\|ACATTGCCAGGGATTTCCAAT | >MI0000079\|hsa-mir-23a&WithFlank&19\|-\|13836581\|13836665 | 54 | 73 | 20(+1U) | ACATTGCCAGGGATTTCCAAT | BT20 | 57 | 53 | 60 | 6 | 16 | 37.8 | 45.5 | 19.5 | 6.0 | 18.6 |
| >MI0000079\|hsa-mir-23a&WithFlank&19\|-\|13836581\|13836665@54.73.20(+1U)&[MIMAT0000078&hsa-miR-23a-3p&offsets\|+3\|+2(+1U);m-31&19\|-\|13836595\|13836615&offsets\|+3\|+2(+1U)]\|ACATTGCCAGGGATTTCCAAT | >MI0000079\|hsa-mir-23a&WithFlank&19\|-\|13836581\|13836665 | 54 | 73 | 20(+1U) | ACATTGCCAGGGATTTCCAAT | MB231 | 210 | 131 | 281 | 39 | 189 | 187.7 | 115.3 | 279.2 | 37.4 | 233.5 |
| >MI0000079\|hsa-mir-23a&WithFlank&19\|-\|13836581\|13836665@54.73.20(+2U)&[MIMAT0000078&hsa-miR-23a-3p&offsets\|+3\|+2(+2U);m-31&19\|-\|13836595\|13836615&offsets\|+3\|+2(+2U)]\|ACATTGCCAGGGATTTCCAATT | >MI0000079\|hsa-mir-23a&WithFlank&19\|-\|13836581\|13836665 | 54 | 73 | 20(+2U) | ACATTGCCAGGGATTTCCAATT | MB231 | 26 | 17 | 32 | 2 | 19 | 22.8 | 14.6 | 30.8 | 2.2 | 21.5 |
| >MI0000079\|hsa-mir-23a&WithFlank&19\|-\|13836581\|13836665@54.73.20&[MIMAT0000078&hsa-miR-23a-3p&offsets\|+3\|+2;m-31&19\|-\|13836595\|13836615&offsets\|+3\|+2]\|ACATTGCCAGGGATTTCCAA | >MI0000079\|hsa-mir-23a&WithFlank&19\|-\|13836581\|13836665 | 54 | 73 | 20 | ACATTGCCAGGGATTTCCAA | BT20 | 88 | 87 | 133 | 7 | 19 | 70.4 | 49.6 | 26.5 | 5.6 | 8.3 |
| >MI0000079\|hsa-mir-23a&WithFlank&19\|-\|13836581\|13836665@54.73.20&[MIMAT0000078&hsa-miR-23a-3p&offsets\|+3\|+2;m-31&19\|-\|13836595\|13836615&offsets\|+3\|+2]\|ACATTGCCAGGGATTTCCAA | >MI0000079\|hsa-mir-23a&WithFlank&19\|-\|13836581\|13836665 | 54 | 73 | 20 | ACATTGCCAGGGATTTCCAA | MB231 | 145 | 150 | 201 | 35 | 72 | 141.9 | 148.8 | 223.2 | 37.3 | 90.3 |
| >MI0000079\|hsa-mir-23a&WithFlank&19\|-\|13836581\|13836665@54.74.21&[MIMAT0000078&hsa-miR-23a-3p&offsets\|+3\|+3;m-31&19\|-\|13836595\|13836615&offsets\|+3\|+3]\|ACATTGCCAGGGATTTCCAAC | >MI0000079\|hsa-mir-23a&WithFlank&19\|-\|13836581\|13836665 | 54 | 74 | 21 | ACATTGCCAGGGATTTCCAAC | BT20 | 11 | 10 | 13 | 2 | 5 | 5.9 | 6.9 | 6.8 | 1.9 | 4.7 |
| >MI0000079\|hsa-mir-23a&WithFlank&19\|-\|13836581\|13836665@54.74.21&[MIMAT0000078&hsa-miR-23a-3p&offsets\|+3\|+3;m-31&19\|-\|13836595\|13836615&offsets\|+3\|+3]\|ACATTGCCAGGGATTTCCAAC | >MI0000079\|hsa-mir-23a&WithFlank&19\|-\|13836581\|13836665 | 54 | 74 | 21 | ACATTGCCAGGGATTTCCAAC | MB231 | 21 | 14 | 31 | 7 | 27 | 18.9 | 11.8 | 31.4 | 6.9 | 32.6 |
| >MI0000081\|hsa-mir-24-2&WithFlank&19\|-\|13836281\|13836365@18.41.24&[MIMAT0004497&hsa-miR-24-2-5p&offsets\|-1\|+1;m-226&19\|-\|13836326\|13836348&offsets\|0\|+1]\|GTGCCTACTGAGCTGAAACACAGT | >MI0000081\|hsa-mir-24-2&WithFlank&19\|-\|13836281\|13836365 | 18 | 41 | 24 | GTGCCTACTGAGCTGAAACACAGT | BT20 | 13 | 11 | 4 | 2 | 5 | 8.4 | 9.0 | 2.1 | 1.6 | 1.4 |
| >MI0000081\|hsa-mir-24-2&WithFlank&19\|-\|13836281\|13836365@18.41.24&[MIMAT0004497&hsa-miR-24-2-5p&offsets\|-1\|+1;m-226&19\|-\|13836326\|13836348&offsets\|0\|+1]\|GTGCCTACTGAGCTGAAACACAGT | >MI0000081\|hsa-mir-24-2&WithFlank&19\|-\|13836281\|13836365 | 18 | 41 | 24 | GTGCCTACTGAGCTGAAACACAGT | MB231 | 23 | 19 | 4 | 18 | 22 | 15.8 | 1.1 | 2.4 | 13.4 | 14.2 |
| >MI0000081\|hsa-mir-24-2&WithFlank&19\|-\|13836281\|13836365@56.74.19;MI0000080\|hsa-mir-24-1&WithFlank&9\|+\|95086015\|95086094@50.68.19&[MIMAT0000080&hsa-miR-24-3p&offsets\|0\|-3;m-36&19\|-\|13836288\|13836310&offsets\|0\|-4];[MIMAT0000080_1&hsa-miR-24-3p&offsets\|0\|-3;m-35&9\|+\|95086064\|95086086&offsets\|0\|-4]\|TGGCTCAGTTCAGCAGGAA | >MI0000081\|hsa-mir-24-2&WithFlank&19\|-\|13836281\|13836365 | 56 | 74 | 19 | TGGCTCAGTTCAGCAGGAA | BT20 | 10 | 7 | 4 | 5 | 2 | 9.1 | 3.8 | 2.1 | 4.2 | 1.9 |
| >MI0000081\|hsa-mir-24-2&WithFlank&19\|-\|13836281\|13836365@56.74.19(+1A);MI0000080\|hsa-mir-24-1&WithFlank&9\|+\|95086015\|95086094@50.68.19(+1A)&[MIMAT0000080&hsa-miR-24-3p&offsets\|0\|-3(+1A);m-36&19\|-\|13836288\|13836310&offsets\|0\|-4(+1A)];[MIMAT0000080_1&hsa-miR-24-3p&offsets\|0\|-3(+1A);m-35&9\|+\|95086064\|95086086&offsets\|0\|-4(+1A)]\|TGGCTCAGTTCAGCAGGAAA | >MI0000081\|hsa-mir-24-2&WithFlank&19\|-\|13836281\|13836365 | 56 | 74 | 19(+1A) | TGGCTCAGTTCAGCAGGAAA | BT20 | 20 | 10 | 9 | 23 | 6 | 16.9 | 6.2 | 6.3 | 18.8 | 1.0 |
| >MI0000081\|hsa-mir-24-2&WithFlank&19\|-\|13836281\|13836365@56.74.19(+1U);MI0000080\|hsa-mir-24-1&WithFlank&9\|+\|95086015\|95086094@50.68.19(+1U)&[MIMAT0000080&hsa-miR-24-3p&offsets\|0\|-3(+1U);m-36&19\|-\|13836288\|13836310&offsets\|0\|-4(+1U)];[MIMAT0000080_1&hsa-miR-24-3p&offsets\|0\|-3(+1U);m-35&9\|+\|95086064\|95086086&offsets\|0\|-4(+1U)]\|TGGCTCAGTTCAGCAGGAAT | >MI0000081\|hsa-mir-24-2&WithFlank&19\|-\|13836281\|13836365 | 56 | 74 | 19(+1U) | TGGCTCAGTTCAGCAGGAAT | BT20 | 91 | 58 | 57 | 188 | 42 | 44.5 | 38.2 | 37.6 | 159.4 | 9.5 |
| >MI0000081\|hsa-mir-24-2&WithFlank&19\|-\|13836281\|13836365@56.74.19(+1U);MI0000080\|hsa-mir-24-1&WithFlank&9\|+\|95086015\|95086094@50.68.19(+1U)&[MIMAT0000080&hsa-miR-24-3p&offsets\|0\|-3(+1U);m-36&19\|-\|13836288\|13836310&offsets\|0\|-4(+1U)];[MIMAT0000080_1&hsa-miR-24-3p&offsets\|0\|-3(+1U);m-35&9\|+\|95086064\|95086086&offsets\|0\|-4(+1U)]\|TGGCTCAGTTCAGCAGGAAT | >MI0000081\|hsa-mir-24-2&WithFlank&19\|-\|13836281\|13836365 | 56 | 74 | 19(+1U) | TGGCTCAGTTCAGCAGGAAT | MB231 | 37 | 29 | 11 | 330 | 64 | 17.6 | 7.6 | 7.2 | 230.7 | 36.9 |
| >MI0000081\|hsa-mir-24-2&WithFlank&19\|-\|13836281\|13836365@56.75.20;MI0000080\|hsa-mir-24-1&WithFlank&9\|+\|95086015\|95086094@50.69.20&[MIMAT0000080&hsa-miR-24-3p&offsets\|0\|-2;m-36&19\|-\|13836288\|13836310&offsets\|0\|-3];[MIMAT0000080_1&hsa-miR-24-3p&offsets\|0\|-2;m-35&9\|+\|95086064\|95086086&offsets\|0\|-3]\|TGGCTCAGTTCAGCAGGAAC | >MI0000081\|hsa-mir-24-2&WithFlank&19\|-\|13836281\|13836365 | 56 | 75 | 20 | TGGCTCAGTTCAGCAGGAAC | BT20 | 710 | 557 | 503 | 1824 | 523 | 380.2 | 248.8 | 383.2 | 1320.1 | 84.8 |
| >MI0000081\|hsa-mir-24-2&WithFlank&19\|-\|13836281\|13836365@56.75.20;MI0000080\|hsa-mir-24-1&WithFlank&9\|+\|95086015\|95086094@50.69.20&[MIMAT0000080&hsa-miR-24-3p&offsets\|0\|-2;m-36&19\|-\|13836288\|13836310&offsets\|0\|-3];[MIMAT0000080_1&hsa-miR-24-3p&offsets\|0\|-2;m-35&9\|+\|95086064\|95086086&offsets\|0\|-3]\|TGGCTCAGTTCAGCAGGAAC | >MI0000081\|hsa-mir-24-2&WithFlank&19\|-\|13836281\|13836365 | 56 | 75 | 20 | TGGCTCAGTTCAGCAGGAAC | MB231 | 526 | 563 | 165 | 2646 | 786 | 212.5 | 177.7 | 109.8 | 1290.2 | 307.2 |
| >MI0000081\|hsa-mir-24-2&WithFlank&19\|-\|13836281\|13836365@56.75.20;MI0000080\|hsa-mir-24-1&WithFlank&9\|+\|95086015\|95086094@50.69.20&[MIMAT0000080&hsa-miR-24-3p&offsets\|0\|-2;m-36&19\|-\|13836288\|13836310&offsets\|0\|-3];[MIMAT0000080_1&hsa-miR-24-3p&offsets\|0\|-2;m-35&9\|+\|95086064\|95086086&offsets\|0\|-3]\|TGGCTCAGTTCAGCAGGAAC | >MI0000081\|hsa-mir-24-2&WithFlank&19\|-\|13836281\|13836365 | 56 | 75 | 20 | TGGCTCAGTTCAGCAGGAAC | MB468 | 71 | 37 | 24 | 173 | 27 | 34.6 | 13.5 | 3.9 | 55.0 | 6.0 |
| >MI0000081\|hsa-mir-24-2&WithFlank&19\|-\|13836281\|13836365@56.75.20(+1G);MI0000080\|hsa-mir-24-1&WithFlank&9\|+\|95086015\|95086094@50.69.20(+1G)&[MIMAT0000080&hsa-miR-24-3p&offsets\|0\|-2(+1G);m-36&19\|-\|13836288\|13836310&offsets\|0\|-3(+1G)];[MIMAT0000080_1&hsa-miR-24-3p&offsets\|0\|-2(+1G);m-35&9\|+\|95086064\|95086086&offsets\|0\|-3(+1G)]\|TGGCTCAGTTCAGCAGGAACG | >MI0000081\|hsa-mir-24-2&WithFlank&19\|-\|13836281\|13836365 | 56 | 75 | 20(+1G) | TGGCTCAGTTCAGCAGGAACG | BT20 | 16 | 4 | 3 | 11 | 3 | 13.5 | 1.6 | 1.3 | 10.2 | 0.6 |
| >MI0000081\|hsa-mir-24-2&WithFlank&19\|-\|13836281\|13836365@56.76.21;MI0000080\|hsa-mir-24-1&WithFlank&9\|+\|95086015\|95086094@50.70.21&[MIMAT0000080&hsa-miR-24-3p&offsets\|0\|-1;m-36&19\|-\|13836288\|13836310&offsets\|0\|-2];[MIMAT0000080_1&hsa-miR-24-3p&offsets\|0\|-1;m-35&9\|+\|95086064\|95086086&offsets\|0\|-2]\|TGGCTCAGTTCAGCAGGAACA | >MI0000081\|hsa-mir-24-2&WithFlank&19\|-\|13836281\|13836365 | 56 | 76 | 21 | TGGCTCAGTTCAGCAGGAACA | BT20 | 208 | 121 | 100 | 411 | 167 | 109.3 | 56.4 | 61.0 | 226.7 | 28.9 |
| >MI0000081\|hsa-mir-24-2&WithFlank&19\|-\|13836281\|13836365@56.76.21;MI0000080\|hsa-mir-24-1&WithFlank&9\|+\|95086015\|95086094@50.70.21&[MIMAT0000080&hsa-miR-24-3p&offsets\|0\|-1;m-36&19\|-\|13836288\|13836310&offsets\|0\|-2];[MIMAT0000080_1&hsa-miR-24-3p&offsets\|0\|-1;m-35&9\|+\|95086064\|95086086&offsets\|0\|-2]\|TGGCTCAGTTCAGCAGGAACA | >MI0000081\|hsa-mir-24-2&WithFlank&19\|-\|13836281\|13836365 | 56 | 76 | 21 | TGGCTCAGTTCAGCAGGAACA | MB231 | 154 | 126 | 30 | 559 | 179 | 108.6 | 15.8 | 17.3 | 296.3 | 43.4 |
| >MI0000081\|hsa-mir-24-2&WithFlank&19\|-\|13836281\|13836365@56.76.21;MI0000080\|hsa-mir-24-1&WithFlank&9\|+\|95086015\|95086094@50.70.21&[MIMAT0000080&hsa-miR-24-3p&offsets\|0\|-1;m-36&19\|-\|13836288\|13836310&offsets\|0\|-2];[MIMAT0000080_1&hsa-miR-24-3p&offsets\|0\|-1;m-35&9\|+\|95086064\|95086086&offsets\|0\|-2]\|TGGCTCAGTTCAGCAGGAACA | >MI0000081\|hsa-mir-24-2&WithFlank&19\|-\|13836281\|13836365 | 56 | 76 | 21 | TGGCTCAGTTCAGCAGGAACA | MB468 | 37 | 13 | 9 | 52 | 14 | 17.9 | 5.1 | 2.0 | 16.2 | 2.8 |
| >MI0000081\|hsa-mir-24-2&WithFlank&19\|-\|13836281\|13836365@56.76.21(+1A);MI0000080\|hsa-mir-24-1&WithFlank&9\|+\|95086015\|95086094@50.70.21(+1A)&[MIMAT0000080&hsa-miR-24-3p&offsets\|0\|-1(+1A);m-36&19\|-\|13836288\|13836310&offsets\|0\|-2(+1A)];[MIMAT0000080_1&hsa-miR-24-3p&offsets\|0\|-1(+1A);m-35&9\|+\|95086064\|95086086&offsets\|0\|-2(+1A)]\|TGGCTCAGTTCAGCAGGAACAA | >MI0000081\|hsa-mir-24-2&WithFlank&19\|-\|13836281\|13836365 | 56 | 76 | 21(+1A) | TGGCTCAGTTCAGCAGGAACAA | BT20 | 93 | 60 | 59 | 319 | 49 | 53.2 | 28.6 | 29.3 | 248.9 | 12.2 |
| >MI0000081\|hsa-mir-24-2&WithFlank&19\|-\|13836281\|13836365@56.76.21(+1A);MI0000080\|hsa-mir-24-1&WithFlank&9\|+\|95086015\|95086094@50.70.21(+1A)&[MIMAT0000080&hsa-miR-24-3p&offsets\|0\|-1(+1A);m-36&19\|-\|13836288\|13836310&offsets\|0\|-2(+1A)];[MIMAT0000080_1&hsa-miR-24-3p&offsets\|0\|-1(+1A);m-35&9\|+\|95086064\|95086086&offsets\|0\|-2(+1A)]\|TGGCTCAGTTCAGCAGGAACAA | >MI0000081\|hsa-mir-24-2&WithFlank&19\|-\|13836281\|13836365 | 56 | 76 | 21(+1A) | TGGCTCAGTTCAGCAGGAACAA | MB231 | 55 | 49 | 14 | 538 | 80 | 37.2 | 9.8 | 7.0 | 310.6 | 29.5 |

<table>
<tr><td>>MI0000081|hsa-mir-24-2&WithFlank&19|-|13836281|13836365@56.77.22;MI0000080|hsa-mir-24-1&WithFlank&9|+|95086015|95086094@50.71.22&[MIMAT0000080&hsa-miR-24-3p&offsets|0|0;m-36&19|-|13836288|13836310&offsets|0|-1];[MIMAT0000080_1&hsa-miR-24-3p&offsets|0|0;m-35&9|+|95086064|95086086&offsets|0|-1]|TGGCTCAGTTCAGCAGGAACAG</td><td>>MI0000081|hsa-mir-24-2&WithFlank&19|-|13836281|13836365</td><td>56</td><td>77</td><td>22</td><td>TGGCTCAGTTCAGCAGGAACAG</td><td>BT20</td><td>536</td><td>472</td><td>360</td><td>2692</td><td>460</td><td>253.8</td><td>200.9</td><td>223.9</td><td>1888.7</td><td>89.6</td></tr>
<tr><td>>MI0000081|hsa-mir-24-2&WithFlank&19|-|13836281|13836365@56.77.22;MI0000080|hsa-mir-24-1&WithFlank&9|+|95086015|95086094@50.71.22&[MIMAT0000080&hsa-miR-24-3p&offsets|0|0;m-36&19|-|13836288|13836310&offsets|0|-1];[MIMAT0000080_1&hsa-miR-24-3p&offsets|0|0;m-35&9|+|95086064|95086086&offsets|0|-1]|TGGCTCAGTTCAGCAGGAACAG</td><td>>MI0000081|hsa-mir-24-2&WithFlank&19|-|13836281|13836365</td><td>56</td><td>77</td><td>22</td><td>TGGCTCAGTTCAGCAGGAACAG</td><td>MB231</td><td>572</td><td>637</td><td>127</td><td>3527</td><td>678</td><td>385.5</td><td>96.9</td><td>42.4</td><td>1487.7</td><td>91.2</td></tr>
<tr><td>>MI0000081|hsa-mir-24-2&WithFlank&19|-|13836281|13836365@56.77.22;MI0000080|hsa-mir-24-1&WithFlank&9|+|95086015|95086094@50.71.22&[MIMAT0000080&hsa-miR-24-3p&offsets|0|0;m-36&19|-|13836288|13836310&offsets|0|-1];[MIMAT0000080_1&hsa-miR-24-3p&offsets|0|0;m-35&9|+|95086064|95086086&offsets|0|-1]|TGGCTCAGTTCAGCAGGAACAG</td><td>>MI0000081|hsa-mir-24-2&WithFlank&19|-|13836281|13836365</td><td>56</td><td>77</td><td>22</td><td>TGGCTCAGTTCAGCAGGAACAG</td><td>MB468</td><td>97</td><td>55</td><td>33</td><td>214</td><td>37</td><td>49.6</td><td>27.5</td><td>4.6</td><td>67.8</td><td>7.5</td></tr>
<tr><td>>MI0000081|hsa-mir-24-2&WithFlank&19|-|13836281|13836365@56.77.22(+1A);MI0000080|hsa-mir-24-1&WithFlank&9|+|95086015|95086094@50.71.22(+1A)&[MIMAT0000080&hsa-miR-24-3p&offsets|0|0(+1A);m-36&19|-|13836288|13836310&offsets|0|-1(+1A)];[MIMAT0000080_1&hsa-miR-24-3p&offsets|0|0(+1A);m-35&9|+|95086064|95086086&offsets|0|-1(+1A)]|TGGCTCAGTTCAGCAGGAACAGA</td><td>>MI0000081|hsa-mir-24-2&WithFlank&19|-|13836281|13836365</td><td>56</td><td>77</td><td>22(+1A)</td><td>TGGCTCAGTTCAGCAGGAACAGA</td><td>BT20</td><td>67</td><td>41</td><td>22</td><td>17</td><td>17</td><td>71.4</td><td>22.4</td><td>11.5</td><td>9.6</td><td>2.6</td></tr>
<tr><td>>MI0000081|hsa-mir-24-2&WithFlank&19|-|13836281|13836365@56.77.22(+1A);MI0000080|hsa-mir-24-1&WithFlank&9|+|95086015|95086094@50.71.22(+1A)&[MIMAT0000080&hsa-miR-24-3p&offsets|0|0(+1A);m-36&19|-|13836288|13836310&offsets|0|-1(+1A)];[MIMAT0000080_1&hsa-miR-24-3p&offsets|0|0(+1A);m-35&9|+|95086064|95086086&offsets|0|-1(+1A)]|TGGCTCAGTTCAGCAGGAACAGA</td><td>>MI0000081|hsa-mir-24-2&WithFlank&19|-|13836281|13836365</td><td>56</td><td>77</td><td>22(+1A)</td><td>TGGCTCAGTTCAGCAGGAACAGA</td><td>MB231</td><td>14</td><td>15</td><td>3</td><td>61</td><td>24</td><td>1.7</td><td>5.1</td><td>2.5</td><td>28.0</td><td>18.8</td></tr>
<tr><td>>MI0000081|hsa-mir-24-2&WithFlank&19|-|13836281|13836365@56.77.22(+1C);MI0000080|hsa-mir-24-1&WithFlank&9|+|95086015|95086094@50.71.22(+1C)&[MIMAT0000080&hsa-miR-24-3p&offsets|0|0(+1C);m-36&19|-|13836288|13836310&offsets|0|-1(+1C)];[MIMAT0000080_1&hsa-miR-24-3p&offsets|0|0(+1C);m-35&9|+|95086064|95086086&offsets|0|-1(+1C)]|TGGCTCAGTTCAGCAGGAACAGC</td><td>>MI0000081|hsa-mir-24-2&WithFlank&19|-|13836281|13836365</td><td>56</td><td>77</td><td>22(+1C)</td><td>TGGCTCAGTTCAGCAGGAACAGC</td><td>BT20</td><td>65</td><td>54</td><td>21</td><td>12</td><td>18</td><td>52.4</td><td>47.8</td><td>12.0</td><td>10.6</td><td>8.9</td></tr>
<tr><td>>MI0000081|hsa-mir-24-2&WithFlank&19|-|13836281|13836365@56.77.22(+1C);MI0000080|hsa-mir-24-1&WithFlank&9|+|95086015|95086094@50.71.22(+1C)&[MIMAT0000080&hsa-miR-24-3p&offsets|0|0(+1C);m-36&19|-|13836288|13836310&offsets|0|-1(+1C)];[MIMAT0000080_1&hsa-miR-24-3p&offsets|0|0(+1C);m-35&9|+|95086064|95086086&offsets|0|-1(+1C)]|TGGCTCAGTTCAGCAGGAACAGC</td><td>>MI0000081|hsa-mir-24-2&WithFlank&19|-|13836281|13836365</td><td>56</td><td>77</td><td>22(+1C)</td><td>TGGCTCAGTTCAGCAGGAACAGC</td><td>MB231</td><td>36</td><td>39</td><td>7</td><td>158</td><td>79</td><td>7.3</td><td>23.8</td><td>5.6</td><td>92.6</td><td>77.1</td></tr>
<tr><td>>MI0000081|hsa-mir-24-2&WithFlank&19|-|13836281|13836365@56.77.22(+1U);MI0000080|hsa-mir-24-1&WithFlank&9|+|95086015|95086094@50.71.22(+1U)&[MIMAT0000080&hsa-miR-24-3p&offsets|0|0(+1U);m-36&19|-|13836288|13836310&offsets|0|-1(+1U)];[MIMAT0000080_1&hsa-miR-24-3p&offsets|0|0(+1U);m-35&9|+|95086064|95086086&offsets|0|-1(+1U)]|TGGCTCAGTTCAGCAGGAACAGT</td><td>>MI0000081|hsa-mir-24-2&WithFlank&19|-|13836281|13836365</td><td>56</td><td>77</td><td>22(+1U)</td><td>TGGCTCAGTTCAGCAGGAACAGT</td><td>BT20</td><td>515</td><td>377</td><td>169</td><td>92</td><td>129</td><td>410.1</td><td>301.8</td><td>98.9</td><td>61.5</td><td>60.6</td></tr>
<tr><td>>MI0000081|hsa-mir-24-2&WithFlank&19|-|13836281|13836365@56.77.22(+1U);MI0000080|hsa-mir-24-1&WithFlank&9|+|95086015|95086094@50.71.22(+1U)&[MIMAT0000080&hsa-miR-24-3p&offsets|0|0(+1U);m-36&19|-|13836288|13836310&offsets|0|-1(+1U)];[MIMAT0000080_1&hsa-miR-24-3p&offsets|0|0(+1U);m-35&9|+|95086064|95086086&offsets|0|-1(+1U)]|TGGCTCAGTTCAGCAGGAACAGT</td><td>>MI0000081|hsa-mir-24-2&WithFlank&19|-|13836281|13836365</td><td>56</td><td>77</td><td>22(+1U)</td><td>TGGCTCAGTTCAGCAGGAACAGT</td><td>MB231</td><td>328</td><td>354</td><td>66</td><td>1023</td><td>574</td><td>50.8</td><td>168.4</td><td>52.8</td><td>484.0</td><td>489.8</td></tr>
<tr><td>>MI0000081|hsa-mir-24-2&WithFlank&19|-|13836281|13836365@56.77.22(+1U);MI0000080|hsa-mir-24-1&WithFlank&9|+|95086015|95086094@50.71.22(+1U)&[MIMAT0000080&hsa-miR-24-3p&offsets|0|0(+1U);m-36&19|-|13836288|13836310&offsets|0|-1(+1U)];[MIMAT0000080_1&hsa-miR-24-3p&offsets|0|0(+1U);m-35&9|+|95086064|95086086&offsets|0|-1(+1U)]|TGGCTCAGTTCAGCAGGAACAGT</td><td>>MI0000081|hsa-mir-24-2&WithFlank&19|-|13836281|13836365</td><td>56</td><td>77</td><td>22(+1U)</td><td>TGGCTCAGTTCAGCAGGAACAGT</td><td>MB468</td><td>29</td><td>13</td><td>8</td><td>89</td><td>11</td><td>10.4</td><td>2.4</td><td>2.6</td><td>87.5</td><td>3.5</td></tr>
<tr><td>>MI0000081|hsa-mir-24-2&WithFlank&19|-|13836281|13836365@56.77.22(+2U);MI0000080|hsa-mir-24-1&WithFlank&9|+|95086015|95086094@50.71.22(+2U)&[MIMAT0000080&hsa-miR-24-3p&offsets|0|0(+2U);m-36&19|-|13836288|13836310&offsets|0|-1(+2U)];[MIMAT0000080_1&hsa-miR-24-3p&offsets|0|0(+2U);m-35&9|+|95086064|95086086&offsets|0|-1(+2U)]|TGGCTCAGTTCAGCAGGAACAGTT</td><td>>MI0000081|hsa-mir-24-2&WithFlank&19|-|13836281|13836365</td><td>56</td><td>77</td><td>22(+2U)</td><td>TGGCTCAGTTCAGCAGGAACAGTT</td><td>BT20</td><td>10</td><td>6</td><td>2</td><td>0</td><td>1</td><td>9.9</td><td>5.9</td><td>1.2</td><td>0.5</td><td>1.5</td></tr>
<tr><td>>MI0000081|hsa-mir-24-2&WithFlank&19|-|13836281|13836365@56.78.23;MI0000080|hsa-mir-24-1&WithFlank&9|+|95086015|95086094@50.72.23&[MIMAT0000080&hsa-miR-24-3p&offsets|0|+1;m-36&19|-|13836288|13836310&offsets|0|0];[MIMAT0000080_1&hsa-miR-24-3p&offsets|0|+1;m-35&9|+|95086064|95086086&offsets|0|0]|TGGCTCAGTTCAGCAGGAACAGG</td><td>>MI0000081|hsa-mir-24-2&WithFlank&19|-|13836281|13836365</td><td>56</td><td>78</td><td>23</td><td>TGGCTCAGTTCAGCAGGAACAGG</td><td>BT20</td><td>45</td><td>13</td><td>5</td><td>2</td><td>2</td><td>55.9</td><td>7.4</td><td>1.5</td><td>2.6</td><td>1.6</td></tr>
<tr><td>>MI0000081|hsa-mir-24-2&WithFlank&19|-|13836281|13836365@56.78.23;MI0000080|hsa-mir-24-1&WithFlank&9|+|95086015|95086094@50.72.23&[MIMAT0000080&hsa-miR-24-3p&offsets|0|+1;m-36&19|-|13836288|13836310&offsets|0|0];[MIMAT0000080_1&hsa-miR-24-3p&offsets|0|+1;m-35&9|+|95086064|95086086&offsets|0|0]|TGGCTCAGTTCAGCAGGAACAGG</td><td>>MI0000081|hsa-mir-24-2&WithFlank&19|-|13836281|13836365</td><td>56</td><td>78</td><td>23</td><td>TGGCTCAGTTCAGCAGGAACAGG</td><td>MB231</td><td>16</td><td>8</td><td>2</td><td>17</td><td>7</td><td>12.2</td><td>4.9</td><td>1.2</td><td>8.8</td><td>6.8</td></tr>
<tr><td>>MI0000082|hsa-mir-25&WithFlank&7|-|100093554|100093649@58.77.20(+1A)&[MIMAT0000081&hsa-miR-25-3p&offsets|0|-2(+1A);m-23&7|-|100093571|100093592&offsets|0|-2(+1A)]|CATTGCACTTGTCTCGGTCTA</td><td>>MI0000082|hsa-mir-25&WithFlank&7|-|100093554|100093649</td><td>58</td><td>77</td><td>20(+1A)</td><td>CATTGCACTTGTCTCGGTCTA</td><td>BT20</td><td>44</td><td>29</td><td>49</td><td>51</td><td>27</td><td>24.8</td><td>10.7</td><td>11.7</td><td>33.2</td><td>11.3</td></tr>
<tr><td>>MI0000082|hsa-mir-25&WithFlank&7|-|100093554|100093649@58.77.20(+1A)&[MIMAT0000081&hsa-miR-25-3p&offsets|0|-2(+1A);m-23&7|-|100093571|100093592&offsets|0|-2(+1A)]|CATTGCACTTGTCTCGGTCTA</td><td>>MI0000082|hsa-mir-25&WithFlank&7|-|100093554|100093649</td><td>58</td><td>77</td><td>20(+1A)</td><td>CATTGCACTTGTCTCGGTCTA</td><td>MB231</td><td>25</td><td>29</td><td>17</td><td>94</td><td>35</td><td>7.7</td><td>3.2</td><td>10.6</td><td>41.5</td><td>5.9</td></tr>
<tr><td>>MI0000082|hsa-mir-25&WithFlank&7|-|100093554|100093649@58.77.20&[MIMAT0000081&hsa-miR-25-3p&offsets|0|-2;m-23&7|-|100093571|100093592&offsets|0|-2]|CATTGCACTTGTCTCGGTCT</td><td>>MI0000082|hsa-mir-25&WithFlank&7|-|100093554|100093649</td><td>58</td><td>77</td><td>20</td><td>CATTGCACTTGTCTCGGTCT</td><td>BT20</td><td>22</td><td>14</td><td>25</td><td>21</td><td>10</td><td>11.5</td><td>5.2</td><td>10.3</td><td>15.6</td><td>5.1</td></tr>
<tr><td>>MI0000082|hsa-mir-25&WithFlank&7|-|100093554|100093649@58.77.20&[MIMAT0000081&hsa-miR-25-3p&offsets|0|-2;m-23&7|-|100093571|100093592&offsets|0|-2]|CATTGCACTTGTCTCGGTCT</td><td>>MI0000082|hsa-mir-25&WithFlank&7|-|100093554|100093649</td><td>58</td><td>77</td><td>20</td><td>CATTGCACTTGTCTCGGTCT</td><td>MB231</td><td>13</td><td>13</td><td>9</td><td>31</td><td>18</td><td>1.6</td><td>4.1</td><td>7.3</td><td>17.9</td><td>8.0</td></tr>
<tr><td>>MI0000082|hsa-mir-25&WithFlank&7|-|100093554|100093649@58.78.21(+1C)&[MIMAT0000081&hsa-miR-25-3p&offsets|0|-1(+1C);m-23&7|-|100093571|100093592&offsets|0|-1(+1C)]|CATTGCACTTGTCTCGGTCTGC</td><td>>MI0000082|hsa-mir-25&WithFlank&7|-|100093554|100093649</td><td>58</td><td>78</td><td>21(+1C)</td><td>CATTGCACTTGTCTCGGTCTGC</td><td>BT20</td><td>24</td><td>17</td><td>26</td><td>25</td><td>13</td><td>7.2</td><td>7.9</td><td>7.6</td><td>20.0</td><td>9.0</td></tr>
<tr><td>>MI0000082|hsa-mir-25&WithFlank&7|-|100093554|100093649@58.78.21(+1C)&[MIMAT0000081&hsa-miR-25-3p&offsets|0|-1(+1C);m-23&7|-|100093571|100093592&offsets|0|-1(+1C)]|CATTGCACTTGTCTCGGTCTGC</td><td>>MI0000082|hsa-mir-25&WithFlank&7|-|100093554|100093649</td><td>58</td><td>78</td><td>21(+1C)</td><td>CATTGCACTTGTCTCGGTCTGC</td><td>MB231</td><td>13</td><td>11</td><td>7</td><td>30</td><td>15</td><td>4.9</td><td>3.8</td><td>4.5</td><td>18.6</td><td>9.6</td></tr>
<tr><td>>MI0000082|hsa-mir-25&WithFlank&7|-|100093554|100093649@58.78.21(+1G)&[MIMAT0000081&hsa-miR-25-3p&offsets|0|-1(+1G);m-23&7|-|100093571|100093592&offsets|0|-1(+1G)]|CATTGCACTTGTCTCGGTCTGG</td><td>>MI0000082|hsa-mir-25&WithFlank&7|-|100093554|100093649</td><td>58</td><td>78</td><td>21(+1G)</td><td>CATTGCACTTGTCTCGGTCTGG</td><td>BT20</td><td>121</td><td>50</td><td>81</td><td>48</td><td>20</td><td>65.6</td><td>26.0</td><td>27.5</td><td>44.0</td><td>10.0</td></tr>
<tr><td>>MI0000082|hsa-mir-25&WithFlank&7|-|100093554|100093649@58.78.21(+1G)&[MIMAT0000081&hsa-miR-25-3p&offsets|0|-1(+1G);m-23&7|-|100093571|100093592&offsets|0|-1(+1G)]|CATTGCACTTGTCTCGGTCTGG</td><td>>MI0000082|hsa-mir-25&WithFlank&7|-|100093554|100093649</td><td>58</td><td>78</td><td>21(+1G)</td><td>CATTGCACTTGTCTCGGTCTGG</td><td>MB231</td><td>78</td><td>49</td><td>17</td><td>38</td><td>19</td><td>52.0</td><td>8.1</td><td>10.7</td><td>22.1</td><td>8.4</td></tr>
<tr><td>>MI0000082|hsa-mir-25&WithFlank&7|-|100093554|100093649@58.78.21(+1U)&[MIMAT0000081&hsa-miR-25-3p&offsets|0|-1(+1U);m-23&7|-|100093571|100093592&offsets|0|-1(+1U)]|CATTGCACTTGTCTCGGTCTGT</td><td>>MI0000082|hsa-mir-25&WithFlank&7|-|100093554|100093649</td><td>58</td><td>78</td><td>21(+1U)</td><td>CATTGCACTTGTCTCGGTCTGT</td><td>BT20</td><td>191</td><td>117</td><td>212</td><td>185</td><td>97</td><td>63.2</td><td>54.6</td><td>81.8</td><td>135.2</td><td>61.6</td></tr>
<tr><td>>MI0000082|hsa-mir-25&WithFlank&7|-|100093554|100093649@58.78.21(+1U)&[MIMAT0000081&hsa-miR-25-3p&offsets|0|-1(+1U);m-23&7|-|100093571|100093592&offsets|0|-1(+1U)]|CATTGCACTTGTCTCGGTCTGT</td><td>>MI0000082|hsa-mir-25&WithFlank&7|-|100093554|100093649</td><td>58</td><td>78</td><td>21(+1U)</td><td>CATTGCACTTGTCTCGGTCTGT</td><td>MB231</td><td>114</td><td>95</td><td>62</td><td>183</td><td>111</td><td>34.5</td><td>21.1</td><td>34.8</td><td>89.8</td><td>62.6</td></tr>
<tr><td>>MI0000082|hsa-mir-25&WithFlank&7|-|100093554|100093649@58.78.21&[MIMAT0000081&hsa-miR-25-3p&offsets|0|-1;m-23&7|-|100093571|100093592&offsets|0|-1]|CATTGCACTTGTCTCGGTCTG</td><td>>MI0000082|hsa-mir-25&WithFlank&7|-|100093554|100093649</td><td>58</td><td>78</td><td>21</td><td>CATTGCACTTGTCTCGGTCTG</td><td>BT20</td><td>237</td><td>205</td><td>278</td><td>413</td><td>216</td><td>110.8</td><td>53.6</td><td>114.9</td><td>266.5</td><td>73.6</td></tr>
</table>

<table>
<tr><td>>MI0000082|hsa-mir-25&WithFlank&7|-|100093554|100093649@58.78.21&[MIMAT0000081&hsa-miR-25-3p&offsets|0|-1;m-23&7|-|100093571|100093592&offsets|0|-1]|CATTGCACTTGTCTCGGTCTG</td><td>>MI0000082|hsa-mir-25&WithFlank&7|-|100093554|100093649</td><td>58</td><td>78</td><td>21</td><td>CATTGCACTTGTCTCGGTCTG</td><td>MB231</td><td>229</td><td>310</td><td>152</td><td>570</td><td>269</td><td>80.2</td><td>26.7</td><td>60.6</td><td>136.3</td><td>14.2</td></tr>
<tr><td>>MI0000082|hsa-mir-25&WithFlank&7|-|100093554|100093649@58.78.21&[MIMAT0000081&hsa-miR-25-3p&offsets|0|-1;m-23&7|-|100093571|100093592&offsets|0|-1]|CATTGCACTTGTCTCGGTCTG</td><td>>MI0000082|hsa-mir-25&WithFlank&7|-|100093554|100093649</td><td>58</td><td>78</td><td>21</td><td>CATTGCACTTGTCTCGGTCTG</td><td>MB468</td><td>14</td><td>26</td><td>9</td><td>25</td><td>7</td><td>3.8</td><td>7.1</td><td>5.1</td><td>6.7</td><td>2.2</td></tr>
<tr><td>>MI0000082|hsa-mir-25&WithFlank&7|-|100093554|100093649@58.79.22(+1A)&[MIMAT0000081&hsa-miR-25-3p&offsets|0|0(+1A);m-23&7|-|100093571|100093592&offsets|0|0(+1A)]|CATTGCACTTGTCTCGGTCTGAA</td><td>>MI0000082|hsa-mir-25&WithFlank&7|-|100093554|100093649</td><td>58</td><td>79</td><td>22(+1A)</td><td>CATTGCACTTGTCTCGGTCTGAA</td><td>BT20</td><td>70</td><td>46</td><td>123</td><td>24</td><td>33</td><td>43.6</td><td>19.2</td><td>42.4</td><td>8.8</td><td>11.6</td></tr>
<tr><td>>MI0000082|hsa-mir-25&WithFlank&7|-|100093554|100093649@58.79.22(+1A)&[MIMAT0000081&hsa-miR-25-3p&offsets|0|0(+1A);m-23&7|-|100093571|100093592&offsets|0|0(+1A)]|CATTGCACTTGTCTCGGTCTGAA</td><td>>MI0000082|hsa-mir-25&WithFlank&7|-|100093554|100093649</td><td>58</td><td>79</td><td>22(+1A)</td><td>CATTGCACTTGTCTCGGTCTGAA</td><td>MB231</td><td>37</td><td>66</td><td>25</td><td>28</td><td>23</td><td>9.8</td><td>18.6</td><td>9.6</td><td>14.0</td><td>9.0</td></tr>
<tr><td>>MI0000082|hsa-mir-25&WithFlank&7|-|100093554|100093649@58.79.22(+1U)&[MIMAT0000081&hsa-miR-25-3p&offsets|0|0(+1U);m-23&7|-|100093571|100093592&offsets|0|0(+1U)]|CATTGCACTTGTCTCGGTCTGAT</td><td>>MI0000082|hsa-mir-25&WithFlank&7|-|100093554|100093649</td><td>58</td><td>79</td><td>22(+1U)</td><td>CATTGCACTTGTCTCGGTCTGAT</td><td>BT20</td><td>47</td><td>28</td><td>70</td><td>9</td><td>21</td><td>23.5</td><td>15.0</td><td>27.3</td><td>1.9</td><td>9.9</td></tr>
<tr><td>>MI0000082|hsa-mir-25&WithFlank&7|-|100093554|100093649@58.79.22(+1U)&[MIMAT0000081&hsa-miR-25-3p&offsets|0|0(+1U);m-23&7|-|100093571|100093592&offsets|0|0(+1U)]|CATTGCACTTGTCTCGGTCTGAT</td><td>>MI0000082|hsa-mir-25&WithFlank&7|-|100093554|100093649</td><td>58</td><td>79</td><td>22(+1U)</td><td>CATTGCACTTGTCTCGGTCTGAT</td><td>MB231</td><td>72</td><td>54</td><td>48</td><td>42</td><td>50</td><td>11.2</td><td>19.1</td><td>28.6</td><td>21.2</td><td>25.9</td></tr>
<tr><td>>MI0000082|hsa-mir-25&WithFlank&7|-|100093554|100093649@58.79.22(+2A)&[MIMAT0000081&hsa-miR-25-3p&offsets|0|0(+2A);m-23&7|-|100093571|100093592&offsets|0|0(+2A)]|CATTGCACTTGTCTCGGTCTGAAA</td><td>>MI0000082|hsa-mir-25&WithFlank&7|-|100093554|100093649</td><td>58</td><td>79</td><td>22(+2A)</td><td>CATTGCACTTGTCTCGGTCTGAAA</td><td>BT20</td><td>31</td><td>17</td><td>40</td><td>7</td><td>20</td><td>30.9</td><td>5.5</td><td>13.8</td><td>7.7</td><td>1.1</td></tr>
<tr><td>>MI0000082|hsa-mir-25&WithFlank&7|-|100093554|100093649@58.79.22(+2A)&[MIMAT0000081&hsa-miR-25-3p&offsets|0|0(+2A);m-23&7|-|100093571|100093592&offsets|0|0(+2A)]|CATTGCACTTGTCTCGGTCTGAAA</td><td>>MI0000082|hsa-mir-25&WithFlank&7|-|100093554|100093649</td><td>58</td><td>79</td><td>22(+2A)</td><td>CATTGCACTTGTCTCGGTCTGAAA</td><td>MB231</td><td>16</td><td>18</td><td>11</td><td>5</td><td>13</td><td>9.5</td><td>3.4</td><td>2.8</td><td>3.0</td><td>1.0</td></tr>
<tr><td>>MI0000082|hsa-mir-25&WithFlank&7|-|100093554|100093649@58.79.22&[MIMAT0000081&hsa-miR-25-3p&offsets|0|0;m-23&7|-|100093571|100093592&offsets|0|0]|CATTGCACTTGTCTCGGTCTGA</td><td>>MI0000082|hsa-mir-25&WithFlank&7|-|100093554|100093649</td><td>58</td><td>79</td><td>22</td><td>CATTGCACTTGTCTCGGTCTGA</td><td>BT20</td><td>2055</td><td>1657</td><td>3145</td><td>2472</td><td>1588</td><td>1222.7</td><td>715.6</td><td>1361.7</td><td>1682.4</td><td>866.6</td></tr>
<tr><td>>MI0000082|hsa-mir-25&WithFlank&7|-|100093554|100093649@58.79.22&[MIMAT0000081&hsa-miR-25-3p&offsets|0|0;m-23&7|-|100093571|100093592&offsets|0|0]|CATTGCACTTGTCTCGGTCTGA</td><td>>MI0000082|hsa-mir-25&WithFlank&7|-|100093554|100093649</td><td>58</td><td>79</td><td>22</td><td>CATTGCACTTGTCTCGGTCTGA</td><td>MB231</td><td>1657</td><td>2719</td><td>1021</td><td>2132</td><td>1444</td><td>411.1</td><td>540.2</td><td>414.9</td><td>851.0</td><td>329.6</td></tr>
<tr><td>>MI0000082|hsa-mir-25&WithFlank&7|-|100093554|100093649@58.79.22&[MIMAT0000081&hsa-miR-25-3p&offsets|0|0;m-23&7|-|100093571|100093592&offsets|0|0]|CATTGCACTTGTCTCGGTCTGA</td><td>>MI0000082|hsa-mir-25&WithFlank&7|-|100093554|100093649</td><td>58</td><td>79</td><td>22</td><td>CATTGCACTTGTCTCGGTCTGA</td><td>MB468</td><td>109</td><td>207</td><td>52</td><td>124</td><td>36</td><td>28.7</td><td>72.9</td><td>19.4</td><td>30.4</td><td>9.9</td></tr>
<tr><td>>MI0000084|hsa-mir-26b&WithFlank&2|+|218402640|218402728@18.37.20(+1C)&[MIMAT0000083&hsa-miR-26b-5p&offsets|0|-1(+1C);m-53&2|+|218402657|218402678&offsets|0|-2(+1C)]|TTCAAGTAATTCAGGATAGGC</td><td>>MI0000084|hsa-mir-26b&WithFlank&2|+|218402640|218402728</td><td>18</td><td>37</td><td>20(+1C)</td><td>TTCAAGTAATTCAGGATAGGC</td><td>BT20</td><td>14</td><td>9</td><td>9</td><td>5</td><td>1</td><td>13.1</td><td>3.8</td><td>6.1</td><td>5.9</td><td>0.5</td></tr>
<tr><td>>MI0000084|hsa-mir-26b&WithFlank&2|+|218402640|218402728@18.37.20&[MIMAT0000083&hsa-miR-26b-5p&offsets|0|-1;m-53&2|+|218402657|218402678&offsets|0|-2]|TTCAAGTAATTCAGGATAGG</td><td>>MI0000084|hsa-mir-26b&WithFlank&2|+|218402640|218402728</td><td>18</td><td>37</td><td>20</td><td>TTCAAGTAATTCAGGATAGG</td><td>BT20</td><td>11</td><td>4</td><td>8</td><td>2</td><td>1</td><td>10.3</td><td>2.1</td><td>6.7</td><td>2.7</td><td>0.5</td></tr>
<tr><td>>MI0000084|hsa-mir-26b&WithFlank&2|+|218402640|218402728@18.38.21(+1A)&[MIMAT0000083&hsa-miR-26b-5p&offsets|0|0(+1A);m-53&2|+|218402657|218402678&offsets|0|-1(+1A)]|TTCAAGTAATTCAGGATAGGTA</td><td>>MI0000084|hsa-mir-26b&WithFlank&2|+|218402640|218402728</td><td>18</td><td>38</td><td>21(+1A)</td><td>TTCAAGTAATTCAGGATAGGTA</td><td>BT20</td><td>56</td><td>15</td><td>24</td><td>3</td><td>2</td><td>75.9</td><td>5.8</td><td>21.3</td><td>2.8</td><td>1.5</td></tr>
<tr><td>>MI0000084|hsa-mir-26b&WithFlank&2|+|218402640|218402728@18.38.21(+1C)&[MIMAT0000083&hsa-miR-26b-5p&offsets|0|0(+1C);m-53&2|+|218402657|218402678&offsets|0|-1(+1C)]|TTCAAGTAATTCAGGATAGGTC</td><td>>MI0000084|hsa-mir-26b&WithFlank&2|+|218402640|218402728</td><td>18</td><td>38</td><td>21(+1C)</td><td>TTCAAGTAATTCAGGATAGGTC</td><td>BT20</td><td>96</td><td>60</td><td>53</td><td>11</td><td>9</td><td>84.1</td><td>27.9</td><td>31.9</td><td>12.4</td><td>2.2</td></tr>
<tr><td>>MI0000084|hsa-mir-26b&WithFlank&2|+|218402640|218402728@18.38.21(+1C)&[MIMAT0000083&hsa-miR-26b-5p&offsets|0|0(+1C);m-53&2|+|218402657|218402678&offsets|0|-1(+1C)]|TTCAAGTAATTCAGGATAGGTC</td><td>>MI0000084|hsa-mir-26b&WithFlank&2|+|218402640|218402728</td><td>18</td><td>38</td><td>21(+1C)</td><td>TTCAAGTAATTCAGGATAGGTC</td><td>MB231</td><td>30</td><td>27</td><td>8</td><td>87</td><td>48</td><td>5.1</td><td>14.7</td><td>5.8</td><td>57.1</td><td>40.3</td></tr>
<tr><td>>MI0000084|hsa-mir-26b&WithFlank&2|+|218402640|218402728@18.38.21(+1G)&[MIMAT0000083&hsa-miR-26b-5p&offsets|0|0(+1G);m-53&2|+|218402657|218402678&offsets|0|-1(+1G)]|TTCAAGTAATTCAGGATAGGTG</td><td>>MI0000084|hsa-mir-26b&WithFlank&2|+|218402640|218402728</td><td>18</td><td>38</td><td>21(+1G)</td><td>TTCAAGTAATTCAGGATAGGTG</td><td>BT20</td><td>49</td><td>7</td><td>8</td><td>1</td><td>0</td><td>67.1</td><td>3.7</td><td>6.4</td><td>1.5</td><td>0.3</td></tr>
<tr><td>>MI0000084|hsa-mir-26b&WithFlank&2|+|218402640|218402728@18.38.21(+1G)&[MIMAT0000083&hsa-miR-26b-5p&offsets|0|0(+1G);m-53&2|+|218402657|218402678&offsets|0|-1(+1G)]|TTCAAGTAATTCAGGATAGGTG</td><td>>MI0000084|hsa-mir-26b&WithFlank&2|+|218402640|218402728</td><td>18</td><td>38</td><td>21(+1G)</td><td>TTCAAGTAATTCAGGATAGGTG</td><td>MB231</td><td>11</td><td>3</td><td>1</td><td>6</td><td>3</td><td>10.0</td><td>2.3</td><td>0.7</td><td>4.3</td><td>2.3</td></tr>
<tr><td>>MI0000084|hsa-mir-26b&WithFlank&2|+|218402640|218402728@18.38.21&[MIMAT0000083&hsa-miR-26b-5p&offsets|0|0;m-53&2|+|218402657|218402678&offsets|0|-1]|TTCAAGTAATTCAGGATAGGT</td><td>>MI0000084|hsa-mir-26b&WithFlank&2|+|218402640|218402728</td><td>18</td><td>38</td><td>21</td><td>TTCAAGTAATTCAGGATAGGT</td><td>BT20</td><td>110</td><td>55</td><td>61</td><td>31</td><td>10</td><td>103.9</td><td>15.0</td><td>40.5</td><td>34.2</td><td>2.7</td></tr>
<tr><td>>MI0000084|hsa-mir-26b&WithFlank&2|+|218402640|218402728@18.38.21&[MIMAT0000083&hsa-miR-26b-5p&offsets|0|0;m-53&2|+|218402657|218402678&offsets|0|-1]|TTCAAGTAATTCAGGATAGGT</td><td>>MI0000084|hsa-mir-26b&WithFlank&2|+|218402640|218402728</td><td>18</td><td>38</td><td>21</td><td>TTCAAGTAATTCAGGATAGGT</td><td>MB231</td><td>35</td><td>36</td><td>7</td><td>149</td><td>41</td><td>10.2</td><td>20.2</td><td>5.0</td><td>90.3</td><td>26.2</td></tr>
<tr><td>>MI0000084|hsa-mir-26b&WithFlank&2|+|218402640|218402728@18.39.22(+1A)&[MIMAT0000083&hsa-miR-26b-5p&offsets|0|+1(+1A);m-53&2|+|218402657|218402678&offsets|0|0(+1A)]|TTCAAGTAATTCAGGATAGGTTA</td><td>>MI0000084|hsa-mir-26b&WithFlank&2|+|218402640|218402728</td><td>18</td><td>39</td><td>22(+1A)</td><td>TTCAAGTAATTCAGGATAGGTTA</td><td>BT20</td><td>20</td><td>11</td><td>5</td><td>6</td><td>2</td><td>21.1</td><td>6.7</td><td>2.8</td><td>6.7</td><td>1.6</td></tr>
<tr><td>>MI0000084|hsa-mir-26b&WithFlank&2|+|218402640|218402728@18.39.22(+1U)&[MIMAT0000083&hsa-miR-26b-5p&offsets|0|+1(+1U);m-53&2|+|218402657|218402678&offsets|0|0(+1U)]|TTCAAGTAATTCAGGATAGGTTT</td><td>>MI0000084|hsa-mir-26b&WithFlank&2|+|218402640|218402728</td><td>18</td><td>39</td><td>22(+1U)</td><td>TTCAAGTAATTCAGGATAGGTTT</td><td>BT20</td><td>27</td><td>15</td><td>5</td><td>12</td><td>4</td><td>24.5</td><td>6.1</td><td>2.4</td><td>11.4</td><td>2.0</td></tr>
<tr><td>>MI0000084|hsa-mir-26b&WithFlank&2|+|218402640|218402728@18.39.22(+1U)&[MIMAT0000083&hsa-miR-26b-5p&offsets|0|+1(+1U);m-53&2|+|218402657|218402678&offsets|0|0(+1U)]|TTCAAGTAATTCAGGATAGGTTT</td><td>>MI0000084|hsa-mir-26b&WithFlank&2|+|218402640|218402728</td><td>18</td><td>39</td><td>22(+1U)</td><td>TTCAAGTAATTCAGGATAGGTTT</td><td>MB231</td><td>18</td><td>14</td><td>1</td><td>94</td><td>41</td><td>3.0</td><td>7.6</td><td>0.9</td><td>39.8</td><td>31.1</td></tr>
<tr><td>>MI0000084|hsa-mir-26b&WithFlank&2|+|218402640|218402728@18.39.22&[MIMAT0000083&hsa-miR-26b-5p&offsets|0|+1;m-53&2|+|218402657|218402678&offsets|0|0]|TTCAAGTAATTCAGGATAGGTT</td><td>>MI0000084|hsa-mir-26b&WithFlank&2|+|218402640|218402728</td><td>18</td><td>39</td><td>22</td><td>TTCAAGTAATTCAGGATAGGTT</td><td>BT20</td><td>703</td><td>435</td><td>387</td><td>66</td><td>73</td><td>626.5</td><td>179.9</td><td>203.3</td><td>64.6</td><td>18.8</td></tr>
<tr><td>>MI0000084|hsa-mir-26b&WithFlank&2|+|218402640|218402728@18.39.22&[MIMAT0000083&hsa-miR-26b-5p&offsets|0|+1;m-53&2|+|218402657|218402678&offsets|0|0]|TTCAAGTAATTCAGGATAGGTT</td><td>>MI0000084|hsa-mir-26b&WithFlank&2|+|218402640|218402728</td><td>18</td><td>39</td><td>22</td><td>TTCAAGTAATTCAGGATAGGTT</td><td>MB231</td><td>291</td><td>259</td><td>75</td><td>517</td><td>355</td><td>59.4</td><td>118.4</td><td>50.1</td><td>270.1</td><td>257.1</td></tr>
<tr><td>>MI0000084|hsa-mir-26b&WithFlank&2|+|218402640|218402728@18.39.22&[MIMAT0000083&hsa-miR-26b-5p&offsets|0|+1;m-53&2|+|218402657|218402678&offsets|0|0]|TTCAAGTAATTCAGGATAGGTT</td><td>>MI0000084|hsa-mir-26b&WithFlank&2|+|218402640|218402728</td><td>18</td><td>39</td><td>22</td><td>TTCAAGTAATTCAGGATAGGTT</td><td>MB468</td><td>40</td><td>17</td><td>18</td><td>45</td><td>13</td><td>17.0</td><td>6.6</td><td>4.9</td><td>38.7</td><td>2.1</td></tr>
<tr><td>>MI0000084|hsa-mir-26b&WithFlank&2|+|218402640|218402728@21.39.19&[MIMAT0000083&hsa-miR-26b-5p&offsets|+3|+1;m-53&2|+|218402657|218402678&offsets|+3|0]|AAGTAATTCAGGATAGGTT</td><td>>MI0000084|hsa-mir-26b&WithFlank&2|+|218402640|218402728</td><td>21</td><td>39</td><td>19</td><td>AAGTAATTCAGGATAGGTT</td><td>BT20</td><td>10</td><td>7</td><td>7</td><td>1</td><td>1</td><td>8.4</td><td>3.8</td><td>3.8</td><td>1.2</td><td>0.9</td></tr>
<tr><td>>MI0000085|hsa-mir-27a&WithFlank&19|-|13836434|13836523@16.36.21&[MIMAT0004501&hsa-miR-27a-5p&offsets|0|-1;m-270&19|-|13836487|13836508&offsets|0|-1]|AGGGCTTAGCTGCTTGTGAGC</td><td>>MI0000085|hsa-mir-27a&WithFlank&19|-|13836434|13836523</td><td>16</td><td>36</td><td>21</td><td>AGGGCTTAGCTGCTTGTGAGC</td><td>MB231</td><td>13</td><td>23</td><td>44</td><td>28</td><td>8</td><td>6.3</td><td>14.9</td><td>20.8</td><td>11.1</td><td>1.2</td></tr>
<tr><td>>MI0000085|hsa-mir-27a&WithFlank&19|-|13836434|13836523@16.37.22&[MIMAT0004501&hsa-miR-27a-5p&offsets|0|0;m-270&19|-|13836487|13836508&offsets|0|0]|AGGGCTTAGCTGCTTGTGAGCA</td><td>>MI0000085|hsa-mir-27a&WithFlank&19|-|13836434|13836523</td><td>16</td><td>37</td><td>22</td><td>AGGGCTTAGCTGCTTGTGAGCA</td><td>BT20</td><td>20</td><td>19</td><td>61</td><td>4</td><td>5</td><td>19.9</td><td>15.9</td><td>40.7</td><td>2.0</td><td>2.0</td></tr>
<tr><td>>MI0000085|hsa-mir-27a&WithFlank&19|-|13836434|13836523@16.37.22&[MIMAT0004501&hsa-miR-27a-5p&offsets|0|0;m-270&19|-|13836487|13836508&offsets|0|0]|AGGGCTTAGCTGCTTGTGAGCA</td><td>>MI0000085|hsa-mir-27a&WithFlank&19|-|13836434|13836523</td><td>16</td><td>37</td><td>22</td><td>AGGGCTTAGCTGCTTGTGAGCA</td><td>MB231</td><td>19</td><td>22</td><td>62</td><td>12</td><td>8</td><td>11.9</td><td>12.3</td><td>31.2</td><td>6.1</td><td>1.1</td></tr>
<tr><td>>MI0000085|hsa-mir-27a&WithFlank&19|-|13836434|13836523@57.74.18;MI0000440|hsa-mir-27b&WithFlank&9|+|95085439|95085547@67.84.18&[MIMAT0000084&hsa-miR-27a-3p&offsets|0|-3;m-43&19|-|13836447|13836467&offsets|0|-3];[MIMAT0000419&hsa-miR-27b-3p&offsets|0|-3;m-39&9|+|95085505|95085525&offsets|0|-3]|TTCACAGTGGCTAAGTTC</td><td>>MI0000085|hsa-mir-27a&WithFlank&19|-|13836434|13836523</td><td>57</td><td>74</td><td>18</td><td>TTCACAGTGGCTAAGTTC</td><td>BT20</td><td>19</td><td>17</td><td>22</td><td>34</td><td>7</td><td>13.6</td><td>10.8</td><td>16.8</td><td>26.9</td><td>2.9</td></tr>
</table>

<table>
<tr><th></th><th></th><th></th><th></th><th></th><th></th><th></th><th></th><th></th><th></th><th></th><th></th><th></th><th></th><th></th><th></th><th></th></tr>
<tr><td>>MI0000085|hsa-mir-27a&WithFlank&19|-|13836434|13836523@57.74.18;MI0000440|hsa-mir-27b&WithFlank&9|+|95085439|95085547@67.84.18&[MIMAT0000084&hsa-miR-27a-3p&offsets|0|-3;m-43&19|-|13836447|13836467&offsets|0|-3];[MIMAT0000419&hsa-miR-27b-3p&offsets|0|-3;m-39&9|+|95085505|95085525&offsets|0|-3]|TTCACAGTGGCTAAGTTC</td><td>>MI0000085|hsa-mir-27a&WithFlank&19|-|13836434|13836523</td><td>57</td><td>74</td><td>18</td><td>TTCACAGTGGCTAAGTTC</td><td>MB231</td><td>26</td><td>34</td><td>13</td><td>141</td><td>45</td><td>2.1</td><td>17.9</td><td>10.1</td><td>47.3</td><td>24.0</td></tr>
<tr><td>>MI0000085|hsa-mir-27a&WithFlank&19|-|13836434|13836523@57.74.18(+1A);MI0000440|hsa-mir-27b&WithFlank&9|+|95085439|95085547@67.84.18(+1A)&[MIMAT0000084&hsa-miR-27a-3p&offsets|0|-3(+1A);m-43&19|-|13836447|13836467&offsets|0|-3(+1A)];[MIMAT0000419&hsa-miR-27b-3p&offsets|0|-3(+1A);m-39&9|+|95085505|95085525&offsets|0|-3(+1A)]|TTCACAGTGGCTAAGTTCA</td><td>>MI0000085|hsa-mir-27a&WithFlank&19|-|13836434|13836523</td><td>57</td><td>74</td><td>18(+1A)</td><td>TTCACAGTGGCTAAGTTCA</td><td>BT20</td><td>15</td><td>6</td><td>12</td><td>10</td><td>3</td><td>10.4</td><td>2.5</td><td>8.2</td><td>8.6</td><td>0.2</td></tr>
<tr><td>>MI0000085|hsa-mir-27a&WithFlank&19|-|13836434|13836523@57.75.19(+1A)&[MIMAT0000084&hsa-miR-27a-3p&offsets|0|-2(+1A);m-43&19|-|13836447|13836467&offsets|0|-2(+1A)]|TTCACAGTGGCTAAGTTCCA</td><td>>MI0000085|hsa-mir-27a&WithFlank&19|-|13836434|13836523</td><td>57</td><td>75</td><td>19(+1A)</td><td>TTCACAGTGGCTAAGTTCCA</td><td>BT20</td><td>91</td><td>45</td><td>50</td><td>66</td><td>22</td><td>57.2</td><td>37.9</td><td>34.7</td><td>51.1</td><td>16.4</td></tr>
<tr><td>>MI0000085|hsa-mir-27a&WithFlank&19|-|13836434|13836523@57.75.19(+1A)&[MIMAT0000084&hsa-miR-27a-3p&offsets|0|-2(+1A);m-43&19|-|13836447|13836467&offsets|0|-2(+1A)]|TTCACAGTGGCTAAGTTCCA</td><td>>MI0000085|hsa-mir-27a&WithFlank&19|-|13836434|13836523</td><td>57</td><td>75</td><td>19(+1A)</td><td>TTCACAGTGGCTAAGTTCCA</td><td>MB231</td><td>110</td><td>109</td><td>29</td><td>394</td><td>183</td><td>20.2</td><td>81.2</td><td>23.5</td><td>308.6</td><td>103.1</td></tr>
<tr><td>>MI0000085|hsa-mir-27a&WithFlank&19|-|13836434|13836523@57.75.19(+1A)&[MIMAT0000084&hsa-miR-27a-3p&offsets|0|-2(+1A);m-43&19|-|13836447|13836467&offsets|0|-2(+1A)]|TTCACAGTGGCTAAGTTCCA</td><td>>MI0000085|hsa-mir-27a&WithFlank&19|-|13836434|13836523</td><td>57</td><td>75</td><td>19(+1A)</td><td>TTCACAGTGGCTAAGTTCCA</td><td>MB468</td><td>13</td><td>3</td><td>3</td><td>18</td><td>6</td><td>8.1</td><td>1.2</td><td>1.3</td><td>2.0</td><td>1.7</td></tr>
<tr><td>>MI0000085|hsa-mir-27a&WithFlank&19|-|13836434|13836523@57.75.19(+1U)&[MIMAT0000084&hsa-miR-27a-3p&offsets|0|-2(+1U);m-43&19|-|13836447|13836467&offsets|0|-2(+1U)]|TTCACAGTGGCTAAGTTCCT</td><td>>MI0000085|hsa-mir-27a&WithFlank&19|-|13836434|13836523</td><td>57</td><td>75</td><td>19(+1U)</td><td>TTCACAGTGGCTAAGTTCCT</td><td>BT20</td><td>15</td><td>6</td><td>6</td><td>10</td><td>3</td><td>9.7</td><td>5.1</td><td>4.4</td><td>7.8</td><td>1.4</td></tr>
<tr><td>>MI0000085|hsa-mir-27a&WithFlank&19|-|13836434|13836523@57.75.19(+1U)&[MIMAT0000084&hsa-miR-27a-3p&offsets|0|-2(+1U);m-43&19|-|13836447|13836467&offsets|0|-2(+1U)]|TTCACAGTGGCTAAGTTCCT</td><td>>MI0000085|hsa-mir-27a&WithFlank&19|-|13836434|13836523</td><td>57</td><td>75</td><td>19(+1U)</td><td>TTCACAGTGGCTAAGTTCCT</td><td>MB231</td><td>15</td><td>14</td><td>3</td><td>56</td><td>27</td><td>0.7</td><td>11.7</td><td>2.6</td><td>44.9</td><td>19.8</td></tr>
<tr><td>>MI0000085|hsa-mir-27a&WithFlank&19|-|13836434|13836523@57.75.19(+3U)&[MIMAT0000084&hsa-miR-27a-3p&offsets|0|-2(+3U);m-43&19|-|13836447|13836467&offsets|0|-2(+3U)]|TTCACAGTGGCTAAGTTCCTTT</td><td>>MI0000085|hsa-mir-27a&WithFlank&19|-|13836434|13836523</td><td>57</td><td>75</td><td>19(+3U)</td><td>TTCACAGTGGCTAAGTTCCTTT</td><td>MB231</td><td>13</td><td>8</td><td>2</td><td>19</td><td>12</td><td>7.4</td><td>4.4</td><td>1.4</td><td>9.8</td><td>7.3</td></tr>
<tr><td>>MI0000085|hsa-mir-27a&WithFlank&19|-|13836434|13836523@57.75.19&[MIMAT0000084&hsa-miR-27a-3p&offsets|0|-2;m-43&19|-|13836447|13836467&offsets|0|-2]|TTCACAGTGGCTAAGTTCC</td><td>>MI0000085|hsa-mir-27a&WithFlank&19|-|13836434|13836523</td><td>57</td><td>75</td><td>19</td><td>TTCACAGTGGCTAAGTTCC</td><td>BT20</td><td>29</td><td>19</td><td>23</td><td>33</td><td>9</td><td>15.6</td><td>10.0</td><td>16.0</td><td>23.7</td><td>3.2</td></tr>
<tr><td>>MI0000085|hsa-mir-27a&WithFlank&19|-|13836434|13836523@57.75.19&[MIMAT0000084&hsa-miR-27a-3p&offsets|0|-2;m-43&19|-|13836447|13836467&offsets|0|-2]|TTCACAGTGGCTAAGTTCC</td><td>>MI0000085|hsa-mir-27a&WithFlank&19|-|13836434|13836523</td><td>57</td><td>75</td><td>19</td><td>TTCACAGTGGCTAAGTTCC</td><td>MB231</td><td>30</td><td>37</td><td>11</td><td>109</td><td>41</td><td>5.9</td><td>18.8</td><td>8.3</td><td>62.3</td><td>18.5</td></tr>
<tr><td>>MI0000085|hsa-mir-27a&WithFlank&19|-|13836434|13836523@57.76.20(+1A)&[MIMAT0000084&hsa-miR-27a-3p&offsets|0|-1(+1A);m-43&19|-|13836447|13836467&offsets|0|-1(+1A)]|TTCACAGTGGCTAAGTTCCGA</td><td>>MI0000085|hsa-mir-27a&WithFlank&19|-|13836434|13836523</td><td>57</td><td>76</td><td>20(+1A)</td><td>TTCACAGTGGCTAAGTTCCGA</td><td>BT20</td><td>127</td><td>67</td><td>73</td><td>125</td><td>37</td><td>98.0</td><td>59.3</td><td>69.9</td><td>92.4</td><td>21.9</td></tr>
<tr><td>>MI0000085|hsa-mir-27a&WithFlank&19|-|13836434|13836523@57.76.20(+1A)&[MIMAT0000084&hsa-miR-27a-3p&offsets|0|-1(+1A);m-43&19|-|13836447|13836467&offsets|0|-1(+1A)]|TTCACAGTGGCTAAGTTCCGA</td><td>>MI0000085|hsa-mir-27a&WithFlank&19|-|13836434|13836523</td><td>57</td><td>76</td><td>20(+1A)</td><td>TTCACAGTGGCTAAGTTCCGA</td><td>MB231</td><td>145</td><td>145</td><td>34</td><td>482</td><td>182</td><td>78.2</td><td>71.7</td><td>29.4</td><td>286.5</td><td>68.7</td></tr>
<tr><td>>MI0000085|hsa-mir-27a&WithFlank&19|-|13836434|13836523@57.76.20(+1A)&[MIMAT0000084&hsa-miR-27a-3p&offsets|0|-1(+1A);m-43&19|-|13836447|13836467&offsets|0|-1(+1A)]|TTCACAGTGGCTAAGTTCCGA</td><td>>MI0000085|hsa-mir-27a&WithFlank&19|-|13836434|13836523</td><td>57</td><td>76</td><td>20(+1A)</td><td>TTCACAGTGGCTAAGTTCCGA</td><td>MB468</td><td>26</td><td>6</td><td>5</td><td>41</td><td>10</td><td>17.0</td><td>2.5</td><td>1.1</td><td>4.4</td><td>3.3</td></tr>
<tr><td>>MI0000085|hsa-mir-27a&WithFlank&19|-|13836434|13836523@57.76.20(+1G)&[MIMAT0000084&hsa-miR-27a-3p&offsets|0|-1(+1G);m-43&19|-|13836447|13836467&offsets|0|-1(+1G)]|TTCACAGTGGCTAAGTTCCGG</td><td>>MI0000085|hsa-mir-27a&WithFlank&19|-|13836434|13836523</td><td>57</td><td>76</td><td>20(+1G)</td><td>TTCACAGTGGCTAAGTTCCGG</td><td>BT20</td><td>16</td><td>5</td><td>3</td><td>5</td><td>1</td><td>14.8</td><td>4.4</td><td>1.4</td><td>5.7</td><td>0.5</td></tr>
<tr><td>>MI0000085|hsa-mir-27a&WithFlank&19|-|13836434|13836523@57.76.20(+1G)&[MIMAT0000084&hsa-miR-27a-3p&offsets|0|-1(+1G);m-43&19|-|13836447|13836467&offsets|0|-1(+1G)]|TTCACAGTGGCTAAGTTCCGG</td><td>>MI0000085|hsa-mir-27a&WithFlank&19|-|13836434|13836523</td><td>57</td><td>76</td><td>20(+1G)</td><td>TTCACAGTGGCTAAGTTCCGG</td><td>MB231</td><td>17</td><td>10</td><td>2</td><td>24</td><td>8</td><td>7.6</td><td>7.9</td><td>1.5</td><td>18.8</td><td>5.6</td></tr>
<tr><td>>MI0000085|hsa-mir-27a&WithFlank&19|-|13836434|13836523@57.76.20(+1U)&[MIMAT0000084&hsa-miR-27a-3p&offsets|0|-1(+1U);m-43&19|-|13836447|13836467&offsets|0|-1(+1U)]|TTCACAGTGGCTAAGTTCCGT</td><td>>MI0000085|hsa-mir-27a&WithFlank&19|-|13836434|13836523</td><td>57</td><td>76</td><td>20(+1U)</td><td>TTCACAGTGGCTAAGTTCCGT</td><td>BT20</td><td>189</td><td>120</td><td>78</td><td>172</td><td>51</td><td>107.8</td><td>108.6</td><td>65.1</td><td>167.0</td><td>28.4</td></tr>
<tr><td>>MI0000085|hsa-mir-27a&WithFlank&19|-|13836434|13836523@57.76.20(+1U)&[MIMAT0000084&hsa-miR-27a-3p&offsets|0|-1(+1U);m-43&19|-|13836447|13836467&offsets|0|-1(+1U)]|TTCACAGTGGCTAAGTTCCGT</td><td>>MI0000085|hsa-mir-27a&WithFlank&19|-|13836434|13836523</td><td>57</td><td>76</td><td>20(+1U)</td><td>TTCACAGTGGCTAAGTTCCGT</td><td>MB231</td><td>298</td><td>331</td><td>57</td><td>1222</td><td>455</td><td>118.8</td><td>198.1</td><td>48.4</td><td>709.4</td><td>253.5</td></tr>
<tr><td>>MI0000085|hsa-mir-27a&WithFlank&19|-|13836434|13836523@57.76.20(+1U)&[MIMAT0000084&hsa-miR-27a-3p&offsets|0|-1(+1U);m-43&19|-|13836447|13836467&offsets|0|-1(+1U)]|TTCACAGTGGCTAAGTTCCGT</td><td>>MI0000085|hsa-mir-27a&WithFlank&19|-|13836434|13836523</td><td>57</td><td>76</td><td>20(+1U)</td><td>TTCACAGTGGCTAAGTTCCGT</td><td>MB468</td><td>57</td><td>11</td><td>10</td><td>92</td><td>21</td><td>41.5</td><td>6.1</td><td>2.9</td><td>19.0</td><td>8.5</td></tr>
<tr><td>>MI0000085|hsa-mir-27a&WithFlank&19|-|13836434|13836523@57.76.20(+2U)&[MIMAT0000084&hsa-miR-27a-3p&offsets|0|-1(+2U);m-43&19|-|13836447|13836467&offsets|0|-1(+2U)]|TTCACAGTGGCTAAGTTCCGTT</td><td>>MI0000085|hsa-mir-27a&WithFlank&19|-|13836434|13836523</td><td>57</td><td>76</td><td>20(+2U)</td><td>TTCACAGTGGCTAAGTTCCGTT</td><td>BT20</td><td>81</td><td>39</td><td>29</td><td>38</td><td>15</td><td>58.4</td><td>35.7</td><td>19.7</td><td>36.9</td><td>8.9</td></tr>
<tr><td>>MI0000085|hsa-mir-27a&WithFlank&19|-|13836434|13836523@57.76.20(+2U)&[MIMAT0000084&hsa-miR-27a-3p&offsets|0|-1(+2U);m-43&19|-|13836447|13836467&offsets|0|-1(+2U)]|TTCACAGTGGCTAAGTTCCGTT</td><td>>MI0000085|hsa-mir-27a&WithFlank&19|-|13836434|13836523</td><td>57</td><td>76</td><td>20(+2U)</td><td>TTCACAGTGGCTAAGTTCCGTT</td><td>MB231</td><td>124</td><td>106</td><td>24</td><td>386</td><td>163</td><td>37.6</td><td>62.5</td><td>18.3</td><td>133.0</td><td>86.8</td></tr>
<tr><td>>MI0000085|hsa-mir-27a&WithFlank&19|-|13836434|13836523@57.76.20(+2U)&[MIMAT0000084&hsa-miR-27a-3p&offsets|0|-1(+2U);m-43&19|-|13836447|13836467&offsets|0|-1(+2U)]|TTCACAGTGGCTAAGTTCCGTT</td><td>>MI0000085|hsa-mir-27a&WithFlank&19|-|13836434|13836523</td><td>57</td><td>76</td><td>20(+2U)</td><td>TTCACAGTGGCTAAGTTCCGTT</td><td>MB468</td><td>28</td><td>4</td><td>5</td><td>48</td><td>9</td><td>20.4</td><td>1.9</td><td>1.3</td><td>26.1</td><td>3.0</td></tr>
<tr><td>>MI0000085|hsa-mir-27a&WithFlank&19|-|13836434|13836523@57.76.20(+3U)&[MIMAT0000084&hsa-miR-27a-3p&offsets|0|-1(+3U);m-43&19|-|13836447|13836467&offsets|0|-1(+3U)]|TTCACAGTGGCTAAGTTCCGTTT</td><td>>MI0000085|hsa-mir-27a&WithFlank&19|-|13836434|13836523</td><td>57</td><td>76</td><td>20(+3U)</td><td>TTCACAGTGGCTAAGTTCCGTTT</td><td>BT20</td><td>21</td><td>13</td><td>8</td><td>8</td><td>4</td><td>14.6</td><td>13.6</td><td>7.5</td><td>7.6</td><td>2.6</td></tr>
<tr><td>>MI0000085|hsa-mir-27a&WithFlank&19|-|13836434|13836523@57.76.20(+3U)&[MIMAT0000084&hsa-miR-27a-3p&offsets|0|-1(+3U);m-43&19|-|13836447|13836467&offsets|0|-1(+3U)]|TTCACAGTGGCTAAGTTCCGTTT</td><td>>MI0000085|hsa-mir-27a&WithFlank&19|-|13836434|13836523</td><td>57</td><td>76</td><td>20(+3U)</td><td>TTCACAGTGGCTAAGTTCCGTTT</td><td>MB231</td><td>58</td><td>48</td><td>10</td><td>211</td><td>97</td><td>17.4</td><td>24.8</td><td>8.7</td><td>103.1</td><td>68.2</td></tr>
<tr><td>>MI0000085|hsa-mir-27a&WithFlank&19|-|13836434|13836523@57.76.20&[MIMAT0000084&hsa-miR-27a-3p&offsets|0|-1;m-43&19|-|13836447|13836467&offsets|0|-1]|TTCACAGTGGCTAAGTTCCG</td><td>>MI0000085|hsa-mir-27a&WithFlank&19|-|13836434|13836523</td><td>57</td><td>76</td><td>20</td><td>TTCACAGTGGCTAAGTTCCG</td><td>BT20</td><td>359</td><td>201</td><td>230</td><td>380</td><td>128</td><td>251.3</td><td>136.1</td><td>223.3</td><td>311.4</td><td>85.4</td></tr>
<tr><td>>MI0000085|hsa-mir-27a&WithFlank&19|-|13836434|13836523@57.76.20&[MIMAT0000084&hsa-miR-27a-3p&offsets|0|-1;m-43&19|-|13836447|13836467&offsets|0|-1]|TTCACAGTGGCTAAGTTCCG</td><td>>MI0000085|hsa-mir-27a&WithFlank&19|-|13836434|13836523</td><td>57</td><td>76</td><td>20</td><td>TTCACAGTGGCTAAGTTCCG</td><td>MB231</td><td>689</td><td>805</td><td>166</td><td>1606</td><td>918</td><td>172.1</td><td>537.3</td><td>130.8</td><td>1072.4</td><td>332.5</td></tr>
<tr><td>>MI0000085|hsa-mir-27a&WithFlank&19|-|13836434|13836523@57.76.20&[MIMAT0000084&hsa-miR-27a-3p&offsets|0|-1;m-43&19|-|13836447|13836467&offsets|0|-1]|TTCACAGTGGCTAAGTTCCG</td><td>>MI0000085|hsa-mir-27a&WithFlank&19|-|13836434|13836523</td><td>57</td><td>76</td><td>20</td><td>TTCACAGTGGCTAAGTTCCG</td><td>MB468</td><td>100</td><td>24</td><td>20</td><td>115</td><td>39</td><td>72.6</td><td>11.4</td><td>5.1</td><td>33.9</td><td>16.9</td></tr>
<tr><td>>MI0000085|hsa-mir-27a&WithFlank&19|-|13836434|13836523@57.77.21(+1A)&[MIMAT0000084&hsa-miR-27a-3p&offsets|0|0(+1A);m-43&19|-|13836447|13836467&offsets|0|0(+1A)]|TTCACAGTGGCTAAGTTCCGCA</td><td>>MI0000085|hsa-mir-27a&WithFlank&19|-|13836434|13836523</td><td>57</td><td>77</td><td>21(+1A)</td><td>TTCACAGTGGCTAAGTTCCGCA</td><td>BT20</td><td>14</td><td>3</td><td>2</td><td>2</td><td>1</td><td>18.3</td><td>4.2</td><td>0.4</td><td>3.1</td><td>0.3</td></tr>
<tr><td>>MI0000085|hsa-mir-27a&WithFlank&19|-|13836434|13836523@57.77.21(+1U)&[MIMAT0000084&hsa-miR-27a-3p&offsets|0|0(+1U);m-43&19|-|13836447|13836467&offsets|0|0(+1U)]|TTCACAGTGGCTAAGTTCCGCT</td><td>>MI0000085|hsa-mir-27a&WithFlank&19|-|13836434|13836523</td><td>57</td><td>77</td><td>21(+1U)</td><td>TTCACAGTGGCTAAGTTCCGCT</td><td>BT20</td><td>13</td><td>4</td><td>2</td><td>3</td><td>1</td><td>14.1</td><td>6.9</td><td>1.2</td><td>3.0</td><td>1.0</td></tr>
<tr><td>>MI0000085|hsa-mir-27a&WithFlank&19|-|13836434|13836523@57.77.21(+1U)&[MIMAT0000084&hsa-miR-27a-3p&offsets|0|0(+1U);m-43&19|-|13836447|13836467&offsets|0|0(+1U)]|TTCACAGTGGCTAAGTTCCGCT</td><td>>MI0000085|hsa-mir-27a&WithFlank&19|-|13836434|13836523</td><td>57</td><td>77</td><td>21(+1U)</td><td>TTCACAGTGGCTAAGTTCCGCT</td><td>MB231</td><td>18</td><td>8</td><td>1</td><td>74</td><td>26</td><td>10.9</td><td>5.8</td><td>0.8</td><td>47.6</td><td>28.9</td></tr>
<tr><td>>MI0000085|hsa-mir-27a&WithFlank&19|-|13836434|13836523@57.77.21&[MIMAT0000084&hsa-miR-27a-3p&offsets|0|0;m-43&19|-|13836447|13836467&offsets|0|0]|TTCACAGTGGCTAAGTTCCGC</td><td>>MI0000085|hsa-mir-27a&WithFlank&19|-|13836434|13836523</td><td>57</td><td>77</td><td>21</td><td>TTCACAGTGGCTAAGTTCCGC</td><td>BT20</td><td>120</td><td>64</td><td>33</td><td>196</td><td>19</td><td>96.7</td><td>68.0</td><td>20.7</td><td>203.4</td><td>9.7</td></tr>
<tr><td>>MI0000085|hsa-mir-27a&WithFlank&19|-|13836434|13836523@57.77.21&[MIMAT0000084&hsa-miR-27a-3p&offsets|0|0;m-43&19|-|13836447|13836467&offsets|0|0]|TTCACAGTGGCTAAGTTCCGC</td><td>>MI0000085|hsa-mir-27a&WithFlank&19|-|13836434|13836523</td><td>57</td><td>77</td><td>21</td><td>TTCACAGTGGCTAAGTTCCGC</td><td>MB231</td><td>113</td><td>88</td><td>19</td><td>1152</td><td>188</td><td>26.8</td><td>53.6</td><td>15.9</td><td>425.4</td><td>159.8</td></tr>
<tr><td>>MI0000085|hsa-mir-27a&WithFlank&19|-|13836434|13836523@57.77.21&[MIMAT0000084&hsa-miR-27a-3p&offsets|0|0;m-43&19|-|13836447|13836467&offsets|0|0]|TTCACAGTGGCTAAGTTCCGC</td><td>>MI0000085|hsa-mir-27a&WithFlank&19|-|13836434|13836523</td><td>57</td><td>77</td><td>21</td><td>TTCACAGTGGCTAAGTTCCGC</td><td>MB468</td><td>20</td><td>2</td><td>3</td><td>99</td><td>6</td><td>12.1</td><td>0.9</td><td>1.4</td><td>99.3</td><td>1.2</td></tr>
<tr><td>>MI0000086|hsa-mir-28&WithFlank&3|+|188688775|188688872@20.40.21&[MIMAT0000085&hsa-miR-28-5p&offsets|0|-1;m-133&3|+|188688794|188688815&offsets|0|-1]|AAGGAGCTCACAGTCTATTGA</td><td>>MI0000086|hsa-mir-28&WithFlank&3|+|188688775|188688872</td><td>20</td><td>40</td><td>21</td><td>AAGGAGCTCACAGTCTATTGA</td><td>BT20</td><td>18</td><td>17</td><td>18</td><td>31</td><td>8</td><td>9.1</td><td>4.6</td><td>5.8</td><td>22.3</td><td>1.9</td></tr>
</table>

<table>
<tr><td>>MI0000086|hsa-mir-28&WithFlank&3|+|188688775|188688872@20.41.22(+1A)&[MIMAT0000085&hsa-miR-28-5p&offsets|0|0(+1A);m-133&3|+|188688794|188688815&offsets|0|0(+1A)]|AAGGAGCTCACAGTCTATTGAGA</td><td>>MI0000086|hsa-mir-28&WithFlank&3|+|188688775|188688872</td><td>20</td><td>41</td><td>22(+1A)</td><td>AAGGAGCTCACAGTCTATTGAGA</td><td>BT20</td><td>22</td><td>19</td><td>14</td><td>12</td><td>9</td><td>16.7</td><td>8.9</td><td>6.5</td><td>8.4</td><td>2.0</td></tr>
<tr><td>>MI0000086|hsa-mir-28&WithFlank&3|+|188688775|188688872@20.41.22&[MIMAT0000085&hsa-miR-28-5p&offsets|0|0;m-133&3|+|188688794|188688815&offsets|0|0]|AAGGAGCTCACAGTCTATTGAG</td><td>>MI0000086|hsa-mir-28&WithFlank&3|+|188688775|188688872</td><td>20</td><td>41</td><td>22</td><td>AAGGAGCTCACAGTCTATTGAG</td><td>BT20</td><td>36</td><td>40</td><td>31</td><td>46</td><td>18</td><td>22.8</td><td>17.9</td><td>18.0</td><td>30.5</td><td>5.2</td></tr>
<tr><td>>MI0000086|hsa-mir-28&WithFlank&3|+|188688775|188688872@20.41.22&[MIMAT0000085&hsa-miR-28-5p&offsets|0|0;m-133&3|+|188688794|188688815&offsets|0|0]|AAGGAGCTCACAGTCTATTGAG</td><td>>MI0000086|hsa-mir-28&WithFlank&3|+|188688775|188688872</td><td>20</td><td>41</td><td>22</td><td>AAGGAGCTCACAGTCTATTGAG</td><td>MB231</td><td>12</td><td>15</td><td>6</td><td>90</td><td>17</td><td>3.0</td><td>4.4</td><td>4.5</td><td>45.0</td><td>5.8</td></tr>
<tr><td>>MI0000086|hsa-mir-28&WithFlank&3|+|188688775|188688872@60.79.20(+1A)&[MIMAT0004502&hsa-miR-28-3p&offsets|0|-2(+1A);m-28&3|+|188688834|188688855&offsets|0|-2(+1A)]|CACTAGATTGTGAGCTCCTGA</td><td>>MI0000086|hsa-mir-28&WithFlank&3|+|188688775|188688872</td><td>60</td><td>79</td><td>20(+1A)</td><td>CACTAGATTGTGAGCTCCTGA</td><td>BT20</td><td>20</td><td>7</td><td>40</td><td>16</td><td>4</td><td>18.0</td><td>1.9</td><td>26.1</td><td>11.4</td><td>1.0</td></tr>
<tr><td>>MI0000086|hsa-mir-28&WithFlank&3|+|188688775|188688872@60.80.21(+1G)&[MIMAT0004502&hsa-miR-28-3p&offsets|0|-1(+1G);m-28&3|+|188688834|188688855&offsets|0|-1(+1G)]|CACTAGATTGTGAGCTCCTGGG</td><td>>MI0000086|hsa-mir-28&WithFlank&3|+|188688775|188688872</td><td>60</td><td>80</td><td>21(+1G)</td><td>CACTAGATTGTGAGCTCCTGGG</td><td>BT20</td><td>44</td><td>7</td><td>46</td><td>10</td><td>2</td><td>39.9</td><td>1.4</td><td>29.9</td><td>11.3</td><td>1.1</td></tr>
<tr><td>>MI0000086|hsa-mir-28&WithFlank&3|+|188688775|188688872@60.80.21(+1G)&[MIMAT0004502&hsa-miR-28-3p&offsets|0|-1(+1G);m-28&3|+|188688834|188688855&offsets|0|-1(+1G)]|CACTAGATTGTGAGCTCCTGGG</td><td>>MI0000086|hsa-mir-28&WithFlank&3|+|188688775|188688872</td><td>60</td><td>80</td><td>21(+1G)</td><td>CACTAGATTGTGAGCTCCTGGG</td><td>MB231</td><td>14</td><td>4</td><td>4</td><td>6</td><td>3</td><td>11.4</td><td>1.5</td><td>3.2</td><td>2.6</td><td>1.5</td></tr>
<tr><td>>MI0000086|hsa-mir-28&WithFlank&3|+|188688775|188688872@60.80.21(+1U)&[MIMAT0004502&hsa-miR-28-3p&offsets|0|-1(+1U);m-28&3|+|188688834|188688855&offsets|0|-1(+1U)]|CACTAGATTGTGAGCTCCTGGT</td><td>>MI0000086|hsa-mir-28&WithFlank&3|+|188688775|188688872</td><td>60</td><td>80</td><td>21(+1U)</td><td>CACTAGATTGTGAGCTCCTGGT</td><td>BT20</td><td>36</td><td>10</td><td>57</td><td>17</td><td>8</td><td>31.6</td><td>2.7</td><td>36.1</td><td>13.1</td><td>2.3</td></tr>
<tr><td>>MI0000086|hsa-mir-28&WithFlank&3|+|188688775|188688872@60.80.21(+1U)&[MIMAT0004502&hsa-miR-28-3p&offsets|0|-1(+1U);m-28&3|+|188688834|188688855&offsets|0|-1(+1U)]|CACTAGATTGTGAGCTCCTGGT</td><td>>MI0000086|hsa-mir-28&WithFlank&3|+|188688775|188688872</td><td>60</td><td>80</td><td>21(+1U)</td><td>CACTAGATTGTGAGCTCCTGGT</td><td>MB231</td><td>16</td><td>12</td><td>12</td><td>20</td><td>14</td><td>6.6</td><td>3.7</td><td>8.6</td><td>4.2</td><td>9.3</td></tr>
<tr><td>>MI0000086|hsa-mir-28&WithFlank&3|+|188688775|188688872@60.80.21&[MIMAT0004502&hsa-miR-28-3p&offsets|0|-1;m-28&3|+|188688834|188688855&offsets|0|-1]|CACTAGATTGTGAGCTCCTGG</td><td>>MI0000086|hsa-mir-28&WithFlank&3|+|188688775|188688872</td><td>60</td><td>80</td><td>21</td><td>CACTAGATTGTGAGCTCCTGG</td><td>BT20</td><td>93</td><td>44</td><td>188</td><td>111</td><td>32</td><td>43.8</td><td>11.9</td><td>76.8</td><td>59.8</td><td>7.5</td></tr>
<tr><td>>MI0000086|hsa-mir-28&WithFlank&3|+|188688775|188688872@60.80.21&[MIMAT0004502&hsa-miR-28-3p&offsets|0|-1;m-28&3|+|188688834|188688855&offsets|0|-1]|CACTAGATTGTGAGCTCCTGG</td><td>>MI0000086|hsa-mir-28&WithFlank&3|+|188688775|188688872</td><td>60</td><td>80</td><td>21</td><td>CACTAGATTGTGAGCTCCTGG</td><td>MB231</td><td>55</td><td>48</td><td>49</td><td>122</td><td>37</td><td>9.5</td><td>11.8</td><td>28.1</td><td>16.0</td><td>10.9</td></tr>
<tr><td>>MI0000086|hsa-mir-28&WithFlank&3|+|188688775|188688872@60.81.22(+1A)&[MIMAT0004502&hsa-miR-28-3p&offsets|0|0(+1A);m-28&3|+|188688834|188688855&offsets|0|0(+1A)]|CACTAGATTGTGAGCTCCTGGAA</td><td>>MI0000086|hsa-mir-28&WithFlank&3|+|188688775|188688872</td><td>60</td><td>81</td><td>22(+1A)</td><td>CACTAGATTGTGAGCTCCTGGAA</td><td>BT20</td><td>91</td><td>49</td><td>238</td><td>68</td><td>39</td><td>46.7</td><td>12.2</td><td>98.7</td><td>43.9</td><td>13.3</td></tr>
<tr><td>>MI0000086|hsa-mir-28&WithFlank&3|+|188688775|188688872@60.81.22(+1A)&[MIMAT0004502&hsa-miR-28-3p&offsets|0|0(+1A);m-28&3|+|188688834|188688855&offsets|0|0(+1A)]|CACTAGATTGTGAGCTCCTGGAA</td><td>>MI0000086|hsa-mir-28&WithFlank&3|+|188688775|188688872</td><td>60</td><td>81</td><td>22(+1A)</td><td>CACTAGATTGTGAGCTCCTGGAA</td><td>MB231</td><td>32</td><td>27</td><td>31</td><td>43</td><td>24</td><td>3.6</td><td>6.7</td><td>21.2</td><td>10.6</td><td>11.9</td></tr>
<tr><td>>MI0000086|hsa-mir-28&WithFlank&3|+|188688775|188688872@60.81.22(+1U)&[MIMAT0004502&hsa-miR-28-3p&offsets|0|0(+1U);m-28&3|+|188688834|188688855&offsets|0|0(+1U)]|CACTAGATTGTGAGCTCCTGGAT</td><td>>MI0000086|hsa-mir-28&WithFlank&3|+|188688775|188688872</td><td>60</td><td>81</td><td>22(+1U)</td><td>CACTAGATTGTGAGCTCCTGGAT</td><td>BT20</td><td>29</td><td>19</td><td>50</td><td>18</td><td>20</td><td>14.0</td><td>5.8</td><td>25.3</td><td>13.5</td><td>3.6</td></tr>
<tr><td>>MI0000086|hsa-mir-28&WithFlank&3|+|188688775|188688872@60.81.22(+1U)&[MIMAT0004502&hsa-miR-28-3p&offsets|0|0(+1U);m-28&3|+|188688834|188688855&offsets|0|0(+1U)]|CACTAGATTGTGAGCTCCTGGAT</td><td>>MI0000086|hsa-mir-28&WithFlank&3|+|188688775|188688872</td><td>60</td><td>81</td><td>22(+1U)</td><td>CACTAGATTGTGAGCTCCTGGAT</td><td>MB231</td><td>27</td><td>24</td><td>18</td><td>36</td><td>32</td><td>8.1</td><td>7.9</td><td>9.8</td><td>14.0</td><td>18.9</td></tr>
<tr><td>>MI0000086|hsa-mir-28&WithFlank&3|+|188688775|188688872@60.81.22(+2A)&[MIMAT0004502&hsa-miR-28-3p&offsets|0|0(+2A);m-28&3|+|188688834|188688855&offsets|0|0(+2A)]|CACTAGATTGTGAGCTCCTGGAAA</td><td>>MI0000086|hsa-mir-28&WithFlank&3|+|188688775|188688872</td><td>60</td><td>81</td><td>22(+2A)</td><td>CACTAGATTGTGAGCTCCTGGAAA</td><td>BT20</td><td>15</td><td>8</td><td>26</td><td>10</td><td>9</td><td>9.2</td><td>3.7</td><td>11.5</td><td>4.8</td><td>1.5</td></tr>
<tr><td>>MI0000086|hsa-mir-28&WithFlank&3|+|188688775|188688872@60.81.22&[MIMAT0004502&hsa-miR-28-3p&offsets|0|0;m-28&3|+|188688834|188688855&offsets|0|0]|CACTAGATTGTGAGCTCCTGGA</td><td>>MI0000086|hsa-mir-28&WithFlank&3|+|188688775|188688872</td><td>60</td><td>81</td><td>22</td><td>CACTAGATTGTGAGCTCCTGGA</td><td>BT20</td><td>491</td><td>249</td><td>1493</td><td>472</td><td>209</td><td>184.3</td><td>40.8</td><td>659.3</td><td>297.8</td><td>77.0</td></tr>
<tr><td>>MI0000086|hsa-mir-28&WithFlank&3|+|188688775|188688872@60.81.22&[MIMAT0004502&hsa-miR-28-3p&offsets|0|0;m-28&3|+|188688834|188688855&offsets|0|0]|CACTAGATTGTGAGCTCCTGGA</td><td>>MI0000086|hsa-mir-28&WithFlank&3|+|188688775|188688872</td><td>60</td><td>81</td><td>22</td><td>CACTAGATTGTGAGCTCCTGGA</td><td>MB231</td><td>244</td><td>206</td><td>258</td><td>356</td><td>172</td><td>35.5</td><td>30.8</td><td>164.0</td><td>65.8</td><td>73.5</td></tr>
<tr><td>>MI0000086|hsa-mir-28&WithFlank&3|+|188688775|188688872@60.82.23(+1U)&[MIMAT0004502&hsa-miR-28-3p&offsets|0|+1(+1U);m-28&3|+|188688834|188688855&offsets|0|+1(+1U)]|CACTAGATTGTGAGCTCCTGGAGT</td><td>>MI0000086|hsa-mir-28&WithFlank&3|+|188688775|188688872</td><td>60</td><td>82</td><td>23(+1U)</td><td>CACTAGATTGTGAGCTCCTGGAGT</td><td>BT20</td><td>20</td><td>6</td><td>19</td><td>39</td><td>4</td><td>20.3</td><td>3.2</td><td>7.2</td><td>41.9</td><td>1.2</td></tr>
<tr><td>>MI0000086|hsa-mir-28&WithFlank&3|+|188688775|188688872@60.82.23(+1U)&[MIMAT0004502&hsa-miR-28-3p&offsets|0|+1(+1U);m-28&3|+|188688834|188688855&offsets|0|+1(+1U)]|CACTAGATTGTGAGCTCCTGGAGT</td><td>>MI0000086|hsa-mir-28&WithFlank&3|+|188688775|188688872</td><td>60</td><td>82</td><td>23(+1U)</td><td>CACTAGATTGTGAGCTCCTGGAGT</td><td>MB231</td><td>11</td><td>5</td><td>5</td><td>39</td><td>11</td><td>5.0</td><td>2.0</td><td>5.0</td><td>13.1</td><td>9.3</td></tr>
<tr><td>>MI0000086|hsa-mir-28&WithFlank&3|+|188688775|188688872@60.82.23&[MIMAT0004502&hsa-miR-28-3p&offsets|0|+1;m-28&3|+|188688834|188688855&offsets|0|+1]|CACTAGATTGTGAGCTCCTGGAG</td><td>>MI0000086|hsa-mir-28&WithFlank&3|+|188688775|188688872</td><td>60</td><td>82</td><td>23</td><td>CACTAGATTGTGAGCTCCTGGAG</td><td>BT20</td><td>43</td><td>24</td><td>94</td><td>36</td><td>20</td><td>27.6</td><td>13.8</td><td>52.3</td><td>23.5</td><td>9.9</td></tr>
<tr><td>>MI0000086|hsa-mir-28&WithFlank&3|+|188688775|188688872@60.82.23&[MIMAT0004502&hsa-miR-28-3p&offsets|0|+1;m-28&3|+|188688834|188688855&offsets|0|+1]|CACTAGATTGTGAGCTCCTGGAG</td><td>>MI0000086|hsa-mir-28&WithFlank&3|+|188688775|188688872</td><td>60</td><td>82</td><td>23</td><td>CACTAGATTGTGAGCTCCTGGAG</td><td>MB231</td><td>15</td><td>11</td><td>12</td><td>21</td><td>11</td><td>2.9</td><td>2.1</td><td>7.5</td><td>2.8</td><td>4.2</td></tr>
<tr><td>>MI0000086|hsa-mir-28&WithFlank&3|+|188688775|188688872@61.82.22(+1U)&[MIMAT0004502&hsa-miR-28-3p&offsets|+1|+1(+1U);m-28&3|+|188688834|188688855&offsets|+1|+1(+1U)]|ACTAGATTGTGAGCTCCTGGAGT</td><td>>MI0000086|hsa-mir-28&WithFlank&3|+|188688775|188688872</td><td>61</td><td>82</td><td>22(+1U)</td><td>ACTAGATTGTGAGCTCCTGGAGT</td><td>BT20</td><td>17</td><td>9</td><td>32</td><td>42</td><td>7</td><td>9.2</td><td>4.0</td><td>11.9</td><td>35.1</td><td>1.5</td></tr>
<tr><td>>MI0000086|hsa-mir-28&WithFlank&3|+|188688775|188688872@61.82.22(+1U)&[MIMAT0004502&hsa-miR-28-3p&offsets|+1|+1(+1U);m-28&3|+|188688834|188688855&offsets|+1|+1(+1U)]|ACTAGATTGTGAGCTCCTGGAGT</td><td>>MI0000086|hsa-mir-28&WithFlank&3|+|188688775|188688872</td><td>61</td><td>82</td><td>22(+1U)</td><td>ACTAGATTGTGAGCTCCTGGAGT</td><td>MB231</td><td>12</td><td>10</td><td>8</td><td>37</td><td>9</td><td>3.3</td><td>3.5</td><td>5.1</td><td>7.8</td><td>3.5</td></tr>
<tr><td>>MI0000086|hsa-mir-28&WithFlank&3|+|188688775|188688872@61.82.22&[MIMAT0004502&hsa-miR-28-3p&offsets|+1|+1;m-28&3|+|188688834|188688855&offsets|+1|+1]|ACTAGATTGTGAGCTCCTGGAG</td><td>>MI0000086|hsa-mir-28&WithFlank&3|+|188688775|188688872</td><td>61</td><td>82</td><td>22</td><td>ACTAGATTGTGAGCTCCTGGAG</td><td>BT20</td><td>22</td><td>19</td><td>66</td><td>36</td><td>12</td><td>8.7</td><td>10.9</td><td>32.2</td><td>19.2</td><td>4.1</td></tr>
<tr><td>>MI0000086|hsa-mir-28&WithFlank&3|+|188688775|188688872@62.82.21(+1U)&[MIMAT0004502&hsa-miR-28-3p&offsets|+2|+1(+1U);m-28&3|+|188688834|188688855&offsets|+2|+1(+1U)]|CTAGATTGTGAGCTCCTGGAGT</td><td>>MI0000086|hsa-mir-28&WithFlank&3|+|188688775|188688872</td><td>62</td><td>82</td><td>21(+1U)</td><td>CTAGATTGTGAGCTCCTGGAGT</td><td>BT20</td><td>21</td><td>12</td><td>28</td><td>68</td><td>9</td><td>9.0</td><td>2.9</td><td>10.2</td><td>54.9</td><td>1.3</td></tr>
<tr><td>>MI0000086|hsa-mir-28&WithFlank&3|+|188688775|188688872@62.82.21(+1U)&[MIMAT0004502&hsa-miR-28-3p&offsets|+2|+1(+1U);m-28&3|+|188688834|188688855&offsets|+2|+1(+1U)]|CTAGATTGTGAGCTCCTGGAGT</td><td>>MI0000086|hsa-mir-28&WithFlank&3|+|188688775|188688872</td><td>62</td><td>82</td><td>21(+1U)</td><td>CTAGATTGTGAGCTCCTGGAGT</td><td>MB231</td><td>11</td><td>9</td><td>6</td><td>33</td><td>7</td><td>3.2</td><td>3.1</td><td>3.4</td><td>9.4</td><td>3.1</td></tr>
<tr><td>>MI0000086|hsa-mir-28&WithFlank&3|+|188688775|188688872@62.82.21&[MIMAT0004502&hsa-miR-28-3p&offsets|+2|+1;m-28&3|+|188688834|188688855&offsets|+2|+1]|CTAGATTGTGAGCTCCTGGAG</td><td>>MI0000086|hsa-mir-28&WithFlank&3|+|188688775|188688872</td><td>62</td><td>82</td><td>21</td><td>CTAGATTGTGAGCTCCTGGAG</td><td>BT20</td><td>13</td><td>9</td><td>25</td><td>29</td><td>6</td><td>6.5</td><td>3.4</td><td>12.6</td><td>22.4</td><td>2.4</td></tr>
</table>

| | | | | | | | | | | | | | | | | |
|---|---|---|---|---|---|---|---|---|---|---|---|---|---|---|---|---|
| >MI0000087\|hsa-mir-29a&WithFlank&7\|-\|130876741\|130876816@47.67.21(+1A)&[MIMAT0000086&hsa-miR-29a-3p&offsets\|-1\|-2(+1A);m-26&7\|-\|130876749\|130876769&offsets\|-1\|-1(+1A)]\|CTAGCACCATCTGAAATCGGTA | >MI0000087\|hsa-mir-29a&WithFlank&7\|-\|130876741\|130876816 | 47 | 67 | 21(+1A) | CTAGCACCATCTGAAATCGGTA | BT20 | 33 | 18 | 21 | 33 | 10 | 19.5 | 16.2 | 11.9 | 27.6 | 5.3 |
| >MI0000087\|hsa-mir-29a&WithFlank&7\|-\|130876741\|130876816@47.67.21(+1A)&[MIMAT0000086&hsa-miR-29a-3p&offsets\|-1\|-2(+1A);m-26&7\|-\|130876749\|130876769&offsets\|-1\|-1(+1A)]\|CTAGCACCATCTGAAATCGGTA | >MI0000087\|hsa-mir-29a&WithFlank&7\|-\|130876741\|130876816 | 47 | 67 | 21(+1A) | CTAGCACCATCTGAAATCGGTA | MB231 | 16 | 16 | 5 | 90 | 24 | 6.3 | 6.3 | 4.4 | 54.3 | 15.3 |
| >MI0000087\|hsa-mir-29a&WithFlank&7\|-\|130876741\|130876816@47.67.21(+1C)&[MIMAT0000086&hsa-miR-29a-3p&offsets\|-1\|-2(+1C);m-26&7\|-\|130876749\|130876769&offsets\|-1\|-1(+1C)]\|CTAGCACCATCTGAAATCGGTC | >MI0000087\|hsa-mir-29a&WithFlank&7\|-\|130876741\|130876816 | 47 | 67 | 21(+1C) | CTAGCACCATCTGAAATCGGTC | BT20 | 93 | 87 | 61 | 143 | 56 | 85.8 | 81.2 | 53.8 | 116.5 | 35.2 |
| >MI0000087\|hsa-mir-29a&WithFlank&7\|-\|130876741\|130876816@47.67.21(+1C)&[MIMAT0000086&hsa-miR-29a-3p&offsets\|-1\|-2(+1C);m-26&7\|-\|130876749\|130876769&offsets\|-1\|-1(+1C)]\|CTAGCACCATCTGAAATCGGTC | >MI0000087\|hsa-mir-29a&WithFlank&7\|-\|130876741\|130876816 | 47 | 67 | 21(+1C) | CTAGCACCATCTGAAATCGGTC | MB231 | 62 | 76 | 15 | 383 | 114 | 32.5 | 33.5 | 12.1 | 237.5 | 73.0 |
| >MI0000087\|hsa-mir-29a&WithFlank&7\|-\|130876741\|130876816@47.67.21(+1G)&[MIMAT0000086&hsa-miR-29a-3p&offsets\|-1\|-2(+1G);m-26&7\|-\|130876749\|130876769&offsets\|-1\|-1(+1G)]\|CTAGCACCATCTGAAATCGGTG | >MI0000087\|hsa-mir-29a&WithFlank&7\|-\|130876741\|130876816 | 47 | 67 | 21(+1G) | CTAGCACCATCTGAAATCGGTG | BT20 | 24 | 10 | 7 | 14 | 3 | 12.1 | 8.8 | 3.3 | 13.9 | 1.1 |
| >MI0000087\|hsa-mir-29a&WithFlank&7\|-\|130876741\|130876816@47.67.21(+1G)&[MIMAT0000086&hsa-miR-29a-3p&offsets\|-1\|-2(+1G);m-26&7\|-\|130876749\|130876769&offsets\|-1\|-1(+1G)]\|CTAGCACCATCTGAAATCGGTG | >MI0000087\|hsa-mir-29a&WithFlank&7\|-\|130876741\|130876816 | 47 | 67 | 21(+1G) | CTAGCACCATCTGAAATCGGTG | MB231 | 15 | 10 | 2 | 27 | 6 | 6.5 | 6.4 | 1.6 | 18.7 | 5.4 |
| >MI0000087\|hsa-mir-29a&WithFlank&7\|-\|130876741\|130876816@47.67.21&[MIMAT0000086&hsa-miR-29a-3p&offsets\|-1\|-2;m-26&7\|-\|130876749\|130876769&offsets\|-1\|-1]\|CTAGCACCATCTGAAATCGGT | >MI0000087\|hsa-mir-29a&WithFlank&7\|-\|130876741\|130876816 | 47 | 67 | 21 | CTAGCACCATCTGAAATCGGT | BT20 | 13 | 9 | 11 | 12 | 3 | 7.4 | 8.9 | 7.4 | 12.5 | 2.1 |
| >MI0000087\|hsa-mir-29a&WithFlank&7\|-\|130876741\|130876816@47.68.22(+1U)&[MIMAT0000086&hsa-miR-29a-3p&offsets\|-1\|-1(+1U);m-26&7\|-\|130876749\|130876769&offsets\|-1\|0(+1U)]\|CTAGCACCATCTGAAATCGGTTT | >MI0000087\|hsa-mir-29a&WithFlank&7\|-\|130876741\|130876816 | 47 | 68 | 22(+1U) | CTAGCACCATCTGAAATCGGTTT | BT20 | 49 | 46 | 30 | 185 | 26 | 50.0 | 55.1 | 34.2 | 175.9 | 23.4 |
| >MI0000087\|hsa-mir-29a&WithFlank&7\|-\|130876741\|130876816@47.68.22(+1U)&[MIMAT0000086&hsa-miR-29a-3p&offsets\|-1\|-1(+1U);m-26&7\|-\|130876749\|130876769&offsets\|-1\|0(+1U)]\|CTAGCACCATCTGAAATCGGTTT | >MI0000087\|hsa-mir-29a&WithFlank&7\|-\|130876741\|130876816 | 47 | 68 | 22(+1U) | CTAGCACCATCTGAAATCGGTTT | MB231 | 58 | 70 | 13 | 372 | 92 | 26.8 | 34.8 | 10.6 | 182.6 | 61.7 |
| >MI0000087\|hsa-mir-29a&WithFlank&7\|-\|130876741\|130876816@47.68.22&[MIMAT0000086&hsa-miR-29a-3p&offsets\|-1\|-1;m-26&7\|-\|130876749\|130876769&offsets\|-1\|0]\|CTAGCACCATCTGAAATCGGTT | >MI0000087\|hsa-mir-29a&WithFlank&7\|-\|130876741\|130876816 | 47 | 68 | 22 | CTAGCACCATCTGAAATCGGTT | BT20 | 762 | 620 | 510 | 1090 | 433 | 745.6 | 542.3 | 492.9 | 845.8 | 259.9 |
| >MI0000087\|hsa-mir-29a&WithFlank&7\|-\|130876741\|130876816@47.68.22&[MIMAT0000086&hsa-miR-29a-3p&offsets\|-1\|-1;m-26&7\|-\|130876749\|130876769&offsets\|-1\|0]\|CTAGCACCATCTGAAATCGGTT | >MI0000087\|hsa-mir-29a&WithFlank&7\|-\|130876741\|130876816 | 47 | 68 | 22 | CTAGCACCATCTGAAATCGGTT | MB231 | 591 | 770 | 150 | 2396 | 864 | 324.6 | 308.3 | 117.7 | 1153.7 | 424.3 |
| >MI0000087\|hsa-mir-29a&WithFlank&7\|-\|130876741\|130876816@47.68.22&[MIMAT0000086&hsa-miR-29a-3p&offsets\|-1\|-1;m-26&7\|-\|130876749\|130876769&offsets\|-1\|0]\|CTAGCACCATCTGAAATCGGTT | >MI0000087\|hsa-mir-29a&WithFlank&7\|-\|130876741\|130876816 | 47 | 68 | 22 | CTAGCACCATCTGAAATCGGTT | MB468 | 11 | 4 | 3 | 24 | 4 | 4.8 | 1.6 | 0.8 | 3.9 | 0.8 |
| >MI0000087\|hsa-mir-29a&WithFlank&7\|-\|130876741\|130876816@47.69.23&[MIMAT0000086&hsa-miR-29a-3p&offsets\|-1\|0;m-26&7\|-\|130876749\|130876769&offsets\|-1\|+1]\|CTAGCACCATCTGAAATCGGTTA | >MI0000087\|hsa-mir-29a&WithFlank&7\|-\|130876741\|130876816 | 47 | 69 | 23 | CTAGCACCATCTGAAATCGGTTA | BT20 | 60 | 55 | 34 | 225 | 25 | 31.6 | 63.6 | 34.1 | 192.2 | 22.9 |
| >MI0000087\|hsa-mir-29a&WithFlank&7\|-\|130876741\|130876816@47.69.23&[MIMAT0000086&hsa-miR-29a-3p&offsets\|-1\|0;m-26&7\|-\|130876749\|130876769&offsets\|-1\|+1]\|CTAGCACCATCTGAAATCGGTTA | >MI0000087\|hsa-mir-29a&WithFlank&7\|-\|130876741\|130876816 | 47 | 69 | 23 | CTAGCACCATCTGAAATCGGTTA | MB231 | 86 | 61 | 14 | 493 | 139 | 31.4 | 22.0 | 13.2 | 166.0 | 132.9 |
| >MI0000087\|hsa-mir-29a&WithFlank&7\|-\|130876741\|130876816@47.70.24&[MIMAT0000086&hsa-miR-29a-3p&offsets\|-1\|+1;m-26&7\|-\|130876749\|130876769&offsets\|-1\|+2]\|CTAGCACCATCTGAAATCGGTTAT | >MI0000087\|hsa-mir-29a&WithFlank&7\|-\|130876741\|130876816 | 47 | 70 | 24 | CTAGCACCATCTGAAATCGGTTAT | MB231 | 11 | 13 | 2 | 62 | 15 | 6.5 | 6.3 | 1.2 | 33.6 | 11.4 |
| >MI0000087\|hsa-mir-29a&WithFlank&7\|-\|130876741\|130876816@48.66.19&[MIMAT0000086&hsa-miR-29a-3p&offsets\|0\|-3;m-26&7\|-\|130876749\|130876769&offsets\|0\|-2]\|TAGCACCATCTGAAATCGG | >MI0000087\|hsa-mir-29a&WithFlank&7\|-\|130876741\|130876816 | 48 | 66 | 19 | TAGCACCATCTGAAATCGG | MB231 | 10 | 10 | 6 | 23 | 10 | 1.5 | 4.2 | 4.6 | 14.7 | 4.1 |
| >MI0000087\|hsa-mir-29a&WithFlank&7\|-\|130876741\|130876816@48.67.20(+1A)&[MIMAT0000086&hsa-miR-29a-3p&offsets\|0\|-2(+1A);m-26&7\|-\|130876749\|130876769&offsets\|0\|-1(+1A)]\|TAGCACCATCTGAAATCGGTA | >MI0000087\|hsa-mir-29a&WithFlank&7\|-\|130876741\|130876816 | 48 | 67 | 20(+1A) | TAGCACCATCTGAAATCGGTA | BT20 | 44 | 16 | 14 | 40 | 3 | 51.1 | 21.5 | 8.3 | 37.2 | 3.1 |
| >MI0000087\|hsa-mir-29a&WithFlank&7\|-\|130876741\|130876816@48.67.20(+1A)&[MIMAT0000086&hsa-miR-29a-3p&offsets\|0\|-2(+1A);m-26&7\|-\|130876749\|130876769&offsets\|0\|-1(+1A)]\|TAGCACCATCTGAAATCGGTA | >MI0000087\|hsa-mir-29a&WithFlank&7\|-\|130876741\|130876816 | 48 | 67 | 20(+1A) | TAGCACCATCTGAAATCGGTA | MB231 | 16 | 7 | 4 | 121 | 22 | 2.0 | 4.0 | 4.7 | 69.0 | 23.9 |
| >MI0000087\|hsa-mir-29a&WithFlank&7\|-\|130876741\|130876816@48.67.20(+1C)&[MIMAT0000086&hsa-miR-29a-3p&offsets\|0\|-2(+1C);m-26&7\|-\|130876749\|130876769&offsets\|0\|-1(+1C)]\|TAGCACCATCTGAAATCGGTC | >MI0000087\|hsa-mir-29a&WithFlank&7\|-\|130876741\|130876816 | 48 | 67 | 20(+1C) | TAGCACCATCTGAAATCGGTC | BT20 | 80 | 76 | 39 | 173 | 17 | 54.1 | 98.4 | 25.9 | 159.9 | 14.5 |
| >MI0000087\|hsa-mir-29a&WithFlank&7\|-\|130876741\|130876816@48.67.20(+1C)&[MIMAT0000086&hsa-miR-29a-3p&offsets\|0\|-2(+1C);m-26&7\|-\|130876749\|130876769&offsets\|0\|-1(+1C)]\|TAGCACCATCTGAAATCGGTC | >MI0000087\|hsa-mir-29a&WithFlank&7\|-\|130876741\|130876816 | 48 | 67 | 20(+1C) | TAGCACCATCTGAAATCGGTC | MB231 | 60 | 39 | 13 | 520 | 102 | 16.2 | 19.1 | 12.3 | 293.3 | 109.7 |
| >MI0000087\|hsa-mir-29a&WithFlank&7\|-\|130876741\|130876816@48.67.20(+1G)&[MIMAT0000086&hsa-miR-29a-3p&offsets\|0\|-2(+1G);m-26&7\|-\|130876749\|130876769&offsets\|0\|-1(+1G)]\|TAGCACCATCTGAAATCGGTG | >MI0000087\|hsa-mir-29a&WithFlank&7\|-\|130876741\|130876816 | 48 | 67 | 20(+1G) | TAGCACCATCTGAAATCGGTG | BT20 | 38 | 9 | 5 | 16 | 1 | 49.2 | 12.3 | 2.8 | 19.7 | 0.6 |
| >MI0000087\|hsa-mir-29a&WithFlank&7\|-\|130876741\|130876816@48.67.20(+1G)&[MIMAT0000086&hsa-miR-29a-3p&offsets\|0\|-2(+1G);m-26&7\|-\|130876749\|130876769&offsets\|0\|-1(+1G)]\|TAGCACCATCTGAAATCGGTG | >MI0000087\|hsa-mir-29a&WithFlank&7\|-\|130876741\|130876816 | 48 | 67 | 20(+1G) | TAGCACCATCTGAAATCGGTG | MB231 | 18 | 5 | 1 | 35 | 6 | 13.3 | 3.0 | 1.4 | 19.2 | 6.4 |
| >MI0000087\|hsa-mir-29a&WithFlank&7\|-\|130876741\|130876816@48.67.20&[MIMAT0000086&hsa-miR-29a-3p&offsets\|0\|-2;m-26&7\|-\|130876749\|130876769&offsets\|0\|-1]\|TAGCACCATCTGAAATCGGT | >MI0000087\|hsa-mir-29a&WithFlank&7\|-\|130876741\|130876816 | 48 | 67 | 20 | TAGCACCATCTGAAATCGGT | BT20 | 28 | 19 | 17 | 35 | 4 | 19.4 | 24.1 | 12.9 | 29.5 | 3.9 |
| >MI0000087\|hsa-mir-29a&WithFlank&7\|-\|130876741\|130876816@48.67.20&[MIMAT0000086&hsa-miR-29a-3p&offsets\|0\|-2;m-26&7\|-\|130876749\|130876769&offsets\|0\|-1]\|TAGCACCATCTGAAATCGGT | >MI0000087\|hsa-mir-29a&WithFlank&7\|-\|130876741\|130876816 | 48 | 67 | 20 | TAGCACCATCTGAAATCGGT | MB231 | 21 | 16 | 7 | 138 | 29 | 1.5 | 8.0 | 6.0 | 64.9 | 27.1 |
| >MI0000087\|hsa-mir-29a&WithFlank&7\|-\|130876741\|130876816@48.68.21(+1G)&[MIMAT0000086&hsa-miR-29a-3p&offsets\|0\|-1(+1G);m-26&7\|-\|130876749\|130876769&offsets\|0\|0(+1G)]\|TAGCACCATCTGAAATCGGTTG | >MI0000087\|hsa-mir-29a&WithFlank&7\|-\|130876741\|130876816 | 48 | 68 | 21(+1G) | TAGCACCATCTGAAATCGGTTG | BT20 | 22 | 6 | 2 | 19 | 1 | 28.3 | 7.5 | 1.3 | 20.2 | 0.8 |
| >MI0000087\|hsa-mir-29a&WithFlank&7\|-\|130876741\|130876816@48.68.21(+1U)&[MIMAT0000086&hsa-miR-29a-3p&offsets\|0\|-1(+1U);m-26&7\|-\|130876749\|130876769&offsets\|0\|0(+1U)]\|TAGCACCATCTGAAATCGGTTT | >MI0000087\|hsa-mir-29a&WithFlank&7\|-\|130876741\|130876816 | 48 | 68 | 21(+1U) | TAGCACCATCTGAAATCGGTTT | BT20 | 28 | 21 | 9 | 90 | 5 | 22.5 | 29.7 | 7.2 | 77.7 | 6.5 |
| >MI0000087\|hsa-mir-29a&WithFlank&7\|-\|130876741\|130876816@48.68.21(+1U)&[MIMAT0000086&hsa-miR-29a-3p&offsets\|0\|-1(+1U);m-26&7\|-\|130876749\|130876769&offsets\|0\|0(+1U)]\|TAGCACCATCTGAAATCGGTTT | >MI0000087\|hsa-mir-29a&WithFlank&7\|-\|130876741\|130876816 | 48 | 68 | 21(+1U) | TAGCACCATCTGAAATCGGTTT | MB231 | 24 | 16 | 4 | 205 | 37 | 3.4 | 8.9 | 2.7 | 92.7 | 38.5 |
| >MI0000087\|hsa-mir-29a&WithFlank&7\|-\|130876741\|130876816@48.68.21&[MIMAT0000086&hsa-miR-29a-3p&offsets\|0\|-1;m-26&7\|-\|130876749\|130876769&offsets\|0\|0]\|TAGCACCATCTGAAATCGGTT | >MI0000087\|hsa-mir-29a&WithFlank&7\|-\|130876741\|130876816 | 48 | 68 | 21 | TAGCACCATCTGAAATCGGTT | BT20 | 621 | 542 | 302 | 1270 | 126 | 418.3 | 692.5 | 231.8 | 1085.8 | 99.7 |
| >MI0000087\|hsa-mir-29a&WithFlank&7\|-\|130876741\|130876816@48.68.21&[MIMAT0000086&hsa-miR-29a-3p&offsets\|0\|-1;m-26&7\|-\|130876749\|130876769&offsets\|0\|0]\|TAGCACCATCTGAAATCGGTT | >MI0000087\|hsa-mir-29a&WithFlank&7\|-\|130876741\|130876816 | 48 | 68 | 21 | TAGCACCATCTGAAATCGGTT | MB231 | 581 | 381 | 127 | 3284 | 751 | 154.7 | 155.7 | 121.3 | 1372.2 | 710.4 |
| >MI0000087\|hsa-mir-29a&WithFlank&7\|-\|130876741\|130876816@48.69.22(+1A)&[MIMAT0000086&hsa-miR-29a-3p&offsets\|0\|0(+1A);m-26&7\|-\|130876749\|130876769&offsets\|0\|+1(+1A)]\|TAGCACCATCTGAAATCGGTTAA | >MI0000087\|hsa-mir-29a&WithFlank&7\|-\|130876741\|130876816 | 48 | 69 | 22(+1A) | TAGCACCATCTGAAATCGGTTAA | BT20 | 17 | 17 | 6 | 36 | 4 | 12.4 | 21.3 | 6.0 | 32.9 | 4.1 |
| >MI0000087\|hsa-mir-29a&WithFlank&7\|-\|130876741\|130876816@48.69.22&[MIMAT0000086&hsa-miR-29a-3p&offsets\|0\|0;m-26&7\|-\|130876749\|130876769&offsets\|0\|+1]\|TAGCACCATCTGAAATCGGTTA | >MI0000087\|hsa-mir-29a&WithFlank&7\|-\|130876741\|130876816 | 48 | 69 | 22 | TAGCACCATCTGAAATCGGTTA | BT20 | 587 | 459 | 199 | 2140 | 170 | 398.4 | 594.3 | 197.2 | 1842.2 | 157.1 |
| >MI0000087\|hsa-mir-29a&WithFlank&7\|-\|130876741\|130876816@48.69.22&[MIMAT0000086&hsa-miR-29a-3p&offsets\|0\|0;m-26&7\|-\|130876749\|130876769&offsets\|0\|+1]\|TAGCACCATCTGAAATCGGTTA | >MI0000087\|hsa-mir-29a&WithFlank&7\|-\|130876741\|130876816 | 48 | 69 | 22 | TAGCACCATCTGAAATCGGTTA | MB231 | 556 | 316 | 61 | 3599 | 875 | 198.3 | 112.9 | 53.4 | 969.7 | 868.7 |
| >MI0000087\|hsa-mir-29a&WithFlank&7\|-\|130876741\|130876816@48.70.23&[MIMAT0000086&hsa-miR-29a-3p&offsets\|0\|+1;m-26&7\|-\|130876749\|130876769&offsets\|0\|+2]\|TAGCACCATCTGAAATCGGTTAT | >MI0000087\|hsa-mir-29a&WithFlank&7\|-\|130876741\|130876816 | 48 | 70 | 23 | TAGCACCATCTGAAATCGGTTAT | MB231 | 10 | 7 | 2 | 78 | 18 | 0.5 | 3.5 | 1.3 | 29.9 | 21.3 |
| >MI0000088\|hsa-mir-30a&WithFlank&6\|-\|71403545\|71403627@12.31.20&[MIMAT0000087&hsa-miR-30a-5p&offsets\|0\|-2;m-15&6\|-\|71403595\|71403616&offsets\|0\|-2]\|TGTAAACATCCTCGACTGGA | >MI0000088\|hsa-mir-30a&WithFlank&6\|-\|71403545\|71403627 | 12 | 31 | 20 | TGTAAACATCCTCGACTGGA | BT20 | 106 | 69 | 33 | 338 | 27 | 57.7 | 88.8 | 23.1 | 314.1 | 9.4 |

<table>
<tr><th></th><th></th><th></th><th></th><th></th><th></th><th></th><th></th><th></th><th></th><th></th><th></th><th></th><th></th><th></th><th></th><th></th></tr>
<tr><td>>MI0000088|hsa-mir-30a&WithFlank&6|-|71403545|71403627@12.31.20&[MIMAT0000087&hsa-miR-30a-5p&offsets|0|-2;m-15&6|-|71403595|71403616&offsets|0|-2]|TGTAAACATCCTCGACTGGA</td><td>>MI0000088|hsa-mir-30a&WithFlank&6|-|71403545|71403627</td><td>12</td><td>31</td><td>20</td><td>TGTAAACATCCTCGACTGGA</td><td>MB231</td><td>139</td><td>146</td><td>23</td><td>1181</td><td>158</td><td>27.5</td><td>110.2</td><td>18.7</td><td>285.8</td><td>103.2</td></tr>
<tr><td>>MI0000088|hsa-mir-30a&WithFlank&6|-|71403545|71403627@12.32.21(+1A)&[MIMAT0000087&hsa-miR-30a-5p&offsets|0|-1(+1A);m-15&6|-|71403595|71403616&offsets|0|-1(+1A)]|TGTAAACATCCTCGACTGGAAA</td><td>>MI0000088|hsa-mir-30a&WithFlank&6|-|71403545|71403627</td><td>12</td><td>32</td><td>21(+1A)</td><td>TGTAAACATCCTCGACTGGAAA</td><td>BT20</td><td>99</td><td>16</td><td>11</td><td>23</td><td>4</td><td>141.5</td><td>17.5</td><td>6.3</td><td>18.6</td><td>0.9</td></tr>
<tr><td>>MI0000088|hsa-mir-30a&WithFlank&6|-|71403545|71403627@12.32.21(+1A)&[MIMAT0000087&hsa-miR-30a-5p&offsets|0|-1(+1A);m-15&6|-|71403595|71403616&offsets|0|-1(+1A)]|TGTAAACATCCTCGACTGGAAA</td><td>>MI0000088|hsa-mir-30a&WithFlank&6|-|71403545|71403627</td><td>12</td><td>32</td><td>21(+1A)</td><td>TGTAAACATCCTCGACTGGAAA</td><td>MB231</td><td>17</td><td>10</td><td>2</td><td>96</td><td>16</td><td>4.2</td><td>7.5</td><td>1.6</td><td>37.9</td><td>15.4</td></tr>
<tr><td>>MI0000088|hsa-mir-30a&WithFlank&6|-|71403545|71403627@12.32.21&[MIMAT0000087&hsa-miR-30a-5p&offsets|0|-1;m-15&6|-|71403595|71403616&offsets|0|-1]|TGTAAACATCCTCGACTGGAA</td><td>>MI0000088|hsa-mir-30a&WithFlank&6|-|71403545|71403627</td><td>12</td><td>32</td><td>21</td><td>TGTAAACATCCTCGACTGGAA</td><td>BT20</td><td>91</td><td>32</td><td>14</td><td>41</td><td>10</td><td>99.2</td><td>37.9</td><td>6.8</td><td>33.5</td><td>2.6</td></tr>
<tr><td>>MI0000088|hsa-mir-30a&WithFlank&6|-|71403545|71403627@12.32.21&[MIMAT0000087&hsa-miR-30a-5p&offsets|0|-1;m-15&6|-|71403595|71403616&offsets|0|-1]|TGTAAACATCCTCGACTGGAA</td><td>>MI0000088|hsa-mir-30a&WithFlank&6|-|71403545|71403627</td><td>12</td><td>32</td><td>21</td><td>TGTAAACATCCTCGACTGGAA</td><td>MB231</td><td>53</td><td>51</td><td>6</td><td>195</td><td>54</td><td>20.0</td><td>40.2</td><td>3.8</td><td>45.6</td><td>30.9</td></tr>
<tr><td>>MI0000088|hsa-mir-30a&WithFlank&6|-|71403545|71403627@12.33.22(+1A)&[MIMAT0000087&hsa-miR-30a-5p&offsets|0|0(+1A);m-15&6|-|71403595|71403616&offsets|0|0(+1A)]|TGTAAACATCCTCGACTGGAAGA</td><td>>MI0000088|hsa-mir-30a&WithFlank&6|-|71403545|71403627</td><td>12</td><td>33</td><td>22(+1A)</td><td>TGTAAACATCCTCGACTGGAAGA</td><td>BT20</td><td>329</td><td>67</td><td>48</td><td>123</td><td>22</td><td>423.5</td><td>70.4</td><td>26.8</td><td>108.3</td><td>2.4</td></tr>
<tr><td>>MI0000088|hsa-mir-30a&WithFlank&6|-|71403545|71403627@12.33.22(+1A)&[MIMAT0000087&hsa-miR-30a-5p&offsets|0|0(+1A);m-15&6|-|71403595|71403616&offsets|0|0(+1A)]|TGTAAACATCCTCGACTGGAAGA</td><td>>MI0000088|hsa-mir-30a&WithFlank&6|-|71403545|71403627</td><td>12</td><td>33</td><td>22(+1A)</td><td>TGTAAACATCCTCGACTGGAAGA</td><td>MB231</td><td>105</td><td>60</td><td>13</td><td>363</td><td>92</td><td>14.2</td><td>38.3</td><td>10.9</td><td>33.0</td><td>78.7</td></tr>
<tr><td>>MI0000088|hsa-mir-30a&WithFlank&6|-|71403545|71403627@12.33.22(+1G)&[MIMAT0000087&hsa-miR-30a-5p&offsets|0|0(+1G);m-15&6|-|71403595|71403616&offsets|0|0(+1G)]|TGTAAACATCCTCGACTGGAAGG</td><td>>MI0000088|hsa-mir-30a&WithFlank&6|-|71403545|71403627</td><td>12</td><td>33</td><td>22(+1G)</td><td>TGTAAACATCCTCGACTGGAAGG</td><td>BT20</td><td>65</td><td>5</td><td>3</td><td>7</td><td>0</td><td>93.9</td><td>5.4</td><td>2.2</td><td>8.9</td><td>0.4</td></tr>
<tr><td>>MI0000088|hsa-mir-30a&WithFlank&6|-|71403545|71403627@12.33.22(+1G)&[MIMAT0000087&hsa-miR-30a-5p&offsets|0|0(+1G);m-15&6|-|71403595|71403616&offsets|0|0(+1G)]|TGTAAACATCCTCGACTGGAAGG</td><td>>MI0000088|hsa-mir-30a&WithFlank&6|-|71403545|71403627</td><td>12</td><td>33</td><td>22(+1G)</td><td>TGTAAACATCCTCGACTGGAAGG</td><td>MB231</td><td>14</td><td>4</td><td>1</td><td>15</td><td>4</td><td>12.9</td><td>4.1</td><td>0.5</td><td>5.9</td><td>4.7</td></tr>
<tr><td>>MI0000088|hsa-mir-30a&WithFlank&6|-|71403545|71403627@12.33.22(+1U)&[MIMAT0000087&hsa-miR-30a-5p&offsets|0|0(+1U);m-15&6|-|71403595|71403616&offsets|0|0(+1U)]|TGTAAACATCCTCGACTGGAAGT</td><td>>MI0000088|hsa-mir-30a&WithFlank&6|-|71403545|71403627</td><td>12</td><td>33</td><td>22(+1U)</td><td>TGTAAACATCCTCGACTGGAAGT</td><td>BT20</td><td>1051</td><td>310</td><td>147</td><td>568</td><td>88</td><td>1188.9</td><td>342.7</td><td>69.1</td><td>545.2</td><td>11.6</td></tr>
<tr><td>>MI0000088|hsa-mir-30a&WithFlank&6|-|71403545|71403627@12.33.22(+1U)&[MIMAT0000087&hsa-miR-30a-5p&offsets|0|0(+1U);m-15&6|-|71403595|71403616&offsets|0|0(+1U)]|TGTAAACATCCTCGACTGGAAGT</td><td>>MI0000088|hsa-mir-30a&WithFlank&6|-|71403545|71403627</td><td>12</td><td>33</td><td>22(+1U)</td><td>TGTAAACATCCTCGACTGGAAGT</td><td>MB231</td><td>462</td><td>251</td><td>44</td><td>1568</td><td>438</td><td>54.2</td><td>161.7</td><td>32.4</td><td>19.7</td><td>417.5</td></tr>
<tr><td>>MI0000088|hsa-mir-30a&WithFlank&6|-|71403545|71403627@12.33.22(+1U)&[MIMAT0000087&hsa-miR-30a-5p&offsets|0|0(+1U);m-15&6|-|71403595|71403616&offsets|0|0(+1U)]|TGTAAACATCCTCGACTGGAAGT</td><td>>MI0000088|hsa-mir-30a&WithFlank&6|-|71403545|71403627</td><td>12</td><td>33</td><td>22(+1U)</td><td>TGTAAACATCCTCGACTGGAAGT</td><td>MB468</td><td>13</td><td>3</td><td>3</td><td>66</td><td>4</td><td>2.9</td><td>2.2</td><td>3.4</td><td>62.5</td><td>3.5</td></tr>
<tr><td>>MI0000088|hsa-mir-30a&WithFlank&6|-|71403545|71403627@12.33.22(+2U)&[MIMAT0000087&hsa-miR-30a-5p&offsets|0|0(+2U);m-15&6|-|71403595|71403616&offsets|0|0(+2U)]|TGTAAACATCCTCGACTGGAAGTT</td><td>>MI0000088|hsa-mir-30a&WithFlank&6|-|71403545|71403627</td><td>12</td><td>33</td><td>22(+2U)</td><td>TGTAAACATCCTCGACTGGAAGTT</td><td>BT20</td><td>62</td><td>6</td><td>4</td><td>7</td><td>1</td><td>90.6</td><td>8.5</td><td>2.0</td><td>6.1</td><td>0.6</td></tr>
<tr><td>>MI0000088|hsa-mir-30a&WithFlank&6|-|71403545|71403627@12.33.22(+2U)&[MIMAT0000087&hsa-miR-30a-5p&offsets|0|0(+2U);m-15&6|-|71403595|71403616&offsets|0|0(+2U)]|TGTAAACATCCTCGACTGGAAGTT</td><td>>MI0000088|hsa-mir-30a&WithFlank&6|-|71403545|71403627</td><td>12</td><td>33</td><td>22(+2U)</td><td>TGTAAACATCCTCGACTGGAAGTT</td><td>MB231</td><td>30</td><td>12</td><td>2</td><td>38</td><td>17</td><td>9.5</td><td>8.4</td><td>1.3</td><td>4.4</td><td>20.4</td></tr>
<tr><td>>MI0000088|hsa-mir-30a&WithFlank&6|-|71403545|71403627@12.33.22&[MIMAT0000087&hsa-miR-30a-5p&offsets|0|0;m-15&6|-|71403595|71403616&offsets|0|0]|TGTAAACATCCTCGACTGGAAG</td><td>>MI0000088|hsa-mir-30a&WithFlank&6|-|71403545|71403627</td><td>12</td><td>33</td><td>22</td><td>TGTAAACATCCTCGACTGGAAG</td><td>BT20</td><td>542</td><td>161</td><td>83</td><td>292</td><td>45</td><td>632.8</td><td>169.4</td><td>17.4</td><td>240.6</td><td>14.5</td></tr>
<tr><td>>MI0000088|hsa-mir-30a&WithFlank&6|-|71403545|71403627@12.33.22&[MIMAT0000087&hsa-miR-30a-5p&offsets|0|0;m-15&6|-|71403595|71403616&offsets|0|0]|TGTAAACATCCTCGACTGGAAG</td><td>>MI0000088|hsa-mir-30a&WithFlank&6|-|71403545|71403627</td><td>12</td><td>33</td><td>22</td><td>TGTAAACATCCTCGACTGGAAG</td><td>MB231</td><td>283</td><td>194</td><td>29</td><td>1196</td><td>200</td><td>47.7</td><td>133.0</td><td>21.2</td><td>835.0</td><td>150.8</td></tr>
<tr><td>>MI0000088|hsa-mir-30a&WithFlank&6|-|71403545|71403627@12.33.22&[MIMAT0000087&hsa-miR-30a-5p&offsets|0|0;m-15&6|-|71403595|71403616&offsets|0|0]|TGTAAACATCCTCGACTGGAAG</td><td>>MI0000088|hsa-mir-30a&WithFlank&6|-|71403545|71403627</td><td>12</td><td>33</td><td>22</td><td>TGTAAACATCCTCGACTGGAAG</td><td>MB468</td><td>12</td><td>3</td><td>2</td><td>88</td><td>3</td><td>3.5</td><td>1.9</td><td>1.6</td><td>49.3</td><td>0.7</td></tr>
<tr><td>>MI0000088|hsa-mir-30a&WithFlank&6|-|71403545|71403627@12.34.23(+1A)&[MIMAT0000087&hsa-miR-30a-5p&offsets|0|+1(+1A);m-15&6|-|71403595|71403616&offsets|0|+1(+1A)]|TGTAAACATCCTCGACTGGAAGCA</td><td>>MI0000088|hsa-mir-30a&WithFlank&6|-|71403545|71403627</td><td>12</td><td>34</td><td>23(+1A)</td><td>TGTAAACATCCTCGACTGGAAGCA</td><td>BT20</td><td>1050</td><td>244</td><td>126</td><td>520</td><td>74</td><td>1237.5</td><td>327.0</td><td>50.3</td><td>435.4</td><td>19.5</td></tr>
<tr><td>>MI0000088|hsa-mir-30a&WithFlank&6|-|71403545|71403627@12.34.23(+1A)&[MIMAT0000087&hsa-miR-30a-5p&offsets|0|+1(+1A);m-15&6|-|71403595|71403616&offsets|0|+1(+1A)]|TGTAAACATCCTCGACTGGAAGCA</td><td>>MI0000088|hsa-mir-30a&WithFlank&6|-|71403545|71403627</td><td>12</td><td>34</td><td>23(+1A)</td><td>TGTAAACATCCTCGACTGGAAGCA</td><td>MB231</td><td>783</td><td>358</td><td>77</td><td>3089</td><td>761</td><td>177.3</td><td>200.8</td><td>40.4</td><td>352.7</td><td>778.1</td></tr>
<tr><td>>MI0000088|hsa-mir-30a&WithFlank&6|-|71403545|71403627@12.34.23(+1A)&[MIMAT0000087&hsa-miR-30a-5p&offsets|0|+1(+1A);m-15&6|-|71403595|71403616&offsets|0|+1(+1A)]|TGTAAACATCCTCGACTGGAAGCA</td><td>>MI0000088|hsa-mir-30a&WithFlank&6|-|71403545|71403627</td><td>12</td><td>34</td><td>23(+1A)</td><td>TGTAAACATCCTCGACTGGAAGCA</td><td>MB468</td><td>20</td><td>3</td><td>4</td><td>86</td><td>6</td><td>7.7</td><td>1.7</td><td>4.1</td><td>67.7</td><td>2.7</td></tr>
<tr><td>>MI0000088|hsa-mir-30a&WithFlank&6|-|71403545|71403627@12.34.23(+1C)&[MIMAT0000087&hsa-miR-30a-5p&offsets|0|+1(+1C);m-15&6|-|71403595|71403616&offsets|0|+1(+1C)]|TGTAAACATCCTCGACTGGAAGCC</td><td>>MI0000088|hsa-mir-30a&WithFlank&6|-|71403545|71403627</td><td>12</td><td>34</td><td>23(+1C)</td><td>TGTAAACATCCTCGACTGGAAGCC</td><td>BT20</td><td>952</td><td>409</td><td>145</td><td>789</td><td>122</td><td>798.1</td><td>557.7</td><td>71.6</td><td>653.9</td><td>46.8</td></tr>
<tr><td>>MI0000088|hsa-mir-30a&WithFlank&6|-|71403545|71403627@12.34.23(+1C)&[MIMAT0000087&hsa-miR-30a-5p&offsets|0|+1(+1C);m-15&6|-|71403595|71403616&offsets|0|+1(+1C)]|TGTAAACATCCTCGACTGGAAGCC</td><td>>MI0000088|hsa-mir-30a&WithFlank&6|-|71403545|71403627</td><td>12</td><td>34</td><td>23(+1C)</td><td>TGTAAACATCCTCGACTGGAAGCC</td><td>MB231</td><td>1213</td><td>669</td><td>114</td><td>4585</td><td>1285</td><td>276.0</td><td>366.7</td><td>43.5</td><td>372.3</td><td>1358.7</td></tr>
<tr><td>>MI0000088|hsa-mir-30a&WithFlank&6|-|71403545|71403627@12.34.23(+1C)&[MIMAT0000087&hsa-miR-30a-5p&offsets|0|+1(+1C);m-15&6|-|71403595|71403616&offsets|0|+1(+1C)]|TGTAAACATCCTCGACTGGAAGCC</td><td>>MI0000088|hsa-mir-30a&WithFlank&6|-|71403545|71403627</td><td>12</td><td>34</td><td>23(+1C)</td><td>TGTAAACATCCTCGACTGGAAGCC</td><td>MB468</td><td>28</td><td>6</td><td>6</td><td>114</td><td>8</td><td>9.1</td><td>4.9</td><td>4.7</td><td>67.7</td><td>2.1</td></tr>
<tr><td>>MI0000088|hsa-mir-30a&WithFlank&6|-|71403545|71403627@12.34.23(+1G)&[MIMAT0000087&hsa-miR-30a-5p&offsets|0|+1(+1G);m-15&6|-|71403595|71403616&offsets|0|+1(+1G)]|TGTAAACATCCTCGACTGGAAGCG</td><td>>MI0000088|hsa-mir-30a&WithFlank&6|-|71403545|71403627</td><td>12</td><td>34</td><td>23(+1G)</td><td>TGTAAACATCCTCGACTGGAAGCG</td><td>BT20</td><td>758</td><td>90</td><td>34</td><td>136</td><td>14</td><td>931.9</td><td>120.3</td><td>6.7</td><td>138.3</td><td>4.1</td></tr>
<tr><td>>MI0000088|hsa-mir-30a&WithFlank&6|-|71403545|71403627@12.34.23(+1G)&[MIMAT0000087&hsa-miR-30a-5p&offsets|0|+1(+1G);m-15&6|-|71403595|71403616&offsets|0|+1(+1G)]|TGTAAACATCCTCGACTGGAAGCG</td><td>>MI0000088|hsa-mir-30a&WithFlank&6|-|71403545|71403627</td><td>12</td><td>34</td><td>23(+1G)</td><td>TGTAAACATCCTCGACTGGAAGCG</td><td>MB231</td><td>606</td><td>155</td><td>21</td><td>611</td><td>154</td><td>436.2</td><td>103.7</td><td>13.1</td><td>121.8</td><td>164.3</td></tr>
<tr><td>>MI0000088|hsa-mir-30a&WithFlank&6|-|71403545|71403627@12.34.23(+1G)&[MIMAT0000087&hsa-miR-30a-5p&offsets|0|+1(+1G);m-15&6|-|71403595|71403616&offsets|0|+1(+1G)]|TGTAAACATCCTCGACTGGAAGCG</td><td>>MI0000088|hsa-mir-30a&WithFlank&6|-|71403545|71403627</td><td>12</td><td>34</td><td>23(+1G)</td><td>TGTAAACATCCTCGACTGGAAGCG</td><td>MB468</td><td>17</td><td>2</td><td>4</td><td>35</td><td>2</td><td>5.8</td><td>1.3</td><td>4.2</td><td>27.9</td><td>0.9</td></tr>
<tr><td>>MI0000088|hsa-mir-30a&WithFlank&6|-|71403545|71403627@12.34.23&[MIMAT0000087&hsa-miR-30a-5p&offsets|0|+1;m-15&6|-|71403595|71403616&offsets|0|+1]|TGTAAACATCCTCGACTGGAAGC</td><td>>MI0000088|hsa-mir-30a&WithFlank&6|-|71403545|71403627</td><td>12</td><td>34</td><td>23</td><td>TGTAAACATCCTCGACTGGAAGC</td><td>BT20</td><td>2269</td><td>911</td><td>458</td><td>1840</td><td>345</td><td>2025.1</td><td>890.6</td><td>165.8</td><td>1460.7</td><td>39.3</td></tr>
<tr><td>>MI0000088|hsa-mir-30a&WithFlank&6|-|71403545|71403627@12.34.23&[MIMAT0000087&hsa-miR-30a-5p&offsets|0|+1;m-15&6|-|71403595|71403616&offsets|0|+1]|TGTAAACATCCTCGACTGGAAGC</td><td>>MI0000088|hsa-mir-30a&WithFlank&6|-|71403545|71403627</td><td>12</td><td>34</td><td>23</td><td>TGTAAACATCCTCGACTGGAAGC</td><td>MB231</td><td>1948</td><td>1386</td><td>218</td><td>5141</td><td>1523</td><td>164.8</td><td>803.8</td><td>158.7</td><td>2476.1</td><td>1120.0</td></tr>
<tr><td>>MI0000088|hsa-mir-30a&WithFlank&6|-|71403545|71403627@12.34.23&[MIMAT0000087&hsa-miR-30a-5p&offsets|0|+1;m-15&6|-|71403595|71403616&offsets|0|+1]|TGTAAACATCCTCGACTGGAAGC</td><td>>MI0000088|hsa-mir-30a&WithFlank&6|-|71403545|71403627</td><td>12</td><td>34</td><td>23</td><td>TGTAAACATCCTCGACTGGAAGC</td><td>MB468</td><td>66</td><td>19</td><td>15</td><td>269</td><td>19</td><td>19.1</td><td>10.9</td><td>7.3</td><td>181.7</td><td>4.5</td></tr>
<tr><td>>MI0000088|hsa-mir-30a&WithFlank&6|-|71403545|71403627@12.35.24(+1A)&[MIMAT0000087&hsa-miR-30a-5p&offsets|0|+2(+1A);m-15&6|-|71403595|71403616&offsets|0|+2(+1A)]|TGTAAACATCCTCGACTGGAAGCTA</td><td>>MI0000088|hsa-mir-30a&WithFlank&6|-|71403545|71403627</td><td>12</td><td>35</td><td>24(+1A)</td><td>TGTAAACATCCTCGACTGGAAGCTA</td><td>BT20</td><td>40</td><td>11</td><td>5</td><td>16</td><td>3</td><td>42.7</td><td>15.1</td><td>3.4</td><td>14.2</td><td>0.9</td></tr>
<tr><td>>MI0000088|hsa-mir-30a&WithFlank&6|-|71403545|71403627@12.35.24(+1A)&[MIMAT0000087&hsa-miR-30a-5p&offsets|0|+2(+1A);m-15&6|-|71403595|71403616&offsets|0|+2(+1A)]|TGTAAACATCCTCGACTGGAAGCTA</td><td>>MI0000088|hsa-mir-30a&WithFlank&6|-|71403545|71403627</td><td>12</td><td>35</td><td>24(+1A)</td><td>TGTAAACATCCTCGACTGGAAGCTA</td><td>MB231</td><td>59</td><td>24</td><td>4</td><td>172</td><td>55</td><td>11.8</td><td>13.1</td><td>2.5</td><td>36.9</td><td>57.1</td></tr>
<tr><td>>MI0000088|hsa-mir-30a&WithFlank&6|-|71403545|71403627@12.35.24(+1C)&[MIMAT0000087&hsa-miR-30a-5p&offsets|0|+2(+1C);m-15&6|-|71403595|71403616&offsets|0|+2(+1C)]|TGTAAACATCCTCGACTGGAAGCTC</td><td>>MI0000088|hsa-mir-30a&WithFlank&6|-|71403545|71403627</td><td>12</td><td>35</td><td>24(+1C)</td><td>TGTAAACATCCTCGACTGGAAGCTC</td><td>BT20</td><td>19</td><td>9</td><td>3</td><td>25</td><td>2</td><td>17.2</td><td>13.2</td><td>1.9</td><td>21.4</td><td>1.2</td></tr>
<tr><td>>MI0000088|hsa-mir-30a&WithFlank&6|-|71403545|71403627@12.35.24(+1C)&[MIMAT0000087&hsa-miR-30a-5p&offsets|0|+2(+1C);m-15&6|-|71403595|71403616&offsets|0|+2(+1C)]|TGTAAACATCCTCGACTGGAAGCTC</td><td>>MI0000088|hsa-mir-30a&WithFlank&6|-|71403545|71403627</td><td>12</td><td>35</td><td>24(+1C)</td><td>TGTAAACATCCTCGACTGGAAGCTC</td><td>MB231</td><td>55</td><td>28</td><td>5</td><td>328</td><td>65</td><td>15.2</td><td>16.3</td><td>2.3</td><td>32.9</td><td>72.9</td></tr>
<tr><td>>MI0000088|hsa-mir-30a&WithFlank&6|-|71403545|71403627@12.35.24(+1U)&[MIMAT0000087&hsa-miR-30a-5p&offsets|0|+2(+1U);m-15&6|-|71403595|71403616&offsets|0|+2(+1U)]|TGTAAACATCCTCGACTGGAAGCTT</td><td>>MI0000088|hsa-mir-30a&WithFlank&6|-|71403545|71403627</td><td>12</td><td>35</td><td>24(+1U)</td><td>TGTAAACATCCTCGACTGGAAGCTT</td><td>BT20</td><td>140</td><td>61</td><td>23</td><td>197</td><td>17</td><td>126.9</td><td>89.0</td><td>15.2</td><td>183.6</td><td>10.1</td></tr>
</table>

<table>
<tr><th></th><th></th><th></th><th></th><th></th><th></th><th></th><th></th><th></th><th></th><th></th><th></th><th></th><th></th><th></th><th></th><th></th></tr>
<tr><td>>MI0000088|hsa-mir-30a&WithFlank&6|-|71403545|71403627@12.35.24(+1U)&[MIMAT0000087&hsa-miR-30a-5p&offsets|0|+2(+1U);m-15&6|-|71403595|71403616&offsets|0|+2(+1U)]|TGTAAACATCCTCGACTGGAAGCTT</td><td>>MI0000088|hsa-mir-30a&WithFlank&6|-|71403545|71403627</td><td>12</td><td>35</td><td>24(+1U)</td><td>TGTAAACATCCTCGACTGGAAGCTT</td><td>MB231</td><td>503</td><td>265</td><td>45</td><td>2323</td><td>465</td><td>136.4</td><td>138.8</td><td>19.1</td><td>843.6</td><td>457.7</td></tr>
<tr><td>>MI0000088|hsa-mir-30a&WithFlank&6|-|71403545|71403627@12.35.24(+2U)&[MIMAT0000087&hsa-miR-30a-5p&offsets|0|+2(+2U);m-15&6|-|71403595|71403616&offsets|0|+2(+2U)]|TGTAAACATCCTCGACTGGAAGCTTT</td><td>>MI0000088|hsa-mir-30a&WithFlank&6|-|71403545|71403627</td><td>12</td><td>35</td><td>24(+2U)</td><td>TGTAAACATCCTCGACTGGAAGCTTT</td><td>BT20</td><td>17</td><td>8</td><td>4</td><td>5</td><td>4</td><td>11.4</td><td>10.8</td><td>2.6</td><td>4.1</td><td>2.2</td></tr>
<tr><td>>MI0000088|hsa-mir-30a&WithFlank&6|-|71403545|71403627@12.35.24(+2U)&[MIMAT0000087&hsa-miR-30a-5p&offsets|0|+2(+2U);m-15&6|-|71403595|71403616&offsets|0|+2(+2U)]|TGTAAACATCCTCGACTGGAAGCTTT</td><td>>MI0000088|hsa-mir-30a&WithFlank&6|-|71403545|71403627</td><td>12</td><td>35</td><td>24(+2U)</td><td>TGTAAACATCCTCGACTGGAAGCTTT</td><td>MB231</td><td>166</td><td>101</td><td>16</td><td>335</td><td>157</td><td>59.3</td><td>57.6</td><td>7.9</td><td>115.6</td><td>136.6</td></tr>
<tr><td>>MI0000088|hsa-mir-30a&WithFlank&6|-|71403545|71403627@12.35.24(+3U)&[MIMAT0000087&hsa-miR-30a-5p&offsets|0|+2(+3U);m-15&6|-|71403595|71403616&offsets|0|+2(+3U)]|TGTAAACATCCTCGACTGGAAGCTTTT</td><td>>MI0000088|hsa-mir-30a&WithFlank&6|-|71403545|71403627</td><td>12</td><td>35</td><td>24(+3U)</td><td>TGTAAACATCCTCGACTGGAAGCTTTT</td><td>MB231</td><td>20</td><td>11</td><td>3</td><td>30</td><td>24</td><td>2.1</td><td>6.4</td><td>1.5</td><td>13.5</td><td>24.8</td></tr>
<tr><td>>MI0000088|hsa-mir-30a&WithFlank&6|-|71403545|71403627@12.35.24&[MIMAT0000087&hsa-miR-30a-5p&offsets|0|+2;m-15&6|-|71403595|71403616&offsets|0|+2]|TGTAAACATCCTCGACTGGAAGCT</td><td>>MI0000088|hsa-mir-30a&WithFlank&6|-|71403545|71403627</td><td>12</td><td>35</td><td>24</td><td>TGTAAACATCCTCGACTGGAAGCT</td><td>BT20</td><td>6010</td><td>2214</td><td>956</td><td>5102</td><td>775</td><td>5002.7</td><td>2939.6</td><td>538.6</td><td>4163.1</td><td>237.0</td></tr>
<tr><td>>MI0000088|hsa-mir-30a&WithFlank&6|-|71403545|71403627@12.35.24&[MIMAT0000087&hsa-miR-30a-5p&offsets|0|+2;m-15&6|-|71403595|71403616&offsets|0|+2]|TGTAAACATCCTCGACTGGAAGCT</td><td>>MI0000088|hsa-mir-30a&WithFlank&6|-|71403545|71403627</td><td>12</td><td>35</td><td>24</td><td>TGTAAACATCCTCGACTGGAAGCT</td><td>MB231</td><td>9108</td><td>4954</td><td>886</td><td>27816</td><td>7602</td><td>2639.6</td><td>2328.6</td><td>333.4</td><td>6501.9</td><td>7001.8</td></tr>
<tr><td>>MI0000088|hsa-mir-30a&WithFlank&6|-|71403545|71403627@12.35.24&[MIMAT0000087&hsa-miR-30a-5p&offsets|0|+2;m-15&6|-|71403595|71403616&offsets|0|+2]|TGTAAACATCCTCGACTGGAAGCT</td><td>>MI0000088|hsa-mir-30a&WithFlank&6|-|71403545|71403627</td><td>12</td><td>35</td><td>24</td><td>TGTAAACATCCTCGACTGGAAGCT</td><td>MB468</td><td>212</td><td>50</td><td>39</td><td>752</td><td>54</td><td>74.7</td><td>41.4</td><td>26.9</td><td>396.6</td><td>14.1</td></tr>
<tr><td>>MI0000088|hsa-mir-30a&WithFlank&6|-|71403545|71403627@12.36.25&[MIMAT0000087&hsa-miR-30a-5p&offsets|0|+3;m-15&6|-|71403595|71403616&offsets|0|+3]|TGTAAACATCCTCGACTGGAAGCTG</td><td>>MI0000088|hsa-mir-30a&WithFlank&6|-|71403545|71403627</td><td>12</td><td>36</td><td>25</td><td>TGTAAACATCCTCGACTGGAAGCTG</td><td>BT20</td><td>17</td><td>2</td><td>1</td><td>4</td><td>1</td><td>20.6</td><td>3.4</td><td>1.0</td><td>3.4</td><td>0.6</td></tr>
<tr><td>>MI0000088|hsa-mir-30a&WithFlank&6|-|71403545|71403627@12.36.25&[MIMAT0000087&hsa-miR-30a-5p&offsets|0|+3;m-15&6|-|71403595|71403616&offsets|0|+3]|TGTAAACATCCTCGACTGGAAGCTG</td><td>>MI0000088|hsa-mir-30a&WithFlank&6|-|71403545|71403627</td><td>12</td><td>36</td><td>25</td><td>TGTAAACATCCTCGACTGGAAGCTG</td><td>MB231</td><td>25</td><td>8</td><td>1</td><td>38</td><td>10</td><td>13.3</td><td>4.9</td><td>0.9</td><td>2.0</td><td>7.8</td></tr>
<tr><td>>MI0000088|hsa-mir-30a&WithFlank&6|-|71403545|71403627@13.34.22(+1A)&[MIMAT0000087&hsa-miR-30a-5p&offsets|+1|+1(+1A);m-15&6|-|71403595|71403616&offsets|+1|+1(+1A)]|GTAAACATCCTCGACTGGAAGCA</td><td>>MI0000088|hsa-mir-30a&WithFlank&6|-|71403545|71403627</td><td>13</td><td>34</td><td>22(+1A)</td><td>GTAAACATCCTCGACTGGAAGCA</td><td>BT20</td><td>19</td><td>5</td><td>3</td><td>17</td><td>2</td><td>22.3</td><td>6.1</td><td>0.9</td><td>15.5</td><td>0.1</td></tr>
<tr><td>>MI0000088|hsa-mir-30a&WithFlank&6|-|71403545|71403627@13.34.22(+1C)&[MIMAT0000087&hsa-miR-30a-5p&offsets|+1|+1(+1C);m-15&6|-|71403595|71403616&offsets|+1|+1(+1C)]|GTAAACATCCTCGACTGGAAGCC</td><td>>MI0000088|hsa-mir-30a&WithFlank&6|-|71403545|71403627</td><td>13</td><td>34</td><td>22(+1C)</td><td>GTAAACATCCTCGACTGGAAGCC</td><td>BT20</td><td>19</td><td>8</td><td>3</td><td>27</td><td>2</td><td>15.1</td><td>9.8</td><td>1.1</td><td>22.8</td><td>0.4</td></tr>
<tr><td>>MI0000088|hsa-mir-30a&WithFlank&6|-|71403545|71403627@13.34.22(+1C)&[MIMAT0000087&hsa-miR-30a-5p&offsets|+1|+1(+1C);m-15&6|-|71403595|71403616&offsets|+1|+1(+1C)]|GTAAACATCCTCGACTGGAAGCC</td><td>>MI0000088|hsa-mir-30a&WithFlank&6|-|71403545|71403627</td><td>13</td><td>34</td><td>22(+1C)</td><td>GTAAACATCCTCGACTGGAAGCC</td><td>MB231</td><td>15</td><td>10</td><td>2</td><td>66</td><td>15</td><td>3.1</td><td>4.5</td><td>1.0</td><td>5.8</td><td>15.0</td></tr>
<tr><td>>MI0000088|hsa-mir-30a&WithFlank&6|-|71403545|71403627@13.34.22(+1G)&[MIMAT0000087&hsa-miR-30a-5p&offsets|+1|+1(+1G);m-15&6|-|71403595|71403616&offsets|+1|+1(+1G)]|GTAAACATCCTCGACTGGAAGCG</td><td>>MI0000088|hsa-mir-30a&WithFlank&6|-|71403545|71403627</td><td>13</td><td>34</td><td>22(+1G)</td><td>GTAAACATCCTCGACTGGAAGCG</td><td>BT20</td><td>15</td><td>2</td><td>1</td><td>4</td><td>0</td><td>18.3</td><td>2.4</td><td>0.2</td><td>5.5</td><td>0.2</td></tr>
<tr><td>>MI0000088|hsa-mir-30a&WithFlank&6|-|71403545|71403627@13.35.23(+1U)&[MIMAT0000087&hsa-miR-30a-5p&offsets|+1|+2(+1U);m-15&6|-|71403595|71403616&offsets|+1|+2(+1U)]|GTAAACATCCTCGACTGGAAGCTT</td><td>>MI0000088|hsa-mir-30a&WithFlank&6|-|71403545|71403627</td><td>13</td><td>35</td><td>23(+1U)</td><td>GTAAACATCCTCGACTGGAAGCTT</td><td>MB231</td><td>20</td><td>14</td><td>3</td><td>100</td><td>16</td><td>3.1</td><td>5.9</td><td>1.1</td><td>37.2</td><td>11.2</td></tr>
<tr><td>>MI0000088|hsa-mir-30a&WithFlank&6|-|71403545|71403627@13.35.23&[MIMAT0000087&hsa-miR-30a-5p&offsets|+1|+2;m-15&6|-|71403595|71403616&offsets|+1|+2]|GTAAACATCCTCGACTGGAAGCT</td><td>>MI0000088|hsa-mir-30a&WithFlank&6|-|71403545|71403627</td><td>13</td><td>35</td><td>23</td><td>GTAAACATCCTCGACTGGAAGCT</td><td>BT20</td><td>117</td><td>43</td><td>20</td><td>167</td><td>17</td><td>89.1</td><td>51.3</td><td>10.1</td><td>134.2</td><td>2.0</td></tr>
<tr><td>>MI0000088|hsa-mir-30a&WithFlank&6|-|71403545|71403627@13.35.23&[MIMAT0000087&hsa-miR-30a-5p&offsets|+1|+2;m-15&6|-|71403595|71403616&offsets|+1|+2]|GTAAACATCCTCGACTGGAAGCT</td><td>>MI0000088|hsa-mir-30a&WithFlank&6|-|71403545|71403627</td><td>13</td><td>35</td><td>23</td><td>GTAAACATCCTCGACTGGAAGCT</td><td>MB231</td><td>116</td><td>75</td><td>16</td><td>406</td><td>88</td><td>30.1</td><td>28.8</td><td>7.7</td><td>75.0</td><td>73.7</td></tr>
<tr><td>>MI0000088|hsa-mir-30a&WithFlank&6|-|71403545|71403627@53.70.18(+1U)&[MIMAT0000088&hsa-miR-30a-3p&offsets|0|-4(+1U);m-38&6|-|71403554|71403575&offsets|0|-4(+1U)]|CTTTCAGTCGGATGTTTGT</td><td>>MI0000088|hsa-mir-30a&WithFlank&6|-|71403545|71403627</td><td>53</td><td>70</td><td>18(+1U)</td><td>CTTTCAGTCGGATGTTTGT</td><td>BT20</td><td>12</td><td>8</td><td>14</td><td>6</td><td>4</td><td>10.6</td><td>6.5</td><td>5.1</td><td>4.9</td><td>1.2</td></tr>
<tr><td>>MI0000088|hsa-mir-30a&WithFlank&6|-|71403545|71403627@53.70.18(+1U)&[MIMAT0000088&hsa-miR-30a-3p&offsets|0|-4(+1U);m-38&6|-|71403554|71403575&offsets|0|-4(+1U)]|CTTTCAGTCGGATGTTTGT</td><td>>MI0000088|hsa-mir-30a&WithFlank&6|-|71403545|71403627</td><td>53</td><td>70</td><td>18(+1U)</td><td>CTTTCAGTCGGATGTTTGT</td><td>MB231</td><td>20</td><td>24</td><td>19</td><td>70</td><td>20</td><td>9.8</td><td>1.0</td><td>7.6</td><td>43.7</td><td>8.3</td></tr>
<tr><td>>MI0000088|hsa-mir-30a&WithFlank&6|-|71403545|71403627@53.71.19&[MIMAT0000088&hsa-miR-30a-3p&offsets|0|-3;m-38&6|-|71403554|71403575&offsets|0|-3]|CTTTCAGTCGGATGTTTGC</td><td>>MI0000088|hsa-mir-30a&WithFlank&6|-|71403545|71403627</td><td>53</td><td>71</td><td>19</td><td>CTTTCAGTCGGATGTTTGC</td><td>BT20</td><td>37</td><td>32</td><td>53</td><td>25</td><td>19</td><td>44.0</td><td>19.9</td><td>31.7</td><td>8.3</td><td>6.3</td></tr>
<tr><td>>MI0000088|hsa-mir-30a&WithFlank&6|-|71403545|71403627@53.71.19&[MIMAT0000088&hsa-miR-30a-3p&offsets|0|-3;m-38&6|-|71403554|71403575&offsets|0|-3]|CTTTCAGTCGGATGTTTGC</td><td>>MI0000088|hsa-mir-30a&WithFlank&6|-|71403545|71403627</td><td>53</td><td>71</td><td>19</td><td>CTTTCAGTCGGATGTTTGC</td><td>MB231</td><td>86</td><td>161</td><td>102</td><td>228</td><td>86</td><td>50.3</td><td>34.5</td><td>38.3</td><td>85.2</td><td>25.3</td></tr>
<tr><td>>MI0000088|hsa-mir-30a&WithFlank&6|-|71403545|71403627@53.73.21(+1A)&[MIMAT0000088&hsa-miR-30a-3p&offsets|0|-1(+1A);m-38&6|-|71403554|71403575&offsets|0|-1(+1A)]|CTTTCAGTCGGATGTTTGCAGA</td><td>>MI0000088|hsa-mir-30a&WithFlank&6|-|71403545|71403627</td><td>53</td><td>73</td><td>21(+1A)</td><td>CTTTCAGTCGGATGTTTGCAGA</td><td>BT20</td><td>23</td><td>15</td><td>38</td><td>4</td><td>5</td><td>16.3</td><td>10.3</td><td>16.8</td><td>3.6</td><td>1.3</td></tr>
<tr><td>>MI0000088|hsa-mir-30a&WithFlank&6|-|71403545|71403627@53.73.21(+1A)&[MIMAT0000088&hsa-miR-30a-3p&offsets|0|-1(+1A);m-38&6|-|71403554|71403575&offsets|0|-1(+1A)]|CTTTCAGTCGGATGTTTGCAGA</td><td>>MI0000088|hsa-mir-30a&WithFlank&6|-|71403545|71403627</td><td>53</td><td>73</td><td>21(+1A)</td><td>CTTTCAGTCGGATGTTTGCAGA</td><td>MB231</td><td>28</td><td>35</td><td>32</td><td>58</td><td>31</td><td>5.8</td><td>3.6</td><td>24.1</td><td>22.0</td><td>14.5</td></tr>
<tr><td>>MI0000088|hsa-mir-30a&WithFlank&6|-|71403545|71403627@53.73.21(+1G)&[MIMAT0000088&hsa-miR-30a-3p&offsets|0|-1(+1G);m-38&6|-|71403554|71403575&offsets|0|-1(+1G)]|CTTTCAGTCGGATGTTTGCAGG</td><td>>MI0000088|hsa-mir-30a&WithFlank&6|-|71403545|71403627</td><td>53</td><td>73</td><td>21(+1G)</td><td>CTTTCAGTCGGATGTTTGCAGG</td><td>BT20</td><td>12</td><td>4</td><td>6</td><td>1</td><td>0</td><td>11.5</td><td>2.6</td><td>2.6</td><td>1.1</td><td>0.3</td></tr>
<tr><td>>MI0000088|hsa-mir-30a&WithFlank&6|-|71403545|71403627@53.73.21(+1G)&[MIMAT0000088&hsa-miR-30a-3p&offsets|0|-1(+1G);m-38&6|-|71403554|71403575&offsets|0|-1(+1G)]|CTTTCAGTCGGATGTTTGCAGG</td><td>>MI0000088|hsa-mir-30a&WithFlank&6|-|71403545|71403627</td><td>53</td><td>73</td><td>21(+1G)</td><td>CTTTCAGTCGGATGTTTGCAGG</td><td>MB231</td><td>13</td><td>8</td><td>4</td><td>7</td><td>4</td><td>7.6</td><td>3.5</td><td>2.9</td><td>2.5</td><td>2.6</td></tr>
<tr><td>>MI0000088|hsa-mir-30a&WithFlank&6|-|71403545|71403627@53.73.21(+1U)&[MIMAT0000088&hsa-miR-30a-3p&offsets|0|-1(+1U);m-38&6|-|71403554|71403575&offsets|0|-1(+1U)]|CTTTCAGTCGGATGTTTGCAGT</td><td>>MI0000088|hsa-mir-30a&WithFlank&6|-|71403545|71403627</td><td>53</td><td>73</td><td>21(+1U)</td><td>CTTTCAGTCGGATGTTTGCAGT</td><td>BT20</td><td>201</td><td>199</td><td>273</td><td>48</td><td>72</td><td>95.1</td><td>133.5</td><td>50.7</td><td>31.7</td><td>31.0</td></tr>
<tr><td>>MI0000088|hsa-mir-30a&WithFlank&6|-|71403545|71403627@53.73.21(+1U)&[MIMAT0000088&hsa-miR-30a-3p&offsets|0|-1(+1U);m-38&6|-|71403554|71403575&offsets|0|-1(+1U)]|CTTTCAGTCGGATGTTTGCAGT</td><td>>MI0000088|hsa-mir-30a&WithFlank&6|-|71403545|71403627</td><td>53</td><td>73</td><td>21(+1U)</td><td>CTTTCAGTCGGATGTTTGCAGT</td><td>MB231</td><td>360</td><td>547</td><td>309</td><td>600</td><td>433</td><td>100.2</td><td>80.9</td><td>173.2</td><td>119.7</td><td>159.8</td></tr>
<tr><td>>MI0000088|hsa-mir-30a&WithFlank&6|-|71403545|71403627@53.73.21(+2U)&[MIMAT0000088&hsa-miR-30a-3p&offsets|0|-1(+2U);m-38&6|-|71403554|71403575&offsets|0|-1(+2U)]|CTTTCAGTCGGATGTTTGCAGTT</td><td>>MI0000088|hsa-mir-30a&WithFlank&6|-|71403545|71403627</td><td>53</td><td>73</td><td>21(+2U)</td><td>CTTTCAGTCGGATGTTTGCAGTT</td><td>MB231</td><td>27</td><td>43</td><td>29</td><td>54</td><td>25</td><td>8.1</td><td>8.9</td><td>11.0</td><td>15.8</td><td>6.8</td></tr>
<tr><td>>MI0000088|hsa-mir-30a&WithFlank&6|-|71403545|71403627@53.73.21&[MIMAT0000088&hsa-miR-30a-3p&offsets|0|-1;m-38&6|-|71403554|71403575&offsets|0|-1]|CTTTCAGTCGGATGTTTGCAG</td><td>>MI0000088|hsa-mir-30a&WithFlank&6|-|71403545|71403627</td><td>53</td><td>73</td><td>21</td><td>CTTTCAGTCGGATGTTTGCAG</td><td>BT20</td><td>24</td><td>25</td><td>37</td><td>25</td><td>12</td><td>12.9</td><td>13.2</td><td>20.0</td><td>17.8</td><td>3.2</td></tr>
<tr><td>>MI0000088|hsa-mir-30a&WithFlank&6|-|71403545|71403627@53.73.21&[MIMAT0000088&hsa-miR-30a-3p&offsets|0|-1;m-38&6|-|71403554|71403575&offsets|0|-1]|CTTTCAGTCGGATGTTTGCAG</td><td>>MI0000088|hsa-mir-30a&WithFlank&6|-|71403545|71403627</td><td>53</td><td>73</td><td>21</td><td>CTTTCAGTCGGATGTTTGCAG</td><td>MB231</td><td>53</td><td>84</td><td>46</td><td>186</td><td>45</td><td>8.7</td><td>21.5</td><td>17.0</td><td>50.3</td><td>12.6</td></tr>
<tr><td>>MI0000088|hsa-mir-30a&WithFlank&6|-|71403545|71403627@53.74.22(+1A)&[MIMAT0000088&hsa-miR-30a-3p&offsets|0|0(+1A);m-38&6|-|71403554|71403575&offsets|0|0(+1A)]|CTTTCAGTCGGATGTTTGCAGCA</td><td>>MI0000088|hsa-mir-30a&WithFlank&6|-|71403545|71403627</td><td>53</td><td>74</td><td>22(+1A)</td><td>CTTTCAGTCGGATGTTTGCAGCA</td><td>MB231</td><td>13</td><td>18</td><td>20</td><td>11</td><td>11</td><td>3.9</td><td>5.9</td><td>12.2</td><td>2.9</td><td>3.5</td></tr>
<tr><td>>MI0000088|hsa-mir-30a&WithFlank&6|-|71403545|71403627@53.74.22&[MIMAT0000088&hsa-miR-30a-3p&offsets|0|0;m-38&6|-|71403554|71403575&offsets|0|0]|CTTTCAGTCGGATGTTTGCAGC</td><td>>MI0000088|hsa-mir-30a&WithFlank&6|-|71403545|71403627</td><td>53</td><td>74</td><td>22</td><td>CTTTCAGTCGGATGTTTGCAGC</td><td>BT20</td><td>205</td><td>222</td><td>393</td><td>59</td><td>87</td><td>117.8</td><td>138.1</td><td>85.3</td><td>30.2</td><td>37.1</td></tr>
<tr><td>>MI0000088|hsa-mir-30a&WithFlank&6|-|71403545|71403627@53.74.22&[MIMAT0000088&hsa-miR-30a-3p&offsets|0|0;m-38&6|-|71403554|71403575&offsets|0|0]|CTTTCAGTCGGATGTTTGCAGC</td><td>>MI0000088|hsa-mir-30a&WithFlank&6|-|71403545|71403627</td><td>53</td><td>74</td><td>22</td><td>CTTTCAGTCGGATGTTTGCAGC</td><td>MB231</td><td>507</td><td>849</td><td>629</td><td>772</td><td>536</td><td>179.6</td><td>166.3</td><td>342.1</td><td>193.0</td><td>142.0</td></tr>
<tr><td>>MI0000088|hsa-mir-30a&WithFlank&6|-|71403545|71403627@53.74.22&[MIMAT0000088&hsa-miR-30a-3p&offsets|0|0;m-38&6|-|71403554|71403575&offsets|0|0]|CTTTCAGTCGGATGTTTGCAGC</td><td>>MI0000088|hsa-mir-30a&WithFlank&6|-|71403545|71403627</td><td>53</td><td>74</td><td>22</td><td>CTTTCAGTCGGATGTTTGCAGC</td><td>MB468</td><td>15</td><td>36</td><td>14</td><td>48</td><td>8</td><td>4.0</td><td>11.0</td><td>5.5</td><td>45.2</td><td>3.6</td></tr>
</table>

<table>
<tr><th></th><th></th><th></th><th></th><th></th><th></th><th></th><th></th><th></th><th></th><th></th><th></th><th></th><th></th><th></th><th></th><th></th></tr>
<tr><td>>MI0000088|hsa-mir-30a&WithFlank&6|-|71403545|71403627@53.75.23&[MIMAT0000088&hsa-miR-30a-3p&offsets|0|+1;m-38&6|-|71403554|71403575&offsets|0|+1]|CTTTCAGTCGGATGTTTGCAGCT</td><td>>MI0000088|hsa-mir-30a&WithFlank&6|-|71403545|71403627</td><td>53</td><td>75</td><td>23</td><td>CTTTCAGTCGGATGTTTGCAGCT</td><td>MB231</td><td>13</td><td>21</td><td>17</td><td>21</td><td>14</td><td>4.5</td><td>7.6</td><td>10.4</td><td>6.3</td><td>5.6</td></tr>
<tr><td>>MI0000088|hsa-mir-30a&WithFlank&6|-|71403545|71403627@54.71.18&[MIMAT0000088&hsa-miR-30a-3p&offsets|+1|-3;m-38&6|-|71403554|71403575&offsets|+1|-3]|TTTCAGTCGGATGTTTGC</td><td>>MI0000088|hsa-mir-30a&WithFlank&6|-|71403545|71403627</td><td>54</td><td>71</td><td>18</td><td>TTTCAGTCGGATGTTTGC</td><td>BT20</td><td>21</td><td>13</td><td>25</td><td>16</td><td>10</td><td>26.4</td><td>8.0</td><td>17.8</td><td>7.0</td><td>2.9</td></tr>
<tr><td>>MI0000088|hsa-mir-30a&WithFlank&6|-|71403545|71403627@54.71.18&[MIMAT0000088&hsa-miR-30a-3p&offsets|+1|-3;m-38&6|-|71403554|71403575&offsets|+1|-3]|TTTCAGTCGGATGTTTGC</td><td>>MI0000088|hsa-mir-30a&WithFlank&6|-|71403545|71403627</td><td>54</td><td>71</td><td>18</td><td>TTTCAGTCGGATGTTTGC</td><td>MB231</td><td>40</td><td>63</td><td>33</td><td>93</td><td>36</td><td>22.9</td><td>14.3</td><td>10.9</td><td>51.7</td><td>11.6</td></tr>
<tr><td>>MI0000088|hsa-mir-30a&WithFlank&6|-|71403545|71403627@54.73.20(+1A)&[MIMAT0000088&hsa-miR-30a-3p&offsets|+1|-1(+1A);m-38&6|-|71403554|71403575&offsets|+1|-1(+1A)]|TTTCAGTCGGATGTTTGCAGA</td><td>>MI0000088|hsa-mir-30a&WithFlank&6|-|71403545|71403627</td><td>54</td><td>73</td><td>20(+1A)</td><td>TTTCAGTCGGATGTTTGCAGA</td><td>MB231</td><td>11</td><td>11</td><td>9</td><td>23</td><td>12</td><td>3.1</td><td>1.5</td><td>6.2</td><td>9.5</td><td>6.9</td></tr>
<tr><td>>MI0000088|hsa-mir-30a&WithFlank&6|-|71403545|71403627@54.73.20(+1U)&[MIMAT0000088&hsa-miR-30a-3p&offsets|+1|-1(+1U);m-38&6|-|71403554|71403575&offsets|+1|-1(+1U)]|TTTCAGTCGGATGTTTGCAGT</td><td>>MI0000088|hsa-mir-30a&WithFlank&6|-|71403545|71403627</td><td>54</td><td>73</td><td>20(+1U)</td><td>TTTCAGTCGGATGTTTGCAGT</td><td>BT20</td><td>99</td><td>82</td><td>115</td><td>34</td><td>35</td><td>44.7</td><td>57.2</td><td>25.9</td><td>31.1</td><td>14.5</td></tr>
<tr><td>>MI0000088|hsa-mir-30a&WithFlank&6|-|71403545|71403627@54.73.20(+1U)&[MIMAT0000088&hsa-miR-30a-3p&offsets|+1|-1(+1U);m-38&6|-|71403554|71403575&offsets|+1|-1(+1U)]|TTTCAGTCGGATGTTTGCAGT</td><td>>MI0000088|hsa-mir-30a&WithFlank&6|-|71403545|71403627</td><td>54</td><td>73</td><td>20(+1U)</td><td>TTTCAGTCGGATGTTTGCAGT</td><td>MB231</td><td>187</td><td>234</td><td>110</td><td>271</td><td>209</td><td>59.4</td><td>45.3</td><td>60.7</td><td>75.9</td><td>101.8</td></tr>
<tr><td>>MI0000088|hsa-mir-30a&WithFlank&6|-|71403545|71403627@54.73.20(+2U)&[MIMAT0000088&hsa-miR-30a-3p&offsets|+1|-1(+2U);m-38&6|-|71403554|71403575&offsets|+1|-1(+2U)]|TTTCAGTCGGATGTTTGCAGTT</td><td>>MI0000088|hsa-mir-30a&WithFlank&6|-|71403545|71403627</td><td>54</td><td>73</td><td>20(+2U)</td><td>TTTCAGTCGGATGTTTGCAGTT</td><td>BT20</td><td>13</td><td>11</td><td>20</td><td>5</td><td>5</td><td>5.6</td><td>7.2</td><td>7.2</td><td>4.7</td><td>1.4</td></tr>
<tr><td>>MI0000088|hsa-mir-30a&WithFlank&6|-|71403545|71403627@54.73.20(+2U)&[MIMAT0000088&hsa-miR-30a-3p&offsets|+1|-1(+2U);m-38&6|-|71403554|71403575&offsets|+1|-1(+2U)]|TTTCAGTCGGATGTTTGCAGTT</td><td>>MI0000088|hsa-mir-30a&WithFlank&6|-|71403545|71403627</td><td>54</td><td>73</td><td>20(+2U)</td><td>TTTCAGTCGGATGTTTGCAGTT</td><td>MB231</td><td>24</td><td>32</td><td>17</td><td>45</td><td>20</td><td>4.9</td><td>11.3</td><td>9.3</td><td>14.6</td><td>8.5</td></tr>
<tr><td>>MI0000088|hsa-mir-30a&WithFlank&6|-|71403545|71403627@54.73.20&[MIMAT0000088&hsa-miR-30a-3p&offsets|+1|-1;m-38&6|-|71403554|71403575&offsets|+1|-1]|TTTCAGTCGGATGTTTGCAG</td><td>>MI0000088|hsa-mir-30a&WithFlank&6|-|71403545|71403627</td><td>54</td><td>73</td><td>20</td><td>TTTCAGTCGGATGTTTGCAG</td><td>MB231</td><td>14</td><td>19</td><td>9</td><td>48</td><td>11</td><td>2.5</td><td>6.1</td><td>3.0</td><td>14.9</td><td>3.9</td></tr>
<tr><td>>MI0000088|hsa-mir-30a&WithFlank&6|-|71403545|71403627@54.74.21(+1A)&[MIMAT0000088&hsa-miR-30a-3p&offsets|+1|0(+1A);m-38&6|-|71403554|71403575&offsets|+1|0(+1A)]|TTTCAGTCGGATGTTTGCAGCA</td><td>>MI0000088|hsa-mir-30a&WithFlank&6|-|71403545|71403627</td><td>54</td><td>74</td><td>21(+1A)</td><td>TTTCAGTCGGATGTTTGCAGCA</td><td>MB231</td><td>11</td><td>12</td><td>10</td><td>15</td><td>10</td><td>4.0</td><td>4.9</td><td>7.7</td><td>8.7</td><td>5.4</td></tr>
<tr><td>>MI0000088|hsa-mir-30a&WithFlank&6|-|71403545|71403627@54.74.21&[MIMAT0000088&hsa-miR-30a-3p&offsets|+1|0;m-38&6|-|71403554|71403575&offsets|+1|0]|TTTCAGTCGGATGTTTGCAGC</td><td>>MI0000088|hsa-mir-30a&WithFlank&6|-|71403545|71403627</td><td>54</td><td>74</td><td>21</td><td>TTTCAGTCGGATGTTTGCAGC</td><td>BT20</td><td>88</td><td>78</td><td>139</td><td>37</td><td>39</td><td>46.7</td><td>45.2</td><td>35.7</td><td>28.6</td><td>14.3</td></tr>
<tr><td>>MI0000088|hsa-mir-30a&WithFlank&6|-|71403545|71403627@54.74.21&[MIMAT0000088&hsa-miR-30a-3p&offsets|+1|0;m-38&6|-|71403554|71403575&offsets|+1|0]|TTTCAGTCGGATGTTTGCAGC</td><td>>MI0000088|hsa-mir-30a&WithFlank&6|-|71403545|71403627</td><td>54</td><td>74</td><td>21</td><td>TTTCAGTCGGATGTTTGCAGC</td><td>MB231</td><td>188</td><td>261</td><td>162</td><td>296</td><td>193</td><td>66.6</td><td>60.3</td><td>89.7</td><td>94.9</td><td>78.9</td></tr>
<tr><td>>MI0000088|hsa-mir-30a&WithFlank&6|-|71403545|71403627@54.75.22&[MIMAT0000088&hsa-miR-30a-3p&offsets|+1|+1;m-38&6|-|71403554|71403575&offsets|+1|+1]|TTTCAGTCGGATGTTTGCAGCT</td><td>>MI0000088|hsa-mir-30a&WithFlank&6|-|71403545|71403627</td><td>54</td><td>75</td><td>22</td><td>TTTCAGTCGGATGTTTGCAGCT</td><td>BT20</td><td>15</td><td>13</td><td>25</td><td>3</td><td>8</td><td>5.7</td><td>8.3</td><td>8.4</td><td>3.4</td><td>2.3</td></tr>
<tr><td>>MI0000088|hsa-mir-30a&WithFlank&6|-|71403545|71403627@54.75.22&[MIMAT0000088&hsa-miR-30a-3p&offsets|+1|+1;m-38&6|-|71403554|71403575&offsets|+1|+1]|TTTCAGTCGGATGTTTGCAGCT</td><td>>MI0000088|hsa-mir-30a&WithFlank&6|-|71403545|71403627</td><td>54</td><td>75</td><td>22</td><td>TTTCAGTCGGATGTTTGCAGCT</td><td>MB231</td><td>41</td><td>53</td><td>32</td><td>68</td><td>47</td><td>21.1</td><td>25.2</td><td>21.5</td><td>30.8</td><td>22.6</td></tr>
<tr><td>>MI0000089|hsa-mir-31&WithFlank&9|-|21512109|21512191@14.34.21(+1A)&[MIMAT0000089&hsa-miR-31-5p&offsets|0|0(+1A);m-360&9|-|21512157|21512178&offsets|0|-1(+1A)]|AGGCAAGATGCTGGCATAGCTA</td><td>>MI0000089|hsa-mir-31&WithFlank&9|-|21512109|21512191</td><td>14</td><td>34</td><td>21(+1A)</td><td>AGGCAAGATGCTGGCATAGCTA</td><td>BT20</td><td>15</td><td>10</td><td>7</td><td>6</td><td>3</td><td>12.5</td><td>9.4</td><td>2.9</td><td>5.1</td><td>2.8</td></tr>
<tr><td>>MI0000089|hsa-mir-31&WithFlank&9|-|21512109|21512191@14.34.21&[MIMAT0000089&hsa-miR-31-5p&offsets|0|0;m-360&9|-|21512157|21512178&offsets|0|-1]|AGGCAAGATGCTGGCATAGCT</td><td>>MI0000089|hsa-mir-31&WithFlank&9|-|21512109|21512191</td><td>14</td><td>34</td><td>21</td><td>AGGCAAGATGCTGGCATAGCT</td><td>BT20</td><td>38</td><td>33</td><td>13</td><td>6</td><td>8</td><td>30.0</td><td>30.1</td><td>5.9</td><td>4.2</td><td>8.4</td></tr>
<tr><td>>MI0000089|hsa-mir-31&WithFlank&9|-|21512109|21512191@14.35.22(+1A)&[MIMAT0000089&hsa-miR-31-5p&offsets|0|+1(+1A);m-360&9|-|21512157|21512178&offsets|0|0(+1A)]|AGGCAAGATGCTGGCATAGCTGA</td><td>>MI0000089|hsa-mir-31&WithFlank&9|-|21512109|21512191</td><td>14</td><td>35</td><td>22(+1A)</td><td>AGGCAAGATGCTGGCATAGCTGA</td><td>BT20</td><td>24</td><td>19</td><td>20</td><td>21</td><td>11</td><td>13.2</td><td>14.7</td><td>15.6</td><td>17.9</td><td>8.8</td></tr>
<tr><td>>MI0000089|hsa-mir-31&WithFlank&9|-|21512109|21512191@14.35.22(+1C)&[MIMAT0000089&hsa-miR-31-5p&offsets|0|+1(+1C);m-360&9|-|21512157|21512178&offsets|0|0(+1C)]|AGGCAAGATGCTGGCATAGCTGC</td><td>>MI0000089|hsa-mir-31&WithFlank&9|-|21512109|21512191</td><td>14</td><td>35</td><td>22(+1C)</td><td>AGGCAAGATGCTGGCATAGCTGC</td><td>BT20</td><td>17</td><td>16</td><td>11</td><td>15</td><td>6</td><td>6.0</td><td>17.4</td><td>7.4</td><td>12.1</td><td>7.3</td></tr>
<tr><td>>MI0000089|hsa-mir-31&WithFlank&9|-|21512109|21512191@14.35.22&[MIMAT0000089&hsa-miR-31-5p&offsets|0|+1;m-360&9|-|21512157|21512178&offsets|0|0]|AGGCAAGATGCTGGCATAGCTG</td><td>>MI0000089|hsa-mir-31&WithFlank&9|-|21512109|21512191</td><td>14</td><td>35</td><td>22</td><td>AGGCAAGATGCTGGCATAGCTG</td><td>BT20</td><td>64</td><td>65</td><td>40</td><td>46</td><td>22</td><td>31.0</td><td>52.8</td><td>28.9</td><td>42.5</td><td>15.2</td></tr>
<tr><td>>MI0000089|hsa-mir-31&WithFlank&9|-|21512109|21512191@14.36.23&[MIMAT0000089&hsa-miR-31-5p&offsets|0|+2;m-360&9|-|21512157|21512178&offsets|0|+1]|AGGCAAGATGCTGGCATAGCTGT</td><td>>MI0000089|hsa-mir-31&WithFlank&9|-|21512109|21512191</td><td>14</td><td>36</td><td>23</td><td>AGGCAAGATGCTGGCATAGCTGT</td><td>BT20</td><td>120</td><td>97</td><td>82</td><td>112</td><td>43</td><td>51.7</td><td>98.6</td><td>62.9</td><td>97.0</td><td>47.6</td></tr>
<tr><td>>MI0000089|hsa-mir-31&WithFlank&9|-|21512109|21512191@50.72.23&[MIMAT0004504&hsa-miR-31-3p&offsets|0|+1;m-521&9|-|21512120|21512142&offsets|0|0]|TGCTATGCCAACATATTGCCATC</td><td>>MI0000089|hsa-mir-31&WithFlank&9|-|21512109|21512191</td><td>50</td><td>72</td><td>23</td><td>TGCTATGCCAACATATTGCCATC</td><td>BT20</td><td>18</td><td>9</td><td>3</td><td>8</td><td>1</td><td>23.7</td><td>7.1</td><td>0.7</td><td>11.6</td><td>0.8</td></tr>
<tr><td>>MI0000090|hsa-mir-32&WithFlank&9|-|109046223|109046304@12.33.22&[MIMAT0000090&hsa-miR-32-5p&offsets|0|0;m-208&9|-|109046273|109046293&offsets|0|+1]|TATTGCACATTACTAAGTTGCA</td><td>>MI0000090|hsa-mir-32&WithFlank&9|-|109046223|109046304</td><td>12</td><td>33</td><td>22</td><td>TATTGCACATTACTAAGTTGCA</td><td>BT20</td><td>31</td><td>14</td><td>3</td><td>10</td><td>6</td><td>23.1</td><td>20.2</td><td>2.9</td><td>13.5</td><td>2.3</td></tr>
<tr><td>>MI0000090|hsa-mir-32&WithFlank&9|-|109046223|109046304@12.33.22&[MIMAT0000090&hsa-miR-32-5p&offsets|0|0;m-208&9|-|109046273|109046293&offsets|0|+1]|TATTGCACATTACTAAGTTGCA</td><td>>MI0000090|hsa-mir-32&WithFlank&9|-|109046223|109046304</td><td>12</td><td>33</td><td>22</td><td>TATTGCACATTACTAAGTTGCA</td><td>MB231</td><td>57</td><td>13</td><td>1</td><td>65</td><td>27</td><td>67.0</td><td>7.8</td><td>0.2</td><td>7.2</td><td>20.3</td></tr>
<tr><td>>MI0000093|hsa-mir-92a-1&WithFlank&13|+|91351308|91351397@54.73.20;MI0000094|hsa-mir-92a-2&WithFlank&X|-|134169532|134169618@54.73.20&[MIMAT0000092&hsa-miR-92a-3p&offsets|0|-2;m-9&13|+|91351361|91351382&offsets|0|-2];[MIMAT0000092_1&hsa-miR-92a-3p&offsets|0|-2;m-11&X|-|134169544|134169565&offsets|0|-2]|TATTGCACTTGTCCCGGCCT</td><td>>MI0000093|hsa-mir-92a-1&WithFlank&13|+|91351308|91351397</td><td>54</td><td>73</td><td>20</td><td>TATTGCACTTGTCCCGGCCT</td><td>MB231</td><td>13</td><td>13</td><td>13</td><td>24</td><td>18</td><td>2.6</td><td>1.6</td><td>6.4</td><td>13.2</td><td>9.1</td></tr>
<tr><td>>MI0000093|hsa-mir-92a-1&WithFlank&13|+|91351308|91351397@54.74.21;MI0000094|hsa-mir-92a-2&WithFlank&X|-|134169532|134169618@54.74.21&[MIMAT0000092&hsa-miR-92a-3p&offsets|0|-1;m-9&13|+|91351361|91351382&offsets|0|-1];[MIMAT0000092_1&hsa-miR-92a-3p&offsets|0|-1;m-11&X|-|134169544|134169565&offsets|0|-1]|TATTGCACTTGTCCCGGCCTG</td><td>>MI0000093|hsa-mir-92a-1&WithFlank&13|+|91351308|91351397</td><td>54</td><td>74</td><td>21</td><td>TATTGCACTTGTCCCGGCCTG</td><td>BT20</td><td>36</td><td>15</td><td>28</td><td>29</td><td>11</td><td>22.9</td><td>6.8</td><td>0.8</td><td>31.4</td><td>4.9</td></tr>
<tr><td>>MI0000093|hsa-mir-92a-1&WithFlank&13|+|91351308|91351397@54.74.21;MI0000094|hsa-mir-92a-2&WithFlank&X|-|134169532|134169618@54.74.21&[MIMAT0000092&hsa-miR-92a-3p&offsets|0|-1;m-9&13|+|91351361|91351382&offsets|0|-1];[MIMAT0000092_1&hsa-miR-92a-3p&offsets|0|-1;m-11&X|-|134169544|134169565&offsets|0|-1]|TATTGCACTTGTCCCGGCCTG</td><td>>MI0000093|hsa-mir-92a-1&WithFlank&13|+|91351308|91351397</td><td>54</td><td>74</td><td>21</td><td>TATTGCACTTGTCCCGGCCTG</td><td>MB231</td><td>57</td><td>59</td><td>40</td><td>103</td><td>50</td><td>2.7</td><td>13.0</td><td>21.8</td><td>72.6</td><td>7.3</td></tr>
<tr><td>>MI0000093|hsa-mir-92a-1&WithFlank&13|+|91351308|91351397@54.74.21(+1A);MI0000094|hsa-mir-92a-2&WithFlank&X|-|134169532|134169618@54.74.21(+1A)&[MIMAT0000092&hsa-miR-92a-3p&offsets|0|-1(+1A);m-9&13|+|91351361|91351382&offsets|0|-1(+1A)];[MIMAT0000092_1&hsa-miR-92a-3p&offsets|0|-1(+1A);m-11&X|-|134169544|134169565&offsets|0|-1(+1A)]|TATTGCACTTGTCCCGGCCTGA</td><td>>MI0000093|hsa-mir-92a-1&WithFlank&13|+|91351308|91351397</td><td>54</td><td>74</td><td>21(+1A)</td><td>TATTGCACTTGTCCCGGCCTGA</td><td>BT20</td><td>114</td><td>60</td><td>138</td><td>51</td><td>37</td><td>62.8</td><td>36.9</td><td>45.0</td><td>43.4</td><td>16.8</td></tr>
<tr><td>>MI0000093|hsa-mir-92a-1&WithFlank&13|+|91351308|91351397@54.74.21(+1A);MI0000094|hsa-mir-92a-2&WithFlank&X|-|134169532|134169618@54.74.21(+1A)&[MIMAT0000092&hsa-miR-92a-3p&offsets|0|-1(+1A);m-9&13|+|91351361|91351382&offsets|0|-1(+1A)];[MIMAT0000092_1&hsa-miR-92a-3p&offsets|0|-1(+1A);m-11&X|-|134169544|134169565&offsets|0|-1(+1A)]|TATTGCACTTGTCCCGGCCTGA</td><td>>MI0000093|hsa-mir-92a-1&WithFlank&13|+|91351308|91351397</td><td>54</td><td>74</td><td>21(+1A)</td><td>TATTGCACTTGTCCCGGCCTGA</td><td>MB231</td><td>101</td><td>106</td><td>92</td><td>191</td><td>152</td><td>21.3</td><td>14.7</td><td>59.5</td><td>123.2</td><td>86.0</td></tr>
<tr><td>>MI0000093|hsa-mir-92a-1&WithFlank&13|+|91351308|91351397@54.74.21(+1C);MI0000094|hsa-mir-92a-2&WithFlank&X|-|134169532|134169618@54.74.21(+1C)&[MIMAT0000092&hsa-miR-92a-3p&offsets|0|-1(+1C);m-9&13|+|91351361|91351382&offsets|0|-1(+1C)];[MIMAT0000092_1&hsa-miR-92a-3p&offsets|0|-1(+1C);m-11&X|-|134169544|134169565&offsets|0|-1(+1C)]|TATTGCACTTGTCCCGGCCTGC</td><td>>MI0000093|hsa-mir-92a-1&WithFlank&13|+|91351308|91351397</td><td>54</td><td>74</td><td>21(+1C)</td><td>TATTGCACTTGTCCCGGCCTGC</td><td>BT20</td><td>275</td><td>246</td><td>334</td><td>193</td><td>165</td><td>91.5</td><td>177.9</td><td>45.5</td><td>154.9</td><td>96.5</td></tr>
</table>

<table>
<tr><td>>MI0000093|hsa-mir-92a-1&WithFlank&13|+|91351308|91351397@54.74.21(+1C);MI0000094|hsa-mir-92a-2&WithFlank&X|-|134169532|134169618@54.74.21(+1C)&[MIMAT0000092&hsa-miR-92a-3p&offsets|0|-1(+1C);m-9&13|+|91351361|91351382&offsets|0|-1(+1C)];[MIMAT0000092_1&hsa-miR-92a-3p&offsets|0|-1(+1C);m-11&X|-|134169544|134169565&offsets|0|-1(+1C)]|TATTGCACTTGTCCCGGCCTGC</td><td>>MI0000093|hsa-mir-92a-1&WithFlank&13|+|91351308|91351397</td><td>54</td><td>74</td><td>21(+1C)</td><td>TATTGCACTTGTCCCGGCCTGC</td><td>MB231</td><td>369</td><td>426</td><td>309</td><td>728</td><td>643</td><td>114.3</td><td>107.6</td><td>178.2</td><td>461.1</td><td>383.9</td></tr>
<tr><td>>MI0000093|hsa-mir-92a-1&WithFlank&13|+|91351308|91351397@54.74.21(+1C);MI0000094|hsa-mir-92a-2&WithFlank&X|-|134169532|134169618@54.74.21(+1C)&[MIMAT0000092&hsa-miR-92a-3p&offsets|0|-1(+1C);m-9&13|+|91351361|91351382&offsets|0|-1(+1C)];[MIMAT0000092_1&hsa-miR-92a-3p&offsets|0|-1(+1C);m-11&X|-|134169544|134169565&offsets|0|-1(+1C)]|TATTGCACTTGTCCCGGCCTGC</td><td>>MI0000093|hsa-mir-92a-1&WithFlank&13|+|91351308|91351397</td><td>54</td><td>74</td><td>21(+1C)</td><td>TATTGCACTTGTCCCGGCCTGC</td><td>MB468</td><td>19</td><td>49</td><td>10</td><td>44</td><td>13</td><td>6.2</td><td>13.1</td><td>4.3</td><td>12.3</td><td>3.7</td></tr>
<tr><td>>MI0000093|hsa-mir-92a-1&WithFlank&13|+|91351308|91351397@54.74.21(+1G);MI0000094|hsa-mir-92a-2&WithFlank&X|-|134169532|134169618@54.74.21(+1G)&[MIMAT0000092&hsa-miR-92a-3p&offsets|0|-1(+1G);m-9&13|+|91351361|91351382&offsets|0|-1(+1G)];[MIMAT0000092_1&hsa-miR-92a-3p&offsets|0|-1(+1G);m-11&X|-|134169544|134169565&offsets|0|-1(+1G)]|TATTGCACTTGTCCCGGCCTGG</td><td>>MI0000093|hsa-mir-92a-1&WithFlank&13|+|91351308|91351397</td><td>54</td><td>74</td><td>21(+1G)</td><td>TATTGCACTTGTCCCGGCCTGG</td><td>BT20</td><td>119</td><td>43</td><td>66</td><td>26</td><td>11</td><td>81.6</td><td>27.4</td><td>30.4</td><td>27.8</td><td>6.5</td></tr>
<tr><td>>MI0000093|hsa-mir-92a-1&WithFlank&13|+|91351308|91351397@54.74.21(+1G);MI0000094|hsa-mir-92a-2&WithFlank&X|-|134169532|134169618@54.74.21(+1G)&[MIMAT0000092&hsa-miR-92a-3p&offsets|0|-1(+1G);m-9&13|+|91351361|91351382&offsets|0|-1(+1G)];[MIMAT0000092_1&hsa-miR-92a-3p&offsets|0|-1(+1G);m-11&X|-|134169544|134169565&offsets|0|-1(+1G)]|TATTGCACTTGTCCCGGCCTGG</td><td>>MI0000093|hsa-mir-92a-1&WithFlank&13|+|91351308|91351397</td><td>54</td><td>74</td><td>21(+1G)</td><td>TATTGCACTTGTCCCGGCCTGG</td><td>MB231</td><td>134</td><td>70</td><td>49</td><td>74</td><td>57</td><td>80.1</td><td>23.8</td><td>36.1</td><td>59.2</td><td>40.0</td></tr>
<tr><td>>MI0000093|hsa-mir-92a-1&WithFlank&13|+|91351308|91351397@54.75.22;MI0000094|hsa-mir-92a-2&WithFlank&X|-|134169532|134169618@54.75.22&[MIMAT0000092&hsa-miR-92a-3p&offsets|0|0;m-9&13|+|91351361|91351382&offsets|0|0];[MIMAT0000092_1&hsa-miR-92a-3p&offsets|0|0;m-11&X|-|134169544|134169565&offsets|0|0]|TATTGCACTTGTCCCGGCCTGT</td><td>>MI0000093|hsa-mir-92a-1&WithFlank&13|+|91351308|91351397</td><td>54</td><td>75</td><td>22</td><td>TATTGCACTTGTCCCGGCCTGT</td><td>BT20</td><td>1890</td><td>1534</td><td>2441</td><td>1343</td><td>1130</td><td>759.6</td><td>925.6</td><td>465.3</td><td>1026.5</td><td>593.3</td></tr>
<tr><td>>MI0000093|hsa-mir-92a-1&WithFlank&13|+|91351308|91351397@54.75.22;MI0000094|hsa-mir-92a-2&WithFlank&X|-|134169532|134169618@54.75.22&[MIMAT0000092&hsa-miR-92a-3p&offsets|0|0;m-9&13|+|91351361|91351382&offsets|0|0];[MIMAT0000092_1&hsa-miR-92a-3p&offsets|0|0;m-11&X|-|134169544|134169565&offsets|0|0]|TATTGCACTTGTCCCGGCCTGT</td><td>>MI0000093|hsa-mir-92a-1&WithFlank&13|+|91351308|91351397</td><td>54</td><td>75</td><td>22</td><td>TATTGCACTTGTCCCGGCCTGT</td><td>MB231</td><td>3041</td><td>3761</td><td>2707</td><td>4233</td><td>4455</td><td>948.3</td><td>962.1</td><td>1518.2</td><td>2092.9</td><td>1932.7</td></tr>
<tr><td>>MI0000093|hsa-mir-92a-1&WithFlank&13|+|91351308|91351397@54.75.22;MI0000094|hsa-mir-92a-2&WithFlank&X|-|134169532|134169618@54.75.22&[MIMAT0000092&hsa-miR-92a-3p&offsets|0|0;m-9&13|+|91351361|91351382&offsets|0|0];[MIMAT0000092_1&hsa-miR-92a-3p&offsets|0|0;m-11&X|-|134169544|134169565&offsets|0|0]|TATTGCACTTGTCCCGGCCTGT</td><td>>MI0000093|hsa-mir-92a-1&WithFlank&13|+|91351308|91351397</td><td>54</td><td>75</td><td>22</td><td>TATTGCACTTGTCCCGGCCTGT</td><td>MB468</td><td>155</td><td>415</td><td>84</td><td>315</td><td>105</td><td>47.4</td><td>122.2</td><td>37.7</td><td>77.2</td><td>34.2</td></tr>
<tr><td>>MI0000093|hsa-mir-92a-1&WithFlank&13|+|91351308|91351397@54.75.22(+1A);MI0000094|hsa-mir-92a-2&WithFlank&X|-|134169532|134169618@54.75.22(+1A)&[MIMAT0000092&hsa-miR-92a-3p&offsets|0|0(+1A);m-9&13|+|91351361|91351382&offsets|0|0(+1A)];[MIMAT0000092_1&hsa-miR-92a-3p&offsets|0|0(+1A);m-11&X|-|134169544|134169565&offsets|0|0(+1A)]|TATTGCACTTGTCCCGGCCTGTA</td><td>>MI0000093|hsa-mir-92a-1&WithFlank&13|+|91351308|91351397</td><td>54</td><td>75</td><td>22(+1A)</td><td>TATTGCACTTGTCCCGGCCTGTA</td><td>BT20</td><td>437</td><td>216</td><td>539</td><td>90</td><td>300</td><td>264.1</td><td>73.3</td><td>187.1</td><td>42.4</td><td>115.3</td></tr>
<tr><td>>MI0000093|hsa-mir-92a-1&WithFlank&13|+|91351308|91351397@54.75.22(+1A);MI0000094|hsa-mir-92a-2&WithFlank&X|-|134169532|134169618@54.75.22(+1A)&[MIMAT0000092&hsa-miR-92a-3p&offsets|0|0(+1A);m-9&13|+|91351361|91351382&offsets|0|0(+1A)];[MIMAT0000092_1&hsa-miR-92a-3p&offsets|0|0(+1A);m-11&X|-|134169544|134169565&offsets|0|0(+1A)]|TATTGCACTTGTCCCGGCCTGTA</td><td>>MI0000093|hsa-mir-92a-1&WithFlank&13|+|91351308|91351397</td><td>54</td><td>75</td><td>22(+1A)</td><td>TATTGCACTTGTCCCGGCCTGTA</td><td>MB231</td><td>434</td><td>422</td><td>366</td><td>294</td><td>470</td><td>170.4</td><td>119.4</td><td>114.1</td><td>60.5</td><td>157.2</td></tr>
<tr><td>>MI0000093|hsa-mir-92a-1&WithFlank&13|+|91351308|91351397@54.75.22(+1A);MI0000094|hsa-mir-92a-2&WithFlank&X|-|134169532|134169618@54.75.22(+1A)&[MIMAT0000092&hsa-miR-92a-3p&offsets|0|0(+1A);m-9&13|+|91351361|91351382&offsets|0|0(+1A)];[MIMAT0000092_1&hsa-miR-92a-3p&offsets|0|0(+1A);m-11&X|-|134169544|134169565&offsets|0|0(+1A)]|TATTGCACTTGTCCCGGCCTGTA</td><td>>MI0000093|hsa-mir-92a-1&WithFlank&13|+|91351308|91351397</td><td>54</td><td>75</td><td>22(+1A)</td><td>TATTGCACTTGTCCCGGCCTGTA</td><td>MB468</td><td>43</td><td>105</td><td>24</td><td>40</td><td>25</td><td>11.8</td><td>28.6</td><td>16.7</td><td>10.1</td><td>7.2</td></tr>
<tr><td>>MI0000093|hsa-mir-92a-1&WithFlank&13|+|91351308|91351397@54.75.22(+1C);MI0000094|hsa-mir-92a-2&WithFlank&X|-|134169532|134169618@54.75.22(+1C)&[MIMAT0000092&hsa-miR-92a-3p&offsets|0|0(+1C);m-9&13|+|91351361|91351382&offsets|0|0(+1C)];[MIMAT0000092_1&hsa-miR-92a-3p&offsets|0|0(+1C);m-11&X|-|134169544|134169565&offsets|0|0(+1C)]|TATTGCACTTGTCCCGGCCTGTC</td><td>>MI0000093|hsa-mir-92a-1&WithFlank&13|+|91351308|91351397</td><td>54</td><td>75</td><td>22(+1C)</td><td>TATTGCACTTGTCCCGGCCTGTC</td><td>BT20</td><td>18</td><td>11</td><td>20</td><td>5</td><td>10</td><td>9.6</td><td>7.0</td><td>2.2</td><td>3.1</td><td>5.5</td></tr>
<tr><td>>MI0000093|hsa-mir-92a-1&WithFlank&13|+|91351308|91351397@54.75.22(+1C);MI0000094|hsa-mir-92a-2&WithFlank&X|-|134169532|134169618@54.75.22(+1C)&[MIMAT0000092&hsa-miR-92a-3p&offsets|0|0(+1C);m-9&13|+|91351361|91351382&offsets|0|0(+1C)];[MIMAT0000092_1&hsa-miR-92a-3p&offsets|0|0(+1C);m-11&X|-|134169544|134169565&offsets|0|0(+1C)]|TATTGCACTTGTCCCGGCCTGTC</td><td>>MI0000093|hsa-mir-92a-1&WithFlank&13|+|91351308|91351397</td><td>54</td><td>75</td><td>22(+1C)</td><td>TATTGCACTTGTCCCGGCCTGTC</td><td>MB231</td><td>42</td><td>32</td><td>30</td><td>42</td><td>57</td><td>10.3</td><td>5.5</td><td>14.7</td><td>24.1</td><td>32.8</td></tr>
<tr><td>>MI0000093|hsa-mir-92a-1&WithFlank&13|+|91351308|91351397@54.75.22(+2A);MI0000094|hsa-mir-92a-2&WithFlank&X|-|134169532|134169618@54.75.22(+2A)&[MIMAT0000092&hsa-miR-92a-3p&offsets|0|0(+2A);m-9&13|+|91351361|91351382&offsets|0|0(+2A)];[MIMAT0000092_1&hsa-miR-92a-3p&offsets|0|0(+2A);m-11&X|-|134169544|134169565&offsets|0|0(+2A)]|TATTGCACTTGTCCCGGCCTGTAA</td><td>>MI0000093|hsa-mir-92a-1&WithFlank&13|+|91351308|91351397</td><td>54</td><td>75</td><td>22(+2A)</td><td>TATTGCACTTGTCCCGGCCTGTAA</td><td>BT20</td><td>31</td><td>19</td><td>31</td><td>2</td><td>18</td><td>24.4</td><td>15.6</td><td>13.4</td><td>1.5</td><td>9.4</td></tr>
<tr><td>>MI0000093|hsa-mir-92a-1&WithFlank&13|+|91351308|91351397@54.75.22(+2A);MI0000094|hsa-mir-92a-2&WithFlank&X|-|134169532|134169618@54.75.22(+2A)&[MIMAT0000092&hsa-miR-92a-3p&offsets|0|0(+2A);m-9&13|+|91351361|91351382&offsets|0|0(+2A)];[MIMAT0000092_1&hsa-miR-92a-3p&offsets|0|0(+2A);m-11&X|-|134169544|134169565&offsets|0|0(+2A)]|TATTGCACTTGTCCCGGCCTGTAA</td><td>>MI0000093|hsa-mir-92a-1&WithFlank&13|+|91351308|91351397</td><td>54</td><td>75</td><td>22(+2A)</td><td>TATTGCACTTGTCCCGGCCTGTAA</td><td>MB231</td><td>34</td><td>41</td><td>21</td><td>11</td><td>35</td><td>15.3</td><td>13.1</td><td>5.5</td><td>4.1</td><td>14.1</td></tr>
<tr><td>>MI0000093|hsa-mir-92a-1&WithFlank&13|+|91351308|91351397@54.76.23(+1U)&[MIMAT0000092&hsa-miR-92a-3p&offsets|0|+1(+1U);m-9&13|+|91351361|91351382&offsets|0|+1(+1U)]|TATTGCACTTGTCCCGGCCTGTTT</td><td>>MI0000093|hsa-mir-92a-1&WithFlank&13|+|91351308|91351397</td><td>54</td><td>76</td><td>23(+1U)</td><td>TATTGCACTTGTCCCGGCCTGTTT</td><td>MB231</td><td>28</td><td>21</td><td>20</td><td>21</td><td>32</td><td>10.3</td><td>4.2</td><td>4.3</td><td>10.8</td><td>10.8</td></tr>
<tr><td>>MI0000093|hsa-mir-92a-1&WithFlank&13|+|91351308|91351397@54.76.23&[MIMAT0000092&hsa-miR-92a-3p&offsets|0|+1;m-9&13|+|91351361|91351382&offsets|0|+1]|TATTGCACTTGTCCCGGCCTGTT</td><td>>MI0000093|hsa-mir-92a-1&WithFlank&13|+|91351308|91351397</td><td>54</td><td>76</td><td>23</td><td>TATTGCACTTGTCCCGGCCTGTT</td><td>BT20</td><td>128</td><td>73</td><td>158</td><td>31</td><td>70</td><td>80.1</td><td>44.9</td><td>28.4</td><td>22.0</td><td>39.9</td></tr>
<tr><td>>MI0000093|hsa-mir-92a-1&WithFlank&13|+|91351308|91351397@54.76.23&[MIMAT0000092&hsa-miR-92a-3p&offsets|0|+1;m-9&13|+|91351361|91351382&offsets|0|+1]|TATTGCACTTGTCCCGGCCTGTT</td><td>>MI0000093|hsa-mir-92a-1&WithFlank&13|+|91351308|91351397</td><td>54</td><td>76</td><td>23</td><td>TATTGCACTTGTCCCGGCCTGTT</td><td>MB231</td><td>353</td><td>299</td><td>271</td><td>247</td><td>400</td><td>100.1</td><td>71.5</td><td>115.3</td><td>114.9</td><td>156.7</td></tr>
<tr><td>>MI0000093|hsa-mir-92a-1&WithFlank&13|+|91351308|91351397@54.76.23&[MIMAT0000092&hsa-miR-92a-3p&offsets|0|+1;m-9&13|+|91351361|91351382&offsets|0|+1]|TATTGCACTTGTCCCGGCCTGTT</td><td>>MI0000093|hsa-mir-92a-1&WithFlank&13|+|91351308|91351397</td><td>54</td><td>76</td><td>23</td><td>TATTGCACTTGTCCCGGCCTGTT</td><td>MB468</td><td>17</td><td>29</td><td>8</td><td>14</td><td>9</td><td>6.1</td><td>7.3</td><td>3.5</td><td>5.1</td><td>3.2</td></tr>
<tr><td>>MI0000093|hsa-mir-92a-1&WithFlank&13|+|91351308|91351397@55.75.21;MI0000094|hsa-mir-92a-2&WithFlank&X|-|134169532|134169618@55.75.21&[MIMAT0000092&hsa-miR-92a-3p&offsets|+1|0;m-9&13|+|91351361|91351382&offsets|+1|0];[MIMAT0000092_1&hsa-miR-92a-3p&offsets|+1|0;m-11&X|-|134169544|134169565&offsets|+1|0]|ATTGCACTTGTCCCGGCCTGT</td><td>>MI0000093|hsa-mir-92a-1&WithFlank&13|+|91351308|91351397</td><td>55</td><td>75</td><td>21</td><td>ATTGCACTTGTCCCGGCCTGT</td><td>BT20</td><td>22</td><td>22</td><td>35</td><td>30</td><td>27</td><td>11.5</td><td>7.3</td><td>4.3</td><td>20.0</td><td>14.4</td></tr>
<tr><td>>MI0000093|hsa-mir-92a-1&WithFlank&13|+|91351308|91351397@55.75.21;MI0000094|hsa-mir-92a-2&WithFlank&X|-|134169532|134169618@55.75.21&[MIMAT0000092&hsa-miR-92a-3p&offsets|+1|0;m-9&13|+|91351361|91351382&offsets|+1|0];[MIMAT0000092_1&hsa-miR-92a-3p&offsets|+1|0;m-11&X|-|134169544|134169565&offsets|+1|0]|ATTGCACTTGTCCCGGCCTGT</td><td>>MI0000093|hsa-mir-92a-1&WithFlank&13|+|91351308|91351397</td><td>55</td><td>75</td><td>21</td><td>ATTGCACTTGTCCCGGCCTGT</td><td>MB231</td><td>37</td><td>54</td><td>39</td><td>67</td><td>61</td><td>11.5</td><td>14.3</td><td>27.4</td><td>23.1</td><td>14.0</td></tr>
<tr><td>>MI0000094|hsa-mir-92a-2&WithFlank&X|-|134169532|134169618@54.76.23&[MIMAT0000092_1&hsa-miR-92a-3p&offsets|0|+1;m-11&X|-|134169544|134169565&offsets|0|+1]|TATTGCACTTGTCCCGGCCTGTG</td><td>>MI0000094|hsa-mir-92a-2&WithFlank&X|-|134169532|134169618</td><td>54</td><td>76</td><td>23</td><td>TATTGCACTTGTCCCGGCCTGTG</td><td>BT20</td><td>26</td><td>7</td><td>16</td><td>3</td><td>4</td><td>15.6</td><td>4.4</td><td>4.1</td><td>1.9</td><td>1.6</td></tr>
</table>

<table>
<tr><td>>MI0000094|hsa-mir-92a-2&WithFlank&X|-|134169532|134169618@54.76.23&[MIMAT0000092_1&hsa-miR-92a-3p&offsets|0|+1;m-11&X|-|134169544|134169565&offsets|0|+1]|TATTGCACTTGTCCCGGCCTGTG</td><td>>MI0000094|hsa-mir-92a-2&WithFlank&X|-|134169532|134169618</td><td>54</td><td>76</td><td>23</td><td>TATTGCACTTGTCCCGGCCTGTG</td><td>MB231</td><td>26</td><td>12</td><td>10</td><td>11</td><td>11</td><td>16.5</td><td>1.9</td><td>5.1</td><td>6.4</td><td>6.4</td></tr>
<tr><td>>MI0000095|hsa-mir-93&WithFlank&7|-|100093762|100093853@17.37.21&[MIMAT0000093&hsa-miR-93-5p&offsets|0|-2;m-29&7|-|100093815|100093837&offsets|0|-2]|CAAAGTGCTGTTCGTGCAGGT</td><td>>MI0000095|hsa-mir-93&WithFlank&7|-|100093762|100093853</td><td>17</td><td>37</td><td>21</td><td>CAAAGTGCTGTTCGTGCAGGT</td><td>BT20</td><td>48</td><td>53</td><td>35</td><td>152</td><td>17</td><td>26.1</td><td>50.5</td><td>23.4</td><td>142.6</td><td>10.8</td></tr>
<tr><td>>MI0000095|hsa-mir-93&WithFlank&7|-|100093762|100093853@17.37.21&[MIMAT0000093&hsa-miR-93-5p&offsets|0|-2;m-29&7|-|100093815|100093837&offsets|0|-2]|CAAAGTGCTGTTCGTGCAGGT</td><td>>MI0000095|hsa-mir-93&WithFlank&7|-|100093762|100093853</td><td>17</td><td>37</td><td>21</td><td>CAAAGTGCTGTTCGTGCAGGT</td><td>MB231</td><td>48</td><td>53</td><td>17</td><td>414</td><td>80</td><td>15.7</td><td>27.0</td><td>16.6</td><td>156.0</td><td>59.4</td></tr>
<tr><td>>MI0000095|hsa-mir-93&WithFlank&7|-|100093762|100093853@17.38.22(+1A)&[MIMAT0000093&hsa-miR-93-5p&offsets|0|-1(+1A);m-29&7|-|100093815|100093837&offsets|0|-1(+1A)]|CAAAGTGCTGTTCGTGCAGGTAA</td><td>>MI0000095|hsa-mir-93&WithFlank&7|-|100093762|100093853</td><td>17</td><td>38</td><td>22(+1A)</td><td>CAAAGTGCTGTTCGTGCAGGTAA</td><td>BT20</td><td>93</td><td>64</td><td>78</td><td>159</td><td>17</td><td>67.7</td><td>59.5</td><td>36.5</td><td>136.7</td><td>10.7</td></tr>
<tr><td>>MI0000095|hsa-mir-93&WithFlank&7|-|100093762|100093853@17.38.22(+1A)&[MIMAT0000093&hsa-miR-93-5p&offsets|0|-1(+1A);m-29&7|-|100093815|100093837&offsets|0|-1(+1A)]|CAAAGTGCTGTTCGTGCAGGTAA</td><td>>MI0000095|hsa-mir-93&WithFlank&7|-|100093762|100093853</td><td>17</td><td>38</td><td>22(+1A)</td><td>CAAAGTGCTGTTCGTGCAGGTAA</td><td>MB231</td><td>46</td><td>33</td><td>18</td><td>362</td><td>59</td><td>3.9</td><td>9.6</td><td>17.2</td><td>189.7</td><td>43.2</td></tr>
<tr><td>>MI0000095|hsa-mir-93&WithFlank&7|-|100093762|100093853@17.38.22&[MIMAT0000093&hsa-miR-93-5p&offsets|0|-1;m-29&7|-|100093815|100093837&offsets|0|-1]|CAAAGTGCTGTTCGTGCAGGTA</td><td>>MI0000095|hsa-mir-93&WithFlank&7|-|100093762|100093853</td><td>17</td><td>38</td><td>22</td><td>CAAAGTGCTGTTCGTGCAGGTA</td><td>BT20</td><td>38</td><td>27</td><td>30</td><td>31</td><td>9</td><td>31.2</td><td>20.6</td><td>19.8</td><td>26.7</td><td>6.5</td></tr>
<tr><td>>MI0000095|hsa-mir-93&WithFlank&7|-|100093762|100093853@17.38.22&[MIMAT0000093&hsa-miR-93-5p&offsets|0|-1;m-29&7|-|100093815|100093837&offsets|0|-1]|CAAAGTGCTGTTCGTGCAGGTA</td><td>>MI0000095|hsa-mir-93&WithFlank&7|-|100093762|100093853</td><td>17</td><td>38</td><td>22</td><td>CAAAGTGCTGTTCGTGCAGGTA</td><td>MB231</td><td>28</td><td>24</td><td>14</td><td>141</td><td>48</td><td>9.2</td><td>11.1</td><td>13.9</td><td>75.3</td><td>37.8</td></tr>
<tr><td>>MI0000095|hsa-mir-93&WithFlank&7|-|100093762|100093853@17.39.23(+1A)&[MIMAT0000093&hsa-miR-93-5p&offsets|0|0(+1A);m-29&7|-|100093815|100093837&offsets|0|0(+1A)]|CAAAGTGCTGTTCGTGCAGGTAGA</td><td>>MI0000095|hsa-mir-93&WithFlank&7|-|100093762|100093853</td><td>17</td><td>39</td><td>23(+1A)</td><td>CAAAGTGCTGTTCGTGCAGGTAGA</td><td>BT20</td><td>126</td><td>87</td><td>149</td><td>29</td><td>46</td><td>59.5</td><td>64.4</td><td>52.0</td><td>18.7</td><td>33.5</td></tr>
<tr><td>>MI0000095|hsa-mir-93&WithFlank&7|-|100093762|100093853@17.39.23(+1A)&[MIMAT0000093&hsa-miR-93-5p&offsets|0|0(+1A);m-29&7|-|100093815|100093837&offsets|0|0(+1A)]|CAAAGTGCTGTTCGTGCAGGTAGA</td><td>>MI0000095|hsa-mir-93&WithFlank&7|-|100093762|100093853</td><td>17</td><td>39</td><td>23(+1A)</td><td>CAAAGTGCTGTTCGTGCAGGTAGA</td><td>MB231</td><td>105</td><td>76</td><td>45</td><td>74</td><td>81</td><td>24.4</td><td>34.2</td><td>46.4</td><td>27.4</td><td>39.7</td></tr>
<tr><td>>MI0000095|hsa-mir-93&WithFlank&7|-|100093762|100093853@17.39.23(+1A)&[MIMAT0000093&hsa-miR-93-5p&offsets|0|0(+1A);m-29&7|-|100093815|100093837&offsets|0|0(+1A)]|CAAAGTGCTGTTCGTGCAGGTAGA</td><td>>MI0000095|hsa-mir-93&WithFlank&7|-|100093762|100093853</td><td>17</td><td>39</td><td>23(+1A)</td><td>CAAAGTGCTGTTCGTGCAGGTAGA</td><td>MB468</td><td>11</td><td>7</td><td>5</td><td>10</td><td>4</td><td>3.7</td><td>1.6</td><td>1.2</td><td>4.2</td><td>1.4</td></tr>
<tr><td>>MI0000095|hsa-mir-93&WithFlank&7|-|100093762|100093853@17.39.23(+1C)&[MIMAT0000093&hsa-miR-93-5p&offsets|0|0(+1C);m-29&7|-|100093815|100093837&offsets|0|0(+1C)]|CAAAGTGCTGTTCGTGCAGGTAGC</td><td>>MI0000095|hsa-mir-93&WithFlank&7|-|100093762|100093853</td><td>17</td><td>39</td><td>23(+1C)</td><td>CAAAGTGCTGTTCGTGCAGGTAGC</td><td>BT20</td><td>13</td><td>11</td><td>11</td><td>2</td><td>5</td><td>5.7</td><td>9.4</td><td>5.4</td><td>1.1</td><td>4.6</td></tr>
<tr><td>>MI0000095|hsa-mir-93&WithFlank&7|-|100093762|100093853@17.39.23(+1C)&[MIMAT0000093&hsa-miR-93-5p&offsets|0|0(+1C);m-29&7|-|100093815|100093837&offsets|0|0(+1C)]|CAAAGTGCTGTTCGTGCAGGTAGC</td><td>>MI0000095|hsa-mir-93&WithFlank&7|-|100093762|100093853</td><td>17</td><td>39</td><td>23(+1C)</td><td>CAAAGTGCTGTTCGTGCAGGTAGC</td><td>MB231</td><td>20</td><td>15</td><td>6</td><td>17</td><td>23</td><td>6.7</td><td>7.3</td><td>5.8</td><td>11.4</td><td>20.4</td></tr>
<tr><td>>MI0000095|hsa-mir-93&WithFlank&7|-|100093762|100093853@17.39.23(+1G)&[MIMAT0000093&hsa-miR-93-5p&offsets|0|0(+1G);m-29&7|-|100093815|100093837&offsets|0|0(+1G)]|CAAAGTGCTGTTCGTGCAGGTAGG</td><td>>MI0000095|hsa-mir-93&WithFlank&7|-|100093762|100093853</td><td>17</td><td>39</td><td>23(+1G)</td><td>CAAAGTGCTGTTCGTGCAGGTAGG</td><td>BT20</td><td>14</td><td>5</td><td>8</td><td>1</td><td>1</td><td>10.7</td><td>3.4</td><td>1.8</td><td>0.7</td><td>1.2</td></tr>
<tr><td>>MI0000095|hsa-mir-93&WithFlank&7|-|100093762|100093853@17.39.23(+1G)&[MIMAT0000093&hsa-miR-93-5p&offsets|0|0(+1G);m-29&7|-|100093815|100093837&offsets|0|0(+1G)]|CAAAGTGCTGTTCGTGCAGGTAGG</td><td>>MI0000095|hsa-mir-93&WithFlank&7|-|100093762|100093853</td><td>17</td><td>39</td><td>23(+1G)</td><td>CAAAGTGCTGTTCGTGCAGGTAGG</td><td>MB231</td><td>14</td><td>4</td><td>2</td><td>3</td><td>4</td><td>8.8</td><td>2.2</td><td>2.2</td><td>1.2</td><td>3.0</td></tr>
<tr><td>>MI0000095|hsa-mir-93&WithFlank&7|-|100093762|100093853@17.39.23&[MIMAT0000093&hsa-miR-93-5p&offsets|0|0;m-29&7|-|100093815|100093837&offsets|0|0]|CAAAGTGCTGTTCGTGCAGGTAG</td><td>>MI0000095|hsa-mir-93&WithFlank&7|-|100093762|100093853</td><td>17</td><td>39</td><td>23</td><td>CAAAGTGCTGTTCGTGCAGGTAG</td><td>BT20</td><td>535</td><td>515</td><td>515</td><td>1548</td><td>192</td><td>236.3</td><td>428.3</td><td>290.6</td><td>1485.2</td><td>127.8</td></tr>
<tr><td>>MI0000095|hsa-mir-93&WithFlank&7|-|100093762|100093853@17.39.23&[MIMAT0000093&hsa-miR-93-5p&offsets|0|0;m-29&7|-|100093815|100093837&offsets|0|0]|CAAAGTGCTGTTCGTGCAGGTAG</td><td>>MI0000095|hsa-mir-93&WithFlank&7|-|100093762|100093853</td><td>17</td><td>39</td><td>23</td><td>CAAAGTGCTGTTCGTGCAGGTAG</td><td>MB231</td><td>508</td><td>481</td><td>168</td><td>2582</td><td>511</td><td>47.6</td><td>161.5</td><td>147.7</td><td>855.2</td><td>245.0</td></tr>
<tr><td>>MI0000095|hsa-mir-93&WithFlank&7|-|100093762|100093853@17.39.23&[MIMAT0000093&hsa-miR-93-5p&offsets|0|0;m-29&7|-|100093815|100093837&offsets|0|0]|CAAAGTGCTGTTCGTGCAGGTAG</td><td>>MI0000095|hsa-mir-93&WithFlank&7|-|100093762|100093853</td><td>17</td><td>39</td><td>23</td><td>CAAAGTGCTGTTCGTGCAGGTAG</td><td>MB468</td><td>66</td><td>29</td><td>24</td><td>158</td><td>18</td><td>36.5</td><td>5.3</td><td>5.5</td><td>54.4</td><td>4.1</td></tr>
<tr><td>>MI0000095|hsa-mir-93&WithFlank&7|-|100093762|100093853@17.40.24&[MIMAT0000093&hsa-miR-93-5p&offsets|0|+1;m-29&7|-|100093815|100093837&offsets|0|+1]|CAAAGTGCTGTTCGTGCAGGTAGT</td><td>>MI0000095|hsa-mir-93&WithFlank&7|-|100093762|100093853</td><td>17</td><td>40</td><td>24</td><td>CAAAGTGCTGTTCGTGCAGGTAGT</td><td>BT20</td><td>102</td><td>74</td><td>84</td><td>13</td><td>32</td><td>44.6</td><td>66.1</td><td>48.4</td><td>8.8</td><td>31.3</td></tr>
<tr><td>>MI0000095|hsa-mir-93&WithFlank&7|-|100093762|100093853@17.40.24&[MIMAT0000093&hsa-miR-93-5p&offsets|0|+1;m-29&7|-|100093815|100093837&offsets|0|+1]|CAAAGTGCTGTTCGTGCAGGTAGT</td><td>>MI0000095|hsa-mir-93&WithFlank&7|-|100093762|100093853</td><td>17</td><td>40</td><td>24</td><td>CAAAGTGCTGTTCGTGCAGGTAGT</td><td>MB231</td><td>174</td><td>144</td><td>56</td><td>111</td><td>176</td><td>47.3</td><td>64.3</td><td>51.9</td><td>57.8</td><td>123.4</td></tr>
<tr><td>>MI0000095|hsa-mir-93&WithFlank&7|-|100093762|100093853@17.40.24&[MIMAT0000093&hsa-miR-93-5p&offsets|0|+1;m-29&7|-|100093815|100093837&offsets|0|+1]|CAAAGTGCTGTTCGTGCAGGTAGT</td><td>>MI0000095|hsa-mir-93&WithFlank&7|-|100093762|100093853</td><td>17</td><td>40</td><td>24</td><td>CAAAGTGCTGTTCGTGCAGGTAGT</td><td>MB468</td><td>15</td><td>8</td><td>5</td><td>8</td><td>4</td><td>7.0</td><td>3.3</td><td>1.0</td><td>6.2</td><td>0.9</td></tr>
<tr><td>>MI0000098|hsa-mir-96&WithFlank&7|-|129774686|129774775@15.35.21(+1U)&[MIMAT0000095&hsa-miR-96-5p&offsets|0|-2(+1U);m-248&7|-|129774739|129774761&offsets|0|-2(+1U)]|TTTGGCACTAGCACATTTTTGT</td><td>>MI0000098|hsa-mir-96&WithFlank&7|-|129774686|129774775</td><td>15</td><td>35</td><td>21(+1U)</td><td>TTTGGCACTAGCACATTTTTGT</td><td>BT20</td><td>11</td><td>4</td><td>3</td><td>2</td><td>1</td><td>14.1</td><td>5.1</td><td>1.6</td><td>2.3</td><td>0.7</td></tr>
<tr><td>>MI0000098|hsa-mir-96&WithFlank&7|-|129774686|129774775@15.36.22(+1A)&[MIMAT0000095&hsa-miR-96-5p&offsets|0|-1(+1A);m-248&7|-|129774739|129774761&offsets|0|-1(+1A)]|TTTGGCACTAGCACATTTTTGCA</td><td>>MI0000098|hsa-mir-96&WithFlank&7|-|129774686|129774775</td><td>15</td><td>36</td><td>22(+1A)</td><td>TTTGGCACTAGCACATTTTTGCA</td><td>BT20</td><td>121</td><td>41</td><td>23</td><td>8</td><td>11</td><td>141.7</td><td>34.8</td><td>12.4</td><td>9.5</td><td>2.1</td></tr>
<tr><td>>MI0000098|hsa-mir-96&WithFlank&7|-|129774686|129774775@15.36.22(+1C)&[MIMAT0000095&hsa-miR-96-5p&offsets|0|-1(+1C);m-248&7|-|129774739|129774761&offsets|0|-1(+1C)]|TTTGGCACTAGCACATTTTTGCC</td><td>>MI0000098|hsa-mir-96&WithFlank&7|-|129774686|129774775</td><td>15</td><td>36</td><td>22(+1C)</td><td>TTTGGCACTAGCACATTTTTGCC</td><td>BT20</td><td>104</td><td>63</td><td>24</td><td>12</td><td>19</td><td>86.9</td><td>56.3</td><td>12.6</td><td>14.0</td><td>4.5</td></tr>
<tr><td>>MI0000098|hsa-mir-96&WithFlank&7|-|129774686|129774775@15.36.22(+1C)&[MIMAT0000095&hsa-miR-96-5p&offsets|0|-1(+1C);m-248&7|-|129774739|129774761&offsets|0|-1(+1C)]|TTTGGCACTAGCACATTTTTGCC</td><td>>MI0000098|hsa-mir-96&WithFlank&7|-|129774686|129774775</td><td>15</td><td>36</td><td>22(+1C)</td><td>TTTGGCACTAGCACATTTTTGCC</td><td>MB231</td><td>11</td><td>7</td><td>1</td><td>12</td><td>12</td><td>5.1</td><td>2.3</td><td>0.1</td><td>2.8</td><td>11.1</td></tr>
<tr><td>>MI0000098|hsa-mir-96&WithFlank&7|-|129774686|129774775@15.36.22(+1G)&[MIMAT0000095&hsa-miR-96-5p&offsets|0|-1(+1G);m-248&7|-|129774739|129774761&offsets|0|-1(+1G)]|TTTGGCACTAGCACATTTTTGCG</td><td>>MI0000098|hsa-mir-96&WithFlank&7|-|129774686|129774775</td><td>15</td><td>36</td><td>22(+1G)</td><td>TTTGGCACTAGCACATTTTTGCG</td><td>BT20</td><td>87</td><td>15</td><td>6</td><td>2</td><td>1</td><td>109.7</td><td>12.6</td><td>3.1</td><td>3.7</td><td>1.0</td></tr>
<tr><td>>MI0000098|hsa-mir-96&WithFlank&7|-|129774686|129774775@15.36.22&[MIMAT0000095&hsa-miR-96-5p&offsets|0|-1;m-248&7|-|129774739|129774761&offsets|0|-1]|TTTGGCACTAGCACATTTTTGC</td><td>>MI0000098|hsa-mir-96&WithFlank&7|-|129774686|129774775</td><td>15</td><td>36</td><td>22</td><td>TTTGGCACTAGCACATTTTTGC</td><td>BT20</td><td>19</td><td>12</td><td>8</td><td>9</td><td>2</td><td>20.4</td><td>11.7</td><td>5.4</td><td>7.6</td><td>2.5</td></tr>
<tr><td>>MI0000098|hsa-mir-96&WithFlank&7|-|129774686|129774775@15.37.23&[MIMAT0000095&hsa-miR-96-5p&offsets|0|0;m-248&7|-|129774739|129774761&offsets|0|0]|TTTGGCACTAGCACATTTTTGCT</td><td>>MI0000098|hsa-mir-96&WithFlank&7|-|129774686|129774775</td><td>15</td><td>37</td><td>23</td><td>TTTGGCACTAGCACATTTTTGCT</td><td>BT20</td><td>663</td><td>373</td><td>160</td><td>75</td><td>123</td><td>534.3</td><td>303.0</td><td>83.6</td><td>78.2</td><td>20.5</td></tr>
<tr><td>>MI0000098|hsa-mir-96&WithFlank&7|-|129774686|129774775@15.37.23&[MIMAT0000095&hsa-miR-96-5p&offsets|0|0;m-248&7|-|129774739|129774761&offsets|0|0]|TTTGGCACTAGCACATTTTTGCT</td><td>>MI0000098|hsa-mir-96&WithFlank&7|-|129774686|129774775</td><td>15</td><td>37</td><td>23</td><td>TTTGGCACTAGCACATTTTTGCT</td><td>MB231</td><td>79</td><td>53</td><td>6</td><td>76</td><td>70</td><td>37.3</td><td>15.0</td><td>1.6</td><td>27.8</td><td>61.8</td></tr>
<tr><td>>MI0000098|hsa-mir-96&WithFlank&7|-|129774686|129774775@15.37.23&[MIMAT0000095&hsa-miR-96-5p&offsets|0|0;m-248&7|-|129774739|129774761&offsets|0|0]|TTTGGCACTAGCACATTTTTGCT</td><td>>MI0000098|hsa-mir-96&WithFlank&7|-|129774686|129774775</td><td>15</td><td>37</td><td>23</td><td>TTTGGCACTAGCACATTTTTGCT</td><td>MB468</td><td>23</td><td>5</td><td>5</td><td>42</td><td>5</td><td>10.6</td><td>3.4</td><td>1.0</td><td>38.0</td><td>0.5</td></tr>
<tr><td>>MI0000100|hsa-mir-98&WithFlank&X|-|53556217|53556347@28.48.21(+1A)&[MIMAT0000096&hsa-miR-98-5p&offsets|0|-1(+1A);m-143&X|-|53556299|53556320&offsets|0|-1(+1A)]|TGAGGTAGTAAGTTGTATTGTA</td><td>>MI0000100|hsa-mir-98&WithFlank&X|-|53556217|53556347</td><td>28</td><td>48</td><td>21(+1A)</td><td>TGAGGTAGTAAGTTGTATTGTA</td><td>BT20</td><td>17</td><td>5</td><td>12</td><td>2</td><td>1</td><td>17.4</td><td>1.5</td><td>10.7</td><td>2.3</td><td>0.4</td></tr>
<tr><td>>MI0000100|hsa-mir-98&WithFlank&X|-|53556217|53556347@28.48.21(+1C)&[MIMAT0000096&hsa-miR-98-5p&offsets|0|-1(+1C);m-143&X|-|53556299|53556320&offsets|0|-1(+1C)]|TGAGGTAGTAAGTTGTATTGTC</td><td>>MI0000100|hsa-mir-98&WithFlank&X|-|53556217|53556347</td><td>28</td><td>48</td><td>21(+1C)</td><td>TGAGGTAGTAAGTTGTATTGTC</td><td>BT20</td><td>33</td><td>24</td><td>33</td><td>10</td><td>9</td><td>18.5</td><td>5.9</td><td>21.0</td><td>10.2</td><td>2.7</td></tr>
<tr><td>>MI0000100|hsa-mir-98&WithFlank&X|-|53556217|53556347@28.48.21(+1C)&[MIMAT0000096&hsa-miR-98-5p&offsets|0|-1(+1C);m-143&X|-|53556299|53556320&offsets|0|-1(+1C)]|TGAGGTAGTAAGTTGTATTGTC</td><td>>MI0000100|hsa-mir-98&WithFlank&X|-|53556217|53556347</td><td>28</td><td>48</td><td>21(+1C)</td><td>TGAGGTAGTAAGTTGTATTGTC</td><td>MB231</td><td>26</td><td>22</td><td>6</td><td>61</td><td>24</td><td>6.6</td><td>6.8</td><td>3.2</td><td>13.4</td><td>11.4</td></tr>
<tr><td>>MI0000100|hsa-mir-98&WithFlank&X|-|53556217|53556347@28.48.21(+1G)&[MIMAT0000096&hsa-miR-98-5p&offsets|0|-1(+1G);m-143&X|-|53556299|53556320&offsets|0|-1(+1G)]|TGAGGTAGTAAGTTGTATTGTG</td><td>>MI0000100|hsa-mir-98&WithFlank&X|-|53556217|53556347</td><td>28</td><td>48</td><td>21(+1G)</td><td>TGAGGTAGTAAGTTGTATTGTG</td><td>BT20</td><td>15</td><td>3</td><td>5</td><td>1</td><td>0</td><td>17.0</td><td>1.2</td><td>4.2</td><td>1.6</td><td>0.2</td></tr>
<tr><td>>MI0000100|hsa-mir-98&WithFlank&X|-|53556217|53556347@28.48.21&[MIMAT0000096&hsa-miR-98-5p&offsets|0|-1;m-143&X|-|53556299|53556320&offsets|0|-1]|TGAGGTAGTAAGTTGTATTGT</td><td>>MI0000100|hsa-mir-98&WithFlank&X|-|53556217|53556347</td><td>28</td><td>48</td><td>21</td><td>TGAGGTAGTAAGTTGTATTGT</td><td>BT20</td><td>15</td><td>11</td><td>11</td><td>10</td><td>5</td><td>7.6</td><td>4.9</td><td>7.6</td><td>9.3</td><td>2.3</td></tr>
</table>

<table>
<tr><th></th><th></th><th></th><th></th><th></th><th></th><th></th><th></th><th></th><th></th><th></th><th></th><th></th><th></th><th></th><th></th><th></th></tr>
<tr><td>>MI0000100|hsa-mir-98&WithFlank&X|-|53556217|53556347@28.48.21&[MIMAT0000096&hsa-miR-98-5p&offsets|0|-1;m-143&X|-|53556299|53556320&offsets|0|-1]|TGAGGTAGTAAGTTGTATTGT</td><td>>MI0000100|hsa-mir-98&WithFlank&X|-|53556217|53556347</td><td>28</td><td>48</td><td>21</td><td>TGAGGTAGTAAGTTGTATTGT</td><td>MB231</td><td>18</td><td>17</td><td>3</td><td>40</td><td>13</td><td>4.4</td><td>0.8</td><td>1.9</td><td>1.8</td><td>4.8</td></tr>
<tr><td>>MI0000100|hsa-mir-98&WithFlank&X|-|53556217|53556347@28.49.22&[MIMAT0000096&hsa-miR-98-5p&offsets|0|0;m-143&X|-|53556299|53556320&offsets|0|0]|TGAGGTAGTAAGTTGTATTGTT</td><td>>MI0000100|hsa-mir-98&WithFlank&X|-|53556217|53556347</td><td>28</td><td>49</td><td>22</td><td>TGAGGTAGTAAGTTGTATTGTT</td><td>BT20</td><td>234</td><td>155</td><td>242</td><td>65</td><td>64</td><td>126.7</td><td>29.2</td><td>153.2</td><td>50.8</td><td>21.3</td></tr>
<tr><td>>MI0000100|hsa-mir-98&WithFlank&X|-|53556217|53556347@28.49.22&[MIMAT0000096&hsa-miR-98-5p&offsets|0|0;m-143&X|-|53556299|53556320&offsets|0|0]|TGAGGTAGTAAGTTGTATTGTT</td><td>>MI0000100|hsa-mir-98&WithFlank&X|-|53556217|53556347</td><td>28</td><td>49</td><td>22</td><td>TGAGGTAGTAAGTTGTATTGTT</td><td>MB231</td><td>223</td><td>206</td><td>55</td><td>404</td><td>174</td><td>56.3</td><td>37.6</td><td>26.9</td><td>55.9</td><td>61.5</td></tr>
<tr><td>>MI0000100|hsa-mir-98&WithFlank&X|-|53556217|53556347@28.49.22&[MIMAT0000096&hsa-miR-98-5p&offsets|0|0;m-143&X|-|53556299|53556320&offsets|0|0]|TGAGGTAGTAAGTTGTATTGTT</td><td>>MI0000100|hsa-mir-98&WithFlank&X|-|53556217|53556347</td><td>28</td><td>49</td><td>22</td><td>TGAGGTAGTAAGTTGTATTGTT</td><td>MB468</td><td>57</td><td>41</td><td>33</td><td>71</td><td>10</td><td>28.1</td><td>14.4</td><td>10.0</td><td>36.5</td><td>3.2</td></tr>
<tr><td>>MI0000101|hsa-mir-99a&WithFlank&21|+|16539083|16539175@19.38.20(+1C)&[MIMAT0000097&hsa-miR-99a-5p&offsets|0|-2(+1C);m-64&21|+|16539101|16539121&offsets|0|-1(+1C)]|AACCCGTAGATCCGATCTTGC</td><td>>MI0000101|hsa-mir-99a&WithFlank&21|+|16539083|16539175</td><td>19</td><td>38</td><td>20(+1C)</td><td>AACCCGTAGATCCGATCTTGC</td><td>MB468</td><td>15</td><td>16</td><td>10</td><td>29</td><td>8</td><td>3.9</td><td>6.5</td><td>2.2</td><td>9.2</td><td>0.8</td></tr>
<tr><td>>MI0000101|hsa-mir-99a&WithFlank&21|+|16539083|16539175@19.39.21&[MIMAT0000097&hsa-miR-99a-5p&offsets|0|-1;m-64&21|+|16539101|16539121&offsets|0|0]|AACCCGTAGATCCGATCTTGT</td><td>>MI0000101|hsa-mir-99a&WithFlank&21|+|16539083|16539175</td><td>19</td><td>39</td><td>21</td><td>AACCCGTAGATCCGATCTTGT</td><td>MB231</td><td>56</td><td>95</td><td>29</td><td>97</td><td>79</td><td>17.4</td><td>14.7</td><td>18.7</td><td>31.2</td><td>22.1</td></tr>
<tr><td>>MI0000101|hsa-mir-99a&WithFlank&21|+|16539083|16539175@19.39.21&[MIMAT0000097&hsa-miR-99a-5p&offsets|0|-1;m-64&21|+|16539101|16539121&offsets|0|0]|AACCCGTAGATCCGATCTTGT</td><td>>MI0000101|hsa-mir-99a&WithFlank&21|+|16539083|16539175</td><td>19</td><td>39</td><td>21</td><td>AACCCGTAGATCCGATCTTGT</td><td>MB468</td><td>124</td><td>150</td><td>86</td><td>221</td><td>63</td><td>42.5</td><td>76.7</td><td>31.6</td><td>81.4</td><td>9.1</td></tr>
<tr><td>>MI0000101|hsa-mir-99a&WithFlank&21|+|16539083|16539175@19.40.22(+1A)&[MIMAT0000097&hsa-miR-99a-5p&offsets|0|0(+1A);m-64&21|+|16539101|16539121&offsets|0|+1(+1A)]|AACCCGTAGATCCGATCTTGTGA</td><td>>MI0000101|hsa-mir-99a&WithFlank&21|+|16539083|16539175</td><td>19</td><td>40</td><td>22(+1A)</td><td>AACCCGTAGATCCGATCTTGTGA</td><td>MB231</td><td>18</td><td>29</td><td>10</td><td>21</td><td>25</td><td>6.2</td><td>6.5</td><td>5.9</td><td>6.2</td><td>9.9</td></tr>
<tr><td>>MI0000101|hsa-mir-99a&WithFlank&21|+|16539083|16539175@19.40.22(+1A)&[MIMAT0000097&hsa-miR-99a-5p&offsets|0|0(+1A);m-64&21|+|16539101|16539121&offsets|0|+1(+1A)]|AACCCGTAGATCCGATCTTGTGA</td><td>>MI0000101|hsa-mir-99a&WithFlank&21|+|16539083|16539175</td><td>19</td><td>40</td><td>22(+1A)</td><td>AACCCGTAGATCCGATCTTGTGA</td><td>MB468</td><td>70</td><td>75</td><td>55</td><td>69</td><td>30</td><td>22.7</td><td>30.1</td><td>18.7</td><td>31.3</td><td>9.3</td></tr>
<tr><td>>MI0000101|hsa-mir-99a&WithFlank&21|+|16539083|16539175@19.40.22(+1U)&[MIMAT0000097&hsa-miR-99a-5p&offsets|0|0(+1U);m-64&21|+|16539101|16539121&offsets|0|+1(+1U)]|AACCCGTAGATCCGATCTTGTGT</td><td>>MI0000101|hsa-mir-99a&WithFlank&21|+|16539083|16539175</td><td>19</td><td>40</td><td>22(+1U)</td><td>AACCCGTAGATCCGATCTTGTGT</td><td>MB468</td><td>12</td><td>9</td><td>9</td><td>14</td><td>5</td><td>3.4</td><td>3.1</td><td>1.9</td><td>8.8</td><td>1.6</td></tr>
<tr><td>>MI0000101|hsa-mir-99a&WithFlank&21|+|16539083|16539175@19.40.22&[MIMAT0000097&hsa-miR-99a-5p&offsets|0|0;m-64&21|+|16539101|16539121&offsets|0|+1]|AACCCGTAGATCCGATCTTGTG</td><td>>MI0000101|hsa-mir-99a&WithFlank&21|+|16539083|16539175</td><td>19</td><td>40</td><td>22</td><td>AACCCGTAGATCCGATCTTGTG</td><td>MB231</td><td>81</td><td>88</td><td>39</td><td>87</td><td>79</td><td>14.6</td><td>30.5</td><td>30.8</td><td>34.7</td><td>40.5</td></tr>
<tr><td>>MI0000101|hsa-mir-99a&WithFlank&21|+|16539083|16539175@19.40.22&[MIMAT0000097&hsa-miR-99a-5p&offsets|0|0;m-64&21|+|16539101|16539121&offsets|0|+1]|AACCCGTAGATCCGATCTTGTG</td><td>>MI0000101|hsa-mir-99a&WithFlank&21|+|16539083|16539175</td><td>19</td><td>40</td><td>22</td><td>AACCCGTAGATCCGATCTTGTG</td><td>MB468</td><td>98</td><td>67</td><td>58</td><td>110</td><td>40</td><td>37.5</td><td>20.1</td><td>9.8</td><td>62.6</td><td>16.7</td></tr>
<tr><td>>MI0000102|hsa-mir-100&WithFlank&11|-|122152223|122152314@19.36.18&[MIMAT0000098&hsa-miR-100-5p&offsets|0|-4;m-27&11|-|122152276|122152296&offsets|0|-3]|AACCCGTAGATCCGAACT</td><td>>MI0000102|hsa-mir-100&WithFlank&11|-|122152223|122152314</td><td>19</td><td>36</td><td>18</td><td>AACCCGTAGATCCGAACT</td><td>MB231</td><td>21</td><td>22</td><td>13</td><td>49</td><td>25</td><td>1.6</td><td>7.0</td><td>5.3</td><td>11.5</td><td>8.8</td></tr>
<tr><td>>MI0000102|hsa-mir-100&WithFlank&11|-|122152223|122152314@19.37.19(+1A)&[MIMAT0000098&hsa-miR-100-5p&offsets|0|-3(+1A);m-27&11|-|122152276|122152296&offsets|0|-2(+1A)]|AACCCGTAGATCCGAACTTA</td><td>>MI0000102|hsa-mir-100&WithFlank&11|-|122152223|122152314</td><td>19</td><td>37</td><td>19(+1A)</td><td>AACCCGTAGATCCGAACTTA</td><td>MB231</td><td>95</td><td>123</td><td>115</td><td>558</td><td>147</td><td>52.3</td><td>84.1</td><td>119.9</td><td>445.6</td><td>92.9</td></tr>
<tr><td>>MI0000102|hsa-mir-100&WithFlank&11|-|122152223|122152314@19.37.19&[MIMAT0000098&hsa-miR-100-5p&offsets|0|-3;m-27&11|-|122152276|122152296&offsets|0|-2]|AACCCGTAGATCCGAACTT</td><td>>MI0000102|hsa-mir-100&WithFlank&11|-|122152223|122152314</td><td>19</td><td>37</td><td>19</td><td>AACCCGTAGATCCGAACTT</td><td>MB231</td><td>19</td><td>19</td><td>11</td><td>61</td><td>24</td><td>7.0</td><td>6.2</td><td>8.0</td><td>22.3</td><td>13.4</td></tr>
<tr><td>>MI0000102|hsa-mir-100&WithFlank&11|-|122152223|122152314@19.38.20(+1A)&[MIMAT0000098&hsa-miR-100-5p&offsets|0|-2(+1A);m-27&11|-|122152276|122152296&offsets|0|-1(+1A)]|AACCCGTAGATCCGAACTTGA</td><td>>MI0000102|hsa-mir-100&WithFlank&11|-|122152223|122152314</td><td>19</td><td>38</td><td>20(+1A)</td><td>AACCCGTAGATCCGAACTTGA</td><td>MB231</td><td>241</td><td>214</td><td>221</td><td>751</td><td>250</td><td>70.9</td><td>59.4</td><td>209.4</td><td>360.8</td><td>145.6</td></tr>
<tr><td>>MI0000102|hsa-mir-100&WithFlank&11|-|122152223|122152314@19.38.20(+1C)&[MIMAT0000098&hsa-miR-100-5p&offsets|0|-2(+1C);m-27&11|-|122152276|122152296&offsets|0|-1(+1C)]|AACCCGTAGATCCGAACTTGC</td><td>>MI0000102|hsa-mir-100&WithFlank&11|-|122152223|122152314</td><td>19</td><td>38</td><td>20(+1C)</td><td>AACCCGTAGATCCGAACTTGC</td><td>BT20</td><td>21</td><td>19</td><td>78</td><td>23</td><td>5</td><td>17.8</td><td>17.8</td><td>101.0</td><td>31.1</td><td>4.9</td></tr>
<tr><td>>MI0000102|hsa-mir-100&WithFlank&11|-|122152223|122152314@19.38.20(+1C)&[MIMAT0000098&hsa-miR-100-5p&offsets|0|-2(+1C);m-27&11|-|122152276|122152296&offsets|0|-1(+1C)]|AACCCGTAGATCCGAACTTGC</td><td>>MI0000102|hsa-mir-100&WithFlank&11|-|122152223|122152314</td><td>19</td><td>38</td><td>20(+1C)</td><td>AACCCGTAGATCCGAACTTGC</td><td>MB231</td><td>1063</td><td>1303</td><td>855</td><td>3077</td><td>1263</td><td>397.7</td><td>508.5</td><td>718.0</td><td>1288.2</td><td>773.0</td></tr>
<tr><td>>MI0000102|hsa-mir-100&WithFlank&11|-|122152223|122152314@19.38.20(+1G)&[MIMAT0000098&hsa-miR-100-5p&offsets|0|-2(+1G);m-27&11|-|122152276|122152296&offsets|0|-1(+1G)]|AACCCGTAGATCCGAACTTGG</td><td>>MI0000102|hsa-mir-100&WithFlank&11|-|122152223|122152314</td><td>19</td><td>38</td><td>20(+1G)</td><td>AACCCGTAGATCCGAACTTGG</td><td>MB231</td><td>476</td><td>198</td><td>133</td><td>277</td><td>99</td><td>398.8</td><td>102.3</td><td>122.2</td><td>137.4</td><td>69.0</td></tr>
<tr><td>>MI0000102|hsa-mir-100&WithFlank&11|-|122152223|122152314@19.38.20(+2G)&[MIMAT0000098&hsa-miR-100-5p&offsets|0|-2(+2G);m-27&11|-|122152276|122152296&offsets|0|-1(+2G)]|AACCCGTAGATCCGAACTTGGG</td><td>>MI0000102|hsa-mir-100&WithFlank&11|-|122152223|122152314</td><td>19</td><td>38</td><td>20(+2G)</td><td>AACCCGTAGATCCGAACTTGGG</td><td>MB231</td><td>114</td><td>71</td><td>36</td><td>54</td><td>59</td><td>72.0</td><td>33.6</td><td>29.3</td><td>12.8</td><td>47.0</td></tr>
<tr><td>>MI0000102|hsa-mir-100&WithFlank&11|-|122152223|122152314@19.38.20&[MIMAT0000098&hsa-miR-100-5p&offsets|0|-2;m-27&11|-|122152276|122152296&offsets|0|-1]|AACCCGTAGATCCGAACTTG</td><td>>MI0000102|hsa-mir-100&WithFlank&11|-|122152223|122152314</td><td>19</td><td>38</td><td>20</td><td>AACCCGTAGATCCGAACTTG</td><td>BT20</td><td>31</td><td>25</td><td>85</td><td>52</td><td>15</td><td>45.1</td><td>23.1</td><td>75.5</td><td>72.4</td><td>13.0</td></tr>
<tr><td>>MI0000102|hsa-mir-100&WithFlank&11|-|122152223|122152314@19.38.20&[MIMAT0000098&hsa-miR-100-5p&offsets|0|-2;m-27&11|-|122152276|122152296&offsets|0|-1]|AACCCGTAGATCCGAACTTG</td><td>>MI0000102|hsa-mir-100&WithFlank&11|-|122152223|122152314</td><td>19</td><td>38</td><td>20</td><td>AACCCGTAGATCCGAACTTG</td><td>MB231</td><td>1034</td><td>1819</td><td>1048</td><td>3781</td><td>1421</td><td>556.2</td><td>1326.6</td><td>1005.4</td><td>2816.0</td><td>811.6</td></tr>
<tr><td>>MI0000102|hsa-mir-100&WithFlank&11|-|122152223|122152314@19.39.21(+1A)&[MIMAT0000098&hsa-miR-100-5p&offsets|0|-1(+1A);m-27&11|-|122152276|122152296&offsets|0|0(+1A)]|AACCCGTAGATCCGAACTTGTA</td><td>>MI0000102|hsa-mir-100&WithFlank&11|-|122152223|122152314</td><td>19</td><td>39</td><td>21(+1A)</td><td>AACCCGTAGATCCGAACTTGTA</td><td>BT20</td><td>163</td><td>70</td><td>631</td><td>30</td><td>24</td><td>146.6</td><td>61.0</td><td>918.0</td><td>30.6</td><td>21.0</td></tr>
<tr><td>>MI0000102|hsa-mir-100&WithFlank&11|-|122152223|122152314@19.39.21(+1A)&[MIMAT0000098&hsa-miR-100-5p&offsets|0|-1(+1A);m-27&11|-|122152276|122152296&offsets|0|0(+1A)]|AACCCGTAGATCCGAACTTGTA</td><td>>MI0000102|hsa-mir-100&WithFlank&11|-|122152223|122152314</td><td>19</td><td>39</td><td>21(+1A)</td><td>AACCCGTAGATCCGAACTTGTA</td><td>MB231</td><td>4929</td><td>3729</td><td>3086</td><td>8060</td><td>5327</td><td>1441.1</td><td>623.0</td><td>2398.4</td><td>2651.3</td><td>3409.1</td></tr>
<tr><td>>MI0000102|hsa-mir-100&WithFlank&11|-|122152223|122152314@19.39.21(+1C)&[MIMAT0000098&hsa-miR-100-5p&offsets|0|-1(+1C);m-27&11|-|122152276|122152296&offsets|0|0(+1C)]|AACCCGTAGATCCGAACTTGTC</td><td>>MI0000102|hsa-mir-100&WithFlank&11|-|122152223|122152314</td><td>19</td><td>39</td><td>21(+1C)</td><td>AACCCGTAGATCCGAACTTGTC</td><td>MB231</td><td>153</td><td>117</td><td>69</td><td>223</td><td>150</td><td>69.3</td><td>44.4</td><td>48.3</td><td>89.4</td><td>110.3</td></tr>
<tr><td>>MI0000102|hsa-mir-100&WithFlank&11|-|122152223|122152314@19.39.21(+1U)&[MIMAT0000098&hsa-miR-100-5p&offsets|0|-1(+1U);m-27&11|-|122152276|122152296&offsets|0|0(+1U)]|AACCCGTAGATCCGAACTTGTT</td><td>>MI0000102|hsa-mir-100&WithFlank&11|-|122152223|122152314</td><td>19</td><td>39</td><td>21(+1U)</td><td>AACCCGTAGATCCGAACTTGTT</td><td>BT20</td><td>16</td><td>7</td><td>57</td><td>3</td><td>2</td><td>14.5</td><td>5.9</td><td>84.9</td><td>2.9</td><td>1.6</td></tr>
<tr><td>>MI0000102|hsa-mir-100&WithFlank&11|-|122152223|122152314@19.39.21(+1U)&[MIMAT0000098&hsa-miR-100-5p&offsets|0|-1(+1U);m-27&11|-|122152276|122152296&offsets|0|0(+1U)]|AACCCGTAGATCCGAACTTGTT</td><td>>MI0000102|hsa-mir-100&WithFlank&11|-|122152223|122152314</td><td>19</td><td>39</td><td>21(+1U)</td><td>AACCCGTAGATCCGAACTTGTT</td><td>MB231</td><td>452</td><td>349</td><td>222</td><td>723</td><td>413</td><td>195.3</td><td>107.9</td><td>153.3</td><td>305.5</td><td>251.7</td></tr>
<tr><td>>MI0000102|hsa-mir-100&WithFlank&11|-|122152223|122152314@19.39.21&[MIMAT0000098&hsa-miR-100-5p&offsets|0|-1;m-27&11|-|122152276|122152296&offsets|0|0]|AACCCGTAGATCCGAACTTGT</td><td>>MI0000102|hsa-mir-100&WithFlank&11|-|122152223|122152314</td><td>19</td><td>39</td><td>21</td><td>AACCCGTAGATCCGAACTTGT</td><td>BT20</td><td>158</td><td>117</td><td>523</td><td>196</td><td>38</td><td>132.1</td><td>110.7</td><td>625.2</td><td>280.6</td><td>36.1</td></tr>
<tr><td>>MI0000102|hsa-mir-100&WithFlank&11|-|122152223|122152314@19.39.21&[MIMAT0000098&hsa-miR-100-5p&offsets|0|-1;m-27&11|-|122152276|122152296&offsets|0|0]|AACCCGTAGATCCGAACTTGT</td><td>>MI0000102|hsa-mir-100&WithFlank&11|-|122152223|122152314</td><td>19</td><td>39</td><td>21</td><td>AACCCGTAGATCCGAACTTGT</td><td>MB231</td><td>9160</td><td>11345</td><td>7610</td><td>19037</td><td>8933</td><td>3133.3</td><td>3850.0</td><td>6497.4</td><td>4702.8</td><td>4170.4</td></tr>
<tr><td>>MI0000102|hsa-mir-100&WithFlank&11|-|122152223|122152314@19.40.22(+1A)&[MIMAT0000098&hsa-miR-100-5p&offsets|0|0(+1A);m-27&11|-|122152276|122152296&offsets|0|+1(+1A)]|AACCCGTAGATCCGAACTTGTGA</td><td>>MI0000102|hsa-mir-100&WithFlank&11|-|122152223|122152314</td><td>19</td><td>40</td><td>22(+1A)</td><td>AACCCGTAGATCCGAACTTGTGA</td><td>BT20</td><td>51</td><td>40</td><td>94</td><td>23</td><td>20</td><td>52.2</td><td>37.4</td><td>92.2</td><td>20.4</td><td>17.2</td></tr>
<tr><td>>MI0000102|hsa-mir-100&WithFlank&11|-|122152223|122152314@19.40.22(+1A)&[MIMAT0000098&hsa-miR-100-5p&offsets|0|0(+1A);m-27&11|-|122152276|122152296&offsets|0|+1(+1A)]|AACCCGTAGATCCGAACTTGTGA</td><td>>MI0000102|hsa-mir-100&WithFlank&11|-|122152223|122152314</td><td>19</td><td>40</td><td>22(+1A)</td><td>AACCCGTAGATCCGAACTTGTGA</td><td>MB231</td><td>1901</td><td>2922</td><td>1077</td><td>3038</td><td>2304</td><td>82.1</td><td>788.5</td><td>598.9</td><td>433.1</td><td>875.1</td></tr>
<tr><td>>MI0000102|hsa-mir-100&WithFlank&11|-|122152223|122152314@19.40.22(+1C)&[MIMAT0000098&hsa-miR-100-5p&offsets|0|0(+1C);m-27&11|-|122152276|122152296&offsets|0|+1(+1C)]|AACCCGTAGATCCGAACTTGTGC</td><td>>MI0000102|hsa-mir-100&WithFlank&11|-|122152223|122152314</td><td>19</td><td>40</td><td>22(+1C)</td><td>AACCCGTAGATCCGAACTTGTGC</td><td>MB231</td><td>396</td><td>410</td><td>257</td><td>665</td><td>466</td><td>132.8</td><td>127.5</td><td>170.7</td><td>179.6</td><td>329.6</td></tr>
</table>

| | | | | | | | | | | | | | | | | |
|---|---|---|---|---|---|---|---|---|---|---|---|---|---|---|---|---|
| >MI0000102\|hsa-mir-100&WithFlank&11\|-\|122152223\|122152314@19.40.22(+1U)&[MIMAT0000098&hsa-miR-100-5p&offsets\|0\|0(+1U);m-27&11\|-\|122152276\|122152296&offsets\|0\|+1(+1U)]\|AACCCGTAGATCCGAACTTGTGT | >MI0000102\|hsa-mir-100&WithFlank&11\|-\|122152223\|122152314 | 19 | 40 | 22(+1U) | AACCCGTAGATCCGAACTTGTGT | BT20 | 39 | 31 | 188 | 13 | 12 | 35.5 | 29.3 | 250.1 | 11.6 | 13.3 |
| >MI0000102\|hsa-mir-100&WithFlank&11\|-\|122152223\|122152314@19.40.22(+1U)&[MIMAT0000098&hsa-miR-100-5p&offsets\|0\|0(+1U);m-27&11\|-\|122152276\|122152296&offsets\|0\|+1(+1U)]\|AACCCGTAGATCCGAACTTGTGT | >MI0000102\|hsa-mir-100&WithFlank&11\|-\|122152223\|122152314 | 19 | 40 | 22(+1U) | AACCCGTAGATCCGAACTTGTGT | MB231 | 3377 | 3608 | 2216 | 4391 | 3264 | 1150.9 | 1150.1 | 1553.5 | 824.4 | 1936.4 |
| >MI0000102\|hsa-mir-100&WithFlank&11\|-\|122152223\|122152314@19.40.22(+2A)&[MIMAT0000098&hsa-miR-100-5p&offsets\|0\|0(+2A);m-27&11\|-\|122152276\|122152296&offsets\|0\|+1(+2A)]\|AACCCGTAGATCCGAACTTGTGAA | >MI0000102\|hsa-mir-100&WithFlank&11\|-\|122152223\|122152314 | 19 | 40 | 22(+2A) | AACCCGTAGATCCGAACTTGTGAA | MB231 | 84 | 129 | 47 | 103 | 108 | 27.1 | 25.6 | 21.5 | 26.9 | 18.7 |
| >MI0000102\|hsa-mir-100&WithFlank&11\|-\|122152223\|122152314@19.40.22(+2U)&[MIMAT0000098&hsa-miR-100-5p&offsets\|0\|0(+2U);m-27&11\|-\|122152276\|122152296&offsets\|0\|+1(+2U)]\|AACCCGTAGATCCGAACTTGTGTT | >MI0000102\|hsa-mir-100&WithFlank&11\|-\|122152223\|122152314 | 19 | 40 | 22(+2U) | AACCCGTAGATCCGAACTTGTGTT | MB231 | 37 | 45 | 20 | 27 | 51 | 6.0 | 6.9 | 12.9 | 8.9 | 25.3 |
| >MI0000102\|hsa-mir-100&WithFlank&11\|-\|122152223\|122152314@19.40.22(+3A)&[MIMAT0000098&hsa-miR-100-5p&offsets\|0\|0(+3A);m-27&11\|-\|122152276\|122152296&offsets\|0\|+1(+3A)]\|AACCCGTAGATCCGAACTTGTGAAA | >MI0000102\|hsa-mir-100&WithFlank&11\|-\|122152223\|122152314 | 19 | 40 | 22(+3A) | AACCCGTAGATCCGAACTTGTGAAA | MB231 | 14 | 19 | 11 | 5 | 17 | 5.9 | 4.2 | 3.6 | 3.8 | 3.4 |
| >MI0000102\|hsa-mir-100&WithFlank&11\|-\|122152223\|122152314@19.40.22&[MIMAT0000098&hsa-miR-100-5p&offsets\|0\|0;m-27&11\|-\|122152276\|122152296&offsets\|0\|+1]\|AACCCGTAGATCCGAACTTGTG | >MI0000102\|hsa-mir-100&WithFlank&11\|-\|122152223\|122152314 | 19 | 40 | 22 | AACCCGTAGATCCGAACTTGTG | BT20 | 880 | 504 | 2787 | 319 | 233 | 787.5 | 423.6 | 3422.2 | 328.8 | 200.7 |
| >MI0000102\|hsa-mir-100&WithFlank&11\|-\|122152223\|122152314@19.40.22&[MIMAT0000098&hsa-miR-100-5p&offsets\|0\|0;m-27&11\|-\|122152276\|122152296&offsets\|0\|+1]\|AACCCGTAGATCCGAACTTGTG | >MI0000102\|hsa-mir-100&WithFlank&11\|-\|122152223\|122152314 | 19 | 40 | 22 | AACCCGTAGATCCGAACTTGTG | MB231 | 47270 | 46193 | 26121 | 57641 | 41964 | 11838.8 | 11689.4 | 16336.9 | 17514.5 | 15974.4 |
| >MI0000102\|hsa-mir-100&WithFlank&11\|-\|122152223\|122152314@19.40.22&[MIMAT0000098&hsa-miR-100-5p&offsets\|0\|0;m-27&11\|-\|122152276\|122152296&offsets\|0\|+1]\|AACCCGTAGATCCGAACTTGTG | >MI0000102\|hsa-mir-100&WithFlank&11\|-\|122152223\|122152314 | 19 | 40 | 22 | AACCCGTAGATCCGAACTTGTG | MB468 | 15 | 1 | 2 | 18 | 5 | 27.0 | 0.8 | 1.4 | 26.3 | 10.3 |
| >MI0000102\|hsa-mir-100&WithFlank&11\|-\|122152223\|122152314@19.41.23&[MIMAT0000098&hsa-miR-100-5p&offsets\|0\|+1;m-27&11\|-\|122152276\|122152296&offsets\|0\|+2]\|AACCCGTAGATCCGAACTTGTGG | >MI0000102\|hsa-mir-100&WithFlank&11\|-\|122152223\|122152314 | 19 | 41 | 23 | AACCCGTAGATCCGAACTTGTGG | MB231 | 332 | 186 | 71 | 221 | 136 | 191.5 | 66.4 | 46.2 | 27.2 | 53.7 |
| >MI0000102\|hsa-mir-100&WithFlank&11\|-\|122152223\|122152314@20.39.20(+1A)&[MIMAT0000098&hsa-miR-100-5p&offsets\|+1\|-1(+1A);m-27&11\|-\|122152276\|122152296&offsets\|+1\|0(+1A)]\|ACCCGTAGATCCGAACTTGTA | >MI0000102\|hsa-mir-100&WithFlank&11\|-\|122152223\|122152314 | 20 | 39 | 20(+1A) | ACCCGTAGATCCGAACTTGTA | MB231 | 10 | 8 | 9 | 18 | 8 | 5.4 | 1.9 | 6.6 | 7.7 | 6.4 |
| >MI0000102\|hsa-mir-100&WithFlank&11\|-\|122152223\|122152314@20.39.20&[MIMAT0000098&hsa-miR-100-5p&offsets\|+1\|-1;m-27&11\|-\|122152276\|122152296&offsets\|+1\|0]\|ACCCGTAGATCCGAACTTGT | >MI0000102\|hsa-mir-100&WithFlank&11\|-\|122152223\|122152314 | 20 | 39 | 20 | ACCCGTAGATCCGAACTTGT | MB231 | 16 | 23 | 15 | 31 | 15 | 5.6 | 10.5 | 11.9 | 11.5 | 6.5 |
| >MI0000102\|hsa-mir-100&WithFlank&11\|-\|122152223\|122152314@20.40.21&[MIMAT0000098&hsa-miR-100-5p&offsets\|+1\|0;m-27&11\|-\|122152276\|122152296&offsets\|+1\|+1]\|ACCCGTAGATCCGAACTTGTG | >MI0000102\|hsa-mir-100&WithFlank&11\|-\|122152223\|122152314 | 20 | 40 | 21 | ACCCGTAGATCCGAACTTGTG | MB231 | 98 | 91 | 71 | 121 | 68 | 45.5 | 31.2 | 48.8 | 25.3 | 29.1 |
| >MI0000102\|hsa-mir-100&WithFlank&11\|-\|122152223\|122152314@21.40.20&[MIMAT0000098&hsa-miR-100-5p&offsets\|+2\|0;m-27&11\|-\|122152276\|122152296&offsets\|+2\|+1]\|CCCGTAGATCCGAACTTGTG | >MI0000102\|hsa-mir-100&WithFlank&11\|-\|122152223\|122152314 | 21 | 40 | 20 | CCCGTAGATCCGAACTTGTG | MB231 | 14 | 8 | 13 | 5 | 4 | 12.9 | 4.3 | 8.9 | 2.4 | 1.2 |
| >MI0000102\|hsa-mir-100&WithFlank&11\|-\|122152223\|122152314@22.40.19&[MIMAT0000098&hsa-miR-100-5p&offsets\|+3\|0;m-27&11\|-\|122152276\|122152296&offsets\|+3\|+1]\|CCGTAGATCCGAACTTGTG | >MI0000102\|hsa-mir-100&WithFlank&11\|-\|122152223\|122152314 | 22 | 40 | 19 | CCGTAGATCCGAACTTGTG | MB231 | 47 | 42 | 29 | 40 | 32 | 39.0 | 34.8 | 25.7 | 35.5 | 29.3 |
| >MI0000103\|hsa-mir-101-1&WithFlank&1\|-\|65058428\|65058514@52.70.19;MI0000739\|hsa-mir-101-2&WithFlank&9\|+\|4850291\|4850381@54.72.19&[MIMAT0000099&hsa-miR-101-3p&offsets\|-1\|-3;m-17&1\|-\|65058443\|65058462&offsets\|-1\|-2];[MIMAT0000099_1&hsa-miR-101-3p&offsets\|-1\|-3;m-16&9\|+\|4850345\|4850364&offsets\|-1\|-2]\|GTACAGTACTGTGATAACT | >MI0000103\|hsa-mir-101-1&WithFlank&1\|-\|65058428\|65058514 | 52 | 70 | 19 | GTACAGTACTGTGATAACT | BT20 | 15 | 5 | 3 | 6 | 1 | 16.3 | 3.5 | 1.6 | 6.9 | 0.7 |
| >MI0000103\|hsa-mir-101-1&WithFlank&1\|-\|65058428\|65058514@52.71.20;MI0000739\|hsa-mir-101-2&WithFlank&9\|+\|4850291\|4850381@54.73.20&[MIMAT0000099&hsa-miR-101-3p&offsets\|-1\|-2;m-17&1\|-\|65058443\|65058462&offsets\|-1\|-1];[MIMAT0000099_1&hsa-miR-101-3p&offsets\|-1\|-2;m-16&9\|+\|4850345\|4850364&offsets\|-1\|-1]\|GTACAGTACTGTGATAACTG | >MI0000103\|hsa-mir-101-1&WithFlank&1\|-\|65058428\|65058514 | 52 | 71 | 20 | GTACAGTACTGTGATAACTG | BT20 | 15 | 5 | 2 | 5 | 1 | 17.0 | 5.4 | 1.1 | 4.4 | 0.0 |
| >MI0000103\|hsa-mir-101-1&WithFlank&1\|-\|65058428\|65058514@52.71.20(+1G);MI0000739\|hsa-mir-101-2&WithFlank&9\|+\|4850291\|4850381@54.73.20(+1G)&[MIMAT0000099&hsa-miR-101-3p&offsets\|-1\|-2(+1G);m-17&1\|-\|65058443\|65058462&offsets\|-1\|-1(+1G)];[MIMAT0000099_1&hsa-miR-101-3p&offsets\|-1\|-2(+1G);m-16&9\|+\|4850345\|4850364&offsets\|-1\|-1(+1G)]\|GTACAGTACTGTGATAACTGG | >MI0000103\|hsa-mir-101-1&WithFlank&1\|-\|65058428\|65058514 | 52 | 71 | 20(+1G) | GTACAGTACTGTGATAACTGG | BT20 | 21 | 1 | 0 | 1 | 0 | 31.5 | 1.9 | 0.4 | 1.3 | 0.0 |
| >MI0000103\|hsa-mir-101-1&WithFlank&1\|-\|65058428\|65058514@52.71.20(+1U);MI0000739\|hsa-mir-101-2&WithFlank&9\|+\|4850291\|4850381@54.73.20(+1U)&[MIMAT0000099&hsa-miR-101-3p&offsets\|-1\|-2(+1U);m-17&1\|-\|65058443\|65058462&offsets\|-1\|-1(+1U)];[MIMAT0000099_1&hsa-miR-101-3p&offsets\|-1\|-2(+1U);m-16&9\|+\|4850345\|4850364&offsets\|-1\|-1(+1U)]\|GTACAGTACTGTGATAACTGT | >MI0000103\|hsa-mir-101-1&WithFlank&1\|-\|65058428\|65058514 | 52 | 71 | 20(+1U) | GTACAGTACTGTGATAACTGT | BT20 | 14 | 1 | 0 | 0 | 0 | 21.7 | 1.4 | 0.3 | 0.6 | 0.0 |
| >MI0000103\|hsa-mir-101-1&WithFlank&1\|-\|65058428\|65058514@52.72.21;MI0000739\|hsa-mir-101-2&WithFlank&9\|+\|4850291\|4850381@54.74.21&[MIMAT0000099&hsa-miR-101-3p&offsets\|-1\|-1;m-17&1\|-\|65058443\|65058462&offsets\|-1\|0];[MIMAT0000099_1&hsa-miR-101-3p&offsets\|-1\|-1;m-16&9\|+\|4850345\|4850364&offsets\|-1\|0]\|GTACAGTACTGTGATAACTGA | >MI0000103\|hsa-mir-101-1&WithFlank&1\|-\|65058428\|65058514 | 52 | 72 | 21 | GTACAGTACTGTGATAACTGA | BT20 | 222 | 48 | 12 | 33 | 6 | 310.6 | 53.3 | 4.7 | 40.3 | 0.6 |
| >MI0000103\|hsa-mir-101-1&WithFlank&1\|-\|65058428\|65058514@52.72.21;MI0000739\|hsa-mir-101-2&WithFlank&9\|+\|4850291\|4850381@54.74.21&[MIMAT0000099&hsa-miR-101-3p&offsets\|-1\|-1;m-17&1\|-\|65058443\|65058462&offsets\|-1\|0];[MIMAT0000099_1&hsa-miR-101-3p&offsets\|-1\|-1;m-16&9\|+\|4850345\|4850364&offsets\|-1\|0]\|GTACAGTACTGTGATAACTGA | >MI0000103\|hsa-mir-101-1&WithFlank&1\|-\|65058428\|65058514 | 52 | 72 | 21 | GTACAGTACTGTGATAACTGA | MB231 | 35 | 27 | 2 | 65 | 27 | 9.3 | 16.2 | 0.7 | 19.4 | 28.1 |
| >MI0000103\|hsa-mir-101-1&WithFlank&1\|-\|65058428\|65058514@52.72.21(+1G);MI0000739\|hsa-mir-101-2&WithFlank&9\|+\|4850291\|4850381@54.74.21(+1G)&[MIMAT0000099&hsa-miR-101-3p&offsets\|-1\|-1(+1G);m-17&1\|-\|65058443\|65058462&offsets\|-1\|0(+1G)];[MIMAT0000099_1&hsa-miR-101-3p&offsets\|-1\|-1(+1G);m-16&9\|+\|4850345\|4850364&offsets\|-1\|0(+1G)]\|GTACAGTACTGTGATAACTGAG | >MI0000103\|hsa-mir-101-1&WithFlank&1\|-\|65058428\|65058514 | 52 | 72 | 21(+1G) | GTACAGTACTGTGATAACTGAG | BT20 | 23 | 1 | 0 | 1 | 0 | 33.8 | 1.4 | 0.6 | 2.2 | 0.0 |
| >MI0000103\|hsa-mir-101-1&WithFlank&1\|-\|65058428\|65058514@52.72.21(+1U);MI0000739\|hsa-mir-101-2&WithFlank&9\|+\|4850291\|4850381@54.74.21(+1U)&[MIMAT0000099&hsa-miR-101-3p&offsets\|-1\|-1(+1U);m-17&1\|-\|65058443\|65058462&offsets\|-1\|0(+1U)];[MIMAT0000099_1&hsa-miR-101-3p&offsets\|-1\|-1(+1U);m-16&9\|+\|4850345\|4850364&offsets\|-1\|0(+1U)]\|GTACAGTACTGTGATAACTGAT | >MI0000103\|hsa-mir-101-1&WithFlank&1\|-\|65058428\|65058514 | 52 | 72 | 21(+1U) | GTACAGTACTGTGATAACTGAT | BT20 | 26 | 7 | 3 | 3 | 0 | 33.5 | 7.6 | 1.0 | 5.0 | 0.4 |
| >MI0000103\|hsa-mir-101-1&WithFlank&1\|-\|65058428\|65058514@52.73.22;MI0000739\|hsa-mir-101-2&WithFlank&9\|+\|4850291\|4850381@54.75.22&[MIMAT0000099&hsa-miR-101-3p&offsets\|-1\|0;m-17&1\|-\|65058443\|65058462&offsets\|-1\|+1];[MIMAT0000099_1&hsa-miR-101-3p&offsets\|-1\|0;m-16&9\|+\|4850345\|4850364&offsets\|-1\|+1]\|GTACAGTACTGTGATAACTGAA | >MI0000103\|hsa-mir-101-1&WithFlank&1\|-\|65058428\|65058514 | 52 | 73 | 22 | GTACAGTACTGTGATAACTGAA | BT20 | 312 | 72 | 23 | 65 | 15 | 411.4 | 53.2 | 12.8 | 87.0 | 2.6 |
| >MI0000103\|hsa-mir-101-1&WithFlank&1\|-\|65058428\|65058514@52.73.22;MI0000739\|hsa-mir-101-2&WithFlank&9\|+\|4850291\|4850381@54.75.22&[MIMAT0000099&hsa-miR-101-3p&offsets\|-1\|0;m-17&1\|-\|65058443\|65058462&offsets\|-1\|+1];[MIMAT0000099_1&hsa-miR-101-3p&offsets\|-1\|0;m-16&9\|+\|4850345\|4850364&offsets\|-1\|+1]\|GTACAGTACTGTGATAACTGAA | >MI0000103\|hsa-mir-101-1&WithFlank&1\|-\|65058428\|65058514 | 52 | 73 | 22 | GTACAGTACTGTGATAACTGAA | MB231 | 93 | 70 | 4 | 193 | 74 | 21.2 | 47.5 | 2.2 | 34.3 | 71.2 |
| >MI0000103\|hsa-mir-101-1&WithFlank&1\|-\|65058428\|65058514@52.73.22;MI0000739\|hsa-mir-101-2&WithFlank&9\|+\|4850291\|4850381@54.75.22&[MIMAT0000099&hsa-miR-101-3p&offsets\|-1\|0;m-17&1\|-\|65058443\|65058462&offsets\|-1\|+1];[MIMAT0000099_1&hsa-miR-101-3p&offsets\|-1\|0;m-16&9\|+\|4850345\|4850364&offsets\|-1\|+1]\|GTACAGTACTGTGATAACTGAA | >MI0000103\|hsa-mir-101-1&WithFlank&1\|-\|65058428\|65058514 | 52 | 73 | 22 | GTACAGTACTGTGATAACTGAA | MB468 | 12 | 2 | 3 | 45 | 2 | 5.4 | 1.2 | 1.3 | 32.9 | 0.7 |
| >MI0000103\|hsa-mir-101-1&WithFlank&1\|-\|65058428\|65058514@52.73.22(+1A);MI0000739\|hsa-mir-101-2&WithFlank&9\|+\|4850291\|4850381@54.75.22(+1A)&[MIMAT0000099&hsa-miR-101-3p&offsets\|-1\|0(+1A);m-17&1\|-\|65058443\|65058462&offsets\|-1\|+1(+1A)];[MIMAT0000099_1&hsa-miR-101-3p&offsets\|-1\|0(+1A);m-16&9\|+\|4850345\|4850364&offsets\|-1\|+1(+1A)]\|GTACAGTACTGTGATAACTGAAA | >MI0000103\|hsa-mir-101-1&WithFlank&1\|-\|65058428\|65058514 | 52 | 73 | 22(+1A) | GTACAGTACTGTGATAACTGAAA | BT20 | 118 | 21 | 7 | 6 | 4 | 170.5 | 18.3 | 4.4 | 7.7 | 1.6 |

<table>
<tr><th></th><th></th><th></th><th></th><th></th><th></th><th></th><th></th><th></th><th></th><th></th><th></th><th></th><th></th><th></th><th></th><th></th></tr>
<tr><td>>MI0000103|hsa-mir-101-1&WithFlank&1|-|65058428|65058514@52.73.22(+1A);MI0000739|hsa-mir-101-2&WithFlank&9|+|4850291|4850381@54.75.22(+1A)&[MIMAT0000099&hsa-miR-101-3p&offsets|-1|0(+1A);m-17&1|-|65058443|65058462&offsets|-1|+1(+1A)];[MIMAT0000099_1&hsa-miR-101-3p&offsets|-1|0(+1A);m-16&9|+|4850345|4850364&offsets|-1|+1(+1A)]|GTACAGTACTGTGATAACTGAAA</td><td>>MI0000103|hsa-mir-101-1&WithFlank&1|-|65058428|65058514</td><td>52</td><td>73</td><td>22(+1A)</td><td>GTACAGTACTGTGATAACTGAAA</td><td>MB231</td><td>17</td><td>13</td><td>1</td><td>21</td><td>17</td><td>7.3</td><td>7.1</td><td>0.4</td><td>7.5</td><td>17.8</td></tr>
<tr><td>>MI0000103|hsa-mir-101-1&WithFlank&1|-|65058428|65058514@52.73.22(+1U);MI0000739|hsa-mir-101-2&WithFlank&9|+|4850291|4850381@54.75.22(+1U)&[MIMAT0000099&hsa-miR-101-3p&offsets|-1|0(+1U);m-17&1|-|65058443|65058462&offsets|-1|+1(+1U)];[MIMAT0000099_1&hsa-miR-101-3p&offsets|-1|0(+1U);m-16&9|+|4850345|4850364&offsets|-1|+1(+1U)]|GTACAGTACTGTGATAACTGAAT</td><td>>MI0000103|hsa-mir-101-1&WithFlank&1|-|65058428|65058514</td><td>52</td><td>73</td><td>22(+1U)</td><td>GTACAGTACTGTGATAACTGAAT</td><td>BT20</td><td>14</td><td>5</td><td>1</td><td>1</td><td>0</td><td>18.5</td><td>6.5</td><td>0.4</td><td>0.9</td><td>0.3</td></tr>
<tr><td>>MI0000103|hsa-mir-101-1&WithFlank&1|-|65058428|65058514@52.74.23;MI0000739|hsa-mir-101-2&WithFlank&9|+|4850291|4850381@54.76.23&[MIMAT0000099&hsa-miR-101-3p&offsets|-1|+1;m-17&1|-|65058443|65058462&offsets|-1|+2];[MIMAT0000099_1&hsa-miR-101-3p&offsets|-1|+1;m-16&9|+|4850345|4850364&offsets|-1|+2]|GTACAGTACTGTGATAACTGAAG</td><td>>MI0000103|hsa-mir-101-1&WithFlank&1|-|65058428|65058514</td><td>52</td><td>74</td><td>23</td><td>GTACAGTACTGTGATAACTGAAG</td><td>BT20</td><td>11</td><td>1</td><td>1</td><td>0</td><td>0</td><td>15.6</td><td>2.0</td><td>0.5</td><td>0.6</td><td>0.0</td></tr>
<tr><td>>MI0000103|hsa-mir-101-1&WithFlank&1|-|65058428|65058514@53.72.20;MI0000739|hsa-mir-101-2&WithFlank&9|+|4850291|4850381@55.74.20&[MIMAT0000099&hsa-miR-101-3p&offsets|0|-1;m-17&1|-|65058443|65058462&offsets|0|0];[MIMAT0000099_1&hsa-miR-101-3p&offsets|0|-1;m-16&9|+|4850345|4850364&offsets|0|0]|TACAGTACTGTGATAACTGA</td><td>>MI0000103|hsa-mir-101-1&WithFlank&1|-|65058428|65058514</td><td>53</td><td>72</td><td>20</td><td>TACAGTACTGTGATAACTGA</td><td>BT20</td><td>122</td><td>58</td><td>30</td><td>99</td><td>18</td><td>74.6</td><td>47.5</td><td>20.6</td><td>103.4</td><td>5.1</td></tr>
<tr><td>>MI0000103|hsa-mir-101-1&WithFlank&1|-|65058428|65058514@53.72.20;MI0000739|hsa-mir-101-2&WithFlank&9|+|4850291|4850381@55.74.20&[MIMAT0000099&hsa-miR-101-3p&offsets|0|-1;m-17&1|-|65058443|65058462&offsets|0|0];[MIMAT0000099_1&hsa-miR-101-3p&offsets|0|-1;m-16&9|+|4850345|4850364&offsets|0|0]|TACAGTACTGTGATAACTGA</td><td>>MI0000103|hsa-mir-101-1&WithFlank&1|-|65058428|65058514</td><td>53</td><td>72</td><td>20</td><td>TACAGTACTGTGATAACTGA</td><td>MB231</td><td>62</td><td>66</td><td>5</td><td>196</td><td>48</td><td>9.7</td><td>46.9</td><td>2.0</td><td>90.4</td><td>29.4</td></tr>
<tr><td>>MI0000103|hsa-mir-101-1&WithFlank&1|-|65058428|65058514@53.72.20(+2U);MI0000739|hsa-mir-101-2&WithFlank&9|+|4850291|4850381@55.74.20(+2U)&[MIMAT0000099&hsa-miR-101-3p&offsets|0|-1(+2U);m-17&1|-|65058443|65058462&offsets|0|0(+2U)];[MIMAT0000099_1&hsa-miR-101-3p&offsets|0|-1(+2U);m-16&9|+|4850345|4850364&offsets|0|0(+2U)]|TACAGTACTGTGATAACTGATT</td><td>>MI0000103|hsa-mir-101-1&WithFlank&1|-|65058428|65058514</td><td>53</td><td>72</td><td>20(+2U)</td><td>TACAGTACTGTGATAACTGATT</td><td>BT20</td><td>25</td><td>7</td><td>3</td><td>2</td><td>0</td><td>36.0</td><td>10.1</td><td>0.7</td><td>2.7</td><td>0.4</td></tr>
<tr><td>>MI0000103|hsa-mir-101-1&WithFlank&1|-|65058428|65058514@53.72.20(+2U);MI0000739|hsa-mir-101-2&WithFlank&9|+|4850291|4850381@55.74.20(+2U)&[MIMAT0000099&hsa-miR-101-3p&offsets|0|-1(+2U);m-17&1|-|65058443|65058462&offsets|0|0(+2U)];[MIMAT0000099_1&hsa-miR-101-3p&offsets|0|-1(+2U);m-16&9|+|4850345|4850364&offsets|0|0(+2U)]|TACAGTACTGTGATAACTGATT</td><td>>MI0000103|hsa-mir-101-1&WithFlank&1|-|65058428|65058514</td><td>53</td><td>72</td><td>20(+2U)</td><td>TACAGTACTGTGATAACTGATT</td><td>MB231</td><td>12</td><td>6</td><td>1</td><td>11</td><td>4</td><td>8.3</td><td>5.2</td><td>0.4</td><td>3.3</td><td>6.2</td></tr>
<tr><td>>MI0000103|hsa-mir-101-1&WithFlank&1|-|65058428|65058514@53.73.21;MI0000739|hsa-mir-101-2&WithFlank&9|+|4850291|4850381@55.75.21&[MIMAT0000099&hsa-miR-101-3p&offsets|0|0;m-17&1|-|65058443|65058462&offsets|0|+1];[MIMAT0000099_1&hsa-miR-101-3p&offsets|0|0;m-16&9|+|4850345|4850364&offsets|0|+1]|TACAGTACTGTGATAACTGAA</td><td>>MI0000103|hsa-mir-101-1&WithFlank&1|-|65058428|65058514</td><td>53</td><td>73</td><td>21</td><td>TACAGTACTGTGATAACTGAA</td><td>BT20</td><td>108</td><td>27</td><td>13</td><td>32</td><td>4</td><td>132.5</td><td>25.9</td><td>4.8</td><td>39.7</td><td>0.8</td></tr>
<tr><td>>MI0000103|hsa-mir-101-1&WithFlank&1|-|65058428|65058514@53.73.21;MI0000739|hsa-mir-101-2&WithFlank&9|+|4850291|4850381@55.75.21&[MIMAT0000099&hsa-miR-101-3p&offsets|0|0;m-17&1|-|65058443|65058462&offsets|0|+1];[MIMAT0000099_1&hsa-miR-101-3p&offsets|0|0;m-16&9|+|4850345|4850364&offsets|0|+1]|TACAGTACTGTGATAACTGAA</td><td>>MI0000103|hsa-mir-101-1&WithFlank&1|-|65058428|65058514</td><td>53</td><td>73</td><td>21</td><td>TACAGTACTGTGATAACTGAA</td><td>MB231</td><td>36</td><td>27</td><td>1</td><td>68</td><td>21</td><td>11.0</td><td>22.7</td><td>1.0</td><td>5.6</td><td>19.0</td></tr>
<tr><td>>MI0000103|hsa-mir-101-1&WithFlank&1|-|65058428|65058514@53.73.21(+1A);MI0000739|hsa-mir-101-2&WithFlank&9|+|4850291|4850381@55.75.21(+1A)&[MIMAT0000099&hsa-miR-101-3p&offsets|0|0(+1A);m-17&1|-|65058443|65058462&offsets|0|+1(+1A)];[MIMAT0000099_1&hsa-miR-101-3p&offsets|0|0(+1A);m-16&9|+|4850345|4850364&offsets|0|+1(+1A)]|TACAGTACTGTGATAACTGAAA</td><td>>MI0000103|hsa-mir-101-1&WithFlank&1|-|65058428|65058514</td><td>53</td><td>73</td><td>21(+1A)</td><td>TACAGTACTGTGATAACTGAAA</td><td>BT20</td><td>184</td><td>26</td><td>11</td><td>13</td><td>3</td><td>275.4</td><td>25.9</td><td>8.5</td><td>17.8</td><td>0.8</td></tr>
<tr><td>>MI0000103|hsa-mir-101-1&WithFlank&1|-|65058428|65058514@53.73.21(+1A);MI0000739|hsa-mir-101-2&WithFlank&9|+|4850291|4850381@55.75.21(+1A)&[MIMAT0000099&hsa-miR-101-3p&offsets|0|0(+1A);m-17&1|-|65058443|65058462&offsets|0|+1(+1A)];[MIMAT0000099_1&hsa-miR-101-3p&offsets|0|0(+1A);m-16&9|+|4850345|4850364&offsets|0|+1(+1A)]|TACAGTACTGTGATAACTGAAA</td><td>>MI0000103|hsa-mir-101-1&WithFlank&1|-|65058428|65058514</td><td>53</td><td>73</td><td>21(+1A)</td><td>TACAGTACTGTGATAACTGAAA</td><td>MB231</td><td>17</td><td>13</td><td>1</td><td>28</td><td>13</td><td>8.0</td><td>9.7</td><td>0.4</td><td>2.1</td><td>14.9</td></tr>
<tr><td>>MI0000103|hsa-mir-101-1&WithFlank&1|-|65058428|65058514@53.73.21(+1U);MI0000739|hsa-mir-101-2&WithFlank&9|+|4850291|4850381@55.75.21(+1U)&[MIMAT0000099&hsa-miR-101-3p&offsets|0|0(+1U);m-17&1|-|65058443|65058462&offsets|0|+1(+1U)];[MIMAT0000099_1&hsa-miR-101-3p&offsets|0|0(+1U);m-16&9|+|4850345|4850364&offsets|0|+1(+1U)]|TACAGTACTGTGATAACTGAAT</td><td>>MI0000103|hsa-mir-101-1&WithFlank&1|-|65058428|65058514</td><td>53</td><td>73</td><td>21(+1U)</td><td>TACAGTACTGTGATAACTGAAT</td><td>BT20</td><td>57</td><td>26</td><td>4</td><td>5</td><td>1</td><td>71.6</td><td>38.7</td><td>1.3</td><td>5.7</td><td>0.9</td></tr>
<tr><td>>MI0000103|hsa-mir-101-1&WithFlank&1|-|65058428|65058514@53.73.21(+1U);MI0000739|hsa-mir-101-2&WithFlank&9|+|4850291|4850381@55.75.21(+1U)&[MIMAT0000099&hsa-miR-101-3p&offsets|0|0(+1U);m-17&1|-|65058443|65058462&offsets|0|+1(+1U)];[MIMAT0000099_1&hsa-miR-101-3p&offsets|0|0(+1U);m-16&9|+|4850345|4850364&offsets|0|+1(+1U)]|TACAGTACTGTGATAACTGAAT</td><td>>MI0000103|hsa-mir-101-1&WithFlank&1|-|65058428|65058514</td><td>53</td><td>73</td><td>21(+1U)</td><td>TACAGTACTGTGATAACTGAAT</td><td>MB231</td><td>43</td><td>46</td><td>2</td><td>49</td><td>28</td><td>25.1</td><td>39.3</td><td>1.2</td><td>21.2</td><td>34.7</td></tr>
<tr><td>>MI0000103|hsa-mir-101-1&WithFlank&1|-|65058428|65058514@53.74.22;MI0000739|hsa-mir-101-2&WithFlank&9|+|4850291|4850381@55.76.22&[MIMAT0000099&hsa-miR-101-3p&offsets|0|+1;m-17&1|-|65058443|65058462&offsets|0|+2];[MIMAT0000099_1&hsa-miR-101-3p&offsets|0|+1;m-16&9|+|4850345|4850364&offsets|0|+2]|TACAGTACTGTGATAACTGAAG</td><td>>MI0000103|hsa-mir-101-1&WithFlank&1|-|65058428|65058514</td><td>53</td><td>74</td><td>22</td><td>TACAGTACTGTGATAACTGAAG</td><td>BT20</td><td>596</td><td>129</td><td>42</td><td>84</td><td>14</td><td>812.5</td><td>122.7</td><td>19.7</td><td>107.3</td><td>2.7</td></tr>
<tr><td>>MI0000103|hsa-mir-101-1&WithFlank&1|-|65058428|65058514@53.74.22;MI0000739|hsa-mir-101-2&WithFlank&9|+|4850291|4850381@55.76.22&[MIMAT0000099&hsa-miR-101-3p&offsets|0|+1;m-17&1|-|65058443|65058462&offsets|0|+2];[MIMAT0000099_1&hsa-miR-101-3p&offsets|0|+1;m-16&9|+|4850345|4850364&offsets|0|+2]|TACAGTACTGTGATAACTGAAG</td><td>>MI0000103|hsa-mir-101-1&WithFlank&1|-|65058428|65058514</td><td>53</td><td>74</td><td>22</td><td>TACAGTACTGTGATAACTGAAG</td><td>MB231</td><td>109</td><td>97</td><td>4</td><td>157</td><td>69</td><td>61.7</td><td>67.1</td><td>2.3</td><td>11.7</td><td>75.6</td></tr>
<tr><td>>MI0000103|hsa-mir-101-1&WithFlank&1|-|65058428|65058514@53.74.22(+1U);MI0000739|hsa-mir-101-2&WithFlank&9|+|4850291|4850381@55.76.22(+1U)&[MIMAT0000099&hsa-miR-101-3p&offsets|0|+1(+1U);m-17&1|-|65058443|65058462&offsets|0|+2(+1U)];[MIMAT0000099_1&hsa-miR-101-3p&offsets|0|+1(+1U);m-16&9|+|4850345|4850364&offsets|0|+2(+1U)]|TACAGTACTGTGATAACTGAAGT</td><td>>MI0000103|hsa-mir-101-1&WithFlank&1|-|65058428|65058514</td><td>53</td><td>74</td><td>22(+1U)</td><td>TACAGTACTGTGATAACTGAAGT</td><td>BT20</td><td>58</td><td>16</td><td>7</td><td>8</td><td>3</td><td>66.7</td><td>16.2</td><td>1.8</td><td>11.2</td><td>1.1</td></tr>
<tr><td>>MI0000103|hsa-mir-101-1&WithFlank&1|-|65058428|65058514@53.74.22(+1U);MI0000739|hsa-mir-101-2&WithFlank&9|+|4850291|4850381@55.76.22(+1U)&[MIMAT0000099&hsa-miR-101-3p&offsets|0|+1(+1U);m-17&1|-|65058443|65058462&offsets|0|+2(+1U)];[MIMAT0000099_1&hsa-miR-101-3p&offsets|0|+1(+1U);m-16&9|+|4850345|4850364&offsets|0|+2(+1U)]|TACAGTACTGTGATAACTGAAGT</td><td>>MI0000103|hsa-mir-101-1&WithFlank&1|-|65058428|65058514</td><td>53</td><td>74</td><td>22(+1U)</td><td>TACAGTACTGTGATAACTGAAGT</td><td>MB231</td><td>37</td><td>32</td><td>2</td><td>50</td><td>28</td><td>22.1</td><td>25.0</td><td>0.7</td><td>12.2</td><td>30.6</td></tr>
<tr><td>>MI0000107|hsa-mir-29b-2&WithFlank&1|-|207802437|207802529@58.79.22(+1C);MI0000105|hsa-mir-29b-1&WithFlank&7|-|130877453|130877545@57.78.22(+1C)&[MIMAT0000100&hsa-miR-29b-3p&offsets|0|-1(+1C);m-81&1|-|207802450|207802472&offsets|0|-1(+1C)];[MIMAT0000100_1&hsa-miR-29b-3p&offsets|0|-1(+1C);m-82&7|-|130877467|130877489&offsets|0|-1(+1C)]|TAGCACCATTTGAAATCAGTGTC</td><td>>MI0000107|hsa-mir-29b-2&WithFlank&1|-|207802437|207802529</td><td>58</td><td>79</td><td>22(+1C)</td><td>TAGCACCATTTGAAATCAGTGTC</td><td>BT20</td><td>17</td><td>14</td><td>0</td><td>5</td><td>0</td><td>14.7</td><td>21.6</td><td>0.3</td><td>8.5</td><td>0.6</td></tr>
<tr><td>>MI0000107|hsa-mir-29b-2&WithFlank&1|-|207802437|207802529@58.80.23;MI0000105|hsa-mir-29b-1&WithFlank&7|-|130877453|130877545@57.79.23&[MIMAT0000100&hsa-miR-29b-3p&offsets|0|0;m-81&1|-|207802450|207802472&offsets|0|0];[MIMAT0000100_1&hsa-miR-29b-3p&offsets|0|0;m-82&7|-|130877467|130877489&offsets|0|0]|TAGCACCATTTGAAATCAGTGTT</td><td>>MI0000107|hsa-mir-29b-2&WithFlank&1|-|207802437|207802529</td><td>58</td><td>80</td><td>23</td><td>TAGCACCATTTGAAATCAGTGTT</td><td>BT20</td><td>123</td><td>107</td><td>4</td><td>35</td><td>6</td><td>108.0</td><td>157.4</td><td>1.0</td><td>49.9</td><td>1.5</td></tr>
</table>

<table>
<tr><td>>MI0000107|hsa-mir-29b-2&WithFlank&1|-|207802437|207802529@58.80.23;MI0000105|hsa-mir-29b-1&WithFlank&7|-|130877453|130877545@57.79.23&[MIMAT0000100&hsa-miR-29b-3p&offsets|0|0;m-81&1|-|207802450|207802472&offsets|0|0];[MIMAT0000100_1&hsa-miR-29b-3p&offsets|0|0;m-82&7|-|130877467|130877489&offsets|0|0]|TAGCACCATTTGAAATCAGTGTT</td><td>>MI0000107|hsa-mir-29b-2&WithFlank&1|-|207802437|207802529</td><td>58</td><td>80</td><td>23</td><td>TAGCACCATTTGAAATCAGTGTT</td><td>MB231</td><td>54</td><td>43</td><td>1</td><td>143</td><td>34</td><td>43.9</td><td>25.0</td><td>0.8</td><td>96.6</td><td>35.7</td></tr>
<tr><td>>MI0000108|hsa-mir-103a-2&WithFlank&20|+|3917488|3917577@54.75.22;MI0000109|hsa-mir-103a-1&WithFlank&5|-|168560890|168560979@54.75.22&[MIMAT0000101&hsa-miR-103a-3p&offsets|0|-1;m-12&20|+|3917541|3917563&offsets|0|-1];[MIMAT0000101_1&hsa-miR-103a-3p&offsets|0|-1;m-13&5|-|168560904|168560926&offsets|0|-1;m-44220&5|-|168560908|168560926&offsets|0|+3]|AGCAGCATTGTACAGGGCTATG</td><td>>MI0000108|hsa-mir-103a-2&WithFlank&20|+|3917488|3917577</td><td>54</td><td>75</td><td>22</td><td>AGCAGCATTGTACAGGGCTATG</td><td>BT20</td><td>437</td><td>293</td><td>247</td><td>122</td><td>214</td><td>224.8</td><td>203.7</td><td>182.9</td><td>93.0</td><td>230.2</td></tr>
<tr><td>>MI0000108|hsa-mir-103a-2&WithFlank&20|+|3917488|3917577@54.75.22;MI0000109|hsa-mir-103a-1&WithFlank&5|-|168560890|168560979@54.75.22&[MIMAT0000101&hsa-miR-103a-3p&offsets|0|-1;m-12&20|+|3917541|3917563&offsets|0|-1];[MIMAT0000101_1&hsa-miR-103a-3p&offsets|0|-1;m-13&5|-|168560904|168560926&offsets|0|-1;m-44220&5|-|168560908|168560926&offsets|0|+3]|AGCAGCATTGTACAGGGCTATG</td><td>>MI0000108|hsa-mir-103a-2&WithFlank&20|+|3917488|3917577</td><td>54</td><td>75</td><td>22</td><td>AGCAGCATTGTACAGGGCTATG</td><td>MB231</td><td>353</td><td>294</td><td>84</td><td>773</td><td>410</td><td>86.2</td><td>82.1</td><td>68.9</td><td>85.2</td><td>190.9</td></tr>
<tr><td>>MI0000108|hsa-mir-103a-2&WithFlank&20|+|3917488|3917577@54.75.22;MI0000109|hsa-mir-103a-1&WithFlank&5|-|168560890|168560979@54.75.22&[MIMAT0000101&hsa-miR-103a-3p&offsets|0|-1;m-12&20|+|3917541|3917563&offsets|0|-1];[MIMAT0000101_1&hsa-miR-103a-3p&offsets|0|-1;m-13&5|-|168560904|168560926&offsets|0|-1;m-44220&5|-|168560908|168560926&offsets|0|+3]|AGCAGCATTGTACAGGGCTATG</td><td>>MI0000108|hsa-mir-103a-2&WithFlank&20|+|3917488|3917577</td><td>54</td><td>75</td><td>22</td><td>AGCAGCATTGTACAGGGCTATG</td><td>MB468</td><td>25</td><td>10</td><td>7</td><td>65</td><td>9</td><td>8.4</td><td>2.9</td><td>2.4</td><td>56.0</td><td>2.9</td></tr>
<tr><td>>MI0000108|hsa-mir-103a-2&WithFlank&20|+|3917488|3917577@54.75.22(+1C);MI0000109|hsa-mir-103a-1&WithFlank&5|-|168560890|168560979@54.75.22(+1C)&[MIMAT0000101&hsa-miR-103a-3p&offsets|0|-1(+1C);m-12&20|+|3917541|3917563&offsets|0|-1(+1C)];[MIMAT0000101_1&hsa-miR-103a-3p&offsets|0|-1(+1C);m-13&5|-|168560904|168560926&offsets|0|-1(+1C);m-44220&5|-|168560908|168560926&offsets|0|+3(+1C)]|AGCAGCATTGTACAGGGCTATGC</td><td>>MI0000108|hsa-mir-103a-2&WithFlank&20|+|3917488|3917577</td><td>54</td><td>75</td><td>22(+1C)</td><td>AGCAGCATTGTACAGGGCTATGC</td><td>BT20</td><td>12</td><td>9</td><td>6</td><td>2</td><td>3</td><td>7.6</td><td>8.2</td><td>2.8</td><td>2.4</td><td>3.0</td></tr>
<tr><td>>MI0000108|hsa-mir-103a-2&WithFlank&20|+|3917488|3917577@54.75.22(+1G);MI0000109|hsa-mir-103a-1&WithFlank&5|-|168560890|168560979@54.75.22(+1G)&[MIMAT0000101&hsa-miR-103a-3p&offsets|0|-1(+1G);m-12&20|+|3917541|3917563&offsets|0|-1(+1G)];[MIMAT0000101_1&hsa-miR-103a-3p&offsets|0|-1(+1G);m-13&5|-|168560904|168560926&offsets|0|-1(+1G);m-44220&5|-|168560908|168560926&offsets|0|+3(+1G)]|AGCAGCATTGTACAGGGCTATGG</td><td>>MI0000108|hsa-mir-103a-2&WithFlank&20|+|3917488|3917577</td><td>54</td><td>75</td><td>22(+1G)</td><td>AGCAGCATTGTACAGGGCTATGG</td><td>BT20</td><td>135</td><td>36</td><td>30</td><td>8</td><td>8</td><td>117.3</td><td>19.2</td><td>4.1</td><td>9.1</td><td>3.3</td></tr>
<tr><td>>MI0000108|hsa-mir-103a-2&WithFlank&20|+|3917488|3917577@54.75.22(+1G);MI0000109|hsa-mir-103a-1&WithFlank&5|-|168560890|168560979@54.75.22(+1G)&[MIMAT0000101&hsa-miR-103a-3p&offsets|0|-1(+1G);m-12&20|+|3917541|3917563&offsets|0|-1(+1G)];[MIMAT0000101_1&hsa-miR-103a-3p&offsets|0|-1(+1G);m-13&5|-|168560904|168560926&offsets|0|-1(+1G);m-44220&5|-|168560908|168560926&offsets|0|+3(+1G)]|AGCAGCATTGTACAGGGCTATGG</td><td>>MI0000108|hsa-mir-103a-2&WithFlank&20|+|3917488|3917577</td><td>54</td><td>75</td><td>22(+1G)</td><td>AGCAGCATTGTACAGGGCTATGG</td><td>MB231</td><td>61</td><td>22</td><td>7</td><td>48</td><td>25</td><td>40.0</td><td>4.7</td><td>5.7</td><td>9.3</td><td>20.5</td></tr>
<tr><td>>MI0000108|hsa-mir-103a-2&WithFlank&20|+|3917488|3917577@54.75.22(+1U);MI0000109|hsa-mir-103a-1&WithFlank&5|-|168560890|168560979@54.75.22(+1U)&[MIMAT0000101&hsa-miR-103a-3p&offsets|0|-1(+1U);m-12&20|+|3917541|3917563&offsets|0|-1(+1U)];[MIMAT0000101_1&hsa-miR-103a-3p&offsets|0|-1(+1U);m-13&5|-|168560904|168560926&offsets|0|-1(+1U);m-44220&5|-|168560908|168560926&offsets|0|+3(+1U)]|AGCAGCATTGTACAGGGCTATGT</td><td>>MI0000108|hsa-mir-103a-2&WithFlank&20|+|3917488|3917577</td><td>54</td><td>75</td><td>22(+1U)</td><td>AGCAGCATTGTACAGGGCTATGT</td><td>BT20</td><td>118</td><td>64</td><td>52</td><td>16</td><td>23</td><td>83.9</td><td>52.1</td><td>27.1</td><td>14.1</td><td>18.3</td></tr>
<tr><td>>MI0000108|hsa-mir-103a-2&WithFlank&20|+|3917488|3917577@54.75.22(+1U);MI0000109|hsa-mir-103a-1&WithFlank&5|-|168560890|168560979@54.75.22(+1U)&[MIMAT0000101&hsa-miR-103a-3p&offsets|0|-1(+1U);m-12&20|+|3917541|3917563&offsets|0|-1(+1U)];[MIMAT0000101_1&hsa-miR-103a-3p&offsets|0|-1(+1U);m-13&5|-|168560904|168560926&offsets|0|-1(+1U);m-44220&5|-|168560908|168560926&offsets|0|+3(+1U)]|AGCAGCATTGTACAGGGCTATGT</td><td>>MI0000108|hsa-mir-103a-2&WithFlank&20|+|3917488|3917577</td><td>54</td><td>75</td><td>22(+1U)</td><td>AGCAGCATTGTACAGGGCTATGT</td><td>MB231</td><td>69</td><td>58</td><td>21</td><td>208</td><td>107</td><td>10.5</td><td>20.4</td><td>18.1</td><td>27.2</td><td>94.9</td></tr>
<tr><td>>MI0000108|hsa-mir-103a-2&WithFlank&20|+|3917488|3917577@54.75.22(+2U);MI0000109|hsa-mir-103a-1&WithFlank&5|-|168560890|168560979@54.75.22(+2U)&[MIMAT0000101&hsa-miR-103a-3p&offsets|0|-1(+2U);m-12&20|+|3917541|3917563&offsets|0|-1(+2U)];[MIMAT0000101_1&hsa-miR-103a-3p&offsets|0|-1(+2U);m-13&5|-|168560904|168560926&offsets|0|-1(+2U);m-44220&5|-|168560908|168560926&offsets|0|+3(+2U)]|AGCAGCATTGTACAGGGCTATGTT</td><td>>MI0000108|hsa-mir-103a-2&WithFlank&20|+|3917488|3917577</td><td>54</td><td>75</td><td>22(+2U)</td><td>AGCAGCATTGTACAGGGCTATGTT</td><td>MB231</td><td>13</td><td>8</td><td>4</td><td>10</td><td>14</td><td>2.2</td><td>3.2</td><td>2.9</td><td>4.4</td><td>6.9</td></tr>
<tr><td>>MI0000108|hsa-mir-103a-2&WithFlank&20|+|3917488|3917577@54.76.23;MI0000109|hsa-mir-103a-1&WithFlank&5|-|168560890|168560979@54.76.23&[MIMAT0000101&hsa-miR-103a-3p&offsets|0|0;m-12&20|+|3917541|3917563&offsets|0|0];[MIMAT0000101_1&hsa-miR-103a-3p&offsets|0|0;m-13&5|-|168560904|168560926&offsets|0|0;m-44220&5|-|168560908|168560926&offsets|0|+4]|AGCAGCATTGTACAGGGCTATGA</td><td>>MI0000108|hsa-mir-103a-2&WithFlank&20|+|3917488|3917577</td><td>54</td><td>76</td><td>23</td><td>AGCAGCATTGTACAGGGCTATGA</td><td>BT20</td><td>1878</td><td>1310</td><td>1249</td><td>365</td><td>625</td><td>702.5</td><td>608.7</td><td>566.2</td><td>257.0</td><td>339.0</td></tr>
<tr><td>>MI0000108|hsa-mir-103a-2&WithFlank&20|+|3917488|3917577@54.76.23;MI0000109|hsa-mir-103a-1&WithFlank&5|-|168560890|168560979@54.76.23&[MIMAT0000101&hsa-miR-103a-3p&offsets|0|0;m-12&20|+|3917541|3917563&offsets|0|0];[MIMAT0000101_1&hsa-miR-103a-3p&offsets|0|0;m-13&5|-|168560904|168560926&offsets|0|0;m-44220&5|-|168560908|168560926&offsets|0|+4]|AGCAGCATTGTACAGGGCTATGA</td><td>>MI0000108|hsa-mir-103a-2&WithFlank&20|+|3917488|3917577</td><td>54</td><td>76</td><td>23</td><td>AGCAGCATTGTACAGGGCTATGA</td><td>MB231</td><td>1390</td><td>1278</td><td>377</td><td>3242</td><td>1892</td><td>284.1</td><td>324.6</td><td>297.8</td><td>1627.1</td><td>1229.8</td></tr>
<tr><td>>MI0000108|hsa-mir-103a-2&WithFlank&20|+|3917488|3917577@54.76.23;MI0000109|hsa-mir-103a-1&WithFlank&5|-|168560890|168560979@54.76.23&[MIMAT0000101&hsa-miR-103a-3p&offsets|0|0;m-12&20|+|3917541|3917563&offsets|0|0];[MIMAT0000101_1&hsa-miR-103a-3p&offsets|0|0;m-13&5|-|168560904|168560926&offsets|0|0;m-44220&5|-|168560908|168560926&offsets|0|+4]|AGCAGCATTGTACAGGGCTATGA</td><td>>MI0000108|hsa-mir-103a-2&WithFlank&20|+|3917488|3917577</td><td>54</td><td>76</td><td>23</td><td>AGCAGCATTGTACAGGGCTATGA</td><td>MB468</td><td>81</td><td>57</td><td>27</td><td>396</td><td>33</td><td>26.3</td><td>22.5</td><td>12.0</td><td>339.4</td><td>15.4</td></tr>
<tr><td>>MI0000108|hsa-mir-103a-2&WithFlank&20|+|3917488|3917577@54.76.23(+1C);MI0000109|hsa-mir-103a-1&WithFlank&5|-|168560890|168560979@54.76.23(+1C)&[MIMAT0000101&hsa-miR-103a-3p&offsets|0|0(+1C);m-12&20|+|3917541|3917563&offsets|0|0(+1C)];[MIMAT0000101_1&hsa-miR-103a-3p&offsets|0|0(+1C);m-13&5|-|168560904|168560926&offsets|0|0(+1C);m-44220&5|-|168560908|168560926&offsets|0|+4(+1C)]|AGCAGCATTGTACAGGGCTATGAC</td><td>>MI0000108|hsa-mir-103a-2&WithFlank&20|+|3917488|3917577</td><td>54</td><td>76</td><td>23(+1C)</td><td>AGCAGCATTGTACAGGGCTATGAC</td><td>BT20</td><td>21</td><td>15</td><td>14</td><td>1</td><td>7</td><td>8.9</td><td>8.4</td><td>4.5</td><td>0.5</td><td>3.6</td></tr>
<tr><td>>MI0000108|hsa-mir-103a-2&WithFlank&20|+|3917488|3917577@54.76.23(+1C);MI0000109|hsa-mir-103a-1&WithFlank&5|-|168560890|168560979@54.76.23(+1C)&[MIMAT0000101&hsa-miR-103a-3p&offsets|0|0(+1C);m-12&20|+|3917541|3917563&offsets|0|0(+1C)];[MIMAT0000101_1&hsa-miR-103a-3p&offsets|0|0(+1C);m-13&5|-|168560904|168560926&offsets|0|0(+1C);m-44220&5|-|168560908|168560926&offsets|0|+4(+1C)]|AGCAGCATTGTACAGGGCTATGAC</td><td>>MI0000108|hsa-mir-103a-2&WithFlank&20|+|3917488|3917577</td><td>54</td><td>76</td><td>23(+1C)</td><td>AGCAGCATTGTACAGGGCTATGAC</td><td>MB231</td><td>19</td><td>14</td><td>7</td><td>11</td><td>36</td><td>2.4</td><td>2.9</td><td>5.1</td><td>3.9</td><td>32.1</td></tr>
<tr><td>>MI0000108|hsa-mir-103a-2&WithFlank&20|+|3917488|3917577@54.76.23(+1G);MI0000109|hsa-mir-103a-1&WithFlank&5|-|168560890|168560979@54.76.23(+1G)&[MIMAT0000101&hsa-miR-103a-3p&offsets|0|0(+1G);m-12&20|+|3917541|3917563&offsets|0|0(+1G)];[MIMAT0000101_1&hsa-miR-103a-3p&offsets|0|0(+1G);m-13&5|-|168560904|168560926&offsets|0|0(+1G);m-44220&5|-|168560908|168560926&offsets|0|+4(+1G)]|AGCAGCATTGTACAGGGCTATGAG</td><td>>MI0000108|hsa-mir-103a-2&WithFlank&20|+|3917488|3917577</td><td>54</td><td>76</td><td>23(+1G)</td><td>AGCAGCATTGTACAGGGCTATGAG</td><td>BT20</td><td>16</td><td>6</td><td>9</td><td>1</td><td>2</td><td>11.9</td><td>1.5</td><td>1.5</td><td>0.7</td><td>0.9</td></tr>
<tr><td>>MI0000108|hsa-mir-103a-2&WithFlank&20|+|3917488|3917577@54.76.23(+1U);MI0000109|hsa-mir-103a-1&WithFlank&5|-|168560890|168560979@54.76.23(+1U)&[MIMAT0000101&hsa-miR-103a-3p&offsets|0|0(+1U);m-12&20|+|3917541|3917563&offsets|0|0(+1U)];[MIMAT0000101_1&hsa-miR-103a-3p&offsets|0|0(+1U);m-13&5|-|168560904|168560926&offsets|0|0(+1U);m-44220&5|-|168560908|168560926&offsets|0|+4(+1U)]|AGCAGCATTGTACAGGGCTATGAT</td><td>>MI0000108|hsa-mir-103a-2&WithFlank&20|+|3917488|3917577</td><td>54</td><td>76</td><td>23(+1U)</td><td>AGCAGCATTGTACAGGGCTATGAT</td><td>BT20</td><td>158</td><td>102</td><td>101</td><td>7</td><td>49</td><td>78.3</td><td>55.7</td><td>22.8</td><td>2.2</td><td>32.3</td></tr>
<tr><td>>MI0000108|hsa-mir-103a-2&WithFlank&20|+|3917488|3917577@54.76.23(+1U);MI0000109|hsa-mir-103a-1&WithFlank&5|-|168560890|168560979@54.76.23(+1U)&[MIMAT0000101&hsa-miR-103a-3p&offsets|0|0(+1U);m-12&20|+|3917541|3917563&offsets|0|0(+1U)];[MIMAT0000101_1&hsa-miR-103a-3p&offsets|0|0(+1U);m-13&5|-|168560904|168560926&offsets|0|0(+1U);m-44220&5|-|168560908|168560926&offsets|0|+4(+1U)]|AGCAGCATTGTACAGGGCTATGAT</td><td>>MI0000108|hsa-mir-103a-2&WithFlank&20|+|3917488|3917577</td><td>54</td><td>76</td><td>23(+1U)</td><td>AGCAGCATTGTACAGGGCTATGAT</td><td>MB231</td><td>185</td><td>149</td><td>65</td><td>78</td><td>283</td><td>27.7</td><td>43.2</td><td>50.2</td><td>39.3</td><td>215.8</td></tr>
</table>

<table>
<tr><td>>MI0000108|hsa-mir-103a-2&WithFlank&20|+|3917488|3917577@54.76.23(+2U);MI0000109|hsa-mir-103a-1&WithFlank&5|-|168560890|168560979@54.76.23(+2U)&[MIMAT0000101&hsa-miR-103a-3p&offsets|0|0(+2U);m-12&20|+|3917541|3917563&offsets|0|0(+2U)];[MIMAT0000101_1&hsa-miR-103a-3p&offsets|0|0(+2U);m-13&5|-|168560904|168560926&offsets|0|0(+2U);m-44220&5|-|168560908|168560926&offsets|0|+4(+2U)]|AGCAGCATTGTACAGGGCTATGATT</td><td>>MI0000108|hsa-mir-103a-2&WithFlank&20|+|3917488|3917577</td><td>54</td><td>76</td><td>23(+2U)</td><td>AGCAGCATTGTACAGGGCTATGATT</td><td>BT20</td><td>21</td><td>11</td><td>21</td><td>3</td><td>7</td><td>10.5</td><td>6.8</td><td>8.5</td><td>0.6</td><td>4.4</td></tr>
<tr><td>>MI0000108|hsa-mir-103a-2&WithFlank&20|+|3917488|3917577@54.76.23(+2U);MI0000109|hsa-mir-103a-1&WithFlank&5|-|168560890|168560979@54.76.23(+2U)&[MIMAT0000101&hsa-miR-103a-3p&offsets|0|0(+2U);m-12&20|+|3917541|3917563&offsets|0|0(+2U)];[MIMAT0000101_1&hsa-miR-103a-3p&offsets|0|0(+2U);m-13&5|-|168560904|168560926&offsets|0|0(+2U);m-44220&5|-|168560908|168560926&offsets|0|+4(+2U)]|AGCAGCATTGTACAGGGCTATGATT</td><td>>MI0000108|hsa-mir-103a-2&WithFlank&20|+|3917488|3917577</td><td>54</td><td>76</td><td>23(+2U)</td><td>AGCAGCATTGTACAGGGCTATGATT</td><td>MB231</td><td>70</td><td>43</td><td>34</td><td>34</td><td>61</td><td>12.8</td><td>11.5</td><td>25.1</td><td>10.3</td><td>40.7</td></tr>
<tr><td>>MI0000108|hsa-mir-103a-2&WithFlank&20|+|3917488|3917577@54.77.24;MI0000109|hsa-mir-103a-1&WithFlank&5|-|168560890|168560979@54.77.24&[MIMAT0000101&hsa-miR-103a-3p&offsets|0|+1;m-12&20|+|3917541|3917563&offsets|0|+1];[MIMAT0000101_1&hsa-miR-103a-3p&offsets|0|+1;m-13&5|-|168560904|168560926&offsets|0|+1;m-44220&5|-|168560908|168560926&offsets|0|+5]|AGCAGCATTGTACAGGGCTATGAA</td><td>>MI0000108|hsa-mir-103a-2&WithFlank&20|+|3917488|3917577</td><td>54</td><td>77</td><td>24</td><td>AGCAGCATTGTACAGGGCTATGAA</td><td>BT20</td><td>81</td><td>54</td><td>45</td><td>4</td><td>15</td><td>58.8</td><td>27.0</td><td>8.5</td><td>2.5</td><td>3.9</td></tr>
<tr><td>>MI0000108|hsa-mir-103a-2&WithFlank&20|+|3917488|3917577@54.77.24;MI0000109|hsa-mir-103a-1&WithFlank&5|-|168560890|168560979@54.77.24&[MIMAT0000101&hsa-miR-103a-3p&offsets|0|+1;m-12&20|+|3917541|3917563&offsets|0|+1];[MIMAT0000101_1&hsa-miR-103a-3p&offsets|0|+1;m-13&5|-|168560904|168560926&offsets|0|+1;m-44220&5|-|168560908|168560926&offsets|0|+5]|AGCAGCATTGTACAGGGCTATGAA</td><td>>MI0000108|hsa-mir-103a-2&WithFlank&20|+|3917488|3917577</td><td>54</td><td>77</td><td>24</td><td>AGCAGCATTGTACAGGGCTATGAA</td><td>MB231</td><td>42</td><td>44</td><td>14</td><td>26</td><td>41</td><td>5.7</td><td>15.5</td><td>11.7</td><td>16.0</td><td>27.8</td></tr>
<tr><td>>MI0000108|hsa-mir-103a-2&WithFlank&20|+|3917488|3917577@54.77.24(+1U);MI0000109|hsa-mir-103a-1&WithFlank&5|-|168560890|168560979@54.77.24(+1U)&[MIMAT0000101&hsa-miR-103a-3p&offsets|0|+1(+1U);m-12&20|+|3917541|3917563&offsets|0|+1(+1U)];[MIMAT0000101_1&hsa-miR-103a-3p&offsets|0|+1(+1U);m-13&5|-|168560904|168560926&offsets|0|+1(+1U);m-44220&5|-|168560908|168560926&offsets|0|+5(+1U)]|AGCAGCATTGTACAGGGCTATGAAT</td><td>>MI0000108|hsa-mir-103a-2&WithFlank&20|+|3917488|3917577</td><td>54</td><td>77</td><td>24(+1U)</td><td>AGCAGCATTGTACAGGGCTATGAAT</td><td>BT20</td><td>17</td><td>10</td><td>10</td><td>0</td><td>2</td><td>12.9</td><td>5.6</td><td>3.0</td><td>0.8</td><td>3.9</td></tr>
<tr><td>>MI0000108|hsa-mir-103a-2&WithFlank&20|+|3917488|3917577@54.77.24(+1U);MI0000109|hsa-mir-103a-1&WithFlank&5|-|168560890|168560979@54.77.24(+1U)&[MIMAT0000101&hsa-miR-103a-3p&offsets|0|+1(+1U);m-12&20|+|3917541|3917563&offsets|0|+1(+1U)];[MIMAT0000101_1&hsa-miR-103a-3p&offsets|0|+1(+1U);m-13&5|-|168560904|168560926&offsets|0|+1(+1U);m-44220&5|-|168560908|168560926&offsets|0|+5(+1U)]|AGCAGCATTGTACAGGGCTATGAAT</td><td>>MI0000108|hsa-mir-103a-2&WithFlank&20|+|3917488|3917577</td><td>54</td><td>77</td><td>24(+1U)</td><td>AGCAGCATTGTACAGGGCTATGAAT</td><td>MB231</td><td>29</td><td>22</td><td>8</td><td>9</td><td>37</td><td>6.6</td><td>8.4</td><td>7.1</td><td>6.7</td><td>28.6</td></tr>
<tr><td>>MI0000108|hsa-mir-103a-2&WithFlank&20|+|3917488|3917577@54.78.25&[MIMAT0000101&hsa-miR-103a-3p&offsets|0|+2;m-12&20|+|3917541|3917563&offsets|0|+2]|AGCAGCATTGTACAGGGCTATGAAA</td><td>>MI0000108|hsa-mir-103a-2&WithFlank&20|+|3917488|3917577</td><td>54</td><td>78</td><td>25</td><td>AGCAGCATTGTACAGGGCTATGAAA</td><td>BT20</td><td>15</td><td>11</td><td>9</td><td>0</td><td>1</td><td>13.4</td><td>6.7</td><td>4.1</td><td>0.6</td><td>2.0</td></tr>
<tr><td>>MI0000108|hsa-mir-103a-2&WithFlank&20|+|3917488|3917577@56.76.21;MI0000109|hsa-mir-103a-1&WithFlank&5|-|168560890|168560979@56.76.21&[MIMAT0000101&hsa-miR-103a-3p&offsets|+2|0;m-12&20|+|3917541|3917563&offsets|+2|0];[MIMAT0000101_1&hsa-miR-103a-3p&offsets|+2|0;m-13&5|-|168560904|168560926&offsets|+2|0;m-44220&5|-|168560908|168560926&offsets|+2|+4]|CAGCATTGTACAGGGCTATGA</td><td>>MI0000108|hsa-mir-103a-2&WithFlank&20|+|3917488|3917577</td><td>56</td><td>76</td><td>21</td><td>CAGCATTGTACAGGGCTATGA</td><td>BT20</td><td>18</td><td>9</td><td>5</td><td>7</td><td>4</td><td>11.2</td><td>4.4</td><td>3.8</td><td>5.1</td><td>2.5</td></tr>
<tr><td>>MI0000108|hsa-mir-103a-2&WithFlank&20|+|3917488|3917577@56.76.21;MI0000109|hsa-mir-103a-1&WithFlank&5|-|168560890|168560979@56.76.21&[MIMAT0000101&hsa-miR-103a-3p&offsets|+2|0;m-12&20|+|3917541|3917563&offsets|+2|0];[MIMAT0000101_1&hsa-miR-103a-3p&offsets|+2|0;m-13&5|-|168560904|168560926&offsets|+2|0;m-44220&5|-|168560908|168560926&offsets|+2|+4]|CAGCATTGTACAGGGCTATGA</td><td>>MI0000108|hsa-mir-103a-2&WithFlank&20|+|3917488|3917577</td><td>56</td><td>76</td><td>21</td><td>CAGCATTGTACAGGGCTATGA</td><td>MB231</td><td>16</td><td>12</td><td>3</td><td>29</td><td>18</td><td>1.4</td><td>3.1</td><td>1.8</td><td>17.5</td><td>11.7</td></tr>
<tr><td>>MI0000111|hsa-mir-105-1&WithFlank&X|-|152392213|152392305@19.40.22;MI0000112|hsa-mir-105-2&WithFlank&X|-|152394406|152394498@19.40.22&[MIMAT0000102&hsa-miR-105-5p&offsets|0|-1;m-735&X|-|152392266|152392287&offsets|0|0];[MIMAT0000102_1&hsa-miR-105-5p&offsets|0|-1;m-735&X|-|152394459|152394480&offsets|0|0]|TCAAATGCTCAGACTCCTGTGG</td><td>>MI0000111|hsa-mir-105-1&WithFlank&X|-|152392213|152392305</td><td>19</td><td>40</td><td>22</td><td>TCAAATGCTCAGACTCCTGTGG</td><td>BT20</td><td>19</td><td>16</td><td>28</td><td>28</td><td>15</td><td>21.6</td><td>4.9</td><td>17.9</td><td>23.8</td><td>2.5</td></tr>
<tr><td>>MI0000111|hsa-mir-105-1&WithFlank&X|-|152392213|152392305@19.41.23;MI0000112|hsa-mir-105-2&WithFlank&X|-|152394406|152394498@19.41.23&[MIMAT0000102&hsa-miR-105-5p&offsets|0|0;m-735&X|-|152392266|152392287&offsets|0|+1];[MIMAT0000102_1&hsa-miR-105-5p&offsets|0|0;m-735&X|-|152394459|152394480&offsets|0|+1]|TCAAATGCTCAGACTCCTGTGGT</td><td>>MI0000111|hsa-mir-105-1&WithFlank&X|-|152392213|152392305</td><td>19</td><td>41</td><td>23</td><td>TCAAATGCTCAGACTCCTGTGGT</td><td>BT20</td><td>12</td><td>10</td><td>12</td><td>8</td><td>5</td><td>6.8</td><td>7.0</td><td>5.7</td><td>6.7</td><td>3.5</td></tr>
<tr><td>>MI0000114|hsa-mir-107&WithFlank&10|-|89592741|89592833@56.76.21;MI0000108|hsa-mir-103a-2&WithFlank&20|+|3917488|3917577@54.74.21;MI0000109|hsa-mir-103a-1&WithFlank&5|-|168560890|168560979@54.74.21&[MIMAT0000104&hsa-miR-107&offsets|0|-2;m-78&10|-|89592758|89592778&offsets|0|0];[MIMAT0000101&hsa-miR-103a-3p&offsets|0|-2;m-12&20|+|3917541|3917563&offsets|0|-2];[MIMAT0000101_1&hsa-miR-103a-3p&offsets|0|-2;m-13&5|-|168560904|168560926&offsets|0|-2;m-44220&5|-|168560908|168560926&offsets|0|+2]|AGCAGCATTGTACAGGGCTAT</td><td>>MI0000114|hsa-mir-107&WithFlank&10|-|89592741|89592833</td><td>56</td><td>76</td><td>21</td><td>AGCAGCATTGTACAGGGCTAT</td><td>BT20</td><td>51</td><td>49</td><td>27</td><td>82</td><td>11</td><td>32.1</td><td>55.1</td><td>20.1</td><td>68.5</td><td>10.1</td></tr>
<tr><td>>MI0000114|hsa-mir-107&WithFlank&10|-|89592741|89592833@56.76.21;MI0000108|hsa-mir-103a-2&WithFlank&20|+|3917488|3917577@54.74.21;MI0000109|hsa-mir-103a-1&WithFlank&5|-|168560890|168560979@54.74.21&[MIMAT0000104&hsa-miR-107&offsets|0|-2;m-78&10|-|89592758|89592778&offsets|0|0];[MIMAT0000101&hsa-miR-103a-3p&offsets|0|-2;m-12&20|+|3917541|3917563&offsets|0|-2];[MIMAT0000101_1&hsa-miR-103a-3p&offsets|0|-2;m-13&5|-|168560904|168560926&offsets|0|-2;m-44220&5|-|168560908|168560926&offsets|0|+2]|AGCAGCATTGTACAGGGCTAT</td><td>>MI0000114|hsa-mir-107&WithFlank&10|-|89592741|89592833</td><td>56</td><td>76</td><td>21</td><td>AGCAGCATTGTACAGGGCTAT</td><td>MB231</td><td>40</td><td>39</td><td>13</td><td>279</td><td>71</td><td>5.7</td><td>19.7</td><td>11.4</td><td>75.7</td><td>68.3</td></tr>
<tr><td>>MI0000114|hsa-mir-107&WithFlank&10|-|89592741|89592833@56.76.21(+1A);MI0000108|hsa-mir-103a-2&WithFlank&20|+|3917488|3917577@54.74.21(+1A);MI0000109|hsa-mir-103a-1&WithFlank&5|-|168560890|168560979@54.74.21(+1A)&[MIMAT0000104&hsa-miR-107&offsets|0|-2(+1A);m-78&10|-|89592758|89592778&offsets|0|0(+1A)];[MIMAT0000101&hsa-miR-103a-3p&offsets|0|-2(+1A);m-12&20|+|3917541|3917563&offsets|0|-2(+1A)];[MIMAT0000101_1&hsa-miR-103a-3p&offsets|0|-2(+1A);m-13&5|-|168560904|168560926&offsets|0|-2(+1A);m-44220&5|-|168560908|168560926&offsets|0|+2(+1A)]|AGCAGCATTGTACAGGGCTATA</td><td>>MI0000114|hsa-mir-107&WithFlank&10|-|89592741|89592833</td><td>56</td><td>76</td><td>21(+1A)</td><td>AGCAGCATTGTACAGGGCTATA</td><td>BT20</td><td>74</td><td>37</td><td>36</td><td>15</td><td>19</td><td>49.7</td><td>32.8</td><td>16.7</td><td>12.9</td><td>19.1</td></tr>
<tr><td>>MI0000114|hsa-mir-107&WithFlank&10|-|89592741|89592833@56.76.21(+1A);MI0000108|hsa-mir-103a-2&WithFlank&20|+|3917488|3917577@54.74.21(+1A);MI0000109|hsa-mir-103a-1&WithFlank&5|-|168560890|168560979@54.74.21(+1A)&[MIMAT0000104&hsa-miR-107&offsets|0|-2(+1A);m-78&10|-|89592758|89592778&offsets|0|0(+1A)];[MIMAT0000101&hsa-miR-103a-3p&offsets|0|-2(+1A);m-12&20|+|3917541|3917563&offsets|0|-2(+1A)];[MIMAT0000101_1&hsa-miR-103a-3p&offsets|0|-2(+1A);m-13&5|-|168560904|168560926&offsets|0|-2(+1A);m-44220&5|-|168560908|168560926&offsets|0|+2(+1A)]|AGCAGCATTGTACAGGGCTATA</td><td>>MI0000114|hsa-mir-107&WithFlank&10|-|89592741|89592833</td><td>56</td><td>76</td><td>21(+1A)</td><td>AGCAGCATTGTACAGGGCTATA</td><td>MB231</td><td>32</td><td>21</td><td>9</td><td>108</td><td>46</td><td>7.1</td><td>4.9</td><td>9.0</td><td>32.0</td><td>33.0</td></tr>
<tr><td>>MI0000114|hsa-mir-107&WithFlank&10|-|89592741|89592833@56.76.21(+1U);MI0000108|hsa-mir-103a-2&WithFlank&20|+|3917488|3917577@54.74.21(+1U);MI0000109|hsa-mir-103a-1&WithFlank&5|-|168560890|168560979@54.74.21(+1U)&[MIMAT0000104&hsa-miR-107&offsets|0|-2(+1U);m-78&10|-|89592758|89592778&offsets|0|0(+1U)];[MIMAT0000101&hsa-miR-103a-3p&offsets|0|-2(+1U);m-12&20|+|3917541|3917563&offsets|0|-2(+1U)];[MIMAT0000101_1&hsa-miR-103a-3p&offsets|0|-2(+1U);m-13&5|-|168560904|168560926&offsets|0|-2(+1U);m-44220&5|-|168560908|168560926&offsets|0|+2(+1U)]|AGCAGCATTGTACAGGGCTATT</td><td>>MI0000114|hsa-mir-107&WithFlank&10|-|89592741|89592833</td><td>56</td><td>76</td><td>21(+1U)</td><td>AGCAGCATTGTACAGGGCTATT</td><td>BT20</td><td>16</td><td>12</td><td>8</td><td>3</td><td>3</td><td>11.4</td><td>14.3</td><td>4.6</td><td>3.0</td><td>3.8</td></tr>
</table>

<table>
<tr><td>>MI0000114|hsa-mir-107&WithFlank&10|-|89592741|89592833@56.76.21(+1U);MI0000108|hsa-mir-103a-2&WithFlank&20|+|3917488|3917577@54.74.21(+1U);MI0000109|hsa-mir-103a-1&WithFlank&5|-|168560890|168560979@54.74.21(+1U)&[MIMAT0000104&hsa-miR-107&offsets|0|-2(+1U);m-78&10|-|89592758|89592778&offsets|0|0(+1U)];[MIMAT0000101&hsa-miR-103a-3p&offsets|0|-2(+1U);m-12&20|+|3917541|3917563&offsets|0|-2(+1U)];[MIMAT0000101_1&hsa-miR-103a-3p&offsets|0|-2(+1U);m-13&5|-|168560904|168560926&offsets|0|-2(+1U);m-44220&5|-|168560908|168560926&offsets|0|+2(+1U)]|AGCAGCATTGTACAGGGCTATT</td><td>>MI0000114|hsa-mir-107&WithFlank&10|-|89592741|89592833</td><td>56</td><td>76</td><td>21(+1U)</td><td>AGCAGCATTGTACAGGGCTATT</td><td>MB231</td><td>12</td><td>10</td><td>4</td><td>47</td><td>19</td><td>1.4</td><td>3.9</td><td>3.4</td><td>15.0</td><td>16.8</td></tr>
<tr><td>>MI0000114|hsa-mir-107&WithFlank&10|-|89592741|89592833@56.77.22&[MIMAT0000104&hsa-miR-107&offsets|0|-1;m-78&10|-|89592758|89592778&offsets|0|+1]|AGCAGCATTGTACAGGGCTATC</td><td>>MI0000114|hsa-mir-107&WithFlank&10|-|89592741|89592833</td><td>56</td><td>77</td><td>22</td><td>AGCAGCATTGTACAGGGCTATC</td><td>BT20</td><td>27</td><td>25</td><td>23</td><td>8</td><td>8</td><td>13.7</td><td>26.8</td><td>17.4</td><td>6.7</td><td>7.9</td></tr>
<tr><td>>MI0000114|hsa-mir-107&WithFlank&10|-|89592741|89592833@56.77.22&[MIMAT0000104&hsa-miR-107&offsets|0|-1;m-78&10|-|89592758|89592778&offsets|0|+1]|AGCAGCATTGTACAGGGCTATC</td><td>>MI0000114|hsa-mir-107&WithFlank&10|-|89592741|89592833</td><td>56</td><td>77</td><td>22</td><td>AGCAGCATTGTACAGGGCTATC</td><td>MB231</td><td>37</td><td>30</td><td>16</td><td>129</td><td>50</td><td>4.9</td><td>11.8</td><td>15.6</td><td>23.0</td><td>39.1</td></tr>
<tr><td>>MI0000114|hsa-mir-107&WithFlank&10|-|89592741|89592833@56.78.23&[MIMAT0000104&hsa-miR-107&offsets|0|0;m-78&10|-|89592758|89592778&offsets|0|+2]|AGCAGCATTGTACAGGGCTATCA</td><td>>MI0000114|hsa-mir-107&WithFlank&10|-|89592741|89592833</td><td>56</td><td>78</td><td>23</td><td>AGCAGCATTGTACAGGGCTATCA</td><td>BT20</td><td>17</td><td>9</td><td>25</td><td>1</td><td>2</td><td>10.3</td><td>7.0</td><td>16.5</td><td>0.9</td><td>2.1</td></tr>
<tr><td>>MI0000114|hsa-mir-107&WithFlank&10|-|89592741|89592833@56.78.23&[MIMAT0000104&hsa-miR-107&offsets|0|0;m-78&10|-|89592758|89592778&offsets|0|+2]|AGCAGCATTGTACAGGGCTATCA</td><td>>MI0000114|hsa-mir-107&WithFlank&10|-|89592741|89592833</td><td>56</td><td>78</td><td>23</td><td>AGCAGCATTGTACAGGGCTATCA</td><td>MB231</td><td>14</td><td>8</td><td>11</td><td>13</td><td>11</td><td>2.4</td><td>2.1</td><td>10.4</td><td>3.1</td><td>8.0</td></tr>
<tr><td>>MI0000115|hsa-mir-16-2&WithFlank&3|+|160404739|160404831@58.79.22&[MIMAT0004518&hsa-miR-16-2-3p&offsets|-1|-1;m-252&3|+|160404796|160404816&offsets|0|+1]|ACCAATATTACTGTGCTGCTTT</td><td>>MI0000115|hsa-mir-16-2&WithFlank&3|+|160404739|160404831</td><td>58</td><td>79</td><td>22</td><td>ACCAATATTACTGTGCTGCTTT</td><td>BT20</td><td>58</td><td>42</td><td>70</td><td>27</td><td>28</td><td>56.3</td><td>40.1</td><td>73.8</td><td>12.7</td><td>22.0</td></tr>
<tr><td>>MI0000115|hsa-mir-16-2&WithFlank&3|+|160404739|160404831@58.79.22&[MIMAT0004518&hsa-miR-16-2-3p&offsets|-1|-1;m-252&3|+|160404796|160404816&offsets|0|+1]|ACCAATATTACTGTGCTGCTTT</td><td>>MI0000115|hsa-mir-16-2&WithFlank&3|+|160404739|160404831</td><td>58</td><td>79</td><td>22</td><td>ACCAATATTACTGTGCTGCTTT</td><td>MB231</td><td>38</td><td>47</td><td>24</td><td>97</td><td>74</td><td>8.0</td><td>23.7</td><td>17.2</td><td>63.5</td><td>34.9</td></tr>
<tr><td>>MI0000115|hsa-mir-16-2&WithFlank&3|+|160404739|160404831@58.80.23&[MIMAT0004518&hsa-miR-16-2-3p&offsets|-1|0;m-252&3|+|160404796|160404816&offsets|0|+2]|ACCAATATTACTGTGCTGCTTTA</td><td>>MI0000115|hsa-mir-16-2&WithFlank&3|+|160404739|160404831</td><td>58</td><td>80</td><td>23</td><td>ACCAATATTACTGTGCTGCTTTA</td><td>BT20</td><td>12</td><td>6</td><td>17</td><td>1</td><td>4</td><td>15.3</td><td>6.1</td><td>20.3</td><td>1.3</td><td>4.2</td></tr>
<tr><td>>MI0000115|hsa-mir-16-2&WithFlank&3|+|160404739|160404831@59.80.22&[MIMAT0004518&hsa-miR-16-2-3p&offsets|0|0;m-252&3|+|160404796|160404816&offsets|+1|+2]|CCAATATTACTGTGCTGCTTTA</td><td>>MI0000115|hsa-mir-16-2&WithFlank&3|+|160404739|160404831</td><td>59</td><td>80</td><td>22</td><td>CCAATATTACTGTGCTGCTTTA</td><td>BT20</td><td>15</td><td>6</td><td>22</td><td>5</td><td>6</td><td>18.6</td><td>5.6</td><td>24.4</td><td>2.3</td><td>3.8</td></tr>
<tr><td>>MI0000234|hsa-mir-192&WithFlank&11|-|64891131|64891252@30.49.20(+1A)&[MIMAT0000222&hsa-miR-192-5p&offsets|0|-1(+1A);m-59&11|-|64891203|64891223&offsets|0|-1(+1A)]|CTGACCTATGAATTGACAGCA</td><td>>MI0000234|hsa-mir-192&WithFlank&11|-|64891131|64891252</td><td>30</td><td>49</td><td>20(+1A)</td><td>CTGACCTATGAATTGACAGCA</td><td>BT20</td><td>19</td><td>7</td><td>8</td><td>12</td><td>3</td><td>11.4</td><td>5.1</td><td>3.0</td><td>11.6</td><td>1.1</td></tr>
<tr><td>>MI0000234|hsa-mir-192&WithFlank&11|-|64891131|64891252@30.49.20(+1U)&[MIMAT0000222&hsa-miR-192-5p&offsets|0|-1(+1U);m-59&11|-|64891203|64891223&offsets|0|-1(+1U)]|CTGACCTATGAATTGACAGCT</td><td>>MI0000234|hsa-mir-192&WithFlank&11|-|64891131|64891252</td><td>30</td><td>49</td><td>20(+1U)</td><td>CTGACCTATGAATTGACAGCT</td><td>BT20</td><td>48</td><td>22</td><td>18</td><td>37</td><td>9</td><td>18.4</td><td>16.7</td><td>10.9</td><td>35.3</td><td>4.2</td></tr>
<tr><td>>MI0000234|hsa-mir-192&WithFlank&11|-|64891131|64891252@30.49.20(+1U)&[MIMAT0000222&hsa-miR-192-5p&offsets|0|-1(+1U);m-59&11|-|64891203|64891223&offsets|0|-1(+1U)]|CTGACCTATGAATTGACAGCT</td><td>>MI0000234|hsa-mir-192&WithFlank&11|-|64891131|64891252</td><td>30</td><td>49</td><td>20(+1U)</td><td>CTGACCTATGAATTGACAGCT</td><td>MB231</td><td>21</td><td>16</td><td>3</td><td>82</td><td>26</td><td>6.2</td><td>5.9</td><td>1.7</td><td>27.4</td><td>24.6</td></tr>
<tr><td>>MI0000234|hsa-mir-192&WithFlank&11|-|64891131|64891252@30.49.20&[MIMAT0000222&hsa-miR-192-5p&offsets|0|-1;m-59&11|-|64891203|64891223&offsets|0|-1]|CTGACCTATGAATTGACAGC</td><td>>MI0000234|hsa-mir-192&WithFlank&11|-|64891131|64891252</td><td>30</td><td>49</td><td>20</td><td>CTGACCTATGAATTGACAGC</td><td>BT20</td><td>13</td><td>6</td><td>5</td><td>10</td><td>2</td><td>8.2</td><td>3.1</td><td>2.2</td><td>9.3</td><td>1.2</td></tr>
<tr><td>>MI0000234|hsa-mir-192&WithFlank&11|-|64891131|64891252@30.50.21&[MIMAT0000222&hsa-miR-192-5p&offsets|0|0;m-59&11|-|64891203|64891223&offsets|0|0]|CTGACCTATGAATTGACAGCC</td><td>>MI0000234|hsa-mir-192&WithFlank&11|-|64891131|64891252</td><td>30</td><td>50</td><td>21</td><td>CTGACCTATGAATTGACAGCC</td><td>BT20</td><td>75</td><td>37</td><td>35</td><td>65</td><td>20</td><td>35.9</td><td>23.3</td><td>25.7</td><td>50.2</td><td>9.4</td></tr>
<tr><td>>MI0000234|hsa-mir-192&WithFlank&11|-|64891131|64891252@30.50.21&[MIMAT0000222&hsa-miR-192-5p&offsets|0|0;m-59&11|-|64891203|64891223&offsets|0|0]|CTGACCTATGAATTGACAGCC</td><td>>MI0000234|hsa-mir-192&WithFlank&11|-|64891131|64891252</td><td>30</td><td>50</td><td>21</td><td>CTGACCTATGAATTGACAGCC</td><td>MB231</td><td>53</td><td>46</td><td>8</td><td>132</td><td>50</td><td>15.9</td><td>17.8</td><td>4.2</td><td>26.8</td><td>34.9</td></tr>
<tr><td>>MI0000234|hsa-mir-192&WithFlank&11|-|64891131|64891252@30.51.22(+1U)&[MIMAT0000222&hsa-miR-192-5p&offsets|0|+1(+1U);m-59&11|-|64891203|64891223&offsets|0|+1(+1U)]|CTGACCTATGAATTGACAGCCAT</td><td>>MI0000234|hsa-mir-192&WithFlank&11|-|64891131|64891252</td><td>30</td><td>51</td><td>22(+1U)</td><td>CTGACCTATGAATTGACAGCCAT</td><td>BT20</td><td>33</td><td>17</td><td>15</td><td>18</td><td>13</td><td>28.3</td><td>9.7</td><td>9.5</td><td>10.1</td><td>4.9</td></tr>
<tr><td>>MI0000234|hsa-mir-192&WithFlank&11|-|64891131|64891252@30.51.22(+1U)&[MIMAT0000222&hsa-miR-192-5p&offsets|0|+1(+1U);m-59&11|-|64891203|64891223&offsets|0|+1(+1U)]|CTGACCTATGAATTGACAGCCAT</td><td>>MI0000234|hsa-mir-192&WithFlank&11|-|64891131|64891252</td><td>30</td><td>51</td><td>22(+1U)</td><td>CTGACCTATGAATTGACAGCCAT</td><td>MB231</td><td>54</td><td>50</td><td>11</td><td>97</td><td>52</td><td>21.6</td><td>18.5</td><td>3.2</td><td>13.0</td><td>23.0</td></tr>
<tr><td>>MI0000234|hsa-mir-192&WithFlank&11|-|64891131|64891252@30.51.22&[MIMAT0000222&hsa-miR-192-5p&offsets|0|+1;m-59&11|-|64891203|64891223&offsets|0|+1]|CTGACCTATGAATTGACAGCCA</td><td>>MI0000234|hsa-mir-192&WithFlank&11|-|64891131|64891252</td><td>30</td><td>51</td><td>22</td><td>CTGACCTATGAATTGACAGCCA</td><td>BT20</td><td>16</td><td>6</td><td>6</td><td>6</td><td>3</td><td>7.3</td><td>4.2</td><td>2.7</td><td>7.0</td><td>1.5</td></tr>
<tr><td>>MI0000234|hsa-mir-192&WithFlank&11|-|64891131|64891252@30.51.22&[MIMAT0000222&hsa-miR-192-5p&offsets|0|+1;m-59&11|-|64891203|64891223&offsets|0|+1]|CTGACCTATGAATTGACAGCCA</td><td>>MI0000234|hsa-mir-192&WithFlank&11|-|64891131|64891252</td><td>30</td><td>51</td><td>22</td><td>CTGACCTATGAATTGACAGCCA</td><td>MB231</td><td>11</td><td>7</td><td>1</td><td>19</td><td>9</td><td>3.3</td><td>2.3</td><td>1.0</td><td>4.5</td><td>9.5</td></tr>
<tr><td>>MI0000234|hsa-mir-192&WithFlank&11|-|64891131|64891252@31.51.21(+1U)&[MIMAT0000222&hsa-miR-192-5p&offsets|+1|+1(+1U);m-59&11|-|64891203|64891223&offsets|+1|+1(+1U)]|TGACCTATGAATTGACAGCCAT</td><td>>MI0000234|hsa-mir-192&WithFlank&11|-|64891131|64891252</td><td>31</td><td>51</td><td>21(+1U)</td><td>TGACCTATGAATTGACAGCCAT</td><td>MB231</td><td>14</td><td>12</td><td>2</td><td>40</td><td>12</td><td>7.1</td><td>4.1</td><td>0.6</td><td>7.4</td><td>6.6</td></tr>
<tr><td>>MI0000234|hsa-mir-192&WithFlank&11|-|64891131|64891252@31.52.22&[MIMAT0000222&hsa-miR-192-5p&offsets|+1|+2;m-59&11|-|64891203|64891223&offsets|+1|+2]|TGACCTATGAATTGACAGCCAG</td><td>>MI0000234|hsa-mir-192&WithFlank&11|-|64891131|64891252</td><td>31</td><td>52</td><td>22</td><td>TGACCTATGAATTGACAGCCAG</td><td>BT20</td><td>11</td><td>4</td><td>6</td><td>10</td><td>4</td><td>5.1</td><td>1.7</td><td>4.4</td><td>8.5</td><td>0.1</td></tr>
<tr><td>>MI0000234|hsa-mir-192&WithFlank&11|-|64891131|64891252@31.53.23&[MIMAT0000222&hsa-miR-192-5p&offsets|+1|+3;m-59&11|-|64891203|64891223&offsets|+1|+3]|TGACCTATGAATTGACAGCCAGT</td><td>>MI0000234|hsa-mir-192&WithFlank&11|-|64891131|64891252</td><td>31</td><td>53</td><td>23</td><td>TGACCTATGAATTGACAGCCAGT</td><td>BT20</td><td>16</td><td>6</td><td>8</td><td>15</td><td>6</td><td>10.7</td><td>4.0</td><td>5.9</td><td>13.1</td><td>2.9</td></tr>
<tr><td>>MI0000234|hsa-mir-192&WithFlank&11|-|64891131|64891252@31.53.23&[MIMAT0000222&hsa-miR-192-5p&offsets|+1|+3;m-59&11|-|64891203|64891223&offsets|+1|+3]|TGACCTATGAATTGACAGCCAGT</td><td>>MI0000234|hsa-mir-192&WithFlank&11|-|64891131|64891252</td><td>31</td><td>53</td><td>23</td><td>TGACCTATGAATTGACAGCCAGT</td><td>MB231</td><td>28</td><td>24</td><td>5</td><td>83</td><td>25</td><td>5.4</td><td>12.1</td><td>3.1</td><td>18.5</td><td>12.6</td></tr>
<tr><td>>MI0000239|hsa-mir-197&WithFlank&1|+|109598887|109598973@54.72.19&[MIMAT0000227&hsa-miR-197-3p&offsets|0|-3;m-99&1|+|109598940|109598961&offsets|0|-3]|TTCACCACCTTCTCCACCC</td><td>>MI0000239|hsa-mir-197&WithFlank&1|+|109598887|109598973</td><td>54</td><td>72</td><td>19</td><td>TTCACCACCTTCTCCACCC</td><td>BT20</td><td>10</td><td>18</td><td>27</td><td>4</td><td>16</td><td>4.4</td><td>5.3</td><td>0.2</td><td>2.0</td><td>1.9</td></tr>
<tr><td>>MI0000239|hsa-mir-197&WithFlank&1|+|109598887|109598973@54.74.21(+1A)&[MIMAT0000227&hsa-miR-197-3p&offsets|0|-1(+1A);m-99&1|+|109598940|109598961&offsets|0|-1(+1A)]|TTCACCACCTTCTCCACCCAGA</td><td>>MI0000239|hsa-mir-197&WithFlank&1|+|109598887|109598973</td><td>54</td><td>74</td><td>21(+1A)</td><td>TTCACCACCTTCTCCACCCAGA</td><td>BT20</td><td>48</td><td>35</td><td>95</td><td>5</td><td>23</td><td>52.2</td><td>8.9</td><td>62.8</td><td>3.1</td><td>10.7</td></tr>
<tr><td>>MI0000239|hsa-mir-197&WithFlank&1|+|109598887|109598973@54.74.21(+1U)&[MIMAT0000227&hsa-miR-197-3p&offsets|0|-1(+1U);m-99&1|+|109598940|109598961&offsets|0|-1(+1U)]|TTCACCACCTTCTCCACCCAGT</td><td>>MI0000239|hsa-mir-197&WithFlank&1|+|109598887|109598973</td><td>54</td><td>74</td><td>21(+1U)</td><td>TTCACCACCTTCTCCACCCAGT</td><td>BT20</td><td>155</td><td>149</td><td>274</td><td>19</td><td>93</td><td>137.8</td><td>38.5</td><td>143.2</td><td>12.5</td><td>36.0</td></tr>
<tr><td>>MI0000239|hsa-mir-197&WithFlank&1|+|109598887|109598973@54.74.21(+1U)&[MIMAT0000227&hsa-miR-197-3p&offsets|0|-1(+1U);m-99&1|+|109598940|109598961&offsets|0|-1(+1U)]|TTCACCACCTTCTCCACCCAGT</td><td>>MI0000239|hsa-mir-197&WithFlank&1|+|109598887|109598973</td><td>54</td><td>74</td><td>21(+1U)</td><td>TTCACCACCTTCTCCACCCAGT</td><td>MB231</td><td>11</td><td>13</td><td>19</td><td>6</td><td>26</td><td>4.1</td><td>2.3</td><td>8.6</td><td>4.0</td><td>16.0</td></tr>
<tr><td>>MI0000239|hsa-mir-197&WithFlank&1|+|109598887|109598973@54.74.21&[MIMAT0000227&hsa-miR-197-3p&offsets|0|-1;m-99&1|+|109598940|109598961&offsets|0|-1]|TTCACCACCTTCTCCACCCAG</td><td>>MI0000239|hsa-mir-197&WithFlank&1|+|109598887|109598973</td><td>54</td><td>74</td><td>21</td><td>TTCACCACCTTCTCCACCCAG</td><td>BT20</td><td>15</td><td>21</td><td>38</td><td>9</td><td>15</td><td>10.6</td><td>8.2</td><td>16.4</td><td>5.8</td><td>5.1</td></tr>
<tr><td>>MI0000239|hsa-mir-197&WithFlank&1|+|109598887|109598973@54.75.22&[MIMAT0000227&hsa-miR-197-3p&offsets|0|0;m-99&1|+|109598940|109598961&offsets|0|0]|TTCACCACCTTCTCCACCCAGC</td><td>>MI0000239|hsa-mir-197&WithFlank&1|+|109598887|109598973</td><td>54</td><td>75</td><td>22</td><td>TTCACCACCTTCTCCACCCAGC</td><td>BT20</td><td>359</td><td>525</td><td>880</td><td>70</td><td>396</td><td>205.1</td><td>122.5</td><td>249.7</td><td>33.2</td><td>119.2</td></tr>
</table>

| | | | | | | | | | | | | | | | |
|---|---|---|---|---|---|---|---|---|---|---|---|---|---|---|---|
| >MI0000239\|hsa-mir-197&WithFlank&1\|+\|109598887\|109598973@54.75.22&[MIMAT0000227&hsa-miR-197-3p&offsets\|0\|0;m-99&1\|+\|109598940\|109598961&offsets\|0\|0]\|TTCACCACCTTCTCCACCCAGC | >MI0000239\|hsa-mir-197&WithFlank&1\|+\|109598887\|109598973 | 54 | 75 | 22 | TTCACCACCTTCTCCACCCAGC | MB231 | 47 | 68 | 93 | 19 | 93 | 19.8 | 17.6 | 44.8 | 6.1 | 37.1 |
| >MI0000239\|hsa-mir-197&WithFlank&1\|+\|109598887\|109598973@54.76.23&[MIMAT0000227&hsa-miR-197-3p&offsets\|0\|+1;m-99&1\|+\|109598940\|109598961&offsets\|0\|+1]\|TTCACCACCTTCTCCACCCAGCA | >MI0000239\|hsa-mir-197&WithFlank&1\|+\|109598887\|109598973 | 54 | 76 | 23 | TTCACCACCTTCTCCACCCAGCA | BT20 | 13 | 13 | 23 | 0 | 0 | 11.0 | 3.2 | 8.9 | 0.0 | 0.0 |
| >MI0000253\|hsa-mir-148a&WithFlank&7\|-\|25949913\|25949992@12.33.22&[MIMAT0004549&hsa-miR-148a-5p&offsets\|0\|0;m-189&7\|-\|25949960\|25949981&offsets\|0\|0]\|AAAGTTCTGAGACACTCCGACT | >MI0000253\|hsa-mir-148a&WithFlank&7\|-\|25949913\|25949992 | 12 | 33 | 22 | AAAGTTCTGAGACACTCCGACT | BT20 | 42 | 36 | 40 | 28 | 12 | 18.8 | 22.9 | 13.0 | 16.6 | 4.7 |
| >MI0000253\|hsa-mir-148a&WithFlank&7\|-\|25949913\|25949992@50.69.20(+2U)&[MIMAT0000243&hsa-miR-148a-3p&offsets\|0\|-2(+2U);m-7&7\|-\|25949922\|25949943&offsets\|0\|-2(+2U)]\|TCAGTGCACTACAGAACTTTTT | >MI0000253\|hsa-mir-148a&WithFlank&7\|-\|25949913\|25949992 | 50 | 69 | 20(+2U) | TCAGTGCACTACAGAACTTTTT | BT20 | 11 | 3 | 5 | 8 | 1 | 9.3 | 0.0 | 2.6 | 10.5 | 0.6 |
| >MI0000253\|hsa-mir-148a&WithFlank&7\|-\|25949913\|25949992@50.70.21(+1A)&[MIMAT0000243&hsa-miR-148a-3p&offsets\|0\|-1(+1A);m-7&7\|-\|25949922\|25949943&offsets\|0\|-1(+1A)]\|TCAGTGCACTACAGAACTTTGA | >MI0000253\|hsa-mir-148a&WithFlank&7\|-\|25949913\|25949992 | 50 | 70 | 21(+1A) | TCAGTGCACTACAGAACTTTGA | BT20 | 709 | 217 | 328 | 1143 | 102 | 290.7 | 82.6 | 100.6 | 1601.4 | 27.2 |
| >MI0000253\|hsa-mir-148a&WithFlank&7\|-\|25949913\|25949992@50.70.21(+1A)&[MIMAT0000243&hsa-miR-148a-3p&offsets\|0\|-1(+1A);m-7&7\|-\|25949922\|25949943&offsets\|0\|-1(+1A)]\|TCAGTGCACTACAGAACTTTGA | >MI0000253\|hsa-mir-148a&WithFlank&7\|-\|25949913\|25949992 | 50 | 70 | 21(+1A) | TCAGTGCACTACAGAACTTTGA | MB231 | 18 | 9 | 5 | 62 | 12 | 9.9 | 1.6 | 2.7 | 23.8 | 9.7 |
| >MI0000253\|hsa-mir-148a&WithFlank&7\|-\|25949913\|25949992@50.70.21(+1C)&[MIMAT0000243&hsa-miR-148a-3p&offsets\|0\|-1(+1C);m-7&7\|-\|25949922\|25949943&offsets\|0\|-1(+1C)]\|TCAGTGCACTACAGAACTTTGC | >MI0000253\|hsa-mir-148a&WithFlank&7\|-\|25949913\|25949992 | 50 | 70 | 21(+1C) | TCAGTGCACTACAGAACTTTGC | BT20 | 2133 | 1248 | 1021 | 5124 | 658 | 344.5 | 516.3 | 113.3 | 6722.3 | 300.8 |
| >MI0000253\|hsa-mir-148a&WithFlank&7\|-\|25949913\|25949992@50.70.21(+1C)&[MIMAT0000243&hsa-miR-148a-3p&offsets\|0\|-1(+1C);m-7&7\|-\|25949922\|25949943&offsets\|0\|-1(+1C)]\|TCAGTGCACTACAGAACTTTGC | >MI0000253\|hsa-mir-148a&WithFlank&7\|-\|25949913\|25949992 | 50 | 70 | 21(+1C) | TCAGTGCACTACAGAACTTTGC | MB231 | 88 | 62 | 21 | 277 | 71 | 54.9 | 19.4 | 10.3 | 85.2 | 64.6 |
| >MI0000253\|hsa-mir-148a&WithFlank&7\|-\|25949913\|25949992@50.70.21(+1C)&[MIMAT0000243&hsa-miR-148a-3p&offsets\|0\|-1(+1C);m-7&7\|-\|25949922\|25949943&offsets\|0\|-1(+1C)]\|TCAGTGCACTACAGAACTTTGC | >MI0000253\|hsa-mir-148a&WithFlank&7\|-\|25949913\|25949992 | 50 | 70 | 21(+1C) | TCAGTGCACTACAGAACTTTGC | MB468 | 43 | 43 | 22 | 247 | 17 | 6.4 | 39.7 | 11.3 | 90.7 | 7.1 |
| >MI0000253\|hsa-mir-148a&WithFlank&7\|-\|25949913\|25949992@50.70.21(+1G)&[MIMAT0000243&hsa-miR-148a-3p&offsets\|0\|-1(+1G);m-7&7\|-\|25949922\|25949943&offsets\|0\|-1(+1G)]\|TCAGTGCACTACAGAACTTTGG | >MI0000253\|hsa-mir-148a&WithFlank&7\|-\|25949913\|25949992 | 50 | 70 | 21(+1G) | TCAGTGCACTACAGAACTTTGG | BT20 | 1101 | 215 | 209 | 792 | 45 | 703.6 | 66.2 | 71.9 | 1218.4 | 17.2 |
| >MI0000253\|hsa-mir-148a&WithFlank&7\|-\|25949913\|25949992@50.70.21(+1G)&[MIMAT0000243&hsa-miR-148a-3p&offsets\|0\|-1(+1G);m-7&7\|-\|25949922\|25949943&offsets\|0\|-1(+1G)]\|TCAGTGCACTACAGAACTTTGG | >MI0000253\|hsa-mir-148a&WithFlank&7\|-\|25949913\|25949992 | 50 | 70 | 21(+1G) | TCAGTGCACTACAGAACTTTGG | MB231 | 34 | 9 | 3 | 24 | 6 | 23.5 | 3.6 | 2.2 | 2.3 | 5.4 |
| >MI0000253\|hsa-mir-148a&WithFlank&7\|-\|25949913\|25949992@50.70.21(+1G)&[MIMAT0000243&hsa-miR-148a-3p&offsets\|0\|-1(+1G);m-7&7\|-\|25949922\|25949943&offsets\|0\|-1(+1G)]\|TCAGTGCACTACAGAACTTTGG | >MI0000253\|hsa-mir-148a&WithFlank&7\|-\|25949913\|25949992 | 50 | 70 | 21(+1G) | TCAGTGCACTACAGAACTTTGG | MB468 | 18 | 7 | 9 | 52 | 3 | 8.8 | 7.4 | 6.8 | 32.4 | 2.1 |
| >MI0000253\|hsa-mir-148a&WithFlank&7\|-\|25949913\|25949992@50.70.21&[MIMAT0000243&hsa-miR-148a-3p&offsets\|0\|-1;m-7&7\|-\|25949922\|25949943&offsets\|0\|-1]\|TCAGTGCACTACAGAACTTTG | >MI0000253\|hsa-mir-148a&WithFlank&7\|-\|25949913\|25949992 | 50 | 70 | 21 | TCAGTGCACTACAGAACTTTG | BT20 | 52 | 23 | 14 | 35 | 4 | 46.7 | 15.8 | 5.5 | 38.0 | 2.2 |
| >MI0000253\|hsa-mir-148a&WithFlank&7\|-\|25949913\|25949992@50.71.22(+1A)&[MIMAT0000243&hsa-miR-148a-3p&offsets\|0\|0(+1A);m-7&7\|-\|25949922\|25949943&offsets\|0\|0(+1A)]\|TCAGTGCACTACAGAACTTTGTA | >MI0000253\|hsa-mir-148a&WithFlank&7\|-\|25949913\|25949992 | 50 | 71 | 22(+1A) | TCAGTGCACTACAGAACTTTGTA | BT20 | 51 | 15 | 16 | 10 | 6 | 34.0 | 9.0 | 3.2 | 12.4 | 1.6 |
| >MI0000253\|hsa-mir-148a&WithFlank&7\|-\|25949913\|25949992@50.71.22(+1U)&[MIMAT0000243&hsa-miR-148a-3p&offsets\|0\|0(+1U);m-7&7\|-\|25949922\|25949943&offsets\|0\|0(+1U)]\|TCAGTGCACTACAGAACTTTGTT | >MI0000253\|hsa-mir-148a&WithFlank&7\|-\|25949913\|25949992 | 50 | 71 | 22(+1U) | TCAGTGCACTACAGAACTTTGTT | BT20 | 110 | 43 | 38 | 20 | 13 | 58.1 | 31.8 | 8.7 | 25.9 | 7.5 |
| >MI0000253\|hsa-mir-148a&WithFlank&7\|-\|25949913\|25949992@50.71.22&[MIMAT0000243&hsa-miR-148a-3p&offsets\|0\|0;m-7&7\|-\|25949922\|25949943&offsets\|0\|0]\|TCAGTGCACTACAGAACTTTGT | >MI0000253\|hsa-mir-148a&WithFlank&7\|-\|25949913\|25949992 | 50 | 71 | 22 | TCAGTGCACTACAGAACTTTGT | BT20 | 16515 | 8830 | 8189 | 34466 | 5027 | 2733.0 | 2655.4 | 1726.0 | 41081.6 | 1989.3 |
| >MI0000253\|hsa-mir-148a&WithFlank&7\|-\|25949913\|25949992@50.71.22&[MIMAT0000243&hsa-miR-148a-3p&offsets\|0\|0;m-7&7\|-\|25949922\|25949943&offsets\|0\|0]\|TCAGTGCACTACAGAACTTTGT | >MI0000253\|hsa-mir-148a&WithFlank&7\|-\|25949913\|25949992 | 50 | 71 | 22 | TCAGTGCACTACAGAACTTTGT | MB231 | 829 | 611 | 221 | 2038 | 533 | 511.1 | 252.0 | 119.2 | 844.1 | 410.0 |
| >MI0000253\|hsa-mir-148a&WithFlank&7\|-\|25949913\|25949992@50.71.22&[MIMAT0000243&hsa-miR-148a-3p&offsets\|0\|0;m-7&7\|-\|25949922\|25949943&offsets\|0\|0]\|TCAGTGCACTACAGAACTTTGT | >MI0000253\|hsa-mir-148a&WithFlank&7\|-\|25949913\|25949992 | 50 | 71 | 22 | TCAGTGCACTACAGAACTTTGT | MB468 | 394 | 427 | 198 | 1969 | 133 | 37.2 | 388.3 | 129.1 | 660.1 | 49.0 |
| >MI0000253\|hsa-mir-148a&WithFlank&7\|-\|25949913\|25949992@50.72.23(+1A)&[MIMAT0000243&hsa-miR-148a-3p&offsets\|0\|+1(+1A);m-7&7\|-\|25949922\|25949943&offsets\|0\|+1(+1A)]\|TCAGTGCACTACAGAACTTTGTCA | >MI0000253\|hsa-mir-148a&WithFlank&7\|-\|25949913\|25949992 | 50 | 72 | 23(+1A) | TCAGTGCACTACAGAACTTTGTCA | BT20 | 18 | 6 | 7 | 2 | 3 | 11.2 | 3.9 | 2.0 | 1.7 | 1.5 |
| >MI0000253\|hsa-mir-148a&WithFlank&7\|-\|25949913\|25949992@50.72.23&[MIMAT0000243&hsa-miR-148a-3p&offsets\|0\|+1;m-7&7\|-\|25949922\|25949943&offsets\|0\|+1]\|TCAGTGCACTACAGAACTTTGTC | >MI0000253\|hsa-mir-148a&WithFlank&7\|-\|25949913\|25949992 | 50 | 72 | 23 | TCAGTGCACTACAGAACTTTGTC | BT20 | 254 | 124 | 127 | 78 | 57 | 83.4 | 62.3 | 50.8 | 89.0 | 28.6 |
| >MI0000253\|hsa-mir-148a&WithFlank&7\|-\|25949913\|25949992@50.73.24&[MIMAT0000243&hsa-miR-148a-3p&offsets\|0\|+2;m-7&7\|-\|25949922\|25949943&offsets\|0\|+2]\|TCAGTGCACTACAGAACTTTGTCT | >MI0000253\|hsa-mir-148a&WithFlank&7\|-\|25949913\|25949992 | 50 | 73 | 24 | TCAGTGCACTACAGAACTTTGTCT | BT20 | 48 | 22 | 17 | 5 | 11 | 20.1 | 15.2 | 10.9 | 5.4 | 6.0 |
| >MI0000254\|hsa-mir-30c-2&WithFlank&6\|-\|71376954\|71377037@53.74.22&[MIMAT0004550&hsa-miR-30c-2-3p&offsets\|0\|0;m-206&6\|-\|71376964\|71376985&offsets\|0\|0]\|CTGGGAGAAGGCTGTTTACTCT | >MI0000254\|hsa-mir-30c-2&WithFlank&6\|-\|71376954\|71377037 | 53 | 74 | 22 | CTGGGAGAAGGCTGTTTACTCT | BT20 | 43 | 39 | 101 | 10 | 22 | 16.0 | 20.3 | 29.3 | 2.2 | 13.2 |
| >MI0000254\|hsa-mir-30c-2&WithFlank&6\|-\|71376954\|71377037@53.74.22&[MIMAT0004550&hsa-miR-30c-2-3p&offsets\|0\|0;m-206&6\|-\|71376964\|71376985&offsets\|0\|0]\|CTGGGAGAAGGCTGTTTACTCT | >MI0000254\|hsa-mir-30c-2&WithFlank&6\|-\|71376954\|71377037 | 53 | 74 | 22 | CTGGGAGAAGGCTGTTTACTCT | MB231 | 73 | 67 | 88 | 39 | 52 | 22.7 | 18.0 | 60.3 | 2.5 | 21.6 |
| >MI0000254\|hsa-mir-30c-2&WithFlank&6\|-\|71376954\|71377037@53.75.23&[MIMAT0004550&hsa-miR-30c-2-3p&offsets\|0\|+1;m-206&6\|-\|71376964\|71376985&offsets\|0\|+1]\|CTGGGAGAAGGCTGTTTACTCTT | >MI0000254\|hsa-mir-30c-2&WithFlank&6\|-\|71376954\|71377037 | 53 | 75 | 23 | CTGGGAGAAGGCTGTTTACTCTT | MB231 | 19 | 19 | 10 | 9 | 13 | 2.7 | 5.8 | 3.2 | 1.4 | 3.5 |
| >MI0000255\|hsa-mir-30d&WithFlank&8\|-\|134804870\|134804951@12.31.20&[MIMAT0000245&hsa-miR-30d-5p&offsets\|0\|-2;m-21&8\|-\|134804919\|134804940&offsets\|0\|-2]\|TGTAAACATCCCCGACTGGA | >MI0000255\|hsa-mir-30d&WithFlank&8\|-\|134804870\|134804951 | 12 | 31 | 20 | TGTAAACATCCCCGACTGGA | BT20 | 115 | 87 | 108 | 223 | 65 | 77.7 | 47.4 | 33.5 | 163.6 | 13.7 |
| >MI0000255\|hsa-mir-30d&WithFlank&8\|-\|134804870\|134804951@12.31.20&[MIMAT0000245&hsa-miR-30d-5p&offsets\|0\|-2;m-21&8\|-\|134804919\|134804940&offsets\|0\|-2]\|TGTAAACATCCCCGACTGGA | >MI0000255\|hsa-mir-30d&WithFlank&8\|-\|134804870\|134804951 | 12 | 31 | 20 | TGTAAACATCCCCGACTGGA | MB231 | 44 | 82 | 25 | 104 | 62 | 8.6 | 58.1 | 22.9 | 50.1 | 26.7 |
| >MI0000255\|hsa-mir-30d&WithFlank&8\|-\|134804870\|134804951@12.32.21(+1A)&[MIMAT0000245&hsa-miR-30d-5p&offsets\|0\|-1(+1A);m-21&8\|-\|134804919\|134804940&offsets\|0\|-1(+1A)]\|TGTAAACATCCCCGACTGGAAA | >MI0000255\|hsa-mir-30d&WithFlank&8\|-\|134804870\|134804951 | 12 | 32 | 21(+1A) | TGTAAACATCCCCGACTGGAAA | BT20 | 87 | 35 | 51 | 11 | 24 | 75.7 | 17.1 | 18.3 | 7.1 | 4.1 |
| >MI0000255\|hsa-mir-30d&WithFlank&8\|-\|134804870\|134804951@12.32.21&[MIMAT0000245&hsa-miR-30d-5p&offsets\|0\|-1;m-21&8\|-\|134804919\|134804940&offsets\|0\|-1]\|TGTAAACATCCCCGACTGGAA | >MI0000255\|hsa-mir-30d&WithFlank&8\|-\|134804870\|134804951 | 12 | 32 | 21 | TGTAAACATCCCCGACTGGAA | BT20 | 79 | 53 | 64 | 23 | 32 | 39.6 | 20.6 | 6.8 | 17.6 | 0.7 |
| >MI0000255\|hsa-mir-30d&WithFlank&8\|-\|134804870\|134804951@12.32.21&[MIMAT0000245&hsa-miR-30d-5p&offsets\|0\|-1;m-21&8\|-\|134804919\|134804940&offsets\|0\|-1]\|TGTAAACATCCCCGACTGGAA | >MI0000255\|hsa-mir-30d&WithFlank&8\|-\|134804870\|134804951 | 12 | 32 | 21 | TGTAAACATCCCCGACTGGAA | MB231 | 15 | 20 | 6 | 12 | 19 | 2.6 | 11.2 | 4.6 | 3.1 | 6.8 |
| >MI0000255\|hsa-mir-30d&WithFlank&8\|-\|134804870\|134804951@12.33.22(+1A)&[MIMAT0000245&hsa-miR-30d-5p&offsets\|0\|0(+1A);m-21&8\|-\|134804919\|134804940&offsets\|0\|0(+1A)]\|TGTAAACATCCCCGACTGGAAGA | >MI0000255\|hsa-mir-30d&WithFlank&8\|-\|134804870\|134804951 | 12 | 33 | 22(+1A) | TGTAAACATCCCCGACTGGAAGA | BT20 | 213 | 111 | 181 | 60 | 100 | 77.7 | 39.1 | 30.6 | 41.9 | 45.8 |
| >MI0000255\|hsa-mir-30d&WithFlank&8\|-\|134804870\|134804951@12.33.22(+1A)&[MIMAT0000245&hsa-miR-30d-5p&offsets\|0\|0(+1A);m-21&8\|-\|134804919\|134804940&offsets\|0\|0(+1A)]\|TGTAAACATCCCCGACTGGAAGA | >MI0000255\|hsa-mir-30d&WithFlank&8\|-\|134804870\|134804951 | 12 | 33 | 22(+1A) | TGTAAACATCCCCGACTGGAAGA | MB231 | 46 | 38 | 16 | 28 | 45 | 16.1 | 12.0 | 12.1 | 5.0 | 12.2 |
| >MI0000255\|hsa-mir-30d&WithFlank&8\|-\|134804870\|134804951@12.33.22(+1G)&[MIMAT0000245&hsa-miR-30d-5p&offsets\|0\|0(+1G);m-21&8\|-\|134804919\|134804940&offsets\|0\|0(+1G)]\|TGTAAACATCCCCGACTGGAAGG | >MI0000255\|hsa-mir-30d&WithFlank&8\|-\|134804870\|134804951 | 12 | 33 | 22(+1G) | TGTAAACATCCCCGACTGGAAGG | BT20 | 30 | 8 | 11 | 3 | 3 | 22.6 | 1.7 | 5.1 | 4.0 | 1.5 |
| >MI0000255\|hsa-mir-30d&WithFlank&8\|-\|134804870\|134804951@12.33.22(+1U)&[MIMAT0000245&hsa-miR-30d-5p&offsets\|0\|0(+1U);m-21&8\|-\|134804919\|134804940&offsets\|0\|0(+1U)]\|TGTAAACATCCCCGACTGGAAGT | >MI0000255\|hsa-mir-30d&WithFlank&8\|-\|134804870\|134804951 | 12 | 33 | 22(+1U) | TGTAAACATCCCCGACTGGAAGT | BT20 | 804 | 527 | 635 | 278 | 485 | 233.6 | 182.1 | 16.7 | 203.9 | 204.6 |

| | | | | | | | | | | | | | | | | |
|---|---|---|---|---|---|---|---|---|---|---|---|---|---|---|---|---|
| >MI0000255\|hsa-mir-30d&WithFlank&8\|-\|134804870\|134804951@12.33.22(+1U)&[MIMAT0000245&hsa-miR-30d-5p&offsets\|0\|0(+1U);m-21&8\|-\|134804919\|134804940&offsets\|0\|0(+1U)]\|TGTAAACATCCCCGACTGGAAGT | >MI0000255\|hsa-mir-30d&WithFlank&8\|-\|134804870\|134804951 | 12 | 33 | 22(+1U) | TGTAAACATCCCCGACTGGAAGT | MB231 | 214 | 179 | 60 | 126 | 216 | 67.0 | 61.7 | 39.1 | 33.6 | 76.6 |
| >MI0000255\|hsa-mir-30d&WithFlank&8\|-\|134804870\|134804951@12.33.22(+2U)&[MIMAT0000245&hsa-miR-30d-5p&offsets\|0\|0(+2U);m-21&8\|-\|134804919\|134804940&offsets\|0\|0(+2U)]\|TGTAAACATCCCCGACTGGAAGTT | >MI0000255\|hsa-mir-30d&WithFlank&8\|-\|134804870\|134804951 | 12 | 33 | 22(+2U) | TGTAAACATCCCCGACTGGAAGTT | BT20 | 42 | 9 | 16 | 2 | 3 | 48.9 | 2.7 | 9.3 | 1.0 | 0.8 |
| >MI0000255\|hsa-mir-30d&WithFlank&8\|-\|134804870\|134804951@12.33.22(+2U)&[MIMAT0000245&hsa-miR-30d-5p&offsets\|0\|0(+2U);m-21&8\|-\|134804919\|134804940&offsets\|0\|0(+2U)]\|TGTAAACATCCCCGACTGGAAGTT | >MI0000255\|hsa-mir-30d&WithFlank&8\|-\|134804870\|134804951 | 12 | 33 | 22(+2U) | TGTAAACATCCCCGACTGGAAGTT | MB231 | 11 | 7 | 2 | 2 | 6 | 5.8 | 3.6 | 1.8 | 0.3 | 5.3 |
| >MI0000255\|hsa-mir-30d&WithFlank&8\|-\|134804870\|134804951@12.33.22&[MIMAT0000245&hsa-miR-30d-5p&offsets\|0\|0;m-21&8\|-\|134804919\|134804940&offsets\|0\|0]\|TGTAAACATCCCCGACTGGAAG | >MI0000255\|hsa-mir-30d&WithFlank&8\|-\|134804870\|134804951 | 12 | 33 | 22 | TGTAAACATCCCCGACTGGAAG | BT20 | 471 | 278 | 280 | 105 | 154 | 300.2 | 108.2 | 67.1 | 54.3 | 17.4 |
| >MI0000255\|hsa-mir-30d&WithFlank&8\|-\|134804870\|134804951@12.33.22&[MIMAT0000245&hsa-miR-30d-5p&offsets\|0\|0;m-21&8\|-\|134804919\|134804940&offsets\|0\|0]\|TGTAAACATCCCCGACTGGAAG | >MI0000255\|hsa-mir-30d&WithFlank&8\|-\|134804870\|134804951 | 12 | 33 | 22 | TGTAAACATCCCCGACTGGAAG | MB231 | 106 | 110 | 28 | 92 | 99 | 19.6 | 64.3 | 22.2 | 40.6 | 31.7 |
| >MI0000255\|hsa-mir-30d&WithFlank&8\|-\|134804870\|134804951@12.34.23(+1A)&[MIMAT0000245&hsa-miR-30d-5p&offsets\|0\|+1(+1A);m-21&8\|-\|134804919\|134804940&offsets\|0\|+1(+1A)]\|TGTAAACATCCCCGACTGGAAGCA | >MI0000255\|hsa-mir-30d&WithFlank&8\|-\|134804870\|134804951 | 12 | 34 | 23(+1A) | TGTAAACATCCCCGACTGGAAGCA | BT20 | 813 | 326 | 464 | 144 | 203 | 531.8 | 148.3 | 95.8 | 92.0 | 86.4 |
| >MI0000255\|hsa-mir-30d&WithFlank&8\|-\|134804870\|134804951@12.34.23(+1A)&[MIMAT0000245&hsa-miR-30d-5p&offsets\|0\|+1(+1A);m-21&8\|-\|134804919\|134804940&offsets\|0\|+1(+1A)]\|TGTAAACATCCCCGACTGGAAGCA | >MI0000255\|hsa-mir-30d&WithFlank&8\|-\|134804870\|134804951 | 12 | 34 | 23(+1A) | TGTAAACATCCCCGACTGGAAGCA | MB231 | 258 | 169 | 74 | 192 | 257 | 24.3 | 58.9 | 57.2 | 44.1 | 186.4 |
| >MI0000255\|hsa-mir-30d&WithFlank&8\|-\|134804870\|134804951@12.34.23(+1C)&[MIMAT0000245&hsa-miR-30d-5p&offsets\|0\|+1(+1C);m-21&8\|-\|134804919\|134804940&offsets\|0\|+1(+1C)]\|TGTAAACATCCCCGACTGGAAGCC | >MI0000255\|hsa-mir-30d&WithFlank&8\|-\|134804870\|134804951 | 12 | 34 | 23(+1C) | TGTAAACATCCCCGACTGGAAGCC | BT20 | 885 | 540 | 538 | 237 | 355 | 197.9 | 237.5 | 129.5 | 132.0 | 133.5 |
| >MI0000255\|hsa-mir-30d&WithFlank&8\|-\|134804870\|134804951@12.34.23(+1C)&[MIMAT0000245&hsa-miR-30d-5p&offsets\|0\|+1(+1C);m-21&8\|-\|134804919\|134804940&offsets\|0\|+1(+1C)]\|TGTAAACATCCCCGACTGGAAGCC | >MI0000255\|hsa-mir-30d&WithFlank&8\|-\|134804870\|134804951 | 12 | 34 | 23(+1C) | TGTAAACATCCCCGACTGGAAGCC | MB231 | 429 | 343 | 114 | 301 | 458 | 70.0 | 141.4 | 74.2 | 57.7 | 338.6 |
| >MI0000255\|hsa-mir-30d&WithFlank&8\|-\|134804870\|134804951@12.34.23(+1G)&[MIMAT0000245&hsa-miR-30d-5p&offsets\|0\|+1(+1G);m-21&8\|-\|134804919\|134804940&offsets\|0\|+1(+1G)]\|TGTAAACATCCCCGACTGGAAGCG | >MI0000255\|hsa-mir-30d&WithFlank&8\|-\|134804870\|134804951 | 12 | 34 | 23(+1G) | TGTAAACATCCCCGACTGGAAGCG | BT20 | 579 | 119 | 139 | 39 | 37 | 466.2 | 40.0 | 68.1 | 36.7 | 15.1 |
| >MI0000255\|hsa-mir-30d&WithFlank&8\|-\|134804870\|134804951@12.34.23(+1G)&[MIMAT0000245&hsa-miR-30d-5p&offsets\|0\|+1(+1G);m-21&8\|-\|134804919\|134804940&offsets\|0\|+1(+1G)]\|TGTAAACATCCCCGACTGGAAGCG | >MI0000255\|hsa-mir-30d&WithFlank&8\|-\|134804870\|134804951 | 12 | 34 | 23(+1G) | TGTAAACATCCCCGACTGGAAGCG | MB231 | 223 | 72 | 22 | 38 | 52 | 171.3 | 38.0 | 17.0 | 12.1 | 41.0 |
| >MI0000255\|hsa-mir-30d&WithFlank&8\|-\|134804870\|134804951@12.34.23&[MIMAT0000245&hsa-miR-30d-5p&offsets\|0\|+1;m-21&8\|-\|134804919\|134804940&offsets\|0\|+1]\|TGTAAACATCCCCGACTGGAAGC | >MI0000255\|hsa-mir-30d&WithFlank&8\|-\|134804870\|134804951 | 12 | 34 | 23 | TGTAAACATCCCCGACTGGAAGC | BT20 | 2234 | 1712 | 2107 | 904 | 1834 | 1265.2 | 356.1 | 676.5 | 329.3 | 513.3 |
| >MI0000255\|hsa-mir-30d&WithFlank&8\|-\|134804870\|134804951@12.34.23&[MIMAT0000245&hsa-miR-30d-5p&offsets\|0\|+1;m-21&8\|-\|134804919\|134804940&offsets\|0\|+1]\|TGTAAACATCCCCGACTGGAAGC | >MI0000255\|hsa-mir-30d&WithFlank&8\|-\|134804870\|134804951 | 12 | 34 | 23 | TGTAAACATCCCCGACTGGAAGC | MB231 | 919 | 1014 | 297 | 365 | 832 | 342.6 | 382.2 | 194.8 | 46.8 | 98.2 |
| >MI0000255\|hsa-mir-30d&WithFlank&8\|-\|134804870\|134804951@12.34.23&[MIMAT0000245&hsa-miR-30d-5p&offsets\|0\|+1;m-21&8\|-\|134804919\|134804940&offsets\|0\|+1]\|TGTAAACATCCCCGACTGGAAGC | >MI0000255\|hsa-mir-30d&WithFlank&8\|-\|134804870\|134804951 | 12 | 34 | 23 | TGTAAACATCCCCGACTGGAAGC | MB468 | 40 | 45 | 17 | 36 | 22 | 12.9 | 10.6 | 10.1 | 21.6 | 10.2 |
| >MI0000255\|hsa-mir-30d&WithFlank&8\|-\|134804870\|134804951@12.35.24(+1A)&[MIMAT0000245&hsa-miR-30d-5p&offsets\|0\|+2(+1A);m-21&8\|-\|134804919\|134804940&offsets\|0\|+2(+1A)]\|TGTAAACATCCCCGACTGGAAGCTA | >MI0000255\|hsa-mir-30d&WithFlank&8\|-\|134804870\|134804951 | 12 | 35 | 24(+1A) | TGTAAACATCCCCGACTGGAAGCTA | BT20 | 30 | 13 | 18 | 4 | 10 | 17.2 | 7.7 | 5.1 | 1.6 | 4.0 |
| >MI0000255\|hsa-mir-30d&WithFlank&8\|-\|134804870\|134804951@12.35.24(+1A)&[MIMAT0000245&hsa-miR-30d-5p&offsets\|0\|+2(+1A);m-21&8\|-\|134804919\|134804940&offsets\|0\|+2(+1A)]\|TGTAAACATCCCCGACTGGAAGCTA | >MI0000255\|hsa-mir-30d&WithFlank&8\|-\|134804870\|134804951 | 12 | 35 | 24(+1A) | TGTAAACATCCCCGACTGGAAGCTA | MB231 | 19 | 11 | 5 | 10 | 18 | 1.1 | 3.8 | 3.6 | 2.0 | 12.5 |
| >MI0000255\|hsa-mir-30d&WithFlank&8\|-\|134804870\|134804951@12.35.24(+1C)&[MIMAT0000245&hsa-miR-30d-5p&offsets\|0\|+2(+1C);m-21&8\|-\|134804919\|134804940&offsets\|0\|+2(+1C)]\|TGTAAACATCCCCGACTGGAAGCTC | >MI0000255\|hsa-mir-30d&WithFlank&8\|-\|134804870\|134804951 | 12 | 35 | 24(+1C) | TGTAAACATCCCCGACTGGAAGCTC | BT20 | 20 | 12 | 12 | 7 | 9 | 6.7 | 6.6 | 2.8 | 4.6 | 4.4 |
| >MI0000255\|hsa-mir-30d&WithFlank&8\|-\|134804870\|134804951@12.35.24(+1C)&[MIMAT0000245&hsa-miR-30d-5p&offsets\|0\|+2(+1C);m-21&8\|-\|134804919\|134804940&offsets\|0\|+2(+1C)]\|TGTAAACATCCCCGACTGGAAGCTC | >MI0000255\|hsa-mir-30d&WithFlank&8\|-\|134804870\|134804951 | 12 | 35 | 24(+1C) | TGTAAACATCCCCGACTGGAAGCTC | MB231 | 21 | 16 | 6 | 23 | 26 | 4.0 | 6.6 | 4.0 | 5.9 | 21.3 |
| >MI0000255\|hsa-mir-30d&WithFlank&8\|-\|134804870\|134804951@12.35.24(+1U)&[MIMAT0000245&hsa-miR-30d-5p&offsets\|0\|+2(+1U);m-21&8\|-\|134804919\|134804940&offsets\|0\|+2(+1U)]\|TGTAAACATCCCCGACTGGAAGCTT | >MI0000255\|hsa-mir-30d&WithFlank&8\|-\|134804870\|134804951 | 12 | 35 | 24(+1U) | TGTAAACATCCCCGACTGGAAGCTT | BT20 | 136 | 79 | 85 | 53 | 56 | 46.3 | 41.6 | 28.5 | 37.5 | 28.3 |
| >MI0000255\|hsa-mir-30d&WithFlank&8\|-\|134804870\|134804951@12.35.24(+1U)&[MIMAT0000245&hsa-miR-30d-5p&offsets\|0\|+2(+1U);m-21&8\|-\|134804919\|134804940&offsets\|0\|+2(+1U)]\|TGTAAACATCCCCGACTGGAAGCTT | >MI0000255\|hsa-mir-30d&WithFlank&8\|-\|134804870\|134804951 | 12 | 35 | 24(+1U) | TGTAAACATCCCCGACTGGAAGCTT | MB231 | 185 | 140 | 47 | 153 | 180 | 20.4 | 50.0 | 27.3 | 33.0 | 118.8 |
| >MI0000255\|hsa-mir-30d&WithFlank&8\|-\|134804870\|134804951@12.35.24(+2U)&[MIMAT0000245&hsa-miR-30d-5p&offsets\|0\|+2(+2U);m-21&8\|-\|134804919\|134804940&offsets\|0\|+2(+2U)]\|TGTAAACATCCCCGACTGGAAGCTTT | >MI0000255\|hsa-mir-30d&WithFlank&8\|-\|134804870\|134804951 | 12 | 35 | 24(+2U) | TGTAAACATCCCCGACTGGAAGCTTT | BT20 | 16 | 9 | 15 | 2 | 6 | 6.6 | 5.8 | 2.8 | 1.4 | 4.2 |
| >MI0000255\|hsa-mir-30d&WithFlank&8\|-\|134804870\|134804951@12.35.24(+2U)&[MIMAT0000245&hsa-miR-30d-5p&offsets\|0\|+2(+2U);m-21&8\|-\|134804919\|134804940&offsets\|0\|+2(+2U)]\|TGTAAACATCCCCGACTGGAAGCTTT | >MI0000255\|hsa-mir-30d&WithFlank&8\|-\|134804870\|134804951 | 12 | 35 | 24(+2U) | TGTAAACATCCCCGACTGGAAGCTTT | MB231 | 46 | 35 | 16 | 16 | 43 | 9.5 | 15.6 | 10.1 | 5.0 | 30.5 |
| >MI0000255\|hsa-mir-30d&WithFlank&8\|-\|134804870\|134804951@12.35.24(+3U)&[MIMAT0000245&hsa-miR-30d-5p&offsets\|0\|+2(+3U);m-21&8\|-\|134804919\|134804940&offsets\|0\|+2(+3U)]\|TGTAAACATCCCCGACTGGAAGCTTTT | >MI0000255\|hsa-mir-30d&WithFlank&8\|-\|134804870\|134804951 | 12 | 35 | 24(+3U) | TGTAAACATCCCCGACTGGAAGCTTTT | MB231 | 25 | 22 | 9 | 3 | 26 | 7.3 | 11.1 | 4.9 | 1.0 | 10.6 |
| >MI0000255\|hsa-mir-30d&WithFlank&8\|-\|134804870\|134804951@12.35.24(+4U)&[MIMAT0000245&hsa-miR-30d-5p&offsets\|0\|+2(+4U);m-21&8\|-\|134804919\|134804940&offsets\|0\|+2(+4U)]\|TGTAAACATCCCCGACTGGAAGCTTTTT | >MI0000255\|hsa-mir-30d&WithFlank&8\|-\|134804870\|134804951 | 12 | 35 | 24(+4U) | TGTAAACATCCCCGACTGGAAGCTTTTT | MB231 | 14 | 12 | 6 | 1 | 12 | 5.5 | 5.8 | 2.6 | 0.7 | 3.7 |
| >MI0000255\|hsa-mir-30d&WithFlank&8\|-\|134804870\|134804951@12.35.24&[MIMAT0000245&hsa-miR-30d-5p&offsets\|0\|+2;m-21&8\|-\|134804919\|134804940&offsets\|0\|+2]\|TGTAAACATCCCCGACTGGAAGCT | >MI0000255\|hsa-mir-30d&WithFlank&8\|-\|134804870\|134804951 | 12 | 35 | 24 | TGTAAACATCCCCGACTGGAAGCT | BT20 | 5508 | 3109 | 3567 | 1493 | 2189 | 1288.4 | 1124.5 | 1137.9 | 654.2 | 696.6 |
| >MI0000255\|hsa-mir-30d&WithFlank&8\|-\|134804870\|134804951@12.35.24&[MIMAT0000245&hsa-miR-30d-5p&offsets\|0\|+2;m-21&8\|-\|134804919\|134804940&offsets\|0\|+2]\|TGTAAACATCCCCGACTGGAAGCT | >MI0000255\|hsa-mir-30d&WithFlank&8\|-\|134804870\|134804951 | 12 | 35 | 24 | TGTAAACATCCCCGACTGGAAGCT | MB231 | 3058 | 2472 | 840 | 1703 | 2703 | 314.4 | 896.8 | 553.1 | 206.8 | 1558.1 |
| >MI0000255\|hsa-mir-30d&WithFlank&8\|-\|134804870\|134804951@12.35.24&[MIMAT0000245&hsa-miR-30d-5p&offsets\|0\|+2;m-21&8\|-\|134804919\|134804940&offsets\|0\|+2]\|TGTAAACATCCCCGACTGGAAGCT | >MI0000255\|hsa-mir-30d&WithFlank&8\|-\|134804870\|134804951 | 12 | 35 | 24 | TGTAAACATCCCCGACTGGAAGCT | MB468 | 72 | 64 | 27 | 87 | 31 | 13.6 | 21.3 | 16.3 | 41.2 | 12.8 |
| >MI0000255\|hsa-mir-30d&WithFlank&8\|-\|134804870\|134804951@12.36.25&[MIMAT0000245&hsa-miR-30d-5p&offsets\|0\|+3;m-21&8\|-\|134804919\|134804940&offsets\|0\|+3]\|TGTAAACATCCCCGACTGGAAGCTG | >MI0000255\|hsa-mir-30d&WithFlank&8\|-\|134804870\|134804951 | 12 | 36 | 25 | TGTAAACATCCCCGACTGGAAGCTG | BT20 | 12 | 3 | 4 | 1 | 1 | 10.1 | 1.4 | 1.2 | 0.7 | 1.1 |

<table>
<tr><td>>MI0000261|hsa-mir-139&WithFlank&11|-|72615057|72615136@13.35.23&[MIMAT0000250&hsa-miR-139-5p&offsets|0|0;m-186&11|-|72615102|72615124&offsets|0|0]|TCTACAGTGCACGTGTCTCCAGT</td><td>>MI0000261|hsa-mir-139&WithFlank&11|-|72615057|72615136</td><td>13</td><td>35</td><td>23</td><td>TCTACAGTGCACGTGTCTCCAGT</td><td>BT20</td><td>21</td><td>22</td><td>42</td><td>23</td><td>11</td><td>13.7</td><td>19.2</td><td>5.3</td><td>18.2</td><td>7.1</td></tr>
<tr><td>>MI0000261|hsa-mir-139&WithFlank&11|-|72615057|72615136@13.35.23&[MIMAT0000250&hsa-miR-139-5p&offsets|0|0;m-186&11|-|72615102|72615124&offsets|0|0]|TCTACAGTGCACGTGTCTCCAGT</td><td>>MI0000261|hsa-mir-139&WithFlank&11|-|72615057|72615136</td><td>13</td><td>35</td><td>23</td><td>TCTACAGTGCACGTGTCTCCAGT</td><td>MB231</td><td>81</td><td>95</td><td>69</td><td>143</td><td>121</td><td>47.5</td><td>39.6</td><td>41.6</td><td>39.8</td><td>64.0</td></tr>
<tr><td>>MI0000263|hsa-mir-7-1&WithFlank&9|-|83969742|83969863@30.54.25&[MIMAT0000252_2&hsa-miR-7-5p&offsets|0|+1;m-401&9|-|83969811|83969834&offsets|0|+1]|TGGAAGACTAGTGATTTTGTTGTTT</td><td>>MI0000263|hsa-mir-7-1&WithFlank&9|-|83969742|83969863</td><td>30</td><td>54</td><td>25</td><td>TGGAAGACTAGTGATTTTGTTGTTT</td><td>BT20</td><td>84</td><td>45</td><td>53</td><td>12</td><td>28</td><td>50.1</td><td>28.6</td><td>42.5</td><td>8.6</td><td>13.8</td></tr>
<tr><td>>MI0000263|hsa-mir-7-1&WithFlank&9|-|83969742|83969863@30.54.25&[MIMAT0000252_2&hsa-miR-7-5p&offsets|0|+1;m-401&9|-|83969811|83969834&offsets|0|+1]|TGGAAGACTAGTGATTTTGTTGTTT</td><td>>MI0000263|hsa-mir-7-1&WithFlank&9|-|83969742|83969863</td><td>30</td><td>54</td><td>25</td><td>TGGAAGACTAGTGATTTTGTTGTTT</td><td>MB231</td><td>129</td><td>79</td><td>12</td><td>221</td><td>109</td><td>4.5</td><td>35.3</td><td>6.2</td><td>33.2</td><td>51.6</td></tr>
<tr><td>>MI0000263|hsa-mir-7-1&WithFlank&9|-|83969742|83969863@30.55.26&[MIMAT0000252_2&hsa-miR-7-5p&offsets|0|+2;m-401&9|-|83969811|83969834&offsets|0|+2]|TGGAAGACTAGTGATTTTGTTGTTTT</td><td>>MI0000263|hsa-mir-7-1&WithFlank&9|-|83969742|83969863</td><td>30</td><td>55</td><td>26</td><td>TGGAAGACTAGTGATTTTGTTGTTTT</td><td>MB231</td><td>18</td><td>12</td><td>2</td><td>13</td><td>16</td><td>3.3</td><td>4.5</td><td>1.1</td><td>6.2</td><td>8.9</td></tr>
<tr><td>>MI0000264|hsa-mir-7-2&WithFlank&15|+|88611819|88611940@38.57.20;MI0000265|hsa-mir-7-3&WithFlank&19|+|4770664|4770785@37.56.20;MI0000263|hsa-mir-7-1&WithFlank&9|-|83969742|83969863@30.49.20&[MIMAT0000252&hsa-miR-7-5p&offsets|0|-4;m-400&15|+|88611856|88611879&offsets|0|-4];[MIMAT0000252_1&hsa-miR-7-5p&offsets|0|-4;m-402&19|+|4770700|4770723&offsets|0|-4];[MIMAT0000252_2&hsa-miR-7-5p&offsets|0|-4;m-401&9|-|83969811|83969834&offsets|0|-4]|TGGAAGACTAGTGATTTTGT</td><td>>MI0000264|hsa-mir-7-2&WithFlank&15|+|88611819|88611940</td><td>38</td><td>57</td><td>20</td><td>TGGAAGACTAGTGATTTTGT</td><td>BT20</td><td>16</td><td>8</td><td>7</td><td>7</td><td>3</td><td>10.8</td><td>4.5</td><td>5.0</td><td>6.8</td><td>1.0</td></tr>
<tr><td>>MI0000264|hsa-mir-7-2&WithFlank&15|+|88611819|88611940@38.57.20(+1C);MI0000265|hsa-mir-7-3&WithFlank&19|+|4770664|4770785@37.56.20(+1C);MI0000263|hsa-mir-7-1&WithFlank&9|-|83969742|83969863@30.49.20(+1C)&[MIMAT0000252&hsa-miR-7-5p&offsets|0|-4(+1C);m-400&15|+|88611856|88611879&offsets|0|-4(+1C)];[MIMAT0000252_1&hsa-miR-7-5p&offsets|0|-4(+1C);m-402&19|+|4770700|4770723&offsets|0|-4(+1C)];[MIMAT0000252_2&hsa-miR-7-5p&offsets|0|-4(+1C);m-401&9|-|83969811|83969834&offsets|0|-4(+1C)]|TGGAAGACTAGTGATTTTGTC</td><td>>MI0000264|hsa-mir-7-2&WithFlank&15|+|88611819|88611940</td><td>38</td><td>57</td><td>20(+1C)</td><td>TGGAAGACTAGTGATTTTGTC</td><td>BT20</td><td>15</td><td>9</td><td>7</td><td>6</td><td>3</td><td>11.1</td><td>7.0</td><td>7.0</td><td>7.4</td><td>0.6</td></tr>
<tr><td>>MI0000264|hsa-mir-7-2&WithFlank&15|+|88611819|88611940@38.57.20(+1C);MI0000265|hsa-mir-7-3&WithFlank&19|+|4770664|4770785@37.56.20(+1C);MI0000263|hsa-mir-7-1&WithFlank&9|-|83969742|83969863@30.49.20(+1C)&[MIMAT0000252&hsa-miR-7-5p&offsets|0|-4(+1C);m-400&15|+|88611856|88611879&offsets|0|-4(+1C)];[MIMAT0000252_1&hsa-miR-7-5p&offsets|0|-4(+1C);m-402&19|+|4770700|4770723&offsets|0|-4(+1C)];[MIMAT0000252_2&hsa-miR-7-5p&offsets|0|-4(+1C);m-401&9|-|83969811|83969834&offsets|0|-4(+1C)]|TGGAAGACTAGTGATTTTGTC</td><td>>MI0000264|hsa-mir-7-2&WithFlank&15|+|88611819|88611940</td><td>38</td><td>57</td><td>20(+1C)</td><td>TGGAAGACTAGTGATTTTGTC</td><td>MB231</td><td>11</td><td>8</td><td>1</td><td>32</td><td>11</td><td>3.2</td><td>1.9</td><td>0.6</td><td>6.0</td><td>7.6</td></tr>
<tr><td>>MI0000264|hsa-mir-7-2&WithFlank&15|+|88611819|88611940@38.58.21;MI0000265|hsa-mir-7-3&WithFlank&19|+|4770664|4770785@37.57.21;MI0000263|hsa-mir-7-1&WithFlank&9|-|83969742|83969863@30.50.21&[MIMAT0000252&hsa-miR-7-5p&offsets|0|-3;m-400&15|+|88611856|88611879&offsets|0|-3];[MIMAT0000252_1&hsa-miR-7-5p&offsets|0|-3;m-402&19|+|4770700|4770723&offsets|0|-3];[MIMAT0000252_2&hsa-miR-7-5p&offsets|0|-3;m-401&9|-|83969811|83969834&offsets|0|-3]|TGGAAGACTAGTGATTTTGTT</td><td>>MI0000264|hsa-mir-7-2&WithFlank&15|+|88611819|88611940</td><td>38</td><td>58</td><td>21</td><td>TGGAAGACTAGTGATTTTGTT</td><td>BT20</td><td>104</td><td>62</td><td>49</td><td>37</td><td>24</td><td>77.4</td><td>45.8</td><td>38.3</td><td>40.2</td><td>6.7</td></tr>
<tr><td>>MI0000264|hsa-mir-7-2&WithFlank&15|+|88611819|88611940@38.58.21;MI0000265|hsa-mir-7-3&WithFlank&19|+|4770664|4770785@37.57.21;MI0000263|hsa-mir-7-1&WithFlank&9|-|83969742|83969863@30.50.21&[MIMAT0000252&hsa-miR-7-5p&offsets|0|-3;m-400&15|+|88611856|88611879&offsets|0|-3];[MIMAT0000252_1&hsa-miR-7-5p&offsets|0|-3;m-402&19|+|4770700|4770723&offsets|0|-3];[MIMAT0000252_2&hsa-miR-7-5p&offsets|0|-3;m-401&9|-|83969811|83969834&offsets|0|-3]|TGGAAGACTAGTGATTTTGTT</td><td>>MI0000264|hsa-mir-7-2&WithFlank&15|+|88611819|88611940</td><td>38</td><td>58</td><td>21</td><td>TGGAAGACTAGTGATTTTGTT</td><td>MB231</td><td>95</td><td>69</td><td>10</td><td>221</td><td>79</td><td>20.4</td><td>15.0</td><td>4.4</td><td>74.3</td><td>44.0</td></tr>
<tr><td>>MI0000264|hsa-mir-7-2&WithFlank&15|+|88611819|88611940@38.58.21(+1A);MI0000265|hsa-mir-7-3&WithFlank&19|+|4770664|4770785@37.57.21(+1A);MI0000263|hsa-mir-7-1&WithFlank&9|-|83969742|83969863@30.50.21(+1A)&[MIMAT0000252&hsa-miR-7-5p&offsets|0|-3(+1A);m-400&15|+|88611856|88611879&offsets|0|-3(+1A)];[MIMAT0000252_1&hsa-miR-7-5p&offsets|0|-3(+1A);m-402&19|+|4770700|4770723&offsets|0|-3(+1A)];[MIMAT0000252_2&hsa-miR-7-5p&offsets|0|-3(+1A);m-401&9|-|83969811|83969834&offsets|0|-3(+1A)]|TGGAAGACTAGTGATTTTGTTA</td><td>>MI0000264|hsa-mir-7-2&WithFlank&15|+|88611819|88611940</td><td>38</td><td>58</td><td>21(+1A)</td><td>TGGAAGACTAGTGATTTTGTTA</td><td>BT20</td><td>26</td><td>8</td><td>11</td><td>7</td><td>3</td><td>32.1</td><td>6.0</td><td>13.4</td><td>7.6</td><td>0.4</td></tr>
<tr><td>>MI0000264|hsa-mir-7-2&WithFlank&15|+|88611819|88611940@38.59.22;MI0000265|hsa-mir-7-3&WithFlank&19|+|4770664|4770785@37.58.22;MI0000263|hsa-mir-7-1&WithFlank&9|-|83969742|83969863@30.51.22&[MIMAT0000252&hsa-miR-7-5p&offsets|0|-2;m-400&15|+|88611856|88611879&offsets|0|-2];[MIMAT0000252_1&hsa-miR-7-5p&offsets|0|-2;m-402&19|+|4770700|4770723&offsets|0|-2];[MIMAT0000252_2&hsa-miR-7-5p&offsets|0|-2;m-401&9|-|83969811|83969834&offsets|0|-2]|TGGAAGACTAGTGATTTTGTTG</td><td>>MI0000264|hsa-mir-7-2&WithFlank&15|+|88611819|88611940</td><td>38</td><td>59</td><td>22</td><td>TGGAAGACTAGTGATTTTGTTG</td><td>BT20</td><td>109</td><td>61</td><td>46</td><td>53</td><td>23</td><td>90.4</td><td>49.2</td><td>41.6</td><td>42.8</td><td>9.3</td></tr>
<tr><td>>MI0000264|hsa-mir-7-2&WithFlank&15|+|88611819|88611940@38.59.22;MI0000265|hsa-mir-7-3&WithFlank&19|+|4770664|4770785@37.58.22;MI0000263|hsa-mir-7-1&WithFlank&9|-|83969742|83969863@30.51.22&[MIMAT0000252&hsa-miR-7-5p&offsets|0|-2;m-400&15|+|88611856|88611879&offsets|0|-2];[MIMAT0000252_1&hsa-miR-7-5p&offsets|0|-2;m-402&19|+|4770700|4770723&offsets|0|-2];[MIMAT0000252_2&hsa-miR-7-5p&offsets|0|-2;m-401&9|-|83969811|83969834&offsets|0|-2]|TGGAAGACTAGTGATTTTGTTG</td><td>>MI0000264|hsa-mir-7-2&WithFlank&15|+|88611819|88611940</td><td>38</td><td>59</td><td>22</td><td>TGGAAGACTAGTGATTTTGTTG</td><td>MB231</td><td>83</td><td>53</td><td>7</td><td>199</td><td>60</td><td>11.2</td><td>17.0</td><td>4.1</td><td>42.3</td><td>25.2</td></tr>
<tr><td>>MI0000264|hsa-mir-7-2&WithFlank&15|+|88611819|88611940@38.59.22(+1A);MI0000265|hsa-mir-7-3&WithFlank&19|+|4770664|4770785@37.58.22(+1A);MI0000263|hsa-mir-7-1&WithFlank&9|-|83969742|83969863@30.51.22(+1A)&[MIMAT0000252&hsa-miR-7-5p&offsets|0|-2(+1A);m-400&15|+|88611856|88611879&offsets|0|-2(+1A)];[MIMAT0000252_1&hsa-miR-7-5p&offsets|0|-2(+1A);m-402&19|+|4770700|4770723&offsets|0|-2(+1A)];[MIMAT0000252_2&hsa-miR-7-5p&offsets|0|-2(+1A);m-401&9|-|83969811|83969834&offsets|0|-2(+1A)]|TGGAAGACTAGTGATTTTGTTGA</td><td>>MI0000264|hsa-mir-7-2&WithFlank&15|+|88611819|88611940</td><td>38</td><td>59</td><td>22(+1A)</td><td>TGGAAGACTAGTGATTTTGTTGA</td><td>BT20</td><td>82</td><td>16</td><td>37</td><td>17</td><td>5</td><td>104.7</td><td>10.8</td><td>49.0</td><td>22.5</td><td>1.2</td></tr>
<tr><td>>MI0000264|hsa-mir-7-2&WithFlank&15|+|88611819|88611940@38.59.22(+1A);MI0000265|hsa-mir-7-3&WithFlank&19|+|4770664|4770785@37.58.22(+1A);MI0000263|hsa-mir-7-1&WithFlank&9|-|83969742|83969863@30.51.22(+1A)&[MIMAT0000252&hsa-miR-7-5p&offsets|0|-2(+1A);m-400&15|+|88611856|88611879&offsets|0|-2(+1A)];[MIMAT0000252_1&hsa-miR-7-5p&offsets|0|-2(+1A);m-402&19|+|4770700|4770723&offsets|0|-2(+1A)];[MIMAT0000252_2&hsa-miR-7-5p&offsets|0|-2(+1A);m-401&9|-|83969811|83969834&offsets|0|-2(+1A)]|TGGAAGACTAGTGATTTTGTTGA</td><td>>MI0000264|hsa-mir-7-2&WithFlank&15|+|88611819|88611940</td><td>38</td><td>59</td><td>22(+1A)</td><td>TGGAAGACTAGTGATTTTGTTGA</td><td>MB231</td><td>16</td><td>7</td><td>2</td><td>38</td><td>11</td><td>3.6</td><td>3.0</td><td>1.2</td><td>4.4</td><td>8.1</td></tr>
<tr><td>>MI0000264|hsa-mir-7-2&WithFlank&15|+|88611819|88611940@38.59.22(+1C);MI0000265|hsa-mir-7-3&WithFlank&19|+|4770664|4770785@37.58.22(+1C);MI0000263|hsa-mir-7-1&WithFlank&9|-|83969742|83969863@30.51.22(+1C)&[MIMAT0000252&hsa-miR-7-5p&offsets|0|-2(+1C);m-400&15|+|88611856|88611879&offsets|0|-2(+1C)];[MIMAT0000252_1&hsa-miR-7-5p&offsets|0|-2(+1C);m-402&19|+|4770700|4770723&offsets|0|-2(+1C)];[MIMAT0000252_2&hsa-miR-7-5p&offsets|0|-2(+1C);m-401&9|-|83969811|83969834&offsets|0|-2(+1C)]|TGGAAGACTAGTGATTTTGTTGC</td><td>>MI0000264|hsa-mir-7-2&WithFlank&15|+|88611819|88611940</td><td>38</td><td>59</td><td>22(+1C)</td><td>TGGAAGACTAGTGATTTTGTTGC</td><td>BT20</td><td>150</td><td>73</td><td>77</td><td>64</td><td>24</td><td>133.1</td><td>43.4</td><td>80.5</td><td>78.2</td><td>11.4</td></tr>
<tr><td>>MI0000264|hsa-mir-7-2&WithFlank&15|+|88611819|88611940@38.59.22(+1C);MI0000265|hsa-mir-7-3&WithFlank&19|+|4770664|4770785@37.58.22(+1C);MI0000263|hsa-mir-7-1&WithFlank&9|-|83969742|83969863@30.51.22(+1C)&[MIMAT0000252&hsa-miR-7-5p&offsets|0|-2(+1C);m-400&15|+|88611856|88611879&offsets|0|-2(+1C)];[MIMAT0000252_1&hsa-miR-7-5p&offsets|0|-2(+1C);m-402&19|+|4770700|4770723&offsets|0|-2(+1C)];[MIMAT0000252_2&hsa-miR-7-5p&offsets|0|-2(+1C);m-401&9|-|83969811|83969834&offsets|0|-2(+1C)]|TGGAAGACTAGTGATTTTGTTGC</td><td>>MI0000264|hsa-mir-7-2&WithFlank&15|+|88611819|88611940</td><td>38</td><td>59</td><td>22(+1C)</td><td>TGGAAGACTAGTGATTTTGTTGC</td><td>MB231</td><td>61</td><td>35</td><td>6</td><td>144</td><td>51</td><td>13.8</td><td>16.9</td><td>3.6</td><td>16.7</td><td>35.7</td></tr>
</table>

<table>
<tr><th></th><th></th><th></th><th></th><th></th><th></th><th></th><th></th><th></th><th></th><th></th><th></th><th></th><th></th><th></th><th></th><th></th></tr>
<tr><td>>MI0000264|hsa-mir-7-2&WithFlank&15|+|88611819|88611940@38.59.22(+1G);MI0000265|hsa-mir-7-3&WithFlank&19|+|4770664|4770785@37.58.22(+1G);MI0000263|hsa-mir-7-1&WithFlank&9|-|83969742|83969863@30.51.22(+1G)&[MIMAT0000252&hsa-miR-7-5p&offsets|0|-2(+1G);m-400&15|+|88611856|88611879&offsets|0|-2(+1G)];[MIMAT0000252_1&hsa-miR-7-5p&offsets|0|-2(+1G);m-402&19|+|4770700|4770723&offsets|0|-2(+1G)];[MIMAT0000252_2&hsa-miR-7-5p&offsets|0|-2(+1G);m-401&9|-|83969811|83969834&offsets|0|-2(+1G)]|TGGAAGACTAGTGATTTTGTTGG</td><td>>MI0000264|hsa-mir-7-2&WithFlank&15|+|88611819|88611940</td><td>38</td><td>59</td><td>22(+1G)</td><td>TGGAAGACTAGTGATTTTGTTGG</td><td>BT20</td><td>104</td><td>11</td><td>18</td><td>10</td><td>2</td><td>135.2</td><td>3.6</td><td>21.9</td><td>14.5</td><td>0.4</td></tr>
<tr><td>>MI0000264|hsa-mir-7-2&WithFlank&15|+|88611819|88611940@38.59.22(+1G);MI0000265|hsa-mir-7-3&WithFlank&19|+|4770664|4770785@37.58.22(+1G);MI0000263|hsa-mir-7-1&WithFlank&9|-|83969742|83969863@30.51.22(+1G)&[MIMAT0000252&hsa-miR-7-5p&offsets|0|-2(+1G);m-400&15|+|88611856|88611879&offsets|0|-2(+1G)];[MIMAT0000252_1&hsa-miR-7-5p&offsets|0|-2(+1G);m-402&19|+|4770700|4770723&offsets|0|-2(+1G)];[MIMAT0000252_2&hsa-miR-7-5p&offsets|0|-2(+1G);m-401&9|-|83969811|83969834&offsets|0|-2(+1G)]|TGGAAGACTAGTGATTTTGTTGG</td><td>>MI0000264|hsa-mir-7-2&WithFlank&15|+|88611819|88611940</td><td>38</td><td>59</td><td>22(+1G)</td><td>TGGAAGACTAGTGATTTTGTTGG</td><td>MB231</td><td>29</td><td>6</td><td>1</td><td>13</td><td>4</td><td>28.2</td><td>4.6</td><td>0.7</td><td>4.1</td><td>3.5</td></tr>
<tr><td>>MI0000264|hsa-mir-7-2&WithFlank&15|+|88611819|88611940@38.59.22(+2C);MI0000265|hsa-mir-7-3&WithFlank&19|+|4770664|4770785@37.58.22(+2C);MI0000263|hsa-mir-7-1&WithFlank&9|-|83969742|83969863@30.51.22(+2C)&[MIMAT0000252&hsa-miR-7-5p&offsets|0|-2(+2C);m-400&15|+|88611856|88611879&offsets|0|-2(+2C)];[MIMAT0000252_1&hsa-miR-7-5p&offsets|0|-2(+2C);m-402&19|+|4770700|4770723&offsets|0|-2(+2C)];[MIMAT0000252_2&hsa-miR-7-5p&offsets|0|-2(+2C);m-401&9|-|83969811|83969834&offsets|0|-2(+2C)]|TGGAAGACTAGTGATTTTGTTGCC</td><td>>MI0000264|hsa-mir-7-2&WithFlank&15|+|88611819|88611940</td><td>38</td><td>59</td><td>22(+2C)</td><td>TGGAAGACTAGTGATTTTGTTGCC</td><td>BT20</td><td>18</td><td>3</td><td>4</td><td>0</td><td>0</td><td>24.8</td><td>1.9</td><td>5.6</td><td>0.6</td><td>0.5</td></tr>
<tr><td>>MI0000264|hsa-mir-7-2&WithFlank&15|+|88611819|88611940@38.60.23;MI0000265|hsa-mir-7-3&WithFlank&19|+|4770664|4770785@37.59.23;MI0000263|hsa-mir-7-1&WithFlank&9|-|83969742|83969863@30.52.23&[MIMAT0000252&hsa-miR-7-5p&offsets|0|-1;m-400&15|+|88611856|88611879&offsets|0|-1];[MIMAT0000252_1&hsa-miR-7-5p&offsets|0|-1;m-402&19|+|4770700|4770723&offsets|0|-1];[MIMAT0000252_2&hsa-miR-7-5p&offsets|0|-1;m-401&9|-|83969811|83969834&offsets|0|-1]|TGGAAGACTAGTGATTTTGTTGT</td><td>>MI0000264|hsa-mir-7-2&WithFlank&15|+|88611819|88611940</td><td>38</td><td>60</td><td>23</td><td>TGGAAGACTAGTGATTTTGTTGT</td><td>BT20</td><td>1112</td><td>486</td><td>548</td><td>417</td><td>177</td><td>977.4</td><td>283.0</td><td>511.9</td><td>453.0</td><td>75.0</td></tr>
<tr><td>>MI0000264|hsa-mir-7-2&WithFlank&15|+|88611819|88611940@38.60.23;MI0000265|hsa-mir-7-3&WithFlank&19|+|4770664|4770785@37.59.23;MI0000263|hsa-mir-7-1&WithFlank&9|-|83969742|83969863@30.52.23&[MIMAT0000252&hsa-miR-7-5p&offsets|0|-1;m-400&15|+|88611856|88611879&offsets|0|-1];[MIMAT0000252_1&hsa-miR-7-5p&offsets|0|-1;m-402&19|+|4770700|4770723&offsets|0|-1];[MIMAT0000252_2&hsa-miR-7-5p&offsets|0|-1;m-401&9|-|83969811|83969834&offsets|0|-1]|TGGAAGACTAGTGATTTTGTTGT</td><td>>MI0000264|hsa-mir-7-2&WithFlank&15|+|88611819|88611940</td><td>38</td><td>60</td><td>23</td><td>TGGAAGACTAGTGATTTTGTTGT</td><td>MB231</td><td>534</td><td>314</td><td>55</td><td>977</td><td>371</td><td>95.9</td><td>115.1</td><td>28.9</td><td>197.7</td><td>199.9</td></tr>
<tr><td>>MI0000264|hsa-mir-7-2&WithFlank&15|+|88611819|88611940@38.60.23;MI0000265|hsa-mir-7-3&WithFlank&19|+|4770664|4770785@37.59.23;MI0000263|hsa-mir-7-1&WithFlank&9|-|83969742|83969863@30.52.23&[MIMAT0000252&hsa-miR-7-5p&offsets|0|-1;m-400&15|+|88611856|88611879&offsets|0|-1];[MIMAT0000252_1&hsa-miR-7-5p&offsets|0|-1;m-402&19|+|4770700|4770723&offsets|0|-1];[MIMAT0000252_2&hsa-miR-7-5p&offsets|0|-1;m-401&9|-|83969811|83969834&offsets|0|-1]|TGGAAGACTAGTGATTTTGTTGT</td><td>>MI0000264|hsa-mir-7-2&WithFlank&15|+|88611819|88611940</td><td>38</td><td>60</td><td>23</td><td>TGGAAGACTAGTGATTTTGTTGT</td><td>MB468</td><td>25</td><td>11</td><td>9</td><td>42</td><td>5</td><td>14.3</td><td>5.4</td><td>3.4</td><td>11.1</td><td>2.0</td></tr>
<tr><td>>MI0000264|hsa-mir-7-2&WithFlank&15|+|88611819|88611940@38.60.23(+1A);MI0000265|hsa-mir-7-3&WithFlank&19|+|4770664|4770785@37.59.23(+1A);MI0000263|hsa-mir-7-1&WithFlank&9|-|83969742|83969863@30.52.23(+1A)&[MIMAT0000252&hsa-miR-7-5p&offsets|0|-1(+1A);m-400&15|+|88611856|88611879&offsets|0|-1(+1A)];[MIMAT0000252_1&hsa-miR-7-5p&offsets|0|-1(+1A);m-402&19|+|4770700|4770723&offsets|0|-1(+1A)];[MIMAT0000252_2&hsa-miR-7-5p&offsets|0|-1(+1A);m-401&9|-|83969811|83969834&offsets|0|-1(+1A)]|TGGAAGACTAGTGATTTTGTTGTA</td><td>>MI0000264|hsa-mir-7-2&WithFlank&15|+|88611819|88611940</td><td>38</td><td>60</td><td>23(+1A)</td><td>TGGAAGACTAGTGATTTTGTTGTA</td><td>BT20</td><td>487</td><td>126</td><td>316</td><td>29</td><td>66</td><td>554.2</td><td>102.9</td><td>361.5</td><td>22.5</td><td>21.5</td></tr>
<tr><td>>MI0000264|hsa-mir-7-2&WithFlank&15|+|88611819|88611940@38.60.23(+1A);MI0000265|hsa-mir-7-3&WithFlank&19|+|4770664|4770785@37.59.23(+1A);MI0000263|hsa-mir-7-1&WithFlank&9|-|83969742|83969863@30.52.23(+1A)&[MIMAT0000252&hsa-miR-7-5p&offsets|0|-1(+1A);m-400&15|+|88611856|88611879&offsets|0|-1(+1A)];[MIMAT0000252_1&hsa-miR-7-5p&offsets|0|-1(+1A);m-402&19|+|4770700|4770723&offsets|0|-1(+1A)];[MIMAT0000252_2&hsa-miR-7-5p&offsets|0|-1(+1A);m-401&9|-|83969811|83969834&offsets|0|-1(+1A)]|TGGAAGACTAGTGATTTTGTTGTA</td><td>>MI0000264|hsa-mir-7-2&WithFlank&15|+|88611819|88611940</td><td>38</td><td>60</td><td>23(+1A)</td><td>TGGAAGACTAGTGATTTTGTTGTA</td><td>MB231</td><td>112</td><td>52</td><td>16</td><td>141</td><td>77</td><td>16.9</td><td>24.5</td><td>7.0</td><td>18.4</td><td>30.4</td></tr>
<tr><td>>MI0000264|hsa-mir-7-2&WithFlank&15|+|88611819|88611940@38.60.23(+1C);MI0000265|hsa-mir-7-3&WithFlank&19|+|4770664|4770785@37.59.23(+1C);MI0000263|hsa-mir-7-1&WithFlank&9|-|83969742|83969863@30.52.23(+1C)&[MIMAT0000252&hsa-miR-7-5p&offsets|0|-1(+1C);m-400&15|+|88611856|88611879&offsets|0|-1(+1C)];[MIMAT0000252_1&hsa-miR-7-5p&offsets|0|-1(+1C);m-402&19|+|4770700|4770723&offsets|0|-1(+1C)];[MIMAT0000252_2&hsa-miR-7-5p&offsets|0|-1(+1C);m-401&9|-|83969811|83969834&offsets|0|-1(+1C)]|TGGAAGACTAGTGATTTTGTTGTC</td><td>>MI0000264|hsa-mir-7-2&WithFlank&15|+|88611819|88611940</td><td>38</td><td>60</td><td>23(+1C)</td><td>TGGAAGACTAGTGATTTTGTTGTC</td><td>BT20</td><td>971</td><td>598</td><td>737</td><td>145</td><td>352</td><td>582.4</td><td>494.0</td><td>641.1</td><td>101.9</td><td>103.8</td></tr>
<tr><td>>MI0000264|hsa-mir-7-2&WithFlank&15|+|88611819|88611940@38.60.23(+1C);MI0000265|hsa-mir-7-3&WithFlank&19|+|4770664|4770785@37.59.23(+1C);MI0000263|hsa-mir-7-1&WithFlank&9|-|83969742|83969863@30.52.23(+1C)&[MIMAT0000252&hsa-miR-7-5p&offsets|0|-1(+1C);m-400&15|+|88611856|88611879&offsets|0|-1(+1C)];[MIMAT0000252_1&hsa-miR-7-5p&offsets|0|-1(+1C);m-402&19|+|4770700|4770723&offsets|0|-1(+1C)];[MIMAT0000252_2&hsa-miR-7-5p&offsets|0|-1(+1C);m-401&9|-|83969811|83969834&offsets|0|-1(+1C)]|TGGAAGACTAGTGATTTTGTTGTC</td><td>>MI0000264|hsa-mir-7-2&WithFlank&15|+|88611819|88611940</td><td>38</td><td>60</td><td>23(+1C)</td><td>TGGAAGACTAGTGATTTTGTTGTC</td><td>MB231</td><td>436</td><td>278</td><td>59</td><td>588</td><td>372</td><td>17.8</td><td>147.4</td><td>13.9</td><td>75.3</td><td>159.1</td></tr>
<tr><td>>MI0000264|hsa-mir-7-2&WithFlank&15|+|88611819|88611940@38.60.23(+1C);MI0000265|hsa-mir-7-3&WithFlank&19|+|4770664|4770785@37.59.23(+1C);MI0000263|hsa-mir-7-1&WithFlank&9|-|83969742|83969863@30.52.23(+1C)&[MIMAT0000252&hsa-miR-7-5p&offsets|0|-1(+1C);m-400&15|+|88611856|88611879&offsets|0|-1(+1C)];[MIMAT0000252_1&hsa-miR-7-5p&offsets|0|-1(+1C);m-402&19|+|4770700|4770723&offsets|0|-1(+1C)];[MIMAT0000252_2&hsa-miR-7-5p&offsets|0|-1(+1C);m-401&9|-|83969811|83969834&offsets|0|-1(+1C)]|TGGAAGACTAGTGATTTTGTTGTC</td><td>>MI0000264|hsa-mir-7-2&WithFlank&15|+|88611819|88611940</td><td>38</td><td>60</td><td>23(+1C)</td><td>TGGAAGACTAGTGATTTTGTTGTC</td><td>MB468</td><td>28</td><td>11</td><td>13</td><td>26</td><td>7</td><td>13.0</td><td>3.9</td><td>4.1</td><td>21.0</td><td>4.4</td></tr>
<tr><td>>MI0000264|hsa-mir-7-2&WithFlank&15|+|88611819|88611940@38.60.23(+1G);MI0000265|hsa-mir-7-3&WithFlank&19|+|4770664|4770785@37.59.23(+1G);MI0000263|hsa-mir-7-1&WithFlank&9|-|83969742|83969863@30.52.23(+1G)&[MIMAT0000252&hsa-miR-7-5p&offsets|0|-1(+1G);m-400&15|+|88611856|88611879&offsets|0|-1(+1G)];[MIMAT0000252_1&hsa-miR-7-5p&offsets|0|-1(+1G);m-402&19|+|4770700|4770723&offsets|0|-1(+1G)];[MIMAT0000252_2&hsa-miR-7-5p&offsets|0|-1(+1G);m-401&9|-|83969811|83969834&offsets|0|-1(+1G)]|TGGAAGACTAGTGATTTTGTTGTG</td><td>>MI0000264|hsa-mir-7-2&WithFlank&15|+|88611819|88611940</td><td>38</td><td>60</td><td>23(+1G)</td><td>TGGAAGACTAGTGATTTTGTTGTG</td><td>BT20</td><td>459</td><td>65</td><td>117</td><td>14</td><td>18</td><td>552.9</td><td>36.1</td><td>131.1</td><td>15.2</td><td>4.2</td></tr>
<tr><td>>MI0000264|hsa-mir-7-2&WithFlank&15|+|88611819|88611940@38.60.23(+1G);MI0000265|hsa-mir-7-3&WithFlank&19|+|4770664|4770785@37.59.23(+1G);MI0000263|hsa-mir-7-1&WithFlank&9|-|83969742|83969863@30.52.23(+1G)&[MIMAT0000252&hsa-miR-7-5p&offsets|0|-1(+1G);m-400&15|+|88611856|88611879&offsets|0|-1(+1G)];[MIMAT0000252_1&hsa-miR-7-5p&offsets|0|-1(+1G);m-402&19|+|4770700|4770723&offsets|0|-1(+1G)];[MIMAT0000252_2&hsa-miR-7-5p&offsets|0|-1(+1G);m-401&9|-|83969811|83969834&offsets|0|-1(+1G)]|TGGAAGACTAGTGATTTTGTTGTG</td><td>>MI0000264|hsa-mir-7-2&WithFlank&15|+|88611819|88611940</td><td>38</td><td>60</td><td>23(+1G)</td><td>TGGAAGACTAGTGATTTTGTTGTG</td><td>MB231</td><td>146</td><td>33</td><td>6</td><td>42</td><td>22</td><td>129.8</td><td>23.8</td><td>3.7</td><td>11.8</td><td>11.5</td></tr>
</table>

<table>
<tr><td>>MI0000264|hsa-mir-7-2&WithFlank&15|+|88611819|88611940@38.61.24;MI0000265|hsa-mir-7-3&WithFlank&19|+|4770664|4770785@37.60.24;MI0000263|hsa-mir-7-1&WithFlank&9|-|83969742|83969863@30.53.24&[MIMAT0000252&hsa-miR-7-5p&offsets|0|0;m-400&15|+|88611856|88611879&offsets|0|0];[MIMAT0000252_1&hsa-miR-7-5p&offsets|0|0;m-402&19|+|4770700|4770723&offsets|0|0];[MIMAT0000252_2&hsa-miR-7-5p&offsets|0|0;m-401&9|-|83969811|83969834&offsets|0|0]|TGGAAGACTAGTGATTTTGTTGTT</td><td>>MI0000264|hsa-mir-7-2&WithFlank&15|+|88611819|88611940</td><td>38</td><td>61</td><td>24</td><td>TGGAAGACTAGTGATTTTGTTGTT</td><td>BT20</td><td>6911</td><td>4104</td><td>5143</td><td>998</td><td>2684</td><td>4051.5</td><td>3231.4</td><td>3826.7</td><td>579.4</td><td>983.4</td></tr>
<tr><td>>MI0000264|hsa-mir-7-2&WithFlank&15|+|88611819|88611940@38.61.24;MI0000265|hsa-mir-7-3&WithFlank&19|+|4770664|4770785@37.60.24;MI0000263|hsa-mir-7-1&WithFlank&9|-|83969742|83969863@30.53.24&[MIMAT0000252&hsa-miR-7-5p&offsets|0|0;m-400&15|+|88611856|88611879&offsets|0|0];[MIMAT0000252_1&hsa-miR-7-5p&offsets|0|0;m-402&19|+|4770700|4770723&offsets|0|0];[MIMAT0000252_2&hsa-miR-7-5p&offsets|0|0;m-401&9|-|83969811|83969834&offsets|0|0]|TGGAAGACTAGTGATTTTGTTGTT</td><td>>MI0000264|hsa-mir-7-2&WithFlank&15|+|88611819|88611940</td><td>38</td><td>61</td><td>24</td><td>TGGAAGACTAGTGATTTTGTTGTT</td><td>MB231</td><td>3921</td><td>2559</td><td>531</td><td>3929</td><td>2747</td><td>211.5</td><td>1078.7</td><td>60.3</td><td>704.2</td><td>908.7</td></tr>
<tr><td>>MI0000264|hsa-mir-7-2&WithFlank&15|+|88611819|88611940@38.61.24;MI0000265|hsa-mir-7-3&WithFlank&19|+|4770664|4770785@37.60.24;MI0000263|hsa-mir-7-1&WithFlank&9|-|83969742|83969863@30.53.24&[MIMAT0000252&hsa-miR-7-5p&offsets|0|0;m-400&15|+|88611856|88611879&offsets|0|0];[MIMAT0000252_1&hsa-miR-7-5p&offsets|0|0;m-402&19|+|4770700|4770723&offsets|0|0];[MIMAT0000252_2&hsa-miR-7-5p&offsets|0|0;m-401&9|-|83969811|83969834&offsets|0|0]|TGGAAGACTAGTGATTTTGTTGTT</td><td>>MI0000264|hsa-mir-7-2&WithFlank&15|+|88611819|88611940</td><td>38</td><td>61</td><td>24</td><td>TGGAAGACTAGTGATTTTGTTGTT</td><td>MB468</td><td>256</td><td>112</td><td>116</td><td>213</td><td>58</td><td>140.6</td><td>55.8</td><td>18.3</td><td>175.1</td><td>47.1</td></tr>
<tr><td>>MI0000264|hsa-mir-7-2&WithFlank&15|+|88611819|88611940@38.61.24(+1A);MI0000265|hsa-mir-7-3&WithFlank&19|+|4770664|4770785@37.60.24(+1A);MI0000263|hsa-mir-7-1&WithFlank&9|-|83969742|83969863@30.53.24(+1A)&[MIMAT0000252&hsa-miR-7-5p&offsets|0|0(+1A);m-400&15|+|88611856|88611879&offsets|0|0(+1A)];[MIMAT0000252_1&hsa-miR-7-5p&offsets|0|0(+1A);m-402&19|+|4770700|4770723&offsets|0|0(+1A)];[MIMAT0000252_2&hsa-miR-7-5p&offsets|0|0(+1A);m-401&9|-|83969811|83969834&offsets|0|0(+1A)]|TGGAAGACTAGTGATTTTGTTGTTA</td><td>>MI0000264|hsa-mir-7-2&WithFlank&15|+|88611819|88611940</td><td>38</td><td>61</td><td>24(+1A)</td><td>TGGAAGACTAGTGATTTTGTTGTTA</td><td>BT20</td><td>74</td><td>30</td><td>48</td><td>6</td><td>18</td><td>51.2</td><td>14.9</td><td>33.7</td><td>3.5</td><td>6.3</td></tr>
<tr><td>>MI0000264|hsa-mir-7-2&WithFlank&15|+|88611819|88611940@38.61.24(+1A);MI0000265|hsa-mir-7-3&WithFlank&19|+|4770664|4770785@37.60.24(+1A);MI0000263|hsa-mir-7-1&WithFlank&9|-|83969742|83969863@30.53.24(+1A)&[MIMAT0000252&hsa-miR-7-5p&offsets|0|0(+1A);m-400&15|+|88611856|88611879&offsets|0|0(+1A)];[MIMAT0000252_1&hsa-miR-7-5p&offsets|0|0(+1A);m-402&19|+|4770700|4770723&offsets|0|0(+1A)];[MIMAT0000252_2&hsa-miR-7-5p&offsets|0|0(+1A);m-401&9|-|83969811|83969834&offsets|0|0(+1A)]|TGGAAGACTAGTGATTTTGTTGTTA</td><td>>MI0000264|hsa-mir-7-2&WithFlank&15|+|88611819|88611940</td><td>38</td><td>61</td><td>24(+1A)</td><td>TGGAAGACTAGTGATTTTGTTGTTA</td><td>MB231</td><td>46</td><td>23</td><td>5</td><td>54</td><td>33</td><td>0.7</td><td>6.2</td><td>2.4</td><td>11.2</td><td>18.5</td></tr>
<tr><td>>MI0000265|hsa-mir-7-3&WithFlank&19|+|4770664|4770785@37.61.25&[MIMAT0000252_1&hsa-miR-7-5p&offsets|0|+1;m-402&19|+|4770700|4770723&offsets|0|+1]|TGGAAGACTAGTGATTTTGTTGTTC</td><td>>MI0000265|hsa-mir-7-3&WithFlank&19|+|4770664|4770785</td><td>37</td><td>61</td><td>25</td><td>TGGAAGACTAGTGATTTTGTTGTTC</td><td>BT20</td><td>11</td><td>6</td><td>7</td><td>1</td><td>4</td><td>6.3</td><td>4.3</td><td>6.3</td><td>0.9</td><td>1.7</td></tr>
<tr><td>>MI0000265|hsa-mir-7-3&WithFlank&19|+|4770664|4770785@37.61.25&[MIMAT0000252_1&hsa-miR-7-5p&offsets|0|+1;m-402&19|+|4770700|4770723&offsets|0|+1]|TGGAAGACTAGTGATTTTGTTGTTC</td><td>>MI0000265|hsa-mir-7-3&WithFlank&19|+|4770664|4770785</td><td>37</td><td>61</td><td>25</td><td>TGGAAGACTAGTGATTTTGTTGTTC</td><td>MB231</td><td>12</td><td>8</td><td>1</td><td>32</td><td>13</td><td>0.2</td><td>3.8</td><td>0.6</td><td>6.3</td><td>9.0</td></tr>
<tr><td>>MI0000266|hsa-mir-10a&WithFlank&17|-|48579832|48579953@28.46.19&[MIMAT0000253&hsa-miR-10a-5p&offsets|0|-4;m-18&17|-|48579905|48579926&offsets|0|-3]|TACCCTGTAGATCCGAATT</td><td>>MI0000266|hsa-mir-10a&WithFlank&17|-|48579832|48579953</td><td>28</td><td>46</td><td>19</td><td>TACCCTGTAGATCCGAATT</td><td>BT20</td><td>10</td><td>5</td><td>13</td><td>6</td><td>2</td><td>8.5</td><td>3.0</td><td>5.4</td><td>6.4</td><td>1.3</td></tr>
<tr><td>>MI0000266|hsa-mir-10a&WithFlank&17|-|48579832|48579953@28.48.21(+1A)&[MIMAT0000253&hsa-miR-10a-5p&offsets|0|-2(+1A);m-18&17|-|48579905|48579926&offsets|0|-1(+1A)]|TACCCTGTAGATCCGAATTTGA</td><td>>MI0000266|hsa-mir-10a&WithFlank&17|-|48579832|48579953</td><td>28</td><td>48</td><td>21(+1A)</td><td>TACCCTGTAGATCCGAATTTGA</td><td>BT20</td><td>54</td><td>16</td><td>77</td><td>33</td><td>7</td><td>37.6</td><td>6.4</td><td>42.7</td><td>30.2</td><td>1.6</td></tr>
<tr><td>>MI0000266|hsa-mir-10a&WithFlank&17|-|48579832|48579953@28.48.21(+1A)&[MIMAT0000253&hsa-miR-10a-5p&offsets|0|-2(+1A);m-18&17|-|48579905|48579926&offsets|0|-1(+1A)]|TACCCTGTAGATCCGAATTTGA</td><td>>MI0000266|hsa-mir-10a&WithFlank&17|-|48579832|48579953</td><td>28</td><td>48</td><td>21(+1A)</td><td>TACCCTGTAGATCCGAATTTGA</td><td>MB231</td><td>10</td><td>8</td><td>8</td><td>25</td><td>12</td><td>1.6</td><td>2.1</td><td>4.1</td><td>6.5</td><td>5.2</td></tr>
<tr><td>>MI0000266|hsa-mir-10a&WithFlank&17|-|48579832|48579953@28.48.21(+1C)&[MIMAT0000253&hsa-miR-10a-5p&offsets|0|-2(+1C);m-18&17|-|48579905|48579926&offsets|0|-1(+1C)]|TACCCTGTAGATCCGAATTTGC</td><td>>MI0000266|hsa-mir-10a&WithFlank&17|-|48579832|48579953</td><td>28</td><td>48</td><td>21(+1C)</td><td>TACCCTGTAGATCCGAATTTGC</td><td>BT20</td><td>147</td><td>89</td><td>219</td><td>149</td><td>42</td><td>55.0</td><td>41.2</td><td>45.5</td><td>122.1</td><td>16.4</td></tr>
<tr><td>>MI0000266|hsa-mir-10a&WithFlank&17|-|48579832|48579953@28.48.21(+1C)&[MIMAT0000253&hsa-miR-10a-5p&offsets|0|-2(+1C);m-18&17|-|48579905|48579926&offsets|0|-1(+1C)]|TACCCTGTAGATCCGAATTTGC</td><td>>MI0000266|hsa-mir-10a&WithFlank&17|-|48579832|48579953</td><td>28</td><td>48</td><td>21(+1C)</td><td>TACCCTGTAGATCCGAATTTGC</td><td>MB231</td><td>49</td><td>54</td><td>34</td><td>105</td><td>63</td><td>13.9</td><td>11.6</td><td>15.5</td><td>21.5</td><td>25.6</td></tr>
<tr><td>>MI0000266|hsa-mir-10a&WithFlank&17|-|48579832|48579953@28.48.21(+1G)&[MIMAT0000253&hsa-miR-10a-5p&offsets|0|-2(+1G);m-18&17|-|48579905|48579926&offsets|0|-1(+1G)]|TACCCTGTAGATCCGAATTTGG</td><td>>MI0000266|hsa-mir-10a&WithFlank&17|-|48579832|48579953</td><td>28</td><td>48</td><td>21(+1G)</td><td>TACCCTGTAGATCCGAATTTGG</td><td>BT20</td><td>71</td><td>15</td><td>42</td><td>18</td><td>3</td><td>60.4</td><td>7.0</td><td>17.4</td><td>21.7</td><td>0.6</td></tr>
<tr><td>>MI0000266|hsa-mir-10a&WithFlank&17|-|48579832|48579953@28.48.21(+1G)&[MIMAT0000253&hsa-miR-10a-5p&offsets|0|-2(+1G);m-18&17|-|48579905|48579926&offsets|0|-1(+1G)]|TACCCTGTAGATCCGAATTTGG</td><td>>MI0000266|hsa-mir-10a&WithFlank&17|-|48579832|48579953</td><td>28</td><td>48</td><td>21(+1G)</td><td>TACCCTGTAGATCCGAATTTGG</td><td>MB231</td><td>17</td><td>8</td><td>4</td><td>9</td><td>4</td><td>10.9</td><td>3.0</td><td>3.0</td><td>3.3</td><td>2.2</td></tr>
<tr><td>>MI0000266|hsa-mir-10a&WithFlank&17|-|48579832|48579953@28.48.21&[MIMAT0000253&hsa-miR-10a-5p&offsets|0|-2;m-18&17|-|48579905|48579926&offsets|0|-1]|TACCCTGTAGATCCGAATTTG</td><td>>MI0000266|hsa-mir-10a&WithFlank&17|-|48579832|48579953</td><td>28</td><td>48</td><td>21</td><td>TACCCTGTAGATCCGAATTTG</td><td>BT20</td><td>29</td><td>16</td><td>48</td><td>35</td><td>11</td><td>20.9</td><td>10.0</td><td>28.8</td><td>27.7</td><td>6.0</td></tr>
<tr><td>>MI0000266|hsa-mir-10a&WithFlank&17|-|48579832|48579953@28.48.21&[MIMAT0000253&hsa-miR-10a-5p&offsets|0|-2;m-18&17|-|48579905|48579926&offsets|0|-1]|TACCCTGTAGATCCGAATTTG</td><td>>MI0000266|hsa-mir-10a&WithFlank&17|-|48579832|48579953</td><td>28</td><td>48</td><td>21</td><td>TACCCTGTAGATCCGAATTTG</td><td>MB231</td><td>11</td><td>12</td><td>9</td><td>27</td><td>10</td><td>3.4</td><td>7.1</td><td>6.4</td><td>12.1</td><td>3.6</td></tr>
<tr><td>>MI0000266|hsa-mir-10a&WithFlank&17|-|48579832|48579953@28.49.22(+1A)&[MIMAT0000253&hsa-miR-10a-5p&offsets|0|-1(+1A);m-18&17|-|48579905|48579926&offsets|0|0(+1A)]|TACCCTGTAGATCCGAATTTGTA</td><td>>MI0000266|hsa-mir-10a&WithFlank&17|-|48579832|48579953</td><td>28</td><td>49</td><td>22(+1A)</td><td>TACCCTGTAGATCCGAATTTGTA</td><td>BT20</td><td>155</td><td>54</td><td>169</td><td>59</td><td>34</td><td>85.9</td><td>19.7</td><td>22.1</td><td>61.4</td><td>22.9</td></tr>
<tr><td>>MI0000266|hsa-mir-10a&WithFlank&17|-|48579832|48579953@28.49.22(+1A)&[MIMAT0000253&hsa-miR-10a-5p&offsets|0|-1(+1A);m-18&17|-|48579905|48579926&offsets|0|0(+1A)]|TACCCTGTAGATCCGAATTTGTA</td><td>>MI0000266|hsa-mir-10a&WithFlank&17|-|48579832|48579953</td><td>28</td><td>49</td><td>22(+1A)</td><td>TACCCTGTAGATCCGAATTTGTA</td><td>MB231</td><td>34</td><td>21</td><td>18</td><td>48</td><td>32</td><td>3.9</td><td>4.1</td><td>8.6</td><td>8.8</td><td>17.5</td></tr>
<tr><td>>MI0000266|hsa-mir-10a&WithFlank&17|-|48579832|48579953@28.49.22(+1U)&[MIMAT0000253&hsa-miR-10a-5p&offsets|0|-1(+1U);m-18&17|-|48579905|48579926&offsets|0|0(+1U)]|TACCCTGTAGATCCGAATTTGTT</td><td>>MI0000266|hsa-mir-10a&WithFlank&17|-|48579832|48579953</td><td>28</td><td>49</td><td>22(+1U)</td><td>TACCCTGTAGATCCGAATTTGTT</td><td>BT20</td><td>18</td><td>7</td><td>18</td><td>7</td><td>3</td><td>10.2</td><td>2.6</td><td>3.4</td><td>7.4</td><td>2.2</td></tr>
<tr><td>>MI0000266|hsa-mir-10a&WithFlank&17|-|48579832|48579953@28.49.22&[MIMAT0000253&hsa-miR-10a-5p&offsets|0|-1;m-18&17|-|48579905|48579926&offsets|0|0]|TACCCTGTAGATCCGAATTTGT</td><td>>MI0000266|hsa-mir-10a&WithFlank&17|-|48579832|48579953</td><td>28</td><td>49</td><td>22</td><td>TACCCTGTAGATCCGAATTTGT</td><td>BT20</td><td>1103</td><td>586</td><td>1639</td><td>1037</td><td>301</td><td>444.2</td><td>213.9</td><td>180.5</td><td>762.5</td><td>101.3</td></tr>
<tr><td>>MI0000266|hsa-mir-10a&WithFlank&17|-|48579832|48579953@28.49.22&[MIMAT0000253&hsa-miR-10a-5p&offsets|0|-1;m-18&17|-|48579905|48579926&offsets|0|0]|TACCCTGTAGATCCGAATTTGT</td><td>>MI0000266|hsa-mir-10a&WithFlank&17|-|48579832|48579953</td><td>28</td><td>49</td><td>22</td><td>TACCCTGTAGATCCGAATTTGT</td><td>MB231</td><td>437</td><td>480</td><td>304</td><td>671</td><td>468</td><td>110.0</td><td>62.0</td><td>137.3</td><td>50.4</td><td>153.1</td></tr>
<tr><td>>MI0000266|hsa-mir-10a&WithFlank&17|-|48579832|48579953@28.50.23(+1A)&[MIMAT0000253&hsa-miR-10a-5p&offsets|0|0(+1A);m-18&17|-|48579905|48579926&offsets|0|+1(+1A)]|TACCCTGTAGATCCGAATTTGTGA</td><td>>MI0000266|hsa-mir-10a&WithFlank&17|-|48579832|48579953</td><td>28</td><td>50</td><td>23(+1A)</td><td>TACCCTGTAGATCCGAATTTGTGA</td><td>BT20</td><td>84</td><td>37</td><td>84</td><td>23</td><td>37</td><td>60.4</td><td>12.8</td><td>57.6</td><td>8.1</td><td>21.7</td></tr>
<tr><td>>MI0000266|hsa-mir-10a&WithFlank&17|-|48579832|48579953@28.50.23(+1A)&[MIMAT0000253&hsa-miR-10a-5p&offsets|0|0(+1A);m-18&17|-|48579905|48579926&offsets|0|+1(+1A)]|TACCCTGTAGATCCGAATTTGTGA</td><td>>MI0000266|hsa-mir-10a&WithFlank&17|-|48579832|48579953</td><td>28</td><td>50</td><td>23(+1A)</td><td>TACCCTGTAGATCCGAATTTGTGA</td><td>MB231</td><td>34</td><td>33</td><td>15</td><td>40</td><td>35</td><td>14.6</td><td>1.4</td><td>3.3</td><td>18.1</td><td>9.4</td></tr>
<tr><td>>MI0000266|hsa-mir-10a&WithFlank&17|-|48579832|48579953@28.50.23&[MIMAT0000253&hsa-miR-10a-5p&offsets|0|0;m-18&17|-|48579905|48579926&offsets|0|+1]|TACCCTGTAGATCCGAATTTGTG</td><td>>MI0000266|hsa-mir-10a&WithFlank&17|-|48579832|48579953</td><td>28</td><td>50</td><td>23</td><td>TACCCTGTAGATCCGAATTTGTG</td><td>BT20</td><td>790</td><td>385</td><td>989</td><td>451</td><td>297</td><td>226.0</td><td>107.4</td><td>521.0</td><td>386.7</td><td>200.7</td></tr>
<tr><td>>MI0000266|hsa-mir-10a&WithFlank&17|-|48579832|48579953@28.50.23&[MIMAT0000253&hsa-miR-10a-5p&offsets|0|0;m-18&17|-|48579905|48579926&offsets|0|+1]|TACCCTGTAGATCCGAATTTGTG</td><td>>MI0000266|hsa-mir-10a&WithFlank&17|-|48579832|48579953</td><td>28</td><td>50</td><td>23</td><td>TACCCTGTAGATCCGAATTTGTG</td><td>MB231</td><td>304</td><td>235</td><td>145</td><td>328</td><td>240</td><td>42.4</td><td>21.1</td><td>50.0</td><td>98.4</td><td>76.0</td></tr>
<tr><td>>MI0000266|hsa-mir-10a&WithFlank&17|-|48579832|48579953@28.51.24&[MIMAT0000253&hsa-miR-10a-5p&offsets|0|+1;m-18&17|-|48579905|48579926&offsets|0|+2]|TACCCTGTAGATCCGAATTTGTGT</td><td>>MI0000266|hsa-mir-10a&WithFlank&17|-|48579832|48579953</td><td>28</td><td>51</td><td>24</td><td>TACCCTGTAGATCCGAATTTGTGT</td><td>BT20</td><td>71</td><td>35</td><td>94</td><td>14</td><td>25</td><td>26.0</td><td>12.7</td><td>35.4</td><td>8.7</td><td>19.0</td></tr>
<tr><td>>MI0000266|hsa-mir-10a&WithFlank&17|-|48579832|48579953@28.51.24&[MIMAT0000253&hsa-miR-10a-5p&offsets|0|+1;m-18&17|-|48579905|48579926&offsets|0|+2]|TACCCTGTAGATCCGAATTTGTGT</td><td>>MI0000266|hsa-mir-10a&WithFlank&17|-|48579832|48579953</td><td>28</td><td>51</td><td>24</td><td>TACCCTGTAGATCCGAATTTGTGT</td><td>MB231</td><td>50</td><td>37</td><td>27</td><td>40</td><td>46</td><td>7.9</td><td>8.7</td><td>12.3</td><td>13.1</td><td>22.7</td></tr>
</table>

<table>
<tr><th></th><th></th><th></th><th></th><th></th><th></th><th></th><th></th><th></th><th></th><th></th><th></th><th></th><th></th><th></th><th></th><th></th></tr>
<tr><td>>MI0000266|hsa-mir-10a&WithFlank&17|-|48579832|48579953@29.48.20(+1C)&[MIMAT0000253&hsa-miR-10a-5p&offsets|+1|-2(+1C);m-18&17|-|48579905|48579926&offsets|+1|-1(+1C)]|ACCCTGTAGATCCGAATTTGC</td><td>>MI0000266|hsa-mir-10a&WithFlank&17|-|48579832|48579953</td><td>29</td><td>48</td><td>20(+1C)</td><td>ACCCTGTAGATCCGAATTTGC</td><td>BT20</td><td>18</td><td>12</td><td>29</td><td>16</td><td>7</td><td>8.2</td><td>5.3</td><td>6.6</td><td>10.4</td><td>2.7</td></tr>
<tr><td>>MI0000266|hsa-mir-10a&WithFlank&17|-|48579832|48579953@29.49.21(+1A)&[MIMAT0000253&hsa-miR-10a-5p&offsets|+1|-1(+1A);m-18&17|-|48579905|48579926&offsets|+1|0(+1A)]|ACCCTGTAGATCCGAATTTGTA</td><td>>MI0000266|hsa-mir-10a&WithFlank&17|-|48579832|48579953</td><td>29</td><td>49</td><td>21(+1A)</td><td>ACCCTGTAGATCCGAATTTGTA</td><td>BT20</td><td>101</td><td>39</td><td>118</td><td>23</td><td>24</td><td>69.7</td><td>9.3</td><td>4.7</td><td>18.6</td><td>13.3</td></tr>
<tr><td>>MI0000266|hsa-mir-10a&WithFlank&17|-|48579832|48579953@29.49.21(+1A)&[MIMAT0000253&hsa-miR-10a-5p&offsets|+1|-1(+1A);m-18&17|-|48579905|48579926&offsets|+1|0(+1A)]|ACCCTGTAGATCCGAATTTGTA</td><td>>MI0000266|hsa-mir-10a&WithFlank&17|-|48579832|48579953</td><td>29</td><td>49</td><td>21(+1A)</td><td>ACCCTGTAGATCCGAATTTGTA</td><td>MB231</td><td>26</td><td>18</td><td>15</td><td>33</td><td>25</td><td>3.2</td><td>2.9</td><td>7.6</td><td>5.7</td><td>11.9</td></tr>
<tr><td>>MI0000266|hsa-mir-10a&WithFlank&17|-|48579832|48579953@29.49.21(+1U)&[MIMAT0000253&hsa-miR-10a-5p&offsets|+1|-1(+1U);m-18&17|-|48579905|48579926&offsets|+1|0(+1U)]|ACCCTGTAGATCCGAATTTGTT</td><td>>MI0000266|hsa-mir-10a&WithFlank&17|-|48579832|48579953</td><td>29</td><td>49</td><td>21(+1U)</td><td>ACCCTGTAGATCCGAATTTGTT</td><td>BT20</td><td>11</td><td>4</td><td>11</td><td>3</td><td>2</td><td>7.1</td><td>1.4</td><td>2.0</td><td>2.1</td><td>1.2</td></tr>
<tr><td>>MI0000266|hsa-mir-10a&WithFlank&17|-|48579832|48579953@29.49.21&[MIMAT0000253&hsa-miR-10a-5p&offsets|+1|-1;m-18&17|-|48579905|48579926&offsets|+1|0]|ACCCTGTAGATCCGAATTTGT</td><td>>MI0000266|hsa-mir-10a&WithFlank&17|-|48579832|48579953</td><td>29</td><td>49</td><td>21</td><td>ACCCTGTAGATCCGAATTTGT</td><td>BT20</td><td>134</td><td>77</td><td>214</td><td>111</td><td>47</td><td>66.1</td><td>28.6</td><td>37.1</td><td>68.2</td><td>16.1</td></tr>
<tr><td>>MI0000266|hsa-mir-10a&WithFlank&17|-|48579832|48579953@29.49.21&[MIMAT0000253&hsa-miR-10a-5p&offsets|+1|-1;m-18&17|-|48579905|48579926&offsets|+1|0]|ACCCTGTAGATCCGAATTTGT</td><td>>MI0000266|hsa-mir-10a&WithFlank&17|-|48579832|48579953</td><td>29</td><td>49</td><td>21</td><td>ACCCTGTAGATCCGAATTTGT</td><td>MB231</td><td>62</td><td>77</td><td>49</td><td>100</td><td>73</td><td>21.6</td><td>3.6</td><td>21.8</td><td>8.7</td><td>9.2</td></tr>
<tr><td>>MI0000266|hsa-mir-10a&WithFlank&17|-|48579832|48579953@29.50.22(+1A)&[MIMAT0000253&hsa-miR-10a-5p&offsets|+1|0(+1A);m-18&17|-|48579905|48579926&offsets|+1|+1(+1A)]|ACCCTGTAGATCCGAATTTGTGA</td><td>>MI0000266|hsa-mir-10a&WithFlank&17|-|48579832|48579953</td><td>29</td><td>50</td><td>22(+1A)</td><td>ACCCTGTAGATCCGAATTTGTGA</td><td>BT20</td><td>75</td><td>37</td><td>77</td><td>19</td><td>39</td><td>42.4</td><td>9.8</td><td>34.3</td><td>5.0</td><td>16.6</td></tr>
<tr><td>>MI0000266|hsa-mir-10a&WithFlank&17|-|48579832|48579953@29.50.22(+1A)&[MIMAT0000253&hsa-miR-10a-5p&offsets|+1|0(+1A);m-18&17|-|48579905|48579926&offsets|+1|+1(+1A)]|ACCCTGTAGATCCGAATTTGTGA</td><td>>MI0000266|hsa-mir-10a&WithFlank&17|-|48579832|48579953</td><td>29</td><td>50</td><td>22(+1A)</td><td>ACCCTGTAGATCCGAATTTGTGA</td><td>MB231</td><td>28</td><td>30</td><td>13</td><td>32</td><td>32</td><td>12.5</td><td>2.4</td><td>4.0</td><td>17.1</td><td>6.6</td></tr>
<tr><td>>MI0000266|hsa-mir-10a&WithFlank&17|-|48579832|48579953@29.50.22&[MIMAT0000253&hsa-miR-10a-5p&offsets|+1|0;m-18&17|-|48579905|48579926&offsets|+1|+1]|ACCCTGTAGATCCGAATTTGTG</td><td>>MI0000266|hsa-mir-10a&WithFlank&17|-|48579832|48579953</td><td>29</td><td>50</td><td>22</td><td>ACCCTGTAGATCCGAATTTGTG</td><td>BT20</td><td>484</td><td>265</td><td>654</td><td>170</td><td>202</td><td>161.1</td><td>18.9</td><td>236.7</td><td>101.5</td><td>113.5</td></tr>
<tr><td>>MI0000266|hsa-mir-10a&WithFlank&17|-|48579832|48579953@29.50.22&[MIMAT0000253&hsa-miR-10a-5p&offsets|+1|0;m-18&17|-|48579905|48579926&offsets|+1|+1]|ACCCTGTAGATCCGAATTTGTG</td><td>>MI0000266|hsa-mir-10a&WithFlank&17|-|48579832|48579953</td><td>29</td><td>50</td><td>22</td><td>ACCCTGTAGATCCGAATTTGTG</td><td>MB231</td><td>234</td><td>206</td><td>125</td><td>232</td><td>187</td><td>29.7</td><td>13.8</td><td>44.0</td><td>95.7</td><td>51.2</td></tr>
<tr><td>>MI0000266|hsa-mir-10a&WithFlank&17|-|48579832|48579953@29.51.23&[MIMAT0000253&hsa-miR-10a-5p&offsets|+1|+1;m-18&17|-|48579905|48579926&offsets|+1|+2]|ACCCTGTAGATCCGAATTTGTGT</td><td>>MI0000266|hsa-mir-10a&WithFlank&17|-|48579832|48579953</td><td>29</td><td>51</td><td>23</td><td>ACCCTGTAGATCCGAATTTGTGT</td><td>BT20</td><td>71</td><td>37</td><td>91</td><td>12</td><td>24</td><td>28.4</td><td>6.8</td><td>14.3</td><td>3.7</td><td>12.9</td></tr>
<tr><td>>MI0000266|hsa-mir-10a&WithFlank&17|-|48579832|48579953@29.51.23&[MIMAT0000253&hsa-miR-10a-5p&offsets|+1|+1;m-18&17|-|48579905|48579926&offsets|+1|+2]|ACCCTGTAGATCCGAATTTGTGT</td><td>>MI0000266|hsa-mir-10a&WithFlank&17|-|48579832|48579953</td><td>29</td><td>51</td><td>23</td><td>ACCCTGTAGATCCGAATTTGTGT</td><td>MB231</td><td>45</td><td>41</td><td>26</td><td>37</td><td>45</td><td>8.4</td><td>10.3</td><td>14.5</td><td>13.4</td><td>21.1</td></tr>
<tr><td>>MI0000266|hsa-mir-10a&WithFlank&17|-|48579832|48579953@69.89.21&[MIMAT0004555&hsa-miR-10a-3p&offsets|0|-1;m-468&17|-|48579865|48579885&offsets|0|0]|CAAATTCGTATCTAGGGGAAT</td><td>>MI0000266|hsa-mir-10a&WithFlank&17|-|48579832|48579953</td><td>69</td><td>89</td><td>21</td><td>CAAATTCGTATCTAGGGGAAT</td><td>BT20</td><td>21</td><td>20</td><td>23</td><td>27</td><td>9</td><td>12.6</td><td>16.0</td><td>26.0</td><td>25.4</td><td>7.5</td></tr>
<tr><td>>MI0000268|hsa-mir-34a&WithFlank&1|-|9151662|9151783@28.49.22&[MIMAT0000255&hsa-miR-34a-5p&offsets|0|0;m-119&1|-|9151735|9151756&offsets|0|0]|TGGCAGTGTCTTAGCTGGTTGT</td><td>>MI0000268|hsa-mir-34a&WithFlank&1|-|9151662|9151783</td><td>28</td><td>49</td><td>22</td><td>TGGCAGTGTCTTAGCTGGTTGT</td><td>BT20</td><td>17</td><td>3</td><td>2</td><td>34</td><td>2</td><td>20.2</td><td>2.8</td><td>1.2</td><td>31.8</td><td>1.5</td></tr>
<tr><td>>MI0000268|hsa-mir-34a&WithFlank&1|-|9151662|9151783@28.50.23&[MIMAT0000255&hsa-miR-34a-5p&offsets|0|+1;m-119&1|-|9151735|9151756&offsets|0|+1]|TGGCAGTGTCTTAGCTGGTTGTT</td><td>>MI0000268|hsa-mir-34a&WithFlank&1|-|9151662|9151783</td><td>28</td><td>50</td><td>23</td><td>TGGCAGTGTCTTAGCTGGTTGTT</td><td>BT20</td><td>22</td><td>7</td><td>2</td><td>12</td><td>4</td><td>24.1</td><td>6.7</td><td>0.3</td><td>12.9</td><td>1.2</td></tr>
<tr><td>>MI0000269|hsa-mir-181a-2&WithFlank&9|+|124692436|124692557@83.103.21(+1U)&[MIMAT0004558&hsa-miR-181a-2-3p&offsets|0|-1(+1U);m-71&9|+|124692518|124692539&offsets|0|-1(+1U)]|ACCACTGACCGTTGACTGTACT</td><td>>MI0000269|hsa-mir-181a-2&WithFlank&9|+|124692436|124692557</td><td>83</td><td>103</td><td>21(+1U)</td><td>ACCACTGACCGTTGACTGTACT</td><td>BT20</td><td>18</td><td>9</td><td>40</td><td>4</td><td>4</td><td>11.6</td><td>4.7</td><td>12.2</td><td>2.4</td><td>1.8</td></tr>
<tr><td>>MI0000269|hsa-mir-181a-2&WithFlank&9|+|124692436|124692557@83.103.21(+1U)&[MIMAT0004558&hsa-miR-181a-2-3p&offsets|0|-1(+1U);m-71&9|+|124692518|124692539&offsets|0|-1(+1U)]|ACCACTGACCGTTGACTGTACT</td><td>>MI0000269|hsa-mir-181a-2&WithFlank&9|+|124692436|124692557</td><td>83</td><td>103</td><td>21(+1U)</td><td>ACCACTGACCGTTGACTGTACT</td><td>MB231</td><td>22</td><td>21</td><td>31</td><td>24</td><td>27</td><td>3.0</td><td>4.3</td><td>12.9</td><td>8.7</td><td>13.0</td></tr>
<tr><td>>MI0000269|hsa-mir-181a-2&WithFlank&9|+|124692436|124692557@83.104.22&[MIMAT0004558&hsa-miR-181a-2-3p&offsets|0|0;m-71&9|+|124692518|124692539&offsets|0|0]|ACCACTGACCGTTGACTGTACC</td><td>>MI0000269|hsa-mir-181a-2&WithFlank&9|+|124692436|124692557</td><td>83</td><td>104</td><td>22</td><td>ACCACTGACCGTTGACTGTACC</td><td>BT20</td><td>34</td><td>20</td><td>94</td><td>10</td><td>12</td><td>23.6</td><td>8.2</td><td>24.9</td><td>6.7</td><td>6.7</td></tr>
<tr><td>>MI0000269|hsa-mir-181a-2&WithFlank&9|+|124692436|124692557@83.104.22&[MIMAT0004558&hsa-miR-181a-2-3p&offsets|0|0;m-71&9|+|124692518|124692539&offsets|0|0]|ACCACTGACCGTTGACTGTACC</td><td>>MI0000269|hsa-mir-181a-2&WithFlank&9|+|124692436|124692557</td><td>83</td><td>104</td><td>22</td><td>ACCACTGACCGTTGACTGTACC</td><td>MB231</td><td>63</td><td>76</td><td>94</td><td>50</td><td>70</td><td>4.2</td><td>23.3</td><td>38.6</td><td>11.6</td><td>20.0</td></tr>
<tr><td>>MI0000270|hsa-mir-181b-1&WithFlank&1|-|198858867|198858988@42.61.20;MI0000683|hsa-mir-181b-2&WithFlank&9|+|124693704|124693804@22.41.20&[MIMAT0000257&hsa-miR-181b-5p&offsets|0|-3;m-73&1|-|198858925|198858947&offsets|0|-3];[MIMAT0000257_1&hsa-miR-181b-5p&offsets|0|-3;m-72&9|+|124693725|124693747&offsets|0|-3]|AACATTCATTGCTGTCGGTG</td><td>>MI0000270|hsa-mir-181b-1&WithFlank&1|-|198858867|198858988</td><td>42</td><td>61</td><td>20</td><td>AACATTCATTGCTGTCGGTG</td><td>MB231</td><td>12</td><td>15</td><td>6</td><td>34</td><td>11</td><td>2.3</td><td>8.1</td><td>5.2</td><td>11.5</td><td>4.3</td></tr>
<tr><td>>MI0000270|hsa-mir-181b-1&WithFlank&1|-|198858867|198858988@42.62.21;MI0000683|hsa-mir-181b-2&WithFlank&9|+|124693704|124693804@22.42.21&[MIMAT0000257&hsa-miR-181b-5p&offsets|0|-2;m-73&1|-|198858925|198858947&offsets|0|-2];[MIMAT0000257_1&hsa-miR-181b-5p&offsets|0|-2;m-72&9|+|124693725|124693747&offsets|0|-2]|AACATTCATTGCTGTCGGTGG</td><td>>MI0000270|hsa-mir-181b-1&WithFlank&1|-|198858867|198858988</td><td>42</td><td>62</td><td>21</td><td>AACATTCATTGCTGTCGGTGG</td><td>MB231</td><td>23</td><td>28</td><td>8</td><td>30</td><td>25</td><td>5.1</td><td>11.3</td><td>5.8</td><td>12.6</td><td>6.4</td></tr>
<tr><td>>MI0000270|hsa-mir-181b-1&WithFlank&1|-|198858867|198858988@42.63.22;MI0000683|hsa-mir-181b-2&WithFlank&9|+|124693704|124693804@22.43.22&[MIMAT0000257&hsa-miR-181b-5p&offsets|0|-1;m-73&1|-|198858925|198858947&offsets|0|-1];[MIMAT0000257_1&hsa-miR-181b-5p&offsets|0|-1;m-72&9|+|124693725|124693747&offsets|0|-1]|AACATTCATTGCTGTCGGTGGG</td><td>>MI0000270|hsa-mir-181b-1&WithFlank&1|-|198858867|198858988</td><td>42</td><td>63</td><td>22</td><td>AACATTCATTGCTGTCGGTGGG</td><td>MB231</td><td>21</td><td>24</td><td>13</td><td>20</td><td>17</td><td>5.1</td><td>7.7</td><td>9.1</td><td>11.0</td><td>4.3</td></tr>
<tr><td>>MI0000270|hsa-mir-181b-1&WithFlank&1|-|198858867|198858988@42.63.22(+1A);MI0000683|hsa-mir-181b-2&WithFlank&9|+|124693704|124693804@22.43.22(+1A)&[MIMAT0000257&hsa-miR-181b-5p&offsets|0|-1(+1A);m-73&1|-|198858925|198858947&offsets|0|-1(+1A)];[MIMAT0000257_1&hsa-miR-181b-5p&offsets|0|-1(+1A);m-72&9|+|124693725|124693747&offsets|0|-1(+1A)]|AACATTCATTGCTGTCGGTGGGA</td><td>>MI0000270|hsa-mir-181b-1&WithFlank&1|-|198858867|198858988</td><td>42</td><td>63</td><td>22(+1A)</td><td>AACATTCATTGCTGTCGGTGGGA</td><td>MB231</td><td>14</td><td>14</td><td>10</td><td>19</td><td>11</td><td>2.2</td><td>4.1</td><td>7.0</td><td>8.2</td><td>5.1</td></tr>
<tr><td>>MI0000270|hsa-mir-181b-1&WithFlank&1|-|198858867|198858988@42.63.22(+1C);MI0000683|hsa-mir-181b-2&WithFlank&9|+|124693704|124693804@22.43.22(+1C)&[MIMAT0000257&hsa-miR-181b-5p&offsets|0|-1(+1C);m-73&1|-|198858925|198858947&offsets|0|-1(+1C)];[MIMAT0000257_1&hsa-miR-181b-5p&offsets|0|-1(+1C);m-72&9|+|124693725|124693747&offsets|0|-1(+1C)]|AACATTCATTGCTGTCGGTGGGC</td><td>>MI0000270|hsa-mir-181b-1&WithFlank&1|-|198858867|198858988</td><td>42</td><td>63</td><td>22(+1C)</td><td>AACATTCATTGCTGTCGGTGGGC</td><td>MB231</td><td>18</td><td>19</td><td>8</td><td>36</td><td>21</td><td>3.3</td><td>7.9</td><td>5.1</td><td>18.1</td><td>13.8</td></tr>
<tr><td>>MI0000270|hsa-mir-181b-1&WithFlank&1|-|198858867|198858988@42.64.23;MI0000683|hsa-mir-181b-2&WithFlank&9|+|124693704|124693804@22.44.23&[MIMAT0000257&hsa-miR-181b-5p&offsets|0|0;m-73&1|-|198858925|198858947&offsets|0|0];[MIMAT0000257_1&hsa-miR-181b-5p&offsets|0|0;m-72&9|+|124693725|124693747&offsets|0|0]|AACATTCATTGCTGTCGGTGGGT</td><td>>MI0000270|hsa-mir-181b-1&WithFlank&1|-|198858867|198858988</td><td>42</td><td>64</td><td>23</td><td>AACATTCATTGCTGTCGGTGGGT</td><td>BT20</td><td>55</td><td>51</td><td>75</td><td>41</td><td>37</td><td>31.6</td><td>26.1</td><td>39.3</td><td>28.1</td><td>15.1</td></tr>
<tr><td>>MI0000270|hsa-mir-181b-1&WithFlank&1|-|198858867|198858988@42.64.23;MI0000683|hsa-mir-181b-2&WithFlank&9|+|124693704|124693804@22.44.23&[MIMAT0000257&hsa-miR-181b-5p&offsets|0|0;m-73&1|-|198858925|198858947&offsets|0|0];[MIMAT0000257_1&hsa-miR-181b-5p&offsets|0|0;m-72&9|+|124693725|124693747&offsets|0|0]|AACATTCATTGCTGTCGGTGGGT</td><td>>MI0000270|hsa-mir-181b-1&WithFlank&1|-|198858867|198858988</td><td>42</td><td>64</td><td>23</td><td>AACATTCATTGCTGTCGGTGGGT</td><td>MB231</td><td>147</td><td>157</td><td>64</td><td>213</td><td>146</td><td>22.8</td><td>67.4</td><td>44.4</td><td>76.5</td><td>73.3</td></tr>
<tr><td>>MI0000270|hsa-mir-181b-1&WithFlank&1|-|198858867|198858988@42.64.23;MI0000683|hsa-mir-181b-2&WithFlank&9|+|124693704|124693804@22.44.23&[MIMAT0000257&hsa-miR-181b-5p&offsets|0|0;m-73&1|-|198858925|198858947&offsets|0|0];[MIMAT0000257_1&hsa-miR-181b-5p&offsets|0|0;m-72&9|+|124693725|124693747&offsets|0|0]|AACATTCATTGCTGTCGGTGGGT</td><td>>MI0000270|hsa-mir-181b-1&WithFlank&1|-|198858867|198858988</td><td>42</td><td>64</td><td>23</td><td>AACATTCATTGCTGTCGGTGGGT</td><td>MB468</td><td>11</td><td>7</td><td>7</td><td>15</td><td>3</td><td>6.1</td><td>5.1</td><td>1.2</td><td>9.0</td><td>0.6</td></tr>
</table>

<table>
<tr><td></td><td></td><td></td><td></td><td></td><td></td><td></td><td></td><td></td><td></td><td></td><td></td><td></td><td></td><td></td><td></td><td></td></tr>
<tr><td>>MI0000270|hsa-mir-181b-1&WithFlank&1|-|198858867|198858988@42.64.23(+1C);MI0000683|hsa-mir-181b-2&WithFlank&9|+|124693704|124693804@22.44.23(+1C)&[MIMAT0000257&hsa-miR-181b-5p&offsets|0|0(+1C);m-73&1|-|198858925|198858947&offsets|0|0(+1C)];[MIMAT0000257_1&hsa-miR-181b-5p&offsets|0|0(+1C);m-72&9|+|124693725|124693747&offsets|0|0(+1C)]|AACATTCATTGCTGTCGGTGGGTC</td><td>>MI0000270|hsa-mir-181b-1&WithFlank&1|-|198858867|198858988</td><td>42</td><td>64</td><td>23(+1C)</td><td>AACATTCATTGCTGTCGGTGGGTC</td><td>MB231</td><td>11</td><td>12</td><td>3</td><td>61</td><td>24</td><td>1.8</td><td>6.6</td><td>2.3</td><td>28.6</td><td>20.6</td></tr>
<tr><td>>MI0000270|hsa-mir-181b-1&WithFlank&1|-|198858867|198858988@42.65.24;MI0000683|hsa-mir-181b-2&WithFlank&9|+|124693704|124693804@22.45.24&[MIMAT0000257&hsa-miR-181b-5p&offsets|0|+1;m-73&1|-|198858925|198858947&offsets|0|+1];[MIMAT0000257_1&hsa-miR-181b-5p&offsets|0|+1;m-72&9|+|124693725|124693747&offsets|0|+1]|AACATTCATTGCTGTCGGTGGGTT</td><td>>MI0000270|hsa-mir-181b-1&WithFlank&1|-|198858867|198858988</td><td>42</td><td>65</td><td>24</td><td>AACATTCATTGCTGTCGGTGGGTT</td><td>BT20</td><td>54</td><td>45</td><td>32</td><td>66</td><td>28</td><td>41.7</td><td>32.3</td><td>9.3</td><td>56.9</td><td>18.3</td></tr>
<tr><td>>MI0000270|hsa-mir-181b-1&WithFlank&1|-|198858867|198858988@42.65.24;MI0000683|hsa-mir-181b-2&WithFlank&9|+|124693704|124693804@22.45.24&[MIMAT0000257&hsa-miR-181b-5p&offsets|0|+1;m-73&1|-|198858925|198858947&offsets|0|+1];[MIMAT0000257_1&hsa-miR-181b-5p&offsets|0|+1;m-72&9|+|124693725|124693747&offsets|0|+1]|AACATTCATTGCTGTCGGTGGGTT</td><td>>MI0000270|hsa-mir-181b-1&WithFlank&1|-|198858867|198858988</td><td>42</td><td>65</td><td>24</td><td>AACATTCATTGCTGTCGGTGGGTT</td><td>MB231</td><td>91</td><td>99</td><td>29</td><td>368</td><td>161</td><td>8.7</td><td>54.7</td><td>23.1</td><td>107.8</td><td>110.5</td></tr>
<tr><td>>MI0000272|hsa-mir-182&WithFlank&7|-|129770377|129770498@29.49.21(+1G)&[MIMAT0000259&hsa-miR-182-5p&offsets|0|-3(+1G);m-22&7|-|129770449|129770470&offsets|0|-1(+1G)]|TTTGGCAATGGTAGAACTCACG</td><td>>MI0000272|hsa-mir-182&WithFlank&7|-|129770377|129770498</td><td>29</td><td>49</td><td>21(+1G)</td><td>TTTGGCAATGGTAGAACTCACG</td><td>BT20</td><td>27</td><td>5</td><td>10</td><td>2</td><td>1</td><td>33.0</td><td>1.2</td><td>7.4</td><td>3.6</td><td>0.6</td></tr>
<tr><td>>MI0000272|hsa-mir-182&WithFlank&7|-|129770377|129770498@29.49.21&[MIMAT0000259&hsa-miR-182-5p&offsets|0|-3;m-22&7|-|129770449|129770470&offsets|0|-1]|TTTGGCAATGGTAGAACTCAC</td><td>>MI0000272|hsa-mir-182&WithFlank&7|-|129770377|129770498</td><td>29</td><td>49</td><td>21</td><td>TTTGGCAATGGTAGAACTCAC</td><td>BT20</td><td>36</td><td>28</td><td>54</td><td>67</td><td>9</td><td>21.5</td><td>10.4</td><td>21.8</td><td>64.0</td><td>0.5</td></tr>
<tr><td>>MI0000272|hsa-mir-182&WithFlank&7|-|129770377|129770498@29.50.22(+1A)&[MIMAT0000259&hsa-miR-182-5p&offsets|0|-2(+1A);m-22&7|-|129770449|129770470&offsets|0|0(+1A)]|TTTGGCAATGGTAGAACTCACAA</td><td>>MI0000272|hsa-mir-182&WithFlank&7|-|129770377|129770498</td><td>29</td><td>50</td><td>22(+1A)</td><td>TTTGGCAATGGTAGAACTCACAA</td><td>BT20</td><td>12</td><td>6</td><td>12</td><td>2</td><td>1</td><td>12.3</td><td>2.0</td><td>8.1</td><td>3.3</td><td>1.2</td></tr>
<tr><td>>MI0000272|hsa-mir-182&WithFlank&7|-|129770377|129770498@29.50.22(+1U)&[MIMAT0000259&hsa-miR-182-5p&offsets|0|-2(+1U);m-22&7|-|129770449|129770470&offsets|0|0(+1U)]|TTTGGCAATGGTAGAACTCACAT</td><td>>MI0000272|hsa-mir-182&WithFlank&7|-|129770377|129770498</td><td>29</td><td>50</td><td>22(+1U)</td><td>TTTGGCAATGGTAGAACTCACAT</td><td>BT20</td><td>45</td><td>34</td><td>47</td><td>14</td><td>10</td><td>30.9</td><td>14.2</td><td>28.9</td><td>13.7</td><td>3.2</td></tr>
<tr><td>>MI0000272|hsa-mir-182&WithFlank&7|-|129770377|129770498@29.50.22(+2U)&[MIMAT0000259&hsa-miR-182-5p&offsets|0|-2(+2U);m-22&7|-|129770449|129770470&offsets|0|0(+2U)]|TTTGGCAATGGTAGAACTCACATT</td><td>>MI0000272|hsa-mir-182&WithFlank&7|-|129770377|129770498</td><td>29</td><td>50</td><td>22(+2U)</td><td>TTTGGCAATGGTAGAACTCACATT</td><td>BT20</td><td>23</td><td>6</td><td>17</td><td>0</td><td>0</td><td>29.4</td><td>0.8</td><td>16.2</td><td>0.7</td><td>0.7</td></tr>
<tr><td>>MI0000272|hsa-mir-182&WithFlank&7|-|129770377|129770498@29.50.22&[MIMAT0000259&hsa-miR-182-5p&offsets|0|-2;m-22&7|-|129770449|129770470&offsets|0|0]|TTTGGCAATGGTAGAACTCACA</td><td>>MI0000272|hsa-mir-182&WithFlank&7|-|129770377|129770498</td><td>29</td><td>50</td><td>22</td><td>TTTGGCAATGGTAGAACTCACA</td><td>BT20</td><td>282</td><td>173</td><td>319</td><td>91</td><td>64</td><td>230.5</td><td>50.7</td><td>143.8</td><td>77.4</td><td>1.5</td></tr>
<tr><td>>MI0000272|hsa-mir-182&WithFlank&7|-|129770377|129770498@29.50.22&[MIMAT0000259&hsa-miR-182-5p&offsets|0|-2;m-22&7|-|129770449|129770470&offsets|0|0]|TTTGGCAATGGTAGAACTCACA</td><td>>MI0000272|hsa-mir-182&WithFlank&7|-|129770377|129770498</td><td>29</td><td>50</td><td>22</td><td>TTTGGCAATGGTAGAACTCACA</td><td>MB231</td><td>19</td><td>22</td><td>11</td><td>48</td><td>18</td><td>3.4</td><td>1.2</td><td>6.9</td><td>8.1</td><td>6.1</td></tr>
<tr><td>>MI0000272|hsa-mir-182&WithFlank&7|-|129770377|129770498@29.51.23(+1A)&[MIMAT0000259&hsa-miR-182-5p&offsets|0|-1(+1A);m-22&7|-|129770449|129770470&offsets|0|+1(+1A)]|TTTGGCAATGGTAGAACTCACACA</td><td>>MI0000272|hsa-mir-182&WithFlank&7|-|129770377|129770498</td><td>29</td><td>51</td><td>23(+1A)</td><td>TTTGGCAATGGTAGAACTCACACA</td><td>BT20</td><td>413</td><td>186</td><td>570</td><td>40</td><td>63</td><td>384.3</td><td>50.2</td><td>447.3</td><td>27.9</td><td>18.1</td></tr>
<tr><td>>MI0000272|hsa-mir-182&WithFlank&7|-|129770377|129770498@29.51.23(+1A)&[MIMAT0000259&hsa-miR-182-5p&offsets|0|-1(+1A);m-22&7|-|129770449|129770470&offsets|0|+1(+1A)]|TTTGGCAATGGTAGAACTCACACA</td><td>>MI0000272|hsa-mir-182&WithFlank&7|-|129770377|129770498</td><td>29</td><td>51</td><td>23(+1A)</td><td>TTTGGCAATGGTAGAACTCACACA</td><td>MB231</td><td>20</td><td>15</td><td>10</td><td>13</td><td>17</td><td>2.8</td><td>1.9</td><td>5.7</td><td>5.8</td><td>10.9</td></tr>
<tr><td>>MI0000272|hsa-mir-182&WithFlank&7|-|129770377|129770498@29.51.23(+1C)&[MIMAT0000259&hsa-miR-182-5p&offsets|0|-1(+1C);m-22&7|-|129770449|129770470&offsets|0|+1(+1C)]|TTTGGCAATGGTAGAACTCACACC</td><td>>MI0000272|hsa-mir-182&WithFlank&7|-|129770377|129770498</td><td>29</td><td>51</td><td>23(+1C)</td><td>TTTGGCAATGGTAGAACTCACACC</td><td>BT20</td><td>524</td><td>412</td><td>771</td><td>89</td><td>150</td><td>256.7</td><td>118.1</td><td>358.1</td><td>55.4</td><td>44.6</td></tr>
<tr><td>>MI0000272|hsa-mir-182&WithFlank&7|-|129770377|129770498@29.51.23(+1C)&[MIMAT0000259&hsa-miR-182-5p&offsets|0|-1(+1C);m-22&7|-|129770449|129770470&offsets|0|+1(+1C)]|TTTGGCAATGGTAGAACTCACACC</td><td>>MI0000272|hsa-mir-182&WithFlank&7|-|129770377|129770498</td><td>29</td><td>51</td><td>23(+1C)</td><td>TTTGGCAATGGTAGAACTCACACC</td><td>MB231</td><td>39</td><td>36</td><td>20</td><td>27</td><td>38</td><td>9.2</td><td>8.3</td><td>8.4</td><td>9.9</td><td>25.2</td></tr>
<tr><td>>MI0000272|hsa-mir-182&WithFlank&7|-|129770377|129770498@29.51.23(+1G)&[MIMAT0000259&hsa-miR-182-5p&offsets|0|-1(+1G);m-22&7|-|129770449|129770470&offsets|0|+1(+1G)]|TTTGGCAATGGTAGAACTCACACG</td><td>>MI0000272|hsa-mir-182&WithFlank&7|-|129770377|129770498</td><td>29</td><td>51</td><td>23(+1G)</td><td>TTTGGCAATGGTAGAACTCACACG</td><td>BT20</td><td>349</td><td>85</td><td>187</td><td>14</td><td>14</td><td>353.5</td><td>19.5</td><td>127.1</td><td>13.8</td><td>4.7</td></tr>
<tr><td>>MI0000272|hsa-mir-182&WithFlank&7|-|129770377|129770498@29.51.23(+1G)&[MIMAT0000259&hsa-miR-182-5p&offsets|0|-1(+1G);m-22&7|-|129770449|129770470&offsets|0|+1(+1G)]|TTTGGCAATGGTAGAACTCACACG</td><td>>MI0000272|hsa-mir-182&WithFlank&7|-|129770377|129770498</td><td>29</td><td>51</td><td>23(+1G)</td><td>TTTGGCAATGGTAGAACTCACACG</td><td>MB231</td><td>20</td><td>7</td><td>3</td><td>3</td><td>4</td><td>17.0</td><td>2.0</td><td>2.4</td><td>0.9</td><td>3.2</td></tr>
<tr><td>>MI0000272|hsa-mir-182&WithFlank&7|-|129770377|129770498@29.51.23&[MIMAT0000259&hsa-miR-182-5p&offsets|0|-1;m-22&7|-|129770449|129770470&offsets|0|+1]|TTTGGCAATGGTAGAACTCACAC</td><td>>MI0000272|hsa-mir-182&WithFlank&7|-|129770377|129770498</td><td>29</td><td>51</td><td>23</td><td>TTTGGCAATGGTAGAACTCACAC</td><td>BT20</td><td>178</td><td>169</td><td>211</td><td>61</td><td>58</td><td>87.9</td><td>36.9</td><td>84.9</td><td>48.1</td><td>11.6</td></tr>
<tr><td>>MI0000272|hsa-mir-182&WithFlank&7|-|129770377|129770498@29.51.23&[MIMAT0000259&hsa-miR-182-5p&offsets|0|-1;m-22&7|-|129770449|129770470&offsets|0|+1]|TTTGGCAATGGTAGAACTCACAC</td><td>>MI0000272|hsa-mir-182&WithFlank&7|-|129770377|129770498</td><td>29</td><td>51</td><td>23</td><td>TTTGGCAATGGTAGAACTCACAC</td><td>MB231</td><td>12</td><td>16</td><td>7</td><td>24</td><td>12</td><td>2.0</td><td>4.3</td><td>4.0</td><td>7.2</td><td>4.9</td></tr>
<tr><td>>MI0000272|hsa-mir-182&WithFlank&7|-|129770377|129770498@29.52.24(+1A)&[MIMAT0000259&hsa-miR-182-5p&offsets|0|0(+1A);m-22&7|-|129770449|129770470&offsets|0|+2(+1A)]|TTTGGCAATGGTAGAACTCACACTA</td><td>>MI0000272|hsa-mir-182&WithFlank&7|-|129770377|129770498</td><td>29</td><td>52</td><td>24(+1A)</td><td>TTTGGCAATGGTAGAACTCACACTA</td><td>BT20</td><td>138</td><td>70</td><td>138</td><td>12</td><td>34</td><td>99.9</td><td>21.6</td><td>37.8</td><td>4.3</td><td>11.1</td></tr>
<tr><td>>MI0000272|hsa-mir-182&WithFlank&7|-|129770377|129770498@29.52.24(+1A)&[MIMAT0000259&hsa-miR-182-5p&offsets|0|0(+1A);m-22&7|-|129770449|129770470&offsets|0|+2(+1A)]|TTTGGCAATGGTAGAACTCACACTA</td><td>>MI0000272|hsa-mir-182&WithFlank&7|-|129770377|129770498</td><td>29</td><td>52</td><td>24(+1A)</td><td>TTTGGCAATGGTAGAACTCACACTA</td><td>MB231</td><td>12</td><td>8</td><td>7</td><td>8</td><td>13</td><td>0.7</td><td>0.4</td><td>3.5</td><td>3.2</td><td>7.9</td></tr>
<tr><td>>MI0000272|hsa-mir-182&WithFlank&7|-|129770377|129770498@29.52.24(+1U)&[MIMAT0000259&hsa-miR-182-5p&offsets|0|0(+1U);m-22&7|-|129770449|129770470&offsets|0|+2(+1U)]|TTTGGCAATGGTAGAACTCACACTT</td><td>>MI0000272|hsa-mir-182&WithFlank&7|-|129770377|129770498</td><td>29</td><td>52</td><td>24(+1U)</td><td>TTTGGCAATGGTAGAACTCACACTT</td><td>BT20</td><td>25</td><td>16</td><td>23</td><td>3</td><td>7</td><td>14.7</td><td>4.4</td><td>7.5</td><td>1.6</td><td>1.9</td></tr>
<tr><td>>MI0000272|hsa-mir-182&WithFlank&7|-|129770377|129770498@29.52.24&[MIMAT0000259&hsa-miR-182-5p&offsets|0|0;m-22&7|-|129770449|129770470&offsets|0|+2]|TTTGGCAATGGTAGAACTCACACT</td><td>>MI0000272|hsa-mir-182&WithFlank&7|-|129770377|129770498</td><td>29</td><td>52</td><td>24</td><td>TTTGGCAATGGTAGAACTCACACT</td><td>BT20</td><td>3666</td><td>2733</td><td>5379</td><td>594</td><td>1032</td><td>1811.7</td><td>467.4</td><td>1861.5</td><td>253.8</td><td>233.1</td></tr>
<tr><td>>MI0000272|hsa-mir-182&WithFlank&7|-|129770377|129770498@29.52.24&[MIMAT0000259&hsa-miR-182-5p&offsets|0|0;m-22&7|-|129770449|129770470&offsets|0|+2]|TTTGGCAATGGTAGAACTCACACT</td><td>>MI0000272|hsa-mir-182&WithFlank&7|-|129770377|129770498</td><td>29</td><td>52</td><td>24</td><td>TTTGGCAATGGTAGAACTCACACT</td><td>MB231</td><td>332</td><td>322</td><td>173</td><td>170</td><td>265</td><td>63.3</td><td>73.7</td><td>68.7</td><td>58.5</td><td>142.7</td></tr>
<tr><td>>MI0000272|hsa-mir-182&WithFlank&7|-|129770377|129770498@29.52.24&[MIMAT0000259&hsa-miR-182-5p&offsets|0|0;m-22&7|-|129770449|129770470&offsets|0|+2]|TTTGGCAATGGTAGAACTCACACT</td><td>>MI0000272|hsa-mir-182&WithFlank&7|-|129770377|129770498</td><td>29</td><td>52</td><td>24</td><td>TTTGGCAATGGTAGAACTCACACT</td><td>MB468</td><td>70</td><td>86</td><td>69</td><td>70</td><td>19</td><td>24.7</td><td>13.9</td><td>27.0</td><td>77.4</td><td>8.2</td></tr>
<tr><td>>MI0000272|hsa-mir-182&WithFlank&7|-|129770377|129770498@29.53.25(+1A)&[MIMAT0000259&hsa-miR-182-5p&offsets|0|+1(+1A);m-22&7|-|129770449|129770470&offsets|0|+3(+1A)]|TTTGGCAATGGTAGAACTCACACTGA</td><td>>MI0000272|hsa-mir-182&WithFlank&7|-|129770377|129770498</td><td>29</td><td>53</td><td>25(+1A)</td><td>TTTGGCAATGGTAGAACTCACACTGA</td><td>BT20</td><td>85</td><td>52</td><td>76</td><td>11</td><td>25</td><td>73.8</td><td>18.8</td><td>34.5</td><td>3.5</td><td>7.0</td></tr>
<tr><td>>MI0000272|hsa-mir-182&WithFlank&7|-|129770377|129770498@29.53.25(+1C)&[MIMAT0000259&hsa-miR-182-5p&offsets|0|+1(+1C);m-22&7|-|129770449|129770470&offsets|0|+3(+1C)]|TTTGGCAATGGTAGAACTCACACTGC</td><td>>MI0000272|hsa-mir-182&WithFlank&7|-|129770377|129770498</td><td>29</td><td>53</td><td>25(+1C)</td><td>TTTGGCAATGGTAGAACTCACACTGC</td><td>BT20</td><td>19</td><td>14</td><td>23</td><td>4</td><td>8</td><td>6.8</td><td>9.0</td><td>8.8</td><td>1.9</td><td>3.6</td></tr>
<tr><td>>MI0000272|hsa-mir-182&WithFlank&7|-|129770377|129770498@29.53.25(+1U)&[MIMAT0000259&hsa-miR-182-5p&offsets|0|+1(+1U);m-22&7|-|129770449|129770470&offsets|0|+3(+1U)]|TTTGGCAATGGTAGAACTCACACTGT</td><td>>MI0000272|hsa-mir-182&WithFlank&7|-|129770377|129770498</td><td>29</td><td>53</td><td>25(+1U)</td><td>TTTGGCAATGGTAGAACTCACACTGT</td><td>BT20</td><td>127</td><td>79</td><td>158</td><td>18</td><td>43</td><td>48.2</td><td>40.9</td><td>59.7</td><td>9.0</td><td>20.5</td></tr>
<tr><td>>MI0000272|hsa-mir-182&WithFlank&7|-|129770377|129770498@29.53.25(+1U)&[MIMAT0000259&hsa-miR-182-5p&offsets|0|+1(+1U);m-22&7|-|129770449|129770470&offsets|0|+3(+1U)]|TTTGGCAATGGTAGAACTCACACTGT</td><td>>MI0000272|hsa-mir-182&WithFlank&7|-|129770377|129770498</td><td>29</td><td>53</td><td>25(+1U)</td><td>TTTGGCAATGGTAGAACTCACACTGT</td><td>MB231</td><td>46</td><td>29</td><td>19</td><td>24</td><td>39</td><td>17.9</td><td>10.4</td><td>10.1</td><td>13.9</td><td>28.2</td></tr>
</table>

<table>
<tr><td></td><td></td><td></td><td></td><td></td><td></td><td></td><td></td><td></td><td></td><td></td><td></td><td></td><td></td><td></td><td></td><td></td></tr>
<tr><td>>MI0000272|hsa-mir-182&WithFlank&7|-|129770377|129770498@29.53.25&[MIMAT0000259&hsa-miR-182-5p&offsets|0|+1;m-22&7|-|129770449|129770470&offsets|0|+3]|TTTGGCAATGGTAGAACTCACACTG</td><td>>MI0000272|hsa-mir-182&WithFlank&7|-|129770377|129770498</td><td>29</td><td>53</td><td>25</td><td>TTTGGCAATGGTAGAACTCACACTG</td><td>BT20</td><td>650</td><td>489</td><td>722</td><td>100</td><td>276</td><td>235.5</td><td>78.4</td><td>177.9</td><td>9.3</td><td>77.4</td></tr>
<tr><td>>MI0000272|hsa-mir-182&WithFlank&7|-|129770377|129770498@29.53.25&[MIMAT0000259&hsa-miR-182-5p&offsets|0|+1;m-22&7|-|129770449|129770470&offsets|0|+3]|TTTGGCAATGGTAGAACTCACACTG</td><td>>MI0000272|hsa-mir-182&WithFlank&7|-|129770377|129770498</td><td>29</td><td>53</td><td>25</td><td>TTTGGCAATGGTAGAACTCACACTG</td><td>MB231</td><td>109</td><td>90</td><td>55</td><td>55</td><td>92</td><td>15.9</td><td>12.4</td><td>18.8</td><td>24.4</td><td>35.0</td></tr>
<tr><td>>MI0000272|hsa-mir-182&WithFlank&7|-|129770377|129770498@29.53.25&[MIMAT0000259&hsa-miR-182-5p&offsets|0|+1;m-22&7|-|129770449|129770470&offsets|0|+3]|TTTGGCAATGGTAGAACTCACACTG</td><td>>MI0000272|hsa-mir-182&WithFlank&7|-|129770377|129770498</td><td>29</td><td>53</td><td>25</td><td>TTTGGCAATGGTAGAACTCACACTG</td><td>MB468</td><td>16</td><td>19</td><td>11</td><td>18</td><td>5</td><td>6.8</td><td>4.6</td><td>4.8</td><td>19.8</td><td>2.0</td></tr>
<tr><td>>MI0000272|hsa-mir-182&WithFlank&7|-|129770377|129770498@29.54.26(+1A)&[MIMAT0000259&hsa-miR-182-5p&offsets|0|+2(+1A);m-22&7|-|129770449|129770470&offsets|0|+4(+1A)]|TTTGGCAATGGTAGAACTCACACTGGA</td><td>>MI0000272|hsa-mir-182&WithFlank&7|-|129770377|129770498</td><td>29</td><td>54</td><td>26(+1A)</td><td>TTTGGCAATGGTAGAACTCACACTGGA</td><td>BT20</td><td>11</td><td>9</td><td>14</td><td>4</td><td>5</td><td>5.9</td><td>2.5</td><td>2.7</td><td>1.5</td><td>0.8</td></tr>
<tr><td>>MI0000272|hsa-mir-182&WithFlank&7|-|129770377|129770498@29.54.26(+1U)&[MIMAT0000259&hsa-miR-182-5p&offsets|0|+2(+1U);m-22&7|-|129770449|129770470&offsets|0|+4(+1U)]|TTTGGCAATGGTAGAACTCACACTGGT</td><td>>MI0000272|hsa-mir-182&WithFlank&7|-|129770377|129770498</td><td>29</td><td>54</td><td>26(+1U)</td><td>TTTGGCAATGGTAGAACTCACACTGGT</td><td>BT20</td><td>38</td><td>30</td><td>45</td><td>14</td><td>21</td><td>16.8</td><td>9.5</td><td>12.0</td><td>7.3</td><td>8.4</td></tr>
<tr><td>>MI0000272|hsa-mir-182&WithFlank&7|-|129770377|129770498@29.54.26(+1U)&[MIMAT0000259&hsa-miR-182-5p&offsets|0|+2(+1U);m-22&7|-|129770449|129770470&offsets|0|+4(+1U)]|TTTGGCAATGGTAGAACTCACACTGGT</td><td>>MI0000272|hsa-mir-182&WithFlank&7|-|129770377|129770498</td><td>29</td><td>54</td><td>26(+1U)</td><td>TTTGGCAATGGTAGAACTCACACTGGT</td><td>MB231</td><td>13</td><td>10</td><td>7</td><td>12</td><td>19</td><td>2.7</td><td>3.5</td><td>2.4</td><td>7.2</td><td>12.6</td></tr>
<tr><td>>MI0000272|hsa-mir-182&WithFlank&7|-|129770377|129770498@29.54.26&[MIMAT0000259&hsa-miR-182-5p&offsets|0|+2;m-22&7|-|129770449|129770470&offsets|0|+4]|TTTGGCAATGGTAGAACTCACACTGG</td><td>>MI0000272|hsa-mir-182&WithFlank&7|-|129770377|129770498</td><td>29</td><td>54</td><td>26</td><td>TTTGGCAATGGTAGAACTCACACTGG</td><td>BT20</td><td>310</td><td>258</td><td>295</td><td>70</td><td>148</td><td>158.0</td><td>74.4</td><td>87.2</td><td>11.0</td><td>25.1</td></tr>
<tr><td>>MI0000272|hsa-mir-182&WithFlank&7|-|129770377|129770498@29.54.26&[MIMAT0000259&hsa-miR-182-5p&offsets|0|+2;m-22&7|-|129770449|129770470&offsets|0|+4]|TTTGGCAATGGTAGAACTCACACTGG</td><td>>MI0000272|hsa-mir-182&WithFlank&7|-|129770377|129770498</td><td>29</td><td>54</td><td>26</td><td>TTTGGCAATGGTAGAACTCACACTGG</td><td>MB231</td><td>50</td><td>44</td><td>21</td><td>34</td><td>55</td><td>10.9</td><td>4.9</td><td>4.0</td><td>20.4</td><td>20.6</td></tr>
<tr><td>>MI0000273|hsa-mir-183&WithFlank&7|-|129774899|129775020@33.53.21(+1A)&[MIMAT0000261&hsa-miR-183-5p&offsets|0|-1(+1A);m-41&7|-|129774966|129774987&offsets|-1|-2(+1A)]|TATGGCACTGGTAGAATTCACA</td><td>>MI0000273|hsa-mir-183&WithFlank&7|-|129774899|129775020</td><td>33</td><td>53</td><td>21(+1A)</td><td>TATGGCACTGGTAGAATTCACA</td><td>BT20</td><td>80</td><td>31</td><td>66</td><td>12</td><td>19</td><td>55.9</td><td>11.8</td><td>19.5</td><td>7.5</td><td>3.4</td></tr>
<tr><td>>MI0000273|hsa-mir-183&WithFlank&7|-|129774899|129775020@33.53.21(+1C)&[MIMAT0000261&hsa-miR-183-5p&offsets|0|-1(+1C);m-41&7|-|129774966|129774987&offsets|-1|-2(+1C)]|TATGGCACTGGTAGAATTCACC</td><td>>MI0000273|hsa-mir-183&WithFlank&7|-|129774899|129775020</td><td>33</td><td>53</td><td>21(+1C)</td><td>TATGGCACTGGTAGAATTCACC</td><td>BT20</td><td>134</td><td>88</td><td>118</td><td>31</td><td>56</td><td>51.1</td><td>33.7</td><td>25.9</td><td>17.1</td><td>15.0</td></tr>
<tr><td>>MI0000273|hsa-mir-183&WithFlank&7|-|129774899|129775020@33.53.21(+1C)&[MIMAT0000261&hsa-miR-183-5p&offsets|0|-1(+1C);m-41&7|-|129774966|129774987&offsets|-1|-2(+1C)]|TATGGCACTGGTAGAATTCACC</td><td>>MI0000273|hsa-mir-183&WithFlank&7|-|129774899|129775020</td><td>33</td><td>53</td><td>21(+1C)</td><td>TATGGCACTGGTAGAATTCACC</td><td>MB231</td><td>21</td><td>21</td><td>8</td><td>23</td><td>21</td><td>7.3</td><td>2.2</td><td>2.7</td><td>8.7</td><td>7.7</td></tr>
<tr><td>>MI0000273|hsa-mir-183&WithFlank&7|-|129774899|129775020@33.53.21(+1G)&[MIMAT0000261&hsa-miR-183-5p&offsets|0|-1(+1G);m-41&7|-|129774966|129774987&offsets|-1|-2(+1G)]|TATGGCACTGGTAGAATTCACG</td><td>>MI0000273|hsa-mir-183&WithFlank&7|-|129774899|129775020</td><td>33</td><td>53</td><td>21(+1G)</td><td>TATGGCACTGGTAGAATTCACG</td><td>BT20</td><td>67</td><td>16</td><td>25</td><td>4</td><td>5</td><td>56.1</td><td>4.8</td><td>5.2</td><td>4.2</td><td>0.8</td></tr>
<tr><td>>MI0000273|hsa-mir-183&WithFlank&7|-|129774899|129775020@33.53.21&[MIMAT0000261&hsa-miR-183-5p&offsets|0|-1;m-41&7|-|129774966|129774987&offsets|-1|-2]|TATGGCACTGGTAGAATTCAC</td><td>>MI0000273|hsa-mir-183&WithFlank&7|-|129774899|129775020</td><td>33</td><td>53</td><td>21</td><td>TATGGCACTGGTAGAATTCAC</td><td>BT20</td><td>13</td><td>12</td><td>21</td><td>27</td><td>8</td><td>7.9</td><td>6.3</td><td>9.5</td><td>19.4</td><td>2.1</td></tr>
<tr><td>>MI0000273|hsa-mir-183&WithFlank&7|-|129774899|129775020@33.54.22(+1A)&[MIMAT0000261&hsa-miR-183-5p&offsets|0|0(+1A);m-41&7|-|129774966|129774987&offsets|-1|-1(+1A)]|TATGGCACTGGTAGAATTCACTA</td><td>>MI0000273|hsa-mir-183&WithFlank&7|-|129774899|129775020</td><td>33</td><td>54</td><td>22(+1A)</td><td>TATGGCACTGGTAGAATTCACTA</td><td>BT20</td><td>53</td><td>28</td><td>45</td><td>26</td><td>16</td><td>35.2</td><td>10.6</td><td>6.0</td><td>23.0</td><td>4.5</td></tr>
<tr><td>>MI0000273|hsa-mir-183&WithFlank&7|-|129774899|129775020@33.54.22&[MIMAT0000261&hsa-miR-183-5p&offsets|0|0;m-41&7|-|129774966|129774987&offsets|-1|-1]|TATGGCACTGGTAGAATTCACT</td><td>>MI0000273|hsa-mir-183&WithFlank&7|-|129774899|129775020</td><td>33</td><td>54</td><td>22</td><td>TATGGCACTGGTAGAATTCACT</td><td>BT20</td><td>948</td><td>597</td><td>894</td><td>211</td><td>401</td><td>385.8</td><td>181.2</td><td>232.0</td><td>75.9</td><td>97.8</td></tr>
<tr><td>>MI0000273|hsa-mir-183&WithFlank&7|-|129774899|129775020@33.54.22&[MIMAT0000261&hsa-miR-183-5p&offsets|0|0;m-41&7|-|129774966|129774987&offsets|-1|-1]|TATGGCACTGGTAGAATTCACT</td><td>>MI0000273|hsa-mir-183&WithFlank&7|-|129774899|129775020</td><td>33</td><td>54</td><td>22</td><td>TATGGCACTGGTAGAATTCACT</td><td>MB231</td><td>176</td><td>195</td><td>73</td><td>145</td><td>146</td><td>61.7</td><td>34.4</td><td>25.2</td><td>34.0</td><td>42.5</td></tr>
<tr><td>>MI0000273|hsa-mir-183&WithFlank&7|-|129774899|129775020@33.54.22&[MIMAT0000261&hsa-miR-183-5p&offsets|0|0;m-41&7|-|129774966|129774987&offsets|-1|-1]|TATGGCACTGGTAGAATTCACT</td><td>>MI0000273|hsa-mir-183&WithFlank&7|-|129774899|129775020</td><td>33</td><td>54</td><td>22</td><td>TATGGCACTGGTAGAATTCACT</td><td>MB468</td><td>37</td><td>59</td><td>21</td><td>44</td><td>13</td><td>14.0</td><td>20.1</td><td>9.7</td><td>20.2</td><td>8.1</td></tr>
<tr><td>>MI0000273|hsa-mir-183&WithFlank&7|-|129774899|129775020@33.55.23&[MIMAT0000261&hsa-miR-183-5p&offsets|0|+1;m-41&7|-|129774966|129774987&offsets|-1|0]|TATGGCACTGGTAGAATTCACTG</td><td>>MI0000273|hsa-mir-183&WithFlank&7|-|129774899|129775020</td><td>33</td><td>55</td><td>23</td><td>TATGGCACTGGTAGAATTCACTG</td><td>BT20</td><td>256</td><td>189</td><td>245</td><td>187</td><td>134</td><td>104.9</td><td>43.2</td><td>73.0</td><td>122.9</td><td>34.5</td></tr>
<tr><td>>MI0000273|hsa-mir-183&WithFlank&7|-|129774899|129775020@33.55.23&[MIMAT0000261&hsa-miR-183-5p&offsets|0|+1;m-41&7|-|129774966|129774987&offsets|-1|0]|TATGGCACTGGTAGAATTCACTG</td><td>>MI0000273|hsa-mir-183&WithFlank&7|-|129774899|129775020</td><td>33</td><td>55</td><td>23</td><td>TATGGCACTGGTAGAATTCACTG</td><td>MB231</td><td>37</td><td>41</td><td>14</td><td>80</td><td>31</td><td>11.2</td><td>8.1</td><td>3.9</td><td>18.7</td><td>6.2</td></tr>
<tr><td>>MI0000273|hsa-mir-183&WithFlank&7|-|129774899|129775020@33.56.24&[MIMAT0000261&hsa-miR-183-5p&offsets|0|+2;m-41&7|-|129774966|129774987&offsets|-1|+1]|TATGGCACTGGTAGAATTCACTGT</td><td>>MI0000273|hsa-mir-183&WithFlank&7|-|129774899|129775020</td><td>33</td><td>56</td><td>24</td><td>TATGGCACTGGTAGAATTCACTGT</td><td>BT20</td><td>44</td><td>26</td><td>79</td><td>9</td><td>14</td><td>18.1</td><td>11.7</td><td>21.5</td><td>6.8</td><td>6.6</td></tr>
<tr><td>>MI0000273|hsa-mir-183&WithFlank&7|-|129774899|129775020@33.56.24&[MIMAT0000261&hsa-miR-183-5p&offsets|0|+2;m-41&7|-|129774966|129774987&offsets|-1|+1]|TATGGCACTGGTAGAATTCACTGT</td><td>>MI0000273|hsa-mir-183&WithFlank&7|-|129774899|129775020</td><td>33</td><td>56</td><td>24</td><td>TATGGCACTGGTAGAATTCACTGT</td><td>MB231</td><td>17</td><td>13</td><td>10</td><td>12</td><td>9</td><td>4.7</td><td>4.6</td><td>6.1</td><td>3.5</td><td>5.1</td></tr>
<tr><td>>MI0000273|hsa-mir-183&WithFlank&7|-|129774899|129775020@34.53.20(+1A)&[MIMAT0000261&hsa-miR-183-5p&offsets|+1|-1(+1A);m-41&7|-|129774966|129774987&offsets|0|-2(+1A)]|ATGGCACTGGTAGAATTCACA</td><td>>MI0000273|hsa-mir-183&WithFlank&7|-|129774899|129775020</td><td>34</td><td>53</td><td>20(+1A)</td><td>ATGGCACTGGTAGAATTCACA</td><td>BT20</td><td>21</td><td>9</td><td>20</td><td>5</td><td>8</td><td>13.9</td><td>3.0</td><td>6.8</td><td>4.5</td><td>1.6</td></tr>
<tr><td>>MI0000273|hsa-mir-183&WithFlank&7|-|129774899|129775020@34.53.20(+1C)&[MIMAT0000261&hsa-miR-183-5p&offsets|+1|-1(+1C);m-41&7|-|129774966|129774987&offsets|0|-2(+1C)]|ATGGCACTGGTAGAATTCACC</td><td>>MI0000273|hsa-mir-183&WithFlank&7|-|129774899|129775020</td><td>34</td><td>53</td><td>20(+1C)</td><td>ATGGCACTGGTAGAATTCACC</td><td>BT20</td><td>37</td><td>27</td><td>36</td><td>14</td><td>25</td><td>13.0</td><td>8.2</td><td>5.2</td><td>8.9</td><td>2.8</td></tr>
<tr><td>>MI0000273|hsa-mir-183&WithFlank&7|-|129774899|129775020@34.53.20(+1G)&[MIMAT0000261&hsa-miR-183-5p&offsets|+1|-1(+1G);m-41&7|-|129774966|129774987&offsets|0|-2(+1G)]|ATGGCACTGGTAGAATTCACG</td><td>>MI0000273|hsa-mir-183&WithFlank&7|-|129774899|129775020</td><td>34</td><td>53</td><td>20(+1G)</td><td>ATGGCACTGGTAGAATTCACG</td><td>BT20</td><td>18</td><td>5</td><td>7</td><td>2</td><td>1</td><td>14.6</td><td>0.5</td><td>2.2</td><td>2.2</td><td>1.1</td></tr>
<tr><td>>MI0000273|hsa-mir-183&WithFlank&7|-|129774899|129775020@34.54.21(+1A)&[MIMAT0000261&hsa-miR-183-5p&offsets|+1|0(+1A);m-41&7|-|129774966|129774987&offsets|0|-1(+1A)]|ATGGCACTGGTAGAATTCACTA</td><td>>MI0000273|hsa-mir-183&WithFlank&7|-|129774899|129775020</td><td>34</td><td>54</td><td>21(+1A)</td><td>ATGGCACTGGTAGAATTCACTA</td><td>BT20</td><td>42</td><td>24</td><td>42</td><td>26</td><td>20</td><td>27.1</td><td>7.0</td><td>11.7</td><td>23.3</td><td>5.2</td></tr>
<tr><td>>MI0000273|hsa-mir-183&WithFlank&7|-|129774899|129775020@34.54.21&[MIMAT0000261&hsa-miR-183-5p&offsets|+1|0;m-41&7|-|129774966|129774987&offsets|0|-1]|ATGGCACTGGTAGAATTCACT</td><td>>MI0000273|hsa-mir-183&WithFlank&7|-|129774899|129775020</td><td>34</td><td>54</td><td>21</td><td>ATGGCACTGGTAGAATTCACT</td><td>BT20</td><td>258</td><td>179</td><td>259</td><td>93</td><td>179</td><td>92.1</td><td>35.2</td><td>44.4</td><td>50.9</td><td>5.9</td></tr>
<tr><td>>MI0000273|hsa-mir-183&WithFlank&7|-|129774899|129775020@34.54.21&[MIMAT0000261&hsa-miR-183-5p&offsets|+1|0;m-41&7|-|129774966|129774987&offsets|0|-1]|ATGGCACTGGTAGAATTCACT</td><td>>MI0000273|hsa-mir-183&WithFlank&7|-|129774899|129775020</td><td>34</td><td>54</td><td>21</td><td>ATGGCACTGGTAGAATTCACT</td><td>MB231</td><td>33</td><td>31</td><td>17</td><td>36</td><td>30</td><td>10.8</td><td>5.4</td><td>6.8</td><td>7.1</td><td>8.8</td></tr>
<tr><td>>MI0000273|hsa-mir-183&WithFlank&7|-|129774899|129775020@34.55.22&[MIMAT0000261&hsa-miR-183-5p&offsets|+1|+1;m-41&7|-|129774966|129774987&offsets|0|0]|ATGGCACTGGTAGAATTCACTG</td><td>>MI0000273|hsa-mir-183&WithFlank&7|-|129774899|129775020</td><td>34</td><td>55</td><td>22</td><td>ATGGCACTGGTAGAATTCACTG</td><td>BT20</td><td>204</td><td>174</td><td>221</td><td>177</td><td>169</td><td>62.2</td><td>46.7</td><td>27.6</td><td>123.0</td><td>19.2</td></tr>
<tr><td>>MI0000273|hsa-mir-183&WithFlank&7|-|129774899|129775020@34.55.22&[MIMAT0000261&hsa-miR-183-5p&offsets|+1|+1;m-41&7|-|129774966|129774987&offsets|0|0]|ATGGCACTGGTAGAATTCACTG</td><td>>MI0000273|hsa-mir-183&WithFlank&7|-|129774899|129775020</td><td>34</td><td>55</td><td>22</td><td>ATGGCACTGGTAGAATTCACTG</td><td>MB231</td><td>25</td><td>25</td><td>12</td><td>57</td><td>22</td><td>7.4</td><td>4.1</td><td>5.0</td><td>14.2</td><td>2.1</td></tr>
<tr><td>>MI0000273|hsa-mir-183&WithFlank&7|-|129774899|129775020@34.56.23&[MIMAT0000261&hsa-miR-183-5p&offsets|+1|+2;m-41&7|-|129774966|129774987&offsets|0|+1]|ATGGCACTGGTAGAATTCACTGT</td><td>>MI0000273|hsa-mir-183&WithFlank&7|-|129774899|129775020</td><td>34</td><td>56</td><td>23</td><td>ATGGCACTGGTAGAATTCACTGT</td><td>BT20</td><td>59</td><td>42</td><td>124</td><td>32</td><td>29</td><td>18.6</td><td>14.1</td><td>20.0</td><td>25.1</td><td>4.3</td></tr>
<tr><td>>MI0000273|hsa-mir-183&WithFlank&7|-|129774899|129775020@34.56.23&[MIMAT0000261&hsa-miR-183-5p&offsets|+1|+2;m-41&7|-|129774966|129774987&offsets|0|+1]|ATGGCACTGGTAGAATTCACTGT</td><td>>MI0000273|hsa-mir-183&WithFlank&7|-|129774899|129775020</td><td>34</td><td>56</td><td>23</td><td>ATGGCACTGGTAGAATTCACTGT</td><td>MB231</td><td>15</td><td>10</td><td>10</td><td>14</td><td>8</td><td>4.1</td><td>2.5</td><td>4.9</td><td>2.4</td><td>2.1</td></tr>
<tr><td>>MI0000279|hsa-mir-196a-2&WithFlank&12|+|53991732|53991853@31.50.20(+1A);MI0000238|hsa-mir-196a-1&WithFlank&17|-|48632484|48632565@13.32.20(+1A)&[MIMAT0000226&hsa-miR-196a-5p&offsets|0|-2(+1A);m-261&12|+|53991762|53991783&offsets|0|-2(+1A)];[MIMAT0000226_1&hsa-miR-196a-5p&offsets|0|-2(+1A);m-268&17|-|48632532|48632553&offsets|0|-2(+1A)]|TAGGTAGTTTCATGTTGTTGA</td><td>>MI0000279|hsa-mir-196a-2&WithFlank&12|+|53991732|53991853</td><td>31</td><td>50</td><td>20(+1A)</td><td>TAGGTAGTTTCATGTTGTTGA</td><td>BT20</td><td>13</td><td>9</td><td>18</td><td>6</td><td>2</td><td>12.3</td><td>4.2</td><td>13.1</td><td>6.6</td><td>0.2</td></tr>
</table>

| | | | | | | | | | | | | | | | | |
|---|---|---|---|---|---|---|---|---|---|---|---|---|---|---|---|---|
| >MI0000279\|hsa-mir-196a-2&WithFlank&12\|+\|53991732\|53991853@31.51.21;MI0000238\|hsa-mir-196a-1&WithFlank&17\|-\|48632484\|48632565@13.33.21&[MIMAT0000226&hsa-miR-196a-5p&offsets\|0\|-1;m-261&12\|+\|53991762\|53991783&offsets\|0\|-1];[MIMAT0000226_1&hsa-miR-196a-5p&offsets\|0\|-1;m-268&17\|-\|48632532\|48632553&offsets\|0\|-1]\|TAGGTAGTTTCATGTTGTTGG | >MI0000279\|hsa-mir-196a-2&WithFlank&12\|+\|53991732\|53991853 | 31 | 51 | 21 | TAGGTAGTTTCATGTTGTTGG | BT20 | 54 | 52 | 92 | 39 | 14 | 28.4 | 13.3 | 59.2 | 36.0 | 1.4 |
| >MI0000279\|hsa-mir-196a-2&WithFlank&12\|+\|53991732\|53991853@31.51.21(+1A);MI0000238\|hsa-mir-196a-1&WithFlank&17\|-\|48632484\|48632565@13.33.21(+1A)&[MIMAT0000226&hsa-miR-196a-5p&offsets\|0\|-1(+1A);m-261&12\|+\|53991762\|53991783&offsets\|0\|-1(+1A)];[MIMAT0000226_1&hsa-miR-196a-5p&offsets\|0\|-1(+1A);m-268&17\|-\|48632532\|48632553&offsets\|0\|-1(+1A)]\|TAGGTAGTTTCATGTTGTTGGA | >MI0000279\|hsa-mir-196a-2&WithFlank&12\|+\|53991732\|53991853 | 31 | 51 | 21(+1A) | TAGGTAGTTTCATGTTGTTGGA | BT20 | 43 | 36 | 62 | 38 | 8 | 38.6 | 14.7 | 35.7 | 42.4 | 2.7 |
| >MI0000279\|hsa-mir-196a-2&WithFlank&12\|+\|53991732\|53991853@31.51.21(+1U);MI0000238\|hsa-mir-196a-1&WithFlank&17\|-\|48632484\|48632565@13.33.21(+1U)&[MIMAT0000226&hsa-miR-196a-5p&offsets\|0\|-1(+1U);m-261&12\|+\|53991762\|53991783&offsets\|0\|-1(+1U)];[MIMAT0000226_1&hsa-miR-196a-5p&offsets\|0\|-1(+1U);m-268&17\|-\|48632532\|48632553&offsets\|0\|-1(+1U)]\|TAGGTAGTTTCATGTTGTTGGT | >MI0000279\|hsa-mir-196a-2&WithFlank&12\|+\|53991732\|53991853 | 31 | 51 | 21(+1U) | TAGGTAGTTTCATGTTGTTGGT | BT20 | 12 | 8 | 13 | 11 | 2 | 11.1 | 3.9 | 7.3 | 13.1 | 0.9 |
| >MI0000279\|hsa-mir-196a-2&WithFlank&12\|+\|53991732\|53991853@31.52.22;MI0000238\|hsa-mir-196a-1&WithFlank&17\|-\|48632484\|48632565@13.34.22&[MIMAT0000226&hsa-miR-196a-5p&offsets\|0\|0;m-261&12\|+\|53991762\|53991783&offsets\|0\|0];[MIMAT0000226_1&hsa-miR-196a-5p&offsets\|0\|0;m-268&17\|-\|48632532\|48632553&offsets\|0\|0]\|TAGGTAGTTTCATGTTGTTGGG | >MI0000279\|hsa-mir-196a-2&WithFlank&12\|+\|53991732\|53991853 | 31 | 52 | 22 | TAGGTAGTTTCATGTTGTTGGG | BT20 | 185 | 215 | 309 | 281 | 59 | 92.3 | 70.2 | 147.4 | 259.7 | 18.9 |
| >MI0000279\|hsa-mir-196a-2&WithFlank&12\|+\|53991732\|53991853@31.52.22(+1U);MI0000238\|hsa-mir-196a-1&WithFlank&17\|-\|48632484\|48632565@13.34.22(+1U)&[MIMAT0000226&hsa-miR-196a-5p&offsets\|0\|0(+1U);m-261&12\|+\|53991762\|53991783&offsets\|0\|0(+1U)];[MIMAT0000226_1&hsa-miR-196a-5p&offsets\|0\|0(+1U);m-268&17\|-\|48632532\|48632553&offsets\|0\|0(+1U)]\|TAGGTAGTTTCATGTTGTTGGGT | >MI0000279\|hsa-mir-196a-2&WithFlank&12\|+\|53991732\|53991853 | 31 | 52 | 22(+1U) | TAGGTAGTTTCATGTTGTTGGGT | BT20 | 28 | 29 | 29 | 38 | 7 | 15.2 | 17.5 | 16.5 | 30.1 | 2.6 |
| >MI0000279\|hsa-mir-196a-2&WithFlank&12\|+\|53991732\|53991853@31.53.23&[MIMAT0000226&hsa-miR-196a-5p&offsets\|0\|+1;m-261&12\|+\|53991762\|53991783&offsets\|0\|+1]\|TAGGTAGTTTCATGTTGTTGGGA | >MI0000279\|hsa-mir-196a-2&WithFlank&12\|+\|53991732\|53991853 | 31 | 53 | 23 | TAGGTAGTTTCATGTTGTTGGGA | BT20 | 81 | 83 | 129 | 84 | 20 | 53.2 | 31.0 | 55.9 | 77.9 | 3.2 |
| >MI0000283\|hsa-mir-203a&WithFlank&14\|+\|104117399\|104117520@71.90.20&[MIMAT0000264&hsa-miR-203a-3p&offsets\|0\|-2;m-79&14\|+\|104117469\|104117490&offsets\|0\|-2]\|GTGAAATGTTTAGGACCACT | >MI0000283\|hsa-mir-203a&WithFlank&14\|+\|104117399\|104117520 | 71 | 90 | 20 | GTGAAATGTTTAGGACCACT | BT20 | 41 | 29 | 38 | 31 | 12 | 17.4 | 17.5 | 17.1 | 23.9 | 4.4 |
| >MI0000283\|hsa-mir-203a&WithFlank&14\|+\|104117399\|104117520@71.91.21(+1A)&[MIMAT0000264&hsa-miR-203a-3p&offsets\|0\|-1(+1A);m-79&14\|+\|104117469\|104117490&offsets\|0\|-1(+1A)]\|GTGAAATGTTTAGGACCACTAA | >MI0000283\|hsa-mir-203a&WithFlank&14\|+\|104117399\|104117520 | 71 | 91 | 21(+1A) | GTGAAATGTTTAGGACCACTAA | BT20 | 24 | 9 | 20 | 7 | 3 | 21.4 | 4.5 | 5.8 | 7.6 | 1.1 |
| >MI0000283\|hsa-mir-203a&WithFlank&14\|+\|104117399\|104117520@71.92.22(+1U)&[MIMAT0000264&hsa-miR-203a-3p&offsets\|0\|0(+1U);m-79&14\|+\|104117469\|104117490&offsets\|0\|0(+1U)]\|GTGAAATGTTTAGGACCACTAGT | >MI0000283\|hsa-mir-203a&WithFlank&14\|+\|104117399\|104117520 | 71 | 92 | 22(+1U) | GTGAAATGTTTAGGACCACTAGT | BT20 | 13 | 8 | 11 | 4 | 4 | 5.6 | 4.7 | 3.7 | 2.6 | 2.8 |
| >MI0000283\|hsa-mir-203a&WithFlank&14\|+\|104117399\|104117520@71.92.22&[MIMAT0000264&hsa-miR-203a-3p&offsets\|0\|0;m-79&14\|+\|104117469\|104117490&offsets\|0\|0]\|GTGAAATGTTTAGGACCACTAG | >MI0000283\|hsa-mir-203a&WithFlank&14\|+\|104117399\|104117520 | 71 | 92 | 22 | GTGAAATGTTTAGGACCACTAG | BT20 | 129 | 73 | 125 | 52 | 24 | 62.1 | 21.7 | 8.2 | 46.6 | 10.5 |
| >MI0000283\|hsa-mir-203a&WithFlank&14\|+\|104117399\|104117520@71.93.23&[MIMAT0000264&hsa-miR-203a-3p&offsets\|0\|+1;m-79&14\|+\|104117469\|104117490&offsets\|0\|+1]\|GTGAAATGTTTAGGACCACTAGA | >MI0000283\|hsa-mir-203a&WithFlank&14\|+\|104117399\|104117520 | 71 | 93 | 23 | GTGAAATGTTTAGGACCACTAGA | BT20 | 28 | 16 | 26 | 6 | 6 | 20.0 | 5.2 | 4.3 | 3.3 | 1.5 |
| >MI0000283\|hsa-mir-203a&WithFlank&14\|+\|104117399\|104117520@72.90.19&[MIMAT0000264&hsa-miR-203a-3p&offsets\|+1\|-2;m-79&14\|+\|104117469\|104117490&offsets\|+1\|-2]\|TGAAATGTTTAGGACCACT | >MI0000283\|hsa-mir-203a&WithFlank&14\|+\|104117399\|104117520 | 72 | 90 | 19 | TGAAATGTTTAGGACCACT | BT20 | 34 | 26 | 44 | 24 | 12 | 11.3 | 8.9 | 8.7 | 17.3 | 2.4 |
| >MI0000283\|hsa-mir-203a&WithFlank&14\|+\|104117399\|104117520@72.91.20(+1A)&[MIMAT0000264&hsa-miR-203a-3p&offsets\|+1\|-1(+1A);m-79&14\|+\|104117469\|104117490&offsets\|+1\|-1(+1A)]\|TGAAATGTTTAGGACCACTAA | >MI0000283\|hsa-mir-203a&WithFlank&14\|+\|104117399\|104117520 | 72 | 91 | 20(+1A) | TGAAATGTTTAGGACCACTAA | BT20 | 22 | 10 | 24 | 7 | 3 | 21.8 | 1.1 | 10.8 | 7.6 | 0.7 |
| >MI0000283\|hsa-mir-203a&WithFlank&14\|+\|104117399\|104117520@72.92.21(+1U)&[MIMAT0000264&hsa-miR-203a-3p&offsets\|+1\|0(+1U);m-79&14\|+\|104117469\|104117490&offsets\|+1\|0(+1U)]\|TGAAATGTTTAGGACCACTAGT | >MI0000283\|hsa-mir-203a&WithFlank&14\|+\|104117399\|104117520 | 72 | 92 | 21(+1U) | TGAAATGTTTAGGACCACTAGT | BT20 | 72 | 45 | 61 | 15 | 20 | 23.0 | 17.7 | 13.0 | 10.2 | 8.4 |
| >MI0000283\|hsa-mir-203a&WithFlank&14\|+\|104117399\|104117520@72.92.21&[MIMAT0000264&hsa-miR-203a-3p&offsets\|+1\|0;m-79&14\|+\|104117469\|104117490&offsets\|+1\|0]\|TGAAATGTTTAGGACCACTAG | >MI0000283\|hsa-mir-203a&WithFlank&14\|+\|104117399\|104117520 | 72 | 92 | 21 | TGAAATGTTTAGGACCACTAG | BT20 | 117 | 80 | 143 | 48 | 25 | 65.7 | 5.6 | 16.4 | 40.1 | 8.0 |
| >MI0000283\|hsa-mir-203a&WithFlank&14\|+\|104117399\|104117520@72.93.22&[MIMAT0000264&hsa-miR-203a-3p&offsets\|+1\|+1;m-79&14\|+\|104117469\|104117490&offsets\|+1\|+1]\|TGAAATGTTTAGGACCACTAGA | >MI0000283\|hsa-mir-203a&WithFlank&14\|+\|104117399\|104117520 | 72 | 93 | 22 | TGAAATGTTTAGGACCACTAGA | BT20 | 57 | 34 | 44 | 14 | 12 | 44.7 | 7.0 | 8.7 | 12.3 | 4.0 |
| >MI0000285\|hsa-mir-205&WithFlank&1\|+\|209432127\|209432248@40.59.20(+1A)&[MIMAT0000266&hsa-miR-205-5p&offsets\|0\|-2(+1A);m-351&1\|+\|209432166\|209432187&offsets\|0\|-2(+1A)]\|TCCTTCATTCCACCGGAGTCA | >MI0000285\|hsa-mir-205&WithFlank&1\|+\|209432127\|209432248 | 40 | 59 | 20(+1A) | TCCTTCATTCCACCGGAGTCA | BT20 | 68 | 44 | 54 | 39 | 17 | 64.1 | 44.3 | 5.2 | 33.3 | 15.2 |
| >MI0000285\|hsa-mir-205&WithFlank&1\|+\|209432127\|209432248@40.59.20(+1C)&[MIMAT0000266&hsa-miR-205-5p&offsets\|0\|-2(+1C);m-351&1\|+\|209432166\|209432187&offsets\|0\|-2(+1C)]\|TCCTTCATTCCACCGGAGTCC | >MI0000285\|hsa-mir-205&WithFlank&1\|+\|209432127\|209432248 | 40 | 59 | 20(+1C) | TCCTTCATTCCACCGGAGTCC | BT20 | 61 | 66 | 56 | 54 | 27 | 32.9 | 68.3 | 16.9 | 46.0 | 25.7 |
| >MI0000285\|hsa-mir-205&WithFlank&1\|+\|209432127\|209432248@40.59.20(+1G)&[MIMAT0000266&hsa-miR-205-5p&offsets\|0\|-2(+1G);m-351&1\|+\|209432166\|209432187&offsets\|0\|-2(+1G)]\|TCCTTCATTCCACCGGAGTCG | >MI0000285\|hsa-mir-205&WithFlank&1\|+\|209432127\|209432248 | 40 | 59 | 20(+1G) | TCCTTCATTCCACCGGAGTCG | BT20 | 45 | 16 | 14 | 9 | 2 | 49.4 | 14.2 | 6.0 | 7.8 | 2.9 |
| >MI0000285\|hsa-mir-205&WithFlank&1\|+\|209432127\|209432248@40.59.20&[MIMAT0000266&hsa-miR-205-5p&offsets\|0\|-2;m-351&1\|+\|209432166\|209432187&offsets\|0\|-2]\|TCCTTCATTCCACCGGAGTC | >MI0000285\|hsa-mir-205&WithFlank&1\|+\|209432127\|209432248 | 40 | 59 | 20 | TCCTTCATTCCACCGGAGTC | BT20 | 27 | 32 | 33 | 34 | 18 | 26.4 | 36.4 | 31.3 | 40.8 | 18.1 |
| >MI0000285\|hsa-mir-205&WithFlank&1\|+\|209432127\|209432248@40.60.21(+1A)&[MIMAT0000266&hsa-miR-205-5p&offsets\|0\|-1(+1A);m-351&1\|+\|209432166\|209432187&offsets\|0\|-1(+1A)]\|TCCTTCATTCCACCGGAGTCTA | >MI0000285\|hsa-mir-205&WithFlank&1\|+\|209432127\|209432248 | 40 | 60 | 21(+1A) | TCCTTCATTCCACCGGAGTCTA | BT20 | 319 | 292 | 507 | 326 | 164 | 177.7 | 337.8 | 315.7 | 277.2 | 182.7 |
| >MI0000285\|hsa-mir-205&WithFlank&1\|+\|209432127\|209432248@40.60.21(+1A)&[MIMAT0000266&hsa-miR-205-5p&offsets\|0\|-1(+1A);m-351&1\|+\|209432166\|209432187&offsets\|0\|-1(+1A)]\|TCCTTCATTCCACCGGAGTCTA | >MI0000285\|hsa-mir-205&WithFlank&1\|+\|209432127\|209432248 | 40 | 60 | 21(+1A) | TCCTTCATTCCACCGGAGTCTA | MB468 | 13 | 5 | 7 | 52 | 9 | 4.7 | 1.9 | 3.2 | 33.5 | 5.3 |
| >MI0000285\|hsa-mir-205&WithFlank&1\|+\|209432127\|209432248@40.60.21(+1C)&[MIMAT0000266&hsa-miR-205-5p&offsets\|0\|-1(+1C);m-351&1\|+\|209432166\|209432187&offsets\|0\|-1(+1C)]\|TCCTTCATTCCACCGGAGTCTC | >MI0000285\|hsa-mir-205&WithFlank&1\|+\|209432127\|209432248 | 40 | 60 | 21(+1C) | TCCTTCATTCCACCGGAGTCTC | BT20 | 12 | 12 | 15 | 14 | 7 | 6.8 | 13.0 | 10.9 | 11.2 | 6.4 |
| >MI0000285\|hsa-mir-205&WithFlank&1\|+\|209432127\|209432248@40.60.21(+1U)&[MIMAT0000266&hsa-miR-205-5p&offsets\|0\|-1(+1U);m-351&1\|+\|209432166\|209432187&offsets\|0\|-1(+1U)]\|TCCTTCATTCCACCGGAGTCTT | >MI0000285\|hsa-mir-205&WithFlank&1\|+\|209432127\|209432248 | 40 | 60 | 21(+1U) | TCCTTCATTCCACCGGAGTCTT | BT20 | 60 | 60 | 85 | 69 | 29 | 34.1 | 69.6 | 61.5 | 58.0 | 30.3 |

<table>
<tr><th></th><th></th><th></th><th></th><th></th><th></th><th></th><th></th><th></th><th></th><th></th><th></th><th></th><th></th><th></th><th></th><th></th></tr>
<tr><td>>MI0000285|hsa-mir-205&WithFlank&1|+|209432127|209432248@40.60.21&[MIMAT0000266&hsa-miR-205-5p&offsets|0|-1;m-351&1|+|209432166|209432187&offsets|0|-1]|TCCTTCATTCCACCGGAGTCT</td><td>>MI0000285|hsa-mir-205&WithFlank&1|+|209432127|209432248</td><td>40</td><td>60</td><td>21</td><td>TCCTTCATTCCACCGGAGTCT</td><td>BT20</td><td>392</td><td>378</td><td>390</td><td>365</td><td>171</td><td>210.6</td><td>360.2</td><td>149.0</td><td>336.5</td><td>164.8</td></tr>
<tr><td>>MI0000285|hsa-mir-205&WithFlank&1|+|209432127|209432248@40.60.21&[MIMAT0000266&hsa-miR-205-5p&offsets|0|-1;m-351&1|+|209432166|209432187&offsets|0|-1]|TCCTTCATTCCACCGGAGTCT</td><td>>MI0000285|hsa-mir-205&WithFlank&1|+|209432127|209432248</td><td>40</td><td>60</td><td>21</td><td>TCCTTCATTCCACCGGAGTCT</td><td>MB468</td><td>18</td><td>10</td><td>8</td><td>70</td><td>12</td><td>6.5</td><td>4.5</td><td>1.7</td><td>50.4</td><td>7.9</td></tr>
<tr><td>>MI0000285|hsa-mir-205&WithFlank&1|+|209432127|209432248@40.61.22(+1A)&[MIMAT0000266&hsa-miR-205-5p&offsets|0|0(+1A);m-351&1|+|209432166|209432187&offsets|0|0(+1A)]|TCCTTCATTCCACCGGAGTCTGA</td><td>>MI0000285|hsa-mir-205&WithFlank&1|+|209432127|209432248</td><td>40</td><td>61</td><td>22(+1A)</td><td>TCCTTCATTCCACCGGAGTCTGA</td><td>BT20</td><td>2887</td><td>2042</td><td>4692</td><td>204</td><td>2074</td><td>3876.5</td><td>2011.4</td><td>4365.1</td><td>68.9</td><td>1878.8</td></tr>
<tr><td>>MI0000285|hsa-mir-205&WithFlank&1|+|209432127|209432248@40.61.22(+1A)&[MIMAT0000266&hsa-miR-205-5p&offsets|0|0(+1A);m-351&1|+|209432166|209432187&offsets|0|0(+1A)]|TCCTTCATTCCACCGGAGTCTGA</td><td>>MI0000285|hsa-mir-205&WithFlank&1|+|209432127|209432248</td><td>40</td><td>61</td><td>22(+1A)</td><td>TCCTTCATTCCACCGGAGTCTGA</td><td>MB468</td><td>199</td><td>188</td><td>113</td><td>168</td><td>128</td><td>98.7</td><td>101.8</td><td>18.1</td><td>37.1</td><td>18.3</td></tr>
<tr><td>>MI0000285|hsa-mir-205&WithFlank&1|+|209432127|209432248@40.61.22(+1C)&[MIMAT0000266&hsa-miR-205-5p&offsets|0|0(+1C);m-351&1|+|209432166|209432187&offsets|0|0(+1C)]|TCCTTCATTCCACCGGAGTCTGC</td><td>>MI0000285|hsa-mir-205&WithFlank&1|+|209432127|209432248</td><td>40</td><td>61</td><td>22(+1C)</td><td>TCCTTCATTCCACCGGAGTCTGC</td><td>BT20</td><td>355</td><td>417</td><td>604</td><td>46</td><td>273</td><td>192.8</td><td>457.3</td><td>401.2</td><td>30.8</td><td>316.1</td></tr>
<tr><td>>MI0000285|hsa-mir-205&WithFlank&1|+|209432127|209432248@40.61.22(+1C)&[MIMAT0000266&hsa-miR-205-5p&offsets|0|0(+1C);m-351&1|+|209432166|209432187&offsets|0|0(+1C)]|TCCTTCATTCCACCGGAGTCTGC</td><td>>MI0000285|hsa-mir-205&WithFlank&1|+|209432127|209432248</td><td>40</td><td>61</td><td>22(+1C)</td><td>TCCTTCATTCCACCGGAGTCTGC</td><td>MB468</td><td>17</td><td>9</td><td>10</td><td>24</td><td>8</td><td>6.9</td><td>1.9</td><td>3.5</td><td>24.5</td><td>5.1</td></tr>
<tr><td>>MI0000285|hsa-mir-205&WithFlank&1|+|209432127|209432248@40.61.22(+1G)&[MIMAT0000266&hsa-miR-205-5p&offsets|0|0(+1G);m-351&1|+|209432166|209432187&offsets|0|0(+1G)]|TCCTTCATTCCACCGGAGTCTGG</td><td>>MI0000285|hsa-mir-205&WithFlank&1|+|209432127|209432248</td><td>40</td><td>61</td><td>22(+1G)</td><td>TCCTTCATTCCACCGGAGTCTGG</td><td>BT20</td><td>217</td><td>130</td><td>200</td><td>10</td><td>44</td><td>37.9</td><td>131.2</td><td>81.6</td><td>7.1</td><td>42.4</td></tr>
<tr><td>>MI0000285|hsa-mir-205&WithFlank&1|+|209432127|209432248@40.61.22(+1G)&[MIMAT0000266&hsa-miR-205-5p&offsets|0|0(+1G);m-351&1|+|209432166|209432187&offsets|0|0(+1G)]|TCCTTCATTCCACCGGAGTCTGG</td><td>>MI0000285|hsa-mir-205&WithFlank&1|+|209432127|209432248</td><td>40</td><td>61</td><td>22(+1G)</td><td>TCCTTCATTCCACCGGAGTCTGG</td><td>MB468</td><td>14</td><td>5</td><td>10</td><td>10</td><td>5</td><td>3.5</td><td>2.2</td><td>6.0</td><td>7.7</td><td>2.7</td></tr>
<tr><td>>MI0000285|hsa-mir-205&WithFlank&1|+|209432127|209432248@40.61.22(+2A)&[MIMAT0000266&hsa-miR-205-5p&offsets|0|0(+2A);m-351&1|+|209432166|209432187&offsets|0|0(+2A)]|TCCTTCATTCCACCGGAGTCTGAA</td><td>>MI0000285|hsa-mir-205&WithFlank&1|+|209432127|209432248</td><td>40</td><td>61</td><td>22(+2A)</td><td>TCCTTCATTCCACCGGAGTCTGAA</td><td>BT20</td><td>76</td><td>66</td><td>188</td><td>4</td><td>46</td><td>68.6</td><td>62.8</td><td>154.6</td><td>1.7</td><td>42.3</td></tr>
<tr><td>>MI0000285|hsa-mir-205&WithFlank&1|+|209432127|209432248@40.61.22(+3A)&[MIMAT0000266&hsa-miR-205-5p&offsets|0|0(+3A);m-351&1|+|209432166|209432187&offsets|0|0(+3A)]|TCCTTCATTCCACCGGAGTCTGAAA</td><td>>MI0000285|hsa-mir-205&WithFlank&1|+|209432127|209432248</td><td>40</td><td>61</td><td>22(+3A)</td><td>TCCTTCATTCCACCGGAGTCTGAAA</td><td>BT20</td><td>13</td><td>13</td><td>30</td><td>1</td><td>6</td><td>9.6</td><td>12.7</td><td>23.5</td><td>0.9</td><td>8.0</td></tr>
<tr><td>>MI0000285|hsa-mir-205&WithFlank&1|+|209432127|209432248@40.61.22&[MIMAT0000266&hsa-miR-205-5p&offsets|0|0;m-351&1|+|209432166|209432187&offsets|0|0]|TCCTTCATTCCACCGGAGTCTG</td><td>>MI0000285|hsa-mir-205&WithFlank&1|+|209432127|209432248</td><td>40</td><td>61</td><td>22</td><td>TCCTTCATTCCACCGGAGTCTG</td><td>BT20</td><td>1741</td><td>1774</td><td>3052</td><td>2444</td><td>1268</td><td>1377.3</td><td>1871.6</td><td>2918.4</td><td>2377.7</td><td>1424.2</td></tr>
<tr><td>>MI0000285|hsa-mir-205&WithFlank&1|+|209432127|209432248@40.61.22&[MIMAT0000266&hsa-miR-205-5p&offsets|0|0;m-351&1|+|209432166|209432187&offsets|0|0]|TCCTTCATTCCACCGGAGTCTG</td><td>>MI0000285|hsa-mir-205&WithFlank&1|+|209432127|209432248</td><td>40</td><td>61</td><td>22</td><td>TCCTTCATTCCACCGGAGTCTG</td><td>MB468</td><td>138</td><td>67</td><td>62</td><td>408</td><td>72</td><td>64.9</td><td>23.6</td><td>6.9</td><td>273.6</td><td>41.8</td></tr>
<tr><td>>MI0000285|hsa-mir-205&WithFlank&1|+|209432127|209432248@40.62.23(+1A)&[MIMAT0000266&hsa-miR-205-5p&offsets|0|+1(+1A);m-351&1|+|209432166|209432187&offsets|0|+1(+1A)]|TCCTTCATTCCACCGGAGTCTGTA</td><td>>MI0000285|hsa-mir-205&WithFlank&1|+|209432127|209432248</td><td>40</td><td>62</td><td>23(+1A)</td><td>TCCTTCATTCCACCGGAGTCTGTA</td><td>BT20</td><td>119</td><td>86</td><td>218</td><td>5</td><td>69</td><td>119.7</td><td>95.4</td><td>166.6</td><td>2.7</td><td>80.4</td></tr>
<tr><td>>MI0000285|hsa-mir-205&WithFlank&1|+|209432127|209432248@40.62.23(+1U)&[MIMAT0000266&hsa-miR-205-5p&offsets|0|+1(+1U);m-351&1|+|209432166|209432187&offsets|0|+1(+1U)]|TCCTTCATTCCACCGGAGTCTGTT</td><td>>MI0000285|hsa-mir-205&WithFlank&1|+|209432127|209432248</td><td>40</td><td>62</td><td>23(+1U)</td><td>TCCTTCATTCCACCGGAGTCTGTT</td><td>BT20</td><td>72</td><td>70</td><td>118</td><td>10</td><td>48</td><td>61.5</td><td>80.7</td><td>104.7</td><td>7.0</td><td>57.0</td></tr>
<tr><td>>MI0000285|hsa-mir-205&WithFlank&1|+|209432127|209432248@40.62.23&[MIMAT0000266&hsa-miR-205-5p&offsets|0|+1;m-351&1|+|209432166|209432187&offsets|0|+1]|TCCTTCATTCCACCGGAGTCTGT</td><td>>MI0000285|hsa-mir-205&WithFlank&1|+|209432127|209432248</td><td>40</td><td>62</td><td>23</td><td>TCCTTCATTCCACCGGAGTCTGT</td><td>BT20</td><td>2552</td><td>2605</td><td>4661</td><td>314</td><td>1848</td><td>1523.1</td><td>2683.8</td><td>3287.3</td><td>202.3</td><td>2107.4</td></tr>
<tr><td>>MI0000285|hsa-mir-205&WithFlank&1|+|209432127|209432248@40.62.23&[MIMAT0000266&hsa-miR-205-5p&offsets|0|+1;m-351&1|+|209432166|209432187&offsets|0|+1]|TCCTTCATTCCACCGGAGTCTGT</td><td>>MI0000285|hsa-mir-205&WithFlank&1|+|209432127|209432248</td><td>40</td><td>62</td><td>23</td><td>TCCTTCATTCCACCGGAGTCTGT</td><td>MB468</td><td>136</td><td>80</td><td>78</td><td>181</td><td>60</td><td>56.5</td><td>19.1</td><td>13.2</td><td>183.2</td><td>34.9</td></tr>
<tr><td>>MI0000285|hsa-mir-205&WithFlank&1|+|209432127|209432248@40.63.24&[MIMAT0000266&hsa-miR-205-5p&offsets|0|+2;m-351&1|+|209432166|209432187&offsets|0|+2]|TCCTTCATTCCACCGGAGTCTGTC</td><td>>MI0000285|hsa-mir-205&WithFlank&1|+|209432127|209432248</td><td>40</td><td>63</td><td>24</td><td>TCCTTCATTCCACCGGAGTCTGTC</td><td>BT20</td><td>13</td><td>15</td><td>22</td><td>2</td><td>9</td><td>11.2</td><td>17.1</td><td>18.9</td><td>1.1</td><td>10.3</td></tr>
<tr><td>>MI0000286|hsa-mir-210&WithFlank&11|-|568083|568204@71.93.23&[MIMAT0000267&hsa-miR-210-3p&offsets|-1|0;m-102&11|-|568112|568133&offsets|-1|0]|ACTGTGCGTGTGACAGCGGCTGA</td><td>>MI0000286|hsa-mir-210&WithFlank&11|-|568083|568204</td><td>71</td><td>93</td><td>23</td><td>ACTGTGCGTGTGACAGCGGCTGA</td><td>BT20</td><td>11</td><td>1</td><td>2</td><td>4</td><td>1</td><td>15.8</td><td>1.0</td><td>1.6</td><td>5.6</td><td>0.4</td></tr>
<tr><td>>MI0000286|hsa-mir-210&WithFlank&11|-|568083|568204@72.91.20&[MIMAT0000267&hsa-miR-210-3p&offsets|0|-2;m-102&11|-|568112|568133&offsets|0|-2]|CTGTGCGTGTGACAGCGGCT</td><td>>MI0000286|hsa-mir-210&WithFlank&11|-|568083|568204</td><td>72</td><td>91</td><td>20</td><td>CTGTGCGTGTGACAGCGGCT</td><td>BT20</td><td>14</td><td>2</td><td>2</td><td>13</td><td>1</td><td>21.3</td><td>3.7</td><td>3.0</td><td>19.6</td><td>0.9</td></tr>
<tr><td>>MI0000286|hsa-mir-210&WithFlank&11|-|568083|568204@72.92.21(+1C)&[MIMAT0000267&hsa-miR-210-3p&offsets|0|-1(+1C);m-102&11|-|568112|568133&offsets|0|-1(+1C)]|CTGTGCGTGTGACAGCGGCTGC</td><td>>MI0000286|hsa-mir-210&WithFlank&11|-|568083|568204</td><td>72</td><td>92</td><td>21(+1C)</td><td>CTGTGCGTGTGACAGCGGCTGC</td><td>BT20</td><td>14</td><td>1</td><td>2</td><td>10</td><td>1</td><td>19.2</td><td>1.2</td><td>1.4</td><td>15.5</td><td>0.5</td></tr>
<tr><td>>MI0000286|hsa-mir-210&WithFlank&11|-|568083|568204@72.92.21(+1G)&[MIMAT0000267&hsa-miR-210-3p&offsets|0|-1(+1G);m-102&11|-|568112|568133&offsets|0|-1(+1G)]|CTGTGCGTGTGACAGCGGCTGG</td><td>>MI0000286|hsa-mir-210&WithFlank&11|-|568083|568204</td><td>72</td><td>92</td><td>21(+1G)</td><td>CTGTGCGTGTGACAGCGGCTGG</td><td>BT20</td><td>43</td><td>1</td><td>1</td><td>7</td><td>0</td><td>64.3</td><td>0.6</td><td>1.4</td><td>10.9</td><td>0.4</td></tr>
<tr><td>>MI0000286|hsa-mir-210&WithFlank&11|-|568083|568204@72.92.21(+1U)&[MIMAT0000267&hsa-miR-210-3p&offsets|0|-1(+1U);m-102&11|-|568112|568133&offsets|0|-1(+1U)]|CTGTGCGTGTGACAGCGGCTGT</td><td>>MI0000286|hsa-mir-210&WithFlank&11|-|568083|568204</td><td>72</td><td>92</td><td>21(+1U)</td><td>CTGTGCGTGTGACAGCGGCTGT</td><td>BT20</td><td>105</td><td>9</td><td>12</td><td>59</td><td>8</td><td>151.8</td><td>6.9</td><td>9.3</td><td>87.9</td><td>3.8</td></tr>
<tr><td>>MI0000286|hsa-mir-210&WithFlank&11|-|568083|568204@72.92.21&[MIMAT0000267&hsa-miR-210-3p&offsets|0|-1;m-102&11|-|568112|568133&offsets|0|-1]|CTGTGCGTGTGACAGCGGCTG</td><td>>MI0000286|hsa-mir-210&WithFlank&11|-|568083|568204</td><td>72</td><td>92</td><td>21</td><td>CTGTGCGTGTGACAGCGGCTG</td><td>BT20</td><td>17</td><td>2</td><td>2</td><td>11</td><td>0</td><td>24.6</td><td>1.9</td><td>1.1</td><td>15.5</td><td>0.7</td></tr>
<tr><td>>MI0000286|hsa-mir-210&WithFlank&11|-|568083|568204@72.93.22(+1A)&[MIMAT0000267&hsa-miR-210-3p&offsets|0|0(+1A);m-102&11|-|568112|568133&offsets|0|0(+1A)]|CTGTGCGTGTGACAGCGGCTGAA</td><td>>MI0000286|hsa-mir-210&WithFlank&11|-|568083|568204</td><td>72</td><td>93</td><td>22(+1A)</td><td>CTGTGCGTGTGACAGCGGCTGAA</td><td>BT20</td><td>240</td><td>44</td><td>40</td><td>107</td><td>39</td><td>335.0</td><td>44.4</td><td>36.4</td><td>137.7</td><td>29.1</td></tr>
<tr><td>>MI0000286|hsa-mir-210&WithFlank&11|-|568083|568204@72.93.22(+1A)&[MIMAT0000267&hsa-miR-210-3p&offsets|0|0(+1A);m-102&11|-|568112|568133&offsets|0|0(+1A)]|CTGTGCGTGTGACAGCGGCTGAA</td><td>>MI0000286|hsa-mir-210&WithFlank&11|-|568083|568204</td><td>72</td><td>93</td><td>22(+1A)</td><td>CTGTGCGTGTGACAGCGGCTGAA</td><td>MB231</td><td>10</td><td>6</td><td>2</td><td>47</td><td>16</td><td>1.1</td><td>3.3</td><td>1.7</td><td>21.2</td><td>15.0</td></tr>
<tr><td>>MI0000286|hsa-mir-210&WithFlank&11|-|568083|568204@72.93.22(+1C)&[MIMAT0000267&hsa-miR-210-3p&offsets|0|0(+1C);m-102&11|-|568112|568133&offsets|0|0(+1C)]|CTGTGCGTGTGACAGCGGCTGAC</td><td>>MI0000286|hsa-mir-210&WithFlank&11|-|568083|568204</td><td>72</td><td>93</td><td>22(+1C)</td><td>CTGTGCGTGTGACAGCGGCTGAC</td><td>BT20</td><td>28</td><td>6</td><td>6</td><td>16</td><td>6</td><td>36.8</td><td>5.8</td><td>5.7</td><td>21.6</td><td>3.6</td></tr>
<tr><td>>MI0000286|hsa-mir-210&WithFlank&11|-|568083|568204@72.93.22(+1G)&[MIMAT0000267&hsa-miR-210-3p&offsets|0|0(+1G);m-102&11|-|568112|568133&offsets|0|0(+1G)]|CTGTGCGTGTGACAGCGGCTGAG</td><td>>MI0000286|hsa-mir-210&WithFlank&11|-|568083|568204</td><td>72</td><td>93</td><td>22(+1G)</td><td>CTGTGCGTGTGACAGCGGCTGAG</td><td>BT20</td><td>35</td><td>2</td><td>2</td><td>5</td><td>1</td><td>53.3</td><td>1.8</td><td>2.4</td><td>9.0</td><td>1.0</td></tr>
<tr><td>>MI0000286|hsa-mir-210&WithFlank&11|-|568083|568204@72.93.22(+2A)&[MIMAT0000267&hsa-miR-210-3p&offsets|0|0(+2A);m-102&11|-|568112|568133&offsets|0|0(+2A)]|CTGTGCGTGTGACAGCGGCTGAAA</td><td>>MI0000286|hsa-mir-210&WithFlank&11|-|568083|568204</td><td>72</td><td>93</td><td>22(+2A)</td><td>CTGTGCGTGTGACAGCGGCTGAAA</td><td>BT20</td><td>228</td><td>185</td><td>118</td><td>216</td><td>213</td><td>178.7</td><td>231.1</td><td>108.7</td><td>186.8</td><td>188.1</td></tr>
<tr><td>>MI0000286|hsa-mir-210&WithFlank&11|-|568083|568204@72.93.22(+2A)&[MIMAT0000267&hsa-miR-210-3p&offsets|0|0(+2A);m-102&11|-|568112|568133&offsets|0|0(+2A)]|CTGTGCGTGTGACAGCGGCTGAAA</td><td>>MI0000286|hsa-mir-210&WithFlank&11|-|568083|568204</td><td>72</td><td>93</td><td>22(+2A)</td><td>CTGTGCGTGTGACAGCGGCTGAAA</td><td>MB231</td><td>33</td><td>25</td><td>6</td><td>139</td><td>48</td><td>11.7</td><td>14.2</td><td>4.3</td><td>60.5</td><td>26.2</td></tr>
<tr><td>>MI0000286|hsa-mir-210&WithFlank&11|-|568083|568204@72.93.22(+3A)&[MIMAT0000267&hsa-miR-210-3p&offsets|0|0(+3A);m-102&11|-|568112|568133&offsets|0|0(+3A)]|CTGTGCGTGTGACAGCGGCTGAAAA</td><td>>MI0000286|hsa-mir-210&WithFlank&11|-|568083|568204</td><td>72</td><td>93</td><td>22(+3A)</td><td>CTGTGCGTGTGACAGCGGCTGAAAA</td><td>BT20</td><td>16</td><td>19</td><td>13</td><td>17</td><td>19</td><td>11.6</td><td>25.4</td><td>15.8</td><td>11.7</td><td>20.7</td></tr>
</table>

<table>
<tr><td>>MI0000286|hsa-mir-210&WithFlank&11|-|568083|568204@72.93.22&[MIMAT0000267&hsa-miR-210-3p&offsets|0|0;m-102&11|-|568112|568133&offsets|0|0]|CTGTGCGTGTGACAGCGGCTGA</td><td>>MI0000286|hsa-mir-210&WithFlank&11|-|568083|568204</td><td>72</td><td>93</td><td>22</td><td>CTGTGCGTGTGACAGCGGCTGA</td><td>BT20</td><td>331</td><td>41</td><td>42</td><td>204</td><td>30</td><td>478.4</td><td>32.0</td><td>36.0</td><td>289.9</td><td>19.2</td></tr>
<tr><td>>MI0000286|hsa-mir-210&WithFlank&11|-|568083|568204@72.93.22&[MIMAT0000267&hsa-miR-210-3p&offsets|0|0;m-102&11|-|568112|568133&offsets|0|0]|CTGTGCGTGTGACAGCGGCTGA</td><td>>MI0000286|hsa-mir-210&WithFlank&11|-|568083|568204</td><td>72</td><td>93</td><td>22</td><td>CTGTGCGTGTGACAGCGGCTGA</td><td>MB231</td><td>14</td><td>7</td><td>2</td><td>93</td><td>28</td><td>1.4</td><td>3.9</td><td>1.9</td><td>29.5</td><td>31.3</td></tr>
<tr><td>>MI0000286|hsa-mir-210&WithFlank&11|-|568083|568204@72.94.23(+1A)&[MIMAT0000267&hsa-miR-210-3p&offsets|0|+1(+1A);m-102&11|-|568112|568133&offsets|0|+1(+1A)]|CTGTGCGTGTGACAGCGGCTGATA</td><td>>MI0000286|hsa-mir-210&WithFlank&11|-|568083|568204</td><td>72</td><td>94</td><td>23(+1A)</td><td>CTGTGCGTGTGACAGCGGCTGATA</td><td>BT20</td><td>12</td><td>1</td><td>2</td><td>1</td><td>0</td><td>18.9</td><td>1.1</td><td>1.5</td><td>2.2</td><td>0.3</td></tr>
<tr><td>>MI0000286|hsa-mir-210&WithFlank&11|-|568083|568204@72.94.23(+1U)&[MIMAT0000267&hsa-miR-210-3p&offsets|0|+1(+1U);m-102&11|-|568112|568133&offsets|0|+1(+1U)]|CTGTGCGTGTGACAGCGGCTGATT</td><td>>MI0000286|hsa-mir-210&WithFlank&11|-|568083|568204</td><td>72</td><td>94</td><td>23(+1U)</td><td>CTGTGCGTGTGACAGCGGCTGATT</td><td>BT20</td><td>38</td><td>4</td><td>3</td><td>7</td><td>3</td><td>57.9</td><td>3.5</td><td>3.6</td><td>9.5</td><td>1.9</td></tr>
<tr><td>>MI0000286|hsa-mir-210&WithFlank&11|-|568083|568204@72.94.23&[MIMAT0000267&hsa-miR-210-3p&offsets|0|+1;m-102&11|-|568112|568133&offsets|0|+1]|CTGTGCGTGTGACAGCGGCTGAT</td><td>>MI0000286|hsa-mir-210&WithFlank&11|-|568083|568204</td><td>72</td><td>94</td><td>23</td><td>CTGTGCGTGTGACAGCGGCTGAT</td><td>BT20</td><td>149</td><td>21</td><td>18</td><td>44</td><td>17</td><td>212.1</td><td>17.5</td><td>17.1</td><td>59.1</td><td>9.5</td></tr>
<tr><td>>MI0000286|hsa-mir-210&WithFlank&11|-|568083|568204@72.94.23&[MIMAT0000267&hsa-miR-210-3p&offsets|0|+1;m-102&11|-|568112|568133&offsets|0|+1]|CTGTGCGTGTGACAGCGGCTGAT</td><td>>MI0000286|hsa-mir-210&WithFlank&11|-|568083|568204</td><td>72</td><td>94</td><td>23</td><td>CTGTGCGTGTGACAGCGGCTGAT</td><td>MB231</td><td>11</td><td>6</td><td>2</td><td>55</td><td>22</td><td>2.3</td><td>3.9</td><td>1.6</td><td>29.4</td><td>23.5</td></tr>
<tr><td>>MI0000289|hsa-mir-181a-1&WithFlank&1|-|198859038|198859159@30.49.20;MI0000269|hsa-mir-181a-2&WithFlank&9|+|124692436|124692557@45.64.20&[MIMAT0000256&hsa-miR-181a-5p&offsets|0|-3;m-32&1|-|198859108|198859130&offsets|0|-3];[MIMAT0000256_1&hsa-miR-181a-5p&offsets|0|-3;m-32&9|+|124692480|124692502&offsets|0|-3]|AACATTCAACGCTGTCGGTG</td><td>>MI0000289|hsa-mir-181a-1&WithFlank&1|-|198859038|198859159</td><td>30</td><td>49</td><td>20</td><td>AACATTCAACGCTGTCGGTG</td><td>MB231</td><td>27</td><td>34</td><td>8</td><td>90</td><td>40</td><td>3.9</td><td>21.5</td><td>5.0</td><td>51.2</td><td>17.6</td></tr>
<tr><td>>MI0000289|hsa-mir-181a-1&WithFlank&1|-|198859038|198859159@30.50.21;MI0000269|hsa-mir-181a-2&WithFlank&9|+|124692436|124692557@45.65.21&[MIMAT0000256&hsa-miR-181a-5p&offsets|0|-2;m-32&1|-|198859108|198859130&offsets|0|-2];[MIMAT0000256_1&hsa-miR-181a-5p&offsets|0|-2;m-32&9|+|124692480|124692502&offsets|0|-2]|AACATTCAACGCTGTCGGTGA</td><td>>MI0000289|hsa-mir-181a-1&WithFlank&1|-|198859038|198859159</td><td>30</td><td>50</td><td>21</td><td>AACATTCAACGCTGTCGGTGA</td><td>MB231</td><td>26</td><td>30</td><td>6</td><td>57</td><td>26</td><td>6.8</td><td>15.6</td><td>4.0</td><td>32.7</td><td>10.6</td></tr>
<tr><td>>MI0000289|hsa-mir-181a-1&WithFlank&1|-|198859038|198859159@30.51.22;MI0000269|hsa-mir-181a-2&WithFlank&9|+|124692436|124692557@45.66.22&[MIMAT0000256&hsa-miR-181a-5p&offsets|0|-1;m-32&1|-|198859108|198859130&offsets|0|-1];[MIMAT0000256_1&hsa-miR-181a-5p&offsets|0|-1;m-32&9|+|124692480|124692502&offsets|0|-1]|AACATTCAACGCTGTCGGTGAG</td><td>>MI0000289|hsa-mir-181a-1&WithFlank&1|-|198859038|198859159</td><td>30</td><td>51</td><td>22</td><td>AACATTCAACGCTGTCGGTGAG</td><td>MB231</td><td>28</td><td>30</td><td>15</td><td>68</td><td>31</td><td>5.7</td><td>17.6</td><td>12.4</td><td>39.7</td><td>11.3</td></tr>
<tr><td>>MI0000289|hsa-mir-181a-1&WithFlank&1|-|198859038|198859159@30.51.22(+1A);MI0000269|hsa-mir-181a-2&WithFlank&9|+|124692436|124692557@45.66.22(+1A)&[MIMAT0000256&hsa-miR-181a-5p&offsets|0|-1(+1A);m-32&1|-|198859108|198859130&offsets|0|-1(+1A)];[MIMAT0000256_1&hsa-miR-181a-5p&offsets|0|-1(+1A);m-32&9|+|124692480|124692502&offsets|0|-1(+1A)]|AACATTCAACGCTGTCGGTGAGA</td><td>>MI0000289|hsa-mir-181a-1&WithFlank&1|-|198859038|198859159</td><td>30</td><td>51</td><td>22(+1A)</td><td>AACATTCAACGCTGTCGGTGAGA</td><td>MB231</td><td>18</td><td>17</td><td>7</td><td>58</td><td>19</td><td>1.1</td><td>9.4</td><td>5.3</td><td>31.5</td><td>8.3</td></tr>
<tr><td>>MI0000289|hsa-mir-181a-1&WithFlank&1|-|198859038|198859159@30.51.22(+1C);MI0000269|hsa-mir-181a-2&WithFlank&9|+|124692436|124692557@45.66.22(+1C)&[MIMAT0000256&hsa-miR-181a-5p&offsets|0|-1(+1C);m-32&1|-|198859108|198859130&offsets|0|-1(+1C)];[MIMAT0000256_1&hsa-miR-181a-5p&offsets|0|-1(+1C);m-32&9|+|124692480|124692502&offsets|0|-1(+1C)]|AACATTCAACGCTGTCGGTGAGC</td><td>>MI0000289|hsa-mir-181a-1&WithFlank&1|-|198859038|198859159</td><td>30</td><td>51</td><td>22(+1C)</td><td>AACATTCAACGCTGTCGGTGAGC</td><td>BT20</td><td>10</td><td>11</td><td>9</td><td>13</td><td>6</td><td>4.7</td><td>7.1</td><td>2.9</td><td>11.3</td><td>2.2</td></tr>
<tr><td>>MI0000289|hsa-mir-181a-1&WithFlank&1|-|198859038|198859159@30.51.22(+1C);MI0000269|hsa-mir-181a-2&WithFlank&9|+|124692436|124692557@45.66.22(+1C)&[MIMAT0000256&hsa-miR-181a-5p&offsets|0|-1(+1C);m-32&1|-|198859108|198859130&offsets|0|-1(+1C)];[MIMAT0000256_1&hsa-miR-181a-5p&offsets|0|-1(+1C);m-32&9|+|124692480|124692502&offsets|0|-1(+1C)]|AACATTCAACGCTGTCGGTGAGC</td><td>>MI0000289|hsa-mir-181a-1&WithFlank&1|-|198859038|198859159</td><td>30</td><td>51</td><td>22(+1C)</td><td>AACATTCAACGCTGTCGGTGAGC</td><td>MB231</td><td>26</td><td>31</td><td>8</td><td>204</td><td>53</td><td>6.0</td><td>21.6</td><td>6.3</td><td>117.8</td><td>44.5</td></tr>
<tr><td>>MI0000289|hsa-mir-181a-1&WithFlank&1|-|198859038|198859159@30.51.22(+1G);MI0000269|hsa-mir-181a-2&WithFlank&9|+|124692436|124692557@45.66.22(+1G)&[MIMAT0000256&hsa-miR-181a-5p&offsets|0|-1(+1G);m-32&1|-|198859108|198859130&offsets|0|-1(+1G)];[MIMAT0000256_1&hsa-miR-181a-5p&offsets|0|-1(+1G);m-32&9|+|124692480|124692502&offsets|0|-1(+1G)]|AACATTCAACGCTGTCGGTGAGG</td><td>>MI0000289|hsa-mir-181a-1&WithFlank&1|-|198859038|198859159</td><td>30</td><td>51</td><td>22(+1G)</td><td>AACATTCAACGCTGTCGGTGAGG</td><td>MB231</td><td>14</td><td>6</td><td>2</td><td>21</td><td>5</td><td>12.3</td><td>4.6</td><td>1.3</td><td>15.3</td><td>4.2</td></tr>
<tr><td>>MI0000289|hsa-mir-181a-1&WithFlank&1|-|198859038|198859159@30.52.23;MI0000269|hsa-mir-181a-2&WithFlank&9|+|124692436|124692557@45.67.23&[MIMAT0000256&hsa-miR-181a-5p&offsets|0|0;m-32&1|-|198859108|198859130&offsets|0|0];[MIMAT0000256_1&hsa-miR-181a-5p&offsets|0|0;m-32&9|+|124692480|124692502&offsets|0|0]|AACATTCAACGCTGTCGGTGAGT</td><td>>MI0000289|hsa-mir-181a-1&WithFlank&1|-|198859038|198859159</td><td>30</td><td>52</td><td>23</td><td>AACATTCAACGCTGTCGGTGAGT</td><td>BT20</td><td>79</td><td>70</td><td>67</td><td>92</td><td>44</td><td>41.0</td><td>42.3</td><td>17.9</td><td>70.8</td><td>17.8</td></tr>
<tr><td>>MI0000289|hsa-mir-181a-1&WithFlank&1|-|198859038|198859159@30.52.23;MI0000269|hsa-mir-181a-2&WithFlank&9|+|124692436|124692557@45.67.23&[MIMAT0000256&hsa-miR-181a-5p&offsets|0|0;m-32&1|-|198859108|198859130&offsets|0|0];[MIMAT0000256_1&hsa-miR-181a-5p&offsets|0|0;m-32&9|+|124692480|124692502&offsets|0|0]|AACATTCAACGCTGTCGGTGAGT</td><td>>MI0000289|hsa-mir-181a-1&WithFlank&1|-|198859038|198859159</td><td>30</td><td>52</td><td>23</td><td>AACATTCAACGCTGTCGGTGAGT</td><td>MB231</td><td>241</td><td>283</td><td>74</td><td>1271</td><td>383</td><td>46.0</td><td>192.9</td><td>57.6</td><td>571.3</td><td>273.9</td></tr>
<tr><td>>MI0000289|hsa-mir-181a-1&WithFlank&1|-|198859038|198859159@30.52.23;MI0000269|hsa-mir-181a-2&WithFlank&9|+|124692436|124692557@45.67.23&[MIMAT0000256&hsa-miR-181a-5p&offsets|0|0;m-32&1|-|198859108|198859130&offsets|0|0];[MIMAT0000256_1&hsa-miR-181a-5p&offsets|0|0;m-32&9|+|124692480|124692502&offsets|0|0]|AACATTCAACGCTGTCGGTGAGT</td><td>>MI0000289|hsa-mir-181a-1&WithFlank&1|-|198859038|198859159</td><td>30</td><td>52</td><td>23</td><td>AACATTCAACGCTGTCGGTGAGT</td><td>MB468</td><td>22</td><td>7</td><td>9</td><td>69</td><td>7</td><td>11.9</td><td>4.6</td><td>1.3</td><td>47.0</td><td>0.9</td></tr>
<tr><td>>MI0000289|hsa-mir-181a-1&WithFlank&1|-|198859038|198859159@30.53.24;MI0000269|hsa-mir-181a-2&WithFlank&9|+|124692436|124692557@45.68.24&[MIMAT0000256&hsa-miR-181a-5p&offsets|0|+1;m-32&1|-|198859108|198859130&offsets|0|+1];[MIMAT0000256_1&hsa-miR-181a-5p&offsets|0|+1;m-32&9|+|124692480|124692502&offsets|0|+1]|AACATTCAACGCTGTCGGTGAGTT</td><td>>MI0000289|hsa-mir-181a-1&WithFlank&1|-|198859038|198859159</td><td>30</td><td>53</td><td>24</td><td>AACATTCAACGCTGTCGGTGAGTT</td><td>BT20</td><td>20</td><td>11</td><td>8</td><td>32</td><td>5</td><td>16.3</td><td>10.2</td><td>0.8</td><td>32.7</td><td>4.7</td></tr>
<tr><td>>MI0000289|hsa-mir-181a-1&WithFlank&1|-|198859038|198859159@30.53.24;MI0000269|hsa-mir-181a-2&WithFlank&9|+|124692436|124692557@45.68.24&[MIMAT0000256&hsa-miR-181a-5p&offsets|0|+1;m-32&1|-|198859108|198859130&offsets|0|+1];[MIMAT0000256_1&hsa-miR-181a-5p&offsets|0|+1;m-32&9|+|124692480|124692502&offsets|0|+1]|AACATTCAACGCTGTCGGTGAGTT</td><td>>MI0000289|hsa-mir-181a-1&WithFlank&1|-|198859038|198859159</td><td>30</td><td>53</td><td>24</td><td>AACATTCAACGCTGTCGGTGAGTT</td><td>MB231</td><td>81</td><td>53</td><td>13</td><td>598</td><td>162</td><td>21.1</td><td>36.8</td><td>11.6</td><td>175.5</td><td>192.0</td></tr>
<tr><td>>MI0000289|hsa-mir-181a-1&WithFlank&1|-|198859038|198859159@30.54.25;MI0000269|hsa-mir-181a-2&WithFlank&9|+|124692436|124692557@45.69.25&[MIMAT0000256&hsa-miR-181a-5p&offsets|0|+2;m-32&1|-|198859108|198859130&offsets|0|+2];[MIMAT0000256_1&hsa-miR-181a-5p&offsets|0|+2;m-32&9|+|124692480|124692502&offsets|0|+2]|AACATTCAACGCTGTCGGTGAGTTT</td><td>>MI0000289|hsa-mir-181a-1&WithFlank&1|-|198859038|198859159</td><td>30</td><td>54</td><td>25</td><td>AACATTCAACGCTGTCGGTGAGTTT</td><td>BT20</td><td>17</td><td>12</td><td>9</td><td>18</td><td>6</td><td>14.6</td><td>11.8</td><td>2.5</td><td>15.1</td><td>4.5</td></tr>
<tr><td>>MI0000289|hsa-mir-181a-1&WithFlank&1|-|198859038|198859159@30.54.25;MI0000269|hsa-mir-181a-2&WithFlank&9|+|124692436|124692557@45.69.25&[MIMAT0000256&hsa-miR-181a-5p&offsets|0|+2;m-32&1|-|198859108|198859130&offsets|0|+2];[MIMAT0000256_1&hsa-miR-181a-5p&offsets|0|+2;m-32&9|+|124692480|124692502&offsets|0|+2]|AACATTCAACGCTGTCGGTGAGTTT</td><td>>MI0000289|hsa-mir-181a-1&WithFlank&1|-|198859038|198859159</td><td>30</td><td>54</td><td>25</td><td>AACATTCAACGCTGTCGGTGAGTTT</td><td>MB231</td><td>88</td><td>67</td><td>17</td><td>483</td><td>194</td><td>7.6</td><td>44.8</td><td>14.2</td><td>134.8</td><td>215.0</td></tr>
<tr><td>>MI0000289|hsa-mir-181a-1&WithFlank&1|-|198859038|198859159@30.54.25(+1U);MI0000269|hsa-mir-181a-2&WithFlank&9|+|124692436|124692557@45.69.25(+1U)&[MIMAT0000256&hsa-miR-181a-5p&offsets|0|+2(+1U);m-32&1|-|198859108|198859130&offsets|0|+2(+1U)];[MIMAT0000256_1&hsa-miR-181a-5p&offsets|0|+2(+1U);m-32&9|+|124692480|124692502&offsets|0|+2(+1U)]|AACATTCAACGCTGTCGGTGAGTTTT</td><td>>MI0000289|hsa-mir-181a-1&WithFlank&1|-|198859038|198859159</td><td>30</td><td>54</td><td>25(+1U)</td><td>AACATTCAACGCTGTCGGTGAGTTTT</td><td>MB231</td><td>33</td><td>24</td><td>4</td><td>92</td><td>58</td><td>5.7</td><td>16.1</td><td>3.1</td><td>21.2</td><td>63.4</td></tr>
<tr><td>>MI0000289|hsa-mir-181a-1&WithFlank&1|-|198859038|198859159@70.90.21(+1A)&[MIMAT0000270&hsa-miR-181a-3p&offsets|0|-1(+1A);m-125&1|-|198859069|198859090&offsets|0|-1(+1A)]|ACCATCGACCGTTGATTGTACA</td><td>>MI0000289|hsa-mir-181a-1&WithFlank&1|-|198859038|198859159</td><td>70</td><td>90</td><td>21(+1A)</td><td>ACCATCGACCGTTGATTGTACA</td><td>MB231</td><td>11</td><td>4</td><td>13</td><td>12</td><td>10</td><td>5.1</td><td>1.2</td><td>9.8</td><td>2.3</td><td>7.6</td></tr>
<tr><td>>MI0000289|hsa-mir-181a-1&WithFlank&1|-|198859038|198859159@70.90.21(+1U)&[MIMAT0000270&hsa-miR-181a-3p&offsets|0|-1(+1U);m-125&1|-|198859069|198859090&offsets|0|-1(+1U)]|ACCATCGACCGTTGATTGTACT</td><td>>MI0000289|hsa-mir-181a-1&WithFlank&1|-|198859038|198859159</td><td>70</td><td>90</td><td>21(+1U)</td><td>ACCATCGACCGTTGATTGTACT</td><td>MB231</td><td>30</td><td>13</td><td>28</td><td>32</td><td>26</td><td>12.1</td><td>4.0</td><td>17.4</td><td>8.8</td><td>20.7</td></tr>
</table>

| | | | | | | | | | | | | | | | | |
|---|---|---|---|---|---|---|---|---|---|---|---|---|---|---|---|---|
| >MI0000289\|hsa-mir-181a-1&WithFlank&1\|-\|198859038\|198859159@70.91.22&[MIMAT0000270&hsa-miR-181a-3p&offsets\|0\|0;m-125&1\|-\|198859069\|198859090&offsets\|0\|0]\|ACCATCGACCGTTGATTGTACC | >MI0000289\|hsa-mir-181a-1&WithFlank&1\|-\|198859038\|198859159 | 70 | 91 | 22 | ACCATCGACCGTTGATTGTACC | MB231 | 83 | 49 | 90 | 60 | 64 | 31.3 | 15.9 | 56.1 | 18.2 | 34.9 |
| >MI0000294\|hsa-mir-218-1&WithFlank&4\|+\|20528269\|20528390@31.51.21;MI0000295\|hsa-mir-218-2&WithFlank&5\|-\|168768140\|168768261@31.51.21&[MIMAT0000275&hsa-miR-218-5p&offsets\|0\|0;m-185&4\|+\|20528299\|20528319&offsets\|0\|0];[MIMAT0000275_1&hsa-miR-218-5p&offsets\|0\|0;m-187&5\|-\|168768211\|168768231&offsets\|0\|0]\|TTGTGCTTGATCTAACCATGT | >MI0000294\|hsa-mir-218-1&WithFlank&4\|+\|20528269\|20528390 | 31 | 51 | 21 | TTGTGCTTGATCTAACCATGT | MB231 | 31 | 25 | 4 | 66 | 24 | 10.6 | 12.0 | 1.6 | 27.1 | 15.1 |
| >MI0000294\|hsa-mir-218-1&WithFlank&4\|+\|20528269\|20528390@31.52.22;MI0000295\|hsa-mir-218-2&WithFlank&5\|-\|168768140\|168768261@31.52.22&[MIMAT0000275&hsa-miR-218-5p&offsets\|0\|+1;m-185&4\|+\|20528299\|20528319&offsets\|0\|+1];[MIMAT0000275_1&hsa-miR-218-5p&offsets\|0\|+1;m-187&5\|-\|168768211\|168768231&offsets\|0\|+1]\|TTGTGCTTGATCTAACCATGTG | >MI0000294\|hsa-mir-218-1&WithFlank&4\|+\|20528269\|20528390 | 31 | 52 | 22 | TTGTGCTTGATCTAACCATGTG | MB231 | 16 | 14 | 2 | 22 | 9 | 3.8 | 7.1 | 1.3 | 10.5 | 4.0 |
| >MI0000294\|hsa-mir-218-1&WithFlank&4\|+\|20528269\|20528390@31.52.22(+1U);MI0000295\|hsa-mir-218-2&WithFlank&5\|-\|168768140\|168768261@31.52.22(+1U)&[MIMAT0000275&hsa-miR-218-5p&offsets\|0\|+1(+1U);m-185&4\|+\|20528299\|20528319&offsets\|0\|+1(+1U)];[MIMAT0000275_1&hsa-miR-218-5p&offsets\|0\|+1(+1U);m-187&5\|-\|168768211\|168768231&offsets\|0\|+1(+1U)]\|TTGTGCTTGATCTAACCATGTGT | >MI0000294\|hsa-mir-218-1&WithFlank&4\|+\|20528269\|20528390 | 31 | 52 | 22(+1U) | TTGTGCTTGATCTAACCATGTGT | MB231 | 13 | 12 | 2 | 23 | 8 | 3.4 | 4.9 | 0.5 | 10.4 | 3.7 |
| >MI0000298\|hsa-mir-221&WithFlank&X\|-\|45746151\|45746272@31.52.22&[MIMAT0004568&hsa-miR-221-5p&offsets\|0\|0;m-405&X\|-\|45746220\|45746242&offsets\|0\|-1]\|ACCTGGCATACAATGTAGATTT | >MI0000298\|hsa-mir-221&WithFlank&X\|-\|45746151\|45746272 | 31 | 52 | 22 | ACCTGGCATACAATGTAGATTT | MB231 | 12 | 8 | 2 | 3 | 10 | 4.7 | 3.0 | 1.8 | 1.7 | 5.4 |
| >MI0000298\|hsa-mir-221&WithFlank&X\|-\|45746151\|45746272@31.53.23(+1C)&[MIMAT0004568&hsa-miR-221-5p&offsets\|0\|+1(+1C);m-405&X\|-\|45746220\|45746242&offsets\|0\|0(+1C)]\|ACCTGGCATACAATGTAGATTTCC | >MI0000298\|hsa-mir-221&WithFlank&X\|-\|45746151\|45746272 | 31 | 53 | 23(+1C) | ACCTGGCATACAATGTAGATTTCC | MB231 | 15 | 9 | 8 | 6 | 17 | 3.3 | 1.3 | 4.5 | 0.9 | 10.6 |
| >MI0000298\|hsa-mir-221&WithFlank&X\|-\|45746151\|45746272@31.53.23&[MIMAT0004568&hsa-miR-221-5p&offsets\|0\|+1;m-405&X\|-\|45746220\|45746242&offsets\|0\|0]\|ACCTGGCATACAATGTAGATTTC | >MI0000298\|hsa-mir-221&WithFlank&X\|-\|45746151\|45746272 | 31 | 53 | 23 | ACCTGGCATACAATGTAGATTTC | BT20 | 15 | 9 | 13 | 12 | 7 | 8.1 | 4.0 | 8.1 | 11.2 | 5.1 |
| >MI0000298\|hsa-mir-221&WithFlank&X\|-\|45746151\|45746272@31.53.23&[MIMAT0004568&hsa-miR-221-5p&offsets\|0\|+1;m-405&X\|-\|45746220\|45746242&offsets\|0\|0]\|ACCTGGCATACAATGTAGATTTC | >MI0000298\|hsa-mir-221&WithFlank&X\|-\|45746151\|45746272 | 31 | 53 | 23 | ACCTGGCATACAATGTAGATTTC | MB231 | 43 | 32 | 20 | 29 | 36 | 9.4 | 8.8 | 13.0 | 6.1 | 10.7 |
| >MI0000298\|hsa-mir-221&WithFlank&X\|-\|45746151\|45746272@31.54.24&[MIMAT0004568&hsa-miR-221-5p&offsets\|0\|+2;m-405&X\|-\|45746220\|45746242&offsets\|0\|+1]\|ACCTGGCATACAATGTAGATTTCT | >MI0000298\|hsa-mir-221&WithFlank&X\|-\|45746151\|45746272 | 31 | 54 | 24 | ACCTGGCATACAATGTAGATTTCT | BT20 | 28 | 14 | 25 | 2 | 13 | 12.5 | 8.2 | 10.8 | 1.6 | 13.6 |
| >MI0000298\|hsa-mir-221&WithFlank&X\|-\|45746151\|45746272@31.54.24&[MIMAT0004568&hsa-miR-221-5p&offsets\|0\|+2;m-405&X\|-\|45746220\|45746242&offsets\|0\|+1]\|ACCTGGCATACAATGTAGATTTCT | >MI0000298\|hsa-mir-221&WithFlank&X\|-\|45746151\|45746272 | 31 | 54 | 24 | ACCTGGCATACAATGTAGATTTCT | MB231 | 141 | 90 | 72 | 33 | 118 | 32.5 | 24.8 | 43.7 | 2.1 | 60.6 |
| >MI0000298\|hsa-mir-221&WithFlank&X\|-\|45746151\|45746272@31.55.25(+1C)&[MIMAT0004568&hsa-miR-221-5p&offsets\|0\|+3(+1C);m-405&X\|-\|45746220\|45746242&offsets\|0\|+2(+1C)]\|ACCTGGCATACAATGTAGATTTCTGC | >MI0000298\|hsa-mir-221&WithFlank&X\|-\|45746151\|45746272 | 31 | 55 | 25(+1C) | ACCTGGCATACAATGTAGATTTCTGC | MB231 | 17 | 11 | 13 | 2 | 13 | 3.5 | 1.6 | 4.5 | 0.3 | 4.8 |
| >MI0000298\|hsa-mir-221&WithFlank&X\|-\|45746151\|45746272@31.55.25&[MIMAT0004568&hsa-miR-221-5p&offsets\|0\|+3;m-405&X\|-\|45746220\|45746242&offsets\|0\|+2]\|ACCTGGCATACAATGTAGATTTCTG | >MI0000298\|hsa-mir-221&WithFlank&X\|-\|45746151\|45746272 | 31 | 55 | 25 | ACCTGGCATACAATGTAGATTTCTG | MB231 | 24 | 16 | 18 | 3 | 14 | 4.8 | 4.7 | 10.8 | 0.6 | 2.5 |
| >MI0000298\|hsa-mir-221&WithFlank&X\|-\|45746151\|45746272@31.56.26(+1U)&[MIMAT0004568&hsa-miR-221-5p&offsets\|0\|+4(+1U);m-405&X\|-\|45746220\|45746242&offsets\|0\|+3(+1U)]\|ACCTGGCATACAATGTAGATTTCTGTT | >MI0000298\|hsa-mir-221&WithFlank&X\|-\|45746151\|45746272 | 31 | 56 | 26(+1U) | ACCTGGCATACAATGTAGATTTCTGTT | MB231 | 11 | 7 | 12 | 1 | 6 | 2.7 | 1.3 | 3.7 | 0.8 | 5.0 |
| >MI0000298\|hsa-mir-221&WithFlank&X\|-\|45746151\|45746272@31.56.26&[MIMAT0004568&hsa-miR-221-5p&offsets\|0\|+4;m-405&X\|-\|45746220\|45746242&offsets\|0\|+3]\|ACCTGGCATACAATGTAGATTTCTGT | >MI0000298\|hsa-mir-221&WithFlank&X\|-\|45746151\|45746272 | 31 | 56 | 26 | ACCTGGCATACAATGTAGATTTCTGT | BT20 | 21 | 13 | 48 | 2 | 7 | 13.7 | 8.3 | 33.0 | 1.4 | 6.4 |
| >MI0000298\|hsa-mir-221&WithFlank&X\|-\|45746151\|45746272@31.56.26&[MIMAT0004568&hsa-miR-221-5p&offsets\|0\|+4;m-405&X\|-\|45746220\|45746242&offsets\|0\|+3]\|ACCTGGCATACAATGTAGATTTCTGT | >MI0000298\|hsa-mir-221&WithFlank&X\|-\|45746151\|45746272 | 31 | 56 | 26 | ACCTGGCATACAATGTAGATTTCTGT | MB231 | 146 | 106 | 126 | 12 | 98 | 21.0 | 23.4 | 36.7 | 4.1 | 19.4 |
| >MI0000298\|hsa-mir-221&WithFlank&X\|-\|45746151\|45746272@71.90.20(+1C)&[MIMAT0000278&hsa-miR-221-3p&offsets\|0\|-3(+1C);m-91&X\|-\|45746181\|45746202&offsets\|0\|-2(+1C)]\|AGCTACATTGTCTGCTGGGTC | >MI0000298\|hsa-mir-221&WithFlank&X\|-\|45746151\|45746272 | 71 | 90 | 20(+1C) | AGCTACATTGTCTGCTGGGTC | MB231 | 10 | 11 | 5 | 95 | 20 | 1.3 | 6.6 | 3.7 | 44.9 | 14.1 |
| >MI0000298\|hsa-mir-221&WithFlank&X\|-\|45746151\|45746272@71.90.20&[MIMAT0000278&hsa-miR-221-3p&offsets\|0\|-3;m-91&X\|-\|45746181\|45746202&offsets\|0\|-2]\|AGCTACATTGTCTGCTGGGT | >MI0000298\|hsa-mir-221&WithFlank&X\|-\|45746151\|45746272 | 71 | 90 | 20 | AGCTACATTGTCTGCTGGGT | BT20 | 11 | 11 | 20 | 39 | 8 | 4.8 | 5.6 | 7.9 | 37.9 | 5.2 |
| >MI0000298\|hsa-mir-221&WithFlank&X\|-\|45746151\|45746272@71.90.20&[MIMAT0000278&hsa-miR-221-3p&offsets\|0\|-3;m-91&X\|-\|45746181\|45746202&offsets\|0\|-2]\|AGCTACATTGTCTGCTGGGT | >MI0000298\|hsa-mir-221&WithFlank&X\|-\|45746151\|45746272 | 71 | 90 | 20 | AGCTACATTGTCTGCTGGGT | MB231 | 30 | 30 | 15 | 112 | 43 | 6.1 | 11.6 | 9.0 | 35.8 | 14.8 |
| >MI0000298\|hsa-mir-221&WithFlank&X\|-\|45746151\|45746272@71.91.21(+1A)&[MIMAT0000278&hsa-miR-221-3p&offsets\|0\|-2(+1A);m-91&X\|-\|45746181\|45746202&offsets\|0\|-1(+1A)]\|AGCTACATTGTCTGCTGGGTTA | >MI0000298\|hsa-mir-221&WithFlank&X\|-\|45746151\|45746272 | 71 | 91 | 21(+1A) | AGCTACATTGTCTGCTGGGTTA | BT20 | 19 | 10 | 19 | 34 | 9 | 12.7 | 6.8 | 9.8 | 29.1 | 7.3 |
| >MI0000298\|hsa-mir-221&WithFlank&X\|-\|45746151\|45746272@71.91.21(+1A)&[MIMAT0000278&hsa-miR-221-3p&offsets\|0\|-2(+1A);m-91&X\|-\|45746181\|45746202&offsets\|0\|-1(+1A)]\|AGCTACATTGTCTGCTGGGTTA | >MI0000298\|hsa-mir-221&WithFlank&X\|-\|45746151\|45746272 | 71 | 91 | 21(+1A) | AGCTACATTGTCTGCTGGGTTA | MB231 | 36 | 28 | 14 | 183 | 61 | 2.8 | 10.8 | 13.4 | 81.0 | 41.4 |
| >MI0000298\|hsa-mir-221&WithFlank&X\|-\|45746151\|45746272@71.91.21(+1C)&[MIMAT0000278&hsa-miR-221-3p&offsets\|0\|-2(+1C);m-91&X\|-\|45746181\|45746202&offsets\|0\|-1(+1C)]\|AGCTACATTGTCTGCTGGGTTC | >MI0000298\|hsa-mir-221&WithFlank&X\|-\|45746151\|45746272 | 71 | 91 | 21(+1C) | AGCTACATTGTCTGCTGGGTTC | BT20 | 48 | 33 | 50 | 130 | 37 | 21.8 | 21.7 | 25.0 | 105.8 | 33.5 |
| >MI0000298\|hsa-mir-221&WithFlank&X\|-\|45746151\|45746272@71.91.21(+1C)&[MIMAT0000278&hsa-miR-221-3p&offsets\|0\|-2(+1C);m-91&X\|-\|45746181\|45746202&offsets\|0\|-1(+1C)]\|AGCTACATTGTCTGCTGGGTTC | >MI0000298\|hsa-mir-221&WithFlank&X\|-\|45746151\|45746272 | 71 | 91 | 21(+1C) | AGCTACATTGTCTGCTGGGTTC | MB231 | 144 | 120 | 49 | 734 | 276 | 29.1 | 49.8 | 39.2 | 343.4 | 207.6 |
| >MI0000298\|hsa-mir-221&WithFlank&X\|-\|45746151\|45746272@71.91.21(+1G)&[MIMAT0000278&hsa-miR-221-3p&offsets\|0\|-2(+1G);m-91&X\|-\|45746181\|45746202&offsets\|0\|-1(+1G)]\|AGCTACATTGTCTGCTGGGTTG | >MI0000298\|hsa-mir-221&WithFlank&X\|-\|45746151\|45746272 | 71 | 91 | 21(+1G) | AGCTACATTGTCTGCTGGGTTG | BT20 | 12 | 3 | 6 | 9 | 1 | 10.5 | 2.3 | 3.0 | 7.5 | 1.6 |
| >MI0000298\|hsa-mir-221&WithFlank&X\|-\|45746151\|45746272@71.91.21(+1G)&[MIMAT0000278&hsa-miR-221-3p&offsets\|0\|-2(+1G);m-91&X\|-\|45746181\|45746202&offsets\|0\|-1(+1G)]\|AGCTACATTGTCTGCTGGGTTG | >MI0000298\|hsa-mir-221&WithFlank&X\|-\|45746151\|45746272 | 71 | 91 | 21(+1G) | AGCTACATTGTCTGCTGGGTTG | MB231 | 30 | 11 | 4 | 39 | 12 | 21.4 | 5.2 | 3.1 | 19.2 | 9.5 |
| >MI0000298\|hsa-mir-221&WithFlank&X\|-\|45746151\|45746272@71.91.21(+2C)&[MIMAT0000278&hsa-miR-221-3p&offsets\|0\|-2(+2C);m-91&X\|-\|45746181\|45746202&offsets\|0\|-1(+2C)]\|AGCTACATTGTCTGCTGGGTTCC | >MI0000298\|hsa-mir-221&WithFlank&X\|-\|45746151\|45746272 | 71 | 91 | 21(+2C) | AGCTACATTGTCTGCTGGGTTCC | MB231 | 10 | 6 | 5 | 15 | 11 | 2.9 | 2.3 | 3.7 | 2.3 | 9.1 |
| >MI0000298\|hsa-mir-221&WithFlank&X\|-\|45746151\|45746272@71.91.21&[MIMAT0000278&hsa-miR-221-3p&offsets\|0\|-2;m-91&X\|-\|45746181\|45746202&offsets\|0\|-1]\|AGCTACATTGTCTGCTGGGTT | >MI0000298\|hsa-mir-221&WithFlank&X\|-\|45746151\|45746272 | 71 | 91 | 21 | AGCTACATTGTCTGCTGGGTT | BT20 | 31 | 32 | 35 | 135 | 25 | 14.9 | 19.1 | 13.0 | 126.7 | 17.1 |
| >MI0000298\|hsa-mir-221&WithFlank&X\|-\|45746151\|45746272@71.91.21&[MIMAT0000278&hsa-miR-221-3p&offsets\|0\|-2;m-91&X\|-\|45746181\|45746202&offsets\|0\|-1]\|AGCTACATTGTCTGCTGGGTT | >MI0000298\|hsa-mir-221&WithFlank&X\|-\|45746151\|45746272 | 71 | 91 | 21 | AGCTACATTGTCTGCTGGGTT | MB231 | 89 | 103 | 39 | 597 | 147 | 12.4 | 47.4 | 28.2 | 141.4 | 64.3 |
| >MI0000298\|hsa-mir-221&WithFlank&X\|-\|45746151\|45746272@71.92.22(+1A)&[MIMAT0000278&hsa-miR-221-3p&offsets\|0\|-1(+1A);m-91&X\|-\|45746181\|45746202&offsets\|0\|0(+1A)]\|AGCTACATTGTCTGCTGGGTTTA | >MI0000298\|hsa-mir-221&WithFlank&X\|-\|45746151\|45746272 | 71 | 92 | 22(+1A) | AGCTACATTGTCTGCTGGGTTTA | BT20 | 23 | 14 | 41 | 32 | 12 | 14.1 | 8.3 | 20.1 | 26.9 | 6.8 |
| >MI0000298\|hsa-mir-221&WithFlank&X\|-\|45746151\|45746272@71.92.22(+1A)&[MIMAT0000278&hsa-miR-221-3p&offsets\|0\|-1(+1A);m-91&X\|-\|45746181\|45746202&offsets\|0\|0(+1A)]\|AGCTACATTGTCTGCTGGGTTTA | >MI0000298\|hsa-mir-221&WithFlank&X\|-\|45746151\|45746272 | 71 | 92 | 22(+1A) | AGCTACATTGTCTGCTGGGTTTA | MB231 | 57 | 39 | 34 | 214 | 66 | 15.1 | 6.1 | 29.2 | 83.6 | 25.8 |
| >MI0000298\|hsa-mir-221&WithFlank&X\|-\|45746151\|45746272@71.92.22(+1U)&[MIMAT0000278&hsa-miR-221-3p&offsets\|0\|-1(+1U);m-91&X\|-\|45746181\|45746202&offsets\|0\|0(+1U)]\|AGCTACATTGTCTGCTGGGTTTT | >MI0000298\|hsa-mir-221&WithFlank&X\|-\|45746151\|45746272 | 71 | 92 | 22(+1U) | AGCTACATTGTCTGCTGGGTTTT | BT20 | 67 | 47 | 96 | 141 | 47 | 39.8 | 31.9 | 51.5 | 118.8 | 35.0 |
| >MI0000298\|hsa-mir-221&WithFlank&X\|-\|45746151\|45746272@71.92.22(+1U)&[MIMAT0000278&hsa-miR-221-3p&offsets\|0\|-1(+1U);m-91&X\|-\|45746181\|45746202&offsets\|0\|0(+1U)]\|AGCTACATTGTCTGCTGGGTTTT | >MI0000298\|hsa-mir-221&WithFlank&X\|-\|45746151\|45746272 | 71 | 92 | 22(+1U) | AGCTACATTGTCTGCTGGGTTTT | MB231 | 272 | 208 | 103 | 1037 | 376 | 51.3 | 60.4 | 75.6 | 334.8 | 172.0 |
| >MI0000298\|hsa-mir-221&WithFlank&X\|-\|45746151\|45746272@71.92.22(+2U)&[MIMAT0000278&hsa-miR-221-3p&offsets\|0\|-1(+2U);m-91&X\|-\|45746181\|45746202&offsets\|0\|0(+2U)]\|AGCTACATTGTCTGCTGGGTTTTT | >MI0000298\|hsa-mir-221&WithFlank&X\|-\|45746151\|45746272 | 71 | 92 | 22(+2U) | AGCTACATTGTCTGCTGGGTTTTT | MB231 | 29 | 22 | 10 | 65 | 39 | 1.1 | 9.2 | 7.3 | 6.9 | 18.7 |

<table>
<tr><th></th><th></th><th></th><th></th><th></th><th></th><th></th><th></th><th></th><th></th><th></th><th></th><th></th><th></th><th></th><th></th><th></th></tr>
<tr><td>>MI0000298|hsa-mir-221&WithFlank&X|-|45746151|45746272@71.92.22&[MIMAT0000278&hsa-miR-221-3p&offsets|0|-1;m-91&X|-|45746181|45746202&offsets|0|0]|AGCTACATTGTCTGCTGGGTTT</td><td>>MI0000298|hsa-mir-221&WithFlank&X|-|45746151|45746272</td><td>71</td><td>92</td><td>22</td><td>AGCTACATTGTCTGCTGGGTTT</td><td>BT20</td><td>341</td><td>245</td><td>369</td><td>963</td><td>292</td><td>189.3</td><td>146.2</td><td>212.1</td><td>871.1</td><td>275.7</td></tr>
<tr><td>>MI0000298|hsa-mir-221&WithFlank&X|-|45746151|45746272@71.92.22&[MIMAT0000278&hsa-miR-221-3p&offsets|0|-1;m-91&X|-|45746181|45746202&offsets|0|0]|AGCTACATTGTCTGCTGGGTTT</td><td>>MI0000298|hsa-mir-221&WithFlank&X|-|45746151|45746272</td><td>71</td><td>92</td><td>22</td><td>AGCTACATTGTCTGCTGGGTTT</td><td>MB231</td><td>1244</td><td>1093</td><td>404</td><td>4389</td><td>1962</td><td>209.3</td><td>393.8</td><td>323.1</td><td>1096.4</td><td>1158.2</td></tr>
<tr><td>>MI0000298|hsa-mir-221&WithFlank&X|-|45746151|45746272@71.92.22&[MIMAT0000278&hsa-miR-221-3p&offsets|0|-1;m-91&X|-|45746181|45746202&offsets|0|0]|AGCTACATTGTCTGCTGGGTTT</td><td>>MI0000298|hsa-mir-221&WithFlank&X|-|45746151|45746272</td><td>71</td><td>92</td><td>22</td><td>AGCTACATTGTCTGCTGGGTTT</td><td>MB468</td><td>31</td><td>11</td><td>16</td><td>91</td><td>14</td><td>15.1</td><td>4.5</td><td>2.5</td><td>38.8</td><td>1.5</td></tr>
<tr><td>>MI0000298|hsa-mir-221&WithFlank&X|-|45746151|45746272@71.93.23(+1C)&[MIMAT0000278&hsa-miR-221-3p&offsets|0|0(+1C);m-91&X|-|45746181|45746202&offsets|0|+1(+1C)]|AGCTACATTGTCTGCTGGGTTTCC</td><td>>MI0000298|hsa-mir-221&WithFlank&X|-|45746151|45746272</td><td>71</td><td>93</td><td>23(+1C)</td><td>AGCTACATTGTCTGCTGGGTTTCC</td><td>BT20</td><td>25</td><td>14</td><td>44</td><td>9</td><td>17</td><td>11.7</td><td>4.3</td><td>14.1</td><td>3.8</td><td>9.1</td></tr>
<tr><td>>MI0000298|hsa-mir-221&WithFlank&X|-|45746151|45746272@71.93.23(+1C)&[MIMAT0000278&hsa-miR-221-3p&offsets|0|0(+1C);m-91&X|-|45746181|45746202&offsets|0|+1(+1C)]|AGCTACATTGTCTGCTGGGTTTCC</td><td>>MI0000298|hsa-mir-221&WithFlank&X|-|45746151|45746272</td><td>71</td><td>93</td><td>23(+1C)</td><td>AGCTACATTGTCTGCTGGGTTTCC</td><td>MB231</td><td>121</td><td>70</td><td>67</td><td>125</td><td>102</td><td>15.2</td><td>9.0</td><td>37.3</td><td>57.7</td><td>57.6</td></tr>
<tr><td>>MI0000298|hsa-mir-221&WithFlank&X|-|45746151|45746272@71.93.23(+1G)&[MIMAT0000278&hsa-miR-221-3p&offsets|0|0(+1G);m-91&X|-|45746181|45746202&offsets|0|+1(+1G)]|AGCTACATTGTCTGCTGGGTTTCG</td><td>>MI0000298|hsa-mir-221&WithFlank&X|-|45746151|45746272</td><td>71</td><td>93</td><td>23(+1G)</td><td>AGCTACATTGTCTGCTGGGTTTCG</td><td>BT20</td><td>16</td><td>5</td><td>18</td><td>3</td><td>4</td><td>5.0</td><td>1.5</td><td>4.5</td><td>0.8</td><td>2.0</td></tr>
<tr><td>>MI0000298|hsa-mir-221&WithFlank&X|-|45746151|45746272@71.93.23(+1G)&[MIMAT0000278&hsa-miR-221-3p&offsets|0|0(+1G);m-91&X|-|45746181|45746202&offsets|0|+1(+1G)]|AGCTACATTGTCTGCTGGGTTTCG</td><td>>MI0000298|hsa-mir-221&WithFlank&X|-|45746151|45746272</td><td>71</td><td>93</td><td>23(+1G)</td><td>AGCTACATTGTCTGCTGGGTTTCG</td><td>MB231</td><td>65</td><td>22</td><td>20</td><td>22</td><td>17</td><td>37.6</td><td>3.6</td><td>11.2</td><td>7.7</td><td>5.3</td></tr>
<tr><td>>MI0000298|hsa-mir-221&WithFlank&X|-|45746151|45746272@71.93.23(+1U)&[MIMAT0000278&hsa-miR-221-3p&offsets|0|0(+1U);m-91&X|-|45746181|45746202&offsets|0|+1(+1U)]|AGCTACATTGTCTGCTGGGTTTCT</td><td>>MI0000298|hsa-mir-221&WithFlank&X|-|45746151|45746272</td><td>71</td><td>93</td><td>23(+1U)</td><td>AGCTACATTGTCTGCTGGGTTTCT</td><td>BT20</td><td>164</td><td>93</td><td>302</td><td>56</td><td>107</td><td>82.8</td><td>21.8</td><td>84.1</td><td>22.2</td><td>54.6</td></tr>
<tr><td>>MI0000298|hsa-mir-221&WithFlank&X|-|45746151|45746272@71.93.23(+1U)&[MIMAT0000278&hsa-miR-221-3p&offsets|0|0(+1U);m-91&X|-|45746181|45746202&offsets|0|+1(+1U)]|AGCTACATTGTCTGCTGGGTTTCT</td><td>>MI0000298|hsa-mir-221&WithFlank&X|-|45746151|45746272</td><td>71</td><td>93</td><td>23(+1U)</td><td>AGCTACATTGTCTGCTGGGTTTCT</td><td>MB231</td><td>956</td><td>604</td><td>523</td><td>697</td><td>668</td><td>119.1</td><td>125.3</td><td>270.0</td><td>223.5</td><td>283.0</td></tr>
<tr><td>>MI0000298|hsa-mir-221&WithFlank&X|-|45746151|45746272@71.93.23(+1U)&[MIMAT0000278&hsa-miR-221-3p&offsets|0|0(+1U);m-91&X|-|45746181|45746202&offsets|0|+1(+1U)]|AGCTACATTGTCTGCTGGGTTTCT</td><td>>MI0000298|hsa-mir-221&WithFlank&X|-|45746151|45746272</td><td>71</td><td>93</td><td>23(+1U)</td><td>AGCTACATTGTCTGCTGGGTTTCT</td><td>MB468</td><td>25</td><td>18</td><td>18</td><td>29</td><td>10</td><td>7.7</td><td>4.5</td><td>8.3</td><td>8.6</td><td>3.3</td></tr>
<tr><td>>MI0000298|hsa-mir-221&WithFlank&X|-|45746151|45746272@71.93.23(+2U)&[MIMAT0000278&hsa-miR-221-3p&offsets|0|0(+2U);m-91&X|-|45746181|45746202&offsets|0|+1(+2U)]|AGCTACATTGTCTGCTGGGTTTCTT</td><td>>MI0000298|hsa-mir-221&WithFlank&X|-|45746151|45746272</td><td>71</td><td>93</td><td>23(+2U)</td><td>AGCTACATTGTCTGCTGGGTTTCTT</td><td>MB231</td><td>14</td><td>10</td><td>12</td><td>5</td><td>10</td><td>5.1</td><td>2.6</td><td>7.2</td><td>2.0</td><td>2.8</td></tr>
<tr><td>>MI0000298|hsa-mir-221&WithFlank&X|-|45746151|45746272@71.93.23&[MIMAT0000278&hsa-miR-221-3p&offsets|0|0;m-91&X|-|45746181|45746202&offsets|0|+1]|AGCTACATTGTCTGCTGGGTTTC</td><td>>MI0000298|hsa-mir-221&WithFlank&X|-|45746151|45746272</td><td>71</td><td>93</td><td>23</td><td>AGCTACATTGTCTGCTGGGTTTC</td><td>BT20</td><td>422</td><td>365</td><td>894</td><td>990</td><td>429</td><td>341.4</td><td>114.8</td><td>410.2</td><td>850.3</td><td>234.4</td></tr>
<tr><td>>MI0000298|hsa-mir-221&WithFlank&X|-|45746151|45746272@71.93.23&[MIMAT0000278&hsa-miR-221-3p&offsets|0|0;m-91&X|-|45746181|45746202&offsets|0|+1]|AGCTACATTGTCTGCTGGGTTTC</td><td>>MI0000298|hsa-mir-221&WithFlank&X|-|45746151|45746272</td><td>71</td><td>93</td><td>23</td><td>AGCTACATTGTCTGCTGGGTTTC</td><td>MB231</td><td>2125</td><td>1901</td><td>1180</td><td>5584</td><td>2196</td><td>888.6</td><td>453.0</td><td>844.7</td><td>818.1</td><td>462.9</td></tr>
<tr><td>>MI0000298|hsa-mir-221&WithFlank&X|-|45746151|45746272@71.93.23&[MIMAT0000278&hsa-miR-221-3p&offsets|0|0;m-91&X|-|45746181|45746202&offsets|0|+1]|AGCTACATTGTCTGCTGGGTTTC</td><td>>MI0000298|hsa-mir-221&WithFlank&X|-|45746151|45746272</td><td>71</td><td>93</td><td>23</td><td>AGCTACATTGTCTGCTGGGTTTC</td><td>MB468</td><td>94</td><td>59</td><td>83</td><td>233</td><td>38</td><td>40.1</td><td>27.3</td><td>30.8</td><td>142.2</td><td>11.9</td></tr>
<tr><td>>MI0000298|hsa-mir-221&WithFlank&X|-|45746151|45746272@71.94.24&[MIMAT0000278&hsa-miR-221-3p&offsets|0|+1;m-91&X|-|45746181|45746202&offsets|0|+2]|AGCTACATTGTCTGCTGGGTTTCA</td><td>>MI0000298|hsa-mir-221&WithFlank&X|-|45746151|45746272</td><td>71</td><td>94</td><td>24</td><td>AGCTACATTGTCTGCTGGGTTTCA</td><td>BT20</td><td>45</td><td>19</td><td>79</td><td>12</td><td>25</td><td>30.1</td><td>8.1</td><td>48.6</td><td>6.5</td><td>14.8</td></tr>
<tr><td>>MI0000298|hsa-mir-221&WithFlank&X|-|45746151|45746272@71.94.24&[MIMAT0000278&hsa-miR-221-3p&offsets|0|+1;m-91&X|-|45746181|45746202&offsets|0|+2]|AGCTACATTGTCTGCTGGGTTTCA</td><td>>MI0000298|hsa-mir-221&WithFlank&X|-|45746151|45746272</td><td>71</td><td>94</td><td>24</td><td>AGCTACATTGTCTGCTGGGTTTCA</td><td>MB231</td><td>186</td><td>109</td><td>91</td><td>194</td><td>161</td><td>55.1</td><td>21.7</td><td>59.2</td><td>44.3</td><td>71.0</td></tr>
<tr><td>>MI0000298|hsa-mir-221&WithFlank&X|-|45746151|45746272@71.94.24&[MIMAT0000278&hsa-miR-221-3p&offsets|0|+1;m-91&X|-|45746181|45746202&offsets|0|+2]|AGCTACATTGTCTGCTGGGTTTCA</td><td>>MI0000298|hsa-mir-221&WithFlank&X|-|45746151|45746272</td><td>71</td><td>94</td><td>24</td><td>AGCTACATTGTCTGCTGGGTTTCA</td><td>MB468</td><td>10</td><td>5</td><td>5</td><td>14</td><td>5</td><td>3.6</td><td>2.5</td><td>1.1</td><td>9.0</td><td>1.2</td></tr>
<tr><td>>MI0000299|hsa-mir-222&WithFlank&X|-|45747009|45747130@75.100.26&[MIMAT0000279&hsa-miR-222-3p&offsets|0|+5;m-140&X|-|45747033|45747056&offsets|0|+2]|AGCTACATCTGGCTACTGGGTCTCTG</td><td>>MI0000299|hsa-mir-222&WithFlank&X|-|45747009|45747130</td><td>75</td><td>100</td><td>26</td><td>AGCTACATCTGGCTACTGGGTCTCTG</td><td>MB231</td><td>11</td><td>3</td><td>3</td><td>1</td><td>2</td><td>5.8</td><td>0.7</td><td>1.8</td><td>0.7</td><td>2.3</td></tr>
<tr><td>>MI0000299|hsa-mir-222&WithFlank&X|-|45747009|45747130@75.94.20(+1C)&[MIMAT0000279&hsa-miR-222-3p&offsets|0|-1(+1C);m-140&X|-|45747033|45747056&offsets|0|-4(+1C)]|AGCTACATCTGGCTACTGGGC</td><td>>MI0000299|hsa-mir-222&WithFlank&X|-|45747009|45747130</td><td>75</td><td>94</td><td>20(+1C)</td><td>AGCTACATCTGGCTACTGGGC</td><td>MB231</td><td>10</td><td>14</td><td>5</td><td>121</td><td>27</td><td>3.4</td><td>6.3</td><td>2.8</td><td>44.0</td><td>16.0</td></tr>
<tr><td>>MI0000299|hsa-mir-222&WithFlank&X|-|45747009|45747130@75.95.21(+1U)&[MIMAT0000279&hsa-miR-222-3p&offsets|0|0(+1U);m-140&X|-|45747033|45747056&offsets|0|-3(+1U)]|AGCTACATCTGGCTACTGGGTT</td><td>>MI0000299|hsa-mir-222&WithFlank&X|-|45747009|45747130</td><td>75</td><td>95</td><td>21(+1U)</td><td>AGCTACATCTGGCTACTGGGTT</td><td>MB231</td><td>12</td><td>15</td><td>9</td><td>48</td><td>23</td><td>3.5</td><td>4.2</td><td>4.0</td><td>26.4</td><td>9.7</td></tr>
<tr><td>>MI0000299|hsa-mir-222&WithFlank&X|-|45747009|45747130@75.95.21&[MIMAT0000279&hsa-miR-222-3p&offsets|0|0;m-140&X|-|45747033|45747056&offsets|0|-3]|AGCTACATCTGGCTACTGGGT</td><td>>MI0000299|hsa-mir-222&WithFlank&X|-|45747009|45747130</td><td>75</td><td>95</td><td>21</td><td>AGCTACATCTGGCTACTGGGT</td><td>BT20</td><td>51</td><td>58</td><td>50</td><td>272</td><td>45</td><td>28.7</td><td>29.1</td><td>18.8</td><td>214.5</td><td>17.9</td></tr>
<tr><td>>MI0000299|hsa-mir-222&WithFlank&X|-|45747009|45747130@75.95.21&[MIMAT0000279&hsa-miR-222-3p&offsets|0|0;m-140&X|-|45747033|45747056&offsets|0|-3]|AGCTACATCTGGCTACTGGGT</td><td>>MI0000299|hsa-mir-222&WithFlank&X|-|45747009|45747130</td><td>75</td><td>95</td><td>21</td><td>AGCTACATCTGGCTACTGGGT</td><td>MB231</td><td>86</td><td>120</td><td>41</td><td>752</td><td>188</td><td>29.1</td><td>36.7</td><td>23.2</td><td>99.2</td><td>78.4</td></tr>
<tr><td>>MI0000299|hsa-mir-222&WithFlank&X|-|45747009|45747130@75.96.22(+1A)&[MIMAT0000279&hsa-miR-222-3p&offsets|0|+1(+1A);m-140&X|-|45747033|45747056&offsets|0|-2(+1A)]|AGCTACATCTGGCTACTGGGTCA</td><td>>MI0000299|hsa-mir-222&WithFlank&X|-|45747009|45747130</td><td>75</td><td>96</td><td>22(+1A)</td><td>AGCTACATCTGGCTACTGGGTCA</td><td>BT20</td><td>26</td><td>15</td><td>21</td><td>17</td><td>17</td><td>17.0</td><td>9.7</td><td>7.3</td><td>13.0</td><td>7.7</td></tr>
<tr><td>>MI0000299|hsa-mir-222&WithFlank&X|-|45747009|45747130@75.96.22(+1A)&[MIMAT0000279&hsa-miR-222-3p&offsets|0|+1(+1A);m-140&X|-|45747033|45747056&offsets|0|-2(+1A)]|AGCTACATCTGGCTACTGGGTCA</td><td>>MI0000299|hsa-mir-222&WithFlank&X|-|45747009|45747130</td><td>75</td><td>96</td><td>22(+1A)</td><td>AGCTACATCTGGCTACTGGGTCA</td><td>MB231</td><td>38</td><td>31</td><td>20</td><td>111</td><td>67</td><td>6.8</td><td>9.3</td><td>14.5</td><td>69.6</td><td>43.8</td></tr>
<tr><td>>MI0000299|hsa-mir-222&WithFlank&X|-|45747009|45747130@75.96.22(+1C)&[MIMAT0000279&hsa-miR-222-3p&offsets|0|+1(+1C);m-140&X|-|45747033|45747056&offsets|0|-2(+1C)]|AGCTACATCTGGCTACTGGGTCC</td><td>>MI0000299|hsa-mir-222&WithFlank&X|-|45747009|45747130</td><td>75</td><td>96</td><td>22(+1C)</td><td>AGCTACATCTGGCTACTGGGTCC</td><td>BT20</td><td>22</td><td>19</td><td>21</td><td>21</td><td>22</td><td>10.1</td><td>12.1</td><td>9.1</td><td>15.8</td><td>13.2</td></tr>
<tr><td>>MI0000299|hsa-mir-222&WithFlank&X|-|45747009|45747130@75.96.22(+1C)&[MIMAT0000279&hsa-miR-222-3p&offsets|0|+1(+1C);m-140&X|-|45747033|45747056&offsets|0|-2(+1C)]|AGCTACATCTGGCTACTGGGTCC</td><td>>MI0000299|hsa-mir-222&WithFlank&X|-|45747009|45747130</td><td>75</td><td>96</td><td>22(+1C)</td><td>AGCTACATCTGGCTACTGGGTCC</td><td>MB231</td><td>51</td><td>43</td><td>24</td><td>132</td><td>90</td><td>11.7</td><td>15.2</td><td>16.6</td><td>82.8</td><td>61.3</td></tr>
<tr><td>>MI0000299|hsa-mir-222&WithFlank&X|-|45747009|45747130@75.96.22(+1G)&[MIMAT0000279&hsa-miR-222-3p&offsets|0|+1(+1G);m-140&X|-|45747033|45747056&offsets|0|-2(+1G)]|AGCTACATCTGGCTACTGGGTCG</td><td>>MI0000299|hsa-mir-222&WithFlank&X|-|45747009|45747130</td><td>75</td><td>96</td><td>22(+1G)</td><td>AGCTACATCTGGCTACTGGGTCG</td><td>BT20</td><td>14</td><td>5</td><td>5</td><td>3</td><td>3</td><td>11.5</td><td>3.2</td><td>0.5</td><td>3.3</td><td>1.5</td></tr>
<tr><td>>MI0000299|hsa-mir-222&WithFlank&X|-|45747009|45747130@75.96.22(+1G)&[MIMAT0000279&hsa-miR-222-3p&offsets|0|+1(+1G);m-140&X|-|45747033|45747056&offsets|0|-2(+1G)]|AGCTACATCTGGCTACTGGGTCG</td><td>>MI0000299|hsa-mir-222&WithFlank&X|-|45747009|45747130</td><td>75</td><td>96</td><td>22(+1G)</td><td>AGCTACATCTGGCTACTGGGTCG</td><td>MB231</td><td>24</td><td>10</td><td>5</td><td>18</td><td>11</td><td>15.6</td><td>4.5</td><td>4.0</td><td>12.2</td><td>7.4</td></tr>
<tr><td>>MI0000299|hsa-mir-222&WithFlank&X|-|45747009|45747130@75.96.22&[MIMAT0000279&hsa-miR-222-3p&offsets|0|+1;m-140&X|-|45747033|45747056&offsets|0|-2]|AGCTACATCTGGCTACTGGGTC</td><td>>MI0000299|hsa-mir-222&WithFlank&X|-|45747009|45747130</td><td>75</td><td>96</td><td>22</td><td>AGCTACATCTGGCTACTGGGTC</td><td>BT20</td><td>31</td><td>31</td><td>41</td><td>59</td><td>34</td><td>16.5</td><td>14.3</td><td>11.2</td><td>43.4</td><td>16.8</td></tr>
<tr><td>>MI0000299|hsa-mir-222&WithFlank&X|-|45747009|45747130@75.96.22&[MIMAT0000279&hsa-miR-222-3p&offsets|0|+1;m-140&X|-|45747033|45747056&offsets|0|-2]|AGCTACATCTGGCTACTGGGTC</td><td>>MI0000299|hsa-mir-222&WithFlank&X|-|45747009|45747130</td><td>75</td><td>96</td><td>22</td><td>AGCTACATCTGGCTACTGGGTC</td><td>MB231</td><td>83</td><td>107</td><td>62</td><td>175</td><td>124</td><td>40.4</td><td>18.9</td><td>28.9</td><td>69.9</td><td>22.7</td></tr>
<tr><td>>MI0000299|hsa-mir-222&WithFlank&X|-|45747009|45747130@75.97.23(+1A)&[MIMAT0000279&hsa-miR-222-3p&offsets|0|+2(+1A);m-140&X|-|45747033|45747056&offsets|0|-1(+1A)]|AGCTACATCTGGCTACTGGGTCTA</td><td>>MI0000299|hsa-mir-222&WithFlank&X|-|45747009|45747130</td><td>75</td><td>97</td><td>23(+1A)</td><td>AGCTACATCTGGCTACTGGGTCTA</td><td>BT20</td><td>29</td><td>14</td><td>35</td><td>14</td><td>17</td><td>18.4</td><td>9.5</td><td>6.8</td><td>9.9</td><td>11.3</td></tr>
<tr><td>>MI0000299|hsa-mir-222&WithFlank&X|-|45747009|45747130@75.97.23(+1A)&[MIMAT0000279&hsa-miR-222-3p&offsets|0|+2(+1A);m-140&X|-|45747033|45747056&offsets|0|-1(+1A)]|AGCTACATCTGGCTACTGGGTCTA</td><td>>MI0000299|hsa-mir-222&WithFlank&X|-|45747009|45747130</td><td>75</td><td>97</td><td>23(+1A)</td><td>AGCTACATCTGGCTACTGGGTCTA</td><td>MB231</td><td>79</td><td>45</td><td>66</td><td>66</td><td>78</td><td>45.5</td><td>11.2</td><td>34.0</td><td>37.0</td><td>31.1</td></tr>
<tr><td>>MI0000299|hsa-mir-222&WithFlank&X|-|45747009|45747130@75.97.23(+1U)&[MIMAT0000279&hsa-miR-222-3p&offsets|0|+2(+1U);m-140&X|-|45747033|45747056&offsets|0|-1(+1U)]|AGCTACATCTGGCTACTGGGTCTT</td><td>>MI0000299|hsa-mir-222&WithFlank&X|-|45747009|45747130</td><td>75</td><td>97</td><td>23(+1U)</td><td>AGCTACATCTGGCTACTGGGTCTT</td><td>BT20</td><td>78</td><td>44</td><td>80</td><td>45</td><td>59</td><td>61.6</td><td>34.8</td><td>40.6</td><td>31.7</td><td>44.1</td></tr>
<tr><td>>MI0000299|hsa-mir-222&WithFlank&X|-|45747009|45747130@75.97.23(+1U)&[MIMAT0000279&hsa-miR-222-3p&offsets|0|+2(+1U);m-140&X|-|45747033|45747056&offsets|0|-1(+1U)]|AGCTACATCTGGCTACTGGGTCTT</td><td>>MI0000299|hsa-mir-222&WithFlank&X|-|45747009|45747130</td><td>75</td><td>97</td><td>23(+1U)</td><td>AGCTACATCTGGCTACTGGGTCTT</td><td>MB231</td><td>238</td><td>146</td><td>141</td><td>224</td><td>276</td><td>138.5</td><td>15.8</td><td>45.1</td><td>112.1</td><td>99.4</td></tr>
</table>

| | | | | | | | | | | | | | | | | |
|---|---|---|---|---|---|---|---|---|---|---|---|---|---|---|---|---|
| >MI0000299\|hsa-mir-222&WithFlank&X\|-\|45747009\|45747130@75.97.23(+2U)&[MIMAT0000279&hsa-miR-222-3p&offsets\|0\|+2(+2U);m-140&X\|-\|45747033\|45747056&offsets\|0\|-1(+2U)]\|AGCTACATCTGGCTACTGGGTCTTT | >MI0000299\|hsa-mir-222&WithFlank&X\|-\|45747009\|45747130 | 75 | 97 | 23(+2U) | AGCTACATCTGGCTACTGGGTCTTT | BT20 | 17 | 8 | 17 | 3 | 11 | 16.2 | 7.4 | 14.5 | 2.4 | 12.4 |
| >MI0000299\|hsa-mir-222&WithFlank&X\|-\|45747009\|45747130@75.97.23(+2U)&[MIMAT0000279&hsa-miR-222-3p&offsets\|0\|+2(+2U);m-140&X\|-\|45747033\|45747056&offsets\|0\|-1(+2U)]\|AGCTACATCTGGCTACTGGGTCTTT | >MI0000299\|hsa-mir-222&WithFlank&X\|-\|45747009\|45747130 | 75 | 97 | 23(+2U) | AGCTACATCTGGCTACTGGGTCTTT | MB231 | 74 | 57 | 36 | 33 | 80 | 29.5 | 16.8 | 22.5 | 14.6 | 30.7 |
| >MI0000299\|hsa-mir-222&WithFlank&X\|-\|45747009\|45747130@75.97.23(+3U)&[MIMAT0000279&hsa-miR-222-3p&offsets\|0\|+2(+3U);m-140&X\|-\|45747033\|45747056&offsets\|0\|-1(+3U)]\|AGCTACATCTGGCTACTGGGTCTTTT | >MI0000299\|hsa-mir-222&WithFlank&X\|-\|45747009\|45747130 | 75 | 97 | 23(+3U) | AGCTACATCTGGCTACTGGGTCTTTT | MB231 | 19 | 10 | 6 | 10 | 23 | 10.5 | 1.4 | 3.3 | 3.3 | 12.2 |
| >MI0000299\|hsa-mir-222&WithFlank&X\|-\|45747009\|45747130@75.97.23&[MIMAT0000279&hsa-miR-222-3p&offsets\|0\|+2;m-140&X\|-\|45747033\|45747056&offsets\|0\|-1]\|AGCTACATCTGGCTACTGGGTCT | >MI0000299\|hsa-mir-222&WithFlank&X\|-\|45747009\|45747130 | 75 | 97 | 23 | AGCTACATCTGGCTACTGGGTCT | BT20 | 141 | 108 | 137 | 134 | 143 | 69.1 | 65.2 | 79.2 | 99.6 | 80.2 |
| >MI0000299\|hsa-mir-222&WithFlank&X\|-\|45747009\|45747130@75.97.23&[MIMAT0000279&hsa-miR-222-3p&offsets\|0\|+2;m-140&X\|-\|45747033\|45747056&offsets\|0\|-1]\|AGCTACATCTGGCTACTGGGTCT | >MI0000299\|hsa-mir-222&WithFlank&X\|-\|45747009\|45747130 | 75 | 97 | 23 | AGCTACATCTGGCTACTGGGTCT | MB231 | 355 | 328 | 177 | 714 | 542 | 108.3 | 77.5 | 113.4 | 367.0 | 250.8 |
| >MI0000299\|hsa-mir-222&WithFlank&X\|-\|45747009\|45747130@75.97.23&[MIMAT0000279&hsa-miR-222-3p&offsets\|0\|+2;m-140&X\|-\|45747033\|45747056&offsets\|0\|-1]\|AGCTACATCTGGCTACTGGGTCT | >MI0000299\|hsa-mir-222&WithFlank&X\|-\|45747009\|45747130 | 75 | 97 | 23 | AGCTACATCTGGCTACTGGGTCT | MB468 | 10 | 13 | 4 | 17 | 7 | 6.1 | 9.6 | 0.9 | 11.1 | 1.0 |
| >MI0000299\|hsa-mir-222&WithFlank&X\|-\|45747009\|45747130@75.98.24(+1A)&[MIMAT0000279&hsa-miR-222-3p&offsets\|0\|+3(+1A);m-140&X\|-\|45747033\|45747056&offsets\|0\|0(+1A)]\|AGCTACATCTGGCTACTGGGTCTCA | >MI0000299\|hsa-mir-222&WithFlank&X\|-\|45747009\|45747130 | 75 | 98 | 24(+1A) | AGCTACATCTGGCTACTGGGTCTCA | BT20 | 88 | 39 | 118 | 19 | 49 | 54.2 | 31.7 | 13.3 | 11.7 | 35.9 |
| >MI0000299\|hsa-mir-222&WithFlank&X\|-\|45747009\|45747130@75.98.24(+1A)&[MIMAT0000279&hsa-miR-222-3p&offsets\|0\|+3(+1A);m-140&X\|-\|45747033\|45747056&offsets\|0\|0(+1A)]\|AGCTACATCTGGCTACTGGGTCTCA | >MI0000299\|hsa-mir-222&WithFlank&X\|-\|45747009\|45747130 | 75 | 98 | 24(+1A) | AGCTACATCTGGCTACTGGGTCTCA | MB231 | 313 | 171 | 217 | 148 | 292 | 167.5 | 37.2 | 97.1 | 56.3 | 119.9 |
| >MI0000299\|hsa-mir-222&WithFlank&X\|-\|45747009\|45747130@75.98.24(+1C)&[MIMAT0000279&hsa-miR-222-3p&offsets\|0\|+3(+1C);m-140&X\|-\|45747033\|45747056&offsets\|0\|0(+1C)]\|AGCTACATCTGGCTACTGGGTCTCC | >MI0000299\|hsa-mir-222&WithFlank&X\|-\|45747009\|45747130 | 75 | 98 | 24(+1C) | AGCTACATCTGGCTACTGGGTCTCC | BT20 | 113 | 60 | 156 | 33 | 99 | 88.8 | 40.9 | 54.1 | 14.1 | 71.1 |
| >MI0000299\|hsa-mir-222&WithFlank&X\|-\|45747009\|45747130@75.98.24(+1C)&[MIMAT0000279&hsa-miR-222-3p&offsets\|0\|+3(+1C);m-140&X\|-\|45747033\|45747056&offsets\|0\|0(+1C)]\|AGCTACATCTGGCTACTGGGTCTCC | >MI0000299\|hsa-mir-222&WithFlank&X\|-\|45747009\|45747130 | 75 | 98 | 24(+1C) | AGCTACATCTGGCTACTGGGTCTCC | MB231 | 565 | 354 | 385 | 239 | 582 | 373.3 | 56.4 | 81.7 | 79.1 | 162.1 |
| >MI0000299\|hsa-mir-222&WithFlank&X\|-\|45747009\|45747130@75.98.24(+1G)&[MIMAT0000279&hsa-miR-222-3p&offsets\|0\|+3(+1G);m-140&X\|-\|45747033\|45747056&offsets\|0\|0(+1G)]\|AGCTACATCTGGCTACTGGGTCTCG | >MI0000299\|hsa-mir-222&WithFlank&X\|-\|45747009\|45747130 | 75 | 98 | 24(+1G) | AGCTACATCTGGCTACTGGGTCTCG | BT20 | 42 | 12 | 33 | 5 | 9 | 11.4 | 8.6 | 12.6 | 1.9 | 6.6 |
| >MI0000299\|hsa-mir-222&WithFlank&X\|-\|45747009\|45747130@75.98.24(+1G)&[MIMAT0000279&hsa-miR-222-3p&offsets\|0\|+3(+1G);m-140&X\|-\|45747033\|45747056&offsets\|0\|0(+1G)]\|AGCTACATCTGGCTACTGGGTCTCG | >MI0000299\|hsa-mir-222&WithFlank&X\|-\|45747009\|45747130 | 75 | 98 | 24(+1G) | AGCTACATCTGGCTACTGGGTCTCG | MB231 | 153 | 45 | 49 | 28 | 45 | 71.9 | 4.0 | 25.9 | 11.0 | 20.9 |
| >MI0000299\|hsa-mir-222&WithFlank&X\|-\|45747009\|45747130@75.98.24&[MIMAT0000279&hsa-miR-222-3p&offsets\|0\|+3;m-140&X\|-\|45747033\|45747056&offsets\|0\|0]\|AGCTACATCTGGCTACTGGGTCTC | >MI0000299\|hsa-mir-222&WithFlank&X\|-\|45747009\|45747130 | 75 | 98 | 24 | AGCTACATCTGGCTACTGGGTCTC | BT20 | 304 | 184 | 398 | 210 | 305 | 307.6 | 106.9 | 224.4 | 160.7 | 231.6 |
| >MI0000299\|hsa-mir-222&WithFlank&X\|-\|45747009\|45747130@75.98.24&[MIMAT0000279&hsa-miR-222-3p&offsets\|0\|+3;m-140&X\|-\|45747033\|45747056&offsets\|0\|0]\|AGCTACATCTGGCTACTGGGTCTC | >MI0000299\|hsa-mir-222&WithFlank&X\|-\|45747009\|45747130 | 75 | 98 | 24 | AGCTACATCTGGCTACTGGGTCTC | MB231 | 1383 | 988 | 1075 | 698 | 1215 | 953.9 | 229.3 | 345.6 | 269.8 | 305.8 |
| >MI0000299\|hsa-mir-222&WithFlank&X\|-\|45747009\|45747130@75.98.24&[MIMAT0000279&hsa-miR-222-3p&offsets\|0\|+3;m-140&X\|-\|45747033\|45747056&offsets\|0\|0]\|AGCTACATCTGGCTACTGGGTCTC | >MI0000299\|hsa-mir-222&WithFlank&X\|-\|45747009\|45747130 | 75 | 98 | 24 | AGCTACATCTGGCTACTGGGTCTC | MB468 | 21 | 34 | 13 | 23 | 14 | 8.8 | 20.3 | 7.3 | 11.8 | 4.8 |
| >MI0000299\|hsa-mir-222&WithFlank&X\|-\|45747009\|45747130@75.99.25(+1A)&[MIMAT0000279&hsa-miR-222-3p&offsets\|0\|+4(+1A);m-140&X\|-\|45747033\|45747056&offsets\|0\|+1(+1A)]\|AGCTACATCTGGCTACTGGGTCTCTA | >MI0000299\|hsa-mir-222&WithFlank&X\|-\|45747009\|45747130 | 75 | 99 | 25(+1A) | AGCTACATCTGGCTACTGGGTCTCTA | MB231 | 25 | 12 | 20 | 5 | 19 | 8.3 | 3.8 | 11.3 | 2.6 | 9.8 |
| >MI0000299\|hsa-mir-222&WithFlank&X\|-\|45747009\|45747130@75.99.25(+1C)&[MIMAT0000279&hsa-miR-222-3p&offsets\|0\|+4(+1C);m-140&X\|-\|45747033\|45747056&offsets\|0\|+1(+1C)]\|AGCTACATCTGGCTACTGGGTCTCTC | >MI0000299\|hsa-mir-222&WithFlank&X\|-\|45747009\|45747130 | 75 | 99 | 25(+1C) | AGCTACATCTGGCTACTGGGTCTCTC | MB231 | 39 | 18 | 25 | 10 | 38 | 16.7 | 5.6 | 9.4 | 5.9 | 22.5 |
| >MI0000299\|hsa-mir-222&WithFlank&X\|-\|45747009\|45747130@75.99.25(+1U)&[MIMAT0000279&hsa-miR-222-3p&offsets\|0\|+4(+1U);m-140&X\|-\|45747033\|45747056&offsets\|0\|+1(+1U)]\|AGCTACATCTGGCTACTGGGTCTCTT | >MI0000299\|hsa-mir-222&WithFlank&X\|-\|45747009\|45747130 | 75 | 99 | 25(+1U) | AGCTACATCTGGCTACTGGGTCTCTT | BT20 | 62 | 24 | 81 | 12 | 41 | 63.3 | 18.2 | 33.3 | 9.6 | 39.6 |
| >MI0000299\|hsa-mir-222&WithFlank&X\|-\|45747009\|45747130@75.99.25(+1U)&[MIMAT0000279&hsa-miR-222-3p&offsets\|0\|+4(+1U);m-140&X\|-\|45747033\|45747056&offsets\|0\|+1(+1U)]\|AGCTACATCTGGCTACTGGGTCTCTT | >MI0000299\|hsa-mir-222&WithFlank&X\|-\|45747009\|45747130 | 75 | 99 | 25(+1U) | AGCTACATCTGGCTACTGGGTCTCTT | MB231 | 345 | 185 | 223 | 64 | 289 | 134.1 | 64.8 | 75.3 | 24.6 | 133.0 |
| >MI0000299\|hsa-mir-222&WithFlank&X\|-\|45747009\|45747130@75.99.25(+2U)&[MIMAT0000279&hsa-miR-222-3p&offsets\|0\|+4(+2U);m-140&X\|-\|45747033\|45747056&offsets\|0\|+1(+2U)]\|AGCTACATCTGGCTACTGGGTCTCTTT | >MI0000299\|hsa-mir-222&WithFlank&X\|-\|45747009\|45747130 | 75 | 99 | 25(+2U) | AGCTACATCTGGCTACTGGGTCTCTTT | BT20 | 27 | 11 | 27 | 4 | 16 | 30.8 | 11.1 | 13.0 | 2.7 | 18.7 |
| >MI0000299\|hsa-mir-222&WithFlank&X\|-\|45747009\|45747130@75.99.25(+2U)&[MIMAT0000279&hsa-miR-222-3p&offsets\|0\|+4(+2U);m-140&X\|-\|45747033\|45747056&offsets\|0\|+1(+2U)]\|AGCTACATCTGGCTACTGGGTCTCTTT | >MI0000299\|hsa-mir-222&WithFlank&X\|-\|45747009\|45747130 | 75 | 99 | 25(+2U) | AGCTACATCTGGCTACTGGGTCTCTTT | MB231 | 189 | 117 | 142 | 34 | 164 | 93.5 | 49.9 | 61.4 | 13.3 | 75.2 |
| >MI0000299\|hsa-mir-222&WithFlank&X\|-\|45747009\|45747130@75.99.25(+3U)&[MIMAT0000279&hsa-miR-222-3p&offsets\|0\|+4(+3U);m-140&X\|-\|45747033\|45747056&offsets\|0\|+1(+3U)]\|AGCTACATCTGGCTACTGGGTCTCTTTT | >MI0000299\|hsa-mir-222&WithFlank&X\|-\|45747009\|45747130 | 75 | 99 | 25(+3U) | AGCTACATCTGGCTACTGGGTCTCTTTT | BT20 | 16 | 5 | 17 | 1 | 9 | 19.5 | 3.5 | 8.1 | 1.3 | 12.5 |
| >MI0000299\|hsa-mir-222&WithFlank&X\|-\|45747009\|45747130@75.99.25(+3U)&[MIMAT0000279&hsa-miR-222-3p&offsets\|0\|+4(+3U);m-140&X\|-\|45747033\|45747056&offsets\|0\|+1(+3U)]\|AGCTACATCTGGCTACTGGGTCTCTTTT | >MI0000299\|hsa-mir-222&WithFlank&X\|-\|45747009\|45747130 | 75 | 99 | 25(+3U) | AGCTACATCTGGCTACTGGGTCTCTTTT | MB231 | 162 | 83 | 103 | 19 | 157 | 71.0 | 38.7 | 34.4 | 8.8 | 80.9 |
| >MI0000299\|hsa-mir-222&WithFlank&X\|-\|45747009\|45747130@75.99.25(+4U)&[MIMAT0000279&hsa-miR-222-3p&offsets\|0\|+4(+4U);m-140&X\|-\|45747033\|45747056&offsets\|0\|+1(+4U)]\|AGCTACATCTGGCTACTGGGTCTCTTTTT | >MI0000299\|hsa-mir-222&WithFlank&X\|-\|45747009\|45747130 | 75 | 99 | 25(+4U) | AGCTACATCTGGCTACTGGGTCTCTTTTT | MB231 | 32 | 18 | 26 | 3 | 31 | 14.6 | 7.0 | 11.9 | 2.3 | 16.9 |
| >MI0000299\|hsa-mir-222&WithFlank&X\|-\|45747009\|45747130@75.99.25&[MIMAT0000279&hsa-miR-222-3p&offsets\|0\|+4;m-140&X\|-\|45747033\|45747056&offsets\|0\|+1]\|AGCTACATCTGGCTACTGGGTCTCT | >MI0000299\|hsa-mir-222&WithFlank&X\|-\|45747009\|45747130 | 75 | 99 | 25 | AGCTACATCTGGCTACTGGGTCTCT | BT20 | 560 | 268 | 810 | 184 | 399 | 395.4 | 175.3 | 295.9 | 116.5 | 278.8 |
| >MI0000299\|hsa-mir-222&WithFlank&X\|-\|45747009\|45747130@75.99.25&[MIMAT0000279&hsa-miR-222-3p&offsets\|0\|+4;m-140&X\|-\|45747033\|45747056&offsets\|0\|+1]\|AGCTACATCTGGCTACTGGGTCTCT | >MI0000299\|hsa-mir-222&WithFlank&X\|-\|45747009\|45747130 | 75 | 99 | 25 | AGCTACATCTGGCTACTGGGTCTCT | MB231 | 2804 | 1665 | 2065 | 1205 | 2330 | 1345.0 | 412.1 | 659.2 | 201.7 | 811.3 |
| >MI0000299\|hsa-mir-222&WithFlank&X\|-\|45747009\|45747130@75.99.25&[MIMAT0000279&hsa-miR-222-3p&offsets\|0\|+4;m-140&X\|-\|45747033\|45747056&offsets\|0\|+1]\|AGCTACATCTGGCTACTGGGTCTCT | >MI0000299\|hsa-mir-222&WithFlank&X\|-\|45747009\|45747130 | 75 | 99 | 25 | AGCTACATCTGGCTACTGGGTCTCT | MB468 | 31 | 46 | 19 | 30 | 19 | 9.7 | 27.7 | 10.2 | 14.8 | 6.7 |
| >MI0000301\|hsa-mir-224&WithFlank&X\|-\|151958572\|151958664@14.33.20(+1C)&[MIMAT0000281&hsa-miR-224-5p&offsets\|+1\|-4(+1C);m-238&X\|-\|151958630\|151958651&offsets\|0\|-2(+1C)]\|CAAGTCACTAGTGGTTCCGTC | >MI0000301\|hsa-mir-224&WithFlank&X\|-\|151958572\|151958664 | 14 | 33 | 20(+1C) | CAAGTCACTAGTGGTTCCGTC | BT20 | 12 | 9 | 10 | 6 | 6 | 6.7 | 4.7 | 4.4 | 4.2 | 3.7 |
| >MI0000301\|hsa-mir-224&WithFlank&X\|-\|151958572\|151958664@14.34.21(+1A)&[MIMAT0000281&hsa-miR-224-5p&offsets\|+1\|-3(+1A);m-238&X\|-\|151958630\|151958651&offsets\|0\|-1(+1A)]\|CAAGTCACTAGTGGTTCCGTTA | >MI0000301\|hsa-mir-224&WithFlank&X\|-\|151958572\|151958664 | 14 | 34 | 21(+1A) | CAAGTCACTAGTGGTTCCGTTA | BT20 | 25 | 11 | 26 | 11 | 6 | 10.3 | 6.4 | 4.8 | 8.4 | 3.2 |
| >MI0000301\|hsa-mir-224&WithFlank&X\|-\|151958572\|151958664@14.34.21(+1C)&[MIMAT0000281&hsa-miR-224-5p&offsets\|+1\|-3(+1C);m-238&X\|-\|151958630\|151958651&offsets\|0\|-1(+1C)]\|CAAGTCACTAGTGGTTCCGTTC | >MI0000301\|hsa-mir-224&WithFlank&X\|-\|151958572\|151958664 | 14 | 34 | 21(+1C) | CAAGTCACTAGTGGTTCCGTTC | BT20 | 75 | 55 | 79 | 60 | 32 | 22.2 | 33.9 | 27.7 | 47.1 | 17.1 |
| >MI0000301\|hsa-mir-224&WithFlank&X\|-\|151958572\|151958664@14.34.21(+1G)&[MIMAT0000281&hsa-miR-224-5p&offsets\|+1\|-3(+1G);m-238&X\|-\|151958630\|151958651&offsets\|0\|-1(+1G)]\|CAAGTCACTAGTGGTTCCGTTG | >MI0000301\|hsa-mir-224&WithFlank&X\|-\|151958572\|151958664 | 14 | 34 | 21(+1G) | CAAGTCACTAGTGGTTCCGTTG | BT20 | 20 | 5 | 8 | 4 | 1 | 12.0 | 3.3 | 1.0 | 4.0 | 1.0 |
| >MI0000301\|hsa-mir-224&WithFlank&X\|-\|151958572\|151958664@14.34.21&[MIMAT0000281&hsa-miR-224-5p&offsets\|+1\|-3;m-238&X\|-\|151958630\|151958651&offsets\|0\|-1]\|CAAGTCACTAGTGGTTCCGTT | >MI0000301\|hsa-mir-224&WithFlank&X\|-\|151958572\|151958664 | 14 | 34 | 21 | CAAGTCACTAGTGGTTCCGTT | BT20 | 91 | 62 | 81 | 47 | 45 | 55.2 | 29.1 | 51.0 | 29.3 | 25.7 |
| >MI0000301\|hsa-mir-224&WithFlank&X\|-\|151958572\|151958664@14.34.21&[MIMAT0000281&hsa-miR-224-5p&offsets\|+1\|-3;m-238&X\|-\|151958630\|151958651&offsets\|0\|-1]\|CAAGTCACTAGTGGTTCCGTT | >MI0000301\|hsa-mir-224&WithFlank&X\|-\|151958572\|151958664 | 14 | 34 | 21 | CAAGTCACTAGTGGTTCCGTT | MB231 | 16 | 19 | 9 | 12 | 15 | 0.8 | 5.4 | 3.0 | 5.6 | 5.2 |
| >MI0000301\|hsa-mir-224&WithFlank&X\|-\|151958572\|151958664@14.35.22(+1U)&[MIMAT0000281&hsa-miR-224-5p&offsets\|+1\|-2(+1U);m-238&X\|-\|151958630\|151958651&offsets\|0\|0(+1U)]\|CAAGTCACTAGTGGTTCCGTTTT | >MI0000301\|hsa-mir-224&WithFlank&X\|-\|151958572\|151958664 | 14 | 35 | 22(+1U) | CAAGTCACTAGTGGTTCCGTTTT | BT20 | 15 | 10 | 14 | 5 | 6 | 5.7 | 5.0 | 3.8 | 3.1 | 3.0 |

<table>
<tr><td>>MI0000301|hsa-mir-224&WithFlank&X|-|151958572|151958664@14.35.22&[MIMAT0000281&hsa-miR-224-5p&offsets|+1|-2;m-238&X|-|151958630|151958651&offsets|0|0]|CAAGTCACTAGTGGTTCCGTTT</td><td>>MI0000301|hsa-mir-224&WithFlank&X|-|151958572|151958664</td><td>14</td><td>35</td><td>22</td><td>CAAGTCACTAGTGGTTCCGTTT</td><td>BT20</td><td>652</td><td>469</td><td>756</td><td>524</td><td>300</td><td>257.9</td><td>265.6</td><td>366.7</td><td>373.4</td><td>172.3</td></tr>
<tr><td>>MI0000301|hsa-mir-224&WithFlank&X|-|151958572|151958664@14.35.22&[MIMAT0000281&hsa-miR-224-5p&offsets|+1|-2;m-238&X|-|151958630|151958651&offsets|0|0]|CAAGTCACTAGTGGTTCCGTTT</td><td>>MI0000301|hsa-mir-224&WithFlank&X|-|151958572|151958664</td><td>14</td><td>35</td><td>22</td><td>CAAGTCACTAGTGGTTCCGTTT</td><td>MB231</td><td>96</td><td>95</td><td>60</td><td>74</td><td>88</td><td>15.4</td><td>43.7</td><td>38.8</td><td>37.7</td><td>37.0</td></tr>
<tr><td>>MI0000301|hsa-mir-224&WithFlank&X|-|151958572|151958664@14.36.23(+1A)&[MIMAT0000281&hsa-miR-224-5p&offsets|+1|-1(+1A);m-238&X|-|151958630|151958651&offsets|0|+1(+1A)]|CAAGTCACTAGTGGTTCCGTTTAA</td><td>>MI0000301|hsa-mir-224&WithFlank&X|-|151958572|151958664</td><td>14</td><td>36</td><td>23(+1A)</td><td>CAAGTCACTAGTGGTTCCGTTTAA</td><td>BT20</td><td>23</td><td>11</td><td>24</td><td>3</td><td>9</td><td>14.9</td><td>7.6</td><td>9.8</td><td>1.1</td><td>4.0</td></tr>
<tr><td>>MI0000301|hsa-mir-224&WithFlank&X|-|151958572|151958664@14.36.23(+1U)&[MIMAT0000281&hsa-miR-224-5p&offsets|+1|-1(+1U);m-238&X|-|151958630|151958651&offsets|0|+1(+1U)]|CAAGTCACTAGTGGTTCCGTTTAT</td><td>>MI0000301|hsa-mir-224&WithFlank&X|-|151958572|151958664</td><td>14</td><td>36</td><td>23(+1U)</td><td>CAAGTCACTAGTGGTTCCGTTTAT</td><td>BT20</td><td>10</td><td>6</td><td>9</td><td>2</td><td>4</td><td>11.8</td><td>3.7</td><td>6.1</td><td>0.6</td><td>2.5</td></tr>
<tr><td>>MI0000301|hsa-mir-224&WithFlank&X|-|151958572|151958664@14.36.23&[MIMAT0000281&hsa-miR-224-5p&offsets|+1|-1;m-238&X|-|151958630|151958651&offsets|0|+1]|CAAGTCACTAGTGGTTCCGTTTA</td><td>>MI0000301|hsa-mir-224&WithFlank&X|-|151958572|151958664</td><td>14</td><td>36</td><td>23</td><td>CAAGTCACTAGTGGTTCCGTTTA</td><td>BT20</td><td>115</td><td>79</td><td>131</td><td>19</td><td>44</td><td>77.6</td><td>65.8</td><td>67.8</td><td>13.3</td><td>27.1</td></tr>
<tr><td>>MI0000301|hsa-mir-224&WithFlank&X|-|151958572|151958664@14.36.23&[MIMAT0000281&hsa-miR-224-5p&offsets|+1|-1;m-238&X|-|151958630|151958651&offsets|0|+1]|CAAGTCACTAGTGGTTCCGTTTA</td><td>>MI0000301|hsa-mir-224&WithFlank&X|-|151958572|151958664</td><td>14</td><td>36</td><td>23</td><td>CAAGTCACTAGTGGTTCCGTTTA</td><td>MB231</td><td>18</td><td>19</td><td>12</td><td>9</td><td>22</td><td>1.7</td><td>8.9</td><td>8.9</td><td>5.3</td><td>9.4</td></tr>
<tr><td>>MI0000301|hsa-mir-224&WithFlank&X|-|151958572|151958664@14.37.24&[MIMAT0000281&hsa-miR-224-5p&offsets|+1|0;m-238&X|-|151958630|151958651&offsets|0|+2]|CAAGTCACTAGTGGTTCCGTTTAG</td><td>>MI0000301|hsa-mir-224&WithFlank&X|-|151958572|151958664</td><td>14</td><td>37</td><td>24</td><td>CAAGTCACTAGTGGTTCCGTTTAG</td><td>BT20</td><td>190</td><td>134</td><td>226</td><td>47</td><td>139</td><td>159.0</td><td>52.3</td><td>156.0</td><td>23.2</td><td>59.2</td></tr>
<tr><td>>MI0000301|hsa-mir-224&WithFlank&X|-|151958572|151958664@14.37.24&[MIMAT0000281&hsa-miR-224-5p&offsets|+1|0;m-238&X|-|151958630|151958651&offsets|0|+2]|CAAGTCACTAGTGGTTCCGTTTAG</td><td>>MI0000301|hsa-mir-224&WithFlank&X|-|151958572|151958664</td><td>14</td><td>37</td><td>24</td><td>CAAGTCACTAGTGGTTCCGTTTAG</td><td>MB231</td><td>84</td><td>91</td><td>47</td><td>33</td><td>89</td><td>18.9</td><td>38.3</td><td>29.6</td><td>17.8</td><td>27.6</td></tr>
<tr><td>>MI0000301|hsa-mir-224&WithFlank&X|-|151958572|151958664@14.38.25&[MIMAT0000281&hsa-miR-224-5p&offsets|+1|+1;m-238&X|-|151958630|151958651&offsets|0|+3]|CAAGTCACTAGTGGTTCCGTTTAGT</td><td>>MI0000301|hsa-mir-224&WithFlank&X|-|151958572|151958664</td><td>14</td><td>38</td><td>25</td><td>CAAGTCACTAGTGGTTCCGTTTAGT</td><td>BT20</td><td>32</td><td>25</td><td>50</td><td>2</td><td>15</td><td>32.7</td><td>22.6</td><td>46.7</td><td>1.8</td><td>19.8</td></tr>
<tr><td>>MI0000301|hsa-mir-224&WithFlank&X|-|151958572|151958664@14.38.25&[MIMAT0000281&hsa-miR-224-5p&offsets|+1|+1;m-238&X|-|151958630|151958651&offsets|0|+3]|CAAGTCACTAGTGGTTCCGTTTAGT</td><td>>MI0000301|hsa-mir-224&WithFlank&X|-|151958572|151958664</td><td>14</td><td>38</td><td>25</td><td>CAAGTCACTAGTGGTTCCGTTTAGT</td><td>MB231</td><td>28</td><td>32</td><td>20</td><td>2</td><td>32</td><td>4.4</td><td>15.5</td><td>13.5</td><td>1.8</td><td>31.2</td></tr>
<tr><td>>MI0000342|hsa-mir-200b&WithFlank&1|+|1167098|1167204@63.83.21&[MIMAT0000318&hsa-miR-200b-3p&offsets|0|-1;m-93&1|+|1167160|1167182&offsets|0|-2]|TAATACTGCCTGGTAATGATG</td><td>>MI0000342|hsa-mir-200b&WithFlank&1|+|1167098|1167204</td><td>63</td><td>83</td><td>21</td><td>TAATACTGCCTGGTAATGATG</td><td>MB468</td><td>39</td><td>20</td><td>19</td><td>76</td><td>16</td><td>21.9</td><td>7.9</td><td>1.2</td><td>20.5</td><td>4.0</td></tr>
<tr><td>>MI0000342|hsa-mir-200b&WithFlank&1|+|1167098|1167204@63.84.22(+1U)&[MIMAT0000318&hsa-miR-200b-3p&offsets|0|0(+1U);m-93&1|+|1167160|1167182&offsets|0|-1(+1U)]|TAATACTGCCTGGTAATGATGAT</td><td>>MI0000342|hsa-mir-200b&WithFlank&1|+|1167098|1167204</td><td>63</td><td>84</td><td>22(+1U)</td><td>TAATACTGCCTGGTAATGATGAT</td><td>MB468</td><td>25</td><td>24</td><td>13</td><td>18</td><td>12</td><td>5.6</td><td>10.9</td><td>5.2</td><td>8.2</td><td>8.8</td></tr>
<tr><td>>MI0000342|hsa-mir-200b&WithFlank&1|+|1167098|1167204@63.84.22&[MIMAT0000318&hsa-miR-200b-3p&offsets|0|0;m-93&1|+|1167160|1167182&offsets|0|-1]|TAATACTGCCTGGTAATGATGA</td><td>>MI0000342|hsa-mir-200b&WithFlank&1|+|1167098|1167204</td><td>63</td><td>84</td><td>22</td><td>TAATACTGCCTGGTAATGATGA</td><td>MB468</td><td>93</td><td>63</td><td>37</td><td>191</td><td>50</td><td>44.1</td><td>19.6</td><td>2.0</td><td>52.2</td><td>25.9</td></tr>
<tr><td>>MI0000342|hsa-mir-200b&WithFlank&1|+|1167098|1167204@63.85.23(+1A)&[MIMAT0000318&hsa-miR-200b-3p&offsets|0|+1(+1A);m-93&1|+|1167160|1167182&offsets|0|0(+1A)]|TAATACTGCCTGGTAATGATGACA</td><td>>MI0000342|hsa-mir-200b&WithFlank&1|+|1167098|1167204</td><td>63</td><td>85</td><td>23(+1A)</td><td>TAATACTGCCTGGTAATGATGACA</td><td>MB468</td><td>87</td><td>43</td><td>39</td><td>37</td><td>31</td><td>32.5</td><td>11.2</td><td>7.6</td><td>9.7</td><td>12.3</td></tr>
<tr><td>>MI0000342|hsa-mir-200b&WithFlank&1|+|1167098|1167204@63.85.23(+1C)&[MIMAT0000318&hsa-miR-200b-3p&offsets|0|+1(+1C);m-93&1|+|1167160|1167182&offsets|0|0(+1C)]|TAATACTGCCTGGTAATGATGACC</td><td>>MI0000342|hsa-mir-200b&WithFlank&1|+|1167098|1167204</td><td>63</td><td>85</td><td>23(+1C)</td><td>TAATACTGCCTGGTAATGATGACC</td><td>MB468</td><td>25</td><td>21</td><td>13</td><td>11</td><td>11</td><td>8.7</td><td>3.2</td><td>1.9</td><td>1.9</td><td>4.0</td></tr>
<tr><td>>MI0000342|hsa-mir-200b&WithFlank&1|+|1167098|1167204@63.85.23(+1U)&[MIMAT0000318&hsa-miR-200b-3p&offsets|0|+1(+1U);m-93&1|+|1167160|1167182&offsets|0|0(+1U)]|TAATACTGCCTGGTAATGATGACT</td><td>>MI0000342|hsa-mir-200b&WithFlank&1|+|1167098|1167204</td><td>63</td><td>85</td><td>23(+1U)</td><td>TAATACTGCCTGGTAATGATGACT</td><td>MB468</td><td>146</td><td>117</td><td>69</td><td>48</td><td>63</td><td>51.0</td><td>34.5</td><td>10.1</td><td>11.3</td><td>32.5</td></tr>
<tr><td>>MI0000342|hsa-mir-200b&WithFlank&1|+|1167098|1167204@63.85.23&[MIMAT0000318&hsa-miR-200b-3p&offsets|0|+1;m-93&1|+|1167160|1167182&offsets|0|0]|TAATACTGCCTGGTAATGATGAC</td><td>>MI0000342|hsa-mir-200b&WithFlank&1|+|1167098|1167204</td><td>63</td><td>85</td><td>23</td><td>TAATACTGCCTGGTAATGATGAC</td><td>BT20</td><td>17</td><td>10</td><td>77</td><td>4</td><td>9</td><td>12.3</td><td>2.6</td><td>44.2</td><td>2.1</td><td>2.5</td></tr>
<tr><td>>MI0000342|hsa-mir-200b&WithFlank&1|+|1167098|1167204@63.85.23&[MIMAT0000318&hsa-miR-200b-3p&offsets|0|+1;m-93&1|+|1167160|1167182&offsets|0|0]|TAATACTGCCTGGTAATGATGAC</td><td>>MI0000342|hsa-mir-200b&WithFlank&1|+|1167098|1167204</td><td>63</td><td>85</td><td>23</td><td>TAATACTGCCTGGTAATGATGAC</td><td>MB468</td><td>288</td><td>313</td><td>148</td><td>153</td><td>101</td><td>129.9</td><td>114.5</td><td>48.3</td><td>19.4</td><td>33.5</td></tr>
<tr><td>>MI0000342|hsa-mir-200b&WithFlank&1|+|1167098|1167204@63.86.24&[MIMAT0000318&hsa-miR-200b-3p&offsets|0|+2;m-93&1|+|1167160|1167182&offsets|0|+1]|TAATACTGCCTGGTAATGATGACG</td><td>>MI0000342|hsa-mir-200b&WithFlank&1|+|1167098|1167204</td><td>63</td><td>86</td><td>24</td><td>TAATACTGCCTGGTAATGATGACG</td><td>MB468</td><td>17</td><td>8</td><td>9</td><td>5</td><td>4</td><td>4.3</td><td>1.2</td><td>4.7</td><td>0.8</td><td>1.8</td></tr>
<tr><td>>MI0000342|hsa-mir-200b&WithFlank&1|+|1167098|1167204@64.85.22(+1U)&[MIMAT0000318&hsa-miR-200b-3p&offsets|+1|+1(+1U);m-93&1|+|1167160|1167182&offsets|+1|0(+1U)]|AATACTGCCTGGTAATGATGACT</td><td>>MI0000342|hsa-mir-200b&WithFlank&1|+|1167098|1167204</td><td>64</td><td>85</td><td>22(+1U)</td><td>AATACTGCCTGGTAATGATGACT</td><td>MB468</td><td>27</td><td>30</td><td>20</td><td>21</td><td>13</td><td>9.3</td><td>17.5</td><td>3.3</td><td>6.7</td><td>7.1</td></tr>
<tr><td>>MI0000433|hsa-let-7g&WithFlank&3|-|52268272|52268367@10.32.23&[MIMAT0000414&hsa-let-7g-5p&offsets|-1|0;m-58&3|-|52268336|52268357&offsets|-1|0]|CTGAGGTAGTAGTTTGTACAGTT</td><td>>MI0000433|hsa-let-7g&WithFlank&3|-|52268272|52268367</td><td>10</td><td>32</td><td>23</td><td>CTGAGGTAGTAGTTTGTACAGTT</td><td>BT20</td><td>27</td><td>20</td><td>18</td><td>5</td><td>13</td><td>12.7</td><td>7.1</td><td>16.9</td><td>2.2</td><td>5.3</td></tr>
<tr><td>>MI0000433|hsa-let-7g&WithFlank&3|-|52268272|52268367@10.32.23&[MIMAT0000414&hsa-let-7g-5p&offsets|-1|0;m-58&3|-|52268336|52268357&offsets|-1|0]|CTGAGGTAGTAGTTTGTACAGTT</td><td>>MI0000433|hsa-let-7g&WithFlank&3|-|52268272|52268367</td><td>10</td><td>32</td><td>23</td><td>CTGAGGTAGTAGTTTGTACAGTT</td><td>MB231</td><td>24</td><td>23</td><td>5</td><td>13</td><td>12</td><td>4.2</td><td>9.2</td><td>2.4</td><td>6.7</td><td>4.3</td></tr>
<tr><td>>MI0000433|hsa-let-7g&WithFlank&3|-|52268272|52268367@11.28.18&[MIMAT0000414&hsa-let-7g-5p&offsets|0|-4;m-58&3|-|52268336|52268357&offsets|0|-4]|TGAGGTAGTAGTTTGTAC</td><td>>MI0000433|hsa-let-7g&WithFlank&3|-|52268272|52268367</td><td>11</td><td>28</td><td>18</td><td>TGAGGTAGTAGTTTGTAC</td><td>BT20</td><td>146</td><td>159</td><td>162</td><td>63</td><td>79</td><td>158.5</td><td>84.6</td><td>191.1</td><td>36.5</td><td>11.9</td></tr>
<tr><td>>MI0000433|hsa-let-7g&WithFlank&3|-|52268272|52268367@11.28.18&[MIMAT0000414&hsa-let-7g-5p&offsets|0|-4;m-58&3|-|52268336|52268357&offsets|0|-4]|TGAGGTAGTAGTTTGTAC</td><td>>MI0000433|hsa-let-7g&WithFlank&3|-|52268272|52268367</td><td>11</td><td>28</td><td>18</td><td>TGAGGTAGTAGTTTGTAC</td><td>MB231</td><td>137</td><td>209</td><td>47</td><td>262</td><td>78</td><td>51.3</td><td>114.1</td><td>33.4</td><td>200.9</td><td>44.1</td></tr>
<tr><td>>MI0000433|hsa-let-7g&WithFlank&3|-|52268272|52268367@11.29.19&[MIMAT0000414&hsa-let-7g-5p&offsets|0|-3;m-58&3|-|52268336|52268357&offsets|0|-3]|TGAGGTAGTAGTTTGTACA</td><td>>MI0000433|hsa-let-7g&WithFlank&3|-|52268272|52268367</td><td>11</td><td>29</td><td>19</td><td>TGAGGTAGTAGTTTGTACA</td><td>BT20</td><td>14</td><td>13</td><td>25</td><td>3</td><td>5</td><td>5.3</td><td>1.5</td><td>11.4</td><td>2.2</td><td>0.3</td></tr>
<tr><td>>MI0000433|hsa-let-7g&WithFlank&3|-|52268272|52268367@11.30.20(+1C)&[MIMAT0000414&hsa-let-7g-5p&offsets|0|-2(+1C);m-58&3|-|52268336|52268357&offsets|0|-2(+1C)]|TGAGGTAGTAGTTTGTACAGC</td><td>>MI0000433|hsa-let-7g&WithFlank&3|-|52268272|52268367</td><td>11</td><td>30</td><td>20(+1C)</td><td>TGAGGTAGTAGTTTGTACAGC</td><td>BT20</td><td>36</td><td>36</td><td>29</td><td>19</td><td>13</td><td>25.5</td><td>23.0</td><td>26.5</td><td>22.2</td><td>4.5</td></tr>
<tr><td>>MI0000433|hsa-let-7g&WithFlank&3|-|52268272|52268367@11.30.20(+1C)&[MIMAT0000414&hsa-let-7g-5p&offsets|0|-2(+1C);m-58&3|-|52268336|52268357&offsets|0|-2(+1C)]|TGAGGTAGTAGTTTGTACAGC</td><td>>MI0000433|hsa-let-7g&WithFlank&3|-|52268272|52268367</td><td>11</td><td>30</td><td>20(+1C)</td><td>TGAGGTAGTAGTTTGTACAGC</td><td>MB231</td><td>27</td><td>33</td><td>9</td><td>47</td><td>12</td><td>3.9</td><td>13.5</td><td>5.7</td><td>37.5</td><td>5.3</td></tr>
<tr><td>>MI0000433|hsa-let-7g&WithFlank&3|-|52268272|52268367@11.30.20(+1G)&[MIMAT0000414&hsa-let-7g-5p&offsets|0|-2(+1G);m-58&3|-|52268336|52268357&offsets|0|-2(+1G)]|TGAGGTAGTAGTTTGTACAGG</td><td>>MI0000433|hsa-let-7g&WithFlank&3|-|52268272|52268367</td><td>11</td><td>30</td><td>20(+1G)</td><td>TGAGGTAGTAGTTTGTACAGG</td><td>BT20</td><td>13</td><td>6</td><td>5</td><td>3</td><td>1</td><td>7.0</td><td>3.8</td><td>1.8</td><td>4.3</td><td>0.7</td></tr>
<tr><td>>MI0000433|hsa-let-7g&WithFlank&3|-|52268272|52268367@11.30.20(+1G)&[MIMAT0000414&hsa-let-7g-5p&offsets|0|-2(+1G);m-58&3|-|52268336|52268357&offsets|0|-2(+1G)]|TGAGGTAGTAGTTTGTACAGG</td><td>>MI0000433|hsa-let-7g&WithFlank&3|-|52268272|52268367</td><td>11</td><td>30</td><td>20(+1G)</td><td>TGAGGTAGTAGTTTGTACAGG</td><td>MB231</td><td>12</td><td>5</td><td>1</td><td>4</td><td>1</td><td>9.2</td><td>2.8</td><td>1.2</td><td>3.5</td><td>0.7</td></tr>
<tr><td>>MI0000433|hsa-let-7g&WithFlank&3|-|52268272|52268367@11.31.21(+1A)&[MIMAT0000414&hsa-let-7g-5p&offsets|0|-1(+1A);m-58&3|-|52268336|52268357&offsets|0|-1(+1A)]|TGAGGTAGTAGTTTGTACAGTA</td><td>>MI0000433|hsa-let-7g&WithFlank&3|-|52268272|52268367</td><td>11</td><td>31</td><td>21(+1A)</td><td>TGAGGTAGTAGTTTGTACAGTA</td><td>BT20</td><td>259</td><td>151</td><td>162</td><td>38</td><td>71</td><td>114.0</td><td>71.2</td><td>115.2</td><td>42.3</td><td>19.3</td></tr>
<tr><td>>MI0000433|hsa-let-7g&WithFlank&3|-|52268272|52268367@11.31.21(+1A)&[MIMAT0000414&hsa-let-7g-5p&offsets|0|-1(+1A);m-58&3|-|52268336|52268357&offsets|0|-1(+1A)]|TGAGGTAGTAGTTTGTACAGTA</td><td>>MI0000433|hsa-let-7g&WithFlank&3|-|52268272|52268367</td><td>11</td><td>31</td><td>21(+1A)</td><td>TGAGGTAGTAGTTTGTACAGTA</td><td>MB231</td><td>160</td><td>121</td><td>35</td><td>179</td><td>64</td><td>12.1</td><td>24.9</td><td>21.4</td><td>139.2</td><td>21.7</td></tr>
<tr><td>>MI0000433|hsa-let-7g&WithFlank&3|-|52268272|52268367@11.31.21(+1C)&[MIMAT0000414&hsa-let-7g-5p&offsets|0|-1(+1C);m-58&3|-|52268336|52268357&offsets|0|-1(+1C)]|TGAGGTAGTAGTTTGTACAGTC</td><td>>MI0000433|hsa-let-7g&WithFlank&3|-|52268272|52268367</td><td>11</td><td>31</td><td>21(+1C)</td><td>TGAGGTAGTAGTTTGTACAGTC</td><td>BT20</td><td>751</td><td>696</td><td>510</td><td>161</td><td>352</td><td>470.1</td><td>349.8</td><td>487.8</td><td>173.8</td><td>96.0</td></tr>
<tr><td>>MI0000433|hsa-let-7g&WithFlank&3|-|52268272|52268367@11.31.21(+1C)&[MIMAT0000414&hsa-let-7g-5p&offsets|0|-1(+1C);m-58&3|-|52268336|52268357&offsets|0|-1(+1C)]|TGAGGTAGTAGTTTGTACAGTC</td><td>>MI0000433|hsa-let-7g&WithFlank&3|-|52268272|52268367</td><td>11</td><td>31</td><td>21(+1C)</td><td>TGAGGTAGTAGTTTGTACAGTC</td><td>MB231</td><td>640</td><td>631</td><td>127</td><td>721</td><td>312</td><td>121.9</td><td>185.4</td><td>74.4</td><td>554.8</td><td>132.5</td></tr>
</table>

<table>
<tr><td>>MI0000433|hsa-let-7g&WithFlank&3|-|52268272|52268367@11.31.21(+1C)&[MIMAT0000414&hsa-let-7g-5p&offsets|0|-1(+1C);m-58&3|-|52268336|52268357&offsets|0|-1(+1C)]|TGAGGTAGTAGTTTGTACAGTC</td><td>>MI0000433|hsa-let-7g&WithFlank&3|-|52268272|52268367</td><td>11</td><td>31</td><td>21(+1C)</td><td>TGAGGTAGTAGTTTGTACAGTC</td><td>MB468</td><td>39</td><td>33</td><td>14</td><td>17</td><td>8</td><td>11.9</td><td>7.1</td><td>5.9</td><td>5.7</td><td>3.2</td></tr>
<tr><td>>MI0000433|hsa-let-7g&WithFlank&3|-|52268272|52268367@11.31.21(+1G)&[MIMAT0000414&hsa-let-7g-5p&offsets|0|-1(+1G);m-58&3|-|52268336|52268357&offsets|0|-1(+1G)]|TGAGGTAGTAGTTTGTACAGTG</td><td>>MI0000433|hsa-let-7g&WithFlank&3|-|52268272|52268367</td><td>11</td><td>31</td><td>21(+1G)</td><td>TGAGGTAGTAGTTTGTACAGTG</td><td>BT20</td><td>184</td><td>76</td><td>52</td><td>14</td><td>15</td><td>83.6</td><td>36.7</td><td>27.9</td><td>19.3</td><td>5.9</td></tr>
<tr><td>>MI0000433|hsa-let-7g&WithFlank&3|-|52268272|52268367@11.31.21(+1G)&[MIMAT0000414&hsa-let-7g-5p&offsets|0|-1(+1G);m-58&3|-|52268336|52268357&offsets|0|-1(+1G)]|TGAGGTAGTAGTTTGTACAGTG</td><td>>MI0000433|hsa-let-7g&WithFlank&3|-|52268272|52268367</td><td>11</td><td>31</td><td>21(+1G)</td><td>TGAGGTAGTAGTTTGTACAGTG</td><td>MB231</td><td>156</td><td>60</td><td>12</td><td>44</td><td>15</td><td>109.5</td><td>22.8</td><td>9.5</td><td>32.3</td><td>7.0</td></tr>
<tr><td>>MI0000433|hsa-let-7g&WithFlank&3|-|52268272|52268367@11.31.21&[MIMAT0000414&hsa-let-7g-5p&offsets|0|-1;m-58&3|-|52268336|52268357&offsets|0|-1]|TGAGGTAGTAGTTTGTACAGT</td><td>>MI0000433|hsa-let-7g&WithFlank&3|-|52268272|52268367</td><td>11</td><td>31</td><td>21</td><td>TGAGGTAGTAGTTTGTACAGT</td><td>BT20</td><td>291</td><td>268</td><td>248</td><td>140</td><td>108</td><td>221.5</td><td>147.8</td><td>240.3</td><td>130.7</td><td>35.4</td></tr>
<tr><td>>MI0000433|hsa-let-7g&WithFlank&3|-|52268272|52268367@11.31.21&[MIMAT0000414&hsa-let-7g-5p&offsets|0|-1;m-58&3|-|52268336|52268357&offsets|0|-1]|TGAGGTAGTAGTTTGTACAGT</td><td>>MI0000433|hsa-let-7g&WithFlank&3|-|52268272|52268367</td><td>11</td><td>31</td><td>21</td><td>TGAGGTAGTAGTTTGTACAGT</td><td>MB231</td><td>280</td><td>323</td><td>86</td><td>303</td><td>100</td><td>31.3</td><td>118.8</td><td>50.3</td><td>239.8</td><td>46.4</td></tr>
<tr><td>>MI0000433|hsa-let-7g&WithFlank&3|-|52268272|52268367@11.31.21&[MIMAT0000414&hsa-let-7g-5p&offsets|0|-1;m-58&3|-|52268336|52268357&offsets|0|-1]|TGAGGTAGTAGTTTGTACAGT</td><td>>MI0000433|hsa-let-7g&WithFlank&3|-|52268272|52268367</td><td>11</td><td>31</td><td>21</td><td>TGAGGTAGTAGTTTGTACAGT</td><td>MB468</td><td>14</td><td>13</td><td>6</td><td>8</td><td>3</td><td>6.1</td><td>3.3</td><td>1.8</td><td>4.6</td><td>1.4</td></tr>
<tr><td>>MI0000433|hsa-let-7g&WithFlank&3|-|52268272|52268367@11.32.22(+1A)&[MIMAT0000414&hsa-let-7g-5p&offsets|0|0(+1A);m-58&3|-|52268336|52268357&offsets|0|0(+1A)]|TGAGGTAGTAGTTTGTACAGTTA</td><td>>MI0000433|hsa-let-7g&WithFlank&3|-|52268272|52268367</td><td>11</td><td>32</td><td>22(+1A)</td><td>TGAGGTAGTAGTTTGTACAGTTA</td><td>BT20</td><td>98</td><td>78</td><td>96</td><td>155</td><td>44</td><td>78.6</td><td>41.7</td><td>107.9</td><td>147.5</td><td>6.6</td></tr>
<tr><td>>MI0000433|hsa-let-7g&WithFlank&3|-|52268272|52268367@11.32.22(+1A)&[MIMAT0000414&hsa-let-7g-5p&offsets|0|0(+1A);m-58&3|-|52268336|52268357&offsets|0|0(+1A)]|TGAGGTAGTAGTTTGTACAGTTA</td><td>>MI0000433|hsa-let-7g&WithFlank&3|-|52268272|52268367</td><td>11</td><td>32</td><td>22(+1A)</td><td>TGAGGTAGTAGTTTGTACAGTTA</td><td>MB231</td><td>70</td><td>61</td><td>16</td><td>159</td><td>26</td><td>14.8</td><td>16.9</td><td>9.8</td><td>110.6</td><td>6.9</td></tr>
<tr><td>>MI0000433|hsa-let-7g&WithFlank&3|-|52268272|52268367@11.32.22(+1C)&[MIMAT0000414&hsa-let-7g-5p&offsets|0|0(+1C);m-58&3|-|52268336|52268357&offsets|0|0(+1C)]|TGAGGTAGTAGTTTGTACAGTTC</td><td>>MI0000433|hsa-let-7g&WithFlank&3|-|52268272|52268367</td><td>11</td><td>32</td><td>22(+1C)</td><td>TGAGGTAGTAGTTTGTACAGTTC</td><td>MB231</td><td>10</td><td>10</td><td>3</td><td>39</td><td>6</td><td>0.3</td><td>4.0</td><td>2.2</td><td>28.4</td><td>2.2</td></tr>
<tr><td>>MI0000433|hsa-let-7g&WithFlank&3|-|52268272|52268367@11.32.22&[MIMAT0000414&hsa-let-7g-5p&offsets|0|0;m-58&3|-|52268336|52268357&offsets|0|0]|TGAGGTAGTAGTTTGTACAGTT</td><td>>MI0000433|hsa-let-7g&WithFlank&3|-|52268272|52268367</td><td>11</td><td>32</td><td>22</td><td>TGAGGTAGTAGTTTGTACAGTT</td><td>BT20</td><td>5440</td><td>4773</td><td>3974</td><td>1044</td><td>2554</td><td>3898.9</td><td>1965.7</td><td>4176.1</td><td>897.3</td><td>599.6</td></tr>
<tr><td>>MI0000433|hsa-let-7g&WithFlank&3|-|52268272|52268367@11.32.22&[MIMAT0000414&hsa-let-7g-5p&offsets|0|0;m-58&3|-|52268336|52268357&offsets|0|0]|TGAGGTAGTAGTTTGTACAGTT</td><td>>MI0000433|hsa-let-7g&WithFlank&3|-|52268272|52268367</td><td>11</td><td>32</td><td>22</td><td>TGAGGTAGTAGTTTGTACAGTT</td><td>MB231</td><td>5681</td><td>5776</td><td>1095</td><td>4101</td><td>2275</td><td>1091.4</td><td>1700.9</td><td>553.5</td><td>2897.1</td><td>819.8</td></tr>
<tr><td>>MI0000433|hsa-let-7g&WithFlank&3|-|52268272|52268367@11.32.22&[MIMAT0000414&hsa-let-7g-5p&offsets|0|0;m-58&3|-|52268336|52268357&offsets|0|0]|TGAGGTAGTAGTTTGTACAGTT</td><td>>MI0000433|hsa-let-7g&WithFlank&3|-|52268272|52268367</td><td>11</td><td>32</td><td>22</td><td>TGAGGTAGTAGTTTGTACAGTT</td><td>MB468</td><td>353</td><td>318</td><td>114</td><td>131</td><td>67</td><td>138.0</td><td>79.6</td><td>25.4</td><td>53.0</td><td>34.1</td></tr>
<tr><td>>MI0000433|hsa-let-7g&WithFlank&3|-|52268272|52268367@11.33.23&[MIMAT0000414&hsa-let-7g-5p&offsets|0|+1;m-58&3|-|52268336|52268357&offsets|0|+1]|TGAGGTAGTAGTTTGTACAGTTT</td><td>>MI0000433|hsa-let-7g&WithFlank&3|-|52268272|52268367</td><td>11</td><td>33</td><td>23</td><td>TGAGGTAGTAGTTTGTACAGTTT</td><td>BT20</td><td>73</td><td>65</td><td>62</td><td>81</td><td>32</td><td>65.4</td><td>41.9</td><td>68.7</td><td>76.3</td><td>15.8</td></tr>
<tr><td>>MI0000433|hsa-let-7g&WithFlank&3|-|52268272|52268367@11.33.23&[MIMAT0000414&hsa-let-7g-5p&offsets|0|+1;m-58&3|-|52268336|52268357&offsets|0|+1]|TGAGGTAGTAGTTTGTACAGTTT</td><td>>MI0000433|hsa-let-7g&WithFlank&3|-|52268272|52268367</td><td>11</td><td>33</td><td>23</td><td>TGAGGTAGTAGTTTGTACAGTTT</td><td>MB231</td><td>113</td><td>117</td><td>31</td><td>233</td><td>45</td><td>8.1</td><td>41.3</td><td>18.5</td><td>169.5</td><td>21.2</td></tr>
<tr><td>>MI0000434|hsa-let-7i&WithFlank&12|+|62603680|62603775@12.31.20(+1A)&[MIMAT0000415&hsa-let-7i-5p&offsets|0|-2(+1A);m-49&12|+|62603691|62603712&offsets|0|-2(+1A)]|TGAGGTAGTAGTTTGTGCTGA</td><td>>MI0000434|hsa-let-7i&WithFlank&12|+|62603680|62603775</td><td>12</td><td>31</td><td>20(+1A)</td><td>TGAGGTAGTAGTTTGTGCTGA</td><td>BT20</td><td>12</td><td>5</td><td>8</td><td>9</td><td>2</td><td>9.5</td><td>2.0</td><td>5.0</td><td>11.6</td><td>0.8</td></tr>
<tr><td>>MI0000434|hsa-let-7i&WithFlank&12|+|62603680|62603775@12.31.20(+1A)&[MIMAT0000415&hsa-let-7i-5p&offsets|0|-2(+1A);m-49&12|+|62603691|62603712&offsets|0|-2(+1A)]|TGAGGTAGTAGTTTGTGCTGA</td><td>>MI0000434|hsa-let-7i&WithFlank&12|+|62603680|62603775</td><td>12</td><td>31</td><td>20(+1A)</td><td>TGAGGTAGTAGTTTGTGCTGA</td><td>MB231</td><td>13</td><td>9</td><td>4</td><td>28</td><td>3</td><td>1.9</td><td>5.0</td><td>2.3</td><td>19.6</td><td>1.5</td></tr>
<tr><td>>MI0000434|hsa-let-7i&WithFlank&12|+|62603680|62603775@12.31.20(+1C)&[MIMAT0000415&hsa-let-7i-5p&offsets|0|-2(+1C);m-49&12|+|62603691|62603712&offsets|0|-2(+1C)]|TGAGGTAGTAGTTTGTGCTGC</td><td>>MI0000434|hsa-let-7i&WithFlank&12|+|62603680|62603775</td><td>12</td><td>31</td><td>20(+1C)</td><td>TGAGGTAGTAGTTTGTGCTGC</td><td>BT20</td><td>30</td><td>25</td><td>24</td><td>40</td><td>12</td><td>9.8</td><td>9.0</td><td>10.9</td><td>46.9</td><td>1.3</td></tr>
<tr><td>>MI0000434|hsa-let-7i&WithFlank&12|+|62603680|62603775@12.31.20(+1C)&[MIMAT0000415&hsa-let-7i-5p&offsets|0|-2(+1C);m-49&12|+|62603691|62603712&offsets|0|-2(+1C)]|TGAGGTAGTAGTTTGTGCTGC</td><td>>MI0000434|hsa-let-7i&WithFlank&12|+|62603680|62603775</td><td>12</td><td>31</td><td>20(+1C)</td><td>TGAGGTAGTAGTTTGTGCTGC</td><td>MB231</td><td>53</td><td>49</td><td>16</td><td>115</td><td>17</td><td>7.8</td><td>29.1</td><td>8.8</td><td>77.4</td><td>9.2</td></tr>
<tr><td>>MI0000434|hsa-let-7i&WithFlank&12|+|62603680|62603775@12.31.20(+1G)&[MIMAT0000415&hsa-let-7i-5p&offsets|0|-2(+1G);m-49&12|+|62603691|62603712&offsets|0|-2(+1G)]|TGAGGTAGTAGTTTGTGCTGG</td><td>>MI0000434|hsa-let-7i&WithFlank&12|+|62603680|62603775</td><td>12</td><td>31</td><td>20(+1G)</td><td>TGAGGTAGTAGTTTGTGCTGG</td><td>BT20</td><td>16</td><td>4</td><td>5</td><td>6</td><td>1</td><td>14.4</td><td>0.4</td><td>2.2</td><td>8.5</td><td>0.1</td></tr>
<tr><td>>MI0000434|hsa-let-7i&WithFlank&12|+|62603680|62603775@12.31.20(+1G)&[MIMAT0000415&hsa-let-7i-5p&offsets|0|-2(+1G);m-49&12|+|62603691|62603712&offsets|0|-2(+1G)]|TGAGGTAGTAGTTTGTGCTGG</td><td>>MI0000434|hsa-let-7i&WithFlank&12|+|62603680|62603775</td><td>12</td><td>31</td><td>20(+1G)</td><td>TGAGGTAGTAGTTTGTGCTGG</td><td>MB231</td><td>24</td><td>9</td><td>2</td><td>11</td><td>1</td><td>19.8</td><td>6.6</td><td>1.8</td><td>9.2</td><td>1.2</td></tr>
<tr><td>>MI0000434|hsa-let-7i&WithFlank&12|+|62603680|62603775@12.31.20(+2C)&[MIMAT0000415&hsa-let-7i-5p&offsets|0|-2(+2C);m-49&12|+|62603691|62603712&offsets|0|-2(+2C)]|TGAGGTAGTAGTTTGTGCTGCC</td><td>>MI0000434|hsa-let-7i&WithFlank&12|+|62603680|62603775</td><td>12</td><td>31</td><td>20(+2C)</td><td>TGAGGTAGTAGTTTGTGCTGCC</td><td>BT20</td><td>11</td><td>3</td><td>3</td><td>3</td><td>1</td><td>11.7</td><td>0.9</td><td>1.2</td><td>4.2</td><td>1.0</td></tr>
<tr><td>>MI0000434|hsa-let-7i&WithFlank&12|+|62603680|62603775@12.31.20(+2C)&[MIMAT0000415&hsa-let-7i-5p&offsets|0|-2(+2C);m-49&12|+|62603691|62603712&offsets|0|-2(+2C)]|TGAGGTAGTAGTTTGTGCTGCC</td><td>>MI0000434|hsa-let-7i&WithFlank&12|+|62603680|62603775</td><td>12</td><td>31</td><td>20(+2C)</td><td>TGAGGTAGTAGTTTGTGCTGCC</td><td>MB231</td><td>14</td><td>6</td><td>2</td><td>7</td><td>4</td><td>8.5</td><td>4.4</td><td>1.2</td><td>4.4</td><td>4.1</td></tr>
<tr><td>>MI0000434|hsa-let-7i&WithFlank&12|+|62603680|62603775@12.31.20&[MIMAT0000415&hsa-let-7i-5p&offsets|0|-2;m-49&12|+|62603691|62603712&offsets|0|-2]|TGAGGTAGTAGTTTGTGCTG</td><td>>MI0000434|hsa-let-7i&WithFlank&12|+|62603680|62603775</td><td>12</td><td>31</td><td>20</td><td>TGAGGTAGTAGTTTGTGCTG</td><td>BT20</td><td>15</td><td>11</td><td>9</td><td>16</td><td>6</td><td>7.7</td><td>4.8</td><td>5.1</td><td>12.8</td><td>2.5</td></tr>
<tr><td>>MI0000434|hsa-let-7i&WithFlank&12|+|62603680|62603775@12.31.20&[MIMAT0000415&hsa-let-7i-5p&offsets|0|-2;m-49&12|+|62603691|62603712&offsets|0|-2]|TGAGGTAGTAGTTTGTGCTG</td><td>>MI0000434|hsa-let-7i&WithFlank&12|+|62603680|62603775</td><td>12</td><td>31</td><td>20</td><td>TGAGGTAGTAGTTTGTGCTG</td><td>MB231</td><td>30</td><td>29</td><td>7</td><td>45</td><td>10</td><td>3.9</td><td>15.5</td><td>4.7</td><td>24.2</td><td>6.0</td></tr>
<tr><td>>MI0000434|hsa-let-7i&WithFlank&12|+|62603680|62603775@12.32.21(+1A)&[MIMAT0000415&hsa-let-7i-5p&offsets|0|-1(+1A);m-49&12|+|62603691|62603712&offsets|0|-1(+1A)]|TGAGGTAGTAGTTTGTGCTGTA</td><td>>MI0000434|hsa-let-7i&WithFlank&12|+|62603680|62603775</td><td>12</td><td>32</td><td>21(+1A)</td><td>TGAGGTAGTAGTTTGTGCTGTA</td><td>BT20</td><td>330</td><td>153</td><td>213</td><td>217</td><td>97</td><td>224.1</td><td>59.5</td><td>98.0</td><td>278.3</td><td>38.2</td></tr>
<tr><td>>MI0000434|hsa-let-7i&WithFlank&12|+|62603680|62603775@12.32.21(+1A)&[MIMAT0000415&hsa-let-7i-5p&offsets|0|-1(+1A);m-49&12|+|62603691|62603712&offsets|0|-1(+1A)]|TGAGGTAGTAGTTTGTGCTGTA</td><td>>MI0000434|hsa-let-7i&WithFlank&12|+|62603680|62603775</td><td>12</td><td>32</td><td>21(+1A)</td><td>TGAGGTAGTAGTTTGTGCTGTA</td><td>MB231</td><td>379</td><td>267</td><td>98</td><td>709</td><td>106</td><td>9.9</td><td>127.9</td><td>49.8</td><td>399.3</td><td>48.4</td></tr>
<tr><td>>MI0000434|hsa-let-7i&WithFlank&12|+|62603680|62603775@12.32.21(+1A)&[MIMAT0000415&hsa-let-7i-5p&offsets|0|-1(+1A);m-49&12|+|62603691|62603712&offsets|0|-1(+1A)]|TGAGGTAGTAGTTTGTGCTGTA</td><td>>MI0000434|hsa-let-7i&WithFlank&12|+|62603680|62603775</td><td>12</td><td>32</td><td>21(+1A)</td><td>TGAGGTAGTAGTTTGTGCTGTA</td><td>MB468</td><td>11</td><td>6</td><td>6</td><td>15</td><td>2</td><td>3.0</td><td>1.5</td><td>4.1</td><td>9.8</td><td>1.4</td></tr>
<tr><td>>MI0000434|hsa-let-7i&WithFlank&12|+|62603680|62603775@12.32.21(+1C)&[MIMAT0000415&hsa-let-7i-5p&offsets|0|-1(+1C);m-49&12|+|62603691|62603712&offsets|0|-1(+1C)]|TGAGGTAGTAGTTTGTGCTGTC</td><td>>MI0000434|hsa-let-7i&WithFlank&12|+|62603680|62603775</td><td>12</td><td>32</td><td>21(+1C)</td><td>TGAGGTAGTAGTTTGTGCTGTC</td><td>BT20</td><td>675</td><td>571</td><td>493</td><td>792</td><td>393</td><td>164.8</td><td>199.7</td><td>213.1</td><td>963.8</td><td>111.7</td></tr>
<tr><td>>MI0000434|hsa-let-7i&WithFlank&12|+|62603680|62603775@12.32.21(+1C)&[MIMAT0000415&hsa-let-7i-5p&offsets|0|-1(+1C);m-49&12|+|62603691|62603712&offsets|0|-1(+1C)]|TGAGGTAGTAGTTTGTGCTGTC</td><td>>MI0000434|hsa-let-7i&WithFlank&12|+|62603680|62603775</td><td>12</td><td>32</td><td>21(+1C)</td><td>TGAGGTAGTAGTTTGTGCTGTC</td><td>MB231</td><td>1266</td><td>1196</td><td>292</td><td>2430</td><td>431</td><td>201.6</td><td>600.2</td><td>118.9</td><td>1324.9</td><td>229.3</td></tr>
</table>

| | | | | | | | | | | | | | | | | |
|---|---|---|---|---|---|---|---|---|---|---|---|---|---|---|---|---|
| >MI0000434\|hsa-let-7i&WithFlank&12\|+\|62603680\|62603775@12.32.21(+1C)&[MIMAT0000415&hsa-let-7i-5p&offsets\|0\|-1(+1C);m-49&12\|+\|62603691\|62603712&offsets\|0\|-1(+1C)]\|TGAGGTAGTAGTTTGTGCTGTC | >MI0000434\|hsa-let-7i&WithFlank&12\|+\|62603680\|62603775 | 12 | 32 | 21(+1C) | TGAGGTAGTAGTTTGTGCTGTC | MB468 | 46 | 31 | 19 | 53 | 10 | 18.9 | 10.9 | 7.7 | 24.3 | 10.7 |
| >MI0000434\|hsa-let-7i&WithFlank&12\|+\|62603680\|62603775@12.32.21(+1G)&[MIMAT0000415&hsa-let-7i-5p&offsets\|0\|-1(+1G);m-49&12\|+\|62603691\|62603712&offsets\|0\|-1(+1G)]\|TGAGGTAGTAGTTTGTGCTGTG | >MI0000434\|hsa-let-7i&WithFlank&12\|+\|62603680\|62603775 | 12 | 32 | 21(+1G) | TGAGGTAGTAGTTTGTGCTGTG | BT20 | 280 | 72 | 74 | 91 | 22 | 222.5 | 6.5 | 26.5 | 131.6 | 8.5 |
| >MI0000434\|hsa-let-7i&WithFlank&12\|+\|62603680\|62603775@12.32.21(+1G)&[MIMAT0000415&hsa-let-7i-5p&offsets\|0\|-1(+1G);m-49&12\|+\|62603691\|62603712&offsets\|0\|-1(+1G)]\|TGAGGTAGTAGTTTGTGCTGTG | >MI0000434\|hsa-let-7i&WithFlank&12\|+\|62603680\|62603775 | 12 | 32 | 21(+1G) | TGAGGTAGTAGTTTGTGCTGTG | MB231 | 409 | 155 | 36 | 191 | 29 | 303.2 | 97.8 | 24.8 | 115.9 | 17.1 |
| >MI0000434\|hsa-let-7i&WithFlank&12\|+\|62603680\|62603775@12.32.21(+1G)&[MIMAT0000415&hsa-let-7i-5p&offsets\|0\|-1(+1G);m-49&12\|+\|62603691\|62603712&offsets\|0\|-1(+1G)]\|TGAGGTAGTAGTTTGTGCTGTG | >MI0000434\|hsa-let-7i&WithFlank&12\|+\|62603680\|62603775 | 12 | 32 | 21(+1G) | TGAGGTAGTAGTTTGTGCTGTG | MB468 | 14 | 3 | 7 | 8 | 1 | 4.4 | 0.8 | 5.6 | 4.5 | 1.0 |
| >MI0000434\|hsa-let-7i&WithFlank&12\|+\|62603680\|62603775@12.32.21&[MIMAT0000415&hsa-let-7i-5p&offsets\|0\|-1;m-49&12\|+\|62603691\|62603712&offsets\|0\|-1]\|TGAGGTAGTAGTTTGTGCTGT | >MI0000434\|hsa-let-7i&WithFlank&12\|+\|62603680\|62603775 | 12 | 32 | 21 | TGAGGTAGTAGTTTGTGCTGT | BT20 | 232 | 167 | 184 | 270 | 88 | 95.0 | 51.0 | 82.9 | 267.7 | 22.1 |
| >MI0000434\|hsa-let-7i&WithFlank&12\|+\|62603680\|62603775@12.32.21&[MIMAT0000415&hsa-let-7i-5p&offsets\|0\|-1;m-49&12\|+\|62603691\|62603712&offsets\|0\|-1]\|TGAGGTAGTAGTTTGTGCTGT | >MI0000434\|hsa-let-7i&WithFlank&12\|+\|62603680\|62603775 | 12 | 32 | 21 | TGAGGTAGTAGTTTGTGCTGT | MB231 | 479 | 437 | 142 | 688 | 130 | 35.4 | 223.9 | 70.5 | 381.8 | 66.4 |
| >MI0000434\|hsa-let-7i&WithFlank&12\|+\|62603680\|62603775@12.32.21&[MIMAT0000415&hsa-let-7i-5p&offsets\|0\|-1;m-49&12\|+\|62603691\|62603712&offsets\|0\|-1]\|TGAGGTAGTAGTTTGTGCTGT | >MI0000434\|hsa-let-7i&WithFlank&12\|+\|62603680\|62603775 | 12 | 32 | 21 | TGAGGTAGTAGTTTGTGCTGT | MB468 | 14 | 8 | 6 | 14 | 2 | 6.7 | 2.8 | 1.7 | 8.4 | 1.8 |
| >MI0000434\|hsa-let-7i&WithFlank&12\|+\|62603680\|62603775@12.33.22(+1A)&[MIMAT0000415&hsa-let-7i-5p&offsets\|0\|0(+1A);m-49&12\|+\|62603691\|62603712&offsets\|0\|0(+1A)]\|TGAGGTAGTAGTTTGTGCTGTTA | >MI0000434\|hsa-let-7i&WithFlank&12\|+\|62603680\|62603775 | 12 | 33 | 22(+1A) | TGAGGTAGTAGTTTGTGCTGTTA | BT20 | 288 | 191 | 179 | 728 | 149 | 129.8 | 60.5 | 79.4 | 690.5 | 26.5 |
| >MI0000434\|hsa-let-7i&WithFlank&12\|+\|62603680\|62603775@12.33.22(+1A)&[MIMAT0000415&hsa-let-7i-5p&offsets\|0\|0(+1A);m-49&12\|+\|62603691\|62603712&offsets\|0\|0(+1A)]\|TGAGGTAGTAGTTTGTGCTGTTA | >MI0000434\|hsa-let-7i&WithFlank&12\|+\|62603680\|62603775 | 12 | 33 | 22(+1A) | TGAGGTAGTAGTTTGTGCTGTTA | MB231 | 331 | 280 | 64 | 821 | 113 | 53.5 | 104.1 | 36.9 | 248.2 | 42.4 |
| >MI0000434\|hsa-let-7i&WithFlank&12\|+\|62603680\|62603775@12.33.22(+1A)&[MIMAT0000415&hsa-let-7i-5p&offsets\|0\|0(+1A);m-49&12\|+\|62603691\|62603712&offsets\|0\|0(+1A)]\|TGAGGTAGTAGTTTGTGCTGTTA | >MI0000434\|hsa-let-7i&WithFlank&12\|+\|62603680\|62603775 | 12 | 33 | 22(+1A) | TGAGGTAGTAGTTTGTGCTGTTA | MB468 | 24 | 12 | 7 | 51 | 5 | 11.3 | 3.4 | 1.9 | 25.8 | 3.9 |
| >MI0000434\|hsa-let-7i&WithFlank&12\|+\|62603680\|62603775@12.33.22(+1C)&[MIMAT0000415&hsa-let-7i-5p&offsets\|0\|0(+1C);m-49&12\|+\|62603691\|62603712&offsets\|0\|0(+1C)]\|TGAGGTAGTAGTTTGTGCTGTTC | >MI0000434\|hsa-let-7i&WithFlank&12\|+\|62603680\|62603775 | 12 | 33 | 22(+1C) | TGAGGTAGTAGTTTGTGCTGTTC | BT20 | 25 | 19 | 16 | 70 | 12 | 9.2 | 7.6 | 6.0 | 91.9 | 1.2 |
| >MI0000434\|hsa-let-7i&WithFlank&12\|+\|62603680\|62603775@12.33.22(+1C)&[MIMAT0000415&hsa-let-7i-5p&offsets\|0\|0(+1C);m-49&12\|+\|62603691\|62603712&offsets\|0\|0(+1C)]\|TGAGGTAGTAGTTTGTGCTGTTC | >MI0000434\|hsa-let-7i&WithFlank&12\|+\|62603680\|62603775 | 12 | 33 | 22(+1C) | TGAGGTAGTAGTTTGTGCTGTTC | MB231 | 56 | 45 | 14 | 175 | 20 | 1.7 | 25.9 | 7.8 | 93.6 | 11.2 |
| >MI0000434\|hsa-let-7i&WithFlank&12\|+\|62603680\|62603775@12.33.22(+1U)&[MIMAT0000415&hsa-let-7i-5p&offsets\|0\|0(+1U);m-49&12\|+\|62603691\|62603712&offsets\|0\|0(+1U)]\|TGAGGTAGTAGTTTGTGCTGTTT | >MI0000434\|hsa-let-7i&WithFlank&12\|+\|62603680\|62603775 | 12 | 33 | 22(+1U) | TGAGGTAGTAGTTTGTGCTGTTT | BT20 | 191 | 146 | 114 | 503 | 99 | 85.8 | 45.6 | 59.6 | 593.0 | 7.2 |
| >MI0000434\|hsa-let-7i&WithFlank&12\|+\|62603680\|62603775@12.33.22(+1U)&[MIMAT0000415&hsa-let-7i-5p&offsets\|0\|0(+1U);m-49&12\|+\|62603691\|62603712&offsets\|0\|0(+1U)]\|TGAGGTAGTAGTTTGTGCTGTTT | >MI0000434\|hsa-let-7i&WithFlank&12\|+\|62603680\|62603775 | 12 | 33 | 22(+1U) | TGAGGTAGTAGTTTGTGCTGTTT | MB231 | 530 | 421 | 119 | 1156 | 156 | 40.7 | 217.0 | 58.9 | 437.7 | 89.4 |
| >MI0000434\|hsa-let-7i&WithFlank&12\|+\|62603680\|62603775@12.33.22(+1U)&[MIMAT0000415&hsa-let-7i-5p&offsets\|0\|0(+1U);m-49&12\|+\|62603691\|62603712&offsets\|0\|0(+1U)]\|TGAGGTAGTAGTTTGTGCTGTTT | >MI0000434\|hsa-let-7i&WithFlank&12\|+\|62603680\|62603775 | 12 | 33 | 22(+1U) | TGAGGTAGTAGTTTGTGCTGTTT | MB468 | 16 | 8 | 5 | 19 | 3 | 7.5 | 1.7 | 1.3 | 10.0 | 2.5 |
| >MI0000434\|hsa-let-7i&WithFlank&12\|+\|62603680\|62603775@12.33.22(+2U)&[MIMAT0000415&hsa-let-7i-5p&offsets\|0\|0(+2U);m-49&12\|+\|62603691\|62603712&offsets\|0\|0(+2U)]\|TGAGGTAGTAGTTTGTGCTGTTTT | >MI0000434\|hsa-let-7i&WithFlank&12\|+\|62603680\|62603775 | 12 | 33 | 22(+2U) | TGAGGTAGTAGTTTGTGCTGTTTT | MB231 | 19 | 13 | 3 | 32 | 9 | 2.9 | 5.2 | 2.0 | 13.8 | 3.9 |
| >MI0000434\|hsa-let-7i&WithFlank&12\|+\|62603680\|62603775@12.33.22&[MIMAT0000415&hsa-let-7i-5p&offsets\|0\|0;m-49&12\|+\|62603691\|62603712&offsets\|0\|0]\|TGAGGTAGTAGTTTGTGCTGTT | >MI0000434\|hsa-let-7i&WithFlank&12\|+\|62603680\|62603775 | 12 | 33 | 22 | TGAGGTAGTAGTTTGTGCTGTT | BT20 | 5025 | 4182 | 3990 | 5360 | 3121 | 1676.6 | 1250.6 | 2041.2 | 5539.2 | 975.5 |
| >MI0000434\|hsa-let-7i&WithFlank&12\|+\|62603680\|62603775@12.33.22&[MIMAT0000415&hsa-let-7i-5p&offsets\|0\|0;m-49&12\|+\|62603691\|62603712&offsets\|0\|0]\|TGAGGTAGTAGTTTGTGCTGTT | >MI0000434\|hsa-let-7i&WithFlank&12\|+\|62603680\|62603775 | 12 | 33 | 22 | TGAGGTAGTAGTTTGTGCTGTT | MB231 | 11786 | 11589 | 2785 | 15493 | 3376 | 2053.7 | 5068.6 | 892.3 | 5783.5 | 1817.2 |
| >MI0000434\|hsa-let-7i&WithFlank&12\|+\|62603680\|62603775@12.33.22&[MIMAT0000415&hsa-let-7i-5p&offsets\|0\|0;m-49&12\|+\|62603691\|62603712&offsets\|0\|0]\|TGAGGTAGTAGTTTGTGCTGTT | >MI0000434\|hsa-let-7i&WithFlank&12\|+\|62603680\|62603775 | 12 | 33 | 22 | TGAGGTAGTAGTTTGTGCTGTT | MB468 | 437 | 334 | 170 | 455 | 95 | 201.9 | 148.5 | 20.8 | 234.5 | 113.1 |
| >MI0000434\|hsa-let-7i&WithFlank&12\|+\|62603680\|62603775@12.34.23&[MIMAT0000415&hsa-let-7i-5p&offsets\|0\|+1;m-49&12\|+\|62603691\|62603712&offsets\|0\|+1]\|TGAGGTAGTAGTTTGTGCTGTTG | >MI0000434\|hsa-let-7i&WithFlank&12\|+\|62603680\|62603775 | 12 | 34 | 23 | TGAGGTAGTAGTTTGTGCTGTTG | BT20 | 22 | 12 | 12 | 38 | 7 | 11.1 | 4.2 | 3.6 | 38.7 | 1.3 |
| >MI0000434\|hsa-let-7i&WithFlank&12\|+\|62603680\|62603775@12.34.23&[MIMAT0000415&hsa-let-7i-5p&offsets\|0\|+1;m-49&12\|+\|62603691\|62603712&offsets\|0\|+1]\|TGAGGTAGTAGTTTGTGCTGTTG | >MI0000434\|hsa-let-7i&WithFlank&12\|+\|62603680\|62603775 | 12 | 34 | 23 | TGAGGTAGTAGTTTGTGCTGTTG | MB231 | 26 | 15 | 5 | 37 | 4 | 12.5 | 6.6 | 2.9 | 14.7 | 2.3 |
| >MI0000438\|hsa-mir-15b&WithFlank&3\|+\|160404582\|160404691@26.46.21(+1G)&[MIMAT0000417&hsa-miR-15b-5p&offsets\|0\|-1(+1G);m-110&3\|+\|160404607\|160404628&offsets\|0\|-1(+1G)]\|TAGCAGCACATCATGGTTTACG | >MI0000438\|hsa-mir-15b&WithFlank&3\|+\|160404582\|160404691 | 26 | 46 | 21(+1G) | TAGCAGCACATCATGGTTTACG | BT20 | 16 | 6 | 2 | 0 | 0 | 20.1 | 5.8 | 0.6 | 0.0 | 0.0 |
| >MI0000438\|hsa-mir-15b&WithFlank&3\|+\|160404582\|160404691@26.46.21&[MIMAT0000417&hsa-miR-15b-5p&offsets\|0\|-1;m-110&3\|+\|160404607\|160404628&offsets\|0\|-1]\|TAGCAGCACATCATGGTTTAC | >MI0000438\|hsa-mir-15b&WithFlank&3\|+\|160404582\|160404691 | 26 | 46 | 21 | TAGCAGCACATCATGGTTTAC | BT20 | 27 | 40 | 14 | 29 | 6 | 17.9 | 37.6 | 7.4 | 30.1 | 4.2 |
| >MI0000438\|hsa-mir-15b&WithFlank&3\|+\|160404582\|160404691@26.46.21&[MIMAT0000417&hsa-miR-15b-5p&offsets\|0\|-1;m-110&3\|+\|160404607\|160404628&offsets\|0\|-1]\|TAGCAGCACATCATGGTTTAC | >MI0000438\|hsa-mir-15b&WithFlank&3\|+\|160404582\|160404691 | 26 | 46 | 21 | TAGCAGCACATCATGGTTTAC | MB231 | 14 | 24 | 5 | 11 | 16 | 2.3 | 13.5 | 4.4 | 6.2 | 10.0 |
| >MI0000438\|hsa-mir-15b&WithFlank&3\|+\|160404582\|160404691@26.47.22&[MIMAT0000417&hsa-miR-15b-5p&offsets\|0\|0;m-110&3\|+\|160404607\|160404628&offsets\|0\|0]\|TAGCAGCACATCATGGTTTACA | >MI0000438\|hsa-mir-15b&WithFlank&3\|+\|160404582\|160404691 | 26 | 47 | 22 | TAGCAGCACATCATGGTTTACA | BT20 | 174 | 183 | 61 | 18 | 28 | 147.0 | 161.1 | 20.2 | 15.8 | 17.1 |
| >MI0000438\|hsa-mir-15b&WithFlank&3\|+\|160404582\|160404691@26.47.22&[MIMAT0000417&hsa-miR-15b-5p&offsets\|0\|0;m-110&3\|+\|160404607\|160404628&offsets\|0\|0]\|TAGCAGCACATCATGGTTTACA | >MI0000438\|hsa-mir-15b&WithFlank&3\|+\|160404582\|160404691 | 26 | 47 | 22 | TAGCAGCACATCATGGTTTACA | MB231 | 72 | 117 | 17 | 26 | 70 | 14.6 | 56.7 | 14.3 | 16.7 | 47.6 |
| >MI0000438\|hsa-mir-15b&WithFlank&3\|+\|160404582\|160404691@64.84.21&[MIMAT0004586&hsa-miR-15b-3p&offsets\|0\|-1;m-204&3\|+\|160404645\|160404665&offsets\|0\|0]\|CGAATCATTATTTGCTGCTCT | >MI0000438\|hsa-mir-15b&WithFlank&3\|+\|160404582\|160404691 | 64 | 84 | 21 | CGAATCATTATTTGCTGCTCT | BT20 | 21 | 19 | 19 | 25 | 6 | 12.5 | 13.5 | 11.2 | 20.7 | 2.4 |
| >MI0000438\|hsa-mir-15b&WithFlank&3\|+\|160404582\|160404691@64.84.21&[MIMAT0004586&hsa-miR-15b-3p&offsets\|0\|-1;m-204&3\|+\|160404645\|160404665&offsets\|0\|0]\|CGAATCATTATTTGCTGCTCT | >MI0000438\|hsa-mir-15b&WithFlank&3\|+\|160404582\|160404691 | 64 | 84 | 21 | CGAATCATTATTTGCTGCTCT | MB231 | 13 | 15 | 6 | 51 | 20 | 2.7 | 5.3 | 3.5 | 31.1 | 14.1 |
| >MI0000439\|hsa-mir-23b&WithFlank&9\|+\|95085202\|95085310@64.86.23(+1U)&[MIMAT0000418&hsa-miR-23b-3p&offsets\|0\|0(+1U);m-42&9\|+\|95085265\|95085285&offsets\|0\|+2(+1U)]\|ATCACATTGCCAGGGATTACCACT | >MI0000439\|hsa-mir-23b&WithFlank&9\|+\|95085202\|95085310 | 64 | 86 | 23(+1U) | ATCACATTGCCAGGGATTACCACT | MB231 | 75 | 50 | 20 | 19 | 75 | 129.1 | 85.9 | 34.3 | 32.8 | 129.5 |

<table>
<tr><td>>MI0000439|hsa-mir-23b&WithFlank&9|+|95085202|95085310@64.86.23(+1U)&[MIMAT0000418&hsa-miR-23b-3p&offsets|0|0(+1U);m-42&9|+|95085265|95085285&offsets|0|+2(+1U)]|ATCACATTGCCAGGGATTACCACT</td><td>>MI0000439|hsa-mir-23b&WithFlank&9|+|95085202|95085310</td><td>64</td><td>86</td><td>23(+1U)</td><td>ATCACATTGCCAGGGATTACCACT</td><td>MB468</td><td>11</td><td>7</td><td>8</td><td>3</td><td>3</td><td>5.2</td><td>2.6</td><td>2.8</td><td>3.1</td><td>0.9</td></tr>
<tr><td>>MI0000439|hsa-mir-23b&WithFlank&9|+|95085202|95085310@64.86.23&[MIMAT0000418&hsa-miR-23b-3p&offsets|0|0;m-42&9|+|95085265|95085285&offsets|0|+2]|ATCACATTGCCAGGGATTACCAC</td><td>>MI0000439|hsa-mir-23b&WithFlank&9|+|95085202|95085310</td><td>64</td><td>86</td><td>23</td><td>ATCACATTGCCAGGGATTACCAC</td><td>MB231</td><td>27</td><td>12</td><td>6</td><td>4</td><td>16</td><td>46.2</td><td>20.6</td><td>10.4</td><td>6.2</td><td>28.2</td></tr>
<tr><td>>MI0000439|hsa-mir-23b&WithFlank&9|+|95085202|95085310@65.86.22(+1U)&[MIMAT0000418&hsa-miR-23b-3p&offsets|+1|0(+1U);m-42&9|+|95085265|95085285&offsets|+1|+2(+1U)]|TCACATTGCCAGGGATTACCACT</td><td>>MI0000439|hsa-mir-23b&WithFlank&9|+|95085202|95085310</td><td>65</td><td>86</td><td>22(+1U)</td><td>TCACATTGCCAGGGATTACCACT</td><td>BT20</td><td>19</td><td>14</td><td>17</td><td>6</td><td>11</td><td>9.4</td><td>4.7</td><td>6.8</td><td>5.4</td><td>7.3</td></tr>
<tr><td>>MI0000439|hsa-mir-23b&WithFlank&9|+|95085202|95085310@65.86.22(+1U)&[MIMAT0000418&hsa-miR-23b-3p&offsets|+1|0(+1U);m-42&9|+|95085265|95085285&offsets|+1|+2(+1U)]|TCACATTGCCAGGGATTACCACT</td><td>>MI0000439|hsa-mir-23b&WithFlank&9|+|95085202|95085310</td><td>65</td><td>86</td><td>22(+1U)</td><td>TCACATTGCCAGGGATTACCACT</td><td>MB231</td><td>17</td><td>14</td><td>14</td><td>11</td><td>22</td><td>5.1</td><td>4.9</td><td>10.9</td><td>7.2</td><td>10.9</td></tr>
<tr><td>>MI0000439|hsa-mir-23b&WithFlank&9|+|95085202|95085310@67.84.18&[MIMAT0000418&hsa-miR-23b-3p&offsets|+3|-2;m-42&9|+|95085265|95085285&offsets|+3|0]|ACATTGCCAGGGATTACC</td><td>>MI0000439|hsa-mir-23b&WithFlank&9|+|95085202|95085310</td><td>67</td><td>84</td><td>18</td><td>ACATTGCCAGGGATTACC</td><td>BT20</td><td>72</td><td>59</td><td>115</td><td>20</td><td>26</td><td>53.4</td><td>10.9</td><td>24.9</td><td>11.9</td><td>7.5</td></tr>
<tr><td>>MI0000439|hsa-mir-23b&WithFlank&9|+|95085202|95085310@67.84.18&[MIMAT0000418&hsa-miR-23b-3p&offsets|+3|-2;m-42&9|+|95085265|95085285&offsets|+3|0]|ACATTGCCAGGGATTACC</td><td>>MI0000439|hsa-mir-23b&WithFlank&9|+|95085202|95085310</td><td>67</td><td>84</td><td>18</td><td>ACATTGCCAGGGATTACC</td><td>MB231</td><td>29</td><td>27</td><td>64</td><td>24</td><td>32</td><td>25.6</td><td>23.5</td><td>64.2</td><td>22.2</td><td>31.8</td></tr>
<tr><td>>MI0000439|hsa-mir-23b&WithFlank&9|+|95085202|95085310@67.85.19(+1A)&[MIMAT0000418&hsa-miR-23b-3p&offsets|+3|-1(+1A);m-42&9|+|95085265|95085285&offsets|+3|+1(+1A)]|ACATTGCCAGGGATTACCAA</td><td>>MI0000439|hsa-mir-23b&WithFlank&9|+|95085202|95085310</td><td>67</td><td>85</td><td>19(+1A)</td><td>ACATTGCCAGGGATTACCAA</td><td>BT20</td><td>11</td><td>4</td><td>17</td><td>0</td><td>1</td><td>12.7</td><td>1.5</td><td>11.2</td><td>0.7</td><td>1.0</td></tr>
<tr><td>>MI0000439|hsa-mir-23b&WithFlank&9|+|95085202|95085310@67.85.19(+1U)&[MIMAT0000418&hsa-miR-23b-3p&offsets|+3|-1(+1U);m-42&9|+|95085265|95085285&offsets|+3|+1(+1U)]|ACATTGCCAGGGATTACCAT</td><td>>MI0000439|hsa-mir-23b&WithFlank&9|+|95085202|95085310</td><td>67</td><td>85</td><td>19(+1U)</td><td>ACATTGCCAGGGATTACCAT</td><td>BT20</td><td>52</td><td>33</td><td>86</td><td>4</td><td>10</td><td>43.1</td><td>11.1</td><td>34.2</td><td>4.6</td><td>3.5</td></tr>
<tr><td>>MI0000439|hsa-mir-23b&WithFlank&9|+|95085202|95085310@67.85.19(+1U)&[MIMAT0000418&hsa-miR-23b-3p&offsets|+3|-1(+1U);m-42&9|+|95085265|95085285&offsets|+3|+1(+1U)]|ACATTGCCAGGGATTACCAT</td><td>>MI0000439|hsa-mir-23b&WithFlank&9|+|95085202|95085310</td><td>67</td><td>85</td><td>19(+1U)</td><td>ACATTGCCAGGGATTACCAT</td><td>MB231</td><td>17</td><td>11</td><td>38</td><td>7</td><td>14</td><td>16.1</td><td>9.7</td><td>37.5</td><td>6.5</td><td>15.4</td></tr>
<tr><td>>MI0000439|hsa-mir-23b&WithFlank&9|+|95085202|95085310@67.85.19(+2U)&[MIMAT0000418&hsa-miR-23b-3p&offsets|+3|-1(+2U);m-42&9|+|95085265|95085285&offsets|+3|+1(+2U)]|ACATTGCCAGGGATTACCATT</td><td>>MI0000439|hsa-mir-23b&WithFlank&9|+|95085202|95085310</td><td>67</td><td>85</td><td>19(+2U)</td><td>ACATTGCCAGGGATTACCATT</td><td>BT20</td><td>30</td><td>20</td><td>31</td><td>2</td><td>7</td><td>20.0</td><td>13.1</td><td>2.5</td><td>1.5</td><td>4.6</td></tr>
<tr><td>>MI0000439|hsa-mir-23b&WithFlank&9|+|95085202|95085310@67.85.19(+2U)&[MIMAT0000418&hsa-miR-23b-3p&offsets|+3|-1(+2U);m-42&9|+|95085265|95085285&offsets|+3|+1(+2U)]|ACATTGCCAGGGATTACCATT</td><td>>MI0000439|hsa-mir-23b&WithFlank&9|+|95085202|95085310</td><td>67</td><td>85</td><td>19(+2U)</td><td>ACATTGCCAGGGATTACCATT</td><td>MB231</td><td>31</td><td>20</td><td>40</td><td>9</td><td>22</td><td>28.5</td><td>18.6</td><td>38.9</td><td>9.3</td><td>26.9</td></tr>
<tr><td>>MI0000439|hsa-mir-23b&WithFlank&9|+|95085202|95085310@67.85.19&[MIMAT0000418&hsa-miR-23b-3p&offsets|+3|-1;m-42&9|+|95085265|95085285&offsets|+3|+1]|ACATTGCCAGGGATTACCA</td><td>>MI0000439|hsa-mir-23b&WithFlank&9|+|95085202|95085310</td><td>67</td><td>85</td><td>19</td><td>ACATTGCCAGGGATTACCA</td><td>BT20</td><td>52</td><td>35</td><td>56</td><td>4</td><td>15</td><td>49.7</td><td>10.5</td><td>11.7</td><td>2.5</td><td>3.6</td></tr>
<tr><td>>MI0000439|hsa-mir-23b&WithFlank&9|+|95085202|95085310@67.85.19&[MIMAT0000418&hsa-miR-23b-3p&offsets|+3|-1;m-42&9|+|95085265|95085285&offsets|+3|+1]|ACATTGCCAGGGATTACCA</td><td>>MI0000439|hsa-mir-23b&WithFlank&9|+|95085202|95085310</td><td>67</td><td>85</td><td>19</td><td>ACATTGCCAGGGATTACCA</td><td>MB231</td><td>14</td><td>11</td><td>28</td><td>6</td><td>15</td><td>12.0</td><td>10.1</td><td>29.4</td><td>6.5</td><td>17.4</td></tr>
<tr><td>>MI0000439|hsa-mir-23b&WithFlank&9|+|95085202|95085310@67.86.20(+1A)&[MIMAT0000418&hsa-miR-23b-3p&offsets|+3|0(+1A);m-42&9|+|95085265|95085285&offsets|+3|+2(+1A)]|ACATTGCCAGGGATTACCACA</td><td>>MI0000439|hsa-mir-23b&WithFlank&9|+|95085202|95085310</td><td>67</td><td>86</td><td>20(+1A)</td><td>ACATTGCCAGGGATTACCACA</td><td>BT20</td><td>59</td><td>27</td><td>53</td><td>9</td><td>19</td><td>52.9</td><td>9.8</td><td>15.4</td><td>8.5</td><td>9.7</td></tr>
<tr><td>>MI0000439|hsa-mir-23b&WithFlank&9|+|95085202|95085310@67.86.20(+1A)&[MIMAT0000418&hsa-miR-23b-3p&offsets|+3|0(+1A);m-42&9|+|95085265|95085285&offsets|+3|+2(+1A)]|ACATTGCCAGGGATTACCACA</td><td>>MI0000439|hsa-mir-23b&WithFlank&9|+|95085202|95085310</td><td>67</td><td>86</td><td>20(+1A)</td><td>ACATTGCCAGGGATTACCACA</td><td>MB231</td><td>21</td><td>16</td><td>28</td><td>13</td><td>29</td><td>18.8</td><td>14.0</td><td>28.0</td><td>12.7</td><td>31.6</td></tr>
<tr><td>>MI0000439|hsa-mir-23b&WithFlank&9|+|95085202|95085310@67.86.20(+1C)&[MIMAT0000418&hsa-miR-23b-3p&offsets|+3|0(+1C);m-42&9|+|95085265|95085285&offsets|+3|+2(+1C)]|ACATTGCCAGGGATTACCACC</td><td>>MI0000439|hsa-mir-23b&WithFlank&9|+|95085202|95085310</td><td>67</td><td>86</td><td>20(+1C)</td><td>ACATTGCCAGGGATTACCACC</td><td>BT20</td><td>54</td><td>42</td><td>62</td><td>17</td><td>34</td><td>26.0</td><td>16.1</td><td>18.6</td><td>15.9</td><td>22.6</td></tr>
<tr><td>>MI0000439|hsa-mir-23b&WithFlank&9|+|95085202|95085310@67.86.20(+1C)&[MIMAT0000418&hsa-miR-23b-3p&offsets|+3|0(+1C);m-42&9|+|95085265|95085285&offsets|+3|+2(+1C)]|ACATTGCCAGGGATTACCACC</td><td>>MI0000439|hsa-mir-23b&WithFlank&9|+|95085202|95085310</td><td>67</td><td>86</td><td>20(+1C)</td><td>ACATTGCCAGGGATTACCACC</td><td>MB231</td><td>29</td><td>22</td><td>37</td><td>23</td><td>47</td><td>26.5</td><td>19.2</td><td>35.4</td><td>20.7</td><td>50.6</td></tr>
<tr><td>>MI0000439|hsa-mir-23b&WithFlank&9|+|95085202|95085310@67.86.20(+1U)&[MIMAT0000418&hsa-miR-23b-3p&offsets|+3|0(+1U);m-42&9|+|95085265|95085285&offsets|+3|+2(+1U)]|ACATTGCCAGGGATTACCACT</td><td>>MI0000439|hsa-mir-23b&WithFlank&9|+|95085202|95085310</td><td>67</td><td>86</td><td>20(+1U)</td><td>ACATTGCCAGGGATTACCACT</td><td>BT20</td><td>352</td><td>249</td><td>411</td><td>105</td><td>216</td><td>174.4</td><td>73.6</td><td>152.8</td><td>78.9</td><td>129.4</td></tr>
<tr><td>>MI0000439|hsa-mir-23b&WithFlank&9|+|95085202|95085310@67.86.20(+1U)&[MIMAT0000418&hsa-miR-23b-3p&offsets|+3|0(+1U);m-42&9|+|95085265|95085285&offsets|+3|+2(+1U)]|ACATTGCCAGGGATTACCACT</td><td>>MI0000439|hsa-mir-23b&WithFlank&9|+|95085202|95085310</td><td>67</td><td>86</td><td>20(+1U)</td><td>ACATTGCCAGGGATTACCACT</td><td>MB231</td><td>221</td><td>188</td><td>297</td><td>127</td><td>299</td><td>199.3</td><td>167.8</td><td>291.9</td><td>121.5</td><td>305.2</td></tr>
<tr><td>>MI0000439|hsa-mir-23b&WithFlank&9|+|95085202|95085310@67.86.20&[MIMAT0000418&hsa-miR-23b-3p&offsets|+3|0;m-42&9|+|95085265|95085285&offsets|+3|+2]|ACATTGCCAGGGATTACCAC</td><td>>MI0000439|hsa-mir-23b&WithFlank&9|+|95085202|95085310</td><td>67</td><td>86</td><td>20</td><td>ACATTGCCAGGGATTACCAC</td><td>BT20</td><td>143</td><td>108</td><td>305</td><td>16</td><td>42</td><td>99.9</td><td>14.9</td><td>96.8</td><td>9.2</td><td>12.0</td></tr>
<tr><td>>MI0000439|hsa-mir-23b&WithFlank&9|+|95085202|95085310@67.86.20&[MIMAT0000418&hsa-miR-23b-3p&offsets|+3|0;m-42&9|+|95085265|95085285&offsets|+3|+2]|ACATTGCCAGGGATTACCAC</td><td>>MI0000439|hsa-mir-23b&WithFlank&9|+|95085202|95085310</td><td>67</td><td>86</td><td>20</td><td>ACATTGCCAGGGATTACCAC</td><td>MB231</td><td>69</td><td>49</td><td>170</td><td>15</td><td>40</td><td>63.9</td><td>42.6</td><td>173.2</td><td>14.3</td><td>39.0</td></tr>
<tr><td>>MI0000439|hsa-mir-23b&WithFlank&9|+|95085202|95085310@67.87.21&[MIMAT0000418&hsa-miR-23b-3p&offsets|+3|+1;m-42&9|+|95085265|95085285&offsets|+3|+3]|ACATTGCCAGGGATTACCACG</td><td>>MI0000439|hsa-mir-23b&WithFlank&9|+|95085202|95085310</td><td>67</td><td>87</td><td>21</td><td>ACATTGCCAGGGATTACCACG</td><td>BT20</td><td>38</td><td>9</td><td>15</td><td>3</td><td>4</td><td>37.5</td><td>3.1</td><td>7.4</td><td>4.1</td><td>1.8</td></tr>
<tr><td>>MI0000439|hsa-mir-23b&WithFlank&9|+|95085202|95085310@67.87.21&[MIMAT0000418&hsa-miR-23b-3p&offsets|+3|+1;m-42&9|+|95085265|95085285&offsets|+3|+3]|ACATTGCCAGGGATTACCACG</td><td>>MI0000439|hsa-mir-23b&WithFlank&9|+|95085202|95085310</td><td>67</td><td>87</td><td>21</td><td>ACATTGCCAGGGATTACCACG</td><td>MB231</td><td>18</td><td>5</td><td>8</td><td>3</td><td>5</td><td>18.2</td><td>4.2</td><td>7.1</td><td>2.8</td><td>5.6</td></tr>
<tr><td>>MI0000440|hsa-mir-27b&WithFlank&9|+|95085439|95085547@66.87.22&[MIMAT0000419&hsa-miR-27b-3p&offsets|-1|0;m-39&9|+|95085505|95085525&offsets|-1|0]|GTTCACAGTGGCTAAGTTCTGC</td><td>>MI0000440|hsa-mir-27b&WithFlank&9|+|95085439|95085547</td><td>66</td><td>87</td><td>22</td><td>GTTCACAGTGGCTAAGTTCTGC</td><td>BT20</td><td>29</td><td>24</td><td>43</td><td>11</td><td>15</td><td>11.2</td><td>4.5</td><td>11.1</td><td>3.0</td><td>2.1</td></tr>
<tr><td>>MI0000440|hsa-mir-27b&WithFlank&9|+|95085439|95085547@66.87.22&[MIMAT0000419&hsa-miR-27b-3p&offsets|-1|0;m-39&9|+|95085505|95085525&offsets|-1|0]|GTTCACAGTGGCTAAGTTCTGC</td><td>>MI0000440|hsa-mir-27b&WithFlank&9|+|95085439|95085547</td><td>66</td><td>87</td><td>22</td><td>GTTCACAGTGGCTAAGTTCTGC</td><td>MB231</td><td>18</td><td>16</td><td>7</td><td>9</td><td>11</td><td>7.0</td><td>4.4</td><td>4.6</td><td>4.2</td><td>2.8</td></tr>
<tr><td>>MI0000440|hsa-mir-27b&WithFlank&9|+|95085439|95085547@67.85.19(+1A)&[MIMAT0000419&hsa-miR-27b-3p&offsets|0|-2(+1A);m-39&9|+|95085505|95085525&offsets|0|-2(+1A)]|TTCACAGTGGCTAAGTTCTA</td><td>>MI0000440|hsa-mir-27b&WithFlank&9|+|95085439|95085547</td><td>67</td><td>85</td><td>19(+1A)</td><td>TTCACAGTGGCTAAGTTCTA</td><td>BT20</td><td>125</td><td>39</td><td>101</td><td>46</td><td>14</td><td>126.7</td><td>12.9</td><td>78.3</td><td>49.2</td><td>2.4</td></tr>
<tr><td>>MI0000440|hsa-mir-27b&WithFlank&9|+|95085439|95085547@67.85.19(+1A)&[MIMAT0000419&hsa-miR-27b-3p&offsets|0|-2(+1A);m-39&9|+|95085505|95085525&offsets|0|-2(+1A)]|TTCACAGTGGCTAAGTTCTA</td><td>>MI0000440|hsa-mir-27b&WithFlank&9|+|95085439|95085547</td><td>67</td><td>85</td><td>19(+1A)</td><td>TTCACAGTGGCTAAGTTCTA</td><td>MB231</td><td>33</td><td>22</td><td>16</td><td>72</td><td>34</td><td>10.9</td><td>11.9</td><td>13.5</td><td>38.1</td><td>21.8</td></tr>
<tr><td>>MI0000440|hsa-mir-27b&WithFlank&9|+|95085439|95085547@67.85.19(+1U)&[MIMAT0000419&hsa-miR-27b-3p&offsets|0|-2(+1U);m-39&9|+|95085505|95085525&offsets|0|-2(+1U)]|TTCACAGTGGCTAAGTTCTT</td><td>>MI0000440|hsa-mir-27b&WithFlank&9|+|95085439|95085547</td><td>67</td><td>85</td><td>19(+1U)</td><td>TTCACAGTGGCTAAGTTCTT</td><td>BT20</td><td>13</td><td>4</td><td>9</td><td>5</td><td>2</td><td>10.6</td><td>1.3</td><td>6.9</td><td>5.3</td><td>0.3</td></tr>
</table>

<table>
<tr><td>>MI0000440|hsa-mir-27b&WithFlank&9|+|95085439|95085547@67.85.19(+2U)&[MIMAT0000419&hsa-miR-27b-3p&offsets|0|-2(+2U);m-39&9|+|95085505|95085525&offsets|0|-2(+2U)]|TTCACAGTGGCTAAGTTCTTT</td><td>>MI0000440|hsa-mir-27b&WithFlank&9|+|95085439|95085547</td><td>67</td><td>85</td><td>19(+2U)</td><td>TTCACAGTGGCTAAGTTCTTT</td><td>BT20</td><td>15</td><td>6</td><td>10</td><td>6</td><td>3</td><td>11.8</td><td>3.5</td><td>7.8</td><td>4.1</td><td>2.0</td></tr>
<tr><td>>MI0000440|hsa-mir-27b&WithFlank&9|+|95085439|95085547@67.85.19&[MIMAT0000419&hsa-miR-27b-3p&offsets|0|-2;m-39&9|+|95085505|95085525&offsets|0|-2]|TTCACAGTGGCTAAGTTCT</td><td>>MI0000440|hsa-mir-27b&WithFlank&9|+|95085439|95085547</td><td>67</td><td>85</td><td>19</td><td>TTCACAGTGGCTAAGTTCT</td><td>BT20</td><td>116</td><td>71</td><td>103</td><td>100</td><td>40</td><td>70.2</td><td>20.9</td><td>66.6</td><td>63.9</td><td>7.3</td></tr>
<tr><td>>MI0000440|hsa-mir-27b&WithFlank&9|+|95085439|95085547@67.85.19&[MIMAT0000419&hsa-miR-27b-3p&offsets|0|-2;m-39&9|+|95085505|95085525&offsets|0|-2]|TTCACAGTGGCTAAGTTCT</td><td>>MI0000440|hsa-mir-27b&WithFlank&9|+|95085439|95085547</td><td>67</td><td>85</td><td>19</td><td>TTCACAGTGGCTAAGTTCT</td><td>MB231</td><td>87</td><td>99</td><td>32</td><td>175</td><td>91</td><td>18.3</td><td>32.4</td><td>13.8</td><td>102.4</td><td>35.2</td></tr>
<tr><td>>MI0000440|hsa-mir-27b&WithFlank&9|+|95085439|95085547@67.86.20(+1A)&[MIMAT0000419&hsa-miR-27b-3p&offsets|0|-1(+1A);m-39&9|+|95085505|95085525&offsets|0|-1(+1A)]|TTCACAGTGGCTAAGTTCTGA</td><td>>MI0000440|hsa-mir-27b&WithFlank&9|+|95085439|95085547</td><td>67</td><td>86</td><td>20(+1A)</td><td>TTCACAGTGGCTAAGTTCTGA</td><td>BT20</td><td>498</td><td>144</td><td>264</td><td>97</td><td>59</td><td>435.4</td><td>46.0</td><td>133.2</td><td>105.8</td><td>5.8</td></tr>
<tr><td>>MI0000440|hsa-mir-27b&WithFlank&9|+|95085439|95085547@67.86.20(+1A)&[MIMAT0000419&hsa-miR-27b-3p&offsets|0|-1(+1A);m-39&9|+|95085505|95085525&offsets|0|-1(+1A)]|TTCACAGTGGCTAAGTTCTGA</td><td>>MI0000440|hsa-mir-27b&WithFlank&9|+|95085439|95085547</td><td>67</td><td>86</td><td>20(+1A)</td><td>TTCACAGTGGCTAAGTTCTGA</td><td>MB231</td><td>163</td><td>81</td><td>39</td><td>211</td><td>169</td><td>27.6</td><td>33.7</td><td>30.6</td><td>118.9</td><td>124.8</td></tr>
<tr><td>>MI0000440|hsa-mir-27b&WithFlank&9|+|95085439|95085547@67.86.20(+1G)&[MIMAT0000419&hsa-miR-27b-3p&offsets|0|-1(+1G);m-39&9|+|95085505|95085525&offsets|0|-1(+1G)]|TTCACAGTGGCTAAGTTCTGG</td><td>>MI0000440|hsa-mir-27b&WithFlank&9|+|95085439|95085547</td><td>67</td><td>86</td><td>20(+1G)</td><td>TTCACAGTGGCTAAGTTCTGG</td><td>BT20</td><td>87</td><td>10</td><td>14</td><td>5</td><td>1</td><td>98.2</td><td>4.2</td><td>9.7</td><td>7.5</td><td>1.0</td></tr>
<tr><td>>MI0000440|hsa-mir-27b&WithFlank&9|+|95085439|95085547@67.86.20(+1G)&[MIMAT0000419&hsa-miR-27b-3p&offsets|0|-1(+1G);m-39&9|+|95085505|95085525&offsets|0|-1(+1G)]|TTCACAGTGGCTAAGTTCTGG</td><td>>MI0000440|hsa-mir-27b&WithFlank&9|+|95085439|95085547</td><td>67</td><td>86</td><td>20(+1G)</td><td>TTCACAGTGGCTAAGTTCTGG</td><td>MB231</td><td>20</td><td>5</td><td>2</td><td>8</td><td>6</td><td>15.0</td><td>2.8</td><td>1.3</td><td>6.2</td><td>6.2</td></tr>
<tr><td>>MI0000440|hsa-mir-27b&WithFlank&9|+|95085439|95085547@67.86.20(+1U)&[MIMAT0000419&hsa-miR-27b-3p&offsets|0|-1(+1U);m-39&9|+|95085505|95085525&offsets|0|-1(+1U)]|TTCACAGTGGCTAAGTTCTGT</td><td>>MI0000440|hsa-mir-27b&WithFlank&9|+|95085439|95085547</td><td>67</td><td>86</td><td>20(+1U)</td><td>TTCACAGTGGCTAAGTTCTGT</td><td>BT20</td><td>1500</td><td>561</td><td>682</td><td>329</td><td>230</td><td>942.4</td><td>251.8</td><td>236.2</td><td>375.5</td><td>40.3</td></tr>
<tr><td>>MI0000440|hsa-mir-27b&WithFlank&9|+|95085439|95085547@67.86.20(+1U)&[MIMAT0000419&hsa-miR-27b-3p&offsets|0|-1(+1U);m-39&9|+|95085505|95085525&offsets|0|-1(+1U)]|TTCACAGTGGCTAAGTTCTGT</td><td>>MI0000440|hsa-mir-27b&WithFlank&9|+|95085439|95085547</td><td>67</td><td>86</td><td>20(+1U)</td><td>TTCACAGTGGCTAAGTTCTGT</td><td>MB231</td><td>629</td><td>289</td><td>103</td><td>737</td><td>681</td><td>110.4</td><td>111.9</td><td>73.3</td><td>448.9</td><td>577.1</td></tr>
<tr><td>>MI0000440|hsa-mir-27b&WithFlank&9|+|95085439|95085547@67.86.20(+1U)&[MIMAT0000419&hsa-miR-27b-3p&offsets|0|-1(+1U);m-39&9|+|95085505|95085525&offsets|0|-1(+1U)]|TTCACAGTGGCTAAGTTCTGT</td><td>>MI0000440|hsa-mir-27b&WithFlank&9|+|95085439|95085547</td><td>67</td><td>86</td><td>20(+1U)</td><td>TTCACAGTGGCTAAGTTCTGT</td><td>MB468</td><td>24</td><td>5</td><td>7</td><td>24</td><td>7</td><td>10.3</td><td>2.4</td><td>4.0</td><td>4.1</td><td>3.7</td></tr>
<tr><td>>MI0000440|hsa-mir-27b&WithFlank&9|+|95085439|95085547@67.86.20(+2U)&[MIMAT0000419&hsa-miR-27b-3p&offsets|0|-1(+2U);m-39&9|+|95085505|95085525&offsets|0|-1(+2U)]|TTCACAGTGGCTAAGTTCTGTT</td><td>>MI0000440|hsa-mir-27b&WithFlank&9|+|95085439|95085547</td><td>67</td><td>86</td><td>20(+2U)</td><td>TTCACAGTGGCTAAGTTCTGTT</td><td>BT20</td><td>12</td><td>6</td><td>4</td><td>2</td><td>1</td><td>9.0</td><td>4.4</td><td>1.8</td><td>2.5</td><td>1.1</td></tr>
<tr><td>>MI0000440|hsa-mir-27b&WithFlank&9|+|95085439|95085547@67.86.20&[MIMAT0000419&hsa-miR-27b-3p&offsets|0|-1;m-39&9|+|95085505|95085525&offsets|0|-1]|TTCACAGTGGCTAAGTTCTG</td><td>>MI0000440|hsa-mir-27b&WithFlank&9|+|95085439|95085547</td><td>67</td><td>86</td><td>20</td><td>TTCACAGTGGCTAAGTTCTG</td><td>BT20</td><td>638</td><td>340</td><td>554</td><td>368</td><td>140</td><td>370.9</td><td>84.4</td><td>226.5</td><td>332.7</td><td>10.7</td></tr>
<tr><td>>MI0000440|hsa-mir-27b&WithFlank&9|+|95085439|95085547@67.86.20&[MIMAT0000419&hsa-miR-27b-3p&offsets|0|-1;m-39&9|+|95085505|95085525&offsets|0|-1]|TTCACAGTGGCTAAGTTCTG</td><td>>MI0000440|hsa-mir-27b&WithFlank&9|+|95085439|95085547</td><td>67</td><td>86</td><td>20</td><td>TTCACAGTGGCTAAGTTCTG</td><td>MB231</td><td>362</td><td>304</td><td>141</td><td>497</td><td>302</td><td>116.9</td><td>139.8</td><td>105.2</td><td>151.3</td><td>132.8</td></tr>
<tr><td>>MI0000440|hsa-mir-27b&WithFlank&9|+|95085439|95085547@67.86.20&[MIMAT0000419&hsa-miR-27b-3p&offsets|0|-1;m-39&9|+|95085505|95085525&offsets|0|-1]|TTCACAGTGGCTAAGTTCTG</td><td>>MI0000440|hsa-mir-27b&WithFlank&9|+|95085439|95085547</td><td>67</td><td>86</td><td>20</td><td>TTCACAGTGGCTAAGTTCTG</td><td>MB468</td><td>16</td><td>7</td><td>8</td><td>28</td><td>4</td><td>7.6</td><td>3.3</td><td>1.2</td><td>3.8</td><td>0.5</td></tr>
<tr><td>>MI0000440|hsa-mir-27b&WithFlank&9|+|95085439|95085547@67.87.21(+1C)&[MIMAT0000419&hsa-miR-27b-3p&offsets|0|0(+1C);m-39&9|+|95085505|95085525&offsets|0|0(+1C)]|TTCACAGTGGCTAAGTTCTGCC</td><td>>MI0000440|hsa-mir-27b&WithFlank&9|+|95085439|95085547</td><td>67</td><td>87</td><td>21(+1C)</td><td>TTCACAGTGGCTAAGTTCTGCC</td><td>BT20</td><td>19</td><td>9</td><td>5</td><td>4</td><td>3</td><td>13.0</td><td>10.6</td><td>2.7</td><td>4.7</td><td>3.3</td></tr>
<tr><td>>MI0000440|hsa-mir-27b&WithFlank&9|+|95085439|95085547@67.87.21(+1C)&[MIMAT0000419&hsa-miR-27b-3p&offsets|0|0(+1C);m-39&9|+|95085505|95085525&offsets|0|0(+1C)]|TTCACAGTGGCTAAGTTCTGCC</td><td>>MI0000440|hsa-mir-27b&WithFlank&9|+|95085439|95085547</td><td>67</td><td>87</td><td>21(+1C)</td><td>TTCACAGTGGCTAAGTTCTGCC</td><td>MB231</td><td>10</td><td>6</td><td>1</td><td>27</td><td>18</td><td>1.7</td><td>4.0</td><td>0.9</td><td>16.8</td><td>18.3</td></tr>
<tr><td>>MI0000440|hsa-mir-27b&WithFlank&9|+|95085439|95085547@67.87.21(+1G)&[MIMAT0000419&hsa-miR-27b-3p&offsets|0|0(+1G);m-39&9|+|95085505|95085525&offsets|0|0(+1G)]|TTCACAGTGGCTAAGTTCTGCG</td><td>>MI0000440|hsa-mir-27b&WithFlank&9|+|95085439|95085547</td><td>67</td><td>87</td><td>21(+1G)</td><td>TTCACAGTGGCTAAGTTCTGCG</td><td>BT20</td><td>67</td><td>12</td><td>16</td><td>2</td><td>3</td><td>59.2</td><td>5.2</td><td>5.6</td><td>2.5</td><td>2.5</td></tr>
<tr><td>>MI0000440|hsa-mir-27b&WithFlank&9|+|95085439|95085547@67.87.21(+1G)&[MIMAT0000419&hsa-miR-27b-3p&offsets|0|0(+1G);m-39&9|+|95085505|95085525&offsets|0|0(+1G)]|TTCACAGTGGCTAAGTTCTGCG</td><td>>MI0000440|hsa-mir-27b&WithFlank&9|+|95085439|95085547</td><td>67</td><td>87</td><td>21(+1G)</td><td>TTCACAGTGGCTAAGTTCTGCG</td><td>MB231</td><td>30</td><td>10</td><td>2</td><td>7</td><td>8</td><td>22.8</td><td>5.9</td><td>1.5</td><td>5.1</td><td>6.2</td></tr>
<tr><td>>MI0000440|hsa-mir-27b&WithFlank&9|+|95085439|95085547@67.87.21(+1U)&[MIMAT0000419&hsa-miR-27b-3p&offsets|0|0(+1U);m-39&9|+|95085505|95085525&offsets|0|0(+1U)]|TTCACAGTGGCTAAGTTCTGCT</td><td>>MI0000440|hsa-mir-27b&WithFlank&9|+|95085439|95085547</td><td>67</td><td>87</td><td>21(+1U)</td><td>TTCACAGTGGCTAAGTTCTGCT</td><td>BT20</td><td>123</td><td>56</td><td>33</td><td>26</td><td>26</td><td>79.2</td><td>58.0</td><td>23.2</td><td>25.0</td><td>10.8</td></tr>
<tr><td>>MI0000440|hsa-mir-27b&WithFlank&9|+|95085439|95085547@67.87.21(+1U)&[MIMAT0000419&hsa-miR-27b-3p&offsets|0|0(+1U);m-39&9|+|95085505|95085525&offsets|0|0(+1U)]|TTCACAGTGGCTAAGTTCTGCT</td><td>>MI0000440|hsa-mir-27b&WithFlank&9|+|95085439|95085547</td><td>67</td><td>87</td><td>21(+1U)</td><td>TTCACAGTGGCTAAGTTCTGCT</td><td>MB231</td><td>79</td><td>50</td><td>8</td><td>149</td><td>114</td><td>25.2</td><td>26.6</td><td>4.4</td><td>76.4</td><td>100.7</td></tr>
<tr><td>>MI0000440|hsa-mir-27b&WithFlank&9|+|95085439|95085547@67.87.21(+2U)&[MIMAT0000419&hsa-miR-27b-3p&offsets|0|0(+2U);m-39&9|+|95085505|95085525&offsets|0|0(+2U)]|TTCACAGTGGCTAAGTTCTGCTT</td><td>>MI0000440|hsa-mir-27b&WithFlank&9|+|95085439|95085547</td><td>67</td><td>87</td><td>21(+2U)</td><td>TTCACAGTGGCTAAGTTCTGCTT</td><td>BT20</td><td>35</td><td>23</td><td>12</td><td>13</td><td>16</td><td>24.9</td><td>23.0</td><td>11.9</td><td>6.6</td><td>7.9</td></tr>
<tr><td>>MI0000440|hsa-mir-27b&WithFlank&9|+|95085439|95085547@67.87.21(+2U)&[MIMAT0000419&hsa-miR-27b-3p&offsets|0|0(+2U);m-39&9|+|95085505|95085525&offsets|0|0(+2U)]|TTCACAGTGGCTAAGTTCTGCTT</td><td>>MI0000440|hsa-mir-27b&WithFlank&9|+|95085439|95085547</td><td>67</td><td>87</td><td>21(+2U)</td><td>TTCACAGTGGCTAAGTTCTGCTT</td><td>MB231</td><td>36</td><td>20</td><td>4</td><td>52</td><td>41</td><td>30.1</td><td>5.7</td><td>0.7</td><td>27.4</td><td>22.7</td></tr>
<tr><td>>MI0000440|hsa-mir-27b&WithFlank&9|+|95085439|95085547@67.87.21&[MIMAT0000419&hsa-miR-27b-3p&offsets|0|0;m-39&9|+|95085505|95085525&offsets|0|0]|TTCACAGTGGCTAAGTTCTGC</td><td>>MI0000440|hsa-mir-27b&WithFlank&9|+|95085439|95085547</td><td>67</td><td>87</td><td>21</td><td>TTCACAGTGGCTAAGTTCTGC</td><td>BT20</td><td>4129</td><td>2228</td><td>2611</td><td>1251</td><td>1180</td><td>1066.8</td><td>541.6</td><td>510.7</td><td>1064.0</td><td>266.8</td></tr>
<tr><td>>MI0000440|hsa-mir-27b&WithFlank&9|+|95085439|95085547@67.87.21&[MIMAT0000419&hsa-miR-27b-3p&offsets|0|0;m-39&9|+|95085505|95085525&offsets|0|0]|TTCACAGTGGCTAAGTTCTGC</td><td>>MI0000440|hsa-mir-27b&WithFlank&9|+|95085439|95085547</td><td>67</td><td>87</td><td>21</td><td>TTCACAGTGGCTAAGTTCTGC</td><td>MB231</td><td>3053</td><td>1920</td><td>594</td><td>2574</td><td>2832</td><td>491.1</td><td>706.9</td><td>408.9</td><td>1021.5</td><td>1527.9</td></tr>
<tr><td>>MI0000440|hsa-mir-27b&WithFlank&9|+|95085439|95085547@67.87.21&[MIMAT0000419&hsa-miR-27b-3p&offsets|0|0;m-39&9|+|95085505|95085525&offsets|0|0]|TTCACAGTGGCTAAGTTCTGC</td><td>>MI0000440|hsa-mir-27b&WithFlank&9|+|95085439|95085547</td><td>67</td><td>87</td><td>21</td><td>TTCACAGTGGCTAAGTTCTGC</td><td>MB468</td><td>151</td><td>43</td><td>45</td><td>138</td><td>36</td><td>75.6</td><td>20.5</td><td>9.5</td><td>31.3</td><td>7.4</td></tr>
<tr><td>>MI0000440|hsa-mir-27b&WithFlank&9|+|95085439|95085547@67.88.22(+1A)&[MIMAT0000419&hsa-miR-27b-3p&offsets|0|+1(+1A);m-39&9|+|95085505|95085525&offsets|0|+1(+1A)]|TTCACAGTGGCTAAGTTCTGCAA</td><td>>MI0000440|hsa-mir-27b&WithFlank&9|+|95085439|95085547</td><td>67</td><td>88</td><td>22(+1A)</td><td>TTCACAGTGGCTAAGTTCTGCAA</td><td>BT20</td><td>41</td><td>22</td><td>32</td><td>6</td><td>20</td><td>17.9</td><td>4.7</td><td>11.7</td><td>1.0</td><td>2.3</td></tr>
<tr><td>>MI0000440|hsa-mir-27b&WithFlank&9|+|95085439|95085547@67.88.22(+1A)&[MIMAT0000419&hsa-miR-27b-3p&offsets|0|+1(+1A);m-39&9|+|95085505|95085525&offsets|0|+1(+1A)]|TTCACAGTGGCTAAGTTCTGCAA</td><td>>MI0000440|hsa-mir-27b&WithFlank&9|+|95085439|95085547</td><td>67</td><td>88</td><td>22(+1A)</td><td>TTCACAGTGGCTAAGTTCTGCAA</td><td>MB231</td><td>20</td><td>19</td><td>5</td><td>14</td><td>21</td><td>2.2</td><td>6.2</td><td>3.2</td><td>8.2</td><td>8.1</td></tr>
</table>

<table>
<tr><th></th><th></th><th></th><th></th><th></th><th></th><th></th><th></th><th></th><th></th><th></th><th></th><th></th><th></th><th></th><th></th><th></th></tr>
<tr><td>>MI0000440|hsa-mir-27b&WithFlank&9|+|95085439|95085547@67.88.22(+1U)&[MIMAT0000419&hsa-miR-27b-3p&offsets|0|+1(+1U);m-39&9|+|95085505|95085525&offsets|0|+1(+1U)]|TTCACAGTGGCTAAGTTCTGCAT</td><td>>MI0000440|hsa-mir-27b&WithFlank&9|+|95085439|95085547</td><td>67</td><td>88</td><td>22(+1U)</td><td>TTCACAGTGGCTAAGTTCTGCAT</td><td>BT20</td><td>28</td><td>16</td><td>21</td><td>4</td><td>9</td><td>8.6</td><td>6.2</td><td>3.7</td><td>1.7</td><td>3.4</td></tr>
<tr><td>>MI0000440|hsa-mir-27b&WithFlank&9|+|95085439|95085547@67.88.22(+1U)&[MIMAT0000419&hsa-miR-27b-3p&offsets|0|+1(+1U);m-39&9|+|95085505|95085525&offsets|0|+1(+1U)]|TTCACAGTGGCTAAGTTCTGCAT</td><td>>MI0000440|hsa-mir-27b&WithFlank&9|+|95085439|95085547</td><td>67</td><td>88</td><td>22(+1U)</td><td>TTCACAGTGGCTAAGTTCTGCAT</td><td>MB231</td><td>22</td><td>14</td><td>5</td><td>12</td><td>22</td><td>3.1</td><td>5.8</td><td>3.3</td><td>4.7</td><td>11.1</td></tr>
<tr><td>>MI0000440|hsa-mir-27b&WithFlank&9|+|95085439|95085547@67.88.22(+2U)&[MIMAT0000419&hsa-miR-27b-3p&offsets|0|+1(+2U);m-39&9|+|95085505|95085525&offsets|0|+1(+2U)]|TTCACAGTGGCTAAGTTCTGCATT</td><td>>MI0000440|hsa-mir-27b&WithFlank&9|+|95085439|95085547</td><td>67</td><td>88</td><td>22(+2U)</td><td>TTCACAGTGGCTAAGTTCTGCATT</td><td>MB231</td><td>12</td><td>10</td><td>3</td><td>3</td><td>10</td><td>2.5</td><td>2.5</td><td>0.9</td><td>0.6</td><td>3.9</td></tr>
<tr><td>>MI0000440|hsa-mir-27b&WithFlank&9|+|95085439|95085547@67.88.22&[MIMAT0000419&hsa-miR-27b-3p&offsets|0|+1;m-39&9|+|95085505|95085525&offsets|0|+1]|TTCACAGTGGCTAAGTTCTGCA</td><td>>MI0000440|hsa-mir-27b&WithFlank&9|+|95085439|95085547</td><td>67</td><td>88</td><td>22</td><td>TTCACAGTGGCTAAGTTCTGCA</td><td>BT20</td><td>494</td><td>224</td><td>389</td><td>33</td><td>151</td><td>194.7</td><td>28.5</td><td>15.3</td><td>19.9</td><td>14.3</td></tr>
<tr><td>>MI0000440|hsa-mir-27b&WithFlank&9|+|95085439|95085547@67.88.22&[MIMAT0000419&hsa-miR-27b-3p&offsets|0|+1;m-39&9|+|95085505|95085525&offsets|0|+1]|TTCACAGTGGCTAAGTTCTGCA</td><td>>MI0000440|hsa-mir-27b&WithFlank&9|+|95085439|95085547</td><td>67</td><td>88</td><td>22</td><td>TTCACAGTGGCTAAGTTCTGCA</td><td>MB231</td><td>345</td><td>312</td><td>70</td><td>103</td><td>240</td><td>48.6</td><td>119.5</td><td>46.0</td><td>43.4</td><td>113.1</td></tr>
<tr><td>>MI0000440|hsa-mir-27b&WithFlank&9|+|95085439|95085547@67.88.22&[MIMAT0000419&hsa-miR-27b-3p&offsets|0|+1;m-39&9|+|95085505|95085525&offsets|0|+1]|TTCACAGTGGCTAAGTTCTGCA</td><td>>MI0000440|hsa-mir-27b&WithFlank&9|+|95085439|95085547</td><td>67</td><td>88</td><td>22</td><td>TTCACAGTGGCTAAGTTCTGCA</td><td>MB468</td><td>25</td><td>10</td><td>8</td><td>8</td><td>5</td><td>13.4</td><td>4.6</td><td>1.8</td><td>2.7</td><td>0.9</td></tr>
<tr><td>>MI0000440|hsa-mir-27b&WithFlank&9|+|95085439|95085547@70.87.18&[MIMAT0000419&hsa-miR-27b-3p&offsets|+3|0;m-39&9|+|95085505|95085525&offsets|+3|0]|ACAGTGGCTAAGTTCTGC</td><td>>MI0000440|hsa-mir-27b&WithFlank&9|+|95085439|95085547</td><td>70</td><td>87</td><td>18</td><td>ACAGTGGCTAAGTTCTGC</td><td>BT20</td><td>69</td><td>40</td><td>45</td><td>23</td><td>22</td><td>15.9</td><td>18.0</td><td>9.5</td><td>18.4</td><td>6.4</td></tr>
<tr><td>>MI0000440|hsa-mir-27b&WithFlank&9|+|95085439|95085547@70.87.18&[MIMAT0000419&hsa-miR-27b-3p&offsets|+3|0;m-39&9|+|95085505|95085525&offsets|+3|0]|ACAGTGGCTAAGTTCTGC</td><td>>MI0000440|hsa-mir-27b&WithFlank&9|+|95085439|95085547</td><td>70</td><td>87</td><td>18</td><td>ACAGTGGCTAAGTTCTGC</td><td>MB231</td><td>22</td><td>17</td><td>6</td><td>23</td><td>27</td><td>19.9</td><td>14.6</td><td>5.7</td><td>20.3</td><td>27.8</td></tr>
<tr><td>>MI0000441|hsa-mir-30b&WithFlank&8|-|134800514|134800613@23.42.20(+1U)&[MIMAT0000420&hsa-miR-30b-5p&offsets|0|-2(+1U);m-76&8|-|134800570|134800591&offsets|0|-2(+1U)]|TGTAAACATCCTACACTCAGT</td><td>>MI0000441|hsa-mir-30b&WithFlank&8|-|134800514|134800613</td><td>23</td><td>42</td><td>20(+1U)</td><td>TGTAAACATCCTACACTCAGT</td><td>BT20</td><td>13</td><td>4</td><td>1</td><td>3</td><td>0</td><td>19.3</td><td>4.5</td><td>0.6</td><td>4.0</td><td>0.3</td></tr>
<tr><td>>MI0000441|hsa-mir-30b&WithFlank&8|-|134800514|134800613@23.42.20(+2U)&[MIMAT0000420&hsa-miR-30b-5p&offsets|0|-2(+2U);m-76&8|-|134800570|134800591&offsets|0|-2(+2U)]|TGTAAACATCCTACACTCAGTT</td><td>>MI0000441|hsa-mir-30b&WithFlank&8|-|134800514|134800613</td><td>23</td><td>42</td><td>20(+2U)</td><td>TGTAAACATCCTACACTCAGTT</td><td>BT20</td><td>15</td><td>1</td><td>1</td><td>0</td><td>0</td><td>24.5</td><td>1.4</td><td>0.9</td><td>0.0</td><td>0.0</td></tr>
<tr><td>>MI0000441|hsa-mir-30b&WithFlank&8|-|134800514|134800613@23.43.21(+1A)&[MIMAT0000420&hsa-miR-30b-5p&offsets|0|-1(+1A);m-76&8|-|134800570|134800591&offsets|0|-1(+1A)]|TGTAAACATCCTACACTCAGCA</td><td>>MI0000441|hsa-mir-30b&WithFlank&8|-|134800514|134800613</td><td>23</td><td>43</td><td>21(+1A)</td><td>TGTAAACATCCTACACTCAGCA</td><td>BT20</td><td>209</td><td>39</td><td>21</td><td>3</td><td>3</td><td>306.8</td><td>37.0</td><td>16.7</td><td>4.0</td><td>3.6</td></tr>
<tr><td>>MI0000441|hsa-mir-30b&WithFlank&8|-|134800514|134800613@23.43.21(+1A)&[MIMAT0000420&hsa-miR-30b-5p&offsets|0|-1(+1A);m-76&8|-|134800570|134800591&offsets|0|-1(+1A)]|TGTAAACATCCTACACTCAGCA</td><td>>MI0000441|hsa-mir-30b&WithFlank&8|-|134800514|134800613</td><td>23</td><td>43</td><td>21(+1A)</td><td>TGTAAACATCCTACACTCAGCA</td><td>MB231</td><td>23</td><td>13</td><td>4</td><td>20</td><td>52</td><td>14.3</td><td>8.7</td><td>3.6</td><td>10.3</td><td>71.9</td></tr>
<tr><td>>MI0000441|hsa-mir-30b&WithFlank&8|-|134800514|134800613@23.43.21(+1C)&[MIMAT0000420&hsa-miR-30b-5p&offsets|0|-1(+1C);m-76&8|-|134800570|134800591&offsets|0|-1(+1C)]|TGTAAACATCCTACACTCAGCC</td><td>>MI0000441|hsa-mir-30b&WithFlank&8|-|134800514|134800613</td><td>23</td><td>43</td><td>21(+1C)</td><td>TGTAAACATCCTACACTCAGCC</td><td>BT20</td><td>163</td><td>62</td><td>21</td><td>5</td><td>7</td><td>205.4</td><td>62.6</td><td>9.4</td><td>5.7</td><td>7.3</td></tr>
<tr><td>>MI0000441|hsa-mir-30b&WithFlank&8|-|134800514|134800613@23.43.21(+1C)&[MIMAT0000420&hsa-miR-30b-5p&offsets|0|-1(+1C);m-76&8|-|134800570|134800591&offsets|0|-1(+1C)]|TGTAAACATCCTACACTCAGCC</td><td>>MI0000441|hsa-mir-30b&WithFlank&8|-|134800514|134800613</td><td>23</td><td>43</td><td>21(+1C)</td><td>TGTAAACATCCTACACTCAGCC</td><td>MB231</td><td>34</td><td>24</td><td>5</td><td>27</td><td>80</td><td>19.3</td><td>14.9</td><td>4.0</td><td>14.0</td><td>111.8</td></tr>
<tr><td>>MI0000441|hsa-mir-30b&WithFlank&8|-|134800514|134800613@23.43.21(+1G)&[MIMAT0000420&hsa-miR-30b-5p&offsets|0|-1(+1G);m-76&8|-|134800570|134800591&offsets|0|-1(+1G)]|TGTAAACATCCTACACTCAGCG</td><td>>MI0000441|hsa-mir-30b&WithFlank&8|-|134800514|134800613</td><td>23</td><td>43</td><td>21(+1G)</td><td>TGTAAACATCCTACACTCAGCG</td><td>BT20</td><td>159</td><td>14</td><td>6</td><td>1</td><td>0</td><td>228.6</td><td>13.7</td><td>3.8</td><td>1.3</td><td>0.3</td></tr>
<tr><td>>MI0000441|hsa-mir-30b&WithFlank&8|-|134800514|134800613@23.43.21(+1G)&[MIMAT0000420&hsa-miR-30b-5p&offsets|0|-1(+1G);m-76&8|-|134800570|134800591&offsets|0|-1(+1G)]|TGTAAACATCCTACACTCAGCG</td><td>>MI0000441|hsa-mir-30b&WithFlank&8|-|134800514|134800613</td><td>23</td><td>43</td><td>21(+1G)</td><td>TGTAAACATCCTACACTCAGCG</td><td>MB231</td><td>24</td><td>6</td><td>1</td><td>4</td><td>10</td><td>22.7</td><td>4.9</td><td>1.0</td><td>1.8</td><td>14.4</td></tr>
<tr><td>>MI0000441|hsa-mir-30b&WithFlank&8|-|134800514|134800613@23.43.21&[MIMAT0000420&hsa-miR-30b-5p&offsets|0|-1;m-76&8|-|134800570|134800591&offsets|0|-1]|TGTAAACATCCTACACTCAGC</td><td>>MI0000441|hsa-mir-30b&WithFlank&8|-|134800514|134800613</td><td>23</td><td>43</td><td>21</td><td>TGTAAACATCCTACACTCAGC</td><td>BT20</td><td>30</td><td>14</td><td>4</td><td>9</td><td>2</td><td>37.5</td><td>14.6</td><td>2.6</td><td>8.9</td><td>1.1</td></tr>
<tr><td>>MI0000441|hsa-mir-30b&WithFlank&8|-|134800514|134800613@23.44.22(+1A)&[MIMAT0000420&hsa-miR-30b-5p&offsets|0|0(+1A);m-76&8|-|134800570|134800591&offsets|0|0(+1A)]|TGTAAACATCCTACACTCAGCTA</td><td>>MI0000441|hsa-mir-30b&WithFlank&8|-|134800514|134800613</td><td>23</td><td>44</td><td>22(+1A)</td><td>TGTAAACATCCTACACTCAGCTA</td><td>BT20</td><td>13</td><td>4</td><td>3</td><td>0</td><td>0</td><td>15.6</td><td>2.7</td><td>1.2</td><td>0.5</td><td>0.9</td></tr>
<tr><td>>MI0000441|hsa-mir-30b&WithFlank&8|-|134800514|134800613@23.44.22&[MIMAT0000420&hsa-miR-30b-5p&offsets|0|0;m-76&8|-|134800570|134800591&offsets|0|0]|TGTAAACATCCTACACTCAGCT</td><td>>MI0000441|hsa-mir-30b&WithFlank&8|-|134800514|134800613</td><td>23</td><td>44</td><td>22</td><td>TGTAAACATCCTACACTCAGCT</td><td>BT20</td><td>1049</td><td>356</td><td>136</td><td>31</td><td>41</td><td>1333.1</td><td>325.4</td><td>48.1</td><td>27.7</td><td>35.4</td></tr>
<tr><td>>MI0000441|hsa-mir-30b&WithFlank&8|-|134800514|134800613@23.44.22&[MIMAT0000420&hsa-miR-30b-5p&offsets|0|0;m-76&8|-|134800570|134800591&offsets|0|0]|TGTAAACATCCTACACTCAGCT</td><td>>MI0000441|hsa-mir-30b&WithFlank&8|-|134800514|134800613</td><td>23</td><td>44</td><td>22</td><td>TGTAAACATCCTACACTCAGCT</td><td>MB231</td><td>250</td><td>185</td><td>39</td><td>160</td><td>460</td><td>142.2</td><td>97.1</td><td>33.7</td><td>55.9</td><td>609.8</td></tr>
<tr><td>>MI0000446|hsa-mir-125b-1&WithFlank&11|-|122099751|122099850@21.39.19;MI0000470|hsa-mir-125b-2&WithFlank&21|+|16590231|16590331@23.41.19&[MIMAT0000423&hsa-miR-125b-5p&offsets|0|-3;m-67&11|-|122099809|122099830&offsets|0|-3];[MIMAT0000423_1&hsa-miR-125b-5p&offsets|0|-3;m-68&21|+|16590253|16590274&offsets|0|-3]|TCCCTGAGACCCTAACTTG</td><td>>MI0000446|hsa-mir-125b-1&WithFlank&11|-|122099751|122099850</td><td>21</td><td>39</td><td>19</td><td>TCCCTGAGACCCTAACTTG</td><td>MB231</td><td>54</td><td>80</td><td>69</td><td>49</td><td>78</td><td>17.2</td><td>56.3</td><td>66.0</td><td>38.1</td><td>48.5</td></tr>
<tr><td>>MI0000446|hsa-mir-125b-1&WithFlank&11|-|122099751|122099850@21.40.20;MI0000470|hsa-mir-125b-2&WithFlank&21|+|16590231|16590331@23.42.20&[MIMAT0000423&hsa-miR-125b-5p&offsets|0|-2;m-67&11|-|122099809|122099830&offsets|0|-2];[MIMAT0000423_1&hsa-miR-125b-5p&offsets|0|-2;m-68&21|+|16590253|16590274&offsets|0|-2]|TCCCTGAGACCCTAACTTGT</td><td>>MI0000446|hsa-mir-125b-1&WithFlank&11|-|122099751|122099850</td><td>21</td><td>40</td><td>20</td><td>TCCCTGAGACCCTAACTTGT</td><td>MB231</td><td>81</td><td>127</td><td>102</td><td>65</td><td>131</td><td>9.9</td><td>50.9</td><td>74.8</td><td>42.6</td><td>64.7</td></tr>
<tr><td>>MI0000446|hsa-mir-125b-1&WithFlank&11|-|122099751|122099850@21.40.20(+1A);MI0000470|hsa-mir-125b-2&WithFlank&21|+|16590231|16590331@23.42.20(+1A)&[MIMAT0000423&hsa-miR-125b-5p&offsets|0|-2(+1A);m-67&11|-|122099809|122099830&offsets|0|-2(+1A)];[MIMAT0000423_1&hsa-miR-125b-5p&offsets|0|-2(+1A);m-68&21|+|16590253|16590274&offsets|0|-2(+1A)]|TCCCTGAGACCCTAACTTGTA</td><td>>MI0000446|hsa-mir-125b-1&WithFlank&11|-|122099751|122099850</td><td>21</td><td>40</td><td>20(+1A)</td><td>TCCCTGAGACCCTAACTTGTA</td><td>MB231</td><td>47</td><td>44</td><td>56</td><td>93</td><td>77</td><td>1.4</td><td>13.8</td><td>49.6</td><td>55.7</td><td>61.1</td></tr>
<tr><td>>MI0000446|hsa-mir-125b-1&WithFlank&11|-|122099751|122099850@21.41.21;MI0000470|hsa-mir-125b-2&WithFlank&21|+|16590231|16590331@23.43.21&[MIMAT0000423&hsa-miR-125b-5p&offsets|0|-1;m-67&11|-|122099809|122099830&offsets|0|-1];[MIMAT0000423_1&hsa-miR-125b-5p&offsets|0|-1;m-68&21|+|16590253|16590274&offsets|0|-1]|TCCCTGAGACCCTAACTTGTG</td><td>>MI0000446|hsa-mir-125b-1&WithFlank&11|-|122099751|122099850</td><td>21</td><td>41</td><td>21</td><td>TCCCTGAGACCCTAACTTGTG</td><td>BT20</td><td>16</td><td>8</td><td>42</td><td>5</td><td>6</td><td>20.7</td><td>10.2</td><td>34.9</td><td>8.0</td><td>8.5</td></tr>
<tr><td>>MI0000446|hsa-mir-125b-1&WithFlank&11|-|122099751|122099850@21.41.21;MI0000470|hsa-mir-125b-2&WithFlank&21|+|16590231|16590331@23.43.21&[MIMAT0000423&hsa-miR-125b-5p&offsets|0|-1;m-67&11|-|122099809|122099830&offsets|0|-1];[MIMAT0000423_1&hsa-miR-125b-5p&offsets|0|-1;m-68&21|+|16590253|16590274&offsets|0|-1]|TCCCTGAGACCCTAACTTGTG</td><td>>MI0000446|hsa-mir-125b-1&WithFlank&11|-|122099751|122099850</td><td>21</td><td>41</td><td>21</td><td>TCCCTGAGACCCTAACTTGTG</td><td>MB231</td><td>451</td><td>552</td><td>462</td><td>583</td><td>570</td><td>16.8</td><td>208.9</td><td>345.9</td><td>246.3</td><td>311.5</td></tr>
<tr><td>>MI0000446|hsa-mir-125b-1&WithFlank&11|-|122099751|122099850@21.41.21;MI0000470|hsa-mir-125b-2&WithFlank&21|+|16590231|16590331@23.43.21&[MIMAT0000423&hsa-miR-125b-5p&offsets|0|-1;m-67&11|-|122099809|122099830&offsets|0|-1];[MIMAT0000423_1&hsa-miR-125b-5p&offsets|0|-1;m-68&21|+|16590253|16590274&offsets|0|-1]|TCCCTGAGACCCTAACTTGTG</td><td>>MI0000446|hsa-mir-125b-1&WithFlank&11|-|122099751|122099850</td><td>21</td><td>41</td><td>21</td><td>TCCCTGAGACCCTAACTTGTG</td><td>MB468</td><td>11</td><td>7</td><td>5</td><td>13</td><td>4</td><td>5.9</td><td>1.5</td><td>1.2</td><td>8.9</td><td>1.9</td></tr>
</table>

<table>
<tr><td></td><td></td><td></td><td></td><td></td><td></td><td></td><td></td><td></td><td></td><td></td><td></td><td></td><td></td><td></td><td></td><td></td></tr>
<tr><td>>MI0000446|hsa-mir-125b-1&WithFlank&11|-|122099751|122099850@21.41.21(+1G);MI0000470|hsa-mir-125b-2&WithFlank&21|+|16590231|16590331@23.43.21(+1G)&[MIMAT0000423&hsa-miR-125b-5p&offsets|0|-1(+1G);m-67&11|-|122099809|122099830&offsets|0|-1(+1G)];[MIMAT0000423_1&hsa-miR-125b-5p&offsets|0|-1(+1G);m-68&21|+|16590253|16590274&offsets|0|-1(+1G)]|TCCCTGAGACCCTAACTTGTGG</td><td>>MI0000446|hsa-mir-125b-1&WithFlank&11|-|122099751|122099850</td><td>21</td><td>41</td><td>21(+1G)</td><td>TCCCTGAGACCCTAACTTGTGG</td><td>MB231</td><td>34</td><td>22</td><td>10</td><td>29</td><td>19</td><td>25.2</td><td>11.8</td><td>7.8</td><td>15.9</td><td>17.6</td></tr>
<tr><td>>MI0000446|hsa-mir-125b-1&WithFlank&11|-|122099751|122099850@21.41.21(+1U);MI0000470|hsa-mir-125b-2&WithFlank&21|+|16590231|16590331@23.43.21(+1U)&[MIMAT0000423&hsa-miR-125b-5p&offsets|0|-1(+1U);m-67&11|-|122099809|122099830&offsets|0|-1(+1U)];[MIMAT0000423_1&hsa-miR-125b-5p&offsets|0|-1(+1U);m-68&21|+|16590253|16590274&offsets|0|-1(+1U)]|TCCCTGAGACCCTAACTTGTGT</td><td>>MI0000446|hsa-mir-125b-1&WithFlank&11|-|122099751|122099850</td><td>21</td><td>41</td><td>21(+1U)</td><td>TCCCTGAGACCCTAACTTGTGT</td><td>MB231</td><td>48</td><td>52</td><td>34</td><td>52</td><td>80</td><td>4.3</td><td>19.1</td><td>24.9</td><td>28.4</td><td>66.8</td></tr>
<tr><td>>MI0000446|hsa-mir-125b-1&WithFlank&11|-|122099751|122099850@21.42.22;MI0000470|hsa-mir-125b-2&WithFlank&21|+|16590231|16590331@23.44.22&[MIMAT0000423&hsa-miR-125b-5p&offsets|0|0;m-67&11|-|122099809|122099830&offsets|0|0];[MIMAT0000423_1&hsa-miR-125b-5p&offsets|0|0;m-68&21|+|16590253|16590274&offsets|0|0]|TCCCTGAGACCCTAACTTGTGA</td><td>>MI0000446|hsa-mir-125b-1&WithFlank&11|-|122099751|122099850</td><td>21</td><td>42</td><td>22</td><td>TCCCTGAGACCCTAACTTGTGA</td><td>BT20</td><td>20</td><td>25</td><td>48</td><td>20</td><td>11</td><td>22.8</td><td>34.7</td><td>40.2</td><td>33.3</td><td>15.3</td></tr>
<tr><td>>MI0000446|hsa-mir-125b-1&WithFlank&11|-|122099751|122099850@21.42.22;MI0000470|hsa-mir-125b-2&WithFlank&21|+|16590231|16590331@23.44.22&[MIMAT0000423&hsa-miR-125b-5p&offsets|0|0;m-67&11|-|122099809|122099830&offsets|0|0];[MIMAT0000423_1&hsa-miR-125b-5p&offsets|0|0;m-68&21|+|16590253|16590274&offsets|0|0]|TCCCTGAGACCCTAACTTGTGA</td><td>>MI0000446|hsa-mir-125b-1&WithFlank&11|-|122099751|122099850</td><td>21</td><td>42</td><td>22</td><td>TCCCTGAGACCCTAACTTGTGA</td><td>MB231</td><td>730</td><td>1120</td><td>519</td><td>1703</td><td>1330</td><td>143.0</td><td>471.3</td><td>410.1</td><td>593.4</td><td>931.4</td></tr>
<tr><td>>MI0000446|hsa-mir-125b-1&WithFlank&11|-|122099751|122099850@21.42.22(+1A);MI0000470|hsa-mir-125b-2&WithFlank&21|+|16590231|16590331@23.44.22(+1A)&[MIMAT0000423&hsa-miR-125b-5p&offsets|0|0(+1A);m-67&11|-|122099809|122099830&offsets|0|0(+1A)];[MIMAT0000423_1&hsa-miR-125b-5p&offsets|0|0(+1A);m-68&21|+|16590253|16590274&offsets|0|0(+1A)]|TCCCTGAGACCCTAACTTGTGAA</td><td>>MI0000446|hsa-mir-125b-1&WithFlank&11|-|122099751|122099850</td><td>21</td><td>42</td><td>22(+1A)</td><td>TCCCTGAGACCCTAACTTGTGAA</td><td>MB231</td><td>248</td><td>248</td><td>183</td><td>104</td><td>319</td><td>56.0</td><td>91.8</td><td>167.9</td><td>21.6</td><td>252.9</td></tr>
<tr><td>>MI0000446|hsa-mir-125b-1&WithFlank&11|-|122099751|122099850@21.42.22(+1C);MI0000470|hsa-mir-125b-2&WithFlank&21|+|16590231|16590331@23.44.22(+1C)&[MIMAT0000423&hsa-miR-125b-5p&offsets|0|0(+1C);m-67&11|-|122099809|122099830&offsets|0|0(+1C)];[MIMAT0000423_1&hsa-miR-125b-5p&offsets|0|0(+1C);m-68&21|+|16590253|16590274&offsets|0|0(+1C)]|TCCCTGAGACCCTAACTTGTGAC</td><td>>MI0000446|hsa-mir-125b-1&WithFlank&11|-|122099751|122099850</td><td>21</td><td>42</td><td>22(+1C)</td><td>TCCCTGAGACCCTAACTTGTGAC</td><td>MB231</td><td>17</td><td>15</td><td>13</td><td>8</td><td>24</td><td>6.0</td><td>5.8</td><td>10.4</td><td>3.4</td><td>21.5</td></tr>
<tr><td>>MI0000446|hsa-mir-125b-1&WithFlank&11|-|122099751|122099850@21.42.22(+2A);MI0000470|hsa-mir-125b-2&WithFlank&21|+|16590231|16590331@23.44.22(+2A)&[MIMAT0000423&hsa-miR-125b-5p&offsets|0|0(+2A);m-67&11|-|122099809|122099830&offsets|0|0(+2A)];[MIMAT0000423_1&hsa-miR-125b-5p&offsets|0|0(+2A);m-68&21|+|16590253|16590274&offsets|0|0(+2A)]|TCCCTGAGACCCTAACTTGTGAAA</td><td>>MI0000446|hsa-mir-125b-1&WithFlank&11|-|122099751|122099850</td><td>21</td><td>42</td><td>22(+2A)</td><td>TCCCTGAGACCCTAACTTGTGAAA</td><td>MB231</td><td>49</td><td>42</td><td>36</td><td>9</td><td>59</td><td>3.6</td><td>13.6</td><td>28.7</td><td>0.8</td><td>39.5</td></tr>
<tr><td>>MI0000446|hsa-mir-125b-1&WithFlank&11|-|122099751|122099850@21.43.23(+1U)&[MIMAT0000423&hsa-miR-125b-5p&offsets|0|+1(+1U);m-67&11|-|122099809|122099830&offsets|0|+1(+1U)]|TCCCTGAGACCCTAACTTGTGATT</td><td>>MI0000446|hsa-mir-125b-1&WithFlank&11|-|122099751|122099850</td><td>21</td><td>43</td><td>23(+1U)</td><td>TCCCTGAGACCCTAACTTGTGATT</td><td>MB231</td><td>18</td><td>16</td><td>11</td><td>6</td><td>24</td><td>4.1</td><td>7.5</td><td>8.6</td><td>2.5</td><td>17.7</td></tr>
<tr><td>>MI0000446|hsa-mir-125b-1&WithFlank&11|-|122099751|122099850@21.43.23&[MIMAT0000423&hsa-miR-125b-5p&offsets|0|+1;m-67&11|-|122099809|122099830&offsets|0|+1]|TCCCTGAGACCCTAACTTGTGAT</td><td>>MI0000446|hsa-mir-125b-1&WithFlank&11|-|122099751|122099850</td><td>21</td><td>43</td><td>23</td><td>TCCCTGAGACCCTAACTTGTGAT</td><td>MB231</td><td>123</td><td>98</td><td>77</td><td>35</td><td>150</td><td>45.3</td><td>38.9</td><td>63.3</td><td>14.6</td><td>131.4</td></tr>
<tr><td>>MI0000446|hsa-mir-125b-1&WithFlank&11|-|122099751|122099850@22.42.21;MI0000470|hsa-mir-125b-2&WithFlank&21|+|16590231|16590331@24.44.21&[MIMAT0000423&hsa-miR-125b-5p&offsets|+1|0;m-67&11|-|122099809|122099830&offsets|+1|0];[MIMAT0000423_1&hsa-miR-125b-5p&offsets|+1|0;m-68&21|+|16590253|16590274&offsets|+1|0]|CCCTGAGACCCTAACTTGTGA</td><td>>MI0000446|hsa-mir-125b-1&WithFlank&11|-|122099751|122099850</td><td>22</td><td>42</td><td>21</td><td>CCCTGAGACCCTAACTTGTGA</td><td>MB231</td><td>11</td><td>18</td><td>7</td><td>8</td><td>17</td><td>1.3</td><td>5.8</td><td>5.3</td><td>1.0</td><td>9.9</td></tr>
<tr><td>>MI0000446|hsa-mir-125b-1&WithFlank&11|-|122099751|122099850@61.81.21&[MIMAT0004592&hsa-miR-125b-1-3p&offsets|0|-1;m-485&11|-|122099769|122099790&offsets|0|-1]|ACGGGTTAGGCTCTTGGGAGC</td><td>>MI0000446|hsa-mir-125b-1&WithFlank&11|-|122099751|122099850</td><td>61</td><td>81</td><td>21</td><td>ACGGGTTAGGCTCTTGGGAGC</td><td>MB231</td><td>13</td><td>12</td><td>50</td><td>35</td><td>9</td><td>8.8</td><td>2.5</td><td>22.4</td><td>14.8</td><td>2.5</td></tr>
<tr><td>>MI0000446|hsa-mir-125b-1&WithFlank&11|-|122099751|122099850@61.82.22&[MIMAT0004592&hsa-miR-125b-1-3p&offsets|0|0;m-485&11|-|122099769|122099790&offsets|0|0]|ACGGGTTAGGCTCTTGGGAGCT</td><td>>MI0000446|hsa-mir-125b-1&WithFlank&11|-|122099751|122099850</td><td>61</td><td>82</td><td>22</td><td>ACGGGTTAGGCTCTTGGGAGCT</td><td>MB231</td><td>40</td><td>35</td><td>52</td><td>105</td><td>32</td><td>32.5</td><td>7.2</td><td>26.8</td><td>45.8</td><td>6.0</td></tr>
<tr><td>>MI0000447|hsa-mir-128-1&WithFlank&2|+|135665391|135665484@56.74.19;MI0000727|hsa-mir-128-2&WithFlank&3|+|35744470|35744565@58.76.19&[MIMAT0000424&hsa-miR-128-3p&offsets|0|-2;m-108&2|+|135665446|135665466&offsets|0|-2];[MIMAT0000424_1&hsa-miR-128-3p&offsets|0|-2;m-117&3|+|35744527|35744547&offsets|0|-2]|TCACAGTGAACCGGTCTCT</td><td>>MI0000447|hsa-mir-128-1&WithFlank&2|+|135665391|135665484</td><td>56</td><td>74</td><td>19</td><td>TCACAGTGAACCGGTCTCT</td><td>BT20</td><td>11</td><td>7</td><td>8</td><td>4</td><td>5</td><td>6.9</td><td>4.3</td><td>3.8</td><td>1.8</td><td>0.8</td></tr>
<tr><td>>MI0000447|hsa-mir-128-1&WithFlank&2|+|135665391|135665484@56.75.20;MI0000727|hsa-mir-128-2&WithFlank&3|+|35744470|35744565@58.77.20&[MIMAT0000424&hsa-miR-128-3p&offsets|0|-1;m-108&2|+|135665446|135665466&offsets|0|-1];[MIMAT0000424_1&hsa-miR-128-3p&offsets|0|-1;m-117&3|+|35744527|35744547&offsets|0|-1]|TCACAGTGAACCGGTCTCTT</td><td>>MI0000447|hsa-mir-128-1&WithFlank&2|+|135665391|135665484</td><td>56</td><td>75</td><td>20</td><td>TCACAGTGAACCGGTCTCTT</td><td>BT20</td><td>68</td><td>35</td><td>49</td><td>35</td><td>33</td><td>45.2</td><td>11.8</td><td>16.5</td><td>21.8</td><td>7.6</td></tr>
<tr><td>>MI0000447|hsa-mir-128-1&WithFlank&2|+|135665391|135665484@56.75.20;MI0000727|hsa-mir-128-2&WithFlank&3|+|35744470|35744565@58.77.20&[MIMAT0000424&hsa-miR-128-3p&offsets|0|-1;m-108&2|+|135665446|135665466&offsets|0|-1];[MIMAT0000424_1&hsa-miR-128-3p&offsets|0|-1;m-117&3|+|35744527|35744547&offsets|0|-1]|TCACAGTGAACCGGTCTCTT</td><td>>MI0000447|hsa-mir-128-1&WithFlank&2|+|135665391|135665484</td><td>56</td><td>75</td><td>20</td><td>TCACAGTGAACCGGTCTCTT</td><td>MB231</td><td>49</td><td>57</td><td>15</td><td>26</td><td>45</td><td>35.8</td><td>18.3</td><td>5.0</td><td>12.5</td><td>8.9</td></tr>
<tr><td>>MI0000447|hsa-mir-128-1&WithFlank&2|+|135665391|135665484@56.75.20(+1A);MI0000727|hsa-mir-128-2&WithFlank&3|+|35744470|35744565@58.77.20(+1A)&[MIMAT0000424&hsa-miR-128-3p&offsets|0|-1(+1A);m-108&2|+|135665446|135665466&offsets|0|-1(+1A)];[MIMAT0000424_1&hsa-miR-128-3p&offsets|0|-1(+1A);m-117&3|+|35744527|35744547&offsets|0|-1(+1A)]|TCACAGTGAACCGGTCTCTTA</td><td>>MI0000447|hsa-mir-128-1&WithFlank&2|+|135665391|135665484</td><td>56</td><td>75</td><td>20(+1A)</td><td>TCACAGTGAACCGGTCTCTTA</td><td>BT20</td><td>29</td><td>10</td><td>18</td><td>8</td><td>8</td><td>17.3</td><td>4.1</td><td>3.3</td><td>3.3</td><td>1.7</td></tr>
<tr><td>>MI0000447|hsa-mir-128-1&WithFlank&2|+|135665391|135665484@56.75.20(+1C);MI0000727|hsa-mir-128-2&WithFlank&3|+|35744470|35744565@58.77.20(+1C)&[MIMAT0000424&hsa-miR-128-3p&offsets|0|-1(+1C);m-108&2|+|135665446|135665466&offsets|0|-1(+1C)];[MIMAT0000424_1&hsa-miR-128-3p&offsets|0|-1(+1C);m-117&3|+|35744527|35744547&offsets|0|-1(+1C)]|TCACAGTGAACCGGTCTCTTC</td><td>>MI0000447|hsa-mir-128-1&WithFlank&2|+|135665391|135665484</td><td>56</td><td>75</td><td>20(+1C)</td><td>TCACAGTGAACCGGTCTCTTC</td><td>BT20</td><td>83</td><td>46</td><td>54</td><td>45</td><td>42</td><td>44.4</td><td>18.1</td><td>10.9</td><td>24.6</td><td>6.3</td></tr>
<tr><td>>MI0000447|hsa-mir-128-1&WithFlank&2|+|135665391|135665484@56.75.20(+1C);MI0000727|hsa-mir-128-2&WithFlank&3|+|35744470|35744565@58.77.20(+1C)&[MIMAT0000424&hsa-miR-128-3p&offsets|0|-1(+1C);m-108&2|+|135665446|135665466&offsets|0|-1(+1C)];[MIMAT0000424_1&hsa-miR-128-3p&offsets|0|-1(+1C);m-117&3|+|35744527|35744547&offsets|0|-1(+1C)]|TCACAGTGAACCGGTCTCTTC</td><td>>MI0000447|hsa-mir-128-1&WithFlank&2|+|135665391|135665484</td><td>56</td><td>75</td><td>20(+1C)</td><td>TCACAGTGAACCGGTCTCTTC</td><td>MB231</td><td>29</td><td>30</td><td>9</td><td>23</td><td>34</td><td>20.8</td><td>9.3</td><td>4.9</td><td>13.2</td><td>12.3</td></tr>
<tr><td>>MI0000447|hsa-mir-128-1&WithFlank&2|+|135665391|135665484@56.75.20(+1G);MI0000727|hsa-mir-128-2&WithFlank&3|+|35744470|35744565@58.77.20(+1G)&[MIMAT0000424&hsa-miR-128-3p&offsets|0|-1(+1G);m-108&2|+|135665446|135665466&offsets|0|-1(+1G)];[MIMAT0000424_1&hsa-miR-128-3p&offsets|0|-1(+1G);m-117&3|+|35744527|35744547&offsets|0|-1(+1G)]|TCACAGTGAACCGGTCTCTTG</td><td>>MI0000447|hsa-mir-128-1&WithFlank&2|+|135665391|135665484</td><td>56</td><td>75</td><td>20(+1G)</td><td>TCACAGTGAACCGGTCTCTTG</td><td>BT20</td><td>17</td><td>3</td><td>5</td><td>2</td><td>2</td><td>13.8</td><td>1.4</td><td>1.0</td><td>1.5</td><td>0.2</td></tr>
</table>

<table>
<tr><th></th><th></th><th></th><th></th><th></th><th></th><th></th><th></th><th></th><th></th><th></th><th></th><th></th><th></th><th></th><th></th><th></th></tr>
<tr><td>>MI0000447|hsa-mir-128-1&WithFlank&2|+|135665391|135665484@56.75.20(+2A);MI0000727|hsa-mir-128-2&WithFlank&3|+|35744470|35744565@58.77.20(+2A)&[MIMAT0000424&hsa-miR-128-3p&offsets|0|-1(+2A);m-108&2|+|135665446|135665466&offsets|0|-1(+2A)];[MIMAT0000424_1&hsa-miR-128-3p&offsets|0|-1(+2A);m-117&3|+|35744527|35744547&offsets|0|-1(+2A)]|TCACAGTGAACCGGTCTCTTAA</td><td>>MI0000447|hsa-mir-128-1&WithFlank&2|+|135665391|135665484</td><td>56</td><td>75</td><td>20(+2A)</td><td>TCACAGTGAACCGGTCTCTTAA</td><td>BT20</td><td>14</td><td>8</td><td>12</td><td>8</td><td>6</td><td>14.8</td><td>3.7</td><td>4.2</td><td>5.1</td><td>1.5</td></tr>
<tr><td>>MI0000447|hsa-mir-128-1&WithFlank&2|+|135665391|135665484@56.76.21;MI0000727|hsa-mir-128-2&WithFlank&3|+|35744470|35744565@58.78.21&[MIMAT0000424&hsa-miR-128-3p&offsets|0|0;m-108&2|+|135665446|135665466&offsets|0|0];[MIMAT0000424_1&hsa-miR-128-3p&offsets|0|0;m-117&3|+|35744527|35744547&offsets|0|0]|TCACAGTGAACCGGTCTCTTT</td><td>>MI0000447|hsa-mir-128-1&WithFlank&2|+|135665391|135665484</td><td>56</td><td>76</td><td>21</td><td>TCACAGTGAACCGGTCTCTTT</td><td>BT20</td><td>698</td><td>379</td><td>479</td><td>362</td><td>364</td><td>432.9</td><td>116.0</td><td>141.2</td><td>176.8</td><td>61.5</td></tr>
<tr><td>>MI0000447|hsa-mir-128-1&WithFlank&2|+|135665391|135665484@56.76.21;MI0000727|hsa-mir-128-2&WithFlank&3|+|35744470|35744565@58.78.21&[MIMAT0000424&hsa-miR-128-3p&offsets|0|0;m-108&2|+|135665446|135665466&offsets|0|0];[MIMAT0000424_1&hsa-miR-128-3p&offsets|0|0;m-117&3|+|35744527|35744547&offsets|0|0]|TCACAGTGAACCGGTCTCTTT</td><td>>MI0000447|hsa-mir-128-1&WithFlank&2|+|135665391|135665484</td><td>56</td><td>76</td><td>21</td><td>TCACAGTGAACCGGTCTCTTT</td><td>MB231</td><td>311</td><td>336</td><td>95</td><td>150</td><td>284</td><td>240.6</td><td>94.0</td><td>37.3</td><td>54.6</td><td>71.2</td></tr>
<tr><td>>MI0000447|hsa-mir-128-1&WithFlank&2|+|135665391|135665484@56.76.21;MI0000727|hsa-mir-128-2&WithFlank&3|+|35744470|35744565@58.78.21&[MIMAT0000424&hsa-miR-128-3p&offsets|0|0;m-108&2|+|135665446|135665466&offsets|0|0];[MIMAT0000424_1&hsa-miR-128-3p&offsets|0|0;m-117&3|+|35744527|35744547&offsets|0|0]|TCACAGTGAACCGGTCTCTTT</td><td>>MI0000447|hsa-mir-128-1&WithFlank&2|+|135665391|135665484</td><td>56</td><td>76</td><td>21</td><td>TCACAGTGAACCGGTCTCTTT</td><td>MB468</td><td>79</td><td>88</td><td>25</td><td>74</td><td>45</td><td>33.5</td><td>57.6</td><td>4.3</td><td>44.0</td><td>19.4</td></tr>
<tr><td>>MI0000447|hsa-mir-128-1&WithFlank&2|+|135665391|135665484@56.76.21(+1A);MI0000727|hsa-mir-128-2&WithFlank&3|+|35744470|35744565@58.78.21(+1A)&[MIMAT0000424&hsa-miR-128-3p&offsets|0|0(+1A);m-108&2|+|135665446|135665466&offsets|0|0(+1A)];[MIMAT0000424_1&hsa-miR-128-3p&offsets|0|0(+1A);m-117&3|+|35744527|35744547&offsets|0|0(+1A)]|TCACAGTGAACCGGTCTCTTTA</td><td>>MI0000447|hsa-mir-128-1&WithFlank&2|+|135665391|135665484</td><td>56</td><td>76</td><td>21(+1A)</td><td>TCACAGTGAACCGGTCTCTTTA</td><td>BT20</td><td>116</td><td>54</td><td>79</td><td>31</td><td>52</td><td>95.2</td><td>19.2</td><td>25.7</td><td>6.6</td><td>12.5</td></tr>
<tr><td>>MI0000447|hsa-mir-128-1&WithFlank&2|+|135665391|135665484@56.76.21(+1A);MI0000727|hsa-mir-128-2&WithFlank&3|+|35744470|35744565@58.78.21(+1A)&[MIMAT0000424&hsa-miR-128-3p&offsets|0|0(+1A);m-108&2|+|135665446|135665466&offsets|0|0(+1A)];[MIMAT0000424_1&hsa-miR-128-3p&offsets|0|0(+1A);m-117&3|+|35744527|35744547&offsets|0|0(+1A)]|TCACAGTGAACCGGTCTCTTTA</td><td>>MI0000447|hsa-mir-128-1&WithFlank&2|+|135665391|135665484</td><td>56</td><td>76</td><td>21(+1A)</td><td>TCACAGTGAACCGGTCTCTTTA</td><td>MB231</td><td>33</td><td>36</td><td>11</td><td>13</td><td>29</td><td>27.5</td><td>8.2</td><td>3.4</td><td>6.4</td><td>8.3</td></tr>
<tr><td>>MI0000447|hsa-mir-128-1&WithFlank&2|+|135665391|135665484@56.76.21(+1A);MI0000727|hsa-mir-128-2&WithFlank&3|+|35744470|35744565@58.78.21(+1A)&[MIMAT0000424&hsa-miR-128-3p&offsets|0|0(+1A);m-108&2|+|135665446|135665466&offsets|0|0(+1A)];[MIMAT0000424_1&hsa-miR-128-3p&offsets|0|0(+1A);m-117&3|+|35744527|35744547&offsets|0|0(+1A)]|TCACAGTGAACCGGTCTCTTTA</td><td>>MI0000447|hsa-mir-128-1&WithFlank&2|+|135665391|135665484</td><td>56</td><td>76</td><td>21(+1A)</td><td>TCACAGTGAACCGGTCTCTTTA</td><td>MB468</td><td>15</td><td>18</td><td>6</td><td>10</td><td>7</td><td>5.4</td><td>9.7</td><td>2.5</td><td>5.4</td><td>2.8</td></tr>
<tr><td>>MI0000447|hsa-mir-128-1&WithFlank&2|+|135665391|135665484@56.76.21(+2A);MI0000727|hsa-mir-128-2&WithFlank&3|+|35744470|35744565@58.78.21(+2A)&[MIMAT0000424&hsa-miR-128-3p&offsets|0|0(+2A);m-108&2|+|135665446|135665466&offsets|0|0(+2A)];[MIMAT0000424_1&hsa-miR-128-3p&offsets|0|0(+2A);m-117&3|+|35744527|35744547&offsets|0|0(+2A)]|TCACAGTGAACCGGTCTCTTTAA</td><td>>MI0000447|hsa-mir-128-1&WithFlank&2|+|135665391|135665484</td><td>56</td><td>76</td><td>21(+2A)</td><td>TCACAGTGAACCGGTCTCTTTAA</td><td>BT20</td><td>53</td><td>29</td><td>42</td><td>21</td><td>24</td><td>52.7</td><td>15.2</td><td>16.4</td><td>9.6</td><td>8.4</td></tr>
<tr><td>>MI0000447|hsa-mir-128-1&WithFlank&2|+|135665391|135665484@56.76.21(+2A);MI0000727|hsa-mir-128-2&WithFlank&3|+|35744470|35744565@58.78.21(+2A)&[MIMAT0000424&hsa-miR-128-3p&offsets|0|0(+2A);m-108&2|+|135665446|135665466&offsets|0|0(+2A)];[MIMAT0000424_1&hsa-miR-128-3p&offsets|0|0(+2A);m-117&3|+|35744527|35744547&offsets|0|0(+2A)]|TCACAGTGAACCGGTCTCTTTAA</td><td>>MI0000447|hsa-mir-128-1&WithFlank&2|+|135665391|135665484</td><td>56</td><td>76</td><td>21(+2A)</td><td>TCACAGTGAACCGGTCTCTTTAA</td><td>MB231</td><td>22</td><td>23</td><td>8</td><td>9</td><td>20</td><td>16.2</td><td>7.7</td><td>2.8</td><td>2.3</td><td>5.1</td></tr>
<tr><td>>MI0000447|hsa-mir-128-1&WithFlank&2|+|135665391|135665484@56.77.22&[MIMAT0000424&hsa-miR-128-3p&offsets|0|+1;m-108&2|+|135665446|135665466&offsets|0|+1]|TCACAGTGAACCGGTCTCTTTT</td><td>>MI0000447|hsa-mir-128-1&WithFlank&2|+|135665391|135665484</td><td>56</td><td>77</td><td>22</td><td>TCACAGTGAACCGGTCTCTTTT</td><td>BT20</td><td>179</td><td>93</td><td>140</td><td>64</td><td>90</td><td>128.8</td><td>23.9</td><td>28.8</td><td>24.3</td><td>17.9</td></tr>
<tr><td>>MI0000447|hsa-mir-128-1&WithFlank&2|+|135665391|135665484@56.77.22&[MIMAT0000424&hsa-miR-128-3p&offsets|0|+1;m-108&2|+|135665446|135665466&offsets|0|+1]|TCACAGTGAACCGGTCTCTTTT</td><td>>MI0000447|hsa-mir-128-1&WithFlank&2|+|135665391|135665484</td><td>56</td><td>77</td><td>22</td><td>TCACAGTGAACCGGTCTCTTTT</td><td>MB231</td><td>134</td><td>127</td><td>41</td><td>48</td><td>121</td><td>104.8</td><td>37.4</td><td>12.2</td><td>22.8</td><td>38.9</td></tr>
<tr><td>>MI0000447|hsa-mir-128-1&WithFlank&2|+|135665391|135665484@56.77.22&[MIMAT0000424&hsa-miR-128-3p&offsets|0|+1;m-108&2|+|135665446|135665466&offsets|0|+1]|TCACAGTGAACCGGTCTCTTTT</td><td>>MI0000447|hsa-mir-128-1&WithFlank&2|+|135665391|135665484</td><td>56</td><td>77</td><td>22</td><td>TCACAGTGAACCGGTCTCTTTT</td><td>MB468</td><td>25</td><td>28</td><td>9</td><td>21</td><td>13</td><td>9.8</td><td>17.8</td><td>1.4</td><td>11.7</td><td>5.1</td></tr>
<tr><td>>MI0000447|hsa-mir-128-1&WithFlank&2|+|135665391|135665484@56.78.23&[MIMAT0000424&hsa-miR-128-3p&offsets|0|+2;m-108&2|+|135665446|135665466&offsets|0|+2]|TCACAGTGAACCGGTCTCTTTTT</td><td>>MI0000447|hsa-mir-128-1&WithFlank&2|+|135665391|135665484</td><td>56</td><td>78</td><td>23</td><td>TCACAGTGAACCGGTCTCTTTTT</td><td>BT20</td><td>14</td><td>7</td><td>13</td><td>5</td><td>7</td><td>11.4</td><td>2.7</td><td>3.2</td><td>0.8</td><td>2.0</td></tr>
<tr><td>>MI0000447|hsa-mir-128-1&WithFlank&2|+|135665391|135665484@56.78.23&[MIMAT0000424&hsa-miR-128-3p&offsets|0|+2;m-108&2|+|135665446|135665466&offsets|0|+2]|TCACAGTGAACCGGTCTCTTTTT</td><td>>MI0000447|hsa-mir-128-1&WithFlank&2|+|135665391|135665484</td><td>56</td><td>78</td><td>23</td><td>TCACAGTGAACCGGTCTCTTTTT</td><td>MB231</td><td>16</td><td>15</td><td>6</td><td>5</td><td>14</td><td>11.9</td><td>4.4</td><td>0.7</td><td>2.5</td><td>3.1</td></tr>
<tr><td>>MI0000448|hsa-mir-130a&WithFlank&11|+|57641192|57641292@61.82.22&[MIMAT0000425&hsa-miR-130a-3p&offsets|0|0;m-139&11|+|57641252|57641273&offsets|0|0]|CAGTGCAATGTTAAAAGGGCAT</td><td>>MI0000448|hsa-mir-130a&WithFlank&11|+|57641192|57641292</td><td>61</td><td>82</td><td>22</td><td>CAGTGCAATGTTAAAAGGGCAT</td><td>BT20</td><td>11</td><td>4</td><td>1</td><td>1</td><td>0</td><td>16.6</td><td>4.1</td><td>0.6</td><td>0.8</td><td>0.0</td></tr>
<tr><td>>MI0000448|hsa-mir-130a&WithFlank&11|+|57641192|57641292@61.83.23&[MIMAT0000425&hsa-miR-130a-3p&offsets|0|+1;m-139&11|+|57641252|57641273&offsets|0|+1]|CAGTGCAATGTTAAAAGGGCATT</td><td>>MI0000448|hsa-mir-130a&WithFlank&11|+|57641192|57641292</td><td>61</td><td>83</td><td>23</td><td>CAGTGCAATGTTAAAAGGGCATT</td><td>BT20</td><td>14</td><td>7</td><td>2</td><td>1</td><td>1</td><td>14.1</td><td>8.2</td><td>0.7</td><td>1.8</td><td>1.1</td></tr>
<tr><td>>MI0000448|hsa-mir-130a&WithFlank&11|+|57641192|57641292@61.83.23&[MIMAT0000425&hsa-miR-130a-3p&offsets|0|+1;m-139&11|+|57641252|57641273&offsets|0|+1]|CAGTGCAATGTTAAAAGGGCATT</td><td>>MI0000448|hsa-mir-130a&WithFlank&11|+|57641192|57641292</td><td>61</td><td>83</td><td>23</td><td>CAGTGCAATGTTAAAAGGGCATT</td><td>MB231</td><td>16</td><td>17</td><td>3</td><td>18</td><td>18</td><td>3.8</td><td>9.4</td><td>3.4</td><td>5.4</td><td>18.7</td></tr>
<tr><td>>MI0000449|hsa-mir-132&WithFlank&17|-|2049902|2050014@29.50.22&[MIMAT0004594&hsa-miR-132-5p&offsets|0|0;m-414&17|-|2049965|2049986&offsets|0|0]|ACCGTGGCTTTCGATTGTTACT</td><td>>MI0000449|hsa-mir-132&WithFlank&17|-|2049902|2050014</td><td>29</td><td>50</td><td>22</td><td>ACCGTGGCTTTCGATTGTTACT</td><td>BT20</td><td>11</td><td>11</td><td>9</td><td>2</td><td>14</td><td>5.3</td><td>5.4</td><td>5.8</td><td>1.0</td><td>7.0</td></tr>
<tr><td>>MI0000449|hsa-mir-132&WithFlank&17|-|2049902|2050014@65.86.22(+1U)&[MIMAT0000426&hsa-miR-132-3p&offsets|0|0(+1U);m-138&17|-|2049929|2049950&offsets|0|0(+1U)]|TAACAGTCTACAGCCATGGTCGT</td><td>>MI0000449|hsa-mir-132&WithFlank&17|-|2049902|2050014</td><td>65</td><td>86</td><td>22(+1U)</td><td>TAACAGTCTACAGCCATGGTCGT</td><td>BT20</td><td>14</td><td>15</td><td>33</td><td>21</td><td>12</td><td>13.8</td><td>9.5</td><td>16.7</td><td>16.3</td><td>6.0</td></tr>
<tr><td>>MI0000449|hsa-mir-132&WithFlank&17|-|2049902|2050014@65.86.22&[MIMAT0000426&hsa-miR-132-3p&offsets|0|0;m-138&17|-|2049929|2049950&offsets|0|0]|TAACAGTCTACAGCCATGGTCG</td><td>>MI0000449|hsa-mir-132&WithFlank&17|-|2049902|2050014</td><td>65</td><td>86</td><td>22</td><td>TAACAGTCTACAGCCATGGTCG</td><td>BT20</td><td>17</td><td>24</td><td>47</td><td>54</td><td>8</td><td>3.7</td><td>16.1</td><td>39.7</td><td>46.3</td><td>4.5</td></tr>
<tr><td>>MI0000456|hsa-mir-140&WithFlank&16|+|69933075|69933186@29.50.22&[MIMAT0000431&hsa-miR-140-5p&offsets|0|0;m-173&16|+|69933103|69933124&offsets|0|0]|CAGTGGTTTTACCCTATGGTAG</td><td>>MI0000456|hsa-mir-140&WithFlank&16|+|69933075|69933186</td><td>29</td><td>50</td><td>22</td><td>CAGTGGTTTTACCCTATGGTAG</td><td>BT20</td><td>23</td><td>17</td><td>8</td><td>53</td><td>10</td><td>10.0</td><td>13.5</td><td>7.4</td><td>42.0</td><td>1.5</td></tr>
<tr><td>>MI0000456|hsa-mir-140&WithFlank&16|+|69933075|69933186@29.50.22&[MIMAT0000431&hsa-miR-140-5p&offsets|0|0;m-173&16|+|69933103|69933124&offsets|0|0]|CAGTGGTTTTACCCTATGGTAG</td><td>>MI0000456|hsa-mir-140&WithFlank&16|+|69933075|69933186</td><td>29</td><td>50</td><td>22</td><td>CAGTGGTTTTACCCTATGGTAG</td><td>MB231</td><td>22</td><td>18</td><td>2</td><td>121</td><td>21</td><td>12.4</td><td>11.7</td><td>1.2</td><td>69.7</td><td>11.8</td></tr>
<tr><td>>MI0000456|hsa-mir-140&WithFlank&16|+|69933075|69933186@68.88.21&[MIMAT0004597&hsa-miR-140-3p&offsets|0|0;m-45&16|+|69933143|69933164&offsets|-1|-2]|TACCACAGGGTAGAACCACGG</td><td>>MI0000456|hsa-mir-140&WithFlank&16|+|69933075|69933186</td><td>68</td><td>88</td><td>21</td><td>TACCACAGGGTAGAACCACGG</td><td>BT20</td><td>15</td><td>7</td><td>20</td><td>16</td><td>3</td><td>7.8</td><td>3.5</td><td>0.7</td><td>13.2</td><td>0.5</td></tr>
</table>

<table>
<tr><td>>MI0000456|hsa-mir-140&WithFlank&16|+|69933075|69933186@68.88.21&[MIMAT0004597&hsa-miR-140-3p&offsets|0|0;m-45&16|+|69933143|69933164&offsets|-1|-2]|TACCACAGGGTAGAACCACGG</td><td>>MI0000456|hsa-mir-140&WithFlank&16|+|69933075|69933186</td><td>68</td><td>88</td><td>21</td><td>TACCACAGGGTAGAACCACGG</td><td>MB231</td><td>10</td><td>9</td><td>9</td><td>33</td><td>8</td><td>2.4</td><td>3.1</td><td>9.0</td><td>3.1</td><td>3.8</td></tr>
<tr><td>>MI0000456|hsa-mir-140&WithFlank&16|+|69933075|69933186@68.89.22&[MIMAT0004597&hsa-miR-140-3p&offsets|0|+1;m-45&16|+|69933143|69933164&offsets|-1|-1]|TACCACAGGGTAGAACCACGGA</td><td>>MI0000456|hsa-mir-140&WithFlank&16|+|69933075|69933186</td><td>68</td><td>89</td><td>22</td><td>TACCACAGGGTAGAACCACGGA</td><td>BT20</td><td>60</td><td>27</td><td>94</td><td>29</td><td>7</td><td>47.9</td><td>15.3</td><td>27.3</td><td>27.0</td><td>1.7</td></tr>
<tr><td>>MI0000456|hsa-mir-140&WithFlank&16|+|69933075|69933186@68.89.22&[MIMAT0004597&hsa-miR-140-3p&offsets|0|+1;m-45&16|+|69933143|69933164&offsets|-1|-1]|TACCACAGGGTAGAACCACGGA</td><td>>MI0000456|hsa-mir-140&WithFlank&16|+|69933075|69933186</td><td>68</td><td>89</td><td>22</td><td>TACCACAGGGTAGAACCACGGA</td><td>MB231</td><td>34</td><td>21</td><td>38</td><td>88</td><td>28</td><td>13.4</td><td>7.7</td><td>43.1</td><td>24.0</td><td>21.1</td></tr>
<tr><td>>MI0000456|hsa-mir-140&WithFlank&16|+|69933075|69933186@68.90.23(+1U)&[MIMAT0004597&hsa-miR-140-3p&offsets|0|+2(+1U);m-45&16|+|69933143|69933164&offsets|-1|0(+1U)]|TACCACAGGGTAGAACCACGGACT</td><td>>MI0000456|hsa-mir-140&WithFlank&16|+|69933075|69933186</td><td>68</td><td>90</td><td>23(+1U)</td><td>TACCACAGGGTAGAACCACGGACT</td><td>MB231</td><td>16</td><td>11</td><td>16</td><td>38</td><td>13</td><td>5.4</td><td>4.4</td><td>13.7</td><td>11.1</td><td>6.8</td></tr>
<tr><td>>MI0000456|hsa-mir-140&WithFlank&16|+|69933075|69933186@68.90.23&[MIMAT0004597&hsa-miR-140-3p&offsets|0|+2;m-45&16|+|69933143|69933164&offsets|-1|0]|TACCACAGGGTAGAACCACGGAC</td><td>>MI0000456|hsa-mir-140&WithFlank&16|+|69933075|69933186</td><td>68</td><td>90</td><td>23</td><td>TACCACAGGGTAGAACCACGGAC</td><td>BT20</td><td>29</td><td>20</td><td>62</td><td>14</td><td>7</td><td>10.3</td><td>9.3</td><td>4.7</td><td>10.8</td><td>2.4</td></tr>
<tr><td>>MI0000456|hsa-mir-140&WithFlank&16|+|69933075|69933186@68.90.23&[MIMAT0004597&hsa-miR-140-3p&offsets|0|+2;m-45&16|+|69933143|69933164&offsets|-1|0]|TACCACAGGGTAGAACCACGGAC</td><td>>MI0000456|hsa-mir-140&WithFlank&16|+|69933075|69933186</td><td>68</td><td>90</td><td>23</td><td>TACCACAGGGTAGAACCACGGAC</td><td>MB231</td><td>29</td><td>20</td><td>36</td><td>44</td><td>24</td><td>13.2</td><td>11.6</td><td>39.2</td><td>17.3</td><td>16.7</td></tr>
<tr><td>>MI0000456|hsa-mir-140&WithFlank&16|+|69933075|69933186@68.91.24&[MIMAT0004597&hsa-miR-140-3p&offsets|0|+3;m-45&16|+|69933143|69933164&offsets|-1|+1]|TACCACAGGGTAGAACCACGGACA</td><td>>MI0000456|hsa-mir-140&WithFlank&16|+|69933075|69933186</td><td>68</td><td>91</td><td>24</td><td>TACCACAGGGTAGAACCACGGACA</td><td>BT20</td><td>19</td><td>10</td><td>35</td><td>14</td><td>3</td><td>10.8</td><td>3.9</td><td>6.1</td><td>11.1</td><td>1.2</td></tr>
<tr><td>>MI0000456|hsa-mir-140&WithFlank&16|+|69933075|69933186@68.91.24&[MIMAT0004597&hsa-miR-140-3p&offsets|0|+3;m-45&16|+|69933143|69933164&offsets|-1|+1]|TACCACAGGGTAGAACCACGGACA</td><td>>MI0000456|hsa-mir-140&WithFlank&16|+|69933075|69933186</td><td>68</td><td>91</td><td>24</td><td>TACCACAGGGTAGAACCACGGACA</td><td>MB231</td><td>12</td><td>7</td><td>10</td><td>22</td><td>8</td><td>3.6</td><td>1.9</td><td>9.6</td><td>0.5</td><td>3.8</td></tr>
<tr><td>>MI0000456|hsa-mir-140&WithFlank&16|+|69933075|69933186@69.88.20&[MIMAT0004597&hsa-miR-140-3p&offsets|+1|0;m-45&16|+|69933143|69933164&offsets|0|-2]|ACCACAGGGTAGAACCACGG</td><td>>MI0000456|hsa-mir-140&WithFlank&16|+|69933075|69933186</td><td>69</td><td>88</td><td>20</td><td>ACCACAGGGTAGAACCACGG</td><td>BT20</td><td>16</td><td>8</td><td>12</td><td>30</td><td>3</td><td>10.2</td><td>5.3</td><td>1.7</td><td>23.5</td><td>0.6</td></tr>
<tr><td>>MI0000456|hsa-mir-140&WithFlank&16|+|69933075|69933186@69.88.20&[MIMAT0004597&hsa-miR-140-3p&offsets|+1|0;m-45&16|+|69933143|69933164&offsets|0|-2]|ACCACAGGGTAGAACCACGG</td><td>>MI0000456|hsa-mir-140&WithFlank&16|+|69933075|69933186</td><td>69</td><td>88</td><td>20</td><td>ACCACAGGGTAGAACCACGG</td><td>MB231</td><td>15</td><td>12</td><td>8</td><td>74</td><td>11</td><td>3.2</td><td>4.6</td><td>8.6</td><td>7.0</td><td>5.8</td></tr>
<tr><td>>MI0000456|hsa-mir-140&WithFlank&16|+|69933075|69933186@69.89.21(+1A)&[MIMAT0004597&hsa-miR-140-3p&offsets|+1|+1(+1A);m-45&16|+|69933143|69933164&offsets|0|-1(+1A)]|ACCACAGGGTAGAACCACGGAA</td><td>>MI0000456|hsa-mir-140&WithFlank&16|+|69933075|69933186</td><td>69</td><td>89</td><td>21(+1A)</td><td>ACCACAGGGTAGAACCACGGAA</td><td>BT20</td><td>15</td><td>7</td><td>14</td><td>8</td><td>2</td><td>14.2</td><td>4.5</td><td>4.1</td><td>7.2</td><td>0.7</td></tr>
<tr><td>>MI0000456|hsa-mir-140&WithFlank&16|+|69933075|69933186@69.89.21(+1U)&[MIMAT0004597&hsa-miR-140-3p&offsets|+1|+1(+1U);m-45&16|+|69933143|69933164&offsets|0|-1(+1U)]|ACCACAGGGTAGAACCACGGAT</td><td>>MI0000456|hsa-mir-140&WithFlank&16|+|69933075|69933186</td><td>69</td><td>89</td><td>21(+1U)</td><td>ACCACAGGGTAGAACCACGGAT</td><td>BT20</td><td>21</td><td>11</td><td>22</td><td>14</td><td>2</td><td>17.1</td><td>7.8</td><td>9.8</td><td>16.4</td><td>0.8</td></tr>
<tr><td>>MI0000456|hsa-mir-140&WithFlank&16|+|69933075|69933186@69.89.21(+1U)&[MIMAT0004597&hsa-miR-140-3p&offsets|+1|+1(+1U);m-45&16|+|69933143|69933164&offsets|0|-1(+1U)]|ACCACAGGGTAGAACCACGGAT</td><td>>MI0000456|hsa-mir-140&WithFlank&16|+|69933075|69933186</td><td>69</td><td>89</td><td>21(+1U)</td><td>ACCACAGGGTAGAACCACGGAT</td><td>MB231</td><td>12</td><td>8</td><td>9</td><td>41</td><td>12</td><td>5.9</td><td>3.5</td><td>10.9</td><td>12.2</td><td>12.0</td></tr>
<tr><td>>MI0000456|hsa-mir-140&WithFlank&16|+|69933075|69933186@69.89.21&[MIMAT0004597&hsa-miR-140-3p&offsets|+1|+1;m-45&16|+|69933143|69933164&offsets|0|-1]|ACCACAGGGTAGAACCACGGA</td><td>>MI0000456|hsa-mir-140&WithFlank&16|+|69933075|69933186</td><td>69</td><td>89</td><td>21</td><td>ACCACAGGGTAGAACCACGGA</td><td>BT20</td><td>90</td><td>40</td><td>78</td><td>74</td><td>11</td><td>81.7</td><td>24.6</td><td>23.1</td><td>67.2</td><td>1.6</td></tr>
<tr><td>>MI0000456|hsa-mir-140&WithFlank&16|+|69933075|69933186@69.89.21&[MIMAT0004597&hsa-miR-140-3p&offsets|+1|+1;m-45&16|+|69933143|69933164&offsets|0|-1]|ACCACAGGGTAGAACCACGGA</td><td>>MI0000456|hsa-mir-140&WithFlank&16|+|69933075|69933186</td><td>69</td><td>89</td><td>21</td><td>ACCACAGGGTAGAACCACGGA</td><td>MB231</td><td>61</td><td>38</td><td>44</td><td>182</td><td>47</td><td>25.0</td><td>14.9</td><td>48.4</td><td>68.3</td><td>39.1</td></tr>
<tr><td>>MI0000456|hsa-mir-140&WithFlank&16|+|69933075|69933186@69.90.22(+1G)&[MIMAT0004597&hsa-miR-140-3p&offsets|+1|+2(+1G);m-45&16|+|69933143|69933164&offsets|0|0(+1G)]|ACCACAGGGTAGAACCACGGACG</td><td>>MI0000456|hsa-mir-140&WithFlank&16|+|69933075|69933186</td><td>69</td><td>90</td><td>22(+1G)</td><td>ACCACAGGGTAGAACCACGGACG</td><td>BT20</td><td>12</td><td>3</td><td>8</td><td>10</td><td>1</td><td>10.2</td><td>1.2</td><td>3.2</td><td>7.4</td><td>0.0</td></tr>
<tr><td>>MI0000456|hsa-mir-140&WithFlank&16|+|69933075|69933186@69.90.22(+1U)&[MIMAT0004597&hsa-miR-140-3p&offsets|+1|+2(+1U);m-45&16|+|69933143|69933164&offsets|0|0(+1U)]|ACCACAGGGTAGAACCACGGACT</td><td>>MI0000456|hsa-mir-140&WithFlank&16|+|69933075|69933186</td><td>69</td><td>90</td><td>22(+1U)</td><td>ACCACAGGGTAGAACCACGGACT</td><td>BT20</td><td>37</td><td>22</td><td>46</td><td>62</td><td>6</td><td>19.1</td><td>14.2</td><td>8.3</td><td>50.2</td><td>3.3</td></tr>
<tr><td>>MI0000456|hsa-mir-140&WithFlank&16|+|69933075|69933186@69.90.22(+1U)&[MIMAT0004597&hsa-miR-140-3p&offsets|+1|+2(+1U);m-45&16|+|69933143|69933164&offsets|0|0(+1U)]|ACCACAGGGTAGAACCACGGACT</td><td>>MI0000456|hsa-mir-140&WithFlank&16|+|69933075|69933186</td><td>69</td><td>90</td><td>22(+1U)</td><td>ACCACAGGGTAGAACCACGGACT</td><td>MB231</td><td>54</td><td>33</td><td>40</td><td>147</td><td>41</td><td>22.7</td><td>14.9</td><td>34.5</td><td>2.8</td><td>31.7</td></tr>
<tr><td>>MI0000456|hsa-mir-140&WithFlank&16|+|69933075|69933186@69.90.22&[MIMAT0004597&hsa-miR-140-3p&offsets|+1|+2;m-45&16|+|69933143|69933164&offsets|0|0]|ACCACAGGGTAGAACCACGGAC</td><td>>MI0000456|hsa-mir-140&WithFlank&16|+|69933075|69933186</td><td>69</td><td>90</td><td>22</td><td>ACCACAGGGTAGAACCACGGAC</td><td>BT20</td><td>85</td><td>57</td><td>118</td><td>84</td><td>17</td><td>40.6</td><td>31.2</td><td>21.2</td><td>74.0</td><td>5.4</td></tr>
<tr><td>>MI0000456|hsa-mir-140&WithFlank&16|+|69933075|69933186@69.90.22&[MIMAT0004597&hsa-miR-140-3p&offsets|+1|+2;m-45&16|+|69933143|69933164&offsets|0|0]|ACCACAGGGTAGAACCACGGAC</td><td>>MI0000456|hsa-mir-140&WithFlank&16|+|69933075|69933186</td><td>69</td><td>90</td><td>22</td><td>ACCACAGGGTAGAACCACGGAC</td><td>MB231</td><td>62</td><td>49</td><td>57</td><td>169</td><td>55</td><td>28.2</td><td>27.5</td><td>63.5</td><td>28.5</td><td>44.0</td></tr>
<tr><td>>MI0000456|hsa-mir-140&WithFlank&16|+|69933075|69933186@69.91.23&[MIMAT0004597&hsa-miR-140-3p&offsets|+1|+3;m-45&16|+|69933143|69933164&offsets|0|+1]|ACCACAGGGTAGAACCACGGACA</td><td>>MI0000456|hsa-mir-140&WithFlank&16|+|69933075|69933186</td><td>69</td><td>91</td><td>23</td><td>ACCACAGGGTAGAACCACGGACA</td><td>BT20</td><td>60</td><td>30</td><td>75</td><td>62</td><td>9</td><td>40.2</td><td>14.1</td><td>21.2</td><td>51.5</td><td>2.1</td></tr>
<tr><td>>MI0000456|hsa-mir-140&WithFlank&16|+|69933075|69933186@69.91.23&[MIMAT0004597&hsa-miR-140-3p&offsets|+1|+3;m-45&16|+|69933143|69933164&offsets|0|+1]|ACCACAGGGTAGAACCACGGACA</td><td>>MI0000456|hsa-mir-140&WithFlank&16|+|69933075|69933186</td><td>69</td><td>91</td><td>23</td><td>ACCACAGGGTAGAACCACGGACA</td><td>MB231</td><td>25</td><td>15</td><td>18</td><td>58</td><td>18</td><td>8.9</td><td>5.0</td><td>17.7</td><td>12.0</td><td>14.8</td></tr>
<tr><td>>MI0000457|hsa-mir-141&WithFlank&12|+|6964091|6964197@23.44.22&[MIMAT0004598&hsa-miR-141-5p&offsets|0|0;m-221&12|+|6964113|6964133&offsets|0|+1]|CATCTTCCAGTACAGTGTTGGA</td><td>>MI0000457|hsa-mir-141&WithFlank&12|+|6964091|6964197</td><td>23</td><td>44</td><td>22</td><td>CATCTTCCAGTACAGTGTTGGA</td><td>BT20</td><td>29</td><td>13</td><td>12</td><td>29</td><td>4</td><td>21.8</td><td>13.3</td><td>9.4</td><td>25.6</td><td>2.7</td></tr>
<tr><td>>MI0000457|hsa-mir-141&WithFlank&12|+|6964091|6964197@65.84.20(+1A)&[MIMAT0000432&hsa-miR-141-3p&offsets|0|-2(+1A);m-134&12|+|6964155|6964175&offsets|0|-1(+1A)]|TAACACTGTCTGGTAAAGATA</td><td>>MI0000457|hsa-mir-141&WithFlank&12|+|6964091|6964197</td><td>65</td><td>84</td><td>20(+1A)</td><td>TAACACTGTCTGGTAAAGATA</td><td>BT20</td><td>17</td><td>5</td><td>5</td><td>11</td><td>1</td><td>20.7</td><td>5.3</td><td>4.1</td><td>10.2</td><td>0.5</td></tr>
<tr><td>>MI0000457|hsa-mir-141&WithFlank&12|+|6964091|6964197@65.85.21&[MIMAT0000432&hsa-miR-141-3p&offsets|0|-1;m-134&12|+|6964155|6964175&offsets|0|0]|TAACACTGTCTGGTAAAGATG</td><td>>MI0000457|hsa-mir-141&WithFlank&12|+|6964091|6964197</td><td>65</td><td>85</td><td>21</td><td>TAACACTGTCTGGTAAAGATG</td><td>BT20</td><td>76</td><td>35</td><td>23</td><td>94</td><td>8</td><td>61.9</td><td>36.5</td><td>16.0</td><td>76.9</td><td>2.9</td></tr>
<tr><td>>MI0000457|hsa-mir-141&WithFlank&12|+|6964091|6964197@65.86.22&[MIMAT0000432&hsa-miR-141-3p&offsets|0|0;m-134&12|+|6964155|6964175&offsets|0|+1]|TAACACTGTCTGGTAAAGATGG</td><td>>MI0000457|hsa-mir-141&WithFlank&12|+|6964091|6964197</td><td>65</td><td>86</td><td>22</td><td>TAACACTGTCTGGTAAAGATGG</td><td>BT20</td><td>17</td><td>5</td><td>1</td><td>13</td><td>0</td><td>23.0</td><td>8.1</td><td>0.6</td><td>11.6</td><td>0.2</td></tr>
<tr><td>>MI0000457|hsa-mir-141&WithFlank&12|+|6964091|6964197@65.87.23&[MIMAT0000432&hsa-miR-141-3p&offsets|0|+1;m-134&12|+|6964155|6964175&offsets|0|+2]|TAACACTGTCTGGTAAAGATGGC</td><td>>MI0000457|hsa-mir-141&WithFlank&12|+|6964091|6964197</td><td>65</td><td>87</td><td>23</td><td>TAACACTGTCTGGTAAAGATGGC</td><td>BT20</td><td>13</td><td>11</td><td>3</td><td>8</td><td>1</td><td>11.7</td><td>18.0</td><td>1.8</td><td>6.9</td><td>1.1</td></tr>
</table>

| | | | | | | | | | | | | | | | | |
|---|---|---|---|---|---|---|---|---|---|---|---|---|---|---|---|---|
| >MI0000462\|hsa-mir-152&WithFlank&17\|-\|48037155\|48037253@60.79.20(+1A)&[MIMAT0000438&hsa-miR-152-3p&offsets\|0\|-1(+1A);m-97&17\|-\|48037174\|48037194&offsets\|0\|-1(+1A)]\|TCAGTGCATGACAGAACTTGA | >MI0000462\|hsa-mir-152&WithFlank&17\|-\|48037155\|48037253 | 60 | 79 | 20(+1A) | TCAGTGCATGACAGAACTTGA | BT20 | 21 | 9 | 10 | 9 | 1 | 24.6 | 6.1 | 8.2 | 9.2 | 0.5 |
| >MI0000462\|hsa-mir-152&WithFlank&17\|-\|48037155\|48037253@60.80.21(+1A)&[MIMAT0000438&hsa-miR-152-3p&offsets\|0\|0(+1A);m-97&17\|-\|48037174\|48037194&offsets\|0\|0(+1A)]\|TCAGTGCATGACAGAACTTGGA | >MI0000462\|hsa-mir-152&WithFlank&17\|-\|48037155\|48037253 | 60 | 80 | 21(+1A) | TCAGTGCATGACAGAACTTGGA | BT20 | 14 | 8 | 12 | 13 | 4 | 9.8 | 4.1 | 8.2 | 13.5 | 0.7 |
| >MI0000462\|hsa-mir-152&WithFlank&17\|-\|48037155\|48037253@60.80.21(+1U)&[MIMAT0000438&hsa-miR-152-3p&offsets\|0\|0(+1U);m-97&17\|-\|48037174\|48037194&offsets\|0\|0(+1U)]\|TCAGTGCATGACAGAACTTGGT | >MI0000462\|hsa-mir-152&WithFlank&17\|-\|48037155\|48037253 | 60 | 80 | 21(+1U) | TCAGTGCATGACAGAACTTGGT | BT20 | 18 | 10 | 19 | 15 | 5 | 9.3 | 5.9 | 10.9 | 14.7 | 1.7 |
| >MI0000462\|hsa-mir-152&WithFlank&17\|-\|48037155\|48037253@60.80.21(+1U)&[MIMAT0000438&hsa-miR-152-3p&offsets\|0\|0(+1U);m-97&17\|-\|48037174\|48037194&offsets\|0\|0(+1U)]\|TCAGTGCATGACAGAACTTGGT | >MI0000462\|hsa-mir-152&WithFlank&17\|-\|48037155\|48037253 | 60 | 80 | 21(+1U) | TCAGTGCATGACAGAACTTGGT | MB231 | 18 | 14 | 8 | 67 | 16 | 6.5 | 4.6 | 6.6 | 16.4 | 13.3 |
| >MI0000462\|hsa-mir-152&WithFlank&17\|-\|48037155\|48037253@60.80.21&[MIMAT0000438&hsa-miR-152-3p&offsets\|0\|0;m-97&17\|-\|48037174\|48037194&offsets\|0\|0]\|TCAGTGCATGACAGAACTTGG | >MI0000462\|hsa-mir-152&WithFlank&17\|-\|48037155\|48037253 | 60 | 80 | 21 | TCAGTGCATGACAGAACTTGG | BT20 | 82 | 58 | 46 | 65 | 10 | 65.4 | 33.4 | 28.1 | 56.5 | 3.3 |
| >MI0000462\|hsa-mir-152&WithFlank&17\|-\|48037155\|48037253@60.80.21&[MIMAT0000438&hsa-miR-152-3p&offsets\|0\|0;m-97&17\|-\|48037174\|48037194&offsets\|0\|0]\|TCAGTGCATGACAGAACTTGG | >MI0000462\|hsa-mir-152&WithFlank&17\|-\|48037155\|48037253 | 60 | 80 | 21 | TCAGTGCATGACAGAACTTGG | MB231 | 37 | 38 | 11 | 131 | 30 | 9.5 | 11.3 | 11.0 | 40.7 | 26.3 |
| >MI0000462\|hsa-mir-152&WithFlank&17\|-\|48037155\|48037253@60.81.22(+1A)&[MIMAT0000438&hsa-miR-152-3p&offsets\|0\|+1(+1A);m-97&17\|-\|48037174\|48037194&offsets\|0\|+1(+1A)]\|TCAGTGCATGACAGAACTTGGGA | >MI0000462\|hsa-mir-152&WithFlank&17\|-\|48037155\|48037253 | 60 | 81 | 22(+1A) | TCAGTGCATGACAGAACTTGGGA | BT20 | 15 | 9 | 26 | 13 | 6 | 7.8 | 4.0 | 6.5 | 11.3 | 1.7 |
| >MI0000462\|hsa-mir-152&WithFlank&17\|-\|48037155\|48037253@60.81.22(+1U)&[MIMAT0000438&hsa-miR-152-3p&offsets\|0\|+1(+1U);m-97&17\|-\|48037174\|48037194&offsets\|0\|+1(+1U)]\|TCAGTGCATGACAGAACTTGGGT | >MI0000462\|hsa-mir-152&WithFlank&17\|-\|48037155\|48037253 | 60 | 81 | 22(+1U) | TCAGTGCATGACAGAACTTGGGT | BT20 | 26 | 15 | 36 | 23 | 12 | 15.9 | 6.5 | 15.8 | 18.1 | 9.4 |
| >MI0000462\|hsa-mir-152&WithFlank&17\|-\|48037155\|48037253@60.81.22(+1U)&[MIMAT0000438&hsa-miR-152-3p&offsets\|0\|+1(+1U);m-97&17\|-\|48037174\|48037194&offsets\|0\|+1(+1U)]\|TCAGTGCATGACAGAACTTGGGT | >MI0000462\|hsa-mir-152&WithFlank&17\|-\|48037155\|48037253 | 60 | 81 | 22(+1U) | TCAGTGCATGACAGAACTTGGGT | MB231 | 24 | 16 | 14 | 54 | 20 | 7.6 | 4.3 | 10.1 | 22.6 | 14.2 |
| >MI0000462\|hsa-mir-152&WithFlank&17\|-\|48037155\|48037253@60.81.22&[MIMAT0000438&hsa-miR-152-3p&offsets\|0\|+1;m-97&17\|-\|48037174\|48037194&offsets\|0\|+1]\|TCAGTGCATGACAGAACTTGGG | >MI0000462\|hsa-mir-152&WithFlank&17\|-\|48037155\|48037253 | 60 | 81 | 22 | TCAGTGCATGACAGAACTTGGG | BT20 | 44 | 37 | 45 | 76 | 25 | 17.1 | 14.8 | 21.0 | 62.9 | 1.4 |
| >MI0000462\|hsa-mir-152&WithFlank&17\|-\|48037155\|48037253@60.81.22&[MIMAT0000438&hsa-miR-152-3p&offsets\|0\|+1;m-97&17\|-\|48037174\|48037194&offsets\|0\|+1]\|TCAGTGCATGACAGAACTTGGG | >MI0000462\|hsa-mir-152&WithFlank&17\|-\|48037155\|48037253 | 60 | 81 | 22 | TCAGTGCATGACAGAACTTGGG | MB231 | 25 | 25 | 14 | 91 | 19 | 5.0 | 4.8 | 8.6 | 32.1 | 6.5 |
| >MI0000465\|hsa-mir-191&WithFlank&3\|-\|49020612\|49020715@22.39.18&[MIMAT0000440&hsa-miR-191-5p&offsets\|0\|-5;m-48&3\|-\|49020672\|49020694&offsets\|0\|-5]\|CAACGGAATCCCAAAAGC | >MI0000465\|hsa-mir-191&WithFlank&3\|-\|49020612\|49020715 | 22 | 39 | 18 | CAACGGAATCCCAAAAGC | BT20 | 11 | 11 | 16 | 18 | 11 | 12.8 | 6.0 | 13.8 | 18.0 | 5.6 |
| >MI0000465\|hsa-mir-191&WithFlank&3\|-\|49020612\|49020715@22.42.21(+1A)&[MIMAT0000440&hsa-miR-191-5p&offsets\|0\|-2(+1A);m-48&3\|-\|49020672\|49020694&offsets\|0\|-2(+1A)]\|CAACGGAATCCCAAAAGCAGCA | >MI0000465\|hsa-mir-191&WithFlank&3\|-\|49020612\|49020715 | 22 | 42 | 21(+1A) | CAACGGAATCCCAAAAGCAGCA | BT20 | 60 | 46 | 65 | 25 | 23 | 40.7 | 22.1 | 26.4 | 24.0 | 14.1 |
| >MI0000465\|hsa-mir-191&WithFlank&3\|-\|49020612\|49020715@22.42.21(+1A)&[MIMAT0000440&hsa-miR-191-5p&offsets\|0\|-2(+1A);m-48&3\|-\|49020672\|49020694&offsets\|0\|-2(+1A)]\|CAACGGAATCCCAAAAGCAGCA | >MI0000465\|hsa-mir-191&WithFlank&3\|-\|49020612\|49020715 | 22 | 42 | 21(+1A) | CAACGGAATCCCAAAAGCAGCA | MB231 | 19 | 23 | 11 | 69 | 32 | 3.3 | 6.2 | 9.3 | 40.6 | 18.2 |
| >MI0000465\|hsa-mir-191&WithFlank&3\|-\|49020612\|49020715@22.42.21(+1C)&[MIMAT0000440&hsa-miR-191-5p&offsets\|0\|-2(+1C);m-48&3\|-\|49020672\|49020694&offsets\|0\|-2(+1C)]\|CAACGGAATCCCAAAAGCAGCC | >MI0000465\|hsa-mir-191&WithFlank&3\|-\|49020612\|49020715 | 22 | 42 | 21(+1C) | CAACGGAATCCCAAAAGCAGCC | BT20 | 68 | 82 | 77 | 42 | 45 | 27.4 | 43.9 | 31.7 | 36.2 | 31.3 |
| >MI0000465\|hsa-mir-191&WithFlank&3\|-\|49020612\|49020715@22.42.21(+1C)&[MIMAT0000440&hsa-miR-191-5p&offsets\|0\|-2(+1C);m-48&3\|-\|49020672\|49020694&offsets\|0\|-2(+1C)]\|CAACGGAATCCCAAAAGCAGCC | >MI0000465\|hsa-mir-191&WithFlank&3\|-\|49020612\|49020715 | 22 | 42 | 21(+1C) | CAACGGAATCCCAAAAGCAGCC | MB231 | 31 | 45 | 17 | 108 | 55 | 8.3 | 16.4 | 12.6 | 57.7 | 29.4 |
| >MI0000465\|hsa-mir-191&WithFlank&3\|-\|49020612\|49020715@22.42.21(+1G)&[MIMAT0000440&hsa-miR-191-5p&offsets\|0\|-2(+1G);m-48&3\|-\|49020672\|49020694&offsets\|0\|-2(+1G)]\|CAACGGAATCCCAAAAGCAGCG | >MI0000465\|hsa-mir-191&WithFlank&3\|-\|49020612\|49020715 | 22 | 42 | 21(+1G) | CAACGGAATCCCAAAAGCAGCG | BT20 | 41 | 19 | 19 | 8 | 5 | 35.6 | 8.9 | 4.5 | 8.6 | 3.2 |
| >MI0000465\|hsa-mir-191&WithFlank&3\|-\|49020612\|49020715@22.42.21(+1G)&[MIMAT0000440&hsa-miR-191-5p&offsets\|0\|-2(+1G);m-48&3\|-\|49020672\|49020694&offsets\|0\|-2(+1G)]\|CAACGGAATCCCAAAAGCAGCG | >MI0000465\|hsa-mir-191&WithFlank&3\|-\|49020612\|49020715 | 22 | 42 | 21(+1G) | CAACGGAATCCCAAAAGCAGCG | MB231 | 14 | 10 | 4 | 13 | 6 | 9.0 | 4.4 | 2.9 | 6.6 | 4.2 |
| >MI0000465\|hsa-mir-191&WithFlank&3\|-\|49020612\|49020715@22.42.21&[MIMAT0000440&hsa-miR-191-5p&offsets\|0\|-2;m-48&3\|-\|49020672\|49020694&offsets\|0\|-2]\|CAACGGAATCCCAAAAGCAGC | >MI0000465\|hsa-mir-191&WithFlank&3\|-\|49020612\|49020715 | 22 | 42 | 21 | CAACGGAATCCCAAAAGCAGC | BT20 | 31 | 40 | 47 | 35 | 18 | 18.8 | 17.4 | 22.4 | 34.2 | 11.4 |
| >MI0000465\|hsa-mir-191&WithFlank&3\|-\|49020612\|49020715@22.42.21&[MIMAT0000440&hsa-miR-191-5p&offsets\|0\|-2;m-48&3\|-\|49020672\|49020694&offsets\|0\|-2]\|CAACGGAATCCCAAAAGCAGC | >MI0000465\|hsa-mir-191&WithFlank&3\|-\|49020612\|49020715 | 22 | 42 | 21 | CAACGGAATCCCAAAAGCAGC | MB231 | 16 | 36 | 13 | 53 | 23 | 3.7 | 13.7 | 10.5 | 23.6 | 4.9 |
| >MI0000465\|hsa-mir-191&WithFlank&3\|-\|49020612\|49020715@22.43.22(+1A)&[MIMAT0000440&hsa-miR-191-5p&offsets\|0\|-1(+1A);m-48&3\|-\|49020672\|49020694&offsets\|0\|-1(+1A)]\|CAACGGAATCCCAAAAGCAGCTA | >MI0000465\|hsa-mir-191&WithFlank&3\|-\|49020612\|49020715 | 22 | 43 | 22(+1A) | CAACGGAATCCCAAAAGCAGCTA | BT20 | 94 | 86 | 144 | 106 | 67 | 44.5 | 42.1 | 50.1 | 78.0 | 42.5 |
| >MI0000465\|hsa-mir-191&WithFlank&3\|-\|49020612\|49020715@22.43.22(+1A)&[MIMAT0000440&hsa-miR-191-5p&offsets\|0\|-1(+1A);m-48&3\|-\|49020672\|49020694&offsets\|0\|-1(+1A)]\|CAACGGAATCCCAAAAGCAGCTA | >MI0000465\|hsa-mir-191&WithFlank&3\|-\|49020612\|49020715 | 22 | 43 | 22(+1A) | CAACGGAATCCCAAAAGCAGCTA | MB231 | 40 | 42 | 36 | 197 | 63 | 9.0 | 12.2 | 33.8 | 115.9 | 32.8 |
| >MI0000465\|hsa-mir-191&WithFlank&3\|-\|49020612\|49020715@22.43.22(+1U)&[MIMAT0000440&hsa-miR-191-5p&offsets\|0\|-1(+1U);m-48&3\|-\|49020672\|49020694&offsets\|0\|-1(+1U)]\|CAACGGAATCCCAAAAGCAGCTT | >MI0000465\|hsa-mir-191&WithFlank&3\|-\|49020612\|49020715 | 22 | 43 | 22(+1U) | CAACGGAATCCCAAAAGCAGCTT | BT20 | 20 | 22 | 28 | 24 | 17 | 10.4 | 10.7 | 15.2 | 19.1 | 11.1 |
| >MI0000465\|hsa-mir-191&WithFlank&3\|-\|49020612\|49020715@22.43.22(+1U)&[MIMAT0000440&hsa-miR-191-5p&offsets\|0\|-1(+1U);m-48&3\|-\|49020672\|49020694&offsets\|0\|-1(+1U)]\|CAACGGAATCCCAAAAGCAGCTT | >MI0000465\|hsa-mir-191&WithFlank&3\|-\|49020612\|49020715 | 22 | 43 | 22(+1U) | CAACGGAATCCCAAAAGCAGCTT | MB231 | 13 | 17 | 9 | 46 | 19 | 1.7 | 6.6 | 7.2 | 27.5 | 8.0 |
| >MI0000465\|hsa-mir-191&WithFlank&3\|-\|49020612\|49020715@22.43.22&[MIMAT0000440&hsa-miR-191-5p&offsets\|0\|-1;m-48&3\|-\|49020672\|49020694&offsets\|0\|-1]\|CAACGGAATCCCAAAAGCAGCT | >MI0000465\|hsa-mir-191&WithFlank&3\|-\|49020612\|49020715 | 22 | 43 | 22 | CAACGGAATCCCAAAAGCAGCT | BT20 | 456 | 502 | 536 | 269 | 291 | 204.9 | 203.4 | 246.0 | 205.5 | 180.2 |
| >MI0000465\|hsa-mir-191&WithFlank&3\|-\|49020612\|49020715@22.43.22&[MIMAT0000440&hsa-miR-191-5p&offsets\|0\|-1;m-48&3\|-\|49020672\|49020694&offsets\|0\|-1]\|CAACGGAATCCCAAAAGCAGCT | >MI0000465\|hsa-mir-191&WithFlank&3\|-\|49020612\|49020715 | 22 | 43 | 22 | CAACGGAATCCCAAAAGCAGCT | MB231 | 238 | 380 | 139 | 626 | 364 | 66.5 | 139.8 | 104.5 | 273.5 | 151.3 |
| >MI0000465\|hsa-mir-191&WithFlank&3\|-\|49020612\|49020715@22.43.22&[MIMAT0000440&hsa-miR-191-5p&offsets\|0\|-1;m-48&3\|-\|49020672\|49020694&offsets\|0\|-1]\|CAACGGAATCCCAAAAGCAGCT | >MI0000465\|hsa-mir-191&WithFlank&3\|-\|49020612\|49020715 | 22 | 43 | 22 | CAACGGAATCCCAAAAGCAGCT | MB468 | 25 | 31 | 11 | 59 | 9 | 12.0 | 11.3 | 0.7 | 48.4 | 2.4 |
| >MI0000465\|hsa-mir-191&WithFlank&3\|-\|49020612\|49020715@22.44.23(+1A)&[MIMAT0000440&hsa-miR-191-5p&offsets\|0\|0(+1A);m-48&3\|-\|49020672\|49020694&offsets\|0\|0(+1A)]\|CAACGGAATCCCAAAAGCAGCTGA | >MI0000465\|hsa-mir-191&WithFlank&3\|-\|49020612\|49020715 | 22 | 44 | 23(+1A) | CAACGGAATCCCAAAAGCAGCTGA | BT20 | 22 | 20 | 26 | 22 | 20 | 11.4 | 14.6 | 17.4 | 16.8 | 17.5 |
| >MI0000465\|hsa-mir-191&WithFlank&3\|-\|49020612\|49020715@22.44.23(+1A)&[MIMAT0000440&hsa-miR-191-5p&offsets\|0\|0(+1A);m-48&3\|-\|49020672\|49020694&offsets\|0\|0(+1A)]\|CAACGGAATCCCAAAAGCAGCTGA | >MI0000465\|hsa-mir-191&WithFlank&3\|-\|49020612\|49020715 | 22 | 44 | 23(+1A) | CAACGGAATCCCAAAAGCAGCTGA | MB231 | 22 | 24 | 13 | 52 | 32 | 3.4 | 6.7 | 9.6 | 16.1 | 16.7 |
| >MI0000465\|hsa-mir-191&WithFlank&3\|-\|49020612\|49020715@22.44.23(+1C)&[MIMAT0000440&hsa-miR-191-5p&offsets\|0\|0(+1C);m-48&3\|-\|49020672\|49020694&offsets\|0\|0(+1C)]\|CAACGGAATCCCAAAAGCAGCTGC | >MI0000465\|hsa-mir-191&WithFlank&3\|-\|49020612\|49020715 | 22 | 44 | 23(+1C) | CAACGGAATCCCAAAAGCAGCTGC | BT20 | 15 | 25 | 18 | 25 | 11 | 7.0 | 25.5 | 12.3 | 20.8 | 12.1 |
| >MI0000465\|hsa-mir-191&WithFlank&3\|-\|49020612\|49020715@22.44.23(+1C)&[MIMAT0000440&hsa-miR-191-5p&offsets\|0\|0(+1C);m-48&3\|-\|49020672\|49020694&offsets\|0\|0(+1C)]\|CAACGGAATCCCAAAAGCAGCTGC | >MI0000465\|hsa-mir-191&WithFlank&3\|-\|49020612\|49020715 | 22 | 44 | 23(+1C) | CAACGGAATCCCAAAAGCAGCTGC | MB231 | 23 | 25 | 13 | 114 | 45 | 3.1 | 14.5 | 11.3 | 66.3 | 44.3 |
| >MI0000465\|hsa-mir-191&WithFlank&3\|-\|49020612\|49020715@22.44.23(+1G)&[MIMAT0000440&hsa-miR-191-5p&offsets\|0\|0(+1G);m-48&3\|-\|49020672\|49020694&offsets\|0\|0(+1G)]\|CAACGGAATCCCAAAAGCAGCTGG | >MI0000465\|hsa-mir-191&WithFlank&3\|-\|49020612\|49020715 | 22 | 44 | 23(+1G) | CAACGGAATCCCAAAAGCAGCTGG | MB231 | 10 | 5 | 2 | 12 | 5 | 8.2 | 2.6 | 2.0 | 6.6 | 5.0 |
| >MI0000465\|hsa-mir-191&WithFlank&3\|-\|49020612\|49020715@22.44.23&[MIMAT0000440&hsa-miR-191-5p&offsets\|0\|0;m-48&3\|-\|49020672\|49020694&offsets\|0\|0]\|CAACGGAATCCCAAAAGCAGCTG | >MI0000465\|hsa-mir-191&WithFlank&3\|-\|49020612\|49020715 | 22 | 44 | 23 | CAACGGAATCCCAAAAGCAGCTG | BT20 | 581 | 653 | 936 | 959 | 639 | 397.9 | 227.7 | 694.8 | 773.8 | 401.2 |
| >MI0000465\|hsa-mir-191&WithFlank&3\|-\|49020612\|49020715@22.44.23&[MIMAT0000440&hsa-miR-191-5p&offsets\|0\|0;m-48&3\|-\|49020672\|49020694&offsets\|0\|0]\|CAACGGAATCCCAAAAGCAGCTG | >MI0000465\|hsa-mir-191&WithFlank&3\|-\|49020612\|49020715 | 22 | 44 | 23 | CAACGGAATCCCAAAAGCAGCTG | MB231 | 394 | 540 | 306 | 1280 | 518 | 107.7 | 204.4 | 263.8 | 562.1 | 150.2 |

<table>
<tr><th></th><th></th><th></th><th></th><th></th><th></th><th></th><th></th><th></th><th></th><th></th><th></th><th></th><th></th><th></th><th></th><th></th></tr>
<tr><td>>MI0000465|hsa-mir-191&WithFlank&3|-|49020612|49020715@22.44.23&[MIMAT0000440&hsa-miR-191-5p&offsets|0|0;m-48&3|-|49020672|49020694&offsets|0|0]|CAACGGAATCCCAAAAGCAGCTG</td><td>>MI0000465|hsa-mir-191&WithFlank&3|-|49020612|49020715</td><td>22</td><td>44</td><td>23</td><td>CAACGGAATCCCAAAAGCAGCTG</td><td>MB468</td><td>44</td><td>50</td><td>20</td><td>91</td><td>16</td><td>23.5</td><td>25.7</td><td>1.9</td><td>40.7</td><td>3.2</td></tr>
<tr><td>>MI0000465|hsa-mir-191&WithFlank&3|-|49020612|49020715@22.45.24&[MIMAT0000440&hsa-miR-191-5p&offsets|0|+1;m-48&3|-|49020672|49020694&offsets|0|+1]|CAACGGAATCCCAAAAGCAGCTGT</td><td>>MI0000465|hsa-mir-191&WithFlank&3|-|49020612|49020715</td><td>22</td><td>45</td><td>24</td><td>CAACGGAATCCCAAAAGCAGCTGT</td><td>BT20</td><td>109</td><td>162</td><td>131</td><td>173</td><td>72</td><td>50.9</td><td>159.5</td><td>99.1</td><td>148.6</td><td>80.2</td></tr>
<tr><td>>MI0000465|hsa-mir-191&WithFlank&3|-|49020612|49020715@22.45.24&[MIMAT0000440&hsa-miR-191-5p&offsets|0|+1;m-48&3|-|49020672|49020694&offsets|0|+1]|CAACGGAATCCCAAAAGCAGCTGT</td><td>>MI0000465|hsa-mir-191&WithFlank&3|-|49020612|49020715</td><td>22</td><td>45</td><td>24</td><td>CAACGGAATCCCAAAAGCAGCTGT</td><td>MB231</td><td>192</td><td>228</td><td>107</td><td>692</td><td>314</td><td>28.5</td><td>133.5</td><td>108.4</td><td>343.6</td><td>267.3</td></tr>
<tr><td>>MI0000465|hsa-mir-191&WithFlank&3|-|49020612|49020715@23.44.22&[MIMAT0000440&hsa-miR-191-5p&offsets|+1|0;m-48&3|-|49020672|49020694&offsets|+1|0]|AACGGAATCCCAAAAGCAGCTG</td><td>>MI0000465|hsa-mir-191&WithFlank&3|-|49020612|49020715</td><td>23</td><td>44</td><td>22</td><td>AACGGAATCCCAAAAGCAGCTG</td><td>BT20</td><td>12</td><td>10</td><td>23</td><td>24</td><td>10</td><td>12.5</td><td>2.0</td><td>18.1</td><td>19.9</td><td>6.1</td></tr>
<tr><td>>MI0000465|hsa-mir-191&WithFlank&3|-|49020612|49020715@23.44.22&[MIMAT0000440&hsa-miR-191-5p&offsets|+1|0;m-48&3|-|49020672|49020694&offsets|+1|0]|AACGGAATCCCAAAAGCAGCTG</td><td>>MI0000465|hsa-mir-191&WithFlank&3|-|49020612|49020715</td><td>23</td><td>44</td><td>22</td><td>AACGGAATCCCAAAAGCAGCTG</td><td>MB231</td><td>11</td><td>17</td><td>9</td><td>41</td><td>14</td><td>3.4</td><td>2.5</td><td>5.6</td><td>13.5</td><td>1.6</td></tr>
<tr><td>>MI0000466|hsa-mir-9-1&WithFlank&1|-|156420335|156420435@22.42.21;MI0000468|hsa-mir-9-3&WithFlank&15|+|89368011|89368112@22.42.21;MI0000467|hsa-mir-9-2&WithFlank&5|-|88666847|88666945@22.42.21&[MIMAT0000441&hsa-miR-9-5p&offsets|0|-2;m-104&1|-|156420392|156420414&offsets|0|-2];[MIMAT0000441_1&hsa-miR-9-5p&offsets|0|-2;m-105&15|+|89368032|89368054&offsets|0|-2];[MIMAT0000441_2&hsa-miR-9-5p&offsets|0|-2;m-104&5|-|88666902|88666924&offsets|0|-2]|TCTTTGGTTATCTAGCTGTAT</td><td>>MI0000466|hsa-mir-9-1&WithFlank&1|-|156420335|156420435</td><td>22</td><td>42</td><td>21</td><td>TCTTTGGTTATCTAGCTGTAT</td><td>BT20</td><td>14</td><td>6</td><td>5</td><td>3</td><td>2</td><td>6.5</td><td>5.9</td><td>3.4</td><td>2.5</td><td>2.3</td></tr>
<tr><td>>MI0000466|hsa-mir-9-1&WithFlank&1|-|156420335|156420435@22.42.21(+1A);MI0000468|hsa-mir-9-3&WithFlank&15|+|89368011|89368112@22.42.21(+1A);MI0000467|hsa-mir-9-2&WithFlank&5|-|88666847|88666945@22.42.21(+1A)&[MIMAT0000441&hsa-miR-9-5p&offsets|0|-2(+1A);m-104&1|-|156420392|156420414&offsets|0|-2(+1A)];[MIMAT0000441_1&hsa-miR-9-5p&offsets|0|-2(+1A);m-105&15|+|89368032|89368054&offsets|0|-2(+1A)];[MIMAT0000441_2&hsa-miR-9-5p&offsets|0|-2(+1A);m-104&5|-|88666902|88666924&offsets|0|-2(+1A)]|TCTTTGGTTATCTAGCTGTATA</td><td>>MI0000466|hsa-mir-9-1&WithFlank&1|-|156420335|156420435</td><td>22</td><td>42</td><td>21(+1A)</td><td>TCTTTGGTTATCTAGCTGTATA</td><td>BT20</td><td>26</td><td>6</td><td>9</td><td>1</td><td>2</td><td>26.5</td><td>4.6</td><td>3.4</td><td>1.5</td><td>1.4</td></tr>
<tr><td>>MI0000466|hsa-mir-9-1&WithFlank&1|-|156420335|156420435@22.43.22;MI0000468|hsa-mir-9-3&WithFlank&15|+|89368011|89368112@22.43.22;MI0000467|hsa-mir-9-2&WithFlank&5|-|88666847|88666945@22.43.22&[MIMAT0000441&hsa-miR-9-5p&offsets|0|-1;m-104&1|-|156420392|156420414&offsets|0|-1];[MIMAT0000441_1&hsa-miR-9-5p&offsets|0|-1;m-105&15|+|89368032|89368054&offsets|0|-1];[MIMAT0000441_2&hsa-miR-9-5p&offsets|0|-1;m-104&5|-|88666902|88666924&offsets|0|-1]|TCTTTGGTTATCTAGCTGTATG</td><td>>MI0000466|hsa-mir-9-1&WithFlank&1|-|156420335|156420435</td><td>22</td><td>43</td><td>22</td><td>TCTTTGGTTATCTAGCTGTATG</td><td>BT20</td><td>162</td><td>63</td><td>71</td><td>12</td><td>35</td><td>89.4</td><td>37.5</td><td>37.3</td><td>8.0</td><td>7.5</td></tr>
<tr><td>>MI0000466|hsa-mir-9-1&WithFlank&1|-|156420335|156420435@22.43.22;MI0000468|hsa-mir-9-3&WithFlank&15|+|89368011|89368112@22.43.22;MI0000467|hsa-mir-9-2&WithFlank&5|-|88666847|88666945@22.43.22&[MIMAT0000441&hsa-miR-9-5p&offsets|0|-1;m-104&1|-|156420392|156420414&offsets|0|-1];[MIMAT0000441_1&hsa-miR-9-5p&offsets|0|-1;m-105&15|+|89368032|89368054&offsets|0|-1];[MIMAT0000441_2&hsa-miR-9-5p&offsets|0|-1;m-104&5|-|88666902|88666924&offsets|0|-1]|TCTTTGGTTATCTAGCTGTATG</td><td>>MI0000466|hsa-mir-9-1&WithFlank&1|-|156420335|156420435</td><td>22</td><td>43</td><td>22</td><td>TCTTTGGTTATCTAGCTGTATG</td><td>MB231</td><td>24</td><td>16</td><td>5</td><td>10</td><td>15</td><td>4.7</td><td>10.8</td><td>3.9</td><td>3.0</td><td>8.7</td></tr>
<tr><td>>MI0000466|hsa-mir-9-1&WithFlank&1|-|156420335|156420435@22.43.22(+1G);MI0000468|hsa-mir-9-3&WithFlank&15|+|89368011|89368112@22.43.22(+1G);MI0000467|hsa-mir-9-2&WithFlank&5|-|88666847|88666945@22.43.22(+1G)&[MIMAT0000441&hsa-miR-9-5p&offsets|0|-1(+1G);m-104&1|-|156420392|156420414&offsets|0|-1(+1G)];[MIMAT0000441_1&hsa-miR-9-5p&offsets|0|-1(+1G);m-105&15|+|89368032|89368054&offsets|0|-1(+1G)];[MIMAT0000441_2&hsa-miR-9-5p&offsets|0|-1(+1G);m-104&5|-|88666902|88666924&offsets|0|-1(+1G)]|TCTTTGGTTATCTAGCTGTATGG</td><td>>MI0000466|hsa-mir-9-1&WithFlank&1|-|156420335|156420435</td><td>22</td><td>43</td><td>22(+1G)</td><td>TCTTTGGTTATCTAGCTGTATGG</td><td>BT20</td><td>16</td><td>3</td><td>2</td><td>0</td><td>0</td><td>18.9</td><td>2.2</td><td>0.6</td><td>0.7</td><td>0.5</td></tr>
<tr><td>>MI0000466|hsa-mir-9-1&WithFlank&1|-|156420335|156420435@22.43.22(+1U);MI0000468|hsa-mir-9-3&WithFlank&15|+|89368011|89368112@22.43.22(+1U);MI0000467|hsa-mir-9-2&WithFlank&5|-|88666847|88666945@22.43.22(+1U)&[MIMAT0000441&hsa-miR-9-5p&offsets|0|-1(+1U);m-104&1|-|156420392|156420414&offsets|0|-1(+1U)];[MIMAT0000441_1&hsa-miR-9-5p&offsets|0|-1(+1U);m-105&15|+|89368032|89368054&offsets|0|-1(+1U)];[MIMAT0000441_2&hsa-miR-9-5p&offsets|0|-1(+1U);m-104&5|-|88666902|88666924&offsets|0|-1(+1U)]|TCTTTGGTTATCTAGCTGTATGT</td><td>>MI0000466|hsa-mir-9-1&WithFlank&1|-|156420335|156420435</td><td>22</td><td>43</td><td>22(+1U)</td><td>TCTTTGGTTATCTAGCTGTATGT</td><td>BT20</td><td>16</td><td>6</td><td>3</td><td>1</td><td>1</td><td>15.0</td><td>4.8</td><td>2.1</td><td>1.0</td><td>1.0</td></tr>
<tr><td>>MI0000466|hsa-mir-9-1&WithFlank&1|-|156420335|156420435@22.44.23;MI0000468|hsa-mir-9-3&WithFlank&15|+|89368011|89368112@22.44.23;MI0000467|hsa-mir-9-2&WithFlank&5|-|88666847|88666945@22.44.23&[MIMAT0000441&hsa-miR-9-5p&offsets|0|0;m-104&1|-|156420392|156420414&offsets|0|0];[MIMAT0000441_1&hsa-miR-9-5p&offsets|0|0;m-105&15|+|89368032|89368054&offsets|0|0];[MIMAT0000441_2&hsa-miR-9-5p&offsets|0|0;m-104&5|-|88666902|88666924&offsets|0|0]|TCTTTGGTTATCTAGCTGTATGA</td><td>>MI0000466|hsa-mir-9-1&WithFlank&1|-|156420335|156420435</td><td>22</td><td>44</td><td>23</td><td>TCTTTGGTTATCTAGCTGTATGA</td><td>BT20</td><td>208</td><td>102</td><td>63</td><td>23</td><td>52</td><td>137.9</td><td>82.1</td><td>36.2</td><td>14.9</td><td>8.4</td></tr>
<tr><td>>MI0000466|hsa-mir-9-1&WithFlank&1|-|156420335|156420435@22.44.23;MI0000468|hsa-mir-9-3&WithFlank&15|+|89368011|89368112@22.44.23;MI0000467|hsa-mir-9-2&WithFlank&5|-|88666847|88666945@22.44.23&[MIMAT0000441&hsa-miR-9-5p&offsets|0|0;m-104&1|-|156420392|156420414&offsets|0|0];[MIMAT0000441_1&hsa-miR-9-5p&offsets|0|0;m-105&15|+|89368032|89368054&offsets|0|0];[MIMAT0000441_2&hsa-miR-9-5p&offsets|0|0;m-104&5|-|88666902|88666924&offsets|0|0]|TCTTTGGTTATCTAGCTGTATGA</td><td>>MI0000466|hsa-mir-9-1&WithFlank&1|-|156420335|156420435</td><td>22</td><td>44</td><td>23</td><td>TCTTTGGTTATCTAGCTGTATGA</td><td>MB231</td><td>20</td><td>20</td><td>3</td><td>26</td><td>18</td><td>2.6</td><td>11.9</td><td>2.3</td><td>3.7</td><td>10.2</td></tr>
<tr><td>>MI0000469|hsa-mir-125a&WithFlank&19|+|51693248|51693345@21.41.21(+1A)&[MIMAT0000443&hsa-miR-125a-5p&offsets|0|-3(+1A);m-63&19|+|51693268|51693289&offsets|0|-1(+1A)]|TCCCTGAGACCCTTTAACCTGA</td><td>>MI0000469|hsa-mir-125a&WithFlank&19|+|51693248|51693345</td><td>21</td><td>41</td><td>21(+1A)</td><td>TCCCTGAGACCCTTTAACCTGA</td><td>BT20</td><td>12</td><td>6</td><td>15</td><td>1</td><td>3</td><td>7.6</td><td>3.2</td><td>5.8</td><td>0.5</td><td>3.1</td></tr>
<tr><td>>MI0000469|hsa-mir-125a&WithFlank&19|+|51693248|51693345@21.41.21(+1C)&[MIMAT0000443&hsa-miR-125a-5p&offsets|0|-3(+1C);m-63&19|+|51693268|51693289&offsets|0|-1(+1C)]|TCCCTGAGACCCTTTAACCTGC</td><td>>MI0000469|hsa-mir-125a&WithFlank&19|+|51693248|51693345</td><td>21</td><td>41</td><td>21(+1C)</td><td>TCCCTGAGACCCTTTAACCTGC</td><td>BT20</td><td>38</td><td>36</td><td>48</td><td>4</td><td>30</td><td>29.1</td><td>20.1</td><td>20.6</td><td>1.5</td><td>14.9</td></tr>
<tr><td>>MI0000469|hsa-mir-125a&WithFlank&19|+|51693248|51693345@21.41.21(+1G)&[MIMAT0000443&hsa-miR-125a-5p&offsets|0|-3(+1G);m-63&19|+|51693268|51693289&offsets|0|-1(+1G)]|TCCCTGAGACCCTTTAACCTGG</td><td>>MI0000469|hsa-mir-125a&WithFlank&19|+|51693248|51693345</td><td>21</td><td>41</td><td>21(+1G)</td><td>TCCCTGAGACCCTTTAACCTGG</td><td>BT20</td><td>13</td><td>6</td><td>8</td><td>0</td><td>0</td><td>8.8</td><td>2.6</td><td>1.6</td><td>0.0</td><td>0.0</td></tr>
<tr><td>>MI0000469|hsa-mir-125a&WithFlank&19|+|51693248|51693345@21.41.21&[MIMAT0000443&hsa-miR-125a-5p&offsets|0|-3;m-63&19|+|51693268|51693289&offsets|0|-1]|TCCCTGAGACCCTTTAACCTG</td><td>>MI0000469|hsa-mir-125a&WithFlank&19|+|51693248|51693345</td><td>21</td><td>41</td><td>21</td><td>TCCCTGAGACCCTTTAACCTG</td><td>BT20</td><td>33</td><td>15</td><td>34</td><td>6</td><td>16</td><td>41.0</td><td>8.5</td><td>37.9</td><td>6.4</td><td>19.4</td></tr>
<tr><td>>MI0000469|hsa-mir-125a&WithFlank&19|+|51693248|51693345@21.42.22(+1A)&[MIMAT0000443&hsa-miR-125a-5p&offsets|0|-2(+1A);m-63&19|+|51693268|51693289&offsets|0|0(+1A)]|TCCCTGAGACCCTTTAACCTGTA</td><td>>MI0000469|hsa-mir-125a&WithFlank&19|+|51693248|51693345</td><td>21</td><td>42</td><td>22(+1A)</td><td>TCCCTGAGACCCTTTAACCTGTA</td><td>BT20</td><td>226</td><td>124</td><td>264</td><td>20</td><td>148</td><td>139.2</td><td>48.2</td><td>136.8</td><td>2.1</td><td>78.3</td></tr>
<tr><td>>MI0000469|hsa-mir-125a&WithFlank&19|+|51693248|51693345@21.42.22(+1A)&[MIMAT0000443&hsa-miR-125a-5p&offsets|0|-2(+1A);m-63&19|+|51693268|51693289&offsets|0|0(+1A)]|TCCCTGAGACCCTTTAACCTGTA</td><td>>MI0000469|hsa-mir-125a&WithFlank&19|+|51693248|51693345</td><td>21</td><td>42</td><td>22(+1A)</td><td>TCCCTGAGACCCTTTAACCTGTA</td><td>MB231</td><td>46</td><td>40</td><td>36</td><td>18</td><td>71</td><td>11.1</td><td>12.3</td><td>25.6</td><td>8.6</td><td>41.7</td></tr>
<tr><td>>MI0000469|hsa-mir-125a&WithFlank&19|+|51693248|51693345@21.42.22(+1U)&[MIMAT0000443&hsa-miR-125a-5p&offsets|0|-2(+1U);m-63&19|+|51693268|51693289&offsets|0|0(+1U)]|TCCCTGAGACCCTTTAACCTGTT</td><td>>MI0000469|hsa-mir-125a&WithFlank&19|+|51693248|51693345</td><td>21</td><td>42</td><td>22(+1U)</td><td>TCCCTGAGACCCTTTAACCTGTT</td><td>BT20</td><td>23</td><td>14</td><td>24</td><td>2</td><td>15</td><td>14.4</td><td>6.1</td><td>12.4</td><td>0.5</td><td>7.1</td></tr>
<tr><td>>MI0000469|hsa-mir-125a&WithFlank&19|+|51693248|51693345@21.42.22&[MIMAT0000443&hsa-miR-125a-5p&offsets|0|-2;m-63&19|+|51693268|51693289&offsets|0|0]|TCCCTGAGACCCTTTAACCTGT</td><td>>MI0000469|hsa-mir-125a&WithFlank&19|+|51693248|51693345</td><td>21</td><td>42</td><td>22</td><td>TCCCTGAGACCCTTTAACCTGT</td><td>BT20</td><td>300</td><td>244</td><td>384</td><td>31</td><td>228</td><td>241.5</td><td>112.2</td><td>182.7</td><td>17.0</td><td>93.2</td></tr>
<tr><td>>MI0000469|hsa-mir-125a&WithFlank&19|+|51693248|51693345@21.42.22&[MIMAT0000443&hsa-miR-125a-5p&offsets|0|-2;m-63&19|+|51693268|51693289&offsets|0|0]|TCCCTGAGACCCTTTAACCTGT</td><td>>MI0000469|hsa-mir-125a&WithFlank&19|+|51693248|51693345</td><td>21</td><td>42</td><td>22</td><td>TCCCTGAGACCCTTTAACCTGT</td><td>MB231</td><td>87</td><td>103</td><td>90</td><td>18</td><td>98</td><td>37.9</td><td>33.3</td><td>64.7</td><td>9.8</td><td>35.0</td></tr>
</table>

<table>
<tr><td>>MI0000469|hsa-mir-125a&WithFlank&19|+|51693248|51693345@21.42.22&[MIMAT0000443&hsa-miR-125a-5p&offsets|0|-2;m-63&19|+|51693268|51693289&offsets|0|0]|TCCCTGAGACCCTTTAACCTGT</td><td>>MI0000469|hsa-mir-125a&WithFlank&19|+|51693248|51693345</td><td>21</td><td>42</td><td>22</td><td>TCCCTGAGACCCTTTAACCTGT</td><td>MB468</td><td>11</td><td>24</td><td>4</td><td>4</td><td>7</td><td>4.7</td><td>12.3</td><td>1.3</td><td>1.3</td><td>1.1</td></tr>
<tr><td>>MI0000469|hsa-mir-125a&WithFlank&19|+|51693248|51693345@21.43.23(+1G)&[MIMAT0000443&hsa-miR-125a-5p&offsets|0|-1(+1G);m-63&19|+|51693268|51693289&offsets|0|+1(+1G)]|TCCCTGAGACCCTTTAACCTGTGG</td><td>>MI0000469|hsa-mir-125a&WithFlank&19|+|51693248|51693345</td><td>21</td><td>43</td><td>23(+1G)</td><td>TCCCTGAGACCCTTTAACCTGTGG</td><td>BT20</td><td>14</td><td>7</td><td>9</td><td>0</td><td>1</td><td>10.4</td><td>5.9</td><td>2.3</td><td>0.3</td><td>0.9</td></tr>
<tr><td>>MI0000469|hsa-mir-125a&WithFlank&19|+|51693248|51693345@21.43.23(+1U)&[MIMAT0000443&hsa-miR-125a-5p&offsets|0|-1(+1U);m-63&19|+|51693268|51693289&offsets|0|+1(+1U)]|TCCCTGAGACCCTTTAACCTGTGT</td><td>>MI0000469|hsa-mir-125a&WithFlank&19|+|51693248|51693345</td><td>21</td><td>43</td><td>23(+1U)</td><td>TCCCTGAGACCCTTTAACCTGTGT</td><td>BT20</td><td>72</td><td>42</td><td>74</td><td>12</td><td>58</td><td>69.6</td><td>23.2</td><td>60.5</td><td>4.9</td><td>43.8</td></tr>
<tr><td>>MI0000469|hsa-mir-125a&WithFlank&19|+|51693248|51693345@21.43.23(+1U)&[MIMAT0000443&hsa-miR-125a-5p&offsets|0|-1(+1U);m-63&19|+|51693268|51693289&offsets|0|+1(+1U)]|TCCCTGAGACCCTTTAACCTGTGT</td><td>>MI0000469|hsa-mir-125a&WithFlank&19|+|51693248|51693345</td><td>21</td><td>43</td><td>23(+1U)</td><td>TCCCTGAGACCCTTTAACCTGTGT</td><td>MB231</td><td>42</td><td>44</td><td>28</td><td>18</td><td>56</td><td>17.5</td><td>13.5</td><td>18.1</td><td>10.4</td><td>31.6</td></tr>
<tr><td>>MI0000469|hsa-mir-125a&WithFlank&19|+|51693248|51693345@21.43.23&[MIMAT0000443&hsa-miR-125a-5p&offsets|0|-1;m-63&19|+|51693268|51693289&offsets|0|+1]|TCCCTGAGACCCTTTAACCTGTG</td><td>>MI0000469|hsa-mir-125a&WithFlank&19|+|51693248|51693345</td><td>21</td><td>43</td><td>23</td><td>TCCCTGAGACCCTTTAACCTGTG</td><td>BT20</td><td>1491</td><td>1014</td><td>1881</td><td>211</td><td>1501</td><td>1400.6</td><td>257.8</td><td>1676.2</td><td>73.1</td><td>795.1</td></tr>
<tr><td>>MI0000469|hsa-mir-125a&WithFlank&19|+|51693248|51693345@21.43.23&[MIMAT0000443&hsa-miR-125a-5p&offsets|0|-1;m-63&19|+|51693268|51693289&offsets|0|+1]|TCCCTGAGACCCTTTAACCTGTG</td><td>>MI0000469|hsa-mir-125a&WithFlank&19|+|51693248|51693345</td><td>21</td><td>43</td><td>23</td><td>TCCCTGAGACCCTTTAACCTGTG</td><td>MB231</td><td>468</td><td>526</td><td>320</td><td>123</td><td>599</td><td>134.8</td><td>212.6</td><td>210.3</td><td>40.5</td><td>206.1</td></tr>
<tr><td>>MI0000469|hsa-mir-125a&WithFlank&19|+|51693248|51693345@21.43.23&[MIMAT0000443&hsa-miR-125a-5p&offsets|0|-1;m-63&19|+|51693268|51693289&offsets|0|+1]|TCCCTGAGACCCTTTAACCTGTG</td><td>>MI0000469|hsa-mir-125a&WithFlank&19|+|51693248|51693345</td><td>21</td><td>43</td><td>23</td><td>TCCCTGAGACCCTTTAACCTGTG</td><td>MB468</td><td>68</td><td>82</td><td>25</td><td>29</td><td>32</td><td>36.5</td><td>28.2</td><td>4.6</td><td>16.6</td><td>10.0</td></tr>
<tr><td>>MI0000469|hsa-mir-125a&WithFlank&19|+|51693248|51693345@21.44.24&[MIMAT0000443&hsa-miR-125a-5p&offsets|0|0;m-63&19|+|51693268|51693289&offsets|0|+2]|TCCCTGAGACCCTTTAACCTGTGA</td><td>>MI0000469|hsa-mir-125a&WithFlank&19|+|51693248|51693345</td><td>21</td><td>44</td><td>24</td><td>TCCCTGAGACCCTTTAACCTGTGA</td><td>BT20</td><td>237</td><td>257</td><td>338</td><td>25</td><td>206</td><td>171.1</td><td>171.2</td><td>208.5</td><td>11.8</td><td>121.3</td></tr>
<tr><td>>MI0000469|hsa-mir-125a&WithFlank&19|+|51693248|51693345@21.44.24&[MIMAT0000443&hsa-miR-125a-5p&offsets|0|0;m-63&19|+|51693268|51693289&offsets|0|+2]|TCCCTGAGACCCTTTAACCTGTGA</td><td>>MI0000469|hsa-mir-125a&WithFlank&19|+|51693248|51693345</td><td>21</td><td>44</td><td>24</td><td>TCCCTGAGACCCTTTAACCTGTGA</td><td>MB231</td><td>119</td><td>137</td><td>78</td><td>32</td><td>180</td><td>56.3</td><td>31.3</td><td>48.9</td><td>7.0</td><td>119.1</td></tr>
<tr><td>>MI0000469|hsa-mir-125a&WithFlank&19|+|51693248|51693345@21.44.24&[MIMAT0000443&hsa-miR-125a-5p&offsets|0|0;m-63&19|+|51693268|51693289&offsets|0|+2]|TCCCTGAGACCCTTTAACCTGTGA</td><td>>MI0000469|hsa-mir-125a&WithFlank&19|+|51693248|51693345</td><td>21</td><td>44</td><td>24</td><td>TCCCTGAGACCCTTTAACCTGTGA</td><td>MB468</td><td>16</td><td>17</td><td>7</td><td>7</td><td>8</td><td>5.3</td><td>5.4</td><td>1.7</td><td>3.8</td><td>1.4</td></tr>
<tr><td>>MI0000470|hsa-mir-125b-2&WithFlank&21|+|16590231|16590331@23.45.23&[MIMAT0000423_1&hsa-miR-125b-5p&offsets|0|+1;m-68&21|+|16590253|16590274&offsets|0|+1]|TCCCTGAGACCCTAACTTGTGAG</td><td>>MI0000470|hsa-mir-125b-2&WithFlank&21|+|16590231|16590331</td><td>23</td><td>45</td><td>23</td><td>TCCCTGAGACCCTAACTTGTGAG</td><td>MB231</td><td>17</td><td>6</td><td>7</td><td>3</td><td>6</td><td>13.9</td><td>2.3</td><td>5.8</td><td>0.5</td><td>5.1</td></tr>
<tr><td>>MI0000471|hsa-mir-126&WithFlank&9|+|136670596|136670692@21.41.21&[MIMAT0000444&hsa-miR-126-5p&offsets|0|0;m-114&9|+|136670616|136670636&offsets|0|0]|CATTATTACTTTTGGTACGCG</td><td>>MI0000471|hsa-mir-126&WithFlank&9|+|136670596|136670692</td><td>21</td><td>41</td><td>21</td><td>CATTATTACTTTTGGTACGCG</td><td>BT20</td><td>20</td><td>18</td><td>13</td><td>7</td><td>2</td><td>15.1</td><td>21.9</td><td>8.3</td><td>5.2</td><td>1.6</td></tr>
<tr><td>>MI0000471|hsa-mir-126&WithFlank&9|+|136670596|136670692@57.78.22(+1A)&[MIMAT0000445&hsa-miR-126-3p&offsets|-1|-1(+1A);m-40&9|+|136670653|136670674&offsets|-1|-1(+1A)]|CTCGTACCGTGAGTAATAATGCA</td><td>>MI0000471|hsa-mir-126&WithFlank&9|+|136670596|136670692</td><td>57</td><td>78</td><td>22(+1A)</td><td>CTCGTACCGTGAGTAATAATGCA</td><td>BT20</td><td>11</td><td>7</td><td>5</td><td>2</td><td>7</td><td>6.3</td><td>5.6</td><td>4.5</td><td>0.6</td><td>2.3</td></tr>
<tr><td>>MI0000471|hsa-mir-126&WithFlank&9|+|136670596|136670692@57.78.22&[MIMAT0000445&hsa-miR-126-3p&offsets|-1|-1;m-40&9|+|136670653|136670674&offsets|-1|-1]|CTCGTACCGTGAGTAATAATGC</td><td>>MI0000471|hsa-mir-126&WithFlank&9|+|136670596|136670692</td><td>57</td><td>78</td><td>22</td><td>CTCGTACCGTGAGTAATAATGC</td><td>BT20</td><td>11</td><td>4</td><td>3</td><td>2</td><td>3</td><td>4.6</td><td>2.9</td><td>2.6</td><td>1.7</td><td>0.4</td></tr>
<tr><td>>MI0000471|hsa-mir-126&WithFlank&9|+|136670596|136670692@57.79.23&[MIMAT0000445&hsa-miR-126-3p&offsets|-1|0;m-40&9|+|136670653|136670674&offsets|-1|0]|CTCGTACCGTGAGTAATAATGCG</td><td>>MI0000471|hsa-mir-126&WithFlank&9|+|136670596|136670692</td><td>57</td><td>79</td><td>23</td><td>CTCGTACCGTGAGTAATAATGCG</td><td>BT20</td><td>47</td><td>27</td><td>25</td><td>13</td><td>35</td><td>42.1</td><td>18.9</td><td>29.8</td><td>0.8</td><td>9.5</td></tr>
<tr><td>>MI0000471|hsa-mir-126&WithFlank&9|+|136670596|136670692@57.79.23&[MIMAT0000445&hsa-miR-126-3p&offsets|-1|0;m-40&9|+|136670653|136670674&offsets|-1|0]|CTCGTACCGTGAGTAATAATGCG</td><td>>MI0000471|hsa-mir-126&WithFlank&9|+|136670596|136670692</td><td>57</td><td>79</td><td>23</td><td>CTCGTACCGTGAGTAATAATGCG</td><td>MB231</td><td>31</td><td>28</td><td>6</td><td>35</td><td>31</td><td>22.5</td><td>5.1</td><td>2.0</td><td>7.3</td><td>2.9</td></tr>
<tr><td>>MI0000471|hsa-mir-126&WithFlank&9|+|136670596|136670692@58.77.20(+1A)&[MIMAT0000445&hsa-miR-126-3p&offsets|0|-2(+1A);m-40&9|+|136670653|136670674&offsets|0|-2(+1A)]|TCGTACCGTGAGTAATAATGA</td><td>>MI0000471|hsa-mir-126&WithFlank&9|+|136670596|136670692</td><td>58</td><td>77</td><td>20(+1A)</td><td>TCGTACCGTGAGTAATAATGA</td><td>BT20</td><td>15</td><td>6</td><td>10</td><td>7</td><td>4</td><td>9.6</td><td>5.3</td><td>9.4</td><td>5.5</td><td>1.4</td></tr>
<tr><td>>MI0000471|hsa-mir-126&WithFlank&9|+|136670596|136670692@58.77.20(+1U)&[MIMAT0000445&hsa-miR-126-3p&offsets|0|-2(+1U);m-40&9|+|136670653|136670674&offsets|0|-2(+1U)]|TCGTACCGTGAGTAATAATGT</td><td>>MI0000471|hsa-mir-126&WithFlank&9|+|136670596|136670692</td><td>58</td><td>77</td><td>20(+1U)</td><td>TCGTACCGTGAGTAATAATGT</td><td>BT20</td><td>60</td><td>34</td><td>38</td><td>34</td><td>18</td><td>40.6</td><td>33.5</td><td>43.4</td><td>29.2</td><td>7.1</td></tr>
<tr><td>>MI0000471|hsa-mir-126&WithFlank&9|+|136670596|136670692@58.77.20(+1U)&[MIMAT0000445&hsa-miR-126-3p&offsets|0|-2(+1U);m-40&9|+|136670653|136670674&offsets|0|-2(+1U)]|TCGTACCGTGAGTAATAATGT</td><td>>MI0000471|hsa-mir-126&WithFlank&9|+|136670596|136670692</td><td>58</td><td>77</td><td>20(+1U)</td><td>TCGTACCGTGAGTAATAATGT</td><td>MB231</td><td>35</td><td>30</td><td>7</td><td>147</td><td>37</td><td>18.8</td><td>7.4</td><td>0.7</td><td>86.1</td><td>14.9</td></tr>
<tr><td>>MI0000471|hsa-mir-126&WithFlank&9|+|136670596|136670692@58.77.20&[MIMAT0000445&hsa-miR-126-3p&offsets|0|-2;m-40&9|+|136670653|136670674&offsets|0|-2]|TCGTACCGTGAGTAATAATG</td><td>>MI0000471|hsa-mir-126&WithFlank&9|+|136670596|136670692</td><td>58</td><td>77</td><td>20</td><td>TCGTACCGTGAGTAATAATG</td><td>BT20</td><td>40</td><td>24</td><td>32</td><td>22</td><td>18</td><td>43.6</td><td>17.0</td><td>42.9</td><td>13.0</td><td>4.9</td></tr>
<tr><td>>MI0000471|hsa-mir-126&WithFlank&9|+|136670596|136670692@58.77.20&[MIMAT0000445&hsa-miR-126-3p&offsets|0|-2;m-40&9|+|136670653|136670674&offsets|0|-2]|TCGTACCGTGAGTAATAATG</td><td>>MI0000471|hsa-mir-126&WithFlank&9|+|136670596|136670692</td><td>58</td><td>77</td><td>20</td><td>TCGTACCGTGAGTAATAATG</td><td>MB231</td><td>15</td><td>20</td><td>4</td><td>35</td><td>16</td><td>4.6</td><td>15.9</td><td>3.1</td><td>23.2</td><td>9.3</td></tr>
<tr><td>>MI0000471|hsa-mir-126&WithFlank&9|+|136670596|136670692@58.78.21(+1A)&[MIMAT0000445&hsa-miR-126-3p&offsets|0|-1(+1A);m-40&9|+|136670653|136670674&offsets|0|-1(+1A)]|TCGTACCGTGAGTAATAATGCA</td><td>>MI0000471|hsa-mir-126&WithFlank&9|+|136670596|136670692</td><td>58</td><td>78</td><td>21(+1A)</td><td>TCGTACCGTGAGTAATAATGCA</td><td>BT20</td><td>72</td><td>54</td><td>59</td><td>43</td><td>27</td><td>47.7</td><td>49.8</td><td>59.6</td><td>36.1</td><td>12.3</td></tr>
<tr><td>>MI0000471|hsa-mir-126&WithFlank&9|+|136670596|136670692@58.78.21(+1A)&[MIMAT0000445&hsa-miR-126-3p&offsets|0|-1(+1A);m-40&9|+|136670653|136670674&offsets|0|-1(+1A)]|TCGTACCGTGAGTAATAATGCA</td><td>>MI0000471|hsa-mir-126&WithFlank&9|+|136670596|136670692</td><td>58</td><td>78</td><td>21(+1A)</td><td>TCGTACCGTGAGTAATAATGCA</td><td>MB231</td><td>30</td><td>25</td><td>5</td><td>149</td><td>39</td><td>15.6</td><td>11.1</td><td>2.6</td><td>89.1</td><td>22.5</td></tr>
<tr><td>>MI0000471|hsa-mir-126&WithFlank&9|+|136670596|136670692@58.78.21(+1U)&[MIMAT0000445&hsa-miR-126-3p&offsets|0|-1(+1U);m-40&9|+|136670653|136670674&offsets|0|-1(+1U)]|TCGTACCGTGAGTAATAATGCT</td><td>>MI0000471|hsa-mir-126&WithFlank&9|+|136670596|136670692</td><td>58</td><td>78</td><td>21(+1U)</td><td>TCGTACCGTGAGTAATAATGCT</td><td>BT20</td><td>11</td><td>7</td><td>7</td><td>6</td><td>4</td><td>6.8</td><td>5.9</td><td>7.8</td><td>5.4</td><td>1.8</td></tr>
<tr><td>>MI0000471|hsa-mir-126&WithFlank&9|+|136670596|136670692@58.78.21&[MIMAT0000445&hsa-miR-126-3p&offsets|0|-1;m-40&9|+|136670653|136670674&offsets|0|-1]|TCGTACCGTGAGTAATAATGC</td><td>>MI0000471|hsa-mir-126&WithFlank&9|+|136670596|136670692</td><td>58</td><td>78</td><td>21</td><td>TCGTACCGTGAGTAATAATGC</td><td>BT20</td><td>169</td><td>105</td><td>136</td><td>115</td><td>77</td><td>155.4</td><td>86.9</td><td>175.2</td><td>66.2</td><td>33.9</td></tr>
<tr><td>>MI0000471|hsa-mir-126&WithFlank&9|+|136670596|136670692@58.78.21&[MIMAT0000445&hsa-miR-126-3p&offsets|0|-1;m-40&9|+|136670653|136670674&offsets|0|-1]|TCGTACCGTGAGTAATAATGC</td><td>>MI0000471|hsa-mir-126&WithFlank&9|+|136670596|136670692</td><td>58</td><td>78</td><td>21</td><td>TCGTACCGTGAGTAATAATGC</td><td>MB231</td><td>153</td><td>161</td><td>35</td><td>421</td><td>140</td><td>89.7</td><td>23.5</td><td>3.2</td><td>151.2</td><td>28.9</td></tr>
<tr><td>>MI0000471|hsa-mir-126&WithFlank&9|+|136670596|136670692@58.79.22&[MIMAT0000445&hsa-miR-126-3p&offsets|0|0;m-40&9|+|136670653|136670674&offsets|0|0]|TCGTACCGTGAGTAATAATGCG</td><td>>MI0000471|hsa-mir-126&WithFlank&9|+|136670596|136670692</td><td>58</td><td>79</td><td>22</td><td>TCGTACCGTGAGTAATAATGCG</td><td>BT20</td><td>285</td><td>216</td><td>269</td><td>224</td><td>145</td><td>269.0</td><td>164.9</td><td>349.6</td><td>149.9</td><td>43.9</td></tr>
</table>

| | | | | | | | | | | | | | | | | |
|---|---|---|---|---|---|---|---|---|---|---|---|---|---|---|---|---|
| >MI0000471\|hsa-mir-126&WithFlank&9\|+\|136670596\|136670692@58.79.22&[MIMAT0000445&hsa-miR-126-3p&offsets\|0\|0;m-40&9\|+\|136670653\|136670674&offsets\|0\|0]\|TCGTACCGTGAGTAATAATGCG | >MI0000471\|hsa-mir-126&WithFlank&9\|+\|136670596\|136670692 | 58 | 79 | 22 | TCGTACCGTGAGTAATAATGCG | MB231 | 182 | 170 | 28 | 591 | 176 | 110.0 | 62.6 | 12.3 | 290.6 | 52.0 |
| >MI0000471\|hsa-mir-126&WithFlank&9\|+\|136670596\|136670692@59.77.19(+1U)&[MIMAT0000445&hsa-miR-126-3p&offsets\|+1\|-2(+1U);m-40&9\|+\|136670653\|136670674&offsets\|+1\|-2(+1U)]\|CGTACCGTGAGTAATAATGT | >MI0000471\|hsa-mir-126&WithFlank&9\|+\|136670596\|136670692 | 59 | 77 | 19(+1U) | CGTACCGTGAGTAATAATGT | BT20 | 11 | 8 | 14 | 14 | 4 | 9.9 | 8.2 | 16.6 | 10.2 | 1.7 |
| >MI0000471\|hsa-mir-126&WithFlank&9\|+\|136670596\|136670692@59.78.20&[MIMAT0000445&hsa-miR-126-3p&offsets\|+1\|-1;m-40&9\|+\|136670653\|136670674&offsets\|+1\|-1]\|CGTACCGTGAGTAATAATGC | >MI0000471\|hsa-mir-126&WithFlank&9\|+\|136670596\|136670692 | 59 | 78 | 20 | CGTACCGTGAGTAATAATGC | BT20 | 36 | 25 | 50 | 46 | 15 | 41.2 | 23.5 | 65.7 | 37.7 | 6.9 |
| >MI0000471\|hsa-mir-126&WithFlank&9\|+\|136670596\|136670692@59.78.20&[MIMAT0000445&hsa-miR-126-3p&offsets\|+1\|-1;m-40&9\|+\|136670653\|136670674&offsets\|+1\|-1]\|CGTACCGTGAGTAATAATGC | >MI0000471\|hsa-mir-126&WithFlank&9\|+\|136670596\|136670692 | 59 | 78 | 20 | CGTACCGTGAGTAATAATGC | MB231 | 25 | 26 | 7 | 92 | 24 | 18.8 | 3.1 | 1.0 | 45.5 | 3.1 |
| >MI0000471\|hsa-mir-126&WithFlank&9\|+\|136670596\|136670692@59.79.21&[MIMAT0000445&hsa-miR-126-3p&offsets\|+1\|0;m-40&9\|+\|136670653\|136670674&offsets\|+1\|0]\|CGTACCGTGAGTAATAATGCG | >MI0000471\|hsa-mir-126&WithFlank&9\|+\|136670596\|136670692 | 59 | 79 | 21 | CGTACCGTGAGTAATAATGCG | BT20 | 19 | 16 | 24 | 32 | 8 | 15.5 | 11.2 | 28.7 | 24.8 | 2.1 |
| >MI0000476\|hsa-mir-138-1&WithFlank&3\|+\|44114206\|44114316@29.52.24&[MIMAT0000430_1&hsa-miR-138-5p&offsets\|0\|+1;m-497&3\|+\|44114234\|44114256&offsets\|0\|+1]\|AGCTGGTGTTGTGAATCAGGCCGT | >MI0000476\|hsa-mir-138-1&WithFlank&3\|+\|44114206\|44114316 | 29 | 52 | 24 | AGCTGGTGTTGTGAATCAGGCCGT | MB231 | 16 | 9 | 7 | 112 | 14 | 5.6 | 6.7 | 8.5 | 32.5 | 10.6 |
| >MI0000477\|hsa-mir-146a&WithFlank&5\|+\|160485346\|160485456@27.46.20(+1C)&[MIMAT0000449&hsa-miR-146a-5p&offsets\|0\|-2(+1C);m-129&5\|+\|160485372\|160485393&offsets\|0\|-2(+1C)]\|TGAGAACTGAATTCCATGGGC | >MI0000477\|hsa-mir-146a&WithFlank&5\|+\|160485346\|160485456 | 27 | 46 | 20(+1C) | TGAGAACTGAATTCCATGGGC | MB231 | 13 | 19 | 6 | 177 | 18 | 10.5 | 20.3 | 4.9 | 166.8 | 13.9 |
| >MI0000477\|hsa-mir-146a&WithFlank&5\|+\|160485346\|160485456@27.47.21(+1A)&[MIMAT0000449&hsa-miR-146a-5p&offsets\|0\|-1(+1A);m-129&5\|+\|160485372\|160485393&offsets\|0\|-1(+1A)]\|TGAGAACTGAATTCCATGGGTA | >MI0000477\|hsa-mir-146a&WithFlank&5\|+\|160485346\|160485456 | 27 | 47 | 21(+1A) | TGAGAACTGAATTCCATGGGTA | MB231 | 32 | 46 | 20 | 593 | 48 | 27.3 | 54.3 | 17.4 | 569.2 | 37.9 |
| >MI0000477\|hsa-mir-146a&WithFlank&5\|+\|160485346\|160485456@27.47.21(+1C)&[MIMAT0000449&hsa-miR-146a-5p&offsets\|0\|-1(+1C);m-129&5\|+\|160485372\|160485393&offsets\|0\|-1(+1C)]\|TGAGAACTGAATTCCATGGGTC | >MI0000477\|hsa-mir-146a&WithFlank&5\|+\|160485346\|160485456 | 27 | 47 | 21(+1C) | TGAGAACTGAATTCCATGGGTC | MB231 | 92 | 189 | 55 | 2096 | 191 | 69.6 | 218.9 | 47.1 | 2057.2 | 149.5 |
| >MI0000477\|hsa-mir-146a&WithFlank&5\|+\|160485346\|160485456@27.47.21(+1G)&[MIMAT0000449&hsa-miR-146a-5p&offsets\|0\|-1(+1G);m-129&5\|+\|160485372\|160485393&offsets\|0\|-1(+1G)]\|TGAGAACTGAATTCCATGGGTG | >MI0000477\|hsa-mir-146a&WithFlank&5\|+\|160485346\|160485456 | 27 | 47 | 21(+1G) | TGAGAACTGAATTCCATGGGTG | MB231 | 46 | 25 | 7 | 168 | 12 | 58.3 | 31.5 | 6.5 | 189.8 | 9.4 |
| >MI0000477\|hsa-mir-146a&WithFlank&5\|+\|160485346\|160485456@27.47.21&[MIMAT0000449&hsa-miR-146a-5p&offsets\|0\|-1;m-129&5\|+\|160485372\|160485393&offsets\|0\|-1]\|TGAGAACTGAATTCCATGGGT | >MI0000477\|hsa-mir-146a&WithFlank&5\|+\|160485346\|160485456 | 27 | 47 | 21 | TGAGAACTGAATTCCATGGGT | MB231 | 117 | 169 | 61 | 1064 | 141 | 96.0 | 172.8 | 47.7 | 883.9 | 103.5 |
| >MI0000477\|hsa-mir-146a&WithFlank&5\|+\|160485346\|160485456@27.48.22(+1A)&[MIMAT0000449&hsa-miR-146a-5p&offsets\|0\|0(+1A);m-129&5\|+\|160485372\|160485393&offsets\|0\|0(+1A)]\|TGAGAACTGAATTCCATGGGTTA | >MI0000477\|hsa-mir-146a&WithFlank&5\|+\|160485346\|160485456 | 27 | 48 | 22(+1A) | TGAGAACTGAATTCCATGGGTTA | MB231 | 20 | 42 | 7 | 234 | 34 | 17.5 | 48.1 | 6.5 | 224.1 | 25.3 |
| >MI0000477\|hsa-mir-146a&WithFlank&5\|+\|160485346\|160485456@27.48.22(+1U)&[MIMAT0000449&hsa-miR-146a-5p&offsets\|0\|0(+1U);m-129&5\|+\|160485372\|160485393&offsets\|0\|0(+1U)]\|TGAGAACTGAATTCCATGGGTTT | >MI0000477\|hsa-mir-146a&WithFlank&5\|+\|160485346\|160485456 | 27 | 48 | 22(+1U) | TGAGAACTGAATTCCATGGGTTT | MB231 | 35 | 82 | 10 | 878 | 78 | 26.1 | 94.8 | 8.5 | 756.4 | 59.4 |
| >MI0000477\|hsa-mir-146a&WithFlank&5\|+\|160485346\|160485456@27.48.22&[MIMAT0000449&hsa-miR-146a-5p&offsets\|0\|0;m-129&5\|+\|160485372\|160485393&offsets\|0\|0]\|TGAGAACTGAATTCCATGGGTT | >MI0000477\|hsa-mir-146a&WithFlank&5\|+\|160485346\|160485456 | 27 | 48 | 22 | TGAGAACTGAATTCCATGGGTT | MB231 | 837 | 1648 | 473 | 11936 | 1369 | 628.3 | 1766.2 | 392.9 | 10350.7 | 1009.7 |
| >MI0000477\|hsa-mir-146a&WithFlank&5\|+\|160485346\|160485456@27.49.23&[MIMAT0000449&hsa-miR-146a-5p&offsets\|0\|+1;m-129&5\|+\|160485372\|160485393&offsets\|0\|+1]\|TGAGAACTGAATTCCATGGGTTG | >MI0000477\|hsa-mir-146a&WithFlank&5\|+\|160485346\|160485456 | 27 | 49 | 23 | TGAGAACTGAATTCCATGGGTTG | MB231 | 72 | 131 | 33 | 583 | 124 | 68.4 | 155.9 | 30.3 | 508.0 | 127.2 |
| >MI0000477\|hsa-mir-146a&WithFlank&5\|+\|160485346\|160485456@27.50.24&[MIMAT0000449&hsa-miR-146a-5p&offsets\|0\|+2;m-129&5\|+\|160485372\|160485393&offsets\|0\|+2]\|TGAGAACTGAATTCCATGGGTTGT | >MI0000477\|hsa-mir-146a&WithFlank&5\|+\|160485346\|160485456 | 27 | 50 | 24 | TGAGAACTGAATTCCATGGGTTGT | MB231 | 24 | 33 | 10 | 146 | 32 | 19.3 | 32.4 | 8.5 | 98.3 | 22.7 |
| >MI0000478\|hsa-mir-149&WithFlank&2\|+\|240455995\|240456095@21.40.20&[MIMAT0000450&hsa-miR-149-5p&offsets\|0\|-3;m-195&2\|+\|240456015\|240456037&offsets\|0\|-3]\|TCTGGCTCCGTGTCTTCACT | >MI0000478\|hsa-mir-149&WithFlank&2\|+\|240455995\|240456095 | 21 | 40 | 20 | TCTGGCTCCGTGTCTTCACT | BT20 | 24 | 10 | 10 | 6 | 6 | 27.1 | 6.5 | 4.9 | 5.2 | 1.0 |
| >MI0000478\|hsa-mir-149&WithFlank&2\|+\|240455995\|240456095@21.41.21(+1A)&[MIMAT0000450&hsa-miR-149-5p&offsets\|0\|-2(+1A);m-195&2\|+\|240456015\|240456037&offsets\|0\|-2(+1A)]\|TCTGGCTCCGTGTCTTCACTCA | >MI0000478\|hsa-mir-149&WithFlank&2\|+\|240455995\|240456095 | 21 | 41 | 21(+1A) | TCTGGCTCCGTGTCTTCACTCA | BT20 | 18 | 12 | 28 | 9 | 8 | 18.0 | 9.5 | 20.9 | 6.5 | 2.3 |
| >MI0000478\|hsa-mir-149&WithFlank&2\|+\|240455995\|240456095@21.41.21(+1U)&[MIMAT0000450&hsa-miR-149-5p&offsets\|0\|-2(+1U);m-195&2\|+\|240456015\|240456037&offsets\|0\|-2(+1U)]\|TCTGGCTCCGTGTCTTCACTCT | >MI0000478\|hsa-mir-149&WithFlank&2\|+\|240455995\|240456095 | 21 | 41 | 21(+1U) | TCTGGCTCCGTGTCTTCACTCT | BT20 | 36 | 31 | 59 | 25 | 22 | 25.5 | 23.2 | 41.1 | 17.7 | 8.0 |
| >MI0000478\|hsa-mir-149&WithFlank&2\|+\|240455995\|240456095@21.41.21&[MIMAT0000450&hsa-miR-149-5p&offsets\|0\|-2;m-195&2\|+\|240456015\|240456037&offsets\|0\|-2]\|TCTGGCTCCGTGTCTTCACTC | >MI0000478\|hsa-mir-149&WithFlank&2\|+\|240455995\|240456095 | 21 | 41 | 21 | TCTGGCTCCGTGTCTTCACTC | BT20 | 32 | 19 | 26 | 7 | 17 | 18.2 | 10.5 | 8.1 | 5.0 | 1.4 |
| >MI0000478\|hsa-mir-149&WithFlank&2\|+\|240455995\|240456095@21.42.22(+1A)&[MIMAT0000450&hsa-miR-149-5p&offsets\|0\|-1(+1A);m-195&2\|+\|240456015\|240456037&offsets\|0\|-1(+1A)]\|TCTGGCTCCGTGTCTTCACTCCA | >MI0000478\|hsa-mir-149&WithFlank&2\|+\|240455995\|240456095 | 21 | 42 | 22(+1A) | TCTGGCTCCGTGTCTTCACTCCA | BT20 | 28 | 23 | 45 | 10 | 16 | 25.3 | 18.0 | 33.3 | 6.2 | 9.8 |
| >MI0000478\|hsa-mir-149&WithFlank&2\|+\|240455995\|240456095@21.42.22(+1U)&[MIMAT0000450&hsa-miR-149-5p&offsets\|0\|-1(+1U);m-195&2\|+\|240456015\|240456037&offsets\|0\|-1(+1U)]\|TCTGGCTCCGTGTCTTCACTCCT | >MI0000478\|hsa-mir-149&WithFlank&2\|+\|240455995\|240456095 | 21 | 42 | 22(+1U) | TCTGGCTCCGTGTCTTCACTCCT | BT20 | 58 | 65 | 88 | 29 | 49 | 38.0 | 52.0 | 56.5 | 18.6 | 27.7 |
| >MI0000478\|hsa-mir-149&WithFlank&2\|+\|240455995\|240456095@21.42.22&[MIMAT0000450&hsa-miR-149-5p&offsets\|0\|-1;m-195&2\|+\|240456015\|240456037&offsets\|0\|-1]\|TCTGGCTCCGTGTCTTCACTCC | >MI0000478\|hsa-mir-149&WithFlank&2\|+\|240455995\|240456095 | 21 | 42 | 22 | TCTGGCTCCGTGTCTTCACTCC | BT20 | 61 | 71 | 109 | 56 | 58 | 30.3 | 50.5 | 40.4 | 23.7 | 19.1 |
| >MI0000478\|hsa-mir-149&WithFlank&2\|+\|240455995\|240456095@21.43.23(+1A)&[MIMAT0000450&hsa-miR-149-5p&offsets\|0\|0(+1A);m-195&2\|+\|240456015\|240456037&offsets\|0\|0(+1A)]\|TCTGGCTCCGTGTCTTCACTCCCA | >MI0000478\|hsa-mir-149&WithFlank&2\|+\|240455995\|240456095 | 21 | 43 | 23(+1A) | TCTGGCTCCGTGTCTTCACTCCCA | BT20 | 10 | 7 | 16 | 3 | 5 | 10.5 | 5.0 | 10.2 | 1.1 | 2.1 |
| >MI0000478\|hsa-mir-149&WithFlank&2\|+\|240455995\|240456095@21.43.23&[MIMAT0000450&hsa-miR-149-5p&offsets\|0\|0;m-195&2\|+\|240456015\|240456037&offsets\|0\|0]\|TCTGGCTCCGTGTCTTCACTCCC | >MI0000478\|hsa-mir-149&WithFlank&2\|+\|240455995\|240456095 | 21 | 43 | 23 | TCTGGCTCCGTGTCTTCACTCCC | BT20 | 82 | 116 | 147 | 56 | 100 | 35.6 | 91.3 | 64.6 | 20.7 | 50.2 |
| >MI0000482\|hsa-mir-185&WithFlank&22\|+\|20033133\|20033226@21.41.21&[MIMAT0000455&hsa-miR-185-5p&offsets\|0\|-1;m-145&22\|+\|20033153\|20033174&offsets\|0\|-1]\|TGGAGAGAAAGGCAGTTCCTG | >MI0000482\|hsa-mir-185&WithFlank&22\|+\|20033133\|20033226 | 21 | 41 | 21 | TGGAGAGAAAGGCAGTTCCTG | BT20 | 11 | 7 | 13 | 24 | 8 | 7.2 | 2.4 | 10.3 | 19.1 | 3.2 |
| >MI0000482\|hsa-mir-185&WithFlank&22\|+\|20033133\|20033226@21.42.22(+1A)&[MIMAT0000455&hsa-miR-185-5p&offsets\|0\|0(+1A);m-145&22\|+\|20033153\|20033174&offsets\|0\|0(+1A)]\|TGGAGAGAAAGGCAGTTCCTGAA | >MI0000482\|hsa-mir-185&WithFlank&22\|+\|20033133\|20033226 | 21 | 42 | 22(+1A) | TGGAGAGAAAGGCAGTTCCTGAA | BT20 | 16 | 15 | 23 | 3 | 10 | 6.1 | 5.8 | 10.5 | 3.0 | 0.2 |
| >MI0000482\|hsa-mir-185&WithFlank&22\|+\|20033133\|20033226@21.42.22&[MIMAT0000455&hsa-miR-185-5p&offsets\|0\|0;m-145&22\|+\|20033153\|20033174&offsets\|0\|0]\|TGGAGAGAAAGGCAGTTCCTGA | >MI0000482\|hsa-mir-185&WithFlank&22\|+\|20033133\|20033226 | 21 | 42 | 22 | TGGAGAGAAAGGCAGTTCCTGA | BT20 | 86 | 82 | 126 | 120 | 69 | 47.7 | 27.7 | 90.8 | 131.6 | 4.7 |

<table>
<tr><td></td><td></td><td></td><td></td><td></td><td></td><td></td><td></td><td></td><td></td><td></td><td></td><td></td><td></td><td></td><td></td><td></td></tr>
<tr><td>>MI0000482|hsa-mir-185&WithFlank&22|+|20033133|20033226@21.42.22&[MIMAT0000455&hsa-miR-185-5p&offsets|0|0;m-145&22|+|20033153|20033174&offsets|0|0]|TGGAGAGAAAGGCAGTTCCTGA</td><td>>MI0000482|hsa-mir-185&WithFlank&22|+|20033133|20033226</td><td>21</td><td>42</td><td>22</td><td>TGGAGAGAAAGGCAGTTCCTGA</td><td>MB231</td><td>38</td><td>35</td><td>8</td><td>91</td><td>32</td><td>26.7</td><td>2.9</td><td>3.1</td><td>17.6</td><td>7.8</td></tr>
<tr><td>>MI0000482|hsa-mir-185&WithFlank&22|+|20033133|20033226@21.43.23&[MIMAT0000455&hsa-miR-185-5p&offsets|0|+1;m-145&22|+|20033153|20033174&offsets|0|+1]|TGGAGAGAAAGGCAGTTCCTGAT</td><td>>MI0000482|hsa-mir-185&WithFlank&22|+|20033133|20033226</td><td>21</td><td>43</td><td>23</td><td>TGGAGAGAAAGGCAGTTCCTGAT</td><td>BT20</td><td>22</td><td>18</td><td>29</td><td>2</td><td>15</td><td>9.4</td><td>6.4</td><td>17.7</td><td>1.3</td><td>3.7</td></tr>
<tr><td>>MI0000483|hsa-mir-186&WithFlank&1|-|71067625|71067722@21.40.20&[MIMAT0000456&hsa-miR-186-5p&offsets|0|-2;m-84&1|-|71067681|71067702&offsets|0|-2]|CAAAGAATTCTCCTTTTGGG</td><td>>MI0000483|hsa-mir-186&WithFlank&1|-|71067625|71067722</td><td>21</td><td>40</td><td>20</td><td>CAAAGAATTCTCCTTTTGGG</td><td>BT20</td><td>10</td><td>12</td><td>13</td><td>10</td><td>6</td><td>3.8</td><td>5.3</td><td>8.9</td><td>5.9</td><td>2.1</td></tr>
<tr><td>>MI0000483|hsa-mir-186&WithFlank&1|-|71067625|71067722@21.40.20&[MIMAT0000456&hsa-miR-186-5p&offsets|0|-2;m-84&1|-|71067681|71067702&offsets|0|-2]|CAAAGAATTCTCCTTTTGGG</td><td>>MI0000483|hsa-mir-186&WithFlank&1|-|71067625|71067722</td><td>21</td><td>40</td><td>20</td><td>CAAAGAATTCTCCTTTTGGG</td><td>MB231</td><td>12</td><td>17</td><td>3</td><td>30</td><td>12</td><td>4.4</td><td>5.5</td><td>1.2</td><td>7.8</td><td>2.4</td></tr>
<tr><td>>MI0000483|hsa-mir-186&WithFlank&1|-|71067625|71067722@21.41.21(+1A)&[MIMAT0000456&hsa-miR-186-5p&offsets|0|-1(+1A);m-84&1|-|71067681|71067702&offsets|0|-1(+1A)]|CAAAGAATTCTCCTTTTGGGCA</td><td>>MI0000483|hsa-mir-186&WithFlank&1|-|71067625|71067722</td><td>21</td><td>41</td><td>21(+1A)</td><td>CAAAGAATTCTCCTTTTGGGCA</td><td>BT20</td><td>25</td><td>14</td><td>15</td><td>5</td><td>5</td><td>24.1</td><td>8.0</td><td>11.0</td><td>3.1</td><td>0.6</td></tr>
<tr><td>>MI0000483|hsa-mir-186&WithFlank&1|-|71067625|71067722@21.41.21(+1C)&[MIMAT0000456&hsa-miR-186-5p&offsets|0|-1(+1C);m-84&1|-|71067681|71067702&offsets|0|-1(+1C)]|CAAAGAATTCTCCTTTTGGGCC</td><td>>MI0000483|hsa-mir-186&WithFlank&1|-|71067625|71067722</td><td>21</td><td>41</td><td>21(+1C)</td><td>CAAAGAATTCTCCTTTTGGGCC</td><td>BT20</td><td>18</td><td>16</td><td>12</td><td>5</td><td>7</td><td>9.7</td><td>9.4</td><td>7.6</td><td>3.8</td><td>1.1</td></tr>
<tr><td>>MI0000483|hsa-mir-186&WithFlank&1|-|71067625|71067722@21.41.21(+1G)&[MIMAT0000456&hsa-miR-186-5p&offsets|0|-1(+1G);m-84&1|-|71067681|71067702&offsets|0|-1(+1G)]|CAAAGAATTCTCCTTTTGGGCG</td><td>>MI0000483|hsa-mir-186&WithFlank&1|-|71067625|71067722</td><td>21</td><td>41</td><td>21(+1G)</td><td>CAAAGAATTCTCCTTTTGGGCG</td><td>BT20</td><td>15</td><td>4</td><td>3</td><td>1</td><td>0</td><td>17.1</td><td>1.9</td><td>2.4</td><td>1.0</td><td>0.4</td></tr>
<tr><td>>MI0000483|hsa-mir-186&WithFlank&1|-|71067625|71067722@21.41.21&[MIMAT0000456&hsa-miR-186-5p&offsets|0|-1;m-84&1|-|71067681|71067702&offsets|0|-1]|CAAAGAATTCTCCTTTTGGGC</td><td>>MI0000483|hsa-mir-186&WithFlank&1|-|71067625|71067722</td><td>21</td><td>41</td><td>21</td><td>CAAAGAATTCTCCTTTTGGGC</td><td>BT20</td><td>21</td><td>18</td><td>26</td><td>9</td><td>6</td><td>9.5</td><td>8.7</td><td>19.4</td><td>2.7</td><td>1.1</td></tr>
<tr><td>>MI0000483|hsa-mir-186&WithFlank&1|-|71067625|71067722@21.42.22(+1A)&[MIMAT0000456&hsa-miR-186-5p&offsets|0|0(+1A);m-84&1|-|71067681|71067702&offsets|0|0(+1A)]|CAAAGAATTCTCCTTTTGGGCTA</td><td>>MI0000483|hsa-mir-186&WithFlank&1|-|71067625|71067722</td><td>21</td><td>42</td><td>22(+1A)</td><td>CAAAGAATTCTCCTTTTGGGCTA</td><td>BT20</td><td>10</td><td>5</td><td>3</td><td>2</td><td>1</td><td>13.1</td><td>3.9</td><td>1.4</td><td>1.6</td><td>0.9</td></tr>
<tr><td>>MI0000483|hsa-mir-186&WithFlank&1|-|71067625|71067722@21.42.22(+1C)&[MIMAT0000456&hsa-miR-186-5p&offsets|0|0(+1C);m-84&1|-|71067681|71067702&offsets|0|0(+1C)]|CAAAGAATTCTCCTTTTGGGCTC</td><td>>MI0000483|hsa-mir-186&WithFlank&1|-|71067625|71067722</td><td>21</td><td>42</td><td>22(+1C)</td><td>CAAAGAATTCTCCTTTTGGGCTC</td><td>BT20</td><td>12</td><td>10</td><td>4</td><td>5</td><td>2</td><td>11.3</td><td>9.5</td><td>1.5</td><td>3.7</td><td>1.4</td></tr>
<tr><td>>MI0000483|hsa-mir-186&WithFlank&1|-|71067625|71067722@21.42.22&[MIMAT0000456&hsa-miR-186-5p&offsets|0|0;m-84&1|-|71067681|71067702&offsets|0|0]|CAAAGAATTCTCCTTTTGGGCT</td><td>>MI0000483|hsa-mir-186&WithFlank&1|-|71067625|71067722</td><td>21</td><td>42</td><td>22</td><td>CAAAGAATTCTCCTTTTGGGCT</td><td>BT20</td><td>122</td><td>105</td><td>80</td><td>41</td><td>47</td><td>65.5</td><td>48.8</td><td>41.4</td><td>22.2</td><td>10.2</td></tr>
<tr><td>>MI0000483|hsa-mir-186&WithFlank&1|-|71067625|71067722@21.42.22&[MIMAT0000456&hsa-miR-186-5p&offsets|0|0;m-84&1|-|71067681|71067702&offsets|0|0]|CAAAGAATTCTCCTTTTGGGCT</td><td>>MI0000483|hsa-mir-186&WithFlank&1|-|71067625|71067722</td><td>21</td><td>42</td><td>22</td><td>CAAAGAATTCTCCTTTTGGGCT</td><td>MB231</td><td>46</td><td>57</td><td>11</td><td>140</td><td>81</td><td>15.3</td><td>33.8</td><td>8.0</td><td>28.3</td><td>45.6</td></tr>
<tr><td>>MI0000483|hsa-mir-186&WithFlank&1|-|71067625|71067722@21.43.23&[MIMAT0000456&hsa-miR-186-5p&offsets|0|+1;m-84&1|-|71067681|71067702&offsets|0|+1]|CAAAGAATTCTCCTTTTGGGCTT</td><td>>MI0000483|hsa-mir-186&WithFlank&1|-|71067625|71067722</td><td>21</td><td>43</td><td>23</td><td>CAAAGAATTCTCCTTTTGGGCTT</td><td>BT20</td><td>79</td><td>70</td><td>28</td><td>31</td><td>15</td><td>73.6</td><td>62.2</td><td>13.8</td><td>21.0</td><td>8.3</td></tr>
<tr><td>>MI0000483|hsa-mir-186&WithFlank&1|-|71067625|71067722@21.43.23&[MIMAT0000456&hsa-miR-186-5p&offsets|0|+1;m-84&1|-|71067681|71067702&offsets|0|+1]|CAAAGAATTCTCCTTTTGGGCTT</td><td>>MI0000483|hsa-mir-186&WithFlank&1|-|71067625|71067722</td><td>21</td><td>43</td><td>23</td><td>CAAAGAATTCTCCTTTTGGGCTT</td><td>MB231</td><td>20</td><td>24</td><td>4</td><td>156</td><td>44</td><td>6.2</td><td>12.6</td><td>3.3</td><td>46.7</td><td>34.8</td></tr>
<tr><td>>MI0000483|hsa-mir-186&WithFlank&1|-|71067625|71067722@21.44.24&[MIMAT0000456&hsa-miR-186-5p&offsets|0|+2;m-84&1|-|71067681|71067702&offsets|0|+2]|CAAAGAATTCTCCTTTTGGGCTTT</td><td>>MI0000483|hsa-mir-186&WithFlank&1|-|71067625|71067722</td><td>21</td><td>44</td><td>24</td><td>CAAAGAATTCTCCTTTTGGGCTTT</td><td>BT20</td><td>16</td><td>15</td><td>7</td><td>17</td><td>2</td><td>15.3</td><td>16.1</td><td>4.2</td><td>17.1</td><td>2.7</td></tr>
<tr><td>>MI0000483|hsa-mir-186&WithFlank&1|-|71067625|71067722@21.44.24&[MIMAT0000456&hsa-miR-186-5p&offsets|0|+2;m-84&1|-|71067681|71067702&offsets|0|+2]|CAAAGAATTCTCCTTTTGGGCTTT</td><td>>MI0000483|hsa-mir-186&WithFlank&1|-|71067625|71067722</td><td>21</td><td>44</td><td>24</td><td>CAAAGAATTCTCCTTTTGGGCTTT</td><td>MB231</td><td>12</td><td>15</td><td>2</td><td>186</td><td>28</td><td>1.9</td><td>7.2</td><td>1.6</td><td>77.4</td><td>29.0</td></tr>
<tr><td>>MI0000487|hsa-mir-193a&WithFlank&17|+|31559990|31560089@27.47.21&[MIMAT0004614&hsa-miR-193a-5p&offsets|0|-1;m-123&17|+|31560016|31560037&offsets|0|-1]|TGGGTCTTTGCGGGCGAGATG</td><td>>MI0000487|hsa-mir-193a&WithFlank&17|+|31559990|31560089</td><td>27</td><td>47</td><td>21</td><td>TGGGTCTTTGCGGGCGAGATG</td><td>BT20</td><td>14</td><td>11</td><td>24</td><td>82</td><td>11</td><td>12.6</td><td>7.3</td><td>12.5</td><td>100.3</td><td>5.1</td></tr>
<tr><td>>MI0000487|hsa-mir-193a&WithFlank&17|+|31559990|31560089@27.48.22&[MIMAT0004614&hsa-miR-193a-5p&offsets|0|0;m-123&17|+|31560016|31560037&offsets|0|0]|TGGGTCTTTGCGGGCGAGATGA</td><td>>MI0000487|hsa-mir-193a&WithFlank&17|+|31559990|31560089</td><td>27</td><td>48</td><td>22</td><td>TGGGTCTTTGCGGGCGAGATGA</td><td>BT20</td><td>22</td><td>24</td><td>29</td><td>82</td><td>18</td><td>22.4</td><td>25.2</td><td>22.0</td><td>96.6</td><td>9.7</td></tr>
<tr><td>>MI0000488|hsa-mir-194-1&WithFlank&1|-|220118151|220118247@21.42.22;MI0000732|hsa-mir-194-2&WithFlank&11|-|64891349|64891445@21.42.22&[MIMAT0000460&hsa-miR-194-5p&offsets|0|0;m-131&1|-|220118206|220118227&offsets|0|0];[MIMAT0000460_1&hsa-miR-194-5p&offsets|0|0;m-126&11|-|64891404|64891425&offsets|0|0]|TGTAACAGCAACTCCATGTGGA</td><td>>MI0000488|hsa-mir-194-1&WithFlank&1|-|220118151|220118247</td><td>21</td><td>42</td><td>22</td><td>TGTAACAGCAACTCCATGTGGA</td><td>BT20</td><td>33</td><td>22</td><td>23</td><td>22</td><td>5</td><td>22.4</td><td>23.2</td><td>7.7</td><td>18.3</td><td>4.8</td></tr>
<tr><td>>MI0000489|hsa-mir-195&WithFlank&17|-|7017609|7017707@21.42.22&[MIMAT0000461&hsa-miR-195-5p&offsets|0|+1;m-191&17|-|7017666|7017687&offsets|0|0]|TAGCAGCACAGAAATATTGGCA</td><td>>MI0000489|hsa-mir-195&WithFlank&17|-|7017609|7017707</td><td>21</td><td>42</td><td>22</td><td>TAGCAGCACAGAAATATTGGCA</td><td>BT20</td><td>23</td><td>21</td><td>8</td><td>3</td><td>1</td><td>23.9</td><td>22.9</td><td>5.1</td><td>2.8</td><td>1.1</td></tr>
<tr><td>>MI0000542|hsa-mir-320a&WithFlank&8|-|22244960|22245043@48.68.21(+1C)&[MIMAT0000510&hsa-miR-320a-3p&offsets|0|-1(+1C);m-60&8|-|22244975|22244996&offsets|0|-1(+1C)]|AAAAGCTGGGTTGAGAGGGCGC</td><td>>MI0000542|hsa-mir-320a&WithFlank&8|-|22244960|22245043</td><td>48</td><td>68</td><td>21(+1C)</td><td>AAAAGCTGGGTTGAGAGGGCGC</td><td>BT20</td><td>37</td><td>30</td><td>203</td><td>392</td><td>12</td><td>16.2</td><td>21.1</td><td>49.6</td><td>279.5</td><td>7.4</td></tr>
<tr><td>>MI0000542|hsa-mir-320a&WithFlank&8|-|22244960|22245043@48.68.21(+1G)&[MIMAT0000510&hsa-miR-320a-3p&offsets|0|-1(+1G);m-60&8|-|22244975|22244996&offsets|0|-1(+1G)]|AAAAGCTGGGTTGAGAGGGCGG</td><td>>MI0000542|hsa-mir-320a&WithFlank&8|-|22244960|22245043</td><td>48</td><td>68</td><td>21(+1G)</td><td>AAAAGCTGGGTTGAGAGGGCGG</td><td>BT20</td><td>69</td><td>23</td><td>118</td><td>145</td><td>4</td><td>64.5</td><td>15.0</td><td>55.9</td><td>114.9</td><td>1.3</td></tr>
<tr><td>>MI0000542|hsa-mir-320a&WithFlank&8|-|22244960|22245043@48.68.21(+1U)&[MIMAT0000510&hsa-miR-320a-3p&offsets|0|-1(+1U);m-60&8|-|22244975|22244996&offsets|0|-1(+1U)]|AAAAGCTGGGTTGAGAGGGCGT</td><td>>MI0000542|hsa-mir-320a&WithFlank&8|-|22244960|22245043</td><td>48</td><td>68</td><td>21(+1U)</td><td>AAAAGCTGGGTTGAGAGGGCGT</td><td>BT20</td><td>289</td><td>197</td><td>1530</td><td>2863</td><td>83</td><td>136.3</td><td>112.6</td><td>382.7</td><td>2155.1</td><td>44.9</td></tr>
<tr><td>>MI0000542|hsa-mir-320a&WithFlank&8|-|22244960|22245043@48.68.21(+1U)&[MIMAT0000510&hsa-miR-320a-3p&offsets|0|-1(+1U);m-60&8|-|22244975|22244996&offsets|0|-1(+1U)]|AAAAGCTGGGTTGAGAGGGCGT</td><td>>MI0000542|hsa-mir-320a&WithFlank&8|-|22244960|22245043</td><td>48</td><td>68</td><td>21(+1U)</td><td>AAAAGCTGGGTTGAGAGGGCGT</td><td>MB231</td><td>58</td><td>75</td><td>75</td><td>416</td><td>23</td><td>16.5</td><td>38.5</td><td>41.0</td><td>192.9</td><td>8.9</td></tr>
<tr><td>>MI0000542|hsa-mir-320a&WithFlank&8|-|22244960|22245043@48.68.21(+2U)&[MIMAT0000510&hsa-miR-320a-3p&offsets|0|-1(+2U);m-60&8|-|22244975|22244996&offsets|0|-1(+2U)]|AAAAGCTGGGTTGAGAGGGCGTT</td><td>>MI0000542|hsa-mir-320a&WithFlank&8|-|22244960|22245043</td><td>48</td><td>68</td><td>21(+2U)</td><td>AAAAGCTGGGTTGAGAGGGCGTT</td><td>BT20</td><td>167</td><td>118</td><td>644</td><td>401</td><td>67</td><td>80.0</td><td>56.8</td><td>157.9</td><td>237.1</td><td>21.8</td></tr>
<tr><td>>MI0000542|hsa-mir-320a&WithFlank&8|-|22244960|22245043@48.68.21(+2U)&[MIMAT0000510&hsa-miR-320a-3p&offsets|0|-1(+2U);m-60&8|-|22244975|22244996&offsets|0|-1(+2U)]|AAAAGCTGGGTTGAGAGGGCGTT</td><td>>MI0000542|hsa-mir-320a&WithFlank&8|-|22244960|22245043</td><td>48</td><td>68</td><td>21(+2U)</td><td>AAAAGCTGGGTTGAGAGGGCGTT</td><td>MB231</td><td>24</td><td>25</td><td>26</td><td>79</td><td>13</td><td>7.1</td><td>13.6</td><td>14.2</td><td>44.4</td><td>5.4</td></tr>
<tr><td>>MI0000542|hsa-mir-320a&WithFlank&8|-|22244960|22245043@48.68.21(+3U)&[MIMAT0000510&hsa-miR-320a-3p&offsets|0|-1(+3U);m-60&8|-|22244975|22244996&offsets|0|-1(+3U)]|AAAAGCTGGGTTGAGAGGGCGTTT</td><td>>MI0000542|hsa-mir-320a&WithFlank&8|-|22244960|22245043</td><td>48</td><td>68</td><td>21(+3U)</td><td>AAAAGCTGGGTTGAGAGGGCGTTT</td><td>BT20</td><td>45</td><td>33</td><td>300</td><td>109</td><td>13</td><td>22.0</td><td>26.3</td><td>209.6</td><td>97.3</td><td>9.7</td></tr>
<tr><td>>MI0000542|hsa-mir-320a&WithFlank&8|-|22244960|22245043@48.68.21&[MIMAT0000510&hsa-miR-320a-3p&offsets|0|-1;m-60&8|-|22244975|22244996&offsets|0|-1]|AAAAGCTGGGTTGAGAGGGCG</td><td>>MI0000542|hsa-mir-320a&WithFlank&8|-|22244960|22245043</td><td>48</td><td>68</td><td>21</td><td>AAAAGCTGGGTTGAGAGGGCG</td><td>BT20</td><td>238</td><td>190</td><td>868</td><td>1574</td><td>136</td><td>122.3</td><td>55.9</td><td>130.3</td><td>997.9</td><td>17.7</td></tr>
<tr><td>>MI0000542|hsa-mir-320a&WithFlank&8|-|22244960|22245043@48.68.21&[MIMAT0000510&hsa-miR-320a-3p&offsets|0|-1;m-60&8|-|22244975|22244996&offsets|0|-1]|AAAAGCTGGGTTGAGAGGGCG</td><td>>MI0000542|hsa-mir-320a&WithFlank&8|-|22244960|22245043</td><td>48</td><td>68</td><td>21</td><td>AAAAGCTGGGTTGAGAGGGCG</td><td>MB231</td><td>34</td><td>49</td><td>46</td><td>157</td><td>39</td><td>8.1</td><td>20.1</td><td>26.7</td><td>73.9</td><td>9.9</td></tr>
<tr><td>>MI0000542|hsa-mir-320a&WithFlank&8|-|22244960|22245043@48.69.22(+1C)&[MIMAT0000510&hsa-miR-320a-3p&offsets|0|0(+1C);m-60&8|-|22244975|22244996&offsets|0|0(+1C)]|AAAAGCTGGGTTGAGAGGGCGAC</td><td>>MI0000542|hsa-mir-320a&WithFlank&8|-|22244960|22245043</td><td>48</td><td>69</td><td>22(+1C)</td><td>AAAAGCTGGGTTGAGAGGGCGAC</td><td>BT20</td><td>69</td><td>58</td><td>218</td><td>100</td><td>20</td><td>33.4</td><td>48.6</td><td>66.0</td><td>54.6</td><td>10.4</td></tr>
<tr><td>>MI0000542|hsa-mir-320a&WithFlank&8|-|22244960|22245043@48.69.22(+1C)&[MIMAT0000510&hsa-miR-320a-3p&offsets|0|0(+1C);m-60&8|-|22244975|22244996&offsets|0|0(+1C)]|AAAAGCTGGGTTGAGAGGGCGAC</td><td>>MI0000542|hsa-mir-320a&WithFlank&8|-|22244960|22245043</td><td>48</td><td>69</td><td>22(+1C)</td><td>AAAAGCTGGGTTGAGAGGGCGAC</td><td>MB231</td><td>12</td><td>15</td><td>9</td><td>36</td><td>5</td><td>6.1</td><td>10.5</td><td>6.4</td><td>24.2</td><td>3.7</td></tr>
<tr><td>>MI0000542|hsa-mir-320a&WithFlank&8|-|22244960|22245043@48.69.22(+1G)&[MIMAT0000510&hsa-miR-320a-3p&offsets|0|0(+1G);m-60&8|-|22244975|22244996&offsets|0|0(+1G)]|AAAAGCTGGGTTGAGAGGGCGAG</td><td>>MI0000542|hsa-mir-320a&WithFlank&8|-|22244960|22245043</td><td>48</td><td>69</td><td>22(+1G)</td><td>AAAAGCTGGGTTGAGAGGGCGAG</td><td>BT20</td><td>61</td><td>23</td><td>112</td><td>51</td><td>8</td><td>46.1</td><td>13.5</td><td>40.7</td><td>29.2</td><td>2.6</td></tr>
<tr><td>>MI0000542|hsa-mir-320a&WithFlank&8|-|22244960|22245043@48.69.22(+1U)&[MIMAT0000510&hsa-miR-320a-3p&offsets|0|0(+1U);m-60&8|-|22244975|22244996&offsets|0|0(+1U)]|AAAAGCTGGGTTGAGAGGGCGAT</td><td>>MI0000542|hsa-mir-320a&WithFlank&8|-|22244960|22245043</td><td>48</td><td>69</td><td>22(+1U)</td><td>AAAAGCTGGGTTGAGAGGGCGAT</td><td>BT20</td><td>542</td><td>406</td><td>1736</td><td>727</td><td>158</td><td>283.3</td><td>314.0</td><td>448.2</td><td>446.4</td><td>84.6</td></tr>
</table>

<table>
<tr><td></td><td></td><td></td><td></td><td></td><td></td><td></td><td></td><td></td><td></td><td></td><td></td><td></td><td></td><td></td><td></td><td></td><td></td></tr>
<tr><td>>MI0000542|hsa-mir-320a&WithFlank&8|-|22244960|22245043@48.69.22(+1U)&[MIMAT0000510&hsa-miR-320a-3p&offsets|0|0(+1U);m-60&8|-|22244975|22244996&offsets|0|0(+1U)]|AAAAGCTGGGTTGAGAGGGCGAT</td><td>>MI0000542|hsa-mir-320a&WithFlank&8|-|22244960|22245043</td><td>48</td><td>69</td><td>22(+1U)</td><td>AAAAGCTGGGTTGAGAGGGCGAT</td><td>MB231</td><td>115</td><td>167</td><td>92</td><td>220</td><td>46</td><td>57.2</td><td>137.0</td><td>65.7</td><td>129.7</td><td>27.1</td><td></td></tr>
<tr><td>>MI0000542|hsa-mir-320a&WithFlank&8|-|22244960|22245043@48.69.22(+2U)&[MIMAT0000510&hsa-miR-320a-3p&offsets|0|0(+2U);m-60&8|-|22244975|22244996&offsets|0|0(+2U)]|AAAAGCTGGGTTGAGAGGGCGATT</td><td>>MI0000542|hsa-mir-320a&WithFlank&8|-|22244960|22245043</td><td>48</td><td>69</td><td>22(+2U)</td><td>AAAAGCTGGGTTGAGAGGGCGATT</td><td>BT20</td><td>203</td><td>130</td><td>1007</td><td>152</td><td>28</td><td>107.8</td><td>110.7</td><td>526.6</td><td>126.0</td><td>23.0</td><td></td></tr>
<tr><td>>MI0000542|hsa-mir-320a&WithFlank&8|-|22244960|22245043@48.69.22(+2U)&[MIMAT0000510&hsa-miR-320a-3p&offsets|0|0(+2U);m-60&8|-|22244975|22244996&offsets|0|0(+2U)]|AAAAGCTGGGTTGAGAGGGCGATT</td><td>>MI0000542|hsa-mir-320a&WithFlank&8|-|22244960|22245043</td><td>48</td><td>69</td><td>22(+2U)</td><td>AAAAGCTGGGTTGAGAGGGCGATT</td><td>MB231</td><td>55</td><td>54</td><td>56</td><td>83</td><td>13</td><td>18.9</td><td>30.5</td><td>32.5</td><td>50.2</td><td>9.0</td><td></td></tr>
<tr><td>>MI0000542|hsa-mir-320a&WithFlank&8|-|22244960|22245043@48.69.22(+3U)&[MIMAT0000510&hsa-miR-320a-3p&offsets|0|0(+3U);m-60&8|-|22244975|22244996&offsets|0|0(+3U)]|AAAAGCTGGGTTGAGAGGGCGATTT</td><td>>MI0000542|hsa-mir-320a&WithFlank&8|-|22244960|22245043</td><td>48</td><td>69</td><td>22(+3U)</td><td>AAAAGCTGGGTTGAGAGGGCGATTT</td><td>BT20</td><td>16</td><td>11</td><td>40</td><td>12</td><td>4</td><td>8.6</td><td>9.6</td><td>13.8</td><td>9.4</td><td>5.2</td><td></td></tr>
<tr><td>>MI0000542|hsa-mir-320a&WithFlank&8|-|22244960|22245043@48.69.22&[MIMAT0000510&hsa-miR-320a-3p&offsets|0|0;m-60&8|-|22244975|22244996&offsets|0|0]|AAAAGCTGGGTTGAGAGGGCGA</td><td>>MI0000542|hsa-mir-320a&WithFlank&8|-|22244960|22245043</td><td>48</td><td>69</td><td>22</td><td>AAAAGCTGGGTTGAGAGGGCGA</td><td>BT20</td><td>861</td><td>654</td><td>2923</td><td>6128</td><td>295</td><td>475.0</td><td>362.6</td><td>329.1</td><td>3892.0</td><td>71.6</td><td></td></tr>
<tr><td>>MI0000542|hsa-mir-320a&WithFlank&8|-|22244960|22245043@48.69.22&[MIMAT0000510&hsa-miR-320a-3p&offsets|0|0;m-60&8|-|22244975|22244996&offsets|0|0]|AAAAGCTGGGTTGAGAGGGCGA</td><td>>MI0000542|hsa-mir-320a&WithFlank&8|-|22244960|22245043</td><td>48</td><td>69</td><td>22</td><td>AAAAGCTGGGTTGAGAGGGCGA</td><td>MB231</td><td>109</td><td>172</td><td>103</td><td>766</td><td>61</td><td>28.9</td><td>108.2</td><td>78.3</td><td>363.6</td><td>22.0</td><td></td></tr>
<tr><td>>MI0000542|hsa-mir-320a&WithFlank&8|-|22244960|22245043@48.70.23(+1C)&[MIMAT0000510&hsa-miR-320a-3p&offsets|0|+1(+1C);m-60&8|-|22244975|22244996&offsets|0|+1(+1C)]|AAAAGCTGGGTTGAGAGGGCGAAC</td><td>>MI0000542|hsa-mir-320a&WithFlank&8|-|22244960|22245043</td><td>48</td><td>70</td><td>23(+1C)</td><td>AAAAGCTGGGTTGAGAGGGCGAAC</td><td>BT20</td><td>47</td><td>33</td><td>173</td><td>92</td><td>21</td><td>24.7</td><td>21.5</td><td>42.6</td><td>44.8</td><td>6.5</td><td></td></tr>
<tr><td>>MI0000542|hsa-mir-320a&WithFlank&8|-|22244960|22245043@48.70.23(+1C)&[MIMAT0000510&hsa-miR-320a-3p&offsets|0|+1(+1C);m-60&8|-|22244975|22244996&offsets|0|+1(+1C)]|AAAAGCTGGGTTGAGAGGGCGAAC</td><td>>MI0000542|hsa-mir-320a&WithFlank&8|-|22244960|22245043</td><td>48</td><td>70</td><td>23(+1C)</td><td>AAAAGCTGGGTTGAGAGGGCGAAC</td><td>MB231</td><td>10</td><td>8</td><td>12</td><td>33</td><td>5</td><td>2.5</td><td>3.3</td><td>6.6</td><td>18.2</td><td>2.7</td><td></td></tr>
<tr><td>>MI0000542|hsa-mir-320a&WithFlank&8|-|22244960|22245043@48.70.23(+1G)&[MIMAT0000510&hsa-miR-320a-3p&offsets|0|+1(+1G);m-60&8|-|22244975|22244996&offsets|0|+1(+1G)]|AAAAGCTGGGTTGAGAGGGCGAAG</td><td>>MI0000542|hsa-mir-320a&WithFlank&8|-|22244960|22245043</td><td>48</td><td>70</td><td>23(+1G)</td><td>AAAAGCTGGGTTGAGAGGGCGAAG</td><td>BT20</td><td>43</td><td>24</td><td>108</td><td>73</td><td>12</td><td>23.4</td><td>11.4</td><td>26.8</td><td>29.9</td><td>2.4</td><td></td></tr>
<tr><td>>MI0000542|hsa-mir-320a&WithFlank&8|-|22244960|22245043@48.70.23(+1U)&[MIMAT0000510&hsa-miR-320a-3p&offsets|0|+1(+1U);m-60&8|-|22244975|22244996&offsets|0|+1(+1U)]|AAAAGCTGGGTTGAGAGGGCGAAT</td><td>>MI0000542|hsa-mir-320a&WithFlank&8|-|22244960|22245043</td><td>48</td><td>70</td><td>23(+1U)</td><td>AAAAGCTGGGTTGAGAGGGCGAAT</td><td>BT20</td><td>353</td><td>224</td><td>1230</td><td>628</td><td>142</td><td>205.1</td><td>142.8</td><td>133.7</td><td>306.3</td><td>57.8</td><td></td></tr>
<tr><td>>MI0000542|hsa-mir-320a&WithFlank&8|-|22244960|22245043@48.70.23(+1U)&[MIMAT0000510&hsa-miR-320a-3p&offsets|0|+1(+1U);m-60&8|-|22244975|22244996&offsets|0|+1(+1U)]|AAAAGCTGGGTTGAGAGGGCGAAT</td><td>>MI0000542|hsa-mir-320a&WithFlank&8|-|22244960|22245043</td><td>48</td><td>70</td><td>23(+1U)</td><td>AAAAGCTGGGTTGAGAGGGCGAAT</td><td>MB231</td><td>107</td><td>91</td><td>121</td><td>214</td><td>41</td><td>19.7</td><td>47.8</td><td>66.3</td><td>87.6</td><td>18.0</td><td></td></tr>
<tr><td>>MI0000542|hsa-mir-320a&WithFlank&8|-|22244960|22245043@48.70.23(+2U)&[MIMAT0000510&hsa-miR-320a-3p&offsets|0|+1(+2U);m-60&8|-|22244975|22244996&offsets|0|+1(+2U)]|AAAAGCTGGGTTGAGAGGGCGAATT</td><td>>MI0000542|hsa-mir-320a&WithFlank&8|-|22244960|22245043</td><td>48</td><td>70</td><td>23(+2U)</td><td>AAAAGCTGGGTTGAGAGGGCGAATT</td><td>BT20</td><td>32</td><td>21</td><td>82</td><td>40</td><td>8</td><td>17.8</td><td>16.8</td><td>2.9</td><td>26.3</td><td>5.8</td><td></td></tr>
<tr><td>>MI0000542|hsa-mir-320a&WithFlank&8|-|22244960|22245043@48.70.23(+2U)&[MIMAT0000510&hsa-miR-320a-3p&offsets|0|+1(+2U);m-60&8|-|22244975|22244996&offsets|0|+1(+2U)]|AAAAGCTGGGTTGAGAGGGCGAATT</td><td>>MI0000542|hsa-mir-320a&WithFlank&8|-|22244960|22245043</td><td>48</td><td>70</td><td>23(+2U)</td><td>AAAAGCTGGGTTGAGAGGGCGAATT</td><td>MB231</td><td>12</td><td>12</td><td>8</td><td>19</td><td>4</td><td>2.4</td><td>8.5</td><td>3.5</td><td>9.9</td><td>1.9</td><td></td></tr>
<tr><td>>MI0000542|hsa-mir-320a&WithFlank&8|-|22244960|22245043@48.70.23&[MIMAT0000510&hsa-miR-320a-3p&offsets|0|+1;m-60&8|-|22244975|22244996&offsets|0|+1]|AAAAGCTGGGTTGAGAGGGCGAA</td><td>>MI0000542|hsa-mir-320a&WithFlank&8|-|22244960|22245043</td><td>48</td><td>70</td><td>23</td><td>AAAAGCTGGGTTGAGAGGGCGAA</td><td>BT20</td><td>456</td><td>348</td><td>1435</td><td>865</td><td>175</td><td>262.5</td><td>158.1</td><td>275.1</td><td>325.4</td><td>23.5</td><td></td></tr>
<tr><td>>MI0000542|hsa-mir-320a&WithFlank&8|-|22244960|22245043@48.70.23&[MIMAT0000510&hsa-miR-320a-3p&offsets|0|+1;m-60&8|-|22244975|22244996&offsets|0|+1]|AAAAGCTGGGTTGAGAGGGCGAA</td><td>>MI0000542|hsa-mir-320a&WithFlank&8|-|22244960|22245043</td><td>48</td><td>70</td><td>23</td><td>AAAAGCTGGGTTGAGAGGGCGAA</td><td>MB231</td><td>78</td><td>134</td><td>73</td><td>199</td><td>39</td><td>15.8</td><td>73.9</td><td>43.0</td><td>110.6</td><td>13.3</td><td></td></tr>
<tr><td>>MI0000542|hsa-mir-320a&WithFlank&8|-|22244960|22245043@48.71.24(+1U)&[MIMAT0000510&hsa-miR-320a-3p&offsets|0|+2(+1U);m-60&8|-|22244975|22244996&offsets|0|+2(+1U)]|AAAAGCTGGGTTGAGAGGGCGAAAT</td><td>>MI0000542|hsa-mir-320a&WithFlank&8|-|22244960|22245043</td><td>48</td><td>71</td><td>24(+1U)</td><td>AAAAGCTGGGTTGAGAGGGCGAAAT</td><td>BT20</td><td>40</td><td>26</td><td>95</td><td>79</td><td>15</td><td>22.8</td><td>19.8</td><td>17.1</td><td>34.4</td><td>7.2</td><td></td></tr>
<tr><td>>MI0000542|hsa-mir-320a&WithFlank&8|-|22244960|22245043@48.71.24&[MIMAT0000510&hsa-miR-320a-3p&offsets|0|+2;m-60&8|-|22244975|22244996&offsets|0|+2]|AAAAGCTGGGTTGAGAGGGCGAAA</td><td>>MI0000542|hsa-mir-320a&WithFlank&8|-|22244960|22245043</td><td>48</td><td>71</td><td>24</td><td>AAAAGCTGGGTTGAGAGGGCGAAA</td><td>BT20</td><td>148</td><td>106</td><td>592</td><td>191</td><td>45</td><td>91.6</td><td>61.0</td><td>160.9</td><td>37.0</td><td>9.3</td><td></td></tr>
<tr><td>>MI0000542|hsa-mir-320a&WithFlank&8|-|22244960|22245043@48.71.24&[MIMAT0000510&hsa-miR-320a-3p&offsets|0|+2;m-60&8|-|22244975|22244996&offsets|0|+2]|AAAAGCTGGGTTGAGAGGGCGAAA</td><td>>MI0000542|hsa-mir-320a&WithFlank&8|-|22244960|22245043</td><td>48</td><td>71</td><td>24</td><td>AAAAGCTGGGTTGAGAGGGCGAAA</td><td>MB231</td><td>21</td><td>26</td><td>24</td><td>42</td><td>9</td><td>4.6</td><td>15.2</td><td>15.8</td><td>21.9</td><td>3.9</td><td></td></tr>
<tr><td>>MI0000542|hsa-mir-320a&WithFlank&8|-|22244960|22245043@48.72.25&[MIMAT0000510&hsa-miR-320a-3p&offsets|0|+3;m-60&8|-|22244975|22244996&offsets|0|+3]|AAAAGCTGGGTTGAGAGGGCGAAAA</td><td>>MI0000542|hsa-mir-320a&WithFlank&8|-|22244960|22245043</td><td>48</td><td>72</td><td>25</td><td>AAAAGCTGGGTTGAGAGGGCGAAAA</td><td>BT20</td><td>13</td><td>10</td><td>41</td><td>21</td><td>3</td><td>9.6</td><td>7.2</td><td>18.8</td><td>3.5</td><td>1.3</td><td></td></tr>
<tr><td>>MI0000542|hsa-mir-320a&WithFlank&8|-|22244960|22245043@49.68.20(+1U)&[MIMAT0000510&hsa-miR-320a-3p&offsets|+1|-1(+1U);m-60&8|-|22244975|22244996&offsets|+1|-1(+1U)]|AAAGCTGGGTTGAGAGGGCGT</td><td>>MI0000542|hsa-mir-320a&WithFlank&8|-|22244960|22245043</td><td>49</td><td>68</td><td>20(+1U)</td><td>AAAGCTGGGTTGAGAGGGCGT</td><td>BT20</td><td>15</td><td>10</td><td>43</td><td>137</td><td>5</td><td>7.3</td><td>7.0</td><td>12.6</td><td>114.6</td><td>3.4</td><td></td></tr>
<tr><td>>MI0000542|hsa-mir-320a&WithFlank&8|-|22244960|22245043@49.68.20(+2U)&[MIMAT0000510&hsa-miR-320a-3p&offsets|+1|-1(+2U);m-60&8|-|22244975|22244996&offsets|+1|-1(+2U)]|AAAGCTGGGTTGAGAGGGCGTT</td><td>>MI0000542|hsa-mir-320a&WithFlank&8|-|22244960|22245043</td><td>49</td><td>68</td><td>20(+2U)</td><td>AAAGCTGGGTTGAGAGGGCGTT</td><td>BT20</td><td>12</td><td>8</td><td>27</td><td>43</td><td>5</td><td>5.9</td><td>3.9</td><td>6.8</td><td>30.1</td><td>1.6</td><td></td></tr>
<tr><td>>MI0000542|hsa-mir-320a&WithFlank&8|-|22244960|22245043@49.68.20&[MIMAT0000510&hsa-miR-320a-3p&offsets|+1|-1;m-60&8|-|22244975|22244996&offsets|+1|-1]|AAAGCTGGGTTGAGAGGGCG</td><td>>MI0000542|hsa-mir-320a&WithFlank&8|-|22244960|22245043</td><td>49</td><td>68</td><td>20</td><td>AAAGCTGGGTTGAGAGGGCG</td><td>BT20</td><td>15</td><td>11</td><td>42</td><td>81</td><td>8</td><td>7.5</td><td>2.8</td><td>6.5</td><td>54.8</td><td>1.0</td><td></td></tr>
<tr><td>>MI0000542|hsa-mir-320a&WithFlank&8|-|22244960|22245043@49.69.21(+1U)&[MIMAT0000510&hsa-miR-320a-3p&offsets|+1|0(+1U);m-60&8|-|22244975|22244996&offsets|+1|0(+1U)]|AAAGCTGGGTTGAGAGGGCGAT</td><td>>MI0000542|hsa-mir-320a&WithFlank&8|-|22244960|22245043</td><td>49</td><td>69</td><td>21(+1U)</td><td>AAAGCTGGGTTGAGAGGGCGAT</td><td>BT20</td><td>33</td><td>24</td><td>56</td><td>62</td><td>11</td><td>15.2</td><td>19.4</td><td>12.1</td><td>42.8</td><td>6.5</td><td></td></tr>
<tr><td>>MI0000542|hsa-mir-320a&WithFlank&8|-|22244960|22245043@49.69.21(+2U)&[MIMAT0000510&hsa-miR-320a-3p&offsets|+1|0(+2U);m-60&8|-|22244975|22244996&offsets|+1|0(+2U)]|AAAGCTGGGTTGAGAGGGCGATT</td><td>>MI0000542|hsa-mir-320a&WithFlank&8|-|22244960|22245043</td><td>49</td><td>69</td><td>21(+2U)</td><td>AAAGCTGGGTTGAGAGGGCGATT</td><td>BT20</td><td>13</td><td>7</td><td>26</td><td>19</td><td>2</td><td>5.8</td><td>6.9</td><td>8.2</td><td>13.7</td><td>2.4</td><td></td></tr>
<tr><td>>MI0000542|hsa-mir-320a&WithFlank&8|-|22244960|22245043@49.69.21&[MIMAT0000510&hsa-miR-320a-3p&offsets|+1|0;m-60&8|-|22244975|22244996&offsets|+1|0]|AAAGCTGGGTTGAGAGGGCGA</td><td>>MI0000542|hsa-mir-320a&WithFlank&8|-|22244960|22245043</td><td>49</td><td>69</td><td>21</td><td>AAAGCTGGGTTGAGAGGGCGA</td><td>BT20</td><td>46</td><td>31</td><td>74</td><td>263</td><td>18</td><td>24.6</td><td>17.4</td><td>5.3</td><td>162.1</td><td>2.9</td><td></td></tr>
<tr><td>>MI0000542|hsa-mir-320a&WithFlank&8|-|22244960|22245043@49.70.22(+1U)&[MIMAT0000510&hsa-miR-320a-3p&offsets|+1|+1(+1U);m-60&8|-|22244975|22244996&offsets|+1|+1(+1U)]|AAAGCTGGGTTGAGAGGGCGAAT</td><td>>MI0000542|hsa-mir-320a&WithFlank&8|-|22244960|22245043</td><td>49</td><td>70</td><td>22(+1U)</td><td>AAAGCTGGGTTGAGAGGGCGAAT</td><td>BT20</td><td>32</td><td>19</td><td>61</td><td>82</td><td>15</td><td>16.2</td><td>11.7</td><td>3.6</td><td>41.1</td><td>5.3</td><td></td></tr>
<tr><td>>MI0000542|hsa-mir-320a&WithFlank&8|-|22244960|22245043@49.70.22&[MIMAT0000510&hsa-miR-320a-3p&offsets|+1|+1;m-60&8|-|22244975|22244996&offsets|+1|+1]|AAAGCTGGGTTGAGAGGGCGAA</td><td>>MI0000542|hsa-mir-320a&WithFlank&8|-|22244960|22245043</td><td>49</td><td>70</td><td>22</td><td>AAAGCTGGGTTGAGAGGGCGAA</td><td>BT20</td><td>30</td><td>21</td><td>54</td><td>80</td><td>13</td><td>17.4</td><td>7.6</td><td>6.7</td><td>38.2</td><td>2.2</td><td></td></tr>
<tr><td>>MI0000542|hsa-mir-320a&WithFlank&8|-|22244960|22245043@49.71.23&[MIMAT0000510&hsa-miR-320a-3p&offsets|+1|+2;m-60&8|-|22244975|22244996&offsets|+1|+2]|AAAGCTGGGTTGAGAGGGCGAAA</td><td>>MI0000542|hsa-mir-320a&WithFlank&8|-|22244960|22245043</td><td>49</td><td>71</td><td>23</td><td>AAAGCTGGGTTGAGAGGGCGAAA</td><td>BT20</td><td>16</td><td>10</td><td>32</td><td>27</td><td>5</td><td>9.2</td><td>5.4</td><td>5.8</td><td>8.9</td><td>2.0</td><td></td></tr>
<tr><td>>MI0000542|hsa-mir-320a&WithFlank&8|-|22244960|22245043@50.70.21(+1U)&[MIMAT0000510&hsa-miR-320a-3p&offsets|+2|+1(+1U);m-60&8|-|22244975|22244996&offsets|+2|+1(+1U)]|AAGCTGGGTTGAGAGGGCGAAT</td><td>>MI0000542|hsa-mir-320a&WithFlank&8|-|22244960|22245043</td><td>50</td><td>70</td><td>21(+1U)</td><td>AAGCTGGGTTGAGAGGGCGAAT</td><td>BT20</td><td>11</td><td>6</td><td>17</td><td>46</td><td>6</td><td>6.4</td><td>3.6</td><td>2.2</td><td>33.3</td><td>2.6</td><td></td></tr>
<tr><td>>MI0000650|hsa-mir-200c&WithFlank&12|+|6963693|6963772@49.70.22&[MIMAT0000617&hsa-miR-200c-3p&offsets|-1|-2;m-54&12|+|6963742|6963764&offsets|-1|-2]|CTAATACTGCCGGGTAATGATG</td><td>>MI0000650|hsa-mir-200c&WithFlank&12|+|6963693|6963772</td><td>49</td><td>70</td><td>22</td><td>CTAATACTGCCGGGTAATGATG</td><td>BT20</td><td>20</td><td>21</td><td>27</td><td>17</td><td>13</td><td>14.0</td><td>4.6</td><td>4.0</td><td>11.9</td><td>3.5</td><td></td></tr>
</table>

<table>
<tr><td>>MI0000650|hsa-mir-200c&WithFlank&12|+|6963693|6963772@49.71.23&[MIMAT0000617&hsa-miR-200c-3p&offsets|-1|-1;m-54&12|+|6963742|6963764&offsets|-1|-1]|CTAATACTGCCGGGTAATGATGG</td><td>>MI0000650|hsa-mir-200c&WithFlank&12|+|6963693|6963772</td><td>49</td><td>71</td><td>23</td><td>CTAATACTGCCGGGTAATGATGG</td><td>BT20</td><td>30</td><td>28</td><td>35</td><td>14</td><td>13</td><td>16.3</td><td>14.2</td><td>10.4</td><td>9.3</td><td>3.7</td></tr>
<tr><td>>MI0000650|hsa-mir-200c&WithFlank&12|+|6963693|6963772@49.72.24&[MIMAT0000617&hsa-miR-200c-3p&offsets|-1|0;m-54&12|+|6963742|6963764&offsets|-1|0]|CTAATACTGCCGGGTAATGATGGA</td><td>>MI0000650|hsa-mir-200c&WithFlank&12|+|6963693|6963772</td><td>49</td><td>72</td><td>24</td><td>CTAATACTGCCGGGTAATGATGGA</td><td>BT20</td><td>27</td><td>25</td><td>34</td><td>8</td><td>21</td><td>16.7</td><td>12.4</td><td>18.7</td><td>1.7</td><td>4.4</td></tr>
<tr><td>>MI0000650|hsa-mir-200c&WithFlank&12|+|6963693|6963772@50.68.19(+1C)&[MIMAT0000617&hsa-miR-200c-3p&offsets|0|-4(+1C);m-54&12|+|6963742|6963764&offsets|0|-4(+1C)]|TAATACTGCCGGGTAATGAC</td><td>>MI0000650|hsa-mir-200c&WithFlank&12|+|6963693|6963772</td><td>50</td><td>68</td><td>19(+1C)</td><td>TAATACTGCCGGGTAATGAC</td><td>BT20</td><td>12</td><td>16</td><td>13</td><td>13</td><td>10</td><td>9.5</td><td>16.6</td><td>8.8</td><td>6.8</td><td>3.8</td></tr>
<tr><td>>MI0000650|hsa-mir-200c&WithFlank&12|+|6963693|6963772@50.69.20(+1A)&[MIMAT0000617&hsa-miR-200c-3p&offsets|0|-3(+1A);m-54&12|+|6963742|6963764&offsets|0|-3(+1A)]|TAATACTGCCGGGTAATGATA</td><td>>MI0000650|hsa-mir-200c&WithFlank&12|+|6963693|6963772</td><td>50</td><td>69</td><td>20(+1A)</td><td>TAATACTGCCGGGTAATGATA</td><td>BT20</td><td>35</td><td>23</td><td>40</td><td>36</td><td>16</td><td>26.2</td><td>15.0</td><td>11.6</td><td>24.9</td><td>3.9</td></tr>
<tr><td>>MI0000650|hsa-mir-200c&WithFlank&12|+|6963693|6963772@50.69.20&[MIMAT0000617&hsa-miR-200c-3p&offsets|0|-3;m-54&12|+|6963742|6963764&offsets|0|-3]|TAATACTGCCGGGTAATGAT</td><td>>MI0000650|hsa-mir-200c&WithFlank&12|+|6963693|6963772</td><td>50</td><td>69</td><td>20</td><td>TAATACTGCCGGGTAATGAT</td><td>BT20</td><td>96</td><td>117</td><td>106</td><td>103</td><td>84</td><td>78.2</td><td>111.5</td><td>82.2</td><td>61.8</td><td>21.8</td></tr>
<tr><td>>MI0000650|hsa-mir-200c&WithFlank&12|+|6963693|6963772@50.70.21(+1A)&[MIMAT0000617&hsa-miR-200c-3p&offsets|0|-2(+1A);m-54&12|+|6963742|6963764&offsets|0|-2(+1A)]|TAATACTGCCGGGTAATGATGA</td><td>>MI0000650|hsa-mir-200c&WithFlank&12|+|6963693|6963772</td><td>50</td><td>70</td><td>21(+1A)</td><td>TAATACTGCCGGGTAATGATGA</td><td>BT20</td><td>267</td><td>180</td><td>232</td><td>224</td><td>73</td><td>219.2</td><td>136.1</td><td>57.7</td><td>179.2</td><td>34.3</td></tr>
<tr><td>>MI0000650|hsa-mir-200c&WithFlank&12|+|6963693|6963772@50.70.21(+1C)&[MIMAT0000617&hsa-miR-200c-3p&offsets|0|-2(+1C);m-54&12|+|6963742|6963764&offsets|0|-2(+1C)]|TAATACTGCCGGGTAATGATGC</td><td>>MI0000650|hsa-mir-200c&WithFlank&12|+|6963693|6963772</td><td>50</td><td>70</td><td>21(+1C)</td><td>TAATACTGCCGGGTAATGATGC</td><td>BT20</td><td>13</td><td>9</td><td>9</td><td>13</td><td>3</td><td>10.2</td><td>7.3</td><td>3.4</td><td>11.1</td><td>1.2</td></tr>
<tr><td>>MI0000650|hsa-mir-200c&WithFlank&12|+|6963693|6963772@50.70.21(+1U)&[MIMAT0000617&hsa-miR-200c-3p&offsets|0|-2(+1U);m-54&12|+|6963742|6963764&offsets|0|-2(+1U)]|TAATACTGCCGGGTAATGATGT</td><td>>MI0000650|hsa-mir-200c&WithFlank&12|+|6963693|6963772</td><td>50</td><td>70</td><td>21(+1U)</td><td>TAATACTGCCGGGTAATGATGT</td><td>BT20</td><td>59</td><td>34</td><td>38</td><td>44</td><td>14</td><td>54.6</td><td>24.3</td><td>14.8</td><td>35.6</td><td>6.5</td></tr>
<tr><td>>MI0000650|hsa-mir-200c&WithFlank&12|+|6963693|6963772@50.70.21(+2A)&[MIMAT0000617&hsa-miR-200c-3p&offsets|0|-2(+2A);m-54&12|+|6963742|6963764&offsets|0|-2(+2A)]|TAATACTGCCGGGTAATGATGAA</td><td>>MI0000650|hsa-mir-200c&WithFlank&12|+|6963693|6963772</td><td>50</td><td>70</td><td>21(+2A)</td><td>TAATACTGCCGGGTAATGATGAA</td><td>BT20</td><td>25</td><td>10</td><td>7</td><td>3</td><td>4</td><td>29.4</td><td>7.1</td><td>2.3</td><td>2.1</td><td>1.4</td></tr>
<tr><td>>MI0000650|hsa-mir-200c&WithFlank&12|+|6963693|6963772@50.70.21&[MIMAT0000617&hsa-miR-200c-3p&offsets|0|-2;m-54&12|+|6963742|6963764&offsets|0|-2]|TAATACTGCCGGGTAATGATG</td><td>>MI0000650|hsa-mir-200c&WithFlank&12|+|6963693|6963772</td><td>50</td><td>70</td><td>21</td><td>TAATACTGCCGGGTAATGATG</td><td>BT20</td><td>186</td><td>171</td><td>252</td><td>321</td><td>149</td><td>115.4</td><td>89.5</td><td>135.9</td><td>198.0</td><td>27.1</td></tr>
<tr><td>>MI0000650|hsa-mir-200c&WithFlank&12|+|6963693|6963772@50.71.22(+1C)&[MIMAT0000617&hsa-miR-200c-3p&offsets|0|-1(+1C);m-54&12|+|6963742|6963764&offsets|0|-1(+1C)]|TAATACTGCCGGGTAATGATGGC</td><td>>MI0000650|hsa-mir-200c&WithFlank&12|+|6963693|6963772</td><td>50</td><td>71</td><td>22(+1C)</td><td>TAATACTGCCGGGTAATGATGGC</td><td>BT20</td><td>42</td><td>28</td><td>19</td><td>22</td><td>11</td><td>33.5</td><td>23.4</td><td>7.0</td><td>20.5</td><td>4.5</td></tr>
<tr><td>>MI0000650|hsa-mir-200c&WithFlank&12|+|6963693|6963772@50.71.22(+1G)&[MIMAT0000617&hsa-miR-200c-3p&offsets|0|-1(+1G);m-54&12|+|6963742|6963764&offsets|0|-1(+1G)]|TAATACTGCCGGGTAATGATGGG</td><td>>MI0000650|hsa-mir-200c&WithFlank&12|+|6963693|6963772</td><td>50</td><td>71</td><td>22(+1G)</td><td>TAATACTGCCGGGTAATGATGGG</td><td>BT20</td><td>495</td><td>259</td><td>199</td><td>120</td><td>84</td><td>343.1</td><td>179.3</td><td>51.4</td><td>106.4</td><td>29.5</td></tr>
<tr><td>>MI0000650|hsa-mir-200c&WithFlank&12|+|6963693|6963772@50.71.22(+1U)&[MIMAT0000617&hsa-miR-200c-3p&offsets|0|-1(+1U);m-54&12|+|6963742|6963764&offsets|0|-1(+1U)]|TAATACTGCCGGGTAATGATGGT</td><td>>MI0000650|hsa-mir-200c&WithFlank&12|+|6963693|6963772</td><td>50</td><td>71</td><td>22(+1U)</td><td>TAATACTGCCGGGTAATGATGGT</td><td>BT20</td><td>381</td><td>212</td><td>175</td><td>167</td><td>98</td><td>300.1</td><td>159.5</td><td>71.5</td><td>133.5</td><td>40.0</td></tr>
<tr><td>>MI0000650|hsa-mir-200c&WithFlank&12|+|6963693|6963772@50.71.22(+2U)&[MIMAT0000617&hsa-miR-200c-3p&offsets|0|-1(+2U);m-54&12|+|6963742|6963764&offsets|0|-1(+2U)]|TAATACTGCCGGGTAATGATGGTT</td><td>>MI0000650|hsa-mir-200c&WithFlank&12|+|6963693|6963772</td><td>50</td><td>71</td><td>22(+2U)</td><td>TAATACTGCCGGGTAATGATGGTT</td><td>BT20</td><td>11</td><td>9</td><td>5</td><td>23</td><td>4</td><td>7.4</td><td>11.0</td><td>2.4</td><td>22.8</td><td>3.6</td></tr>
<tr><td>>MI0000650|hsa-mir-200c&WithFlank&12|+|6963693|6963772@50.71.22&[MIMAT0000617&hsa-miR-200c-3p&offsets|0|-1;m-54&12|+|6963742|6963764&offsets|0|-1]|TAATACTGCCGGGTAATGATGG</td><td>>MI0000650|hsa-mir-200c&WithFlank&12|+|6963693|6963772</td><td>50</td><td>71</td><td>22</td><td>TAATACTGCCGGGTAATGATGG</td><td>BT20</td><td>1261</td><td>1095</td><td>1318</td><td>1711</td><td>548</td><td>628.0</td><td>686.4</td><td>742.3</td><td>1257.0</td><td>231.8</td></tr>
<tr><td>>MI0000650|hsa-mir-200c&WithFlank&12|+|6963693|6963772@50.71.22&[MIMAT0000617&hsa-miR-200c-3p&offsets|0|-1;m-54&12|+|6963742|6963764&offsets|0|-1]|TAATACTGCCGGGTAATGATGG</td><td>>MI0000650|hsa-mir-200c&WithFlank&12|+|6963693|6963772</td><td>50</td><td>71</td><td>22</td><td>TAATACTGCCGGGTAATGATGG</td><td>MB468</td><td>26</td><td>21</td><td>12</td><td>43</td><td>9</td><td>11.9</td><td>3.7</td><td>3.9</td><td>16.5</td><td>3.1</td></tr>
<tr><td>>MI0000650|hsa-mir-200c&WithFlank&12|+|6963693|6963772@50.72.23(+1A)&[MIMAT0000617&hsa-miR-200c-3p&offsets|0|0(+1A);m-54&12|+|6963742|6963764&offsets|0|0(+1A)]|TAATACTGCCGGGTAATGATGGAA</td><td>>MI0000650|hsa-mir-200c&WithFlank&12|+|6963693|6963772</td><td>50</td><td>72</td><td>23(+1A)</td><td>TAATACTGCCGGGTAATGATGGAA</td><td>BT20</td><td>825</td><td>733</td><td>983</td><td>239</td><td>568</td><td>488.9</td><td>338.6</td><td>287.8</td><td>65.6</td><td>40.0</td></tr>
<tr><td>>MI0000650|hsa-mir-200c&WithFlank&12|+|6963693|6963772@50.72.23(+1A)&[MIMAT0000617&hsa-miR-200c-3p&offsets|0|0(+1A);m-54&12|+|6963742|6963764&offsets|0|0(+1A)]|TAATACTGCCGGGTAATGATGGAA</td><td>>MI0000650|hsa-mir-200c&WithFlank&12|+|6963693|6963772</td><td>50</td><td>72</td><td>23(+1A)</td><td>TAATACTGCCGGGTAATGATGGAA</td><td>MB468</td><td>13</td><td>17</td><td>6</td><td>10</td><td>4</td><td>4.4</td><td>4.7</td><td>2.7</td><td>4.4</td><td>1.8</td></tr>
<tr><td>>MI0000650|hsa-mir-200c&WithFlank&12|+|6963693|6963772@50.72.23(+1C)&[MIMAT0000617&hsa-miR-200c-3p&offsets|0|0(+1C);m-54&12|+|6963742|6963764&offsets|0|0(+1C)]|TAATACTGCCGGGTAATGATGGAC</td><td>>MI0000650|hsa-mir-200c&WithFlank&12|+|6963693|6963772</td><td>50</td><td>72</td><td>23(+1C)</td><td>TAATACTGCCGGGTAATGATGGAC</td><td>BT20</td><td>89</td><td>70</td><td>97</td><td>22</td><td>53</td><td>41.2</td><td>33.7</td><td>22.2</td><td>14.4</td><td>20.0</td></tr>
<tr><td>>MI0000650|hsa-mir-200c&WithFlank&12|+|6963693|6963772@50.72.23(+1U)&[MIMAT0000617&hsa-miR-200c-3p&offsets|0|0(+1U);m-54&12|+|6963742|6963764&offsets|0|0(+1U)]|TAATACTGCCGGGTAATGATGGAT</td><td>>MI0000650|hsa-mir-200c&WithFlank&12|+|6963693|6963772</td><td>50</td><td>72</td><td>23(+1U)</td><td>TAATACTGCCGGGTAATGATGGAT</td><td>BT20</td><td>599</td><td>445</td><td>651</td><td>143</td><td>357</td><td>357.8</td><td>219.7</td><td>235.8</td><td>70.4</td><td>153.7</td></tr>
<tr><td>>MI0000650|hsa-mir-200c&WithFlank&12|+|6963693|6963772@50.72.23(+1U)&[MIMAT0000617&hsa-miR-200c-3p&offsets|0|0(+1U);m-54&12|+|6963742|6963764&offsets|0|0(+1U)]|TAATACTGCCGGGTAATGATGGAT</td><td>>MI0000650|hsa-mir-200c&WithFlank&12|+|6963693|6963772</td><td>50</td><td>72</td><td>23(+1U)</td><td>TAATACTGCCGGGTAATGATGGAT</td><td>MB468</td><td>14</td><td>15</td><td>6</td><td>9</td><td>4</td><td>4.5</td><td>3.9</td><td>2.3</td><td>3.3</td><td>1.4</td></tr>
<tr><td>>MI0000650|hsa-mir-200c&WithFlank&12|+|6963693|6963772@50.72.23(+2A)&[MIMAT0000617&hsa-miR-200c-3p&offsets|0|0(+2A);m-54&12|+|6963742|6963764&offsets|0|0(+2A)]|TAATACTGCCGGGTAATGATGGAAA</td><td>>MI0000650|hsa-mir-200c&WithFlank&12|+|6963693|6963772</td><td>50</td><td>72</td><td>23(+2A)</td><td>TAATACTGCCGGGTAATGATGGAAA</td><td>BT20</td><td>56</td><td>46</td><td>76</td><td>9</td><td>37</td><td>38.5</td><td>23.1</td><td>28.2</td><td>2.8</td><td>6.5</td></tr>
<tr><td>>MI0000650|hsa-mir-200c&WithFlank&12|+|6963693|6963772@50.72.23(+2U)&[MIMAT0000617&hsa-miR-200c-3p&offsets|0|0(+2U);m-54&12|+|6963742|6963764&offsets|0|0(+2U)]|TAATACTGCCGGGTAATGATGGATT</td><td>>MI0000650|hsa-mir-200c&WithFlank&12|+|6963693|6963772</td><td>50</td><td>72</td><td>23(+2U)</td><td>TAATACTGCCGGGTAATGATGGATT</td><td>BT20</td><td>24</td><td>20</td><td>33</td><td>6</td><td>17</td><td>22.4</td><td>15.0</td><td>27.1</td><td>1.8</td><td>9.4</td></tr>
<tr><td>>MI0000650|hsa-mir-200c&WithFlank&12|+|6963693|6963772@50.72.23&[MIMAT0000617&hsa-miR-200c-3p&offsets|0|0;m-54&12|+|6963742|6963764&offsets|0|0]|TAATACTGCCGGGTAATGATGGA</td><td>>MI0000650|hsa-mir-200c&WithFlank&12|+|6963693|6963772</td><td>50</td><td>72</td><td>23</td><td>TAATACTGCCGGGTAATGATGGA</td><td>BT20</td><td>8675</td><td>8641</td><td>8590</td><td>6154</td><td>6469</td><td>5751.8</td><td>5296.0</td><td>6055.5</td><td>3480.2</td><td>2210.6</td></tr>
<tr><td>>MI0000650|hsa-mir-200c&WithFlank&12|+|6963693|6963772@50.72.23&[MIMAT0000617&hsa-miR-200c-3p&offsets|0|0;m-54&12|+|6963742|6963764&offsets|0|0]|TAATACTGCCGGGTAATGATGGA</td><td>>MI0000650|hsa-mir-200c&WithFlank&12|+|6963693|6963772</td><td>50</td><td>72</td><td>23</td><td>TAATACTGCCGGGTAATGATGGA</td><td>MB468</td><td>185</td><td>173</td><td>71</td><td>167</td><td>66</td><td>89.2</td><td>48.2</td><td>20.4</td><td>45.6</td><td>20.0</td></tr>
<tr><td>>MI0000650|hsa-mir-200c&WithFlank&12|+|6963693|6963772@50.73.24&[MIMAT0000617&hsa-miR-200c-3p&offsets|0|+1;m-54&12|+|6963742|6963764&offsets|0|+1]|TAATACTGCCGGGTAATGATGGAG</td><td>>MI0000650|hsa-mir-200c&WithFlank&12|+|6963693|6963772</td><td>50</td><td>73</td><td>24</td><td>TAATACTGCCGGGTAATGATGGAG</td><td>BT20</td><td>65</td><td>30</td><td>39</td><td>20</td><td>12</td><td>43.1</td><td>19.6</td><td>6.5</td><td>15.6</td><td>3.9</td></tr>
<tr><td>>MI0000683|hsa-mir-181b-2&WithFlank&9|+|124693704|124693804@22.46.25(+1U)&[MIMAT0000257_1&hsa-miR-181b-5p&offsets|0|+2(+1U);m-72&9|+|124693725|124693747&offsets|0|+2(+1U)]|AACATTCATTGCTGTCGGTGGGTTTT</td><td>>MI0000683|hsa-mir-181b-2&WithFlank&9|+|124693704|124693804</td><td>22</td><td>46</td><td>25(+1U)</td><td>AACATTCATTGCTGTCGGTGGGTTTT</td><td>MB231</td><td>23</td><td>18</td><td>5</td><td>50</td><td>39</td><td>3.0</td><td>10.5</td><td>4.0</td><td>19.2</td><td>35.3</td></tr>
</table>

<table>
<tr><th></th><th></th><th></th><th></th><th></th><th></th><th></th><th></th><th></th><th></th><th></th><th></th><th></th><th></th><th></th><th></th><th></th></tr>
<tr><td>>MI0000683|hsa-mir-181b-2&WithFlank&9|+|124693704|124693804@22.46.25&[MIMAT0000257_1&hsa-miR-181b-5p&offsets|0|+2;m-72&9|+|124693725|124693747&offsets|0|+2]|AACATTCATTGCTGTCGGTGGGTTT</td><td>>MI0000683|hsa-mir-181b-2&WithFlank&9|+|124693704|124693804</td><td>22</td><td>46</td><td>25</td><td>AACATTCATTGCTGTCGGTGGGTTT</td><td>BT20</td><td>15</td><td>13</td><td>9</td><td>9</td><td>10</td><td>9.6</td><td>11.1</td><td>4.3</td><td>7.9</td><td>9.0</td></tr>
<tr><td>>MI0000683|hsa-mir-181b-2&WithFlank&9|+|124693704|124693804@22.46.25&[MIMAT0000257_1&hsa-miR-181b-5p&offsets|0|+2;m-72&9|+|124693725|124693747&offsets|0|+2]|AACATTCATTGCTGTCGGTGGGTTT</td><td>>MI0000683|hsa-mir-181b-2&WithFlank&9|+|124693704|124693804</td><td>22</td><td>46</td><td>25</td><td>AACATTCATTGCTGTCGGTGGGTTT</td><td>MB231</td><td>42</td><td>33</td><td>9</td><td>105</td><td>86</td><td>5.1</td><td>17.9</td><td>7.9</td><td>33.5</td><td>62.0</td></tr>
<tr><td>>MI0000727|hsa-mir-128-2&WithFlank&3|+|35744470|35744565@58.79.22&[MIMAT0000424_1&hsa-miR-128-3p&offsets|0|+1;m-117&3|+|35744527|35744547&offsets|0|+1]|TCACAGTGAACCGGTCTCTTTC</td><td>>MI0000727|hsa-mir-128-2&WithFlank&3|+|35744470|35744565</td><td>58</td><td>79</td><td>22</td><td>TCACAGTGAACCGGTCTCTTTC</td><td>BT20</td><td>18</td><td>8</td><td>15</td><td>7</td><td>9</td><td>12.0</td><td>2.3</td><td>2.4</td><td>2.7</td><td>1.2</td></tr>
<tr><td>>MI0000727|hsa-mir-128-2&WithFlank&3|+|35744470|35744565@58.79.22&[MIMAT0000424_1&hsa-miR-128-3p&offsets|0|+1;m-117&3|+|35744527|35744547&offsets|0|+1]|TCACAGTGAACCGGTCTCTTTC</td><td>>MI0000727|hsa-mir-128-2&WithFlank&3|+|35744470|35744565</td><td>58</td><td>79</td><td>22</td><td>TCACAGTGAACCGGTCTCTTTC</td><td>MB231</td><td>11</td><td>9</td><td>3</td><td>6</td><td>11</td><td>8.5</td><td>1.0</td><td>0.7</td><td>3.1</td><td>4.3</td></tr>
<tr><td>>MI0000732|hsa-mir-194-2&WithFlank&11|-|64891349|64891445@21.43.23&[MIMAT0000460_1&hsa-miR-194-5p&offsets|0|+1;m-126&11|-|64891404|64891425&offsets|0|+1]|TGTAACAGCAACTCCATGTGGAA</td><td>>MI0000732|hsa-mir-194-2&WithFlank&11|-|64891349|64891445</td><td>21</td><td>43</td><td>23</td><td>TGTAACAGCAACTCCATGTGGAA</td><td>BT20</td><td>13</td><td>10</td><td>8</td><td>5</td><td>2</td><td>7.3</td><td>11.1</td><td>5.6</td><td>4.1</td><td>2.6</td></tr>
<tr><td>>MI0000734|hsa-mir-106b&WithFlank&7|-|100093987|100094080@18.37.20(+1C)&[MIMAT0000680&hsa-miR-106b-5p&offsets|0|-1(+1C);m-103&7|-|100094043|100094063&offsets|0|-1(+1C)]|TAAAGTGCTGACAGTGCAGAC</td><td>>MI0000734|hsa-mir-106b&WithFlank&7|-|100093987|100094080</td><td>18</td><td>37</td><td>20(+1C)</td><td>TAAAGTGCTGACAGTGCAGAC</td><td>BT20</td><td>21</td><td>16</td><td>2</td><td>1</td><td>1</td><td>27.1</td><td>20.8</td><td>1.1</td><td>1.2</td><td>0.7</td></tr>
<tr><td>>MI0000734|hsa-mir-106b&WithFlank&7|-|100093987|100094080@18.37.20&[MIMAT0000680&hsa-miR-106b-5p&offsets|0|-1;m-103&7|-|100094043|100094063&offsets|0|-1]|TAAAGTGCTGACAGTGCAGA</td><td>>MI0000734|hsa-mir-106b&WithFlank&7|-|100093987|100094080</td><td>18</td><td>37</td><td>20</td><td>TAAAGTGCTGACAGTGCAGA</td><td>BT20</td><td>26</td><td>15</td><td>4</td><td>4</td><td>1</td><td>30.6</td><td>16.9</td><td>2.4</td><td>3.4</td><td>1.3</td></tr>
<tr><td>>MI0000734|hsa-mir-106b&WithFlank&7|-|100093987|100094080@18.38.21(+1G)&[MIMAT0000680&hsa-miR-106b-5p&offsets|0|0(+1G);m-103&7|-|100094043|100094063&offsets|0|0(+1G)]|TAAAGTGCTGACAGTGCAGATG</td><td>>MI0000734|hsa-mir-106b&WithFlank&7|-|100093987|100094080</td><td>18</td><td>38</td><td>21(+1G)</td><td>TAAAGTGCTGACAGTGCAGATG</td><td>BT20</td><td>20</td><td>4</td><td>1</td><td>2</td><td>0</td><td>29.4</td><td>3.5</td><td>0.3</td><td>2.6</td><td>0.2</td></tr>
<tr><td>>MI0000734|hsa-mir-106b&WithFlank&7|-|100093987|100094080@18.38.21(+1U)&[MIMAT0000680&hsa-miR-106b-5p&offsets|0|0(+1U);m-103&7|-|100094043|100094063&offsets|0|0(+1U)]|TAAAGTGCTGACAGTGCAGATT</td><td>>MI0000734|hsa-mir-106b&WithFlank&7|-|100093987|100094080</td><td>18</td><td>38</td><td>21(+1U)</td><td>TAAAGTGCTGACAGTGCAGATT</td><td>BT20</td><td>17</td><td>7</td><td>2</td><td>2</td><td>0</td><td>24.7</td><td>7.7</td><td>0.8</td><td>1.9</td><td>0.4</td></tr>
<tr><td>>MI0000734|hsa-mir-106b&WithFlank&7|-|100093987|100094080@18.38.21&[MIMAT0000680&hsa-miR-106b-5p&offsets|0|0;m-103&7|-|100094043|100094063&offsets|0|0]|TAAAGTGCTGACAGTGCAGAT</td><td>>MI0000734|hsa-mir-106b&WithFlank&7|-|100093987|100094080</td><td>18</td><td>38</td><td>21</td><td>TAAAGTGCTGACAGTGCAGAT</td><td>BT20</td><td>151</td><td>108</td><td>13</td><td>7</td><td>5</td><td>196.5</td><td>135.7</td><td>6.9</td><td>7.5</td><td>5.3</td></tr>
<tr><td>>MI0000734|hsa-mir-106b&WithFlank&7|-|100093987|100094080@18.38.21&[MIMAT0000680&hsa-miR-106b-5p&offsets|0|0;m-103&7|-|100094043|100094063&offsets|0|0]|TAAAGTGCTGACAGTGCAGAT</td><td>>MI0000734|hsa-mir-106b&WithFlank&7|-|100093987|100094080</td><td>18</td><td>38</td><td>21</td><td>TAAAGTGCTGACAGTGCAGAT</td><td>MB231</td><td>40</td><td>53</td><td>3</td><td>81</td><td>67</td><td>10.8</td><td>30.4</td><td>2.4</td><td>12.6</td><td>80.8</td></tr>
<tr><td>>MI0000734|hsa-mir-106b&WithFlank&7|-|100093987|100094080@18.39.22&[MIMAT0000680&hsa-miR-106b-5p&offsets|0|+1;m-103&7|-|100094043|100094063&offsets|0|+1]|TAAAGTGCTGACAGTGCAGATA</td><td>>MI0000734|hsa-mir-106b&WithFlank&7|-|100093987|100094080</td><td>18</td><td>39</td><td>22</td><td>TAAAGTGCTGACAGTGCAGATA</td><td>BT20</td><td>278</td><td>132</td><td>26</td><td>27</td><td>14</td><td>368.9</td><td>121.7</td><td>10.7</td><td>34.1</td><td>3.4</td></tr>
<tr><td>>MI0000734|hsa-mir-106b&WithFlank&7|-|100093987|100094080@18.39.22&[MIMAT0000680&hsa-miR-106b-5p&offsets|0|+1;m-103&7|-|100094043|100094063&offsets|0|+1]|TAAAGTGCTGACAGTGCAGATA</td><td>>MI0000734|hsa-mir-106b&WithFlank&7|-|100093987|100094080</td><td>18</td><td>39</td><td>22</td><td>TAAAGTGCTGACAGTGCAGATA</td><td>MB231</td><td>56</td><td>69</td><td>5</td><td>175</td><td>91</td><td>9.2</td><td>40.6</td><td>5.1</td><td>10.6</td><td>99.1</td></tr>
<tr><td>>MI0000734|hsa-mir-106b&WithFlank&7|-|100093987|100094080@56.76.21(+1A)&[MIMAT0004672&hsa-miR-106b-3p&offsets|-2|-3(+1A);m-87&7|-|100094003|100094023&offsets|-2|-2(+1A)]|TACCGCACTGTGGGTACTTGCA</td><td>>MI0000734|hsa-mir-106b&WithFlank&7|-|100093987|100094080</td><td>56</td><td>76</td><td>21(+1A)</td><td>TACCGCACTGTGGGTACTTGCA</td><td>BT20</td><td>11</td><td>5</td><td>10</td><td>3</td><td>2</td><td>7.3</td><td>3.9</td><td>2.0</td><td>2.5</td><td>1.8</td></tr>
<tr><td>>MI0000734|hsa-mir-106b&WithFlank&7|-|100093987|100094080@56.76.21(+1C)&[MIMAT0004672&hsa-miR-106b-3p&offsets|-2|-3(+1C);m-87&7|-|100094003|100094023&offsets|-2|-2(+1C)]|TACCGCACTGTGGGTACTTGCC</td><td>>MI0000734|hsa-mir-106b&WithFlank&7|-|100093987|100094080</td><td>56</td><td>76</td><td>21(+1C)</td><td>TACCGCACTGTGGGTACTTGCC</td><td>BT20</td><td>11</td><td>8</td><td>10</td><td>6</td><td>4</td><td>2.8</td><td>7.1</td><td>2.9</td><td>4.1</td><td>3.6</td></tr>
<tr><td>>MI0000734|hsa-mir-106b&WithFlank&7|-|100093987|100094080@56.77.22&[MIMAT0004672&hsa-miR-106b-3p&offsets|-2|-2;m-87&7|-|100094003|100094023&offsets|-2|-1]|TACCGCACTGTGGGTACTTGCT</td><td>>MI0000734|hsa-mir-106b&WithFlank&7|-|100093987|100094080</td><td>56</td><td>77</td><td>22</td><td>TACCGCACTGTGGGTACTTGCT</td><td>BT20</td><td>72</td><td>47</td><td>80</td><td>40</td><td>25</td><td>24.5</td><td>38.1</td><td>28.0</td><td>31.8</td><td>24.4</td></tr>
<tr><td>>MI0000734|hsa-mir-106b&WithFlank&7|-|100093987|100094080@56.77.22&[MIMAT0004672&hsa-miR-106b-3p&offsets|-2|-2;m-87&7|-|100094003|100094023&offsets|-2|-1]|TACCGCACTGTGGGTACTTGCT</td><td>>MI0000734|hsa-mir-106b&WithFlank&7|-|100093987|100094080</td><td>56</td><td>77</td><td>22</td><td>TACCGCACTGTGGGTACTTGCT</td><td>MB231</td><td>60</td><td>45</td><td>31</td><td>55</td><td>45</td><td>27.5</td><td>21.1</td><td>23.1</td><td>28.9</td><td>25.8</td></tr>
<tr><td>>MI0000734|hsa-mir-106b&WithFlank&7|-|100093987|100094080@58.76.19&[MIMAT0004672&hsa-miR-106b-3p&offsets|0|-3;m-87&7|-|100094003|100094023&offsets|0|-2]|CCGCACTGTGGGTACTTGC</td><td>>MI0000734|hsa-mir-106b&WithFlank&7|-|100093987|100094080</td><td>58</td><td>76</td><td>19</td><td>CCGCACTGTGGGTACTTGC</td><td>BT20</td><td>11</td><td>10</td><td>21</td><td>25</td><td>9</td><td>8.8</td><td>4.5</td><td>6.4</td><td>18.9</td><td>1.9</td></tr>
<tr><td>>MI0000734|hsa-mir-106b&WithFlank&7|-|100093987|100094080@58.77.20(+1A)&[MIMAT0004672&hsa-miR-106b-3p&offsets|0|-2(+1A);m-87&7|-|100094003|100094023&offsets|0|-1(+1A)]|CCGCACTGTGGGTACTTGCTA</td><td>>MI0000734|hsa-mir-106b&WithFlank&7|-|100093987|100094080</td><td>58</td><td>77</td><td>20(+1A)</td><td>CCGCACTGTGGGTACTTGCTA</td><td>BT20</td><td>21</td><td>12</td><td>44</td><td>18</td><td>8</td><td>13.1</td><td>4.4</td><td>17.5</td><td>9.9</td><td>3.2</td></tr>
<tr><td>>MI0000734|hsa-mir-106b&WithFlank&7|-|100093987|100094080@58.77.20&[MIMAT0004672&hsa-miR-106b-3p&offsets|0|-2;m-87&7|-|100094003|100094023&offsets|0|-1]|CCGCACTGTGGGTACTTGCT</td><td>>MI0000734|hsa-mir-106b&WithFlank&7|-|100093987|100094080</td><td>58</td><td>77</td><td>20</td><td>CCGCACTGTGGGTACTTGCT</td><td>BT20</td><td>63</td><td>46</td><td>96</td><td>73</td><td>38</td><td>30.2</td><td>19.9</td><td>28.5</td><td>61.0</td><td>20.9</td></tr>
<tr><td>>MI0000734|hsa-mir-106b&WithFlank&7|-|100093987|100094080@58.77.20&[MIMAT0004672&hsa-miR-106b-3p&offsets|0|-2;m-87&7|-|100094003|100094023&offsets|0|-1]|CCGCACTGTGGGTACTTGCT</td><td>>MI0000734|hsa-mir-106b&WithFlank&7|-|100093987|100094080</td><td>58</td><td>77</td><td>20</td><td>CCGCACTGTGGGTACTTGCT</td><td>MB231</td><td>48</td><td>52</td><td>45</td><td>55</td><td>36</td><td>13.6</td><td>18.5</td><td>24.0</td><td>27.0</td><td>7.3</td></tr>
<tr><td>>MI0000734|hsa-mir-106b&WithFlank&7|-|100093987|100094080@58.78.21(+1A)&[MIMAT0004672&hsa-miR-106b-3p&offsets|0|-1(+1A);m-87&7|-|100094003|100094023&offsets|0|0(+1A)]|CCGCACTGTGGGTACTTGCTGA</td><td>>MI0000734|hsa-mir-106b&WithFlank&7|-|100093987|100094080</td><td>58</td><td>78</td><td>21(+1A)</td><td>CCGCACTGTGGGTACTTGCTGA</td><td>BT20</td><td>26</td><td>16</td><td>48</td><td>30</td><td>10</td><td>8.6</td><td>6.0</td><td>19.8</td><td>22.9</td><td>3.8</td></tr>
<tr><td>>MI0000734|hsa-mir-106b&WithFlank&7|-|100093987|100094080@58.78.21(+1A)&[MIMAT0004672&hsa-miR-106b-3p&offsets|0|-1(+1A);m-87&7|-|100094003|100094023&offsets|0|0(+1A)]|CCGCACTGTGGGTACTTGCTGA</td><td>>MI0000734|hsa-mir-106b&WithFlank&7|-|100093987|100094080</td><td>58</td><td>78</td><td>21(+1A)</td><td>CCGCACTGTGGGTACTTGCTGA</td><td>MB231</td><td>15</td><td>16</td><td>15</td><td>18</td><td>10</td><td>3.9</td><td>2.2</td><td>6.8</td><td>10.7</td><td>0.3</td></tr>
<tr><td>>MI0000734|hsa-mir-106b&WithFlank&7|-|100093987|100094080@58.78.21(+1U)&[MIMAT0004672&hsa-miR-106b-3p&offsets|0|-1(+1U);m-87&7|-|100094003|100094023&offsets|0|0(+1U)]|CCGCACTGTGGGTACTTGCTGT</td><td>>MI0000734|hsa-mir-106b&WithFlank&7|-|100093987|100094080</td><td>58</td><td>78</td><td>21(+1U)</td><td>CCGCACTGTGGGTACTTGCTGT</td><td>BT20</td><td>98</td><td>70</td><td>156</td><td>119</td><td>48</td><td>29.7</td><td>31.0</td><td>56.8</td><td>76.9</td><td>22.2</td></tr>
<tr><td>>MI0000734|hsa-mir-106b&WithFlank&7|-|100093987|100094080@58.78.21(+1U)&[MIMAT0004672&hsa-miR-106b-3p&offsets|0|-1(+1U);m-87&7|-|100094003|100094023&offsets|0|0(+1U)]|CCGCACTGTGGGTACTTGCTGT</td><td>>MI0000734|hsa-mir-106b&WithFlank&7|-|100093987|100094080</td><td>58</td><td>78</td><td>21(+1U)</td><td>CCGCACTGTGGGTACTTGCTGT</td><td>MB231</td><td>73</td><td>79</td><td>64</td><td>84</td><td>47</td><td>18.9</td><td>15.9</td><td>21.4</td><td>44.7</td><td>8.2</td></tr>
<tr><td>>MI0000734|hsa-mir-106b&WithFlank&7|-|100093987|100094080@58.78.21&[MIMAT0004672&hsa-miR-106b-3p&offsets|0|-1;m-87&7|-|100094003|100094023&offsets|0|0]|CCGCACTGTGGGTACTTGCTG</td><td>>MI0000734|hsa-mir-106b&WithFlank&7|-|100093987|100094080</td><td>58</td><td>78</td><td>21</td><td>CCGCACTGTGGGTACTTGCTG</td><td>BT20</td><td>114</td><td>93</td><td>259</td><td>166</td><td>77</td><td>27.7</td><td>23.8</td><td>76.0</td><td>108.9</td><td>21.2</td></tr>
<tr><td>>MI0000734|hsa-mir-106b&WithFlank&7|-|100093987|100094080@58.78.21&[MIMAT0004672&hsa-miR-106b-3p&offsets|0|-1;m-87&7|-|100094003|100094023&offsets|0|0]|CCGCACTGTGGGTACTTGCTG</td><td>>MI0000734|hsa-mir-106b&WithFlank&7|-|100093987|100094080</td><td>58</td><td>78</td><td>21</td><td>CCGCACTGTGGGTACTTGCTG</td><td>MB231</td><td>90</td><td>92</td><td>113</td><td>113</td><td>53</td><td>20.5</td><td>23.1</td><td>59.8</td><td>36.0</td><td>5.8</td></tr>
<tr><td>>MI0000734|hsa-mir-106b&WithFlank&7|-|100093987|100094080@58.79.22(+1A)&[MIMAT0004672&hsa-miR-106b-3p&offsets|0|0(+1A);m-87&7|-|100094003|100094023&offsets|0|+1(+1A)]|CCGCACTGTGGGTACTTGCTGCA</td><td>>MI0000734|hsa-mir-106b&WithFlank&7|-|100093987|100094080</td><td>58</td><td>79</td><td>22(+1A)</td><td>CCGCACTGTGGGTACTTGCTGCA</td><td>BT20</td><td>10</td><td>7</td><td>20</td><td>4</td><td>6</td><td>3.5</td><td>1.0</td><td>9.4</td><td>0.4</td><td>1.3</td></tr>
<tr><td>>MI0000734|hsa-mir-106b&WithFlank&7|-|100093987|100094080@58.79.22&[MIMAT0004672&hsa-miR-106b-3p&offsets|0|0;m-87&7|-|100094003|100094023&offsets|0|+1]|CCGCACTGTGGGTACTTGCTGC</td><td>>MI0000734|hsa-mir-106b&WithFlank&7|-|100093987|100094080</td><td>58</td><td>79</td><td>22</td><td>CCGCACTGTGGGTACTTGCTGC</td><td>BT20</td><td>250</td><td>206</td><td>515</td><td>414</td><td>171</td><td>147.9</td><td>57.1</td><td>285.2</td><td>315.1</td><td>84.4</td></tr>
<tr><td>>MI0000734|hsa-mir-106b&WithFlank&7|-|100093987|100094080@58.79.22&[MIMAT0004672&hsa-miR-106b-3p&offsets|0|0;m-87&7|-|100094003|100094023&offsets|0|+1]|CCGCACTGTGGGTACTTGCTGC</td><td>>MI0000734|hsa-mir-106b&WithFlank&7|-|100093987|100094080</td><td>58</td><td>79</td><td>22</td><td>CCGCACTGTGGGTACTTGCTGC</td><td>MB231</td><td>245</td><td>347</td><td>254</td><td>195</td><td>155</td><td>88.4</td><td>88.9</td><td>86.2</td><td>68.1</td><td>23.4</td></tr>
<tr><td>>MI0000734|hsa-mir-106b&WithFlank&7|-|100093987|100094080@58.79.22&[MIMAT0004672&hsa-miR-106b-3p&offsets|0|0;m-87&7|-|100094003|100094023&offsets|0|+1]|CCGCACTGTGGGTACTTGCTGC</td><td>>MI0000734|hsa-mir-106b&WithFlank&7|-|100093987|100094080</td><td>58</td><td>79</td><td>22</td><td>CCGCACTGTGGGTACTTGCTGC</td><td>MB468</td><td>15</td><td>29</td><td>14</td><td>18</td><td>5</td><td>6.0</td><td>8.3</td><td>7.9</td><td>18.0</td><td>1.6</td></tr>
<tr><td>>MI0000734|hsa-mir-106b&WithFlank&7|-|100093987|100094080@58.80.23&[MIMAT0004672&hsa-miR-106b-3p&offsets|0|+1;m-87&7|-|100094003|100094023&offsets|0|+2]|CCGCACTGTGGGTACTTGCTGCT</td><td>>MI0000734|hsa-mir-106b&WithFlank&7|-|100093987|100094080</td><td>58</td><td>80</td><td>23</td><td>CCGCACTGTGGGTACTTGCTGCT</td><td>MB231</td><td>10</td><td>11</td><td>9</td><td>9</td><td>6</td><td>2.9</td><td>3.2</td><td>1.5</td><td>2.7</td><td>0.8</td></tr>
</table>

<table>
<tr><th></th><th></th><th></th><th></th><th></th><th></th><th></th><th></th><th></th><th></th><th></th><th></th><th></th><th></th><th></th><th></th><th></th></tr>
<tr><td>>MI0000736|hsa-mir-30c-1&WithFlank&1|+|40757278|40757378@23.42.20;MI0000254|hsa-mir-30c-2&WithFlank&6|-|71376954|71377037@13.32.20&[MIMAT0000244&hsa-miR-30c-5p&offsets|0|-3;m-57&1|+|40757300|40757323&offsets|0|-4];[MIMAT0000244_1&hsa-miR-30c-5p&offsets|0|-3;m-56&6|-|71377002|71377025&offsets|0|-4]|TGTAAACATCCTACACTCTC</td><td>>MI0000736|hsa-mir-30c-1&WithFlank&1|+|40757278|40757378</td><td>23</td><td>42</td><td>20</td><td>TGTAAACATCCTACACTCTC</td><td>BT20</td><td>44</td><td>42</td><td>40</td><td>18</td><td>18</td><td>53.1</td><td>48.9</td><td>49.1</td><td>12.8</td><td>17.1</td></tr>
<tr><td>>MI0000736|hsa-mir-30c-1&WithFlank&1|+|40757278|40757378@23.42.20;MI0000254|hsa-mir-30c-2&WithFlank&6|-|71376954|71377037@13.32.20&[MIMAT0000244&hsa-miR-30c-5p&offsets|0|-3;m-57&1|+|40757300|40757323&offsets|0|-4];[MIMAT0000244_1&hsa-miR-30c-5p&offsets|0|-3;m-56&6|-|71377002|71377025&offsets|0|-4]|TGTAAACATCCTACACTCTC</td><td>>MI0000736|hsa-mir-30c-1&WithFlank&1|+|40757278|40757378</td><td>23</td><td>42</td><td>20</td><td>TGTAAACATCCTACACTCTC</td><td>MB231</td><td>66</td><td>101</td><td>23</td><td>84</td><td>134</td><td>21.4</td><td>44.2</td><td>17.0</td><td>57.9</td><td>74.7</td></tr>
<tr><td>>MI0000736|hsa-mir-30c-1&WithFlank&1|+|40757278|40757378@23.44.22(+1A);MI0000254|hsa-mir-30c-2&WithFlank&6|-|71376954|71377037@13.34.22(+1A)&[MIMAT0000244&hsa-miR-30c-5p&offsets|0|-1(+1A);m-57&1|+|40757300|40757323&offsets|0|-2(+1A)];[MIMAT0000244_1&hsa-miR-30c-5p&offsets|0|-1(+1A);m-56&6|-|71377002|71377025&offsets|0|-2(+1A)]|TGTAAACATCCTACACTCTCAGA</td><td>>MI0000736|hsa-mir-30c-1&WithFlank&1|+|40757278|40757378</td><td>23</td><td>44</td><td>22(+1A)</td><td>TGTAAACATCCTACACTCTCAGA</td><td>BT20</td><td>74</td><td>32</td><td>42</td><td>8</td><td>11</td><td>53.6</td><td>28.2</td><td>18.9</td><td>6.6</td><td>7.2</td></tr>
<tr><td>>MI0000736|hsa-mir-30c-1&WithFlank&1|+|40757278|40757378@23.44.22(+1A);MI0000254|hsa-mir-30c-2&WithFlank&6|-|71376954|71377037@13.34.22(+1A)&[MIMAT0000244&hsa-miR-30c-5p&offsets|0|-1(+1A);m-57&1|+|40757300|40757323&offsets|0|-2(+1A)];[MIMAT0000244_1&hsa-miR-30c-5p&offsets|0|-1(+1A);m-56&6|-|71377002|71377025&offsets|0|-2(+1A)]|TGTAAACATCCTACACTCTCAGA</td><td>>MI0000736|hsa-mir-30c-1&WithFlank&1|+|40757278|40757378</td><td>23</td><td>44</td><td>22(+1A)</td><td>TGTAAACATCCTACACTCTCAGA</td><td>MB231</td><td>83</td><td>73</td><td>22</td><td>46</td><td>144</td><td>12.3</td><td>13.2</td><td>15.3</td><td>37.9</td><td>115.5</td></tr>
<tr><td>>MI0000736|hsa-mir-30c-1&WithFlank&1|+|40757278|40757378@23.44.22(+1G);MI0000254|hsa-mir-30c-2&WithFlank&6|-|71376954|71377037@13.34.22(+1G)&[MIMAT0000244&hsa-miR-30c-5p&offsets|0|-1(+1G);m-57&1|+|40757300|40757323&offsets|0|-2(+1G)];[MIMAT0000244_1&hsa-miR-30c-5p&offsets|0|-1(+1G);m-56&6|-|71377002|71377025&offsets|0|-2(+1G)]|TGTAAACATCCTACACTCTCAGG</td><td>>MI0000736|hsa-mir-30c-1&WithFlank&1|+|40757278|40757378</td><td>23</td><td>44</td><td>22(+1G)</td><td>TGTAAACATCCTACACTCTCAGG</td><td>BT20</td><td>11</td><td>3</td><td>2</td><td>0</td><td>0</td><td>12.8</td><td>2.3</td><td>0.5</td><td>0.0</td><td>0.0</td></tr>
<tr><td>>MI0000736|hsa-mir-30c-1&WithFlank&1|+|40757278|40757378@23.44.22(+1G);MI0000254|hsa-mir-30c-2&WithFlank&6|-|71376954|71377037@13.34.22(+1G)&[MIMAT0000244&hsa-miR-30c-5p&offsets|0|-1(+1G);m-57&1|+|40757300|40757323&offsets|0|-2(+1G)];[MIMAT0000244_1&hsa-miR-30c-5p&offsets|0|-1(+1G);m-56&6|-|71377002|71377025&offsets|0|-2(+1G)]|TGTAAACATCCTACACTCTCAGG</td><td>>MI0000736|hsa-mir-30c-1&WithFlank&1|+|40757278|40757378</td><td>23</td><td>44</td><td>22(+1G)</td><td>TGTAAACATCCTACACTCTCAGG</td><td>MB231</td><td>11</td><td>5</td><td>1</td><td>2</td><td>7</td><td>9.0</td><td>2.9</td><td>0.9</td><td>2.1</td><td>9.1</td></tr>
<tr><td>>MI0000736|hsa-mir-30c-1&WithFlank&1|+|40757278|40757378@23.44.22(+1U);MI0000254|hsa-mir-30c-2&WithFlank&6|-|71376954|71377037@13.34.22(+1U)&[MIMAT0000244&hsa-miR-30c-5p&offsets|0|-1(+1U);m-57&1|+|40757300|40757323&offsets|0|-2(+1U)];[MIMAT0000244_1&hsa-miR-30c-5p&offsets|0|-1(+1U);m-56&6|-|71377002|71377025&offsets|0|-2(+1U)]|TGTAAACATCCTACACTCTCAGT</td><td>>MI0000736|hsa-mir-30c-1&WithFlank&1|+|40757278|40757378</td><td>23</td><td>44</td><td>22(+1U)</td><td>TGTAAACATCCTACACTCTCAGT</td><td>BT20</td><td>269</td><td>154</td><td>147</td><td>28</td><td>51</td><td>159.6</td><td>141.7</td><td>86.7</td><td>23.0</td><td>37.2</td></tr>
<tr><td>>MI0000736|hsa-mir-30c-1&WithFlank&1|+|40757278|40757378@23.44.22(+1U);MI0000254|hsa-mir-30c-2&WithFlank&6|-|71376954|71377037@13.34.22(+1U)&[MIMAT0000244&hsa-miR-30c-5p&offsets|0|-1(+1U);m-57&1|+|40757300|40757323&offsets|0|-2(+1U)];[MIMAT0000244_1&hsa-miR-30c-5p&offsets|0|-1(+1U);m-56&6|-|71377002|71377025&offsets|0|-2(+1U)]|TGTAAACATCCTACACTCTCAGT</td><td>>MI0000736|hsa-mir-30c-1&WithFlank&1|+|40757278|40757378</td><td>23</td><td>44</td><td>22(+1U)</td><td>TGTAAACATCCTACACTCTCAGT</td><td>MB231</td><td>380</td><td>339</td><td>81</td><td>201</td><td>689</td><td>35.2</td><td>68.1</td><td>50.1</td><td>163.4</td><td>620.2</td></tr>
<tr><td>>MI0000736|hsa-mir-30c-1&WithFlank&1|+|40757278|40757378@23.44.22(+1U);MI0000254|hsa-mir-30c-2&WithFlank&6|-|71376954|71377037@13.34.22(+1U)&[MIMAT0000244&hsa-miR-30c-5p&offsets|0|-1(+1U);m-57&1|+|40757300|40757323&offsets|0|-2(+1U)];[MIMAT0000244_1&hsa-miR-30c-5p&offsets|0|-1(+1U);m-56&6|-|71377002|71377025&offsets|0|-2(+1U)]|TGTAAACATCCTACACTCTCAGT</td><td>>MI0000736|hsa-mir-30c-1&WithFlank&1|+|40757278|40757378</td><td>23</td><td>44</td><td>22(+1U)</td><td>TGTAAACATCCTACACTCTCAGT</td><td>MB468</td><td>19</td><td>18</td><td>5</td><td>6</td><td>9</td><td>5.0</td><td>6.3</td><td>2.4</td><td>3.0</td><td>7.1</td></tr>
<tr><td>>MI0000736|hsa-mir-30c-1&WithFlank&1|+|40757278|40757378@23.45.23;MI0000254|hsa-mir-30c-2&WithFlank&6|-|71376954|71377037@13.35.23&[MIMAT0000244&hsa-miR-30c-5p&offsets|0|0;m-57&1|+|40757300|40757323&offsets|0|-1];[MIMAT0000244_1&hsa-miR-30c-5p&offsets|0|0;m-56&6|-|71377002|71377025&offsets|0|-1]|TGTAAACATCCTACACTCTCAGC</td><td>>MI0000736|hsa-mir-30c-1&WithFlank&1|+|40757278|40757378</td><td>23</td><td>45</td><td>23</td><td>TGTAAACATCCTACACTCTCAGC</td><td>BT20</td><td>744</td><td>491</td><td>533</td><td>99</td><td>221</td><td>513.7</td><td>365.2</td><td>413.6</td><td>58.9</td><td>174.7</td></tr>
<tr><td>>MI0000736|hsa-mir-30c-1&WithFlank&1|+|40757278|40757378@23.45.23;MI0000254|hsa-mir-30c-2&WithFlank&6|-|71376954|71377037@13.35.23&[MIMAT0000244&hsa-miR-30c-5p&offsets|0|0;m-57&1|+|40757300|40757323&offsets|0|-1];[MIMAT0000244_1&hsa-miR-30c-5p&offsets|0|0;m-56&6|-|71377002|71377025&offsets|0|-1]|TGTAAACATCCTACACTCTCAGC</td><td>>MI0000736|hsa-mir-30c-1&WithFlank&1|+|40757278|40757378</td><td>23</td><td>45</td><td>23</td><td>TGTAAACATCCTACACTCTCAGC</td><td>MB231</td><td>1602</td><td>1884</td><td>398</td><td>510</td><td>2396</td><td>303.0</td><td>550.6</td><td>233.6</td><td>342.8</td><td>1517.8</td></tr>
<tr><td>>MI0000736|hsa-mir-30c-1&WithFlank&1|+|40757278|40757378@23.45.23;MI0000254|hsa-mir-30c-2&WithFlank&6|-|71376954|71377037@13.35.23&[MIMAT0000244&hsa-miR-30c-5p&offsets|0|0;m-57&1|+|40757300|40757323&offsets|0|-1];[MIMAT0000244_1&hsa-miR-30c-5p&offsets|0|0;m-56&6|-|71377002|71377025&offsets|0|-1]|TGTAAACATCCTACACTCTCAGC</td><td>>MI0000736|hsa-mir-30c-1&WithFlank&1|+|40757278|40757378</td><td>23</td><td>45</td><td>23</td><td>TGTAAACATCCTACACTCTCAGC</td><td>MB468</td><td>103</td><td>112</td><td>27</td><td>29</td><td>40</td><td>44.7</td><td>14.8</td><td>7.1</td><td>11.5</td><td>15.0</td></tr>
<tr><td>>MI0000736|hsa-mir-30c-1&WithFlank&1|+|40757278|40757378@23.45.23(+1A);MI0000254|hsa-mir-30c-2&WithFlank&6|-|71376954|71377037@13.35.23(+1A)&[MIMAT0000244&hsa-miR-30c-5p&offsets|0|0(+1A);m-57&1|+|40757300|40757323&offsets|0|-1(+1A)];[MIMAT0000244_1&hsa-miR-30c-5p&offsets|0|0(+1A);m-56&6|-|71377002|71377025&offsets|0|-1(+1A)]|TGTAAACATCCTACACTCTCAGCA</td><td>>MI0000736|hsa-mir-30c-1&WithFlank&1|+|40757278|40757378</td><td>23</td><td>45</td><td>23(+1A)</td><td>TGTAAACATCCTACACTCTCAGCA</td><td>BT20</td><td>156</td><td>102</td><td>76</td><td>18</td><td>32</td><td>131.2</td><td>112.2</td><td>42.6</td><td>14.9</td><td>33.0</td></tr>
<tr><td>>MI0000736|hsa-mir-30c-1&WithFlank&1|+|40757278|40757378@23.45.23(+1A);MI0000254|hsa-mir-30c-2&WithFlank&6|-|71376954|71377037@13.35.23(+1A)&[MIMAT0000244&hsa-miR-30c-5p&offsets|0|0(+1A);m-57&1|+|40757300|40757323&offsets|0|-1(+1A)];[MIMAT0000244_1&hsa-miR-30c-5p&offsets|0|0(+1A);m-56&6|-|71377002|71377025&offsets|0|-1(+1A)]|TGTAAACATCCTACACTCTCAGCA</td><td>>MI0000736|hsa-mir-30c-1&WithFlank&1|+|40757278|40757378</td><td>23</td><td>45</td><td>23(+1A)</td><td>TGTAAACATCCTACACTCTCAGCA</td><td>MB231</td><td>118</td><td>119</td><td>40</td><td>106</td><td>283</td><td>7.2</td><td>27.9</td><td>29.8</td><td>86.8</td><td>250.2</td></tr>
<tr><td>>MI0000736|hsa-mir-30c-1&WithFlank&1|+|40757278|40757378@23.45.23(+1A);MI0000254|hsa-mir-30c-2&WithFlank&6|-|71376954|71377037@13.35.23(+1A)&[MIMAT0000244&hsa-miR-30c-5p&offsets|0|0(+1A);m-57&1|+|40757300|40757323&offsets|0|-1(+1A)];[MIMAT0000244_1&hsa-miR-30c-5p&offsets|0|0(+1A);m-56&6|-|71377002|71377025&offsets|0|-1(+1A)]|TGTAAACATCCTACACTCTCAGCA</td><td>>MI0000736|hsa-mir-30c-1&WithFlank&1|+|40757278|40757378</td><td>23</td><td>45</td><td>23(+1A)</td><td>TGTAAACATCCTACACTCTCAGCA</td><td>MB468</td><td>11</td><td>10</td><td>4</td><td>5</td><td>6</td><td>3.0</td><td>2.0</td><td>2.3</td><td>4.9</td><td>3.4</td></tr>
<tr><td>>MI0000736|hsa-mir-30c-1&WithFlank&1|+|40757278|40757378@23.45.23(+1C);MI0000254|hsa-mir-30c-2&WithFlank&6|-|71376954|71377037@13.35.23(+1C)&[MIMAT0000244&hsa-miR-30c-5p&offsets|0|0(+1C);m-57&1|+|40757300|40757323&offsets|0|-1(+1C)];[MIMAT0000244_1&hsa-miR-30c-5p&offsets|0|0(+1C);m-56&6|-|71377002|71377025&offsets|0|-1(+1C)]|TGTAAACATCCTACACTCTCAGCC</td><td>>MI0000736|hsa-mir-30c-1&WithFlank&1|+|40757278|40757378</td><td>23</td><td>45</td><td>23(+1C)</td><td>TGTAAACATCCTACACTCTCAGCC</td><td>BT20</td><td>138</td><td>145</td><td>80</td><td>26</td><td>51</td><td>83.0</td><td>165.7</td><td>62.9</td><td>21.2</td><td>55.9</td></tr>
<tr><td>>MI0000736|hsa-mir-30c-1&WithFlank&1|+|40757278|40757378@23.45.23(+1C);MI0000254|hsa-mir-30c-2&WithFlank&6|-|71376954|71377037@13.35.23(+1C)&[MIMAT0000244&hsa-miR-30c-5p&offsets|0|0(+1C);m-57&1|+|40757300|40757323&offsets|0|-1(+1C)];[MIMAT0000244_1&hsa-miR-30c-5p&offsets|0|0(+1C);m-56&6|-|71377002|71377025&offsets|0|-1(+1C)]|TGTAAACATCCTACACTCTCAGCC</td><td>>MI0000736|hsa-mir-30c-1&WithFlank&1|+|40757278|40757378</td><td>23</td><td>45</td><td>23(+1C)</td><td>TGTAAACATCCTACACTCTCAGCC</td><td>MB231</td><td>157</td><td>183</td><td>44</td><td>142</td><td>402</td><td>25.6</td><td>47.9</td><td>29.9</td><td>119.9</td><td>364.5</td></tr>
<tr><td>>MI0000736|hsa-mir-30c-1&WithFlank&1|+|40757278|40757378@23.45.23(+1C);MI0000254|hsa-mir-30c-2&WithFlank&6|-|71376954|71377037@13.35.23(+1C)&[MIMAT0000244&hsa-miR-30c-5p&offsets|0|0(+1C);m-57&1|+|40757300|40757323&offsets|0|-1(+1C)];[MIMAT0000244_1&hsa-miR-30c-5p&offsets|0|0(+1C);m-56&6|-|71377002|71377025&offsets|0|-1(+1C)]|TGTAAACATCCTACACTCTCAGCC</td><td>>MI0000736|hsa-mir-30c-1&WithFlank&1|+|40757278|40757378</td><td>23</td><td>45</td><td>23(+1C)</td><td>TGTAAACATCCTACACTCTCAGCC</td><td>MB468</td><td>17</td><td>18</td><td>6</td><td>8</td><td>9</td><td>6.2</td><td>0.6</td><td>2.2</td><td>7.5</td><td>3.3</td></tr>
</table>

<table>
<tr><td>>MI0000736|hsa-mir-30c-1&WithFlank&1|+|40757278|40757378@23.45.23(+1G);MI0000254|hsa-mir-30c-2&WithFlank&6|-|71376954|71377037@13.35.23(+1G)&[MIMAT0000244&hsa-miR-30c-5p&offsets|0|0(+1G);m-57&1|+|40757300|40757323&offsets|0|-1(+1G)];[MIMAT0000244_1&hsa-miR-30c-5p&offsets|0|0(+1G);m-56&6|-|71377002|71377025&offsets|0|-1(+1G)]|TGTAAACATCCTACACTCTCAGCG</td><td>>MI0000736|hsa-mir-30c-1&WithFlank&1|+|40757278|40757378</td><td>23</td><td>45</td><td>23(+1G)</td><td>TGTAAACATCCTACACTCTCAGCG</td><td>BT20</td><td>93</td><td>37</td><td>19</td><td>5</td><td>6</td><td>102.7</td><td>36.7</td><td>7.1</td><td>5.0</td><td>5.9</td></tr>
<tr><td>>MI0000736|hsa-mir-30c-1&WithFlank&1|+|40757278|40757378@23.45.23(+1G);MI0000254|hsa-mir-30c-2&WithFlank&6|-|71376954|71377037@13.35.23(+1G)&[MIMAT0000244&hsa-miR-30c-5p&offsets|0|0(+1G);m-57&1|+|40757300|40757323&offsets|0|-1(+1G)];[MIMAT0000244_1&hsa-miR-30c-5p&offsets|0|0(+1G);m-56&6|-|71377002|71377025&offsets|0|-1(+1G)]|TGTAAACATCCTACACTCTCAGCG</td><td>>MI0000736|hsa-mir-30c-1&WithFlank&1|+|40757278|40757378</td><td>23</td><td>45</td><td>23(+1G)</td><td>TGTAAACATCCTACACTCTCAGCG</td><td>MB231</td><td>88</td><td>44</td><td>10</td><td>20</td><td>52</td><td>62.5</td><td>18.5</td><td>7.7</td><td>13.6</td><td>51.9</td></tr>
<tr><td>>MI0000736|hsa-mir-30c-1&WithFlank&1|+|40757278|40757378@23.45.23(+1G);MI0000254|hsa-mir-30c-2&WithFlank&6|-|71376954|71377037@13.35.23(+1G)&[MIMAT0000244&hsa-miR-30c-5p&offsets|0|0(+1G);m-57&1|+|40757300|40757323&offsets|0|-1(+1G)];[MIMAT0000244_1&hsa-miR-30c-5p&offsets|0|0(+1G);m-56&6|-|71377002|71377025&offsets|0|-1(+1G)]|TGTAAACATCCTACACTCTCAGCG</td><td>>MI0000736|hsa-mir-30c-1&WithFlank&1|+|40757278|40757378</td><td>23</td><td>45</td><td>23(+1G)</td><td>TGTAAACATCCTACACTCTCAGCG</td><td>MB468</td><td>10</td><td>5</td><td>4</td><td>2</td><td>2</td><td>2.0</td><td>1.8</td><td>2.0</td><td>2.6</td><td>1.9</td></tr>
<tr><td>>MI0000736|hsa-mir-30c-1&WithFlank&1|+|40757278|40757378@23.46.24;MI0000254|hsa-mir-30c-2&WithFlank&6|-|71376954|71377037@13.36.24&[MIMAT0000244&hsa-miR-30c-5p&offsets|0|+1;m-57&1|+|40757300|40757323&offsets|0|0];[MIMAT0000244_1&hsa-miR-30c-5p&offsets|0|+1;m-56&6|-|71377002|71377025&offsets|0|0]|TGTAAACATCCTACACTCTCAGCT</td><td>>MI0000736|hsa-mir-30c-1&WithFlank&1|+|40757278|40757378</td><td>23</td><td>46</td><td>24</td><td>TGTAAACATCCTACACTCTCAGCT</td><td>BT20</td><td>912</td><td>830</td><td>566</td><td>169</td><td>309</td><td>550.4</td><td>863.0</td><td>491.6</td><td>123.4</td><td>325.4</td></tr>
<tr><td>>MI0000736|hsa-mir-30c-1&WithFlank&1|+|40757278|40757378@23.46.24;MI0000254|hsa-mir-30c-2&WithFlank&6|-|71376954|71377037@13.36.24&[MIMAT0000244&hsa-miR-30c-5p&offsets|0|+1;m-57&1|+|40757300|40757323&offsets|0|0];[MIMAT0000244_1&hsa-miR-30c-5p&offsets|0|+1;m-56&6|-|71377002|71377025&offsets|0|0]|TGTAAACATCCTACACTCTCAGCT</td><td>>MI0000736|hsa-mir-30c-1&WithFlank&1|+|40757278|40757378</td><td>23</td><td>46</td><td>24</td><td>TGTAAACATCCTACACTCTCAGCT</td><td>MB231</td><td>1191</td><td>1441</td><td>355</td><td>818</td><td>2515</td><td>227.5</td><td>337.1</td><td>225.6</td><td>616.3</td><td>1890.7</td></tr>
<tr><td>>MI0000736|hsa-mir-30c-1&WithFlank&1|+|40757278|40757378@23.46.24;MI0000254|hsa-mir-30c-2&WithFlank&6|-|71376954|71377037@13.36.24&[MIMAT0000244&hsa-miR-30c-5p&offsets|0|+1;m-57&1|+|40757300|40757323&offsets|0|0];[MIMAT0000244_1&hsa-miR-30c-5p&offsets|0|+1;m-56&6|-|71377002|71377025&offsets|0|0]|TGTAAACATCCTACACTCTCAGCT</td><td>>MI0000736|hsa-mir-30c-1&WithFlank&1|+|40757278|40757378</td><td>23</td><td>46</td><td>24</td><td>TGTAAACATCCTACACTCTCAGCT</td><td>MB468</td><td>138</td><td>147</td><td>41</td><td>52</td><td>62</td><td>56.2</td><td>17.0</td><td>14.2</td><td>51.4</td><td>16.7</td></tr>
<tr><td>>MI0000736|hsa-mir-30c-1&WithFlank&1|+|40757278|40757378@23.46.24(+1A);MI0000254|hsa-mir-30c-2&WithFlank&6|-|71376954|71377037@13.36.24(+1A)&[MIMAT0000244&hsa-miR-30c-5p&offsets|0|+1(+1A);m-57&1|+|40757300|40757323&offsets|0|0(+1A)];[MIMAT0000244_1&hsa-miR-30c-5p&offsets|0|+1(+1A);m-56&6|-|71377002|71377025&offsets|0|0(+1A)]|TGTAAACATCCTACACTCTCAGCTA</td><td>>MI0000736|hsa-mir-30c-1&WithFlank&1|+|40757278|40757378</td><td>23</td><td>46</td><td>24(+1A)</td><td>TGTAAACATCCTACACTCTCAGCTA</td><td>BT20</td><td>13</td><td>7</td><td>11</td><td>1</td><td>3</td><td>10.4</td><td>8.2</td><td>10.6</td><td>1.4</td><td>4.3</td></tr>
<tr><td>>MI0000736|hsa-mir-30c-1&WithFlank&1|+|40757278|40757378@23.46.24(+1A);MI0000254|hsa-mir-30c-2&WithFlank&6|-|71376954|71377037@13.36.24(+1A)&[MIMAT0000244&hsa-miR-30c-5p&offsets|0|+1(+1A);m-57&1|+|40757300|40757323&offsets|0|0(+1A)];[MIMAT0000244_1&hsa-miR-30c-5p&offsets|0|+1(+1A);m-56&6|-|71377002|71377025&offsets|0|0(+1A)]|TGTAAACATCCTACACTCTCAGCTA</td><td>>MI0000736|hsa-mir-30c-1&WithFlank&1|+|40757278|40757378</td><td>23</td><td>46</td><td>24(+1A)</td><td>TGTAAACATCCTACACTCTCAGCTA</td><td>MB231</td><td>22</td><td>15</td><td>5</td><td>6</td><td>26</td><td>6.9</td><td>3.7</td><td>2.5</td><td>5.6</td><td>16.7</td></tr>
<tr><td>>MI0000736|hsa-mir-30c-1&WithFlank&1|+|40757278|40757378@23.46.24(+1U);MI0000254|hsa-mir-30c-2&WithFlank&6|-|71376954|71377037@13.36.24(+1U)&[MIMAT0000244&hsa-miR-30c-5p&offsets|0|+1(+1U);m-57&1|+|40757300|40757323&offsets|0|0(+1U)];[MIMAT0000244_1&hsa-miR-30c-5p&offsets|0|+1(+1U);m-56&6|-|71377002|71377025&offsets|0|0(+1U)]|TGTAAACATCCTACACTCTCAGCTT</td><td>>MI0000736|hsa-mir-30c-1&WithFlank&1|+|40757278|40757378</td><td>23</td><td>46</td><td>24(+1U)</td><td>TGTAAACATCCTACACTCTCAGCTT</td><td>BT20</td><td>17</td><td>15</td><td>12</td><td>7</td><td>4</td><td>11.5</td><td>16.8</td><td>11.2</td><td>5.1</td><td>5.9</td></tr>
<tr><td>>MI0000736|hsa-mir-30c-1&WithFlank&1|+|40757278|40757378@23.46.24(+1U);MI0000254|hsa-mir-30c-2&WithFlank&6|-|71376954|71377037@13.36.24(+1U)&[MIMAT0000244&hsa-miR-30c-5p&offsets|0|+1(+1U);m-57&1|+|40757300|40757323&offsets|0|0(+1U)];[MIMAT0000244_1&hsa-miR-30c-5p&offsets|0|+1(+1U);m-56&6|-|71377002|71377025&offsets|0|0(+1U)]|TGTAAACATCCTACACTCTCAGCTT</td><td>>MI0000736|hsa-mir-30c-1&WithFlank&1|+|40757278|40757378</td><td>23</td><td>46</td><td>24(+1U)</td><td>TGTAAACATCCTACACTCTCAGCTT</td><td>MB231</td><td>66</td><td>65</td><td>16</td><td>46</td><td>105</td><td>13.0</td><td>18.4</td><td>6.6</td><td>36.7</td><td>86.8</td></tr>
<tr><td>>MI0000736|hsa-mir-30c-1&WithFlank&1|+|40757278|40757378@23.46.24(+2U);MI0000254|hsa-mir-30c-2&WithFlank&6|-|71376954|71377037@13.36.24(+2U)&[MIMAT0000244&hsa-miR-30c-5p&offsets|0|+1(+2U);m-57&1|+|40757300|40757323&offsets|0|0(+2U)];[MIMAT0000244_1&hsa-miR-30c-5p&offsets|0|+1(+2U);m-56&6|-|71377002|71377025&offsets|0|0(+2U)]|TGTAAACATCCTACACTCTCAGCTTT</td><td>>MI0000736|hsa-mir-30c-1&WithFlank&1|+|40757278|40757378</td><td>23</td><td>46</td><td>24(+2U)</td><td>TGTAAACATCCTACACTCTCAGCTTT</td><td>MB231</td><td>21</td><td>20</td><td>7</td><td>17</td><td>39</td><td>6.0</td><td>6.7</td><td>4.2</td><td>13.7</td><td>34.9</td></tr>
<tr><td>>MI0000736|hsa-mir-30c-1&WithFlank&1|+|40757278|40757378@23.48.26;MI0000254|hsa-mir-30c-2&WithFlank&6|-|71376954|71377037@13.38.26&[MIMAT0000244&hsa-miR-30c-5p&offsets|0|+3;m-57&1|+|40757300|40757323&offsets|0|+2];[MIMAT0000244_1&hsa-miR-30c-5p&offsets|0|+3;m-56&6|-|71377002|71377025&offsets|0|+2]|TGTAAACATCCTACACTCTCAGCTGT</td><td>>MI0000736|hsa-mir-30c-1&WithFlank&1|+|40757278|40757378</td><td>23</td><td>48</td><td>26</td><td>TGTAAACATCCTACACTCTCAGCTGT</td><td>MB231</td><td>17</td><td>18</td><td>7</td><td>10</td><td>28</td><td>12.8</td><td>5.4</td><td>3.0</td><td>7.0</td><td>13.9</td></tr>
<tr><td>>MI0000737|hsa-mir-200a&WithFlank&1|+|1167857|1167958@22.43.22&[MIMAT0001620&hsa-miR-200a-5p&offsets|0|0;m-154&1|+|1167878|1167899&offsets|0|0]|CATCTTACCGGACAGTGCTGGA</td><td>>MI0000737|hsa-mir-200a&WithFlank&1|+|1167857|1167958</td><td>22</td><td>43</td><td>22</td><td>CATCTTACCGGACAGTGCTGGA</td><td>MB468</td><td>21</td><td>16</td><td>12</td><td>61</td><td>7</td><td>6.0</td><td>7.6</td><td>5.8</td><td>19.5</td><td>1.8</td></tr>
<tr><td>>MI0000737|hsa-mir-200a&WithFlank&1|+|1167857|1167958@60.80.21&[MIMAT0000682&hsa-miR-200a-3p&offsets|0|-1;m-142&1|+|1167916|1167938&offsets|0|-2]|TAACACTGTCTGGTAACGATG</td><td>>MI0000737|hsa-mir-200a&WithFlank&1|+|1167857|1167958</td><td>60</td><td>80</td><td>21</td><td>TAACACTGTCTGGTAACGATG</td><td>MB468</td><td>11</td><td>6</td><td>5</td><td>26</td><td>3</td><td>3.4</td><td>2.2</td><td>0.9</td><td>8.4</td><td>1.2</td></tr>
<tr><td>>MI0000737|hsa-mir-200a&WithFlank&1|+|1167857|1167958@60.81.22(+1C)&[MIMAT0000682&hsa-miR-200a-3p&offsets|0|0(+1C);m-142&1|+|1167916|1167938&offsets|0|-1(+1C)]|TAACACTGTCTGGTAACGATGTC</td><td>>MI0000737|hsa-mir-200a&WithFlank&1|+|1167857|1167958</td><td>60</td><td>81</td><td>22(+1C)</td><td>TAACACTGTCTGGTAACGATGTC</td><td>MB468</td><td>20</td><td>2</td><td>3</td><td>56</td><td>5</td><td>13.6</td><td>0.7</td><td>2.3</td><td>32.3</td><td>2.7</td></tr>
<tr><td>>MI0000737|hsa-mir-200a&WithFlank&1|+|1167857|1167958@60.81.22&[MIMAT0000682&hsa-miR-200a-3p&offsets|0|0;m-142&1|+|1167916|1167938&offsets|0|-1]|TAACACTGTCTGGTAACGATGT</td><td>>MI0000737|hsa-mir-200a&WithFlank&1|+|1167857|1167958</td><td>60</td><td>81</td><td>22</td><td>TAACACTGTCTGGTAACGATGT</td><td>MB468</td><td>35</td><td>6</td><td>7</td><td>87</td><td>8</td><td>20.2</td><td>1.2</td><td>3.0</td><td>30.2</td><td>0.5</td></tr>
<tr><td>>MI0000737|hsa-mir-200a&WithFlank&1|+|1167857|1167958@60.82.23(+1U)&[MIMAT0000682&hsa-miR-200a-3p&offsets|0|+1(+1U);m-142&1|+|1167916|1167938&offsets|0|0(+1U)]|TAACACTGTCTGGTAACGATGTTT</td><td>>MI0000737|hsa-mir-200a&WithFlank&1|+|1167857|1167958</td><td>60</td><td>82</td><td>23(+1U)</td><td>TAACACTGTCTGGTAACGATGTTT</td><td>MB468</td><td>15</td><td>1</td><td>1</td><td>63</td><td>4</td><td>12.4</td><td>0.4</td><td>0.8</td><td>39.4</td><td>1.0</td></tr>
<tr><td>>MI0000737|hsa-mir-200a&WithFlank&1|+|1167857|1167958@60.82.23&[MIMAT0000682&hsa-miR-200a-3p&offsets|0|+1;m-142&1|+|1167916|1167938&offsets|0|0]|TAACACTGTCTGGTAACGATGTT</td><td>>MI0000737|hsa-mir-200a&WithFlank&1|+|1167857|1167958</td><td>60</td><td>82</td><td>23</td><td>TAACACTGTCTGGTAACGATGTT</td><td>MB468</td><td>195</td><td>17</td><td>24</td><td>417</td><td>42</td><td>143.0</td><td>4.1</td><td>13.6</td><td>235.3</td><td>15.2</td></tr>
<tr><td>>MI0000739|hsa-mir-101-2&WithFlank&9|+|4850291|4850381@55.77.23&[MIMAT0000099_1&hsa-miR-101-3p&offsets|0|+2;m-16&9|+|4850345|4850364&offsets|0|+3]|TACAGTACTGTGATAACTGAAGA</td><td>>MI0000739|hsa-mir-101-2&WithFlank&9|+|4850291|4850381</td><td>55</td><td>77</td><td>23</td><td>TACAGTACTGTGATAACTGAAGA</td><td>BT20</td><td>40</td><td>9</td><td>4</td><td>8</td><td>2</td><td>51.4</td><td>4.6</td><td>2.3</td><td>10.3</td><td>0.9</td></tr>
<tr><td>>MI0000743|hsa-mir-34c&WithFlank&11|+|111513433|111513521@19.40.22(+1U)&[MIMAT0000686&hsa-miR-34c-5p&offsets|0|-1(+1U);m-336&11|+|111513451|111513473&offsets|0|-1(+1U)]|AGGCAGTGTAGTTAGCTGATTGT</td><td>>MI0000743|hsa-mir-34c&WithFlank&11|+|111513433|111513521</td><td>19</td><td>40</td><td>22(+1U)</td><td>AGGCAGTGTAGTTAGCTGATTGT</td><td>MB231</td><td>14</td><td>3</td><td>3</td><td>38</td><td>16</td><td>5.2</td><td>0.3</td><td>2.3</td><td>15.0</td><td>16.4</td></tr>
<tr><td>>MI0000743|hsa-mir-34c&WithFlank&11|+|111513433|111513521@19.41.23&[MIMAT0000686&hsa-miR-34c-5p&offsets|0|0;m-336&11|+|111513451|111513473&offsets|0|0]|AGGCAGTGTAGTTAGCTGATTGC</td><td>>MI0000743|hsa-mir-34c&WithFlank&11|+|111513433|111513521</td><td>19</td><td>41</td><td>23</td><td>AGGCAGTGTAGTTAGCTGATTGC</td><td>MB231</td><td>56</td><td>18</td><td>14</td><td>135</td><td>47</td><td>25.8</td><td>2.2</td><td>11.6</td><td>109.2</td><td>38.2</td></tr>
<tr><td>>MI0000746|hsa-mir-99b&WithFlank&19|+|51692606|51692687@13.31.19(+1A)&[MIMAT0000689&hsa-miR-99b-5p&offsets|0|-3(+1A);m-10&19|+|51692618|51692639&offsets|0|-3(+1A)]|CACCCGTAGAACCGACCTTA</td><td>>MI0000746|hsa-mir-99b&WithFlank&19|+|51692606|51692687</td><td>13</td><td>31</td><td>19(+1A)</td><td>CACCCGTAGAACCGACCTTA</td><td>BT20</td><td>15</td><td>11</td><td>32</td><td>16</td><td>7</td><td>8.8</td><td>6.1</td><td>14.7</td><td>9.5</td><td>1.7</td></tr>
<tr><td>>MI0000746|hsa-mir-99b&WithFlank&19|+|51692606|51692687@13.31.19&[MIMAT0000689&hsa-miR-99b-5p&offsets|0|-3;m-10&19|+|51692618|51692639&offsets|0|-3]|CACCCGTAGAACCGACCTT</td><td>>MI0000746|hsa-mir-99b&WithFlank&19|+|51692606|51692687</td><td>13</td><td>31</td><td>19</td><td>CACCCGTAGAACCGACCTT</td><td>BT20</td><td>10</td><td>9</td><td>17</td><td>17</td><td>5</td><td>4.5</td><td>4.3</td><td>2.8</td><td>13.4</td><td>0.4</td></tr>
</table>

| | | | | | | | | | | | | | | | | |
|---|---|---|---|---|---|---|---|---|---|---|---|---|---|---|---|---|
| >MI0000746\|hsa-mir-99b&WithFlank&19\|+\|51692606\|51692687@13.32.20(+1A)&[MIMAT0000689&hsa-miR-99b-5p&offsets\|0\|-2(+1A);m-10&19\|+\|51692618\|51692639&offsets\|0\|-2(+1A)]\|CACCCGTAGAACCGACCTTGA | >MI0000746\|hsa-mir-99b&WithFlank&19\|+\|51692606\|51692687 | 13 | 32 | 20(+1A) | CACCCGTAGAACCGACCTTGA | BT20 | 73 | 33 | 152 | 35 | 16 | 62.9 | 16.9 | 96.0 | 20.0 | 1.4 |
| >MI0000746\|hsa-mir-99b&WithFlank&19\|+\|51692606\|51692687@13.32.20(+1A)&[MIMAT0000689&hsa-miR-99b-5p&offsets\|0\|-2(+1A);m-10&19\|+\|51692618\|51692639&offsets\|0\|-2(+1A)]\|CACCCGTAGAACCGACCTTGA | >MI0000746\|hsa-mir-99b&WithFlank&19\|+\|51692606\|51692687 | 13 | 32 | 20(+1A) | CACCCGTAGAACCGACCTTGA | MB231 | 16 | 21 | 20 | 24 | 20 | 5.8 | 1.6 | 11.3 | 10.7 | 5.7 |
| >MI0000746\|hsa-mir-99b&WithFlank&19\|+\|51692606\|51692687@13.32.20(+1G)&[MIMAT0000689&hsa-miR-99b-5p&offsets\|0\|-2(+1G);m-10&19\|+\|51692618\|51692639&offsets\|0\|-2(+1G)]\|CACCCGTAGAACCGACCTTGG | >MI0000746\|hsa-mir-99b&WithFlank&19\|+\|51692606\|51692687 | 13 | 32 | 20(+1G) | CACCCGTAGAACCGACCTTGG | BT20 | 53 | 12 | 50 | 8 | 2 | 58.4 | 3.8 | 31.9 | 8.0 | 0.5 |
| >MI0000746\|hsa-mir-99b&WithFlank&19\|+\|51692606\|51692687@13.32.20(+1U)&[MIMAT0000689&hsa-miR-99b-5p&offsets\|0\|-2(+1U);m-10&19\|+\|51692618\|51692639&offsets\|0\|-2(+1U)]\|CACCCGTAGAACCGACCTTGT | >MI0000746\|hsa-mir-99b&WithFlank&19\|+\|51692606\|51692687 | 13 | 32 | 20(+1U) | CACCCGTAGAACCGACCTTGT | BT20 | 695 | 502 | 1681 | 537 | 228 | 410.5 | 196.8 | 597.1 | 284.8 | 38.8 |
| >MI0000746\|hsa-mir-99b&WithFlank&19\|+\|51692606\|51692687@13.32.20(+1U)&[MIMAT0000689&hsa-miR-99b-5p&offsets\|0\|-2(+1U);m-10&19\|+\|51692618\|51692639&offsets\|0\|-2(+1U)]\|CACCCGTAGAACCGACCTTGT | >MI0000746\|hsa-mir-99b&WithFlank&19\|+\|51692606\|51692687 | 13 | 32 | 20(+1U) | CACCCGTAGAACCGACCTTGT | MB231 | 60 | 79 | 64 | 89 | 78 | 17.3 | 10.4 | 26.5 | 40.9 | 23.1 |
| >MI0000746\|hsa-mir-99b&WithFlank&19\|+\|51692606\|51692687@13.32.20(+2G)&[MIMAT0000689&hsa-miR-99b-5p&offsets\|0\|-2(+2G);m-10&19\|+\|51692618\|51692639&offsets\|0\|-2(+2G)]\|CACCCGTAGAACCGACCTTGGG | >MI0000746\|hsa-mir-99b&WithFlank&19\|+\|51692606\|51692687 | 13 | 32 | 20(+2G) | CACCCGTAGAACCGACCTTGGG | BT20 | 28 | 6 | 19 | 3 | 2 | 38.3 | 2.3 | 13.7 | 2.8 | 0.7 |
| >MI0000746\|hsa-mir-99b&WithFlank&19\|+\|51692606\|51692687@13.32.20(+2U)&[MIMAT0000689&hsa-miR-99b-5p&offsets\|0\|-2(+2U);m-10&19\|+\|51692618\|51692639&offsets\|0\|-2(+2U)]\|CACCCGTAGAACCGACCTTGTT | >MI0000746\|hsa-mir-99b&WithFlank&19\|+\|51692606\|51692687 | 13 | 32 | 20(+2U) | CACCCGTAGAACCGACCTTGTT | BT20 | 118 | 43 | 143 | 29 | 19 | 137.2 | 29.5 | 119.9 | 15.6 | 6.0 |
| >MI0000746\|hsa-mir-99b&WithFlank&19\|+\|51692606\|51692687@13.32.20&[MIMAT0000689&hsa-miR-99b-5p&offsets\|0\|-2;m-10&19\|+\|51692618\|51692639&offsets\|0\|-2]\|CACCCGTAGAACCGACCTTG | >MI0000746\|hsa-mir-99b&WithFlank&19\|+\|51692606\|51692687 | 13 | 32 | 20 | CACCCGTAGAACCGACCTTG | BT20 | 98 | 102 | 207 | 174 | 72 | 73.6 | 55.8 | 56.2 | 131.0 | 14.0 |
| >MI0000746\|hsa-mir-99b&WithFlank&19\|+\|51692606\|51692687@13.32.20&[MIMAT0000689&hsa-miR-99b-5p&offsets\|0\|-2;m-10&19\|+\|51692618\|51692639&offsets\|0\|-2]\|CACCCGTAGAACCGACCTTG | >MI0000746\|hsa-mir-99b&WithFlank&19\|+\|51692606\|51692687 | 13 | 32 | 20 | CACCCGTAGAACCGACCTTG | MB231 | 20 | 34 | 37 | 34 | 22 | 5.6 | 18.7 | 28.6 | 16.9 | 5.5 |
| >MI0000746\|hsa-mir-99b&WithFlank&19\|+\|51692606\|51692687@13.33.21(+1A)&[MIMAT0000689&hsa-miR-99b-5p&offsets\|0\|-1(+1A);m-10&19\|+\|51692618\|51692639&offsets\|0\|-1(+1A)]\|CACCCGTAGAACCGACCTTGCA | >MI0000746\|hsa-mir-99b&WithFlank&19\|+\|51692606\|51692687 | 13 | 33 | 21(+1A) | CACCCGTAGAACCGACCTTGCA | BT20 | 596 | 248 | 1240 | 103 | 126 | 540.3 | 120.1 | 1057.4 | 25.8 | 36.9 |
| >MI0000746\|hsa-mir-99b&WithFlank&19\|+\|51692606\|51692687@13.33.21(+1A)&[MIMAT0000689&hsa-miR-99b-5p&offsets\|0\|-1(+1A);m-10&19\|+\|51692618\|51692639&offsets\|0\|-1(+1A)]\|CACCCGTAGAACCGACCTTGCA | >MI0000746\|hsa-mir-99b&WithFlank&19\|+\|51692606\|51692687 | 13 | 33 | 21(+1A) | CACCCGTAGAACCGACCTTGCA | MB231 | 211 | 172 | 216 | 182 | 196 | 29.2 | 25.8 | 135.9 | 51.1 | 94.5 |
| >MI0000746\|hsa-mir-99b&WithFlank&19\|+\|51692606\|51692687@13.33.21(+1A)&[MIMAT0000689&hsa-miR-99b-5p&offsets\|0\|-1(+1A);m-10&19\|+\|51692618\|51692639&offsets\|0\|-1(+1A)]\|CACCCGTAGAACCGACCTTGCA | >MI0000746\|hsa-mir-99b&WithFlank&19\|+\|51692606\|51692687 | 13 | 33 | 21(+1A) | CACCCGTAGAACCGACCTTGCA | MB468 | 13 | 17 | 12 | 18 | 5 | 3.6 | 4.4 | 5.0 | 6.9 | 1.3 |
| >MI0000746\|hsa-mir-99b&WithFlank&19\|+\|51692606\|51692687@13.33.21(+1C)&[MIMAT0000689&hsa-miR-99b-5p&offsets\|0\|-1(+1C);m-10&19\|+\|51692618\|51692639&offsets\|0\|-1(+1C)]\|CACCCGTAGAACCGACCTTGCC | >MI0000746\|hsa-mir-99b&WithFlank&19\|+\|51692606\|51692687 | 13 | 33 | 21(+1C) | CACCCGTAGAACCGACCTTGCC | BT20 | 29 | 10 | 45 | 4 | 5 | 28.2 | 4.7 | 42.5 | 1.9 | 0.3 |
| >MI0000746\|hsa-mir-99b&WithFlank&19\|+\|51692606\|51692687@13.33.21(+1C)&[MIMAT0000689&hsa-miR-99b-5p&offsets\|0\|-1(+1C);m-10&19\|+\|51692618\|51692639&offsets\|0\|-1(+1C)]\|CACCCGTAGAACCGACCTTGCC | >MI0000746\|hsa-mir-99b&WithFlank&19\|+\|51692606\|51692687 | 13 | 33 | 21(+1C) | CACCCGTAGAACCGACCTTGCC | MB231 | 11 | 7 | 8 | 9 | 9 | 2.8 | 0.6 | 4.5 | 2.8 | 5.7 |
| >MI0000746\|hsa-mir-99b&WithFlank&19\|+\|51692606\|51692687@13.33.21(+1U)&[MIMAT0000689&hsa-miR-99b-5p&offsets\|0\|-1(+1U);m-10&19\|+\|51692618\|51692639&offsets\|0\|-1(+1U)]\|CACCCGTAGAACCGACCTTGCT | >MI0000746\|hsa-mir-99b&WithFlank&19\|+\|51692606\|51692687 | 13 | 33 | 21(+1U) | CACCCGTAGAACCGACCTTGCT | BT20 | 98 | 30 | 129 | 12 | 15 | 102.5 | 15.6 | 118.8 | 5.4 | 3.5 |
| >MI0000746\|hsa-mir-99b&WithFlank&19\|+\|51692606\|51692687@13.33.21(+1U)&[MIMAT0000689&hsa-miR-99b-5p&offsets\|0\|-1(+1U);m-10&19\|+\|51692618\|51692639&offsets\|0\|-1(+1U)]\|CACCCGTAGAACCGACCTTGCT | >MI0000746\|hsa-mir-99b&WithFlank&19\|+\|51692606\|51692687 | 13 | 33 | 21(+1U) | CACCCGTAGAACCGACCTTGCT | MB231 | 30 | 20 | 21 | 22 | 29 | 11.0 | 2.9 | 10.8 | 8.4 | 19.8 |
| >MI0000746\|hsa-mir-99b&WithFlank&19\|+\|51692606\|51692687@13.33.21&[MIMAT0000689&hsa-miR-99b-5p&offsets\|0\|-1;m-10&19\|+\|51692618\|51692639&offsets\|0\|-1]\|CACCCGTAGAACCGACCTTGC | >MI0000746\|hsa-mir-99b&WithFlank&19\|+\|51692606\|51692687 | 13 | 33 | 21 | CACCCGTAGAACCGACCTTGC | BT20 | 443 | 377 | 976 | 479 | 255 | 256.7 | 165.9 | 117.0 | 349.6 | 75.4 |
| >MI0000746\|hsa-mir-99b&WithFlank&19\|+\|51692606\|51692687@13.33.21&[MIMAT0000689&hsa-miR-99b-5p&offsets\|0\|-1;m-10&19\|+\|51692618\|51692639&offsets\|0\|-1]\|CACCCGTAGAACCGACCTTGC | >MI0000746\|hsa-mir-99b&WithFlank&19\|+\|51692606\|51692687 | 13 | 33 | 21 | CACCCGTAGAACCGACCTTGC | MB231 | 297 | 544 | 360 | 306 | 364 | 119.8 | 133.6 | 135.0 | 53.0 | 74.2 |
| >MI0000746\|hsa-mir-99b&WithFlank&19\|+\|51692606\|51692687@13.33.21&[MIMAT0000689&hsa-miR-99b-5p&offsets\|0\|-1;m-10&19\|+\|51692618\|51692639&offsets\|0\|-1]\|CACCCGTAGAACCGACCTTGC | >MI0000746\|hsa-mir-99b&WithFlank&19\|+\|51692606\|51692687 | 13 | 33 | 21 | CACCCGTAGAACCGACCTTGC | MB468 | 18 | 58 | 16 | 32 | 12 | 6.0 | 26.7 | 12.0 | 10.3 | 2.6 |
| >MI0000746\|hsa-mir-99b&WithFlank&19\|+\|51692606\|51692687@13.34.22(+1A)&[MIMAT0000689&hsa-miR-99b-5p&offsets\|0\|0(+1A);m-10&19\|+\|51692618\|51692639&offsets\|0\|0(+1A)]\|CACCCGTAGAACCGACCTTGCGA | >MI0000746\|hsa-mir-99b&WithFlank&19\|+\|51692606\|51692687 | 13 | 34 | 22(+1A) | CACCCGTAGAACCGACCTTGCGA | BT20 | 88 | 67 | 194 | 28 | 41 | 48.5 | 28.1 | 59.0 | 10.2 | 10.8 |
| >MI0000746\|hsa-mir-99b&WithFlank&19\|+\|51692606\|51692687@13.34.22(+1A)&[MIMAT0000689&hsa-miR-99b-5p&offsets\|0\|0(+1A);m-10&19\|+\|51692618\|51692639&offsets\|0\|0(+1A)]\|CACCCGTAGAACCGACCTTGCGA | >MI0000746\|hsa-mir-99b&WithFlank&19\|+\|51692606\|51692687 | 13 | 34 | 22(+1A) | CACCCGTAGAACCGACCTTGCGA | MB231 | 33 | 36 | 38 | 26 | 34 | 2.9 | 10.4 | 23.6 | 4.9 | 14.3 |
| >MI0000746\|hsa-mir-99b&WithFlank&19\|+\|51692606\|51692687@13.34.22(+1C)&[MIMAT0000689&hsa-miR-99b-5p&offsets\|0\|0(+1C);m-10&19\|+\|51692618\|51692639&offsets\|0\|0(+1C)]\|CACCCGTAGAACCGACCTTGCGC | >MI0000746\|hsa-mir-99b&WithFlank&19\|+\|51692606\|51692687 | 13 | 34 | 22(+1C) | CACCCGTAGAACCGACCTTGCGC | BT20 | 16 | 15 | 30 | 6 | 8 | 7.1 | 7.9 | 5.8 | 1.5 | 3.5 |
| >MI0000746\|hsa-mir-99b&WithFlank&19\|+\|51692606\|51692687@13.34.22(+1C)&[MIMAT0000689&hsa-miR-99b-5p&offsets\|0\|0(+1C);m-10&19\|+\|51692618\|51692639&offsets\|0\|0(+1C)]\|CACCCGTAGAACCGACCTTGCGC | >MI0000746\|hsa-mir-99b&WithFlank&19\|+\|51692606\|51692687 | 13 | 34 | 22(+1C) | CACCCGTAGAACCGACCTTGCGC | MB231 | 10 | 13 | 12 | 13 | 14 | 1.7 | 4.8 | 8.1 | 6.6 | 9.0 |
| >MI0000746\|hsa-mir-99b&WithFlank&19\|+\|51692606\|51692687@13.34.22(+1U)&[MIMAT0000689&hsa-miR-99b-5p&offsets\|0\|0(+1U);m-10&19\|+\|51692618\|51692639&offsets\|0\|0(+1U)]\|CACCCGTAGAACCGACCTTGCGT | >MI0000746\|hsa-mir-99b&WithFlank&19\|+\|51692606\|51692687 | 13 | 34 | 22(+1U) | CACCCGTAGAACCGACCTTGCGT | BT20 | 110 | 101 | 221 | 40 | 59 | 49.7 | 46.2 | 19.2 | 7.4 | 23.9 |
| >MI0000746\|hsa-mir-99b&WithFlank&19\|+\|51692606\|51692687@13.34.22(+1U)&[MIMAT0000689&hsa-miR-99b-5p&offsets\|0\|0(+1U);m-10&19\|+\|51692618\|51692639&offsets\|0\|0(+1U)]\|CACCCGTAGAACCGACCTTGCGT | >MI0000746\|hsa-mir-99b&WithFlank&19\|+\|51692606\|51692687 | 13 | 34 | 22(+1U) | CACCCGTAGAACCGACCTTGCGT | MB231 | 92 | 109 | 112 | 81 | 99 | 17.4 | 42.6 | 74.0 | 25.7 | 45.0 |
| >MI0000746\|hsa-mir-99b&WithFlank&19\|+\|51692606\|51692687@13.34.22&[MIMAT0000689&hsa-miR-99b-5p&offsets\|0\|0;m-10&19\|+\|51692618\|51692639&offsets\|0\|0]\|CACCCGTAGAACCGACCTTGCG | >MI0000746\|hsa-mir-99b&WithFlank&19\|+\|51692606\|51692687 | 13 | 34 | 22 | CACCCGTAGAACCGACCTTGCG | BT20 | 1914 | 1134 | 4088 | 603 | 718 | 1124.4 | 512.9 | 2194.8 | 70.7 | 132.4 |
| >MI0000746\|hsa-mir-99b&WithFlank&19\|+\|51692606\|51692687@13.34.22&[MIMAT0000689&hsa-miR-99b-5p&offsets\|0\|0;m-10&19\|+\|51692618\|51692639&offsets\|0\|0]\|CACCCGTAGAACCGACCTTGCG | >MI0000746\|hsa-mir-99b&WithFlank&19\|+\|51692606\|51692687 | 13 | 34 | 22 | CACCCGTAGAACCGACCTTGCG | MB231 | 1204 | 1204 | 1233 | 797 | 945 | 76.9 | 306.0 | 583.8 | 53.0 | 195.6 |

| | | | | | | | | | | | | | | | |
|---|---|---|---|---|---|---|---|---|---|---|---|---|---|---|---|
| >MI0000746\|hsa-mir-99b&WithFlank&19\|+\|51692606\|51692687@13.34.22&[MIMAT0000689&hsa-miR-99b-5p&offsets\|0\|0;m-10&19\|+\|51692618\|51692639&offsets\|0\|0]\|CACCCGTAGAACCGACCTTGCG | >MI0000746\|hsa-mir-99b&WithFlank&19\|+\|51692606\|51692687 | 13 | 34 | 22 | CACCCGTAGAACCGACCTTGCG | MB468 | 90 | 133 | 79 | 108 | 34 | 36.1 | 53.7 | 25.4 | 48.9 | 9.1 |
| >MI0000746\|hsa-mir-99b&WithFlank&19\|+\|51692606\|51692687@13.35.23&[MIMAT0000689&hsa-miR-99b-5p&offsets\|0\|+1;m-10&19\|+\|51692618\|51692639&offsets\|0\|+1]\|CACCCGTAGAACCGACCTTGCGG | >MI0000746\|hsa-mir-99b&WithFlank&19\|+\|51692606\|51692687 | 13 | 35 | 23 | CACCCGTAGAACCGACCTTGCGG | BT20 | 14 | 5 | 11 | 1 | 1 | 14.0 | 1.8 | 5.0 | 0.9 | 0.5 |
| >MI0000746\|hsa-mir-99b&WithFlank&19\|+\|51692606\|51692687@14.34.21&[MIMAT0000689&hsa-miR-99b-5p&offsets\|+1\|0;m-10&19\|+\|51692618\|51692639&offsets\|+1\|0]\|ACCCGTAGAACCGACCTTGCG | >MI0000746\|hsa-mir-99b&WithFlank&19\|+\|51692606\|51692687 | 14 | 34 | 21 | ACCCGTAGAACCGACCTTGCG | BT20 | 16 | 12 | 30 | 5 | 5 | 9.2 | 6.9 | 17.4 | 0.5 | 1.2 |
| >MI0000746\|hsa-mir-99b&WithFlank&19\|+\|51692606\|51692687@51.69.19&[MIMAT0004678&hsa-miR-99b-3p&offsets\|0\|-3;m-151&19\|+\|51692656\|51692678&offsets\|0\|-4]\|CAAGCTCGTGTCTGTGGGT | >MI0000746\|hsa-mir-99b&WithFlank&19\|+\|51692606\|51692687 | 51 | 69 | 19 | CAAGCTCGTGTCTGTGGGT | BT20 | 19 | 7 | 16 | 23 | 3 | 19.5 | 4.0 | 8.3 | 18.4 | 1.4 |
| >MI0000746\|hsa-mir-99b&WithFlank&19\|+\|51692606\|51692687@51.70.20(+1A)&[MIMAT0004678&hsa-miR-99b-3p&offsets\|0\|-2(+1A);m-151&19\|+\|51692656\|51692678&offsets\|0\|-3(+1A)]\|CAAGCTCGTGTCTGTGGGTCA | >MI0000746\|hsa-mir-99b&WithFlank&19\|+\|51692606\|51692687 | 51 | 70 | 20(+1A) | CAAGCTCGTGTCTGTGGGTCA | BT20 | 13 | 5 | 14 | 10 | 2 | 13.4 | 2.4 | 7.8 | 6.7 | 0.3 |
| >MI0000746\|hsa-mir-99b&WithFlank&19\|+\|51692606\|51692687@51.70.20(+1U)&[MIMAT0004678&hsa-miR-99b-3p&offsets\|0\|-2(+1U);m-151&19\|+\|51692656\|51692678&offsets\|0\|-3(+1U)]\|CAAGCTCGTGTCTGTGGGTCT | >MI0000746\|hsa-mir-99b&WithFlank&19\|+\|51692606\|51692687 | 51 | 70 | 20(+1U) | CAAGCTCGTGTCTGTGGGTCT | BT20 | 26 | 12 | 28 | 31 | 6 | 19.7 | 7.3 | 12.8 | 22.1 | 1.8 |
| >MI0000746\|hsa-mir-99b&WithFlank&19\|+\|51692606\|51692687@51.70.20&[MIMAT0004678&hsa-miR-99b-3p&offsets\|0\|-2;m-151&19\|+\|51692656\|51692678&offsets\|0\|-3]\|CAAGCTCGTGTCTGTGGGTC | >MI0000746\|hsa-mir-99b&WithFlank&19\|+\|51692606\|51692687 | 51 | 70 | 20 | CAAGCTCGTGTCTGTGGGTC | BT20 | 23 | 10 | 39 | 31 | 4 | 17.8 | 4.8 | 17.4 | 20.9 | 1.6 |
| >MI0000746\|hsa-mir-99b&WithFlank&19\|+\|51692606\|51692687@51.71.21(+1A)&[MIMAT0004678&hsa-miR-99b-3p&offsets\|0\|-1(+1A);m-151&19\|+\|51692656\|51692678&offsets\|0\|-2(+1A)]\|CAAGCTCGTGTCTGTGGGTCCA | >MI0000746\|hsa-mir-99b&WithFlank&19\|+\|51692606\|51692687 | 51 | 71 | 21(+1A) | CAAGCTCGTGTCTGTGGGTCCA | BT20 | 35 | 21 | 43 | 20 | 16 | 25.2 | 18.9 | 22.5 | 7.4 | 6.9 |
| >MI0000746\|hsa-mir-99b&WithFlank&19\|+\|51692606\|51692687@51.71.21(+1U)&[MIMAT0004678&hsa-miR-99b-3p&offsets\|0\|-1(+1U);m-151&19\|+\|51692656\|51692678&offsets\|0\|-2(+1U)]\|CAAGCTCGTGTCTGTGGGTCCT | >MI0000746\|hsa-mir-99b&WithFlank&19\|+\|51692606\|51692687 | 51 | 71 | 21(+1U) | CAAGCTCGTGTCTGTGGGTCCT | BT20 | 11 | 7 | 10 | 8 | 5 | 6.5 | 6.6 | 4.6 | 1.2 | 2.3 |
| >MI0000746\|hsa-mir-99b&WithFlank&19\|+\|51692606\|51692687@51.71.21&[MIMAT0004678&hsa-miR-99b-3p&offsets\|0\|-1;m-151&19\|+\|51692656\|51692678&offsets\|0\|-2]\|CAAGCTCGTGTCTGTGGGTCC | >MI0000746\|hsa-mir-99b&WithFlank&19\|+\|51692606\|51692687 | 51 | 71 | 21 | CAAGCTCGTGTCTGTGGGTCC | BT20 | 32 | 18 | 40 | 44 | 11 | 17.5 | 9.9 | 9.4 | 30.5 | 4.0 |
| >MI0000746\|hsa-mir-99b&WithFlank&19\|+\|51692606\|51692687@51.71.21&[MIMAT0004678&hsa-miR-99b-3p&offsets\|0\|-1;m-151&19\|+\|51692656\|51692678&offsets\|0\|-2]\|CAAGCTCGTGTCTGTGGGTCC | >MI0000746\|hsa-mir-99b&WithFlank&19\|+\|51692606\|51692687 | 51 | 71 | 21 | CAAGCTCGTGTCTGTGGGTCC | MB231 | 10 | 7 | 5 | 8 | 6 | 2.9 | 3.6 | 3.1 | 2.6 | 1.6 |
| >MI0000746\|hsa-mir-99b&WithFlank&19\|+\|51692606\|51692687@51.72.22(+1A)&[MIMAT0004678&hsa-miR-99b-3p&offsets\|0\|0(+1A);m-151&19\|+\|51692656\|51692678&offsets\|0\|-1(+1A)]\|CAAGCTCGTGTCTGTGGGTCCGA | >MI0000746\|hsa-mir-99b&WithFlank&19\|+\|51692606\|51692687 | 51 | 72 | 22(+1A) | CAAGCTCGTGTCTGTGGGTCCGA | BT20 | 49 | 27 | 95 | 46 | 19 | 22.3 | 12.3 | 39.3 | 24.2 | 2.7 |
| >MI0000746\|hsa-mir-99b&WithFlank&19\|+\|51692606\|51692687@51.72.22(+1C)&[MIMAT0004678&hsa-miR-99b-3p&offsets\|0\|0(+1C);m-151&19\|+\|51692656\|51692678&offsets\|0\|-1(+1C)]\|CAAGCTCGTGTCTGTGGGTCCGC | >MI0000746\|hsa-mir-99b&WithFlank&19\|+\|51692606\|51692687 | 51 | 72 | 22(+1C) | CAAGCTCGTGTCTGTGGGTCCGC | BT20 | 11 | 6 | 19 | 6 | 5 | 0.8 | 1.8 | 6.0 | 1.4 | 1.2 |
| >MI0000746\|hsa-mir-99b&WithFlank&19\|+\|51692606\|51692687@51.72.22&[MIMAT0004678&hsa-miR-99b-3p&offsets\|0\|0;m-151&19\|+\|51692656\|51692678&offsets\|0\|-1]\|CAAGCTCGTGTCTGTGGGTCCG | >MI0000746\|hsa-mir-99b&WithFlank&19\|+\|51692606\|51692687 | 51 | 72 | 22 | CAAGCTCGTGTCTGTGGGTCCG | BT20 | 100 | 83 | 128 | 109 | 80 | 36.5 | 78.0 | 33.1 | 53.9 | 42.8 |
| >MI0000746\|hsa-mir-99b&WithFlank&19\|+\|51692606\|51692687@51.72.22&[MIMAT0004678&hsa-miR-99b-3p&offsets\|0\|0;m-151&19\|+\|51692656\|51692678&offsets\|0\|-1]\|CAAGCTCGTGTCTGTGGGTCCG | >MI0000746\|hsa-mir-99b&WithFlank&19\|+\|51692606\|51692687 | 51 | 72 | 22 | CAAGCTCGTGTCTGTGGGTCCG | MB231 | 36 | 34 | 23 | 18 | 23 | 15.7 | 5.8 | 7.5 | 4.7 | 8.7 |
| >MI0000746\|hsa-mir-99b&WithFlank&19\|+\|51692606\|51692687@51.73.23(+1U)&[MIMAT0004678&hsa-miR-99b-3p&offsets\|0\|+1(+1U);m-151&19\|+\|51692656\|51692678&offsets\|0\|0(+1U)]\|CAAGCTCGTGTCTGTGGGTCCGTT | >MI0000746\|hsa-mir-99b&WithFlank&19\|+\|51692606\|51692687 | 51 | 73 | 23(+1U) | CAAGCTCGTGTCTGTGGGTCCGTT | BT20 | 18 | 11 | 25 | 10 | 8 | 4.6 | 3.9 | 7.3 | 2.6 | 1.6 |
| >MI0000746\|hsa-mir-99b&WithFlank&19\|+\|51692606\|51692687@51.73.23&[MIMAT0004678&hsa-miR-99b-3p&offsets\|0\|+1;m-151&19\|+\|51692656\|51692678&offsets\|0\|0]\|CAAGCTCGTGTCTGTGGGTCCGT | >MI0000746\|hsa-mir-99b&WithFlank&19\|+\|51692606\|51692687 | 51 | 73 | 23 | CAAGCTCGTGTCTGTGGGTCCGT | BT20 | 79 | 40 | 131 | 39 | 31 | 11.1 | 9.9 | 53.5 | 12.6 | 5.3 |
| >MI0000746\|hsa-mir-99b&WithFlank&19\|+\|51692606\|51692687@51.73.23&[MIMAT0004678&hsa-miR-99b-3p&offsets\|0\|+1;m-151&19\|+\|51692656\|51692678&offsets\|0\|0]\|CAAGCTCGTGTCTGTGGGTCCGT | >MI0000746\|hsa-mir-99b&WithFlank&19\|+\|51692606\|51692687 | 51 | 73 | 23 | CAAGCTCGTGTCTGTGGGTCCGT | MB231 | 31 | 23 | 18 | 22 | 17 | 5.7 | 11.8 | 10.0 | 8.3 | 7.1 |
| >MI0000748\|hsa-mir-130b&WithFlank&22\|+\|21653298\|21653391@19.40.22&[MIMAT0004680&hsa-miR-130b-5p&offsets\|0\|+1;m-234&22\|+\|21653316\|21653337&offsets\|0\|0]\|ACTCTTTCCCTGTTGCACTACT | >MI0000748\|hsa-mir-130b&WithFlank&22\|+\|21653298\|21653391 | 19 | 40 | 22 | ACTCTTTCCCTGTTGCACTACT | BT20 | 37 | 21 | 67 | 0 | 0 | 37.7 | 13.0 | 28.7 | 0.0 | 0.0 |
| >MI0000748\|hsa-mir-130b&WithFlank&22\|+\|21653298\|21653391@19.40.22&[MIMAT0004680&hsa-miR-130b-5p&offsets\|0\|+1;m-234&22\|+\|21653316\|21653337&offsets\|0\|0]\|ACTCTTTCCCTGTTGCACTACT | >MI0000748\|hsa-mir-130b&WithFlank&22\|+\|21653298\|21653391 | 19 | 40 | 22 | ACTCTTTCCCTGTTGCACTACT | MB231 | 50 | 60 | 67 | 2 | 55 | 12.3 | 22.9 | 32.2 | 1.7 | 18.2 |
| >MI0000748\|hsa-mir-130b&WithFlank&22\|+\|21653298\|21653391@57.78.22&[MIMAT0000691&hsa-miR-130b-3p&offsets\|0\|0;m-205&22\|+\|21653354\|21653375&offsets\|0\|0]\|CAGTGCAATGATGAAAGGGCAT | >MI0000748\|hsa-mir-130b&WithFlank&22\|+\|21653298\|21653391 | 57 | 78 | 22 | CAGTGCAATGATGAAAGGGCAT | BT20 | 18 | 14 | 2 | 2 | 1 | 21.8 | 15.2 | 0.6 | 2.1 | 1.4 |
| >MI0000749\|hsa-mir-30e&WithFlank&1\|+\|40754349\|40754452@23.44.22(+1U)&[MIMAT0000692&hsa-miR-30e-5p&offsets\|0\|0(+1U);m-24&1\|+\|40754371\|40754394&offsets\|0\|-2(+1U)]\|TGTAAACATCCTTGACTGGAAGT | >MI0000749\|hsa-mir-30e&WithFlank&1\|+\|40754349\|40754452 | 23 | 44 | 22(+1U) | TGTAAACATCCTTGACTGGAAGT | BT20 | 15 | 6 | 1 | 8 | 1 | 19.8 | 8.1 | 0.5 | 9.0 | 0.4 |
| >MI0000749\|hsa-mir-30e&WithFlank&1\|+\|40754349\|40754452@23.44.22&[MIMAT0000692&hsa-miR-30e-5p&offsets\|0\|0;m-24&1\|+\|40754371\|40754394&offsets\|0\|-2]\|TGTAAACATCCTTGACTGGAAG | >MI0000749\|hsa-mir-30e&WithFlank&1\|+\|40754349\|40754452 | 23 | 44 | 22 | TGTAAACATCCTTGACTGGAAG | BT20 | 23 | 7 | 2 | 25 | 1 | 30.3 | 9.5 | 0.7 | 21.9 | 1.1 |
| >MI0000749\|hsa-mir-30e&WithFlank&1\|+\|40754349\|40754452@23.45.23(+1A)&[MIMAT0000692&hsa-miR-30e-5p&offsets\|0\|+1(+1A);m-24&1\|+\|40754371\|40754394&offsets\|0\|-1(+1A)]\|TGTAAACATCCTTGACTGGAAGCA | >MI0000749\|hsa-mir-30e&WithFlank&1\|+\|40754349\|40754452 | 23 | 45 | 23(+1A) | TGTAAACATCCTTGACTGGAAGCA | BT20 | 72 | 28 | 10 | 82 | 6 | 82.0 | 40.7 | 5.4 | 78.3 | 1.4 |
| >MI0000749\|hsa-mir-30e&WithFlank&1\|+\|40754349\|40754452@23.45.23(+1A)&[MIMAT0000692&hsa-miR-30e-5p&offsets\|0\|+1(+1A);m-24&1\|+\|40754371\|40754394&offsets\|0\|-1(+1A)]\|TGTAAACATCCTTGACTGGAAGCA | >MI0000749\|hsa-mir-30e&WithFlank&1\|+\|40754349\|40754452 | 23 | 45 | 23(+1A) | TGTAAACATCCTTGACTGGAAGCA | MB231 | 25 | 15 | 2 | 200 | 37 | 12.9 | 10.7 | 1.4 | 95.7 | 41.2 |
| >MI0000749\|hsa-mir-30e&WithFlank&1\|+\|40754349\|40754452@23.45.23(+1C)&[MIMAT0000692&hsa-miR-30e-5p&offsets\|0\|+1(+1C);m-24&1\|+\|40754371\|40754394&offsets\|0\|-1(+1C)]\|TGTAAACATCCTTGACTGGAAGCC | >MI0000749\|hsa-mir-30e&WithFlank&1\|+\|40754349\|40754452 | 23 | 45 | 23(+1C) | TGTAAACATCCTTGACTGGAAGCC | BT20 | 69 | 47 | 12 | 129 | 11 | 55.4 | 71.3 | 9.0 | 120.7 | 5.0 |
| >MI0000749\|hsa-mir-30e&WithFlank&1\|+\|40754349\|40754452@23.45.23(+1C)&[MIMAT0000692&hsa-miR-30e-5p&offsets\|0\|+1(+1C);m-24&1\|+\|40754371\|40754394&offsets\|0\|-1(+1C)]\|TGTAAACATCCTTGACTGGAAGCC | >MI0000749\|hsa-mir-30e&WithFlank&1\|+\|40754349\|40754452 | 23 | 45 | 23(+1C) | TGTAAACATCCTTGACTGGAAGCC | MB231 | 40 | 29 | 3 | 303 | 63 | 21.2 | 22.7 | 1.2 | 138.4 | 72.3 |
| >MI0000749\|hsa-mir-30e&WithFlank&1\|+\|40754349\|40754452@23.45.23(+1C)&[MIMAT0000692&hsa-miR-30e-5p&offsets\|0\|+1(+1C);m-24&1\|+\|40754371\|40754394&offsets\|0\|-1(+1C)]\|TGTAAACATCCTTGACTGGAAGCC | >MI0000749\|hsa-mir-30e&WithFlank&1\|+\|40754349\|40754452 | 23 | 45 | 23(+1C) | TGTAAACATCCTTGACTGGAAGCC | MB468 | 16 | 1 | 2 | 53 | 4 | 10.3 | 0.5 | 1.1 | 28.5 | 1.0 |

<table>
<tr><td></td><td></td><td></td><td></td><td></td><td></td><td></td><td></td><td></td><td></td><td></td><td></td><td></td><td></td><td></td><td></td><td></td></tr>
<tr><td>>MI0000749|hsa-mir-30e&WithFlank&1|+|40754349|40754452@23.45.23(+1G)&[MIMAT0000692&hsa-miR-30e-5p&offsets|0|+1(+1G);m-24&1|+|40754371|40754394&offsets|0|-1(+1G)]|TGTAAACATCCTTGACTGGAAGCG</td><td>>MI0000749|hsa-mir-30e&WithFlank&1|+|40754349|40754452</td><td>23</td><td>45</td><td>23(+1G)</td><td>TGTAAACATCCTTGACTGGAAGCG</td><td>BT20</td><td>51</td><td>10</td><td>3</td><td>23</td><td>1</td><td>62.8</td><td>15.6</td><td>1.0</td><td>26.3</td><td>0.4</td></tr>
<tr><td>>MI0000749|hsa-mir-30e&WithFlank&1|+|40754349|40754452@23.45.23(+1G)&[MIMAT0000692&hsa-miR-30e-5p&offsets|0|+1(+1G);m-24&1|+|40754371|40754394&offsets|0|-1(+1G)]|TGTAAACATCCTTGACTGGAAGCG</td><td>>MI0000749|hsa-mir-30e&WithFlank&1|+|40754349|40754452</td><td>23</td><td>45</td><td>23(+1G)</td><td>TGTAAACATCCTTGACTGGAAGCG</td><td>MB231</td><td>17</td><td>7</td><td>1</td><td>39</td><td>7</td><td>10.2</td><td>6.3</td><td>0.3</td><td>20.7</td><td>8.3</td></tr>
<tr><td>>MI0000749|hsa-mir-30e&WithFlank&1|+|40754349|40754452@23.45.23&[MIMAT0000692&hsa-miR-30e-5p&offsets|0|+1;m-24&1|+|40754371|40754394&offsets|0|-1]|TGTAAACATCCTTGACTGGAAGC</td><td>>MI0000749|hsa-mir-30e&WithFlank&1|+|40754349|40754452</td><td>23</td><td>45</td><td>23</td><td>TGTAAACATCCTTGACTGGAAGC</td><td>BT20</td><td>30</td><td>17</td><td>4</td><td>26</td><td>3</td><td>34.7</td><td>23.1</td><td>2.0</td><td>23.7</td><td>0.7</td></tr>
<tr><td>>MI0000749|hsa-mir-30e&WithFlank&1|+|40754349|40754452@23.46.24(+1U)&[MIMAT0000692&hsa-miR-30e-5p&offsets|0|+2(+1U);m-24&1|+|40754371|40754394&offsets|0|0(+1U)]|TGTAAACATCCTTGACTGGAAGCTT</td><td>>MI0000749|hsa-mir-30e&WithFlank&1|+|40754349|40754452</td><td>23</td><td>46</td><td>24(+1U)</td><td>TGTAAACATCCTTGACTGGAAGCTT</td><td>BT20</td><td>13</td><td>9</td><td>2</td><td>30</td><td>2</td><td>9.3</td><td>13.6</td><td>2.0</td><td>26.2</td><td>2.0</td></tr>
<tr><td>>MI0000749|hsa-mir-30e&WithFlank&1|+|40754349|40754452@23.46.24(+1U)&[MIMAT0000692&hsa-miR-30e-5p&offsets|0|+2(+1U);m-24&1|+|40754371|40754394&offsets|0|0(+1U)]|TGTAAACATCCTTGACTGGAAGCTT</td><td>>MI0000749|hsa-mir-30e&WithFlank&1|+|40754349|40754452</td><td>23</td><td>46</td><td>24(+1U)</td><td>TGTAAACATCCTTGACTGGAAGCTT</td><td>MB231</td><td>26</td><td>20</td><td>2</td><td>144</td><td>32</td><td>11.9</td><td>15.2</td><td>1.4</td><td>47.8</td><td>32.9</td></tr>
<tr><td>>MI0000749|hsa-mir-30e&WithFlank&1|+|40754349|40754452@23.46.24(+2U)&[MIMAT0000692&hsa-miR-30e-5p&offsets|0|+2(+2U);m-24&1|+|40754371|40754394&offsets|0|0(+2U)]|TGTAAACATCCTTGACTGGAAGCTTT</td><td>>MI0000749|hsa-mir-30e&WithFlank&1|+|40754349|40754452</td><td>23</td><td>46</td><td>24(+2U)</td><td>TGTAAACATCCTTGACTGGAAGCTTT</td><td>MB231</td><td>13</td><td>11</td><td>1</td><td>34</td><td>16</td><td>6.3</td><td>7.6</td><td>0.8</td><td>14.0</td><td>15.9</td></tr>
<tr><td>>MI0000749|hsa-mir-30e&WithFlank&1|+|40754349|40754452@23.46.24&[MIMAT0000692&hsa-miR-30e-5p&offsets|0|+2;m-24&1|+|40754371|40754394&offsets|0|0]|TGTAAACATCCTTGACTGGAAGCT</td><td>>MI0000749|hsa-mir-30e&WithFlank&1|+|40754349|40754452</td><td>23</td><td>46</td><td>24</td><td>TGTAAACATCCTTGACTGGAAGCT</td><td>BT20</td><td>434</td><td>266</td><td>82</td><td>808</td><td>72</td><td>334.2</td><td>393.4</td><td>64.3</td><td>695.5</td><td>23.0</td></tr>
<tr><td>>MI0000749|hsa-mir-30e&WithFlank&1|+|40754349|40754452@23.46.24&[MIMAT0000692&hsa-miR-30e-5p&offsets|0|+2;m-24&1|+|40754371|40754394&offsets|0|0]|TGTAAACATCCTTGACTGGAAGCT</td><td>>MI0000749|hsa-mir-30e&WithFlank&1|+|40754349|40754452</td><td>23</td><td>46</td><td>24</td><td>TGTAAACATCCTTGACTGGAAGCT</td><td>MB231</td><td>314</td><td>217</td><td>21</td><td>1707</td><td>369</td><td>195.2</td><td>146.3</td><td>10.4</td><td>542.3</td><td>378.9</td></tr>
<tr><td>>MI0000749|hsa-mir-30e&WithFlank&1|+|40754349|40754452@23.46.24&[MIMAT0000692&hsa-miR-30e-5p&offsets|0|+2;m-24&1|+|40754371|40754394&offsets|0|0]|TGTAAACATCCTTGACTGGAAGCT</td><td>>MI0000749|hsa-mir-30e&WithFlank&1|+|40754349|40754452</td><td>23</td><td>46</td><td>24</td><td>TGTAAACATCCTTGACTGGAAGCT</td><td>MB468</td><td>119</td><td>12</td><td>13</td><td>349</td><td>26</td><td>77.0</td><td>4.6</td><td>7.7</td><td>158.5</td><td>7.2</td></tr>
<tr><td>>MI0000749|hsa-mir-30e&WithFlank&1|+|40754349|40754452@24.46.23&[MIMAT0000692&hsa-miR-30e-5p&offsets|+1|+2;m-24&1|+|40754371|40754394&offsets|+1|0]|GTAAACATCCTTGACTGGAAGCT</td><td>>MI0000749|hsa-mir-30e&WithFlank&1|+|40754349|40754452</td><td>24</td><td>46</td><td>23</td><td>GTAAACATCCTTGACTGGAAGCT</td><td>BT20</td><td>32</td><td>18</td><td>6</td><td>90</td><td>5</td><td>20.8</td><td>24.0</td><td>4.7</td><td>79.9</td><td>0.5</td></tr>
<tr><td>>MI0000749|hsa-mir-30e&WithFlank&1|+|40754349|40754452@24.46.23&[MIMAT0000692&hsa-miR-30e-5p&offsets|+1|+2;m-24&1|+|40754371|40754394&offsets|+1|0]|GTAAACATCCTTGACTGGAAGCT</td><td>>MI0000749|hsa-mir-30e&WithFlank&1|+|40754349|40754452</td><td>24</td><td>46</td><td>23</td><td>GTAAACATCCTTGACTGGAAGCT</td><td>MB231</td><td>12</td><td>10</td><td>1</td><td>84</td><td>14</td><td>6.6</td><td>5.9</td><td>1.0</td><td>34.5</td><td>13.4</td></tr>
<tr><td>>MI0000749|hsa-mir-30e&WithFlank&1|+|40754349|40754452@65.83.19&[MIMAT0000693&hsa-miR-30e-3p&offsets|0|-3;m-25&1|+|40754413|40754434&offsets|0|-3]|CTTTCAGTCGGATGTTTAC</td><td>>MI0000749|hsa-mir-30e&WithFlank&1|+|40754349|40754452</td><td>65</td><td>83</td><td>19</td><td>CTTTCAGTCGGATGTTTAC</td><td>BT20</td><td>30</td><td>21</td><td>35</td><td>11</td><td>17</td><td>34.9</td><td>10.7</td><td>24.6</td><td>3.3</td><td>4.7</td></tr>
<tr><td>>MI0000749|hsa-mir-30e&WithFlank&1|+|40754349|40754452@65.83.19&[MIMAT0000693&hsa-miR-30e-3p&offsets|0|-3;m-25&1|+|40754413|40754434&offsets|0|-3]|CTTTCAGTCGGATGTTTAC</td><td>>MI0000749|hsa-mir-30e&WithFlank&1|+|40754349|40754452</td><td>65</td><td>83</td><td>19</td><td>CTTTCAGTCGGATGTTTAC</td><td>MB231</td><td>30</td><td>60</td><td>26</td><td>49</td><td>36</td><td>15.4</td><td>19.8</td><td>10.0</td><td>16.9</td><td>13.9</td></tr>
<tr><td>>MI0000749|hsa-mir-30e&WithFlank&1|+|40754349|40754452@65.85.21(+1A)&[MIMAT0000693&hsa-miR-30e-3p&offsets|0|-1(+1A);m-25&1|+|40754413|40754434&offsets|0|-1(+1A)]|CTTTCAGTCGGATGTTTACAGA</td><td>>MI0000749|hsa-mir-30e&WithFlank&1|+|40754349|40754452</td><td>65</td><td>85</td><td>21(+1A)</td><td>CTTTCAGTCGGATGTTTACAGA</td><td>BT20</td><td>14</td><td>7</td><td>12</td><td>2</td><td>2</td><td>12.2</td><td>4.0</td><td>4.1</td><td>1.6</td><td>0.4</td></tr>
<tr><td>>MI0000749|hsa-mir-30e&WithFlank&1|+|40754349|40754452@65.85.21(+1G)&[MIMAT0000693&hsa-miR-30e-3p&offsets|0|-1(+1G);m-25&1|+|40754413|40754434&offsets|0|-1(+1G)]|CTTTCAGTCGGATGTTTACAGG</td><td>>MI0000749|hsa-mir-30e&WithFlank&1|+|40754349|40754452</td><td>65</td><td>85</td><td>21(+1G)</td><td>CTTTCAGTCGGATGTTTACAGG</td><td>BT20</td><td>10</td><td>3</td><td>3</td><td>0</td><td>0</td><td>11.1</td><td>2.2</td><td>1.4</td><td>0.7</td><td>0.2</td></tr>
<tr><td>>MI0000749|hsa-mir-30e&WithFlank&1|+|40754349|40754452@65.85.21(+1U)&[MIMAT0000693&hsa-miR-30e-3p&offsets|0|-1(+1U);m-25&1|+|40754413|40754434&offsets|0|-1(+1U)]|CTTTCAGTCGGATGTTTACAGT</td><td>>MI0000749|hsa-mir-30e&WithFlank&1|+|40754349|40754452</td><td>65</td><td>85</td><td>21(+1U)</td><td>CTTTCAGTCGGATGTTTACAGT</td><td>BT20</td><td>153</td><td>122</td><td>98</td><td>24</td><td>35</td><td>83.1</td><td>72.2</td><td>6.7</td><td>18.9</td><td>17.7</td></tr>
<tr><td>>MI0000749|hsa-mir-30e&WithFlank&1|+|40754349|40754452@65.85.21(+1U)&[MIMAT0000693&hsa-miR-30e-3p&offsets|0|-1(+1U);m-25&1|+|40754413|40754434&offsets|0|-1(+1U)]|CTTTCAGTCGGATGTTTACAGT</td><td>>MI0000749|hsa-mir-30e&WithFlank&1|+|40754349|40754452</td><td>65</td><td>85</td><td>21(+1U)</td><td>CTTTCAGTCGGATGTTTACAGT</td><td>MB231</td><td>120</td><td>183</td><td>54</td><td>140</td><td>124</td><td>35.8</td><td>59.8</td><td>38.7</td><td>28.9</td><td>54.1</td></tr>
<tr><td>>MI0000749|hsa-mir-30e&WithFlank&1|+|40754349|40754452@65.85.21(+1U)&[MIMAT0000693&hsa-miR-30e-3p&offsets|0|-1(+1U);m-25&1|+|40754413|40754434&offsets|0|-1(+1U)]|CTTTCAGTCGGATGTTTACAGT</td><td>>MI0000749|hsa-mir-30e&WithFlank&1|+|40754349|40754452</td><td>65</td><td>85</td><td>21(+1U)</td><td>CTTTCAGTCGGATGTTTACAGT</td><td>MB468</td><td>15</td><td>26</td><td>6</td><td>50</td><td>10</td><td>0.9</td><td>8.7</td><td>2.9</td><td>44.6</td><td>2.9</td></tr>
<tr><td>>MI0000749|hsa-mir-30e&WithFlank&1|+|40754349|40754452@65.85.21(+2U)&[MIMAT0000693&hsa-miR-30e-3p&offsets|0|-1(+2U);m-25&1|+|40754413|40754434&offsets|0|-1(+2U)]|CTTTCAGTCGGATGTTTACAGTT</td><td>>MI0000749|hsa-mir-30e&WithFlank&1|+|40754349|40754452</td><td>65</td><td>85</td><td>21(+2U)</td><td>CTTTCAGTCGGATGTTTACAGTT</td><td>BT20</td><td>24</td><td>16</td><td>30</td><td>4</td><td>8</td><td>15.4</td><td>8.2</td><td>12.4</td><td>2.1</td><td>2.8</td></tr>
<tr><td>>MI0000749|hsa-mir-30e&WithFlank&1|+|40754349|40754452@65.85.21(+2U)&[MIMAT0000693&hsa-miR-30e-3p&offsets|0|-1(+2U);m-25&1|+|40754413|40754434&offsets|0|-1(+2U)]|CTTTCAGTCGGATGTTTACAGTT</td><td>>MI0000749|hsa-mir-30e&WithFlank&1|+|40754349|40754452</td><td>65</td><td>85</td><td>21(+2U)</td><td>CTTTCAGTCGGATGTTTACAGTT</td><td>MB231</td><td>39</td><td>51</td><td>22</td><td>36</td><td>31</td><td>10.8</td><td>14.9</td><td>8.2</td><td>6.3</td><td>10.8</td></tr>
<tr><td>>MI0000749|hsa-mir-30e&WithFlank&1|+|40754349|40754452@65.85.21&[MIMAT0000693&hsa-miR-30e-3p&offsets|0|-1;m-25&1|+|40754413|40754434&offsets|0|-1]|CTTTCAGTCGGATGTTTACAG</td><td>>MI0000749|hsa-mir-30e&WithFlank&1|+|40754349|40754452</td><td>65</td><td>85</td><td>21</td><td>CTTTCAGTCGGATGTTTACAG</td><td>BT20</td><td>17</td><td>12</td><td>18</td><td>3</td><td>4</td><td>11.2</td><td>3.9</td><td>6.1</td><td>1.6</td><td>0.3</td></tr>
<tr><td>>MI0000749|hsa-mir-30e&WithFlank&1|+|40754349|40754452@65.85.21&[MIMAT0000693&hsa-miR-30e-3p&offsets|0|-1;m-25&1|+|40754413|40754434&offsets|0|-1]|CTTTCAGTCGGATGTTTACAG</td><td>>MI0000749|hsa-mir-30e&WithFlank&1|+|40754349|40754452</td><td>65</td><td>85</td><td>21</td><td>CTTTCAGTCGGATGTTTACAG</td><td>MB231</td><td>13</td><td>18</td><td>8</td><td>20</td><td>9</td><td>4.4</td><td>8.3</td><td>5.1</td><td>3.5</td><td>2.9</td></tr>
<tr><td>>MI0000749|hsa-mir-30e&WithFlank&1|+|40754349|40754452@65.86.22&[MIMAT0000693&hsa-miR-30e-3p&offsets|0|0;m-25&1|+|40754413|40754434&offsets|0|0]|CTTTCAGTCGGATGTTTACAGC</td><td>>MI0000749|hsa-mir-30e&WithFlank&1|+|40754349|40754452</td><td>65</td><td>86</td><td>22</td><td>CTTTCAGTCGGATGTTTACAGC</td><td>BT20</td><td>78</td><td>65</td><td>92</td><td>17</td><td>25</td><td>32.3</td><td>34.6</td><td>16.4</td><td>10.4</td><td>11.0</td></tr>
<tr><td>>MI0000749|hsa-mir-30e&WithFlank&1|+|40754349|40754452@65.86.22&[MIMAT0000693&hsa-miR-30e-3p&offsets|0|0;m-25&1|+|40754413|40754434&offsets|0|0]|CTTTCAGTCGGATGTTTACAGC</td><td>>MI0000749|hsa-mir-30e&WithFlank&1|+|40754349|40754452</td><td>65</td><td>86</td><td>22</td><td>CTTTCAGTCGGATGTTTACAGC</td><td>MB231</td><td>92</td><td>142</td><td>66</td><td>105</td><td>90</td><td>23.0</td><td>35.6</td><td>34.2</td><td>13.1</td><td>20.2</td></tr>
<tr><td>>MI0000749|hsa-mir-30e&WithFlank&1|+|40754349|40754452@65.86.22&[MIMAT0000693&hsa-miR-30e-3p&offsets|0|0;m-25&1|+|40754413|40754434&offsets|0|0]|CTTTCAGTCGGATGTTTACAGC</td><td>>MI0000749|hsa-mir-30e&WithFlank&1|+|40754349|40754452</td><td>65</td><td>86</td><td>22</td><td>CTTTCAGTCGGATGTTTACAGC</td><td>MB468</td><td>21</td><td>34</td><td>13</td><td>60</td><td>10</td><td>5.1</td><td>9.9</td><td>5.4</td><td>62.1</td><td>4.1</td></tr>
<tr><td>>MI0000750|hsa-mir-26a-2&WithFlank&12|-|57824603|57824698@20.39.20;MI0000083|hsa-mir-26a-1&WithFlank&3|+|37969398|37969486@16.35.20&[MIMAT0000082&hsa-miR-26a-5p&offsets|0|-2;m-33&12|-|57824658|57824679&offsets|0|-2];[MIMAT0000082_1&hsa-miR-26a-5p&offsets|0|-2;m-34&3|+|37969413|37969434&offsets|0|-2]|TTCAAGTAATCCAGGATAGG</td><td>>MI0000750|hsa-mir-26a-2&WithFlank&12|-|57824603|57824698</td><td>20</td><td>39</td><td>20</td><td>TTCAAGTAATCCAGGATAGG</td><td>BT20</td><td>40</td><td>16</td><td>41</td><td>3</td><td>2</td><td>38.6</td><td>4.3</td><td>19.8</td><td>3.2</td><td>1.4</td></tr>
<tr><td>>MI0000750|hsa-mir-26a-2&WithFlank&12|-|57824603|57824698@20.39.20;MI0000083|hsa-mir-26a-1&WithFlank&3|+|37969398|37969486@16.35.20&[MIMAT0000082&hsa-miR-26a-5p&offsets|0|-2;m-33&12|-|57824658|57824679&offsets|0|-2];[MIMAT0000082_1&hsa-miR-26a-5p&offsets|0|-2;m-34&3|+|37969413|37969434&offsets|0|-2]|TTCAAGTAATCCAGGATAGG</td><td>>MI0000750|hsa-mir-26a-2&WithFlank&12|-|57824603|57824698</td><td>20</td><td>39</td><td>20</td><td>TTCAAGTAATCCAGGATAGG</td><td>MB231</td><td>36</td><td>27</td><td>20</td><td>34</td><td>27</td><td>16.0</td><td>10.4</td><td>16.3</td><td>17.1</td><td>16.0</td></tr>
<tr><td>>MI0000750|hsa-mir-26a-2&WithFlank&12|-|57824603|57824698@20.39.20(+1U);MI0000083|hsa-mir-26a-1&WithFlank&3|+|37969398|37969486@16.35.20(+1U)&[MIMAT0000082&hsa-miR-26a-5p&offsets|0|-2(+1U);m-33&12|-|57824658|57824679&offsets|0|-2(+1U)];[MIMAT0000082_1&hsa-miR-26a-5p&offsets|0|-2(+1U);m-34&3|+|37969413|37969434&offsets|0|-2(+1U)]|TTCAAGTAATCCAGGATAGGT</td><td>>MI0000750|hsa-mir-26a-2&WithFlank&12|-|57824603|57824698</td><td>20</td><td>39</td><td>20(+1U)</td><td>TTCAAGTAATCCAGGATAGGT</td><td>BT20</td><td>25</td><td>20</td><td>21</td><td>3</td><td>3</td><td>20.1</td><td>15.4</td><td>10.0</td><td>3.0</td><td>2.5</td></tr>
</table>

<table>
<tr><th></th><th></th><th></th><th></th><th></th><th></th><th></th><th></th><th></th><th></th><th></th><th></th><th></th><th></th><th></th><th></th><th></th></tr>
<tr><td>>MI0000750|hsa-mir-26a-2&WithFlank&12|-|57824603|57824698@20.39.20(+1U);MI0000083|hsa-mir-26a-1&WithFlank&3|+|37969398|37969486@16.35.20(+1U)&[MIMAT0000082&hsa-miR-26a-5p&offsets|0|-2(+1U);m-33&12|-|57824658|57824679&offsets|0|-2(+1U)];[MIMAT0000082_1&hsa-miR-26a-5p&offsets|0|-2(+1U);m-34&3|+|37969413|37969434&offsets|0|-2(+1U)]|TTCAAGTAATCCAGGATAGGT</td><td>>MI0000750|hsa-mir-26a-2&WithFlank&12|-|57824603|57824698</td><td>20</td><td>39</td><td>20(+1U)</td><td>TTCAAGTAATCCAGGATAGGT</td><td>MB231</td><td>14</td><td>16</td><td>8</td><td>49</td><td>23</td><td>3.9</td><td>8.2</td><td>7.2</td><td>33.4</td><td>21.4</td></tr>
<tr><td>>MI0000750|hsa-mir-26a-2&WithFlank&12|-|57824603|57824698@20.39.20(+2U);MI0000083|hsa-mir-26a-1&WithFlank&3|+|37969398|37969486@16.35.20(+2U)&[MIMAT0000082&hsa-miR-26a-5p&offsets|0|-2(+2U);m-33&12|-|57824658|57824679&offsets|0|-2(+2U)];[MIMAT0000082_1&hsa-miR-26a-5p&offsets|0|-2(+2U);m-34&3|+|37969413|37969434&offsets|0|-2(+2U)]|TTCAAGTAATCCAGGATAGGTT</td><td>>MI0000750|hsa-mir-26a-2&WithFlank&12|-|57824603|57824698</td><td>20</td><td>39</td><td>20(+2U)</td><td>TTCAAGTAATCCAGGATAGGTT</td><td>BT20</td><td>22</td><td>10</td><td>16</td><td>1</td><td>1</td><td>18.8</td><td>5.2</td><td>7.0</td><td>1.5</td><td>1.2</td></tr>
<tr><td>>MI0000750|hsa-mir-26a-2&WithFlank&12|-|57824603|57824698@20.39.20(+2U);MI0000083|hsa-mir-26a-1&WithFlank&3|+|37969398|37969486@16.35.20(+2U)&[MIMAT0000082&hsa-miR-26a-5p&offsets|0|-2(+2U);m-33&12|-|57824658|57824679&offsets|0|-2(+2U)];[MIMAT0000082_1&hsa-miR-26a-5p&offsets|0|-2(+2U);m-34&3|+|37969413|37969434&offsets|0|-2(+2U)]|TTCAAGTAATCCAGGATAGGTT</td><td>>MI0000750|hsa-mir-26a-2&WithFlank&12|-|57824603|57824698</td><td>20</td><td>39</td><td>20(+2U)</td><td>TTCAAGTAATCCAGGATAGGTT</td><td>MB231</td><td>19</td><td>14</td><td>8</td><td>11</td><td>17</td><td>8.4</td><td>8.7</td><td>7.1</td><td>6.8</td><td>15.8</td></tr>
<tr><td>>MI0000750|hsa-mir-26a-2&WithFlank&12|-|57824603|57824698@20.40.21;MI0000083|hsa-mir-26a-1&WithFlank&3|+|37969398|37969486@16.36.21&[MIMAT0000082&hsa-miR-26a-5p&offsets|0|-1;m-33&12|-|57824658|57824679&offsets|0|-1];[MIMAT0000082_1&hsa-miR-26a-5p&offsets|0|-1;m-34&3|+|37969413|37969434&offsets|0|-1]|TTCAAGTAATCCAGGATAGGC</td><td>>MI0000750|hsa-mir-26a-2&WithFlank&12|-|57824603|57824698</td><td>20</td><td>40</td><td>21</td><td>TTCAAGTAATCCAGGATAGGC</td><td>BT20</td><td>61</td><td>59</td><td>67</td><td>10</td><td>14</td><td>32.0</td><td>36.8</td><td>21.5</td><td>7.4</td><td>8.2</td></tr>
<tr><td>>MI0000750|hsa-mir-26a-2&WithFlank&12|-|57824603|57824698@20.40.21;MI0000083|hsa-mir-26a-1&WithFlank&3|+|37969398|37969486@16.36.21&[MIMAT0000082&hsa-miR-26a-5p&offsets|0|-1;m-33&12|-|57824658|57824679&offsets|0|-1];[MIMAT0000082_1&hsa-miR-26a-5p&offsets|0|-1;m-34&3|+|37969413|37969434&offsets|0|-1]|TTCAAGTAATCCAGGATAGGC</td><td>>MI0000750|hsa-mir-26a-2&WithFlank&12|-|57824603|57824698</td><td>20</td><td>40</td><td>21</td><td>TTCAAGTAATCCAGGATAGGC</td><td>MB231</td><td>54</td><td>81</td><td>38</td><td>139</td><td>80</td><td>14.3</td><td>36.6</td><td>35.1</td><td>72.3</td><td>50.7</td></tr>
<tr><td>>MI0000750|hsa-mir-26a-2&WithFlank&12|-|57824603|57824698@20.40.21(+1A);MI0000083|hsa-mir-26a-1&WithFlank&3|+|37969398|37969486@16.36.21(+1A)&[MIMAT0000082&hsa-miR-26a-5p&offsets|0|-1(+1A);m-33&12|-|57824658|57824679&offsets|0|-1(+1A)];[MIMAT0000082_1&hsa-miR-26a-5p&offsets|0|-1(+1A);m-34&3|+|37969413|37969434&offsets|0|-1(+1A)]|TTCAAGTAATCCAGGATAGGCA</td><td>>MI0000750|hsa-mir-26a-2&WithFlank&12|-|57824603|57824698</td><td>20</td><td>40</td><td>21(+1A)</td><td>TTCAAGTAATCCAGGATAGGCA</td><td>BT20</td><td>383</td><td>240</td><td>408</td><td>81</td><td>172</td><td>183.9</td><td>126.2</td><td>178.3</td><td>51.4</td><td>83.8</td></tr>
<tr><td>>MI0000750|hsa-mir-26a-2&WithFlank&12|-|57824603|57824698@20.40.21(+1A);MI0000083|hsa-mir-26a-1&WithFlank&3|+|37969398|37969486@16.36.21(+1A)&[MIMAT0000082&hsa-miR-26a-5p&offsets|0|-1(+1A);m-33&12|-|57824658|57824679&offsets|0|-1(+1A)];[MIMAT0000082_1&hsa-miR-26a-5p&offsets|0|-1(+1A);m-34&3|+|37969413|37969434&offsets|0|-1(+1A)]|TTCAAGTAATCCAGGATAGGCA</td><td>>MI0000750|hsa-mir-26a-2&WithFlank&12|-|57824603|57824698</td><td>20</td><td>40</td><td>21(+1A)</td><td>TTCAAGTAATCCAGGATAGGCA</td><td>MB231</td><td>348</td><td>288</td><td>177</td><td>860</td><td>619</td><td>44.4</td><td>131.0</td><td>145.1</td><td>598.2</td><td>358.1</td></tr>
<tr><td>>MI0000750|hsa-mir-26a-2&WithFlank&12|-|57824603|57824698@20.40.21(+1A);MI0000083|hsa-mir-26a-1&WithFlank&3|+|37969398|37969486@16.36.21(+1A)&[MIMAT0000082&hsa-miR-26a-5p&offsets|0|-1(+1A);m-33&12|-|57824658|57824679&offsets|0|-1(+1A)];[MIMAT0000082_1&hsa-miR-26a-5p&offsets|0|-1(+1A);m-34&3|+|37969413|37969434&offsets|0|-1(+1A)]|TTCAAGTAATCCAGGATAGGCA</td><td>>MI0000750|hsa-mir-26a-2&WithFlank&12|-|57824603|57824698</td><td>20</td><td>40</td><td>21(+1A)</td><td>TTCAAGTAATCCAGGATAGGCA</td><td>MB468</td><td>24</td><td>14</td><td>12</td><td>28</td><td>13</td><td>6.8</td><td>4.5</td><td>5.9</td><td>15.6</td><td>4.7</td></tr>
<tr><td>>MI0000750|hsa-mir-26a-2&WithFlank&12|-|57824603|57824698@20.40.21(+1C);MI0000083|hsa-mir-26a-1&WithFlank&3|+|37969398|37969486@16.36.21(+1C)&[MIMAT0000082&hsa-miR-26a-5p&offsets|0|-1(+1C);m-33&12|-|57824658|57824679&offsets|0|-1(+1C)];[MIMAT0000082_1&hsa-miR-26a-5p&offsets|0|-1(+1C);m-34&3|+|37969413|37969434&offsets|0|-1(+1C)]|TTCAAGTAATCCAGGATAGGCC</td><td>>MI0000750|hsa-mir-26a-2&WithFlank&12|-|57824603|57824698</td><td>20</td><td>40</td><td>21(+1C)</td><td>TTCAAGTAATCCAGGATAGGCC</td><td>BT20</td><td>453</td><td>387</td><td>489</td><td>130</td><td>301</td><td>297.9</td><td>224.6</td><td>339.7</td><td>69.4</td><td>169.7</td></tr>
<tr><td>>MI0000750|hsa-mir-26a-2&WithFlank&12|-|57824603|57824698@20.40.21(+1C);MI0000083|hsa-mir-26a-1&WithFlank&3|+|37969398|37969486@16.36.21(+1C)&[MIMAT0000082&hsa-miR-26a-5p&offsets|0|-1(+1C);m-33&12|-|57824658|57824679&offsets|0|-1(+1C)];[MIMAT0000082_1&hsa-miR-26a-5p&offsets|0|-1(+1C);m-34&3|+|37969413|37969434&offsets|0|-1(+1C)]|TTCAAGTAATCCAGGATAGGCC</td><td>>MI0000750|hsa-mir-26a-2&WithFlank&12|-|57824603|57824698</td><td>20</td><td>40</td><td>21(+1C)</td><td>TTCAAGTAATCCAGGATAGGCC</td><td>MB231</td><td>535</td><td>519</td><td>241</td><td>1242</td><td>1020</td><td>120.7</td><td>225.4</td><td>177.8</td><td>841.8</td><td>575.3</td></tr>
<tr><td>>MI0000750|hsa-mir-26a-2&WithFlank&12|-|57824603|57824698@20.40.21(+1C);MI0000083|hsa-mir-26a-1&WithFlank&3|+|37969398|37969486@16.36.21(+1C)&[MIMAT0000082&hsa-miR-26a-5p&offsets|0|-1(+1C);m-33&12|-|57824658|57824679&offsets|0|-1(+1C)];[MIMAT0000082_1&hsa-miR-26a-5p&offsets|0|-1(+1C);m-34&3|+|37969413|37969434&offsets|0|-1(+1C)]|TTCAAGTAATCCAGGATAGGCC</td><td>>MI0000750|hsa-mir-26a-2&WithFlank&12|-|57824603|57824698</td><td>20</td><td>40</td><td>21(+1C)</td><td>TTCAAGTAATCCAGGATAGGCC</td><td>MB468</td><td>46</td><td>30</td><td>22</td><td>45</td><td>24</td><td>14.1</td><td>10.7</td><td>6.9</td><td>25.3</td><td>6.0</td></tr>
<tr><td>>MI0000750|hsa-mir-26a-2&WithFlank&12|-|57824603|57824698@20.40.21(+1G);MI0000083|hsa-mir-26a-1&WithFlank&3|+|37969398|37969486@16.36.21(+1G)&[MIMAT0000082&hsa-miR-26a-5p&offsets|0|-1(+1G);m-33&12|-|57824658|57824679&offsets|0|-1(+1G)];[MIMAT0000082_1&hsa-miR-26a-5p&offsets|0|-1(+1G);m-34&3|+|37969413|37969434&offsets|0|-1(+1G)]|TTCAAGTAATCCAGGATAGGCG</td><td>>MI0000750|hsa-mir-26a-2&WithFlank&12|-|57824603|57824698</td><td>20</td><td>40</td><td>21(+1G)</td><td>TTCAAGTAATCCAGGATAGGCG</td><td>BT20</td><td>229</td><td>94</td><td>111</td><td>22</td><td>33</td><td>113.1</td><td>46.1</td><td>27.5</td><td>21.4</td><td>16.9</td></tr>
<tr><td>>MI0000750|hsa-mir-26a-2&WithFlank&12|-|57824603|57824698@20.40.21(+1G);MI0000083|hsa-mir-26a-1&WithFlank&3|+|37969398|37969486@16.36.21(+1G)&[MIMAT0000082&hsa-miR-26a-5p&offsets|0|-1(+1G);m-33&12|-|57824658|57824679&offsets|0|-1(+1G)];[MIMAT0000082_1&hsa-miR-26a-5p&offsets|0|-1(+1G);m-34&3|+|37969413|37969434&offsets|0|-1(+1G)]|TTCAAGTAATCCAGGATAGGCG</td><td>>MI0000750|hsa-mir-26a-2&WithFlank&12|-|57824603|57824698</td><td>20</td><td>40</td><td>21(+1G)</td><td>TTCAAGTAATCCAGGATAGGCG</td><td>MB231</td><td>277</td><td>124</td><td>54</td><td>170</td><td>125</td><td>175.9</td><td>74.2</td><td>43.5</td><td>119.3</td><td>81.6</td></tr>
<tr><td>>MI0000750|hsa-mir-26a-2&WithFlank&12|-|57824603|57824698@20.40.21(+1G);MI0000083|hsa-mir-26a-1&WithFlank&3|+|37969398|37969486@16.36.21(+1G)&[MIMAT0000082&hsa-miR-26a-5p&offsets|0|-1(+1G);m-33&12|-|57824658|57824679&offsets|0|-1(+1G)];[MIMAT0000082_1&hsa-miR-26a-5p&offsets|0|-1(+1G);m-34&3|+|37969413|37969434&offsets|0|-1(+1G)]|TTCAAGTAATCCAGGATAGGCG</td><td>>MI0000750|hsa-mir-26a-2&WithFlank&12|-|57824603|57824698</td><td>20</td><td>40</td><td>21(+1G)</td><td>TTCAAGTAATCCAGGATAGGCG</td><td>MB468</td><td>25</td><td>6</td><td>13</td><td>12</td><td>6</td><td>6.1</td><td>1.9</td><td>7.1</td><td>8.9</td><td>1.8</td></tr>
<tr><td>>MI0000750|hsa-mir-26a-2&WithFlank&12|-|57824603|57824698@20.41.22;MI0000083|hsa-mir-26a-1&WithFlank&3|+|37969398|37969486@16.37.22&[MIMAT0000082&hsa-miR-26a-5p&offsets|0|0;m-33&12|-|57824658|57824679&offsets|0|0];[MIMAT0000082_1&hsa-miR-26a-5p&offsets|0|0;m-34&3|+|37969413|37969434&offsets|0|0]|TTCAAGTAATCCAGGATAGGCT</td><td>>MI0000750|hsa-mir-26a-2&WithFlank&12|-|57824603|57824698</td><td>20</td><td>41</td><td>22</td><td>TTCAAGTAATCCAGGATAGGCT</td><td>BT20</td><td>3156</td><td>2409</td><td>3519</td><td>893</td><td>2022</td><td>2262.6</td><td>1120.3</td><td>2630.0</td><td>378.9</td><td>974.7</td></tr>
<tr><td>>MI0000750|hsa-mir-26a-2&WithFlank&12|-|57824603|57824698@20.41.22;MI0000083|hsa-mir-26a-1&WithFlank&3|+|37969398|37969486@16.37.22&[MIMAT0000082&hsa-miR-26a-5p&offsets|0|0;m-33&12|-|57824658|57824679&offsets|0|0];[MIMAT0000082_1&hsa-miR-26a-5p&offsets|0|0;m-34&3|+|37969413|37969434&offsets|0|0]|TTCAAGTAATCCAGGATAGGCT</td><td>>MI0000750|hsa-mir-26a-2&WithFlank&12|-|57824603|57824698</td><td>20</td><td>41</td><td>22</td><td>TTCAAGTAATCCAGGATAGGCT</td><td>MB231</td><td>4301</td><td>4350</td><td>2005</td><td>7284</td><td>6846</td><td>1109.1</td><td>1674.6</td><td>1488.3</td><td>4423.3</td><td>2818.9</td></tr>
<tr><td>>MI0000750|hsa-mir-26a-2&WithFlank&12|-|57824603|57824698@20.41.22;MI0000083|hsa-mir-26a-1&WithFlank&3|+|37969398|37969486@16.37.22&[MIMAT0000082&hsa-miR-26a-5p&offsets|0|0;m-33&12|-|57824658|57824679&offsets|0|0];[MIMAT0000082_1&hsa-miR-26a-5p&offsets|0|0;m-34&3|+|37969413|37969434&offsets|0|0]|TTCAAGTAATCCAGGATAGGCT</td><td>>MI0000750|hsa-mir-26a-2&WithFlank&12|-|57824603|57824698</td><td>20</td><td>41</td><td>22</td><td>TTCAAGTAATCCAGGATAGGCT</td><td>MB468</td><td>367</td><td>255</td><td>160</td><td>316</td><td>181</td><td>150.3</td><td>104.9</td><td>26.3</td><td>181.5</td><td>55.1</td></tr>
<tr><td>>MI0000750|hsa-mir-26a-2&WithFlank&12|-|57824603|57824698@20.41.22(+1A);MI0000083|hsa-mir-26a-1&WithFlank&3|+|37969398|37969486@16.37.22(+1A)&[MIMAT0000082&hsa-miR-26a-5p&offsets|0|0(+1A);m-33&12|-|57824658|57824679&offsets|0|0(+1A)];[MIMAT0000082_1&hsa-miR-26a-5p&offsets|0|0(+1A);m-34&3|+|37969413|37969434&offsets|0|0(+1A)]|TTCAAGTAATCCAGGATAGGCTA</td><td>>MI0000750|hsa-mir-26a-2&WithFlank&12|-|57824603|57824698</td><td>20</td><td>41</td><td>22(+1A)</td><td>TTCAAGTAATCCAGGATAGGCTA</td><td>BT20</td><td>70</td><td>46</td><td>66</td><td>39</td><td>27</td><td>37.0</td><td>25.9</td><td>36.8</td><td>30.8</td><td>23.4</td></tr>
</table>

<table>
<tr><td>>MI0000750|hsa-mir-26a-2&WithFlank&12|-|57824603|57824698@20.41.22(+1A);MI0000083|hsa-mir-26a-1&WithFlank&3|+|37969398|37969486@16.37.22(+1A)&[MIMAT0000082&hsa-miR-26a-5p&offsets|0|0(+1A);m-33&12|-|57824658|57824679&offsets|0|0(+1A)];[MIMAT0000082_1&hsa-miR-26a-5p&offsets|0|0(+1A);m-34&3|+|37969413|37969434&offsets|0|0(+1A)]|TTCAAGTAATCCAGGATAGGCTA</td><td>>MI0000750|hsa-mir-26a-2&WithFlank&12|-|57824603|57824698</td><td>20</td><td>41</td><td>22(+1A)</td><td>TTCAAGTAATCCAGGATAGGCTA</td><td>MB231</td><td>52</td><td>36</td><td>16</td><td>111</td><td>63</td><td>6.4</td><td>12.1</td><td>13.3</td><td>61.7</td><td>35.9</td></tr>
<tr><td>>MI0000750|hsa-mir-26a-2&WithFlank&12|-|57824603|57824698@20.41.22(+1C);MI0000083|hsa-mir-26a-1&WithFlank&3|+|37969398|37969486@16.37.22(+1C)&[MIMAT0000082&hsa-miR-26a-5p&offsets|0|0(+1C);m-33&12|-|57824658|57824679&offsets|0|0(+1C)];[MIMAT0000082_1&hsa-miR-26a-5p&offsets|0|0(+1C);m-34&3|+|37969413|37969434&offsets|0|0(+1C)]|TTCAAGTAATCCAGGATAGGCTC</td><td>>MI0000750|hsa-mir-26a-2&WithFlank&12|-|57824603|57824698</td><td>20</td><td>41</td><td>22(+1C)</td><td>TTCAAGTAATCCAGGATAGGCTC</td><td>BT20</td><td>28</td><td>21</td><td>25</td><td>15</td><td>8</td><td>10.8</td><td>9.4</td><td>12.2</td><td>13.5</td><td>5.4</td></tr>
<tr><td>>MI0000750|hsa-mir-26a-2&WithFlank&12|-|57824603|57824698@20.41.22(+1C);MI0000083|hsa-mir-26a-1&WithFlank&3|+|37969398|37969486@16.37.22(+1C)&[MIMAT0000082&hsa-miR-26a-5p&offsets|0|0(+1C);m-33&12|-|57824658|57824679&offsets|0|0(+1C)];[MIMAT0000082_1&hsa-miR-26a-5p&offsets|0|0(+1C);m-34&3|+|37969413|37969434&offsets|0|0(+1C)]|TTCAAGTAATCCAGGATAGGCTC</td><td>>MI0000750|hsa-mir-26a-2&WithFlank&12|-|57824603|57824698</td><td>20</td><td>41</td><td>22(+1C)</td><td>TTCAAGTAATCCAGGATAGGCTC</td><td>MB231</td><td>33</td><td>23</td><td>8</td><td>82</td><td>44</td><td>11.1</td><td>12.6</td><td>5.9</td><td>54.3</td><td>36.3</td></tr>
<tr><td>>MI0000750|hsa-mir-26a-2&WithFlank&12|-|57824603|57824698@20.41.22(+1U);MI0000083|hsa-mir-26a-1&WithFlank&3|+|37969398|37969486@16.37.22(+1U)&[MIMAT0000082&hsa-miR-26a-5p&offsets|0|0(+1U);m-33&12|-|57824658|57824679&offsets|0|0(+1U)];[MIMAT0000082_1&hsa-miR-26a-5p&offsets|0|0(+1U);m-34&3|+|37969413|37969434&offsets|0|0(+1U)]|TTCAAGTAATCCAGGATAGGCTT</td><td>>MI0000750|hsa-mir-26a-2&WithFlank&12|-|57824603|57824698</td><td>20</td><td>41</td><td>22(+1U)</td><td>TTCAAGTAATCCAGGATAGGCTT</td><td>BT20</td><td>191</td><td>132</td><td>177</td><td>99</td><td>53</td><td>82.0</td><td>50.6</td><td>87.0</td><td>79.7</td><td>33.7</td></tr>
<tr><td>>MI0000750|hsa-mir-26a-2&WithFlank&12|-|57824603|57824698@20.41.22(+1U);MI0000083|hsa-mir-26a-1&WithFlank&3|+|37969398|37969486@16.37.22(+1U)&[MIMAT0000082&hsa-miR-26a-5p&offsets|0|0(+1U);m-33&12|-|57824658|57824679&offsets|0|0(+1U)];[MIMAT0000082_1&hsa-miR-26a-5p&offsets|0|0(+1U);m-34&3|+|37969413|37969434&offsets|0|0(+1U)]|TTCAAGTAATCCAGGATAGGCTT</td><td>>MI0000750|hsa-mir-26a-2&WithFlank&12|-|57824603|57824698</td><td>20</td><td>41</td><td>22(+1U)</td><td>TTCAAGTAATCCAGGATAGGCTT</td><td>MB231</td><td>291</td><td>202</td><td>62</td><td>487</td><td>302</td><td>99.3</td><td>113.2</td><td>45.5</td><td>279.2</td><td>223.2</td></tr>
<tr><td>>MI0000750|hsa-mir-26a-2&WithFlank&12|-|57824603|57824698@20.41.22(+1U);MI0000083|hsa-mir-26a-1&WithFlank&3|+|37969398|37969486@16.37.22(+1U)&[MIMAT0000082&hsa-miR-26a-5p&offsets|0|0(+1U);m-33&12|-|57824658|57824679&offsets|0|0(+1U)];[MIMAT0000082_1&hsa-miR-26a-5p&offsets|0|0(+1U);m-34&3|+|37969413|37969434&offsets|0|0(+1U)]|TTCAAGTAATCCAGGATAGGCTT</td><td>>MI0000750|hsa-mir-26a-2&WithFlank&12|-|57824603|57824698</td><td>20</td><td>41</td><td>22(+1U)</td><td>TTCAAGTAATCCAGGATAGGCTT</td><td>MB468</td><td>12</td><td>5</td><td>5</td><td>11</td><td>3</td><td>4.8</td><td>3.4</td><td>2.2</td><td>5.8</td><td>1.1</td></tr>
<tr><td>>MI0000750|hsa-mir-26a-2&WithFlank&12|-|57824603|57824698@20.41.22(+2U);MI0000083|hsa-mir-26a-1&WithFlank&3|+|37969398|37969486@16.37.22(+2U)&[MIMAT0000082&hsa-miR-26a-5p&offsets|0|0(+2U);m-33&12|-|57824658|57824679&offsets|0|0(+2U)];[MIMAT0000082_1&hsa-miR-26a-5p&offsets|0|0(+2U);m-34&3|+|37969413|37969434&offsets|0|0(+2U)]|TTCAAGTAATCCAGGATAGGCTTT</td><td>>MI0000750|hsa-mir-26a-2&WithFlank&12|-|57824603|57824698</td><td>20</td><td>41</td><td>22(+2U)</td><td>TTCAAGTAATCCAGGATAGGCTTT</td><td>BT20</td><td>82</td><td>56</td><td>81</td><td>47</td><td>30</td><td>39.0</td><td>24.4</td><td>44.7</td><td>38.5</td><td>24.6</td></tr>
<tr><td>>MI0000750|hsa-mir-26a-2&WithFlank&12|-|57824603|57824698@20.41.22(+2U);MI0000083|hsa-mir-26a-1&WithFlank&3|+|37969398|37969486@16.37.22(+2U)&[MIMAT0000082&hsa-miR-26a-5p&offsets|0|0(+2U);m-33&12|-|57824658|57824679&offsets|0|0(+2U)];[MIMAT0000082_1&hsa-miR-26a-5p&offsets|0|0(+2U);m-34&3|+|37969413|37969434&offsets|0|0(+2U)]|TTCAAGTAATCCAGGATAGGCTTT</td><td>>MI0000750|hsa-mir-26a-2&WithFlank&12|-|57824603|57824698</td><td>20</td><td>41</td><td>22(+2U)</td><td>TTCAAGTAATCCAGGATAGGCTTT</td><td>MB231</td><td>267</td><td>148</td><td>54</td><td>286</td><td>277</td><td>76.7</td><td>61.2</td><td>36.3</td><td>123.3</td><td>192.3</td></tr>
<tr><td>>MI0000750|hsa-mir-26a-2&WithFlank&12|-|57824603|57824698@20.41.22(+3U);MI0000083|hsa-mir-26a-1&WithFlank&3|+|37969398|37969486@16.37.22(+3U)&[MIMAT0000082&hsa-miR-26a-5p&offsets|0|0(+3U);m-33&12|-|57824658|57824679&offsets|0|0(+3U)];[MIMAT0000082_1&hsa-miR-26a-5p&offsets|0|0(+3U);m-34&3|+|37969413|37969434&offsets|0|0(+3U)]|TTCAAGTAATCCAGGATAGGCTTTT</td><td>>MI0000750|hsa-mir-26a-2&WithFlank&12|-|57824603|57824698</td><td>20</td><td>41</td><td>22(+3U)</td><td>TTCAAGTAATCCAGGATAGGCTTTT</td><td>MB231</td><td>12</td><td>8</td><td>3</td><td>23</td><td>21</td><td>1.3</td><td>4.8</td><td>2.3</td><td>11.0</td><td>15.8</td></tr>
<tr><td>>MI0000750|hsa-mir-26a-2&WithFlank&12|-|57824603|57824698@20.42.23;MI0000083|hsa-mir-26a-1&WithFlank&3|+|37969398|37969486@16.38.23&[MIMAT0000082&hsa-miR-26a-5p&offsets|0|+1;m-33&12|-|57824658|57824679&offsets|0|+1];[MIMAT0000082_1&hsa-miR-26a-5p&offsets|0|+1;m-34&3|+|37969413|37969434&offsets|0|+1]|TTCAAGTAATCCAGGATAGGCTG</td><td>>MI0000750|hsa-mir-26a-2&WithFlank&12|-|57824603|57824698</td><td>20</td><td>42</td><td>23</td><td>TTCAAGTAATCCAGGATAGGCTG</td><td>BT20</td><td>25</td><td>13</td><td>14</td><td>21</td><td>6</td><td>12.7</td><td>3.8</td><td>6.8</td><td>19.0</td><td>2.9</td></tr>
<tr><td>>MI0000750|hsa-mir-26a-2&WithFlank&12|-|57824603|57824698@20.42.23;MI0000083|hsa-mir-26a-1&WithFlank&3|+|37969398|37969486@16.38.23&[MIMAT0000082&hsa-miR-26a-5p&offsets|0|+1;m-33&12|-|57824658|57824679&offsets|0|+1];[MIMAT0000082_1&hsa-miR-26a-5p&offsets|0|+1;m-34&3|+|37969413|37969434&offsets|0|+1]|TTCAAGTAATCCAGGATAGGCTG</td><td>>MI0000750|hsa-mir-26a-2&WithFlank&12|-|57824603|57824698</td><td>20</td><td>42</td><td>23</td><td>TTCAAGTAATCCAGGATAGGCTG</td><td>MB231</td><td>23</td><td>12</td><td>3</td><td>42</td><td>13</td><td>9.5</td><td>6.4</td><td>2.9</td><td>18.1</td><td>4.6</td></tr>
<tr><td>>MI0000750|hsa-mir-26a-2&WithFlank&12|-|57824603|57824698@20.43.24;MI0000083|hsa-mir-26a-1&WithFlank&3|+|37969398|37969486@16.39.24&[MIMAT0000082&hsa-miR-26a-5p&offsets|0|+2;m-33&12|-|57824658|57824679&offsets|0|+2];[MIMAT0000082_1&hsa-miR-26a-5p&offsets|0|+2;m-34&3|+|37969413|37969434&offsets|0|+2]|TTCAAGTAATCCAGGATAGGCTGT</td><td>>MI0000750|hsa-mir-26a-2&WithFlank&12|-|57824603|57824698</td><td>20</td><td>43</td><td>24</td><td>TTCAAGTAATCCAGGATAGGCTGT</td><td>BT20</td><td>11</td><td>7</td><td>8</td><td>7</td><td>3</td><td>4.1</td><td>2.5</td><td>4.2</td><td>7.3</td><td>2.7</td></tr>
<tr><td>>MI0000750|hsa-mir-26a-2&WithFlank&12|-|57824603|57824698@20.43.24;MI0000083|hsa-mir-26a-1&WithFlank&3|+|37969398|37969486@16.39.24&[MIMAT0000082&hsa-miR-26a-5p&offsets|0|+2;m-33&12|-|57824658|57824679&offsets|0|+2];[MIMAT0000082_1&hsa-miR-26a-5p&offsets|0|+2;m-34&3|+|37969413|37969434&offsets|0|+2]|TTCAAGTAATCCAGGATAGGCTGT</td><td>>MI0000750|hsa-mir-26a-2&WithFlank&12|-|57824603|57824698</td><td>20</td><td>43</td><td>24</td><td>TTCAAGTAATCCAGGATAGGCTGT</td><td>MB231</td><td>19</td><td>9</td><td>3</td><td>20</td><td>14</td><td>4.0</td><td>3.6</td><td>2.1</td><td>6.8</td><td>10.5</td></tr>
<tr><td>>MI0000750|hsa-mir-26a-2&WithFlank&12|-|57824603|57824698@23.41.19;MI0000083|hsa-mir-26a-1&WithFlank&3|+|37969398|37969486@19.37.19&[MIMAT0000082&hsa-miR-26a-5p&offsets|+3|0;m-33&12|-|57824658|57824679&offsets|+3|0];[MIMAT0000082_1&hsa-miR-26a-5p&offsets|+3|0;m-34&3|+|37969413|37969434&offsets|+3|0]|AAGTAATCCAGGATAGGCT</td><td>>MI0000750|hsa-mir-26a-2&WithFlank&12|-|57824603|57824698</td><td>23</td><td>41</td><td>19</td><td>AAGTAATCCAGGATAGGCT</td><td>BT20</td><td>52</td><td>40</td><td>55</td><td>14</td><td>32</td><td>44.7</td><td>24.5</td><td>41.8</td><td>6.7</td><td>16.1</td></tr>
<tr><td>>MI0000750|hsa-mir-26a-2&WithFlank&12|-|57824603|57824698@23.41.19;MI0000083|hsa-mir-26a-1&WithFlank&3|+|37969398|37969486@19.37.19&[MIMAT0000082&hsa-miR-26a-5p&offsets|+3|0;m-33&12|-|57824658|57824679&offsets|+3|0];[MIMAT0000082_1&hsa-miR-26a-5p&offsets|+3|0;m-34&3|+|37969413|37969434&offsets|+3|0]|AAGTAATCCAGGATAGGCT</td><td>>MI0000750|hsa-mir-26a-2&WithFlank&12|-|57824603|57824698</td><td>23</td><td>41</td><td>19</td><td>AAGTAATCCAGGATAGGCT</td><td>MB231</td><td>32</td><td>50</td><td>25</td><td>85</td><td>82</td><td>28.1</td><td>43.4</td><td>23.3</td><td>74.5</td><td>73.8</td></tr>
<tr><td>>MI0000760|hsa-mir-361&WithFlank&X|-|85903630|85903713@12.33.22&[MIMAT0000703&hsa-miR-361-5p&offsets|0|0;m-89&X|-|85903681|85903702&offsets|0|0]|TTATCAGAATCTCCAGGGGTAC</td><td>>MI0000760|hsa-mir-361&WithFlank&X|-|85903630|85903713</td><td>12</td><td>33</td><td>22</td><td>TTATCAGAATCTCCAGGGGTAC</td><td>BT20</td><td>15</td><td>18</td><td>18</td><td>18</td><td>7</td><td>7.6</td><td>14.2</td><td>12.4</td><td>16.9</td><td>6.1</td></tr>
<tr><td>>MI0000760|hsa-mir-361&WithFlank&X|-|85903630|85903713@12.33.22&[MIMAT0000703&hsa-miR-361-5p&offsets|0|0;m-89&X|-|85903681|85903702&offsets|0|0]|TTATCAGAATCTCCAGGGGTAC</td><td>>MI0000760|hsa-mir-361&WithFlank&X|-|85903630|85903713</td><td>12</td><td>33</td><td>22</td><td>TTATCAGAATCTCCAGGGGTAC</td><td>MB231</td><td>14</td><td>16</td><td>12</td><td>26</td><td>23</td><td>1.4</td><td>6.2</td><td>10.5</td><td>12.2</td><td>16.4</td></tr>
<tr><td>>MI0000760|hsa-mir-361&WithFlank&X|-|85903630|85903713@51.73.23&[MIMAT0004682&hsa-miR-361-3p&offsets|0|0;m-124&X|-|85903642|85903663&offsets|0|+1]|TCCCCCAGGTGTGATTCTGATTT</td><td>>MI0000760|hsa-mir-361&WithFlank&X|-|85903630|85903713</td><td>51</td><td>73</td><td>23</td><td>TCCCCCAGGTGTGATTCTGATTT</td><td>BT20</td><td>22</td><td>10</td><td>42</td><td>5</td><td>6</td><td>7.8</td><td>3.4</td><td>6.9</td><td>2.0</td><td>4.3</td></tr>
<tr><td>>MI0000760|hsa-mir-361&WithFlank&X|-|85903630|85903713@51.73.23&[MIMAT0004682&hsa-miR-361-3p&offsets|0|0;m-124&X|-|85903642|85903663&offsets|0|+1]|TCCCCCAGGTGTGATTCTGATTT</td><td>>MI0000760|hsa-mir-361&WithFlank&X|-|85903630|85903713</td><td>51</td><td>73</td><td>23</td><td>TCCCCCAGGTGTGATTCTGATTT</td><td>MB231</td><td>15</td><td>9</td><td>13</td><td>7</td><td>12</td><td>3.3</td><td>1.9</td><td>11.1</td><td>2.1</td><td>6.0</td></tr>
<tr><td>>MI0000760|hsa-mir-361&WithFlank&X|-|85903630|85903713@51.74.24&[MIMAT0004682&hsa-miR-361-3p&offsets|0|+1;m-124&X|-|85903642|85903663&offsets|0|+2]|TCCCCCAGGTGTGATTCTGATTTG</td><td>>MI0000760|hsa-mir-361&WithFlank&X|-|85903630|85903713</td><td>51</td><td>74</td><td>24</td><td>TCCCCCAGGTGTGATTCTGATTTG</td><td>BT20</td><td>15</td><td>8</td><td>30</td><td>5</td><td>6</td><td>7.4</td><td>2.1</td><td>9.9</td><td>2.3</td><td>3.4</td></tr>
<tr><td>>MI0000760|hsa-mir-361&WithFlank&X|-|85903630|85903713@51.74.24&[MIMAT0004682&hsa-miR-361-3p&offsets|0|+1;m-124&X|-|85903642|85903663&offsets|0|+2]|TCCCCCAGGTGTGATTCTGATTTG</td><td>>MI0000760|hsa-mir-361&WithFlank&X|-|85903630|85903713</td><td>51</td><td>74</td><td>24</td><td>TCCCCCAGGTGTGATTCTGATTTG</td><td>MB231</td><td>23</td><td>17</td><td>17</td><td>6</td><td>19</td><td>0.7</td><td>0.9</td><td>12.8</td><td>2.3</td><td>5.9</td></tr>
<tr><td>>MI0000760|hsa-mir-361&WithFlank&X|-|85903630|85903713@51.75.25&[MIMAT0004682&hsa-miR-361-3p&offsets|0|+2;m-124&X|-|85903642|85903663&offsets|0|+3]|TCCCCCAGGTGTGATTCTGATTTGC</td><td>>MI0000760|hsa-mir-361&WithFlank&X|-|85903630|85903713</td><td>51</td><td>75</td><td>25</td><td>TCCCCCAGGTGTGATTCTGATTTGC</td><td>MB231</td><td>10</td><td>11</td><td>12</td><td>3</td><td>7</td><td>5.8</td><td>1.5</td><td>3.6</td><td>0.3</td><td>1.6</td></tr>
</table>

<table>
<tr><td>>MI0000760|hsa-mir-361&WithFlank&X|-|85903630|85903713@52.75.24&[MIMAT0004682&hsa-miR-361-3p&offsets|+1|+2;m-124&X|-|85903642|85903663&offsets|+1|+3]|CCCCCAGGTGTGATTCTGATTTGC</td><td>>MI0000760|hsa-mir-361&WithFlank&X|-|85903630|85903713</td><td>52</td><td>75</td><td>24</td><td>CCCCCAGGTGTGATTCTGATTTGC</td><td>BT20</td><td>14</td><td>5</td><td>29</td><td>7</td><td>8</td><td>11.1</td><td>1.2</td><td>12.6</td><td>4.5</td><td>4.1</td></tr>
<tr><td>>MI0000760|hsa-mir-361&WithFlank&X|-|85903630|85903713@52.75.24&[MIMAT0004682&hsa-miR-361-3p&offsets|+1|+2;m-124&X|-|85903642|85903663&offsets|+1|+3]|CCCCCAGGTGTGATTCTGATTTGC</td><td>>MI0000760|hsa-mir-361&WithFlank&X|-|85903630|85903713</td><td>52</td><td>75</td><td>24</td><td>CCCCCAGGTGTGATTCTGATTTGC</td><td>MB231</td><td>18</td><td>25</td><td>22</td><td>10</td><td>16</td><td>10.6</td><td>6.3</td><td>8.0</td><td>0.5</td><td>1.8</td></tr>
<tr><td>>MI0000762|hsa-mir-362&WithFlank&X|+|50008958|50009034@11.34.24&[MIMAT0000705&hsa-miR-362-5p&offsets|0|0;m-267&X|+|50008968|50008991&offsets|0|0]|AATCCTTGGAACCTAGGTGTGAGT</td><td>>MI0000762|hsa-mir-362&WithFlank&X|+|50008958|50009034</td><td>11</td><td>34</td><td>24</td><td>AATCCTTGGAACCTAGGTGTGAGT</td><td>BT20</td><td>21</td><td>11</td><td>18</td><td>4</td><td>6</td><td>5.7</td><td>6.5</td><td>3.7</td><td>2.6</td><td>4.9</td></tr>
<tr><td>>MI0000764|hsa-mir-363&WithFlank&X|-|134169372|134169458@56.76.21&[MIMAT0000707&hsa-miR-363-3p&offsets|0|-1;m-178&X|-|134169381|134169403&offsets|0|-2]|AATTGCACGGTATCCATCTGT</td><td>>MI0000764|hsa-mir-363&WithFlank&X|-|134169372|134169458</td><td>56</td><td>76</td><td>21</td><td>AATTGCACGGTATCCATCTGT</td><td>BT20</td><td>13</td><td>9</td><td>9</td><td>15</td><td>3</td><td>7.5</td><td>8.2</td><td>3.1</td><td>14.5</td><td>1.0</td></tr>
<tr><td>>MI0000764|hsa-mir-363&WithFlank&X|-|134169372|134169458@56.77.22&[MIMAT0000707&hsa-miR-363-3p&offsets|0|0;m-178&X|-|134169381|134169403&offsets|0|-1]|AATTGCACGGTATCCATCTGTA</td><td>>MI0000764|hsa-mir-363&WithFlank&X|-|134169372|134169458</td><td>56</td><td>77</td><td>22</td><td>AATTGCACGGTATCCATCTGTA</td><td>BT20</td><td>113</td><td>74</td><td>52</td><td>108</td><td>66</td><td>98.0</td><td>61.0</td><td>44.5</td><td>65.5</td><td>17.6</td></tr>
<tr><td>>MI0000764|hsa-mir-363&WithFlank&X|-|134169372|134169458@56.77.22&[MIMAT0000707&hsa-miR-363-3p&offsets|0|0;m-178&X|-|134169381|134169403&offsets|0|-1]|AATTGCACGGTATCCATCTGTA</td><td>>MI0000764|hsa-mir-363&WithFlank&X|-|134169372|134169458</td><td>56</td><td>77</td><td>22</td><td>AATTGCACGGTATCCATCTGTA</td><td>MB468</td><td>17</td><td>6</td><td>4</td><td>29</td><td>6</td><td>12.0</td><td>2.9</td><td>0.3</td><td>6.3</td><td>2.4</td></tr>
<tr><td>>MI0000767|hsa-mir-365a&WithFlank&16|+|14309279|14309377@22.44.23&[MIMAT0009199&hsa-miR-365a-5p&offsets|0|0;m-595&16|+|14309300|14309322&offsets|0|0]|AGGGACTTTTGGGGGCAGATGTG</td><td>>MI0000767|hsa-mir-365a&WithFlank&16|+|14309279|14309377</td><td>22</td><td>44</td><td>23</td><td>AGGGACTTTTGGGGGCAGATGTG</td><td>BT20</td><td>12</td><td>7</td><td>15</td><td>8</td><td>2</td><td>12.9</td><td>7.4</td><td>6.4</td><td>3.3</td><td>0.4</td></tr>
<tr><td>>MI0000767|hsa-mir-365a&WithFlank&16|+|14309279|14309377@22.45.24&[MIMAT0009199&hsa-miR-365a-5p&offsets|0|+1;m-595&16|+|14309300|14309322&offsets|0|+1]|AGGGACTTTTGGGGGCAGATGTGT</td><td>>MI0000767|hsa-mir-365a&WithFlank&16|+|14309279|14309377</td><td>22</td><td>45</td><td>24</td><td>AGGGACTTTTGGGGGCAGATGTGT</td><td>BT20</td><td>13</td><td>5</td><td>9</td><td>17</td><td>4</td><td>14.0</td><td>4.0</td><td>3.3</td><td>12.1</td><td>0.8</td></tr>
<tr><td>>MI0000767|hsa-mir-365a&WithFlank&16|+|14309279|14309377@62.81.20;MI0000769|hsa-mir-365b&WithFlank&17|+|31575405|31575527@74.93.20&[MIMAT0000710&hsa-miR-365a-3p&offsets|0|-2;m-176&16|+|14309340|14309361&offsets|0|-2];[MIMAT0022834&hsa-miR-365b-3p&offsets|0|-2;m-175&17|+|31575478|31575499&offsets|0|-2]|TAATGCCCCTAAAAATCCTT</td><td>>MI0000767|hsa-mir-365a&WithFlank&16|+|14309279|14309377</td><td>62</td><td>81</td><td>20</td><td>TAATGCCCCTAAAAATCCTT</td><td>BT20</td><td>16</td><td>19</td><td>16</td><td>4</td><td>13</td><td>14.3</td><td>12.2</td><td>6.0</td><td>3.1</td><td>2.6</td></tr>
<tr><td>>MI0000767|hsa-mir-365a&WithFlank&16|+|14309279|14309377@62.81.20(+2U);MI0000769|hsa-mir-365b&WithFlank&17|+|31575405|31575527@74.93.20(+2U)&[MIMAT0000710&hsa-miR-365a-3p&offsets|0|-2(+2U);m-176&16|+|14309340|14309361&offsets|0|-2(+2U)];[MIMAT0022834&hsa-miR-365b-3p&offsets|0|-2(+2U);m-175&17|+|31575478|31575499&offsets|0|-2(+2U)]|TAATGCCCCTAAAAATCCTTTT</td><td>>MI0000767|hsa-mir-365a&WithFlank&16|+|14309279|14309377</td><td>62</td><td>81</td><td>20(+2U)</td><td>TAATGCCCCTAAAAATCCTTTT</td><td>BT20</td><td>23</td><td>16</td><td>29</td><td>10</td><td>11</td><td>20.5</td><td>8.1</td><td>13.8</td><td>7.3</td><td>2.1</td></tr>
<tr><td>>MI0000767|hsa-mir-365a&WithFlank&16|+|14309279|14309377@62.82.21;MI0000769|hsa-mir-365b&WithFlank&17|+|31575405|31575527@74.94.21&[MIMAT0000710&hsa-miR-365a-3p&offsets|0|-1;m-176&16|+|14309340|14309361&offsets|0|-1];[MIMAT0022834&hsa-miR-365b-3p&offsets|0|-1;m-175&17|+|31575478|31575499&offsets|0|-1]|TAATGCCCCTAAAAATCCTTA</td><td>>MI0000767|hsa-mir-365a&WithFlank&16|+|14309279|14309377</td><td>62</td><td>82</td><td>21</td><td>TAATGCCCCTAAAAATCCTTA</td><td>BT20</td><td>32</td><td>25</td><td>31</td><td>5</td><td>20</td><td>34.3</td><td>12.5</td><td>14.9</td><td>3.7</td><td>5.2</td></tr>
<tr><td>>MI0000767|hsa-mir-365a&WithFlank&16|+|14309279|14309377@62.82.21(+1C);MI0000769|hsa-mir-365b&WithFlank&17|+|31575405|31575527@74.94.21(+1C)&[MIMAT0000710&hsa-miR-365a-3p&offsets|0|-1(+1C);m-176&16|+|14309340|14309361&offsets|0|-1(+1C)];[MIMAT0022834&hsa-miR-365b-3p&offsets|0|-1(+1C);m-175&17|+|31575478|31575499&offsets|0|-1(+1C)]|TAATGCCCCTAAAAATCCTTAC</td><td>>MI0000767|hsa-mir-365a&WithFlank&16|+|14309279|14309377</td><td>62</td><td>82</td><td>21(+1C)</td><td>TAATGCCCCTAAAAATCCTTAC</td><td>BT20</td><td>16</td><td>12</td><td>14</td><td>3</td><td>8</td><td>16.9</td><td>5.3</td><td>4.3</td><td>2.0</td><td>0.9</td></tr>
<tr><td>>MI0000767|hsa-mir-365a&WithFlank&16|+|14309279|14309377@62.83.22;MI0000769|hsa-mir-365b&WithFlank&17|+|31575405|31575527@74.95.22&[MIMAT0000710&hsa-miR-365a-3p&offsets|0|0;m-176&16|+|14309340|14309361&offsets|0|0];[MIMAT0022834&hsa-miR-365b-3p&offsets|0|0;m-175&17|+|31575478|31575499&offsets|0|0]|TAATGCCCCTAAAAATCCTTAT</td><td>>MI0000767|hsa-mir-365a&WithFlank&16|+|14309279|14309377</td><td>62</td><td>83</td><td>22</td><td>TAATGCCCCTAAAAATCCTTAT</td><td>BT20</td><td>140</td><td>104</td><td>110</td><td>23</td><td>66</td><td>137.6</td><td>36.4</td><td>23.4</td><td>14.6</td><td>8.0</td></tr>
<tr><td>>MI0000767|hsa-mir-365a&WithFlank&16|+|14309279|14309377@62.84.23;MI0000769|hsa-mir-365b&WithFlank&17|+|31575405|31575527@74.96.23&[MIMAT0000710&hsa-miR-365a-3p&offsets|0|+1;m-176&16|+|14309340|14309361&offsets|0|+1];[MIMAT0022834&hsa-miR-365b-3p&offsets|0|+1;m-175&17|+|31575478|31575499&offsets|0|+1]|TAATGCCCCTAAAAATCCTTATT</td><td>>MI0000767|hsa-mir-365a&WithFlank&16|+|14309279|14309377</td><td>62</td><td>84</td><td>23</td><td>TAATGCCCCTAAAAATCCTTATT</td><td>BT20</td><td>50</td><td>32</td><td>42</td><td>0</td><td>7</td><td>47.1</td><td>9.3</td><td>9.1</td><td>0.3</td><td>12.3</td></tr>
<tr><td>>MI0000782|hsa-mir-374a&WithFlank&X|-|74287280|74287363@18.38.21(+1A)&[MIMAT0000727&hsa-miR-374a-5p&offsets|0|-1(+1A);m-184&X|-|74287325|74287346&offsets|0|-1(+1A)]|TTATAATACAACCTGATAAGTA</td><td>>MI0000782|hsa-mir-374a&WithFlank&X|-|74287280|74287363</td><td>18</td><td>38</td><td>21(+1A)</td><td>TTATAATACAACCTGATAAGTA</td><td>BT20</td><td>25</td><td>3</td><td>4</td><td>0</td><td>0</td><td>38.3</td><td>4.0</td><td>3.0</td><td>0.0</td><td>0.0</td></tr>
<tr><td>>MI0000782|hsa-mir-374a&WithFlank&X|-|74287280|74287363@18.38.21&[MIMAT0000727&hsa-miR-374a-5p&offsets|0|-1;m-184&X|-|74287325|74287346&offsets|0|-1]|TTATAATACAACCTGATAAGT</td><td>>MI0000782|hsa-mir-374a&WithFlank&X|-|74287280|74287363</td><td>18</td><td>38</td><td>21</td><td>TTATAATACAACCTGATAAGT</td><td>BT20</td><td>40</td><td>17</td><td>6</td><td>3</td><td>1</td><td>48.6</td><td>20.6</td><td>5.1</td><td>4.4</td><td>1.4</td></tr>
<tr><td>>MI0000782|hsa-mir-374a&WithFlank&X|-|74287280|74287363@18.38.21&[MIMAT0000727&hsa-miR-374a-5p&offsets|0|-1;m-184&X|-|74287325|74287346&offsets|0|-1]|TTATAATACAACCTGATAAGT</td><td>>MI0000782|hsa-mir-374a&WithFlank&X|-|74287280|74287363</td><td>18</td><td>38</td><td>21</td><td>TTATAATACAACCTGATAAGT</td><td>MB231</td><td>12</td><td>14</td><td>1</td><td>42</td><td>19</td><td>2.2</td><td>9.4</td><td>1.1</td><td>25.0</td><td>21.2</td></tr>
<tr><td>>MI0000782|hsa-mir-374a&WithFlank&X|-|74287280|74287363@18.39.22&[MIMAT0000727&hsa-miR-374a-5p&offsets|0|0;m-184&X|-|74287325|74287346&offsets|0|0]|TTATAATACAACCTGATAAGTG</td><td>>MI0000782|hsa-mir-374a&WithFlank&X|-|74287280|74287363</td><td>18</td><td>39</td><td>22</td><td>TTATAATACAACCTGATAAGTG</td><td>BT20</td><td>89</td><td>17</td><td>17</td><td>3</td><td>2</td><td>122.8</td><td>19.1</td><td>8.8</td><td>2.4</td><td>0.9</td></tr>
<tr><td>>MI0000782|hsa-mir-374a&WithFlank&X|-|74287280|74287363@18.39.22&[MIMAT0000727&hsa-miR-374a-5p&offsets|0|0;m-184&X|-|74287325|74287346&offsets|0|0]|TTATAATACAACCTGATAAGTG</td><td>>MI0000782|hsa-mir-374a&WithFlank&X|-|74287280|74287363</td><td>18</td><td>39</td><td>22</td><td>TTATAATACAACCTGATAAGTG</td><td>MB231</td><td>33</td><td>19</td><td>6</td><td>42</td><td>25</td><td>10.8</td><td>11.9</td><td>5.8</td><td>18.1</td><td>25.8</td></tr>
<tr><td>>MI0000782|hsa-mir-374a&WithFlank&X|-|74287280|74287363@48.68.21(+1C)&[MIMAT0004688&hsa-miR-374a-3p&offsets|0|-1(+1C);m-74&X|-|74287295|74287316&offsets|0|-1(+1C)]|CTTATCAGATTGTATTGTAATC</td><td>>MI0000782|hsa-mir-374a&WithFlank&X|-|74287280|74287363</td><td>48</td><td>68</td><td>21(+1C)</td><td>CTTATCAGATTGTATTGTAATC</td><td>BT20</td><td>13</td><td>1</td><td>1</td><td>0</td><td>0</td><td>19.8</td><td>2.5</td><td>0.7</td><td>0.0</td><td>0.0</td></tr>
<tr><td>>MI0000782|hsa-mir-374a&WithFlank&X|-|74287280|74287363@48.69.22&[MIMAT0004688&hsa-miR-374a-3p&offsets|0|0;m-74&X|-|74287295|74287316&offsets|0|0]|CTTATCAGATTGTATTGTAATT</td><td>>MI0000782|hsa-mir-374a&WithFlank&X|-|74287280|74287363</td><td>48</td><td>69</td><td>22</td><td>CTTATCAGATTGTATTGTAATT</td><td>BT20</td><td>113</td><td>15</td><td>5</td><td>0</td><td>0</td><td>172.4</td><td>15.6</td><td>3.7</td><td>0.7</td><td>0.0</td></tr>
<tr><td>>MI0000782|hsa-mir-374a&WithFlank&X|-|74287280|74287363@48.69.22&[MIMAT0004688&hsa-miR-374a-3p&offsets|0|0;m-74&X|-|74287295|74287316&offsets|0|0]|CTTATCAGATTGTATTGTAATT</td><td>>MI0000782|hsa-mir-374a&WithFlank&X|-|74287280|74287363</td><td>48</td><td>69</td><td>22</td><td>CTTATCAGATTGTATTGTAATT</td><td>MB231</td><td>30</td><td>14</td><td>1</td><td>17</td><td>12</td><td>16.0</td><td>12.2</td><td>0.4</td><td>5.8</td><td>13.6</td></tr>
<tr><td>>MI0000802|hsa-mir-340&WithFlank&5|-|180015297|180015403@22.42.21(+1A)&[MIMAT0004692&hsa-miR-340-5p&offsets|0|-1(+1A);m-220&5|-|180015361|180015382&offsets|0|-1(+1A)]|TTATAAAGCAATGAGACTGATA</td><td>>MI0000802|hsa-mir-340&WithFlank&5|-|180015297|180015403</td><td>22</td><td>42</td><td>21(+1A)</td><td>TTATAAAGCAATGAGACTGATA</td><td>BT20</td><td>22</td><td>5</td><td>2</td><td>2</td><td>0</td><td>33.0</td><td>3.7</td><td>2.0</td><td>3.9</td><td>0.2</td></tr>
<tr><td>>MI0000802|hsa-mir-340&WithFlank&5|-|180015297|180015403@22.42.21(+1C)&[MIMAT0004692&hsa-miR-340-5p&offsets|0|-1(+1C);m-220&5|-|180015361|180015382&offsets|0|-1(+1C)]|TTATAAAGCAATGAGACTGATC</td><td>>MI0000802|hsa-mir-340&WithFlank&5|-|180015297|180015403</td><td>22</td><td>42</td><td>21(+1C)</td><td>TTATAAAGCAATGAGACTGATC</td><td>BT20</td><td>49</td><td>19</td><td>6</td><td>14</td><td>1</td><td>60.8</td><td>16.8</td><td>5.5</td><td>23.4</td><td>1.2</td></tr>
<tr><td>>MI0000802|hsa-mir-340&WithFlank&5|-|180015297|180015403@22.42.21(+1G)&[MIMAT0004692&hsa-miR-340-5p&offsets|0|-1(+1G);m-220&5|-|180015361|180015382&offsets|0|-1(+1G)]|TTATAAAGCAATGAGACTGATG</td><td>>MI0000802|hsa-mir-340&WithFlank&5|-|180015297|180015403</td><td>22</td><td>42</td><td>21(+1G)</td><td>TTATAAAGCAATGAGACTGATG</td><td>BT20</td><td>24</td><td>2</td><td>1</td><td>2</td><td>0</td><td>36.3</td><td>1.3</td><td>0.8</td><td>3.3</td><td>0.2</td></tr>
<tr><td>>MI0000802|hsa-mir-340&WithFlank&5|-|180015297|180015403@22.42.21&[MIMAT0004692&hsa-miR-340-5p&offsets|0|-1;m-220&5|-|180015361|180015382&offsets|0|-1]|TTATAAAGCAATGAGACTGAT</td><td>>MI0000802|hsa-mir-340&WithFlank&5|-|180015297|180015403</td><td>22</td><td>42</td><td>21</td><td>TTATAAAGCAATGAGACTGAT</td><td>BT20</td><td>46</td><td>18</td><td>5</td><td>6</td><td>2</td><td>54.8</td><td>11.9</td><td>3.9</td><td>8.0</td><td>1.2</td></tr>
<tr><td>>MI0000802|hsa-mir-340&WithFlank&5|-|180015297|180015403@22.43.22&[MIMAT0004692&hsa-miR-340-5p&offsets|0|0;m-220&5|-|180015361|180015382&offsets|0|0]|TTATAAAGCAATGAGACTGATT</td><td>>MI0000802|hsa-mir-340&WithFlank&5|-|180015297|180015403</td><td>22</td><td>43</td><td>22</td><td>TTATAAAGCAATGAGACTGATT</td><td>BT20</td><td>486</td><td>197</td><td>62</td><td>116</td><td>25</td><td>580.2</td><td>151.4</td><td>41.4</td><td>176.0</td><td>7.9</td></tr>
<tr><td>>MI0000802|hsa-mir-340&WithFlank&5|-|180015297|180015403@22.43.22&[MIMAT0004692&hsa-miR-340-5p&offsets|0|0;m-220&5|-|180015361|180015382&offsets|0|0]|TTATAAAGCAATGAGACTGATT</td><td>>MI0000802|hsa-mir-340&WithFlank&5|-|180015297|180015403</td><td>22</td><td>43</td><td>22</td><td>TTATAAAGCAATGAGACTGATT</td><td>MB231</td><td>91</td><td>84</td><td>4</td><td>207</td><td>75</td><td>18.6</td><td>55.9</td><td>3.0</td><td>22.2</td><td>78.2</td></tr>
<tr><td>>MI0000802|hsa-mir-340&WithFlank&5|-|180015297|180015403@22.43.22&[MIMAT0004692&hsa-miR-340-5p&offsets|0|0;m-220&5|-|180015361|180015382&offsets|0|0]|TTATAAAGCAATGAGACTGATT</td><td>>MI0000802|hsa-mir-340&WithFlank&5|-|180015297|180015403</td><td>22</td><td>43</td><td>22</td><td>TTATAAAGCAATGAGACTGATT</td><td>MB468</td><td>21</td><td>6</td><td>4</td><td>53</td><td>4</td><td>9.5</td><td>4.1</td><td>1.7</td><td>35.5</td><td>1.4</td></tr>
</table>

| | | | | | | | | | | | | | | | |
|---|---|---|---|---|---|---|---|---|---|---|---|---|---|---|---|
| >MI0000803\|hsa-mir-330&WithFlank&19\|-\|45638988\|45639093@63.85.23&[MIMAT0000751&hsa-miR-330-3p&offsets\|0\|0;m-347&19\|-\|45639009\|45639030&offsets\|-1\|0]\|GCAAAGCACACGGCCTGCAGAGA | >MI0000803\|hsa-mir-330&WithFlank&19\|-\|45638988\|45639093 | 63 | 85 | 23 | GCAAAGCACACGGCCTGCAGAGA | BT20 | 11 | 11 | 18 | 11 | 6 | 4.6 | 5.6 | 4.8 | 7.8 | 2.8 |
| >MI0000803\|hsa-mir-330&WithFlank&19\|-\|45638988\|45639093@63.86.24(+1U)&[MIMAT0000751&hsa-miR-330-3p&offsets\|0\|+1(+1U);m-347&19\|-\|45639009\|45639030&offsets\|-1\|+1(+1U)]\|GCAAAGCACACGGCCTGCAGAGAGT | >MI0000803\|hsa-mir-330&WithFlank&19\|-\|45638988\|45639093 | 63 | 86 | 24(+1U) | GCAAAGCACACGGCCTGCAGAGAGT | BT20 | 14 | 7 | 29 | 14 | 13 | 15.2 | 2.5 | 9.1 | 9.3 | 5.6 |
| >MI0000803\|hsa-mir-330&WithFlank&19\|-\|45638988\|45639093@63.86.24(+1U)&[MIMAT0000751&hsa-miR-330-3p&offsets\|0\|+1(+1U);m-347&19\|-\|45639009\|45639030&offsets\|-1\|+1(+1U)]\|GCAAAGCACACGGCCTGCAGAGAGT | >MI0000803\|hsa-mir-330&WithFlank&19\|-\|45638988\|45639093 | 63 | 86 | 24(+1U) | GCAAAGCACACGGCCTGCAGAGAGT | MB231 | 42 | 28 | 35 | 31 | 29 | 30.7 | 10.5 | 11.5 | 8.4 | 7.9 |
| >MI0000803\|hsa-mir-330&WithFlank&19\|-\|45638988\|45639093@64.86.23(+1U)&[MIMAT0000751&hsa-miR-330-3p&offsets\|+1\|+1(+1U);m-347&19\|-\|45639009\|45639030&offsets\|0\|+1(+1U)]\|CAAAGCACACGGCCTGCAGAGAGT | >MI0000803\|hsa-mir-330&WithFlank&19\|-\|45638988\|45639093 | 64 | 86 | 23(+1U) | CAAAGCACACGGCCTGCAGAGAGT | MB231 | 13 | 11 | 9 | 20 | 12 | 1.3 | 4.2 | 4.5 | 4.9 | 5.3 |
| >MI0000804\|hsa-mir-328&WithFlank&16\|-\|67202315\|67202401@54.75.22&[MIMAT0000752&hsa-miR-328-3p&offsets\|0\|0;m-160&16\|-\|67202327\|67202348&offsets\|0\|0]\|CTGGCCCTCTCTGCCCTTCCGT | >MI0000804\|hsa-mir-328&WithFlank&16\|-\|67202315\|67202401 | 54 | 75 | 22 | CTGGCCCTCTCTGCCCTTCCGT | BT20 | 49 | 33 | 87 | 13 | 38 | 39.2 | 8.4 | 25.9 | 7.6 | 21.0 |
| >MI0000805\|hsa-mir-342&WithFlank&14\|+\|100109649\|100109759@67.90.24&[MIMAT0000753&hsa-miR-342-3p&offsets\|0\|+1;m-128&14\|+\|100109715\|100109737&offsets\|0\|+1]\|TCTCACACAGAAATCGCACCCGTC | >MI0000805\|hsa-mir-342&WithFlank&14\|+\|100109649\|100109759 | 67 | 90 | 24 | TCTCACACAGAAATCGCACCCGTC | BT20 | 16 | 24 | 52 | 2 | 8 | 10.9 | 9.9 | 24.4 | 1.5 | 0.4 |
| >MI0000805\|hsa-mir-342&WithFlank&14\|+\|100109649\|100109759@67.90.24&[MIMAT0000753&hsa-miR-342-3p&offsets\|0\|+1;m-128&14\|+\|100109715\|100109737&offsets\|0\|+1]\|TCTCACACAGAAATCGCACCCGTC | >MI0000805\|hsa-mir-342&WithFlank&14\|+\|100109649\|100109759 | 67 | 90 | 24 | TCTCACACAGAAATCGCACCCGTC | MB231 | 10 | 16 | 17 | 7 | 20 | 3.1 | 3.0 | 10.3 | 3.2 | 9.8 |
| >MI0000809\|hsa-mir-151a&WithFlank&8\|-\|140732558\|140732659@51.71.21&[MIMAT0000757&hsa-miR-151a-3p&offsets\|-2\|-2;m-30&8\|-\|140732587\|140732607&offsets\|-2\|-2]\|TACTAGACTGAAGCTCCTTGA | >MI0000809\|hsa-mir-151a&WithFlank&8\|-\|140732558\|140732659 | 51 | 71 | 21 | TACTAGACTGAAGCTCCTTGA | BT20 | 12 | 4 | 9 | 14 | 7 | 7.9 | 1.9 | 4.1 | 10.4 | 2.7 |
| >MI0000809\|hsa-mir-151a&WithFlank&8\|-\|140732558\|140732659@51.72.22(+1A)&[MIMAT0000757&hsa-miR-151a-3p&offsets\|-2\|-1(+1A);m-30&8\|-\|140732587\|140732607&offsets\|-2\|-1(+1A)]\|TACTAGACTGAAGCTCCTTGAGA | >MI0000809\|hsa-mir-151a&WithFlank&8\|-\|140732558\|140732659 | 51 | 72 | 22(+1A) | TACTAGACTGAAGCTCCTTGAGA | BT20 | 52 | 20 | 72 | 25 | 23 | 16.4 | 5.0 | 7.9 | 12.9 | 9.0 |
| >MI0000809\|hsa-mir-151a&WithFlank&8\|-\|140732558\|140732659@51.72.22(+1A)&[MIMAT0000757&hsa-miR-151a-3p&offsets\|-2\|-1(+1A);m-30&8\|-\|140732587\|140732607&offsets\|-2\|-1(+1A)]\|TACTAGACTGAAGCTCCTTGAGA | >MI0000809\|hsa-mir-151a&WithFlank&8\|-\|140732558\|140732659 | 51 | 72 | 22(+1A) | TACTAGACTGAAGCTCCTTGAGA | MB231 | 29 | 20 | 15 | 40 | 30 | 10.1 | 0.8 | 6.3 | 6.7 | 12.0 |
| >MI0000809\|hsa-mir-151a&WithFlank&8\|-\|140732558\|140732659@51.72.22(+1U)&[MIMAT0000757&hsa-miR-151a-3p&offsets\|-2\|-1(+1U);m-30&8\|-\|140732587\|140732607&offsets\|-2\|-1(+1U)]\|TACTAGACTGAAGCTCCTTGAGT | >MI0000809\|hsa-mir-151a&WithFlank&8\|-\|140732558\|140732659 | 51 | 72 | 22(+1U) | TACTAGACTGAAGCTCCTTGAGT | BT20 | 32 | 12 | 28 | 16 | 13 | 10.6 | 4.4 | 4.3 | 9.8 | 3.4 |
| >MI0000809\|hsa-mir-151a&WithFlank&8\|-\|140732558\|140732659@51.72.22(+1U)&[MIMAT0000757&hsa-miR-151a-3p&offsets\|-2\|-1(+1U);m-30&8\|-\|140732587\|140732607&offsets\|-2\|-1(+1U)]\|TACTAGACTGAAGCTCCTTGAGT | >MI0000809\|hsa-mir-151a&WithFlank&8\|-\|140732558\|140732659 | 51 | 72 | 22(+1U) | TACTAGACTGAAGCTCCTTGAGT | MB231 | 37 | 30 | 12 | 50 | 40 | 7.3 | 12.7 | 4.3 | 15.8 | 21.4 |
| >MI0000809\|hsa-mir-151a&WithFlank&8\|-\|140732558\|140732659@51.72.22&[MIMAT0000757&hsa-miR-151a-3p&offsets\|-2\|-1;m-30&8\|-\|140732587\|140732607&offsets\|-2\|-1]\|TACTAGACTGAAGCTCCTTGAG | >MI0000809\|hsa-mir-151a&WithFlank&8\|-\|140732558\|140732659 | 51 | 72 | 22 | TACTAGACTGAAGCTCCTTGAG | BT20 | 53 | 20 | 52 | 65 | 26 | 19.8 | 3.9 | 12.8 | 47.5 | 7.2 |
| >MI0000809\|hsa-mir-151a&WithFlank&8\|-\|140732558\|140732659@51.72.22&[MIMAT0000757&hsa-miR-151a-3p&offsets\|-2\|-1;m-30&8\|-\|140732587\|140732607&offsets\|-2\|-1]\|TACTAGACTGAAGCTCCTTGAG | >MI0000809\|hsa-mir-151a&WithFlank&8\|-\|140732558\|140732659 | 51 | 72 | 22 | TACTAGACTGAAGCTCCTTGAG | MB231 | 33 | 30 | 11 | 58 | 31 | 5.8 | 7.3 | 1.9 | 11.0 | 10.7 |
| >MI0000809\|hsa-mir-151a&WithFlank&8\|-\|140732558\|140732659@51.73.23(+1U)&[MIMAT0000757&hsa-miR-151a-3p&offsets\|-2\|0(+1U);m-30&8\|-\|140732587\|140732607&offsets\|-2\|0(+1U)]\|TACTAGACTGAAGCTCCTTGAGGT | >MI0000809\|hsa-mir-151a&WithFlank&8\|-\|140732558\|140732659 | 51 | 73 | 23(+1U) | TACTAGACTGAAGCTCCTTGAGGT | BT20 | 15 | 7 | 22 | 4 | 9 | 8.5 | 2.4 | 9.9 | 2.0 | 4.4 |
| >MI0000809\|hsa-mir-151a&WithFlank&8\|-\|140732558\|140732659@51.73.23(+1U)&[MIMAT0000757&hsa-miR-151a-3p&offsets\|-2\|0(+1U);m-30&8\|-\|140732587\|140732607&offsets\|-2\|0(+1U)]\|TACTAGACTGAAGCTCCTTGAGGT | >MI0000809\|hsa-mir-151a&WithFlank&8\|-\|140732558\|140732659 | 51 | 73 | 23(+1U) | TACTAGACTGAAGCTCCTTGAGGT | MB231 | 28 | 21 | 14 | 18 | 28 | 7.5 | 3.3 | 2.2 | 2.5 | 11.4 |
| >MI0000809\|hsa-mir-151a&WithFlank&8\|-\|140732558\|140732659@51.73.23&[MIMAT0000757&hsa-miR-151a-3p&offsets\|-2\|0;m-30&8\|-\|140732587\|140732607&offsets\|-2\|0]\|TACTAGACTGAAGCTCCTTGAGG | >MI0000809\|hsa-mir-151a&WithFlank&8\|-\|140732558\|140732659 | 51 | 73 | 23 | TACTAGACTGAAGCTCCTTGAGG | BT20 | 264 | 119 | 377 | 180 | 160 | 151.1 | 21.9 | 90.9 | 80.6 | 55.5 |
| >MI0000809\|hsa-mir-151a&WithFlank&8\|-\|140732558\|140732659@51.73.23&[MIMAT0000757&hsa-miR-151a-3p&offsets\|-2\|0;m-30&8\|-\|140732587\|140732607&offsets\|-2\|0]\|TACTAGACTGAAGCTCCTTGAGG | >MI0000809\|hsa-mir-151a&WithFlank&8\|-\|140732558\|140732659 | 51 | 73 | 23 | TACTAGACTGAAGCTCCTTGAGG | MB231 | 241 | 196 | 139 | 273 | 207 | 96.6 | 16.4 | 83.7 | 101.5 | 14.6 |
| >MI0000809\|hsa-mir-151a&WithFlank&8\|-\|140732558\|140732659@53.74.22(+1A)&[MIMAT0000757&hsa-miR-151a-3p&offsets\|0\|+1(+1A);m-30&8\|-\|140732587\|140732607&offsets\|0\|+1(+1A)]\|CTAGACTGAAGCTCCTTGAGGAA | >MI0000809\|hsa-mir-151a&WithFlank&8\|-\|140732558\|140732659 | 53 | 74 | 22(+1A) | CTAGACTGAAGCTCCTTGAGGAA | BT20 | 288 | 218 | 414 | 346 | 319 | 174.2 | 101.7 | 164.0 | 70.4 | 28.5 |
| >MI0000809\|hsa-mir-151a&WithFlank&8\|-\|140732558\|140732659@53.74.22(+1A)&[MIMAT0000757&hsa-miR-151a-3p&offsets\|0\|+1(+1A);m-30&8\|-\|140732587\|140732607&offsets\|0\|+1(+1A)]\|CTAGACTGAAGCTCCTTGAGGAA | >MI0000809\|hsa-mir-151a&WithFlank&8\|-\|140732558\|140732659 | 53 | 74 | 22(+1A) | CTAGACTGAAGCTCCTTGAGGAA | MB231 | 133 | 121 | 64 | 218 | 123 | 100.7 | 8.9 | 14.8 | 22.6 | 4.7 |
| >MI0000809\|hsa-mir-151a&WithFlank&8\|-\|140732558\|140732659@53.74.22(+1G)&[MIMAT0000757&hsa-miR-151a-3p&offsets\|0\|+1(+1G);m-30&8\|-\|140732587\|140732607&offsets\|0\|+1(+1G)]\|CTAGACTGAAGCTCCTTGAGGAG | >MI0000809\|hsa-mir-151a&WithFlank&8\|-\|140732558\|140732659 | 53 | 74 | 22(+1G) | CTAGACTGAAGCTCCTTGAGGAG | BT20 | 18 | 7 | 11 | 12 | 4 | 9.4 | 1.1 | 3.9 | 7.1 | 0.9 |
| >MI0000809\|hsa-mir-151a&WithFlank&8\|-\|140732558\|140732659@53.74.22(+1U)&[MIMAT0000757&hsa-miR-151a-3p&offsets\|0\|+1(+1U);m-30&8\|-\|140732587\|140732607&offsets\|0\|+1(+1U)]\|CTAGACTGAAGCTCCTTGAGGAT | >MI0000809\|hsa-mir-151a&WithFlank&8\|-\|140732558\|140732659 | 53 | 74 | 22(+1U) | CTAGACTGAAGCTCCTTGAGGAT | BT20 | 83 | 56 | 90 | 72 | 77 | 39.9 | 15.7 | 29.9 | 27.6 | 28.1 |
| >MI0000809\|hsa-mir-151a&WithFlank&8\|-\|140732558\|140732659@53.74.22(+1U)&[MIMAT0000757&hsa-miR-151a-3p&offsets\|0\|+1(+1U);m-30&8\|-\|140732587\|140732607&offsets\|0\|+1(+1U)]\|CTAGACTGAAGCTCCTTGAGGAT | >MI0000809\|hsa-mir-151a&WithFlank&8\|-\|140732558\|140732659 | 53 | 74 | 22(+1U) | CTAGACTGAAGCTCCTTGAGGAT | MB231 | 75 | 76 | 28 | 136 | 86 | 14.0 | 22.3 | 1.4 | 9.8 | 28.5 |
| >MI0000809\|hsa-mir-151a&WithFlank&8\|-\|140732558\|140732659@53.74.22(+2A)&[MIMAT0000757&hsa-miR-151a-3p&offsets\|0\|+1(+2A);m-30&8\|-\|140732587\|140732607&offsets\|0\|+1(+2A)]\|CTAGACTGAAGCTCCTTGAGGAAA | >MI0000809\|hsa-mir-151a&WithFlank&8\|-\|140732558\|140732659 | 53 | 74 | 22(+2A) | CTAGACTGAAGCTCCTTGAGGAAA | BT20 | 52 | 37 | 76 | 55 | 49 | 27.4 | 18.4 | 20.5 | 15.8 | 5.1 |
| >MI0000809\|hsa-mir-151a&WithFlank&8\|-\|140732558\|140732659@53.74.22(+2A)&[MIMAT0000757&hsa-miR-151a-3p&offsets\|0\|+1(+2A);m-30&8\|-\|140732587\|140732607&offsets\|0\|+1(+2A)]\|CTAGACTGAAGCTCCTTGAGGAAA | >MI0000809\|hsa-mir-151a&WithFlank&8\|-\|140732558\|140732659 | 53 | 74 | 22(+2A) | CTAGACTGAAGCTCCTTGAGGAAA | MB231 | 24 | 19 | 10 | 32 | 20 | 18.6 | 1.0 | 2.1 | 7.4 | 3.4 |
| >MI0000809\|hsa-mir-151a&WithFlank&8\|-\|140732558\|140732659@53.74.22(+2U)&[MIMAT0000757&hsa-miR-151a-3p&offsets\|0\|+1(+2U);m-30&8\|-\|140732587\|140732607&offsets\|0\|+1(+2U)]\|CTAGACTGAAGCTCCTTGAGGATT | >MI0000809\|hsa-mir-151a&WithFlank&8\|-\|140732558\|140732659 | 53 | 74 | 22(+2U) | CTAGACTGAAGCTCCTTGAGGATT | BT20 | 26 | 15 | 29 | 38 | 15 | 10.3 | 4.3 | 7.1 | 26.5 | 7.8 |
| >MI0000809\|hsa-mir-151a&WithFlank&8\|-\|140732558\|140732659@53.74.22(+2U)&[MIMAT0000757&hsa-miR-151a-3p&offsets\|0\|+1(+2U);m-30&8\|-\|140732587\|140732607&offsets\|0\|+1(+2U)]\|CTAGACTGAAGCTCCTTGAGGATT | >MI0000809\|hsa-mir-151a&WithFlank&8\|-\|140732558\|140732659 | 53 | 74 | 22(+2U) | CTAGACTGAAGCTCCTTGAGGATT | MB231 | 40 | 38 | 17 | 79 | 36 | 2.7 | 11.9 | 3.1 | 2.2 | 12.3 |
| >MI0000809\|hsa-mir-151a&WithFlank&8\|-\|140732558\|140732659@53.74.22&[MIMAT0000757&hsa-miR-151a-3p&offsets\|0\|+1;m-30&8\|-\|140732587\|140732607&offsets\|0\|+1]\|CTAGACTGAAGCTCCTTGAGGA | >MI0000809\|hsa-mir-151a&WithFlank&8\|-\|140732558\|140732659 | 53 | 74 | 22 | CTAGACTGAAGCTCCTTGAGGA | BT20 | 896 | 606 | 1277 | 862 | 785 | 357.9 | 182.3 | 527.8 | 261.3 | 168.8 |
| >MI0000809\|hsa-mir-151a&WithFlank&8\|-\|140732558\|140732659@53.74.22&[MIMAT0000757&hsa-miR-151a-3p&offsets\|0\|+1;m-30&8\|-\|140732587\|140732607&offsets\|0\|+1]\|CTAGACTGAAGCTCCTTGAGGA | >MI0000809\|hsa-mir-151a&WithFlank&8\|-\|140732558\|140732659 | 53 | 74 | 22 | CTAGACTGAAGCTCCTTGAGGA | MB231 | 560 | 514 | 267 | 812 | 506 | 347.0 | 22.6 | 75.6 | 76.9 | 44.1 |

<table>
<tr><td>>MI0000809|hsa-mir-151a&WithFlank&8|-|140732558|140732659@53.74.22&[MIMAT0000757&hsa-miR-151a-3p&offsets|0|+1;m-30&8|-|140732587|140732607&offsets|0|+1]|CTAGACTGAAGCTCCTTGAGGA</td><td>>MI0000809|hsa-mir-151a&WithFlank&8|-|140732558|140732659</td><td>53</td><td>74</td><td>22</td><td>CTAGACTGAAGCTCCTTGAGGA</td><td>MB468</td><td>16</td><td>25</td><td>10</td><td>33</td><td>8</td><td>4.1</td><td>5.6</td><td>6.9</td><td>11.1</td><td>3.6</td></tr>
<tr><td>>MI0000809|hsa-mir-151a&WithFlank&8|-|140732558|140732659@53.75.23(+1U)&[MIMAT0000757&hsa-miR-151a-3p&offsets|0|+2(+1U);m-30&8|-|140732587|140732607&offsets|0|+2(+1U)]|CTAGACTGAAGCTCCTTGAGGACT</td><td>>MI0000809|hsa-mir-151a&WithFlank&8|-|140732558|140732659</td><td>53</td><td>75</td><td>23(+1U)</td><td>CTAGACTGAAGCTCCTTGAGGACT</td><td>BT20</td><td>13</td><td>7</td><td>9</td><td>13</td><td>5</td><td>7.1</td><td>3.8</td><td>1.6</td><td>7.7</td><td>0.3</td></tr>
<tr><td>>MI0000809|hsa-mir-151a&WithFlank&8|-|140732558|140732659@53.75.23&[MIMAT0000757&hsa-miR-151a-3p&offsets|0|+2;m-30&8|-|140732587|140732607&offsets|0|+2]|CTAGACTGAAGCTCCTTGAGGAC</td><td>>MI0000809|hsa-mir-151a&WithFlank&8|-|140732558|140732659</td><td>53</td><td>75</td><td>23</td><td>CTAGACTGAAGCTCCTTGAGGAC</td><td>BT20</td><td>15</td><td>11</td><td>16</td><td>12</td><td>13</td><td>6.7</td><td>3.3</td><td>3.7</td><td>5.0</td><td>4.5</td></tr>
<tr><td>>MI0000809|hsa-mir-151a&WithFlank&8|-|140732558|140732659@53.75.23&[MIMAT0000757&hsa-miR-151a-3p&offsets|0|+2;m-30&8|-|140732587|140732607&offsets|0|+2]|CTAGACTGAAGCTCCTTGAGGAC</td><td>>MI0000809|hsa-mir-151a&WithFlank&8|-|140732558|140732659</td><td>53</td><td>75</td><td>23</td><td>CTAGACTGAAGCTCCTTGAGGAC</td><td>MB231</td><td>10</td><td>9</td><td>3</td><td>22</td><td>13</td><td>2.7</td><td>2.7</td><td>0.6</td><td>6.5</td><td>5.5</td></tr>
<tr><td>>MI0000810|hsa-mir-135b&WithFlank&1|-|205448296|205448404@22.43.22&[MIMAT0000758&hsa-miR-135b-5p&offsets|0|-1;m-321&1|-|205448361|205448383&offsets|0|-1]|TATGGCTTTTCATTCCTATGTG</td><td>>MI0000810|hsa-mir-135b&WithFlank&1|-|205448296|205448404</td><td>22</td><td>43</td><td>22</td><td>TATGGCTTTTCATTCCTATGTG</td><td>BT20</td><td>18</td><td>22</td><td>5</td><td>12</td><td>5</td><td>17.4</td><td>24.3</td><td>7.4</td><td>9.9</td><td>4.2</td></tr>
<tr><td>>MI0000810|hsa-mir-135b&WithFlank&1|-|205448296|205448404@22.44.23&[MIMAT0000758&hsa-miR-135b-5p&offsets|0|0;m-321&1|-|205448361|205448383&offsets|0|0]|TATGGCTTTTCATTCCTATGTGA</td><td>>MI0000810|hsa-mir-135b&WithFlank&1|-|205448296|205448404</td><td>22</td><td>44</td><td>23</td><td>TATGGCTTTTCATTCCTATGTGA</td><td>BT20</td><td>111</td><td>144</td><td>31</td><td>109</td><td>30</td><td>144.0</td><td>219.5</td><td>49.7</td><td>116.1</td><td>34.2</td></tr>
<tr><td>>MI0000810|hsa-mir-135b&WithFlank&1|-|205448296|205448404@22.44.23&[MIMAT0000758&hsa-miR-135b-5p&offsets|0|0;m-321&1|-|205448361|205448383&offsets|0|0]|TATGGCTTTTCATTCCTATGTGA</td><td>>MI0000810|hsa-mir-135b&WithFlank&1|-|205448296|205448404</td><td>22</td><td>44</td><td>23</td><td>TATGGCTTTTCATTCCTATGTGA</td><td>MB231</td><td>28</td><td>37</td><td>1</td><td>140</td><td>44</td><td>19.9</td><td>19.8</td><td>0.5</td><td>40.8</td><td>31.1</td></tr>
<tr><td>>MI0000810|hsa-mir-135b&WithFlank&1|-|205448296|205448404@22.44.23&[MIMAT0000758&hsa-miR-135b-5p&offsets|0|0;m-321&1|-|205448361|205448383&offsets|0|0]|TATGGCTTTTCATTCCTATGTGA</td><td>>MI0000810|hsa-mir-135b&WithFlank&1|-|205448296|205448404</td><td>22</td><td>44</td><td>23</td><td>TATGGCTTTTCATTCCTATGTGA</td><td>MB468</td><td>17</td><td>3</td><td>3</td><td>27</td><td>4</td><td>10.7</td><td>0.6</td><td>1.0</td><td>2.9</td><td>1.2</td></tr>
<tr><td>>MI0000811|hsa-mir-148b&WithFlank&12|+|54337210|54337320@69.89.21(+1A)&[MIMAT0000759&hsa-miR-148b-3p&offsets|0|-1(+1A);m-77&12|+|54337278|54337299&offsets|0|-1(+1A)]|TCAGTGCATCACAGAACTTTGA</td><td>>MI0000811|hsa-mir-148b&WithFlank&12|+|54337210|54337320</td><td>69</td><td>89</td><td>21(+1A)</td><td>TCAGTGCATCACAGAACTTTGA</td><td>BT20</td><td>41</td><td>7</td><td>14</td><td>23</td><td>1</td><td>48.2</td><td>4.3</td><td>5.9</td><td>32.1</td><td>0.6</td></tr>
<tr><td>>MI0000811|hsa-mir-148b&WithFlank&12|+|54337210|54337320@69.89.21(+1C)&[MIMAT0000759&hsa-miR-148b-3p&offsets|0|-1(+1C);m-77&12|+|54337278|54337299&offsets|0|-1(+1C)]|TCAGTGCATCACAGAACTTTGC</td><td>>MI0000811|hsa-mir-148b&WithFlank&12|+|54337210|54337320</td><td>69</td><td>89</td><td>21(+1C)</td><td>TCAGTGCATCACAGAACTTTGC</td><td>BT20</td><td>89</td><td>39</td><td>42</td><td>98</td><td>10</td><td>57.9</td><td>25.6</td><td>5.9</td><td>132.5</td><td>4.6</td></tr>
<tr><td>>MI0000811|hsa-mir-148b&WithFlank&12|+|54337210|54337320@69.89.21(+1C)&[MIMAT0000759&hsa-miR-148b-3p&offsets|0|-1(+1C);m-77&12|+|54337278|54337299&offsets|0|-1(+1C)]|TCAGTGCATCACAGAACTTTGC</td><td>>MI0000811|hsa-mir-148b&WithFlank&12|+|54337210|54337320</td><td>69</td><td>89</td><td>21(+1C)</td><td>TCAGTGCATCACAGAACTTTGC</td><td>MB231</td><td>48</td><td>24</td><td>20</td><td>233</td><td>43</td><td>24.0</td><td>10.7</td><td>18.7</td><td>52.9</td><td>46.8</td></tr>
<tr><td>>MI0000811|hsa-mir-148b&WithFlank&12|+|54337210|54337320@69.89.21(+1G)&[MIMAT0000759&hsa-miR-148b-3p&offsets|0|-1(+1G);m-77&12|+|54337278|54337299&offsets|0|-1(+1G)]|TCAGTGCATCACAGAACTTTGG</td><td>>MI0000811|hsa-mir-148b&WithFlank&12|+|54337210|54337320</td><td>69</td><td>89</td><td>21(+1G)</td><td>TCAGTGCATCACAGAACTTTGG</td><td>BT20</td><td>59</td><td>7</td><td>10</td><td>15</td><td>1</td><td>64.9</td><td>4.4</td><td>4.8</td><td>23.8</td><td>0.6</td></tr>
<tr><td>>MI0000811|hsa-mir-148b&WithFlank&12|+|54337210|54337320@69.89.21(+1G)&[MIMAT0000759&hsa-miR-148b-3p&offsets|0|-1(+1G);m-77&12|+|54337278|54337299&offsets|0|-1(+1G)]|TCAGTGCATCACAGAACTTTGG</td><td>>MI0000811|hsa-mir-148b&WithFlank&12|+|54337210|54337320</td><td>69</td><td>89</td><td>21(+1G)</td><td>TCAGTGCATCACAGAACTTTGG</td><td>MB231</td><td>24</td><td>4</td><td>3</td><td>21</td><td>3</td><td>21.6</td><td>2.3</td><td>3.0</td><td>7.0</td><td>3.9</td></tr>
<tr><td>>MI0000811|hsa-mir-148b&WithFlank&12|+|54337210|54337320@69.89.21&[MIMAT0000759&hsa-miR-148b-3p&offsets|0|-1;m-77&12|+|54337278|54337299&offsets|0|-1]|TCAGTGCATCACAGAACTTTG</td><td>>MI0000811|hsa-mir-148b&WithFlank&12|+|54337210|54337320</td><td>69</td><td>89</td><td>21</td><td>TCAGTGCATCACAGAACTTTG</td><td>BT20</td><td>50</td><td>20</td><td>15</td><td>44</td><td>8</td><td>40.9</td><td>12.6</td><td>13.3</td><td>36.6</td><td>3.3</td></tr>
<tr><td>>MI0000811|hsa-mir-148b&WithFlank&12|+|54337210|54337320@69.89.21&[MIMAT0000759&hsa-miR-148b-3p&offsets|0|-1;m-77&12|+|54337278|54337299&offsets|0|-1]|TCAGTGCATCACAGAACTTTG</td><td>>MI0000811|hsa-mir-148b&WithFlank&12|+|54337210|54337320</td><td>69</td><td>89</td><td>21</td><td>TCAGTGCATCACAGAACTTTG</td><td>MB231</td><td>26</td><td>27</td><td>7</td><td>104</td><td>29</td><td>11.8</td><td>19.4</td><td>6.4</td><td>54.7</td><td>19.0</td></tr>
<tr><td>>MI0000811|hsa-mir-148b&WithFlank&12|+|54337210|54337320@69.90.22(+1A)&[MIMAT0000759&hsa-miR-148b-3p&offsets|0|0(+1A);m-77&12|+|54337278|54337299&offsets|0|0(+1A)]|TCAGTGCATCACAGAACTTTGTA</td><td>>MI0000811|hsa-mir-148b&WithFlank&12|+|54337210|54337320</td><td>69</td><td>90</td><td>22(+1A)</td><td>TCAGTGCATCACAGAACTTTGTA</td><td>BT20</td><td>42</td><td>12</td><td>16</td><td>5</td><td>5</td><td>30.8</td><td>6.6</td><td>1.8</td><td>4.2</td><td>1.0</td></tr>
<tr><td>>MI0000811|hsa-mir-148b&WithFlank&12|+|54337210|54337320@69.90.22(+1A)&[MIMAT0000759&hsa-miR-148b-3p&offsets|0|0(+1A);m-77&12|+|54337278|54337299&offsets|0|0(+1A)]|TCAGTGCATCACAGAACTTTGTA</td><td>>MI0000811|hsa-mir-148b&WithFlank&12|+|54337210|54337320</td><td>69</td><td>90</td><td>22(+1A)</td><td>TCAGTGCATCACAGAACTTTGTA</td><td>MB231</td><td>18</td><td>8</td><td>6</td><td>18</td><td>15</td><td>5.2</td><td>3.3</td><td>5.8</td><td>6.9</td><td>11.9</td></tr>
<tr><td>>MI0000811|hsa-mir-148b&WithFlank&12|+|54337210|54337320@69.90.22(+1G)&[MIMAT0000759&hsa-miR-148b-3p&offsets|0|0(+1G);m-77&12|+|54337278|54337299&offsets|0|0(+1G)]|TCAGTGCATCACAGAACTTTGTG</td><td>>MI0000811|hsa-mir-148b&WithFlank&12|+|54337210|54337320</td><td>69</td><td>90</td><td>22(+1G)</td><td>TCAGTGCATCACAGAACTTTGTG</td><td>BT20</td><td>26</td><td>3</td><td>5</td><td>1</td><td>0</td><td>25.9</td><td>1.4</td><td>1.9</td><td>1.8</td><td>0.6</td></tr>
<tr><td>>MI0000811|hsa-mir-148b&WithFlank&12|+|54337210|54337320@69.90.22(+1G)&[MIMAT0000759&hsa-miR-148b-3p&offsets|0|0(+1G);m-77&12|+|54337278|54337299&offsets|0|0(+1G)]|TCAGTGCATCACAGAACTTTGTG</td><td>>MI0000811|hsa-mir-148b&WithFlank&12|+|54337210|54337320</td><td>69</td><td>90</td><td>22(+1G)</td><td>TCAGTGCATCACAGAACTTTGTG</td><td>MB231</td><td>16</td><td>3</td><td>2</td><td>4</td><td>3</td><td>13.8</td><td>1.1</td><td>1.6</td><td>1.2</td><td>2.2</td></tr>
<tr><td>>MI0000811|hsa-mir-148b&WithFlank&12|+|54337210|54337320@69.90.22(+1U)&[MIMAT0000759&hsa-miR-148b-3p&offsets|0|0(+1U);m-77&12|+|54337278|54337299&offsets|0|0(+1U)]|TCAGTGCATCACAGAACTTTGTT</td><td>>MI0000811|hsa-mir-148b&WithFlank&12|+|54337210|54337320</td><td>69</td><td>90</td><td>22(+1U)</td><td>TCAGTGCATCACAGAACTTTGTT</td><td>BT20</td><td>411</td><td>171</td><td>236</td><td>35</td><td>53</td><td>171.9</td><td>100.4</td><td>58.0</td><td>42.9</td><td>24.2</td></tr>
<tr><td>>MI0000811|hsa-mir-148b&WithFlank&12|+|54337210|54337320@69.90.22(+1U)&[MIMAT0000759&hsa-miR-148b-3p&offsets|0|0(+1U);m-77&12|+|54337278|54337299&offsets|0|0(+1U)]|TCAGTGCATCACAGAACTTTGTT</td><td>>MI0000811|hsa-mir-148b&WithFlank&12|+|54337210|54337320</td><td>69</td><td>90</td><td>22(+1U)</td><td>TCAGTGCATCACAGAACTTTGTT</td><td>MB231</td><td>419</td><td>184</td><td>160</td><td>260</td><td>273</td><td>215.5</td><td>76.7</td><td>140.2</td><td>60.7</td><td>252.1</td></tr>
<tr><td>>MI0000811|hsa-mir-148b&WithFlank&12|+|54337210|54337320@69.90.22(+1U)&[MIMAT0000759&hsa-miR-148b-3p&offsets|0|0(+1U);m-77&12|+|54337278|54337299&offsets|0|0(+1U)]|TCAGTGCATCACAGAACTTTGTT</td><td>>MI0000811|hsa-mir-148b&WithFlank&12|+|54337210|54337320</td><td>69</td><td>90</td><td>22(+1U)</td><td>TCAGTGCATCACAGAACTTTGTT</td><td>MB468</td><td>17</td><td>6</td><td>9</td><td>25</td><td>4</td><td>5.1</td><td>3.2</td><td>2.8</td><td>17.9</td><td>1.6</td></tr>
<tr><td>>MI0000811|hsa-mir-148b&WithFlank&12|+|54337210|54337320@69.90.22(+2U)&[MIMAT0000759&hsa-miR-148b-3p&offsets|0|0(+2U);m-77&12|+|54337278|54337299&offsets|0|0(+2U)]|TCAGTGCATCACAGAACTTTGTTT</td><td>>MI0000811|hsa-mir-148b&WithFlank&12|+|54337210|54337320</td><td>69</td><td>90</td><td>22(+2U)</td><td>TCAGTGCATCACAGAACTTTGTTT</td><td>BT20</td><td>23</td><td>12</td><td>8</td><td>4</td><td>5</td><td>10.8</td><td>10.0</td><td>3.3</td><td>5.1</td><td>2.7</td></tr>
<tr><td>>MI0000811|hsa-mir-148b&WithFlank&12|+|54337210|54337320@69.90.22(+2U)&[MIMAT0000759&hsa-miR-148b-3p&offsets|0|0(+2U);m-77&12|+|54337278|54337299&offsets|0|0(+2U)]|TCAGTGCATCACAGAACTTTGTTT</td><td>>MI0000811|hsa-mir-148b&WithFlank&12|+|54337210|54337320</td><td>69</td><td>90</td><td>22(+2U)</td><td>TCAGTGCATCACAGAACTTTGTTT</td><td>MB231</td><td>29</td><td>18</td><td>6</td><td>44</td><td>33</td><td>10.1</td><td>8.6</td><td>4.5</td><td>9.2</td><td>30.8</td></tr>
<tr><td>>MI0000811|hsa-mir-148b&WithFlank&12|+|54337210|54337320@69.90.22(+3U)&[MIMAT0000759&hsa-miR-148b-3p&offsets|0|0(+3U);m-77&12|+|54337278|54337299&offsets|0|0(+3U)]|TCAGTGCATCACAGAACTTTGTTTT</td><td>>MI0000811|hsa-mir-148b&WithFlank&12|+|54337210|54337320</td><td>69</td><td>90</td><td>22(+3U)</td><td>TCAGTGCATCACAGAACTTTGTTTT</td><td>MB231</td><td>18</td><td>11</td><td>3</td><td>21</td><td>18</td><td>3.6</td><td>3.4</td><td>1.3</td><td>3.8</td><td>6.9</td></tr>
<tr><td>>MI0000811|hsa-mir-148b&WithFlank&12|+|54337210|54337320@69.90.22&[MIMAT0000759&hsa-miR-148b-3p&offsets|0|0;m-77&12|+|54337278|54337299&offsets|0|0]|TCAGTGCATCACAGAACTTTGT</td><td>>MI0000811|hsa-mir-148b&WithFlank&12|+|54337210|54337320</td><td>69</td><td>90</td><td>22</td><td>TCAGTGCATCACAGAACTTTGT</td><td>BT20</td><td>685</td><td>271</td><td>333</td><td>657</td><td>73</td><td>457.5</td><td>145.7</td><td>72.9</td><td>830.6</td><td>27.6</td></tr>
<tr><td>>MI0000811|hsa-mir-148b&WithFlank&12|+|54337210|54337320@69.90.22&[MIMAT0000759&hsa-miR-148b-3p&offsets|0|0;m-77&12|+|54337278|54337299&offsets|0|0]|TCAGTGCATCACAGAACTTTGT</td><td>>MI0000811|hsa-mir-148b&WithFlank&12|+|54337210|54337320</td><td>69</td><td>90</td><td>22</td><td>TCAGTGCATCACAGAACTTTGT</td><td>MB231</td><td>449</td><td>230</td><td>195</td><td>1597</td><td>318</td><td>213.9</td><td>99.4</td><td>193.6</td><td>215.7</td><td>316.8</td></tr>
<tr><td>>MI0000811|hsa-mir-148b&WithFlank&12|+|54337210|54337320@69.90.22&[MIMAT0000759&hsa-miR-148b-3p&offsets|0|0;m-77&12|+|54337278|54337299&offsets|0|0]|TCAGTGCATCACAGAACTTTGT</td><td>>MI0000811|hsa-mir-148b&WithFlank&12|+|54337210|54337320</td><td>69</td><td>90</td><td>22</td><td>TCAGTGCATCACAGAACTTTGT</td><td>MB468</td><td>22</td><td>8</td><td>10</td><td>108</td><td>5</td><td>7.8</td><td>3.5</td><td>2.3</td><td>64.6</td><td>2.1</td></tr>
</table>

| | | | | | | | | | | | | | | | | |
|---|---|---|---|---|---|---|---|---|---|---|---|---|---|---|---|---|
| >MI0000811\|hsa-mir-148b&WithFlank&12\|+\|54337210\|54337320@69.91.23&[MIMAT0000759&hsa-miR-148b-3p&offsets\|0\|+1;m-77&12\|+\|54337278\|54337299&offsets\|0\|+1]\|TCAGTGCATCACAGAACTTTGTC | >MI0000811\|hsa-mir-148b&WithFlank&12\|+\|54337210\|54337320 | 69 | 91 | 23 | TCAGTGCATCACAGAACTTTGTC | BT20 | 57 | 25 | 31 | 6 | 8 | 24.3 | 16.0 | 3.7 | 8.8 | 3.5 |
| >MI0000811\|hsa-mir-148b&WithFlank&12\|+\|54337210\|54337320@69.91.23&[MIMAT0000759&hsa-miR-148b-3p&offsets\|0\|+1;m-77&12\|+\|54337278\|54337299&offsets\|0\|+1]\|TCAGTGCATCACAGAACTTTGTC | >MI0000811\|hsa-mir-148b&WithFlank&12\|+\|54337210\|54337320 | 69 | 91 | 23 | TCAGTGCATCACAGAACTTTGTC | MB231 | 46 | 19 | 18 | 40 | 39 | 22.8 | 8.6 | 15.0 | 12.9 | 40.7 |
| >MI0000812\|hsa-mir-331&WithFlank&12\|+\|95308414\|95308519@67.86.20&[MIMAT0000760&hsa-miR-331-3p&offsets\|0\|-1;m-192&12\|+\|95308480\|95308501&offsets\|0\|-2]\|GCCCCTGGGCCTATCCTAGA | >MI0000812\|hsa-mir-331&WithFlank&12\|+\|95308414\|95308519 | 67 | 86 | 20 | GCCCCTGGGCCTATCCTAGA | BT20 | 18 | 15 | 10 | 1 | 4 | 20.4 | 8.6 | 6.0 | 0.6 | 0.4 |
| >MI0000812\|hsa-mir-331&WithFlank&12\|+\|95308414\|95308519@67.87.21(+1U)&[MIMAT0000760&hsa-miR-331-3p&offsets\|0\|0(+1U);m-192&12\|+\|95308480\|95308501&offsets\|0\|-1(+1U)]\|GCCCCTGGGCCTATCCTAGAAT | >MI0000812\|hsa-mir-331&WithFlank&12\|+\|95308414\|95308519 | 67 | 87 | 21(+1U) | GCCCCTGGGCCTATCCTAGAAT | BT20 | 38 | 51 | 27 | 1 | 8 | 15.0 | 31.4 | 6.9 | 2.0 | 13.2 |
| >MI0000812\|hsa-mir-331&WithFlank&12\|+\|95308414\|95308519@67.87.21(+1U)&[MIMAT0000760&hsa-miR-331-3p&offsets\|0\|0(+1U);m-192&12\|+\|95308480\|95308501&offsets\|0\|-1(+1U)]\|GCCCCTGGGCCTATCCTAGAAT | >MI0000812\|hsa-mir-331&WithFlank&12\|+\|95308414\|95308519 | 67 | 87 | 21(+1U) | GCCCCTGGGCCTATCCTAGAAT | MB231 | 24 | 41 | 8 | 5 | 46 | 8.8 | 26.1 | 5.7 | 1.3 | 28.7 |
| >MI0000812\|hsa-mir-331&WithFlank&12\|+\|95308414\|95308519@67.87.21&[MIMAT0000760&hsa-miR-331-3p&offsets\|0\|0;m-192&12\|+\|95308480\|95308501&offsets\|0\|-1]\|GCCCCTGGGCCTATCCTAGAA | >MI0000812\|hsa-mir-331&WithFlank&12\|+\|95308414\|95308519 | 67 | 87 | 21 | GCCCCTGGGCCTATCCTAGAA | BT20 | 11 | 9 | 4 | 0 | 0 | 13.3 | 5.0 | 3.0 | 0.0 | 0.0 |
| >MI0000815\|hsa-mir-339&WithFlank&7\|-\|1022927\|1023032@21.43.23&[MIMAT0000764&hsa-miR-339-5p&offsets\|0\|0;m-163&7\|-\|1022990\|1023012&offsets\|0\|0]\|TCCCTGTCCTCCAGGAGCTCACG | >MI0000815\|hsa-mir-339&WithFlank&7\|-\|1022927\|1023032 | 21 | 43 | 23 | TCCCTGTCCTCCAGGAGCTCACG | BT20 | 49 | 36 | 82 | 3 | 20 | 23.9 | 31.4 | 31.3 | 1.0 | 16.9 |
| >MI0000815\|hsa-mir-339&WithFlank&7\|-\|1022927\|1023032@21.43.23&[MIMAT0000764&hsa-miR-339-5p&offsets\|0\|0;m-163&7\|-\|1022990\|1023012&offsets\|0\|0]\|TCCCTGTCCTCCAGGAGCTCACG | >MI0000815\|hsa-mir-339&WithFlank&7\|-\|1022927\|1023032 | 21 | 43 | 23 | TCCCTGTCCTCCAGGAGCTCACG | MB231 | 25 | 19 | 28 | 2 | 28 | 4.8 | 9.3 | 27.2 | 0.3 | 16.6 |
| >MI0000815\|hsa-mir-339&WithFlank&7\|-\|1022927\|1023032@21.44.24&[MIMAT0000764&hsa-miR-339-5p&offsets\|0\|+1;m-163&7\|-\|1022990\|1023012&offsets\|0\|+1]\|TCCCTGTCCTCCAGGAGCTCACGT | >MI0000815\|hsa-mir-339&WithFlank&7\|-\|1022927\|1023032 | 21 | 44 | 24 | TCCCTGTCCTCCAGGAGCTCACGT | MB231 | 11 | 10 | 9 | 1 | 17 | 2.0 | 5.7 | 8.0 | 0.7 | 11.8 |
| >MI0000815\|hsa-mir-339&WithFlank&7\|-\|1022927\|1023032@56.76.21&[MIMAT0004702&hsa-miR-339-3p&offsets\|0\|-2;m-235&7\|-\|1022957\|1022977&offsets\|0\|0]\|TGAGCGCCTCGACGACAGAGC | >MI0000815\|hsa-mir-339&WithFlank&7\|-\|1022927\|1023032 | 56 | 76 | 21 | TGAGCGCCTCGACGACAGAGC | BT20 | 19 | 15 | 20 | 85 | 14 | 17.8 | 8.2 | 9.4 | 70.1 | 4.9 |
| >MI0000815\|hsa-mir-339&WithFlank&7\|-\|1022927\|1023032@56.77.22&[MIMAT0004702&hsa-miR-339-3p&offsets\|0\|-1;m-235&7\|-\|1022957\|1022977&offsets\|0\|+1]\|TGAGCGCCTCGACGACAGAGCC | >MI0000815\|hsa-mir-339&WithFlank&7\|-\|1022927\|1023032 | 56 | 77 | 22 | TGAGCGCCTCGACGACAGAGCC | BT20 | 10 | 9 | 10 | 19 | 6 | 6.7 | 6.4 | 4.8 | 19.2 | 3.9 |
| >MI0000816\|hsa-mir-335&WithFlank&7\|+\|130496105\|130496210@22.43.22&[MIMAT0000765&hsa-miR-335-5p&offsets\|0\|-1;m-249&7\|+\|130496126\|130496147&offsets\|0\|0]\|TCAAGAGCAATAACGAAAAATG | >MI0000816\|hsa-mir-335&WithFlank&7\|+\|130496105\|130496210 | 22 | 43 | 22 | TCAAGAGCAATAACGAAAAATG | BT20 | 11 | 10 | 7 | 5 | 1 | 7.1 | 10.1 | 6.3 | 4.5 | 0.6 |
| >MI0000816\|hsa-mir-335&WithFlank&7\|+\|130496105\|130496210@57.79.23&[MIMAT0004703&hsa-miR-335-3p&offsets\|-1\|0;m-159&7\|+\|130496162\|130496183&offsets\|-1\|0]\|GTTTTTCATTATTGCTCCTGACC | >MI0000816\|hsa-mir-335&WithFlank&7\|+\|130496105\|130496210 | 57 | 79 | 23 | GTTTTTCATTATTGCTCCTGACC | MB231 | 14 | 10 | 12 | 3 | 15 | 5.1 | 0.7 | 2.5 | 0.7 | 6.2 |
| >MI0000816\|hsa-mir-335&WithFlank&7\|+\|130496105\|130496210@58.78.21(+1A)&[MIMAT0004703&hsa-miR-335-3p&offsets\|0\|-1(+1A);m-159&7\|+\|130496162\|130496183&offsets\|0\|-1(+1A)]\|TTTTTCATTATTGCTCCTGACA | >MI0000816\|hsa-mir-335&WithFlank&7\|+\|130496105\|130496210 | 58 | 78 | 21(+1A) | TTTTTCATTATTGCTCCTGACA | BT20 | 25 | 8 | 24 | 3 | 4 | 18.6 | 9.1 | 14.2 | 3.2 | 4.3 |
| >MI0000816\|hsa-mir-335&WithFlank&7\|+\|130496105\|130496210@58.78.21(+1A)&[MIMAT0004703&hsa-miR-335-3p&offsets\|0\|-1(+1A);m-159&7\|+\|130496162\|130496183&offsets\|0\|-1(+1A)]\|TTTTTCATTATTGCTCCTGACA | >MI0000816\|hsa-mir-335&WithFlank&7\|+\|130496105\|130496210 | 58 | 78 | 21(+1A) | TTTTTCATTATTGCTCCTGACA | MB231 | 23 | 15 | 19 | 14 | 30 | 1.6 | 2.1 | 14.9 | 8.4 | 26.0 |
| >MI0000816\|hsa-mir-335&WithFlank&7\|+\|130496105\|130496210@58.78.21(+1U)&[MIMAT0004703&hsa-miR-335-3p&offsets\|0\|-1(+1U);m-159&7\|+\|130496162\|130496183&offsets\|0\|-1(+1U)]\|TTTTTCATTATTGCTCCTGACT | >MI0000816\|hsa-mir-335&WithFlank&7\|+\|130496105\|130496210 | 58 | 78 | 21(+1U) | TTTTTCATTATTGCTCCTGACT | BT20 | 49 | 21 | 50 | 8 | 8 | 36.1 | 25.1 | 37.1 | 9.0 | 9.3 |
| >MI0000816\|hsa-mir-335&WithFlank&7\|+\|130496105\|130496210@58.78.21(+1U)&[MIMAT0004703&hsa-miR-335-3p&offsets\|0\|-1(+1U);m-159&7\|+\|130496162\|130496183&offsets\|0\|-1(+1U)]\|TTTTTCATTATTGCTCCTGACT | >MI0000816\|hsa-mir-335&WithFlank&7\|+\|130496105\|130496210 | 58 | 78 | 21(+1U) | TTTTTCATTATTGCTCCTGACT | MB231 | 55 | 34 | 43 | 36 | 73 | 3.6 | 6.4 | 26.7 | 21.5 | 58.2 |
| >MI0000816\|hsa-mir-335&WithFlank&7\|+\|130496105\|130496210@58.78.21&[MIMAT0004703&hsa-miR-335-3p&offsets\|0\|-1;m-159&7\|+\|130496162\|130496183&offsets\|0\|-1]\|TTTTTCATTATTGCTCCTGAC | >MI0000816\|hsa-mir-335&WithFlank&7\|+\|130496105\|130496210 | 58 | 78 | 21 | TTTTTCATTATTGCTCCTGAC | BT20 | 15 | 7 | 19 | 5 | 2 | 10.0 | 8.0 | 14.5 | 5.2 | 1.7 |
| >MI0000816\|hsa-mir-335&WithFlank&7\|+\|130496105\|130496210@58.78.21&[MIMAT0004703&hsa-miR-335-3p&offsets\|0\|-1;m-159&7\|+\|130496162\|130496183&offsets\|0\|-1]\|TTTTTCATTATTGCTCCTGAC | >MI0000816\|hsa-mir-335&WithFlank&7\|+\|130496105\|130496210 | 58 | 78 | 21 | TTTTTCATTATTGCTCCTGAC | MB231 | 29 | 18 | 19 | 18 | 24 | 7.5 | 4.5 | 14.9 | 9.1 | 18.7 |
| >MI0000816\|hsa-mir-335&WithFlank&7\|+\|130496105\|130496210@58.79.22&[MIMAT0004703&hsa-miR-335-3p&offsets\|0\|0;m-159&7\|+\|130496162\|130496183&offsets\|0\|0]\|TTTTTCATTATTGCTCCTGACC | >MI0000816\|hsa-mir-335&WithFlank&7\|+\|130496105\|130496210 | 58 | 79 | 22 | TTTTTCATTATTGCTCCTGACC | BT20 | 96 | 43 | 109 | 20 | 24 | 83.4 | 45.4 | 91.9 | 24.7 | 28.0 |
| >MI0000816\|hsa-mir-335&WithFlank&7\|+\|130496105\|130496210@58.79.22&[MIMAT0004703&hsa-miR-335-3p&offsets\|0\|0;m-159&7\|+\|130496162\|130496183&offsets\|0\|0]\|TTTTTCATTATTGCTCCTGACC | >MI0000816\|hsa-mir-335&WithFlank&7\|+\|130496105\|130496210 | 58 | 79 | 22 | TTTTTCATTATTGCTCCTGACC | MB231 | 153 | 120 | 142 | 62 | 167 | 16.2 | 10.1 | 84.5 | 24.0 | 106.7 |
| >MI0000825\|hsa-mir-345&WithFlank&14\|+\|100307853\|100307962@24.45.22(+1U)&[MIMAT0000772&hsa-miR-345-5p&offsets\|0\|0(+1U);m-193&14\|+\|100307876\|100307896&offsets\|0\|+1(+1U)]\|GCTGACTCCTAGTCCAGGGCTCT | >MI0000825\|hsa-mir-345&WithFlank&14\|+\|100307853\|100307962 | 24 | 45 | 22(+1U) | GCTGACTCCTAGTCCAGGGCTCT | MB231 | 13 | 9 | 5 | 22 | 19 | 3.8 | 3.8 | 4.0 | 10.6 | 14.3 |
| >MI0000825\|hsa-mir-345&WithFlank&14\|+\|100307853\|100307962@24.45.22&[MIMAT0000772&hsa-miR-345-5p&offsets\|0\|0;m-193&14\|+\|100307876\|100307896&offsets\|0\|+1]\|GCTGACTCCTAGTCCAGGGCTC | >MI0000825\|hsa-mir-345&WithFlank&14\|+\|100307853\|100307962 | 24 | 45 | 22 | GCTGACTCCTAGTCCAGGGCTC | MB231 | 18 | 16 | 10 | 57 | 17 | 13.1 | 4.7 | 5.6 | 11.1 | 4.4 |
| >MI0001445\|hsa-mir-423&WithFlank&17\|+\|30117073\|30117178@23.43.21&[MIMAT0004748&hsa-miR-423-5p&offsets\|0\|-2;m-141&17\|+\|30117095\|30117117&offsets\|0\|-2]\|TGAGGGGCAGAGAGCGAGACT | >MI0001445\|hsa-mir-423&WithFlank&17\|+\|30117073\|30117178 | 23 | 43 | 21 | TGAGGGGCAGAGAGCGAGACT | BT20 | 42 | 29 | 140 | 704 | 17 | 14.2 | 7.5 | 46.8 | 472.7 | 4.7 |
| >MI0001445\|hsa-mir-423&WithFlank&17\|+\|30117073\|30117178@23.43.21&[MIMAT0004748&hsa-miR-423-5p&offsets\|0\|-2;m-141&17\|+\|30117095\|30117117&offsets\|0\|-2]\|TGAGGGGCAGAGAGCGAGACT | >MI0001445\|hsa-mir-423&WithFlank&17\|+\|30117073\|30117178 | 23 | 43 | 21 | TGAGGGGCAGAGAGCGAGACT | MB231 | 41 | 39 | 76 | 277 | 14 | 12.3 | 13.6 | 37.3 | 56.2 | 4.5 |
| >MI0001445\|hsa-mir-423&WithFlank&17\|+\|30117073\|30117178@23.44.22(+1C)&[MIMAT0004748&hsa-miR-423-5p&offsets\|0\|-1(+1C);m-141&17\|+\|30117095\|30117117&offsets\|0\|-1(+1C)]\|TGAGGGGCAGAGAGCGAGACTTC | >MI0001445\|hsa-mir-423&WithFlank&17\|+\|30117073\|30117178 | 23 | 44 | 22(+1C) | TGAGGGGCAGAGAGCGAGACTTC | BT20 | 27 | 20 | 66 | 645 | 13 | 8.3 | 9.9 | 18.0 | 536.8 | 8.6 |
| >MI0001445\|hsa-mir-423&WithFlank&17\|+\|30117073\|30117178@23.44.22(+1C)&[MIMAT0004748&hsa-miR-423-5p&offsets\|0\|-1(+1C);m-141&17\|+\|30117095\|30117117&offsets\|0\|-1(+1C)]\|TGAGGGGCAGAGAGCGAGACTTC | >MI0001445\|hsa-mir-423&WithFlank&17\|+\|30117073\|30117178 | 23 | 44 | 22(+1C) | TGAGGGGCAGAGAGCGAGACTTC | MB231 | 15 | 12 | 21 | 205 | 6 | 6.5 | 3.2 | 9.3 | 105.9 | 2.1 |
| >MI0001445\|hsa-mir-423&WithFlank&17\|+\|30117073\|30117178@23.44.22&[MIMAT0004748&hsa-miR-423-5p&offsets\|0\|-1;m-141&17\|+\|30117095\|30117117&offsets\|0\|-1]\|TGAGGGGCAGAGAGCGAGACTT | >MI0001445\|hsa-mir-423&WithFlank&17\|+\|30117073\|30117178 | 23 | 44 | 22 | TGAGGGGCAGAGAGCGAGACTT | BT20 | 28 | 18 | 92 | 419 | 15 | 16.7 | 7.9 | 18.5 | 265.6 | 6.3 |
| >MI0001445\|hsa-mir-423&WithFlank&17\|+\|30117073\|30117178@23.44.22&[MIMAT0004748&hsa-miR-423-5p&offsets\|0\|-1;m-141&17\|+\|30117095\|30117117&offsets\|0\|-1]\|TGAGGGGCAGAGAGCGAGACTT | >MI0001445\|hsa-mir-423&WithFlank&17\|+\|30117073\|30117178 | 23 | 44 | 22 | TGAGGGGCAGAGAGCGAGACTT | MB231 | 25 | 22 | 43 | 184 | 8 | 6.1 | 9.7 | 22.4 | 89.0 | 1.5 |

<table>
<tr><th></th><th></th><th></th><th></th><th></th><th></th><th></th><th></th><th></th><th></th><th></th><th></th><th></th><th></th><th></th><th></th><th></th></tr>
<tr><td>>MI0001445|hsa-mir-423&WithFlank&17|+|30117073|30117178@23.45.23&[MIMAT0004748&hsa-miR-423-5p&offsets|0|0;m-141&17|+|30117095|30117117&offsets|0|0]|TGAGGGGCAGAGAGCGAGACTTT</td><td>>MI0001445|hsa-mir-423&WithFlank&17|+|30117073|30117178</td><td>23</td><td>45</td><td>23</td><td>TGAGGGGCAGAGAGCGAGACTTT</td><td>BT20</td><td>234</td><td>174</td><td>619</td><td>5510</td><td>123</td><td>89.9</td><td>77.4</td><td>198.2</td><td>3782.3</td><td>61.2</td></tr>
<tr><td>>MI0001445|hsa-mir-423&WithFlank&17|+|30117073|30117178@23.45.23&[MIMAT0004748&hsa-miR-423-5p&offsets|0|0;m-141&17|+|30117095|30117117&offsets|0|0]|TGAGGGGCAGAGAGCGAGACTTT</td><td>>MI0001445|hsa-mir-423&WithFlank&17|+|30117073|30117178</td><td>23</td><td>45</td><td>23</td><td>TGAGGGGCAGAGAGCGAGACTTT</td><td>MB231</td><td>171</td><td>152</td><td>247</td><td>1466</td><td>56</td><td>69.0</td><td>55.4</td><td>104.6</td><td>459.9</td><td>17.7</td></tr>
<tr><td>>MI0001445|hsa-mir-423&WithFlank&17|+|30117073|30117178@23.46.24&[MIMAT0004748&hsa-miR-423-5p&offsets|0|+1;m-141&17|+|30117095|30117117&offsets|0|+1]|TGAGGGGCAGAGAGCGAGACTTTT</td><td>>MI0001445|hsa-mir-423&WithFlank&17|+|30117073|30117178</td><td>23</td><td>46</td><td>24</td><td>TGAGGGGCAGAGAGCGAGACTTTT</td><td>BT20</td><td>54</td><td>37</td><td>142</td><td>467</td><td>27</td><td>17.9</td><td>16.1</td><td>41.4</td><td>336.7</td><td>10.3</td></tr>
<tr><td>>MI0001445|hsa-mir-423&WithFlank&17|+|30117073|30117178@23.46.24&[MIMAT0004748&hsa-miR-423-5p&offsets|0|+1;m-141&17|+|30117095|30117117&offsets|0|+1]|TGAGGGGCAGAGAGCGAGACTTTT</td><td>>MI0001445|hsa-mir-423&WithFlank&17|+|30117073|30117178</td><td>23</td><td>46</td><td>24</td><td>TGAGGGGCAGAGAGCGAGACTTTT</td><td>MB231</td><td>45</td><td>35</td><td>56</td><td>232</td><td>16</td><td>11.5</td><td>12.7</td><td>24.9</td><td>65.4</td><td>5.9</td></tr>
<tr><td>>MI0001445|hsa-mir-423&WithFlank&17|+|30117073|30117178@58.78.21&[MIMAT0001340&hsa-miR-423-3p&offsets|-1|-3;m-100&17|+|30117131|30117153&offsets|-1|-3]|AAGCTCGGTCTGAGGCCCCTC</td><td>>MI0001445|hsa-mir-423&WithFlank&17|+|30117073|30117178</td><td>58</td><td>78</td><td>21</td><td>AAGCTCGGTCTGAGGCCCCTC</td><td>BT20</td><td>46</td><td>36</td><td>54</td><td>54</td><td>81</td><td>45.8</td><td>16.6</td><td>19.2</td><td>29.7</td><td>13.3</td></tr>
<tr><td>>MI0001445|hsa-mir-423&WithFlank&17|+|30117073|30117178@58.78.21&[MIMAT0001340&hsa-miR-423-3p&offsets|-1|-3;m-100&17|+|30117131|30117153&offsets|-1|-3]|AAGCTCGGTCTGAGGCCCCTC</td><td>>MI0001445|hsa-mir-423&WithFlank&17|+|30117073|30117178</td><td>58</td><td>78</td><td>21</td><td>AAGCTCGGTCTGAGGCCCCTC</td><td>MB231</td><td>38</td><td>35</td><td>21</td><td>20</td><td>35</td><td>31.9</td><td>12.6</td><td>9.2</td><td>9.5</td><td>6.5</td></tr>
<tr><td>>MI0001445|hsa-mir-423&WithFlank&17|+|30117073|30117178@58.79.22&[MIMAT0001340&hsa-miR-423-3p&offsets|-1|-2;m-100&17|+|30117131|30117153&offsets|-1|-2]|AAGCTCGGTCTGAGGCCCCTCA</td><td>>MI0001445|hsa-mir-423&WithFlank&17|+|30117073|30117178</td><td>58</td><td>79</td><td>22</td><td>AAGCTCGGTCTGAGGCCCCTCA</td><td>BT20</td><td>25</td><td>12</td><td>29</td><td>15</td><td>21</td><td>20.2</td><td>5.6</td><td>6.5</td><td>5.7</td><td>5.1</td></tr>
<tr><td>>MI0001445|hsa-mir-423&WithFlank&17|+|30117073|30117178@58.79.22&[MIMAT0001340&hsa-miR-423-3p&offsets|-1|-2;m-100&17|+|30117131|30117153&offsets|-1|-2]|AAGCTCGGTCTGAGGCCCCTCA</td><td>>MI0001445|hsa-mir-423&WithFlank&17|+|30117073|30117178</td><td>58</td><td>79</td><td>22</td><td>AAGCTCGGTCTGAGGCCCCTCA</td><td>MB231</td><td>13</td><td>8</td><td>5</td><td>6</td><td>9</td><td>9.5</td><td>2.5</td><td>1.9</td><td>1.3</td><td>1.3</td></tr>
<tr><td>>MI0001445|hsa-mir-423&WithFlank&17|+|30117073|30117178@58.80.23(+1C)&[MIMAT0001340&hsa-miR-423-3p&offsets|-1|-1(+1C);m-100&17|+|30117131|30117153&offsets|-1|-1(+1C)]|AAGCTCGGTCTGAGGCCCCTCAGC</td><td>>MI0001445|hsa-mir-423&WithFlank&17|+|30117073|30117178</td><td>58</td><td>80</td><td>23(+1C)</td><td>AAGCTCGGTCTGAGGCCCCTCAGC</td><td>BT20</td><td>21</td><td>19</td><td>25</td><td>19</td><td>34</td><td>6.8</td><td>12.3</td><td>5.0</td><td>13.8</td><td>14.1</td></tr>
<tr><td>>MI0001445|hsa-mir-423&WithFlank&17|+|30117073|30117178@58.80.23(+1C)&[MIMAT0001340&hsa-miR-423-3p&offsets|-1|-1(+1C);m-100&17|+|30117131|30117153&offsets|-1|-1(+1C)]|AAGCTCGGTCTGAGGCCCCTCAGC</td><td>>MI0001445|hsa-mir-423&WithFlank&17|+|30117073|30117178</td><td>58</td><td>80</td><td>23(+1C)</td><td>AAGCTCGGTCTGAGGCCCCTCAGC</td><td>MB231</td><td>16</td><td>9</td><td>7</td><td>11</td><td>27</td><td>7.0</td><td>3.0</td><td>4.4</td><td>7.1</td><td>20.4</td></tr>
<tr><td>>MI0001445|hsa-mir-423&WithFlank&17|+|30117073|30117178@58.80.23&[MIMAT0001340&hsa-miR-423-3p&offsets|-1|-1;m-100&17|+|30117131|30117153&offsets|-1|-1]|AAGCTCGGTCTGAGGCCCCTCAG</td><td>>MI0001445|hsa-mir-423&WithFlank&17|+|30117073|30117178</td><td>58</td><td>80</td><td>23</td><td>AAGCTCGGTCTGAGGCCCCTCAG</td><td>BT20</td><td>11</td><td>9</td><td>16</td><td>11</td><td>20</td><td>10.3</td><td>3.8</td><td>8.4</td><td>3.4</td><td>2.7</td></tr>
<tr><td>>MI0001445|hsa-mir-423&WithFlank&17|+|30117073|30117178@58.80.23&[MIMAT0001340&hsa-miR-423-3p&offsets|-1|-1;m-100&17|+|30117131|30117153&offsets|-1|-1]|AAGCTCGGTCTGAGGCCCCTCAG</td><td>>MI0001445|hsa-mir-423&WithFlank&17|+|30117073|30117178</td><td>58</td><td>80</td><td>23</td><td>AAGCTCGGTCTGAGGCCCCTCAG</td><td>MB231</td><td>14</td><td>11</td><td>6</td><td>8</td><td>13</td><td>7.1</td><td>5.1</td><td>3.1</td><td>4.4</td><td>2.8</td></tr>
<tr><td>>MI0001445|hsa-mir-423&WithFlank&17|+|30117073|30117178@58.81.24(+1U)&[MIMAT0001340&hsa-miR-423-3p&offsets|-1|0(+1U);m-100&17|+|30117131|30117153&offsets|-1|0(+1U)]|AAGCTCGGTCTGAGGCCCCTCAGTT</td><td>>MI0001445|hsa-mir-423&WithFlank&17|+|30117073|30117178</td><td>58</td><td>81</td><td>24(+1U)</td><td>AAGCTCGGTCTGAGGCCCCTCAGTT</td><td>BT20</td><td>14</td><td>8</td><td>13</td><td>3</td><td>20</td><td>8.0</td><td>4.8</td><td>5.3</td><td>0.7</td><td>7.1</td></tr>
<tr><td>>MI0001445|hsa-mir-423&WithFlank&17|+|30117073|30117178@58.81.24(+1U)&[MIMAT0001340&hsa-miR-423-3p&offsets|-1|0(+1U);m-100&17|+|30117131|30117153&offsets|-1|0(+1U)]|AAGCTCGGTCTGAGGCCCCTCAGTT</td><td>>MI0001445|hsa-mir-423&WithFlank&17|+|30117073|30117178</td><td>58</td><td>81</td><td>24(+1U)</td><td>AAGCTCGGTCTGAGGCCCCTCAGTT</td><td>MB231</td><td>19</td><td>11</td><td>6</td><td>3</td><td>19</td><td>7.7</td><td>7.2</td><td>3.0</td><td>1.7</td><td>8.9</td></tr>
<tr><td>>MI0001445|hsa-mir-423&WithFlank&17|+|30117073|30117178@58.81.24&[MIMAT0001340&hsa-miR-423-3p&offsets|-1|0;m-100&17|+|30117131|30117153&offsets|-1|0]|AAGCTCGGTCTGAGGCCCCTCAGT</td><td>>MI0001445|hsa-mir-423&WithFlank&17|+|30117073|30117178</td><td>58</td><td>81</td><td>24</td><td>AAGCTCGGTCTGAGGCCCCTCAGT</td><td>BT20</td><td>161</td><td>142</td><td>203</td><td>136</td><td>259</td><td>59.2</td><td>75.3</td><td>62.7</td><td>75.8</td><td>98.3</td></tr>
<tr><td>>MI0001445|hsa-mir-423&WithFlank&17|+|30117073|30117178@58.81.24&[MIMAT0001340&hsa-miR-423-3p&offsets|-1|0;m-100&17|+|30117131|30117153&offsets|-1|0]|AAGCTCGGTCTGAGGCCCCTCAGT</td><td>>MI0001445|hsa-mir-423&WithFlank&17|+|30117073|30117178</td><td>58</td><td>81</td><td>24</td><td>AAGCTCGGTCTGAGGCCCCTCAGT</td><td>MB231</td><td>152</td><td>91</td><td>65</td><td>76</td><td>199</td><td>60.6</td><td>36.7</td><td>43.2</td><td>37.7</td><td>120.4</td></tr>
<tr><td>>MI0001445|hsa-mir-423&WithFlank&17|+|30117073|30117178@59.77.19(+1A)&[MIMAT0001340&hsa-miR-423-3p&offsets|0|-4(+1A);m-100&17|+|30117131|30117153&offsets|0|-4(+1A)]|AGCTCGGTCTGAGGCCCCTA</td><td>>MI0001445|hsa-mir-423&WithFlank&17|+|30117073|30117178</td><td>59</td><td>77</td><td>19(+1A)</td><td>AGCTCGGTCTGAGGCCCCTA</td><td>BT20</td><td>10</td><td>8</td><td>15</td><td>9</td><td>16</td><td>8.1</td><td>5.1</td><td>2.1</td><td>4.6</td><td>5.4</td></tr>
<tr><td>>MI0001445|hsa-mir-423&WithFlank&17|+|30117073|30117178@59.77.19(+1U)&[MIMAT0001340&hsa-miR-423-3p&offsets|0|-4(+1U);m-100&17|+|30117131|30117153&offsets|0|-4(+1U)]|AGCTCGGTCTGAGGCCCCTT</td><td>>MI0001445|hsa-mir-423&WithFlank&17|+|30117073|30117178</td><td>59</td><td>77</td><td>19(+1U)</td><td>AGCTCGGTCTGAGGCCCCTT</td><td>BT20</td><td>28</td><td>25</td><td>32</td><td>28</td><td>46</td><td>26.2</td><td>15.3</td><td>10.1</td><td>12.2</td><td>14.1</td></tr>
<tr><td>>MI0001445|hsa-mir-423&WithFlank&17|+|30117073|30117178@59.77.19(+1U)&[MIMAT0001340&hsa-miR-423-3p&offsets|0|-4(+1U);m-100&17|+|30117131|30117153&offsets|0|-4(+1U)]|AGCTCGGTCTGAGGCCCCTT</td><td>>MI0001445|hsa-mir-423&WithFlank&17|+|30117073|30117178</td><td>59</td><td>77</td><td>19(+1U)</td><td>AGCTCGGTCTGAGGCCCCTT</td><td>MB231</td><td>20</td><td>16</td><td>9</td><td>21</td><td>20</td><td>14.7</td><td>5.7</td><td>2.1</td><td>13.3</td><td>4.4</td></tr>
<tr><td>>MI0001445|hsa-mir-423&WithFlank&17|+|30117073|30117178@59.78.20&[MIMAT0001340&hsa-miR-423-3p&offsets|0|-3;m-100&17|+|30117131|30117153&offsets|0|-3]|AGCTCGGTCTGAGGCCCCTC</td><td>>MI0001445|hsa-mir-423&WithFlank&17|+|30117073|30117178</td><td>59</td><td>78</td><td>20</td><td>AGCTCGGTCTGAGGCCCCTC</td><td>BT20</td><td>169</td><td>150</td><td>211</td><td>181</td><td>345</td><td>189.0</td><td>68.6</td><td>114.0</td><td>100.1</td><td>74.0</td></tr>
<tr><td>>MI0001445|hsa-mir-423&WithFlank&17|+|30117073|30117178@59.78.20&[MIMAT0001340&hsa-miR-423-3p&offsets|0|-3;m-100&17|+|30117131|30117153&offsets|0|-3]|AGCTCGGTCTGAGGCCCCTC</td><td>>MI0001445|hsa-mir-423&WithFlank&17|+|30117073|30117178</td><td>59</td><td>78</td><td>20</td><td>AGCTCGGTCTGAGGCCCCTC</td><td>MB231</td><td>170</td><td>178</td><td>93</td><td>99</td><td>173</td><td>137.6</td><td>78.5</td><td>26.8</td><td>52.7</td><td>42.9</td></tr>
<tr><td>>MI0001445|hsa-mir-423&WithFlank&17|+|30117073|30117178@59.79.21&[MIMAT0001340&hsa-miR-423-3p&offsets|0|-2;m-100&17|+|30117131|30117153&offsets|0|-2]|AGCTCGGTCTGAGGCCCCTCA</td><td>>MI0001445|hsa-mir-423&WithFlank&17|+|30117073|30117178</td><td>59</td><td>79</td><td>21</td><td>AGCTCGGTCTGAGGCCCCTCA</td><td>BT20</td><td>48</td><td>25</td><td>62</td><td>38</td><td>48</td><td>53.1</td><td>12.1</td><td>19.1</td><td>17.7</td><td>10.6</td></tr>
<tr><td>>MI0001445|hsa-mir-423&WithFlank&17|+|30117073|30117178@59.79.21&[MIMAT0001340&hsa-miR-423-3p&offsets|0|-2;m-100&17|+|30117131|30117153&offsets|0|-2]|AGCTCGGTCTGAGGCCCCTCA</td><td>>MI0001445|hsa-mir-423&WithFlank&17|+|30117073|30117178</td><td>59</td><td>79</td><td>21</td><td>AGCTCGGTCTGAGGCCCCTCA</td><td>MB231</td><td>40</td><td>29</td><td>19</td><td>19</td><td>31</td><td>31.5</td><td>11.2</td><td>10.2</td><td>9.9</td><td>1.8</td></tr>
<tr><td>>MI0001445|hsa-mir-423&WithFlank&17|+|30117073|30117178@59.80.22(+1A)&[MIMAT0001340&hsa-miR-423-3p&offsets|0|-1(+1A);m-100&17|+|30117131|30117153&offsets|0|-1(+1A)]|AGCTCGGTCTGAGGCCCCTCAGA</td><td>>MI0001445|hsa-mir-423&WithFlank&17|+|30117073|30117178</td><td>59</td><td>80</td><td>22(+1A)</td><td>AGCTCGGTCTGAGGCCCCTCAGA</td><td>BT20</td><td>47</td><td>38</td><td>70</td><td>51</td><td>62</td><td>14.5</td><td>20.1</td><td>11.3</td><td>27.3</td><td>17.2</td></tr>
<tr><td>>MI0001445|hsa-mir-423&WithFlank&17|+|30117073|30117178@59.80.22(+1A)&[MIMAT0001340&hsa-miR-423-3p&offsets|0|-1(+1A);m-100&17|+|30117131|30117153&offsets|0|-1(+1A)]|AGCTCGGTCTGAGGCCCCTCAGA</td><td>>MI0001445|hsa-mir-423&WithFlank&17|+|30117073|30117178</td><td>59</td><td>80</td><td>22(+1A)</td><td>AGCTCGGTCTGAGGCCCCTCAGA</td><td>MB231</td><td>34</td><td>22</td><td>16</td><td>31</td><td>41</td><td>15.1</td><td>8.9</td><td>10.8</td><td>18.3</td><td>20.9</td></tr>
<tr><td>>MI0001445|hsa-mir-423&WithFlank&17|+|30117073|30117178@59.80.22(+1C)&[MIMAT0001340&hsa-miR-423-3p&offsets|0|-1(+1C);m-100&17|+|30117131|30117153&offsets|0|-1(+1C)]|AGCTCGGTCTGAGGCCCCTCAGC</td><td>>MI0001445|hsa-mir-423&WithFlank&17|+|30117073|30117178</td><td>59</td><td>80</td><td>22(+1C)</td><td>AGCTCGGTCTGAGGCCCCTCAGC</td><td>BT20</td><td>126</td><td>139</td><td>165</td><td>113</td><td>207</td><td>39.5</td><td>87.8</td><td>33.9</td><td>81.9</td><td>88.5</td></tr>
</table>

<table>
<tr><td>>MI0001445|hsa-mir-423&WithFlank&17|+|30117073|30117178@59.80.22(+1C)&[MIMAT0001340&hsa-miR-423-3p&offsets|0|-1(+1C);m-100&17|+|30117131|30117153&offsets|0|-1(+1C)]|AGCTCGGTCTGAGGCCCCTCAGC</td><td>>MI0001445|hsa-mir-423&WithFlank&17|+|30117073|30117178</td><td>59</td><td>80</td><td>22(+1C)</td><td>AGCTCGGTCTGAGGCCCCTCAGC</td><td>MB231</td><td>118</td><td>73</td><td>53</td><td>99</td><td>175</td><td>47.7</td><td>33.5</td><td>38.8</td><td>61.2</td><td>127.3</td></tr>
<tr><td>>MI0001445|hsa-mir-423&WithFlank&17|+|30117073|30117178@59.80.22(+1G)&[MIMAT0001340&hsa-miR-423-3p&offsets|0|-1(+1G);m-100&17|+|30117131|30117153&offsets|0|-1(+1G)]|AGCTCGGTCTGAGGCCCCTCAGG</td><td>>MI0001445|hsa-mir-423&WithFlank&17|+|30117073|30117178</td><td>59</td><td>80</td><td>22(+1G)</td><td>AGCTCGGTCTGAGGCCCCTCAGG</td><td>BT20</td><td>58</td><td>23</td><td>31</td><td>14</td><td>14</td><td>32.7</td><td>12.5</td><td>7.3</td><td>15.7</td><td>6.0</td></tr>
<tr><td>>MI0001445|hsa-mir-423&WithFlank&17|+|30117073|30117178@59.80.22(+1G)&[MIMAT0001340&hsa-miR-423-3p&offsets|0|-1(+1G);m-100&17|+|30117131|30117153&offsets|0|-1(+1G)]|AGCTCGGTCTGAGGCCCCTCAGG</td><td>>MI0001445|hsa-mir-423&WithFlank&17|+|30117073|30117178</td><td>59</td><td>80</td><td>22(+1G)</td><td>AGCTCGGTCTGAGGCCCCTCAGG</td><td>MB231</td><td>42</td><td>11</td><td>9</td><td>10</td><td>16</td><td>22.3</td><td>6.2</td><td>7.4</td><td>5.9</td><td>13.2</td></tr>
<tr><td>>MI0001445|hsa-mir-423&WithFlank&17|+|30117073|30117178@59.80.22&[MIMAT0001340&hsa-miR-423-3p&offsets|0|-1;m-100&17|+|30117131|30117153&offsets|0|-1]|AGCTCGGTCTGAGGCCCCTCAG</td><td>>MI0001445|hsa-mir-423&WithFlank&17|+|30117073|30117178</td><td>59</td><td>80</td><td>22</td><td>AGCTCGGTCTGAGGCCCCTCAG</td><td>BT20</td><td>53</td><td>41</td><td>83</td><td>62</td><td>84</td><td>56.5</td><td>19.6</td><td>53.7</td><td>29.6</td><td>19.4</td></tr>
<tr><td>>MI0001445|hsa-mir-423&WithFlank&17|+|30117073|30117178@59.80.22&[MIMAT0001340&hsa-miR-423-3p&offsets|0|-1;m-100&17|+|30117131|30117153&offsets|0|-1]|AGCTCGGTCTGAGGCCCCTCAG</td><td>>MI0001445|hsa-mir-423&WithFlank&17|+|30117073|30117178</td><td>59</td><td>80</td><td>22</td><td>AGCTCGGTCTGAGGCCCCTCAG</td><td>MB231</td><td>76</td><td>57</td><td>41</td><td>51</td><td>72</td><td>32.7</td><td>29.6</td><td>27.4</td><td>30.7</td><td>18.3</td></tr>
<tr><td>>MI0001445|hsa-mir-423&WithFlank&17|+|30117073|30117178@59.81.23(+1A)&[MIMAT0001340&hsa-miR-423-3p&offsets|0|0(+1A);m-100&17|+|30117131|30117153&offsets|0|0(+1A)]|AGCTCGGTCTGAGGCCCCTCAGTA</td><td>>MI0001445|hsa-mir-423&WithFlank&17|+|30117073|30117178</td><td>59</td><td>81</td><td>23(+1A)</td><td>AGCTCGGTCTGAGGCCCCTCAGTA</td><td>BT20</td><td>66</td><td>47</td><td>71</td><td>9</td><td>95</td><td>54.0</td><td>27.4</td><td>35.5</td><td>1.7</td><td>38.1</td></tr>
<tr><td>>MI0001445|hsa-mir-423&WithFlank&17|+|30117073|30117178@59.81.23(+1A)&[MIMAT0001340&hsa-miR-423-3p&offsets|0|0(+1A);m-100&17|+|30117131|30117153&offsets|0|0(+1A)]|AGCTCGGTCTGAGGCCCCTCAGTA</td><td>>MI0001445|hsa-mir-423&WithFlank&17|+|30117073|30117178</td><td>59</td><td>81</td><td>23(+1A)</td><td>AGCTCGGTCTGAGGCCCCTCAGTA</td><td>MB231</td><td>44</td><td>33</td><td>15</td><td>6</td><td>41</td><td>22.6</td><td>19.2</td><td>9.0</td><td>3.4</td><td>13.4</td></tr>
<tr><td>>MI0001445|hsa-mir-423&WithFlank&17|+|30117073|30117178@59.81.23(+1U)&[MIMAT0001340&hsa-miR-423-3p&offsets|0|0(+1U);m-100&17|+|30117131|30117153&offsets|0|0(+1U)]|AGCTCGGTCTGAGGCCCCTCAGTT</td><td>>MI0001445|hsa-mir-423&WithFlank&17|+|30117073|30117178</td><td>59</td><td>81</td><td>23(+1U)</td><td>AGCTCGGTCTGAGGCCCCTCAGTT</td><td>BT20</td><td>83</td><td>61</td><td>89</td><td>22</td><td>121</td><td>43.9</td><td>33.6</td><td>33.4</td><td>6.8</td><td>44.0</td></tr>
<tr><td>>MI0001445|hsa-mir-423&WithFlank&17|+|30117073|30117178@59.81.23(+1U)&[MIMAT0001340&hsa-miR-423-3p&offsets|0|0(+1U);m-100&17|+|30117131|30117153&offsets|0|0(+1U)]|AGCTCGGTCTGAGGCCCCTCAGTT</td><td>>MI0001445|hsa-mir-423&WithFlank&17|+|30117073|30117178</td><td>59</td><td>81</td><td>23(+1U)</td><td>AGCTCGGTCTGAGGCCCCTCAGTT</td><td>MB231</td><td>114</td><td>68</td><td>39</td><td>24</td><td>112</td><td>41.0</td><td>35.8</td><td>22.9</td><td>11.5</td><td>61.1</td></tr>
<tr><td>>MI0001445|hsa-mir-423&WithFlank&17|+|30117073|30117178@59.81.23(+2U)&[MIMAT0001340&hsa-miR-423-3p&offsets|0|0(+2U);m-100&17|+|30117131|30117153&offsets|0|0(+2U)]|AGCTCGGTCTGAGGCCCCTCAGTTT</td><td>>MI0001445|hsa-mir-423&WithFlank&17|+|30117073|30117178</td><td>59</td><td>81</td><td>23(+2U)</td><td>AGCTCGGTCTGAGGCCCCTCAGTTT</td><td>MB231</td><td>11</td><td>8</td><td>4</td><td>3</td><td>12</td><td>5.2</td><td>3.8</td><td>1.7</td><td>0.9</td><td>4.9</td></tr>
<tr><td>>MI0001445|hsa-mir-423&WithFlank&17|+|30117073|30117178@59.81.23&[MIMAT0001340&hsa-miR-423-3p&offsets|0|0;m-100&17|+|30117131|30117153&offsets|0|0]|AGCTCGGTCTGAGGCCCCTCAGT</td><td>>MI0001445|hsa-mir-423&WithFlank&17|+|30117073|30117178</td><td>59</td><td>81</td><td>23</td><td>AGCTCGGTCTGAGGCCCCTCAGT</td><td>BT20</td><td>984</td><td>968</td><td>1335</td><td>838</td><td>1571</td><td>375.4</td><td>539.2</td><td>410.3</td><td>494.3</td><td>583.2</td></tr>
<tr><td>>MI0001445|hsa-mir-423&WithFlank&17|+|30117073|30117178@59.81.23&[MIMAT0001340&hsa-miR-423-3p&offsets|0|0;m-100&17|+|30117131|30117153&offsets|0|0]|AGCTCGGTCTGAGGCCCCTCAGT</td><td>>MI0001445|hsa-mir-423&WithFlank&17|+|30117073|30117178</td><td>59</td><td>81</td><td>23</td><td>AGCTCGGTCTGAGGCCCCTCAGT</td><td>MB231</td><td>1092</td><td>681</td><td>487</td><td>635</td><td>1298</td><td>427.5</td><td>327.8</td><td>354.1</td><td>335.7</td><td>750.9</td></tr>
<tr><td>>MI0001445|hsa-mir-423&WithFlank&17|+|30117073|30117178@59.81.23&[MIMAT0001340&hsa-miR-423-3p&offsets|0|0;m-100&17|+|30117131|30117153&offsets|0|0]|AGCTCGGTCTGAGGCCCCTCAGT</td><td>>MI0001445|hsa-mir-423&WithFlank&17|+|30117073|30117178</td><td>59</td><td>81</td><td>23</td><td>AGCTCGGTCTGAGGCCCCTCAGT</td><td>MB468</td><td>32</td><td>27</td><td>13</td><td>26</td><td>22</td><td>10.1</td><td>15.7</td><td>3.7</td><td>6.6</td><td>6.0</td></tr>
<tr><td>>MI0001445|hsa-mir-423&WithFlank&17|+|30117073|30117178@59.82.24&[MIMAT0001340&hsa-miR-423-3p&offsets|0|+1;m-100&17|+|30117131|30117153&offsets|0|+1]|AGCTCGGTCTGAGGCCCCTCAGTC</td><td>>MI0001445|hsa-mir-423&WithFlank&17|+|30117073|30117178</td><td>59</td><td>82</td><td>24</td><td>AGCTCGGTCTGAGGCCCCTCAGTC</td><td>BT20</td><td>12</td><td>9</td><td>12</td><td>4</td><td>17</td><td>5.5</td><td>6.0</td><td>3.7</td><td>1.1</td><td>7.1</td></tr>
<tr><td>>MI0001445|hsa-mir-423&WithFlank&17|+|30117073|30117178@59.82.24&[MIMAT0001340&hsa-miR-423-3p&offsets|0|+1;m-100&17|+|30117131|30117153&offsets|0|+1]|AGCTCGGTCTGAGGCCCCTCAGTC</td><td>>MI0001445|hsa-mir-423&WithFlank&17|+|30117073|30117178</td><td>59</td><td>82</td><td>24</td><td>AGCTCGGTCTGAGGCCCCTCAGTC</td><td>MB231</td><td>13</td><td>8</td><td>5</td><td>5</td><td>17</td><td>4.9</td><td>3.9</td><td>3.1</td><td>3.1</td><td>11.4</td></tr>
<tr><td>>MI0001446|hsa-mir-424&WithFlank&X|-|134546608|134546717@17.37.21&[MIMAT0001341&hsa-miR-424-5p&offsets|0|-1;m-152&X|-|134546681|134546701&offsets|0|0]|CAGCAGCAATTCATGTTTTGA</td><td>>MI0001446|hsa-mir-424&WithFlank&X|-|134546608|134546717</td><td>17</td><td>37</td><td>21</td><td>CAGCAGCAATTCATGTTTTGA</td><td>BT20</td><td>59</td><td>22</td><td>2</td><td>18</td><td>2</td><td>83.9</td><td>28.2</td><td>2.3</td><td>18.6</td><td>2.3</td></tr>
<tr><td>>MI0001446|hsa-mir-424&WithFlank&X|-|134546608|134546717@17.38.22(+1A)&[MIMAT0001341&hsa-miR-424-5p&offsets|0|0(+1A);m-152&X|-|134546681|134546701&offsets|0|+1(+1A)]|CAGCAGCAATTCATGTTTTGAAA</td><td>>MI0001446|hsa-mir-424&WithFlank&X|-|134546608|134546717</td><td>17</td><td>38</td><td>22(+1A)</td><td>CAGCAGCAATTCATGTTTTGAAA</td><td>BT20</td><td>16</td><td>5</td><td>1</td><td>0</td><td>0</td><td>23.5</td><td>5.8</td><td>0.5</td><td>0.0</td><td>0.0</td></tr>
<tr><td>>MI0001446|hsa-mir-424&WithFlank&X|-|134546608|134546717@17.38.22&[MIMAT0001341&hsa-miR-424-5p&offsets|0|0;m-152&X|-|134546681|134546701&offsets|0|+1]|CAGCAGCAATTCATGTTTTGAA</td><td>>MI0001446|hsa-mir-424&WithFlank&X|-|134546608|134546717</td><td>17</td><td>38</td><td>22</td><td>CAGCAGCAATTCATGTTTTGAA</td><td>BT20</td><td>26</td><td>12</td><td>1</td><td>2</td><td>1</td><td>36.1</td><td>15.4</td><td>1.1</td><td>2.6</td><td>0.7</td></tr>
<tr><td>>MI0001446|hsa-mir-424&WithFlank&X|-|134546608|134546717@54.74.21&[MIMAT0004749&hsa-miR-424-3p&offsets|0|0;m-305&X|-|134546644|134546664&offsets|0|0]|CAAAACGTGAGGCGCTGCTAT</td><td>>MI0001446|hsa-mir-424&WithFlank&X|-|134546608|134546717</td><td>54</td><td>74</td><td>21</td><td>CAAAACGTGAGGCGCTGCTAT</td><td>BT20</td><td>48</td><td>26</td><td>63</td><td>29</td><td>24</td><td>41.4</td><td>21.3</td><td>37.2</td><td>19.8</td><td>20.8</td></tr>
<tr><td>>MI0001446|hsa-mir-424&WithFlank&X|-|134546608|134546717@54.74.21&[MIMAT0004749&hsa-miR-424-3p&offsets|0|0;m-305&X|-|134546644|134546664&offsets|0|0]|CAAAACGTGAGGCGCTGCTAT</td><td>>MI0001446|hsa-mir-424&WithFlank&X|-|134546608|134546717</td><td>54</td><td>74</td><td>21</td><td>CAAAACGTGAGGCGCTGCTAT</td><td>MB231</td><td>27</td><td>21</td><td>14</td><td>14</td><td>12</td><td>7.5</td><td>10.3</td><td>10.0</td><td>10.3</td><td>6.0</td></tr>
<tr><td>>MI0001446|hsa-mir-424&WithFlank&X|-|134546608|134546717@54.75.22&[MIMAT0004749&hsa-miR-424-3p&offsets|0|+1;m-305&X|-|134546644|134546664&offsets|0|+1]|CAAAACGTGAGGCGCTGCTATA</td><td>>MI0001446|hsa-mir-424&WithFlank&X|-|134546608|134546717</td><td>54</td><td>75</td><td>22</td><td>CAAAACGTGAGGCGCTGCTATA</td><td>BT20</td><td>18</td><td>8</td><td>27</td><td>3</td><td>6</td><td>12.1</td><td>6.2</td><td>10.4</td><td>1.4</td><td>4.7</td></tr>
<tr><td>>MI0001446|hsa-mir-424&WithFlank&X|-|134546608|134546717@54.75.22&[MIMAT0004749&hsa-miR-424-3p&offsets|0|+1;m-305&X|-|134546644|134546664&offsets|0|+1]|CAAAACGTGAGGCGCTGCTATA</td><td>>MI0001446|hsa-mir-424&WithFlank&X|-|134546608|134546717</td><td>54</td><td>75</td><td>22</td><td>CAAAACGTGAGGCGCTGCTATA</td><td>MB231</td><td>14</td><td>10</td><td>7</td><td>4</td><td>6</td><td>4.2</td><td>5.1</td><td>5.3</td><td>2.2</td><td>3.2</td></tr>
<tr><td>>MI0001448|hsa-mir-425&WithFlank&3|-|49020142|49020240@20.39.20&[MIMAT0003393&hsa-miR-425-5p&offsets|0|-3;m-95&3|-|49020200|49020221&offsets|0|-2]|AATGACACGATCACTCCCGT</td><td>>MI0001448|hsa-mir-425&WithFlank&3|-|49020142|49020240</td><td>20</td><td>39</td><td>20</td><td>AATGACACGATCACTCCCGT</td><td>BT20</td><td>14</td><td>15</td><td>18</td><td>9</td><td>8</td><td>20.5</td><td>13.7</td><td>17.3</td><td>11.1</td><td>6.6</td></tr>
<tr><td>>MI0001448|hsa-mir-425&WithFlank&3|-|49020142|49020240@20.41.22&[MIMAT0003393&hsa-miR-425-5p&offsets|0|-1;m-95&3|-|49020200|49020221&offsets|0|0]|AATGACACGATCACTCCCGTTG</td><td>>MI0001448|hsa-mir-425&WithFlank&3|-|49020142|49020240</td><td>20</td><td>41</td><td>22</td><td>AATGACACGATCACTCCCGTTG</td><td>BT20</td><td>15</td><td>12</td><td>27</td><td>21</td><td>3</td><td>9.0</td><td>7.1</td><td>7.7</td><td>17.0</td><td>1.6</td></tr>
<tr><td>>MI0001448|hsa-mir-425&WithFlank&3|-|49020142|49020240@20.42.23&[MIMAT0003393&hsa-miR-425-5p&offsets|0|0;m-95&3|-|49020200|49020221&offsets|0|+1]|AATGACACGATCACTCCCGTTGA</td><td>>MI0001448|hsa-mir-425&WithFlank&3|-|49020142|49020240</td><td>20</td><td>42</td><td>23</td><td>AATGACACGATCACTCCCGTTGA</td><td>BT20</td><td>16</td><td>15</td><td>28</td><td>10</td><td>4</td><td>9.0</td><td>12.3</td><td>5.3</td><td>8.9</td><td>3.3</td></tr>
<tr><td>>MI0001448|hsa-mir-425&WithFlank&3|-|49020142|49020240@20.43.24(+1A)&[MIMAT0003393&hsa-miR-425-5p&offsets|0|+1(+1A);m-95&3|-|49020200|49020221&offsets|0|+2(+1A)]|AATGACACGATCACTCCCGTTGAGA</td><td>>MI0001448|hsa-mir-425&WithFlank&3|-|49020142|49020240</td><td>20</td><td>43</td><td>24(+1A)</td><td>AATGACACGATCACTCCCGTTGAGA</td><td>BT20</td><td>15</td><td>7</td><td>19</td><td>1</td><td>2</td><td>14.9</td><td>3.9</td><td>15.6</td><td>0.5</td><td>2.0</td></tr>
<tr><td>>MI0001448|hsa-mir-425&WithFlank&3|-|49020142|49020240@20.43.24(+1C)&[MIMAT0003393&hsa-miR-425-5p&offsets|0|+1(+1C);m-95&3|-|49020200|49020221&offsets|0|+2(+1C)]|AATGACACGATCACTCCCGTTGAGC</td><td>>MI0001448|hsa-mir-425&WithFlank&3|-|49020142|49020240</td><td>20</td><td>43</td><td>24(+1C)</td><td>AATGACACGATCACTCCCGTTGAGC</td><td>BT20</td><td>32</td><td>25</td><td>46</td><td>3</td><td>7</td><td>22.3</td><td>20.6</td><td>32.2</td><td>1.9</td><td>5.9</td></tr>
<tr><td>>MI0001448|hsa-mir-425&WithFlank&3|-|49020142|49020240@20.43.24(+1C)&[MIMAT0003393&hsa-miR-425-5p&offsets|0|+1(+1C);m-95&3|-|49020200|49020221&offsets|0|+2(+1C)]|AATGACACGATCACTCCCGTTGAGC</td><td>>MI0001448|hsa-mir-425&WithFlank&3|-|49020142|49020240</td><td>20</td><td>43</td><td>24(+1C)</td><td>AATGACACGATCACTCCCGTTGAGC</td><td>MB231</td><td>32</td><td>29</td><td>12</td><td>19</td><td>42</td><td>8.3</td><td>9.1</td><td>7.8</td><td>6.0</td><td>36.7</td></tr>
<tr><td>>MI0001448|hsa-mir-425&WithFlank&3|-|49020142|49020240@20.43.24(+1G)&[MIMAT0003393&hsa-miR-425-5p&offsets|0|+1(+1G);m-95&3|-|49020200|49020221&offsets|0|+2(+1G)]|AATGACACGATCACTCCCGTTGAGG</td><td>>MI0001448|hsa-mir-425&WithFlank&3|-|49020142|49020240</td><td>20</td><td>43</td><td>24(+1G)</td><td>AATGACACGATCACTCCCGTTGAGG</td><td>BT20</td><td>20</td><td>5</td><td>11</td><td>0</td><td>0</td><td>24.6</td><td>3.9</td><td>9.3</td><td>0.0</td><td>0.0</td></tr>
</table>

| | | | | | | | | | | | | | | | | |
|---|---|---|---|---|---|---|---|---|---|---|---|---|---|---|---|---|
| >MI0001448\|hsa-mir-425&WithFlank&3\|-\|49020142\|49020240@20.43.24(+1G)&[MIMAT0003393&hsa-miR-425-5p&offsets\|0\|+1(+1G);m-95&3\|-\|49020200\|49020221&offsets\|0\|+2(+1G)]\|AATGACACGATCACTCCCGTTGAGG | >MI0001448\|hsa-mir-425&WithFlank&3\|-\|49020142\|49020240 | 20 | 43 | 24(+1G) | AATGACACGATCACTCCCGTTGAGG | MB231 | 14 | 5 | 2 | 2 | 4 | 9.9 | 2.2 | 1.8 | 0.4 | 3.1 |
| >MI0001448\|hsa-mir-425&WithFlank&3\|-\|49020142\|49020240@20.43.24&[MIMAT0003393&hsa-miR-425-5p&offsets\|0\|+1;m-95&3\|-\|49020200\|49020221&offsets\|0\|+2]\|AATGACACGATCACTCCCGTTGAG | >MI0001448\|hsa-mir-425&WithFlank&3\|-\|49020142\|49020240 | 20 | 43 | 24 | AATGACACGATCACTCCCGTTGAG | BT20 | 44 | 54 | 87 | 10 | 24 | 23.6 | 29.0 | 17.0 | 6.9 | 18.8 |
| >MI0001448\|hsa-mir-425&WithFlank&3\|-\|49020142\|49020240@20.43.24&[MIMAT0003393&hsa-miR-425-5p&offsets\|0\|+1;m-95&3\|-\|49020200\|49020221&offsets\|0\|+2]\|AATGACACGATCACTCCCGTTGAG | >MI0001448\|hsa-mir-425&WithFlank&3\|-\|49020142\|49020240 | 20 | 43 | 24 | AATGACACGATCACTCCCGTTGAG | MB231 | 69 | 89 | 41 | 53 | 80 | 17.9 | 30.7 | 22.0 | 16.9 | 38.0 |
| >MI0001448\|hsa-mir-425&WithFlank&3\|-\|49020142\|49020240@20.44.25&[MIMAT0003393&hsa-miR-425-5p&offsets\|0\|+2;m-95&3\|-\|49020200\|49020221&offsets\|0\|+3]\|AATGACACGATCACTCCCGTTGAGT | >MI0001448\|hsa-mir-425&WithFlank&3\|-\|49020142\|49020240 | 20 | 44 | 25 | AATGACACGATCACTCCCGTTGAGT | BT20 | 245 | 174 | 348 | 27 | 51 | 176.0 | 125.8 | 196.0 | 14.7 | 39.2 |
| >MI0001448\|hsa-mir-425&WithFlank&3\|-\|49020142\|49020240@20.44.25&[MIMAT0003393&hsa-miR-425-5p&offsets\|0\|+2;m-95&3\|-\|49020200\|49020221&offsets\|0\|+3]\|AATGACACGATCACTCCCGTTGAGT | >MI0001448\|hsa-mir-425&WithFlank&3\|-\|49020142\|49020240 | 20 | 44 | 25 | AATGACACGATCACTCCCGTTGAGT | MB231 | 299 | 285 | 117 | 142 | 321 | 79.5 | 95.8 | 74.8 | 68.0 | 233.5 |
| >MI0001448\|hsa-mir-425&WithFlank&3\|-\|49020142\|49020240@21.44.24&[MIMAT0003393&hsa-miR-425-5p&offsets\|+1\|+2;m-95&3\|-\|49020200\|49020221&offsets\|+1\|+3]\|ATGACACGATCACTCCCGTTGAGT | >MI0001448\|hsa-mir-425&WithFlank&3\|-\|49020142\|49020240 | 21 | 44 | 24 | ATGACACGATCACTCCCGTTGAGT | BT20 | 27 | 16 | 39 | 5 | 4 | 26.0 | 10.2 | 25.4 | 3.5 | 2.9 |
| >MI0001448\|hsa-mir-425&WithFlank&3\|-\|49020142\|49020240@21.44.24&[MIMAT0003393&hsa-miR-425-5p&offsets\|+1\|+2;m-95&3\|-\|49020200\|49020221&offsets\|+1\|+3]\|ATGACACGATCACTCCCGTTGAGT | >MI0001448\|hsa-mir-425&WithFlank&3\|-\|49020142\|49020240 | 21 | 44 | 24 | ATGACACGATCACTCCCGTTGAGT | MB231 | 14 | 14 | 6 | 16 | 15 | 3.0 | 5.1 | 4.2 | 6.2 | 10.7 |
| >MI0001519\|hsa-mir-20b&WithFlank&X\|-\|134169803\|134169883@12.34.23&[MIMAT0001413&hsa-miR-20b-5p&offsets\|0\|0;m-241&X\|-\|134169850\|134169872&offsets\|0\|0]\|CAAAGTGCTCATAGTGCAGGTAG | >MI0001519\|hsa-mir-20b&WithFlank&X\|-\|134169803\|134169883 | 12 | 34 | 23 | CAAAGTGCTCATAGTGCAGGTAG | BT20 | 14 | 10 | 6 | 14 | 3 | 8.9 | 8.9 | 6.1 | 12.9 | 2.1 |
| >MI0001641\|hsa-mir-429&WithFlank&1\|+\|1168999\|1169093@57.77.21(+1C)&[MIMAT0001536&hsa-miR-429&offsets\|0\|-1(+1C);m-207&1\|+\|1169055\|1169076&offsets\|0\|-1(+1C)]\|TAATACTGTCTGGTAAAACCGC | >MI0001641\|hsa-mir-429&WithFlank&1\|+\|1168999\|1169093 | 57 | 77 | 21(+1C) | TAATACTGTCTGGTAAAACCGC | MB468 | 12 | 2 | 3 | 16 | 3 | 6.1 | 0.5 | 1.9 | 1.2 | 0.3 |
| >MI0001641\|hsa-mir-429&WithFlank&1\|+\|1168999\|1169093@57.78.22&[MIMAT0001536&hsa-miR-429&offsets\|0\|0;m-207&1\|+\|1169055\|1169076&offsets\|0\|0]\|TAATACTGTCTGGTAAAACCGT | >MI0001641\|hsa-mir-429&WithFlank&1\|+\|1168999\|1169093 | 57 | 78 | 22 | TAATACTGTCTGGTAAAACCGT | MB468 | 105 | 17 | 24 | 125 | 24 | 62.4 | 7.0 | 11.1 | 5.8 | 4.0 |
| >MI0001652\|hsa-mir-450a-1&WithFlank&X\|-\|134540335\|134540437@24.43.20;MI0003187\|hsa-mir-450a-2&WithFlank&X\|-\|134540502\|134540613@29.48.20&[MIMAT0001545&hsa-miR-450a-5p&offsets\|0\|-2;m-288&X\|-\|134540393\|134540414&offsets\|0\|-2];[MIMAT0001545_1&hsa-miR-450a-5p&offsets\|0\|-2;m-287&X\|-\|134540564\|134540585&offsets\|0\|-2]\|TTTTGCGATGTGTTCCTAAT | >MI0001652\|hsa-mir-450a-1&WithFlank&X\|-\|134540335\|134540437 | 24 | 43 | 20 | TTTTGCGATGTGTTCCTAAT | BT20 | 19 | 10 | 11 | 1 | 4 | 13.2 | 6.1 | 6.5 | 0.8 | 2.2 |
| >MI0001652\|hsa-mir-450a-1&WithFlank&X\|-\|134540335\|134540437@24.45.22;MI0003187\|hsa-mir-450a-2&WithFlank&X\|-\|134540502\|134540613@29.50.22&[MIMAT0001545&hsa-miR-450a-5p&offsets\|0\|0;m-288&X\|-\|134540393\|134540414&offsets\|0\|0];[MIMAT0001545_1&hsa-miR-450a-5p&offsets\|0\|0;m-287&X\|-\|134540564\|134540585&offsets\|0\|0]\|TTTTGCGATGTGTTCCTAATAT | >MI0001652\|hsa-mir-450a-1&WithFlank&X\|-\|134540335\|134540437 | 24 | 45 | 22 | TTTTGCGATGTGTTCCTAATAT | BT20 | 23 | 7 | 5 | 2 | 2 | 29.3 | 4.4 | 3.0 | 1.1 | 1.8 |
| >MI0001733\|hsa-mir-452&WithFlank&X\|-\|151959622\|151959718@20.41.22(+1A)&[MIMAT0001635&hsa-miR-452-5p&offsets\|0\|0(+1A);m-188&X\|-\|151959677\|151959699&offsets\|0\|-1(+1A)]\|AACTGTTTGCAGAGGAAACTGAA | >MI0001733\|hsa-mir-452&WithFlank&X\|-\|151959622\|151959718 | 20 | 41 | 22(+1A) | AACTGTTTGCAGAGGAAACTGAA | BT20 | 13 | 7 | 9 | 7 | 3 | 5.6 | 1.1 | 3.9 | 8.5 | 2.5 |
| >MI0001733\|hsa-mir-452&WithFlank&X\|-\|151959622\|151959718@20.41.22&[MIMAT0001635&hsa-miR-452-5p&offsets\|0\|0;m-188&X\|-\|151959677\|151959699&offsets\|0\|-1]\|AACTGTTTGCAGAGGAAACTGA | >MI0001733\|hsa-mir-452&WithFlank&X\|-\|151959622\|151959718 | 20 | 41 | 22 | AACTGTTTGCAGAGGAAACTGA | BT20 | 36 | 24 | 12 | 33 | 8 | 17.5 | 5.8 | 1.0 | 43.6 | 3.3 |
| >MI0001733\|hsa-mir-452&WithFlank&X\|-\|151959622\|151959718@20.42.23(+1U)&[MIMAT0001635&hsa-miR-452-5p&offsets\|0\|+1(+1U);m-188&X\|-\|151959677\|151959699&offsets\|0\|0(+1U)]\|AACTGTTTGCAGAGGAAACTGAGT | >MI0001733\|hsa-mir-452&WithFlank&X\|-\|151959622\|151959718 | 20 | 42 | 23(+1U) | AACTGTTTGCAGAGGAAACTGAGT | BT20 | 20 | 13 | 16 | 4 | 5 | 4.3 | 1.5 | 2.4 | 5.1 | 1.5 |
| >MI0001733\|hsa-mir-452&WithFlank&X\|-\|151959622\|151959718@20.42.23&[MIMAT0001635&hsa-miR-452-5p&offsets\|0\|+1;m-188&X\|-\|151959677\|151959699&offsets\|0\|0]\|AACTGTTTGCAGAGGAAACTGAG | >MI0001733\|hsa-mir-452&WithFlank&X\|-\|151959622\|151959718 | 20 | 42 | 23 | AACTGTTTGCAGAGGAAACTGAG | BT20 | 82 | 64 | 64 | 52 | 38 | 5.2 | 8.9 | 2.2 | 57.6 | 19.9 |
| >MI0001733\|hsa-mir-452&WithFlank&X\|-\|151959622\|151959718@20.43.24&[MIMAT0001635&hsa-miR-452-5p&offsets\|0\|+2;m-188&X\|-\|151959677\|151959699&offsets\|0\|+1]\|AACTGTTTGCAGAGGAAACTGAGA | >MI0001733\|hsa-mir-452&WithFlank&X\|-\|151959622\|151959718 | 20 | 43 | 24 | AACTGTTTGCAGAGGAAACTGAGA | BT20 | 75 | 61 | 61 | 11 | 26 | 26.1 | 20.0 | 17.7 | 10.0 | 4.7 |
| >MI0001733\|hsa-mir-452&WithFlank&X\|-\|151959622\|151959718@20.43.24&[MIMAT0001635&hsa-miR-452-5p&offsets\|0\|+2;m-188&X\|-\|151959677\|151959699&offsets\|0\|+1]\|AACTGTTTGCAGAGGAAACTGAGA | >MI0001733\|hsa-mir-452&WithFlank&X\|-\|151959622\|151959718 | 20 | 43 | 24 | AACTGTTTGCAGAGGAAACTGAGA | MB231 | 11 | 10 | 3 | 15 | 11 | 1.8 | 1.6 | 3.1 | 6.0 | 6.3 |
| >MI0001733\|hsa-mir-452&WithFlank&X\|-\|151959622\|151959718@23.43.21&[MIMAT0001635&hsa-miR-452-5p&offsets\|+3\|+2;m-188&X\|-\|151959677\|151959699&offsets\|+3\|+1]\|TGTTTGCAGAGGAAACTGAGA | >MI0001733\|hsa-mir-452&WithFlank&X\|-\|151959622\|151959718 | 23 | 43 | 21 | TGTTTGCAGAGGAAACTGAGA | BT20 | 49 | 35 | 43 | 48 | 21 | 2.0 | 2.5 | 17.5 | 56.5 | 8.5 |
| >MI0001733\|hsa-mir-452&WithFlank&X\|-\|151959622\|151959718@23.44.22&[MIMAT0001635&hsa-miR-452-5p&offsets\|+3\|+3;m-188&X\|-\|151959677\|151959699&offsets\|+3\|+2]\|TGTTTGCAGAGGAAACTGAGAC | >MI0001733\|hsa-mir-452&WithFlank&X\|-\|151959622\|151959718 | 23 | 44 | 22 | TGTTTGCAGAGGAAACTGAGAC | BT20 | 29 | 22 | 19 | 127 | 18 | 18.4 | 11.2 | 16.3 | 124.1 | 6.0 |
| >MI0001733\|hsa-mir-452&WithFlank&X\|-\|151959622\|151959718@59.80.22&[MIMAT0001636&hsa-miR-452-3p&offsets\|-5\|-5;m-457&X\|-\|151959639\|151959660&offsets\|0\|0]\|TCAGTCTCATCTGCAAAGAAGT | >MI0001733\|hsa-mir-452&WithFlank&X\|-\|151959622\|151959718 | 59 | 80 | 22 | TCAGTCTCATCTGCAAAGAAGT | BT20 | 21 | 13 | 8 | 28 | 3 | 12.7 | 9.2 | 3.7 | 27.8 | 0.8 |
| >MI0002468\|hsa-mir-484&WithFlank&16\|+\|15643288\|15643378@14.33.20(+2U)&[MIMAT0002174&hsa-miR-484&offsets\|0\|-2(+2U);m-127&16\|+\|15643301\|15643321&offsets\|0\|-1(+2U)]\|TCAGGCTCAGTCCCCTCCCGTT | >MI0002468\|hsa-mir-484&WithFlank&16\|+\|15643288\|15643378 | 14 | 33 | 20(+2U) | TCAGGCTCAGTCCCCTCCCGTT | BT20 | 13 | 8 | 12 | 5 | 7 | 7.7 | 5.7 | 2.2 | 2.7 | 3.6 |
| >MI0002468\|hsa-mir-484&WithFlank&16\|+\|15643288\|15643378@14.35.22(+1U)&[MIMAT0002174&hsa-miR-484&offsets\|0\|0(+1U);m-127&16\|+\|15643301\|15643321&offsets\|0\|+1(+1U)]\|TCAGGCTCAGTCCCCTCCCGATT | >MI0002468\|hsa-mir-484&WithFlank&16\|+\|15643288\|15643378 | 14 | 35 | 22(+1U) | TCAGGCTCAGTCCCCTCCCGATT | BT20 | 29 | 17 | 28 | 6 | 19 | 18.2 | 12.8 | 3.1 | 3.9 | 17.0 |
| >MI0002468\|hsa-mir-484&WithFlank&16\|+\|15643288\|15643378@14.35.22(+1U)&[MIMAT0002174&hsa-miR-484&offsets\|0\|0(+1U);m-127&16\|+\|15643301\|15643321&offsets\|0\|+1(+1U)]\|TCAGGCTCAGTCCCCTCCCGATT | >MI0002468\|hsa-mir-484&WithFlank&16\|+\|15643288\|15643378 | 14 | 35 | 22(+1U) | TCAGGCTCAGTCCCCTCCCGATT | MB231 | 30 | 23 | 14 | 16 | 44 | 12.7 | 8.7 | 7.0 | 5.6 | 26.5 |
| >MI0002468\|hsa-mir-484&WithFlank&16\|+\|15643288\|15643378@14.35.22&[MIMAT0002174&hsa-miR-484&offsets\|0\|0;m-127&16\|+\|15643301\|15643321&offsets\|0\|+1]\|TCAGGCTCAGTCCCCTCCCGAT | >MI0002468\|hsa-mir-484&WithFlank&16\|+\|15643288\|15643378 | 14 | 35 | 22 | TCAGGCTCAGTCCCCTCCCGAT | BT20 | 76 | 43 | 70 | 6 | 52 | 52.1 | 38.8 | 28.6 | 4.5 | 50.5 |
| >MI0002468\|hsa-mir-484&WithFlank&16\|+\|15643288\|15643378@14.35.22&[MIMAT0002174&hsa-miR-484&offsets\|0\|0;m-127&16\|+\|15643301\|15643321&offsets\|0\|+1]\|TCAGGCTCAGTCCCCTCCCGAT | >MI0002468\|hsa-mir-484&WithFlank&16\|+\|15643288\|15643378 | 14 | 35 | 22 | TCAGGCTCAGTCCCCTCCCGAT | MB231 | 72 | 43 | 30 | 14 | 102 | 37.5 | 11.4 | 20.0 | 7.6 | 71.4 |
| >MI0002468\|hsa-mir-484&WithFlank&16\|+\|15643288\|15643378@14.36.23&[MIMAT0002174&hsa-miR-484&offsets\|0\|+1;m-127&16\|+\|15643301\|15643321&offsets\|0\|+2]\|TCAGGCTCAGTCCCCTCCCGATA | >MI0002468\|hsa-mir-484&WithFlank&16\|+\|15643288\|15643378 | 14 | 36 | 23 | TCAGGCTCAGTCCCCTCCCGATA | BT20 | 13 | 7 | 12 | 1 | 6 | 8.0 | 3.8 | 0.6 | 0.9 | 6.1 |
| >MI0002470\|hsa-mir-486-1&WithFlank&8\|-\|41660435\|41660514@10.31.22;MI0023622\|hsa-mir-486-2&WithFlank&8\|+\|41660438\|41660513@7.28.22&[MIMAT0002177&hsa-miR-486-5p&offsets\|0\|0;m-107&8\|-\|41660484\|41660505&offsets\|0\|0];[MIMAT0002177_1&hsa-miR-486-5p&offsets\|0\|0;TJU_CMC_MD2.ID03138.5p-miR&offsets\|0\|-2;m-106&8\|+\|41660444\|41660465&offsets\|0\|0]\|TCCTGTACTGAGCTGCCCCGAG | >MI0002470\|hsa-mir-486-1&WithFlank&8\|-\|41660435\|41660514 | 10 | 31 | 22 | TCCTGTACTGAGCTGCCCCGAG | MB231 | 12 | 8 | 13 | 19 | 7 | 2.6 | 2.9 | 5.4 | 0.9 | 2.1 |
| >MI0002470\|hsa-mir-486-1&WithFlank&8\|-\|41660435\|41660514@10.31.22(+1U);MI0023622\|hsa-mir-486-2&WithFlank&8\|+\|41660438\|41660513@7.28.22(+1U)&[MIMAT0002177&hsa-miR-486-5p&offsets\|0\|0(+1U);m-107&8\|-\|41660484\|41660505&offsets\|0\|0(+1U)];[MIMAT0002177_1&hsa-miR-486-5p&offsets\|0\|0(+1U);TJU_CMC_MD2.ID03138.5p-miR&offsets\|0\|-2(+1U);m-106&8\|+\|41660444\|41660465&offsets\|0\|0(+1U)]\|TCCTGTACTGAGCTGCCCCGAGT | >MI0002470\|hsa-mir-486-1&WithFlank&8\|-\|41660435\|41660514 | 10 | 31 | 22(+1U) | TCCTGTACTGAGCTGCCCCGAGT | MB231 | 17 | 15 | 22 | 36 | 16 | 4.2 | 5.6 | 11.8 | 14.9 | 5.8 |

| | | | | | | | | | | | | | | | | |
|---|---|---|---|---|---|---|---|---|---|---|---|---|---|---|---|---|
| >MI0003129\|hsa-mir-146b&WithFlank&10\|+\|102436506\|102436590@15.38.24&[MIMAT0002809&hsa-miR-146b-5p&offsets\|0\|+1;m-70&10\|+\|102436520\|102436543&offsets\|0\|0]\|TGAGAACTGAATTCCATAGGCTGT | >MI0003129\|hsa-mir-146b&WithFlank&10\|+\|102436506\|102436590 | 15 | 38 | 24 | TGAGAACTGAATTCCATAGGCTGT | BT20 | 21 | 17 | 20 | 23 | 11 | 18.4 | 11.6 | 16.2 | 17.9 | 6.7 |
| >MI0003129\|hsa-mir-146b&WithFlank&10\|+\|102436506\|102436590@15.38.24&[MIMAT0002809&hsa-miR-146b-5p&offsets\|0\|+1;m-70&10\|+\|102436520\|102436543&offsets\|0\|0]\|TGAGAACTGAATTCCATAGGCTGT | >MI0003129\|hsa-mir-146b&WithFlank&10\|+\|102436506\|102436590 | 15 | 38 | 24 | TGAGAACTGAATTCCATAGGCTGT | MB231 | 16 | 17 | 5 | 66 | 16 | 3.1 | 6.0 | 3.5 | 12.8 | 9.2 |
| >MI0003137\|hsa-mir-193b&WithFlank&16\|+\|14303961\|14304055@57.76.20(+1U)&[MIMAT0002819&hsa-miR-193b-3p&offsets\|0\|-2(+1U);m-166&16\|+\|14304017\|14304038&offsets\|0\|-2(+1U)]\|AACTGGCCCTCAAAGTCCCGT | >MI0003137\|hsa-mir-193b&WithFlank&16\|+\|14303961\|14304055 | 57 | 76 | 20(+1U) | AACTGGCCCTCAAAGTCCCGT | BT20 | 10 | 4 | 2 | 30 | 0 | 15.1 | 2.3 | 1.3 | 28.9 | 0.4 |
| >MI0003137\|hsa-mir-193b&WithFlank&16\|+\|14303961\|14304055@57.77.21(+1A)&[MIMAT0002819&hsa-miR-193b-3p&offsets\|0\|-1(+1A);m-166&16\|+\|14304017\|14304038&offsets\|0\|-1(+1A)]\|AACTGGCCCTCAAAGTCCCGCA | >MI0003137\|hsa-mir-193b&WithFlank&16\|+\|14303961\|14304055 | 57 | 77 | 21(+1A) | AACTGGCCCTCAAAGTCCCGCA | BT20 | 49 | 31 | 21 | 47 | 9 | 67.3 | 20.3 | 10.5 | 50.6 | 1.4 |
| >MI0003137\|hsa-mir-193b&WithFlank&16\|+\|14303961\|14304055@57.77.21(+1C)&[MIMAT0002819&hsa-miR-193b-3p&offsets\|0\|-1(+1C);m-166&16\|+\|14304017\|14304038&offsets\|0\|-1(+1C)]\|AACTGGCCCTCAAAGTCCCGCC | >MI0003137\|hsa-mir-193b&WithFlank&16\|+\|14303961\|14304055 | 57 | 77 | 21(+1C) | AACTGGCCCTCAAAGTCCCGCC | BT20 | 43 | 54 | 23 | 78 | 17 | 46.6 | 34.6 | 7.3 | 78.2 | 3.0 |
| >MI0003137\|hsa-mir-193b&WithFlank&16\|+\|14303961\|14304055@57.77.21(+1G)&[MIMAT0002819&hsa-miR-193b-3p&offsets\|0\|-1(+1G);m-166&16\|+\|14304017\|14304038&offsets\|0\|-1(+1G)]\|AACTGGCCCTCAAAGTCCCGCG | >MI0003137\|hsa-mir-193b&WithFlank&16\|+\|14303961\|14304055 | 57 | 77 | 21(+1G) | AACTGGCCCTCAAAGTCCCGCG | BT20 | 36 | 11 | 6 | 14 | 1 | 50.2 | 5.5 | 4.1 | 18.9 | 0.4 |
| >MI0003137\|hsa-mir-193b&WithFlank&16\|+\|14303961\|14304055@57.77.21&[MIMAT0002819&hsa-miR-193b-3p&offsets\|0\|-1;m-166&16\|+\|14304017\|14304038&offsets\|0\|-1]\|AACTGGCCCTCAAAGTCCCGC | >MI0003137\|hsa-mir-193b&WithFlank&16\|+\|14303961\|14304055 | 57 | 77 | 21 | AACTGGCCCTCAAAGTCCCGC | BT20 | 20 | 12 | 7 | 91 | 3 | 27.5 | 7.0 | 3.1 | 69.2 | 0.5 |
| >MI0003137\|hsa-mir-193b&WithFlank&16\|+\|14303961\|14304055@57.78.22(+1A)&[MIMAT0002819&hsa-miR-193b-3p&offsets\|0\|0(+1A);m-166&16\|+\|14304017\|14304038&offsets\|0\|0(+1A)]\|AACTGGCCCTCAAAGTCCCGCTA | >MI0003137\|hsa-mir-193b&WithFlank&16\|+\|14303961\|14304055 | 57 | 78 | 22(+1A) | AACTGGCCCTCAAAGTCCCGCTA | BT20 | 19 | 13 | 11 | 16 | 4 | 25.1 | 6.3 | 5.4 | 13.8 | 1.0 |
| >MI0003137\|hsa-mir-193b&WithFlank&16\|+\|14303961\|14304055@57.78.22&[MIMAT0002819&hsa-miR-193b-3p&offsets\|0\|0;m-166&16\|+\|14304017\|14304038&offsets\|0\|0]\|AACTGGCCCTCAAAGTCCCGCT | >MI0003137\|hsa-mir-193b&WithFlank&16\|+\|14303961\|14304055 | 57 | 78 | 22 | AACTGGCCCTCAAAGTCCCGCT | BT20 | 269 | 310 | 156 | 475 | 101 | 295.3 | 193.2 | 54.4 | 412.5 | 21.2 |
| >MI0003137\|hsa-mir-193b&WithFlank&16\|+\|14303961\|14304055@57.79.23&[MIMAT0002819&hsa-miR-193b-3p&offsets\|0\|+1;m-166&16\|+\|14304017\|14304038&offsets\|0\|+1]\|AACTGGCCCTCAAAGTCCCGCTT | >MI0003137\|hsa-mir-193b&WithFlank&16\|+\|14303961\|14304055 | 57 | 79 | 23 | AACTGGCCCTCAAAGTCCCGCTT | BT20 | 23 | 14 | 11 | 122 | 3 | 31.2 | 7.6 | 7.0 | 119.1 | 0.8 |
| >MI0003139\|hsa-mir-181d&WithFlank&19\|+\|13874869\|13875017@42.64.23&[MIMAT0002821&hsa-miR-181d-5p&offsets\|0\|0;m-194&19\|+\|13874910\|13874932&offsets\|0\|0]\|AACATTCATTGTTGTCGGTGGGT | >MI0003139\|hsa-mir-181d&WithFlank&19\|+\|13874869\|13875017 | 42 | 64 | 23 | AACATTCATTGTTGTCGGTGGGT | BT20 | 13 | 7 | 9 | 8 | 6 | 3.8 | 4.2 | 2.7 | 5.6 | 2.2 |
| >MI0003188\|hsa-mir-503&WithFlank&X\|-\|134546322\|134546404@12.31.20(+1A)&[MIMAT0002874&hsa-miR-503-5p&offsets\|0\|-3(+1A);m-297&X\|-\|134546371\|134546393&offsets\|0\|-3(+1A)]\|TAGCAGCGGGAACAGTTCTGA | >MI0003188\|hsa-mir-503&WithFlank&X\|-\|134546322\|134546404 | 12 | 31 | 20(+1A) | TAGCAGCGGGAACAGTTCTGA | BT20 | 14 | 9 | 16 | 26 | 8 | 6.1 | 3.8 | 7.3 | 21.3 | 1.3 |
| >MI0003188\|hsa-mir-503&WithFlank&X\|-\|134546322\|134546404@12.31.20&[MIMAT0002874&hsa-miR-503-5p&offsets\|0\|-3;m-297&X\|-\|134546371\|134546393&offsets\|0\|-3]\|TAGCAGCGGGAACAGTTCTG | >MI0003188\|hsa-mir-503&WithFlank&X\|-\|134546322\|134546404 | 12 | 31 | 20 | TAGCAGCGGGAACAGTTCTG | BT20 | 22 | 17 | 26 | 63 | 19 | 11.4 | 6.0 | 13.2 | 46.3 | 2.4 |
| >MI0003188\|hsa-mir-503&WithFlank&X\|-\|134546322\|134546404@12.31.20&[MIMAT0002874&hsa-miR-503-5p&offsets\|0\|-3;m-297&X\|-\|134546371\|134546393&offsets\|0\|-3]\|TAGCAGCGGGAACAGTTCTG | >MI0003188\|hsa-mir-503&WithFlank&X\|-\|134546322\|134546404 | 12 | 31 | 20 | TAGCAGCGGGAACAGTTCTG | MB231 | 12 | 11 | 6 | 36 | 14 | 4.5 | 2.9 | 4.2 | 9.6 | 2.7 |
| >MI0003188\|hsa-mir-503&WithFlank&X\|-\|134546322\|134546404@12.33.22&[MIMAT0002874&hsa-miR-503-5p&offsets\|0\|-1;m-297&X\|-\|134546371\|134546393&offsets\|0\|-1]\|TAGCAGCGGGAACAGTTCTGCA | >MI0003188\|hsa-mir-503&WithFlank&X\|-\|134546322\|134546404 | 12 | 33 | 22 | TAGCAGCGGGAACAGTTCTGCA | BT20 | 12 | 4 | 2 | 2 | 1 | 15.2 | 3.8 | 1.1 | 1.7 | 0.8 |
| >MI0003190\|hsa-mir-505&WithFlank&X\|-\|139924142\|139924237@57.80.24&[MIMAT0002876&hsa-miR-505-3p&offsets\|0\|+2;m-196&X\|-\|139924160\|139924181&offsets\|0\|+2]\|CGTCAACACTTGCTGGTTTCCTCT | >MI0003190\|hsa-mir-505&WithFlank&X\|-\|139924142\|139924237 | 57 | 80 | 24 | CGTCAACACTTGCTGGTTTCCTCT | BT20 | 10 | 11 | 10 | 1 | 5 | 5.0 | 4.9 | 2.0 | 0.7 | 6.1 |
| >MI0003205\|hsa-mir-532&WithFlank&X\|+\|50003142\|50003244@26.46.21(+1A)&[MIMAT0002888&hsa-miR-532-5p&offsets\|0\|-1(+1A);m-65&X\|+\|50003167\|50003188&offsets\|0\|-1(+1A)]\|CATGCCTTGAGTGTAGGACCGA | >MI0003205\|hsa-mir-532&WithFlank&X\|+\|50003142\|50003244 | 26 | 46 | 21(+1A) | CATGCCTTGAGTGTAGGACCGA | BT20 | 10 | 4 | 13 | 17 | 3 | 3.8 | 0.9 | 5.0 | 17.0 | 1.1 |
| >MI0003205\|hsa-mir-532&WithFlank&X\|+\|50003142\|50003244@26.46.21(+1C)&[MIMAT0002888&hsa-miR-532-5p&offsets\|0\|-1(+1C);m-65&X\|+\|50003167\|50003188&offsets\|0\|-1(+1C)]\|CATGCCTTGAGTGTAGGACCGC | >MI0003205\|hsa-mir-532&WithFlank&X\|+\|50003142\|50003244 | 26 | 46 | 21(+1C) | CATGCCTTGAGTGTAGGACCGC | BT20 | 24 | 17 | 31 | 58 | 11 | 3.5 | 4.1 | 13.0 | 54.0 | 3.7 |
| >MI0003205\|hsa-mir-532&WithFlank&X\|+\|50003142\|50003244@26.46.21(+1C)&[MIMAT0002888&hsa-miR-532-5p&offsets\|0\|-1(+1C);m-65&X\|+\|50003167\|50003188&offsets\|0\|-1(+1C)]\|CATGCCTTGAGTGTAGGACCGC | >MI0003205\|hsa-mir-532&WithFlank&X\|+\|50003142\|50003244 | 26 | 46 | 21(+1C) | CATGCCTTGAGTGTAGGACCGC | MB231 | 15 | 15 | 5 | 48 | 13 | 5.2 | 3.3 | 0.7 | 15.1 | 4.7 |
| >MI0003205\|hsa-mir-532&WithFlank&X\|+\|50003142\|50003244@26.46.21(+1G)&[MIMAT0002888&hsa-miR-532-5p&offsets\|0\|-1(+1G);m-65&X\|+\|50003167\|50003188&offsets\|0\|-1(+1G)]\|CATGCCTTGAGTGTAGGACCGG | >MI0003205\|hsa-mir-532&WithFlank&X\|+\|50003142\|50003244 | 26 | 46 | 21(+1G) | CATGCCTTGAGTGTAGGACCGG | BT20 | 13 | 3 | 7 | 8 | 1 | 8.7 | 1.0 | 5.3 | 10.9 | 0.6 |
| >MI0003205\|hsa-mir-532&WithFlank&X\|+\|50003142\|50003244@26.47.22&[MIMAT0002888&hsa-miR-532-5p&offsets\|0\|0;m-65&X\|+\|50003167\|50003188&offsets\|0\|0]\|CATGCCTTGAGTGTAGGACCGT | >MI0003205\|hsa-mir-532&WithFlank&X\|+\|50003142\|50003244 | 26 | 47 | 22 | CATGCCTTGAGTGTAGGACCGT | BT20 | 181 | 110 | 240 | 393 | 85 | 22.1 | 20.3 | 116.1 | 309.3 | 27.3 |
| >MI0003205\|hsa-mir-532&WithFlank&X\|+\|50003142\|50003244@26.47.22&[MIMAT0002888&hsa-miR-532-5p&offsets\|0\|0;m-65&X\|+\|50003167\|50003188&offsets\|0\|0]\|CATGCCTTGAGTGTAGGACCGT | >MI0003205\|hsa-mir-532&WithFlank&X\|+\|50003142\|50003244 | 26 | 47 | 22 | CATGCCTTGAGTGTAGGACCGT | MB231 | 140 | 140 | 49 | 320 | 103 | 55.4 | 25.7 | 12.7 | 54.4 | 25.3 |
| >MI0003205\|hsa-mir-532&WithFlank&X\|+\|50003142\|50003244@26.47.22&[MIMAT0002888&hsa-miR-532-5p&offsets\|0\|0;m-65&X\|+\|50003167\|50003188&offsets\|0\|0]\|CATGCCTTGAGTGTAGGACCGT | >MI0003205\|hsa-mir-532&WithFlank&X\|+\|50003142\|50003244 | 26 | 47 | 22 | CATGCCTTGAGTGTAGGACCGT | MB468 | 21 | 19 | 13 | 54 | 8 | 4.4 | 6.5 | 5.3 | 26.4 | 3.6 |
| >MI0003513\|hsa-mir-455&WithFlank&9\|+\|114209428\|114209535@22.43.22&[MIMAT0003150&hsa-miR-455-5p&offsets\|0\|0;m-232&9\|+\|114209449\|114209470&offsets\|0\|0]\|TATGTGCCTTTGGACTACATCG | >MI0003513\|hsa-mir-455&WithFlank&9\|+\|114209428\|114209535 | 22 | 43 | 22 | TATGTGCCTTTGGACTACATCG | MB231 | 15 | 7 | 1 | 37 | 10 | 3.8 | 3.9 | 1.0 | 5.5 | 9.1 |
| >MI0003513\|hsa-mir-455&WithFlank&9\|+\|114209428\|114209535@22.44.23&[MIMAT0003150&hsa-miR-455-5p&offsets\|0\|+1;m-232&9\|+\|114209449\|114209470&offsets\|0\|+1]\|TATGTGCCTTTGGACTACATCGT | >MI0003513\|hsa-mir-455&WithFlank&9\|+\|114209428\|114209535 | 22 | 44 | 23 | TATGTGCCTTTGGACTACATCGT | MB231 | 81 | 54 | 11 | 131 | 71 | 41.4 | 16.3 | 3.9 | 24.0 | 38.9 |
| >MI0003513\|hsa-mir-455&WithFlank&9\|+\|114209428\|114209535@22.45.24&[MIMAT0003150&hsa-miR-455-5p&offsets\|0\|+2;m-232&9\|+\|114209449\|114209470&offsets\|0\|+2]\|TATGTGCCTTTGGACTACATCGTG | >MI0003513\|hsa-mir-455&WithFlank&9\|+\|114209428\|114209535 | 22 | 45 | 24 | TATGTGCCTTTGGACTACATCGTG | MB231 | 18 | 12 | 3 | 13 | 14 | 3.7 | 3.7 | 1.6 | 3.9 | 5.1 |
| >MI0003513\|hsa-mir-455&WithFlank&9\|+\|114209428\|114209535@60.80.21(+1C)&[MIMAT0004784&hsa-miR-455-3p&offsets\|0\|0(+1C);m-135&9\|+\|114209487\|114209508&offsets\|0\|-1(+1C)]\|GCAGTCCATGGGCATATACACC | >MI0003513\|hsa-mir-455&WithFlank&9\|+\|114209428\|114209535 | 60 | 80 | 21(+1C) | GCAGTCCATGGGCATATACACC | MB231 | 16 | 19 | 11 | 10 | 16 | 7.4 | 1.2 | 3.9 | 2.9 | 2.3 |
| >MI0003513\|hsa-mir-455&WithFlank&9\|+\|114209428\|114209535@60.81.22&[MIMAT0004784&hsa-miR-455-3p&offsets\|0\|+1;m-135&9\|+\|114209487\|114209508&offsets\|0\|0]\|GCAGTCCATGGGCATATACACT | >MI0003513\|hsa-mir-455&WithFlank&9\|+\|114209428\|114209535 | 60 | 81 | 22 | GCAGTCCATGGGCATATACACT | MB231 | 17 | 23 | 14 | 12 | 22 | 4.1 | 5.7 | 8.6 | 7.8 | 9.3 |

| | | | | | | | | | | | | | | | | |
|---|---|---|---|---|---|---|---|---|---|---|---|---|---|---|---|---|
| >MI0003560\|hsa-mir-92b&WithFlank&1\|+\|155195171\|155195278@67.86.20(+1U)&[MIMAT0003218&hsa-miR-92b-3p&offsets\|0\|-2(+1U);m-115&1\|+\|155195237\|155195258&offsets\|0\|-2(+1U)]\|TATTGCACTCGTCCCGGCCTT | >MI0003560\|hsa-mir-92b&WithFlank&1\|+\|155195171\|155195278 | 67 | 86 | 20(+1U) | TATTGCACTCGTCCCGGCCTT | BT20 | 21 | 17 | 25 | 29 | 10 | 14.4 | 14.3 | 5.5 | 33.4 | 6.7 |
| >MI0003560\|hsa-mir-92b&WithFlank&1\|+\|155195171\|155195278@67.86.20(+1U)&[MIMAT0003218&hsa-miR-92b-3p&offsets\|0\|-2(+1U);m-115&1\|+\|155195237\|155195258&offsets\|0\|-2(+1U)]\|TATTGCACTCGTCCCGGCCTT | >MI0003560\|hsa-mir-92b&WithFlank&1\|+\|155195171\|155195278 | 67 | 86 | 20(+1U) | TATTGCACTCGTCCCGGCCTT | MB231 | 13 | 13 | 16 | 33 | 16 | 1.9 | 3.6 | 7.4 | 20.8 | 9.7 |
| >MI0003560\|hsa-mir-92b&WithFlank&1\|+\|155195171\|155195278@67.86.20&[MIMAT0003218&hsa-miR-92b-3p&offsets\|0\|-2;m-115&1\|+\|155195237\|155195258&offsets\|0\|-2]\|TATTGCACTCGTCCCGGCCT | >MI0003560\|hsa-mir-92b&WithFlank&1\|+\|155195171\|155195278 | 67 | 86 | 20 | TATTGCACTCGTCCCGGCCT | BT20 | 25 | 20 | 36 | 28 | 13 | 15.8 | 15.7 | 5.5 | 30.5 | 9.6 |
| >MI0003560\|hsa-mir-92b&WithFlank&1\|+\|155195171\|155195278@67.86.20&[MIMAT0003218&hsa-miR-92b-3p&offsets\|0\|-2;m-115&1\|+\|155195237\|155195258&offsets\|0\|-2]\|TATTGCACTCGTCCCGGCCT | >MI0003560\|hsa-mir-92b&WithFlank&1\|+\|155195171\|155195278 | 67 | 86 | 20 | TATTGCACTCGTCCCGGCCT | MB231 | 20 | 22 | 29 | 36 | 21 | 2.2 | 4.7 | 13.4 | 22.8 | 9.0 |
| >MI0003560\|hsa-mir-92b&WithFlank&1\|+\|155195171\|155195278@67.87.21(+1A)&[MIMAT0003218&hsa-miR-92b-3p&offsets\|0\|-1(+1A);m-115&1\|+\|155195237\|155195258&offsets\|0\|-1(+1A)]\|TATTGCACTCGTCCCGGCCTCA | >MI0003560\|hsa-mir-92b&WithFlank&1\|+\|155195171\|155195278 | 67 | 87 | 21(+1A) | TATTGCACTCGTCCCGGCCTCA | BT20 | 91 | 54 | 125 | 68 | 39 | 57.3 | 46.1 | 19.6 | 73.4 | 28.4 |
| >MI0003560\|hsa-mir-92b&WithFlank&1\|+\|155195171\|155195278@67.87.21(+1A)&[MIMAT0003218&hsa-miR-92b-3p&offsets\|0\|-1(+1A);m-115&1\|+\|155195237\|155195258&offsets\|0\|-1(+1A)]\|TATTGCACTCGTCCCGGCCTCA | >MI0003560\|hsa-mir-92b&WithFlank&1\|+\|155195171\|155195278 | 67 | 87 | 21(+1A) | TATTGCACTCGTCCCGGCCTCA | MB231 | 30 | 28 | 38 | 67 | 41 | 7.2 | 4.6 | 18.2 | 49.8 | 24.6 |
| >MI0003560\|hsa-mir-92b&WithFlank&1\|+\|155195171\|155195278@67.87.21(+1G)&[MIMAT0003218&hsa-miR-92b-3p&offsets\|0\|-1(+1G);m-115&1\|+\|155195237\|155195258&offsets\|0\|-1(+1G)]\|TATTGCACTCGTCCCGGCCTCG | >MI0003560\|hsa-mir-92b&WithFlank&1\|+\|155195171\|155195278 | 67 | 87 | 21(+1G) | TATTGCACTCGTCCCGGCCTCG | BT20 | 15 | 5 | 11 | 4 | 2 | 13.3 | 5.3 | 3.0 | 3.8 | 2.3 |
| >MI0003560\|hsa-mir-92b&WithFlank&1\|+\|155195171\|155195278@67.87.21(+1U)&[MIMAT0003218&hsa-miR-92b-3p&offsets\|0\|-1(+1U);m-115&1\|+\|155195237\|155195258&offsets\|0\|-1(+1U)]\|TATTGCACTCGTCCCGGCCTCT | >MI0003560\|hsa-mir-92b&WithFlank&1\|+\|155195171\|155195278 | 67 | 87 | 21(+1U) | TATTGCACTCGTCCCGGCCTCT | BT20 | 190 | 137 | 253 | 207 | 96 | 115.1 | 127.5 | 44.8 | 227.2 | 67.7 |
| >MI0003560\|hsa-mir-92b&WithFlank&1\|+\|155195171\|155195278@67.87.21(+1U)&[MIMAT0003218&hsa-miR-92b-3p&offsets\|0\|-1(+1U);m-115&1\|+\|155195237\|155195258&offsets\|0\|-1(+1U)]\|TATTGCACTCGTCCCGGCCTCT | >MI0003560\|hsa-mir-92b&WithFlank&1\|+\|155195171\|155195278 | 67 | 87 | 21(+1U) | TATTGCACTCGTCCCGGCCTCT | MB231 | 77 | 71 | 96 | 194 | 104 | 22.2 | 12.6 | 44.3 | 142.8 | 61.6 |
| >MI0003560\|hsa-mir-92b&WithFlank&1\|+\|155195171\|155195278@67.87.21&[MIMAT0003218&hsa-miR-92b-3p&offsets\|0\|-1;m-115&1\|+\|155195237\|155195258&offsets\|0\|-1]\|TATTGCACTCGTCCCGGCCTC | >MI0003560\|hsa-mir-92b&WithFlank&1\|+\|155195171\|155195278 | 67 | 87 | 21 | TATTGCACTCGTCCCGGCCTC | BT20 | 58 | 46 | 76 | 114 | 35 | 47.5 | 35.5 | 22.8 | 150.8 | 27.0 |
| >MI0003560\|hsa-mir-92b&WithFlank&1\|+\|155195171\|155195278@67.87.21&[MIMAT0003218&hsa-miR-92b-3p&offsets\|0\|-1;m-115&1\|+\|155195237\|155195258&offsets\|0\|-1]\|TATTGCACTCGTCCCGGCCTC | >MI0003560\|hsa-mir-92b&WithFlank&1\|+\|155195171\|155195278 | 67 | 87 | 21 | TATTGCACTCGTCCCGGCCTC | MB231 | 34 | 41 | 54 | 71 | 42 | 6.7 | 12.3 | 24.5 | 43.0 | 19.1 |
| >MI0003560\|hsa-mir-92b&WithFlank&1\|+\|155195171\|155195278@67.88.22(+1A)&[MIMAT0003218&hsa-miR-92b-3p&offsets\|0\|0(+1A);m-115&1\|+\|155195237\|155195258&offsets\|0\|0(+1A)]\|TATTGCACTCGTCCCGGCCTCCA | >MI0003560\|hsa-mir-92b&WithFlank&1\|+\|155195171\|155195278 | 67 | 88 | 22(+1A) | TATTGCACTCGTCCCGGCCTCCA | BT20 | 22 | 14 | 33 | 6 | 13 | 13.4 | 11.6 | 3.5 | 7.6 | 10.4 |
| >MI0003560\|hsa-mir-92b&WithFlank&1\|+\|155195171\|155195278@67.88.22(+1U)&[MIMAT0003218&hsa-miR-92b-3p&offsets\|0\|0(+1U);m-115&1\|+\|155195237\|155195258&offsets\|0\|0(+1U)]\|TATTGCACTCGTCCCGGCCTCCT | >MI0003560\|hsa-mir-92b&WithFlank&1\|+\|155195171\|155195278 | 67 | 88 | 22(+1U) | TATTGCACTCGTCCCGGCCTCCT | BT20 | 33 | 25 | 47 | 21 | 23 | 21.4 | 24.6 | 14.8 | 26.0 | 20.5 |
| >MI0003560\|hsa-mir-92b&WithFlank&1\|+\|155195171\|155195278@67.88.22(+1U)&[MIMAT0003218&hsa-miR-92b-3p&offsets\|0\|0(+1U);m-115&1\|+\|155195237\|155195258&offsets\|0\|0(+1U)]\|TATTGCACTCGTCCCGGCCTCCT | >MI0003560\|hsa-mir-92b&WithFlank&1\|+\|155195171\|155195278 | 67 | 88 | 22(+1U) | TATTGCACTCGTCCCGGCCTCCT | MB231 | 23 | 18 | 24 | 24 | 25 | 7.5 | 3.3 | 7.6 | 16.4 | 13.6 |
| >MI0003560\|hsa-mir-92b&WithFlank&1\|+\|155195171\|155195278@67.88.22(+2U)&[MIMAT0003218&hsa-miR-92b-3p&offsets\|0\|0(+2U);m-115&1\|+\|155195237\|155195258&offsets\|0\|0(+2U)]\|TATTGCACTCGTCCCGGCCTCCTT | >MI0003560\|hsa-mir-92b&WithFlank&1\|+\|155195171\|155195278 | 67 | 88 | 22(+2U) | TATTGCACTCGTCCCGGCCTCCTT | BT20 | 14 | 9 | 15 | 9 | 8 | 8.7 | 8.5 | 4.6 | 9.7 | 6.6 |
| >MI0003560\|hsa-mir-92b&WithFlank&1\|+\|155195171\|155195278@67.88.22(+3U)&[MIMAT0003218&hsa-miR-92b-3p&offsets\|0\|0(+3U);m-115&1\|+\|155195237\|155195258&offsets\|0\|0(+3U)]\|TATTGCACTCGTCCCGGCCTCCTTT | >MI0003560\|hsa-mir-92b&WithFlank&1\|+\|155195171\|155195278 | 67 | 88 | 22(+3U) | TATTGCACTCGTCCCGGCCTCCTTT | BT20 | 11 | 7 | 14 | 4 | 8 | 8.0 | 6.7 | 4.6 | 4.6 | 8.3 |
| >MI0003560\|hsa-mir-92b&WithFlank&1\|+\|155195171\|155195278@67.88.22(+3U)&[MIMAT0003218&hsa-miR-92b-3p&offsets\|0\|0(+3U);m-115&1\|+\|155195237\|155195258&offsets\|0\|0(+3U)]\|TATTGCACTCGTCCCGGCCTCCTTT | >MI0003560\|hsa-mir-92b&WithFlank&1\|+\|155195171\|155195278 | 67 | 88 | 22(+3U) | TATTGCACTCGTCCCGGCCTCCTTT | MB231 | 12 | 8 | 14 | 7 | 16 | 1.6 | 3.4 | 7.8 | 4.7 | 8.9 |
| >MI0003560\|hsa-mir-92b&WithFlank&1\|+\|155195171\|155195278@67.88.22&[MIMAT0003218&hsa-miR-92b-3p&offsets\|0\|0;m-115&1\|+\|155195237\|155195258&offsets\|0\|0]\|TATTGCACTCGTCCCGGCCTCC | >MI0003560\|hsa-mir-92b&WithFlank&1\|+\|155195171\|155195278 | 67 | 88 | 22 | TATTGCACTCGTCCCGGCCTCC | BT20 | 276 | 210 | 411 | 427 | 183 | 196.8 | 152.7 | 103.6 | 526.0 | 129.6 |
| >MI0003560\|hsa-mir-92b&WithFlank&1\|+\|155195171\|155195278@67.88.22&[MIMAT0003218&hsa-miR-92b-3p&offsets\|0\|0;m-115&1\|+\|155195237\|155195258&offsets\|0\|0]\|TATTGCACTCGTCCCGGCCTCC | >MI0003560\|hsa-mir-92b&WithFlank&1\|+\|155195171\|155195278 | 67 | 88 | 22 | TATTGCACTCGTCCCGGCCTCC | MB231 | 139 | 142 | 170 | 242 | 168 | 49.5 | 31.6 | 71.1 | 159.1 | 70.0 |
| >MI0003581\|hsa-mir-574&WithFlank&4\|+\|38868026\|38868133@67.88.22&[MIMAT0003239&hsa-miR-574-3p&offsets\|0\|0;m-149&4\|+\|38868092\|38868113&offsets\|0\|0]\|CACGCTCATGCACACACCCACA | >MI0003581\|hsa-mir-574&WithFlank&4\|+\|38868026\|38868133 | 67 | 88 | 22 | CACGCTCATGCACACACCCACA | BT20 | 32 | 28 | 141 | 15 | 11 | 23.6 | 11.9 | 80.7 | 14.6 | 3.3 |
| >MI0003581\|hsa-mir-574&WithFlank&4\|+\|38868026\|38868133@67.88.22&[MIMAT0003239&hsa-miR-574-3p&offsets\|0\|0;m-149&4\|+\|38868092\|38868113&offsets\|0\|0]\|CACGCTCATGCACACACCCACA | >MI0003581\|hsa-mir-574&WithFlank&4\|+\|38868026\|38868133 | 67 | 88 | 22 | CACGCTCATGCACACACCCACA | MB231 | 40 | 44 | 178 | 76 | 66 | 14.0 | 15.5 | 188.4 | 30.5 | 34.4 |
| >MI0003591\|hsa-mir-584&WithFlank&5\|-\|149062307\|149062415@22.40.19&[MIMAT0003249&hsa-miR-584-5p&offsets\|0\|-3;m-167&5\|-\|149062374\|149062394&offsets\|0\|-2]\|TTATGGTTTGCCTGGGACT | >MI0003591\|hsa-mir-584&WithFlank&5\|-\|149062307\|149062415 | 22 | 40 | 19 | TTATGGTTTGCCTGGGACT | BT20 | 38 | 20 | 25 | 104 | 23 | 11.8 | 3.0 | 7.1 | 79.5 | 5.0 |
| >MI0003591\|hsa-mir-584&WithFlank&5\|-\|149062307\|149062415@22.40.19&[MIMAT0003249&hsa-miR-584-5p&offsets\|0\|-3;m-167&5\|-\|149062374\|149062394&offsets\|0\|-2]\|TTATGGTTTGCCTGGGACT | >MI0003591\|hsa-mir-584&WithFlank&5\|-\|149062307\|149062415 | 22 | 40 | 19 | TTATGGTTTGCCTGGGACT | MB231 | 37 | 30 | 12 | 74 | 36 | 9.9 | 4.9 | 3.4 | 25.1 | 12.3 |
| >MI0003591\|hsa-mir-584&WithFlank&5\|-\|149062307\|149062415@22.41.20(+1G)&[MIMAT0003249&hsa-miR-584-5p&offsets\|0\|-2(+1G);m-167&5\|-\|149062374\|149062394&offsets\|0\|-1(+1G)]\|TTATGGTTTGCCTGGGACTGG | >MI0003591\|hsa-mir-584&WithFlank&5\|-\|149062307\|149062415 | 22 | 41 | 20(+1G) | TTATGGTTTGCCTGGGACTGG | BT20 | 13 | 3 | 3 | 13 | 2 | 9.6 | 0.7 | 0.1 | 13.6 | 0.5 |
| >MI0003591\|hsa-mir-584&WithFlank&5\|-\|149062307\|149062415@22.41.20(+1U)&[MIMAT0003249&hsa-miR-584-5p&offsets\|0\|-2(+1U);m-167&5\|-\|149062374\|149062394&offsets\|0\|-1(+1U)]\|TTATGGTTTGCCTGGGACTGT | >MI0003591\|hsa-mir-584&WithFlank&5\|-\|149062307\|149062415 | 22 | 41 | 20(+1U) | TTATGGTTTGCCTGGGACTGT | BT20 | 46 | 26 | 31 | 114 | 35 | 10.0 | 6.1 | 10.3 | 89.5 | 10.1 |
| >MI0003591\|hsa-mir-584&WithFlank&5\|-\|149062307\|149062415@22.41.20(+1U)&[MIMAT0003249&hsa-miR-584-5p&offsets\|0\|-2(+1U);m-167&5\|-\|149062374\|149062394&offsets\|0\|-1(+1U)]\|TTATGGTTTGCCTGGGACTGT | >MI0003591\|hsa-mir-584&WithFlank&5\|-\|149062307\|149062415 | 22 | 41 | 20(+1U) | TTATGGTTTGCCTGGGACTGT | MB231 | 42 | 31 | 13 | 93 | 38 | 19.6 | 14.9 | 9.1 | 49.7 | 28.9 |
| >MI0003591\|hsa-mir-584&WithFlank&5\|-\|149062307\|149062415@22.41.20&[MIMAT0003249&hsa-miR-584-5p&offsets\|0\|-2;m-167&5\|-\|149062374\|149062394&offsets\|0\|-1]\|TTATGGTTTGCCTGGGACTG | >MI0003591\|hsa-mir-584&WithFlank&5\|-\|149062307\|149062415 | 22 | 41 | 20 | TTATGGTTTGCCTGGGACTG | BT20 | 22 | 14 | 13 | 79 | 23 | 7.1 | 1.1 | 7.0 | 49.5 | 4.1 |
| >MI0003591\|hsa-mir-584&WithFlank&5\|-\|149062307\|149062415@22.41.20&[MIMAT0003249&hsa-miR-584-5p&offsets\|0\|-2;m-167&5\|-\|149062374\|149062394&offsets\|0\|-1]\|TTATGGTTTGCCTGGGACTG | >MI0003591\|hsa-mir-584&WithFlank&5\|-\|149062307\|149062415 | 22 | 41 | 20 | TTATGGTTTGCCTGGGACTG | MB231 | 21 | 18 | 6 | 55 | 19 | 7.7 | 1.6 | 1.4 | 9.2 | 1.9 |

| | | | | | | | | | | | | | | | | |
|---|---|---|---|---|---|---|---|---|---|---|---|---|---|---|---|---|
| >MI0003591\|hsa-mir-584&WithFlank&5\|-\|149062307\|149062415@22.42.21(+1A)&[MIMAT0003249&hsa-miR-584-5p&offsets\|0\|-1(+1A);m-167&5\|-\|149062374\|149062394&offsets\|0\|0(+1A)]\|TTATGGTTTGCCTGGGACTGAA | >MI0003591\|hsa-mir-584&WithFlank&5\|-\|149062307\|149062415 | 22 | 42 | 21(+1A) | TTATGGTTTGCCTGGGACTGAA | BT20 | 16 | 11 | 10 | 14 | 16 | 6.0 | 2.7 | 2.5 | 9.8 | 3.9 |
| >MI0003591\|hsa-mir-584&WithFlank&5\|-\|149062307\|149062415@22.42.21(+1U)&[MIMAT0003249&hsa-miR-584-5p&offsets\|0\|-1(+1U);m-167&5\|-\|149062374\|149062394&offsets\|0\|0(+1U)]\|TTATGGTTTGCCTGGGACTGAT | >MI0003591\|hsa-mir-584&WithFlank&5\|-\|149062307\|149062415 | 22 | 42 | 21(+1U) | TTATGGTTTGCCTGGGACTGAT | BT20 | 21 | 11 | 14 | 6 | 18 | 5.9 | 3.9 | 5.4 | 4.5 | 8.6 |
| >MI0003591\|hsa-mir-584&WithFlank&5\|-\|149062307\|149062415@22.42.21(+1U)&[MIMAT0003249&hsa-miR-584-5p&offsets\|0\|-1(+1U);m-167&5\|-\|149062374\|149062394&offsets\|0\|0(+1U)]\|TTATGGTTTGCCTGGGACTGAT | >MI0003591\|hsa-mir-584&WithFlank&5\|-\|149062307\|149062415 | 22 | 42 | 21(+1U) | TTATGGTTTGCCTGGGACTGAT | MB231 | 24 | 16 | 8 | 19 | 25 | 6.7 | 5.4 | 4.0 | 3.7 | 14.3 |
| >MI0003591\|hsa-mir-584&WithFlank&5\|-\|149062307\|149062415@22.42.21(+2A)&[MIMAT0003249&hsa-miR-584-5p&offsets\|0\|-1(+2A);m-167&5\|-\|149062374\|149062394&offsets\|0\|0(+2A)]\|TTATGGTTTGCCTGGGACTGAAA | >MI0003591\|hsa-mir-584&WithFlank&5\|-\|149062307\|149062415 | 22 | 42 | 21(+2A) | TTATGGTTTGCCTGGGACTGAAA | BT20 | 13 | 8 | 7 | 6 | 13 | 4.5 | 2.3 | 3.3 | 3.3 | 2.3 |
| >MI0003591\|hsa-mir-584&WithFlank&5\|-\|149062307\|149062415@22.42.21(+2U)&[MIMAT0003249&hsa-miR-584-5p&offsets\|0\|-1(+2U);m-167&5\|-\|149062374\|149062394&offsets\|0\|0(+2U)]\|TTATGGTTTGCCTGGGACTGATT | >MI0003591\|hsa-mir-584&WithFlank&5\|-\|149062307\|149062415 | 22 | 42 | 21(+2U) | TTATGGTTTGCCTGGGACTGATT | MB231 | 10 | 6 | 3 | 16 | 9 | 3.0 | 3.3 | 1.2 | 2.5 | 5.0 |
| >MI0003591\|hsa-mir-584&WithFlank&5\|-\|149062307\|149062415@22.42.21&[MIMAT0003249&hsa-miR-584-5p&offsets\|0\|-1;m-167&5\|-\|149062374\|149062394&offsets\|0\|0]\|TTATGGTTTGCCTGGGACTGA | >MI0003591\|hsa-mir-584&WithFlank&5\|-\|149062307\|149062415 | 22 | 42 | 21 | TTATGGTTTGCCTGGGACTGA | BT20 | 128 | 85 | 74 | 476 | 122 | 46.4 | 21.7 | 34.3 | 354.5 | 18.9 |
| >MI0003591\|hsa-mir-584&WithFlank&5\|-\|149062307\|149062415@22.42.21&[MIMAT0003249&hsa-miR-584-5p&offsets\|0\|-1;m-167&5\|-\|149062374\|149062394&offsets\|0\|0]\|TTATGGTTTGCCTGGGACTGA | >MI0003591\|hsa-mir-584&WithFlank&5\|-\|149062307\|149062415 | 22 | 42 | 21 | TTATGGTTTGCCTGGGACTGA | MB231 | 95 | 83 | 22 | 263 | 108 | 24.5 | 19.4 | 8.1 | 23.3 | 39.9 |
| >MI0003591\|hsa-mir-584&WithFlank&5\|-\|149062307\|149062415@22.43.22(+1U)&[MIMAT0003249&hsa-miR-584-5p&offsets\|0\|0(+1U);m-167&5\|-\|149062374\|149062394&offsets\|0\|+1(+1U)]\|TTATGGTTTGCCTGGGACTGAGT | >MI0003591\|hsa-mir-584&WithFlank&5\|-\|149062307\|149062415 | 22 | 43 | 22(+1U) | TTATGGTTTGCCTGGGACTGAGT | BT20 | 10 | 6 | 8 | 11 | 7 | 4.2 | 2.2 | 4.3 | 6.2 | 5.0 |
| >MI0003591\|hsa-mir-584&WithFlank&5\|-\|149062307\|149062415@22.43.22(+1U)&[MIMAT0003249&hsa-miR-584-5p&offsets\|0\|0(+1U);m-167&5\|-\|149062374\|149062394&offsets\|0\|+1(+1U)]\|TTATGGTTTGCCTGGGACTGAGT | >MI0003591\|hsa-mir-584&WithFlank&5\|-\|149062307\|149062415 | 22 | 43 | 22(+1U) | TTATGGTTTGCCTGGGACTGAGT | MB231 | 13 | 8 | 6 | 22 | 12 | 3.8 | 1.8 | 1.2 | 7.5 | 6.7 |
| >MI0003591\|hsa-mir-584&WithFlank&5\|-\|149062307\|149062415@22.43.22&[MIMAT0003249&hsa-miR-584-5p&offsets\|0\|0;m-167&5\|-\|149062374\|149062394&offsets\|0\|+1]\|TTATGGTTTGCCTGGGACTGAG | >MI0003591\|hsa-mir-584&WithFlank&5\|-\|149062307\|149062415 | 22 | 43 | 22 | TTATGGTTTGCCTGGGACTGAG | BT20 | 13 | 8 | 9 | 19 | 13 | 3.5 | 1.8 | 3.4 | 10.4 | 3.0 |
| >MI0003599\|hsa-mir-589&WithFlank&7\|-\|5495813\|5495923@30.50.21&[MIMAT0004799&hsa-miR-589-5p&offsets\|0\|-1;m-162&7\|-\|5495874\|5495894&offsets\|0\|0]\|TGAGAACCACGTCTGCTCTGA | >MI0003599\|hsa-mir-589&WithFlank&7\|-\|5495813\|5495923 | 30 | 50 | 21 | TGAGAACCACGTCTGCTCTGA | BT20 | 18 | 16 | 17 | 30 | 11 | 10.0 | 6.1 | 6.6 | 23.3 | 5.2 |
| >MI0003599\|hsa-mir-589&WithFlank&7\|-\|5495813\|5495923@30.50.21&[MIMAT0004799&hsa-miR-589-5p&offsets\|0\|-1;m-162&7\|-\|5495874\|5495894&offsets\|0\|0]\|TGAGAACCACGTCTGCTCTGA | >MI0003599\|hsa-mir-589&WithFlank&7\|-\|5495813\|5495923 | 30 | 50 | 21 | TGAGAACCACGTCTGCTCTGA | MB231 | 17 | 27 | 10 | 48 | 28 | 4.5 | 9.3 | 8.4 | 8.7 | 9.8 |
| >MI0003599\|hsa-mir-589&WithFlank&7\|-\|5495813\|5495923@30.51.22(+1U)&[MIMAT0004799&hsa-miR-589-5p&offsets\|0\|0(+1U);m-162&7\|-\|5495874\|5495894&offsets\|0\|+1(+1U)]\|TGAGAACCACGTCTGCTCTGAGT | >MI0003599\|hsa-mir-589&WithFlank&7\|-\|5495813\|5495923 | 30 | 51 | 22(+1U) | TGAGAACCACGTCTGCTCTGAGT | BT20 | 10 | 9 | 21 | 6 | 7 | 4.1 | 2.3 | 11.3 | 4.7 | 4.1 |
| >MI0003599\|hsa-mir-589&WithFlank&7\|-\|5495813\|5495923@30.52.23&[MIMAT0004799&hsa-miR-589-5p&offsets\|0\|+1;m-162&7\|-\|5495874\|5495894&offsets\|0\|+2]\|TGAGAACCACGTCTGCTCTGAGC | >MI0003599\|hsa-mir-589&WithFlank&7\|-\|5495813\|5495923 | 30 | 52 | 23 | TGAGAACCACGTCTGCTCTGAGC | BT20 | 37 | 38 | 92 | 28 | 33 | 25.5 | 4.3 | 61.7 | 12.1 | 17.5 |
| >MI0003599\|hsa-mir-589&WithFlank&7\|-\|5495813\|5495923@30.52.23&[MIMAT0004799&hsa-miR-589-5p&offsets\|0\|+1;m-162&7\|-\|5495874\|5495894&offsets\|0\|+2]\|TGAGAACCACGTCTGCTCTGAGC | >MI0003599\|hsa-mir-589&WithFlank&7\|-\|5495813\|5495923 | 30 | 52 | 23 | TGAGAACCACGTCTGCTCTGAGC | MB231 | 26 | 40 | 37 | 22 | 21 | 13.3 | 25.1 | 21.8 | 7.3 | 3.8 |
| >MI0003602\|hsa-mir-590&WithFlank&7\|+\|74191192\|74191300@62.82.21&[MIMAT0004801&hsa-miR-590-3p&offsets\|0\|0;m-332&7\|+\|74191253\|74191273&offsets\|0\|0]\|TAATTTTATGTATAAGCTAGT | >MI0003602\|hsa-mir-590&WithFlank&7\|+\|74191192\|74191300 | 62 | 82 | 21 | TAATTTTATGTATAAGCTAGT | BT20 | 31 | 8 | 2 | 0 | 0 | 47.6 | 10.6 | 1.6 | 0.0 | 0.0 |
| >MI0003628\|hsa-mir-615&WithFlank&12\|+\|54033944\|54034051@67.85.19&[MIMAT0003283&hsa-miR-615-3p&offsets\|0\|-3;m-495&12\|+\|54034010\|54034030&offsets\|0\|-2]\|TCCGAGCCTGGGTCTCCCT | >MI0003628\|hsa-mir-615&WithFlank&12\|+\|54033944\|54034051 | 67 | 85 | 19 | TCCGAGCCTGGGTCTCCCT | BT20 | 17 | 14 | 13 | 5 | 21 | 17.1 | 6.5 | 4.7 | 1.6 | 6.2 |
| >MI0003628\|hsa-mir-615&WithFlank&12\|+\|54033944\|54034051@67.86.20(+1A)&[MIMAT0003283&hsa-miR-615-3p&offsets\|0\|-2(+1A);m-495&12\|+\|54034010\|54034030&offsets\|0\|-1(+1A)]\|TCCGAGCCTGGGTCTCCCTCA | >MI0003628\|hsa-mir-615&WithFlank&12\|+\|54033944\|54034051 | 67 | 86 | 20(+1A) | TCCGAGCCTGGGTCTCCCTCA | BT20 | 17 | 13 | 18 | 2 | 20 | 14.8 | 4.9 | 1.6 | 0.5 | 1.6 |
| >MI0003628\|hsa-mir-615&WithFlank&12\|+\|54033944\|54034051@67.86.20(+1C)&[MIMAT0003283&hsa-miR-615-3p&offsets\|0\|-2(+1C);m-495&12\|+\|54034010\|54034030&offsets\|0\|-1(+1C)]\|TCCGAGCCTGGGTCTCCCTCC | >MI0003628\|hsa-mir-615&WithFlank&12\|+\|54033944\|54034051 | 67 | 86 | 20(+1C) | TCCGAGCCTGGGTCTCCCTCC | BT20 | 17 | 15 | 16 | 3 | 27 | 14.7 | 5.0 | 3.0 | 0.7 | 3.1 |
| >MI0003628\|hsa-mir-615&WithFlank&12\|+\|54033944\|54034051@67.86.20&[MIMAT0003283&hsa-miR-615-3p&offsets\|0\|-2;m-495&12\|+\|54034010\|54034030&offsets\|0\|-1]\|TCCGAGCCTGGGTCTCCCTC | >MI0003628\|hsa-mir-615&WithFlank&12\|+\|54033944\|54034051 | 67 | 86 | 20 | TCCGAGCCTGGGTCTCCCTC | BT20 | 31 | 30 | 35 | 7 | 46 | 31.2 | 11.6 | 16.4 | 1.4 | 7.7 |
| >MI0003628\|hsa-mir-615&WithFlank&12\|+\|54033944\|54034051@67.87.21(+1A)&[MIMAT0003283&hsa-miR-615-3p&offsets\|0\|-1(+1A);m-495&12\|+\|54034010\|54034030&offsets\|0\|0(+1A)]\|TCCGAGCCTGGGTCTCCCTCTA | >MI0003628\|hsa-mir-615&WithFlank&12\|+\|54033944\|54034051 | 67 | 87 | 21(+1A) | TCCGAGCCTGGGTCTCCCTCTA | BT20 | 10 | 5 | 12 | 1 | 9 | 8.9 | 2.5 | 1.0 | 0.2 | 0.9 |
| >MI0003628\|hsa-mir-615&WithFlank&12\|+\|54033944\|54034051@67.87.21&[MIMAT0003283&hsa-miR-615-3p&offsets\|0\|-1;m-495&12\|+\|54034010\|54034030&offsets\|0\|0]\|TCCGAGCCTGGGTCTCCCTCT | >MI0003628\|hsa-mir-615&WithFlank&12\|+\|54033944\|54034051 | 67 | 87 | 21 | TCCGAGCCTGGGTCTCCCTCT | BT20 | 119 | 97 | 115 | 22 | 189 | 114.2 | 37.2 | 30.1 | 2.7 | 11.0 |
| >MI0003628\|hsa-mir-615&WithFlank&12\|+\|54033944\|54034051@67.88.22&[MIMAT0003283&hsa-miR-615-3p&offsets\|0\|0;m-495&12\|+\|54034010\|54034030&offsets\|0\|+1]\|TCCGAGCCTGGGTCTCCCTCTT | >MI0003628\|hsa-mir-615&WithFlank&12\|+\|54033944\|54034051 | 67 | 88 | 22 | TCCGAGCCTGGGTCTCCCTCTT | BT20 | 76 | 64 | 96 | 24 | 111 | 65.9 | 26.3 | 13.6 | 8.6 | 10.4 |
| >MI0003643\|hsa-mir-629&WithFlank&15\|-\|70079366\|70079474@28.48.21(+1C)&[MIMAT0004810&hsa-miR-629-5p&offsets\|0\|0(+1C);m-150&15\|-\|70079427\|70079447&offsets\|0\|0(+1C)]\|TGGGTTTACGTTGGGAGAACTC | >MI0003643\|hsa-mir-629&WithFlank&15\|-\|70079366\|70079474 | 28 | 48 | 21(+1C) | TGGGTTTACGTTGGGAGAACTC | BT20 | 15 | 13 | 22 | 133 | 13 | 4.0 | 6.9 | 7.8 | 122.5 | 3.1 |
| >MI0003643\|hsa-mir-629&WithFlank&15\|-\|70079366\|70079474@28.48.21&[MIMAT0004810&hsa-miR-629-5p&offsets\|0\|0;m-150&15\|-\|70079427\|70079447&offsets\|0\|0]\|TGGGTTTACGTTGGGAGAACT | >MI0003643\|hsa-mir-629&WithFlank&15\|-\|70079366\|70079474 | 28 | 48 | 21 | TGGGTTTACGTTGGGAGAACT | BT20 | 29 | 31 | 42 | 401 | 24 | 11.6 | 14.5 | 15.4 | 321.7 | 1.6 |
| >MI0003643\|hsa-mir-629&WithFlank&15\|-\|70079366\|70079474@28.49.22&[MIMAT0004810&hsa-miR-629-5p&offsets\|0\|+1;m-150&15\|-\|70079427\|70079447&offsets\|0\|+1]\|TGGGTTTACGTTGGGAGAACTT | >MI0003643\|hsa-mir-629&WithFlank&15\|-\|70079366\|70079474 | 28 | 49 | 22 | TGGGTTTACGTTGGGAGAACTT | BT20 | 119 | 96 | 168 | 935 | 98 | 37.0 | 45.1 | 37.9 | 762.2 | 15.2 |
| >MI0003643\|hsa-mir-629&WithFlank&15\|-\|70079366\|70079474@28.49.22&[MIMAT0004810&hsa-miR-629-5p&offsets\|0\|+1;m-150&15\|-\|70079427\|70079447&offsets\|0\|+1]\|TGGGTTTACGTTGGGAGAACTT | >MI0003643\|hsa-mir-629&WithFlank&15\|-\|70079366\|70079474 | 28 | 49 | 22 | TGGGTTTACGTTGGGAGAACTT | MB231 | 31 | 19 | 27 | 292 | 31 | 7.8 | 7.0 | 13.8 | 51.5 | 10.0 |
| >MI0003666\|hsa-mir-651&WithFlank&X\|+\|8126959\|8127067@22.43.22&[MIMAT0003321&hsa-miR-651-5p&offsets\|0\|0;m-399&X\|+\|8126980\|8127001&offsets\|0\|0]\|TTTAGGATAAGCTTGACTTTTG | >MI0003666\|hsa-mir-651&WithFlank&X\|+\|8126959\|8127067 | 22 | 43 | 22 | TTTAGGATAAGCTTGACTTTTG | BT20 | 28 | 9 | 12 | 5 | 1 | 28.0 | 0.9 | 7.8 | 6.7 | 1.0 |
| >MI0003666\|hsa-mir-651&WithFlank&X\|+\|8126959\|8127067@22.43.22&[MIMAT0003321&hsa-miR-651-5p&offsets\|0\|0;m-399&X\|+\|8126980\|8127001&offsets\|0\|0]\|TTTAGGATAAGCTTGACTTTTG | >MI0003666\|hsa-mir-651&WithFlank&X\|+\|8126959\|8127067 | 22 | 43 | 22 | TTTAGGATAAGCTTGACTTTTG | MB231 | 17 | 10 | 2 | 14 | 7 | 8.3 | 3.6 | 1.3 | 7.4 | 4.3 |
| >MI0003667\|hsa-mir-652&WithFlank&X\|+\|110055323\|110055432@67.87.21(+1U)&[MIMAT0003322&hsa-miR-652-3p&offsets\|0\|0(+1U);m-148&X\|+\|110055389\|110055410&offsets\|0\|-1(+1U)]\|AATGGCGCCACTAGGGTTGTGT | >MI0003667\|hsa-mir-652&WithFlank&X\|+\|110055323\|110055432 | 67 | 87 | 21(+1U) | AATGGCGCCACTAGGGTTGTGT | BT20 | 45 | 31 | 71 | 131 | 11 | 32.1 | 12.6 | 34.3 | 126.2 | 3.8 |
| >MI0003667\|hsa-mir-652&WithFlank&X\|+\|110055323\|110055432@67.87.21&[MIMAT0003322&hsa-miR-652-3p&offsets\|0\|0;m-148&X\|+\|110055389\|110055410&offsets\|0\|-1]\|AATGGCGCCACTAGGGTTGTG | >MI0003667\|hsa-mir-652&WithFlank&X\|+\|110055323\|110055432 | 67 | 87 | 21 | AATGGCGCCACTAGGGTTGTG | BT20 | 13 | 12 | 21 | 104 | 7 | 7.2 | 5.2 | 6.3 | 101.5 | 0.7 |
| >MI0003674\|hsa-mir-653&WithFlank&7\|-\|93482754\|93482861@22.43.22&[MIMAT0003328&hsa-miR-653-5p&offsets\|+3\|+4;m-368&7\|-\|93482819\|93482840&offsets\|0\|0]\|TTGAAACAATCTCTACTGAACC | >MI0003674\|hsa-mir-653&WithFlank&7\|-\|93482754\|93482861 | 22 | 43 | 22 | TTGAAACAATCTCTACTGAACC | MB231 | 12 | 10 | 6 | 14 | 11 | 1.2 | 3.8 | 5.0 | 10.7 | 5.1 |

| | | | | | | | | | | | | | | | |
|---|---|---|---|---|---|---|---|---|---|---|---|---|---|---|---|
| >MI0003684\|hsa-mir-660&WithFlank&X\|+\|50013235\|50013343@22.44.23&[MIMAT0003338&hsa-miR-660-5p&offsets\|0\|+1;m-157&X\|+\|50013256\|50013278&offsets\|0\|0]\|TACCCATTGCATATCGGAGTTGT | >MI0003684\|hsa-mir-660&WithFlank&X\|+\|50013235\|50013343 | 22 | 44 | 23 | TACCCATTGCATATCGGAGTTGT | BT20 | 18 | 9 | 9 | 23 | 2 | 14.0 | 7.4 | 3.7 | 25.0 | 2.3 |
| >MI0003684\|hsa-mir-660&WithFlank&X\|+\|50013235\|50013343@22.44.23&[MIMAT0003338&hsa-miR-660-5p&offsets\|0\|+1;m-157&X\|+\|50013256\|50013278&offsets\|0\|0]\|TACCCATTGCATATCGGAGTTGT | >MI0003684\|hsa-mir-660&WithFlank&X\|+\|50013235\|50013343 | 22 | 44 | 23 | TACCCATTGCATATCGGAGTTGT | MB231 | 15 | 10 | 3 | 56 | 18 | 1.7 | 3.4 | 2.4 | 5.0 | 12.8 |
| >MI0003686\|hsa-mir-542&WithFlank&X\|-\|134541335\|134541443@58.78.21&[MIMAT0003389&hsa-miR-542-3p&offsets\|-1\|-2;m-136&X\|-\|134541364\|134541385&offsets\|-1\|-2]\|TTGTGACAGATTGATAACTGA | >MI0003686\|hsa-mir-542&WithFlank&X\|-\|134541335\|134541443 | 58 | 78 | 21 | TTGTGACAGATTGATAACTGA | BT20 | 12 | 2 | 1 | 1 | 0 | 16.9 | 1.1 | 1.0 | 1.9 | 0.0 |
| >MI0003686\|hsa-mir-542&WithFlank&X\|-\|134541335\|134541443@58.79.22&[MIMAT0003389&hsa-miR-542-3p&offsets\|-1\|-1;m-136&X\|-\|134541364\|134541385&offsets\|-1\|-1]\|TTGTGACAGATTGATAACTGAA | >MI0003686\|hsa-mir-542&WithFlank&X\|-\|134541335\|134541443 | 58 | 79 | 22 | TTGTGACAGATTGATAACTGAA | BT20 | 11 | 2 | 1 | 0 | 0 | 16.1 | 1.1 | 1.1 | 0.6 | 0.0 |
| >MI0003686\|hsa-mir-542&WithFlank&X\|-\|134541335\|134541443@58.80.23&[MIMAT0003389&hsa-miR-542-3p&offsets\|-1\|0;m-136&X\|-\|134541364\|134541385&offsets\|-1\|0]\|TTGTGACAGATTGATAACTGAAA | >MI0003686\|hsa-mir-542&WithFlank&X\|-\|134541335\|134541443 | 58 | 80 | 23 | TTGTGACAGATTGATAACTGAAA | BT20 | 12 | 2 | 1 | 0 | 0 | 17.4 | 1.0 | 1.1 | 0.0 | 0.0 |
| >MI0003686\|hsa-mir-542&WithFlank&X\|-\|134541335\|134541443@59.78.20&[MIMAT0003389&hsa-miR-542-3p&offsets\|0\|-2;m-136&X\|-\|134541364\|134541385&offsets\|0\|-2]\|TGTGACAGATTGATAACTGA | >MI0003686\|hsa-mir-542&WithFlank&X\|-\|134541335\|134541443 | 59 | 78 | 20 | TGTGACAGATTGATAACTGA | BT20 | 17 | 4 | 3 | 3 | 0 | 23.6 | 2.2 | 2.1 | 4.7 | 0.4 |
| >MI0003686\|hsa-mir-542&WithFlank&X\|-\|134541335\|134541443@59.79.21&[MIMAT0003389&hsa-miR-542-3p&offsets\|0\|-1;m-136&X\|-\|134541364\|134541385&offsets\|0\|-1]\|TGTGACAGATTGATAACTGAA | >MI0003686\|hsa-mir-542&WithFlank&X\|-\|134541335\|134541443 | 59 | 79 | 21 | TGTGACAGATTGATAACTGAA | BT20 | 22 | 4 | 4 | 2 | 0 | 31.4 | 2.2 | 3.3 | 1.4 | 0.2 |
| >MI0003686\|hsa-mir-542&WithFlank&X\|-\|134541335\|134541443@59.80.22&[MIMAT0003389&hsa-miR-542-3p&offsets\|0\|0;m-136&X\|-\|134541364\|134541385&offsets\|0\|0]\|TGTGACAGATTGATAACTGAAA | >MI0003686\|hsa-mir-542&WithFlank&X\|-\|134541335\|134541443 | 59 | 80 | 22 | TGTGACAGATTGATAACTGAAA | BT20 | 69 | 13 | 9 | 3 | 1 | 101.1 | 7.0 | 9.4 | 5.0 | 1.2 |
| >MI0003760\|hsa-mir-671&WithFlank&7\|+\|151238415\|151238544@35.58.24&[MIMAT0003880&hsa-miR-671-5p&offsets\|0\|+1;m-316&7\|+\|151238449\|151238472&offsets\|0\|0]\|AGGAAGCCCTGGAGGGGCTGGAGG | >MI0003760\|hsa-mir-671&WithFlank&7\|+\|151238415\|151238544 | 35 | 58 | 24 | AGGAAGCCCTGGAGGGGCTGGAGG | BT20 | 16 | 10 | 15 | 30 | 20 | 2.2 | 7.1 | 7.4 | 24.3 | 5.3 |
| >MI0003760\|hsa-mir-671&WithFlank&7\|+\|151238415\|151238544@74.94.21&[MIMAT0004819&hsa-miR-671-3p&offsets\|0\|0;m-323&7\|+\|151238488\|151238508&offsets\|0\|0]\|TCCGGTTCTCAGGGCTCCACC | >MI0003760\|hsa-mir-671&WithFlank&7\|+\|151238415\|151238544 | 74 | 94 | 21 | TCCGGTTCTCAGGGCTCCACC | BT20 | 11 | 6 | 17 | 3 | 8 | 11.5 | 1.7 | 7.9 | 2.9 | 4.8 |
| >MI0003763\|hsa-mir-767&WithFlank&X\|-\|152393415\|152393535@33.56.24&[MIMAT0003882&hsa-miR-767-5p&offsets\|0\|+1;m-788&X\|-\|152393481\|152393503&offsets\|0\|+1]\|TGCACCATGGTTGTCTGAGCATGC | >MI0003763\|hsa-mir-767&WithFlank&X\|-\|152393415\|152393535 | 33 | 56 | 24 | TGCACCATGGTTGTCTGAGCATGC | BT20 | 16 | 17 | 41 | 6 | 15 | 16.9 | 9.1 | 28.4 | 4.5 | 12.6 |
| >MI0003776\|hsa-mir-320b-1&WithFlank&1\|+\|116671740\|116671823@48.66.19;MI0003839\|hsa-mir-320b-2&WithFlank&1\|-\|224257034\|224257116@47.65.19;MI0003778\|hsa-mir-320c-1&WithFlank&18\|+\|21683512\|21683595@48.66.19;MI0008191\|hsa-mir-320c-2&WithFlank&18\|+\|24321669\|24321752@48.66.19;MI0000542\|hsa-mir-320a&WithFlank&8\|-\|22244960\|22245043@48.66.19&[MIMAT0005792&hsa-miR-320b&offsets\|0\|-3;m-183&1\|+\|116671787\|116671806&offsets\|0\|-1];[MIMAT0005792_1&hsa-miR-320b&offsets\|0\|-3;m-183&1\|-\|224257051\|224257070&offsets\|0\|-1];[MIMAT0005793&hsa-miR-320c&offsets\|0\|-1;m-222&18\|+\|21683559\|21683578&offsets\|0\|-1];[MIMAT0005793_1&hsa-miR-320c&offsets\|0\|-1;m-219&18\|+\|24321716\|24321735&offsets\|0\|-1];[MIMAT0000510&hsa-miR-320a-3p&offsets\|0\|-3;m-60&8\|-\|22244975\|22244996&offsets\|0\|-3]\|AAAAGCTGGGTTGAGAGGG | >MI0003776\|hsa-mir-320b-1&WithFlank&1\|+\|116671740\|116671823 | 48 | 66 | 19 | AAAAGCTGGGTTGAGAGGG | BT20 | 17 | 18 | 32 | 172 | 10 | 8.3 | 6.4 | 6.5 | 156.6 | 1.9 |
| >MI0003776\|hsa-mir-320b-1&WithFlank&1\|+\|116671740\|116671823@48.66.19(+1A);MI0003839\|hsa-mir-320b-2&WithFlank&1\|-\|224257034\|224257116@47.65.19(+1A);MI0003778\|hsa-mir-320c-1&WithFlank&18\|+\|21683512\|21683595@48.66.19(+1A);MI0008191\|hsa-mir-320c-2&WithFlank&18\|+\|24321669\|24321752@48.66.19(+1A);MI0000542\|hsa-mir-320a&WithFlank&8\|-\|22244960\|22245043@48.66.19(+1A)&[MIMAT0005792&hsa-miR-320b&offsets\|0\|-3(+1A);m-183&1\|+\|116671787\|116671806&offsets\|0\|-1(+1A)];[MIMAT0005792_1&hsa-miR-320b&offsets\|0\|-3(+1A);m-183&1\|-\|224257051\|224257070&offsets\|0\|-1(+1A)];[MIMAT0005793&hsa-miR-320c&offsets\|0\|-1(+1A);m-222&18\|+\|21683559\|21683578&offsets\|0\|-1(+1A)];[MIMAT0005793_1&hsa-miR-320c&offsets\|0\|-1(+1A);m-219&18\|+\|24321716\|24321735&offsets\|0\|-1(+1A)];[MIMAT0000510&hsa-miR-320a-3p&offsets\|0\|-3(+1A);m-60&8\|-\|22244975\|22244996&offsets\|0\|-3(+1A)]\|AAAAGCTGGGTTGAGAGGGA | >MI0003776\|hsa-mir-320b-1&WithFlank&1\|+\|116671740\|116671823 | 48 | 66 | 19(+1A) | AAAAGCTGGGTTGAGAGGGA | BT20 | 17 | 14 | 50 | 121 | 7 | 9.4 | 6.9 | 23.5 | 84.8 | 1.6 |
| >MI0003776\|hsa-mir-320b-1&WithFlank&1\|+\|116671740\|116671823@48.67.20;MI0003839\|hsa-mir-320b-2&WithFlank&1\|-\|224257034\|224257116@47.66.20;MI0000542\|hsa-mir-320a&WithFlank&8\|-\|22244960\|22245043@48.67.20&[MIMAT0005792&hsa-miR-320b&offsets\|0\|-2;m-183&1\|+\|116671787\|116671806&offsets\|0\|0];[MIMAT0005792_1&hsa-miR-320b&offsets\|0\|-2;m-183&1\|-\|224257051\|224257070&offsets\|0\|0];[MIMAT0000510&hsa-miR-320a-3p&offsets\|0\|-2;m-60&8\|-\|22244975\|22244996&offsets\|0\|-2]\|AAAAGCTGGGTTGAGAGGGC | >MI0003776\|hsa-mir-320b-1&WithFlank&1\|+\|116671740\|116671823 | 48 | 67 | 20 | AAAAGCTGGGTTGAGAGGGC | BT20 | 78 | 67 | 248 | 850 | 45 | 59.5 | 22.6 | 34.8 | 549.5 | 9.2 |
| >MI0003776\|hsa-mir-320b-1&WithFlank&1\|+\|116671740\|116671823@48.67.20;MI0003839\|hsa-mir-320b-2&WithFlank&1\|-\|224257034\|224257116@47.66.20;MI0000542\|hsa-mir-320a&WithFlank&8\|-\|22244960\|22245043@48.67.20&[MIMAT0005792&hsa-miR-320b&offsets\|0\|-2;m-183&1\|+\|116671787\|116671806&offsets\|0\|0];[MIMAT0005792_1&hsa-miR-320b&offsets\|0\|-2;m-183&1\|-\|224257051\|224257070&offsets\|0\|0];[MIMAT0000510&hsa-miR-320a-3p&offsets\|0\|-2;m-60&8\|-\|22244975\|22244996&offsets\|0\|-2]\|AAAAGCTGGGTTGAGAGGGC | >MI0003776\|hsa-mir-320b-1&WithFlank&1\|+\|116671740\|116671823 | 48 | 67 | 20 | AAAAGCTGGGTTGAGAGGGC | MB231 | 11 | 12 | 15 | 68 | 5 | 3.9 | 3.2 | 3.1 | 25.5 | 0.7 |
| >MI0003776\|hsa-mir-320b-1&WithFlank&1\|+\|116671740\|116671823@48.67.20(+1U);MI0003839\|hsa-mir-320b-2&WithFlank&1\|-\|224257034\|224257116@47.66.20(+1U);MI0000542\|hsa-mir-320a&WithFlank&8\|-\|22244960\|22245043@48.67.20(+1U)&[MIMAT0005792&hsa-miR-320b&offsets\|0\|-2(+1U);m-183&1\|+\|116671787\|116671806&offsets\|0\|0(+1U)];[MIMAT0005792_1&hsa-miR-320b&offsets\|0\|-2(+1U);m-183&1\|-\|224257051\|224257070&offsets\|0\|0(+1U)];[MIMAT0000510&hsa-miR-320a-3p&offsets\|0\|-2(+1U);m-60&8\|-\|22244975\|22244996&offsets\|0\|-2(+1U)]\|AAAAGCTGGGTTGAGAGGGCT | >MI0003776\|hsa-mir-320b-1&WithFlank&1\|+\|116671740\|116671823 | 48 | 67 | 20(+1U) | AAAAGCTGGGTTGAGAGGGCT | BT20 | 24 | 15 | 68 | 128 | 10 | 13.2 | 5.4 | 24.6 | 89.2 | 2.6 |
| >MI0003776\|hsa-mir-320b-1&WithFlank&1\|+\|116671740\|116671823@48.68.21;MI0003839\|hsa-mir-320b-2&WithFlank&1\|-\|224257034\|224257116@47.67.21&[MIMAT0005792&hsa-miR-320b&offsets\|0\|-1;m-183&1\|+\|116671787\|116671806&offsets\|0\|+1];[MIMAT0005792_1&hsa-miR-320b&offsets\|0\|-1;m-183&1\|-\|224257051\|224257070&offsets\|0\|+1]\|AAAAGCTGGGTTGAGAGGGCA | >MI0003776\|hsa-mir-320b-1&WithFlank&1\|+\|116671740\|116671823 | 48 | 68 | 21 | AAAAGCTGGGTTGAGAGGGCA | BT20 | 72 | 48 | 273 | 318 | 29 | 46.3 | 20.7 | 125.4 | 193.1 | 8.8 |
| >MI0003778\|hsa-mir-320c-1&WithFlank&18\|+\|21683512\|21683595@48.67.20;MI0008191\|hsa-mir-320c-2&WithFlank&18\|+\|24321669\|24321752@48.67.20&[MIMAT0005793&hsa-miR-320c&offsets\|0\|0;m-222&18\|+\|21683559\|21683578&offsets\|0\|0];[MIMAT0005793_1&hsa-miR-320c&offsets\|0\|0;m-219&18\|+\|24321716\|24321735&offsets\|0\|0]\|AAAAGCTGGGTTGAGAGGGT | >MI0003778\|hsa-mir-320c-1&WithFlank&18\|+\|21683512\|21683595 | 48 | 67 | 20 | AAAAGCTGGGTTGAGAGGGT | BT20 | 33 | 27 | 98 | 282 | 13 | 16.0 | 14.7 | 38.4 | 189.6 | 4.0 |
| >MI0003780\|hsa-mir-1296&WithFlank&10\|-\|63372951\|63373054@22.42.21(+1U)&[MIMAT0005794&hsa-miR-1296-5p&offsets\|0\|-1(+1U);m-304&10\|-\|63373012\|63373033&offsets\|0\|-1(+1U)]\|TTAGGGCCCTGGCTCCATCTCT | >MI0003780\|hsa-mir-1296&WithFlank&10\|-\|63372951\|63373054 | 22 | 42 | 21(+1U) | TTAGGGCCCTGGCTCCATCTCT | BT20 | 12 | 5 | 9 | 10 | 2 | 13.4 | 3.5 | 5.0 | 7.0 | 0.7 |
| >MI0003780\|hsa-mir-1296&WithFlank&10\|-\|63372951\|63373054@22.43.22&[MIMAT0005794&hsa-miR-1296-5p&offsets\|0\|0;m-304&10\|-\|63373012\|63373033&offsets\|0\|0]\|TTAGGGCCCTGGCTCCATCTCC | >MI0003780\|hsa-mir-1296&WithFlank&10\|-\|63372951\|63373054 | 22 | 43 | 22 | TTAGGGCCCTGGCTCCATCTCC | BT20 | 21 | 13 | 21 | 24 | 7 | 18.6 | 6.5 | 7.0 | 18.0 | 3.2 |
| >MI0003820\|hsa-mir-454&WithFlank&17\|-\|59137752\|59137878@70.92.23&[MIMAT0003885&hsa-miR-454-3p&offsets\|0\|0;m-272&17\|-\|59137787\|59137809&offsets\|0\|0]\|TAGTGCAATATTGCTTATAGGGT | >MI0003820\|hsa-mir-454&WithFlank&17\|-\|59137752\|59137878 | 70 | 92 | 23 | TAGTGCAATATTGCTTATAGGGT | BT20 | 35 | 19 | 18 | 6 | 2 | 38.4 | 14.6 | 8.5 | 5.4 | 2.0 |
| >MI0003820\|hsa-mir-454&WithFlank&17\|-\|59137752\|59137878@70.93.24&[MIMAT0003885&hsa-miR-454-3p&offsets\|0\|+1;m-272&17\|-\|59137787\|59137809&offsets\|0\|+1]\|TAGTGCAATATTGCTTATAGGGTT | >MI0003820\|hsa-mir-454&WithFlank&17\|-\|59137752\|59137878 | 70 | 93 | 24 | TAGTGCAATATTGCTTATAGGGTT | BT20 | 42 | 34 | 18 | 4 | 4 | 38.7 | 29.3 | 7.9 | 3.4 | 4.2 |

| | | | | | | | | | | | | | | | | |
|---|---|---|---|---|---|---|---|---|---|---|---|---|---|---|---|---|
| >MI0003820\|hsa-mir-454&WithFlank&17\|-\|59137752\|59137878@70.93.24&[MIMAT0003885&hsa-miR-454-3p&offsets\|0\|+1;m-272&17\|-\|59137787\|59137809&offsets\|0\|+1]\|TAGTGCAATATTGCTTATAGGGTT | >MI0003820\|hsa-mir-454&WithFlank&17\|-\|59137752\|59137878 | 70 | 93 | 24 | TAGTGCAATATTGCTTATAGGGTT | MB231 | 25 | 28 | 6 | 76 | 45 | 6.5 | 11.2 | 5.9 | 22.4 | 32.7 |
| >MI0003820\|hsa-mir-454&WithFlank&17\|-\|59137752\|59137878@70.94.25&[MIMAT0003885&hsa-miR-454-3p&offsets\|0\|+2;m-272&17\|-\|59137787\|59137809&offsets\|0\|+2]\|TAGTGCAATATTGCTTATAGGGTTT | >MI0003820\|hsa-mir-454&WithFlank&17\|-\|59137752\|59137878 | 70 | 94 | 25 | TAGTGCAATATTGCTTATAGGGTTT | BT20 | 36 | 23 | 10 | 2 | 3 | 35.7 | 22.2 | 5.4 | 1.5 | 2.9 |
| >MI0003820\|hsa-mir-454&WithFlank&17\|-\|59137752\|59137878@70.94.25&[MIMAT0003885&hsa-miR-454-3p&offsets\|0\|+2;m-272&17\|-\|59137787\|59137809&offsets\|0\|+2]\|TAGTGCAATATTGCTTATAGGGTTT | >MI0003820\|hsa-mir-454&WithFlank&17\|-\|59137752\|59137878 | 70 | 94 | 25 | TAGTGCAATATTGCTTATAGGGTTT | MB231 | 35 | 31 | 5 | 76 | 75 | 9.3 | 12.9 | 4.1 | 19.8 | 58.6 |
| >MI0003834\|hsa-mir-769&WithFlank&19\|+\|46018926\|46019055@36.55.20(+1U)&[MIMAT0003886&hsa-miR-769-5p&offsets\|0\|-2(+1U);m-216&19\|+\|46018961\|46018982&offsets\|0\|-2(+1U)]\|TGAGACCTCTGGGTTCTGAGT | >MI0003834\|hsa-mir-769&WithFlank&19\|+\|46018926\|46019055 | 36 | 55 | 20(+1U) | TGAGACCTCTGGGTTCTGAGT | BT20 | 14 | 3 | 4 | 7 | 1 | 16.4 | 1.6 | 2.4 | 9.2 | 0.7 |
| >MI0003834\|hsa-mir-769&WithFlank&19\|+\|46018926\|46019055@36.56.21(+1A)&[MIMAT0003886&hsa-miR-769-5p&offsets\|0\|-1(+1A);m-216&19\|+\|46018961\|46018982&offsets\|0\|-1(+1A)]\|TGAGACCTCTGGGTTCTGAGCA | >MI0003834\|hsa-mir-769&WithFlank&19\|+\|46018926\|46019055 | 36 | 56 | 21(+1A) | TGAGACCTCTGGGTTCTGAGCA | BT20 | 27 | 6 | 12 | 11 | 2 | 33.2 | 3.1 | 10.7 | 12.8 | 0.6 |
| >MI0003834\|hsa-mir-769&WithFlank&19\|+\|46018926\|46019055@36.56.21(+1C)&[MIMAT0003886&hsa-miR-769-5p&offsets\|0\|-1(+1C);m-216&19\|+\|46018961\|46018982&offsets\|0\|-1(+1C)]\|TGAGACCTCTGGGTTCTGAGCC | >MI0003834\|hsa-mir-769&WithFlank&19\|+\|46018926\|46019055 | 36 | 56 | 21(+1C) | TGAGACCTCTGGGTTCTGAGCC | BT20 | 22 | 9 | 11 | 15 | 4 | 20.9 | 4.1 | 6.8 | 16.8 | 0.7 |
| >MI0003834\|hsa-mir-769&WithFlank&19\|+\|46018926\|46019055@36.56.21(+1G)&[MIMAT0003886&hsa-miR-769-5p&offsets\|0\|-1(+1G);m-216&19\|+\|46018961\|46018982&offsets\|0\|-1(+1G)]\|TGAGACCTCTGGGTTCTGAGCG | >MI0003834\|hsa-mir-769&WithFlank&19\|+\|46018926\|46019055 | 36 | 56 | 21(+1G) | TGAGACCTCTGGGTTCTGAGCG | BT20 | 21 | 2 | 3 | 3 | 0 | 25.0 | 0.4 | 2.4 | 4.2 | 0.3 |
| >MI0003834\|hsa-mir-769&WithFlank&19\|+\|46018926\|46019055@36.56.21&[MIMAT0003886&hsa-miR-769-5p&offsets\|0\|-1;m-216&19\|+\|46018961\|46018982&offsets\|0\|-1]\|TGAGACCTCTGGGTTCTGAGC | >MI0003834\|hsa-mir-769&WithFlank&19\|+\|46018926\|46019055 | 36 | 56 | 21 | TGAGACCTCTGGGTTCTGAGC | BT20 | 34 | 14 | 14 | 25 | 5 | 32.2 | 6.2 | 5.9 | 26.1 | 0.3 |
| >MI0003834\|hsa-mir-769&WithFlank&19\|+\|46018926\|46019055@36.56.21&[MIMAT0003886&hsa-miR-769-5p&offsets\|0\|-1;m-216&19\|+\|46018961\|46018982&offsets\|0\|-1]\|TGAGACCTCTGGGTTCTGAGC | >MI0003834\|hsa-mir-769&WithFlank&19\|+\|46018926\|46019055 | 36 | 56 | 21 | TGAGACCTCTGGGTTCTGAGC | MB231 | 17 | 13 | 6 | 48 | 15 | 5.7 | 6.7 | 5.3 | 10.9 | 10.6 |
| >MI0003834\|hsa-mir-769&WithFlank&19\|+\|46018926\|46019055@36.57.22(+1U)&[MIMAT0003886&hsa-miR-769-5p&offsets\|0\|0(+1U);m-216&19\|+\|46018961\|46018982&offsets\|0\|0(+1U)]\|TGAGACCTCTGGGTTCTGAGCTT | >MI0003834\|hsa-mir-769&WithFlank&19\|+\|46018926\|46019055 | 36 | 57 | 22(+1U) | TGAGACCTCTGGGTTCTGAGCTT | BT20 | 13 | 6 | 8 | 24 | 2 | 12.6 | 2.3 | 4.8 | 26.8 | 0.5 |
| >MI0003834\|hsa-mir-769&WithFlank&19\|+\|46018926\|46019055@36.57.22(+2U)&[MIMAT0003886&hsa-miR-769-5p&offsets\|0\|0(+2U);m-216&19\|+\|46018961\|46018982&offsets\|0\|0(+2U)]\|TGAGACCTCTGGGTTCTGAGCTTT | >MI0003834\|hsa-mir-769&WithFlank&19\|+\|46018926\|46019055 | 36 | 57 | 22(+2U) | TGAGACCTCTGGGTTCTGAGCTTT | BT20 | 11 | 5 | 8 | 23 | 1 | 10.3 | 1.9 | 4.6 | 22.4 | 1.2 |
| >MI0003834\|hsa-mir-769&WithFlank&19\|+\|46018926\|46019055@36.57.22&[MIMAT0003886&hsa-miR-769-5p&offsets\|0\|0;m-216&19\|+\|46018961\|46018982&offsets\|0\|0]\|TGAGACCTCTGGGTTCTGAGCT | >MI0003834\|hsa-mir-769&WithFlank&19\|+\|46018926\|46019055 | 36 | 57 | 22 | TGAGACCTCTGGGTTCTGAGCT | BT20 | 139 | 53 | 71 | 98 | 25 | 134.3 | 21.7 | 36.7 | 98.6 | 4.0 |
| >MI0003834\|hsa-mir-769&WithFlank&19\|+\|46018926\|46019055@36.57.22&[MIMAT0003886&hsa-miR-769-5p&offsets\|0\|0;m-216&19\|+\|46018961\|46018982&offsets\|0\|0]\|TGAGACCTCTGGGTTCTGAGCT | >MI0003834\|hsa-mir-769&WithFlank&19\|+\|46018926\|46019055 | 36 | 57 | 22 | TGAGACCTCTGGGTTCTGAGCT | MB231 | 66 | 43 | 25 | 160 | 60 | 26.4 | 22.9 | 24.7 | 40.1 | 48.7 |
| >MI0005531\|hsa-mir-450b&WithFlank&X\|-\|134540179\|134540268@17.37.21&[MIMAT0004909&hsa-miR-450b-5p&offsets\|0\|-1;m-266&X\|-\|134540232\|134540252&offsets\|0\|0]\|TTTTGCAATATGTTCCTGAAT | >MI0005531\|hsa-mir-450b&WithFlank&X\|-\|134540179\|134540268 | 17 | 37 | 21 | TTTTGCAATATGTTCCTGAAT | BT20 | 72 | 23 | 31 | 10 | 3 | 92.0 | 8.9 | 23.8 | 13.2 | 0.4 |
| >MI0005531\|hsa-mir-450b&WithFlank&X\|-\|134540179\|134540268@17.37.21&[MIMAT0004909&hsa-miR-450b-5p&offsets\|0\|-1;m-266&X\|-\|134540232\|134540252&offsets\|0\|0]\|TTTTGCAATATGTTCCTGAAT | >MI0005531\|hsa-mir-450b&WithFlank&X\|-\|134540179\|134540268 | 17 | 37 | 21 | TTTTGCAATATGTTCCTGAAT | MB231 | 13 | 9 | 3 | 14 | 8 | 3.9 | 2.7 | 1.6 | 1.7 | 7.7 |
| >MI0005531\|hsa-mir-450b&WithFlank&X\|-\|134540179\|134540268@17.38.22&[MIMAT0004909&hsa-miR-450b-5p&offsets\|0\|0;m-266&X\|-\|134540232\|134540252&offsets\|0\|+1]\|TTTTGCAATATGTTCCTGAATA | >MI0005531\|hsa-mir-450b&WithFlank&X\|-\|134540179\|134540268 | 17 | 38 | 22 | TTTTGCAATATGTTCCTGAATA | BT20 | 31 | 10 | 15 | 3 | 3 | 34.5 | 3.8 | 9.1 | 2.1 | 0.9 |
| >MI0005559\|hsa-mir-744&WithFlank&17\|+\|12081893\|12082002@17.36.20(+1U)&[MIMAT0004945&hsa-miR-744-5p&offsets\|0\|-2(+1U);m-172&17\|+\|12081909\|12081929&offsets\|0\|-1(+1U)]\|TGCGGGGCTAGGGCTAACAGT | >MI0005559\|hsa-mir-744&WithFlank&17\|+\|12081893\|12082002 | 17 | 36 | 20(+1U) | TGCGGGGCTAGGGCTAACAGT | BT20 | 23 | 19 | 40 | 51 | 35 | 11.6 | 10.2 | 2.3 | 30.0 | 11.7 |
| >MI0005559\|hsa-mir-744&WithFlank&17\|+\|12081893\|12082002@17.37.21&[MIMAT0004945&hsa-miR-744-5p&offsets\|0\|-1;m-172&17\|+\|12081909\|12081929&offsets\|0\|0]\|TGCGGGGCTAGGGCTAACAGC | >MI0005559\|hsa-mir-744&WithFlank&17\|+\|12081893\|12082002 | 17 | 37 | 21 | TGCGGGGCTAGGGCTAACAGC | BT20 | 95 | 83 | 182 | 229 | 184 | 72.0 | 38.1 | 56.0 | 54.0 | 47.5 |
| >MI0005559\|hsa-mir-744&WithFlank&17\|+\|12081893\|12082002@17.37.21&[MIMAT0004945&hsa-miR-744-5p&offsets\|0\|-1;m-172&17\|+\|12081909\|12081929&offsets\|0\|0]\|TGCGGGGCTAGGGCTAACAGC | >MI0005559\|hsa-mir-744&WithFlank&17\|+\|12081893\|12082002 | 17 | 37 | 21 | TGCGGGGCTAGGGCTAACAGC | MB231 | 55 | 45 | 61 | 71 | 30 | 54.3 | 14.3 | 27.9 | 45.3 | 4.1 |
| >MI0005559\|hsa-mir-744&WithFlank&17\|+\|12081893\|12082002@17.38.22&[MIMAT0004945&hsa-miR-744-5p&offsets\|0\|0;m-172&17\|+\|12081909\|12081929&offsets\|0\|+1]\|TGCGGGGCTAGGGCTAACAGCA | >MI0005559\|hsa-mir-744&WithFlank&17\|+\|12081893\|12082002 | 17 | 38 | 22 | TGCGGGGCTAGGGCTAACAGCA | BT20 | 42 | 34 | 49 | 45 | 49 | 18.8 | 17.0 | 6.6 | 29.7 | 3.1 |
| >MI0005559\|hsa-mir-744&WithFlank&17\|+\|12081893\|12082002@17.38.22&[MIMAT0004945&hsa-miR-744-5p&offsets\|0\|0;m-172&17\|+\|12081909\|12081929&offsets\|0\|+1]\|TGCGGGGCTAGGGCTAACAGCA | >MI0005559\|hsa-mir-744&WithFlank&17\|+\|12081893\|12082002 | 17 | 38 | 22 | TGCGGGGCTAGGGCTAACAGCA | MB231 | 21 | 13 | 10 | 39 | 16 | 17.6 | 3.3 | 5.7 | 24.8 | 4.2 |
| >MI0005566\|hsa-mir-374b&WithFlank&X\|-\|74218541\|74218624@17.36.20(+1C)&[MIMAT0004955&hsa-miR-374b-5p&offsets\|0\|-2(+1C);m-153&X\|-\|74218587\|74218608&offsets\|0\|-2(+1C)]\|ATATAATACAACCTGCTAAGC | >MI0005566\|hsa-mir-374b&WithFlank&X\|-\|74218541\|74218624 | 17 | 36 | 20(+1C) | ATATAATACAACCTGCTAAGC | BT20 | 11 | 4 | 5 | 1 | 2 | 11.6 | 2.3 | 1.9 | 1.2 | 2.2 |
| >MI0005566\|hsa-mir-374b&WithFlank&X\|-\|74218541\|74218624@17.37.21(+1A)&[MIMAT0004955&hsa-miR-374b-5p&offsets\|0\|-1(+1A);m-153&X\|-\|74218587\|74218608&offsets\|0\|-1(+1A)]\|ATATAATACAACCTGCTAAGTA | >MI0005566\|hsa-mir-374b&WithFlank&X\|-\|74218541\|74218624 | 17 | 37 | 21(+1A) | ATATAATACAACCTGCTAAGTA | BT20 | 22 | 5 | 5 | 1 | 2 | 30.8 | 3.0 | 2.7 | 1.5 | 2.0 |
| >MI0005566\|hsa-mir-374b&WithFlank&X\|-\|74218541\|74218624@17.37.21&[MIMAT0004955&hsa-miR-374b-5p&offsets\|0\|-1;m-153&X\|-\|74218587\|74218608&offsets\|0\|-1]\|ATATAATACAACCTGCTAAGT | >MI0005566\|hsa-mir-374b&WithFlank&X\|-\|74218541\|74218624 | 17 | 37 | 21 | ATATAATACAACCTGCTAAGT | BT20 | 88 | 33 | 35 | 7 | 19 | 95.9 | 15.7 | 10.7 | 6.7 | 15.0 |
| >MI0005566\|hsa-mir-374b&WithFlank&X\|-\|74218541\|74218624@17.37.21&[MIMAT0004955&hsa-miR-374b-5p&offsets\|0\|-1;m-153&X\|-\|74218587\|74218608&offsets\|0\|-1]\|ATATAATACAACCTGCTAAGT | >MI0005566\|hsa-mir-374b&WithFlank&X\|-\|74218541\|74218624 | 17 | 37 | 21 | ATATAATACAACCTGCTAAGT | MB231 | 40 | 44 | 8 | 44 | 80 | 19.7 | 18.9 | 6.8 | 27.4 | 63.8 |
| >MI0005566\|hsa-mir-374b&WithFlank&X\|-\|74218541\|74218624@17.38.22&[MIMAT0004955&hsa-miR-374b-5p&offsets\|0\|0;m-153&X\|-\|74218587\|74218608&offsets\|0\|0]\|ATATAATACAACCTGCTAAGTG | >MI0005566\|hsa-mir-374b&WithFlank&X\|-\|74218541\|74218624 | 17 | 38 | 22 | ATATAATACAACCTGCTAAGTG | BT20 | 79 | 35 | 27 | 11 | 22 | 85.8 | 13.2 | 5.4 | 10.2 | 13.3 |
| >MI0005566\|hsa-mir-374b&WithFlank&X\|-\|74218541\|74218624@17.38.22&[MIMAT0004955&hsa-miR-374b-5p&offsets\|0\|0;m-153&X\|-\|74218587\|74218608&offsets\|0\|0]\|ATATAATACAACCTGCTAAGTG | >MI0005566\|hsa-mir-374b&WithFlank&X\|-\|74218541\|74218624 | 17 | 38 | 22 | ATATAATACAACCTGCTAAGTG | MB231 | 52 | 54 | 9 | 87 | 118 | 25.0 | 29.8 | 8.2 | 46.1 | 87.3 |
| >MI0005566\|hsa-mir-374b&WithFlank&X\|-\|74218541\|74218624@17.39.23&[MIMAT0004955&hsa-miR-374b-5p&offsets\|0\|+1;m-153&X\|-\|74218587\|74218608&offsets\|0\|+1]\|ATATAATACAACCTGCTAAGTGT | >MI0005566\|hsa-mir-374b&WithFlank&X\|-\|74218541\|74218624 | 17 | 39 | 23 | ATATAATACAACCTGCTAAGTGT | BT20 | 11 | 4 | 3 | 2 | 2 | 12.0 | 3.5 | 0.6 | 1.2 | 1.2 |
| >MI0005566\|hsa-mir-374b&WithFlank&X\|-\|74218541\|74218624@17.39.23&[MIMAT0004955&hsa-miR-374b-5p&offsets\|0\|+1;m-153&X\|-\|74218587\|74218608&offsets\|0\|+1]\|ATATAATACAACCTGCTAAGTGT | >MI0005566\|hsa-mir-374b&WithFlank&X\|-\|74218541\|74218624 | 17 | 39 | 23 | ATATAATACAACCTGCTAAGTGT | MB231 | 15 | 16 | 1 | 42 | 28 | 8.0 | 11.3 | 1.3 | 20.6 | 24.3 |
| >MI0005756\|hsa-mir-934&WithFlank&X\|+\|136550872\|136550966@21.40.20(+1A)&[MIMAT0004977&hsa-miR-934&offsets\|0\|-2(+1A);m-943&X\|+\|136550892\|136550913&offsets\|0\|-2(+1A)]\|TGTCTACTACTGGAGACACTA | >MI0005756\|hsa-mir-934&WithFlank&X\|+\|136550872\|136550966 | 21 | 40 | 20(+1A) | TGTCTACTACTGGAGACACTA | BT20 | 66 | 14 | 10 | 61 | 11 | 83.2 | 9.6 | 5.8 | 87.9 | 3.1 |
| >MI0005756\|hsa-mir-934&WithFlank&X\|+\|136550872\|136550966@21.40.20&[MIMAT0004977&hsa-miR-934&offsets\|0\|-2;m-943&X\|+\|136550892\|136550913&offsets\|0\|-2]\|TGTCTACTACTGGAGACACT | >MI0005756\|hsa-mir-934&WithFlank&X\|+\|136550872\|136550966 | 21 | 40 | 20 | TGTCTACTACTGGAGACACT | BT20 | 25 | 13 | 4 | 15 | 7 | 27.1 | 15.0 | 2.6 | 16.0 | 5.4 |

<table>
<tr><td>>MI0005756|hsa-mir-934&WithFlank&X|+|136550872|136550966@21.41.21(+1A)&[MIMAT0004977&hsa-miR-934&offsets|0|-1(+1A);m-943&X|+|136550892|136550913&offsets|0|-1(+1A)]|TGTCTACTACTGGAGACACTGA</td><td>>MI0005756|hsa-mir-934&WithFlank&X|+|136550872|136550966</td><td>21</td><td>41</td><td>21(+1A)</td><td>TGTCTACTACTGGAGACACTGA</td><td>BT20</td><td>158</td><td>27</td><td>12</td><td>21</td><td>11</td><td>242.4</td><td>26.8</td><td>4.6</td><td>28.0</td><td>9.4</td></tr>
<tr><td>>MI0005756|hsa-mir-934&WithFlank&X|+|136550872|136550966@21.41.21(+1U)&[MIMAT0004977&hsa-miR-934&offsets|0|-1(+1U);m-943&X|+|136550892|136550913&offsets|0|-1(+1U)]|TGTCTACTACTGGAGACACTGT</td><td>>MI0005756|hsa-mir-934&WithFlank&X|+|136550872|136550966</td><td>21</td><td>41</td><td>21(+1U)</td><td>TGTCTACTACTGGAGACACTGT</td><td>BT20</td><td>53</td><td>14</td><td>7</td><td>14</td><td>9</td><td>65.2</td><td>16.1</td><td>4.0</td><td>15.0</td><td>9.6</td></tr>
<tr><td>>MI0005756|hsa-mir-934&WithFlank&X|+|136550872|136550966@21.41.21&[MIMAT0004977&hsa-miR-934&offsets|0|-1;m-943&X|+|136550892|136550913&offsets|0|-1]|TGTCTACTACTGGAGACACTG</td><td>>MI0005756|hsa-mir-934&WithFlank&X|+|136550872|136550966</td><td>21</td><td>41</td><td>21</td><td>TGTCTACTACTGGAGACACTG</td><td>BT20</td><td>283</td><td>100</td><td>51</td><td>407</td><td>93</td><td>265.3</td><td>57.5</td><td>14.8</td><td>523.9</td><td>27.2</td></tr>
<tr><td>>MI0005756|hsa-mir-934&WithFlank&X|+|136550872|136550966@21.42.22(+1A)&[MIMAT0004977&hsa-miR-934&offsets|0|0(+1A);m-943&X|+|136550892|136550913&offsets|0|0(+1A)]|TGTCTACTACTGGAGACACTGGA</td><td>>MI0005756|hsa-mir-934&WithFlank&X|+|136550872|136550966</td><td>21</td><td>42</td><td>22(+1A)</td><td>TGTCTACTACTGGAGACACTGGA</td><td>BT20</td><td>330</td><td>109</td><td>81</td><td>40</td><td>48</td><td>422.1</td><td>79.1</td><td>20.9</td><td>42.2</td><td>32.0</td></tr>
<tr><td>>MI0005756|hsa-mir-934&WithFlank&X|+|136550872|136550966@21.42.22(+1C)&[MIMAT0004977&hsa-miR-934&offsets|0|0(+1C);m-943&X|+|136550892|136550913&offsets|0|0(+1C)]|TGTCTACTACTGGAGACACTGGC</td><td>>MI0005756|hsa-mir-934&WithFlank&X|+|136550872|136550966</td><td>21</td><td>42</td><td>22(+1C)</td><td>TGTCTACTACTGGAGACACTGGC</td><td>BT20</td><td>18</td><td>13</td><td>12</td><td>9</td><td>13</td><td>10.2</td><td>12.6</td><td>8.3</td><td>7.1</td><td>13.7</td></tr>
<tr><td>>MI0005756|hsa-mir-934&WithFlank&X|+|136550872|136550966@21.42.22(+1G)&[MIMAT0004977&hsa-miR-934&offsets|0|0(+1G);m-943&X|+|136550892|136550913&offsets|0|0(+1G)]|TGTCTACTACTGGAGACACTGGG</td><td>>MI0005756|hsa-mir-934&WithFlank&X|+|136550872|136550966</td><td>21</td><td>42</td><td>22(+1G)</td><td>TGTCTACTACTGGAGACACTGGG</td><td>BT20</td><td>39</td><td>6</td><td>4</td><td>2</td><td>1</td><td>51.0</td><td>5.0</td><td>1.8</td><td>2.4</td><td>1.5</td></tr>
<tr><td>>MI0005756|hsa-mir-934&WithFlank&X|+|136550872|136550966@21.42.22(+2A)&[MIMAT0004977&hsa-miR-934&offsets|0|0(+2A);m-943&X|+|136550892|136550913&offsets|0|0(+2A)]|TGTCTACTACTGGAGACACTGGAA</td><td>>MI0005756|hsa-mir-934&WithFlank&X|+|136550872|136550966</td><td>21</td><td>42</td><td>22(+2A)</td><td>TGTCTACTACTGGAGACACTGGAA</td><td>BT20</td><td>78</td><td>37</td><td>26</td><td>7</td><td>12</td><td>99.1</td><td>27.4</td><td>10.6</td><td>6.3</td><td>8.0</td></tr>
<tr><td>>MI0005756|hsa-mir-934&WithFlank&X|+|136550872|136550966@21.42.22(+3A)&[MIMAT0004977&hsa-miR-934&offsets|0|0(+3A);m-943&X|+|136550892|136550913&offsets|0|0(+3A)]|TGTCTACTACTGGAGACACTGGAAA</td><td>>MI0005756|hsa-mir-934&WithFlank&X|+|136550872|136550966</td><td>21</td><td>42</td><td>22(+3A)</td><td>TGTCTACTACTGGAGACACTGGAAA</td><td>BT20</td><td>18</td><td>9</td><td>6</td><td>1</td><td>2</td><td>23.2</td><td>5.9</td><td>3.8</td><td>0.8</td><td>2.4</td></tr>
<tr><td>>MI0005756|hsa-mir-934&WithFlank&X|+|136550872|136550966@21.42.22&[MIMAT0004977&hsa-miR-934&offsets|0|0;m-943&X|+|136550892|136550913&offsets|0|0]|TGTCTACTACTGGAGACACTGG</td><td>>MI0005756|hsa-mir-934&WithFlank&X|+|136550872|136550966</td><td>21</td><td>42</td><td>22</td><td>TGTCTACTACTGGAGACACTGG</td><td>BT20</td><td>539</td><td>146</td><td>60</td><td>123</td><td>79</td><td>735.5</td><td>128.8</td><td>14.9</td><td>148.9</td><td>67.2</td></tr>
<tr><td>>MI0005756|hsa-mir-934&WithFlank&X|+|136550872|136550966@21.43.23(+1U)&[MIMAT0004977&hsa-miR-934&offsets|0|+1(+1U);m-943&X|+|136550892|136550913&offsets|0|+1(+1U)]|TGTCTACTACTGGAGACACTGGTT</td><td>>MI0005756|hsa-mir-934&WithFlank&X|+|136550872|136550966</td><td>21</td><td>43</td><td>23(+1U)</td><td>TGTCTACTACTGGAGACACTGGTT</td><td>BT20</td><td>11</td><td>6</td><td>4</td><td>7</td><td>7</td><td>8.6</td><td>7.2</td><td>3.1</td><td>5.9</td><td>9.9</td></tr>
<tr><td>>MI0005756|hsa-mir-934&WithFlank&X|+|136550872|136550966@21.43.23&[MIMAT0004977&hsa-miR-934&offsets|0|+1;m-943&X|+|136550892|136550913&offsets|0|+1]|TGTCTACTACTGGAGACACTGGT</td><td>>MI0005756|hsa-mir-934&WithFlank&X|+|136550872|136550966</td><td>21</td><td>43</td><td>23</td><td>TGTCTACTACTGGAGACACTGGT</td><td>BT20</td><td>142</td><td>86</td><td>98</td><td>67</td><td>91</td><td>79.1</td><td>78.3</td><td>75.5</td><td>49.5</td><td>98.1</td></tr>
<tr><td>>MI0006273|hsa-mir-1180&WithFlank&17|-|19344500|19344580@47.66.20&[MIMAT0005825&hsa-miR-1180-3p&offsets|0|-2;m-214&17|-|19344513|19344534&offsets|0|-2]|TTTCCGGCTCGCGTGGGTGT</td><td>>MI0006273|hsa-mir-1180&WithFlank&17|-|19344500|19344580</td><td>47</td><td>66</td><td>20</td><td>TTTCCGGCTCGCGTGGGTGT</td><td>BT20</td><td>19</td><td>18</td><td>53</td><td>36</td><td>16</td><td>1.3</td><td>1.8</td><td>28.5</td><td>22.6</td><td>7.6</td></tr>
<tr><td>>MI0006273|hsa-mir-1180&WithFlank&17|-|19344500|19344580@47.67.21(+1C)&[MIMAT0005825&hsa-miR-1180-3p&offsets|0|-1(+1C);m-214&17|-|19344513|19344534&offsets|0|-1(+1C)]|TTTCCGGCTCGCGTGGGTGTGC</td><td>>MI0006273|hsa-mir-1180&WithFlank&17|-|19344500|19344580</td><td>47</td><td>67</td><td>21(+1C)</td><td>TTTCCGGCTCGCGTGGGTGTGC</td><td>BT20</td><td>12</td><td>11</td><td>29</td><td>19</td><td>9</td><td>1.5</td><td>2.4</td><td>13.6</td><td>14.6</td><td>4.3</td></tr>
<tr><td>>MI0006273|hsa-mir-1180&WithFlank&17|-|19344500|19344580@47.68.22&[MIMAT0005825&hsa-miR-1180-3p&offsets|0|0;m-214&17|-|19344513|19344534&offsets|0|0]|TTTCCGGCTCGCGTGGGTGTGT</td><td>>MI0006273|hsa-mir-1180&WithFlank&17|-|19344500|19344580</td><td>47</td><td>68</td><td>22</td><td>TTTCCGGCTCGCGTGGGTGTGT</td><td>BT20</td><td>82</td><td>65</td><td>197</td><td>128</td><td>60</td><td>16.3</td><td>11.5</td><td>88.7</td><td>78.6</td><td>24.8</td></tr>
<tr><td>>MI0006273|hsa-mir-1180&WithFlank&17|-|19344500|19344580@47.68.22&[MIMAT0005825&hsa-miR-1180-3p&offsets|0|0;m-214&17|-|19344513|19344534&offsets|0|0]|TTTCCGGCTCGCGTGGGTGTGT</td><td>>MI0006273|hsa-mir-1180&WithFlank&17|-|19344500|19344580</td><td>47</td><td>68</td><td>22</td><td>TTTCCGGCTCGCGTGGGTGTGT</td><td>MB231</td><td>26</td><td>24</td><td>21</td><td>43</td><td>21</td><td>14.9</td><td>2.4</td><td>11.0</td><td>15.7</td><td>6.9</td></tr>
<tr><td>>MI0006444|hsa-mir-1307&WithFlank&10|-|103394247|103394407@86.107.22(+1A)&[MIMAT0005951&hsa-miR-1307-3p&offsets|0|0(+1A);m-66&10|-|103394300|103394322&offsets|0|-1(+1A)]|ACTCGGCGTGGCGTCGGTCGTGA</td><td>>MI0006444|hsa-mir-1307&WithFlank&10|-|103394247|103394407</td><td>86</td><td>107</td><td>22(+1A)</td><td>ACTCGGCGTGGCGTCGGTCGTGA</td><td>BT20</td><td>34</td><td>19</td><td>67</td><td>17</td><td>27</td><td>17.4</td><td>8.8</td><td>11.6</td><td>5.3</td><td>11.5</td></tr>
<tr><td>>MI0006444|hsa-mir-1307&WithFlank&10|-|103394247|103394407@86.107.22(+1A)&[MIMAT0005951&hsa-miR-1307-3p&offsets|0|0(+1A);m-66&10|-|103394300|103394322&offsets|0|-1(+1A)]|ACTCGGCGTGGCGTCGGTCGTGA</td><td>>MI0006444|hsa-mir-1307&WithFlank&10|-|103394247|103394407</td><td>86</td><td>107</td><td>22(+1A)</td><td>ACTCGGCGTGGCGTCGGTCGTGA</td><td>MB231</td><td>35</td><td>24</td><td>24</td><td>21</td><td>27</td><td>17.6</td><td>3.4</td><td>9.0</td><td>13.4</td><td>7.4</td></tr>
<tr><td>>MI0006444|hsa-mir-1307&WithFlank&10|-|103394247|103394407@86.107.22(+1U)&[MIMAT0005951&hsa-miR-1307-3p&offsets|0|0(+1U);m-66&10|-|103394300|103394322&offsets|0|-1(+1U)]|ACTCGGCGTGGCGTCGGTCGTGT</td><td>>MI0006444|hsa-mir-1307&WithFlank&10|-|103394247|103394407</td><td>86</td><td>107</td><td>22(+1U)</td><td>ACTCGGCGTGGCGTCGGTCGTGT</td><td>BT20</td><td>39</td><td>23</td><td>62</td><td>20</td><td>29</td><td>21.3</td><td>12.3</td><td>25.6</td><td>11.3</td><td>18.6</td></tr>
<tr><td>>MI0006444|hsa-mir-1307&WithFlank&10|-|103394247|103394407@86.107.22(+1U)&[MIMAT0005951&hsa-miR-1307-3p&offsets|0|0(+1U);m-66&10|-|103394300|103394322&offsets|0|-1(+1U)]|ACTCGGCGTGGCGTCGGTCGTGT</td><td>>MI0006444|hsa-mir-1307&WithFlank&10|-|103394247|103394407</td><td>86</td><td>107</td><td>22(+1U)</td><td>ACTCGGCGTGGCGTCGGTCGTGT</td><td>MB231</td><td>61</td><td>44</td><td>33</td><td>40</td><td>59</td><td>20.5</td><td>15.8</td><td>16.6</td><td>16.1</td><td>36.3</td></tr>
<tr><td>>MI0006444|hsa-mir-1307&WithFlank&10|-|103394247|103394407@86.107.22&[MIMAT0005951&hsa-miR-1307-3p&offsets|0|0;m-66&10|-|103394300|103394322&offsets|0|-1]|ACTCGGCGTGGCGTCGGTCGTG</td><td>>MI0006444|hsa-mir-1307&WithFlank&10|-|103394247|103394407</td><td>86</td><td>107</td><td>22</td><td>ACTCGGCGTGGCGTCGGTCGTG</td><td>BT20</td><td>25</td><td>17</td><td>39</td><td>20</td><td>24</td><td>19.5</td><td>7.8</td><td>20.4</td><td>8.5</td><td>8.2</td></tr>
<tr><td>>MI0006444|hsa-mir-1307&WithFlank&10|-|103394247|103394407@86.107.22&[MIMAT0005951&hsa-miR-1307-3p&offsets|0|0;m-66&10|-|103394300|103394322&offsets|0|-1]|ACTCGGCGTGGCGTCGGTCGTG</td><td>>MI0006444|hsa-mir-1307&WithFlank&10|-|103394247|103394407</td><td>86</td><td>107</td><td>22</td><td>ACTCGGCGTGGCGTCGGTCGTG</td><td>MB231</td><td>54</td><td>42</td><td>27</td><td>33</td><td>42</td><td>35.6</td><td>11.4</td><td>9.3</td><td>14.2</td><td>3.9</td></tr>
<tr><td>>MI0006444|hsa-mir-1307&WithFlank&10|-|103394247|103394407@86.108.23(+1A)&[MIMAT0005951&hsa-miR-1307-3p&offsets|0|+1(+1A);m-66&10|-|103394300|103394322&offsets|0|0(+1A)]|ACTCGGCGTGGCGTCGGTCGTGGA</td><td>>MI0006444|hsa-mir-1307&WithFlank&10|-|103394247|103394407</td><td>86</td><td>108</td><td>23(+1A)</td><td>ACTCGGCGTGGCGTCGGTCGTGGA</td><td>BT20</td><td>14</td><td>9</td><td>26</td><td>5</td><td>15</td><td>12.5</td><td>4.6</td><td>15.6</td><td>2.3</td><td>7.4</td></tr>
<tr><td>>MI0006444|hsa-mir-1307&WithFlank&10|-|103394247|103394407@86.108.23(+1A)&[MIMAT0005951&hsa-miR-1307-3p&offsets|0|+1(+1A);m-66&10|-|103394300|103394322&offsets|0|0(+1A)]|ACTCGGCGTGGCGTCGGTCGTGGA</td><td>>MI0006444|hsa-mir-1307&WithFlank&10|-|103394247|103394407</td><td>86</td><td>108</td><td>23(+1A)</td><td>ACTCGGCGTGGCGTCGGTCGTGGA</td><td>MB231</td><td>11</td><td>9</td><td>12</td><td>3</td><td>9</td><td>7.9</td><td>3.2</td><td>5.7</td><td>1.8</td><td>1.5</td></tr>
<tr><td>>MI0006444|hsa-mir-1307&WithFlank&10|-|103394247|103394407@86.108.23&[MIMAT0005951&hsa-miR-1307-3p&offsets|0|+1;m-66&10|-|103394300|103394322&offsets|0|0]|ACTCGGCGTGGCGTCGGTCGTGG</td><td>>MI0006444|hsa-mir-1307&WithFlank&10|-|103394247|103394407</td><td>86</td><td>108</td><td>23</td><td>ACTCGGCGTGGCGTCGGTCGTGG</td><td>BT20</td><td>131</td><td>80</td><td>264</td><td>94</td><td>124</td><td>85.0</td><td>29.1</td><td>97.2</td><td>50.7</td><td>50.9</td></tr>
<tr><td>>MI0006444|hsa-mir-1307&WithFlank&10|-|103394247|103394407@86.108.23&[MIMAT0005951&hsa-miR-1307-3p&offsets|0|+1;m-66&10|-|103394300|103394322&offsets|0|0]|ACTCGGCGTGGCGTCGGTCGTGG</td><td>>MI0006444|hsa-mir-1307&WithFlank&10|-|103394247|103394407</td><td>86</td><td>108</td><td>23</td><td>ACTCGGCGTGGCGTCGGTCGTGG</td><td>MB231</td><td>216</td><td>153</td><td>136</td><td>93</td><td>135</td><td>109.9</td><td>29.2</td><td>57.7</td><td>41.4</td><td>5.1</td></tr>
<tr><td>>MI0006444|hsa-mir-1307&WithFlank&10|-|103394247|103394407@86.109.24&[MIMAT0005951&hsa-miR-1307-3p&offsets|0|+2;m-66&10|-|103394300|103394322&offsets|0|+1]|ACTCGGCGTGGCGTCGGTCGTGGT</td><td>>MI0006444|hsa-mir-1307&WithFlank&10|-|103394247|103394407</td><td>86</td><td>109</td><td>24</td><td>ACTCGGCGTGGCGTCGGTCGTGGT</td><td>BT20</td><td>17</td><td>12</td><td>33</td><td>3</td><td>18</td><td>15.4</td><td>7.7</td><td>18.0</td><td>2.9</td><td>12.9</td></tr>
<tr><td>>MI0006444|hsa-mir-1307&WithFlank&10|-|103394247|103394407@86.109.24&[MIMAT0005951&hsa-miR-1307-3p&offsets|0|+2;m-66&10|-|103394300|103394322&offsets|0|+1]|ACTCGGCGTGGCGTCGGTCGTGGT</td><td>>MI0006444|hsa-mir-1307&WithFlank&10|-|103394247|103394407</td><td>86</td><td>109</td><td>24</td><td>ACTCGGCGTGGCGTCGGTCGTGGT</td><td>MB231</td><td>22</td><td>17</td><td>15</td><td>7</td><td>25</td><td>10.1</td><td>7.0</td><td>8.6</td><td>4.0</td><td>12.8</td></tr>
<tr><td>>MI0006444|hsa-mir-1307&WithFlank&10|-|103394247|103394407@87.107.21(+1A)&[MIMAT0005951&hsa-miR-1307-3p&offsets|+1|0(+1A);m-66&10|-|103394300|103394322&offsets|+1|-1(+1A)]|CTCGGCGTGGCGTCGGTCGTGA</td><td>>MI0006444|hsa-mir-1307&WithFlank&10|-|103394247|103394407</td><td>87</td><td>107</td><td>21(+1A)</td><td>CTCGGCGTGGCGTCGGTCGTGA</td><td>BT20</td><td>11</td><td>8</td><td>19</td><td>11</td><td>10</td><td>8.8</td><td>4.0</td><td>5.8</td><td>3.9</td><td>2.8</td></tr>
</table>

<table>
<tr><th></th><th></th><th></th><th></th><th></th><th></th><th></th><th></th><th></th><th></th><th></th><th></th><th></th><th></th><th></th><th></th><th></th></tr>
<tr><td>>MI0006444|hsa-mir-1307&WithFlank&10|-|103394247|103394407@87.107.21(+1A)&[MIMAT0005951&hsa-miR-1307-3p&offsets|+1|0(+1A);m-66&10|-|103394300|103394322&offsets|+1|-1(+1A)]|CTCGGCGTGGCGTCGGTCGTGA</td><td>>MI0006444|hsa-mir-1307&WithFlank&10|-|103394247|103394407</td><td>87</td><td>107</td><td>21(+1A)</td><td>CTCGGCGTGGCGTCGGTCGTGA</td><td>MB231</td><td>12</td><td>11</td><td>9</td><td>13</td><td>8</td><td>6.9</td><td>0.4</td><td>2.4</td><td>7.4</td><td>0.8</td></tr>
<tr><td>>MI0006444|hsa-mir-1307&WithFlank&10|-|103394247|103394407@87.107.21(+1U)&[MIMAT0005951&hsa-miR-1307-3p&offsets|+1|0(+1U);m-66&10|-|103394300|103394322&offsets|+1|-1(+1U)]|CTCGGCGTGGCGTCGGTCGTGT</td><td>>MI0006444|hsa-mir-1307&WithFlank&10|-|103394247|103394407</td><td>87</td><td>107</td><td>21(+1U)</td><td>CTCGGCGTGGCGTCGGTCGTGT</td><td>BT20</td><td>14</td><td>8</td><td>18</td><td>14</td><td>10</td><td>12.0</td><td>4.1</td><td>7.6</td><td>9.0</td><td>5.3</td></tr>
<tr><td>>MI0006444|hsa-mir-1307&WithFlank&10|-|103394247|103394407@87.107.21(+1U)&[MIMAT0005951&hsa-miR-1307-3p&offsets|+1|0(+1U);m-66&10|-|103394300|103394322&offsets|+1|-1(+1U)]|CTCGGCGTGGCGTCGGTCGTGT</td><td>>MI0006444|hsa-mir-1307&WithFlank&10|-|103394247|103394407</td><td>87</td><td>107</td><td>21(+1U)</td><td>CTCGGCGTGGCGTCGGTCGTGT</td><td>MB231</td><td>22</td><td>19</td><td>13</td><td>17</td><td>13</td><td>11.4</td><td>1.9</td><td>4.9</td><td>4.5</td><td>2.4</td></tr>
<tr><td>>MI0006444|hsa-mir-1307&WithFlank&10|-|103394247|103394407@87.107.21&[MIMAT0005951&hsa-miR-1307-3p&offsets|+1|0;m-66&10|-|103394300|103394322&offsets|+1|-1]|CTCGGCGTGGCGTCGGTCGTG</td><td>>MI0006444|hsa-mir-1307&WithFlank&10|-|103394247|103394407</td><td>87</td><td>107</td><td>21</td><td>CTCGGCGTGGCGTCGGTCGTG</td><td>MB231</td><td>12</td><td>14</td><td>6</td><td>20</td><td>11</td><td>8.2</td><td>5.3</td><td>2.4</td><td>8.4</td><td>1.8</td></tr>
<tr><td>>MI0006444|hsa-mir-1307&WithFlank&10|-|103394247|103394407@87.108.22&[MIMAT0005951&hsa-miR-1307-3p&offsets|+1|+1;m-66&10|-|103394300|103394322&offsets|+1|0]|CTCGGCGTGGCGTCGGTCGTGG</td><td>>MI0006444|hsa-mir-1307&WithFlank&10|-|103394247|103394407</td><td>87</td><td>108</td><td>22</td><td>CTCGGCGTGGCGTCGGTCGTGG</td><td>BT20</td><td>49</td><td>37</td><td>81</td><td>67</td><td>55</td><td>43.1</td><td>16.5</td><td>44.2</td><td>34.7</td><td>12.6</td></tr>
<tr><td>>MI0006444|hsa-mir-1307&WithFlank&10|-|103394247|103394407@87.108.22&[MIMAT0005951&hsa-miR-1307-3p&offsets|+1|+1;m-66&10|-|103394300|103394322&offsets|+1|0]|CTCGGCGTGGCGTCGGTCGTGG</td><td>>MI0006444|hsa-mir-1307&WithFlank&10|-|103394247|103394407</td><td>87</td><td>108</td><td>22</td><td>CTCGGCGTGGCGTCGGTCGTGG</td><td>MB231</td><td>75</td><td>86</td><td>56</td><td>57</td><td>47</td><td>49.3</td><td>8.9</td><td>15.4</td><td>19.9</td><td>8.5</td></tr>
<tr><td>>MI0006444|hsa-mir-1307&WithFlank&10|-|103394247|103394407@87.109.23&[MIMAT0005951&hsa-miR-1307-3p&offsets|+1|+2;m-66&10|-|103394300|103394322&offsets|+1|+1]|CTCGGCGTGGCGTCGGTCGTGGT</td><td>>MI0006444|hsa-mir-1307&WithFlank&10|-|103394247|103394407</td><td>87</td><td>109</td><td>23</td><td>CTCGGCGTGGCGTCGGTCGTGGT</td><td>MB231</td><td>13</td><td>13</td><td>14</td><td>7</td><td>9</td><td>6.2</td><td>1.7</td><td>4.3</td><td>2.0</td><td>2.4</td></tr>
<tr><td>>MI0015866|hsa-mir-4326&WithFlank&20|+|63286802|63286872@17.39.23&[MIMAT0016888&hsa-miR-4326&offsets|0|+3;m-339&20|+|63286818|63286841&offsets|0|-1]|TGTTCCTCTGTCTCCCAGACTCT</td><td>>MI0015866|hsa-mir-4326&WithFlank&20|+|63286802|63286872</td><td>17</td><td>39</td><td>23</td><td>TGTTCCTCTGTCTCCCAGACTCT</td><td>BT20</td><td>16</td><td>7</td><td>22</td><td>7</td><td>9</td><td>9.9</td><td>3.4</td><td>3.9</td><td>3.8</td><td>4.0</td></tr>
<tr><td>>MI0015866|hsa-mir-4326&WithFlank&20|+|63286802|63286872@17.39.23&[MIMAT0016888&hsa-miR-4326&offsets|0|+3;m-339&20|+|63286818|63286841&offsets|0|-1]|TGTTCCTCTGTCTCCCAGACTCT</td><td>>MI0015866|hsa-mir-4326&WithFlank&20|+|63286802|63286872</td><td>17</td><td>39</td><td>23</td><td>TGTTCCTCTGTCTCCCAGACTCT</td><td>MB231</td><td>11</td><td>11</td><td>11</td><td>9</td><td>15</td><td>5.2</td><td>2.8</td><td>3.9</td><td>3.4</td><td>5.1</td></tr>
<tr><td>>MI0015866|hsa-mir-4326&WithFlank&20|+|63286802|63286872@17.40.24(+1U)&[MIMAT0016888&hsa-miR-4326&offsets|0|+4(+1U);m-339&20|+|63286818|63286841&offsets|0|0(+1U)]|TGTTCCTCTGTCTCCCAGACTCTGT</td><td>>MI0015866|hsa-mir-4326&WithFlank&20|+|63286802|63286872</td><td>17</td><td>40</td><td>24(+1U)</td><td>TGTTCCTCTGTCTCCCAGACTCTGT</td><td>MB231</td><td>11</td><td>8</td><td>7</td><td>11</td><td>12</td><td>1.4</td><td>3.1</td><td>2.9</td><td>2.9</td><td>8.0</td></tr>
<tr><td>>MI0015866|hsa-mir-4326&WithFlank&20|+|63286802|63286872@17.40.24&[MIMAT0016888&hsa-miR-4326&offsets|0|+4;m-339&20|+|63286818|63286841&offsets|0|0]|TGTTCCTCTGTCTCCCAGACTCTG</td><td>>MI0015866|hsa-mir-4326&WithFlank&20|+|63286802|63286872</td><td>17</td><td>40</td><td>24</td><td>TGTTCCTCTGTCTCCCAGACTCTG</td><td>BT20</td><td>22</td><td>14</td><td>40</td><td>31</td><td>12</td><td>12.3</td><td>11.0</td><td>12.8</td><td>12.5</td><td>1.7</td></tr>
<tr><td>>MI0015866|hsa-mir-4326&WithFlank&20|+|63286802|63286872@17.40.24&[MIMAT0016888&hsa-miR-4326&offsets|0|+4;m-339&20|+|63286818|63286841&offsets|0|0]|TGTTCCTCTGTCTCCCAGACTCTG</td><td>>MI0015866|hsa-mir-4326&WithFlank&20|+|63286802|63286872</td><td>17</td><td>40</td><td>24</td><td>TGTTCCTCTGTCTCCCAGACTCTG</td><td>MB231</td><td>16</td><td>14</td><td>13</td><td>43</td><td>15</td><td>3.9</td><td>3.4</td><td>6.7</td><td>20.8</td><td>4.5</td></tr>
<tr><td>>MI0016003|hsa-mir-3613&WithFlank&13|-|49996409|49996507@22.43.22&[MIMAT0017990&hsa-miR-3613-5p&offsets|0|0;m-259&13|-|49996465|49996486&offsets|0|0]|TGTTGTACTTTTTTTTTTGTTC</td><td>>MI0016003|hsa-mir-3613&WithFlank&13|-|49996409|49996507</td><td>22</td><td>43</td><td>22</td><td>TGTTGTACTTTTTTTTTTGTTC</td><td>BT20</td><td>20</td><td>23</td><td>13</td><td>15</td><td>4</td><td>15.2</td><td>17.6</td><td>10.9</td><td>22.5</td><td>1.6</td></tr>
<tr><td>>MI0016005|hsa-mir-3615&WithFlank&17|+|74748607|74748705@57.78.22&[MIMAT0017994&hsa-miR-3615&offsets|0|+1;m-271&17|+|74748663|74748684&offsets|0|0]|TCTCTCGGCTCCTCGCGGCTCG</td><td>>MI0016005|hsa-mir-3615&WithFlank&17|+|74748607|74748705</td><td>57</td><td>78</td><td>22</td><td>TCTCTCGGCTCCTCGCGGCTCG</td><td>BT20</td><td>13</td><td>9</td><td>54</td><td>2</td><td>6</td><td>15.2</td><td>4.6</td><td>10.4</td><td>1.7</td><td>7.0</td></tr>
<tr><td>>MI0016005|hsa-mir-3615&WithFlank&17|+|74748607|74748705@57.78.22&[MIMAT0017994&hsa-miR-3615&offsets|0|+1;m-271&17|+|74748663|74748684&offsets|0|0]|TCTCTCGGCTCCTCGCGGCTCG</td><td>>MI0016005|hsa-mir-3615&WithFlank&17|+|74748607|74748705</td><td>57</td><td>78</td><td>22</td><td>TCTCTCGGCTCCTCGCGGCTCG</td><td>MB231</td><td>27</td><td>45</td><td>61</td><td>5</td><td>30</td><td>23.9</td><td>15.1</td><td>38.5</td><td>0.6</td><td>12.0</td></tr>
<tr><td>>MI0016005|hsa-mir-3615&WithFlank&17|+|74748607|74748705@57.79.23&[MIMAT0017994&hsa-miR-3615&offsets|0|+2;m-271&17|+|74748663|74748684&offsets|0|+1]|TCTCTCGGCTCCTCGCGGCTCGC</td><td>>MI0016005|hsa-mir-3615&WithFlank&17|+|74748607|74748705</td><td>57</td><td>79</td><td>23</td><td>TCTCTCGGCTCCTCGCGGCTCGC</td><td>MB231</td><td>13</td><td>14</td><td>16</td><td>2</td><td>15</td><td>9.4</td><td>4.5</td><td>6.7</td><td>0.6</td><td>4.9</td></tr>
<tr><td>>MI0016746|hsa-mir-548o-2&WithFlank&20|+|38516557|38516638@51.72.22;MI0006402|hsa-mir-548o&WithFlank&7|-|102405736|102405861@93.114.22&[MIMAT0005919&hsa-miR-548o-3p&offsets|0|0;m-431&20|+|38516607|38516628&offsets|0|0];[MIMAT0005919_1&hsa-miR-548o-3p&offsets|0|0;m-430&7|-|102405748|102405769&offsets|0|0]|CCAAAACTGCAGTTACTTTTGC</td><td>>MI0016746|hsa-mir-548o-2&WithFlank&20|+|38516557|38516638</td><td>51</td><td>72</td><td>22</td><td>CCAAAACTGCAGTTACTTTTGC</td><td>BT20</td><td>32</td><td>19</td><td>75</td><td>36</td><td>14</td><td>14.1</td><td>7.9</td><td>27.6</td><td>25.8</td><td>9.8</td></tr>
<tr><td>>MI0016746|hsa-mir-548o-2&WithFlank&20|+|38516557|38516638@51.72.22;MI0006402|hsa-mir-548o&WithFlank&7|-|102405736|102405861@93.114.22&[MIMAT0005919&hsa-miR-548o-3p&offsets|0|0;m-431&20|+|38516607|38516628&offsets|0|0];[MIMAT0005919_1&hsa-miR-548o-3p&offsets|0|0;m-430&7|-|102405748|102405769&offsets|0|0]|CCAAAACTGCAGTTACTTTTGC</td><td>>MI0016746|hsa-mir-548o-2&WithFlank&20|+|38516557|38516638</td><td>51</td><td>72</td><td>22</td><td>CCAAAACTGCAGTTACTTTTGC</td><td>MB231</td><td>16</td><td>19</td><td>19</td><td>63</td><td>17</td><td>5.7</td><td>4.5</td><td>10.7</td><td>43.2</td><td>5.6</td></tr>
<tr><td>>MI0016887|hsa-mir-4521&WithFlank&17|+|8186939|8187010@10.34.25&[MIMAT0019058&hsa-miR-4521&offsets|0|+3;m-765&17|+|8186948|8186969&offsets|0|+3]|GCTAAGGAAGTCCTGTGCTCAGTTT</td><td>>MI0016887|hsa-mir-4521&WithFlank&17|+|8186939|8187010</td><td>10</td><td>34</td><td>25</td><td>GCTAAGGAAGTCCTGTGCTCAGTTT</td><td>BT20</td><td>59</td><td>36</td><td>115</td><td>4</td><td>2</td><td>53.3</td><td>45.8</td><td>86.2</td><td>5.6</td><td>3.4</td></tr>
<tr><td>>MI0016887|hsa-mir-4521&WithFlank&17|+|8186939|8187010@10.34.25&[MIMAT0019058&hsa-miR-4521&offsets|0|+3;m-765&17|+|8186948|8186969&offsets|0|+3]|GCTAAGGAAGTCCTGTGCTCAGTTT</td><td>>MI0016887|hsa-mir-4521&WithFlank&17|+|8186939|8187010</td><td>10</td><td>34</td><td>25</td><td>GCTAAGGAAGTCCTGTGCTCAGTTT</td><td>MB231</td><td>19</td><td>12</td><td>31</td><td>7</td><td>5</td><td>3.2</td><td>1.9</td><td>27.0</td><td>3.8</td><td>3.0</td></tr>
<tr><td>>MI0016887|hsa-mir-4521&WithFlank&17|+|8186939|8187010@10.35.26&[MIMAT0019058&hsa-miR-4521&offsets|0|+4;m-765&17|+|8186948|8186969&offsets|0|+4]|GCTAAGGAAGTCCTGTGCTCAGTTTT</td><td>>MI0016887|hsa-mir-4521&WithFlank&17|+|8186939|8187010</td><td>10</td><td>35</td><td>26</td><td>GCTAAGGAAGTCCTGTGCTCAGTTTT</td><td>BT20</td><td>58</td><td>27</td><td>75</td><td>2</td><td>2</td><td>62.4</td><td>31.9</td><td>63.8</td><td>3.0</td><td>2.6</td></tr>
<tr><td>>MI0016887|hsa-mir-4521&WithFlank&17|+|8186939|8187010@10.35.26&[MIMAT0019058&hsa-miR-4521&offsets|0|+4;m-765&17|+|8186948|8186969&offsets|0|+4]|GCTAAGGAAGTCCTGTGCTCAGTTTT</td><td>>MI0016887|hsa-mir-4521&WithFlank&17|+|8186939|8187010</td><td>10</td><td>35</td><td>26</td><td>GCTAAGGAAGTCCTGTGCTCAGTTTT</td><td>MB231</td><td>33</td><td>18</td><td>28</td><td>11</td><td>8</td><td>10.3</td><td>3.7</td><td>19.3</td><td>8.1</td><td>4.2</td></tr>
<tr><td>>MI0016887|hsa-mir-4521&WithFlank&17|+|8186939|8187010@12.34.23&[MIMAT0019058&hsa-miR-4521&offsets|+2|+3;m-765&17|+|8186948|8186969&offsets|+2|+3]|TAAGGAAGTCCTGTGCTCAGTTT</td><td>>MI0016887|hsa-mir-4521&WithFlank&17|+|8186939|8187010</td><td>12</td><td>34</td><td>23</td><td>TAAGGAAGTCCTGTGCTCAGTTT</td><td>BT20</td><td>11</td><td>7</td><td>16</td><td>0</td><td>0</td><td>10.1</td><td>7.6</td><td>9.1</td><td>0.0</td><td>0.0</td></tr>
<tr><td>>MI0016887|hsa-mir-4521&WithFlank&17|+|8186939|8187010@12.35.24&[MIMAT0019058&hsa-miR-4521&offsets|+2|+4;m-765&17|+|8186948|8186969&offsets|+2|+4]|TAAGGAAGTCCTGTGCTCAGTTTT</td><td>>MI0016887|hsa-mir-4521&WithFlank&17|+|8186939|8187010</td><td>12</td><td>35</td><td>24</td><td>TAAGGAAGTCCTGTGCTCAGTTTT</td><td>BT20</td><td>12</td><td>6</td><td>14</td><td>0</td><td>0</td><td>8.8</td><td>5.6</td><td>9.0</td><td>0.0</td><td>0.0</td></tr>
<tr><td>>MI0016887|hsa-mir-4521&WithFlank&17|+|8186939|8187010@13.34.22&[MIMAT0019058&hsa-miR-4521&offsets|+3|+3;m-765&17|+|8186948|8186969&offsets|+3|+3]|AAGGAAGTCCTGTGCTCAGTTT</td><td>>MI0016887|hsa-mir-4521&WithFlank&17|+|8186939|8187010</td><td>13</td><td>34</td><td>22</td><td>AAGGAAGTCCTGTGCTCAGTTT</td><td>BT20</td><td>23</td><td>26</td><td>26</td><td>3</td><td>2</td><td>21.1</td><td>25.4</td><td>16.4</td><td>2.8</td><td>2.4</td></tr>
<tr><td>>MI0016887|hsa-mir-4521&WithFlank&17|+|8186939|8187010@13.35.23&[MIMAT0019058&hsa-miR-4521&offsets|+3|+4;m-765&17|+|8186948|8186969&offsets|+3|+4]|AAGGAAGTCCTGTGCTCAGTTTT</td><td>>MI0016887|hsa-mir-4521&WithFlank&17|+|8186939|8187010</td><td>13</td><td>35</td><td>23</td><td>AAGGAAGTCCTGTGCTCAGTTTT</td><td>BT20</td><td>23</td><td>21</td><td>25</td><td>1</td><td>2</td><td>19.7</td><td>18.4</td><td>18.2</td><td>1.7</td><td>3.2</td></tr>
</table>

| | | | | | | | | | | | | | | | | |
|---|---|---|---|---|---|---|---|---|---|---|---|---|---|---|---|---|
| >MI0016887\|hsa-mir-4521&WithFlank&17\|+\|8186939\|8187010@14.34.21&[MIMAT0019058&hsa-miR-4521&offsets\|+4\|+3;m-765&17\|+\|8186948\|8186969&offsets\|+4\|+3]\|AGGAAGTCCTGTGCTCAGTTT | >MI0016887\|hsa-mir-4521&WithFlank&17\|+\|8186939\|8187010 | 14 | 34 | 21 | AGGAAGTCCTGTGCTCAGTTT | BT20 | 15 | 12 | 26 | 1 | 1 | 12.5 | 11.6 | 13.5 | 2.3 | 1.2 |
| >MI0016887\|hsa-mir-4521&WithFlank&17\|+\|8186939\|8187010@14.35.22&[MIMAT0019058&hsa-miR-4521&offsets\|+4\|+4;m-765&17\|+\|8186948\|8186969&offsets\|+4\|+4]\|AGGAAGTCCTGTGCTCAGTTTT | >MI0016887\|hsa-mir-4521&WithFlank&17\|+\|8186939\|8187010 | 14 | 35 | 22 | AGGAAGTCCTGTGCTCAGTTTT | BT20 | 14 | 10 | 18 | 1 | 1 | 9.7 | 7.9 | 11.5 | 1.6 | 1.1 |
| >MI0017308\|hsa-mir-4677&WithFlank&1\|+\|243346170\|243346261@56.77.22&[MIMAT0019761&hsa-miR-4677-3p&offsets\|0\|0;m-295&1\|+\|243346225\|243346246&offsets\|0\|0]\|TCTGTGAGACCAAAGAACTACT | >MI0017308\|hsa-mir-4677&WithFlank&1\|+\|243346170\|243346261 | 56 | 77 | 22 | TCTGTGAGACCAAAGAACTACT | BT20 | 11 | 5 | 4 | 1 | 1 | 8.4 | 1.8 | 0.4 | 0.7 | 0.8 |
| >MI0025242\|hsa-mir-7706&WithFlank&15\|+\|85380590\|85380668@50.71.22(+1U)&[MIMAT0030021&hsa-miR-7706&offsets\|0\|-2(+1U);m-390&15\|+\|85380639\|85380662&offsets\|0\|-2(+1U)]\|TGAAGCGCCTGTGCTCTGCCGAT | >MI0025242\|hsa-mir-7706&WithFlank&15\|+\|85380590\|85380668 | 50 | 71 | 22(+1U) | TGAAGCGCCTGTGCTCTGCCGAT | BT20 | 10 | 6 | 11 | 3 | 7 | 2.5 | 1.0 | 8.1 | 1.0 | 0.1 |
| >MI0025242\|hsa-mir-7706&WithFlank&15\|+\|85380590\|85380668@50.71.22&[MIMAT0030021&hsa-miR-7706&offsets\|0\|-2;m-390&15\|+\|85380639\|85380662&offsets\|0\|-2]\|TGAAGCGCCTGTGCTCTGCCGA | >MI0025242\|hsa-mir-7706&WithFlank&15\|+\|85380590\|85380668 | 50 | 71 | 22 | TGAAGCGCCTGTGCTCTGCCGA | BT20 | 15 | 11 | 22 | 14 | 12 | 4.8 | 2.9 | 16.9 | 3.9 | 2.2 |
| >MI0025242\|hsa-mir-7706&WithFlank&15\|+\|85380590\|85380668@50.72.23(+1U)&[MIMAT0030021&hsa-miR-7706&offsets\|0\|-1(+1U);m-390&15\|+\|85380639\|85380662&offsets\|0\|-1(+1U)]\|TGAAGCGCCTGTGCTCTGCCGAGT | >MI0025242\|hsa-mir-7706&WithFlank&15\|+\|85380590\|85380668 | 50 | 72 | 23(+1U) | TGAAGCGCCTGTGCTCTGCCGAGT | BT20 | 11 | 7 | 16 | 3 | 5 | 2.6 | 1.4 | 15.2 | 1.5 | 1.0 |
| >MI0025242\|hsa-mir-7706&WithFlank&15\|+\|85380590\|85380668@50.72.23&[MIMAT0030021&hsa-miR-7706&offsets\|0\|-1;m-390&15\|+\|85380639\|85380662&offsets\|0\|-1]\|TGAAGCGCCTGTGCTCTGCCGAG | >MI0025242\|hsa-mir-7706&WithFlank&15\|+\|85380590\|85380668 | 50 | 72 | 23 | TGAAGCGCCTGTGCTCTGCCGAG | BT20 | 14 | 10 | 22 | 10 | 9 | 3.7 | 1.5 | 17.8 | 1.3 | 1.1 |
| >MI0025750\|hsa-mir-7974&WithFlank&19\|-\|11495538\|11495628@62.85.24&[MIMAT0031177&hsa-miR-7974&offsets\|0\|0;m-867&19\|-\|11495544\|11495567&offsets\|0\|0]\|AGGCTGTGATGCTCTCCTGAGCCC | >MI0025750\|hsa-mir-7974&WithFlank&19\|-\|11495538\|11495628 | 62 | 85 | 24 | AGGCTGTGATGCTCTCCTGAGCCC | BT20 | 10 | 12 | 9 | 0 | 1 | 8.4 | 11.1 | 8.6 | 0.3 | 1.0 |
| >MI0025750\|hsa-mir-7974&WithFlank&19\|-\|11495538\|11495628@62.85.24&[MIMAT0031177&hsa-miR-7974&offsets\|0\|0;m-867&19\|-\|11495544\|11495567&offsets\|0\|0]\|AGGCTGTGATGCTCTCCTGAGCCC | >MI0025750\|hsa-mir-7974&WithFlank&19\|-\|11495538\|11495628 | 62 | 85 | 24 | AGGCTGTGATGCTCTCCTGAGCCC | MB231 | 15 | 26 | 8 | 0 | 0 | 6.6 | 3.3 | 4.5 | 0.0 | 0.0 |
| >MI0025750\|hsa-mir-7974&WithFlank&19\|-\|11495538\|11495628@62.85.24&[MIMAT0031177&hsa-miR-7974&offsets\|0\|0;m-867&19\|-\|11495544\|11495567&offsets\|0\|0]\|AGGCTGTGATGCTCTCCTGAGCCC | >MI0025750\|hsa-mir-7974&WithFlank&19\|-\|11495538\|11495628 | 62 | 85 | 24 | AGGCTGTGATGCTCTCCTGAGCCC | MB468 | 13 | 35 | 1 | 0 | 1 | 3.9 | 26.8 | 1.4 | 0.3 | 1.0 |
| >MI0033423\|hsa-mir-10399&WithFlank&7\|-\|139044094\|139044163@43.63.21(+1U)&[MIMAT0041630&hsa-miR-10399-3p&offsets\|-1\|-1(+1U);TJU_CMC_MD2.ID03061.3p-miR&offsets\|+3\|+2(+1U);m-2657&7\|-\|139044103\|139044124&offsets\|+3\|+2(+1U)]\|TCTCTCGGACAAGCTGTAGGTT | >MI0033423\|hsa-mir-10399&WithFlank&7\|-\|139044094\|139044163 | 43 | 63 | 21(+1U) | TCTCTCGGACAAGCTGTAGGTT | MB231 | 13 | 16 | 13 | 31 | 12 | 1.8 | 2.9 | 6.6 | 1.8 | 1.6 |
| >multi-am_trna2_LeuTAA_X_-_55207755_55207829@1.18.18,trna4_LeuTAA_20_-_55933428_55933502@1.18.18,trna60_LeuTAA_1_-_238106956_238107030@1.18.18,trna9_LeuTAA_2_+_117781417_117781491@1.18.18,trnaMT_LeuTAA_MT_+_3230_3304@1.18.18\|GTTAAGATGGCAGAGCCC | >multi-am_trna2_LeuTAA_X_-_55207755_55207829 | 1 | 18 | 18 | GTTAAGATGGCAGAGCCC | BT20 | 19 | 14 | 17 | 20 | 8 | 15.8 | 8.4 | 10.7 | 29.6 | 8.9 |
| >multi-am_trnaMT_ArgTCG_MT_+_10405_10469@28.48.21\|ATTTCGACTCATTAAATTATG | >multi-am_trnaMT_ArgTCG_MT_+_10405_10469 | 28 | 48 | 21 | ATTTCGACTCATTAAATTATG | MB468 | 88 | 179 | 57 | 33 | 63 | 36.8 | 268.8 | 58.7 | 20.7 | 57.7 |
| >multi-am_trnaMT_ArgTCG_MT_+_10405_10469@29.48.20\|TTTCGACTCATTAAATTATG | >multi-am_trnaMT_ArgTCG_MT_+_10405_10469 | 29 | 48 | 20 | TTTCGACTCATTAAATTATG | MB468 | 18 | 51 | 19 | 9 | 9 | 10.9 | 81.2 | 18.9 | 7.1 | 7.7 |
| >multi-am_trnaMT_ArgTCG_MT_+_10405_10469@31.48.18\|TCGACTCATTAAATTATG | >multi-am_trnaMT_ArgTCG_MT_+_10405_10469 | 31 | 48 | 18 | TCGACTCATTAAATTATG | MB468 | 18 | 12 | 17 | 10 | 19 | 5.6 | 13.0 | 14.8 | 6.0 | 28.8 |
| >multi-am_trnaMT_GluTTC_MT_-_14674_14742@27.48.22,trnalookalike8_GluTTC_5_-_93905172_93905240@27.48.22\|GTTTTTCATATCATTGGTCGTG | >multi-am_trnaMT_GluTTC_MT_-_14674_14742 | 27 | 48 | 22 | GTTTTTCATATCATTGGTCGTG | MB468 | 15 | 12 | 13 | 9 | 4 | 8.3 | 7.4 | 7.0 | 3.8 | 1.2 |
| >multi-am_trnaMT_GluTTC_MT_-_14674_14742@27.52.26,trnalookalike8_GluTTC_5_-_93905172_93905240@27.52.26\|GTTTTTCATATCATTGGTCGTGGTTG | >multi-am_trnaMT_GluTTC_MT_-_14674_14742 | 27 | 52 | 26 | GTTTTTCATATCATTGGTCGTGGTTG | MB468 | 38 | 16 | 24 | 19 | 14 | 20.6 | 8.0 | 23.0 | 6.6 | 10.2 |
| >multi-am_trnaMT_GluTTC_MT_-_14674_14742@27.54.28,trnalookalike8_GluTTC_5_-_93905172_93905240@27.54.28\|GTTTTTCATATCATTGGTCGTGGTTGTA | >multi-am_trnaMT_GluTTC_MT_-_14674_14742 | 27 | 54 | 28 | GTTTTTCATATCATTGGTCGTGGTTGTA | MB468 | 11 | 7 | 7 | 12 | 4 | 5.8 | 8.0 | 7.1 | 12.4 | 1.7 |
| >multi-am_trnaMT_GluTTC_MT_-_14674_14742@27.55.29,trnalookalike8_GluTTC_5_-_93905172_93905240@27.55.29\|GTTTTTCATATCATTGGTCGTGGTTGTAG | >multi-am_trnaMT_GluTTC_MT_-_14674_14742 | 27 | 55 | 29 | GTTTTTCATATCATTGGTCGTGGTTGTAG | MB468 | 12 | 9 | 7 | 10 | 5 | 8.0 | 11.1 | 7.2 | 6.8 | 1.7 |
| >multi-am_trnaMT_GluTTC_MT_-_14674_14742@41.58.18,trnalookalike8_GluTTC_5_-_93905172_93905240@41.58.18\|TGGTCGTGGTTGTAGTCC | >multi-am_trnaMT_GluTTC_MT_-_14674_14742 | 41 | 58 | 18 | TGGTCGTGGTTGTAGTCC | MB231 | 13 | 9 | 12 | 17 | 10 | 10.1 | 3.9 | 2.3 | 14.1 | 8.4 |
| >multi-am_trnaMT_GlyTCC_MT_+_9991_10058@28.48.21\|ACTTCCAATTAACTAGTTTTG | >multi-am_trnaMT_GlyTCC_MT_+_9991_10058 | 28 | 48 | 21 | ACTTCCAATTAACTAGTTTTG | MB468 | 15 | 7 | 9 | 9 | 7 | 8.3 | 9.2 | 9.4 | 5.1 | 6.9 |
| >multi-am_trnaMT_HisGTG_MT_+_12138_12206@18.38.21\|AAAACATCAGATTGTGAATCT | >multi-am_trnaMT_HisGTG_MT_+_12138_12206 | 18 | 38 | 21 | AAAACATCAGATTGTGAATCT | BT20 | 18 | 6 | 6 | 16 | 6 | 4.9 | 6.4 | 1.9 | 13.2 | 5.7 |
| >multi-am_trnaMT_HisGTG_MT_+_12138_12206@18.51.34\|AAAACATCAGATTGTGAATCTGACAACAGAGGCT | >multi-am_trnaMT_HisGTG_MT_+_12138_12206 | 18 | 51 | 34 | AAAACATCAGATTGTGAATCTGACAACAGAGGCT | BT20 | 12 | 5 | 2 | 74 | 36 | 5.4 | 3.6 | 0.3 | 56.4 | 27.9 |
| >multi-am_trnaMT_HisGTG_MT_+_12138_12206@19.38.20\|AAACATCAGATTGTGAATCT | >multi-am_trnaMT_HisGTG_MT_+_12138_12206 | 19 | 38 | 20 | AAACATCAGATTGTGAATCT | BT20 | 15 | 5 | 5 | 11 | 5 | 4.6 | 6.3 | 2.5 | 8.9 | 5.0 |
| >multi-am_trnaMT_HisGTG_MT_+_12138_12206@19.51.33\|AAACATCAGATTGTGAATCTGACAACAGAGGCT | >multi-am_trnaMT_HisGTG_MT_+_12138_12206 | 19 | 51 | 33 | AAACATCAGATTGTGAATCTGACAACAGAGGCT | BT20 | 21 | 18 | 7 | 82 | 41 | 13.9 | 18.4 | 5.8 | 68.5 | 30.0 |
| >multi-am_trnaMT_HisGTG_MT_+_12138_12206@20.51.32\|AACATCAGATTGTGAATCTGACAACAGAGGCT | >multi-am_trnaMT_HisGTG_MT_+_12138_12206 | 20 | 51 | 32 | AACATCAGATTGTGAATCTGACAACAGAGGCT | BT20 | 14 | 15 | 6 | 70 | 35 | 11.5 | 17.8 | 5.1 | 64.3 | 24.0 |
| >multi-am_trnaMT_HisGTG_MT_+_12138_12206@26.43.18\|AGATTGTGAATCTGACAA | >multi-am_trnaMT_HisGTG_MT_+_12138_12206 | 26 | 43 | 18 | AGATTGTGAATCTGACAA | BT20 | 12 | 2 | 0 | 2 | 9 | 16.3 | 1.7 | 0.1 | 1.5 | 6.6 |
| >multi-am_trnaMT_HisGTG_MT_+_12138_12206@26.44.19\|AGATTGTGAATCTGACAAC | >multi-am_trnaMT_HisGTG_MT_+_12138_12206 | 26 | 44 | 19 | AGATTGTGAATCTGACAAC | BT20 | 25 | 5 | 0 | 16 | 39 | 24.0 | 4.7 | 0.3 | 12.7 | 23.5 |
| >multi-am_trnaMT_HisGTG_MT_+_12138_12206@26.45.20\|AGATTGTGAATCTGACAACA | >multi-am_trnaMT_HisGTG_MT_+_12138_12206 | 26 | 45 | 20 | AGATTGTGAATCTGACAACA | BT20 | 51 | 6 | 0 | 12 | 27 | 67.3 | 3.8 | 0.2 | 11.1 | 13.9 |
| >multi-am_trnaMT_HisGTG_MT_+_12138_12206@28.45.18\|ATTGTGAATCTGACAACA | >multi-am_trnaMT_HisGTG_MT_+_12138_12206 | 28 | 45 | 18 | ATTGTGAATCTGACAACA | BT20 | 11 | 1 | 4 | 3 | 1 | 8.9 | 1.3 | 1.8 | 2.9 | 1.7 |
| >multi-am_trnaMT_IleGAT_MT_+_4263_4331@1.32.32\|AGAAATATGTCTGATAAAAGAGTTACTTTGAT | >multi-am_trnaMT_IleGAT_MT_+_4263_4331 | 1 | 32 | 32 | AGAAATATGTCTGATAAAAGAGTTACTTTGAT | MB231 | 18 | 1 | 0 | 6 | 16 | 22.8 | 1.0 | 0.0 | 0.8 | 21.4 |
| >multi-am_trnaMT_LysTTT_MT_+_8295_8364@27.52.26,trnalookalike7_LysTTT_1_+_568844_568913@27.52.26\|CTTTTAAGTTAAAGATTAAGAGAACC | >multi-am_trnaMT_LysTTT_MT_+_8295_8364 | 27 | 52 | 26 | CTTTTAAGTTAAAGATTAAGAGAACC | BT20 | 16 | 5 | 7 | 7 | 3 | 11.6 | 7.5 | 9.8 | 6.5 | 3.8 |
| >multi-am_trnaMT_LysTTT_MT_+_8295_8364@28.52.25,trnalookalike7_LysTTT_1_+_568844_568913@28.52.25\|TTTTAAGTTAAAGATTAAGAGAACC | >multi-am_trnaMT_LysTTT_MT_+_8295_8364 | 28 | 52 | 25 | TTTTAAGTTAAAGATTAAGAGAACC | BT20 | 28 | 9 | 26 | 8 | 4 | 15.9 | 8.0 | 20.9 | 6.9 | 4.5 |
| >multi-am_trnaMT_LysTTT_MT_+_8295_8364@28.52.25,trnalookalike7_LysTTT_1_+_568844_568913@28.52.25\|TTTTAAGTTAAAGATTAAGAGAACC | >multi-am_trnaMT_LysTTT_MT_+_8295_8364 | 28 | 52 | 25 | TTTTAAGTTAAAGATTAAGAGAACC | MB231 | 14 | 10 | 5 | 20 | 5 | 15.6 | 10.2 | 4.7 | 16.5 | 2.1 |
| >multi-am_trnaMT_LysTTT_MT_+_8295_8364@29.52.24,trnalookalike7_LysTTT_1_+_568844_568913@29.52.24\|TTTAAGTTAAAGATTAAGAGAACC | >multi-am_trnaMT_LysTTT_MT_+_8295_8364 | 29 | 52 | 24 | TTTAAGTTAAAGATTAAGAGAACC | BT20 | 25 | 9 | 27 | 8 | 4 | 13.5 | 5.1 | 17.3 | 7.6 | 3.7 |
| >multi-am_trnaMT_LysTTT_MT_+_8295_8364@30.52.23,trnalookalike7_LysTTT_1_+_568844_568913@30.52.23\|TTAAGTTAAAGATTAAGAGAACC | >multi-am_trnaMT_LysTTT_MT_+_8295_8364 | 30 | 52 | 23 | TTAAGTTAAAGATTAAGAGAACC | BT20 | 25 | 8 | 30 | 6 | 2 | 14.5 | 5.0 | 15.2 | 6.1 | 2.9 |
| >multi-am_trnaMT_LysTTT_MT_+_8295_8364@31.52.22,trnalookalike7_LysTTT_1_+_568844_568913@31.52.22\|TAAGTTAAAGATTAAGAGAACC | >multi-am_trnaMT_LysTTT_MT_+_8295_8364 | 31 | 52 | 22 | TAAGTTAAAGATTAAGAGAACC | BT20 | 13 | 7 | 15 | 3 | 1 | 13.4 | 6.8 | 13.7 | 2.5 | 1.3 |
| >multi-am_trnaMT_LysTTT_MT_+_8295_8364@32.52.21,trnalookalike7_LysTTT_1_+_568844_568913@32.52.21\|AAGTTAAAGATTAAGAGAACC | >multi-am_trnaMT_LysTTT_MT_+_8295_8364 | 32 | 52 | 21 | AAGTTAAAGATTAAGAGAACC | BT20 | 14 | 6 | 18 | 3 | 1 | 15.9 | 6.1 | 18.3 | 2.5 | 1.9 |
| >multi-am_trnaMT_LysTTT_MT_+_8295_8364@33.52.20,trnalookalike7_LysTTT_1_+_568844_568913@33.52.20\|AGTTAAAGATTAAGAGAACC | >multi-am_trnaMT_LysTTT_MT_+_8295_8364 | 33 | 52 | 20 | AGTTAAAGATTAAGAGAACC | BT20 | 17 | 6 | 20 | 5 | 2 | 17.0 | 4.4 | 16.9 | 4.1 | 1.8 |
| >multi-am_trnaMT_LysTTT_MT_+_8295_8364@34.52.19,trnalookalike7_LysTTT_1_+_568844_568913@34.52.19\|GTTAAAGATTAAGAGAACC | >multi-am_trnaMT_LysTTT_MT_+_8295_8364 | 34 | 52 | 19 | GTTAAAGATTAAGAGAACC | BT20 | 29 | 10 | 38 | 10 | 4 | 25.7 | 5.0 | 24.2 | 6.9 | 3.5 |

| | | | | | | | | | | | | | | | | |
|---|---|---|---|---|---|---|---|---|---|---|---|---|---|---|---|---|
| >multi-am_trnaMT_LysTTT_MT_+_8295_8364@35.52.18,trnalookalike7_LysTTT_1_+_568844_568913@35.52.18\|TTAAAGATTAAGAGAACC | >multi-am_trnaMT_LysTTT_MT_+_8295_8364 | 35 | 52 | 18 | TTAAAGATTAAGAGAACC | BT20 | 11 | 2 | 13 | 1 | 1 | 10.4 | 2.4 | 6.8 | 1.3 | 0.7 |
| >multi-am_trnaMT_MetCAT_MT_+_4402_4469@1.18.18\|AGTAAGGTCAGCTAAATA | >multi-am_trnaMT_MetCAT_MT_+_4402_4469 | 1 | 18 | 18 | AGTAAGGTCAGCTAAATA | BT20 | 18 | 11 | 6 | 22 | 6 | 10.3 | 6.4 | 2.8 | 28.3 | 8.0 |
| >multi-am_trnaMT_MetCAT_MT_+_4402_4469@1.21.21\|AGTAAGGTCAGCTAAATAAGC | >multi-am_trnaMT_MetCAT_MT_+_4402_4469 | 1 | 21 | 21 | AGTAAGGTCAGCTAAATAAGC | BT20 | 28 | 14 | 14 | 68 | 34 | 15.8 | 10.4 | 10.6 | 72.5 | 46.6 |
| >multi-am_trnaMT_MetCAT_MT_+_4402_4469@1.21.21\|AGTAAGGTCAGCTAAATAAGC | >multi-am_trnaMT_MetCAT_MT_+_4402_4469 | 1 | 21 | 21 | AGTAAGGTCAGCTAAATAAGC | MB231 | 27 | 36 | 5 | 378 | 221 | 9.5 | 9.2 | 3.2 | 178.3 | 104.8 |
| >multi-am_trnaMT_MetCAT_MT_+_4402_4469@1.22.22\|AGTAAGGTCAGCTAAATAAGCT | >multi-am_trnaMT_MetCAT_MT_+_4402_4469 | 1 | 22 | 22 | AGTAAGGTCAGCTAAATAAGCT | BT20 | 62 | 7 | 3 | 35 | 20 | 36.0 | 2.6 | 1.0 | 29.5 | 20.2 |
| >multi-am_trnaMT_MetCAT_MT_+_4402_4469@1.22.22\|AGTAAGGTCAGCTAAATAAGCT | >multi-am_trnaMT_MetCAT_MT_+_4402_4469 | 1 | 22 | 22 | AGTAAGGTCAGCTAAATAAGCT | MB231 | 17 | 11 | 1 | 98 | 57 | 9.5 | 5.8 | 0.4 | 44.4 | 30.4 |
| >multi-am_trnaMT_MetCAT_MT_+_4402_4469@1.47.47\|AGTAAGGTCAGCTAAATAAGCTATCGGGCCCATACCCCGAAAATGTT | >multi-am_trnaMT_MetCAT_MT_+_4402_4469 | 1 | 47 | 47 | AGTAAGGTCAGCTAAATAAGCTATCGGGCCCATACCCCGAAAATGTT | BT20 | 25 | 11 | 6 | 128 | 21 | 35.5 | 7.0 | 9.7 | 161.5 | 27.9 |
| >multi-am_trnaMT_MetCAT_MT_+_4402_4469@1.47.47\|AGTAAGGTCAGCTAAATAAGCTATCGGGCCCATACCCCGAAAATGTT | >multi-am_trnaMT_MetCAT_MT_+_4402_4469 | 1 | 47 | 47 | AGTAAGGTCAGCTAAATAAGCTATCGGGCCCATACCCCGAAAATGTT | MB231 | 25 | 11 | 5 | 21 | 41 | 22.4 | 13.6 | 4.5 | 30.7 | 32.3 |
| >multi-am_trnaMT_PheGAA_MT_+_577_647@40.57.18\|TGTTTAGACGGGCTCACA | >multi-am_trnaMT_PheGAA_MT_+_577_647 | 40 | 57 | 18 | TGTTTAGACGGGCTCACA | BT20 | 40 | 14 | 28 | 39 | 64 | 8.5 | 4.5 | 23.7 | 19.4 | 23.4 |
| >multi-am_trnaMT_PheGAA_MT_+_577_647@40.57.18\|TGTTTAGACGGGCTCACA | >multi-am_trnaMT_PheGAA_MT_+_577_647 | 40 | 57 | 18 | TGTTTAGACGGGCTCACA | MB231 | 22 | 22 | 12 | 17 | 62 | 8.0 | 5.9 | 0.7 | 1.3 | 12.5 |
| >multi-am_trnaMT_PheGAA_MT_+_577_647@40.58.19\|TGTTTAGACGGGCTCACAT | >multi-am_trnaMT_PheGAA_MT_+_577_647 | 40 | 58 | 19 | TGTTTAGACGGGCTCACAT | BT20 | 14 | 3 | 6 | 15 | 28 | 9.2 | 0.5 | 4.9 | 9.0 | 13.5 |
| >multi-am_trnaMT_PheGAA_MT_+_577_647@40.58.19\|TGTTTAGACGGGCTCACAT | >multi-am_trnaMT_PheGAA_MT_+_577_647 | 40 | 58 | 19 | TGTTTAGACGGGCTCACAT | MB231 | 10 | 12 | 4 | 12 | 39 | 3.9 | 3.9 | 1.1 | 5.6 | 14.7 |
| >multi-am_trnaMT_PheGAA_MT_+_577_647@40.59.20\|TGTTTAGACGGGCTCACATC | >multi-am_trnaMT_PheGAA_MT_+_577_647 | 40 | 59 | 20 | TGTTTAGACGGGCTCACATC | BT20 | 16 | 3 | 19 | 20 | 15 | 8.1 | 2.3 | 19.9 | 22.0 | 14.5 |
| >multi-am_trnaMT_PheGAA_MT_+_577_647@40.62.23\|TGTTTAGACGGGCTCACATCACC | >multi-am_trnaMT_PheGAA_MT_+_577_647 | 40 | 62 | 23 | TGTTTAGACGGGCTCACATCACC | BT20 | 15 | 5 | 18 | 26 | 24 | 9.3 | 3.2 | 16.4 | 25.2 | 25.0 |
| >multi-am_trnaMT_PheGAA_MT_+_577_647@40.62.23\|TGTTTAGACGGGCTCACATCACC | >multi-am_trnaMT_PheGAA_MT_+_577_647 | 40 | 62 | 23 | TGTTTAGACGGGCTCACATCACC | MB231 | 14 | 10 | 10 | 11 | 26 | 6.4 | 1.6 | 5.2 | 8.9 | 16.4 |
| >multi-am_trnaMT_PheGAA_MT_+_577_647@41.58.18\|GTTTAGACGGGCTCACAT | >multi-am_trnaMT_PheGAA_MT_+_577_647 | 41 | 58 | 18 | GTTTAGACGGGCTCACAT | BT20 | 35 | 10 | 14 | 47 | 93 | 25.6 | 2.5 | 9.2 | 24.3 | 52.7 |
| >multi-am_trnaMT_PheGAA_MT_+_577_647@41.58.18\|GTTTAGACGGGCTCACAT | >multi-am_trnaMT_PheGAA_MT_+_577_647 | 41 | 58 | 18 | GTTTAGACGGGCTCACAT | MB231 | 20 | 28 | 8 | 35 | 113 | 1.6 | 8.1 | 1.5 | 16.6 | 46.9 |
| >multi-am_trnaMT_PheGAA_MT_+_577_647@41.59.19\|GTTTAGACGGGCTCACATC | >multi-am_trnaMT_PheGAA_MT_+_577_647 | 41 | 59 | 19 | GTTTAGACGGGCTCACATC | BT20 | 30 | 8 | 34 | 53 | 40 | 19.3 | 3.2 | 26.0 | 57.3 | 42.4 |
| >multi-am_trnaMT_PheGAA_MT_+_577_647@41.59.19\|GTTTAGACGGGCTCACATC | >multi-am_trnaMT_PheGAA_MT_+_577_647 | 41 | 59 | 19 | GTTTAGACGGGCTCACATC | MB231 | 15 | 10 | 16 | 16 | 22 | 9.3 | 2.3 | 6.3 | 9.9 | 9.8 |
| >multi-am_trnaMT_PheGAA_MT_+_577_647@41.60.20\|GTTTAGACGGGCTCACATCA | >multi-am_trnaMT_PheGAA_MT_+_577_647 | 41 | 60 | 20 | GTTTAGACGGGCTCACATCA | BT20 | 11 | 4 | 10 | 17 | 17 | 7.3 | 3.5 | 9.0 | 14.7 | 16.2 |
| >multi-am_trnaMT_PheGAA_MT_+_577_647@41.62.22\|GTTTAGACGGGCTCACATCACC | >multi-am_trnaMT_PheGAA_MT_+_577_647 | 41 | 62 | 22 | GTTTAGACGGGCTCACATCACC | BT20 | 31 | 13 | 54 | 65 | 57 | 21.6 | 7.6 | 44.3 | 73.5 | 69.6 |
| >multi-am_trnaMT_PheGAA_MT_+_577_647@41.62.22\|GTTTAGACGGGCTCACATCACC | >multi-am_trnaMT_PheGAA_MT_+_577_647 | 41 | 62 | 22 | GTTTAGACGGGCTCACATCACC | MB231 | 26 | 23 | 27 | 25 | 66 | 11.8 | 5.6 | 12.7 | 20.0 | 40.8 |
| >multi-am_trnaMT_PheGAA_MT_+_577_647@42.59.18\|TTTAGACGGGCTCACATC | >multi-am_trnaMT_PheGAA_MT_+_577_647 | 42 | 59 | 18 | TTTAGACGGGCTCACATC | BT20 | 11 | 2 | 11 | 26 | 16 | 7.5 | 1.6 | 8.4 | 29.2 | 20.9 |
| >multi-am_trnaMT_PheGAA_MT_+_577_647@42.62.21\|TTTAGACGGGCTCACATCACC | >multi-am_trnaMT_PheGAA_MT_+_577_647 | 42 | 62 | 21 | TTTAGACGGGCTCACATCACC | BT20 | 18 | 5 | 30 | 30 | 24 | 12.5 | 3.7 | 23.5 | 37.6 | 30.7 |
| >multi-am_trnaMT_PheGAA_MT_+_577_647@42.62.21\|TTTAGACGGGCTCACATCACC | >multi-am_trnaMT_PheGAA_MT_+_577_647 | 42 | 62 | 21 | TTTAGACGGGCTCACATCACC | MB231 | 13 | 8 | 11 | 8 | 22 | 7.9 | 1.2 | 5.4 | 6.8 | 10.4 |
| >multi-am_trnaMT_PheGAA_MT_+_577_647@43.62.20\|TTAGACGGGCTCACATCACC | >multi-am_trnaMT_PheGAA_MT_+_577_647 | 43 | 62 | 20 | TTAGACGGGCTCACATCACC | BT20 | 10 | 4 | 19 | 15 | 13 | 7.6 | 2.7 | 16.0 | 18.0 | 16.8 |
| >multi-am_trnaMT_PheGAA_MT_+_577_647@44.62.19\|TAGACGGGCTCACATCACC | >multi-am_trnaMT_PheGAA_MT_+_577_647 | 44 | 62 | 19 | TAGACGGGCTCACATCACC | BT20 | 39 | 15 | 60 | 56 | 54 | 31.6 | 10.2 | 46.9 | 63.3 | 68.0 |
| >multi-am_trnaMT_PheGAA_MT_+_577_647@44.62.19\|TAGACGGGCTCACATCACC | >multi-am_trnaMT_PheGAA_MT_+_577_647 | 44 | 62 | 19 | TAGACGGGCTCACATCACC | MB231 | 26 | 19 | 20 | 15 | 51 | 16.4 | 4.4 | 11.1 | 9.2 | 26.4 |
| >multi-am_trnaMT_PheGAA_MT_+_577_647@45.62.18\|AGACGGGCTCACATCACC | >multi-am_trnaMT_PheGAA_MT_+_577_647 | 45 | 62 | 18 | AGACGGGCTCACATCACC | BT20 | 22 | 12 | 34 | 25 | 26 | 21.4 | 9.1 | 17.8 | 25.2 | 26.1 |
| >multi-am_trnaMT_PheGAA_MT_+_577_647@45.62.18\|AGACGGGCTCACATCACC | >multi-am_trnaMT_PheGAA_MT_+_577_647 | 45 | 62 | 18 | AGACGGGCTCACATCACC | MB468 | 10 | 23 | 7 | 5 | 9 | 8.3 | 30.1 | 7.7 | 1.2 | 2.9 |
| >multi-am_trnaMT_ProTGG_MT_-_15956_16023@52.70.19\|AAAGACTTTTTCTCTGACC | >multi-am_trnaMT_ProTGG_MT_-_15956_16023 | 52 | 70 | 19 | AAAGACTTTTTCTCTGACC | BT20 | 112 | 10 | 14 | 42 | 37 | 69.8 | 0.5 | 8.8 | 29.1 | 26.3 |
| >multi-am_trnaMT_ProTGG_MT_-_15956_16023@52.70.19\|AAAGACTTTTTCTCTGACC | >multi-am_trnaMT_ProTGG_MT_-_15956_16023 | 52 | 70 | 19 | AAAGACTTTTTCTCTGACC | MB231 | 123 | 36 | 13 | 211 | 66 | 113.7 | 20.4 | 9.0 | 209.9 | 22.6 |
| >multi-am_trnaMT_ProTGG_MT_-_15956_16023@52.71.20\|AAAGACTTTTTCTCTGACCA | >multi-am_trnaMT_ProTGG_MT_-_15956_16023 | 52 | 71 | 20 | AAAGACTTTTTCTCTGACCA | BT20 | 355 | 40 | 67 | 55 | 121 | 193.0 | 19.6 | 12.3 | 33.7 | 113.7 |
| >multi-am_trnaMT_ProTGG_MT_-_15956_16023@52.71.20\|AAAGACTTTTTCTCTGACCA | >multi-am_trnaMT_ProTGG_MT_-_15956_16023 | 52 | 71 | 20 | AAAGACTTTTTCTCTGACCA | MB231 | 311 | 97 | 38 | 353 | 195 | 292.7 | 48.4 | 26.1 | 400.4 | 133.8 |
| >multi-am_trnaMT_ProTGG_MT_-_15956_16023@53.70.18\|AAGACTTTTTCTCTGACC | >multi-am_trnaMT_ProTGG_MT_-_15956_16023 | 53 | 70 | 18 | AAGACTTTTTCTCTGACC | BT20 | 88 | 9 | 26 | 27 | 42 | 39.7 | 0.7 | 18.8 | 14.0 | 42.0 |
| >multi-am_trnaMT_ProTGG_MT_-_15956_16023@53.70.18\|AAGACTTTTTCTCTGACC | >multi-am_trnaMT_ProTGG_MT_-_15956_16023 | 53 | 70 | 18 | AAGACTTTTTCTCTGACC | MB231 | 99 | 26 | 17 | 156 | 65 | 89.0 | 12.8 | 10.9 | 148.3 | 28.3 |
| >multi-am_trnaMT_ProTGG_MT_-_15956_16023@53.71.19\|AAGACTTTTTCTCTGACCA | >multi-am_trnaMT_ProTGG_MT_-_15956_16023 | 53 | 71 | 19 | AAGACTTTTTCTCTGACCA | BT20 | 266 | 39 | 97 | 43 | 181 | 59.8 | 20.2 | 32.9 | 19.0 | 230.0 |
| >multi-am_trnaMT_ProTGG_MT_-_15956_16023@53.71.19\|AAGACTTTTTCTCTGACCA | >multi-am_trnaMT_ProTGG_MT_-_15956_16023 | 53 | 71 | 19 | AAGACTTTTTCTCTGACCA | MB231 | 261 | 87 | 52 | 274 | 229 | 183.0 | 22.9 | 24.4 | 298.1 | 180.5 |
| >multi-am_trnaMT_ProTGG_MT_-_15956_16023@53.71.19\|AAGACTTTTTCTCTGACCA | >multi-am_trnaMT_ProTGG_MT_-_15956_16023 | 53 | 71 | 19 | AAGACTTTTTCTCTGACCA | MB468 | 54 | 31 | 36 | 56 | 156 | 15.0 | 41.1 | 39.5 | 32.4 | 167.5 |
| >multi-am_trnaMT_ProTGG_MT_-_15956_16023@54.71.18\|AGACTTTTTCTCTGACCA | >multi-am_trnaMT_ProTGG_MT_-_15956_16023 | 54 | 71 | 18 | AGACTTTTTCTCTGACCA | BT20 | 383 | 44 | 126 | 36 | 104 | 123.7 | 19.8 | 23.1 | 15.3 | 110.9 |
| >multi-am_trnaMT_ProTGG_MT_-_15956_16023@54.71.18\|AGACTTTTTCTCTGACCA | >multi-am_trnaMT_ProTGG_MT_-_15956_16023 | 54 | 71 | 18 | AGACTTTTTCTCTGACCA | MB231 | 328 | 93 | 72 | 187 | 153 | 220.8 | 29.7 | 35.3 | 195.5 | 117.2 |
| >multi-am_trnaMT_ProTGG_MT_-_15956_16023@54.71.18\|AGACTTTTTCTCTGACCA | >multi-am_trnaMT_ProTGG_MT_-_15956_16023 | 54 | 71 | 18 | AGACTTTTTCTCTGACCA | MB468 | 22 | 11 | 14 | 25 | 71 | 4.8 | 14.4 | 12.8 | 16.8 | 71.1 |
| >multi-am_trnaMT_SerGCT_MT_+_12207_12265@12.31.20\|CAAGAACTGCTAACTCATGC | >multi-am_trnaMT_SerGCT_MT_+_12207_12265 | 12 | 31 | 20 | CAAGAACTGCTAACTCATGC | BT20 | 13 | 3 | 12 | 11 | 9 | 4.2 | 0.8 | 6.9 | 7.1 | 7.6 |
| >multi-am_trnaMT_SerGCT_MT_+_12207_12265@12.32.21\|CAAGAACTGCTAACTCATGCC | >multi-am_trnaMT_SerGCT_MT_+_12207_12265 | 12 | 32 | 21 | CAAGAACTGCTAACTCATGCC | BT20 | 17 | 3 | 19 | 9 | 9 | 2.8 | 1.2 | 11.9 | 7.1 | 9.8 |
| >multi-am_trnaMT_SerGCT_MT_+_12207_12265@12.32.21\|CAAGAACTGCTAACTCATGCC | >multi-am_trnaMT_SerGCT_MT_+_12207_12265 | 12 | 32 | 21 | CAAGAACTGCTAACTCATGCC | MB231 | 10 | 2 | 2 | 5 | 5 | 10.2 | 0.9 | 0.5 | 3.1 | 1.0 |
| >multi-am_trnaMT_SerGCT_MT_+_12207_12265@12.38.27\|CAAGAACTGCTAACTCATGCCCCCATG | >multi-am_trnaMT_SerGCT_MT_+_12207_12265 | 12 | 38 | 27 | CAAGAACTGCTAACTCATGCCCCCATG | MB468 | 20 | 100 | 20 | 10 | 42 | 17.5 | 111.5 | 24.6 | 4.3 | 15.5 |
| >multi-am_trnaMT_SerGCT_MT_+_12207_12265@13.31.19\|AAGAACTGCTAACTCATGC | >multi-am_trnaMT_SerGCT_MT_+_12207_12265 | 13 | 31 | 19 | AAGAACTGCTAACTCATGC | BT20 | 13 | 4 | 12 | 16 | 7 | 5.3 | 0.9 | 7.3 | 11.8 | 4.2 |
| >multi-am_trnaMT_SerGCT_MT_+_12207_12265@13.32.20\|AAGAACTGCTAACTCATGCC | >multi-am_trnaMT_SerGCT_MT_+_12207_12265 | 13 | 32 | 20 | AAGAACTGCTAACTCATGCC | BT20 | 12 | 3 | 16 | 8 | 6 | 2.8 | 1.0 | 10.7 | 6.6 | 6.1 |
| >multi-am_trnaMT_SerGCT_MT_+_12207_12265@13.38.26\|AAGAACTGCTAACTCATGCCCCCATG | >multi-am_trnaMT_SerGCT_MT_+_12207_12265 | 13 | 38 | 26 | AAGAACTGCTAACTCATGCCCCCATG | MB468 | 15 | 76 | 19 | 10 | 18 | 13.0 | 79.5 | 23.8 | 7.5 | 11.3 |
| >multi-am_trnaMT_SerGCT_MT_+_12207_12265@14.31.18\|AGAACTGCTAACTCATGC | >multi-am_trnaMT_SerGCT_MT_+_12207_12265 | 14 | 31 | 18 | AGAACTGCTAACTCATGC | BT20 | 32 | 9 | 29 | 22 | 9 | 18.2 | 3.2 | 19.5 | 20.5 | 2.8 |
| >multi-am_trnaMT_SerGCT_MT_+_12207_12265@14.31.18\|AGAACTGCTAACTCATGC | >multi-am_trnaMT_SerGCT_MT_+_12207_12265 | 14 | 31 | 18 | AGAACTGCTAACTCATGC | MB231 | 23 | 7 | 5 | 34 | 8 | 19.1 | 3.9 | 1.6 | 27.1 | 2.6 |
| >multi-am_trnaMT_SerGCT_MT_+_12207_12265@14.32.19\|AGAACTGCTAACTCATGCC | >multi-am_trnaMT_SerGCT_MT_+_12207_12265 | 14 | 32 | 19 | AGAACTGCTAACTCATGCC | BT20 | 22 | 6 | 35 | 8 | 7 | 6.3 | 2.3 | 24.7 | 5.9 | 5.7 |
| >multi-am_trnaMT_SerGCT_MT_+_12207_12265@14.32.19\|AGAACTGCTAACTCATGCC | >multi-am_trnaMT_SerGCT_MT_+_12207_12265 | 14 | 32 | 19 | AGAACTGCTAACTCATGCC | MB231 | 20 | 4 | 4 | 9 | 6 | 22.9 | 2.9 | 1.9 | 7.3 | 2.6 |
| >multi-am_trnaMT_SerGCT_MT_+_12207_12265@16.38.23\|AACTGCTAACTCATGCCCCCATG | >multi-am_trnaMT_SerGCT_MT_+_12207_12265 | 16 | 38 | 23 | AACTGCTAACTCATGCCCCCATG | MB468 | 22 | 57 | 13 | 21 | 21 | 16.2 | 50.6 | 14.1 | 15.1 | 11.1 |
| >multi-am_trnaMT_TyrGTA_MT_-_5826_5891@1.32.32\|GGTAAAATGGCTGAGTGAAGCATTGGACTGTA | >multi-am_trnaMT_TyrGTA_MT_-_5826_5891 | 1 | 32 | 32 | GGTAAAATGGCTGAGTGAAGCATTGGACTGTA | BT20 | 38 | 3 | 1 | 20 | 14 | 29.3 | 0.7 | 0.6 | 27.6 | 20.8 |
| >multi-am_trnaMT_TyrGTA_MT_-_5826_5891@1.32.32\|GGTAAAATGGCTGAGTGAAGCATTGGACTGTA | >multi-am_trnaMT_TyrGTA_MT_-_5826_5891 | 1 | 32 | 32 | GGTAAAATGGCTGAGTGAAGCATTGGACTGTA | MB231 | 64 | 10 | 2 | 43 | 123 | 48.9 | 11.0 | 3.6 | 44.2 | 135.0 |
| >multi-am_trnaMT_TyrGTA_MT_-_5826_5891@1.33.33\|GGTAAAATGGCTGAGTGAAGCATTGGACTGTAA | >multi-am_trnaMT_TyrGTA_MT_-_5826_5891 | 1 | 33 | 33 | GGTAAAATGGCTGAGTGAAGCATTGGACTGTAA | MB231 | 12 | 2 | 1 | 16 | 26 | 7.6 | 2.0 | 0.7 | 19.5 | 24.1 |
| >multi-am_trnaMT_ValTAC_MT_+_1602_1670@28.47.20\|AACTTACACTTAGGAGATTT | >multi-am_trnaMT_ValTAC_MT_+_1602_1670 | 28 | 47 | 20 | AACTTACACTTAGGAGATTT | BT20 | 17 | 11 | 4 | 1 | 0 | 14.5 | 14.0 | 1.3 | 1.0 | 0.5 |
| >multi-am_trnaMT_ValTAC_MT_+_1602_1670@28.48.21\|AACTTACACTTAGGAGATTTC | >multi-am_trnaMT_ValTAC_MT_+_1602_1670 | 28 | 48 | 21 | AACTTACACTTAGGAGATTTC | BT20 | 12 | 12 | 8 | 8 | 2 | 4.0 | 13.1 | 2.7 | 8.1 | 1.9 |
| >multi-am_trnaMT_ValTAC_MT_+_1602_1670@28.52.25\|AACTTACACTTAGGAGATTTCAACT | >multi-am_trnaMT_ValTAC_MT_+_1602_1670 | 28 | 52 | 25 | AACTTACACTTAGGAGATTTCAACT | BT20 | 13 | 7 | 17 | 6 | 4 | 1.2 | 5.9 | 6.8 | 7.7 | 5.1 |
| >multi-am_trnaMT_ValTAC_MT_+_1602_1670@28.57.30\|AACTTACACTTAGGAGATTTCAACTTAACT | >multi-am_trnaMT_ValTAC_MT_+_1602_1670 | 28 | 57 | 30 | AACTTACACTTAGGAGATTTCAACTTAACT | BT20 | 11 | 2 | 5 | 26 | 4 | 4.2 | 2.3 | 2.3 | 25.0 | 5.0 |
| >multi-am_trnaMT_ValTAC_MT_+_1602_1670@29.52.24\|ACTTACACTTAGGAGATTTCAACT | >multi-am_trnaMT_ValTAC_MT_+_1602_1670 | 29 | 52 | 24 | ACTTACACTTAGGAGATTTCAACT | BT20 | 14 | 6 | 23 | 6 | 6 | 10.4 | 5.4 | 20.1 | 8.4 | 8.4 |
| >multi-am_trnaMT_ValTAC_MT_+_1602_1670@33.52.20\|ACACTTAGGAGATTTCAACT | >multi-am_trnaMT_ValTAC_MT_+_1602_1670 | 33 | 52 | 20 | ACACTTAGGAGATTTCAACT | BT20 | 11 | 6 | 18 | 3 | 3 | 11.3 | 5.4 | 16.5 | 3.0 | 3.9 |
| >multi-am_trnaMT_ValTAC_MT_+_1602_1670@36.62.27\|CTTAGGAGATTTCAACTTAACTTGACC | >multi-am_trnaMT_ValTAC_MT_+_1602_1670 | 36 | 62 | 27 | CTTAGGAGATTTCAACTTAACTTGACC | BT20 | 13 | 2 | 6 | 39 | 18 | 10.5 | 2.3 | 7.1 | 55.0 | 26.6 |
| >multi-am_trnaMT_ValTAC_MT_+_1602_1670@38.57.20\|TAGGAGATTTCAACTTAACT | >multi-am_trnaMT_ValTAC_MT_+_1602_1670 | 38 | 57 | 20 | TAGGAGATTTCAACTTAACT | BT20 | 19 | 12 | 13 | 25 | 6 | 8.6 | 15.1 | 13.1 | 25.1 | 6.7 |
| >multi-am_trnaMT_ValTAC_MT_+_1602_1670@38.62.25\|TAGGAGATTTCAACTTAACTTGACC | >multi-am_trnaMT_ValTAC_MT_+_1602_1670 | 38 | 62 | 25 | TAGGAGATTTCAACTTAACTTGACC | BT20 | 21 | 3 | 11 | 55 | 31 | 19.2 | 2.6 | 14.7 | 83.6 | 45.8 |
| >multi-am_trnaMT_ValTAC_MT_+_1602_1670@44.68.25\|ATTTCAACTTAACTTGACCGCTCTG | >multi-am_trnaMT_ValTAC_MT_+_1602_1670 | 44 | 68 | 25 | ATTTCAACTTAACTTGACCGCTCTG | MB468 | 16 | 67 | 26 | 5 | 22 | 15.7 | 93.8 | 26.0 | 3.1 | 27.8 |
| >multi-am_trnaMT_ValTAC_MT_+_1602_1670@48.65.18\|CAACTTAACTTGACCGCT | >multi-am_trnaMT_ValTAC_MT_+_1602_1670 | 48 | 65 | 18 | CAACTTAACTTGACCGCT | BT20 | 11 | 6 | 21 | 3 | 9 | 4.8 | 6.8 | 18.9 | 3.3 | 12.8 |
| >multi-am_trnaMT_ValTAC_MT_+_1602_1670@48.66.19\|CAACTTAACTTGACCGCTC | >multi-am_trnaMT_ValTAC_MT_+_1602_1670 | 48 | 66 | 19 | CAACTTAACTTGACCGCTC | BT20 | 15 | 12 | 48 | 6 | 9 | 12.9 | 15.4 | 46.8 | 7.3 | 12.5 |
| >multi-am_trnaMT_ValTAC_MT_+_1602_1670@48.67.20\|CAACTTAACTTGACCGCTCT | >multi-am_trnaMT_ValTAC_MT_+_1602_1670 | 48 | 67 | 20 | CAACTTAACTTGACCGCTCT | BT20 | 255 | 230 | 784 | 13 | 105 | 253.6 | 282.6 | 791.4 | 16.1 | 137.3 |
| >multi-am_trnaMT_ValTAC_MT_+_1602_1670@48.67.20\|CAACTTAACTTGACCGCTCT | >multi-am_trnaMT_ValTAC_MT_+_1602_1670 | 48 | 67 | 20 | CAACTTAACTTGACCGCTCT | MB231 | 85 | 66 | 132 | 4 | 51 | 32.0 | 17.8 | 46.7 | 4.1 | 14.1 |
| >multi-am_trnaMT_ValTAC_MT_+_1602_1670@48.68.21\|CAACTTAACTTGACCGCTCTG | >multi-am_trnaMT_ValTAC_MT_+_1602_1670 | 48 | 68 | 21 | CAACTTAACTTGACCGCTCTG | BT20 | 18 | 9 | 40 | 8 | 16 | 14.6 | 6.1 | 23.3 | 9.7 | 14.9 |
| >multi-am_trnaMT_ValTAC_MT_+_1602_1670@48.68.21\|CAACTTAACTTGACCGCTCTG | >multi-am_trnaMT_ValTAC_MT_+_1602_1670 | 48 | 68 | 21 | CAACTTAACTTGACCGCTCTG | MB468 | 12 | 54 | 12 | 8 | 31 | 10.8 | 76.0 | 12.0 | 2.5 | 36.7 |
| >multi-am_trnaMT_ValTAC_MT_+_1602_1670@49.66.18\|AACTTAACTTGACCGCTC | >multi-am_trnaMT_ValTAC_MT_+_1602_1670 | 49 | 66 | 18 | AACTTAACTTGACCGCTC | BT20 | 13 | 5 | 22 | 4 | 5 | 13.8 | 6.0 | 23.9 | 4.8 | 6.3 |
| >multi-am_trnaMT_ValTAC_MT_+_1602_1670@49.67.19\|AACTTAACTTGACCGCTCT | >multi-am_trnaMT_ValTAC_MT_+_1602_1670 | 49 | 67 | 19 | AACTTAACTTGACCGCTCT | BT20 | 166 | 82 | 302 | 11 | 61 | 182.7 | 92.5 | 303.2 | 12.2 | 69.7 |
| >multi-am_trnaMT_ValTAC_MT_+_1602_1670@49.67.19\|AACTTAACTTGACCGCTCT | >multi-am_trnaMT_ValTAC_MT_+_1602_1670 | 49 | 67 | 19 | AACTTAACTTGACCGCTCT | MB231 | 26 | 21 | 21 | 3 | 14 | 15.1 | 5.8 | 6.2 | 2.5 | 1.7 |
| >multi-am_trnaMT_ValTAC_MT_+_1602_1670@49.68.20\|AACTTAACTTGACCGCTCTG | >multi-am_trnaMT_ValTAC_MT_+_1602_1670 | 49 | 68 | 20 | AACTTAACTTGACCGCTCTG | BT20 | 11 | 4 | 15 | 15 | 16 | 9.0 | 1.8 | 9.5 | 14.5 | 13.5 |
| >multi-am_trnaMT_ValTAC_MT_+_1602_1670@49.68.20\|AACTTAACTTGACCGCTCTG | >multi-am_trnaMT_ValTAC_MT_+_1602_1670 | 49 | 68 | 20 | AACTTAACTTGACCGCTCTG | MB468 | 53 | 250 | 40 | 26 | 41 | 47.4 | 355.7 | 40.6 | 9.5 | 36.6 |
| >multi-am_trnaMT_ValTAC_MT_+_1602_1670@50.67.18\|ACTTAACTTGACCGCTCT | >multi-am_trnaMT_ValTAC_MT_+_1602_1670 | 50 | 67 | 18 | ACTTAACTTGACCGCTCT | BT20 | 190 | 83 | 351 | 13 | 73 | 216.8 | 101.9 | 378.0 | 16.8 | 91.4 |
| >multi-am_trnaMT_ValTAC_MT_+_1602_1670@50.67.18\|ACTTAACTTGACCGCTCT | >multi-am_trnaMT_ValTAC_MT_+_1602_1670 | 50 | 67 | 18 | ACTTAACTTGACCGCTCT | MB231 | 41 | 39 | 34 | 5 | 22 | 20.9 | 21.7 | 2.6 | 5.6 | 2.0 |
| >multi-am_trnaMT_ValTAC_MT_+_1602_1670@50.67.18\|ACTTAACTTGACCGCTCT | >multi-am_trnaMT_ValTAC_MT_+_1602_1670 | 50 | 67 | 18 | ACTTAACTTGACCGCTCT | MB468 | 22 | 42 | 28 | 5 | 9 | 23.5 | 54.9 | 32.6 | 2.6 | 3.7 |
| >multi-am_trnaMT_ValTAC_MT_+_1602_1670@50.68.19\|ACTTAACTTGACCGCTCTG | >multi-am_trnaMT_ValTAC_MT_+_1602_1670 | 50 | 68 | 19 | ACTTAACTTGACCGCTCTG | BT20 | 14 | 6 | 27 | 17 | 17 | 11.6 | 2.8 | 20.7 | 10.0 | 13.8 |
| >multi-am_trnaMT_ValTAC_MT_+_1602_1670@50.68.19\|ACTTAACTTGACCGCTCTG | >multi-am_trnaMT_ValTAC_MT_+_1602_1670 | 50 | 68 | 19 | ACTTAACTTGACCGCTCTG | MB468 | 399 | 1768 | 340 | 205 | 186 | 366.5 | 2421.3 | 376.1 | 93.2 | 83.5 |
| >multi-am_trnaMT_ValTAC_MT_+_1602_1670@51.68.18\|CTTAACTTGACCGCTCTG | >multi-am_trnaMT_ValTAC_MT_+_1602_1670 | 51 | 68 | 18 | CTTAACTTGACCGCTCTG | MB468 | 1785 | 3832 | 1943 | 753 | 941 | 1621.0 | 5560.1 | 1962.8 | 556.1 | 514.0 |
| >multi-am_trnaMT_ValTAC_MT_+_1602_1670@51.69.19\|CTTAACTTGACCGCTCTGA | >multi-am_trnaMT_ValTAC_MT_+_1602_1670 | 51 | 69 | 19 | CTTAACTTGACCGCTCTGA | MB468 | 16 | 34 | 16 | 9 | 10 | 13.5 | 39.9 | 14.6 | 6.0 | 6.6 |
| >multi-hg38_MT_+_1621_3279_RNR2_16S_50ntFlanks@1000.1017.18\|GGGACCTGTATGAATGGC | >multi-hg38_MT_+_1621_3279_RNR2_16S_50ntFlanks | 1000 | 1017 | 18 | GGGACCTGTATGAATGGC | MB231 | 24 | 4 | 2 | 35 | 14 | 33.3 | 4.4 | 1.1 | 31.2 | 12.4 |

| | | | | | | | | | | | | | | | | |
|---|---|---|---|---|---|---|---|---|---|---|---|---|---|---|---|---|
| >multi-hg38_MT_+_1621_3279_RNR2_16S_50ntFlanks@1095.1120.26\|AGACGAGAAGACCCTATGGAGCTTTA | >multi-hg38_MT_+_1621_3279_RNR2_16S_50ntFlanks | 1095 | 1120 | 26 | AGACGAGAAGACCCTATGGAGCTTTA | MB231 | 13 | 9 | 1 | 49 | 17 | 13.8 | 8.8 | 0.7 | 36.0 | 9.5 |
| >multi-hg38_MT_+_1621_3279_RNR2_16S_50ntFlanks@1349.1382.34\|ACAATAGGGTTTACGACCTCGATGTTGGATCAGG | >multi-hg38_MT_+_1621_3279_RNR2_16S_50ntFlanks | 1349 | 1382 | 34 | ACAATAGGGTTTACGACCTCGATGTTGGATCAGG | MB231 | 11 | 4 | 0 | 27 | 10 | 13.7 | 4.7 | 0.7 | 20.2 | 6.8 |
| >multi-hg38_MT_+_1621_3279_RNR2_16S_50ntFlanks@1349.1385.37\|ACAATAGGGTTTACGACCTCGATGTTGGATCAGGACA | >multi-hg38_MT_+_1621_3279_RNR2_16S_50ntFlanks | 1349 | 1385 | 37 | ACAATAGGGTTTACGACCTCGATGTTGGATCAGGACA | MB231 | 25 | 6 | 2 | 33 | 17 | 33.6 | 4.7 | 1.7 | 21.7 | 10.9 |
| >multi-hg38_MT_+_1621_3279_RNR2_16S_50ntFlanks@1349.1388.40\|ACAATAGGGTTTACGACCTCGATGTTGGATCAGGACATCC | >multi-hg38_MT_+_1621_3279_RNR2_16S_50ntFlanks | 1349 | 1388 | 40 | ACAATAGGGTTTACGACCTCGATGTTGGATCAGGACATCC | BT20 | 15 | 3 | 1 | 24 | 13 | 9.0 | 2.8 | 0.7 | 25.1 | 8.4 |
| >multi-hg38_MT_+_1621_3279_RNR2_16S_50ntFlanks@1349.1388.40\|ACAATAGGGTTTACGACCTCGATGTTGGATCAGGACATCC | >multi-hg38_MT_+_1621_3279_RNR2_16S_50ntFlanks | 1349 | 1388 | 40 | ACAATAGGGTTTACGACCTCGATGTTGGATCAGGACATCC | MB231 | 88 | 17 | 6 | 61 | 51 | 119.6 | 10.5 | 4.8 | 42.3 | 20.5 |
| >multi-hg38_MT_+_1621_3279_RNR2_16S_50ntFlanks@1351.1388.38\|AATAGGGTTTACGACCTCGATGTTGGATCAGGACATCC | >multi-hg38_MT_+_1621_3279_RNR2_16S_50ntFlanks | 1351 | 1388 | 38 | AATAGGGTTTACGACCTCGATGTTGGATCAGGACATCC | MB231 | 12 | 3 | 1 | 10 | 7 | 15.3 | 1.9 | 1.1 | 7.3 | 3.6 |
| >multi-hg38_MT_+_1621_3279_RNR2_16S_50ntFlanks@1352.1388.37\|ATAGGGTTTACGACCTCGATGTTGGATCAGGACATCC | >multi-hg38_MT_+_1621_3279_RNR2_16S_50ntFlanks | 1352 | 1388 | 37 | ATAGGGTTTACGACCTCGATGTTGGATCAGGACATCC | MB231 | 17 | 4 | 1 | 13 | 9 | 23.7 | 3.6 | 1.1 | 9.9 | 4.9 |
| >multi-hg38_MT_+_1621_3279_RNR2_16S_50ntFlanks@1356.1388.33\|GGTTTACGACCTCGATGTTGGATCAGGACATCC | >multi-hg38_MT_+_1621_3279_RNR2_16S_50ntFlanks | 1356 | 1388 | 33 | GGTTTACGACCTCGATGTTGGATCAGGACATCC | MB231 | 12 | 4 | 2 | 24 | 8 | 16.5 | 5.6 | 1.7 | 21.3 | 3.6 |
| >multi-hg38_MT_+_1621_3279_RNR2_16S_50ntFlanks@1357.1388.32\|GTTTACGACCTCGATGTTGGATCAGGACATCC | >multi-hg38_MT_+_1621_3279_RNR2_16S_50ntFlanks | 1357 | 1388 | 32 | GTTTACGACCTCGATGTTGGATCAGGACATCC | MB231 | 14 | 8 | 1 | 25 | 15 | 16.8 | 8.2 | 1.7 | 18.8 | 9.5 |
| >multi-hg38_MT_+_1621_3279_RNR2_16S_50ntFlanks@1362.1388.27\|CGACCTCGATGTTGGATCAGGACATCC | >multi-hg38_MT_+_1621_3279_RNR2_16S_50ntFlanks | 1362 | 1388 | 27 | CGACCTCGATGTTGGATCAGGACATCC | MB231 | 27 | 7 | 3 | 17 | 10 | 33.9 | 7.4 | 1.6 | 14.8 | 9.6 |
| >multi-hg38_MT_+_1621_3279_RNR2_16S_50ntFlanks@1363.1388.26\|GACCTCGATGTTGGATCAGGACATCC | >multi-hg38_MT_+_1621_3279_RNR2_16S_50ntFlanks | 1363 | 1388 | 26 | GACCTCGATGTTGGATCAGGACATCC | MB231 | 25 | 9 | 2 | 62 | 19 | 29.0 | 11.7 | 1.2 | 51.4 | 11.7 |
| >multi-hg38_MT_+_1621_3279_RNR2_16S_50ntFlanks@1367.1385.19\|TCGATGTTGGATCAGGACA | >multi-hg38_MT_+_1621_3279_RNR2_16S_50ntFlanks | 1367 | 1385 | 19 | TCGATGTTGGATCAGGACA | MB231 | 18 | 4 | 2 | 38 | 11 | 25.3 | 4.9 | 0.7 | 32.3 | 10.4 |
| >multi-hg38_MT_+_1621_3279_RNR2_16S_50ntFlanks@1370.1387.18\|ATGTTGGATCAGGACATC | >multi-hg38_MT_+_1621_3279_RNR2_16S_50ntFlanks | 1370 | 1387 | 18 | ATGTTGGATCAGGACATC | MB231 | 14 | 2 | 0 | 13 | 4 | 22.0 | 2.8 | 0.5 | 10.6 | 2.0 |
| >multi-hg38_MT_+_1621_3279_RNR2_16S_50ntFlanks@1370.1388.19\|ATGTTGGATCAGGACATCC | >multi-hg38_MT_+_1621_3279_RNR2_16S_50ntFlanks | 1370 | 1388 | 19 | ATGTTGGATCAGGACATCC | BT20 | 23 | 2 | 1 | 21 | 10 | 10.0 | 2.1 | 0.4 | 19.3 | 6.8 |
| >multi-hg38_MT_+_1621_3279_RNR2_16S_50ntFlanks@1370.1388.19\|ATGTTGGATCAGGACATCC | >multi-hg38_MT_+_1621_3279_RNR2_16S_50ntFlanks | 1370 | 1388 | 19 | ATGTTGGATCAGGACATCC | MB231 | 44 | 9 | 2 | 80 | 20 | 57.7 | 13.4 | 2.0 | 66.1 | 14.3 |
| >multi-hg38_MT_+_1621_3279_RNR2_16S_50ntFlanks@1370.1389.20\|ATGTTGGATCAGGACATCCC | >multi-hg38_MT_+_1621_3279_RNR2_16S_50ntFlanks | 1370 | 1389 | 20 | ATGTTGGATCAGGACATCCC | BT20 | 10 | 0 | 0 | 7 | 2 | 8.6 | 0.3 | 0.2 | 6.3 | 1.1 |
| >multi-hg38_MT_+_1621_3279_RNR2_16S_50ntFlanks@1370.1389.20\|ATGTTGGATCAGGACATCCC | >multi-hg38_MT_+_1621_3279_RNR2_16S_50ntFlanks | 1370 | 1389 | 20 | ATGTTGGATCAGGACATCCC | MB231 | 19 | 2 | 1 | 25 | 4 | 28.6 | 2.6 | 0.8 | 19.4 | 1.9 |
| >multi-hg38_MT_+_1621_3279_RNR2_16S_50ntFlanks@1404.1421.18\|TATTAAAGGTTCGTTTGT | >multi-hg38_MT_+_1621_3279_RNR2_16S_50ntFlanks | 1404 | 1421 | 18 | TATTAAAGGTTCGTTTGT | BT20 | 20 | 1 | 2 | 2 | 4 | 16.8 | 0.7 | 2.8 | 2.4 | 6.3 |
| >multi-hg38_MT_+_1621_3279_RNR2_16S_50ntFlanks@1404.1421.18\|TATTAAAGGTTCGTTTGT | >multi-hg38_MT_+_1621_3279_RNR2_16S_50ntFlanks | 1404 | 1421 | 18 | TATTAAAGGTTCGTTTGT | MB231 | 19 | 4 | 1 | 47 | 36 | 6.4 | 2.9 | 0.8 | 30.3 | 27.3 |
| >multi-hg38_MT_+_1621_3279_RNR2_16S_50ntFlanks@1416.1446.31\|GTTTGTTCAACGATTAAAGTCCTACGTGATC | >multi-hg38_MT_+_1621_3279_RNR2_16S_50ntFlanks | 1416 | 1446 | 31 | GTTTGTTCAACGATTAAAGTCCTACGTGATC | MB231 | 10 | 6 | 1 | 26 | 12 | 7.6 | 5.0 | 0.6 | 17.9 | 6.9 |
| >multi-hg38_MT_+_1621_3279_RNR2_16S_50ntFlanks@1427.1451.25\|GATTAAAGTCCTACGTGATCTGAGT | >multi-hg38_MT_+_1621_3279_RNR2_16S_50ntFlanks | 1427 | 1451 | 25 | GATTAAAGTCCTACGTGATCTGAGT | MB231 | 14 | 4 | 2 | 52 | 16 | 18.1 | 3.3 | 0.8 | 46.2 | 4.8 |
| >multi-hg38_MT_+_1621_3279_RNR2_16S_50ntFlanks@1427.1458.32\|GATTAAAGTCCTACGTGATCTGAGTTCAGACC | >multi-hg38_MT_+_1621_3279_RNR2_16S_50ntFlanks | 1427 | 1458 | 32 | GATTAAAGTCCTACGTGATCTGAGTTCAGACC | MB231 | 24 | 8 | 3 | 62 | 31 | 21.4 | 6.7 | 1.5 | 41.7 | 15.0 |
| >multi-hg38_MT_+_1621_3279_RNR2_16S_50ntFlanks@1428.1451.24\|ATTAAAGTCCTACGTGATCTGAGT | >multi-hg38_MT_+_1621_3279_RNR2_16S_50ntFlanks | 1428 | 1451 | 24 | ATTAAAGTCCTACGTGATCTGAGT | MB231 | 10 | 3 | 1 | 24 | 8 | 14.3 | 2.6 | 0.6 | 22.2 | 1.2 |
| >multi-hg38_MT_+_1621_3279_RNR2_16S_50ntFlanks@1428.1452.25\|ATTAAAGTCCTACGTGATCTGAGTT | >multi-hg38_MT_+_1621_3279_RNR2_16S_50ntFlanks | 1428 | 1452 | 25 | ATTAAAGTCCTACGTGATCTGAGTT | MB231 | 18 | 7 | 1 | 40 | 6 | 26.5 | 9.0 | 1.4 | 47.4 | 2.6 |
| >multi-hg38_MT_+_1621_3279_RNR2_16S_50ntFlanks@1428.1453.26\|ATTAAAGTCCTACGTGATCTGAGTTC | >multi-hg38_MT_+_1621_3279_RNR2_16S_50ntFlanks | 1428 | 1453 | 26 | ATTAAAGTCCTACGTGATCTGAGTTC | MB231 | 15 | 6 | 2 | 50 | 9 | 20.2 | 6.2 | 1.2 | 41.6 | 4.3 |
| >multi-hg38_MT_+_1621_3279_RNR2_16S_50ntFlanks@1428.1457.30\|ATTAAAGTCCTACGTGATCTGAGTTCAGAC | >multi-hg38_MT_+_1621_3279_RNR2_16S_50ntFlanks | 1428 | 1457 | 30 | ATTAAAGTCCTACGTGATCTGAGTTCAGAC | MB231 | 12 | 5 | 3 | 25 | 11 | 8.4 | 3.2 | 0.8 | 20.7 | 7.6 |
| >multi-hg38_MT_+_1621_3279_RNR2_16S_50ntFlanks@1428.1458.31\|ATTAAAGTCCTACGTGATCTGAGTTCAGACC | >multi-hg38_MT_+_1621_3279_RNR2_16S_50ntFlanks | 1428 | 1458 | 31 | ATTAAAGTCCTACGTGATCTGAGTTCAGACC | BT20 | 18 | 2 | 1 | 31 | 20 | 8.9 | 1.0 | 0.5 | 36.6 | 22.9 |
| >multi-hg38_MT_+_1621_3279_RNR2_16S_50ntFlanks@1428.1458.31\|ATTAAAGTCCTACGTGATCTGAGTTCAGACC | >multi-hg38_MT_+_1621_3279_RNR2_16S_50ntFlanks | 1428 | 1458 | 31 | ATTAAAGTCCTACGTGATCTGAGTTCAGACC | MB231 | 47 | 20 | 6 | 58 | 36 | 44.6 | 15.4 | 4.4 | 42.2 | 18.7 |
| >multi-hg38_MT_+_1621_3279_RNR2_16S_50ntFlanks@1430.1458.29\|TAAAGTCCTACGTGATCTGAGTTCAGACC | >multi-hg38_MT_+_1621_3279_RNR2_16S_50ntFlanks | 1430 | 1458 | 29 | TAAAGTCCTACGTGATCTGAGTTCAGACC | MB231 | 16 | 3 | 2 | 15 | 9 | 17.1 | 3.2 | 1.4 | 12.9 | 5.8 |
| >multi-hg38_MT_+_1621_3279_RNR2_16S_50ntFlanks@1431.1451.21\|AAAGTCCTACGTGATCTGAGT | >multi-hg38_MT_+_1621_3279_RNR2_16S_50ntFlanks | 1431 | 1451 | 21 | AAAGTCCTACGTGATCTGAGT | MB231 | 16 | 3 | 1 | 30 | 15 | 22.3 | 3.5 | 0.7 | 21.6 | 4.4 |
| >multi-hg38_MT_+_1621_3279_RNR2_16S_50ntFlanks@1431.1452.22\|AAAGTCCTACGTGATCTGAGTT | >multi-hg38_MT_+_1621_3279_RNR2_16S_50ntFlanks | 1431 | 1452 | 22 | AAAGTCCTACGTGATCTGAGTT | MB231 | 12 | 2 | 1 | 21 | 4 | 17.8 | 3.4 | 0.9 | 23.0 | 1.1 |
| >multi-hg38_MT_+_1621_3279_RNR2_16S_50ntFlanks@1431.1453.23\|AAAGTCCTACGTGATCTGAGTTC | >multi-hg38_MT_+_1621_3279_RNR2_16S_50ntFlanks | 1431 | 1453 | 23 | AAAGTCCTACGTGATCTGAGTTC | MB231 | 12 | 4 | 1 | 46 | 7 | 15.5 | 4.3 | 0.8 | 37.3 | 3.8 |
| >multi-hg38_MT_+_1621_3279_RNR2_16S_50ntFlanks@1431.1458.28\|AAAGTCCTACGTGATCTGAGTTCAGACC | >multi-hg38_MT_+_1621_3279_RNR2_16S_50ntFlanks | 1431 | 1458 | 28 | AAAGTCCTACGTGATCTGAGTTCAGACC | MB231 | 14 | 4 | 1 | 27 | 11 | 14.8 | 4.0 | 1.0 | 25.4 | 4.4 |
| >multi-hg38_MT_+_1621_3279_RNR2_16S_50ntFlanks@1432.1451.20\|AAGTCCTACGTGATCTGAGT | >multi-hg38_MT_+_1621_3279_RNR2_16S_50ntFlanks | 1432 | 1451 | 20 | AAGTCCTACGTGATCTGAGT | MB231 | 11 | 2 | 1 | 12 | 6 | 17.0 | 2.5 | 1.4 | 12.5 | 2.5 |
| >multi-hg38_MT_+_1621_3279_RNR2_16S_50ntFlanks@1432.1458.27\|AAGTCCTACGTGATCTGAGTTCAGACC | >multi-hg38_MT_+_1621_3279_RNR2_16S_50ntFlanks | 1432 | 1458 | 27 | AAGTCCTACGTGATCTGAGTTCAGACC | MB231 | 17 | 4 | 1 | 30 | 14 | 19.7 | 5.4 | 1.0 | 28.6 | 5.1 |
| >multi-hg38_MT_+_1621_3279_RNR2_16S_50ntFlanks@1437.1457.21\|CTACGTGATCTGAGTTCAGAC | >multi-hg38_MT_+_1621_3279_RNR2_16S_50ntFlanks | 1437 | 1457 | 21 | CTACGTGATCTGAGTTCAGAC | MB231 | 11 | 2 | 1 | 14 | 3 | 9.8 | 2.0 | 0.7 | 9.9 | 2.2 |

| | | | | | | | | | | | | | | | | |
|---|---|---|---|---|---|---|---|---|---|---|---|---|---|---|---|---|
| >multi-hg38_MT_+_1621_3279_RNR2_16S_50ntFlanks@1438.1455.18\|TACGTGATCTGAGTTCAG | >multi-hg38_MT_+_1621_3279_RNR2_16S_50ntFlanks | 1438 | 1455 | 18 | TACGTGATCTGAGTTCAG | MB231 | 15 | 4 | 1 | 11 | 5 | 15.6 | 3.2 | 0.4 | 9.4 | 4.0 |
| >multi-hg38_MT_+_1621_3279_RNR2_16S_50ntFlanks@1438.1457.20\|TACGTGATCTGAGTTCAGAC | >multi-hg38_MT_+_1621_3279_RNR2_16S_50ntFlanks | 1438 | 1457 | 20 | TACGTGATCTGAGTTCAGAC | BT20 | 13 | 0 | 0 | 13 | 5 | 4.6 | 0.3 | 0.4 | 10.6 | 3.7 |
| >multi-hg38_MT_+_1621_3279_RNR2_16S_50ntFlanks@1438.1457.20\|TACGTGATCTGAGTTCAGAC | >multi-hg38_MT_+_1621_3279_RNR2_16S_50ntFlanks | 1438 | 1457 | 20 | TACGTGATCTGAGTTCAGAC | MB231 | 36 | 7 | 1 | 56 | 13 | 35.1 | 7.2 | 0.8 | 43.7 | 6.7 |
| >multi-hg38_MT_+_1621_3279_RNR2_16S_50ntFlanks@1438.1458.21\|TACGTGATCTGAGTTCAGACC | >multi-hg38_MT_+_1621_3279_RNR2_16S_50ntFlanks | 1438 | 1458 | 21 | TACGTGATCTGAGTTCAGACC | MB231 | 22 | 3 | 1 | 32 | 7 | 21.6 | 4.1 | 0.4 | 26.1 | 4.9 |
| >multi-hg38_MT_+_1621_3279_RNR2_16S_50ntFlanks@1439.1457.19\|ACGTGATCTGAGTTCAGAC | >multi-hg38_MT_+_1621_3279_RNR2_16S_50ntFlanks | 1439 | 1457 | 19 | ACGTGATCTGAGTTCAGAC | MB231 | 13 | 3 | 1 | 37 | 7 | 12.7 | 2.8 | 0.3 | 29.1 | 2.8 |
| >multi-hg38_MT_+_1621_3279_RNR2_16S_50ntFlanks@322.340.19\|CACACCCGTCTATGTAGCA | >multi-hg38_MT_+_1621_3279_RNR2_16S_50ntFlanks | 322 | 340 | 19 | CACACCCGTCTATGTAGCA | MB231 | 24 | 4 | 5 | 8 | 10 | 28.7 | 2.4 | 2.1 | 6.7 | 4.8 |
| >multi-hg38_MT_+_1621_3279_RNR2_16S_50ntFlanks@459.492.34\|CTTGTAAATTTAACTGTTAGTCCAAAGAGGAACA | >multi-hg38_MT_+_1621_3279_RNR2_16S_50ntFlanks | 459 | 492 | 34 | CTTGTAAATTTAACTGTTAGTCCAAAGAGGAACA | BT20 | 24 | 2 | 0 | 25 | 23 | 8.0 | 1.3 | 0.5 | 27.6 | 11.0 |
| >multi-hg38_MT_+_1621_3279_RNR2_16S_50ntFlanks@459.492.34\|CTTGTAAATTTAACTGTTAGTCCAAAGAGGAACA | >multi-hg38_MT_+_1621_3279_RNR2_16S_50ntFlanks | 459 | 492 | 34 | CTTGTAAATTTAACTGTTAGTCCAAAGAGGAACA | MB231 | 66 | 17 | 2 | 36 | 48 | 36.8 | 12.3 | 0.8 | 15.9 | 25.3 |
| >multi-hg38_MT_+_1621_3279_RNR2_16S_50ntFlanks@460.491.32\|TTGTAAATTTAACTGTTAGTCCAAAGAGGAAC | >multi-hg38_MT_+_1621_3279_RNR2_16S_50ntFlanks | 460 | 491 | 32 | TTGTAAATTTAACTGTTAGTCCAAAGAGGAAC | MB231 | 12 | 6 | 1 | 11 | 15 | 7.9 | 6.1 | 0.3 | 6.4 | 15.4 |
| >multi-hg38_MT_+_1621_3279_RNR2_16S_50ntFlanks@460.492.33\|TTGTAAATTTAACTGTTAGTCCAAAGAGGAACA | >multi-hg38_MT_+_1621_3279_RNR2_16S_50ntFlanks | 460 | 492 | 33 | TTGTAAATTTAACTGTTAGTCCAAAGAGGAACA | BT20 | 19 | 1 | 0 | 6 | 7 | 14.4 | 0.9 | 0.2 | 6.7 | 3.8 |
| >multi-hg38_MT_+_1621_3279_RNR2_16S_50ntFlanks@460.492.33\|TTGTAAATTTAACTGTTAGTCCAAAGAGGAACA | >multi-hg38_MT_+_1621_3279_RNR2_16S_50ntFlanks | 460 | 492 | 33 | TTGTAAATTTAACTGTTAGTCCAAAGAGGAACA | MB231 | 46 | 14 | 1 | 13 | 23 | 40.5 | 12.7 | 0.8 | 5.4 | 7.4 |
| >multi-hg38_MT_+_1621_3279_RNR2_16S_50ntFlanks@461.492.32\|TGTAAATTTAACTGTTAGTCCAAAGAGGAACA | >multi-hg38_MT_+_1621_3279_RNR2_16S_50ntFlanks | 461 | 492 | 32 | TGTAAATTTAACTGTTAGTCCAAAGAGGAACA | MB231 | 12 | 3 | 0 | 3 | 4 | 9.3 | 2.3 | 0.3 | 1.8 | 1.1 |
| >multi-hg38_MT_+_1621_3279_RNR2_16S_50ntFlanks@465.492.28\|AATTTAACTGTTAGTCCAAAGAGGAACA | >multi-hg38_MT_+_1621_3279_RNR2_16S_50ntFlanks | 465 | 492 | 28 | AATTTAACTGTTAGTCCAAAGAGGAACA | MB231 | 13 | 5 | 0 | 7 | 7 | 16.9 | 5.7 | 0.4 | 4.4 | 2.0 |
| >multi-hg38_MT_+_1621_3279_RNR2_16S_50ntFlanks@486.506.21\|AGGAACAGCTCTTTGGACACT | >multi-hg38_MT_+_1621_3279_RNR2_16S_50ntFlanks | 486 | 506 | 21 | AGGAACAGCTCTTTGGACACT | MB231 | 22 | 5 | 6 | 31 | 13 | 29.4 | 4.7 | 6.7 | 27.1 | 8.5 |
| >multi-hg38_MT_+_1621_3279_RNR2_16S_50ntFlanks@486.517.32\|AGGAACAGCTCTTTGGACACTAGGAAAAAACC | >multi-hg38_MT_+_1621_3279_RNR2_16S_50ntFlanks | 486 | 517 | 32 | AGGAACAGCTCTTTGGACACTAGGAAAAAACC | MB231 | 10 | 2 | 7 | 20 | 14 | 7.0 | 1.1 | 2.4 | 16.0 | 9.8 |
| >multi-hg38_MT_+_1621_3279_RNR2_16S_50ntFlanks@486.518.33\|AGGAACAGCTCTTTGGACACTAGGAAAAAACCT | >multi-hg38_MT_+_1621_3279_RNR2_16S_50ntFlanks | 486 | 518 | 33 | AGGAACAGCTCTTTGGACACTAGGAAAAAACCT | MB231 | 16 | 4 | 3 | 32 | 16 | 14.7 | 1.9 | 0.6 | 32.8 | 6.2 |
| >multi-hg38_MT_+_1621_3279_RNR2_16S_50ntFlanks@488.506.19\|GAACAGCTCTTTGGACACT | >multi-hg38_MT_+_1621_3279_RNR2_16S_50ntFlanks | 488 | 506 | 19 | GAACAGCTCTTTGGACACT | MB231 | 10 | 2 | 1 | 3 | 4 | 13.1 | 2.9 | 0.9 | 3.0 | 3.6 |
| >multi-hg38_MT_+_1621_3279_RNR2_16S_50ntFlanks@488.518.31\|GAACAGCTCTTTGGACACTAGGAAAAAACCT | >multi-hg38_MT_+_1621_3279_RNR2_16S_50ntFlanks | 488 | 518 | 31 | GAACAGCTCTTTGGACACTAGGAAAAAACCT | MB231 | 12 | 3 | 3 | 14 | 17 | 8.0 | 1.4 | 1.0 | 12.0 | 6.8 |
| >multi-hg38_MT_+_1621_3279_RNR2_16S_50ntFlanks@493.517.25\|GCTCTTTGGACACTAGGAAAAAACC | >multi-hg38_MT_+_1621_3279_RNR2_16S_50ntFlanks | 493 | 517 | 25 | GCTCTTTGGACACTAGGAAAAAACC | MB231 | 40 | 22 | 19 | 24 | 64 | 24.2 | 15.5 | 8.5 | 13.6 | 30.4 |
| >multi-hg38_MT_+_1621_3279_RNR2_16S_50ntFlanks@493.518.26\|GCTCTTTGGACACTAGGAAAAAACCT | >multi-hg38_MT_+_1621_3279_RNR2_16S_50ntFlanks | 493 | 518 | 26 | GCTCTTTGGACACTAGGAAAAAACCT | BT20 | 13 | 5 | 1 | 18 | 28 | 1.1 | 4.6 | 0.2 | 11.2 | 23.5 |
| >multi-hg38_MT_+_1621_3279_RNR2_16S_50ntFlanks@493.518.26\|GCTCTTTGGACACTAGGAAAAAACCT | >multi-hg38_MT_+_1621_3279_RNR2_16S_50ntFlanks | 493 | 518 | 26 | GCTCTTTGGACACTAGGAAAAAACCT | MB231 | 59 | 24 | 7 | 35 | 69 | 44.3 | 15.9 | 2.7 | 21.4 | 15.8 |
| >multi-hg38_MT_+_1621_3279_RNR2_16S_50ntFlanks@51.70.20\|GCTAAACCTAGCCCCAAACC | >multi-hg38_MT_+_1621_3279_RNR2_16S_50ntFlanks | 51 | 70 | 20 | GCTAAACCTAGCCCCAAACC | BT20 | 11 | 4 | 0 | 65 | 18 | 10.9 | 3.2 | 0.1 | 94.5 | 23.8 |
| >multi-hg38_MT_+_1621_3279_RNR2_16S_50ntFlanks@51.70.20\|GCTAAACCTAGCCCCAAACC | >multi-hg38_MT_+_1621_3279_RNR2_16S_50ntFlanks | 51 | 70 | 20 | GCTAAACCTAGCCCCAAACC | MB231 | 12 | 9 | 1 | 310 | 116 | 6.4 | 2.2 | 0.5 | 320.0 | 95.3 |
| >multi-hg38_MT_+_1621_3279_RNR2_16S_50ntFlanks@51.71.21\|GCTAAACCTAGCCCCAAACCC | >multi-hg38_MT_+_1621_3279_RNR2_16S_50ntFlanks | 51 | 71 | 21 | GCTAAACCTAGCCCCAAACCC | BT20 | 39 | 27 | 1 | 197 | 135 | 24.6 | 5.5 | 0.5 | 203.3 | 118.8 |
| >multi-hg38_MT_+_1621_3279_RNR2_16S_50ntFlanks@51.71.21\|GCTAAACCTAGCCCCAAACCC | >multi-hg38_MT_+_1621_3279_RNR2_16S_50ntFlanks | 51 | 71 | 21 | GCTAAACCTAGCCCCAAACCC | MB231 | 22 | 22 | 2 | 410 | 311 | 6.4 | 4.7 | 1.3 | 248.2 | 237.1 |
| >multi-hg38_MT_+_1621_3279_RNR2_16S_50ntFlanks@51.72.22\|GCTAAACCTAGCCCCAAACCCA | >multi-hg38_MT_+_1621_3279_RNR2_16S_50ntFlanks | 51 | 72 | 22 | GCTAAACCTAGCCCCAAACCCA | BT20 | 45 | 26 | 2 | 36 | 156 | 34.1 | 4.1 | 0.4 | 32.2 | 129.5 |
| >multi-hg38_MT_+_1621_3279_RNR2_16S_50ntFlanks@51.72.22\|GCTAAACCTAGCCCCAAACCCA | >multi-hg38_MT_+_1621_3279_RNR2_16S_50ntFlanks | 51 | 72 | 22 | GCTAAACCTAGCCCCAAACCCA | MB231 | 19 | 17 | 2 | 130 | 287 | 3.3 | 4.1 | 1.0 | 109.9 | 195.8 |
| >multi-hg38_MT_+_1621_3279_RNR2_16S_50ntFlanks@51.73.23\|GCTAAACCTAGCCCCAAACCCAC | >multi-hg38_MT_+_1621_3279_RNR2_16S_50ntFlanks | 51 | 73 | 23 | GCTAAACCTAGCCCCAAACCCAC | BT20 | 19 | 15 | 8 | 42 | 87 | 11.1 | 7.0 | 12.4 | 38.7 | 80.1 |
| >multi-hg38_MT_+_1621_3279_RNR2_16S_50ntFlanks@51.74.24\|GCTAAACCTAGCCCCAAACCCACT | >multi-hg38_MT_+_1621_3279_RNR2_16S_50ntFlanks | 51 | 74 | 24 | GCTAAACCTAGCCCCAAACCCACT | BT20 | 80 | 57 | 7 | 368 | 332 | 34.0 | 10.6 | 3.0 | 395.7 | 353.6 |
| >multi-hg38_MT_+_1621_3279_RNR2_16S_50ntFlanks@51.74.24\|GCTAAACCTAGCCCCAAACCCACT | >multi-hg38_MT_+_1621_3279_RNR2_16S_50ntFlanks | 51 | 74 | 24 | GCTAAACCTAGCCCCAAACCCACT | MB231 | 48 | 51 | 7 | 368 | 573 | 7.6 | 17.4 | 4.2 | 226.2 | 335.5 |
| >multi-hg38_MT_+_1621_3279_RNR2_16S_50ntFlanks@51.77.27\|GCTAAACCTAGCCCCAAACCCACTCCA | >multi-hg38_MT_+_1621_3279_RNR2_16S_50ntFlanks | 51 | 77 | 27 | GCTAAACCTAGCCCCAAACCCACTCCA | BT20 | 10 | 2 | 3 | 9 | 17 | 5.2 | 0.2 | 0.8 | 9.9 | 19.2 |
| >multi-hg38_MT_+_1621_3279_RNR2_16S_50ntFlanks@51.77.27\|GCTAAACCTAGCCCCAAACCCACTCCA | >multi-hg38_MT_+_1621_3279_RNR2_16S_50ntFlanks | 51 | 77 | 27 | GCTAAACCTAGCCCCAAACCCACTCCA | MB231 | 11 | 4 | 3 | 15 | 28 | 1.7 | 1.4 | 1.8 | 12.8 | 17.2 |
| >multi-hg38_MT_+_1621_3279_RNR2_16S_50ntFlanks@51.79.29\|GCTAAACCTAGCCCCAAACCCACTCCACC | >multi-hg38_MT_+_1621_3279_RNR2_16S_50ntFlanks | 51 | 79 | 29 | GCTAAACCTAGCCCCAAACCCACTCCACC | BT20 | 32 | 32 | 32 | 612 | 331 | 17.4 | 8.3 | 46.7 | 873.4 | 342.0 |
| >multi-hg38_MT_+_1621_3279_RNR2_16S_50ntFlanks@51.79.29\|GCTAAACCTAGCCCCAAACCCACTCCACC | >multi-hg38_MT_+_1621_3279_RNR2_16S_50ntFlanks | 51 | 79 | 29 | GCTAAACCTAGCCCCAAACCCACTCCACC | MB231 | 28 | 24 | 11 | 218 | 229 | 10.6 | 10.4 | 5.8 | 219.0 | 178.3 |
| >multi-hg38_MT_+_1621_3279_RNR2_16S_50ntFlanks@51.80.30\|GCTAAACCTAGCCCCAAACCCACTCCACCT | >multi-hg38_MT_+_1621_3279_RNR2_16S_50ntFlanks | 51 | 80 | 30 | GCTAAACCTAGCCCCAAACCCACTCCACCT | BT20 | 13 | 11 | 8 | 85 | 90 | 4.7 | 1.4 | 11.9 | 108.4 | 96.8 |
| >multi-hg38_MT_+_1621_3279_RNR2_16S_50ntFlanks@51.81.31\|GCTAAACCTAGCCCCAAACCCACTCCACCTT | >multi-hg38_MT_+_1621_3279_RNR2_16S_50ntFlanks | 51 | 81 | 31 | GCTAAACCTAGCCCCAAACCCACTCCACCTT | BT20 | 17 | 40 | 4 | 208 | 140 | 9.2 | 38.1 | 2.9 | 273.5 | 140.8 |
| >multi-hg38_MT_+_1621_3279_RNR2_16S_50ntFlanks@51.82.32\|GCTAAACCTAGCCCCAAACCCACTCCACCTTA | >multi-hg38_MT_+_1621_3279_RNR2_16S_50ntFlanks | 51 | 82 | 32 | GCTAAACCTAGCCCCAAACCCACTCCACCTTA | BT20 | 10 | 9 | 2 | 702 | 97 | 5.8 | 1.5 | 1.5 | 977.8 | 101.1 |
| >multi-hg38_MT_+_1621_3279_RNR2_16S_50ntFlanks@51.84.34\|GCTAAACCTAGCCCCAAACCCACTCCACCTTACT | >multi-hg38_MT_+_1621_3279_RNR2_16S_50ntFlanks | 51 | 84 | 34 | GCTAAACCTAGCCCCAAACCCACTCCACCTTACT | BT20 | 42 | 36 | 6 | 764 | 429 | 14.5 | 15.5 | 1.5 | 931.5 | 389.9 |

| | | | | | | | | | | | | | | | | |
|---|---|---|---|---|---|---|---|---|---|---|---|---|---|---|---|---|
| >multi-hg38_MT_+_1621_3279_RNR2_16S_50ntFlanks@51.84.34\|GCTAAACCTAGCCCCAAACCCACTCCACCTTACT | >multi-hg38_MT_+_1621_3279_RNR2_16S_50ntFlanks | 51 | 84 | 34 | GCTAAACCTAGCCCCAAACCCACTCCACCTTACT | MB231 | 26 | 25 | 8 | 244 | 257 | 16.3 | 18.3 | 4.7 | 323.8 | 288.1 |
| >multi-hg38_MT_+_1621_3279_RNR2_16S_50ntFlanks@51.85.35\|GCTAAACCTAGCCCCAAACCCACTCCACCTTACTA | >multi-hg38_MT_+_1621_3279_RNR2_16S_50ntFlanks | 51 | 85 | 35 | GCTAAACCTAGCCCCAAACCCACTCCACCTTACTA | MB231 | 12 | 9 | 5 | 61 | 107 | 6.2 | 7.0 | 3.6 | 72.3 | 106.5 |
| >multi-hg38_MT_+_1621_3279_RNR2_16S_50ntFlanks@51.87.37\|GCTAAACCTAGCCCCAAACCCACTCCACCTTACTACC | >multi-hg38_MT_+_1621_3279_RNR2_16S_50ntFlanks | 51 | 87 | 37 | GCTAAACCTAGCCCCAAACCCACTCCACCTTACTACC | BT20 | 39 | 18 | 6 | 1183 | 175 | 23.7 | 6.1 | 1.8 | 1607.3 | 154.0 |
| >multi-hg38_MT_+_1621_3279_RNR2_16S_50ntFlanks@51.87.37\|GCTAAACCTAGCCCCAAACCCACTCCACCTTACTACC | >multi-hg38_MT_+_1621_3279_RNR2_16S_50ntFlanks | 51 | 87 | 37 | GCTAAACCTAGCCCCAAACCCACTCCACCTTACTACC | MB231 | 60 | 51 | 16 | 400 | 387 | 24.2 | 35.7 | 7.8 | 304.3 | 352.0 |
| >multi-hg38_MT_+_1621_3279_RNR2_16S_50ntFlanks@51.88.38\|GCTAAACCTAGCCCCAAACCCACTCCACCTTACTACCA | >multi-hg38_MT_+_1621_3279_RNR2_16S_50ntFlanks | 51 | 88 | 38 | GCTAAACCTAGCCCCAAACCCACTCCACCTTACTACCA | BT20 | 26 | 17 | 3 | 276 | 127 | 16.2 | 8.7 | 1.5 | 325.2 | 114.2 |
| >multi-hg38_MT_+_1621_3279_RNR2_16S_50ntFlanks@51.88.38\|GCTAAACCTAGCCCCAAACCCACTCCACCTTACTACCA | >multi-hg38_MT_+_1621_3279_RNR2_16S_50ntFlanks | 51 | 88 | 38 | GCTAAACCTAGCCCCAAACCCACTCCACCTTACTACCA | MB231 | 62 | 73 | 15 | 765 | 628 | 34.8 | 60.5 | 9.5 | 813.9 | 589.8 |
| >multi-hg38_MT_+_1621_3279_RNR2_16S_50ntFlanks@939.958.20\|TGTTTAACGGCCGCGGTACC | >multi-hg38_MT_+_1621_3279_RNR2_16S_50ntFlanks | 939 | 958 | 20 | TGTTTAACGGCCGCGGTACC | MB231 | 17 | 4 | 4 | 4 | 8 | 19.4 | 3.4 | 2.5 | 5.1 | 5.9 |
| >multi-hg38_MT_+_1621_3279_RNR2_16S_50ntFlanks@939.959.21\|TGTTTAACGGCCGCGGTACCC | >multi-hg38_MT_+_1621_3279_RNR2_16S_50ntFlanks | 939 | 959 | 21 | TGTTTAACGGCCGCGGTACCC | MB231 | 10 | 2 | 2 | 1 | 4 | 10.7 | 1.9 | 1.3 | 1.8 | 4.1 |
| >multi-hg38_MT_+_1621_3279_RNR2_16S_50ntFlanks@940.958.19\|GTTTAACGGCCGCGGTACC | >multi-hg38_MT_+_1621_3279_RNR2_16S_50ntFlanks | 940 | 958 | 19 | GTTTAACGGCCGCGGTACC | MB231 | 14 | 5 | 3 | 10 | 16 | 15.9 | 4.1 | 2.3 | 10.5 | 9.1 |
| >multi-hg38_MT_+_1621_3279_RNR2_16S_50ntFlanks@941.958.18\|TTTAACGGCCGCGGTACC | >multi-hg38_MT_+_1621_3279_RNR2_16S_50ntFlanks | 941 | 958 | 18 | TTTAACGGCCGCGGTACC | MB231 | 17 | 4 | 4 | 6 | 12 | 21.4 | 3.3 | 3.1 | 7.1 | 7.9 |
| >multi-hg38_MT_+_1621_3279_RNR2_16S_50ntFlanks@941.959.19\|TTTAACGGCCGCGGTACCC | >multi-hg38_MT_+_1621_3279_RNR2_16S_50ntFlanks | 941 | 959 | 19 | TTTAACGGCCGCGGTACCC | MB231 | 10 | 2 | 2 | 2 | 6 | 11.9 | 1.7 | 1.5 | 2.9 | 6.1 |
| >multi-hg38_MT_+_1621_3279_RNR2_16S_50ntFlanks@999.1017.19\|AGGGACCTGTATGAATGGC | >multi-hg38_MT_+_1621_3279_RNR2_16S_50ntFlanks | 999 | 1017 | 19 | AGGGACCTGTATGAATGGC | MB231 | 11 | 1 | 1 | 17 | 6 | 14.4 | 2.0 | 0.4 | 14.2 | 5.3 |
| >multi-hg38_MT_+_598_1651_RNR1_12S_50ntFlanks@367.404.38\|CCAATAAAGCTAAAACTCACCTGAGTTGTAAAAAACTC | >multi-hg38_MT_+_598_1651_RNR1_12S_50ntFlanks | 367 | 404 | 38 | CCAATAAAGCTAAAACTCACCTGAGTTGTAAAAAACTC | BT20 | 12 | 1 | 1 | 29 | 32 | 9.2 | 0.9 | 0.6 | 42.4 | 21.7 |
| >multi-hg38_MT_+_598_1651_RNR1_12S_50ntFlanks@367.404.38\|CCAATAAAGCTAAAACTCACCTGAGTTGTAAAAAACTC | >multi-hg38_MT_+_598_1651_RNR1_12S_50ntFlanks | 367 | 404 | 38 | CCAATAAAGCTAAAACTCACCTGAGTTGTAAAAAACTC | MB231 | 12 | 5 | 3 | 20 | 72 | 3.0 | 3.3 | 1.6 | 25.2 | 76.8 |
| >multi-hg38_MT_+_598_1651_RNR1_12S_50ntFlanks@368.404.37\|CAATAAAGCTAAAACTCACCTGAGTTGTAAAAAACTC | >multi-hg38_MT_+_598_1651_RNR1_12S_50ntFlanks | 368 | 404 | 37 | CAATAAAGCTAAAACTCACCTGAGTTGTAAAAAACTC | BT20 | 20 | 2 | 1 | 41 | 44 | 6.2 | 0.6 | 0.4 | 54.3 | 26.6 |
| >multi-hg38_MT_+_598_1651_RNR1_12S_50ntFlanks@368.404.37\|CAATAAAGCTAAAACTCACCTGAGTTGTAAAAAACTC | >multi-hg38_MT_+_598_1651_RNR1_12S_50ntFlanks | 368 | 404 | 37 | CAATAAAGCTAAAACTCACCTGAGTTGTAAAAAACTC | MB231 | 15 | 7 | 3 | 28 | 68 | 2.2 | 1.6 | 0.5 | 27.6 | 68.9 |
| >multi-hg38_MT_+_598_1651_RNR1_12S_50ntFlanks@369.404.36\|AATAAAGCTAAAACTCACCTGAGTTGTAAAAAACTC | >multi-hg38_MT_+_598_1651_RNR1_12S_50ntFlanks | 369 | 404 | 36 | AATAAAGCTAAAACTCACCTGAGTTGTAAAAAACTC | BT20 | 28 | 4 | 1 | 52 | 58 | 7.0 | 2.7 | 0.5 | 61.5 | 32.2 |
| >multi-hg38_MT_+_598_1651_RNR1_12S_50ntFlanks@369.404.36\|AATAAAGCTAAAACTCACCTGAGTTGTAAAAAACTC | >multi-hg38_MT_+_598_1651_RNR1_12S_50ntFlanks | 369 | 404 | 36 | AATAAAGCTAAAACTCACCTGAGTTGTAAAAAACTC | MB231 | 16 | 9 | 2 | 37 | 66 | 7.6 | 2.4 | 0.3 | 28.9 | 52.9 |
| >multi-hg38_MT_+_598_1651_RNR1_12S_50ntFlanks@370.404.35\|ATAAAGCTAAAACTCACCTGAGTTGTAAAAAACTC | >multi-hg38_MT_+_598_1651_RNR1_12S_50ntFlanks | 370 | 404 | 35 | ATAAAGCTAAAACTCACCTGAGTTGTAAAAAACTC | BT20 | 15 | 1 | 1 | 25 | 39 | 5.2 | 1.7 | 0.3 | 23.7 | 20.9 |
| >multi-hg38_MT_+_598_1651_RNR1_12S_50ntFlanks@370.404.35\|ATAAAGCTAAAACTCACCTGAGTTGTAAAAAACTC | >multi-hg38_MT_+_598_1651_RNR1_12S_50ntFlanks | 370 | 404 | 35 | ATAAAGCTAAAACTCACCTGAGTTGTAAAAAACTC | MB231 | 11 | 5 | 2 | 31 | 67 | 4.7 | 0.3 | 1.5 | 26.3 | 44.9 |
| >multi-hg38_MT_+_598_1651_RNR1_12S_50ntFlanks@424.452.29\|TACGAAAGTGGCTTTAACATATCTGAACA | >multi-hg38_MT_+_598_1651_RNR1_12S_50ntFlanks | 424 | 452 | 29 | TACGAAAGTGGCTTTAACATATCTGAACA | MB231 | 10 | 6 | 2 | 8 | 10 | 13.4 | 7.5 | 1.9 | 9.1 | 8.8 |
| >multi-hg38_MT_+_598_1651_RNR1_12S_50ntFlanks@424.455.32\|TACGAAAGTGGCTTTAACATATCTGAACACAC | >multi-hg38_MT_+_598_1651_RNR1_12S_50ntFlanks | 424 | 455 | 32 | TACGAAAGTGGCTTTAACATATCTGAACACAC | MB231 | 10 | 7 | 2 | 14 | 18 | 9.9 | 7.3 | 1.2 | 10.9 | 15.1 |
| >multi-hg38_MT_+_598_1651_RNR1_12S_50ntFlanks@424.456.33\|TACGAAAGTGGCTTTAACATATCTGAACACACA | >multi-hg38_MT_+_598_1651_RNR1_12S_50ntFlanks | 424 | 456 | 33 | TACGAAAGTGGCTTTAACATATCTGAACACACA | MB231 | 18 | 5 | 1 | 8 | 10 | 24.0 | 5.7 | 0.8 | 7.2 | 5.5 |
| >multi-hg38_MT_+_598_1651_RNR1_12S_50ntFlanks@424.462.39\|TACGAAAGTGGCTTTAACATATCTGAACACACAATAGCT | >multi-hg38_MT_+_598_1651_RNR1_12S_50ntFlanks | 424 | 462 | 39 | TACGAAAGTGGCTTTAACATATCTGAACACACAATAGCT | MB231 | 12 | 2 | 1 | 3 | 6 | 16.4 | 2.0 | 1.2 | 1.9 | 1.4 |
| >multi-hg38_MT_+_598_1651_RNR1_12S_50ntFlanks@425.452.28\|ACGAAAGTGGCTTTAACATATCTGAACA | >multi-hg38_MT_+_598_1651_RNR1_12S_50ntFlanks | 425 | 452 | 28 | ACGAAAGTGGCTTTAACATATCTGAACA | BT20 | 11 | 4 | 1 | 23 | 21 | 1.8 | 0.5 | 0.1 | 27.3 | 23.0 |
| >multi-hg38_MT_+_598_1651_RNR1_12S_50ntFlanks@425.452.28\|ACGAAAGTGGCTTTAACATATCTGAACA | >multi-hg38_MT_+_598_1651_RNR1_12S_50ntFlanks | 425 | 452 | 28 | ACGAAAGTGGCTTTAACATATCTGAACA | MB231 | 11 | 8 | 2 | 13 | 13 | 14.3 | 10.5 | 1.6 | 14.2 | 12.3 |
| >multi-hg38_MT_+_598_1651_RNR1_12S_50ntFlanks@425.455.31\|ACGAAAGTGGCTTTAACATATCTGAACACAC | >multi-hg38_MT_+_598_1651_RNR1_12S_50ntFlanks | 425 | 455 | 31 | ACGAAAGTGGCTTTAACATATCTGAACACAC | MB231 | 10 | 8 | 2 | 20 | 20 | 10.3 | 8.9 | 1.1 | 15.7 | 17.5 |
| >multi-hg38_MT_+_598_1651_RNR1_12S_50ntFlanks@425.456.32\|ACGAAAGTGGCTTTAACATATCTGAACACACA | >multi-hg38_MT_+_598_1651_RNR1_12S_50ntFlanks | 425 | 456 | 32 | ACGAAAGTGGCTTTAACATATCTGAACACACA | BT20 | 14 | 4 | 0 | 19 | 13 | 7.0 | 1.0 | 0.3 | 22.9 | 14.6 |
| >multi-hg38_MT_+_598_1651_RNR1_12S_50ntFlanks@425.456.32\|ACGAAAGTGGCTTTAACATATCTGAACACACA | >multi-hg38_MT_+_598_1651_RNR1_12S_50ntFlanks | 425 | 456 | 32 | ACGAAAGTGGCTTTAACATATCTGAACACACA | MB231 | 18 | 6 | 1 | 14 | 11 | 25.9 | 6.2 | 0.6 | 10.5 | 6.5 |
| >multi-hg38_MT_+_598_1651_RNR1_12S_50ntFlanks@425.462.38\|ACGAAAGTGGCTTTAACATATCTGAACACACAATAGCT | >multi-hg38_MT_+_598_1651_RNR1_12S_50ntFlanks | 425 | 462 | 38 | ACGAAAGTGGCTTTAACATATCTGAACACACAATAGCT | MB231 | 11 | 3 | 1 | 3 | 7 | 16.6 | 2.2 | 1.1 | 2.2 | 1.8 |
| >multi-hg38_MT_+_598_1651_RNR1_12S_50ntFlanks@427.468.42\|GAAAGTGGCTTTAACATATCTGAACACACAATAGCTAAGACC | >multi-hg38_MT_+_598_1651_RNR1_12S_50ntFlanks | 427 | 468 | 42 | GAAAGTGGCTTTAACATATCTGAACACACAATAGCTAAGACC | MB231 | 11 | 2 | 3 | 10 | 17 | 12.9 | 0.8 | 2.2 | 10.0 | 8.5 |
| >multi-hg38_MT_+_598_1651_RNR1_12S_50ntFlanks@428.448.21\|AAAGTGGCTTTAACATATCTG | >multi-hg38_MT_+_598_1651_RNR1_12S_50ntFlanks | 428 | 448 | 21 | AAAGTGGCTTTAACATATCTG | MB231 | 11 | 1 | 1 | 3 | 1 | 16.8 | 1.6 | 1.6 | 3.6 | 1.5 |
| >multi-hg38_MT_+_598_1651_RNR1_12S_50ntFlanks@428.452.25\|AAAGTGGCTTTAACATATCTGAACA | >multi-hg38_MT_+_598_1651_RNR1_12S_50ntFlanks | 428 | 452 | 25 | AAAGTGGCTTTAACATATCTGAACA | BT20 | 16 | 2 | 1 | 31 | 16 | 1.4 | 1.1 | 0.1 | 22.0 | 13.0 |
| >multi-hg38_MT_+_598_1651_RNR1_12S_50ntFlanks@428.452.25\|AAAGTGGCTTTAACATATCTGAACA | >multi-hg38_MT_+_598_1651_RNR1_12S_50ntFlanks | 428 | 452 | 25 | AAAGTGGCTTTAACATATCTGAACA | MB231 | 26 | 10 | 3 | 50 | 21 | 32.1 | 11.4 | 3.3 | 65.7 | 15.5 |
| >multi-hg38_MT_+_598_1651_RNR1_12S_50ntFlanks@428.453.26\|AAAGTGGCTTTAACATATCTGAACAC | >multi-hg38_MT_+_598_1651_RNR1_12S_50ntFlanks | 428 | 453 | 26 | AAAGTGGCTTTAACATATCTGAACAC | MB231 | 12 | 7 | 1 | 42 | 13 | 12.0 | 7.0 | 1.4 | 39.7 | 8.6 |
| >multi-hg38_MT_+_598_1651_RNR1_12S_50ntFlanks@428.454.27\|AAAGTGGCTTTAACATATCTGAACACA | >multi-hg38_MT_+_598_1651_RNR1_12S_50ntFlanks | 428 | 454 | 27 | AAAGTGGCTTTAACATATCTGAACACA | BT20 | 11 | 1 | 0 | 8 | 4 | 8.7 | 0.7 | 0.0 | 6.3 | 3.7 |

| | | | | | | | | | | | | | | | | |
|---|---|---|---|---|---|---|---|---|---|---|---|---|---|---|---|---|
| >multi-hg38_MT_+_598_1651_RNR1_12S_50ntFlanks@428.454.27\|AAAGTGGCTTTAACATATCTGAACACA | >multi-hg38_MT_+_598_1651_RNR1_12S_50ntFlanks | 428 | 454 | 27 | AAAGTGGCTTTAACATATCTGAACACA | MB231 | 14 | 5 | 0 | 24 | 8 | 19.2 | 6.3 | 0.6 | 23.8 | 5.3 |
| >multi-hg38_MT_+_598_1651_RNR1_12S_50ntFlanks@428.455.28\|AAAGTGGCTTTAACATATCTGAACACAC | >multi-hg38_MT_+_598_1651_RNR1_12S_50ntFlanks | 428 | 455 | 28 | AAAGTGGCTTTAACATATCTGAACACAC | BT20 | 18 | 2 | 1 | 37 | 20 | 6.0 | 2.0 | 0.1 | 28.0 | 18.0 |
| >multi-hg38_MT_+_598_1651_RNR1_12S_50ntFlanks@428.455.28\|AAAGTGGCTTTAACATATCTGAACACAC | >multi-hg38_MT_+_598_1651_RNR1_12S_50ntFlanks | 428 | 455 | 28 | AAAGTGGCTTTAACATATCTGAACACAC | MB231 | 42 | 18 | 3 | 77 | 33 | 46.7 | 19.4 | 2.7 | 63.7 | 19.9 |
| >multi-hg38_MT_+_598_1651_RNR1_12S_50ntFlanks@428.456.29\|AAAGTGGCTTTAACATATCTGAACACACA | >multi-hg38_MT_+_598_1651_RNR1_12S_50ntFlanks | 428 | 456 | 29 | AAAGTGGCTTTAACATATCTGAACACACA | BT20 | 30 | 2 | 0 | 19 | 12 | 25.0 | 1.0 | 0.3 | 15.5 | 10.1 |
| >multi-hg38_MT_+_598_1651_RNR1_12S_50ntFlanks@428.456.29\|AAAGTGGCTTTAACATATCTGAACACACA | >multi-hg38_MT_+_598_1651_RNR1_12S_50ntFlanks | 428 | 456 | 29 | AAAGTGGCTTTAACATATCTGAACACACA | MB231 | 51 | 11 | 1 | 41 | 21 | 71.4 | 12.4 | 1.2 | 35.8 | 8.4 |
| >multi-hg38_MT_+_598_1651_RNR1_12S_50ntFlanks@428.462.35\|AAAGTGGCTTTAACATATCTGAACACACAATAGCT | >multi-hg38_MT_+_598_1651_RNR1_12S_50ntFlanks | 428 | 462 | 35 | AAAGTGGCTTTAACATATCTGAACACACAATAGCT | BT20 | 11 | 1 | 1 | 9 | 9 | 6.9 | 0.8 | 0.5 | 8.6 | 9.3 |
| >multi-hg38_MT_+_598_1651_RNR1_12S_50ntFlanks@428.462.35\|AAAGTGGCTTTAACATATCTGAACACACAATAGCT | >multi-hg38_MT_+_598_1651_RNR1_12S_50ntFlanks | 428 | 462 | 35 | AAAGTGGCTTTAACATATCTGAACACACAATAGCT | MB231 | 44 | 6 | 4 | 23 | 22 | 62.7 | 6.5 | 4.1 | 21.0 | 9.4 |
| >multi-hg38_MT_+_598_1651_RNR1_12S_50ntFlanks@428.465.38\|AAAGTGGCTTTAACATATCTGAACACACAATAGCTAAG | >multi-hg38_MT_+_598_1651_RNR1_12S_50ntFlanks | 428 | 465 | 38 | AAAGTGGCTTTAACATATCTGAACACACAATAGCTAAG | MB231 | 12 | 2 | 1 | 20 | 18 | 13.6 | 2.0 | 1.0 | 12.7 | 12.3 |
| >multi-hg38_MT_+_598_1651_RNR1_12S_50ntFlanks@428.468.41\|AAAGTGGCTTTAACATATCTGAACACACAATAGCTAAGACC | >multi-hg38_MT_+_598_1651_RNR1_12S_50ntFlanks | 428 | 468 | 41 | AAAGTGGCTTTAACATATCTGAACACACAATAGCTAAGACC | BT20 | 14 | 3 | 1 | 60 | 38 | 5.8 | 3.9 | 0.9 | 66.5 | 10.0 |
| >multi-hg38_MT_+_598_1651_RNR1_12S_50ntFlanks@428.468.41\|AAAGTGGCTTTAACATATCTGAACACACAATAGCTAAGACC | >multi-hg38_MT_+_598_1651_RNR1_12S_50ntFlanks | 428 | 468 | 41 | AAAGTGGCTTTAACATATCTGAACACACAATAGCTAAGACC | MB231 | 76 | 13 | 13 | 62 | 86 | 92.5 | 9.2 | 11.2 | 46.6 | 53.3 |
| >multi-hg38_MT_+_598_1651_RNR1_12S_50ntFlanks@428.469.42\|AAAGTGGCTTTAACATATCTGAACACACAATAGCTAAGACCC | >multi-hg38_MT_+_598_1651_RNR1_12S_50ntFlanks | 428 | 469 | 42 | AAAGTGGCTTTAACATATCTGAACACACAATAGCTAAGACCC | BT20 | 12 | 3 | 0 | 45 | 30 | 6.5 | 3.7 | 0.4 | 47.8 | 4.6 |
| >multi-hg38_MT_+_598_1651_RNR1_12S_50ntFlanks@428.469.42\|AAAGTGGCTTTAACATATCTGAACACACAATAGCTAAGACCC | >multi-hg38_MT_+_598_1651_RNR1_12S_50ntFlanks | 428 | 469 | 42 | AAAGTGGCTTTAACATATCTGAACACACAATAGCTAAGACCC | MB231 | 58 | 10 | 11 | 37 | 55 | 67.0 | 5.5 | 9.8 | 30.6 | 19.3 |
| >multi-hg38_MT_+_598_1651_RNR1_12S_50ntFlanks@428.470.43\|AAAGTGGCTTTAACATATCTGAACACACAATAGCTAAGACCCA | >multi-hg38_MT_+_598_1651_RNR1_12S_50ntFlanks | 428 | 470 | 43 | AAAGTGGCTTTAACATATCTGAACACACAATAGCTAAGACCCA | MB231 | 15 | 2 | 2 | 5 | 11 | 19.0 | 0.8 | 2.2 | 5.1 | 1.7 |
| >multi-hg38_MT_+_598_1651_RNR1_12S_50ntFlanks@430.468.39\|AGTGGCTTTAACATATCTGAACACACAATAGCTAAGACC | >multi-hg38_MT_+_598_1651_RNR1_12S_50ntFlanks | 430 | 468 | 39 | AGTGGCTTTAACATATCTGAACACACAATAGCTAAGACC | MB231 | 16 | 3 | 2 | 14 | 17 | 18.6 | 2.8 | 1.8 | 9.6 | 12.5 |
| >multi-hg38_MT_+_598_1651_RNR1_12S_50ntFlanks@430.469.40\|AGTGGCTTTAACATATCTGAACACACAATAGCTAAGACCC | >multi-hg38_MT_+_598_1651_RNR1_12S_50ntFlanks | 430 | 469 | 40 | AGTGGCTTTAACATATCTGAACACACAATAGCTAAGACCC | MB231 | 12 | 3 | 1 | 8 | 10 | 15.3 | 2.0 | 1.5 | 5.2 | 5.2 |
| >multi-hg38_MT_+_598_1651_RNR1_12S_50ntFlanks@439.462.24\|AACATATCTGAACACACAATAGCT | >multi-hg38_MT_+_598_1651_RNR1_12S_50ntFlanks | 439 | 462 | 24 | AACATATCTGAACACACAATAGCT | MB231 | 11 | 1 | 0 | 8 | 2 | 16.3 | 1.3 | 0.8 | 10.2 | 2.1 |
| >multi-hg38_MT_+_598_1651_RNR1_12S_50ntFlanks@439.468.30\|AACATATCTGAACACACAATAGCTAAGACC | >multi-hg38_MT_+_598_1651_RNR1_12S_50ntFlanks | 439 | 468 | 30 | AACATATCTGAACACACAATAGCTAAGACC | MB231 | 13 | 2 | 1 | 20 | 6 | 17.3 | 2.8 | 1.7 | 16.3 | 4.2 |
| >multi-hg38_MT_+_598_1651_RNR1_12S_50ntFlanks@439.469.31\|AACATATCTGAACACACAATAGCTAAGACCC | >multi-hg38_MT_+_598_1651_RNR1_12S_50ntFlanks | 439 | 469 | 31 | AACATATCTGAACACACAATAGCTAAGACCC | MB231 | 11 | 2 | 3 | 20 | 9 | 12.1 | 2.7 | 3.2 | 14.8 | 5.9 |
| >multi-hg38_MT_+_598_1651_RNR1_12S_50ntFlanks@440.468.29\|ACATATCTGAACACACAATAGCTAAGACC | >multi-hg38_MT_+_598_1651_RNR1_12S_50ntFlanks | 440 | 468 | 29 | ACATATCTGAACACACAATAGCTAAGACC | MB231 | 11 | 2 | 1 | 18 | 5 | 15.7 | 2.4 | 0.8 | 15.0 | 3.1 |
| >multi-hg38_MT_+_598_1651_RNR1_12S_50ntFlanks@440.469.30\|ACATATCTGAACACACAATAGCTAAGACCC | >multi-hg38_MT_+_598_1651_RNR1_12S_50ntFlanks | 440 | 469 | 30 | ACATATCTGAACACACAATAGCTAAGACCC | MB231 | 10 | 2 | 2 | 18 | 9 | 11.9 | 2.9 | 2.2 | 14.1 | 5.5 |
| >multi-hg38_MT_+_598_1651_RNR1_12S_50ntFlanks@442.468.27\|ATATCTGAACACACAATAGCTAAGACC | >multi-hg38_MT_+_598_1651_RNR1_12S_50ntFlanks | 442 | 468 | 27 | ATATCTGAACACACAATAGCTAAGACC | MB231 | 12 | 2 | 0 | 22 | 4 | 16.7 | 3.1 | 0.4 | 22.9 | 2.8 |
| >multi-hg38_MT_+_598_1651_RNR1_12S_50ntFlanks@442.469.28\|ATATCTGAACACACAATAGCTAAGACCC | >multi-hg38_MT_+_598_1651_RNR1_12S_50ntFlanks | 442 | 469 | 28 | ATATCTGAACACACAATAGCTAAGACCC | MB231 | 12 | 3 | 1 | 21 | 7 | 15.3 | 3.6 | 1.6 | 20.8 | 4.3 |
| >multi-hg38_MT_+_598_1651_RNR1_12S_50ntFlanks@442.480.39\|ATATCTGAACACACAATAGCTAAGACCCAAACTGGGATT | >multi-hg38_MT_+_598_1651_RNR1_12S_50ntFlanks | 442 | 480 | 39 | ATATCTGAACACACAATAGCTAAGACCCAAACTGGGATT | MB231 | 11 | 3 | 3 | 49 | 37 | 6.3 | 1.7 | 2.5 | 61.7 | 24.5 |
| >multi-hg38_MT_+_598_1651_RNR1_12S_50ntFlanks@443.462.20\|TATCTGAACACACAATAGCT | >multi-hg38_MT_+_598_1651_RNR1_12S_50ntFlanks | 443 | 462 | 20 | TATCTGAACACACAATAGCT | MB231 | 16 | 1 | 0 | 5 | 1 | 24.5 | 1.7 | 0.4 | 6.1 | 1.0 |
| >multi-hg38_MT_+_598_1651_RNR1_12S_50ntFlanks@443.468.26\|TATCTGAACACACAATAGCTAAGACC | >multi-hg38_MT_+_598_1651_RNR1_12S_50ntFlanks | 443 | 468 | 26 | TATCTGAACACACAATAGCTAAGACC | MB231 | 21 | 4 | 1 | 39 | 5 | 29.1 | 4.0 | 1.3 | 49.7 | 2.6 |
| >multi-hg38_MT_+_598_1651_RNR1_12S_50ntFlanks@443.469.27\|TATCTGAACACACAATAGCTAAGACCC | >multi-hg38_MT_+_598_1651_RNR1_12S_50ntFlanks | 443 | 469 | 27 | TATCTGAACACACAATAGCTAAGACCC | MB231 | 25 | 5 | 4 | 38 | 10 | 29.9 | 5.8 | 4.0 | 44.9 | 6.4 |
| >multi-hg38_MT_+_598_1651_RNR1_12S_50ntFlanks@443.480.38\|TATCTGAACACACAATAGCTAAGACCCAAACTGGGATT | >multi-hg38_MT_+_598_1651_RNR1_12S_50ntFlanks | 443 | 480 | 38 | TATCTGAACACACAATAGCTAAGACCCAAACTGGGATT | MB231 | 13 | 6 | 3 | 47 | 61 | 2.7 | 1.0 | 2.0 | 57.0 | 20.9 |
| >multi-hg38_MT_+_598_1651_RNR1_12S_50ntFlanks@52.74.23\|ATAGGTTTGGTCCTAGCCTTTCT | >multi-hg38_MT_+_598_1651_RNR1_12S_50ntFlanks | 52 | 74 | 23 | ATAGGTTTGGTCCTAGCCTTTCT | BT20 | 11 | 6 | 0 | 404 | 23 | 11.1 | 2.9 | 0.2 | 532.5 | 29.8 |
| >multi-hg38_MT_+_598_1651_RNR1_12S_50ntFlanks@52.77.26\|ATAGGTTTGGTCCTAGCCTTTCTATT | >multi-hg38_MT_+_598_1651_RNR1_12S_50ntFlanks | 52 | 77 | 26 | ATAGGTTTGGTCCTAGCCTTTCTATT | BT20 | 37 | 16 | 3 | 3511 | 301 | 48.4 | 11.3 | 2.8 | 5229.8 | 364.3 |
| >multi-hg38_MT_+_598_1651_RNR1_12S_50ntFlanks@52.77.26\|ATAGGTTTGGTCCTAGCCTTTCTATT | >multi-hg38_MT_+_598_1651_RNR1_12S_50ntFlanks | 52 | 77 | 26 | ATAGGTTTGGTCCTAGCCTTTCTATT | MB231 | 11 | 11 | 2 | 656 | 183 | 7.2 | 4.9 | 0.3 | 650.7 | 80.0 |
| >multi-hg38_MT_+_598_1651_RNR1_12S_50ntFlanks@52.83.32\|ATAGGTTTGGTCCTAGCCTTTCTATTAGCTCT | >multi-hg38_MT_+_598_1651_RNR1_12S_50ntFlanks | 52 | 83 | 32 | ATAGGTTTGGTCCTAGCCTTTCTATTAGCTCT | BT20 | 66 | 14 | 8 | 336 | 89 | 31.5 | 4.4 | 2.0 | 448.3 | 123.2 |
| >multi-hg38_MT_+_598_1651_RNR1_12S_50ntFlanks@52.83.32\|ATAGGTTTGGTCCTAGCCTTTCTATTAGCTCT | >multi-hg38_MT_+_598_1651_RNR1_12S_50ntFlanks | 52 | 83 | 32 | ATAGGTTTGGTCCTAGCCTTTCTATTAGCTCT | MB231 | 60 | 15 | 11 | 367 | 169 | 15.0 | 7.6 | 11.1 | 272.7 | 78.9 |
| >multi-hg38_MT_+_598_1651_RNR1_12S_50ntFlanks@52.84.33\|ATAGGTTTGGTCCTAGCCTTTCTATTAGCTCTT | >multi-hg38_MT_+_598_1651_RNR1_12S_50ntFlanks | 52 | 84 | 33 | ATAGGTTTGGTCCTAGCCTTTCTATTAGCTCTT | BT20 | 22 | 20 | 3 | 1224 | 83 | 7.4 | 10.0 | 1.0 | 1574.4 | 87.2 |
| >multi-hg38_MT_+_598_1651_RNR1_12S_50ntFlanks@52.87.36\|ATAGGTTTGGTCCTAGCCTTTCTATTAGCTCTTAGT | >multi-hg38_MT_+_598_1651_RNR1_12S_50ntFlanks | 52 | 87 | 36 | ATAGGTTTGGTCCTAGCCTTTCTATTAGCTCTTAGT | BT20 | 41 | 27 | 4 | 999 | 170 | 33.5 | 17.4 | 1.1 | 1497.5 | 208.2 |
| >multi-hg38_MT_+_598_1651_RNR1_12S_50ntFlanks@52.87.36\|ATAGGTTTGGTCCTAGCCTTTCTATTAGCTCTTAGT | >multi-hg38_MT_+_598_1651_RNR1_12S_50ntFlanks | 52 | 87 | 36 | ATAGGTTTGGTCCTAGCCTTTCTATTAGCTCTTAGT | MB231 | 19 | 12 | 4 | 292 | 142 | 19.7 | 4.8 | 1.9 | 272.9 | 113.6 |

| | | | | | | | | | | | | | | | | |
|---|---|---|---|---|---|---|---|---|---|---|---|---|---|---|---|---|
| >multi-hg38_MT_+_598_1651_RNR1_12S_50ntFlanks@52.88.37\|ATAGGTTTGGTCCTAGCCTTTCTATTAGCTCTTAGTA | >multi-hg38_MT_+_598_1651_RNR1_12S_50ntFlanks | 52 | 88 | 37 | ATAGGTTTGGTCCTAGCCTTTCTATTAGCTCTTAGTA | BT20 | 95 | 52 | 9 | 11415 | 697 | 122.2 | 31.7 | 4.7 | 18187.1 | 697.1 |
| >multi-hg38_MT_+_598_1651_RNR1_12S_50ntFlanks@52.88.37\|ATAGGTTTGGTCCTAGCCTTTCTATTAGCTCTTAGTA | >multi-hg38_MT_+_598_1651_RNR1_12S_50ntFlanks | 52 | 88 | 37 | ATAGGTTTGGTCCTAGCCTTTCTATTAGCTCTTAGTA | MB231 | 50 | 40 | 13 | 7004 | 601 | 37.9 | 6.5 | 8.4 | 2188.7 | 476.2 |
| >multi-hg38_MT_+_598_1651_RNR1_12S_50ntFlanks@52.93.42\|ATAGGTTTGGTCCTAGCCTTTCTATTAGCTCTTAGTAAGATT | >multi-hg38_MT_+_598_1651_RNR1_12S_50ntFlanks | 52 | 93 | 42 | ATAGGTTTGGTCCTAGCCTTTCTATTAGCTCTTAGTAAGATT | BT20 | 17 | 11 | 2 | 207 | 22 | 20.2 | 6.2 | 2.8 | 273.8 | 11.2 |
| >multi-hg38_MT_+_598_1651_RNR1_12S_50ntFlanks@606.633.28\|CCTGTTCTGTAATCGATAAACCCCGATC | >multi-hg38_MT_+_598_1651_RNR1_12S_50ntFlanks | 606 | 633 | 28 | CCTGTTCTGTAATCGATAAACCCCGATC | MB231 | 24 | 9 | 15 | 7 | 17 | 26.6 | 7.2 | 5.3 | 8.1 | 15.7 |
| >multi-hg38_MT_+_598_1651_RNR1_12S_50ntFlanks@607.633.27\|CTGTTCTGTAATCGATAAACCCCGATC | >multi-hg38_MT_+_598_1651_RNR1_12S_50ntFlanks | 607 | 633 | 27 | CTGTTCTGTAATCGATAAACCCCGATC | MB231 | 12 | 4 | 5 | 11 | 4 | 13.3 | 3.7 | 2.0 | 13.4 | 3.9 |
| >multi-hg38_MT_+_598_1651_RNR1_12S_50ntFlanks@608.633.26\|TGTTCTGTAATCGATAAACCCCGATC | >multi-hg38_MT_+_598_1651_RNR1_12S_50ntFlanks | 608 | 633 | 26 | TGTTCTGTAATCGATAAACCCCGATC | BT20 | 10 | 1 | 2 | 14 | 7 | 2.9 | 1.2 | 0.3 | 13.5 | 5.6 |
| >multi-hg38_MT_+_598_1651_RNR1_12S_50ntFlanks@608.633.26\|TGTTCTGTAATCGATAAACCCCGATC | >multi-hg38_MT_+_598_1651_RNR1_12S_50ntFlanks | 608 | 633 | 26 | TGTTCTGTAATCGATAAACCCCGATC | MB231 | 32 | 10 | 10 | 36 | 11 | 39.7 | 9.7 | 4.8 | 43.3 | 7.9 |
| >multi-hg38_MT_+_598_1651_RNR1_12S_50ntFlanks@634.665.32\|AACCTCACCACCTCTTGCTCAGCCTATATACC | >multi-hg38_MT_+_598_1651_RNR1_12S_50ntFlanks | 634 | 665 | 32 | AACCTCACCACCTCTTGCTCAGCCTATATACC | MB231 | 14 | 3 | 2 | 21 | 21 | 17.5 | 2.9 | 1.1 | 12.1 | 12.7 |
| >multi-hg38_MT_+_598_1651_RNR1_12S_50ntFlanks@716.744.29\|ACGTAAAGACGTTAGGTCAAGGTGTAGCC | >multi-hg38_MT_+_598_1651_RNR1_12S_50ntFlanks | 716 | 744 | 29 | ACGTAAAGACGTTAGGTCAAGGTGTAGCC | MB231 | 13 | 3 | 3 | 17 | 9 | 15.9 | 3.9 | 2.3 | 21.2 | 7.8 |
| >multi-hg38_MT_+_598_1651_RNR1_12S_50ntFlanks@717.744.28\|CGTAAAGACGTTAGGTCAAGGTGTAGCC | >multi-hg38_MT_+_598_1651_RNR1_12S_50ntFlanks | 717 | 744 | 28 | CGTAAAGACGTTAGGTCAAGGTGTAGCC | MB231 | 14 | 4 | 3 | 17 | 9 | 16.7 | 4.2 | 3.1 | 21.2 | 8.8 |
| >multi-hg38_MT_+_598_1651_RNR1_12S_50ntFlanks@718.744.27\|GTAAAGACGTTAGGTCAAGGTGTAGCC | >multi-hg38_MT_+_598_1651_RNR1_12S_50ntFlanks | 718 | 744 | 27 | GTAAAGACGTTAGGTCAAGGTGTAGCC | MB231 | 19 | 6 | 4 | 39 | 17 | 22.8 | 6.9 | 3.7 | 43.4 | 13.8 |
| >multi-hg38_MT_+_598_1651_RNR1_12S_50ntFlanks@719.744.26\|TAAAGACGTTAGGTCAAGGTGTAGCC | >multi-hg38_MT_+_598_1651_RNR1_12S_50ntFlanks | 719 | 744 | 26 | TAAAGACGTTAGGTCAAGGTGTAGCC | MB231 | 44 | 13 | 8 | 62 | 33 | 51.4 | 15.4 | 7.5 | 70.2 | 27.3 |
| >multi-hg38_MT_+_598_1651_RNR1_12S_50ntFlanks@719.745.27\|TAAAGACGTTAGGTCAAGGTGTAGCCC | >multi-hg38_MT_+_598_1651_RNR1_12S_50ntFlanks | 719 | 745 | 27 | TAAAGACGTTAGGTCAAGGTGTAGCCC | MB231 | 20 | 6 | 2 | 20 | 14 | 24.8 | 6.8 | 2.0 | 24.8 | 12.6 |
| >multi-hg38_MT_+_598_1651_RNR1_12S_50ntFlanks@726.744.19\|GTTAGGTCAAGGTGTAGCC | >multi-hg38_MT_+_598_1651_RNR1_12S_50ntFlanks | 726 | 744 | 19 | GTTAGGTCAAGGTGTAGCC | MB231 | 12 | 2 | 2 | 12 | 5 | 16.1 | 3.5 | 1.6 | 14.4 | 4.9 |
| >multi-hg38_MT_+_598_1651_RNR1_12S_50ntFlanks@903.929.27\|CAAGTATACTTCAAAGGACATTTAACT | >multi-hg38_MT_+_598_1651_RNR1_12S_50ntFlanks | 903 | 929 | 27 | CAAGTATACTTCAAAGGACATTTAACT | MB231 | 13 | 1 | 1 | 19 | 4 | 17.1 | 1.4 | 0.7 | 20.7 | 1.2 |
| >multi-hg38_MT_+_598_1651_RNR1_12S_50ntFlanks@904.929.26\|AAGTATACTTCAAAGGACATTTAACT | >multi-hg38_MT_+_598_1651_RNR1_12S_50ntFlanks | 904 | 929 | 26 | AAGTATACTTCAAAGGACATTTAACT | BT20 | 11 | 1 | 1 | 6 | 5 | 5.3 | 0.6 | 0.3 | 5.0 | 6.5 |
| >multi-hg38_MT_+_598_1651_RNR1_12S_50ntFlanks@904.929.26\|AAGTATACTTCAAAGGACATTTAACT | >multi-hg38_MT_+_598_1651_RNR1_12S_50ntFlanks | 904 | 929 | 26 | AAGTATACTTCAAAGGACATTTAACT | MB231 | 21 | 4 | 1 | 51 | 7 | 27.6 | 3.9 | 1.0 | 57.8 | 1.6 |
| >multi-hg38_MT_+_598_1651_RNR1_12S_50ntFlanks@905.929.25\|AGTATACTTCAAAGGACATTTAACT | >multi-hg38_MT_+_598_1651_RNR1_12S_50ntFlanks | 905 | 929 | 25 | AGTATACTTCAAAGGACATTTAACT | BT20 | 42 | 2 | 3 | 40 | 21 | 6.6 | 1.2 | 1.3 | 31.1 | 25.0 |
| >multi-hg38_MT_+_598_1651_RNR1_12S_50ntFlanks@905.929.25\|AGTATACTTCAAAGGACATTTAACT | >multi-hg38_MT_+_598_1651_RNR1_12S_50ntFlanks | 905 | 929 | 25 | AGTATACTTCAAAGGACATTTAACT | MB231 | 78 | 13 | 4 | 418 | 34 | 91.5 | 15.1 | 4.5 | 567.2 | 12.6 |
| >multi-hg38_MT_+_598_1651_RNR1_12S_50ntFlanks@905.935.31\|AGTATACTTCAAAGGACATTTAACTAAAACC | >multi-hg38_MT_+_598_1651_RNR1_12S_50ntFlanks | 905 | 935 | 31 | AGTATACTTCAAAGGACATTTAACTAAAACC | BT20 | 11 | 0 | 1 | 12 | 3 | 2.5 | 0.5 | 0.1 | 16.3 | 4.5 |
| >multi-hg38_MT_+_598_1651_RNR1_12S_50ntFlanks@905.935.31\|AGTATACTTCAAAGGACATTTAACTAAAACC | >multi-hg38_MT_+_598_1651_RNR1_12S_50ntFlanks | 905 | 935 | 31 | AGTATACTTCAAAGGACATTTAACTAAAACC | MB231 | 27 | 4 | 8 | 31 | 8 | 31.1 | 3.6 | 5.8 | 23.7 | 4.5 |
| >multi-hg38_MT_+_598_1651_RNR1_12S_50ntFlanks@905.936.32\|AGTATACTTCAAAGGACATTTAACTAAAACCC | >multi-hg38_MT_+_598_1651_RNR1_12S_50ntFlanks | 905 | 936 | 32 | AGTATACTTCAAAGGACATTTAACTAAAACCC | MB231 | 10 | 1 | 1 | 4 | 4 | 13.1 | 1.1 | 1.7 | 2.8 | 1.4 |
| >multi-hg38_MT_+_598_1651_RNR1_12S_50ntFlanks@915.935.21\|AAAGGACATTTAACTAAAACC | >multi-hg38_MT_+_598_1651_RNR1_12S_50ntFlanks | 915 | 935 | 21 | AAAGGACATTTAACTAAAACC | MB231 | 10 | 2 | 2 | 26 | 4 | 12.2 | 1.9 | 1.8 | 23.8 | 2.1 |
| >multi-hg38_MT_+_598_1651_RNR1_12S_50ntFlanks@916.935.20\|AAGGACATTTAACTAAAACC | >multi-hg38_MT_+_598_1651_RNR1_12S_50ntFlanks | 916 | 935 | 20 | AAGGACATTTAACTAAAACC | BT20 | 19 | 2 | 3 | 14 | 4 | 7.9 | 0.6 | 1.0 | 14.1 | 3.8 |
| >multi-hg38_MT_+_598_1651_RNR1_12S_50ntFlanks@916.935.20\|AAGGACATTTAACTAAAACC | >multi-hg38_MT_+_598_1651_RNR1_12S_50ntFlanks | 916 | 935 | 20 | AAGGACATTTAACTAAAACC | MB231 | 32 | 5 | 4 | 72 | 11 | 37.8 | 4.7 | 3.2 | 71.4 | 6.2 |
| >multi-hg38_MT_+_598_1651_RNR1_12S_50ntFlanks@916.936.21\|AAGGACATTTAACTAAAACCC | >multi-hg38_MT_+_598_1651_RNR1_12S_50ntFlanks | 916 | 936 | 21 | AAGGACATTTAACTAAAACCC | BT20 | 12 | 1 | 1 | 6 | 3 | 3.8 | 0.5 | 0.3 | 5.3 | 4.0 |
| >multi-hg38_MT_+_598_1651_RNR1_12S_50ntFlanks@916.936.21\|AAGGACATTTAACTAAAACCC | >multi-hg38_MT_+_598_1651_RNR1_12S_50ntFlanks | 916 | 936 | 21 | AAGGACATTTAACTAAAACCC | MB231 | 20 | 1 | 1 | 17 | 6 | 27.9 | 1.9 | 1.5 | 14.2 | 2.7 |
| >multi-hg38_MT_+_598_1651_RNR1_12S_50ntFlanks@917.935.19\|AGGACATTTAACTAAAACC | >multi-hg38_MT_+_598_1651_RNR1_12S_50ntFlanks | 917 | 935 | 19 | AGGACATTTAACTAAAACC | MB231 | 14 | 3 | 1 | 52 | 5 | 14.8 | 3.6 | 1.1 | 39.5 | 3.1 |
| >multi-hg38_MT_+_598_1651_RNR1_12S_50ntFlanks@965.1004.40\|GTAACATGGTAAGTGTACTGGAAAGTGCACTTGGACGAAC | >multi-hg38_MT_+_598_1651_RNR1_12S_50ntFlanks | 965 | 1004 | 40 | GTAACATGGTAAGTGTACTGGAAAGTGCACTTGGACGAAC | MB231 | 13 | 5 | 2 | 12 | 14 | 13.0 | 2.8 | 1.6 | 11.0 | 7.4 |
| >multi-hg38_MT_+_598_1651_RNR1_12S_50ntFlanks@966.1004.39\|TAACATGGTAAGTGTACTGGAAAGTGCACTTGGACGAAC | >multi-hg38_MT_+_598_1651_RNR1_12S_50ntFlanks | 966 | 1004 | 39 | TAACATGGTAAGTGTACTGGAAAGTGCACTTGGACGAAC | MB231 | 30 | 11 | 6 | 15 | 27 | 27.3 | 4.2 | 3.4 | 19.0 | 22.0 |
| >multi-hg38_MT_+_598_1651_RNR1_12S_50ntFlanks@967.1004.38\|AACATGGTAAGTGTACTGGAAAGTGCACTTGGACGAAC | >multi-hg38_MT_+_598_1651_RNR1_12S_50ntFlanks | 967 | 1004 | 38 | AACATGGTAAGTGTACTGGAAAGTGCACTTGGACGAAC | MB231 | 15 | 7 | 3 | 15 | 20 | 10.6 | 3.6 | 1.8 | 18.1 | 17.4 |
| >multi-hg38_MT_+_598_1651_RNR1_12S_50ntFlanks@968.1004.37\|ACATGGTAAGTGTACTGGAAAGTGCACTTGGACGAAC | >multi-hg38_MT_+_598_1651_RNR1_12S_50ntFlanks | 968 | 1004 | 37 | ACATGGTAAGTGTACTGGAAAGTGCACTTGGACGAAC | MB231 | 24 | 10 | 3 | 19 | 29 | 20.3 | 6.1 | 1.5 | 22.7 | 19.8 |
| >multi-hg38_MT_+_598_1651_RNR1_12S_50ntFlanks@970.1004.35\|ATGGTAAGTGTACTGGAAAGTGCACTTGGACGAAC | >multi-hg38_MT_+_598_1651_RNR1_12S_50ntFlanks | 970 | 1004 | 35 | ATGGTAAGTGTACTGGAAAGTGCACTTGGACGAAC | MB231 | 16 | 9 | 3 | 29 | 31 | 4.1 | 4.8 | 1.0 | 34.7 | 19.5 |
| >multi-hg38_MT_+_598_1651_RNR1_12S_50ntFlanks@971.1004.34\|TGGTAAGTGTACTGGAAAGTGCACTTGGACGAAC | >multi-hg38_MT_+_598_1651_RNR1_12S_50ntFlanks | 971 | 1004 | 34 | TGGTAAGTGTACTGGAAAGTGCACTTGGACGAAC | MB231 | 21 | 12 | 4 | 26 | 22 | 9.6 | 5.9 | 1.8 | 32.4 | 14.6 |
| >multi-hg38_MT_+_598_1651_RNR1_12S_50ntFlanks@972.1004.33\|GGTAAGTGTACTGGAAAGTGCACTTGGACGAAC | >multi-hg38_MT_+_598_1651_RNR1_12S_50ntFlanks | 972 | 1004 | 33 | GGTAAGTGTACTGGAAAGTGCACTTGGACGAAC | MB231 | 18 | 8 | 2 | 39 | 42 | 4.2 | 4.9 | 1.1 | 38.6 | 21.6 |
| >multi-trnaMT_AsnGTT_MT_-_5657_5729@1.40.40,trnalookalike5_AsnGTT_1_-_566207_566279@1.40.40\|TAGATTGAAGCCAGTTGATTAGGGTGCTTAGCTGTTAACT | >multi-trnaMT_AsnGTT_MT_-_5657_5729 | 1 | 40 | 40 | TAGATTGAAGCCAGTTGATTAGGGTGCTTAGCTGTTAACT | MB231 | 14 | 2 | 3 | 15 | 14 | 11.2 | 1.1 | 2.1 | 18.5 | 9.7 |
| >multi-trnaMT_GlnTTG_MT_-_4329_4400@1.29.29,trnalookalike2_GlnTTG_1_-_564879_564950@1.29.29\|TAGGATGGGGTGTGATAGGTGGCACGGAG | >multi-trnaMT_GlnTTG_MT_-_4329_4400 | 1 | 29 | 29 | TAGGATGGGGTGTGATAGGTGGCACGGAG | BT20 | 11 | 2 | 2 | 64 | 26 | 5.8 | 0.7 | 0.7 | 91.3 | 34.0 |

| | | | | | | | | | | | | | | | | |
|---|---|---|---|---|---|---|---|---|---|---|---|---|---|---|---|---|
| >multi-trnaMT_GlnTTG_MT_-_4329_4400@1.30.30,trnalookalike2_GlnTTG_1_-_564879_564950@1.30.30\|TAGGATGGGGTGTGATAGGTGGCACGGAGA | >multi-trnaMT_GlnTTG_MT_-_4329_4400 | 1 | 30 | 30 | TAGGATGGGGTGTGATAGGTGGCACGGAGA | BT20 | 28 | 7 | 11 | 139 | 46 | 14.4 | 2.1 | 6.1 | 190.7 | 55.4 |
| >multi-trnaMT_GlnTTG_MT_-_4329_4400@1.33.33,trnalookalike2_GlnTTG_1_-_564879_564950@1.33.33\|TAGGATGGGGTGTGATAGGTGGCACGGAGAATT | >multi-trnaMT_GlnTTG_MT_-_4329_4400 | 1 | 33 | 33 | TAGGATGGGGTGTGATAGGTGGCACGGAGAATT | BT20 | 28 | 15 | 48 | 56 | 27 | 23.1 | 8.1 | 63.1 | 73.7 | 38.5 |
| >multi-trnaMT_GluTTC_MT_-_14674_14742@13.61.49,trnalookalike8_GluTTC_5_-_93905172_93905240@13.61.49\|GAAATACAACGATGGTTTTTCATATCATTGGTCGTGGTTGTAGTCCGTG | >multi-trnaMT_GluTTC_MT_-_14674_14742 | 13 | 61 | 49 | GAAATACAACGATGGTTTTTCATATCATTGGTCGTGGTTGTAGTCCGTG | MB231 | 10 | 2 | 0 | 2 | 2 | 14.4 | 2.8 | 0.2 | 3.1 | 4.1 |
| >multi-trnaMT_GluTTC_MT_-_14674_14742@20.58.39,trnalookalike8_GluTTC_5_-_93905172_93905240@20.58.39\|AACGATGGTTTTTCATATCATTGGTCGTGGTTGTAGTCC | >multi-trnaMT_GluTTC_MT_-_14674_14742 | 20 | 58 | 39 | AACGATGGTTTTTCATATCATTGGTCGTGGTTGTAGTCC | MB231 | 10 | 4 | 2 | 5 | 9 | 11.1 | 2.6 | 1.5 | 2.5 | 7.6 |
| >multi-trnaMT_GluTTC_MT_-_14674_14742@21.58.38,trnalookalike8_GluTTC_5_-_93905172_93905240@21.58.38\|ACGATGGTTTTTCATATCATTGGTCGTGGTTGTAGTCC | >multi-trnaMT_GluTTC_MT_-_14674_14742 | 21 | 58 | 38 | ACGATGGTTTTTCATATCATTGGTCGTGGTTGTAGTCC | MB231 | 18 | 6 | 3 | 12 | 16 | 21.6 | 5.4 | 2.8 | 6.2 | 14.1 |
| >multi-trnaMT_GluTTC_MT_-_14674_14742@34.57.24,trnalookalike8_GluTTC_5_-_93905172_93905240@34.57.24\|ATATCATTGGTCGTGGTTGTAGTC | >multi-trnaMT_GluTTC_MT_-_14674_14742 | 34 | 57 | 24 | ATATCATTGGTCGTGGTTGTAGTC | MB231 | 10 | 2 | 0 | 19 | 8 | 10.6 | 1.8 | 0.4 | 14.9 | 2.6 |
| >multi-trnaMT_GluTTC_MT_-_14674_14742@38.66.29,trnalookalike8_GluTTC_5_-_93905172_93905240@38.66.29\|CATTGGTCGTGGTTGTAGTCCGTGCGAGA | >multi-trnaMT_GluTTC_MT_-_14674_14742 | 38 | 66 | 29 | CATTGGTCGTGGTTGTAGTCCGTGCGAGA | BT20 | 11 | 9 | 1 | 24 | 33 | 5.8 | 8.1 | 1.2 | 18.5 | 23.7 |
| >multi-trnaMT_GluTTC_MT_-_14674_14742@39.57.19,trnalookalike8_GluTTC_5_-_93905172_93905240@39.57.19\|ATTGGTCGTGGTTGTAGTC | >multi-trnaMT_GluTTC_MT_-_14674_14742 | 39 | 57 | 19 | ATTGGTCGTGGTTGTAGTC | BT20 | 15 | 4 | 14 | 50 | 22 | 5.9 | 0.1 | 2.2 | 38.3 | 12.4 |
| >multi-trnaMT_GluTTC_MT_-_14674_14742@39.57.19,trnalookalike8_GluTTC_5_-_93905172_93905240@39.57.19\|ATTGGTCGTGGTTGTAGTC | >multi-trnaMT_GluTTC_MT_-_14674_14742 | 39 | 57 | 19 | ATTGGTCGTGGTTGTAGTC | MB231 | 17 | 7 | 12 | 42 | 17 | 18.6 | 5.2 | 2.9 | 34.5 | 11.4 |
| >multi-trnaMT_GluTTC_MT_-_14674_14742@39.58.20,trnalookalike8_GluTTC_5_-_93905172_93905240@39.58.20\|ATTGGTCGTGGTTGTAGTCC | >multi-trnaMT_GluTTC_MT_-_14674_14742 | 39 | 58 | 20 | ATTGGTCGTGGTTGTAGTCC | BT20 | 27 | 9 | 20 | 91 | 50 | 6.3 | 4.1 | 4.1 | 40.5 | 5.5 |
| >multi-trnaMT_GluTTC_MT_-_14674_14742@39.58.20,trnalookalike8_GluTTC_5_-_93905172_93905240@39.58.20\|ATTGGTCGTGGTTGTAGTCC | >multi-trnaMT_GluTTC_MT_-_14674_14742 | 39 | 58 | 20 | ATTGGTCGTGGTTGTAGTCC | MB231 | 33 | 18 | 24 | 51 | 27 | 36.4 | 14.8 | 8.2 | 51.3 | 29.5 |
| >multi-trnaMT_GluTTC_MT_-_14674_14742@39.61.23,trnalookalike8_GluTTC_5_-_93905172_93905240@39.61.23\|ATTGGTCGTGGTTGTAGTCCGTG | >multi-trnaMT_GluTTC_MT_-_14674_14742 | 39 | 61 | 23 | ATTGGTCGTGGTTGTAGTCCGTG | BT20 | 16 | 7 | 4 | 66 | 37 | 2.8 | 4.2 | 4.2 | 52.3 | 19.0 |
| >multi-trnaMT_GluTTC_MT_-_14674_14742@39.61.23,trnalookalike8_GluTTC_5_-_93905172_93905240@39.61.23\|ATTGGTCGTGGTTGTAGTCCGTG | >multi-trnaMT_GluTTC_MT_-_14674_14742 | 39 | 61 | 23 | ATTGGTCGTGGTTGTAGTCCGTG | MB231 | 19 | 12 | 4 | 31 | 16 | 21.9 | 9.8 | 2.9 | 26.6 | 14.3 |
| >multi-trnaMT_GluTTC_MT_-_14674_14742@39.63.25,trnalookalike8_GluTTC_5_-_93905172_93905240@39.63.25\|ATTGGTCGTGGTTGTAGTCCGTGCG | >multi-trnaMT_GluTTC_MT_-_14674_14742 | 39 | 63 | 25 | ATTGGTCGTGGTTGTAGTCCGTGCG | BT20 | 16 | 7 | 3 | 69 | 48 | 6.2 | 3.3 | 2.6 | 59.7 | 37.1 |
| >multi-trnaMT_GluTTC_MT_-_14674_14742@39.63.25,trnalookalike8_GluTTC_5_-_93905172_93905240@39.63.25\|ATTGGTCGTGGTTGTAGTCCGTGCG | >multi-trnaMT_GluTTC_MT_-_14674_14742 | 39 | 63 | 25 | ATTGGTCGTGGTTGTAGTCCGTGCG | MB231 | 17 | 13 | 4 | 28 | 17 | 19.5 | 8.2 | 2.7 | 24.0 | 15.2 |
| >multi-trnaMT_GluTTC_MT_-_14674_14742@39.65.27,trnalookalike8_GluTTC_5_-_93905172_93905240@39.65.27\|ATTGGTCGTGGTTGTAGTCCGTGCGAG | >multi-trnaMT_GluTTC_MT_-_14674_14742 | 39 | 65 | 27 | ATTGGTCGTGGTTGTAGTCCGTGCGAG | BT20 | 12 | 5 | 1 | 34 | 38 | 2.9 | 1.4 | 0.8 | 33.1 | 38.1 |
| >multi-trnaMT_GluTTC_MT_-_14674_14742@39.65.27,trnalookalike8_GluTTC_5_-_93905172_93905240@39.65.27\|ATTGGTCGTGGTTGTAGTCCGTGCGAG | >multi-trnaMT_GluTTC_MT_-_14674_14742 | 39 | 65 | 27 | ATTGGTCGTGGTTGTAGTCCGTGCGAG | MB231 | 12 | 7 | 2 | 20 | 12 | 12.6 | 4.4 | 1.5 | 13.2 | 9.8 |
| >multi-trnaMT_GluTTC_MT_-_14674_14742@39.66.28,trnalookalike8_GluTTC_5_-_93905172_93905240@39.66.28\|ATTGGTCGTGGTTGTAGTCCGTGCGAGA | >multi-trnaMT_GluTTC_MT_-_14674_14742 | 39 | 66 | 28 | ATTGGTCGTGGTTGTAGTCCGTGCGAGA | BT20 | 17 | 12 | 5 | 62 | 50 | 5.9 | 8.8 | 6.0 | 28.9 | 25.7 |
| >multi-trnaMT_GluTTC_MT_-_14674_14742@39.66.28,trnalookalike8_GluTTC_5_-_93905172_93905240@39.66.28\|ATTGGTCGTGGTTGTAGTCCGTGCGAGA | >multi-trnaMT_GluTTC_MT_-_14674_14742 | 39 | 66 | 28 | ATTGGTCGTGGTTGTAGTCCGTGCGAGA | MB231 | 11 | 17 | 6 | 24 | 16 | 7.0 | 11.6 | 5.4 | 17.9 | 11.8 |
| >multi-trnaMT_GluTTC_MT_-_14674_14742@40.57.18,trnalookalike8_GluTTC_5_-_93905172_93905240@40.57.18\|TTGGTCGTGGTTGTAGTC | >multi-trnaMT_GluTTC_MT_-_14674_14742 | 40 | 57 | 18 | TTGGTCGTGGTTGTAGTC | BT20 | 13 | 3 | 15 | 28 | 17 | 3.7 | 0.8 | 1.8 | 16.8 | 7.9 |
| >multi-trnaMT_GluTTC_MT_-_14674_14742@40.57.18,trnalookalike8_GluTTC_5_-_93905172_93905240@40.57.18\|TTGGTCGTGGTTGTAGTC | >multi-trnaMT_GluTTC_MT_-_14674_14742 | 40 | 57 | 18 | TTGGTCGTGGTTGTAGTC | MB231 | 18 | 7 | 15 | 30 | 14 | 19.7 | 4.7 | 2.9 | 27.6 | 9.2 |
| >multi-trnaMT_GluTTC_MT_-_14674_14742@40.58.19,trnalookalike8_GluTTC_5_-_93905172_93905240@40.58.19\|TTGGTCGTGGTTGTAGTCC | >multi-trnaMT_GluTTC_MT_-_14674_14742 | 40 | 58 | 19 | TTGGTCGTGGTTGTAGTCC | BT20 | 29 | 8 | 27 | 60 | 43 | 5.0 | 2.4 | 4.9 | 22.7 | 4.7 |
| >multi-trnaMT_GluTTC_MT_-_14674_14742@40.58.19,trnalookalike8_GluTTC_5_-_93905172_93905240@40.58.19\|TTGGTCGTGGTTGTAGTCC | >multi-trnaMT_GluTTC_MT_-_14674_14742 | 40 | 58 | 19 | TTGGTCGTGGTTGTAGTCC | MB231 | 45 | 22 | 34 | 47 | 27 | 50.6 | 17.7 | 10.7 | 51.9 | 29.6 |
| >multi-trnaMT_GluTTC_MT_-_14674_14742@40.59.20,trnalookalike8_GluTTC_5_-_93905172_93905240@40.59.20\|TTGGTCGTGGTTGTAGTCCG | >multi-trnaMT_GluTTC_MT_-_14674_14742 | 40 | 59 | 20 | TTGGTCGTGGTTGTAGTCCG | MB231 | 10 | 5 | 3 | 9 | 7 | 12.5 | 4.5 | 1.6 | 9.2 | 6.8 |
| >multi-trnaMT_GluTTC_MT_-_14674_14742@40.61.22,trnalookalike8_GluTTC_5_-_93905172_93905240@40.61.22\|TTGGTCGTGGTTGTAGTCCGTG | >multi-trnaMT_GluTTC_MT_-_14674_14742 | 40 | 61 | 22 | TTGGTCGTGGTTGTAGTCCGTG | BT20 | 11 | 4 | 2 | 42 | 24 | 1.0 | 2.6 | 1.2 | 30.5 | 14.1 |
| >multi-trnaMT_GluTTC_MT_-_14674_14742@40.61.22,trnalookalike8_GluTTC_5_-_93905172_93905240@40.61.22\|TTGGTCGTGGTTGTAGTCCGTG | >multi-trnaMT_GluTTC_MT_-_14674_14742 | 40 | 61 | 22 | TTGGTCGTGGTTGTAGTCCGTG | MB231 | 16 | 9 | 3 | 26 | 13 | 19.2 | 7.9 | 1.5 | 23.4 | 13.0 |
| >multi-trnaMT_GluTTC_MT_-_14674_14742@40.63.24,trnalookalike8_GluTTC_5_-_93905172_93905240@40.63.24\|TTGGTCGTGGTTGTAGTCCGTGCG | >multi-trnaMT_GluTTC_MT_-_14674_14742 | 40 | 63 | 24 | TTGGTCGTGGTTGTAGTCCGTGCG | BT20 | 14 | 6 | 2 | 50 | 36 | 4.4 | 3.7 | 1.4 | 44.0 | 29.9 |
| >multi-trnaMT_GluTTC_MT_-_14674_14742@40.63.24,trnalookalike8_GluTTC_5_-_93905172_93905240@40.63.24\|TTGGTCGTGGTTGTAGTCCGTGCG | >multi-trnaMT_GluTTC_MT_-_14674_14742 | 40 | 63 | 24 | TTGGTCGTGGTTGTAGTCCGTGCG | MB231 | 19 | 12 | 4 | 27 | 15 | 24.4 | 8.3 | 1.9 | 24.6 | 15.3 |
| >multi-trnaMT_GluTTC_MT_-_14674_14742@40.65.26,trnalookalike8_GluTTC_5_-_93905172_93905240@40.65.26\|TTGGTCGTGGTTGTAGTCCGTGCGAG | >multi-trnaMT_GluTTC_MT_-_14674_14742 | 40 | 65 | 26 | TTGGTCGTGGTTGTAGTCCGTGCGAG | MB231 | 11 | 7 | 2 | 17 | 10 | 13.3 | 4.8 | 1.2 | 13.3 | 8.1 |
| >multi-trnaMT_GluTTC_MT_-_14674_14742@40.66.27,trnalookalike8_GluTTC_5_-_93905172_93905240@40.66.27\|TTGGTCGTGGTTGTAGTCCGTGCGAGA | >multi-trnaMT_GluTTC_MT_-_14674_14742 | 40 | 66 | 27 | TTGGTCGTGGTTGTAGTCCGTGCGAGA | BT20 | 21 | 16 | 4 | 53 | 49 | 11.2 | 14.4 | 4.1 | 31.4 | 19.6 |
| >multi-trnaMT_GluTTC_MT_-_14674_14742@40.66.27,trnalookalike8_GluTTC_5_-_93905172_93905240@40.66.27\|TTGGTCGTGGTTGTAGTCCGTGCGAGA | >multi-trnaMT_GluTTC_MT_-_14674_14742 | 40 | 66 | 27 | TTGGTCGTGGTTGTAGTCCGTGCGAGA | MB231 | 16 | 21 | 5 | 29 | 21 | 11.6 | 11.2 | 3.4 | 23.4 | 16.6 |
| >multi-trnaMT_LysTTT_MT_+_8295_8364@1.20.20,trnalookalike7_LysTTT_1_+_568844_568913@1.20.20\|CACTGTAAAGCTAACTTAGC | >multi-trnaMT_LysTTT_MT_+_8295_8364 | 1 | 20 | 20 | CACTGTAAAGCTAACTTAGC | BT20 | 18 | 4 | 6 | 21 | 7 | 17.6 | 2.3 | 4.0 | 28.5 | 9.7 |
| >multi-trnaMT_LysTTT_MT_+_8295_8364@1.21.21,trnalookalike7_LysTTT_1_+_568844_568913@1.21.21\|CACTGTAAAGCTAACTTAGCA | >multi-trnaMT_LysTTT_MT_+_8295_8364 | 1 | 21 | 21 | CACTGTAAAGCTAACTTAGCA | BT20 | 14 | 2 | 3 | 4 | 1 | 17.8 | 2.3 | 1.2 | 4.8 | 2.5 |
| >multi-trnaMT_LysTTT_MT_+_8295_8364@1.23.23,trnalookalike7_LysTTT_1_+_568844_568913@1.23.23\|CACTGTAAAGCTAACTTAGCATT | >multi-trnaMT_LysTTT_MT_+_8295_8364 | 1 | 23 | 23 | CACTGTAAAGCTAACTTAGCATT | BT20 | 11 | 8 | 3 | 60 | 6 | 12.0 | 6.9 | 2.6 | 75.6 | 7.2 |
| >multi-trnaMT_LysTTT_MT_+_8295_8364@1.26.26,trnalookalike7_LysTTT_1_+_568844_568913@1.26.26\|CACTGTAAAGCTAACTTAGCATTAAC | >multi-trnaMT_LysTTT_MT_+_8295_8364 | 1 | 26 | 26 | CACTGTAAAGCTAACTTAGCATTAAC | BT20 | 15 | 3 | 1 | 44 | 20 | 12.3 | 2.1 | 1.3 | 56.9 | 28.2 |
| >multi-trnaMT_LysTTT_MT_+_8295_8364@1.26.26,trnalookalike7_LysTTT_1_+_568844_568913@1.26.26\|CACTGTAAAGCTAACTTAGCATTAAC | >multi-trnaMT_LysTTT_MT_+_8295_8364 | 1 | 26 | 26 | CACTGTAAAGCTAACTTAGCATTAAC | MB231 | 27 | 10 | 1 | 115 | 40 | 16.9 | 5.4 | 0.4 | 124.3 | 19.3 |
| >multi-trnaMT_LysTTT_MT_+_8295_8364@1.36.36,trnalookalike7_LysTTT_1_+_568844_568913@1.36.36\|CACTGTAAAGCTAACTTAGCATTAACCTTTTAAGTT | >multi-trnaMT_LysTTT_MT_+_8295_8364 | 1 | 36 | 36 | CACTGTAAAGCTAACTTAGCATTAACCTTTTAAGTT | BT20 | 18 | 8 | 3 | 63 | 29 | 13.1 | 7.0 | 1.2 | 95.3 | 45.1 |
| >multi-trnaMT_LysTTT_MT_+_8295_8364@1.36.36,trnalookalike7_LysTTT_1_+_568844_568913@1.36.36\|CACTGTAAAGCTAACTTAGCATTAACCTTTTAAGTT | >multi-trnaMT_LysTTT_MT_+_8295_8364 | 1 | 36 | 36 | CACTGTAAAGCTAACTTAGCATTAACCTTTTAAGTT | MB231 | 28 | 14 | 4 | 56 | 42 | 17.1 | 6.9 | 1.1 | 55.4 | 16.1 |
| >multi-trnaMT_LysTTT_MT_+_8295_8364@1.38.38,trnalookalike7_LysTTT_1_+_568844_568913@1.38.38\|CACTGTAAAGCTAACTTAGCATTAACCTTTTAAGTTAA | >multi-trnaMT_LysTTT_MT_+_8295_8364 | 1 | 38 | 38 | CACTGTAAAGCTAACTTAGCATTAACCTTTTAAGTTAA | BT20 | 12 | 6 | 2 | 79 | 31 | 11.0 | 6.3 | 1.2 | 115.6 | 31.8 |
| >multi-trnaMT_LysTTT_MT_+_8295_8364@1.43.43,trnalookalike7_LysTTT_1_+_568844_568913@1.43.43\|CACTGTAAAGCTAACTTAGCATTAACCTTTTAAGTTAAAGATT | >multi-trnaMT_LysTTT_MT_+_8295_8364 | 1 | 43 | 43 | CACTGTAAAGCTAACTTAGCATTAACCTTTTAAGTTAAAGATT | MB231 | 11 | 4 | 3 | 15 | 23 | 7.2 | 3.4 | 0.4 | 19.9 | 21.0 |
| >multi-trnaMT_LysTTT_MT_+_8295_8364@1.47.47,trnalookalike7_LysTTT_1_+_568844_568913@1.47.47\|CACTGTAAAGCTAACTTAGCATTAACCTTTTAAGTTAAAGATTAAGA | >multi-trnaMT_LysTTT_MT_+_8295_8364 | 1 | 47 | 47 | CACTGTAAAGCTAACTTAGCATTAACCTTTTAAGTTAAAGATTAAGA | MB231 | 20 | 4 | 1 | 7 | 28 | 26.3 | 5.3 | 1.3 | 10.1 | 26.1 |
| >multi-trnaMT_SerTGA_MT_-_7446_7514@1.26.26,trnalookalike6_SerTGA_1_-_567997_568065@1.26.26\|GAAAAAGTCATGGAGGCCATGGGGTT | >multi-trnaMT_SerTGA_MT_-_7446_7514 | 1 | 26 | 26 | GAAAAAGTCATGGAGGCCATGGGGTT | BT20 | 19 | 56 | 9 | 361 | 34 | 8.1 | 31.9 | 4.1 | 513.1 | 47.3 |
| >trna1_GlnCTG_6_+_18836402_18836473@58.75.18,trna3_GlnCTG_17_+_8023070_8023141@58.75.18,trna42_GlnCTG_6_+_27263212_27263283@58.75.18,trna49_GlnCTG_6_+_27487308_27487379@58.75.18,trna7_GlnCTG_15_-_66161400_66161471@58.75.18,trna99_GlnCTG_6_-_28909378_28909449@58.75.18\|ATCTCGGTGGAACCTCCA | >trna1_GlnCTG_6_+_18836402_18836473 | 58 | 75 | 18 | ATCTCGGTGGAACCTCCA | MB231 | 15 | 15 | 49 | 2 | 10 | 26.7 | 25.3 | 85.0 | 3.9 | 17.3 |

| | | | | | | | | | | | | | | | | |
|---|---|---|---|---|---|---|---|---|---|---|---|---|---|---|---|---|
| >trna10_AlaCGC_6_+_26553731_26553802@58.75.18,trna119_AlaCGC_6_-_28641613_28641684@58.75.18,trna13_AlaCGC_2_+_157257281_157257352@58.75.18,trna13_AlaTGC_12_-_125406301_125406372@58.75.18,trna66_AlaTGC_6_+_28611222_28611293@58.75.18,trna8_AlaTGC_12_+_125424512_125424583@58.75.18,trna8_AlaTGC_5_+_180633868_180633939@58.75.18\|TCCCCGGCATCTCCACCA | >trna10_AlaCGC_6_+_26553731_26553802 | 58 | 75 | 18 | TCCCCGGCATCTCCACCA | BT20 | 108 | 27 | 880 | 1 | 17 | 66.1 | 7.8 | 395.2 | 0.7 | 19.7 |
| >trna10_AlaCGC_6_+_26553731_26553802@58.75.18,trna119_AlaCGC_6_-_28641613_28641684@58.75.18,trna13_AlaCGC_2_+_157257281_157257352@58.75.18,trna13_AlaTGC_12_-_125406301_125406372@58.75.18,trna66_AlaTGC_6_+_28611222_28611293@58.75.18,trna8_AlaTGC_12_+_125424512_125424583@58.75.18,trna8_AlaTGC_5_+_180633868_180633939@58.75.18\|TCCCCGGCATCTCCACCA | >trna10_AlaCGC_6_+_26553731_26553802 | 58 | 75 | 18 | TCCCCGGCATCTCCACCA | MB231 | 89 | 42 | 346 | 2 | 44 | 22.7 | 19.3 | 109.6 | 0.8 | 25.5 |
| >trna10_AspGTC_12_-_125424193_125424264@1.29.29,trna12_AspGTC_12_-_125411891_125411962@1.29.29,trna38_AspGTC_17_-_8125556_8125627@1.29.29,trna45_AspGTC_6_+_27447453_27447524@1.29.29,trna48_AspGTC_6_+_27471523_27471594@1.29.29,trna4_AspGTC_12_+_96429799_96429870@1.29.29,trna69_AspGTC_1_-_161440205_161440276@1.29.29,trna72_AspGTC_1_-_161432824_161432895@1.29.29,trna75_AspGTC_1_-_161425414_161425485@1.29.29,trna78_AspGTC_1_-_161418033_161418104@1.29.29,trna81_AspGTC_1_-_161410615_161410686@1.29.29\|TCCTCGTTAGTATAGTGGTGAGTATCCCC | >trna10_AspGTC_12_-_125424193_125424264 | 1 | 29 | 29 | TCCTCGTTAGTATAGTGGTGAGTATCCCC | MB468 | 26 | 9 | 46 | 16 | 10 | 10.7 | 2.8 | 14.5 | 5.8 | 7.6 |
| >trna10_AspGTC_12_-_125424193_125424264@1.30.30,trna12_AspGTC_12_-_125411891_125411962@1.30.30,trna38_AspGTC_17_-_8125556_8125627@1.30.30,trna45_AspGTC_6_+_27447453_27447524@1.30.30,trna48_AspGTC_6_+_27471523_27471594@1.30.30,trna4_AspGTC_12_+_96429799_96429870@1.30.30,trna69_AspGTC_1_-_161440205_161440276@1.30.30,trna72_AspGTC_1_-_161432824_161432895@1.30.30,trna75_AspGTC_1_-_161425414_161425485@1.30.30,trna78_AspGTC_1_-_161418033_161418104@1.30.30,trna81_AspGTC_1_-_161410615_161410686@1.30.30\|TCCTCGTTAGTATAGTGGTGAGTATCCCCG | >trna10_AspGTC_12_-_125424193_125424264 | 1 | 30 | 30 | TCCTCGTTAGTATAGTGGTGAGTATCCCCG | MB468 | 49 | 43 | 77 | 16 | 16 | 16.8 | 11.2 | 13.7 | 6.2 | 7.5 |
| >trna10_AspGTC_12_-_125424193_125424264@1.31.31,trna12_AspGTC_12_-_125411891_125411962@1.31.31,trna38_AspGTC_17_-_8125556_8125627@1.31.31,trna45_AspGTC_6_+_27447453_27447524@1.31.31,trna48_AspGTC_6_+_27471523_27471594@1.31.31,trna4_AspGTC_12_+_96429799_96429870@1.31.31,trna69_AspGTC_1_-_161440205_161440276@1.31.31,trna72_AspGTC_1_-_161432824_161432895@1.31.31,trna75_AspGTC_1_-_161425414_161425485@1.31.31,trna78_AspGTC_1_-_161418033_161418104@1.31.31,trna81_AspGTC_1_-_161410615_161410686@1.31.31\|TCCTCGTTAGTATAGTGGTGAGTATCCCCGC | >trna10_AspGTC_12_-_125424193_125424264 | 1 | 31 | 31 | TCCTCGTTAGTATAGTGGTGAGTATCCCCGC | MB468 | 19 | 13 | 28 | 6 | 4 | 12.2 | 4.3 | 7.7 | 4.7 | 4.1 |
| >trna10_AspGTC_12_-_125424193_125424264@1.34.34,trna12_AspGTC_12_-_125411891_125411962@1.34.34,trna38_AspGTC_17_-_8125556_8125627@1.34.34,trna45_AspGTC_6_+_27447453_27447524@1.34.34,trna48_AspGTC_6_+_27471523_27471594@1.34.34,trna4_AspGTC_12_+_96429799_96429870@1.34.34,trna69_AspGTC_1_-_161440205_161440276@1.34.34,trna72_AspGTC_1_-_161432824_161432895@1.34.34,trna75_AspGTC_1_-_161425414_161425485@1.34.34,trna78_AspGTC_1_-_161418033_161418104@1.34.34,trna81_AspGTC_1_-_161410615_161410686@1.34.34\|TCCTCGTTAGTATAGTGGTGAGTATCCCCGCCTG | >trna10_AspGTC_12_-_125424193_125424264 | 1 | 34 | 34 | TCCTCGTTAGTATAGTGGTGAGTATCCCCGCCTG | MB231 | 31 | 13 | 44 | 12 | 29 | 22.6 | 13.9 | 34.0 | 7.2 | 19.5 |
| >trna10_AspGTC_12_-_125424193_125424264@1.36.36,trna12_AspGTC_12_-_125411891_125411962@1.36.36,trna38_AspGTC_17_-_8125556_8125627@1.36.36,trna45_AspGTC_6_+_27447453_27447524@1.36.36,trna48_AspGTC_6_+_27471523_27471594@1.36.36,trna4_AspGTC_12_+_96429799_96429870@1.36.36,trna69_AspGTC_1_-_161440205_161440276@1.36.36,trna72_AspGTC_1_-_161432824_161432895@1.36.36,trna75_AspGTC_1_-_161425414_161425485@1.36.36,trna78_AspGTC_1_-_161418033_161418104@1.36.36,trna81_AspGTC_1_-_161410615_161410686@1.36.36\|TCCTCGTTAGTATAGTGGTGAGTATCCCCGCCTGTC | >trna10_AspGTC_12_-_125424193_125424264 | 1 | 36 | 36 | TCCTCGTTAGTATAGTGGTGAGTATCCCCGCCTGTC | MB231 | 11 | 13 | 42 | 7 | 9 | 10.5 | 17.7 | 44.4 | 7.1 | 6.7 |
| >trna10_AspGTC_12_-_125424193_125424264@31.72.42,trna12_AspGTC_12_-_125411891_125411962@31.72.42,trna38_AspGTC_17_-_8125556_8125627@31.72.42,trna45_AspGTC_6_+_27447453_27447524@31.72.42,trna48_AspGTC_6_+_27471523_27471594@31.72.42,trna4_AspGTC_12_+_96429799_96429870@31.72.42,trna69_AspGTC_1_-_161440205_161440276@31.72.42,trna72_AspGTC_1_-_161432824_161432895@31.72.42,trna75_AspGTC_1_-_161425414_161425485@31.72.42,trna78_AspGTC_1_-_161418033_161418104@31.72.42,trna81_AspGTC_1_-_161410615_161410686@31.72.42\|CCTGTCACGCGGGAGACCGGGGTTCGATTCCCCGACGGGGAG | >trna10_AspGTC_12_-_125424193_125424264 | 31 | 72 | 42 | CCTGTCACGCGGGAGACCGGGGTTCGATTCCCCGACGGGGAG | MB468 | 29 | 50 | 135 | 7 | 47 | 19.8 | 45.6 | 68.7 | 5.6 | 60.1 |
| >trna10_AspGTC_12_-_125424193_125424264@32.70.39,trna12_AspGTC_12_-_125411891_125411962@32.70.39,trna144_AspGTC_6_-_27551236_27551307@32.70.39,trna38_AspGTC_17_-_8125556_8125627@32.70.39,trna45_AspGTC_6_+_27447453_27447524@32.70.39,trna48_AspGTC_6_+_27471523_27471594@32.70.39,trna4_AspGTC_12_+_96429799_96429870@32.70.39,trna69_AspGTC_1_-_161440205_161440276@32.70.39,trna72_AspGTC_1_-_161432824_161432895@32.70.39,trna75_AspGTC_1_-_161425414_161425485@32.70.39,trna78_AspGTC_1_-_161418033_161418104@32.70.39,trna81_AspGTC_1_-_161410615_161410686@32.70.39\|CTGTCACGCGGGAGACCGGGGTTCGATTCCCCGACGGGG | >trna10_AspGTC_12_-_125424193_125424264 | 32 | 70 | 39 | CTGTCACGCGGGAGACCGGGGTTCGATTCCCCGACGGGG | MB468 | 13 | 36 | 46 | 21 | 12 | 3.9 | 20.4 | 15.0 | 16.4 | 13.1 |
| >trna10_AspGTC_12_-_125424193_125424264@32.72.41,trna12_AspGTC_12_-_125411891_125411962@32.72.41,trna144_AspGTC_6_-_27551236_27551307@32.72.41,trna38_AspGTC_17_-_8125556_8125627@32.72.41,trna45_AspGTC_6_+_27447453_27447524@32.72.41,trna48_AspGTC_6_+_27471523_27471594@32.72.41,trna4_AspGTC_12_+_96429799_96429870@32.72.41,trna69_AspGTC_1_-_161440205_161440276@32.72.41,trna72_AspGTC_1_-_161432824_161432895@32.72.41,trna75_AspGTC_1_-_161425414_161425485@32.72.41,trna78_AspGTC_1_-_161418033_161418104@32.72.41,trna81_AspGTC_1_-_161410615_161410686@32.72.41\|CTGTCACGCGGGAGACCGGGGTTCGATTCCCCGACGGGGAG | >trna10_AspGTC_12_-_125424193_125424264 | 32 | 72 | 41 | CTGTCACGCGGGAGACCGGGGTTCGATTCCCCGACGGGGAG | MB468 | 37 | 106 | 159 | 19 | 28 | 15.6 | 89.9 | 66.9 | 17.9 | 32.9 |
| >trna10_AspGTC_12_-_125424193_125424264@35.73.39,trna12_AspGTC_12_-_125411891_125411962@35.73.39,trna144_AspGTC_6_-_27551236_27551307@35.73.39,trna38_AspGTC_17_-_8125556_8125627@35.73.39,trna45_AspGTC_6_+_27447453_27447524@35.73.39,trna48_AspGTC_6_+_27471523_27471594@35.73.39,trna4_AspGTC_12_+_96429799_96429870@35.73.39,trna69_AspGTC_1_-_161440205_161440276@35.73.39,trna72_AspGTC_1_-_161432824_161432895@35.73.39,trna75_AspGTC_1_-_161425414_161425485@35.73.39,trna78_AspGTC_1_-_161418033_161418104@35.73.39,trna81_AspGTC_1_-_161410615_161410686@35.73.39\|TCACGCGGGAGACCGGGGTTCGATTCCCCGACGGGGAGC | >trna10_AspGTC_12_-_125424193_125424264 | 35 | 73 | 39 | TCACGCGGGAGACCGGGGTTCGATTCCCCGACGGGGAGC | MB231 | 21 | 13 | 217 | 19 | 11 | 20.3 | 12.4 | 179.6 | 13.7 | 7.9 |

| | | | | | | | | | | | | | | | | |
|---|---|---|---|---|---|---|---|---|---|---|---|---|---|---|---|---|
| >trna10_AspGTC_12_-_125424193_125424264@35.74.40,trna12_AspGTC_12_-_125411891_125411962@35.74.40,trna144_AspGTC_6_-_27551236_27551307@35.74.40,trna38_AspGTC_17_-_8125556_8125627@35.74.40,trna45_AspGTC_6_+_27447453_27447524@35.74.40,trna48_AspGTC_6_+_27471523_27471594@35.74.40,trna4_AspGTC_12_+_96429799_96429870@35.74.40,trna69_AspGTC_1_-_161440205_161440276@35.74.40,trna72_AspGTC_1_-_161432824_161432895@35.74.40,trna75_AspGTC_1_-_161425414_161425485@35.74.40,trna78_AspGTC_1_-_161418033_161418104@35.74.40,trna81_AspGTC_1_-_161410615_161410686@35.74.40\|TCACGCGGGAGACCGGGGTTCGATTCCCCGACGGGGAGCC | >trna10_AspGTC_12_-_125424193_125424264 | 35 | 74 | 40 | TCACGCGGGAGACCGGGGTTCGATTCCCCGACGGGGAGCC | MB231 | 33 | 28 | 533 | 19 | 17 | 28.2 | 23.3 | 586.1 | 17.9 | 17.8 |
| >trna10_AspGTC_12_-_125424193_125424264@35.75.41,trna12_AspGTC_12_-_125411891_125411962@35.75.41,trna144_AspGTC_6_-_27551236_27551307@35.75.41,trna38_AspGTC_17_-_8125556_8125627@35.75.41,trna45_AspGTC_6_+_27447453_27447524@35.75.41,trna48_AspGTC_6_+_27471523_27471594@35.75.41,trna4_AspGTC_12_+_96429799_96429870@35.75.41,trna69_AspGTC_1_-_161440205_161440276@35.75.41,trna72_AspGTC_1_-_161432824_161432895@35.75.41,trna75_AspGTC_1_-_161425414_161425485@35.75.41,trna78_AspGTC_1_-_161418033_161418104@35.75.41,trna81_AspGTC_1_-_161410615_161410686@35.75.41\|TCACGCGGGAGACCGGGGTTCGATTCCCCGACGGGGAGCCA | >trna10_AspGTC_12_-_125424193_125424264 | 35 | 75 | 41 | TCACGCGGGAGACCGGGGTTCGATTCCCCGACGGGGAGCCA | MB231 | 83 | 100 | 1638 | 49 | 30 | 74.7 | 83.3 | 2119.6 | 57.3 | 33.3 |
| >trna10_AspGTC_12_-_125424193_125424264@36.73.38,trna12_AspGTC_12_-_125411891_125411962@36.73.38,trna144_AspGTC_6_-_27551236_27551307@36.73.38,trna38_AspGTC_17_-_8125556_8125627@36.73.38,trna45_AspGTC_6_+_27447453_27447524@36.73.38,trna48_AspGTC_6_+_27471523_27471594@36.73.38,trna4_AspGTC_12_+_96429799_96429870@36.73.38,trna69_AspGTC_1_-_161440205_161440276@36.73.38,trna72_AspGTC_1_-_161432824_161432895@36.73.38,trna75_AspGTC_1_-_161425414_161425485@36.73.38,trna78_AspGTC_1_-_161418033_161418104@36.73.38,trna81_AspGTC_1_-_161410615_161410686@36.73.38\|CACGCGGGAGACCGGGGTTCGATTCCCCGACGGGGAGC | >trna10_AspGTC_12_-_125424193_125424264 | 36 | 73 | 38 | CACGCGGGAGACCGGGGTTCGATTCCCCGACGGGGAGC | MB231 | 14 | 13 | 232 | 13 | 7 | 11.6 | 16.7 | 252.3 | 8.9 | 3.3 |
| >trna10_AspGTC_12_-_125424193_125424264@36.74.39,trna12_AspGTC_12_-_125411891_125411962@36.74.39,trna144_AspGTC_6_-_27551236_27551307@36.74.39,trna38_AspGTC_17_-_8125556_8125627@36.74.39,trna45_AspGTC_6_+_27447453_27447524@36.74.39,trna48_AspGTC_6_+_27471523_27471594@36.74.39,trna4_AspGTC_12_+_96429799_96429870@36.74.39,trna69_AspGTC_1_-_161440205_161440276@36.74.39,trna72_AspGTC_1_-_161432824_161432895@36.74.39,trna75_AspGTC_1_-_161425414_161425485@36.74.39,trna78_AspGTC_1_-_161418033_161418104@36.74.39,trna81_AspGTC_1_-_161410615_161410686@36.74.39\|CACGCGGGAGACCGGGGTTCGATTCCCCGACGGGGAGCC | >trna10_AspGTC_12_-_125424193_125424264 | 36 | 74 | 39 | CACGCGGGAGACCGGGGTTCGATTCCCCGACGGGGAGCC | MB231 | 21 | 17 | 465 | 14 | 9 | 17.3 | 20.6 | 591.7 | 9.1 | 3.9 |
| >trna10_AspGTC_12_-_125424193_125424264@36.75.40,trna12_AspGTC_12_-_125411891_125411962@36.75.40,trna144_AspGTC_6_-_27551236_27551307@36.75.40,trna38_AspGTC_17_-_8125556_8125627@36.75.40,trna45_AspGTC_6_+_27447453_27447524@36.75.40,trna48_AspGTC_6_+_27471523_27471594@36.75.40,trna4_AspGTC_12_+_96429799_96429870@36.75.40,trna69_AspGTC_1_-_161440205_161440276@36.75.40,trna72_AspGTC_1_-_161432824_161432895@36.75.40,trna75_AspGTC_1_-_161425414_161425485@36.75.40,trna78_AspGTC_1_-_161418033_161418104@36.75.40,trna81_AspGTC_1_-_161410615_161410686@36.75.40\|CACGCGGGAGACCGGGGTTCGATTCCCCGACGGGGAGCCA | >trna10_AspGTC_12_-_125424193_125424264 | 36 | 75 | 40 | CACGCGGGAGACCGGGGTTCGATTCCCCGACGGGGAGCCA | MB231 | 29 | 25 | 914 | 24 | 11 | 20.8 | 24.2 | 1290.9 | 19.4 | 6.9 |
| >trna10_AspGTC_12_-_125424193_125424264@37.72.36,trna12_AspGTC_12_-_125411891_125411962@37.72.36,trna144_AspGTC_6_-_27551236_27551307@37.72.36,trna38_AspGTC_17_-_8125556_8125627@37.72.36,trna45_AspGTC_6_+_27447453_27447524@37.72.36,trna48_AspGTC_6_+_27471523_27471594@37.72.36,trna4_AspGTC_12_+_96429799_96429870@37.72.36,trna69_AspGTC_1_-_161440205_161440276@37.72.36,trna72_AspGTC_1_-_161432824_161432895@37.72.36,trna75_AspGTC_1_-_161425414_161425485@37.72.36,trna78_AspGTC_1_-_161418033_161418104@37.72.36,trna81_AspGTC_1_-_161410615_161410686@37.72.36\|ACGCGGGAGACCGGGGTTCGATTCCCCGACGGGGAG | >trna10_AspGTC_12_-_125424193_125424264 | 37 | 72 | 36 | ACGCGGGAGACCGGGGTTCGATTCCCCGACGGGGAG | MB468 | 11 | 19 | 58 | 42 | 8 | 3.2 | 5.4 | 18.1 | 37.6 | 6.0 |
| >trna10_AspGTC_12_-_125424193_125424264@37.73.37,trna12_AspGTC_12_-_125411891_125411962@37.73.37,trna144_AspGTC_6_-_27551236_27551307@37.73.37,trna38_AspGTC_17_-_8125556_8125627@37.73.37,trna45_AspGTC_6_+_27447453_27447524@37.73.37,trna48_AspGTC_6_+_27471523_27471594@37.73.37,trna4_AspGTC_12_+_96429799_96429870@37.73.37,trna69_AspGTC_1_-_161440205_161440276@37.73.37,trna72_AspGTC_1_-_161432824_161432895@37.73.37,trna75_AspGTC_1_-_161425414_161425485@37.73.37,trna78_AspGTC_1_-_161418033_161418104@37.73.37,trna81_AspGTC_1_-_161410615_161410686@37.73.37\|ACGCGGGAGACCGGGGTTCGATTCCCCGACGGGGAGC | >trna10_AspGTC_12_-_125424193_125424264 | 37 | 73 | 37 | ACGCGGGAGACCGGGGTTCGATTCCCCGACGGGGAGC | MB231 | 48 | 67 | 1040 | 31 | 7 | 59.4 | 101.6 | 1369.3 | 30.5 | 4.3 |
| >trna10_AspGTC_12_-_125424193_125424264@37.74.38,trna12_AspGTC_12_-_125411891_125411962@37.74.38,trna144_AspGTC_6_-_27551236_27551307@37.74.38,trna38_AspGTC_17_-_8125556_8125627@37.74.38,trna45_AspGTC_6_+_27447453_27447524@37.74.38,trna48_AspGTC_6_+_27471523_27471594@37.74.38,trna4_AspGTC_12_+_96429799_96429870@37.74.38,trna69_AspGTC_1_-_161440205_161440276@37.74.38,trna72_AspGTC_1_-_161432824_161432895@37.74.38,trna75_AspGTC_1_-_161425414_161425485@37.74.38,trna78_AspGTC_1_-_161418033_161418104@37.74.38,trna81_AspGTC_1_-_161410615_161410686@37.74.38\|ACGCGGGAGACCGGGGTTCGATTCCCCGACGGGGAGCC | >trna10_AspGTC_12_-_125424193_125424264 | 37 | 74 | 38 | ACGCGGGAGACCGGGGTTCGATTCCCCGACGGGGAGCC | MB231 | 44 | 60 | 1071 | 26 | 9 | 55.9 | 89.3 | 1454.0 | 27.8 | 7.5 |
| >trna10_AspGTC_12_-_125424193_125424264@37.75.39,trna12_AspGTC_12_-_125411891_125411962@37.75.39,trna144_AspGTC_6_-_27551236_27551307@37.75.39,trna38_AspGTC_17_-_8125556_8125627@37.75.39,trna45_AspGTC_6_+_27447453_27447524@37.75.39,trna48_AspGTC_6_+_27471523_27471594@37.75.39,trna4_AspGTC_12_+_96429799_96429870@37.75.39,trna69_AspGTC_1_-_161440205_161440276@37.75.39,trna72_AspGTC_1_-_161432824_161432895@37.75.39,trna75_AspGTC_1_-_161425414_161425485@37.75.39,trna78_AspGTC_1_-_161418033_161418104@37.75.39,trna81_AspGTC_1_-_161410615_161410686@37.75.39\|ACGCGGGAGACCGGGGTTCGATTCCCCGACGGGGAGCCA | >trna10_AspGTC_12_-_125424193_125424264 | 37 | 75 | 39 | ACGCGGGAGACCGGGGTTCGATTCCCCGACGGGGAGCCA | MB231 | 30 | 54 | 926 | 17 | 7 | 35.2 | 76.6 | 1378.2 | 14.9 | 3.5 |

| | | | | | | | | | | | | | | | |
|---|---|---|---|---|---|---|---|---|---|---|---|---|---|---|---|
| >trna10_AspGTC_12_-_125424193_125424264@39.74.36,trna12_AspGTC_12_-_125411891_125411962@39.74.36,trna144_AspGTC_6_-_27551236_27551307@39.74.36,trna38_AspGTC_17_-_8125556_8125627@39.74.36,trna45_AspGTC_6_+_27447453_27447524@39.74.36,trna48_AspGTC_6_+_27471523_27471594@39.74.36,trna4_AspGTC_12_+_96429799_96429870@39.74.36,trna69_AspGTC_1_-_161440205_161440276@39.74.36,trna72_AspGTC_1_-_161432824_161432895@39.74.36,trna75_AspGTC_1_-_161425414_161425485@39.74.36,trna78_AspGTC_1_-_161418033_161418104@39.74.36,trna81_AspGTC_1_-_161410615_161410686@39.74.36\|GCGGGAGACCGGGGTTCGATTCCCCGACGGGGAGCC | >trna10_AspGTC_12_-_125424193_125424264 | 39 | 74 | 36 | GCGGGAGACCGGGGTTCGATTCCCCGACGGGGAGCC | MB231 | 11 | 3 | 112 | 4 | 2 | 6.6 | 3.1 | 107.3 | 2.5 | 1.6 |
| >trna10_AspGTC_12_-_125424193_125424264@39.75.37,trna12_AspGTC_12_-_125411891_125411962@39.75.37,trna144_AspGTC_6_-_27551236_27551307@39.75.37,trna38_AspGTC_17_-_8125556_8125627@39.75.37,trna45_AspGTC_6_+_27447453_27447524@39.75.37,trna48_AspGTC_6_+_27471523_27471594@39.75.37,trna4_AspGTC_12_+_96429799_96429870@39.75.37,trna69_AspGTC_1_-_161440205_161440276@39.75.37,trna72_AspGTC_1_-_161432824_161432895@39.75.37,trna75_AspGTC_1_-_161425414_161425485@39.75.37,trna78_AspGTC_1_-_161418033_161418104@39.75.37,trna81_AspGTC_1_-_161410615_161410686@39.75.37\|GCGGGAGACCGGGGTTCGATTCCCCGACGGGGAGCCA | >trna10_AspGTC_12_-_125424193_125424264 | 39 | 75 | 37 | GCGGGAGACCGGGGTTCGATTCCCCGACGGGGAGCCA | BT20 | 30 | 3 | 127 | 7 | 1 | 50.4 | 5.5 | 173.0 | 9.5 | 2.0 |
| >trna10_AspGTC_12_-_125424193_125424264@39.75.37,trna12_AspGTC_12_-_125411891_125411962@39.75.37,trna144_AspGTC_6_-_27551236_27551307@39.75.37,trna38_AspGTC_17_-_8125556_8125627@39.75.37,trna45_AspGTC_6_+_27447453_27447524@39.75.37,trna48_AspGTC_6_+_27471523_27471594@39.75.37,trna4_AspGTC_12_+_96429799_96429870@39.75.37,trna69_AspGTC_1_-_161440205_161440276@39.75.37,trna72_AspGTC_1_-_161432824_161432895@39.75.37,trna75_AspGTC_1_-_161425414_161425485@39.75.37,trna78_AspGTC_1_-_161418033_161418104@39.75.37,trna81_AspGTC_1_-_161410615_161410686@39.75.37\|GCGGGAGACCGGGGTTCGATTCCCCGACGGGGAGCCA | >trna10_AspGTC_12_-_125424193_125424264 | 39 | 75 | 37 | GCGGGAGACCGGGGTTCGATTCCCCGACGGGGAGCCA | MB231 | 19 | 6 | 191 | 7 | 2 | 13.2 | 4.9 | 238.2 | 4.8 | 1.7 |
| >trna10_GluCTC_3_-_103879524_103879595@16.42.27,trna116_GluCTC_1_-_145399233_145399304@16.42.27,trna59_GluCTC_1_+_249168447_249168518@16.42.27,trna71_GluCTC_1_-_161439189_161439260@16.42.27,trna74_GluCTC_1_-_161431809_161431880@16.42.27,trna77_GluCTC_1_-_161424398_161424469@16.42.27,trna77_GluCTC_6_+_28949976_28950047@16.42.27,trna80_GluCTC_1_-_161417018_161417089@16.42.27,trna87_GluCTC_6_-_126101393_126101464@16.42.27\|TGGTTAGGATTCGGCGCTCTCACCGCC | >trna10_GluCTC_3_-_103879524_103879595 | 16 | 42 | 27 | TGGTTAGGATTCGGCGCTCTCACCGCC | BT20 | 22 | 0 | 77 | 27 | 6 | 27.7 | 0.4 | 91.8 | 41.3 | 7.8 |
| >trna10_GluCTC_3_-_103879524_103879595@16.42.27,trna116_GluCTC_1_-_145399233_145399304@16.42.27,trna59_GluCTC_1_+_249168447_249168518@16.42.27,trna71_GluCTC_1_-_161439189_161439260@16.42.27,trna74_GluCTC_1_-_161431809_161431880@16.42.27,trna77_GluCTC_1_-_161424398_161424469@16.42.27,trna77_GluCTC_6_+_28949976_28950047@16.42.27,trna80_GluCTC_1_-_161417018_161417089@16.42.27,trna87_GluCTC_6_-_126101393_126101464@16.42.27\|TGGTTAGGATTCGGCGCTCTCACCGCC | >trna10_GluCTC_3_-_103879524_103879595 | 16 | 42 | 27 | TGGTTAGGATTCGGCGCTCTCACCGCC | MB231 | 11 | 1 | 27 | 14 | 5 | 3.8 | 0.8 | 17.5 | 4.8 | 1.8 |
| >trna10_GlyTCC_17_+_8124866_8124937@1.29.29\|GCGTTGGTGGTATAGTGGTAAGCATAGCT | >trna10_GlyTCC_17_+_8124866_8124937 | 1 | 29 | 29 | GCGTTGGTGGTATAGTGGTAAGCATAGCT | MB468 | 13 | 13 | 30 | 7 | 4 | 7.3 | 14.8 | 14.5 | 1.9 | 3.3 |
| >trna10_GlyTCC_17_+_8124866_8124937@1.30.30\|GCGTTGGTGGTATAGTGGTAAGCATAGCTG | >trna10_GlyTCC_17_+_8124866_8124937 | 1 | 30 | 30 | GCGTTGGTGGTATAGTGGTAAGCATAGCTG | MB468 | 27 | 19 | 82 | 24 | 15 | 15.1 | 13.3 | 25.0 | 9.6 | 15.0 |
| >trna10_LysCTT_16_+_3241501_3241573@1.26.26,trna119_LysCTT_1_-_145395522_145395594@1.26.26,trna11_LysCTT_5_-_180648979_180649051@1.26.26,trna13_LysCTT_14_-_58706613_58706685@1.26.26,trna13_LysCTT_6_+_26556774_26556846@1.26.26,trna2_LysCTT_15_+_79152904_79152976@1.26.26,trna32_LysCTT_16_-_3207406_3207478@1.26.26,trna7_LysCTT_16_+_3225692_3225764@1.26.26,trna9_LysCTT_5_+_180634755_180634827@1.26.26\|GCCCGGCTAGCTCAGTCGGTAGAGCA | >trna10_LysCTT_16_+_3241501_3241573 | 1 | 26 | 26 | GCCCGGCTAGCTCAGTCGGTAGAGCA | MB468 | 10 | 1 | 8 | 1 | 3 | 1.3 | 0.4 | 3.8 | 1.0 | 2.0 |
| >trna10_LysCTT_16_+_3241501_3241573@1.28.28,trna119_LysCTT_1_-_145395522_145395594@1.28.28,trna11_LysCTT_5_-_180648979_180649051@1.28.28,trna13_LysCTT_14_-_58706613_58706685@1.28.28,trna13_LysCTT_6_+_26556774_26556846@1.28.28,trna2_LysCTT_15_+_79152904_79152976@1.28.28,trna32_LysCTT_16_-_3207406_3207478@1.28.28,trna7_LysCTT_16_+_3225692_3225764@1.28.28,trna9_LysCTT_5_+_180634755_180634827@1.28.28\|GCCCGGCTAGCTCAGTCGGTAGAGCATG | >trna10_LysCTT_16_+_3241501_3241573 | 1 | 28 | 28 | GCCCGGCTAGCTCAGTCGGTAGAGCATG | MB468 | 403 | 136 | 447 | 68 | 155 | 142.2 | 83.8 | 227.7 | 32.3 | 30.9 |
| >trna10_LysCTT_16_+_3241501_3241573@1.29.29,trna13_LysCTT_14_-_58706613_58706685@1.29.29,trna2_LysCTT_15_+_79152904_79152976@1.29.29\|GCCCGGCTAGCTCAGTCGGTAGAGCATGG | >trna10_LysCTT_16_+_3241501_3241573 | 1 | 29 | 29 | GCCCGGCTAGCTCAGTCGGTAGAGCATGG | MB468 | 71 | 22 | 86 | 19 | 16 | 25.5 | 9.7 | 21.6 | 6.8 | 6.6 |
| >trna10_LysCTT_16_+_3241501_3241573@1.30.30,trna13_LysCTT_14_-_58706613_58706685@1.30.30,trna2_LysCTT_15_+_79152904_79152976@1.30.30\|GCCCGGCTAGCTCAGTCGGTAGAGCATGGG | >trna10_LysCTT_16_+_3241501_3241573 | 1 | 30 | 30 | GCCCGGCTAGCTCAGTCGGTAGAGCATGGG | MB468 | 21 | 4 | 48 | 5 | 5 | 10.7 | 1.0 | 26.9 | 4.8 | 3.5 |
| >trna10_LysCTT_16_+_3241501_3241573@1.33.33,trna13_LysCTT_14_-_58706613_58706685@1.33.33,trna2_LysCTT_15_+_79152904_79152976@1.33.33\|GCCCGGCTAGCTCAGTCGGTAGAGCATGGGACT | >trna10_LysCTT_16_+_3241501_3241573 | 1 | 33 | 33 | GCCCGGCTAGCTCAGTCGGTAGAGCATGGGACT | BT20 | 11 | 1 | 39 | 20 | 3 | 6.9 | 1.0 | 21.4 | 33.2 | 5.6 |
| >trna10_LysCTT_16_+_3241501_3241573@1.33.33,trna13_LysCTT_14_-_58706613_58706685@1.33.33,trna2_LysCTT_15_+_79152904_79152976@1.33.33\|GCCCGGCTAGCTCAGTCGGTAGAGCATGGGACT | >trna10_LysCTT_16_+_3241501_3241573 | 1 | 33 | 33 | GCCCGGCTAGCTCAGTCGGTAGAGCATGGGACT | MB231 | 17 | 7 | 38 | 33 | 15 | 16.5 | 9.5 | 50.6 | 23.7 | 18.2 |
| >trna10_LysCTT_16_+_3241501_3241573@1.34.34,trna13_LysCTT_14_-_58706613_58706685@1.34.34,trna2_LysCTT_15_+_79152904_79152976@1.34.34\|GCCCGGCTAGCTCAGTCGGTAGAGCATGGGACTC | >trna10_LysCTT_16_+_3241501_3241573 | 1 | 34 | 34 | GCCCGGCTAGCTCAGTCGGTAGAGCATGGGACTC | BT20 | 111 | 9 | 77 | 334 | 37 | 115.9 | 5.3 | 46.6 | 530.1 | 57.9 |
| >trna10_LysCTT_16_+_3241501_3241573@1.34.34,trna13_LysCTT_14_-_58706613_58706685@1.34.34,trna2_LysCTT_15_+_79152904_79152976@1.34.34\|GCCCGGCTAGCTCAGTCGGTAGAGCATGGGACTC | >trna10_LysCTT_16_+_3241501_3241573 | 1 | 34 | 34 | GCCCGGCTAGCTCAGTCGGTAGAGCATGGGACTC | MB231 | 99 | 31 | 144 | 952 | 180 | 101.9 | 41.7 | 207.9 | 823.9 | 150.9 |
| >trna10_LysCTT_16_+_3241501_3241573@1.35.35,trna13_LysCTT_14_-_58706613_58706685@1.35.35,trna2_LysCTT_15_+_79152904_79152976@1.35.35\|GCCCGGCTAGCTCAGTCGGTAGAGCATGGGACTCT | >trna10_LysCTT_16_+_3241501_3241573 | 1 | 35 | 35 | GCCCGGCTAGCTCAGTCGGTAGAGCATGGGACTCT | BT20 | 8737 | 189 | 11251 | 416 | 136 | 12017.1 | 121.5 | 10321.4 | 563.3 | 179.1 |
| >trna10_LysCTT_16_+_3241501_3241573@1.35.35,trna13_LysCTT_14_-_58706613_58706685@1.35.35,trna2_LysCTT_15_+_79152904_79152976@1.35.35\|GCCCGGCTAGCTCAGTCGGTAGAGCATGGGACTCT | >trna10_LysCTT_16_+_3241501_3241573 | 1 | 35 | 35 | GCCCGGCTAGCTCAGTCGGTAGAGCATGGGACTCT | MB231 | 323 | 60 | 698 | 206 | 359 | 191.5 | 52.0 | 671.5 | 83.9 | 337.8 |
| >trna10_LysCTT_16_+_3241501_3241573@1.36.36,trna13_LysCTT_14_-_58706613_58706685@1.36.36,trna2_LysCTT_15_+_79152904_79152976@1.36.36\|GCCCGGCTAGCTCAGTCGGTAGAGCATGGGACTCTT | >trna10_LysCTT_16_+_3241501_3241573 | 1 | 36 | 36 | GCCCGGCTAGCTCAGTCGGTAGAGCATGGGACTCTT | BT20 | 337 | 16 | 421 | 488 | 22 | 405.9 | 8.1 | 306.5 | 673.4 | 27.6 |
| >trna10_LysCTT_16_+_3241501_3241573@1.36.36,trna13_LysCTT_14_-_58706613_58706685@1.36.36,trna2_LysCTT_15_+_79152904_79152976@1.36.36\|GCCCGGCTAGCTCAGTCGGTAGAGCATGGGACTCTT | >trna10_LysCTT_16_+_3241501_3241573 | 1 | 36 | 36 | GCCCGGCTAGCTCAGTCGGTAGAGCATGGGACTCTT | MB231 | 55 | 25 | 31 | 302 | 260 | 25.9 | 17.9 | 19.9 | 171.6 | 264.5 |
| >trna10_LysCTT_16_+_3241501_3241573@14.31.18,trna13_LysCTT_14_-_58706613_58706685@14.31.18,trna2_LysCTT_15_+_79152904_79152976@14.31.18,trna4_LysCTT_18_-_43669271_43669343@14.31.18\|AGTCGGTAGAGCATGGGA | >trna10_LysCTT_16_+_3241501_3241573 | 14 | 31 | 18 | AGTCGGTAGAGCATGGGA | MB468 | 99 | 92 | 86 | 29 | 4 | 56.2 | 50.3 | 26.0 | 15.1 | 1.0 |
| >trna10_LysCTT_16_+_3241501_3241573@3.33.31,trna13_LysCTT_14_-_58706613_58706685@3.33.31,trna2_LysCTT_15_+_79152904_79152976@3.33.31\|CCGGCTAGCTCAGTCGGTAGAGCATGGGACT | >trna10_LysCTT_16_+_3241501_3241573 | 3 | 33 | 31 | CCGGCTAGCTCAGTCGGTAGAGCATGGGACT | MB231 | 27 | 38 | 24 | 38 | 110 | 14.0 | 32.3 | 22.0 | 18.5 | 38.9 |
| >trna10_LysCTT_16_+_3241501_3241573@3.35.33,trna13_LysCTT_14_-_58706613_58706685@3.35.33,trna2_LysCTT_15_+_79152904_79152976@3.35.33\|CCGGCTAGCTCAGTCGGTAGAGCATGGGACTCT | >trna10_LysCTT_16_+_3241501_3241573 | 3 | 35 | 33 | CCGGCTAGCTCAGTCGGTAGAGCATGGGACTCT | MB231 | 20 | 16 | 21 | 49 | 127 | 6.9 | 7.7 | 14.4 | 20.5 | 68.7 |

| | | | | | | | | | | | | | | | | |
|---|---|---|---|---|---|---|---|---|---|---|---|---|---|---|---|---|
| >trna10_LysCTT_16_+_3241501_3241573@4.33.30,trna13_LysCTT_14_-_58706613_58706685@4.33.30,trna2_LysCTT_15_+_79152904_79152976@4.33.30\|CGGCTAGCTCAGTCGGTAGAGCATGGGACT | >trna10_LysCTT_16_+_3241501_3241573 | 4 | 33 | 30 | CGGCTAGCTCAGTCGGTAGAGCATGGGACT | MB231 | 28 | 53 | 10 | 53 | 125 | 20.6 | 48.5 | 9.9 | 31.2 | 72.8 |
| >trna10_LysCTT_16_+_3241501_3241573@4.35.32,trna13_LysCTT_14_-_58706613_58706685@4.35.32,trna2_LysCTT_15_+_79152904_79152976@4.35.32\|CGGCTAGCTCAGTCGGTAGAGCATGGGACTCT | >trna10_LysCTT_16_+_3241501_3241573 | 4 | 35 | 32 | CGGCTAGCTCAGTCGGTAGAGCATGGGACTCT | MB231 | 20 | 18 | 3 | 27 | 109 | 9.9 | 12.1 | 3.4 | 15.1 | 57.7 |
| >trna10_MetCAT_8_-_124169470_124169542@31.51.21,trna22_MetCAT_16_-_87417628_87417700@31.51.21\|TCTCATAATCTGAAGGTCGTG | >trna10_MetCAT_8_-_124169470_124169542 | 31 | 51 | 21 | TCTCATAATCTGAAGGTCGTG | MB468 | 117 | 30 | 91 | 53 | 22 | 33.2 | 17.2 | 33.6 | 29.1 | 8.7 |
| >trna10_MetCAT_8_-_124169470_124169542@32.51.20,trna22_MetCAT_16_-_87417628_87417700@32.51.20\|CTCATAATCTGAAGGTCGTG | >trna10_MetCAT_8_-_124169470_124169542 | 32 | 51 | 20 | CTCATAATCTGAAGGTCGTG | MB468 | 12 | 3 | 11 | 5 | 2 | 6.1 | 1.5 | 4.7 | 3.1 | 0.5 |
| >trna10_MetCAT_8_-_124169470_124169542@34.51.18,trna22_MetCAT_16_-_87417628_87417700@34.51.18\|CATAATCTGAAGGTCGTG | >trna10_MetCAT_8_-_124169470_124169542 | 34 | 51 | 18 | CATAATCTGAAGGTCGTG | MB468 | 17 | 3 | 16 | 4 | 2 | 5.8 | 2.4 | 5.9 | 3.8 | 0.7 |
| >trna10_ValCAC_5_-_180649395_180649467@35.54.20,trna12_ValAAC_5_-_180645270_180645342@35.54.20,trna132_ValAAC_6_-_27721179_27721251@35.54.20,trna13_ValCAC_19_-_4724647_4724719@35.54.20,trna152_ValCAC_6_-_27248049_27248121@35.54.20,trna157_ValCAC_6_-_27173867_27173939@35.54.20,trna15_ValAAC_5_-_180615416_180615488@35.54.20,trna18_ValCAC_5_-_180529253_180529325@35.54.20,trna2_ValAAC_3_+_169490018_169490090@35.54.20,trna2_ValCAC_5_+_180524070_180524142@35.54.20,trna37_ValAAC_6_+_27203288_27203360@35.54.20,trna4_ValAAC_5_+_180591154_180591226@35.54.20,trna5_ValAAC_5_+_180596610_180596682@35.54.20,trna6_ValCAC_5_+_180600650_180600722@35.54.20,trna85_ValCAC_1_-_161369490_161369562@35.54.20,trna98_ValCAC_1_-_149298555_149298627@35.54.20,trna9_ValCAC_6_+_26538282_26538354@35.54.20\|ACACGCGAAAGGTCCCCGGT | >trna10_ValCAC_5_-_180649395_180649467 | 35 | 54 | 20 | ACACGCGAAAGGTCCCCGGT | MB468 | 11 | 4 | 16 | 45 | 8 | 5.4 | 1.1 | 8.8 | 45.2 | 4.6 |
| >trna101_AlaAGC_6_-_28831462_28831533@58.75.18,trna102_AlaAGC_6_-_28806221_28806292@58.75.18,trna110_AlaTGC_6_-_28757547_28757618@58.75.18,trna113_AlaTGC_6_-_28726141_28726212@58.75.18,trna65_AlaAGC_6_+_28574933_28575004@58.75.18\|TCCCCGGCACCTCCACCA | >trna101_AlaAGC_6_-_28831462_28831533 | 58 | 75 | 18 | TCCCCGGCACCTCCACCA | BT20 | 98 | 45 | 714 | 1 | 30 | 71.5 | 8.0 | 206.2 | 1.4 | 33.1 |
| >trna101_AlaAGC_6_-_28831462_28831533@58.75.18,trna102_AlaAGC_6_-_28806221_28806292@58.75.18,trna110_AlaTGC_6_-_28757547_28757618@58.75.18,trna113_AlaTGC_6_-_28726141_28726212@58.75.18,trna65_AlaAGC_6_+_28574933_28575004@58.75.18\|TCCCCGGCACCTCCACCA | >trna101_AlaAGC_6_-_28831462_28831533 | 58 | 75 | 18 | TCCCCGGCACCTCCACCA | MB231 | 77 | 57 | 489 | 2 | 70 | 41.5 | 31.0 | 218.0 | 1.2 | 46.4 |
| >trna103_AsnGTT_1_-_149230570_149230643@33.53.21,trna107_AsnGTT_1_-_148760356_148760429@33.53.21,trna108_AsnGTT_1_-_148598314_148598387@33.53.21,trna1_AsnGTT_19_+_1383562_1383635@33.53.21,trna26_AsnGTT_1_+_148248115_148248188@33.53.21,trna31_AsnGTT_17_-_36908034_36908107@33.53.21,trna47_AsnGTT_1_+_161510031_161510104@33.53.21,trna4_AsnGTT_10_-_22518438_22518511@33.53.21,trna7_AsnGTT_13_-_31248101_31248174@33.53.21,trna83_AsnGTT_1_-_161397867_161397940@33.53.21\|CTGTTAACCGAAAGGTTGGTG | >trna103_AsnGTT_1_-_149230570_149230643 | 33 | 53 | 21 | CTGTTAACCGAAAGGTTGGTG | MB468 | 11 | 0 | 8 | 7 | 1 | 10.4 | 0.1 | 12.8 | 6.5 | 0.5 |
| >trna106_HisGTG_1_-_149155828_149155899@13.31.19,trna111_HisGTG_1_-_147774845_147774916@13.31.19,trna118_HisGTG_1_-_145396881_145396952@13.31.19,trna16_HisGTG_1_+_146544773_146544844@13.31.19,trna1_HisGTG_15_+_45493349_45493420@13.31.19,trna21_HisGTG_1_+_147753471_147753542@13.31.19,trna33_HisGTG_6_+_27125906_27125977@13.31.19,trna7_HisGTG_9_-_14433938_14434009@13.31.19,trna8_HisGTG_15_-_45492611_45492682@13.31.19,trna9_HisGTG_15_-_45490804_45490875@13.31.19\|TAGTGGTTAGTACTCTGCG | >trna106_HisGTG_1_-_149155828_149155899 | 13 | 31 | 19 | TAGTGGTTAGTACTCTGCG | MB468 | 14 | 3 | 15 | 7 | 9 | 1.2 | 2.3 | 7.9 | 3.0 | 10.8 |
| >trna108_AlaAGC_6_-_28763741_28763812@58.75.18\|TCCCCAGTACCTCCACCA | >trna108_AlaAGC_6_-_28763741_28763812 | 58 | 75 | 18 | TCCCCAGTACCTCCACCA | BT20 | 24 | 20 | 145 | 0 | 0 | 13.8 | 11.2 | 60.3 | 0.0 | 0.0 |
| >trna108_AlaAGC_6_-_28763741_28763812@58.75.18\|TCCCCAGTACCTCCACCA | >trna108_AlaAGC_6_-_28763741_28763812 | 58 | 75 | 18 | TCCCCAGTACCTCCACCA | MB231 | 14 | 24 | 87 | 0 | 8 | 7.3 | 16.2 | 54.6 | 0.5 | 13.5 |
| >trna11_GluTTC_15_-_26327381_26327452@1.27.27,trna3_GluTTC_13_-_45492062_45492133@1.27.27\|TCCCACATGGTCTAGCGGTTAGGATTC | >trna11_GluTTC_15_-_26327381_26327452 | 1 | 27 | 27 | TCCCACATGGTCTAGCGGTTAGGATTC | MB468 | 26 | 7 | 5 | 1 | 1 | 17.0 | 3.9 | 1.6 | 0.5 | 0.4 |
| >trna11_GluTTC_15_-_26327381_26327452@1.28.28,trna3_GluTTC_13_-_45492062_45492133@1.28.28\|TCCCACATGGTCTAGCGGTTAGGATTCC | >trna11_GluTTC_15_-_26327381_26327452 | 1 | 28 | 28 | TCCCACATGGTCTAGCGGTTAGGATTCC | MB468 | 30 | 10 | 23 | 2 | 3 | 22.7 | 6.6 | 12.8 | 1.3 | 1.5 |
| >trna11_GluTTC_15_-_26327381_26327452@1.29.29,trna3_GluTTC_13_-_45492062_45492133@1.29.29\|TCCCACATGGTCTAGCGGTTAGGATTCCT | >trna11_GluTTC_15_-_26327381_26327452 | 1 | 29 | 29 | TCCCACATGGTCTAGCGGTTAGGATTCCT | BT20 | 18 | 2 | 129 | 24 | 8 | 19.2 | 1.0 | 107.9 | 34.2 | 12.9 |
| >trna11_GluTTC_15_-_26327381_26327452@1.29.29,trna3_GluTTC_13_-_45492062_45492133@1.29.29\|TCCCACATGGTCTAGCGGTTAGGATTCCT | >trna11_GluTTC_15_-_26327381_26327452 | 1 | 29 | 29 | TCCCACATGGTCTAGCGGTTAGGATTCCT | MB231 | 12 | 3 | 34 | 2 | 9 | 13.2 | 1.8 | 40.8 | 1.0 | 1.7 |
| >trna11_GluTTC_15_-_26327381_26327452@1.29.29,trna3_GluTTC_13_-_45492062_45492133@1.29.29\|TCCCACATGGTCTAGCGGTTAGGATTCCT | >trna11_GluTTC_15_-_26327381_26327452 | 1 | 29 | 29 | TCCCACATGGTCTAGCGGTTAGGATTCCT | MB468 | 273 | 116 | 371 | 65 | 122 | 66.7 | 92.3 | 200.4 | 55.9 | 36.1 |
| >trna11_GluTTC_15_-_26327381_26327452@1.30.30,trna3_GluTTC_13_-_45492062_45492133@1.30.30\|TCCCACATGGTCTAGCGGTTAGGATTCCTG | >trna11_GluTTC_15_-_26327381_26327452 | 1 | 30 | 30 | TCCCACATGGTCTAGCGGTTAGGATTCCTG | MB468 | 180 | 67 | 190 | 98 | 59 | 40.2 | 25.8 | 90.7 | 51.5 | 29.0 |
| >trna11_GluTTC_15_-_26327381_26327452@1.31.31,trna3_GluTTC_13_-_45492062_45492133@1.31.31\|TCCCACATGGTCTAGCGGTTAGGATTCCTGG | >trna11_GluTTC_15_-_26327381_26327452 | 1 | 31 | 31 | TCCCACATGGTCTAGCGGTTAGGATTCCTGG | MB231 | 25 | 11 | 30 | 24 | 33 | 13.4 | 12.1 | 31.9 | 10.4 | 17.3 |
| >trna11_GluTTC_15_-_26327381_26327452@1.31.31,trna3_GluTTC_13_-_45492062_45492133@1.31.31\|TCCCACATGGTCTAGCGGTTAGGATTCCTGG | >trna11_GluTTC_15_-_26327381_26327452 | 1 | 31 | 31 | TCCCACATGGTCTAGCGGTTAGGATTCCTGG | MB468 | 128 | 39 | 190 | 35 | 25 | 30.2 | 10.1 | 92.1 | 13.3 | 14.6 |
| >trna11_GluTTC_15_-_26327381_26327452@1.32.32,trna3_GluTTC_13_-_45492062_45492133@1.32.32\|TCCCACATGGTCTAGCGGTTAGGATTCCTGGT | >trna11_GluTTC_15_-_26327381_26327452 | 1 | 32 | 32 | TCCCACATGGTCTAGCGGTTAGGATTCCTGGT | BT20 | 15 | 2 | 18 | 13 | 19 | 15.7 | 0.6 | 8.8 | 18.9 | 30.9 |
| >trna11_GluTTC_15_-_26327381_26327452@1.32.32,trna3_GluTTC_13_-_45492062_45492133@1.32.32\|TCCCACATGGTCTAGCGGTTAGGATTCCTGGT | >trna11_GluTTC_15_-_26327381_26327452 | 1 | 32 | 32 | TCCCACATGGTCTAGCGGTTAGGATTCCTGGT | MB231 | 25 | 9 | 36 | 27 | 42 | 11.2 | 9.6 | 32.2 | 6.1 | 26.4 |
| >trna11_GluTTC_15_-_26327381_26327452@1.33.33,trna3_GluTTC_13_-_45492062_45492133@1.33.33\|TCCCACATGGTCTAGCGGTTAGGATTCCTGGTT | >trna11_GluTTC_15_-_26327381_26327452 | 1 | 33 | 33 | TCCCACATGGTCTAGCGGTTAGGATTCCTGGTT | BT20 | 238 | 39 | 346 | 75 | 138 | 247.5 | 7.9 | 117.1 | 101.4 | 205.3 |
| >trna11_GluTTC_15_-_26327381_26327452@1.33.33,trna3_GluTTC_13_-_45492062_45492133@1.33.33\|TCCCACATGGTCTAGCGGTTAGGATTCCTGGTT | >trna11_GluTTC_15_-_26327381_26327452 | 1 | 33 | 33 | TCCCACATGGTCTAGCGGTTAGGATTCCTGGTT | MB231 | 267 | 108 | 333 | 144 | 304 | 85.7 | 98.6 | 180.5 | 21.2 | 145.6 |
| >trna11_GluTTC_15_-_26327381_26327452@1.34.34,trna3_GluTTC_13_-_45492062_45492133@1.34.34\|TCCCACATGGTCTAGCGGTTAGGATTCCTGGTTT | >trna11_GluTTC_15_-_26327381_26327452 | 1 | 34 | 34 | TCCCACATGGTCTAGCGGTTAGGATTCCTGGTTT | BT20 | 20 | 5 | 25 | 5 | 11 | 11.5 | 1.5 | 14.4 | 5.2 | 12.3 |
| >trna11_GluTTC_15_-_26327381_26327452@1.34.34,trna3_GluTTC_13_-_45492062_45492133@1.34.34\|TCCCACATGGTCTAGCGGTTAGGATTCCTGGTTT | >trna11_GluTTC_15_-_26327381_26327452 | 1 | 34 | 34 | TCCCACATGGTCTAGCGGTTAGGATTCCTGGTTT | MB231 | 12 | 6 | 18 | 6 | 14 | 6.2 | 6.4 | 17.2 | 3.2 | 9.3 |
| >trna11_GluTTC_15_-_26327381_26327452@1.35.35,trna3_GluTTC_13_-_45492062_45492133@1.35.35\|TCCCACATGGTCTAGCGGTTAGGATTCCTGGTTTT | >trna11_GluTTC_15_-_26327381_26327452 | 1 | 35 | 35 | TCCCACATGGTCTAGCGGTTAGGATTCCTGGTTTT | BT20 | 11 | 3 | 16 | 6 | 11 | 2.8 | 0.6 | 4.7 | 7.9 | 14.4 |
| >trna11_GluTTC_15_-_26327381_26327452@1.36.36,trna3_GluTTC_13_-_45492062_45492133@1.36.36\|TCCCACATGGTCTAGCGGTTAGGATTCCTGGTTTTC | >trna11_GluTTC_15_-_26327381_26327452 | 1 | 36 | 36 | TCCCACATGGTCTAGCGGTTAGGATTCCTGGTTTTC | MB231 | 20 | 10 | 74 | 4 | 34 | 14.3 | 9.8 | 79.0 | 3.1 | 14.0 |
| >trna111_HisGTG_1_-_147774845_147774916@-1A.23.24,trna118_HisGTG_1_-_145396881_145396952@-1A.23.24,trna16_HisGTG_1_+_146544773_146544844@-1A.23.24,trna1_HisGTG_15_+_45493349_45493420@-1A.23.24,trna21_HisGTG_1_+_147753471_147753542@-1A.23.24,trna33_HisGTG_6_+_27125906_27125977@-1A.23.24,trna7_HisGTG_9_-_14433938_14434009@-1A.23.24,trna8_HisGTG_15_-_45492611_45492682@-1A.23.24,trna9_HisGTG_15_-_45490804_45490875@-1A.23.24\|AGCCGTGATCGTATAGTGGTTAGT | >trna111_HisGTG_1_-_147774845_147774916 | -1A | 23 | 24 | AGCCGTGATCGTATAGTGGTTAGT | MB231 | 30 | 37 | 8 | 13 | 40 | 9.6 | 20.4 | 6.4 | 7.6 | 18.2 |
| >trna111_HisGTG_1_-_147774845_147774916@-1A.26.27,trna118_HisGTG_1_-_145396881_145396952@-1A.26.27,trna16_HisGTG_1_+_146544773_146544844@-1A.26.27,trna1_HisGTG_15_+_45493349_45493420@-1A.26.27,trna21_HisGTG_1_+_147753471_147753542@-1A.26.27,trna33_HisGTG_6_+_27125906_27125977@-1A.26.27,trna7_HisGTG_9_-_14433938_14434009@-1A.26.27,trna8_HisGTG_15_-_45492611_45492682@-1A.26.27,trna9_HisGTG_15_-_45490804_45490875@-1A.26.27\|AGCCGTGATCGTATAGTGGTTAGTACT | >trna111_HisGTG_1_-_147774845_147774916 | -1A | 26 | 27 | AGCCGTGATCGTATAGTGGTTAGTACT | MB231 | 28 | 31 | 24 | 6 | 28 | 18.4 | 17.5 | 21.1 | 3.9 | 11.1 |

| | | | | | | | | | | | | | | | |
|---|---|---|---|---|---|---|---|---|---|---|---|---|---|---|---|
| >trna111_HisGTG_1_-_147774845_147774916@-1A.29.30,trna118_HisGTG_1_-_145396881_145396952@-1A.29.30,trna16_HisGTG_1_+_146544773_146544844@-1A.29.30,trna1_HisGTG_15_+_45493349_45493420@-1A.29.30,trna21_HisGTG_1_+_147753471_147753542@-1A.29.30,trna33_HisGTG_6_+_27125906_27125977@-1A.29.30,trna7_HisGTG_9_-_14433938_14434009@-1A.29.30,trna8_HisGTG_15_-_45492611_45492682@-1A.29.30,trna9_HisGTG_15_-_45490804_45490875@-1A.29.30\|AGCCGTGATCGTATAGTGGTTAGTACTCTG | >trna111_HisGTG_1_-_147774845_147774916 | -1A | 29 | 30 | AGCCGTGATCGTATAGTGGTTAGTACTCTG | MB468 | 41 | 62 | 98 | 18 | 14 | 22.0 | 32.2 | 85.9 | 8.7 | 9.1 |
| >trna111_HisGTG_1_-_147774845_147774916@-1A.31.32,trna118_HisGTG_1_-_145396881_145396952@-1A.31.32,trna16_HisGTG_1_+_146544773_146544844@-1A.31.32,trna1_HisGTG_15_+_45493349_45493420@-1A.31.32,trna21_HisGTG_1_+_147753471_147753542@-1A.31.32,trna33_HisGTG_6_+_27125906_27125977@-1A.31.32,trna7_HisGTG_9_-_14433938_14434009@-1A.31.32,trna8_HisGTG_15_-_45492611_45492682@-1A.31.32,trna9_HisGTG_15_-_45490804_45490875@-1A.31.32\|AGCCGTGATCGTATAGTGGTTAGTACTCTGCG | >trna111_HisGTG_1_-_147774845_147774916 | -1A | 31 | 32 | AGCCGTGATCGTATAGTGGTTAGTACTCTGCG | MB468 | 98 | 129 | 284 | 61 | 24 | 45.7 | 56.4 | 306.5 | 5.6 | 19.4 |
| >trna111_HisGTG_1_-_147774845_147774916@-1A.33.34,trna118_HisGTG_1_-_145396881_145396952@-1A.33.34,trna16_HisGTG_1_+_146544773_146544844@-1A.33.34,trna1_HisGTG_15_+_45493349_45493420@-1A.33.34,trna21_HisGTG_1_+_147753471_147753542@-1A.33.34,trna33_HisGTG_6_+_27125906_27125977@-1A.33.34,trna7_HisGTG_9_-_14433938_14434009@-1A.33.34,trna8_HisGTG_15_-_45492611_45492682@-1A.33.34,trna9_HisGTG_15_-_45490804_45490875@-1A.33.34\|AGCCGTGATCGTATAGTGGTTAGTACTCTGCGTT | >trna111_HisGTG_1_-_147774845_147774916 | -1A | 33 | 34 | AGCCGTGATCGTATAGTGGTTAGTACTCTGCGTT | MB231 | 16 | 18 | 34 | 15 | 12 | 11.8 | 22.9 | 26.2 | 9.6 | 8.5 |
| >trna111_HisGTG_1_-_147774845_147774916@-1A.34.35,trna118_HisGTG_1_-_145396881_145396952@-1A.34.35,trna16_HisGTG_1_+_146544773_146544844@-1A.34.35,trna1_HisGTG_15_+_45493349_45493420@-1A.34.35,trna21_HisGTG_1_+_147753471_147753542@-1A.34.35,trna33_HisGTG_6_+_27125906_27125977@-1A.34.35,trna7_HisGTG_9_-_14433938_14434009@-1A.34.35,trna8_HisGTG_15_-_45492611_45492682@-1A.34.35,trna9_HisGTG_15_-_45490804_45490875@-1A.34.35\|AGCCGTGATCGTATAGTGGTTAGTACTCTGCGTTG | >trna111_HisGTG_1_-_147774845_147774916 | -1A | 34 | 35 | AGCCGTGATCGTATAGTGGTTAGTACTCTGCGTTG | BT20 | 11 | 1 | 10 | 4 | 4 | 16.9 | 0.7 | 13.0 | 7.1 | 6.9 |
| >trna111_HisGTG_1_-_147774845_147774916@-1A.34.35,trna118_HisGTG_1_-_145396881_145396952@-1A.34.35,trna16_HisGTG_1_+_146544773_146544844@-1A.34.35,trna1_HisGTG_15_+_45493349_45493420@-1A.34.35,trna21_HisGTG_1_+_147753471_147753542@-1A.34.35,trna33_HisGTG_6_+_27125906_27125977@-1A.34.35,trna7_HisGTG_9_-_14433938_14434009@-1A.34.35,trna8_HisGTG_15_-_45492611_45492682@-1A.34.35,trna9_HisGTG_15_-_45490804_45490875@-1A.34.35\|AGCCGTGATCGTATAGTGGTTAGTACTCTGCGTTG | >trna111_HisGTG_1_-_147774845_147774916 | -1A | 34 | 35 | AGCCGTGATCGTATAGTGGTTAGTACTCTGCGTTG | MB231 | 16 | 11 | 48 | 6 | 9 | 12.7 | 11.1 | 35.8 | 2.8 | 7.0 |
| >trna111_HisGTG_1_-_147774845_147774916@-1G.30.31,trna118_HisGTG_1_-_145396881_145396952@-1G.30.31,trna16_HisGTG_1_+_146544773_146544844@-1G.30.31,trna1_HisGTG_15_+_45493349_45493420@-1G.30.31,trna21_HisGTG_1_+_147753471_147753542@-1G.30.31,trna33_HisGTG_6_+_27125906_27125977@-1G.30.31,trna7_HisGTG_9_-_14433938_14434009@-1G.30.31,trna8_HisGTG_15_-_45492611_45492682@-1G.30.31,trna9_HisGTG_15_-_45490804_45490875@-1G.30.31\|GGCCGTGATCGTATAGTGGTTAGTACTCTGC | >trna111_HisGTG_1_-_147774845_147774916 | -1G | 30 | 31 | GGCCGTGATCGTATAGTGGTTAGTACTCTGC | BT20 | 14 | 5 | 42 | 22 | 6 | 21.5 | 5.2 | 51.9 | 23.0 | 6.3 |
| >trna111_HisGTG_1_-_147774845_147774916@-1G.30.31,trna118_HisGTG_1_-_145396881_145396952@-1G.30.31,trna16_HisGTG_1_+_146544773_146544844@-1G.30.31,trna1_HisGTG_15_+_45493349_45493420@-1G.30.31,trna21_HisGTG_1_+_147753471_147753542@-1G.30.31,trna33_HisGTG_6_+_27125906_27125977@-1G.30.31,trna7_HisGTG_9_-_14433938_14434009@-1G.30.31,trna8_HisGTG_15_-_45492611_45492682@-1G.30.31,trna9_HisGTG_15_-_45490804_45490875@-1G.30.31\|GGCCGTGATCGTATAGTGGTTAGTACTCTGC | >trna111_HisGTG_1_-_147774845_147774916 | -1G | 30 | 31 | GGCCGTGATCGTATAGTGGTTAGTACTCTGC | MB231 | 89 | 85 | 364 | 233 | 95 | 72.3 | 54.4 | 238.1 | 161.1 | 29.2 |
| >trna111_HisGTG_1_-_147774845_147774916@-1G.30.31,trna118_HisGTG_1_-_145396881_145396952@-1G.30.31,trna16_HisGTG_1_+_146544773_146544844@-1G.30.31,trna1_HisGTG_15_+_45493349_45493420@-1G.30.31,trna21_HisGTG_1_+_147753471_147753542@-1G.30.31,trna33_HisGTG_6_+_27125906_27125977@-1G.30.31,trna7_HisGTG_9_-_14433938_14434009@-1G.30.31,trna8_HisGTG_15_-_45492611_45492682@-1G.30.31,trna9_HisGTG_15_-_45490804_45490875@-1G.30.31\|GGCCGTGATCGTATAGTGGTTAGTACTCTGC | >trna111_HisGTG_1_-_147774845_147774916 | -1G | 30 | 31 | GGCCGTGATCGTATAGTGGTTAGTACTCTGC | MB468 | 69 | 136 | 160 | 35 | 17 | 37.8 | 55.4 | 109.2 | 9.9 | 9.0 |
| >trna111_HisGTG_1_-_147774845_147774916@-1G.31.32,trna118_HisGTG_1_-_145396881_145396952@-1G.31.32,trna16_HisGTG_1_+_146544773_146544844@-1G.31.32,trna1_HisGTG_15_+_45493349_45493420@-1G.31.32,trna21_HisGTG_1_+_147753471_147753542@-1G.31.32,trna33_HisGTG_6_+_27125906_27125977@-1G.31.32,trna7_HisGTG_9_-_14433938_14434009@-1G.31.32,trna8_HisGTG_15_-_45492611_45492682@-1G.31.32,trna9_HisGTG_15_-_45490804_45490875@-1G.31.32\|GGCCGTGATCGTATAGTGGTTAGTACTCTGCG | >trna111_HisGTG_1_-_147774845_147774916 | -1G | 31 | 32 | GGCCGTGATCGTATAGTGGTTAGTACTCTGCG | BT20 | 45 | 9 | 91 | 66 | 14 | 69.6 | 9.4 | 121.2 | 76.6 | 19.6 |
| >trna111_HisGTG_1_-_147774845_147774916@-1G.31.32,trna118_HisGTG_1_-_145396881_145396952@-1G.31.32,trna16_HisGTG_1_+_146544773_146544844@-1G.31.32,trna1_HisGTG_15_+_45493349_45493420@-1G.31.32,trna21_HisGTG_1_+_147753471_147753542@-1G.31.32,trna33_HisGTG_6_+_27125906_27125977@-1G.31.32,trna7_HisGTG_9_-_14433938_14434009@-1G.31.32,trna8_HisGTG_15_-_45492611_45492682@-1G.31.32,trna9_HisGTG_15_-_45490804_45490875@-1G.31.32\|GGCCGTGATCGTATAGTGGTTAGTACTCTGCG | >trna111_HisGTG_1_-_147774845_147774916 | -1G | 31 | 32 | GGCCGTGATCGTATAGTGGTTAGTACTCTGCG | MB231 | 293 | 237 | 1334 | 1090 | 397 | 252.0 | 195.7 | 1509.6 | 887.8 | 310.6 |
| >trna111_HisGTG_1_-_147774845_147774916@-1G.31.32,trna118_HisGTG_1_-_145396881_145396952@-1G.31.32,trna16_HisGTG_1_+_146544773_146544844@-1G.31.32,trna1_HisGTG_15_+_45493349_45493420@-1G.31.32,trna21_HisGTG_1_+_147753471_147753542@-1G.31.32,trna33_HisGTG_6_+_27125906_27125977@-1G.31.32,trna7_HisGTG_9_-_14433938_14434009@-1G.31.32,trna8_HisGTG_15_-_45492611_45492682@-1G.31.32,trna9_HisGTG_15_-_45490804_45490875@-1G.31.32\|GGCCGTGATCGTATAGTGGTTAGTACTCTGCG | >trna111_HisGTG_1_-_147774845_147774916 | -1G | 31 | 32 | GGCCGTGATCGTATAGTGGTTAGTACTCTGCG | MB468 | 1032 | 1875 | 2308 | 875 | 286 | 377.4 | 537.1 | 2063.2 | 135.9 | 115.9 |
| >trna111_HisGTG_1_-_147774845_147774916@-1G.32.33,trna118_HisGTG_1_-_145396881_145396952@-1G.32.33,trna16_HisGTG_1_+_146544773_146544844@-1G.32.33,trna1_HisGTG_15_+_45493349_45493420@-1G.32.33,trna21_HisGTG_1_+_147753471_147753542@-1G.32.33,trna33_HisGTG_6_+_27125906_27125977@-1G.32.33,trna7_HisGTG_9_-_14433938_14434009@-1G.32.33,trna8_HisGTG_15_-_45492611_45492682@-1G.32.33,trna9_HisGTG_15_-_45490804_45490875@-1G.32.33\|GGCCGTGATCGTATAGTGGTTAGTACTCTGCGT | >trna111_HisGTG_1_-_147774845_147774916 | -1G | 32 | 33 | GGCCGTGATCGTATAGTGGTTAGTACTCTGCGT | BT20 | 26 | 3 | 21 | 34 | 20 | 40.7 | 2.5 | 27.4 | 52.0 | 29.2 |

| | | | | | | | | | | | | | | | |
|---|---|---|---|---|---|---|---|---|---|---|---|---|---|---|---|
| >trna111_HisGTG_1_-_147774845_147774916@-1G.32.33,trna118_HisGTG_1_-_145396881_145396952@-1G.32.33,trna16_HisGTG_1_+_146544773_146544844@-1G.32.33,trna1_HisGTG_15_+_45493349_45493420@-1G.32.33,trna21_HisGTG_1_+_147753471_147753542@-1G.32.33,trna33_HisGTG_6_+_27125906_27125977@-1G.32.33,trna7_HisGTG_9_-_14433938_14434009@-1G.32.33,trna8_HisGTG_15_-_45492611_45492682@-1G.32.33,trna9_HisGTG_15_-_45490804_45490875@-1G.32.33\|GGCCGTGATCGTATAGTGGTTAGTACTCTGCGT | >trna111_HisGTG_1_-_147774845_147774916 | -1G | 32 | 33 | GGCCGTGATCGTATAGTGGTTAGTACTCTGCGT | MB231 | 120 | 138 | 252 | 328 | 159 | 94.0 | 157.8 | 151.4 | 254.3 | 104.8 |
| >trna111_HisGTG_1_-_147774845_147774916@-1G.33.34,trna118_HisGTG_1_-_145396881_145396952@-1G.33.34,trna16_HisGTG_1_+_146544773_146544844@-1G.33.34,trna1_HisGTG_15_+_45493349_45493420@-1G.33.34,trna21_HisGTG_1_+_147753471_147753542@-1G.33.34,trna33_HisGTG_6_+_27125906_27125977@-1G.33.34,trna7_HisGTG_9_-_14433938_14434009@-1G.33.34,trna8_HisGTG_15_-_45492611_45492682@-1G.33.34,trna9_HisGTG_15_-_45490804_45490875@-1G.33.34\|GGCCGTGATCGTATAGTGGTTAGTACTCTGCGTT | >trna111_HisGTG_1_-_147774845_147774916 | -1G | 33 | 34 | GGCCGTGATCGTATAGTGGTTAGTACTCTGCGTT | BT20 | 207 | 67 | 293 | 712 | 232 | 300.6 | 62.7 | 331.6 | 1178.3 | 386.3 |
| >trna111_HisGTG_1_-_147774845_147774916@-1G.33.34,trna118_HisGTG_1_-_145396881_145396952@-1G.33.34,trna16_HisGTG_1_+_146544773_146544844@-1G.33.34,trna1_HisGTG_15_+_45493349_45493420@-1G.33.34,trna21_HisGTG_1_+_147753471_147753542@-1G.33.34,trna33_HisGTG_6_+_27125906_27125977@-1G.33.34,trna7_HisGTG_9_-_14433938_14434009@-1G.33.34,trna8_HisGTG_15_-_45492611_45492682@-1G.33.34,trna9_HisGTG_15_-_45490804_45490875@-1G.33.34\|GGCCGTGATCGTATAGTGGTTAGTACTCTGCGTT | >trna111_HisGTG_1_-_147774845_147774916 | -1G | 33 | 34 | GGCCGTGATCGTATAGTGGTTAGTACTCTGCGTT | MB231 | 1338 | 1843 | 3216 | 1811 | 1175 | 929.7 | 2364.2 | 2105.3 | 1246.4 | 760.7 |
| >trna111_HisGTG_1_-_147774845_147774916@-1G.34.35,trna118_HisGTG_1_-_145396881_145396952@-1G.34.35,trna16_HisGTG_1_+_146544773_146544844@-1G.34.35,trna1_HisGTG_15_+_45493349_45493420@-1G.34.35,trna21_HisGTG_1_+_147753471_147753542@-1G.34.35,trna33_HisGTG_6_+_27125906_27125977@-1G.34.35,trna7_HisGTG_9_-_14433938_14434009@-1G.34.35,trna8_HisGTG_15_-_45492611_45492682@-1G.34.35,trna9_HisGTG_15_-_45490804_45490875@-1G.34.35\|GGCCGTGATCGTATAGTGGTTAGTACTCTGCGTTG | >trna111_HisGTG_1_-_147774845_147774916 | -1G | 34 | 35 | GGCCGTGATCGTATAGTGGTTAGTACTCTGCGTTG | BT20 | 150 | 13 | 108 | 80 | 110 | 242.5 | 10.1 | 153.0 | 131.9 | 179.1 |
| >trna111_HisGTG_1_-_147774845_147774916@-1G.34.35,trna118_HisGTG_1_-_145396881_145396952@-1G.34.35,trna16_HisGTG_1_+_146544773_146544844@-1G.34.35,trna1_HisGTG_15_+_45493349_45493420@-1G.34.35,trna21_HisGTG_1_+_147753471_147753542@-1G.34.35,trna33_HisGTG_6_+_27125906_27125977@-1G.34.35,trna7_HisGTG_9_-_14433938_14434009@-1G.34.35,trna8_HisGTG_15_-_45492611_45492682@-1G.34.35,trna9_HisGTG_15_-_45490804_45490875@-1G.34.35\|GGCCGTGATCGTATAGTGGTTAGTACTCTGCGTTG | >trna111_HisGTG_1_-_147774845_147774916 | -1G | 34 | 35 | GGCCGTGATCGTATAGTGGTTAGTACTCTGCGTTG | MB231 | 898 | 508 | 2619 | 452 | 609 | 941.1 | 445.3 | 2860.0 | 268.5 | 524.0 |
| >trna111_HisGTG_1_-_147774845_147774916@-1G.34.35,trna118_HisGTG_1_-_145396881_145396952@-1G.34.35,trna16_HisGTG_1_+_146544773_146544844@-1G.34.35,trna1_HisGTG_15_+_45493349_45493420@-1G.34.35,trna21_HisGTG_1_+_147753471_147753542@-1G.34.35,trna33_HisGTG_6_+_27125906_27125977@-1G.34.35,trna7_HisGTG_9_-_14433938_14434009@-1G.34.35,trna8_HisGTG_15_-_45492611_45492682@-1G.34.35,trna9_HisGTG_15_-_45490804_45490875@-1G.34.35\|GGCCGTGATCGTATAGTGGTTAGTACTCTGCGTTG | >trna111_HisGTG_1_-_147774845_147774916 | -1G | 34 | 35 | GGCCGTGATCGTATAGTGGTTAGTACTCTGCGTTG | MB468 | 14 | 19 | 1 | 4 | 6 | 10.8 | 22.5 | 1.0 | 5.0 | 4.3 |
| >trna111_HisGTG_1_-_147774845_147774916@-1G.35.36,trna118_HisGTG_1_-_145396881_145396952@-1G.35.36,trna16_HisGTG_1_+_146544773_146544844@-1G.35.36,trna1_HisGTG_15_+_45493349_45493420@-1G.35.36,trna21_HisGTG_1_+_147753471_147753542@-1G.35.36,trna33_HisGTG_6_+_27125906_27125977@-1G.35.36,trna7_HisGTG_9_-_14433938_14434009@-1G.35.36,trna8_HisGTG_15_-_45492611_45492682@-1G.35.36,trna9_HisGTG_15_-_45490804_45490875@-1G.35.36\|GGCCGTGATCGTATAGTGGTTAGTACTCTGCGTTGT | >trna111_HisGTG_1_-_147774845_147774916 | -1G | 35 | 36 | GGCCGTGATCGTATAGTGGTTAGTACTCTGCGTTGT | BT20 | 15 | 15 | 30 | 9 | 9 | 2.3 | 10.6 | 37.6 | 12.3 | 8.7 |
| >trna111_HisGTG_1_-_147774845_147774916@-1G.35.36,trna118_HisGTG_1_-_145396881_145396952@-1G.35.36,trna16_HisGTG_1_+_146544773_146544844@-1G.35.36,trna1_HisGTG_15_+_45493349_45493420@-1G.35.36,trna21_HisGTG_1_+_147753471_147753542@-1G.35.36,trna33_HisGTG_6_+_27125906_27125977@-1G.35.36,trna7_HisGTG_9_-_14433938_14434009@-1G.35.36,trna8_HisGTG_15_-_45492611_45492682@-1G.35.36,trna9_HisGTG_15_-_45490804_45490875@-1G.35.36\|GGCCGTGATCGTATAGTGGTTAGTACTCTGCGTTGT | >trna111_HisGTG_1_-_147774845_147774916 | -1G | 35 | 36 | GGCCGTGATCGTATAGTGGTTAGTACTCTGCGTTGT | MB231 | 76 | 94 | 201 | 54 | 74 | 62.4 | 120.2 | 170.7 | 49.5 | 43.8 |
| >trna111_HisGTG_1_-_147774845_147774916@-1T.22.23,trna118_HisGTG_1_-_145396881_145396952@-1T.22.23,trna16_HisGTG_1_+_146544773_146544844@-1T.22.23,trna1_HisGTG_15_+_45493349_45493420@-1T.22.23,trna21_HisGTG_1_+_147753471_147753542@-1T.22.23,trna33_HisGTG_6_+_27125906_27125977@-1T.22.23,trna7_HisGTG_9_-_14433938_14434009@-1T.22.23,trna8_HisGTG_15_-_45492611_45492682@-1T.22.23,trna9_HisGTG_15_-_45490804_45490875@-1T.22.23\|TGCCGTGATCGTATAGTGGTTAG | >trna111_HisGTG_1_-_147774845_147774916 | -1T | 22 | 23 | TGCCGTGATCGTATAGTGGTTAG | MB231 | 13 | 22 | 5 | 31 | 16 | 2.7 | 13.2 | 3.5 | 19.1 | 8.8 |
| >trna111_HisGTG_1_-_147774845_147774916@-1T.23.24,trna118_HisGTG_1_-_145396881_145396952@-1T.23.24,trna16_HisGTG_1_+_146544773_146544844@-1T.23.24,trna1_HisGTG_15_+_45493349_45493420@-1T.23.24,trna21_HisGTG_1_+_147753471_147753542@-1T.23.24,trna33_HisGTG_6_+_27125906_27125977@-1T.23.24,trna7_HisGTG_9_-_14433938_14434009@-1T.23.24,trna8_HisGTG_15_-_45492611_45492682@-1T.23.24,trna9_HisGTG_15_-_45490804_45490875@-1T.23.24\|TGCCGTGATCGTATAGTGGTTAGT | >trna111_HisGTG_1_-_147774845_147774916 | -1T | 23 | 24 | TGCCGTGATCGTATAGTGGTTAGT | BT20 | 11 | 10 | 9 | 2 | 11 | 3.7 | 3.5 | 4.1 | 1.1 | 6.4 |
| >trna111_HisGTG_1_-_147774845_147774916@-1T.23.24,trna118_HisGTG_1_-_145396881_145396952@-1T.23.24,trna16_HisGTG_1_+_146544773_146544844@-1T.23.24,trna1_HisGTG_15_+_45493349_45493420@-1T.23.24,trna21_HisGTG_1_+_147753471_147753542@-1T.23.24,trna33_HisGTG_6_+_27125906_27125977@-1T.23.24,trna7_HisGTG_9_-_14433938_14434009@-1T.23.24,trna8_HisGTG_15_-_45492611_45492682@-1T.23.24,trna9_HisGTG_15_-_45490804_45490875@-1T.23.24\|TGCCGTGATCGTATAGTGGTTAGT | >trna111_HisGTG_1_-_147774845_147774916 | -1T | 23 | 24 | TGCCGTGATCGTATAGTGGTTAGT | MB231 | 257 | 329 | 75 | 98 | 308 | 95.5 | 174.5 | 51.4 | 65.3 | 121.5 |
| >trna111_HisGTG_1_-_147774845_147774916@-1T.24.25,trna118_HisGTG_1_-_145396881_145396952@-1T.24.25,trna16_HisGTG_1_+_146544773_146544844@-1T.24.25,trna1_HisGTG_15_+_45493349_45493420@-1T.24.25,trna21_HisGTG_1_+_147753471_147753542@-1T.24.25,trna33_HisGTG_6_+_27125906_27125977@-1T.24.25,trna7_HisGTG_9_-_14433938_14434009@-1T.24.25,trna8_HisGTG_15_-_45492611_45492682@-1T.24.25,trna9_HisGTG_15_-_45490804_45490875@-1T.24.25\|TGCCGTGATCGTATAGTGGTTAGTA | >trna111_HisGTG_1_-_147774845_147774916 | -1T | 24 | 25 | TGCCGTGATCGTATAGTGGTTAGTA | MB231 | 13 | 17 | 4 | 5 | 13 | 7.9 | 6.4 | 1.4 | 3.6 | 2.9 |

| | | | | | | | | | | | | | | | |
|---|---|---|---|---|---|---|---|---|---|---|---|---|---|---|---|
| >trna111_HisGTG_1_-_147774845_147774916@-1T.25.26,trna118_HisGTG_1_-_145396881_145396952@-1T.25.26,trna16_HisGTG_1_+_146544773_146544844@-1T.25.26,trna1_HisGTG_15_+_45493349_45493420@-1T.25.26,trna21_HisGTG_1_+_147753471_147753542@-1T.25.26,trna33_HisGTG_6_+_27125906_27125977@-1T.25.26,trna7_HisGTG_9_-_14433938_14434009@-1T.25.26,trna8_HisGTG_15_-_45492611_45492682@-1T.25.26,trna9_HisGTG_15_-_45490804_45490875@-1T.25.26\|TGCCGTGATCGTATAGTGGTTAGTAC | >trna111_HisGTG_1_-_147774845_147774916 | -1T | 25 | 26 | TGCCGTGATCGTATAGTGGTTAGTAC | MB231 | 23 | 29 | 8 | 6 | 26 | 15.6 | 13.8 | 2.9 | 4.4 | 2.2 |
| >trna111_HisGTG_1_-_147774845_147774916@-1T.26.27,trna118_HisGTG_1_-_145396881_145396952@-1T.26.27,trna16_HisGTG_1_+_146544773_146544844@-1T.26.27,trna1_HisGTG_15_+_45493349_45493420@-1T.26.27,trna21_HisGTG_1_+_147753471_147753542@-1T.26.27,trna33_HisGTG_6_+_27125906_27125977@-1T.26.27,trna7_HisGTG_9_-_14433938_14434009@-1T.26.27,trna8_HisGTG_15_-_45492611_45492682@-1T.26.27,trna9_HisGTG_15_-_45490804_45490875@-1T.26.27\|TGCCGTGATCGTATAGTGGTTAGTACT | >trna111_HisGTG_1_-_147774845_147774916 | -1T | 26 | 27 | TGCCGTGATCGTATAGTGGTTAGTACT | BT20 | 37 | 25 | 45 | 20 | 27 | 29.5 | 25.8 | 42.3 | 16.9 | 30.4 |
| >trna111_HisGTG_1_-_147774845_147774916@-1T.26.27,trna118_HisGTG_1_-_145396881_145396952@-1T.26.27,trna16_HisGTG_1_+_146544773_146544844@-1T.26.27,trna1_HisGTG_15_+_45493349_45493420@-1T.26.27,trna21_HisGTG_1_+_147753471_147753542@-1T.26.27,trna33_HisGTG_6_+_27125906_27125977@-1T.26.27,trna7_HisGTG_9_-_14433938_14434009@-1T.26.27,trna8_HisGTG_15_-_45492611_45492682@-1T.26.27,trna9_HisGTG_15_-_45490804_45490875@-1T.26.27\|TGCCGTGATCGTATAGTGGTTAGTACT | >trna111_HisGTG_1_-_147774845_147774916 | -1T | 26 | 27 | TGCCGTGATCGTATAGTGGTTAGTACT | MB231 | 230 | 236 | 142 | 36 | 217 | 153.8 | 139.8 | 103.1 | 24.2 | 79.1 |
| >trna111_HisGTG_1_-_147774845_147774916@-1T.28.29,trna118_HisGTG_1_-_145396881_145396952@-1T.28.29,trna16_HisGTG_1_+_146544773_146544844@-1T.28.29,trna1_HisGTG_15_+_45493349_45493420@-1T.28.29,trna21_HisGTG_1_+_147753471_147753542@-1T.28.29,trna33_HisGTG_6_+_27125906_27125977@-1T.28.29,trna7_HisGTG_9_-_14433938_14434009@-1T.28.29,trna8_HisGTG_15_-_45492611_45492682@-1T.28.29,trna9_HisGTG_15_-_45490804_45490875@-1T.28.29\|TGCCGTGATCGTATAGTGGTTAGTACTCT | >trna111_HisGTG_1_-_147774845_147774916 | -1T | 28 | 29 | TGCCGTGATCGTATAGTGGTTAGTACTCT | MB231 | 31 | 38 | 30 | 8 | 36 | 18.6 | 18.9 | 15.1 | 4.2 | 6.8 |
| >trna111_HisGTG_1_-_147774845_147774916@-1T.28.29,trna118_HisGTG_1_-_145396881_145396952@-1T.28.29,trna16_HisGTG_1_+_146544773_146544844@-1T.28.29,trna1_HisGTG_15_+_45493349_45493420@-1T.28.29,trna21_HisGTG_1_+_147753471_147753542@-1T.28.29,trna33_HisGTG_6_+_27125906_27125977@-1T.28.29,trna7_HisGTG_9_-_14433938_14434009@-1T.28.29,trna8_HisGTG_15_-_45492611_45492682@-1T.28.29,trna9_HisGTG_15_-_45490804_45490875@-1T.28.29\|TGCCGTGATCGTATAGTGGTTAGTACTCT | >trna111_HisGTG_1_-_147774845_147774916 | -1T | 28 | 29 | TGCCGTGATCGTATAGTGGTTAGTACTCT | MB468 | 56 | 76 | 99 | 7 | 6 | 11.3 | 49.8 | 32.5 | 1.5 | 3.0 |
| >trna111_HisGTG_1_-_147774845_147774916@-1T.29.30,trna118_HisGTG_1_-_145396881_145396952@-1T.29.30,trna16_HisGTG_1_+_146544773_146544844@-1T.29.30,trna1_HisGTG_15_+_45493349_45493420@-1T.29.30,trna21_HisGTG_1_+_147753471_147753542@-1T.29.30,trna33_HisGTG_6_+_27125906_27125977@-1T.29.30,trna7_HisGTG_9_-_14433938_14434009@-1T.29.30,trna8_HisGTG_15_-_45492611_45492682@-1T.29.30,trna9_HisGTG_15_-_45490804_45490875@-1T.29.30\|TGCCGTGATCGTATAGTGGTTAGTACTCTG | >trna111_HisGTG_1_-_147774845_147774916 | -1T | 29 | 30 | TGCCGTGATCGTATAGTGGTTAGTACTCTG | MB468 | 328 | 457 | 930 | 166 | 118 | 155.2 | 232.1 | 656.2 | 90.0 | 69.9 |
| >trna111_HisGTG_1_-_147774845_147774916@-1T.30.31,trna118_HisGTG_1_-_145396881_145396952@-1T.30.31,trna16_HisGTG_1_+_146544773_146544844@-1T.30.31,trna1_HisGTG_15_+_45493349_45493420@-1T.30.31,trna21_HisGTG_1_+_147753471_147753542@-1T.30.31,trna33_HisGTG_6_+_27125906_27125977@-1T.30.31,trna7_HisGTG_9_-_14433938_14434009@-1T.30.31,trna8_HisGTG_15_-_45492611_45492682@-1T.30.31,trna9_HisGTG_15_-_45490804_45490875@-1T.30.31\|TGCCGTGATCGTATAGTGGTTAGTACTCTGC | >trna111_HisGTG_1_-_147774845_147774916 | -1T | 30 | 31 | TGCCGTGATCGTATAGTGGTTAGTACTCTGC | MB231 | 12 | 11 | 33 | 10 | 12 | 7.4 | 6.3 | 15.3 | 4.4 | 1.2 |
| >trna111_HisGTG_1_-_147774845_147774916@-1T.30.31,trna118_HisGTG_1_-_145396881_145396952@-1T.30.31,trna16_HisGTG_1_+_146544773_146544844@-1T.30.31,trna1_HisGTG_15_+_45493349_45493420@-1T.30.31,trna21_HisGTG_1_+_147753471_147753542@-1T.30.31,trna33_HisGTG_6_+_27125906_27125977@-1T.30.31,trna7_HisGTG_9_-_14433938_14434009@-1T.30.31,trna8_HisGTG_15_-_45492611_45492682@-1T.30.31,trna9_HisGTG_15_-_45490804_45490875@-1T.30.31\|TGCCGTGATCGTATAGTGGTTAGTACTCTGC | >trna111_HisGTG_1_-_147774845_147774916 | -1T | 30 | 31 | TGCCGTGATCGTATAGTGGTTAGTACTCTGC | MB468 | 37 | 45 | 110 | 15 | 7 | 25.6 | 15.3 | 71.4 | 2.4 | 5.7 |
| >trna111_HisGTG_1_-_147774845_147774916@-1T.31.32,trna118_HisGTG_1_-_145396881_145396952@-1T.31.32,trna16_HisGTG_1_+_146544773_146544844@-1T.31.32,trna1_HisGTG_15_+_45493349_45493420@-1T.31.32,trna21_HisGTG_1_+_147753471_147753542@-1T.31.32,trna33_HisGTG_6_+_27125906_27125977@-1T.31.32,trna7_HisGTG_9_-_14433938_14434009@-1T.31.32,trna8_HisGTG_15_-_45492611_45492682@-1T.31.32,trna9_HisGTG_15_-_45490804_45490875@-1T.31.32\|TGCCGTGATCGTATAGTGGTTAGTACTCTGCG | >trna111_HisGTG_1_-_147774845_147774916 | -1T | 31 | 32 | TGCCGTGATCGTATAGTGGTTAGTACTCTGCG | BT20 | 13 | 3 | 55 | 10 | 3 | 15.3 | 1.7 | 59.0 | 10.1 | 2.9 |
| >trna111_HisGTG_1_-_147774845_147774916@-1T.31.32,trna118_HisGTG_1_-_145396881_145396952@-1T.31.32,trna16_HisGTG_1_+_146544773_146544844@-1T.31.32,trna1_HisGTG_15_+_45493349_45493420@-1T.31.32,trna21_HisGTG_1_+_147753471_147753542@-1T.31.32,trna33_HisGTG_6_+_27125906_27125977@-1T.31.32,trna7_HisGTG_9_-_14433938_14434009@-1T.31.32,trna8_HisGTG_15_-_45492611_45492682@-1T.31.32,trna9_HisGTG_15_-_45490804_45490875@-1T.31.32\|TGCCGTGATCGTATAGTGGTTAGTACTCTGCG | >trna111_HisGTG_1_-_147774845_147774916 | -1T | 31 | 32 | TGCCGTGATCGTATAGTGGTTAGTACTCTGCG | MB231 | 32 | 21 | 126 | 47 | 31 | 20.8 | 14.0 | 95.3 | 38.8 | 17.4 |
| >trna111_HisGTG_1_-_147774845_147774916@-1T.31.32,trna118_HisGTG_1_-_145396881_145396952@-1T.31.32,trna16_HisGTG_1_+_146544773_146544844@-1T.31.32,trna1_HisGTG_15_+_45493349_45493420@-1T.31.32,trna21_HisGTG_1_+_147753471_147753542@-1T.31.32,trna33_HisGTG_6_+_27125906_27125977@-1T.31.32,trna7_HisGTG_9_-_14433938_14434009@-1T.31.32,trna8_HisGTG_15_-_45492611_45492682@-1T.31.32,trna9_HisGTG_15_-_45490804_45490875@-1T.31.32\|TGCCGTGATCGTATAGTGGTTAGTACTCTGCG | >trna111_HisGTG_1_-_147774845_147774916 | -1T | 31 | 32 | TGCCGTGATCGTATAGTGGTTAGTACTCTGCG | MB468 | 605 | 680 | 2005 | 448 | 175 | 366.2 | 264.7 | 1914.2 | 68.9 | 138.6 |
| >trna111_HisGTG_1_-_147774845_147774916@-1T.32.33,trna118_HisGTG_1_-_145396881_145396952@-1T.32.33,trna16_HisGTG_1_+_146544773_146544844@-1T.32.33,trna1_HisGTG_15_+_45493349_45493420@-1T.32.33,trna21_HisGTG_1_+_147753471_147753542@-1T.32.33,trna33_HisGTG_6_+_27125906_27125977@-1T.32.33,trna7_HisGTG_9_-_14433938_14434009@-1T.32.33,trna8_HisGTG_15_-_45492611_45492682@-1T.32.33,trna9_HisGTG_15_-_45490804_45490875@-1T.32.33\|TGCCGTGATCGTATAGTGGTTAGTACTCTGCGT | >trna111_HisGTG_1_-_147774845_147774916 | -1T | 32 | 33 | TGCCGTGATCGTATAGTGGTTAGTACTCTGCGT | MB231 | 14 | 17 | 29 | 20 | 18 | 9.5 | 17.4 | 20.5 | 12.4 | 8.5 |

| | | | | | | | | | | | | | | | | |
|---|---|---|---|---|---|---|---|---|---|---|---|---|---|---|---|---|
| >trna111_HisGTG_1_-_147774845_147774916@-1T.33.34,trna118_HisGTG_1_-_145396881_145396952@-1T.33.34,trna16_HisGTG_1_+_146544773_146544844@-1T.33.34,trna1_HisGTG_15_+_45493349_45493420@-1T.33.34,trna21_HisGTG_1_+_147753471_147753542@-1T.33.34,trna33_HisGTG_6_+_27125906_27125977@-1T.33.34,trna7_HisGTG_9_-_14433938_14434009@-1T.33.34,trna8_HisGTG_15_-_45492611_45492682@-1T.33.34,trna9_HisGTG_15_-_45490804_45490875@-1T.33.34\|TGCCGTGATCGTATAGTGGTTAGTACTCTGCGTT | >trna111_HisGTG_1_-_147774845_147774916 | -1T | 33 | 34 | TGCCGTGATCGTATAGTGGTTAGTACTCTGCGTT | BT20 | 43 | 18 | 102 | 122 | 36 | 42.5 | 12.0 | 59.5 | 200.8 | 56.0 |
| >trna111_HisGTG_1_-_147774845_147774916@-1T.33.34,trna118_HisGTG_1_-_145396881_145396952@-1T.33.34,trna16_HisGTG_1_+_146544773_146544844@-1T.33.34,trna1_HisGTG_15_+_45493349_45493420@-1T.33.34,trna21_HisGTG_1_+_147753471_147753542@-1T.33.34,trna33_HisGTG_6_+_27125906_27125977@-1T.33.34,trna7_HisGTG_9_-_14433938_14434009@-1T.33.34,trna8_HisGTG_15_-_45492611_45492682@-1T.33.34,trna9_HisGTG_15_-_45490804_45490875@-1T.33.34\|TGCCGTGATCGTATAGTGGTTAGTACTCTGCGTT | >trna111_HisGTG_1_-_147774845_147774916 | -1T | 33 | 34 | TGCCGTGATCGTATAGTGGTTAGTACTCTGCGTT | MB231 | 88 | 104 | 184 | 73 | 62 | 82.1 | 136.4 | 163.4 | 46.2 | 43.1 |
| >trna111_HisGTG_1_-_147774845_147774916@-1T.34.35,trna118_HisGTG_1_-_145396881_145396952@-1T.34.35,trna16_HisGTG_1_+_146544773_146544844@-1T.34.35,trna1_HisGTG_15_+_45493349_45493420@-1T.34.35,trna21_HisGTG_1_+_147753471_147753542@-1T.34.35,trna33_HisGTG_6_+_27125906_27125977@-1T.34.35,trna7_HisGTG_9_-_14433938_14434009@-1T.34.35,trna8_HisGTG_15_-_45492611_45492682@-1T.34.35,trna9_HisGTG_15_-_45490804_45490875@-1T.34.35\|TGCCGTGATCGTATAGTGGTTAGTACTCTGCGTTG | >trna111_HisGTG_1_-_147774845_147774916 | -1T | 34 | 35 | TGCCGTGATCGTATAGTGGTTAGTACTCTGCGTTG | BT20 | 41 | 5 | 51 | 15 | 18 | 58.7 | 2.4 | 53.0 | 24.5 | 28.6 |
| >trna111_HisGTG_1_-_147774845_147774916@-1T.34.35,trna118_HisGTG_1_-_145396881_145396952@-1T.34.35,trna16_HisGTG_1_+_146544773_146544844@-1T.34.35,trna1_HisGTG_15_+_45493349_45493420@-1T.34.35,trna21_HisGTG_1_+_147753471_147753542@-1T.34.35,trna33_HisGTG_6_+_27125906_27125977@-1T.34.35,trna7_HisGTG_9_-_14433938_14434009@-1T.34.35,trna8_HisGTG_15_-_45492611_45492682@-1T.34.35,trna9_HisGTG_15_-_45490804_45490875@-1T.34.35\|TGCCGTGATCGTATAGTGGTTAGTACTCTGCGTTG | >trna111_HisGTG_1_-_147774845_147774916 | -1T | 34 | 35 | TGCCGTGATCGTATAGTGGTTAGTACTCTGCGTTG | MB231 | 80 | 49 | 235 | 31 | 54 | 60.8 | 45.3 | 168.1 | 18.3 | 38.9 |
| >trna111_HisGTG_1_-_147774845_147774916@-1T.35.36,trna118_HisGTG_1_-_145396881_145396952@-1T.35.36,trna16_HisGTG_1_+_146544773_146544844@-1T.35.36,trna1_HisGTG_15_+_45493349_45493420@-1T.35.36,trna21_HisGTG_1_+_147753471_147753542@-1T.35.36,trna33_HisGTG_6_+_27125906_27125977@-1T.35.36,trna7_HisGTG_9_-_14433938_14434009@-1T.35.36,trna8_HisGTG_15_-_45492611_45492682@-1T.35.36,trna9_HisGTG_15_-_45490804_45490875@-1T.35.36\|TGCCGTGATCGTATAGTGGTTAGTACTCTGCGTTGT | >trna111_HisGTG_1_-_147774845_147774916 | -1T | 35 | 36 | TGCCGTGATCGTATAGTGGTTAGTACTCTGCGTTGT | MB231 | 12 | 13 | 27 | 5 | 13 | 9.0 | 13.1 | 27.9 | 3.5 | 6.9 |
| >trna111_HisGTG_1_-_147774845_147774916@1.30.30,trna118_HisGTG_1_-_145396881_145396952@1.30.30,trna16_HisGTG_1_+_146544773_146544844@1.30.30,trna1_HisGTG_15_+_45493349_45493420@1.30.30,trna21_HisGTG_1_+_147753471_147753542@1.30.30,trna33_HisGTG_6_+_27125906_27125977@1.30.30,trna7_HisGTG_9_-_14433938_14434009@1.30.30,trna8_HisGTG_15_-_45492611_45492682@1.30.30,trna9_HisGTG_15_-_45490804_45490875@1.30.30\|GCCGTGATCGTATAGTGGTTAGTACTCTGC | >trna111_HisGTG_1_-_147774845_147774916 | 1 | 30 | 30 | GCCGTGATCGTATAGTGGTTAGTACTCTGC | BT20 | 10 | 7 | 49 | 9 | 10 | 6.8 | 7.2 | 24.8 | 4.6 | 5.1 |
| >trna111_HisGTG_1_-_147774845_147774916@1.30.30,trna118_HisGTG_1_-_145396881_145396952@1.30.30,trna16_HisGTG_1_+_146544773_146544844@1.30.30,trna1_HisGTG_15_+_45493349_45493420@1.30.30,trna21_HisGTG_1_+_147753471_147753542@1.30.30,trna33_HisGTG_6_+_27125906_27125977@1.30.30,trna7_HisGTG_9_-_14433938_14434009@1.30.30,trna8_HisGTG_15_-_45492611_45492682@1.30.30,trna9_HisGTG_15_-_45490804_45490875@1.30.30\|GCCGTGATCGTATAGTGGTTAGTACTCTGC | >trna111_HisGTG_1_-_147774845_147774916 | 1 | 30 | 30 | GCCGTGATCGTATAGTGGTTAGTACTCTGC | MB231 | 36 | 27 | 141 | 93 | 51 | 29.5 | 28.6 | 116.1 | 64.6 | 28.6 |
| >trna111_HisGTG_1_-_147774845_147774916@1.30.30,trna118_HisGTG_1_-_145396881_145396952@1.30.30,trna16_HisGTG_1_+_146544773_146544844@1.30.30,trna1_HisGTG_15_+_45493349_45493420@1.30.30,trna21_HisGTG_1_+_147753471_147753542@1.30.30,trna33_HisGTG_6_+_27125906_27125977@1.30.30,trna7_HisGTG_9_-_14433938_14434009@1.30.30,trna8_HisGTG_15_-_45492611_45492682@1.30.30,trna9_HisGTG_15_-_45490804_45490875@1.30.30\|GCCGTGATCGTATAGTGGTTAGTACTCTGC | >trna111_HisGTG_1_-_147774845_147774916 | 1 | 30 | 30 | GCCGTGATCGTATAGTGGTTAGTACTCTGC | MB468 | 56 | 62 | 183 | 17 | 8 | 37.8 | 20.1 | 125.7 | 1.2 | 6.3 |
| >trna111_HisGTG_1_-_147774845_147774916@1.31.31,trna118_HisGTG_1_-_145396881_145396952@1.31.31,trna16_HisGTG_1_+_146544773_146544844@1.31.31,trna1_HisGTG_15_+_45493349_45493420@1.31.31,trna21_HisGTG_1_+_147753471_147753542@1.31.31,trna33_HisGTG_6_+_27125906_27125977@1.31.31,trna7_HisGTG_9_-_14433938_14434009@1.31.31,trna8_HisGTG_15_-_45492611_45492682@1.31.31,trna9_HisGTG_15_-_45490804_45490875@1.31.31\|GCCGTGATCGTATAGTGGTTAGTACTCTGCG | >trna111_HisGTG_1_-_147774845_147774916 | 1 | 31 | 31 | GCCGTGATCGTATAGTGGTTAGTACTCTGCG | BT20 | 33 | 20 | 116 | 23 | 13 | 18.6 | 21.4 | 41.1 | 19.1 | 3.6 |
| >trna111_HisGTG_1_-_147774845_147774916@1.31.31,trna118_HisGTG_1_-_145396881_145396952@1.31.31,trna16_HisGTG_1_+_146544773_146544844@1.31.31,trna1_HisGTG_15_+_45493349_45493420@1.31.31,trna21_HisGTG_1_+_147753471_147753542@1.31.31,trna33_HisGTG_6_+_27125906_27125977@1.31.31,trna7_HisGTG_9_-_14433938_14434009@1.31.31,trna8_HisGTG_15_-_45492611_45492682@1.31.31,trna9_HisGTG_15_-_45490804_45490875@1.31.31\|GCCGTGATCGTATAGTGGTTAGTACTCTGCG | >trna111_HisGTG_1_-_147774845_147774916 | 1 | 31 | 31 | GCCGTGATCGTATAGTGGTTAGTACTCTGCG | MB231 | 80 | 57 | 410 | 198 | 102 | 59.5 | 54.6 | 301.6 | 119.1 | 64.1 |
| >trna111_HisGTG_1_-_147774845_147774916@1.31.31,trna118_HisGTG_1_-_145396881_145396952@1.31.31,trna16_HisGTG_1_+_146544773_146544844@1.31.31,trna1_HisGTG_15_+_45493349_45493420@1.31.31,trna21_HisGTG_1_+_147753471_147753542@1.31.31,trna33_HisGTG_6_+_27125906_27125977@1.31.31,trna7_HisGTG_9_-_14433938_14434009@1.31.31,trna8_HisGTG_15_-_45492611_45492682@1.31.31,trna9_HisGTG_15_-_45490804_45490875@1.31.31\|GCCGTGATCGTATAGTGGTTAGTACTCTGCG | >trna111_HisGTG_1_-_147774845_147774916 | 1 | 31 | 31 | GCCGTGATCGTATAGTGGTTAGTACTCTGCG | MB468 | 935 | 964 | 3460 | 452 | 132 | 556.7 | 430.0 | 3642.8 | 103.0 | 98.5 |
| >trna111_HisGTG_1_-_147774845_147774916@1.32.32,trna118_HisGTG_1_-_145396881_145396952@1.32.32,trna16_HisGTG_1_+_146544773_146544844@1.32.32,trna1_HisGTG_15_+_45493349_45493420@1.32.32,trna21_HisGTG_1_+_147753471_147753542@1.32.32,trna33_HisGTG_6_+_27125906_27125977@1.32.32,trna7_HisGTG_9_-_14433938_14434009@1.32.32,trna8_HisGTG_15_-_45492611_45492682@1.32.32,trna9_HisGTG_15_-_45490804_45490875@1.32.32\|GCCGTGATCGTATAGTGGTTAGTACTCTGCGT | >trna111_HisGTG_1_-_147774845_147774916 | 1 | 32 | 32 | GCCGTGATCGTATAGTGGTTAGTACTCTGCGT | BT20 | 19 | 15 | 32 | 9 | 24 | 18.1 | 17.6 | 17.6 | 9.3 | 23.1 |
| >trna111_HisGTG_1_-_147774845_147774916@1.32.32,trna118_HisGTG_1_-_145396881_145396952@1.32.32,trna16_HisGTG_1_+_146544773_146544844@1.32.32,trna1_HisGTG_15_+_45493349_45493420@1.32.32,trna21_HisGTG_1_+_147753471_147753542@1.32.32,trna33_HisGTG_6_+_27125906_27125977@1.32.32,trna7_HisGTG_9_-_14433938_14434009@1.32.32,trna8_HisGTG_15_-_45492611_45492682@1.32.32,trna9_HisGTG_15_-_45490804_45490875@1.32.32\|GCCGTGATCGTATAGTGGTTAGTACTCTGCGT | >trna111_HisGTG_1_-_147774845_147774916 | 1 | 32 | 32 | GCCGTGATCGTATAGTGGTTAGTACTCTGCGT | MB231 | 16 | 18 | 45 | 33 | 20 | 8.4 | 15.8 | 22.2 | 21.4 | 7.8 |

| | | | | | | | | | | | | | | | |
|---|---|---|---|---|---|---|---|---|---|---|---|---|---|---|---|
| >trna111_HisGTG_1_-_147774845_147774916@1.33.33,trna118_HisGTG_1_-_145396881_145396952@1.33.33,trna16_HisGTG_1_+_146544773_146544844@1.33.33,trna1_HisGTG_15_+_45493349_45493420@1.33.33,trna21_HisGTG_1_+_147753471_147753542@1.33.33,trna33_HisGTG_6_+_27125906_27125977@1.33.33,trna7_HisGTG_9_-_14433938_14434009@1.33.33,trna8_HisGTG_15_-_45492611_45492682@1.33.33,trna9_HisGTG_15_-_45490804_45490875@1.33.33\|GCCGTGATCGTATAGTGGTTAGTACTCTGCGTT | >trna111_HisGTG_1_-_147774845_147774916 | 1 | 33 | 33 | GCCGTGATCGTATAGTGGTTAGTACTCTGCGTT | BT20 | 198 | 197 | 776 | 173 | 61 | 159.3 | 221.5 | 900.2 | 205.3 | 60.5 |
| >trna111_HisGTG_1_-_147774845_147774916@1.33.33,trna118_HisGTG_1_-_145396881_145396952@1.33.33,trna16_HisGTG_1_+_146544773_146544844@1.33.33,trna1_HisGTG_15_+_45493349_45493420@1.33.33,trna21_HisGTG_1_+_147753471_147753542@1.33.33,trna33_HisGTG_6_+_27125906_27125977@1.33.33,trna7_HisGTG_9_-_14433938_14434009@1.33.33,trna8_HisGTG_15_-_45492611_45492682@1.33.33,trna9_HisGTG_15_-_45490804_45490875@1.33.33\|GCCGTGATCGTATAGTGGTTAGTACTCTGCGTT | >trna111_HisGTG_1_-_147774845_147774916 | 1 | 33 | 33 | GCCGTGATCGTATAGTGGTTAGTACTCTGCGTT | MB231 | 80 | 67 | 265 | 78 | 46 | 49.9 | 60.3 | 170.1 | 29.3 | 22.3 |
| >trna111_HisGTG_1_-_147774845_147774916@1.34.34,trna118_HisGTG_1_-_145396881_145396952@1.34.34,trna16_HisGTG_1_+_146544773_146544844@1.34.34,trna1_HisGTG_15_+_45493349_45493420@1.34.34,trna21_HisGTG_1_+_147753471_147753542@1.34.34,trna33_HisGTG_6_+_27125906_27125977@1.34.34,trna7_HisGTG_9_-_14433938_14434009@1.34.34,trna8_HisGTG_15_-_45492611_45492682@1.34.34,trna9_HisGTG_15_-_45490804_45490875@1.34.34\|GCCGTGATCGTATAGTGGTTAGTACTCTGCGTTG | >trna111_HisGTG_1_-_147774845_147774916 | 1 | 34 | 34 | GCCGTGATCGTATAGTGGTTAGTACTCTGCGTTG | BT20 | 91 | 34 | 179 | 26 | 122 | 66.9 | 20.4 | 86.4 | 28.8 | 143.9 |
| >trna111_HisGTG_1_-_147774845_147774916@1.34.34,trna118_HisGTG_1_-_145396881_145396952@1.34.34,trna16_HisGTG_1_+_146544773_146544844@1.34.34,trna1_HisGTG_15_+_45493349_45493420@1.34.34,trna21_HisGTG_1_+_147753471_147753542@1.34.34,trna33_HisGTG_6_+_27125906_27125977@1.34.34,trna7_HisGTG_9_-_14433938_14434009@1.34.34,trna8_HisGTG_15_-_45492611_45492682@1.34.34,trna9_HisGTG_15_-_45490804_45490875@1.34.34\|GCCGTGATCGTATAGTGGTTAGTACTCTGCGTTG | >trna111_HisGTG_1_-_147774845_147774916 | 1 | 34 | 34 | GCCGTGATCGTATAGTGGTTAGTACTCTGCGTTG | MB231 | 115 | 70 | 449 | 60 | 73 | 100.5 | 57.2 | 398.6 | 34.6 | 39.0 |
| >trna111_HisGTG_1_-_147774845_147774916@1.34.34,trna118_HisGTG_1_-_145396881_145396952@1.34.34,trna16_HisGTG_1_+_146544773_146544844@1.34.34,trna1_HisGTG_15_+_45493349_45493420@1.34.34,trna21_HisGTG_1_+_147753471_147753542@1.34.34,trna33_HisGTG_6_+_27125906_27125977@1.34.34,trna7_HisGTG_9_-_14433938_14434009@1.34.34,trna8_HisGTG_15_-_45492611_45492682@1.34.34,trna9_HisGTG_15_-_45490804_45490875@1.34.34\|GCCGTGATCGTATAGTGGTTAGTACTCTGCGTTG | >trna111_HisGTG_1_-_147774845_147774916 | 1 | 34 | 34 | GCCGTGATCGTATAGTGGTTAGTACTCTGCGTTG | MB468 | 14 | 9 | 3 | 3 | 3 | 10.6 | 12.4 | 1.8 | 2.8 | 1.3 |
| >trna111_HisGTG_1_-_147774845_147774916@1.35.35,trna118_HisGTG_1_-_145396881_145396952@1.35.35,trna16_HisGTG_1_+_146544773_146544844@1.35.35,trna1_HisGTG_15_+_45493349_45493420@1.35.35,trna21_HisGTG_1_+_147753471_147753542@1.35.35,trna33_HisGTG_6_+_27125906_27125977@1.35.35,trna7_HisGTG_9_-_14433938_14434009@1.35.35,trna8_HisGTG_15_-_45492611_45492682@1.35.35,trna9_HisGTG_15_-_45490804_45490875@1.35.35\|GCCGTGATCGTATAGTGGTTAGTACTCTGCGTTGT | >trna111_HisGTG_1_-_147774845_147774916 | 1 | 35 | 35 | GCCGTGATCGTATAGTGGTTAGTACTCTGCGTTGT | BT20 | 45 | 48 | 194 | 5 | 4 | 49.7 | 52.5 | 301.8 | 3.8 | 2.1 |
| >trna111_HisGTG_1_-_147774845_147774916@2.30.29,trna118_HisGTG_1_-_145396881_145396952@2.30.29,trna16_HisGTG_1_+_146544773_146544844@2.30.29,trna1_HisGTG_15_+_45493349_45493420@2.30.29,trna21_HisGTG_1_+_147753471_147753542@2.30.29,trna33_HisGTG_6_+_27125906_27125977@2.30.29,trna7_HisGTG_9_-_14433938_14434009@2.30.29,trna8_HisGTG_15_-_45492611_45492682@2.30.29,trna9_HisGTG_15_-_45490804_45490875@2.30.29\|CCGTGATCGTATAGTGGTTAGTACTCTGC | >trna111_HisGTG_1_-_147774845_147774916 | 2 | 30 | 29 | CCGTGATCGTATAGTGGTTAGTACTCTGC | BT20 | 21 | 1 | 137 | 31 | 5 | 26.5 | 2.0 | 133.5 | 42.2 | 6.0 |
| >trna111_HisGTG_1_-_147774845_147774916@2.31.30,trna118_HisGTG_1_-_145396881_145396952@2.31.30,trna16_HisGTG_1_+_146544773_146544844@2.31.30,trna1_HisGTG_15_+_45493349_45493420@2.31.30,trna21_HisGTG_1_+_147753471_147753542@2.31.30,trna33_HisGTG_6_+_27125906_27125977@2.31.30,trna7_HisGTG_9_-_14433938_14434009@2.31.30,trna8_HisGTG_15_-_45492611_45492682@2.31.30,trna9_HisGTG_15_-_45490804_45490875@2.31.30\|CCGTGATCGTATAGTGGTTAGTACTCTGCG | >trna111_HisGTG_1_-_147774845_147774916 | 2 | 31 | 30 | CCGTGATCGTATAGTGGTTAGTACTCTGCG | BT20 | 10 | 1 | 49 | 19 | 3 | 8.2 | 1.1 | 39.5 | 25.5 | 3.8 |
| >trna111_HisGTG_1_-_147774845_147774916@3.31.29,trna118_HisGTG_1_-_145396881_145396952@3.31.29,trna16_HisGTG_1_+_146544773_146544844@3.31.29,trna1_HisGTG_15_+_45493349_45493420@3.31.29,trna21_HisGTG_1_+_147753471_147753542@3.31.29,trna33_HisGTG_6_+_27125906_27125977@3.31.29,trna7_HisGTG_9_-_14433938_14434009@3.31.29,trna8_HisGTG_15_-_45492611_45492682@3.31.29,trna9_HisGTG_15_-_45490804_45490875@3.31.29\|CGTGATCGTATAGTGGTTAGTACTCTGCG | >trna111_HisGTG_1_-_147774845_147774916 | 3 | 31 | 29 | CGTGATCGTATAGTGGTTAGTACTCTGCG | MB468 | 12 | 8 | 26 | 11 | 7 | 5.9 | 7.1 | 16.6 | 5.5 | 3.0 |
| >trna112_GlnCTG_1_-_147737382_147737453@30.50.21,trna131_GlnCTG_6_-_27759135_27759206@30.50.21,trna146_GlnCTG_6_-_27515531_27515602@30.50.21,trna15_GlnCTG_1_+_145963304_145963375@30.50.21,trna19_GlnCTG_1_+_147505038_147505109@30.50.21,trna1_GlnCTG_6_+_18836402_18836473@30.50.21,trna28_GlnCTG_1_+_149186125_149186196@30.50.21,trna3_GlnCTG_17_+_8023070_8023141@30.50.21,trna49_GlnCTG_6_+_27487308_27487379@30.50.21,trna7_GlnCTG_15_-_66161400_66161471@30.50.21,trna99_GlnCTG_6_-_28909378_28909449@30.50.21\|GACTCTGAATCCAGCGATCCG | >trna112_GlnCTG_1_-_147737382_147737453 | 30 | 50 | 21 | GACTCTGAATCCAGCGATCCG | MB468 | 48 | 29 | 23 | 16 | 28 | 27.1 | 20.7 | 12.7 | 6.4 | 16.1 |
| >trna112_GlnCTG_1_-_147737382_147737453@31.50.20,trna131_GlnCTG_6_-_27759135_27759206@31.50.20,trna146_GlnCTG_6_-_27515531_27515602@31.50.20,trna15_GlnCTG_1_+_145963304_145963375@31.50.20,trna19_GlnCTG_1_+_147505038_147505109@31.50.20,trna1_GlnCTG_6_+_18836402_18836473@31.50.20,trna28_GlnCTG_1_+_149186125_149186196@31.50.20,trna3_GlnCTG_17_+_8023070_8023141@31.50.20,trna49_GlnCTG_6_+_27487308_27487379@31.50.20,trna7_GlnCTG_15_-_66161400_66161471@31.50.20,trna99_GlnCTG_6_-_28909378_28909449@31.50.20\|ACTCTGAATCCAGCGATCCG | >trna112_GlnCTG_1_-_147737382_147737453 | 31 | 50 | 20 | ACTCTGAATCCAGCGATCCG | MB468 | 23 | 9 | 12 | 7 | 12 | 13.9 | 7.5 | 3.1 | 3.4 | 8.6 |
| >trna116_GluCTC_1_-_145399233_145399304@1.31.31,trna59_GluCTC_1_+_249168447_249168518@1.31.31,trna71_GluCTC_1_-_161439189_161439260@1.31.31,trna74_GluCTC_1_-_161431809_161431880@1.31.31,trna77_GluCTC_1_-_161424398_161424469@1.31.31,trna77_GluCTC_6_+_28949976_28950047@1.31.31,trna80_GluCTC_1_-_161417018_161417089@1.31.31,trna87_GluCTC_6_-_126101393_126101464@1.31.31\|TCCCTGGTGGTCTAGTGGTTAGGATTCGGCG | >trna116_GluCTC_1_-_145399233_145399304 | 1 | 31 | 31 | TCCCTGGTGGTCTAGTGGTTAGGATTCGGCG | BT20 | 59 | 37 | 352 | 420 | 150 | 47.8 | 43.7 | 217.4 | 685.1 | 160.2 |
| >trna116_GluCTC_1_-_145399233_145399304@1.31.31,trna59_GluCTC_1_+_249168447_249168518@1.31.31,trna71_GluCTC_1_-_161439189_161439260@1.31.31,trna74_GluCTC_1_-_161431809_161431880@1.31.31,trna77_GluCTC_1_-_161424398_161424469@1.31.31,trna77_GluCTC_6_+_28949976_28950047@1.31.31,trna80_GluCTC_1_-_161417018_161417089@1.31.31,trna87_GluCTC_6_-_126101393_126101464@1.31.31\|TCCCTGGTGGTCTAGTGGTTAGGATTCGGCG | >trna116_GluCTC_1_-_145399233_145399304 | 1 | 31 | 31 | TCCCTGGTGGTCTAGTGGTTAGGATTCGGCG | MB231 | 251 | 134 | 595 | 133 | 162 | 182.3 | 126.7 | 387.6 | 73.9 | 42.4 |
| >trna116_GluCTC_1_-_145399233_145399304@1.31.31,trna59_GluCTC_1_+_249168447_249168518@1.31.31,trna71_GluCTC_1_-_161439189_161439260@1.31.31,trna74_GluCTC_1_-_161431809_161431880@1.31.31,trna77_GluCTC_1_-_161424398_161424469@1.31.31,trna77_GluCTC_6_+_28949976_28950047@1.31.31,trna80_GluCTC_1_-_161417018_161417089@1.31.31,trna87_GluCTC_6_-_126101393_126101464@1.31.31\|TCCCTGGTGGTCTAGTGGTTAGGATTCGGCG | >trna116_GluCTC_1_-_145399233_145399304 | 1 | 31 | 31 | TCCCTGGTGGTCTAGTGGTTAGGATTCGGCG | MB468 | 10579 | 9603 | 23061 | 1168 | 915 | 5239.1 | 4076.4 | 13373.9 | 757.5 | 573.3 |

| | | | | | | | | | | | | | | | | |
|---|---|---|---|---|---|---|---|---|---|---|---|---|---|---|---|---|
| >trna116_GluCTC_1_-_145399233_145399304@1.32.32,trna59_GluCTC_1_+_249168447_249168518@1.32.32,trna71_GluCTC_1_-_161439189_161439260@1.32.32,trna74_GluCTC_1_-_161431809_161431880@1.32.32,trna77_GluCTC_1_-_161424398_161424469@1.32.32,trna77_GluCTC_6_+_28949976_28950047@1.32.32,trna80_GluCTC_1_-_161417018_161417089@1.32.32,trna87_GluCTC_6_-_126101393_126101464@1.32.32\|TCCCTGGTGGTCTAGTGGTTAGGATTCGGCGC | >trna116_GluCTC_1_-_145399233_145399304 | 1 | 32 | 32 | TCCCTGGTGGTCTAGTGGTTAGGATTCGGCGC | BT20 | 191 | 44 | 324 | 72 | 226 | 226.5 | 9.5 | 102.1 | 83.9 | 262.8 |
| >trna116_GluCTC_1_-_145399233_145399304@1.32.32,trna59_GluCTC_1_+_249168447_249168518@1.32.32,trna71_GluCTC_1_-_161439189_161439260@1.32.32,trna74_GluCTC_1_-_161431809_161431880@1.32.32,trna77_GluCTC_1_-_161424398_161424469@1.32.32,trna77_GluCTC_6_+_28949976_28950047@1.32.32,trna80_GluCTC_1_-_161417018_161417089@1.32.32,trna87_GluCTC_6_-_126101393_126101464@1.32.32\|TCCCTGGTGGTCTAGTGGTTAGGATTCGGCGC | >trna116_GluCTC_1_-_145399233_145399304 | 1 | 32 | 32 | TCCCTGGTGGTCTAGTGGTTAGGATTCGGCGC | MB231 | 94 | 88 | 313 | 89 | 109 | 48.7 | 99.5 | 156.0 | 34.6 | 40.1 |
| >trna116_GluCTC_1_-_145399233_145399304@1.33.33,trna59_GluCTC_1_+_249168447_249168518@1.33.33,trna71_GluCTC_1_-_161439189_161439260@1.33.33,trna74_GluCTC_1_-_161431809_161431880@1.33.33,trna77_GluCTC_1_-_161424398_161424469@1.33.33,trna77_GluCTC_6_+_28949976_28950047@1.33.33,trna80_GluCTC_1_-_161417018_161417089@1.33.33,trna87_GluCTC_6_-_126101393_126101464@1.33.33\|TCCCTGGTGGTCTAGTGGTTAGGATTCGGCGCT | >trna116_GluCTC_1_-_145399233_145399304 | 1 | 33 | 33 | TCCCTGGTGGTCTAGTGGTTAGGATTCGGCGCT | BT20 | 1409 | 642 | 6927 | 595 | 1838 | 1332.8 | 545.6 | 7609.2 | 680.9 | 1753.6 |
| >trna116_GluCTC_1_-_145399233_145399304@1.33.33,trna59_GluCTC_1_+_249168447_249168518@1.33.33,trna71_GluCTC_1_-_161439189_161439260@1.33.33,trna74_GluCTC_1_-_161431809_161431880@1.33.33,trna77_GluCTC_1_-_161424398_161424469@1.33.33,trna77_GluCTC_6_+_28949976_28950047@1.33.33,trna80_GluCTC_1_-_161417018_161417089@1.33.33,trna87_GluCTC_6_-_126101393_126101464@1.33.33\|TCCCTGGTGGTCTAGTGGTTAGGATTCGGCGCT | >trna116_GluCTC_1_-_145399233_145399304 | 1 | 33 | 33 | TCCCTGGTGGTCTAGTGGTTAGGATTCGGCGCT | MB231 | 1357 | 2368 | 6388 | 365 | 653 | 737.2 | 3070.7 | 3093.0 | 244.9 | 140.2 |
| >trna116_GluCTC_1_-_145399233_145399304@1.33.33,trna59_GluCTC_1_+_249168447_249168518@1.33.33,trna71_GluCTC_1_-_161439189_161439260@1.33.33,trna74_GluCTC_1_-_161431809_161431880@1.33.33,trna77_GluCTC_1_-_161424398_161424469@1.33.33,trna77_GluCTC_6_+_28949976_28950047@1.33.33,trna80_GluCTC_1_-_161417018_161417089@1.33.33,trna87_GluCTC_6_-_126101393_126101464@1.33.33\|TCCCTGGTGGTCTAGTGGTTAGGATTCGGCGCT | >trna116_GluCTC_1_-_145399233_145399304 | 1 | 33 | 33 | TCCCTGGTGGTCTAGTGGTTAGGATTCGGCGCT | MB468 | 26 | 74 | 74 | 22 | 23 | 8.4 | 38.3 | 68.5 | 21.3 | 18.0 |
| >trna116_GluCTC_1_-_145399233_145399304@1.34.34,trna59_GluCTC_1_+_249168447_249168518@1.34.34,trna71_GluCTC_1_-_161439189_161439260@1.34.34,trna74_GluCTC_1_-_161431809_161431880@1.34.34,trna77_GluCTC_1_-_161424398_161424469@1.34.34,trna77_GluCTC_6_+_28949976_28950047@1.34.34,trna80_GluCTC_1_-_161417018_161417089@1.34.34,trna87_GluCTC_6_-_126101393_126101464@1.34.34\|TCCCTGGTGGTCTAGTGGTTAGGATTCGGCGCTC | >trna116_GluCTC_1_-_145399233_145399304 | 1 | 34 | 34 | TCCCTGGTGGTCTAGTGGTTAGGATTCGGCGCTC | BT20 | 408 | 151 | 665 | 176 | 546 | 499.4 | 80.9 | 242.5 | 207.5 | 463.2 |
| >trna116_GluCTC_1_-_145399233_145399304@1.34.34,trna59_GluCTC_1_+_249168447_249168518@1.34.34,trna71_GluCTC_1_-_161439189_161439260@1.34.34,trna74_GluCTC_1_-_161431809_161431880@1.34.34,trna77_GluCTC_1_-_161424398_161424469@1.34.34,trna77_GluCTC_6_+_28949976_28950047@1.34.34,trna80_GluCTC_1_-_161417018_161417089@1.34.34,trna87_GluCTC_6_-_126101393_126101464@1.34.34\|TCCCTGGTGGTCTAGTGGTTAGGATTCGGCGCTC | >trna116_GluCTC_1_-_145399233_145399304 | 1 | 34 | 34 | TCCCTGGTGGTCTAGTGGTTAGGATTCGGCGCTC | MB231 | 145 | 115 | 558 | 205 | 259 | 91.8 | 104.4 | 156.1 | 100.1 | 162.4 |
| >trna116_GluCTC_1_-_145399233_145399304@1.35.35,trna59_GluCTC_1_+_249168447_249168518@1.35.35,trna71_GluCTC_1_-_161439189_161439260@1.35.35,trna74_GluCTC_1_-_161431809_161431880@1.35.35,trna77_GluCTC_1_-_161424398_161424469@1.35.35,trna77_GluCTC_6_+_28949976_28950047@1.35.35,trna80_GluCTC_1_-_161417018_161417089@1.35.35,trna87_GluCTC_6_-_126101393_126101464@1.35.35\|TCCCTGGTGGTCTAGTGGTTAGGATTCGGCGCTCT | >trna116_GluCTC_1_-_145399233_145399304 | 1 | 35 | 35 | TCCCTGGTGGTCTAGTGGTTAGGATTCGGCGCTCT | BT20 | 358 | 215 | 2139 | 123 | 338 | 221.0 | 257.9 | 2846.7 | 154.0 | 238.1 |
| >trna116_GluCTC_1_-_145399233_145399304@1.35.35,trna59_GluCTC_1_+_249168447_249168518@1.35.35,trna71_GluCTC_1_-_161439189_161439260@1.35.35,trna74_GluCTC_1_-_161431809_161431880@1.35.35,trna77_GluCTC_1_-_161424398_161424469@1.35.35,trna77_GluCTC_6_+_28949976_28950047@1.35.35,trna80_GluCTC_1_-_161417018_161417089@1.35.35,trna87_GluCTC_6_-_126101393_126101464@1.35.35\|TCCCTGGTGGTCTAGTGGTTAGGATTCGGCGCTCT | >trna116_GluCTC_1_-_145399233_145399304 | 1 | 35 | 35 | TCCCTGGTGGTCTAGTGGTTAGGATTCGGCGCTCT | MB231 | 144 | 128 | 605 | 80 | 132 | 57.6 | 130.0 | 256.0 | 46.4 | 34.1 |
| >trna116_GluCTC_1_-_145399233_145399304@1.36.36,trna59_GluCTC_1_+_249168447_249168518@1.36.36,trna71_GluCTC_1_-_161439189_161439260@1.36.36,trna74_GluCTC_1_-_161431809_161431880@1.36.36,trna77_GluCTC_1_-_161424398_161424469@1.36.36,trna77_GluCTC_6_+_28949976_28950047@1.36.36,trna80_GluCTC_1_-_161417018_161417089@1.36.36,trna87_GluCTC_6_-_126101393_126101464@1.36.36\|TCCCTGGTGGTCTAGTGGTTAGGATTCGGCGCTCTC | >trna116_GluCTC_1_-_145399233_145399304 | 1 | 36 | 36 | TCCCTGGTGGTCTAGTGGTTAGGATTCGGCGCTCTC | BT20 | 134 | 56 | 327 | 82 | 139 | 91.6 | 24.6 | 156.6 | 104.4 | 98.7 |
| >trna116_GluCTC_1_-_145399233_145399304@1.36.36,trna59_GluCTC_1_+_249168447_249168518@1.36.36,trna71_GluCTC_1_-_161439189_161439260@1.36.36,trna74_GluCTC_1_-_161431809_161431880@1.36.36,trna77_GluCTC_1_-_161424398_161424469@1.36.36,trna77_GluCTC_6_+_28949976_28950047@1.36.36,trna80_GluCTC_1_-_161417018_161417089@1.36.36,trna87_GluCTC_6_-_126101393_126101464@1.36.36\|TCCCTGGTGGTCTAGTGGTTAGGATTCGGCGCTCTC | >trna116_GluCTC_1_-_145399233_145399304 | 1 | 36 | 36 | TCCCTGGTGGTCTAGTGGTTAGGATTCGGCGCTCTC | MB231 | 138 | 162 | 951 | 85 | 161 | 102.0 | 190.8 | 791.7 | 73.0 | 48.0 |
| >trna116_GluCTC_1_-_145399233_145399304@1.39.39,trna59_GluCTC_1_+_249168447_249168518@1.39.39,trna71_GluCTC_1_-_161439189_161439260@1.39.39,trna74_GluCTC_1_-_161431809_161431880@1.39.39,trna77_GluCTC_1_-_161424398_161424469@1.39.39,trna77_GluCTC_6_+_28949976_28950047@1.39.39,trna80_GluCTC_1_-_161417018_161417089@1.39.39,trna87_GluCTC_6_-_126101393_126101464@1.39.39\|TCCCTGGTGGTCTAGTGGTTAGGATTCGGCGCTCTCACC | >trna116_GluCTC_1_-_145399233_145399304 | 1 | 39 | 39 | TCCCTGGTGGTCTAGTGGTTAGGATTCGGCGCTCTCACC | BT20 | 89 | 47 | 420 | 58 | 313 | 107.2 | 38.0 | 16.7 | 62.5 | 240.0 |
| >trna116_GluCTC_1_-_145399233_145399304@1.39.39,trna59_GluCTC_1_+_249168447_249168518@1.39.39,trna71_GluCTC_1_-_161439189_161439260@1.39.39,trna74_GluCTC_1_-_161431809_161431880@1.39.39,trna77_GluCTC_1_-_161424398_161424469@1.39.39,trna77_GluCTC_6_+_28949976_28950047@1.39.39,trna80_GluCTC_1_-_161417018_161417089@1.39.39,trna87_GluCTC_6_-_126101393_126101464@1.39.39\|TCCCTGGTGGTCTAGTGGTTAGGATTCGGCGCTCTCACC | >trna116_GluCTC_1_-_145399233_145399304 | 1 | 39 | 39 | TCCCTGGTGGTCTAGTGGTTAGGATTCGGCGCTCTCACC | MB231 | 140 | 153 | 1732 | 15 | 75 | 176.7 | 162.0 | 1666.0 | 8.5 | 55.7 |
| >trna116_GluCTC_1_-_145399233_145399304@1.42.42,trna59_GluCTC_1_+_249168447_249168518@1.42.42,trna71_GluCTC_1_-_161439189_161439260@1.42.42,trna74_GluCTC_1_-_161431809_161431880@1.42.42,trna77_GluCTC_1_-_161424398_161424469@1.42.42,trna77_GluCTC_6_+_28949976_28950047@1.42.42,trna80_GluCTC_1_-_161417018_161417089@1.42.42,trna87_GluCTC_6_-_126101393_126101464@1.42.42\|TCCCTGGTGGTCTAGTGGTTAGGATTCGGCGCTCTCACCGCC | >trna116_GluCTC_1_-_145399233_145399304 | 1 | 42 | 42 | TCCCTGGTGGTCTAGTGGTTAGGATTCGGCGCTCTCACCGCC | BT20 | 10 | 2 | 72 | 5 | 3 | 15.3 | 0.4 | 49.1 | 7.0 | 4.0 |

| | | | | | | | | | | | | | | | | |
|---|---|---|---|---|---|---|---|---|---|---|---|---|---|---|---|---|
| >trna116_GluCTC_1_-_145399233_145399304@13.31.19,trna25_GluCTC_2_-_71273488_71273560@13.31.19,trna59_GluCTC_1_+_249168447_249168518@13.31.19,trna71_GluCTC_1_-_161439189_161439260@13.31.19,trna74_GluCTC_1_-_161431809_161431880@13.31.19,trna77_GluCTC_1_-_161424398_161424469@13.31.19,trna77_GluCTC_6_+_28949976_28950047@13.31.19,trna80_GluCTC_1_-_161417018_161417089@13.31.19,trna87_GluCTC_6_-_126101393_126101464@13.31.19\|TAGTGGTTAGGATTCGGCG | >trna116_GluCTC_1_-_145399233_145399304 | 13 | 31 | 19 | TAGTGGTTAGGATTCGGCG | MB468 | 11 | 10 | 22 | 3 | 4 | 4.6 | 5.2 | 11.8 | 0.8 | 1.9 |
| >trna116_GluCTC_1_-_145399233_145399304@2.31.30,trna59_GluCTC_1_+_249168447_249168518@2.31.30,trna71_GluCTC_1_-_161439189_161439260@2.31.30,trna74_GluCTC_1_-_161431809_161431880@2.31.30,trna77_GluCTC_1_-_161424398_161424469@2.31.30,trna77_GluCTC_6_+_28949976_28950047@2.31.30,trna80_GluCTC_1_-_161417018_161417089@2.31.30,trna87_GluCTC_6_-_126101393_126101464@2.31.30\|CCCTGGTGGTCTAGTGGTTAGGATTCGGCG | >trna116_GluCTC_1_-_145399233_145399304 | 2 | 31 | 30 | CCCTGGTGGTCTAGTGGTTAGGATTCGGCG | MB468 | 24 | 25 | 38 | 3 | 3 | 8.0 | 8.8 | 18.0 | 1.6 | 2.3 |
| >trna116_GluCTC_1_-_145399233_145399304@3.33.31,trna59_GluCTC_1_+_249168447_249168518@3.33.31,trna71_GluCTC_1_-_161439189_161439260@3.33.31,trna74_GluCTC_1_-_161431809_161431880@3.33.31,trna77_GluCTC_1_-_161424398_161424469@3.33.31,trna77_GluCTC_6_+_28949976_28950047@3.33.31,trna80_GluCTC_1_-_161417018_161417089@3.33.31,trna87_GluCTC_6_-_126101393_126101464@3.33.31\|CCTGGTGGTCTAGTGGTTAGGATTCGGCGCT | >trna116_GluCTC_1_-_145399233_145399304 | 3 | 33 | 31 | CCTGGTGGTCTAGTGGTTAGGATTCGGCGCT | MB231 | 13 | 22 | 23 | 12 | 42 | 10.0 | 10.6 | 7.9 | 8.6 | 7.6 |
| >trna116_GluCTC_1_-_145399233_145399304@32.69.38,trna59_GluCTC_1_+_249168447_249168518@32.69.38,trna71_GluCTC_1_-_161439189_161439260@32.69.38,trna74_GluCTC_1_-_161431809_161431880@32.69.38,trna77_GluCTC_1_-_161424398_161424469@32.69.38,trna77_GluCTC_6_+_28949976_28950047@32.69.38,trna80_GluCTC_1_-_161417018_161417089@32.69.38,trna87_GluCTC_6_-_126101393_126101464@32.69.38\|CTCTCACCGCCGCGGCCCGGGTTCGATTCCCGGTCAGG | >trna116_GluCTC_1_-_145399233_145399304 | 32 | 69 | 38 | CTCTCACCGCCGCGGCCCGGGTTCGATTCCCGGTCAGG | MB468 | 19 | 58 | 50 | 1 | 5 | 9.5 | 35.2 | 34.3 | 1.1 | 6.4 |
| >trna116_GluCTC_1_-_145399233_145399304@32.70.39,trna71_GluCTC_1_-_161439189_161439260@32.70.39,trna74_GluCTC_1_-_161431809_161431880@32.70.39,trna77_GluCTC_1_-_161424398_161424469@32.70.39,trna77_GluCTC_6_+_28949976_28950047@32.70.39,trna80_GluCTC_1_-_161417018_161417089@32.70.39,trna87_GluCTC_6_-_126101393_126101464@32.70.39\|CTCTCACCGCCGCGGCCCGGGTTCGATTCCCGGTCAGGG | >trna116_GluCTC_1_-_145399233_145399304 | 32 | 70 | 39 | CTCTCACCGCCGCGGCCCGGGTTCGATTCCCGGTCAGGG | MB468 | 132 | 373 | 571 | 13 | 117 | 68.0 | 153.7 | 298.2 | 10.5 | 125.1 |
| >trna116_GluCTC_1_-_145399233_145399304@32.71.40,trna71_GluCTC_1_-_161439189_161439260@32.71.40,trna74_GluCTC_1_-_161431809_161431880@32.71.40,trna77_GluCTC_1_-_161424398_161424469@32.71.40,trna77_GluCTC_6_+_28949976_28950047@32.71.40,trna80_GluCTC_1_-_161417018_161417089@32.71.40,trna87_GluCTC_6_-_126101393_126101464@32.71.40\|CTCTCACCGCCGCGGCCCGGGTTCGATTCCCGGTCAGGGA | >trna116_GluCTC_1_-_145399233_145399304 | 32 | 71 | 40 | CTCTCACCGCCGCGGCCCGGGTTCGATTCCCGGTCAGGGA | MB468 | 38 | 61 | 160 | 6 | 37 | 23.5 | 21.8 | 70.5 | 4.3 | 40.1 |
| >trna116_GluCTC_1_-_145399233_145399304@32.72.41,trna71_GluCTC_1_-_161439189_161439260@32.72.41,trna74_GluCTC_1_-_161431809_161431880@32.72.41,trna77_GluCTC_1_-_161424398_161424469@32.72.41,trna77_GluCTC_6_+_28949976_28950047@32.72.41,trna80_GluCTC_1_-_161417018_161417089@32.72.41,trna87_GluCTC_6_-_126101393_126101464@32.72.41\|CTCTCACCGCCGCGGCCCGGGTTCGATTCCCGGTCAGGGAA | >trna116_GluCTC_1_-_145399233_145399304 | 32 | 72 | 41 | CTCTCACCGCCGCGGCCCGGGTTCGATTCCCGGTCAGGGAA | MB468 | 12 | 21 | 64 | 1 | 13 | 5.5 | 9.5 | 33.0 | 0.7 | 13.5 |
| >trna116_GluCTC_1_-_145399233_145399304@32.73.42,trna71_GluCTC_1_-_161439189_161439260@32.73.42,trna74_GluCTC_1_-_161431809_161431880@32.73.42,trna77_GluCTC_1_-_161424398_161424469@32.73.42,trna77_GluCTC_6_+_28949976_28950047@32.73.42,trna80_GluCTC_1_-_161417018_161417089@32.73.42,trna87_GluCTC_6_-_126101393_126101464@32.73.42\|CTCTCACCGCCGCGGCCCGGGTTCGATTCCCGGTCAGGGAAC | >trna116_GluCTC_1_-_145399233_145399304 | 32 | 73 | 42 | CTCTCACCGCCGCGGCCCGGGTTCGATTCCCGGTCAGGGAAC | MB468 | 23 | 55 | 140 | 1 | 1 | 17.4 | 24.1 | 103.2 | 1.8 | 1.8 |
| >trna116_GluCTC_1_-_145399233_145399304@32.75.44,trna71_GluCTC_1_-_161439189_161439260@32.75.44,trna74_GluCTC_1_-_161431809_161431880@32.75.44,trna77_GluCTC_1_-_161424398_161424469@32.75.44,trna77_GluCTC_6_+_28949976_28950047@32.75.44,trna80_GluCTC_1_-_161417018_161417089@32.75.44,trna87_GluCTC_6_-_126101393_126101464@32.75.44\|CTCTCACCGCCGCGGCCCGGGTTCGATTCCCGGTCAGGGAACCA | >trna116_GluCTC_1_-_145399233_145399304 | 32 | 75 | 44 | CTCTCACCGCCGCGGCCCGGGTTCGATTCCCGGTCAGGGAACCA | MB468 | 16 | 92 | 89 | 0 | 2 | 11.1 | 73.0 | 47.2 | 0.3 | 3.0 |
| >trna116_GluCTC_1_-_145399233_145399304@35.74.40,trna134_GluTTC_1_-_16861774_16861845@35.74.40,trna71_GluCTC_1_-_161439189_161439260@35.74.40,trna74_GluCTC_1_-_161431809_161431880@35.74.40,trna77_GluCTC_1_-_161424398_161424469@35.74.40,trna77_GluCTC_6_+_28949976_28950047@35.74.40,trna80_GluCTC_1_-_161417018_161417089@35.74.40,trna84_GluTTC_1_-_161391883_161391954@35.74.40,trna87_GluCTC_6_-_126101393_126101464@35.74.40\|TCACCGCCGCGGCCCGGGTTCGATTCCCGGTCAGGGAACC | >trna116_GluCTC_1_-_145399233_145399304 | 35 | 74 | 40 | TCACCGCCGCGGCCCGGGTTCGATTCCCGGTCAGGGAACC | MB231 | 13 | 5 | 88 | 0 | 0 | 9.6 | 3.8 | 84.0 | 0.0 | 0.0 |
| >trna116_GluCTC_1_-_145399233_145399304@35.75.41,trna134_GluTTC_1_-_16861774_16861845@35.75.41,trna71_GluCTC_1_-_161439189_161439260@35.75.41,trna74_GluCTC_1_-_161431809_161431880@35.75.41,trna77_GluCTC_1_-_161424398_161424469@35.75.41,trna77_GluCTC_6_+_28949976_28950047@35.75.41,trna80_GluCTC_1_-_161417018_161417089@35.75.41,trna84_GluTTC_1_-_161391883_161391954@35.75.41,trna87_GluCTC_6_-_126101393_126101464@35.75.41\|TCACCGCCGCGGCCCGGGTTCGATTCCCGGTCAGGGAACCA | >trna116_GluCTC_1_-_145399233_145399304 | 35 | 75 | 41 | TCACCGCCGCGGCCCGGGTTCGATTCCCGGTCAGGGAACCA | MB231 | 19 | 8 | 160 | 0 | 0 | 17.2 | 5.7 | 177.0 | 0.0 | 0.0 |
| >trna116_GluCTC_1_-_145399233_145399304@36.74.39,trna134_GluTTC_1_-_16861774_16861845@36.74.39,trna71_GluCTC_1_-_161439189_161439260@36.74.39,trna74_GluCTC_1_-_161431809_161431880@36.74.39,trna77_GluCTC_1_-_161424398_161424469@36.74.39,trna77_GluCTC_6_+_28949976_28950047@36.74.39,trna80_GluCTC_1_-_161417018_161417089@36.74.39,trna84_GluTTC_1_-_161391883_161391954@36.74.39,trna87_GluCTC_6_-_126101393_126101464@36.74.39\|CACCGCCGCGGCCCGGGTTCGATTCCCGGTCAGGGAACC | >trna116_GluCTC_1_-_145399233_145399304 | 36 | 74 | 39 | CACCGCCGCGGCCCGGGTTCGATTCCCGGTCAGGGAACC | MB231 | 15 | 8 | 118 | 1 | 3 | 7.2 | 8.7 | 115.1 | 1.0 | 4.6 |
| >trna116_GluCTC_1_-_145399233_145399304@36.75.40,trna134_GluTTC_1_-_16861774_16861845@36.75.40,trna71_GluCTC_1_-_161439189_161439260@36.75.40,trna74_GluCTC_1_-_161431809_161431880@36.75.40,trna77_GluCTC_1_-_161424398_161424469@36.75.40,trna77_GluCTC_6_+_28949976_28950047@36.75.40,trna80_GluCTC_1_-_161417018_161417089@36.75.40,trna84_GluTTC_1_-_161391883_161391954@36.75.40,trna87_GluCTC_6_-_126101393_126101464@36.75.40\|CACCGCCGCGGCCCGGGTTCGATTCCCGGTCAGGGAACCA | >trna116_GluCTC_1_-_145399233_145399304 | 36 | 75 | 40 | CACCGCCGCGGCCCGGGTTCGATTCCCGGTCAGGGAACCA | MB231 | 17 | 11 | 164 | 0 | 1 | 6.7 | 11.2 | 182.9 | 0.3 | 1.6 |

| | | | | | | | | | | | | | | | | |
|---|---|---|---|---|---|---|---|---|---|---|---|---|---|---|---|---|
| >trna116_GluCTC_1_-_145399233_145399304@37.73.37,trna134_GluTTC_1_-_16861774_16861845@37.73.37,trna71_GluCTC_1_-_161439189_161439260@37.73.37,trna74_GluCTC_1_-_161431809_161431880@37.73.37,trna77_GluCTC_1_-_161424398_161424469@37.73.37,trna77_GluCTC_6_+_28949976_28950047@37.73.37,trna80_GluCTC_1_-_161417018_161417089@37.73.37,trna84_GluTTC_1_-_161391883_161391954@37.73.37,trna87_GluCTC_6_-_126101393_126101464@37.73.37\|ACCGCCGCGGCCCGGGTTCGATTCCCGGTCAGGGAAC | >trna116_GluCTC_1_-_145399233_145399304 | 37 | 73 | 37 | ACCGCCGCGGCCCGGGTTCGATTCCCGGTCAGGGAAC | MB231 | 24 | 48 | 110 | 3 | 7 | 24.4 | 71.6 | 142.0 | 3.7 | 8.9 |
| >trna116_GluCTC_1_-_145399233_145399304@37.74.38,trna134_GluTTC_1_-_16861774_16861845@37.74.38,trna71_GluCTC_1_-_161439189_161439260@37.74.38,trna74_GluCTC_1_-_161431809_161431880@37.74.38,trna77_GluCTC_1_-_161424398_161424469@37.74.38,trna77_GluCTC_6_+_28949976_28950047@37.74.38,trna80_GluCTC_1_-_161417018_161417089@37.74.38,trna84_GluTTC_1_-_161391883_161391954@37.74.38,trna87_GluCTC_6_-_126101393_126101464@37.74.38\|ACCGCCGCGGCCCGGGTTCGATTCCCGGTCAGGGAACC | >trna116_GluCTC_1_-_145399233_145399304 | 37 | 74 | 38 | ACCGCCGCGGCCCGGGTTCGATTCCCGGTCAGGGAACC | MB231 | 102 | 236 | 658 | 4 | 31 | 101.2 | 347.1 | 741.8 | 4.4 | 32.6 |
| >trna116_GluCTC_1_-_145399233_145399304@37.75.39,trna134_GluTTC_1_-_16861774_16861845@37.75.39,trna71_GluCTC_1_-_161439189_161439260@37.75.39,trna74_GluCTC_1_-_161431809_161431880@37.75.39,trna77_GluCTC_1_-_161424398_161424469@37.75.39,trna77_GluCTC_6_+_28949976_28950047@37.75.39,trna80_GluCTC_1_-_161417018_161417089@37.75.39,trna84_GluTTC_1_-_161391883_161391954@37.75.39,trna87_GluCTC_6_-_126101393_126101464@37.75.39\|ACCGCCGCGGCCCGGGTTCGATTCCCGGTCAGGGAACCA | >trna116_GluCTC_1_-_145399233_145399304 | 37 | 75 | 39 | ACCGCCGCGGCCCGGGTTCGATTCCCGGTCAGGGAACCA | MB231 | 98 | 324 | 824 | 2 | 24 | 102.7 | 490.8 | 1061.2 | 2.4 | 26.9 |
| >trna116_GluCTC_1_-_145399233_145399304@39.74.36,trna134_GluTTC_1_-_16861774_16861845@39.74.36,trna71_GluCTC_1_-_161439189_161439260@39.74.36,trna74_GluCTC_1_-_161431809_161431880@39.74.36,trna77_GluCTC_1_-_161424398_161424469@39.74.36,trna77_GluCTC_6_+_28949976_28950047@39.74.36,trna80_GluCTC_1_-_161417018_161417089@39.74.36,trna84_GluTTC_1_-_161391883_161391954@39.74.36,trna87_GluCTC_6_-_126101393_126101464@39.74.36\|CGCCGCGGCCCGGGTTCGATTCCCGGTCAGGGAACC | >trna116_GluCTC_1_-_145399233_145399304 | 39 | 74 | 36 | CGCCGCGGCCCGGGTTCGATTCCCGGTCAGGGAACC | MB231 | 11 | 11 | 42 | 4 | 9 | 6.6 | 11.3 | 47.2 | 2.8 | 6.6 |
| >trna119_LysCTT_1_-_145395522_145395594@1.29.29,trna11_LysCTT_5_-_180648979_180649051@1.29.29,trna13_LysCTT_6_+_26556774_26556846@1.29.29,trna32_LysCTT_16_-_3207406_3207478@1.29.29,trna7_LysCTT_16_+_3225692_3225764@1.29.29,trna9_LysCTT_5_+_180634755_180634827@1.29.29\|GCCCGGCTAGCTCAGTCGGTAGAGCATGA | >trna119_LysCTT_1_-_145395522_145395594 | 1 | 29 | 29 | GCCCGGCTAGCTCAGTCGGTAGAGCATGA | MB468 | 217 | 68 | 209 | 34 | 47 | 71.9 | 19.8 | 73.4 | 6.7 | 21.7 |
| >trna119_LysCTT_1_-_145395522_145395594@1.30.30,trna11_LysCTT_5_-_180648979_180649051@1.30.30,trna13_LysCTT_6_+_26556774_26556846@1.30.30,trna32_LysCTT_16_-_3207406_3207478@1.30.30,trna7_LysCTT_16_+_3225692_3225764@1.30.30,trna9_LysCTT_5_+_180634755_180634827@1.30.30\|GCCCGGCTAGCTCAGTCGGTAGAGCATGAG | >trna119_LysCTT_1_-_145395522_145395594 | 1 | 30 | 30 | GCCCGGCTAGCTCAGTCGGTAGAGCATGAG | MB468 | 1687 | 709 | 2952 | 215 | 525 | 876.1 | 254.7 | 1738.3 | 37.5 | 265.0 |
| >trna119_LysCTT_1_-_145395522_145395594@1.31.31,trna11_LysCTT_5_-_180648979_180649051@1.31.31,trna13_LysCTT_6_+_26556774_26556846@1.31.31,trna32_LysCTT_16_-_3207406_3207478@1.31.31,trna7_LysCTT_16_+_3225692_3225764@1.31.31,trna9_LysCTT_5_+_180634755_180634827@1.31.31\|GCCCGGCTAGCTCAGTCGGTAGAGCATGAGA | >trna119_LysCTT_1_-_145395522_145395594 | 1 | 31 | 31 | GCCCGGCTAGCTCAGTCGGTAGAGCATGAGA | MB231 | 20 | 12 | 22 | 9 | 32 | 12.5 | 9.0 | 21.4 | 6.4 | 16.1 |
| >trna119_LysCTT_1_-_145395522_145395594@1.31.31,trna11_LysCTT_5_-_180648979_180649051@1.31.31,trna13_LysCTT_6_+_26556774_26556846@1.31.31,trna32_LysCTT_16_-_3207406_3207478@1.31.31,trna7_LysCTT_16_+_3225692_3225764@1.31.31,trna9_LysCTT_5_+_180634755_180634827@1.31.31\|GCCCGGCTAGCTCAGTCGGTAGAGCATGAGA | >trna119_LysCTT_1_-_145395522_145395594 | 1 | 31 | 31 | GCCCGGCTAGCTCAGTCGGTAGAGCATGAGA | MB468 | 597 | 167 | 1015 | 86 | 110 | 155.7 | 115.6 | 132.2 | 27.4 | 44.0 |
| >trna119_LysCTT_1_-_145395522_145395594@1.32.32,trna11_LysCTT_5_-_180648979_180649051@1.32.32,trna13_LysCTT_6_+_26556774_26556846@1.32.32,trna32_LysCTT_16_-_3207406_3207478@1.32.32,trna7_LysCTT_16_+_3225692_3225764@1.32.32,trna9_LysCTT_5_+_180634755_180634827@1.32.32\|GCCCGGCTAGCTCAGTCGGTAGAGCATGAGAC | >trna119_LysCTT_1_-_145395522_145395594 | 1 | 32 | 32 | GCCCGGCTAGCTCAGTCGGTAGAGCATGAGAC | BT20 | 19 | 5 | 43 | 10 | 32 | 17.8 | 3.2 | 25.0 | 11.7 | 48.7 |
| >trna119_LysCTT_1_-_145395522_145395594@1.32.32,trna11_LysCTT_5_-_180648979_180649051@1.32.32,trna13_LysCTT_6_+_26556774_26556846@1.32.32,trna32_LysCTT_16_-_3207406_3207478@1.32.32,trna7_LysCTT_16_+_3225692_3225764@1.32.32,trna9_LysCTT_5_+_180634755_180634827@1.32.32\|GCCCGGCTAGCTCAGTCGGTAGAGCATGAGAC | >trna119_LysCTT_1_-_145395522_145395594 | 1 | 32 | 32 | GCCCGGCTAGCTCAGTCGGTAGAGCATGAGAC | MB231 | 97 | 65 | 196 | 66 | 168 | 48.9 | 74.4 | 224.9 | 17.1 | 110.6 |
| >trna119_LysCTT_1_-_145395522_145395594@1.32.32,trna11_LysCTT_5_-_180648979_180649051@1.32.32,trna13_LysCTT_6_+_26556774_26556846@1.32.32,trna32_LysCTT_16_-_3207406_3207478@1.32.32,trna7_LysCTT_16_+_3225692_3225764@1.32.32,trna9_LysCTT_5_+_180634755_180634827@1.32.32\|GCCCGGCTAGCTCAGTCGGTAGAGCATGAGAC | >trna119_LysCTT_1_-_145395522_145395594 | 1 | 32 | 32 | GCCCGGCTAGCTCAGTCGGTAGAGCATGAGAC | MB468 | 47 | 18 | 97 | 14 | 12 | 12.6 | 13.7 | 13.9 | 4.9 | 5.6 |
| >trna119_LysCTT_1_-_145395522_145395594@1.33.33,trna11_LysCTT_5_-_180648979_180649051@1.33.33,trna13_LysCTT_6_+_26556774_26556846@1.33.33,trna7_LysCTT_16_+_3225692_3225764@1.33.33,trna9_LysCTT_5_+_180634755_180634827@1.33.33\|GCCCGGCTAGCTCAGTCGGTAGAGCATGAGACT | >trna119_LysCTT_1_-_145395522_145395594 | 1 | 33 | 33 | GCCCGGCTAGCTCAGTCGGTAGAGCATGAGACT | BT20 | 68 | 23 | 221 | 111 | 108 | 29.7 | 10.7 | 134.6 | 178.9 | 169.6 |
| >trna119_LysCTT_1_-_145395522_145395594@1.33.33,trna11_LysCTT_5_-_180648979_180649051@1.33.33,trna13_LysCTT_6_+_26556774_26556846@1.33.33,trna7_LysCTT_16_+_3225692_3225764@1.33.33,trna9_LysCTT_5_+_180634755_180634827@1.33.33\|GCCCGGCTAGCTCAGTCGGTAGAGCATGAGACT | >trna119_LysCTT_1_-_145395522_145395594 | 1 | 33 | 33 | GCCCGGCTAGCTCAGTCGGTAGAGCATGAGACT | MB231 | 675 | 660 | 1602 | 264 | 477 | 617.9 | 912.2 | 1799.7 | 205.1 | 311.4 |
| >trna119_LysCTT_1_-_145395522_145395594@1.33.33,trna11_LysCTT_5_-_180648979_180649051@1.33.33,trna13_LysCTT_6_+_26556774_26556846@1.33.33,trna7_LysCTT_16_+_3225692_3225764@1.33.33,trna9_LysCTT_5_+_180634755_180634827@1.33.33\|GCCCGGCTAGCTCAGTCGGTAGAGCATGAGACT | >trna119_LysCTT_1_-_145395522_145395594 | 1 | 33 | 33 | GCCCGGCTAGCTCAGTCGGTAGAGCATGAGACT | MB468 | 11 | 5 | 18 | 9 | 7 | 2.3 | 1.6 | 11.4 | 5.6 | 2.5 |
| >trna119_LysCTT_1_-_145395522_145395594@1.34.34,trna11_LysCTT_5_-_180648979_180649051@1.34.34,trna13_LysCTT_6_+_26556774_26556846@1.34.34,trna7_LysCTT_16_+_3225692_3225764@1.34.34,trna9_LysCTT_5_+_180634755_180634827@1.34.34\|GCCCGGCTAGCTCAGTCGGTAGAGCATGAGACTC | >trna119_LysCTT_1_-_145395522_145395594 | 1 | 34 | 34 | GCCCGGCTAGCTCAGTCGGTAGAGCATGAGACTC | BT20 | 93 | 19 | 134 | 48 | 65 | 61.5 | 6.6 | 50.4 | 77.3 | 101.6 |
| >trna119_LysCTT_1_-_145395522_145395594@1.34.34,trna11_LysCTT_5_-_180648979_180649051@1.34.34,trna13_LysCTT_6_+_26556774_26556846@1.34.34,trna7_LysCTT_16_+_3225692_3225764@1.34.34,trna9_LysCTT_5_+_180634755_180634827@1.34.34\|GCCCGGCTAGCTCAGTCGGTAGAGCATGAGACTC | >trna119_LysCTT_1_-_145395522_145395594 | 1 | 34 | 34 | GCCCGGCTAGCTCAGTCGGTAGAGCATGAGACTC | MB231 | 226 | 271 | 576 | 144 | 317 | 199.8 | 364.8 | 743.5 | 72.6 | 217.1 |
| >trna119_LysCTT_1_-_145395522_145395594@1.35.35,trna11_LysCTT_5_-_180648979_180649051@1.35.35,trna13_LysCTT_6_+_26556774_26556846@1.35.35,trna7_LysCTT_16_+_3225692_3225764@1.35.35,trna9_LysCTT_5_+_180634755_180634827@1.35.35\|GCCCGGCTAGCTCAGTCGGTAGAGCATGAGACTCT | >trna119_LysCTT_1_-_145395522_145395594 | 1 | 35 | 35 | GCCCGGCTAGCTCAGTCGGTAGAGCATGAGACTCT | BT20 | 812 | 53 | 1627 | 79 | 86 | 638.3 | 31.0 | 1381.4 | 115.0 | 124.9 |
| >trna119_LysCTT_1_-_145395522_145395594@1.35.35,trna11_LysCTT_5_-_180648979_180649051@1.35.35,trna13_LysCTT_6_+_26556774_26556846@1.35.35,trna7_LysCTT_16_+_3225692_3225764@1.35.35,trna9_LysCTT_5_+_180634755_180634827@1.35.35\|GCCCGGCTAGCTCAGTCGGTAGAGCATGAGACTCT | >trna119_LysCTT_1_-_145395522_145395594 | 1 | 35 | 35 | GCCCGGCTAGCTCAGTCGGTAGAGCATGAGACTCT | MB231 | 116 | 60 | 233 | 136 | 229 | 30.4 | 43.6 | 163.0 | 51.8 | 135.2 |
| >trna119_LysCTT_1_-_145395522_145395594@1.36.36,trna11_LysCTT_5_-_180648979_180649051@1.36.36,trna13_LysCTT_6_+_26556774_26556846@1.36.36,trna7_LysCTT_16_+_3225692_3225764@1.36.36,trna9_LysCTT_5_+_180634755_180634827@1.36.36\|GCCCGGCTAGCTCAGTCGGTAGAGCATGAGACTCTT | >trna119_LysCTT_1_-_145395522_145395594 | 1 | 36 | 36 | GCCCGGCTAGCTCAGTCGGTAGAGCATGAGACTCTT | BT20 | 94 | 14 | 249 | 45 | 10 | 74.9 | 13.6 | 214.1 | 64.4 | 12.4 |

| | | | | | | | | | | | | | | | |
|---|---|---|---|---|---|---|---|---|---|---|---|---|---|---|---|
| >trna119_LysCTT_1_-_145395522_145395594@1.36.36,trna11_LysCTT_5_-_180648979_180649051@1.36.36,trna13_LysCTT_6_+_26556774_26556846@1.36.36,trna7_LysCTT_16_+_3225692_3225764@1.36.36,trna9_LysCTT_5_+_180634755_180634827@1.36.36\|GCCCGGCTAGCTCAGTCGGTAGAGCATGAGACTCTT | >trna119_LysCTT_1_-_145395522_145395594 | 1 | 36 | 36 | GCCCGGCTAGCTCAGTCGGTAGAGCATGAGACTCTT | MB231 | 42 | 30 | 35 | 47 | 102 | 9.8 | 15.1 | 23.6 | 18.2 | 78.2 |
| >trna119_LysCTT_1_-_145395522_145395594@2.33.32,trna11_LysCTT_5_-_180648979_180649051@2.33.32,trna13_LysCTT_6_+_26556774_26556846@2.33.32,trna7_LysCTT_16_+_3225692_3225764@2.33.32,trna9_LysCTT_5_+_180634755_180634827@2.33.32\|CCCGGCTAGCTCAGTCGGTAGAGCATGAGACT | >trna119_LysCTT_1_-_145395522_145395594 | 2 | 33 | 32 | CCCGGCTAGCTCAGTCGGTAGAGCATGAGACT | MB231 | 17 | 42 | 15 | 7 | 23 | 14.3 | 51.9 | 18.4 | 2.3 | 7.7 |
| >trna119_LysCTT_1_-_145395522_145395594@3.33.31,trna11_LysCTT_5_-_180648979_180649051@3.33.31,trna13_LysCTT_6_+_26556774_26556846@3.33.31,trna7_LysCTT_16_+_3225692_3225764@3.33.31,trna9_LysCTT_5_+_180634755_180634827@3.33.31\|CCGGCTAGCTCAGTCGGTAGAGCATGAGACT | >trna119_LysCTT_1_-_145395522_145395594 | 3 | 33 | 31 | CCGGCTAGCTCAGTCGGTAGAGCATGAGACT | MB231 | 34 | 75 | 33 | 29 | 79 | 22.6 | 85.0 | 33.1 | 15.0 | 32.6 |
| >trna119_LysCTT_1_-_145395522_145395594@3.34.32,trna11_LysCTT_5_-_180648979_180649051@3.34.32,trna13_LysCTT_6_+_26556774_26556846@3.34.32,trna7_LysCTT_16_+_3225692_3225764@3.34.32,trna9_LysCTT_5_+_180634755_180634827@3.34.32\|CCGGCTAGCTCAGTCGGTAGAGCATGAGACTC | >trna119_LysCTT_1_-_145395522_145395594 | 3 | 34 | 32 | CCGGCTAGCTCAGTCGGTAGAGCATGAGACTC | MB231 | 11 | 18 | 5 | 13 | 39 | 5.6 | 8.8 | 4.8 | 11.8 | 17.9 |
| >trna119_LysCTT_1_-_145395522_145395594@4.33.30,trna11_LysCTT_5_-_180648979_180649051@4.33.30,trna13_LysCTT_6_+_26556774_26556846@4.33.30,trna7_LysCTT_16_+_3225692_3225764@4.33.30,trna9_LysCTT_5_+_180634755_180634827@4.33.30\|CGGCTAGCTCAGTCGGTAGAGCATGAGACT | >trna119_LysCTT_1_-_145395522_145395594 | 4 | 33 | 30 | CGGCTAGCTCAGTCGGTAGAGCATGAGACT | MB231 | 32 | 80 | 16 | 54 | 160 | 21.2 | 85.4 | 15.4 | 33.0 | 120.0 |
| >trna119_LysCTT_1_-_145395522_145395594@4.34.31,trna11_LysCTT_5_-_180648979_180649051@4.34.31,trna13_LysCTT_6_+_26556774_26556846@4.34.31,trna7_LysCTT_16_+_3225692_3225764@4.34.31,trna9_LysCTT_5_+_180634755_180634827@4.34.31\|CGGCTAGCTCAGTCGGTAGAGCATGAGACTC | >trna119_LysCTT_1_-_145395522_145395594 | 4 | 34 | 31 | CGGCTAGCTCAGTCGGTAGAGCATGAGACTC | MB231 | 11 | 21 | 2 | 17 | 45 | 5.2 | 15.8 | 2.2 | 10.1 | 18.8 |
| >trna119_LysCTT_1_-_145395522_145395594@4.35.32,trna11_LysCTT_5_-_180648979_180649051@4.35.32,trna13_LysCTT_6_+_26556774_26556846@4.35.32,trna7_LysCTT_16_+_3225692_3225764@4.35.32,trna9_LysCTT_5_+_180634755_180634827@4.35.32\|CGGCTAGCTCAGTCGGTAGAGCATGAGACTCT | >trna119_LysCTT_1_-_145395522_145395594 | 4 | 35 | 32 | CGGCTAGCTCAGTCGGTAGAGCATGAGACTCT | MB231 | 60 | 64 | 9 | 71 | 331 | 26.5 | 33.2 | 7.9 | 27.8 | 118.8 |
| >trna119_LysCTT_1_-_145395522_145395594@4.36.33,trna11_LysCTT_5_-_180648979_180649051@4.36.33,trna13_LysCTT_6_+_26556774_26556846@4.36.33,trna7_LysCTT_16_+_3225692_3225764@4.36.33,trna9_LysCTT_5_+_180634755_180634827@4.36.33\|CGGCTAGCTCAGTCGGTAGAGCATGAGACTCTT | >trna119_LysCTT_1_-_145395522_145395594 | 4 | 36 | 33 | CGGCTAGCTCAGTCGGTAGAGCATGAGACTCTT | MB231 | 33 | 29 | 4 | 61 | 122 | 20.4 | 23.3 | 4.8 | 32.2 | 107.3 |
| >trna12_ArgCCT_16_+_3243918_3243990@34.51.18,trna18_ArgCCT_17_+_73030001_73030073@34.51.18,trna21_ArgCCT_17_-_73030526_73030598@34.51.18,trna2_ArgCCT_16_+_3202901_3202973@34.51.18,trna3_ArgCCT_7_+_139025446_139025518@34.51.18\|CCTAAGCCAGGGATTGTG | >trna12_ArgCCT_16_+_3243918_3243990 | 34 | 51 | 18 | CCTAAGCCAGGGATTGTG | MB468 | 13 | 4 | 10 | 3 | 2 | 5.0 | 2.6 | 6.0 | 0.7 | 0.4 |
| >trna12_ArgCCT_16_+_3243918_3243990@35.52.18,trna18_ArgCCT_17_+_73030001_73030073@35.52.18,trna21_ArgCCT_17_-_73030526_73030598@35.52.18,trna2_ArgCCT_16_+_3202901_3202973@35.52.18,trna3_ArgCCT_7_+_139025446_139025518@35.52.18\|CTAAGCCAGGGATTGTGG | >trna12_ArgCCT_16_+_3243918_3243990 | 35 | 52 | 18 | CTAAGCCAGGGATTGTGG | MB468 | 12 | 5 | 9 | 5 | 1 | 4.9 | 2.6 | 2.9 | 2.6 | 0.6 |
| >trna12_ArgCCT_16_+_3243918_3243990@35.54.20,trna18_ArgCCT_17_+_73030001_73030073@35.54.20,trna21_ArgCCT_17_-_73030526_73030598@35.54.20,trna2_ArgCCT_16_+_3202901_3202973@35.54.20,trna3_ArgCCT_7_+_139025446_139025518@35.54.20\|CTAAGCCAGGGATTGTGGGT | >trna12_ArgCCT_16_+_3243918_3243990 | 35 | 54 | 20 | CTAAGCCAGGGATTGTGGGT | BT20 | 11 | 1 | 54 | 6 | 2 | 8.5 | 0.8 | 39.4 | 7.2 | 2.4 |
| >trna12_ArgCCT_16_+_3243918_3243990@35.54.20,trna18_ArgCCT_17_+_73030001_73030073@35.54.20,trna21_ArgCCT_17_-_73030526_73030598@35.54.20,trna2_ArgCCT_16_+_3202901_3202973@35.54.20,trna3_ArgCCT_7_+_139025446_139025518@35.54.20\|CTAAGCCAGGGATTGTGGGT | >trna12_ArgCCT_16_+_3243918_3243990 | 35 | 54 | 20 | CTAAGCCAGGGATTGTGGGT | MB231 | 17 | 1 | 40 | 16 | 3 | 6.0 | 1.4 | 29.9 | 5.6 | 1.3 |
| >trna12_ArgCCT_16_+_3243918_3243990@35.54.20,trna18_ArgCCT_17_+_73030001_73030073@35.54.20,trna21_ArgCCT_17_-_73030526_73030598@35.54.20,trna2_ArgCCT_16_+_3202901_3202973@35.54.20,trna3_ArgCCT_7_+_139025446_139025518@35.54.20\|CTAAGCCAGGGATTGTGGGT | >trna12_ArgCCT_16_+_3243918_3243990 | 35 | 54 | 20 | CTAAGCCAGGGATTGTGGGT | MB468 | 27 | 7 | 31 | 7 | 1 | 12.8 | 3.5 | 7.7 | 2.8 | 0.2 |
| >trna12_ArgCCT_16_+_3243918_3243990@36.54.19,trna18_ArgCCT_17_+_73030001_73030073@36.54.19,trna21_ArgCCT_17_-_73030526_73030598@36.54.19,trna2_ArgCCT_16_+_3202901_3202973@36.54.19,trna3_ArgCCT_7_+_139025446_139025518@36.54.19\|TAAGCCAGGGATTGTGGGT | >trna12_ArgCCT_16_+_3243918_3243990 | 36 | 54 | 19 | TAAGCCAGGGATTGTGGGT | MB468 | 23 | 6 | 26 | 5 | 1 | 9.8 | 2.9 | 3.2 | 1.6 | 0.3 |
| >trna12_TrpCCA_17_+_19411494_19411565@30.52.23,trna168_TrpCCA_6_-_26331672_26331743@30.52.23,trna170_TrpCCA_6_-_26319330_26319401@30.52.23,trna1_TrpCCA_7_+_99067307_99067378@30.52.23,trna39_TrpCCA_17_-_8124187_8124258@30.52.23,trna6_TrpCCA_17_+_8089676_8089747@30.52.23\|ACTCCAGATCAGAAGGTTGCGTG | >trna12_TrpCCA_17_+_19411494_19411565 | 30 | 52 | 23 | ACTCCAGATCAGAAGGTTGCGTG | MB468 | 30 | 38 | 12 | 8 | 8 | 13.3 | 29.0 | 2.4 | 3.2 | 2.9 |
| >trna126_LeuAAG_6_-_28446400_28446481@68.85.18,trna16_LeuAAG_16_+_22308461_22308542@68.85.18,trna16_LeuAAG_5_-_180601044_180601125@68.85.18,trna19_LeuAAG_5_-_180524474_180524555@68.85.18,trna1_LeuAAG_14_+_21078291_21078372@68.85.18,trna27_LeuTAG_16_-_22207032_22207113@68.85.18,trna3_LeuAAG_5_+_180528840_180528921@68.85.18,trna42_LeuTAG_17_-_8023632_8023713@68.85.18,trna78_LeuAAG_6_+_28956779_28956860@68.85.18,trna7_LeuAAG_5_+_180614701_180614782@68.85.18,trna98_LeuAAG_6_-_28911399_28911480@68.85.18\|ATCCCACCGCTGCCACCA | >trna126_LeuAAG_6_-_28446400_28446481 | 68 | 85 | 18 | ATCCCACCGCTGCCACCA | MB231 | 12 | 15 | 110 | 2 | 11 | 20.5 | 25.8 | 190.4 | 3.0 | 19.3 |
| >trna128_GlyGCC_6_-_27870686_27870756@19.38.20,trna19_GlyGCC_16_+_70823410_70823480@19.38.20,trna19_GlyGCC_2_-_157257659_157257729@19.38.20,trna24_GlyGCC_16_-_70812942_70813012@19.38.20,trna25_GlyGCC_16_-_70812114_70812184@19.38.20,trna2_GlyGCC_21_-_18827107_18827177@19.38.20,trna35_GlyGCC_1_+_161413094_161413164@19.38.20,trna37_GlyGCC_1_+_161420467_161420537@19.38.20,trna39_GlyGCC_1_+_161427898_161427968@19.38.20,trna41_GlyGCC_1_+_161435258_161435328@19.38.20,trna5_GlyGCC_17_+_8029064_8029134@19.38.20,trna68_GlyGCC_1_-_161493637_161493707@19.38.20\|TAGAATTCTCGCCTGCCACG | >trna128_GlyGCC_6_-_27870686_27870756 | 19 | 38 | 20 | TAGAATTCTCGCCTGCCACG | BT20 | 26 | 3 | 149 | 14 | 8 | 29.5 | 1.5 | 136.2 | 19.5 | 9.4 |
| >trna129_MetCAT_6_-_27870271_27870342@31.50.20,trna142_MetCAT_6_-_27560600_27560671@31.50.20,trna150_MetCAT_6_-_27300764_27300835@31.50.20,trna169_MetCAT_6_-_26330529_26330600@31.50.20,trna171_MetCAT_6_-_26313352_26313423@31.50.20,trna20_MetCAT_17_-_80452597_80452668@31.50.20,trna2_MetCAT_6_+_26286754_26286825@31.50.20,trna32_MetCAT_1_+_153643726_153643797@31.50.20,trna61_MetCAT_6_+_27745664_27745735@31.50.20\|CCCATAACCCAGAGGTCGAT | >trna129_MetCAT_6_-_27870271_27870342 | 31 | 50 | 20 | CCCATAACCCAGAGGTCGAT | MB468 | 12 | 4 | 4 | 5 | 2 | 5.1 | 3.0 | 1.8 | 1.9 | 1.6 |
| >trna129_MetCAT_6_-_27870271_27870342@31.51.21,trna142_MetCAT_6_-_27560600_27560671@31.51.21,trna150_MetCAT_6_-_27300764_27300835@31.51.21,trna169_MetCAT_6_-_26330529_26330600@31.51.21,trna171_MetCAT_6_-_26313352_26313423@31.51.21,trna20_MetCAT_17_-_80452597_80452668@31.51.21,trna2_MetCAT_6_+_26286754_26286825@31.51.21,trna32_MetCAT_1_+_153643726_153643797@31.51.21,trna61_MetCAT_6_+_27745664_27745735@31.51.21\|CCCATAACCCAGAGGTCGATG | >trna129_MetCAT_6_-_27870271_27870342 | 31 | 51 | 21 | CCCATAACCCAGAGGTCGATG | MB468 | 151 | 44 | 55 | 96 | 28 | 80.0 | 31.7 | 28.9 | 51.2 | 22.1 |

| | | | | | | | | | | | | | | | | |
|---|---|---|---|---|---|---|---|---|---|---|---|---|---|---|---|---|
| >trna129_MetCAT_6_-_27870271_27870342@32.51.20,trna142_MetCAT_6_-_27560600_27560671@32.51.20,trna150_MetCAT_6_-_27300764_27300835@32.51.20,trna169_MetCAT_6_-_26330529_26330600@32.51.20,trna171_MetCAT_6_-_26313352_26313423@32.51.20,trna20_MetCAT_17_-_80452597_80452668@32.51.20,trna2_MetCAT_6_+_26286754_26286825@32.51.20,trna32_MetCAT_1_+_153643726_153643797@32.51.20,trna61_MetCAT_6_+_27745664_27745735@32.51.20\|CCATAACCCAGAGGTCGATG | >trna129_MetCAT_6_-_27870271_27870342 | 32 | 51 | 20 | CCATAACCCAGAGGTCGATG | MB468 | 28 | 4 | 8 | 31 | 5 | 18.4 | 2.4 | 5.4 | 25.5 | 2.7 |
| >trna13_ValCAC_19_-_4724647_4724719@1.34.34\|GTTTCCGTAGTGTAGCGGTTATCACATTCGCCTC | >trna13_ValCAC_19_-_4724647_4724719 | 1 | 34 | 34 | GTTTCCGTAGTGTAGCGGTTATCACATTCGCCTC | MB231 | 19 | 20 | 48 | 14 | 14 | 17.7 | 24.3 | 48.2 | 9.9 | 7.9 |
| >trna130_GlnTTG_6_-_27763640_27763711@1.29.29,trna173_GlnTTG_6_-_26311975_26312046@1.29.29,trna174_GlnTTG_6_-_26311424_26311495@1.29.29\|GGCCCCATGGTGTAATGGTTAGCACTCTG | >trna130_GlnTTG_6_-_27763640_27763711 | 1 | 29 | 29 | GGCCCCATGGTGTAATGGTTAGCACTCTG | MB468 | 85 | 61 | 153 | 6 | 14 | 73.0 | 23.4 | 130.4 | 5.0 | 10.3 |
| >trna134_GluTTC_1_-_16861774_16861845@1.31.31,trna5_GluTTC_1_+_17199078_17199149@1.31.31,trna84_GluTTC_1_-_161391883_161391954@1.31.31,trna94_GluTTC_1_-_149664355_149664427@1.31.31\|TCCCTGGTGGTCTAGTGGCTAGGATTCGGCG | >trna134_GluTTC_1_-_16861774_16861845 | 1 | 31 | 31 | TCCCTGGTGGTCTAGTGGCTAGGATTCGGCG | MB468 | 171 | 421 | 459 | 28 | 25 | 88.7 | 280.3 | 276.5 | 16.4 | 16.3 |
| >trna134_GluTTC_1_-_16861774_16861845@1.33.33,trna5_GluTTC_1_+_17199078_17199149@1.33.33,trna84_GluTTC_1_-_161391883_161391954@1.33.33,trna94_GluTTC_1_-_149664355_149664427@1.33.33\|TCCCTGGTGGTCTAGTGGCTAGGATTCGGCGCT | >trna134_GluTTC_1_-_16861774_16861845 | 1 | 33 | 33 | TCCCTGGTGGTCTAGTGGCTAGGATTCGGCGCT | BT20 | 101 | 27 | 378 | 25 | 63 | 104.0 | 12.4 | 186.8 | 27.1 | 60.7 |
| >trna134_GluTTC_1_-_16861774_16861845@1.34.34,trna5_GluTTC_1_+_17199078_17199149@1.34.34,trna84_GluTTC_1_-_161391883_161391954@1.34.34,trna94_GluTTC_1_-_149664355_149664427@1.34.34\|TCCCTGGTGGTCTAGTGGCTAGGATTCGGCGCTT | >trna134_GluTTC_1_-_16861774_16861845 | 1 | 34 | 34 | TCCCTGGTGGTCTAGTGGCTAGGATTCGGCGCTT | BT20 | 13 | 4 | 30 | 3 | 6 | 11.6 | 1.7 | 12.1 | 3.9 | 8.3 |
| >trna139_ValAAC_6_-_27618707_27618779@36.56.21\|CACGCGAAAGGTCCCTGGATC | >trna139_ValAAC_6_-_27618707_27618779 | 36 | 56 | 21 | CACGCGAAAGGTCCCTGGATC | BT20 | 11 | 8 | 24 | 28 | 7 | 12.5 | 4.2 | 14.5 | 31.4 | 5.1 |
| >trna14_LysTTT_11_-_59327808_59327880@1.24.24,trna2_LysTTT_17_+_8022473_8022545@1.24.24,trna54_LysTTT_1_+_204475655_204475727@1.24.24,trna5_LysTTT_11_+_59323902_59323974@1.24.24,trna62_LysTTT_1_-_204476158_204476230@1.24.24,trna76_LysTTT_6_+_28918806_28918878@1.24.24\|GCCCGGATAGCTCAGTCGGTAGAG | >trna14_LysTTT_11_-_59327808_59327880 | 1 | 24 | 24 | GCCCGGATAGCTCAGTCGGTAGAG | MB468 | 55 | 21 | 19 | 5 | 24 | 18.3 | 13.3 | 5.1 | 1.8 | 9.9 |
| >trna14_LysTTT_11_-_59327808_59327880@1.28.28,trna2_LysTTT_17_+_8022473_8022545@1.28.28,trna54_LysTTT_1_+_204475655_204475727@1.28.28,trna5_LysTTT_11_+_59323902_59323974@1.28.28,trna62_LysTTT_1_-_204476158_204476230@1.28.28,trna76_LysTTT_6_+_28918806_28918878@1.28.28\|GCCCGGATAGCTCAGTCGGTAGAGCATC | >trna14_LysTTT_11_-_59327808_59327880 | 1 | 28 | 28 | GCCCGGATAGCTCAGTCGGTAGAGCATC | MB468 | 21 | 2 | 19 | 7 | 3 | 11.6 | 0.7 | 17.0 | 2.8 | 2.0 |
| >trna14_LysTTT_11_-_59327808_59327880@1.29.29,trna2_LysTTT_17_+_8022473_8022545@1.29.29,trna54_LysTTT_1_+_204475655_204475727@1.29.29,trna5_LysTTT_11_+_59323902_59323974@1.29.29,trna62_LysTTT_1_-_204476158_204476230@1.29.29,trna76_LysTTT_6_+_28918806_28918878@1.29.29\|GCCCGGATAGCTCAGTCGGTAGAGCATCA | >trna14_LysTTT_11_-_59327808_59327880 | 1 | 29 | 29 | GCCCGGATAGCTCAGTCGGTAGAGCATCA | MB468 | 26 | 1 | 9 | 3 | 2 | 27.6 | 0.6 | 3.8 | 1.0 | 1.3 |
| >trna14_LysTTT_11_-_59327808_59327880@1.30.30,trna2_LysTTT_17_+_8022473_8022545@1.30.30,trna54_LysTTT_1_+_204475655_204475727@1.30.30,trna5_LysTTT_11_+_59323902_59323974@1.30.30,trna62_LysTTT_1_-_204476158_204476230@1.30.30,trna76_LysTTT_6_+_28918806_28918878@1.30.30\|GCCCGGATAGCTCAGTCGGTAGAGCATCAG | >trna14_LysTTT_11_-_59327808_59327880 | 1 | 30 | 30 | GCCCGGATAGCTCAGTCGGTAGAGCATCAG | MB468 | 106 | 4 | 87 | 29 | 11 | 56.0 | 1.2 | 61.7 | 15.3 | 4.2 |
| >trna14_LysTTT_11_-_59327808_59327880@1.31.31,trna2_LysTTT_17_+_8022473_8022545@1.31.31,trna54_LysTTT_1_+_204475655_204475727@1.31.31,trna5_LysTTT_11_+_59323902_59323974@1.31.31,trna62_LysTTT_1_-_204476158_204476230@1.31.31,trna76_LysTTT_6_+_28918806_28918878@1.31.31\|GCCCGGATAGCTCAGTCGGTAGAGCATCAGA | >trna14_LysTTT_11_-_59327808_59327880 | 1 | 31 | 31 | GCCCGGATAGCTCAGTCGGTAGAGCATCAGA | MB468 | 28 | 1 | 16 | 20 | 3 | 10.6 | 0.6 | 7.5 | 14.5 | 1.7 |
| >trna14_LysTTT_11_-_59327808_59327880@1.32.32,trna2_LysTTT_17_+_8022473_8022545@1.32.32,trna54_LysTTT_1_+_204475655_204475727@1.32.32,trna5_LysTTT_11_+_59323902_59323974@1.32.32,trna62_LysTTT_1_-_204476158_204476230@1.32.32,trna76_LysTTT_6_+_28918806_28918878@1.32.32\|GCCCGGATAGCTCAGTCGGTAGAGCATCAGAC | >trna14_LysTTT_11_-_59327808_59327880 | 1 | 32 | 32 | GCCCGGATAGCTCAGTCGGTAGAGCATCAGAC | MB231 | 13 | 3 | 5 | 43 | 14 | 8.5 | 3.7 | 4.1 | 9.0 | 10.5 |
| >trna14_LysTTT_11_-_59327808_59327880@1.33.33,trna2_LysTTT_17_+_8022473_8022545@1.33.33,trna54_LysTTT_1_+_204475655_204475727@1.33.33,trna5_LysTTT_11_+_59323902_59323974@1.33.33,trna62_LysTTT_1_-_204476158_204476230@1.33.33,trna76_LysTTT_6_+_28918806_28918878@1.33.33\|GCCCGGATAGCTCAGTCGGTAGAGCATCAGACT | >trna14_LysTTT_11_-_59327808_59327880 | 1 | 33 | 33 | GCCCGGATAGCTCAGTCGGTAGAGCATCAGACT | BT20 | 739 | 34 | 133 | 203 | 68 | 814.0 | 22.4 | 55.4 | 292.4 | 93.7 |
| >trna14_LysTTT_11_-_59327808_59327880@1.33.33,trna2_LysTTT_17_+_8022473_8022545@1.33.33,trna54_LysTTT_1_+_204475655_204475727@1.33.33,trna5_LysTTT_11_+_59323902_59323974@1.33.33,trna62_LysTTT_1_-_204476158_204476230@1.33.33,trna76_LysTTT_6_+_28918806_28918878@1.33.33\|GCCCGGATAGCTCAGTCGGTAGAGCATCAGACT | >trna14_LysTTT_11_-_59327808_59327880 | 1 | 33 | 33 | GCCCGGATAGCTCAGTCGGTAGAGCATCAGACT | MB231 | 661 | 135 | 146 | 1167 | 620 | 365.5 | 112.0 | 97.5 | 592.0 | 491.8 |
| >trna14_LysTTT_11_-_59327808_59327880@1.34.34,trna2_LysTTT_17_+_8022473_8022545@1.34.34,trna54_LysTTT_1_+_204475655_204475727@1.34.34,trna5_LysTTT_11_+_59323902_59323974@1.34.34,trna62_LysTTT_1_-_204476158_204476230@1.34.34,trna76_LysTTT_6_+_28918806_28918878@1.34.34\|GCCCGGATAGCTCAGTCGGTAGAGCATCAGACTT | >trna14_LysTTT_11_-_59327808_59327880 | 1 | 34 | 34 | GCCCGGATAGCTCAGTCGGTAGAGCATCAGACTT | BT20 | 13 | 1 | 5 | 4 | 2 | 13.4 | 0.2 | 2.7 | 6.2 | 2.7 |
| >trna14_LysTTT_11_-_59327808_59327880@1.35.35,trna2_LysTTT_17_+_8022473_8022545@1.35.35,trna54_LysTTT_1_+_204475655_204475727@1.35.35,trna5_LysTTT_11_+_59323902_59323974@1.35.35,trna62_LysTTT_1_-_204476158_204476230@1.35.35,trna76_LysTTT_6_+_28918806_28918878@1.35.35\|GCCCGGATAGCTCAGTCGGTAGAGCATCAGACTTT | >trna14_LysTTT_11_-_59327808_59327880 | 1 | 35 | 35 | GCCCGGATAGCTCAGTCGGTAGAGCATCAGACTTT | BT20 | 54 | 6 | 16 | 7 | 2 | 70.1 | 2.2 | 9.4 | 8.8 | 3.2 |
| >trna14_LysTTT_11_-_59327808_59327880@1.36.36,trna2_LysTTT_17_+_8022473_8022545@1.36.36,trna54_LysTTT_1_+_204475655_204475727@1.36.36,trna5_LysTTT_11_+_59323902_59323974@1.36.36,trna62_LysTTT_1_-_204476158_204476230@1.36.36,trna76_LysTTT_6_+_28918806_28918878@1.36.36\|GCCCGGATAGCTCAGTCGGTAGAGCATCAGACTTTT | >trna14_LysTTT_11_-_59327808_59327880 | 1 | 36 | 36 | GCCCGGATAGCTCAGTCGGTAGAGCATCAGACTTTT | BT20 | 65 | 9 | 24 | 6 | 2 | 82.0 | 3.4 | 14.3 | 8.4 | 3.7 |
| >trna143_LysTTT_6_-_27559593_27559665@13.30.18,trna149_LysTTT_6_-_27302769_27302841@13.30.18,trna14_LysTTT_11_-_59327808_59327880@13.30.18,trna2_LysTTT_17_+_8022473_8022545@13.30.18,trna54_LysTTT_1_+_204475655_204475727@13.30.18,trna5_LysTTT_11_+_59323902_59323974@13.30.18,trna62_LysTTT_1_-_204476158_204476230@13.30.18,trna76_LysTTT_6_+_28918806_28918878@13.30.18\|CAGTCGGTAGAGCATCAG | >trna143_LysTTT_6_-_27559593_27559665 | 13 | 30 | 18 | CAGTCGGTAGAGCATCAG | MB468 | 87 | 24 | 70 | 17 | 8 | 47.8 | 15.7 | 35.3 | 7.5 | 2.3 |

| | | | | | | | | | | | | | | | | |
|---|---|---|---|---|---|---|---|---|---|---|---|---|---|---|---|---|
| >trna143_LysTTT_6_-_27559593_27559665@14.31.18,trna149_LysTTT_6_-_27302769_27302841@14.31.18,trna14_LysTTT_11_-_59327808_59327880@14.31.18,trna2_LysTTT_17_+_8022473_8022545@14.31.18,trna54_LysTTT_1_+_204475655_204475727@14.31.18,trna5_LysTTT_11_+_59323902_59323974@14.31.18,trna62_LysTTT_1_-_204476158_204476230@14.31.18,trna76_LysTTT_6_+_28918806_28918878@14.31.18\|AGTCGGTAGAGCATCAGA | >trna143_LysTTT_6_-_27559593_27559665 | 14 | 31 | 18 | AGTCGGTAGAGCATCAGA | MB468 | 40 | 14 | 27 | 14 | 2 | 29.2 | 9.8 | 16.9 | 7.8 | 0.6 |
| >trna146_GlnCTG_6_-_27515531_27515602@1.30.30,trna1_GlnCTG_6_+_18836402_18836473@1.30.30,trna3_GlnCTG_17_+_8023070_8023141@1.30.30,trna42_GlnCTG_6_+_27263212_27263283@1.30.30,trna49_GlnCTG_6_+_27487308_27487379@1.30.30,trna7_GlnCTG_15_-_66161400_66161471@1.30.30,trna99_GlnCTG_6_-_28909378_28909449@1.30.30\|GGTTCCATGGTGTAATGGTTAGCACTCTGG | >trna146_GlnCTG_6_-_27515531_27515602 | 1 | 30 | 30 | GGTTCCATGGTGTAATGGTTAGCACTCTGG | MB468 | 18 | 2 | 25 | 14 | 2 | 21.0 | 0.7 | 22.1 | 14.5 | 2.8 |
| >trna15_ValAAC_5_-_180615416_180615488@1.23.23\|GTTTCCGTAGTGTAGTGGTCATC | >trna15_ValAAC_5_-_180615416_180615488 | 1 | 23 | 23 | GTTTCCGTAGTGTAGTGGTCATC | BT20 | 23 | 6 | 29 | 58 | 40 | 33.9 | 4.2 | 38.3 | 61.2 | 58.6 |
| >trna15_ValAAC_5_-_180615416_180615488@1.23.23\|GTTTCCGTAGTGTAGTGGTCATC | >trna15_ValAAC_5_-_180615416_180615488 | 1 | 23 | 23 | GTTTCCGTAGTGTAGTGGTCATC | MB468 | 37 | 14 | 36 | 10 | 7 | 9.0 | 4.7 | 10.8 | 9.7 | 1.2 |
| >trna15_ValAAC_5_-_180615416_180615488@1.29.29\|GTTTCCGTAGTGTAGTGGTCATCACGTTC | >trna15_ValAAC_5_-_180615416_180615488 | 1 | 29 | 29 | GTTTCCGTAGTGTAGTGGTCATCACGTTC | MB468 | 259 | 39 | 173 | 273 | 45 | 229.6 | 17.1 | 82.0 | 126.3 | 41.5 |
| >trna15_ValAAC_5_-_180615416_180615488@1.32.32\|GTTTCCGTAGTGTAGTGGTCATCACGTTCGCC | >trna15_ValAAC_5_-_180615416_180615488 | 1 | 32 | 32 | GTTTCCGTAGTGTAGTGGTCATCACGTTCGCC | BT20 | 18 | 3 | 24 | 14 | 22 | 20.8 | 0.9 | 20.8 | 20.1 | 35.7 |
| >trna15_ValAAC_5_-_180615416_180615488@1.32.32\|GTTTCCGTAGTGTAGTGGTCATCACGTTCGCC | >trna15_ValAAC_5_-_180615416_180615488 | 1 | 32 | 32 | GTTTCCGTAGTGTAGTGGTCATCACGTTCGCC | MB231 | 24 | 13 | 55 | 29 | 43 | 16.1 | 13.9 | 69.6 | 18.1 | 27.3 |
| >trna152_ValCAC_6_-_27248049_27248121@1.23.23\|GCTTCTGTAGTGTAGTGGTTATC | >trna152_ValCAC_6_-_27248049_27248121 | 1 | 23 | 23 | GCTTCTGTAGTGTAGTGGTTATC | BT20 | 11 | 2 | 15 | 45 | 14 | 16.1 | 0.0 | 13.0 | 55.8 | 18.1 |
| >trna152_ValCAC_6_-_27248049_27248121@1.23.23\|GCTTCTGTAGTGTAGTGGTTATC | >trna152_ValCAC_6_-_27248049_27248121 | 1 | 23 | 23 | GCTTCTGTAGTGTAGTGGTTATC | MB231 | 12 | 13 | 22 | 8 | 18 | 8.0 | 1.0 | 8.4 | 6.3 | 9.6 |
| >trna152_ValCAC_6_-_27248049_27248121@1.23.23\|GCTTCTGTAGTGTAGTGGTTATC | >trna152_ValCAC_6_-_27248049_27248121 | 1 | 23 | 23 | GCTTCTGTAGTGTAGTGGTTATC | MB468 | 20 | 11 | 38 | 7 | 4 | 5.2 | 6.3 | 11.0 | 8.3 | 2.8 |
| >trna152_ValCAC_6_-_27248049_27248121@1.24.24\|GCTTCTGTAGTGTAGTGGTTATCA | >trna152_ValCAC_6_-_27248049_27248121 | 1 | 24 | 24 | GCTTCTGTAGTGTAGTGGTTATCA | MB468 | 13 | 18 | 24 | 10 | 3 | 8.4 | 19.3 | 14.9 | 14.1 | 3.7 |
| >trna152_ValCAC_6_-_27248049_27248121@1.28.28\|GCTTCTGTAGTGTAGTGGTTATCACGTT | >trna152_ValCAC_6_-_27248049_27248121 | 1 | 28 | 28 | GCTTCTGTAGTGTAGTGGTTATCACGTT | MB468 | 26 | 5 | 20 | 2 | 2 | 18.7 | 1.6 | 5.5 | 0.7 | 1.7 |
| >trna152_ValCAC_6_-_27248049_27248121@1.29.29\|GCTTCTGTAGTGTAGTGGTTATCACGTTC | >trna152_ValCAC_6_-_27248049_27248121 | 1 | 29 | 29 | GCTTCTGTAGTGTAGTGGTTATCACGTTC | MB468 | 1255 | 459 | 2214 | 487 | 171 | 1098.2 | 250.1 | 1034.0 | 311.5 | 129.2 |
| >trna152_ValCAC_6_-_27248049_27248121@1.30.30\|GCTTCTGTAGTGTAGTGGTTATCACGTTCG | >trna152_ValCAC_6_-_27248049_27248121 | 1 | 30 | 30 | GCTTCTGTAGTGTAGTGGTTATCACGTTCG | MB468 | 20 | 6 | 32 | 3 | 2 | 20.0 | 1.4 | 23.5 | 0.8 | 2.5 |
| >trna152_ValCAC_6_-_27248049_27248121@1.31.31\|GCTTCTGTAGTGTAGTGGTTATCACGTTCGC | >trna152_ValCAC_6_-_27248049_27248121 | 1 | 31 | 31 | GCTTCTGTAGTGTAGTGGTTATCACGTTCGC | MB468 | 11 | 3 | 20 | 16 | 1 | 9.5 | 1.0 | 12.8 | 15.0 | 1.4 |
| >trna152_ValCAC_6_-_27248049_27248121@1.32.32\|GCTTCTGTAGTGTAGTGGTTATCACGTTCGCC | >trna152_ValCAC_6_-_27248049_27248121 | 1 | 32 | 32 | GCTTCTGTAGTGTAGTGGTTATCACGTTCGCC | BT20 | 19 | 4 | 40 | 48 | 26 | 14.8 | 2.6 | 11.7 | 53.5 | 33.8 |
| >trna152_ValCAC_6_-_27248049_27248121@1.32.32\|GCTTCTGTAGTGTAGTGGTTATCACGTTCGCC | >trna152_ValCAC_6_-_27248049_27248121 | 1 | 32 | 32 | GCTTCTGTAGTGTAGTGGTTATCACGTTCGCC | MB231 | 112 | 79 | 322 | 166 | 116 | 85.5 | 95.0 | 399.4 | 117.0 | 54.4 |
| >trna152_ValCAC_6_-_27248049_27248121@1.33.33\|GCTTCTGTAGTGTAGTGGTTATCACGTTCGCCT | >trna152_ValCAC_6_-_27248049_27248121 | 1 | 33 | 33 | GCTTCTGTAGTGTAGTGGTTATCACGTTCGCCT | BT20 | 11 | 3 | 25 | 39 | 19 | 4.0 | 1.2 | 6.5 | 54.6 | 28.4 |
| >trna152_ValCAC_6_-_27248049_27248121@1.33.33\|GCTTCTGTAGTGTAGTGGTTATCACGTTCGCCT | >trna152_ValCAC_6_-_27248049_27248121 | 1 | 33 | 33 | GCTTCTGTAGTGTAGTGGTTATCACGTTCGCCT | MB231 | 73 | 55 | 273 | 97 | 55 | 73.7 | 66.7 | 298.8 | 90.3 | 38.3 |
| >trna152_ValCAC_6_-_27248049_27248121@1.34.34\|GCTTCTGTAGTGTAGTGGTTATCACGTTCGCCTC | >trna152_ValCAC_6_-_27248049_27248121 | 1 | 34 | 34 | GCTTCTGTAGTGTAGTGGTTATCACGTTCGCCTC | BT20 | 25 | 6 | 45 | 113 | 50 | 31.4 | 1.0 | 32.5 | 177.4 | 72.0 |
| >trna152_ValCAC_6_-_27248049_27248121@1.34.34\|GCTTCTGTAGTGTAGTGGTTATCACGTTCGCCTC | >trna152_ValCAC_6_-_27248049_27248121 | 1 | 34 | 34 | GCTTCTGTAGTGTAGTGGTTATCACGTTCGCCTC | MB231 | 144 | 260 | 604 | 119 | 84 | 150.4 | 352.2 | 608.1 | 112.3 | 47.0 |
| >trna152_ValCAC_6_-_27248049_27248121@1.36.36\|GCTTCTGTAGTGTAGTGGTTATCACGTTCGCCTCAC | >trna152_ValCAC_6_-_27248049_27248121 | 1 | 36 | 36 | GCTTCTGTAGTGTAGTGGTTATCACGTTCGCCTCAC | MB231 | 21 | 36 | 92 | 13 | 44 | 10.4 | 33.9 | 70.9 | 10.3 | 7.8 |
| >trna16_GlnTTG_17_+_47269890_47269961@1.29.29,trna64_GlnTTG_6_+_28557156_28557227@1.29.29,trna84_GlnTTG_6_+_145503859_145503930@1.29.29\|GGTCCCATGGTGTAATGGTTAGCACTCTG | >trna16_GlnTTG_17_+_47269890_47269961 | 1 | 29 | 29 | GGTCCCATGGTGTAATGGTTAGCACTCTG | MB468 | 266 | 73 | 423 | 62 | 49 | 316.2 | 32.2 | 463.6 | 43.5 | 58.5 |
| >trna16_TyrGTA_14_-_21131351_21131444@59.76.18,trna17_TyrGTA_14_-_21128117_21128210@59.76.18,trna19_TyrGTA_14_-_21121258_21121351@59.76.18,trna2_TyrGTA_2_+_27273650_27273738@59.76.18,trna4_TyrGTA_8_+_67025602_67025694@59.76.18,trna5_TyrGTA_14_+_21151432_21151520@59.76.18,trna5_TyrGTA_8_+_67026223_67026311@59.76.18\|TTCCGGCTCGAAGGACCA | >trna16_TyrGTA_14_-_21131351_21131444 | 59 | 76 | 18 | TTCCGGCTCGAAGGACCA | BT20 | 138 | 59 | 159 | 110 | 57 | 189.6 | 76.8 | 77.0 | 157.5 | 70.6 |
| >trna16_TyrGTA_14_-_21131351_21131444@59.76.18,trna17_TyrGTA_14_-_21128117_21128210@59.76.18,trna19_TyrGTA_14_-_21121258_21121351@59.76.18,trna2_TyrGTA_2_+_27273650_27273738@59.76.18,trna4_TyrGTA_8_+_67025602_67025694@59.76.18,trna5_TyrGTA_14_+_21151432_21151520@59.76.18,trna5_TyrGTA_8_+_67026223_67026311@59.76.18\|TTCCGGCTCGAAGGACCA | >trna16_TyrGTA_14_-_21131351_21131444 | 59 | 76 | 18 | TTCCGGCTCGAAGGACCA | MB231 | 35 | 23 | 32 | 37 | 23 | 18.9 | 8.6 | 15.1 | 14.8 | 11.3 |
| >trna16_ValTAC_11_-_59318460_59318532@1.30.30\|GGTTCCATAGTGTAGCGGTTATCACGTCTG | >trna16_ValTAC_11_-_59318460_59318532 | 1 | 30 | 30 | GGTTCCATAGTGTAGCGGTTATCACGTCTG | MB468 | 15 | 7 | 25 | 2 | 2 | 14.7 | 4.8 | 9.3 | 1.3 | 1.9 |
| >trna16_ValTAC_11_-_59318460_59318532@1.34.34\|GGTTCCATAGTGTAGCGGTTATCACGTCTGCTTT | >trna16_ValTAC_11_-_59318460_59318532 | 1 | 34 | 34 | GGTTCCATAGTGTAGCGGTTATCACGTCTGCTTT | MB231 | 12 | 9 | 15 | 24 | 21 | 12.2 | 9.9 | 15.1 | 25.0 | 18.3 |
| >trna17_ValTAC_11_-_59318102_59318174@1.30.30,trna4_ValTAC_X_-_18693029_18693101@1.30.30\|GGTTCCATAGTGTAGTGGTTATCACGTCTG | >trna17_ValTAC_11_-_59318102_59318174 | 1 | 30 | 30 | GGTTCCATAGTGTAGTGGTTATCACGTCTG | MB468 | 43 | 16 | 68 | 6 | 6 | 44.2 | 9.3 | 40.8 | 3.9 | 5.2 |
| >trna17_ValTAC_11_-_59318102_59318174@1.34.34,trna4_ValTAC_X_-_18693029_18693101@1.34.34\|GGTTCCATAGTGTAGTGGTTATCACGTCTGCTTT | >trna17_ValTAC_11_-_59318102_59318174 | 1 | 34 | 34 | GGTTCCATAGTGTAGTGGTTATCACGTCTGCTTT | BT20 | 13 | 3 | 7 | 27 | 64 | 13.2 | 3.2 | 5.8 | 45.9 | 109.8 |
| >trna17_ValTAC_11_-_59318102_59318174@1.34.34,trna4_ValTAC_X_-_18693029_18693101@1.34.34\|GGTTCCATAGTGTAGTGGTTATCACGTCTGCTTT | >trna17_ValTAC_11_-_59318102_59318174 | 1 | 34 | 34 | GGTTCCATAGTGTAGTGGTTATCACGTCTGCTTT | MB231 | 11 | 11 | 18 | 16 | 22 | 10.1 | 11.4 | 17.3 | 16.7 | 17.0 |
| >trna2_ArgCCT_16_+_3202901_3202973@13.30.18,trna3_ArgCCT_7_+_139025446_139025518@13.30.18\|TAATGGATAAGGCATTGG | >trna2_ArgCCT_16_+_3202901_3202973 | 13 | 30 | 18 | TAATGGATAAGGCATTGG | MB468 | 15 | 7 | 8 | 2 | 1 | 9.2 | 4.9 | 2.2 | 0.6 | 0.3 |
| >trna2_GlyGCC_21_-_18827107_18827177@1.28.28,trna35_GlyGCC_1_+_161413094_161413164@1.28.28,trna37_GlyGCC_1_+_161420467_161420537@1.28.28,trna39_GlyGCC_1_+_161427898_161427968@1.28.28,trna41_GlyGCC_1_+_161435258_161435328@1.28.28\|GCATGGGTGGTTCAGTGGTAGAATTCTC | >trna2_GlyGCC_21_-_18827107_18827177 | 1 | 28 | 28 | GCATGGGTGGTTCAGTGGTAGAATTCTC | BT20 | 75 | 55 | 344 | 307 | 123 | 88.2 | 84.0 | 342.9 | 58.9 | 168.5 |
| >trna2_GlyGCC_21_-_18827107_18827177@1.28.28,trna35_GlyGCC_1_+_161413094_161413164@1.28.28,trna37_GlyGCC_1_+_161420467_161420537@1.28.28,trna39_GlyGCC_1_+_161427898_161427968@1.28.28,trna41_GlyGCC_1_+_161435258_161435328@1.28.28\|GCATGGGTGGTTCAGTGGTAGAATTCTC | >trna2_GlyGCC_21_-_18827107_18827177 | 1 | 28 | 28 | GCATGGGTGGTTCAGTGGTAGAATTCTC | MB468 | 1265 | 1607 | 3703 | 1121 | 222 | 487.1 | 1181.2 | 1634.1 | 718.2 | 168.2 |
| >trna2_GlyGCC_21_-_18827107_18827177@1.29.29,trna35_GlyGCC_1_+_161413094_161413164@1.29.29,trna37_GlyGCC_1_+_161420467_161420537@1.29.29,trna39_GlyGCC_1_+_161427898_161427968@1.29.29,trna41_GlyGCC_1_+_161435258_161435328@1.29.29\|GCATGGGTGGTTCAGTGGTAGAATTCTCG | >trna2_GlyGCC_21_-_18827107_18827177 | 1 | 29 | 29 | GCATGGGTGGTTCAGTGGTAGAATTCTCG | BT20 | 19 | 12 | 100 | 56 | 19 | 14.1 | 15.8 | 66.1 | 24.7 | 18.0 |
| >trna2_GlyGCC_21_-_18827107_18827177@1.29.29,trna35_GlyGCC_1_+_161413094_161413164@1.29.29,trna37_GlyGCC_1_+_161420467_161420537@1.29.29,trna39_GlyGCC_1_+_161427898_161427968@1.29.29,trna41_GlyGCC_1_+_161435258_161435328@1.29.29\|GCATGGGTGGTTCAGTGGTAGAATTCTCG | >trna2_GlyGCC_21_-_18827107_18827177 | 1 | 29 | 29 | GCATGGGTGGTTCAGTGGTAGAATTCTCG | MB468 | 1350 | 1105 | 3570 | 851 | 227 | 582.2 | 361.7 | 1540.3 | 627.5 | 185.1 |
| >trna2_GlyGCC_21_-_18827107_18827177@1.30.30,trna35_GlyGCC_1_+_161413094_161413164@1.30.30,trna37_GlyGCC_1_+_161420467_161420537@1.30.30,trna39_GlyGCC_1_+_161427898_161427968@1.30.30,trna41_GlyGCC_1_+_161435258_161435328@1.30.30\|GCATGGGTGGTTCAGTGGTAGAATTCTCGC | >trna2_GlyGCC_21_-_18827107_18827177 | 1 | 30 | 30 | GCATGGGTGGTTCAGTGGTAGAATTCTCGC | BT20 | 44 | 13 | 160 | 255 | 44 | 33.8 | 6.8 | 87.3 | 170.2 | 51.0 |
| >trna2_GlyGCC_21_-_18827107_18827177@1.30.30,trna35_GlyGCC_1_+_161413094_161413164@1.30.30,trna37_GlyGCC_1_+_161420467_161420537@1.30.30,trna39_GlyGCC_1_+_161427898_161427968@1.30.30,trna41_GlyGCC_1_+_161435258_161435328@1.30.30\|GCATGGGTGGTTCAGTGGTAGAATTCTCGC | >trna2_GlyGCC_21_-_18827107_18827177 | 1 | 30 | 30 | GCATGGGTGGTTCAGTGGTAGAATTCTCGC | MB468 | 480 | 279 | 1335 | 432 | 75 | 241.2 | 57.5 | 742.7 | 303.6 | 63.5 |

| | | | | | | | | | | | | | | | |
|---|---|---|---|---|---|---|---|---|---|---|---|---|---|---|---|
| >trna2_GlyGCC_21_-_18827107_18827177@1.31.31,trna35_GlyGCC_1_+_161413094_161413164@1.31.31,trna37_GlyGCC_1_+_161420467_161420537@1.31.31,trna39_GlyGCC_1_+_161427898_161427968@1.31.31,trna41_GlyGCC_1_+_161435258_161435328@1.31.31\|GCATGGGTGGTTCAGTGGTAGAATTCTCGCC | >trna2_GlyGCC_21_-_18827107_18827177 | 1 | 31 | 31 | GCATGGGTGGTTCAGTGGTAGAATTCTCGCC | BT20 | 1948 | 247 | 2550 | 4913 | 1911 | 1892.7 | 90.8 | 989.6 | 6414.3 | 2615.6 |
| >trna2_GlyGCC_21_-_18827107_18827177@1.31.31,trna35_GlyGCC_1_+_161413094_161413164@1.31.31,trna37_GlyGCC_1_+_161420467_161420537@1.31.31,trna39_GlyGCC_1_+_161427898_161427968@1.31.31,trna41_GlyGCC_1_+_161435258_161435328@1.31.31\|GCATGGGTGGTTCAGTGGTAGAATTCTCGCC | >trna2_GlyGCC_21_-_18827107_18827177 | 1 | 31 | 31 | GCATGGGTGGTTCAGTGGTAGAATTCTCGCC | MB231 | 24 | 9 | 51 | 60 | 21 | 16.5 | 9.4 | 36.2 | 40.5 | 18.4 |
| >trna2_GlyGCC_21_-_18827107_18827177@1.31.31,trna35_GlyGCC_1_+_161413094_161413164@1.31.31,trna37_GlyGCC_1_+_161420467_161420537@1.31.31,trna39_GlyGCC_1_+_161427898_161427968@1.31.31,trna41_GlyGCC_1_+_161435258_161435328@1.31.31\|GCATGGGTGGTTCAGTGGTAGAATTCTCGCC | >trna2_GlyGCC_21_-_18827107_18827177 | 1 | 31 | 31 | GCATGGGTGGTTCAGTGGTAGAATTCTCGCC | MB468 | 25 | 18 | 80 | 25 | 9 | 11.3 | 14.9 | 19.5 | 13.6 | 5.3 |
| >trna2_GlyGCC_21_-_18827107_18827177@1.32.32,trna35_GlyGCC_1_+_161413094_161413164@1.32.32,trna37_GlyGCC_1_+_161420467_161420537@1.32.32,trna39_GlyGCC_1_+_161427898_161427968@1.32.32,trna41_GlyGCC_1_+_161435258_161435328@1.32.32\|GCATGGGTGGTTCAGTGGTAGAATTCTCGCCT | >trna2_GlyGCC_21_-_18827107_18827177 | 1 | 32 | 32 | GCATGGGTGGTTCAGTGGTAGAATTCTCGCCT | BT20 | 432 | 89 | 484 | 2327 | 682 | 167.5 | 39.4 | 218.0 | 3695.3 | 1087.0 |
| >trna2_GlyGCC_21_-_18827107_18827177@1.32.32,trna35_GlyGCC_1_+_161413094_161413164@1.32.32,trna37_GlyGCC_1_+_161420467_161420537@1.32.32,trna39_GlyGCC_1_+_161427898_161427968@1.32.32,trna41_GlyGCC_1_+_161435258_161435328@1.32.32\|GCATGGGTGGTTCAGTGGTAGAATTCTCGCCT | >trna2_GlyGCC_21_-_18827107_18827177 | 1 | 32 | 32 | GCATGGGTGGTTCAGTGGTAGAATTCTCGCCT | MB231 | 15 | 5 | 20 | 34 | 13 | 9.2 | 5.3 | 20.4 | 18.5 | 8.0 |
| >trna2_GlyGCC_21_-_18827107_18827177@1.33.33,trna35_GlyGCC_1_+_161413094_161413164@1.33.33,trna37_GlyGCC_1_+_161420467_161420537@1.33.33,trna39_GlyGCC_1_+_161427898_161427968@1.33.33,trna41_GlyGCC_1_+_161435258_161435328@1.33.33\|GCATGGGTGGTTCAGTGGTAGAATTCTCGCCTG | >trna2_GlyGCC_21_-_18827107_18827177 | 1 | 33 | 33 | GCATGGGTGGTTCAGTGGTAGAATTCTCGCCTG | BT20 | 61 | 16 | 85 | 110 | 35 | 43.8 | 17.2 | 29.9 | 97.0 | 30.3 |
| >trna2_GlyGCC_21_-_18827107_18827177@1.35.35,trna35_GlyGCC_1_+_161413094_161413164@1.35.35,trna37_GlyGCC_1_+_161420467_161420537@1.35.35,trna39_GlyGCC_1_+_161427898_161427968@1.35.35,trna41_GlyGCC_1_+_161435258_161435328@1.35.35\|GCATGGGTGGTTCAGTGGTAGAATTCTCGCCTGCC | >trna2_GlyGCC_21_-_18827107_18827177 | 1 | 35 | 35 | GCATGGGTGGTTCAGTGGTAGAATTCTCGCCTGCC | BT20 | 21 | 1 | 16 | 39 | 3 | 30.2 | 0.5 | 8.0 | 36.4 | 3.2 |
| >trna2_LeuTAA_4_-_156384978_156385052@1.18.18\|GTTAAGATGGCAGAGCCT | >trna2_LeuTAA_4_-_156384978_156385052 | 1 | 18 | 18 | GTTAAGATGGCAGAGCCT | BT20 | 11 | 9 | 9 | 9 | 4 | 7.7 | 7.1 | 6.4 | 11.9 | 3.8 |
| >trna20_GluTTC_2_-_131094701_131094772@1.29.29,trna5_GluTTC_13_-_41634874_41634945@1.29.29\|TCCCATATGGTCTAGCGGTTAGGATTCCT | >trna20_GluTTC_2_-_131094701_131094772 | 1 | 29 | 29 | TCCCATATGGTCTAGCGGTTAGGATTCCT | MB468 | 34 | 7 | 27 | 11 | 13 | 18.5 | 3.7 | 11.2 | 9.9 | 3.9 |
| >trna20_GluTTC_2_-_131094701_131094772@1.30.30,trna5_GluTTC_13_-_41634874_41634945@1.30.30\|TCCCATATGGTCTAGCGGTTAGGATTCCTG | >trna20_GluTTC_2_-_131094701_131094772 | 1 | 30 | 30 | TCCCATATGGTCTAGCGGTTAGGATTCCTG | MB468 | 28 | 6 | 27 | 32 | 19 | 7.2 | 2.3 | 13.4 | 23.5 | 4.3 |
| >trna20_GluTTC_2_-_131094701_131094772@1.31.31,trna5_GluTTC_13_-_41634874_41634945@1.31.31\|TCCCATATGGTCTAGCGGTTAGGATTCCTGG | >trna20_GluTTC_2_-_131094701_131094772 | 1 | 31 | 31 | TCCCATATGGTCTAGCGGTTAGGATTCCTGG | MB231 | 17 | 6 | 13 | 35 | 19 | 10.8 | 5.5 | 12.2 | 23.0 | 11.1 |
| >trna20_GluTTC_2_-_131094701_131094772@1.31.31,trna5_GluTTC_13_-_41634874_41634945@1.31.31\|TCCCATATGGTCTAGCGGTTAGGATTCCTGG | >trna20_GluTTC_2_-_131094701_131094772 | 1 | 31 | 31 | TCCCATATGGTCTAGCGGTTAGGATTCCTGG | MB468 | 28 | 7 | 38 | 13 | 9 | 3.2 | 2.2 | 16.0 | 7.8 | 6.2 |
| >trna20_GluTTC_2_-_131094701_131094772@1.32.32,trna5_GluTTC_13_-_41634874_41634945@1.32.32\|TCCCATATGGTCTAGCGGTTAGGATTCCTGGT | >trna20_GluTTC_2_-_131094701_131094772 | 1 | 32 | 32 | TCCCATATGGTCTAGCGGTTAGGATTCCTGGT | MB231 | 12 | 3 | 13 | 30 | 19 | 6.0 | 3.2 | 11.4 | 12.8 | 13.1 |
| >trna20_GluTTC_2_-_131094701_131094772@1.33.33,trna5_GluTTC_13_-_41634874_41634945@1.33.33\|TCCCATATGGTCTAGCGGTTAGGATTCCTGGTT | >trna20_GluTTC_2_-_131094701_131094772 | 1 | 33 | 33 | TCCCATATGGTCTAGCGGTTAGGATTCCTGGTT | BT20 | 29 | 5 | 43 | 29 | 12 | 18.5 | 0.7 | 12.0 | 43.5 | 18.4 |
| >trna20_GluTTC_2_-_131094701_131094772@1.33.33,trna5_GluTTC_13_-_41634874_41634945@1.33.33\|TCCCATATGGTCTAGCGGTTAGGATTCCTGGTT | >trna20_GluTTC_2_-_131094701_131094772 | 1 | 33 | 33 | TCCCATATGGTCTAGCGGTTAGGATTCCTGGTT | MB231 | 120 | 37 | 126 | 182 | 138 | 43.5 | 34.6 | 64.6 | 95.5 | 89.3 |
| >trna20_GluTTC_2_-_131094701_131094772@1.36.36,trna5_GluTTC_13_-_41634874_41634945@1.36.36\|TCCCATATGGTCTAGCGGTTAGGATTCCTGGTTTTC | >trna20_GluTTC_2_-_131094701_131094772 | 1 | 36 | 36 | TCCCATATGGTCTAGCGGTTAGGATTCCTGGTTTTC | MB231 | 15 | 6 | 41 | 7 | 25 | 11.0 | 6.3 | 44.5 | 5.9 | 13.2 |
| >trna23_ArgCCG_17_-_66016013_66016085@14.31.18\|AATGGATAAGGCATCAGC | >trna23_ArgCCG_17_-_66016013_66016085 | 14 | 31 | 18 | AATGGATAAGGCATCAGC | MB468 | 20 | 9 | 10 | 3 | 1 | 9.7 | 6.3 | 5.5 | 1.2 | 0.4 |
| >trna27_GlyCCC_2_-_70476123_70476193@1.32.32,trna34_GlyCCC_16_-_686736_686806@1.32.32\|GCGCCGCTGGTGTAGTGGTATCATGCAAGATT | >trna27_GlyCCC_2_-_70476123_70476193 | 1 | 32 | 32 | GCGCCGCTGGTGTAGTGGTATCATGCAAGATT | MB231 | 10 | 5 | 21 | 11 | 10 | 9.7 | 6.0 | 27.2 | 4.7 | 11.1 |
| >trna32_LysCTT_16_-_3207406_3207478@1.33.33\|GCCCGGCTAGCTCAGTCGGTAGAGCATGAGACC | >trna32_LysCTT_16_-_3207406_3207478 | 1 | 33 | 33 | GCCCGGCTAGCTCAGTCGGTAGAGCATGAGACC | BT20 | 11 | 3 | 32 | 15 | 17 | 4.7 | 0.9 | 17.3 | 23.6 | 25.8 |
| >trna32_LysCTT_16_-_3207406_3207478@1.33.33\|GCCCGGCTAGCTCAGTCGGTAGAGCATGAGACC | >trna32_LysCTT_16_-_3207406_3207478 | 1 | 33 | 33 | GCCCGGCTAGCTCAGTCGGTAGAGCATGAGACC | MB231 | 89 | 75 | 208 | 46 | 74 | 85.4 | 99.5 | 229.2 | 37.6 | 55.7 |
| >trna32_LysCTT_16_-_3207406_3207478@1.35.35\|GCCCGGCTAGCTCAGTCGGTAGAGCATGAGACCCT | >trna32_LysCTT_16_-_3207406_3207478 | 1 | 35 | 35 | GCCCGGCTAGCTCAGTCGGTAGAGCATGAGACCCT | BT20 | 10 | 0 | 16 | 0 | 0 | 7.8 | 0.8 | 2.8 | 0.0 | 0.0 |
| >trna37_ValAAC_6_+_27203288_27203360@1.29.29\|GTTTCCGTAGTGTAGTGGTTATCACGTTT | >trna37_ValAAC_6_+_27203288_27203360 | 1 | 29 | 29 | GTTTCCGTAGTGTAGTGGTTATCACGTTT | MB468 | 127 | 18 | 107 | 73 | 28 | 91.9 | 8.0 | 90.4 | 27.1 | 21.3 |
| >trna5_AspGTC_12_+_98897281_98897352@35.75.41\|TCACGCGGGAGACCGGGGTTCAATTCCCCGACGGGGAGCCA | >trna5_AspGTC_12_+_98897281_98897352 | 35 | 75 | 41 | TCACGCGGGAGACCGGGGTTCAATTCCCCGACGGGGAGCCA | MB231 | 15 | 16 | 298 | 7 | 4 | 13.6 | 14.2 | 348.1 | 9.3 | 5.7 |
| >trna59_GluCTC_1_+_249168447_249168518@32.70.39\|CTCTCACCGCCGCGGCCCGGGTTCGATTCCCGGTCAGGA | >trna59_GluCTC_1_+_249168447_249168518 | 32 | 70 | 39 | CTCTCACCGCCGCGGCCCGGGTTCGATTCCCGGTCAGGA | MB468 | 11 | 29 | 48 | 1 | 5 | 5.7 | 10.5 | 20.2 | 0.8 | 6.9 |
| >trna6_TrpCCA_12_+_98898030_98898101@30.52.23\|ACTCCAGATCAGAAGGCTGCGTG | >trna6_TrpCCA_12_+_98898030_98898101 | 30 | 52 | 23 | ACTCCAGATCAGAAGGCTGCGTG | MB468 | 33 | 31 | 10 | 9 | 11 | 12.3 | 21.5 | 2.9 | 4.0 | 2.1 |
| >trna9_ArgTCT_1_+_94313129_94313213@1.29.29\|GGCTCCGTGGCGCAATGGATAGCGCATTG | >trna9_ArgTCT_1_+_94313129_94313213 | 1 | 29 | 29 | GGCTCCGTGGCGCAATGGATAGCGCATTG | MB468 | 17 | 2 | 29 | 5 | 3 | 12.0 | 0.6 | 12.6 | 1.6 | 2.4 |
| >trnaMT_CysGCA_MT_-_5761_5826@47.69.23\|CTTCAAACCTGCCGGGGCTTCCA | >trnaMT_CysGCA_MT_-_5761_5826 | 47 | 69 | 23 | CTTCAAACCTGCCGGGGCTTCCA | MB468 | 19 | 18 | 14 | 4 | 25 | 8.4 | 11.0 | 13.7 | 1.7 | 14.2 |
| >trnaMT_GlnTTG_MT_-_4329_4400@57.75.19,trnalookalike2_GlnTTG_1_-_564879_564950@57.75.19\|ATTCTCATAGTCCTAGCCA | >trnaMT_GlnTTG_MT_-_4329_4400 | 57 | 75 | 19 | ATTCTCATAGTCCTAGCCA | MB231 | 11 | 3 | 2 | 21 | 15 | 10.7 | 1.9 | 0.8 | 21.9 | 10.7 |
| >trnaMT_GluTTC_MT_-_14674_14742@25.72.48,trnalookalike8_GluTTC_5_-_93905172_93905240@25.72.48\|TGGTTTTTCATATCATTGGTCGTGGTTGTAGTCCGTGCGAGAATACCA | >trnaMT_GluTTC_MT_-_14674_14742 | 25 | 72 | 48 | TGGTTTTTCATATCATTGGTCGTGGTTGTAGTCCGTGCGAGAATACCA | MB231 | 36 | 18 | 18 | 6 | 28 | 35.0 | 24.4 | 12.5 | 10.3 | 28.2 |
| >trnaMT_GluTTC_MT_-_14674_14742@26.72.47,trnalookalike8_GluTTC_5_-_93905172_93905240@26.72.47\|GGTTTTTCATATCATTGGTCGTGGTTGTAGTCCGTGCGAGAATACCA | >trnaMT_GluTTC_MT_-_14674_14742 | 26 | 72 | 47 | GGTTTTTCATATCATTGGTCGTGGTTGTAGTCCGTGCGAGAATACCA | BT20 | 11 | 1 | 1 | 16 | 3 | 17.7 | 1.3 | 1.3 | 17.0 | 3.5 |
| >trnaMT_GluTTC_MT_-_14674_14742@26.72.47,trnalookalike8_GluTTC_5_-_93905172_93905240@26.72.47\|GGTTTTTCATATCATTGGTCGTGGTTGTAGTCCGTGCGAGAATACCA | >trnaMT_GluTTC_MT_-_14674_14742 | 26 | 72 | 47 | GGTTTTTCATATCATTGGTCGTGGTTGTAGTCCGTGCGAGAATACCA | MB231 | 12 | 5 | 3 | 17 | 30 | 11.2 | 8.8 | 1.5 | 28.4 | 33.2 |
| >trnaMT_GluTTC_MT_-_14674_14742@31.72.42,trnalookalike8_GluTTC_5_-_93905172_93905240@31.72.42\|TTCATATCATTGGTCGTGGTTGTAGTCCGTGCGAGAATACCA | >trnaMT_GluTTC_MT_-_14674_14742 | 31 | 72 | 42 | TTCATATCATTGGTCGTGGTTGTAGTCCGTGCGAGAATACCA | MB231 | 41 | 61 | 96 | 86 | 130 | 18.4 | 53.4 | 60.6 | 112.9 | 142.7 |
| >trnaMT_GluTTC_MT_-_14674_14742@32.72.41,trnalookalike8_GluTTC_5_-_93905172_93905240@32.72.41\|TCATATCATTGGTCGTGGTTGTAGTCCGTGCGAGAATACCA | >trnaMT_GluTTC_MT_-_14674_14742 | 32 | 72 | 41 | TCATATCATTGGTCGTGGTTGTAGTCCGTGCGAGAATACCA | BT20 | 24 | 6 | 8 | 46 | 39 | 16.1 | 1.0 | 4.8 | 45.6 | 19.6 |
| >trnaMT_GluTTC_MT_-_14674_14742@32.72.41,trnalookalike8_GluTTC_5_-_93905172_93905240@32.72.41\|TCATATCATTGGTCGTGGTTGTAGTCCGTGCGAGAATACCA | >trnaMT_GluTTC_MT_-_14674_14742 | 32 | 72 | 41 | TCATATCATTGGTCGTGGTTGTAGTCCGTGCGAGAATACCA | MB231 | 28 | 28 | 37 | 56 | 53 | 9.4 | 14.3 | 20.8 | 55.6 | 34.4 |
| >trnaMT_GluTTC_MT_-_14674_14742@33.72.40,trnalookalike8_GluTTC_5_-_93905172_93905240@33.72.40\|CATATCATTGGTCGTGGTTGTAGTCCGTGCGAGAATACCA | >trnaMT_GluTTC_MT_-_14674_14742 | 33 | 72 | 40 | CATATCATTGGTCGTGGTTGTAGTCCGTGCGAGAATACCA | BT20 | 22 | 6 | 13 | 46 | 41 | 14.2 | 2.5 | 13.9 | 48.2 | 24.8 |
| >trnaMT_GluTTC_MT_-_14674_14742@33.72.40,trnalookalike8_GluTTC_5_-_93905172_93905240@33.72.40\|CATATCATTGGTCGTGGTTGTAGTCCGTGCGAGAATACCA | >trnaMT_GluTTC_MT_-_14674_14742 | 33 | 72 | 40 | CATATCATTGGTCGTGGTTGTAGTCCGTGCGAGAATACCA | MB231 | 39 | 23 | 30 | 55 | 52 | 19.0 | 11.1 | 12.7 | 59.8 | 38.5 |
| >trnaMT_GluTTC_MT_-_14674_14742@34.71.38,trnalookalike8_GluTTC_5_-_93905172_93905240@34.71.38\|ATATCATTGGTCGTGGTTGTAGTCCGTGCGAGAATACC | >trnaMT_GluTTC_MT_-_14674_14742 | 34 | 71 | 38 | ATATCATTGGTCGTGGTTGTAGTCCGTGCGAGAATACC | MB231 | 33 | 21 | 29 | 100 | 51 | 15.9 | 11.5 | 27.7 | 120.1 | 24.9 |
| >trnaMT_GluTTC_MT_-_14674_14742@34.72.39,trnalookalike8_GluTTC_5_-_93905172_93905240@34.72.39\|ATATCATTGGTCGTGGTTGTAGTCCGTGCGAGAATACCA | >trnaMT_GluTTC_MT_-_14674_14742 | 34 | 72 | 39 | ATATCATTGGTCGTGGTTGTAGTCCGTGCGAGAATACCA | BT20 | 106 | 23 | 21 | 243 | 204 | 68.3 | 12.2 | 5.6 | 269.2 | 141.5 |
| >trnaMT_GluTTC_MT_-_14674_14742@34.72.39,trnalookalike8_GluTTC_5_-_93905172_93905240@34.72.39\|ATATCATTGGTCGTGGTTGTAGTCCGTGCGAGAATACCA | >trnaMT_GluTTC_MT_-_14674_14742 | 34 | 72 | 39 | ATATCATTGGTCGTGGTTGTAGTCCGTGCGAGAATACCA | MB231 | 196 | 150 | 121 | 557 | 345 | 67.5 | 72.3 | 118.3 | 684.7 | 238.0 |
| >trnaMT_GluTTC_MT_-_14674_14742@35.71.37,trnalookalike8_GluTTC_5_-_93905172_93905240@35.71.37\|TATCATTGGTCGTGGTTGTAGTCCGTGCGAGAATACC | >trnaMT_GluTTC_MT_-_14674_14742 | 35 | 71 | 37 | TATCATTGGTCGTGGTTGTAGTCCGTGCGAGAATACC | MB231 | 28 | 30 | 72 | 43 | 33 | 5.5 | 18.1 | 36.4 | 40.2 | 14.1 |

| | | | | | | | | | | | | | | | | |
|---|---|---|---|---|---|---|---|---|---|---|---|---|---|---|---|---|
| >trnaMT_GluTTC_MT_-_14674_14742@35.72.38,trnalookalike8_GluTTC_5_-_93905172_93905240@35.72.38\|TATCATTGGTCGTGGTTGTAGTCCGTGCGAGAATACCA | >trnaMT_GluTTC_MT_-_14674_14742 | 35 | 72 | 38 | TATCATTGGTCGTGGTTGTAGTCCGTGCGAGAATACCA | BT20 | 91 | 21 | 43 | 211 | 176 | 81.9 | 10.1 | 44.9 | 219.7 | 134.2 |
| >trnaMT_GluTTC_MT_-_14674_14742@35.72.38,trnalookalike8_GluTTC_5_-_93905172_93905240@35.72.38\|TATCATTGGTCGTGGTTGTAGTCCGTGCGAGAATACCA | >trnaMT_GluTTC_MT_-_14674_14742 | 35 | 72 | 38 | TATCATTGGTCGTGGTTGTAGTCCGTGCGAGAATACCA | MB231 | 237 | 367 | 488 | 334 | 276 | 111.1 | 244.2 | 336.6 | 302.8 | 177.7 |
| >trnaMT_GluTTC_MT_-_14674_14742@36.72.37,trnalookalike8_GluTTC_5_-_93905172_93905240@36.72.37\|ATCATTGGTCGTGGTTGTAGTCCGTGCGAGAATACCA | >trnaMT_GluTTC_MT_-_14674_14742 | 36 | 72 | 37 | ATCATTGGTCGTGGTTGTAGTCCGTGCGAGAATACCA | MB231 | 46 | 109 | 65 | 125 | 71 | 79.8 | 188.8 | 111.8 | 216.2 | 123.4 |
| >trnaMT_GluTTC_MT_-_14674_14742@37.71.35,trnalookalike8_GluTTC_5_-_93905172_93905240@37.71.35\|TCATTGGTCGTGGTTGTAGTCCGTGCGAGAATACC | >trnaMT_GluTTC_MT_-_14674_14742 | 37 | 71 | 35 | TCATTGGTCGTGGTTGTAGTCCGTGCGAGAATACC | MB231 | 17 | 47 | 48 | 17 | 14 | 5.0 | 45.2 | 28.5 | 15.7 | 7.4 |
| >trnaMT_GluTTC_MT_-_14674_14742@37.72.36,trnalookalike8_GluTTC_5_-_93905172_93905240@37.72.36\|TCATTGGTCGTGGTTGTAGTCCGTGCGAGAATACCA | >trnaMT_GluTTC_MT_-_14674_14742 | 37 | 72 | 36 | TCATTGGTCGTGGTTGTAGTCCGTGCGAGAATACCA | BT20 | 45 | 29 | 104 | 106 | 94 | 21.0 | 25.7 | 137.2 | 66.5 | 49.0 |
| >trnaMT_GluTTC_MT_-_14674_14742@37.72.36,trnalookalike8_GluTTC_5_-_93905172_93905240@37.72.36\|TCATTGGTCGTGGTTGTAGTCCGTGCGAGAATACCA | >trnaMT_GluTTC_MT_-_14674_14742 | 37 | 72 | 36 | TCATTGGTCGTGGTTGTAGTCCGTGCGAGAATACCA | MB231 | 135 | 481 | 291 | 127 | 119 | 45.5 | 397.3 | 196.5 | 107.5 | 72.9 |
| >trnaMT_GluTTC_MT_-_14674_14742@38.71.34,trnalookalike8_GluTTC_5_-_93905172_93905240@38.71.34\|CATTGGTCGTGGTTGTAGTCCGTGCGAGAATACC | >trnaMT_GluTTC_MT_-_14674_14742 | 38 | 71 | 34 | CATTGGTCGTGGTTGTAGTCCGTGCGAGAATACC | BT20 | 12 | 4 | 3 | 36 | 36 | 9.7 | 1.4 | 1.6 | 30.2 | 22.4 |
| >trnaMT_GluTTC_MT_-_14674_14742@38.71.34,trnalookalike8_GluTTC_5_-_93905172_93905240@38.71.34\|CATTGGTCGTGGTTGTAGTCCGTGCGAGAATACC | >trnaMT_GluTTC_MT_-_14674_14742 | 38 | 71 | 34 | CATTGGTCGTGGTTGTAGTCCGTGCGAGAATACC | MB231 | 22 | 22 | 15 | 39 | 33 | 7.4 | 10.5 | 5.9 | 27.1 | 16.3 |
| >trnaMT_GluTTC_MT_-_14674_14742@38.72.35,trnalookalike8_GluTTC_5_-_93905172_93905240@38.72.35\|CATTGGTCGTGGTTGTAGTCCGTGCGAGAATACCA | >trnaMT_GluTTC_MT_-_14674_14742 | 38 | 72 | 35 | CATTGGTCGTGGTTGTAGTCCGTGCGAGAATACCA | BT20 | 160 | 48 | 44 | 573 | 457 | 132.0 | 19.7 | 34.4 | 385.4 | 205.4 |
| >trnaMT_GluTTC_MT_-_14674_14742@38.72.35,trnalookalike8_GluTTC_5_-_93905172_93905240@38.72.35\|CATTGGTCGTGGTTGTAGTCCGTGCGAGAATACCA | >trnaMT_GluTTC_MT_-_14674_14742 | 38 | 72 | 35 | CATTGGTCGTGGTTGTAGTCCGTGCGAGAATACCA | MB231 | 257 | 276 | 129 | 474 | 370 | 113.5 | 108.7 | 43.5 | 334.3 | 222.1 |
| >trnaMT_GluTTC_MT_-_14674_14742@39.71.33,trnalookalike8_GluTTC_5_-_93905172_93905240@39.71.33\|ATTGGTCGTGGTTGTAGTCCGTGCGAGAATACC | >trnaMT_GluTTC_MT_-_14674_14742 | 39 | 71 | 33 | ATTGGTCGTGGTTGTAGTCCGTGCGAGAATACC | BT20 | 38 | 15 | 14 | 151 | 97 | 27.5 | 7.4 | 15.3 | 137.9 | 58.6 |
| >trnaMT_GluTTC_MT_-_14674_14742@39.71.33,trnalookalike8_GluTTC_5_-_93905172_93905240@39.71.33\|ATTGGTCGTGGTTGTAGTCCGTGCGAGAATACC | >trnaMT_GluTTC_MT_-_14674_14742 | 39 | 71 | 33 | ATTGGTCGTGGTTGTAGTCCGTGCGAGAATACC | MB231 | 89 | 132 | 98 | 189 | 113 | 28.7 | 116.2 | 110.4 | 96.5 | 58.8 |
| >trnaMT_GluTTC_MT_-_14674_14742@39.72.34,trnalookalike8_GluTTC_5_-_93905172_93905240@39.72.34\|ATTGGTCGTGGTTGTAGTCCGTGCGAGAATACCA | >trnaMT_GluTTC_MT_-_14674_14742 | 39 | 72 | 34 | ATTGGTCGTGGTTGTAGTCCGTGCGAGAATACCA | BT20 | 559 | 198 | 209 | 2092 | 1249 | 457.9 | 104.6 | 234.1 | 1699.7 | 731.1 |
| >trnaMT_GluTTC_MT_-_14674_14742@39.72.34,trnalookalike8_GluTTC_5_-_93905172_93905240@39.72.34\|ATTGGTCGTGGTTGTAGTCCGTGCGAGAATACCA | >trnaMT_GluTTC_MT_-_14674_14742 | 39 | 72 | 34 | ATTGGTCGTGGTTGTAGTCCGTGCGAGAATACCA | MB231 | 819 | 1291 | 528 | 2009 | 989 | 205.5 | 940.9 | 553.2 | 1316.9 | 369.5 |
| >trnaMT_GluTTC_MT_-_14674_14742@39.72.34,trnalookalike8_GluTTC_5_-_93905172_93905240@39.72.34\|ATTGGTCGTGGTTGTAGTCCGTGCGAGAATACCA | >trnaMT_GluTTC_MT_-_14674_14742 | 39 | 72 | 34 | ATTGGTCGTGGTTGTAGTCCGTGCGAGAATACCA | MB468 | 11 | 43 | 5 | 145 | 46 | 10.0 | 25.3 | 4.9 | 95.3 | 26.9 |
| >trnaMT_GluTTC_MT_-_14674_14742@40.71.32,trnalookalike8_GluTTC_5_-_93905172_93905240@40.71.32\|TTGGTCGTGGTTGTAGTCCGTGCGAGAATACC | >trnaMT_GluTTC_MT_-_14674_14742 | 40 | 71 | 32 | TTGGTCGTGGTTGTAGTCCGTGCGAGAATACC | BT20 | 24 | 11 | 6 | 91 | 67 | 11.7 | 5.9 | 4.4 | 58.0 | 28.2 |
| >trnaMT_GluTTC_MT_-_14674_14742@40.71.32,trnalookalike8_GluTTC_5_-_93905172_93905240@40.71.32\|TTGGTCGTGGTTGTAGTCCGTGCGAGAATACC | >trnaMT_GluTTC_MT_-_14674_14742 | 40 | 71 | 32 | TTGGTCGTGGTTGTAGTCCGTGCGAGAATACC | MB231 | 81 | 81 | 72 | 143 | 88 | 41.9 | 19.6 | 39.8 | 88.2 | 52.3 |
| >trnaMT_GluTTC_MT_-_14674_14742@40.72.33,trnalookalike8_GluTTC_5_-_93905172_93905240@40.72.33\|TTGGTCGTGGTTGTAGTCCGTGCGAGAATACCA | >trnaMT_GluTTC_MT_-_14674_14742 | 40 | 72 | 33 | TTGGTCGTGGTTGTAGTCCGTGCGAGAATACCA | BT20 | 456 | 146 | 99 | 1776 | 1148 | 328.7 | 86.5 | 88.6 | 1168.4 | 438.2 |
| >trnaMT_GluTTC_MT_-_14674_14742@40.72.33,trnalookalike8_GluTTC_5_-_93905172_93905240@40.72.33\|TTGGTCGTGGTTGTAGTCCGTGCGAGAATACCA | >trnaMT_GluTTC_MT_-_14674_14742 | 40 | 72 | 33 | TTGGTCGTGGTTGTAGTCCGTGCGAGAATACCA | MB231 | 918 | 1134 | 724 | 1894 | 985 | 443.3 | 473.2 | 526.9 | 1209.1 | 545.5 |
| >trnaMT_GluTTC_MT_-_14674_14742@40.72.33,trnalookalike8_GluTTC_5_-_93905172_93905240@40.72.33\|TTGGTCGTGGTTGTAGTCCGTGCGAGAATACCA | >trnaMT_GluTTC_MT_-_14674_14742 | 40 | 72 | 33 | TTGGTCGTGGTTGTAGTCCGTGCGAGAATACCA | MB468 | 21 | 146 | 10 | 172 | 55 | 10.2 | 136.2 | 7.6 | 111.5 | 29.4 |
| >trnaMT_GluTTC_MT_-_14674_14742@41.71.31,trnalookalike8_GluTTC_5_-_93905172_93905240@41.71.31\|TGGTCGTGGTTGTAGTCCGTGCGAGAATACC | >trnaMT_GluTTC_MT_-_14674_14742 | 41 | 71 | 31 | TGGTCGTGGTTGTAGTCCGTGCGAGAATACC | MB231 | 20 | 28 | 32 | 32 | 21 | 8.3 | 12.6 | 21.1 | 21.6 | 11.3 |
| >trnaMT_GluTTC_MT_-_14674_14742@41.72.32,trnalookalike8_GluTTC_5_-_93905172_93905240@41.72.32\|TGGTCGTGGTTGTAGTCCGTGCGAGAATACCA | >trnaMT_GluTTC_MT_-_14674_14742 | 41 | 72 | 32 | TGGTCGTGGTTGTAGTCCGTGCGAGAATACCA | BT20 | 97 | 42 | 41 | 536 | 346 | 84.0 | 20.1 | 42.8 | 295.2 | 95.0 |
| >trnaMT_GluTTC_MT_-_14674_14742@41.72.32,trnalookalike8_GluTTC_5_-_93905172_93905240@41.72.32\|TGGTCGTGGTTGTAGTCCGTGCGAGAATACCA | >trnaMT_GluTTC_MT_-_14674_14742 | 41 | 72 | 32 | TGGTCGTGGTTGTAGTCCGTGCGAGAATACCA | MB231 | 260 | 469 | 426 | 469 | 256 | 159.4 | 255.9 | 385.7 | 346.7 | 146.2 |
| >trnaMT_GluTTC_MT_-_14674_14742@44.72.29,trnalookalike8_GluTTC_5_-_93905172_93905240@44.72.29\|TCGTGGTTGTAGTCCGTGCGAGAATACCA | >trnaMT_GluTTC_MT_-_14674_14742 | 44 | 72 | 29 | TCGTGGTTGTAGTCCGTGCGAGAATACCA | MB231 | 32 | 52 | 53 | 232 | 137 | 10.7 | 32.7 | 51.8 | 265.8 | 102.5 |
| >trnaMT_GluTTC_MT_-_14674_14742@45.72.28,trnalookalike8_GluTTC_5_-_93905172_93905240@45.72.28\|CGTGGTTGTAGTCCGTGCGAGAATACCA | >trnaMT_GluTTC_MT_-_14674_14742 | 45 | 72 | 28 | CGTGGTTGTAGTCCGTGCGAGAATACCA | BT20 | 16 | 7 | 12 | 38 | 36 | 3.9 | 2.4 | 14.4 | 26.3 | 28.5 |
| >trnaMT_GluTTC_MT_-_14674_14742@45.72.28,trnalookalike8_GluTTC_5_-_93905172_93905240@45.72.28\|CGTGGTTGTAGTCCGTGCGAGAATACCA | >trnaMT_GluTTC_MT_-_14674_14742 | 45 | 72 | 28 | CGTGGTTGTAGTCCGTGCGAGAATACCA | MB231 | 33 | 41 | 36 | 47 | 40 | 6.1 | 18.2 | 31.8 | 35.4 | 19.9 |
| >trnaMT_GluTTC_MT_-_14674_14742@46.71.26,trnalookalike8_GluTTC_5_-_93905172_93905240@46.71.26\|GTGGTTGTAGTCCGTGCGAGAATACC | >trnaMT_GluTTC_MT_-_14674_14742 | 46 | 71 | 26 | GTGGTTGTAGTCCGTGCGAGAATACC | MB231 | 12 | 11 | 10 | 33 | 14 | 9.4 | 3.0 | 5.4 | 20.4 | 6.0 |
| >trnaMT_GluTTC_MT_-_14674_14742@46.72.27,trnalookalike8_GluTTC_5_-_93905172_93905240@46.72.27\|GTGGTTGTAGTCCGTGCGAGAATACCA | >trnaMT_GluTTC_MT_-_14674_14742 | 46 | 72 | 27 | GTGGTTGTAGTCCGTGCGAGAATACCA | BT20 | 23 | 8 | 8 | 67 | 59 | 9.2 | 1.7 | 6.2 | 73.1 | 67.2 |
| >trnaMT_GluTTC_MT_-_14674_14742@46.72.27,trnalookalike8_GluTTC_5_-_93905172_93905240@46.72.27\|GTGGTTGTAGTCCGTGCGAGAATACCA | >trnaMT_GluTTC_MT_-_14674_14742 | 46 | 72 | 27 | GTGGTTGTAGTCCGTGCGAGAATACCA | MB231 | 42 | 33 | 25 | 103 | 55 | 27.4 | 6.0 | 18.6 | 62.0 | 25.5 |
| >trnaMT_GluTTC_MT_-_14674_14742@53.72.20,trnalookalike8_GluTTC_5_-_93905172_93905240@53.72.20\|TAGTCCGTGCGAGAATACCA | >trnaMT_GluTTC_MT_-_14674_14742 | 53 | 72 | 20 | TAGTCCGTGCGAGAATACCA | MB468 | 16 | 17 | 9 | 6 | 52 | 8.5 | 17.9 | 8.5 | 3.1 | 39.7 |
| >trnaMT_HisGTG_MT_+_12138_12206@28.72.45\|ATTGTGAATCTGACAACAGAGGCTTACGACCCCTTATTTACCCCA | >trnaMT_HisGTG_MT_+_12138_12206 | 28 | 72 | 45 | ATTGTGAATCTGACAACAGAGGCTTACGACCCCTTATTTACCCCA | BT20 | 12 | 0 | 1 | 10 | 3 | 19.3 | 0.8 | 1.1 | 9.7 | 2.9 |
| >trnaMT_HisGTG_MT_+_12138_12206@28.72.45\|ATTGTGAATCTGACAACAGAGGCTTACGACCCCTTATTTACCCCA | >trnaMT_HisGTG_MT_+_12138_12206 | 28 | 72 | 45 | ATTGTGAATCTGACAACAGAGGCTTACGACCCCTTATTTACCCCA | MB231 | 14 | 6 | 5 | 5 | 15 | 9.4 | 9.6 | 2.1 | 8.3 | 26.6 |
| >trnaMT_HisGTG_MT_+_12138_12206@32.61.30\|TGAATCTGACAACAGAGGCTTACGACCCCT | >trnaMT_HisGTG_MT_+_12138_12206 | 32 | 61 | 30 | TGAATCTGACAACAGAGGCTTACGACCCCT | BT20 | 13 | 9 | 4 | 8 | 32 | 10.6 | 9.3 | 5.2 | 7.6 | 36.2 |
| >trnaMT_HisGTG_MT_+_12138_12206@32.63.32\|TGAATCTGACAACAGAGGCTTACGACCCCTTA | >trnaMT_HisGTG_MT_+_12138_12206 | 32 | 63 | 32 | TGAATCTGACAACAGAGGCTTACGACCCCTTA | BT20 | 34 | 10 | 25 | 75 | 94 | 39.5 | 9.5 | 40.2 | 94.5 | 135.4 |
| >trnaMT_HisGTG_MT_+_12138_12206@32.63.32\|TGAATCTGACAACAGAGGCTTACGACCCCTTA | >trnaMT_HisGTG_MT_+_12138_12206 | 32 | 63 | 32 | TGAATCTGACAACAGAGGCTTACGACCCCTTA | MB231 | 31 | 7 | 2 | 88 | 95 | 20.6 | 5.5 | 3.0 | 89.8 | 74.3 |
| >trnaMT_HisGTG_MT_+_12138_12206@32.67.36\|TGAATCTGACAACAGAGGCTTACGACCCCTTATTTA | >trnaMT_HisGTG_MT_+_12138_12206 | 32 | 67 | 36 | TGAATCTGACAACAGAGGCTTACGACCCCTTATTTA | BT20 | 13 | 1 | 1 | 16 | 29 | 20.5 | 0.9 | 0.9 | 24.4 | 44.0 |
| >trnaMT_HisGTG_MT_+_12138_12206@32.67.36\|TGAATCTGACAACAGAGGCTTACGACCCCTTATTTA | >trnaMT_HisGTG_MT_+_12138_12206 | 32 | 67 | 36 | TGAATCTGACAACAGAGGCTTACGACCCCTTATTTA | MB231 | 15 | 3 | 1 | 23 | 55 | 14.2 | 2.3 | 0.6 | 24.1 | 54.0 |
| >trnaMT_HisGTG_MT_+_12138_12206@32.69.38\|TGAATCTGACAACAGAGGCTTACGACCCCTTATTTACC | >trnaMT_HisGTG_MT_+_12138_12206 | 32 | 69 | 38 | TGAATCTGACAACAGAGGCTTACGACCCCTTATTTACC | BT20 | 22 | 3 | 3 | 21 | 108 | 35.6 | 2.4 | 2.8 | 30.2 | 174.8 |
| >trnaMT_HisGTG_MT_+_12138_12206@32.69.38\|TGAATCTGACAACAGAGGCTTACGACCCCTTATTTACC | >trnaMT_HisGTG_MT_+_12138_12206 | 32 | 69 | 38 | TGAATCTGACAACAGAGGCTTACGACCCCTTATTTACC | MB231 | 36 | 26 | 4 | 67 | 216 | 19.5 | 17.9 | 2.8 | 58.9 | 159.4 |
| >trnaMT_HisGTG_MT_+_12138_12206@32.70.39\|TGAATCTGACAACAGAGGCTTACGACCCCTTATTTACCC | >trnaMT_HisGTG_MT_+_12138_12206 | 32 | 70 | 39 | TGAATCTGACAACAGAGGCTTACGACCCCTTATTTACCC | BT20 | 42 | 3 | 4 | 117 | 147 | 69.6 | 2.8 | 5.0 | 163.7 | 228.3 |
| >trnaMT_HisGTG_MT_+_12138_12206@32.70.39\|TGAATCTGACAACAGAGGCTTACGACCCCTTATTTACCC | >trnaMT_HisGTG_MT_+_12138_12206 | 32 | 70 | 39 | TGAATCTGACAACAGAGGCTTACGACCCCTTATTTACCC | MB231 | 115 | 77 | 27 | 324 | 528 | 46.5 | 50.4 | 21.0 | 334.4 | 365.0 |
| >trnaMT_HisGTG_MT_+_12138_12206@32.71.40\|TGAATCTGACAACAGAGGCTTACGACCCCTTATTTACCCC | >trnaMT_HisGTG_MT_+_12138_12206 | 32 | 71 | 40 | TGAATCTGACAACAGAGGCTTACGACCCCTTATTTACCCC | BT20 | 74 | 6 | 9 | 674 | 292 | 124.1 | 8.2 | 10.8 | 1027.1 | 447.8 |
| >trnaMT_HisGTG_MT_+_12138_12206@32.71.40\|TGAATCTGACAACAGAGGCTTACGACCCCTTATTTACCCC | >trnaMT_HisGTG_MT_+_12138_12206 | 32 | 71 | 40 | TGAATCTGACAACAGAGGCTTACGACCCCTTATTTACCCC | MB231 | 193 | 120 | 44 | 846 | 1268 | 100.6 | 101.4 | 29.1 | 989.4 | 1167.0 |
| >trnaMT_HisGTG_MT_+_12138_12206@32.72.41\|TGAATCTGACAACAGAGGCTTACGACCCCTTATTTACCCCA | >trnaMT_HisGTG_MT_+_12138_12206 | 32 | 72 | 41 | TGAATCTGACAACAGAGGCTTACGACCCCTTATTTACCCCA | BT20 | 27 | 4 | 4 | 255 | 120 | 43.7 | 6.0 | 4.3 | 392.1 | 183.7 |
| >trnaMT_HisGTG_MT_+_12138_12206@32.72.41\|TGAATCTGACAACAGAGGCTTACGACCCCTTATTTACCCCA | >trnaMT_HisGTG_MT_+_12138_12206 | 32 | 72 | 41 | TGAATCTGACAACAGAGGCTTACGACCCCTTATTTACCCCA | MB231 | 73 | 36 | 26 | 211 | 541 | 50.3 | 41.2 | 19.6 | 267.8 | 514.9 |
| >trnaMT_LeuTAA_MT_+_3230_3304@1.25.25\|GTTAAGATGGCAGAGCCCGGTAATC | >trnaMT_LeuTAA_MT_+_3230_3304 | 1 | 25 | 25 | GTTAAGATGGCAGAGCCCGGTAATC | BT20 | 13 | 4 | 8 | 66 | 23 | 17.3 | 2.1 | 6.6 | 104.1 | 32.1 |
| >trnaMT_LeuTAA_MT_+_3230_3304@1.30.30\|GTTAAGATGGCAGAGCCCGGTAATCGCATA | >trnaMT_LeuTAA_MT_+_3230_3304 | 1 | 30 | 30 | GTTAAGATGGCAGAGCCCGGTAATCGCATA | BT20 | 35 | 24 | 7 | 352 | 74 | 40.5 | 10.0 | 3.2 | 564.4 | 105.3 |
| >trnaMT_LeuTAA_MT_+_3230_3304@1.30.30\|GTTAAGATGGCAGAGCCCGGTAATCGCATA | >trnaMT_LeuTAA_MT_+_3230_3304 | 1 | 30 | 30 | GTTAAGATGGCAGAGCCCGGTAATCGCATA | MB231 | 19 | 6 | 5 | 45 | 45 | 14.1 | 2.8 | 3.4 | 17.8 | 11.9 |
| >trnaMT_LeuTAA_MT_+_3230_3304@1.37.37\|GTTAAGATGGCAGAGCCCGGTAATCGCATAAAACTTA | >trnaMT_LeuTAA_MT_+_3230_3304 | 1 | 37 | 37 | GTTAAGATGGCAGAGCCCGGTAATCGCATAAAACTTA | MB231 | 13 | 3 | 8 | 9 | 25 | 7.7 | 2.3 | 10.1 | 3.7 | 16.6 |
| >trnaMT_LeuTAA_MT_+_3230_3304@16.44.29\|CCCGGTAATCGCATAAAACTTAAAACTTT | >trnaMT_LeuTAA_MT_+_3230_3304 | 16 | 44 | 29 | CCCGGTAATCGCATAAAACTTAAAACTTT | MB468 | 57 | 181 | 100 | 81 | 87 | 30.9 | 101.4 | 65.4 | 81.8 | 38.4 |
| >trnaMT_LeuTAA_MT_+_3230_3304@18.42.25\|CGGTAATCGCATAAAACTTAAAACT | >trnaMT_LeuTAA_MT_+_3230_3304 | 18 | 42 | 25 | CGGTAATCGCATAAAACTTAAAACT | BT20 | 19 | 6 | 24 | 4 | 2 | 14.0 | 3.6 | 10.1 | 3.5 | 2.8 |
| >trnaMT_LeuTAA_MT_+_3230_3304@18.44.27\|CGGTAATCGCATAAAACTTAAAACTTT | >trnaMT_LeuTAA_MT_+_3230_3304 | 18 | 44 | 27 | CGGTAATCGCATAAAACTTAAAACTTT | BT20 | 14 | 4 | 29 | 13 | 11 | 6.9 | 1.7 | 15.5 | 14.6 | 12.0 |
| >trnaMT_LeuTAA_MT_+_3230_3304@18.44.27\|CGGTAATCGCATAAAACTTAAAACTTT | >trnaMT_LeuTAA_MT_+_3230_3304 | 18 | 44 | 27 | CGGTAATCGCATAAAACTTAAAACTTT | MB468 | 22 | 44 | 32 | 47 | 43 | 11.2 | 52.3 | 28.0 | 38.6 | 33.7 |
| >trnaMT_LeuTAA_MT_+_3230_3304@19.42.24\|GGTAATCGCATAAAACTTAAAACT | >trnaMT_LeuTAA_MT_+_3230_3304 | 19 | 42 | 24 | GGTAATCGCATAAAACTTAAAACT | BT20 | 11 | 6 | 18 | 2 | 1 | 3.7 | 4.6 | 10.0 | 2.3 | 1.2 |

| | | | | | | | | | | | | | | | | |
|---|---|---|---|---|---|---|---|---|---|---|---|---|---|---|---|---|
| >trnaMT_MetCAT_MT_+_4402_4469@34.71.38\|ACCCCGAAAATGTTGGTTATACCCTTCCCGTACTACCA | >trnaMT_MetCAT_MT_+_4402_4469 | 34 | 71 | 38 | ACCCCGAAAATGTTGGTTATACCCTTCCCGTACTACCA | MB231 | 13 | 26 | 29 | 1 | 4 | 7.1 | 20.5 | 26.8 | 1.4 | 7.4 |
| >trnaMT_MetCAT_MT_+_4402_4469@35.71.37\|CCCCGAAAATGTTGGTTATACCCTTCCCGTACTACCA | >trnaMT_MetCAT_MT_+_4402_4469 | 35 | 71 | 37 | CCCCGAAAATGTTGGTTATACCCTTCCCGTACTACCA | MB231 | 10 | 29 | 16 | 1 | 6 | 3.5 | 20.2 | 7.1 | 1.3 | 6.9 |
| >trnaMT_MetCAT_MT_+_4402_4469@36.71.36\|CCCGAAAATGTTGGTTATACCCTTCCCGTACTACCA | >trnaMT_MetCAT_MT_+_4402_4469 | 36 | 71 | 36 | CCCGAAAATGTTGGTTATACCCTTCCCGTACTACCA | MB231 | 14 | 37 | 24 | 4 | 12 | 7.0 | 22.8 | 11.8 | 4.1 | 9.6 |
| >trnaMT_MetCAT_MT_+_4402_4469@37.71.35\|CCGAAAATGTTGGTTATACCCTTCCCGTACTACCA | >trnaMT_MetCAT_MT_+_4402_4469 | 37 | 71 | 35 | CCGAAAATGTTGGTTATACCCTTCCCGTACTACCA | BT20 | 21 | 6 | 10 | 64 | 35 | 11.1 | 3.2 | 4.4 | 83.0 | 24.8 |
| >trnaMT_MetCAT_MT_+_4402_4469@37.71.35\|CCGAAAATGTTGGTTATACCCTTCCCGTACTACCA | >trnaMT_MetCAT_MT_+_4402_4469 | 37 | 71 | 35 | CCGAAAATGTTGGTTATACCCTTCCCGTACTACCA | MB231 | 22 | 33 | 34 | 36 | 21 | 1.8 | 16.3 | 23.1 | 41.1 | 15.2 |
| >trnaMT_MetCAT_MT_+_4402_4469@38.71.34\|CGAAAATGTTGGTTATACCCTTCCCGTACTACCA | >trnaMT_MetCAT_MT_+_4402_4469 | 38 | 71 | 34 | CGAAAATGTTGGTTATACCCTTCCCGTACTACCA | MB231 | 10 | 14 | 5 | 32 | 31 | 1.1 | 5.9 | 2.1 | 17.2 | 6.9 |
| >trnaMT_MetCAT_MT_+_4402_4469@40.71.32\|AAAATGTTGGTTATACCCTTCCCGTACTACCA | >trnaMT_MetCAT_MT_+_4402_4469 | 40 | 71 | 32 | AAAATGTTGGTTATACCCTTCCCGTACTACCA | BT20 | 10 | 1 | 3 | 26 | 10 | 1.9 | 1.1 | 1.2 | 28.1 | 5.1 |
| >trnaMT_MetCAT_MT_+_4402_4469@46.71.26\|TTGGTTATACCCTTCCCGTACTACCA | >trnaMT_MetCAT_MT_+_4402_4469 | 46 | 71 | 26 | TTGGTTATACCCTTCCCGTACTACCA | BT20 | 11 | 0 | 3 | 3 | 3 | 6.1 | 0.4 | 3.3 | 1.9 | 2.4 |
| >trnaMT_MetCAT_MT_+_4402_4469@46.71.26\|TTGGTTATACCCTTCCCGTACTACCA | >trnaMT_MetCAT_MT_+_4402_4469 | 46 | 71 | 26 | TTGGTTATACCCTTCCCGTACTACCA | MB231 | 18 | 7 | 13 | 3 | 3 | 8.3 | 8.3 | 15.1 | 2.9 | 1.6 |
| >trnaMT_MetCAT_MT_+_4402_4469@50.71.22\|TTATACCCTTCCCGTACTACCA | >trnaMT_MetCAT_MT_+_4402_4469 | 50 | 71 | 22 | TTATACCCTTCCCGTACTACCA | MB231 | 10 | 17 | 29 | 2 | 5 | 2.6 | 11.6 | 19.3 | 3.0 | 6.4 |
| >trnaMT_MetCAT_MT_+_4402_4469@52.70.19\|ATACCCTTCCCGTACTACC | >trnaMT_MetCAT_MT_+_4402_4469 | 52 | 70 | 19 | ATACCCTTCCCGTACTACC | MB231 | 21 | 28 | 66 | 0 | 1 | 20.1 | 33.8 | 86.0 | 0.6 | 2.5 |
| >trnaMT_MetCAT_MT_+_4402_4469@52.71.20\|ATACCCTTCCCGTACTACCA | >trnaMT_MetCAT_MT_+_4402_4469 | 52 | 71 | 20 | ATACCCTTCCCGTACTACCA | BT20 | 10 | 7 | 38 | 1 | 5 | 3.0 | 6.4 | 46.0 | 1.5 | 9.2 |
| >trnaMT_MetCAT_MT_+_4402_4469@52.71.20\|ATACCCTTCCCGTACTACCA | >trnaMT_MetCAT_MT_+_4402_4469 | 52 | 71 | 20 | ATACCCTTCCCGTACTACCA | MB231 | 30 | 71 | 97 | 3 | 17 | 23.9 | 82.6 | 106.3 | 2.8 | 8.4 |
| >trnaMT_PheGAA_MT_+_577_647@1.18.18\|GTTTATGTAGCTTACCTC | >trnaMT_PheGAA_MT_+_577_647 | 1 | 18 | 18 | GTTTATGTAGCTTACCTC | BT20 | 30 | 3 | 2 | 14 | 24 | 17.1 | 1.3 | 2.0 | 21.1 | 34.0 |
| >trnaMT_PheGAA_MT_+_577_647@1.18.18\|GTTTATGTAGCTTACCTC | >trnaMT_PheGAA_MT_+_577_647 | 1 | 18 | 18 | GTTTATGTAGCTTACCTC | MB231 | 63 | 26 | 5 | 77 | 49 | 55.0 | 27.1 | 3.1 | 47.1 | 13.7 |
| >trnaMT_PheGAA_MT_+_577_647@1.19.19\|GTTTATGTAGCTTACCTCC | >trnaMT_PheGAA_MT_+_577_647 | 1 | 19 | 19 | GTTTATGTAGCTTACCTCC | BT20 | 43 | 7 | 12 | 15 | 36 | 24.9 | 2.2 | 8.4 | 21.8 | 53.2 |
| >trnaMT_PheGAA_MT_+_577_647@1.19.19\|GTTTATGTAGCTTACCTCC | >trnaMT_PheGAA_MT_+_577_647 | 1 | 19 | 19 | GTTTATGTAGCTTACCTCC | MB231 | 84 | 41 | 16 | 55 | 79 | 75.7 | 34.1 | 8.9 | 36.2 | 33.7 |
| >trnaMT_PheGAA_MT_+_577_647@1.20.20\|GTTTATGTAGCTTACCTCCT | >trnaMT_PheGAA_MT_+_577_647 | 1 | 20 | 20 | GTTTATGTAGCTTACCTCCT | BT20 | 12 | 3 | 3 | 14 | 14 | 11.0 | 1.9 | 3.0 | 22.9 | 21.1 |
| >trnaMT_PheGAA_MT_+_577_647@1.20.20\|GTTTATGTAGCTTACCTCCT | >trnaMT_PheGAA_MT_+_577_647 | 1 | 20 | 20 | GTTTATGTAGCTTACCTCCT | MB231 | 27 | 19 | 5 | 26 | 32 | 21.6 | 16.5 | 4.0 | 17.1 | 12.5 |
| >trnaMT_PheGAA_MT_+_577_647@1.34.34\|GTTTATGTAGCTTACCTCCTCAAAGCAATACACT | >trnaMT_PheGAA_MT_+_577_647 | 1 | 34 | 34 | GTTTATGTAGCTTACCTCCTCAAAGCAATACACT | BT20 | 26 | 7 | 4 | 30 | 49 | 19.2 | 6.0 | 2.2 | 48.3 | 68.7 |
| >trnaMT_PheGAA_MT_+_577_647@1.34.34\|GTTTATGTAGCTTACCTCCTCAAAGCAATACACT | >trnaMT_PheGAA_MT_+_577_647 | 1 | 34 | 34 | GTTTATGTAGCTTACCTCCTCAAAGCAATACACT | MB231 | 32 | 11 | 1 | 58 | 126 | 25.3 | 4.5 | 0.8 | 21.0 | 123.5 |
| >trnaMT_PheGAA_MT_+_577_647@1.35.35\|GTTTATGTAGCTTACCTCCTCAAAGCAATACACTG | >trnaMT_PheGAA_MT_+_577_647 | 1 | 35 | 35 | GTTTATGTAGCTTACCTCCTCAAAGCAATACACTG | MB231 | 17 | 7 | 1 | 30 | 58 | 16.4 | 2.2 | 0.3 | 11.6 | 57.8 |
| >trnaMT_PheGAA_MT_+_577_647@35.53.19\|GAAAATGTTTAGACGGGCT | >trnaMT_PheGAA_MT_+_577_647 | 35 | 53 | 19 | GAAAATGTTTAGACGGGCT | BT20 | 24 | 9 | 12 | 64 | 24 | 8.1 | 5.1 | 9.4 | 59.4 | 6.2 |
| >trnaMT_PheGAA_MT_+_577_647@35.54.20\|GAAAATGTTTAGACGGGCTC | >trnaMT_PheGAA_MT_+_577_647 | 35 | 54 | 20 | GAAAATGTTTAGACGGGCTC | BT20 | 28 | 19 | 12 | 61 | 48 | 9.0 | 13.9 | 6.8 | 40.1 | 11.8 |
| >trnaMT_PheGAA_MT_+_577_647@35.54.20\|GAAAATGTTTAGACGGGCTC | >trnaMT_PheGAA_MT_+_577_647 | 35 | 54 | 20 | GAAAATGTTTAGACGGGCTC | MB231 | 34 | 20 | 9 | 54 | 54 | 19.5 | 12.4 | 6.1 | 42.0 | 35.6 |
| >trnaMT_PheGAA_MT_+_577_647@35.55.21\|GAAAATGTTTAGACGGGCTCA | >trnaMT_PheGAA_MT_+_577_647 | 35 | 55 | 21 | GAAAATGTTTAGACGGGCTCA | BT20 | 48 | 22 | 15 | 151 | 93 | 22.2 | 23.4 | 15.4 | 116.6 | 13.7 |
| >trnaMT_PheGAA_MT_+_577_647@35.55.21\|GAAAATGTTTAGACGGGCTCA | >trnaMT_PheGAA_MT_+_577_647 | 35 | 55 | 21 | GAAAATGTTTAGACGGGCTCA | MB231 | 14 | 9 | 3 | 45 | 27 | 6.4 | 3.7 | 1.2 | 20.7 | 15.7 |
| >trnaMT_PheGAA_MT_+_577_647@35.56.22\|GAAAATGTTTAGACGGGCTCAC | >trnaMT_PheGAA_MT_+_577_647 | 35 | 56 | 22 | GAAAATGTTTAGACGGGCTCAC | BT20 | 45 | 23 | 16 | 90 | 81 | 28.1 | 13.4 | 10.6 | 76.1 | 66.2 |
| >trnaMT_PheGAA_MT_+_577_647@35.56.22\|GAAAATGTTTAGACGGGCTCAC | >trnaMT_PheGAA_MT_+_577_647 | 35 | 56 | 22 | GAAAATGTTTAGACGGGCTCAC | MB231 | 17 | 13 | 11 | 35 | 37 | 4.4 | 6.4 | 5.0 | 7.2 | 3.5 |
| >trnaMT_PheGAA_MT_+_577_647@35.57.23\|GAAAATGTTTAGACGGGCTCACA | >trnaMT_PheGAA_MT_+_577_647 | 35 | 57 | 23 | GAAAATGTTTAGACGGGCTCACA | BT20 | 150 | 103 | 36 | 146 | 148 | 60.5 | 82.1 | 41.3 | 99.8 | 58.8 |
| >trnaMT_PheGAA_MT_+_577_647@35.57.23\|GAAAATGTTTAGACGGGCTCACA | >trnaMT_PheGAA_MT_+_577_647 | 35 | 57 | 23 | GAAAATGTTTAGACGGGCTCACA | MB231 | 21 | 14 | 5 | 68 | 72 | 7.3 | 4.9 | 2.2 | 26.6 | 18.9 |
| >trnaMT_PheGAA_MT_+_577_647@35.59.25\|GAAAATGTTTAGACGGGCTCACATC | >trnaMT_PheGAA_MT_+_577_647 | 35 | 59 | 25 | GAAAATGTTTAGACGGGCTCACATC | BT20 | 29 | 14 | 5 | 49 | 52 | 17.5 | 13.5 | 4.5 | 44.1 | 40.2 |
| >trnaMT_PheGAA_MT_+_577_647@35.59.25\|GAAAATGTTTAGACGGGCTCACATC | >trnaMT_PheGAA_MT_+_577_647 | 35 | 59 | 25 | GAAAATGTTTAGACGGGCTCACATC | MB231 | 13 | 7 | 2 | 20 | 26 | 6.0 | 1.8 | 0.9 | 10.9 | 10.2 |
| >trnaMT_PheGAA_MT_+_577_647@35.60.26\|GAAAATGTTTAGACGGGCTCACATCA | >trnaMT_PheGAA_MT_+_577_647 | 35 | 60 | 26 | GAAAATGTTTAGACGGGCTCACATCA | BT20 | 27 | 19 | 6 | 39 | 53 | 24.8 | 28.0 | 8.1 | 25.9 | 32.9 |
| >trnaMT_PheGAA_MT_+_577_647@35.62.28\|GAAAATGTTTAGACGGGCTCACATCACC | >trnaMT_PheGAA_MT_+_577_647 | 35 | 62 | 28 | GAAAATGTTTAGACGGGCTCACATCACC | BT20 | 27 | 29 | 8 | 63 | 50 | 24.2 | 42.5 | 10.1 | 52.0 | 35.6 |
| >trnaMT_PheGAA_MT_+_577_647@35.62.28\|GAAAATGTTTAGACGGGCTCACATCACC | >trnaMT_PheGAA_MT_+_577_647 | 35 | 62 | 28 | GAAAATGTTTAGACGGGCTCACATCACC | MB231 | 14 | 12 | 2 | 37 | 63 | 7.9 | 3.4 | 0.9 | 37.8 | 41.7 |
| >trnaMT_PheGAA_MT_+_577_647@36.53.18\|AAAATGTTTAGACGGGCT | >trnaMT_PheGAA_MT_+_577_647 | 36 | 53 | 18 | AAAATGTTTAGACGGGCT | BT20 | 13 | 4 | 6 | 22 | 10 | 3.4 | 1.9 | 5.7 | 20.5 | 4.1 |
| >trnaMT_PheGAA_MT_+_577_647@36.54.19\|AAAATGTTTAGACGGGCTC | >trnaMT_PheGAA_MT_+_577_647 | 36 | 54 | 19 | AAAATGTTTAGACGGGCTC | BT20 | 29 | 11 | 16 | 39 | 25 | 10.9 | 6.2 | 17.0 | 26.9 | 5.7 |
| >trnaMT_PheGAA_MT_+_577_647@36.54.19\|AAAATGTTTAGACGGGCTC | >trnaMT_PheGAA_MT_+_577_647 | 36 | 54 | 19 | AAAATGTTTAGACGGGCTC | MB231 | 18 | 9 | 7 | 18 | 21 | 10.4 | 4.3 | 3.3 | 10.4 | 13.2 |
| >trnaMT_PheGAA_MT_+_577_647@36.55.20\|AAAATGTTTAGACGGGCTCA | >trnaMT_PheGAA_MT_+_577_647 | 36 | 55 | 20 | AAAATGTTTAGACGGGCTCA | BT20 | 33 | 11 | 13 | 66 | 38 | 14.1 | 10.6 | 16.0 | 56.4 | 5.6 |
| >trnaMT_PheGAA_MT_+_577_647@36.56.21\|AAAATGTTTAGACGGGCTCAC | >trnaMT_PheGAA_MT_+_577_647 | 36 | 56 | 21 | AAAATGTTTAGACGGGCTCAC | BT20 | 32 | 11 | 11 | 46 | 44 | 18.5 | 4.0 | 9.0 | 34.4 | 26.4 |
| >trnaMT_PheGAA_MT_+_577_647@36.56.21\|AAAATGTTTAGACGGGCTCAC | >trnaMT_PheGAA_MT_+_577_647 | 36 | 56 | 21 | AAAATGTTTAGACGGGCTCAC | MB231 | 10 | 7 | 6 | 15 | 21 | 2.6 | 2.6 | 2.8 | 2.0 | 6.7 |
| >trnaMT_PheGAA_MT_+_577_647@36.57.22\|AAAATGTTTAGACGGGCTCACA | >trnaMT_PheGAA_MT_+_577_647 | 36 | 57 | 22 | AAAATGTTTAGACGGGCTCACA | BT20 | 90 | 31 | 19 | 65 | 64 | 44.2 | 26.7 | 24.8 | 38.9 | 22.6 |
| >trnaMT_PheGAA_MT_+_577_647@36.57.22\|AAAATGTTTAGACGGGCTCACA | >trnaMT_PheGAA_MT_+_577_647 | 36 | 57 | 22 | AAAATGTTTAGACGGGCTCACA | MB231 | 12 | 7 | 2 | 24 | 29 | 3.6 | 1.3 | 0.6 | 7.9 | 6.2 |
| >trnaMT_PheGAA_MT_+_577_647@36.59.24\|AAAATGTTTAGACGGGCTCACATC | >trnaMT_PheGAA_MT_+_577_647 | 36 | 59 | 24 | AAAATGTTTAGACGGGCTCACATC | BT20 | 17 | 6 | 3 | 21 | 27 | 9.5 | 4.5 | 2.4 | 20.2 | 21.8 |
| >trnaMT_PheGAA_MT_+_577_647@36.60.25\|AAAATGTTTAGACGGGCTCACATCA | >trnaMT_PheGAA_MT_+_577_647 | 36 | 60 | 25 | AAAATGTTTAGACGGGCTCACATCA | BT20 | 13 | 7 | 3 | 11 | 22 | 12.2 | 10.6 | 3.7 | 8.4 | 12.7 |
| >trnaMT_PheGAA_MT_+_577_647@36.62.27\|AAAATGTTTAGACGGGCTCACATCACC | >trnaMT_PheGAA_MT_+_577_647 | 36 | 62 | 27 | AAAATGTTTAGACGGGCTCACATCACC | BT20 | 22 | 19 | 8 | 40 | 41 | 18.9 | 27.4 | 10.1 | 32.6 | 31.9 |
| >trnaMT_PheGAA_MT_+_577_647@36.62.27\|AAAATGTTTAGACGGGCTCACATCACC | >trnaMT_PheGAA_MT_+_577_647 | 36 | 62 | 27 | AAAATGTTTAGACGGGCTCACATCACC | MB231 | 14 | 11 | 2 | 24 | 52 | 6.1 | 4.0 | 0.4 | 23.4 | 34.7 |
| >trnaMT_PheGAA_MT_+_577_647@37.54.18\|AAATGTTTAGACGGGCTC | >trnaMT_PheGAA_MT_+_577_647 | 37 | 54 | 18 | AAATGTTTAGACGGGCTC | BT20 | 13 | 5 | 10 | 13 | 10 | 7.3 | 3.4 | 10.4 | 10.1 | 3.4 |
| >trnaMT_PheGAA_MT_+_577_647@37.56.20\|AAATGTTTAGACGGGCTCAC | >trnaMT_PheGAA_MT_+_577_647 | 37 | 56 | 20 | AAATGTTTAGACGGGCTCAC | BT20 | 17 | 5 | 6 | 20 | 31 | 12.4 | 1.7 | 3.0 | 13.9 | 9.6 |
| >trnaMT_PheGAA_MT_+_577_647@37.57.21\|AAATGTTTAGACGGGCTCACA | >trnaMT_PheGAA_MT_+_577_647 | 37 | 57 | 21 | AAATGTTTAGACGGGCTCACA | BT20 | 12 | 3 | 3 | 9 | 11 | 2.6 | 2.6 | 3.4 | 6.2 | 5.6 |
| >trnaMT_PheGAA_MT_+_577_647@38.55.18\|AATGTTTAGACGGGCTCA | >trnaMT_PheGAA_MT_+_577_647 | 38 | 55 | 18 | AATGTTTAGACGGGCTCA | BT20 | 19 | 4 | 4 | 18 | 12 | 8.4 | 2.4 | 6.3 | 13.0 | 1.0 |
| >trnaMT_PheGAA_MT_+_577_647@38.56.19\|AATGTTTAGACGGGCTCAC | >trnaMT_PheGAA_MT_+_577_647 | 38 | 56 | 19 | AATGTTTAGACGGGCTCAC | BT20 | 30 | 7 | 8 | 33 | 45 | 22.3 | 1.0 | 3.9 | 18.1 | 12.2 |
| >trnaMT_PheGAA_MT_+_577_647@38.56.19\|AATGTTTAGACGGGCTCAC | >trnaMT_PheGAA_MT_+_577_647 | 38 | 56 | 19 | AATGTTTAGACGGGCTCAC | MB231 | 11 | 9 | 4 | 10 | 29 | 2.2 | 1.7 | 1.5 | 1.3 | 9.2 |
| >trnaMT_PheGAA_MT_+_577_647@38.57.20\|AATGTTTAGACGGGCTCACA | >trnaMT_PheGAA_MT_+_577_647 | 38 | 57 | 20 | AATGTTTAGACGGGCTCACA | BT20 | 41 | 9 | 5 | 17 | 19 | 13.5 | 4.7 | 4.1 | 10.8 | 6.3 |
| >trnaMT_PheGAA_MT_+_577_647@38.59.22\|AATGTTTAGACGGGCTCACATC | >trnaMT_PheGAA_MT_+_577_647 | 38 | 59 | 22 | AATGTTTAGACGGGCTCACATC | BT20 | 15 | 3 | 2 | 12 | 15 | 7.6 | 0.5 | 1.2 | 13.6 | 10.7 |
| >trnaMT_PheGAA_MT_+_577_647@38.62.25\|AATGTTTAGACGGGCTCACATCACC | >trnaMT_PheGAA_MT_+_577_647 | 38 | 62 | 25 | AATGTTTAGACGGGCTCACATCACC | BT20 | 16 | 7 | 3 | 24 | 20 | 12.1 | 9.2 | 2.4 | 21.7 | 14.3 |
| >trnaMT_PheGAA_MT_+_577_647@39.56.18\|ATGTTTAGACGGGCTCAC | >trnaMT_PheGAA_MT_+_577_647 | 39 | 56 | 18 | ATGTTTAGACGGGCTCAC | BT20 | 175 | 46 | 63 | 284 | 436 | 135.0 | 9.0 | 30.6 | 130.2 | 112.7 |
| >trnaMT_PheGAA_MT_+_577_647@39.56.18\|ATGTTTAGACGGGCTCAC | >trnaMT_PheGAA_MT_+_577_647 | 39 | 56 | 18 | ATGTTTAGACGGGCTCAC | MB231 | 90 | 116 | 50 | 123 | 356 | 48.2 | 35.1 | 23.6 | 27.3 | 99.5 |
| >trnaMT_PheGAA_MT_+_577_647@39.57.19\|ATGTTTAGACGGGCTCACA | >trnaMT_PheGAA_MT_+_577_647 | 39 | 57 | 19 | ATGTTTAGACGGGCTCACA | BT20 | 75 | 19 | 30 | 49 | 72 | 19.2 | 9.5 | 30.8 | 27.8 | 33.4 |
| >trnaMT_PheGAA_MT_+_577_647@39.57.19\|ATGTTTAGACGGGCTCACA | >trnaMT_PheGAA_MT_+_577_647 | 39 | 57 | 19 | ATGTTTAGACGGGCTCACA | MB231 | 23 | 18 | 8 | 31 | 64 | 6.8 | 2.7 | 2.8 | 7.2 | 18.1 |
| >trnaMT_PheGAA_MT_+_577_647@39.59.21\|ATGTTTAGACGGGCTCACATC | >trnaMT_PheGAA_MT_+_577_647 | 39 | 59 | 21 | ATGTTTAGACGGGCTCACATC | BT20 | 24 | 5 | 10 | 34 | 39 | 13.8 | 1.7 | 8.4 | 32.8 | 34.2 |
| >trnaMT_PheGAA_MT_+_577_647@39.59.21\|ATGTTTAGACGGGCTCACATC | >trnaMT_PheGAA_MT_+_577_647 | 39 | 59 | 21 | ATGTTTAGACGGGCTCACATC | MB231 | 14 | 10 | 5 | 12 | 22 | 5.6 | 1.4 | 0.4 | 7.8 | 4.9 |
| >trnaMT_PheGAA_MT_+_577_647@39.62.24\|ATGTTTAGACGGGCTCACATCACC | >trnaMT_PheGAA_MT_+_577_647 | 39 | 62 | 24 | ATGTTTAGACGGGCTCACATCACC | BT20 | 19 | 9 | 12 | 42 | 41 | 12.2 | 9.5 | 9.9 | 43.9 | 38.6 |
| >trnaMT_PheGAA_MT_+_577_647@39.62.24\|ATGTTTAGACGGGCTCACATCACC | >trnaMT_PheGAA_MT_+_577_647 | 39 | 62 | 24 | ATGTTTAGACGGGCTCACATCACC | MB231 | 18 | 16 | 7 | 23 | 59 | 7.3 | 5.4 | 1.8 | 22.8 | 28.6 |
| >trnaMT_PheGAA_MT_+_577_647@42.74.33\|TTTAGACGGGCTCACATCACCCCATAAACACCA | >trnaMT_PheGAA_MT_+_577_647 | 42 | 74 | 33 | TTTAGACGGGCTCACATCACCCCATAAACACCA | BT20 | 20 | 11 | 3 | 92 | 27 | 8.1 | 4.9 | 0.7 | 98.0 | 15.8 |
| >trnaMT_PheGAA_MT_+_577_647@42.74.33\|TTTAGACGGGCTCACATCACCCCATAAACACCA | >trnaMT_PheGAA_MT_+_577_647 | 42 | 74 | 33 | TTTAGACGGGCTCACATCACCCCATAAACACCA | MB231 | 11 | 10 | 1 | 43 | 38 | 3.6 | 3.3 | 0.6 | 20.8 | 18.9 |
| >trnaMT_PheGAA_MT_+_577_647@43.74.32\|TTAGACGGGCTCACATCACCCCATAAACACCA | >trnaMT_PheGAA_MT_+_577_647 | 43 | 74 | 32 | TTAGACGGGCTCACATCACCCCATAAACACCA | BT20 | 23 | 18 | 12 | 48 | 16 | 0.6 | 10.6 | 4.5 | 64.8 | 17.1 |
| >trnaMT_PheGAA_MT_+_577_647@43.74.32\|TTAGACGGGCTCACATCACCCCATAAACACCA | >trnaMT_PheGAA_MT_+_577_647 | 43 | 74 | 32 | TTAGACGGGCTCACATCACCCCATAAACACCA | MB231 | 23 | 21 | 5 | 36 | 20 | 8.9 | 5.0 | 0.4 | 20.9 | 6.1 |
| >trnaMT_PheGAA_MT_+_577_647@44.73.30\|TAGACGGGCTCACATCACCCCATAAACACC | >trnaMT_PheGAA_MT_+_577_647 | 44 | 73 | 30 | TAGACGGGCTCACATCACCCCATAAACACC | MB231 | 11 | 10 | 2 | 4 | 6 | 10.6 | 3.4 | 0.5 | 1.6 | 1.0 |
| >trnaMT_PheGAA_MT_+_577_647@44.74.31\|TAGACGGGCTCACATCACCCCATAAACACCA | >trnaMT_PheGAA_MT_+_577_647 | 44 | 74 | 31 | TAGACGGGCTCACATCACCCCATAAACACCA | BT20 | 41 | 35 | 12 | 28 | 23 | 5.9 | 21.2 | 6.6 | 28.0 | 15.6 |
| >trnaMT_PheGAA_MT_+_577_647@44.74.31\|TAGACGGGCTCACATCACCCCATAAACACCA | >trnaMT_PheGAA_MT_+_577_647 | 44 | 74 | 31 | TAGACGGGCTCACATCACCCCATAAACACCA | MB231 | 37 | 29 | 5 | 26 | 23 | 25.2 | 7.5 | 0.8 | 13.4 | 2.2 |
| >trnaMT_PheGAA_MT_+_577_647@46.74.29\|GACGGGCTCACATCACCCCATAAACACCA | >trnaMT_PheGAA_MT_+_577_647 | 46 | 74 | 29 | GACGGGCTCACATCACCCCATAAACACCA | BT20 | 12 | 13 | 4 | 14 | 13 | 2.9 | 6.5 | 1.9 | 16.1 | 11.4 |
| >trnaMT_PheGAA_MT_+_577_647@46.74.29\|GACGGGCTCACATCACCCCATAAACACCA | >trnaMT_PheGAA_MT_+_577_647 | 46 | 74 | 29 | GACGGGCTCACATCACCCCATAAACACCA | MB231 | 15 | 16 | 2 | 15 | 13 | 8.1 | 0.3 | 0.1 | 8.5 | 0.4 |
| >trnaMT_PheGAA_MT_+_577_647@47.74.28\|ACGGGCTCACATCACCCCATAAACACCA | >trnaMT_PheGAA_MT_+_577_647 | 47 | 74 | 28 | ACGGGCTCACATCACCCCATAAACACCA | BT20 | 11 | 10 | 4 | 5 | 6 | 3.0 | 6.2 | 2.3 | 4.5 | 4.4 |
| >trnaMT_PheGAA_MT_+_577_647@54.74.21\|CACATCACCCCATAAACACCA | >trnaMT_PheGAA_MT_+_577_647 | 54 | 74 | 21 | CACATCACCCCATAAACACCA | BT20 | 13 | 7 | 13 | 4 | 19 | 2.9 | 3.4 | 9.0 | 4.4 | 20.4 |
| >trnaMT_PheGAA_MT_+_577_647@54.74.21\|CACATCACCCCATAAACACCA | >trnaMT_PheGAA_MT_+_577_647 | 54 | 74 | 21 | CACATCACCCCATAAACACCA | MB468 | 13 | 35 | 7 | 12 | 30 | 8.4 | 48.5 | 9.4 | 6.8 | 15.0 |
| >trnaMT_PheGAA_MT_+_577_647@56.74.19\|CATCACCCCATAAACACCA | >trnaMT_PheGAA_MT_+_577_647 | 56 | 74 | 19 | CATCACCCCATAAACACCA | BT20 | 47 | 21 | 22 | 5 | 46 | 7.8 | 14.2 | 22.4 | 4.7 | 43.0 |
| >trnaMT_PheGAA_MT_+_577_647@56.74.19\|CATCACCCCATAAACACCA | >trnaMT_PheGAA_MT_+_577_647 | 56 | 74 | 19 | CATCACCCCATAAACACCA | MB231 | 30 | 19 | 22 | 12 | 54 | 13.1 | 4.2 | 16.0 | 8.8 | 18.4 |
| >trnaMT_ProTGG_MT_-_15956_16023@1.32.32\|CAGAGAATAGTTTAAATTAGAATCTTAGCTTT | >trnaMT_ProTGG_MT_-_15956_16023 | 1 | 32 | 32 | CAGAGAATAGTTTAAATTAGAATCTTAGCTTT | BT20 | 22 | 50 | 3 | 10 | 3 | 25.0 | 75.6 | 2.9 | 13.6 | 3.9 |
| >trnaMT_ProTGG_MT_-_15956_16023@1.32.32\|CAGAGAATAGTTTAAATTAGAATCTTAGCTTT | >trnaMT_ProTGG_MT_-_15956_16023 | 1 | 32 | 32 | CAGAGAATAGTTTAAATTAGAATCTTAGCTTT | MB231 | 13 | 3 | 1 | 30 | 7 | 9.4 | 2.8 | 0.4 | 10.9 | 3.7 |
| >trnaMT_ProTGG_MT_-_15956_16023@40.58.19\|AATGGTGGAGTTAAAGACT | >trnaMT_ProTGG_MT_-_15956_16023 | 40 | 58 | 19 | AATGGTGGAGTTAAAGACT | BT20 | 15 | 4 | 9 | 47 | 38 | 6.9 | 2.6 | 6.7 | 34.1 | 28.3 |
| >trnaMT_ProTGG_MT_-_15956_16023@40.58.19\|AATGGTGGAGTTAAAGACT | >trnaMT_ProTGG_MT_-_15956_16023 | 40 | 58 | 19 | AATGGTGGAGTTAAAGACT | MB231 | 37 | 9 | 9 | 100 | 32 | 39.2 | 8.0 | 7.5 | 103.7 | 20.6 |
| >trnaMT_ProTGG_MT_-_15956_16023@41.67.27\|ATGGTGGAGTTAAAGACTTTTTCTCTG | >trnaMT_ProTGG_MT_-_15956_16023 | 41 | 67 | 27 | ATGGTGGAGTTAAAGACTTTTTCTCTG | MB468 | 12 | 8 | 11 | 15 | 11 | 8.0 | 10.1 | 12.3 | 6.8 | 11.4 |
| >trnaMT_ProTGG_MT_-_15956_16023@44.70.27\|GTGGAGTTAAAGACTTTTTCTCTGACC | >trnaMT_ProTGG_MT_-_15956_16023 | 44 | 70 | 27 | GTGGAGTTAAAGACTTTTTCTCTGACC | MB231 | 17 | 10 | 8 | 74 | 24 | 8.9 | 0.5 | 1.6 | 74.6 | 9.5 |
| >trnaMT_ProTGG_MT_-_15956_16023@50.70.21\|TTAAAGACTTTTTCTCTGACC | >trnaMT_ProTGG_MT_-_15956_16023 | 50 | 70 | 21 | TTAAAGACTTTTTCTCTGACC | MB231 | 16 | 14 | 5 | 23 | 11 | 3.9 | 4.8 | 4.5 | 21.7 | 4.2 |
| >trnaMT_ProTGG_MT_-_15956_16023@50.71.22\|TTAAAGACTTTTTCTCTGACCA | >trnaMT_ProTGG_MT_-_15956_16023 | 50 | 71 | 22 | TTAAAGACTTTTTCTCTGACCA | BT20 | 33 | 4 | 7 | 8 | 19 | 20.5 | 0.6 | 2.6 | 4.4 | 19.2 |
| >trnaMT_ProTGG_MT_-_15956_16023@50.71.22\|TTAAAGACTTTTTCTCTGACCA | >trnaMT_ProTGG_MT_-_15956_16023 | 50 | 71 | 22 | TTAAAGACTTTTTCTCTGACCA | MB231 | 50 | 49 | 20 | 37 | 32 | 9.0 | 16.6 | 19.2 | 39.8 | 21.5 |
| >trnaMT_ProTGG_MT_-_15956_16023@51.70.20\|TAAAGACTTTTTCTCTGACC | >trnaMT_ProTGG_MT_-_15956_16023 | 51 | 70 | 20 | TAAAGACTTTTTCTCTGACC | MB231 | 13 | 11 | 4 | 20 | 9 | 4.9 | 4.8 | 3.9 | 21.8 | 3.2 |
| >trnaMT_ProTGG_MT_-_15956_16023@51.71.21\|TAAAGACTTTTTCTCTGACCA | >trnaMT_ProTGG_MT_-_15956_16023 | 51 | 71 | 21 | TAAAGACTTTTTCTCTGACCA | BT20 | 29 | 4 | 7 | 10 | 17 | 14.4 | 1.2 | 4.5 | 4.9 | 17.0 |
| >trnaMT_ProTGG_MT_-_15956_16023@51.71.21\|TAAAGACTTTTTCTCTGACCA | >trnaMT_ProTGG_MT_-_15956_16023 | 51 | 71 | 21 | TAAAGACTTTTTCTCTGACCA | MB231 | 39 | 35 | 16 | 30 | 27 | 9.8 | 9.2 | 14.6 | 32.2 | 17.8 |

| | | | | | | | | | | | | | | | | |
|---|---|---|---|---|---|---|---|---|---|---|---|---|---|---|---|---|
| >trnaMT_SerGCT_MT_+_12207_12265@1.18.18\|GAGAAAGCTCACAAGAAC | >trnaMT_SerGCT_MT_+_12207_12265 | 1 | 18 | 18 | GAGAAAGCTCACAAGAAC | BT20 | 1640 | 1532 | 138 | 738 | 366 | 2494.5 | 2483.7 | 171.6 | 243.2 | 434.4 |
| >trnaMT_SerGCT_MT_+_12207_12265@1.18.18\|GAGAAAGCTCACAAGAAC | >trnaMT_SerGCT_MT_+_12207_12265 | 1 | 18 | 18 | GAGAAAGCTCACAAGAAC | MB231 | 236 | 300 | 33 | 11740 | 4505 | 178.4 | 58.5 | 45.7 | 8647.0 | 3821.6 |
| >trnaMT_SerGCT_MT_+_12207_12265@1.19.19\|GAGAAAGCTCACAAGAACT | >trnaMT_SerGCT_MT_+_12207_12265 | 1 | 19 | 19 | GAGAAAGCTCACAAGAACT | BT20 | 372 | 284 | 52 | 1111 | 328 | 96.8 | 155.2 | 14.5 | 964.9 | 419.1 |
| >trnaMT_SerGCT_MT_+_12207_12265@1.19.19\|GAGAAAGCTCACAAGAACT | >trnaMT_SerGCT_MT_+_12207_12265 | 1 | 19 | 19 | GAGAAAGCTCACAAGAACT | MB231 | 259 | 76 | 14 | 1516 | 330 | 176.2 | 37.0 | 2.8 | 847.2 | 78.0 |
| >trnaMT_SerGCT_MT_+_12207_12265@1.19.19\|GAGAAAGCTCACAAGAACT | >trnaMT_SerGCT_MT_+_12207_12265 | 1 | 19 | 19 | GAGAAAGCTCACAAGAACT | MB468 | 10 | 4 | 1 | 338 | 62 | 8.7 | 4.1 | 1.3 | 180.2 | 51.6 |
| >trnaMT_SerGCT_MT_+_12207_12265@1.20.20\|GAGAAAGCTCACAAGAACTG | >trnaMT_SerGCT_MT_+_12207_12265 | 1 | 20 | 20 | GAGAAAGCTCACAAGAACTG | BT20 | 41 | 8 | 6 | 213 | 68 | 21.4 | 0.9 | 2.9 | 185.3 | 87.6 |
| >trnaMT_SerGCT_MT_+_12207_12265@1.20.20\|GAGAAAGCTCACAAGAACTG | >trnaMT_SerGCT_MT_+_12207_12265 | 1 | 20 | 20 | GAGAAAGCTCACAAGAACTG | MB231 | 30 | 11 | 3 | 272 | 83 | 20.8 | 9.9 | 2.0 | 173.2 | 40.7 |
| >trnaMT_SerGCT_MT_+_12207_12265@1.20.20\|GAGAAAGCTCACAAGAACTG | >trnaMT_SerGCT_MT_+_12207_12265 | 1 | 20 | 20 | GAGAAAGCTCACAAGAACTG | MB468 | 15 | 4 | 4 | 125 | 80 | 6.8 | 4.9 | 3.4 | 93.6 | 111.6 |
| >trnaMT_SerGCT_MT_+_12207_12265@1.21.21\|GAGAAAGCTCACAAGAACTGC | >trnaMT_SerGCT_MT_+_12207_12265 | 1 | 21 | 21 | GAGAAAGCTCACAAGAACTGC | BT20 | 47 | 22 | 18 | 199 | 185 | 23.3 | 23.6 | 16.5 | 192.3 | 278.7 |
| >trnaMT_SerGCT_MT_+_12207_12265@1.21.21\|GAGAAAGCTCACAAGAACTGC | >trnaMT_SerGCT_MT_+_12207_12265 | 1 | 21 | 21 | GAGAAAGCTCACAAGAACTGC | MB231 | 47 | 22 | 4 | 1270 | 694 | 8.1 | 7.0 | 2.2 | 429.2 | 628.6 |
| >trnaMT_SerGCT_MT_+_12207_12265@1.22.22\|GAGAAAGCTCACAAGAACTGCT | >trnaMT_SerGCT_MT_+_12207_12265 | 1 | 22 | 22 | GAGAAAGCTCACAAGAACTGCT | BT20 | 261 | 134 | 41 | 1503 | 764 | 99.1 | 98.0 | 17.7 | 1516.4 | 1104.5 |
| >trnaMT_SerGCT_MT_+_12207_12265@1.22.22\|GAGAAAGCTCACAAGAACTGCT | >trnaMT_SerGCT_MT_+_12207_12265 | 1 | 22 | 22 | GAGAAAGCTCACAAGAACTGCT | MB231 | 209 | 45 | 12 | 2011 | 370 | 180.8 | 34.7 | 6.6 | 1849.5 | 56.1 |
| >trnaMT_SerGCT_MT_+_12207_12265@1.22.22\|GAGAAAGCTCACAAGAACTGCT | >trnaMT_SerGCT_MT_+_12207_12265 | 1 | 22 | 22 | GAGAAAGCTCACAAGAACTGCT | MB468 | 14 | 3 | 2 | 258 | 72 | 6.1 | 2.9 | 1.5 | 130.5 | 64.1 |
| >trnaMT_SerGCT_MT_+_12207_12265@1.23.23\|GAGAAAGCTCACAAGAACTGCTA | >trnaMT_SerGCT_MT_+_12207_12265 | 1 | 23 | 23 | GAGAAAGCTCACAAGAACTGCTA | BT20 | 26 | 5 | 10 | 179 | 89 | 11.6 | 1.9 | 8.1 | 224.5 | 135.2 |
| >trnaMT_SerGCT_MT_+_12207_12265@1.23.23\|GAGAAAGCTCACAAGAACTGCTA | >trnaMT_SerGCT_MT_+_12207_12265 | 1 | 23 | 23 | GAGAAAGCTCACAAGAACTGCTA | MB231 | 14 | 4 | 1 | 183 | 44 | 7.2 | 2.0 | 0.8 | 126.6 | 7.9 |
| >trnaMT_SerGCT_MT_+_12207_12265@1.24.24\|GAGAAAGCTCACAAGAACTGCTAA | >trnaMT_SerGCT_MT_+_12207_12265 | 1 | 24 | 24 | GAGAAAGCTCACAAGAACTGCTAA | BT20 | 15 | 2 | 4 | 132 | 99 | 11.6 | 0.4 | 2.8 | 191.5 | 158.3 |
| >trnaMT_SerGCT_MT_+_12207_12265@1.25.25\|GAGAAAGCTCACAAGAACTGCTAAC | >trnaMT_SerGCT_MT_+_12207_12265 | 1 | 25 | 25 | GAGAAAGCTCACAAGAACTGCTAAC | BT20 | 502 | 238 | 132 | 1059 | 2517 | 275.5 | 293.9 | 80.7 | 950.4 | 3833.8 |
| >trnaMT_SerGCT_MT_+_12207_12265@1.25.25\|GAGAAAGCTCACAAGAACTGCTAAC | >trnaMT_SerGCT_MT_+_12207_12265 | 1 | 25 | 25 | GAGAAAGCTCACAAGAACTGCTAAC | MB231 | 271 | 151 | 30 | 4870 | 3646 | 11.8 | 32.1 | 14.7 | 4437.4 | 2465.5 |
| >trnaMT_SerGCT_MT_+_12207_12265@1.25.25\|GAGAAAGCTCACAAGAACTGCTAAC | >trnaMT_SerGCT_MT_+_12207_12265 | 1 | 25 | 25 | GAGAAAGCTCACAAGAACTGCTAAC | MB468 | 15 | 6 | 2 | 339 | 113 | 11.4 | 6.6 | 2.1 | 203.8 | 57.3 |
| >trnaMT_SerGCT_MT_+_12207_12265@1.26.26\|GAGAAAGCTCACAAGAACTGCTAACT | >trnaMT_SerGCT_MT_+_12207_12265 | 1 | 26 | 26 | GAGAAAGCTCACAAGAACTGCTAACT | BT20 | 349 | 84 | 42 | 1950 | 939 | 257.5 | 19.9 | 13.4 | 2505.0 | 1359.7 |
| >trnaMT_SerGCT_MT_+_12207_12265@1.26.26\|GAGAAAGCTCACAAGAACTGCTAACT | >trnaMT_SerGCT_MT_+_12207_12265 | 1 | 26 | 26 | GAGAAAGCTCACAAGAACTGCTAACT | MB231 | 266 | 99 | 19 | 2994 | 568 | 146.9 | 41.6 | 3.7 | 2400.3 | 288.3 |
| >trnaMT_SerGCT_MT_+_12207_12265@1.26.26\|GAGAAAGCTCACAAGAACTGCTAACT | >trnaMT_SerGCT_MT_+_12207_12265 | 1 | 26 | 26 | GAGAAAGCTCACAAGAACTGCTAACT | MB468 | 23 | 9 | 8 | 273 | 74 | 17.8 | 6.3 | 9.1 | 195.2 | 52.0 |
| >trnaMT_SerGCT_MT_+_12207_12265@1.38.38\|GAGAAAGCTCACAAGAACTGCTAACTCATGCC CCCATG | >trnaMT_SerGCT_MT_+_12207_12265 | 1 | 38 | 38 | GAGAAAGCTCACAAGAACTGCTAACTCATGCCCCCATG | BT20 | 20 | 16 | 8 | 6 | 54 | 15.3 | 8.5 | 6.7 | 8.0 | 69.2 |
| >trnaMT_SerGCT_MT_+_12207_12265@1.38.38\|GAGAAAGCTCACAAGAACTGCTAACTCATGCC CCCATG | >trnaMT_SerGCT_MT_+_12207_12265 | 1 | 38 | 38 | GAGAAAGCTCACAAGAACTGCTAACTCATGCCCCCATG | MB231 | 22 | 16 | 12 | 5 | 134 | 12.3 | 9.2 | 4.2 | 2.6 | 99.0 |
| >trnaMT_SerGCT_MT_+_12207_12265@1.48.48\|GAGAAAGCTCACAAGAACTGCTAACTCATGCC CCCATGTCTAACAACA | >trnaMT_SerGCT_MT_+_12207_12265 | 1 | 48 | 48 | GAGAAAGCTCACAAGAACTGCTAACTCATGCCCCCATGTCTAACAACA | BT20 | 11 | 5 | 0 | 2 | 2 | 19.2 | 8.7 | 0.8 | 1.9 | 2.0 |
| >trnaMT_SerGCT_MT_+_12207_12265@1.48.48\|GAGAAAGCTCACAAGAACTGCTAACTCATGCC CCCATGTCTAACAACA | >trnaMT_SerGCT_MT_+_12207_12265 | 1 | 48 | 48 | GAGAAAGCTCACAAGAACTGCTAACTCATGCCCCCATGTCTAACAACA | MB231 | 16 | 7 | 10 | 7 | 103 | 22.0 | 11.5 | 8.5 | 10.6 | 94.6 |
| >trnaMT_SerGCT_MT_+_12207_12265@10.38.29\|CACAAGAACTGCTAACTCATGCCCCCATG | >trnaMT_SerGCT_MT_+_12207_12265 | 10 | 38 | 29 | CACAAGAACTGCTAACTCATGCCCCCATG | MB468 | 22 | 104 | 24 | 11 | 45 | 16.8 | 95.0 | 30.2 | 7.6 | 28.5 |
| >trnaMT_SerGCT_MT_+_12207_12265@13.62.50\|AAGAACTGCTAACTCATGCCCCCATGTCTAA CAACATGGCTTTCTCACCA | >trnaMT_SerGCT_MT_+_12207_12265 | 13 | 62 | 50 | AAGAACTGCTAACTCATGCCCCCATGTCTAACAACATGGCTTTCTCACCA | BT20 | 17 | 1 | 1 | 70 | 13 | 22.8 | 1.8 | 1.4 | 60.2 | 12.6 |
| >trnaMT_SerGCT_MT_+_12207_12265@13.62.50\|AAGAACTGCTAACTCATGCCCCCATGTCTAA CAACATGGCTTTCTCACCA | >trnaMT_SerGCT_MT_+_12207_12265 | 13 | 62 | 50 | AAGAACTGCTAACTCATGCCCCCATGTCTAACAACATGGCTTTCTCACCA | MB231 | 21 | 5 | 4 | 13 | 39 | 23.4 | 8.5 | 1.8 | 22.5 | 67.7 |
| >trnaMT_SerGCT_MT_+_12207_12265@14.45.32\|AGAACTGCTAACTCATGCCCCCATGTCTAAC A | >trnaMT_SerGCT_MT_+_12207_12265 | 14 | 45 | 32 | AGAACTGCTAACTCATGCCCCCATGTCTAACA | MB231 | 15 | 6 | 5 | 15 | 19 | 15.5 | 4.9 | 2.8 | 10.1 | 10.9 |
| >trnaMT_SerGCT_MT_+_12207_12265@14.48.35\|AGAACTGCTAACTCATGCCCCCATGTCTAAC AACA | >trnaMT_SerGCT_MT_+_12207_12265 | 14 | 48 | 35 | AGAACTGCTAACTCATGCCCCCATGTCTAACAACA | MB231 | 15 | 5 | 6 | 40 | 32 | 9.3 | 0.8 | 1.6 | 45.8 | 24.3 |
| >trnaMT_SerGCT_MT_+_12207_12265@14.61.48\|AGAACTGCTAACTCATGCCCCCATGTCTAAC AACATGGCTTTCTCACC | >trnaMT_SerGCT_MT_+_12207_12265 | 14 | 61 | 48 | AGAACTGCTAACTCATGCCCCCATGTCTAACAACATGGCTTTCTCACC | MB231 | 10 | 3 | 3 | 6 | 13 | 4.0 | 4.9 | 0.6 | 10.0 | 23.2 |
| >trnaMT_SerGCT_MT_+_12207_12265@14.62.49\|AGAACTGCTAACTCATGCCCCCATGTCTAAC AACATGGCTTTCTCACCA | >trnaMT_SerGCT_MT_+_12207_12265 | 14 | 62 | 49 | AGAACTGCTAACTCATGCCCCCATGTCTAACAACATGGCTTTCTCACCA | BT20 | 20 | 2 | 2 | 50 | 40 | 25.8 | 2.7 | 2.9 | 42.9 | 30.7 |
| >trnaMT_SerGCT_MT_+_12207_12265@14.62.49\|AGAACTGCTAACTCATGCCCCCATGTCTAAC AACATGGCTTTCTCACCA | >trnaMT_SerGCT_MT_+_12207_12265 | 14 | 62 | 49 | AGAACTGCTAACTCATGCCCCCATGTCTAACAACATGGCTTTCTCACCA | MB231 | 34 | 10 | 10 | 13 | 52 | 20.6 | 15.7 | 3.1 | 22.5 | 86.4 |
| >trnaMT_SerGCT_MT_+_12207_12265@15.62.48\|GAACTGCTAACTCATGCCCCCATGTCTAACA ACATGGCTTTCTCACCA | >trnaMT_SerGCT_MT_+_12207_12265 | 15 | 62 | 48 | GAACTGCTAACTCATGCCCCCATGTCTAACAACATGGCTTTCTCACCA | MB231 | 16 | 6 | 7 | 13 | 60 | 3.9 | 9.4 | 1.9 | 21.8 | 104.0 |
| >trnaMT_SerGCT_MT_+_12207_12265@16.62.47\|AACTGCTAACTCATGCCCCCATGTCTAACAA CATGGCTTTCTCACCA | >trnaMT_SerGCT_MT_+_12207_12265 | 16 | 62 | 47 | AACTGCTAACTCATGCCCCCATGTCTAACAACATGGCTTTCTCACCA | MB231 | 10 | 4 | 6 | 7 | 29 | 1.1 | 6.3 | 1.2 | 11.9 | 50.6 |
| >trnaMT_SerGCT_MT_+_12207_12265@23.62.40\|AACTCATGCCCCCATGTCTAACAACATGGCT TTCTCACCA | >trnaMT_SerGCT_MT_+_12207_12265 | 23 | 62 | 40 | AACTCATGCCCCCATGTCTAACAACATGGCTTTCTCACCA | BT20 | 10 | 2 | 1 | 6 | 25 | 4.5 | 0.1 | 0.4 | 6.9 | 20.4 |
| >trnaMT_SerGCT_MT_+_12207_12265@27.62.36\|CATGCCCCCATGTCTAACAACATGGCTTTCT CACCA | >trnaMT_SerGCT_MT_+_12207_12265 | 27 | 62 | 36 | CATGCCCCCATGTCTAACAACATGGCTTTCTCACCA | BT20 | 16 | 3 | 2 | 9 | 33 | 6.0 | 0.5 | 0.6 | 10.2 | 41.8 |
| >trnaMT_SerGCT_MT_+_12207_12265@27.62.36\|CATGCCCCCATGTCTAACAACATGGCTTTCT CACCA | >trnaMT_SerGCT_MT_+_12207_12265 | 27 | 62 | 36 | CATGCCCCCATGTCTAACAACATGGCTTTCTCACCA | MB231 | 13 | 3 | 3 | 13 | 20 | 9.8 | 0.4 | 1.4 | 12.6 | 10.8 |
| >trnaMT_SerGCT_MT_+_12207_12265@28.62.35\|ATGCCCCCATGTCTAACAACATGGCTTTCTC ACCA | >trnaMT_SerGCT_MT_+_12207_12265 | 28 | 62 | 35 | ATGCCCCCATGTCTAACAACATGGCTTTCTCACCA | BT20 | 13 | 2 | 1 | 6 | 21 | 5.6 | 0.1 | 0.2 | 7.5 | 25.4 |
| >trnaMT_SerGCT_MT_+_12207_12265@8.30.23\|CTCACAAGAACTGCTAACTCATG | >trnaMT_SerGCT_MT_+_12207_12265 | 8 | 30 | 23 | CTCACAAGAACTGCTAACTCATG | MB468 | 13 | 33 | 15 | 8 | 12 | 8.3 | 37.0 | 13.4 | 7.9 | 10.6 |
| >trnaMT_SerGCT_MT_+_12207_12265@8.31.24\|CTCACAAGAACTGCTAACTCATGC | >trnaMT_SerGCT_MT_+_12207_12265 | 8 | 31 | 24 | CTCACAAGAACTGCTAACTCATGC | MB468 | 11 | 33 | 17 | 7 | 5 | 7.8 | 41.9 | 17.4 | 7.8 | 4.3 |
| >trnaMT_SerGCT_MT_+_12207_12265@8.38.31\|CTCACAAGAACTGCTAACTCATGCCCCCATG | >trnaMT_SerGCT_MT_+_12207_12265 | 8 | 38 | 31 | CTCACAAGAACTGCTAACTCATGCCCCCATG | MB468 | 32 | 139 | 36 | 9 | 56 | 23.9 | 120.2 | 44.1 | 9.4 | 39.5 |
| >trnaMT_ThrTGT_MT_+_15888_15953@31.69.39\|TTGTAAACCGGAGATGAAAACCTTTTTCCAAG GACACCA | >trnaMT_ThrTGT_MT_+_15888_15953 | 31 | 69 | 39 | TTGTAAACCGGAGATGAAAACCTTTTTCCAAGGACACCA | BT20 | 30 | 5 | 1 | 69 | 25 | 22.3 | 3.3 | 1.0 | 74.9 | 30.5 |
| >trnaMT_ThrTGT_MT_+_15888_15953@31.69.39\|TTGTAAACCGGAGATGAAAACCTTTTTCCAAG GACACCA | >trnaMT_ThrTGT_MT_+_15888_15953 | 31 | 69 | 39 | TTGTAAACCGGAGATGAAAACCTTTTTCCAAGGACACCA | MB231 | 24 | 7 | 2 | 41 | 22 | 20.2 | 1.3 | 1.8 | 43.4 | 15.1 |
| >trnaMT_ThrTGT_MT_+_15888_15953@34.69.36\|TAAACCGGAGATGAAAACCTTTTTCCAAGGAC ACCA | >trnaMT_ThrTGT_MT_+_15888_15953 | 34 | 69 | 36 | TAAACCGGAGATGAAAACCTTTTTCCAAGGACACCA | MB231 | 10 | 5 | 7 | 23 | 14 | 0.9 | 2.2 | 6.6 | 20.5 | 10.5 |
| >trnaMT_ThrTGT_MT_+_15888_15953@37.69.33\|ACCGGAGATGAAAACCTTTTTCCAAGGACAC CA | >trnaMT_ThrTGT_MT_+_15888_15953 | 37 | 69 | 33 | ACCGGAGATGAAAACCTTTTTCCAAGGACACCA | BT20 | 16 | 3 | 1 | 60 | 25 | 3.9 | 0.7 | 0.3 | 50.3 | 30.0 |
| >trnaMT_ThrTGT_MT_+_15888_15953@37.69.33\|ACCGGAGATGAAAACCTTTTTCCAAGGACAC CA | >trnaMT_ThrTGT_MT_+_15888_15953 | 37 | 69 | 33 | ACCGGAGATGAAAACCTTTTTCCAAGGACACCA | MB231 | 14 | 5 | 2 | 69 | 20 | 10.3 | 1.0 | 0.6 | 59.3 | 11.3 |
| >trnaMT_TrpTCA_MT_+_5512_5579@53.71.19,trnalookalike3_TrpTCA_1_+_566062_566129@5 3.71.19\|CAATACTTAATTTCTGCCA | >trnaMT_TrpTCA_MT_+_5512_5579 | 53 | 71 | 19 | CAATACTTAATTTCTGCCA | MB468 | 18 | 10 | 7 | 46 | 32 | 6.7 | 15.3 | 8.2 | 23.7 | 26.2 |
| >trnaMT_ValTAC_MT_+_1602_1670@28.71.44\|AACTTACACTTAGGAGATTTCAACTTAACTTGAC CGCTCTGACC | >trnaMT_ValTAC_MT_+_1602_1670 | 28 | 71 | 44 | AACTTACACTTAGGAGATTTCAACTTAACTTGACCGCTCTGACC | MB231 | 16 | 5 | 5 | 17 | 21 | 17.5 | 4.6 | 3.6 | 20.9 | 16.2 |
| >trnaMT_ValTAC_MT_+_1602_1670@28.72.45\|AACTTACACTTAGGAGATTTCAACTTAACTTGAC CGCTCTGACCA | >trnaMT_ValTAC_MT_+_1602_1670 | 28 | 72 | 45 | AACTTACACTTAGGAGATTTCAACTTAACTTGACCGCTCTGACCA | BT20 | 17 | 3 | 2 | 95 | 45 | 13.3 | 2.1 | 1.5 | 125.0 | 7.4 |
| >trnaMT_ValTAC_MT_+_1602_1670@28.72.45\|AACTTACACTTAGGAGATTTCAACTTAACTTGAC CGCTCTGACCA | >trnaMT_ValTAC_MT_+_1602_1670 | 28 | 72 | 45 | AACTTACACTTAGGAGATTTCAACTTAACTTGACCGCTCTGACCA | MB231 | 20 | 28 | 9 | 32 | 106 | 5.4 | 39.3 | 4.9 | 44.4 | 110.4 |
| >trnaMT_ValTAC_MT_+_1602_1670@41.71.31\|GAGATTTCAACTTAACTTGACCGCTCTGACC | >trnaMT_ValTAC_MT_+_1602_1670 | 41 | 71 | 31 | GAGATTTCAACTTAACTTGACCGCTCTGACC | MB231 | 14 | 12 | 6 | 55 | 65 | 8.1 | 2.1 | 3.4 | 51.0 | 40.8 |
| >trnaMT_ValTAC_MT_+_1602_1670@41.72.32\|GAGATTTCAACTTAACTTGACCGCTCTGACCA | >trnaMT_ValTAC_MT_+_1602_1670 | 41 | 72 | 32 | GAGATTTCAACTTAACTTGACCGCTCTGACCA | BT20 | 12 | 8 | 4 | 31 | 41 | 5.7 | 5.6 | 2.8 | 38.0 | 53.7 |
| >trnaMT_ValTAC_MT_+_1602_1670@41.72.32\|GAGATTTCAACTTAACTTGACCGCTCTGACCA | >trnaMT_ValTAC_MT_+_1602_1670 | 41 | 72 | 32 | GAGATTTCAACTTAACTTGACCGCTCTGACCA | MB231 | 13 | 24 | 6 | 39 | 84 | 4.7 | 7.7 | 3.4 | 30.7 | 60.7 |
| >trnaMT_ValTAC_MT_+_1602_1670@42.71.30\|AGATTTCAACTTAACTTGACCGCTCTGACC | >trnaMT_ValTAC_MT_+_1602_1670 | 42 | 71 | 30 | AGATTTCAACTTAACTTGACCGCTCTGACC | BT20 | 19 | 12 | 6 | 80 | 211 | 3.7 | 11.5 | 5.4 | 52.2 | 254.0 |
| >trnaMT_ValTAC_MT_+_1602_1670@42.71.30\|AGATTTCAACTTAACTTGACCGCTCTGACC | >trnaMT_ValTAC_MT_+_1602_1670 | 42 | 71 | 30 | AGATTTCAACTTAACTTGACCGCTCTGACC | MB231 | 26 | 76 | 6 | 441 | 807 | 7.3 | 10.4 | 1.8 | 340.0 | 477.3 |
| >trnaMT_ValTAC_MT_+_1602_1670@42.72.31\|AGATTTCAACTTAACTTGACCGCTCTGACCA | >trnaMT_ValTAC_MT_+_1602_1670 | 42 | 72 | 31 | AGATTTCAACTTAACTTGACCGCTCTGACCA | BT20 | 50 | 53 | 12 | 134 | 782 | 28.8 | 28.5 | 8.6 | 112.3 | 1063.1 |
| >trnaMT_ValTAC_MT_+_1602_1670@42.72.31\|AGATTTCAACTTAACTTGACCGCTCTGACCA | >trnaMT_ValTAC_MT_+_1602_1670 | 42 | 72 | 31 | AGATTTCAACTTAACTTGACCGCTCTGACCA | MB231 | 74 | 283 | 12 | 318 | 1740 | 27.4 | 103.9 | 6.3 | 208.4 | 1263.6 |
| >trnaMT_ValTAC_MT_+_1602_1670@43.72.30\|GATTTCAACTTAACTTGACCGCTCTGACCA | >trnaMT_ValTAC_MT_+_1602_1670 | 43 | 72 | 30 | GATTTCAACTTAACTTGACCGCTCTGACCA | BT20 | 12 | 8 | 5 | 27 | 157 | 8.1 | 4.8 | 4.3 | 22.9 | 210.2 |
| >trnaMT_ValTAC_MT_+_1602_1670@44.72.29\|ATTTCAACTTAACTTGACCGCTCTGACCA | >trnaMT_ValTAC_MT_+_1602_1670 | 44 | 72 | 29 | ATTTCAACTTAACTTGACCGCTCTGACCA | BT20 | 12 | 12 | 7 | 33 | 237 | 6.8 | 8.0 | 7.4 | 17.4 | 303.9 |
| >trnaMT_ValTAC_MT_+_1602_1670@45.72.28\|TTTCAACTTAACTTGACCGCTCTGACCA | >trnaMT_ValTAC_MT_+_1602_1670 | 45 | 72 | 28 | TTTCAACTTAACTTGACCGCTCTGACCA | MB231 | 13 | 47 | 17 | 4 | 56 | 12.3 | 34.9 | 14.0 | 2.7 | 44.5 |
| >trnaMT_ValTAC_MT_+_1602_1670@46.72.27\|TTCAACTTAACTTGACCGCTCTGACCA | >trnaMT_ValTAC_MT_+_1602_1670 | 46 | 72 | 27 | TTCAACTTAACTTGACCGCTCTGACCA | BT20 | 20 | 9 | 17 | 13 | 81 | 14.8 | 6.1 | 17.4 | 9.7 | 86.5 |
| >trnaMT_ValTAC_MT_+_1602_1670@46.72.27\|TTCAACTTAACTTGACCGCTCTGACCA | >trnaMT_ValTAC_MT_+_1602_1670 | 46 | 72 | 27 | TTCAACTTAACTTGACCGCTCTGACCA | MB231 | 14 | 52 | 15 | 6 | 58 | 11.4 | 37.5 | 10.0 | 2.7 | 41.3 |
| >trnaMT_ValTAC_MT_+_1602_1670@47.72.26\|TCAACTTAACTTGACCGCTCTGACCA | >trnaMT_ValTAC_MT_+_1602_1670 | 47 | 72 | 26 | TCAACTTAACTTGACCGCTCTGACCA | BT20 | 15 | 8 | 16 | 12 | 76 | 11.3 | 5.7 | 17.0 | 8.1 | 67.8 |
| >trnaMT_ValTAC_MT_+_1602_1670@47.72.26\|TCAACTTAACTTGACCGCTCTGACCA | >trnaMT_ValTAC_MT_+_1602_1670 | 47 | 72 | 26 | TCAACTTAACTTGACCGCTCTGACCA | MB231 | 15 | 67 | 10 | 5 | 57 | 16.0 | 57.8 | 7.3 | 4.8 | 47.3 |
| >trnaMT_ValTAC_MT_+_1602_1670@48.70.23\|CAACTTAACTTGACCGCTCTGAC | >trnaMT_ValTAC_MT_+_1602_1670 | 48 | 70 | 23 | CAACTTAACTTGACCGCTCTGAC | BT20 | 16 | 7 | 35 | 10 | 15 | 14.2 | 5.9 | 20.9 | 13.7 | 19.6 |
| >trnaMT_ValTAC_MT_+_1602_1670@48.71.24\|CAACTTAACTTGACCGCTCTGACC | >trnaMT_ValTAC_MT_+_1602_1670 | 48 | 71 | 24 | CAACTTAACTTGACCGCTCTGACC | BT20 | 36 | 12 | 86 | 26 | 48 | 22.5 | 9.4 | 110.3 | 20.3 | 46.7 |
| >trnaMT_ValTAC_MT_+_1602_1670@48.71.24\|CAACTTAACTTGACCGCTCTGACC | >trnaMT_ValTAC_MT_+_1602_1670 | 48 | 71 | 24 | CAACTTAACTTGACCGCTCTGACC | MB231 | 38 | 30 | 37 | 32 | 51 | 27.2 | 11.3 | 17.4 | 21.6 | 30.6 |
| >trnaMT_ValTAC_MT_+_1602_1670@48.72.25\|CAACTTAACTTGACCGCTCTGACCA | >trnaMT_ValTAC_MT_+_1602_1670 | 48 | 72 | 25 | CAACTTAACTTGACCGCTCTGACCA | BT20 | 127 | 66 | 132 | 52 | 193 | 116.2 | 42.0 | 93.1 | 58.6 | 236.4 |
| >trnaMT_ValTAC_MT_+_1602_1670@48.72.25\|CAACTTAACTTGACCGCTCTGACCA | >trnaMT_ValTAC_MT_+_1602_1670 | 48 | 72 | 25 | CAACTTAACTTGACCGCTCTGACCA | MB231 | 45 | 106 | 32 | 22 | 114 | 28.6 | 76.0 | 15.9 | 15.9 | 106.9 |
| >trnaMT_ValTAC_MT_+_1602_1670@49.71.23\|AACTTAACTTGACCGCTCTGACC | >trnaMT_ValTAC_MT_+_1602_1670 | 49 | 71 | 23 | AACTTAACTTGACCGCTCTGACC | BT20 | 21 | 8 | 39 | 25 | 50 | 16.7 | 7.7 | 53.1 | 17.9 | 41.5 |

| | | | | | | | | | | | | | | | | |
|---|---|---|---|---|---|---|---|---|---|---|---|---|---|---|---|---|
| >trnaMT_ValTAC_MT_+_1602_1670@49.71.23\|AACTTAACTTGACCGCTCTGACC | >trnaMT_ValTAC_MT_+_1602_1670 | 49 | 71 | 23 | AACTTAACTTGACCGCTCTGACC | MB231 | 13 | 13 | 10 | 16 | 29 | 10.5 | 3.0 | 1.6 | 9.5 | 11.4 |
| >trnaMT_ValTAC_MT_+_1602_1670@49.72.24\|AACTTAACTTGACCGCTCTGACCA | >trnaMT_ValTAC_MT_+_1602_1670 | 49 | 72 | 24 | AACTTAACTTGACCGCTCTGACCA | BT20 | 82 | 39 | 68 | 69 | 201 | 76.6 | 22.6 | 54.2 | 64.1 | 219.0 |
| >trnaMT_ValTAC_MT_+_1602_1670@49.72.24\|AACTTAACTTGACCGCTCTGACCA | >trnaMT_ValTAC_MT_+_1602_1670 | 49 | 72 | 24 | AACTTAACTTGACCGCTCTGACCA | MB231 | 17 | 48 | 9 | 20 | 77 | 9.2 | 23.4 | 1.6 | 13.3 | 50.7 |
| >trnaMT_ValTAC_MT_+_1602_1670@50.70.21\|ACTTAACTTGACCGCTCTGAC | >trnaMT_ValTAC_MT_+_1602_1670 | 50 | 70 | 21 | ACTTAACTTGACCGCTCTGAC | BT20 | 15 | 4 | 21 | 14 | 15 | 12.9 | 2.9 | 11.4 | 16.9 | 16.8 |
| >trnaMT_ValTAC_MT_+_1602_1670@50.71.22\|ACTTAACTTGACCGCTCTGACC | >trnaMT_ValTAC_MT_+_1602_1670 | 50 | 71 | 22 | ACTTAACTTGACCGCTCTGACC | BT20 | 33 | 10 | 61 | 33 | 65 | 19.6 | 8.5 | 79.1 | 24.2 | 58.5 |
| >trnaMT_ValTAC_MT_+_1602_1670@50.71.22\|ACTTAACTTGACCGCTCTGACC | >trnaMT_ValTAC_MT_+_1602_1670 | 50 | 71 | 22 | ACTTAACTTGACCGCTCTGACC | MB231 | 20 | 21 | 14 | 24 | 42 | 15.1 | 5.0 | 3.8 | 12.8 | 11.7 |
| >trnaMT_ValTAC_MT_+_1602_1670@50.72.23\|ACTTAACTTGACCGCTCTGACCA | >trnaMT_ValTAC_MT_+_1602_1670 | 50 | 72 | 23 | ACTTAACTTGACCGCTCTGACCA | BT20 | 141 | 60 | 135 | 99 | 354 | 137.5 | 35.2 | 101.3 | 101.5 | 441.0 |
| >trnaMT_ValTAC_MT_+_1602_1670@50.72.23\|ACTTAACTTGACCGCTCTGACCA | >trnaMT_ValTAC_MT_+_1602_1670 | 50 | 72 | 23 | ACTTAACTTGACCGCTCTGACCA | MB231 | 32 | 94 | 21 | 33 | 124 | 17.5 | 50.6 | 10.7 | 24.9 | 80.7 |
| >trnaMT_ValTAC_MT_+_1602_1670@50.72.23\|ACTTAACTTGACCGCTCTGACCA | >trnaMT_ValTAC_MT_+_1602_1670 | 50 | 72 | 23 | ACTTAACTTGACCGCTCTGACCA | MB468 | 16 | 161 | 3 | 24 | 81 | 6.5 | 137.6 | 3.6 | 15.0 | 99.2 |
| >trnaMT_ValTAC_MT_+_1602_1670@51.71.21\|CTTAACTTGACCGCTCTGACC | >trnaMT_ValTAC_MT_+_1602_1670 | 51 | 71 | 21 | CTTAACTTGACCGCTCTGACC | MB468 | 11 | 62 | 5 | 34 | 75 | 4.8 | 97.7 | 5.3 | 14.8 | 79.4 |
| >trnaMT_ValTAC_MT_+_1602_1670@51.72.22\|CTTAACTTGACCGCTCTGACCA | >trnaMT_ValTAC_MT_+_1602_1670 | 51 | 72 | 22 | CTTAACTTGACCGCTCTGACCA | BT20 | 19 | 8 | 25 | 14 | 53 | 17.1 | 3.6 | 17.8 | 6.2 | 48.0 |
| >trnaMT_ValTAC_MT_+_1602_1670@51.72.22\|CTTAACTTGACCGCTCTGACCA | >trnaMT_ValTAC_MT_+_1602_1670 | 51 | 72 | 22 | CTTAACTTGACCGCTCTGACCA | MB468 | 69 | 401 | 15 | 97 | 313 | 28.1 | 515.0 | 13.4 | 58.3 | 362.6 |
| >trnaMT_ValTAC_MT_+_1602_1670@52.72.21\|TTAACTTGACCGCTCTGACCA | >trnaMT_ValTAC_MT_+_1602_1670 | 52 | 72 | 21 | TTAACTTGACCGCTCTGACCA | BT20 | 44 | 14 | 29 | 24 | 94 | 37.6 | 9.8 | 24.4 | 15.4 | 93.7 |
| >trnaMT_ValTAC_MT_+_1602_1670@52.72.21\|TTAACTTGACCGCTCTGACCA | >trnaMT_ValTAC_MT_+_1602_1670 | 52 | 72 | 21 | TTAACTTGACCGCTCTGACCA | MB231 | 14 | 55 | 6 | 11 | 59 | 9.1 | 33.5 | 2.9 | 7.5 | 32.5 |
| >trnaMT_ValTAC_MT_+_1602_1670@53.71.19\|TAACTTGACCGCTCTGACC | >trnaMT_ValTAC_MT_+_1602_1670 | 53 | 71 | 19 | TAACTTGACCGCTCTGACC | BT20 | 27 | 9 | 48 | 17 | 32 | 23.2 | 10.6 | 72.9 | 12.9 | 21.0 |
| >trnaMT_ValTAC_MT_+_1602_1670@53.71.19\|TAACTTGACCGCTCTGACC | >trnaMT_ValTAC_MT_+_1602_1670 | 53 | 71 | 19 | TAACTTGACCGCTCTGACC | MB231 | 21 | 25 | 12 | 18 | 32 | 16.0 | 6.5 | 4.8 | 12.2 | 14.4 |
| >trnaMT_ValTAC_MT_+_1602_1670@53.72.20\|TAACTTGACCGCTCTGACCA | >trnaMT_ValTAC_MT_+_1602_1670 | 53 | 72 | 20 | TAACTTGACCGCTCTGACCA | BT20 | 121 | 56 | 95 | 48 | 183 | 100.3 | 43.8 | 100.6 | 38.0 | 167.1 |
| >trnaMT_ValTAC_MT_+_1602_1670@53.72.20\|TAACTTGACCGCTCTGACCA | >trnaMT_ValTAC_MT_+_1602_1670 | 53 | 72 | 20 | TAACTTGACCGCTCTGACCA | MB231 | 34 | 121 | 12 | 19 | 94 | 20.9 | 64.0 | 6.9 | 14.0 | 56.7 |
| >trnaMT_ValTAC_MT_+_1602_1670@54.71.18\|AACTTGACCGCTCTGACC | >trnaMT_ValTAC_MT_+_1602_1670 | 54 | 71 | 18 | AACTTGACCGCTCTGACC | BT20 | 31 | 19 | 76 | 20 | 40 | 27.4 | 18.1 | 116.3 | 14.6 | 27.2 |
| >trnaMT_ValTAC_MT_+_1602_1670@54.71.18\|AACTTGACCGCTCTGACC | >trnaMT_ValTAC_MT_+_1602_1670 | 54 | 71 | 18 | AACTTGACCGCTCTGACC | MB231 | 23 | 50 | 19 | 20 | 34 | 8.7 | 30.8 | 11.7 | 8.2 | 5.8 |
| >trnaMT_ValTAC_MT_+_1602_1670@54.72.19\|AACTTGACCGCTCTGACCA | >trnaMT_ValTAC_MT_+_1602_1670 | 54 | 72 | 19 | AACTTGACCGCTCTGACCA | BT20 | 113 | 116 | 131 | 59 | 190 | 86.2 | 67.9 | 152.9 | 30.9 | 125.9 |
| >trnaMT_ValTAC_MT_+_1602_1670@54.72.19\|AACTTGACCGCTCTGACCA | >trnaMT_ValTAC_MT_+_1602_1670 | 54 | 72 | 19 | AACTTGACCGCTCTGACCA | MB231 | 32 | 166 | 18 | 25 | 98 | 5.8 | 45.4 | 7.3 | 15.4 | 43.8 |
| >trnaMT_ValTAC_MT_+_1602_1670@55.72.18\|ACTTGACCGCTCTGACCA | >trnaMT_ValTAC_MT_+_1602_1670 | 55 | 72 | 18 | ACTTGACCGCTCTGACCA | BT20 | 155 | 102 | 147 | 65 | 252 | 135.7 | 54.7 | 146.1 | 35.8 | 174.2 |
| >trnaMT_ValTAC_MT_+_1602_1670@55.72.18\|ACTTGACCGCTCTGACCA | >trnaMT_ValTAC_MT_+_1602_1670 | 55 | 72 | 18 | ACTTGACCGCTCTGACCA | MB231 | 40 | 165 | 18 | 26 | 156 | 13.4 | 29.3 | 8.7 | 18.2 | 64.6 |
