## Supplementary material for "Single-nucleotide Differences and Cell Type Decide the Subcellular Localization of miRNA Isoforms (isomiRs), tRNA-derived Fragments (tRFs) and rRNA-derived Fragments (rRFs)": Supp. Table S3

**Supplemental Table S3:** Data table of the top 10% most abundant short RNA from each cell line based on average abundance of the cell line replicates. The annotation "am" is used to indicate abiguous short RNA, those that can be mapped to a noncoding RNA gene as well as somewhere else on the geneome. The annotation "multi" referrs to short RNA that map to locations in both the mitochondrial and nuclear genomes. The rRF are flanked by 50 nucleotdies upstream and downstream and their coordiantes reflect this. The coordiantes for the isomiR and rRF are from the hg38 human genome assembly while the coordintes for the tRF are from the hg19 human genome assembly. Columns H-J indicate whether or not a given short RNA is present in the top 10% of the short RNA in one of the three cell lines and is indicated by blue (TRUE) or white (FALSE) sorted from most abundant to least abundant.

<table>
<tr><th>Full Label</th><th>Coordinates</th><th>Start</th><th>End</th><th>Length</th><th>Sequence</th><th>RNA</th><th>BT20</th><th>MB231</th><th>MB468</th></tr>
<tr><td>>MI0000077|hsa-mir-21&WithFlank&17|+|59841260|59841343@14.36.23&[MIMAT0000076&hsa-miR-21-5p&offsets|0|+1;m-1&17|+|59841273|59841294&offsets|0|+1]|TAGCTTATCAGACTGATGTTGAC</td><td>>MI0000077|hsa-mir-21&WithFlank&17|+|59841260|59841343</td><td>14</td><td>36</td><td>23</td><td>TAGCTTATCAGACTGATGTTGAC</td><td>isomiR</td><td>TRUE</td><td>TRUE</td><td>TRUE</td></tr>
<tr><td>>MI0000253|hsa-mir-148a&WithFlank&7|-|25949913|25949992@50.71.22&[MIMAT0000243&hsa-miR-148a-3p&offsets|0|0;m-7&7|-|25949922|25949943&offsets|0|0]|TCAGTGCACTACAGAACTTTGT</td><td>>MI0000253|hsa-mir-148a&WithFlank&7|-|25949913|25949992</td><td>50</td><td>71</td><td>22</td><td>TCAGTGCACTACAGAACTTTGT</td><td>isomiR</td><td>TRUE</td><td>TRUE</td><td>TRUE</td></tr>
<tr><td>>MI0000077|hsa-mir-21&WithFlank&17|+|59841260|59841343@14.35.22&[MIMAT0000076&hsa-miR-21-5p&offsets|0|0;m-1&17|+|59841273|59841294&offsets|0|0]|TAGCTTATCAGACTGATGTTGA</td><td>>MI0000077|hsa-mir-21&WithFlank&17|+|59841260|59841343</td><td>14</td><td>35</td><td>22</td><td>TAGCTTATCAGACTGATGTTGA</td><td>isomiR</td><td>TRUE</td><td>TRUE</td><td>TRUE</td></tr>
<tr><td>>MI0000077|hsa-mir-21&WithFlank&17|+|59841260|59841343@14.35.22(+1U)&[MIMAT0000076&hsa-miR-21-5p&offsets|0|0(+1U);m-1&17|+|59841273|59841294&offsets|0|0(+1U)]|TAGCTTATCAGACTGATGTTGAT</td><td>>MI0000077|hsa-mir-21&WithFlank&17|+|59841260|59841343</td><td>14</td><td>35</td><td>22(+1U)</td><td>TAGCTTATCAGACTGATGTTGAT</td><td>isomiR</td><td>TRUE</td><td>TRUE</td><td>FALSE</td></tr>
<tr><td>>MI0000077|hsa-mir-21&WithFlank&17|+|59841260|59841343@14.37.24&[MIMAT0000076&hsa-miR-21-5p&offsets|0|+2;m-1&17|+|59841273|59841294&offsets|0|+2]|TAGCTTATCAGACTGATGTTGACT</td><td>>MI0000077|hsa-mir-21&WithFlank&17|+|59841260|59841343</td><td>14</td><td>37</td><td>24</td><td>TAGCTTATCAGACTGATGTTGACT</td><td>isomiR</td><td>TRUE</td><td>TRUE</td><td>TRUE</td></tr>
<tr><td>>MI0000077|hsa-mir-21&WithFlank&17|+|59841260|59841343@14.36.23(+1A)&[MIMAT0000076&hsa-miR-21-5p&offsets|0|+1(+1A);m-1&17|+|59841273|59841294&offsets|0|+1(+1A)]|TAGCTTATCAGACTGATGTTGACA</td><td>>MI0000077|hsa-mir-21&WithFlank&17|+|59841260|59841343</td><td>14</td><td>36</td><td>23(+1A)</td><td>TAGCTTATCAGACTGATGTTGACA</td><td>isomiR</td><td>TRUE</td><td>TRUE</td><td>TRUE</td></tr>
<tr><td>>trna10_LysCTT_16_+_3241501_3241573@1.35.35,trna13_LysCTT_14_-_58706613_58706685@1.35.35,trna2_LysCTT_15_+_79152904_79152976@1.35.35|GCCCGGCTAGCTCAGTCGGTAGAGCATGGGACTCT</td><td>>trna10_LysCTT_16_+_3241501_3241573</td><td>1</td><td>35</td><td>35</td><td>GCCCGGCTAGCTCAGTCGGTAGAGCATGGGACTCT</td><td>tRF</td><td>TRUE</td><td>TRUE</td><td>FALSE</td></tr>
<tr><td>>MI0000650|hsa-mir-200c&WithFlank&12|+|6963693|6963772@50.72.23&[MIMAT0000617&hsa-miR-200c-3p&offsets|0|0;m-54&12|+|6963742|6963764&offsets|0|0]|TAATACTGCCGGGTAATGATGGA</td><td>>MI0000650|hsa-mir-200c&WithFlank&12|+|6963693|6963772</td><td>50</td><td>72</td><td>23</td><td>TAATACTGCCGGGTAATGATGGA</td><td>isomiR</td><td>TRUE</td><td>FALSE</td><td>FALSE</td></tr>
<tr><td>>MI0000067|hsa-let-7f-1&WithFlank&9|+|94176341|94176439@13.34.22;MI0000068|hsa-let-7f-2&WithFlank&X|-|53557186|53557280@14.35.22&[MIMAT0000067&hsa-let-7f-5p&offsets|0|0;m-20&9|+|94176353|94176374&offsets|0|0];[MIMAT0000067_1&hsa-let-7f-5p&offsets|0|0;m-19&X|-|53557246|53557267&offsets|0|0]|TGAGGTAGTAGATTGTATAGTT</td><td>>MI0000067|hsa-let-7f-1&WithFlank&9|+|94176341|94176439</td><td>13</td><td>34</td><td>22</td><td>TGAGGTAGTAGATTGTATAGTT</td><td>isomiR</td><td>TRUE</td><td>TRUE</td><td>TRUE</td></tr>
<tr><td>>MI0000264|hsa-mir-7-2&WithFlank&15|+|88611819|88611940@38.61.24;MI0000265|hsa-mir-7-3&WithFlank&19|+|4770664|4770785@37.60.24;MI0000263|hsa-mir-7-1&WithFlank&9|-|83969742|83969863@30.53.24&[MIMAT0000252&hsa-miR-7-5p&offsets|0|0;m-400&15|+|88611856|88611879&offsets|0|0];[MIMAT0000252_1&hsa-miR-7-5p&offsets|0|0;m-402&19|+|4770700|4770723&offsets|0|0];[MIMAT0000252_2&hsa-miR-7-5p&offsets|0|0;m-401&9|-|83969811|83969834&offsets|0|0]|TGGAAGACTAGTGATTTTGTTGTT</td><td>>MI0000264|hsa-mir-7-2&WithFlank&15|+|88611819|88611940</td><td>38</td><td>61</td><td>24</td><td>TGGAAGACTAGTGATTTTGTTGTT</td><td>isomiR</td><td>TRUE</td><td>TRUE</td><td>TRUE</td></tr>
<tr><td>>MI0000088|hsa-mir-30a&WithFlank&6|-|71403545|71403627@12.35.24&[MIMAT0000087&hsa-miR-30a-5p&offsets|0|+2;m-15&6|-|71403595|71403616&offsets|0|+2]|TGTAAACATCCTCGACTGGAAGCT</td><td>>MI0000088|hsa-mir-30a&WithFlank&6|-|71403545|71403627</td><td>12</td><td>35</td><td>24</td><td>TGTAAACATCCTCGACTGGAAGCT</td><td>isomiR</td><td>TRUE</td><td>TRUE</td><td>TRUE</td></tr>
<tr><td>>MI0000255|hsa-mir-30d&WithFlank&8|-|134804870|134804951@12.35.24&[MIMAT0000245&hsa-miR-30d-5p&offsets|0|+2;m-21&8|-|134804919|134804940&offsets|0|+2]|TGTAAACATCCCCGACTGGAAGCT</td><td>>MI0000255|hsa-mir-30d&WithFlank&8|-|134804870|134804951</td><td>12</td><td>35</td><td>24</td><td>TGTAAACATCCCCGACTGGAAGCT</td><td>isomiR</td><td>TRUE</td><td>TRUE</td><td>FALSE</td></tr>
<tr><td>>MI0000433|hsa-let-7g&WithFlank&3|-|52268272|52268367@11.32.22&[MIMAT0000414&hsa-let-7g-5p&offsets|0|0;m-58&3|-|52268336|52268357&offsets|0|0]|TGAGGTAGTAGTTTGTACAGTT</td><td>>MI0000433|hsa-let-7g&WithFlank&3|-|52268272|52268367</td><td>11</td><td>32</td><td>22</td><td>TGAGGTAGTAGTTTGTACAGTT</td><td>isomiR</td><td>TRUE</td><td>TRUE</td><td>TRUE</td></tr>
<tr><td>>MI0000434|hsa-let-7i&WithFlank&12|+|62603680|62603775@12.33.22&[MIMAT0000415&hsa-let-7i-5p&offsets|0|0;m-49&12|+|62603691|62603712&offsets|0|0]|TGAGGTAGTAGTTTGTGCTGTT</td><td>>MI0000434|hsa-let-7i&WithFlank&12|+|62603680|62603775</td><td>12</td><td>33</td><td>22</td><td>TGAGGTAGTAGTTTGTGCTGTT</td><td>isomiR</td><td>TRUE</td><td>TRUE</td><td>TRUE</td></tr>
<tr><td>>MI0000061|hsa-let-7a-2&WithFlank&11|-|122146516|122146599@11.32.22;MI0000062|hsa-let-7a-3&WithFlank&22|+|46112743|46112828@10.31.22;MI0000060|hsa-let-7a-1&WithFlank&9|+|94175951|94176042@12.33.22&[MIMAT0000062&hsa-let-7a-5p&offsets|0|0;m-6&11|-|122146568|122146589&offsets|0|0];[MIMAT0000062_1&hsa-let-7a-5p&offsets|0|0;m-5&22|+|46112752|46112773&offsets|0|0];[MIMAT0000062_2&hsa-let-7a-5p&offsets|0|0;m-4&9|+|94175962|94175983&offsets|0|0]|TGAGGTAGTAGGTTGTATAGTT</td><td>>MI0000061|hsa-let-7a-2&WithFlank&11|-|122146516|122146599</td><td>11</td><td>32</td><td>22</td><td>TGAGGTAGTAGGTTGTATAGTT</td><td>isomiR</td><td>TRUE</td><td>TRUE</td><td>TRUE</td></tr>
<tr><td>>am_MI0000809|hsa-mir-151a&WithFlank&8|-|140732558|140732659@53.73.21&[MIMAT0000757&hsa-miR-151a-3p&offsets|0|0;m-30&8|-|140732587|140732607&offsets|0|0]|CTAGACTGAAGCTCCTTGAGG</td><td>>am_MI0000809|hsa-mir-151a&WithFlank&8|-|140732558|140732659</td><td>53</td><td>73</td><td>21</td><td>CTAGACTGAAGCTCCTTGAGG</td><td>isomiR</td><td>TRUE</td><td>TRUE</td><td>FALSE</td></tr>
<tr><td>>MI0000440|hsa-mir-27b&WithFlank&9|+|95085439|95085547@67.87.21&[MIMAT0000419&hsa-miR-27b-3p&offsets|0|0;m-39&9|+|95085505|95085525&offsets|0|0]|TTCACAGTGGCTAAGTTCTGC</td><td>>MI0000440|hsa-mir-27b&WithFlank&9|+|95085439|95085547</td><td>67</td><td>87</td><td>21</td><td>TTCACAGTGGCTAAGTTCTGC</td><td>isomiR</td><td>TRUE</td><td>TRUE</td><td>FALSE</td></tr>
<tr><td>>hg38_21_+_8433172_8446622_RNA45SN1_45S_50ntFlanks@7975.8011.37|CGCGACCTCAGATCAGACGTGGCGACCCGCTGAATTT</td><td>>hg38_21_+_8433172_8446622_RNA45SN1_45S_50ntFlanks</td><td>7975</td><td>8011</td><td>37</td><td>CGCGACCTCAGATCAGACGTGGCGACCCGCTGAATTT</td><td>rRF</td><td>TRUE</td><td>TRUE</td><td>FALSE</td></tr>
<tr><td>>MI0000272|hsa-mir-182&WithFlank&7|-|129770377|129770498@29.52.24&[MIMAT0000259&hsa-miR-182-5p&offsets|0|0;m-22&7|-|129770449|129770470&offsets|0|+2]|TTTGGCAATGGTAGAACTCACACT</td><td>>MI0000272|hsa-mir-182&WithFlank&7|-|129770377|129770498</td><td>29</td><td>52</td><td>24</td><td>TTTGGCAATGGTAGAACTCACACT</td><td>isomiR</td><td>TRUE</td><td>TRUE</td><td>FALSE</td></tr>
<tr><td>>hg38_21_+_8433172_8446622_RNA45SN1_45S_50ntFlanks@7975.8010.36|CGCGACCTCAGATCAGACGTGGCGACCCGCTGAATT</td><td>>hg38_21_+_8433172_8446622_RNA45SN1_45S_50ntFlanks</td><td>7975</td><td>8010</td><td>36</td><td>CGCGACCTCAGATCAGACGTGGCGACCCGCTGAATT</td><td>rRF</td><td>TRUE</td><td>TRUE</td><td>FALSE</td></tr>
<tr><td>>MI0000750|hsa-mir-26a-2&WithFlank&12|-|57824603|57824698@20.41.22;MI0000083|hsa-mir-26a-1&WithFlank&3|+|37969398|37969486@16.37.22&[MIMAT0000082&hsa-miR-26a-5p&offsets|0|0;m-33&12|-|57824658|57824679&offsets|0|0];[MIMAT0000082_1&hsa-miR-26a-5p&offsets|0|0;m-34&3|+|37969413|37969434&offsets|0|0]|TTCAAGTAATCCAGGATAGGCT</td><td>>MI0000750|hsa-mir-26a-2&WithFlank&12|-|57824603|57824698</td><td>20</td><td>41</td><td>22</td><td>TTCAAGTAATCCAGGATAGGCT</td><td>isomiR</td><td>TRUE</td><td>TRUE</td><td>TRUE</td></tr>
<tr><td>>MI0000077|hsa-mir-21&WithFlank&17|+|59841260|59841343@14.35.22(+1A)&[MIMAT0000076&hsa-miR-21-5p&offsets|0|0(+1A);m-1&17|+|59841273|59841294&offsets|0|0(+1A)]|TAGCTTATCAGACTGATGTTGAA</td><td>>MI0000077|hsa-mir-21&WithFlank&17|+|59841260|59841343</td><td>14</td><td>35</td><td>22(+1A)</td><td>TAGCTTATCAGACTGATGTTGAA</td><td>isomiR</td><td>TRUE</td><td>TRUE</td><td>FALSE</td></tr>
<tr><td>>MI0000285|hsa-mir-205&WithFlank&1|+|209432127|209432248@40.61.22(+1A)&[MIMAT0000266&hsa-miR-205-5p&offsets|0|0(+1A);m-351&1|+|209432166|209432187&offsets|0|0(+1A)]|TCCTTCATTCCACCGGAGTCTGA</td><td>>MI0000285|hsa-mir-205&WithFlank&1|+|209432127|209432248</td><td>40</td><td>61</td><td>22(+1A)</td><td>TCCTTCATTCCACCGGAGTCTGA</td><td>isomiR</td><td>TRUE</td><td>FALSE</td><td>TRUE</td></tr>
</table>

<table>
<tr><th></th><th></th><th></th><th></th><th></th><th></th><th></th><th></th><th></th><th></th></tr>
<tr><td>>hg38_21_+_8433172_8446622_RNA45SN1_45S_50ntFlanks@12995.13039.45|CTCGCTGCGATCTATTGAAAGTCAGCCCTCGACACAAGGGTTTGT</td><td>>hg38_21_+_8433172_8446622_RNA45SN1_45S_50ntFlanks</td><td>12995</td><td>13039</td><td>45</td><td>CTCGCTGCGATCTATTGAAAGTCAGCCCTCGACACAAGGGTTTGT</td><td>rRF</td><td>TRUE</td><td>TRUE</td><td>FALSE</td></tr>
<tr><td>>MI0000285|hsa-mir-205&WithFlank&1|+|209432127|209432248@40.62.23&[MIMAT0000266&hsa-miR-205-5p&offsets|0|+1;m-351&1|+|209432166|209432187&offsets|0|+1]|TCCTTCATTCCACCGGAGTCTGT</td><td>>MI0000285|hsa-mir-205&WithFlank&1|+|209432127|209432248</td><td>40</td><td>62</td><td>23</td><td>TCCTTCATTCCACCGGAGTCTGT</td><td>isomiR</td><td>TRUE</td><td>FALSE</td><td>FALSE</td></tr>
<tr><td>>MI0000077|hsa-mir-21&WithFlank&17|+|59841260|59841343@14.36.23(+1C)&[MIMAT0000076&hsa-miR-21-5p&offsets|0|+1(+1C);m-1&17|+|59841273|59841294&offsets|0|+1(+1C)]|TAGCTTATCAGACTGATGTTGACC</td><td>>MI0000077|hsa-mir-21&WithFlank&17|+|59841260|59841343</td><td>14</td><td>36</td><td>23(+1C)</td><td>TAGCTTATCAGACTGATGTTGACC</td><td>isomiR</td><td>TRUE</td><td>TRUE</td><td>FALSE</td></tr>
<tr><td>>MI0000063|hsa-let-7b&WithFlank&22|+|46113680|46113774@12.33.22&[MIMAT0000063&hsa-let-7b-5p&offsets|0|0;m-14&22|+|46113691|46113712&offsets|0|0]|TGAGGTAGTAGGTTGTGTGGTT</td><td>>MI0000063|hsa-let-7b&WithFlank&22|+|46113680|46113774</td><td>12</td><td>33</td><td>22</td><td>TGAGGTAGTAGGTTGTGTGGTT</td><td>isomiR</td><td>TRUE</td><td>TRUE</td><td>FALSE</td></tr>
<tr><td>>MI0000088|hsa-mir-30a&WithFlank&6|-|71403545|71403627@12.34.23&[MIMAT0000087&hsa-miR-30a-5p&offsets|0|+1;m-15&6|-|71403595|71403616&offsets|0|+1]|TGTAAACATCCTCGACTGGAAGC</td><td>>MI0000088|hsa-mir-30a&WithFlank&6|-|71403545|71403627</td><td>12</td><td>34</td><td>23</td><td>TGTAAACATCCTCGACTGGAAGC</td><td>isomiR</td><td>TRUE</td><td>TRUE</td><td>FALSE</td></tr>
<tr><td>>hg38_21_+_8433172_8446622_RNA45SN1_45S_50ntFlanks@7975.8009.35|CGCGACCTCAGATCAGACGTGGCGACCCGCTGAAT</td><td>>hg38_21_+_8433172_8446622_RNA45SN1_45S_50ntFlanks</td><td>7975</td><td>8009</td><td>35</td><td>CGCGACCTCAGATCAGACGTGGCGACCCGCTGAAT</td><td>rRF</td><td>TRUE</td><td>TRUE</td><td>FALSE</td></tr>
<tr><td>>MI0000255|hsa-mir-30d&WithFlank&8|-|134804870|134804951@12.34.23&[MIMAT0000245&hsa-miR-30d-5p&offsets|0|+1;m-21&8|-|134804919|134804940&offsets|0|+1]|TGTAAACATCCCCGACTGGAAGC</td><td>>MI0000255|hsa-mir-30d&WithFlank&8|-|134804870|134804951</td><td>12</td><td>34</td><td>23</td><td>TGTAAACATCCCCGACTGGAAGC</td><td>isomiR</td><td>TRUE</td><td>TRUE</td><td>FALSE</td></tr>
<tr><td>>MI0000253|hsa-mir-148a&WithFlank&7|-|25949913|25949992@50.70.21(+1C)&[MIMAT0000243&hsa-miR-148a-3p&offsets|0|-1(+1C);m-7&7|-|25949922|25949943&offsets|0|-1(+1C)]|TCAGTGCACTACAGAACTTTGC</td><td>>MI0000253|hsa-mir-148a&WithFlank&7|-|25949913|25949992</td><td>50</td><td>70</td><td>21(+1C)</td><td>TCAGTGCACTACAGAACTTTGC</td><td>isomiR</td><td>TRUE</td><td>FALSE</td><td>FALSE</td></tr>
<tr><td>>MI0000082|hsa-mir-25&WithFlank&7|-|100093554|100093649@58.79.22&[MIMAT0000081&hsa-miR-25-3p&offsets|0|0;m-23&7|-|100093571|100093592&offsets|0|0]|CATTGCACTTGTCTCGGTCTGA</td><td>>MI0000082|hsa-mir-25&WithFlank&7|-|100093554|100093649</td><td>58</td><td>79</td><td>22</td><td>CATTGCACTTGTCTCGGTCTGA</td><td>isomiR</td><td>TRUE</td><td>TRUE</td><td>FALSE</td></tr>
<tr><td>>hg38_21_+_8433172_8446622_RNA45SN1_45S_50ntFlanks@6646.6665.20|TCGTACGACTCTTAGCGGTG</td><td>>hg38_21_+_8433172_8446622_RNA45SN1_45S_50ntFlanks</td><td>6646</td><td>6665</td><td>20</td><td>TCGTACGACTCTTAGCGGTG</td><td>rRF</td><td>TRUE</td><td>TRUE</td><td>FALSE</td></tr>
<tr><td>>MI0000077|hsa-mir-21&WithFlank&17|+|59841260|59841343@14.36.23(+1G)&[MIMAT0000076&hsa-miR-21-5p&offsets|0|+1(+1G);m-1&17|+|59841273|59841294&offsets|0|+1(+1G)]|TAGCTTATCAGACTGATGTTGACG</td><td>>MI0000077|hsa-mir-21&WithFlank&17|+|59841260|59841343</td><td>14</td><td>36</td><td>23(+1G)</td><td>TAGCTTATCAGACTGATGTTGACG</td><td>isomiR</td><td>TRUE</td><td>TRUE</td><td>FALSE</td></tr>
<tr><td>>trna2_GlyGCC_21_-_18827107_18827177@1.31.31,trna35_GlyGCC_1_+_161413094_161413164@1.31.31,trna37_GlyGCC_1_+_161420467_161420537@1.31.31,trna39_GlyGCC_1_+_161427898_161427968@1.31.31,trna41_GlyGCC_1_+_161435258_161435328@1.31.31|GCATGGGTGGTTCAGTGGTAGAATTCTCGCC</td><td>>trna2_GlyGCC_21_-_18827107_18827177</td><td>1</td><td>31</td><td>31</td><td>GCATGGGTGGTTCAGTGGTAGAATTCTCGCC</td><td>tRF</td><td>TRUE</td><td>FALSE</td><td>FALSE</td></tr>
<tr><td>>hg38_21_+_8433172_8446622_RNA45SN1_45S_50ntFlanks@7975.8005.31|CGCGACCTCAGATCAGACGTGGCGACCCGCT</td><td>>hg38_21_+_8433172_8446622_RNA45SN1_45S_50ntFlanks</td><td>7975</td><td>8005</td><td>31</td><td>CGCGACCTCAGATCAGACGTGGCGACCCGCT</td><td>rRF</td><td>TRUE</td><td>TRUE</td><td>FALSE</td></tr>
<tr><td>>MI0000746|hsa-mir-99b&WithFlank&19|+|51692606|51692687@13.34.22&[MIMAT0000689&hsa-miR-99b-5p&offsets|0|0;m-10&19|+|51692618|51692639&offsets|0|0]|CACCCGTAGAACCGACCTTGCG</td><td>>MI0000746|hsa-mir-99b&WithFlank&19|+|51692606|51692687</td><td>13</td><td>34</td><td>22</td><td>CACCCGTAGAACCGACCTTGCG</td><td>isomiR</td><td>TRUE</td><td>TRUE</td><td>FALSE</td></tr>
<tr><td>>hg38_21_+_8433172_8446622_RNA45SN1_45S_50ntFlanks@7975.7994.20|CGCGACCTCAGATCAGACGT</td><td>>hg38_21_+_8433172_8446622_RNA45SN1_45S_50ntFlanks</td><td>7975</td><td>7994</td><td>20</td><td>CGCGACCTCAGATCAGACGT</td><td>rRF</td><td>TRUE</td><td>TRUE</td><td>TRUE</td></tr>
<tr><td>>MI0000093|hsa-mir-92a-1&WithFlank&13|+|91351308|91351397@54.75.22;MI0000094|hsa-mir-92a-2&WithFlank&X|-|134169532|134169618@54.75.22&[MIMAT0000092&hsa-miR-92a-3p&offsets|0|0;m-9&13|+|91351361|91351382&offsets|0|0];[MIMAT0000092_1&hsa-miR-92a-3p&offsets|0|0;m-11&X|-|134169544|134169565&offsets|0|0]|TATTGCACTTGTCCCGGCCTGT</td><td>>MI0000093|hsa-mir-92a-1&WithFlank&13|+|91351308|91351397</td><td>54</td><td>75</td><td>22</td><td>TATTGCACTTGTCCCGGCCTGT</td><td>isomiR</td><td>TRUE</td><td>TRUE</td><td>FALSE</td></tr>
<tr><td>>MI0000108|hsa-mir-103a-2&WithFlank&20|+|3917488|3917577@54.76.23;MI0000109|hsa-mir-103a-1&WithFlank&5|-|168560890|168560979@54.76.23&[MIMAT0000101&hsa-miR-103a-3p&offsets|0|0;m-12&20|+|3917541|3917563&offsets|0|0];[MIMAT0000101_1&hsa-miR-103a-3p&offsets|0|0;m-13&5|-|168560904|168560926&offsets|0|0;m-44220&5|-|168560908|168560926&offsets|0|+4]|AGCAGCATTGTACAGGGCTATGA</td><td>>MI0000108|hsa-mir-103a-2&WithFlank&20|+|3917488|3917577</td><td>54</td><td>76</td><td>23</td><td>AGCAGCATTGTACAGGGCTATGA</td><td>isomiR</td><td>TRUE</td><td>TRUE</td><td>FALSE</td></tr>
<tr><td>>hg38_21_+_8433172_8446622_RNA45SN1_45S_50ntFlanks@12997.13039.43|CGCTGCGATCTATTGAAAGTCAGCCCTCGACACAAGGGTTTGT</td><td>>hg38_21_+_8433172_8446622_RNA45SN1_45S_50ntFlanks</td><td>12997</td><td>13039</td><td>43</td><td>CGCTGCGATCTATTGAAAGTCAGCCCTCGACACAAGGGTTTGT</td><td>rRF</td><td>TRUE</td><td>TRUE</td><td>FALSE</td></tr>
<tr><td>>hg38_21_+_8433172_8446622_RNA45SN1_45S_50ntFlanks@12996.13039.44|TCGCTGCGATCTATTGAAAGTCAGCCCTCGACACAAGGGTTTGT</td><td>>hg38_21_+_8433172_8446622_RNA45SN1_45S_50ntFlanks</td><td>12996</td><td>13039</td><td>44</td><td>TCGCTGCGATCTATTGAAAGTCAGCCCTCGACACAAGGGTTTGT</td><td>rRF</td><td>TRUE</td><td>TRUE</td><td>FALSE</td></tr>
<tr><td>>MI0000285|hsa-mir-205&WithFlank&1|+|209432127|209432248@40.61.22&[MIMAT0000266&hsa-miR-205-5p&offsets|0|0;m-351&1|+|209432166|209432187&offsets|0|0]|TCCTTCATTCCACCGGAGTCTG</td><td>>MI0000285|hsa-mir-205&WithFlank&1|+|209432127|209432248</td><td>40</td><td>61</td><td>22</td><td>TCCTTCATTCCACCGGAGTCTG</td><td>isomiR</td><td>TRUE</td><td>FALSE</td><td>FALSE</td></tr>
<tr><td>>trnaMT_SerGCT_MT_+_12207_12265@1.18.18|GAGAAAGCTCACAAGAAC</td><td>>trnaMT_SerGCT_MT_+_12207_12265</td><td>1</td><td>18</td><td>18</td><td>GAGAAAGCTCACAAGAAC</td><td>tRF</td><td>TRUE</td><td>TRUE</td><td>FALSE</td></tr>
<tr><td>>MI0000440|hsa-mir-27b&WithFlank&9|+|95085439|95085547@67.86.20(+1U)&[MIMAT0000419&hsa-miR-27b-3p&offsets|0|-1(+1U);m-39&9|+|95085505|95085525&offsets|0|-1(+1U)]|TTCACAGTGGCTAAGTTCTGT</td><td>>MI0000440|hsa-mir-27b&WithFlank&9|+|95085439|95085547</td><td>67</td><td>86</td><td>20(+1U)</td><td>TTCACAGTGGCTAAGTTCTGT</td><td>isomiR</td><td>TRUE</td><td>TRUE</td><td>FALSE</td></tr>
<tr><td>>MI0000469|hsa-mir-125a&WithFlank&19|+|51693248|51693345@21.43.23&[MIMAT0000443&hsa-miR-125a-5p&offsets|0|-1;m-63&19|+|51693268|51693289&offsets|0|+1]|TCCCTGAGACCCTTTAACCTGTG</td><td>>MI0000469|hsa-mir-125a&WithFlank&19|+|51693248|51693345</td><td>21</td><td>43</td><td>23</td><td>TCCCTGAGACCCTTTAACCTGTG</td><td>isomiR</td><td>TRUE</td><td>TRUE</td><td>FALSE</td></tr>
<tr><td>>trna116_GluCTC_1_-_145399233_145399304@1.33.33,trna59_GluCTC_1_+_249168447_249168518@1.33.33,trna71_GluCTC_1_-_161439189_161439260@1.33.33,trna74_GluCTC_1_-_161431809_161431880@1.33.33,trna77_GluCTC_1_-_161424398_161424469@1.33.33,trna77_GluCTC_6_+_28949976_28950047@1.33.33,trna80_GluCTC_1_-_161417018_161417089@1.33.33,trna87_GluCTC_6_-_126101393_126101464@1.33.33|TCCCTGGTGGTCTAGTGGTTAGGATTCGGCGCT</td><td>>trna116_GluCTC_1_-_145399233_145399304</td><td>1</td><td>33</td><td>33</td><td>TCCCTGGTGGTCTAGTGGTTAGGATTCGGCGCT</td><td>tRF</td><td>TRUE</td><td>TRUE</td><td>FALSE</td></tr>
<tr><td>>hg38_1_-_228634819_228635039_RNA5S12_5S_50ntFlanks@134.170.37|TGGGAGACCGCCTGGGAATACCGGGTGCTGTAGGCTT</td><td>>hg38_1_-_228634819_228635039_RNA5S12_5S_50ntFlanks</td><td>134</td><td>170</td><td>37</td><td>TGGGAGACCGCCTGGGAATACCGGGTGCTGTAGGCTT</td><td>rRF</td><td>TRUE</td><td>TRUE</td><td>FALSE</td></tr>
<tr><td>>hg38_1_-_228634819_228635039_RNA5S12_5S_50ntFlanks@153.170.18|ACCGGGTGCTGTAGGCTT</td><td>>hg38_1_-_228634819_228635039_RNA5S12_5S_50ntFlanks</td><td>153</td><td>170</td><td>18</td><td>ACCGGGTGCTGTAGGCTT</td><td>rRF</td><td>TRUE</td><td>TRUE</td><td>FALSE</td></tr>
<tr><td>>MI0000650|hsa-mir-200c&WithFlank&12|+|6963693|6963772@50.71.22&[MIMAT0000617&hsa-miR-200c-3p&offsets|0|-1;m-54&12|+|6963742|6963764&offsets|0|-1]|TAATACTGCCGGGTAATGATGG</td><td>>MI0000650|hsa-mir-200c&WithFlank&12|+|6963693|6963772</td><td>50</td><td>71</td><td>22</td><td>TAATACTGCCGGGTAATGATGG</td><td>isomiR</td><td>TRUE</td><td>FALSE</td><td>FALSE</td></tr>
</table>

| | | | | | | | | | |
|---|---|---|---|---|---|---|---|---|---|
| >MI0000077\|hsa-mir-21&WithFlank&17\|+\|59841260\|59841343@14.34.21&[MIMAT0000076&hsa-miR-21-5p&offsets\|0\|-1;m-1&17\|+\|59841273\|59841294&offsets\|0\|-1]\|TAGCTTATCAGACTGATGTTG | >MI0000077\|hsa-mir-21&WithFlank&17\|+\|59841260\|59841343 | 14 | 34 | 21 | TAGCTTATCAGACTGATGTTG | isomiR | TRUE | TRUE | FALSE |
| >am_trna128_GlyGCC_6_-_27870686_27870756@1.31.31,trna133_GlyCCC_1_-_16872434_16872504@1.31.31,trna18_GlyGCC_16_+_70822597_70822667@1.31.31,trna19_GlyGCC_16_+_70823410_70823480@1.31.31,trna19_GlyGCC_2_-_157257659_157257729@1.31.31,trna24_GlyGCC_16_-_70812942_70813012@1.31.31,trna25_GlyGCC_16_-_70812114_70812184@1.31.31,trna4_GlyCCC_1_+_17188416_17188486@1.31.31,trna5_GlyGCC_17_+_8029064_8029134@1.31.31,trna68_GlyGCC_1_-_161493637_161493707@1.31.31\|GCATTGGTGGTTCAGTGGTAGAATTCTCGCC | >am_trna128_GlyGCC_6_-_27870686_27870756 | 1 | 31 | 31 | GCATTGGTGGTTCAGTGGTAGAATTCTCGCC | tRF | TRUE | TRUE | FALSE |
| >MI0000067\|hsa-let-7f-1&WithFlank&9\|+\|94176341\|94176439@13.33.21(+1C);MI0000068\|hsa-let-7f-2&WithFlank&X\|-\|53557186\|53557280@14.34.21(+1C)&[MIMAT0000067&hsa-let-7f-5p&offsets\|0\|-1(+1C);m-20&9\|+\|94176353\|94176374&offsets\|0\|-1(+1C)];[MIMAT0000067_1&hsa-let-7f-5p&offsets\|0\|-1(+1C);m-19&X\|-\|53557246\|53557267&offsets\|0\|-1(+1C)]\|TGAGGTAGTAGATTGTATAGTC | >MI0000067\|hsa-let-7f-1&WithFlank&9\|+\|94176341\|94176439 | 13 | 33 | 21(+1C) | TGAGGTAGTAGATTGTATAGTC | isomiR | TRUE | TRUE | FALSE |
| >MI0000264\|hsa-mir-7-2&WithFlank&15\|+\|88611819\|88611940@38.60.23;MI0000265\|hsa-mir-7-3&WithFlank&19\|+\|4770664\|4770785@37.59.23;MI0000263\|hsa-mir-7-1&WithFlank&9\|-\|83969742\|83969863@30.52.23&[MIMAT0000252&hsa-miR-7-5p&offsets\|0\|-1;m-400&15\|+\|88611856\|88611879&offsets\|0\|-1];[MIMAT0000252_1&hsa-miR-7-5p&offsets\|0\|-1;m-402&19\|+\|4770700\|4770723&offsets\|0\|-1];[MIMAT0000252_2&hsa-miR-7-5p&offsets\|0\|-1;m-401&9\|-\|83969811\|83969834&offsets\|0\|-1]\|TGGAAGACTAGTGATTTTGTTGT | >MI0000264\|hsa-mir-7-2&WithFlank&15\|+\|88611819\|88611940 | 38 | 60 | 23 | TGGAAGACTAGTGATTTTGTTGT | isomiR | TRUE | TRUE | FALSE |
| >MI0000266\|hsa-mir-10a&WithFlank&17\|-\|48579832\|48579953@28.49.22&[MIMAT0000253&hsa-miR-10a-5p&offsets\|0\|-1;m-18&17\|-\|48579905\|48579926&offsets\|0\|0]\|TACCCTGTAGATCCGAATTTGT | >MI0000266\|hsa-mir-10a&WithFlank&17\|-\|48579832\|48579953 | 28 | 49 | 22 | TACCCTGTAGATCCGAATTTGT | isomiR | TRUE | TRUE | FALSE |
| >MI0000253\|hsa-mir-148a&WithFlank&7\|-\|25949913\|25949992@50.70.21(+1G)&[MIMAT0000243&hsa-miR-148a-3p&offsets\|0\|-1(+1G);m-7&7\|-\|25949922\|25949943&offsets\|0\|-1(+1G)]\|TCAGTGCACTACAGAACTTTGG | >MI0000253\|hsa-mir-148a&WithFlank&7\|-\|25949913\|25949992 | 50 | 70 | 21(+1G) | TCAGTGCACTACAGAACTTTGG | isomiR | TRUE | FALSE | FALSE |
| >hg38_21_+_8433172_8446622_RNA45SN1_45S_50ntFlanks@6646.6664.19\|TCGTACGACTCTTAGCGGT | >hg38_21_+_8433172_8446622_RNA45SN1_45S_50ntFlanks | 6646 | 6664 | 19 | TCGTACGACTCTTAGCGGT | rRF | TRUE | TRUE | FALSE |
| >MI0000088\|hsa-mir-30a&WithFlank&6\|-\|71403545\|71403627@12.33.22(+1U)&[MIMAT0000087&hsa-miR-30a-5p&offsets\|0\|0(+1U);m-15&6\|-\|71403595\|71403616&offsets\|0\|0(+1U)]\|TGTAAACATCCTCGACTGGAAGT | >MI0000088\|hsa-mir-30a&WithFlank&6\|-\|71403545\|71403627 | 12 | 33 | 22(+1U) | TGTAAACATCCTCGACTGGAAGT | isomiR | TRUE | TRUE | FALSE |
| >MI0000088\|hsa-mir-30a&WithFlank&6\|-\|71403545\|71403627@12.34.23(+1A)&[MIMAT0000087&hsa-miR-30a-5p&offsets\|0\|+1(+1A);m-15&6\|-\|71403595\|71403616&offsets\|0\|+1(+1A)]\|TGTAAACATCCTCGACTGGAAGCA | >MI0000088\|hsa-mir-30a&WithFlank&6\|-\|71403545\|71403627 | 12 | 34 | 23(+1A) | TGTAAACATCCTCGACTGGAAGCA | isomiR | TRUE | TRUE | FALSE |
| >MI0000441\|hsa-mir-30b&WithFlank&8\|-\|134800514\|134800613@23.44.22&[MIMAT0000420&hsa-miR-30b-5p&offsets\|0\|0;m-76&8\|-\|134800570\|134800591&offsets\|0\|0]\|TGTAAACATCCTACACTCAGCT | >MI0000441\|hsa-mir-30b&WithFlank&8\|-\|134800514\|134800613 | 23 | 44 | 22 | TGTAAACATCCTACACTCAGCT | isomiR | TRUE | TRUE | FALSE |
| >MI0000077\|hsa-mir-21&WithFlank&17\|+\|59841260\|59841343@14.34.21(+1G)&[MIMAT0000076&hsa-miR-21-5p&offsets\|0\|-1(+1G);m-1&17\|+\|59841273\|59841294&offsets\|0\|-1(+1G)]\|TAGCTTATCAGACTGATGTTGG | >MI0000077\|hsa-mir-21&WithFlank&17\|+\|59841260\|59841343 | 14 | 34 | 21(+1G) | TAGCTTATCAGACTGATGTTGG | isomiR | TRUE | TRUE | FALSE |
| >MI0001445\|hsa-mir-423&WithFlank&17\|+\|30117073\|30117178@59.81.23&[MIMAT0001340&hsa-miR-423-3p&offsets\|0\|0;m-100&17\|+\|30117131\|30117153&offsets\|0\|0]\|AGCTCGGTCTGAGGCCCCTCAGT | >MI0001445\|hsa-mir-423&WithFlank&17\|+\|30117073\|30117178 | 59 | 81 | 23 | AGCTCGGTCTGAGGCCCCTCAGT | isomiR | TRUE | TRUE | FALSE |
| >MI0000264\|hsa-mir-7-2&WithFlank&15\|+\|88611819\|88611940@38.60.23(+1C);MI0000265\|hsa-mir-7-3&WithFlank&19\|+\|4770664\|4770785@37.59.23(+1C);MI0000263\|hsa-mir-7-1&WithFlank&9\|-\|83969742\|83969863@30.52.23(+1C)&[MIMAT0000252&hsa-miR-7-5p&offsets\|0\|-1(+1C);m-400&15\|+\|88611856\|88611879&offsets\|0\|-1(+1C)];[MIMAT0000252_1&hsa-miR-7-5p&offsets\|0\|-1(+1C);m-402&19\|+\|4770700\|4770723&offsets\|0\|-1(+1C)];[MIMAT0000252_2&hsa-miR-7-5p&offsets\|0\|-1(+1C);m-401&9\|-\|83969811\|83969834&offsets\|0\|-1(+1C)]\|TGGAAGACTAGTGATTTTGTTGTC | >MI0000264\|hsa-mir-7-2&WithFlank&15\|+\|88611819\|88611940 | 38 | 60 | 23(+1C) | TGGAAGACTAGTGATTTTGTTGTC | isomiR | TRUE | TRUE | FALSE |
| >hg38_21_+_8433172_8446622_RNA45SN1_45S_50ntFlanks@6646.6669.24\|TCGTACGACTCTTAGCGGTGGATC | >hg38_21_+_8433172_8446622_RNA45SN1_45S_50ntFlanks | 6646 | 6669 | 24 | TCGTACGACTCTTAGCGGTGGATC | rRF | TRUE | TRUE | TRUE |
| >MI0000088\|hsa-mir-30a&WithFlank&6\|-\|71403545\|71403627@12.34.23(+1C)&[MIMAT0000087&hsa-miR-30a-5p&offsets\|0\|+1(+1C);m-15&6\|-\|71403595\|71403616&offsets\|0\|+1(+1C)]\|TGTAAACATCCTCGACTGGAAGCC | >MI0000088\|hsa-mir-30a&WithFlank&6\|-\|71403545\|71403627 | 12 | 34 | 23(+1C) | TGTAAACATCCTCGACTGGAAGCC | isomiR | TRUE | TRUE | FALSE |
| >hg38_21_+_8433172_8446622_RNA45SN1_45S_50ntFlanks@475.505.31\|CTTCGTGATCGATGTGGTGACGTCGTGCTCT | >hg38_21_+_8433172_8446622_RNA45SN1_45S_50ntFlanks | 475 | 505 | 31 | CTTCGTGATCGATGTGGTGACGTCGTGCTCT | rRF | TRUE | TRUE | FALSE |
| >MI0000273\|hsa-mir-183&WithFlank&7\|-\|129774899\|129775020@33.54.22&[MIMAT0000261&hsa-miR-183-5p&offsets\|0\|0;m-41&7\|-\|129774966\|129774987&offsets\|-1\|-1]\|TATGGCACTGGTAGAATTCACT | >MI0000273\|hsa-mir-183&WithFlank&7\|-\|129774899\|129775020 | 33 | 54 | 22 | TATGGCACTGGTAGAATTCACT | isomiR | TRUE | FALSE | FALSE |
| >hg38_1_-_228634819_228635039_RNA5S12_5S_50ntFlanks@51.89.39\|GTCTACGGCCATACCACCCTGAACGCGCCCGATCTCGTC | >hg38_1_-_228634819_228635039_RNA5S12_5S_50ntFlanks | 51 | 89 | 39 | GTCTACGGCCATACCACCCTGAACGCGCCCGATCTCGTC | rRF | TRUE | TRUE | FALSE |
| >MI0000736\|hsa-mir-30c-1&WithFlank&1\|+\|40757278\|40757378@23.46.24;MI0000254\|hsa-mir-30c-2&WithFlank&6\|-\|71376954\|71377037@13.36.24&[MIMAT0000244&hsa-miR-30c-5p&offsets\|0\|+1;m-57&1\|+\|40757300\|40757323&offsets\|0\|0];[MIMAT0000244_1&hsa-miR-30c-5p&offsets\|0\|+1;m-56&6\|-\|71377002\|71377025&offsets\|0\|0]\|TGTAAACATCCTACACTCTCAGCT | >MI0000736\|hsa-mir-30c-1&WithFlank&1\|+\|40757278\|40757378 | 23 | 46 | 24 | TGTAAACATCCTACACTCTCAGCT | isomiR | TRUE | TRUE | FALSE |
| >MI0000809\|hsa-mir-151a&WithFlank&8\|-\|140732558\|140732659@53.74.22&[MIMAT0000757&hsa-miR-151a-3p&offsets\|0\|+1;m-30&8\|-\|140732587\|140732607&offsets\|0\|+1]\|CTAGACTGAAGCTCCTTGAGGA | >MI0000809\|hsa-mir-151a&WithFlank&8\|-\|140732558\|140732659 | 53 | 74 | 22 | CTAGACTGAAGCTCCTTGAGGA | isomiR | TRUE | TRUE | FALSE |
| >MI0000255\|hsa-mir-30d&WithFlank&8\|-\|134804870\|134804951@12.34.23(+1C)&[MIMAT0000245&hsa-miR-30d-5p&offsets\|0\|+1(+1C);m-21&8\|-\|134804919\|134804940&offsets\|0\|+1(+1C)]\|TGTAAACATCCCCGACTGGAAGCC | >MI0000255\|hsa-mir-30d&WithFlank&8\|-\|134804870\|134804951 | 12 | 34 | 23(+1C) | TGTAAACATCCCCGACTGGAAGCC | isomiR | TRUE | TRUE | FALSE |
| >MI0000102\|hsa-mir-100&WithFlank&11\|-\|122152223\|122152314@19.40.22&[MIMAT0000098&hsa-miR-100-5p&offsets\|0\|0;m-27&11\|-\|122152276\|122152296&offsets\|0\|+1]\|AACCCGTAGATCCGAACTTGTG | >MI0000102\|hsa-mir-100&WithFlank&11\|-\|122152223\|122152314 | 19 | 40 | 22 | AACCCGTAGATCCGAACTTGTG | isomiR | TRUE | TRUE | FALSE |
| >MI0000542\|hsa-mir-320a&WithFlank&8\|-\|22244960\|22245043@48.69.22&[MIMAT0000510&hsa-miR-320a-3p&offsets\|0\|0;m-60&8\|-\|22244975\|22244996&offsets\|0\|0]\|AAAAGCTGGGTTGAGAGGGCGA | >MI0000542\|hsa-mir-320a&WithFlank&8\|-\|22244960\|22245043 | 48 | 69 | 22 | AAAAGCTGGGTTGAGAGGGCGA | isomiR | TRUE | FALSE | FALSE |

| | | | | | | | | | |
|---|---|---|---|---|---|---|---|---|---|
| >MI0000650\|hsa-mir-200c&WithFlank&12\|+\|6963693\|6963772@50.72.23(+1A)&[MIMAT0000617&hsa-miR-200c-3p&offsets\|0\|0(+1A);m-54&12\|+\|6963742\|6963764&offsets\|0\|0(+1A)]\|TAATACTGCCGGGTAATGATGGAA | >MI0000650\|hsa-mir-200c&WithFlank&12\|+\|6963693\|6963772 | 50 | 72 | 23(+1A) | TAATACTGCCGGGTAATGATGGAA | isomiR | TRUE | FALSE | FALSE |
| >hg38_1_-_228634819_228635039_RNA5S12_5S_50ntFlanks@153.171.19\|ACCGGGTGCTGTAGGCTTT | >hg38_1_-_228634819_228635039_RNA5S12_5S_50ntFlanks | 153 | 171 | 19 | ACCGGGTGCTGTAGGCTTT | rRF | TRUE | TRUE | FALSE |
| >MI0000255\|hsa-mir-30d&WithFlank&8\|-\|134804870\|134804951@12.34.23(+1A)&[MIMAT0000245&hsa-miR-30d-5p&offsets\|0\|+1(+1A);m-21&8\|-\|134804919\|134804940&offsets\|0\|+1(+1A)]\|TGTAAACATCCCCGACTGGAAGCA | >MI0000255\|hsa-mir-30d&WithFlank&8\|-\|134804870\|134804951 | 12 | 34 | 23(+1A) | TGTAAACATCCCCGACTGGAAGCA | isomiR | TRUE | TRUE | FALSE |
| >trna119_LysCTT_1_-_145395522_145395594@1.35.35,trna11_LysCTT_5_-_180648979_180649051@1.35.35,trna13_LysCTT_6_+_26556774_26556846@1.35.35,trna7_LysCTT_16_+_3225692_3225764@1.35.35,trna9_LysCTT_5_+_180634755_180634827@1.35.35\|GCCCGGCTAGCTCAGTCGGTAGAGCATGAGACTCT | >trna119_LysCTT_1_-_145395522_145395594 | 1 | 35 | 35 | GCCCGGCTAGCTCAGTCGGTAGAGCATGAGACTCT | tRF | TRUE | FALSE | FALSE |
| >hg38_21_+_8433172_8446622_RNA45SN1_45S_50ntFlanks@485.505.21\|GATGTGGTGACGTCGTGCTCT | >hg38_21_+_8433172_8446622_RNA45SN1_45S_50ntFlanks | 485 | 505 | 21 | GATGTGGTGACGTCGTGCTCT | rRF | TRUE | TRUE | FALSE |
| >MI0000255\|hsa-mir-30d&WithFlank&8\|-\|134804870\|134804951@12.33.22(+1U)&[MIMAT0000245&hsa-miR-30d-5p&offsets\|0\|0(+1U);m-21&8\|-\|134804919\|134804940&offsets\|0\|0(+1U)]\|TGTAAACATCCCCGACTGGAAGT | >MI0000255\|hsa-mir-30d&WithFlank&8\|-\|134804870\|134804951 | 12 | 33 | 22(+1U) | TGTAAACATCCCCGACTGGAAGT | isomiR | TRUE | TRUE | FALSE |
| >am_MI0000809\|hsa-mir-151a&WithFlank&8\|-\|140732558\|140732659@53.72.20(+1A)&[MIMAT0000757&hsa-miR-151a-3p&offsets\|0\|-1(+1A);m-30&8\|-\|140732587\|140732607&offsets\|0\|-1(+1A)]\|CTAGACTGAAGCTCCTTGAGA | >am_MI0000809\|hsa-mir-151a&WithFlank&8\|-\|140732558\|140732659 | 53 | 72 | 20(+1A) | CTAGACTGAAGCTCCTTGAGA | isomiR | TRUE | TRUE | FALSE |
| >MI0000266\|hsa-mir-10a&WithFlank&17\|-\|48579832\|48579953@28.50.23&[MIMAT0000253&hsa-miR-10a-5p&offsets\|0\|0;m-18&17\|-\|48579905\|48579926&offsets\|0\|+1]\|TACCCTGTAGATCCGAATTTGTG | >MI0000266\|hsa-mir-10a&WithFlank&17\|-\|48579832\|48579953 | 28 | 50 | 23 | TACCCTGTAGATCCGAATTTGTG | isomiR | TRUE | TRUE | FALSE |
| >MI0000087\|hsa-mir-29a&WithFlank&7\|-\|130876741\|130876816@47.68.22&[MIMAT0000086&hsa-miR-29a-3p&offsets\|-1\|-1;m-26&7\|-\|130876749\|130876769&offsets\|-1\|0]\|CTAGCACCATCTGAAATCGGTT | >MI0000087\|hsa-mir-29a&WithFlank&7\|-\|130876741\|130876816 | 47 | 68 | 22 | CTAGCACCATCTGAAATCGGTT | isomiR | TRUE | TRUE | FALSE |
| >MI0000066\|hsa-let-7e&WithFlank&19\|+\|51692780\|51692870@14.35.22&[MIMAT0000066&hsa-let-7e-5p&offsets\|0\|0;m-50&19\|+\|51692793\|51692814&offsets\|0\|0]\|TGAGGTAGGAGGTTGTATAGTT | >MI0000066\|hsa-let-7e&WithFlank&19\|+\|51692780\|51692870 | 14 | 35 | 22 | TGAGGTAGGAGGTTGTATAGTT | isomiR | TRUE | TRUE | FALSE |
| >MI0000088\|hsa-mir-30a&WithFlank&6\|-\|71403545\|71403627@12.34.23(+1G)&[MIMAT0000087&hsa-miR-30a-5p&offsets\|0\|+1(+1G);m-15&6\|-\|71403595\|71403616&offsets\|0\|+1(+1G)]\|TGTAAACATCCTCGACTGGAAGCG | >MI0000088\|hsa-mir-30a&WithFlank&6\|-\|71403545\|71403627 | 12 | 34 | 23(+1G) | TGTAAACATCCTCGACTGGAAGCG | isomiR | TRUE | TRUE | FALSE |
| >MI0000433\|hsa-let-7g&WithFlank&3\|-\|52268272\|52268367@11.31.21(+1C)&[MIMAT0000414&hsa-let-7g-5p&offsets\|0\|-1(+1C);m-58&3\|-\|52268336\|52268357&offsets\|0\|-1(+1C)]\|TGAGGTAGTAGTTTGTACAGTC | >MI0000433\|hsa-let-7g&WithFlank&3\|-\|52268272\|52268367 | 11 | 31 | 21(+1C) | TGAGGTAGTAGTTTGTACAGTC | isomiR | TRUE | TRUE | FALSE |
| >MI0000736\|hsa-mir-30c-1&WithFlank&1\|+\|40757278\|40757378@23.45.23;MI0000254\|hsa-mir-30c-2&WithFlank&6\|-\|71376954\|71377037@13.35.23&[MIMAT0000244&hsa-miR-30c-5p&offsets\|0\|0;m-57&1\|+\|40757300\|40757323&offsets\|0\|-1];[MIMAT0000244_1&hsa-miR-30c-5p&offsets\|0\|0;m-56&6\|-\|71377002\|71377025&offsets\|0\|-1]\|TGTAAACATCCTACACTCTCAGC | >MI0000736\|hsa-mir-30c-1&WithFlank&1\|+\|40757278\|40757378 | 23 | 45 | 23 | TGTAAACATCCTACACTCTCAGC | isomiR | TRUE | TRUE | FALSE |
| >trna14_LysTTT_11_-_59327808_59327880@1.33.33,trna2_LysTTT_17_+_8022473_8022545@1.33.33,trna54_LysTTT_1_+_204475655_204475727@1.33.33,trna5_LysTTT_11_+_59323902_59323974@1.33.33,trna62_LysTTT_1_-_204476158_204476230@1.33.33,trna76_LysTTT_6_+_28918806_28918878@1.33.33\|GCCCGGATAGCTCAGTCGGTAGAGCATCAGACT | >trna14_LysTTT_11_-_59327808_59327880 | 1 | 33 | 33 | GCCCGGATAGCTCAGTCGGTAGAGCATCAGACT | tRF | TRUE | TRUE | FALSE |
| >MI0000081\|hsa-mir-24-2&WithFlank&19\|-\|13836281\|13836365@56.75.20;MI0000080\|hsa-mir-24-1&WithFlank&9\|+\|95086015\|95086094@50.69.20&[MIMAT0000080&hsa-miR-24-3p&offsets\|0\|-2;m-36&19\|-\|13836288\|13836310&offsets\|0\|-3];[MIMAT0000080_1&hsa-miR-24-3p&offsets\|0\|-2;m-35&9\|+\|95086064\|95086086&offsets\|0\|-3]\|TGGCTCAGTTCAGCAGGAAC | >MI0000081\|hsa-mir-24-2&WithFlank&19\|-\|13836281\|13836365 | 56 | 75 | 20 | TGGCTCAGTTCAGCAGGAAC | isomiR | TRUE | TRUE | FALSE |
| >MI0000253\|hsa-mir-148a&WithFlank&7\|-\|25949913\|25949992@50.70.21(+1A)&[MIMAT0000243&hsa-miR-148a-3p&offsets\|0\|-1(+1A);m-7&7\|-\|25949922\|25949943&offsets\|0\|-1(+1A)]\|TCAGTGCACTACAGAACTTTGA | >MI0000253\|hsa-mir-148a&WithFlank&7\|-\|25949913\|25949992 | 50 | 70 | 21(+1A) | TCAGTGCACTACAGAACTTTGA | isomiR | TRUE | FALSE | FALSE |
| >MI0000077\|hsa-mir-21&WithFlank&17\|+\|59841260\|59841343@14.34.21(+1U)&[MIMAT0000076&hsa-miR-21-5p&offsets\|0\|-1(+1U);m-1&17\|+\|59841273\|59841294&offsets\|0\|-1(+1U)]\|TAGCTTATCAGACTGATGTTGT | >MI0000077\|hsa-mir-21&WithFlank&17\|+\|59841260\|59841343 | 14 | 34 | 21(+1U) | TAGCTTATCAGACTGATGTTGT | isomiR | TRUE | FALSE | FALSE |
| >MI0000084\|hsa-mir-26b&WithFlank&2\|+\|218402640\|218402728@18.39.22&[MIMAT0000083&hsa-miR-26b-5p&offsets\|0\|+1;m-53&2\|+\|218402657\|218402678&offsets\|0\|0]\|TTCAAGTAATTCAGGATAGGTT | >MI0000084\|hsa-mir-26b&WithFlank&2\|+\|218402640\|218402728 | 18 | 39 | 22 | TTCAAGTAATTCAGGATAGGTT | isomiR | TRUE | TRUE | FALSE |
| >MI0000447\|hsa-mir-128-1&WithFlank&2\|+\|135665391\|135665484@56.76.21;MI0000727\|hsa-mir-128-2&WithFlank&3\|+\|35744470\|35744565@58.78.21&[MIMAT0000424&hsa-miR-128-3p&offsets\|0\|0;m-108&2\|+\|135665446\|135665466&offsets\|0\|0];[MIMAT0000424_1&hsa-miR-128-3p&offsets\|0\|0;m-117&3\|+\|35744527\|35744547&offsets\|0\|0]\|TCACAGTGAACCGGTCTCTTT | >MI0000447\|hsa-mir-128-1&WithFlank&2\|+\|135665391\|135665484 | 56 | 76 | 21 | TCACAGTGAACCGGTCTCTTT | isomiR | TRUE | TRUE | FALSE |
| >MI0000746\|hsa-mir-99b&WithFlank&19\|+\|51692606\|51692687@13.32.20(+1U)&[MIMAT0000689&hsa-miR-99b-5p&offsets\|0\|-2(+1U);m-10&19\|+\|51692618\|51692639&offsets\|0\|-2(+1U)]\|CACCCGTAGAACCGACCTTGT | >MI0000746\|hsa-mir-99b&WithFlank&19\|+\|51692606\|51692687 | 13 | 32 | 20(+1U) | CACCCGTAGAACCGACCTTGT | isomiR | TRUE | FALSE | FALSE |
| >MI0000811\|hsa-mir-148b&WithFlank&12\|+\|54337210\|54337320@69.90.22&[MIMAT0000759&hsa-miR-148b-3p&offsets\|0\|0;m-77&12\|+\|54337278\|54337299&offsets\|0\|0]\|TCAGTGCATCACAGAACTTTGT | >MI0000811\|hsa-mir-148b&WithFlank&12\|+\|54337210\|54337320 | 69 | 90 | 22 | TCAGTGCATCACAGAACTTTGT | isomiR | TRUE | TRUE | FALSE |
| >MI0000078\|hsa-mir-22&WithFlank&17\|-\|1713897\|1713993@59.80.22&[MIMAT0000077&hsa-miR-22-3p&offsets\|0\|0;m-2&17\|-\|1713914\|1713935&offsets\|0\|0]\|AAGCTGCCAGTTGAAGAACTGT | >MI0000078\|hsa-mir-22&WithFlank&17\|-\|1713897\|1713993 | 59 | 80 | 22 | AAGCTGCCAGTTGAAGAACTGT | isomiR | TRUE | TRUE | FALSE |
| >MI0000434\|hsa-let-7i&WithFlank&12\|+\|62603680\|62603775@12.32.21(+1C)&[MIMAT0000415&hsa-let-7i-5p&offsets\|0\|-1(+1C);m-49&12\|+\|62603691\|62603712&offsets\|0\|-1(+1C)]\|TGAGGTAGTAGTTTGTGCTGTC | >MI0000434\|hsa-let-7i&WithFlank&12\|+\|62603680\|62603775 | 12 | 32 | 21(+1C) | TGAGGTAGTAGTTTGTGCTGTC | isomiR | TRUE | TRUE | FALSE |
| >hg38_21_+_8433172_8446622_RNA45SN1_45S_50ntFlanks@7975.8008.34\|CGCGACCTCAGATCAGACGTGGCGACCCGCTGAA | >hg38_21_+_8433172_8446622_RNA45SN1_45S_50ntFlanks | 7975 | 8008 | 34 | CGCGACCTCAGATCAGACGTGGCGACCCGCTGAA | rRF | TRUE | TRUE | TRUE |

<table>
<tr><td>>MI0000061|hsa-let-7a-2&WithFlank&11|-|122146516|122146599@11.31.21(+1C);MI0000062|hsa-let-7a-3&WithFlank&22|+|46112743|46112828@10.30.21(+1C);MI0000060|hsa-let-7a-1&WithFlank&9|+|94175951|94176042@12.32.21(+1C)&[MIMAT0000062&hsa-let-7a-5p&offsets|0|-1(+1C);m-6&11|-|122146568|122146589&offsets|0|-1(+1C)];[MIMAT0000062_1&hsa-let-7a-5p&offsets|0|-1(+1C);m-5&22|+|46112752|46112773&offsets|0|-1(+1C)];[MIMAT0000062_2&hsa-let-7a-5p&offsets|0|-1(+1C);m-4&9|+|94175962|94175983&offsets|0|-1(+1C)]|TGAGGTAGTAGGTTGTATAGTC</td><td>>MI0000061|hsa-let-7a-2&WithFlank&11|-|122146516|122146599</td><td>11</td><td>31</td><td>21(+1C)</td><td>TGAGGTAGTAGGTTGTATAGTC</td><td>isomiR</td><td>TRUE</td><td>TRUE</td><td>FALSE</td></tr>
<tr><td>>MI0000098|hsa-mir-96&WithFlank&7|-|129774686|129774775@15.37.23&[MIMAT0000095&hsa-miR-96-5p&offsets|0|0;m-248&7|-|129774739|129774761&offsets|0|0]|TTTGGCACTAGCACATTTTTGCT</td><td>>MI0000098|hsa-mir-96&WithFlank&7|-|129774686|129774775</td><td>15</td><td>37</td><td>23</td><td>TTTGGCACTAGCACATTTTTGCT</td><td>isomiR</td><td>TRUE</td><td>FALSE</td><td>FALSE</td></tr>
<tr><td>>MI0000067|hsa-let-7f-1&WithFlank&9|+|94176341|94176439@13.33.21;MI0000068|hsa-let-7f-2&WithFlank&X|-|53557186|53557280@14.34.21&[MIMAT0000067&hsa-let-7f-5p&offsets|0|-1;m-20&9|+|94176353|94176374&offsets|0|-1];[MIMAT0000067_1&hsa-let-7f-5p&offsets|0|-1;m-19&X|-|53557246|53557267&offsets|0|-1]|TGAGGTAGTAGATTGTATAGT</td><td>>MI0000067|hsa-let-7f-1&WithFlank&9|+|94176341|94176439</td><td>13</td><td>33</td><td>21</td><td>TGAGGTAGTAGATTGTATAGT</td><td>isomiR</td><td>TRUE</td><td>TRUE</td><td>FALSE</td></tr>
<tr><td>>hg38_21_+_8433172_8446622_RNA45SN1_45S_50ntFlanks@7975.7993.19|CGCGACCTCAGATCAGACG</td><td>>hg38_21_+_8433172_8446622_RNA45SN1_45S_50ntFlanks</td><td>7975</td><td>7993</td><td>19</td><td>CGCGACCTCAGATCAGACG</td><td>rRF</td><td>TRUE</td><td>TRUE</td><td>FALSE</td></tr>
<tr><td>>MI0000301|hsa-mir-224&WithFlank&X|-|151958572|151958664@14.35.22&[MIMAT0000281&hsa-miR-224-5p&offsets|+1|-2;m-238&X|-|151958630|151958651&offsets|0|0]|CAAGTCACTAGTGGTTCCGTTT</td><td>>MI0000301|hsa-mir-224&WithFlank&X|-|151958572|151958664</td><td>14</td><td>35</td><td>22</td><td>CAAGTCACTAGTGGTTCCGTTT</td><td>isomiR</td><td>TRUE</td><td>FALSE</td><td>FALSE</td></tr>
<tr><td>>MI0000272|hsa-mir-182&WithFlank&7|-|129770377|129770498@29.53.25&[MIMAT0000259&hsa-miR-182-5p&offsets|0|+1;m-22&7|-|129770449|129770470&offsets|0|+3]|TTTGGCAATGGTAGAACTCACACTG</td><td>>MI0000272|hsa-mir-182&WithFlank&7|-|129770377|129770498</td><td>29</td><td>53</td><td>25</td><td>TTTGGCAATGGTAGAACTCACACTG</td><td>isomiR</td><td>TRUE</td><td>FALSE</td><td>FALSE</td></tr>
<tr><td>>hg38_21_+_8433172_8446622_RNA45SN1_45S_50ntFlanks@5492.5516.25|AGAAGACGGTCGAACTTGACTATCT</td><td>>hg38_21_+_8433172_8446622_RNA45SN1_45S_50ntFlanks</td><td>5492</td><td>5516</td><td>25</td><td>AGAAGACGGTCGAACTTGACTATCT</td><td>rRF</td><td>TRUE</td><td>FALSE</td><td>FALSE</td></tr>
<tr><td>>MI0000440|hsa-mir-27b&WithFlank&9|+|95085439|95085547@67.86.20&[MIMAT0000419&hsa-miR-27b-3p&offsets|0|-1;m-39&9|+|95085505|95085525&offsets|0|-1]|TTCACAGTGGCTAAGTTCTG</td><td>>MI0000440|hsa-mir-27b&WithFlank&9|+|95085439|95085547</td><td>67</td><td>86</td><td>20</td><td>TTCACAGTGGCTAAGTTCTG</td><td>isomiR</td><td>TRUE</td><td>TRUE</td><td>FALSE</td></tr>
<tr><td>>MI0000076|hsa-mir-20a&WithFlank&13|+|91351059|91351141@14.36.23&[MIMAT0000075&hsa-miR-20a-5p&offsets|0|0;m-96&13|+|91351072|91351094&offsets|0|0]|TAAAGTGCTTATAGTGCAGGTAG</td><td>>MI0000076|hsa-mir-20a&WithFlank&13|+|91351059|91351141</td><td>14</td><td>36</td><td>23</td><td>TAAAGTGCTTATAGTGCAGGTAG</td><td>isomiR</td><td>TRUE</td><td>TRUE</td><td>FALSE</td></tr>
<tr><td>>MI0000087|hsa-mir-29a&WithFlank&7|-|130876741|130876816@48.68.21&[MIMAT0000086&hsa-miR-29a-3p&offsets|0|-1;m-26&7|-|130876749|130876769&offsets|0|0]|TAGCACCATCTGAAATCGGTT</td><td>>MI0000087|hsa-mir-29a&WithFlank&7|-|130876741|130876816</td><td>48</td><td>68</td><td>21</td><td>TAGCACCATCTGAAATCGGTT</td><td>isomiR</td><td>TRUE</td><td>TRUE</td><td>FALSE</td></tr>
<tr><td>>am_MI0000786|hsa-mir-378a&WithFlank&5|+|149732819|149732896@49.70.22&[MIMAT0000732&hsa-miR-378a-3p&offsets|0|0;m-44&5|+|149732867|149732888&offsets|0|0]|ACTGGACTTGGAGTCAGAAGGC</td><td>>am_MI0000786|hsa-mir-378a&WithFlank&5|+|149732819|149732896</td><td>49</td><td>70</td><td>22</td><td>ACTGGACTTGGAGTCAGAAGGC</td><td>isomiR</td><td>TRUE</td><td>TRUE</td><td>FALSE</td></tr>
<tr><td>>MI0000070|hsa-mir-16-1&WithFlank&13|-|50048967|50049067@20.41.22;MI0000115|hsa-mir-16-2&WithFlank&3|+|160404739|160404831@16.37.22&[MIMAT0000069&hsa-miR-16-5p&offsets|0|0;m-62&13|-|50049027|50049048&offsets|0|0];[MIMAT0000069_1&hsa-miR-16-5p&offsets|0|0;m-61&3|+|160404754|160404775&offsets|0|0]|TAGCAGCACGTAAATATTGGCG</td><td>>MI0000070|hsa-mir-16-1&WithFlank&13|-|50048967|50049067</td><td>20</td><td>41</td><td>22</td><td>TAGCAGCACGTAAATATTGGCG</td><td>isomiR</td><td>TRUE</td><td>FALSE</td><td>FALSE</td></tr>
<tr><td>>MI0000650|hsa-mir-200c&WithFlank&12|+|6963693|6963772@50.72.23(+1U)&[MIMAT0000617&hsa-miR-200c-3p&offsets|0|0(+1U);m-54&12|+|6963742|6963764&offsets|0|0(+1U)]|TAATACTGCCGGGTAATGATGGAT</td><td>>MI0000650|hsa-mir-200c&WithFlank&12|+|6963693|6963772</td><td>50</td><td>72</td><td>23(+1U)</td><td>TAATACTGCCGGGTAATGATGGAT</td><td>isomiR</td><td>TRUE</td><td>FALSE</td><td>FALSE</td></tr>
<tr><td>>MI0000746|hsa-mir-99b&WithFlank&19|+|51692606|51692687@13.33.21(+1A)&[MIMAT0000689&hsa-miR-99b-5p&offsets|0|-1(+1A);m-10&19|+|51692618|51692639&offsets|0|-1(+1A)]|CACCCGTAGAACCGACCTTGCA</td><td>>MI0000746|hsa-mir-99b&WithFlank&19|+|51692606|51692687</td><td>13</td><td>33</td><td>21(+1A)</td><td>CACCCGTAGAACCGACCTTGCA</td><td>isomiR</td><td>TRUE</td><td>TRUE</td><td>FALSE</td></tr>
<tr><td>>MI0000103|hsa-mir-101-1&WithFlank&1|-|65058428|65058514@53.74.22;MI0000739|hsa-mir-101-2&WithFlank&9|+|4850291|4850381@55.76.22&[MIMAT0000099&hsa-miR-101-3p&offsets|0|+1;m-17&1|-|65058443|65058462&offsets|0|+2];[MIMAT0000099_1&hsa-miR-101-3p&offsets|0|+1;m-16&9|+|4850345|4850364&offsets|0|+2]|TACAGTACTGTGATAACTGAAG</td><td>>MI0000103|hsa-mir-101-1&WithFlank&1|-|65058428|65058514</td><td>53</td><td>74</td><td>22</td><td>TACAGTACTGTGATAACTGAAG</td><td>isomiR</td><td>TRUE</td><td>FALSE</td><td>FALSE</td></tr>
<tr><td>>MI0000065|hsa-let-7d&WithFlank&9|+|94178828|94178926@14.35.22&[MIMAT0000065&hsa-let-7d-5p&offsets|0|0;m-90&9|+|94178841|94178862&offsets|0|0]|AGAGGTAGTAGGTTGCATAGTT</td><td>>MI0000065|hsa-let-7d&WithFlank&9|+|94178828|94178926</td><td>14</td><td>35</td><td>22</td><td>AGAGGTAGTAGGTTGCATAGTT</td><td>isomiR</td><td>TRUE</td><td>TRUE</td><td>FALSE</td></tr>
<tr><td>>MI0000087|hsa-mir-29a&WithFlank&7|-|130876741|130876816@48.69.22&[MIMAT0000086&hsa-miR-29a-3p&offsets|0|0;m-26&7|-|130876749|130876769&offsets|0|+1]|TAGCACCATCTGAAATCGGTTA</td><td>>MI0000087|hsa-mir-29a&WithFlank&7|-|130876741|130876816</td><td>48</td><td>69</td><td>22</td><td>TAGCACCATCTGAAATCGGTTA</td><td>isomiR</td><td>TRUE</td><td>TRUE</td><td>FALSE</td></tr>
<tr><td>>MI0000465|hsa-mir-191&WithFlank&3|-|49020612|49020715@22.44.23&[MIMAT0000440&hsa-miR-191-5p&offsets|0|0;m-48&3|-|49020672|49020694&offsets|0|0]|CAACGGAATCCCAAAAGCAGCTG</td><td>>MI0000465|hsa-mir-191&WithFlank&3|-|49020612|49020715</td><td>22</td><td>44</td><td>23</td><td>CAACGGAATCCCAAAAGCAGCTG</td><td>isomiR</td><td>TRUE</td><td>TRUE</td><td>FALSE</td></tr>
<tr><td>>MI0000255|hsa-mir-30d&WithFlank&8|-|134804870|134804951@12.34.23(+1G)&[MIMAT0000245&hsa-miR-30d-5p&offsets|0|+1(+1G);m-21&8|-|134804919|134804940&offsets|0|+1(+1G)]|TGTAAACATCCCCGACTGGAAGCG</td><td>>MI0000255|hsa-mir-30d&WithFlank&8|-|134804870|134804951</td><td>12</td><td>34</td><td>23(+1G)</td><td>TGTAAACATCCCCGACTGGAAGCG</td><td>isomiR</td><td>TRUE</td><td>TRUE</td><td>FALSE</td></tr>
<tr><td>>hg38_1_-_228634819_228635039_RNA5S12_5S_50ntFlanks@132.170.39|GATGGGAGACCGCCTGGGAATACCGGGTGCTGTAGGCTT</td><td>>hg38_1_-_228634819_228635039_RNA5S12_5S_50ntFlanks</td><td>132</td><td>170</td><td>39</td><td>GATGGGAGACCGCCTGGGAATACCGGGTGCTGTAGGCTT</td><td>rRF</td><td>TRUE</td><td>TRUE</td><td>TRUE</td></tr>
<tr><td>>am_MI0005763|hsa-mir-941-1&WithFlank&20|+|63919443|63919526@53.75.23;MI0005764|hsa-mir-941-2&WithFlank&20|+|63919499|63919582@53.75.23;MI0005765|hsa-mir-941-3&WithFlank&20|+|63919555|63919638@53.75.23;MI0005766|hsa-mir-941-4&WithFlank&20|+|63919750|63919833@53.75.23;MI0031520|hsa-mir-941-5&WithFlank&20|+|63919862|63919945@53.75.23&[MIMAT0004984&hsa-miR-941&offsets|0|0;m-263&20|+|63919495|63919517&offsets|0|0];[MIMAT0004984_1&hsa-miR-941&offsets|0|0;m-263&20|+|63919551|63919573&offsets|0|0];[MIMAT0004984_2&hsa-miR-941&offsets|0|0;m-263&20|+|63919607|63919629&offsets|0|0];[MIMAT0004984_3&hsa-miR-941&offsets|0|0;m-263&20|+|63919802|63919824&offsets|0|0];[MIMAT0004984_4&hsa-miR-941&offsets|0|0;m-263&20|+|63919914|63919936&offsets|0|0]|CACCCGGCTGTGTGCACATGTGC</td><td>>am_MI0005763|hsa-mir-941-1&WithFlank&20|+|63919443|63919526</td><td>53</td><td>75</td><td>23</td><td>CACCCGGCTGTGTGCACATGTGC</td><td>isomiR</td><td>TRUE</td><td>TRUE</td><td>FALSE</td></tr>
<tr><td>>MI0000299|hsa-mir-222&WithFlank&X|-|45747009|45747130@75.99.25&[MIMAT0000279&hsa-miR-222-3p&offsets|0|+4;m-140&X|-|45747033|45747056&offsets|0|+1]|AGCTACATCTGGCTACTGGGTCTCT</td><td>>MI0000299|hsa-mir-222&WithFlank&X|-|45747009|45747130</td><td>75</td><td>99</td><td>25</td><td>AGCTACATCTGGCTACTGGGTCTCT</td><td>isomiR</td><td>TRUE</td><td>TRUE</td><td>FALSE</td></tr>
<tr><td>>trnaMT_GluTTC_MT_-_14674_14742@39.72.34,trnalookalike8_GluTTC_5_-_93905172_93905240@39.72.34|ATTGGTCGTGGTTGTAGTCCGTGCGAGAATACCA</td><td>>trnaMT_GluTTC_MT_-_14674_14742</td><td>39</td><td>72</td><td>34</td><td>ATTGGTCGTGGTTGTAGTCCGTGCGAGAATACCA</td><td>tRF</td><td>TRUE</td><td>TRUE</td><td>FALSE</td></tr>
</table>

<table>
<tr><th></th><th></th><th></th><th></th><th></th><th></th><th></th><th></th><th></th><th></th></tr>
<tr><td>>am_MI0000809|hsa-mir-151a&WithFlank&8|-|140732558|140732659@53.73.21(+1U)&[MIMAT0000757&hsa-miR-151a-3p&offsets|0|0(+1U);m-30&8|-|140732587|140732607&offsets|0|0(+1U)]|CTAGACTGAAGCTCCTTGAGGT</td><td>>am_MI0000809|hsa-mir-151a&WithFlank&8|-|140732558|140732659</td><td>53</td><td>73</td><td>21(+1U)</td><td>CTAGACTGAAGCTCCTTGAGGT</td><td>isomiR</td><td>TRUE</td><td>TRUE</td><td>FALSE</td></tr>
<tr><td>>MI0000088|hsa-mir-30a&WithFlank&6|-|71403545|71403627@12.33.22&[MIMAT0000087&hsa-miR-30a-5p&offsets|0|0;m-15&6|-|71403595|71403616&offsets|0|0]|TGTAAACATCCTCGACTGGAAG</td><td>>MI0000088|hsa-mir-30a&WithFlank&6|-|71403545|71403627</td><td>12</td><td>33</td><td>22</td><td>TGTAAACATCCTCGACTGGAAG</td><td>isomiR</td><td>TRUE</td><td>TRUE</td><td>FALSE</td></tr>
<tr><td>>MI0000542|hsa-mir-320a&WithFlank&8|-|22244960|22245043@48.69.22(+1U)&[MIMAT0000510&hsa-miR-320a-3p&offsets|0|0(+1U);m-60&8|-|22244975|22244996&offsets|0|0(+1U)]|AAAAGCTGGGTTGAGAGGGCGAT</td><td>>MI0000542|hsa-mir-320a&WithFlank&8|-|22244960|22245043</td><td>48</td><td>69</td><td>22(+1U)</td><td>AAAAGCTGGGTTGAGAGGGCGAT</td><td>isomiR</td><td>TRUE</td><td>FALSE</td><td>FALSE</td></tr>
<tr><td>>hg38_21_+_8433172_8446622_RNA45SN1_45S_50ntFlanks@5491.5516.26|GAGAAGACGGTCGAACTTGACTATCT</td><td>>hg38_21_+_8433172_8446622_RNA45SN1_45S_50ntFlanks</td><td>5491</td><td>5516</td><td>26</td><td>GAGAAGACGGTCGAACTTGACTATCT</td><td>rRF</td><td>TRUE</td><td>FALSE</td><td>FALSE</td></tr>
<tr><td>>MI0005756|hsa-mir-934&WithFlank&X|+|136550872|136550966@21.42.22&[MIMAT0004977&hsa-miR-934&offsets|0|0;m-943&X|+|136550892|136550913&offsets|0|0]|TGTCTACTACTGGAGACACTGG</td><td>>MI0005756|hsa-mir-934&WithFlank&X|+|136550872|136550966</td><td>21</td><td>42</td><td>22</td><td>TGTCTACTACTGGAGACACTGG</td><td>isomiR</td><td>TRUE</td><td>FALSE</td><td>FALSE</td></tr>
<tr><td>>hg38_21_+_8433172_8446622_RNA45SN1_45S_50ntFlanks@7975.7997.23|CGCGACCTCAGATCAGACGTGGC</td><td>>hg38_21_+_8433172_8446622_RNA45SN1_45S_50ntFlanks</td><td>7975</td><td>7997</td><td>23</td><td>CGCGACCTCAGATCAGACGTGGC</td><td>rRF</td><td>TRUE</td><td>FALSE</td><td>FALSE</td></tr>
<tr><td>>MI0000081|hsa-mir-24-2&WithFlank&19|-|13836281|13836365@56.77.22;MI0000080|hsa-mir-24-1&WithFlank&9|+|95086015|95086094@50.71.22&[MIMAT0000080&hsa-miR-24-3p&offsets|0|0;m-36&19|-|13836288|13836310&offsets|0|-1];[MIMAT0000080_1&hsa-miR-24-3p&offsets|0|0;m-35&9|+|95086064|95086086&offsets|0|-1]|TGGCTCAGTTCAGCAGGAACAG</td><td>>MI0000081|hsa-mir-24-2&WithFlank&19|-|13836281|13836365</td><td>56</td><td>77</td><td>22</td><td>TGGCTCAGTTCAGCAGGAACAG</td><td>isomiR</td><td>TRUE</td><td>TRUE</td><td>FALSE</td></tr>
<tr><td>>MI0000095|hsa-mir-93&WithFlank&7|-|100093762|100093853@17.39.23&[MIMAT0000093&hsa-miR-93-5p&offsets|0|0;m-29&7|-|100093815|100093837&offsets|0|0]|CAAAGTGCTGTTCGTGCAGGTAG</td><td>>MI0000095|hsa-mir-93&WithFlank&7|-|100093762|100093853</td><td>17</td><td>39</td><td>23</td><td>CAAAGTGCTGTTCGTGCAGGTAG</td><td>isomiR</td><td>TRUE</td><td>TRUE</td><td>FALSE</td></tr>
<tr><td>>MI0000272|hsa-mir-182&WithFlank&7|-|129770377|129770498@29.51.23(+1C)&[MIMAT0000259&hsa-miR-182-5p&offsets|0|-1(+1C);m-22&7|-|129770449|129770470&offsets|0|+1(+1C)]|TTTGGCAATGGTAGAACTCACACC</td><td>>MI0000272|hsa-mir-182&WithFlank&7|-|129770377|129770498</td><td>29</td><td>51</td><td>23(+1C)</td><td>TTTGGCAATGGTAGAACTCACACC</td><td>isomiR</td><td>TRUE</td><td>FALSE</td><td>FALSE</td></tr>
<tr><td>>hg38_21_+_8433172_8446622_RNA45SN1_45S_50ntFlanks@7975.8006.32|CGCGACCTCAGATCAGACGTGGCGACCCGCTG</td><td>>hg38_21_+_8433172_8446622_RNA45SN1_45S_50ntFlanks</td><td>7975</td><td>8006</td><td>32</td><td>CGCGACCTCAGATCAGACGTGGCGACCCGCTG</td><td>rRF</td><td>TRUE</td><td>FALSE</td><td>TRUE</td></tr>
<tr><td>>MI0000081|hsa-mir-24-2&WithFlank&19|-|13836281|13836365@56.77.22(+1U);MI0000080|hsa-mir-24-1&WithFlank&9|+|95086015|95086094@50.71.22(+1U)&[MIMAT0000080&hsa-miR-24-3p&offsets|0|0(+1U);m-36&19|-|13836288|13836310&offsets|0|-1(+1U)];[MIMAT0000080_1&hsa-miR-24-3p&offsets|0|0(+1U);m-35&9|+|95086064|95086086&offsets|0|-1(+1U)]|TGGCTCAGTTCAGCAGGAACAGT</td><td>>MI0000081|hsa-mir-24-2&WithFlank&19|-|13836281|13836365</td><td>56</td><td>77</td><td>22(+1U)</td><td>TGGCTCAGTTCAGCAGGAACAGT</td><td>isomiR</td><td>TRUE</td><td>TRUE</td><td>FALSE</td></tr>
<tr><td>>hg38_21_+_8433172_8446622_RNA45SN1_45S_50ntFlanks@6646.6663.18|TCGTACGACTCTTAGCGG</td><td>>hg38_21_+_8433172_8446622_RNA45SN1_45S_50ntFlanks</td><td>6646</td><td>6663</td><td>18</td><td>TCGTACGACTCTTAGCGG</td><td>rRF</td><td>TRUE</td><td>FALSE</td><td>TRUE</td></tr>
<tr><td>>trnaMT_SerGCT_MT_+_12207_12265@1.25.25|GAGAAAGCTCACAAGAACTGCTAAC</td><td>>trnaMT_SerGCT_MT_+_12207_12265</td><td>1</td><td>25</td><td>25</td><td>GAGAAAGCTCACAAGAACTGCTAAC</td><td>tRF</td><td>TRUE</td><td>TRUE</td><td>FALSE</td></tr>
<tr><td>>MI0000440|hsa-mir-27b&WithFlank&9|+|95085439|95085547@67.86.20(+1A)&[MIMAT0000419&hsa-miR-27b-3p&offsets|0|-1(+1A);m-39&9|+|95085505|95085525&offsets|0|-1(+1A)]|TTCACAGTGGCTAAGTTCTGA</td><td>>MI0000440|hsa-mir-27b&WithFlank&9|+|95085439|95085547</td><td>67</td><td>86</td><td>20(+1A)</td><td>TTCACAGTGGCTAAGTTCTGA</td><td>isomiR</td><td>TRUE</td><td>FALSE</td><td>FALSE</td></tr>
<tr><td>>hg38_21_+_8433172_8446622_RNA45SN1_45S_50ntFlanks@475.498.24|CTTCGTGATCGATGTGGTGACGTC</td><td>>hg38_21_+_8433172_8446622_RNA45SN1_45S_50ntFlanks</td><td>475</td><td>498</td><td>24</td><td>CTTCGTGATCGATGTGGTGACGTC</td><td>rRF</td><td>TRUE</td><td>TRUE</td><td>FALSE</td></tr>
<tr><td>>MI0000650|hsa-mir-200c&WithFlank&12|+|6963693|6963772@50.71.22(+1G)&[MIMAT0000617&hsa-miR-200c-3p&offsets|0|-1(+1G);m-54&12|+|6963742|6963764&offsets|0|-1(+1G)]|TAATACTGCCGGGTAATGATGGG</td><td>>MI0000650|hsa-mir-200c&WithFlank&12|+|6963693|6963772</td><td>50</td><td>71</td><td>22(+1G)</td><td>TAATACTGCCGGGTAATGATGGG</td><td>isomiR</td><td>TRUE</td><td>FALSE</td><td>FALSE</td></tr>
<tr><td>>MI0000440|hsa-mir-27b&WithFlank&9|+|95085439|95085547@67.88.22&[MIMAT0000419&hsa-miR-27b-3p&offsets|0|+1;m-39&9|+|95085505|95085525&offsets|0|+1]|TTCACAGTGGCTAAGTTCTGCA</td><td>>MI0000440|hsa-mir-27b&WithFlank&9|+|95085439|95085547</td><td>67</td><td>88</td><td>22</td><td>TTCACAGTGGCTAAGTTCTGCA</td><td>isomiR</td><td>TRUE</td><td>TRUE</td><td>FALSE</td></tr>
<tr><td>>MI0000086|hsa-mir-28&WithFlank&3|+|188688775|188688872@60.81.22&[MIMAT0004502&hsa-miR-28-3p&offsets|0|0;m-28&3|+|188688834|188688855&offsets|0|0]|CACTAGATTGTGAGCTCCTGGA</td><td>>MI0000086|hsa-mir-28&WithFlank&3|+|188688775|188688872</td><td>60</td><td>81</td><td>22</td><td>CACTAGATTGTGAGCTCCTGGA</td><td>isomiR</td><td>TRUE</td><td>TRUE</td><td>FALSE</td></tr>
<tr><td>>MI0000077|hsa-mir-21&WithFlank&17|+|59841260|59841343@14.35.22(+1G)&[MIMAT0000076&hsa-miR-21-5p&offsets|0|0(+1G);m-1&17|+|59841273|59841294&offsets|0|0(+1G)]|TAGCTTATCAGACTGATGTTGAG</td><td>>MI0000077|hsa-mir-21&WithFlank&17|+|59841260|59841343</td><td>14</td><td>35</td><td>22(+1G)</td><td>TAGCTTATCAGACTGATGTTGAG</td><td>isomiR</td><td>TRUE</td><td>TRUE</td><td>FALSE</td></tr>
<tr><td>>MI0000264|hsa-mir-7-2&WithFlank&15|+|88611819|88611940@38.60.23(+1A);MI0000265|hsa-mir-7-3&WithFlank&19|+|4770664|4770785@37.59.23(+1A);MI0000263|hsa-mir-7-1&WithFlank&9|-|83969742|83969863@30.52.23(+1A)&[MIMAT0000252&hsa-miR-7-5p&offsets|0|-1(+1A);m-400&15|+|88611856|88611879&offsets|0|-1(+1A)];[MIMAT0000252_1&hsa-miR-7-5p&offsets|0|-1(+1A);m-402&19|+|4770700|4770723&offsets|0|-1(+1A)];[MIMAT0000252_2&hsa-miR-7-5p&offsets|0|-1(+1A);m-401&9|-|83969811|83969834&offsets|0|-1(+1A)]|TGGAAGACTAGTGATTTTGTTGTA</td><td>>MI0000264|hsa-mir-7-2&WithFlank&15|+|88611819|88611940</td><td>38</td><td>60</td><td>23(+1A)</td><td>TGGAAGACTAGTGATTTTGTTGTA</td><td>isomiR</td><td>TRUE</td><td>FALSE</td><td>FALSE</td></tr>
<tr><td>>hg38_1_-_228634819_228635039_RNA5S12_5S_50ntFlanks@140.170.31|ACCGCCTGGGAATACCGGGTGCTGTAGGCTT</td><td>>hg38_1_-_228634819_228635039_RNA5S12_5S_50ntFlanks</td><td>140</td><td>170</td><td>31</td><td>ACCGCCTGGGAATACCGGGTGCTGTAGGCTT</td><td>rRF</td><td>TRUE</td><td>TRUE</td><td>FALSE</td></tr>
<tr><td>>MI0000802|hsa-mir-340&WithFlank&5|-|180015297|180015403@22.43.22&[MIMAT0004692&hsa-miR-340-5p&offsets|0|0;m-220&5|-|180015361|180015382&offsets|0|0]|TTATAAAGCAATGAGACTGATT</td><td>>MI0000802|hsa-mir-340&WithFlank&5|-|180015297|180015403</td><td>22</td><td>43</td><td>22</td><td>TTATAAAGCAATGAGACTGATT</td><td>isomiR</td><td>TRUE</td><td>FALSE</td><td>FALSE</td></tr>
<tr><td>>MI0000266|hsa-mir-10a&WithFlank&17|-|48579832|48579953@29.50.22&[MIMAT0000253&hsa-miR-10a-5p&offsets|+1|0;m-18&17|-|48579905|48579926&offsets|+1|+1]|ACCCTGTAGATCCGAATTTGTG</td><td>>MI0000266|hsa-mir-10a&WithFlank&17|-|48579832|48579953</td><td>29</td><td>50</td><td>22</td><td>ACCCTGTAGATCCGAATTTGTG</td><td>isomiR</td><td>TRUE</td><td>TRUE</td><td>FALSE</td></tr>
<tr><td>>MI0000255|hsa-mir-30d&WithFlank&8|-|134804870|134804951@12.33.22&[MIMAT0000245&hsa-miR-30d-5p&offsets|0|0;m-21&8|-|134804919|134804940&offsets|0|0]|TGTAAACATCCCCGACTGGAAG</td><td>>MI0000255|hsa-mir-30d&WithFlank&8|-|134804870|134804951</td><td>12</td><td>33</td><td>22</td><td>TGTAAACATCCCCGACTGGAAG</td><td>isomiR</td><td>TRUE</td><td>FALSE</td><td>FALSE</td></tr>
<tr><td>>hg38_1_-_228634819_228635039_RNA5S12_5S_50ntFlanks@135.170.36|GGGAGACCGCCTGGGAATACCGGGTGCTGTAGGCTT</td><td>>hg38_1_-_228634819_228635039_RNA5S12_5S_50ntFlanks</td><td>135</td><td>170</td><td>36</td><td>GGGAGACCGCCTGGGAATACCGGGTGCTGTAGGCTT</td><td>rRF</td><td>TRUE</td><td>TRUE</td><td>FALSE</td></tr>
</table>

| | | | | | | | | | |
|---|---|---|---|---|---|---|---|---|---|
| >am_trna128_GlyGCC_6_-_27870686_27870756@1.32.32,trna133_GlyCCC_1_-_16872434_16872504@1.32.32,trna18_GlyGCC_16_+_70822597_70822667@1.32.32,trna19_GlyGCC_16_+_70823410_70823480@1.32.32,trna19_GlyGCC_2_-_157257659_157257729@1.32.32,trna24_GlyGCC_16_-_70812942_70813012@1.32.32,trna25_GlyGCC_16_-_70812114_70812184@1.32.32,trna4_GlyCCC_1_+_17188416_17188486@1.32.32,trna5_GlyGCC_17_+_8029064_8029134@1.32.32,trna68_GlyGCC_1_-_161493637_161493707@1.32.32\|GCATTGGTGGTTCAGTGGTAGAATTCTCGCCT | >am_trna128_GlyGCC_6_-_27870686_27870756 | 1 | 32 | 32 | GCATTGGTGGTTCAGTGGTAGAATTCTCGCCT | tRF | TRUE | TRUE | FALSE |
| >MI0000264\|hsa-mir-7-2&WithFlank&15\|+\|88611819\|88611940@38.60.23(+1G);MI0000265\|hsa-mir-7-3&WithFlank&19\|+\|4770664\|4770785@37.59.23(+1G);MI0000263\|hsa-mir-7-1&WithFlank&9\|-\|83969742\|83969863@30.52.23(+1G)&[MIMAT0000252&hsa-miR-7-5p&offsets\|0\|-1(+1G);m-400&15\|+\|88611856\|88611879&offsets\|0\|-1(+1G)];[MIMAT0000252_1&hsa-miR-7-5p&offsets\|0\|-1(+1G);m-402&19\|+\|4770700\|4770723&offsets\|0\|-1(+1G)];[MIMAT0000252_2&hsa-miR-7-5p&offsets\|0\|-1(+1G);m-401&9\|-\|83969811\|83969834&offsets\|0\|-1(+1G)]\|TGGAAGACTAGTGATTTTGTTGTG | >MI0000264\|hsa-mir-7-2&WithFlank&15\|+\|88611819\|88611940 | 38 | 60 | 23(+1G) | TGGAAGACTAGTGATTTTGTTGTG | isomiR | TRUE | FALSE | FALSE |
| >hg38_21_+_8433172_8446622_RNA45SN1_45S_50ntFlanks@3705.3725.21\|TACCTGGTTGATCCTGCCAGT | >hg38_21_+_8433172_8446622_RNA45SN1_45S_50ntFlanks | 3705 | 3725 | 21 | TACCTGGTTGATCCTGCCAGT | rRF | TRUE | FALSE | FALSE |
| >MI0000465\|hsa-mir-191&WithFlank&3\|-\|49020612\|49020715@22.43.22&[MIMAT0000440&hsa-miR-191-5p&offsets\|0\|-1;m-48&3\|-\|49020672\|49020694&offsets\|0\|-1]\|CAACGGAATCCCAAAAGCAGCT | >MI0000465\|hsa-mir-191&WithFlank&3\|-\|49020612\|49020715 | 22 | 43 | 22 | CAACGGAATCCCAAAAGCAGCT | isomiR | TRUE | TRUE | FALSE |
| >MI0000542\|hsa-mir-320a&WithFlank&8\|-\|22244960\|22245043@48.70.23&[MIMAT0000510&hsa-miR-320a-3p&offsets\|0\|+1;m-60&8\|-\|22244975\|22244996&offsets\|0\|+1]\|AAAAGCTGGGTTGAGAGGGCGAA | >MI0000542\|hsa-mir-320a&WithFlank&8\|-\|22244960\|22245043 | 48 | 70 | 23 | AAAAGCTGGGTTGAGAGGGCGAA | isomiR | TRUE | FALSE | FALSE |
| >trnaMT_GluTTC_MT_-_14674_14742@40.72.33,trnalookalike8_GluTTC_5_-_93905172_93905240@40.72.33\|TTGGTCGTGGTTGTAGTCCGTGCGAGAATACCA | >trnaMT_GluTTC_MT_-_14674_14742 | 40 | 72 | 33 | TTGGTCGTGGTTGTAGTCCGTGCGAGAATACCA | tRF | TRUE | TRUE | FALSE |
| >MI0000061\|hsa-let-7a-2&WithFlank&11\|-\|122146516\|122146599@11.31.21;MI0000062\|hsa-let-7a-3&WithFlank&22\|+\|46112743\|46112828@10.30.21;MI0000060\|hsa-let-7a-1&WithFlank&9\|+\|94175951\|94176042@12.32.21&[MIMAT0000062&hsa-let-7a-5p&offsets\|0\|-1;m-6&11\|-\|122146568\|122146589&offsets\|0\|-1];[MIMAT0000062_1&hsa-let-7a-5p&offsets\|0\|-1;m-5&22\|+\|46112752\|46112773&offsets\|0\|-1];[MIMAT0000062_2&hsa-let-7a-5p&offsets\|0\|-1;m-4&9\|+\|94175962\|94175983&offsets\|0\|-1]\|TGAGGTAGTAGGTTGTATAGT | >MI0000061\|hsa-let-7a-2&WithFlank&11\|-\|122146516\|122146599 | 11 | 31 | 21 | TGAGGTAGTAGGTTGTATAGT | isomiR | TRUE | TRUE | FALSE |
| >MI0000750\|hsa-mir-26a-2&WithFlank&12\|-\|57824603\|57824698@20.40.21(+1C);MI0000083\|hsa-mir-26a-1&WithFlank&3\|+\|37969398\|37969486@16.36.21(+1C)&[MIMAT0000082&hsa-miR-26a-5p&offsets\|0\|-1(+1C);m-33&12\|-\|57824658\|57824679&offsets\|0\|-1(+1C)];[MIMAT0000082_1&hsa-miR-26a-5p&offsets\|0\|-1(+1C);m-34&3\|+\|37969413\|37969434&offsets\|0\|-1(+1C)]\|TTCAAGTAATCCAGGATAGGCC | >MI0000750\|hsa-mir-26a-2&WithFlank&12\|-\|57824603\|57824698 | 20 | 40 | 21(+1C) | TTCAAGTAATCCAGGATAGGCC | isomiR | TRUE | TRUE | FALSE |
| >hg38_21_+_8433172_8446622_RNA45SN1_45S_50ntFlanks@6646.6666.21\|TCGTACGACTCTTAGCGGTGG | >hg38_21_+_8433172_8446622_RNA45SN1_45S_50ntFlanks | 6646 | 6666 | 21 | TCGTACGACTCTTAGCGGTGG | rRF | TRUE | FALSE | TRUE |
| >MI0000746\|hsa-mir-99b&WithFlank&19\|+\|51692606\|51692687@13.33.21&[MIMAT0000689&hsa-miR-99b-5p&offsets\|0\|-1;m-10&19\|+\|51692618\|51692639&offsets\|0\|-1]\|CACCCGTAGAACCGACCTTGC | >MI0000746\|hsa-mir-99b&WithFlank&19\|+\|51692606\|51692687 | 13 | 33 | 21 | CACCCGTAGAACCGACCTTGC | isomiR | TRUE | TRUE | FALSE |
| >hg38_21_+_8433172_8446622_RNA45SN1_45S_50ntFlanks@475.504.30\|CTTCGTGATCGATGTGGTGACGTCGTGCTC | >hg38_21_+_8433172_8446622_RNA45SN1_45S_50ntFlanks | 475 | 504 | 30 | CTTCGTGATCGATGTGGTGACGTCGTGCTC | rRF | TRUE | TRUE | FALSE |
| >MI0000108\|hsa-mir-103a-2&WithFlank&20\|+\|3917488\|3917577@54.75.22;MI0000109\|hsa-mir-103a-1&WithFlank&5\|-\|168560890\|168560979@54.75.22&[MIMAT0000101&hsa-miR-103a-3p&offsets\|0\|-1;m-12&20\|+\|3917541\|3917563&offsets\|0\|-1];[MIMAT0000101_1&hsa-miR-103a-3p&offsets\|0\|-1;m-13&5\|-\|168560904\|168560926&offsets\|0\|-1;m-44220&5\|-\|168560908\|168560926&offsets\|0\|+3]\|AGCAGCATTGTACAGGGCTATG | >MI0000108\|hsa-mir-103a-2&WithFlank&20\|+\|3917488\|3917577 | 54 | 75 | 22 | AGCAGCATTGTACAGGGCTATG | isomiR | TRUE | TRUE | FALSE |
| >MI0000093\|hsa-mir-92a-1&WithFlank&13\|+\|91351308\|91351397@54.75.22(+1A);MI0000094\|hsa-mir-92a-2&WithFlank&X\|-\|134169532\|134169618@54.75.22(+1A)&[MIMAT0000092&hsa-miR-92a-3p&offsets\|0\|0(+1A);m-9&13\|+\|91351361\|91351382&offsets\|0\|0(+1A)];[MIMAT0000092_1&hsa-miR-92a-3p&offsets\|0\|0(+1A);m-11&X\|-\|134169544\|134169565&offsets\|0\|0(+1A)]\|TATTGCACTTGTCCCGGCCTGTA | >MI0000093\|hsa-mir-92a-1&WithFlank&13\|+\|91351308\|91351397 | 54 | 75 | 22(+1A) | TATTGCACTTGTCCCGGCCTGTA | isomiR | TRUE | TRUE | FALSE |
| >MI0000749\|hsa-mir-30e&WithFlank&1\|+\|40754349\|40754452@23.46.24&[MIMAT0000692&hsa-miR-30e-5p&offsets\|0\|+2;m-24&1\|+\|40754371\|40754394&offsets\|0\|0]\|TGTAAACATCCTTGACTGGAAGCT | >MI0000749\|hsa-mir-30e&WithFlank&1\|+\|40754349\|40754452 | 23 | 46 | 24 | TGTAAACATCCTTGACTGGAAGCT | isomiR | TRUE | TRUE | FALSE |
| >trna2_GlyGCC_21_-_18827107_18827177@1.32.32,trna35_GlyGCC_1_+_161413094_161413164@1.32.32,trna37_GlyGCC_1_+_161420467_161420537@1.32.32,trna39_GlyGCC_1_+_161427898_161427968@1.32.32,trna41_GlyGCC_1_+_161435258_161435328@1.32.32\|GCATGGGTGGTTCAGTGGTAGAATTCTCGCCT | >trna2_GlyGCC_21_-_18827107_18827177 | 1 | 32 | 32 | GCATGGGTGGTTCAGTGGTAGAATTCTCGCCT | tRF | TRUE | FALSE | FALSE |
| >hg38_1_-_228634819_228635039_RNA5S12_5S_50ntFlanks@134.171.38\|TGGGAGACCGCCTGGGAATACCGGGTGCTGTAGGCTTT | >hg38_1_-_228634819_228635039_RNA5S12_5S_50ntFlanks | 134 | 171 | 38 | TGGGAGACCGCCTGGGAATACCGGGTGCTGTAGGCTTT | rRF | TRUE | TRUE | FALSE |
| >hg38_21_+_8433172_8446622_RNA45SN1_45S_50ntFlanks@7975.7992.18\|CGCGACCTCAGATCAGAC | >hg38_21_+_8433172_8446622_RNA45SN1_45S_50ntFlanks | 7975 | 7992 | 18 | CGCGACCTCAGATCAGAC | rRF | TRUE | FALSE | FALSE |
| >MI0000298\|hsa-mir-221&WithFlank&X\|-\|45746151\|45746272@71.93.23&[MIMAT0000278&hsa-miR-221-3p&offsets\|0\|0;m-91&X\|-\|45746181\|45746202&offsets\|0\|+1]\|AGCTACATTGTCTGCTGGGTTTC | >MI0000298\|hsa-mir-221&WithFlank&X\|-\|45746151\|45746272 | 71 | 93 | 23 | AGCTACATTGTCTGCTGGGTTTC | isomiR | TRUE | TRUE | FALSE |
| >hg38_21_+_8433172_8446622_RNA45SN1_45S_50ntFlanks@6646.6667.22\|TCGTACGACTCTTAGCGGTGGA | >hg38_21_+_8433172_8446622_RNA45SN1_45S_50ntFlanks | 6646 | 6667 | 22 | TCGTACGACTCTTAGCGGTGGA | rRF | TRUE | FALSE | FALSE |
| >MI0000272\|hsa-mir-182&WithFlank&7\|-\|129770377\|129770498@29.51.23(+1A)&[MIMAT0000259&hsa-miR-182-5p&offsets\|0\|-1(+1A);m-22&7\|-\|129770449\|129770470&offsets\|0\|+1(+1A)]\|TTTGGCAATGGTAGAACTCACACA | >MI0000272\|hsa-mir-182&WithFlank&7\|-\|129770377\|129770498 | 29 | 51 | 23(+1A) | TTTGGCAATGGTAGAACTCACACA | isomiR | TRUE | FALSE | FALSE |
| >hg38_21_+_8433172_8446622_RNA45SN1_45S_50ntFlanks@6646.6668.23\|TCGTACGACTCTTAGCGGTGGAT | >hg38_21_+_8433172_8446622_RNA45SN1_45S_50ntFlanks | 6646 | 6668 | 23 | TCGTACGACTCTTAGCGGTGGAT | rRF | TRUE | FALSE | FALSE |
| >MI0000811\|hsa-mir-148b&WithFlank&12\|+\|54337210\|54337320@69.90.22(+1U)&[MIMAT0000759&hsa-miR-148b-3p&offsets\|0\|0(+1U);m-77&12\|+\|54337278\|54337299&offsets\|0\|0(+1U)]\|TCAGTGCATCACAGAACTTTGTT | >MI0000811\|hsa-mir-148b&WithFlank&12\|+\|54337210\|54337320 | 69 | 90 | 22(+1U) | TCAGTGCATCACAGAACTTTGTT | isomiR | TRUE | TRUE | FALSE |

<table>
<tr><td>>trna116_GluCTC_1_-_145399233_145399304@1.34.34,trna59_GluCTC_1_+_249168447_249168518@1.34.34,trna71_GluCTC_1_-_161439189_161439260@1.34.34,trna74_GluCTC_1_-_161431809_161431880@1.34.34,trna77_GluCTC_1_-_161424398_161424469@1.34.34,trna77_GluCTC_6_+_28949976_28950047@1.34.34,trna80_GluCTC_1_-_161417018_161417089@1.34.34,trna87_GluCTC_6_-_126101393_126101464@1.34.34|TCCCTGGTGGTCTAGTGGTTAGGATTCGGCGCTC</td><td>>trna116_GluCTC_1_-_145399233_145399304</td><td>1</td><td>34</td><td>34</td><td>TCCCTGGTGGTCTAGTGGTTAGGATTCGGCGCTC</td><td>tRF</td><td>TRUE</td><td>FALSE</td><td>FALSE</td></tr>
<tr><td>>hg38_21_+_8433172_8446622_RNA45SN1_45S_50ntFlanks@6645.6665.21|CTCGTACGACTCTTAGCGGTG</td><td>>hg38_21_+_8433172_8446622_RNA45SN1_45S_50ntFlanks</td><td>6645</td><td>6665</td><td>21</td><td>CTCGTACGACTCTTAGCGGTG</td><td>rRF</td><td>TRUE</td><td>FALSE</td><td>FALSE</td></tr>
<tr><td>>hg38_21_+_8433172_8446622_RNA45SN1_45S_50ntFlanks@475.506.32|CTTCGTGATCGATGTGGTGACGTCGTGCTCTC</td><td>>hg38_21_+_8433172_8446622_RNA45SN1_45S_50ntFlanks</td><td>475</td><td>506</td><td>32</td><td>CTTCGTGATCGATGTGGTGACGTCGTGCTCTC</td><td>rRF</td><td>TRUE</td><td>TRUE</td><td>FALSE</td></tr>
<tr><td>>MI0000285|hsa-mir-205&WithFlank&1|+|209432127|209432248@40.60.21&[MIMAT0000266&hsa-miR-205-5p&offsets|0|-1;m-351&1|+|209432166|209432187&offsets|0|-1]|TCCTTCATTCCACCGGAGTCT</td><td>>MI0000285|hsa-mir-205&WithFlank&1|+|209432127|209432248</td><td>40</td><td>60</td><td>21</td><td>TCCTTCATTCCACCGGAGTCT</td><td>isomiR</td><td>TRUE</td><td>FALSE</td><td>FALSE</td></tr>
<tr><td>>hg38_21_+_8433172_8446622_RNA45SN1_45S_50ntFlanks@475.497.23|CTTCGTGATCGATGTGGTGACGT</td><td>>hg38_21_+_8433172_8446622_RNA45SN1_45S_50ntFlanks</td><td>475</td><td>497</td><td>23</td><td>CTTCGTGATCGATGTGGTGACGT</td><td>rRF</td><td>TRUE</td><td>TRUE</td><td>FALSE</td></tr>
<tr><td>>hg38_21_+_8433172_8446622_RNA45SN1_45S_50ntFlanks@6651.6668.18|CGACTCTTAGCGGTGGAT</td><td>>hg38_21_+_8433172_8446622_RNA45SN1_45S_50ntFlanks</td><td>6651</td><td>6668</td><td>18</td><td>CGACTCTTAGCGGTGGAT</td><td>rRF</td><td>TRUE</td><td>FALSE</td><td>FALSE</td></tr>
<tr><td>>hg38_21_+_8433172_8446622_RNA45SN1_45S_50ntFlanks@7974.8011.38|ACGCGACCTCAGATCAGACGTGGCGACCCGCTGAATTT</td><td>>hg38_21_+_8433172_8446622_RNA45SN1_45S_50ntFlanks</td><td>7974</td><td>8011</td><td>38</td><td>ACGCGACCTCAGATCAGACGTGGCGACCCGCTGAATTT</td><td>rRF</td><td>TRUE</td><td>TRUE</td><td>FALSE</td></tr>
<tr><td>>MI0000750|hsa-mir-26a-2&WithFlank&12|-|57824603|57824698@20.40.21(+1A);MI0000083|hsa-mir-26a-1&WithFlank&3|+|37969398|37969486@16.36.21(+1A)&[MIMAT0000082&hsa-miR-26a-5p&offsets|0|-1(+1A);m-33&12|-|57824658|57824679&offsets|0|-1(+1A)];[MIMAT0000082_1&hsa-miR-26a-5p&offsets|0|-1(+1A);m-34&3|+|37969413|37969434&offsets|0|-1(+1A)]|TTCAAGTAATCCAGGATAGGCA</td><td>>MI0000750|hsa-mir-26a-2&WithFlank&12|-|57824603|57824698</td><td>20</td><td>40</td><td>21(+1A)</td><td>TTCAAGTAATCCAGGATAGGCA</td><td>isomiR</td><td>TRUE</td><td>TRUE</td><td>FALSE</td></tr>
<tr><td>>multi-am_trnaMT_ProTGG_MT_-_15956_16023@54.71.18|AGACTTTTTCTCTGACCA</td><td>>multi-am_trnaMT_ProTGG_MT_-_15956_16023</td><td>54</td><td>71</td><td>18</td><td>AGACTTTTTCTCTGACCA</td><td>tRF</td><td>TRUE</td><td>TRUE</td><td>FALSE</td></tr>
<tr><td>>MI0000650|hsa-mir-200c&WithFlank&12|+|6963693|6963772@50.71.22(+1U)&[MIMAT0000617&hsa-miR-200c-3p&offsets|0|-1(+1U);m-54&12|+|6963742|6963764&offsets|0|-1(+1U)]|TAATACTGCCGGGTAATGATGGT</td><td>>MI0000650|hsa-mir-200c&WithFlank&12|+|6963693|6963772</td><td>50</td><td>71</td><td>22(+1U)</td><td>TAATACTGCCGGGTAATGATGGT</td><td>isomiR</td><td>TRUE</td><td>FALSE</td><td>FALSE</td></tr>
<tr><td>>MI0000067|hsa-let-7f-1&WithFlank&9|+|94176341|94176439@13.33.21(+1A);MI0000068|hsa-let-7f-2&WithFlank&X|-|53557186|53557280@14.34.21(+1A)&[MIMAT0000067&hsa-let-7f-5p&offsets|0|-1(+1A);m-20&9|+|94176353|94176374&offsets|0|-1(+1A)];[MIMAT0000067_1&hsa-let-7f-5p&offsets|0|-1(+1A);m-19&X|-|53557246|53557267&offsets|0|-1(+1A)]|TGAGGTAGTAGATTGTATAGTA</td><td>>MI0000067|hsa-let-7f-1&WithFlank&9|+|94176341|94176439</td><td>13</td><td>33</td><td>21(+1A)</td><td>TGAGGTAGTAGATTGTATAGTA</td><td>isomiR</td><td>TRUE</td><td>FALSE</td><td>FALSE</td></tr>
<tr><td>>am_MI0000809|hsa-mir-151a&WithFlank&8|-|140732558|140732659@53.72.20(+1U)&[MIMAT0000757&hsa-miR-151a-3p&offsets|0|-1(+1U);m-30&8|-|140732587|140732607&offsets|0|-1(+1U)]|CTAGACTGAAGCTCCTTGAGT</td><td>>am_MI0000809|hsa-mir-151a&WithFlank&8|-|140732558|140732659</td><td>53</td><td>72</td><td>20(+1U)</td><td>CTAGACTGAAGCTCCTTGAGT</td><td>isomiR</td><td>TRUE</td><td>TRUE</td><td>FALSE</td></tr>
<tr><td>>trnaMT_SerGCT_MT_+_12207_12265@1.19.19|GAGAAAGCTCACAAGAACT</td><td>>trnaMT_SerGCT_MT_+_12207_12265</td><td>1</td><td>19</td><td>19</td><td>GAGAAAGCTCACAAGAACT</td><td>tRF</td><td>TRUE</td><td>TRUE</td><td>FALSE</td></tr>
<tr><td>>MI0000063|hsa-let-7b&WithFlank&22|+|46113680|46113774@12.34.23&[MIMAT0000063&hsa-let-7b-5p&offsets|0|+1;m-14&22|+|46113691|46113712&offsets|0|+1]|TGAGGTAGTAGGTTGTGTGGTTT</td><td>>MI0000063|hsa-let-7b&WithFlank&22|+|46113680|46113774</td><td>12</td><td>34</td><td>23</td><td>TGAGGTAGTAGGTTGTGTGGTTT</td><td>isomiR</td><td>TRUE</td><td>FALSE</td><td>FALSE</td></tr>
<tr><td>>MI0000239|hsa-mir-197&WithFlank&1|+|109598887|109598973@54.75.22&[MIMAT0000227&hsa-miR-197-3p&offsets|0|0;m-99&1|+|109598940|109598961&offsets|0|0]|TTCACCACCTTCTCCACCCAGC</td><td>>MI0000239|hsa-mir-197&WithFlank&1|+|109598887|109598973</td><td>54</td><td>75</td><td>22</td><td>TTCACCACCTTCTCCACCCAGC</td><td>isomiR</td><td>TRUE</td><td>FALSE</td><td>FALSE</td></tr>
<tr><td>>MI0000079|hsa-mir-23a&WithFlank&19|-|13836581|13836665@54.72.19&[MIMAT0000078&hsa-miR-23a-3p&offsets|+3|+1;m-31&19|-|13836595|13836615&offsets|+3|+1]|ACATTGCCAGGGATTTCCA</td><td>>MI0000079|hsa-mir-23a&WithFlank&19|-|13836581|13836665</td><td>54</td><td>72</td><td>19</td><td>ACATTGCCAGGGATTTCCA</td><td>isomiR</td><td>TRUE</td><td>TRUE</td><td>FALSE</td></tr>
<tr><td>>MI0000085|hsa-mir-27a&WithFlank&19|-|13836434|13836523@57.76.20&[MIMAT0000084&hsa-miR-27a-3p&offsets|0|-1;m-43&19|-|13836447|13836467&offsets|0|-1]|TTCACAGTGGCTAAGTTCCG</td><td>>MI0000085|hsa-mir-27a&WithFlank&19|-|13836434|13836523</td><td>57</td><td>76</td><td>20</td><td>TTCACAGTGGCTAAGTTCCG</td><td>isomiR</td><td>TRUE</td><td>TRUE</td><td>FALSE</td></tr>
<tr><td>>trna116_GluCTC_1_-_145399233_145399304@1.35.35,trna59_GluCTC_1_+_249168447_249168518@1.35.35,trna71_GluCTC_1_-_161439189_161439260@1.35.35,trna74_GluCTC_1_-_161431809_161431880@1.35.35,trna77_GluCTC_1_-_161424398_161424469@1.35.35,trna77_GluCTC_6_+_28949976_28950047@1.35.35,trna80_GluCTC_1_-_161417018_161417089@1.35.35,trna87_GluCTC_6_-_126101393_126101464@1.35.35|TCCCTGGTGGTCTAGTGGTTAGGATTCGGCGCTCT</td><td>>trna116_GluCTC_1_-_145399233_145399304</td><td>1</td><td>35</td><td>35</td><td>TCCCTGGTGGTCTAGTGGTTAGGATTCGGCGCTCT</td><td>tRF</td><td>TRUE</td><td>FALSE</td><td>FALSE</td></tr>
<tr><td>>MI0000070|hsa-mir-16-1&WithFlank&13|-|50048967|50049067@20.42.23;MI0000115|hsa-mir-16-2&WithFlank&3|+|160404739|160404831@16.38.23&[MIMAT0000069&hsa-miR-16-5p&offsets|0|+1;m-62&13|-|50049027|50049048&offsets|0|+1];[MIMAT0000069_1&hsa-miR-16-5p&offsets|0|+1;m-61&3|+|160404754|160404775&offsets|0|+1]|TAGCAGCACGTAAATATTGGCGT</td><td>>MI0000070|hsa-mir-16-1&WithFlank&13|-|50048967|50049067</td><td>20</td><td>42</td><td>23</td><td>TAGCAGCACGTAAATATTGGCGT</td><td>isomiR</td><td>TRUE</td><td>FALSE</td><td>FALSE</td></tr>
<tr><td>>MI0000285|hsa-mir-205&WithFlank&1|+|209432127|209432248@40.61.22(+1C)&[MIMAT0000266&hsa-miR-205-5p&offsets|0|0(+1C);m-351&1|+|209432166|209432187&offsets|0|0(+1C)]|TCCTTCATTCCACCGGAGTCTGC</td><td>>MI0000285|hsa-mir-205&WithFlank&1|+|209432127|209432248</td><td>40</td><td>61</td><td>22(+1C)</td><td>TCCTTCATTCCACCGGAGTCTGC</td><td>isomiR</td><td>TRUE</td><td>FALSE</td><td>FALSE</td></tr>
<tr><td>>hg38_21_+_8433172_8446622_RNA45SN1_45S_50ntFlanks@7975.8015.41|CGCGACCTCAGATCAGACGTGGCGACCCGCTGAATTTAAGC</td><td>>hg38_21_+_8433172_8446622_RNA45SN1_45S_50ntFlanks</td><td>7975</td><td>8015</td><td>41</td><td>CGCGACCTCAGATCAGACGTGGCGACCCGCTGAATTTAAGC</td><td>rRF</td><td>TRUE</td><td>TRUE</td><td>FALSE</td></tr>
<tr><td>>multi-am_trnaMT_ProTGG_MT_-_15956_16023@52.71.20|AAAGACTTTTTCTCTGACCA</td><td>>multi-am_trnaMT_ProTGG_MT_-_15956_16023</td><td>52</td><td>71</td><td>20</td><td>AAAGACTTTTTCTCTGACCA</td><td>tRF</td><td>TRUE</td><td>TRUE</td><td>FALSE</td></tr>
<tr><td>>MI0000542|hsa-mir-320a&WithFlank&8|-|22244960|22245043@48.70.23(+1U)&[MIMAT0000510&hsa-miR-320a-3p&offsets|0|+1(+1U);m-60&8|-|22244975|22244996&offsets|0|+1(+1U)]|AAAAGCTGGGTTGAGAGGGCGAAT</td><td>>MI0000542|hsa-mir-320a&WithFlank&8|-|22244960|22245043</td><td>48</td><td>70</td><td>23(+1U)</td><td>AAAAGCTGGGTTGAGAGGGCGAAT</td><td>isomiR</td><td>TRUE</td><td>FALSE</td><td>FALSE</td></tr>
<tr><td>>MI0000439|hsa-mir-23b&WithFlank&9|+|95085202|95085310@67.86.20(+1U)&[MIMAT0000418&hsa-miR-23b-3p&offsets|+3|0(+1U);m-42&9|+|95085265|95085285&offsets|+3|+2(+1U)]|ACATTGCCAGGGATTACCACT</td><td>>MI0000439|hsa-mir-23b&WithFlank&9|+|95085202|95085310</td><td>67</td><td>86</td><td>20(+1U)</td><td>ACATTGCCAGGGATTACCACT</td><td>isomiR</td><td>TRUE</td><td>TRUE</td><td>FALSE</td></tr>
<tr><td>>hg38_21_+_8433172_8446622_RNA45SN1_45S_50ntFlanks@7975.8012.38|CGCGACCTCAGATCAGACGTGGCGACCCGCTGAATTTA</td><td>>hg38_21_+_8433172_8446622_RNA45SN1_45S_50ntFlanks</td><td>7975</td><td>8012</td><td>38</td><td>CGCGACCTCAGATCAGACGTGGCGACCCGCTGAATTTA</td><td>rRF</td><td>TRUE</td><td>TRUE</td><td>TRUE</td></tr>
<tr><td>>trnaMT_SerGCT_MT_+_12207_12265@1.26.26|GAGAAAGCTCACAAGAACTGCTAACT</td><td>>trnaMT_SerGCT_MT_+_12207_12265</td><td>1</td><td>26</td><td>26</td><td>GAGAAAGCTCACAAGAACTGCTAACT</td><td>tRF</td><td>TRUE</td><td>TRUE</td><td>FALSE</td></tr>
</table>

| | | | | | | | | | |
|---|---|---|---|---|---|---|---|---|---|
| >MI0000272\|hsa-mir-182&WithFlank&7\|-\|129770377\|129770498@29.51.23(+1G)&[MIMAT0000259&hsa-miR-182-5p&offsets\|0\|-1(+1G);m-22&7\|-\|129770449\|129770470&offsets\|0\|+1(+1G)]\|TTTGGCAATGGTAGAACTCACACG | >MI0000272\|hsa-mir-182&WithFlank&7\|-\|129770377\|129770498 | 29 | 51 | 23(+1G) | TTTGGCAATGGTAGAACTCACACG | isomiR | TRUE | FALSE | FALSE |
| >MI0000298\|hsa-mir-221&WithFlank&X\|-\|45746151\|45746272@71.92.22&[MIMAT0000278&hsa-miR-221-3p&offsets\|0\|-1;m-91&X\|-\|45746181\|45746202&offsets\|0\|0]\|AGCTACATTGTCTGCTGGGTTT | >MI0000298\|hsa-mir-221&WithFlank&X\|-\|45746151\|45746272 | 71 | 92 | 22 | AGCTACATTGTCTGCTGGGTTT | isomiR | TRUE | TRUE | FALSE |
| >MI0000063\|hsa-let-7b&WithFlank&22\|+\|46113680\|46113774@12.32.21(+1C)&[MIMAT0000063&hsa-let-7b-5p&offsets\|0\|-1(+1C);m-14&22\|+\|46113691\|46113712&offsets\|0\|-1(+1C)]\|TGAGGTAGTAGGTTGTGTGGTC | >MI0000063\|hsa-let-7b&WithFlank&22\|+\|46113680\|46113774 | 12 | 32 | 21(+1C) | TGAGGTAGTAGGTTGTGTGGTC | isomiR | TRUE | FALSE | FALSE |
| >trna10_LysCTT_16_+_3241501_3241573@1.36.36,trna13_LysCTT_14_-_58706613_58706685@1.36.36,trna2_LysCTT_15_+_79152904_79152976@1.36.36\|GCCCGGCTAGCTCAGTCGGTAGAGCATGGGACTCTT | >trna10_LysCTT_16_+_3241501_3241573 | 1 | 36 | 36 | GCCCGGCTAGCTCAGTCGGTAGAGCATGGGACTCTT | tRF | TRUE | FALSE | FALSE |
| >MI0000286\|hsa-mir-210&WithFlank&11\|-\|568083\|568204@72.93.22&[MIMAT0000267&hsa-miR-210-3p&offsets\|0\|0;m-102&11\|-\|568112\|568133&offsets\|0\|0]\|CTGTGCGTGTGACAGCGGCTGA | >MI0000286\|hsa-mir-210&WithFlank&11\|-\|568083\|568204 | 72 | 93 | 22 | CTGTGCGTGTGACAGCGGCTGA | isomiR | TRUE | FALSE | FALSE |
| >MI0005756\|hsa-mir-934&WithFlank&X\|+\|136550872\|136550966@21.42.22(+1A)&[MIMAT0004977&hsa-miR-934&offsets\|0\|0(+1A);m-943&X\|+\|136550892\|136550913&offsets\|0\|0(+1A)]\|TGTCTACTACTGGAGACACTGGA | >MI0005756\|hsa-mir-934&WithFlank&X\|+\|136550872\|136550966 | 21 | 42 | 22(+1A) | TGTCTACTACTGGAGACACTGGA | isomiR | TRUE | FALSE | FALSE |
| >MI0000434\|hsa-let-7i&WithFlank&12\|+\|62603680\|62603775@12.32.21(+1A)&[MIMAT0000415&hsa-let-7i-5p&offsets\|0\|-1(+1A);m-49&12\|+\|62603691\|62603712&offsets\|0\|-1(+1A)]\|TGAGGTAGTAGTTTGTGCTGTA | >MI0000434\|hsa-let-7i&WithFlank&12\|+\|62603680\|62603775 | 12 | 32 | 21(+1A) | TGAGGTAGTAGTTTGTGCTGTA | isomiR | TRUE | TRUE | FALSE |
| >MI0000088\|hsa-mir-30a&WithFlank&6\|-\|71403545\|71403627@12.33.22(+1A)&[MIMAT0000087&hsa-miR-30a-5p&offsets\|0\|0(+1A);m-15&6\|-\|71403595\|71403616&offsets\|0\|0(+1A)]\|TGTAAACATCCTCGACTGGAAGA | >MI0000088\|hsa-mir-30a&WithFlank&6\|-\|71403545\|71403627 | 12 | 33 | 22(+1A) | TGTAAACATCCTCGACTGGAAGA | isomiR | TRUE | FALSE | FALSE |
| >hg38_21_+_8433172_8446622_RNA45SN1_45S_50ntFlanks@3705.3722.18\|TACCTGGTTGATCCTGCC | >hg38_21_+_8433172_8446622_RNA45SN1_45S_50ntFlanks | 3705 | 3722 | 18 | TACCTGGTTGATCCTGCC | rRF | TRUE | FALSE | FALSE |
| >hg38_21_+_8433172_8446622_RNA45SN1_45S_50ntFlanks@7974.8010.37\|ACGCGACCTCAGATCAGACGTGGCGACCCGCTGAATT | >hg38_21_+_8433172_8446622_RNA45SN1_45S_50ntFlanks | 7974 | 8010 | 37 | ACGCGACCTCAGATCAGACGTGGCGACCCGCTGAATT | rRF | TRUE | TRUE | FALSE |
| >hg38_1_-_228634819_228635039_RNA5S12_5S_50ntFlanks@135.171.37\|GGGAGACCGCCTGGGAATACCGGGTGCTGTAGGCTTT | >hg38_1_-_228634819_228635039_RNA5S12_5S_50ntFlanks | 135 | 171 | 37 | GGGAGACCGCCTGGGAATACCGGGTGCTGTAGGCTTT | rRF | TRUE | TRUE | FALSE |
| >MI0000067\|hsa-let-7f-1&WithFlank&9\|+\|94176341\|94176439@13.33.21(+1G);MI0000068\|hsa-let-7f-2&WithFlank&X\|-\|53557186\|53557280@14.34.21(+1G)&[MIMAT0000067&hsa-let-7f-5p&offsets\|0\|-1(+1G);m-20&9\|+\|94176353\|94176374&offsets\|0\|-1(+1G)];[MIMAT0000067_1&hsa-let-7f-5p&offsets\|0\|-1(+1G);m-19&X\|-\|53557246\|53557267&offsets\|0\|-1(+1G)]\|TGAGGTAGTAGATTGTATAGTG | >MI0000067\|hsa-let-7f-1&WithFlank&9\|+\|94176341\|94176439 | 13 | 33 | 21(+1G) | TGAGGTAGTAGATTGTATAGTG | isomiR | TRUE | FALSE | FALSE |
| >hg38_21_+_8433172_8446622_RNA45SN1_45S_50ntFlanks@6646.6672.27\|TCGTACGACTCTTAGCGGTGGATCACT | >hg38_21_+_8433172_8446622_RNA45SN1_45S_50ntFlanks | 6646 | 6672 | 27 | TCGTACGACTCTTAGCGGTGGATCACT | rRF | TRUE | FALSE | FALSE |
| >MI0000285\|hsa-mir-205&WithFlank&1\|+\|209432127\|209432248@40.60.21(+1A)&[MIMAT0000266&hsa-miR-205-5p&offsets\|0\|-1(+1A);m-351&1\|+\|209432166\|209432187&offsets\|0\|-1(+1A)]\|TCCTTCATTCCACCGGAGTCTA | >MI0000285\|hsa-mir-205&WithFlank&1\|+\|209432127\|209432248 | 40 | 60 | 21(+1A) | TCCTTCATTCCACCGGAGTCTA | isomiR | TRUE | FALSE | FALSE |
| >hg38_21_+_8433172_8446622_RNA45SN1_45S_50ntFlanks@3705.3748.44\|TACCTGGTTGATCCTGCCAGTAGCATATGCTTGTCTCAAAGATT | >hg38_21_+_8433172_8446622_RNA45SN1_45S_50ntFlanks | 3705 | 3748 | 44 | TACCTGGTTGATCCTGCCAGTAGCATATGCTTGTCTCAAAGATT | rRF | TRUE | FALSE | FALSE |
| >hg38_1_-_228634819_228635039_RNA5S12_5S_50ntFlanks@133.170.38\|ATGGGAGACCGCCTGGGAATACCGGGTGCTGTAGGCTT | >hg38_1_-_228634819_228635039_RNA5S12_5S_50ntFlanks | 133 | 170 | 38 | ATGGGAGACCGCCTGGGAATACCGGGTGCTGTAGGCTT | rRF | TRUE | TRUE | FALSE |
| >MI0000103\|hsa-mir-101-1&WithFlank&1\|-\|65058428\|65058514@52.73.22;MI0000739\|hsa-mir-101-2&WithFlank&9\|+\|4850291\|4850381@54.75.22&[MIMAT0000099&hsa-miR-101-3p&offsets\|-1\|0;m-17&1\|-\|65058443\|65058462&offsets\|-1\|+1];[MIMAT0000099_1&hsa-miR-101-3p&offsets\|-1\|0;m-16&9\|+\|4850345\|4850364&offsets\|-1\|+1]\|GTACAGTACTGTGATAACTGAA | >MI0000103\|hsa-mir-101-1&WithFlank&1\|-\|65058428\|65058514 | 52 | 73 | 22 | GTACAGTACTGTGATAACTGAA | isomiR | TRUE | FALSE | FALSE |
| >hg38_21_+_8433172_8446622_RNA45SN1_45S_50ntFlanks@476.505.30\|TTCGTGATCGATGTGGTGACGTCGTGCTCT | >hg38_21_+_8433172_8446622_RNA45SN1_45S_50ntFlanks | 476 | 505 | 30 | TTCGTGATCGATGTGGTGACGTCGTGCTCT | rRF | TRUE | TRUE | FALSE |
| >MI0000272\|hsa-mir-182&WithFlank&7\|-\|129770377\|129770498@29.54.26&[MIMAT0000259&hsa-miR-182-5p&offsets\|0\|+2;m-22&7\|-\|129770449\|129770470&offsets\|0\|+4]\|TTTGGCAATGGTAGAACTCACACTGG | >MI0000272\|hsa-mir-182&WithFlank&7\|-\|129770377\|129770498 | 29 | 54 | 26 | TTTGGCAATGGTAGAACTCACACTGG | isomiR | TRUE | FALSE | FALSE |
| >MI0000299\|hsa-mir-222&WithFlank&X\|-\|45747009\|45747130@75.98.24&[MIMAT0000279&hsa-miR-222-3p&offsets\|0\|+3;m-140&X\|-\|45747033\|45747056&offsets\|0\|0]\|AGCTACATCTGGCTACTGGGTCTC | >MI0000299\|hsa-mir-222&WithFlank&X\|-\|45747009\|45747130 | 75 | 98 | 24 | AGCTACATCTGGCTACTGGGTCTC | isomiR | TRUE | TRUE | FALSE |
| >MI0000071\|hsa-mir-17&WithFlank&13\|+\|91350599\|91350694@20.42.23&[MIMAT0000070&hsa-miR-17-5p&offsets\|0\|0;m-88&13\|+\|91350618\|91350640&offsets\|0\|0]\|CAAAGTGCTTACAGTGCAGGTAG | >MI0000071\|hsa-mir-17&WithFlank&13\|+\|91350599\|91350694 | 20 | 42 | 23 | CAAAGTGCTTACAGTGCAGGTAG | isomiR | TRUE | TRUE | FALSE |
| >MI0000469\|hsa-mir-125a&WithFlank&19\|+\|51693248\|51693345@21.42.22&[MIMAT0000443&hsa-miR-125a-5p&offsets\|0\|-2;m-63&19\|+\|51693268\|51693289&offsets\|0\|0]\|TCCCTGAGACCCTTTAACCTGT | >MI0000469\|hsa-mir-125a&WithFlank&19\|+\|51693248\|51693345 | 21 | 42 | 22 | TCCCTGAGACCCTTTAACCTGT | isomiR | TRUE | FALSE | FALSE |
| >hg38_1_-_228634819_228635039_RNA5S12_5S_50ntFlanks@134.169.36\|TGGGAGACCGCCTGGGAATACCGGGTGCTGTAGGCT | >hg38_1_-_228634819_228635039_RNA5S12_5S_50ntFlanks | 134 | 169 | 36 | TGGGAGACCGCCTGGGAATACCGGGTGCTGTAGGCT | rRF | TRUE | TRUE | FALSE |
| >hg38_21_+_8433172_8446622_RNA45SN1_45S_50ntFlanks@485.506.22\|GATGTGGTGACGTCGTGCTCTC | >hg38_21_+_8433172_8446622_RNA45SN1_45S_50ntFlanks | 485 | 506 | 22 | GATGTGGTGACGTCGTGCTCTC | rRF | TRUE | TRUE | FALSE |
| >MI0000433\|hsa-let-7g&WithFlank&3\|-\|52268272\|52268367@11.31.21&[MIMAT0000414&hsa-let-7g-5p&offsets\|0\|-1;m-58&3\|-\|52268336\|52268357&offsets\|0\|-1]\|TGAGGTAGTAGTTTGTACAGT | >MI0000433\|hsa-let-7g&WithFlank&3\|-\|52268272\|52268367 | 11 | 31 | 21 | TGAGGTAGTAGTTTGTACAGT | isomiR | TRUE | TRUE | FALSE |
| >MI0000542\|hsa-mir-320a&WithFlank&8\|-\|22244960\|22245043@48.68.21(+1U)&[MIMAT0000510&hsa-miR-320a-3p&offsets\|0\|-1(+1U);m-60&8\|-\|22244975\|22244996&offsets\|0\|-1(+1U)]\|AAAAGCTGGGTTGAGAGGGCGT | >MI0000542\|hsa-mir-320a&WithFlank&8\|-\|22244960\|22245043 | 48 | 68 | 21(+1U) | AAAAGCTGGGTTGAGAGGGCGT | isomiR | TRUE | FALSE | FALSE |
| >hg38_21_+_8433172_8446622_RNA45SN1_45S_50ntFlanks@6646.6685.40\|TCGTACGACTCTTAGCGGTGGATCACTCGGCTCGTGCGTC | >hg38_21_+_8433172_8446622_RNA45SN1_45S_50ntFlanks | 6646 | 6685 | 40 | TCGTACGACTCTTAGCGGTGGATCACTCGGCTCGTGCGTC | rRF | TRUE | FALSE | FALSE |
| >MI0000809\|hsa-mir-151a&WithFlank&8\|-\|140732558\|140732659@53.74.22(+1A)&[MIMAT0000757&hsa-miR-151a-3p&offsets\|0\|+1(+1A);m-30&8\|-\|140732587\|140732607&offsets\|0\|+1(+1A)]\|CTAGACTGAAGCTCCTTGAGGAA | >MI0000809\|hsa-mir-151a&WithFlank&8\|-\|140732558\|140732659 | 53 | 74 | 22(+1A) | CTAGACTGAAGCTCCTTGAGGAA | isomiR | TRUE | FALSE | FALSE |

<table>
<tr><td>>hg38_21_+_8433172_8446622_RNA45SN1_45S_50ntFlanks@7975.8007.33|CGCGACCTCAGATCAGACGTGGCGACCCGCTGA</td><td>>hg38_21_+_8433172_8446622_RNA45SN1_45S_50ntFlanks</td><td>7975</td><td>8007</td><td>33</td><td>CGCGACCTCAGATCAGACGTGGCGACCCGCTGA</td><td>rRF</td><td>TRUE</td><td>FALSE</td><td>FALSE</td></tr>
<tr><td>>MI0000434|hsa-let-7i&WithFlank&12|+|62603680|62603775@12.33.22(+1A)&[MIMAT0000415&hsa-let-7i-5p&offsets|0|0(+1A);m-49&12|+|62603691|62603712&offsets|0|0(+1A)]|TGAGGTAGTAGTTTGTGCTGTTA</td><td>>MI0000434|hsa-let-7i&WithFlank&12|+|62603680|62603775</td><td>12</td><td>33</td><td>22(+1A)</td><td>TGAGGTAGTAGTTTGTGCTGTTA</td><td>isomiR</td><td>TRUE</td><td>TRUE</td><td>FALSE</td></tr>
<tr><td>>MI0000471|hsa-mir-126&WithFlank&9|+|136670596|136670692@58.79.22&[MIMAT0000445&hsa-miR-126-3p&offsets|0|0;m-40&9|+|136670653|136670674&offsets|0|0]|TCGTACCGTGAGTAATAATGCG</td><td>>MI0000471|hsa-mir-126&WithFlank&9|+|136670596|136670692</td><td>58</td><td>79</td><td>22</td><td>TCGTACCGTGAGTAATAATGCG</td><td>isomiR</td><td>TRUE</td><td>FALSE</td><td>FALSE</td></tr>
<tr><td>>hg38_21_+_8433172_8446622_RNA45SN1_45S_50ntFlanks@12998.13039.42|GCTGCGATCTATTGAAAGTCAGCCCTCGACACAAGGGTTTGT</td><td>>hg38_21_+_8433172_8446622_RNA45SN1_45S_50ntFlanks</td><td>12998</td><td>13039</td><td>42</td><td>GCTGCGATCTATTGAAAGTCAGCCCTCGACACAAGGGTTTGT</td><td>rRF</td><td>TRUE</td><td>TRUE</td><td>FALSE</td></tr>
<tr><td>>MI0005756|hsa-mir-934&WithFlank&X|+|136550872|136550966@21.41.21&[MIMAT0004977&hsa-miR-934&offsets|0|-1;m-943&X|+|136550892|136550913&offsets|0|-1]|TGTCTACTACTGGAGACACTG</td><td>>MI0005756|hsa-mir-934&WithFlank&X|+|136550872|136550966</td><td>21</td><td>41</td><td>21</td><td>TGTCTACTACTGGAGACACTG</td><td>isomiR</td><td>TRUE</td><td>FALSE</td><td>FALSE</td></tr>
<tr><td>>MI0000272|hsa-mir-182&WithFlank&7|-|129770377|129770498@29.50.22&[MIMAT0000259&hsa-miR-182-5p&offsets|0|-2;m-22&7|-|129770449|129770470&offsets|0|0]|TTTGGCAATGGTAGAACTCACA</td><td>>MI0000272|hsa-mir-182&WithFlank&7|-|129770377|129770498</td><td>29</td><td>50</td><td>22</td><td>TTTGGCAATGGTAGAACTCACA</td><td>isomiR</td><td>TRUE</td><td>FALSE</td><td>FALSE</td></tr>
<tr><td>>MI0000063|hsa-let-7b&WithFlank&22|+|46113680|46113774@12.32.21&[MIMAT0000063&hsa-let-7b-5p&offsets|0|-1;m-14&22|+|46113691|46113712&offsets|0|-1]|TGAGGTAGTAGGTTGTGTGGT</td><td>>MI0000063|hsa-let-7b&WithFlank&22|+|46113680|46113774</td><td>12</td><td>32</td><td>21</td><td>TGAGGTAGTAGGTTGTGTGGT</td><td>isomiR</td><td>TRUE</td><td>FALSE</td><td>FALSE</td></tr>
<tr><td>>hg38_21_+_8433172_8446622_RNA45SN1_45S_50ntFlanks@7975.7995.21|CGCGACCTCAGATCAGACGTG</td><td>>hg38_21_+_8433172_8446622_RNA45SN1_45S_50ntFlanks</td><td>7975</td><td>7995</td><td>21</td><td>CGCGACCTCAGATCAGACGTG</td><td>rRF</td><td>TRUE</td><td>FALSE</td><td>TRUE</td></tr>
<tr><td>>MI0000434|hsa-let-7i&WithFlank&12|+|62603680|62603775@12.32.21(+1G)&[MIMAT0000415&hsa-let-7i-5p&offsets|0|-1(+1G);m-49&12|+|62603691|62603712&offsets|0|-1(+1G)]|TGAGGTAGTAGTTTGTGCTGTG</td><td>>MI0000434|hsa-let-7i&WithFlank&12|+|62603680|62603775</td><td>12</td><td>32</td><td>21(+1G)</td><td>TGAGGTAGTAGTTTGTGCTGTG</td><td>isomiR</td><td>TRUE</td><td>TRUE</td><td>FALSE</td></tr>
<tr><td>>hg38_21_+_8433172_8446622_RNA45SN1_45S_50ntFlanks@6647.6665.19|CGTACGACTCTTAGCGGTG</td><td>>hg38_21_+_8433172_8446622_RNA45SN1_45S_50ntFlanks</td><td>6647</td><td>6665</td><td>19</td><td>CGTACGACTCTTAGCGGTG</td><td>rRF</td><td>TRUE</td><td>FALSE</td><td>FALSE</td></tr>
<tr><td>>MI0000734|hsa-mir-106b&WithFlank&7|-|100093987|100094080@18.39.22&[MIMAT0000680&hsa-miR-106b-5p&offsets|0|+1;m-103&7|-|100094043|100094063&offsets|0|+1]|TAAAGTGCTGACAGTGCAGATA</td><td>>MI0000734|hsa-mir-106b&WithFlank&7|-|100093987|100094080</td><td>18</td><td>39</td><td>22</td><td>TAAAGTGCTGACAGTGCAGATA</td><td>isomiR</td><td>TRUE</td><td>FALSE</td><td>FALSE</td></tr>
<tr><td>>MI0003560|hsa-mir-92b&WithFlank&1|+|155195171|155195278@67.88.22&[MIMAT0003218&hsa-miR-92b-3p&offsets|0|0;m-115&1|+|155195237|155195258&offsets|0|0]|TATTGCACTCGTCCCGGCCTCC</td><td>>MI0003560|hsa-mir-92b&WithFlank&1|+|155195171|155195278</td><td>67</td><td>88</td><td>22</td><td>TATTGCACTCGTCCCGGCCTCC</td><td>isomiR</td><td>TRUE</td><td>FALSE</td><td>FALSE</td></tr>
<tr><td>>hg38_21_+_8433172_8446622_RNA45SN1_45S_50ntFlanks@6651.6669.19|CGACTCTTAGCGGTGGATC</td><td>>hg38_21_+_8433172_8446622_RNA45SN1_45S_50ntFlanks</td><td>6651</td><td>6669</td><td>19</td><td>CGACTCTTAGCGGTGGATC</td><td>rRF</td><td>TRUE</td><td>FALSE</td><td>TRUE</td></tr>
<tr><td>>MI0000093|hsa-mir-92a-1&WithFlank&13|+|91351308|91351397@54.74.21(+1C);MI0000094|hsa-mir-92a-2&WithFlank&X|-|134169532|134169618@54.74.21(+1C)&[MIMAT0000092&hsa-miR-92a-3p&offsets|0|-1(+1C);m-9&13|+|91351361|91351382&offsets|0|-1(+1C)];[MIMAT0000092_1&hsa-miR-92a-3p&offsets|0|-1(+1C);m-11&X|-|134169544|134169565&offsets|0|-1(+1C)]|TATTGCACTTGTCCCGGCCTGC</td><td>>MI0000093|hsa-mir-92a-1&WithFlank&13|+|91351308|91351397</td><td>54</td><td>74</td><td>21(+1C)</td><td>TATTGCACTTGTCCCGGCCTGC</td><td>isomiR</td><td>TRUE</td><td>TRUE</td><td>FALSE</td></tr>
<tr><td>>MI0000736|hsa-mir-30c-1&WithFlank&1|+|40757278|40757378@23.44.22(+1U);MI0000254|hsa-mir-30c-2&WithFlank&6|-|71376954|71377037@13.34.22(+1U)&[MIMAT0000244&hsa-miR-30c-5p&offsets|0|-1(+1U);m-57&1|+|40757300|40757323&offsets|0|-2(+1U)];[MIMAT0000244_1&hsa-miR-30c-5p&offsets|0|-1(+1U);m-56&6|-|71377002|71377025&offsets|0|-2(+1U)]|TGTAAACATCCTACACTCTCAGT</td><td>>MI0000736|hsa-mir-30c-1&WithFlank&1|+|40757278|40757378</td><td>23</td><td>44</td><td>22(+1U)</td><td>TGTAAACATCCTACACTCTCAGT</td><td>isomiR</td><td>TRUE</td><td>TRUE</td><td>FALSE</td></tr>
<tr><td>>MI0003137|hsa-mir-193b&WithFlank&16|+|14303961|14304055@57.78.22&[MIMAT0002819&hsa-miR-193b-3p&offsets|0|0;m-166&16|+|14304017|14304038&offsets|0|0]|AACTGGCCCTCAAAGTCCCGCT</td><td>>MI0003137|hsa-mir-193b&WithFlank&16|+|14303961|14304055</td><td>57</td><td>78</td><td>22</td><td>AACTGGCCCTCAAAGTCCCGCT</td><td>isomiR</td><td>TRUE</td><td>FALSE</td><td>FALSE</td></tr>
<tr><td>>MI0000650|hsa-mir-200c&WithFlank&12|+|6963693|6963772@50.70.21(+1A)&[MIMAT0000617&hsa-miR-200c-3p&offsets|0|-2(+1A);m-54&12|+|6963742|6963764&offsets|0|-2(+1A)]|TAATACTGCCGGGTAATGATGA</td><td>>MI0000650|hsa-mir-200c&WithFlank&12|+|6963693|6963772</td><td>50</td><td>70</td><td>21(+1A)</td><td>TAATACTGCCGGGTAATGATGA</td><td>isomiR</td><td>TRUE</td><td>FALSE</td><td>FALSE</td></tr>
<tr><td>>multi-am_trnaMT_ProTGG_MT_-_15956_16023@53.71.19|AAGACTTTTTCTCTGACCA</td><td>>multi-am_trnaMT_ProTGG_MT_-_15956_16023</td><td>53</td><td>71</td><td>19</td><td>AAGACTTTTTCTCTGACCA</td><td>tRF</td><td>TRUE</td><td>TRUE</td><td>FALSE</td></tr>
<tr><td>>MI0000809|hsa-mir-151a&WithFlank&8|-|140732558|140732659@51.73.23&[MIMAT0000757&hsa-miR-151a-3p&offsets|-2|0;m-30&8|-|140732587|140732607&offsets|-2|0]|TACTAGACTGAAGCTCCTTGAGG</td><td>>MI0000809|hsa-mir-151a&WithFlank&8|-|140732558|140732659</td><td>51</td><td>73</td><td>23</td><td>TACTAGACTGAAGCTCCTTGAGG</td><td>isomiR</td><td>TRUE</td><td>TRUE</td><td>FALSE</td></tr>
<tr><td>>trnaMT_SerGCT_MT_+_12207_12265@1.22.22|GAGAAAGCTCACAAGAACTGCT</td><td>>trnaMT_SerGCT_MT_+_12207_12265</td><td>1</td><td>22</td><td>22</td><td>GAGAAAGCTCACAAGAACTGCT</td><td>tRF</td><td>TRUE</td><td>TRUE</td><td>FALSE</td></tr>
<tr><td>>MI0000433|hsa-let-7g&WithFlank&3|-|52268272|52268367@11.31.21(+1A)&[MIMAT0000414&hsa-let-7g-5p&offsets|0|-1(+1A);m-58&3|-|52268336|52268357&offsets|0|-1(+1A)]|TGAGGTAGTAGTTTGTACAGTA</td><td>>MI0000433|hsa-let-7g&WithFlank&3|-|52268272|52268367</td><td>11</td><td>31</td><td>21(+1A)</td><td>TGAGGTAGTAGTTTGTACAGTA</td><td>isomiR</td><td>TRUE</td><td>FALSE</td><td>FALSE</td></tr>
<tr><td>>MI0000273|hsa-mir-183&WithFlank&7|-|129774899|129775020@34.54.21&[MIMAT0000261&hsa-miR-183-5p&offsets|+1|0;m-41&7|-|129774966|129774987&offsets|0|-1]|ATGGCACTGGTAGAATTCACT</td><td>>MI0000273|hsa-mir-183&WithFlank&7|-|129774899|129775020</td><td>34</td><td>54</td><td>21</td><td>ATGGCACTGGTAGAATTCACT</td><td>isomiR</td><td>TRUE</td><td>FALSE</td><td>FALSE</td></tr>
<tr><td>>MI0000273|hsa-mir-183&WithFlank&7|-|129774899|129775020@33.55.23&[MIMAT0000261&hsa-miR-183-5p&offsets|0|+1;m-41&7|-|129774966|129774987&offsets|-1|0]|TATGGCACTGGTAGAATTCACTG</td><td>>MI0000273|hsa-mir-183&WithFlank&7|-|129774899|129775020</td><td>33</td><td>55</td><td>23</td><td>TATGGCACTGGTAGAATTCACTG</td><td>isomiR</td><td>TRUE</td><td>FALSE</td><td>FALSE</td></tr>
<tr><td>>MI0000102|hsa-mir-100&WithFlank&11|-|122152223|122152314@19.39.21&[MIMAT0000098&hsa-miR-100-5p&offsets|0|-1;m-27&11|-|122152276|122152296&offsets|0|0]|AACCCGTAGATCCGAACTTGT</td><td>>MI0000102|hsa-mir-100&WithFlank&11|-|122152223|122152314</td><td>19</td><td>39</td><td>21</td><td>AACCCGTAGATCCGAACTTGT</td><td>isomiR</td><td>FALSE</td><td>TRUE</td><td>FALSE</td></tr>
<tr><td>>MI0000102|hsa-mir-100&WithFlank&11|-|122152223|122152314@19.39.21(+1A)&[MIMAT0000098&hsa-miR-100-5p&offsets|0|-1(+1A);m-27&11|-|122152276|122152296&offsets|0|0(+1A)]|AACCCGTAGATCCGAACTTGTA</td><td>>MI0000102|hsa-mir-100&WithFlank&11|-|122152223|122152314</td><td>19</td><td>39</td><td>21(+1A)</td><td>AACCCGTAGATCCGAACTTGTA</td><td>isomiR</td><td>FALSE</td><td>TRUE</td><td>FALSE</td></tr>
<tr><td>>MI0000102|hsa-mir-100&WithFlank&11|-|122152223|122152314@19.40.22(+1U)&[MIMAT0000098&hsa-miR-100-5p&offsets|0|0(+1U);m-27&11|-|122152276|122152296&offsets|0|+1(+1U)]|AACCCGTAGATCCGAACTTGTGT</td><td>>MI0000102|hsa-mir-100&WithFlank&11|-|122152223|122152314</td><td>19</td><td>40</td><td>22(+1U)</td><td>AACCCGTAGATCCGAACTTGTGT</td><td>isomiR</td><td>FALSE</td><td>TRUE</td><td>FALSE</td></tr>
<tr><td>>MI0000102|hsa-mir-100&WithFlank&11|-|122152223|122152314@19.40.22(+1A)&[MIMAT0000098&hsa-miR-100-5p&offsets|0|0(+1A);m-27&11|-|122152276|122152296&offsets|0|+1(+1A)]|AACCCGTAGATCCGAACTTGTGA</td><td>>MI0000102|hsa-mir-100&WithFlank&11|-|122152223|122152314</td><td>19</td><td>40</td><td>22(+1A)</td><td>AACCCGTAGATCCGAACTTGTGA</td><td>isomiR</td><td>FALSE</td><td>TRUE</td><td>FALSE</td></tr>
<tr><td>>hg38_21_+_8433172_8446622_RNA45SN1_45S_50ntFlanks@12994.13039.46|CCTCGCTGCGATCTATTGAAAGTCAGCCCTCGACACAAGGGTTTGT</td><td>>hg38_21_+_8433172_8446622_RNA45SN1_45S_50ntFlanks</td><td>12994</td><td>13039</td><td>46</td><td>CCTCGCTGCGATCTATTGAAAGTCAGCCCTCGACACAAGGGTTTGT</td><td>rRF</td><td>FALSE</td><td>TRUE</td><td>FALSE</td></tr>
</table>

<table>
<tr><td>>trna111_HisGTG_1_-_147774845_147774916@-1G.33.34,trna118_HisGTG_1_-_145396881_145396952@-1G.33.34,trna16_HisGTG_1_+_146544773_146544844@-1G.33.34,trna1_HisGTG_15_+_45493349_45493420@-1G.33.34,trna21_HisGTG_1_+_147753471_147753542@-1G.33.34,trna33_HisGTG_6_+_27125906_27125977@-1G.33.34,trna7_HisGTG_9_-_14433938_14434009@-1G.33.34,trna8_HisGTG_15_-_45492611_45492682@-1G.33.34,trna9_HisGTG_15_-_45490804_45490875@-1G.33.34|GGCCGTGATCGTATAGTGGTTAGTACTCTGCGTT</td><td>>trna111_HisGTG_1_-_147774845_147774916</td><td>-1G</td><td>33</td><td>34</td><td>GGCCGTGATCGTATAGTGGTTAGTACTCTGCGTT</td><td>tRF</td><td>FALSE</td><td>TRUE</td><td>FALSE</td></tr>
<tr><td>>MI0000102|hsa-mir-100&WithFlank&11|-|122152223|122152314@19.38.20(+1C)&[MIMAT0000098&hsa-miR-100-5p&offsets|0|-2(+1C);m-27&11|-|122152276|122152296&offsets|0|-1(+1C)]|AACCCGTAGATCCGAACTTGC</td><td>>MI0000102|hsa-mir-100&WithFlank&11|-|122152223|122152314</td><td>19</td><td>38</td><td>20(+1C)</td><td>AACCCGTAGATCCGAACTTGC</td><td>isomiR</td><td>FALSE</td><td>TRUE</td><td>FALSE</td></tr>
<tr><td>>MI0000102|hsa-mir-100&WithFlank&11|-|122152223|122152314@19.38.20&[MIMAT0000098&hsa-miR-100-5p&offsets|0|-2;m-27&11|-|122152276|122152296&offsets|0|-1]|AACCCGTAGATCCGAACTTG</td><td>>MI0000102|hsa-mir-100&WithFlank&11|-|122152223|122152314</td><td>19</td><td>38</td><td>20</td><td>AACCCGTAGATCCGAACTTG</td><td>isomiR</td><td>FALSE</td><td>TRUE</td><td>FALSE</td></tr>
<tr><td>>MI0000298|hsa-mir-221&WithFlank&X|-|45746151|45746272@71.93.23(+1U)&[MIMAT0000278&hsa-miR-221-3p&offsets|0|0(+1U);m-91&X|-|45746181|45746202&offsets|0|+1(+1U)]|AGCTACATTGTCTGCTGGGTTTCT</td><td>>MI0000298|hsa-mir-221&WithFlank&X|-|45746151|45746272</td><td>71</td><td>93</td><td>23(+1U)</td><td>AGCTACATTGTCTGCTGGGTTTCT</td><td>isomiR</td><td>FALSE</td><td>TRUE</td><td>FALSE</td></tr>
<tr><td>>trna111_HisGTG_1_-_147774845_147774916@-1G.34.35,trna118_HisGTG_1_-_145396881_145396952@-1G.34.35,trna16_HisGTG_1_+_146544773_146544844@-1G.34.35,trna1_HisGTG_15_+_45493349_45493420@-1G.34.35,trna21_HisGTG_1_+_147753471_147753542@-1G.34.35,trna33_HisGTG_6_+_27125906_27125977@-1G.34.35,trna7_HisGTG_9_-_14433938_14434009@-1G.34.35,trna8_HisGTG_15_-_45492611_45492682@-1G.34.35,trna9_HisGTG_15_-_45490804_45490875@-1G.34.35|GGCCGTGATCGTATAGTGGTTAGTACTCTGCGTTG</td><td>>trna111_HisGTG_1_-_147774845_147774916</td><td>-1G</td><td>34</td><td>35</td><td>GGCCGTGATCGTATAGTGGTTAGTACTCTGCGTTG</td><td>tRF</td><td>FALSE</td><td>TRUE</td><td>FALSE</td></tr>
<tr><td>>MI0000477|hsa-mir-146a&WithFlank&5|+|160485346|160485456@27.48.22&[MIMAT0000449&hsa-miR-146a-5p&offsets|0|0;m-129&5|+|160485372|160485393&offsets|0|0]|TGAGAACTGAATTCCATGGGTT</td><td>>MI0000477|hsa-mir-146a&WithFlank&5|+|160485346|160485456</td><td>27</td><td>48</td><td>22</td><td>TGAGAACTGAATTCCATGGGTT</td><td>isomiR</td><td>FALSE</td><td>TRUE</td><td>FALSE</td></tr>
<tr><td>>MI0000446|hsa-mir-125b-1&WithFlank&11|-|122099751|122099850@21.42.22;MI0000470|hsa-mir-125b-2&WithFlank&21|+|16590231|16590331@23.44.22&[MIMAT0000423&hsa-miR-125b-5p&offsets|0|0;m-67&11|-|122099809|122099830&offsets|0|0];[MIMAT0000423_1&hsa-miR-125b-5p&offsets|0|0;m-68&21|+|16590253|16590274&offsets|0|0]|TCCCTGAGACCCTAACTTGTGA</td><td>>MI0000446|hsa-mir-125b-1&WithFlank&11|-|122099751|122099850</td><td>21</td><td>42</td><td>22</td><td>TCCCTGAGACCCTAACTTGTGA</td><td>isomiR</td><td>FALSE</td><td>TRUE</td><td>FALSE</td></tr>
<tr><td>>hg38_21_+_8433172_8446622_RNA45SN1_45S_50ntFlanks@6651.6696.46|CGACTCTTAGCGGTGGATCACTCGGCTCGTGCGTCGATGAAGAACG</td><td>>hg38_21_+_8433172_8446622_RNA45SN1_45S_50ntFlanks</td><td>6651</td><td>6696</td><td>46</td><td>CGACTCTTAGCGGTGGATCACTCGGCTCGTGCGTCGATGAAGAACG</td><td>rRF</td><td>FALSE</td><td>TRUE</td><td>FALSE</td></tr>
<tr><td>>trna119_LysCTT_1_-_145395522_145395594@1.33.33,trna11_LysCTT_5_-_180648979_180649051@1.33.33,trna13_LysCTT_6_+_26556774_26556846@1.33.33,trna7_LysCTT_16_+_3225692_3225764@1.33.33,trna9_LysCTT_5_+_180634755_180634827@1.33.33|GCCCGGCTAGCTCAGTCGGTAGAGCATGAGACT</td><td>>trna119_LysCTT_1_-_145395522_145395594</td><td>1</td><td>33</td><td>33</td><td>GCCCGGCTAGCTCAGTCGGTAGAGCATGAGACT</td><td>tRF</td><td>FALSE</td><td>TRUE</td><td>FALSE</td></tr>
<tr><td>>hg38_21_+_8433172_8446622_RNA45SN1_45S_50ntFlanks@7975.8016.42|CGCGACCTCAGATCAGACGTGGCGACCCGCTGAATTTAAGCA</td><td>>hg38_21_+_8433172_8446622_RNA45SN1_45S_50ntFlanks</td><td>7975</td><td>8016</td><td>42</td><td>CGCGACCTCAGATCAGACGTGGCGACCCGCTGAATTTAAGCA</td><td>rRF</td><td>FALSE</td><td>TRUE</td><td>FALSE</td></tr>
<tr><td>>MI0000299|hsa-mir-222&WithFlank&X|-|45747009|45747130@75.98.24(+1C)&[MIMAT0000279&hsa-miR-222-3p&offsets|0|+3(+1C);m-140&X|-|45747033|45747056&offsets|0|0(+1C)]|AGCTACATCTGGCTACTGGGTCTCC</td><td>>MI0000299|hsa-mir-222&WithFlank&X|-|45747009|45747130</td><td>75</td><td>98</td><td>24(+1C)</td><td>AGCTACATCTGGCTACTGGGTCTCC</td><td>isomiR</td><td>FALSE</td><td>TRUE</td><td>FALSE</td></tr>
<tr><td>>MI0000434|hsa-let-7i&WithFlank&12|+|62603680|62603775@12.33.22(+1U)&[MIMAT0000415&hsa-let-7i-5p&offsets|0|0(+1U);m-49&12|+|62603691|62603712&offsets|0|0(+1U)]|TGAGGTAGTAGTTTGTGCTGTTT</td><td>>MI0000434|hsa-let-7i&WithFlank&12|+|62603680|62603775</td><td>12</td><td>33</td><td>22(+1U)</td><td>TGAGGTAGTAGTTTGTGCTGTTT</td><td>isomiR</td><td>FALSE</td><td>TRUE</td><td>FALSE</td></tr>
<tr><td>>hg38_21_+_8433172_8446622_RNA45SN1_45S_50ntFlanks@12999.13039.41|CTGCGATCTATTGAAAGTCAGCCCTCGACACAAGGGTTTGT</td><td>>hg38_21_+_8433172_8446622_RNA45SN1_45S_50ntFlanks</td><td>12999</td><td>13039</td><td>41</td><td>CTGCGATCTATTGAAAGTCAGCCCTCGACACAAGGGTTTGT</td><td>rRF</td><td>FALSE</td><td>TRUE</td><td>FALSE</td></tr>
<tr><td>>MI0000088|hsa-mir-30a&WithFlank&6|-|71403545|71403627@53.74.22&[MIMAT0000088&hsa-miR-30a-3p&offsets|0|0;m-38&6|-|71403554|71403575&offsets|0|0]|CTTTCAGTCGGATGTTTGCAGC</td><td>>MI0000088|hsa-mir-30a&WithFlank&6|-|71403545|71403627</td><td>53</td><td>74</td><td>22</td><td>CTTTCAGTCGGATGTTTGCAGC</td><td>isomiR</td><td>FALSE</td><td>TRUE</td><td>FALSE</td></tr>
<tr><td>>MI0000088|hsa-mir-30a&WithFlank&6|-|71403545|71403627@12.35.24(+1U)&[MIMAT0000087&hsa-miR-30a-5p&offsets|0|+2(+1U);m-15&6|-|71403595|71403616&offsets|0|+2(+1U)]|TGTAAACATCCTCGACTGGAAGCTT</td><td>>MI0000088|hsa-mir-30a&WithFlank&6|-|71403545|71403627</td><td>12</td><td>35</td><td>24(+1U)</td><td>TGTAAACATCCTCGACTGGAAGCTT</td><td>isomiR</td><td>FALSE</td><td>TRUE</td><td>FALSE</td></tr>
<tr><td>>MI0000434|hsa-let-7i&WithFlank&12|+|62603680|62603775@12.32.21&[MIMAT0000415&hsa-let-7i-5p&offsets|0|-1;m-49&12|+|62603691|62603712&offsets|0|-1]|TGAGGTAGTAGTTTGTGCTGT</td><td>>MI0000434|hsa-let-7i&WithFlank&12|+|62603680|62603775</td><td>12</td><td>32</td><td>21</td><td>TGAGGTAGTAGTTTGTGCTGT</td><td>isomiR</td><td>FALSE</td><td>TRUE</td><td>FALSE</td></tr>
<tr><td>>MI0000102|hsa-mir-100&WithFlank&11|-|122152223|122152314@19.38.20(+1G)&[MIMAT0000098&hsa-miR-100-5p&offsets|0|-2(+1G);m-27&11|-|122152276|122152296&offsets|0|-1(+1G)]|AACCCGTAGATCCGAACTTGG</td><td>>MI0000102|hsa-mir-100&WithFlank&11|-|122152223|122152314</td><td>19</td><td>38</td><td>20(+1G)</td><td>AACCCGTAGATCCGAACTTGG</td><td>isomiR</td><td>FALSE</td><td>TRUE</td><td>FALSE</td></tr>
<tr><td>>hg38_21_+_8433172_8446622_RNA45SN1_45S_50ntFlanks@6651.6681.31|CGACTCTTAGCGGTGGATCACTCGGCTCGTG</td><td>>hg38_21_+_8433172_8446622_RNA45SN1_45S_50ntFlanks</td><td>6651</td><td>6681</td><td>31</td><td>CGACTCTTAGCGGTGGATCACTCGGCTCGTG</td><td>rRF</td><td>FALSE</td><td>TRUE</td><td>TRUE</td></tr>
<tr><td>>MI0000102|hsa-mir-100&WithFlank&11|-|122152223|122152314@19.39.21(+1U)&[MIMAT0000098&hsa-miR-100-5p&offsets|0|-1(+1U);m-27&11|-|122152276|122152296&offsets|0|0(+1U)]|AACCCGTAGATCCGAACTTGTT</td><td>>MI0000102|hsa-mir-100&WithFlank&11|-|122152223|122152314</td><td>19</td><td>39</td><td>21(+1U)</td><td>AACCCGTAGATCCGAACTTGTT</td><td>isomiR</td><td>FALSE</td><td>TRUE</td><td>FALSE</td></tr>
<tr><td>>MI0000446|hsa-mir-125b-1&WithFlank&11|-|122099751|122099850@21.41.21;MI0000470|hsa-mir-125b-2&WithFlank&21|+|16590231|16590331@23.43.21&[MIMAT0000423&hsa-miR-125b-5p&offsets|0|-1;m-67&11|-|122099809|122099830&offsets|0|-1];[MIMAT0000423_1&hsa-miR-125b-5p&offsets|0|-1;m-68&21|+|16590253|16590274&offsets|0|-1]|TCCCTGAGACCCTAACTTGTG</td><td>>MI0000446|hsa-mir-125b-1&WithFlank&11|-|122099751|122099850</td><td>21</td><td>41</td><td>21</td><td>TCCCTGAGACCCTAACTTGTG</td><td>isomiR</td><td>FALSE</td><td>TRUE</td><td>FALSE</td></tr>
<tr><td>>am_trna128_GlyGCC_6_-_27870686_27870756@1.35.35,trna18_GlyGCC_16_+_70822597_70822667@1.35.35,trna19_GlyGCC_16_+_70823410_70823480@1.35.35,trna19_GlyGCC_2_-_157257659_157257729@1.35.35,trna24_GlyGCC_16_-_70812942_70813012@1.35.35,trna25_GlyGCC_16_-_70812114_70812184@1.35.35,trna5_GlyGCC_17_+_8029064_8029134@1.35.35,trna68_GlyGCC_1_-_161493637_161493707@1.35.35|GCATTGGTGGTTCAGTGGTAGAATTCTCGCCTGCC</td><td>>am_trna128_GlyGCC_6_-_27870686_27870756</td><td>1</td><td>35</td><td>35</td><td>GCATTGGTGGTTCAGTGGTAGAATTCTCGCCTGCC</td><td>tRF</td><td>FALSE</td><td>TRUE</td><td>FALSE</td></tr>
<tr><td>>hg38_1_-_228634819_228635039_RNA5S12_5S_50ntFlanks@144.171.28|CCTGGGAATACCGGGTGCTGTAGGCTTT</td><td>>hg38_1_-_228634819_228635039_RNA5S12_5S_50ntFlanks</td><td>144</td><td>171</td><td>28</td><td>CCTGGGAATACCGGGTGCTGTAGGCTTT</td><td>rRF</td><td>FALSE</td><td>TRUE</td><td>FALSE</td></tr>
</table>

| | | | | | | | | | |
|---|---|---|---|---|---|---|---|---|---|
| >hg38_21_+_8433172_8446622_RNA45SN1_45S_50ntFlanks@7975.8013.39\|CGCGACCTCAGATCAGACGTGGCGACCCGCTGAATTTAA | >hg38_21_+_8433172_8446622_RNA45SN1_45S_50ntFlanks | 7975 | 8013 | 39 | CGCGACCTCAGATCAGACGTGGCGACCCGCTGAATTTAA | rRF | FALSE | TRUE | FALSE |
| >MI0000102\|hsa-mir-100&WithFlank&11\|-\|122152223\|122152314@19.40.22(+1C)&[MIMAT0000098&hsa-miR-100-5p&offsets\|0\|0(+1C);m-27&11\|-\|122152276\|122152296&offsets\|0\|+1(+1C)]\|AACCCGTAGATCCGAACTTGTGC | >MI0000102\|hsa-mir-100&WithFlank&11\|-\|122152223\|122152314 | 19 | 40 | 22(+1C) | AACCCGTAGATCCGAACTTGTGC | isomiR | FALSE | TRUE | FALSE |
| >hg38_21_+_8433172_8446622_RNA45SN1_45S_50ntFlanks@475.496.22\|CTTCGTGATCGATGTGGTGACG | >hg38_21_+_8433172_8446622_RNA45SN1_45S_50ntFlanks | 475 | 496 | 22 | CTTCGTGATCGATGTGGTGACG | rRF | FALSE | TRUE | FALSE |
| >MI0000088\|hsa-mir-30a&WithFlank&6\|-\|71403545\|71403627@53.73.21(+1U)&[MIMAT0000088&hsa-miR-30a-3p&offsets\|0\|-1(+1U);m-38&6\|-\|71403554\|71403575&offsets\|0\|-1(+1U)]\|CTTTCAGTCGGATGTTTGCAGT | >MI0000088\|hsa-mir-30a&WithFlank&6\|-\|71403545\|71403627 | 53 | 73 | 21(+1U) | CTTTCAGTCGGATGTTTGCAGT | isomiR | FALSE | TRUE | FALSE |
| >MI0000299\|hsa-mir-222&WithFlank&X\|-\|45747009\|45747130@75.97.23&[MIMAT0000279&hsa-miR-222-3p&offsets\|0\|+2;m-140&X\|-\|45747033\|45747056&offsets\|0\|-1]\|AGCTACATCTGGCTACTGGGTCT | >MI0000299\|hsa-mir-222&WithFlank&X\|-\|45747009\|45747130 | 75 | 97 | 23 | AGCTACATCTGGCTACTGGGTCT | isomiR | FALSE | TRUE | FALSE |
| >MI0000093\|hsa-mir-92a-1&WithFlank&13\|+\|91351308\|91351397@54.76.23&[MIMAT0000092&hsa-miR-92a-3p&offsets\|0\|+1;m-9&13\|+\|91351361\|91351382&offsets\|0\|+1]\|TATTGCACTTGTCCCGGCCTGTT | >MI0000093\|hsa-mir-92a-1&WithFlank&13\|+\|91351308\|91351397 | 54 | 76 | 23 | TATTGCACTTGTCCCGGCCTGTT | isomiR | FALSE | TRUE | FALSE |
| >MI0000299\|hsa-mir-222&WithFlank&X\|-\|45747009\|45747130@75.99.25(+1U)&[MIMAT0000279&hsa-miR-222-3p&offsets\|0\|+4(+1U);m-140&X\|-\|45747033\|45747056&offsets\|0\|+1(+1U)]\|AGCTACATCTGGCTACTGGGTCTCTT | >MI0000299\|hsa-mir-222&WithFlank&X\|-\|45747009\|45747130 | 75 | 99 | 25(+1U) | AGCTACATCTGGCTACTGGGTCTCTT | isomiR | FALSE | TRUE | FALSE |
| >MI0000102\|hsa-mir-100&WithFlank&11\|-\|122152223\|122152314@19.41.23&[MIMAT0000098&hsa-miR-100-5p&offsets\|0\|+1;m-27&11\|-\|122152276\|122152296&offsets\|0\|+2]\|AACCCGTAGATCCGAACTTGTGG | >MI0000102\|hsa-mir-100&WithFlank&11\|-\|122152223\|122152314 | 19 | 41 | 23 | AACCCGTAGATCCGAACTTGTGG | isomiR | FALSE | TRUE | FALSE |
| >hg38_21_+_8433172_8446622_RNA45SN1_45S_50ntFlanks@6651.6699.49\|CGACTCTTAGCGGTGGATCACTCGGCTCGTGCGTCGATGAAGAACGCAG | >hg38_21_+_8433172_8446622_RNA45SN1_45S_50ntFlanks | 6651 | 6699 | 49 | CGACTCTTAGCGGTGGATCACTCGGCTCGTGCGTCGATGAAGAACGCAG | rRF | FALSE | TRUE | TRUE |
| >hg38_21_+_8433172_8446622_RNA45SN1_45S_50ntFlanks@485.532.48\|GATGTGGTGACGTCGTGCTCTCCCGGGCCGGGTCCGAGCCGCGACGGG | >hg38_21_+_8433172_8446622_RNA45SN1_45S_50ntFlanks | 485 | 532 | 48 | GATGTGGTGACGTCGTGCTCTCCCGGGCCGGGTCCGAGCCGCGACGGG | rRF | FALSE | TRUE | FALSE |
| >hg38_21_+_8433172_8446622_RNA45SN1_45S_50ntFlanks@7974.8009.36\|ACGCGACCTCAGATCAGACGTGGCGACCCGCTGAAT | >hg38_21_+_8433172_8446622_RNA45SN1_45S_50ntFlanks | 7974 | 8009 | 36 | ACGCGACCTCAGATCAGACGTGGCGACCCGCTGAAT | rRF | FALSE | TRUE | FALSE |
| >MI0000299\|hsa-mir-222&WithFlank&X\|-\|45747009\|45747130@75.98.24(+1A)&[MIMAT0000279&hsa-miR-222-3p&offsets\|0\|+3(+1A);m-140&X\|-\|45747033\|45747056&offsets\|0\|0(+1A)]\|AGCTACATCTGGCTACTGGGTCTCA | >MI0000299\|hsa-mir-222&WithFlank&X\|-\|45747009\|45747130 | 75 | 98 | 24(+1A) | AGCTACATCTGGCTACTGGGTCTCA | isomiR | FALSE | TRUE | FALSE |
| >hg38_21_+_8433172_8446622_RNA45SN1_45S_50ntFlanks@6651.6695.45\|CGACTCTTAGCGGTGGATCACTCGGCTCGTGCGTCGATGAAGAAC | >hg38_21_+_8433172_8446622_RNA45SN1_45S_50ntFlanks | 6651 | 6695 | 45 | CGACTCTTAGCGGTGGATCACTCGGCTCGTGCGTCGATGAAGAAC | rRF | FALSE | TRUE | FALSE |
| >hg38_21_+_8433172_8446622_RNA45SN1_45S_50ntFlanks@482.505.24\|ATCGATGTGGTGACGTCGTGCTCT | >hg38_21_+_8433172_8446622_RNA45SN1_45S_50ntFlanks | 482 | 505 | 24 | ATCGATGTGGTGACGTCGTGCTCT | rRF | FALSE | TRUE | FALSE |
| >MI0001448\|hsa-mir-425&WithFlank&3\|-\|49020142\|49020240@20.44.25&[MIMAT0003393&hsa-miR-425-5p&offsets\|0\|+2;m-95&3\|-\|49020200\|49020221&offsets\|0\|+3]\|AATGACACGATCACTCCCGTTGAGT | >MI0001448\|hsa-mir-425&WithFlank&3\|-\|49020142\|49020240 | 20 | 44 | 25 | AATGACACGATCACTCCCGTTGAGT | isomiR | FALSE | TRUE | FALSE |
| >MI0000085\|hsa-mir-27a&WithFlank&19\|-\|13836434\|13836523@57.76.20(+1U)&[MIMAT0000084&hsa-miR-27a-3p&offsets\|0\|-1(+1U);m-43&19\|-\|13836447\|13836467&offsets\|0\|-1(+1U)]\|TTCACAGTGGCTAAGTTCCGT | >MI0000085\|hsa-mir-27a&WithFlank&19\|-\|13836434\|13836523 | 57 | 76 | 20(+1U) | TTCACAGTGGCTAAGTTCCGT | isomiR | FALSE | TRUE | FALSE |
| >hg38_1_-_228634819_228635039_RNA5S12_5S_50ntFlanks@140.171.32\|ACCGCCTGGGAATACCGGGTGCTGTAGGCTTT | >hg38_1_-_228634819_228635039_RNA5S12_5S_50ntFlanks | 140 | 171 | 32 | ACCGCCTGGGAATACCGGGTGCTGTAGGCTTT | rRF | FALSE | TRUE | FALSE |
| >trna111_HisGTG_1_-_147774845_147774916@-1G.31.32,trna118_HisGTG_1_-_145396881_145396952@-1G.31.32,trna16_HisGTG_1_+_146544773_146544844@-1G.31.32,trna1_HisGTG_15_+_45493349_45493420@-1G.31.32,trna21_HisGTG_1_+_147753471_147753542@-1G.31.32,trna33_HisGTG_6_+_27125906_27125977@-1G.31.32,trna7_HisGTG_9_-_14433938_14434009@-1G.31.32,trna8_HisGTG_15_-_45492611_45492682@-1G.31.32,trna9_HisGTG_15_-_45490804_45490875@-1G.31.32\|GGCCGTGATCGTATAGTGGTTAGTACTCTGCG | >trna111_HisGTG_1_-_147774845_147774916 | -1G | 31 | 32 | GGCCGTGATCGTATAGTGGTTAGTACTCTGCG | tRF | FALSE | TRUE | TRUE |
| >hg38_21_+_8433172_8446622_RNA45SN1_45S_50ntFlanks@5472.5516.45\|ACGGCCCTGGCGGAGCGCTGAGAAGACGGTCGAACTTGACTATCT | >hg38_21_+_8433172_8446622_RNA45SN1_45S_50ntFlanks | 5472 | 5516 | 45 | ACGGCCCTGGCGGAGCGCTGAGAAGACGGTCGAACTTGACTATCT | rRF | FALSE | TRUE | FALSE |
| >am_trna128_GlyGCC_6_-_27870686_27870756@1.30.30,trna133_GlyCCC_1_-_16872434_16872504@1.30.30,trna18_GlyGCC_16_+_70822597_70822667@1.30.30,trna19_GlyGCC_16_+_70823410_70823480@1.30.30,trna19_GlyGCC_2_-_157257659_157257729@1.30.30,trna24_GlyGCC_16_-_70812942_70813012@1.30.30,trna25_GlyGCC_16_-_70812114_70812184@1.30.30,trna4_GlyCCC_1_+_17188416_17188486@1.30.30,trna5_GlyGCC_17_+_8029064_8029134@1.30.30,trna68_GlyGCC_1_-_161493637_161493707@1.30.30\|GCATTGGTGGTTCAGTGGTAGAATTCTCGC | >am_trna128_GlyGCC_6_-_27870686_27870756 | 1 | 30 | 30 | GCATTGGTGGTTCAGTGGTAGAATTCTCGC | tRF | FALSE | TRUE | TRUE |
| >MI0000750\|hsa-mir-26a-2&WithFlank&12\|-\|57824603\|57824698@20.41.22(+1U);MI0000083\|hsa-mir-26a-1&WithFlank&3\|+\|37969398\|37969486@16.37.22(+1U)&[MIMAT0000082&hsa-miR-26a-5p&offsets\|0\|0(+1U);m-33&12\|-\|57824658\|57824679&offsets\|0\|0(+1U)];[MIMAT0000082_1&hsa-miR-26a-5p&offsets\|0\|0(+1U);m-34&3\|+\|37969413\|37969434&offsets\|0\|0(+1U)]\|TTCAAGTAATCCAGGATAGGCTT | >MI0000750\|hsa-mir-26a-2&WithFlank&12\|-\|57824603\|57824698 | 20 | 41 | 22(+1U) | TTCAAGTAATCCAGGATAGGCTT | isomiR | FALSE | TRUE | FALSE |
| >hg38_21_+_8433172_8446622_RNA45SN1_45S_50ntFlanks@6651.6683.33\|CGACTCTTAGCGGTGGATCACTCGGCTCGTGCG | >hg38_21_+_8433172_8446622_RNA45SN1_45S_50ntFlanks | 6651 | 6683 | 33 | CGACTCTTAGCGGTGGATCACTCGGCTCGTGCG | rRF | FALSE | TRUE | FALSE |
| >MI0000750\|hsa-mir-26a-2&WithFlank&12\|-\|57824603\|57824698@20.40.21(+1G);MI0000083\|hsa-mir-26a-1&WithFlank&3\|+\|37969398\|37969486@16.36.21(+1G)&[MIMAT0000082&hsa-miR-26a-5p&offsets\|0\|-1(+1G);m-33&12\|-\|57824658\|57824679&offsets\|0\|-1(+1G)];[MIMAT0000082_1&hsa-miR-26a-5p&offsets\|0\|-1(+1G);m-34&3\|+\|37969413\|37969434&offsets\|0\|-1(+1G)]\|TTCAAGTAATCCAGGATAGGCG | >MI0000750\|hsa-mir-26a-2&WithFlank&12\|-\|57824603\|57824698 | 20 | 40 | 21(+1G) | TTCAAGTAATCCAGGATAGGCG | isomiR | FALSE | TRUE | FALSE |
| >hg38_21_+_8433172_8446622_RNA45SN1_45S_50ntFlanks@476.506.31\|TTCGTGATCGATGTGGTGACGTCGTGCTCTC | >hg38_21_+_8433172_8446622_RNA45SN1_45S_50ntFlanks | 476 | 506 | 31 | TTCGTGATCGATGTGGTGACGTCGTGCTCTC | rRF | FALSE | TRUE | FALSE |
| >MI0000298\|hsa-mir-221&WithFlank&X\|-\|45746151\|45746272@71.92.22(+1U)&[MIMAT0000278&hsa-miR-221-3p&offsets\|0\|-1(+1U);m-91&X\|-\|45746181\|45746202&offsets\|0\|0(+1U)]\|AGCTACATTGTCTGCTGGGTTTT | >MI0000298\|hsa-mir-221&WithFlank&X\|-\|45746151\|45746272 | 71 | 92 | 22(+1U) | AGCTACATTGTCTGCTGGGTTTT | isomiR | FALSE | TRUE | FALSE |
| >hg38_1_-_228634819_228635039_RNA5S12_5S_50ntFlanks@132.171.40\|GATGGGAGACCGCCTGGGAATACCGGGTGCTGTAGGCTTT | >hg38_1_-_228634819_228635039_RNA5S12_5S_50ntFlanks | 132 | 171 | 40 | GATGGGAGACCGCCTGGGAATACCGGGTGCTGTAGGCTTT | rRF | FALSE | TRUE | FALSE |

| | | | | | | | | | |
|---|---|---|---|---|---|---|---|---|---|
| >MI0000750\|hsa-mir-26a-2&WithFlank&12\|-\|57824603\|57824698@20.41.22(+2U);MI0000083\|hsa-mir-26a-1&WithFlank&3\|+\|37969398\|37969486@16.37.22(+2U)&[MIMAT0000082&hsa-miR-26a-5p&offsets\|0\|0(+2U);m-33&12\|-\|57824658\|57824679&offsets\|0\|0(+2U)];[MIMAT0000082_1&hsa-miR-26a-5p&offsets\|0\|0(+2U);m-34&3\|+\|37969413\|37969434&offsets\|0\|0(+2U)]\|TTCAAGTAATCCAGGATAGGCTTT | >MI0000750\|hsa-mir-26a-2&WithFlank&12\|-\|57824603\|57824698 | 20 | 41 | 22(+2U) | TTCAAGTAATCCAGGATAGGCTTT | isomiR | FALSE | TRUE | FALSE |
| >trna11_GluTTC_15_-_26327381_26327452@1.33.33,trna3_GluTTC_13_-_45492062_45492133@1.33.33\|TCCCACATGGTCTAGCGGTTAGGATTCCTGGTT | >trna11_GluTTC_15_-_26327381_26327452 | 1 | 33 | 33 | TCCCACATGGTCTAGCGGTTAGGATTCCTGGTT | tRF | FALSE | TRUE | FALSE |
| >hg38_21_+_8433172_8446622_RNA45SN1_45S_50ntFlanks@11567.11584.18\|AGCAGCCGACTTAGAACT | >hg38_21_+_8433172_8446622_RNA45SN1_45S_50ntFlanks | 11567 | 11584 | 18 | AGCAGCCGACTTAGAACT | rRF | FALSE | TRUE | FALSE |
| >am_trna128_GlyGCC_6_-_27870686_27870756@1.33.33,trna18_GlyGCC_16_+_70822597_70822667@1.33.33,trna19_GlyGCC_16_+_70823410_70823480@1.33.33,trna19_GlyGCC_2_-_157257659_157257729@1.33.33,trna24_GlyGCC_16_-_70812942_70813012@1.33.33,trna25_GlyGCC_16_-_70812114_70812184@1.33.33,trna5_GlyGCC_17_+_8029064_8029134@1.33.33,trna68_GlyGCC_1_-_161493637_161493707@1.33.33\|GCATTGGTGGTTCAGTGGTAGAATTCTCGCCTG | >am_trna128_GlyGCC_6_-_27870686_27870756 | 1 | 33 | 33 | GCATTGGTGGTTCAGTGGTAGAATTCTCGCCTG | tRF | FALSE | TRUE | FALSE |
| >hg38_21_+_8433172_8446622_RNA45SN1_45S_50ntFlanks@12995.13038.44\|CTCGCTGCGATCTATTGAAAGTCAGCCCTCGACACAAGGGTTTG | >hg38_21_+_8433172_8446622_RNA45SN1_45S_50ntFlanks | 12995 | 13038 | 44 | CTCGCTGCGATCTATTGAAAGTCAGCCCTCGACACAAGGGTTTG | rRF | FALSE | TRUE | FALSE |
| >hg38_21_+_8433172_8446622_RNA45SN1_45S_50ntFlanks@6646.6681.36\|TCGTACGACTCTTAGCGGTGGATCACTCGGCTCGTG | >hg38_21_+_8433172_8446622_RNA45SN1_45S_50ntFlanks | 6646 | 6681 | 36 | TCGTACGACTCTTAGCGGTGGATCACTCGGCTCGTG | rRF | FALSE | TRUE | TRUE |
| >hg38_21_+_8433172_8446622_RNA45SN1_45S_50ntFlanks@6788.6806.19\|CGCCTGTCTGAGCGTCGCT | >hg38_21_+_8433172_8446622_RNA45SN1_45S_50ntFlanks | 6788 | 6806 | 19 | CGCCTGTCTGAGCGTCGCT | rRF | FALSE | TRUE | FALSE |
| >trnaMT_GluTTC_MT_-_14674_14742@41.72.32,trnalookalike8_GluTTC_5_-_93905172_93905240@41.72.32\|TGGTCGTGGTTGTAGTCCGTGCGAGAATACCA | >trnaMT_GluTTC_MT_-_14674_14742 | 41 | 72 | 32 | TGGTCGTGGTTGTAGTCCGTGCGAGAATACCA | tRF | FALSE | TRUE | FALSE |
| >trna111_HisGTG_1_-_147774845_147774916@-1T.23.24,trna118_HisGTG_1_-_145396881_145396952@-1T.23.24,trna16_HisGTG_1_+_146544773_146544844@-1T.23.24,trna1_HisGTG_15_+_45493349_45493420@-1T.23.24,trna21_HisGTG_1_+_147753471_147753542@-1T.23.24,trna33_HisGTG_6_+_27125906_27125977@-1T.23.24,trna7_HisGTG_9_-_14433938_14434009@-1T.23.24,trna8_HisGTG_15_-_45492611_45492682@-1T.23.24,trna9_HisGTG_15_-_45490804_45490875@-1T.23.24\|TGCCGTGATCGTATAGTGGTTAGT | >trna111_HisGTG_1_-_147774845_147774916 | -1T | 23 | 24 | TGCCGTGATCGTATAGTGGTTAGT | tRF | FALSE | TRUE | FALSE |
| >MI0000071\|hsa-mir-17&WithFlank&13\|+\|91350599\|91350694@20.42.23(+1A)&[MIMAT0000070&hsa-miR-17-5p&offsets\|0\|0(+1A);m-88&13\|+\|91350618\|91350640&offsets\|0\|0(+1A)]\|CAAAGTGCTTACAGTGCAGGTAGA | >MI0000071\|hsa-mir-17&WithFlank&13\|+\|91350599\|91350694 | 20 | 42 | 23(+1A) | CAAAGTGCTTACAGTGCAGGTAGA | isomiR | FALSE | TRUE | FALSE |
| >trnaMT_GluTTC_MT_-_14674_14742@38.72.35,trnalookalike8_GluTTC_5_-_93905172_93905240@38.72.35\|CATTGGTCGTGGTTGTAGTCCGTGCGAGAATACCA | >trnaMT_GluTTC_MT_-_14674_14742 | 38 | 72 | 35 | CATTGGTCGTGGTTGTAGTCCGTGCGAGAATACCA | tRF | FALSE | TRUE | FALSE |
| >am_trna10_ValCAC_5_-_180649395_180649467@1.32.32,trna12_ValAAC_5_-_180645270_180645342@1.32.32,trna132_ValAAC_6_-_27721179_27721251@1.32.32,trna136_ValAAC_6_-_27648885_27648957@1.32.32,trna139_ValAAC_6_-_27618707_27618779@1.32.32,trna18_ValCAC_5_-_180529253_180529325@1.32.32,trna2_ValAAC_3_+_169490018_169490090@1.32.32,trna2_ValCAC_5_+_180524070_180524142@1.32.32,trna4_ValAAC_5_+_180591154_180591226@1.32.32,trna5_ValAAC_5_+_180596610_180596682@1.32.32,trna6_ValCAC_5_+_180600650_180600722@1.32.32,trna85_ValCAC_1_-_161369490_161369562@1.32.32,trna90_ValCAC_1_-_149684088_149684161@1.32.32,trna98_ValCAC_1_-_149298555_149298627@1.32.32,trna9_ValCAC_6_+_26538282_26538354@1.32.32\|GTTTCCGTAGTGTAGTGGTTATCACGTTCGCC | >am_trna10_ValCAC_5_-_180649395_180649467 | 1 | 32 | 32 | GTTTCCGTAGTGTAGTGGTTATCACGTTCGCC | tRF | FALSE | TRUE | FALSE |
| >hg38_21_+_8433172_8446622_RNA45SN1_45S_50ntFlanks@6651.6692.42\|CGACTCTTAGCGGTGGATCACTCGGCTCGTGCGTCGATGAAG | >hg38_21_+_8433172_8446622_RNA45SN1_45S_50ntFlanks | 6651 | 6692 | 42 | CGACTCTTAGCGGTGGATCACTCGGCTCGTGCGTCGATGAAG | rRF | FALSE | TRUE | FALSE |
| >hg38_21_+_8433172_8446622_RNA45SN1_45S_50ntFlanks@6651.6689.39\|CGACTCTTAGCGGTGGATCACTCGGCTCGTGCGTCGATG | >hg38_21_+_8433172_8446622_RNA45SN1_45S_50ntFlanks | 6651 | 6689 | 39 | CGACTCTTAGCGGTGGATCACTCGGCTCGTGCGTCGATG | rRF | FALSE | TRUE | TRUE |
| >trna116_GluCTC_1_-_145399233_145399304@1.31.31,trna59_GluCTC_1_+_249168447_249168518@1.31.31,trna71_GluCTC_1_-_161439189_161439260@1.31.31,trna74_GluCTC_1_-_161431809_161431880@1.31.31,trna77_GluCTC_1_-_161424398_161424469@1.31.31,trna77_GluCTC_6_+_28949976_28950047@1.31.31,trna80_GluCTC_1_-_161417018_161417089@1.31.31,trna87_GluCTC_6_-_126101393_126101464@1.31.31\|TCCCTGGTGGTCTAGTGGTTAGGATTCGGCG | >trna116_GluCTC_1_-_145399233_145399304 | 1 | 31 | 31 | TCCCTGGTGGTCTAGTGGTTAGGATTCGGCG | tRF | FALSE | TRUE | TRUE |
| >hg38_21_+_8433172_8446622_RNA45SN1_45S_50ntFlanks@475.516.42\|CTTCGTGATCGATGTGGTGACGTCGTGCTCTCCCGGGCCGGG | >hg38_21_+_8433172_8446622_RNA45SN1_45S_50ntFlanks | 475 | 516 | 42 | CTTCGTGATCGATGTGGTGACGTCGTGCTCTCCCGGGCCGGG | rRF | FALSE | TRUE | FALSE |
| >hg38_1_-_228634819_228635039_RNA5S12_5S_50ntFlanks@144.170.27\|CCTGGGAATACCGGGTGCTGTAGGCTT | >hg38_1_-_228634819_228635039_RNA5S12_5S_50ntFlanks | 144 | 170 | 27 | CCTGGGAATACCGGGTGCTGTAGGCTT | rRF | FALSE | TRUE | FALSE |
| >MI0000446\|hsa-mir-125b-1&WithFlank&11\|-\|122099751\|122099850@21.42.22(+1A);MI0000470\|hsa-mir-125b-2&WithFlank&21\|+\|16590231\|16590331@23.44.22(+1A)&[MIMAT0000423&hsa-miR-125b-5p&offsets\|0\|0(+1A);m-67&11\|-\|122099809\|122099830&offsets\|0\|0(+1A)];[MIMAT0000423_1&hsa-miR-125b-5p&offsets\|0\|0(+1A);m-68&21\|+\|16590253\|16590274&offsets\|0\|0(+1A)]\|TCCCTGAGACCCTAACTTGTGAA | >MI0000446\|hsa-mir-125b-1&WithFlank&11\|-\|122099751\|122099850 | 21 | 42 | 22(+1A) | TCCCTGAGACCCTAACTTGTGAA | isomiR | FALSE | TRUE | FALSE |
| >hg38_21_+_8433172_8446622_RNA45SN1_45S_50ntFlanks@6651.6698.48\|CGACTCTTAGCGGTGGATCACTCGGCTCGTGCGTCGATGAAGAACGCA | >hg38_21_+_8433172_8446622_RNA45SN1_45S_50ntFlanks | 6651 | 6698 | 48 | CGACTCTTAGCGGTGGATCACTCGGCTCGTGCGTCGATGAAGAACGCA | rRF | FALSE | TRUE | FALSE |
| >MI0000734\|hsa-mir-106b&WithFlank&7\|-\|100093987\|100094080@58.79.22&[MIMAT0004672&hsa-miR-106b-3p&offsets\|0\|0;m-87&7\|-\|100094003\|100094023&offsets\|0\|+1]\|CCGCACTGTGGGTACTTGCTGC | >MI0000734\|hsa-mir-106b&WithFlank&7\|-\|100093987\|100094080 | 58 | 79 | 22 | CCGCACTGTGGGTACTTGCTGC | isomiR | FALSE | TRUE | FALSE |
| >MI0000076\|hsa-mir-20a&WithFlank&13\|+\|91351059\|91351141@14.36.23(+1A)&[MIMAT0000075&hsa-miR-20a-5p&offsets\|0\|0(+1A);m-96&13\|+\|91351072\|91351094&offsets\|0\|0(+1A)]\|TAAAGTGCTTATAGTGCAGGTAGA | >MI0000076\|hsa-mir-20a&WithFlank&13\|+\|91351059\|91351141 | 14 | 36 | 23(+1A) | TAAAGTGCTTATAGTGCAGGTAGA | isomiR | FALSE | TRUE | FALSE |

<table>
<tr><td></td><td></td><td></td><td></td><td></td><td></td><td></td><td></td><td></td><td></td></tr>
<tr><td>>MI0000289|hsa-mir-181a-1&WithFlank&1|-|198859038|198859159@30.52.23;MI0000269|hsa-mir-181a-2&WithFlank&9|+|124692436|124692557@45.67.23&[MIMAT0000256&hsa-miR-181a-5p&offsets|0|0;m-32&1|-|198859108|198859130&offsets|0|0];[MIMAT0000256_1&hsa-miR-181a-5p&offsets|0|0;m-32&9|+|124692480|124692502&offsets|0|0]|AACATTCAACGCTGTCGGTGAGT</td><td>>MI0000289|hsa-mir-181a-1&WithFlank&1|-|198859038|198859159</td><td>30</td><td>52</td><td>23</td><td>AACATTCAACGCTGTCGGTGAGT</td><td>isomiR</td><td>FALSE</td><td>TRUE</td><td>FALSE</td></tr>
<tr><td>>MI0000102|hsa-mir-100&WithFlank&11|-|122152223|122152314@19.38.20(+1A)&[MIMAT0000098&hsa-miR-100-5p&offsets|0|-2(+1A);m-27&11|-|122152276|122152296&offsets|0|-1(+1A)]|AACCCGTAGATCCGAACTTGA</td><td>>MI0000102|hsa-mir-100&WithFlank&11|-|122152223|122152314</td><td>19</td><td>38</td><td>20(+1A)</td><td>AACCCGTAGATCCGAACTTGA</td><td>isomiR</td><td>FALSE</td><td>TRUE</td><td>FALSE</td></tr>
<tr><td>>hg38_21_+_8433172_8446622_RNA45SN1_45S_50ntFlanks@12997.13014.18|CGCTGCGATCTATTGAAA</td><td>>hg38_21_+_8433172_8446622_RNA45SN1_45S_50ntFlanks</td><td>12997</td><td>13014</td><td>18</td><td>CGCTGCGATCTATTGAAA</td><td>rRF</td><td>FALSE</td><td>TRUE</td><td>FALSE</td></tr>
<tr><td>>hg38_21_+_8433172_8446622_RNA45SN1_45S_50ntFlanks@475.510.36|CTTCGTGATCGATGTGGTGACGTCGTGCTCTCCCGG</td><td>>hg38_21_+_8433172_8446622_RNA45SN1_45S_50ntFlanks</td><td>475</td><td>510</td><td>36</td><td>CTTCGTGATCGATGTGGTGACGTCGTGCTCTCCCGG</td><td>rRF</td><td>FALSE</td><td>TRUE</td><td>FALSE</td></tr>
<tr><td>>MI0000299|hsa-mir-222&WithFlank&X|-|45747009|45747130@75.97.23(+1U)&[MIMAT0000279&hsa-miR-222-3p&offsets|0|+2(+1U);m-140&X|-|45747033|45747056&offsets|0|-1(+1U)]|AGCTACATCTGGCTACTGGGTCTT</td><td>>MI0000299|hsa-mir-222&WithFlank&X|-|45747009|45747130</td><td>75</td><td>97</td><td>23(+1U)</td><td>AGCTACATCTGGCTACTGGGTCTT</td><td>isomiR</td><td>FALSE</td><td>TRUE</td><td>FALSE</td></tr>
<tr><td>>trnaMT_GluTTC_MT_-_14674_14742@35.72.38,trnalookalike8_GluTTC_5_-_93905172_93905240@35.72.38|TATCATTGGTCGTGGTTGTAGTCCGTGCGAGAATACCA</td><td>>trnaMT_GluTTC_MT_-_14674_14742</td><td>35</td><td>72</td><td>38</td><td>TATCATTGGTCGTGGTTGTAGTCCGTGCGAGAATACCA</td><td>tRF</td><td>FALSE</td><td>TRUE</td><td>FALSE</td></tr>
<tr><td>>trna111_HisGTG_1_-_147774845_147774916@-1T.26.27,trna118_HisGTG_1_-_145396881_145396952@-1T.26.27,trna16_HisGTG_1_+_146544773_146544844@-1T.26.27,trna1_HisGTG_15_+_45493349_45493420@-1T.26.27,trna21_HisGTG_1_+_147753471_147753542@-1T.26.27,trna33_HisGTG_6_+_27125906_27125977@-1T.26.27,trna7_HisGTG_9_-_14433938_14434009@-1T.26.27,trna8_HisGTG_15_-_45492611_45492682@-1T.26.27,trna9_HisGTG_15_-_45490804_45490875@-1T.26.27|TGCCGTGATCGTATAGTGGTTAGTACT</td><td>>trna111_HisGTG_1_-_147774845_147774916</td><td>-1T</td><td>26</td><td>27</td><td>TGCCGTGATCGTATAGTGGTTAGTACT</td><td>tRF</td><td>FALSE</td><td>TRUE</td><td>FALSE</td></tr>
<tr><td>>MI0000082|hsa-mir-25&WithFlank&7|-|100093554|100093649@58.78.21&[MIMAT0000081&hsa-miR-25-3p&offsets|0|-1;m-23&7|-|100093571|100093592&offsets|0|-1]|CATTGCACTTGTCTCGGTCTG</td><td>>MI0000082|hsa-mir-25&WithFlank&7|-|100093554|100093649</td><td>58</td><td>78</td><td>21</td><td>CATTGCACTTGTCTCGGTCTG</td><td>isomiR</td><td>FALSE</td><td>TRUE</td><td>FALSE</td></tr>
<tr><td>>hg38_21_+_8433172_8446622_RNA45SN1_45S_50ntFlanks@9554.9571.18|CGTAGCGGTCCTGACGTG</td><td>>hg38_21_+_8433172_8446622_RNA45SN1_45S_50ntFlanks</td><td>9554</td><td>9571</td><td>18</td><td>CGTAGCGGTCCTGACGTG</td><td>rRF</td><td>FALSE</td><td>TRUE</td><td>TRUE</td></tr>
<tr><td>>trna119_LysCTT_1_-_145395522_145395594@1.34.34,trna11_LysCTT_5_-_180648979_180649051@1.34.34,trna13_LysCTT_6_+_26556774_26556846@1.34.34,trna7_LysCTT_16_+_3225692_3225764@1.34.34,trna9_LysCTT_5_+_180634755_180634827@1.34.34|GCCCGGCTAGCTCAGTCGGTAGAGCATGAGACTC</td><td>>trna119_LysCTT_1_-_145395522_145395594</td><td>1</td><td>34</td><td>34</td><td>GCCCGGCTAGCTCAGTCGGTAGAGCATGAGACTC</td><td>tRF</td><td>FALSE</td><td>TRUE</td><td>FALSE</td></tr>
<tr><td>>MI0000079|hsa-mir-23a&WithFlank&19|-|13836581|13836665@54.71.18&[MIMAT0000078&hsa-miR-23a-3p&offsets|+3|0;m-31&19|-|13836595|13836615&offsets|+3|0]|ACATTGCCAGGGATTTCC</td><td>>MI0000079|hsa-mir-23a&WithFlank&19|-|13836581|13836665</td><td>54</td><td>71</td><td>18</td><td>ACATTGCCAGGGATTTCC</td><td>isomiR</td><td>FALSE</td><td>TRUE</td><td>FALSE</td></tr>
<tr><td>>MI0000100|hsa-mir-98&WithFlank&X|-|53556217|53556347@28.49.22&[MIMAT0000096&hsa-miR-98-5p&offsets|0|0;m-143&X|-|53556299|53556320&offsets|0|0]|TGAGGTAGTAAGTTGTATTGTT</td><td>>MI0000100|hsa-mir-98&WithFlank&X|-|53556217|53556347</td><td>28</td><td>49</td><td>22</td><td>TGAGGTAGTAAGTTGTATTGTT</td><td>isomiR</td><td>FALSE</td><td>TRUE</td><td>FALSE</td></tr>
<tr><td>>hg38_21_+_8433172_8446622_RNA45SN1_45S_50ntFlanks@12419.12437.19|GATGTCGGCTCTTCCTATC</td><td>>hg38_21_+_8433172_8446622_RNA45SN1_45S_50ntFlanks</td><td>12419</td><td>12437</td><td>19</td><td>GATGTCGGCTCTTCCTATC</td><td>rRF</td><td>FALSE</td><td>TRUE</td><td>FALSE</td></tr>
<tr><td>>hg38_21_+_8433172_8446622_RNA45SN1_45S_50ntFlanks@12997.13038.42|CGCTGCGATCTATTGAAAGTCAGCCCTCGACACAAGGGTTTG</td><td>>hg38_21_+_8433172_8446622_RNA45SN1_45S_50ntFlanks</td><td>12997</td><td>13038</td><td>42</td><td>CGCTGCGATCTATTGAAAGTCAGCCCTCGACACAAGGGTTTG</td><td>rRF</td><td>FALSE</td><td>TRUE</td><td>FALSE</td></tr>
<tr><td>>hg38_21_+_8433172_8446622_RNA45SN1_45S_50ntFlanks@7974.8008.35|ACGCGACCTCAGATCAGACGTGGCGACCCGCTGAA</td><td>>hg38_21_+_8433172_8446622_RNA45SN1_45S_50ntFlanks</td><td>7974</td><td>8008</td><td>35</td><td>ACGCGACCTCAGATCAGACGTGGCGACCCGCTGAA</td><td>rRF</td><td>FALSE</td><td>TRUE</td><td>FALSE</td></tr>
<tr><td>>MI0006444|hsa-mir-1307&WithFlank&10|-|103394247|103394407@86.108.23&[MIMAT0005951&hsa-miR-1307-3p&offsets|0|+1;m-66&10|-|103394300|103394322&offsets|0|0]|ACTCGGCGTGGCGTCGGTCGTGG</td><td>>MI0006444|hsa-mir-1307&WithFlank&10|-|103394247|103394407</td><td>86</td><td>108</td><td>23</td><td>ACTCGGCGTGGCGTCGGTCGTGG</td><td>isomiR</td><td>FALSE</td><td>TRUE</td><td>FALSE</td></tr>
<tr><td>>hg38_21_+_8433172_8446622_RNA45SN1_45S_50ntFlanks@7975.8014.40|CGCGACCTCAGATCAGACGTGGCGACCCGCTGAATTTAAG</td><td>>hg38_21_+_8433172_8446622_RNA45SN1_45S_50ntFlanks</td><td>7975</td><td>8014</td><td>40</td><td>CGCGACCTCAGATCAGACGTGGCGACCCGCTGAATTTAAG</td><td>rRF</td><td>FALSE</td><td>TRUE</td><td>FALSE</td></tr>
<tr><td>>hg38_21_+_8433172_8446622_RNA45SN1_45S_50ntFlanks@7989.8011.23|AGACGTGGCGACCCGCTGAATTT</td><td>>hg38_21_+_8433172_8446622_RNA45SN1_45S_50ntFlanks</td><td>7989</td><td>8011</td><td>23</td><td>AGACGTGGCGACCCGCTGAATTT</td><td>rRF</td><td>FALSE</td><td>TRUE</td><td>FALSE</td></tr>
<tr><td>>hg38_21_+_8433172_8446622_RNA45SN1_45S_50ntFlanks@476.504.29|TTCGTGATCGATGTGGTGACGTCGTGCTC</td><td>>hg38_21_+_8433172_8446622_RNA45SN1_45S_50ntFlanks</td><td>476</td><td>504</td><td>29</td><td>TTCGTGATCGATGTGGTGACGTCGTGCTC</td><td>rRF</td><td>FALSE</td><td>TRUE</td><td>FALSE</td></tr>
<tr><td>>MI0000079|hsa-mir-23a&WithFlank&19|-|13836581|13836665@54.73.20(+1U)&[MIMAT0000078&hsa-miR-23a-3p&offsets|+3|+2(+1U);m-31&19|-|13836595|13836615&offsets|+3|+2(+1U)]|ACATTGCCAGGGATTTCCAAT</td><td>>MI0000079|hsa-mir-23a&WithFlank&19|-|13836581|13836665</td><td>54</td><td>73</td><td>20(+1U)</td><td>ACATTGCCAGGGATTTCCAAT</td><td>isomiR</td><td>FALSE</td><td>TRUE</td><td>FALSE</td></tr>
<tr><td>>hg38_21_+_8433172_8446622_RNA45SN1_45S_50ntFlanks@6651.6697.47|CGACTCTTAGCGGTGGATCACTCGGCTCGTGCGTCGATGAAGAACGC</td><td>>hg38_21_+_8433172_8446622_RNA45SN1_45S_50ntFlanks</td><td>6651</td><td>6697</td><td>47</td><td>CGACTCTTAGCGGTGGATCACTCGGCTCGTGCGTCGATGAAGAACGC</td><td>rRF</td><td>FALSE</td><td>TRUE</td><td>FALSE</td></tr>
<tr><td>>hg38_1_-_228634819_228635039_RNA5S12_5S_50ntFlanks@133.171.39|ATGGGAGACCGCCTGGGAATACCGGGTGCTGTAGGCTTT</td><td>>hg38_1_-_228634819_228635039_RNA5S12_5S_50ntFlanks</td><td>133</td><td>171</td><td>39</td><td>ATGGGAGACCGCCTGGGAATACCGGGTGCTGTAGGCTTT</td><td>rRF</td><td>FALSE</td><td>TRUE</td><td>FALSE</td></tr>
<tr><td>>trnaMT_GluTTC_MT_-_14674_14742@34.72.39,trnalookalike8_GluTTC_5_-_93905172_93905240@34.72.39|ATATCATTGGTCGTGGTTGTAGTCCGTGCGAGAATACCA</td><td>>trnaMT_GluTTC_MT_-_14674_14742</td><td>34</td><td>72</td><td>39</td><td>ATATCATTGGTCGTGGTTGTAGTCCGTGCGAGAATACCA</td><td>tRF</td><td>FALSE</td><td>TRUE</td><td>FALSE</td></tr>
<tr><td>>hg38_21_+_8433172_8446622_RNA45SN1_45S_50ntFlanks@12416.12448.33|TTCGATGTCGGCTCTTCCTATCATTGTGAAGCA</td><td>>hg38_21_+_8433172_8446622_RNA45SN1_45S_50ntFlanks</td><td>12416</td><td>12448</td><td>33</td><td>TTCGATGTCGGCTCTTCCTATCATTGTGAAGCA</td><td>rRF</td><td>FALSE</td><td>TRUE</td><td>FALSE</td></tr>
<tr><td>>trnaMT_HisGTG_MT_+_12138_12206@32.71.40|TGAATCTGACAACAGAGGCTTACGACCCCTTATTTACCCC</td><td>>trnaMT_HisGTG_MT_+_12138_12206</td><td>32</td><td>71</td><td>40</td><td>TGAATCTGACAACAGAGGCTTACGACCCCTTATTTACCCC</td><td>tRF</td><td>FALSE</td><td>TRUE</td><td>FALSE</td></tr>
<tr><td>>MI0000465|hsa-mir-191&WithFlank&3|-|49020612|49020715@22.45.24&[MIMAT0000440&hsa-miR-191-5p&offsets|0|+1;m-48&3|-|49020672|49020694&offsets|0|+1]|CAACGGAATCCCAAAAGCAGCTGT</td><td>>MI0000465|hsa-mir-191&WithFlank&3|-|49020612|49020715</td><td>22</td><td>45</td><td>24</td><td>CAACGGAATCCCAAAAGCAGCTGT</td><td>isomiR</td><td>FALSE</td><td>TRUE</td><td>FALSE</td></tr>
<tr><td>>MI0000299|hsa-mir-222&WithFlank&X|-|45747009|45747130@75.99.25(+2U)&[MIMAT0000279&hsa-miR-222-3p&offsets|0|+4(+2U);m-140&X|-|45747033|45747056&offsets|0|+1(+2U)]|AGCTACATCTGGCTACTGGGTCTCTTT</td><td>>MI0000299|hsa-mir-222&WithFlank&X|-|45747009|45747130</td><td>75</td><td>99</td><td>25(+2U)</td><td>AGCTACATCTGGCTACTGGGTCTCTTT</td><td>isomiR</td><td>FALSE</td><td>TRUE</td><td>FALSE</td></tr>
<tr><td>>hg38_21_+_8433172_8446622_RNA45SN1_45S_50ntFlanks@10678.10710.33|CGGTTCCGGCGGCGTCCGGTGAGCTCTCGCTGG</td><td>>hg38_21_+_8433172_8446622_RNA45SN1_45S_50ntFlanks</td><td>10678</td><td>10710</td><td>33</td><td>CGGTTCCGGCGGCGTCCGGTGAGCTCTCGCTGG</td><td>rRF</td><td>FALSE</td><td>FALSE</td><td>TRUE</td></tr>
<tr><td>>hg38_21_+_8433172_8446622_RNA45SN1_45S_50ntFlanks@9534.9553.20|AGGAAACTCTGGTGGAGGTC</td><td>>hg38_21_+_8433172_8446622_RNA45SN1_45S_50ntFlanks</td><td>9534</td><td>9553</td><td>20</td><td>AGGAAACTCTGGTGGAGGTC</td><td>rRF</td><td>FALSE</td><td>FALSE</td><td>TRUE</td></tr>
<tr><td>>hg38_21_+_8433172_8446622_RNA45SN1_45S_50ntFlanks@11729.11751.23|TAACTATGACTCTCTTAAGGTAG</td><td>>hg38_21_+_8433172_8446622_RNA45SN1_45S_50ntFlanks</td><td>11729</td><td>11751</td><td>23</td><td>TAACTATGACTCTCTTAAGGTAG</td><td>rRF</td><td>FALSE</td><td>FALSE</td><td>TRUE</td></tr>
<tr><td>>hg38_21_+_8433172_8446622_RNA45SN1_45S_50ntFlanks@8357.8379.23|AAAAGAACTTTGAAGAGAGAGTT</td><td>>hg38_21_+_8433172_8446622_RNA45SN1_45S_50ntFlanks</td><td>8357</td><td>8379</td><td>23</td><td>AAAAGAACTTTGAAGAGAGAGTT</td><td>rRF</td><td>FALSE</td><td>FALSE</td><td>TRUE</td></tr>
<tr><td>>hg38_21_+_8433172_8446622_RNA45SN1_45S_50ntFlanks@12456.12487.32|CCAAGCGTTGGATTGTTCACCCACTAATAGGG</td><td>>hg38_21_+_8433172_8446622_RNA45SN1_45S_50ntFlanks</td><td>12456</td><td>12487</td><td>32</td><td>CCAAGCGTTGGATTGTTCACCCACTAATAGGG</td><td>rRF</td><td>FALSE</td><td>FALSE</td><td>TRUE</td></tr>
</table>

| | | | | | | | | | |
|---|---|---|---|---|---|---|---|---|---|
| >hg38_21_+_8433172_8446622_RNA45SN1_45S_50ntFlanks@12456.12476.21\|CCAAGCGTTGGATTGTTCACC | >hg38_21_+_8433172_8446622_RNA45SN1_45S_50ntFlanks | 12456 | 12476 | 21 | CCAAGCGTTGGATTGTTCACC | rRF | FALSE | FALSE | TRUE |
| >hg38_21_+_8433172_8446622_RNA45SN1_45S_50ntFlanks@10841.10865.25\|CGTAACTTCGGGATAAGGATTGGCT | >hg38_21_+_8433172_8446622_RNA45SN1_45S_50ntFlanks | 10841 | 10865 | 25 | CGTAACTTCGGGATAAGGATTGGCT | rRF | FALSE | FALSE | TRUE |
| >am_trna128_GlyGCC_6_-_27870686_27870756@1.28.28,trna133_GlyCCC_1_-_16872434_16872504@1.28.28,trna18_GlyGCC_16_+_70822597_70822667@1.28.28,trna19_GlyGCC_16_+_70823410_70823480@1.28.28,trna19_GlyGCC_2_-_157257659_157257729@1.28.28,trna24_GlyGCC_16_-_70812942_70813012@1.28.28,trna25_GlyGCC_16_-_70812114_70812184@1.28.28,trna4_GlyCCC_1_+_17188416_17188486@1.28.28,trna5_GlyGCC_17_+_8029064_8029134@1.28.28,trna68_GlyGCC_1_-_161493637_161493707@1.28.28\|GCATTGGTGGTTCAGTGGTAGAATTCTC | >am_trna128_GlyGCC_6_-_27870686_27870756 | 1 | 28 | 28 | GCATTGGTGGTTCAGTGGTAGAATTCTC | tRF | FALSE | FALSE | TRUE |
| >hg38_21_+_8433172_8446622_RNA45SN1_45S_50ntFlanks@10679.10710.32\|GGTTCCGGCGGCGTCCGGTGAGCTCTCGCTGG | >hg38_21_+_8433172_8446622_RNA45SN1_45S_50ntFlanks | 10679 | 10710 | 32 | GGTTCCGGCGGCGTCCGGTGAGCTCTCGCTGG | rRF | FALSE | FALSE | TRUE |
| >hg38_21_+_8433172_8446622_RNA45SN1_45S_50ntFlanks@10678.10711.34\|CGGTTCCGGCGGCGTCCGGTGAGCTCTCGCTGGC | >hg38_21_+_8433172_8446622_RNA45SN1_45S_50ntFlanks | 10678 | 10711 | 34 | CGGTTCCGGCGGCGTCCGGTGAGCTCTCGCTGGC | rRF | FALSE | FALSE | TRUE |
| >hg38_21_+_8433172_8446622_RNA45SN1_45S_50ntFlanks@10351.10368.18\|CGAGAACTTTGAAGGCCG | >hg38_21_+_8433172_8446622_RNA45SN1_45S_50ntFlanks | 10351 | 10368 | 18 | CGAGAACTTTGAAGGCCG | rRF | FALSE | FALSE | TRUE |
| >am_trna116_GluCTC_1_-_145399233_145399304@1.29.29,trna59_GluCTC_1_+_249168447_249168518@1.29.29,trna71_GluCTC_1_-_161439189_161439260@1.29.29,trna74_GluCTC_1_-_161431809_161431880@1.29.29,trna77_GluCTC_1_-_161424398_161424469@1.29.29,trna77_GluCTC_6_+_28949976_28950047@1.29.29,trna80_GluCTC_1_-_161417018_161417089@1.29.29,trna87_GluCTC_6_-_126101393_126101464@1.29.29\|TCCCTGGTGGTCTAGTGGTTAGGATTCGG | >am_trna116_GluCTC_1_-_145399233_145399304 | 1 | 29 | 29 | TCCCTGGTGGTCTAGTGGTTAGGATTCGG | tRF | FALSE | FALSE | TRUE |
| >hg38_21_+_8433172_8446622_RNA45SN1_45S_50ntFlanks@6699.6720.22\|GCTAGCTGCGAGAATTAATGTG | >hg38_21_+_8433172_8446622_RNA45SN1_45S_50ntFlanks | 6699 | 6720 | 22 | GCTAGCTGCGAGAATTAATGTG | rRF | FALSE | FALSE | TRUE |
| >hg38_21_+_8433172_8446622_RNA45SN1_45S_50ntFlanks@11783.11804.22\|CATGAATGGATGAACGAGATTC | >hg38_21_+_8433172_8446622_RNA45SN1_45S_50ntFlanks | 11783 | 11804 | 22 | CATGAATGGATGAACGAGATTC | rRF | FALSE | FALSE | TRUE |
| >am_trna128_GlyGCC_6_-_27870686_27870756@1.29.29,trna133_GlyCCC_1_-_16872434_16872504@1.29.29,trna18_GlyGCC_16_+_70822597_70822667@1.29.29,trna19_GlyGCC_16_+_70823410_70823480@1.29.29,trna19_GlyGCC_2_-_157257659_157257729@1.29.29,trna24_GlyGCC_16_-_70812942_70813012@1.29.29,trna25_GlyGCC_16_-_70812114_70812184@1.29.29,trna4_GlyCCC_1_+_17188416_17188486@1.29.29,trna5_GlyGCC_17_+_8029064_8029134@1.29.29,trna68_GlyGCC_1_-_161493637_161493707@1.29.29\|GCATTGGTGGTTCAGTGGTAGAATTCTCG | >am_trna128_GlyGCC_6_-_27870686_27870756 | 1 | 29 | 29 | GCATTGGTGGTTCAGTGGTAGAATTCTCG | tRF | FALSE | FALSE | TRUE |
| >hg38_21_+_8433172_8446622_RNA45SN1_45S_50ntFlanks@12456.12475.20\|CCAAGCGTTGGATTGTTCAC | >hg38_21_+_8433172_8446622_RNA45SN1_45S_50ntFlanks | 12456 | 12475 | 20 | CCAAGCGTTGGATTGTTCAC | rRF | FALSE | FALSE | TRUE |
| >hg38_21_+_8433172_8446622_RNA45SN1_45S_50ntFlanks@10678.10709.32\|CGGTTCCGGCGGCGTCCGGTGAGCTCTCGCTG | >hg38_21_+_8433172_8446622_RNA45SN1_45S_50ntFlanks | 10678 | 10709 | 32 | CGGTTCCGGCGGCGTCCGGTGAGCTCTCGCTG | rRF | FALSE | FALSE | TRUE |
| >hg38_21_+_8433172_8446622_RNA45SN1_45S_50ntFlanks@12437.12455.19\|CATTGTGAAGCAGAATTCA | >hg38_21_+_8433172_8446622_RNA45SN1_45S_50ntFlanks | 12437 | 12455 | 19 | CATTGTGAAGCAGAATTCA | rRF | FALSE | FALSE | TRUE |
| >hg38_21_+_8433172_8446622_RNA45SN1_45S_50ntFlanks@8356.8379.24\|GAAAAGAACTTTGAAGAGAGAGTT | >hg38_21_+_8433172_8446622_RNA45SN1_45S_50ntFlanks | 8356 | 8379 | 24 | GAAAAGAACTTTGAAGAGAGAGTT | rRF | FALSE | FALSE | TRUE |
| >hg38_21_+_8433172_8446622_RNA45SN1_45S_50ntFlanks@6721.6744.24\|AATTGCAGGACACATTGATCATCG | >hg38_21_+_8433172_8446622_RNA45SN1_45S_50ntFlanks | 6721 | 6744 | 24 | AATTGCAGGACACATTGATCATCG | rRF | FALSE | FALSE | TRUE |
| >hg38_21_+_8433172_8446622_RNA45SN1_45S_50ntFlanks@8357.8378.22\|AAAAGAACTTTGAAGAGAGAGT | >hg38_21_+_8433172_8446622_RNA45SN1_45S_50ntFlanks | 8357 | 8378 | 22 | AAAAGAACTTTGAAGAGAGAGT | rRF | FALSE | FALSE | TRUE |
| >hg38_21_+_8433172_8446622_RNA45SN1_45S_50ntFlanks@10678.10706.29\|CGGTTCCGGCGGCGTCCGGTGAGCTCTCG | >hg38_21_+_8433172_8446622_RNA45SN1_45S_50ntFlanks | 10678 | 10706 | 29 | CGGTTCCGGCGGCGTCCGGTGAGCTCTCG | rRF | FALSE | FALSE | TRUE |
| >am_trna116_GluCTC_1_-_145399233_145399304@1.28.28,trna59_GluCTC_1_+_249168447_249168518@1.28.28,trna71_GluCTC_1_-_161439189_161439260@1.28.28,trna74_GluCTC_1_-_161431809_161431880@1.28.28,trna77_GluCTC_1_-_161424398_161424469@1.28.28,trna77_GluCTC_6_+_28949976_28950047@1.28.28,trna80_GluCTC_1_-_161417018_161417089@1.28.28,trna87_GluCTC_6_-_126101393_126101464@1.28.28\|TCCCTGGTGGTCTAGTGGTTAGGATTCG | >am_trna116_GluCTC_1_-_145399233_145399304 | 1 | 28 | 28 | TCCCTGGTGGTCTAGTGGTTAGGATTCG | tRF | FALSE | FALSE | TRUE |
| >hg38_21_+_8433172_8446622_RNA45SN1_45S_50ntFlanks@10680.10710.31\|GTTCCGGCGGCGTCCGGTGAGCTCTCGCTGG | >hg38_21_+_8433172_8446622_RNA45SN1_45S_50ntFlanks | 10680 | 10710 | 31 | GTTCCGGCGGCGTCCGGTGAGCTCTCGCTGG | rRF | FALSE | FALSE | TRUE |
| >am_trna10_ValCAC_5_-_180649395_180649467@1.29.29,trna12_ValAAC_5_-_180645270_180645342@1.29.29,trna132_ValAAC_6_-_27721179_27721251@1.29.29,trna136_ValAAC_6_-_27648885_27648957@1.29.29,trna139_ValAAC_6_-_27618707_27618779@1.29.29,trna18_ValCAC_5_-_180529253_180529325@1.29.29,trna2_ValAAC_3_+_169490018_169490090@1.29.29,trna2_ValCAC_5_+_180524070_180524142@1.29.29,trna4_ValAAC_5_+_180591154_180591226@1.29.29,trna5_ValAAC_5_+_180596610_180596682@1.29.29,trna6_ValCAC_5_+_180600650_180600722@1.29.29,trna85_ValCAC_1_-_161369490_161369562@1.29.29,trna90_ValCAC_1_-_149684088_149684161@1.29.29,trna98_ValCAC_1_-_149298555_149298627@1.29.29,trna9_ValCAC_6_+_26538282_26538354@1.29.29\|GTTTCCGTAGTGTAGTGGTTATCACGTTC | >am_trna10_ValCAC_5_-_180649395_180649467 | 1 | 29 | 29 | GTTTCCGTAGTGTAGTGGTTATCACGTTC | tRF | FALSE | FALSE | TRUE |
| >hg38_21_+_8433172_8446622_RNA45SN1_45S_50ntFlanks@10680.10708.29\|GTTCCGGCGGCGTCCGGTGAGCTCTCGCT | >hg38_21_+_8433172_8446622_RNA45SN1_45S_50ntFlanks | 10680 | 10708 | 29 | GTTCCGGCGGCGTCCGGTGAGCTCTCGCT | rRF | FALSE | FALSE | TRUE |
| >multi-am_trnaMT_ValTAC_MT_+_1602_1670@51.68.18\|CTTAACTTGACCGCTCTG | >multi-am_trnaMT_ValTAC_MT_+_1602_1670 | 51 | 68 | 18 | CTTAACTTGACCGCTCTG | tRF | FALSE | FALSE | TRUE |
| >trna119_LysCTT_1_-_145395522_145395594@1.30.30,trna11_LysCTT_5_-_180648979_180649051@1.30.30,trna13_LysCTT_6_+_26556774_26556846@1.30.30,trna32_LysCTT_16_-_3207406_3207478@1.30.30,trna7_LysCTT_16_+_3225692_3225764@1.30.30,trna9_LysCTT_5_+_180634755_180634827@1.30.30\|GCCCGGCTAGCTCAGTCGGTAGAGCATGAG | >trna119_LysCTT_1_-_145395522_145395594 | 1 | 30 | 30 | GCCCGGCTAGCTCAGTCGGTAGAGCATGAG | tRF | FALSE | FALSE | TRUE |
| >hg38_21_+_8433172_8446622_RNA45SN1_45S_50ntFlanks@9564.9596.33\|CTGACGTGCAAATCGGTCGTCCGACCTGGGTAT | >hg38_21_+_8433172_8446622_RNA45SN1_45S_50ntFlanks | 9564 | 9596 | 33 | CTGACGTGCAAATCGGTCGTCCGACCTGGGTAT | rRF | FALSE | FALSE | TRUE |
| >hg38_21_+_8433172_8446622_RNA45SN1_45S_50ntFlanks@10351.10383.33\|CGAGAACTTTGAAGGCCGAAGTGGAGAAGGGTT | >hg38_21_+_8433172_8446622_RNA45SN1_45S_50ntFlanks | 10351 | 10383 | 33 | CGAGAACTTTGAAGGCCGAAGTGGAGAAGGGTT | rRF | FALSE | FALSE | TRUE |

| | | | | | | | | | |
|---|---|---|---|---|---|---|---|---|---|
| >hg38_21_+_8433172_8446622_RNA45SN1_45S_50ntFlanks@12418.12436.19\|CGATGTCGGCTCTTCCTAT | >hg38_21_+_8433172_8446622_RNA45SN1_45S_50ntFlanks | 12418 | 12436 | 19 | CGATGTCGGCTCTTCCTAT | rRF | FALSE | FALSE | TRUE |
| >hg38_21_+_8433172_8446622_RNA45SN1_45S_50ntFlanks@6700.6720.21\|CTAGCTGCGAGAATTAATGTG | >hg38_21_+_8433172_8446622_RNA45SN1_45S_50ntFlanks | 6700 | 6720 | 21 | CTAGCTGCGAGAATTAATGTG | rRF | FALSE | FALSE | TRUE |
| >hg38_21_+_8433172_8446622_RNA45SN1_45S_50ntFlanks@6698.6720.23\|AGCTAGCTGCGAGAATTAATGTG | >hg38_21_+_8433172_8446622_RNA45SN1_45S_50ntFlanks | 6698 | 6720 | 23 | AGCTAGCTGCGAGAATTAATGTG | rRF | FALSE | FALSE | TRUE |
| >hg38_21_+_8433172_8446622_RNA45SN1_45S_50ntFlanks@12456.12486.31\|CCAAGCGTTGGATTGTTCACCCACTAATAGG | >hg38_21_+_8433172_8446622_RNA45SN1_45S_50ntFlanks | 12456 | 12486 | 31 | CCAAGCGTTGGATTGTTCACCCACTAATAGG | rRF | FALSE | FALSE | TRUE |
| >hg38_21_+_8433172_8446622_RNA45SN1_45S_50ntFlanks@10411.10429.19\|CAGTCGGTCCTGAGAGATG | >hg38_21_+_8433172_8446622_RNA45SN1_45S_50ntFlanks | 10411 | 10429 | 19 | CAGTCGGTCCTGAGAGATG | rRF | FALSE | FALSE | TRUE |
| >hg38_21_+_8433172_8446622_RNA45SN1_45S_50ntFlanks@12633.12654.22\|CGAAGCTACCATCTGTGGGATT | >hg38_21_+_8433172_8446622_RNA45SN1_45S_50ntFlanks | 12633 | 12654 | 22 | CGAAGCTACCATCTGTGGGATT | rRF | FALSE | FALSE | TRUE |
| >hg38_21_+_8433172_8446622_RNA45SN1_45S_50ntFlanks@12456.12478.23\|CCAAGCGTTGGATTGTTCACCCA | >hg38_21_+_8433172_8446622_RNA45SN1_45S_50ntFlanks | 12456 | 12478 | 23 | CCAAGCGTTGGATTGTTCACCCA | rRF | FALSE | FALSE | TRUE |
| >hg38_21_+_8433172_8446622_RNA45SN1_45S_50ntFlanks@6699.6718.20\|GCTAGCTGCGAGAATTAATG | >hg38_21_+_8433172_8446622_RNA45SN1_45S_50ntFlanks | 6699 | 6718 | 20 | GCTAGCTGCGAGAATTAATG | rRF | FALSE | FALSE | TRUE |
| >hg38_21_+_8433172_8446622_RNA45SN1_45S_50ntFlanks@9554.9572.19\|CGTAGCGGTCCTGACGTGC | >hg38_21_+_8433172_8446622_RNA45SN1_45S_50ntFlanks | 9554 | 9572 | 19 | CGTAGCGGTCCTGACGTGC | rRF | FALSE | FALSE | TRUE |
| >trna2_GlyGCC_21_-_18827107_18827177@1.29.29,trna35_GlyGCC_1_+_161413094_161413164@1.29.29,trna37_GlyGCC_1_+_161420467_161420537@1.29.29,trna39_GlyGCC_1_+_161427898_161427968@1.29.29,trna41_GlyGCC_1_+_161435258_161435328@1.29.29\|GCATGGGTGGTTCAGTGGTAGAATTCTCG | >trna2_GlyGCC_21_-_18827107_18827177 | 1 | 29 | 29 | GCATGGGTGGTTCAGTGGTAGAATTCTCG | tRF | FALSE | FALSE | TRUE |
| >am_trna146_GlnCTG_6_-_27515531_27515602@1.29.29,trna1_GlnCTG_6_+_18836402_18836473@1.29.29,trna3_GlnCTG_17_+_8023070_8023141@1.29.29,trna42_GlnCTG_6_+_27263212_27263283@1.29.29,trna49_GlnCTG_6_+_27487308_27487379@1.29.29,trna7_GlnCTG_15_-_66161400_66161471@1.29.29,trna99_GlnCTG_6_-_28909378_28909449@1.29.29\|GGTTCCATGGTGTAATGGTTAGCACTCTG | >am_trna146_GlnCTG_6_-_27515531_27515602 | 1 | 29 | 29 | GGTTCCATGGTGTAATGGTTAGCACTCTG | tRF | FALSE | FALSE | TRUE |
| >hg38_21_+_8433172_8446622_RNA45SN1_45S_50ntFlanks@11641.11660.20\|CGGCGGGTGTTGACGCGATG | >hg38_21_+_8433172_8446622_RNA45SN1_45S_50ntFlanks | 11641 | 11660 | 20 | CGGCGGGTGTTGACGCGATG | rRF | FALSE | FALSE | TRUE |
| >hg38_21_+_8433172_8446622_RNA45SN1_45S_50ntFlanks@10678.10708.31\|CGGTTCCGGCGGCGTCCGGTGAGCTCTCGCT | >hg38_21_+_8433172_8446622_RNA45SN1_45S_50ntFlanks | 10678 | 10708 | 31 | CGGTTCCGGCGGCGTCCGGTGAGCTCTCGCT | rRF | FALSE | FALSE | TRUE |
| >trna2_GlyGCC_21_-_18827107_18827177@1.28.28,trna35_GlyGCC_1_+_161413094_161413164@1.28.28,trna37_GlyGCC_1_+_161420467_161420537@1.28.28,trna39_GlyGCC_1_+_161427898_161427968@1.28.28,trna41_GlyGCC_1_+_161435258_161435328@1.28.28\|GCATGGGTGGTTCAGTGGTAGAATTCTC | >trna2_GlyGCC_21_-_18827107_18827177 | 1 | 28 | 28 | GCATGGGTGGTTCAGTGGTAGAATTCTC | tRF | FALSE | FALSE | TRUE |
| >trna152_ValCAC_6_-_27248049_27248121@1.29.29\|GCTTCTGTAGTGTAGTGGTTATCACGTTC | >trna152_ValCAC_6_-_27248049_27248121 | 1 | 29 | 29 | GCTTCTGTAGTGTAGTGGTTATCACGTTC | tRF | FALSE | FALSE | TRUE |
| >hg38_21_+_8433172_8446622_RNA45SN1_45S_50ntFlanks@11209.11232.24\|CGGCGACTCTGGACGCGAGCCGGG | >hg38_21_+_8433172_8446622_RNA45SN1_45S_50ntFlanks | 11209 | 11232 | 24 | CGGCGACTCTGGACGCGAGCCGGG | rRF | FALSE | FALSE | TRUE |
| >hg38_21_+_8433172_8446622_RNA45SN1_45S_50ntFlanks@12456.12484.29\|CCAAGCGTTGGATTGTTCACCCACTAATA | >hg38_21_+_8433172_8446622_RNA45SN1_45S_50ntFlanks | 12456 | 12484 | 29 | CCAAGCGTTGGATTGTTCACCCACTAATA | rRF | FALSE | FALSE | TRUE |
| >hg38_21_+_8433172_8446622_RNA45SN1_45S_50ntFlanks@9554.9573.20\|CGTAGCGGTCCTGACGTGCA | >hg38_21_+_8433172_8446622_RNA45SN1_45S_50ntFlanks | 9554 | 9573 | 20 | CGTAGCGGTCCTGACGTGCA | rRF | FALSE | FALSE | TRUE |
| >hg38_21_+_8433172_8446622_RNA45SN1_45S_50ntFlanks@11785.11804.20\|TGAATGGATGAACGAGATTC | >hg38_21_+_8433172_8446622_RNA45SN1_45S_50ntFlanks | 11785 | 11804 | 20 | TGAATGGATGAACGAGATTC | rRF | FALSE | FALSE | TRUE |
| >hg38_21_+_8433172_8446622_RNA45SN1_45S_50ntFlanks@6719.6744.26\|TGAATTGCAGGACACATTGATCATCG | >hg38_21_+_8433172_8446622_RNA45SN1_45S_50ntFlanks | 6719 | 6744 | 26 | TGAATTGCAGGACACATTGATCATCG | rRF | FALSE | FALSE | TRUE |
| >am_trna116_GluCTC_1_-_145399233_145399304@1.30.30,trna59_GluCTC_1_+_249168447_249168518@1.30.30,trna71_GluCTC_1_-_161439189_161439260@1.30.30,trna74_GluCTC_1_-_161431809_161431880@1.30.30,trna77_GluCTC_1_-_161424398_161424469@1.30.30,trna77_GluCTC_6_+_28949976_28950047@1.30.30,trna80_GluCTC_1_-_161417018_161417089@1.30.30,trna87_GluCTC_6_-_126101393_126101464@1.30.30\|TCCCTGGTGGTCTAGTGGTTAGGATTCGGC | >am_trna116_GluCTC_1_-_145399233_145399304 | 1 | 30 | 30 | TCCCTGGTGGTCTAGTGGTTAGGATTCGGC | tRF | FALSE | FALSE | TRUE |
| >hg38_21_+_8433172_8446622_RNA45SN1_45S_50ntFlanks@10350.10368.19\|ACGAGAACTTTGAAGGCCG | >hg38_21_+_8433172_8446622_RNA45SN1_45S_50ntFlanks | 10350 | 10368 | 19 | ACGAGAACTTTGAAGGCCG | rRF | FALSE | FALSE | TRUE |
| >hg38_21_+_8433172_8446622_RNA45SN1_45S_50ntFlanks@10679.10709.31\|GGTTCCGGCGGCGTCCGGTGAGCTCTCGCTG | >hg38_21_+_8433172_8446622_RNA45SN1_45S_50ntFlanks | 10679 | 10709 | 31 | GGTTCCGGCGGCGTCCGGTGAGCTCTCGCTG | rRF | FALSE | FALSE | TRUE |
| >hg38_21_+_8433172_8446622_RNA45SN1_45S_50ntFlanks@5444.5466.23\|CTCGGATCGGCCCCGCCGGGGTC | >hg38_21_+_8433172_8446622_RNA45SN1_45S_50ntFlanks | 5444 | 5466 | 23 | CTCGGATCGGCCCCGCCGGGGTC | rRF | FALSE | FALSE | TRUE |
| >hg38_21_+_8433172_8446622_RNA45SN1_45S_50ntFlanks@6721.6740.20\|AATTGCAGGACACATTGATC | >hg38_21_+_8433172_8446622_RNA45SN1_45S_50ntFlanks | 6721 | 6740 | 20 | AATTGCAGGACACATTGATC | rRF | FALSE | FALSE | TRUE |
| >hg38_21_+_8433172_8446622_RNA45SN1_45S_50ntFlanks@9534.9563.30\|AGGAAACTCTGGTGGAGGTCCGTAGCGGTC | >hg38_21_+_8433172_8446622_RNA45SN1_45S_50ntFlanks | 9534 | 9563 | 30 | AGGAAACTCTGGTGGAGGTCCGTAGCGGTC | rRF | FALSE | FALSE | TRUE |
| >hg38_21_+_8433172_8446622_RNA45SN1_45S_50ntFlanks@12655.12675.21\|ATGACTGAACGCCTCTAAGTC | >hg38_21_+_8433172_8446622_RNA45SN1_45S_50ntFlanks | 12655 | 12675 | 21 | ATGACTGAACGCCTCTAAGTC | rRF | FALSE | FALSE | TRUE |
| >hg38_21_+_8433172_8446622_RNA45SN1_45S_50ntFlanks@12973.12990.18\|TCGTACGTAGCAGAGCAG | >hg38_21_+_8433172_8446622_RNA45SN1_45S_50ntFlanks | 12973 | 12990 | 18 | TCGTACGTAGCAGAGCAG | rRF | FALSE | FALSE | TRUE |
| >hg38_21_+_8433172_8446622_RNA45SN1_45S_50ntFlanks@11783.11803.21\|CATGAATGGATGAACGAGATT | >hg38_21_+_8433172_8446622_RNA45SN1_45S_50ntFlanks | 11783 | 11803 | 21 | CATGAATGGATGAACGAGATT | rRF | FALSE | FALSE | TRUE |
| >hg38_21_+_8433172_8446622_RNA45SN1_45S_50ntFlanks@12418.12435.18\|CGATGTCGGCTCTTCCTA | >hg38_21_+_8433172_8446622_RNA45SN1_45S_50ntFlanks | 12418 | 12435 | 18 | CGATGTCGGCTCTTCCTA | rRF | FALSE | FALSE | TRUE |
| >hg38_21_+_8433172_8446622_RNA45SN1_45S_50ntFlanks@10680.10706.27\|GTTCCGGCGGCGTCCGGTGAGCTCTCG | >hg38_21_+_8433172_8446622_RNA45SN1_45S_50ntFlanks | 10680 | 10706 | 27 | GTTCCGGCGGCGTCCGGTGAGCTCTCG | rRF | FALSE | FALSE | TRUE |
| >hg38_21_+_8433172_8446622_RNA45SN1_45S_50ntFlanks@12626.12654.29\|AATGGGGCGAAGCTACCATCTGTGGGATT | >hg38_21_+_8433172_8446622_RNA45SN1_45S_50ntFlanks | 12626 | 12654 | 29 | AATGGGGCGAAGCTACCATCTGTGGGATT | rRF | FALSE | FALSE | TRUE |
| >hg38_21_+_8433172_8446622_RNA45SN1_45S_50ntFlanks@12633.12653.21\|CGAAGCTACCATCTGTGGGAT | >hg38_21_+_8433172_8446622_RNA45SN1_45S_50ntFlanks | 12633 | 12653 | 21 | CGAAGCTACCATCTGTGGGAT | rRF | FALSE | FALSE | TRUE |
| >hg38_21_+_8433172_8446622_RNA45SN1_45S_50ntFlanks@12635.12654.20\|AAGCTACCATCTGTGGGATT | >hg38_21_+_8433172_8446622_RNA45SN1_45S_50ntFlanks | 12635 | 12654 | 20 | AAGCTACCATCTGTGGGATT | rRF | FALSE | FALSE | TRUE |
| >hg38_21_+_8433172_8446622_RNA45SN1_45S_50ntFlanks@10680.10711.32\|GTTCCGGCGGCGTCCGGTGAGCTCTCGCTGGC | >hg38_21_+_8433172_8446622_RNA45SN1_45S_50ntFlanks | 10680 | 10711 | 32 | GTTCCGGCGGCGTCCGGTGAGCTCTCGCTGGC | rRF | FALSE | FALSE | TRUE |

| | | | | | | | | | |
|---|---|---|---|---|---|---|---|---|---|
| >hg38_21_+_8433172_8446622_RNA45SN1_45S_50ntFlanks@10392.10410.19\|ACAGCAGTTGAACATGGGT | >hg38_21_+_8433172_8446622_RNA45SN1_45S_50ntFlanks | 10392 | 10410 | 19 | ACAGCAGTTGAACATGGGT | rRF | FALSE | FALSE | TRUE |
| >trna111_HisGTG_1_-_147774845_147774916@1.31.31,trna118_HisGTG_1_-_145396881_145396952@1.31.31,trna16_HisGTG_1_+_146544773_146544844@1.31.31,trna1_HisGTG_15_+_45493349_45493420@1.31.31,trna21_HisGTG_1_+_147753471_147753542@1.31.31,trna33_HisGTG_6_+_27125906_27125977@1.31.31,trna7_HisGTG_9_-_14433938_14434009@1.31.31,trna8_HisGTG_15_-_45492611_45492682@1.31.31,trna9_HisGTG_15_-_45490804_45490875@1.31.31\|GCCGTGATCGTATAGTGGTTAGTACTCTGCG | >trna111_HisGTG_1_-_147774845_147774916 | 1 | 31 | 31 | GCCGTGATCGTATAGTGGTTAGTACTCTGCG | tRF | FALSE | FALSE | TRUE |
| >hg38_21_+_8433172_8446622_RNA45SN1_45S_50ntFlanks@11787.11804.18\|AATGGATGAACGAGATTC | >hg38_21_+_8433172_8446622_RNA45SN1_45S_50ntFlanks | 11787 | 11804 | 18 | AATGGATGAACGAGATTC | rRF | FALSE | FALSE | TRUE |
| >hg38_21_+_8433172_8446622_RNA45SN1_45S_50ntFlanks@12405.12431.27\|CTTTTTGATCCTTCGATGTCGGCTCTT | >hg38_21_+_8433172_8446622_RNA45SN1_45S_50ntFlanks | 12405 | 12431 | 27 | CTTTTTGATCCTTCGATGTCGGCTCTT | rRF | FALSE | FALSE | TRUE |
| >hg38_21_+_8433172_8446622_RNA45SN1_45S_50ntFlanks@9572.9597.26\|CAAATCGGTCGTCCGACCTGGGTATA | >hg38_21_+_8433172_8446622_RNA45SN1_45S_50ntFlanks | 9572 | 9597 | 26 | CAAATCGGTCGTCCGACCTGGGTATA | rRF | FALSE | FALSE | TRUE |
| >hg38_21_+_8433172_8446622_RNA45SN1_45S_50ntFlanks@4759.4780.22\|ATCAGATACCGTCGTAGTTCCG | >hg38_21_+_8433172_8446622_RNA45SN1_45S_50ntFlanks | 4759 | 4780 | 22 | ATCAGATACCGTCGTAGTTCCG | rRF | FALSE | FALSE | TRUE |
| >hg38_21_+_8433172_8446622_RNA45SN1_45S_50ntFlanks@11572.11591.20\|CCGACTTAGAACTGGTGCGG | >hg38_21_+_8433172_8446622_RNA45SN1_45S_50ntFlanks | 11572 | 11591 | 20 | CCGACTTAGAACTGGTGCGG | rRF | FALSE | FALSE | TRUE |
| >hg38_21_+_8433172_8446622_RNA45SN1_45S_50ntFlanks@9572.9596.25\|CAAATCGGTCGTCCGACCTGGGTAT | >hg38_21_+_8433172_8446622_RNA45SN1_45S_50ntFlanks | 9572 | 9596 | 25 | CAAATCGGTCGTCCGACCTGGGTAT | rRF | FALSE | FALSE | TRUE |
| >hg38_21_+_8433172_8446622_RNA45SN1_45S_50ntFlanks@12456.12477.22\|CCAAGCGTTGGATTGTTCACCC | >hg38_21_+_8433172_8446622_RNA45SN1_45S_50ntFlanks | 12456 | 12477 | 22 | CCAAGCGTTGGATTGTTCACCC | rRF | FALSE | FALSE | TRUE |
| >hg38_21_+_8433172_8446622_RNA45SN1_45S_50ntFlanks@10684.10710.27\|CGGCGGCGTCCGGTGAGCTCTCGCTGG | >hg38_21_+_8433172_8446622_RNA45SN1_45S_50ntFlanks | 10684 | 10710 | 27 | CGGCGGCGTCCGGTGAGCTCTCGCTGG | rRF | FALSE | FALSE | TRUE |
| >hg38_21_+_8433172_8446622_RNA45SN1_45S_50ntFlanks@9534.9571.38\|AGGAAACTCTGGTGGAGGTCCGTAGCGGTCCTGACGTG | >hg38_21_+_8433172_8446622_RNA45SN1_45S_50ntFlanks | 9534 | 9571 | 38 | AGGAAACTCTGGTGGAGGTCCGTAGCGGTCCTGACGTG | rRF | FALSE | FALSE | TRUE |
| >hg38_21_+_8433172_8446622_RNA45SN1_45S_50ntFlanks@10411.10430.20\|CAGTCGGTCCTGAGAGATGG | >hg38_21_+_8433172_8446622_RNA45SN1_45S_50ntFlanks | 10411 | 10430 | 20 | CAGTCGGTCCTGAGAGATGG | rRF | FALSE | FALSE | TRUE |
| >hg38_21_+_8433172_8446622_RNA45SN1_45S_50ntFlanks@12008.12032.25\|CCGCCGGTGAAATACCACTACTCTG | >hg38_21_+_8433172_8446622_RNA45SN1_45S_50ntFlanks | 12008 | 12032 | 25 | CCGCCGGTGAAATACCACTACTCTG | rRF | FALSE | FALSE | TRUE |
| >hg38_21_+_8433172_8446622_RNA45SN1_45S_50ntFlanks@9545.9571.27\|GTGGAGGTCCGTAGCGGTCCTGACGTG | >hg38_21_+_8433172_8446622_RNA45SN1_45S_50ntFlanks | 9545 | 9571 | 27 | GTGGAGGTCCGTAGCGGTCCTGACGTG | rRF | FALSE | FALSE | TRUE |
| >hg38_21_+_8433172_8446622_RNA45SN1_45S_50ntFlanks@12973.12991.19\|TCGTACGTAGCAGAGCAGC | >hg38_21_+_8433172_8446622_RNA45SN1_45S_50ntFlanks | 12973 | 12991 | 19 | TCGTACGTAGCAGAGCAGC | rRF | FALSE | FALSE | TRUE |
| >hg38_21_+_8433172_8446622_RNA45SN1_45S_50ntFlanks@12605.12625.21\|TATGTGCTTGGCTGAGGAGCC | >hg38_21_+_8433172_8446622_RNA45SN1_45S_50ntFlanks | 12605 | 12625 | 21 | TATGTGCTTGGCTGAGGAGCC | rRF | FALSE | FALSE | TRUE |
| >hg38_21_+_8433172_8446622_RNA45SN1_45S_50ntFlanks@10678.10712.35\|CGGTTCCGGCGGCGTCCGGTGAGCTCTCGCTGGCC | >hg38_21_+_8433172_8446622_RNA45SN1_45S_50ntFlanks | 10678 | 10712 | 35 | CGGTTCCGGCGGCGTCCGGTGAGCTCTCGCTGGCC | rRF | FALSE | FALSE | TRUE |
| >hg38_21_+_8433172_8446622_RNA45SN1_45S_50ntFlanks@12627.12654.28\|ATGGGGCGAAGCTACCATCTGTGGGATT | >hg38_21_+_8433172_8446622_RNA45SN1_45S_50ntFlanks | 12627 | 12654 | 28 | ATGGGGCGAAGCTACCATCTGTGGGATT | rRF | FALSE | FALSE | TRUE |
| >hg38_21_+_8433172_8446622_RNA45SN1_45S_50ntFlanks@12991.13013.23\|CTCCCTCGCTGCGATCTATTGAA | >hg38_21_+_8433172_8446622_RNA45SN1_45S_50ntFlanks | 12991 | 13013 | 23 | CTCCCTCGCTGCGATCTATTGAA | rRF | FALSE | FALSE | TRUE |
| >hg38_21_+_8433172_8446622_RNA45SN1_45S_50ntFlanks@11323.11344.22\|CGTCTCGTCGCGCGCGCGTCCG | >hg38_21_+_8433172_8446622_RNA45SN1_45S_50ntFlanks | 11323 | 11344 | 22 | CGTCTCGTCGCGCGCGCGTCCG | rRF | FALSE | FALSE | TRUE |
| >hg38_21_+_8433172_8446622_RNA45SN1_45S_50ntFlanks@7983.8006.24\|CAGATCAGACGTGGCGACCCGCTG | >hg38_21_+_8433172_8446622_RNA45SN1_45S_50ntFlanks | 7983 | 8006 | 24 | CAGATCAGACGTGGCGACCCGCTG | rRF | FALSE | FALSE | TRUE |
| >hg38_21_+_8433172_8446622_RNA45SN1_45S_50ntFlanks@10786.10818.33\|AACAGCCTCTGGCATGTTGGAACAATGTAGGTA | >hg38_21_+_8433172_8446622_RNA45SN1_45S_50ntFlanks | 10786 | 10818 | 33 | AACAGCCTCTGGCATGTTGGAACAATGTAGGTA | rRF | FALSE | FALSE | TRUE |
| >hg38_21_+_8433172_8446622_RNA45SN1_45S_50ntFlanks@11572.11601.30\|CCGACTTAGAACTGGTGCGGACCAGGGGAA | >hg38_21_+_8433172_8446622_RNA45SN1_45S_50ntFlanks | 11572 | 11601 | 30 | CCGACTTAGAACTGGTGCGGACCAGGGGAA | rRF | FALSE | FALSE | TRUE |
| >hg38_21_+_8433172_8446622_RNA45SN1_45S_50ntFlanks@12405.12434.30\|CTTTTTGATCCTTCGATGTCGGCTCTTCCT | >hg38_21_+_8433172_8446622_RNA45SN1_45S_50ntFlanks | 12405 | 12434 | 30 | CTTTTTGATCCTTCGATGTCGGCTCTTCCT | rRF | FALSE | FALSE | TRUE |
| >hg38_21_+_8433172_8446622_RNA45SN1_45S_50ntFlanks@6700.6718.19\|CTAGCTGCGAGAATTAATG | >hg38_21_+_8433172_8446622_RNA45SN1_45S_50ntFlanks | 6700 | 6718 | 19 | CTAGCTGCGAGAATTAATG | rRF | FALSE | FALSE | TRUE |
| >hg38_21_+_8433172_8446622_RNA45SN1_45S_50ntFlanks@6719.6743.25\|TGAATTGCAGGACACATTGATCATC | >hg38_21_+_8433172_8446622_RNA45SN1_45S_50ntFlanks | 6719 | 6743 | 25 | TGAATTGCAGGACACATTGATCATC | rRF | FALSE | FALSE | TRUE |
| >hg38_21_+_8433172_8446622_RNA45SN1_45S_50ntFlanks@12456.12488.33\|CCAAGCGTTGGATTGTTCACCCACTAATAGGGA | >hg38_21_+_8433172_8446622_RNA45SN1_45S_50ntFlanks | 12456 | 12488 | 33 | CCAAGCGTTGGATTGTTCACCCACTAATAGGGA | rRF | FALSE | FALSE | TRUE |
| >hg38_21_+_8433172_8446622_RNA45SN1_45S_50ntFlanks@12456.12473.18\|CCAAGCGTTGGATTGTTC | >hg38_21_+_8433172_8446622_RNA45SN1_45S_50ntFlanks | 12456 | 12473 | 18 | CCAAGCGTTGGATTGTTC | rRF | FALSE | FALSE | TRUE |
| >hg38_21_+_8433172_8446622_RNA45SN1_45S_50ntFlanks@10841.10868.28\|CGTAACTTCGGGATAAGGATTGGCTCTA | >hg38_21_+_8433172_8446622_RNA45SN1_45S_50ntFlanks | 10841 | 10868 | 28 | CGTAACTTCGGGATAAGGATTGGCTCTA | rRF | FALSE | FALSE | TRUE |
| >hg38_21_+_8433172_8446622_RNA45SN1_45S_50ntFlanks@7991.8008.18\|ACGTGGCGACCCGCTGAA | >hg38_21_+_8433172_8446622_RNA45SN1_45S_50ntFlanks | 7991 | 8008 | 18 | ACGTGGCGACCCGCTGAA | rRF | FALSE | FALSE | TRUE |
| >hg38_21_+_8433172_8446622_RNA45SN1_45S_50ntFlanks@6721.6743.23\|AATTGCAGGACACATTGATCATC | >hg38_21_+_8433172_8446622_RNA45SN1_45S_50ntFlanks | 6721 | 6743 | 23 | AATTGCAGGACACATTGATCATC | rRF | FALSE | FALSE | TRUE |
| >hg38_21_+_8433172_8446622_RNA45SN1_45S_50ntFlanks@11208.11232.25\|GCGGCGACTCTGGACGCGAGCCGGG | >hg38_21_+_8433172_8446622_RNA45SN1_45S_50ntFlanks | 11208 | 11232 | 25 | GCGGCGACTCTGGACGCGAGCCGGG | rRF | FALSE | FALSE | TRUE |
| >hg38_21_+_8433172_8446622_RNA45SN1_45S_50ntFlanks@10410.10429.20\|TCAGTCGGTCCTGAGAGATG | >hg38_21_+_8433172_8446622_RNA45SN1_45S_50ntFlanks | 10410 | 10429 | 20 | TCAGTCGGTCCTGAGAGATG | rRF | FALSE | FALSE | TRUE |
| >hg38_21_+_8433172_8446622_RNA45SN1_45S_50ntFlanks@11729.11749.21\|TAACTATGACTCTCTTAAGGT | >hg38_21_+_8433172_8446622_RNA45SN1_45S_50ntFlanks | 11729 | 11749 | 21 | TAACTATGACTCTCTTAAGGT | rRF | FALSE | FALSE | TRUE |
| >hg38_21_+_8433172_8446622_RNA45SN1_45S_50ntFlanks@13002.13027.26\|CGATCTATTGAAAGTCAGCCCTCGAC | >hg38_21_+_8433172_8446622_RNA45SN1_45S_50ntFlanks | 13002 | 13027 | 26 | CGATCTATTGAAAGTCAGCCCTCGAC | rRF | FALSE | FALSE | TRUE |
| >hg38_21_+_8433172_8446622_RNA45SN1_45S_50ntFlanks@10679.10711.33\|GGTTCCGGCGGCGTCCGGTGAGCTCTCGCTGGC | >hg38_21_+_8433172_8446622_RNA45SN1_45S_50ntFlanks | 10679 | 10711 | 33 | GGTTCCGGCGGCGTCCGGTGAGCTCTCGCTGGC | rRF | FALSE | FALSE | TRUE |
| >hg38_21_+_8433172_8446622_RNA45SN1_45S_50ntFlanks@9533.9553.21\|GAGGAAACTCTGGTGGAGGTC | >hg38_21_+_8433172_8446622_RNA45SN1_45S_50ntFlanks | 9533 | 9553 | 21 | GAGGAAACTCTGGTGGAGGTC | rRF | FALSE | FALSE | TRUE |
| >hg38_21_+_8433172_8446622_RNA45SN1_45S_50ntFlanks@10351.10377.27\|CGAGAACTTTGAAGGCCGAAGTGGAGA | >hg38_21_+_8433172_8446622_RNA45SN1_45S_50ntFlanks | 10351 | 10377 | 27 | CGAGAACTTTGAAGGCCGAAGTGGAGA | rRF | FALSE | FALSE | TRUE |
| >hg38_21_+_8433172_8446622_RNA45SN1_45S_50ntFlanks@11784.11804.21\|ATGAATGGATGAACGAGATTC | >hg38_21_+_8433172_8446622_RNA45SN1_45S_50ntFlanks | 11784 | 11804 | 21 | ATGAATGGATGAACGAGATTC | rRF | FALSE | FALSE | TRUE |
| >hg38_21_+_8433172_8446622_RNA45SN1_45S_50ntFlanks@12934.12956.23\|CTAAACCATTCGTAGACGACCTG | >hg38_21_+_8433172_8446622_RNA45SN1_45S_50ntFlanks | 12934 | 12956 | 23 | CTAAACCATTCGTAGACGACCTG | rRF | FALSE | FALSE | TRUE |

| | | | | | | | | | |
|---|---|---|---|---|---|---|---|---|---|
| >hg38_21_+_8433172_8446622_RNA45SN1_45S_50ntFlanks@12405.12432.28\|CTTTTTGATCCTTCGATGTCGGCTCTTC | >hg38_21_+_8433172_8446622_RNA45SN1_45S_50ntFlanks | 12405 | 12432 | 28 | CTTTTTGATCCTTCGATGTCGGCTCTTC | rRF | FALSE | FALSE | TRUE |
| >hg38_21_+_8433172_8446622_RNA45SN1_45S_50ntFlanks@10683.10706.24\|CCGGCGGCGTCCGGTGAGCTCTCG | >hg38_21_+_8433172_8446622_RNA45SN1_45S_50ntFlanks | 10683 | 10706 | 24 | CCGGCGGCGTCCGGTGAGCTCTCG | rRF | FALSE | FALSE | TRUE |
| >hg38_21_+_8433172_8446622_RNA45SN1_45S_50ntFlanks@10351.10388.38\|CGAGAACTTTGAAGGCCGAAGTGGAGAAGGGTTCCATG | >hg38_21_+_8433172_8446622_RNA45SN1_45S_50ntFlanks | 10351 | 10388 | 38 | CGAGAACTTTGAAGGCCGAAGTGGAGAAGGGTTCCATG | rRF | FALSE | FALSE | TRUE |
| >hg38_21_+_8433172_8446622_RNA45SN1_45S_50ntFlanks@12405.12436.32\|CTTTTTGATCCTTCGATGTCGGCTCTTCCTAT | >hg38_21_+_8433172_8446622_RNA45SN1_45S_50ntFlanks | 12405 | 12436 | 32 | CTTTTTGATCCTTCGATGTCGGCTCTTCCTAT | rRF | FALSE | FALSE | TRUE |
| >hg38_21_+_8433172_8446622_RNA45SN1_45S_50ntFlanks@10391.10410.20\|AACAGCAGTTGAACATGGGT | >hg38_21_+_8433172_8446622_RNA45SN1_45S_50ntFlanks | 10391 | 10410 | 20 | AACAGCAGTTGAACATGGGT | rRF | FALSE | FALSE | TRUE |
| >hg38_21_+_8433172_8446622_RNA45SN1_45S_50ntFlanks@10350.10383.34\|ACGAGAACTTTGAAGGCCGAAGTGGAGAAGGGTT | >hg38_21_+_8433172_8446622_RNA45SN1_45S_50ntFlanks | 10350 | 10383 | 34 | ACGAGAACTTTGAAGGCCGAAGTGGAGAAGGGTT | rRF | FALSE | FALSE | TRUE |
| >hg38_21_+_8433172_8446622_RNA45SN1_45S_50ntFlanks@9574.9597.24\|AATCGGTCGTCCGACCTGGGTATA | >hg38_21_+_8433172_8446622_RNA45SN1_45S_50ntFlanks | 9574 | 9597 | 24 | AATCGGTCGTCCGACCTGGGTATA | rRF | FALSE | FALSE | TRUE |
| >hg38_21_+_8433172_8446622_RNA45SN1_45S_50ntFlanks@11316.11344.29\|CACCCCACGTCTCGTCGCGCGCGCGTCCG | >hg38_21_+_8433172_8446622_RNA45SN1_45S_50ntFlanks | 11316 | 11344 | 29 | CACCCCACGTCTCGTCGCGCGCGCGTCCG | rRF | FALSE | FALSE | TRUE |
| >hg38_21_+_8433172_8446622_RNA45SN1_45S_50ntFlanks@12488.12509.22\|AACGTGAGCTGGGTTTAGACCG | >hg38_21_+_8433172_8446622_RNA45SN1_45S_50ntFlanks | 12488 | 12509 | 22 | AACGTGAGCTGGGTTTAGACCG | rRF | FALSE | FALSE | TRUE |
| >hg38_21_+_8433172_8446622_RNA45SN1_45S_50ntFlanks@10679.10708.30\|GGTTCCGGCGGCGTCCGGTGAGCTCTCGCT | >hg38_21_+_8433172_8446622_RNA45SN1_45S_50ntFlanks | 10679 | 10708 | 30 | GGTTCCGGCGGCGTCCGGTGAGCTCTCGCT | rRF | FALSE | FALSE | TRUE |
| >hg38_21_+_8433172_8446622_RNA45SN1_45S_50ntFlanks@12456.12474.19\|CCAAGCGTTGGATTGTTCA | >hg38_21_+_8433172_8446622_RNA45SN1_45S_50ntFlanks | 12456 | 12474 | 19 | CCAAGCGTTGGATTGTTCA | rRF | FALSE | FALSE | TRUE |
| >hg38_21_+_8433172_8446622_RNA45SN1_45S_50ntFlanks@12974.12991.18\|CGTACGTAGCAGAGCAGC | >hg38_21_+_8433172_8446622_RNA45SN1_45S_50ntFlanks | 12974 | 12991 | 18 | CGTACGTAGCAGAGCAGC | rRF | FALSE | FALSE | TRUE |
| >hg38_21_+_8433172_8446622_RNA45SN1_45S_50ntFlanks@12438.12455.18\|ATTGTGAAGCAGAATTCA | >hg38_21_+_8433172_8446622_RNA45SN1_45S_50ntFlanks | 12438 | 12455 | 18 | ATTGTGAAGCAGAATTCA | rRF | FALSE | FALSE | TRUE |
| >hg38_21_+_8433172_8446622_RNA45SN1_45S_50ntFlanks@12457.12476.20\|CAAGCGTTGGATTGTTCACC | >hg38_21_+_8433172_8446622_RNA45SN1_45S_50ntFlanks | 12457 | 12476 | 20 | CAAGCGTTGGATTGTTCACC | rRF | FALSE | FALSE | TRUE |
| >hg38_21_+_8433172_8446622_RNA45SN1_45S_50ntFlanks@12936.12956.21\|AAACCATTCGTAGACGACCTG | >hg38_21_+_8433172_8446622_RNA45SN1_45S_50ntFlanks | 12936 | 12956 | 21 | AAACCATTCGTAGACGACCTG | rRF | FALSE | FALSE | TRUE |
| >hg38_21_+_8433172_8446622_RNA45SN1_45S_50ntFlanks@8357.8375.19\|AAAAGAACTTTGAAGAGAG | >hg38_21_+_8433172_8446622_RNA45SN1_45S_50ntFlanks | 8357 | 8375 | 19 | AAAAGAACTTTGAAGAGAG | rRF | FALSE | FALSE | TRUE |
| >hg38_21_+_8433172_8446622_RNA45SN1_45S_50ntFlanks@9565.9596.32\|TGACGTGCAAATCGGTCGTCCGACCTGGGTAT | >hg38_21_+_8433172_8446622_RNA45SN1_45S_50ntFlanks | 9565 | 9596 | 32 | TGACGTGCAAATCGGTCGTCCGACCTGGGTAT | rRF | FALSE | FALSE | TRUE |
| >hg38_21_+_8433172_8446622_RNA45SN1_45S_50ntFlanks@6698.6718.21\|AGCTAGCTGCGAGAATTAATG | >hg38_21_+_8433172_8446622_RNA45SN1_45S_50ntFlanks | 6698 | 6718 | 21 | AGCTAGCTGCGAGAATTAATG | rRF | FALSE | FALSE | TRUE |
| >hg38_21_+_8433172_8446622_RNA45SN1_45S_50ntFlanks@11999.12032.34\|CCCTGCGGGCCGCCGGTGAAATACCACTACTCTG | >hg38_21_+_8433172_8446622_RNA45SN1_45S_50ntFlanks | 11999 | 12032 | 34 | CCCTGCGGGCCGCCGGTGAAATACCACTACTCTG | rRF | FALSE | FALSE | TRUE |
| >trna111_HisGTG_1_-_147774845_147774916@-1T.31.32,trna118_HisGTG_1_-_145396881_145396952@-1T.31.32,trna16_HisGTG_1_+_146544773_146544844@-1T.31.32,trna1_HisGTG_15_+_45493349_45493420@-1T.31.32,trna21_HisGTG_1_+_147753471_147753542@-1T.31.32,trna33_HisGTG_6_+_27125906_27125977@-1T.31.32,trna7_HisGTG_9_-_14433938_14434009@-1T.31.32,trna8_HisGTG_15_-_45492611_45492682@-1T.31.32,trna9_HisGTG_15_-_45490804_45490875@-1T.31.32\|TGCCGTGATCGTATAGTGGTTAGTACTCTGCG | >trna111_HisGTG_1_-_147774845_147774916 | -1T | 31 | 32 | TGCCGTGATCGTATAGTGGTTAGTACTCTGCG | tRF | FALSE | FALSE | TRUE |
| >hg38_21_+_8433172_8446622_RNA45SN1_45S_50ntFlanks@9564.9597.34\|CTGACGTGCAAATCGGTCGTCCGACCTGGGTATA | >hg38_21_+_8433172_8446622_RNA45SN1_45S_50ntFlanks | 9564 | 9597 | 34 | CTGACGTGCAAATCGGTCGTCCGACCTGGGTATA | rRF | FALSE | FALSE | TRUE |
| >trna119_LysCTT_1_-_145395522_145395594@1.31.31,trna11_LysCTT_5_-_180648979_180649051@1.31.31,trna13_LysCTT_6_+_26556774_26556846@1.31.31,trna32_LysCTT_16_-_3207406_3207478@1.31.31,trna7_LysCTT_16_+_3225692_3225764@1.31.31,trna9_LysCTT_5_+_180634755_180634827@1.31.31\|GCCCGGCTAGCTCAGTCGGTAGAGCATGAGA | >trna119_LysCTT_1_-_145395522_145395594 | 1 | 31 | 31 | GCCCGGCTAGCTCAGTCGGTAGAGCATGAGA | tRF | FALSE | FALSE | TRUE |
| >hg38_21_+_8433172_8446622_RNA45SN1_45S_50ntFlanks@10845.10865.21\|ACTTCGGGATAAGGATTGGCT | >hg38_21_+_8433172_8446622_RNA45SN1_45S_50ntFlanks | 10845 | 10865 | 21 | ACTTCGGGATAAGGATTGGCT | rRF | FALSE | FALSE | TRUE |
| >hg38_21_+_8433172_8446622_RNA45SN1_45S_50ntFlanks@10690.10708.19\|CGTCCGGTGAGCTCTCGCT | >hg38_21_+_8433172_8446622_RNA45SN1_45S_50ntFlanks | 10690 | 10708 | 19 | CGTCCGGTGAGCTCTCGCT | rRF | FALSE | FALSE | TRUE |
| >hg38_21_+_8433172_8446622_RNA45SN1_45S_50ntFlanks@9574.9596.23\|AATCGGTCGTCCGACCTGGGTAT | >hg38_21_+_8433172_8446622_RNA45SN1_45S_50ntFlanks | 9574 | 9596 | 23 | AATCGGTCGTCCGACCTGGGTAT | rRF | FALSE | FALSE | TRUE |
| >hg38_21_+_8433172_8446622_RNA45SN1_45S_50ntFlanks@12417.12436.20\|TCGATGTCGGCTCTTCCTAT | >hg38_21_+_8433172_8446622_RNA45SN1_45S_50ntFlanks | 12417 | 12436 | 20 | TCGATGTCGGCTCTTCCTAT | rRF | FALSE | FALSE | TRUE |
| >hg38_21_+_8433172_8446622_RNA45SN1_45S_50ntFlanks@11661.11685.25\|TGATTTCTGCCCAGTGCTCTGAATG | >hg38_21_+_8433172_8446622_RNA45SN1_45S_50ntFlanks | 11661 | 11685 | 25 | TGATTTCTGCCCAGTGCTCTGAATG | rRF | FALSE | FALSE | TRUE |
| >hg38_21_+_8433172_8446622_RNA45SN1_45S_50ntFlanks@10680.10709.30\|GTTCCGGCGGCGTCCGGTGAGCTCTCGCTG | >hg38_21_+_8433172_8446622_RNA45SN1_45S_50ntFlanks | 10680 | 10709 | 30 | GTTCCGGCGGCGTCCGGTGAGCTCTCGCTG | rRF | FALSE | FALSE | TRUE |
| >hg38_21_+_8433172_8446622_RNA45SN1_45S_50ntFlanks@10684.10706.23\|CGGCGGCGTCCGGTGAGCTCTCG | >hg38_21_+_8433172_8446622_RNA45SN1_45S_50ntFlanks | 10684 | 10706 | 23 | CGGCGGCGTCCGGTGAGCTCTCG | rRF | FALSE | FALSE | TRUE |
| >hg38_21_+_8433172_8446622_RNA45SN1_45S_50ntFlanks@6719.6736.18\|TGAATTGCAGGACACATT | >hg38_21_+_8433172_8446622_RNA45SN1_45S_50ntFlanks | 6719 | 6736 | 18 | TGAATTGCAGGACACATT | rRF | FALSE | FALSE | TRUE |
| >hg38_21_+_8433172_8446622_RNA45SN1_45S_50ntFlanks@11874.11893.20\|AGAAGACCCTGTTGAGCTTG | >hg38_21_+_8433172_8446622_RNA45SN1_45S_50ntFlanks | 11874 | 11893 | 20 | AGAAGACCCTGTTGAGCTTG | rRF | FALSE | FALSE | TRUE |
| >hg38_21_+_8433172_8446622_RNA45SN1_45S_50ntFlanks@4724.4758.35\|ATTAATCAAGAACGAAAGTCGGAGGTTCGAAGACG | >hg38_21_+_8433172_8446622_RNA45SN1_45S_50ntFlanks | 4724 | 4758 | 35 | ATTAATCAAGAACGAAAGTCGGAGGTTCGAAGACG | rRF | FALSE | FALSE | TRUE |
| >hg38_21_+_8433172_8446622_RNA45SN1_45S_50ntFlanks@10687.10710.24\|CGGCGTCCGGTGAGCTCTCGCTGG | >hg38_21_+_8433172_8446622_RNA45SN1_45S_50ntFlanks | 10687 | 10710 | 24 | CGGCGTCCGGTGAGCTCTCGCTGG | rRF | FALSE | FALSE | TRUE |
| >hg38_21_+_8433172_8446622_RNA45SN1_45S_50ntFlanks@12605.12626.22\|TATGTGCTTGGCTGAGGAGCCA | >hg38_21_+_8433172_8446622_RNA45SN1_45S_50ntFlanks | 12605 | 12626 | 22 | TATGTGCTTGGCTGAGGAGCCA | rRF | FALSE | FALSE | TRUE |
| >hg38_21_+_8433172_8446622_RNA45SN1_45S_50ntFlanks@10679.10706.28\|GGTTCCGGCGGCGTCCGGTGAGCTCTCG | >hg38_21_+_8433172_8446622_RNA45SN1_45S_50ntFlanks | 10679 | 10706 | 28 | GGTTCCGGCGGCGTCCGGTGAGCTCTCG | rRF | FALSE | FALSE | TRUE |
| >hg38_21_+_8433172_8446622_RNA45SN1_45S_50ntFlanks@8356.8378.23\|GAAAAGAACTTTGAAGAGAGAGT | >hg38_21_+_8433172_8446622_RNA45SN1_45S_50ntFlanks | 8356 | 8378 | 23 | GAAAAGAACTTTGAAGAGAGAGT | rRF | FALSE | FALSE | TRUE |
| >hg38_21_+_8433172_8446622_RNA45SN1_45S_50ntFlanks@8357.8377.21\|AAAAGAACTTTGAAGAGAGAG | >hg38_21_+_8433172_8446622_RNA45SN1_45S_50ntFlanks | 8357 | 8377 | 21 | AAAAGAACTTTGAAGAGAGAG | rRF | FALSE | FALSE | TRUE |
| >hg38_21_+_8433172_8446622_RNA45SN1_45S_50ntFlanks@7979.8006.28\|ACCTCAGATCAGACGTGGCGACCCGCTG | >hg38_21_+_8433172_8446622_RNA45SN1_45S_50ntFlanks | 7979 | 8006 | 28 | ACCTCAGATCAGACGTGGCGACCCGCTG | rRF | FALSE | FALSE | TRUE |

<table>
<tr><th></th><th></th><th></th><th></th><th></th><th></th><th></th><th></th><th></th><th></th></tr>
<tr><td>>hg38_21_+_8433172_8446622_RNA45SN1_45S_50ntFlanks@11322.11344.23|ACGTCTCGTCGCGCGCGCGTCCG</td><td>>hg38_21_+_8433172_8446622_RNA45SN1_45S_50ntFlanks</td><td>11322</td><td>11344</td><td>23</td><td>ACGTCTCGTCGCGCGCGCGTCCG</td><td>rRF</td><td>FALSE</td><td>FALSE</td><td>TRUE</td></tr>
<tr><td>>hg38_21_+_8433172_8446622_RNA45SN1_45S_50ntFlanks@9504.9526.23|GTGAACTATGCCTGGGCAGGGCG</td><td>>hg38_21_+_8433172_8446622_RNA45SN1_45S_50ntFlanks</td><td>9504</td><td>9526</td><td>23</td><td>GTGAACTATGCCTGGGCAGGGCG</td><td>rRF</td><td>FALSE</td><td>FALSE</td><td>TRUE</td></tr>
<tr><td>>hg38_21_+_8433172_8446622_RNA45SN1_45S_50ntFlanks@10351.10390.40|CGAGAACTTTGAAGGCCGAAGTGGAGAAGGGTTCCATGTG</td><td>>hg38_21_+_8433172_8446622_RNA45SN1_45S_50ntFlanks</td><td>10351</td><td>10390</td><td>40</td><td>CGAGAACTTTGAAGGCCGAAGTGGAGAAGGGTTCCATGTG</td><td>rRF</td><td>FALSE</td><td>FALSE</td><td>TRUE</td></tr>
<tr><td>>am_trna111_HisGTG_1_-_147774845_147774916@1.29.29,trna118_HisGTG_1_-_145396881_145396952@1.29.29,trna16_HisGTG_1_+_146544773_146544844@1.29.29,trna1_HisGTG_15_+_45493349_45493420@1.29.29,trna21_HisGTG_1_+_147753471_147753542@1.29.29,trna33_HisGTG_6_+_27125906_27125977@1.29.29,trna7_HisGTG_9_-_14433938_14434009@1.29.29,trna8_HisGTG_15_-_45492611_45492682@1.29.29,trna9_HisGTG_15_-_45490804_45490875@1.29.29|GCCGTGATCGTATAGTGGTTAGTACTCTG</td><td>>am_trna111_HisGTG_1_-_147774845_147774916</td><td>1</td><td>29</td><td>29</td><td>GCCGTGATCGTATAGTGGTTAGTACTCTG</td><td>tRF</td><td>FALSE</td><td>FALSE</td><td>TRUE</td></tr>
<tr><td>>hg38_21_+_8433172_8446622_RNA45SN1_45S_50ntFlanks@12420.12437.18|ATGTCGGCTCTTCCTATC</td><td>>hg38_21_+_8433172_8446622_RNA45SN1_45S_50ntFlanks</td><td>12420</td><td>12437</td><td>18</td><td>ATGTCGGCTCTTCCTATC</td><td>rRF</td><td>FALSE</td><td>FALSE</td><td>TRUE</td></tr>
<tr><td>>hg38_21_+_8433172_8446622_RNA45SN1_45S_50ntFlanks@12436.12455.20|TCATTGTGAAGCAGAATTCA</td><td>>hg38_21_+_8433172_8446622_RNA45SN1_45S_50ntFlanks</td><td>12436</td><td>12455</td><td>20</td><td>TCATTGTGAAGCAGAATTCA</td><td>rRF</td><td>FALSE</td><td>FALSE</td><td>TRUE</td></tr>
<tr><td>>hg38_21_+_8433172_8446622_RNA45SN1_45S_50ntFlanks@9567.9596.30|ACGTGCAAATCGGTCGTCCGACCTGGGTAT</td><td>>hg38_21_+_8433172_8446622_RNA45SN1_45S_50ntFlanks</td><td>9567</td><td>9596</td><td>30</td><td>ACGTGCAAATCGGTCGTCCGACCTGGGTAT</td><td>rRF</td><td>FALSE</td><td>FALSE</td><td>TRUE</td></tr>
<tr><td>>hg38_21_+_8433172_8446622_RNA45SN1_45S_50ntFlanks@11873.11893.21|AAGAAGACCCTGTTGAGCTTG</td><td>>hg38_21_+_8433172_8446622_RNA45SN1_45S_50ntFlanks</td><td>11873</td><td>11893</td><td>21</td><td>AAGAAGACCCTGTTGAGCTTG</td><td>rRF</td><td>FALSE</td><td>FALSE</td><td>TRUE</td></tr>
<tr><td>>hg38_21_+_8433172_8446622_RNA45SN1_45S_50ntFlanks@10410.10430.21|TCAGTCGGTCCTGAGAGATGG</td><td>>hg38_21_+_8433172_8446622_RNA45SN1_45S_50ntFlanks</td><td>10410</td><td>10430</td><td>21</td><td>TCAGTCGGTCCTGAGAGATGG</td><td>rRF</td><td>FALSE</td><td>FALSE</td><td>TRUE</td></tr>
<tr><td>>hg38_21_+_8433172_8446622_RNA45SN1_45S_50ntFlanks@12655.12674.20|ATGACTGAACGCCTCTAAGT</td><td>>hg38_21_+_8433172_8446622_RNA45SN1_45S_50ntFlanks</td><td>12655</td><td>12674</td><td>20</td><td>ATGACTGAACGCCTCTAAGT</td><td>rRF</td><td>FALSE</td><td>FALSE</td><td>TRUE</td></tr>
<tr><td>>hg38_21_+_8433172_8446622_RNA45SN1_45S_50ntFlanks@9534.9551.18|AGGAAACTCTGGTGGAGG</td><td>>hg38_21_+_8433172_8446622_RNA45SN1_45S_50ntFlanks</td><td>9534</td><td>9551</td><td>18</td><td>AGGAAACTCTGGTGGAGG</td><td>rRF</td><td>FALSE</td><td>FALSE</td><td>TRUE</td></tr>
<tr><td>>hg38_21_+_8433172_8446622_RNA45SN1_45S_50ntFlanks@11572.11603.32|CCGACTTAGAACTGGTGCGGACCAGGGGAATC</td><td>>hg38_21_+_8433172_8446622_RNA45SN1_45S_50ntFlanks</td><td>11572</td><td>11603</td><td>32</td><td>CCGACTTAGAACTGGTGCGGACCAGGGGAATC</td><td>rRF</td><td>FALSE</td><td>FALSE</td><td>TRUE</td></tr>
<tr><td>>hg38_21_+_8433172_8446622_RNA45SN1_45S_50ntFlanks@6719.6740.22|TGAATTGCAGGACACATTGATC</td><td>>hg38_21_+_8433172_8446622_RNA45SN1_45S_50ntFlanks</td><td>6719</td><td>6740</td><td>22</td><td>TGAATTGCAGGACACATTGATC</td><td>rRF</td><td>FALSE</td><td>FALSE</td><td>TRUE</td></tr>
<tr><td>>hg38_21_+_8433172_8446622_RNA45SN1_45S_50ntFlanks@11575.11603.29|ACTTAGAACTGGTGCGGACCAGGGGAATC</td><td>>hg38_21_+_8433172_8446622_RNA45SN1_45S_50ntFlanks</td><td>11575</td><td>11603</td><td>29</td><td>ACTTAGAACTGGTGCGGACCAGGGGAATC</td><td>rRF</td><td>FALSE</td><td>FALSE</td><td>TRUE</td></tr>
<tr><td>>am_trna111_HisGTG_1_-_147774845_147774916@-1G.29.30,trna118_HisGTG_1_-_145396881_145396952@-1G.29.30,trna16_HisGTG_1_+_146544773_146544844@-1G.29.30,trna1_HisGTG_15_+_45493349_45493420@-1G.29.30,trna21_HisGTG_1_+_147753471_147753542@-1G.29.30,trna33_HisGTG_6_+_27125906_27125977@-1G.29.30,trna7_HisGTG_9_-_14433938_14434009@-1G.29.30,trna8_HisGTG_15_-_45492611_45492682@-1G.29.30,trna9_HisGTG_15_-_45490804_45490875@-1G.29.30|GGCCGTGATCGTATAGTGGTTAGTACTCTG</td><td>>am_trna111_HisGTG_1_-_147774845_147774916</td><td>-1G</td><td>29</td><td>30</td><td>GGCCGTGATCGTATAGTGGTTAGTACTCTG</td><td>tRF</td><td>FALSE</td><td>FALSE</td><td>TRUE</td></tr>
<tr><td>>hg38_21_+_8433172_8446622_RNA45SN1_45S_50ntFlanks@12626.12653.28|AATGGGGCGAAGCTACCATCTGTGGGAT</td><td>>hg38_21_+_8433172_8446622_RNA45SN1_45S_50ntFlanks</td><td>12626</td><td>12653</td><td>28</td><td>AATGGGGCGAAGCTACCATCTGTGGGAT</td><td>rRF</td><td>FALSE</td><td>FALSE</td><td>TRUE</td></tr>
<tr><td>>hg38_21_+_8433172_8446622_RNA45SN1_45S_50ntFlanks@12625.12654.30|CAATGGGGCGAAGCTACCATCTGTGGGATT</td><td>>hg38_21_+_8433172_8446622_RNA45SN1_45S_50ntFlanks</td><td>12625</td><td>12654</td><td>30</td><td>CAATGGGGCGAAGCTACCATCTGTGGGATT</td><td>rRF</td><td>FALSE</td><td>FALSE</td><td>TRUE</td></tr>
<tr><td>>hg38_21_+_8433172_8446622_RNA45SN1_45S_50ntFlanks@12420.12446.27|ATGTCGGCTCTTCCTATCATTGTGAAG</td><td>>hg38_21_+_8433172_8446622_RNA45SN1_45S_50ntFlanks</td><td>12420</td><td>12446</td><td>27</td><td>ATGTCGGCTCTTCCTATCATTGTGAAG</td><td>rRF</td><td>FALSE</td><td>FALSE</td><td>TRUE</td></tr>
<tr><td>>hg38_21_+_8433172_8446622_RNA45SN1_45S_50ntFlanks@6719.6737.19|TGAATTGCAGGACACATTG</td><td>>hg38_21_+_8433172_8446622_RNA45SN1_45S_50ntFlanks</td><td>6719</td><td>6737</td><td>19</td><td>TGAATTGCAGGACACATTG</td><td>rRF</td><td>FALSE</td><td>FALSE</td><td>TRUE</td></tr>
<tr><td>>hg38_21_+_8433172_8446622_RNA45SN1_45S_50ntFlanks@12456.12485.30|CCAAGCGTTGGATTGTTCACCCACTAATAG</td><td>>hg38_21_+_8433172_8446622_RNA45SN1_45S_50ntFlanks</td><td>12456</td><td>12485</td><td>30</td><td>CCAAGCGTTGGATTGTTCACCCACTAATAG</td><td>rRF</td><td>FALSE</td><td>FALSE</td><td>TRUE</td></tr>
<tr><td>>trna2_GlyGCC_21_-_18827107_18827177@1.30.30,trna35_GlyGCC_1_+_161413094_161413164@1.30.30,trna37_GlyGCC_1_+_161420467_161420537@1.30.30,trna39_GlyGCC_1_+_161427898_161427968@1.30.30,trna41_GlyGCC_1_+_161435258_161435328@1.30.30|GCATGGGTGGTTCAGTGGTAGAATTCTCGC</td><td>>trna2_GlyGCC_21_-_18827107_18827177</td><td>1</td><td>30</td><td>30</td><td>GCATGGGTGGTTCAGTGGTAGAATTCTCGC</td><td>tRF</td><td>FALSE</td><td>FALSE</td><td>TRUE</td></tr>
<tr><td>>hg38_21_+_8433172_8446622_RNA45SN1_45S_50ntFlanks@11581.11603.23|AACTGGTGCGGACCAGGGGAATC</td><td>>hg38_21_+_8433172_8446622_RNA45SN1_45S_50ntFlanks</td><td>11581</td><td>11603</td><td>23</td><td>AACTGGTGCGGACCAGGGGAATC</td><td>rRF</td><td>FALSE</td><td>FALSE</td><td>TRUE</td></tr>
<tr><td>>hg38_21_+_8433172_8446622_RNA45SN1_45S_50ntFlanks@7980.8006.27|CCTCAGATCAGACGTGGCGACCCGCTG</td><td>>hg38_21_+_8433172_8446622_RNA45SN1_45S_50ntFlanks</td><td>7980</td><td>8006</td><td>27</td><td>CCTCAGATCAGACGTGGCGACCCGCTG</td><td>rRF</td><td>FALSE</td><td>FALSE</td><td>TRUE</td></tr>
<tr><td>>hg38_21_+_8433172_8446622_RNA45SN1_45S_50ntFlanks@12937.12956.20|AACCATTCGTAGACGACCTG</td><td>>hg38_21_+_8433172_8446622_RNA45SN1_45S_50ntFlanks</td><td>12937</td><td>12956</td><td>20</td><td>AACCATTCGTAGACGACCTG</td><td>rRF</td><td>FALSE</td><td>FALSE</td><td>TRUE</td></tr>
<tr><td>>hg38_21_+_8433172_8446622_RNA45SN1_45S_50ntFlanks@9568.9596.29|CGTGCAAATCGGTCGTCCGACCTGGGTAT</td><td>>hg38_21_+_8433172_8446622_RNA45SN1_45S_50ntFlanks</td><td>9568</td><td>9596</td><td>29</td><td>CGTGCAAATCGGTCGTCCGACCTGGGTAT</td><td>rRF</td><td>FALSE</td><td>FALSE</td><td>TRUE</td></tr>
<tr><td>>hg38_21_+_8433172_8446622_RNA45SN1_45S_50ntFlanks@5446.5466.21|CGGATCGGCCCCGCCGGGGTC</td><td>>hg38_21_+_8433172_8446622_RNA45SN1_45S_50ntFlanks</td><td>5446</td><td>5466</td><td>21</td><td>CGGATCGGCCCCGCCGGGGTC</td><td>rRF</td><td>FALSE</td><td>FALSE</td><td>TRUE</td></tr>
<tr><td>>hg38_21_+_8433172_8446622_RNA45SN1_45S_50ntFlanks@12624.12654.31|CCAATGGGGCGAAGCTACCATCTGTGGGATT</td><td>>hg38_21_+_8433172_8446622_RNA45SN1_45S_50ntFlanks</td><td>12624</td><td>12654</td><td>31</td><td>CCAATGGGGCGAAGCTACCATCTGTGGGATT</td><td>rRF</td><td>FALSE</td><td>FALSE</td><td>TRUE</td></tr>
<tr><td>>hg38_21_+_8433172_8446622_RNA45SN1_45S_50ntFlanks@12418.12443.26|CGATGTCGGCTCTTCCTATCATTGTG</td><td>>hg38_21_+_8433172_8446622_RNA45SN1_45S_50ntFlanks</td><td>12418</td><td>12443</td><td>26</td><td>CGATGTCGGCTCTTCCTATCATTGTG</td><td>rRF</td><td>FALSE</td><td>FALSE</td><td>TRUE</td></tr>
<tr><td>>hg38_21_+_8433172_8446622_RNA45SN1_45S_50ntFlanks@11323.11342.20|CGTCTCGTCGCGCGCGCGTC</td><td>>hg38_21_+_8433172_8446622_RNA45SN1_45S_50ntFlanks</td><td>11323</td><td>11342</td><td>20</td><td>CGTCTCGTCGCGCGCGCGTC</td><td>rRF</td><td>FALSE</td><td>FALSE</td><td>TRUE</td></tr>
<tr><td>>hg38_21_+_8433172_8446622_RNA45SN1_45S_50ntFlanks@11641.11665.25|CGGCGGGTGTTGACGCGATGTGATT</td><td>>hg38_21_+_8433172_8446622_RNA45SN1_45S_50ntFlanks</td><td>11641</td><td>11665</td><td>25</td><td>CGGCGGGTGTTGACGCGATGTGATT</td><td>rRF</td><td>FALSE</td><td>FALSE</td><td>TRUE</td></tr>
<tr><td>>hg38_21_+_8433172_8446622_RNA45SN1_45S_50ntFlanks@11661.11681.21|TGATTTCTGCCCAGTGCTCTG</td><td>>hg38_21_+_8433172_8446622_RNA45SN1_45S_50ntFlanks</td><td>11661</td><td>11681</td><td>21</td><td>TGATTTCTGCCCAGTGCTCTG</td><td>rRF</td><td>FALSE</td><td>FALSE</td><td>TRUE</td></tr>
<tr><td>>hg38_21_+_8433172_8446622_RNA45SN1_45S_50ntFlanks@12635.12653.19|AAGCTACCATCTGTGGGAT</td><td>>hg38_21_+_8433172_8446622_RNA45SN1_45S_50ntFlanks</td><td>12635</td><td>12653</td><td>19</td><td>AAGCTACCATCTGTGGGAT</td><td>rRF</td><td>FALSE</td><td>FALSE</td><td>TRUE</td></tr>
<tr><td>>hg38_21_+_8433172_8446622_RNA45SN1_45S_50ntFlanks@4760.4780.21|TCAGATACCGTCGTAGTTCCG</td><td>>hg38_21_+_8433172_8446622_RNA45SN1_45S_50ntFlanks</td><td>4760</td><td>4780</td><td>21</td><td>TCAGATACCGTCGTAGTTCCG</td><td>rRF</td><td>FALSE</td><td>FALSE</td><td>TRUE</td></tr>
<tr><td>>hg38_21_+_8433172_8446622_RNA45SN1_45S_50ntFlanks@9568.9597.30|CGTGCAAATCGGTCGTCCGACCTGGGTATA</td><td>>hg38_21_+_8433172_8446622_RNA45SN1_45S_50ntFlanks</td><td>9568</td><td>9597</td><td>30</td><td>CGTGCAAATCGGTCGTCCGACCTGGGTATA</td><td>rRF</td><td>FALSE</td><td>FALSE</td><td>TRUE</td></tr>
<tr><td>>hg38_21_+_8433172_8446622_RNA45SN1_45S_50ntFlanks@6652.6681.30|GACTCTTAGCGGTGGATCACTCGGCTCGTG</td><td>>hg38_21_+_8433172_8446622_RNA45SN1_45S_50ntFlanks</td><td>6652</td><td>6681</td><td>30</td><td>GACTCTTAGCGGTGGATCACTCGGCTCGTG</td><td>rRF</td><td>FALSE</td><td>FALSE</td><td>TRUE</td></tr>
</table>

| | | | | | | | | | |
|---|---|---|---|---|---|---|---|---|---|
| >hg38_21_+_8433172_8446622_RNA45SN1_45S_50ntFlanks@12627.12653.27\|ATGGGGCGAAGCTACCATCTGTGGGAT | >hg38_21_+_8433172_8446622_RNA45SN1_45S_50ntFlanks | 12627 | 12653 | 27 | ATGGGGCGAAGCTACCATCTGTGGGAT | rRF | FALSE | FALSE | TRUE |
| >hg38_21_+_8433172_8446622_RNA45SN1_45S_50ntFlanks@10841.10862.22\|CGTAACTTCGGGATAAGGATTG | >hg38_21_+_8433172_8446622_RNA45SN1_45S_50ntFlanks | 10841 | 10862 | 22 | CGTAACTTCGGGATAAGGATTG | rRF | FALSE | FALSE | TRUE |
| >hg38_21_+_8433172_8446622_RNA45SN1_45S_50ntFlanks@12418.12437.20\|CGATGTCGGCTCTTCCTATC | >hg38_21_+_8433172_8446622_RNA45SN1_45S_50ntFlanks | 12418 | 12437 | 20 | CGATGTCGGCTCTTCCTATC | rRF | FALSE | FALSE | TRUE |
| >hg38_21_+_8433172_8446622_RNA45SN1_45S_50ntFlanks@6651.6679.29\|CGACTCTTAGCGGTGGATCACTCGGCTCG | >hg38_21_+_8433172_8446622_RNA45SN1_45S_50ntFlanks | 6651 | 6679 | 29 | CGACTCTTAGCGGTGGATCACTCGGCTCG | rRF | FALSE | FALSE | TRUE |
| >hg38_21_+_8433172_8446622_RNA45SN1_45S_50ntFlanks@9567.9597.31\|ACGTGCAAATCGGTCGTCCGACCTGGGTATA | >hg38_21_+_8433172_8446622_RNA45SN1_45S_50ntFlanks | 9567 | 9597 | 31 | ACGTGCAAATCGGTCGTCCGACCTGGGTATA | rRF | FALSE | FALSE | TRUE |
| >hg38_21_+_8433172_8446622_RNA45SN1_45S_50ntFlanks@6745.6770.26\|ACACTTCGAACGCACTTGCGGCCCCG | >hg38_21_+_8433172_8446622_RNA45SN1_45S_50ntFlanks | 6745 | 6770 | 26 | ACACTTCGAACGCACTTGCGGCCCCG | rRF | FALSE | FALSE | TRUE |
| >hg38_21_+_8433172_8446622_RNA45SN1_45S_50ntFlanks@11572.11602.31\|CCGACTTAGAACTGGTGCGGACCAGGGGAAT | >hg38_21_+_8433172_8446622_RNA45SN1_45S_50ntFlanks | 11572 | 11602 | 31 | CCGACTTAGAACTGGTGCGGACCAGGGGAAT | rRF | FALSE | FALSE | TRUE |
| >hg38_21_+_8433172_8446622_RNA45SN1_45S_50ntFlanks@11317.11344.28\|ACCCCACGTCTCGTCGCGCGCGCGTCCG | >hg38_21_+_8433172_8446622_RNA45SN1_45S_50ntFlanks | 11317 | 11344 | 28 | ACCCCACGTCTCGTCGCGCGCGCGTCCG | rRF | FALSE | FALSE | TRUE |
| >hg38_21_+_8433172_8446622_RNA45SN1_45S_50ntFlanks@10393.10410.18\|CAGCAGTTGAACATGGGT | >hg38_21_+_8433172_8446622_RNA45SN1_45S_50ntFlanks | 10393 | 10410 | 18 | CAGCAGTTGAACATGGGT | rRF | FALSE | FALSE | TRUE |
| >hg38_21_+_8433172_8446622_RNA45SN1_45S_50ntFlanks@11321.11344.24\|CACGTCTCGTCGCGCGCGCGTCCG | >hg38_21_+_8433172_8446622_RNA45SN1_45S_50ntFlanks | 11321 | 11344 | 24 | CACGTCTCGTCGCGCGCGCGTCCG | rRF | FALSE | FALSE | TRUE |
| >hg38_21_+_8433172_8446622_RNA45SN1_45S_50ntFlanks@11729.11747.19\|TAACTATGACTCTCTTAAG | >hg38_21_+_8433172_8446622_RNA45SN1_45S_50ntFlanks | 11729 | 11747 | 19 | TAACTATGACTCTCTTAAG | rRF | FALSE | FALSE | TRUE |
| >hg38_21_+_8433172_8446622_RNA45SN1_45S_50ntFlanks@12457.12478.22\|CAAGCGTTGGATTGTTCACCCA | >hg38_21_+_8433172_8446622_RNA45SN1_45S_50ntFlanks | 12457 | 12478 | 22 | CAAGCGTTGGATTGTTCACCCA | rRF | FALSE | FALSE | TRUE |
| >am_trna116_GluCTC_1_-_145399233_145399304@1.27.27,trna59_GluCTC_1_+_249168447_249168518@1.27.27,trna71_GluCTC_1_-_161439189_161439260@1.27.27,trna74_GluCTC_1_-_161431809_161431880@1.27.27,trna77_GluCTC_1_-_161424398_161424469@1.27.27,trna77_GluCTC_6_+_28949976_28950047@1.27.27,trna80_GluCTC_1_-_161417018_161417089@1.27.27,trna87_GluCTC_6_-_126101393_126101464@1.27.27\|TCCCTGGTGGTCTAGTGGTTAGGATTC | >am_trna116_GluCTC_1_-_145399233_145399304 | 1 | 27 | 27 | TCCCTGGTGGTCTAGTGGTTAGGATTC | tRF | FALSE | FALSE | TRUE |
| >hg38_21_+_8433172_8446622_RNA45SN1_45S_50ntFlanks@12405.12426.22\|CTTTTTGATCCTTCGATGTCGG | >hg38_21_+_8433172_8446622_RNA45SN1_45S_50ntFlanks | 12405 | 12426 | 22 | CTTTTTGATCCTTCGATGTCGG | rRF | FALSE | FALSE | TRUE |
| >hg38_21_+_8433172_8446622_RNA45SN1_45S_50ntFlanks@11575.11592.18\|ACTTAGAACTGGTGCGGA | >hg38_21_+_8433172_8446622_RNA45SN1_45S_50ntFlanks | 11575 | 11592 | 18 | ACTTAGAACTGGTGCGGA | rRF | FALSE | FALSE | TRUE |
| >hg38_21_+_8433172_8446622_RNA45SN1_45S_50ntFlanks@12654.12675.22\|TATGACTGAACGCCTCTAAGTC | >hg38_21_+_8433172_8446622_RNA45SN1_45S_50ntFlanks | 12654 | 12675 | 22 | TATGACTGAACGCCTCTAAGTC | rRF | FALSE | FALSE | TRUE |
| >hg38_21_+_8433172_8446622_RNA45SN1_45S_50ntFlanks@11323.11343.21\|CGTCTCGTCGCGCGCGCGTCC | >hg38_21_+_8433172_8446622_RNA45SN1_45S_50ntFlanks | 11323 | 11343 | 21 | CGTCTCGTCGCGCGCGCGTCC | rRF | FALSE | FALSE | TRUE |
| >hg38_21_+_8433172_8446622_RNA45SN1_45S_50ntFlanks@9534.9552.19\|AGGAAACTCTGGTGGAGGT | >hg38_21_+_8433172_8446622_RNA45SN1_45S_50ntFlanks | 9534 | 9552 | 19 | AGGAAACTCTGGTGGAGGT | rRF | FALSE | FALSE | TRUE |
| >hg38_21_+_8433172_8446622_RNA45SN1_45S_50ntFlanks@10349.10368.20\|AACGAGAACTTTGAAGGCCG | >hg38_21_+_8433172_8446622_RNA45SN1_45S_50ntFlanks | 10349 | 10368 | 20 | AACGAGAACTTTGAAGGCCG | rRF | FALSE | FALSE | TRUE |
| >trna10_LysCTT_16_+_3241501_3241573@1.28.28,trna119_LysCTT_1_-_145395522_145395594@1.28.28,trna11_LysCTT_5_-_180648979_180649051@1.28.28,trna13_LysCTT_14_-_58706613_58706685@1.28.28,trna13_LysCTT_6_+_26556774_26556846@1.28.28,trna2_LysCTT_15_+_79152904_79152976@1.28.28,trna32_LysCTT_16_-_3207406_3207478@1.28.28,trna7_LysCTT_16_+_3225692_3225764@1.28.28,trna9_LysCTT_5_+_180634755_180634827@1.28.28\|GCCCGGCTAGCTCAGTCGGTAGAGCATG | >trna10_LysCTT_16_+_3241501_3241573 | 1 | 28 | 28 | GCCCGGCTAGCTCAGTCGGTAGAGCATG | tRF | FALSE | FALSE | TRUE |
| >hg38_21_+_8433172_8446622_RNA45SN1_45S_50ntFlanks@12163.12182.20\|ACTGGGGCGGTACACCTGTC | >hg38_21_+_8433172_8446622_RNA45SN1_45S_50ntFlanks | 12163 | 12182 | 20 | ACTGGGGCGGTACACCTGTC | rRF | FALSE | FALSE | TRUE |
| >hg38_21_+_8433172_8446622_RNA45SN1_45S_50ntFlanks@10794.10818.25\|CTGGCATGTTGGAACAATGTAGGTA | >hg38_21_+_8433172_8446622_RNA45SN1_45S_50ntFlanks | 10794 | 10818 | 25 | CTGGCATGTTGGAACAATGTAGGTA | rRF | FALSE | FALSE | TRUE |
| >hg38_21_+_8433172_8446622_RNA45SN1_45S_50ntFlanks@6652.6689.38\|GACTCTTAGCGGTGGATCACTCGGCTCGTGCGTCGATG | >hg38_21_+_8433172_8446622_RNA45SN1_45S_50ntFlanks | 6652 | 6689 | 38 | GACTCTTAGCGGTGGATCACTCGGCTCGTGCGTCGATG | rRF | FALSE | FALSE | TRUE |
| >multi-am_trnaMT_ValTAC_MT_+_1602_1670@50.68.19\|ACTTAACTTGACCGCTCTG | >multi-am_trnaMT_ValTAC_MT_+_1602_1670 | 50 | 68 | 19 | ACTTAACTTGACCGCTCTG | tRF | FALSE | FALSE | TRUE |
| >hg38_21_+_8433172_8446622_RNA45SN1_45S_50ntFlanks@11785.11803.19\|TGAATGGATGAACGAGATT | >hg38_21_+_8433172_8446622_RNA45SN1_45S_50ntFlanks | 11785 | 11803 | 19 | TGAATGGATGAACGAGATT | rRF | FALSE | FALSE | TRUE |
| >hg38_21_+_8433172_8446622_RNA45SN1_45S_50ntFlanks@10687.10706.20\|CGGCGTCCGGTGAGCTCTCG | >hg38_21_+_8433172_8446622_RNA45SN1_45S_50ntFlanks | 10687 | 10706 | 20 | CGGCGTCCGGTGAGCTCTCG | rRF | FALSE | FALSE | TRUE |
| >hg38_21_+_8433172_8446622_RNA45SN1_45S_50ntFlanks@7986.8006.21\|ATCAGACGTGGCGACCCGCTG | >hg38_21_+_8433172_8446622_RNA45SN1_45S_50ntFlanks | 7986 | 8006 | 21 | ATCAGACGTGGCGACCCGCTG | rRF | FALSE | FALSE | TRUE |
| >hg38_21_+_8433172_8446622_RNA45SN1_45S_50ntFlanks@9572.9595.24\|CAAATCGGTCGTCCGACCTGGGTA | >hg38_21_+_8433172_8446622_RNA45SN1_45S_50ntFlanks | 9572 | 9595 | 24 | CAAATCGGTCGTCCGACCTGGGTA | rRF | FALSE | FALSE | TRUE |
| >hg38_21_+_8433172_8446622_RNA45SN1_45S_50ntFlanks@10819.10840.22\|AGGGAAGTCGGCAAGCCGGATC | >hg38_21_+_8433172_8446622_RNA45SN1_45S_50ntFlanks | 10819 | 10840 | 22 | AGGGAAGTCGGCAAGCCGGATC | rRF | FALSE | FALSE | TRUE |
| >hg38_21_+_8433172_8446622_RNA45SN1_45S_50ntFlanks@13002.13028.27\|CGATCTATTGAAAGTCAGCCCTCGACA | >hg38_21_+_8433172_8446622_RNA45SN1_45S_50ntFlanks | 13002 | 13028 | 27 | CGATCTATTGAAAGTCAGCCCTCGACA | rRF | FALSE | FALSE | TRUE |
| >hg38_21_+_8433172_8446622_RNA45SN1_45S_50ntFlanks@10797.10818.22\|GCATGTTGGAACAATGTAGGTA | >hg38_21_+_8433172_8446622_RNA45SN1_45S_50ntFlanks | 10797 | 10818 | 22 | GCATGTTGGAACAATGTAGGTA | rRF | FALSE | FALSE | TRUE |
| >hg38_21_+_8433172_8446622_RNA45SN1_45S_50ntFlanks@7981.8006.26\|CTCAGATCAGACGTGGCGACCCGCTG | >hg38_21_+_8433172_8446622_RNA45SN1_45S_50ntFlanks | 7981 | 8006 | 26 | CTCAGATCAGACGTGGCGACCCGCTG | rRF | FALSE | FALSE | TRUE |
| >hg38_21_+_8433172_8446622_RNA45SN1_45S_50ntFlanks@6745.6771.27\|ACACTTCGAACGCACTTGCGGCCCCGG | >hg38_21_+_8433172_8446622_RNA45SN1_45S_50ntFlanks | 6745 | 6771 | 27 | ACACTTCGAACGCACTTGCGGCCCCGG | rRF | FALSE | FALSE | TRUE |
| >hg38_21_+_8433172_8446622_RNA45SN1_45S_50ntFlanks@10411.10431.21\|CAGTCGGTCCTGAGAGATGGG | >hg38_21_+_8433172_8446622_RNA45SN1_45S_50ntFlanks | 10411 | 10431 | 21 | CAGTCGGTCCTGAGAGATGGG | rRF | FALSE | FALSE | TRUE |
| >hg38_21_+_8433172_8446622_RNA45SN1_45S_50ntFlanks@10402.10429.28\|AACATGGGTCAGTCGGTCCTGAGAGATG | >hg38_21_+_8433172_8446622_RNA45SN1_45S_50ntFlanks | 10402 | 10429 | 28 | AACATGGGTCAGTCGGTCCTGAGAGATG | rRF | FALSE | FALSE | TRUE |
| >hg38_21_+_8433172_8446622_RNA45SN1_45S_50ntFlanks@11208.11225.18\|GCGGCGACTCTGGACGCG | >hg38_21_+_8433172_8446622_RNA45SN1_45S_50ntFlanks | 11208 | 11225 | 18 | GCGGCGACTCTGGACGCG | rRF | FALSE | FALSE | TRUE |
| >hg38_21_+_8433172_8446622_RNA45SN1_45S_50ntFlanks@12417.12435.19\|TCGATGTCGGCTCTTCCTA | >hg38_21_+_8433172_8446622_RNA45SN1_45S_50ntFlanks | 12417 | 12435 | 19 | TCGATGTCGGCTCTTCCTA | rRF | FALSE | FALSE | TRUE |
| >hg38_21_+_8433172_8446622_RNA45SN1_45S_50ntFlanks@7991.8012.22\|ACGTGGCGACCCGCTGAATTTA | >hg38_21_+_8433172_8446622_RNA45SN1_45S_50ntFlanks | 7991 | 8012 | 22 | ACGTGGCGACCCGCTGAATTTA | rRF | FALSE | FALSE | TRUE |

| | | | | | | | | | |
|---|---|---|---|---|---|---|---|---|---|
| >hg38_21_+_8433172_8446622_RNA45SN1_45S_50ntFlanks@12635.12661.27\|AAGCTACCATCTGTGGGATTATGACTG | >hg38_21_+_8433172_8446622_RNA45SN1_45S_50ntFlanks | 12635 | 12661 | 27 | AAGCTACCATCTGTGGGATTATGACTG | rRF | FALSE | FALSE | TRUE |
| >hg38_21_+_8433172_8446622_RNA45SN1_45S_50ntFlanks@10680.10712.33\|GTTCCGGCGGCGTCCGGTGAGCTCTCGCTGGCC | >hg38_21_+_8433172_8446622_RNA45SN1_45S_50ntFlanks | 10680 | 10712 | 33 | GTTCCGGCGGCGTCCGGTGAGCTCTCGCTGGCC | rRF | FALSE | FALSE | TRUE |
| >hg38_21_+_8433172_8446622_RNA45SN1_45S_50ntFlanks@9573.9597.25\|AAATCGGTCGTCCGACCTGGGTATA | >hg38_21_+_8433172_8446622_RNA45SN1_45S_50ntFlanks | 9573 | 9597 | 25 | AAATCGGTCGTCCGACCTGGGTATA | rRF | FALSE | FALSE | TRUE |
| >hg38_21_+_8433172_8446622_RNA45SN1_45S_50ntFlanks@11783.11806.24\|CATGAATGGATGAACGAGATTCCC | >hg38_21_+_8433172_8446622_RNA45SN1_45S_50ntFlanks | 11783 | 11806 | 24 | CATGAATGGATGAACGAGATTCCC | rRF | FALSE | FALSE | TRUE |
| >hg38_21_+_8433172_8446622_RNA45SN1_45S_50ntFlanks@12405.12435.31\|CTTTTTGATCCTTCGATGTCGGCTCTTCCTA | >hg38_21_+_8433172_8446622_RNA45SN1_45S_50ntFlanks | 12405 | 12435 | 31 | CTTTTTGATCCTTCGATGTCGGCTCTTCCTA | rRF | FALSE | FALSE | TRUE |
| >hg38_21_+_8433172_8446622_RNA45SN1_45S_50ntFlanks@4722.4758.37\|TCATTAATCAAGAACGAAAGTCGGAGGTTCGAAGACG | >hg38_21_+_8433172_8446622_RNA45SN1_45S_50ntFlanks | 4722 | 4758 | 37 | TCATTAATCAAGAACGAAAGTCGGAGGTTCGAAGACG | rRF | FALSE | FALSE | TRUE |
| >hg38_21_+_8433172_8446622_RNA45SN1_45S_50ntFlanks@10687.10709.23\|CGGCGTCCGGTGAGCTCTCGCTG | >hg38_21_+_8433172_8446622_RNA45SN1_45S_50ntFlanks | 10687 | 10709 | 23 | CGGCGTCCGGTGAGCTCTCGCTG | rRF | FALSE | FALSE | TRUE |
| >hg38_21_+_8433172_8446622_RNA45SN1_45S_50ntFlanks@12159.12179.21\|TTTGACTGGGGCGGTACACCT | >hg38_21_+_8433172_8446622_RNA45SN1_45S_50ntFlanks | 12159 | 12179 | 21 | TTTGACTGGGGCGGTACACCT | rRF | FALSE | FALSE | TRUE |
| >hg38_21_+_8433172_8446622_RNA45SN1_45S_50ntFlanks@11729.11765.37\|TAACTATGACTCTCTTAAGGTAGCCAAATGCCTCGTC | >hg38_21_+_8433172_8446622_RNA45SN1_45S_50ntFlanks | 11729 | 11765 | 37 | TAACTATGACTCTCTTAAGGTAGCCAAATGCCTCGTC | rRF | FALSE | FALSE | TRUE |
| >hg38_21_+_8433172_8446622_RNA45SN1_45S_50ntFlanks@11781.11804.24\|CGCATGAATGGATGAACGAGATTC | >hg38_21_+_8433172_8446622_RNA45SN1_45S_50ntFlanks | 11781 | 11804 | 24 | CGCATGAATGGATGAACGAGATTC | rRF | FALSE | FALSE | TRUE |
| >hg38_21_+_8433172_8446622_RNA45SN1_45S_50ntFlanks@10389.10407.19\|TGAACAGCAGTTGAACATG | >hg38_21_+_8433172_8446622_RNA45SN1_45S_50ntFlanks | 10389 | 10407 | 19 | TGAACAGCAGTTGAACATG | rRF | FALSE | FALSE | TRUE |
| >hg38_21_+_8433172_8446622_RNA45SN1_45S_50ntFlanks@12405.12429.25\|CTTTTTGATCCTTCGATGTCGGCTC | >hg38_21_+_8433172_8446622_RNA45SN1_45S_50ntFlanks | 12405 | 12429 | 25 | CTTTTTGATCCTTCGATGTCGGCTC | rRF | FALSE | FALSE | TRUE |
| >hg38_21_+_8433172_8446622_RNA45SN1_45S_50ntFlanks@11576.11603.28\|CTTAGAACTGGTGCGGACCAGGGGAATC | >hg38_21_+_8433172_8446622_RNA45SN1_45S_50ntFlanks | 11576 | 11603 | 28 | CTTAGAACTGGTGCGGACCAGGGGAATC | rRF | FALSE | FALSE | TRUE |
| >hg38_21_+_8433172_8446622_RNA45SN1_45S_50ntFlanks@10841.10864.24\|CGTAACTTCGGGATAAGGATTGGC | >hg38_21_+_8433172_8446622_RNA45SN1_45S_50ntFlanks | 10841 | 10864 | 24 | CGTAACTTCGGGATAAGGATTGGC | rRF | FALSE | FALSE | TRUE |
| >hg38_21_+_8433172_8446622_RNA45SN1_45S_50ntFlanks@4723.4758.36\|CATTAATCAAGAACGAAAGTCGGAGGTTCGAAGACG | >hg38_21_+_8433172_8446622_RNA45SN1_45S_50ntFlanks | 4723 | 4758 | 36 | CATTAATCAAGAACGAAAGTCGGAGGTTCGAAGACG | rRF | FALSE | FALSE | TRUE |
| >hg38_21_+_8433172_8446622_RNA45SN1_45S_50ntFlanks@11729.11748.20\|TAACTATGACTCTCTTAAGG | >hg38_21_+_8433172_8446622_RNA45SN1_45S_50ntFlanks | 11729 | 11748 | 20 | TAACTATGACTCTCTTAAGG | rRF | FALSE | FALSE | TRUE |
| >hg38_21_+_8433172_8446622_RNA45SN1_45S_50ntFlanks@11580.11603.24\|GAACTGGTGCGGACCAGGGGAATC | >hg38_21_+_8433172_8446622_RNA45SN1_45S_50ntFlanks | 11580 | 11603 | 24 | GAACTGGTGCGGACCAGGGGAATC | rRF | FALSE | FALSE | TRUE |
| >hg38_21_+_8433172_8446622_RNA45SN1_45S_50ntFlanks@12991.13015.25\|CTCCCTCGCTGCGATCTATTGAAAG | >hg38_21_+_8433172_8446622_RNA45SN1_45S_50ntFlanks | 12991 | 13015 | 25 | CTCCCTCGCTGCGATCTATTGAAAG | rRF | FALSE | FALSE | TRUE |
| >hg38_21_+_8433172_8446622_RNA45SN1_45S_50ntFlanks@11318.11344.27\|CCCCACGTCTCGTCGCGCGCGCGTCCG | >hg38_21_+_8433172_8446622_RNA45SN1_45S_50ntFlanks | 11318 | 11344 | 27 | CCCCACGTCTCGTCGCGCGCGCGTCCG | rRF | FALSE | FALSE | TRUE |
| >hg38_21_+_8433172_8446622_RNA45SN1_45S_50ntFlanks@12457.12477.21\|CAAGCGTTGGATTGTTCACCC | >hg38_21_+_8433172_8446622_RNA45SN1_45S_50ntFlanks | 12457 | 12477 | 21 | CAAGCGTTGGATTGTTCACCC | rRF | FALSE | FALSE | TRUE |
| >hg38_21_+_8433172_8446622_RNA45SN1_45S_50ntFlanks@6738.6770.33\|ATCATCGACACTTCGAACGCACTTGCGGCCCCG | >hg38_21_+_8433172_8446622_RNA45SN1_45S_50ntFlanks | 6738 | 6770 | 33 | ATCATCGACACTTCGAACGCACTTGCGGCCCCG | rRF | FALSE | FALSE | TRUE |
| >hg38_21_+_8433172_8446622_RNA45SN1_45S_50ntFlanks@7984.8006.23\|AGATCAGACGTGGCGACCCGCTG | >hg38_21_+_8433172_8446622_RNA45SN1_45S_50ntFlanks | 7984 | 8006 | 23 | AGATCAGACGTGGCGACCCGCTG | rRF | FALSE | FALSE | TRUE |
| >hg38_21_+_8433172_8446622_RNA45SN1_45S_50ntFlanks@12415.12432.18\|CTTCGATGTCGGCTCTTC | >hg38_21_+_8433172_8446622_RNA45SN1_45S_50ntFlanks | 12415 | 12432 | 18 | CTTCGATGTCGGCTCTTC | rRF | FALSE | FALSE | TRUE |
| >hg38_21_+_8433172_8446622_RNA45SN1_45S_50ntFlanks@10687.10711.25\|CGGCGTCCGGTGAGCTCTCGCTGGC | >hg38_21_+_8433172_8446622_RNA45SN1_45S_50ntFlanks | 10687 | 10711 | 25 | CGGCGTCCGGTGAGCTCTCGCTGGC | rRF | FALSE | FALSE | TRUE |
| >hg38_21_+_8433172_8446622_RNA45SN1_45S_50ntFlanks@3985.4002.18\|CGGCTTTGGTGACTCTAG | >hg38_21_+_8433172_8446622_RNA45SN1_45S_50ntFlanks | 3985 | 4002 | 18 | CGGCTTTGGTGACTCTAG | rRF | FALSE | FALSE | TRUE |
| >hg38_21_+_8433172_8446622_RNA45SN1_45S_50ntFlanks@9573.9596.24\|AAATCGGTCGTCCGACCTGGGTAT | >hg38_21_+_8433172_8446622_RNA45SN1_45S_50ntFlanks | 9573 | 9596 | 24 | AAATCGGTCGTCCGACCTGGGTAT | rRF | FALSE | FALSE | TRUE |
| >hg38_21_+_8433172_8446622_RNA45SN1_45S_50ntFlanks@12417.12434.18\|TCGATGTCGGCTCTTCCT | >hg38_21_+_8433172_8446622_RNA45SN1_45S_50ntFlanks | 12417 | 12434 | 18 | TCGATGTCGGCTCTTCCT | rRF | FALSE | FALSE | TRUE |
| >hg38_21_+_8433172_8446622_RNA45SN1_45S_50ntFlanks@12076.12095.20\|AGGGGCTCTCGCTTCTGGCG | >hg38_21_+_8433172_8446622_RNA45SN1_45S_50ntFlanks | 12076 | 12095 | 20 | AGGGGCTCTCGCTTCTGGCG | rRF | FALSE | FALSE | TRUE |
| >hg38_21_+_8433172_8446622_RNA45SN1_45S_50ntFlanks@9570.9596.27\|TGCAAATCGGTCGTCCGACCTGGGTAT | >hg38_21_+_8433172_8446622_RNA45SN1_45S_50ntFlanks | 9570 | 9596 | 27 | TGCAAATCGGTCGTCCGACCTGGGTAT | rRF | FALSE | FALSE | TRUE |
| >hg38_21_+_8433172_8446622_RNA45SN1_45S_50ntFlanks@12420.12443.24\|ATGTCGGCTCTTCCTATCATTGTG | >hg38_21_+_8433172_8446622_RNA45SN1_45S_50ntFlanks | 12420 | 12443 | 24 | ATGTCGGCTCTTCCTATCATTGTG | rRF | FALSE | FALSE | TRUE |
| >hg38_21_+_8433172_8446622_RNA45SN1_45S_50ntFlanks@8380.8400.21\|CAAGAGGGCGTGAAACCGTTA | >hg38_21_+_8433172_8446622_RNA45SN1_45S_50ntFlanks | 8380 | 8400 | 21 | CAAGAGGGCGTGAAACCGTTA | rRF | FALSE | FALSE | TRUE |
| >trna111_HisGTG_1_-_147774845_147774916@-1T.29.30,trna118_HisGTG_1_-_145396881_145396952@-1T.29.30,trna16_HisGTG_1_+_146544773_146544844@-1T.29.30,trna1_HisGTG_15_+_45493349_45493420@-1T.29.30,trna21_HisGTG_1_+_147753471_147753542@-1T.29.30,trna33_HisGTG_6_+_27125906_27125977@-1T.29.30,trna7_HisGTG_9_-_14433938_14434009@-1T.29.30,trna8_HisGTG_15_-_45492611_45492682@-1T.29.30,trna9_HisGTG_15_-_45490804_45490875@-1T.29.30\|TGCCGTGATCGTATAGTGGTTAGTACTCTG | >trna111_HisGTG_1_-_147774845_147774916 | -1T | 29 | 30 | TGCCGTGATCGTATAGTGGTTAGTACTCTG | tRF | FALSE | FALSE | TRUE |
| >hg38_21_+_8433172_8446622_RNA45SN1_45S_50ntFlanks@10362.10388.27\|AAGGCCGAAGTGGAGAAGGGTTCCATG | >hg38_21_+_8433172_8446622_RNA45SN1_45S_50ntFlanks | 10362 | 10388 | 27 | AAGGCCGAAGTGGAGAAGGGTTCCATG | rRF | FALSE | FALSE | TRUE |
| >hg38_21_+_8433172_8446622_RNA45SN1_45S_50ntFlanks@6721.6739.19\|AATTGCAGGACACATTGAT | >hg38_21_+_8433172_8446622_RNA45SN1_45S_50ntFlanks | 6721 | 6739 | 19 | AATTGCAGGACACATTGAT | rRF | FALSE | FALSE | TRUE |
| >hg38_21_+_8433172_8446622_RNA45SN1_45S_50ntFlanks@10684.10711.28\|CGGCGGCGTCCGGTGAGCTCTCGCTGGC | >hg38_21_+_8433172_8446622_RNA45SN1_45S_50ntFlanks | 10684 | 10711 | 28 | CGGCGGCGTCCGGTGAGCTCTCGCTGGC | rRF | FALSE | FALSE | TRUE |
| >hg38_21_+_8433172_8446622_RNA45SN1_45S_50ntFlanks@9545.9563.19\|GTGGAGGTCCGTAGCGGTC | >hg38_21_+_8433172_8446622_RNA45SN1_45S_50ntFlanks | 9545 | 9563 | 19 | GTGGAGGTCCGTAGCGGTC | rRF | FALSE | FALSE | TRUE |
| >hg38_21_+_8433172_8446622_RNA45SN1_45S_50ntFlanks@12002.12032.31\|TGCGGGCCGCCGGTGAAATACCACTACTCTG | >hg38_21_+_8433172_8446622_RNA45SN1_45S_50ntFlanks | 12002 | 12032 | 31 | TGCGGGCCGCCGGTGAAATACCACTACTCTG | rRF | FALSE | FALSE | TRUE |
| >hg38_21_+_8433172_8446622_RNA45SN1_45S_50ntFlanks@9570.9597.28\|TGCAAATCGGTCGTCCGACCTGGGTATA | >hg38_21_+_8433172_8446622_RNA45SN1_45S_50ntFlanks | 9570 | 9597 | 28 | TGCAAATCGGTCGTCCGACCTGGGTATA | rRF | FALSE | FALSE | TRUE |
| >hg38_21_+_8433172_8446622_RNA45SN1_45S_50ntFlanks@8105.8143.39\|CGCGGCGGGGCGCGGGACATGTGGCGTACGGAAGACCCG | >hg38_21_+_8433172_8446622_RNA45SN1_45S_50ntFlanks | 8105 | 8143 | 39 | CGCGGCGGGGCGCGGGACATGTGGCGTACGGAAGACCCG | rRF | FALSE | FALSE | TRUE |

| | | | | | | | | | |
|---|---|---|---|---|---|---|---|---|---|
| >hg38_21_+_8433172_8446622_RNA45SN1_45S_50ntFlanks@11235.11254.20\|CTTCCCGTGGATCGCCCCAG | >hg38_21_+_8433172_8446622_RNA45SN1_45S_50ntFlanks | 11235 | 11254 | 20 | CTTCCCGTGGATCGCCCCAG | rRF | FALSE | FALSE | TRUE |
| >hg38_21_+_8433172_8446622_RNA45SN1_45S_50ntFlanks@10798.10818.21\|CATGTTGGAACAATGTAGGTA | >hg38_21_+_8433172_8446622_RNA45SN1_45S_50ntFlanks | 10798 | 10818 | 21 | CATGTTGGAACAATGTAGGTA | rRF | FALSE | FALSE | TRUE |
| >hg38_21_+_8433172_8446622_RNA45SN1_45S_50ntFlanks@8104.8143.40\|CCGCGGCGGGGCGCGGGACATGTGGCGTACGGAAGACCCG | >hg38_21_+_8433172_8446622_RNA45SN1_45S_50ntFlanks | 8104 | 8143 | 40 | CCGCGGCGGGGCGCGGGACATGTGGCGTACGGAAGACCCG | rRF | FALSE | FALSE | TRUE |
| >hg38_21_+_8433172_8446622_RNA45SN1_45S_50ntFlanks@10846.10865.20\|CTTCGGGATAAGGATTGGCT | >hg38_21_+_8433172_8446622_RNA45SN1_45S_50ntFlanks | 10846 | 10865 | 20 | CTTCGGGATAAGGATTGGCT | rRF | FALSE | FALSE | TRUE |
| >hg38_21_+_8433172_8446622_RNA45SN1_45S_50ntFlanks@6738.6771.34\|ATCATCGACACTTCGAACGCACTTGCGGCCCCGG | >hg38_21_+_8433172_8446622_RNA45SN1_45S_50ntFlanks | 6738 | 6771 | 34 | ATCATCGACACTTCGAACGCACTTGCGGCCCCGG | rRF | FALSE | FALSE | TRUE |
| >hg38_21_+_8433172_8446622_RNA45SN1_45S_50ntFlanks@11641.11670.30\|CGGCGGGTGTTGACGCGATGTGATTTCTGC | >hg38_21_+_8433172_8446622_RNA45SN1_45S_50ntFlanks | 11641 | 11670 | 30 | CGGCGGGTGTTGACGCGATGTGATTTCTGC | rRF | FALSE | FALSE | TRUE |
| >hg38_21_+_8433172_8446622_RNA45SN1_45S_50ntFlanks@10411.10428.18\|CAGTCGGTCCTGAGAGAT | >hg38_21_+_8433172_8446622_RNA45SN1_45S_50ntFlanks | 10411 | 10428 | 18 | CAGTCGGTCCTGAGAGAT | rRF | FALSE | FALSE | TRUE |
| >hg38_21_+_8433172_8446622_RNA45SN1_45S_50ntFlanks@11579.11603.25\|AGAACTGGTGCGGACCAGGGGAATC | >hg38_21_+_8433172_8446622_RNA45SN1_45S_50ntFlanks | 11579 | 11603 | 25 | AGAACTGGTGCGGACCAGGGGAATC | rRF | FALSE | FALSE | TRUE |
| >hg38_21_+_8433172_8446622_RNA45SN1_45S_50ntFlanks@12415.12436.22\|CTTCGATGTCGGCTCTTCCTAT | >hg38_21_+_8433172_8446622_RNA45SN1_45S_50ntFlanks | 12415 | 12436 | 22 | CTTCGATGTCGGCTCTTCCTAT | rRF | FALSE | FALSE | TRUE |
| >hg38_21_+_8433172_8446622_RNA45SN1_45S_50ntFlanks@11593.11611.19\|CCAGGGGAATCCGACTGTT | >hg38_21_+_8433172_8446622_RNA45SN1_45S_50ntFlanks | 11593 | 11611 | 19 | CCAGGGGAATCCGACTGTT | rRF | FALSE | FALSE | TRUE |
| >hg38_21_+_8433172_8446622_RNA45SN1_45S_50ntFlanks@11768.11794.27\|CTAATTAGTGACGCGCATGAATGGATG | >hg38_21_+_8433172_8446622_RNA45SN1_45S_50ntFlanks | 11768 | 11794 | 27 | CTAATTAGTGACGCGCATGAATGGATG | rRF | FALSE | FALSE | TRUE |
| >hg38_21_+_8433172_8446622_RNA45SN1_45S_50ntFlanks@10392.10409.18\|ACAGCAGTTGAACATGGG | >hg38_21_+_8433172_8446622_RNA45SN1_45S_50ntFlanks | 10392 | 10409 | 18 | ACAGCAGTTGAACATGGG | rRF | FALSE | FALSE | TRUE |
| >hg38_21_+_8433172_8446622_RNA45SN1_45S_50ntFlanks@12636.12654.19\|AGCTACCATCTGTGGGATT | >hg38_21_+_8433172_8446622_RNA45SN1_45S_50ntFlanks | 12636 | 12654 | 19 | AGCTACCATCTGTGGGATT | rRF | FALSE | FALSE | TRUE |
| >hg38_21_+_8433172_8446622_RNA45SN1_45S_50ntFlanks@12628.12654.27\|TGGGGCGAAGCTACCATCTGTGGGATT | >hg38_21_+_8433172_8446622_RNA45SN1_45S_50ntFlanks | 12628 | 12654 | 27 | TGGGGCGAAGCTACCATCTGTGGGATT | rRF | FALSE | FALSE | TRUE |
| >hg38_21_+_8433172_8446622_RNA45SN1_45S_50ntFlanks@12437.12454.18\|CATTGTGAAGCAGAATTC | >hg38_21_+_8433172_8446622_RNA45SN1_45S_50ntFlanks | 12437 | 12454 | 18 | CATTGTGAAGCAGAATTC | rRF | FALSE | FALSE | TRUE |
| >hg38_21_+_8433172_8446622_RNA45SN1_45S_50ntFlanks@6646.6670.25\|TCGTACGACTCTTAGCGGTGGATCA | >hg38_21_+_8433172_8446622_RNA45SN1_45S_50ntFlanks | 6646 | 6670 | 25 | TCGTACGACTCTTAGCGGTGGATCA | rRF | FALSE | FALSE | TRUE |
| >hg38_21_+_8433172_8446622_RNA45SN1_45S_50ntFlanks@10349.10383.35\|AACGAGAACTTTGAAGGCCGAAGTGGAGAAGGGTT | >hg38_21_+_8433172_8446622_RNA45SN1_45S_50ntFlanks | 10349 | 10383 | 35 | AACGAGAACTTTGAAGGCCGAAGTGGAGAAGGGTT | rRF | FALSE | FALSE | TRUE |
| >hg38_21_+_8433172_8446622_RNA45SN1_45S_50ntFlanks@9738.9758.21\|GAAACGATCTCAACCTATTCT | >hg38_21_+_8433172_8446622_RNA45SN1_45S_50ntFlanks | 9738 | 9758 | 21 | GAAACGATCTCAACCTATTCT | rRF | FALSE | FALSE | TRUE |
| >hg38_21_+_8433172_8446622_RNA45SN1_45S_50ntFlanks@12159.12180.22\|TTTGACTGGGGCGGTACACCTG | >hg38_21_+_8433172_8446622_RNA45SN1_45S_50ntFlanks | 12159 | 12180 | 22 | TTTGACTGGGGCGGTACACCTG | rRF | FALSE | FALSE | TRUE |
| >hg38_21_+_8433172_8446622_RNA45SN1_45S_50ntFlanks@7988.8008.21\|CAGACGTGGCGACCCGCTGAA | >hg38_21_+_8433172_8446622_RNA45SN1_45S_50ntFlanks | 7988 | 8008 | 21 | CAGACGTGGCGACCCGCTGAA | rRF | FALSE | FALSE | TRUE |
| >hg38_21_+_8433172_8446622_RNA45SN1_45S_50ntFlanks@6722.6744.23\|ATTGCAGGACACATTGATCATCG | >hg38_21_+_8433172_8446622_RNA45SN1_45S_50ntFlanks | 6722 | 6744 | 23 | ATTGCAGGACACATTGATCATCG | rRF | FALSE | FALSE | TRUE |
| >hg38_21_+_8433172_8446622_RNA45SN1_45S_50ntFlanks@10788.10818.31\|CAGCCTCTGGCATGTTGGAACAATGTAGGTA | >hg38_21_+_8433172_8446622_RNA45SN1_45S_50ntFlanks | 10788 | 10818 | 31 | CAGCCTCTGGCATGTTGGAACAATGTAGGTA | rRF | FALSE | FALSE | TRUE |
| >hg38_21_+_8433172_8446622_RNA45SN1_45S_50ntFlanks@10458.10478.21\|ATGGCCTCCGTTGCCCTCGGC | >hg38_21_+_8433172_8446622_RNA45SN1_45S_50ntFlanks | 10458 | 10478 | 21 | ATGGCCTCCGTTGCCCTCGGC | rRF | FALSE | FALSE | TRUE |
| >hg38_21_+_8433172_8446622_RNA45SN1_45S_50ntFlanks@4657.4689.33\|CTAGAGGTGAAATTCTTGGACCGGCGCAAGACG | >hg38_21_+_8433172_8446622_RNA45SN1_45S_50ntFlanks | 4657 | 4689 | 33 | CTAGAGGTGAAATTCTTGGACCGGCGCAAGACG | rRF | FALSE | FALSE | TRUE |
| >hg38_21_+_8433172_8446622_RNA45SN1_45S_50ntFlanks@11784.11803.20\|ATGAATGGATGAACGAGATT | >hg38_21_+_8433172_8446622_RNA45SN1_45S_50ntFlanks | 11784 | 11803 | 20 | ATGAATGGATGAACGAGATT | rRF | FALSE | FALSE | TRUE |
| >hg38_21_+_8433172_8446622_RNA45SN1_45S_50ntFlanks@10389.10410.22\|TGAACAGCAGTTGAACATGGGT | >hg38_21_+_8433172_8446622_RNA45SN1_45S_50ntFlanks | 10389 | 10410 | 22 | TGAACAGCAGTTGAACATGGGT | rRF | FALSE | FALSE | TRUE |
| >hg38_21_+_8433172_8446622_RNA45SN1_45S_50ntFlanks@12615.12654.40\|GCTGAGGAGCCAATGGGGCGAAGCTACCATCTGTGGGATT | >hg38_21_+_8433172_8446622_RNA45SN1_45S_50ntFlanks | 12615 | 12654 | 40 | GCTGAGGAGCCAATGGGGCGAAGCTACCATCTGTGGGATT | rRF | FALSE | FALSE | TRUE |
| >hg38_21_+_8433172_8446622_RNA45SN1_45S_50ntFlanks@6671.6699.29\|CTCGGCTCGTGCGTCGATGAAGAACGCAG | >hg38_21_+_8433172_8446622_RNA45SN1_45S_50ntFlanks | 6671 | 6699 | 29 | CTCGGCTCGTGCGTCGATGAAGAACGCAG | rRF | FALSE | FALSE | TRUE |
| >hg38_21_+_8433172_8446622_RNA45SN1_45S_50ntFlanks@12163.12180.18\|ACTGGGGCGGTACACCTG | >hg38_21_+_8433172_8446622_RNA45SN1_45S_50ntFlanks | 12163 | 12180 | 18 | ACTGGGGCGGTACACCTG | rRF | FALSE | FALSE | TRUE |
| >hg38_21_+_8433172_8446622_RNA45SN1_45S_50ntFlanks@11729.11753.25\|TAACTATGACTCTCTTAAGGTAGCC | >hg38_21_+_8433172_8446622_RNA45SN1_45S_50ntFlanks | 11729 | 11753 | 25 | TAACTATGACTCTCTTAAGGTAGCC | rRF | FALSE | FALSE | TRUE |
| >hg38_21_+_8433172_8446622_RNA45SN1_45S_50ntFlanks@12459.12476.18\|AGCGTTGGATTGTTCACC | >hg38_21_+_8433172_8446622_RNA45SN1_45S_50ntFlanks | 12459 | 12476 | 18 | AGCGTTGGATTGTTCACC | rRF | FALSE | FALSE | TRUE |
| >MI0000342\|hsa-mir-200b&WithFlank&1\|+\|1167098\|1167204@63.85.23&[MIMAT0000318&hsa-miR-200b-3p&offsets\|0\|+1;m-93&1\|+\|1167160\|1167182&offsets\|0\|0]\|TAATACTGCCTGGTAATGATGAC | >MI0000342\|hsa-mir-200b&WithFlank&1\|+\|1167098\|1167204 | 63 | 85 | 23 | TAATACTGCCTGGTAATGATGAC | isomiR | FALSE | FALSE | TRUE |
| >hg38_21_+_8433172_8446622_RNA45SN1_45S_50ntFlanks@10408.10429.22\|GGTCAGTCGGTCCTGAGAGATG | >hg38_21_+_8433172_8446622_RNA45SN1_45S_50ntFlanks | 10408 | 10429 | 22 | GGTCAGTCGGTCCTGAGAGATG | rRF | FALSE | FALSE | TRUE |
| >hg38_21_+_8433172_8446622_RNA45SN1_45S_50ntFlanks@6721.6747.27\|AATTGCAGGACACATTGATCATCGACA | >hg38_21_+_8433172_8446622_RNA45SN1_45S_50ntFlanks | 6721 | 6747 | 27 | AATTGCAGGACACATTGATCATCGACA | rRF | FALSE | FALSE | TRUE |
| >hg38_21_+_8433172_8446622_RNA45SN1_45S_50ntFlanks@10391.10409.19\|AACAGCAGTTGAACATGGG | >hg38_21_+_8433172_8446622_RNA45SN1_45S_50ntFlanks | 10391 | 10409 | 19 | AACAGCAGTTGAACATGGG | rRF | FALSE | FALSE | TRUE |
| >hg38_21_+_8433172_8446622_RNA45SN1_45S_50ntFlanks@11569.11601.33\|CAGCCGACTTAGAACTGGTGCGGACCAGGGGAA | >hg38_21_+_8433172_8446622_RNA45SN1_45S_50ntFlanks | 11569 | 11601 | 33 | CAGCCGACTTAGAACTGGTGCGGACCAGGGGAA | rRF | FALSE | FALSE | TRUE |
| >hg38_21_+_8433172_8446622_RNA45SN1_45S_50ntFlanks@11572.11600.29\|CCGACTTAGAACTGGTGCGGACCAGGGGA | >hg38_21_+_8433172_8446622_RNA45SN1_45S_50ntFlanks | 11572 | 11600 | 29 | CCGACTTAGAACTGGTGCGGACCAGGGGA | rRF | FALSE | FALSE | TRUE |
| >hg38_21_+_8433172_8446622_RNA45SN1_45S_50ntFlanks@10351.10384.34\|CGAGAACTTTGAAGGCCGAAGTGGAGAAGGGTTC | >hg38_21_+_8433172_8446622_RNA45SN1_45S_50ntFlanks | 10351 | 10384 | 34 | CGAGAACTTTGAAGGCCGAAGTGGAGAAGGGTTC | rRF | FALSE | FALSE | TRUE |
| >hg38_21_+_8433172_8446622_RNA45SN1_45S_50ntFlanks@11783.11805.23\|CATGAATGGATGAACGAGATTCC | >hg38_21_+_8433172_8446622_RNA45SN1_45S_50ntFlanks | 11783 | 11805 | 23 | CATGAATGGATGAACGAGATTCC | rRF | FALSE | FALSE | TRUE |
| >hg38_21_+_8433172_8446622_RNA45SN1_45S_50ntFlanks@11208.11234.27\|GCGGCGACTCTGGACGCGAGCCGGGCC | >hg38_21_+_8433172_8446622_RNA45SN1_45S_50ntFlanks | 11208 | 11234 | 27 | GCGGCGACTCTGGACGCGAGCCGGGCC | rRF | FALSE | FALSE | TRUE |
| >trna11_GluTTC_15_-_26327381_26327452@1.29.29,trna3_GluTTC_13_-_45492062_45492133@1.29.29\|TCCCACATGGTCTAGCGGTTAGGATTCCT | >trna11_GluTTC_15_-_26327381_26327452 | 1 | 29 | 29 | TCCCACATGGTCTAGCGGTTAGGATTCCT | tRF | FALSE | FALSE | TRUE |
| >hg38_21_+_8433172_8446622_RNA45SN1_45S_50ntFlanks@4731.4758.28\|AAGAACGAAAGTCGGAGGTTCGAAGACG | >hg38_21_+_8433172_8446622_RNA45SN1_45S_50ntFlanks | 4731 | 4758 | 28 | AAGAACGAAAGTCGGAGGTTCGAAGACG | rRF | FALSE | FALSE | TRUE |

| | | | | | | | | | |
|---|---|---|---|---|---|---|---|---|---|
| >hg38_21_+_8433172_8446622_RNA45SN1_45S_50ntFlanks@6721.6746.26\|AATTGCAGGACACATTGATCATCGAC | >hg38_21_+_8433172_8446622_RNA45SN1_45S_50ntFlanks | 6721 | 6746 | 26 | AATTGCAGGACACATTGATCATCGAC | rRF | FALSE | FALSE | TRUE |
| >hg38_21_+_8433172_8446622_RNA45SN1_45S_50ntFlanks@7974.8006.33\|ACGCGACCTCAGATCAGACGTGGCGACCCGCTG | >hg38_21_+_8433172_8446622_RNA45SN1_45S_50ntFlanks | 7974 | 8006 | 33 | ACGCGACCTCAGATCAGACGTGGCGACCCGCTG | rRF | FALSE | FALSE | TRUE |
| >hg38_21_+_8433172_8446622_RNA45SN1_45S_50ntFlanks@10684.10709.26\|CGGCGGCGTCCGGTGAGCTCTCGCTG | >hg38_21_+_8433172_8446622_RNA45SN1_45S_50ntFlanks | 10684 | 10709 | 26 | CGGCGGCGTCCGGTGAGCTCTCGCTG | rRF | FALSE | FALSE | TRUE |
| >hg38_21_+_8433172_8446622_RNA45SN1_45S_50ntFlanks@10350.10388.39\|ACGAGAACTTTGAAGGCCGAAGTGGAGAAGGGTTCCATG | >hg38_21_+_8433172_8446622_RNA45SN1_45S_50ntFlanks | 10350 | 10388 | 39 | ACGAGAACTTTGAAGGCCGAAGTGGAGAAGGGTTCCATG | rRF | FALSE | FALSE | TRUE |
| >hg38_21_+_8433172_8446622_RNA45SN1_45S_50ntFlanks@5445.5466.22\|TCGGATCGGCCCCGCCGGGGTC | >hg38_21_+_8433172_8446622_RNA45SN1_45S_50ntFlanks | 5445 | 5466 | 22 | TCGGATCGGCCCCGCCGGGGTC | rRF | FALSE | FALSE | TRUE |
| >hg38_21_+_8433172_8446622_RNA45SN1_45S_50ntFlanks@10686.10710.25\|GCGGCGTCCGGTGAGCTCTCGCTGG | >hg38_21_+_8433172_8446622_RNA45SN1_45S_50ntFlanks | 10686 | 10710 | 25 | GCGGCGTCCGGTGAGCTCTCGCTGG | rRF | FALSE | FALSE | TRUE |
| >trna16_GlnTTG_17_+_47269890_47269961@1.29.29,trna64_GlnTTG_6_+_28557156_28557227@1.29.29,trna84_GlnTTG_6_+_145503859_145503930@1.29.29\|GGTCCCATGGTGTAATGGTTAGCACTCTG | >trna16_GlnTTG_17_+_47269890_47269961 | 1 | 29 | 29 | GGTCCCATGGTGTAATGGTTAGCACTCTG | tRF | FALSE | FALSE | TRUE |
| >hg38_21_+_8433172_8446622_RNA45SN1_45S_50ntFlanks@9575.9597.23\|ATCGGTCGTCCGACCTGGGTATA | >hg38_21_+_8433172_8446622_RNA45SN1_45S_50ntFlanks | 9575 | 9597 | 23 | ATCGGTCGTCCGACCTGGGTATA | rRF | FALSE | FALSE | TRUE |
| >hg38_21_+_8433172_8446622_RNA45SN1_45S_50ntFlanks@10821.10840.20\|GGAAGTCGGCAAGCCGGATC | >hg38_21_+_8433172_8446622_RNA45SN1_45S_50ntFlanks | 10821 | 10840 | 20 | GGAAGTCGGCAAGCCGGATC | rRF | FALSE | FALSE | TRUE |
| >hg38_21_+_8433172_8446622_RNA45SN1_45S_50ntFlanks@11876.11893.18\|AAGACCCTGTTGAGCTTG | >hg38_21_+_8433172_8446622_RNA45SN1_45S_50ntFlanks | 11876 | 11893 | 18 | AAGACCCTGTTGAGCTTG | rRF | FALSE | FALSE | TRUE |
| >hg38_21_+_8433172_8446622_RNA45SN1_45S_50ntFlanks@3775.3792.18\|GCCGGTACAGTGAAACTG | >hg38_21_+_8433172_8446622_RNA45SN1_45S_50ntFlanks | 3775 | 3792 | 18 | GCCGGTACAGTGAAACTG | rRF | FALSE | FALSE | TRUE |
| >hg38_21_+_8433172_8446622_RNA45SN1_45S_50ntFlanks@10681.10710.30\|TTCCGGCGGCGTCCGGTGAGCTCTCGCTGG | >hg38_21_+_8433172_8446622_RNA45SN1_45S_50ntFlanks | 10681 | 10710 | 30 | TTCCGGCGGCGTCCGGTGAGCTCTCGCTGG | rRF | FALSE | FALSE | TRUE |
| >hg38_21_+_8433172_8446622_RNA45SN1_45S_50ntFlanks@9574.9595.22\|AATCGGTCGTCCGACCTGGGTA | >hg38_21_+_8433172_8446622_RNA45SN1_45S_50ntFlanks | 9574 | 9595 | 22 | AATCGGTCGTCCGACCTGGGTA | rRF | FALSE | FALSE | TRUE |
| >hg38_21_+_8433172_8446622_RNA45SN1_45S_50ntFlanks@11572.11590.19\|CCGACTTAGAACTGGTGCG | >hg38_21_+_8433172_8446622_RNA45SN1_45S_50ntFlanks | 11572 | 11590 | 19 | CCGACTTAGAACTGGTGCG | rRF | FALSE | FALSE | TRUE |
| >hg38_21_+_8433172_8446622_RNA45SN1_45S_50ntFlanks@10458.10477.20\|ATGGCCTCCGTTGCCCTCGG | >hg38_21_+_8433172_8446622_RNA45SN1_45S_50ntFlanks | 10458 | 10477 | 20 | ATGGCCTCCGTTGCCCTCGG | rRF | FALSE | FALSE | TRUE |
| >trna15_ValAAC_5_-_180615416_180615488@1.29.29\|GTTTCCGTAGTGTAGTGGTCATCACGTTC | >trna15_ValAAC_5_-_180615416_180615488 | 1 | 29 | 29 | GTTTCCGTAGTGTAGTGGTCATCACGTTC | tRF | FALSE | FALSE | TRUE |
| >hg38_21_+_8433172_8446622_RNA45SN1_45S_50ntFlanks@6648.6681.34\|GTACGACTCTTAGCGGTGGATCACTCGGCTCGTG | >hg38_21_+_8433172_8446622_RNA45SN1_45S_50ntFlanks | 6648 | 6681 | 34 | GTACGACTCTTAGCGGTGGATCACTCGGCTCGTG | rRF | FALSE | FALSE | TRUE |
| >hg38_21_+_8433172_8446622_RNA45SN1_45S_50ntFlanks@7982.8006.25\|TCAGATCAGACGTGGCGACCCGCTG | >hg38_21_+_8433172_8446622_RNA45SN1_45S_50ntFlanks | 7982 | 8006 | 25 | TCAGATCAGACGTGGCGACCCGCTG | rRF | FALSE | FALSE | TRUE |
| >hg38_21_+_8433172_8446622_RNA45SN1_45S_50ntFlanks@12476.12498.23\|CCACTAATAGGGAACGTGAGCTG | >hg38_21_+_8433172_8446622_RNA45SN1_45S_50ntFlanks | 12476 | 12498 | 23 | CCACTAATAGGGAACGTGAGCTG | rRF | FALSE | FALSE | TRUE |
| >hg38_21_+_8433172_8446622_RNA45SN1_45S_50ntFlanks@12456.12480.25\|CCAAGCGTTGGATTGTTCACCCACT | >hg38_21_+_8433172_8446622_RNA45SN1_45S_50ntFlanks | 12456 | 12480 | 25 | CCAAGCGTTGGATTGTTCACCCACT | rRF | FALSE | FALSE | TRUE |
| >hg38_21_+_8433172_8446622_RNA45SN1_45S_50ntFlanks@11208.11233.26\|GCGGCGACTCTGGACGCGAGCCGGGC | >hg38_21_+_8433172_8446622_RNA45SN1_45S_50ntFlanks | 11208 | 11233 | 26 | GCGGCGACTCTGGACGCGAGCCGGGC | rRF | FALSE | FALSE | TRUE |
| >hg38_21_+_8433172_8446622_RNA45SN1_45S_50ntFlanks@8380.8402.23\|CAAGAGGGCGTGAAACCGTTAAG | >hg38_21_+_8433172_8446622_RNA45SN1_45S_50ntFlanks | 8380 | 8402 | 23 | CAAGAGGGCGTGAAACCGTTAAG | rRF | FALSE | FALSE | TRUE |
| >hg38_21_+_8433172_8446622_RNA45SN1_45S_50ntFlanks@9570.9595.26\|TGCAAATCGGTCGTCCGACCTGGGTA | >hg38_21_+_8433172_8446622_RNA45SN1_45S_50ntFlanks | 9570 | 9595 | 26 | TGCAAATCGGTCGTCCGACCTGGGTA | rRF | FALSE | FALSE | TRUE |
| >hg38_21_+_8433172_8446622_RNA45SN1_45S_50ntFlanks@8198.8215.18\|TGGACGGTGTGAGGCCGG | >hg38_21_+_8433172_8446622_RNA45SN1_45S_50ntFlanks | 8198 | 8215 | 18 | TGGACGGTGTGAGGCCGG | rRF | FALSE | FALSE | TRUE |
| >hg38_21_+_8433172_8446622_RNA45SN1_45S_50ntFlanks@6650.6681.32\|ACGACTCTTAGCGGTGGATCACTCGGCTCGTG | >hg38_21_+_8433172_8446622_RNA45SN1_45S_50ntFlanks | 6650 | 6681 | 32 | ACGACTCTTAGCGGTGGATCACTCGGCTCGTG | rRF | FALSE | FALSE | TRUE |
| >hg38_21_+_8433172_8446622_RNA45SN1_45S_50ntFlanks@9554.9574.21\|CGTAGCGGTCCTGACGTGCAA | >hg38_21_+_8433172_8446622_RNA45SN1_45S_50ntFlanks | 9554 | 9574 | 21 | CGTAGCGGTCCTGACGTGCAA | rRF | FALSE | FALSE | TRUE |
| >hg38_21_+_8433172_8446622_RNA45SN1_45S_50ntFlanks@10841.10863.23\|CGTAACTTCGGGATAAGGATTGG | >hg38_21_+_8433172_8446622_RNA45SN1_45S_50ntFlanks | 10841 | 10863 | 23 | CGTAACTTCGGGATAAGGATTGG | rRF | FALSE | FALSE | TRUE |
| >hg38_21_+_8433172_8446622_RNA45SN1_45S_50ntFlanks@12458.12476.19\|AAGCGTTGGATTGTTCACC | >hg38_21_+_8433172_8446622_RNA45SN1_45S_50ntFlanks | 12458 | 12476 | 19 | AAGCGTTGGATTGTTCACC | rRF | FALSE | FALSE | TRUE |
| >hg38_21_+_8433172_8446622_RNA45SN1_45S_50ntFlanks@12405.12443.39\|CTTTTTGATCCTTCGATGTCGGCTCTTCCTATCATTGTG | >hg38_21_+_8433172_8446622_RNA45SN1_45S_50ntFlanks | 12405 | 12443 | 39 | CTTTTTGATCCTTCGATGTCGGCTCTTCCTATCATTGTG | rRF | FALSE | FALSE | TRUE |
| >hg38_21_+_8433172_8446622_RNA45SN1_45S_50ntFlanks@3874.3898.25\|AATACATGCCGACGGGCGCTGACCC | >hg38_21_+_8433172_8446622_RNA45SN1_45S_50ntFlanks | 3874 | 3898 | 25 | AATACATGCCGACGGGCGCTGACCC | rRF | FALSE | FALSE | TRUE |
| >hg38_21_+_8433172_8446622_RNA45SN1_45S_50ntFlanks@12632.12654.23\|GCGAAGCTACCATCTGTGGGATT | >hg38_21_+_8433172_8446622_RNA45SN1_45S_50ntFlanks | 12632 | 12654 | 23 | GCGAAGCTACCATCTGTGGGATT | rRF | FALSE | FALSE | TRUE |
| >hg38_21_+_8433172_8446622_RNA45SN1_45S_50ntFlanks@10346.10368.23\|TCAAACGAGAACTTTGAAGGCCG | >hg38_21_+_8433172_8446622_RNA45SN1_45S_50ntFlanks | 10346 | 10368 | 23 | TCAAACGAGAACTTTGAAGGCCG | rRF | FALSE | FALSE | TRUE |
| >hg38_21_+_8433172_8446622_RNA45SN1_45S_50ntFlanks@12417.12443.27\|TCGATGTCGGCTCTTCCTATCATTGTG | >hg38_21_+_8433172_8446622_RNA45SN1_45S_50ntFlanks | 12417 | 12443 | 27 | TCGATGTCGGCTCTTCCTATCATTGTG | rRF | FALSE | FALSE | TRUE |
| >hg38_21_+_8433172_8446622_RNA45SN1_45S_50ntFlanks@9566.9596.31\|GACGTGCAAATCGGTCGTCCGACCTGGGTAT | >hg38_21_+_8433172_8446622_RNA45SN1_45S_50ntFlanks | 9566 | 9596 | 31 | GACGTGCAAATCGGTCGTCCGACCTGGGTAT | rRF | FALSE | FALSE | TRUE |
| >hg38_21_+_8433172_8446622_RNA45SN1_45S_50ntFlanks@3875.3898.24\|ATACATGCCGACGGGCGCTGACCC | >hg38_21_+_8433172_8446622_RNA45SN1_45S_50ntFlanks | 3875 | 3898 | 24 | ATACATGCCGACGGGCGCTGACCC | rRF | FALSE | FALSE | TRUE |
| >hg38_21_+_8433172_8446622_RNA45SN1_45S_50ntFlanks@11729.11752.24\|TAACTATGACTCTCTTAAGGTAGC | >hg38_21_+_8433172_8446622_RNA45SN1_45S_50ntFlanks | 11729 | 11752 | 24 | TAACTATGACTCTCTTAAGGTAGC | rRF | FALSE | FALSE | TRUE |
| >hg38_21_+_8433172_8446622_RNA45SN1_45S_50ntFlanks@11583.11611.29\|CTGGTGCGGACCAGGGGAATCCGACTGTT | >hg38_21_+_8433172_8446622_RNA45SN1_45S_50ntFlanks | 11583 | 11611 | 29 | CTGGTGCGGACCAGGGGAATCCGACTGTT | rRF | FALSE | FALSE | TRUE |
| >hg38_21_+_8433172_8446622_RNA45SN1_45S_50ntFlanks@7989.8008.20\|AGACGTGGCGACCCGCTGAA | >hg38_21_+_8433172_8446622_RNA45SN1_45S_50ntFlanks | 7989 | 8008 | 20 | AGACGTGGCGACCCGCTGAA | rRF | FALSE | FALSE | TRUE |
| >hg38_21_+_8433172_8446622_RNA45SN1_45S_50ntFlanks@10350.10377.28\|ACGAGAACTTTGAAGGCCGAAGTGGAGA | >hg38_21_+_8433172_8446622_RNA45SN1_45S_50ntFlanks | 10350 | 10377 | 28 | ACGAGAACTTTGAAGGCCGAAGTGGAGA | rRF | FALSE | FALSE | TRUE |
| >hg38_21_+_8433172_8446622_RNA45SN1_45S_50ntFlanks@12435.12455.21\|ATCATTGTGAAGCAGAATTCA | >hg38_21_+_8433172_8446622_RNA45SN1_45S_50ntFlanks | 12435 | 12455 | 21 | ATCATTGTGAAGCAGAATTCA | rRF | FALSE | FALSE | TRUE |
| >hg38_21_+_8433172_8446622_RNA45SN1_45S_50ntFlanks@8176.8203.28\|CTTCTGATCGAGGCCCAGCCCGTGGACG | >hg38_21_+_8433172_8446622_RNA45SN1_45S_50ntFlanks | 8176 | 8203 | 28 | CTTCTGATCGAGGCCCAGCCCGTGGACG | rRF | FALSE | FALSE | TRUE |
| >hg38_21_+_8433172_8446622_RNA45SN1_45S_50ntFlanks@9739.9758.20\|AAACGATCTCAACCTATTCT | >hg38_21_+_8433172_8446622_RNA45SN1_45S_50ntFlanks | 9739 | 9758 | 20 | AAACGATCTCAACCTATTCT | rRF | FALSE | FALSE | TRUE |

| | | | | | | | | | |
|---|---|---|---|---|---|---|---|---|---|
| >hg38_21_+_8433172_8446622_RNA45SN1_45S_50ntFlanks@10351.10374.24\|CGAGAACTTTGAAGGCCGAAGTGG | >hg38_21_+_8433172_8446622_RNA45SN1_45S_50ntFlanks | 10351 | 10374 | 24 | CGAGAACTTTGAAGGCCGAAGTGG | rRF | FALSE | FALSE | TRUE |
| >hg38_21_+_8433172_8446622_RNA45SN1_45S_50ntFlanks@7989.8006.18\|AGACGTGGCGACCCGCTG | >hg38_21_+_8433172_8446622_RNA45SN1_45S_50ntFlanks | 7989 | 8006 | 18 | AGACGTGGCGACCCGCTG | rRF | FALSE | FALSE | TRUE |
| >hg38_21_+_8433172_8446622_RNA45SN1_45S_50ntFlanks@10840.10865.26\|CCGTAACTTCGGGATAAGGATTGGCT | >hg38_21_+_8433172_8446622_RNA45SN1_45S_50ntFlanks | 10840 | 10865 | 26 | CCGTAACTTCGGGATAAGGATTGGCT | rRF | FALSE | FALSE | TRUE |
| >hg38_21_+_8433172_8446622_RNA45SN1_45S_50ntFlanks@11320.11344.25\|CCACGTCTCGTCGCGCGCGCGTCCG | >hg38_21_+_8433172_8446622_RNA45SN1_45S_50ntFlanks | 11320 | 11344 | 25 | CCACGTCTCGTCGCGCGCGCGTCCG | rRF | FALSE | FALSE | TRUE |
| >hg38_21_+_8433172_8446622_RNA45SN1_45S_50ntFlanks@12324.12346.23\|TTGGGTTTTAAGCAGGAGGTGTC | >hg38_21_+_8433172_8446622_RNA45SN1_45S_50ntFlanks | 12324 | 12346 | 23 | TTGGGTTTTAAGCAGGAGGTGTC | rRF | FALSE | FALSE | TRUE |
| >hg38_21_+_8433172_8446622_RNA45SN1_45S_50ntFlanks@11586.11611.26\|GTGCGGACCAGGGGAATCCGACTGTT | >hg38_21_+_8433172_8446622_RNA45SN1_45S_50ntFlanks | 11586 | 11611 | 26 | GTGCGGACCAGGGGAATCCGACTGTT | rRF | FALSE | FALSE | TRUE |
| >hg38_21_+_8433172_8446622_RNA45SN1_45S_50ntFlanks@12405.12433.29\|CTTTTTGATCCTTCGATGTCGGCTCTTCC | >hg38_21_+_8433172_8446622_RNA45SN1_45S_50ntFlanks | 12405 | 12433 | 29 | CTTTTTGATCCTTCGATGTCGGCTCTTCC | rRF | FALSE | FALSE | TRUE |
| >hg38_21_+_8433172_8446622_RNA45SN1_45S_50ntFlanks@12420.12455.36\|ATGTCGGCTCTTCCTATCATTGTGAAGCAGAATTCA | >hg38_21_+_8433172_8446622_RNA45SN1_45S_50ntFlanks | 12420 | 12455 | 36 | ATGTCGGCTCTTCCTATCATTGTGAAGCAGAATTCA | rRF | FALSE | FALSE | TRUE |
| >hg38_21_+_8433172_8446622_RNA45SN1_45S_50ntFlanks@9575.9596.22\|ATCGGTCGTCCGACCTGGGTAT | >hg38_21_+_8433172_8446622_RNA45SN1_45S_50ntFlanks | 9575 | 9596 | 22 | ATCGGTCGTCCGACCTGGGTAT | rRF | FALSE | FALSE | TRUE |
| >hg38_21_+_8433172_8446622_RNA45SN1_45S_50ntFlanks@12991.13012.22\|CTCCCTCGCTGCGATCTATTGA | >hg38_21_+_8433172_8446622_RNA45SN1_45S_50ntFlanks | 12991 | 13012 | 22 | CTCCCTCGCTGCGATCTATTGA | rRF | FALSE | FALSE | TRUE |
| >hg38_21_+_8433172_8446622_RNA45SN1_45S_50ntFlanks@12457.12487.31\|CAAGCGTTGGATTGTTCACCCACTAATAGGG | >hg38_21_+_8433172_8446622_RNA45SN1_45S_50ntFlanks | 12457 | 12487 | 31 | CAAGCGTTGGATTGTTCACCCACTAATAGGG | rRF | FALSE | FALSE | TRUE |
| >hg38_21_+_8433172_8446622_RNA45SN1_45S_50ntFlanks@4046.4063.18\|CATTCGAACGTCTGCCCT | >hg38_21_+_8433172_8446622_RNA45SN1_45S_50ntFlanks | 4046 | 4063 | 18 | CATTCGAACGTCTGCCCT | rRF | FALSE | FALSE | TRUE |
| >hg38_21_+_8433172_8446622_RNA45SN1_45S_50ntFlanks@10362.10383.22\|AAGGCCGAAGTGGAGAAGGGTT | >hg38_21_+_8433172_8446622_RNA45SN1_45S_50ntFlanks | 10362 | 10383 | 22 | AAGGCCGAAGTGGAGAAGGGTT | rRF | FALSE | FALSE | TRUE |
| >hg38_21_+_8433172_8446622_RNA45SN1_45S_50ntFlanks@11661.11679.19\|TGATTTCTGCCCAGTGCTC | >hg38_21_+_8433172_8446622_RNA45SN1_45S_50ntFlanks | 11661 | 11679 | 19 | TGATTTCTGCCCAGTGCTC | rRF | FALSE | FALSE | TRUE |
| >hg38_21_+_8433172_8446622_RNA45SN1_45S_50ntFlanks@6656.6681.26\|CTTAGCGGTGGATCACTCGGCTCGTG | >hg38_21_+_8433172_8446622_RNA45SN1_45S_50ntFlanks | 6656 | 6681 | 26 | CTTAGCGGTGGATCACTCGGCTCGTG | rRF | FALSE | FALSE | TRUE |
| >hg38_21_+_8433172_8446622_RNA45SN1_45S_50ntFlanks@11569.11603.35\|CAGCCGACTTAGAACTGGTGCGGACCAGGGGAATC | >hg38_21_+_8433172_8446622_RNA45SN1_45S_50ntFlanks | 11569 | 11603 | 35 | CAGCCGACTTAGAACTGGTGCGGACCAGGGGAATC | rRF | FALSE | FALSE | TRUE |
| >hg38_21_+_8433172_8446622_RNA45SN1_45S_50ntFlanks@11768.11792.25\|CTAATTAGTGACGCGCATGAATGGA | >hg38_21_+_8433172_8446622_RNA45SN1_45S_50ntFlanks | 11768 | 11792 | 25 | CTAATTAGTGACGCGCATGAATGGA | rRF | FALSE | FALSE | TRUE |
| >hg38_21_+_8433172_8446622_RNA45SN1_45S_50ntFlanks@11316.11342.27\|CACCCCACGTCTCGTCGCGCGCGCGTC | >hg38_21_+_8433172_8446622_RNA45SN1_45S_50ntFlanks | 11316 | 11342 | 27 | CACCCCACGTCTCGTCGCGCGCGCGTC | rRF | FALSE | FALSE | TRUE |
| >hg38_21_+_8433172_8446622_RNA45SN1_45S_50ntFlanks@12991.13014.24\|CTCCCTCGCTGCGATCTATTGAAA | >hg38_21_+_8433172_8446622_RNA45SN1_45S_50ntFlanks | 12991 | 13014 | 24 | CTCCCTCGCTGCGATCTATTGAAA | rRF | FALSE | FALSE | TRUE |
| >hg38_1_-_228634819_228635039_RNA5S12_5S_50ntFlanks@84.101.18\|CTCGTCTGATCTCGGAAG | >hg38_1_-_228634819_228635039_RNA5S12_5S_50ntFlanks | 84 | 101 | 18 | CTCGTCTGATCTCGGAAG | rRF | FALSE | FALSE | TRUE |
| >hg38_21_+_8433172_8446622_RNA45SN1_45S_50ntFlanks@12073.12095.23\|CCGAGGGGCTCTCGCTTCTGGCG | >hg38_21_+_8433172_8446622_RNA45SN1_45S_50ntFlanks | 12073 | 12095 | 23 | CCGAGGGGCTCTCGCTTCTGGCG | rRF | FALSE | FALSE | TRUE |
| >hg38_21_+_8433172_8446622_RNA45SN1_45S_50ntFlanks@7985.8006.22\|GATCAGACGTGGCGACCCGCTG | >hg38_21_+_8433172_8446622_RNA45SN1_45S_50ntFlanks | 7985 | 8006 | 22 | GATCAGACGTGGCGACCCGCTG | rRF | FALSE | FALSE | TRUE |
| >hg38_21_+_8433172_8446622_RNA45SN1_45S_50ntFlanks@13002.13029.28\|CGATCTATTGAAAGTCAGCCCTCGACAC | >hg38_21_+_8433172_8446622_RNA45SN1_45S_50ntFlanks | 13002 | 13029 | 28 | CGATCTATTGAAAGTCAGCCCTCGACAC | rRF | FALSE | FALSE | TRUE |
| >hg38_21_+_8433172_8446622_RNA45SN1_45S_50ntFlanks@6661.6689.29\|CGGTGGATCACTCGGCTCGTGCGTCGATG | >hg38_21_+_8433172_8446622_RNA45SN1_45S_50ntFlanks | 6661 | 6689 | 29 | CGGTGGATCACTCGGCTCGTGCGTCGATG | rRF | FALSE | FALSE | TRUE |
| >hg38_21_+_8433172_8446622_RNA45SN1_45S_50ntFlanks@5485.5514.30\|AGCGCTGAGAAGACGGTCGAACTTGACTAT | >hg38_21_+_8433172_8446622_RNA45SN1_45S_50ntFlanks | 5485 | 5514 | 30 | AGCGCTGAGAAGACGGTCGAACTTGACTAT | rRF | FALSE | FALSE | TRUE |
| >hg38_21_+_8433172_8446622_RNA45SN1_45S_50ntFlanks@12476.12496.21\|CCACTAATAGGGAACGTGAGC | >hg38_21_+_8433172_8446622_RNA45SN1_45S_50ntFlanks | 12476 | 12496 | 21 | CCACTAATAGGGAACGTGAGC | rRF | FALSE | FALSE | TRUE |
| >hg38_21_+_8433172_8446622_RNA45SN1_45S_50ntFlanks@10681.10706.26\|TTCCGGCGGCGTCCGGTGAGCTCTCG | >hg38_21_+_8433172_8446622_RNA45SN1_45S_50ntFlanks | 10681 | 10706 | 26 | TTCCGGCGGCGTCCGGTGAGCTCTCG | rRF | FALSE | FALSE | TRUE |
| >hg38_21_+_8433172_8446622_RNA45SN1_45S_50ntFlanks@9565.9597.33\|TGACGTGCAAATCGGTCGTCCGACCTGGGTATA | >hg38_21_+_8433172_8446622_RNA45SN1_45S_50ntFlanks | 9565 | 9597 | 33 | TGACGTGCAAATCGGTCGTCCGACCTGGGTATA | rRF | FALSE | FALSE | TRUE |
| >trna119_LysCTT_1_-_145395522_145395594@1.29.29,trna11_LysCTT_5_-_180648979_180649051@1.29.29,trna13_LysCTT_6_+_26556774_26556846@1.29.29,trna32_LysCTT_16_-_3207406_3207478@1.29.29,trna7_LysCTT_16_+_3225692_3225764@1.29.29,trna9_LysCTT_5_+_180634755_180634827@1.29.29\|GCCCGGCTAGCTCAGTCGGTAGAGCATGA | >trna119_LysCTT_1_-_145395522_145395594 | 1 | 29 | 29 | GCCCGGCTAGCTCAGTCGGTAGAGCATGA | tRF | FALSE | FALSE | TRUE |
| >hg38_21_+_8433172_8446622_RNA45SN1_45S_50ntFlanks@11581.11612.32\|AACTGGTGCGGACCAGGGGAATCCGACTGTTT | >hg38_21_+_8433172_8446622_RNA45SN1_45S_50ntFlanks | 11581 | 11612 | 32 | AACTGGTGCGGACCAGGGGAATCCGACTGTTT | rRF | FALSE | FALSE | TRUE |
| >hg38_21_+_8433172_8446622_RNA45SN1_45S_50ntFlanks@9534.9569.36\|AGGAAACTCTGGTGGAGGTCCGTAGCGGTCCTGACG | >hg38_21_+_8433172_8446622_RNA45SN1_45S_50ntFlanks | 9534 | 9569 | 36 | AGGAAACTCTGGTGGAGGTCCGTAGCGGTCCTGACG | rRF | FALSE | FALSE | TRUE |
| >hg38_21_+_8433172_8446622_RNA45SN1_45S_50ntFlanks@4003.4024.22\|ATAACCTCGGGCCGATCGCACG | >hg38_21_+_8433172_8446622_RNA45SN1_45S_50ntFlanks | 4003 | 4024 | 22 | ATAACCTCGGGCCGATCGCACG | rRF | FALSE | FALSE | TRUE |
| >hg38_21_+_8433172_8446622_RNA45SN1_45S_50ntFlanks@12405.12437.33\|CTTTTTGATCCTTCGATGTCGGCTCTTCCTATC | >hg38_21_+_8433172_8446622_RNA45SN1_45S_50ntFlanks | 12405 | 12437 | 33 | CTTTTTGATCCTTCGATGTCGGCTCTTCCTATC | rRF | FALSE | FALSE | TRUE |
| >hg38_21_+_8433172_8446622_RNA45SN1_45S_50ntFlanks@13004.13027.24\|ATCTATTGAAAGTCAGCCCTCGAC | >hg38_21_+_8433172_8446622_RNA45SN1_45S_50ntFlanks | 13004 | 13027 | 24 | ATCTATTGAAAGTCAGCCCTCGAC | rRF | FALSE | FALSE | TRUE |
| >hg38_21_+_8433172_8446622_RNA45SN1_45S_50ntFlanks@5484.5514.31\|GAGCGCTGAGAAGACGGTCGAACTTGACTAT | >hg38_21_+_8433172_8446622_RNA45SN1_45S_50ntFlanks | 5484 | 5514 | 31 | GAGCGCTGAGAAGACGGTCGAACTTGACTAT | rRF | FALSE | FALSE | TRUE |
| >hg38_21_+_8433172_8446622_RNA45SN1_45S_50ntFlanks@10841.10867.27\|CGTAACTTCGGGATAAGGATTGGCTCT | >hg38_21_+_8433172_8446622_RNA45SN1_45S_50ntFlanks | 10841 | 10867 | 27 | CGTAACTTCGGGATAAGGATTGGCTCT | rRF | FALSE | FALSE | TRUE |
| >hg38_21_+_8433172_8446622_RNA45SN1_45S_50ntFlanks@11581.11605.25\|AACTGGTGCGGACCAGGGGAATCCG | >hg38_21_+_8433172_8446622_RNA45SN1_45S_50ntFlanks | 11581 | 11605 | 25 | AACTGGTGCGGACCAGGGGAATCCG | rRF | FALSE | FALSE | TRUE |
| >hg38_21_+_8433172_8446622_RNA45SN1_45S_50ntFlanks@11768.11803.36\|CTAATTAGTGACGCGCATGAATGGATGAACGAGATT | >hg38_21_+_8433172_8446622_RNA45SN1_45S_50ntFlanks | 11768 | 11803 | 36 | CTAATTAGTGACGCGCATGAATGGATGAACGAGATT | rRF | FALSE | FALSE | TRUE |
| >hg38_21_+_8433172_8446622_RNA45SN1_45S_50ntFlanks@12935.12956.22\|TAAACCATTCGTAGACGACCTG | >hg38_21_+_8433172_8446622_RNA45SN1_45S_50ntFlanks | 12935 | 12956 | 22 | TAAACCATTCGTAGACGACCTG | rRF | FALSE | FALSE | TRUE |
| >hg38_21_+_8433172_8446622_RNA45SN1_45S_50ntFlanks@3882.3899.18\|CCGACGGGCGCTGACCCC | >hg38_21_+_8433172_8446622_RNA45SN1_45S_50ntFlanks | 3882 | 3899 | 18 | CCGACGGGCGCTGACCCC | rRF | FALSE | FALSE | TRUE |
| >hg38_21_+_8433172_8446622_RNA45SN1_45S_50ntFlanks@12419.12436.18\|GATGTCGGCTCTTCCTAT | >hg38_21_+_8433172_8446622_RNA45SN1_45S_50ntFlanks | 12419 | 12436 | 18 | GATGTCGGCTCTTCCTAT | rRF | FALSE | FALSE | TRUE |

<table>
<tr><td>>hg38_21_+_8433172_8446622_RNA45SN1_45S_50ntFlanks@4759.4782.24|ATCAGATACCGTCGTAGTTCCGAC</td><td>>hg38_21_+_8433172_8446622_RNA45SN1_45S_50ntFlanks</td><td>4759</td><td>4782</td><td>24</td><td>ATCAGATACCGTCGTAGTTCCGAC</td><td>rRF</td><td>FALSE</td><td>FALSE</td><td>TRUE</td></tr>
<tr><td>>hg38_21_+_8433172_8446622_RNA45SN1_45S_50ntFlanks@11703.11728.26|AATGAAGCGCGGGTAAACGGCGGGAG</td><td>>hg38_21_+_8433172_8446622_RNA45SN1_45S_50ntFlanks</td><td>11703</td><td>11728</td><td>26</td><td>AATGAAGCGCGGGTAAACGGCGGGAG</td><td>rRF</td><td>FALSE</td><td>FALSE</td><td>TRUE</td></tr>
<tr><td>>hg38_21_+_8433172_8446622_RNA45SN1_45S_50ntFlanks@10787.10818.32|ACAGCCTCTGGCATGTTGGAACAATGTAGGTA</td><td>>hg38_21_+_8433172_8446622_RNA45SN1_45S_50ntFlanks</td><td>10787</td><td>10818</td><td>32</td><td>ACAGCCTCTGGCATGTTGGAACAATGTAGGTA</td><td>rRF</td><td>FALSE</td><td>FALSE</td><td>TRUE</td></tr>
<tr><td>>hg38_21_+_8433172_8446622_RNA45SN1_45S_50ntFlanks@11641.11662.22|CGGCGGGTGTTGACGCGATGTG</td><td>>hg38_21_+_8433172_8446622_RNA45SN1_45S_50ntFlanks</td><td>11641</td><td>11662</td><td>22</td><td>CGGCGGGTGTTGACGCGATGTG</td><td>rRF</td><td>FALSE</td><td>FALSE</td><td>TRUE</td></tr>
<tr><td>>hg38_21_+_8433172_8446622_RNA45SN1_45S_50ntFlanks@12007.12032.26|GCCGCCGGTGAAATACCACTACTCTG</td><td>>hg38_21_+_8433172_8446622_RNA45SN1_45S_50ntFlanks</td><td>12007</td><td>12032</td><td>26</td><td>GCCGCCGGTGAAATACCACTACTCTG</td><td>rRF</td><td>FALSE</td><td>FALSE</td><td>TRUE</td></tr>
<tr><td>>hg38_21_+_8433172_8446622_RNA45SN1_45S_50ntFlanks@9567.9595.29|ACGTGCAAATCGGTCGTCCGACCTGGGTA</td><td>>hg38_21_+_8433172_8446622_RNA45SN1_45S_50ntFlanks</td><td>9567</td><td>9595</td><td>29</td><td>ACGTGCAAATCGGTCGTCCGACCTGGGTA</td><td>rRF</td><td>FALSE</td><td>FALSE</td><td>TRUE</td></tr>
<tr><td>>hg38_21_+_8433172_8446622_RNA45SN1_45S_50ntFlanks@11874.11898.25|AGAAGACCCTGTTGAGCTTGACTCT</td><td>>hg38_21_+_8433172_8446622_RNA45SN1_45S_50ntFlanks</td><td>11874</td><td>11898</td><td>25</td><td>AGAAGACCCTGTTGAGCTTGACTCT</td><td>rRF</td><td>FALSE</td><td>FALSE</td><td>TRUE</td></tr>
<tr><td>>hg38_21_+_8433172_8446622_RNA45SN1_45S_50ntFlanks@12456.12483.28|CCAAGCGTTGGATTGTTCACCCACTAAT</td><td>>hg38_21_+_8433172_8446622_RNA45SN1_45S_50ntFlanks</td><td>12456</td><td>12483</td><td>28</td><td>CCAAGCGTTGGATTGTTCACCCACTAAT</td><td>rRF</td><td>FALSE</td><td>FALSE</td><td>TRUE</td></tr>
<tr><td>>hg38_21_+_8433172_8446622_RNA45SN1_45S_50ntFlanks@12415.12434.20|CTTCGATGTCGGCTCTTCCT</td><td>>hg38_21_+_8433172_8446622_RNA45SN1_45S_50ntFlanks</td><td>12415</td><td>12434</td><td>20</td><td>CTTCGATGTCGGCTCTTCCT</td><td>rRF</td><td>FALSE</td><td>FALSE</td><td>TRUE</td></tr>
<tr><td>>hg38_21_+_8433172_8446622_RNA45SN1_45S_50ntFlanks@9568.9595.28|CGTGCAAATCGGTCGTCCGACCTGGGTA</td><td>>hg38_21_+_8433172_8446622_RNA45SN1_45S_50ntFlanks</td><td>9568</td><td>9595</td><td>28</td><td>CGTGCAAATCGGTCGTCCGACCTGGGTA</td><td>rRF</td><td>FALSE</td><td>FALSE</td><td>TRUE</td></tr>
<tr><td>>hg38_21_+_8433172_8446622_RNA45SN1_45S_50ntFlanks@11580.11611.32|GAACTGGTGCGGACCAGGGGAATCCGACTGTT</td><td>>hg38_21_+_8433172_8446622_RNA45SN1_45S_50ntFlanks</td><td>11580</td><td>11611</td><td>32</td><td>GAACTGGTGCGGACCAGGGGAATCCGACTGTT</td><td>rRF</td><td>FALSE</td><td>FALSE</td><td>TRUE</td></tr>
<tr><td>>hg38_21_+_8433172_8446622_RNA45SN1_45S_50ntFlanks@12605.12624.20|TATGTGCTTGGCTGAGGAGC</td><td>>hg38_21_+_8433172_8446622_RNA45SN1_45S_50ntFlanks</td><td>12605</td><td>12624</td><td>20</td><td>TATGTGCTTGGCTGAGGAGC</td><td>rRF</td><td>FALSE</td><td>FALSE</td><td>TRUE</td></tr>
<tr><td>>hg38_21_+_8433172_8446622_RNA45SN1_45S_50ntFlanks@11641.11669.29|CGGCGGGTGTTGACGCGATGTGATTTCTG</td><td>>hg38_21_+_8433172_8446622_RNA45SN1_45S_50ntFlanks</td><td>11641</td><td>11669</td><td>29</td><td>CGGCGGGTGTTGACGCGATGTGATTTCTG</td><td>rRF</td><td>FALSE</td><td>FALSE</td><td>TRUE</td></tr>
<tr><td>>hg38_21_+_8433172_8446622_RNA45SN1_45S_50ntFlanks@11594.11611.18|CAGGGGAATCCGACTGTT</td><td>>hg38_21_+_8433172_8446622_RNA45SN1_45S_50ntFlanks</td><td>11594</td><td>11611</td><td>18</td><td>CAGGGGAATCCGACTGTT</td><td>rRF</td><td>FALSE</td><td>FALSE</td><td>TRUE</td></tr>
<tr><td>>hg38_21_+_8433172_8446622_RNA45SN1_45S_50ntFlanks@12163.12181.19|ACTGGGGCGGTACACCTGT</td><td>>hg38_21_+_8433172_8446622_RNA45SN1_45S_50ntFlanks</td><td>12163</td><td>12181</td><td>19</td><td>ACTGGGGCGGTACACCTGT</td><td>rRF</td><td>FALSE</td><td>FALSE</td><td>TRUE</td></tr>
<tr><td>>hg38_21_+_8433172_8446622_RNA45SN1_45S_50ntFlanks@11581.11611.31|AACTGGTGCGGACCAGGGGAATCCGACTGTT</td><td>>hg38_21_+_8433172_8446622_RNA45SN1_45S_50ntFlanks</td><td>11581</td><td>11611</td><td>31</td><td>AACTGGTGCGGACCAGGGGAATCCGACTGTT</td><td>rRF</td><td>FALSE</td><td>FALSE</td><td>TRUE</td></tr>
<tr><td>>hg38_21_+_8433172_8446622_RNA45SN1_45S_50ntFlanks@3854.3881.28|ACTGTGGTAATTCTAGAGCTAATACATG</td><td>>hg38_21_+_8433172_8446622_RNA45SN1_45S_50ntFlanks</td><td>3854</td><td>3881</td><td>28</td><td>ACTGTGGTAATTCTAGAGCTAATACATG</td><td>rRF</td><td>FALSE</td><td>FALSE</td><td>TRUE</td></tr>
<tr><td>>hg38_21_+_8433172_8446622_RNA45SN1_45S_50ntFlanks@4726.4758.33|TAATCAAGAACGAAAGTCGGAGGTTCGAAGACG</td><td>>hg38_21_+_8433172_8446622_RNA45SN1_45S_50ntFlanks</td><td>4726</td><td>4758</td><td>33</td><td>TAATCAAGAACGAAAGTCGGAGGTTCGAAGACG</td><td>rRF</td><td>FALSE</td><td>FALSE</td><td>TRUE</td></tr>
<tr><td>>hg38_21_+_8433172_8446622_RNA45SN1_45S_50ntFlanks@12633.12661.29|CGAAGCTACCATCTGTGGGATTATGACTG</td><td>>hg38_21_+_8433172_8446622_RNA45SN1_45S_50ntFlanks</td><td>12633</td><td>12661</td><td>29</td><td>CGAAGCTACCATCTGTGGGATTATGACTG</td><td>rRF</td><td>FALSE</td><td>FALSE</td><td>TRUE</td></tr>
<tr><td>>hg38_21_+_8433172_8446622_RNA45SN1_45S_50ntFlanks@11209.11234.26|CGGCGACTCTGGACGCGAGCCGGGCC</td><td>>hg38_21_+_8433172_8446622_RNA45SN1_45S_50ntFlanks</td><td>11209</td><td>11234</td><td>26</td><td>CGGCGACTCTGGACGCGAGCCGGGCC</td><td>rRF</td><td>FALSE</td><td>FALSE</td><td>TRUE</td></tr>
<tr><td>>hg38_21_+_8433172_8446622_RNA45SN1_45S_50ntFlanks@10794.10816.23|CTGGCATGTTGGAACAATGTAGG</td><td>>hg38_21_+_8433172_8446622_RNA45SN1_45S_50ntFlanks</td><td>10794</td><td>10816</td><td>23</td><td>CTGGCATGTTGGAACAATGTAGG</td><td>rRF</td><td>FALSE</td><td>FALSE</td><td>TRUE</td></tr>
<tr><td>>hg38_21_+_8433172_8446622_RNA45SN1_45S_50ntFlanks@6721.6745.25|AATTGCAGGACACATTGATCATCGA</td><td>>hg38_21_+_8433172_8446622_RNA45SN1_45S_50ntFlanks</td><td>6721</td><td>6745</td><td>25</td><td>AATTGCAGGACACATTGATCATCGA</td><td>rRF</td><td>FALSE</td><td>FALSE</td><td>TRUE</td></tr>
<tr><td>>hg38_21_+_8433172_8446622_RNA45SN1_45S_50ntFlanks@7983.8008.26|CAGATCAGACGTGGCGACCCGCTGAA</td><td>>hg38_21_+_8433172_8446622_RNA45SN1_45S_50ntFlanks</td><td>7983</td><td>8008</td><td>26</td><td>CAGATCAGACGTGGCGACCCGCTGAA</td><td>rRF</td><td>FALSE</td><td>FALSE</td><td>TRUE</td></tr>
<tr><td>>hg38_21_+_8433172_8446622_RNA45SN1_45S_50ntFlanks@4773.4793.21|TAGTTCCGACCATAAACGATG</td><td>>hg38_21_+_8433172_8446622_RNA45SN1_45S_50ntFlanks</td><td>4773</td><td>4793</td><td>21</td><td>TAGTTCCGACCATAAACGATG</td><td>rRF</td><td>FALSE</td><td>FALSE</td><td>TRUE</td></tr>
<tr><td>>hg38_21_+_8433172_8446622_RNA45SN1_45S_50ntFlanks@6647.6681.35|CGTACGACTCTTAGCGGTGGATCACTCGGCTCGTG</td><td>>hg38_21_+_8433172_8446622_RNA45SN1_45S_50ntFlanks</td><td>6647</td><td>6681</td><td>35</td><td>CGTACGACTCTTAGCGGTGGATCACTCGGCTCGTG</td><td>rRF</td><td>FALSE</td><td>FALSE</td><td>TRUE</td></tr>
<tr><td>>hg38_21_+_8433172_8446622_RNA45SN1_45S_50ntFlanks@7988.8012.25|CAGACGTGGCGACCCGCTGAATTTA</td><td>>hg38_21_+_8433172_8446622_RNA45SN1_45S_50ntFlanks</td><td>7988</td><td>8012</td><td>25</td><td>CAGACGTGGCGACCCGCTGAATTTA</td><td>rRF</td><td>FALSE</td><td>FALSE</td><td>TRUE</td></tr>
<tr><td>>hg38_21_+_8433172_8446622_RNA45SN1_45S_50ntFlanks@9572.9603.32|CAAATCGGTCGTCCGACCTGGGTATAGGGGCG</td><td>>hg38_21_+_8433172_8446622_RNA45SN1_45S_50ntFlanks</td><td>9572</td><td>9603</td><td>32</td><td>CAAATCGGTCGTCCGACCTGGGTATAGGGGCG</td><td>rRF</td><td>FALSE</td><td>FALSE</td><td>TRUE</td></tr>
<tr><td>>hg38_21_+_8433172_8446622_RNA45SN1_45S_50ntFlanks@9534.9561.28|AGGAAACTCTGGTGGAGGTCCGTAGCGG</td><td>>hg38_21_+_8433172_8446622_RNA45SN1_45S_50ntFlanks</td><td>9534</td><td>9561</td><td>28</td><td>AGGAAACTCTGGTGGAGGTCCGTAGCGG</td><td>rRF</td><td>FALSE</td><td>FALSE</td><td>TRUE</td></tr>
<tr><td>>hg38_21_+_8433172_8446622_RNA45SN1_45S_50ntFlanks@11781.11803.23|CGCATGAATGGATGAACGAGATT</td><td>>hg38_21_+_8433172_8446622_RNA45SN1_45S_50ntFlanks</td><td>11781</td><td>11803</td><td>23</td><td>CGCATGAATGGATGAACGAGATT</td><td>rRF</td><td>FALSE</td><td>FALSE</td><td>TRUE</td></tr>
<tr><td>>hg38_21_+_8433172_8446622_RNA45SN1_45S_50ntFlanks@12001.12032.32|CTGCGGGCCGCCGGTGAAATACCACTACTCTG</td><td>>hg38_21_+_8433172_8446622_RNA45SN1_45S_50ntFlanks</td><td>12001</td><td>12032</td><td>32</td><td>CTGCGGGCCGCCGGTGAAATACCACTACTCTG</td><td>rRF</td><td>FALSE</td><td>FALSE</td><td>TRUE</td></tr>
<tr><td>>hg38_21_+_8433172_8446622_RNA45SN1_45S_50ntFlanks@12420.12441.22|ATGTCGGCTCTTCCTATCATTG</td><td>>hg38_21_+_8433172_8446622_RNA45SN1_45S_50ntFlanks</td><td>12420</td><td>12441</td><td>22</td><td>ATGTCGGCTCTTCCTATCATTG</td><td>rRF</td><td>FALSE</td><td>FALSE</td><td>TRUE</td></tr>
<tr><td>>hg38_21_+_8433172_8446622_RNA45SN1_45S_50ntFlanks@11568.11585.18|GCAGCCGACTTAGAACTG</td><td>>hg38_21_+_8433172_8446622_RNA45SN1_45S_50ntFlanks</td><td>11568</td><td>11585</td><td>18</td><td>GCAGCCGACTTAGAACTG</td><td>rRF</td><td>FALSE</td><td>FALSE</td><td>TRUE</td></tr>
<tr><td>>hg38_21_+_8433172_8446622_RNA45SN1_45S_50ntFlanks@6651.6680.30|CGACTCTTAGCGGTGGATCACTCGGCTCGT</td><td>>hg38_21_+_8433172_8446622_RNA45SN1_45S_50ntFlanks</td><td>6651</td><td>6680</td><td>30</td><td>CGACTCTTAGCGGTGGATCACTCGGCTCGT</td><td>rRF</td><td>FALSE</td><td>FALSE</td><td>TRUE</td></tr>
<tr><td>>MI0000737|hsa-mir-200a&WithFlank&1|+|1167857|1167958@60.82.23&[MIMAT0000682&hsa-miR-200a-3p&offsets|0|+1;m-142&1|+|1167916|1167938&offsets|0|0]|TAACACTGTCTGGTAACGATGTT</td><td>>MI0000737|hsa-mir-200a&WithFlank&1|+|1167857|1167958</td><td>60</td><td>82</td><td>23</td><td>TAACACTGTCTGGTAACGATGTT</td><td>isomiR</td><td>FALSE</td><td>FALSE</td><td>TRUE</td></tr>
<tr><td>>hg38_21_+_8433172_8446622_RNA45SN1_45S_50ntFlanks@10781.10818.38|AGGTGAACAGCCTCTGGCATGTTGGAACAATGTAGGTA</td><td>>hg38_21_+_8433172_8446622_RNA45SN1_45S_50ntFlanks</td><td>10781</td><td>10818</td><td>38</td><td>AGGTGAACAGCCTCTGGCATGTTGGAACAATGTAGGTA</td><td>rRF</td><td>FALSE</td><td>FALSE</td><td>TRUE</td></tr>
<tr><td>>hg38_21_+_8433172_8446622_RNA45SN1_45S_50ntFlanks@4367.4389.23|CAGTTAAAAAGCTCGTAGTTGGA</td><td>>hg38_21_+_8433172_8446622_RNA45SN1_45S_50ntFlanks</td><td>4367</td><td>4389</td><td>23</td><td>CAGTTAAAAAGCTCGTAGTTGGA</td><td>rRF</td><td>FALSE</td><td>FALSE</td><td>TRUE</td></tr>
</table>
