## Supplementary material for "Single-nucleotide Differences and Cell Type Decide the Subcellular Localization of miRNA Isoforms (isomiRs), tRNA-derived Fragments (tRFs) and rRNA-derived Fragments (rRFs)": Supp. Table S4

**Supplemental Table S4:** Data table of the differentially abundant (DA) short RNA for the cell line RNA between the three cell lines with FDR < 0.05. The table is displayed as the output from DESeq2 where MedianFC (median fold change) is presented as raw reads . The annotation "am" is used to indicate abiguous short RNA, those that can be mapped to a noncoding RNA gene as well as somewhere else on the geneome. The annotation "multi" referrs to short RNA that map to locations in both the mitochondrial and nuclear genomes. The rRF are flanked by 50 nucleotdies upstream and downstream and their coordiantes reflect this. The coordiantes for the isomiR and rRF are from the hg38 human genome assembly while the coordintes for the tRF are from the hg19 human genome assembly.

<table>
<tr><th>Full Name</th><th>Short Name</th><th>Start</th><th>End</th><th>Length</th><th>Sequence</th><th>MedianFC</th><th>log2FC</th><th>lfcSE</th><th>stat</th><th>p-value</th><th>adjp-value</th><th>Numerator/Denominator</th></tr>
<tr><td>>am_MI0000809|hsa-mir-151a&WithFlank&8|-|140732558|140732659@53.73.21&[MIMAT0000757&hsa-miR-151a-3p&offsets|0|0;m-30&8|-|140732587|140732607&offsets|0|0]|CTAGACTGAAGCTCCTTGAGG</td><td>>am_MI0000809|hsa-mir-151a&WithFlank&8|-|140732558|140732659</td><td>53</td><td>73</td><td>21</td><td>CTAGACTGAAGCTCCTTGAGG</td><td>152086.2</td><td>8.8</td><td>0.4</td><td>22.0</td><td>4.1E-107</td><td>2.4E-103</td><td>BT20/MB468</td></tr>
<tr><td>>hg38_21_+_8433172_8446622_RNA45SN1_45S_50ntFlanks@6721.6744.24|AATTGCAGGACACATTGATCATCG</td><td>>hg38_21_+_8433172_8446622_RNA45SN1_45S_50ntFlanks</td><td>6721</td><td>6744</td><td>24</td><td>AATTGCAGGACACATTGATCATCG</td><td>20028.6</td><td>-8.6</td><td>0.4</td><td>-20.7</td><td>8.4E-95</td><td>2.5E-91</td><td>BT20/MB468</td></tr>
<tr><td>>trna116_GluCTC_1_-_145399233_145399304@1.35.35,trna59_GluCTC_1_+_249168447_249168518@1.35.35,trna71_GluCTC_1_-_161439189_161439260@1.35.35,trna74_GluCTC_1_-_161431809_161431880@1.35.35,trna77_GluCTC_1_-_161424398_161424469@1.35.35,trna77_GluCTC_6_+_28949976_28950047@1.35.35,trna80_GluCTC_1_-_161417018_161417089@1.35.35,trna87_GluCTC_6_-_126101393_126101464@1.35.35|TCCCTGGTGGTCTAGTGGTTAGGATTCGGCGCTCT</td><td>>trna116_GluCTC_1_-_145399233_145399304</td><td>1</td><td>35</td><td>35</td><td>TCCCTGGTGGTCTAGTGGTTAGGATTCGGCGCTCT</td><td>10385.9</td><td>9.5</td><td>0.5</td><td>20.2</td><td>1.8E-90</td><td>3.6E-87</td><td>BT20/MB468</td></tr>
<tr><td>>hg38_21_+_8433172_8446622_RNA45SN1_45S_50ntFlanks@12456.12486.31|CCAAGCGTTGGATTGTTCACCCACTAATAGG</td><td>>hg38_21_+_8433172_8446622_RNA45SN1_45S_50ntFlanks</td><td>12456</td><td>12486</td><td>31</td><td>CCAAGCGTTGGATTGTTCACCCACTAATAGG</td><td>12047.4</td><td>-8.0</td><td>0.4</td><td>-19.5</td><td>5.2E-85</td><td>7.7E-82</td><td>BT20/MB468</td></tr>
<tr><td>>hg38_21_+_8433172_8446622_RNA45SN1_45S_50ntFlanks@12456.12478.23|CCAAGCGTTGGATTGTTCACCCA</td><td>>hg38_21_+_8433172_8446622_RNA45SN1_45S_50ntFlanks</td><td>12456</td><td>12478</td><td>23</td><td>CCAAGCGTTGGATTGTTCACCCA</td><td>11035.1</td><td>-8.0</td><td>0.4</td><td>-19.5</td><td>8.9E-85</td><td>1.0E-81</td><td>BT20/MB468</td></tr>
<tr><td>>hg38_21_+_8433172_8446622_RNA45SN1_45S_50ntFlanks@10841.10865.25|CGTAACTTCGGGATAAGGATTGGCT</td><td>>hg38_21_+_8433172_8446622_RNA45SN1_45S_50ntFlanks</td><td>10841</td><td>10865</td><td>25</td><td>CGTAACTTCGGGATAAGGATTGGCT</td><td>36363.5</td><td>-8.7</td><td>0.4</td><td>-19.4</td><td>7.9E-84</td><td>7.8E-81</td><td>BT20/MB468</td></tr>
<tr><td>>MI0000253|hsa-mir-148a&WithFlank&7|-|25949913|25949992@50.70.21(+1C)&[MIMAT0000243&hsa-miR-148a-3p&offsets|0|-1(+1C);m-7&7|-|25949922|25949943&offsets|0|-1(+1C)]|TCAGTGCACTACAGAACTTTGC</td><td>>MI0000253|hsa-mir-148a&WithFlank&7|-|25949913|25949992</td><td>50</td><td>70</td><td>21(+1C)</td><td>TCAGTGCACTACAGAACTTTGC</td><td>75765.1</td><td>8.2</td><td>0.4</td><td>19.2</td><td>5.5E-82</td><td>4.6E-79</td><td>BT20/MB468</td></tr>
<tr><td>>MI0000301|hsa-mir-224&WithFlank&X|-|151958572|151958664@14.35.22&[MIMAT0000281&hsa-miR-224-5p&offsets|+1|-2;m-238&X|-|151958630|151958651&offsets|0|0]|CAAGTCACTAGTGGTTCCGTTT</td><td>>MI0000301|hsa-mir-224&WithFlank&X|-|151958572|151958664</td><td>14</td><td>35</td><td>22</td><td>CAAGTCACTAGTGGTTCCGTTT</td><td>23030.8</td><td>12.1</td><td>0.6</td><td>19.1</td><td>1.5E-81</td><td>1.1E-78</td><td>BT20/MB468</td></tr>
<tr><td>>MI0000253|hsa-mir-148a&WithFlank&7|-|25949913|25949992@50.71.22&[MIMAT0000243&hsa-miR-148a-3p&offsets|0|0;m-7&7|-|25949922|25949943&offsets|0|0]|TCAGTGCACTACAGAACTTTGT</td><td>>MI0000253|hsa-mir-148a&WithFlank&7|-|25949913|25949992</td><td>50</td><td>71</td><td>22</td><td>TCAGTGCACTACAGAACTTTGT</td><td>590961.2</td><td>7.9</td><td>0.4</td><td>18.7</td><td>7.9E-78</td><td>4.6E-75</td><td>BT20/MB468</td></tr>
<tr><td>>MI0000255|hsa-mir-30d&WithFlank&8|-|134804870|134804951@12.33.22(+1U)&[MIMAT0000245&hsa-miR-30d-5p&offsets|0|0(+1U);m-21&8|-|134804919|134804940&offsets|0|0(+1U)]|TGTAAACATCCCCGACTGGAAGT</td><td>>MI0000255|hsa-mir-30d&WithFlank&8|-|134804870|134804951</td><td>12</td><td>33</td><td>22(+1U)</td><td>TGTAAACATCCCCGACTGGAAGT</td><td>29657.7</td><td>9.3</td><td>0.5</td><td>18.7</td><td>7.7E-78</td><td>4.6E-75</td><td>BT20/MB468</td></tr>
<tr><td>>hg38_21_+_8433172_8446622_RNA45SN1_45S_50ntFlanks@10679.10710.32|GGTTCCGGCGGCGTCCGGTGAGCTCTCGCTGG</td><td>>hg38_21_+_8433172_8446622_RNA45SN1_45S_50ntFlanks</td><td>10679</td><td>10710</td><td>32</td><td>GGTTCCGGCGGCGTCCGGTGAGCTCTCGCTGG</td><td>27771.3</td><td>-9.0</td><td>0.5</td><td>-18.7</td><td>2.9E-78</td><td>1.7E-74</td><td>MB231/MB468</td></tr>
<tr><td>>am_MI0000809|hsa-mir-151a&WithFlank&8|-|140732558|140732659@53.72.20(+1A)&[MIMAT0000757&hsa-miR-151a-3p&offsets|0|-1(+1A);m-30&8|-|140732587|140732607&offsets|0|-1(+1A)]|CTAGACTGAAGCTCCTTGAGA</td><td>>am_MI0000809|hsa-mir-151a&WithFlank&8|-|140732558|140732659</td><td>53</td><td>72</td><td>20(+1A)</td><td>CTAGACTGAAGCTCCTTGAGA</td><td>31140.1</td><td>9.9</td><td>0.5</td><td>18.1</td><td>2.1E-73</td><td>1.1E-70</td><td>BT20/MB468</td></tr>
<tr><td>>hg38_21_+_8433172_8446622_RNA45SN1_45S_50ntFlanks@10411.10429.19|CAGTCGGTCCTGAGAGATG</td><td>>hg38_21_+_8433172_8446622_RNA45SN1_45S_50ntFlanks</td><td>10411</td><td>10429</td><td>19</td><td>CAGTCGGTCCTGAGAGATG</td><td>11716.5</td><td>-7.2</td><td>0.4</td><td>-18.0</td><td>4.2E-72</td><td>2.1E-69</td><td>BT20/MB468</td></tr>
<tr><td>>hg38_21_+_8433172_8446622_RNA45SN1_45S_50ntFlanks@10391.10410.20|AACAGCAGTTGAACATGGGT</td><td>>hg38_21_+_8433172_8446622_RNA45SN1_45S_50ntFlanks</td><td>10391</td><td>10410</td><td>20</td><td>AACAGCAGTTGAACATGGGT</td><td>5413.9</td><td>-7.9</td><td>0.5</td><td>-17.4</td><td>1.7E-67</td><td>7.7E-65</td><td>BT20/MB468</td></tr>
<tr><td>>trna111_HisGTG_1_-_147774845_147774916@1.33.33,trna118_HisGTG_1_-_145396881_145396952@1.33.33,trna16_HisGTG_1_+_146544773_146544844@1.33.33,trna1_HisGTG_15_+_45493349_45493420@1.33.33,trna21_HisGTG_1_+_147753471_147753542@1.33.33,trna33_HisGTG_6_+_27125906_27125977@1.33.33,trna7_HisGTG_9_-_14433938_14434009@1.33.33,trna8_HisGTG_15_-_45492611_45492682@1.33.33,trna9_HisGTG_15_-_45490804_45490875@1.33.33|GCCGTGATCGTATAGTGGTTAGTACTCTGCGTT</td><td>>trna111_HisGTG_1_-_147774845_147774916</td><td>1</td><td>33</td><td>33</td><td>GCCGTGATCGTATAGTGGTTAGTACTCTGCGTT</td><td>5669.4</td><td>9.0</td><td>0.5</td><td>17.3</td><td>2.1E-67</td><td>8.6E-65</td><td>BT20/MB468</td></tr>
<tr><td>>hg38_21_+_8433172_8446622_RNA45SN1_45S_50ntFlanks@12456.12487.32|CCAAGCGTTGGATTGTTCACCCACTAATAGGG</td><td>>hg38_21_+_8433172_8446622_RNA45SN1_45S_50ntFlanks</td><td>12456</td><td>12487</td><td>32</td><td>CCAAGCGTTGGATTGTTCACCCACTAATAGGG</td><td>44902.3</td><td>-11.2</td><td>0.6</td><td>-17.3</td><td>2.9E-67</td><td>1.1E-64</td><td>BT20/MB468</td></tr>
<tr><td>>hg38_21_+_8433172_8446622_RNA45SN1_45S_50ntFlanks@12456.12476.21|CCAAGCGTTGGATTGTTCACC</td><td>>hg38_21_+_8433172_8446622_RNA45SN1_45S_50ntFlanks</td><td>12456</td><td>12476</td><td>21</td><td>CCAAGCGTTGGATTGTTCACC</td><td>39598.1</td><td>-8.1</td><td>0.5</td><td>-17.2</td><td>1.4E-66</td><td>5.3E-64</td><td>BT20/MB468</td></tr>
<tr><td>>hg38_21_+_8433172_8446622_RNA45SN1_45S_50ntFlanks@6719.6744.26|TGAATTGCAGGACACATTGATCATCG</td><td>>hg38_21_+_8433172_8446622_RNA45SN1_45S_50ntFlanks</td><td>6719</td><td>6744</td><td>26</td><td>TGAATTGCAGGACACATTGATCATCG</td><td>9240.3</td><td>-9.2</td><td>0.5</td><td>-17.2</td><td>4.4E-66</td><td>1.5E-63</td><td>BT20/MB468</td></tr>
<tr><td>>hg38_21_+_8433172_8446622_RNA45SN1_45S_50ntFlanks@12626.12654.29|AATGGGGCGAAGCTACCATCTGTGGGATT</td><td>>hg38_21_+_8433172_8446622_RNA45SN1_45S_50ntFlanks</td><td>12626</td><td>12654</td><td>29</td><td>AATGGGGCGAAGCTACCATCTGTGGGATT</td><td>7850.0</td><td>-7.3</td><td>0.4</td><td>-17.1</td><td>2.6E-65</td><td>8.6E-63</td><td>BT20/MB468</td></tr>
<tr><td>>MI0000650|hsa-mir-200c&WithFlank&12|+|6963693|6963772@50.72.23&[MIMAT0000617&hsa-miR-200c-3p&offsets|0|0;m-54&12|+|6963742|6963764&offsets|0|0]|TAATACTGCCGGGTAATGATGGA</td><td>>MI0000650|hsa-mir-200c&WithFlank&12|+|6963693|6963772</td><td>50</td><td>72</td><td>23</td><td>TAATACTGCCGGGTAATGATGGA</td><td>140065.0</td><td>10.5</td><td>0.6</td><td>17.2</td><td>4.3E-66</td><td>1.3E-62</td><td>BT20/MB231</td></tr>
<tr><td>>am_MI0005763|hsa-mir-941-1&WithFlank&20|+|63919443|63919526@53.74.22(+1U);MI0005764|hsa-mir-941-2&WithFlank&20|+|63919499|63919582@53.74.22(+1U);MI0005765|hsa-mir-941-3&WithFlank&20|+|63919555|63919638@53.74.22(+1U);MI0005766|hsa-mir-941-4&WithFlank&20|+|63919750|63919833@53.74.22(+1U);MI0031520|hsa-mir-941-5&WithFlank&20|+|63919862|63919945@53.74.22(+1U)&[MIMAT0004984&hsa-miR-941&offsets|0|-1(+1U);m-263&20|+|63919495|63919517&offsets|0|-1(+1U)];[MIMAT0004984_1&hsa-miR-941&offsets|0|-1(+1U);m-263&20|+|63919551|63919573&offsets|0|-1(+1U)];[MIMAT0004984_2&hsa-miR-941&offsets|0|-1(+1U);m-263&20|+|63919607|63919629&offsets|0|-1(+1U)];[MIMAT0004984_3&hsa-miR-941&offsets|0|-1(+1U);m-263&20|+|63919802|63919824&offsets|0|-1(+1U)];[MIMAT0004984_4&hsa-miR-941&offsets|0|-1(+1U);m-263&20|+|63919914|63919936&offsets|0|-1(+1U)]|CACCCGGCTGTGTGCACATGTGT</td><td>>am_MI0005763|hsa-mir-941-1&WithFlank&20|+|63919443|63919526</td><td>53</td><td>74</td><td>22(+1U)</td><td>CACCCGGCTGTGTGCACATGTGT</td><td>7588.0</td><td>8.5</td><td>0.5</td><td>17.0</td><td>5.6E-65</td><td>1.7E-62</td><td>BT20/MB468</td></tr>
<tr><td>>hg38_21_+_8433172_8446622_RNA45SN1_45S_50ntFlanks@9534.9553.20|AGGAAACTCTGGTGGAGGTC</td><td>>hg38_21_+_8433172_8446622_RNA45SN1_45S_50ntFlanks</td><td>9534</td><td>9553</td><td>20</td><td>AGGAAACTCTGGTGGAGGTC</td><td>69638.5</td><td>-9.4</td><td>0.6</td><td>-16.9</td><td>1.9E-64</td><td>5.7E-62</td><td>BT20/MB468</td></tr>
<tr><td>>hg38_21_+_8433172_8446622_RNA45SN1_45S_50ntFlanks@4731.4765.35|AAGAACGAAAGTCGGAGGTTCGAAGACGATCAGAT</td><td>>hg38_21_+_8433172_8446622_RNA45SN1_45S_50ntFlanks</td><td>4731</td><td>4765</td><td>35</td><td>AAGAACGAAAGTCGGAGGTTCGAAGACGATCAGAT</td><td>754.8</td><td>6.5</td><td>0.4</td><td>16.9</td><td>2.1E-64</td><td>5.8E-62</td><td>BT20/MB468</td></tr>
<tr><td>>am_MI0000809|hsa-mir-151a&WithFlank&8|-|140732558|140732659@53.72.20(+1U)&[MIMAT0000757&hsa-miR-151a-3p&offsets|0|-1(+1U);m-30&8|-|140732587|140732607&offsets|0|-1(+1U)]|CTAGACTGAAGCTCCTTGAGT</td><td>>am_MI0000809|hsa-mir-151a&WithFlank&8|-|140732558|140732659</td><td>53</td><td>72</td><td>20(+1U)</td><td>CTAGACTGAAGCTCCTTGAGT</td><td>14662.1</td><td>9.9</td><td>0.6</td><td>16.9</td><td>3.1E-64</td><td>8.2E-62</td><td>BT20/MB468</td></tr>
<tr><td>>am_MI0000809|hsa-mir-151a&WithFlank&8|-|140732558|140732659@53.73.21(+1U)&[MIMAT0000757&hsa-miR-151a-3p&offsets|0|0(+1U);m-30&8|-|140732587|140732607&offsets|0|0(+1U)]|CTAGACTGAAGCTCCTTGAGGT</td><td>>am_MI0000809|hsa-mir-151a&WithFlank&8|-|140732558|140732659</td><td>53</td><td>73</td><td>21(+1U)</td><td>CTAGACTGAAGCTCCTTGAGGT</td><td>19907.0</td><td>9.2</td><td>0.5</td><td>16.9</td><td>4.7E-64</td><td>1.2E-61</td><td>BT20/MB468</td></tr>
<tr><td>>hg38_21_+_8433172_8446622_RNA45SN1_45S_50ntFlanks@6700.6718.19|CTAGCTGCGAGAATTAATG</td><td>>hg38_21_+_8433172_8446622_RNA45SN1_45S_50ntFlanks</td><td>6700</td><td>6718</td><td>19</td><td>CTAGCTGCGAGAATTAATG</td><td>6501.0</td><td>-7.9</td><td>0.5</td><td>-16.8</td><td>2.8E-63</td><td>6.7E-61</td><td>BT20/MB468</td></tr>
<tr><td>>hg38_21_+_8433172_8446622_RNA45SN1_45S_50ntFlanks@7975.8009.35|CGCGACCTCAGATCAGACGTGGCGACCCGCTGAAT</td><td>>hg38_21_+_8433172_8446622_RNA45SN1_45S_50ntFlanks</td><td>7975</td><td>8009</td><td>35</td><td>CGCGACCTCAGATCAGACGTGGCGACCCGCTGAAT</td><td>74305.5</td><td>6.7</td><td>0.4</td><td>16.8</td><td>2.8E-63</td><td>6.7E-61</td><td>BT20/MB468</td></tr>
<tr><td>>MI0000650|hsa-mir-200c&WithFlank&12|+|6963693|6963772@50.71.22&[MIMAT0000617&hsa-miR-200c-3p&offsets|0|-1;m-54&12|+|6963742|6963764&offsets|0|-1]|TAATACTGCCGGGTAATGATGG</td><td>>MI0000650|hsa-mir-200c&WithFlank&12|+|6963693|6963772</td><td>50</td><td>71</td><td>22</td><td>TAATACTGCCGGGTAATGATGG</td><td>21473.7</td><td>11.1</td><td>0.7</td><td>16.9</td><td>6.5E-64</td><td>9.7E-61</td><td>BT20/MB231</td></tr>
<tr><td>>hg38_21_+_8433172_8446622_RNA45SN1_45S_50ntFlanks@11783.11804.22|CATGAATGGATGAACGAGATTC</td><td>>hg38_21_+_8433172_8446622_RNA45SN1_45S_50ntFlanks</td><td>11783</td><td>11804</td><td>22</td><td>CATGAATGGATGAACGAGATTC</td><td>24645.2</td><td>-8.4</td><td>0.5</td><td>-16.7</td><td>6.4E-63</td><td>1.4E-60</td><td>BT20/MB468</td></tr>
<tr><td>>hg38_21_+_8433172_8446622_RNA45SN1_45S_50ntFlanks@10678.10710.33|CGGTTCCGGCGGCGTCCGGTGAGCTCTCGCTGG</td><td>>hg38_21_+_8433172_8446622_RNA45SN1_45S_50ntFlanks</td><td>10678</td><td>10710</td><td>33</td><td>CGGTTCCGGCGGCGTCCGGTGAGCTCTCGCTGG</td><td>140001.6</td><td>-8.5</td><td>0.5</td><td>-16.8</td><td>1.1E-63</td><td>3.3E-60</td><td>MB231/MB468</td></tr>
<tr><td>>hg38_21_+_8433172_8446622_RNA45SN1_45S_50ntFlanks@6651.6695.45|CGACTCTTAGCGGTGGATCACTCGGCTCGTGCGTCGATGAAGAAC</td><td>>hg38_21_+_8433172_8446622_RNA45SN1_45S_50ntFlanks</td><td>6651</td><td>6695</td><td>45</td><td>CGACTCTTAGCGGTGGATCACTCGGCTCGTGCGTCGATGAAGAAC</td><td>11574.1</td><td>9.9</td><td>0.6</td><td>16.8</td><td>1.7E-63</td><td>3.5E-60</td><td>MB231/MB468</td></tr>
<tr><td>>hg38_21_+_8433172_8446622_RNA45SN1_45S_50ntFlanks@12456.12484.29|CCAAGCGTTGGATTGTTCACCCACTAATA</td><td>>hg38_21_+_8433172_8446622_RNA45SN1_45S_50ntFlanks</td><td>12456</td><td>12484</td><td>29</td><td>CCAAGCGTTGGATTGTTCACCCACTAATA</td><td>10099.9</td><td>-9.6</td><td>0.6</td><td>-16.6</td><td>4.0E-62</td><td>8.8E-60</td><td>BT20/MB468</td></tr>
<tr><td>>hg38_21_+_8433172_8446622_RNA45SN1_45S_50ntFlanks@3988.4023.36|CTTTGGTGACTCTAGATAACCTCGGGCCGATCGCAC</td><td>>hg38_21_+_8433172_8446622_RNA45SN1_45S_50ntFlanks</td><td>3988</td><td>4023</td><td>36</td><td>CTTTGGTGACTCTAGATAACCTCGGGCCGATCGCAC</td><td>1386.2</td><td>6.5</td><td>0.4</td><td>16.6</td><td>4.5E-62</td><td>9.4E-60</td><td>BT20/MB468</td></tr>
</table>

| | | | | | | | | | | | | |
|---|---|---|---|---|---|---|---|---|---|---|---|---|
| >hg38_21_+_8433172_8446622_RNA45SN1_45S_50ntFlanks@12456.12487.32\|CCAAGCGTTGGATTGTTCACCCACTAATAGGG | >hg38_21_+_8433172_8446622_RNA45SN1_45S_50ntFlanks | 12456 | 12487 | 32 | CCAAGCGTTGGATTGTTCACCCACTAATAGGG | 37787.5 | -9.4 | 0.6 | -16.7 | 1.4E-62 | 2.1E-59 | MB231/MB468 |
| >hg38_21_+_8433172_8446622_RNA45SN1_45S_50ntFlanks@10351.10368.18\|CGAGAACTTTGAAGGCCG | >hg38_21_+_8433172_8446622_RNA45SN1_45S_50ntFlanks | 10351 | 10368 | 18 | CGAGAACTTTGAAGGCCG | 28878.1 | -8.7 | 0.5 | -16.6 | 1.1E-61 | 2.1E-59 | BT20/MB468 |
| >MI0000809\|hsa-mir-151a&WithFlank&8\|-\|140732558\|140732659@51.73.23&[MIMAT0000757&hsa-miR-151a-3p&offsets\|-2\|0;m-30&8\|-\|140732587\|140732607&offsets\|-2\|0]\|TACTAGACTGAAGCTCCTTGAGG | >MI0000809\|hsa-mir-151a&WithFlank&8\|-\|140732558\|140732659 | 51 | 73 | 23 | TACTAGACTGAAGCTCCTTGAGG | 8777.0 | 7.8 | 0.5 | 16.5 | 2.6E-61 | 5.0E-59 | BT20/MB468 |
| >hg38_21_+_8433172_8446622_RNA45SN1_45S_50ntFlanks@7975.8010.36\|CGCGACCTCAGATCAGACGTGGCGACCCGCTGAATT | >hg38_21_+_8433172_8446622_RNA45SN1_45S_50ntFlanks | 7975 | 8010 | 36 | CGCGACCTCAGATCAGACGTGGCGACCCGCTGAATT | 98712.3 | 8.0 | 0.5 | 16.5 | 3.3E-61 | 6.2E-59 | BT20/MB468 |
| >hg38_21_+_8433172_8446622_RNA45SN1_45S_50ntFlanks@9564.9596.33\|CTGACGTGCAAATCGGTCGTCCGACCTGGGTAT | >hg38_21_+_8433172_8446622_RNA45SN1_45S_50ntFlanks | 9564 | 9596 | 33 | CTGACGTGCAAATCGGTCGTCCGACCTGGGTAT | 13021.5 | -8.7 | 0.5 | -16.3 | 6.0E-60 | 1.1E-57 | BT20/MB468 |
| >hg38_21_+_8433172_8446622_RNA45SN1_45S_50ntFlanks@12974.12991.18\|CGTACGTAGCAGAGCAGC | >hg38_21_+_8433172_8446622_RNA45SN1_45S_50ntFlanks | 12974 | 12991 | 18 | CGTACGTAGCAGAGCAGC | 5085.3 | -7.6 | 0.5 | -16.3 | 2.1E-59 | 3.7E-57 | BT20/MB468 |
| >am_MI0000809\|hsa-mir-151a&WithFlank&8\|-\|140732558\|140732659@52.73.22&[MIMAT0000757&hsa-miR-151a-3p&offsets\|-1\|0;m-30&8\|-\|140732587\|140732607&offsets\|-1\|0]\|ACTAGACTGAAGCTCCTTGAGG | >am_MI0000809\|hsa-mir-151a&WithFlank&8\|-\|140732558\|140732659 | 52 | 73 | 22 | ACTAGACTGAAGCTCCTTGAGG | 2761.4 | 7.9 | 0.5 | 16.2 | 3.8E-59 | 6.6E-57 | BT20/MB468 |
| >am_MI0000809\|hsa-mir-151a&WithFlank&8\|-\|140732558\|140732659@53.72.20&[MIMAT0000757&hsa-miR-151a-3p&offsets\|0\|-1;m-30&8\|-\|140732587\|140732607&offsets\|0\|-1]\|CTAGACTGAAGCTCCTTGAG | >am_MI0000809\|hsa-mir-151a&WithFlank&8\|-\|140732558\|140732659 | 53 | 72 | 20 | CTAGACTGAAGCTCCTTGAG | 9785.3 | 9.3 | 0.6 | 16.1 | 3.4E-58 | 5.7E-56 | BT20/MB468 |
| >MI0000255\|hsa-mir-30d&WithFlank&8\|-\|134804870\|134804951@12.34.23&[MIMAT0000245&hsa-miR-30d-5p&offsets\|0\|+1;m-21&8\|-\|134804919\|134804940&offsets\|0\|+1]\|TGTAAACATCCCCGACTGGAAGC | >MI0000255\|hsa-mir-30d&WithFlank&8\|-\|134804870\|134804951 | 12 | 34 | 23 | TGTAAACATCCCCGACTGGAAGC | 74914.7 | 8.3 | 0.5 | 15.8 | 1.9E-56 | 3.1E-54 | BT20/MB468 |
| >am_MI0000809\|hsa-mir-151a&WithFlank&8\|-\|140732558\|140732659@53.73.21(+1C)&[MIMAT0000757&hsa-miR-151a-3p&offsets\|0\|0(+1C);m-30&8\|-\|140732587\|140732607&offsets\|0\|0(+1C)]\|CTAGACTGAAGCTCCTTGAGGC | >am_MI0000809\|hsa-mir-151a&WithFlank&8\|-\|140732558\|140732659 | 53 | 73 | 21(+1C) | CTAGACTGAAGCTCCTTGAGGC | 3045.7 | 9.1 | 0.6 | 15.8 | 6.4E-56 | 1.0E-53 | BT20/MB468 |
| >hg38_21_+_8433172_8446622_RNA45SN1_45S_50ntFlanks@12934.12956.23\|CTAAACCATTCGTAGACGACCTG | >hg38_21_+_8433172_8446622_RNA45SN1_45S_50ntFlanks | 12934 | 12956 | 23 | CTAAACCATTCGTAGACGACCTG | 5660.7 | -9.2 | 0.6 | -15.7 | 2.9E-55 | 4.4E-53 | BT20/MB468 |
| >MI0000650\|hsa-mir-200c&WithFlank&12\|+\|6963693\|6963772@50.72.23(+1A)&[MIMAT0000617&hsa-miR-200c-3p&offsets\|0\|0(+1A);m-54&12\|+\|6963742\|6963764&offsets\|0\|0(+1A)]\|TAATACTGCCGGGTAATGATGGAA | >MI0000650\|hsa-mir-200c&WithFlank&12\|+\|6963693\|6963772 | 50 | 72 | 23(+1A) | TAATACTGCCGGGTAATGATGGAA | 13515.6 | 10.9 | 0.7 | 15.7 | 6.9E-56 | 6.9E-53 | BT20/MB231 |
| >trnaMT_GluTTC_MT_-_14674_14742@35.72.38,trnalookalike8_GluTTC_5_-_93905172_93905240@35.72.38\|TATCATTGGTCGTGGTTGTAGTCCGTGCGAGAATACCA | >trnaMT_GluTTC_MT_-_14674_14742 | 35 | 72 | 38 | TATCATTGGTCGTGGTTGTAGTCCGTGCGAGAATACCA | 11344.8 | 10.2 | 0.6 | 15.7 | 8.2E-56 | 9.9E-53 | MB231/MB468 |
| >hg38_21_+_8433172_8446622_RNA45SN1_45S_50ntFlanks@10411.10430.20\|CAGTCGGTCCTGAGAGATGG | >hg38_21_+_8433172_8446622_RNA45SN1_45S_50ntFlanks | 10411 | 10430 | 20 | CAGTCGGTCCTGAGAGATGG | 7025.5 | -10.0 | 0.6 | -15.5 | 2.3E-54 | 3.4E-52 | BT20/MB468 |
| >hg38_21_+_8433172_8446622_RNA45SN1_45S_50ntFlanks@12605.12626.22\|TATGTGCTTGGCTGAGGAGCCA | >hg38_21_+_8433172_8446622_RNA45SN1_45S_50ntFlanks | 12605 | 12626 | 22 | TATGTGCTTGGCTGAGGAGCCA | 4640.3 | -8.3 | 0.5 | -15.5 | 3.5E-54 | 5.2E-52 | BT20/MB468 |
| >hg38_21_+_8433172_8446622_RNA45SN1_45S_50ntFlanks@10678.10711.34\|CGGTTCCGGCGGCGTCCGGTGAGCTCTCGCTGGC | >hg38_21_+_8433172_8446622_RNA45SN1_45S_50ntFlanks | 10678 | 10711 | 34 | CGGTTCCGGCGGCGTCCGGTGAGCTCTCGCTGGC | 27135.7 | -7.5 | 0.5 | -15.6 | 8.8E-55 | 8.8E-52 | MB231/MB468 |
| >trnaMT_PheGAA_MT_+_577_647@46.74.29\|GACGGGCTCACATCACCCCATAAACACCA | >trnaMT_PheGAA_MT_+_577_647 | 46 | 74 | 29 | GACGGGCTCACATCACCCCATAAACACCA | 418.6 | 6.7 | 0.4 | 15.4 | 1.1E-53 | 1.5E-51 | BT20/MB468 |
| >hg38_21_+_8433172_8446622_RNA45SN1_45S_50ntFlanks@3987.4023.37\|GCTTTGGTGACTCTAGATAACCTCGGGCCGATCGCAC | >hg38_21_+_8433172_8446622_RNA45SN1_45S_50ntFlanks | 3987 | 4023 | 37 | GCTTTGGTGACTCTAGATAACCTCGGGCCGATCGCAC | 683.1 | 7.1 | 0.5 | 15.4 | 1.1E-53 | 1.6E-51 | BT20/MB468 |
| >hg38_21_+_8433172_8446622_RNA45SN1_45S_50ntFlanks@12633.12654.22\|CGAAGCTACCATCTGTGGGATT | >hg38_21_+_8433172_8446622_RNA45SN1_45S_50ntFlanks | 12633 | 12654 | 22 | CGAAGCTACCATCTGTGGGATT | 11462.9 | -8.9 | 0.6 | -15.4 | 2.0E-53 | 2.7E-51 | BT20/MB468 |
| >hg38_21_+_8433172_8446622_RNA45SN1_45S_50ntFlanks@6652.6695.44\|GACTCTTAGCGGTGGATCACTCGGCTCGTGCGTCGATGAAGAAC | >hg38_21_+_8433172_8446622_RNA45SN1_45S_50ntFlanks | 6652 | 6695 | 44 | GACTCTTAGCGGTGGATCACTCGGCTCGTGCGTCGATGAAGAAC | 3035.3 | 10.0 | 0.6 | 15.4 | 1.3E-53 | 1.1E-50 | MB231/MB468 |
| >hg38_21_+_8433172_8446622_RNA45SN1_45S_50ntFlanks@12997.13036.40\|CGCTGCGATCTATTGAAAGTCAGCCCTCGACACAAGGGTT | >hg38_21_+_8433172_8446622_RNA45SN1_45S_50ntFlanks | 12997 | 13036 | 40 | CGCTGCGATCTATTGAAAGTCAGCCCTCGACACAAGGGTT | 5359.4 | 10.0 | 0.7 | 15.4 | 2.1E-53 | 1.6E-50 | MB231/MB468 |
| >MI0000285\|hsa-mir-205&WithFlank&1\|+\|209432127\|209432248@40.62.23&[MIMAT0000266&hsa-miR-205-5p&offsets\|0\|+1;m-351&1\|+\|209432166\|209432187&offsets\|0\|+1]\|TCCTTCATTCCACCGGAGTCTGT | >MI0000285\|hsa-mir-205&WithFlank&1\|+\|209432127\|209432248 | 40 | 62 | 23 | TCCTTCATTCCACCGGAGTCTGT | 41861.6 | 14.4 | 0.9 | 15.3 | 3.8E-53 | 2.8E-50 | BT20/MB231 |
| >MI0000255\|hsa-mir-30d&WithFlank&8\|-\|134804870\|134804951@12.34.23(+1C)&[MIMAT0000245&hsa-miR-30d-5p&offsets\|0\|+1(+1C);m-21&8\|-\|134804919\|134804940&offsets\|0\|+1(+1C)]\|TGTAAACATCCCCGACTGGAAGCC | >MI0000255\|hsa-mir-30d&WithFlank&8\|-\|134804870\|134804951 | 12 | 34 | 23(+1C) | TGTAAACATCCCCGACTGGAAGCC | 36063.9 | 9.2 | 0.6 | 15.2 | 6.7E-52 | 9.0E-50 | BT20/MB468 |
| >trnaMT_ValTAC_MT_+_1602_1670@54.72.19\|AACTTGACCGCTCTGACCA | >trnaMT_ValTAC_MT_+_1602_1670 | 54 | 72 | 19 | AACTTGACCGCTCTGACCA | 3378.5 | 8.4 | 0.6 | 15.1 | 1.1E-51 | 1.5E-49 | BT20/MB468 |
| >am_trna128_GlyGCC_6_-_27870686_27870756@1.32.32,trna133_GlyCCC_1_-_16872434_16872504@1.32.32,trna18_GlyGCC_16_+_70822597_70822667@1.32.32,trna19_GlyGCC_16_+_70823410_70823480@1.32.32,trna19_GlyGCC_2_-_157257659_157257729@1.32.32,trna24_GlyGCC_16_-_70812942_70813012@1.32.32,trna25_GlyGCC_16_-_70812114_70812184@1.32.32,trna4_GlyCCC_1_+_17188416_17188486@1.32.32,trna5_GlyGCC_17_+_8029064_8029134@1.32.32,trna68_GlyGCC_1_-_161493637_161493707@1.32.32\|GCATTGGTGGTTCAGTGGTAGAATTCTCGCCT | >am_trna128_GlyGCC_6_-_27870686_27870756 | 1 | 32 | 32 | GCATTGGTGGTTCAGTGGTAGAATTCTCGCCT | 17231.9 | 8.3 | 0.6 | 15.1 | 1.2E-51 | 1.6E-49 | BT20/MB468 |
| >MI0001445\|hsa-mir-423&WithFlank&17\|+\|30117073\|30117178@23.45.23&[MIMAT0004748&hsa-miR-423-5p&offsets\|0\|0;m-141&17\|+\|30117095\|30117117&offsets\|0\|0]\|TGAGGGGCAGAGAGCGAGACTTT | >MI0001445\|hsa-mir-423&WithFlank&17\|+\|30117073\|30117178 | 23 | 45 | 23 | TGAGGGGCAGAGAGCGAGACTTT | 8086.6 | 8.0 | 0.5 | 15.1 | 1.6E-51 | 2.0E-49 | BT20/MB468 |
| >hg38_21_+_8433172_8446622_RNA45SN1_45S_50ntFlanks@12625.12654.30\|CAATGGGGCGAAGCTACCATCTGTGGGATT | >hg38_21_+_8433172_8446622_RNA45SN1_45S_50ntFlanks | 12625 | 12654 | 30 | CAATGGGGCGAAGCTACCATCTGTGGGATT | 3946.5 | -6.1 | 0.4 | -15.1 | 1.8E-51 | 2.2E-49 | BT20/MB468 |
| >MI0000809\|hsa-mir-151a&WithFlank&8\|-\|140732558\|140732659@53.74.22&[MIMAT0000757&hsa-miR-151a-3p&offsets\|0\|+1;m-30&8\|-\|140732587\|140732607&offsets\|0\|+1]\|CTAGACTGAAGCTCCTTGAGGA | >MI0000809\|hsa-mir-151a&WithFlank&8\|-\|140732558\|140732659 | 53 | 74 | 22 | CTAGACTGAAGCTCCTTGAGGA | 33820.5 | 8.4 | 0.6 | 15.1 | 2.9E-51 | 3.4E-49 | BT20/MB468 |
| >am_MI0000809\|hsa-mir-151a&WithFlank&8\|-\|140732558\|140732659@53.70.18&[MIMAT0000757&hsa-miR-151a-3p&offsets\|0\|-3;m-30&8\|-\|140732587\|140732607&offsets\|0\|-3]\|CTAGACTGAAGCTCCTTG | >am_MI0000809\|hsa-mir-151a&WithFlank&8\|-\|140732558\|140732659 | 53 | 70 | 18 | CTAGACTGAAGCTCCTTG | 3101.0 | 9.6 | 0.6 | 15.1 | 3.1E-51 | 3.6E-49 | BT20/MB468 |
| >MI0000809\|hsa-mir-151a&WithFlank&8\|-\|140732558\|140732659@53.74.22(+1A)&[MIMAT0000757&hsa-miR-151a-3p&offsets\|0\|+1(+1A);m-30&8\|-\|140732587\|140732607&offsets\|0\|+1(+1A)]\|CTAGACTGAAGCTCCTTGAGGAA | >MI0000809\|hsa-mir-151a&WithFlank&8\|-\|140732558\|140732659 | 53 | 74 | 22(+1A) | CTAGACTGAAGCTCCTTGAGGAA | 10047.0 | 8.2 | 0.5 | 15.0 | 4.1E-51 | 4.7E-49 | BT20/MB468 |
| >hg38_21_+_8433172_8446622_RNA45SN1_45S_50ntFlanks@10410.10430.21\|TCAGTCGGTCCTGAGAGATGG | >hg38_21_+_8433172_8446622_RNA45SN1_45S_50ntFlanks | 10410 | 10430 | 21 | TCAGTCGGTCCTGAGAGATGG | 4335.7 | -8.7 | 0.6 | -15.0 | 4.5E-51 | 5.1E-49 | BT20/MB468 |
| >hg38_21_+_8433172_8446622_RNA45SN1_45S_50ntFlanks@12456.12475.20\|CCAAGCGTTGGATTGTTCAC | >hg38_21_+_8433172_8446622_RNA45SN1_45S_50ntFlanks | 12456 | 12475 | 20 | CCAAGCGTTGGATTGTTCAC | 18280.7 | -8.7 | 0.6 | -15.1 | 1.0E-51 | 6.7E-49 | MB231/MB468 |
| >hg38_21_+_8433172_8446622_RNA45SN1_45S_50ntFlanks@6661.6695.35\|CGGTGGATCACTCGGCTCGTGCGTCGATGAAGAAC | >hg38_21_+_8433172_8446622_RNA45SN1_45S_50ntFlanks | 6661 | 6695 | 35 | CGGTGGATCACTCGGCTCGTGCGTCGATGAAGAAC | 1197.3 | 7.1 | 0.5 | 15.0 | 7.8E-51 | 8.6E-49 | BT20/MB468 |
| >trnaMT_GluTTC_MT_-_14674_14742@40.71.32,trnalookalike8_GluTTC_5_-_93905172_93905240@40.71.32\|TTGGTCGTGGTTGTAGTCCGTGCGAGAATACC | >trnaMT_GluTTC_MT_-_14674_14742 | 40 | 71 | 32 | TTGGTCGTGGTTGTAGTCCGTGCGAGAATACC | 3445.4 | 9.4 | 0.6 | 15.0 | 3.6E-51 | 2.1E-48 | MB231/MB468 |
| >hg38_21_+_8433172_8446622_RNA45SN1_45S_50ntFlanks@6719.6737.19\|TGAATTGCAGGACACATTG | >hg38_21_+_8433172_8446622_RNA45SN1_45S_50ntFlanks | 6719 | 6737 | 19 | TGAATTGCAGGACACATTG | 4065.7 | -8.1 | 0.5 | -14.9 | 4.7E-50 | 5.1E-48 | BT20/MB468 |
| >hg38_21_+_8433172_8446622_RNA45SN1_45S_50ntFlanks@9534.9552.19\|AGGAAACTCTGGTGGAGGT | >hg38_21_+_8433172_8446622_RNA45SN1_45S_50ntFlanks | 9534 | 9552 | 19 | AGGAAACTCTGGTGGAGGT | 3345.0 | -6.0 | 0.4 | -14.8 | 1.0E-49 | 1.1E-47 | BT20/MB468 |
| >hg38_21_+_8433172_8446622_RNA45SN1_45S_50ntFlanks@12605.12625.21\|TATGTGCTTGGCTGAGGAGCC | >hg38_21_+_8433172_8446622_RNA45SN1_45S_50ntFlanks | 12605 | 12625 | 21 | TATGTGCTTGGCTGAGGAGCC | 7047.8 | -6.8 | 0.5 | -14.8 | 1.3E-49 | 1.3E-47 | BT20/MB468 |
| >MI0000255\|hsa-mir-30d&WithFlank&8\|-\|134804870\|134804951@12.35.24&[MIMAT0000245&hsa-miR-30d-5p&offsets\|0\|+2;m-21&8\|-\|134804919\|134804940&offsets\|0\|+2]\|TGTAAACATCCCCGACTGGAAGCT | >MI0000255\|hsa-mir-30d&WithFlank&8\|-\|134804870\|134804951 | 12 | 35 | 24 | TGTAAACATCCCCGACTGGAAGCT | 224143.7 | 9.0 | 0.6 | 14.8 | 1.5E-49 | 1.5E-47 | BT20/MB468 |
| >hg38_21_+_8433172_8446622_RNA45SN1_45S_50ntFlanks@6699.6718.20\|GCTAGCTGCGAGAATTAATG | >hg38_21_+_8433172_8446622_RNA45SN1_45S_50ntFlanks | 6699 | 6718 | 20 | GCTAGCTGCGAGAATTAATG | 11362.1 | -7.3 | 0.5 | -14.8 | 1.6E-49 | 1.6E-47 | BT20/MB468 |
| >hg38_21_+_8433172_8446622_RNA45SN1_45S_50ntFlanks@10410.10429.20\|TCAGTCGGTCCTGAGAGATG | >hg38_21_+_8433172_8446622_RNA45SN1_45S_50ntFlanks | 10410 | 10429 | 20 | TCAGTCGGTCCTGAGAGATG | 6237.7 | -6.3 | 0.4 | -14.8 | 2.9E-49 | 2.9E-47 | BT20/MB468 |
| >hg38_21_+_8433172_8446622_RNA45SN1_45S_50ntFlanks@12997.13017.21\|CGCTGCGATCTATTGAAAGTC | >hg38_21_+_8433172_8446622_RNA45SN1_45S_50ntFlanks | 12997 | 13017 | 21 | CGCTGCGATCTATTGAAAGTC | 4307.9 | 7.9 | 0.5 | 14.9 | 5.5E-50 | 3.0E-47 | MB231/MB468 |
| >MI0000809\|hsa-mir-151a&WithFlank&8\|-\|140732558\|140732659@53.74.22(+1U)&[MIMAT0000757&hsa-miR-151a-3p&offsets\|0\|+1(+1U);m-30&8\|-\|140732587\|140732607&offsets\|0\|+1(+1U)]\|CTAGACTGAAGCTCCTTGAGGAT | >MI0000809\|hsa-mir-151a&WithFlank&8\|-\|140732558\|140732659 | 53 | 74 | 22(+1U) | CTAGACTGAAGCTCCTTGAGGAT | 2992.9 | 8.3 | 0.6 | 14.7 | 3.5E-49 | 3.4E-47 | BT20/MB468 |

<table>
<tr><th></th><th></th><th></th><th></th><th></th><th></th><th></th><th></th><th></th><th></th><th></th><th></th><th></th><th></th></tr>
<tr><td>>hg38_21_+_8433172_8446622_RNA45SN1_45S_50ntFlanks@7974.8010.37|ACGCGACCTCAGATCAGACGTGGCGACCCGCTGAATT</td><td>>hg38_21_+_8433172_8446622_RNA45SN1_45S_50ntFlanks</td><td>7974</td><td>8010</td><td>37</td><td>ACGCGACCTCAGATCAGACGTGGCGACCCGCTGAATT</td><td>10198.8</td><td>7.5</td><td>0.5</td><td>14.7</td><td>4.5E-49</td><td>4.3E-47</td><td>BT20/MB468</td></tr>
<tr><td>>MI0000255|hsa-mir-30d&WithFlank&8|-|134804870|134804951@12.33.22(+1A)&[MIMAT0000245&hsa-miR-30d-5p&offsets|0|0(+1A);m-21&8|-|134804919|134804940&offsets|0|0(+1A)]|TGTAAACATCCCCGACTGGAAGA</td><td>>MI0000255|hsa-mir-30d&WithFlank&8|-|134804870|134804951</td><td>12</td><td>33</td><td>22(+1A)</td><td>TGTAAACATCCCCGACTGGAAGA</td><td>8821.5</td><td>9.6</td><td>0.7</td><td>14.7</td><td>6.0E-49</td><td>5.6E-47</td><td>BT20/MB468</td></tr>
<tr><td>>hg38_21_+_8433172_8446622_RNA45SN1_45S_50ntFlanks@10679.10709.31|GGTTCCGGCGGCGTCCGGTGAGCTCTCGCTG</td><td>>hg38_21_+_8433172_8446622_RNA45SN1_45S_50ntFlanks</td><td>10679</td><td>10709</td><td>31</td><td>GGTTCCGGCGGCGTCCGGTGAGCTCTCGCTG</td><td>7001.4</td><td>-7.4</td><td>0.5</td><td>-14.8</td><td>1.3E-49</td><td>6.4E-47</td><td>MB231/MB468</td></tr>
<tr><td>>hg38_21_+_8433172_8446622_RNA45SN1_45S_50ntFlanks@6645.6672.28|CTCGTACGACTCTTAGCGGTGGATCACT</td><td>>hg38_21_+_8433172_8446622_RNA45SN1_45S_50ntFlanks</td><td>6645</td><td>6672</td><td>28</td><td>CTCGTACGACTCTTAGCGGTGGATCACT</td><td>1447.7</td><td>7.1</td><td>0.5</td><td>14.7</td><td>9.0E-49</td><td>8.4E-47</td><td>BT20/MB468</td></tr>
<tr><td>>MI0001445|hsa-mir-423&WithFlank&17|+|30117073|30117178@59.80.22(+1C)&[MIMAT0001340&hsa-miR-423-3p&offsets|0|-1(+1C);m-100&17|+|30117131|30117153&offsets|0|-1(+1C)]|AGCTCGGTCTGAGGCCCCTCAGC</td><td>>MI0001445|hsa-mir-423&WithFlank&17|+|30117073|30117178</td><td>59</td><td>80</td><td>22(+1C)</td><td>AGCTCGGTCTGAGGCCCCTCAGC</td><td>4454.7</td><td>7.7</td><td>0.5</td><td>14.6</td><td>1.4E-48</td><td>1.3E-46</td><td>BT20/MB468</td></tr>
<tr><td>>hg38_21_+_8433172_8446622_RNA45SN1_45S_50ntFlanks@6745.6770.26|ACACTTCGAACGCACTTGCGGCCCCG</td><td>>hg38_21_+_8433172_8446622_RNA45SN1_45S_50ntFlanks</td><td>6745</td><td>6770</td><td>26</td><td>ACACTTCGAACGCACTTGCGGCCCCG</td><td>3445.5</td><td>-8.9</td><td>0.6</td><td>-14.6</td><td>1.5E-48</td><td>1.3E-46</td><td>BT20/MB468</td></tr>
<tr><td>>MI0000299|hsa-mir-222&WithFlank&X|-|45747009|45747130@75.97.23(+1A)&[MIMAT0000279&hsa-miR-222-3p&offsets|0|+2(+1A);m-140&X|-|45747033|45747056&offsets|0|-1(+1A)]|AGCTACATCTGGCTACTGGGTCTA</td><td>>MI0000299|hsa-mir-222&WithFlank&X|-|45747009|45747130</td><td>75</td><td>97</td><td>23(+1A)</td><td>AGCTACATCTGGCTACTGGGTCTA</td><td>4044.5</td><td>9.7</td><td>0.7</td><td>14.7</td><td>3.5E-49</td><td>1.6E-46</td><td>MB231/MB468</td></tr>
<tr><td>>MI0000285|hsa-mir-205&WithFlank&1|+|209432127|209432248@40.61.22&[MIMAT0000266&hsa-miR-205-5p&offsets|0|0;m-351&1|+|209432166|209432187&offsets|0|0]|TCCTTCATTCCACCGGAGTCTG</td><td>>MI0000285|hsa-mir-205&WithFlank&1|+|209432127|209432248</td><td>40</td><td>61</td><td>22</td><td>TCCTTCATTCCACCGGAGTCTG</td><td>27563.1</td><td>14.1</td><td>1.0</td><td>14.7</td><td>5.7E-49</td><td>3.4E-46</td><td>BT20/MB231</td></tr>
<tr><td>>trna2_GlyGCC_21_-_18827107_18827177@1.32.32,trna35_GlyGCC_1_+_161413094_161413164@1.32.32,trna37_GlyGCC_1_+_161420467_161420537@1.32.32,trna39_GlyGCC_1_+_161427898_161427968@1.32.32,trna41_GlyGCC_1_+_161435258_161435328@1.32.32|GCATGGGTGGTTCAGTGGTAGAATTCTCGCCT</td><td>>trna2_GlyGCC_21_-_18827107_18827177</td><td>1</td><td>32</td><td>32</td><td>GCATGGGTGGTTCAGTGGTAGAATTCTCGCCT</td><td>15751.4</td><td>8.9</td><td>0.6</td><td>14.5</td><td>6.8E-48</td><td>6.1E-46</td><td>BT20/MB468</td></tr>
<tr><td>>hg38_21_+_8433172_8446622_RNA45SN1_45S_50ntFlanks@7983.8006.24|CAGATCAGACGTGGCGACCCGCTG</td><td>>hg38_21_+_8433172_8446622_RNA45SN1_45S_50ntFlanks</td><td>7983</td><td>8006</td><td>24</td><td>CAGATCAGACGTGGCGACCCGCTG</td><td>6290.9</td><td>-8.5</td><td>0.6</td><td>-14.5</td><td>1.0E-47</td><td>8.7E-46</td><td>BT20/MB468</td></tr>
<tr><td>>hg38_1_-_228634819_228635039_RNA5S12_5S_50ntFlanks@146.170.25|TGGGAATACCGGGTGCTGTAGGCTT</td><td>>hg38_1_-_228634819_228635039_RNA5S12_5S_50ntFlanks</td><td>146</td><td>170</td><td>25</td><td>TGGGAATACCGGGTGCTGTAGGCTT</td><td>2180.9</td><td>7.2</td><td>0.5</td><td>14.5</td><td>1.2E-47</td><td>1.1E-45</td><td>BT20/MB468</td></tr>
<tr><td>>trnaMT_GluTTC_MT_-_14674_14742@40.72.33,trnalookalike8_GluTTC_5_-_93905172_93905240@40.72.33|TTGGTCGTGGTTGTAGTCCGTGCGAGAATACCA</td><td>>trnaMT_GluTTC_MT_-_14674_14742</td><td>40</td><td>72</td><td>33</td><td>TTGGTCGTGGTTGTAGTCCGTGCGAGAATACCA</td><td>39433.1</td><td>8.6</td><td>0.6</td><td>14.6</td><td>5.6E-48</td><td>2.4E-45</td><td>MB231/MB468</td></tr>
<tr><td>>hg38_21_+_8433172_8446622_RNA45SN1_45S_50ntFlanks@6661.6678.18|CGGTGGATCACTCGGCTC</td><td>>hg38_21_+_8433172_8446622_RNA45SN1_45S_50ntFlanks</td><td>6661</td><td>6678</td><td>18</td><td>CGGTGGATCACTCGGCTC</td><td>1397.9</td><td>6.9</td><td>0.5</td><td>14.4</td><td>5.1E-47</td><td>4.4E-45</td><td>BT20/MB468</td></tr>
<tr><td>>hg38_21_+_8433172_8446622_RNA45SN1_45S_50ntFlanks@12628.12654.27|TGGGGCGAAGCTACCATCTGTGGGATT</td><td>>hg38_21_+_8433172_8446622_RNA45SN1_45S_50ntFlanks</td><td>12628</td><td>12654</td><td>27</td><td>TGGGGCGAAGCTACCATCTGTGGGATT</td><td>2501.3</td><td>-9.1</td><td>0.6</td><td>-14.4</td><td>8.8E-47</td><td>7.3E-45</td><td>BT20/MB468</td></tr>
<tr><td>>hg38_21_+_8433172_8446622_RNA45SN1_45S_50ntFlanks@12627.12654.28|ATGGGGCGAAGCTACCATCTGTGGGATT</td><td>>hg38_21_+_8433172_8446622_RNA45SN1_45S_50ntFlanks</td><td>12627</td><td>12654</td><td>28</td><td>ATGGGGCGAAGCTACCATCTGTGGGATT</td><td>5749.3</td><td>-7.1</td><td>0.5</td><td>-14.5</td><td>1.9E-47</td><td>7.6E-45</td><td>MB231/MB468</td></tr>
<tr><td>>trnaMT_ProTGG_MT_-_15956_16023@50.71.22|TTAAAGACTTTTTCTCTGACCA</td><td>>trnaMT_ProTGG_MT_-_15956_16023</td><td>50</td><td>71</td><td>22</td><td>TTAAAGACTTTTTCTCTGACCA</td><td>2747.9</td><td>9.6</td><td>0.7</td><td>14.5</td><td>2.3E-47</td><td>8.8E-45</td><td>MB231/MB468</td></tr>
<tr><td>>MI0000746|hsa-mir-99b&WithFlank&19|+|51692606|51692687@13.32.20(+1U)&[MIMAT0000689&hsa-miR-99b-5p&offsets|0|-2(+1U);m-10&19|+|51692618|51692639&offsets|0|-2(+1U)]|CACCCGTAGAACCGACCTTGT</td><td>>MI0000746|hsa-mir-99b&WithFlank&19|+|51692606|51692687</td><td>13</td><td>32</td><td>20(+1U)</td><td>CACCCGTAGAACCGACCTTGT</td><td>31669.0</td><td>10.8</td><td>0.8</td><td>14.3</td><td>1.2E-46</td><td>9.6E-45</td><td>BT20/MB468</td></tr>
<tr><td>>hg38_21_+_8433172_8446622_RNA45SN1_45S_50ntFlanks@6719.6740.22|TGAATTGCAGGACACATTGATC</td><td>>hg38_21_+_8433172_8446622_RNA45SN1_45S_50ntFlanks</td><td>6719</td><td>6740</td><td>22</td><td>TGAATTGCAGGACACATTGATC</td><td>4409.5</td><td>-6.9</td><td>0.5</td><td>-14.3</td><td>1.4E-46</td><td>1.2E-44</td><td>BT20/MB468</td></tr>
<tr><td>>am_MI0005763|hsa-mir-941-1&WithFlank&20|+|63919443|63919526@53.75.23;MI0005764|hsa-mir-941-2&WithFlank&20|+|63919499|63919582@53.75.23;MI0005765|hsa-mir-941-3&WithFlank&20|+|63919555|63919638@53.75.23;MI0005766|hsa-mir-941-4&WithFlank&20|+|63919750|63919833@53.75.23;MI0031520|hsa-mir-941-5&WithFlank&20|+|63919862|63919945@53.75.23&[MIMAT0004984&hsa-miR-941&offsets|0|0;m-263&20|+|63919495|63919517&offsets|0|0];[MIMAT0004984_1&hsa-miR-941&offsets|0|0;m-263&20|+|63919551|63919573&offsets|0|0];[MIMAT0004984_2&hsa-miR-941&offsets|0|0;m-263&20|+|63919607|63919629&offsets|0|0];[MIMAT0004984_3&hsa-miR-941&offsets|0|0;m-263&20|+|63919802|63919824&offsets|0|0];[MIMAT0004984_4&hsa-miR-941&offsets|0|0;m-263&20|+|63919914|63919936&offsets|0|0]|CACCCGGCTGTGTGCACATGTGC</td><td>>am_MI0005763|hsa-mir-941-1&WithFlank&20|+|63919443|63919526</td><td>53</td><td>75</td><td>23</td><td>CACCCGGCTGTGTGCACATGTGC</td><td>18217.3</td><td>7.5</td><td>0.5</td><td>14.3</td><td>1.8E-46</td><td>1.5E-44</td><td>BT20/MB468</td></tr>
<tr><td>>MI0000542|hsa-mir-320a&WithFlank&8|-|22244960|22245043@48.68.21(+1U)&[MIMAT0000510&hsa-miR-320a-3p&offsets|0|-1(+1U);m-60&8|-|22244975|22244996&offsets|0|-1(+1U)]|AAAAGCTGGGTTGAGAGGGCGT</td><td>>MI0000542|hsa-mir-320a&WithFlank&8|-|22244960|22245043</td><td>48</td><td>68</td><td>21(+1U)</td><td>AAAAGCTGGGTTGAGAGGGCGT</td><td>11519.7</td><td>9.9</td><td>0.7</td><td>14.3</td><td>2.0E-46</td><td>1.6E-44</td><td>BT20/MB468</td></tr>
<tr><td>>hg38_21_+_8433172_8446622_RNA45SN1_45S_50ntFlanks@12938.12956.19|ACCATTCGTAGACGACCTG</td><td>>hg38_21_+_8433172_8446622_RNA45SN1_45S_50ntFlanks</td><td>12938</td><td>12956</td><td>19</td><td>ACCATTCGTAGACGACCTG</td><td>1493.0</td><td>-5.6</td><td>0.4</td><td>-14.3</td><td>2.4E-46</td><td>1.9E-44</td><td>BT20/MB468</td></tr>
<tr><td>>MI0001445|hsa-mir-423&WithFlank&17|+|30117073|30117178@23.46.24&[MIMAT0004748&hsa-miR-423-5p&offsets|0|+1;m-141&17|+|30117095|30117117&offsets|0|+1]|TGAGGGGCAGAGAGCGAGACTTTT</td><td>>MI0001445|hsa-mir-423&WithFlank&17|+|30117073|30117178</td><td>23</td><td>46</td><td>24</td><td>TGAGGGGCAGAGAGCGAGACTTTT</td><td>1904.7</td><td>8.1</td><td>0.6</td><td>14.3</td><td>2.7E-46</td><td>2.1E-44</td><td>BT20/MB468</td></tr>
<tr><td>>MI0000809|hsa-mir-151a&WithFlank&8|-|140732558|140732659@51.72.22(+1A)&[MIMAT0000757&hsa-miR-151a-3p&offsets|-2|-1(+1A);m-30&8|-|140732587|140732607&offsets|-2|-1(+1A)]|TACTAGACTGAAGCTCCTTGAGA</td><td>>MI0000809|hsa-mir-151a&WithFlank&8|-|140732558|140732659</td><td>51</td><td>72</td><td>22(+1A)</td><td>TACTAGACTGAAGCTCCTTGAGA</td><td>2005.4</td><td>8.7</td><td>0.6</td><td>14.2</td><td>7.2E-46</td><td>5.5E-44</td><td>BT20/MB468</td></tr>
<tr><td>>hg38_21_+_8433172_8446622_RNA45SN1_45S_50ntFlanks@12456.12477.22|CCAAGCGTTGGATTGTTCACCC</td><td>>hg38_21_+_8433172_8446622_RNA45SN1_45S_50ntFlanks</td><td>12456</td><td>12477</td><td>22</td><td>CCAAGCGTTGGATTGTTCACCC</td><td>7434.0</td><td>-6.5</td><td>0.5</td><td>-14.2</td><td>9.3E-46</td><td>7.0E-44</td><td>BT20/MB468</td></tr>
<tr><td>>MI0000650|hsa-mir-200c&WithFlank&12|+|6963693|6963772@50.72.23(+1U)&[MIMAT0000617&hsa-miR-200c-3p&offsets|0|0(+1U);m-54&12|+|6963742|6963764&offsets|0|0(+1U)]|TAATACTGCCGGGTAATGATGGAT</td><td>>MI0000650|hsa-mir-200c&WithFlank&12|+|6963693|6963772</td><td>50</td><td>72</td><td>23(+1U)</td><td>TAATACTGCCGGGTAATGATGGAT</td><td>9810.3</td><td>9.2</td><td>0.6</td><td>14.3</td><td>2.3E-46</td><td>1.2E-43</td><td>BT20/MB231</td></tr>
<tr><td>>MI0001445|hsa-mir-423&WithFlank&17|+|30117073|30117178@58.81.24&[MIMAT0001340&hsa-miR-423-3p&offsets|-1|0;m-100&17|+|30117131|30117153&offsets|-1|0]|AAGCTCGGTCTGAGGCCCCTCAGT</td><td>>MI0001445|hsa-mir-423&WithFlank&17|+|30117073|30117178</td><td>58</td><td>81</td><td>24</td><td>AAGCTCGGTCTGAGGCCCCTCAGT</td><td>5593.9</td><td>7.7</td><td>0.5</td><td>14.2</td><td>1.8E-45</td><td>1.4E-43</td><td>BT20/MB468</td></tr>
<tr><td>>trna11_GluTTC_15_-_26327381_26327452@1.33.33,trna3_GluTTC_13_-_45492062_45492133@1.33.33|TCCCACATGGTCTAGCGGTTAGGATTCCTGGTT</td><td>>trna11_GluTTC_15_-_26327381_26327452</td><td>1</td><td>33</td><td>33</td><td>TCCCACATGGTCTAGCGGTTAGGATTCCTGGTT</td><td>7055.8</td><td>9.9</td><td>0.7</td><td>14.2</td><td>1.9E-45</td><td>1.4E-43</td><td>BT20/MB468</td></tr>
<tr><td>>hg38_21_+_8433172_8446622_RNA45SN1_45S_50ntFlanks@6699.6719.21|GCTAGCTGCGAGAATTAATGT</td><td>>hg38_21_+_8433172_8446622_RNA45SN1_45S_50ntFlanks</td><td>6699</td><td>6719</td><td>21</td><td>GCTAGCTGCGAGAATTAATGT</td><td>1129.6</td><td>-6.0</td><td>0.4</td><td>-14.1</td><td>1.9E-45</td><td>1.4E-43</td><td>BT20/MB468</td></tr>
<tr><td>>hg38_21_+_8433172_8446622_RNA45SN1_45S_50ntFlanks@9534.9551.18|AGGAAACTCTGGTGGAGG</td><td>>hg38_21_+_8433172_8446622_RNA45SN1_45S_50ntFlanks</td><td>9534</td><td>9551</td><td>18</td><td>AGGAAACTCTGGTGGAGG</td><td>4398.6</td><td>-6.9</td><td>0.5</td><td>-14.1</td><td>2.1E-45</td><td>1.5E-43</td><td>BT20/MB468</td></tr>
<tr><td>>MI0000088|hsa-mir-30a&WithFlank&6|-|71403545|71403627@12.35.24(+2U)&[MIMAT0000087&hsa-miR-30a-5p&offsets|0|+2(+2U);m-15&6|-|71403595|71403616&offsets|0|+2(+2U)]|TGTAAACATCCTCGACTGGAAGCTTT</td><td>>MI0000088|hsa-mir-30a&WithFlank&6|-|71403545|71403627</td><td>12</td><td>35</td><td>24(+2U)</td><td>TGTAAACATCCTCGACTGGAAGCTTT</td><td>9522.3</td><td>9.9</td><td>0.7</td><td>14.3</td><td>4.3E-46</td><td>1.5E-43</td><td>MB231/MB468</td></tr>
<tr><td>>trnaMT_ValTAC_MT_+_1602_1670@48.71.24|CAACTTAACTTGACCGCTCTGACC</td><td>>trnaMT_ValTAC_MT_+_1602_1670</td><td>48</td><td>71</td><td>24</td><td>CAACTTAACTTGACCGCTCTGACC</td><td>1051.3</td><td>7.6</td><td>0.5</td><td>14.1</td><td>3.0E-45</td><td>2.1E-43</td><td>BT20/MB468</td></tr>
<tr><td>>hg38_21_+_8433172_8446622_RNA45SN1_45S_50ntFlanks@6646.6665.20|TCGTACGACTCTTAGCGGTG</td><td>>hg38_21_+_8433172_8446622_RNA45SN1_45S_50ntFlanks</td><td>6646</td><td>6665</td><td>20</td><td>TCGTACGACTCTTAGCGGTG</td><td>67948.4</td><td>6.4</td><td>0.5</td><td>14.1</td><td>3.1E-45</td><td>2.2E-43</td><td>BT20/MB468</td></tr>
<tr><td>>hg38_21_+_8433172_8446622_RNA45SN1_45S_50ntFlanks@9570.9595.26|TGCAAATCGGTCGTCCGACCTGGGTA</td><td>>hg38_21_+_8433172_8446622_RNA45SN1_45S_50ntFlanks</td><td>9570</td><td>9595</td><td>26</td><td>TGCAAATCGGTCGTCCGACCTGGGTA</td><td>2037.2</td><td>-6.2</td><td>0.4</td><td>-14.1</td><td>5.5E-45</td><td>3.8E-43</td><td>BT20/MB468</td></tr>
<tr><td>>hg38_21_+_8433172_8446622_RNA45SN1_45S_50ntFlanks@12934.12956.23|CTAAACCATTCGTAGACGACCTG</td><td>>hg38_21_+_8433172_8446622_RNA45SN1_45S_50ntFlanks</td><td>12934</td><td>12956</td><td>23</td><td>CTAAACCATTCGTAGACGACCTG</td><td>4743.1</td><td>-7.3</td><td>0.5</td><td>-14.2</td><td>1.4E-45</td><td>4.6E-43</td><td>MB231/MB468</td></tr>
<tr><td>>hg38_21_+_8433172_8446622_RNA45SN1_45S_50ntFlanks@10392.10410.19|ACAGCAGTTGAACATGGGT</td><td>>hg38_21_+_8433172_8446622_RNA45SN1_45S_50ntFlanks</td><td>10392</td><td>10410</td><td>19</td><td>ACAGCAGTTGAACATGGGT</td><td>7880.2</td><td>-9.0</td><td>0.6</td><td>-14.1</td><td>7.6E-45</td><td>5.2E-43</td><td>BT20/MB468</td></tr>
<tr><td>>trna11_GluTTC_15_-_26327381_26327452@1.33.33,trna3_GluTTC_13_-_45492062_45492133@1.33.33|TCCCACATGGTCTAGCGGTTAGGATTCCTGGTT</td><td>>trna11_GluTTC_15_-_26327381_26327452</td><td>1</td><td>33</td><td>33</td><td>TCCCACATGGTCTAGCGGTTAGGATTCCTGGTT</td><td>18525.4</td><td>11.5</td><td>0.8</td><td>14.2</td><td>1.7E-45</td><td>5.4E-43</td><td>MB231/MB468</td></tr>
<tr><td>>hg38_21_+_8433172_8446622_RNA45SN1_45S_50ntFlanks@12324.12346.23|TTGGGTTTTAAGCAGGAGGTGTC</td><td>>hg38_21_+_8433172_8446622_RNA45SN1_45S_50ntFlanks</td><td>12324</td><td>12346</td><td>23</td><td>TTGGGTTTTAAGCAGGAGGTGTC</td><td>1860.4</td><td>-8.4</td><td>0.6</td><td>-14.0</td><td>8.1E-45</td><td>5.5E-43</td><td>BT20/MB468</td></tr>
<tr><td>>MI0000086|hsa-mir-28&WithFlank&3|+|188688775|188688872@60.81.22&[MIMAT0004502&hsa-miR-28-3p&offsets|0|0;m-28&3|+|188688834|188688855&offsets|0|0]|CACTAGATTGTGAGCTCCTGGA</td><td>>MI0000086|hsa-mir-28&WithFlank&3|+|188688775|188688872</td><td>60</td><td>81</td><td>22</td><td>CACTAGATTGTGAGCTCCTGGA</td><td>21049.0</td><td>10.3</td><td>0.7</td><td>14.0</td><td>9.1E-45</td><td>6.0E-43</td><td>BT20/MB468</td></tr>
<tr><td>>hg38_21_+_8433172_8446622_RNA45SN1_45S_50ntFlanks@9534.9563.30|AGGAAACTCTGGTGGAGGTCCGTAGCGGTC</td><td>>hg38_21_+_8433172_8446622_RNA45SN1_45S_50ntFlanks</td><td>9534</td><td>9563</td><td>30</td><td>AGGAAACTCTGGTGGAGGTCCGTAGCGGTC</td><td>8716.6</td><td>-7.7</td><td>0.6</td><td>-14.0</td><td>9.5E-45</td><td>6.3E-43</td><td>BT20/MB468</td></tr>
<tr><td>>hg38_21_+_8433172_8446622_RNA45SN1_45S_50ntFlanks@12405.12435.31|CTTTTTGATCCTTCGATGTCGGCTCTTCCTA</td><td>>hg38_21_+_8433172_8446622_RNA45SN1_45S_50ntFlanks</td><td>12405</td><td>12435</td><td>31</td><td>CTTTTTGATCCTTCGATGTCGGCTCTTCCTA</td><td>2961.9</td><td>-9.3</td><td>0.7</td><td>-14.0</td><td>1.0E-44</td><td>6.7E-43</td><td>BT20/MB468</td></tr>
</table>

<table>
<tr><td>>hg38_21_+_8433172_8446622_RNA45SN1_45S_50ntFlanks@12437.12455.19|CATTGTGAAGCAGAATTCA</td><td>>hg38_21_+_8433172_8446622_RNA45SN1_45S_50ntFlanks</td><td>12437</td><td>12455</td><td>19</td><td>CATTGTGAAGCAGAATTCA</td><td>21373.3</td><td>-10.9</td><td>0.8</td><td>-14.0</td><td>1.0E-44</td><td>6.7E-43</td><td>BT20/MB468</td></tr>
<tr><td>>hg38_21_+_8433172_8446622_RNA45SN1_45S_50ntFlanks@11703.11728.26|AATGAAGCGCGGGTAAACGGCGGGAG</td><td>>hg38_21_+_8433172_8446622_RNA45SN1_45S_50ntFlanks</td><td>11703</td><td>11728</td><td>26</td><td>AATGAAGCGCGGGTAAACGGCGGGAG</td><td>1687.1</td><td>-8.2</td><td>0.6</td><td>-14.0</td><td>1.8E-44</td><td>1.1E-42</td><td>BT20/MB468</td></tr>
<tr><td>>trnaMT_GluTTC_MT_-_14674_14742@34.72.39,trnalookalike8_GluTTC_5_-_93905172_93905240@34.72.39|ATATCATTGGTCGTGGTTGTAGTCCGTGCGAGAATACCA</td><td>>trnaMT_GluTTC_MT_-_14674_14742</td><td>34</td><td>72</td><td>39</td><td>ATATCATTGGTCGTGGTTGTAGTCCGTGCGAGAATACCA</td><td>9592.5</td><td>9.8</td><td>0.7</td><td>14.1</td><td>4.4E-45</td><td>1.3E-42</td><td>MB231/MB468</td></tr>
<tr><td>>trna111_HisGTG_1_-_147774845_147774916@-1T.23.24,trna118_HisGTG_1_-_145396881_145396952@-1T.23.24,trna16_HisGTG_1_+_146544773_146544844@-1T.23.24,trna1_HisGTG_15_+_45493349_45493420@-1T.23.24,trna21_HisGTG_1_+_147753471_147753542@-1T.23.24,trna33_HisGTG_6_+_27125906_27125977@-1T.23.24,trna7_HisGTG_9_-_14433938_14434009@-1T.23.24,trna8_HisGTG_15_-_45492611_45492682@-1T.23.24,trna9_HisGTG_15_-_45490804_45490875@-1T.23.24|TGCCGTGATCGTATAGTGGTTAGT</td><td>>trna111_HisGTG_1_-_147774845_147774916</td><td>-1T</td><td>23</td><td>24</td><td>TGCCGTGATCGTATAGTGGTTAGT</td><td>18605.9</td><td>11.7</td><td>0.8</td><td>14.1</td><td>4.8E-45</td><td>1.4E-42</td><td>MB231/MB468</td></tr>
<tr><td>>MI0000809|hsa-mir-151a&WithFlank&8|-|140732558|140732659@51.72.22&[MIMAT0000757&hsa-miR-151a-3p&offsets|-2|-1;m-30&8|-|140732587|140732607&offsets|-2|-1]|TACTAGACTGAAGCTCCTTGAG</td><td>>MI0000809|hsa-mir-151a&WithFlank&8|-|140732558|140732659</td><td>51</td><td>72</td><td>22</td><td>TACTAGACTGAAGCTCCTTGAG</td><td>1909.5</td><td>8.5</td><td>0.6</td><td>14.0</td><td>3.0E-44</td><td>1.9E-42</td><td>BT20/MB468</td></tr>
<tr><td>>hg38_21_+_8433172_8446622_RNA45SN1_45S_50ntFlanks@6721.6746.26|AATTGCAGGACACATTGATCATCGAC</td><td>>hg38_21_+_8433172_8446622_RNA45SN1_45S_50ntFlanks</td><td>6721</td><td>6746</td><td>26</td><td>AATTGCAGGACACATTGATCATCGAC</td><td>2222.8</td><td>-6.8</td><td>0.5</td><td>-13.9</td><td>3.4E-44</td><td>2.1E-42</td><td>BT20/MB468</td></tr>
<tr><td>>hg38_21_+_8433172_8446622_RNA45SN1_45S_50ntFlanks@12973.12990.18|TCGTACGTAGCAGAGCAG</td><td>>hg38_21_+_8433172_8446622_RNA45SN1_45S_50ntFlanks</td><td>12973</td><td>12990</td><td>18</td><td>TCGTACGTAGCAGAGCAG</td><td>6963.6</td><td>-8.3</td><td>0.6</td><td>-14.0</td><td>8.7E-45</td><td>2.4E-42</td><td>MB231/MB468</td></tr>
<tr><td>>hg38_21_+_8433172_8446622_RNA45SN1_45S_50ntFlanks@12995.13017.23|CTCGCTGCGATCTATTGAAAGTC</td><td>>hg38_21_+_8433172_8446622_RNA45SN1_45S_50ntFlanks</td><td>12995</td><td>13017</td><td>23</td><td>CTCGCTGCGATCTATTGAAAGTC</td><td>1221.7</td><td>7.0</td><td>0.5</td><td>14.0</td><td>9.5E-45</td><td>2.5E-42</td><td>MB231/MB468</td></tr>
<tr><td>>MI0000299|hsa-mir-222&WithFlank&X|-|45747009|45747130@75.97.23(+1U)&[MIMAT0000279&hsa-miR-222-3p&offsets|0|+2(+1U);m-140&X|-|45747033|45747056&offsets|0|-1(+1U)]|AGCTACATCTGGCTACTGGGTCTT</td><td>>MI0000299|hsa-mir-222&WithFlank&X|-|45747009|45747130</td><td>75</td><td>97</td><td>23(+1U)</td><td>AGCTACATCTGGCTACTGGGTCTT</td><td>12273.3</td><td>9.4</td><td>0.7</td><td>14.0</td><td>1.0E-44</td><td>2.5E-42</td><td>MB231/MB468</td></tr>
<tr><td>>hg38_21_+_8433172_8446622_RNA45SN1_45S_50ntFlanks@12973.12991.19|TCGTACGTAGCAGAGCAGC</td><td>>hg38_21_+_8433172_8446622_RNA45SN1_45S_50ntFlanks</td><td>12973</td><td>12991</td><td>19</td><td>TCGTACGTAGCAGAGCAGC</td><td>6888.7</td><td>-9.1</td><td>0.7</td><td>-13.9</td><td>5.9E-44</td><td>3.7E-42</td><td>BT20/MB468</td></tr>
<tr><td>>hg38_21_+_8433172_8446622_RNA45SN1_45S_50ntFlanks@6721.6740.20|AATTGCAGGACACATTGATC</td><td>>hg38_21_+_8433172_8446622_RNA45SN1_45S_50ntFlanks</td><td>6721</td><td>6740</td><td>20</td><td>AATTGCAGGACACATTGATC</td><td>8798.9</td><td>-7.3</td><td>0.5</td><td>-13.9</td><td>8.7E-44</td><td>5.3E-42</td><td>BT20/MB468</td></tr>
<tr><td>>MI0000299|hsa-mir-222&WithFlank&X|-|45747009|45747130@75.98.24(+1A)&[MIMAT0000279&hsa-miR-222-3p&offsets|0|+3(+1A);m-140&X|-|45747033|45747056&offsets|0|0(+1A)]|AGCTACATCTGGCTACTGGGTCTCA</td><td>>MI0000299|hsa-mir-222&WithFlank&X|-|45747009|45747130</td><td>75</td><td>98</td><td>24(+1A)</td><td>AGCTACATCTGGCTACTGGGTCTCA</td><td>16414.0</td><td>9.4</td><td>0.7</td><td>14.0</td><td>2.3E-44</td><td>5.6E-42</td><td>MB231/MB468</td></tr>
<tr><td>>trna111_HisGTG_1_-_147774845_147774916@-1G.33.34,trna118_HisGTG_1_-_145396881_145396952@-1G.33.34,trna16_HisGTG_1_+_146544773_146544844@-1G.33.34,trna1_HisGTG_15_+_45493349_45493420@-1G.33.34,trna21_HisGTG_1_+_147753471_147753542@-1G.33.34,trna33_HisGTG_6_+_27125906_27125977@-1G.33.34,trna7_HisGTG_9_-_14433938_14434009@-1G.33.34,trna8_HisGTG_15_-_45492611_45492682@-1G.33.34,trna9_HisGTG_15_-_45490804_45490875@-1G.33.34|GGCCGTGATCGTATAGTGGTTAGTACTCTGCGTT</td><td>>trna111_HisGTG_1_-_147774845_147774916</td><td>-1G</td><td>33</td><td>34</td><td>GGCCGTGATCGTATAGTGGTTAGTACTCTGCGTT</td><td>106487.0</td><td>13.5</td><td>1.0</td><td>14.0</td><td>2.5E-44</td><td>5.7E-42</td><td>MB231/MB468</td></tr>
<tr><td>>trnaMT_ValTAC_MT_+_1602_1670@48.71.24|CAACTTAACTTGACCGCTCTGACC</td><td>>trnaMT_ValTAC_MT_+_1602_1670</td><td>48</td><td>71</td><td>24</td><td>CAACTTAACTTGACCGCTCTGACC</td><td>1444.2</td><td>8.3</td><td>0.6</td><td>14.0</td><td>2.6E-44</td><td>5.9E-42</td><td>MB231/MB468</td></tr>
<tr><td>>hg38_21_+_8433172_8446622_RNA45SN1_45S_50ntFlanks@6721.6739.19|AATTGCAGGACACATTGAT</td><td>>hg38_21_+_8433172_8446622_RNA45SN1_45S_50ntFlanks</td><td>6721</td><td>6739</td><td>19</td><td>AATTGCAGGACACATTGAT</td><td>2726.6</td><td>-7.9</td><td>0.6</td><td>-13.9</td><td>9.7E-44</td><td>5.9E-42</td><td>BT20/MB468</td></tr>
<tr><td>>MI0001445|hsa-mir-423&WithFlank&17|+|30117073|30117178@23.43.21&[MIMAT0004748&hsa-miR-423-5p&offsets|0|-2;m-141&17|+|30117095|30117117&offsets|0|-2]|TGAGGGGCAGAGAGCGAGACT</td><td>>MI0001445|hsa-mir-423&WithFlank&17|+|30117073|30117178</td><td>23</td><td>43</td><td>21</td><td>TGAGGGGCAGAGAGCGAGACT</td><td>1513.5</td><td>7.8</td><td>0.6</td><td>13.9</td><td>1.0E-43</td><td>6.1E-42</td><td>BT20/MB468</td></tr>
<tr><td>>MI0000063|hsa-let-7b&WithFlank&22|+|46113680|46113774@12.33.22(+1C)&[MIMAT0000063&hsa-let-7b-5p&offsets|0|0(+1C);m-14&22|+|46113691|46113712&offsets|0|0(+1C)]|TGAGGTAGTAGGTTGTGTGGTTC</td><td>>MI0000063|hsa-let-7b&WithFlank&22|+|46113680|46113774</td><td>12</td><td>33</td><td>22(+1C)</td><td>TGAGGTAGTAGGTTGTGTGGTTC</td><td>1786.3</td><td>8.5</td><td>0.6</td><td>13.9</td><td>1.0E-43</td><td>6.2E-42</td><td>BT20/MB468</td></tr>
<tr><td>>trnaMT_PheGAA_MT_+_577_647@46.74.29|GACGGGCTCACATCACCCCATAAACACCA</td><td>>trnaMT_PheGAA_MT_+_577_647</td><td>46</td><td>74</td><td>29</td><td>GACGGGCTCACATCACCCCATAAACACCA</td><td>706.4</td><td>7.7</td><td>0.6</td><td>13.9</td><td>3.4E-44</td><td>7.3E-42</td><td>MB231/MB468</td></tr>
<tr><td>>hg38_21_+_8433172_8446622_RNA45SN1_45S_50ntFlanks@475.515.41|CTTCGTGATCGATGTGGTGACGTCGTGCTCTCCCGGGCCGG</td><td>>hg38_21_+_8433172_8446622_RNA45SN1_45S_50ntFlanks</td><td>475</td><td>515</td><td>41</td><td>CTTCGTGATCGATGTGGTGACGTCGTGCTCTCCCGGGCCGG</td><td>5487.4</td><td>9.9</td><td>0.7</td><td>13.9</td><td>3.8E-44</td><td>7.7E-42</td><td>MB231/MB468</td></tr>
<tr><td>>trnaMT_LysTTT_MT_+_8295_8364@1.36.36,trnalookalike7_LysTTT_1_+_568844_568913@1.36.36|CACTGTAAAGCTAACTTAGCATTAACCTTTTAAGTT</td><td>>trnaMT_LysTTT_MT_+_8295_8364</td><td>1</td><td>36</td><td>36</td><td>CACTGTAAAGCTAACTTAGCATTAACCTTTTAAGTT</td><td>1177.4</td><td>6.9</td><td>0.5</td><td>13.9</td><td>3.9E-44</td><td>7.7E-42</td><td>MB231/MB468</td></tr>
<tr><td>>MI0000088|hsa-mir-30a&WithFlank&6|-|71403545|71403627@12.35.24(+1U)&[MIMAT0000087&hsa-miR-30a-5p&offsets|0|+2(+1U);m-15&6|-|71403595|71403616&offsets|0|+2(+1U)]|TGTAAACATCCTCGACTGGAAGCTT</td><td>>MI0000088|hsa-mir-30a&WithFlank&6|-|71403545|71403627</td><td>12</td><td>35</td><td>24(+1U)</td><td>TGTAAACATCCTCGACTGGAAGCTT</td><td>27822.6</td><td>9.5</td><td>0.7</td><td>13.9</td><td>4.2E-44</td><td>8.0E-42</td><td>MB231/MB468</td></tr>
<tr><td>>trnaMT_GluTTC_MT_-_14674_14742@38.72.35,trnalookalike8_GluTTC_5_-_93905172_93905240@38.72.35|CATTGGTCGTGGTTGTAGTCCGTGCGAGAATACCA</td><td>>trnaMT_GluTTC_MT_-_14674_14742</td><td>38</td><td>72</td><td>35</td><td>CATTGGTCGTGGTTGTAGTCCGTGCGAGAATACCA</td><td>12061.3</td><td>9.7</td><td>0.7</td><td>13.9</td><td>4.2E-44</td><td>8.0E-42</td><td>MB231/MB468</td></tr>
<tr><td>>MI0000299|hsa-mir-222&WithFlank&X|-|45747009|45747130@75.98.24(+1G)&[MIMAT0000279&hsa-miR-222-3p&offsets|0|+3(+1G);m-140&X|-|45747033|45747056&offsets|0|0(+1G)]|AGCTACATCTGGCTACTGGGTCTCG</td><td>>MI0000299|hsa-mir-222&WithFlank&X|-|45747009|45747130</td><td>75</td><td>98</td><td>24(+1G)</td><td>AGCTACATCTGGCTACTGGGTCTCG</td><td>8681.6</td><td>9.4</td><td>0.7</td><td>13.9</td><td>5.3E-44</td><td>9.7E-42</td><td>MB231/MB468</td></tr>
<tr><td>>hg38_21_+_8433172_8446622_RNA45SN1_45S_50ntFlanks@12476.12498.23|CCACTAATAGGGAACGTGAGCTG</td><td>>hg38_21_+_8433172_8446622_RNA45SN1_45S_50ntFlanks</td><td>12476</td><td>12498</td><td>23</td><td>CCACTAATAGGGAACGTGAGCTG</td><td>2014.1</td><td>-7.1</td><td>0.5</td><td>-13.8</td><td>2.1E-43</td><td>1.3E-41</td><td>BT20/MB468</td></tr>
<tr><td>>MI0000299|hsa-mir-222&WithFlank&X|-|45747009|45747130@75.99.25(+3U)&[MIMAT0000279&hsa-miR-222-3p&offsets|0|+4(+3U);m-140&X|-|45747033|45747056&offsets|0|+1(+3U)]|AGCTACATCTGGCTACTGGGTCTCTTTT</td><td>>MI0000299|hsa-mir-222&WithFlank&X|-|45747009|45747130</td><td>75</td><td>99</td><td>25(+3U)</td><td>AGCTACATCTGGCTACTGGGTCTCTTTT</td><td>9527.5</td><td>10.7</td><td>0.8</td><td>13.8</td><td>1.4E-43</td><td>2.4E-41</td><td>MB231/MB468</td></tr>
<tr><td>>trna134_GluTTC_1_-_16861774_16861845@1.33.33,trna5_GluTTC_1_+_17199078_17199149@1.33.33,trna84_GluTTC_1_-_161391883_161391954@1.33.33,trna94_GluTTC_1_-_149664355_149664427@1.33.33|TCCCTGGTGGTCTAGTGGCTAGGATTCGGCGCT</td><td>>trna134_GluTTC_1_-_16861774_16861845</td><td>1</td><td>33</td><td>33</td><td>TCCCTGGTGGTCTAGTGGCTAGGATTCGGCGCT</td><td>3092.0</td><td>9.6</td><td>0.7</td><td>13.8</td><td>4.7E-43</td><td>2.7E-41</td><td>BT20/MB468</td></tr>
<tr><td>>hg38_21_+_8433172_8446622_RNA45SN1_45S_50ntFlanks@6700.6720.21|CTAGCTGCGAGAATTAATGTG</td><td>>hg38_21_+_8433172_8446622_RNA45SN1_45S_50ntFlanks</td><td>6700</td><td>6720</td><td>21</td><td>CTAGCTGCGAGAATTAATGTG</td><td>12602.4</td><td>-6.1</td><td>0.4</td><td>-13.7</td><td>6.0E-43</td><td>3.4E-41</td><td>BT20/MB468</td></tr>
<tr><td>>MI0000299|hsa-mir-222&WithFlank&X|-|45747009|45747130@75.99.25&[MIMAT0000279&hsa-miR-222-3p&offsets|0|+4;m-140&X|-|45747033|45747056&offsets|0|+1]|AGCTACATCTGGCTACTGGGTCTCT</td><td>>MI0000299|hsa-mir-222&WithFlank&X|-|45747009|45747130</td><td>75</td><td>99</td><td>25</td><td>AGCTACATCTGGCTACTGGGTCTCT</td><td>156206.3</td><td>9.9</td><td>0.7</td><td>13.8</td><td>2.5E-43</td><td>4.3E-41</td><td>MB231/MB468</td></tr>
<tr><td>>hg38_21_+_8433172_8446622_RNA45SN1_45S_50ntFlanks@10411.10428.18|CAGTCGGTCCTGAGAGAT</td><td>>hg38_21_+_8433172_8446622_RNA45SN1_45S_50ntFlanks</td><td>10411</td><td>10428</td><td>18</td><td>CAGTCGGTCCTGAGAGAT</td><td>2545.4</td><td>-6.9</td><td>0.5</td><td>-13.7</td><td>1.3E-42</td><td>7.4E-41</td><td>BT20/MB468</td></tr>
<tr><td>>MI0000063|hsa-let-7b&WithFlank&22|+|46113680|46113774@12.32.21(+1C)&[MIMAT0000063&hsa-let-7b-5p&offsets|0|-1(+1C);m-14&22|+|46113691|46113712&offsets|0|-1(+1C)]|TGAGGTAGTAGGTTGTGTGGTC</td><td>>MI0000063|hsa-let-7b&WithFlank&22|+|46113680|46113774</td><td>12</td><td>32</td><td>21(+1C)</td><td>TGAGGTAGTAGGTTGTGTGGTC</td><td>13231.6</td><td>8.5</td><td>0.6</td><td>13.7</td><td>1.3E-42</td><td>7.4E-41</td><td>BT20/MB468</td></tr>
<tr><td>>hg38_21_+_8433172_8446622_RNA45SN1_45S_50ntFlanks@12456.12484.29|CCAAGCGTTGGATTGTTCACCCACTAATA</td><td>>hg38_21_+_8433172_8446622_RNA45SN1_45S_50ntFlanks</td><td>12456</td><td>12484</td><td>29</td><td>CCAAGCGTTGGATTGTTCACCCACTAATA</td><td>8413.3</td><td>-7.5</td><td>0.5</td><td>-13.8</td><td>4.8E-43</td><td>7.8E-41</td><td>MB231/MB468</td></tr>
<tr><td>>hg38_21_+_8433172_8446622_RNA45SN1_45S_50ntFlanks@3989.4023.35|TTTGGTGACTCTAGATAACCTCGGGCCGATCGCAC</td><td>>hg38_21_+_8433172_8446622_RNA45SN1_45S_50ntFlanks</td><td>3989</td><td>4023</td><td>35</td><td>TTTGGTGACTCTAGATAACCTCGGGCCGATCGCAC</td><td>391.6</td><td>6.2</td><td>0.5</td><td>13.8</td><td>4.8E-43</td><td>7.8E-41</td><td>MB231/MB468</td></tr>
<tr><td>>MI0000446|hsa-mir-125b-1&WithFlank&11|-|122099751|122099850@21.42.22(+1A);MI0000470|hsa-mir-125b-2&WithFlank&21|+|16590231|16590331@23.44.22(+1A)&[MIMAT0000423&hsa-miR-125b-5p&offsets|0|0(+1A);m-67&11|-|122099809|122099830&offsets|0|0(+1A)];[MIMAT0000423_1&hsa-miR-125b-5p&offsets|0|0(+1A);m-68&21|+|16590253|16590274&offsets|0|0(+1A)]|TCCCTGAGACCCTAACTTGTGAA</td><td>>MI0000446|hsa-mir-125b-1&WithFlank&11|-|122099751|122099850</td><td>21</td><td>42</td><td>22(+1A)</td><td>TCCCTGAGACCCTAACTTGTGAA</td><td>15962.5</td><td>10.7</td><td>0.8</td><td>13.7</td><td>5.5E-43</td><td>8.6E-41</td><td>MB231/MB468</td></tr>
<tr><td>>MI0000446|hsa-mir-125b-1&WithFlank&11|-|122099751|122099850@21.42.22(+2A);MI0000470|hsa-mir-125b-2&WithFlank&21|+|16590231|16590331@23.44.22(+2A)&[MIMAT0000423&hsa-miR-125b-5p&offsets|0|0(+2A);m-67&11|-|122099809|122099830&offsets|0|0(+2A)];[MIMAT0000423_1&hsa-miR-125b-5p&offsets|0|0(+2A);m-68&21|+|16590253|16590274&offsets|0|0(+2A)]|TCCCTGAGACCCTAACTTGTGAAA</td><td>>MI0000446|hsa-mir-125b-1&WithFlank&11|-|122099751|122099850</td><td>21</td><td>42</td><td>22(+2A)</td><td>TCCCTGAGACCCTAACTTGTGAAA</td><td>2857.2</td><td>9.9</td><td>0.7</td><td>13.7</td><td>5.7E-43</td><td>8.8E-41</td><td>MB231/MB468</td></tr>
<tr><td>>hg38_21_+_8433172_8446622_RNA45SN1_45S_50ntFlanks@3989.4023.35|TTTGGTGACTCTAGATAACCTCGGGCCGATCGCAC</td><td>>hg38_21_+_8433172_8446622_RNA45SN1_45S_50ntFlanks</td><td>3989</td><td>4023</td><td>35</td><td>TTTGGTGACTCTAGATAACCTCGGGCCGATCGCAC</td><td>292.4</td><td>5.5</td><td>0.4</td><td>13.7</td><td>1.6E-42</td><td>9.0E-41</td><td>BT20/MB468</td></tr>
</table>

| | | | | | | | | | | | | |
|---|---|---|---|---|---|---|---|---|---|---|---|---|
| >am_MI0005763\|hsa-mir-941-1&WithFlank&20\|+\|63919443\|63919526@53.74.22(+1A);MI0005764\|hsa-mir-941-2&WithFlank&20\|+\|63919499\|63919582@53.74.22(+1A);MI0005765\|hsa-mir-941-3&WithFlank&20\|+\|63919555\|63919638@53.74.22(+1A);MI0005766\|hsa-mir-941-4&WithFlank&20\|+\|63919750\|63919833@53.74.22(+1A);MI0031520\|hsa-mir-941-5&WithFlank&20\|+\|63919862\|63919945@53.74.22(+1A)&[MIMAT0004984&hsa-miR-941&offsets\|0\|-1(+1A);m-263&20\|+\|63919495\|63919517&offsets\|0\|-1(+1A)];[MIMAT0004984_1&hsa-miR-941&offsets\|0\|-1(+1A);m-263&20\|+\|63919551\|63919573&offsets\|0\|-1(+1A)];[MIMAT0004984_2&hsa-miR-941&offsets\|0\|-1(+1A);m-263&20\|+\|63919607\|63919629&offsets\|0\|-1(+1A)];[MIMAT0004984_3&hsa-miR-941&offsets\|0\|-1(+1A);m-263&20\|+\|63919802\|63919824&offsets\|0\|-1(+1A)];[MIMAT0004984_4&hsa-miR-941&offsets\|0\|-1(+1A);m-263&20\|+\|63919914\|63919936&offsets\|0\|-1(+1A)]\|CACCCGGCTGTGTGCACATGTGA | >am_MI0005763\|hsa-mir-941-1&WithFlank&20\|+\|63919443\|63919526 | 53 | 74 | 22(+1A) | CACCCGGCTGTGTGCACATGTGA | 2285.9 | 8.7 | 0.6 | 13.6 | 2.1E-42 | 1.2E-40 | BT20/MB468 |
| >hg38_21_+_8433172_8446622_RNA45SN1_45S_50ntFlanks@6646.6672.27\|TCGTACGACTCTTAGCGGTGGATCACT | >hg38_21_+_8433172_8446622_RNA45SN1_45S_50ntFlanks | 6646 | 6672 | 27 | TCGTACGACTCTTAGCGGTGGATCACT | 11337.3 | 7.6 | 0.6 | 13.6 | 2.1E-42 | 1.2E-40 | BT20/MB468 |
| >MI0000446\|hsa-mir-125b-1&WithFlank&11\|-\|122099751\|122099850@21.42.22;MI0000470\|hsa-mir-125b-2&WithFlank&21\|+\|16590231\|16590331@23.44.22&[MIMAT0000423&hsa-miR-125b-5p&offsets\|0\|0;m-67&11\|-\|122099809\|122099830&offsets\|0\|0];[MIMAT0000423_1&hsa-miR-125b-5p&offsets\|0\|0;m-68&21\|+\|16590253\|16590274&offsets\|0\|0]\|TCCCTGAGACCCTAACTTGTGA | >MI0000446\|hsa-mir-125b-1&WithFlank&11\|-\|122099751\|122099850 | 21 | 42 | 22 | TCCCTGAGACCCTAACTTGTGA | 43521.6 | 9.8 | 0.7 | 13.7 | 7.8E-43 | 1.2E-40 | MB231/MB468 |
| >hg38_21_+_8433172_8446622_RNA45SN1_45S_50ntFlanks@476.516.41\|TTCGTGATCGATGTGGTGACGTCGTGCTCTCCCGGGCCGGG | >hg38_21_+_8433172_8446622_RNA45SN1_45S_50ntFlanks | 476 | 516 | 41 | TTCGTGATCGATGTGGTGACGTCGTGCTCTCCCGGGCCGGG | 10270.5 | 10.0 | 0.7 | 13.7 | 8.8E-43 | 1.3E-40 | MB231/MB468 |
| >hg38_21_+_8433172_8446622_RNA45SN1_45S_50ntFlanks@9572.9596.25\|CAAATCGGTCGTCCGACCTGGGTAT | >hg38_21_+_8433172_8446622_RNA45SN1_45S_50ntFlanks | 9572 | 9596 | 25 | CAAATCGGTCGTCCGACCTGGGTAT | 6186.8 | -8.2 | 0.6 | -13.7 | 9.4E-43 | 1.3E-40 | MB231/MB468 |
| >hg38_21_+_8433172_8446622_RNA45SN1_45S_50ntFlanks@6646.6664.19\|TCGTACGACTCTTAGCGGT | >hg38_21_+_8433172_8446622_RNA45SN1_45S_50ntFlanks | 6646 | 6664 | 19 | TCGTACGACTCTTAGCGGT | 37468.1 | 5.9 | 0.4 | 13.6 | 2.6E-42 | 1.4E-40 | BT20/MB468 |
| >hg38_21_+_8433172_8446622_RNA45SN1_45S_50ntFlanks@12001.12032.32\|CTGCGGGCCGCCGGTGAAATACCACTACTCTG | >hg38_21_+_8433172_8446622_RNA45SN1_45S_50ntFlanks | 12001 | 12032 | 32 | CTGCGGGCCGCCGGTGAAATACCACTACTCTG | 1575.5 | -8.0 | 0.6 | -13.6 | 3.0E-42 | 1.6E-40 | BT20/MB468 |
| >am_MI0000809\|hsa-mir-151a&WithFlank&8\|-\|140732558\|140732659@53.73.21(+1U)&[MIMAT0000757&hsa-miR-151a-3p&offsets\|0\|0(+1U);m-30&8\|-\|140732587\|140732607&offsets\|0\|0(+1U)]\|CTAGACTGAAGCTCCTTGAGGT | >am_MI0000809\|hsa-mir-151a&WithFlank&8\|-\|140732558\|140732659 | 53 | 73 | 21(+1U) | CTAGACTGAAGCTCCTTGAGGT | 26587.5 | 9.9 | 0.7 | 13.7 | 1.2E-42 | 1.6E-40 | MB231/MB468 |
| >MI0000102\|hsa-mir-100&WithFlank&11\|-\|122152223\|122152314@19.41.23&[MIMAT0000098&hsa-miR-100-5p&offsets\|0\|+1;m-27&11\|-\|122152276\|122152296&offsets\|0\|+2]\|AACCCGTAGATCCGAACTTGTGG | >MI0000102\|hsa-mir-100&WithFlank&11\|-\|122152223\|122152314 | 19 | 41 | 23 | AACCCGTAGATCCGAACTTGTGG | 18859.3 | 16.6 | 1.2 | 13.7 | 1.3E-42 | 1.8E-40 | MB231/MB468 |
| >hg38_21_+_8433172_8446622_RNA45SN1_45S_50ntFlanks@6653.6696.44\|ACTCTTAGCGGTGGATCACTCGGCTCGTGCGTCGATGAAGAACG | >hg38_21_+_8433172_8446622_RNA45SN1_45S_50ntFlanks | 6653 | 6696 | 44 | ACTCTTAGCGGTGGATCACTCGGCTCGTGCGTCGATGAAGAACG | 656.7 | 7.5 | 0.5 | 13.7 | 2.0E-42 | 2.7E-40 | MB231/MB468 |
| >hg38_1_-_228634819_228635039_RNA5S12_5S_50ntFlanks@135.171.37\|GGGAGACCGCCTGGGAATACCGGGTGCTGTAGGCTTT | >hg38_1_-_228634819_228635039_RNA5S12_5S_50ntFlanks | 135 | 171 | 37 | GGGAGACCGCCTGGGAATACCGGGTGCTGTAGGCTTT | 34224.2 | 9.0 | 0.7 | 13.6 | 2.4E-42 | 3.1E-40 | MB231/MB468 |
| >hg38_21_+_8433172_8446622_RNA45SN1_45S_50ntFlanks@10411.10430.20\|CAGTCGGTCCTGAGAGATGG | >hg38_21_+_8433172_8446622_RNA45SN1_45S_50ntFlanks | 10411 | 10430 | 20 | CAGTCGGTCCTGAGAGATGG | 5949.8 | -8.4 | 0.6 | -13.6 | 3.6E-42 | 4.6E-40 | MB231/MB468 |
| >hg38_21_+_8433172_8446622_RNA45SN1_45S_50ntFlanks@12655.12675.21\|ATGACTGAACGCCTCTAAGTC | >hg38_21_+_8433172_8446622_RNA45SN1_45S_50ntFlanks | 12655 | 12675 | 21 | ATGACTGAACGCCTCTAAGTC | 8576.7 | -8.2 | 0.6 | -13.5 | 8.9E-42 | 4.7E-40 | BT20/MB468 |
| >MI0000542\|hsa-mir-320a&WithFlank&8\|-\|22244960\|22245043@48.69.22(+1U)&[MIMAT0000510&hsa-miR-320a-3p&offsets\|0\|0(+1U);m-60&8\|-\|22244975\|22244996&offsets\|0\|0(+1U)]\|AAAAGCTGGGTTGAGAGGGCGAT | >MI0000542\|hsa-mir-320a&WithFlank&8\|-\|22244960\|22245043 | 48 | 69 | 22(+1U) | AAAAGCTGGGTTGAGAGGGCGAT | 22242.2 | 10.1 | 0.7 | 13.5 | 9.3E-42 | 4.9E-40 | BT20/MB468 |
| >MI0000255\|hsa-mir-30d&WithFlank&8\|-\|134804870\|134804951@12.35.24(+1U)&[MIMAT0000245&hsa-miR-30d-5p&offsets\|0\|+2(+1U);m-21&8\|-\|134804919\|134804940&offsets\|0\|+2(+1U)]\|TGTAAACATCCCCGACTGGAAGCTT | >MI0000255\|hsa-mir-30d&WithFlank&8\|-\|134804870\|134804951 | 12 | 35 | 24(+1U) | TGTAAACATCCCCGACTGGAAGCTT | 11812.7 | 9.7 | 0.7 | 13.6 | 4.7E-42 | 5.9E-40 | MB231/MB468 |
| >hg38_21_+_8433172_8446622_RNA45SN1_45S_50ntFlanks@475.521.47\|CTTCGTGATCGATGTGGTGACGTCGTGCTCTCCCGGGCCGGGTCCGA | >hg38_21_+_8433172_8446622_RNA45SN1_45S_50ntFlanks | 475 | 521 | 47 | CTTCGTGATCGATGTGGTGACGTCGTGCTCTCCCGGGCCGGGTCCGA | 4070.7 | 10.5 | 0.8 | 13.5 | 9.1E-42 | 1.1E-39 | MB231/MB468 |
| >MI0000301\|hsa-mir-224&WithFlank&X\|-\|151958572\|151958664@14.35.22&[MIMAT0000281&hsa-miR-224-5p&offsets\|+1\|-2;m-238&X\|-\|151958630\|151958651&offsets\|0\|0]\|CAAGTCACTAGTGGTTCCGTTT | >MI0000301\|hsa-mir-224&WithFlank&X\|-\|151958572\|151958664 | 14 | 35 | 22 | CAAGTCACTAGTGGTTCCGTTT | 5997.2 | 10.4 | 0.8 | 13.5 | 9.0E-42 | 1.1E-39 | MB231/MB468 |
| >hg38_21_+_8433172_8446622_RNA45SN1_45S_50ntFlanks@11661.11685.25\|TGATTTCTGCCCAGTGCTCTGAATG | >hg38_21_+_8433172_8446622_RNA45SN1_45S_50ntFlanks | 11661 | 11685 | 25 | TGATTTCTGCCCAGTGCTCTGAATG | 4852.9 | -6.6 | 0.5 | -13.5 | 2.5E-41 | 1.3E-39 | BT20/MB468 |
| >trnaMT_SerGCT_MT_+_12207_12265@1.22.22\|GAGAAAGCTCACAAGAACTGCT | >trnaMT_SerGCT_MT_+_12207_12265 | 1 | 22 | 22 | GAGAAAGCTCACAAGAACTGCT | 9192.8 | 6.8 | 0.5 | 13.5 | 2.7E-41 | 1.4E-39 | BT20/MB468 |
| >hg38_21_+_8433172_8446622_RNA45SN1_45S_50ntFlanks@12456.12488.33\|CCAAGCGTTGGATTGTTCACCCACTAATAGGGA | >hg38_21_+_8433172_8446622_RNA45SN1_45S_50ntFlanks | 12456 | 12488 | 33 | CCAAGCGTTGGATTGTTCACCCACTAATAGGGA | 6235.4 | -9.2 | 0.7 | -13.4 | 3.2E-41 | 1.6E-39 | BT20/MB468 |
| >hg38_21_+_8433172_8446622_RNA45SN1_45S_50ntFlanks@7974.8009.36\|ACGCGACCTCAGATCAGACGTGGCGACCCGCTGAAT | >hg38_21_+_8433172_8446622_RNA45SN1_45S_50ntFlanks | 7974 | 8009 | 36 | ACGCGACCTCAGATCAGACGTGGCGACCCGCTGAAT | 8158.3 | 6.0 | 0.4 | 13.4 | 3.2E-41 | 1.6E-39 | BT20/MB468 |
| >hg38_21_+_8433172_8446622_RNA45SN1_45S_50ntFlanks@6647.6664.18\|CGTACGACTCTTAGCGGT | >hg38_21_+_8433172_8446622_RNA45SN1_45S_50ntFlanks | 6647 | 6664 | 18 | CGTACGACTCTTAGCGGT | 4407.0 | 5.8 | 0.4 | 13.4 | 3.7E-41 | 1.9E-39 | BT20/MB468 |
| >MI0003776\|hsa-mir-320b-1&WithFlank&1\|+\|116671740\|116671823@48.67.20;MI0003839\|hsa-mir-320b-2&WithFlank&1\|-\|224257034\|224257116@47.66.20;MI0000542\|hsa-mir-320a&WithFlank&8\|-\|22244960\|22245043@48.67.20&[MIMAT0005792&hsa-miR-320b&offsets\|0\|-2;m-183&1\|+\|116671787\|116671806&offsets\|0\|0];[MIMAT0005792_1&hsa-miR-320b&offsets\|0\|-2;m-183&1\|-\|224257051\|224257070&offsets\|0\|0];[MIMAT0000510&hsa-miR-320a-3p&offsets\|0\|-2;m-60&8\|-\|22244975\|22244996&offsets\|0\|-2]\|AAAAGCTGGGTTGAGAGGGC | >MI0003776\|hsa-mir-320b-1&WithFlank&1\|+\|116671740\|116671823 | 48 | 67 | 20 | AAAAGCTGGGTTGAGAGGGC | 2664.1 | 8.9 | 0.7 | 13.4 | 8.1E-41 | 4.1E-39 | BT20/MB468 |
| >hg38_MT_+_1621_3279_RNR2_16S_50ntFlanks@51.79.29\|GCTAAACCTAGCCCCAAACCCACTCCACC | >hg38_MT_+_1621_3279_RNR2_16S_50ntFlanks | 51 | 79 | 29 | GCTAAACCTAGCCCCAAACCCACTCCACC | 1131.8 | 7.5 | 0.6 | 13.4 | 8.5E-41 | 4.2E-39 | BT20/MB468 |
| >MI0000439\|hsa-mir-23b&WithFlank&9\|+\|95085202\|95085310@67.86.20(+1U)&[MIMAT0000418&hsa-miR-23b-3p&offsets\|+3\|0(+1U);m-42&9\|+\|95085265\|95085285&offsets\|+3\|+2(+1U)]\|ACATTGCCAGGGATTACCACT | >MI0000439\|hsa-mir-23b&WithFlank&9\|+\|95085202\|95085310 | 67 | 86 | 20(+1U) | ACATTGCCAGGGATTACCACT | 15632.9 | 18.6 | 1.4 | 13.4 | 8.6E-41 | 4.2E-39 | BT20/MB468 |
| >MI0000087\|hsa-mir-29a&WithFlank&7\|-\|130876741\|130876816@48.67.20(+1C)&[MIMAT0000086&hsa-miR-29a-3p&offsets\|0\|-2(+1C);m-26&7\|-\|130876749\|130876769&offsets\|0\|-1(+1C)]\|TAGCACCATCTGAAATCGGTC | >MI0000087\|hsa-mir-29a&WithFlank&7\|-\|130876741\|130876816 | 48 | 67 | 20(+1C) | TAGCACCATCTGAAATCGGTC | 3576.0 | 10.1 | 0.8 | 13.4 | 4.8E-41 | 5.7E-39 | MB231/MB468 |
| >hg38_21_+_8433172_8446622_RNA45SN1_45S_50ntFlanks@11768.11790.23\|CTAATTAGTGACGCGCATGAATG | >hg38_21_+_8433172_8446622_RNA45SN1_45S_50ntFlanks | 11768 | 11790 | 23 | CTAATTAGTGACGCGCATGAATG | 1155.7 | -5.8 | 0.4 | -13.3 | 1.3E-40 | 6.1E-39 | BT20/MB468 |
| >trna2_GlyGCC_21_-_18827107_18827177@1.31.31,trna35_GlyGCC_1_+_161413094_161413164@1.31.31,trna37_GlyGCC_1_+_161420467_161420537@1.31.31,trna39_GlyGCC_1_+_161427898_161427968@1.31.31,trna41_GlyGCC_1_+_161435258_161435328@1.31.31\|GCATGGGTGGTTCAGTGGTAGAATTCTCGCC | >trna2_GlyGCC_21_-_18827107_18827177 | 1 | 31 | 31 | GCATGGGTGGTTCAGTGGTAGAATTCTCGCC | 57978.8 | 8.6 | 0.6 | 13.3 | 1.7E-40 | 8.3E-39 | BT20/MB468 |
| >MI0000299\|hsa-mir-222&WithFlank&X\|-\|45747009\|45747130@75.99.25(+2U)&[MIMAT0000279&hsa-miR-222-3p&offsets\|0\|+4(+2U);m-140&X\|-\|45747033\|45747056&offsets\|0\|+1(+2U)]\|AGCTACATCTGGCTACTGGGTCTCTTT | >MI0000299\|hsa-mir-222&WithFlank&X\|-\|45747009\|45747130 | 75 | 99 | 25(+2U) | AGCTACATCTGGCTACTGGGTCTCTTT | 10690.9 | 10.3 | 0.8 | 13.4 | 7.4E-41 | 8.5E-39 | MB231/MB468 |
| >MI0000087\|hsa-mir-29a&WithFlank&7\|-\|130876741\|130876816@47.68.22(+1U)&[MIMAT0000086&hsa-miR-29a-3p&offsets\|-1\|-1(+1U);m-26&7\|-\|130876749\|130876769&offsets\|-1\|0(+1U)]\|CTAGCACCATCTGAAATCGGTTT | >MI0000087\|hsa-mir-29a&WithFlank&7\|-\|130876741\|130876816 | 47 | 68 | 22(+1U) | CTAGCACCATCTGAAATCGGTTT | 3373.8 | 10.1 | 0.8 | 13.4 | 9.8E-41 | 1.1E-38 | MB231/MB468 |
| >hg38_21_+_8433172_8446622_RNA45SN1_45S_50ntFlanks@6651.6676.26\|CGACTCTTAGCGGTGGATCACTCGGC | >hg38_21_+_8433172_8446622_RNA45SN1_45S_50ntFlanks | 6651 | 6676 | 26 | CGACTCTTAGCGGTGGATCACTCGGC | 6287.8 | 6.4 | 0.5 | 13.3 | 2.8E-40 | 1.3E-38 | BT20/MB468 |
| >MI0000734\|hsa-mir-106b&WithFlank&7\|-\|100093987\|100094080@58.78.21(+1U)&[MIMAT0004672&hsa-miR-106b-3p&offsets\|0\|-1(+1U);m-87&7\|-\|100094003\|100094023&offsets\|0\|0(+1U)]\|CCGCACTGTGGGTACTTGCTGT | >MI0000734\|hsa-mir-106b&WithFlank&7\|-\|100093987\|100094080 | 58 | 78 | 21(+1U) | CCGCACTGTGGGTACTTGCTGT | 3644.0 | 7.8 | 0.6 | 13.3 | 2.8E-40 | 1.4E-38 | BT20/MB468 |
| >hg38_21_+_8433172_8446622_RNA45SN1_45S_50ntFlanks@7973.8011.39\|GACGCGACCTCAGATCAGACGTGGCGACCCGCTGAATTT | >hg38_21_+_8433172_8446622_RNA45SN1_45S_50ntFlanks | 7973 | 8011 | 39 | GACGCGACCTCAGATCAGACGTGGCGACCCGCTGAATTT | 3012.6 | 8.3 | 0.6 | 13.3 | 1.2E-40 | 1.4E-38 | MB231/MB468 |
| >MI0000087\|hsa-mir-29a&WithFlank&7\|-\|130876741\|130876816@47.69.23&[MIMAT0000086&hsa-miR-29a-3p&offsets\|-1\|0;m-26&7\|-\|130876749\|130876769&offsets\|-1\|+1]\|CTAGCACCATCTGAAATCGGTTA | >MI0000087\|hsa-mir-29a&WithFlank&7\|-\|130876741\|130876816 | 47 | 69 | 23 | CTAGCACCATCTGAAATCGGTTA | 5010.5 | 9.8 | 0.7 | 13.3 | 1.4E-40 | 1.5E-38 | MB231/MB468 |

<table>
<tr><td>>MI0000542|hsa-mir-320a&WithFlank&8|-|22244960|22245043@48.70.23(+1U)&[MIMAT0000510&hsa-miR-320a-3p&offsets|0|+1(+1U);m-60&8|-|22244975|22244996&offsets|0|+1(+1U)]|AAAAGCTGGGTTGAGAGGGCGAAT</td><td>>MI0000542|hsa-mir-320a&WithFlank&8|-|22244960|22245043</td><td>48</td><td>70</td><td>23(+1U)</td><td>AAAAGCTGGGTTGAGAGGGCGAAT</td><td>13401.7</td><td>9.3</td><td>0.7</td><td>13.3</td><td>3.3E-40</td><td>1.5E-38</td><td>BT20/MB468</td></tr>
<tr><td>>MI0000809|hsa-mir-151a&WithFlank&8|-|140732558|140732659@53.74.22(+1U)&[MIMAT0000757&hsa-miR-151a-3p&offsets|0|+1(+1U);m-30&8|-|140732587|140732607&offsets|0|+1(+1U)]|CTAGACTGAAGCTCCTTGAGGAT</td><td>>MI0000809|hsa-mir-151a&WithFlank&8|-|140732558|140732659</td><td>53</td><td>74</td><td>22(+1U)</td><td>CTAGACTGAAGCTCCTTGAGGAT</td><td>4501.5</td><td>9.1</td><td>0.7</td><td>13.3</td><td>1.5E-40</td><td>1.6E-38</td><td>MB231/MB468</td></tr>
<tr><td>>hg38_21_+_8433172_8446622_RNA45SN1_45S_50ntFlanks@10389.10410.22|TGAACAGCAGTTGAACATGGGT</td><td>>hg38_21_+_8433172_8446622_RNA45SN1_45S_50ntFlanks</td><td>10389</td><td>10410</td><td>22</td><td>TGAACAGCAGTTGAACATGGGT</td><td>2467.3</td><td>-7.0</td><td>0.5</td><td>-13.3</td><td>3.7E-40</td><td>1.7E-38</td><td>BT20/MB468</td></tr>
<tr><td>>MI0001445|hsa-mir-423&WithFlank&17|+|30117073|30117178@59.81.23&[MIMAT0001340&hsa-miR-423-3p&offsets|0|0;m-100&17|+|30117131|30117153&offsets|0|0]|AGCTCGGTCTGAGGCCCCTCAGT</td><td>>MI0001445|hsa-mir-423&WithFlank&17|+|30117073|30117178</td><td>59</td><td>81</td><td>23</td><td>AGCTCGGTCTGAGGCCCCTCAGT</td><td>34704.2</td><td>7.4</td><td>0.6</td><td>13.3</td><td>3.7E-40</td><td>1.7E-38</td><td>BT20/MB468</td></tr>
<tr><td>>MI0000299|hsa-mir-222&WithFlank&X|-|45747009|45747130@75.98.24(+1C)&[MIMAT0000279&hsa-miR-222-3p&offsets|0|+3(+1C);m-140&X|-|45747033|45747056&offsets|0|0(+1C)]|AGCTACATCTGGCTACTGGGTCTCC</td><td>>MI0000299|hsa-mir-222&WithFlank&X|-|45747009|45747130</td><td>75</td><td>98</td><td>24(+1C)</td><td>AGCTACATCTGGCTACTGGGTCTCC</td><td>29160.5</td><td>9.5</td><td>0.7</td><td>13.3</td><td>2.6E-40</td><td>2.8E-38</td><td>MB231/MB468</td></tr>
<tr><td>>hg38_21_+_8433172_8446622_RNA45SN1_45S_50ntFlanks@485.505.21|GATGTGGTGACGTCGTGCTCT</td><td>>hg38_21_+_8433172_8446622_RNA45SN1_45S_50ntFlanks</td><td>485</td><td>505</td><td>21</td><td>GATGTGGTGACGTCGTGCTCT</td><td>40181.2</td><td>19.9</td><td>1.5</td><td>13.2</td><td>6.0E-40</td><td>2.8E-38</td><td>BT20/MB468</td></tr>
<tr><td>>MI0001733|hsa-mir-452&WithFlank&X|-|151959622|151959718@20.42.23&[MIMAT0001635&hsa-miR-452-5p&offsets|0|+1;m-188&X|-|151959677|151959699&offsets|0|0]|AACTGTTTGCAGAGGAAACTGAG</td><td>>MI0001733|hsa-mir-452&WithFlank&X|-|151959622|151959718</td><td>20</td><td>42</td><td>23</td><td>AACTGTTTGCAGAGGAAACTGAG</td><td>3012.0</td><td>16.2</td><td>1.2</td><td>13.2</td><td>6.4E-40</td><td>2.9E-38</td><td>BT20/MB468</td></tr>
<tr><td>>trnaMT_ValTAC_MT_+_1602_1670@50.71.22|ACTTAACTTGACCGCTCTGACC</td><td>>trnaMT_ValTAC_MT_+_1602_1670</td><td>50</td><td>71</td><td>22</td><td>ACTTAACTTGACCGCTCTGACC</td><td>1000.5</td><td>5.8</td><td>0.4</td><td>13.2</td><td>7.3E-40</td><td>3.3E-38</td><td>BT20/MB468</td></tr>
<tr><td>>MI0000299|hsa-mir-222&WithFlank&X|-|45747009|45747130@75.99.25(+1U)&[MIMAT0000279&hsa-miR-222-3p&offsets|0|+4(+1U);m-140&X|-|45747033|45747056&offsets|0|+1(+1U)]|AGCTACATCTGGCTACTGGGTCTCTT</td><td>>MI0000299|hsa-mir-222&WithFlank&X|-|45747009|45747130</td><td>75</td><td>99</td><td>25(+1U)</td><td>AGCTACATCTGGCTACTGGGTCTCTT</td><td>20230.6</td><td>10.2</td><td>0.8</td><td>13.3</td><td>3.3E-40</td><td>3.4E-38</td><td>MB231/MB468</td></tr>
<tr><td>>trna119_LysCTT_1_-_145395522_145395594@1.35.35,trna11_LysCTT_5_-_180648979_180649051@1.35.35,trna13_LysCTT_6_+_26556774_26556846@1.35.35,trna7_LysCTT_16_+_3225692_3225764@1.35.35,trna9_LysCTT_5_+_180634755_180634827@1.35.35|GCCCGGCTAGCTCAGTCGGTAGAGCATGAGACTCT</td><td>>trna119_LysCTT_1_-_145395522_145395594</td><td>1</td><td>35</td><td>35</td><td>GCCCGGCTAGCTCAGTCGGTAGAGCATGAGACTCT</td><td>39491.9</td><td>11.3</td><td>0.9</td><td>13.2</td><td>8.4E-40</td><td>3.8E-38</td><td>BT20/MB468</td></tr>
<tr><td>>MI0000542|hsa-mir-320a&WithFlank&8|-|22244960|22245043@48.68.21(+2U)&[MIMAT0000510&hsa-miR-320a-3p&offsets|0|-1(+2U);m-60&8|-|22244975|22244996&offsets|0|-1(+2U)]|AAAAGCTGGGTTGAGAGGGCGTT</td><td>>MI0000542|hsa-mir-320a&WithFlank&8|-|22244960|22245043</td><td>48</td><td>68</td><td>21(+2U)</td><td>AAAAGCTGGGTTGAGAGGGCGTT</td><td>6761.3</td><td>10.4</td><td>0.8</td><td>13.2</td><td>8.5E-40</td><td>3.8E-38</td><td>BT20/MB468</td></tr>
<tr><td>>hg38_21_+_8433172_8446622_RNA45SN1_45S_50ntFlanks@9568.9596.29|CGTGCAAATCGGTCGTCCGACCTGGGTAT</td><td>>hg38_21_+_8433172_8446622_RNA45SN1_45S_50ntFlanks</td><td>9568</td><td>9596</td><td>29</td><td>CGTGCAAATCGGTCGTCCGACCTGGGTAT</td><td>3201.3</td><td>-7.3</td><td>0.6</td><td>-13.3</td><td>4.0E-40</td><td>4.1E-38</td><td>MB231/MB468</td></tr>
<tr><td>>MI0000086|hsa-mir-28&WithFlank&3|+|188688775|188688872@60.81.22&[MIMAT0004502&hsa-miR-28-3p&offsets|0|0;m-28&3|+|188688834|188688855&offsets|0|0]|CACTAGATTGTGAGCTCCTGGA</td><td>>MI0000086|hsa-mir-28&WithFlank&3|+|188688775|188688872</td><td>60</td><td>81</td><td>22</td><td>CACTAGATTGTGAGCTCCTGGA</td><td>15899.1</td><td>10.2</td><td>0.8</td><td>13.2</td><td>4.6E-40</td><td>4.6E-38</td><td>MB231/MB468</td></tr>
<tr><td>>MI0000746|hsa-mir-99b&WithFlank&19|+|51692606|51692687@51.73.23&[MIMAT0004678&hsa-miR-99b-3p&offsets|0|+1;m-151&19|+|51692656|51692678&offsets|0|0]|CAAGCTCGTGTCTGTGGGTCCGT</td><td>>MI0000746|hsa-mir-99b&WithFlank&19|+|51692606|51692687</td><td>51</td><td>73</td><td>23</td><td>CAAGCTCGTGTCTGTGGGTCCGT</td><td>3064.5</td><td>8.0</td><td>0.6</td><td>13.2</td><td>1.0E-39</td><td>4.7E-38</td><td>BT20/MB468</td></tr>
<tr><td>>hg38_21_+_8433172_8446622_RNA45SN1_45S_50ntFlanks@10391.10410.20|AACAGCAGTTGAACATGGGT</td><td>>hg38_21_+_8433172_8446622_RNA45SN1_45S_50ntFlanks</td><td>10391</td><td>10410</td><td>20</td><td>AACAGCAGTTGAACATGGGT</td><td>4523.3</td><td>-7.1</td><td>0.5</td><td>-13.2</td><td>6.0E-40</td><td>5.9E-38</td><td>MB231/MB468</td></tr>
<tr><td>>hg38_21_+_8433172_8446622_RNA45SN1_45S_50ntFlanks@5472.5490.19|ACGGCCCTGGCGGAGCGCT</td><td>>hg38_21_+_8433172_8446622_RNA45SN1_45S_50ntFlanks</td><td>5472</td><td>5490</td><td>19</td><td>ACGGCCCTGGCGGAGCGCT</td><td>709.1</td><td>7.4</td><td>0.6</td><td>13.2</td><td>6.4E-40</td><td>6.2E-38</td><td>MB231/MB468</td></tr>
<tr><td>>MI0000255|hsa-mir-30d&WithFlank&8|-|134804870|134804951@12.34.23(+1A)&[MIMAT0000245&hsa-miR-30d-5p&offsets|0|+1(+1A);m-21&8|-|134804919|134804940&offsets|0|+1(+1A)]|TGTAAACATCCCCGACTGGAAGCA</td><td>>MI0000255|hsa-mir-30d&WithFlank&8|-|134804870|134804951</td><td>12</td><td>34</td><td>23(+1A)</td><td>TGTAAACATCCCCGACTGGAAGCA</td><td>38083.7</td><td>10.1</td><td>0.8</td><td>13.2</td><td>1.5E-39</td><td>6.4E-38</td><td>BT20/MB468</td></tr>
<tr><td>>MI0003560|hsa-mir-92b&WithFlank&1|+|155195171|155195278@67.87.21(+1U)&[MIMAT0003218&hsa-miR-92b-3p&offsets|0|-1(+1U);m-115&1|+|155195237|155195258&offsets|0|-1(+1U)]|TATTGCACTCGTCCCGGCCTCT</td><td>>MI0003560|hsa-mir-92b&WithFlank&1|+|155195171|155195278</td><td>67</td><td>87</td><td>21(+1U)</td><td>TATTGCACTCGTCCCGGCCTCT</td><td>7541.7</td><td>9.9</td><td>0.8</td><td>13.2</td><td>1.6E-39</td><td>7.0E-38</td><td>BT20/MB468</td></tr>
<tr><td>>hg38_21_+_8433172_8446622_RNA45SN1_45S_50ntFlanks@475.516.42|CTTCGTGATCGATGTGGTGACGTCGTGCTCTCCCGGGCCGGG</td><td>>hg38_21_+_8433172_8446622_RNA45SN1_45S_50ntFlanks</td><td>475</td><td>516</td><td>42</td><td>CTTCGTGATCGATGTGGTGACGTCGTGCTCTCCCGGGCCGGG</td><td>13469.6</td><td>9.7</td><td>0.7</td><td>13.2</td><td>7.8E-40</td><td>7.4E-38</td><td>MB231/MB468</td></tr>
<tr><td>>hg38_21_+_8433172_8446622_RNA45SN1_45S_50ntFlanks@9554.9571.18|CGTAGCGGTCCTGACGTG</td><td>>hg38_21_+_8433172_8446622_RNA45SN1_45S_50ntFlanks</td><td>9554</td><td>9571</td><td>18</td><td>CGTAGCGGTCCTGACGTG</td><td>113975.6</td><td>-5.9</td><td>0.4</td><td>-13.1</td><td>1.7E-39</td><td>7.5E-38</td><td>BT20/MB468</td></tr>
<tr><td>>hg38_21_+_8433172_8446622_RNA45SN1_45S_50ntFlanks@4003.4024.22|ATAACCTCGGGCCGATCGCACG</td><td>>hg38_21_+_8433172_8446622_RNA45SN1_45S_50ntFlanks</td><td>4003</td><td>4024</td><td>22</td><td>ATAACCTCGGGCCGATCGCACG</td><td>1732.4</td><td>-6.6</td><td>0.5</td><td>-13.1</td><td>2.2E-39</td><td>9.5E-38</td><td>BT20/MB468</td></tr>
<tr><td>>hg38_21_+_8433172_8446622_RNA45SN1_45S_50ntFlanks@6647.6672.26|CGTACGACTCTTAGCGGTGGATCACT</td><td>>hg38_21_+_8433172_8446622_RNA45SN1_45S_50ntFlanks</td><td>6647</td><td>6672</td><td>26</td><td>CGTACGACTCTTAGCGGTGGATCACT</td><td>1716.3</td><td>7.5</td><td>0.6</td><td>13.1</td><td>2.3E-39</td><td>9.7E-38</td><td>BT20/MB468</td></tr>
<tr><td>>hg38_21_+_8433172_8446622_RNA45SN1_45S_50ntFlanks@10362.10383.22|AAGGCCGAAGTGGAGAAGGGTT</td><td>>hg38_21_+_8433172_8446622_RNA45SN1_45S_50ntFlanks</td><td>10362</td><td>10383</td><td>22</td><td>AAGGCCGAAGTGGAGAAGGGTT</td><td>1869.1</td><td>-7.0</td><td>0.5</td><td>-13.1</td><td>2.6E-39</td><td>1.1E-37</td><td>BT20/MB468</td></tr>
<tr><td>>MI0000542|hsa-mir-320a&WithFlank&8|-|22244960|22245043@48.69.22&[MIMAT0000510&hsa-miR-320a-3p&offsets|0|0;m-60&8|-|22244975|22244996&offsets|0|0]|AAAAGCTGGGTTGAGAGGGCGA</td><td>>MI0000542|hsa-mir-320a&WithFlank&8|-|22244960|22245043</td><td>48</td><td>69</td><td>22</td><td>AAAAGCTGGGTTGAGAGGGCGA</td><td>36901.5</td><td>10.2</td><td>0.8</td><td>13.1</td><td>2.6E-39</td><td>1.1E-37</td><td>BT20/MB468</td></tr>
<tr><td>>hg38_21_+_8433172_8446622_RNA45SN1_45S_50ntFlanks@6646.6677.32|TCGTACGACTCTTAGCGGTGGATCACTCGGCT</td><td>>hg38_21_+_8433172_8446622_RNA45SN1_45S_50ntFlanks</td><td>6646</td><td>6677</td><td>32</td><td>TCGTACGACTCTTAGCGGTGGATCACTCGGCT</td><td>4671.2</td><td>5.6</td><td>0.4</td><td>13.1</td><td>3.0E-39</td><td>1.2E-37</td><td>BT20/MB468</td></tr>
<tr><td>>MI0000650|hsa-mir-200c&WithFlank&12|+|6963693|6963772@50.72.23(+1C)&[MIMAT0000617&hsa-miR-200c-3p&offsets|0|0(+1C);m-54&12|+|6963742|6963764&offsets|0|0(+1C)]|TAATACTGCCGGGTAATGATGGAC</td><td>>MI0000650|hsa-mir-200c&WithFlank&12|+|6963693|6963772</td><td>50</td><td>72</td><td>23(+1C)</td><td>TAATACTGCCGGGTAATGATGGAC</td><td>3352.2</td><td>8.5</td><td>0.7</td><td>13.1</td><td>3.8E-39</td><td>1.6E-37</td><td>BT20/MB468</td></tr>
<tr><td>>hg38_21_+_8433172_8446622_RNA45SN1_45S_50ntFlanks@10410.10430.21|TCAGTCGGTCCTGAGAGATGG</td><td>>hg38_21_+_8433172_8446622_RNA45SN1_45S_50ntFlanks</td><td>10410</td><td>10430</td><td>21</td><td>TCAGTCGGTCCTGAGAGATGG</td><td>3658.7</td><td>-7.3</td><td>0.6</td><td>-13.1</td><td>2.0E-39</td><td>1.9E-37</td><td>MB231/MB468</td></tr>
<tr><td>>hg38_21_+_8433172_8446622_RNA45SN1_45S_50ntFlanks@10350.10368.19|ACGAGAACTTTGAAGGCCG</td><td>>hg38_21_+_8433172_8446622_RNA45SN1_45S_50ntFlanks</td><td>10350</td><td>10368</td><td>19</td><td>ACGAGAACTTTGAAGGCCG</td><td>8725.2</td><td>-6.6</td><td>0.5</td><td>-13.1</td><td>4.6E-39</td><td>1.9E-37</td><td>BT20/MB468</td></tr>
<tr><td>>trna32_LysCTT_16_-_3207406_3207478@1.33.33|GCCCGGCTAGCTCAGTCGGTAGAGCATGAGACC</td><td>>trna32_LysCTT_16_-_3207406_3207478</td><td>1</td><td>33</td><td>33</td><td>GCCCGGCTAGCTCAGTCGGTAGAGCATGAGACC</td><td>348.9</td><td>5.5</td><td>0.4</td><td>13.1</td><td>5.2E-39</td><td>2.1E-37</td><td>BT20/MB468</td></tr>
<tr><td>>MI0000446|hsa-mir-125b-1&WithFlank&11|-|122099751|122099850@21.40.20(+1A);MI0000470|hsa-mir-125b-2&WithFlank&21|+|16590231|16590331@23.42.20(+1A)&[MIMAT0000423&hsa-miR-125b-5p&offsets|0|-2(+1A);m-67&11|-|122099809|122099830&offsets|0|-2(+1A)];[MIMAT0000423_1&hsa-miR-125b-5p&offsets|0|-2(+1A);m-68&21|+|16590253|16590274&offsets|0|-2(+1A)]|TCCCTGAGACCCTAACTTGTA</td><td>>MI0000446|hsa-mir-125b-1&WithFlank&11|-|122099751|122099850</td><td>21</td><td>40</td><td>20(+1A)</td><td>TCCCTGAGACCCTAACTTGTA</td><td>2803.7</td><td>9.7</td><td>0.7</td><td>13.1</td><td>2.3E-39</td><td>2.1E-37</td><td>MB231/MB468</td></tr>
<tr><td>>hg38_21_+_8433172_8446622_RNA45SN1_45S_50ntFlanks@12996.13017.22|TCGCTGCGATCTATTGAAAGTC</td><td>>hg38_21_+_8433172_8446622_RNA45SN1_45S_50ntFlanks</td><td>12996</td><td>13017</td><td>22</td><td>TCGCTGCGATCTATTGAAAGTC</td><td>2831.4</td><td>8.2</td><td>0.6</td><td>13.1</td><td>2.4E-39</td><td>2.2E-37</td><td>MB231/MB468</td></tr>
<tr><td>>MI0000255|hsa-mir-30d&WithFlank&8|-|134804870|134804951@12.34.23(+1A)&[MIMAT0000245&hsa-miR-30d-5p&offsets|0|+1(+1A);m-21&8|-|134804919|134804940&offsets|0|+1(+1A)]|TGTAAACATCCCCGACTGGAAGCA</td><td>>MI0000255|hsa-mir-30d&WithFlank&8|-|134804870|134804951</td><td>12</td><td>34</td><td>23(+1A)</td><td>TGTAAACATCCCCGACTGGAAGCA</td><td>16148.5</td><td>9.2</td><td>0.7</td><td>13.1</td><td>2.5E-39</td><td>2.2E-37</td><td>MB231/MB468</td></tr>
<tr><td>>trna10_LysCTT_16_+_3241501_3241573@1.35.35,trna13_LysCTT_14_-_58706613_58706685@1.35.35,trna2_LysCTT_15_+_79152904_79152976@1.35.35|GCCCGGCTAGCTCAGTCGGTAGAGCATGGGACTCT</td><td>>trna10_LysCTT_16_+_3241501_3241573</td><td>1</td><td>35</td><td>35</td><td>GCCCGGCTAGCTCAGTCGGTAGAGCATGGGACTCT</td><td>497564.9</td><td>14.6</td><td>1.1</td><td>13.1</td><td>6.3E-39</td><td>2.6E-37</td><td>BT20/MB468</td></tr>
<tr><td>>MI0001445|hsa-mir-423&WithFlank&17|+|30117073|30117178@59.81.23(+1U)&[MIMAT0001340&hsa-miR-423-3p&offsets|0|0(+1U);m-100&17|+|30117131|30117153&offsets|0|0(+1U)]|AGCTCGGTCTGAGGCCCCTCAGTT</td><td>>MI0001445|hsa-mir-423&WithFlank&17|+|30117073|30117178</td><td>59</td><td>81</td><td>23(+1U)</td><td>AGCTCGGTCTGAGGCCCCTCAGTT</td><td>2853.9</td><td>7.2</td><td>0.5</td><td>13.0</td><td>7.3E-39</td><td>3.0E-37</td><td>BT20/MB468</td></tr>
<tr><td>>MI0000086|hsa-mir-28&WithFlank&3|+|188688775|188688872@60.80.21&[MIMAT0004502&hsa-miR-28-3p&offsets|0|-1;m-28&3|+|188688834|188688855&offsets|0|-1]|CACTAGATTGTGAGCTCCTGG</td><td>>MI0000086|hsa-mir-28&WithFlank&3|+|188688775|188688872</td><td>60</td><td>80</td><td>21</td><td>CACTAGATTGTGAGCTCCTGG</td><td>3436.4</td><td>9.7</td><td>0.7</td><td>13.1</td><td>3.4E-39</td><td>3.0E-37</td><td>MB231/MB468</td></tr>
<tr><td>>am_trnaMT_ProTGG_MT_-_15956_16023@54.71.18|AGACTTTTTCTCTGACCA</td><td>>am_trnaMT_ProTGG_MT_-_15956_16023</td><td>54</td><td>71</td><td>18</td><td>AGACTTTTTCTCTGACCA</td><td>14081.6</td><td>7.0</td><td>0.5</td><td>13.1</td><td>3.8E-39</td><td>3.3E-37</td><td>MB231/MB468</td></tr>
<tr><td>>hg38_21_+_8433172_8446622_RNA45SN1_45S_50ntFlanks@10840.10865.26|CCGTAACTTCGGGATAAGGATTGGCT</td><td>>hg38_21_+_8433172_8446622_RNA45SN1_45S_50ntFlanks</td><td>10840</td><td>10865</td><td>26</td><td>CCGTAACTTCGGGATAAGGATTGGCT</td><td>1907.5</td><td>-6.2</td><td>0.5</td><td>-13.0</td><td>8.9E-39</td><td>3.6E-37</td><td>BT20/MB468</td></tr>
<tr><td>>hg38_21_+_8433172_8446622_RNA45SN1_45S_50ntFlanks@10350.10377.28|ACGAGAACTTTGAAGGCCGAAGTGGAGA</td><td>>hg38_21_+_8433172_8446622_RNA45SN1_45S_50ntFlanks</td><td>10350</td><td>10377</td><td>28</td><td>ACGAGAACTTTGAAGGCCGAAGTGGAGA</td><td>1927.6</td><td>-5.4</td><td>0.4</td><td>-13.0</td><td>9.0E-39</td><td>3.6E-37</td><td>BT20/MB468</td></tr>
<tr><td>>hg38_21_+_8433172_8446622_RNA45SN1_45S_50ntFlanks@6647.6667.21|CGTACGACTCTTAGCGGTGGA</td><td>>hg38_21_+_8433172_8446622_RNA45SN1_45S_50ntFlanks</td><td>6647</td><td>6667</td><td>21</td><td>CGTACGACTCTTAGCGGTGGA</td><td>2400.8</td><td>6.3</td><td>0.5</td><td>13.0</td><td>9.4E-39</td><td>3.7E-37</td><td>BT20/MB468</td></tr>
<tr><td>>MI0000087|hsa-mir-29a&WithFlank&7|-|130876741|130876816@48.69.22&[MIMAT0000086&hsa-miR-29a-3p&offsets|0|0;m-26&7|-|130876749|130876769&offsets|0|+1]|TAGCACCATCTGAAATCGGTTA</td><td>>MI0000087|hsa-mir-29a&WithFlank&7|-|130876741|130876816</td><td>48</td><td>69</td><td>22</td><td>TAGCACCATCTGAAATCGGTTA</td><td>30782.7</td><td>9.6</td><td>0.7</td><td>13.1</td><td>4.4E-39</td><td>3.8E-37</td><td>MB231/MB468</td></tr>
<tr><td>>hg38_21_+_8433172_8446622_RNA45SN1_45S_50ntFlanks@6699.6720.22|GCTAGCTGCGAGAATTAATGTG</td><td>>hg38_21_+_8433172_8446622_RNA45SN1_45S_50ntFlanks</td><td>6699</td><td>6720</td><td>22</td><td>GCTAGCTGCGAGAATTAATGTG</td><td>24755.9</td><td>-6.4</td><td>0.5</td><td>-13.0</td><td>9.7E-39</td><td>3.8E-37</td><td>BT20/MB468</td></tr>
</table>

| | | | | | | | | | | | | |
|---|---|---|---|---|---|---|---|---|---|---|---|---|
| >hg38_21_+_8433172_8446622_RNA45SN1_45S_50ntFlanks@12936.12956.21\|AAACCATTCGTAGACGACCTG | >hg38_21_+_8433172_8446622_RNA45SN1_45S_50ntFlanks | 12936 | 12956 | 21 | AAACCATTCGTAGACGACCTG | 4969.8 | -7.1 | 0.5 | -13.0 | 9.9E-39 | 3.9E-37 | BT20/MB468 |
| >MI0003513\|hsa-mir-455&WithFlank&9\|+\|114209428\|114209535@22.44.23&[MIMAT0003150&hsa-miR-455-5p&offsets\|0\|+1;m-232&9\|+\|114209449\|114209470&offsets\|0\|+1]\|TATGTGCCTTTGGACTACATCGT | >MI0003513\|hsa-mir-455&WithFlank&9\|+\|114209428\|114209535 | 22 | 44 | 23 | TATGTGCCTTTGGACTACATCGT | 4291.7 | 17.0 | 1.3 | 13.0 | 6.4E-39 | 5.4E-37 | MB231/MB468 |
| >MI0000301\|hsa-mir-224&WithFlank&X\|-\|151958572\|151958664@14.34.21&[MIMAT0000281&hsa-miR-224-5p&offsets\|+1\|-3;m-238&X\|-\|151958630\|151958651&offsets\|0\|-1]\|CAAGTCACTAGTGGTTCCGTT | >MI0000301\|hsa-mir-224&WithFlank&X\|-\|151958572\|151958664 | 14 | 34 | 21 | CAAGTCACTAGTGGTTCCGTT | 3082.0 | 16.2 | 1.3 | 13.0 | 1.4E-38 | 5.4E-37 | BT20/MB468 |
| >am_MI0000809\|hsa-mir-151a&WithFlank&8\|-\|140732558\|140732659@53.72.20(+1A)&[MIMAT0000757&hsa-miR-151a-3p&offsets\|0\|-1(+1A);m-30&8\|-\|140732587\|140732607&offsets\|0\|-1(+1A)]\|CTAGACTGAAGCTCCTTGAGA | >am_MI0000809\|hsa-mir-151a&WithFlank&8\|-\|140732558\|140732659 | 53 | 72 | 20(+1A) | CTAGACTGAAGCTCCTTGAGA | 17292.1 | 9.3 | 0.7 | 13.0 | 7.9E-39 | 6.5E-37 | MB231/MB468 |
| >hg38_21_+_8433172_8446622_RNA45SN1_45S_50ntFlanks@12997.13039.43\|CGCTGCGATCTATTGAAAGTCAGCCCTCGACACAAGGGTTTGT | >hg38_21_+_8433172_8446622_RNA45SN1_45S_50ntFlanks | 12997 | 13039 | 43 | CGCTGCGATCTATTGAAAGTCAGCCCTCGACACAAGGGTTTGT | 1024536.4 | 12.0 | 0.9 | 13.0 | 7.9E-39 | 6.5E-37 | MB231/MB468 |
| >hg38_MT_+_1621_3279_RNR2_16S_50ntFlanks@1216.1262.47\|CAACCTCCGAGCAGTACATGCTAAGACTTCACCAGTCAAAGCGAACT | >hg38_MT_+_1621_3279_RNR2_16S_50ntFlanks | 1216 | 1262 | 47 | CAACCTCCGAGCAGTACATGCTAAGACTTCACCAGTCAAAGCGAACT | 3277.1 | 14.5 | 1.1 | 13.0 | 8.4E-39 | 6.8E-37 | MB231/MB468 |
| >hg38_21_+_8433172_8446622_RNA45SN1_45S_50ntFlanks@9564.9597.34\|CTGACGTGCAAATCGGTCGTCCGACCTGGGTATA | >hg38_21_+_8433172_8446622_RNA45SN1_45S_50ntFlanks | 9564 | 9597 | 34 | CTGACGTGCAAATCGGTCGTCCGACCTGGGTATA | 4876.4 | -7.0 | 0.5 | -13.0 | 2.2E-38 | 8.3E-37 | BT20/MB468 |
| >MI0000082\|hsa-mir-25&WithFlank&7\|-\|100093554\|100093649@58.78.21(+1U)&[MIMAT0000081&hsa-miR-25-3p&offsets\|0\|-1(+1U);m-23&7\|-\|100093571\|100093592&offsets\|0\|-1(+1U)]\|CATTGCACTTGTCTCGGTCTGT | >MI0000082\|hsa-mir-25&WithFlank&7\|-\|100093554\|100093649 | 58 | 78 | 21(+1U) | CATTGCACTTGTCTCGGTCTGT | 7647.5 | 8.5 | 0.7 | 13.0 | 2.2E-38 | 8.3E-37 | BT20/MB468 |
| >MI0003560\|hsa-mir-92b&WithFlank&1\|+\|155195171\|155195278@67.87.21(+1U)&[MIMAT0003218&hsa-miR-92b-3p&offsets\|0\|-1(+1U);m-115&1\|+\|155195237\|155195258&offsets\|0\|-1(+1U)]\|TATTGCACTCGTCCCGGCCTCT | >MI0003560\|hsa-mir-92b&WithFlank&1\|+\|155195171\|155195278 | 67 | 87 | 21(+1U) | TATTGCACTCGTCCCGGCCTCT | 4673.6 | 9.5 | 0.7 | 13.0 | 1.1E-38 | 8.9E-37 | MB231/MB468 |
| >hg38_1_-_228634819_228635039_RNA5S12_5S_50ntFlanks@137.171.35\|GAGACCGCCTGGGAATACCGGGTGCTGTAGGCTTT | >hg38_1_-_228634819_228635039_RNA5S12_5S_50ntFlanks | 137 | 171 | 35 | GAGACCGCCTGGGAATACCGGGTGCTGTAGGCTTT | 5586.9 | 9.6 | 0.7 | 13.0 | 1.2E-38 | 9.6E-37 | MB231/MB468 |
| >hg38_21_+_8433172_8446622_RNA45SN1_45S_50ntFlanks@6648.6676.29\|GTACGACTCTTAGCGGTGGATCACTCGGC | >hg38_21_+_8433172_8446622_RNA45SN1_45S_50ntFlanks | 6648 | 6676 | 29 | GTACGACTCTTAGCGGTGGATCACTCGGC | 862.4 | 6.5 | 0.5 | 12.9 | 2.8E-38 | 1.1E-36 | BT20/MB468 |
| >MI0000088\|hsa-mir-30a&WithFlank&6\|-\|71403545\|71403627@54.73.20(+1U)&[MIMAT0000088&hsa-miR-30a-3p&offsets\|+1\|-1(+1U);m-38&6\|-\|71403554\|71403575&offsets\|+1\|-1(+1U)]\|TTTCAGTCGGATGTTTGCAGT | >MI0000088\|hsa-mir-30a&WithFlank&6\|-\|71403545\|71403627 | 54 | 73 | 20(+1U) | TTTCAGTCGGATGTTTGCAGT | 13163.6 | 10.2 | 0.8 | 13.0 | 1.4E-38 | 1.1E-36 | MB231/MB468 |
| >MI0000809\|hsa-mir-151a&WithFlank&8\|-\|140732558\|140732659@53.74.22(+2A)&[MIMAT0000757&hsa-miR-151a-3p&offsets\|0\|+1(+2A);m-30&8\|-\|140732587\|140732607&offsets\|0\|+1(+2A)]\|CTAGACTGAAGCTCCTTGAGGAAA | >MI0000809\|hsa-mir-151a&WithFlank&8\|-\|140732558\|140732659 | 53 | 74 | 22(+2A) | CTAGACTGAAGCTCCTTGAGGAAA | 1925.1 | 8.2 | 0.6 | 12.9 | 3.2E-38 | 1.2E-36 | BT20/MB468 |
| >hg38_21_+_8433172_8446622_RNA45SN1_45S_50ntFlanks@12437.12454.18\|CATTGTGAAGCAGAATTC | >hg38_21_+_8433172_8446622_RNA45SN1_45S_50ntFlanks | 12437 | 12454 | 18 | CATTGTGAAGCAGAATTC | 2582.1 | -7.0 | 0.5 | -12.9 | 3.5E-38 | 1.3E-36 | BT20/MB468 |
| >am_trna111_HisGTG_1_-_147774845_147774916@-1G.23.24,trna118_HisGTG_1_-_145396881_145396952@-1G.23.24,trna16_HisGTG_1_+_146544773_146544844@-1G.23.24,trna1_HisGTG_15_+_45493349_45493420@-1G.23.24,trna21_HisGTG_1_+_147753471_147753542@-1G.23.24,trna33_HisGTG_6_+_27125906_27125977@-1G.23.24,trna7_HisGTG_9_-_14433938_14434009@-1G.23.24,trna8_HisGTG_15_-_45492611_45492682@-1G.23.24,trna9_HisGTG_15_-_45490804_45490875@-1G.23.24\|GGCCGTGATCGTATAGTGGTTAGT | >am_trna111_HisGTG_1_-_147774845_147774916 | -1G | 23 | 24 | GGCCGTGATCGTATAGTGGTTAGT | 3692.5 | 10.3 | 0.8 | 13.0 | 2.1E-38 | 1.6E-36 | MB231/MB468 |
| >MI0000088\|hsa-mir-30a&WithFlank&6\|-\|71403545\|71403627@12.35.24(+1C)&[MIMAT0000087&hsa-miR-30a-5p&offsets\|0\|+2(+1C);m-15&6\|-\|71403595\|71403616&offsets\|0\|+2(+1C)]\|TGTAAACATCCTCGACTGGAAGCTC | >MI0000088\|hsa-mir-30a&WithFlank&6\|-\|71403545\|71403627 | 12 | 35 | 24(+1C) | TGTAAACATCCTCGACTGGAAGCTC | 3075.2 | 9.4 | 0.7 | 13.0 | 2.1E-38 | 1.6E-36 | MB231/MB468 |
| >hg38_21_+_8433172_8446622_RNA45SN1_45S_50ntFlanks@10841.10865.25\|CGTAACTTCGGGATAAGGATTGGCT | >hg38_21_+_8433172_8446622_RNA45SN1_45S_50ntFlanks | 10841 | 10865 | 25 | CGTAACTTCGGGATAAGGATTGGCT | 30154.5 | -7.0 | 0.5 | -13.0 | 2.3E-38 | 1.7E-36 | MB231/MB468 |
| >MI0000087\|hsa-mir-29a&WithFlank&7\|-\|130876741\|130876816@48.68.21&[MIMAT0000086&hsa-miR-29a-3p&offsets\|0\|-1;m-26&7\|-\|130876749\|130876769&offsets\|0\|0]\|TAGCACCATCTGAAATCGGTT | >MI0000087\|hsa-mir-29a&WithFlank&7\|-\|130876741\|130876816 | 48 | 68 | 21 | TAGCACCATCTGAAATCGGTT | 34258.2 | 9.7 | 0.7 | 13.0 | 2.3E-38 | 1.7E-36 | MB231/MB468 |
| >MI0000299\|hsa-mir-222&WithFlank&X\|-\|45747009\|45747130@75.99.25(+1C)&[MIMAT0000279&hsa-miR-222-3p&offsets\|0\|+4(+1C);m-140&X\|-\|45747033\|45747056&offsets\|0\|+1(+1C)]\|AGCTACATCTGGCTACTGGGTCTCTC | >MI0000299\|hsa-mir-222&WithFlank&X\|-\|45747009\|45747130 | 75 | 99 | 25(+1C) | AGCTACATCTGGCTACTGGGTCTCTC | 2276.7 | 9.9 | 0.8 | 13.0 | 2.3E-38 | 1.7E-36 | MB231/MB468 |
| >trna14_LysTTT_11_-_59327808_59327880@1.33.33,trna2_LysTTT_17_+_8022473_8022545@1.33.33,trna54_LysTTT_1_+_204475655_204475727@1.33.33,trna5_LysTTT_11_+_59323902_59323974@1.33.33,trna62_LysTTT_1_-_204476158_204476230@1.33.33,trna76_LysTTT_6_+_28918806_28918878@1.33.33\|GCCCGGATAGCTCAGTCGGTAGAGCATCAGACT | >trna14_LysTTT_11_-_59327808_59327880 | 1 | 33 | 33 | GCCCGGATAGCTCAGTCGGTAGAGCATCAGACT | 38120.6 | 10.3 | 0.8 | 13.0 | 2.3E-38 | 1.7E-36 | MB231/MB468 |
| >hg38_21_+_8433172_8446622_RNA45SN1_45S_50ntFlanks@7975.8005.31\|CGCGACCTCAGATCAGACGTGGCGACCCGCT | >hg38_21_+_8433172_8446622_RNA45SN1_45S_50ntFlanks | 7975 | 8005 | 31 | CGCGACCTCAGATCAGACGTGGCGACCCGCT | 62383.8 | 6.1 | 0.5 | 12.9 | 5.0E-38 | 1.9E-36 | BT20/MB468 |
| >hg38_21_+_8433172_8446622_RNA45SN1_45S_50ntFlanks@6721.6745.25\|AATTGCAGGACACATTGATCATCGA | >hg38_21_+_8433172_8446622_RNA45SN1_45S_50ntFlanks | 6721 | 6745 | 25 | AATTGCAGGACACATTGATCATCGA | 1642.4 | -8.4 | 0.7 | -12.9 | 5.1E-38 | 1.9E-36 | BT20/MB468 |
| >MI0000253\|hsa-mir-148a&WithFlank&7\|-\|25949913\|25949992@50.72.23&[MIMAT0000243&hsa-miR-148a-3p&offsets\|0\|+1;m-7&7\|-\|25949922\|25949943&offsets\|0\|+1]\|TCAGTGCACTACAGAACTTTGTC | >MI0000253\|hsa-mir-148a&WithFlank&7\|-\|25949913\|25949992 | 50 | 72 | 23 | TCAGTGCACTACAGAACTTTGTC | 10212.2 | 8.3 | 0.6 | 12.9 | 5.6E-38 | 2.1E-36 | BT20/MB468 |
| >MI0003643\|hsa-mir-629&WithFlank&15\|-\|70079366\|70079474@28.49.22&[MIMAT0004810&hsa-miR-629-5p&offsets\|0\|+1;m-150&15\|-\|70079427\|70079447&offsets\|0\|+1]\|TGGGTTTACGTTGGGAGAACTT | >MI0003643\|hsa-mir-629&WithFlank&15\|-\|70079366\|70079474 | 28 | 49 | 22 | TGGGTTTACGTTGGGAGAACTT | 4764.7 | 16.9 | 1.3 | 12.9 | 6.1E-38 | 2.2E-36 | BT20/MB468 |
| >hg38_21_+_8433172_8446622_RNA45SN1_45S_50ntFlanks@12624.12654.31\|CCAATGGGGCGAAGCTACCATCTGTGGGATT | >hg38_21_+_8433172_8446622_RNA45SN1_45S_50ntFlanks | 12624 | 12654 | 31 | CCAATGGGGCGAAGCTACCATCTGTGGGATT | 3122.3 | -7.5 | 0.6 | -12.9 | 3.4E-38 | 2.5E-36 | MB231/MB468 |
| >MI0000088\|hsa-mir-30a&WithFlank&6\|-\|71403545\|71403627@12.33.22(+1A)&[MIMAT0000087&hsa-miR-30a-5p&offsets\|0\|0(+1A);m-15&6\|-\|71403595\|71403616&offsets\|0\|0(+1A)]\|TGTAAACATCCTCGACTGGAAGA | >MI0000088\|hsa-mir-30a&WithFlank&6\|-\|71403545\|71403627 | 12 | 33 | 22(+1A) | TGTAAACATCCTCGACTGGAAGA | 5924.9 | 8.4 | 0.6 | 12.9 | 3.7E-38 | 2.6E-36 | MB231/MB468 |
| >MI0000255\|hsa-mir-30d&WithFlank&8\|-\|134804870\|134804951@12.35.24(+1U)&[MIMAT0000245&hsa-miR-30d-5p&offsets\|0\|+2(+1U);m-21&8\|-\|134804919\|134804940&offsets\|0\|+2(+1U)]\|TGTAAACATCCCCGACTGGAAGCTT | >MI0000255\|hsa-mir-30d&WithFlank&8\|-\|134804870\|134804951 | 12 | 35 | 24(+1U) | TGTAAACATCCCCGACTGGAAGCTT | 5658.2 | 8.4 | 0.7 | 12.9 | 7.4E-38 | 2.7E-36 | BT20/MB468 |
| >MI0000542\|hsa-mir-320a&WithFlank&8\|-\|22244960\|22245043@48.70.23(+1C)&[MIMAT0000510&hsa-miR-320a-3p&offsets\|0\|+1(+1C);m-60&8\|-\|22244975\|22244996&offsets\|0\|+1(+1C)]\|AAAAGCTGGGTTGAGAGGGCGAAC | >MI0000542\|hsa-mir-320a&WithFlank&8\|-\|22244960\|22245043 | 48 | 70 | 23(+1C) | AAAAGCTGGGTTGAGAGGGCGAAC | 1803.0 | 9.4 | 0.7 | 12.9 | 7.6E-38 | 2.7E-36 | BT20/MB468 |
| >MI0000301\|hsa-mir-224&WithFlank&X\|-\|151958572\|151958664@14.36.23&[MIMAT0000281&hsa-miR-224-5p&offsets\|+1\|-1;m-238&X\|-\|151958630\|151958651&offsets\|0\|+1]\|CAAGTCACTAGTGGTTCCGTTTA | >MI0000301\|hsa-mir-224&WithFlank&X\|-\|151958572\|151958664 | 14 | 36 | 23 | CAAGTCACTAGTGGTTCCGTTTA | 4038.4 | 16.6 | 1.3 | 12.9 | 8.4E-38 | 3.0E-36 | BT20/MB468 |
| >hg38_21_+_8433172_8446622_RNA45SN1_45S_50ntFlanks@485.506.22\|GATGTGGTGACGTCGTGCTCTC | >hg38_21_+_8433172_8446622_RNA45SN1_45S_50ntFlanks | 485 | 506 | 22 | GATGTGGTGACGTCGTGCTCTC | 13301.9 | 18.3 | 1.4 | 12.8 | 8.9E-38 | 3.2E-36 | BT20/MB468 |
| >hg38_21_+_8433172_8446622_RNA45SN1_45S_50ntFlanks@12633.12654.22\|CGAAGCTACCATCTGTGGGATT | >hg38_21_+_8433172_8446622_RNA45SN1_45S_50ntFlanks | 12633 | 12654 | 22 | CGAAGCTACCATCTGTGGGATT | 9566.4 | -8.0 | 0.6 | -12.9 | 4.6E-38 | 3.2E-36 | MB231/MB468 |
| >hg38_21_+_8433172_8446622_RNA45SN1_45S_50ntFlanks@9545.9563.19\|GTGGAGGTCCGTAGCGGTC | >hg38_21_+_8433172_8446622_RNA45SN1_45S_50ntFlanks | 9545 | 9563 | 19 | GTGGAGGTCCGTAGCGGTC | 2684.2 | -8.5 | 0.7 | -12.8 | 9.1E-38 | 3.3E-36 | BT20/MB468 |
| >MI0000809\|hsa-mir-151a&WithFlank&8\|-\|140732558\|140732659@53.74.22&[MIMAT0000757&hsa-miR-151a-3p&offsets\|0\|+1;m-30&8\|-\|140732587\|140732607&offsets\|0\|+1]\|CTAGACTGAAGCTCCTTGAGGA | >MI0000809\|hsa-mir-151a&WithFlank&8\|-\|140732558\|140732659 | 53 | 74 | 22 | CTAGACTGAAGCTCCTTGAGGA | 29060.6 | 8.5 | 0.7 | 12.9 | 4.8E-38 | 3.4E-36 | MB231/MB468 |
| >hg38_1_-_228634819_228635039_RNA5S12_5S_50ntFlanks@147.169.23\|GGGAATACCGGGTGCTGTAGGCT | >hg38_1_-_228634819_228635039_RNA5S12_5S_50ntFlanks | 147 | 169 | 23 | GGGAATACCGGGTGCTGTAGGCT | 810.9 | 5.8 | 0.4 | 12.8 | 9.9E-38 | 3.5E-36 | BT20/MB468 |
| >am_MI0000809\|hsa-mir-151a&WithFlank&8\|-\|140732558\|140732659@53.72.20(+1U)&[MIMAT0000757&hsa-miR-151a-3p&offsets\|0\|-1(+1U);m-30&8\|-\|140732587\|140732607&offsets\|0\|-1(+1U)]\|CTAGACTGAAGCTCCTTGAGT | >am_MI0000809\|hsa-mir-151a&WithFlank&8\|-\|140732558\|140732659 | 53 | 72 | 20(+1U) | CTAGACTGAAGCTCCTTGAGT | 13608.7 | 10.0 | 0.8 | 12.9 | 5.5E-38 | 3.8E-36 | MB231/MB468 |
| >MI0000809\|hsa-mir-151a&WithFlank&8\|-\|140732558\|140732659@51.72.22(+1U)&[MIMAT0000757&hsa-miR-151a-3p&offsets\|-2\|-1(+1U);m-30&8\|-\|140732587\|140732607&offsets\|-2\|-1(+1U)]\|TACTAGACTGAAGCTCCTTGAGT | >MI0000809\|hsa-mir-151a&WithFlank&8\|-\|140732558\|140732659 | 51 | 72 | 22(+1U) | TACTAGACTGAAGCTCCTTGAGT | 1264.5 | 8.1 | 0.6 | 12.8 | 1.1E-37 | 3.9E-36 | BT20/MB468 |

<table>
<tr><th></th><th></th><th></th><th></th><th></th><th></th><th></th><th></th><th></th><th></th><th></th><th></th><th></th></tr>
<tr><td>>hg38_21_+_8433172_8446622_RNA45SN1_45S_50ntFlanks@11668.11692.25|TGCCCAGTGCTCTGAATGTCAAAGT</td><td>>hg38_21_+_8433172_8446622_RNA45SN1_45S_50ntFlanks</td><td>11668</td><td>11692</td><td>25</td><td>TGCCCAGTGCTCTGAATGTCAAAGT</td><td>485.0</td><td>6.8</td><td>0.5</td><td>12.9</td><td>7.5E-38</td><td>5.1E-36</td><td>MB231/MB468</td></tr>
<tr><td>>MI0000102|hsa-mir-100&WithFlank&11|-|122152223|122152314@19.38.20(+1G)&[MIMAT0000098&hsa-miR-100-5p&offsets|0|-2(+1G);m-27&11|-|122152276|122152296&offsets|0|-1(+1G)]|AACCCGTAGATCCGAACTTGG</td><td>>MI0000102|hsa-mir-100&WithFlank&11|-|122152223|122152314</td><td>19</td><td>38</td><td>20(+1G)</td><td>AACCCGTAGATCCGAACTTGG</td><td>28712.7</td><td>17.6</td><td>1.4</td><td>12.8</td><td>9.0E-38</td><td>6.1E-36</td><td>MB231/MB468</td></tr>
<tr><td>>am_MI0005763|hsa-mir-941-1&WithFlank&20|+|63919443|63919526@53.74.22(+1U);MI0005764|hsa-mir-941-2&WithFlank&20|+|63919499|63919582@53.74.22(+1U);MI0005765|hsa-mir-941-3&WithFlank&20|+|63919555|63919638@53.74.22(+1U);MI0005766|hsa-mir-941-4&WithFlank&20|+|63919750|63919833@53.74.22(+1U);MI0031520|hsa-mir-941-5&WithFlank&20|+|63919862|63919945@53.74.22(+1U)&[MIMAT0004984&hsa-miR-941&offsets|0|-1(+1U);m-263&20|+|63919495|63919517&offsets|0|-1(+1U)];[MIMAT0004984_1&hsa-miR-941&offsets|0|-1(+1U);m-263&20|+|63919551|63919573&offsets|0|-1(+1U)];[MIMAT0004984_2&hsa-miR-941&offsets|0|-1(+1U);m-263&20|+|63919607|63919629&offsets|0|-1(+1U)];[MIMAT0004984_3&hsa-miR-941&offsets|0|-1(+1U);m-263&20|+|63919802|63919824&offsets|0|-1(+1U)];[MIMAT0004984_4&hsa-miR-941&offsets|0|-1(+1U);m-263&20|+|63919914|63919936&offsets|0|-1(+1U)]|CACCCGGCTGTGTGCACATGTGT</td><td>>am_MI0005763|hsa-mir-941-1&WithFlank&20|+|63919443|63919526</td><td>53</td><td>74</td><td>22(+1U)</td><td>CACCCGGCTGTGTGCACATGTGT</td><td>10373.3</td><td>9.2</td><td>0.7</td><td>12.8</td><td>9.5E-38</td><td>6.2E-36</td><td>MB231/MB468</td></tr>
<tr><td>>hg38_21_+_8433172_8446622_RNA45SN1_45S_50ntFlanks@12437.12455.19|CATTGTGAAGCAGAATTCA</td><td>>hg38_21_+_8433172_8446622_RNA45SN1_45S_50ntFlanks</td><td>12437</td><td>12455</td><td>19</td><td>CATTGTGAAGCAGAATTCA</td><td>17589.4</td><td>-9.0</td><td>0.7</td><td>-12.8</td><td>9.5E-38</td><td>6.2E-36</td><td>MB231/MB468</td></tr>
<tr><td>>MI0000253|hsa-mir-148a&WithFlank&7|-|25949913|25949992@50.70.21(+1A)&[MIMAT0000243&hsa-miR-148a-3p&offsets|0|-1(+1A);m-7&7|-|25949922|25949943&offsets|0|-1(+1A)]|TCAGTGCACTACAGAACTTTGA</td><td>>MI0000253|hsa-mir-148a&WithFlank&7|-|25949913|25949992</td><td>50</td><td>70</td><td>21(+1A)</td><td>TCAGTGCACTACAGAACTTTGA</td><td>30728.0</td><td>9.4</td><td>0.7</td><td>12.8</td><td>1.8E-37</td><td>6.3E-36</td><td>BT20/MB468</td></tr>
<tr><td>>hg38_21_+_8433172_8446622_RNA45SN1_45S_50ntFlanks@12633.12661.29|CGAAGCTACCATCTGTGGGATTATGACTG</td><td>>hg38_21_+_8433172_8446622_RNA45SN1_45S_50ntFlanks</td><td>12633</td><td>12661</td><td>29</td><td>CGAAGCTACCATCTGTGGGATTATGACTG</td><td>1652.5</td><td>-5.5</td><td>0.4</td><td>-12.8</td><td>1.9E-37</td><td>6.7E-36</td><td>BT20/MB468</td></tr>
<tr><td>>MI0000809|hsa-mir-151a&WithFlank&8|-|140732558|140732659@51.73.23&[MIMAT0000757&hsa-miR-151a-3p&offsets|-2|0;m-30&8|-|140732587|140732607&offsets|-2|0]|TACTAGACTGAAGCTCCTTGAGG</td><td>>MI0000809|hsa-mir-151a&WithFlank&8|-|140732558|140732659</td><td>51</td><td>73</td><td>23</td><td>TACTAGACTGAAGCTCCTTGAGG</td><td>13999.6</td><td>8.8</td><td>0.7</td><td>12.8</td><td>1.1E-37</td><td>7.2E-36</td><td>MB231/MB468</td></tr>
<tr><td>>hg38_21_+_8433172_8446622_RNA45SN1_45S_50ntFlanks@12624.12654.31|CCAATGGGGCGAAGCTACCATCTGTGGGATT</td><td>>hg38_21_+_8433172_8446622_RNA45SN1_45S_50ntFlanks</td><td>12624</td><td>12654</td><td>31</td><td>CCAATGGGGCGAAGCTACCATCTGTGGGATT</td><td>3699.7</td><td>-7.8</td><td>0.6</td><td>-12.8</td><td>2.4E-37</td><td>8.2E-36</td><td>BT20/MB468</td></tr>
<tr><td>>hg38_1_-_228634819_228635039_RNA5S12_5S_50ntFlanks@51.88.38|GTCTACGGCCATACCACCCTGAACGCGCCCGATCTCGT</td><td>>hg38_1_-_228634819_228635039_RNA5S12_5S_50ntFlanks</td><td>51</td><td>88</td><td>38</td><td>GTCTACGGCCATACCACCCTGAACGCGCCCGATCTCGT</td><td>3136.0</td><td>9.3</td><td>0.7</td><td>12.8</td><td>2.5E-37</td><td>8.5E-36</td><td>BT20/MB468</td></tr>
<tr><td>>MI0000063|hsa-let-7b&WithFlank&22|+|46113680|46113774@12.33.22&[MIMAT0000063&hsa-let-7b-5p&offsets|0|0;m-14&22|+|46113691|46113712&offsets|0|0]|TGAGGTAGTAGGTTGTGTGGTT</td><td>>MI0000063|hsa-let-7b&WithFlank&22|+|46113680|46113774</td><td>12</td><td>33</td><td>22</td><td>TGAGGTAGTAGGTTGTGTGGTT</td><td>91586.0</td><td>8.2</td><td>0.6</td><td>12.8</td><td>2.8E-37</td><td>9.5E-36</td><td>BT20/MB468</td></tr>
<tr><td>>hg38_21_+_8433172_8446622_RNA45SN1_45S_50ntFlanks@6645.6678.34|CTCGTACGACTCTTAGCGGTGGATCACTCGGCTC</td><td>>hg38_21_+_8433172_8446622_RNA45SN1_45S_50ntFlanks</td><td>6645</td><td>6678</td><td>34</td><td>CTCGTACGACTCTTAGCGGTGGATCACTCGGCTC</td><td>521.4</td><td>5.1</td><td>0.4</td><td>12.8</td><td>2.9E-37</td><td>9.7E-36</td><td>BT20/MB468</td></tr>
<tr><td>>hg38_21_+_8433172_8446622_RNA45SN1_45S_50ntFlanks@12973.12991.19|TCGTACGTAGCAGAGCAGC</td><td>>hg38_21_+_8433172_8446622_RNA45SN1_45S_50ntFlanks</td><td>12973</td><td>12991</td><td>19</td><td>TCGTACGTAGCAGAGCAGC</td><td>5807.1</td><td>-6.5</td><td>0.5</td><td>-12.8</td><td>1.6E-37</td><td>1.0E-35</td><td>MB231/MB468</td></tr>
<tr><td>>hg38_21_+_8433172_8446622_RNA45SN1_45S_50ntFlanks@12655.12675.21|ATGACTGAACGCCTCTAAGTC</td><td>>hg38_21_+_8433172_8446622_RNA45SN1_45S_50ntFlanks</td><td>12655</td><td>12675</td><td>21</td><td>ATGACTGAACGCCTCTAAGTC</td><td>7062.1</td><td>-8.1</td><td>0.6</td><td>-12.8</td><td>1.7E-37</td><td>1.1E-35</td><td>MB231/MB468</td></tr>
<tr><td>>MI0000088|hsa-mir-30a&WithFlank&6|-|71403545|71403627@12.33.22(+1U)&[MIMAT0000087&hsa-miR-30a-5p&offsets|0|0(+1U);m-15&6|-|71403595|71403616&offsets|0|0(+1U)]|TGTAAACATCCTCGACTGGAAGT</td><td>>MI0000088|hsa-mir-30a&WithFlank&6|-|71403545|71403627</td><td>12</td><td>33</td><td>22(+1U)</td><td>TGTAAACATCCTCGACTGGAAGT</td><td>26550.9</td><td>8.6</td><td>0.7</td><td>12.8</td><td>1.8E-37</td><td>1.1E-35</td><td>MB231/MB468</td></tr>
<tr><td>>MI0000299|hsa-mir-222&WithFlank&X|-|45747009|45747130@75.98.24&[MIMAT0000279&hsa-miR-222-3p&offsets|0|+3;m-140&X|-|45747033|45747056&offsets|0|0]|AGCTACATCTGGCTACTGGGTCTC</td><td>>MI0000299|hsa-mir-222&WithFlank&X|-|45747009|45747130</td><td>75</td><td>98</td><td>24</td><td>AGCTACATCTGGCTACTGGGTCTC</td><td>69992.3</td><td>9.3</td><td>0.7</td><td>12.8</td><td>1.8E-37</td><td>1.1E-35</td><td>MB231/MB468</td></tr>
<tr><td>>hg38_21_+_8433172_8446622_RNA45SN1_45S_50ntFlanks@12405.12431.27|CTTTTTGATCCTTCGATGTCGGCTCTT</td><td>>hg38_21_+_8433172_8446622_RNA45SN1_45S_50ntFlanks</td><td>12405</td><td>12431</td><td>27</td><td>CTTTTTGATCCTTCGATGTCGGCTCTT</td><td>7760.4</td><td>-8.3</td><td>0.6</td><td>-12.7</td><td>3.5E-37</td><td>1.2E-35</td><td>BT20/MB468</td></tr>
<tr><td>>MI0000298|hsa-mir-221&WithFlank&X|-|45746151|45746272@71.92.22(+1U)&[MIMAT0000278&hsa-miR-221-3p&offsets|0|-1(+1U);m-91&X|-|45746181|45746202&offsets|0|0(+1U)]|AGCTACATTGTCTGCTGGGTTTT</td><td>>MI0000298|hsa-mir-221&WithFlank&X|-|45746151|45746272</td><td>71</td><td>92</td><td>22(+1U)</td><td>AGCTACATTGTCTGCTGGGTTTT</td><td>16006.0</td><td>8.8</td><td>0.7</td><td>12.8</td><td>2.4E-37</td><td>1.5E-35</td><td>MB231/MB468</td></tr>
<tr><td>>MI0000255|hsa-mir-30d&WithFlank&8|-|134804870|134804951@12.33.22(+1U)&[MIMAT0000245&hsa-miR-30d-5p&offsets|0|0(+1U);m-21&8|-|134804919|134804940&offsets|0|0(+1U)]|TGTAAACATCCCCGACTGGAAGT</td><td>>MI0000255|hsa-mir-30d&WithFlank&8|-|134804870|134804951</td><td>12</td><td>33</td><td>22(+1U)</td><td>TGTAAACATCCCCGACTGGAAGT</td><td>12159.6</td><td>8.3</td><td>0.7</td><td>12.8</td><td>2.5E-37</td><td>1.5E-35</td><td>MB231/MB468</td></tr>
<tr><td>>MI0000734|hsa-mir-106b&WithFlank&7|-|100093987|100094080@56.77.22&[MIMAT0004672&hsa-miR-106b-3p&offsets|-2|-2;m-87&7|-|100094003|100094023&offsets|-2|-1]|TACCGCACTGTGGGTACTTGCT</td><td>>MI0000734|hsa-mir-106b&WithFlank&7|-|100093987|100094080</td><td>56</td><td>77</td><td>22</td><td>TACCGCACTGTGGGTACTTGCT</td><td>2984.5</td><td>9.5</td><td>0.7</td><td>12.7</td><td>5.8E-37</td><td>1.9E-35</td><td>BT20/MB468</td></tr>
<tr><td>>MI0000086|hsa-mir-28&WithFlank&3|+|188688775|188688872@60.80.21&[MIMAT0004502&hsa-miR-28-3p&offsets|0|-1;m-28&3|+|188688834|188688855&offsets|0|-1]|CACTAGATTGTGAGCTCCTGG</td><td>>MI0000086|hsa-mir-28&WithFlank&3|+|188688775|188688872</td><td>60</td><td>80</td><td>21</td><td>CACTAGATTGTGAGCTCCTGG</td><td>4131.0</td><td>9.7</td><td>0.8</td><td>12.7</td><td>6.0E-37</td><td>2.0E-35</td><td>BT20/MB468</td></tr>
<tr><td>>MI0000253|hsa-mir-148a&WithFlank&7|-|25949913|25949992@50.73.24&[MIMAT0000243&hsa-miR-148a-3p&offsets|0|+2;m-7&7|-|25949922|25949943&offsets|0|+2]|TCAGTGCACTACAGAACTTTGTCT</td><td>>MI0000253|hsa-mir-148a&WithFlank&7|-|25949913|25949992</td><td>50</td><td>73</td><td>24</td><td>TCAGTGCACTACAGAACTTTGTCT</td><td>1939.6</td><td>8.7</td><td>0.7</td><td>12.7</td><td>6.3E-37</td><td>2.1E-35</td><td>BT20/MB468</td></tr>
<tr><td>>am_trna128_GlyGCC_6_-_27870686_27870756@1.31.31,trna133_GlyCCC_1_-_16872434_16872504@1.31.31,trna18_GlyGCC_16_+_70822597_70822667@1.31.31,trna19_GlyGCC_16_+_70823410_70823480@1.31.31,trna19_GlyGCC_2_-_157257659_157257729@1.31.31,trna24_GlyGCC_16_-_70812942_70813012@1.31.31,trna25_GlyGCC_16_-_70812114_70812184@1.31.31,trna4_GlyCCC_1_+_17188416_17188486@1.31.31,trna5_GlyGCC_17_+_8029064_8029134@1.31.31,trna68_GlyGCC_1_-_161493637_161493707@1.31.31|GCATTGGTGGTTCAGTGGTAGAATTCTCGCC</td><td>>am_trna128_GlyGCC_6_-_27870686_27870756</td><td>1</td><td>31</td><td>31</td><td>GCATTGGTGGTTCAGTGGTAGAATTCTCGCC</td><td>38501.9</td><td>6.5</td><td>0.5</td><td>12.7</td><td>6.6E-37</td><td>2.2E-35</td><td>BT20/MB468</td></tr>
<tr><td>>MI0000542|hsa-mir-320a&WithFlank&8|-|22244960|22245043@48.68.21&[MIMAT0000510&hsa-miR-320a-3p&offsets|0|-1;m-60&8|-|22244975|22244996&offsets|0|-1]|AAAAGCTGGGTTGAGAGGGCG</td><td>>MI0000542|hsa-mir-320a&WithFlank&8|-|22244960|22245043</td><td>48</td><td>68</td><td>21</td><td>AAAAGCTGGGTTGAGAGGGCG</td><td>9584.3</td><td>9.1</td><td>0.7</td><td>12.7</td><td>6.6E-37</td><td>2.2E-35</td><td>BT20/MB468</td></tr>
<tr><td>>am_trna128_GlyGCC_6_-_27870686_27870756@1.29.29,trna133_GlyCCC_1_-_16872434_16872504@1.29.29,trna18_GlyGCC_16_+_70822597_70822667@1.29.29,trna19_GlyGCC_16_+_70823410_70823480@1.29.29,trna19_GlyGCC_2_-_157257659_157257729@1.29.29,trna24_GlyGCC_16_-_70812942_70813012@1.29.29,trna25_GlyGCC_16_-_70812114_70812184@1.29.29,trna4_GlyCCC_1_+_17188416_17188486@1.29.29,trna5_GlyGCC_17_+_8029064_8029134@1.29.29,trna68_GlyGCC_1_-_161493637_161493707@1.29.29|GCATTGGTGGTTCAGTGGTAGAATTCTCG</td><td>>am_trna128_GlyGCC_6_-_27870686_27870756</td><td>1</td><td>29</td><td>29</td><td>GCATTGGTGGTTCAGTGGTAGAATTCTCG</td><td>22419.7</td><td>-6.2</td><td>0.5</td><td>-12.7</td><td>6.7E-37</td><td>2.2E-35</td><td>BT20/MB468</td></tr>
<tr><td>>hg38_21_+_8433172_8446622_RNA45SN1_45S_50ntFlanks@9504.9526.23|GTGAACTATGCCTGGGCAGGGCG</td><td>>hg38_21_+_8433172_8446622_RNA45SN1_45S_50ntFlanks</td><td>9504</td><td>9526</td><td>23</td><td>GTGAACTATGCCTGGGCAGGGCG</td><td>4398.0</td><td>-11.0</td><td>0.9</td><td>-12.7</td><td>7.3E-37</td><td>2.4E-35</td><td>BT20/MB468</td></tr>
<tr><td>>hg38_21_+_8433172_8446622_RNA45SN1_45S_50ntFlanks@11669.11691.23|GCCCAGTGCTCTGAATGTCAAAG</td><td>>hg38_21_+_8433172_8446622_RNA45SN1_45S_50ntFlanks</td><td>11669</td><td>11691</td><td>23</td><td>GCCCAGTGCTCTGAATGTCAAAG</td><td>484.0</td><td>6.4</td><td>0.5</td><td>12.7</td><td>4.6E-37</td><td>2.8E-35</td><td>MB231/MB468</td></tr>
<tr><td>>hg38_21_+_8433172_8446622_RNA45SN1_45S_50ntFlanks@12456.12486.31|CCAAGCGTTGGATTGTTCACCCACTAATAGG</td><td>>hg38_21_+_8433172_8446622_RNA45SN1_45S_50ntFlanks</td><td>12456</td><td>12486</td><td>31</td><td>CCAAGCGTTGGATTGTTCACCCACTAATAGG</td><td>10226.6</td><td>-6.3</td><td>0.5</td><td>-12.7</td><td>4.7E-37</td><td>2.8E-35</td><td>MB231/MB468</td></tr>
<tr><td>>trnaMT_PheGAA_MT_+_577_647@36.55.20|AAAATGTTTAGACGGGCTCA</td><td>>trnaMT_PheGAA_MT_+_577_647</td><td>36</td><td>55</td><td>20</td><td>AAAATGTTTAGACGGGCTCA</td><td>1051.3</td><td>14.7</td><td>1.2</td><td>12.7</td><td>8.8E-37</td><td>2.9E-35</td><td>BT20/MB468</td></tr>
<tr><td>>trnaMT_ValTAC_MT_+_1602_1670@42.72.31|AGATTTCAACTTAACTTGACCGCTCTGACCA</td><td>>trnaMT_ValTAC_MT_+_1602_1670</td><td>42</td><td>72</td><td>31</td><td>AGATTTCAACTTAACTTGACCGCTCTGACCA</td><td>3881.9</td><td>8.3</td><td>0.7</td><td>12.7</td><td>4.9E-37</td><td>2.9E-35</td><td>MB231/MB468</td></tr>
<tr><td>>hg38_21_+_8433172_8446622_RNA45SN1_45S_50ntFlanks@12937.12956.20|AACCATTCGTAGACGACCTG</td><td>>hg38_21_+_8433172_8446622_RNA45SN1_45S_50ntFlanks</td><td>12937</td><td>12956</td><td>20</td><td>AACCATTCGTAGACGACCTG</td><td>3749.1</td><td>-6.7</td><td>0.5</td><td>-12.7</td><td>9.2E-37</td><td>2.9E-35</td><td>BT20/MB468</td></tr>
<tr><td>>hg38_21_+_8433172_8446622_RNA45SN1_45S_50ntFlanks@6661.6701.41|CGGTGGATCACTCGGCTCGTGCGTCGATGAAGAACGCAGCT</td><td>>hg38_21_+_8433172_8446622_RNA45SN1_45S_50ntFlanks</td><td>6661</td><td>6701</td><td>41</td><td>CGGTGGATCACTCGGCTCGTGCGTCGATGAAGAACGCAGCT</td><td>1550.3</td><td>7.7</td><td>0.6</td><td>12.7</td><td>9.4E-37</td><td>3.0E-35</td><td>BT20/MB468</td></tr>
<tr><td>>hg38_21_+_8433172_8446622_RNA45SN1_45S_50ntFlanks@11783.11803.21|CATGAATGGATGAACGAGATT</td><td>>hg38_21_+_8433172_8446622_RNA45SN1_45S_50ntFlanks</td><td>11783</td><td>11803</td><td>21</td><td>CATGAATGGATGAACGAGATT</td><td>8540.0</td><td>-6.7</td><td>0.5</td><td>-12.7</td><td>9.9E-37</td><td>3.1E-35</td><td>BT20/MB468</td></tr>
<tr><td>>hg38_21_+_8433172_8446622_RNA45SN1_45S_50ntFlanks@10410.10431.22|TCAGTCGGTCCTGAGAGATGGG</td><td>>hg38_21_+_8433172_8446622_RNA45SN1_45S_50ntFlanks</td><td>10410</td><td>10431</td><td>22</td><td>TCAGTCGGTCCTGAGAGATGGG</td><td>1542.6</td><td>-7.2</td><td>0.6</td><td>-12.7</td><td>1.0E-36</td><td>3.2E-35</td><td>BT20/MB468</td></tr>
</table>

| | | | | | | | | | | | |
|---|---|---|---|---|---|---|---|---|---|---|---|
| >hg38_21_+_8433172_8446622_RNA45SN1_45S_50ntFlanks@9567.9596.30\|ACGTGCAAATCGGTCGTCCGACCTGGGTAT | >hg38_21_+_8433172_8446622_RNA45SN1_45S_50ntFlanks | 9567 | 9596 | 30 | ACGTGCAAATCGGTCGTCCGACCTGGGTAT | 4361.7 | -10.7 | 0.8 | -12.7 | 1.0E-36 | 3.3E-35 | BT20/MB468 |
| >am_MI0000809\|hsa-mir-151a&WithFlank&8\|-\|140732558\|140732659@53.73.21(+1C)&[MIMAT0000757&hsa-miR-151a-3p&offsets\|0\|0(+1C);m-30&8\|-\|140732587\|140732607&offsets\|0\|0(+1C)]\|CTAGACTGAAGCTCCTTGAGGC | >am_MI0000809\|hsa-mir-151a&WithFlank&8\|-\|140732558\|140732659 | 53 | 73 | 21(+1C) | CTAGACTGAAGCTCCTTGAGGC | 3555.0 | 9.5 | 0.8 | 12.7 | 5.7E-37 | 3.3E-35 | MB231/MB468 |
| >MI0016746\|hsa-mir-548o-2&WithFlank&20\|+\|38516557\|38516638@51.72.22;MI0006402\|hsa-mir-548o&WithFlank&7\|-\|102405736\|102405861@93.114.22&[MIMAT0005919&hsa-miR-548o-3p&offsets\|0\|0;m-431&20\|+\|38516607\|38516628&offsets\|0\|0];[MIMAT0005919_1&hsa-miR-548o-3p&offsets\|0\|0;m-430&7\|-\|102405748\|102405769&offsets\|0\|0]\|CCAAAACTGCAGTTACTTTTGC | >MI0016746\|hsa-mir-548o-2&WithFlank&20\|+\|38516557\|38516638 | 51 | 72 | 22 | CCAAAACTGCAGTTACTTTTGC | 1177.6 | 7.6 | 0.6 | 12.6 | 1.2E-36 | 3.7E-35 | BT20/MB468 |
| >hg38_MT_+_1621_3279_RNR2_16S_50ntFlanks@51.79.29\|GCTAAACCTAGCCCCAAACCCACTCCACC | >hg38_MT_+_1621_3279_RNR2_16S_50ntFlanks | 51 | 79 | 29 | GCTAAACCTAGCCCCAAACCCACTCCACC | 1456.1 | 8.1 | 0.6 | 12.7 | 6.5E-37 | 3.7E-35 | MB231/MB468 |
| >trna116_GluCTC_1_-_145399233_145399304@1.36.36,trna59_GluCTC_1_+_249168447_249168518@1.36.36,trna71_GluCTC_1_-_161439189_161439260@1.36.36,trna74_GluCTC_1_-_161431809_161431880@1.36.36,trna77_GluCTC_1_-_161424398_161424469@1.36.36,trna77_GluCTC_6_+_28949976_28950047@1.36.36,trna80_GluCTC_1_-_161417018_161417089@1.36.36,trna87_GluCTC_6_-_126101393_126101464@1.36.36\|TCCCTGGTGGTCTAGTGGTTAGGATTCGGCGCTCTC | >trna116_GluCTC_1_-_145399233_145399304 | 1 | 36 | 36 | TCCCTGGTGGTCTAGTGGTTAGGATTCGGCGCTCTC | 5091.4 | 9.6 | 0.8 | 12.6 | 1.2E-36 | 3.8E-35 | BT20/MB468 |
| >MI0000079\|hsa-mir-23a&WithFlank&19\|-\|13836581\|13836665@54.71.18&[MIMAT0000078&hsa-miR-23a-3p&offsets\|+3\|0;m-31&19\|-\|13836595\|13836615&offsets\|+3\|0]\|ACATTGCCAGGGATTTCC | >MI0000079\|hsa-mir-23a&WithFlank&19\|-\|13836581\|13836665 | 54 | 71 | 18 | ACATTGCCAGGGATTTCC | 9636.5 | 17.9 | 1.4 | 12.6 | 1.3E-36 | 4.0E-35 | BT20/MB468 |
| >hg38_21_+_8433172_8446622_RNA45SN1_45S_50ntFlanks@9739.9758.20\|AAACGATCTCAACCTATTCT | >hg38_21_+_8433172_8446622_RNA45SN1_45S_50ntFlanks | 9739 | 9758 | 20 | AAACGATCTCAACCTATTCT | 1947.2 | -4.8 | 0.4 | -12.6 | 1.3E-36 | 4.0E-35 | BT20/MB468 |
| >hg38_21_+_8433172_8446622_RNA45SN1_45S_50ntFlanks@11641.11660.20\|CGGCGGGTGTTGACGCGATG | >hg38_21_+_8433172_8446622_RNA45SN1_45S_50ntFlanks | 11641 | 11660 | 20 | CGGCGGGTGTTGACGCGATG | 8688.9 | -9.0 | 0.7 | -12.7 | 7.0E-37 | 4.0E-35 | MB231/MB468 |
| >trnaMT_GluTTC_MT_-_14674_14742@41.72.32,trnalookalike8_GluTTC_5_-_93905172_93905240@41.72.32\|TGGTCGTGGTTGTAGTCCGTGCGAGAATACCA | >trnaMT_GluTTC_MT_-_14674_14742 | 41 | 72 | 32 | TGGTCGTGGTTGTAGTCCGTGCGAGAATACCA | 11975.9 | 7.9 | 0.6 | 12.7 | 7.1E-37 | 4.0E-35 | MB231/MB468 |
| >hg38_21_+_8433172_8446622_RNA45SN1_45S_50ntFlanks@476.512.37\|TTCGTGATCGATGTGGTGACGTCGTGCTCTCCCGGGC | >hg38_21_+_8433172_8446622_RNA45SN1_45S_50ntFlanks | 476 | 512 | 37 | TTCGTGATCGATGTGGTGACGTCGTGCTCTCCCGGGC | 1937.3 | 8.9 | 0.7 | 12.7 | 7.6E-37 | 4.3E-35 | MB231/MB468 |
| >trnaMT_PheGAA_MT_+_577_647@1.19.19\|GTTTATGTAGCTTACCTCC | >trnaMT_PheGAA_MT_+_577_647 | 1 | 19 | 19 | GTTTATGTAGCTTACCTCC | 3435.1 | 8.9 | 0.7 | 12.7 | 8.6E-37 | 4.8E-35 | MB231/MB468 |
| >MI0000301\|hsa-mir-224&WithFlank&X\|-\|151958572\|151958664@14.34.21(+1C)&[MIMAT0000281&hsa-miR-224-5p&offsets\|+1\|-3(+1C);m-238&X\|-\|151958630\|151958651&offsets\|0\|-1(+1C)]\|CAAGTCACTAGTGGTTCCGTTC | >MI0000301\|hsa-mir-224&WithFlank&X\|-\|151958572\|151958664 | 14 | 34 | 21(+1C) | CAAGTCACTAGTGGTTCCGTTC | 2690.9 | 14.0 | 1.1 | 12.6 | 1.6E-36 | 5.0E-35 | BT20/MB468 |
| >MI0000102\|hsa-mir-100&WithFlank&11\|-\|122152223\|122152314@20.40.21&[MIMAT0000098&hsa-miR-100-5p&offsets\|+1\|0;m-27&11\|-\|122152276\|122152296&offsets\|+1\|+1]\|ACCCGTAGATCCGAACTTGTG | >MI0000102\|hsa-mir-100&WithFlank&11\|-\|122152223\|122152314 | 20 | 40 | 21 | ACCCGTAGATCCGAACTTGTG | 6232.5 | 17.5 | 1.4 | 12.7 | 9.3E-37 | 5.2E-35 | MB231/MB468 |
| >am_MI0005763\|hsa-mir-941-1&WithFlank&20\|+\|63919443\|63919526@53.75.23(+1U);MI0005764\|hsa-mir-941-2&WithFlank&20\|+\|63919499\|63919582@53.75.23(+1U);MI0005765\|hsa-mir-941-3&WithFlank&20\|+\|63919555\|63919638@53.75.23(+1U);MI0005766\|hsa-mir-941-4&WithFlank&20\|+\|63919750\|63919833@53.75.23(+1U);MI0031520\|hsa-mir-941-5&WithFlank&20\|+\|63919862\|63919945@53.75.23(+1U)&[MIMAT0004984&hsa-miR-941&offsets\|0\|0(+1U);m-263&20\|+\|63919495\|63919517&offsets\|0\|0(+1U)];[MIMAT0004984_1&hsa-miR-941&offsets\|0\|0(+1U);m-263&20\|+\|63919551\|63919573&offsets\|0\|0(+1U)];[MIMAT0004984_2&hsa-miR-941&offsets\|0\|0(+1U);m-263&20\|+\|63919607\|63919629&offsets\|0\|0(+1U)];[MIMAT0004984_3&hsa-miR-941&offsets\|0\|0(+1U);m-263&20\|+\|63919802\|63919824&offsets\|0\|0(+1U)];[MIMAT0004984_4&hsa-miR-941&offsets\|0\|0(+1U);m-263&20\|+\|63919914\|63919936&offsets\|0\|0(+1U)]\|CACCCGGCTGTGTGCACATGTGCT | >am_MI0005763\|hsa-mir-941-1&WithFlank&20\|+\|63919443\|63919526 | 53 | 75 | 23(+1U) | CACCCGGCTGTGTGCACATGTGCT | 841.9 | 7.6 | 0.6 | 12.6 | 1.7E-36 | 5.3E-35 | BT20/MB468 |
| >hg38_21_+_8433172_8446622_RNA45SN1_45S_50ntFlanks@10786.10816.31\|AACAGCCTCTGGCATGTTGGAACAATGTAGG | >hg38_21_+_8433172_8446622_RNA45SN1_45S_50ntFlanks | 10786 | 10816 | 31 | AACAGCCTCTGGCATGTTGGAACAATGTAGG | 956.4 | -5.2 | 0.4 | -12.6 | 2.2E-36 | 6.6E-35 | BT20/MB468 |
| >MI0000088\|hsa-mir-30a&WithFlank&6\|-\|71403545\|71403627@13.35.23&[MIMAT0000087&hsa-miR-30a-5p&offsets\|+1\|+2;m-15&6\|-\|71403595\|71403616&offsets\|+1\|+2]\|GTAAACATCCTCGACTGGAAGCT | >MI0000088\|hsa-mir-30a&WithFlank&6\|-\|71403545\|71403627 | 13 | 35 | 23 | GTAAACATCCTCGACTGGAAGCT | 6680.0 | 8.9 | 0.7 | 12.6 | 1.2E-36 | 6.8E-35 | MB231/MB468 |
| >hg38_21_+_8433172_8446622_RNA45SN1_45S_50ntFlanks@475.517.43\|CTTCGTGATCGATGTGGTGACGTCGTGCTCTCCCGGGCCGGGT | >hg38_21_+_8433172_8446622_RNA45SN1_45S_50ntFlanks | 475 | 517 | 43 | CTTCGTGATCGATGTGGTGACGTCGTGCTCTCCCGGGCCGGGT | 4492.8 | 9.7 | 0.8 | 12.6 | 1.3E-36 | 6.9E-35 | MB231/MB468 |
| >hg38_21_+_8433172_8446622_RNA45SN1_45S_50ntFlanks@7973.8010.38\|GACGCGACCTCAGATCAGACGTGGCGACCCGCTGAATT | >hg38_21_+_8433172_8446622_RNA45SN1_45S_50ntFlanks | 7973 | 8010 | 38 | GACGCGACCTCAGATCAGACGTGGCGACCCGCTGAATT | 1818.5 | 8.4 | 0.7 | 12.6 | 1.4E-36 | 7.6E-35 | MB231/MB468 |
| >hg38_21_+_8433172_8446622_RNA45SN1_45S_50ntFlanks@12605.12626.22\|TATGTGCTTGGCTGAGGAGCCA | >hg38_21_+_8433172_8446622_RNA45SN1_45S_50ntFlanks | 12605 | 12626 | 22 | TATGTGCTTGGCTGAGGAGCCA | 3864.3 | -6.9 | 0.5 | -12.6 | 1.4E-36 | 7.7E-35 | MB231/MB468 |
| >am_MI0000809\|hsa-mir-151a&WithFlank&8\|-\|140732558\|140732659@53.73.21&[MIMAT0000757&hsa-miR-151a-3p&offsets\|0\|0;m-30&8\|-\|140732587\|140732607&offsets\|0\|0]\|CTAGACTGAAGCTCCTTGAGG | >am_MI0000809\|hsa-mir-151a&WithFlank&8\|-\|140732558\|140732659 | 53 | 73 | 21 | CTAGACTGAAGCTCCTTGAGG | 153876.9 | 9.1 | 0.7 | 12.6 | 1.5E-36 | 7.8E-35 | MB231/MB468 |
| >hg38_21_+_8433172_8446622_RNA45SN1_45S_50ntFlanks@10392.10410.19\|ACAGCAGTTGAACATGGGT | >hg38_21_+_8433172_8446622_RNA45SN1_45S_50ntFlanks | 10392 | 10410 | 19 | ACAGCAGTTGAACATGGGT | 6478.1 | -7.6 | 0.6 | -12.6 | 1.7E-36 | 9.0E-35 | MB231/MB468 |
| >hg38_21_+_8433172_8446622_RNA45SN1_45S_50ntFlanks@485.507.23\|GATGTGGTGACGTCGTGCTCTCC | >hg38_21_+_8433172_8446622_RNA45SN1_45S_50ntFlanks | 485 | 507 | 23 | GATGTGGTGACGTCGTGCTCTCC | 6018.2 | 17.2 | 1.4 | 12.6 | 3.1E-36 | 9.4E-35 | BT20/MB468 |
| >hg38_21_+_8433172_8446622_RNA45SN1_45S_50ntFlanks@12635.12659.25\|AAGCTACCATCTGTGGGATTATGAC | >hg38_21_+_8433172_8446622_RNA45SN1_45S_50ntFlanks | 12635 | 12659 | 25 | AAGCTACCATCTGTGGGATTATGAC | 1540.4 | -6.3 | 0.5 | -12.6 | 3.6E-36 | 1.1E-34 | BT20/MB468 |
| >hg38_21_+_8433172_8446622_RNA45SN1_45S_50ntFlanks@6671.6699.29\|CTCGGCTCGTGCGTCGATGAAGAACGCAG | >hg38_21_+_8433172_8446622_RNA45SN1_45S_50ntFlanks | 6671 | 6699 | 29 | CTCGGCTCGTGCGTCGATGAAGAACGCAG | 2362.4 | -6.3 | 0.5 | -12.6 | 3.8E-36 | 1.1E-34 | BT20/MB468 |
| >hg38_21_+_8433172_8446622_RNA45SN1_45S_50ntFlanks@3705.3745.41\|TACCTGGTTGATCCTGCCAGTAGCATATGCTTGTCTCAAAG | >hg38_21_+_8433172_8446622_RNA45SN1_45S_50ntFlanks | 3705 | 3745 | 41 | TACCTGGTTGATCCTGCCAGTAGCATATGCTTGTCTCAAAG | 537.1 | 6.4 | 0.5 | 12.6 | 2.2E-36 | 1.2E-34 | MB231/MB468 |
| >MI0000446\|hsa-mir-125b-1&WithFlank&11\|-\|122099751\|122099850@21.40.20;MI0000470\|hsa-mir-125b-2&WithFlank&21\|+\|16590231\|16590331@23.42.20&[MIMAT0000423&hsa-miR-125b-5p&offsets\|0\|-2;m-67&11\|-\|122099809\|122099830&offsets\|0\|-2];[MIMAT0000423_1&hsa-miR-125b-5p&offsets\|0\|-2;m-68&21\|+\|16590253\|16590274&offsets\|0\|-2]\|TCCCTGAGACCCTAACTTGT | >MI0000446\|hsa-mir-125b-1&WithFlank&11\|-\|122099751\|122099850 | 21 | 40 | 20 | TCCCTGAGACCCTAACTTGT | 4872.9 | 9.7 | 0.8 | 12.6 | 2.3E-36 | 1.2E-34 | MB231/MB468 |
| >MI0000079\|hsa-mir-23a&WithFlank&19\|-\|13836581\|13836665@54.72.19&[MIMAT0000078&hsa-miR-23a-3p&offsets\|+3\|+1;m-31&19\|-\|13836595\|13836615&offsets\|+3\|+1]\|ACATTGCCAGGGATTTCCA | >MI0000079\|hsa-mir-23a&WithFlank&19\|-\|13836581\|13836665 | 54 | 72 | 19 | ACATTGCCAGGGATTTCCA | 17600.4 | 18.8 | 1.5 | 12.5 | 4.6E-36 | 1.4E-34 | BT20/MB468 |
| >hg38_21_+_8433172_8446622_RNA45SN1_45S_50ntFlanks@12625.12653.29\|CAATGGGGCGAAGCTACCATCTGTGGGAT | >hg38_21_+_8433172_8446622_RNA45SN1_45S_50ntFlanks | 12625 | 12653 | 29 | CAATGGGGCGAAGCTACCATCTGTGGGAT | 1446.8 | -8.1 | 0.6 | -12.5 | 5.0E-36 | 1.5E-34 | BT20/MB468 |
| >hg38_21_+_8433172_8446622_RNA45SN1_45S_50ntFlanks@11666.11703.38\|TCTGCCCAGTGCTCTGAATGTCAAAGTGAAGAAATTCA | >hg38_21_+_8433172_8446622_RNA45SN1_45S_50ntFlanks | 11666 | 11703 | 38 | TCTGCCCAGTGCTCTGAATGTCAAAGTGAAGAAATTCA | 346.9 | 5.9 | 0.5 | 12.6 | 3.0E-36 | 1.5E-34 | MB231/MB468 |
| >hg38_21_+_8433172_8446622_RNA45SN1_45S_50ntFlanks@6645.6665.21\|CTCGTACGACTCTTAGCGGTG | >hg38_21_+_8433172_8446622_RNA45SN1_45S_50ntFlanks | 6645 | 6665 | 21 | CTCGTACGACTCTTAGCGGTG | 14489.5 | 6.2 | 0.5 | 12.5 | 5.6E-36 | 1.6E-34 | BT20/MB468 |
| >MI0003560\|hsa-mir-92b&WithFlank&1\|+\|155195171\|155195278@67.88.22&[MIMAT0003218&hsa-miR-92b-3p&offsets\|0\|0;m-115&1\|+\|155195237\|155195258&offsets\|0\|0]\|TATTGCACTCGTCCCGGCCTCC | >MI0003560\|hsa-mir-92b&WithFlank&1\|+\|155195171\|155195278 | 67 | 88 | 22 | TATTGCACTCGTCCCGGCCTCC | 7898.1 | 8.7 | 0.7 | 12.5 | 4.3E-36 | 2.2E-34 | MB231/MB468 |
| >hg38_21_+_8433172_8446622_RNA45SN1_45S_50ntFlanks@9568.9597.30\|CGTGCAAATCGGTCGTCCGACCTGGGTATA | >hg38_21_+_8433172_8446622_RNA45SN1_45S_50ntFlanks | 9568 | 9597 | 30 | CGTGCAAATCGGTCGTCCGACCTGGGTATA | 2988.7 | -7.2 | 0.6 | -12.5 | 4.4E-36 | 2.2E-34 | MB231/MB468 |
| >hg38_21_+_8433172_8446622_RNA45SN1_45S_50ntFlanks@11575.11592.18\|ACTTAGAACTGGTGCGGA | >hg38_21_+_8433172_8446622_RNA45SN1_45S_50ntFlanks | 11575 | 11592 | 18 | ACTTAGAACTGGTGCGGA | 3448.8 | -8.0 | 0.6 | -12.5 | 8.3E-36 | 2.4E-34 | BT20/MB468 |
| >hg38_21_+_8433172_8446622_RNA45SN1_45S_50ntFlanks@12999.13039.41\|CTGCGATCTATTGAAAGTCAGCCCTCGACACAAGGGTTTGT | >hg38_21_+_8433172_8446622_RNA45SN1_45S_50ntFlanks | 12999 | 13039 | 41 | CTGCGATCTATTGAAAGTCAGCCCTCGACACAAGGGTTTGT | 22985.0 | 8.3 | 0.7 | 12.5 | 5.2E-36 | 2.6E-34 | MB231/MB468 |
| >hg38_21_+_8433172_8446622_RNA45SN1_45S_50ntFlanks@10679.10711.33\|GGTTCCGGCGGCGTCCGGTGAGCTCTCGCTGGC | >hg38_21_+_8433172_8446622_RNA45SN1_45S_50ntFlanks | 10679 | 10711 | 33 | GGTTCCGGCGGCGTCCGGTGAGCTCTCGCTGGC | 4971.2 | -7.7 | 0.6 | -12.5 | 5.3E-36 | 2.6E-34 | MB231/MB468 |
| >hg38_21_+_8433172_8446622_RNA45SN1_45S_50ntFlanks@10411.10429.19\|CAGTCGGTCCTGAGAGATG | >hg38_21_+_8433172_8446622_RNA45SN1_45S_50ntFlanks | 10411 | 10429 | 19 | CAGTCGGTCCTGAGAGATG | 9864.9 | -6.6 | 0.5 | -12.5 | 5.4E-36 | 2.6E-34 | MB231/MB468 |
| >hg38_21_+_8433172_8446622_RNA45SN1_45S_50ntFlanks@12416.12457.42\|TTCGATGTCGGCTCTTCCTATCATTGTGAAGCAGAATTCACC | >hg38_21_+_8433172_8446622_RNA45SN1_45S_50ntFlanks | 12416 | 12457 | 42 | TTCGATGTCGGCTCTTCCTATCATTGTGAAGCAGAATTCACC | 1935.7 | 13.4 | 1.1 | 12.5 | 6.9E-36 | 3.3E-34 | MB231/MB468 |

<table>
<tr><td>>MI0000743|hsa-mir-34c&WithFlank&11|+|111513433|111513521@19.41.23&[MIMAT0000686&hsa-miR-34c-5p&offsets|0|0;m-336&11|+|111513451|111513473&offsets|0|0]|AGGCAGTGTAGTTAGCTGATTGC</td><td>>MI0000743|hsa-mir-34c&WithFlank&11|+|111513433|111513521</td><td>19</td><td>41</td><td>23</td><td>AGGCAGTGTAGTTAGCTGATTGC</td><td>3141.4</td><td>16.5</td><td>1.3</td><td>12.5</td><td>6.9E-36</td><td>3.3E-34</td><td>MB231/MB468</td></tr>
<tr><td>>MI0000077|hsa-mir-21&WithFlank&17|+|59841260|59841343@14.35.22(+1U)&[MIMAT0000076&hsa-miR-21-5p&offsets|0|0(+1U);m-1&17|+|59841273|59841294&offsets|0|0(+1U)]|TAGCTTATCAGACTGATGTTGAT</td><td>>MI0000077|hsa-mir-21&WithFlank&17|+|59841260|59841343</td><td>14</td><td>35</td><td>22(+1U)</td><td>TAGCTTATCAGACTGATGTTGAT</td><td>538380.2</td><td>9.0</td><td>0.7</td><td>12.5</td><td>1.2E-35</td><td>3.5E-34</td><td>BT20/MB468</td></tr>
<tr><td>>hg38_21_+_8433172_8446622_RNA45SN1_45S_50ntFlanks@12654.12675.22|TATGACTGAACGCCTCTAAGTC</td><td>>hg38_21_+_8433172_8446622_RNA45SN1_45S_50ntFlanks</td><td>12654</td><td>12675</td><td>22</td><td>TATGACTGAACGCCTCTAAGTC</td><td>3496.3</td><td>-7.1</td><td>0.6</td><td>-12.5</td><td>1.2E-35</td><td>3.6E-34</td><td>BT20/MB468</td></tr>
<tr><td>>MI0000061|hsa-let-7a-2&WithFlank&11|-|122146516|122146599@11.31.21(+1A);MI0000062|hsa-let-7a-3&WithFlank&22|+|46112743|46112828@10.30.21(+1A);MI0000060|hsa-let-7a-1&WithFlank&9|+|94175951|94176042@12.32.21(+1A)&[MIMAT0000062&hsa-let-7a-5p&offsets|0|-1(+1A);m-6&11|-|122146568|122146589&offsets|0|-1(+1A)];[MIMAT0000062_1&hsa-let-7a-5p&offsets|0|-1(+1A);m-5&22|+|46112752|46112773&offsets|0|-1(+1A)];[MIMAT0000062_2&hsa-let-7a-5p&offsets|0|-1(+1A);m-4&9|+|94175962|94175983&offsets|0|-1(+1A)]|TGAGGTAGTAGGTTGTATAGTA</td><td>>MI0000061|hsa-let-7a-2&WithFlank&11|-|122146516|122146599</td><td>11</td><td>31</td><td>21(+1A)</td><td>TGAGGTAGTAGGTTGTATAGTA</td><td>6837.7</td><td>8.4</td><td>0.7</td><td>12.5</td><td>7.7E-36</td><td>3.7E-34</td><td>MB231/MB468</td></tr>
<tr><td>>MI0000255|hsa-mir-30d&WithFlank&8|-|134804870|134804951@12.31.20&[MIMAT0000245&hsa-miR-30d-5p&offsets|0|-2;m-21&8|-|134804919|134804940&offsets|0|-2]|TGTAAACATCCCCGACTGGA</td><td>>MI0000255|hsa-mir-30d&WithFlank&8|-|134804870|134804951</td><td>12</td><td>31</td><td>20</td><td>TGTAAACATCCCCGACTGGA</td><td>4117.5</td><td>8.4</td><td>0.7</td><td>12.5</td><td>1.3E-35</td><td>3.7E-34</td><td>BT20/MB468</td></tr>
<tr><td>>hg38_21_+_8433172_8446622_RNA45SN1_45S_50ntFlanks@12395.12414.20|AGCGACGTCGCTTTTTGATC</td><td>>hg38_21_+_8433172_8446622_RNA45SN1_45S_50ntFlanks</td><td>12395</td><td>12414</td><td>20</td><td>AGCGACGTCGCTTTTTGATC</td><td>1874.6</td><td>6.5</td><td>0.5</td><td>12.5</td><td>8.9E-36</td><td>4.2E-34</td><td>MB231/MB468</td></tr>
<tr><td>>MI0000088|hsa-mir-30a&WithFlank&6|-|71403545|71403627@12.34.23(+1C)&[MIMAT0000087&hsa-miR-30a-5p&offsets|0|+1(+1C);m-15&6|-|71403595|71403616&offsets|0|+1(+1C)]|TGTAAACATCCTCGACTGGAAGCC</td><td>>MI0000088|hsa-mir-30a&WithFlank&6|-|71403545|71403627</td><td>12</td><td>34</td><td>23(+1C)</td><td>TGTAAACATCCTCGACTGGAAGCC</td><td>70084.3</td><td>8.9</td><td>0.7</td><td>12.5</td><td>8.8E-36</td><td>4.2E-34</td><td>MB231/MB468</td></tr>
<tr><td>>hg38_21_+_8433172_8446622_RNA45SN1_45S_50ntFlanks@11575.11593.19|ACTTAGAACTGGTGCGGAC</td><td>>hg38_21_+_8433172_8446622_RNA45SN1_45S_50ntFlanks</td><td>11575</td><td>11593</td><td>19</td><td>ACTTAGAACTGGTGCGGAC</td><td>1371.0</td><td>-6.2</td><td>0.5</td><td>-12.4</td><td>1.5E-35</td><td>4.3E-34</td><td>BT20/MB468</td></tr>
<tr><td>>am_trna10_ValCAC_5_-_180649395_180649467@1.33.33,trna12_ValAAC_5_-_180645270_180645342@1.33.33,trna132_ValAAC_6_-_27721179_27721251@1.33.33,trna136_ValAAC_6_-_27648885_27648957@1.33.33,trna139_ValAAC_6_-_27618707_27618779@1.33.33,trna18_ValCAC_5_-_180529253_180529325@1.33.33,trna2_ValAAC_3_+_169490018_169490090@1.33.33,trna2_ValCAC_5_+_180524070_180524142@1.33.33,trna4_ValAAC_5_+_180591154_180591226@1.33.33,trna5_ValAAC_5_+_180596610_180596682@1.33.33,trna6_ValCAC_5_+_180600650_180600722@1.33.33,trna85_ValCAC_1_-_161369490_161369562@1.33.33,trna90_ValCAC_1_-_149684088_149684161@1.33.33,trna98_ValCAC_1_-_149298555_149298627@1.33.33,trna9_ValCAC_6_+_26538282_26538354@1.33.33|GTTTCCGTAGTGTAGTGGTTATCACGTTCGCCT</td><td>>am_trna10_ValCAC_5_-_180649395_180649467</td><td>1</td><td>33</td><td>33</td><td>GTTTCCGTAGTGTAGTGGTTATCACGTTCGCCT</td><td>1568.6</td><td>8.4</td><td>0.7</td><td>12.4</td><td>1.5E-35</td><td>4.3E-34</td><td>BT20/MB468</td></tr>
<tr><td>>MI0000266|hsa-mir-10a&WithFlank&17|-|48579832|48579953@28.51.24&[MIMAT0000253&hsa-miR-10a-5p&offsets|0|+1;m-18&17|-|48579905|48579926&offsets|0|+2]|TACCCTGTAGATCCGAATTTGTGT</td><td>>MI0000266|hsa-mir-10a&WithFlank&17|-|48579832|48579953</td><td>28</td><td>51</td><td>24</td><td>TACCCTGTAGATCCGAATTTGTGT</td><td>2720.0</td><td>16.1</td><td>1.3</td><td>12.4</td><td>1.5E-35</td><td>4.3E-34</td><td>BT20/MB468</td></tr>
<tr><td>>hg38_21_+_8433172_8446622_RNA45SN1_45S_50ntFlanks@476.517.42|TTCGTGATCGATGTGGTGACGTCGTGCTCTCCCGGGCCGGGT</td><td>>hg38_21_+_8433172_8446622_RNA45SN1_45S_50ntFlanks</td><td>476</td><td>517</td><td>42</td><td>TTCGTGATCGATGTGGTGACGTCGTGCTCTCCCGGGCCGGGT</td><td>4583.6</td><td>9.9</td><td>0.8</td><td>12.5</td><td>9.3E-36</td><td>4.4E-34</td><td>MB231/MB468</td></tr>
<tr><td>>hg38_21_+_8433172_8446622_RNA45SN1_45S_50ntFlanks@11572.11603.32|CCGACTTAGAACTGGTGCGGACCAGGGGAATC</td><td>>hg38_21_+_8433172_8446622_RNA45SN1_45S_50ntFlanks</td><td>11572</td><td>11603</td><td>32</td><td>CCGACTTAGAACTGGTGCGGACCAGGGGAATC</td><td>4435.8</td><td>-6.2</td><td>0.5</td><td>-12.4</td><td>1.7E-35</td><td>4.8E-34</td><td>BT20/MB468</td></tr>
<tr><td>>MI0000088|hsa-mir-30a&WithFlank&6|-|71403545|71403627@12.34.23(+1A)&[MIMAT0000087&hsa-miR-30a-5p&offsets|0|+1(+1A);m-15&6|-|71403595|71403616&offsets|0|+1(+1A)]|TGTAAACATCCTCGACTGGAAGCA</td><td>>MI0000088|hsa-mir-30a&WithFlank&6|-|71403545|71403627</td><td>12</td><td>34</td><td>23(+1A)</td><td>TGTAAACATCCTCGACTGGAAGCA</td><td>42988.4</td><td>8.7</td><td>0.7</td><td>12.5</td><td>1.3E-35</td><td>5.8E-34</td><td>MB231/MB468</td></tr>
<tr><td>>MI0000087|hsa-mir-29a&WithFlank&7|-|130876741|130876816@47.68.22&[MIMAT0000086&hsa-miR-29a-3p&offsets|-1|-1;m-26&7|-|130876749|130876769&offsets|-1|0]|CTAGCACCATCTGAAATCGGTT</td><td>>MI0000087|hsa-mir-29a&WithFlank&7|-|130876741|130876816</td><td>47</td><td>68</td><td>22</td><td>CTAGCACCATCTGAAATCGGTT</td><td>33411.9</td><td>9.2</td><td>0.7</td><td>12.5</td><td>1.3E-35</td><td>6.0E-34</td><td>MB231/MB468</td></tr>
<tr><td>>MI0000255|hsa-mir-30d&WithFlank&8|-|134804870|134804951@12.34.23(+1C)&[MIMAT0000245&hsa-miR-30d-5p&offsets|0|+1(+1C);m-21&8|-|134804919|134804940&offsets|0|+1(+1C)]|TGTAAACATCCCCGACTGGAAGCC</td><td>>MI0000255|hsa-mir-30d&WithFlank&8|-|134804870|134804951</td><td>12</td><td>34</td><td>23(+1C)</td><td>TGTAAACATCCCCGACTGGAAGCC</td><td>28304.4</td><td>9.1</td><td>0.7</td><td>12.4</td><td>1.4E-35</td><td>6.4E-34</td><td>MB231/MB468</td></tr>
<tr><td>>MI0000273|hsa-mir-183&WithFlank&7|-|129774899|129775020@34.53.20(+1C)&[MIMAT0000261&hsa-miR-183-5p&offsets|+1|-1(+1C);m-41&7|-|129774966|129774987&offsets|0|-2(+1C)]|ATGGCACTGGTAGAATTCACC</td><td>>MI0000273|hsa-mir-183&WithFlank&7|-|129774899|129775020</td><td>34</td><td>53</td><td>20(+1C)</td><td>ATGGCACTGGTAGAATTCACC</td><td>1497.6</td><td>8.3</td><td>0.7</td><td>12.4</td><td>2.3E-35</td><td>6.5E-34</td><td>BT20/MB468</td></tr>
<tr><td>>hg38_21_+_8433172_8446622_RNA45SN1_45S_50ntFlanks@6646.6673.28|TCGTACGACTCTTAGCGGTGGATCACTC</td><td>>hg38_21_+_8433172_8446622_RNA45SN1_45S_50ntFlanks</td><td>6646</td><td>6673</td><td>28</td><td>TCGTACGACTCTTAGCGGTGGATCACTC</td><td>1562.7</td><td>6.1</td><td>0.5</td><td>12.4</td><td>2.4E-35</td><td>6.7E-34</td><td>BT20/MB468</td></tr>
<tr><td>>MI0000542|hsa-mir-320a&WithFlank&8|-|22244960|22245043@48.69.22(+2U)&[MIMAT0000510&hsa-miR-320a-3p&offsets|0|0(+2U);m-60&8|-|22244975|22244996&offsets|0|0(+2U)]|AAAAGCTGGGTTGAGAGGGCGATT</td><td>>MI0000542|hsa-mir-320a&WithFlank&8|-|22244960|22245043</td><td>48</td><td>69</td><td>22(+2U)</td><td>AAAAGCTGGGTTGAGAGGGCGATT</td><td>8937.3</td><td>9.6</td><td>0.8</td><td>12.4</td><td>2.4E-35</td><td>6.8E-34</td><td>BT20/MB468</td></tr>
<tr><td>>MI0000446|hsa-mir-125b-1&WithFlank&11|-|122099751|122099850@21.43.23&[MIMAT0000423&hsa-miR-125b-5p&offsets|0|+1;m-67&11|-|122099809|122099830&offsets|0|+1]|TCCCTGAGACCCTAACTTGTGAT</td><td>>MI0000446|hsa-mir-125b-1&WithFlank&11|-|122099751|122099850</td><td>21</td><td>43</td><td>23</td><td>TCCCTGAGACCCTAACTTGTGAT</td><td>8098.5</td><td>15.8</td><td>1.3</td><td>12.4</td><td>1.6E-35</td><td>7.1E-34</td><td>MB231/MB468</td></tr>
<tr><td>>hg38_21_+_8433172_8446622_RNA45SN1_45S_50ntFlanks@11208.11232.25|GCGGCGACTCTGGACGCGAGCCGGG</td><td>>hg38_21_+_8433172_8446622_RNA45SN1_45S_50ntFlanks</td><td>11208</td><td>11232</td><td>25</td><td>GCGGCGACTCTGGACGCGAGCCGGG</td><td>5204.0</td><td>-8.5</td><td>0.7</td><td>-12.4</td><td>1.7E-35</td><td>7.8E-34</td><td>MB231/MB468</td></tr>
<tr><td>>hg38_21_+_8433172_8446622_RNA45SN1_45S_50ntFlanks@11787.11804.18|AATGGATGAACGAGATTC</td><td>>hg38_21_+_8433172_8446622_RNA45SN1_45S_50ntFlanks</td><td>11787</td><td>11804</td><td>18</td><td>AATGGATGAACGAGATTC</td><td>7861.8</td><td>-6.8</td><td>0.5</td><td>-12.4</td><td>3.0E-35</td><td>8.3E-34</td><td>BT20/MB468</td></tr>
<tr><td>>hg38_21_+_8433172_8446622_RNA45SN1_45S_50ntFlanks@12632.12654.23|GCGAAGCTACCATCTGTGGGATT</td><td>>hg38_21_+_8433172_8446622_RNA45SN1_45S_50ntFlanks</td><td>12632</td><td>12654</td><td>23</td><td>GCGAAGCTACCATCTGTGGGATT</td><td>1955.4</td><td>-6.7</td><td>0.5</td><td>-12.4</td><td>3.5E-35</td><td>9.5E-34</td><td>BT20/MB468</td></tr>
<tr><td>>MI0000255|hsa-mir-30d&WithFlank&8|-|134804870|134804951@12.33.22(+1A)&[MIMAT0000245&hsa-miR-30d-5p&offsets|0|0(+1A);m-21&8|-|134804919|134804940&offsets|0|0(+1A)]|TGTAAACATCCCCGACTGGAAGA</td><td>>MI0000255|hsa-mir-30d&WithFlank&8|-|134804870|134804951</td><td>12</td><td>33</td><td>22(+1A)</td><td>TGTAAACATCCCCGACTGGAAGA</td><td>2496.7</td><td>8.1</td><td>0.6</td><td>12.4</td><td>2.3E-35</td><td>1.0E-33</td><td>MB231/MB468</td></tr>
<tr><td>>MI0000440|hsa-mir-27b&WithFlank&9|+|95085439|95085547@70.87.18&[MIMAT0000419&hsa-miR-27b-3p&offsets|+3|0;m-39&9|+|95085505|95085525&offsets|+3|0]|ACAGTGGCTAAGTTCTGC</td><td>>MI0000440|hsa-mir-27b&WithFlank&9|+|95085439|95085547</td><td>70</td><td>87</td><td>18</td><td>ACAGTGGCTAAGTTCTGC</td><td>2748.3</td><td>16.1</td><td>1.3</td><td>12.4</td><td>3.8E-35</td><td>1.0E-33</td><td>BT20/MB468</td></tr>
<tr><td>>hg38_21_+_8433172_8446622_RNA45SN1_45S_50ntFlanks@12405.12426.22|CTTTTTGATCCTTCGATGTCGG</td><td>>hg38_21_+_8433172_8446622_RNA45SN1_45S_50ntFlanks</td><td>12405</td><td>12426</td><td>22</td><td>CTTTTTGATCCTTCGATGTCGG</td><td>3313.4</td><td>-8.6</td><td>0.7</td><td>-12.4</td><td>3.9E-35</td><td>1.1E-33</td><td>BT20/MB468</td></tr>
<tr><td>>MI0001733|hsa-mir-452&WithFlank&X|-|151959622|151959718@23.43.21&[MIMAT0001635&hsa-miR-452-5p&offsets|+3|+2;m-188&X|-|151959677|151959699&offsets|+3|+1]|TGTTTGCAGAGGAAACTGAGA</td><td>>MI0001733|hsa-mir-452&WithFlank&X|-|151959622|151959718</td><td>23</td><td>43</td><td>21</td><td>TGTTTGCAGAGGAAACTGAGA</td><td>1871.7</td><td>15.5</td><td>1.3</td><td>12.4</td><td>4.0E-35</td><td>1.1E-33</td><td>BT20/MB468</td></tr>
<tr><td>>trnaMT_GluTTC_MT_-_14674_14742@40.71.32,trnalookalike8_GluTTC_5_-_93905172_93905240@40.71.32|TTGGTCGTGGTTGTAGTCCGTGCGAGAATACC</td><td>>trnaMT_GluTTC_MT_-_14674_14742</td><td>40</td><td>71</td><td>32</td><td>TTGGTCGTGGTTGTAGTCCGTGCGAGAATACC</td><td>783.9</td><td>7.0</td><td>0.6</td><td>12.4</td><td>4.1E-35</td><td>1.1E-33</td><td>BT20/MB468</td></tr>
<tr><td>>MI0000266|hsa-mir-10a&WithFlank&17|-|48579832|48579953@28.49.22(+1A)&[MIMAT0000253&hsa-miR-10a-5p&offsets|0|-1(+1A);m-18&17|-|48579905|48579926&offsets|0|0(+1A)]|TACCCTGTAGATCCGAATTTGTA</td><td>>MI0000266|hsa-mir-10a&WithFlank&17|-|48579832|48579953</td><td>28</td><td>49</td><td>22(+1A)</td><td>TACCCTGTAGATCCGAATTTGTA</td><td>6997.6</td><td>17.4</td><td>1.4</td><td>12.4</td><td>4.5E-35</td><td>1.2E-33</td><td>BT20/MB468</td></tr>
<tr><td>>MI0000650|hsa-mir-200c&WithFlank&12|+|6963693|6963772@50.71.22(+1U)&[MIMAT0000617&hsa-miR-200c-3p&offsets|0|-1(+1U);m-54&12|+|6963742|6963764&offsets|0|-1(+1U)]|TAATACTGCCGGGTAATGATGGT</td><td>>MI0000650|hsa-mir-200c&WithFlank&12|+|6963693|6963772</td><td>50</td><td>71</td><td>22(+1U)</td><td>TAATACTGCCGGGTAATGATGGT</td><td>6803.6</td><td>11.9</td><td>0.9</td><td>12.6</td><td>3.0E-36</td><td>1.3E-33</td><td>BT20/MB231</td></tr>
<tr><td>>hg38_21_+_8433172_8446622_RNA45SN1_45S_50ntFlanks@6648.6695.48|GTACGACTCTTAGCGGTGGATCACTCGGCTCGTGCGTCGATGAAGAAC</td><td>>hg38_21_+_8433172_8446622_RNA45SN1_45S_50ntFlanks</td><td>6648</td><td>6695</td><td>48</td><td>GTACGACTCTTAGCGGTGGATCACTCGGCTCGTGCGTCGATGAAGAAC</td><td>1772.2</td><td>13.2</td><td>1.1</td><td>12.4</td><td>3.1E-35</td><td>1.4E-33</td><td>MB231/MB468</td></tr>
<tr><td>>hg38_21_+_8433172_8446622_RNA45SN1_45S_50ntFlanks@7974.8005.32|ACGCGACCTCAGATCAGACGTGGCGACCCGCT</td><td>>hg38_21_+_8433172_8446622_RNA45SN1_45S_50ntFlanks</td><td>7974</td><td>8005</td><td>32</td><td>ACGCGACCTCAGATCAGACGTGGCGACCCGCT</td><td>6825.9</td><td>6.0</td><td>0.5</td><td>12.3</td><td>5.7E-35</td><td>1.5E-33</td><td>BT20/MB468</td></tr>
<tr><td>>MI0000255|hsa-mir-30d&WithFlank&8|-|134804870|134804951@12.35.24(+2U)&[MIMAT0000245&hsa-miR-30d-5p&offsets|0|+2(+2U);m-21&8|-|134804919|134804940&offsets|0|+2(+2U)]|TGTAAACATCCCCGACTGGAAGCTTT</td><td>>MI0000255|hsa-mir-30d&WithFlank&8|-|134804870|134804951</td><td>12</td><td>35</td><td>24(+2U)</td><td>TGTAAACATCCCCGACTGGAAGCTTT</td><td>3113.2</td><td>10.1</td><td>0.8</td><td>12.4</td><td>3.6E-35</td><td>1.6E-33</td><td>MB231/MB468</td></tr>
<tr><td>>MI0000809|hsa-mir-151a&WithFlank&8|-|140732558|140732659@51.72.22(+1A)&[MIMAT0000757&hsa-miR-151a-3p&offsets|-2|-1(+1A);m-30&8|-|140732587|140732607&offsets|-2|-1(+1A)]|TACTAGACTGAAGCTCCTTGAGA</td><td>>MI0000809|hsa-mir-151a&WithFlank&8|-|140732558|140732659</td><td>51</td><td>72</td><td>22(+1A)</td><td>TACTAGACTGAAGCTCCTTGAGA</td><td>1671.6</td><td>8.7</td><td>0.7</td><td>12.4</td><td>3.7E-35</td><td>1.6E-33</td><td>MB231/MB468</td></tr>
</table>

<table>
<tr><td>>trnaMT_GluTTC_MT_-_14674_14742@45.72.28,trnalookalike8_GluTTC_5_-_93905172_93905240@45.72.28|CGTGGTTGTAGTCCGTGCGAGAATACCA</td><td>>trnaMT_GluTTC_MT_-_14674_14742</td><td>45</td><td>72</td><td>28</td><td>CGTGGTTGTAGTCCGTGCGAGAATACCA</td><td>1849.1</td><td>8.7</td><td>0.7</td><td>12.4</td><td>3.9E-35</td><td>1.7E-33</td><td>MB231/MB468</td></tr>
<tr><td>>hg38_21_+_8433172_8446622_RNA45SN1_45S_50ntFlanks@11669.11687.19|GCCCAGTGCTCTGAATGTC</td><td>>hg38_21_+_8433172_8446622_RNA45SN1_45S_50ntFlanks</td><td>11669</td><td>11687</td><td>19</td><td>GCCCAGTGCTCTGAATGTC</td><td>2863.0</td><td>7.2</td><td>0.6</td><td>12.4</td><td>4.0E-35</td><td>1.7E-33</td><td>MB231/MB468</td></tr>
<tr><td>>hg38_21_+_8433172_8446622_RNA45SN1_45S_50ntFlanks@6661.6695.35|CGGTGGATCACTCGGCTCGTGCGTCGATGAAGAAC</td><td>>hg38_21_+_8433172_8446622_RNA45SN1_45S_50ntFlanks</td><td>6661</td><td>6695</td><td>35</td><td>CGGTGGATCACTCGGCTCGTGCGTCGATGAAGAAC</td><td>644.9</td><td>6.4</td><td>0.5</td><td>12.4</td><td>4.0E-35</td><td>1.7E-33</td><td>MB231/MB468</td></tr>
<tr><td>>hg38_21_+_8433172_8446622_RNA45SN1_45S_50ntFlanks@6741.6771.31|ATCGACACTTCGAACGCACTTGCGGCCCCGG</td><td>>hg38_21_+_8433172_8446622_RNA45SN1_45S_50ntFlanks</td><td>6741</td><td>6771</td><td>31</td><td>ATCGACACTTCGAACGCACTTGCGGCCCCGG</td><td>1361.6</td><td>-6.9</td><td>0.6</td><td>-12.3</td><td>6.8E-35</td><td>1.8E-33</td><td>BT20/MB468</td></tr>
<tr><td>>MI0000102|hsa-mir-100&WithFlank&11|-|122152223|122152314@19.38.20(+2G)&[MIMAT0000098&hsa-miR-100-5p&offsets|0|-2(+2G);m-27&11|-|122152276|122152296&offsets|0|-1(+2G)]|AACCCGTAGATCCGAACTTGGG</td><td>>MI0000102|hsa-mir-100&WithFlank&11|-|122152223|122152314</td><td>19</td><td>38</td><td>20(+2G)</td><td>AACCCGTAGATCCGAACTTGGG</td><td>7639.5</td><td>17.8</td><td>1.4</td><td>12.4</td><td>4.4E-35</td><td>1.9E-33</td><td>MB231/MB468</td></tr>
<tr><td>>trnaMT_ValTAC_MT_+_1602_1670@49.71.23|AACTTAACTTGACCGCTCTGACC</td><td>>trnaMT_ValTAC_MT_+_1602_1670</td><td>49</td><td>71</td><td>23</td><td>AACTTAACTTGACCGCTCTGACC</td><td>483.5</td><td>6.6</td><td>0.5</td><td>12.4</td><td>4.4E-35</td><td>1.9E-33</td><td>MB231/MB468</td></tr>
<tr><td>>hg38_21_+_8433172_8446622_RNA45SN1_45S_50ntFlanks@10411.10431.21|CAGTCGGTCCTGAGAGATGGG</td><td>>hg38_21_+_8433172_8446622_RNA45SN1_45S_50ntFlanks</td><td>10411</td><td>10431</td><td>21</td><td>CAGTCGGTCCTGAGAGATGGG</td><td>3080.8</td><td>-7.8</td><td>0.6</td><td>-12.3</td><td>7.4E-35</td><td>2.0E-33</td><td>BT20/MB468</td></tr>
<tr><td>>trnaMT_ProTGG_MT_-_15956_16023@44.70.27|GTGGAGTTAAAGACTTTTTCTCTGACC</td><td>>trnaMT_ProTGG_MT_-_15956_16023</td><td>44</td><td>70</td><td>27</td><td>GTGGAGTTAAAGACTTTTTCTCTGACC</td><td>737.9</td><td>10.8</td><td>0.9</td><td>12.4</td><td>4.7E-35</td><td>2.0E-33</td><td>MB231/MB468</td></tr>
<tr><td>>hg38_21_+_8433172_8446622_RNA45SN1_45S_50ntFlanks@12488.12509.22|AACGTGAGCTGGGTTTAGACCG</td><td>>hg38_21_+_8433172_8446622_RNA45SN1_45S_50ntFlanks</td><td>12488</td><td>12509</td><td>22</td><td>AACGTGAGCTGGGTTTAGACCG</td><td>4396.3</td><td>-6.5</td><td>0.5</td><td>-12.3</td><td>5.1E-35</td><td>2.1E-33</td><td>MB231/MB468</td></tr>
<tr><td>>hg38_21_+_8433172_8446622_RNA45SN1_45S_50ntFlanks@9564.9592.29|CTGACGTGCAAATCGGTCGTCCGACCTGG</td><td>>hg38_21_+_8433172_8446622_RNA45SN1_45S_50ntFlanks</td><td>9564</td><td>9592</td><td>29</td><td>CTGACGTGCAAATCGGTCGTCCGACCTGG</td><td>1090.3</td><td>-5.5</td><td>0.5</td><td>-12.3</td><td>1.2E-34</td><td>3.1E-33</td><td>BT20/MB468</td></tr>
<tr><td>>MI0000088|hsa-mir-30a&WithFlank&6|-|71403545|71403627@54.74.21&[MIMAT0000088&hsa-miR-30a-3p&offsets|+1|0;m-38&6|-|71403554|71403575&offsets|+1|0]|TTTCAGTCGGATGTTTGCAGC</td><td>>MI0000088|hsa-mir-30a&WithFlank&6|-|71403545|71403627</td><td>54</td><td>74</td><td>21</td><td>TTTCAGTCGGATGTTTGCAGC</td><td>12795.6</td><td>9.6</td><td>0.8</td><td>12.3</td><td>7.4E-35</td><td>3.1E-33</td><td>MB231/MB468</td></tr>
<tr><td>>hg38_1_-_228634819_228635039_RNA5S12_5S_50ntFlanks@134.171.38|TGGGAGACCGCCTGGGAATACCGGGTGCTGTAGGCTTT</td><td>>hg38_1_-_228634819_228635039_RNA5S12_5S_50ntFlanks</td><td>134</td><td>171</td><td>38</td><td>TGGGAGACCGCCTGGGAATACCGGGTGCTGTAGGCTTT</td><td>123529.1</td><td>10.0</td><td>0.8</td><td>12.3</td><td>7.7E-35</td><td>3.1E-33</td><td>MB231/MB468</td></tr>
<tr><td>>MI0000804|hsa-mir-328&WithFlank&16|-|67202315|67202401@54.75.22&[MIMAT0000752&hsa-miR-328-3p&offsets|0|0;m-160&16|-|67202327|67202348&offsets|0|0]|CTGGCCCTCTCTGCCCTTCCGT</td><td>>MI0000804|hsa-mir-328&WithFlank&16|-|67202315|67202401</td><td>54</td><td>75</td><td>22</td><td>CTGGCCCTCTCTGCCCTTCCGT</td><td>1573.8</td><td>8.1</td><td>0.7</td><td>12.3</td><td>1.3E-34</td><td>3.4E-33</td><td>BT20/MB468</td></tr>
<tr><td>>trna116_GluCTC_1_-_145399233_145399304@1.34.34,trna59_GluCTC_1_+_249168447_249168518@1.34.34,trna71_GluCTC_1_-_161439189_161439260@1.34.34,trna74_GluCTC_1_-_161431809_161431880@1.34.34,trna77_GluCTC_1_-_161424398_161424469@1.34.34,trna77_GluCTC_6_+_28949976_28950047@1.34.34,trna80_GluCTC_1_-_161417018_161417089@1.34.34,trna87_GluCTC_6_-_126101393_126101464@1.34.34|TCCCTGGTGGTCTAGTGGTTAGGATTCGGCGCTC</td><td>>trna116_GluCTC_1_-_145399233_145399304</td><td>1</td><td>34</td><td>34</td><td>TCCCTGGTGGTCTAGTGGTTAGGATTCGGCGCTC</td><td>12191.6</td><td>9.6</td><td>0.8</td><td>12.3</td><td>1.3E-34</td><td>3.4E-33</td><td>BT20/MB468</td></tr>
<tr><td>>hg38_21_+_8433172_8446622_RNA45SN1_45S_50ntFlanks@6722.6744.23|ATTGCAGGACACATTGATCATCG</td><td>>hg38_21_+_8433172_8446622_RNA45SN1_45S_50ntFlanks</td><td>6722</td><td>6744</td><td>23</td><td>ATTGCAGGACACATTGATCATCG</td><td>2507.4</td><td>-6.0</td><td>0.5</td><td>-12.3</td><td>1.5E-34</td><td>3.9E-33</td><td>BT20/MB468</td></tr>
<tr><td>>hg38_21_+_8433172_8446622_RNA45SN1_45S_50ntFlanks@6645.6667.23|CTCGTACGACTCTTAGCGGTGGA</td><td>>hg38_21_+_8433172_8446622_RNA45SN1_45S_50ntFlanks</td><td>6645</td><td>6667</td><td>23</td><td>CTCGTACGACTCTTAGCGGTGGA</td><td>2876.1</td><td>6.3</td><td>0.5</td><td>12.3</td><td>1.5E-34</td><td>3.9E-33</td><td>BT20/MB468</td></tr>
<tr><td>>MI0003591|hsa-mir-584&WithFlank&5|-|149062307|149062415@22.42.21&[MIMAT0003249&hsa-miR-584-5p&offsets|0|-1;m-167&5|-|149062374|149062394&offsets|0|0]|TTATGGTTTGCCTGGGACTGA</td><td>>MI0003591|hsa-mir-584&WithFlank&5|-|149062307|149062415</td><td>22</td><td>42</td><td>21</td><td>TTATGGTTTGCCTGGGACTGA</td><td>5219.5</td><td>8.8</td><td>0.7</td><td>12.3</td><td>1.5E-34</td><td>3.9E-33</td><td>BT20/MB468</td></tr>
<tr><td>>MI0000299|hsa-mir-222&WithFlank&X|-|45747009|45747130@75.97.23(+2U)&[MIMAT0000279&hsa-miR-222-3p&offsets|0|+2(+2U);m-140&X|-|45747033|45747056&offsets|0|-1(+2U)]|AGCTACATCTGGCTACTGGGTCTTT</td><td>>MI0000299|hsa-mir-222&WithFlank&X|-|45747009|45747130</td><td>75</td><td>97</td><td>23(+2U)</td><td>AGCTACATCTGGCTACTGGGTCTTT</td><td>4071.7</td><td>9.5</td><td>0.8</td><td>12.3</td><td>9.8E-35</td><td>4.0E-33</td><td>MB231/MB468</td></tr>
<tr><td>>MI0000809|hsa-mir-151a&WithFlank&8|-|140732558|140732659@51.72.22(+1U)&[MIMAT0000757&hsa-miR-151a-3p&offsets|-2|-1(+1U);m-30&8|-|140732587|140732607&offsets|-2|-1(+1U)]|TACTAGACTGAAGCTCCTTGAGT</td><td>>MI0000809|hsa-mir-151a&WithFlank&8|-|140732558|140732659</td><td>51</td><td>72</td><td>22(+1U)</td><td>TACTAGACTGAAGCTCCTTGAGT</td><td>2488.1</td><td>9.3</td><td>0.8</td><td>12.3</td><td>1.1E-34</td><td>4.5E-33</td><td>MB231/MB468</td></tr>
<tr><td>>MI0003205|hsa-mir-532&WithFlank&X|+|50003142|50003244@26.46.21(+1C)&[MIMAT0002888&hsa-miR-532-5p&offsets|0|-1(+1C);m-65&X|+|50003167|50003188&offsets|0|-1(+1C)]|CATGCCTTGAGTGTAGGACCGC</td><td>>MI0003205|hsa-mir-532&WithFlank&X|+|50003142|50003244</td><td>26</td><td>46</td><td>21(+1C)</td><td>CATGCCTTGAGTGTAGGACCGC</td><td>867.8</td><td>5.8</td><td>0.5</td><td>12.2</td><td>1.9E-34</td><td>4.8E-33</td><td>BT20/MB468</td></tr>
<tr><td>>hg38_21_+_8433172_8446622_RNA45SN1_45S_50ntFlanks@12991.13013.23|CTCCCTCGCTGCGATCTATTGAA</td><td>>hg38_21_+_8433172_8446622_RNA45SN1_45S_50ntFlanks</td><td>12991</td><td>13013</td><td>23</td><td>CTCCCTCGCTGCGATCTATTGAA</td><td>7059.7</td><td>-7.8</td><td>0.6</td><td>-12.2</td><td>2.1E-34</td><td>5.4E-33</td><td>BT20/MB468</td></tr>
<tr><td>>trnaMT_GluTTC_MT_-_14674_14742@37.72.36,trnalookalike8_GluTTC_5_-_93905172_93905240@37.72.36|TCATTGGTCGTGGTTGTAGTCCGTGCGAGAATACCA</td><td>>trnaMT_GluTTC_MT_-_14674_14742</td><td>37</td><td>72</td><td>36</td><td>TCATTGGTCGTGGTTGTAGTCCGTGCGAGAATACCA</td><td>8860.9</td><td>9.6</td><td>0.8</td><td>12.3</td><td>1.4E-34</td><td>5.5E-33</td><td>MB231/MB468</td></tr>
<tr><td>>hg38_21_+_8433172_8446622_RNA45SN1_45S_50ntFlanks@12633.12653.21|CGAAGCTACCATCTGTGGGAT</td><td>>hg38_21_+_8433172_8446622_RNA45SN1_45S_50ntFlanks</td><td>12633</td><td>12653</td><td>21</td><td>CGAAGCTACCATCTGTGGGAT</td><td>6478.1</td><td>-9.8</td><td>0.8</td><td>-12.3</td><td>1.4E-34</td><td>5.7E-33</td><td>MB231/MB468</td></tr>
<tr><td>>MI0000298|hsa-mir-221&WithFlank&X|-|45746151|45746272@31.56.26&[MIMAT0004568&hsa-miR-221-5p&offsets|0|+4;m-405&X|-|45746220|45746242&offsets|0|+3]|ACCTGGCATACAATGTAGATTTCTGT</td><td>>MI0000298|hsa-mir-221&WithFlank&X|-|45746151|45746272</td><td>31</td><td>56</td><td>26</td><td>ACCTGGCATACAATGTAGATTTCTGT</td><td>9159.8</td><td>8.9</td><td>0.7</td><td>12.3</td><td>1.4E-34</td><td>5.7E-33</td><td>MB231/MB468</td></tr>
<tr><td>>MI0000255|hsa-mir-30d&WithFlank&8|-|134804870|134804951@12.35.24&[MIMAT0000245&hsa-miR-30d-5p&offsets|0|+2;m-21&8|-|134804919|134804940&offsets|0|+2]|TGTAAACATCCCCGACTGGAAGCT</td><td>>MI0000255|hsa-mir-30d&WithFlank&8|-|134804870|134804951</td><td>12</td><td>35</td><td>24</td><td>TGTAAACATCCCCGACTGGAAGCT</td><td>194367.8</td><td>9.1</td><td>0.7</td><td>12.3</td><td>1.5E-34</td><td>6.0E-33</td><td>MB231/MB468</td></tr>
<tr><td>>MI0000298|hsa-mir-221&WithFlank&X|-|45746151|45746272@71.93.23(+1C)&[MIMAT0000278&hsa-miR-221-3p&offsets|0|0(+1C);m-91&X|-|45746181|45746202&offsets|0|+1(+1C)]|AGCTACATTGTCTGCTGGGTTTCC</td><td>>MI0000298|hsa-mir-221&WithFlank&X|-|45746151|45746272</td><td>71</td><td>93</td><td>23(+1C)</td><td>AGCTACATTGTCTGCTGGGTTTCC</td><td>7686.5</td><td>8.9</td><td>0.7</td><td>12.3</td><td>1.6E-34</td><td>6.3E-33</td><td>MB231/MB468</td></tr>
<tr><td>>MI0000746|hsa-mir-99b&WithFlank&19|+|51692606|51692687@13.32.20&[MIMAT0000689&hsa-miR-99b-5p&offsets|0|-2;m-10&19|+|51692618|51692639&offsets|0|-2]|CACCCGTAGAACCGACCTTG</td><td>>MI0000746|hsa-mir-99b&WithFlank&19|+|51692606|51692687</td><td>13</td><td>32</td><td>20</td><td>CACCCGTAGAACCGACCTTG</td><td>3515.2</td><td>9.1</td><td>0.7</td><td>12.2</td><td>2.6E-34</td><td>6.7E-33</td><td>BT20/MB468</td></tr>
<tr><td>>hg38_21_+_8433172_8446622_RNA45SN1_45S_50ntFlanks@12997.13037.41|CGCTGCGATCTATTGAAAGTCAGCCCTCGACACAAGGGTTT</td><td>>hg38_21_+_8433172_8446622_RNA45SN1_45S_50ntFlanks</td><td>12997</td><td>13037</td><td>41</td><td>CGCTGCGATCTATTGAAAGTCAGCCCTCGACACAAGGGTTT</td><td>1442.6</td><td>15.4</td><td>1.3</td><td>12.2</td><td>1.8E-34</td><td>6.9E-33</td><td>MB231/MB468</td></tr>
<tr><td>>MI0000088|hsa-mir-30a&WithFlank&6|-|71403545|71403627@53.73.21(+1U)&[MIMAT0000088&hsa-miR-30a-3p&offsets|0|-1(+1U);m-38&6|-|71403554|71403575&offsets|0|-1(+1U)]|CTTTCAGTCGGATGTTTGCAGT</td><td>>MI0000088|hsa-mir-30a&WithFlank&6|-|71403545|71403627</td><td>53</td><td>73</td><td>21(+1U)</td><td>CTTTCAGTCGGATGTTTGCAGT</td><td>23834.8</td><td>9.1</td><td>0.7</td><td>12.2</td><td>2.0E-34</td><td>7.7E-33</td><td>MB231/MB468</td></tr>
<tr><td>>am_trna128_GlyGCC_6_-_27870686_27870756@1.35.35,trna18_GlyGCC_16_+_70822597_70822667@1.35.35,trna19_GlyGCC_16_+_70823410_70823480@1.35.35,trna19_GlyGCC_2_-_157257659_157257729@1.35.35,trna24_GlyGCC_16_-_70812942_70813012@1.35.35,trna25_GlyGCC_16_-_70812114_70812184@1.35.35,trna5_GlyGCC_17_+_8029064_8029134@1.35.35,trna68_GlyGCC_1_-_161493637_161493707@1.35.35|GCATTGGTGGTTCAGTGGTAGAATTCTCGCCTGCC</td><td>>am_trna128_GlyGCC_6_-_27870686_27870756</td><td>1</td><td>35</td><td>35</td><td>GCATTGGTGGTTCAGTGGTAGAATTCTCGCCTGCC</td><td>32730.5</td><td>11.0</td><td>0.9</td><td>12.2</td><td>2.2E-34</td><td>8.3E-33</td><td>MB231/MB468</td></tr>
<tr><td>>MI0000542|hsa-mir-320a&WithFlank&8|-|22244960|22245043@48.70.23&[MIMAT0000510&hsa-miR-320a-3p&offsets|0|+1;m-60&8|-|22244975|22244996&offsets|0|+1]|AAAAGCTGGGTTGAGAGGGCGAA</td><td>>MI0000542|hsa-mir-320a&WithFlank&8|-|22244960|22245043</td><td>48</td><td>70</td><td>23</td><td>AAAAGCTGGGTTGAGAGGGCGAA</td><td>19437.3</td><td>9.8</td><td>0.8</td><td>12.2</td><td>3.5E-34</td><td>8.8E-33</td><td>BT20/MB468</td></tr>
<tr><td>>am_MI0003581|hsa-mir-574&WithFlank&4|+|38868026|38868133@31.55.25&[MIMAT0004795&hsa-miR-574-5p&offsets|0|+2;m-213&4|+|38868056|38868078&offsets|0|+2]|TGAGTGTGTGTGTGTGAGTGTGTGT</td><td>>am_MI0003581|hsa-mir-574&WithFlank&4|+|38868026|38868133</td><td>31</td><td>55</td><td>25</td><td>TGAGTGTGTGTGTGTGAGTGTGTGT</td><td>2120.2</td><td>9.5</td><td>0.8</td><td>12.2</td><td>2.4E-34</td><td>9.0E-33</td><td>MB231/MB468</td></tr>
<tr><td>>hg38_1_-_228634819_228635039_RNA5S12_5S_50ntFlanks@51.89.39|GTCTACGGCCATACCACCCTGAACGCGCCCGATCTCGTC</td><td>>hg38_1_-_228634819_228635039_RNA5S12_5S_50ntFlanks</td><td>51</td><td>89</td><td>39</td><td>GTCTACGGCCATACCACCCTGAACGCGCCCGATCTCGTC</td><td>24994.9</td><td>10.4</td><td>0.9</td><td>12.2</td><td>2.4E-34</td><td>9.0E-33</td><td>MB231/MB468</td></tr>
<tr><td>>MI0000434|hsa-let-7i&WithFlank&12|+|62603680|62603775@12.33.22(+1C)&[MIMAT0000415&hsa-let-7i-5p&offsets|0|0(+1C);m-49&12|+|62603691|62603712&offsets|0|0(+1C)]|TGAGGTAGTAGTTTGTGCTGTTC</td><td>>MI0000434|hsa-let-7i&WithFlank&12|+|62603680|62603775</td><td>12</td><td>33</td><td>22(+1C)</td><td>TGAGGTAGTAGTTTGTGCTGTTC</td><td>3406.1</td><td>8.9</td><td>0.7</td><td>12.2</td><td>2.6E-34</td><td>1.0E-32</td><td>MB231/MB468</td></tr>
<tr><td>>MI0003591|hsa-mir-584&WithFlank&5|-|149062307|149062415@22.42.21&[MIMAT0003249&hsa-miR-584-5p&offsets|0|-1;m-167&5|-|149062374|149062394&offsets|0|0]|TTATGGTTTGCCTGGGACTGA</td><td>>MI0003591|hsa-mir-584&WithFlank&5|-|149062307|149062415</td><td>22</td><td>42</td><td>21</td><td>TTATGGTTTGCCTGGGACTGA</td><td>5872.0</td><td>9.2</td><td>0.8</td><td>12.2</td><td>2.8E-34</td><td>1.0E-32</td><td>MB231/MB468</td></tr>
<tr><td>>MI0000088|hsa-mir-30a&WithFlank&6|-|71403545|71403627@12.32.21(+1A)&[MIMAT0000087&hsa-miR-30a-5p&offsets|0|-1(+1A);m-15&6|-|71403595|71403616&offsets|0|-1(+1A)]|TGTAAACATCCTCGACTGGAAA</td><td>>MI0000088|hsa-mir-30a&WithFlank&6|-|71403545|71403627</td><td>12</td><td>32</td><td>21(+1A)</td><td>TGTAAACATCCTCGACTGGAAA</td><td>918.1</td><td>7.8</td><td>0.6</td><td>12.2</td><td>2.8E-34</td><td>1.1E-32</td><td>MB231/MB468</td></tr>
</table>

<table>
<tr><td>>MI0000650|hsa-mir-200c&WithFlank&12|+|6963693|6963772@50.72.23(+1U)&[MIMAT0000617&hsa-miR-200c-3p&offsets|0|0(+1U);m-54&12|+|6963742|6963764&offsets|0|0(+1U)]|TAATACTGCCGGGTAATGATGGAT</td><td>>MI0000650|hsa-mir-200c&WithFlank&12|+|6963693|6963772</td><td>50</td><td>72</td><td>23(+1U)</td><td>TAATACTGCCGGGTAATGATGGAT</td><td>21676.9</td><td>8.0</td><td>0.7</td><td>12.2</td><td>4.4E-34</td><td>1.1E-32</td><td>BT20/MB468</td></tr>
<tr><td>>hg38_21_+_8433172_8446622_RNA45SN1_45S_50ntFlanks@6646.6676.31|TCGTACGACTCTTAGCGGTGGATCACTCGGC</td><td>>hg38_21_+_8433172_8446622_RNA45SN1_45S_50ntFlanks</td><td>6646</td><td>6676</td><td>31</td><td>TCGTACGACTCTTAGCGGTGGATCACTCGGC</td><td>2756.5</td><td>6.6</td><td>0.5</td><td>12.2</td><td>4.5E-34</td><td>1.1E-32</td><td>BT20/MB468</td></tr>
<tr><td>>hg38_21_+_8433172_8446622_RNA45SN1_45S_50ntFlanks@6721.6747.27|AATTGCAGGACACATTGATCATCGACA</td><td>>hg38_21_+_8433172_8446622_RNA45SN1_45S_50ntFlanks</td><td>6721</td><td>6747</td><td>27</td><td>AATTGCAGGACACATTGATCATCGACA</td><td>2316.5</td><td>-4.5</td><td>0.4</td><td>-12.2</td><td>4.6E-34</td><td>1.2E-32</td><td>BT20/MB468</td></tr>
<tr><td>>am_trna128_GlyGCC_6_-_27870686_27870756@1.32.32,trna133_GlyCCC_1_-_16872434_16872504@1.32.32,trna18_GlyGCC_16_+_70822597_70822667@1.32.32,trna19_GlyGCC_16_+_70823410_70823480@1.32.32,trna19_GlyGCC_2_-_157257659_157257729@1.32.32,trna24_GlyGCC_16_-_70812942_70813012@1.32.32,trna25_GlyGCC_16_-_70812114_70812184@1.32.32,trna4_GlyCCC_1_+_17188416_17188486@1.32.32,trna5_GlyGCC_17_+_8029064_8029134@1.32.32,trna68_GlyGCC_1_-_161493637_161493707@1.32.32|GCATTGGTGGTTCAGTGGTAGAATTCTCGCCT</td><td>>am_trna128_GlyGCC_6_-_27870686_27870756</td><td>1</td><td>32</td><td>32</td><td>GCATTGGTGGTTCAGTGGTAGAATTCTCGCCT</td><td>142940.1</td><td>11.7</td><td>1.0</td><td>12.2</td><td>3.2E-34</td><td>1.2E-32</td><td>MB231/MB468</td></tr>
<tr><td>>MI0000434|hsa-let-7i&WithFlank&12|+|62603680|62603775@12.32.21(+1A)&[MIMAT0000415&hsa-let-7i-5p&offsets|0|-1(+1A);m-49&12|+|62603691|62603712&offsets|0|-1(+1A)]|TGAGGTAGTAGTTTGTGCTGTA</td><td>>MI0000434|hsa-let-7i&WithFlank&12|+|62603680|62603775</td><td>12</td><td>32</td><td>21(+1A)</td><td>TGAGGTAGTAGTTTGTGCTGTA</td><td>23146.9</td><td>8.7</td><td>0.7</td><td>12.2</td><td>3.6E-34</td><td>1.3E-32</td><td>MB231/MB468</td></tr>
<tr><td>>trnaMT_GluTTC_MT_-_14674_14742@37.72.36,trnalookalike8_GluTTC_5_-_93905172_93905240@37.72.36|TCATTGGTCGTGGTTGTAGTCCGTGCGAGAATACCA</td><td>>trnaMT_GluTTC_MT_-_14674_14742</td><td>37</td><td>72</td><td>36</td><td>TCATTGGTCGTGGTTGTAGTCCGTGCGAGAATACCA</td><td>1546.5</td><td>6.8</td><td>0.6</td><td>12.2</td><td>5.5E-34</td><td>1.4E-32</td><td>BT20/MB468</td></tr>
<tr><td>>MI0000102|hsa-mir-100&WithFlank&11|-|122152223|122152314@19.37.19(+1A)&[MIMAT0000098&hsa-miR-100-5p&offsets|0|-3(+1A);m-27&11|-|122152276|122152296&offsets|0|-2(+1A)]|AACCCGTAGATCCGAACTTA</td><td>>MI0000102|hsa-mir-100&WithFlank&11|-|122152223|122152314</td><td>19</td><td>37</td><td>19(+1A)</td><td>AACCCGTAGATCCGAACTTA</td><td>7297.9</td><td>17.7</td><td>1.5</td><td>12.2</td><td>3.8E-34</td><td>1.4E-32</td><td>MB231/MB468</td></tr>
<tr><td>>hg38_21_+_8433172_8446622_RNA45SN1_45S_50ntFlanks@6652.6676.25|GACTCTTAGCGGTGGATCACTCGGC</td><td>>hg38_21_+_8433172_8446622_RNA45SN1_45S_50ntFlanks</td><td>6652</td><td>6676</td><td>25</td><td>GACTCTTAGCGGTGGATCACTCGGC</td><td>613.8</td><td>5.7</td><td>0.5</td><td>12.1</td><td>5.8E-34</td><td>1.4E-32</td><td>BT20/MB468</td></tr>
<tr><td>>MI0000266|hsa-mir-10a&WithFlank&17|-|48579832|48579953@28.51.24&[MIMAT0000253&hsa-miR-10a-5p&offsets|0|+1;m-18&17|-|48579905|48579926&offsets|0|+2]|TACCCTGTAGATCCGAATTTGTGT</td><td>>MI0000266|hsa-mir-10a&WithFlank&17|-|48579832|48579953</td><td>28</td><td>51</td><td>24</td><td>TACCCTGTAGATCCGAATTTGTGT</td><td>3260.3</td><td>16.6</td><td>1.4</td><td>12.2</td><td>4.0E-34</td><td>1.4E-32</td><td>MB231/MB468</td></tr>
<tr><td>>MI0000650|hsa-mir-200c&WithFlank&12|+|6963693|6963772@50.72.23(+1A)&[MIMAT0000617&hsa-miR-200c-3p&offsets|0|0(+1A);m-54&12|+|6963742|6963764&offsets|0|0(+1A)]|TAATACTGCCGGGTAATGATGGAA</td><td>>MI0000650|hsa-mir-200c&WithFlank&12|+|6963693|6963772</td><td>50</td><td>72</td><td>23(+1A)</td><td>TAATACTGCCGGGTAATGATGGAA</td><td>32067.9</td><td>8.7</td><td>0.7</td><td>12.1</td><td>6.2E-34</td><td>1.5E-32</td><td>BT20/MB468</td></tr>
<tr><td>>MI0000477|hsa-mir-146a&WithFlank&5|+|160485346|160485456@27.49.23&[MIMAT0000449&hsa-miR-146a-5p&offsets|0|+1;m-129&5|+|160485372|160485393&offsets|0|+1]|TGAGAACTGAATTCCATGGGTTG</td><td>>MI0000477|hsa-mir-146a&WithFlank&5|+|160485346|160485456</td><td>27</td><td>49</td><td>23</td><td>TGAGAACTGAATTCCATGGGTTG</td><td>3575.6</td><td>16.7</td><td>1.4</td><td>12.2</td><td>4.3E-34</td><td>1.6E-32</td><td>MB231/MB468</td></tr>
<tr><td>>MI0000269|hsa-mir-181a-2&WithFlank&9|+|124692436|124692557@83.104.22&[MIMAT0004558&hsa-miR-181a-2-3p&offsets|0|0;m-71&9|+|124692518|124692539&offsets|0|0]|ACCACTGACCGTTGACTGTACC</td><td>>MI0000269|hsa-mir-181a-2&WithFlank&9|+|124692436|124692557</td><td>83</td><td>104</td><td>22</td><td>ACCACTGACCGTTGACTGTACC</td><td>3961.2</td><td>9.1</td><td>0.7</td><td>12.2</td><td>4.5E-34</td><td>1.6E-32</td><td>MB231/MB468</td></tr>
<tr><td>>hg38_21_+_8433172_8446622_RNA45SN1_45S_50ntFlanks@11539.11584.46|GCGCGGCGCCTCGCCTCGGCCGGCGCCTAGCAGCCGACTTAGAACT</td><td>>hg38_21_+_8433172_8446622_RNA45SN1_45S_50ntFlanks</td><td>11539</td><td>11584</td><td>46</td><td>GCGCGGCGCCTCGCCTCGGCCGGCGCCTAGCAGCCGACTTAGAACT</td><td>988.3</td><td>14.9</td><td>1.2</td><td>12.2</td><td>4.7E-34</td><td>1.7E-32</td><td>MB231/MB468</td></tr>
<tr><td>>hg38_21_+_8433172_8446622_RNA45SN1_45S_50ntFlanks@12416.12458.43|TTCGATGTCGGCTCTTCCTATCATTGTGAAGCAGAATTCACCA</td><td>>hg38_21_+_8433172_8446622_RNA45SN1_45S_50ntFlanks</td><td>12416</td><td>12458</td><td>43</td><td>TTCGATGTCGGCTCTTCCTATCATTGTGAAGCAGAATTCACCA</td><td>592.0</td><td>14.1</td><td>1.2</td><td>12.2</td><td>4.8E-34</td><td>1.7E-32</td><td>MB231/MB468</td></tr>
<tr><td>>hg38_21_+_8433172_8446622_RNA45SN1_45S_50ntFlanks@476.518.43|TTCGTGATCGATGTGGTGACGTCGTGCTCTCCCGGGCCGGGTC</td><td>>hg38_21_+_8433172_8446622_RNA45SN1_45S_50ntFlanks</td><td>476</td><td>518</td><td>43</td><td>TTCGTGATCGATGTGGTGACGTCGTGCTCTCCCGGGCCGGGTC</td><td>3021.0</td><td>9.9</td><td>0.8</td><td>12.2</td><td>4.9E-34</td><td>1.7E-32</td><td>MB231/MB468</td></tr>
<tr><td>>hg38_21_+_8433172_8446622_RNA45SN1_45S_50ntFlanks@9570.9596.27|TGCAAATCGGTCGTCCGACCTGGGTAT</td><td>>hg38_21_+_8433172_8446622_RNA45SN1_45S_50ntFlanks</td><td>9570</td><td>9596</td><td>27</td><td>TGCAAATCGGTCGTCCGACCTGGGTAT</td><td>2783.3</td><td>-7.5</td><td>0.6</td><td>-12.1</td><td>7.6E-34</td><td>1.9E-32</td><td>BT20/MB468</td></tr>
<tr><td>>hg38_21_+_8433172_8446622_RNA45SN1_45S_50ntFlanks@9572.9595.24|CAAATCGGTCGTCCGACCTGGGTA</td><td>>hg38_21_+_8433172_8446622_RNA45SN1_45S_50ntFlanks</td><td>9572</td><td>9595</td><td>24</td><td>CAAATCGGTCGTCCGACCTGGGTA</td><td>3166.9</td><td>-7.0</td><td>0.6</td><td>-12.1</td><td>8.8E-34</td><td>2.2E-32</td><td>BT20/MB468</td></tr>
<tr><td>>hg38_21_+_8433172_8446622_RNA45SN1_45S_50ntFlanks@6651.6696.46|CGACTCTTAGCGGTGGATCACTCGGCTCGTGCGTCGATGAAGAACG</td><td>>hg38_21_+_8433172_8446622_RNA45SN1_45S_50ntFlanks</td><td>6651</td><td>6696</td><td>46</td><td>CGACTCTTAGCGGTGGATCACTCGGCTCGTGCGTCGATGAAGAACG</td><td>33236.4</td><td>7.5</td><td>0.6</td><td>12.1</td><td>6.5E-34</td><td>2.3E-32</td><td>MB231/MB468</td></tr>
<tr><td>>hg38_21_+_8433172_8446622_RNA45SN1_45S_50ntFlanks@12997.13036.40|CGCTGCGATCTATTGAAAGTCAGCCCTCGACACAAGGGTT</td><td>>hg38_21_+_8433172_8446622_RNA45SN1_45S_50ntFlanks</td><td>12997</td><td>13036</td><td>40</td><td>CGCTGCGATCTATTGAAAGTCAGCCCTCGACACAAGGGTT</td><td>2625.7</td><td>8.7</td><td>0.7</td><td>12.1</td><td>9.4E-34</td><td>2.3E-32</td><td>BT20/MB468</td></tr>
<tr><td>>hg38_21_+_8433172_8446622_RNA45SN1_45S_50ntFlanks@6645.6664.20|CTCGTACGACTCTTAGCGGT</td><td>>hg38_21_+_8433172_8446622_RNA45SN1_45S_50ntFlanks</td><td>6645</td><td>6664</td><td>20</td><td>CTCGTACGACTCTTAGCGGT</td><td>4082.8</td><td>5.8</td><td>0.5</td><td>12.1</td><td>9.5E-34</td><td>2.3E-32</td><td>BT20/MB468</td></tr>
<tr><td>>hg38_21_+_8433172_8446622_RNA45SN1_45S_50ntFlanks@11874.11893.20|AGAAGACCCTGTTGAGCTTG</td><td>>hg38_21_+_8433172_8446622_RNA45SN1_45S_50ntFlanks</td><td>11874</td><td>11893</td><td>20</td><td>AGAAGACCCTGTTGAGCTTG</td><td>3926.9</td><td>-7.1</td><td>0.6</td><td>-12.1</td><td>7.1E-34</td><td>2.5E-32</td><td>MB231/MB468</td></tr>
<tr><td>>hg38_21_+_8433172_8446622_RNA45SN1_45S_50ntFlanks@12438.12455.18|ATTGTGAAGCAGAATTCA</td><td>>hg38_21_+_8433172_8446622_RNA45SN1_45S_50ntFlanks</td><td>12438</td><td>12455</td><td>18</td><td>ATTGTGAAGCAGAATTCA</td><td>5288.3</td><td>-8.3</td><td>0.7</td><td>-12.1</td><td>1.1E-33</td><td>2.6E-32</td><td>BT20/MB468</td></tr>
<tr><td>>hg38_21_+_8433172_8446622_RNA45SN1_45S_50ntFlanks@12008.12032.25|CCGCCGGTGAAATACCACTACTCTG</td><td>>hg38_21_+_8433172_8446622_RNA45SN1_45S_50ntFlanks</td><td>12008</td><td>12032</td><td>25</td><td>CCGCCGGTGAAATACCACTACTCTG</td><td>7154.9</td><td>-11.0</td><td>0.9</td><td>-12.1</td><td>1.1E-33</td><td>2.6E-32</td><td>BT20/MB468</td></tr>
<tr><td>>MI0000477|hsa-mir-146a&WithFlank&5|+|160485346|160485456@27.47.21(+1C)&[MIMAT0000449&hsa-miR-146a-5p&offsets|0|-1(+1C);m-129&5|+|160485372|160485393&offsets|0|-1(+1C)]|TGAGAACTGAATTCCATGGGTC</td><td>>MI0000477|hsa-mir-146a&WithFlank&5|+|160485346|160485456</td><td>27</td><td>47</td><td>21(+1C)</td><td>TGAGAACTGAATTCCATGGGTC</td><td>5614.3</td><td>17.4</td><td>1.4</td><td>12.1</td><td>7.8E-34</td><td>2.7E-32</td><td>MB231/MB468</td></tr>
<tr><td>>hg38_21_+_8433172_8446622_RNA45SN1_45S_50ntFlanks@9534.9564.31|AGGAAACTCTGGTGGAGGTCCGTAGCGGTCC</td><td>>hg38_21_+_8433172_8446622_RNA45SN1_45S_50ntFlanks</td><td>9534</td><td>9564</td><td>31</td><td>AGGAAACTCTGGTGGAGGTCCGTAGCGGTCC</td><td>1116.1</td><td>-6.0</td><td>0.5</td><td>-12.1</td><td>1.2E-33</td><td>2.8E-32</td><td>BT20/MB468</td></tr>
<tr><td>>hg38_21_+_8433172_8446622_RNA45SN1_45S_50ntFlanks@7981.8006.26|CTCAGATCAGACGTGGCGACCCGCTG</td><td>>hg38_21_+_8433172_8446622_RNA45SN1_45S_50ntFlanks</td><td>7981</td><td>8006</td><td>26</td><td>CTCAGATCAGACGTGGCGACCCGCTG</td><td>3048.6</td><td>-6.8</td><td>0.6</td><td>-12.1</td><td>1.2E-33</td><td>2.9E-32</td><td>BT20/MB468</td></tr>
<tr><td>>hg38_21_+_8433172_8446622_RNA45SN1_45S_50ntFlanks@6649.6695.47|TACGACTCTTAGCGGTGGATCACTCGGCTCGTGCGTCGATGAAGAAC</td><td>>hg38_21_+_8433172_8446622_RNA45SN1_45S_50ntFlanks</td><td>6649</td><td>6695</td><td>47</td><td>TACGACTCTTAGCGGTGGATCACTCGGCTCGTGCGTCGATGAAGAAC</td><td>654.3</td><td>14.3</td><td>1.2</td><td>12.1</td><td>8.6E-34</td><td>2.9E-32</td><td>MB231/MB468</td></tr>
<tr><td>>MI0003560|hsa-mir-92b&WithFlank&1|+|155195171|155195278@67.88.22&[MIMAT0003218&hsa-miR-92b-3p&offsets|0|0;m-115&1|+|155195237|155195258&offsets|0|0]|TATTGCACTCGTCCCGGCCTCC</td><td>>MI0003560|hsa-mir-92b&WithFlank&1|+|155195171|155195278</td><td>67</td><td>88</td><td>22</td><td>TATTGCACTCGTCCCGGCCTCC</td><td>10282.2</td><td>8.8</td><td>0.7</td><td>12.1</td><td>1.3E-33</td><td>3.1E-32</td><td>BT20/MB468</td></tr>
<tr><td>>hg38_21_+_8433172_8446622_RNA45SN1_45S_50ntFlanks@12627.12654.28|ATGGGGCGAAGCTACCATCTGTGGGATT</td><td>>hg38_21_+_8433172_8446622_RNA45SN1_45S_50ntFlanks</td><td>12627</td><td>12654</td><td>28</td><td>ATGGGGCGAAGCTACCATCTGTGGGATT</td><td>6851.7</td><td>-9.7</td><td>0.8</td><td>-12.1</td><td>1.3E-33</td><td>3.1E-32</td><td>BT20/MB468</td></tr>
<tr><td>>hg38_21_+_8433172_8446622_RNA45SN1_45S_50ntFlanks@12935.12956.22|TAAACCATTCGTAGACGACCTG</td><td>>hg38_21_+_8433172_8446622_RNA45SN1_45S_50ntFlanks</td><td>12935</td><td>12956</td><td>22</td><td>TAAACCATTCGTAGACGACCTG</td><td>1707.9</td><td>-7.3</td><td>0.6</td><td>-12.1</td><td>1.5E-33</td><td>3.5E-32</td><td>BT20/MB468</td></tr>
<tr><td>>hg38_21_+_8433172_8446622_RNA45SN1_45S_50ntFlanks@12995.13037.43|CTCGCTGCGATCTATTGAAAGTCAGCCCTCGACACAAGGGTTT</td><td>>hg38_21_+_8433172_8446622_RNA45SN1_45S_50ntFlanks</td><td>12995</td><td>13037</td><td>43</td><td>CTCGCTGCGATCTATTGAAAGTCAGCCCTCGACACAAGGGTTT</td><td>2931.3</td><td>16.4</td><td>1.4</td><td>12.1</td><td>1.0E-33</td><td>3.6E-32</td><td>MB231/MB468</td></tr>
<tr><td>>hg38_21_+_8433172_8446622_RNA45SN1_45S_50ntFlanks@11668.11687.20|TGCCCAGTGCTCTGAATGTC</td><td>>hg38_21_+_8433172_8446622_RNA45SN1_45S_50ntFlanks</td><td>11668</td><td>11687</td><td>20</td><td>TGCCCAGTGCTCTGAATGTC</td><td>3044.3</td><td>7.4</td><td>0.6</td><td>12.1</td><td>1.1E-33</td><td>3.8E-32</td><td>MB231/MB468</td></tr>
<tr><td>>hg38_21_+_8433172_8446622_RNA45SN1_45S_50ntFlanks@12415.12450.36|CTTCGATGTCGGCTCTTCCTATCATTGTGAAGCAGA</td><td>>hg38_21_+_8433172_8446622_RNA45SN1_45S_50ntFlanks</td><td>12415</td><td>12450</td><td>36</td><td>CTTCGATGTCGGCTCTTCCTATCATTGTGAAGCAGA</td><td>404.2</td><td>5.2</td><td>0.4</td><td>12.1</td><td>1.1E-33</td><td>3.8E-32</td><td>MB231/MB468</td></tr>
<tr><td>>MI0000477|hsa-mir-146a&WithFlank&5|+|160485346|160485456@27.48.22&[MIMAT0000449&hsa-miR-146a-5p&offsets|0|0;m-129&5|+|160485372|160485393&offsets|0|0]|TGAGAACTGAATTCCATGGGTT</td><td>>MI0000477|hsa-mir-146a&WithFlank&5|+|160485346|160485456</td><td>27</td><td>48</td><td>22</td><td>TGAGAACTGAATTCCATGGGTT</td><td>49692.7</td><td>15.8</td><td>1.3</td><td>12.1</td><td>1.1E-33</td><td>3.8E-32</td><td>MB231/MB468</td></tr>
<tr><td>>trnaMT_SerGCT_MT_+_12207_12265@14.48.35|AGAACTGCTAACTCATGCCCCCATGTCTAACAACA</td><td>>trnaMT_SerGCT_MT_+_12207_12265</td><td>14</td><td>48</td><td>35</td><td>AGAACTGCTAACTCATGCCCCCATGTCTAACAACA</td><td>618.8</td><td>14.2</td><td>1.2</td><td>12.1</td><td>1.1E-33</td><td>3.8E-32</td><td>MB231/MB468</td></tr>
<tr><td>>MI0000088|hsa-mir-30a&WithFlank&6|-|71403545|71403627@12.34.23&[MIMAT0000087&hsa-miR-30a-5p&offsets|0|+1;m-15&6|-|71403595|71403616&offsets|0|+1]|TGTAAACATCCTCGACTGGAAGC</td><td>>MI0000088|hsa-mir-30a&WithFlank&6|-|71403545|71403627</td><td>12</td><td>34</td><td>23</td><td>TGTAAACATCCTCGACTGGAAGC</td><td>112363.2</td><td>8.4</td><td>0.7</td><td>12.1</td><td>1.1E-33</td><td>3.8E-32</td><td>MB231/MB468</td></tr>
<tr><td>>hg38_21_+_8433172_8446622_RNA45SN1_45S_50ntFlanks@475.514.40|CTTCGTGATCGATGTGGTGACGTCGTGCTCTCCCGGGCCG</td><td>>hg38_21_+_8433172_8446622_RNA45SN1_45S_50ntFlanks</td><td>475</td><td>514</td><td>40</td><td>CTTCGTGATCGATGTGGTGACGTCGTGCTCTCCCGGGCCG</td><td>4753.1</td><td>8.4</td><td>0.7</td><td>12.1</td><td>1.2E-33</td><td>3.9E-32</td><td>MB231/MB468</td></tr>
<tr><td>>MI0000088|hsa-mir-30a&WithFlank&6|-|71403545|71403627@54.73.20(+1U)&[MIMAT0000088&hsa-miR-30a-3p&offsets|+1|-1(+1U);m-38&6|-|71403554|71403575&offsets|+1|-1(+1U)]|TTTCAGTCGGATGTTTGCAGT</td><td>>MI0000088|hsa-mir-30a&WithFlank&6|-|71403545|71403627</td><td>54</td><td>73</td><td>20(+1U)</td><td>TTTCAGTCGGATGTTTGCAGT</td><td>4034.2</td><td>8.3</td><td>0.7</td><td>12.1</td><td>1.9E-33</td><td>4.5E-32</td><td>BT20/MB468</td></tr>
<tr><td>>hg38_21_+_8433172_8446622_RNA45SN1_45S_50ntFlanks@6698.6718.21|AGCTAGCTGCGAGAATTAATG</td><td>>hg38_21_+_8433172_8446622_RNA45SN1_45S_50ntFlanks</td><td>6698</td><td>6718</td><td>21</td><td>AGCTAGCTGCGAGAATTAATG</td><td>5131.3</td><td>-6.4</td><td>0.5</td><td>-12.1</td><td>1.9E-33</td><td>4.5E-32</td><td>BT20/MB468</td></tr>
<tr><td>>hg38_21_+_8433172_8446622_RNA45SN1_45S_50ntFlanks@12007.12036.30|GCCGCCGGTGAAATACCACTACTCTGATCG</td><td>>hg38_21_+_8433172_8446622_RNA45SN1_45S_50ntFlanks</td><td>12007</td><td>12036</td><td>30</td><td>GCCGCCGGTGAAATACCACTACTCTGATCG</td><td>345.2</td><td>5.7</td><td>0.5</td><td>12.1</td><td>1.4E-33</td><td>4.6E-32</td><td>MB231/MB468</td></tr>
</table>

| | | | | | | | | | | | | |
|---|---|---|---|---|---|---|---|---|---|---|---|---|
| >trna119_LysCTT_1_-_145395522_145395594@1.30.30,trna11_LysCTT_5_-_180648979_180649051@1.30.30,trna13_LysCTT_6_+_26556774_26556846@1.30.30,trna32_LysCTT_16_-_3207406_3207478@1.30.30,trna7_LysCTT_16_+_3225692_3225764@1.30.30,trna9_LysCTT_5_+_180634755_180634827@1.30.30\|GCCCGGCTAGCTCAGTCGGTAGAGCATGAG | >trna119_LysCTT_1_-_145395522_145395594 | 1 | 30 | 30 | GCCCGGCTAGCTCAGTCGGTAGAGCATGAG | 13346.6 | -7.8 | 0.6 | -12.0 | 2.0E-33 | 4.6E-32 | BT20/MB468 |
| >MI0003643\|hsa-mir-629&WithFlank&15\|-\|70079366\|70079474@28.49.22&[MIMAT0004810&hsa-miR-629-5p&offsets\|0\|+1;m-150&15\|-\|70079427\|70079447&offsets\|0\|+1]\|TGGGTTTACGTTGGGAGAACTT | >MI0003643\|hsa-mir-629&WithFlank&15\|-\|70079366\|70079474 | 28 | 49 | 22 | TGGGTTTACGTTGGGAGAACTT | 1693.9 | 15.6 | 1.3 | 12.1 | 1.5E-33 | 4.8E-32 | MB231/MB468 |
| >hg38_MT_+_1621_3279_RNR2_16S_50ntFlanks@51.92.42\|GCTAAACCTAGCCCCAAACCCACTCCACCTTACTACCAGACA | >hg38_MT_+_1621_3279_RNR2_16S_50ntFlanks | 51 | 92 | 42 | GCTAAACCTAGCCCCAAACCCACTCCACCTTACTACCAGACA | 2997.1 | 16.5 | 1.4 | 12.1 | 1.5E-33 | 4.9E-32 | MB231/MB468 |
| >hg38_21_+_8433172_8446622_RNA45SN1_45S_50ntFlanks@4724.4758.35\|ATTAATCAAGAACGAAAGTCGGAGGTTCGAAGACG | >hg38_21_+_8433172_8446622_RNA45SN1_45S_50ntFlanks | 4724 | 4758 | 35 | ATTAATCAAGAACGAAAGTCGGAGGTTCGAAGACG | 4586.9 | -6.2 | 0.5 | -12.0 | 2.2E-33 | 5.0E-32 | BT20/MB468 |
| >MI0000077\|hsa-mir-21&WithFlank&17\|+\|59841260\|59841343@14.36.23(+1C)&[MIMAT0000076&hsa-miR-21-5p&offsets\|0\|+1(+1C);m-1&17\|+\|59841273\|59841294&offsets\|0\|+1(+1C)]\|TAGCTTATCAGACTGATGTTGACC | >MI0000077\|hsa-mir-21&WithFlank&17\|+\|59841260\|59841343 | 14 | 36 | 23(+1C) | TAGCTTATCAGACTGATGTTGACC | 93439.0 | 8.0 | 0.7 | 12.0 | 2.4E-33 | 5.5E-32 | BT20/MB468 |
| >MI0003560\|hsa-mir-92b&WithFlank&1\|+\|155195171\|155195278@67.87.21(+1A)&[MIMAT0003218&hsa-miR-92b-3p&offsets\|0\|-1(+1A);m-115&1\|+\|155195237\|155195258&offsets\|0\|-1(+1A)]\|TATTGCACTCGTCCCGGCCTCA | >MI0003560\|hsa-mir-92b&WithFlank&1\|+\|155195171\|155195278 | 67 | 87 | 21(+1A) | TATTGCACTCGTCCCGGCCTCA | 1764.8 | 9.0 | 0.7 | 12.1 | 1.8E-33 | 5.7E-32 | MB231/MB468 |
| >trnaMT_GluTTC_MT_-_14674_14742@35.71.37,trnalookalike8_GluTTC_5_-_93905172_93905240@35.71.37\|TATCATTGGTCGTGGTTGTAGTCCGTGCGAGAATACC | >trnaMT_GluTTC_MT_-_14674_14742 | 35 | 71 | 37 | TATCATTGGTCGTGGTTGTAGTCCGTGCGAGAATACC | 1483.3 | 15.4 | 1.3 | 12.0 | 2.3E-33 | 7.3E-32 | MB231/MB468 |
| >trnaMT_ValTAC_MT_+_1602_1670@46.71.26\|TTCAACTTAACTTGACCGCTCTGACC | >trnaMT_ValTAC_MT_+_1602_1670 | 46 | 71 | 26 | TTCAACTTAACTTGACCGCTCTGACC | 420.0 | 11.2 | 0.9 | 12.0 | 2.3E-33 | 7.4E-32 | MB231/MB468 |
| >MI0000253\|hsa-mir-148a&WithFlank&7\|-\|25949913\|25949992@50.71.22(+1U)&[MIMAT0000243&hsa-miR-148a-3p&offsets\|0\|0(+1U);m-7&7\|-\|25949922\|25949943&offsets\|0\|0(+1U)]\|TCAGTGCACTACAGAACTTTGTT | >MI0000253\|hsa-mir-148a&WithFlank&7\|-\|25949913\|25949992 | 50 | 71 | 22(+1U) | TCAGTGCACTACAGAACTTTGTT | 4941.4 | 9.1 | 0.8 | 12.0 | 3.5E-33 | 8.0E-32 | BT20/MB468 |
| >MI0000088\|hsa-mir-30a&WithFlank&6\|-\|71403545\|71403627@12.36.25&[MIMAT0000087&hsa-miR-30a-5p&offsets\|0\|+3;m-15&6\|-\|71403595\|71403616&offsets\|0\|+3]\|TGTAAACATCCTCGACTGGAAGCTG | >MI0000088\|hsa-mir-30a&WithFlank&6\|-\|71403545\|71403627 | 12 | 36 | 25 | TGTAAACATCCTCGACTGGAAGCTG | 1363.4 | 9.3 | 0.8 | 12.0 | 2.8E-33 | 8.8E-32 | MB231/MB468 |
| >hg38_21_+_8433172_8446622_RNA45SN1_45S_50ntFlanks@7983.8006.24\|CAGATCAGACGTGGCGACCCGCTG | >hg38_21_+_8433172_8446622_RNA45SN1_45S_50ntFlanks | 7983 | 8006 | 24 | CAGATCAGACGTGGCGACCCGCTG | 5301.1 | -8.0 | 0.7 | -12.0 | 2.8E-33 | 8.9E-32 | MB231/MB468 |
| >MI0000266\|hsa-mir-10a&WithFlank&17\|-\|48579832\|48579953@29.51.23&[MIMAT0000253&hsa-miR-10a-5p&offsets\|+1\|+1;m-18&17\|-\|48579905\|48579926&offsets\|+1\|+2]\|ACCCTGTAGATCCGAATTTGTGT | >MI0000266\|hsa-mir-10a&WithFlank&17\|-\|48579832\|48579953 | 29 | 51 | 23 | ACCCTGTAGATCCGAATTTGTGT | 3001.3 | 16.5 | 1.4 | 12.0 | 2.9E-33 | 9.0E-32 | MB231/MB468 |
| >hg38_21_+_8433172_8446622_RNA45SN1_45S_50ntFlanks@12996.13035.40\|TCGCTGCGATCTATTGAAAGTCAGCCCTCGACACAAGGGT | >hg38_21_+_8433172_8446622_RNA45SN1_45S_50ntFlanks | 12996 | 13035 | 40 | TCGCTGCGATCTATTGAAAGTCAGCCCTCGACACAAGGGT | 403.9 | 11.1 | 0.9 | 12.0 | 2.9E-33 | 9.1E-32 | MB231/MB468 |
| >MI0000088\|hsa-mir-30a&WithFlank&6\|-\|71403545\|71403627@12.35.24&[MIMAT0000087&hsa-miR-30a-5p&offsets\|0\|+2;m-15&6\|-\|71403595\|71403616&offsets\|0\|+2]\|TGTAAACATCCTCGACTGGAAGCT | >MI0000088\|hsa-mir-30a&WithFlank&6\|-\|71403545\|71403627 | 12 | 35 | 24 | TGTAAACATCCTCGACTGGAAGCT | 505795.3 | 8.8 | 0.7 | 12.0 | 3.0E-33 | 9.5E-32 | MB231/MB468 |
| >hg38_21_+_8433172_8446622_RNA45SN1_45S_50ntFlanks@10350.10384.35\|ACGAGAACTTTGAAGGCCGAAGTGGAGAAGGGTTC | >hg38_21_+_8433172_8446622_RNA45SN1_45S_50ntFlanks | 10350 | 10384 | 35 | ACGAGAACTTTGAAGGCCGAAGTGGAGAAGGGTTC | 1180.2 | -5.3 | 0.4 | -12.0 | 4.2E-33 | 9.6E-32 | BT20/MB468 |
| >hg38_21_+_8433172_8446622_RNA45SN1_45S_50ntFlanks@12388.12433.46\|CGTTCATAGCGACGTCGCTTTTTGATCCTTCGATGTCGGCTCTTCC | >hg38_21_+_8433172_8446622_RNA45SN1_45S_50ntFlanks | 12388 | 12433 | 46 | CGTTCATAGCGACGTCGCTTTTTGATCCTTCGATGTCGGCTCTTCC | 592.4 | 14.1 | 1.2 | 12.0 | 3.1E-33 | 9.7E-32 | MB231/MB468 |
| >hg38_21_+_8433172_8446622_RNA45SN1_45S_50ntFlanks@11785.11804.20\|TGAATGGATGAACGAGATTC | >hg38_21_+_8433172_8446622_RNA45SN1_45S_50ntFlanks | 11785 | 11804 | 20 | TGAATGGATGAACGAGATTC | 9957.4 | -7.5 | 0.6 | -12.0 | 4.2E-33 | 9.7E-32 | BT20/MB468 |
| >hg38_21_+_8433172_8446622_RNA45SN1_45S_50ntFlanks@12456.12485.30\|CCAAGCGTTGGATTGTTCACCCACTAATAG | >hg38_21_+_8433172_8446622_RNA45SN1_45S_50ntFlanks | 12456 | 12485 | 30 | CCAAGCGTTGGATTGTTCACCCACTAATAG | 3311.3 | -6.8 | 0.6 | -12.0 | 3.2E-33 | 9.8E-32 | MB231/MB468 |
| >hg38_21_+_8433172_8446622_RNA45SN1_45S_50ntFlanks@7975.8016.42\|CGCGACCTCAGATCAGACGTGGCGACCCGCTGAATTTAAGCA | >hg38_21_+_8433172_8446622_RNA45SN1_45S_50ntFlanks | 7975 | 8016 | 42 | CGCGACCTCAGATCAGACGTGGCGACCCGCTGAATTTAAGCA | 22607.5 | 8.8 | 0.7 | 12.0 | 3.3E-33 | 1.0E-31 | MB231/MB468 |
| >MI0000266\|hsa-mir-10a&WithFlank&17\|-\|48579832\|48579953@29.51.23&[MIMAT0000253&hsa-miR-10a-5p&offsets\|+1\|+1;m-18&17\|-\|48579905\|48579926&offsets\|+1\|+2]\|ACCCTGTAGATCCGAATTTGTGT | >MI0000266\|hsa-mir-10a&WithFlank&17\|-\|48579832\|48579953 | 29 | 51 | 23 | ACCCTGTAGATCCGAATTTGTGT | 2945.2 | 16.2 | 1.3 | 12.0 | 4.5E-33 | 1.0E-31 | BT20/MB468 |
| >am_MI0000809\|hsa-mir-151a&WithFlank&8\|-\|140732558\|140732659@53.72.20&[MIMAT0000757&hsa-miR-151a-3p&offsets\|0\|-1;m-30&8\|-\|140732587\|140732607&offsets\|0\|-1]\|CTAGACTGAAGCTCCTTGAG | >am_MI0000809\|hsa-mir-151a&WithFlank&8\|-\|140732558\|140732659 | 53 | 72 | 20 | CTAGACTGAAGCTCCTTGAG | 5594.9 | 8.8 | 0.7 | 12.0 | 3.3E-33 | 1.0E-31 | MB231/MB468 |
| >MI0000298\|hsa-mir-221&WithFlank&X\|-\|45746151\|45746272@71.93.23(+1U)&[MIMAT0000278&hsa-miR-221-3p&offsets\|0\|0(+1U);m-91&X\|-\|45746181\|45746202&offsets\|0\|+1(+1U)]\|AGCTACATTGTCTGCTGGGTTTCT | >MI0000298\|hsa-mir-221&WithFlank&X\|-\|45746151\|45746272 | 71 | 93 | 23(+1U) | AGCTACATTGTCTGCTGGGTTTCT | 59574.6 | 8.9 | 0.7 | 12.0 | 3.4E-33 | 1.0E-31 | MB231/MB468 |
| >MI0000088\|hsa-mir-30a&WithFlank&6\|-\|71403545\|71403627@53.73.21&[MIMAT0000088&hsa-miR-30a-3p&offsets\|0\|-1;m-38&6\|-\|71403554\|71403575&offsets\|0\|-1]\|CTTTCAGTCGGATGTTTGCAG | >MI0000088\|hsa-mir-30a&WithFlank&6\|-\|71403545\|71403627 | 53 | 73 | 21 | CTTTCAGTCGGATGTTTGCAG | 3501.7 | 9.0 | 0.7 | 12.0 | 3.8E-33 | 1.1E-31 | MB231/MB468 |
| >MI0000746\|hsa-mir-99b&WithFlank&19\|+\|51692606\|51692687@13.32.20(+1A)&[MIMAT0000689&hsa-miR-99b-5p&offsets\|0\|-2(+1A);m-10&19\|+\|51692618\|51692639&offsets\|0\|-2(+1A)]\|CACCCGTAGAACCGACCTTGA | >MI0000746\|hsa-mir-99b&WithFlank&19\|+\|51692606\|51692687 | 13 | 32 | 20(+1A) | CACCCGTAGAACCGACCTTGA | 908.4 | 8.2 | 0.7 | 12.0 | 3.8E-33 | 1.1E-31 | MB231/MB468 |
| >MI0000063\|hsa-let-7b&WithFlank&22\|+\|46113680\|46113774@12.34.23&[MIMAT0000063&hsa-let-7b-5p&offsets\|0\|+1;m-14&22\|+\|46113691\|46113712&offsets\|0\|+1]\|TGAGGTAGTAGGTTGTGTGGTTT | >MI0000063\|hsa-let-7b&WithFlank&22\|+\|46113680\|46113774 | 12 | 34 | 23 | TGAGGTAGTAGGTTGTGTGGTTT | 13303.9 | 8.0 | 0.7 | 12.0 | 5.1E-33 | 1.1E-31 | BT20/MB468 |
| >hg38_21_+_8433172_8446622_RNA45SN1_45S_50ntFlanks@11729.11749.21\|TAACTATGACTCTCTTAAGGT | >hg38_21_+_8433172_8446622_RNA45SN1_45S_50ntFlanks | 11729 | 11749 | 21 | TAACTATGACTCTCTTAAGGT | 6271.0 | -6.6 | 0.6 | -12.0 | 5.3E-33 | 1.2E-31 | BT20/MB468 |
| >MI0000439\|hsa-mir-23b&WithFlank&9\|+\|95085202\|95085310@67.86.20&[MIMAT0000418&hsa-miR-23b-3p&offsets\|+3\|0;m-42&9\|+\|95085265\|95085285&offsets\|+3\|+2]\|ACATTGCCAGGGATTACCAC | >MI0000439\|hsa-mir-23b&WithFlank&9\|+\|95085202\|95085310 | 67 | 86 | 20 | ACATTGCCAGGGATTACCAC | 6767.1 | 17.4 | 1.5 | 12.0 | 5.6E-33 | 1.3E-31 | BT20/MB468 |
| >trna116_GluCTC_1_-_145399233_145399304@1.35.35,trna59_GluCTC_1_+_249168447_249168518@1.35.35,trna71_GluCTC_1_-_161439189_161439260@1.35.35,trna74_GluCTC_1_-_161431809_161431880@1.35.35,trna77_GluCTC_1_-_161424398_161424469@1.35.35,trna77_GluCTC_6_+_28949976_28950047@1.35.35,trna80_GluCTC_1_-_161417018_161417089@1.35.35,trna87_GluCTC_6_-_126101393_126101464@1.35.35\|TCCCTGGTGGTCTAGTGGTTAGGATTCGGCGCTCT | >trna116_GluCTC_1_-_145399233_145399304 | 1 | 35 | 35 | TCCCTGGTGGTCTAGTGGTTAGGATTCGGCGCTCT | 10689.9 | 9.8 | 0.8 | 12.0 | 4.2E-33 | 1.3E-31 | MB231/MB468 |
| >MI0006273\|hsa-mir-1180&WithFlank&17\|-\|19344500\|19344580@47.66.20&[MIMAT0005825&hsa-miR-1180-3p&offsets\|0\|-2;m-214&17\|-\|19344513\|19344534&offsets\|0\|-2]\|TTTCCGGCTCGCGTGGGTGT | >MI0006273\|hsa-mir-1180&WithFlank&17\|-\|19344500\|19344580 | 47 | 66 | 20 | TTTCCGGCTCGCGTGGGTGT | 734.5 | 7.3 | 0.6 | 12.0 | 5.7E-33 | 1.3E-31 | BT20/MB468 |
| >hg38_21_+_8433172_8446622_RNA45SN1_45S_50ntFlanks@12995.13039.45\|CTCGCTGCGATCTATTGAAAGTCAGCCCTCGACACAAGGGTTTGT | >hg38_21_+_8433172_8446622_RNA45SN1_45S_50ntFlanks | 12995 | 13039 | 45 | CTCGCTGCGATCTATTGAAAGTCAGCCCTCGACACAAGGGTTTGT | 2398039.9 | 13.0 | 1.1 | 12.0 | 4.6E-33 | 1.4E-31 | MB231/MB468 |
| >hg38_21_+_8433172_8446622_RNA45SN1_45S_50ntFlanks@12627.12653.27\|ATGGGGCGAAGCTACCATCTGTGGGAT | >hg38_21_+_8433172_8446622_RNA45SN1_45S_50ntFlanks | 12627 | 12653 | 27 | ATGGGGCGAAGCTACCATCTGTGGGAT | 2955.5 | -8.5 | 0.7 | -12.0 | 4.7E-33 | 1.4E-31 | MB231/MB468 |
| >MI0000298\|hsa-mir-221&WithFlank&X\|-\|45746151\|45746272@31.54.24&[MIMAT0004568&hsa-miR-221-5p&offsets\|0\|+2;m-405&X\|-\|45746220\|45746242&offsets\|0\|+1]\|ACCTGGCATACAATGTAGATTTCT | >MI0000298\|hsa-mir-221&WithFlank&X\|-\|45746151\|45746272 | 31 | 54 | 24 | ACCTGGCATACAATGTAGATTTCT | 9597.1 | 9.2 | 0.8 | 12.0 | 4.7E-33 | 1.4E-31 | MB231/MB468 |
| >trna119_LysCTT_1_-_145395522_145395594@1.35.35,trna11_LysCTT_5_-_180648979_180649051@1.35.35,trna13_LysCTT_6_+_26556774_26556846@1.35.35,trna7_LysCTT_16_+_3225692_3225764@1.35.35,trna9_LysCTT_5_+_180634755_180634827@1.35.35\|GCCCGGCTAGCTCAGTCGGTAGAGCATGAGACTCT | >trna119_LysCTT_1_-_145395522_145395594 | 1 | 35 | 35 | GCCCGGCTAGCTCAGTCGGTAGAGCATGAGACTCT | 7800.1 | 9.2 | 0.8 | 12.0 | 5.0E-33 | 1.5E-31 | MB231/MB468 |
| >hg38_21_+_8433172_8446622_RNA45SN1_45S_50ntFlanks@476.511.36\|TTCGTGATCGATGTGGTGACGTCGTGCTCTCCCGGG | >hg38_21_+_8433172_8446622_RNA45SN1_45S_50ntFlanks | 476 | 511 | 36 | TTCGTGATCGATGTGGTGACGTCGTGCTCTCCCGGG | 4474.3 | 8.4 | 0.7 | 12.0 | 5.2E-33 | 1.5E-31 | MB231/MB468 |
| >am_MI0000809\|hsa-mir-151a&WithFlank&8\|-\|140732558\|140732659@52.73.22&[MIMAT0000757&hsa-miR-151a-3p&offsets\|-1\|0;m-30&8\|-\|140732587\|140732607&offsets\|-1\|0]\|ACTAGACTGAAGCTCCTTGAGG | >am_MI0000809\|hsa-mir-151a&WithFlank&8\|-\|140732558\|140732659 | 52 | 73 | 22 | ACTAGACTGAAGCTCCTTGAGG | 3227.9 | 8.4 | 0.7 | 12.0 | 5.3E-33 | 1.5E-31 | MB231/MB468 |
| >hg38_21_+_8433172_8446622_RNA45SN1_45S_50ntFlanks@6650.6696.47\|ACGACTCTTAGCGGTGGATCACTCGGCTCGTGCGTCGATGAAGAACG | >hg38_21_+_8433172_8446622_RNA45SN1_45S_50ntFlanks | 6650 | 6696 | 47 | ACGACTCTTAGCGGTGGATCACTCGGCTCGTGCGTCGATGAAGAACG | 2725.3 | 8.2 | 0.7 | 12.0 | 5.6E-33 | 1.6E-31 | MB231/MB468 |
| >hg38_1_-_228634819_228635039_RNA5S12_5S_50ntFlanks@134.169.36\|TGGGAGACCGCCTGGGAATACCGGGTGCTGTAGGCT | >hg38_1_-_228634819_228635039_RNA5S12_5S_50ntFlanks | 134 | 169 | 36 | TGGGAGACCGCCTGGGAATACCGGGTGCTGTAGGCT | 19182.3 | 8.2 | 0.7 | 12.0 | 5.8E-33 | 1.7E-31 | MB231/MB468 |
| >hg38_21_+_8433172_8446622_RNA45SN1_45S_50ntFlanks@7974.8011.38\|ACGCGACCTCAGATCAGACGTGGCGACCCGCTGAATTT | >hg38_21_+_8433172_8446622_RNA45SN1_45S_50ntFlanks | 7974 | 8011 | 38 | ACGCGACCTCAGATCAGACGTGGCGACCCGCTGAATTT | 19775.4 | 7.7 | 0.6 | 12.0 | 6.0E-33 | 1.7E-31 | MB231/MB468 |

<table>
<tr><td>>am_trnaMT_ProTGG_MT_-_15956_16023@52.71.20|AAAGACTTTTTCTCTGACCA</td><td>>am_trnaMT_ProTGG_MT_-_15956_16023</td><td>52</td><td>71</td><td>20</td><td>AAAGACTTTTTCTCTGACCA</td><td>12728.6</td><td>9.9</td><td>0.8</td><td>12.0</td><td>6.2E-33</td><td>1.8E-31</td><td>MB231/MB468</td></tr>
<tr><td>>MI0000077|hsa-mir-21&WithFlank&17|+|59841260|59841343@14.36.23&[MIMAT0000076&hsa-miR-21-5p&offsets|0|+1;m-1&17|+|59841273|59841294&offsets|0|+1]|TAGCTTATCAGACTGATGTTGAC</td><td>>MI0000077|hsa-mir-21&WithFlank&17|+|59841260|59841343</td><td>14</td><td>36</td><td>23</td><td>TAGCTTATCAGACTGATGTTGAC</td><td>2544052.4</td><td>7.6</td><td>0.6</td><td>11.9</td><td>8.4E-33</td><td>1.9E-31</td><td>BT20/MB468</td></tr>
<tr><td>>am_MI0005763|hsa-mir-941-1&WithFlank&20|+|63919443|63919526@53.74.22(+1A);MI0005764|hsa-mir-941-2&WithFlank&20|+|63919499|63919582@53.74.22(+1A);MI0005765|hsa-mir-941-3&WithFlank&20|+|63919555|63919638@53.74.22(+1A);MI0005766|hsa-mir-941-4&WithFlank&20|+|63919750|63919833@53.74.22(+1A);MI0031520|hsa-mir-941-5&WithFlank&20|+|63919862|63919945@53.74.22(+1A)&[MIMAT0004984&hsa-miR-941&offsets|0|-1(+1A);m-263&20|+|63919495|63919517&offsets|0|-1(+1A)];[MIMAT0004984_1&hsa-miR-941&offsets|0|-1(+1A);m-263&20|+|63919551|63919573&offsets|0|-1(+1A)];[MIMAT0004984_2&hsa-miR-941&offsets|0|-1(+1A);m-263&20|+|63919607|63919629&offsets|0|-1(+1A)];[MIMAT0004984_3&hsa-miR-941&offsets|0|-1(+1A);m-263&20|+|63919802|63919824&offsets|0|-1(+1A)];[MIMAT0004984_4&hsa-miR-941&offsets|0|-1(+1A);m-263&20|+|63919914|63919936&offsets|0|-1(+1A)]|CACCCGGCTGTGTGCACATGTGA</td><td>>am_MI0005763|hsa-mir-941-1&WithFlank&20|+|63919443|63919526</td><td>53</td><td>74</td><td>22(+1A)</td><td>CACCCGGCTGTGTGCACATGTGA</td><td>2186.9</td><td>8.9</td><td>0.7</td><td>11.9</td><td>6.7E-33</td><td>1.9E-31</td><td>MB231/MB468</td></tr>
<tr><td>>hg38_21_+_8433172_8446622_RNA45SN1_45S_50ntFlanks@12419.12462.44|GATGTCGGCTCTTCCTATCATTGTGAAGCAGAATTCACCAAGCG</td><td>>hg38_21_+_8433172_8446622_RNA45SN1_45S_50ntFlanks</td><td>12419</td><td>12462</td><td>44</td><td>GATGTCGGCTCTTCCTATCATTGTGAAGCAGAATTCACCAAGCG</td><td>392.3</td><td>13.5</td><td>1.1</td><td>11.9</td><td>6.7E-33</td><td>1.9E-31</td><td>MB231/MB468</td></tr>
<tr><td>>MI0000466|hsa-mir-9-1&WithFlank&1|-|156420335|156420435@22.43.22;MI0000468|hsa-mir-9-3&WithFlank&15|+|89368011|89368112@22.43.22;MI0000467|hsa-mir-9-2&WithFlank&5|-|88666847|88666945@22.43.22&[MIMAT0000441&hsa-miR-9-5p&offsets|0|-1;m-104&1|-|156420392|156420414&offsets|0|-1];[MIMAT0000441_1&hsa-miR-9-5p&offsets|0|-1;m-105&15|+|89368032|89368054&offsets|0|-1];[MIMAT0000441_2&hsa-miR-9-5p&offsets|0|-1;m-104&5|-|88666902|88666924&offsets|0|-1]|TCTTTGGTTATCTAGCTGTATG</td><td>>MI0000466|hsa-mir-9-1&WithFlank&1|-|156420335|156420435</td><td>22</td><td>43</td><td>22</td><td>TCTTTGGTTATCTAGCTGTATG</td><td>7201.5</td><td>10.3</td><td>0.9</td><td>11.9</td><td>8.6E-33</td><td>1.9E-31</td><td>BT20/MB468</td></tr>
<tr><td>>hg38_21_+_8433172_8446622_RNA45SN1_45S_50ntFlanks@6645.6677.33|CTCGTACGACTCTTAGCGGTGGATCACTCGGCT</td><td>>hg38_21_+_8433172_8446622_RNA45SN1_45S_50ntFlanks</td><td>6645</td><td>6677</td><td>33</td><td>CTCGTACGACTCTTAGCGGTGGATCACTCGGCT</td><td>662.4</td><td>5.3</td><td>0.4</td><td>11.9</td><td>8.6E-33</td><td>1.9E-31</td><td>BT20/MB468</td></tr>
<tr><td>>hg38_21_+_8433172_8446622_RNA45SN1_45S_50ntFlanks@476.510.35|TTCGTGATCGATGTGGTGACGTCGTGCTCTCCCGG</td><td>>hg38_21_+_8433172_8446622_RNA45SN1_45S_50ntFlanks</td><td>476</td><td>510</td><td>35</td><td>TTCGTGATCGATGTGGTGACGTCGTGCTCTCCCGG</td><td>6498.2</td><td>8.2</td><td>0.7</td><td>11.9</td><td>7.2E-33</td><td>2.0E-31</td><td>MB231/MB468</td></tr>
<tr><td>>hg38_21_+_8433172_8446622_RNA45SN1_45S_50ntFlanks@11669.11709.41|GCCCAGTGCTCTGAATGTCAAAGTGAAGAAATTCAATGAAG</td><td>>hg38_21_+_8433172_8446622_RNA45SN1_45S_50ntFlanks</td><td>11669</td><td>11709</td><td>41</td><td>GCCCAGTGCTCTGAATGTCAAAGTGAAGAAATTCAATGAAG</td><td>401.5</td><td>13.6</td><td>1.1</td><td>11.9</td><td>7.3E-33</td><td>2.0E-31</td><td>MB231/MB468</td></tr>
<tr><td>>am_MI0005763|hsa-mir-941-1&WithFlank&20|+|63919443|63919526@53.75.23(+1U);MI0005764|hsa-mir-941-2&WithFlank&20|+|63919499|63919582@53.75.23(+1U);MI0005765|hsa-mir-941-3&WithFlank&20|+|63919555|63919638@53.75.23(+1U);MI0005766|hsa-mir-941-4&WithFlank&20|+|63919750|63919833@53.75.23(+1U);MI0031520|hsa-mir-941-5&WithFlank&20|+|63919862|63919945@53.75.23(+1U)&[MIMAT0004984&hsa-miR-941&offsets|0|0(+1U);m-263&20|+|63919495|63919517&offsets|0|0(+1U)];[MIMAT0004984_1&hsa-miR-941&offsets|0|0(+1U);m-263&20|+|63919551|63919573&offsets|0|0(+1U)];[MIMAT0004984_2&hsa-miR-941&offsets|0|0(+1U);m-263&20|+|63919607|63919629&offsets|0|0(+1U)];[MIMAT0004984_3&hsa-miR-941&offsets|0|0(+1U);m-263&20|+|63919802|63919824&offsets|0|0(+1U)];[MIMAT0004984_4&hsa-miR-941&offsets|0|0(+1U);m-263&20|+|63919914|63919936&offsets|0|0(+1U)]|CACCCGGCTGTGTGCACATGTGCT</td><td>>am_MI0005763|hsa-mir-941-1&WithFlank&20|+|63919443|63919526</td><td>53</td><td>75</td><td>23(+1U)</td><td>CACCCGGCTGTGTGCACATGTGCT</td><td>2104.4</td><td>9.2</td><td>0.8</td><td>11.9</td><td>7.3E-33</td><td>2.0E-31</td><td>MB231/MB468</td></tr>
<tr><td>>MI0000077|hsa-mir-21&WithFlank&17|+|59841260|59841343@15.36.22&[MIMAT0000076&hsa-miR-21-5p&offsets|+1|+1;m-1&17|+|59841273|59841294&offsets|+1|+1]|AGCTTATCAGACTGATGTTGAC</td><td>>MI0000077|hsa-mir-21&WithFlank&17|+|59841260|59841343</td><td>15</td><td>36</td><td>22</td><td>AGCTTATCAGACTGATGTTGAC</td><td>4108.5</td><td>7.8</td><td>0.7</td><td>11.9</td><td>9.4E-33</td><td>2.1E-31</td><td>BT20/MB468</td></tr>
<tr><td>>MI0000087|hsa-mir-29a&WithFlank&7|-|130876741|130876816@47.67.21(+1C)&[MIMAT0000086&hsa-miR-29a-3p&offsets|-1|-2(+1C);m-26&7|-|130876749|130876769&offsets|-1|-1(+1C)]|CTAGCACCATCTGAAATCGGTC</td><td>>MI0000087|hsa-mir-29a&WithFlank&7|-|130876741|130876816</td><td>47</td><td>67</td><td>21(+1C)</td><td>CTAGCACCATCTGAAATCGGTC</td><td>3581.1</td><td>9.2</td><td>0.8</td><td>11.9</td><td>7.5E-33</td><td>2.1E-31</td><td>MB231/MB468</td></tr>
<tr><td>>hg38_21_+_8433172_8446622_RNA45SN1_45S_50ntFlanks@6646.6678.33|TCGTACGACTCTTAGCGGTGGATCACTCGGCTC</td><td>>hg38_21_+_8433172_8446622_RNA45SN1_45S_50ntFlanks</td><td>6646</td><td>6678</td><td>33</td><td>TCGTACGACTCTTAGCGGTGGATCACTCGGCTC</td><td>3138.1</td><td>5.3</td><td>0.4</td><td>11.9</td><td>9.5E-33</td><td>2.1E-31</td><td>BT20/MB468</td></tr>
<tr><td>>MI0000750|hsa-mir-26a-2&WithFlank&12|-|57824603|57824698@20.41.22(+2U);MI0000083|hsa-mir-26a-1&WithFlank&3|+|37969398|37969486@16.37.22(+2U)&[MIMAT0000082&hsa-miR-26a-5p&offsets|0|0(+2U);m-33&12|-|57824658|57824679&offsets|0|0(+2U)];[MIMAT0000082_1&hsa-miR-26a-5p&offsets|0|0(+2U);m-34&3|+|37969413|37969434&offsets|0|0(+2U)]|TTCAAGTAATCCAGGATAGGCTTT</td><td>>MI0000750|hsa-mir-26a-2&WithFlank&12|-|57824603|57824698</td><td>20</td><td>41</td><td>22(+2U)</td><td>TTCAAGTAATCCAGGATAGGCTTT</td><td>18492.8</td><td>9.4</td><td>0.8</td><td>11.9</td><td>8.2E-33</td><td>2.3E-31</td><td>MB231/MB468</td></tr>
<tr><td>>MI0000077|hsa-mir-21&WithFlank&17|+|59841260|59841343@14.36.23(+1C)&[MIMAT0000076&hsa-miR-21-5p&offsets|0|+1(+1C);m-1&17|+|59841273|59841294&offsets|0|+1(+1C)]|TAGCTTATCAGACTGATGTTGACC</td><td>>MI0000077|hsa-mir-21&WithFlank&17|+|59841260|59841343</td><td>14</td><td>36</td><td>23(+1C)</td><td>TAGCTTATCAGACTGATGTTGACC</td><td>158707.6</td><td>9.0</td><td>0.8</td><td>11.9</td><td>8.6E-33</td><td>2.4E-31</td><td>MB231/MB468</td></tr>
<tr><td>>MI0000102|hsa-mir-100&WithFlank&11|-|122152223|122152314@19.40.22(+2U)&[MIMAT0000098&hsa-miR-100-5p&offsets|0|0(+2U);m-27&11|-|122152276|122152296&offsets|0|+1(+2U)]|AACCCGTAGATCCGAACTTGTGTT</td><td>>MI0000102|hsa-mir-100&WithFlank&11|-|122152223|122152314</td><td>19</td><td>40</td><td>22(+2U)</td><td>AACCCGTAGATCCGAACTTGTGTT</td><td>2301.1</td><td>16.1</td><td>1.3</td><td>11.9</td><td>8.7E-33</td><td>2.4E-31</td><td>MB231/MB468</td></tr>
<tr><td>>hg38_21_+_8433172_8446622_RNA45SN1_45S_50ntFlanks@12456.12488.33|CCAAGCGTTGGATTGTTCACCCACTAATAGGGA</td><td>>hg38_21_+_8433172_8446622_RNA45SN1_45S_50ntFlanks</td><td>12456</td><td>12488</td><td>33</td><td>CCAAGCGTTGGATTGTTCACCCACTAATAGGGA</td><td>5261.2</td><td>-7.6</td><td>0.6</td><td>-11.9</td><td>8.9E-33</td><td>2.4E-31</td><td>MB231/MB468</td></tr>
<tr><td>>MI0000734|hsa-mir-106b&WithFlank&7|-|100093987|100094080@56.77.22&[MIMAT0004672&hsa-miR-106b-3p&offsets|-2|-2;m-87&7|-|100094003|100094023&offsets|-2|-1]|TACCGCACTGTGGGTACTTGCT</td><td>>MI0000734|hsa-mir-106b&WithFlank&7|-|100093987|100094080</td><td>56</td><td>77</td><td>22</td><td>TACCGCACTGTGGGTACTTGCT</td><td>4220.4</td><td>10.2</td><td>0.9</td><td>11.9</td><td>9.1E-33</td><td>2.5E-31</td><td>MB231/MB468</td></tr>
<tr><td>>MI0000088|hsa-mir-30a&WithFlank&6|-|71403545|71403627@12.33.22&[MIMAT0000087&hsa-miR-30a-5p&offsets|0|0;m-15&6|-|71403595|71403616&offsets|0|0]|TGTAAACATCCTCGACTGGAAG</td><td>>MI0000088|hsa-mir-30a&WithFlank&6|-|71403545|71403627</td><td>12</td><td>33</td><td>22</td><td>TGTAAACATCCTCGACTGGAAG</td><td>16012.8</td><td>8.0</td><td>0.7</td><td>11.9</td><td>9.6E-33</td><td>2.6E-31</td><td>MB231/MB468</td></tr>
<tr><td>>MI0000261|hsa-mir-139&WithFlank&11|-|72615057|72615136@13.35.23&[MIMAT0000250&hsa-miR-139-5p&offsets|0|0;m-186&11|-|72615102|72615124&offsets|0|0]|TCTACAGTGCACGTGTCTCCAGT</td><td>>MI0000261|hsa-mir-139&WithFlank&11|-|72615057|72615136</td><td>13</td><td>35</td><td>23</td><td>TCTACAGTGCACGTGTCTCCAGT</td><td>4817.2</td><td>14.7</td><td>1.2</td><td>11.9</td><td>9.8E-33</td><td>2.6E-31</td><td>MB231/MB468</td></tr>
<tr><td>>MI0000061|hsa-let-7a-2&WithFlank&11|-|122146516|122146599@11.30.20(+1C);MI0000062|hsa-let-7a-3&WithFlank&22|+|46112743|46112828@10.29.20(+1C);MI0000060|hsa-let-7a-1&WithFlank&9|+|94175951|94176042@12.31.20(+1C)&[MIMAT0000062&hsa-let-7a-5p&offsets|0|-2(+1C);m-6&11|-|122146568|122146589&offsets|0|-2(+1C)];[MIMAT0000062_1&hsa-let-7a-5p&offsets|0|-2(+1C);m-5&22|+|46112752|46112773&offsets|0|-2(+1C)];[MIMAT0000062_2&hsa-let-7a-5p&offsets|0|-2(+1C);m-4&9|+|94175962|94175983&offsets|0|-2(+1C)]|TGAGGTAGTAGGTTGTATAGC</td><td>>MI0000061|hsa-let-7a-2&WithFlank&11|-|122146516|122146599</td><td>11</td><td>30</td><td>20(+1C)</td><td>TGAGGTAGTAGGTTGTATAGC</td><td>3460.7</td><td>8.7</td><td>0.7</td><td>11.9</td><td>1.0E-32</td><td>2.7E-31</td><td>MB231/MB468</td></tr>
<tr><td>>MI0000088|hsa-mir-30a&WithFlank&6|-|71403545|71403627@53.73.21(+1A)&[MIMAT0000088&hsa-miR-30a-3p&offsets|0|-1(+1A);m-38&6|-|71403554|71403575&offsets|0|-1(+1A)]|CTTTCAGTCGGATGTTTGCAGA</td><td>>MI0000088|hsa-mir-30a&WithFlank&6|-|71403545|71403627</td><td>53</td><td>73</td><td>21(+1A)</td><td>CTTTCAGTCGGATGTTTGCAGA</td><td>1789.9</td><td>9.2</td><td>0.8</td><td>11.9</td><td>1.1E-32</td><td>2.8E-31</td><td>MB231/MB468</td></tr>
<tr><td>>hg38_21_+_8433172_8446622_RNA45SN1_45S_50ntFlanks@8357.8379.23|AAAAGAACTTTGAAGAGAGAGTT</td><td>>hg38_21_+_8433172_8446622_RNA45SN1_45S_50ntFlanks</td><td>8357</td><td>8379</td><td>23</td><td>AAAAGAACTTTGAAGAGAGAGTT</td><td>52591.5</td><td>-7.7</td><td>0.6</td><td>-11.9</td><td>1.1E-32</td><td>2.9E-31</td><td>MB231/MB468</td></tr>
<tr><td>>hg38_21_+_8433172_8446622_RNA45SN1_45S_50ntFlanks@474.497.24|TCTTCGTGATCGATGTGGTGACGT</td><td>>hg38_21_+_8433172_8446622_RNA45SN1_45S_50ntFlanks</td><td>474</td><td>497</td><td>24</td><td>TCTTCGTGATCGATGTGGTGACGT</td><td>3067.2</td><td>9.3</td><td>0.8</td><td>11.9</td><td>1.1E-32</td><td>2.9E-31</td><td>MB231/MB468</td></tr>
<tr><td>>hg38_21_+_8433172_8446622_RNA45SN1_45S_50ntFlanks@6647.6695.49|CGTACGACTCTTAGCGGTGGATCACTCGGCTCGTGCGTCGATGAAGAAC</td><td>>hg38_21_+_8433172_8446622_RNA45SN1_45S_50ntFlanks</td><td>6647</td><td>6695</td><td>49</td><td>CGTACGACTCTTAGCGGTGGATCACTCGGCTCGTGCGTCGATGAAGAAC</td><td>817.3</td><td>14.6</td><td>1.2</td><td>11.9</td><td>1.1E-32</td><td>2.9E-31</td><td>MB231/MB468</td></tr>
<tr><td>>am_MI0003581|hsa-mir-574&WithFlank&4|+|38868026|38868133@31.53.23&[MIMAT0004795&hsa-miR-574-5p&offsets|0|0;m-213&4|+|38868056|38868078&offsets|0|0]|TGAGTGTGTGTGTGTGAGTGTGT</td><td>>am_MI0003581|hsa-mir-574&WithFlank&4|+|38868026|38868133</td><td>31</td><td>53</td><td>23</td><td>TGAGTGTGTGTGTGTGAGTGTGT</td><td>2227.4</td><td>8.9</td><td>0.7</td><td>11.9</td><td>1.2E-32</td><td>3.0E-31</td><td>MB231/MB468</td></tr>
<tr><td>>MI0000298|hsa-mir-221&WithFlank&X|-|45746151|45746272@71.92.22(+1A)&[MIMAT0000278&hsa-miR-221-3p&offsets|0|-1(+1A);m-91&X|-|45746181|45746202&offsets|0|0(+1A)]|AGCTACATTGTCTGCTGGGTTTA</td><td>>MI0000298|hsa-mir-221&WithFlank&X|-|45746151|45746272</td><td>71</td><td>92</td><td>22(+1A)</td><td>AGCTACATTGTCTGCTGGGTTTA</td><td>3210.3</td><td>8.0</td><td>0.7</td><td>11.9</td><td>1.2E-32</td><td>3.0E-31</td><td>MB231/MB468</td></tr>
<tr><td>>trnaMT_ValTAC_MT_+_1602_1670@49.71.23|AACTTAACTTGACCGCTCTGACC</td><td>>trnaMT_ValTAC_MT_+_1602_1670</td><td>49</td><td>71</td><td>23</td><td>AACTTAACTTGACCGCTCTGACC</td><td>596.8</td><td>6.6</td><td>0.6</td><td>11.9</td><td>1.4E-32</td><td>3.0E-31</td><td>BT20/MB468</td></tr>
<tr><td>>MI0006273|hsa-mir-1180&WithFlank&17|-|19344500|19344580@47.68.22&[MIMAT0005825&hsa-miR-1180-3p&offsets|0|0;m-214&17|-|19344513|19344534&offsets|0|0]|TTTCCGGCTCGCGTGGGTGTGT</td><td>>MI0006273|hsa-mir-1180&WithFlank&17|-|19344500|19344580</td><td>47</td><td>68</td><td>22</td><td>TTTCCGGCTCGCGTGGGTGTGT</td><td>3186.9</td><td>7.5</td><td>0.6</td><td>11.9</td><td>1.4E-32</td><td>3.1E-31</td><td>BT20/MB468</td></tr>
</table>

<table>
<tr><td>>hg38_21_+_8433172_8446622_RNA45SN1_45S_50ntFlanks@7975.8019.45|CGCGACCTCAGATCAGACGTGGCGACCCGCTGAATTTAAGCATAT</td><td>>hg38_21_+_8433172_8446622_RNA45SN1_45S_50ntFlanks</td><td>7975</td><td>8019</td><td>45</td><td>CGCGACCTCAGATCAGACGTGGCGACCCGCTGAATTTAAGCATAT</td><td>591.6</td><td>14.1</td><td>1.2</td><td>11.9</td><td>1.2E-32</td><td>3.2E-31</td><td>MB231/MB468</td></tr>
<tr><td>>hg38_21_+_8433172_8446622_RNA45SN1_45S_50ntFlanks@12996.13037.42|TCGCTGCGATCTATTGAAAGTCAGCCCTCGACACAAGGGTTT</td><td>>hg38_21_+_8433172_8446622_RNA45SN1_45S_50ntFlanks</td><td>12996</td><td>13037</td><td>42</td><td>TCGCTGCGATCTATTGAAAGTCAGCCCTCGACACAAGGGTTT</td><td>940.4</td><td>14.8</td><td>1.2</td><td>11.9</td><td>1.3E-32</td><td>3.3E-31</td><td>MB231/MB468</td></tr>
<tr><td>>MI0000809|hsa-mir-151a&WithFlank&8|-|140732558|140732659@51.72.22&[MIMAT0000757&hsa-miR-151a-3p&offsets|-2|-1;m-30&8|-|140732587|140732607&offsets|-2|-1]|TACTAGACTGAAGCTCCTTGAG</td><td>>MI0000809|hsa-mir-151a&WithFlank&8|-|140732558|140732659</td><td>51</td><td>72</td><td>22</td><td>TACTAGACTGAAGCTCCTTGAG</td><td>1963.6</td><td>8.8</td><td>0.7</td><td>11.9</td><td>1.4E-32</td><td>3.6E-31</td><td>MB231/MB468</td></tr>
<tr><td>>MI0000088|hsa-mir-30a&WithFlank&6|-|71403545|71403627@12.35.24(+1A)&[MIMAT0000087&hsa-miR-30a-5p&offsets|0|+2(+1A);m-15&6|-|71403595|71403616&offsets|0|+2(+1A)]|TGTAAACATCCTCGACTGGAAGCTA</td><td>>MI0000088|hsa-mir-30a&WithFlank&6|-|71403545|71403627</td><td>12</td><td>35</td><td>24(+1A)</td><td>TGTAAACATCCTCGACTGGAAGCTA</td><td>3236.0</td><td>8.7</td><td>0.7</td><td>11.9</td><td>1.6E-32</td><td>4.0E-31</td><td>MB231/MB468</td></tr>
<tr><td>>hg38_21_+_8433172_8446622_RNA45SN1_45S_50ntFlanks@11665.11687.23|TTCTGCCCAGTGCTCTGAATGTC</td><td>>hg38_21_+_8433172_8446622_RNA45SN1_45S_50ntFlanks</td><td>11665</td><td>11687</td><td>23</td><td>TTCTGCCCAGTGCTCTGAATGTC</td><td>754.1</td><td>7.1</td><td>0.6</td><td>11.9</td><td>1.6E-32</td><td>4.0E-31</td><td>MB231/MB468</td></tr>
<tr><td>>MI0000087|hsa-mir-29a&WithFlank&7|-|130876741|130876816@47.68.22&[MIMAT0000086&hsa-miR-29a-3p&offsets|-1|-1;m-26&7|-|130876749|130876769&offsets|-1|0]|CTAGCACCATCTGAAATCGGTT</td><td>>MI0000087|hsa-mir-29a&WithFlank&7|-|130876741|130876816</td><td>47</td><td>68</td><td>22</td><td>CTAGCACCATCTGAAATCGGTT</td><td>25010.2</td><td>8.5</td><td>0.7</td><td>11.9</td><td>1.9E-32</td><td>4.1E-31</td><td>BT20/MB468</td></tr>
<tr><td>>hg38_21_+_8433172_8446622_RNA45SN1_45S_50ntFlanks@7974.8010.37|ACGCGACCTCAGATCAGACGTGGCGACCCGCTGAATT</td><td>>hg38_21_+_8433172_8446622_RNA45SN1_45S_50ntFlanks</td><td>7974</td><td>8010</td><td>37</td><td>ACGCGACCTCAGATCAGACGTGGCGACCCGCTGAATT</td><td>11208.9</td><td>7.9</td><td>0.7</td><td>11.9</td><td>1.9E-32</td><td>4.7E-31</td><td>MB231/MB468</td></tr>
<tr><td>>hg38_21_+_8433172_8446622_RNA45SN1_45S_50ntFlanks@12996.13039.44|TCGCTGCGATCTATTGAAAGTCAGCCCTCGACACAAGGGTTTGT</td><td>>hg38_21_+_8433172_8446622_RNA45SN1_45S_50ntFlanks</td><td>12996</td><td>13039</td><td>44</td><td>TCGCTGCGATCTATTGAAAGTCAGCCCTCGACACAAGGGTTTGT</td><td>838486.1</td><td>11.3</td><td>1.0</td><td>11.9</td><td>2.0E-32</td><td>5.0E-31</td><td>MB231/MB468</td></tr>
<tr><td>>MI0000746|hsa-mir-99b&WithFlank&19|+|51692606|51692687@13.32.20(+1U)&[MIMAT0000689&hsa-miR-99b-5p&offsets|0|-2(+1U);m-10&19|+|51692618|51692639&offsets|0|-2(+1U)]|CACCCGTAGAACCGACCTTGT</td><td>>MI0000746|hsa-mir-99b&WithFlank&19|+|51692606|51692687</td><td>13</td><td>32</td><td>20(+1U)</td><td>CACCCGTAGAACCGACCTTGT</td><td>3498.4</td><td>7.9</td><td>0.7</td><td>11.9</td><td>2.0E-32</td><td>5.1E-31</td><td>MB231/MB468</td></tr>
<tr><td>>hg38_21_+_8433172_8446622_RNA45SN1_45S_50ntFlanks@11572.11605.34|CCGACTTAGAACTGGTGCGGACCAGGGGAATCCG</td><td>>hg38_21_+_8433172_8446622_RNA45SN1_45S_50ntFlanks</td><td>11572</td><td>11605</td><td>34</td><td>CCGACTTAGAACTGGTGCGGACCAGGGGAATCCG</td><td>1469.2</td><td>-6.1</td><td>0.5</td><td>-11.8</td><td>2.4E-32</td><td>5.2E-31</td><td>BT20/MB468</td></tr>
<tr><td>>hg38_21_+_8433172_8446622_RNA45SN1_45S_50ntFlanks@10393.10410.18|CAGCAGTTGAACATGGGT</td><td>>hg38_21_+_8433172_8446622_RNA45SN1_45S_50ntFlanks</td><td>10393</td><td>10410</td><td>18</td><td>CAGCAGTTGAACATGGGT</td><td>3522.7</td><td>-7.2</td><td>0.6</td><td>-11.8</td><td>2.4E-32</td><td>5.2E-31</td><td>BT20/MB468</td></tr>
<tr><td>>hg38_21_+_8433172_8446622_RNA45SN1_45S_50ntFlanks@10680.10710.31|GTTCCGGCGGCGTCCGGTGAGCTCTCGCTGG</td><td>>hg38_21_+_8433172_8446622_RNA45SN1_45S_50ntFlanks</td><td>10680</td><td>10710</td><td>31</td><td>GTTCCGGCGGCGTCCGGTGAGCTCTCGCTGG</td><td>15352.7</td><td>-7.2</td><td>0.6</td><td>-11.8</td><td>2.2E-32</td><td>5.6E-31</td><td>MB231/MB468</td></tr>
<tr><td>>hg38_21_+_8433172_8446622_RNA45SN1_45S_50ntFlanks@11667.11706.40|CTGCCCAGTGCTCTGAATGTCAAAGTGAAGAAATTCAATG</td><td>>hg38_21_+_8433172_8446622_RNA45SN1_45S_50ntFlanks</td><td>11667</td><td>11706</td><td>40</td><td>CTGCCCAGTGCTCTGAATGTCAAAGTGAAGAAATTCAATG</td><td>351.3</td><td>13.4</td><td>1.1</td><td>11.8</td><td>2.3E-32</td><td>5.7E-31</td><td>MB231/MB468</td></tr>
<tr><td>>hg38_MT_+_598_1651_RNR1_12S_50ntFlanks@428.468.41|AAAGTGGCTTTAACATATCTGAACACACAATAGCTAAGACC</td><td>>hg38_MT_+_598_1651_RNR1_12S_50ntFlanks</td><td>428</td><td>468</td><td>41</td><td>AAAGTGGCTTTAACATATCTGAACACACAATAGCTAAGACC</td><td>3244.3</td><td>16.6</td><td>1.4</td><td>11.8</td><td>2.5E-32</td><td>6.2E-31</td><td>MB231/MB468</td></tr>
<tr><td>>MI0000087|hsa-mir-29a&WithFlank&7|-|130876741|130876816@47.67.21(+1C)&[MIMAT0000086&hsa-miR-29a-3p&offsets|-1|-2(+1C);m-26&7|-|130876749|130876769&offsets|-1|-1(+1C)]|CTAGCACCATCTGAAATCGGTC</td><td>>MI0000087|hsa-mir-29a&WithFlank&7|-|130876741|130876816</td><td>47</td><td>67</td><td>21(+1C)</td><td>CTAGCACCATCTGAAATCGGTC</td><td>3094.7</td><td>8.7</td><td>0.7</td><td>11.8</td><td>3.0E-32</td><td>6.4E-31</td><td>BT20/MB468</td></tr>
<tr><td>>hg38_MT_+_1621_3279_RNR2_16S_50ntFlanks@1217.1262.46|AACCTCCGAGCAGTACATGCTAAGACTTCACCAGTCAAAGCGAACT</td><td>>hg38_MT_+_1621_3279_RNR2_16S_50ntFlanks</td><td>1217</td><td>1262</td><td>46</td><td>AACCTCCGAGCAGTACATGCTAAGACTTCACCAGTCAAAGCGAACT</td><td>523.8</td><td>13.9</td><td>1.2</td><td>11.8</td><td>2.6E-32</td><td>6.4E-31</td><td>MB231/MB468</td></tr>
<tr><td>>hg38_MT_+_598_1651_RNR1_12S_50ntFlanks@370.404.35|ATAAAGCTAAAACTCACCTGAGTTGTAAAAAACTC</td><td>>hg38_MT_+_598_1651_RNR1_12S_50ntFlanks</td><td>370</td><td>404</td><td>35</td><td>ATAAAGCTAAAACTCACCTGAGTTGTAAAAAACTC</td><td>508.6</td><td>13.9</td><td>1.2</td><td>11.8</td><td>2.7E-32</td><td>6.8E-31</td><td>MB231/MB468</td></tr>
<tr><td>>hg38_21_+_8433172_8446622_RNA45SN1_45S_50ntFlanks@9572.9596.25|CAAATCGGTCGTCCGACCTGGGTAT</td><td>>hg38_21_+_8433172_8446622_RNA45SN1_45S_50ntFlanks</td><td>9572</td><td>9596</td><td>25</td><td>CAAATCGGTCGTCCGACCTGGGTAT</td><td>7442.3</td><td>-10.3</td><td>0.9</td><td>-11.8</td><td>3.3E-32</td><td>7.1E-31</td><td>BT20/MB468</td></tr>
<tr><td>>trna111_HisGTG_1_-_147774845_147774916@-1T.23.24,trna118_HisGTG_1_-_145396881_145396952@-1T.23.24,trna16_HisGTG_1_+_146544773_146544844@-1T.23.24,trna1_HisGTG_15_+_45493349_45493420@-1T.23.24,trna21_HisGTG_1_+_147753471_147753542@-1T.23.24,trna33_HisGTG_6_+_27125906_27125977@-1T.23.24,trna7_HisGTG_9_-_14433938_14434009@-1T.23.24,trna8_HisGTG_15_-_45492611_45492682@-1T.23.24,trna9_HisGTG_15_-_45490804_45490875@-1T.23.24|TGCCGTGATCGTATAGTGGTTAGT</td><td>>trna111_HisGTG_1_-_147774845_147774916</td><td>-1T</td><td>23</td><td>24</td><td>TGCCGTGATCGTATAGTGGTTAGT</td><td>370.7</td><td>5.7</td><td>0.5</td><td>11.8</td><td>3.3E-32</td><td>7.1E-31</td><td>BT20/MB468</td></tr>
<tr><td>>hg38_21_+_8433172_8446622_RNA45SN1_45S_50ntFlanks@10678.10712.35|CGGTTCCGGCGGCGTCCGGTGAGCTCTCGCTGGCC</td><td>>hg38_21_+_8433172_8446622_RNA45SN1_45S_50ntFlanks</td><td>10678</td><td>10712</td><td>35</td><td>CGGTTCCGGCGGCGTCCGGTGAGCTCTCGCTGGCC</td><td>5748.8</td><td>-5.7</td><td>0.5</td><td>-11.8</td><td>3.0E-32</td><td>7.3E-31</td><td>MB231/MB468</td></tr>
<tr><td>>hg38_MT_+_1621_3279_RNR2_16S_50ntFlanks@1349.1388.40|ACAATAGGGTTTACGACCTCGATGTTGGATCAGGACATCC</td><td>>hg38_MT_+_1621_3279_RNR2_16S_50ntFlanks</td><td>1349</td><td>1388</td><td>40</td><td>ACAATAGGGTTTACGACCTCGATGTTGGATCAGGACATCC</td><td>3137.5</td><td>16.5</td><td>1.4</td><td>11.8</td><td>3.0E-32</td><td>7.4E-31</td><td>MB231/MB468</td></tr>
<tr><td>>hg38_21_+_8433172_8446622_RNA45SN1_45S_50ntFlanks@11729.11751.23|TAACTATGACTCTCTTAAGGTAG</td><td>>hg38_21_+_8433172_8446622_RNA45SN1_45S_50ntFlanks</td><td>11729</td><td>11751</td><td>23</td><td>TAACTATGACTCTCTTAAGGTAG</td><td>65156.8</td><td>-10.8</td><td>0.9</td><td>-11.8</td><td>3.5E-32</td><td>7.5E-31</td><td>BT20/MB468</td></tr>
<tr><td>>am_trnaMT_HisGTG_MT_+_12138_12206@19.51.33|AAACATCAGATTGTGAATCTGACAACAGAGGCT</td><td>>am_trnaMT_HisGTG_MT_+_12138_12206</td><td>19</td><td>51</td><td>33</td><td>AAACATCAGATTGTGAATCTGACAACAGAGGCT</td><td>632.9</td><td>14.0</td><td>1.2</td><td>11.8</td><td>3.7E-32</td><td>7.8E-31</td><td>BT20/MB468</td></tr>
<tr><td>>am_MI0005763|hsa-mir-941-1&WithFlank&20|+|63919443|63919526@53.75.23;MI0005764|hsa-mir-941-2&WithFlank&20|+|63919499|63919582@53.75.23;MI0005765|hsa-mir-941-3&WithFlank&20|+|63919555|63919638@53.75.23;MI0005766|hsa-mir-941-4&WithFlank&20|+|63919750|63919833@53.75.23;MI0031520|hsa-mir-941-5&WithFlank&20|+|63919862|63919945@53.75.23&[MIMAT0004984&hsa-miR-941&offsets|0|0;m-263&20|+|63919495|63919517&offsets|0|0];[MIMAT0004984_1&hsa-miR-941&offsets|0|0;m-263&20|+|63919551|63919573&offsets|0|0];[MIMAT0004984_2&hsa-miR-941&offsets|0|0;m-263&20|+|63919607|63919629&offsets|0|0];[MIMAT0004984_3&hsa-miR-941&offsets|0|0;m-263&20|+|63919802|63919824&offsets|0|0];[MIMAT0004984_4&hsa-miR-941&offsets|0|0;m-263&20|+|63919914|63919936&offsets|0|0]|CACCCGGCTGTGTGCACATGTGC</td><td>>am_MI0005763|hsa-mir-941-1&WithFlank&20|+|63919443|63919526</td><td>53</td><td>75</td><td>23</td><td>CACCCGGCTGTGTGCACATGTGC</td><td>35873.1</td><td>8.8</td><td>0.7</td><td>11.8</td><td>3.2E-32</td><td>7.9E-31</td><td>MB231/MB468</td></tr>
<tr><td>>MI0001445|hsa-mir-423&WithFlank&17|+|30117073|30117178@23.45.23&[MIMAT0004748&hsa-miR-423-5p&offsets|0|0;m-141&17|+|30117095|30117117&offsets|0|0]|TGAGGGGCAGAGAGCGAGACTTT</td><td>>MI0001445|hsa-mir-423&WithFlank&17|+|30117073|30117178</td><td>23</td><td>45</td><td>23</td><td>TGAGGGGCAGAGAGCGAGACTTT</td><td>10302.4</td><td>8.6</td><td>0.7</td><td>11.8</td><td>3.2E-32</td><td>7.9E-31</td><td>MB231/MB468</td></tr>
<tr><td>>hg38_21_+_8433172_8446622_RNA45SN1_45S_50ntFlanks@4731.4765.35|AAGAACGAAAGTCGGAGGTTCGAAGACGATCAGAT</td><td>>hg38_21_+_8433172_8446622_RNA45SN1_45S_50ntFlanks</td><td>4731</td><td>4765</td><td>35</td><td>AAGAACGAAAGTCGGAGGTTCGAAGACGATCAGAT</td><td>1585.8</td><td>7.9</td><td>0.7</td><td>11.8</td><td>3.5E-32</td><td>8.4E-31</td><td>MB231/MB468</td></tr>
<tr><td>>hg38_21_+_8433172_8446622_RNA45SN1_45S_50ntFlanks@11668.11709.42|TGCCCAGTGCTCTGAATGTCAAAGTGAAGAAATTCAATGAAG</td><td>>hg38_21_+_8433172_8446622_RNA45SN1_45S_50ntFlanks</td><td>11668</td><td>11709</td><td>42</td><td>TGCCCAGTGCTCTGAATGTCAAAGTGAAGAAATTCAATGAAG</td><td>373.2</td><td>13.5</td><td>1.1</td><td>11.8</td><td>3.7E-32</td><td>8.8E-31</td><td>MB231/MB468</td></tr>
<tr><td>>MI0000816|hsa-mir-335&WithFlank&7|+|130496105|130496210@58.78.21(+1U)&[MIMAT0004703&hsa-miR-335-3p&offsets|0|-1(+1U);m-159&7|+|130496162|130496183&offsets|0|-1(+1U)]|TTTTTCATTATTGCTCCTGACT</td><td>>MI0000816|hsa-mir-335&WithFlank&7|+|130496105|130496210</td><td>58</td><td>78</td><td>21(+1U)</td><td>TTTTTCATTATTGCTCCTGACT</td><td>3405.8</td><td>8.4</td><td>0.7</td><td>11.8</td><td>4.0E-32</td><td>9.7E-31</td><td>MB231/MB468</td></tr>
<tr><td>>hg38_21_+_8433172_8446622_RNA45SN1_45S_50ntFlanks@11541.11584.44|GCGGCGCCTCGCCTCGGCCGGCGCCTAGCAGCCGACTTAGAACT</td><td>>hg38_21_+_8433172_8446622_RNA45SN1_45S_50ntFlanks</td><td>11541</td><td>11584</td><td>44</td><td>GCGGCGCCTCGCCTCGGCCGGCGCCTAGCAGCCGACTTAGAACT</td><td>475.0</td><td>13.8</td><td>1.2</td><td>11.8</td><td>4.1E-32</td><td>9.7E-31</td><td>MB231/MB468</td></tr>
<tr><td>>hg38_21_+_8433172_8446622_RNA45SN1_45S_50ntFlanks@475.496.22|CTTCGTGATCGATGTGGTGACG</td><td>>hg38_21_+_8433172_8446622_RNA45SN1_45S_50ntFlanks</td><td>475</td><td>496</td><td>22</td><td>CTTCGTGATCGATGTGGTGACG</td><td>21377.3</td><td>8.1</td><td>0.7</td><td>11.8</td><td>4.2E-32</td><td>9.9E-31</td><td>MB231/MB468</td></tr>
<tr><td>>hg38_21_+_8433172_8446622_RNA45SN1_45S_50ntFlanks@12996.13016.21|TCGCTGCGATCTATTGAAAGT</td><td>>hg38_21_+_8433172_8446622_RNA45SN1_45S_50ntFlanks</td><td>12996</td><td>13016</td><td>21</td><td>TCGCTGCGATCTATTGAAAGT</td><td>1101.5</td><td>7.4</td><td>0.6</td><td>11.8</td><td>4.2E-32</td><td>1.0E-30</td><td>MB231/MB468</td></tr>
<tr><td>>hg38_21_+_8433172_8446622_RNA45SN1_45S_50ntFlanks@475.511.37|CTTCGTGATCGATGTGGTGACGTCGTGCTCTCCCGGG</td><td>>hg38_21_+_8433172_8446622_RNA45SN1_45S_50ntFlanks</td><td>475</td><td>511</td><td>37</td><td>CTTCGTGATCGATGTGGTGACGTCGTGCTCTCCCGGG</td><td>7651.6</td><td>8.0</td><td>0.7</td><td>11.8</td><td>4.2E-32</td><td>1.0E-30</td><td>MB231/MB468</td></tr>
<tr><td>>MI0001445|hsa-mir-423&WithFlank&17|+|30117073|30117178@58.79.22&[MIMAT0001340&hsa-miR-423-3p&offsets|-1|-2;m-100&17|+|30117131|30117153&offsets|-1|-2]|AAGCTCGGTCTGAGGCCCCTCA</td><td>>MI0001445|hsa-mir-423&WithFlank&17|+|30117073|30117178</td><td>58</td><td>79</td><td>22</td><td>AAGCTCGGTCTGAGGCCCCTCA</td><td>853.8</td><td>7.3</td><td>0.6</td><td>11.8</td><td>4.9E-32</td><td>1.0E-30</td><td>BT20/MB468</td></tr>
<tr><td>>MI0000255|hsa-mir-30d&WithFlank&8|-|134804870|134804951@12.33.22&[MIMAT0000245&hsa-miR-30d-5p&offsets|0|0;m-21&8|-|134804919|134804940&offsets|0|0]|TGTAAACATCCCCGACTGGAAG</td><td>>MI0000255|hsa-mir-30d&WithFlank&8|-|134804870|134804951</td><td>12</td><td>33</td><td>22</td><td>TGTAAACATCCCCGACTGGAAG</td><td>21954.6</td><td>9.4</td><td>0.8</td><td>11.8</td><td>5.3E-32</td><td>1.1E-30</td><td>BT20/MB468</td></tr>
<tr><td>>MI0000239|hsa-mir-197&WithFlank&1|+|109598887|109598973@54.74.21(+1U)&[MIMAT0000227&hsa-miR-197-3p&offsets|0|-1(+1U);m-99&1|+|109598940|109598961&offsets|0|-1(+1U)]|TTCACCACCTTCTCCACCCAGT</td><td>>MI0000239|hsa-mir-197&WithFlank&1|+|109598887|109598973</td><td>54</td><td>74</td><td>21(+1U)</td><td>TTCACCACCTTCTCCACCCAGT</td><td>7786.3</td><td>10.3</td><td>0.9</td><td>11.8</td><td>6.0E-32</td><td>1.3E-30</td><td>BT20/MB468</td></tr>
<tr><td>>trnaMT_ValTAC_MT_+_1602_1670@54.72.19|AACTTGACCGCTCTGACCA</td><td>>trnaMT_ValTAC_MT_+_1602_1670</td><td>54</td><td>72</td><td>19</td><td>AACTTGACCGCTCTGACCA</td><td>1726.3</td><td>7.7</td><td>0.7</td><td>11.8</td><td>5.4E-32</td><td>1.3E-30</td><td>MB231/MB468</td></tr>
<tr><td>>MI0000088|hsa-mir-30a&WithFlank&6|-|71403545|71403627@53.73.21(+2U)&[MIMAT0000088&hsa-miR-30a-3p&offsets|0|-1(+2U);m-38&6|-|71403554|71403575&offsets|0|-1(+2U)]|CTTTCAGTCGGATGTTTGCAGTT</td><td>>MI0000088|hsa-mir-30a&WithFlank&6|-|71403545|71403627</td><td>53</td><td>73</td><td>21(+2U)</td><td>CTTTCAGTCGGATGTTTGCAGTT</td><td>1817.1</td><td>9.6</td><td>0.8</td><td>11.8</td><td>5.4E-32</td><td>1.3E-30</td><td>MB231/MB468</td></tr>
</table>

| | | | | | | | | | | | | |
|---|---|---|---|---|---|---|---|---|---|---|---|---|
| >hg38_21_+_8433172_8446622_RNA45SN1_45S_50ntFlanks@4759.4780.22\|ATCAGATACCGTCGTAGTTCCG | >hg38_21_+_8433172_8446622_RNA45SN1_45S_50ntFlanks | 4759 | 4780 | 22 | ATCAGATACCGTCGTAGTTCCG | 7414.9 | -14.8 | 1.3 | -11.8 | 6.6E-32 | 1.4E-30 | BT20/MB468 |
| >hg38_21_+_8433172_8446622_RNA45SN1_45S_50ntFlanks@12419.12459.41\|GATGTCGGCTCTTCCTATCATTGTGAAGCAGAATTCACCAA | >hg38_21_+_8433172_8446622_RNA45SN1_45S_50ntFlanks | 12419 | 12459 | 41 | GATGTCGGCTCTTCCTATCATTGTGAAGCAGAATTCACCAA | 385.4 | 13.5 | 1.1 | 11.8 | 6.2E-32 | 1.4E-30 | MB231/MB468 |
| >hg38_MT_+_598_1651_RNR1_12S_50ntFlanks@968.1004.37\|ACATGGTAAGTGTACTGGAAAGTGCACTTGGACGAAC | >hg38_MT_+_598_1651_RNR1_12S_50ntFlanks | 968 | 1004 | 37 | ACATGGTAAGTGTACTGGAAAGTGCACTTGGACGAAC | 1002.1 | 14.9 | 1.3 | 11.8 | 6.2E-32 | 1.4E-30 | MB231/MB468 |
| >hg38_21_+_8433172_8446622_RNA45SN1_45S_50ntFlanks@12456.12485.30\|CCAAGCGTTGGATTGTTCACCCACTAATAG | >hg38_21_+_8433172_8446622_RNA45SN1_45S_50ntFlanks | 12456 | 12485 | 30 | CCAAGCGTTGGATTGTTCACCCACTAATAG | 3962.7 | -10.5 | 0.9 | -11.8 | 7.0E-32 | 1.5E-30 | BT20/MB468 |
| >MI0000102\|hsa-mir-100&WithFlank&11\|-\|122152223\|122152314@19.38.20(+1C)&[MIMAT0000098&hsa-miR-100-5p&offsets\|0\|-2(+1C);m-27&11\|-\|122152276\|122152296&offsets\|0\|-1(+1C)]\|AACCCGTAGATCCGAACTTGC | >MI0000102\|hsa-mir-100&WithFlank&11\|-\|122152223\|122152314 | 19 | 38 | 20(+1C) | AACCCGTAGATCCGAACTTGC | 76950.5 | 15.6 | 1.3 | 11.8 | 6.4E-32 | 1.5E-30 | MB231/MB468 |
| >hg38_MT_+_1621_3279_RNR2_16S_50ntFlanks@714.740.27\|CATAAGCCTGCGTCAGATTAAAACACT | >hg38_MT_+_1621_3279_RNR2_16S_50ntFlanks | 714 | 740 | 27 | CATAAGCCTGCGTCAGATTAAAACACT | 520.4 | 13.9 | 1.2 | 11.8 | 6.5E-32 | 1.5E-30 | MB231/MB468 |
| >MI0000077\|hsa-mir-21&WithFlank&17\|+\|59841260\|59841343@14.35.22(+1A)&[MIMAT0000076&hsa-miR-21-5p&offsets\|0\|0(+1A);m-1&17\|+\|59841273\|59841294&offsets\|0\|0(+1A)]\|TAGCTTATCAGACTGATGTTGAA | >MI0000077\|hsa-mir-21&WithFlank&17\|+\|59841260\|59841343 | 14 | 35 | 22(+1A) | TAGCTTATCAGACTGATGTTGAA | 70237.7 | 8.5 | 0.7 | 11.8 | 6.5E-32 | 1.5E-30 | MB231/MB468 |
| >hg38_1_-_228634819_228635039_RNA5S12_5S_50ntFlanks@135.170.36\|GGGAGACCGCCTGGGAATACCGGGTGCTGTAGGCTT | >hg38_1_-_228634819_228635039_RNA5S12_5S_50ntFlanks | 135 | 170 | 36 | GGGAGACCGCCTGGGAATACCGGGTGCTGTAGGCTT | 14262.3 | 5.6 | 0.5 | 11.7 | 7.2E-32 | 1.5E-30 | BT20/MB468 |
| >MI0000266\|hsa-mir-10a&WithFlank&17\|-\|48579832\|48579953@28.49.22(+1A)&[MIMAT0000253&hsa-miR-10a-5p&offsets\|0\|-1(+1A);m-18&17\|-\|48579905\|48579926&offsets\|0\|0(+1A)]\|TACCCTGTAGATCCGAATTTGTA | >MI0000266\|hsa-mir-10a&WithFlank&17\|-\|48579832\|48579953 | 28 | 49 | 22(+1A) | TACCCTGTAGATCCGAATTTGTA | 2127.4 | 16.0 | 1.4 | 11.8 | 6.8E-32 | 1.6E-30 | MB231/MB468 |
| >hg38_21_+_8433172_8446622_RNA45SN1_45S_50ntFlanks@6647.6665.19\|CGTACGACTCTTAGCGGTG | >hg38_21_+_8433172_8446622_RNA45SN1_45S_50ntFlanks | 6647 | 6665 | 19 | CGTACGACTCTTAGCGGTG | 9218.3 | 5.8 | 0.5 | 11.7 | 7.8E-32 | 1.6E-30 | BT20/MB468 |
| >hg38_21_+_8433172_8446622_RNA45SN1_45S_50ntFlanks@12451.12476.26\|ATTCACCAAGCGTTGGATTGTTCACC | >hg38_21_+_8433172_8446622_RNA45SN1_45S_50ntFlanks | 12451 | 12476 | 26 | ATTCACCAAGCGTTGGATTGTTCACC | 774.4 | 6.2 | 0.5 | 11.7 | 7.7E-32 | 1.7E-30 | MB231/MB468 |
| >MI0000298\|hsa-mir-221&WithFlank&X\|-\|45746151\|45746272@71.91.21(+1A)&[MIMAT0000278&hsa-miR-221-3p&offsets\|0\|-2(+1A);m-91&X\|-\|45746181\|45746202&offsets\|0\|-1(+1A)]\|AGCTACATTGTCTGCTGGGTTA | >MI0000298\|hsa-mir-221&WithFlank&X\|-\|45746151\|45746272 | 71 | 91 | 21(+1A) | AGCTACATTGTCTGCTGGGTTA | 2285.9 | 8.9 | 0.8 | 11.7 | 7.7E-32 | 1.7E-30 | MB231/MB468 |
| >hg38_21_+_8433172_8446622_RNA45SN1_45S_50ntFlanks@12997.13035.39\|CGCTGCGATCTATTGAAAGTCAGCCCTCGACACAAGGGT | >hg38_21_+_8433172_8446622_RNA45SN1_45S_50ntFlanks | 12997 | 13035 | 39 | CGCTGCGATCTATTGAAAGTCAGCCCTCGACACAAGGGT | 727.8 | 12.4 | 1.1 | 11.7 | 7.8E-32 | 1.8E-30 | MB231/MB468 |
| >hg38_21_+_8433172_8446622_RNA45SN1_45S_50ntFlanks@12973.12990.18\|TCGTACGTAGCAGAGCAG | >hg38_21_+_8433172_8446622_RNA45SN1_45S_50ntFlanks | 12973 | 12990 | 18 | TCGTACGTAGCAGAGCAG | 8385.6 | -15.0 | 1.3 | -11.7 | 8.6E-32 | 1.8E-30 | BT20/MB468 |
| >hg38_21_+_8433172_8446622_RNA45SN1_45S_50ntFlanks@11666.11706.41\|TCTGCCCAGTGCTCTGAATGTCAAAGTGAAGAAATTCAATG | >hg38_21_+_8433172_8446622_RNA45SN1_45S_50ntFlanks | 11666 | 11706 | 41 | TCTGCCCAGTGCTCTGAATGTCAAAGTGAAGAAATTCAATG | 400.8 | 13.6 | 1.2 | 11.7 | 8.7E-32 | 2.0E-30 | MB231/MB468 |
| >trna116_GluCTC_1_-_145399233_145399304@1.36.36,trna59_GluCTC_1_+_249168447_249168518@1.36.36,trna71_GluCTC_1_-_161439189_161439260@1.36.36,trna74_GluCTC_1_-_161431809_161431880@1.36.36,trna77_GluCTC_1_-_161424398_161424469@1.36.36,trna77_GluCTC_6_+_28949976_28950047@1.36.36,trna80_GluCTC_1_-_161417018_161417089@1.36.36,trna87_GluCTC_6_-_126101393_126101464@1.36.36\|TCCCTGGTGGTCTAGTGGTTAGGATTCGGCGCTCTC | >trna116_GluCTC_1_-_145399233_145399304 | 1 | 36 | 36 | TCCCTGGTGGTCTAGTGGTTAGGATTCGGCGCTCTC | 11867.6 | 11.1 | 0.9 | 11.7 | 8.7E-32 | 2.0E-30 | MB231/MB468 |
| >hg38_MT_+_1621_3279_RNR2_16S_50ntFlanks@1175.1215.41\|TGCATTAAAAATTTCGGTTGGGGCGACCTCGGAGCAGAACC | >hg38_MT_+_1621_3279_RNR2_16S_50ntFlanks | 1175 | 1215 | 41 | TGCATTAAAAATTTCGGTTGGGGCGACCTCGGAGCAGAACC | 590.9 | 14.1 | 1.2 | 11.7 | 9.1E-32 | 2.0E-30 | MB231/MB468 |
| >MI0000809\|hsa-mir-151a&WithFlank&8\|-\|140732558\|140732659@51.73.23(+1U)&[MIMAT0000757&hsa-miR-151a-3p&offsets\|-2\|0(+1U);m-30&8\|-\|140732587\|140732607&offsets\|-2\|0(+1U)]\|TACTAGACTGAAGCTCCTTGAGGT | >MI0000809\|hsa-mir-151a&WithFlank&8\|-\|140732558\|140732659 | 51 | 73 | 23(+1U) | TACTAGACTGAAGCTCCTTGAGGT | 1713.1 | 8.8 | 0.8 | 11.7 | 9.3E-32 | 2.1E-30 | MB231/MB468 |
| >trnaMT_PheGAA_MT_+_577_647@36.56.21\|AAAATGTTTAGACGGGCTCAC | >trnaMT_PheGAA_MT_+_577_647 | 36 | 56 | 21 | AAAATGTTTAGACGGGCTCAC | 966.5 | 12.5 | 1.1 | 11.7 | 1.0E-31 | 2.1E-30 | BT20/MB468 |
| >trna116_GluCTC_1_-_145399233_145399304@1.34.34,trna59_GluCTC_1_+_249168447_249168518@1.34.34,trna71_GluCTC_1_-_161439189_161439260@1.34.34,trna74_GluCTC_1_-_161431809_161431880@1.34.34,trna77_GluCTC_1_-_161424398_161424469@1.34.34,trna77_GluCTC_6_+_28949976_28950047@1.34.34,trna80_GluCTC_1_-_161417018_161417089@1.34.34,trna87_GluCTC_6_-_126101393_126101464@1.34.34\|TCCCTGGTGGTCTAGTGGTTAGGATTCGGCGCTC | >trna116_GluCTC_1_-_145399233_145399304 | 1 | 34 | 34 | TCCCTGGTGGTCTAGTGGTTAGGATTCGGCGCTC | 8900.8 | 9.4 | 0.8 | 11.7 | 1.0E-31 | 2.3E-30 | MB231/MB468 |
| >hg38_21_+_8433172_8446622_RNA45SN1_45S_50ntFlanks@476.497.22\|TTCGTGATCGATGTGGTGACGT | >hg38_21_+_8433172_8446622_RNA45SN1_45S_50ntFlanks | 476 | 497 | 22 | TTCGTGATCGATGTGGTGACGT | 4336.2 | 9.0 | 0.8 | 11.7 | 1.0E-31 | 2.3E-30 | MB231/MB468 |
| >MI0000063\|hsa-let-7b&WithFlank&22\|+\|46113680\|46113774@12.32.21(+1A)&[MIMAT0000063&hsa-let-7b-5p&offsets\|0\|-1(+1A);m-14&22\|+\|46113691\|46113712&offsets\|0\|-1(+1A)]\|TGAGGTAGTAGGTTGTGTGGTA | >MI0000063\|hsa-let-7b&WithFlank&22\|+\|46113680\|46113774 | 12 | 32 | 21(+1A) | TGAGGTAGTAGGTTGTGTGGTA | 8176.7 | 9.3 | 0.8 | 11.7 | 1.1E-31 | 2.3E-30 | BT20/MB468 |
| >hg38_21_+_8433172_8446622_RNA45SN1_45S_50ntFlanks@12997.13016.20\|CGCTGCGATCTATTGAAAGT | >hg38_21_+_8433172_8446622_RNA45SN1_45S_50ntFlanks | 12997 | 13016 | 20 | CGCTGCGATCTATTGAAAGT | 1430.7 | 7.6 | 0.6 | 11.7 | 1.1E-31 | 2.4E-30 | MB231/MB468 |
| >hg38_21_+_8433172_8446622_RNA45SN1_45S_50ntFlanks@4727.4765.39\|AATCAAGAACGAAAGTCGGAGGTTCGAAGACGATCAGAT | >hg38_21_+_8433172_8446622_RNA45SN1_45S_50ntFlanks | 4727 | 4765 | 39 | AATCAAGAACGAAAGTCGGAGGTTCGAAGACGATCAGAT | 1153.8 | 8.1 | 0.7 | 11.7 | 1.1E-31 | 2.4E-30 | MB231/MB468 |
| >MI0000102\|hsa-mir-100&WithFlank&11\|-\|122152223\|122152314@19.40.22(+2A)&[MIMAT0000098&hsa-miR-100-5p&offsets\|0\|0(+2A);m-27&11\|-\|122152276\|122152296&offsets\|0\|+1(+2A)]\|AACCCGTAGATCCGAACTTGTGAA | >MI0000102\|hsa-mir-100&WithFlank&11\|-\|122152223\|122152314 | 19 | 40 | 22(+2A) | AACCCGTAGATCCGAACTTGTGAA | 5031.5 | 14.1 | 1.2 | 11.7 | 1.1E-31 | 2.4E-30 | MB231/MB468 |
| >hg38_21_+_8433172_8446622_RNA45SN1_45S_50ntFlanks@12416.12449.34\|TTCGATGTCGGCTCTTCCTATCATTGTGAAGCAG | >hg38_21_+_8433172_8446622_RNA45SN1_45S_50ntFlanks | 12416 | 12449 | 34 | TTCGATGTCGGCTCTTCCTATCATTGTGAAGCAG | 2799.1 | 7.8 | 0.7 | 11.7 | 1.1E-31 | 2.5E-30 | MB231/MB468 |
| >MI0001445\|hsa-mir-423&WithFlank&17\|+\|30117073\|30117178@23.43.21&[MIMAT0004748&hsa-miR-423-5p&offsets\|0\|-2;m-141&17\|+\|30117095\|30117117&offsets\|0\|-2]\|TGAGGGGCAGAGAGCGAGACT | >MI0001445\|hsa-mir-423&WithFlank&17\|+\|30117073\|30117178 | 23 | 43 | 21 | TGAGGGGCAGAGAGCGAGACT | 2664.9 | 8.9 | 0.8 | 11.7 | 1.2E-31 | 2.5E-30 | MB231/MB468 |
| >hg38_21_+_8433172_8446622_RNA45SN1_45S_50ntFlanks@10351.10383.33\|CGAGAACTTTGAAGGCCGAAGTGGAGAAGGGTT | >hg38_21_+_8433172_8446622_RNA45SN1_45S_50ntFlanks | 10351 | 10383 | 33 | CGAGAACTTTGAAGGCCGAAGTGGAGAAGGGTT | 12784.3 | -7.0 | 0.6 | -11.7 | 1.2E-31 | 2.5E-30 | BT20/MB468 |
| >hg38_21_+_8433172_8446622_RNA45SN1_45S_50ntFlanks@6653.6695.43\|ACTCTTAGCGGTGGATCACTCGGCTCGTGCGTCGATGAAGAAC | >hg38_21_+_8433172_8446622_RNA45SN1_45S_50ntFlanks | 6653 | 6695 | 43 | ACTCTTAGCGGTGGATCACTCGGCTCGTGCGTCGATGAAGAAC | 342.3 | 13.3 | 1.1 | 11.7 | 1.2E-31 | 2.6E-30 | MB231/MB468 |
| >hg38_MT_+_1621_3279_RNR2_16S_50ntFlanks@460.491.32\|TTGTAAATTTAACTGTTAGTCCAAAGAGGAAC | >hg38_MT_+_1621_3279_RNR2_16S_50ntFlanks | 460 | 491 | 32 | TTGTAAATTTAACTGTTAGTCCAAAGAGGAAC | 526.5 | 13.9 | 1.2 | 11.7 | 1.2E-31 | 2.7E-30 | MB231/MB468 |
| >hg38_21_+_8433172_8446622_RNA45SN1_45S_50ntFlanks@11669.11707.39\|GCCCAGTGCTCTGAATGTCAAAGTGAAGAAATTCAATGA | >hg38_21_+_8433172_8446622_RNA45SN1_45S_50ntFlanks | 11669 | 11707 | 39 | GCCCAGTGCTCTGAATGTCAAAGTGAAGAAATTCAATGA | 603.6 | 14.1 | 1.2 | 11.7 | 1.2E-31 | 2.7E-30 | MB231/MB468 |
| >hg38_21_+_8433172_8446622_RNA45SN1_45S_50ntFlanks@5472.5500.29\|ACGGCCCTGGCGGAGCGCTGAGAAGACGG | >hg38_21_+_8433172_8446622_RNA45SN1_45S_50ntFlanks | 5472 | 5500 | 29 | ACGGCCCTGGCGGAGCGCTGAGAAGACGG | 489.8 | 5.9 | 0.5 | 11.7 | 1.3E-31 | 2.7E-30 | MB231/MB468 |
| >MI0000299\|hsa-mir-222&WithFlank&X\|-\|45747009\|45747130@75.96.22(+1A)&[MIMAT0000279&hsa-miR-222-3p&offsets\|0\|+1(+1A);m-140&X\|-\|45747033\|45747056&offsets\|0\|-2(+1A)]\|AGCTACATCTGGCTACTGGGTCA | >MI0000299\|hsa-mir-222&WithFlank&X\|-\|45747009\|45747130 | 75 | 96 | 22(+1A) | AGCTACATCTGGCTACTGGGTCA | 2235.7 | 8.8 | 0.8 | 11.7 | 1.3E-31 | 2.8E-30 | MB231/MB468 |
| >hg38_MT_+_1621_3279_RNR2_16S_50ntFlanks@718.740.23\|AGCCTGCGTCAGATTAAAACACT | >hg38_MT_+_1621_3279_RNR2_16S_50ntFlanks | 718 | 740 | 23 | AGCCTGCGTCAGATTAAAACACT | 472.6 | 13.8 | 1.2 | 11.7 | 1.4E-31 | 2.9E-30 | MB231/MB468 |
| >MI0000273\|hsa-mir-183&WithFlank&7\|-\|129774899\|129775020@34.54.21&[MIMAT0000261&hsa-miR-183-5p&offsets\|+1\|0;m-41&7\|-\|129774966\|129774987&offsets\|0\|-1]\|ATGGCACTGGTAGAATTCACT | >MI0000273\|hsa-mir-183&WithFlank&7\|-\|129774899\|129775020 | 34 | 54 | 21 | ATGGCACTGGTAGAATTCACT | 10375.2 | 7.8 | 0.7 | 11.7 | 1.6E-31 | 3.2E-30 | BT20/MB468 |
| >hg38_21_+_8433172_8446622_RNA45SN1_45S_50ntFlanks@11668.11707.40\|TGCCCAGTGCTCTGAATGTCAAAGTGAAGAAATTCAATGA | >hg38_21_+_8433172_8446622_RNA45SN1_45S_50ntFlanks | 11668 | 11707 | 40 | TGCCCAGTGCTCTGAATGTCAAAGTGAAGAAATTCAATGA | 833.4 | 12.6 | 1.1 | 11.7 | 1.6E-31 | 3.4E-30 | MB231/MB468 |
| >hg38_21_+_8433172_8446622_RNA45SN1_45S_50ntFlanks@485.510.26\|GATGTGGTGACGTCGTGCTCTCCCGG | >hg38_21_+_8433172_8446622_RNA45SN1_45S_50ntFlanks | 485 | 510 | 26 | GATGTGGTGACGTCGTGCTCTCCCGG | 2353.1 | 15.8 | 1.4 | 11.7 | 1.7E-31 | 3.5E-30 | BT20/MB468 |
| >hg38_21_+_8433172_8446622_RNA45SN1_45S_50ntFlanks@11539.11583.45\|GCGCGGCGCCTCGCCTCGGCCGGCGCCTAGCAGCCGACTTAGAAC | >hg38_21_+_8433172_8446622_RNA45SN1_45S_50ntFlanks | 11539 | 11583 | 45 | GCGCGGCGCCTCGCCTCGGCCGGCGCCTAGCAGCCGACTTAGAAC | 489.7 | 13.8 | 1.2 | 11.7 | 1.7E-31 | 3.6E-30 | MB231/MB468 |
| >hg38_21_+_8433172_8446622_RNA45SN1_45S_50ntFlanks@10351.10368.18\|CGAGAACTTTGAAGGCCG | >hg38_21_+_8433172_8446622_RNA45SN1_45S_50ntFlanks | 10351 | 10368 | 18 | CGAGAACTTTGAAGGCCG | 24016.1 | -6.8 | 0.6 | -11.7 | 1.8E-31 | 3.8E-30 | MB231/MB468 |
| >hg38_21_+_8433172_8446622_RNA45SN1_45S_50ntFlanks@7973.8010.38\|GACGCGACCTCAGATCAGACGTGGCGACCCGCTGAATT | >hg38_21_+_8433172_8446622_RNA45SN1_45S_50ntFlanks | 7973 | 8010 | 38 | GACGCGACCTCAGATCAGACGTGGCGACCCGCTGAATT | 1456.2 | 7.8 | 0.7 | 11.7 | 1.9E-31 | 3.8E-30 | BT20/MB468 |
| >trnaMT_ThrTGT_MT_+_15888_15953@31.69.39\|TTGTAAACCGGAGATGAAAACCTTTTTCCAAGGACACCA | >trnaMT_ThrTGT_MT_+_15888_15953 | 31 | 69 | 39 | TTGTAAACCGGAGATGAAAACCTTTTTCCAAGGACACCA | 1059.1 | 15.0 | 1.3 | 11.7 | 1.9E-31 | 4.1E-30 | MB231/MB468 |

<table>
<tr><td>>hg38_MT_+_1621_3279_RNR2_16S_50ntFlanks@1370.1414.45|ATGTTGGATCAGGACATCCCGATGGTGCAGCCGCTATTAAAGGTT</td><td>>hg38_MT_+_1621_3279_RNR2_16S_50ntFlanks</td><td>1370</td><td>1414</td><td>45</td><td>ATGTTGGATCAGGACATCCCGATGGTGCAGCCGCTATTAAAGGTT</td><td>593.1</td><td>14.1</td><td>1.2</td><td>11.7</td><td>1.9E-31</td><td>4.1E-30</td><td>MB231/MB468</td></tr>
<tr><td>>MI0000088|hsa-mir-30a&WithFlank&6|-|71403545|71403627@54.71.18&[MIMAT0000088&hsa-miR-30a-3p&offsets|+1|-3;m-38&6|-|71403554|71403575&offsets|+1|-3]|TTTCAGTCGGATGTTTGC</td><td>>MI0000088|hsa-mir-30a&WithFlank&6|-|71403545|71403627</td><td>54</td><td>71</td><td>18</td><td>TTTCAGTCGGATGTTTGC</td><td>2514.3</td><td>9.4</td><td>0.8</td><td>11.7</td><td>2.0E-31</td><td>4.2E-30</td><td>MB231/MB468</td></tr>
<tr><td>>hg38_21_+_8433172_8446622_RNA45SN1_45S_50ntFlanks@10687.10706.20|CGGCGTCCGGTGAGCTCTCG</td><td>>hg38_21_+_8433172_8446622_RNA45SN1_45S_50ntFlanks</td><td>10687</td><td>10706</td><td>20</td><td>CGGCGTCCGGTGAGCTCTCG</td><td>2673.8</td><td>-7.9</td><td>0.7</td><td>-11.7</td><td>2.0E-31</td><td>4.2E-30</td><td>MB231/MB468</td></tr>
<tr><td>>trnaMT_PheGAA_MT_+_577_647@35.57.23|GAAAATGTTTAGACGGGCTCACA</td><td>>trnaMT_PheGAA_MT_+_577_647</td><td>35</td><td>57</td><td>23</td><td>GAAAATGTTTAGACGGGCTCACA</td><td>6271.8</td><td>13.3</td><td>1.1</td><td>11.7</td><td>2.1E-31</td><td>4.3E-30</td><td>BT20/MB468</td></tr>
<tr><td>>MI0000283|hsa-mir-203a&WithFlank&14|+|104117399|104117520@72.92.21(+1U)&[MIMAT0000264&hsa-miR-203a-3p&offsets|+1|0(+1U);m-79&14|+|104117469|104117490&offsets|+1|0(+1U)]|TGAAATGTTTAGGACCACTAGT</td><td>>MI0000283|hsa-mir-203a&WithFlank&14|+|104117399|104117520</td><td>72</td><td>92</td><td>21(+1U)</td><td>TGAAATGTTTAGGACCACTAGT</td><td>2961.0</td><td>9.0</td><td>0.8</td><td>11.6</td><td>2.3E-31</td><td>4.6E-30</td><td>BT20/MB468</td></tr>
<tr><td>>MI0000088|hsa-mir-30a&WithFlank&6|-|71403545|71403627@53.74.22&[MIMAT0000088&hsa-miR-30a-3p&offsets|0|0;m-38&6|-|71403554|71403575&offsets|0|0]|CTTTCAGTCGGATGTTTGCAGC</td><td>>MI0000088|hsa-mir-30a&WithFlank&6|-|71403545|71403627</td><td>53</td><td>74</td><td>22</td><td>CTTTCAGTCGGATGTTTGCAGC</td><td>32427.6</td><td>8.7</td><td>0.7</td><td>11.6</td><td>2.3E-31</td><td>4.8E-30</td><td>MB231/MB468</td></tr>
<tr><td>>hg38_21_+_8433172_8446622_RNA45SN1_45S_50ntFlanks@10350.10376.27|ACGAGAACTTTGAAGGCCGAAGTGGAG</td><td>>hg38_21_+_8433172_8446622_RNA45SN1_45S_50ntFlanks</td><td>10350</td><td>10376</td><td>27</td><td>ACGAGAACTTTGAAGGCCGAAGTGGAG</td><td>1136.9</td><td>-6.1</td><td>0.5</td><td>-11.6</td><td>2.4E-31</td><td>4.8E-30</td><td>BT20/MB468</td></tr>
<tr><td>>trnaMT_PheGAA_MT_+_577_647@35.55.21|GAAAATGTTTAGACGGGCTCA</td><td>>trnaMT_PheGAA_MT_+_577_647</td><td>35</td><td>55</td><td>21</td><td>GAAAATGTTTAGACGGGCTCA</td><td>1504.8</td><td>11.2</td><td>1.0</td><td>11.6</td><td>2.4E-31</td><td>4.8E-30</td><td>BT20/MB468</td></tr>
<tr><td>>hg38_21_+_8433172_8446622_RNA45SN1_45S_50ntFlanks@11320.11344.25|CCACGTCTCGTCGCGCGCGCGTCCG</td><td>>hg38_21_+_8433172_8446622_RNA45SN1_45S_50ntFlanks</td><td>11320</td><td>11344</td><td>25</td><td>CCACGTCTCGTCGCGCGCGCGTCCG</td><td>1582.1</td><td>-7.6</td><td>0.7</td><td>-11.6</td><td>2.3E-31</td><td>4.8E-30</td><td>MB231/MB468</td></tr>
<tr><td>>trnaMT_PheGAA_MT_+_577_647@38.55.18|AATGTTTAGACGGGCTCA</td><td>>trnaMT_PheGAA_MT_+_577_647</td><td>38</td><td>55</td><td>18</td><td>AATGTTTAGACGGGCTCA</td><td>604.3</td><td>11.9</td><td>1.0</td><td>11.6</td><td>2.5E-31</td><td>4.9E-30</td><td>BT20/MB468</td></tr>
<tr><td>>MI0000542|hsa-mir-320a&WithFlank&8|-|22244960|22245043@48.71.24&[MIMAT0000510&hsa-miR-320a-3p&offsets|0|+2;m-60&8|-|22244975|22244996&offsets|0|+2]|AAAAGCTGGGTTGAGAGGGCGAAA</td><td>>MI0000542|hsa-mir-320a&WithFlank&8|-|22244960|22245043</td><td>48</td><td>71</td><td>24</td><td>AAAAGCTGGGTTGAGAGGGCGAAA</td><td>6515.1</td><td>9.9</td><td>0.8</td><td>11.6</td><td>3.0E-31</td><td>5.9E-30</td><td>BT20/MB468</td></tr>
<tr><td>>MI0025242|hsa-mir-7706&WithFlank&15|+|85380590|85380668@50.72.23&[MIMAT0030021&hsa-miR-7706&offsets|0|-1;m-390&15|+|85380639|85380662&offsets|0|-1]|TGAAGCGCCTGTGCTCTGCCGAG</td><td>>MI0025242|hsa-mir-7706&WithFlank&15|+|85380590|85380668</td><td>50</td><td>72</td><td>23</td><td>TGAAGCGCCTGTGCTCTGCCGAG</td><td>526.6</td><td>6.8</td><td>0.6</td><td>11.6</td><td>3.1E-31</td><td>6.2E-30</td><td>BT20/MB468</td></tr>
<tr><td>>MI0000299|hsa-mir-222&WithFlank&X|-|45747009|45747130@75.95.21(+1U)&[MIMAT0000279&hsa-miR-222-3p&offsets|0|0(+1U);m-140&X|-|45747033|45747056&offsets|0|-3(+1U)]|AGCTACATCTGGCTACTGGGTT</td><td>>MI0000299|hsa-mir-222&WithFlank&X|-|45747009|45747130</td><td>75</td><td>95</td><td>21(+1U)</td><td>AGCTACATCTGGCTACTGGGTT</td><td>713.4</td><td>8.2</td><td>0.7</td><td>11.6</td><td>3.0E-31</td><td>6.2E-30</td><td>MB231/MB468</td></tr>
<tr><td>>am_MI0000809|hsa-mir-151a&WithFlank&8|-|140732558|140732659@53.73.21(+1G)&[MIMAT0000757&hsa-miR-151a-3p&offsets|0|0(+1G);m-30&8|-|140732587|140732607&offsets|0|0(+1G)]|CTAGACTGAAGCTCCTTGAGGG</td><td>>am_MI0000809|hsa-mir-151a&WithFlank&8|-|140732558|140732659</td><td>53</td><td>73</td><td>21(+1G)</td><td>CTAGACTGAAGCTCCTTGAGGG</td><td>3314.6</td><td>8.9</td><td>0.8</td><td>11.6</td><td>3.2E-31</td><td>6.3E-30</td><td>BT20/MB468</td></tr>
<tr><td>>hg38_21_+_8433172_8446622_RNA45SN1_45S_50ntFlanks@475.512.38|CTTCGTGATCGATGTGGTGACGTCGTGCTCTCCCGGGC</td><td>>hg38_21_+_8433172_8446622_RNA45SN1_45S_50ntFlanks</td><td>475</td><td>512</td><td>38</td><td>CTTCGTGATCGATGTGGTGACGTCGTGCTCTCCCGGGC</td><td>1277.7</td><td>8.2</td><td>0.7</td><td>11.6</td><td>3.1E-31</td><td>6.4E-30</td><td>MB231/MB468</td></tr>
<tr><td>>hg38_21_+_8433172_8446622_RNA45SN1_45S_50ntFlanks@11729.11748.20|TAACTATGACTCTCTTAAGG</td><td>>hg38_21_+_8433172_8446622_RNA45SN1_45S_50ntFlanks</td><td>11729</td><td>11748</td><td>20</td><td>TAACTATGACTCTCTTAAGG</td><td>2895.7</td><td>-4.5</td><td>0.4</td><td>-11.6</td><td>3.3E-31</td><td>6.5E-30</td><td>BT20/MB468</td></tr>
<tr><td>>hg38_21_+_8433172_8446622_RNA45SN1_45S_50ntFlanks@4453.4473.21|TCGGCGCCCCCTCGATGCTCT</td><td>>hg38_21_+_8433172_8446622_RNA45SN1_45S_50ntFlanks</td><td>4453</td><td>4473</td><td>21</td><td>TCGGCGCCCCCTCGATGCTCT</td><td>811.1</td><td>14.6</td><td>1.3</td><td>11.6</td><td>3.2E-31</td><td>6.6E-30</td><td>MB231/MB468</td></tr>
<tr><td>>hg38_21_+_8433172_8446622_RNA45SN1_45S_50ntFlanks@11666.11697.32|TCTGCCCAGTGCTCTGAATGTCAAAGTGAAGA</td><td>>hg38_21_+_8433172_8446622_RNA45SN1_45S_50ntFlanks</td><td>11666</td><td>11697</td><td>32</td><td>TCTGCCCAGTGCTCTGAATGTCAAAGTGAAGA</td><td>366.6</td><td>5.3</td><td>0.5</td><td>11.6</td><td>3.3E-31</td><td>6.7E-30</td><td>MB231/MB468</td></tr>
<tr><td>>MI0000298|hsa-mir-221&WithFlank&X|-|45746151|45746272@71.93.23(+1G)&[MIMAT0000278&hsa-miR-221-3p&offsets|0|0(+1G);m-91&X|-|45746181|45746202&offsets|0|+1(+1G)]|AGCTACATTGTCTGCTGGGTTTCG</td><td>>MI0000298|hsa-mir-221&WithFlank&X|-|45746151|45746272</td><td>71</td><td>93</td><td>23(+1G)</td><td>AGCTACATTGTCTGCTGGGTTTCG</td><td>3592.6</td><td>8.1</td><td>0.7</td><td>11.6</td><td>3.3E-31</td><td>6.8E-30</td><td>MB231/MB468</td></tr>
<tr><td>>MI0000266|hsa-mir-10a&WithFlank&17|-|48579832|48579953@29.49.21(+1A)&[MIMAT0000253&hsa-miR-10a-5p&offsets|+1|-1(+1A);m-18&17|-|48579905|48579926&offsets|+1|0(+1A)]|ACCCTGTAGATCCGAATTTGTA</td><td>>MI0000266|hsa-mir-10a&WithFlank&17|-|48579832|48579953</td><td>29</td><td>49</td><td>21(+1A)</td><td>ACCCTGTAGATCCGAATTTGTA</td><td>4769.4</td><td>16.9</td><td>1.5</td><td>11.6</td><td>3.5E-31</td><td>6.8E-30</td><td>BT20/MB468</td></tr>
<tr><td>>MI0000063|hsa-let-7b&WithFlank&22|+|46113680|46113774@12.32.21&[MIMAT0000063&hsa-let-7b-5p&offsets|0|-1;m-14&22|+|46113691|46113712&offsets|0|-1]|TGAGGTAGTAGGTTGTGTGGT</td><td>>MI0000063|hsa-let-7b&WithFlank&22|+|46113680|46113774</td><td>12</td><td>32</td><td>21</td><td>TGAGGTAGTAGGTTGTGTGGT</td><td>11366.6</td><td>8.0</td><td>0.7</td><td>11.6</td><td>3.6E-31</td><td>6.9E-30</td><td>BT20/MB468</td></tr>
<tr><td>>hg38_21_+_8433172_8446622_RNA45SN1_45S_50ntFlanks@6745.6770.26|ACACTTCGAACGCACTTGCGGCCCCG</td><td>>hg38_21_+_8433172_8446622_RNA45SN1_45S_50ntFlanks</td><td>6745</td><td>6770</td><td>26</td><td>ACACTTCGAACGCACTTGCGGCCCCG</td><td>2885.3</td><td>-6.2</td><td>0.5</td><td>-11.6</td><td>3.6E-31</td><td>7.2E-30</td><td>MB231/MB468</td></tr>
<tr><td>>MI0000803|hsa-mir-330&WithFlank&19|-|45638988|45639093@63.86.24(+1U)&[MIMAT0000751&hsa-miR-330-3p&offsets|0|+1(+1U);m-347&19|-|45639009|45639030&offsets|-1|+1(+1U)]|GCAAAGCACACGGCCTGCAGAGAGT</td><td>>MI0000803|hsa-mir-330&WithFlank&19|-|45638988|45639093</td><td>63</td><td>86</td><td>24(+1U)</td><td>GCAAAGCACACGGCCTGCAGAGAGT</td><td>2187.5</td><td>8.2</td><td>0.7</td><td>11.6</td><td>3.5E-31</td><td>7.2E-30</td><td>MB231/MB468</td></tr>
<tr><td>>am_trna128_GlyGCC_6_-_27870686_27870756@1.36.36,trna18_GlyGCC_16_+_70822597_70822667@1.36.36,trna19_GlyGCC_16_+_70823410_70823480@1.36.36,trna19_GlyGCC_2_-_157257659_157257729@1.36.36,trna24_GlyGCC_16_-_70812942_70813012@1.36.36,trna25_GlyGCC_16_-_70812114_70812184@1.36.36,trna5_GlyGCC_17_+_8029064_8029134@1.36.36,trna68_GlyGCC_1_-_161493637_161493707@1.36.36|GCATTGGTGGTTCAGTGGTAGAATTCTCGCCTGCCA</td><td>>am_trna128_GlyGCC_6_-_27870686_27870756</td><td>1</td><td>36</td><td>36</td><td>GCATTGGTGGTTCAGTGGTAGAATTCTCGCCTGCCA</td><td>2188.5</td><td>8.0</td><td>0.7</td><td>11.6</td><td>3.6E-31</td><td>7.2E-30</td><td>MB231/MB468</td></tr>
<tr><td>>MI0000089|hsa-mir-31&WithFlank&9|-|21512109|21512191@14.36.23&[MIMAT0000089&hsa-miR-31-5p&offsets|0|+2;m-360&9|-|21512157|21512178&offsets|0|+1]|AGGCAAGATGCTGGCATAGCTGT</td><td>>MI0000089|hsa-mir-31&WithFlank&9|-|21512109|21512191</td><td>14</td><td>36</td><td>23</td><td>AGGCAAGATGCTGGCATAGCTGT</td><td>4934.0</td><td>8.1</td><td>0.7</td><td>11.6</td><td>3.8E-31</td><td>7.4E-30</td><td>BT20/MB468</td></tr>
<tr><td>>hg38_1_-_228634819_228635039_RNA5S12_5S_50ntFlanks@135.171.37|GGGAGACCGCCTGGGAATACCGGGTGCTGTAGGCTTT</td><td>>hg38_1_-_228634819_228635039_RNA5S12_5S_50ntFlanks</td><td>135</td><td>171</td><td>37</td><td>GGGAGACCGCCTGGGAATACCGGGTGCTGTAGGCTTT</td><td>12683.6</td><td>7.4</td><td>0.6</td><td>11.6</td><td>3.8E-31</td><td>7.5E-30</td><td>BT20/MB468</td></tr>
<tr><td>>hg38_1_-_228634819_228635039_RNA5S12_5S_50ntFlanks@147.170.24|GGGAATACCGGGTGCTGTAGGCTT</td><td>>hg38_1_-_228634819_228635039_RNA5S12_5S_50ntFlanks</td><td>147</td><td>170</td><td>24</td><td>GGGAATACCGGGTGCTGTAGGCTT</td><td>2570.8</td><td>5.5</td><td>0.5</td><td>11.6</td><td>3.9E-31</td><td>7.5E-30</td><td>BT20/MB468</td></tr>
<tr><td>>MI0001445|hsa-mir-423&WithFlank&17|+|30117073|30117178@23.46.24&[MIMAT0004748&hsa-miR-423-5p&offsets|0|+1;m-141&17|+|30117095|30117117&offsets|0|+1]|TGAGGGGCAGAGAGCGAGACTTTT</td><td>>MI0001445|hsa-mir-423&WithFlank&17|+|30117073|30117178</td><td>23</td><td>46</td><td>24</td><td>TGAGGGGCAGAGAGCGAGACTTTT</td><td>2972.5</td><td>9.0</td><td>0.8</td><td>11.6</td><td>3.8E-31</td><td>7.7E-30</td><td>MB231/MB468</td></tr>
<tr><td>>MI0000255|hsa-mir-30d&WithFlank&8|-|134804870|134804951@12.34.23&[MIMAT0000245&hsa-miR-30d-5p&offsets|0|+1;m-21&8|-|134804919|134804940&offsets|0|+1]|TGTAAACATCCCCGACTGGAAGC</td><td>>MI0000255|hsa-mir-30d&WithFlank&8|-|134804870|134804951</td><td>12</td><td>34</td><td>23</td><td>TGTAAACATCCCCGACTGGAAGC</td><td>52234.8</td><td>8.0</td><td>0.7</td><td>11.6</td><td>3.9E-31</td><td>7.7E-30</td><td>MB231/MB468</td></tr>
<tr><td>>hg38_21_+_8433172_8446622_RNA45SN1_45S_50ntFlanks@476.505.30|TTCGTGATCGATGTGGTGACGTCGTGCTCT</td><td>>hg38_21_+_8433172_8446622_RNA45SN1_45S_50ntFlanks</td><td>476</td><td>505</td><td>30</td><td>TTCGTGATCGATGTGGTGACGTCGTGCTCT</td><td>39690.0</td><td>9.3</td><td>0.8</td><td>11.6</td><td>4.0E-31</td><td>7.9E-30</td><td>MB231/MB468</td></tr>
<tr><td>>hg38_1_-_228634819_228635039_RNA5S12_5S_50ntFlanks@144.171.28|CCTGGGAATACCGGGTGCTGTAGGCTTT</td><td>>hg38_1_-_228634819_228635039_RNA5S12_5S_50ntFlanks</td><td>144</td><td>171</td><td>28</td><td>CCTGGGAATACCGGGTGCTGTAGGCTTT</td><td>22595.9</td><td>8.5</td><td>0.7</td><td>11.6</td><td>4.1E-31</td><td>8.0E-30</td><td>MB231/MB468</td></tr>
<tr><td>>MI0000542|hsa-mir-320a&WithFlank&8|-|22244960|22245043@49.70.22(+1U)&[MIMAT0000510&hsa-miR-320a-3p&offsets|+1|+1(+1U);m-60&8|-|22244975|22244996&offsets|+1|+1(+1U)]|AAAGCTGGGTTGAGAGGGCGAAT</td><td>>MI0000542|hsa-mir-320a&WithFlank&8|-|22244960|22245043</td><td>49</td><td>70</td><td>22(+1U)</td><td>AAAGCTGGGTTGAGAGGGCGAAT</td><td>1209.5</td><td>8.3</td><td>0.7</td><td>11.6</td><td>4.6E-31</td><td>8.8E-30</td><td>BT20/MB468</td></tr>
<tr><td>>hg38_MT_+_598_1651_RNR1_12S_50ntFlanks@370.404.35|ATAAAGCTAAAACTCACCTGAGTTGTAAAAAACTC</td><td>>hg38_MT_+_598_1651_RNR1_12S_50ntFlanks</td><td>370</td><td>404</td><td>35</td><td>ATAAAGCTAAAACTCACCTGAGTTGTAAAAAACTC</td><td>527.8</td><td>13.7</td><td>1.2</td><td>11.6</td><td>4.7E-31</td><td>9.1E-30</td><td>BT20/MB468</td></tr>
<tr><td>>MI0000077|hsa-mir-21&WithFlank&17|+|59841260|59841343@14.31.18&[MIMAT0000076&hsa-miR-21-5p&offsets|0|-4;m-1&17|+|59841273|59841294&offsets|0|-4]|TAGCTTATCAGACTGATG</td><td>>MI0000077|hsa-mir-21&WithFlank&17|+|59841260|59841343</td><td>14</td><td>31</td><td>18</td><td>TAGCTTATCAGACTGATG</td><td>4946.8</td><td>7.9</td><td>0.7</td><td>11.6</td><td>4.8E-31</td><td>9.3E-30</td><td>BT20/MB468</td></tr>
<tr><td>>hg38_MT_+_598_1651_RNR1_12S_50ntFlanks@428.469.42|AAAGTGGCTTTAACATATCTGAACACACAATAGCTAAGACCC</td><td>>hg38_MT_+_598_1651_RNR1_12S_50ntFlanks</td><td>428</td><td>469</td><td>42</td><td>AAAGTGGCTTTAACATATCTGAACACACAATAGCTAAGACCC</td><td>2563.7</td><td>16.2</td><td>1.4</td><td>11.6</td><td>4.7E-31</td><td>9.4E-30</td><td>MB231/MB468</td></tr>
<tr><td>>MI0003513|hsa-mir-455&WithFlank&9|+|114209428|114209535@22.43.22&[MIMAT0003150&hsa-miR-455-5p&offsets|0|0;m-232&9|+|114209449|114209470&offsets|0|0]|TATGTGCCTTTGGACTACATCG</td><td>>MI0003513|hsa-mir-455&WithFlank&9|+|114209428|114209535</td><td>22</td><td>43</td><td>22</td><td>TATGTGCCTTTGGACTACATCG</td><td>799.4</td><td>14.5</td><td>1.3</td><td>11.6</td><td>4.8E-31</td><td>9.4E-30</td><td>MB231/MB468</td></tr>
<tr><td>>MI0000736|hsa-mir-30c-1&WithFlank&1|+|40757278|40757378@23.44.22(+1A);MI0000254|hsa-mir-30c-2&WithFlank&6|-|71376954|71377037@13.34.22(+1A)&[MIMAT0000244&hsa-miR-30c-5p&offsets|0|-1(+1A);m-57&1|+|40757300|40757323&offsets|0|-2(+1A)];[MIMAT0000244_1&hsa-miR-30c-5p&offsets|0|-1(+1A);m-56&6|-|71377002|71377025&offsets|0|-2(+1A)]|TGTAAACATCCTACACTCTCAGA</td><td>>MI0000736|hsa-mir-30c-1&WithFlank&1|+|40757278|40757378</td><td>23</td><td>44</td><td>22(+1A)</td><td>TGTAAACATCCTACACTCTCAGA</td><td>4802.6</td><td>7.8</td><td>0.7</td><td>11.6</td><td>5.1E-31</td><td>1.0E-29</td><td>MB231/MB468</td></tr>
<tr><td>>hg38_21_+_8433172_8446622_RNA45SN1_45S_50ntFlanks@11668.11693.26|TGCCCAGTGCTCTGAATGTCAAAGTG</td><td>>hg38_21_+_8433172_8446622_RNA45SN1_45S_50ntFlanks</td><td>11668</td><td>11693</td><td>26</td><td>TGCCCAGTGCTCTGAATGTCAAAGTG</td><td>973.8</td><td>7.3</td><td>0.6</td><td>11.6</td><td>5.5E-31</td><td>1.1E-29</td><td>MB231/MB468</td></tr>
<tr><td>>hg38_21_+_8433172_8446622_RNA45SN1_45S_50ntFlanks@11784.11804.21|ATGAATGGATGAACGAGATTC</td><td>>hg38_21_+_8433172_8446622_RNA45SN1_45S_50ntFlanks</td><td>11784</td><td>11804</td><td>21</td><td>ATGAATGGATGAACGAGATTC</td><td>5949.1</td><td>-6.5</td><td>0.6</td><td>-11.6</td><td>5.6E-31</td><td>1.1E-29</td><td>BT20/MB468</td></tr>
</table>

| | | | | | | | | | | | | |
|---|---|---|---|---|---|---|---|---|---|---|---|---|
| >hg38_21_+_8433172_8446622_RNA45SN1_45S_50ntFlanks@11768.11803.36\|CTAATTAGTGACGCGCATGAATGGATGAACGAGATT | >hg38_21_+_8433172_8446622_RNA45SN1_45S_50ntFlanks | 11768 | 11803 | 36 | CTAATTAGTGACGCGCATGAATGGATGAACGAGATT | 1787.8 | -6.6 | 0.6 | -11.6 | 5.7E-31 | 1.1E-29 | BT20/MB468 |
| >hg38_21_+_8433172_8446622_RNA45SN1_45S_50ntFlanks@6646.6667.22\|TCGTACGACTCTTAGCGGTGGA | >hg38_21_+_8433172_8446622_RNA45SN1_45S_50ntFlanks | 6646 | 6667 | 22 | TCGTACGACTCTTAGCGGTGGA | 15778.9 | 6.6 | 0.6 | 11.6 | 5.7E-31 | 1.1E-29 | BT20/MB468 |
| >MI0000750\|hsa-mir-26a-2&WithFlank&12\|-\|57824603\|57824698@23.41.19;MI0000083\|hsa-mir-26a-1&WithFlank&3\|+\|37969398\|37969486@19.37.19&[MIMAT0000082&hsa-miR-26a-5p&offsets\|+3\|0;m-33&12\|-\|57824658\|57824679&offsets\|+3\|0];[MIMAT0000082_1&hsa-miR-26a-5p&offsets\|+3\|0;m-34&3\|+\|37969413\|37969434&offsets\|+3\|0]\|AAGTAATCCAGGATAGGCT | >MI0000750\|hsa-mir-26a-2&WithFlank&12\|-\|57824603\|57824698 | 23 | 41 | 19 | AAGTAATCCAGGATAGGCT | 1738.5 | 15.4 | 1.3 | 11.6 | 5.9E-31 | 1.1E-29 | BT20/MB468 |
| >hg38_21_+_8433172_8446622_RNA45SN1_45S_50ntFlanks@475.495.21\|CTTCGTGATCGATGTGGTGAC | >hg38_21_+_8433172_8446622_RNA45SN1_45S_50ntFlanks | 475 | 495 | 21 | CTTCGTGATCGATGTGGTGAC | 4118.8 | 7.9 | 0.7 | 11.6 | 5.9E-31 | 1.1E-29 | MB231/MB468 |
| >hg38_21_+_8433172_8446622_RNA45SN1_45S_50ntFlanks@12996.13014.19\|TCGCTGCGATCTATTGAAA | >hg38_21_+_8433172_8446622_RNA45SN1_45S_50ntFlanks | 12996 | 13014 | 19 | TCGCTGCGATCTATTGAAA | 3612.9 | 6.1 | 0.5 | 11.6 | 6.0E-31 | 1.2E-29 | MB231/MB468 |
| >hg38_21_+_8433172_8446622_RNA45SN1_45S_50ntFlanks@11209.11232.24\|CGGCGACTCTGGACGCGAGCCGGG | >hg38_21_+_8433172_8446622_RNA45SN1_45S_50ntFlanks | 11209 | 11232 | 24 | CGGCGACTCTGGACGCGAGCCGGG | 8609.8 | -7.2 | 0.6 | -11.6 | 6.2E-31 | 1.2E-29 | MB231/MB468 |
| >MI0000077\|hsa-mir-21&WithFlank&17\|+\|59841260\|59841343@14.39.26(+1U)&[MIMAT0000076&hsa-miR-21-5p&offsets\|0\|+4(+1U);m-1&17\|+\|59841273\|59841294&offsets\|0\|+4(+1U)]\|TAGCTTATCAGACTGATGTTGACTGTT | >MI0000077\|hsa-mir-21&WithFlank&17\|+\|59841260\|59841343 | 14 | 39 | 26(+1U) | TAGCTTATCAGACTGATGTTGACTGTT | 944.1 | 6.6 | 0.6 | 11.6 | 7.0E-31 | 1.3E-29 | BT20/MB468 |
| >hg38_1_-_228634819_228635039_RNA5S12_5S_50ntFlanks@138.171.34\|AGACCGCCTGGGAATACCGGGTGCTGTAGGCTTT | >hg38_1_-_228634819_228635039_RNA5S12_5S_50ntFlanks | 138 | 171 | 34 | AGACCGCCTGGGAATACCGGGTGCTGTAGGCTTT | 3851.9 | 8.9 | 0.8 | 11.6 | 6.8E-31 | 1.3E-29 | MB231/MB468 |
| >hg38_21_+_8433172_8446622_RNA45SN1_45S_50ntFlanks@11669.11693.25\|GCCCAGTGCTCTGAATGTCAAAGTG | >hg38_21_+_8433172_8446622_RNA45SN1_45S_50ntFlanks | 11669 | 11693 | 25 | GCCCAGTGCTCTGAATGTCAAAGTG | 706.7 | 6.9 | 0.6 | 11.6 | 7.0E-31 | 1.4E-29 | MB231/MB468 |
| >MI0000285\|hsa-mir-205&WithFlank&1\|+\|209432127\|209432248@40.61.22(+1C)&[MIMAT0000266&hsa-miR-205-5p&offsets\|0\|0(+1C);m-351&1\|+\|209432166\|209432187&offsets\|0\|0(+1C)]\|TCCTTCATTCCACCGGAGTCTGC | >MI0000285\|hsa-mir-205&WithFlank&1\|+\|209432127\|209432248 | 40 | 61 | 22(+1C) | TCCTTCATTCCACCGGAGTCTGC | 5886.3 | 16.0 | 1.4 | 11.8 | 4.2E-32 | 1.4E-29 | BT20/MB231 |
| >MI0000650\|hsa-mir-200c&WithFlank&12\|+\|6963693\|6963772@50.71.22(+1G)&[MIMAT0000617&hsa-miR-200c-3p&offsets\|0\|-1(+1G);m-54&12\|+\|6963742\|6963764&offsets\|0\|-1(+1G)]\|TAATACTGCCGGGTAATGATGGG | >MI0000650\|hsa-mir-200c&WithFlank&12\|+\|6963693\|6963772 | 50 | 71 | 22(+1G) | TAATACTGCCGGGTAATGATGGG | 9129.0 | 11.2 | 1.0 | 11.8 | 3.9E-32 | 1.4E-29 | BT20/MB231 |
| >MI0000298\|hsa-mir-221&WithFlank&X\|-\|45746151\|45746272@71.91.21(+1G)&[MIMAT0000278&hsa-miR-221-3p&offsets\|0\|-2(+1G);m-91&X\|-\|45746181\|45746202&offsets\|0\|-1(+1G)]\|AGCTACATTGTCTGCTGGGTTG | >MI0000298\|hsa-mir-221&WithFlank&X\|-\|45746151\|45746272 | 71 | 91 | 21(+1G) | AGCTACATTGTCTGCTGGGTTG | 1755.5 | 9.2 | 0.8 | 11.6 | 7.3E-31 | 1.4E-29 | MB231/MB468 |
| >trnaMT_ValTAC_MT_+_1602_1670@41.71.31\|GAGATTTCAACTTAACTTGACCGCTCTGACC | >trnaMT_ValTAC_MT_+_1602_1670 | 41 | 71 | 31 | GAGATTTCAACTTAACTTGACCGCTCTGACC | 684.2 | 7.4 | 0.6 | 11.5 | 7.4E-31 | 1.4E-29 | MB231/MB468 |
| >MI0003560\|hsa-mir-92b&WithFlank&1\|+\|155195171\|155195278@67.87.21(+1A)&[MIMAT0003218&hsa-miR-92b-3p&offsets\|0\|-1(+1A);m-115&1\|+\|155195237\|155195258&offsets\|0\|-1(+1A)]\|TATTGCACTCGTCCCGGCCTCA | >MI0003560\|hsa-mir-92b&WithFlank&1\|+\|155195171\|155195278 | 67 | 87 | 21(+1A) | TATTGCACTCGTCCCGGCCTCA | 3931.4 | 9.8 | 0.9 | 11.5 | 8.0E-31 | 1.5E-29 | BT20/MB468 |
| >hg38_1_-_228634819_228635039_RNA5S12_5S_50ntFlanks@51.88.38\|GTCTACGGCCATACCACCCTGAACGCGCCCGATCTCGT | >hg38_1_-_228634819_228635039_RNA5S12_5S_50ntFlanks | 51 | 88 | 38 | GTCTACGGCCATACCACCCTGAACGCGCCCGATCTCGT | 3115.1 | 9.5 | 0.8 | 11.5 | 9.0E-31 | 1.7E-29 | MB231/MB468 |
| >trnaMT_PheGAA_MT_+_577_647@35.62.28\|GAAAATGTTTAGACGGGCTCACATCACC | >trnaMT_PheGAA_MT_+_577_647 | 35 | 62 | 28 | GAAAATGTTTAGACGGGCTCACATCACC | 670.2 | 10.7 | 0.9 | 11.5 | 9.0E-31 | 1.7E-29 | MB231/MB468 |
| >am_trnaMT_HisGTG_MT_+_12138_12206@18.51.34\|AAAACATCAGATTGTGAATCTGACAACAGAGGCT | >am_trnaMT_HisGTG_MT_+_12138_12206 | 18 | 51 | 34 | AAAACATCAGATTGTGAATCTGACAACAGAGGCT | 378.0 | 13.2 | 1.1 | 11.5 | 9.5E-31 | 1.8E-29 | BT20/MB468 |
| >MI0006444\|hsa-mir-1307&WithFlank&10\|-\|103394247\|103394407@86.107.22(+1U)&[MIMAT0005951&hsa-miR-1307-3p&offsets\|0\|0(+1U);m-66&10\|-\|103394300\|103394322&offsets\|0\|-1(+1U)]\|ACTCGGCGTGGCGTCGGTCGTGT | >MI0006444\|hsa-mir-1307&WithFlank&10\|-\|103394247\|103394407 | 86 | 107 | 22(+1U) | ACTCGGCGTGGCGTCGGTCGTGT | 3712.4 | 8.6 | 0.7 | 11.5 | 1.0E-30 | 1.9E-29 | MB231/MB468 |
| >MI0000294\|hsa-mir-218-1&WithFlank&4\|+\|20528269\|20528390@31.52.22;MI0000295\|hsa-mir-218-2&WithFlank&5\|-\|168768140\|168768261@31.52.22&[MIMAT0000275&hsa-miR-218-5p&offsets\|0\|+1;m-185&4\|+\|20528299\|20528319&offsets\|0\|+1];[MIMAT0000275_1&hsa-miR-218-5p&offsets\|0\|+1;m-187&5\|-\|168768211\|168768231&offsets\|0\|+1]\|TTGTGCTTGATCTAACCATGTG | >MI0000294\|hsa-mir-218-1&WithFlank&4\|+\|20528269\|20528390 | 31 | 52 | 22 | TTGTGCTTGATCTAACCATGTG | 830.1 | 14.6 | 1.3 | 11.5 | 1.1E-30 | 2.0E-29 | MB231/MB468 |
| >hg38_MT_+_1621_3279_RNR2_16S_50ntFlanks@1184.1216.33\|AATTTCGGTTGGGGCGACCTCGGAGCAGAACCC | >hg38_MT_+_1621_3279_RNR2_16S_50ntFlanks | 1184 | 1216 | 33 | AATTTCGGTTGGGGCGACCTCGGAGCAGAACCC | 402.6 | 13.6 | 1.2 | 11.5 | 1.1E-30 | 2.1E-29 | MB231/MB468 |
| >MI0000809\|hsa-mir-151a&WithFlank&8\|-\|140732558\|140732659@53.74.22(+1A)&[MIMAT0000757&hsa-miR-151a-3p&offsets\|0\|+1(+1A);m-30&8\|-\|140732587\|140732607&offsets\|0\|+1(+1A)]\|CTAGACTGAAGCTCCTTGAGGAA | >MI0000809\|hsa-mir-151a&WithFlank&8\|-\|140732558\|140732659 | 53 | 74 | 22(+1A) | CTAGACTGAAGCTCCTTGAGGAA | 6820.2 | 7.9 | 0.7 | 11.5 | 1.1E-30 | 2.1E-29 | MB231/MB468 |
| >MI0000285\|hsa-mir-205&WithFlank&1\|+\|209432127\|209432248@40.60.21&[MIMAT0000266&hsa-miR-205-5p&offsets\|0\|-1;m-351&1\|+\|209432166\|209432187&offsets\|0\|-1]\|TCCTTCATTCCACCGGAGTCT | >MI0000285\|hsa-mir-205&WithFlank&1\|+\|209432127\|209432248 | 40 | 60 | 21 | TCCTTCATTCCACCGGAGTCT | 6906.1 | 16.3 | 1.4 | 11.8 | 7.0E-32 | 2.1E-29 | BT20/MB231 |
| >MI0000255\|hsa-mir-30d&WithFlank&8\|-\|134804870\|134804951@12.33.22&[MIMAT0000245&hsa-miR-30d-5p&offsets\|0\|0;m-21&8\|-\|134804919\|134804940&offsets\|0\|0]\|TGTAAACATCCCCGACTGGAAG | >MI0000255\|hsa-mir-30d&WithFlank&8\|-\|134804870\|134804951 | 12 | 33 | 22 | TGTAAACATCCCCGACTGGAAG | 6156.5 | 7.9 | 0.7 | 11.5 | 1.2E-30 | 2.2E-29 | MB231/MB468 |
| >hg38_21_+_8433172_8446622_RNA45SN1_45S_50ntFlanks@12627.12653.27\|ATGGGGCGAAGCTACCATCTGTGGGAT | >hg38_21_+_8433172_8446622_RNA45SN1_45S_50ntFlanks | 12627 | 12653 | 27 | ATGGGGCGAAGCTACCATCTGTGGGAT | 3586.7 | -10.7 | 0.9 | -11.5 | 1.2E-30 | 2.2E-29 | BT20/MB468 |
| >MI0001733\|hsa-mir-452&WithFlank&X\|-\|151959622\|151959718@20.43.24&[MIMAT0001635&hsa-miR-452-5p&offsets\|0\|+2;m-188&X\|-\|151959677\|151959699&offsets\|0\|+1]\|AACTGTTTGCAGAGGAAACTGAGA | >MI0001733\|hsa-mir-452&WithFlank&X\|-\|151959622\|151959718 | 20 | 43 | 24 | AACTGTTTGCAGAGGAAACTGAGA | 3129.6 | 14.2 | 1.2 | 11.5 | 1.3E-30 | 2.3E-29 | BT20/MB468 |
| >hg38_21_+_8433172_8446622_RNA45SN1_45S_50ntFlanks@12456.12475.20\|CCAAGCGTTGGATTGTTCAC | >hg38_21_+_8433172_8446622_RNA45SN1_45S_50ntFlanks | 12456 | 12475 | 20 | CCAAGCGTTGGATTGTTCAC | 22197.5 | -10.8 | 0.9 | -11.5 | 1.3E-30 | 2.4E-29 | BT20/MB468 |
| >hg38_21_+_8433172_8446622_RNA45SN1_45S_50ntFlanks@475.497.23\|CTTCGTGATCGATGTGGTGACGT | >hg38_21_+_8433172_8446622_RNA45SN1_45S_50ntFlanks | 475 | 497 | 23 | CTTCGTGATCGATGTGGTGACGT | 59049.5 | 9.1 | 0.8 | 11.5 | 1.4E-30 | 2.6E-29 | MB231/MB468 |
| >trnaMT_GluTTC_MT_-_14674_14742@38.72.35,trnalookalike8_GluTTC_5_-_93905172_93905240@38.72.35\|CATTGGTCGTGGTTGTAGTCCGTGCGAGAATACCA | >trnaMT_GluTTC_MT_-_14674_14742 | 38 | 72 | 35 | CATTGGTCGTGGTTGTAGTCCGTGCGAGAATACCA | 5322.2 | 8.3 | 0.7 | 11.5 | 1.4E-30 | 2.6E-29 | BT20/MB468 |
| >MI0001445\|hsa-mir-423&WithFlank&17\|+\|30117073\|30117178@59.81.23(+1U)&[MIMAT0001340&hsa-miR-423-3p&offsets\|0\|0(+1U);m-100&17\|+\|30117131\|30117153&offsets\|0\|0(+1U)]\|AGCTCGGTCTGAGGCCCCTCAGTT | >MI0001445\|hsa-mir-423&WithFlank&17\|+\|30117073\|30117178 | 59 | 81 | 23(+1U) | AGCTCGGTCTGAGGCCCCTCAGTT | 7588.3 | 8.9 | 0.8 | 11.5 | 1.5E-30 | 2.7E-29 | MB231/MB468 |
| >hg38_21_+_8433172_8446622_RNA45SN1_45S_50ntFlanks@10351.10377.27\|CGAGAACTTTGAAGGCCGAAGTGGAGA | >hg38_21_+_8433172_8446622_RNA45SN1_45S_50ntFlanks | 10351 | 10377 | 27 | CGAGAACTTTGAAGGCCGAAGTGGAGA | 4935.7 | -5.7 | 0.5 | -11.5 | 1.5E-30 | 2.8E-29 | MB231/MB468 |
| >MI0001733\|hsa-mir-452&WithFlank&X\|-\|151959622\|151959718@23.44.22&[MIMAT0001635&hsa-miR-452-5p&offsets\|+3\|+3;m-188&X\|-\|151959677\|151959699&offsets\|+3\|+2]\|TGTTTGCAGAGGAAACTGAGAC | >MI0001733\|hsa-mir-452&WithFlank&X\|-\|151959622\|151959718 | 23 | 44 | 22 | TGTTTGCAGAGGAAACTGAGAC | 987.9 | 14.6 | 1.3 | 11.5 | 1.6E-30 | 2.9E-29 | BT20/MB468 |
| >hg38_21_+_8433172_8446622_RNA45SN1_45S_50ntFlanks@5491.5516.26\|GAGAAGACGGTCGAACTTGACTATCT | >hg38_21_+_8433172_8446622_RNA45SN1_45S_50ntFlanks | 5491 | 5516 | 26 | GAGAAGACGGTCGAACTTGACTATCT | 2594.5 | 8.4 | 0.7 | 11.5 | 1.6E-30 | 2.9E-29 | MB231/MB468 |
| >hg38_21_+_8433172_8446622_RNA45SN1_45S_50ntFlanks@12626.12653.28\|AATGGGGCGAAGCTACCATCTGTGGGAT | >hg38_21_+_8433172_8446622_RNA45SN1_45S_50ntFlanks | 12626 | 12653 | 28 | AATGGGGCGAAGCTACCATCTGTGGGAT | 3352.7 | -6.8 | 0.6 | -11.5 | 1.6E-30 | 2.9E-29 | MB231/MB468 |
| >hg38_21_+_8433172_8446622_RNA45SN1_45S_50ntFlanks@10351.10377.27\|CGAGAACTTTGAAGGCCGAAGTGGAGA | >hg38_21_+_8433172_8446622_RNA45SN1_45S_50ntFlanks | 10351 | 10377 | 27 | CGAGAACTTTGAAGGCCGAAGTGGAGA | 5863.8 | -6.7 | 0.6 | -11.5 | 1.7E-30 | 3.0E-29 | BT20/MB468 |
| >MI0000067\|hsa-let-7f-1&WithFlank&9\|+\|94176341\|94176439@13.31.19;MI0000068\|hsa-let-7f-2&WithFlank&X\|-\|53557186\|53557280@14.32.19&[MIMAT0000067&hsa-let-7f-5p&offsets\|0\|-3;m-20&9\|+\|94176353\|94176374&offsets\|0\|-3];[MIMAT0000067_1&hsa-let-7f-5p&offsets\|0\|-3;m-19&X\|-\|53557246\|53557267&offsets\|0\|-3]\|TGAGGTAGTAGATTGTATA | >MI0000067\|hsa-let-7f-1&WithFlank&9\|+\|94176341\|94176439 | 13 | 31 | 19 | TGAGGTAGTAGATTGTATA | 2378.9 | 7.6 | 0.7 | 11.5 | 1.7E-30 | 3.0E-29 | BT20/MB468 |
| >hg38_21_+_8433172_8446622_RNA45SN1_45S_50ntFlanks@6652.6695.44\|GACTCTTAGCGGTGGATCACTCGGCTCGTGCGTCGATGAAGAAC | >hg38_21_+_8433172_8446622_RNA45SN1_45S_50ntFlanks | 6652 | 6695 | 44 | GACTCTTAGCGGTGGATCACTCGGCTCGTGCGTCGATGAAGAAC | 1120.2 | 8.3 | 0.7 | 11.5 | 1.8E-30 | 3.2E-29 | BT20/MB468 |
| >MI0000542\|hsa-mir-320a&WithFlank&8\|-\|22244960\|22245043@48.70.23(+2U)&[MIMAT0000510&hsa-miR-320a-3p&offsets\|0\|+1(+2U);m-60&8\|-\|22244975\|22244996&offsets\|0\|+1(+2U)]\|AAAAGCTGGGTTGAGAGGGCGAATT | >MI0000542\|hsa-mir-320a&WithFlank&8\|-\|22244960\|22245043 | 48 | 70 | 23(+2U) | AAAAGCTGGGTTGAGAGGGCGAATT | 1242.6 | 8.9 | 0.8 | 11.5 | 1.8E-30 | 3.2E-29 | BT20/MB468 |
| >hg38_21_+_8433172_8446622_RNA45SN1_45S_50ntFlanks@6647.6678.32\|CGTACGACTCTTAGCGGTGGATCACTCGGCTC | >hg38_21_+_8433172_8446622_RNA45SN1_45S_50ntFlanks | 6647 | 6678 | 32 | CGTACGACTCTTAGCGGTGGATCACTCGGCTC | 567.2 | 5.2 | 0.5 | 11.5 | 1.8E-30 | 3.3E-29 | BT20/MB468 |

<table>
<tr><th></th><th></th><th></th><th></th><th></th><th></th><th></th><th></th><th></th><th></th><th></th><th></th><th></th></tr>
<tr><td>>hg38_1_-_228634819_228635039_RNA5S12_5S_50ntFlanks@51.89.39|GTCTACGGCCATACCACCCTGAACGCGCCCGATCTCGTC</td><td>>hg38_1_-_228634819_228635039_RNA5S12_5S_50ntFlanks</td><td>51</td><td>89</td><td>39</td><td>GTCTACGGCCATACCACCCTGAACGCGCCCGATCTCGTC</td><td>44400.0</td><td>11.0</td><td>1.0</td><td>11.5</td><td>1.8E-30</td><td>3.3E-29</td><td>BT20/MB468</td></tr>
<tr><td>>hg38_MT_+_1621_3279_RNR2_16S_50ntFlanks@617.642.26|AAACATATAACTGAACTCCTCACACC</td><td>>hg38_MT_+_1621_3279_RNR2_16S_50ntFlanks</td><td>617</td><td>642</td><td>26</td><td>AAACATATAACTGAACTCCTCACACC</td><td>766.2</td><td>14.5</td><td>1.3</td><td>11.5</td><td>1.8E-30</td><td>3.3E-29</td><td>MB231/MB468</td></tr>
<tr><td>>MI0003599|hsa-mir-589&WithFlank&7|-|5495813|5495923@30.51.22(+1U)&[MIMAT0004799&hsa-miR-589-5p&offsets|0|0(+1U);m-162&7|-|5495874|5495894&offsets|0|+1(+1U)]|TGAGAACCACGTCTGCTCTGAGT</td><td>>MI0003599|hsa-mir-589&WithFlank&7|-|5495813|5495923</td><td>30</td><td>51</td><td>22(+1U)</td><td>TGAGAACCACGTCTGCTCTGAGT</td><td>333.5</td><td>13.0</td><td>1.1</td><td>11.5</td><td>1.9E-30</td><td>3.4E-29</td><td>BT20/MB468</td></tr>
<tr><td>>am_MI0000809|hsa-mir-151a&WithFlank&8|-|140732558|140732659@53.70.18&[MIMAT0000757&hsa-miR-151a-3p&offsets|0|-3;m-30&8|-|140732587|140732607&offsets|0|-3]|CTAGACTGAAGCTCCTTG</td><td>>am_MI0000809|hsa-mir-151a&WithFlank&8|-|140732558|140732659</td><td>53</td><td>70</td><td>18</td><td>CTAGACTGAAGCTCCTTG</td><td>2417.5</td><td>9.5</td><td>0.8</td><td>11.5</td><td>1.8E-30</td><td>3.4E-29</td><td>MB231/MB468</td></tr>
<tr><td>>hg38_21_+_8433172_8446622_RNA45SN1_45S_50ntFlanks@12995.13036.42|CTCGCTGCGATCTATTGAAAGTCAGCCCTCGACACAAGGGTT</td><td>>hg38_21_+_8433172_8446622_RNA45SN1_45S_50ntFlanks</td><td>12995</td><td>13036</td><td>42</td><td>CTCGCTGCGATCTATTGAAAGTCAGCCCTCGACACAAGGGTT</td><td>5920.6</td><td>11.2</td><td>1.0</td><td>11.5</td><td>2.0E-30</td><td>3.6E-29</td><td>MB231/MB468</td></tr>
<tr><td>>hg38_21_+_8433172_8446622_RNA45SN1_45S_50ntFlanks@11575.11592.18|ACTTAGAACTGGTGCGGA</td><td>>hg38_21_+_8433172_8446622_RNA45SN1_45S_50ntFlanks</td><td>11575</td><td>11592</td><td>18</td><td>ACTTAGAACTGGTGCGGA</td><td>2837.6</td><td>-7.3</td><td>0.6</td><td>-11.5</td><td>2.0E-30</td><td>3.6E-29</td><td>MB231/MB468</td></tr>
<tr><td>>MI0000253|hsa-mir-148a&WithFlank&7|-|25949913|25949992@12.33.22&[MIMAT0004549&hsa-miR-148a-5p&offsets|0|0;m-189&7|-|25949960|25949981&offsets|0|0]|AAAGTTCTGAGACACTCCGACT</td><td>>MI0000253|hsa-mir-148a&WithFlank&7|-|25949913|25949992</td><td>12</td><td>33</td><td>22</td><td>AAAGTTCTGAGACACTCCGACT</td><td>1526.9</td><td>7.4</td><td>0.6</td><td>11.5</td><td>2.1E-30</td><td>3.7E-29</td><td>BT20/MB468</td></tr>
<tr><td>>MI0000433|hsa-let-7g&WithFlank&3|-|52268272|52268367@11.31.21(+1A)&[MIMAT0000414&hsa-let-7g-5p&offsets|0|-1(+1A);m-58&3|-|52268336|52268357&offsets|0|-1(+1A)]|TGAGGTAGTAGTTTGTACAGTA</td><td>>MI0000433|hsa-let-7g&WithFlank&3|-|52268272|52268367</td><td>11</td><td>31</td><td>21(+1A)</td><td>TGAGGTAGTAGTTTGTACAGTA</td><td>10591.4</td><td>7.7</td><td>0.7</td><td>11.5</td><td>2.1E-30</td><td>3.8E-29</td><td>BT20/MB468</td></tr>
<tr><td>>hg38_21_+_8433172_8446622_RNA45SN1_45S_50ntFlanks@485.509.25|GATGTGGTGACGTCGTGCTCTCCCG</td><td>>hg38_21_+_8433172_8446622_RNA45SN1_45S_50ntFlanks</td><td>485</td><td>509</td><td>25</td><td>GATGTGGTGACGTCGTGCTCTCCCG</td><td>2846.3</td><td>16.1</td><td>1.4</td><td>11.5</td><td>2.2E-30</td><td>3.9E-29</td><td>BT20/MB468</td></tr>
<tr><td>>MI0000272|hsa-mir-182&WithFlank&7|-|129770377|129770498@29.51.23(+1C)&[MIMAT0000259&hsa-miR-182-5p&offsets|0|-1(+1C);m-22&7|-|129770449|129770470&offsets|0|+1(+1C)]|TTTGGCAATGGTAGAACTCACACC</td><td>>MI0000272|hsa-mir-182&WithFlank&7|-|129770377|129770498</td><td>29</td><td>51</td><td>23(+1C)</td><td>TTTGGCAATGGTAGAACTCACACC</td><td>23332.0</td><td>8.8</td><td>0.8</td><td>11.5</td><td>2.2E-30</td><td>3.9E-29</td><td>BT20/MB468</td></tr>
<tr><td>>MI0000266|hsa-mir-10a&WithFlank&17|-|48579832|48579953@29.49.21(+1A)&[MIMAT0000253&hsa-miR-10a-5p&offsets|+1|-1(+1A);m-18&17|-|48579905|48579926&offsets|+1|0(+1A)]|ACCCTGTAGATCCGAATTTGTA</td><td>>MI0000266|hsa-mir-10a&WithFlank&17|-|48579832|48579953</td><td>29</td><td>49</td><td>21(+1A)</td><td>ACCCTGTAGATCCGAATTTGTA</td><td>1642.9</td><td>15.6</td><td>1.4</td><td>11.5</td><td>2.3E-30</td><td>4.2E-29</td><td>MB231/MB468</td></tr>
<tr><td>>MI0001448|hsa-mir-425&WithFlank&3|-|49020142|49020240@20.44.25&[MIMAT0003393&hsa-miR-425-5p&offsets|0|+2;m-95&3|-|49020200|49020221&offsets|0|+3]|AATGACACGATCACTCCCGTTGAGT</td><td>>MI0001448|hsa-mir-425&WithFlank&3|-|49020142|49020240</td><td>20</td><td>44</td><td>25</td><td>AATGACACGATCACTCCCGTTGAGT</td><td>20819.1</td><td>8.8</td><td>0.8</td><td>11.4</td><td>2.4E-30</td><td>4.4E-29</td><td>MB231/MB468</td></tr>
<tr><td>>hg38_MT_+_598_1651_RNR1_12S_50ntFlanks@967.1004.38|AACATGGTAAGTGTACTGGAAAGTGCACTTGGACGAAC</td><td>>hg38_MT_+_598_1651_RNR1_12S_50ntFlanks</td><td>967</td><td>1004</td><td>38</td><td>AACATGGTAAGTGTACTGGAAAGTGCACTTGGACGAAC</td><td>678.6</td><td>14.3</td><td>1.3</td><td>11.4</td><td>2.5E-30</td><td>4.4E-29</td><td>MB231/MB468</td></tr>
<tr><td>>hg38_MT_+_1621_3279_RNR2_16S_50ntFlanks@1184.1215.32|AATTTCGGTTGGGGCGACCTCGGAGCAGAACC</td><td>>hg38_MT_+_1621_3279_RNR2_16S_50ntFlanks</td><td>1184</td><td>1215</td><td>32</td><td>AATTTCGGTTGGGGCGACCTCGGAGCAGAACC</td><td>580.5</td><td>14.1</td><td>1.2</td><td>11.4</td><td>2.5E-30</td><td>4.5E-29</td><td>MB231/MB468</td></tr>
<tr><td>>hg38_MT_+_1621_3279_RNR2_16S_50ntFlanks@1416.1446.31|GTTTGTTCAACGATTAAAGTCCTACGTGATC</td><td>>hg38_MT_+_1621_3279_RNR2_16S_50ntFlanks</td><td>1416</td><td>1446</td><td>31</td><td>GTTTGTTCAACGATTAAAGTCCTACGTGATC</td><td>369.0</td><td>10.4</td><td>0.9</td><td>11.4</td><td>2.5E-30</td><td>4.6E-29</td><td>MB231/MB468</td></tr>
<tr><td>>MI0000108|hsa-mir-103a-2&WithFlank&20|+|3917488|3917577@54.76.23(+2U);MI0000109|hsa-mir-103a-1&WithFlank&5|-|168560890|168560979@54.76.23(+2U)&[MIMAT0000101&hsa-miR-103a-3p&offsets|0|0(+2U);m-12&20|+|3917541|3917563&offsets|0|0(+2U)];[MIMAT0000101_1&hsa-miR-103a-3p&offsets|0|0(+2U);m-13&5|-|168560904|168560926&offsets|0|0(+2U);m-44220&5|-|168560908|168560926&offsets|0|+4(+2U)]|AGCAGCATTGTACAGGGCTATGATT</td><td>>MI0000108|hsa-mir-103a-2&WithFlank&20|+|3917488|3917577</td><td>54</td><td>76</td><td>23(+2U)</td><td>AGCAGCATTGTACAGGGCTATGATT</td><td>4695.6</td><td>8.7</td><td>0.8</td><td>11.4</td><td>2.6E-30</td><td>4.6E-29</td><td>MB231/MB468</td></tr>
<tr><td>>MI0001445|hsa-mir-423&WithFlank&17|+|30117073|30117178@58.81.24&[MIMAT0001340&hsa-miR-423-3p&offsets|-1|0;m-100&17|+|30117131|30117153&offsets|-1|0]|AAGCTCGGTCTGAGGCCCCTCAGT</td><td>>MI0001445|hsa-mir-423&WithFlank&17|+|30117073|30117178</td><td>58</td><td>81</td><td>24</td><td>AAGCTCGGTCTGAGGCCCCTCAGT</td><td>9688.5</td><td>8.7</td><td>0.8</td><td>11.4</td><td>2.6E-30</td><td>4.6E-29</td><td>MB231/MB468</td></tr>
<tr><td>>hg38_21_+_8433172_8446622_RNA45SN1_45S_50ntFlanks@11703.11728.26|AATGAAGCGCGGGTAAACGGCGGGAG</td><td>>hg38_21_+_8433172_8446622_RNA45SN1_45S_50ntFlanks</td><td>11703</td><td>11728</td><td>26</td><td>AATGAAGCGCGGGTAAACGGCGGGAG</td><td>1427.8</td><td>-6.0</td><td>0.5</td><td>-11.4</td><td>2.6E-30</td><td>4.6E-29</td><td>MB231/MB468</td></tr>
<tr><td>>hg38_21_+_8433172_8446622_RNA45SN1_45S_50ntFlanks@475.515.41|CTTCGTGATCGATGTGGTGACGTCGTGCTCTCCCGGGCCGG</td><td>>hg38_21_+_8433172_8446622_RNA45SN1_45S_50ntFlanks</td><td>475</td><td>515</td><td>41</td><td>CTTCGTGATCGATGTGGTGACGTCGTGCTCTCCCGGGCCGG</td><td>1650.1</td><td>7.9</td><td>0.7</td><td>11.4</td><td>2.6E-30</td><td>4.6E-29</td><td>BT20/MB468</td></tr>
<tr><td>>hg38_21_+_8433172_8446622_RNA45SN1_45S_50ntFlanks@10678.10706.29|CGGTTCCGGCGGCGTCCGGTGAGCTCTCG</td><td>>hg38_21_+_8433172_8446622_RNA45SN1_45S_50ntFlanks</td><td>10678</td><td>10706</td><td>29</td><td>CGGTTCCGGCGGCGTCCGGTGAGCTCTCG</td><td>15551.2</td><td>-8.2</td><td>0.7</td><td>-11.4</td><td>2.7E-30</td><td>4.8E-29</td><td>MB231/MB468</td></tr>
<tr><td>>am_MI0005763|hsa-mir-941-1&WithFlank&20|+|63919443|63919526@53.74.22;MI0005764|hsa-mir-941-2&WithFlank&20|+|63919499|63919582@53.74.22;MI0005765|hsa-mir-941-3&WithFlank&20|+|63919555|63919638@53.74.22;MI0005766|hsa-mir-941-4&WithFlank&20|+|63919750|63919833@53.74.22;MI0031520|hsa-mir-941-5&WithFlank&20|+|63919862|63919945@53.74.22&[MIMAT0004984&hsa-miR-941&offsets|0|-1;m-263&20|+|63919495|63919517&offsets|0|-1];[MIMAT0004984_1&hsa-miR-941&offsets|0|-1;m-263&20|+|63919551|63919573&offsets|0|-1];[MIMAT0004984_2&hsa-miR-941&offsets|0|-1;m-263&20|+|63919607|63919629&offsets|0|-1];[MIMAT0004984_3&hsa-miR-941&offsets|0|-1;m-263&20|+|63919802|63919824&offsets|0|-1];[MIMAT0004984_4&hsa-miR-941&offsets|0|-1;m-263&20|+|63919914|63919936&offsets|0|-1]|CACCCGGCTGTGTGCACATGTG</td><td>>am_MI0005763|hsa-mir-941-1&WithFlank&20|+|63919443|63919526</td><td>53</td><td>74</td><td>22</td><td>CACCCGGCTGTGTGCACATGTG</td><td>2286.4</td><td>8.9</td><td>0.8</td><td>11.4</td><td>2.7E-30</td><td>4.9E-29</td><td>MB231/MB468</td></tr>
<tr><td>>trnaMT_GluTTC_MT_-_14674_14742@40.72.33,trnalookalike8_GluTTC_5_-_93905172_93905240@40.72.33|TTGGTCGTGGTTGTAGTCCGTGCGAGAATACCA</td><td>>trnaMT_GluTTC_MT_-_14674_14742</td><td>40</td><td>72</td><td>33</td><td>TTGGTCGTGGTTGTAGTCCGTGCGAGAATACCA</td><td>15051.6</td><td>6.9</td><td>0.6</td><td>11.4</td><td>2.8E-30</td><td>5.0E-29</td><td>BT20/MB468</td></tr>
<tr><td>>trna111_HisGTG_1_-_147774845_147774916@-1G.35.36,trna118_HisGTG_1_-_145396881_145396952@-1G.35.36,trna16_HisGTG_1_+_146544773_146544844@-1G.35.36,trna1_HisGTG_15_+_45493349_45493420@-1G.35.36,trna21_HisGTG_1_+_147753471_147753542@-1G.35.36,trna33_HisGTG_6_+_27125906_27125977@-1G.35.36,trna7_HisGTG_9_-_14433938_14434009@-1G.35.36,trna8_HisGTG_15_-_45492611_45492682@-1G.35.36,trna9_HisGTG_15_-_45490804_45490875@-1G.35.36|GGCCGTGATCGTATAGTGGTTAGTACTCTGCGTTGT</td><td>>trna111_HisGTG_1_-_147774845_147774916</td><td>-1G</td><td>35</td><td>36</td><td>GGCCGTGATCGTATAGTGGTTAGTACTCTGCGTTGT</td><td>6631.6</td><td>17.6</td><td>1.5</td><td>11.4</td><td>3.0E-30</td><td>5.3E-29</td><td>MB231/MB468</td></tr>
<tr><td>>MI0000077|hsa-mir-21&WithFlank&17|+|59841260|59841343@14.37.24&[MIMAT0000076&hsa-miR-21-5p&offsets|0|+2;m-1&17|+|59841273|59841294&offsets|0|+2]|TAGCTTATCAGACTGATGTTGACT</td><td>>MI0000077|hsa-mir-21&WithFlank&17|+|59841260|59841343</td><td>14</td><td>37</td><td>24</td><td>TAGCTTATCAGACTGATGTTGACT</td><td>486091.2</td><td>7.7</td><td>0.7</td><td>11.4</td><td>3.0E-30</td><td>5.3E-29</td><td>BT20/MB468</td></tr>
<tr><td>>MI0000434|hsa-let-7i&WithFlank&12|+|62603680|62603775@12.33.22(+1U)&[MIMAT0000415&hsa-let-7i-5p&offsets|0|0(+1U);m-49&12|+|62603691|62603712&offsets|0|0(+1U)]|TGAGGTAGTAGTTTGTGCTGTTT</td><td>>MI0000434|hsa-let-7i&WithFlank&12|+|62603680|62603775</td><td>12</td><td>33</td><td>22(+1U)</td><td>TGAGGTAGTAGTTTGTGCTGTTT</td><td>31567.9</td><td>8.6</td><td>0.8</td><td>11.4</td><td>3.0E-30</td><td>5.3E-29</td><td>MB231/MB468</td></tr>
<tr><td>>trna111_HisGTG_1_-_147774845_147774916@-1G.35.36,trna118_HisGTG_1_-_145396881_145396952@-1G.35.36,trna16_HisGTG_1_+_146544773_146544844@-1G.35.36,trna1_HisGTG_15_+_45493349_45493420@-1G.35.36,trna21_HisGTG_1_+_147753471_147753542@-1G.35.36,trna33_HisGTG_6_+_27125906_27125977@-1G.35.36,trna7_HisGTG_9_-_14433938_14434009@-1G.35.36,trna8_HisGTG_15_-_45492611_45492682@-1G.35.36,trna9_HisGTG_15_-_45490804_45490875@-1G.35.36|GGCCGTGATCGTATAGTGGTTAGTACTCTGCGTTGT</td><td>>trna111_HisGTG_1_-_147774845_147774916</td><td>-1G</td><td>35</td><td>36</td><td>GGCCGTGATCGTATAGTGGTTAGTACTCTGCGTTGT</td><td>531.1</td><td>13.7</td><td>1.2</td><td>11.4</td><td>3.0E-30</td><td>5.3E-29</td><td>BT20/MB468</td></tr>
<tr><td>>hg38_21_+_8433172_8446622_RNA45SN1_45S_50ntFlanks@10845.10865.21|ACTTCGGGATAAGGATTGGCT</td><td>>hg38_21_+_8433172_8446622_RNA45SN1_45S_50ntFlanks</td><td>10845</td><td>10865</td><td>21</td><td>ACTTCGGGATAAGGATTGGCT</td><td>4121.6</td><td>-6.8</td><td>0.6</td><td>-11.4</td><td>3.1E-30</td><td>5.4E-29</td><td>MB231/MB468</td></tr>
<tr><td>>hg38_1_-_228634819_228635039_RNA5S12_5S_50ntFlanks@144.170.27|CCTGGGAATACCGGGTGCTGTAGGCTT</td><td>>hg38_1_-_228634819_228635039_RNA5S12_5S_50ntFlanks</td><td>144</td><td>170</td><td>27</td><td>CCTGGGAATACCGGGTGCTGTAGGCTT</td><td>14047.2</td><td>8.3</td><td>0.7</td><td>11.4</td><td>3.2E-30</td><td>5.6E-29</td><td>MB231/MB468</td></tr>
<tr><td>>MI0000102|hsa-mir-100&WithFlank&11|-|122152223|122152314@19.36.18&[MIMAT0000098&hsa-miR-100-5p&offsets|0|-4;m-27&11|-|122152276|122152296&offsets|0|-3]|AACCCGTAGATCCGAACT</td><td>>MI0000102|hsa-mir-100&WithFlank&11|-|122152223|122152314</td><td>19</td><td>36</td><td>18</td><td>AACCCGTAGATCCGAACT</td><td>1230.9</td><td>15.2</td><td>1.3</td><td>11.4</td><td>3.2E-30</td><td>5.7E-29</td><td>MB231/MB468</td></tr>
<tr><td>>hg38_21_+_8433172_8446622_RNA45SN1_45S_50ntFlanks@485.508.24|GATGTGGTGACGTCGTGCTCTCCC</td><td>>hg38_21_+_8433172_8446622_RNA45SN1_45S_50ntFlanks</td><td>485</td><td>508</td><td>24</td><td>GATGTGGTGACGTCGTGCTCTCCC</td><td>1974.0</td><td>15.6</td><td>1.4</td><td>11.4</td><td>3.3E-30</td><td>5.7E-29</td><td>BT20/MB468</td></tr>
<tr><td>>hg38_21_+_8433172_8446622_RNA45SN1_45S_50ntFlanks@6786.6806.21|TACGCCTGTCTGAGCGTCGCT</td><td>>hg38_21_+_8433172_8446622_RNA45SN1_45S_50ntFlanks</td><td>6786</td><td>6806</td><td>21</td><td>TACGCCTGTCTGAGCGTCGCT</td><td>4279.6</td><td>7.5</td><td>0.7</td><td>11.4</td><td>3.4E-30</td><td>5.9E-29</td><td>MB231/MB468</td></tr>
<tr><td>>MI0000433|hsa-let-7g&WithFlank&3|-|52268272|52268367@11.31.21(+1A)&[MIMAT0000414&hsa-let-7g-5p&offsets|0|-1(+1A);m-58&3|-|52268336|52268357&offsets|0|-1(+1A)]|TGAGGTAGTAGTTTGTACAGTA</td><td>>MI0000433|hsa-let-7g&WithFlank&3|-|52268272|52268367</td><td>11</td><td>31</td><td>21(+1A)</td><td>TGAGGTAGTAGTTTGTACAGTA</td><td>9811.0</td><td>7.9</td><td>0.7</td><td>11.4</td><td>3.4E-30</td><td>5.9E-29</td><td>MB231/MB468</td></tr>
</table>

<table>
<tr><td>>MI0001445|hsa-mir-423&WithFlank&17|+|30117073|30117178@59.80.22(+1A)&[MIMAT0001340&hsa-miR-423-3p&offsets|0|-1(+1A);m-100&17|+|30117131|30117153&offsets|0|-1(+1A)]|AGCTCGGTCTGAGGCCCCTCAGA</td><td>>MI0001445|hsa-mir-423&WithFlank&17|+|30117073|30117178</td><td>59</td><td>80</td><td>22(+1A)</td><td>AGCTCGGTCTGAGGCCCCTCAGA</td><td>1837.5</td><td>7.4</td><td>0.7</td><td>11.4</td><td>3.6E-30</td><td>6.2E-29</td><td>BT20/MB468</td></tr>
<tr><td>>MI0000439|hsa-mir-23b&WithFlank&9|+|95085202|95085310@67.86.20(+1C)&[MIMAT0000418&hsa-miR-23b-3p&offsets|+3|0(+1C);m-42&9|+|95085265|95085285&offsets|+3|+2(+1C)]|ACATTGCCAGGGATTACCACC</td><td>>MI0000439|hsa-mir-23b&WithFlank&9|+|95085202|95085310</td><td>67</td><td>86</td><td>20(+1C)</td><td>ACATTGCCAGGGATTACCACC</td><td>2386.9</td><td>15.9</td><td>1.4</td><td>11.4</td><td>3.6E-30</td><td>6.3E-29</td><td>BT20/MB468</td></tr>
<tr><td>>hg38_21_+_8433172_8446622_RNA45SN1_45S_50ntFlanks@9568.9596.29|CGTGCAAATCGGTCGTCCGACCTGGGTAT</td><td>>hg38_21_+_8433172_8446622_RNA45SN1_45S_50ntFlanks</td><td>9568</td><td>9596</td><td>29</td><td>CGTGCAAATCGGTCGTCCGACCTGGGTAT</td><td>3825.6</td><td>-10.3</td><td>0.9</td><td>-11.4</td><td>3.8E-30</td><td>6.6E-29</td><td>BT20/MB468</td></tr>
<tr><td>>hg38_MT_+_1621_3279_RNR2_16S_50ntFlanks@461.492.32|TGTAAATTTAACTGTTAGTCCAAAGAGGAACA</td><td>>hg38_MT_+_1621_3279_RNR2_16S_50ntFlanks</td><td>461</td><td>492</td><td>32</td><td>TGTAAATTTAACTGTTAGTCCAAAGAGGAACA</td><td>494.8</td><td>11.4</td><td>1.0</td><td>11.4</td><td>3.9E-30</td><td>6.7E-29</td><td>MB231/MB468</td></tr>
<tr><td>>hg38_21_+_8433172_8446622_RNA45SN1_45S_50ntFlanks@11768.11794.27|CTAATTAGTGACGCGCATGAATGGATG</td><td>>hg38_21_+_8433172_8446622_RNA45SN1_45S_50ntFlanks</td><td>11768</td><td>11794</td><td>27</td><td>CTAATTAGTGACGCGCATGAATGGATG</td><td>2550.0</td><td>-6.0</td><td>0.5</td><td>-11.4</td><td>3.9E-30</td><td>6.7E-29</td><td>BT20/MB468</td></tr>
<tr><td>>hg38_21_+_8433172_8446622_RNA45SN1_45S_50ntFlanks@12995.13035.41|CTCGCTGCGATCTATTGAAAGTCAGCCCTCGACACAAGGGT</td><td>>hg38_21_+_8433172_8446622_RNA45SN1_45S_50ntFlanks</td><td>12995</td><td>13035</td><td>41</td><td>CTCGCTGCGATCTATTGAAAGTCAGCCCTCGACACAAGGGT</td><td>642.1</td><td>14.2</td><td>1.2</td><td>11.4</td><td>4.1E-30</td><td>7.1E-29</td><td>MB231/MB468</td></tr>
<tr><td>>am_trna10_ValCAC_5_-_180649395_180649467@1.33.33,trna12_ValAAC_5_-_180645270_180645342@1.33.33,trna132_ValAAC_6_-_27721179_27721251@1.33.33,trna136_ValAAC_6_-_27648885_27648957@1.33.33,trna139_ValAAC_6_-_27618707_27618779@1.33.33,trna18_ValCAC_5_-_180529253_180529325@1.33.33,trna2_ValAAC_3_+_169490018_169490090@1.33.33,trna2_ValCAC_5_+_180524070_180524142@1.33.33,trna4_ValAAC_5_+_180591154_180591226@1.33.33,trna5_ValAAC_5_+_180596610_180596682@1.33.33,trna6_ValCAC_5_+_180600650_180600722@1.33.33,trna85_ValCAC_1_-_161369490_161369562@1.33.33,trna90_ValCAC_1_-_149684088_149684161@1.33.33,trna98_ValCAC_1_-_149298555_149298627@1.33.33,trna9_ValCAC_6_+_26538282_26538354@1.33.33|GTTTCCGTAGTGTAGTGGTTATCACGTTCGCCT</td><td>>am_trna10_ValCAC_5_-_180649395_180649467</td><td>1</td><td>33</td><td>33</td><td>GTTTCCGTAGTGTAGTGGTTATCACGTTCGCCT</td><td>8308.0</td><td>11.1</td><td>1.0</td><td>11.4</td><td>4.1E-30</td><td>7.1E-29</td><td>MB231/MB468</td></tr>
<tr><td>>MI0000734|hsa-mir-106b&WithFlank&7|-|100093987|100094080@58.78.21&[MIMAT0004672&hsa-miR-106b-3p&offsets|0|-1;m-87&7|-|100094003|100094023&offsets|0|0]|CCGCACTGTGGGTACTTGCTG</td><td>>MI0000734|hsa-mir-106b&WithFlank&7|-|100093987|100094080</td><td>58</td><td>78</td><td>21</td><td>CCGCACTGTGGGTACTTGCTG</td><td>4535.6</td><td>7.8</td><td>0.7</td><td>11.4</td><td>4.4E-30</td><td>7.6E-29</td><td>BT20/MB468</td></tr>
<tr><td>>hg38_21_+_8433172_8446622_RNA45SN1_45S_50ntFlanks@12159.12179.21|TTTGACTGGGGCGGTACACCT</td><td>>hg38_21_+_8433172_8446622_RNA45SN1_45S_50ntFlanks</td><td>12159</td><td>12179</td><td>21</td><td>TTTGACTGGGGCGGTACACCT</td><td>2395.6</td><td>-7.8</td><td>0.7</td><td>-11.4</td><td>4.5E-30</td><td>7.7E-29</td><td>MB231/MB468</td></tr>
<tr><td>>MI0000433|hsa-let-7g&WithFlank&3|-|52268272|52268367@11.31.21(+1C)&[MIMAT0000414&hsa-let-7g-5p&offsets|0|-1(+1C);m-58&3|-|52268336|52268357&offsets|0|-1(+1C)]|TGAGGTAGTAGTTTGTACAGTC</td><td>>MI0000433|hsa-let-7g&WithFlank&3|-|52268272|52268367</td><td>11</td><td>31</td><td>21(+1C)</td><td>TGAGGTAGTAGTTTGTACAGTC</td><td>25579.6</td><td>6.7</td><td>0.6</td><td>11.4</td><td>4.7E-30</td><td>8.2E-29</td><td>BT20/MB468</td></tr>
<tr><td>>hg38_21_+_8433172_8446622_RNA45SN1_45S_50ntFlanks@11881.11904.24|CCTGTTGAGCTTGACTCTAGTCTG</td><td>>hg38_21_+_8433172_8446622_RNA45SN1_45S_50ntFlanks</td><td>11881</td><td>11904</td><td>24</td><td>CCTGTTGAGCTTGACTCTAGTCTG</td><td>1303.6</td><td>-6.8</td><td>0.6</td><td>-11.4</td><td>4.8E-30</td><td>8.2E-29</td><td>MB231/MB468</td></tr>
<tr><td>>MI0001445|hsa-mir-423&WithFlank&17|+|30117073|30117178@59.80.22&[MIMAT0001340&hsa-miR-423-3p&offsets|0|-1;m-100&17|+|30117131|30117153&offsets|0|-1]|AGCTCGGTCTGAGGCCCCTCAG</td><td>>MI0001445|hsa-mir-423&WithFlank&17|+|30117073|30117178</td><td>59</td><td>80</td><td>22</td><td>AGCTCGGTCTGAGGCCCCTCAG</td><td>4417.0</td><td>8.1</td><td>0.7</td><td>11.4</td><td>4.8E-30</td><td>8.2E-29</td><td>MB231/MB468</td></tr>
<tr><td>>MI0000088|hsa-mir-30a&WithFlank&6|-|71403545|71403627@54.73.20(+2U)&[MIMAT0000088&hsa-miR-30a-3p&offsets|+1|-1(+2U);m-38&6|-|71403554|71403575&offsets|+1|-1(+2U)]|TTTCAGTCGGATGTTTGCAGTT</td><td>>MI0000088|hsa-mir-30a&WithFlank&6|-|71403545|71403627</td><td>54</td><td>73</td><td>20(+2U)</td><td>TTTCAGTCGGATGTTTGCAGTT</td><td>1592.1</td><td>9.0</td><td>0.8</td><td>11.4</td><td>5.0E-30</td><td>8.4E-29</td><td>MB231/MB468</td></tr>
<tr><td>>MI0003513|hsa-mir-455&WithFlank&9|+|114209428|114209535@22.45.24&[MIMAT0003150&hsa-miR-455-5p&offsets|0|+2;m-232&9|+|114209449|114209470&offsets|0|+2]|TATGTGCCTTTGGACTACATCGTG</td><td>>MI0003513|hsa-mir-455&WithFlank&9|+|114209428|114209535</td><td>22</td><td>45</td><td>24</td><td>TATGTGCCTTTGGACTACATCGTG</td><td>1059.1</td><td>15.0</td><td>1.3</td><td>11.4</td><td>5.0E-30</td><td>8.4E-29</td><td>MB231/MB468</td></tr>
<tr><td>>hg38_21_+_8433172_8446622_RNA45SN1_45S_50ntFlanks@12488.12509.22|AACGTGAGCTGGGTTTAGACCG</td><td>>hg38_21_+_8433172_8446622_RNA45SN1_45S_50ntFlanks</td><td>12488</td><td>12509</td><td>22</td><td>AACGTGAGCTGGGTTTAGACCG</td><td>5186.5</td><td>-8.0</td><td>0.7</td><td>-11.4</td><td>4.9E-30</td><td>8.5E-29</td><td>BT20/MB468</td></tr>
<tr><td>>MI0001446|hsa-mir-424&WithFlank&X|-|134546608|134546717@54.74.21&[MIMAT0004749&hsa-miR-424-3p&offsets|0|0;m-305&X|-|134546644|134546664&offsets|0|0]|CAAAACGTGAGGCGCTGCTAT</td><td>>MI0001446|hsa-mir-424&WithFlank&X|-|134546608|134546717</td><td>54</td><td>74</td><td>21</td><td>CAAAACGTGAGGCGCTGCTAT</td><td>1622.4</td><td>15.3</td><td>1.3</td><td>11.4</td><td>5.0E-30</td><td>8.5E-29</td><td>BT20/MB468</td></tr>
<tr><td>>trnaMT_ValTAC_MT_+_1602_1670@52.71.20|TTAACTTGACCGCTCTGACC</td><td>>trnaMT_ValTAC_MT_+_1602_1670</td><td>52</td><td>71</td><td>20</td><td>TTAACTTGACCGCTCTGACC</td><td>276.3</td><td>12.8</td><td>1.1</td><td>11.4</td><td>5.1E-30</td><td>8.7E-29</td><td>BT20/MB468</td></tr>
<tr><td>>MI0000067|hsa-let-7f-1&WithFlank&9|+|94176341|94176439@13.33.21(+1A);MI0000068|hsa-let-7f-2&WithFlank&X|-|53557186|53557280@14.34.21(+1A)&[MIMAT0000067&hsa-let-7f-5p&offsets|0|-1(+1A);m-20&9|+|94176353|94176374&offsets|0|-1(+1A)];[MIMAT0000067_1&hsa-let-7f-5p&offsets|0|-1(+1A);m-19&X|-|53557246|53557267&offsets|0|-1(+1A)]|TGAGGTAGTAGATTGTATAGTA</td><td>>MI0000067|hsa-let-7f-1&WithFlank&9|+|94176341|94176439</td><td>13</td><td>33</td><td>21(+1A)</td><td>TGAGGTAGTAGATTGTATAGTA</td><td>8371.7</td><td>7.5</td><td>0.7</td><td>11.4</td><td>5.5E-30</td><td>9.2E-29</td><td>MB231/MB468</td></tr>
<tr><td>>MI0000446|hsa-mir-125b-1&WithFlank&11|-|122099751|122099850@21.41.21;MI0000470|hsa-mir-125b-2&WithFlank&21|+|16590231|16590331@23.43.21&[MIMAT0000423&hsa-miR-125b-5p&offsets|0|-1;m-67&11|-|122099809|122099830&offsets|0|-1];[MIMAT0000423_1&hsa-miR-125b-5p&offsets|0|-1;m-68&21|+|16590253|16590274&offsets|0|-1]|TCCCTGAGACCCTAACTTGTG</td><td>>MI0000446|hsa-mir-125b-1&WithFlank&11|-|122099751|122099850</td><td>21</td><td>41</td><td>21</td><td>TCCCTGAGACCCTAACTTGTG</td><td>27037.7</td><td>9.0</td><td>0.8</td><td>11.4</td><td>5.5E-30</td><td>9.2E-29</td><td>MB231/MB468</td></tr>
<tr><td>>MI0000542|hsa-mir-320a&WithFlank&8|-|22244960|22245043@48.70.23(+1U)&[MIMAT0000510&hsa-miR-320a-3p&offsets|0|+1(+1U);m-60&8|-|22244975|22244996&offsets|0|+1(+1U)]|AAAAGCTGGGTTGAGAGGGCGAAT</td><td>>MI0000542|hsa-mir-320a&WithFlank&8|-|22244960|22245043</td><td>48</td><td>70</td><td>23(+1U)</td><td>AAAAGCTGGGTTGAGAGGGCGAAT</td><td>7028.4</td><td>8.6</td><td>0.8</td><td>11.4</td><td>5.6E-30</td><td>9.5E-29</td><td>MB231/MB468</td></tr>
<tr><td>>MI0000446|hsa-mir-125b-1&WithFlank&11|-|122099751|122099850@21.39.19;MI0000470|hsa-mir-125b-2&WithFlank&21|+|16590231|16590331@23.41.19&[MIMAT0000423&hsa-miR-125b-5p&offsets|0|-3;m-67&11|-|122099809|122099830&offsets|0|-3];[MIMAT0000423_1&hsa-miR-125b-5p&offsets|0|-3;m-68&21|+|16590253|16590274&offsets|0|-3]|TCCCTGAGACCCTAACTTG</td><td>>MI0000446|hsa-mir-125b-1&WithFlank&11|-|122099751|122099850</td><td>21</td><td>39</td><td>19</td><td>TCCCTGAGACCCTAACTTG</td><td>3862.6</td><td>14.3</td><td>1.3</td><td>11.4</td><td>5.7E-30</td><td>9.5E-29</td><td>MB231/MB468</td></tr>
<tr><td>>MI0000088|hsa-mir-30a&WithFlank&6|-|71403545|71403627@53.70.18(+1U)&[MIMAT0000088&hsa-miR-30a-3p&offsets|0|-4(+1U);m-38&6|-|71403554|71403575&offsets|0|-4(+1U)]|CTTTCAGTCGGATGTTTGT</td><td>>MI0000088|hsa-mir-30a&WithFlank&6|-|71403545|71403627</td><td>53</td><td>70</td><td>18(+1U)</td><td>CTTTCAGTCGGATGTTTGT</td><td>1197.9</td><td>8.8</td><td>0.8</td><td>11.4</td><td>5.9E-30</td><td>9.8E-29</td><td>MB231/MB468</td></tr>
<tr><td>>hg38_21_+_8433172_8446622_RNA45SN1_45S_50ntFlanks@12405.12436.32|CTTTTTGATCCTTCGATGTCGGCTCTTCCTAT</td><td>>hg38_21_+_8433172_8446622_RNA45SN1_45S_50ntFlanks</td><td>12405</td><td>12436</td><td>32</td><td>CTTTTTGATCCTTCGATGTCGGCTCTTCCTAT</td><td>4579.9</td><td>-7.9</td><td>0.7</td><td>-11.4</td><td>6.3E-30</td><td>1.0E-28</td><td>MB231/MB468</td></tr>
<tr><td>>MI0000264|hsa-mir-7-2&WithFlank&15|+|88611819|88611940@38.60.23(+1A);MI0000265|hsa-mir-7-3&WithFlank&19|+|4770664|4770785@37.59.23(+1A);MI0000263|hsa-mir-7-1&WithFlank&9|-|83969742|83969863@30.52.23(+1A)&[MIMAT0000252&hsa-miR-7-5p&offsets|0|-1(+1A);m-400&15|+|88611856|88611879&offsets|0|-1(+1A)];[MIMAT0000252_1&hsa-miR-7-5p&offsets|0|-1(+1A);m-402&19|+|4770700|4770723&offsets|0|-1(+1A)];[MIMAT0000252_2&hsa-miR-7-5p&offsets|0|-1(+1A);m-401&9|-|83969811|83969834&offsets|0|-1(+1A)]|TGGAAGACTAGTGATTTTGTTGTA</td><td>>MI0000264|hsa-mir-7-2&WithFlank&15|+|88611819|88611940</td><td>38</td><td>60</td><td>23(+1A)</td><td>TGGAAGACTAGTGATTTTGTTGTA</td><td>6312.5</td><td>7.9</td><td>0.7</td><td>11.4</td><td>6.4E-30</td><td>1.1E-28</td><td>MB231/MB468</td></tr>
<tr><td>>hg38_21_+_8433172_8446622_RNA45SN1_45S_50ntFlanks@475.518.44|CTTCGTGATCGATGTGGTGACGTCGTGCTCTCCCGGGCCGGGTC</td><td>>hg38_21_+_8433172_8446622_RNA45SN1_45S_50ntFlanks</td><td>475</td><td>518</td><td>44</td><td>CTTCGTGATCGATGTGGTGACGTCGTGCTCTCCCGGGCCGGGTC</td><td>3240.2</td><td>9.5</td><td>0.8</td><td>11.4</td><td>6.5E-30</td><td>1.1E-28</td><td>MB231/MB468</td></tr>
<tr><td>>hg38_21_+_8433172_8446622_RNA45SN1_45S_50ntFlanks@7975.8011.37|CGCGACCTCAGATCAGACGTGGCGACCCGCTGAATTT</td><td>>hg38_21_+_8433172_8446622_RNA45SN1_45S_50ntFlanks</td><td>7975</td><td>8011</td><td>37</td><td>CGCGACCTCAGATCAGACGTGGCGACCCGCTGAATTT</td><td>147279.0</td><td>7.8</td><td>0.7</td><td>11.4</td><td>6.6E-30</td><td>1.1E-28</td><td>MB231/MB468</td></tr>
<tr><td>>hg38_MT_+_598_1651_RNR1_12S_50ntFlanks@52.84.33|ATAGGTTTGGTCCTAGCCTTTCTATTAGCTCTT</td><td>>hg38_MT_+_598_1651_RNR1_12S_50ntFlanks</td><td>52</td><td>84</td><td>33</td><td>ATAGGTTTGGTCCTAGCCTTTCTATTAGCTCTT</td><td>487.5</td><td>6.4</td><td>0.6</td><td>11.4</td><td>6.7E-30</td><td>1.1E-28</td><td>MB231/MB468</td></tr>
<tr><td>>hg38_21_+_8433172_8446622_RNA45SN1_45S_50ntFlanks@12626.12654.29|AATGGGGCGAAGCTACCATCTGTGGGATT</td><td>>hg38_21_+_8433172_8446622_RNA45SN1_45S_50ntFlanks</td><td>12626</td><td>12654</td><td>29</td><td>AATGGGGCGAAGCTACCATCTGTGGGATT</td><td>6630.9</td><td>-5.4</td><td>0.5</td><td>-11.4</td><td>6.7E-30</td><td>1.1E-28</td><td>MB231/MB468</td></tr>
<tr><td>>hg38_21_+_8433172_8446622_RNA45SN1_45S_50ntFlanks@6761.6806.46|TGCGGCCCCGGGTTCCTCCCGGGGCTACGCCTGTCTGAGCGTCGCT</td><td>>hg38_21_+_8433172_8446622_RNA45SN1_45S_50ntFlanks</td><td>6761</td><td>6806</td><td>46</td><td>TGCGGCCCCGGGTTCCTCCCGGGGCTACGCCTGTCTGAGCGTCGCT</td><td>1006.1</td><td>14.9</td><td>1.3</td><td>11.4</td><td>6.9E-30</td><td>1.1E-28</td><td>MB231/MB468</td></tr>
<tr><td>>MI0000065|hsa-let-7d&WithFlank&9|+|94178828|94178926@68.89.22&[MIMAT0004484&hsa-let-7d-3p&offsets|0|0;m-98&9|+|94178895|94178916&offsets|0|0]|CTATACGACCTGCTGCCTTTCT</td><td>>MI0000065|hsa-let-7d&WithFlank&9|+|94178828|94178926</td><td>68</td><td>89</td><td>22</td><td>CTATACGACCTGCTGCCTTTCT</td><td>6626.6</td><td>7.7</td><td>0.7</td><td>11.4</td><td>6.8E-30</td><td>1.2E-28</td><td>BT20/MB468</td></tr>
<tr><td>>hg38_21_+_8433172_8446622_RNA45SN1_45S_50ntFlanks@10679.10708.30|GGTTCCGGCGGCGTCCGGTGAGCTCTCGCT</td><td>>hg38_21_+_8433172_8446622_RNA45SN1_45S_50ntFlanks</td><td>10679</td><td>10708</td><td>30</td><td>GGTTCCGGCGGCGTCCGGTGAGCTCTCGCT</td><td>4467.0</td><td>-5.6</td><td>0.5</td><td>-11.4</td><td>7.2E-30</td><td>1.2E-28</td><td>MB231/MB468</td></tr>
<tr><td>>MI0000088|hsa-mir-30a&WithFlank&6|-|71403545|71403627@54.74.21&[MIMAT0000088&hsa-miR-30a-3p&offsets|+1|0;m-38&6|-|71403554|71403575&offsets|+1|0]|TTTCAGTCGGATGTTTGCAGC</td><td>>MI0000088|hsa-mir-30a&WithFlank&6|-|71403545|71403627</td><td>54</td><td>74</td><td>21</td><td>TTTCAGTCGGATGTTTGCAGC</td><td>3389.5</td><td>7.4</td><td>0.6</td><td>11.4</td><td>7.0E-30</td><td>1.2E-28</td><td>BT20/MB468</td></tr>
<tr><td>>hg38_21_+_8433172_8446622_RNA45SN1_45S_50ntFlanks@9569.9610.42|GTGCAAATCGGTCGTCCGACCTGGGTATAGGGGCGAAAGACT</td><td>>hg38_21_+_8433172_8446622_RNA45SN1_45S_50ntFlanks</td><td>9569</td><td>9610</td><td>42</td><td>GTGCAAATCGGTCGTCCGACCTGGGTATAGGGGCGAAAGACT</td><td>321.2</td><td>13.2</td><td>1.2</td><td>11.4</td><td>7.4E-30</td><td>1.2E-28</td><td>MB231/MB468</td></tr>
</table>

| | | | | | | | | | | | | |
|---|---|---|---|---|---|---|---|---|---|---|---|---|
| >hg38_21_+_8433172_8446622_RNA45SN1_45S_50ntFlanks@12163.12182.20\|ACTGGGGCGGTACACCTGTC | >hg38_21_+_8433172_8446622_RNA45SN1_45S_50ntFlanks | 12163 | 12182 | 20 | ACTGGGGCGGTACACCTGTC | 3309.6 | -6.8 | 0.6 | -11.4 | 7.2E-30 | 1.2E-28 | BT20/MB468 |
| >MI0000272\|hsa-mir-182&WithFlank&7\|-\|129770377\|129770498@29.53.25(+1C)&[MIMAT0000259&hsa-miR-182-5p&offsets\|0\|+1(+1C);m-22&7\|-\|129770449\|129770470&offsets\|0\|+3(+1C)]\|TTTGGCAATGGTAGAACTCACACTGC | >MI0000272\|hsa-mir-182&WithFlank&7\|-\|129770377\|129770498 | 29 | 53 | 25(+1C) | TTTGGCAATGGTAGAACTCACACTGC | 790.2 | 8.0 | 0.7 | 11.4 | 7.2E-30 | 1.2E-28 | BT20/MB468 |
| >trnaMT_SerGCT_MT_+_12207_12265@1.22.22\|GAGAAAGCTCACAAGAACTGCT | >trnaMT_SerGCT_MT_+_12207_12265 | 1 | 22 | 22 | GAGAAAGCTCACAAGAACTGCT | 8819.6 | 7.0 | 0.6 | 11.3 | 7.6E-30 | 1.2E-28 | MB231/MB468 |
| >hg38_21_+_8433172_8446622_RNA45SN1_45S_50ntFlanks@475.505.31\|CTTCGTGATCGATGTGGTGACGTCGTGCTCT | >hg38_21_+_8433172_8446622_RNA45SN1_45S_50ntFlanks | 475 | 505 | 31 | CTTCGTGATCGATGTGGTGACGTCGTGCTCT | 40080.8 | 7.5 | 0.7 | 11.3 | 7.8E-30 | 1.3E-28 | BT20/MB468 |
| >MI0000298\|hsa-mir-221&WithFlank&X\|-\|45746151\|45746272@31.55.25&[MIMAT0004568&hsa-miR-221-5p&offsets\|0\|+3;m-405&X\|-\|45746220\|45746242&offsets\|0\|+2]\|ACCTGGCATACAATGTAGATTTCTG | >MI0000298\|hsa-mir-221&WithFlank&X\|-\|45746151\|45746272 | 31 | 55 | 25 | ACCTGGCATACAATGTAGATTTCTG | 1526.5 | 8.7 | 0.8 | 11.3 | 8.3E-30 | 1.3E-28 | MB231/MB468 |
| >MI0000736\|hsa-mir-30c-1&WithFlank&1\|+\|40757278\|40757378@23.44.22(+1U);MI0000254\|hsa-mir-30c-2&WithFlank&6\|-\|71376954\|71377037@13.34.22(+1U)&[MIMAT0000244&hsa-miR-30c-5p&offsets\|0\|-1(+1U);m-57&1\|+\|40757300\|40757323&offsets\|0\|-2(+1U)];[MIMAT0000244_1&hsa-miR-30c-5p&offsets\|0\|-1(+1U);m-56&6\|-\|71377002\|71377025&offsets\|0\|-2(+1U)]\|TGTAAACATCCTACACTCTCAGT | >MI0000736\|hsa-mir-30c-1&WithFlank&1\|+\|40757278\|40757378 | 23 | 44 | 22(+1U) | TGTAAACATCCTACACTCTCAGT | 22748.9 | 7.9 | 0.7 | 11.3 | 8.4E-30 | 1.4E-28 | MB231/MB468 |
| >MI0006444\|hsa-mir-1307&WithFlank&10\|-\|103394247\|103394407@87.107.21(+1U)&[MIMAT0005951&hsa-miR-1307-3p&offsets\|+1\|0(+1U);m-66&10\|-\|103394300\|103394322&offsets\|+1\|-1(+1U)]\|CTCGGCGTGGCGTCGGTCGTGT | >MI0006444\|hsa-mir-1307&WithFlank&10\|-\|103394247\|103394407 | 87 | 107 | 21(+1U) | CTCGGCGTGGCGTCGGTCGTGT | 1216.7 | 8.2 | 0.7 | 11.3 | 8.9E-30 | 1.4E-28 | MB231/MB468 |
| >hg38_21_+_8433172_8446622_RNA45SN1_45S_50ntFlanks@10680.10709.30\|GTTCCGGCGGCGTCCGGTGAGCTCTCGCTG | >hg38_21_+_8433172_8446622_RNA45SN1_45S_50ntFlanks | 10680 | 10709 | 30 | GTTCCGGCGGCGTCCGGTGAGCTCTCGCTG | 4042.4 | -5.6 | 0.5 | -11.3 | 9.2E-30 | 1.5E-28 | MB231/MB468 |
| >hg38_21_+_8433172_8446622_RNA45SN1_45S_50ntFlanks@12657.12674.18\|GACTGAACGCCTCTAAGT | >hg38_21_+_8433172_8446622_RNA45SN1_45S_50ntFlanks | 12657 | 12674 | 18 | GACTGAACGCCTCTAAGT | 502.3 | -5.0 | 0.4 | -11.3 | 9.0E-30 | 1.5E-28 | BT20/MB468 |
| >hg38_21_+_8433172_8446622_RNA45SN1_45S_50ntFlanks@476.507.32\|TTCGTGATCGATGTGGTGACGTCGTGCTCTCC | >hg38_21_+_8433172_8446622_RNA45SN1_45S_50ntFlanks | 476 | 507 | 32 | TTCGTGATCGATGTGGTGACGTCGTGCTCTCC | 6411.2 | 8.2 | 0.7 | 11.3 | 9.5E-30 | 1.5E-28 | MB231/MB468 |
| >MI0000077\|hsa-mir-21&WithFlank&17\|+\|59841260\|59841343@14.35.22(+2U)&[MIMAT0000076&hsa-miR-21-5p&offsets\|0\|0(+2U);m-1&17\|+\|59841273\|59841294&offsets\|0\|0(+2U)]\|TAGCTTATCAGACTGATGTTGATT | >MI0000077\|hsa-mir-21&WithFlank&17\|+\|59841260\|59841343 | 14 | 35 | 22(+2U) | TAGCTTATCAGACTGATGTTGATT | 4090.6 | 9.8 | 0.9 | 11.3 | 9.6E-30 | 1.5E-28 | MB231/MB468 |
| >hg38_MT_+_598_1651_RNR1_12S_50ntFlanks@970.1004.35\|ATGGTAAGTGTACTGGAAAGTGCACTTGGACGAAC | >hg38_MT_+_598_1651_RNR1_12S_50ntFlanks | 970 | 1004 | 35 | ATGGTAAGTGTACTGGAAAGTGCACTTGGACGAAC | 897.1 | 14.7 | 1.3 | 11.3 | 9.8E-30 | 1.6E-28 | MB231/MB468 |
| >MI0000077\|hsa-mir-21&WithFlank&17\|+\|59841260\|59841343@14.31.18&[MIMAT0000076&hsa-miR-21-5p&offsets\|0\|-4;m-1&17\|+\|59841273\|59841294&offsets\|0\|-4]\|TAGCTTATCAGACTGATG | >MI0000077\|hsa-mir-21&WithFlank&17\|+\|59841260\|59841343 | 14 | 31 | 18 | TAGCTTATCAGACTGATG | 1951.2 | 6.8 | 0.6 | 11.3 | 1.0E-29 | 1.6E-28 | MB231/MB468 |
| >MI0001445\|hsa-mir-423&WithFlank&17\|+\|30117073\|30117178@59.80.22(+1C)&[MIMAT0001340&hsa-miR-423-3p&offsets\|0\|-1(+1C);m-100&17\|+\|30117131\|30117153&offsets\|0\|-1(+1C)]\|AGCTCGGTCTGAGGCCCCTCAGC | >MI0001445\|hsa-mir-423&WithFlank&17\|+\|30117073\|30117178 | 59 | 80 | 22(+1C) | AGCTCGGTCTGAGGCCCCTCAGC | 7918.8 | 8.8 | 0.8 | 11.3 | 1.1E-29 | 1.7E-28 | MB231/MB468 |
| >trnaMT_PheGAA_MT_+_577_647@37.55.19\|AAATGTTTAGACGGGCTCA | >trnaMT_PheGAA_MT_+_577_647 | 37 | 55 | 19 | AAATGTTTAGACGGGCTCA | 270.7 | 12.7 | 1.1 | 11.3 | 1.0E-29 | 1.7E-28 | BT20/MB468 |
| >trnaMT_GluTTC_MT_-_14674_14742@39.72.34,trnalookalike8_GluTTC_5_-_93905172_93905240@39.72.34\|ATTGGTCGTGGTTGTAGTCCGTGCGAGAATACCA | >trnaMT_GluTTC_MT_-_14674_14742 | 39 | 72 | 34 | ATTGGTCGTGGTTGTAGTCCGTGCGAGAATACCA | 42006.6 | 9.6 | 0.8 | 11.3 | 1.1E-29 | 1.7E-28 | MB231/MB468 |
| >hg38_21_+_8433172_8446622_RNA45SN1_45S_50ntFlanks@10845.10865.21\|ACTTCGGGATAAGGATTGGCT | >hg38_21_+_8433172_8446622_RNA45SN1_45S_50ntFlanks | 10845 | 10865 | 21 | ACTTCGGGATAAGGATTGGCT | 4975.2 | -8.6 | 0.8 | -11.3 | 1.0E-29 | 1.8E-28 | BT20/MB468 |
| >hg38_21_+_8433172_8446622_RNA45SN1_45S_50ntFlanks@12405.12431.27\|CTTTTTGATCCTTCGATGTCGGCTCTT | >hg38_21_+_8433172_8446622_RNA45SN1_45S_50ntFlanks | 12405 | 12431 | 27 | CTTTTTGATCCTTCGATGTCGGCTCTT | 6426.4 | -7.4 | 0.7 | -11.3 | 1.2E-29 | 1.9E-28 | MB231/MB468 |
| >hg38_21_+_8433172_8446622_RNA45SN1_45S_50ntFlanks@12997.13040.44\|CGCTGCGATCTATTGAAAGTCAGCCCTCGACACAAGGGTTTGTC | >hg38_21_+_8433172_8446622_RNA45SN1_45S_50ntFlanks | 12997 | 13040 | 44 | CGCTGCGATCTATTGAAAGTCAGCCCTCGACACAAGGGTTTGTC | 1965.0 | 15.8 | 1.4 | 11.3 | 1.2E-29 | 1.9E-28 | MB231/MB468 |
| >hg38_21_+_8433172_8446622_RNA45SN1_45S_50ntFlanks@12997.13016.20\|CGCTGCGATCTATTGAAAGT | >hg38_21_+_8433172_8446622_RNA45SN1_45S_50ntFlanks | 12997 | 13016 | 20 | CGCTGCGATCTATTGAAAGT | 385.9 | 5.4 | 0.5 | 11.3 | 1.2E-29 | 1.9E-28 | BT20/MB468 |
| >hg38_21_+_8433172_8446622_RNA45SN1_45S_50ntFlanks@476.509.34\|TTCGTGATCGATGTGGTGACGTCGTGCTCTCCCG | >hg38_21_+_8433172_8446622_RNA45SN1_45S_50ntFlanks | 476 | 509 | 34 | TTCGTGATCGATGTGGTGACGTCGTGCTCTCCCG | 8197.7 | 8.4 | 0.7 | 11.3 | 1.2E-29 | 1.9E-28 | MB231/MB468 |
| >hg38_21_+_8433172_8446622_RNA45SN1_45S_50ntFlanks@11729.11751.23\|TAACTATGACTCTCTTAAGGTAG | >hg38_21_+_8433172_8446622_RNA45SN1_45S_50ntFlanks | 11729 | 11751 | 23 | TAACTATGACTCTCTTAAGGTAG | 54039.7 | -8.0 | 0.7 | -11.3 | 1.2E-29 | 2.0E-28 | MB231/MB468 |
| >trnaMT_ProTGG_MT_-_15956_16023@50.70.21\|TTAAAGACTTTTTCTCTGACC | >trnaMT_ProTGG_MT_-_15956_16023 | 50 | 70 | 21 | TTAAAGACTTTTTCTCTGACC | 901.9 | 14.7 | 1.3 | 11.3 | 1.3E-29 | 2.0E-28 | MB231/MB468 |
| >MI0000734\|hsa-mir-106b&WithFlank&7\|-\|100093987\|100094080@58.77.20&[MIMAT0004672&hsa-miR-106b-3p&offsets\|0\|-2;m-87&7\|-\|100094003\|100094023&offsets\|0\|-1]\|CCGCACTGTGGGTACTTGCT | >MI0000734\|hsa-mir-106b&WithFlank&7\|-\|100093987\|100094080 | 58 | 77 | 20 | CCGCACTGTGGGTACTTGCT | 2370.6 | 8.1 | 0.7 | 11.3 | 1.2E-29 | 2.1E-28 | BT20/MB468 |
| >MI0000272\|hsa-mir-182&WithFlank&7\|-\|129770377\|129770498@29.52.24&[MIMAT0000259&hsa-miR-182-5p&offsets\|0\|0;m-22&7\|-\|129770449\|129770470&offsets\|0\|+2]\|TTTGGCAATGGTAGAACTCACACT | >MI0000272\|hsa-mir-182&WithFlank&7\|-\|129770377\|129770498 | 29 | 52 | 24 | TTTGGCAATGGTAGAACTCACACT | 163020.2 | 8.5 | 0.8 | 11.3 | 1.4E-29 | 2.3E-28 | BT20/MB468 |
| >hg38_21_+_8433172_8446622_RNA45SN1_45S_50ntFlanks@476.512.37\|TTCGTGATCGATGTGGTGACGTCGTGCTCTCCCGGGC | >hg38_21_+_8433172_8446622_RNA45SN1_45S_50ntFlanks | 476 | 512 | 37 | TTCGTGATCGATGTGGTGACGTCGTGCTCTCCCGGGC | 636.7 | 7.0 | 0.6 | 11.3 | 1.4E-29 | 2.3E-28 | BT20/MB468 |
| >MI0000077\|hsa-mir-21&WithFlank&17\|+\|59841260\|59841343@14.35.22(+1U)&[MIMAT0000076&hsa-miR-21-5p&offsets\|0\|0(+1U);m-1&17\|+\|59841273\|59841294&offsets\|0\|0(+1U)]\|TAGCTTATCAGACTGATGTTGAT | >MI0000077\|hsa-mir-21&WithFlank&17\|+\|59841260\|59841343 | 14 | 35 | 22(+1U) | TAGCTTATCAGACTGATGTTGAT | 409193.9 | 8.8 | 0.8 | 11.3 | 1.5E-29 | 2.3E-28 | MB231/MB468 |
| >MI0000477\|hsa-mir-146a&WithFlank&5\|+\|160485346\|160485456@27.48.22(+1U)&[MIMAT0000449&hsa-miR-146a-5p&offsets\|0\|0(+1U);m-129&5\|+\|160485372\|160485393&offsets\|0\|0(+1U)]\|TGAGAACTGAATTCCATGGGTTT | >MI0000477\|hsa-mir-146a&WithFlank&5\|+\|160485346\|160485456 | 27 | 48 | 22(+1U) | TGAGAACTGAATTCCATGGGTTT | 1998.8 | 15.9 | 1.4 | 11.3 | 1.5E-29 | 2.3E-28 | MB231/MB468 |
| >MI0001446\|hsa-mir-424&WithFlank&X\|-\|134546608\|134546717@54.74.21&[MIMAT0004749&hsa-miR-424-3p&offsets\|0\|0;m-305&X\|-\|134546644\|134546664&offsets\|0\|0]\|CAAAACGTGAGGCGCTGCTAT | >MI0001446\|hsa-mir-424&WithFlank&X\|-\|134546608\|134546717 | 54 | 74 | 21 | CAAAACGTGAGGCGCTGCTAT | 1833.1 | 15.7 | 1.4 | 11.3 | 1.6E-29 | 2.4E-28 | MB231/MB468 |
| >hg38_21_+_8433172_8446622_RNA45SN1_45S_50ntFlanks@12628.12654.27\|TGGGGCGAAGCTACCATCTGTGGGATT | >hg38_21_+_8433172_8446622_RNA45SN1_45S_50ntFlanks | 12628 | 12654 | 27 | TGGGGCGAAGCTACCATCTGTGGGATT | 2096.2 | -7.0 | 0.6 | -11.3 | 1.6E-29 | 2.4E-28 | MB231/MB468 |
| >hg38_MT_+_598_1651_RNR1_12S_50ntFlanks@971.1004.34\|TGGTAAGTGTACTGGAAAGTGCACTTGGACGAAC | >hg38_MT_+_598_1651_RNR1_12S_50ntFlanks | 971 | 1004 | 34 | TGGTAAGTGTACTGGAAAGTGCACTTGGACGAAC | 1002.5 | 12.8 | 1.1 | 11.3 | 1.6E-29 | 2.5E-28 | MB231/MB468 |
| >hg38_21_+_8433172_8446622_RNA45SN1_45S_50ntFlanks@6651.6700.50\|CGACTCTTAGCGGTGGATCACTCGGCTCGTGCGTCGATGAAGAACGCAGC | >hg38_21_+_8433172_8446622_RNA45SN1_45S_50ntFlanks | 6651 | 6700 | 50 | CGACTCTTAGCGGTGGATCACTCGGCTCGTGCGTCGATGAAGAACGCAGC | 10241.7 | 11.3 | 1.0 | 11.3 | 1.7E-29 | 2.6E-28 | MB231/MB468 |
| >hg38_21_+_8433172_8446622_RNA45SN1_45S_50ntFlanks@11542.11584.43\|CGGCGCCTCGCCTCGGCCGGCGCCTAGCAGCCGACTTAGAACT | >hg38_21_+_8433172_8446622_RNA45SN1_45S_50ntFlanks | 11542 | 11584 | 43 | CGGCGCCTCGCCTCGGCCGGCGCCTAGCAGCCGACTTAGAACT | 466.5 | 11.0 | 1.0 | 11.3 | 1.7E-29 | 2.6E-28 | MB231/MB468 |
| >hg38_21_+_8433172_8446622_RNA45SN1_45S_50ntFlanks@483.505.23\|TCGATGTGGTGACGTCGTGCTCT | >hg38_21_+_8433172_8446622_RNA45SN1_45S_50ntFlanks | 483 | 505 | 23 | TCGATGTGGTGACGTCGTGCTCT | 2982.4 | 9.2 | 0.8 | 11.3 | 1.8E-29 | 2.8E-28 | MB231/MB468 |
| >MI0000255\|hsa-mir-30d&WithFlank&8\|-\|134804870\|134804951@12.32.21(+1A)&[MIMAT0000245&hsa-miR-30d-5p&offsets\|0\|-1(+1A);m-21&8\|-\|134804919\|134804940&offsets\|0\|-1(+1A)]\|TGTAAACATCCCCGACTGGAAA | >MI0000255\|hsa-mir-30d&WithFlank&8\|-\|134804870\|134804951 | 12 | 32 | 21(+1A) | TGTAAACATCCCCGACTGGAAA | 4323.7 | 10.1 | 0.9 | 11.3 | 1.8E-29 | 3.0E-28 | BT20/MB468 |
| >hg38_21_+_8433172_8446622_RNA45SN1_45S_50ntFlanks@9532.9571.40\|AGAGGAAACTCTGGTGGAGGTCCGTAGCGGTCCTGACGTG | >hg38_21_+_8433172_8446622_RNA45SN1_45S_50ntFlanks | 9532 | 9571 | 40 | AGAGGAAACTCTGGTGGAGGTCCGTAGCGGTCCTGACGTG | 354.3 | 9.3 | 0.8 | 11.3 | 2.0E-29 | 3.0E-28 | MB231/MB468 |
| >hg38_MT_+_1621_3279_RNR2_16S_50ntFlanks@1437.1457.21\|CTACGTGATCTGAGTTCAGAC | >hg38_MT_+_1621_3279_RNR2_16S_50ntFlanks | 1437 | 1457 | 21 | CTACGTGATCTGAGTTCAGAC | 371.9 | 13.5 | 1.2 | 11.3 | 2.0E-29 | 3.0E-28 | MB231/MB468 |
| >hg38_21_+_8433172_8446622_RNA45SN1_45S_50ntFlanks@11781.11804.24\|CGCATGAATGGATGAACGAGATTC | >hg38_21_+_8433172_8446622_RNA45SN1_45S_50ntFlanks | 11781 | 11804 | 24 | CGCATGAATGGATGAACGAGATTC | 2905.3 | -8.0 | 0.7 | -11.3 | 1.8E-29 | 3.0E-28 | BT20/MB468 |
| >hg38_21_+_8433172_8446622_RNA45SN1_45S_50ntFlanks@11999.12032.34\|CCCTGCGGGCCGCCGGTGAAATACCACTACTCTG | >hg38_21_+_8433172_8446622_RNA45SN1_45S_50ntFlanks | 11999 | 12032 | 34 | CCCTGCGGGCCGCCGGTGAAATACCACTACTCTG | 4798.9 | -14.2 | 1.3 | -11.3 | 1.8E-29 | 3.0E-28 | BT20/MB468 |
| >MI0000650\|hsa-mir-200c&WithFlank&12\|+\|6963693\|6963772@50.73.24&[MIMAT0000617&hsa-miR-200c-3p&offsets\|0\|+1;m-54&12\|+\|6963742\|6963764&offsets\|0\|+1]\|TAATACTGCCGGGTAATGATGGAG | >MI0000650\|hsa-mir-200c&WithFlank&12\|+\|6963693\|6963772 | 50 | 73 | 24 | TAATACTGCCGGGTAATGATGGAG | 3064.0 | 9.2 | 0.8 | 11.3 | 1.9E-29 | 3.0E-28 | BT20/MB468 |
| >hg38_21_+_8433172_8446622_RNA45SN1_45S_50ntFlanks@8380.8400.21\|CAAGAGGGCGTGAAACCGTTA | >hg38_21_+_8433172_8446622_RNA45SN1_45S_50ntFlanks | 8380 | 8400 | 21 | CAAGAGGGCGTGAAACCGTTA | 2249.4 | -8.5 | 0.8 | -11.3 | 2.0E-29 | 3.1E-28 | MB231/MB468 |
| >MI0000108\|hsa-mir-103a-2&WithFlank&20\|+\|3917488\|3917577@54.76.23(+1U);MI0000109\|hsa-mir-103a-1&WithFlank&5\|-\|168560890\|168560979@54.76.23(+1U)&[MIMAT0000101&hsa-miR-103a-3p&offsets\|0\|0(+1U);m-12&20\|+\|3917541\|3917563&offsets\|0\|0(+1U)];[MIMAT0000101_1&hsa-miR-103a-3p&offsets\|0\|0(+1U);m-13&5\|-\|168560904\|168560926&offsets\|0\|0(+1U);m-44220&5\|-\|168560908\|168560926&offsets\|0\|+4(+1U)]\|AGCAGCATTGTACAGGGCTATGAT | >MI0000108\|hsa-mir-103a-2&WithFlank&20\|+\|3917488\|3917577 | 54 | 76 | 23(+1U) | AGCAGCATTGTACAGGGCTATGAT | 12028.8 | 8.4 | 0.7 | 11.3 | 2.0E-29 | 3.1E-28 | MB231/MB468 |

<table>
<tr><td>>hg38_21_+_8433172_8446622_RNA45SN1_45S_50ntFlanks@12995.13016.22|CTCGCTGCGATCTATTGAAAGT</td><td>>hg38_21_+_8433172_8446622_RNA45SN1_45S_50ntFlanks</td><td>12995</td><td>13016</td><td>22</td><td>CTCGCTGCGATCTATTGAAAGT</td><td>2034.6</td><td>7.7</td><td>0.7</td><td>11.3</td><td>2.0E-29</td><td>3.1E-28</td><td>MB231/MB468</td></tr>
<tr><td>>trna10_LysCTT_16_+_3241501_3241573@1.35.35,trna13_LysCTT_14_-_58706613_58706685@1.35.35,trna2_LysCTT_15_+_79152904_79152976@1.35.35|GCCCGGCTAGCTCAGTCGGTAGAGCATGGGACTCT</td><td>>trna10_LysCTT_16_+_3241501_3241573</td><td>1</td><td>35</td><td>35</td><td>GCCCGGCTAGCTCAGTCGGTAGAGCATGGGACTCT</td><td>24519.4</td><td>10.5</td><td>0.9</td><td>11.3</td><td>2.1E-29</td><td>3.2E-28</td><td>MB231/MB468</td></tr>
<tr><td>>trnaMT_PheGAA_MT_+_577_647@44.74.31|TAGACGGGCTCACATCACCCCATAAACACCA</td><td>>trnaMT_PheGAA_MT_+_577_647</td><td>44</td><td>74</td><td>31</td><td>TAGACGGGCTCACATCACCCCATAAACACCA</td><td>1485.4</td><td>6.3</td><td>0.6</td><td>11.3</td><td>2.0E-29</td><td>3.2E-28</td><td>BT20/MB468</td></tr>
<tr><td>>MI0006444|hsa-mir-1307&WithFlank&10|-|103394247|103394407@86.107.22&[MIMAT0005951&hsa-miR-1307-3p&offsets|0|0;m-66&10|-|103394300|103394322&offsets|0|-1]|ACTCGGCGTGGCGTCGGTCGTG</td><td>>MI0006444|hsa-mir-1307&WithFlank&10|-|103394247|103394407</td><td>86</td><td>107</td><td>22</td><td>ACTCGGCGTGGCGTCGGTCGTG</td><td>2760.3</td><td>7.6</td><td>0.7</td><td>11.3</td><td>2.2E-29</td><td>3.3E-28</td><td>MB231/MB468</td></tr>
<tr><td>>hg38_21_+_8433172_8446622_RNA45SN1_45S_50ntFlanks@476.504.29|TTCGTGATCGATGTGGTGACGTCGTGCTC</td><td>>hg38_21_+_8433172_8446622_RNA45SN1_45S_50ntFlanks</td><td>476</td><td>504</td><td>29</td><td>TTCGTGATCGATGTGGTGACGTCGTGCTC</td><td>13211.7</td><td>8.7</td><td>0.8</td><td>11.3</td><td>2.2E-29</td><td>3.3E-28</td><td>MB231/MB468</td></tr>
<tr><td>>hg38_21_+_8433172_8446622_RNA45SN1_45S_50ntFlanks@12635.12654.20|AAGCTACCATCTGTGGGATT</td><td>>hg38_21_+_8433172_8446622_RNA45SN1_45S_50ntFlanks</td><td>12635</td><td>12654</td><td>20</td><td>AAGCTACCATCTGTGGGATT</td><td>7958.0</td><td>-9.6</td><td>0.9</td><td>-11.3</td><td>2.1E-29</td><td>3.4E-28</td><td>BT20/MB468</td></tr>
<tr><td>>MI0000087|hsa-mir-29a&WithFlank&7|-|130876741|130876816@48.67.20(+1C)&[MIMAT0000086&hsa-miR-29a-3p&offsets|0|-2(+1C);m-26&7|-|130876749|130876769&offsets|0|-1(+1C)]|TAGCACCATCTGAAATCGGTC</td><td>>MI0000087|hsa-mir-29a&WithFlank&7|-|130876741|130876816</td><td>48</td><td>67</td><td>20(+1C)</td><td>TAGCACCATCTGAAATCGGTC</td><td>3723.1</td><td>9.9</td><td>0.9</td><td>11.3</td><td>2.2E-29</td><td>3.5E-28</td><td>BT20/MB468</td></tr>
<tr><td>>hg38_21_+_8433172_8446622_RNA45SN1_45S_50ntFlanks@7975.8010.36|CGCGACCTCAGATCAGACGTGGCGACCCGCTGAATT</td><td>>hg38_21_+_8433172_8446622_RNA45SN1_45S_50ntFlanks</td><td>7975</td><td>8010</td><td>36</td><td>CGCGACCTCAGATCAGACGTGGCGACCCGCTGAATT</td><td>83313.0</td><td>8.0</td><td>0.7</td><td>11.2</td><td>2.5E-29</td><td>3.7E-28</td><td>MB231/MB468</td></tr>
<tr><td>>MI0000286|hsa-mir-210&WithFlank&11|-|568083|568204@72.93.22(+2A)&[MIMAT0000267&hsa-miR-210-3p&offsets|0|0(+2A);m-102&11|-|568112|568133&offsets|0|0(+2A)]|CTGTGCGTGTGACAGCGGCTGAAA</td><td>>MI0000286|hsa-mir-210&WithFlank&11|-|568083|568204</td><td>72</td><td>93</td><td>22(+2A)</td><td>CTGTGCGTGTGACAGCGGCTGAAA</td><td>11049.5</td><td>9.7</td><td>0.9</td><td>11.3</td><td>2.3E-29</td><td>3.7E-28</td><td>BT20/MB468</td></tr>
<tr><td>>MI0000743|hsa-mir-34c&WithFlank&11|+|111513433|111513521@19.40.22(+1U)&[MIMAT0000686&hsa-miR-34c-5p&offsets|0|-1(+1U);m-336&11|+|111513451|111513473&offsets|0|-1(+1U)]|AGGCAGTGTAGTTAGCTGATTGT</td><td>>MI0000743|hsa-mir-34c&WithFlank&11|+|111513433|111513521</td><td>19</td><td>40</td><td>22(+1U)</td><td>AGGCAGTGTAGTTAGCTGATTGT</td><td>790.7</td><td>14.5</td><td>1.3</td><td>11.2</td><td>2.5E-29</td><td>3.8E-28</td><td>MB231/MB468</td></tr>
<tr><td>>MI0000285|hsa-mir-205&WithFlank&1|+|209432127|209432248@40.60.21(+1U)&[MIMAT0000266&hsa-miR-205-5p&offsets|0|-1(+1U);m-351&1|+|209432166|209432187&offsets|0|-1(+1U)]|TCCTTCATTCCACCGGAGTCTT</td><td>>MI0000285|hsa-mir-205&WithFlank&1|+|209432127|209432248</td><td>40</td><td>60</td><td>21(+1U)</td><td>TCCTTCATTCCACCGGAGTCTT</td><td>2305.8</td><td>7.6</td><td>0.7</td><td>11.2</td><td>2.3E-29</td><td>3.8E-28</td><td>BT20/MB468</td></tr>
<tr><td>>MI0000093|hsa-mir-92a-1&WithFlank&13|+|91351308|91351397@54.74.21(+1A);MI0000094|hsa-mir-92a-2&WithFlank&X|-|134169532|134169618@54.74.21(+1A)&[MIMAT0000092&hsa-miR-92a-3p&offsets|0|-1(+1A);m-9&13|+|91351361|91351382&offsets|0|-1(+1A)];[MIMAT0000092_1&hsa-miR-92a-3p&offsets|0|-1(+1A);m-11&X|-|134169544|134169565&offsets|0|-1(+1A)]|TATTGCACTTGTCCCGGCCTGA</td><td>>MI0000093|hsa-mir-92a-1&WithFlank&13|+|91351308|91351397</td><td>54</td><td>74</td><td>21(+1A)</td><td>TATTGCACTTGTCCCGGCCTGA</td><td>6410.9</td><td>8.0</td><td>0.7</td><td>11.2</td><td>2.7E-29</td><td>4.0E-28</td><td>MB231/MB468</td></tr>
<tr><td>>hg38_21_+_8433172_8446622_RNA45SN1_45S_50ntFlanks@9534.9553.20|AGGAAACTCTGGTGGAGGTC</td><td>>hg38_21_+_8433172_8446622_RNA45SN1_45S_50ntFlanks</td><td>9534</td><td>9553</td><td>20</td><td>AGGAAACTCTGGTGGAGGTC</td><td>57111.3</td><td>-7.1</td><td>0.6</td><td>-11.2</td><td>2.8E-29</td><td>4.1E-28</td><td>MB231/MB468</td></tr>
<tr><td>>MI0000285|hsa-mir-205&WithFlank&1|+|209432127|209432248@40.60.21(+1A)&[MIMAT0000266&hsa-miR-205-5p&offsets|0|-1(+1A);m-351&1|+|209432166|209432187&offsets|0|-1(+1A)]|TCCTTCATTCCACCGGAGTCTA</td><td>>MI0000285|hsa-mir-205&WithFlank&1|+|209432127|209432248</td><td>40</td><td>60</td><td>21(+1A)</td><td>TCCTTCATTCCACCGGAGTCTA</td><td>5294.8</td><td>15.9</td><td>1.4</td><td>11.5</td><td>1.5E-30</td><td>4.1E-28</td><td>BT20/MB231</td></tr>
<tr><td>>hg38_21_+_8433172_8446622_RNA45SN1_45S_50ntFlanks@6652.6696.45|GACTCTTAGCGGTGGATCACTCGGCTCGTGCGTCGATGAAGAACG</td><td>>hg38_21_+_8433172_8446622_RNA45SN1_45S_50ntFlanks</td><td>6652</td><td>6696</td><td>45</td><td>GACTCTTAGCGGTGGATCACTCGGCTCGTGCGTCGATGAAGAACG</td><td>4915.7</td><td>7.3</td><td>0.6</td><td>11.2</td><td>2.8E-29</td><td>4.2E-28</td><td>MB231/MB468</td></tr>
<tr><td>>MI0000255|hsa-mir-30d&WithFlank&8|-|134804870|134804951@12.34.23(+1G)&[MIMAT0000245&hsa-miR-30d-5p&offsets|0|+1(+1G);m-21&8|-|134804919|134804940&offsets|0|+1(+1G)]|TGTAAACATCCCCGACTGGAAGCG</td><td>>MI0000255|hsa-mir-30d&WithFlank&8|-|134804870|134804951</td><td>12</td><td>34</td><td>23(+1G)</td><td>TGTAAACATCCCCGACTGGAAGCG</td><td>27431.4</td><td>9.7</td><td>0.9</td><td>11.2</td><td>2.6E-29</td><td>4.2E-28</td><td>BT20/MB468</td></tr>
<tr><td>>MI0000301|hsa-mir-224&WithFlank&X|-|151958572|151958664@14.34.21&[MIMAT0000281&hsa-miR-224-5p&offsets|+1|-3;m-238&X|-|151958630|151958651&offsets|0|-1]|CAAGTCACTAGTGGTTCCGTT</td><td>>MI0000301|hsa-mir-224&WithFlank&X|-|151958572|151958664</td><td>14</td><td>34</td><td>21</td><td>CAAGTCACTAGTGGTTCCGTT</td><td>948.7</td><td>14.8</td><td>1.3</td><td>11.2</td><td>2.8E-29</td><td>4.2E-28</td><td>MB231/MB468</td></tr>
<tr><td>>hg38_21_+_8433172_8446622_RNA45SN1_45S_50ntFlanks@10678.10709.32|CGGTTCCGGCGGCGTCCGGTGAGCTCTCGCTG</td><td>>hg38_21_+_8433172_8446622_RNA45SN1_45S_50ntFlanks</td><td>10678</td><td>10709</td><td>32</td><td>CGGTTCCGGCGGCGTCCGGTGAGCTCTCGCTG</td><td>17639.5</td><td>-6.5</td><td>0.6</td><td>-11.2</td><td>2.9E-29</td><td>4.3E-28</td><td>MB231/MB468</td></tr>
<tr><td>>hg38_MT_+_1621_3279_RNR2_16S_50ntFlanks@1347.1389.43|CAACAATAGGGTTTACGACCTCGATGTTGGATCAGGACATCCC</td><td>>hg38_MT_+_1621_3279_RNR2_16S_50ntFlanks</td><td>1347</td><td>1389</td><td>43</td><td>CAACAATAGGGTTTACGACCTCGATGTTGGATCAGGACATCCC</td><td>987.5</td><td>14.9</td><td>1.3</td><td>11.2</td><td>3.0E-29</td><td>4.4E-28</td><td>MB231/MB468</td></tr>
<tr><td>>hg38_21_+_8433172_8446622_RNA45SN1_45S_50ntFlanks@13002.13027.26|CGATCTATTGAAAGTCAGCCCTCGAC</td><td>>hg38_21_+_8433172_8446622_RNA45SN1_45S_50ntFlanks</td><td>13002</td><td>13027</td><td>26</td><td>CGATCTATTGAAAGTCAGCCCTCGAC</td><td>6266.0</td><td>-14.6</td><td>1.3</td><td>-11.2</td><td>2.8E-29</td><td>4.5E-28</td><td>BT20/MB468</td></tr>
<tr><td>>MI0000088|hsa-mir-30a&WithFlank&6|-|71403545|71403627@12.32.21&[MIMAT0000087&hsa-miR-30a-5p&offsets|0|-1;m-15&6|-|71403595|71403616&offsets|0|-1]|TGTAAACATCCTCGACTGGAA</td><td>>MI0000088|hsa-mir-30a&WithFlank&6|-|71403545|71403627</td><td>12</td><td>32</td><td>21</td><td>TGTAAACATCCTCGACTGGAA</td><td>2959.1</td><td>7.5</td><td>0.7</td><td>11.2</td><td>3.2E-29</td><td>4.7E-28</td><td>MB231/MB468</td></tr>
<tr><td>>hg38_21_+_8433172_8446622_RNA45SN1_45S_50ntFlanks@11564.11582.19|CCTAGCAGCCGACTTAGAA</td><td>>hg38_21_+_8433172_8446622_RNA45SN1_45S_50ntFlanks</td><td>11564</td><td>11582</td><td>19</td><td>CCTAGCAGCCGACTTAGAA</td><td>1252.6</td><td>-7.3</td><td>0.7</td><td>-11.2</td><td>3.1E-29</td><td>5.0E-28</td><td>BT20/MB468</td></tr>
<tr><td>>trnaMT_PheGAA_MT_+_577_647@36.54.19|AAAATGTTTAGACGGGCTC</td><td>>trnaMT_PheGAA_MT_+_577_647</td><td>36</td><td>54</td><td>19</td><td>AAAATGTTTAGACGGGCTC</td><td>976.0</td><td>10.7</td><td>1.0</td><td>11.2</td><td>3.3E-29</td><td>5.2E-28</td><td>BT20/MB468</td></tr>
<tr><td>>MI0000811|hsa-mir-148b&WithFlank&12|+|54337210|54337320@69.90.22(+3U)&[MIMAT0000759&hsa-miR-148b-3p&offsets|0|0(+3U);m-77&12|+|54337278|54337299&offsets|0|0(+3U)]|TCAGTGCATCACAGAACTTTGTTTT</td><td>>MI0000811|hsa-mir-148b&WithFlank&12|+|54337210|54337320</td><td>69</td><td>90</td><td>22(+3U)</td><td>TCAGTGCATCACAGAACTTTGTTTT</td><td>1137.9</td><td>8.3</td><td>0.7</td><td>11.2</td><td>3.7E-29</td><td>5.4E-28</td><td>MB231/MB468</td></tr>
<tr><td>>MI0000087|hsa-mir-29a&WithFlank&7|-|130876741|130876816@47.68.22(+1U)&[MIMAT0000086&hsa-miR-29a-3p&offsets|-1|-1(+1U);m-26&7|-|130876749|130876769&offsets|-1|0(+1U)]|CTAGCACCATCTGAAATCGGTTT</td><td>>MI0000087|hsa-mir-29a&WithFlank&7|-|130876741|130876816</td><td>47</td><td>68</td><td>22(+1U)</td><td>CTAGCACCATCTGAAATCGGTTT</td><td>1616.6</td><td>8.8</td><td>0.8</td><td>11.2</td><td>3.4E-29</td><td>5.4E-28</td><td>BT20/MB468</td></tr>
<tr><td>>hg38_21_+_8433172_8446622_RNA45SN1_45S_50ntFlanks@9545.9571.27|GTGGAGGTCCGTAGCGGTCCTGACGTG</td><td>>hg38_21_+_8433172_8446622_RNA45SN1_45S_50ntFlanks</td><td>9545</td><td>9571</td><td>27</td><td>GTGGAGGTCCGTAGCGGTCCTGACGTG</td><td>7028.6</td><td>-5.5</td><td>0.5</td><td>-11.2</td><td>3.5E-29</td><td>5.5E-28</td><td>BT20/MB468</td></tr>
<tr><td>>hg38_21_+_8433172_8446622_RNA45SN1_45S_50ntFlanks@8357.8378.22|AAAAGAACTTTGAAGAGAGAGT</td><td>>hg38_21_+_8433172_8446622_RNA45SN1_45S_50ntFlanks</td><td>8357</td><td>8378</td><td>22</td><td>AAAAGAACTTTGAAGAGAGAGT</td><td>19223.4</td><td>-6.4</td><td>0.6</td><td>-11.2</td><td>3.7E-29</td><td>5.8E-28</td><td>BT20/MB468</td></tr>
<tr><td>>hg38_21_+_8433172_8446622_RNA45SN1_45S_50ntFlanks@485.515.31|GATGTGGTGACGTCGTGCTCTCCCGGGCCGG</td><td>>hg38_21_+_8433172_8446622_RNA45SN1_45S_50ntFlanks</td><td>485</td><td>515</td><td>31</td><td>GATGTGGTGACGTCGTGCTCTCCCGGGCCGG</td><td>1743.0</td><td>15.4</td><td>1.4</td><td>11.2</td><td>3.8E-29</td><td>6.0E-28</td><td>BT20/MB468</td></tr>
<tr><td>>MI0000079|hsa-mir-23a&WithFlank&19|-|13836581|13836665@54.73.20&[MIMAT0000078&hsa-miR-23a-3p&offsets|+3|+2;m-31&19|-|13836595|13836615&offsets|+3|+2]|ACATTGCCAGGGATTTCCAA</td><td>>MI0000079|hsa-mir-23a&WithFlank&19|-|13836581|13836665</td><td>54</td><td>73</td><td>20</td><td>ACATTGCCAGGGATTTCCAA</td><td>4303.8</td><td>16.7</td><td>1.5</td><td>11.2</td><td>3.8E-29</td><td>6.0E-28</td><td>BT20/MB468</td></tr>
<tr><td>>MI0000082|hsa-mir-25&WithFlank&7|-|100093554|100093649@58.78.21&[MIMAT0000081&hsa-miR-25-3p&offsets|0|-1;m-23&7|-|100093571|100093592&offsets|0|-1]|CATTGCACTTGTCTCGGTCTG</td><td>>MI0000082|hsa-mir-25&WithFlank&7|-|100093554|100093649</td><td>58</td><td>78</td><td>21</td><td>CATTGCACTTGTCTCGGTCTG</td><td>12958.7</td><td>7.5</td><td>0.7</td><td>11.2</td><td>4.2E-29</td><td>6.1E-28</td><td>MB231/MB468</td></tr>
<tr><td>>hg38_21_+_8433172_8446622_RNA45SN1_45S_50ntFlanks@12324.12346.23|TTGGGTTTTAAGCAGGAGGTGTC</td><td>>hg38_21_+_8433172_8446622_RNA45SN1_45S_50ntFlanks</td><td>12324</td><td>12346</td><td>23</td><td>TTGGGTTTTAAGCAGGAGGTGTC</td><td>1556.2</td><td>-8.0</td><td>0.7</td><td>-11.2</td><td>4.7E-29</td><td>6.9E-28</td><td>MB231/MB468</td></tr>
<tr><td>>MI0000254|hsa-mir-30c-2&WithFlank&6|-|71376954|71377037@53.74.22&[MIMAT0004550&hsa-miR-30c-2-3p&offsets|0|0;m-206&6|-|71376964|71376985&offsets|0|0]|CTGGGAGAAGGCTGTTTACTCT</td><td>>MI0000254|hsa-mir-30c-2&WithFlank&6|-|71376954|71377037</td><td>53</td><td>74</td><td>22</td><td>CTGGGAGAAGGCTGTTTACTCT</td><td>5186.5</td><td>8.8</td><td>0.8</td><td>11.2</td><td>4.8E-29</td><td>6.9E-28</td><td>MB231/MB468</td></tr>
<tr><td>>MI0000446|hsa-mir-125b-1&WithFlank&11|-|122099751|122099850@61.82.22&[MIMAT0004592&hsa-miR-125b-1-3p&offsets|0|0;m-485&11|-|122099769|122099790&offsets|0|0]|ACGGGTTAGGCTCTTGGGAGCT</td><td>>MI0000446|hsa-mir-125b-1&WithFlank&11|-|122099751|122099850</td><td>61</td><td>82</td><td>22</td><td>ACGGGTTAGGCTCTTGGGAGCT</td><td>1979.3</td><td>13.4</td><td>1.2</td><td>11.2</td><td>4.8E-29</td><td>6.9E-28</td><td>MB231/MB468</td></tr>
<tr><td>>hg38_21_+_8433172_8446622_RNA45SN1_45S_50ntFlanks@6786.6807.22|TACGCCTGTCTGAGCGTCGCTT</td><td>>hg38_21_+_8433172_8446622_RNA45SN1_45S_50ntFlanks</td><td>6786</td><td>6807</td><td>22</td><td>TACGCCTGTCTGAGCGTCGCTT</td><td>527.6</td><td>6.9</td><td>0.6</td><td>11.2</td><td>4.9E-29</td><td>7.0E-28</td><td>MB231/MB468</td></tr>
<tr><td>>MI0025242|hsa-mir-7706&WithFlank&15|+|85380590|85380668@50.71.22&[MIMAT0030021&hsa-miR-7706&offsets|0|-2;m-390&15|+|85380639|85380662&offsets|0|-2]|TGAAGCGCCTGTGCTCTGCCGA</td><td>>MI0025242|hsa-mir-7706&WithFlank&15|+|85380590|85380668</td><td>50</td><td>71</td><td>22</td><td>TGAAGCGCCTGTGCTCTGCCGA</td><td>594.7</td><td>6.9</td><td>0.6</td><td>11.2</td><td>4.6E-29</td><td>7.2E-28</td><td>BT20/MB468</td></tr>
<tr><td>>trnaMT_PheGAA_MT_+_577_647@37.54.18|AAATGTTTAGACGGGCTC</td><td>>trnaMT_PheGAA_MT_+_577_647</td><td>37</td><td>54</td><td>18</td><td>AAATGTTTAGACGGGCTC</td><td>375.1</td><td>10.8</td><td>1.0</td><td>11.2</td><td>4.9E-29</td><td>7.6E-28</td><td>BT20/MB468</td></tr>
<tr><td>>MI0000077|hsa-mir-21&WithFlank&17|+|59841260|59841343@14.37.24&[MIMAT0000076&hsa-miR-21-5p&offsets|0|+2;m-1&17|+|59841273|59841294&offsets|0|+2]|TAGCTTATCAGACTGATGTTGACT</td><td>>MI0000077|hsa-mir-21&WithFlank&17|+|59841260|59841343</td><td>14</td><td>37</td><td>24</td><td>TAGCTTATCAGACTGATGTTGACT</td><td>884511.7</td><td>8.8</td><td>0.8</td><td>11.2</td><td>5.4E-29</td><td>7.8E-28</td><td>MB231/MB468</td></tr>
<tr><td>>MI0000299|hsa-mir-222&WithFlank&X|-|45747009|45747130@75.95.21&[MIMAT0000279&hsa-miR-222-3p&offsets|0|0;m-140&X|-|45747033|45747056&offsets|0|-3]|AGCTACATCTGGCTACTGGGT</td><td>>MI0000299|hsa-mir-222&WithFlank&X|-|45747009|45747130</td><td>75</td><td>95</td><td>21</td><td>AGCTACATCTGGCTACTGGGT</td><td>4851.8</td><td>7.7</td><td>0.7</td><td>11.2</td><td>5.4E-29</td><td>7.8E-28</td><td>MB231/MB468</td></tr>
<tr><td>>MI0000439|hsa-mir-23b&WithFlank&9|+|95085202|95085310@67.84.18&[MIMAT0000418&hsa-miR-23b-3p&offsets|+3|-2;m-42&9|+|95085265|95085285&offsets|+3|0]|ACATTGCCAGGGATTACC</td><td>>MI0000439|hsa-mir-23b&WithFlank&9|+|95085202|95085310</td><td>67</td><td>84</td><td>18</td><td>ACATTGCCAGGGATTACC</td><td>3452.5</td><td>16.4</td><td>1.5</td><td>11.2</td><td>5.1E-29</td><td>7.9E-28</td><td>BT20/MB468</td></tr>
</table>

<table>
<tr><td>>MI0000734|hsa-mir-106b&WithFlank&7|-|100093987|100094080@58.78.21(+1A)&[MIMAT0004672&hsa-miR-106b-3p&offsets|0|-1(+1A);m-87&7|-|100094003|100094023&offsets|0|0(+1A)]|CCGCACTGTGGGTACTTGCTGA</td><td>>MI0000734|hsa-mir-106b&WithFlank&7|-|100093987|100094080</td><td>58</td><td>78</td><td>21(+1A)</td><td>CCGCACTGTGGGTACTTGCTGA</td><td>1068.3</td><td>7.8</td><td>0.7</td><td>11.2</td><td>5.1E-29</td><td>7.9E-28</td><td>BT20/MB468</td></tr>
<tr><td>>hg38_21_+_8433172_8446622_RNA45SN1_45S_50ntFlanks@12419.12456.38|GATGTCGGCTCTTCCTATCATTGTGAAGCAGAATTCAC</td><td>>hg38_21_+_8433172_8446622_RNA45SN1_45S_50ntFlanks</td><td>12419</td><td>12456</td><td>38</td><td>GATGTCGGCTCTTCCTATCATTGTGAAGCAGAATTCAC</td><td>823.7</td><td>6.2</td><td>0.6</td><td>11.2</td><td>5.5E-29</td><td>7.9E-28</td><td>MB231/MB468</td></tr>
<tr><td>>MI0000298|hsa-mir-221&WithFlank&X|-|45746151|45746272@71.92.22&[MIMAT0000278&hsa-miR-221-3p&offsets|0|-1;m-91&X|-|45746181|45746202&offsets|0|0]|AGCTACATTGTCTGCTGGGTTT</td><td>>MI0000298|hsa-mir-221&WithFlank&X|-|45746151|45746272</td><td>71</td><td>92</td><td>22</td><td>AGCTACATTGTCTGCTGGGTTT</td><td>80110.4</td><td>9.0</td><td>0.8</td><td>11.2</td><td>5.5E-29</td><td>7.9E-28</td><td>MB231/MB468</td></tr>
<tr><td>>MI0000093|hsa-mir-92a-1&WithFlank&13|+|91351308|91351397@54.75.22(+1C);MI0000094|hsa-mir-92a-2&WithFlank&X|-|134169532|134169618@54.75.22(+1C)&[MIMAT0000092&hsa-miR-92a-3p&offsets|0|0(+1C);m-9&13|+|91351361|91351382&offsets|0|0(+1C)];[MIMAT0000092_1&hsa-miR-92a-3p&offsets|0|0(+1C);m-11&X|-|134169544|134169565&offsets|0|0(+1C)]|TATTGCACTTGTCCCGGCCTGTC</td><td>>MI0000093|hsa-mir-92a-1&WithFlank&13|+|91351308|91351397</td><td>54</td><td>75</td><td>22(+1C)</td><td>TATTGCACTTGTCCCGGCCTGTC</td><td>2580.1</td><td>8.0</td><td>0.7</td><td>11.2</td><td>5.6E-29</td><td>8.0E-28</td><td>MB231/MB468</td></tr>
<tr><td>>MI0000254|hsa-mir-30c-2&WithFlank&6|-|71376954|71377037@53.75.23&[MIMAT0004550&hsa-miR-30c-2-3p&offsets|0|+1;m-206&6|-|71376964|71376985&offsets|0|+1]|CTGGGAGAAGGCTGTTTACTCTT</td><td>>MI0000254|hsa-mir-30c-2&WithFlank&6|-|71376954|71377037</td><td>53</td><td>75</td><td>23</td><td>CTGGGAGAAGGCTGTTTACTCTT</td><td>1192.8</td><td>8.9</td><td>0.8</td><td>11.2</td><td>5.6E-29</td><td>8.0E-28</td><td>MB231/MB468</td></tr>
<tr><td>>MI0025242|hsa-mir-7706&WithFlank&15|+|85380590|85380668@50.72.23(+1U)&[MIMAT0030021&hsa-miR-7706&offsets|0|-1(+1U);m-390&15|+|85380639|85380662&offsets|0|-1(+1U)]|TGAAGCGCCTGTGCTCTGCCGAGT</td><td>>MI0025242|hsa-mir-7706&WithFlank&15|+|85380590|85380668</td><td>50</td><td>72</td><td>23(+1U)</td><td>TGAAGCGCCTGTGCTCTGCCGAGT</td><td>429.6</td><td>6.8</td><td>0.6</td><td>11.2</td><td>5.2E-29</td><td>8.0E-28</td><td>BT20/MB468</td></tr>
<tr><td>>MI0000434|hsa-let-7i&WithFlank&12|+|62603680|62603775@12.32.21&[MIMAT0000415&hsa-let-7i-5p&offsets|0|-1;m-49&12|+|62603691|62603712&offsets|0|-1]|TGAGGTAGTAGTTTGTGCTGT</td><td>>MI0000434|hsa-let-7i&WithFlank&12|+|62603680|62603775</td><td>12</td><td>32</td><td>21</td><td>TGAGGTAGTAGTTTGTGCTGT</td><td>29998.1</td><td>8.7</td><td>0.8</td><td>11.2</td><td>5.6E-29</td><td>8.0E-28</td><td>MB231/MB468</td></tr>
<tr><td>>hg38_21_+_8433172_8446622_RNA45SN1_45S_50ntFlanks@11891.11912.22|TTGACTCTAGTCTGGCACGGTG</td><td>>hg38_21_+_8433172_8446622_RNA45SN1_45S_50ntFlanks</td><td>11891</td><td>11912</td><td>22</td><td>TTGACTCTAGTCTGGCACGGTG</td><td>793.9</td><td>-5.5</td><td>0.5</td><td>-11.2</td><td>5.3E-29</td><td>8.1E-28</td><td>BT20/MB468</td></tr>
<tr><td>>hg38_21_+_8433172_8446622_RNA45SN1_45S_50ntFlanks@475.504.30|CTTCGTGATCGATGTGGTGACGTCGTGCTC</td><td>>hg38_21_+_8433172_8446622_RNA45SN1_45S_50ntFlanks</td><td>475</td><td>504</td><td>30</td><td>CTTCGTGATCGATGTGGTGACGTCGTGCTC</td><td>17624.2</td><td>7.4</td><td>0.7</td><td>11.2</td><td>5.3E-29</td><td>8.2E-28</td><td>BT20/MB468</td></tr>
<tr><td>>hg38_21_+_8433172_8446622_RNA45SN1_45S_50ntFlanks@475.505.31|CTTCGTGATCGATGTGGTGACGTCGTGCTCT</td><td>>hg38_21_+_8433172_8446622_RNA45SN1_45S_50ntFlanks</td><td>475</td><td>505</td><td>31</td><td>CTTCGTGATCGATGTGGTGACGTCGTGCTCT</td><td>102122.1</td><td>9.1</td><td>0.8</td><td>11.2</td><td>5.8E-29</td><td>8.2E-28</td><td>MB231/MB468</td></tr>
<tr><td>>MI0000273|hsa-mir-183&WithFlank&7|-|129774899|129775020@34.54.21(+1A)&[MIMAT0000261&hsa-miR-183-5p&offsets|+1|0(+1A);m-41&7|-|129774966|129774987&offsets|0|-1(+1A)]|ATGGCACTGGTAGAATTCACTA</td><td>>MI0000273|hsa-mir-183&WithFlank&7|-|129774899|129775020</td><td>34</td><td>54</td><td>21(+1A)</td><td>ATGGCACTGGTAGAATTCACTA</td><td>1966.1</td><td>9.2</td><td>0.8</td><td>11.2</td><td>5.4E-29</td><td>8.2E-28</td><td>BT20/MB468</td></tr>
<tr><td>>MI0000748|hsa-mir-130b&WithFlank&22|+|21653298|21653391@19.40.22&[MIMAT0004680&hsa-miR-130b-5p&offsets|0|+1;m-234&22|+|21653316|21653337&offsets|0|0]|ACTCTTTCCCTGTTGCACTACT</td><td>>MI0000748|hsa-mir-130b&WithFlank&22|+|21653298|21653391</td><td>19</td><td>40</td><td>22</td><td>ACTCTTTCCCTGTTGCACTACT</td><td>3052.4</td><td>8.6</td><td>0.8</td><td>11.2</td><td>5.9E-29</td><td>8.4E-28</td><td>MB231/MB468</td></tr>
<tr><td>>hg38_21_+_8433172_8446622_RNA45SN1_45S_50ntFlanks@12415.12452.38|CTTCGATGTCGGCTCTTCCTATCATTGTGAAGCAGAAT</td><td>>hg38_21_+_8433172_8446622_RNA45SN1_45S_50ntFlanks</td><td>12415</td><td>12452</td><td>38</td><td>CTTCGATGTCGGCTCTTCCTATCATTGTGAAGCAGAAT</td><td>458.2</td><td>8.5</td><td>0.8</td><td>11.2</td><td>6.1E-29</td><td>8.6E-28</td><td>MB231/MB468</td></tr>
<tr><td>>hg38_21_+_8433172_8446622_RNA45SN1_45S_50ntFlanks@9534.9551.18|AGGAAACTCTGGTGGAGG</td><td>>hg38_21_+_8433172_8446622_RNA45SN1_45S_50ntFlanks</td><td>9534</td><td>9551</td><td>18</td><td>AGGAAACTCTGGTGGAGG</td><td>3652.8</td><td>-5.9</td><td>0.5</td><td>-11.2</td><td>6.2E-29</td><td>8.8E-28</td><td>MB231/MB468</td></tr>
<tr><td>>MI0000082|hsa-mir-25&WithFlank&7|-|100093554|100093649@58.77.20(+1A)&[MIMAT0000081&hsa-miR-25-3p&offsets|0|-2(+1A);m-23&7|-|100093571|100093592&offsets|0|-2(+1A)]|CATTGCACTTGTCTCGGTCTA</td><td>>MI0000082|hsa-mir-25&WithFlank&7|-|100093554|100093649</td><td>58</td><td>77</td><td>20(+1A)</td><td>CATTGCACTTGTCTCGGTCTA</td><td>1428.5</td><td>7.6</td><td>0.7</td><td>11.2</td><td>6.3E-29</td><td>8.8E-28</td><td>MB231/MB468</td></tr>
<tr><td>>MI0000253|hsa-mir-148a&WithFlank&7|-|25949913|25949992@50.71.22(+1A)&[MIMAT0000243&hsa-miR-148a-3p&offsets|0|0(+1A);m-7&7|-|25949922|25949943&offsets|0|0(+1A)]|TCAGTGCACTACAGAACTTTGTA</td><td>>MI0000253|hsa-mir-148a&WithFlank&7|-|25949913|25949992</td><td>50</td><td>71</td><td>22(+1A)</td><td>TCAGTGCACTACAGAACTTTGTA</td><td>2385.6</td><td>9.1</td><td>0.8</td><td>11.2</td><td>5.8E-29</td><td>8.9E-28</td><td>BT20/MB468</td></tr>
<tr><td>>am_MI0000809|hsa-mir-151a&WithFlank&8|-|140732558|140732659@53.73.21(+1G)&[MIMAT0000757&hsa-miR-151a-3p&offsets|0|0(+1G);m-30&8|-|140732587|140732607&offsets|0|0(+1G)]|CTAGACTGAAGCTCCTTGAGGG</td><td>>am_MI0000809|hsa-mir-151a&WithFlank&8|-|140732558|140732659</td><td>53</td><td>73</td><td>21(+1G)</td><td>CTAGACTGAAGCTCCTTGAGGG</td><td>2164.1</td><td>8.6</td><td>0.8</td><td>11.2</td><td>6.4E-29</td><td>9.0E-28</td><td>MB231/MB468</td></tr>
<tr><td>>MI0000734|hsa-mir-106b&WithFlank&7|-|100093987|100094080@58.78.21(+1U)&[MIMAT0004672&hsa-miR-106b-3p&offsets|0|-1(+1U);m-87&7|-|100094003|100094023&offsets|0|0(+1U)]|CCGCACTGTGGGTACTTGCTGT</td><td>>MI0000734|hsa-mir-106b&WithFlank&7|-|100093987|100094080</td><td>58</td><td>78</td><td>21(+1U)</td><td>CCGCACTGTGGGTACTTGCTGT</td><td>4912.3</td><td>8.5</td><td>0.8</td><td>11.2</td><td>6.4E-29</td><td>9.0E-28</td><td>MB231/MB468</td></tr>
<tr><td>>MI0000301|hsa-mir-224&WithFlank&X|-|151958572|151958664@14.36.23&[MIMAT0000281&hsa-miR-224-5p&offsets|+1|-1;m-238&X|-|151958630|151958651&offsets|0|+1]|CAAGTCACTAGTGGTTCCGTTTA</td><td>>MI0000301|hsa-mir-224&WithFlank&X|-|151958572|151958664</td><td>14</td><td>36</td><td>23</td><td>CAAGTCACTAGTGGTTCCGTTTA</td><td>1143.3</td><td>15.1</td><td>1.4</td><td>11.2</td><td>6.6E-29</td><td>9.2E-28</td><td>MB231/MB468</td></tr>
<tr><td>>hg38_21_+_8433172_8446622_RNA45SN1_45S_50ntFlanks@11667.11687.21|CTGCCCAGTGCTCTGAATGTC</td><td>>hg38_21_+_8433172_8446622_RNA45SN1_45S_50ntFlanks</td><td>11667</td><td>11687</td><td>21</td><td>CTGCCCAGTGCTCTGAATGTC</td><td>947.7</td><td>6.6</td><td>0.6</td><td>11.2</td><td>6.8E-29</td><td>9.4E-28</td><td>MB231/MB468</td></tr>
<tr><td>>trnaMT_HisGTG_MT_+_12138_12206@19.55.37|AAACATCAGATTGTGAATCTGACAACAGAGGCTTACG</td><td>>trnaMT_HisGTG_MT_+_12138_12206</td><td>19</td><td>55</td><td>37</td><td>AAACATCAGATTGTGAATCTGACAACAGAGGCTTACG</td><td>286.5</td><td>12.8</td><td>1.1</td><td>11.2</td><td>6.3E-29</td><td>9.6E-28</td><td>BT20/MB468</td></tr>
<tr><td>>hg38_1_-_228634819_228635039_RNA5S12_5S_50ntFlanks@146.171.26|TGGGAATACCGGGTGCTGTAGGCTTT</td><td>>hg38_1_-_228634819_228635039_RNA5S12_5S_50ntFlanks</td><td>146</td><td>171</td><td>26</td><td>TGGGAATACCGGGTGCTGTAGGCTTT</td><td>1294.6</td><td>7.1</td><td>0.6</td><td>11.2</td><td>6.3E-29</td><td>9.6E-28</td><td>BT20/MB468</td></tr>
<tr><td>>MI0000063|hsa-let-7b&WithFlank&22|+|46113680|46113774@12.33.22(+1A)&[MIMAT0000063&hsa-let-7b-5p&offsets|0|0(+1A);m-14&22|+|46113691|46113712&offsets|0|0(+1A)]|TGAGGTAGTAGGTTGTGTGGTTA</td><td>>MI0000063|hsa-let-7b&WithFlank&22|+|46113680|46113774</td><td>12</td><td>33</td><td>22(+1A)</td><td>TGAGGTAGTAGGTTGTGTGGTTA</td><td>6959.0</td><td>8.1</td><td>0.7</td><td>11.2</td><td>6.5E-29</td><td>9.8E-28</td><td>BT20/MB468</td></tr>
<tr><td>>MI0000466|hsa-mir-9-1&WithFlank&1|-|156420335|156420435@22.44.23;MI0000468|hsa-mir-9-3&WithFlank&15|+|89368011|89368112@22.44.23;MI0000467|hsa-mir-9-2&WithFlank&5|-|88666847|88666945@22.44.23&[MIMAT0000441&hsa-miR-9-5p&offsets|0|0;m-104&1|-|156420392|156420414&offsets|0|0];[MIMAT0000441_1&hsa-miR-9-5p&offsets|0|0;m-105&15|+|89368032|89368054&offsets|0|0];[MIMAT0000441_2&hsa-miR-9-5p&offsets|0|0;m-104&5|-|88666902|88666924&offsets|0|0]|TCTTTGGTTATCTAGCTGTATGA</td><td>>MI0000466|hsa-mir-9-1&WithFlank&1|-|156420335|156420435</td><td>22</td><td>44</td><td>23</td><td>TCTTTGGTTATCTAGCTGTATGA</td><td>9557.2</td><td>10.1</td><td>0.9</td><td>11.2</td><td>6.5E-29</td><td>9.9E-28</td><td>BT20/MB468</td></tr>
<tr><td>>hg38_21_+_8433172_8446622_RNA45SN1_45S_50ntFlanks@12998.13039.42|GCTGCGATCTATTGAAAGTCAGCCCTCGACACAAGGGTTTGT</td><td>>hg38_21_+_8433172_8446622_RNA45SN1_45S_50ntFlanks</td><td>12998</td><td>13039</td><td>42</td><td>GCTGCGATCTATTGAAAGTCAGCCCTCGACACAAGGGTTTGT</td><td>31632.7</td><td>10.1</td><td>0.9</td><td>11.2</td><td>7.1E-29</td><td>9.9E-28</td><td>MB231/MB468</td></tr>
<tr><td>>am_MI0003581|hsa-mir-574&WithFlank&4|+|38868026|38868133@31.54.24&[MIMAT0004795&hsa-miR-574-5p&offsets|0|+1;m-213&4|+|38868056|38868078&offsets|0|+1]|TGAGTGTGTGTGTGTGAGTGTGTG</td><td>>am_MI0003581|hsa-mir-574&WithFlank&4|+|38868026|38868133</td><td>31</td><td>54</td><td>24</td><td>TGAGTGTGTGTGTGTGAGTGTGTG</td><td>1390.8</td><td>8.3</td><td>0.7</td><td>11.1</td><td>7.2E-29</td><td>1.0E-27</td><td>MB231/MB468</td></tr>
<tr><td>>hg38_MT_+_1621_3279_RNR2_16S_50ntFlanks@51.91.41|GCTAAACCTAGCCCCAAACCCACTCCACCTTACTACCAGAC</td><td>>hg38_MT_+_1621_3279_RNR2_16S_50ntFlanks</td><td>51</td><td>91</td><td>41</td><td>GCTAAACCTAGCCCCAAACCCACTCCACCTTACTACCAGAC</td><td>724.1</td><td>14.4</td><td>1.3</td><td>11.1</td><td>7.2E-29</td><td>1.0E-27</td><td>MB231/MB468</td></tr>
<tr><td>>hg38_MT_+_598_1651_RNR1_12S_50ntFlanks@972.1004.33|GGTAAGTGTACTGGAAAGTGCACTTGGACGAAC</td><td>>hg38_MT_+_598_1651_RNR1_12S_50ntFlanks</td><td>972</td><td>1004</td><td>33</td><td>GGTAAGTGTACTGGAAAGTGCACTTGGACGAAC</td><td>1095.0</td><td>15.0</td><td>1.3</td><td>11.1</td><td>7.3E-29</td><td>1.0E-27</td><td>MB231/MB468</td></tr>
<tr><td>>trnaMT_GluTTC_MT_-_14674_14742@32.72.41,trnalookalike8_GluTTC_5_-_93905172_93905240@32.72.41|TCATATCATTGGTCGTGGTTGTAGTCCGTGCGAGAATACCA</td><td>>trnaMT_GluTTC_MT_-_14674_14742</td><td>32</td><td>72</td><td>41</td><td>TCATATCATTGGTCGTGGTTGTAGTCCGTGCGAGAATACCA</td><td>1407.9</td><td>11.2</td><td>1.0</td><td>11.1</td><td>7.3E-29</td><td>1.0E-27</td><td>MB231/MB468</td></tr>
<tr><td>>hg38_21_+_8433172_8446622_RNA45SN1_45S_50ntFlanks@12457.12478.22|CAAGCGTTGGATTGTTCACCCA</td><td>>hg38_21_+_8433172_8446622_RNA45SN1_45S_50ntFlanks</td><td>12457</td><td>12478</td><td>22</td><td>CAAGCGTTGGATTGTTCACCCA</td><td>3557.6</td><td>-6.2</td><td>0.6</td><td>-11.2</td><td>7.0E-29</td><td>1.1E-27</td><td>BT20/MB468</td></tr>
<tr><td>>hg38_1_-_228634819_228635039_RNA5S12_5S_50ntFlanks@139.171.33|GACCGCCTGGGAATACCGGGTGCTGTAGGCTTT</td><td>>hg38_1_-_228634819_228635039_RNA5S12_5S_50ntFlanks</td><td>139</td><td>171</td><td>33</td><td>GACCGCCTGGGAATACCGGGTGCTGTAGGCTTT</td><td>3114.6</td><td>8.6</td><td>0.8</td><td>11.1</td><td>8.0E-29</td><td>1.1E-27</td><td>MB231/MB468</td></tr>
<tr><td>>hg38_21_+_8433172_8446622_RNA45SN1_45S_50ntFlanks@10350.10383.34|ACGAGAACTTTGAAGGCCGAAGTGGAGAAGGGTT</td><td>>hg38_21_+_8433172_8446622_RNA45SN1_45S_50ntFlanks</td><td>10350</td><td>10383</td><td>34</td><td>ACGAGAACTTTGAAGGCCGAAGTGGAGAAGGGTT</td><td>5291.0</td><td>-5.7</td><td>0.5</td><td>-11.1</td><td>7.4E-29</td><td>1.1E-27</td><td>BT20/MB468</td></tr>
<tr><td>>hg38_21_+_8433172_8446622_RNA45SN1_45S_50ntFlanks@476.506.31|TTCGTGATCGATGTGGTGACGTCGTGCTCTC</td><td>>hg38_21_+_8433172_8446622_RNA45SN1_45S_50ntFlanks</td><td>476</td><td>506</td><td>31</td><td>TTCGTGATCGATGTGGTGACGTCGTGCTCTC</td><td>18356.4</td><td>8.6</td><td>0.8</td><td>11.1</td><td>8.4E-29</td><td>1.2E-27</td><td>MB231/MB468</td></tr>
<tr><td>>hg38_21_+_8433172_8446622_RNA45SN1_45S_50ntFlanks@12418.12457.40|CGATGTCGGCTCTTCCTATCATTGTGAAGCAGAATTCACC</td><td>>hg38_21_+_8433172_8446622_RNA45SN1_45S_50ntFlanks</td><td>12418</td><td>12457</td><td>40</td><td>CGATGTCGGCTCTTCCTATCATTGTGAAGCAGAATTCACC</td><td>631.6</td><td>6.2</td><td>0.6</td><td>11.1</td><td>8.5E-29</td><td>1.2E-27</td><td>MB231/MB468</td></tr>
<tr><td>>hg38_21_+_8433172_8446622_RNA45SN1_45S_50ntFlanks@474.505.32|TCTTCGTGATCGATGTGGTGACGTCGTGCTCT</td><td>>hg38_21_+_8433172_8446622_RNA45SN1_45S_50ntFlanks</td><td>474</td><td>505</td><td>32</td><td>TCTTCGTGATCGATGTGGTGACGTCGTGCTCT</td><td>3523.5</td><td>9.1</td><td>0.8</td><td>11.1</td><td>9.3E-29</td><td>1.3E-27</td><td>MB231/MB468</td></tr>
<tr><td>>MI0000082|hsa-mir-25&WithFlank&7|-|100093554|100093649@58.78.21(+1U)&[MIMAT0000081&hsa-miR-25-3p&offsets|0|-1(+1U);m-23&7|-|100093571|100093592&offsets|0|-1(+1U)]|CATTGCACTTGTCTCGGTCTGT</td><td>>MI0000082|hsa-mir-25&WithFlank&7|-|100093554|100093649</td><td>58</td><td>78</td><td>21(+1U)</td><td>CATTGCACTTGTCTCGGTCTGT</td><td>8084.8</td><td>8.8</td><td>0.8</td><td>11.1</td><td>9.4E-29</td><td>1.3E-27</td><td>MB231/MB468</td></tr>
</table>

<table>
<tr><th></th><th></th><th></th><th></th><th></th><th></th><th></th><th></th><th></th><th></th><th></th><th></th><th></th></tr>
<tr><td>>am_trna10_ValCAC_5_-_180649395_180649467@1.32.32,trna12_ValAAC_5_-_180645270_180645342@1.32.32,trna132_ValAAC_6_-_27721179_27721251@1.32.32,trna136_ValAAC_6_-_27648885_27648957@1.32.32,trna139_ValAAC_6_-_27618707_27618779@1.32.32,trna18_ValCAC_5_-_180529253_180529325@1.32.32,trna2_ValAAC_3_+_169490018_169490090@1.32.32,trna2_ValCAC_5_+_180524070_180524142@1.32.32,trna4_ValAAC_5_+_180591154_180591226@1.32.32,trna5_ValAAC_5_+_180596610_180596682@1.32.32,trna6_ValCAC_5_+_180600650_180600722@1.32.32,trna85_ValCAC_1_-_161369490_161369562@1.32.32,trna90_ValCAC_1_-_149684088_149684161@1.32.32,trna98_ValCAC_1_-_149298555_149298627@1.32.32,trna9_ValCAC_6_+_26538282_26538354@1.32.32|GTTTCCGTAGTGTAGTGGTTATCACGTTCGCC</td><td>>am_trna10_ValCAC_5_-_180649395_180649467</td><td>1</td><td>32</td><td>32</td><td>GTTTCCGTAGTGTAGTGGTTATCACGTTCGCC</td><td>19682.5</td><td>10.2</td><td>0.9</td><td>11.1</td><td>9.4E-29</td><td>1.3E-27</td><td>MB231/MB468</td></tr>
<tr><td>>trnaMT_GluTTC_MT_-_14674_14742@29.72.44,trnalookalike8_GluTTC_5_-_93905172_93905240@29.72.44|TTTTCATATCATTGGTCGTGGTTGTAGTCCGTGCGAGAATACCA</td><td>>trnaMT_GluTTC_MT_-_14674_14742</td><td>29</td><td>72</td><td>44</td><td>TTTTCATATCATTGGTCGTGGTTGTAGTCCGTGCGAGAATACCA</td><td>499.0</td><td>13.9</td><td>1.2</td><td>11.1</td><td>9.6E-29</td><td>1.3E-27</td><td>MB231/MB468</td></tr>
<tr><td>>MI0002470|hsa-mir-486-1&WithFlank&8|-|41660435|41660514@10.31.22(+1U);MI0023622|hsa-mir-486-2&WithFlank&8|+|41660438|41660513@7.28.22(+1U)&[MIMAT0002177&hsa-miR-486-5p&offsets|0|0(+1U);m-107&8|-|41660484|41660505&offsets|0|0(+1U)];[MIMAT0002177_1&hsa-miR-486-5p&offsets|0|0(+1U);TJU_CMC_MD2.ID03138.5p-miR&offsets|0|-2(+1U);m-106&8|+|41660444|41660465&offsets|0|0(+1U)]|TCCTGTACTGAGCTGCCCCGAGT</td><td>>MI0002470|hsa-mir-486-1&WithFlank&8|-|41660435|41660514</td><td>10</td><td>31</td><td>22(+1U)</td><td>TCCTGTACTGAGCTGCCCCGAGT</td><td>1005.4</td><td>14.9</td><td>1.3</td><td>11.1</td><td>9.9E-29</td><td>1.3E-27</td><td>MB231/MB468</td></tr>
<tr><td>>trnaMT_SerGCT_MT_+_12207_12265@14.62.49|AGAACTGCTAACTCATGCCCCCATGTCTAACAACATGGCTTTCTCACCA</td><td>>trnaMT_SerGCT_MT_+_12207_12265</td><td>14</td><td>62</td><td>49</td><td>AGAACTGCTAACTCATGCCCCCATGTCTAACAACATGGCTTTCTCACCA</td><td>2792.8</td><td>16.4</td><td>1.5</td><td>11.1</td><td>9.9E-29</td><td>1.3E-27</td><td>MB231/MB468</td></tr>
<tr><td>>MI0006444|hsa-mir-1307&WithFlank&10|-|103394247|103394407@86.108.23&[MIMAT0005951&hsa-miR-1307-3p&offsets|0|+1;m-66&10|-|103394300|103394322&offsets|0|0]|ACTCGGCGTGGCGTCGGTCGTGG</td><td>>MI0006444|hsa-mir-1307&WithFlank&10|-|103394247|103394407</td><td>86</td><td>108</td><td>23</td><td>ACTCGGCGTGGCGTCGGTCGTGG</td><td>11540.3</td><td>7.9</td><td>0.7</td><td>11.1</td><td>1.0E-28</td><td>1.4E-27</td><td>MB231/MB468</td></tr>
<tr><td>>MI0000272|hsa-mir-182&WithFlank&7|-|129770377|129770498@29.51.23(+1A)&[MIMAT0000259&hsa-miR-182-5p&offsets|0|-1(+1A);m-22&7|-|129770449|129770470&offsets|0|+1(+1A)]|TTTGGCAATGGTAGAACTCACACA</td><td>>MI0000272|hsa-mir-182&WithFlank&7|-|129770377|129770498</td><td>29</td><td>51</td><td>23(+1A)</td><td>TTTGGCAATGGTAGAACTCACACA</td><td>20994.6</td><td>10.0</td><td>0.9</td><td>11.1</td><td>9.2E-29</td><td>1.4E-27</td><td>BT20/MB468</td></tr>
<tr><td>>MI0015866|hsa-mir-4326&WithFlank&20|+|63286802|63286872@17.40.24&[MIMAT0016888&hsa-miR-4326&offsets|0|+4;m-339&20|+|63286818|63286841&offsets|0|0]|TGTTCCTCTGTCTCCCAGACTCTG</td><td>>MI0015866|hsa-mir-4326&WithFlank&20|+|63286802|63286872</td><td>17</td><td>40</td><td>24</td><td>TGTTCCTCTGTCTCCCAGACTCTG</td><td>937.1</td><td>14.8</td><td>1.3</td><td>11.1</td><td>1.0E-28</td><td>1.4E-27</td><td>MB231/MB468</td></tr>
<tr><td>>MI0000255|hsa-mir-30d&WithFlank&8|-|134804870|134804951@12.34.23(+1G)&[MIMAT0000245&hsa-miR-30d-5p&offsets|0|+1(+1G);m-21&8|-|134804919|134804940&offsets|0|+1(+1G)]|TGTAAACATCCCCGACTGGAAGCG</td><td>>MI0000255|hsa-mir-30d&WithFlank&8|-|134804870|134804951</td><td>12</td><td>34</td><td>23(+1G)</td><td>TGTAAACATCCCCGACTGGAAGCG</td><td>13114.7</td><td>8.9</td><td>0.8</td><td>11.1</td><td>1.0E-28</td><td>1.4E-27</td><td>MB231/MB468</td></tr>
<tr><td>>MI0003205|hsa-mir-532&WithFlank&X|+|50003142|50003244@26.47.22&[MIMAT0002888&hsa-miR-532-5p&offsets|0|0;m-65&X|+|50003167|50003188&offsets|0|0]|CATGCCTTGAGTGTAGGACCGT</td><td>>MI0003205|hsa-mir-532&WithFlank&X|+|50003142|50003244</td><td>26</td><td>47</td><td>22</td><td>CATGCCTTGAGTGTAGGACCGT</td><td>6704.0</td><td>5.7</td><td>0.5</td><td>11.1</td><td>9.7E-29</td><td>1.5E-27</td><td>BT20/MB468</td></tr>
<tr><td>>MI0000478|hsa-mir-149&WithFlank&2|+|240455995|240456095@21.42.22(+1U)&[MIMAT0000450&hsa-miR-149-5p&offsets|0|-1(+1U);m-195&2|+|240456015|240456037&offsets|0|-1(+1U)]|TCTGGCTCCGTGTCTTCACTCCT</td><td>>MI0000478|hsa-mir-149&WithFlank&2|+|240455995|240456095</td><td>21</td><td>42</td><td>22(+1U)</td><td>TCTGGCTCCGTGTCTTCACTCCT</td><td>2692.3</td><td>16.0</td><td>1.4</td><td>11.1</td><td>9.7E-29</td><td>1.5E-27</td><td>BT20/MB468</td></tr>
<tr><td>>MI0000650|hsa-mir-200c&WithFlank&12|+|6963693|6963772@50.71.22(+1U)&[MIMAT0000617&hsa-miR-200c-3p&offsets|0|-1(+1U);m-54&12|+|6963742|6963764&offsets|0|-1(+1U)]|TAATACTGCCGGGTAATGATGGT</td><td>>MI0000650|hsa-mir-200c&WithFlank&12|+|6963693|6963772</td><td>50</td><td>71</td><td>22(+1U)</td><td>TAATACTGCCGGGTAATGATGGT</td><td>18561.0</td><td>9.5</td><td>0.9</td><td>11.1</td><td>1.0E-28</td><td>1.5E-27</td><td>BT20/MB468</td></tr>
<tr><td>>MI0000746|hsa-mir-99b&WithFlank&19|+|51692606|51692687@13.32.20(+1A)&[MIMAT0000689&hsa-miR-99b-5p&offsets|0|-2(+1A);m-10&19|+|51692618|51692639&offsets|0|-2(+1A)]|CACCCGTAGAACCGACCTTGA</td><td>>MI0000746|hsa-mir-99b&WithFlank&19|+|51692606|51692687</td><td>13</td><td>32</td><td>20(+1A)</td><td>CACCCGTAGAACCGACCTTGA</td><td>3622.4</td><td>9.9</td><td>0.9</td><td>11.1</td><td>1.0E-28</td><td>1.5E-27</td><td>BT20/MB468</td></tr>
<tr><td>>hg38_1_-_228634819_228635039_RNA5S12_5S_50ntFlanks@133.171.39|ATGGGAGACCGCCTGGGAATACCGGGTGCTGTAGGCTTT</td><td>>hg38_1_-_228634819_228635039_RNA5S12_5S_50ntFlanks</td><td>133</td><td>171</td><td>39</td><td>ATGGGAGACCGCCTGGGAATACCGGGTGCTGTAGGCTTT</td><td>10107.3</td><td>7.2</td><td>0.6</td><td>11.1</td><td>1.2E-28</td><td>1.5E-27</td><td>MB231/MB468</td></tr>
<tr><td>>hg38_21_+_8433172_8446622_RNA45SN1_45S_50ntFlanks@485.511.27|GATGTGGTGACGTCGTGCTCTCCCGGG</td><td>>hg38_21_+_8433172_8446622_RNA45SN1_45S_50ntFlanks</td><td>485</td><td>511</td><td>27</td><td>GATGTGGTGACGTCGTGCTCTCCCGGG</td><td>1341.8</td><td>15.0</td><td>1.4</td><td>11.1</td><td>1.1E-28</td><td>1.6E-27</td><td>BT20/MB468</td></tr>
<tr><td>>MI0000746|hsa-mir-99b&WithFlank&19|+|51692606|51692687@13.34.22(+1U)&[MIMAT0000689&hsa-miR-99b-5p&offsets|0|0(+1U);m-10&19|+|51692618|51692639&offsets|0|0(+1U)]|CACCCGTAGAACCGACCTTGCGT</td><td>>MI0000746|hsa-mir-99b&WithFlank&19|+|51692606|51692687</td><td>13</td><td>34</td><td>22(+1U)</td><td>CACCCGTAGAACCGACCTTGCGT</td><td>6147.2</td><td>8.3</td><td>0.7</td><td>11.1</td><td>1.2E-28</td><td>1.6E-27</td><td>MB231/MB468</td></tr>
<tr><td>>hg38_21_+_8433172_8446622_RNA45SN1_45S_50ntFlanks@10684.10706.23|CGGCGGCGTCCGGTGAGCTCTCG</td><td>>hg38_21_+_8433172_8446622_RNA45SN1_45S_50ntFlanks</td><td>10684</td><td>10706</td><td>23</td><td>CGGCGGCGTCCGGTGAGCTCTCG</td><td>3963.9</td><td>-6.9</td><td>0.6</td><td>-11.1</td><td>1.2E-28</td><td>1.6E-27</td><td>MB231/MB468</td></tr>
<tr><td>>MI0000734|hsa-mir-106b&WithFlank&7|-|100093987|100094080@58.78.21&[MIMAT0004672&hsa-miR-106b-3p&offsets|0|-1;m-87&7|-|100094003|100094023&offsets|0|0]|CCGCACTGTGGGTACTTGCTG</td><td>>MI0000734|hsa-mir-106b&WithFlank&7|-|100093987|100094080</td><td>58</td><td>78</td><td>21</td><td>CCGCACTGTGGGTACTTGCTG</td><td>5701.6</td><td>8.4</td><td>0.8</td><td>11.1</td><td>1.2E-28</td><td>1.6E-27</td><td>MB231/MB468</td></tr>
<tr><td>>am_trnaMT_MetCAT_MT_+_4402_4469@1.22.22|AGTAAGGTCAGCTAAATAAGCT</td><td>>am_trnaMT_MetCAT_MT_+_4402_4469</td><td>1</td><td>22</td><td>22</td><td>AGTAAGGTCAGCTAAATAAGCT</td><td>828.2</td><td>6.9</td><td>0.6</td><td>11.1</td><td>1.2E-28</td><td>1.6E-27</td><td>MB231/MB468</td></tr>
<tr><td>>MI0000239|hsa-mir-197&WithFlank&1|+|109598887|109598973@54.75.22&[MIMAT0000227&hsa-miR-197-3p&offsets|0|0;m-99&1|+|109598940|109598961&offsets|0|0]|TTCACCACCTTCTCCACCCAGC</td><td>>MI0000239|hsa-mir-197&WithFlank&1|+|109598887|109598973</td><td>54</td><td>75</td><td>22</td><td>TTCACCACCTTCTCCACCCAGC</td><td>16400.1</td><td>8.9</td><td>0.8</td><td>11.1</td><td>1.1E-28</td><td>1.7E-27</td><td>BT20/MB468</td></tr>
<tr><td>>MI0000061|hsa-let-7a-2&WithFlank&11|-|122146516|122146599@11.30.20(+1C);MI0000062|hsa-let-7a-3&WithFlank&22|+|46112743|46112828@10.29.20(+1C);MI0000060|hsa-let-7a-1&WithFlank&9|+|94175951|94176042@12.31.20(+1C)&[MIMAT0000062&hsa-let-7a-5p&offsets|0|-2(+1C);m-6&11|-|122146568|122146589&offsets|0|-2(+1C)];[MIMAT0000062_1&hsa-let-7a-5p&offsets|0|-2(+1C);m-5&22|+|46112752|46112773&offsets|0|-2(+1C)];[MIMAT0000062_2&hsa-let-7a-5p&offsets|0|-2(+1C);m-4&9|+|94175962|94175983&offsets|0|-2(+1C)]|TGAGGTAGTAGGTTGTATAGC</td><td>>MI0000061|hsa-let-7a-2&WithFlank&11|-|122146516|122146599</td><td>11</td><td>30</td><td>20(+1C)</td><td>TGAGGTAGTAGGTTGTATAGC</td><td>2198.6</td><td>7.7</td><td>0.7</td><td>11.1</td><td>1.2E-28</td><td>1.8E-27</td><td>BT20/MB468</td></tr>
<tr><td>>MI0000449|hsa-mir-132&WithFlank&17|-|2049902|2050014@65.86.22&[MIMAT0000426&hsa-miR-132-3p&offsets|0|0;m-138&17|-|2049929|2049950&offsets|0|0]|TAACAGTCTACAGCCATGGTCG</td><td>>MI0000449|hsa-mir-132&WithFlank&17|-|2049902|2050014</td><td>65</td><td>86</td><td>22</td><td>TAACAGTCTACAGCCATGGTCG</td><td>621.2</td><td>13.9</td><td>1.3</td><td>11.1</td><td>1.3E-28</td><td>1.9E-27</td><td>BT20/MB468</td></tr>
<tr><td>>hg38_21_+_8433172_8446622_RNA45SN1_45S_50ntFlanks@476.495.20|TTCGTGATCGATGTGGTGAC</td><td>>hg38_21_+_8433172_8446622_RNA45SN1_45S_50ntFlanks</td><td>476</td><td>495</td><td>20</td><td>TTCGTGATCGATGTGGTGAC</td><td>4858.1</td><td>8.3</td><td>0.7</td><td>11.1</td><td>1.5E-28</td><td>1.9E-27</td><td>MB231/MB468</td></tr>
<tr><td>>am_trnaMT_ProTGG_MT_-_15956_16023@52.71.20|AAAGACTTTTTCTCTGACCA</td><td>>am_trnaMT_ProTGG_MT_-_15956_16023</td><td>52</td><td>71</td><td>20</td><td>AAAGACTTTTTCTCTGACCA</td><td>15945.8</td><td>10.1</td><td>0.9</td><td>11.1</td><td>1.3E-28</td><td>2.0E-27</td><td>BT20/MB468</td></tr>
<tr><td>>hg38_21_+_8433172_8446622_RNA45SN1_45S_50ntFlanks@6650.6672.23|ACGACTCTTAGCGGTGGATCACT</td><td>>hg38_21_+_8433172_8446622_RNA45SN1_45S_50ntFlanks</td><td>6650</td><td>6672</td><td>23</td><td>ACGACTCTTAGCGGTGGATCACT</td><td>675.6</td><td>6.9</td><td>0.6</td><td>11.1</td><td>1.4E-28</td><td>2.0E-27</td><td>BT20/MB468</td></tr>
<tr><td>>MI0003643|hsa-mir-629&WithFlank&15|-|70079366|70079474@28.48.21&[MIMAT0004810&hsa-miR-629-5p&offsets|0|0;m-150&15|-|70079427|70079447&offsets|0|0]|TGGGTTTACGTTGGGAGAACT</td><td>>MI0003643|hsa-mir-629&WithFlank&15|-|70079366|70079474</td><td>28</td><td>48</td><td>21</td><td>TGGGTTTACGTTGGGAGAACT</td><td>1134.9</td><td>14.8</td><td>1.3</td><td>11.1</td><td>1.4E-28</td><td>2.0E-27</td><td>BT20/MB468</td></tr>
<tr><td>>MI0000462|hsa-mir-152&WithFlank&17|-|48037155|48037253@60.81.22(+1U)&[MIMAT0000438&hsa-miR-152-3p&offsets|0|+1(+1U);m-97&17|-|48037174|48037194&offsets|0|+1(+1U)]|TCAGTGCATGACAGAACTTGGGT</td><td>>MI0000462|hsa-mir-152&WithFlank&17|-|48037155|48037253</td><td>60</td><td>81</td><td>22(+1U)</td><td>TCAGTGCATGACAGAACTTGGGT</td><td>1703.8</td><td>15.6</td><td>1.4</td><td>11.1</td><td>1.5E-28</td><td>2.0E-27</td><td>MB231/MB468</td></tr>
<tr><td>>hg38_21_+_8433172_8446622_RNA45SN1_45S_50ntFlanks@12419.12438.20|GATGTCGGCTCTTCCTATCA</td><td>>hg38_21_+_8433172_8446622_RNA45SN1_45S_50ntFlanks</td><td>12419</td><td>12438</td><td>20</td><td>GATGTCGGCTCTTCCTATCA</td><td>4033.4</td><td>7.3</td><td>0.7</td><td>11.1</td><td>1.6E-28</td><td>2.0E-27</td><td>MB231/MB468</td></tr>
<tr><td>>hg38_21_+_8433172_8446622_RNA45SN1_45S_50ntFlanks@475.510.36|CTTCGTGATCGATGTGGTGACGTCGTGCTCTCCCGG</td><td>>hg38_21_+_8433172_8446622_RNA45SN1_45S_50ntFlanks</td><td>475</td><td>510</td><td>36</td><td>CTTCGTGATCGATGTGGTGACGTCGTGCTCTCCCGG</td><td>12774.1</td><td>7.6</td><td>0.7</td><td>11.1</td><td>1.6E-28</td><td>2.0E-27</td><td>MB231/MB468</td></tr>
<tr><td>>am_trnaMT_ProTGG_MT_-_15956_16023@53.70.18|AAGACTTTTTCTCTGACC</td><td>>am_trnaMT_ProTGG_MT_-_15956_16023</td><td>53</td><td>70</td><td>18</td><td>AAGACTTTTTCTCTGACC</td><td>4268.5</td><td>7.3</td><td>0.7</td><td>11.1</td><td>1.6E-28</td><td>2.1E-27</td><td>MB231/MB468</td></tr>
<tr><td>>MI0000542|hsa-mir-320a&WithFlank&8|-|22244960|22245043@48.68.21(+1U)&[MIMAT0000510&hsa-miR-320a-3p&offsets|0|-1(+1U);m-60&8|-|22244975|22244996&offsets|0|-1(+1U)]|AAAAGCTGGGTTGAGAGGGCGT</td><td>>MI0000542|hsa-mir-320a&WithFlank&8|-|22244960|22245043</td><td>48</td><td>68</td><td>21(+1U)</td><td>AAAAGCTGGGTTGAGAGGGCGT</td><td>4081.2</td><td>8.6</td><td>0.8</td><td>11.1</td><td>1.7E-28</td><td>2.2E-27</td><td>MB231/MB468</td></tr>
<tr><td>>trna116_GluCTC_1_-_145399233_145399304@1.39.39,trna59_GluCTC_1_+_249168447_249168518@1.39.39,trna71_GluCTC_1_-_161439189_161439260@1.39.39,trna74_GluCTC_1_-_161431809_161431880@1.39.39,trna77_GluCTC_1_-_161424398_161424469@1.39.39,trna77_GluCTC_6_+_28949976_28950047@1.39.39,trna80_GluCTC_1_-_161417018_161417089@1.39.39,trna87_GluCTC_6_-_126101393_126101464@1.39.39|TCCCTGGTGGTCTAGTGGTTAGGATTCGGCGCTCTCACC</td><td>>trna116_GluCTC_1_-_145399233_145399304</td><td>1</td><td>39</td><td>39</td><td>TCCCTGGTGGTCTAGTGGTTAGGATTCGGCGCTCTCACC</td><td>4748.9</td><td>8.0</td><td>0.7</td><td>11.1</td><td>1.7E-28</td><td>2.2E-27</td><td>MB231/MB468</td></tr>
</table>

<table>
<tr><td>>MI0006444|hsa-mir-1307&WithFlank&10|-|103394247|103394407@86.107.22(+1A)&[MIMAT0005951&hsa-miR-1307-3p&offsets|0|0(+1A);m-66&10|-|103394300|103394322&offsets|0|-1(+1A)]|ACTCGGCGTGGCGTCGGTCGTGA</td><td>>MI0006444|hsa-mir-1307&WithFlank&10|-|103394247|103394407</td><td>86</td><td>107</td><td>22(+1A)</td><td>ACTCGGCGTGGCGTCGGTCGTGA</td><td>1871.5</td><td>8.0</td><td>0.7</td><td>11.1</td><td>1.7E-28</td><td>2.2E-27</td><td>MB231/MB468</td></tr>
<tr><td>>MI0000088|hsa-mir-30a&WithFlank&6|-|71403545|71403627@12.34.23(+1G)&[MIMAT0000087&hsa-miR-30a-5p&offsets|0|+1(+1G);m-15&6|-|71403595|71403616&offsets|0|+1(+1G)]|TGTAAACATCCTCGACTGGAAGCG</td><td>>MI0000088|hsa-mir-30a&WithFlank&6|-|71403545|71403627</td><td>12</td><td>34</td><td>23(+1G)</td><td>TGTAAACATCCTCGACTGGAAGCG</td><td>33017.5</td><td>8.6</td><td>0.8</td><td>11.1</td><td>1.7E-28</td><td>2.2E-27</td><td>MB231/MB468</td></tr>
<tr><td>>MI0000273|hsa-mir-183&WithFlank&7|-|129774899|129775020@33.53.21(+1C)&[MIMAT0000261&hsa-miR-183-5p&offsets|0|-1(+1C);m-41&7|-|129774966|129774987&offsets|-1|-2(+1C)]|TATGGCACTGGTAGAATTCACC</td><td>>MI0000273|hsa-mir-183&WithFlank&7|-|129774899|129775020</td><td>33</td><td>53</td><td>21(+1C)</td><td>TATGGCACTGGTAGAATTCACC</td><td>5438.0</td><td>7.7</td><td>0.7</td><td>11.1</td><td>1.6E-28</td><td>2.3E-27</td><td>BT20/MB468</td></tr>
<tr><td>>hg38_21_+_8433172_8446622_RNA45SN1_45S_50ntFlanks@6719.6736.18|TGAATTGCAGGACACATT</td><td>>hg38_21_+_8433172_8446622_RNA45SN1_45S_50ntFlanks</td><td>6719</td><td>6736</td><td>18</td><td>TGAATTGCAGGACACATT</td><td>4965.0</td><td>-6.7</td><td>0.6</td><td>-11.1</td><td>1.6E-28</td><td>2.3E-27</td><td>BT20/MB468</td></tr>
<tr><td>>hg38_21_+_8433172_8446622_RNA45SN1_45S_50ntFlanks@11669.11706.38|GCCCAGTGCTCTGAATGTCAAAGTGAAGAAATTCAATG</td><td>>hg38_21_+_8433172_8446622_RNA45SN1_45S_50ntFlanks</td><td>11669</td><td>11706</td><td>38</td><td>GCCCAGTGCTCTGAATGTCAAAGTGAAGAAATTCAATG</td><td>1083.7</td><td>11.9</td><td>1.1</td><td>11.1</td><td>1.8E-28</td><td>2.4E-27</td><td>MB231/MB468</td></tr>
<tr><td>>hg38_MT_+_1621_3279_RNR2_16S_50ntFlanks@51.87.37|GCTAAACCTAGCCCCAAACCCACTCCACCTTACTACC</td><td>>hg38_MT_+_1621_3279_RNR2_16S_50ntFlanks</td><td>51</td><td>87</td><td>37</td><td>GCTAAACCTAGCCCCAAACCCACTCCACCTTACTACC</td><td>3151.0</td><td>10.7</td><td>1.0</td><td>11.1</td><td>1.9E-28</td><td>2.4E-27</td><td>MB231/MB468</td></tr>
<tr><td>>MI0000809|hsa-mir-151a&WithFlank&8|-|140732558|140732659@53.74.22(+2A)&[MIMAT0000757&hsa-miR-151a-3p&offsets|0|+1(+2A);m-30&8|-|140732587|140732607&offsets|0|+1(+2A)]|CTAGACTGAAGCTCCTTGAGGAAA</td><td>>MI0000809|hsa-mir-151a&WithFlank&8|-|140732558|140732659</td><td>53</td><td>74</td><td>22(+2A)</td><td>CTAGACTGAAGCTCCTTGAGGAAA</td><td>1226.8</td><td>7.8</td><td>0.7</td><td>11.1</td><td>1.9E-28</td><td>2.4E-27</td><td>MB231/MB468</td></tr>
<tr><td>>trna119_LysCTT_1_-_145395522_145395594@1.33.33,trna11_LysCTT_5_-_180648979_180649051@1.33.33,trna13_LysCTT_6_+_26556774_26556846@1.33.33,trna7_LysCTT_16_+_3225692_3225764@1.33.33,trna9_LysCTT_5_+_180634755_180634827@1.33.33|GCCCGGCTAGCTCAGTCGGTAGAGCATGAGACT</td><td>>trna119_LysCTT_1_-_145395522_145395594</td><td>1</td><td>33</td><td>33</td><td>GCCCGGCTAGCTCAGTCGGTAGAGCATGAGACT</td><td>2284.0</td><td>5.0</td><td>0.5</td><td>11.1</td><td>1.7E-28</td><td>2.5E-27</td><td>BT20/MB468</td></tr>
<tr><td>>MI0000273|hsa-mir-183&WithFlank&7|-|129774899|129775020@33.53.21(+1A)&[MIMAT0000261&hsa-miR-183-5p&offsets|0|-1(+1A);m-41&7|-|129774966|129774987&offsets|-1|-2(+1A)]|TATGGCACTGGTAGAATTCACA</td><td>>MI0000273|hsa-mir-183&WithFlank&7|-|129774899|129775020</td><td>33</td><td>53</td><td>21(+1A)</td><td>TATGGCACTGGTAGAATTCACA</td><td>3780.7</td><td>9.0</td><td>0.8</td><td>11.1</td><td>1.7E-28</td><td>2.5E-27</td><td>BT20/MB468</td></tr>
<tr><td>>MI0000266|hsa-mir-10a&WithFlank&17|-|48579832|48579953@28.48.21&[MIMAT0000253&hsa-miR-10a-5p&offsets|0|-2;m-18&17|-|48579905|48579926&offsets|0|-1]|TACCCTGTAGATCCGAATTTG</td><td>>MI0000266|hsa-mir-10a&WithFlank&17|-|48579832|48579953</td><td>28</td><td>48</td><td>21</td><td>TACCCTGTAGATCCGAATTTG</td><td>1024.8</td><td>14.6</td><td>1.3</td><td>11.1</td><td>1.8E-28</td><td>2.6E-27</td><td>BT20/MB468</td></tr>
<tr><td>>hg38_21_+_8433172_8446622_RNA45SN1_45S_50ntFlanks@10351.10376.26|CGAGAACTTTGAAGGCCGAAGTGGAG</td><td>>hg38_21_+_8433172_8446622_RNA45SN1_45S_50ntFlanks</td><td>10351</td><td>10376</td><td>26</td><td>CGAGAACTTTGAAGGCCGAAGTGGAG</td><td>1416.0</td><td>-6.3</td><td>0.6</td><td>-11.1</td><td>1.8E-28</td><td>2.6E-27</td><td>BT20/MB468</td></tr>
<tr><td>>hg38_21_+_8433172_8446622_RNA45SN1_45S_50ntFlanks@476.525.50|TTCGTGATCGATGTGGTGACGTCGTGCTCTCCCGGGCCGGGTCCGAGCCG</td><td>>hg38_21_+_8433172_8446622_RNA45SN1_45S_50ntFlanks</td><td>476</td><td>525</td><td>50</td><td>TTCGTGATCGATGTGGTGACGTCGTGCTCTCCCGGGCCGGGTCCGAGCCG</td><td>3266.6</td><td>16.6</td><td>1.5</td><td>11.1</td><td>2.0E-28</td><td>2.6E-27</td><td>MB231/MB468</td></tr>
<tr><td>>hg38_21_+_8433172_8446622_RNA45SN1_45S_50ntFlanks@11581.11605.25|AACTGGTGCGGACCAGGGGAATCCG</td><td>>hg38_21_+_8433172_8446622_RNA45SN1_45S_50ntFlanks</td><td>11581</td><td>11605</td><td>25</td><td>AACTGGTGCGGACCAGGGGAATCCG</td><td>1803.6</td><td>-5.4</td><td>0.5</td><td>-11.1</td><td>1.8E-28</td><td>2.6E-27</td><td>BT20/MB468</td></tr>
<tr><td>>hg38_21_+_8433172_8446622_RNA45SN1_45S_50ntFlanks@483.505.23|TCGATGTGGTGACGTCGTGCTCT</td><td>>hg38_21_+_8433172_8446622_RNA45SN1_45S_50ntFlanks</td><td>483</td><td>505</td><td>23</td><td>TCGATGTGGTGACGTCGTGCTCT</td><td>2008.5</td><td>8.4</td><td>0.8</td><td>11.1</td><td>1.8E-28</td><td>2.6E-27</td><td>BT20/MB468</td></tr>
<tr><td>>MI0000299|hsa-mir-222&WithFlank&X|-|45747009|45747130@75.98.24(+1G)&[MIMAT0000279&hsa-miR-222-3p&offsets|0|+3(+1G);m-140&X|-|45747033|45747056&offsets|0|0(+1G)]|AGCTACATCTGGCTACTGGGTCTCG</td><td>>MI0000299|hsa-mir-222&WithFlank&X|-|45747009|45747130</td><td>75</td><td>98</td><td>24(+1G)</td><td>AGCTACATCTGGCTACTGGGTCTCG</td><td>1741.9</td><td>6.9</td><td>0.6</td><td>11.1</td><td>1.9E-28</td><td>2.7E-27</td><td>BT20/MB468</td></tr>
<tr><td>>MI0000272|hsa-mir-182&WithFlank&7|-|129770377|129770498@29.53.25&[MIMAT0000259&hsa-miR-182-5p&offsets|0|+1;m-22&7|-|129770449|129770470&offsets|0|+3]|TTTGGCAATGGTAGAACTCACACTG</td><td>>MI0000272|hsa-mir-182&WithFlank&7|-|129770377|129770498</td><td>29</td><td>53</td><td>25</td><td>TTTGGCAATGGTAGAACTCACACTG</td><td>26826.5</td><td>8.0</td><td>0.7</td><td>11.1</td><td>2.0E-28</td><td>2.8E-27</td><td>BT20/MB468</td></tr>
<tr><td>>hg38_21_+_8433172_8446622_RNA45SN1_45S_50ntFlanks@12997.13038.42|CGCTGCGATCTATTGAAAGTCAGCCCTCGACACAAGGGTTTG</td><td>>hg38_21_+_8433172_8446622_RNA45SN1_45S_50ntFlanks</td><td>12997</td><td>13038</td><td>42</td><td>CGCTGCGATCTATTGAAAGTCAGCCCTCGACACAAGGGTTTG</td><td>10795.4</td><td>13.0</td><td>1.2</td><td>11.0</td><td>2.4E-28</td><td>3.0E-27</td><td>MB231/MB468</td></tr>
<tr><td>>hg38_21_+_8433172_8446622_RNA45SN1_45S_50ntFlanks@10393.10410.18|CAGCAGTTGAACATGGGT</td><td>>hg38_21_+_8433172_8446622_RNA45SN1_45S_50ntFlanks</td><td>10393</td><td>10410</td><td>18</td><td>CAGCAGTTGAACATGGGT</td><td>2911.1</td><td>-6.3</td><td>0.6</td><td>-11.0</td><td>2.5E-28</td><td>3.2E-27</td><td>MB231/MB468</td></tr>
<tr><td>>trnaMT_TrpTCA_MT_+_5512_5579@18.35.18,trnalookalike3_TrpTCA_1_+_566062_566129@18.35.18|ACAGACCAAGAGCCTTCA</td><td>>trnaMT_TrpTCA_MT_+_5512_5579</td><td>18</td><td>35</td><td>18</td><td>ACAGACCAAGAGCCTTCA</td><td>294.1</td><td>10.4</td><td>0.9</td><td>11.0</td><td>2.3E-28</td><td>3.3E-27</td><td>BT20/MB468</td></tr>
<tr><td>>hg38_21_+_8433172_8446622_RNA45SN1_45S_50ntFlanks@6645.6668.24|CTCGTACGACTCTTAGCGGTGGAT</td><td>>hg38_21_+_8433172_8446622_RNA45SN1_45S_50ntFlanks</td><td>6645</td><td>6668</td><td>24</td><td>CTCGTACGACTCTTAGCGGTGGAT</td><td>2543.7</td><td>5.3</td><td>0.5</td><td>11.0</td><td>2.3E-28</td><td>3.3E-27</td><td>BT20/MB468</td></tr>
<tr><td>>hg38_21_+_8433172_8446622_RNA45SN1_45S_50ntFlanks@12635.12654.20|AAGCTACCATCTGTGGGATT</td><td>>hg38_21_+_8433172_8446622_RNA45SN1_45S_50ntFlanks</td><td>12635</td><td>12654</td><td>20</td><td>AAGCTACCATCTGTGGGATT</td><td>6528.6</td><td>-8.3</td><td>0.8</td><td>-11.0</td><td>2.6E-28</td><td>3.3E-27</td><td>MB231/MB468</td></tr>
<tr><td>>MI0000061|hsa-let-7a-2&WithFlank&11|-|122146516|122146599@11.33.23;MI0000062|hsa-let-7a-3&WithFlank&22|+|46112743|46112828@10.32.23;MI0000060|hsa-let-7a-1&WithFlank&9|+|94175951|94176042@12.34.23&[MIMAT0000062&hsa-let-7a-5p&offsets|0|+1;m-6&11|-|122146568|122146589&offsets|0|+1];[MIMAT0000062_1&hsa-let-7a-5p&offsets|0|+1;m-5&22|+|46112752|46112773&offsets|0|+1];[MIMAT0000062_2&hsa-let-7a-5p&offsets|0|+1;m-4&9|+|94175962|94175983&offsets|0|+1]|TGAGGTAGTAGGTTGTATAGTTT</td><td>>MI0000061|hsa-let-7a-2&WithFlank&11|-|122146516|122146599</td><td>11</td><td>33</td><td>23</td><td>TGAGGTAGTAGGTTGTATAGTTT</td><td>10356.7</td><td>8.4</td><td>0.8</td><td>11.0</td><td>2.6E-28</td><td>3.3E-27</td><td>MB231/MB468</td></tr>
<tr><td>>hg38_21_+_8433172_8446622_RNA45SN1_45S_50ntFlanks@11567.11584.18|AGCAGCCGACTTAGAACT</td><td>>hg38_21_+_8433172_8446622_RNA45SN1_45S_50ntFlanks</td><td>11567</td><td>11584</td><td>18</td><td>AGCAGCCGACTTAGAACT</td><td>9640.5</td><td>7.7</td><td>0.7</td><td>11.0</td><td>2.6E-28</td><td>3.3E-27</td><td>MB231/MB468</td></tr>
<tr><td>>MI0001445|hsa-mir-423&WithFlank&17|+|30117073|30117178@59.81.23(+1A)&[MIMAT0001340&hsa-miR-423-3p&offsets|0|0(+1A);m-100&17|+|30117131|30117153&offsets|0|0(+1A)]|AGCTCGGTCTGAGGCCCCTCAGTA</td><td>>MI0001445|hsa-mir-423&WithFlank&17|+|30117073|30117178</td><td>59</td><td>81</td><td>23(+1A)</td><td>AGCTCGGTCTGAGGCCCCTCAGTA</td><td>2178.2</td><td>6.9</td><td>0.6</td><td>11.0</td><td>2.4E-28</td><td>3.4E-27</td><td>BT20/MB468</td></tr>
<tr><td>>MI0000088|hsa-mir-30a&WithFlank&6|-|71403545|71403627@12.31.20&[MIMAT0000087&hsa-miR-30a-5p&offsets|0|-2;m-15&6|-|71403595|71403616&offsets|0|-2]|TGTAAACATCCTCGACTGGA</td><td>>MI0000088|hsa-mir-30a&WithFlank&6|-|71403545|71403627</td><td>12</td><td>31</td><td>20</td><td>TGTAAACATCCTCGACTGGA</td><td>8316.5</td><td>8.2</td><td>0.7</td><td>11.0</td><td>2.7E-28</td><td>3.5E-27</td><td>MB231/MB468</td></tr>
<tr><td>>hg38_21_+_8433172_8446622_RNA45SN1_45S_50ntFlanks@475.513.39|CTTCGTGATCGATGTGGTGACGTCGTGCTCTCCCGGGCC</td><td>>hg38_21_+_8433172_8446622_RNA45SN1_45S_50ntFlanks</td><td>475</td><td>513</td><td>39</td><td>CTTCGTGATCGATGTGGTGACGTCGTGCTCTCCCGGGCC</td><td>1184.3</td><td>8.5</td><td>0.8</td><td>11.0</td><td>2.7E-28</td><td>3.5E-27</td><td>MB231/MB468</td></tr>
<tr><td>>MI0000082|hsa-mir-25&WithFlank&7|-|100093554|100093649@58.79.22&[MIMAT0000081&hsa-miR-25-3p&offsets|0|0;m-23&7|-|100093571|100093592&offsets|0|0]|CATTGCACTTGTCTCGGTCTGA</td><td>>MI0000082|hsa-mir-25&WithFlank&7|-|100093554|100093649</td><td>58</td><td>79</td><td>22</td><td>CATTGCACTTGTCTCGGTCTGA</td><td>74781.0</td><td>6.8</td><td>0.6</td><td>11.0</td><td>2.6E-28</td><td>3.6E-27</td><td>BT20/MB468</td></tr>
<tr><td>>trnaMT_HisGTG_MT_+_12138_12206@26.61.36|AGATTGTGAATCTGACAACAGAGGCTTACGACCCCT</td><td>>trnaMT_HisGTG_MT_+_12138_12206</td><td>26</td><td>61</td><td>36</td><td>AGATTGTGAATCTGACAACAGAGGCTTACGACCCCT</td><td>339.5</td><td>13.1</td><td>1.2</td><td>11.0</td><td>2.6E-28</td><td>3.6E-27</td><td>BT20/MB468</td></tr>
<tr><td>>trnaMT_PheGAA_MT_+_577_647@47.74.28|ACGGGCTCACATCACCCCATAAACACCA</td><td>>trnaMT_PheGAA_MT_+_577_647</td><td>47</td><td>74</td><td>28</td><td>ACGGGCTCACATCACCCCATAAACACCA</td><td>439.2</td><td>9.6</td><td>0.9</td><td>11.0</td><td>2.9E-28</td><td>3.6E-27</td><td>MB231/MB468</td></tr>
<tr><td>>MI0000298|hsa-mir-221&WithFlank&X|-|45746151|45746272@71.94.24&[MIMAT0000278&hsa-miR-221-3p&offsets|0|+1;m-91&X|-|45746181|45746202&offsets|0|+2]|AGCTACATTGTCTGCTGGGTTTCA</td><td>>MI0000298|hsa-mir-221&WithFlank&X|-|45746151|45746272</td><td>71</td><td>94</td><td>24</td><td>AGCTACATTGTCTGCTGGGTTTCA</td><td>10622.6</td><td>7.7</td><td>0.7</td><td>11.0</td><td>2.9E-28</td><td>3.6E-27</td><td>MB231/MB468</td></tr>
<tr><td>>MI0000067|hsa-let-7f-1&WithFlank&9|+|94176341|94176439@13.35.23&[MIMAT0000067&hsa-let-7f-5p&offsets|0|+1;m-20&9|+|94176353|94176374&offsets|0|+1]|TGAGGTAGTAGATTGTATAGTTG</td><td>>MI0000067|hsa-let-7f-1&WithFlank&9|+|94176341|94176439</td><td>13</td><td>35</td><td>23</td><td>TGAGGTAGTAGATTGTATAGTTG</td><td>506.8</td><td>6.6</td><td>0.6</td><td>11.0</td><td>3.0E-28</td><td>3.8E-27</td><td>MB231/MB468</td></tr>
<tr><td>>MI0000087|hsa-mir-29a&WithFlank&7|-|130876741|130876816@48.68.21(+1U)&[MIMAT0000086&hsa-miR-29a-3p&offsets|0|-1(+1U);m-26&7|-|130876749|130876769&offsets|0|0(+1U)]|TAGCACCATCTGAAATCGGTTT</td><td>>MI0000087|hsa-mir-29a&WithFlank&7|-|130876741|130876816</td><td>48</td><td>68</td><td>21(+1U)</td><td>TAGCACCATCTGAAATCGGTTT</td><td>1377.7</td><td>13.3</td><td>1.2</td><td>11.0</td><td>3.1E-28</td><td>3.8E-27</td><td>MB231/MB468</td></tr>
<tr><td>>hg38_21_+_8433172_8446622_RNA45SN1_45S_50ntFlanks@12476.12498.23|CCACTAATAGGGAACGTGAGCTG</td><td>>hg38_21_+_8433172_8446622_RNA45SN1_45S_50ntFlanks</td><td>12476</td><td>12498</td><td>23</td><td>CCACTAATAGGGAACGTGAGCTG</td><td>1707.8</td><td>-6.8</td><td>0.6</td><td>-11.0</td><td>3.1E-28</td><td>3.9E-27</td><td>MB231/MB468</td></tr>
<tr><td>>MI0000446|hsa-mir-125b-1&WithFlank&11|-|122099751|122099850@21.43.23(+1U)&[MIMAT0000423&hsa-miR-125b-5p&offsets|0|+1(+1U);m-67&11|-|122099809|122099830&offsets|0|+1(+1U)]|TCCCTGAGACCCTAACTTGTGATT</td><td>>MI0000446|hsa-mir-125b-1&WithFlank&11|-|122099751|122099850</td><td>21</td><td>43</td><td>23(+1U)</td><td>TCCCTGAGACCCTAACTTGTGATT</td><td>1193.5</td><td>15.1</td><td>1.4</td><td>11.0</td><td>3.1E-28</td><td>3.9E-27</td><td>MB231/MB468</td></tr>
<tr><td>>MI0000746|hsa-mir-99b&WithFlank&19|+|51692606|51692687@51.72.22(+1A)&[MIMAT0004678&hsa-miR-99b-3p&offsets|0|0(+1A);m-151&19|+|51692656|51692678&offsets|0|-1(+1A)]|CAAGCTCGTGTCTGTGGGTCCGA</td><td>>MI0000746|hsa-mir-99b&WithFlank&19|+|51692606|51692687</td><td>51</td><td>72</td><td>22(+1A)</td><td>CAAGCTCGTGTCTGTGGGTCCGA</td><td>2162.7</td><td>8.3</td><td>0.8</td><td>11.0</td><td>2.9E-28</td><td>4.1E-27</td><td>BT20/MB468</td></tr>
<tr><td>>MI0001448|hsa-mir-425&WithFlank&3|-|49020142|49020240@20.43.24(+1C)&[MIMAT0003393&hsa-miR-425-5p&offsets|0|+1(+1C);m-95&3|-|49020200|49020221&offsets|0|+2(+1C)]|AATGACACGATCACTCCCGTTGAGC</td><td>>MI0001448|hsa-mir-425&WithFlank&3|-|49020142|49020240</td><td>20</td><td>43</td><td>24(+1C)</td><td>AATGACACGATCACTCCCGTTGAGC</td><td>2206.0</td><td>8.6</td><td>0.8</td><td>11.0</td><td>3.3E-28</td><td>4.2E-27</td><td>MB231/MB468</td></tr>
<tr><td>>MI0000285|hsa-mir-205&WithFlank&1|+|209432127|209432248@40.61.22(+1G)&[MIMAT0000266&hsa-miR-205-5p&offsets|0|0(+1G);m-351&1|+|209432166|209432187&offsets|0|0(+1G)]|TCCTTCATTCCACCGGAGTCTGG</td><td>>MI0000285|hsa-mir-205&WithFlank&1|+|209432127|209432248</td><td>40</td><td>61</td><td>22(+1G)</td><td>TCCTTCATTCCACCGGAGTCTGG</td><td>3939.0</td><td>15.4</td><td>1.4</td><td>11.3</td><td>1.7E-29</td><td>4.3E-27</td><td>BT20/MB231</td></tr>
<tr><td>>hg38_21_+_8433172_8446622_RNA45SN1_45S_50ntFlanks@6719.6743.25|TGAATTGCAGGACACATTGATCATC</td><td>>hg38_21_+_8433172_8446622_RNA45SN1_45S_50ntFlanks</td><td>6719</td><td>6743</td><td>25</td><td>TGAATTGCAGGACACATTGATCATC</td><td>6679.4</td><td>-6.9</td><td>0.6</td><td>-11.0</td><td>3.0E-28</td><td>4.3E-27</td><td>BT20/MB468</td></tr>
</table>

<table>
<tr><td>>MI0000108|hsa-mir-103a-2&WithFlank&20|+|3917488|3917577@54.77.24(+1U);MI0000109|hsa-mir-103a-1&WithFlank&5|-|168560890|168560979@54.77.24(+1U)&[MIMAT0000101&hsa-miR-103a-3p&offsets|0|+1(+1U);m-12&20|+|3917541|3917563&offsets|0|+1(+1U)];[MIMAT0000101_1&hsa-miR-103a-3p&offsets|0|+1(+1U);m-13&5|-|168560904|168560926&offsets|0|+1(+1U);m-44220&5|-|168560908|168560926&offsets|0|+5(+1U)]|AGCAGCATTGTACAGGGCTATGAAT</td><td>>MI0000108|hsa-mir-103a-2&WithFlank&20|+|3917488|3917577</td><td>54</td><td>77</td><td>24(+1U)</td><td>AGCAGCATTGTACAGGGCTATGAAT</td><td>1954.3</td><td>8.7</td><td>0.8</td><td>11.0</td><td>3.5E-28</td><td>4.3E-27</td><td>MB231/MB468</td></tr>
<tr><td>>MI0000061|hsa-let-7a-2&WithFlank&11|-|122146516|122146599@11.31.21;MI0000062|hsa-let-7a-3&WithFlank&22|+|46112743|46112828@10.30.21;MI0000060|hsa-let-7a-1&WithFlank&9|+|94175951|94176042@12.32.21&[MIMAT0000062&hsa-let-7a-5p&offsets|0|-1;m-6&11|-|122146568|122146589&offsets|0|-1];[MIMAT0000062_1&hsa-let-7a-5p&offsets|0|-1;m-5&22|+|46112752|46112773&offsets|0|-1];[MIMAT0000062_2&hsa-let-7a-5p&offsets|0|-1;m-4&9|+|94175962|94175983&offsets|0|-1]|TGAGGTAGTAGGTTGTATAGT</td><td>>MI0000061|hsa-let-7a-2&WithFlank&11|-|122146516|122146599</td><td>11</td><td>31</td><td>21</td><td>TGAGGTAGTAGGTTGTATAGT</td><td>33204.9</td><td>8.5</td><td>0.8</td><td>11.0</td><td>3.7E-28</td><td>4.6E-27</td><td>MB231/MB468</td></tr>
<tr><td>>MI0001445|hsa-mir-423&WithFlank&17|+|30117073|30117178@59.81.23&[MIMAT0001340&hsa-miR-423-3p&offsets|0|0;m-100&17|+|30117131|30117153&offsets|0|0]|AGCTCGGTCTGAGGCCCCTCAGT</td><td>>MI0001445|hsa-mir-423&WithFlank&17|+|30117073|30117178</td><td>59</td><td>81</td><td>23</td><td>AGCTCGGTCTGAGGCCCCTCAGT</td><td>72485.4</td><td>8.8</td><td>0.8</td><td>11.0</td><td>3.7E-28</td><td>4.6E-27</td><td>MB231/MB468</td></tr>
<tr><td>>MI0000301|hsa-mir-224&WithFlank&X|-|151958572|151958664@14.37.24&[MIMAT0000281&hsa-miR-224-5p&offsets|+1|0;m-238&X|-|151958630|151958651&offsets|0|+2]|CAAGTCACTAGTGGTTCCGTTTAG</td><td>>MI0000301|hsa-mir-224&WithFlank&X|-|151958572|151958664</td><td>14</td><td>37</td><td>24</td><td>CAAGTCACTAGTGGTTCCGTTTAG</td><td>4598.4</td><td>11.1</td><td>1.0</td><td>11.0</td><td>3.8E-28</td><td>4.7E-27</td><td>MB231/MB468</td></tr>
<tr><td>>MI0003513|hsa-mir-455&WithFlank&9|+|114209428|114209535@60.81.22&[MIMAT0004784&hsa-miR-455-3p&offsets|0|+1;m-135&9|+|114209487|114209508&offsets|0|0]|GCAGTCCATGGGCATATACACT</td><td>>MI0003513|hsa-mir-455&WithFlank&9|+|114209428|114209535</td><td>60</td><td>81</td><td>22</td><td>GCAGTCCATGGGCATATACACT</td><td>1062.3</td><td>15.0</td><td>1.4</td><td>11.0</td><td>3.8E-28</td><td>4.7E-27</td><td>MB231/MB468</td></tr>
<tr><td>>MI0000299|hsa-mir-222&WithFlank&X|-|45747009|45747130@75.96.22&[MIMAT0000279&hsa-miR-222-3p&offsets|0|+1;m-140&X|-|45747033|45747056&offsets|0|-2]|AGCTACATCTGGCTACTGGGTC</td><td>>MI0000299|hsa-mir-222&WithFlank&X|-|45747009|45747130</td><td>75</td><td>96</td><td>22</td><td>AGCTACATCTGGCTACTGGGTC</td><td>4648.7</td><td>8.3</td><td>0.8</td><td>11.0</td><td>3.9E-28</td><td>4.8E-27</td><td>MB231/MB468</td></tr>
<tr><td>>MI0000272|hsa-mir-182&WithFlank&7|-|129770377|129770498@29.52.24(+1A)&[MIMAT0000259&hsa-miR-182-5p&offsets|0|0(+1A);m-22&7|-|129770449|129770470&offsets|0|+2(+1A)]|TTTGGCAATGGTAGAACTCACACTA</td><td>>MI0000272|hsa-mir-182&WithFlank&7|-|129770377|129770498</td><td>29</td><td>52</td><td>24(+1A)</td><td>TTTGGCAATGGTAGAACTCACACTA</td><td>6585.3</td><td>9.4</td><td>0.9</td><td>11.0</td><td>3.5E-28</td><td>4.9E-27</td><td>BT20/MB468</td></tr>
<tr><td>>hg38_21_+_8433172_8446622_RNA45SN1_45S_50ntFlanks@11667.11702.36|CTGCCCAGTGCTCTGAATGTCAAAGTGAAGAAATTC</td><td>>hg38_21_+_8433172_8446622_RNA45SN1_45S_50ntFlanks</td><td>11667</td><td>11702</td><td>36</td><td>CTGCCCAGTGCTCTGAATGTCAAAGTGAAGAAATTC</td><td>679.1</td><td>6.2</td><td>0.6</td><td>11.0</td><td>4.0E-28</td><td>5.0E-27</td><td>MB231/MB468</td></tr>
<tr><td>>MI0000298|hsa-mir-221&WithFlank&X|-|45746151|45746272@71.90.20&[MIMAT0000278&hsa-miR-221-3p&offsets|0|-3;m-91&X|-|45746181|45746202&offsets|0|-2]|AGCTACATTGTCTGCTGGGT</td><td>>MI0000298|hsa-mir-221&WithFlank&X|-|45746151|45746272</td><td>71</td><td>90</td><td>20</td><td>AGCTACATTGTCTGCTGGGT</td><td>1801.1</td><td>8.2</td><td>0.7</td><td>11.0</td><td>4.1E-28</td><td>5.1E-27</td><td>MB231/MB468</td></tr>
<tr><td>>MI0000077|hsa-mir-21&WithFlank&17|+|59841260|59841343@14.34.21(+2C)&[MIMAT0000076&hsa-miR-21-5p&offsets|0|-1(+2C);m-1&17|+|59841273|59841294&offsets|0|-1(+2C)]|TAGCTTATCAGACTGATGTTGCC</td><td>>MI0000077|hsa-mir-21&WithFlank&17|+|59841260|59841343</td><td>14</td><td>34</td><td>21(+2C)</td><td>TAGCTTATCAGACTGATGTTGCC</td><td>11669.8</td><td>9.6</td><td>0.9</td><td>11.0</td><td>4.2E-28</td><td>5.2E-27</td><td>MB231/MB468</td></tr>
<tr><td>>hg38_MT_+_1621_3279_RNR2_16S_50ntFlanks@486.517.32|AGGAACAGCTCTTTGGACACTAGGAAAAAACC</td><td>>hg38_MT_+_1621_3279_RNR2_16S_50ntFlanks</td><td>486</td><td>517</td><td>32</td><td>AGGAACAGCTCTTTGGACACTAGGAAAAAACC</td><td>421.2</td><td>9.3</td><td>0.8</td><td>11.0</td><td>4.3E-28</td><td>5.3E-27</td><td>MB231/MB468</td></tr>
<tr><td>>trna116_GluCTC_1_-_145399233_145399304@1.33.33,trna59_GluCTC_1_+_249168447_249168518@1.33.33,trna71_GluCTC_1_-_161439189_161439260@1.33.33,trna74_GluCTC_1_-_161431809_161431880@1.33.33,trna77_GluCTC_1_-_161424398_161424469@1.33.33,trna77_GluCTC_6_+_28949976_28950047@1.33.33,trna80_GluCTC_1_-_161417018_161417089@1.33.33,trna87_GluCTC_6_-_126101393_126101464@1.33.33|TCCCTGGTGGTCTAGTGGTTAGGATTCGGCGCT</td><td>>trna116_GluCTC_1_-_145399233_145399304</td><td>1</td><td>33</td><td>33</td><td>TCCCTGGTGGTCTAGTGGTTAGGATTCGGCGCT</td><td>102808.6</td><td>9.6</td><td>0.9</td><td>11.0</td><td>4.4E-28</td><td>5.4E-27</td><td>MB231/MB468</td></tr>
<tr><td>>MI0000079|hsa-mir-23a&WithFlank&19|-|13836581|13836665@54.73.20(+1U)&[MIMAT0000078&hsa-miR-23a-3p&offsets|+3|+2(+1U);m-31&19|-|13836595|13836615&offsets|+3|+2(+1U)]|ACATTGCCAGGGATTTCCAAT</td><td>>MI0000079|hsa-mir-23a&WithFlank&19|-|13836581|13836665</td><td>54</td><td>73</td><td>20(+1U)</td><td>ACATTGCCAGGGATTTCCAAT</td><td>2646.4</td><td>16.0</td><td>1.5</td><td>11.0</td><td>4.0E-28</td><td>5.6E-27</td><td>BT20/MB468</td></tr>
<tr><td>>MI0000253|hsa-mir-148a&WithFlank&7|-|25949913|25949992@50.70.21(+1G)&[MIMAT0000243&hsa-miR-148a-3p&offsets|0|-1(+1G);m-7&7|-|25949922|25949943&offsets|0|-1(+1G)]|TCAGTGCACTACAGAACTTTGG</td><td>>MI0000253|hsa-mir-148a&WithFlank&7|-|25949913|25949992</td><td>50</td><td>70</td><td>21(+1G)</td><td>TCAGTGCACTACAGAACTTTGG</td><td>45803.4</td><td>8.7</td><td>0.8</td><td>11.0</td><td>4.2E-28</td><td>5.8E-27</td><td>BT20/MB468</td></tr>
<tr><td>>hg38_21_+_8433172_8446622_RNA45SN1_45S_50ntFlanks@11572.11601.30|CCGACTTAGAACTGGTGCGGACCAGGGGAA</td><td>>hg38_21_+_8433172_8446622_RNA45SN1_45S_50ntFlanks</td><td>11572</td><td>11601</td><td>30</td><td>CCGACTTAGAACTGGTGCGGACCAGGGGAA</td><td>6864.4</td><td>-6.7</td><td>0.6</td><td>-11.0</td><td>4.3E-28</td><td>6.0E-27</td><td>BT20/MB468</td></tr>
<tr><td>>hg38_21_+_8433172_8446622_RNA45SN1_45S_50ntFlanks@4727.4767.41|AATCAAGAACGAAAGTCGGAGGTTCGAAGACGATCAGATAC</td><td>>hg38_21_+_8433172_8446622_RNA45SN1_45S_50ntFlanks</td><td>4727</td><td>4767</td><td>41</td><td>AATCAAGAACGAAAGTCGGAGGTTCGAAGACGATCAGATAC</td><td>1117.2</td><td>15.0</td><td>1.4</td><td>11.0</td><td>4.9E-28</td><td>6.0E-27</td><td>MB231/MB468</td></tr>
<tr><td>>am_trna128_GlyGCC_6_-_27870686_27870756@1.33.33,trna18_GlyGCC_16_+_70822597_70822667@1.33.33,trna19_GlyGCC_16_+_70823410_70823480@1.33.33,trna19_GlyGCC_2_-_157257659_157257729@1.33.33,trna24_GlyGCC_16_-_70812942_70813012@1.33.33,trna25_GlyGCC_16_-_70812114_70812184@1.33.33,trna5_GlyGCC_17_+_8029064_8029134@1.33.33,trna68_GlyGCC_1_-_161493637_161493707@1.33.33|GCATTGGTGGTTCAGTGGTAGAATTCTCGCCTG</td><td>>am_trna128_GlyGCC_6_-_27870686_27870756</td><td>1</td><td>33</td><td>33</td><td>GCATTGGTGGTTCAGTGGTAGAATTCTCGCCTG</td><td>18539.2</td><td>8.7</td><td>0.8</td><td>11.0</td><td>5.0E-28</td><td>6.0E-27</td><td>MB231/MB468</td></tr>
<tr><td>>MI0000434|hsa-let-7i&WithFlank&12|+|62603680|62603775@12.31.20&[MIMAT0000415&hsa-let-7i-5p&offsets|0|-2;m-49&12|+|62603691|62603712&offsets|0|-2]|TGAGGTAGTAGTTTGTGCTG</td><td>>MI0000434|hsa-let-7i&WithFlank&12|+|62603680|62603775</td><td>12</td><td>31</td><td>20</td><td>TGAGGTAGTAGTTTGTGCTG</td><td>1823.2</td><td>8.3</td><td>0.8</td><td>11.0</td><td>5.2E-28</td><td>6.3E-27</td><td>MB231/MB468</td></tr>
<tr><td>>MI0003188|hsa-mir-503&WithFlank&X|-|134546322|134546404@12.31.20&[MIMAT0002874&hsa-miR-503-5p&offsets|0|-3;m-297&X|-|134546371|134546393&offsets|0|-3]|TAGCAGCGGGAACAGTTCTG</td><td>>MI0003188|hsa-mir-503&WithFlank&X|-|134546322|134546404</td><td>12</td><td>31</td><td>20</td><td>TAGCAGCGGGAACAGTTCTG</td><td>828.3</td><td>14.3</td><td>1.3</td><td>11.0</td><td>4.5E-28</td><td>6.3E-27</td><td>BT20/MB468</td></tr>
<tr><td>>hg38_21_+_8433172_8446622_RNA45SN1_45S_50ntFlanks@12633.12653.21|CGAAGCTACCATCTGTGGGAT</td><td>>hg38_21_+_8433172_8446622_RNA45SN1_45S_50ntFlanks</td><td>12633</td><td>12653</td><td>21</td><td>CGAAGCTACCATCTGTGGGAT</td><td>7869.9</td><td>-14.9</td><td>1.4</td><td>-11.0</td><td>4.6E-28</td><td>6.3E-27</td><td>BT20/MB468</td></tr>
<tr><td>>MI0000650|hsa-mir-200c&WithFlank&12|+|6963693|6963772@50.72.23&[MIMAT0000617&hsa-miR-200c-3p&offsets|0|0;m-54&12|+|6963742|6963764&offsets|0|0]|TAATACTGCCGGGTAATGATGGA</td><td>>MI0000650|hsa-mir-200c&WithFlank&12|+|6963693|6963772</td><td>50</td><td>72</td><td>23</td><td>TAATACTGCCGGGTAATGATGGA</td><td>324763.3</td><td>8.1</td><td>0.7</td><td>11.0</td><td>4.6E-28</td><td>6.4E-27</td><td>BT20/MB468</td></tr>
<tr><td>>am_MI0005763|hsa-mir-941-1&WithFlank&20|+|63919443|63919526@53.74.22;MI0005764|hsa-mir-941-2&WithFlank&20|+|63919499|63919582@53.74.22;MI0005765|hsa-mir-941-3&WithFlank&20|+|63919555|63919638@53.74.22;MI0005766|hsa-mir-941-4&WithFlank&20|+|63919750|63919833@53.74.22;MI0031520|hsa-mir-941-5&WithFlank&20|+|63919862|63919945@53.74.22&[MIMAT0004984&hsa-miR-941&offsets|0|-1;m-263&20|+|63919495|63919517&offsets|0|-1];[MIMAT0004984_1&hsa-miR-941&offsets|0|-1;m-263&20|+|63919551|63919573&offsets|0|-1];[MIMAT0004984_2&hsa-miR-941&offsets|0|-1;m-263&20|+|63919607|63919629&offsets|0|-1];[MIMAT0004984_3&hsa-miR-941&offsets|0|-1;m-263&20|+|63919802|63919824&offsets|0|-1];[MIMAT0004984_4&hsa-miR-941&offsets|0|-1;m-263&20|+|63919914|63919936&offsets|0|-1]|CACCCGGCTGTGTGCACATGTG</td><td>>am_MI0005763|hsa-mir-941-1&WithFlank&20|+|63919443|63919526</td><td>53</td><td>74</td><td>22</td><td>CACCCGGCTGTGTGCACATGTG</td><td>1982.8</td><td>8.4</td><td>0.8</td><td>11.0</td><td>4.7E-28</td><td>6.4E-27</td><td>BT20/MB468</td></tr>
<tr><td>>hg38_MT_+_1621_3279_RNR2_16S_50ntFlanks@1348.1385.38|AACAATAGGGTTTACGACCTCGATGTTGGATCAGGACA</td><td>>hg38_MT_+_1621_3279_RNR2_16S_50ntFlanks</td><td>1348</td><td>1385</td><td>38</td><td>AACAATAGGGTTTACGACCTCGATGTTGGATCAGGACA</td><td>1449.5</td><td>15.4</td><td>1.4</td><td>11.0</td><td>5.4E-28</td><td>6.5E-27</td><td>MB231/MB468</td></tr>
<tr><td>>hg38_21_+_8433172_8446622_RNA45SN1_45S_50ntFlanks@9567.9595.29|ACGTGCAAATCGGTCGTCCGACCTGGGTA</td><td>>hg38_21_+_8433172_8446622_RNA45SN1_45S_50ntFlanks</td><td>9567</td><td>9595</td><td>29</td><td>ACGTGCAAATCGGTCGTCCGACCTGGGTA</td><td>1613.1</td><td>-6.5</td><td>0.6</td><td>-11.0</td><td>4.8E-28</td><td>6.6E-27</td><td>BT20/MB468</td></tr>
<tr><td>>MI0003760|hsa-mir-671&WithFlank&7|+|151238415|151238544@35.58.24&[MIMAT0003880&hsa-miR-671-5p&offsets|0|+1;m-316&7|+|151238449|151238472&offsets|0|0]|AGGAAGCCCTGGAGGGGCTGGAGG</td><td>>MI0003760|hsa-mir-671&WithFlank&7|+|151238415|151238544</td><td>35</td><td>58</td><td>24</td><td>AGGAAGCCCTGGAGGGGCTGGAGG</td><td>624.8</td><td>7.1</td><td>0.6</td><td>11.0</td><td>4.8E-28</td><td>6.6E-27</td><td>BT20/MB468</td></tr>
<tr><td>>hg38_21_+_8433172_8446622_RNA45SN1_45S_50ntFlanks@4735.4761.27|ACGAAAGTCGGAGGTTCGAAGACGATC</td><td>>hg38_21_+_8433172_8446622_RNA45SN1_45S_50ntFlanks</td><td>4735</td><td>4761</td><td>27</td><td>ACGAAAGTCGGAGGTTCGAAGACGATC</td><td>1406.3</td><td>7.7</td><td>0.7</td><td>11.0</td><td>5.5E-28</td><td>6.7E-27</td><td>MB231/MB468</td></tr>
<tr><td>>MI0000294|hsa-mir-218-1&WithFlank&4|+|20528269|20528390@31.52.22(+1U);MI0000295|hsa-mir-218-2&WithFlank&5|-|168768140|168768261@31.52.22(+1U)&[MIMAT0000275&hsa-miR-218-5p&offsets|0|+1(+1U);m-185&4|+|20528299|20528319&offsets|0|+1(+1U)];[MIMAT0000275_1&hsa-miR-218-5p&offsets|0|+1(+1U);m-187&5|-|168768211|168768231&offsets|0|+1(+1U)]|TTGTGCTTGATCTAACCATGTGT</td><td>>MI0000294|hsa-mir-218-1&WithFlank&4|+|20528269|20528390</td><td>31</td><td>52</td><td>22(+1U)</td><td>TTGTGCTTGATCTAACCATGTGT</td><td>769.3</td><td>14.5</td><td>1.3</td><td>11.0</td><td>5.6E-28</td><td>6.7E-27</td><td>MB231/MB468</td></tr>
<tr><td>>MI0000108|hsa-mir-103a-2&WithFlank&20|+|3917488|3917577@54.76.23(+1C);MI0000109|hsa-mir-103a-1&WithFlank&5|-|168560890|168560979@54.76.23(+1C)&[MIMAT0000101&hsa-miR-103a-3p&offsets|0|0(+1C);m-12&20|+|3917541|3917563&offsets|0|0(+1C)];[MIMAT0000101_1&hsa-miR-103a-3p&offsets|0|0(+1C);m-13&5|-|168560904|168560926&offsets|0|0(+1C);m-44220&5|-|168560908|168560926&offsets|0|+4(+1C)]|AGCAGCATTGTACAGGGCTATGAC</td><td>>MI0000108|hsa-mir-103a-2&WithFlank&20|+|3917488|3917577</td><td>54</td><td>76</td><td>23(+1C)</td><td>AGCAGCATTGTACAGGGCTATGAC</td><td>1201.4</td><td>8.4</td><td>0.8</td><td>11.0</td><td>5.7E-28</td><td>6.9E-27</td><td>MB231/MB468</td></tr>
</table>

| | | | | | | | | | | | | |
|---|---|---|---|---|---|---|---|---|---|---|---|---|
| >hg38_21_+_8433172_8446622_RNA45SN1_45S_50ntFlanks@10352.10369.18\|GAGAACTTTGAAGGCCGA | >hg38_21_+_8433172_8446622_RNA45SN1_45S_50ntFlanks | 10352 | 10369 | 18 | GAGAACTTTGAAGGCCGA | 1038.0 | 6.8 | 0.6 | 11.0 | 5.8E-28 | 7.0E-27 | MB231/MB468 |
| >MI0001733\|hsa-mir-452&WithFlank&X\|-\|151959622\|151959718@20.41.22&[MIMAT0001635&hsa-miR-452-5p&offsets\|0\|0;m-188&X\|-\|151959677\|151959699&offsets\|0\|-1]\|AACTGTTTGCAGAGGAAACTGA | >MI0001733\|hsa-mir-452&WithFlank&X\|-\|151959622\|151959718 | 20 | 41 | 22 | AACTGTTTGCAGAGGAAACTGA | 1584.8 | 15.3 | 1.4 | 11.0 | 5.2E-28 | 7.1E-27 | BT20/MB468 |
| >MI0000542\|hsa-mir-320a&WithFlank&8\|-\|22244960\|22245043@48.70.23(+1G)&[MIMAT0000510&hsa-miR-320a-3p&offsets\|0\|+1(+1G);m-60&8\|-\|22244975\|22244996&offsets\|0\|+1(+1G)]\|AAAAGCTGGGTTGAGAGGGCGAAG | >MI0000542\|hsa-mir-320a&WithFlank&8\|-\|22244960\|22245043 | 48 | 70 | 23(+1G) | AAAAGCTGGGTTGAGAGGGCGAAG | 1909.5 | 15.5 | 1.4 | 11.0 | 5.5E-28 | 7.5E-27 | BT20/MB468 |
| >MI0000266\|hsa-mir-10a&WithFlank&17\|-\|48579832\|48579953@28.48.21(+1A)&[MIMAT0000253&hsa-miR-10a-5p&offsets\|0\|-2(+1A);m-18&17\|-\|48579905\|48579926&offsets\|0\|-1(+1A)]\|TACCCTGTAGATCCGAATTTGA | >MI0000266\|hsa-mir-10a&WithFlank&17\|-\|48579832\|48579953 | 28 | 48 | 21(+1A) | TACCCTGTAGATCCGAATTTGA | 599.3 | 14.1 | 1.3 | 11.0 | 6.4E-28 | 7.6E-27 | MB231/MB468 |
| >hg38_21_+_8433172_8446622_RNA45SN1_45S_50ntFlanks@12416.12439.24\|TTCGATGTCGGCTCTTCCTATCAT | >hg38_21_+_8433172_8446622_RNA45SN1_45S_50ntFlanks | 12416 | 12439 | 24 | TTCGATGTCGGCTCTTCCTATCAT | 595.5 | 5.5 | 0.5 | 11.0 | 5.7E-28 | 7.7E-27 | BT20/MB468 |
| >hg38_21_+_8433172_8446622_RNA45SN1_45S_50ntFlanks@6651.6684.34\|CGACTCTTAGCGGTGGATCACTCGGCTCGTGCGT | >hg38_21_+_8433172_8446622_RNA45SN1_45S_50ntFlanks | 6651 | 6684 | 34 | CGACTCTTAGCGGTGGATCACTCGGCTCGTGCGT | 8536.6 | 6.6 | 0.6 | 11.0 | 6.6E-28 | 8.0E-27 | MB231/MB468 |
| >MI0000298\|hsa-mir-221&WithFlank&X\|-\|45746151\|45746272@71.91.21(+1C)&[MIMAT0000278&hsa-miR-221-3p&offsets\|0\|-2(+1C);m-91&X\|-\|45746181\|45746202&offsets\|0\|-1(+1C)]\|AGCTACATTGTCTGCTGGGTTC | >MI0000298\|hsa-mir-221&WithFlank&X\|-\|45746151\|45746272 | 71 | 91 | 21(+1C) | AGCTACATTGTCTGCTGGGTTC | 9677.1 | 9.2 | 0.8 | 10.9 | 6.7E-28 | 8.0E-27 | MB231/MB468 |
| >MI0001733\|hsa-mir-452&WithFlank&X\|-\|151959622\|151959718@20.42.23(+1U)&[MIMAT0001635&hsa-miR-452-5p&offsets\|0\|+1(+1U);m-188&X\|-\|151959677\|151959699&offsets\|0\|0(+1U)]\|AACTGTTTGCAGAGGAAACTGAGT | >MI0001733\|hsa-mir-452&WithFlank&X\|-\|151959622\|151959718 | 20 | 42 | 23(+1U) | AACTGTTTGCAGAGGAAACTGAGT | 823.0 | 14.3 | 1.3 | 11.0 | 5.9E-28 | 8.0E-27 | BT20/MB468 |
| >trnaMT_GluTTC_MT_-_14674_14742@31.72.42,trnalookalike8_GluTTC_5_-_93905172_93905240@31.72.42\|TTCATATCATTGGTCGTGGTTGTAGTCCGTGCGAGAATACCA | >trnaMT_GluTTC_MT_-_14674_14742 | 31 | 72 | 42 | TTCATATCATTGGTCGTGGTTGTAGTCCGTGCGAGAATACCA | 2283.2 | 11.8 | 1.1 | 10.9 | 6.9E-28 | 8.2E-27 | MB231/MB468 |
| >MI0000077\|hsa-mir-21&WithFlank&17\|+\|59841260\|59841343@14.34.21(+1U)&[MIMAT0000076&hsa-miR-21-5p&offsets\|0\|-1(+1U);m-1&17\|+\|59841273\|59841294&offsets\|0\|-1(+1U)]\|TAGCTTATCAGACTGATGTTGT | >MI0000077\|hsa-mir-21&WithFlank&17\|+\|59841260\|59841343 | 14 | 34 | 21(+1U) | TAGCTTATCAGACTGATGTTGT | 8604.8 | 8.5 | 0.8 | 10.9 | 6.9E-28 | 8.2E-27 | MB231/MB468 |
| >MI0000299\|hsa-mir-222&WithFlank&X\|-\|45747009\|45747130@75.94.20(+1C)&[MIMAT0000279&hsa-miR-222-3p&offsets\|0\|-1(+1C);m-140&X\|-\|45747033\|45747056&offsets\|0\|-4(+1C)]\|AGCTACATCTGGCTACTGGGC | >MI0000299\|hsa-mir-222&WithFlank&X\|-\|45747009\|45747130 | 75 | 94 | 20(+1C) | AGCTACATCTGGCTACTGGGC | 605.2 | 7.8 | 0.7 | 10.9 | 6.9E-28 | 8.2E-27 | MB231/MB468 |
| >hg38_21_+_8433172_8446622_RNA45SN1_45S_50ntFlanks@12995.13016.22\|CTCGCTGCGATCTATTGAAAGT | >hg38_21_+_8433172_8446622_RNA45SN1_45S_50ntFlanks | 12995 | 13016 | 22 | CTCGCTGCGATCTATTGAAAGT | 862.0 | 6.2 | 0.6 | 11.0 | 6.1E-28 | 8.3E-27 | BT20/MB468 |
| >am_trnaMT_HisGTG_MT_+_12138_12206@20.51.32\|AACATCAGATTGTGAATCTGACAACAGAGGCT | >am_trnaMT_HisGTG_MT_+_12138_12206 | 20 | 51 | 32 | AACATCAGATTGTGAATCTGACAACAGAGGCT | 397.3 | 13.3 | 1.2 | 11.0 | 6.2E-28 | 8.4E-27 | BT20/MB468 |
| >MI0000087\|hsa-mir-29a&WithFlank&7\|-\|130876741\|130876816@47.69.23&[MIMAT0000086&hsa-miR-29a-3p&offsets\|-1\|0;m-26&7\|-\|130876749\|130876769&offsets\|-1\|+1]\|CTAGCACCATCTGAAATCGGTTA | >MI0000087\|hsa-mir-29a&WithFlank&7\|-\|130876741\|130876816 | 47 | 69 | 23 | CTAGCACCATCTGAAATCGGTTA | 2494.0 | 8.5 | 0.8 | 11.0 | 6.5E-28 | 8.8E-27 | BT20/MB468 |
| >hg38_21_+_8433172_8446622_RNA45SN1_45S_50ntFlanks@9554.9572.19\|CGTAGCGGTCCTGACGTGC | >hg38_21_+_8433172_8446622_RNA45SN1_45S_50ntFlanks | 9554 | 9572 | 19 | CGTAGCGGTCCTGACGTGC | 11471.5 | -4.4 | 0.4 | -11.0 | 6.6E-28 | 8.8E-27 | BT20/MB468 |
| >MI0015866\|hsa-mir-4326&WithFlank&20\|+\|63286802\|63286872@17.39.23&[MIMAT0016888&hsa-miR-4326&offsets\|0\|+3;m-339&20\|+\|63286818\|63286841&offsets\|0\|-1]\|TGTTCCTCTGTCTCCCAGACTCT | >MI0015866\|hsa-mir-4326&WithFlank&20\|+\|63286802\|63286872 | 17 | 39 | 23 | TGTTCCTCTGTCTCCCAGACTCT | 622.5 | 14.2 | 1.3 | 10.9 | 7.5E-28 | 8.8E-27 | MB231/MB468 |
| >hg38_21_+_8433172_8446622_RNA45SN1_45S_50ntFlanks@11781.11803.23\|CGCATGAATGGATGAACGAGATT | >hg38_21_+_8433172_8446622_RNA45SN1_45S_50ntFlanks | 11781 | 11803 | 23 | CGCATGAATGGATGAACGAGATT | 1613.6 | -7.2 | 0.7 | -10.9 | 6.9E-28 | 9.3E-27 | BT20/MB468 |
| >MI0000298\|hsa-mir-221&WithFlank&X\|-\|45746151\|45746272@71.91.21&[MIMAT0000278&hsa-miR-221-3p&offsets\|0\|-2;m-91&X\|-\|45746181\|45746202&offsets\|0\|-1]\|AGCTACATTGTCTGCTGGGTT | >MI0000298\|hsa-mir-221&WithFlank&X\|-\|45746151\|45746272 | 71 | 91 | 21 | AGCTACATTGTCTGCTGGGTT | 5649.2 | 8.6 | 0.8 | 10.9 | 8.1E-28 | 9.6E-27 | MB231/MB468 |
| >MI0000301\|hsa-mir-224&WithFlank&X\|-\|151958572\|151958664@14.38.25&[MIMAT0000281&hsa-miR-224-5p&offsets\|+1\|+1;m-238&X\|-\|151958630\|151958651&offsets\|0\|+3]\|CAAGTCACTAGTGGTTCCGTTTAGT | >MI0000301\|hsa-mir-224&WithFlank&X\|-\|151958572\|151958664 | 14 | 38 | 25 | CAAGTCACTAGTGGTTCCGTTTAGT | 1742.6 | 13.6 | 1.2 | 10.9 | 8.2E-28 | 9.7E-27 | MB231/MB468 |
| >MI0000102\|hsa-mir-100&WithFlank&11\|-\|122152223\|122152314@19.37.19&[MIMAT0000098&hsa-miR-100-5p&offsets\|0\|-3;m-27&11\|-\|122152276\|122152296&offsets\|0\|-2]\|AACCCGTAGATCCGAACTT | >MI0000102\|hsa-mir-100&WithFlank&11\|-\|122152223\|122152314 | 19 | 37 | 19 | AACCCGTAGATCCGAACTT | 1224.6 | 15.2 | 1.4 | 10.9 | 8.3E-28 | 9.8E-27 | MB231/MB468 |
| >MI0000462\|hsa-mir-152&WithFlank&17\|-\|48037155\|48037253@60.81.22(+1U)&[MIMAT0000438&hsa-miR-152-3p&offsets\|0\|+1(+1U);m-97&17\|-\|48037174\|48037194&offsets\|0\|+1(+1U)]\|TCAGTGCATGACAGAACTTGGGT | >MI0000462\|hsa-mir-152&WithFlank&17\|-\|48037155\|48037253 | 60 | 81 | 22(+1U) | TCAGTGCATGACAGAACTTGGGT | 951.6 | 14.5 | 1.3 | 10.9 | 7.4E-28 | 9.9E-27 | BT20/MB468 |
| >hg38_21_+_8433172_8446622_RNA45SN1_45S_50ntFlanks@4727.4768.42\|AATCAAGAACGAAAGTCGGAGGTTCGAAGACGATCAGATACC | >hg38_21_+_8433172_8446622_RNA45SN1_45S_50ntFlanks | 4727 | 4768 | 42 | AATCAAGAACGAAAGTCGGAGGTTCGAAGACGATCAGATACC | 655.9 | 14.3 | 1.3 | 10.9 | 8.5E-28 | 1.0E-26 | MB231/MB468 |
| >hg38_21_+_8433172_8446622_RNA45SN1_45S_50ntFlanks@12410.12433.24\|TGATCCTTCGATGTCGGCTCTTCC | >hg38_21_+_8433172_8446622_RNA45SN1_45S_50ntFlanks | 12410 | 12433 | 24 | TGATCCTTCGATGTCGGCTCTTCC | 381.2 | 5.9 | 0.5 | 10.9 | 8.6E-28 | 1.0E-26 | MB231/MB468 |
| >hg38_21_+_8433172_8446622_RNA45SN1_45S_50ntFlanks@11668.11688.21\|TGCCCAGTGCTCTGAATGTCA | >hg38_21_+_8433172_8446622_RNA45SN1_45S_50ntFlanks | 11668 | 11688 | 21 | TGCCCAGTGCTCTGAATGTCA | 1266.5 | 7.3 | 0.7 | 10.9 | 8.6E-28 | 1.0E-26 | MB231/MB468 |
| >MI0000477\|hsa-mir-146a&WithFlank&5\|+\|160485346\|160485456@27.47.21(+1A)&[MIMAT0000449&hsa-miR-146a-5p&offsets\|0\|-1(+1A);m-129&5\|+\|160485372\|160485393&offsets\|0\|-1(+1A)]\|TGAGAACTGAATTCCATGGGTA | >MI0000477\|hsa-mir-146a&WithFlank&5\|+\|160485346\|160485456 | 27 | 47 | 21(+1A) | TGAGAACTGAATTCCATGGGTA | 1774.8 | 15.7 | 1.4 | 10.9 | 8.6E-28 | 1.0E-26 | MB231/MB468 |
| >hg38_MT_+_1621_3279_RNR2_16S_50ntFlanks@617.642.26\|AAACATATAACTGAACTCCTCACACC | >hg38_MT_+_1621_3279_RNR2_16S_50ntFlanks | 617 | 642 | 26 | AAACATATAACTGAACTCCTCACACC | 380.8 | 13.2 | 1.2 | 10.9 | 7.6E-28 | 1.0E-26 | BT20/MB468 |
| >trnaMT_PheGAA_MT_+_577_647@35.55.21\|GAAAATGTTTAGACGGGCTCA | >trnaMT_PheGAA_MT_+_577_647 | 35 | 55 | 21 | GAAAATGTTTAGACGGGCTCA | 626.6 | 10.1 | 0.9 | 10.9 | 8.8E-28 | 1.0E-26 | MB231/MB468 |
| >MI0000255\|hsa-mir-30d&WithFlank&8\|-\|134804870\|134804951@12.32.21&[MIMAT0000245&hsa-miR-30d-5p&offsets\|0\|-1;m-21&8\|-\|134804919\|134804940&offsets\|0\|-1]\|TGTAAACATCCCCGACTGGAA | >MI0000255\|hsa-mir-30d&WithFlank&8\|-\|134804870\|134804951 | 12 | 32 | 21 | TGTAAACATCCCCGACTGGAA | 3459.0 | 8.9 | 0.8 | 10.9 | 7.9E-28 | 1.0E-26 | BT20/MB468 |
| >hg38_21_+_8433172_8446622_RNA45SN1_45S_50ntFlanks@13000.13039.40\|TGCGATCTATTGAAAGTCAGCCCTCGACACAAGGGTTTGT | >hg38_21_+_8433172_8446622_RNA45SN1_45S_50ntFlanks | 13000 | 13039 | 40 | TGCGATCTATTGAAAGTCAGCCCTCGACACAAGGGTTTGT | 1864.2 | 7.8 | 0.7 | 10.9 | 9.0E-28 | 1.1E-26 | MB231/MB468 |
| >hg38_21_+_8433172_8446622_RNA45SN1_45S_50ntFlanks@11566.11584.19\|TAGCAGCCGACTTAGAACT | >hg38_21_+_8433172_8446622_RNA45SN1_45S_50ntFlanks | 11566 | 11584 | 19 | TAGCAGCCGACTTAGAACT | 3250.9 | 8.6 | 0.8 | 10.9 | 9.3E-28 | 1.1E-26 | MB231/MB468 |
| >hg38_21_+_8433172_8446622_RNA45SN1_45S_50ntFlanks@11208.11225.18\|GCGGCGACTCTGGACGCG | >hg38_21_+_8433172_8446622_RNA45SN1_45S_50ntFlanks | 11208 | 11225 | 18 | GCGGCGACTCTGGACGCG | 2599.4 | -8.0 | 0.7 | -10.9 | 1.0E-27 | 1.2E-26 | MB231/MB468 |
| >hg38_21_+_8433172_8446622_RNA45SN1_45S_50ntFlanks@6651.6691.41\|CGACTCTTAGCGGTGGATCACTCGGCTCGTGCGTCGATGAA | >hg38_21_+_8433172_8446622_RNA45SN1_45S_50ntFlanks | 6651 | 6691 | 41 | CGACTCTTAGCGGTGGATCACTCGGCTCGTGCGTCGATGAA | 979.7 | 6.5 | 0.6 | 10.9 | 1.0E-27 | 1.2E-26 | MB231/MB468 |
| >hg38_MT_+_1621_3279_RNR2_16S_50ntFlanks@51.74.24\|GCTAAACCTAGCCCCAAACCCACT | >hg38_MT_+_1621_3279_RNR2_16S_50ntFlanks | 51 | 74 | 24 | GCTAAACCTAGCCCCAAACCCACT | 3466.4 | 7.8 | 0.7 | 10.9 | 8.9E-28 | 1.2E-26 | BT20/MB468 |
| >MI0003513\|hsa-mir-455&WithFlank&9\|+\|114209428\|114209535@60.80.21(+1C)&[MIMAT0004784&hsa-miR-455-3p&offsets\|0\|0(+1C);m-135&9\|+\|114209487\|114209508&offsets\|0\|-1(+1C)]\|GCAGTCCATGGGCATATACACC | >MI0003513\|hsa-mir-455&WithFlank&9\|+\|114209428\|114209535 | 60 | 80 | 21(+1C) | GCAGTCCATGGGCATATACACC | 931.5 | 14.8 | 1.4 | 10.9 | 1.0E-27 | 1.2E-26 | MB231/MB468 |
| >hg38_21_+_8433172_8446622_RNA45SN1_45S_50ntFlanks@12419.12439.21\|GATGTCGGCTCTTCCTATCAT | >hg38_21_+_8433172_8446622_RNA45SN1_45S_50ntFlanks | 12419 | 12439 | 21 | GATGTCGGCTCTTCCTATCAT | 2963.2 | 6.6 | 0.6 | 10.9 | 1.0E-27 | 1.2E-26 | MB231/MB468 |
| >MI0000440\|hsa-mir-27b&WithFlank&9\|+\|95085439\|95085547@67.85.19(+1A)&[MIMAT0000419&hsa-miR-27b-3p&offsets\|0\|-2(+1A);m-39&9\|+\|95085505\|95085525&offsets\|0\|-2(+1A)]\|TTCACAGTGGCTAAGTTCTA | >MI0000440\|hsa-mir-27b&WithFlank&9\|+\|95085439\|95085547 | 67 | 85 | 19(+1A) | TTCACAGTGGCTAAGTTCTA | 2075.1 | 8.5 | 0.8 | 10.9 | 1.1E-27 | 1.2E-26 | MB231/MB468 |
| >MI0000477\|hsa-mir-146a&WithFlank&5\|+\|160485346\|160485456@27.50.24&[MIMAT0000449&hsa-miR-146a-5p&offsets\|0\|+2;m-129&5\|+\|160485372\|160485393&offsets\|0\|+2]\|TGAGAACTGAATTCCATGGGTTGT | >MI0000477\|hsa-mir-146a&WithFlank&5\|+\|160485346\|160485456 | 27 | 50 | 24 | TGAGAACTGAATTCCATGGGTTGT | 1315.3 | 15.3 | 1.4 | 10.9 | 1.1E-27 | 1.2E-26 | MB231/MB468 |
| >hg38_21_+_8433172_8446622_RNA45SN1_45S_50ntFlanks@11873.11893.21\|AAGAAGACCCTGTTGAGCTTG | >hg38_21_+_8433172_8446622_RNA45SN1_45S_50ntFlanks | 11873 | 11893 | 21 | AAGAAGACCCTGTTGAGCTTG | 3689.6 | -6.9 | 0.6 | -10.9 | 1.1E-27 | 1.2E-26 | MB231/MB468 |
| >hg38_21_+_8433172_8446622_RNA45SN1_45S_50ntFlanks@6647.6677.31\|CGTACGACTCTTAGCGGTGGATCACTCGGCT | >hg38_21_+_8433172_8446622_RNA45SN1_45S_50ntFlanks | 6647 | 6677 | 31 | CGTACGACTCTTAGCGGTGGATCACTCGGCT | 774.4 | 5.4 | 0.5 | 10.9 | 9.3E-28 | 1.2E-26 | BT20/MB468 |
| >MI0000088\|hsa-mir-30a&WithFlank&6\|-\|71403545\|71403627@53.71.19&[MIMAT0000088&hsa-miR-30a-3p&offsets\|0\|-3;m-38&6\|-\|71403554\|71403575&offsets\|0\|-3]\|CTTTCAGTCGGATGTTTGC | >MI0000088\|hsa-mir-30a&WithFlank&6\|-\|71403545\|71403627 | 53 | 71 | 19 | CTTTCAGTCGGATGTTTGC | 5141.4 | 8.8 | 0.8 | 10.9 | 1.1E-27 | 1.2E-26 | MB231/MB468 |
| >hg38_21_+_8433172_8446622_RNA45SN1_45S_50ntFlanks@12405.12435.31\|CTTTTTGATCCTTCGATGTCGGCTCTTCCTA | >hg38_21_+_8433172_8446622_RNA45SN1_45S_50ntFlanks | 12405 | 12435 | 31 | CTTTTTGATCCTTCGATGTCGGCTCTTCCTA | 2470.5 | -8.3 | 0.8 | -10.9 | 1.1E-27 | 1.3E-26 | MB231/MB468 |
| >MI0000234\|hsa-mir-192&WithFlank&11\|-\|64891131\|64891252@30.49.20(+1U)&[MIMAT0000222&hsa-miR-192-5p&offsets\|0\|-1(+1U);m-59&11\|-\|64891203\|64891223&offsets\|0\|-1(+1U)]\|CTGACCTATGAATTGACAGCT | >MI0000234\|hsa-mir-192&WithFlank&11\|-\|64891131\|64891252 | 30 | 49 | 20(+1U) | CTGACCTATGAATTGACAGCT | 1954.0 | 7.3 | 0.7 | 10.9 | 9.7E-28 | 1.3E-26 | BT20/MB468 |
| >MI0000086\|hsa-mir-28&WithFlank&3\|+\|188688775\|188688872@60.82.23&[MIMAT0004502&hsa-miR-28-3p&offsets\|0\|+1;m-28&3\|+\|188688834\|188688855&offsets\|0\|+1]\|CACTAGATTGTGAGCTCCTGGAG | >MI0000086\|hsa-mir-28&WithFlank&3\|+\|188688775\|188688872 | 60 | 82 | 23 | CACTAGATTGTGAGCTCCTGGAG | 953.3 | 14.8 | 1.4 | 10.9 | 1.1E-27 | 1.3E-26 | MB231/MB468 |

<table>
<tr><td>>MI0000446|hsa-mir-125b-1&WithFlank&11|-|122099751|122099850@61.81.21&[MIMAT0004592&hsa-miR-125b-1-3p&offsets|0|-1;m-485&11|-|122099769|122099790&offsets|0|-1]|ACGGGTTAGGCTCTTGGGAGC</td><td>>MI0000446|hsa-mir-125b-1&WithFlank&11|-|122099751|122099850</td><td>61</td><td>81</td><td>21</td><td>ACGGGTTAGGCTCTTGGGAGC</td><td>687.2</td><td>14.3</td><td>1.3</td><td>10.9</td><td>1.1E-27</td><td>1.3E-26</td><td>MB231/MB468</td></tr>
<tr><td>>hg38_21_+_8433172_8446622_RNA45SN1_45S_50ntFlanks@6650.6695.46|ACGACTCTTAGCGGTGGATCACTCGGCTCGTGCGTCGATGAAGAAC</td><td>>hg38_21_+_8433172_8446622_RNA45SN1_45S_50ntFlanks</td><td>6650</td><td>6695</td><td>46</td><td>ACGACTCTTAGCGGTGGATCACTCGGCTCGTGCGTCGATGAAGAAC</td><td>1425.4</td><td>10.9</td><td>1.0</td><td>10.9</td><td>1.2E-27</td><td>1.3E-26</td><td>MB231/MB468</td></tr>
<tr><td>>MI0000086|hsa-mir-28&WithFlank&3|+|188688775|188688872@20.41.22&[MIMAT0000085&hsa-miR-28-5p&offsets|0|0;m-133&3|+|188688794|188688815&offsets|0|0]|AAGGAGCTCACAGTCTATTGAG</td><td>>MI0000086|hsa-mir-28&WithFlank&3|+|188688775|188688872</td><td>20</td><td>41</td><td>22</td><td>AAGGAGCTCACAGTCTATTGAG</td><td>667.2</td><td>7.5</td><td>0.7</td><td>10.9</td><td>1.2E-27</td><td>1.4E-26</td><td>MB231/MB468</td></tr>
<tr><td>>MI0000456|hsa-mir-140&WithFlank&16|+|69933075|69933186@29.50.22&[MIMAT0000431&hsa-miR-140-5p&offsets|0|0;m-173&16|+|69933103|69933124&offsets|0|0]|CAGTGGTTTTACCCTATGGTAG</td><td>>MI0000456|hsa-mir-140&WithFlank&16|+|69933075|69933186</td><td>29</td><td>50</td><td>22</td><td>CAGTGGTTTTACCCTATGGTAG</td><td>1089.6</td><td>8.0</td><td>0.7</td><td>10.9</td><td>1.2E-27</td><td>1.4E-26</td><td>MB231/MB468</td></tr>
<tr><td>>MI0003674|hsa-mir-653&WithFlank&7|-|93482754|93482861@22.43.22&[MIMAT0003328&hsa-miR-653-5p&offsets|+3|+4;m-368&7|-|93482819|93482840&offsets|0|0]|TTGAAACAATCTCTACTGAACC</td><td>>MI0003674|hsa-mir-653&WithFlank&7|-|93482754|93482861</td><td>22</td><td>43</td><td>22</td><td>TTGAAACAATCTCTACTGAACC</td><td>788.5</td><td>14.5</td><td>1.3</td><td>10.9</td><td>1.2E-27</td><td>1.4E-26</td><td>MB231/MB468</td></tr>
<tr><td>>hg38_21_+_8433172_8446622_RNA45SN1_45S_50ntFlanks@11661.11681.21|TGATTTCTGCCCAGTGCTCTG</td><td>>hg38_21_+_8433172_8446622_RNA45SN1_45S_50ntFlanks</td><td>11661</td><td>11681</td><td>21</td><td>TGATTTCTGCCCAGTGCTCTG</td><td>3755.8</td><td>-6.2</td><td>0.6</td><td>-10.9</td><td>1.1E-27</td><td>1.4E-26</td><td>BT20/MB468</td></tr>
<tr><td>>hg38_21_+_8433172_8446622_RNA45SN1_45S_50ntFlanks@12456.12476.21|CCAAGCGTTGGATTGTTCACC</td><td>>hg38_21_+_8433172_8446622_RNA45SN1_45S_50ntFlanks</td><td>12456</td><td>12476</td><td>21</td><td>CCAAGCGTTGGATTGTTCACC</td><td>32924.3</td><td>-5.8</td><td>0.5</td><td>-10.9</td><td>1.3E-27</td><td>1.4E-26</td><td>MB231/MB468</td></tr>
<tr><td>>MI0000061|hsa-let-7a-2&WithFlank&11|-|122146516|122146599@11.31.21(+1C);MI0000062|hsa-let-7a-3&WithFlank&22|+|46112743|46112828@10.30.21(+1C);MI0000060|hsa-let-7a-1&WithFlank&9|+|94175951|94176042@12.32.21(+1C)&[MIMAT0000062&hsa-let-7a-5p&offsets|0|-1(+1C);m-6&11|-|122146568|122146589&offsets|0|-1(+1C)];[MIMAT0000062_1&hsa-let-7a-5p&offsets|0|-1(+1C);m-5&22|+|46112752|46112773&offsets|0|-1(+1C)];[MIMAT0000062_2&hsa-let-7a-5p&offsets|0|-1(+1C);m-4&9|+|94175962|94175983&offsets|0|-1(+1C)]|TGAGGTAGTAGGTTGTATAGTC</td><td>>MI0000061|hsa-let-7a-2&WithFlank&11|-|122146516|122146599</td><td>11</td><td>31</td><td>21(+1C)</td><td>TGAGGTAGTAGGTTGTATAGTC</td><td>36616.8</td><td>8.3</td><td>0.8</td><td>10.9</td><td>1.3E-27</td><td>1.4E-26</td><td>MB231/MB468</td></tr>
<tr><td>>MI0000102|hsa-mir-100&WithFlank&11|-|122152223|122152314@19.40.22(+3A)&[MIMAT0000098&hsa-miR-100-5p&offsets|0|0(+3A);m-27&11|-|122152276|122152296&offsets|0|+1(+3A)]|AACCCGTAGATCCGAACTTGTGAAA</td><td>>MI0000102|hsa-mir-100&WithFlank&11|-|122152223|122152314</td><td>19</td><td>40</td><td>22(+3A)</td><td>AACCCGTAGATCCGAACTTGTGAAA</td><td>840.9</td><td>14.6</td><td>1.3</td><td>10.9</td><td>1.4E-27</td><td>1.5E-26</td><td>MB231/MB468</td></tr>
<tr><td>>MI0000077|hsa-mir-21&WithFlank&17|+|59841260|59841343@14.35.22(+1A)&[MIMAT0000076&hsa-miR-21-5p&offsets|0|0(+1A);m-1&17|+|59841273|59841294&offsets|0|0(+1A)]|TAGCTTATCAGACTGATGTTGAA</td><td>>MI0000077|hsa-mir-21&WithFlank&17|+|59841260|59841343</td><td>14</td><td>35</td><td>22(+1A)</td><td>TAGCTTATCAGACTGATGTTGAA</td><td>141975.8</td><td>9.3</td><td>0.9</td><td>10.9</td><td>1.2E-27</td><td>1.6E-26</td><td>BT20/MB468</td></tr>
<tr><td>>hg38_21_+_8433172_8446622_RNA45SN1_45S_50ntFlanks@12625.12667.43|CAATGGGGCGAAGCTACCATCTGTGGGATTATGACTGAACGCC</td><td>>hg38_21_+_8433172_8446622_RNA45SN1_45S_50ntFlanks</td><td>12625</td><td>12667</td><td>43</td><td>CAATGGGGCGAAGCTACCATCTGTGGGATTATGACTGAACGCC</td><td>831.8</td><td>12.1</td><td>1.1</td><td>10.9</td><td>1.4E-27</td><td>1.6E-26</td><td>MB231/MB468</td></tr>
<tr><td>>MI0000650|hsa-mir-200c&WithFlank&12|+|6963693|6963772@50.69.20&[MIMAT0000617&hsa-miR-200c-3p&offsets|0|-3;m-54&12|+|6963742|6963764&offsets|0|-3]|TAATACTGCCGGGTAATGAT</td><td>>MI0000650|hsa-mir-200c&WithFlank&12|+|6963693|6963772</td><td>50</td><td>69</td><td>20</td><td>TAATACTGCCGGGTAATGAT</td><td>3429.1</td><td>9.2</td><td>0.8</td><td>10.9</td><td>1.2E-27</td><td>1.6E-26</td><td>BT20/MB468</td></tr>
<tr><td>>trnaMT_MetCAT_MT_+_4402_4469@37.71.35|CCGAAAATGTTGGTTATACCCTTCCCGTACTACCA</td><td>>trnaMT_MetCAT_MT_+_4402_4469</td><td>37</td><td>71</td><td>35</td><td>CCGAAAATGTTGGTTATACCCTTCCCGTACTACCA</td><td>1308.6</td><td>13.2</td><td>1.2</td><td>10.9</td><td>1.5E-27</td><td>1.6E-26</td><td>MB231/MB468</td></tr>
<tr><td>>trnaMT_GluTTC_MT_-_14674_14742@35.72.38,trnalookalike8_GluTTC_5_-_93905172_93905240@35.72.38|TATCATTGGTCGTGGTTGTAGTCCGTGCGAGAATACCA</td><td>>trnaMT_GluTTC_MT_-_14674_14742</td><td>35</td><td>72</td><td>38</td><td>TATCATTGGTCGTGGTTGTAGTCCGTGCGAGAATACCA</td><td>3112.7</td><td>8.1</td><td>0.7</td><td>10.9</td><td>1.3E-27</td><td>1.6E-26</td><td>BT20/MB468</td></tr>
<tr><td>>MI0000266|hsa-mir-10a&WithFlank&17|-|48579832|48579953@28.48.21(+1A)&[MIMAT0000253&hsa-miR-10a-5p&offsets|0|-2(+1A);m-18&17|-|48579905|48579926&offsets|0|-1(+1A)]|TACCCTGTAGATCCGAATTTGA</td><td>>MI0000266|hsa-mir-10a&WithFlank&17|-|48579832|48579953</td><td>28</td><td>48</td><td>21(+1A)</td><td>TACCCTGTAGATCCGAATTTGA</td><td>2532.8</td><td>16.0</td><td>1.5</td><td>10.9</td><td>1.3E-27</td><td>1.6E-26</td><td>BT20/MB468</td></tr>
<tr><td>>MI0000061|hsa-let-7a-2&WithFlank&11|-|122146516|122146599@11.32.22(+1C);MI0000062|hsa-let-7a-3&WithFlank&22|+|46112743|46112828@10.31.22(+1C);MI0000060|hsa-let-7a-1&WithFlank&9|+|94175951|94176042@12.33.22(+1C)&[MIMAT0000062&hsa-let-7a-5p&offsets|0|0(+1C);m-6&11|-|122146568|122146589&offsets|0|0(+1C)];[MIMAT0000062_1&hsa-let-7a-5p&offsets|0|0(+1C);m-5&22|+|46112752|46112773&offsets|0|0(+1C)];[MIMAT0000062_2&hsa-let-7a-5p&offsets|0|0(+1C);m-4&9|+|94175962|94175983&offsets|0|0(+1C)]|TGAGGTAGTAGGTTGTATAGTTC</td><td>>MI0000061|hsa-let-7a-2&WithFlank&11|-|122146516|122146599</td><td>11</td><td>32</td><td>22(+1C)</td><td>TGAGGTAGTAGGTTGTATAGTTC</td><td>1060.3</td><td>8.5</td><td>0.8</td><td>10.9</td><td>1.6E-27</td><td>1.8E-26</td><td>MB231/MB468</td></tr>
<tr><td>>MI0000086|hsa-mir-28&WithFlank&3|+|188688775|188688872@60.81.22(+1A)&[MIMAT0004502&hsa-miR-28-3p&offsets|0|0(+1A);m-28&3|+|188688834|188688855&offsets|0|0(+1A)]|CACTAGATTGTGAGCTCCTGGAA</td><td>>MI0000086|hsa-mir-28&WithFlank&3|+|188688775|188688872</td><td>60</td><td>81</td><td>22(+1A)</td><td>CACTAGATTGTGAGCTCCTGGAA</td><td>4083.7</td><td>11.3</td><td>1.0</td><td>10.9</td><td>1.5E-27</td><td>1.9E-26</td><td>BT20/MB468</td></tr>
<tr><td>>MI0000266|hsa-mir-10a&WithFlank&17|-|48579832|48579953@69.89.21&[MIMAT0004555&hsa-miR-10a-3p&offsets|0|-1;m-468&17|-|48579865|48579885&offsets|0|0]|CAAATTCGTATCTAGGGGAAT</td><td>>MI0000266|hsa-mir-10a&WithFlank&17|-|48579832|48579953</td><td>69</td><td>89</td><td>21</td><td>CAAATTCGTATCTAGGGGAAT</td><td>782.9</td><td>14.3</td><td>1.3</td><td>10.9</td><td>1.5E-27</td><td>2.0E-26</td><td>BT20/MB468</td></tr>
<tr><td>>hg38_21_+_8433172_8446622_RNA45SN1_45S_50ntFlanks@11575.11603.29|ACTTAGAACTGGTGCGGACCAGGGGAATC</td><td>>hg38_21_+_8433172_8446622_RNA45SN1_45S_50ntFlanks</td><td>11575</td><td>11603</td><td>29</td><td>ACTTAGAACTGGTGCGGACCAGGGGAATC</td><td>4191.4</td><td>-6.4</td><td>0.6</td><td>-10.9</td><td>1.5E-27</td><td>2.0E-26</td><td>BT20/MB468</td></tr>
<tr><td>>MI0000542|hsa-mir-320a&WithFlank&8|-|22244960|22245043@48.71.24(+1U)&[MIMAT0000510&hsa-miR-320a-3p&offsets|0|+2(+1U);m-60&8|-|22244975|22244996&offsets|0|+2(+1U)]|AAAAGCTGGGTTGAGAGGGCGAAAT</td><td>>MI0000542|hsa-mir-320a&WithFlank&8|-|22244960|22245043</td><td>48</td><td>71</td><td>24(+1U)</td><td>AAAAGCTGGGTTGAGAGGGCGAAAT</td><td>1629.8</td><td>9.2</td><td>0.8</td><td>10.9</td><td>1.5E-27</td><td>2.0E-26</td><td>BT20/MB468</td></tr>
<tr><td>>MI0005559|hsa-mir-744&WithFlank&17|+|12081893|12082002@17.37.21&[MIMAT0004945&hsa-miR-744-5p&offsets|0|-1;m-172&17|+|12081909|12081929&offsets|0|0]|TGCGGGGCTAGGGCTAACAGC</td><td>>MI0005559|hsa-mir-744&WithFlank&17|+|12081893|12082002</td><td>17</td><td>37</td><td>21</td><td>TGCGGGGCTAGGGCTAACAGC</td><td>2989.1</td><td>7.3</td><td>0.7</td><td>10.9</td><td>1.5E-27</td><td>2.0E-26</td><td>BT20/MB468</td></tr>
<tr><td>>hg38_MT_+_598_1651_RNR1_12S_50ntFlanks@966.1004.39|TAACATGGTAAGTGTACTGGAAAGTGCACTTGGACGAAC</td><td>>hg38_MT_+_598_1651_RNR1_12S_50ntFlanks</td><td>966</td><td>1004</td><td>39</td><td>TAACATGGTAAGTGTACTGGAAAGTGCACTTGGACGAAC</td><td>1274.0</td><td>12.4</td><td>1.1</td><td>10.9</td><td>1.8E-27</td><td>2.0E-26</td><td>MB231/MB468</td></tr>
<tr><td>>hg38_21_+_8433172_8446622_RNA45SN1_45S_50ntFlanks@8342.8377.36|CGTAAGGGAAAGTTGAAAAGAACTTTGAAGAGAGAG</td><td>>hg38_21_+_8433172_8446622_RNA45SN1_45S_50ntFlanks</td><td>8342</td><td>8377</td><td>36</td><td>CGTAAGGGAAAGTTGAAAAGAACTTTGAAGAGAGAG</td><td>351.2</td><td>13.4</td><td>1.2</td><td>10.9</td><td>1.9E-27</td><td>2.1E-26</td><td>MB231/MB468</td></tr>
<tr><td>>hg38_21_+_8433172_8446622_RNA45SN1_45S_50ntFlanks@12417.12457.41|TCGATGTCGGCTCTTCCTATCATTGTGAAGCAGAATTCACC</td><td>>hg38_21_+_8433172_8446622_RNA45SN1_45S_50ntFlanks</td><td>12417</td><td>12457</td><td>41</td><td>TCGATGTCGGCTCTTCCTATCATTGTGAAGCAGAATTCACC</td><td>623.3</td><td>8.8</td><td>0.8</td><td>10.9</td><td>1.9E-27</td><td>2.1E-26</td><td>MB231/MB468</td></tr>
<tr><td>>MI0000299|hsa-mir-222&WithFlank&X|-|45747009|45747130@75.99.25&[MIMAT0000279&hsa-miR-222-3p&offsets|0|+4;m-140&X|-|45747033|45747056&offsets|0|+1]|AGCTACATCTGGCTACTGGGTCTCT</td><td>>MI0000299|hsa-mir-222&WithFlank&X|-|45747009|45747130</td><td>75</td><td>99</td><td>25</td><td>AGCTACATCTGGCTACTGGGTCTCT</td><td>19163.3</td><td>6.6</td><td>0.6</td><td>10.9</td><td>1.6E-27</td><td>2.1E-26</td><td>BT20/MB468</td></tr>
<tr><td>>MI0000542|hsa-mir-320a&WithFlank&8|-|22244960|22245043@48.69.22(+2U)&[MIMAT0000510&hsa-miR-320a-3p&offsets|0|0(+2U);m-60&8|-|22244975|22244996&offsets|0|0(+2U)]|AAAAGCTGGGTTGAGAGGGCGATT</td><td>>MI0000542|hsa-mir-320a&WithFlank&8|-|22244960|22245043</td><td>48</td><td>69</td><td>22(+2U)</td><td>AAAAGCTGGGTTGAGAGGGCGATT</td><td>3884.9</td><td>8.7</td><td>0.8</td><td>10.9</td><td>2.0E-27</td><td>2.2E-26</td><td>MB231/MB468</td></tr>
<tr><td>>MI0000446|hsa-mir-125b-1&WithFlank&11|-|122099751|122099850@22.42.21;MI0000470|hsa-mir-125b-2&WithFlank&21|+|16590231|16590331@24.44.21&[MIMAT0000423&hsa-miR-125b-5p&offsets|+1|0;m-67&11|-|122099809|122099830&offsets|+1|0];[MIMAT0000423_1&hsa-miR-125b-5p&offsets|+1|0;m-68&21|+|16590253|16590274&offsets|+1|0]|CCCTGAGACCCTAACTTGTGA</td><td>>MI0000446|hsa-mir-125b-1&WithFlank&11|-|122099751|122099850</td><td>22</td><td>42</td><td>21</td><td>CCCTGAGACCCTAACTTGTGA</td><td>710.1</td><td>14.4</td><td>1.3</td><td>10.9</td><td>2.0E-27</td><td>2.2E-26</td><td>MB231/MB468</td></tr>
<tr><td>>MI0000299|hsa-mir-222&WithFlank&X|-|45747009|45747130@75.97.23&[MIMAT0000279&hsa-miR-222-3p&offsets|0|+2;m-140&X|-|45747033|45747056&offsets|0|-1]|AGCTACATCTGGCTACTGGGTCT</td><td>>MI0000299|hsa-mir-222&WithFlank&X|-|45747009|45747130</td><td>75</td><td>97</td><td>23</td><td>AGCTACATCTGGCTACTGGGTCT</td><td>20641.4</td><td>8.6</td><td>0.8</td><td>10.8</td><td>2.2E-27</td><td>2.4E-26</td><td>MB231/MB468</td></tr>
<tr><td>>hg38_21_+_8433172_8446622_RNA45SN1_45S_50ntFlanks@6641.6683.43|CGACCTCGTACGACTCTTAGCGGTGGATCACTCGGCTCGTGCG</td><td>>hg38_21_+_8433172_8446622_RNA45SN1_45S_50ntFlanks</td><td>6641</td><td>6683</td><td>43</td><td>CGACCTCGTACGACTCTTAGCGGTGGATCACTCGGCTCGTGCG</td><td>558.5</td><td>14.0</td><td>1.3</td><td>10.8</td><td>2.2E-27</td><td>2.4E-26</td><td>MB231/MB468</td></tr>
<tr><td>>MI0000434|hsa-let-7i&WithFlank&12|+|62603680|62603775@12.34.23&[MIMAT0000415&hsa-let-7i-5p&offsets|0|+1;m-49&12|+|62603691|62603712&offsets|0|+1]|TGAGGTAGTAGTTTGTGCTGTTG</td><td>>MI0000434|hsa-let-7i&WithFlank&12|+|62603680|62603775</td><td>12</td><td>34</td><td>23</td><td>TGAGGTAGTAGTTTGTGCTGTTG</td><td>1415.6</td><td>7.7</td><td>0.7</td><td>10.8</td><td>2.2E-27</td><td>2.4E-26</td><td>MB231/MB468</td></tr>
<tr><td>>hg38_21_+_8433172_8446622_RNA45SN1_45S_50ntFlanks@475.504.30|CTTCGTGATCGATGTGGTGACGTCGTGCTC</td><td>>hg38_21_+_8433172_8446622_RNA45SN1_45S_50ntFlanks</td><td>475</td><td>504</td><td>30</td><td>CTTCGTGATCGATGTGGTGACGTCGTGCTC</td><td>30318.9</td><td>8.4</td><td>0.8</td><td>10.8</td><td>2.2E-27</td><td>2.4E-26</td><td>MB231/MB468</td></tr>
<tr><td>>MI0000289|hsa-mir-181a-1&WithFlank&1|-|198859038|198859159@30.54.25(+1U);MI0000269|hsa-mir-181a-2&WithFlank&9|+|124692436|124692557@45.69.25(+1U)&[MIMAT0000256&hsa-miR-181a-5p&offsets|0|+2(+1U);m-32&1|-|198859108|198859130&offsets|0|+2(+1U)];[MIMAT0000256_1&hsa-miR-181a-5p&offsets|0|+2(+1U);m-32&9|+|124692480|124692502&offsets|0|+2(+1U)]|AACATTCAACGCTGTCGGTGAGTTTT</td><td>>MI0000289|hsa-mir-181a-1&WithFlank&1|-|198859038|198859159</td><td>30</td><td>54</td><td>25(+1U)</td><td>AACATTCAACGCTGTCGGTGAGTTTT</td><td>2216.9</td><td>8.8</td><td>0.8</td><td>10.8</td><td>2.4E-27</td><td>2.6E-26</td><td>MB231/MB468</td></tr>
<tr><td>>MI0000434|hsa-let-7i&WithFlank&12|+|62603680|62603775@12.32.21(+1C)&[MIMAT0000415&hsa-let-7i-5p&offsets|0|-1(+1C);m-49&12|+|62603691|62603712&offsets|0|-1(+1C)]|TGAGGTAGTAGTTTGTGCTGTC</td><td>>MI0000434|hsa-let-7i&WithFlank&12|+|62603680|62603775</td><td>12</td><td>32</td><td>21(+1C)</td><td>TGAGGTAGTAGTTTGTGCTGTC</td><td>80326.9</td><td>8.4</td><td>0.8</td><td>10.8</td><td>2.4E-27</td><td>2.6E-26</td><td>MB231/MB468</td></tr>
<tr><td>>MI0000446|hsa-mir-125b-1&WithFlank&11|-|122099751|122099850@21.42.22(+1C);MI0000470|hsa-mir-125b-2&WithFlank&21|+|16590231|16590331@23.44.22(+1C)&[MIMAT0000423&hsa-miR-125b-5p&offsets|0|0(+1C);m-67&11|-|122099809|122099830&offsets|0|0(+1C)];[MIMAT0000423_1&hsa-miR-125b-5p&offsets|0|0(+1C);m-68&21|+|16590253|16590274&offsets|0|0(+1C)]|TCCCTGAGACCCTAACTTGTGAC</td><td>>MI0000446|hsa-mir-125b-1&WithFlank&11|-|122099751|122099850</td><td>21</td><td>42</td><td>22(+1C)</td><td>TCCCTGAGACCCTAACTTGTGAC</td><td>1171.8</td><td>15.1</td><td>1.4</td><td>10.8</td><td>2.5E-27</td><td>2.8E-26</td><td>MB231/MB468</td></tr>
</table>

<table>
<tr><td>>hg38_1_-_228634819_228635039_RNA5S12_5S_50ntFlanks@141.171.31|CCGCCTGGGAATACCGGGTGCTGTAGGCTTT</td><td>>hg38_1_-_228634819_228635039_RNA5S12_5S_50ntFlanks</td><td>141</td><td>171</td><td>31</td><td>CCGCCTGGGAATACCGGGTGCTGTAGGCTTT</td><td>7435.8</td><td>7.2</td><td>0.7</td><td>10.8</td><td>2.6E-27</td><td>2.8E-26</td><td>MB231/MB468</td></tr>
<tr><td>>MI0000299|hsa-mir-222&WithFlank&X|-|45747009|45747130@75.97.23(+1A)&[MIMAT0000279&hsa-miR-222-3p&offsets|0|+2(+1A);m-140&X|-|45747033|45747056&offsets|0|-1(+1A)]|AGCTACATCTGGCTACTGGGTCTA</td><td>>MI0000299|hsa-mir-222&WithFlank&X|-|45747009|45747130</td><td>75</td><td>97</td><td>23(+1A)</td><td>AGCTACATCTGGCTACTGGGTCTA</td><td>1100.6</td><td>7.5</td><td>0.7</td><td>10.8</td><td>2.3E-27</td><td>2.9E-26</td><td>BT20/MB468</td></tr>
<tr><td>>hg38_21_+_8433172_8446622_RNA45SN1_45S_50ntFlanks@9534.9563.30|AGGAAACTCTGGTGGAGGTCCGTAGCGGTC</td><td>>hg38_21_+_8433172_8446622_RNA45SN1_45S_50ntFlanks</td><td>9534</td><td>9563</td><td>30</td><td>AGGAAACTCTGGTGGAGGTCCGTAGCGGTC</td><td>7235.2</td><td>-6.0</td><td>0.6</td><td>-10.8</td><td>2.7E-27</td><td>2.9E-26</td><td>MB231/MB468</td></tr>
<tr><td>>MI0000093|hsa-mir-92a-1&WithFlank&13|+|91351308|91351397@54.76.23&[MIMAT0000092&hsa-miR-92a-3p&offsets|0|+1;m-9&13|+|91351361|91351382&offsets|0|+1]|TATTGCACTTGTCCCGGCCTGTT</td><td>>MI0000093|hsa-mir-92a-1&WithFlank&13|+|91351308|91351397</td><td>54</td><td>76</td><td>23</td><td>TATTGCACTTGTCCCGGCCTGTT</td><td>21198.9</td><td>7.9</td><td>0.7</td><td>10.8</td><td>2.7E-27</td><td>3.0E-26</td><td>MB231/MB468</td></tr>
<tr><td>>hg38_1_-_228634819_228635039_RNA5S12_5S_50ntFlanks@135.169.35|GGGAGACCGCCTGGGAATACCGGGTGCTGTAGGCT</td><td>>hg38_1_-_228634819_228635039_RNA5S12_5S_50ntFlanks</td><td>135</td><td>169</td><td>35</td><td>GGGAGACCGCCTGGGAATACCGGGTGCTGTAGGCT</td><td>2369.8</td><td>6.9</td><td>0.6</td><td>10.8</td><td>2.7E-27</td><td>3.0E-26</td><td>MB231/MB468</td></tr>
<tr><td>>hg38_21_+_8433172_8446622_RNA45SN1_45S_50ntFlanks@8342.8376.35|CGTAAGGGAAAGTTGAAAAGAACTTTGAAGAGAGA</td><td>>hg38_21_+_8433172_8446622_RNA45SN1_45S_50ntFlanks</td><td>8342</td><td>8376</td><td>35</td><td>CGTAAGGGAAAGTTGAAAAGAACTTTGAAGAGAGA</td><td>410.7</td><td>10.8</td><td>1.0</td><td>10.8</td><td>2.8E-27</td><td>3.0E-26</td><td>MB231/MB468</td></tr>
<tr><td>>am_MI0000809|hsa-mir-151a&WithFlank&8|-|140732558|140732659@53.71.19&[MIMAT0000757&hsa-miR-151a-3p&offsets|0|-2;m-30&8|-|140732587|140732607&offsets|0|-2]|CTAGACTGAAGCTCCTTGA</td><td>>am_MI0000809|hsa-mir-151a&WithFlank&8|-|140732558|140732659</td><td>53</td><td>71</td><td>19</td><td>CTAGACTGAAGCTCCTTGA</td><td>2541.0</td><td>9.6</td><td>0.9</td><td>10.8</td><td>2.5E-27</td><td>3.2E-26</td><td>BT20/MB468</td></tr>
<tr><td>>hg38_21_+_8433172_8446622_RNA45SN1_45S_50ntFlanks@10350.10374.25|ACGAGAACTTTGAAGGCCGAAGTGG</td><td>>hg38_21_+_8433172_8446622_RNA45SN1_45S_50ntFlanks</td><td>10350</td><td>10374</td><td>25</td><td>ACGAGAACTTTGAAGGCCGAAGTGG</td><td>1477.0</td><td>-5.7</td><td>0.5</td><td>-10.8</td><td>2.5E-27</td><td>3.2E-26</td><td>BT20/MB468</td></tr>
<tr><td>>MI0000734|hsa-mir-106b&WithFlank&7|-|100093987|100094080@58.77.20&[MIMAT0004672&hsa-miR-106b-3p&offsets|0|-2;m-87&7|-|100094003|100094023&offsets|0|-1]|CCGCACTGTGGGTACTTGCT</td><td>>MI0000734|hsa-mir-106b&WithFlank&7|-|100093987|100094080</td><td>58</td><td>77</td><td>20</td><td>CCGCACTGTGGGTACTTGCT</td><td>3387.8</td><td>8.9</td><td>0.8</td><td>10.8</td><td>3.0E-27</td><td>3.3E-26</td><td>MB231/MB468</td></tr>
<tr><td>>trnaMT_PheGAA_MT_+_577_647@35.57.23|GAAAATGTTTAGACGGGCTCACA</td><td>>trnaMT_PheGAA_MT_+_577_647</td><td>35</td><td>57</td><td>23</td><td>GAAAATGTTTAGACGGGCTCACA</td><td>1018.2</td><td>10.8</td><td>1.0</td><td>10.8</td><td>3.1E-27</td><td>3.3E-26</td><td>MB231/MB468</td></tr>
<tr><td>>MI0006444|hsa-mir-1307&WithFlank&10|-|103394247|103394407@87.108.22&[MIMAT0005951&hsa-miR-1307-3p&offsets|+1|+1;m-66&10|-|103394300|103394322&offsets|+1|0]|CTCGGCGTGGCGTCGGTCGTGG</td><td>>MI0006444|hsa-mir-1307&WithFlank&10|-|103394247|103394407</td><td>87</td><td>108</td><td>22</td><td>CTCGGCGTGGCGTCGGTCGTGG</td><td>3781.3</td><td>7.5</td><td>0.7</td><td>10.8</td><td>3.2E-27</td><td>3.4E-26</td><td>MB231/MB468</td></tr>
<tr><td>>MI0000266|hsa-mir-10a&WithFlank&17|-|48579832|48579953@28.48.21(+1G)&[MIMAT0000253&hsa-miR-10a-5p&offsets|0|-2(+1G);m-18&17|-|48579905|48579926&offsets|0|-1(+1G)]|TACCCTGTAGATCCGAATTTGG</td><td>>MI0000266|hsa-mir-10a&WithFlank&17|-|48579832|48579953</td><td>28</td><td>48</td><td>21(+1G)</td><td>TACCCTGTAGATCCGAATTTGG</td><td>3471.6</td><td>16.4</td><td>1.5</td><td>10.8</td><td>2.7E-27</td><td>3.4E-26</td><td>BT20/MB468</td></tr>
<tr><td>>am_trnaMT_ProTGG_MT_-_15956_16023@54.71.18|AGACTTTTTCTCTGACCA</td><td>>am_trnaMT_ProTGG_MT_-_15956_16023</td><td>54</td><td>71</td><td>18</td><td>AGACTTTTTCTCTGACCA</td><td>16266.7</td><td>6.9</td><td>0.6</td><td>10.8</td><td>2.7E-27</td><td>3.4E-26</td><td>BT20/MB468</td></tr>
<tr><td>>MI0000266|hsa-mir-10a&WithFlank&17|-|48579832|48579953@28.48.21(+1G)&[MIMAT0000253&hsa-miR-10a-5p&offsets|0|-2(+1G);m-18&17|-|48579905|48579926&offsets|0|-1(+1G)]|TACCCTGTAGATCCGAATTTGG</td><td>>MI0000266|hsa-mir-10a&WithFlank&17|-|48579832|48579953</td><td>28</td><td>48</td><td>21(+1G)</td><td>TACCCTGTAGATCCGAATTTGG</td><td>1031.7</td><td>14.9</td><td>1.4</td><td>10.8</td><td>3.2E-27</td><td>3.5E-26</td><td>MB231/MB468</td></tr>
<tr><td>>MI0003188|hsa-mir-503&WithFlank&X|-|134546322|134546404@12.31.20&[MIMAT0002874&hsa-miR-503-5p&offsets|0|-3;m-297&X|-|134546371|134546393&offsets|0|-3]|TAGCAGCGGGAACAGTTCTG</td><td>>MI0003188|hsa-mir-503&WithFlank&X|-|134546322|134546404</td><td>12</td><td>31</td><td>20</td><td>TAGCAGCGGGAACAGTTCTG</td><td>764.9</td><td>14.5</td><td>1.3</td><td>10.8</td><td>3.2E-27</td><td>3.5E-26</td><td>MB231/MB468</td></tr>
<tr><td>>MI0000264|hsa-mir-7-2&WithFlank&15|+|88611819|88611940@38.59.22(+1A);MI0000265|hsa-mir-7-3&WithFlank&19|+|4770664|4770785@37.58.22(+1A);MI0000263|hsa-mir-7-1&WithFlank&9|-|83969742|83969863@30.51.22(+1A)&[MIMAT0000252&hsa-miR-7-5p&offsets|0|-2(+1A);m-400&15|+|88611856|88611879&offsets|0|-2(+1A)];[MIMAT0000252_1&hsa-miR-7-5p&offsets|0|-2(+1A);m-402&19|+|4770700|4770723&offsets|0|-2(+1A)];[MIMAT0000252_2&hsa-miR-7-5p&offsets|0|-2(+1A);m-401&9|-|83969811|83969834&offsets|0|-2(+1A)]|TGGAAGACTAGTGATTTTGTTGA</td><td>>MI0000264|hsa-mir-7-2&WithFlank&15|+|88611819|88611940</td><td>38</td><td>59</td><td>22(+1A)</td><td>TGGAAGACTAGTGATTTTGTTGA</td><td>991.1</td><td>8.2</td><td>0.8</td><td>10.8</td><td>3.3E-27</td><td>3.6E-26</td><td>MB231/MB468</td></tr>
<tr><td>>MI0015866|hsa-mir-4326&WithFlank&20|+|63286802|63286872@17.40.24(+1U)&[MIMAT0016888&hsa-miR-4326&offsets|0|+4(+1U);m-339&20|+|63286818|63286841&offsets|0|0(+1U)]|TGTTCCTCTGTCTCCCAGACTCTGT</td><td>>MI0015866|hsa-mir-4326&WithFlank&20|+|63286802|63286872</td><td>17</td><td>40</td><td>24(+1U)</td><td>TGTTCCTCTGTCTCCCAGACTCTGT</td><td>707.7</td><td>14.4</td><td>1.3</td><td>10.8</td><td>3.4E-27</td><td>3.6E-26</td><td>MB231/MB468</td></tr>
<tr><td>>hg38_1_-_228634819_228635039_RNA5S12_5S_50ntFlanks@134.170.37|TGGGAGACCGCCTGGGAATACCGGGTGCTGTAGGCTT</td><td>>hg38_1_-_228634819_228635039_RNA5S12_5S_50ntFlanks</td><td>134</td><td>170</td><td>37</td><td>TGGGAGACCGCCTGGGAATACCGGGTGCTGTAGGCTT</td><td>222943.6</td><td>8.8</td><td>0.8</td><td>10.8</td><td>3.4E-27</td><td>3.7E-26</td><td>MB231/MB468</td></tr>
<tr><td>>trnaMT_HisGTG_MT_+_12138_12206@20.55.36|AACATCAGATTGTGAATCTGACAACAGAGGCTTACG</td><td>>trnaMT_HisGTG_MT_+_12138_12206</td><td>20</td><td>55</td><td>36</td><td>AACATCAGATTGTGAATCTGACAACAGAGGCTTACG</td><td>260.6</td><td>12.7</td><td>1.2</td><td>10.8</td><td>2.9E-27</td><td>3.7E-26</td><td>BT20/MB468</td></tr>
<tr><td>>hg38_21_+_8433172_8446622_RNA45SN1_45S_50ntFlanks@476.505.30|TTCGTGATCGATGTGGTGACGTCGTGCTCT</td><td>>hg38_21_+_8433172_8446622_RNA45SN1_45S_50ntFlanks</td><td>476</td><td>505</td><td>30</td><td>TTCGTGATCGATGTGGTGACGTCGTGCTCT</td><td>13560.3</td><td>7.5</td><td>0.7</td><td>10.8</td><td>3.0E-27</td><td>3.8E-26</td><td>BT20/MB468</td></tr>
<tr><td>>hg38_21_+_8433172_8446622_RNA45SN1_45S_50ntFlanks@12996.13038.43|TCGCTGCGATCTATTGAAAGTCAGCCCTCGACACAAGGGTTTG</td><td>>hg38_21_+_8433172_8446622_RNA45SN1_45S_50ntFlanks</td><td>12996</td><td>13038</td><td>43</td><td>TCGCTGCGATCTATTGAAAGTCAGCCCTCGACACAAGGGTTTG</td><td>7353.6</td><td>12.7</td><td>1.2</td><td>10.8</td><td>3.6E-27</td><td>3.8E-26</td><td>MB231/MB468</td></tr>
<tr><td>>MI0000299|hsa-mir-222&WithFlank&X|-|45747009|45747130@75.100.26&[MIMAT0000279&hsa-miR-222-3p&offsets|0|+5;m-140&X|-|45747033|45747056&offsets|0|+2]|AGCTACATCTGGCTACTGGGTCTCTG</td><td>>MI0000299|hsa-mir-222&WithFlank&X|-|45747009|45747130</td><td>75</td><td>100</td><td>26</td><td>AGCTACATCTGGCTACTGGGTCTCTG</td><td>670.1</td><td>14.3</td><td>1.3</td><td>10.8</td><td>3.7E-27</td><td>3.9E-26</td><td>MB231/MB468</td></tr>
<tr><td>>MI0000086|hsa-mir-28&WithFlank&3|+|188688775|188688872@60.82.23&[MIMAT0004502&hsa-miR-28-3p&offsets|0|+1;m-28&3|+|188688834|188688855&offsets|0|+1]|CACTAGATTGTGAGCTCCTGGAG</td><td>>MI0000086|hsa-mir-28&WithFlank&3|+|188688775|188688872</td><td>60</td><td>82</td><td>23</td><td>CACTAGATTGTGAGCTCCTGGAG</td><td>1985.0</td><td>15.6</td><td>1.4</td><td>10.8</td><td>3.1E-27</td><td>4.0E-26</td><td>BT20/MB468</td></tr>
<tr><td>>hg38_21_+_8433172_8446622_RNA45SN1_45S_50ntFlanks@10841.10863.23|CGTAACTTCGGGATAAGGATTGG</td><td>>hg38_21_+_8433172_8446622_RNA45SN1_45S_50ntFlanks</td><td>10841</td><td>10863</td><td>23</td><td>CGTAACTTCGGGATAAGGATTGG</td><td>2121.9</td><td>-6.9</td><td>0.6</td><td>-10.8</td><td>3.4E-27</td><td>4.3E-26</td><td>BT20/MB468</td></tr>
<tr><td>>MI0000746|hsa-mir-99b&WithFlank&19|+|51692606|51692687@13.33.21&[MIMAT0000689&hsa-miR-99b-5p&offsets|0|-1;m-10&19|+|51692618|51692639&offsets|0|-1]|CACCCGTAGAACCGACCTTGC</td><td>>MI0000746|hsa-mir-99b&WithFlank&19|+|51692606|51692687</td><td>13</td><td>33</td><td>21</td><td>CACCCGTAGAACCGACCTTGC</td><td>17211.3</td><td>7.5</td><td>0.7</td><td>10.8</td><td>4.0E-27</td><td>4.3E-26</td><td>MB231/MB468</td></tr>
<tr><td>>hg38_21_+_8433172_8446622_RNA45SN1_45S_50ntFlanks@9572.9597.26|CAAATCGGTCGTCCGACCTGGGTATA</td><td>>hg38_21_+_8433172_8446622_RNA45SN1_45S_50ntFlanks</td><td>9572</td><td>9597</td><td>26</td><td>CAAATCGGTCGTCCGACCTGGGTATA</td><td>6445.9</td><td>-7.0</td><td>0.7</td><td>-10.8</td><td>4.1E-27</td><td>4.4E-26</td><td>MB231/MB468</td></tr>
<tr><td>>MI0000272|hsa-mir-182&WithFlank&7|-|129770377|129770498@29.54.26&[MIMAT0000259&hsa-miR-182-5p&offsets|0|+2;m-22&7|-|129770449|129770470&offsets|0|+4]|TTTGGCAATGGTAGAACTCACACTGG</td><td>>MI0000272|hsa-mir-182&WithFlank&7|-|129770377|129770498</td><td>29</td><td>54</td><td>26</td><td>TTTGGCAATGGTAGAACTCACACTGG</td><td>13764.9</td><td>8.4</td><td>0.8</td><td>10.8</td><td>3.5E-27</td><td>4.5E-26</td><td>BT20/MB468</td></tr>
<tr><td>>MI0000082|hsa-mir-25&WithFlank&7|-|100093554|100093649@58.79.22(+1U)&[MIMAT0000081&hsa-miR-25-3p&offsets|0|0(+1U);m-23&7|-|100093571|100093592&offsets|0|0(+1U)]|CATTGCACTTGTCTCGGTCTGAT</td><td>>MI0000082|hsa-mir-25&WithFlank&7|-|100093554|100093649</td><td>58</td><td>79</td><td>22(+1U)</td><td>CATTGCACTTGTCTCGGTCTGAT</td><td>4634.5</td><td>8.3</td><td>0.8</td><td>10.8</td><td>4.4E-27</td><td>4.6E-26</td><td>MB231/MB468</td></tr>
<tr><td>>MI0000301|hsa-mir-224&WithFlank&X|-|151958572|151958664@14.35.22(+1U)&[MIMAT0000281&hsa-miR-224-5p&offsets|+1|-2(+1U);m-238&X|-|151958630|151958651&offsets|0|0(+1U)]|CAAGTCACTAGTGGTTCCGTTTT</td><td>>MI0000301|hsa-mir-224&WithFlank&X|-|151958572|151958664</td><td>14</td><td>35</td><td>22(+1U)</td><td>CAAGTCACTAGTGGTTCCGTTTT</td><td>550.1</td><td>13.8</td><td>1.3</td><td>10.8</td><td>3.7E-27</td><td>4.6E-26</td><td>BT20/MB468</td></tr>
<tr><td>>MI0000102|hsa-mir-100&WithFlank&11|-|122152223|122152314@19.39.21&[MIMAT0000098&hsa-miR-100-5p&offsets|0|-1;m-27&11|-|122152276|122152296&offsets|0|0]|AACCCGTAGATCCGAACTTGT</td><td>>MI0000102|hsa-mir-100&WithFlank&11|-|122152223|122152314</td><td>19</td><td>39</td><td>21</td><td>AACCCGTAGATCCGAACTTGT</td><td>649273.7</td><td>15.3</td><td>1.4</td><td>10.8</td><td>4.5E-27</td><td>4.8E-26</td><td>MB231/MB468</td></tr>
<tr><td>>MI0000746|hsa-mir-99b&WithFlank&19|+|51692606|51692687@13.34.22(+1U)&[MIMAT0000689&hsa-miR-99b-5p&offsets|0|0(+1U);m-10&19|+|51692618|51692639&offsets|0|0(+1U)]|CACCCGTAGAACCGACCTTGCGT</td><td>>MI0000746|hsa-mir-99b&WithFlank&19|+|51692606|51692687</td><td>13</td><td>34</td><td>22(+1U)</td><td>CACCCGTAGAACCGACCTTGCGT</td><td>4611.9</td><td>7.6</td><td>0.7</td><td>10.8</td><td>3.9E-27</td><td>4.9E-26</td><td>BT20/MB468</td></tr>
<tr><td>>MI0000462|hsa-mir-152&WithFlank&17|-|48037155|48037253@60.80.21(+1U)&[MIMAT0000438&hsa-miR-152-3p&offsets|0|0(+1U);m-97&17|-|48037174|48037194&offsets|0|0(+1U)]|TCAGTGCATGACAGAACTTGGT</td><td>>MI0000462|hsa-mir-152&WithFlank&17|-|48037155|48037253</td><td>60</td><td>80</td><td>21(+1U)</td><td>TCAGTGCATGACAGAACTTGGT</td><td>1299.4</td><td>15.2</td><td>1.4</td><td>10.8</td><td>5.0E-27</td><td>5.3E-26</td><td>MB231/MB468</td></tr>
<tr><td>>hg38_21_+_8433172_8446622_RNA45SN1_45S_50ntFlanks@7984.8006.23|AGATCAGACGTGGCGACCCGCTG</td><td>>hg38_21_+_8433172_8446622_RNA45SN1_45S_50ntFlanks</td><td>7984</td><td>8006</td><td>23</td><td>AGATCAGACGTGGCGACCCGCTG</td><td>2231.5</td><td>-8.5</td><td>0.8</td><td>-10.8</td><td>5.0E-27</td><td>5.3E-26</td><td>MB231/MB468</td></tr>
<tr><td>>hg38_21_+_8433172_8446622_RNA45SN1_45S_50ntFlanks@476.506.31|TTCGTGATCGATGTGGTGACGTCGTGCTCTC</td><td>>hg38_21_+_8433172_8446622_RNA45SN1_45S_50ntFlanks</td><td>476</td><td>506</td><td>31</td><td>TTCGTGATCGATGTGGTGACGTCGTGCTCTC</td><td>5217.3</td><td>6.5</td><td>0.6</td><td>10.8</td><td>4.3E-27</td><td>5.3E-26</td><td>BT20/MB468</td></tr>
<tr><td>>MI0000272|hsa-mir-182&WithFlank&7|-|129770377|129770498@29.53.25(+1U)&[MIMAT0000259&hsa-miR-182-5p&offsets|0|+1(+1U);m-22&7|-|129770449|129770470&offsets|0|+3(+1U)]|TTTGGCAATGGTAGAACTCACACTGT</td><td>>MI0000272|hsa-mir-182&WithFlank&7|-|129770377|129770498</td><td>29</td><td>53</td><td>25(+1U)</td><td>TTTGGCAATGGTAGAACTCACACTGT</td><td>5293.9</td><td>7.9</td><td>0.7</td><td>10.8</td><td>4.4E-27</td><td>5.5E-26</td><td>BT20/MB468</td></tr>
<tr><td>>MI0000542|hsa-mir-320a&WithFlank&8|-|22244960|22245043@48.70.23(+2U)&[MIMAT0000510&hsa-miR-320a-3p&offsets|0|+1(+2U);m-60&8|-|22244975|22244996&offsets|0|+1(+2U)]|AAAAGCTGGGTTGAGAGGGCGAATT</td><td>>MI0000542|hsa-mir-320a&WithFlank&8|-|22244960|22245043</td><td>48</td><td>70</td><td>23(+2U)</td><td>AAAAGCTGGGTTGAGAGGGCGAATT</td><td>796.8</td><td>8.5</td><td>0.8</td><td>10.8</td><td>5.3E-27</td><td>5.5E-26</td><td>MB231/MB468</td></tr>
</table>

<table>
<tr><td>>trna116_GluCTC_1_-_145399233_145399304@1.32.32,trna59_GluCTC_1_+_249168447_249168518@1.32.32,trna71_GluCTC_1_-_161439189_161439260@1.32.32,trna74_GluCTC_1_-_161431809_161431880@1.32.32,trna77_GluCTC_1_-_161424398_161424469@1.32.32,trna77_GluCTC_6_+_28949976_28950047@1.32.32,trna80_GluCTC_1_-_161417018_161417089@1.32.32,trna87_GluCTC_6_-_126101393_126101464@1.32.32|TCCCTGGTGGTCTAGTGGTTAGGATTCGGCGC</td><td>>trna116_GluCTC_1_-_145399233_145399304</td><td>1</td><td>32</td><td>32</td><td>TCCCTGGTGGTCTAGTGGTTAGGATTCGGCGC</td><td>5530.1</td><td>7.8</td><td>0.7</td><td>10.8</td><td>4.5E-27</td><td>5.6E-26</td><td>BT20/MB468</td></tr>
<tr><td>>MI0000266|hsa-mir-10a&WithFlank&17|-|48579832|48579953@29.48.20(+1C)&[MIMAT0000253&hsa-miR-10a-5p&offsets|+1|-2(+1C);m-18&17|-|48579905|48579926&offsets|+1|-1(+1C)]|ACCCTGTAGATCCGAATTTGC</td><td>>MI0000266|hsa-mir-10a&WithFlank&17|-|48579832|48579953</td><td>29</td><td>48</td><td>20(+1C)</td><td>ACCCTGTAGATCCGAATTTGC</td><td>696.9</td><td>14.1</td><td>1.3</td><td>10.8</td><td>4.5E-27</td><td>5.7E-26</td><td>BT20/MB468</td></tr>
<tr><td>>hg38_21_+_8433172_8446622_RNA45SN1_45S_50ntFlanks@12456.12478.23|CCAAGCGTTGGATTGTTCACCCA</td><td>>hg38_21_+_8433172_8446622_RNA45SN1_45S_50ntFlanks</td><td>12456</td><td>12478</td><td>23</td><td>CCAAGCGTTGGATTGTTCACCCA</td><td>9311.5</td><td>-6.0</td><td>0.6</td><td>-10.8</td><td>5.4E-27</td><td>5.7E-26</td><td>MB231/MB468</td></tr>
<tr><td>>MI0000077|hsa-mir-21&WithFlank&17|+|59841260|59841343@16.36.21&[MIMAT0000076&hsa-miR-21-5p&offsets|+2|+1;m-1&17|+|59841273|59841294&offsets|+2|+1]|GCTTATCAGACTGATGTTGAC</td><td>>MI0000077|hsa-mir-21&WithFlank&17|+|59841260|59841343</td><td>16</td><td>36</td><td>21</td><td>GCTTATCAGACTGATGTTGAC</td><td>926.2</td><td>12.5</td><td>1.2</td><td>10.8</td><td>4.6E-27</td><td>5.7E-26</td><td>BT20/MB468</td></tr>
<tr><td>>MI0000433|hsa-let-7g&WithFlank&3|-|52268272|52268367@11.33.23&[MIMAT0000414&hsa-let-7g-5p&offsets|0|+1;m-58&3|-|52268336|52268357&offsets|0|+1]|TGAGGTAGTAGTTTGTACAGTTT</td><td>>MI0000433|hsa-let-7g&WithFlank&3|-|52268272|52268367</td><td>11</td><td>33</td><td>23</td><td>TGAGGTAGTAGTTTGTACAGTTT</td><td>6735.6</td><td>8.1</td><td>0.7</td><td>10.8</td><td>5.5E-27</td><td>5.8E-26</td><td>MB231/MB468</td></tr>
<tr><td>>hg38_21_+_8433172_8446622_RNA45SN1_45S_50ntFlanks@6648.6696.49|GTACGACTCTTAGCGGTGGATCACTCGGCTCGTGCGTCGATGAAGAACG</td><td>>hg38_21_+_8433172_8446622_RNA45SN1_45S_50ntFlanks</td><td>6648</td><td>6696</td><td>49</td><td>GTACGACTCTTAGCGGTGGATCACTCGGCTCGTGCGTCGATGAAGAACG</td><td>3899.6</td><td>8.6</td><td>0.8</td><td>10.8</td><td>5.6E-27</td><td>5.8E-26</td><td>MB231/MB468</td></tr>
<tr><td>>hg38_21_+_8433172_8446622_RNA45SN1_45S_50ntFlanks@11669.11687.19|GCCCAGTGCTCTGAATGTC</td><td>>hg38_21_+_8433172_8446622_RNA45SN1_45S_50ntFlanks</td><td>11669</td><td>11687</td><td>19</td><td>GCCCAGTGCTCTGAATGTC</td><td>629.6</td><td>4.8</td><td>0.4</td><td>10.8</td><td>4.7E-27</td><td>5.8E-26</td><td>BT20/MB468</td></tr>
<tr><td>>hg38_1_-_228634819_228635039_RNA5S12_5S_50ntFlanks@142.171.30|CGCCTGGGAATACCGGGTGCTGTAGGCTTT</td><td>>hg38_1_-_228634819_228635039_RNA5S12_5S_50ntFlanks</td><td>142</td><td>171</td><td>30</td><td>CGCCTGGGAATACCGGGTGCTGTAGGCTTT</td><td>5517.4</td><td>7.2</td><td>0.7</td><td>10.8</td><td>5.7E-27</td><td>6.0E-26</td><td>MB231/MB468</td></tr>
<tr><td>>MI0000683|hsa-mir-181b-2&WithFlank&9|+|124693704|124693804@22.46.25(+1U)&[MIMAT0000257_1&hsa-miR-181b-5p&offsets|0|+2(+1U);m-72&9|+|124693725|124693747&offsets|0|+2(+1U)]|AACATTCATTGCTGTCGGTGGGTTTT</td><td>>MI0000683|hsa-mir-181b-2&WithFlank&9|+|124693704|124693804</td><td>22</td><td>46</td><td>25(+1U)</td><td>AACATTCATTGCTGTCGGTGGGTTTT</td><td>1518.1</td><td>8.4</td><td>0.8</td><td>10.8</td><td>5.9E-27</td><td>6.2E-26</td><td>MB231/MB468</td></tr>
<tr><td>>hg38_21_+_8433172_8446622_RNA45SN1_45S_50ntFlanks@11593.11611.19|CCAGGGGAATCCGACTGTT</td><td>>hg38_21_+_8433172_8446622_RNA45SN1_45S_50ntFlanks</td><td>11593</td><td>11611</td><td>19</td><td>CCAGGGGAATCCGACTGTT</td><td>2639.0</td><td>-4.9</td><td>0.5</td><td>-10.8</td><td>5.0E-27</td><td>6.2E-26</td><td>BT20/MB468</td></tr>
<tr><td>>hg38_21_+_8433172_8446622_RNA45SN1_45S_50ntFlanks@4760.4780.21|TCAGATACCGTCGTAGTTCCG</td><td>>hg38_21_+_8433172_8446622_RNA45SN1_45S_50ntFlanks</td><td>4760</td><td>4780</td><td>21</td><td>TCAGATACCGTCGTAGTTCCG</td><td>2974.4</td><td>-7.3</td><td>0.7</td><td>-10.7</td><td>6.1E-27</td><td>6.3E-26</td><td>MB231/MB468</td></tr>
<tr><td>>MI0000440|hsa-mir-27b&WithFlank&9|+|95085439|95085547@67.86.20(+1A)&[MIMAT0000419&hsa-miR-27b-3p&offsets|0|-1(+1A);m-39&9|+|95085505|95085525&offsets|0|-1(+1A)]|TTCACAGTGGCTAAGTTCTGA</td><td>>MI0000440|hsa-mir-27b&WithFlank&9|+|95085439|95085547</td><td>67</td><td>86</td><td>20(+1A)</td><td>TTCACAGTGGCTAAGTTCTGA</td><td>10674.5</td><td>8.4</td><td>0.8</td><td>10.7</td><td>6.2E-27</td><td>6.5E-26</td><td>MB231/MB468</td></tr>
<tr><td>>MI0001445|hsa-mir-423&WithFlank&17|+|30117073|30117178@58.79.22&[MIMAT0001340&hsa-miR-423-3p&offsets|-1|-2;m-100&17|+|30117131|30117153&offsets|-1|-2]|AAGCTCGGTCTGAGGCCCCTCA</td><td>>MI0001445|hsa-mir-423&WithFlank&17|+|30117073|30117178</td><td>58</td><td>79</td><td>22</td><td>AAGCTCGGTCTGAGGCCCCTCA</td><td>691.6</td><td>7.3</td><td>0.7</td><td>10.7</td><td>6.4E-27</td><td>6.7E-26</td><td>MB231/MB468</td></tr>
<tr><td>>hg38_21_+_8433172_8446622_RNA45SN1_45S_50ntFlanks@6646.6668.23|TCGTACGACTCTTAGCGGTGGAT</td><td>>hg38_21_+_8433172_8446622_RNA45SN1_45S_50ntFlanks</td><td>6646</td><td>6668</td><td>23</td><td>TCGTACGACTCTTAGCGGTGGAT</td><td>15377.8</td><td>5.5</td><td>0.5</td><td>10.8</td><td>5.4E-27</td><td>6.7E-26</td><td>BT20/MB468</td></tr>
<tr><td>>hg38_21_+_8433172_8446622_RNA45SN1_45S_50ntFlanks@6643.6683.41|ACCTCGTACGACTCTTAGCGGTGGATCACTCGGCTCGTGCG</td><td>>hg38_21_+_8433172_8446622_RNA45SN1_45S_50ntFlanks</td><td>6643</td><td>6683</td><td>41</td><td>ACCTCGTACGACTCTTAGCGGTGGATCACTCGGCTCGTGCG</td><td>930.3</td><td>14.8</td><td>1.4</td><td>10.7</td><td>6.8E-27</td><td>7.0E-26</td><td>MB231/MB468</td></tr>
<tr><td>>MI0000470|hsa-mir-125b-2&WithFlank&21|+|16590231|16590331@23.45.23&[MIMAT0000423_1&hsa-miR-125b-5p&offsets|0|+1;m-68&21|+|16590253|16590274&offsets|0|+1]|TCCCTGAGACCCTAACTTGTGAG</td><td>>MI0000470|hsa-mir-125b-2&WithFlank&21|+|16590231|16590331</td><td>23</td><td>45</td><td>23</td><td>TCCCTGAGACCCTAACTTGTGAG</td><td>869.5</td><td>14.7</td><td>1.4</td><td>10.7</td><td>6.9E-27</td><td>7.2E-26</td><td>MB231/MB468</td></tr>
<tr><td>>hg38_21_+_8433172_8446622_RNA45SN1_45S_50ntFlanks@6646.6683.38|TCGTACGACTCTTAGCGGTGGATCACTCGGCTCGTGCG</td><td>>hg38_21_+_8433172_8446622_RNA45SN1_45S_50ntFlanks</td><td>6646</td><td>6683</td><td>38</td><td>TCGTACGACTCTTAGCGGTGGATCACTCGGCTCGTGCG</td><td>4815.2</td><td>8.6</td><td>0.8</td><td>10.7</td><td>7.1E-27</td><td>7.4E-26</td><td>MB231/MB468</td></tr>
<tr><td>>MI0000746|hsa-mir-99b&WithFlank&19|+|51692606|51692687@13.33.21(+1C)&[MIMAT0000689&hsa-miR-99b-5p&offsets|0|-1(+1C);m-10&19|+|51692618|51692639&offsets|0|-1(+1C)]|CACCCGTAGAACCGACCTTGCC</td><td>>MI0000746|hsa-mir-99b&WithFlank&19|+|51692606|51692687</td><td>13</td><td>33</td><td>21(+1C)</td><td>CACCCGTAGAACCGACCTTGCC</td><td>660.4</td><td>7.5</td><td>0.7</td><td>10.7</td><td>7.4E-27</td><td>7.7E-26</td><td>MB231/MB468</td></tr>
<tr><td>>MI0000061|hsa-let-7a-2&WithFlank&11|-|122146516|122146599@11.31.21(+1G);MI0000062|hsa-let-7a-3&WithFlank&22|+|46112743|46112828@10.30.21(+1G);MI0000060|hsa-let-7a-1&WithFlank&9|+|94175951|94176042@12.32.21(+1G)&[MIMAT0000062&hsa-let-7a-5p&offsets|0|-1(+1G);m-6&11|-|122146568|122146589&offsets|0|-1(+1G)];[MIMAT0000062_1&hsa-let-7a-5p&offsets|0|-1(+1G);m-5&22|+|46112752|46112773&offsets|0|-1(+1G)];[MIMAT0000062_2&hsa-let-7a-5p&offsets|0|-1(+1G);m-4&9|+|94175962|94175983&offsets|0|-1(+1G)]|TGAGGTAGTAGGTTGTATAGTG</td><td>>MI0000061|hsa-let-7a-2&WithFlank&11|-|122146516|122146599</td><td>11</td><td>31</td><td>21(+1G)</td><td>TGAGGTAGTAGGTTGTATAGTG</td><td>7213.7</td><td>8.3</td><td>0.8</td><td>10.7</td><td>7.5E-27</td><td>7.7E-26</td><td>MB231/MB468</td></tr>
<tr><td>>MI0000066|hsa-let-7e&WithFlank&19|+|51692780|51692870@14.34.21(+1C)&[MIMAT0000066&hsa-let-7e-5p&offsets|0|-1(+1C);m-50&19|+|51692793|51692814&offsets|0|-1(+1C)]|TGAGGTAGGAGGTTGTATAGTC</td><td>>MI0000066|hsa-let-7e&WithFlank&19|+|51692780|51692870</td><td>14</td><td>34</td><td>21(+1C)</td><td>TGAGGTAGGAGGTTGTATAGTC</td><td>4224.7</td><td>7.6</td><td>0.7</td><td>10.7</td><td>6.3E-27</td><td>7.8E-26</td><td>BT20/MB468</td></tr>
<tr><td>>hg38_21_+_8433172_8446622_RNA45SN1_45S_50ntFlanks@11575.11603.29|ACTTAGAACTGGTGCGGACCAGGGGAATC</td><td>>hg38_21_+_8433172_8446622_RNA45SN1_45S_50ntFlanks</td><td>11575</td><td>11603</td><td>29</td><td>ACTTAGAACTGGTGCGGACCAGGGGAATC</td><td>3511.1</td><td>-5.9</td><td>0.6</td><td>-10.7</td><td>7.6E-27</td><td>7.8E-26</td><td>MB231/MB468</td></tr>
<tr><td>>MI0000809|hsa-mir-151a&WithFlank&8|-|140732558|140732659@53.74.22(+2U)&[MIMAT0000757&hsa-miR-151a-3p&offsets|0|+1(+2U);m-30&8|-|140732587|140732607&offsets|0|+1(+2U)]|CTAGACTGAAGCTCCTTGAGGATT</td><td>>MI0000809|hsa-mir-151a&WithFlank&8|-|140732558|140732659</td><td>53</td><td>74</td><td>22(+2U)</td><td>CTAGACTGAAGCTCCTTGAGGATT</td><td>2506.7</td><td>10.6</td><td>1.0</td><td>10.7</td><td>7.6E-27</td><td>7.8E-26</td><td>MB231/MB468</td></tr>
<tr><td>>trnaMT_GluTTC_MT_-_14674_14742@38.71.34,trnalookalike8_GluTTC_5_-_93905172_93905240@38.71.34|CATTGGTCGTGGTTGTAGTCCGTGCGAGAATACC</td><td>>trnaMT_GluTTC_MT_-_14674_14742</td><td>38</td><td>71</td><td>34</td><td>CATTGGTCGTGGTTGTAGTCCGTGCGAGAATACC</td><td>1054.8</td><td>11.9</td><td>1.1</td><td>10.7</td><td>7.6E-27</td><td>7.8E-26</td><td>MB231/MB468</td></tr>
<tr><td>>MI0000273|hsa-mir-183&WithFlank&7|-|129774899|129775020@33.54.22&[MIMAT0000261&hsa-miR-183-5p&offsets|0|0;m-41&7|-|129774966|129774987&offsets|-1|-1]|TATGGCACTGGTAGAATTCACT</td><td>>MI0000273|hsa-mir-183&WithFlank&7|-|129774899|129775020</td><td>33</td><td>54</td><td>22</td><td>TATGGCACTGGTAGAATTCACT</td><td>38420.3</td><td>7.4</td><td>0.7</td><td>10.7</td><td>6.4E-27</td><td>7.8E-26</td><td>BT20/MB468</td></tr>
<tr><td>>trnaMT_PheGAA_MT_+_577_647@35.60.26|GAAAATGTTTAGACGGGCTCACATCA</td><td>>trnaMT_PheGAA_MT_+_577_647</td><td>35</td><td>60</td><td>26</td><td>GAAAATGTTTAGACGGGCTCACATCA</td><td>720.7</td><td>14.1</td><td>1.3</td><td>10.7</td><td>6.4E-27</td><td>7.9E-26</td><td>BT20/MB468</td></tr>
<tr><td>>hg38_MT_+_1621_3279_RNR2_16S_50ntFlanks@1438.1477.40|TACGTGATCTGAGTTCAGACCGGAGTAATCCAGGTCGGTT</td><td>>hg38_MT_+_1621_3279_RNR2_16S_50ntFlanks</td><td>1438</td><td>1477</td><td>40</td><td>TACGTGATCTGAGTTCAGACCGGAGTAATCCAGGTCGGTT</td><td>1016.7</td><td>12.8</td><td>1.2</td><td>10.7</td><td>7.9E-27</td><td>8.1E-26</td><td>MB231/MB468</td></tr>
<tr><td>>trnaMT_PheGAA_MT_+_577_647@1.19.19|GTTTATGTAGCTTACCTCC</td><td>>trnaMT_PheGAA_MT_+_577_647</td><td>1</td><td>19</td><td>19</td><td>GTTTATGTAGCTTACCTCC</td><td>1546.7</td><td>7.5</td><td>0.7</td><td>10.7</td><td>7.0E-27</td><td>8.6E-26</td><td>BT20/MB468</td></tr>
<tr><td>>MI0000650|hsa-mir-200c&WithFlank&12|+|6963693|6963772@50.71.22&[MIMAT0000617&hsa-miR-200c-3p&offsets|0|-1;m-54&12|+|6963742|6963764&offsets|0|-1]|TAATACTGCCGGGTAATGATGG</td><td>>MI0000650|hsa-mir-200c&WithFlank&12|+|6963693|6963772</td><td>50</td><td>71</td><td>22</td><td>TAATACTGCCGGGTAATGATGG</td><td>53548.8</td><td>8.4</td><td>0.8</td><td>10.7</td><td>7.0E-27</td><td>8.6E-26</td><td>BT20/MB468</td></tr>
<tr><td>>hg38_21_+_8433172_8446622_RNA45SN1_45S_50ntFlanks@475.506.32|CTTCGTGATCGATGTGGTGACGTCGTGCTCTC</td><td>>hg38_21_+_8433172_8446622_RNA45SN1_45S_50ntFlanks</td><td>475</td><td>506</td><td>32</td><td>CTTCGTGATCGATGTGGTGACGTCGTGCTCTC</td><td>14904.1</td><td>6.4</td><td>0.6</td><td>10.7</td><td>7.1E-27</td><td>8.6E-26</td><td>BT20/MB468</td></tr>
<tr><td>>MI0000760|hsa-mir-361&WithFlank&X|-|85903630|85903713@51.74.24&[MIMAT0004682&hsa-miR-361-3p&offsets|0|+1;m-124&X|-|85903642|85903663&offsets|0|+2]|TCCCCCAGGTGTGATTCTGATTTG</td><td>>MI0000760|hsa-mir-361&WithFlank&X|-|85903630|85903713</td><td>51</td><td>74</td><td>24</td><td>TCCCCCAGGTGTGATTCTGATTTG</td><td>1382.2</td><td>8.0</td><td>0.7</td><td>10.7</td><td>8.5E-27</td><td>8.7E-26</td><td>MB231/MB468</td></tr>
<tr><td>>trnaMT_MetCAT_MT_+_4402_4469@35.71.37|CCCCGAAAATGTTGGTTATACCCTTCCCGTACTACCA</td><td>>trnaMT_MetCAT_MT_+_4402_4469</td><td>35</td><td>71</td><td>37</td><td>CCCCGAAAATGTTGGTTATACCCTTCCCGTACTACCA</td><td>646.8</td><td>14.2</td><td>1.3</td><td>10.7</td><td>8.6E-27</td><td>8.8E-26</td><td>MB231/MB468</td></tr>
<tr><td>>MI0000476|hsa-mir-138-1&WithFlank&3|+|44114206|44114316@29.52.24&[MIMAT0000430_1&hsa-miR-138-5p&offsets|0|+1;m-497&3|+|44114234|44114256&offsets|0|+1]|AGCTGGTGTTGTGAATCAGGCCGT</td><td>>MI0000476|hsa-mir-138-1&WithFlank&3|+|44114206|44114316</td><td>29</td><td>52</td><td>24</td><td>AGCTGGTGTTGTGAATCAGGCCGT</td><td>1185.3</td><td>15.1</td><td>1.4</td><td>10.7</td><td>8.8E-27</td><td>9.0E-26</td><td>MB231/MB468</td></tr>
<tr><td>>trnaMT_PheGAA_MT_+_577_647@36.62.27|AAAATGTTTAGACGGGCTCACATCACC</td><td>>trnaMT_PheGAA_MT_+_577_647</td><td>36</td><td>62</td><td>27</td><td>AAAATGTTTAGACGGGCTCACATCACC</td><td>649.9</td><td>11.2</td><td>1.0</td><td>10.7</td><td>8.9E-27</td><td>9.0E-26</td><td>MB231/MB468</td></tr>
<tr><td>>hg38_21_+_8433172_8446622_RNA45SN1_45S_50ntFlanks@3705.3743.39|TACCTGGTTGATCCTGCCAGTAGCATATGCTTGTCTCAA</td><td>>hg38_21_+_8433172_8446622_RNA45SN1_45S_50ntFlanks</td><td>3705</td><td>3743</td><td>39</td><td>TACCTGGTTGATCCTGCCAGTAGCATATGCTTGTCTCAA</td><td>2154.2</td><td>7.9</td><td>0.7</td><td>10.7</td><td>7.5E-27</td><td>9.1E-26</td><td>BT20/MB468</td></tr>
<tr><td>>MI0000285|hsa-mir-205&WithFlank&1|+|209432127|209432248@40.61.22(+1G)&[MIMAT0000266&hsa-miR-205-5p&offsets|0|0(+1G);m-351&1|+|209432166|209432187&offsets|0|0(+1G)]|TCCTTCATTCCACCGGAGTCTGG</td><td>>MI0000285|hsa-mir-205&WithFlank&1|+|209432127|209432248</td><td>40</td><td>61</td><td>22(+1G)</td><td>TCCTTCATTCCACCGGAGTCTGG</td><td>8803.2</td><td>6.6</td><td>0.6</td><td>10.7</td><td>7.5E-27</td><td>9.2E-26</td><td>BT20/MB468</td></tr>
<tr><td>>MI0000255|hsa-mir-30d&WithFlank&8|-|134804870|134804951@12.31.20&[MIMAT0000245&hsa-miR-30d-5p&offsets|0|-2;m-21&8|-|134804919|134804940&offsets|0|-2]|TGTAAACATCCCCGACTGGA</td><td>>MI0000255|hsa-mir-30d&WithFlank&8|-|134804870|134804951</td><td>12</td><td>31</td><td>20</td><td>TGTAAACATCCCCGACTGGA</td><td>2908.0</td><td>8.2</td><td>0.8</td><td>10.7</td><td>9.1E-27</td><td>9.2E-26</td><td>MB231/MB468</td></tr>
<tr><td>>hg38_21_+_8433172_8446622_RNA45SN1_45S_50ntFlanks@6651.6668.18|CGACTCTTAGCGGTGGAT</td><td>>hg38_21_+_8433172_8446622_RNA45SN1_45S_50ntFlanks</td><td>6651</td><td>6668</td><td>18</td><td>CGACTCTTAGCGGTGGAT</td><td>14331.6</td><td>4.7</td><td>0.4</td><td>10.7</td><td>7.6E-27</td><td>9.3E-26</td><td>BT20/MB468</td></tr>
<tr><td>>hg38_21_+_8433172_8446622_RNA45SN1_45S_50ntFlanks@11641.11665.25|CGGCGGGTGTTGACGCGATGTGATT</td><td>>hg38_21_+_8433172_8446622_RNA45SN1_45S_50ntFlanks</td><td>11641</td><td>11665</td><td>25</td><td>CGGCGGGTGTTGACGCGATGTGATT</td><td>3041.2</td><td>-6.7</td><td>0.6</td><td>-10.7</td><td>9.3E-27</td><td>9.4E-26</td><td>MB231/MB468</td></tr>
<tr><td>>am_trnaMT_ProTGG_MT_-_15956_16023@53.70.18|AAGACTTTTTCTCTGACC</td><td>>am_trnaMT_ProTGG_MT_-_15956_16023</td><td>53</td><td>70</td><td>18</td><td>AAGACTTTTTCTCTGACC</td><td>3322.8</td><td>6.7</td><td>0.6</td><td>10.7</td><td>8.0E-27</td><td>9.6E-26</td><td>BT20/MB468</td></tr>
</table>

<table>
<tr><td>>MI0000301|hsa-mir-224&WithFlank&X|-|151958572|151958664@14.34.21(+1A)&[MIMAT0000281&hsa-miR-224-5p&offsets|+1|-3(+1A);m-238&X|-|151958630|151958651&offsets|0|-1(+1A)]|CAAGTCACTAGTGGTTCCGTTA</td><td>>MI0000301|hsa-mir-224&WithFlank&X|-|151958572|151958664</td><td>14</td><td>34</td><td>21(+1A)</td><td>CAAGTCACTAGTGGTTCCGTTA</td><td>1070.4</td><td>14.7</td><td>1.4</td><td>10.7</td><td>8.2E-27</td><td>9.9E-26</td><td>BT20/MB468</td></tr>
<tr><td>>hg38_21_+_8433172_8446622_RNA45SN1_45S_50ntFlanks@485.514.30|GATGTGGTGACGTCGTGCTCTCCCGGGCCG</td><td>>hg38_21_+_8433172_8446622_RNA45SN1_45S_50ntFlanks</td><td>485</td><td>514</td><td>30</td><td>GATGTGGTGACGTCGTGCTCTCCCGGGCCG</td><td>1428.7</td><td>15.1</td><td>1.4</td><td>10.7</td><td>8.3E-27</td><td>1.0E-25</td><td>BT20/MB468</td></tr>
<tr><td>>MI0000067|hsa-let-7f-1&WithFlank&9|+|94176341|94176439@13.32.20(+1C);MI0000068|hsa-let-7f-2&WithFlank&X|-|53557186|53557280@14.33.20(+1C)&[MIMAT0000067&hsa-let-7f-5p&offsets|0|-2(+1C);m-20&9|+|94176353|94176374&offsets|0|-2(+1C)];[MIMAT0000067_1&hsa-let-7f-5p&offsets|0|-2(+1C);m-19&X|-|53557246|53557267&offsets|0|-2(+1C)]|TGAGGTAGTAGATTGTATAGC</td><td>>MI0000067|hsa-let-7f-1&WithFlank&9|+|94176341|94176439</td><td>13</td><td>32</td><td>20(+1C)</td><td>TGAGGTAGTAGATTGTATAGC</td><td>3172.3</td><td>6.9</td><td>0.6</td><td>10.7</td><td>8.3E-27</td><td>1.0E-25</td><td>BT20/MB468</td></tr>
<tr><td>>trna111_HisGTG_1_-_147774845_147774916@1.33.33,trna118_HisGTG_1_-_145396881_145396952@1.33.33,trna16_HisGTG_1_+_146544773_146544844@1.33.33,trna1_HisGTG_15_+_45493349_45493420@1.33.33,trna21_HisGTG_1_+_147753471_147753542@1.33.33,trna33_HisGTG_6_+_27125906_27125977@1.33.33,trna7_HisGTG_9_-_14433938_14434009@1.33.33,trna8_HisGTG_15_-_45492611_45492682@1.33.33,trna9_HisGTG_15_-_45490804_45490875@1.33.33|GCCGTGATCGTATAGTGGTTAGTACTCTGCGTT</td><td>>trna111_HisGTG_1_-_147774845_147774916</td><td>1</td><td>33</td><td>33</td><td>GCCGTGATCGTATAGTGGTTAGTACTCTGCGTT</td><td>6497.5</td><td>9.4</td><td>0.9</td><td>10.7</td><td>1.0E-26</td><td>1.0E-25</td><td>MB231/MB468</td></tr>
<tr><td>>hg38_21_+_8433172_8446622_RNA45SN1_45S_50ntFlanks@6789.6806.18|GCCTGTCTGAGCGTCGCT</td><td>>hg38_21_+_8433172_8446622_RNA45SN1_45S_50ntFlanks</td><td>6789</td><td>6806</td><td>18</td><td>GCCTGTCTGAGCGTCGCT</td><td>4564.5</td><td>7.4</td><td>0.7</td><td>10.7</td><td>1.0E-26</td><td>1.0E-25</td><td>MB231/MB468</td></tr>
<tr><td>>MI0000102|hsa-mir-100&WithFlank&11|-|122152223|122152314@20.39.20&[MIMAT0000098&hsa-miR-100-5p&offsets|+1|-1;m-27&11|-|122152276|122152296&offsets|+1|0]|ACCCGTAGATCCGAACTTGT</td><td>>MI0000102|hsa-mir-100&WithFlank&11|-|122152223|122152314</td><td>20</td><td>39</td><td>20</td><td>ACCCGTAGATCCGAACTTGT</td><td>1102.9</td><td>15.0</td><td>1.4</td><td>10.7</td><td>1.0E-26</td><td>1.0E-25</td><td>MB231/MB468</td></tr>
<tr><td>>hg38_21_+_8433172_8446622_RNA45SN1_45S_50ntFlanks@6721.6745.25|AATTGCAGGACACATTGATCATCGA</td><td>>hg38_21_+_8433172_8446622_RNA45SN1_45S_50ntFlanks</td><td>6721</td><td>6745</td><td>25</td><td>AATTGCAGGACACATTGATCATCGA</td><td>1373.9</td><td>-6.1</td><td>0.6</td><td>-10.7</td><td>1.0E-26</td><td>1.0E-25</td><td>MB231/MB468</td></tr>
<tr><td>>trna111_HisGTG_1_-_147774845_147774916@-1A.23.24,trna118_HisGTG_1_-_145396881_145396952@-1A.23.24,trna16_HisGTG_1_+_146544773_146544844@-1A.23.24,trna1_HisGTG_15_+_45493349_45493420@-1A.23.24,trna21_HisGTG_1_+_147753471_147753542@-1A.23.24,trna33_HisGTG_6_+_27125906_27125977@-1A.23.24,trna7_HisGTG_9_-_14433938_14434009@-1A.23.24,trna8_HisGTG_15_-_45492611_45492682@-1A.23.24,trna9_HisGTG_15_-_45490804_45490875@-1A.23.24|AGCCGTGATCGTATAGTGGTTAGT</td><td>>trna111_HisGTG_1_-_147774845_147774916</td><td>-1A</td><td>23</td><td>24</td><td>AGCCGTGATCGTATAGTGGTTAGT</td><td>2159.8</td><td>13.9</td><td>1.3</td><td>10.7</td><td>1.0E-26</td><td>1.0E-25</td><td>MB231/MB468</td></tr>
<tr><td>>MI0000077|hsa-mir-21&WithFlank&17|+|59841260|59841343@15.36.22&[MIMAT0000076&hsa-miR-21-5p&offsets|+1|+1;m-1&17|+|59841273|59841294&offsets|+1|+1]|AGCTTATCAGACTGATGTTGAC</td><td>>MI0000077|hsa-mir-21&WithFlank&17|+|59841260|59841343</td><td>15</td><td>36</td><td>22</td><td>AGCTTATCAGACTGATGTTGAC</td><td>4315.4</td><td>8.1</td><td>0.8</td><td>10.7</td><td>1.0E-26</td><td>1.0E-25</td><td>MB231/MB468</td></tr>
<tr><td>>MI0000736|hsa-mir-30c-1&WithFlank&1|+|40757278|40757378@23.46.24(+2U);MI0000254|hsa-mir-30c-2&WithFlank&6|-|71376954|71377037@13.36.24(+2U)&[MIMAT0000244&hsa-miR-30c-5p&offsets|0|+1(+2U);m-57&1|+|40757300|40757323&offsets|0|0(+2U)];[MIMAT0000244_1&hsa-miR-30c-5p&offsets|0|+1(+2U);m-56&6|-|71377002|71377025&offsets|0|0(+2U)]|TGTAAACATCCTACACTCTCAGCTTT</td><td>>MI0000736|hsa-mir-30c-1&WithFlank&1|+|40757278|40757378</td><td>23</td><td>46</td><td>24(+2U)</td><td>TGTAAACATCCTACACTCTCAGCTTT</td><td>1264.1</td><td>8.4</td><td>0.8</td><td>10.7</td><td>1.1E-26</td><td>1.1E-25</td><td>MB231/MB468</td></tr>
<tr><td>>MI0000746|hsa-mir-99b&WithFlank&19|+|51692606|51692687@13.33.21(+1A)&[MIMAT0000689&hsa-miR-99b-5p&offsets|0|-1(+1A);m-10&19|+|51692618|51692639&offsets|0|-1(+1A)]|CACCCGTAGAACCGACCTTGCA</td><td>>MI0000746|hsa-mir-99b&WithFlank&19|+|51692606|51692687</td><td>13</td><td>33</td><td>21(+1A)</td><td>CACCCGTAGAACCGACCTTGCA</td><td>12905.6</td><td>7.6</td><td>0.7</td><td>10.7</td><td>1.1E-26</td><td>1.1E-25</td><td>MB231/MB468</td></tr>
<tr><td>>MI0003643|hsa-mir-629&WithFlank&15|-|70079366|70079474@28.48.21(+1C)&[MIMAT0004810&hsa-miR-629-5p&offsets|0|0(+1C);m-150&15|-|70079427|70079447&offsets|0|0(+1C)]|TGGGTTTACGTTGGGAGAACTC</td><td>>MI0003643|hsa-mir-629&WithFlank&15|-|70079366|70079474</td><td>28</td><td>48</td><td>21(+1C)</td><td>TGGGTTTACGTTGGGAGAACTC</td><td>598.3</td><td>13.9</td><td>1.3</td><td>10.7</td><td>8.8E-27</td><td>1.1E-25</td><td>BT20/MB468</td></tr>
<tr><td>>hg38_21_+_8433172_8446622_RNA45SN1_45S_50ntFlanks@10381.10424.44|GTTCCATGTGAACAGCAGTTGAACATGGGTCAGTCGGTCCTGAG</td><td>>hg38_21_+_8433172_8446622_RNA45SN1_45S_50ntFlanks</td><td>10381</td><td>10424</td><td>44</td><td>GTTCCATGTGAACAGCAGTTGAACATGGGTCAGTCGGTCCTGAG</td><td>805.7</td><td>12.5</td><td>1.2</td><td>10.7</td><td>1.1E-26</td><td>1.1E-25</td><td>MB231/MB468</td></tr>
<tr><td>>hg38_21_+_8433172_8446622_RNA45SN1_45S_50ntFlanks@3872.3898.27|CTAATACATGCCGACGGGCGCTGACCC</td><td>>hg38_21_+_8433172_8446622_RNA45SN1_45S_50ntFlanks</td><td>3872</td><td>3898</td><td>27</td><td>CTAATACATGCCGACGGGCGCTGACCC</td><td>571.4</td><td>-6.5</td><td>0.6</td><td>-10.7</td><td>9.0E-27</td><td>1.1E-25</td><td>BT20/MB468</td></tr>
<tr><td>>hg38_21_+_8433172_8446622_RNA45SN1_45S_50ntFlanks@11661.11685.25|TGATTTCTGCCCAGTGCTCTGAATG</td><td>>hg38_21_+_8433172_8446622_RNA45SN1_45S_50ntFlanks</td><td>11661</td><td>11685</td><td>25</td><td>TGATTTCTGCCCAGTGCTCTGAATG</td><td>4062.9</td><td>-5.6</td><td>0.5</td><td>-10.7</td><td>1.1E-26</td><td>1.1E-25</td><td>MB231/MB468</td></tr>
<tr><td>>am_MI0000809|hsa-mir-151a&WithFlank&8|-|140732558|140732659@53.72.20(+2U)&[MIMAT0000757&hsa-miR-151a-3p&offsets|0|-1(+2U);m-30&8|-|140732587|140732607&offsets|0|-1(+2U)]|CTAGACTGAAGCTCCTTGAGTT</td><td>>am_MI0000809|hsa-mir-151a&WithFlank&8|-|140732558|140732659</td><td>53</td><td>72</td><td>20(+2U)</td><td>CTAGACTGAAGCTCCTTGAGTT</td><td>1161.1</td><td>15.1</td><td>1.4</td><td>10.7</td><td>1.1E-26</td><td>1.1E-25</td><td>MB231/MB468</td></tr>
<tr><td>>am_trnaMT_MetCAT_MT_+_4402_4469@1.22.22|AGTAAGGTCAGCTAAATAAGCT</td><td>>am_trnaMT_MetCAT_MT_+_4402_4469</td><td>1</td><td>22</td><td>22</td><td>AGTAAGGTCAGCTAAATAAGCT</td><td>2848.8</td><td>8.5</td><td>0.8</td><td>10.7</td><td>9.5E-27</td><td>1.1E-25</td><td>BT20/MB468</td></tr>
<tr><td>>MI0003137|hsa-mir-193b&WithFlank&16|+|14303961|14304055@57.78.22&[MIMAT0002819&hsa-miR-193b-3p&offsets|0|0;m-166&16|+|14304017|14304038&offsets|0|0]|AACTGGCCCTCAAAGTCCCGCT</td><td>>MI0003137|hsa-mir-193b&WithFlank&16|+|14303961|14304055</td><td>57</td><td>78</td><td>22</td><td>AACTGGCCCTCAAAGTCCCGCT</td><td>14287.5</td><td>10.5</td><td>1.0</td><td>10.7</td><td>9.6E-27</td><td>1.1E-25</td><td>BT20/MB468</td></tr>
<tr><td>>hg38_21_+_8433172_8446622_RNA45SN1_45S_50ntFlanks@12419.12447.29|GATGTCGGCTCTTCCTATCATTGTGAAGC</td><td>>hg38_21_+_8433172_8446622_RNA45SN1_45S_50ntFlanks</td><td>12419</td><td>12447</td><td>29</td><td>GATGTCGGCTCTTCCTATCATTGTGAAGC</td><td>1869.4</td><td>7.9</td><td>0.7</td><td>10.7</td><td>1.2E-26</td><td>1.2E-25</td><td>MB231/MB468</td></tr>
<tr><td>>MI0000077|hsa-mir-21&WithFlank&17|+|59841260|59841343@15.37.23&[MIMAT0000076&hsa-miR-21-5p&offsets|+1|+2;m-1&17|+|59841273|59841294&offsets|+1|+2]|AGCTTATCAGACTGATGTTGACT</td><td>>MI0000077|hsa-mir-21&WithFlank&17|+|59841260|59841343</td><td>15</td><td>37</td><td>23</td><td>AGCTTATCAGACTGATGTTGACT</td><td>773.5</td><td>7.6</td><td>0.7</td><td>10.7</td><td>1.2E-26</td><td>1.2E-25</td><td>MB231/MB468</td></tr>
<tr><td>>MI0000289|hsa-mir-181a-1&WithFlank&1|-|198859038|198859159@70.91.22&[MIMAT0000270&hsa-miR-181a-3p&offsets|0|0;m-125&1|-|198859069|198859090&offsets|0|0]|ACCATCGACCGTTGATTGTACC</td><td>>MI0000289|hsa-mir-181a-1&WithFlank&1|-|198859038|198859159</td><td>70</td><td>91</td><td>22</td><td>ACCATCGACCGTTGATTGTACC</td><td>5148.7</td><td>8.6</td><td>0.8</td><td>10.7</td><td>1.2E-26</td><td>1.2E-25</td><td>MB231/MB468</td></tr>
<tr><td>>hg38_MT_+_598_1651_RNR1_12S_50ntFlanks@52.84.33|ATAGGTTTGGTCCTAGCCTTTCTATTAGCTCTT</td><td>>hg38_MT_+_598_1651_RNR1_12S_50ntFlanks</td><td>52</td><td>84</td><td>33</td><td>ATAGGTTTGGTCCTAGCCTTTCTATTAGCTCTT</td><td>875.9</td><td>7.0</td><td>0.7</td><td>10.7</td><td>1.0E-26</td><td>1.2E-25</td><td>BT20/MB468</td></tr>
<tr><td>>trnaMT_ValTAC_MT_+_1602_1670@50.71.22|ACTTAACTTGACCGCTCTGACC</td><td>>trnaMT_ValTAC_MT_+_1602_1670</td><td>50</td><td>71</td><td>22</td><td>ACTTAACTTGACCGCTCTGACC</td><td>838.7</td><td>5.8</td><td>0.5</td><td>10.7</td><td>1.2E-26</td><td>1.2E-25</td><td>MB231/MB468</td></tr>
<tr><td>>hg38_21_+_8433172_8446622_RNA45SN1_45S_50ntFlanks@9567.9596.30|ACGTGCAAATCGGTCGTCCGACCTGGGTAT</td><td>>hg38_21_+_8433172_8446622_RNA45SN1_45S_50ntFlanks</td><td>9567</td><td>9596</td><td>30</td><td>ACGTGCAAATCGGTCGTCCGACCTGGGTAT</td><td>3668.3</td><td>-7.4</td><td>0.7</td><td>-10.7</td><td>1.3E-26</td><td>1.3E-25</td><td>MB231/MB468</td></tr>
<tr><td>>hg38_21_+_8433172_8446622_RNA45SN1_45S_50ntFlanks@476.503.28|TTCGTGATCGATGTGGTGACGTCGTGCT</td><td>>hg38_21_+_8433172_8446622_RNA45SN1_45S_50ntFlanks</td><td>476</td><td>503</td><td>28</td><td>TTCGTGATCGATGTGGTGACGTCGTGCT</td><td>8835.7</td><td>7.9</td><td>0.7</td><td>10.7</td><td>1.3E-26</td><td>1.3E-25</td><td>MB231/MB468</td></tr>
<tr><td>>MI0005756|hsa-mir-934&WithFlank&X|+|136550872|136550966@21.41.21&[MIMAT0004977&hsa-miR-934&offsets|0|-1;m-943&X|+|136550892|136550913&offsets|0|-1]|TGTCTACTACTGGAGACACTG</td><td>>MI0005756|hsa-mir-934&WithFlank&X|+|136550872|136550966</td><td>21</td><td>41</td><td>21</td><td>TGTCTACTACTGGAGACACTG</td><td>5177.6</td><td>15.8</td><td>1.4</td><td>11.0</td><td>5.7E-28</td><td>1.3E-25</td><td>BT20/MB231</td></tr>
<tr><td>>hg38_21_+_8433172_8446622_RNA45SN1_45S_50ntFlanks@6745.6771.27|ACACTTCGAACGCACTTGCGGCCCCGG</td><td>>hg38_21_+_8433172_8446622_RNA45SN1_45S_50ntFlanks</td><td>6745</td><td>6771</td><td>27</td><td>ACACTTCGAACGCACTTGCGGCCCCGG</td><td>2608.5</td><td>-6.6</td><td>0.6</td><td>-10.7</td><td>1.4E-26</td><td>1.3E-25</td><td>MB231/MB468</td></tr>
<tr><td>>hg38_21_+_8433172_8446622_RNA45SN1_45S_50ntFlanks@7975.8015.41|CGCGACCTCAGATCAGACGTGGCGACCCGCTGAATTTAAGC</td><td>>hg38_21_+_8433172_8446622_RNA45SN1_45S_50ntFlanks</td><td>7975</td><td>8015</td><td>41</td><td>CGCGACCTCAGATCAGACGTGGCGACCCGCTGAATTTAAGC</td><td>10136.9</td><td>8.6</td><td>0.8</td><td>10.7</td><td>1.4E-26</td><td>1.4E-25</td><td>MB231/MB468</td></tr>
<tr><td>>MI0000811|hsa-mir-148b&WithFlank&12|+|54337210|54337320@69.89.21&[MIMAT0000759&hsa-miR-148b-3p&offsets|0|-1;m-77&12|+|54337278|54337299&offsets|0|-1]|TCAGTGCATCACAGAACTTTG</td><td>>MI0000811|hsa-mir-148b&WithFlank&12|+|54337210|54337320</td><td>69</td><td>89</td><td>21</td><td>TCAGTGCATCACAGAACTTTG</td><td>1839.1</td><td>8.1</td><td>0.8</td><td>10.7</td><td>1.2E-26</td><td>1.4E-25</td><td>BT20/MB468</td></tr>
<tr><td>>hg38_21_+_8433172_8446622_RNA45SN1_45S_50ntFlanks@475.520.46|CTTCGTGATCGATGTGGTGACGTCGTGCTCTCCCGGGCCGGGTCCG</td><td>>hg38_21_+_8433172_8446622_RNA45SN1_45S_50ntFlanks</td><td>475</td><td>520</td><td>46</td><td>CTTCGTGATCGATGTGGTGACGTCGTGCTCTCCCGGGCCGGGTCCG</td><td>2935.6</td><td>11.9</td><td>1.1</td><td>10.7</td><td>1.4E-26</td><td>1.4E-25</td><td>MB231/MB468</td></tr>
<tr><td>>MI0000456|hsa-mir-140&WithFlank&16|+|69933075|69933186@68.90.23(+1U)&[MIMAT0004597&hsa-miR-140-3p&offsets|0|+2(+1U);m-45&16|+|69933143|69933164&offsets|-1|0(+1U)]|TACCACAGGGTAGAACCACGGACT</td><td>>MI0000456|hsa-mir-140&WithFlank&16|+|69933075|69933186</td><td>68</td><td>90</td><td>23(+1U)</td><td>TACCACAGGGTAGAACCACGGACT</td><td>1119.4</td><td>15.0</td><td>1.4</td><td>10.7</td><td>1.5E-26</td><td>1.4E-25</td><td>MB231/MB468</td></tr>
<tr><td>>hg38_21_+_8433172_8446622_RNA45SN1_45S_50ntFlanks@12420.12457.38|ATGTCGGCTCTTCCTATCATTGTGAAGCAGAATTCACC</td><td>>hg38_21_+_8433172_8446622_RNA45SN1_45S_50ntFlanks</td><td>12420</td><td>12457</td><td>38</td><td>ATGTCGGCTCTTCCTATCATTGTGAAGCAGAATTCACC</td><td>1786.9</td><td>6.6</td><td>0.6</td><td>10.7</td><td>1.5E-26</td><td>1.5E-25</td><td>MB231/MB468</td></tr>
<tr><td>>MI0000299|hsa-mir-222&WithFlank&X|-|45747009|45747130@75.96.22(+1G)&[MIMAT0000279&hsa-miR-222-3p&offsets|0|+1(+1G);m-140&X|-|45747033|45747056&offsets|0|-2(+1G)]|AGCTACATCTGGCTACTGGGTCG</td><td>>MI0000299|hsa-mir-222&WithFlank&X|-|45747009|45747130</td><td>75</td><td>96</td><td>22(+1G)</td><td>AGCTACATCTGGCTACTGGGTCG</td><td>1405.2</td><td>8.4</td><td>0.8</td><td>10.7</td><td>1.5E-26</td><td>1.5E-25</td><td>MB231/MB468</td></tr>
<tr><td>>trna116_GluCTC_1_-_145399233_145399304@1.33.33,trna59_GluCTC_1_+_249168447_249168518@1.33.33,trna71_GluCTC_1_-_161439189_161439260@1.33.33,trna74_GluCTC_1_-_161431809_161431880@1.33.33,trna77_GluCTC_1_-_161424398_161424469@1.33.33,trna77_GluCTC_6_+_28949976_28950047@1.33.33,trna80_GluCTC_1_-_161417018_161417089@1.33.33,trna87_GluCTC_6_-_126101393_126101464@1.33.33|TCCCTGGTGGTCTAGTGGTTAGGATTCGGCGCT</td><td>>trna116_GluCTC_1_-_145399233_145399304</td><td>1</td><td>33</td><td>33</td><td>TCCCTGGTGGTCTAGTGGTTAGGATTCGGCGCT</td><td>38267.4</td><td>7.9</td><td>0.7</td><td>10.7</td><td>1.2E-26</td><td>1.5E-25</td><td>BT20/MB468</td></tr>
<tr><td>>MI0000086|hsa-mir-28&WithFlank&3|+|188688775|188688872@61.82.22&[MIMAT0004502&hsa-miR-28-3p&offsets|+1|+1;m-28&3|+|188688834|188688855&offsets|+1|+1]|ACTAGATTGTGAGCTCCTGGAG</td><td>>MI0000086|hsa-mir-28&WithFlank&3|+|188688775|188688872</td><td>61</td><td>82</td><td>22</td><td>ACTAGATTGTGAGCTCCTGGAG</td><td>966.9</td><td>14.6</td><td>1.4</td><td>10.7</td><td>1.3E-26</td><td>1.5E-25</td><td>BT20/MB468</td></tr>
</table>

<table>
<tr><th></th><th></th><th></th><th></th><th></th><th></th><th></th><th></th><th></th><th></th><th></th><th></th><th></th></tr>
<tr><td>>hg38_21_+_8433172_8446622_RNA45SN1_45S_50ntFlanks@13002.13028.27|CGATCTATTGAAAGTCAGCCCTCGACA</td><td>>hg38_21_+_8433172_8446622_RNA45SN1_45S_50ntFlanks</td><td>13002</td><td>13028</td><td>27</td><td>CGATCTATTGAAAGTCAGCCCTCGACA</td><td>3181.2</td><td>-13.6</td><td>1.3</td><td>-10.7</td><td>1.3E-26</td><td>1.6E-25</td><td>BT20/MB468</td></tr>
<tr><td>>MI0000087|hsa-mir-29a&WithFlank&7|-|130876741|130876816@48.66.19&[MIMAT0000086&hsa-miR-29a-3p&offsets|0|-3;m-26&7|-|130876749|130876769&offsets|0|-2]|TAGCACCATCTGAAATCGG</td><td>>MI0000087|hsa-mir-29a&WithFlank&7|-|130876741|130876816</td><td>48</td><td>66</td><td>19</td><td>TAGCACCATCTGAAATCGG</td><td>559.6</td><td>12.0</td><td>1.1</td><td>10.7</td><td>1.6E-26</td><td>1.6E-25</td><td>MB231/MB468</td></tr>
<tr><td>>MI0000746|hsa-mir-99b&WithFlank&19|+|51692606|51692687@13.32.20(+2U)&[MIMAT0000689&hsa-miR-99b-5p&offsets|0|-2(+2U);m-10&19|+|51692618|51692639&offsets|0|-2(+2U)]|CACCCGTAGAACCGACCTTGTT</td><td>>MI0000746|hsa-mir-99b&WithFlank&19|+|51692606|51692687</td><td>13</td><td>32</td><td>20(+2U)</td><td>CACCCGTAGAACCGACCTTGTT</td><td>6368.9</td><td>17.3</td><td>1.6</td><td>10.7</td><td>1.3E-26</td><td>1.6E-25</td><td>BT20/MB468</td></tr>
<tr><td>>MI0000266|hsa-mir-10a&WithFlank&17|-|48579832|48579953@28.48.21&[MIMAT0000253&hsa-miR-10a-5p&offsets|0|-2;m-18&17|-|48579905|48579926&offsets|0|-1]|TACCCTGTAGATCCGAATTTG</td><td>>MI0000266|hsa-mir-10a&WithFlank&17|-|48579832|48579953</td><td>28</td><td>48</td><td>21</td><td>TACCCTGTAGATCCGAATTTG</td><td>688.8</td><td>14.3</td><td>1.3</td><td>10.7</td><td>1.6E-26</td><td>1.6E-25</td><td>MB231/MB468</td></tr>
<tr><td>>MI0000063|hsa-let-7b&WithFlank&22|+|46113680|46113774@12.31.20&[MIMAT0000063&hsa-let-7b-5p&offsets|0|-2;m-14&22|+|46113691|46113712&offsets|0|-2]|TGAGGTAGTAGGTTGTGTGG</td><td>>MI0000063|hsa-let-7b&WithFlank&22|+|46113680|46113774</td><td>12</td><td>31</td><td>20</td><td>TGAGGTAGTAGGTTGTGTGG</td><td>1300.2</td><td>7.7</td><td>0.7</td><td>10.7</td><td>1.4E-26</td><td>1.6E-25</td><td>BT20/MB468</td></tr>
<tr><td>>MI0000439|hsa-mir-23b&WithFlank&9|+|95085202|95085310@67.85.19(+1U)&[MIMAT0000418&hsa-miR-23b-3p&offsets|+3|-1(+1U);m-42&9|+|95085265|95085285&offsets|+3|+1(+1U)]|ACATTGCCAGGGATTACCAT</td><td>>MI0000439|hsa-mir-23b&WithFlank&9|+|95085202|95085310</td><td>67</td><td>85</td><td>19(+1U)</td><td>ACATTGCCAGGGATTACCAT</td><td>2567.8</td><td>16.0</td><td>1.5</td><td>10.7</td><td>1.4E-26</td><td>1.6E-25</td><td>BT20/MB468</td></tr>
<tr><td>>MI0000440|hsa-mir-27b&WithFlank&9|+|95085439|95085547@67.86.20(+1U)&[MIMAT0000419&hsa-miR-27b-3p&offsets|0|-1(+1U);m-39&9|+|95085505|95085525&offsets|0|-1(+1U)]|TTCACAGTGGCTAAGTTCTGT</td><td>>MI0000440|hsa-mir-27b&WithFlank&9|+|95085439|95085547</td><td>67</td><td>86</td><td>20(+1U)</td><td>TTCACAGTGGCTAAGTTCTGT</td><td>69546.5</td><td>8.8</td><td>0.8</td><td>10.7</td><td>1.4E-26</td><td>1.7E-25</td><td>BT20/MB468</td></tr>
<tr><td>>hg38_21_+_8433172_8446622_RNA45SN1_45S_50ntFlanks@11873.11906.34|AAGAAGACCCTGTTGAGCTTGACTCTAGTCTGGC</td><td>>hg38_21_+_8433172_8446622_RNA45SN1_45S_50ntFlanks</td><td>11873</td><td>11906</td><td>34</td><td>AAGAAGACCCTGTTGAGCTTGACTCTAGTCTGGC</td><td>892.7</td><td>-6.4</td><td>0.6</td><td>-10.7</td><td>1.7E-26</td><td>1.7E-25</td><td>MB231/MB468</td></tr>
<tr><td>>hg38_21_+_8433172_8446622_RNA45SN1_45S_50ntFlanks@4734.4768.35|AACGAAAGTCGGAGGTTCGAAGACGATCAGATACC</td><td>>hg38_21_+_8433172_8446622_RNA45SN1_45S_50ntFlanks</td><td>4734</td><td>4768</td><td>35</td><td>AACGAAAGTCGGAGGTTCGAAGACGATCAGATACC</td><td>422.9</td><td>13.6</td><td>1.3</td><td>10.7</td><td>1.7E-26</td><td>1.7E-25</td><td>MB231/MB468</td></tr>
<tr><td>>trnaMT_ProTGG_MT_-_15956_16023@50.71.22|TTAAAGACTTTTTCTCTGACCA</td><td>>trnaMT_ProTGG_MT_-_15956_16023</td><td>50</td><td>71</td><td>22</td><td>TTAAAGACTTTTTCTCTGACCA</td><td>1521.5</td><td>8.5</td><td>0.8</td><td>10.7</td><td>1.5E-26</td><td>1.8E-25</td><td>BT20/MB468</td></tr>
<tr><td>>MI0000102|hsa-mir-100&WithFlank&11|-|122152223|122152314@19.40.22(+1U)&[MIMAT0000098&hsa-miR-100-5p&offsets|0|0(+1U);m-27&11|-|122152276|122152296&offsets|0|+1(+1U)]|AACCCGTAGATCCGAACTTGTGT</td><td>>MI0000102|hsa-mir-100&WithFlank&11|-|122152223|122152314</td><td>19</td><td>40</td><td>22(+1U)</td><td>AACCCGTAGATCCGAACTTGTGT</td><td>233062.3</td><td>15.7</td><td>1.5</td><td>10.6</td><td>2.0E-26</td><td>1.9E-25</td><td>MB231/MB468</td></tr>
<tr><td>>hg38_21_+_8433172_8446622_RNA45SN1_45S_50ntFlanks@12636.12659.24|AGCTACCATCTGTGGGATTATGAC</td><td>>hg38_21_+_8433172_8446622_RNA45SN1_45S_50ntFlanks</td><td>12636</td><td>12659</td><td>24</td><td>AGCTACCATCTGTGGGATTATGAC</td><td>669.7</td><td>-5.9</td><td>0.6</td><td>-10.7</td><td>1.6E-26</td><td>1.9E-25</td><td>BT20/MB468</td></tr>
<tr><td>>MI0000811|hsa-mir-148b&WithFlank&12|+|54337210|54337320@69.91.23&[MIMAT0000759&hsa-miR-148b-3p&offsets|0|+1;m-77&12|+|54337278|54337299&offsets|0|+1]|TCAGTGCATCACAGAACTTTGTC</td><td>>MI0000811|hsa-mir-148b&WithFlank&12|+|54337210|54337320</td><td>69</td><td>91</td><td>23</td><td>TCAGTGCATCACAGAACTTTGTC</td><td>2479.7</td><td>7.8</td><td>0.7</td><td>10.7</td><td>1.6E-26</td><td>1.9E-25</td><td>BT20/MB468</td></tr>
<tr><td>>MI0000439|hsa-mir-23b&WithFlank&9|+|95085202|95085310@67.86.20(+1A)&[MIMAT0000418&hsa-miR-23b-3p&offsets|+3|0(+1A);m-42&9|+|95085265|95085285&offsets|+3|+2(+1A)]|ACATTGCCAGGGATTACCACA</td><td>>MI0000439|hsa-mir-23b&WithFlank&9|+|95085202|95085310</td><td>67</td><td>86</td><td>20(+1A)</td><td>ACATTGCCAGGGATTACCACA</td><td>2977.2</td><td>16.2</td><td>1.5</td><td>10.7</td><td>1.7E-26</td><td>1.9E-25</td><td>BT20/MB468</td></tr>
<tr><td>>MI0000061|hsa-let-7a-2&WithFlank&11|-|122146516|122146599@11.31.21(+1C);MI0000062|hsa-let-7a-3&WithFlank&22|+|46112743|46112828@10.30.21(+1C);MI0000060|hsa-let-7a-1&WithFlank&9|+|94175951|94176042@12.32.21(+1C)&[MIMAT0000062&hsa-let-7a-5p&offsets|0|-1(+1C);m-6&11|-|122146568|122146589&offsets|0|-1(+1C)];[MIMAT0000062_1&hsa-let-7a-5p&offsets|0|-1(+1C);m-5&22|+|46112752|46112773&offsets|0|-1(+1C)];[MIMAT0000062_2&hsa-let-7a-5p&offsets|0|-1(+1C);m-4&9|+|94175962|94175983&offsets|0|-1(+1C)]|TGAGGTAGTAGGTTGTATAGTC</td><td>>MI0000061|hsa-let-7a-2&WithFlank&11|-|122146516|122146599</td><td>11</td><td>31</td><td>21(+1C)</td><td>TGAGGTAGTAGGTTGTATAGTC</td><td>26286.9</td><td>7.5</td><td>0.7</td><td>10.7</td><td>1.7E-26</td><td>2.0E-25</td><td>BT20/MB468</td></tr>
<tr><td>>hg38_21_+_8433172_8446622_RNA45SN1_45S_50ntFlanks@7975.8011.37|CGCGACCTCAGATCAGACGTGGCGACCCGCTGAATTT</td><td>>hg38_21_+_8433172_8446622_RNA45SN1_45S_50ntFlanks</td><td>7975</td><td>8011</td><td>37</td><td>CGCGACCTCAGATCAGACGTGGCGACCCGCTGAATTT</td><td>126311.2</td><td>7.3</td><td>0.7</td><td>10.7</td><td>1.7E-26</td><td>2.0E-25</td><td>BT20/MB468</td></tr>
<tr><td>>MI0003599|hsa-mir-589&WithFlank&7|-|5495813|5495923@30.52.23&[MIMAT0004799&hsa-miR-589-5p&offsets|0|+1;m-162&7|-|5495874|5495894&offsets|0|+2]|TGAGAACCACGTCTGCTCTGAGC</td><td>>MI0003599|hsa-mir-589&WithFlank&7|-|5495813|5495923</td><td>30</td><td>52</td><td>23</td><td>TGAGAACCACGTCTGCTCTGAGC</td><td>1089.8</td><td>10.6</td><td>1.0</td><td>10.7</td><td>1.7E-26</td><td>2.0E-25</td><td>BT20/MB468</td></tr>
<tr><td>>MI0000254|hsa-mir-30c-2&WithFlank&6|-|71376954|71377037@53.74.22&[MIMAT0004550&hsa-miR-30c-2-3p&offsets|0|0;m-206&6|-|71376964|71376985&offsets|0|0]|CTGGGAGAAGGCTGTTTACTCT</td><td>>MI0000254|hsa-mir-30c-2&WithFlank&6|-|71376954|71377037</td><td>53</td><td>74</td><td>22</td><td>CTGGGAGAAGGCTGTTTACTCT</td><td>1698.8</td><td>6.9</td><td>0.6</td><td>10.6</td><td>1.8E-26</td><td>2.1E-25</td><td>BT20/MB468</td></tr>
<tr><td>>hg38_21_+_8433172_8446622_RNA45SN1_45S_50ntFlanks@6647.6668.22|CGTACGACTCTTAGCGGTGGAT</td><td>>hg38_21_+_8433172_8446622_RNA45SN1_45S_50ntFlanks</td><td>6647</td><td>6668</td><td>22</td><td>CGTACGACTCTTAGCGGTGGAT</td><td>2832.6</td><td>5.3</td><td>0.5</td><td>10.6</td><td>1.8E-26</td><td>2.1E-25</td><td>BT20/MB468</td></tr>
<tr><td>>MI0000298|hsa-mir-221&WithFlank&X|-|45746151|45746272@31.53.23&[MIMAT0004568&hsa-miR-221-5p&offsets|0|+1;m-405&X|-|45746220|45746242&offsets|0|0]|ACCTGGCATACAATGTAGATTTC</td><td>>MI0000298|hsa-mir-221&WithFlank&X|-|45746151|45746272</td><td>31</td><td>53</td><td>23</td><td>ACCTGGCATACAATGTAGATTTC</td><td>2825.5</td><td>8.2</td><td>0.8</td><td>10.6</td><td>2.2E-26</td><td>2.2E-25</td><td>MB231/MB468</td></tr>
<tr><td>>hg38_21_+_8433172_8446622_RNA45SN1_45S_50ntFlanks@11666.11691.26|TCTGCCCAGTGCTCTGAATGTCAAAG</td><td>>hg38_21_+_8433172_8446622_RNA45SN1_45S_50ntFlanks</td><td>11666</td><td>11691</td><td>26</td><td>TCTGCCCAGTGCTCTGAATGTCAAAG</td><td>399.5</td><td>4.7</td><td>0.4</td><td>10.6</td><td>2.3E-26</td><td>2.2E-25</td><td>MB231/MB468</td></tr>
<tr><td>>MI0000063|hsa-let-7b&WithFlank&22|+|46113680|46113774@12.34.23(+1U)&[MIMAT0000063&hsa-let-7b-5p&offsets|0|+1(+1U);m-14&22|+|46113691|46113712&offsets|0|+1(+1U)]|TGAGGTAGTAGGTTGTGTGGTTTT</td><td>>MI0000063|hsa-let-7b&WithFlank&22|+|46113680|46113774</td><td>12</td><td>34</td><td>23(+1U)</td><td>TGAGGTAGTAGGTTGTGTGGTTTT</td><td>1232.3</td><td>7.6</td><td>0.7</td><td>10.6</td><td>2.0E-26</td><td>2.3E-25</td><td>BT20/MB468</td></tr>
<tr><td>>hg38_21_+_8433172_8446622_RNA45SN1_45S_50ntFlanks@6651.6695.45|CGACTCTTAGCGGTGGATCACTCGGCTCGTGCGTCGATGAAGAAC</td><td>>hg38_21_+_8433172_8446622_RNA45SN1_45S_50ntFlanks</td><td>6651</td><td>6695</td><td>45</td><td>CGACTCTTAGCGGTGGATCACTCGGCTCGTGCGTCGATGAAGAAC</td><td>5943.2</td><td>8.7</td><td>0.8</td><td>10.6</td><td>2.0E-26</td><td>2.3E-25</td><td>BT20/MB468</td></tr>
<tr><td>>hg38_21_+_8433172_8446622_RNA45SN1_45S_50ntFlanks@7975.8002.28|CGCGACCTCAGATCAGACGTGGCGACCC</td><td>>hg38_21_+_8433172_8446622_RNA45SN1_45S_50ntFlanks</td><td>7975</td><td>8002</td><td>28</td><td>CGCGACCTCAGATCAGACGTGGCGACCC</td><td>3283.9</td><td>6.5</td><td>0.6</td><td>10.6</td><td>2.0E-26</td><td>2.3E-25</td><td>BT20/MB468</td></tr>
<tr><td>>hg38_21_+_8433172_8446622_RNA45SN1_45S_50ntFlanks@8380.8399.20|CAAGAGGGCGTGAAACCGTT</td><td>>hg38_21_+_8433172_8446622_RNA45SN1_45S_50ntFlanks</td><td>8380</td><td>8399</td><td>20</td><td>CAAGAGGGCGTGAAACCGTT</td><td>723.5</td><td>-6.1</td><td>0.6</td><td>-10.6</td><td>2.1E-26</td><td>2.4E-25</td><td>BT20/MB468</td></tr>
<tr><td>>hg38_MT_+_598_1651_RNR1_12S_50ntFlanks@905.935.31|AGTATACTTCAAAGGACATTTAACTAAAACC</td><td>>hg38_MT_+_598_1651_RNR1_12S_50ntFlanks</td><td>905</td><td>935</td><td>31</td><td>AGTATACTTCAAAGGACATTTAACTAAAACC</td><td>981.1</td><td>12.8</td><td>1.2</td><td>10.6</td><td>2.5E-26</td><td>2.4E-25</td><td>MB231/MB468</td></tr>
<tr><td>>trna119_LysCTT_1_-_145395522_145395594@1.34.34,trna11_LysCTT_5_-_180648979_180649051@1.34.34,trna13_LysCTT_6_+_26556774_26556846@1.34.34,trna7_LysCTT_16_+_3225692_3225764@1.34.34,trna9_LysCTT_5_+_180634755_180634827@1.34.34|GCCCGGCTAGCTCAGTCGGTAGAGCATGAGACTC</td><td>>trna119_LysCTT_1_-_145395522_145395594</td><td>1</td><td>34</td><td>34</td><td>GCCCGGCTAGCTCAGTCGGTAGAGCATGAGACTC</td><td>20641.0</td><td>10.5</td><td>1.0</td><td>10.6</td><td>2.5E-26</td><td>2.4E-25</td><td>MB231/MB468</td></tr>
<tr><td>>MI0000082|hsa-mir-25&WithFlank&7|-|100093554|100093649@58.79.22(+1A)&[MIMAT0000081&hsa-miR-25-3p&offsets|0|0(+1A);m-23&7|-|100093571|100093592&offsets|0|0(+1A)]|CATTGCACTTGTCTCGGTCTGAA</td><td>>MI0000082|hsa-mir-25&WithFlank&7|-|100093554|100093649</td><td>58</td><td>79</td><td>22(+1A)</td><td>CATTGCACTTGTCTCGGTCTGAA</td><td>2567.9</td><td>7.0</td><td>0.7</td><td>10.6</td><td>2.1E-26</td><td>2.4E-25</td><td>BT20/MB468</td></tr>
<tr><td>>MI0001445|hsa-mir-423&WithFlank&17|+|30117073|30117178@59.79.21&[MIMAT0001340&hsa-miR-423-3p&offsets|0|-2;m-100&17|+|30117131|30117153&offsets|0|-2]|AGCTCGGTCTGAGGCCCCTCA</td><td>>MI0001445|hsa-mir-423&WithFlank&17|+|30117073|30117178</td><td>59</td><td>79</td><td>21</td><td>AGCTCGGTCTGAGGCCCCTCA</td><td>2110.3</td><td>8.1</td><td>0.8</td><td>10.6</td><td>2.5E-26</td><td>2.4E-25</td><td>MB231/MB468</td></tr>
<tr><td>>MI0000279|hsa-mir-196a-2&WithFlank&12|+|53991732|53991853@31.52.22;MI0000238|hsa-mir-196a-1&WithFlank&17|-|48632484|48632565@13.34.22&[MIMAT0000226&hsa-miR-196a-5p&offsets|0|0;m-261&12|+|53991762|53991783&offsets|0|0];[MIMAT0000226_1&hsa-miR-196a-5p&offsets|0|0;m-268&17|-|48632532|48632553&offsets|0|0]|TAGGTAGTTTCATGTTGTTGGG</td><td>>MI0000279|hsa-mir-196a-2&WithFlank&12|+|53991732|53991853</td><td>31</td><td>52</td><td>22</td><td>TAGGTAGTTTCATGTTGTTGGG</td><td>3226.6</td><td>15.2</td><td>1.4</td><td>10.9</td><td>1.1E-27</td><td>2.5E-25</td><td>BT20/MB231</td></tr>
<tr><td>>hg38_MT_+_1621_3279_RNR2_16S_50ntFlanks@1438.1473.36|TACGTGATCTGAGTTCAGACCGGAGTAATCCAGGTC</td><td>>hg38_MT_+_1621_3279_RNR2_16S_50ntFlanks</td><td>1438</td><td>1473</td><td>36</td><td>TACGTGATCTGAGTTCAGACCGGAGTAATCCAGGTC</td><td>423.4</td><td>9.4</td><td>0.9</td><td>10.6</td><td>2.6E-26</td><td>2.5E-25</td><td>MB231/MB468</td></tr>
<tr><td>>MI0000067|hsa-let-7f-1&WithFlank&9|+|94176341|94176439@13.32.20(+1C);MI0000068|hsa-let-7f-2&WithFlank&X|-|53557186|53557280@14.33.20(+1C)&[MIMAT0000067&hsa-let-7f-5p&offsets|0|-2(+1C);m-20&9|+|94176353|94176374&offsets|0|-2(+1C)];[MIMAT0000067_1&hsa-let-7f-5p&offsets|0|-2(+1C);m-19&X|-|53557246|53557267&offsets|0|-2(+1C)]|TGAGGTAGTAGATTGTATAGC</td><td>>MI0000067|hsa-let-7f-1&WithFlank&9|+|94176341|94176439</td><td>13</td><td>32</td><td>20(+1C)</td><td>TGAGGTAGTAGATTGTATAGC</td><td>4160.1</td><td>7.6</td><td>0.7</td><td>10.6</td><td>2.7E-26</td><td>2.6E-25</td><td>MB231/MB468</td></tr>
<tr><td>>hg38_21_+_8433172_8446622_RNA45SN1_45S_50ntFlanks@11783.11806.24|CATGAATGGATGAACGAGATTCCC</td><td>>hg38_21_+_8433172_8446622_RNA45SN1_45S_50ntFlanks</td><td>11783</td><td>11806</td><td>24</td><td>CATGAATGGATGAACGAGATTCCC</td><td>3115.3</td><td>-6.6</td><td>0.6</td><td>-10.6</td><td>2.3E-26</td><td>2.6E-25</td><td>BT20/MB468</td></tr>
<tr><td>>hg38_21_+_8433172_8446622_RNA45SN1_45S_50ntFlanks@12995.13040.46|CTCGCTGCGATCTATTGAAAGTCAGCCCTCGACACAAGGGTTTGTC</td><td>>hg38_21_+_8433172_8446622_RNA45SN1_45S_50ntFlanks</td><td>12995</td><td>13040</td><td>46</td><td>CTCGCTGCGATCTATTGAAAGTCAGCCCTCGACACAAGGGTTTGTC</td><td>12923.2</td><td>14.4</td><td>1.4</td><td>10.6</td><td>2.8E-26</td><td>2.6E-25</td><td>MB231/MB468</td></tr>
<tr><td>>MI0000286|hsa-mir-210&WithFlank&11|-|568083|568204@72.93.22(+2A)&[MIMAT0000267&hsa-miR-210-3p&offsets|0|0(+2A);m-102&11|-|568112|568133&offsets|0|0(+2A)]|CTGTGCGTGTGACAGCGGCTGAAA</td><td>>MI0000286|hsa-mir-210&WithFlank&11|-|568083|568204</td><td>72</td><td>93</td><td>22(+2A)</td><td>CTGTGCGTGTGACAGCGGCTGAAA</td><td>1849.4</td><td>7.4</td><td>0.7</td><td>10.6</td><td>2.8E-26</td><td>2.7E-25</td><td>MB231/MB468</td></tr>
<tr><td>>hg38_21_+_8433172_8446622_RNA45SN1_45S_50ntFlanks@11999.12032.34|CCCTGCGGGCCGCCGGTGAAATACCACTACTCTG</td><td>>hg38_21_+_8433172_8446622_RNA45SN1_45S_50ntFlanks</td><td>11999</td><td>12032</td><td>34</td><td>CCCTGCGGGCCGCCGGTGAAATACCACTACTCTG</td><td>4051.8</td><td>-9.8</td><td>0.9</td><td>-10.6</td><td>2.8E-26</td><td>2.7E-25</td><td>MB231/MB468</td></tr>
<tr><td>>hg38_1_-_228634819_228635039_RNA5S12_5S_50ntFlanks@146.171.26|TGGGAATACCGGGTGCTGTAGGCTTT</td><td>>hg38_1_-_228634819_228635039_RNA5S12_5S_50ntFlanks</td><td>146</td><td>171</td><td>26</td><td>TGGGAATACCGGGTGCTGTAGGCTTT</td><td>2352.4</td><td>8.2</td><td>0.8</td><td>10.6</td><td>2.9E-26</td><td>2.7E-25</td><td>MB231/MB468</td></tr>
<tr><td>>trnaMT_ValTAC_MT_+_1602_1670@49.70.22|AACTTAACTTGACCGCTCTGAC</td><td>>trnaMT_ValTAC_MT_+_1602_1670</td><td>49</td><td>70</td><td>22</td><td>AACTTAACTTGACCGCTCTGAC</td><td>276.0</td><td>12.8</td><td>1.2</td><td>10.6</td><td>2.4E-26</td><td>2.7E-25</td><td>BT20/MB468</td></tr>
</table>

| | | | | | | | | | | | | |
|---|---|---|---|---|---|---|---|---|---|---|---|---|
| >MI0000434\|hsa-let-7i&WithFlank&12\|+\|62603680\|62603775@12.31.20(+1C)&[MIMAT0000415&hsa-let-7i-5p&offsets\|0\|-2(+1C);m-49&12\|+\|62603691\|62603712&offsets\|0\|-2(+1C)]\|TGAGGTAGTAGTTTGTGCTGC | >MI0000434\|hsa-let-7i&WithFlank&12\|+\|62603680\|62603775 | 12 | 31 | 20(+1C) | TGAGGTAGTAGTTTGTGCTGC | 3450.5 | 8.9 | 0.8 | 10.6 | 3.0E-26 | 2.9E-25 | MB231/MB468 |
| >MI0000093\|hsa-mir-92a-1&WithFlank&13\|+\|91351308\|91351397@54.74.21;MI0000094\|hsa-mir-92a-2&WithFlank&X\|-\|134169532\|134169618@54.74.21&[MIMAT0000092&hsa-miR-92a-3p&offsets\|0\|-1;m-9&13\|+\|91351361\|91351382&offsets\|0\|-1];[MIMAT0000092_1&hsa-miR-92a-3p&offsets\|0\|-1;m-11&X\|-\|134169544\|134169565&offsets\|0\|-1]\|TATTGCACTTGTCCCGGCCTG | >MI0000093\|hsa-mir-92a-1&WithFlank&13\|+\|91351308\|91351397 | 54 | 74 | 21 | TATTGCACTTGTCCCGGCCTG | 3462.3 | 7.8 | 0.7 | 10.6 | 3.1E-26 | 2.9E-25 | MB231/MB468 |
| >hg38_21_+_8433172_8446622_RNA45SN1_45S_50ntFlanks@12655.12674.20\|ATGACTGAACGCCTCTAAGT | >hg38_21_+_8433172_8446622_RNA45SN1_45S_50ntFlanks | 12655 | 12674 | 20 | ATGACTGAACGCCTCTAAGT | 4170.5 | -5.8 | 0.5 | -10.6 | 2.5E-26 | 2.9E-25 | BT20/MB468 |
| >hg38_21_+_8433172_8446622_RNA45SN1_45S_50ntFlanks@11768.11792.25\|CTAATTAGTGACGCGCATGAATGGA | >hg38_21_+_8433172_8446622_RNA45SN1_45S_50ntFlanks | 11768 | 11792 | 25 | CTAATTAGTGACGCGCATGAATGGA | 1840.7 | -9.6 | 0.9 | -10.6 | 2.6E-26 | 2.9E-25 | BT20/MB468 |
| >MI0000102\|hsa-mir-100&WithFlank&11\|-\|122152223\|122152314@21.40.20&[MIMAT0000098&hsa-miR-100-5p&offsets\|+2\|0;m-27&11\|-\|122152276\|122152296&offsets\|+2\|+1]\|CCCGTAGATCCGAACTTGTG | >MI0000102\|hsa-mir-100&WithFlank&11\|-\|122152223\|122152314 | 21 | 40 | 20 | CCCGTAGATCCGAACTTGTG | 709.0 | 14.4 | 1.4 | 10.6 | 3.1E-26 | 2.9E-25 | MB231/MB468 |
| >am_MI0000809\|hsa-mir-151a&WithFlank&8\|-\|140732558\|140732659@53.73.21(+3U)&[MIMAT0000757&hsa-miR-151a-3p&offsets\|0\|0(+3U);m-30&8\|-\|140732587\|140732607&offsets\|0\|0(+3U)]\|CTAGACTGAAGCTCCTTGAGGTTT | >am_MI0000809\|hsa-mir-151a&WithFlank&8\|-\|140732558\|140732659 | 53 | 73 | 21(+3U) | CTAGACTGAAGCTCCTTGAGGTTT | 1810.8 | 10.7 | 1.0 | 10.6 | 3.2E-26 | 3.0E-25 | MB231/MB468 |
| >hg38_21_+_8433172_8446622_RNA45SN1_45S_50ntFlanks@12416.12438.23\|TTCGATGTCGGCTCTTCCTATCA | >hg38_21_+_8433172_8446622_RNA45SN1_45S_50ntFlanks | 12416 | 12438 | 23 | TTCGATGTCGGCTCTTCCTATCA | 2898.0 | 8.0 | 0.8 | 10.6 | 3.3E-26 | 3.1E-25 | MB231/MB468 |
| >MI0001445\|hsa-mir-423&WithFlank&17\|+\|30117073\|30117178@59.78.20&[MIMAT0001340&hsa-miR-423-3p&offsets\|0\|-3;m-100&17\|+\|30117131\|30117153&offsets\|0\|-3]\|AGCTCGGTCTGAGGCCCCTC | >MI0001445\|hsa-mir-423&WithFlank&17\|+\|30117073\|30117178 | 59 | 78 | 20 | AGCTCGGTCTGAGGCCCCTC | 9321.5 | 8.5 | 0.8 | 10.6 | 3.3E-26 | 3.1E-25 | MB231/MB468 |
| >hg38_21_+_8433172_8446622_RNA45SN1_45S_50ntFlanks@476.504.29\|TTCGTGATCGATGTGGTGACGTCGTGCTC | >hg38_21_+_8433172_8446622_RNA45SN1_45S_50ntFlanks | 476 | 504 | 29 | TTCGTGATCGATGTGGTGACGTCGTGCTC | 6317.6 | 7.4 | 0.7 | 10.6 | 2.8E-26 | 3.1E-25 | BT20/MB468 |
| >hg38_21_+_8433172_8446622_RNA45SN1_45S_50ntFlanks@10408.10429.22\|GGTCAGTCGGTCCTGAGAGATG | >hg38_21_+_8433172_8446622_RNA45SN1_45S_50ntFlanks | 10408 | 10429 | 22 | GGTCAGTCGGTCCTGAGAGATG | 2389.7 | -7.5 | 0.7 | -10.6 | 2.9E-26 | 3.2E-25 | BT20/MB468 |
| >hg38_21_+_8433172_8446622_RNA45SN1_45S_50ntFlanks@5057.5081.25\|AGCGATTTGTCTGGTTAATTCCGAT | >hg38_21_+_8433172_8446622_RNA45SN1_45S_50ntFlanks | 5057 | 5081 | 25 | AGCGATTTGTCTGGTTAATTCCGAT | 558.0 | 5.6 | 0.5 | 10.6 | 2.9E-26 | 3.2E-25 | BT20/MB468 |
| >MI0000542\|hsa-mir-320a&WithFlank&8\|-\|22244960\|22245043@48.70.23&[MIMAT0000510&hsa-miR-320a-3p&offsets\|0\|+1;m-60&8\|-\|22244975\|22244996&offsets\|0\|+1]\|AAAAGCTGGGTTGAGAGGGCGAA | >MI0000542\|hsa-mir-320a&WithFlank&8\|-\|22244960\|22245043 | 48 | 70 | 23 | AAAAGCTGGGTTGAGAGGGCGAA | 5107.2 | 8.1 | 0.8 | 10.6 | 3.5E-26 | 3.3E-25 | MB231/MB468 |
| >trnaMT_GluTTC_MT_-_14674_14742@45.72.28,trnalookalike8_GluTTC_5_-_93905172_93905240@45.72.28\|CGTGGTTGTAGTCCGTGCGAGAATACCA | >trnaMT_GluTTC_MT_-_14674_14742 | 45 | 72 | 28 | CGTGGTTGTAGTCCGTGCGAGAATACCA | 662.4 | 7.0 | 0.7 | 10.6 | 3.0E-26 | 3.3E-25 | BT20/MB468 |
| >MI0000465\|hsa-mir-191&WithFlank&3\|-\|49020612\|49020715@22.44.23(+1A)&[MIMAT0000440&hsa-miR-191-5p&offsets\|0\|0(+1A);m-48&3\|-\|49020672\|49020694&offsets\|0\|0(+1A)]\|CAACGGAATCCCAAAAGCAGCTGA | >MI0000465\|hsa-mir-191&WithFlank&3\|-\|49020612\|49020715 | 22 | 44 | 23(+1A) | CAACGGAATCCCAAAAGCAGCTGA | 1343.1 | 7.6 | 0.7 | 10.6 | 3.6E-26 | 3.4E-25 | MB231/MB468 |
| >MI0000478\|hsa-mir-149&WithFlank&2\|+\|240455995\|240456095@21.42.22&[MIMAT0000450&hsa-miR-149-5p&offsets\|0\|-1;m-195&2\|+\|240456015\|240456037&offsets\|0\|-1]\|TCTGGCTCCGTGTCTTCACTCC | >MI0000478\|hsa-mir-149&WithFlank&2\|+\|240455995\|240456095 | 21 | 42 | 22 | TCTGGCTCCGTGTCTTCACTCC | 2719.1 | 13.6 | 1.3 | 10.6 | 3.1E-26 | 3.4E-25 | BT20/MB468 |
| >MI0000102\|hsa-mir-100&WithFlank&11\|-\|122152223\|122152314@20.39.20(+1A)&[MIMAT0000098&hsa-miR-100-5p&offsets\|+1\|-1(+1A);m-27&11\|-\|122152276\|122152296&offsets\|+1\|0(+1A)]\|ACCCGTAGATCCGAACTTGTA | >MI0000102\|hsa-mir-100&WithFlank&11\|-\|122152223\|122152314 | 20 | 39 | 20(+1A) | ACCCGTAGATCCGAACTTGTA | 644.8 | 14.2 | 1.3 | 10.6 | 3.7E-26 | 3.5E-25 | MB231/MB468 |
| >hg38_21_+_8433172_8446622_RNA45SN1_45S_50ntFlanks@7973.8005.33\|GACGCGACCTCAGATCAGACGTGGCGACCCGCT | >hg38_21_+_8433172_8446622_RNA45SN1_45S_50ntFlanks | 7973 | 8005 | 33 | GACGCGACCTCAGATCAGACGTGGCGACCCGCT | 1054.5 | 6.1 | 0.6 | 10.6 | 3.1E-26 | 3.5E-25 | BT20/MB468 |
| >hg38_21_+_8433172_8446622_RNA45SN1_45S_50ntFlanks@10846.10865.20\|CTTCGGGATAAGGATTGGCT | >hg38_21_+_8433172_8446622_RNA45SN1_45S_50ntFlanks | 10846 | 10865 | 20 | CTTCGGGATAAGGATTGGCT | 2218.5 | -6.5 | 0.6 | -10.6 | 3.9E-26 | 3.7E-25 | MB231/MB468 |
| >MI0000542\|hsa-mir-320a&WithFlank&8\|-\|22244960\|22245043@48.70.23(+1C)&[MIMAT0000510&hsa-miR-320a-3p&offsets\|0\|+1(+1C);m-60&8\|-\|22244975\|22244996&offsets\|0\|+1(+1C)]\|AAAAGCTGGGTTGAGAGGGCGAAC | >MI0000542\|hsa-mir-320a&WithFlank&8\|-\|22244960\|22245043 | 48 | 70 | 23(+1C) | AAAAGCTGGGTTGAGAGGGCGAAC | 703.4 | 8.3 | 0.8 | 10.6 | 3.9E-26 | 3.7E-25 | MB231/MB468 |
| >MI0000650\|hsa-mir-200c&WithFlank&12\|+\|6963693\|6963772@50.71.22(+1G)&[MIMAT0000617&hsa-miR-200c-3p&offsets\|0\|-1(+1G);m-54&12\|+\|6963742\|6963764&offsets\|0\|-1(+1G)]\|TAATACTGCCGGGTAATGATGGG | >MI0000650\|hsa-mir-200c&WithFlank&12\|+\|6963693\|6963772 | 50 | 71 | 22(+1G) | TAATACTGCCGGGTAATGATGGG | 23431.2 | 8.7 | 0.8 | 10.6 | 3.4E-26 | 3.7E-25 | BT20/MB468 |
| >MI0000734\|hsa-mir-106b&WithFlank&7\|-\|100093987\|100094080@58.79.22&[MIMAT0004672&hsa-miR-106b-3p&offsets\|0\|0;m-87&7\|-\|100094003\|100094023&offsets\|0\|+1]\|CCGCACTGTGGGTACTTGCTGC | >MI0000734\|hsa-mir-106b&WithFlank&7\|-\|100093987\|100094080 | 58 | 79 | 22 | CCGCACTGTGGGTACTTGCTGC | 8465.6 | 6.5 | 0.6 | 10.6 | 3.3E-26 | 3.7E-25 | BT20/MB468 |
| >MI0000102\|hsa-mir-100&WithFlank&11\|-\|122152223\|122152314@19.38.20&[MIMAT0000098&hsa-miR-100-5p&offsets\|0\|-2;m-27&11\|-\|122152276\|122152296&offsets\|0\|-1]\|AACCCGTAGATCCGAACTTG | >MI0000102\|hsa-mir-100&WithFlank&11\|-\|122152223\|122152314 | 19 | 38 | 20 | AACCCGTAGATCCGAACTTG | 79222.1 | 15.0 | 1.4 | 10.6 | 4.1E-26 | 3.8E-25 | MB231/MB468 |
| >MI0000477\|hsa-mir-146a&WithFlank&5\|+\|160485346\|160485456@27.48.22(+1A)&[MIMAT0000449&hsa-miR-146a-5p&offsets\|0\|0(+1A);m-129&5\|+\|160485372\|160485393&offsets\|0\|0(+1A)]\|TGAGAACTGAATTCCATGGGTTA | >MI0000477\|hsa-mir-146a&WithFlank&5\|+\|160485346\|160485456 | 27 | 48 | 22(+1A) | TGAGAACTGAATTCCATGGGTTA | 1092.5 | 15.0 | 1.4 | 10.6 | 4.1E-26 | 3.8E-25 | MB231/MB468 |
| >hg38_21_+_8433172_8446622_RNA45SN1_45S_50ntFlanks@476.508.33\|TTCGTGATCGATGTGGTGACGTCGTGCTCTCCC | >hg38_21_+_8433172_8446622_RNA45SN1_45S_50ntFlanks | 476 | 508 | 33 | TTCGTGATCGATGTGGTGACGTCGTGCTCTCCC | 1364.3 | 7.9 | 0.8 | 10.6 | 4.5E-26 | 4.2E-25 | MB231/MB468 |
| >hg38_MT_+_598_1651_RNR1_12S_50ntFlanks@368.404.37\|CAATAAAGCTAAAACTCACCTGAGTTGTAAAAAACTC | >hg38_MT_+_598_1651_RNR1_12S_50ntFlanks | 368 | 404 | 37 | CAATAAAGCTAAAACTCACCTGAGTTGTAAAAAACTC | 839.8 | 11.8 | 1.1 | 10.6 | 4.5E-26 | 4.2E-25 | MB231/MB468 |
| >MI0000542\|hsa-mir-320a&WithFlank&8\|-\|22244960\|22245043@48.69.22(+1U)&[MIMAT0000510&hsa-miR-320a-3p&offsets\|0\|0(+1U);m-60&8\|-\|22244975\|22244996&offsets\|0\|0(+1U)]\|AAAAGCTGGGTTGAGAGGGCGAT | >MI0000542\|hsa-mir-320a&WithFlank&8\|-\|22244960\|22245043 | 48 | 69 | 22(+1U) | AAAAGCTGGGTTGAGAGGGCGAT | 8961.2 | 9.0 | 0.9 | 10.6 | 4.6E-26 | 4.2E-25 | MB231/MB468 |
| >MI0000440\|hsa-mir-27b&WithFlank&9\|+\|95085439\|95085547@67.86.20(+1U)&[MIMAT0000419&hsa-miR-27b-3p&offsets\|0\|-1(+1U);m-39&9\|+\|95085505\|95085525&offsets\|0\|-1(+1U)]\|TTCACAGTGGCTAAGTTCTGT | >MI0000440\|hsa-mir-27b&WithFlank&9\|+\|95085439\|95085547 | 67 | 86 | 20(+1U) | TTCACAGTGGCTAAGTTCTGT | 41634.3 | 8.4 | 0.8 | 10.6 | 4.7E-26 | 4.3E-25 | MB231/MB468 |
| >hg38_21_+_8433172_8446622_RNA45SN1_45S_50ntFlanks@4730.4768.39\|CAAGAACGAAAGTCGGAGGTTCGAAGACGATCAGATACC | >hg38_21_+_8433172_8446622_RNA45SN1_45S_50ntFlanks | 4730 | 4768 | 39 | CAAGAACGAAAGTCGGAGGTTCGAAGACGATCAGATACC | 438.9 | 13.7 | 1.3 | 10.6 | 4.7E-26 | 4.4E-25 | MB231/MB468 |
| >hg38_21_+_8433172_8446622_RNA45SN1_45S_50ntFlanks@8194.8214.21\|CCCGTGGACGGTGTGAGGCCG | >hg38_21_+_8433172_8446622_RNA45SN1_45S_50ntFlanks | 8194 | 8214 | 21 | CCCGTGGACGGTGTGAGGCCG | 614.5 | -6.3 | 0.6 | -10.6 | 4.0E-26 | 4.4E-25 | BT20/MB468 |
| >hg38_21_+_8433172_8446622_RNA45SN1_45S_50ntFlanks@476.498.23\|TTCGTGATCGATGTGGTGACGTC | >hg38_21_+_8433172_8446622_RNA45SN1_45S_50ntFlanks | 476 | 498 | 23 | TTCGTGATCGATGTGGTGACGTC | 8750.9 | 8.6 | 0.8 | 10.6 | 4.9E-26 | 4.5E-25 | MB231/MB468 |
| >hg38_21_+_8433172_8446622_RNA45SN1_45S_50ntFlanks@475.509.35\|CTTCGTGATCGATGTGGTGACGTCGTGCTCTCCCG | >hg38_21_+_8433172_8446622_RNA45SN1_45S_50ntFlanks | 475 | 509 | 35 | CTTCGTGATCGATGTGGTGACGTCGTGCTCTCCCG | 8656.1 | 7.7 | 0.7 | 10.6 | 5.0E-26 | 4.5E-25 | MB231/MB468 |
| >trnaMT_PheGAA_MT_+_577_647@35.59.25\|GAAAATGTTTAGACGGGCTCACATC | >trnaMT_PheGAA_MT_+_577_647 | 35 | 59 | 25 | GAAAATGTTTAGACGGGCTCACATC | 563.4 | 9.6 | 0.9 | 10.6 | 4.9E-26 | 4.5E-25 | MB231/MB468 |
| >MI0000434\|hsa-let-7i&WithFlank&12\|+\|62603680\|62603775@13.33.21&[MIMAT0000415&hsa-let-7i-5p&offsets\|+1\|0;m-49&12\|+\|62603691\|62603712&offsets\|+1\|0]\|GAGGTAGTAGTTTGTGCTGTT | >MI0000434\|hsa-let-7i&WithFlank&12\|+\|62603680\|62603775 | 13 | 33 | 21 | GAGGTAGTAGTTTGTGCTGTT | 625.5 | 14.2 | 1.3 | 10.6 | 5.0E-26 | 4.6E-25 | MB231/MB468 |
| >MI0000114\|hsa-mir-107&WithFlank&10\|-\|89592741\|89592833@56.76.21(+1A);MI0000108\|hsa-mir-103a-2&WithFlank&20\|+\|3917488\|3917577@54.74.21(+1A);MI0000109\|hsa-mir-103a-1&WithFlank&5\|-\|168560890\|168560979@54.74.21(+1A)&[MIMAT0000104&hsa-miR-107&offsets\|0\|-2(+1A);m-78&10\|-\|89592758\|89592778&offsets\|0\|0(+1A)];[MIMAT0000101&hsa-miR-103a-3p&offsets\|0\|-2(+1A);m-12&20\|+\|3917541\|3917563&offsets\|0\|-2(+1A)];[MIMAT0000101_1&hsa-miR-103a-3p&offsets\|0\|-2(+1A);m-13&5\|-\|168560904\|168560926&offsets\|0\|-2(+1A);m-44220&5\|-\|168560908\|168560926&offsets\|0\|+2(+1A)]\|AGCAGCATTGTACAGGGCTATA | >MI0000114\|hsa-mir-107&WithFlank&10\|-\|89592741\|89592833 | 56 | 76 | 21(+1A) | AGCAGCATTGTACAGGGCTATA | 1894.4 | 7.4 | 0.7 | 10.5 | 5.1E-26 | 4.7E-25 | MB231/MB468 |
| >hg38_21_+_8433172_8446622_RNA45SN1_45S_50ntFlanks@12616.12654.39\|CTGAGGAGCCAATGGGGCGAAGCTACCATCTGTGGGATT | >hg38_21_+_8433172_8446622_RNA45SN1_45S_50ntFlanks | 12616 | 12654 | 39 | CTGAGGAGCCAATGGGGCGAAGCTACCATCTGTGGGATT | 486.0 | -5.3 | 0.5 | -10.6 | 4.3E-26 | 4.8E-25 | BT20/MB468 |
| >hg38_21_+_8433172_8446622_RNA45SN1_45S_50ntFlanks@485.504.20\|GATGTGGTGACGTCGTGCTC | >hg38_21_+_8433172_8446622_RNA45SN1_45S_50ntFlanks | 485 | 504 | 20 | GATGTGGTGACGTCGTGCTC | 2533.7 | 16.0 | 1.5 | 10.6 | 4.3E-26 | 4.8E-25 | BT20/MB468 |
| >MI0000542\|hsa-mir-320a&WithFlank&8\|-\|22244960\|22245043@48.68.21(+3U)&[MIMAT0000510&hsa-miR-320a-3p&offsets\|0\|-1(+3U);m-60&8\|-\|22244975\|22244996&offsets\|0\|-1(+3U)]\|AAAAGCTGGGTTGAGAGGGCGTTT | >MI0000542\|hsa-mir-320a&WithFlank&8\|-\|22244960\|22245043 | 48 | 68 | 21(+3U) | AAAAGCTGGGTTGAGAGGGCGTTT | 1645.2 | 12.5 | 1.2 | 10.6 | 4.4E-26 | 4.8E-25 | BT20/MB468 |

<table>
<tr><th></th><th></th><th></th><th></th><th></th><th></th><th></th><th></th><th></th><th></th><th></th><th></th><th></th></tr>
<tr><td>>MI0000088|hsa-mir-30a&WithFlank&6|-|71403545|71403627@53.73.21(+1U)&[MIMAT0000088&hsa-miR-30a-3p&offsets|0|-1(+1U);m-38&6|-|71403554|71403575&offsets|0|-1(+1U)]|CTTTCAGTCGGATGTTTGCAGT</td><td>>MI0000088|hsa-mir-30a&WithFlank&6|-|71403545|71403627</td><td>53</td><td>73</td><td>21(+1U)</td><td>CTTTCAGTCGGATGTTTGCAGT</td><td>8428.9</td><td>7.4</td><td>0.7</td><td>10.6</td><td>4.4E-26</td><td>4.9E-25</td><td>BT20/MB468</td></tr>
<tr><td>>am_trna128_GlyGCC_6_-_27870686_27870756@3.31.29,trna133_GlyCCC_1_-_16872434_16872504@3.31.29,trna18_GlyGCC_16_+_70822597_70822667@3.31.29,trna19_GlyGCC_16_+_70823410_70823480@3.31.29,trna19_GlyGCC_2_-_157257659_157257729@3.31.29,trna24_GlyGCC_16_-_70812942_70813012@3.31.29,trna25_GlyGCC_16_-_70812114_70812184@3.31.29,trna4_GlyCCC_1_+_17188416_17188486@3.31.29,trna5_GlyGCC_17_+_8029064_8029134@3.31.29,trna68_GlyGCC_1_-_161493637_161493707@3.31.29|ATTGGTGGTTCAGTGGTAGAATTCTCGCC</td><td>>am_trna128_GlyGCC_6_-_27870686_27870756</td><td>3</td><td>31</td><td>29</td><td>ATTGGTGGTTCAGTGGTAGAATTCTCGCC</td><td>765.6</td><td>14.5</td><td>1.4</td><td>10.5</td><td>5.3E-26</td><td>4.9E-25</td><td>MB231/MB468</td></tr>
<tr><td>>hg38_21_+_8433172_8446622_RNA45SN1_45S_50ntFlanks@11572.11591.20|CCGACTTAGAACTGGTGCGG</td><td>>hg38_21_+_8433172_8446622_RNA45SN1_45S_50ntFlanks</td><td>11572</td><td>11591</td><td>20</td><td>CCGACTTAGAACTGGTGCGG</td><td>7923.3</td><td>-8.1</td><td>0.8</td><td>-10.6</td><td>4.5E-26</td><td>5.0E-25</td><td>BT20/MB468</td></tr>
<tr><td>>MI0000108|hsa-mir-103a-2&WithFlank&20|+|3917488|3917577@54.77.24;MI0000109|hsa-mir-103a-1&WithFlank&5|-|168560890|168560979@54.77.24&[MIMAT0000101&hsa-miR-103a-3p&offsets|0|+1;m-12&20|+|3917541|3917563&offsets|0|+1];[MIMAT0000101_1&hsa-miR-103a-3p&offsets|0|+1;m-13&5|-|168560904|168560926&offsets|0|+1;m-44220&5|-|168560908|168560926&offsets|0|+5]|AGCAGCATTGTACAGGGCTATGAA</td><td>>MI0000108|hsa-mir-103a-2&WithFlank&20|+|3917488|3917577</td><td>54</td><td>77</td><td>24</td><td>AGCAGCATTGTACAGGGCTATGAA</td><td>2759.4</td><td>7.7</td><td>0.7</td><td>10.5</td><td>5.5E-26</td><td>5.0E-25</td><td>MB231/MB468</td></tr>
<tr><td>>hg38_MT_+_1621_3279_RNR2_16S_50ntFlanks@51.74.24|GCTAAACCTAGCCCCAAACCCACT</td><td>>hg38_MT_+_1621_3279_RNR2_16S_50ntFlanks</td><td>51</td><td>74</td><td>24</td><td>GCTAAACCTAGCCCCAAACCCACT</td><td>3115.7</td><td>7.8</td><td>0.7</td><td>10.5</td><td>5.5E-26</td><td>5.0E-25</td><td>MB231/MB468</td></tr>
<tr><td>>MI0000082|hsa-mir-25&WithFlank&7|-|100093554|100093649@58.79.22&[MIMAT0000081&hsa-miR-25-3p&offsets|0|0;m-23&7|-|100093571|100093592&offsets|0|0]|CATTGCACTTGTCTCGGTCTGA</td><td>>MI0000082|hsa-mir-25&WithFlank&7|-|100093554|100093649</td><td>58</td><td>79</td><td>22</td><td>CATTGCACTTGTCTCGGTCTGA</td><td>98566.9</td><td>7.5</td><td>0.7</td><td>10.5</td><td>5.5E-26</td><td>5.0E-25</td><td>MB231/MB468</td></tr>
<tr><td>>MI0000088|hsa-mir-30a&WithFlank&6|-|71403545|71403627@54.73.20(+1A)&[MIMAT0000088&hsa-miR-30a-3p&offsets|+1|-1(+1A);m-38&6|-|71403554|71403575&offsets|+1|-1(+1A)]|TTTCAGTCGGATGTTTGCAGA</td><td>>MI0000088|hsa-mir-30a&WithFlank&6|-|71403545|71403627</td><td>54</td><td>73</td><td>20(+1A)</td><td>TTTCAGTCGGATGTTTGCAGA</td><td>753.2</td><td>14.5</td><td>1.4</td><td>10.5</td><td>5.5E-26</td><td>5.0E-25</td><td>MB231/MB468</td></tr>
<tr><td>>trnaMT_LeuTAA_MT_+_3230_3304@1.30.30|GTTAAGATGGCAGAGCCCGGTAATCGCATA</td><td>>trnaMT_LeuTAA_MT_+_3230_3304</td><td>1</td><td>30</td><td>30</td><td>GTTAAGATGGCAGAGCCCGGTAATCGCATA</td><td>946.7</td><td>7.5</td><td>0.7</td><td>10.5</td><td>5.5E-26</td><td>5.0E-25</td><td>MB231/MB468</td></tr>
<tr><td>>MI0000087|hsa-mir-29a&WithFlank&7|-|130876741|130876816@48.68.21&[MIMAT0000086&hsa-miR-29a-3p&offsets|0|-1;m-26&7|-|130876749|130876769&offsets|0|0]|TAGCACCATCTGAAATCGGTT</td><td>>MI0000087|hsa-mir-29a&WithFlank&7|-|130876741|130876816</td><td>48</td><td>68</td><td>21</td><td>TAGCACCATCTGAAATCGGTT</td><td>28757.8</td><td>9.2</td><td>0.9</td><td>10.6</td><td>4.6E-26</td><td>5.1E-25</td><td>BT20/MB468</td></tr>
<tr><td>>hg38_21_+_8433172_8446622_RNA45SN1_45S_50ntFlanks@3844.3866.23|TACTTGGATAACTGTGGTAATTC</td><td>>hg38_21_+_8433172_8446622_RNA45SN1_45S_50ntFlanks</td><td>3844</td><td>3866</td><td>23</td><td>TACTTGGATAACTGTGGTAATTC</td><td>317.0</td><td>6.5</td><td>0.6</td><td>10.5</td><td>6.1E-26</td><td>5.5E-25</td><td>MB231/MB468</td></tr>
<tr><td>>trnaMT_MetCAT_MT_+_4402_4469@33.71.39|TACCCCGAAAATGTTGGTTATACCCTTCCCGTACTACCA</td><td>>trnaMT_MetCAT_MT_+_4402_4469</td><td>33</td><td>71</td><td>39</td><td>TACCCCGAAAATGTTGGTTATACCCTTCCCGTACTACCA</td><td>636.7</td><td>14.2</td><td>1.4</td><td>10.5</td><td>6.3E-26</td><td>5.7E-25</td><td>MB231/MB468</td></tr>
<tr><td>>MI0000063|hsa-let-7b&WithFlank&22|+|46113680|46113774@12.33.22(+1C)&[MIMAT0000063&hsa-let-7b-5p&offsets|0|0(+1C);m-14&22|+|46113691|46113712&offsets|0|0(+1C)]|TGAGGTAGTAGGTTGTGTGGTTC</td><td>>MI0000063|hsa-let-7b&WithFlank&22|+|46113680|46113774</td><td>12</td><td>33</td><td>22(+1C)</td><td>TGAGGTAGTAGGTTGTGTGGTTC</td><td>662.5</td><td>7.3</td><td>0.7</td><td>10.5</td><td>6.6E-26</td><td>6.0E-25</td><td>MB231/MB468</td></tr>
<tr><td>>trna2_GlyGCC_21_-_18827107_18827177@1.33.33,trna35_GlyGCC_1_+_161413094_161413164@1.33.33,trna37_GlyGCC_1_+_161420467_161420537@1.33.33,trna39_GlyGCC_1_+_161427898_161427968@1.33.33,trna41_GlyGCC_1_+_161435258_161435328@1.33.33|GCATGGGTGGTTCAGTGGTAGAATTCTCGCCTG</td><td>>trna2_GlyGCC_21_-_18827107_18827177</td><td>1</td><td>33</td><td>33</td><td>GCATGGGTGGTTCAGTGGTAGAATTCTCGCCTG</td><td>1933.0</td><td>6.0</td><td>0.6</td><td>10.5</td><td>5.5E-26</td><td>6.0E-25</td><td>BT20/MB468</td></tr>
<tr><td>>hg38_21_+_8433172_8446622_RNA45SN1_45S_50ntFlanks@12419.12451.33|GATGTCGGCTCTTCCTATCATTGTGAAGCAGAA</td><td>>hg38_21_+_8433172_8446622_RNA45SN1_45S_50ntFlanks</td><td>12419</td><td>12451</td><td>33</td><td>GATGTCGGCTCTTCCTATCATTGTGAAGCAGAA</td><td>394.2</td><td>5.2</td><td>0.5</td><td>10.5</td><td>6.7E-26</td><td>6.0E-25</td><td>MB231/MB468</td></tr>
<tr><td>>trnaMT_ValTAC_MT_+_1602_1670@28.72.45|AACTTACACTTAGGAGATTTCAACTTAACTTGACCGCTCTGACCA</td><td>>trnaMT_ValTAC_MT_+_1602_1670</td><td>28</td><td>72</td><td>45</td><td>AACTTACACTTAGGAGATTTCAACTTAACTTGACCGCTCTGACCA</td><td>988.4</td><td>10.9</td><td>1.0</td><td>10.5</td><td>6.9E-26</td><td>6.2E-25</td><td>MB231/MB468</td></tr>
<tr><td>>hg38_21_+_8433172_8446622_RNA45SN1_45S_50ntFlanks@11668.11691.24|TGCCCAGTGCTCTGAATGTCAAAG</td><td>>hg38_21_+_8433172_8446622_RNA45SN1_45S_50ntFlanks</td><td>11668</td><td>11691</td><td>24</td><td>TGCCCAGTGCTCTGAATGTCAAAG</td><td>686.3</td><td>6.3</td><td>0.6</td><td>10.5</td><td>6.9E-26</td><td>6.2E-25</td><td>MB231/MB468</td></tr>
<tr><td>>MI0016746|hsa-mir-548o-2&WithFlank&20|+|38516557|38516638@51.72.22;MI0006402|hsa-mir-548o&WithFlank&7|-|102405736|102405861@93.114.22&[MIMAT0005919&hsa-miR-548o-3p&offsets|0|0;m-431&20|+|38516607|38516628&offsets|0|0];[MIMAT0005919_1&hsa-miR-548o-3p&offsets|0|0;m-430&7|-|102405748|102405769&offsets|0|0]|CCAAAACTGCAGTTACTTTTGC</td><td>>MI0016746|hsa-mir-548o-2&WithFlank&20|+|38516557|38516638</td><td>51</td><td>72</td><td>22</td><td>CCAAAACTGCAGTTACTTTTGC</td><td>951.1</td><td>7.5</td><td>0.7</td><td>10.5</td><td>7.0E-26</td><td>6.3E-25</td><td>MB231/MB468</td></tr>
<tr><td>>MI0000650|hsa-mir-200c&WithFlank&12|+|6963693|6963772@50.70.21(+1A)&[MIMAT0000617&hsa-miR-200c-3p&offsets|0|-2(+1A);m-54&12|+|6963742|6963764&offsets|0|-2(+1A)]|TAATACTGCCGGGTAATGATGA</td><td>>MI0000650|hsa-mir-200c&WithFlank&12|+|6963693|6963772</td><td>50</td><td>70</td><td>21(+1A)</td><td>TAATACTGCCGGGTAATGATGA</td><td>4708.2</td><td>13.7</td><td>1.3</td><td>10.8</td><td>3.2E-27</td><td>6.5E-25</td><td>BT20/MB231</td></tr>
<tr><td>>MI0000465|hsa-mir-191&WithFlank&3|-|49020612|49020715@22.45.24&[MIMAT0000440&hsa-miR-191-5p&offsets|0|+1;m-48&3|-|49020672|49020694&offsets|0|+1]|CAACGGAATCCCAAAAGCAGCTGT</td><td>>MI0000465|hsa-mir-191&WithFlank&3|-|49020612|49020715</td><td>22</td><td>45</td><td>24</td><td>CAACGGAATCCCAAAAGCAGCTGT</td><td>12611.2</td><td>8.2</td><td>0.8</td><td>10.5</td><td>7.3E-26</td><td>6.6E-25</td><td>MB231/MB468</td></tr>
<tr><td>>hg38_21_+_8433172_8446622_RNA45SN1_45S_50ntFlanks@11668.11687.20|TGCCCAGTGCTCTGAATGTC</td><td>>hg38_21_+_8433172_8446622_RNA45SN1_45S_50ntFlanks</td><td>11668</td><td>11687</td><td>20</td><td>TGCCCAGTGCTCTGAATGTC</td><td>717.3</td><td>5.0</td><td>0.5</td><td>10.5</td><td>6.0E-26</td><td>6.6E-25</td><td>BT20/MB468</td></tr>
<tr><td>>hg38_21_+_8433172_8446622_RNA45SN1_45S_50ntFlanks@12405.12441.37|CTTTTTGATCCTTCGATGTCGGCTCTTCCTATCATTG</td><td>>hg38_21_+_8433172_8446622_RNA45SN1_45S_50ntFlanks</td><td>12405</td><td>12441</td><td>37</td><td>CTTTTTGATCCTTCGATGTCGGCTCTTCCTATCATTG</td><td>1108.2</td><td>-5.6</td><td>0.5</td><td>-10.5</td><td>6.0E-26</td><td>6.6E-25</td><td>BT20/MB468</td></tr>
<tr><td>>MI0003778|hsa-mir-320c-1&WithFlank&18|+|21683512|21683595@48.67.20;MI0008191|hsa-mir-320c-2&WithFlank&18|+|24321669|24321752@48.67.20&[MIMAT0005793&hsa-miR-320c&offsets|0|0;m-222&18|+|21683559|21683578&offsets|0|0];[MIMAT0005793_1&hsa-miR-320c&offsets|0|0;m-219&18|+|24321716|24321735&offsets|0|0]|AAAAGCTGGGTTGAGAGGGT</td><td>>MI0003778|hsa-mir-320c-1&WithFlank&18|+|21683512|21683595</td><td>48</td><td>67</td><td>20</td><td>AAAAGCTGGGTTGAGAGGGT</td><td>1297.6</td><td>10.2</td><td>1.0</td><td>10.5</td><td>6.1E-26</td><td>6.7E-25</td><td>BT20/MB468</td></tr>
<tr><td>>MI0000082|hsa-mir-25&WithFlank&7|-|100093554|100093649@58.78.21&[MIMAT0000081&hsa-miR-25-3p&offsets|0|-1;m-23&7|-|100093571|100093592&offsets|0|-1]|CATTGCACTTGTCTCGGTCTG</td><td>>MI0000082|hsa-mir-25&WithFlank&7|-|100093554|100093649</td><td>58</td><td>78</td><td>21</td><td>CATTGCACTTGTCTCGGTCTG</td><td>9165.9</td><td>6.7</td><td>0.6</td><td>10.5</td><td>6.2E-26</td><td>6.7E-25</td><td>BT20/MB468</td></tr>
<tr><td>>hg38_21_+_8433172_8446622_RNA45SN1_45S_50ntFlanks@7975.7997.23|CGCGACCTCAGATCAGACGTGGC</td><td>>hg38_21_+_8433172_8446622_RNA45SN1_45S_50ntFlanks</td><td>7975</td><td>7997</td><td>23</td><td>CGCGACCTCAGATCAGACGTGGC</td><td>16056.3</td><td>6.7</td><td>0.6</td><td>10.5</td><td>6.3E-26</td><td>6.9E-25</td><td>BT20/MB468</td></tr>
<tr><td>>hg38_21_+_8433172_8446622_RNA45SN1_45S_50ntFlanks@12416.12439.24|TTCGATGTCGGCTCTTCCTATCAT</td><td>>hg38_21_+_8433172_8446622_RNA45SN1_45S_50ntFlanks</td><td>12416</td><td>12439</td><td>24</td><td>TTCGATGTCGGCTCTTCCTATCAT</td><td>1828.7</td><td>7.4</td><td>0.7</td><td>10.5</td><td>7.8E-26</td><td>7.0E-25</td><td>MB231/MB468</td></tr>
<tr><td>>hg38_21_+_8433172_8446622_RNA45SN1_45S_50ntFlanks@10846.10865.20|CTTCGGGATAAGGATTGGCT</td><td>>hg38_21_+_8433172_8446622_RNA45SN1_45S_50ntFlanks</td><td>10846</td><td>10865</td><td>20</td><td>CTTCGGGATAAGGATTGGCT</td><td>2692.9</td><td>-8.1</td><td>0.8</td><td>-10.5</td><td>6.5E-26</td><td>7.1E-25</td><td>BT20/MB468</td></tr>
<tr><td>>MI0000086|hsa-mir-28&WithFlank&3|+|188688775|188688872@60.81.22(+1A)&[MIMAT0004502&hsa-miR-28-3p&offsets|0|0(+1A);m-28&3|+|188688834|188688855&offsets|0|0(+1A)]|CACTAGATTGTGAGCTCCTGGAA</td><td>>MI0000086|hsa-mir-28&WithFlank&3|+|188688775|188688872</td><td>60</td><td>81</td><td>22(+1A)</td><td>CACTAGATTGTGAGCTCCTGGAA</td><td>2021.6</td><td>10.5</td><td>1.0</td><td>10.5</td><td>8.3E-26</td><td>7.4E-25</td><td>MB231/MB468</td></tr>
<tr><td>>MI0000077|hsa-mir-21&WithFlank&17|+|59841260|59841343@14.35.22(+1G)&[MIMAT0000076&hsa-miR-21-5p&offsets|0|0(+1G);m-1&17|+|59841273|59841294&offsets|0|0(+1G)]|TAGCTTATCAGACTGATGTTGAG</td><td>>MI0000077|hsa-mir-21&WithFlank&17|+|59841260|59841343</td><td>14</td><td>35</td><td>22(+1G)</td><td>TAGCTTATCAGACTGATGTTGAG</td><td>13208.4</td><td>8.8</td><td>0.8</td><td>10.5</td><td>8.3E-26</td><td>7.4E-25</td><td>MB231/MB468</td></tr>
<tr><td>>MI0000446|hsa-mir-125b-1&WithFlank&11|-|122099751|122099850@21.41.21(+1U);MI0000470|hsa-mir-125b-2&WithFlank&21|+|16590231|16590331@23.43.21(+1U)&[MIMAT0000423&hsa-miR-125b-5p&offsets|0|-1(+1U);m-67&11|-|122099809|122099830&offsets|0|-1(+1U)];[MIMAT0000423_1&hsa-miR-125b-5p&offsets|0|-1(+1U);m-68&21|+|16590253|16590274&offsets|0|-1(+1U)]|TCCCTGAGACCCTAACTTGTGT</td><td>>MI0000446|hsa-mir-125b-1&WithFlank&11|-|122099751|122099850</td><td>21</td><td>41</td><td>21(+1U)</td><td>TCCCTGAGACCCTAACTTGTGT</td><td>2861.4</td><td>10.2</td><td>1.0</td><td>10.5</td><td>8.3E-26</td><td>7.4E-25</td><td>MB231/MB468</td></tr>
<tr><td>>am_trna128_GlyGCC_6_-_27870686_27870756@1.34.34,trna18_GlyGCC_16_+_70822597_70822667@1.34.34,trna19_GlyGCC_16_+_70823410_70823480@1.34.34,trna19_GlyGCC_2_-_157257659_157257729@1.34.34,trna24_GlyGCC_16_-_70812942_70813012@1.34.34,trna25_GlyGCC_16_-_70812114_70812184@1.34.34,trna5_GlyGCC_17_+_8029064_8029134@1.34.34,trna68_GlyGCC_1_-_161493637_161493707@1.34.34|GCATTGGTGGTTCAGTGGTAGAATTCTCGCCTGC</td><td>>am_trna128_GlyGCC_6_-_27870686_27870756</td><td>1</td><td>34</td><td>34</td><td>GCATTGGTGGTTCAGTGGTAGAATTCTCGCCTGC</td><td>14891.1</td><td>10.2</td><td>1.0</td><td>10.5</td><td>8.4E-26</td><td>7.5E-25</td><td>MB231/MB468</td></tr>
<tr><td>>hg38_MT_+_1621_3279_RNR2_16S_50ntFlanks@51.72.22|GCTAAACCTAGCCCCAAACCCA</td><td>>hg38_MT_+_1621_3279_RNR2_16S_50ntFlanks</td><td>51</td><td>72</td><td>22</td><td>GCTAAACCTAGCCCCAAACCCA</td><td>1270.0</td><td>8.0</td><td>0.8</td><td>10.5</td><td>8.5E-26</td><td>7.6E-25</td><td>MB231/MB468</td></tr>
<tr><td>>MI0003560|hsa-mir-92b&WithFlank&1|+|155195171|155195278@67.86.20&[MIMAT0003218&hsa-miR-92b-3p&offsets|0|-2;m-115&1|+|155195237|155195258&offsets|0|-2]|TATTGCACTCGTCCCGGCCT</td><td>>MI0003560|hsa-mir-92b&WithFlank&1|+|155195171|155195278</td><td>67</td><td>86</td><td>20</td><td>TATTGCACTCGTCCCGGCCT</td><td>1282.2</td><td>12.7</td><td>1.2</td><td>10.5</td><td>8.5E-26</td><td>7.6E-25</td><td>MB231/MB468</td></tr>
<tr><td>>hg38_21_+_8433172_8446622_RNA45SN1_45S_50ntFlanks@5476.5516.41|CCCTGGCGGAGCGCTGAGAAGACGGTCGAACTTGACTATCT</td><td>>hg38_21_+_8433172_8446622_RNA45SN1_45S_50ntFlanks</td><td>5476</td><td>5516</td><td>41</td><td>CCCTGGCGGAGCGCTGAGAAGACGGTCGAACTTGACTATCT</td><td>617.3</td><td>14.2</td><td>1.3</td><td>10.5</td><td>8.6E-26</td><td>7.6E-25</td><td>MB231/MB468</td></tr>
</table>

| | | | | | | | | | | | | |
|---|---|---|---|---|---|---|---|---|---|---|---|---|
| >MI0001445\|hsa-mir-423&WithFlank&17\|+\|30117073\|30117178@59.80.22(+1G)&[MIMAT0001340&hsa-miR-423-3p&offsets\|0\|-1(+1G);m-100&17\|+\|30117131\|30117153&offsets\|0\|-1(+1G)]\|AGCTCGGTCTGAGGCCCCTCAGG | >MI0001445\|hsa-mir-423&WithFlank&17\|+\|30117073\|30117178 | 59 | 80 | 22(+1G) | AGCTCGGTCTGAGGCCCCTCAGG | 2468.9 | 8.0 | 0.8 | 10.5 | 7.2E-26 | 7.8E-25 | BT20/MB468 |
| >hg38_21_+_8433172_8446622_RNA45SN1_45S_50ntFlanks@11586.11603.18\|GTGCGGACCAGGGGAATC | >hg38_21_+_8433172_8446622_RNA45SN1_45S_50ntFlanks | 11586 | 11603 | 18 | GTGCGGACCAGGGGAATC | 1194.4 | -7.2 | 0.7 | -10.5 | 7.3E-26 | 7.9E-25 | BT20/MB468 |
| >hg38_21_+_8433172_8446622_RNA45SN1_45S_50ntFlanks@6698.6719.22\|AGCTAGCTGCGAGAATTAATGT | >hg38_21_+_8433172_8446622_RNA45SN1_45S_50ntFlanks | 6698 | 6719 | 22 | AGCTAGCTGCGAGAATTAATGT | 566.3 | -4.6 | 0.4 | -10.5 | 7.4E-26 | 8.0E-25 | BT20/MB468 |
| >hg38_21_+_8433172_8446622_RNA45SN1_45S_50ntFlanks@475.506.32\|CTTCGTGATCGATGTGGTGACGTCGTGCTCTC | >hg38_21_+_8433172_8446622_RNA45SN1_45S_50ntFlanks | 475 | 506 | 32 | CTTCGTGATCGATGTGGTGACGTCGTGCTCTC | 41775.1 | 8.2 | 0.8 | 10.5 | 9.1E-26 | 8.1E-25 | MB231/MB468 |
| >trnaMT_PheGAA_MT_+_577_647@1.18.18\|GTTTATGTAGCTTACCTC | >trnaMT_PheGAA_MT_+_577_647 | 1 | 18 | 18 | GTTTATGTAGCTTACCTC | 2677.2 | 8.2 | 0.8 | 10.5 | 9.4E-26 | 8.3E-25 | MB231/MB468 |
| >hg38_21_+_8433172_8446622_RNA45SN1_45S_50ntFlanks@6646.6695.50\|TCGTACGACTCTTAGCGGTGGATCACTCGGCTCGTGCGTCGATGAAGAAC | >hg38_21_+_8433172_8446622_RNA45SN1_45S_50ntFlanks | 6646 | 6695 | 50 | TCGTACGACTCTTAGCGGTGGATCACTCGGCTCGTGCGTCGATGAAGAAC | 2622.9 | 10.3 | 1.0 | 10.5 | 9.5E-26 | 8.4E-25 | MB231/MB468 |
| >hg38_21_+_8433172_8446622_RNA45SN1_45S_50ntFlanks@6740.6771.32\|CATCGACACTTCGAACGCACTTGCGGCCCCGG | >hg38_21_+_8433172_8446622_RNA45SN1_45S_50ntFlanks | 6740 | 6771 | 32 | CATCGACACTTCGAACGCACTTGCGGCCCCGG | 1395.8 | -6.8 | 0.7 | -10.5 | 7.9E-26 | 8.5E-25 | BT20/MB468 |
| >MI0000650\|hsa-mir-200c&WithFlank&12\|+\|6963693\|6963772@50.70.21(+1A)&[MIMAT0000617&hsa-miR-200c-3p&offsets\|0\|-2(+1A);m-54&12\|+\|6963742\|6963764&offsets\|0\|-2(+1A)]\|TAATACTGCCGGGTAATGATGA | >MI0000650\|hsa-mir-200c&WithFlank&12\|+\|6963693\|6963772 | 50 | 70 | 21(+1A) | TAATACTGCCGGGTAATGATGA | 13080.5 | 9.5 | 0.9 | 10.5 | 8.0E-26 | 8.6E-25 | BT20/MB468 |
| >MI0000298\|hsa-mir-221&WithFlank&X\|-\|45746151\|45746272@71.93.23&[MIMAT0000278&hsa-miR-221-3p&offsets\|0\|0;m-91&X\|-\|45746181\|45746202&offsets\|0\|+1]\|AGCTACATTGTCTGCTGGGTTTC | >MI0000298\|hsa-mir-221&WithFlank&X\|-\|45746151\|45746272 | 71 | 93 | 23 | AGCTACATTGTCTGCTGGGTTTC | 121551.6 | 8.0 | 0.8 | 10.5 | 9.9E-26 | 8.7E-25 | MB231/MB468 |
| >hg38_21_+_8433172_8446622_RNA45SN1_45S_50ntFlanks@476.502.27\|TTCGTGATCGATGTGGTGACGTCGTGC | >hg38_21_+_8433172_8446622_RNA45SN1_45S_50ntFlanks | 476 | 502 | 27 | TTCGTGATCGATGTGGTGACGTCGTGC | 3489.0 | 7.9 | 0.8 | 10.5 | 1.0E-25 | 8.9E-25 | MB231/MB468 |
| >MI0000298\|hsa-mir-221&WithFlank&X\|-\|45746151\|45746272@71.93.23(+2U)&[MIMAT0000278&hsa-miR-221-3p&offsets\|0\|0(+2U);m-91&X\|-\|45746181\|45746202&offsets\|0\|+1(+2U)]\|AGCTACATTGTCTGCTGGGTTTCTT | >MI0000298\|hsa-mir-221&WithFlank&X\|-\|45746151\|45746272 | 71 | 93 | 23(+2U) | AGCTACATTGTCTGCTGGGTTTCTT | 861.7 | 8.3 | 0.8 | 10.5 | 1.0E-25 | 8.9E-25 | MB231/MB468 |
| >hg38_21_+_8433172_8446622_RNA45SN1_45S_50ntFlanks@12419.12450.32\|GATGTCGGCTCTTCCTATCATTGTGAAGCAGA | >hg38_21_+_8433172_8446622_RNA45SN1_45S_50ntFlanks | 12419 | 12450 | 32 | GATGTCGGCTCTTCCTATCATTGTGAAGCAGA | 793.5 | 6.3 | 0.6 | 10.5 | 1.0E-25 | 9.0E-25 | MB231/MB468 |
| >MI0000301\|hsa-mir-224&WithFlank&X\|-\|151958572\|151958664@14.36.23(+1A)&[MIMAT0000281&hsa-miR-224-5p&offsets\|+1\|-1(+1A);m-238&X\|-\|151958630\|151958651&offsets\|0\|+1(+1A)]\|CAAGTCACTAGTGGTTCCGTTTAA | >MI0000301\|hsa-mir-224&WithFlank&X\|-\|151958572\|151958664 | 14 | 36 | 23(+1A) | CAAGTCACTAGTGGTTCCGTTTAA | 871.7 | 14.4 | 1.4 | 10.5 | 8.4E-26 | 9.1E-25 | BT20/MB468 |
| >hg38_21_+_8433172_8446622_RNA45SN1_45S_50ntFlanks@12459.12476.18\|AGCGTTGGATTGTTCACC | >hg38_21_+_8433172_8446622_RNA45SN1_45S_50ntFlanks | 12459 | 12476 | 18 | AGCGTTGGATTGTTCACC | 2462.4 | -5.3 | 0.5 | -10.5 | 8.8E-26 | 9.5E-25 | BT20/MB468 |
| >MI0000285\|hsa-mir-205&WithFlank&1\|+\|209432127\|209432248@40.61.22(+1C)&[MIMAT0000266&hsa-miR-205-5p&offsets\|0\|0(+1C);m-351&1\|+\|209432166\|209432187&offsets\|0\|0(+1C)]\|TCCTTCATTCCACCGGAGTCTGC | >MI0000285\|hsa-mir-205&WithFlank&1\|+\|209432127\|209432248 | 40 | 61 | 22(+1C) | TCCTTCATTCCACCGGAGTCTGC | 12555.1 | 6.9 | 0.7 | 10.5 | 9.1E-26 | 9.8E-25 | BT20/MB468 |
| >hg38_21_+_8433172_8446622_RNA45SN1_45S_50ntFlanks@11874.11906.33\|AGAAGACCCTGTTGAGCTTGACTCTAGTCTGGC | >hg38_21_+_8433172_8446622_RNA45SN1_45S_50ntFlanks | 11874 | 11906 | 33 | AGAAGACCCTGTTGAGCTTGACTCTAGTCTGGC | 1273.7 | -6.6 | 0.6 | -10.5 | 9.3E-26 | 9.9E-25 | BT20/MB468 |
| >hg38_21_+_8433172_8446622_RNA45SN1_45S_50ntFlanks@10680.10708.29\|GTTCCGGCGGCGTCCGGTGAGCTCTCGCT | >hg38_21_+_8433172_8446622_RNA45SN1_45S_50ntFlanks | 10680 | 10708 | 29 | GTTCCGGCGGCGTCCGGTGAGCTCTCGCT | 12637.7 | -5.8 | 0.6 | -10.5 | 1.1E-25 | 9.9E-25 | MB231/MB468 |
| >hg38_21_+_8433172_8446622_RNA45SN1_45S_50ntFlanks@6738.6771.34\|ATCATCGACACTTCGAACGCACTTGCGGCCCCGG | >hg38_21_+_8433172_8446622_RNA45SN1_45S_50ntFlanks | 6738 | 6771 | 34 | ATCATCGACACTTCGAACGCACTTGCGGCCCCGG | 2568.6 | -13.3 | 1.3 | -10.5 | 9.5E-26 | 1.0E-24 | BT20/MB468 |
| >hg38_21_+_8433172_8446622_RNA45SN1_45S_50ntFlanks@12438.12455.18\|ATTGTGAAGCAGAATTCA | >hg38_21_+_8433172_8446622_RNA45SN1_45S_50ntFlanks | 12438 | 12455 | 18 | ATTGTGAAGCAGAATTCA | 4333.8 | -6.9 | 0.7 | -10.5 | 1.2E-25 | 1.0E-24 | MB231/MB468 |
| >trna32_LysCTT_16_-_3207406_3207478@1.33.33\|GCCCGGCTAGCTCAGTCGGTAGAGCATGAGACC | >trna32_LysCTT_16_-_3207406_3207478 | 1 | 33 | 33 | GCCCGGCTAGCTCAGTCGGTAGAGCATGAGACC | 8416.2 | 10.3 | 1.0 | 10.5 | 1.2E-25 | 1.0E-24 | MB231/MB468 |
| >MI0000478\|hsa-mir-149&WithFlank&2\|+\|240455995\|240456095@21.41.21(+1U)&[MIMAT0000450&hsa-miR-149-5p&offsets\|0\|-2(+1U);m-195&2\|+\|240456015\|240456037&offsets\|0\|-2(+1U)]\|TCTGGCTCCGTGTCTTCACTCT | >MI0000478\|hsa-mir-149&WithFlank&2\|+\|240455995\|240456095 | 21 | 41 | 21(+1U) | TCTGGCTCCGTGTCTTCACTCT | 1709.9 | 15.4 | 1.5 | 10.5 | 9.9E-26 | 1.1E-24 | BT20/MB468 |
| >hg38_21_+_8433172_8446622_RNA45SN1_45S_50ntFlanks@11564.11585.22\|CCTAGCAGCCGACTTAGAACTG | >hg38_21_+_8433172_8446622_RNA45SN1_45S_50ntFlanks | 11564 | 11585 | 22 | CCTAGCAGCCGACTTAGAACTG | 1577.9 | -5.1 | 0.5 | -10.5 | 1.0E-25 | 1.1E-24 | BT20/MB468 |
| >MI0000469\|hsa-mir-125a&WithFlank&19\|+\|51693248\|51693345@21.42.22(+1A)&[MIMAT0000443&hsa-miR-125a-5p&offsets\|0\|-2(+1A);m-63&19\|+\|51693268\|51693289&offsets\|0\|0(+1A)]\|TCCCTGAGACCCTTTAACCTGTA | >MI0000469\|hsa-mir-125a&WithFlank&19\|+\|51693248\|51693345 | 21 | 42 | 22(+1A) | TCCCTGAGACCCTTTAACCTGTA | 9097.5 | 8.0 | 0.8 | 10.5 | 1.0E-25 | 1.1E-24 | BT20/MB468 |
| >am_MI0000809\|hsa-mir-151a&WithFlank&8\|-\|140732558\|140732659@53.72.20(+2U)&[MIMAT0000757&hsa-miR-151a-3p&offsets\|0\|-1(+2U);m-30&8\|-\|140732587\|140732607&offsets\|0\|-1(+2U)]\|CTAGACTGAAGCTCCTTGAGTT | >am_MI0000809\|hsa-mir-151a&WithFlank&8\|-\|140732558\|140732659 | 53 | 72 | 20(+2U) | CTAGACTGAAGCTCCTTGAGTT | 602.5 | 13.9 | 1.3 | 10.5 | 1.0E-25 | 1.1E-24 | BT20/MB468 |
| >MI0000273\|hsa-mir-183&WithFlank&7\|-\|129774899\|129775020@33.55.23&[MIMAT0000261&hsa-miR-183-5p&offsets\|0\|+1;m-41&7\|-\|129774966\|129774987&offsets\|-1\|0]\|TATGGCACTGGTAGAATTCACTG | >MI0000273\|hsa-mir-183&WithFlank&7\|-\|129774899\|129775020 | 33 | 55 | 23 | TATGGCACTGGTAGAATTCACTG | 10370.3 | 7.4 | 0.7 | 10.5 | 1.1E-25 | 1.1E-24 | BT20/MB468 |
| >hg38_21_+_8433172_8446622_RNA45SN1_45S_50ntFlanks@11729.11765.37\|TAACTATGACTCTCTTAAGGTAGCCAAATGCCTCGTC | >hg38_21_+_8433172_8446622_RNA45SN1_45S_50ntFlanks | 11729 | 11765 | 37 | TAACTATGACTCTCTTAAGGTAGCCAAATGCCTCGTC | 2978.7 | -7.1 | 0.7 | -10.5 | 1.1E-25 | 1.2E-24 | BT20/MB468 |
| >hg38_21_+_8433172_8446622_RNA45SN1_45S_50ntFlanks@4823.4857.35\|ATGACCCGCCGGGCAGCTTCCGGGAAACCAAAGTC | >hg38_21_+_8433172_8446622_RNA45SN1_45S_50ntFlanks | 4823 | 4857 | 35 | ATGACCCGCCGGGCAGCTTCCGGGAAACCAAAGTC | 328.8 | 9.6 | 0.9 | 10.5 | 1.4E-25 | 1.2E-24 | MB231/MB468 |
| >MI0000114\|hsa-mir-107&WithFlank&10\|-\|89592741\|89592833@56.76.21(+1U);MI0000108\|hsa-mir-103a-2&WithFlank&20\|+\|3917488\|3917577@54.74.21(+1U);MI0000109\|hsa-mir-103a-1&WithFlank&5\|-\|168560890\|168560979@54.74.21(+1U)&[MIMAT0000104&hsa-miR-107&offsets\|0\|-2(+1U);m-78&10\|-\|89592758\|89592778&offsets\|0\|0(+1U)];[MIMAT0000101&hsa-miR-103a-3p&offsets\|0\|-2(+1U);m-12&20\|+\|3917541\|3917563&offsets\|0\|-2(+1U)];[MIMAT0000101_1&hsa-miR-103a-3p&offsets\|0\|-2(+1U);m-13&5\|-\|168560904\|168560926&offsets\|0\|-2(+1U);m-44220&5\|-\|168560908\|168560926&offsets\|0\|+2(+1U)]\|AGCAGCATTGTACAGGGCTATT | >MI0000114\|hsa-mir-107&WithFlank&10\|-\|89592741\|89592833 | 56 | 76 | 21(+1U) | AGCAGCATTGTACAGGGCTATT | 760.2 | 7.9 | 0.8 | 10.5 | 1.4E-25 | 1.2E-24 | MB231/MB468 |
| >hg38_21_+_8433172_8446622_RNA45SN1_45S_50ntFlanks@12419.12454.36\|GATGTCGGCTCTTCCTATCATTGTGAAGCAGAATTC | >hg38_21_+_8433172_8446622_RNA45SN1_45S_50ntFlanks | 12419 | 12454 | 36 | GATGTCGGCTCTTCCTATCATTGTGAAGCAGAATTC | 1284.7 | 7.0 | 0.7 | 10.5 | 1.4E-25 | 1.2E-24 | MB231/MB468 |
| >hg38_21_+_8433172_8446622_RNA45SN1_45S_50ntFlanks@7973.8009.37\|GACGCGACCTCAGATCAGACGTGGCGACCCGCTGAAT | >hg38_21_+_8433172_8446622_RNA45SN1_45S_50ntFlanks | 7973 | 8009 | 37 | GACGCGACCTCAGATCAGACGTGGCGACCCGCTGAAT | 1899.0 | 7.2 | 0.7 | 10.5 | 1.4E-25 | 1.2E-24 | MB231/MB468 |
| >MI0001733\|hsa-mir-452&WithFlank&X\|-\|151959622\|151959718@20.43.24&[MIMAT0001635&hsa-miR-452-5p&offsets\|0\|+2;m-188&X\|-\|151959677\|151959699&offsets\|0\|+1]\|AACTGTTTGCAGAGGAAACTGAGA | >MI0001733\|hsa-mir-452&WithFlank&X\|-\|151959622\|151959718 | 20 | 43 | 24 | AACTGTTTGCAGAGGAAACTGAGA | 620.9 | 12.1 | 1.2 | 10.5 | 1.4E-25 | 1.2E-24 | MB231/MB468 |
| >MI0000067\|hsa-let-7f-1&WithFlank&9\|+\|94176341\|94176439@13.33.21(+1C);MI0000068\|hsa-let-7f-2&WithFlank&X\|-\|53557186\|53557280@14.34.21(+1C)&[MIMAT0000067&hsa-let-7f-5p&offsets\|0\|-1(+1C);m-20&9\|+\|94176353\|94176374&offsets\|0\|-1(+1C)];[MIMAT0000067_1&hsa-let-7f-5p&offsets\|0\|-1(+1C);m-19&X\|-\|53557246\|53557267&offsets\|0\|-1(+1C)]\|TGAGGTAGTAGATTGTATAGTC | >MI0000067\|hsa-let-7f-1&WithFlank&9\|+\|94176341\|94176439 | 13 | 33 | 21(+1C) | TGAGGTAGTAGATTGTATAGTC | 44406.1 | 6.9 | 0.7 | 10.5 | 1.2E-25 | 1.3E-24 | BT20/MB468 |
| >MI0000264\|hsa-mir-7-2&WithFlank&15\|+\|88611819\|88611940@38.58.21;MI0000265\|hsa-mir-7-3&WithFlank&19\|+\|4770664\|4770785@37.57.21;MI0000263\|hsa-mir-7-1&WithFlank&9\|-\|83969742\|83969863@30.50.21&[MIMAT0000252&hsa-miR-7-5p&offsets\|0\|-3;m-400&15\|+\|88611856\|88611879&offsets\|0\|-3];[MIMAT0000252_1&hsa-miR-7-5p&offsets\|0\|-3;m-402&19\|+\|4770700\|4770723&offsets\|0\|-3];[MIMAT0000252_2&hsa-miR-7-5p&offsets\|0\|-3;m-401&9\|-\|83969811\|83969834&offsets\|0\|-3]\|TGGAAGACTAGTGATTTTGTT | >MI0000264\|hsa-mir-7-2&WithFlank&15\|+\|88611819\|88611940 | 38 | 58 | 21 | TGGAAGACTAGTGATTTTGTT | 6418.5 | 8.6 | 0.8 | 10.4 | 1.5E-25 | 1.3E-24 | MB231/MB468 |
| >MI0001445\|hsa-mir-423&WithFlank&17\|+\|30117073\|30117178@59.80.22(+1A)&[MIMAT0001340&hsa-miR-423-3p&offsets\|0\|-1(+1A);m-100&17\|+\|30117131\|30117153&offsets\|0\|-1(+1A)]\|AGCTCGGTCTGAGGCCCCTCAGA | >MI0001445\|hsa-mir-423&WithFlank&17\|+\|30117073\|30117178 | 59 | 80 | 22(+1A) | AGCTCGGTCTGAGGCCCCTCAGA | 2060.8 | 7.9 | 0.8 | 10.4 | 1.5E-25 | 1.3E-24 | MB231/MB468 |
| >hg38_21_+_8433172_8446622_RNA45SN1_45S_50ntFlanks@4733.4768.36\|GAACGAAAGTCGGAGGTTCGAAGACGATCAGATACC | >hg38_21_+_8433172_8446622_RNA45SN1_45S_50ntFlanks | 4733 | 4768 | 36 | GAACGAAAGTCGGAGGTTCGAAGACGATCAGATACC | 687.2 | 14.3 | 1.4 | 10.4 | 1.5E-25 | 1.3E-24 | MB231/MB468 |
| >hg38_21_+_8433172_8446622_RNA45SN1_45S_50ntFlanks@11316.11344.29\|CACCCCACGTCTCGTCGCGCGCGCGTCCG | >hg38_21_+_8433172_8446622_RNA45SN1_45S_50ntFlanks | 11316 | 11344 | 29 | CACCCCACGTCTCGTCGCGCGCGCGTCCG | 4359.7 | -13.7 | 1.3 | -10.4 | 1.5E-25 | 1.3E-24 | MB231/MB468 |

| | | | | | | | | | | | | |
|---|---|---|---|---|---|---|---|---|---|---|---|---|
| >MI0000065\|hsa-let-7d&WithFlank&9\|+\|94178828\|94178926@68.89.22&[MIMAT0004484&hsa-let-7d-3p&offsets\|0\|0;m-98&9\|+\|94178895\|94178916&offsets\|0\|0]\|CTATACGACCTGCTGCCTTTCT | >MI0000065\|hsa-let-7d&WithFlank&9\|+\|94178828\|94178926 | 68 | 89 | 22 | CTATACGACCTGCTGCCTTTCT | 6008.9 | 7.8 | 0.7 | 10.4 | 1.5E-25 | 1.3E-24 | MB231/MB468 |
| >MI0000448\|hsa-mir-130a&WithFlank&11\|+\|57641192\|57641292@61.83.23&[MIMAT0000425&hsa-miR-130a-3p&offsets\|0\|+1;m-139&11\|+\|57641252\|57641273&offsets\|0\|+1]\|CAGTGCAATGTTAAAAGGGCATT | >MI0000448\|hsa-mir-130a&WithFlank&11\|+\|57641192\|57641292 | 61 | 83 | 23 | CAGTGCAATGTTAAAAGGGCATT | 1081.6 | 8.1 | 0.8 | 10.4 | 1.5E-25 | 1.3E-24 | MB231/MB468 |
| >MI0001448\|hsa-mir-425&WithFlank&3\|-\|49020142\|49020240@20.43.24&[MIMAT0003393&hsa-miR-425-5p&offsets\|0\|+1;m-95&3\|-\|49020200\|49020221&offsets\|0\|+2]\|AATGACACGATCACTCCCGTTGAG | >MI0001448\|hsa-mir-425&WithFlank&3\|-\|49020142\|49020240 | 20 | 43 | 24 | AATGACACGATCACTCCCGTTGAG | 4443.3 | 8.1 | 0.8 | 10.4 | 1.5E-25 | 1.3E-24 | MB231/MB468 |
| >MI0000469\|hsa-mir-125a&WithFlank&19\|+\|51693248\|51693345@21.41.21(+1C)&[MIMAT0000443&hsa-miR-125a-5p&offsets\|0\|-3(+1C);m-63&19\|+\|51693268\|51693289&offsets\|0\|-1(+1C)]\|TCCCTGAGACCCTTTAACCTGC | >MI0000469\|hsa-mir-125a&WithFlank&19\|+\|51693248\|51693345 | 21 | 41 | 21(+1C) | TCCCTGAGACCCTTTAACCTGC | 1409.4 | 7.7 | 0.7 | 10.5 | 1.3E-25 | 1.3E-24 | BT20/MB468 |
| >trna119_LysCTT_1_-_145395522_145395594@1.36.36,trna11_LysCTT_5_-_180648979_180649051@1.36.36,trna13_LysCTT_6_+_26556774_26556846@1.36.36,trna7_LysCTT_16_+_3225692_3225764@1.36.36,trna9_LysCTT_5_+_180634755_180634827@1.36.36\|GCCCGGCTAGCTCAGTCGGTAGAGCATGAGACTCTT | >trna119_LysCTT_1_-_145395522_145395594 | 1 | 36 | 36 | GCCCGGCTAGCTCAGTCGGTAGAGCATGAGACTCTT | 2849.0 | 8.4 | 0.8 | 10.4 | 1.6E-25 | 1.3E-24 | MB231/MB468 |
| >MI0000750\|hsa-mir-26a-2&WithFlank&12\|-\|57824603\|57824698@20.40.21(+1A);MI0000083\|hsa-mir-26a-1&WithFlank&3\|+\|37969398\|37969486@16.36.21(+1A)&[MIMAT0000082&hsa-miR-26a-5p&offsets\|0\|-1(+1A);m-33&12\|-\|57824658\|57824679&offsets\|0\|-1(+1A)];[MIMAT0000082_1&hsa-miR-26a-5p&offsets\|0\|-1(+1A);m-34&3\|+\|37969413\|37969434&offsets\|0\|-1(+1A)]\|TTCAAGTAATCCAGGATAGGCA | >MI0000750\|hsa-mir-26a-2&WithFlank&12\|-\|57824603\|57824698 | 20 | 40 | 21(+1A) | TTCAAGTAATCCAGGATAGGCA | 21381.3 | 7.4 | 0.7 | 10.4 | 1.6E-25 | 1.4E-24 | MB231/MB468 |
| >hg38_21_+_8433172_8446622_RNA45SN1_45S_50ntFlanks@8687.8706.20\|CGGCTCCGGGACGGCTGGGA | >hg38_21_+_8433172_8446622_RNA45SN1_45S_50ntFlanks | 8687 | 8706 | 20 | CGGCTCCGGGACGGCTGGGA | 723.0 | 14.4 | 1.4 | 10.4 | 1.8E-25 | 1.5E-24 | MB231/MB468 |
| >hg38_21_+_8433172_8446622_RNA45SN1_45S_50ntFlanks@476.524.49\|TTCGTGATCGATGTGGTGACGTCGTGCTCTCCCGGGCCGGGTCCGAGCC | >hg38_21_+_8433172_8446622_RNA45SN1_45S_50ntFlanks | 476 | 524 | 49 | TTCGTGATCGATGTGGTGACGTCGTGCTCTCCCGGGCCGGGTCCGAGCC | 2410.9 | 14.1 | 1.3 | 10.4 | 1.8E-25 | 1.5E-24 | MB231/MB468 |
| >hg38_21_+_8433172_8446622_RNA45SN1_45S_50ntFlanks@10679.10706.28\|GGTTCCGGCGGCGTCCGGTGAGCTCTCG | >hg38_21_+_8433172_8446622_RNA45SN1_45S_50ntFlanks | 10679 | 10706 | 28 | GGTTCCGGCGGCGTCCGGTGAGCTCTCG | 3925.6 | -7.3 | 0.7 | -10.4 | 1.8E-25 | 1.6E-24 | MB231/MB468 |
| >hg38_21_+_8433172_8446622_RNA45SN1_45S_50ntFlanks@485.521.37\|GATGTGGTGACGTCGTGCTCTCCCGGGCCGGGTCCGA | >hg38_21_+_8433172_8446622_RNA45SN1_45S_50ntFlanks | 485 | 521 | 37 | GATGTGGTGACGTCGTGCTCTCCCGGGCCGGGTCCGA | 2171.3 | 15.7 | 1.5 | 10.4 | 1.5E-25 | 1.6E-24 | BT20/MB468 |
| >hg38_21_+_8433172_8446622_RNA45SN1_45S_50ntFlanks@476.496.21\|TTCGTGATCGATGTGGTGACG | >hg38_21_+_8433172_8446622_RNA45SN1_45S_50ntFlanks | 476 | 496 | 21 | TTCGTGATCGATGTGGTGACG | 3755.1 | 7.8 | 0.8 | 10.4 | 1.9E-25 | 1.6E-24 | MB231/MB468 |
| >MI0000746\|hsa-mir-99b&WithFlank&19\|+\|51692606\|51692687@13.33.21(+1U)&[MIMAT0000689&hsa-miR-99b-5p&offsets\|0\|-1(+1U);m-10&19\|+\|51692618\|51692639&offsets\|0\|-1(+1U)]\|CACCCGTAGAACCGACCTTGCT | >MI0000746\|hsa-mir-99b&WithFlank&19\|+\|51692606\|51692687 | 13 | 33 | 21(+1U) | CACCCGTAGAACCGACCTTGCT | 1873.7 | 8.0 | 0.8 | 10.4 | 1.9E-25 | 1.6E-24 | MB231/MB468 |
| >am_MI0000786\|hsa-mir-378a&WithFlank&5\|+\|149732819\|149732896@49.69.21(+1U)&[MIMAT0000732&hsa-miR-378a-3p&offsets\|0\|-1(+1U);m-44&5\|+\|149732867\|149732888&offsets\|0\|-1(+1U)]\|ACTGGACTTGGAGTCAGAAGGT | >am_MI0000786\|hsa-mir-378a&WithFlank&5\|+\|149732819\|149732896 | 49 | 69 | 21(+1U) | ACTGGACTTGGAGTCAGAAGGT | 8747.6 | 5.9 | 0.6 | 10.4 | 1.5E-25 | 1.6E-24 | BT20/MB468 |
| >MI0000301\|hsa-mir-224&WithFlank&X\|-\|151958572\|151958664@14.37.24&[MIMAT0000281&hsa-miR-224-5p&offsets\|+1\|0;m-238&X\|-\|151958630\|151958651&offsets\|0\|+2]\|CAAGTCACTAGTGGTTCCGTTTAG | >MI0000301\|hsa-mir-224&WithFlank&X\|-\|151958572\|151958664 | 14 | 37 | 24 | CAAGTCACTAGTGGTTCCGTTTAG | 6342.6 | 11.3 | 1.1 | 10.4 | 1.7E-25 | 1.8E-24 | BT20/MB468 |
| >MI0000067\|hsa-let-7f-1&WithFlank&9\|+\|94176341\|94176439@69.89.21(+1U)&[MIMAT0004486&hsa-let-7f-1-3p&offsets\|0\|-1(+1U);m-275&9\|+\|94176409\|94176430&offsets\|0\|-1(+1U)]\|CTATACAATCTATTGCCTTCCT | >MI0000067\|hsa-let-7f-1&WithFlank&9\|+\|94176341\|94176439 | 69 | 89 | 21(+1U) | CTATACAATCTATTGCCTTCCT | 739.5 | 7.7 | 0.7 | 10.4 | 2.1E-25 | 1.8E-24 | MB231/MB468 |
| >MI0000542\|hsa-mir-320a&WithFlank&8\|-\|22244960\|22245043@49.68.20(+1U)&[MIMAT0000510&hsa-miR-320a-3p&offsets\|+1\|-1(+1U);m-60&8\|-\|22244975\|22244996&offsets\|+1\|-1(+1U)]\|AAAGCTGGGTTGAGAGGGCGT | >MI0000542\|hsa-mir-320a&WithFlank&8\|-\|22244960\|22245043 | 49 | 68 | 20(+1U) | AAAGCTGGGTTGAGAGGGCGT | 570.8 | 13.8 | 1.3 | 10.4 | 1.7E-25 | 1.8E-24 | BT20/MB468 |
| >hg38_21_+_8433172_8446622_RNA45SN1_45S_50ntFlanks@7986.8006.21\|ATCAGACGTGGCGACCCGCTG | >hg38_21_+_8433172_8446622_RNA45SN1_45S_50ntFlanks | 7986 | 8006 | 21 | ATCAGACGTGGCGACCCGCTG | 3095.9 | -13.6 | 1.3 | -10.4 | 1.7E-25 | 1.8E-24 | BT20/MB468 |
| >MI0000065\|hsa-let-7d&WithFlank&9\|+\|94178828\|94178926@68.88.21(+1C)&[MIMAT0004484&hsa-let-7d-3p&offsets\|0\|-1(+1C);m-98&9\|+\|94178895\|94178916&offsets\|0\|-1(+1C)]\|CTATACGACCTGCTGCCTTTCC | >MI0000065\|hsa-let-7d&WithFlank&9\|+\|94178828\|94178926 | 68 | 88 | 21(+1C) | CTATACGACCTGCTGCCTTTCC | 1031.8 | 7.8 | 0.8 | 10.4 | 1.7E-25 | 1.8E-24 | BT20/MB468 |
| >MI0000087\|hsa-mir-29a&WithFlank&7\|-\|130876741\|130876816@48.69.22&[MIMAT0000086&hsa-miR-29a-3p&offsets\|0\|0;m-26&7\|-\|130876749\|130876769&offsets\|0\|+1]\|TAGCACCATCTGAAATCGGTTA | >MI0000087\|hsa-mir-29a&WithFlank&7\|-\|130876741\|130876816 | 48 | 69 | 22 | TAGCACCATCTGAAATCGGTTA | 27276.3 | 9.2 | 0.9 | 10.4 | 1.7E-25 | 1.8E-24 | BT20/MB468 |
| >MI0003628\|hsa-mir-615&WithFlank&12\|+\|54033944\|54034051@67.86.20(+1C)&[MIMAT0003283&hsa-miR-615-3p&offsets\|0\|-2(+1C);m-495&12\|+\|54034010\|54034030&offsets\|0\|-1(+1C)]\|TCCGAGCCTGGGTCTCCCTCC | >MI0003628\|hsa-mir-615&WithFlank&12\|+\|54033944\|54034051 | 67 | 86 | 20(+1C) | TCCGAGCCTGGGTCTCCCTCC | 553.5 | 7.0 | 0.7 | 10.4 | 1.7E-25 | 1.8E-24 | BT20/MB468 |
| >hg38_21_+_8433172_8446622_RNA45SN1_45S_50ntFlanks@6721.6740.20\|AATTGCAGGACACATTGATC | >hg38_21_+_8433172_8446622_RNA45SN1_45S_50ntFlanks | 6721 | 6740 | 20 | AATTGCAGGACACATTGATC | 7273.7 | -5.9 | 0.6 | -10.4 | 2.1E-25 | 1.8E-24 | MB231/MB468 |
| >MI0000746\|hsa-mir-99b&WithFlank&19\|+\|51692606\|51692687@13.33.21&[MIMAT0000689&hsa-miR-99b-5p&offsets\|0\|-1;m-10&19\|+\|51692618\|51692639&offsets\|0\|-1]\|CACCCGTAGAACCGACCTTGC | >MI0000746\|hsa-mir-99b&WithFlank&19\|+\|51692606\|51692687 | 13 | 33 | 21 | CACCCGTAGAACCGACCTTGC | 17369.4 | 7.3 | 0.7 | 10.4 | 1.8E-25 | 1.8E-24 | BT20/MB468 |
| >hg38_21_+_8433172_8446622_RNA45SN1_45S_50ntFlanks@3992.4023.32\|GGTGACTCTAGATAACCTCGGGCCGATCGCAC | >hg38_21_+_8433172_8446622_RNA45SN1_45S_50ntFlanks | 3992 | 4023 | 32 | GGTGACTCTAGATAACCTCGGGCCGATCGCAC | 415.1 | 5.6 | 0.5 | 10.4 | 1.8E-25 | 1.8E-24 | BT20/MB468 |
| >MI0000061\|hsa-let-7a-2&WithFlank&11\|-\|122146516\|122146599@11.31.21(+1A);MI0000062\|hsa-let-7a-3&WithFlank&22\|+\|46112743\|46112828@10.30.21(+1A);MI0000060\|hsa-let-7a-1&WithFlank&9\|+\|94175951\|94176042@12.32.21(+1A)&[MIMAT0000062&hsa-let-7a-5p&offsets\|0\|-1(+1A);m-6&11\|-\|122146568\|122146589&offsets\|0\|-1(+1A)];[MIMAT0000062_1&hsa-let-7a-5p&offsets\|0\|-1(+1A);m-5&22\|+\|46112752\|46112773&offsets\|0\|-1(+1A)];[MIMAT0000062_2&hsa-let-7a-5p&offsets\|0\|-1(+1A);m-4&9\|+\|94175962\|94175983&offsets\|0\|-1(+1A)]\|TGAGGTAGTAGGTTGTATAGTA | >MI0000061\|hsa-let-7a-2&WithFlank&11\|-\|122146516\|122146599 | 11 | 31 | 21(+1A) | TGAGGTAGTAGGTTGTATAGTA | 10922.5 | 8.9 | 0.8 | 10.4 | 1.8E-25 | 1.9E-24 | BT20/MB468 |
| >trnaMT_PheGAA_MT_+_577_647@36.59.24\|AAAATGTTTAGACGGGCTCACATC | >trnaMT_PheGAA_MT_+_577_647 | 36 | 59 | 24 | AAAATGTTTAGACGGGCTCACATC | 492.9 | 9.6 | 0.9 | 10.4 | 1.8E-25 | 1.9E-24 | BT20/MB468 |
| >hg38_MT_+_1621_3279_RNR2_16S_50ntFlanks@51.94.44\|GCTAAACCTAGCCCCAAACCCACTCCACCTTACTACCAGACAAC | >hg38_MT_+_1621_3279_RNR2_16S_50ntFlanks | 51 | 94 | 44 | GCTAAACCTAGCCCCAAACCCACTCCACCTTACTACCAGACAAC | 2184.0 | 13.9 | 1.3 | 10.4 | 2.2E-25 | 1.9E-24 | MB231/MB468 |
| >MI0000283\|hsa-mir-203a&WithFlank&14\|+\|104117399\|104117520@72.92.21&[MIMAT0000264&hsa-miR-203a-3p&offsets\|+1\|0;m-79&14\|+\|104117469\|104117490&offsets\|+1\|0]\|TGAAATGTTTAGGACCACTAG | >MI0000283\|hsa-mir-203a&WithFlank&14\|+\|104117399\|104117520 | 72 | 92 | 21 | TGAAATGTTTAGGACCACTAG | 5324.4 | 9.1 | 0.9 | 10.4 | 1.8E-25 | 1.9E-24 | BT20/MB468 |
| >MI0003591\|hsa-mir-584&WithFlank&5\|-\|149062307\|149062415@22.42.21(+2A)&[MIMAT0003249&hsa-miR-584-5p&offsets\|0\|-1(+2A);m-167&5\|-\|149062374\|149062394&offsets\|0\|0(+2A)]\|TTATGGTTTGCCTGGGACTGAAA | >MI0003591\|hsa-mir-584&WithFlank&5\|-\|149062307\|149062415 | 22 | 42 | 21(+2A) | TTATGGTTTGCCTGGGACTGAAA | 505.6 | 13.6 | 1.3 | 10.4 | 1.8E-25 | 1.9E-24 | BT20/MB468 |
| >trnaMT_ThrTGT_MT_+_15888_15953@34.69.36\|TAAACCGGAGATGAAAACCTTTTTCCAAGGACACCA | >trnaMT_ThrTGT_MT_+_15888_15953 | 34 | 69 | 36 | TAAACCGGAGATGAAAACCTTTTTCCAAGGACACCA | 582.4 | 11.3 | 1.1 | 10.4 | 2.3E-25 | 1.9E-24 | MB231/MB468 |
| >MI0000542\|hsa-mir-320a&WithFlank&8\|-\|22244960\|22245043@48.69.22&[MIMAT0000510&hsa-miR-320a-3p&offsets\|0\|0;m-60&8\|-\|22244975\|22244996&offsets\|0\|0]\|AAAAGCTGGGTTGAGAGGGCGA | >MI0000542\|hsa-mir-320a&WithFlank&8\|-\|22244960\|22245043 | 48 | 69 | 22 | AAAAGCTGGGTTGAGAGGGCGA | 7543.3 | 8.2 | 0.8 | 10.4 | 2.3E-25 | 2.0E-24 | MB231/MB468 |
| >MI0001445\|hsa-mir-423&WithFlank&17\|+\|30117073\|30117178@59.80.22(+1G)&[MIMAT0001340&hsa-miR-423-3p&offsets\|0\|-1(+1G);m-100&17\|+\|30117131\|30117153&offsets\|0\|-1(+1G)]\|AGCTCGGTCTGAGGCCCCTCAGG | >MI0001445\|hsa-mir-423&WithFlank&17\|+\|30117073\|30117178 | 59 | 80 | 22(+1G) | AGCTCGGTCTGAGGCCCCTCAGG | 2861.9 | 8.5 | 0.8 | 10.4 | 2.4E-25 | 2.0E-24 | MB231/MB468 |
| >hg38_MT_+_1621_3279_RNR2_16S_50ntFlanks@1438.1477.40\|TACGTGATCTGAGTTCAGACCGGAGTAATCCAGGTCGGTT | >hg38_MT_+_1621_3279_RNR2_16S_50ntFlanks | 1438 | 1477 | 40 | TACGTGATCTGAGTTCAGACCGGAGTAATCCAGGTCGGTT | 454.8 | 11.5 | 1.1 | 10.4 | 2.0E-25 | 2.0E-24 | BT20/MB468 |
| >MI0000078\|hsa-mir-22&WithFlank&17\|-\|1713897\|1713993@59.79.21(+1C)&[MIMAT0000077&hsa-miR-22-3p&offsets\|0\|-1(+1C);m-2&17\|-\|1713914\|1713935&offsets\|0\|-1(+1C)]\|AAGCTGCCAGTTGAAGAACTGC | >MI0000078\|hsa-mir-22&WithFlank&17\|-\|1713897\|1713993 | 59 | 79 | 21(+1C) | AAGCTGCCAGTTGAAGAACTGC | 2644.7 | 8.1 | 0.8 | 10.4 | 2.4E-25 | 2.1E-24 | MB231/MB468 |
| >MI0000108\|hsa-mir-103a-2&WithFlank&20\|+\|3917488\|3917577@54.75.22;MI0000109\|hsa-mir-103a-1&WithFlank&5\|-\|168560890\|168560979@54.75.22&[MIMAT0000101&hsa-miR-103a-3p&offsets\|0\|-1;m-12&20\|+\|3917541\|3917563&offsets\|0\|-1];[MIMAT0000101_1&hsa-miR-103a-3p&offsets\|0\|-1;m-13&5\|-\|168560904\|168560926&offsets\|0\|-1;m-44220&5\|-\|168560908\|168560926&offsets\|0\|+3]\|AGCAGCATTGTACAGGGCTATG | >MI0000108\|hsa-mir-103a-2&WithFlank&20\|+\|3917488\|3917577 | 54 | 75 | 22 | AGCAGCATTGTACAGGGCTATG | 20628.0 | 7.3 | 0.7 | 10.4 | 2.5E-25 | 2.1E-24 | MB231/MB468 |

<table>
<tr><td>>MI0000077|hsa-mir-21&WithFlank&17|+|59841260|59841343@15.36.22(+1A)&[MIMAT0000076&hsa-miR-21-5p&offsets|+1|+1(+1A);m-1&17|+|59841273|59841294&offsets|+1|+1(+1A)]|AGCTTATCAGACTGATGTTGACA</td><td>>MI0000077|hsa-mir-21&WithFlank&17|+|59841260|59841343</td><td>15</td><td>36</td><td>22(+1A)</td><td>AGCTTATCAGACTGATGTTGACA</td><td>1526.3</td><td>8.1</td><td>0.8</td><td>10.4</td><td>2.1E-25</td><td>2.2E-24</td><td>BT20/MB468</td></tr>
<tr><td>>MI0000299|hsa-mir-222&WithFlank&X|-|45747009|45747130@75.96.22(+1C)&[MIMAT0000279&hsa-miR-222-3p&offsets|0|+1(+1C);m-140&X|-|45747033|45747056&offsets|0|-2(+1C)]|AGCTACATCTGGCTACTGGGTCC</td><td>>MI0000299|hsa-mir-222&WithFlank&X|-|45747009|45747130</td><td>75</td><td>96</td><td>22(+1C)</td><td>AGCTACATCTGGCTACTGGGTCC</td><td>3112.9</td><td>8.6</td><td>0.8</td><td>10.4</td><td>2.6E-25</td><td>2.2E-24</td><td>MB231/MB468</td></tr>
<tr><td>>hg38_21_+_8433172_8446622_RNA45SN1_45S_50ntFlanks@12405.12432.28|CTTTTTGATCCTTCGATGTCGGCTCTTC</td><td>>hg38_21_+_8433172_8446622_RNA45SN1_45S_50ntFlanks</td><td>12405</td><td>12432</td><td>28</td><td>CTTTTTGATCCTTCGATGTCGGCTCTTC</td><td>4904.8</td><td>-6.3</td><td>0.6</td><td>-10.4</td><td>2.7E-25</td><td>2.3E-24</td><td>MB231/MB468</td></tr>
<tr><td>>hg38_21_+_8433172_8446622_RNA45SN1_45S_50ntFlanks@8194.8215.22|CCCGTGGACGGTGTGAGGCCGG</td><td>>hg38_21_+_8433172_8446622_RNA45SN1_45S_50ntFlanks</td><td>8194</td><td>8215</td><td>22</td><td>CCCGTGGACGGTGTGAGGCCGG</td><td>1432.1</td><td>-7.9</td><td>0.8</td><td>-10.4</td><td>2.3E-25</td><td>2.3E-24</td><td>BT20/MB468</td></tr>
<tr><td>>hg38_21_+_8433172_8446622_RNA45SN1_45S_50ntFlanks@12633.12657.25|CGAAGCTACCATCTGTGGGATTATG</td><td>>hg38_21_+_8433172_8446622_RNA45SN1_45S_50ntFlanks</td><td>12633</td><td>12657</td><td>25</td><td>CGAAGCTACCATCTGTGGGATTATG</td><td>576.6</td><td>-6.5</td><td>0.6</td><td>-10.4</td><td>2.3E-25</td><td>2.4E-24</td><td>BT20/MB468</td></tr>
<tr><td>>trnaMT_ProTGG_MT_-_15956_16023@51.71.21|TAAAGACTTTTTCTCTGACCA</td><td>>trnaMT_ProTGG_MT_-_15956_16023</td><td>51</td><td>71</td><td>21</td><td>TAAAGACTTTTTCTCTGACCA</td><td>2200.3</td><td>7.8</td><td>0.8</td><td>10.4</td><td>2.8E-25</td><td>2.4E-24</td><td>MB231/MB468</td></tr>
<tr><td>>MI0000108|hsa-mir-103a-2&WithFlank&20|+|3917488|3917577@54.76.23;MI0000109|hsa-mir-103a-1&WithFlank&5|-|168560890|168560979@54.76.23&[MIMAT0000101&hsa-miR-103a-3p&offsets|0|0;m-12&20|+|3917541|3917563&offsets|0|0];[MIMAT0000101_1&hsa-miR-103a-3p&offsets|0|0;m-13&5|-|168560904|168560926&offsets|0|0;m-44220&5|-|168560908|168560926&offsets|0|+4]|AGCAGCATTGTACAGGGCTATGA</td><td>>MI0000108|hsa-mir-103a-2&WithFlank&20|+|3917488|3917577</td><td>54</td><td>76</td><td>23</td><td>AGCAGCATTGTACAGGGCTATGA</td><td>85129.0</td><td>7.7</td><td>0.7</td><td>10.4</td><td>2.8E-25</td><td>2.4E-24</td><td>MB231/MB468</td></tr>
<tr><td>>hg38_21_+_8433172_8446622_RNA45SN1_45S_50ntFlanks@11668.11706.39|TGCCCAGTGCTCTGAATGTCAAAGTGAAGAAATTCAATG</td><td>>hg38_21_+_8433172_8446622_RNA45SN1_45S_50ntFlanks</td><td>11668</td><td>11706</td><td>39</td><td>TGCCCAGTGCTCTGAATGTCAAAGTGAAGAAATTCAATG</td><td>1273.6</td><td>9.4</td><td>0.9</td><td>10.4</td><td>2.8E-25</td><td>2.4E-24</td><td>MB231/MB468</td></tr>
<tr><td>>hg38_21_+_8433172_8446622_RNA45SN1_45S_50ntFlanks@12449.12486.38|GAATTCACCAAGCGTTGGATTGTTCACCCACTAATAGG</td><td>>hg38_21_+_8433172_8446622_RNA45SN1_45S_50ntFlanks</td><td>12449</td><td>12486</td><td>38</td><td>GAATTCACCAAGCGTTGGATTGTTCACCCACTAATAGG</td><td>369.8</td><td>9.1</td><td>0.9</td><td>10.4</td><td>2.9E-25</td><td>2.5E-24</td><td>MB231/MB468</td></tr>
<tr><td>>hg38_21_+_8433172_8446622_RNA45SN1_45S_50ntFlanks@6719.6737.19|TGAATTGCAGGACACATTG</td><td>>hg38_21_+_8433172_8446622_RNA45SN1_45S_50ntFlanks</td><td>6719</td><td>6737</td><td>19</td><td>TGAATTGCAGGACACATTG</td><td>3390.4</td><td>-6.1</td><td>0.6</td><td>-10.4</td><td>3.0E-25</td><td>2.5E-24</td><td>MB231/MB468</td></tr>
<tr><td>>hg38_MT_+_1621_3279_RNR2_16S_50ntFlanks@51.93.43|GCTAAACCTAGCCCCAAACCCACTCCACCTTACTACCAGACAA</td><td>>hg38_MT_+_1621_3279_RNR2_16S_50ntFlanks</td><td>51</td><td>93</td><td>43</td><td>GCTAAACCTAGCCCCAAACCCACTCCACCTTACTACCAGACAA</td><td>644.7</td><td>14.2</td><td>1.4</td><td>10.4</td><td>3.0E-25</td><td>2.5E-24</td><td>MB231/MB468</td></tr>
<tr><td>>MI0000433|hsa-let-7g&WithFlank&3|-|52268272|52268367@11.31.21&[MIMAT0000414&hsa-let-7g-5p&offsets|0|-1;m-58&3|-|52268336|52268357&offsets|0|-1]|TGAGGTAGTAGTTTGTACAGT</td><td>>MI0000433|hsa-let-7g&WithFlank&3|-|52268272|52268367</td><td>11</td><td>31</td><td>21</td><td>TGAGGTAGTAGTTTGTACAGT</td><td>17691.2</td><td>7.9</td><td>0.8</td><td>10.4</td><td>3.1E-25</td><td>2.5E-24</td><td>MB231/MB468</td></tr>
<tr><td>>trnaMT_MetCAT_MT_+_4402_4469@37.71.35|CCGAAAATGTTGGTTATACCCTTCCCGTACTACCA</td><td>>trnaMT_MetCAT_MT_+_4402_4469</td><td>37</td><td>71</td><td>35</td><td>CCGAAAATGTTGGTTATACCCTTCCCGTACTACCA</td><td>745.8</td><td>12.2</td><td>1.2</td><td>10.4</td><td>2.5E-25</td><td>2.6E-24</td><td>BT20/MB468</td></tr>
<tr><td>>trna10_LysCTT_16_+_3241501_3241573@1.36.36,trna13_LysCTT_14_-_58706613_58706685@1.36.36,trna2_LysCTT_15_+_79152904_79152976@1.36.36|GCCCGGCTAGCTCAGTCGGTAGAGCATGGGACTCTT</td><td>>trna10_LysCTT_16_+_3241501_3241573</td><td>1</td><td>36</td><td>36</td><td>GCCCGGCTAGCTCAGTCGGTAGAGCATGGGACTCTT</td><td>3821.8</td><td>8.9</td><td>0.9</td><td>10.4</td><td>3.2E-25</td><td>2.6E-24</td><td>MB231/MB468</td></tr>
<tr><td>>MI0000434|hsa-let-7i&WithFlank&12|+|62603680|62603775@12.32.21(+1G)&[MIMAT0000415&hsa-let-7i-5p&offsets|0|-1(+1G);m-49&12|+|62603691|62603712&offsets|0|-1(+1G)]|TGAGGTAGTAGTTTGTGCTGTG</td><td>>MI0000434|hsa-let-7i&WithFlank&12|+|62603680|62603775</td><td>12</td><td>32</td><td>21(+1G)</td><td>TGAGGTAGTAGTTTGTGCTGTG</td><td>23435.5</td><td>8.3</td><td>0.8</td><td>10.4</td><td>3.3E-25</td><td>2.7E-24</td><td>MB231/MB468</td></tr>
<tr><td>>MI0000266|hsa-mir-10a&WithFlank&17|-|48579832|48579953@29.50.22&[MIMAT0000253&hsa-miR-10a-5p&offsets|+1|0;m-18&17|-|48579905|48579926&offsets|+1|+1]|ACCCTGTAGATCCGAATTTGTG</td><td>>MI0000266|hsa-mir-10a&WithFlank&17|-|48579832|48579953</td><td>29</td><td>50</td><td>22</td><td>ACCCTGTAGATCCGAATTTGTG</td><td>14697.0</td><td>13.2</td><td>1.3</td><td>10.4</td><td>3.3E-25</td><td>2.7E-24</td><td>MB231/MB468</td></tr>
<tr><td>>am_trna128_GlyGCC_6_-_27870686_27870756@1.31.31,trna133_GlyCCC_1_-_16872434_16872504@1.31.31,trna18_GlyGCC_16_+_70822597_70822667@1.31.31,trna19_GlyGCC_16_+_70823410_70823480@1.31.31,trna19_GlyGCC_2_-_157257659_157257729@1.31.31,trna24_GlyGCC_16_-_70812942_70813012@1.31.31,trna25_GlyGCC_16_-_70812114_70812184@1.31.31,trna4_GlyCCC_1_+_17188416_17188486@1.31.31,trna5_GlyGCC_17_+_8029064_8029134@1.31.31,trna68_GlyGCC_1_-_161493637_161493707@1.31.31|GCATTGGTGGTTCAGTGGTAGAATTCTCGCC</td><td>>am_trna128_GlyGCC_6_-_27870686_27870756</td><td>1</td><td>31</td><td>31</td><td>GCATTGGTGGTTCAGTGGTAGAATTCTCGCC</td><td>177657.0</td><td>9.0</td><td>0.9</td><td>10.4</td><td>3.4E-25</td><td>2.8E-24</td><td>MB231/MB468</td></tr>
<tr><td>>MI0000477|hsa-mir-146a&WithFlank&5|+|160485346|160485456@27.47.21(+1G)&[MIMAT0000449&hsa-miR-146a-5p&offsets|0|-1(+1G);m-129&5|+|160485372|160485393&offsets|0|-1(+1G)]|TGAGAACTGAATTCCATGGGTG</td><td>>MI0000477|hsa-mir-146a&WithFlank&5|+|160485346|160485456</td><td>27</td><td>47</td><td>21(+1G)</td><td>TGAGAACTGAATTCCATGGGTG</td><td>2088.4</td><td>15.9</td><td>1.5</td><td>10.4</td><td>3.4E-25</td><td>2.8E-24</td><td>MB231/MB468</td></tr>
<tr><td>>hg38_21_+_8433172_8446622_RNA45SN1_45S_50ntFlanks@474.504.31|TCTTCGTGATCGATGTGGTGACGTCGTGCTC</td><td>>hg38_21_+_8433172_8446622_RNA45SN1_45S_50ntFlanks</td><td>474</td><td>504</td><td>31</td><td>TCTTCGTGATCGATGTGGTGACGTCGTGCTC</td><td>1129.0</td><td>8.3</td><td>0.8</td><td>10.4</td><td>3.6E-25</td><td>3.0E-24</td><td>MB231/MB468</td></tr>
<tr><td>>MI0000067|hsa-let-7f-1&WithFlank&9|+|94176341|94176439@13.33.21;MI0000068|hsa-let-7f-2&WithFlank&X|-|53557186|53557280@14.34.21&[MIMAT0000067&hsa-let-7f-5p&offsets|0|-1;m-20&9|+|94176353|94176374&offsets|0|-1];[MIMAT0000067_1&hsa-let-7f-5p&offsets|0|-1;m-19&X|-|53557246|53557267&offsets|0|-1]|TGAGGTAGTAGATTGTATAGT</td><td>>MI0000067|hsa-let-7f-1&WithFlank&9|+|94176341|94176439</td><td>13</td><td>33</td><td>21</td><td>TGAGGTAGTAGATTGTATAGT</td><td>41387.5</td><td>7.5</td><td>0.7</td><td>10.4</td><td>3.6E-25</td><td>3.0E-24</td><td>MB231/MB468</td></tr>
<tr><td>>MI0000079|hsa-mir-23a&WithFlank&19|-|13836581|13836665@54.72.19(+1U)&[MIMAT0000078&hsa-miR-23a-3p&offsets|+3|+1(+1U);m-31&19|-|13836595|13836615&offsets|+3|+1(+1U)]|ACATTGCCAGGGATTTCCAT</td><td>>MI0000079|hsa-mir-23a&WithFlank&19|-|13836581|13836665</td><td>54</td><td>72</td><td>19(+1U)</td><td>ACATTGCCAGGGATTTCCAT</td><td>1534.7</td><td>15.2</td><td>1.5</td><td>10.4</td><td>3.0E-25</td><td>3.0E-24</td><td>BT20/MB468</td></tr>
<tr><td>>hg38_21_+_8433172_8446622_RNA45SN1_45S_50ntFlanks@6774.6806.33|TCCTCCCGGGGCTACGCCTGTCTGAGCGTCGCT</td><td>>hg38_21_+_8433172_8446622_RNA45SN1_45S_50ntFlanks</td><td>6774</td><td>6806</td><td>33</td><td>TCCTCCCGGGGCTACGCCTGTCTGAGCGTCGCT</td><td>5601.1</td><td>7.8</td><td>0.8</td><td>10.4</td><td>3.0E-25</td><td>3.0E-24</td><td>BT20/MB468</td></tr>
<tr><td>>hg38_21_+_8433172_8446622_RNA45SN1_45S_50ntFlanks@6721.6743.23|AATTGCAGGACACATTGATCATC</td><td>>hg38_21_+_8433172_8446622_RNA45SN1_45S_50ntFlanks</td><td>6721</td><td>6743</td><td>23</td><td>AATTGCAGGACACATTGATCATC</td><td>6536.5</td><td>-6.2</td><td>0.6</td><td>-10.4</td><td>3.1E-25</td><td>3.1E-24</td><td>BT20/MB468</td></tr>
<tr><td>>MI0000108|hsa-mir-103a-2&WithFlank&20|+|3917488|3917577@54.75.22(+1U);MI0000109|hsa-mir-103a-1&WithFlank&5|-|168560890|168560979@54.75.22(+1U)&[MIMAT0000101&hsa-miR-103a-3p&offsets|0|-1(+1U);m-12&20|+|3917541|3917563&offsets|0|-1(+1U)];[MIMAT0000101_1&hsa-miR-103a-3p&offsets|0|-1(+1U);m-13&5|-|168560904|168560926&offsets|0|-1(+1U);m-44220&5|-|168560908|168560926&offsets|0|+3(+1U)]|AGCAGCATTGTACAGGGCTATGT</td><td>>MI0000108|hsa-mir-103a-2&WithFlank&20|+|3917488|3917577</td><td>54</td><td>75</td><td>22(+1U)</td><td>AGCAGCATTGTACAGGGCTATGT</td><td>4543.6</td><td>8.0</td><td>0.8</td><td>10.4</td><td>4.0E-25</td><td>3.3E-24</td><td>MB231/MB468</td></tr>
<tr><td>>MI0000264|hsa-mir-7-2&WithFlank&15|+|88611819|88611940@38.61.24(+1A);MI0000265|hsa-mir-7-3&WithFlank&19|+|4770664|4770785@37.60.24(+1A);MI0000263|hsa-mir-7-1&WithFlank&9|-|83969742|83969863@30.53.24(+1A)&[MIMAT0000252&hsa-miR-7-5p&offsets|0|0(+1A);m-400&15|+|88611856|88611879&offsets|0|0(+1A)];[MIMAT0000252_1&hsa-miR-7-5p&offsets|0|0(+1A);m-402&19|+|4770700|4770723&offsets|0|0(+1A)];[MIMAT0000252_2&hsa-miR-7-5p&offsets|0|0(+1A);m-401&9|-|83969811|83969834&offsets|0|0(+1A)]|TGGAAGACTAGTGATTTTGTTGTTA</td><td>>MI0000264|hsa-mir-7-2&WithFlank&15|+|88611819|88611940</td><td>38</td><td>61</td><td>24(+1A)</td><td>TGGAAGACTAGTGATTTTGTTGTTA</td><td>2777.3</td><td>7.8</td><td>0.8</td><td>10.4</td><td>4.0E-25</td><td>3.3E-24</td><td>MB231/MB468</td></tr>
<tr><td>>MI0000439|hsa-mir-23b&WithFlank&9|+|95085202|95085310@67.85.19(+2U)&[MIMAT0000418&hsa-miR-23b-3p&offsets|+3|-1(+2U);m-42&9|+|95085265|95085285&offsets|+3|+1(+2U)]|ACATTGCCAGGGATTACCATT</td><td>>MI0000439|hsa-mir-23b&WithFlank&9|+|95085202|95085310</td><td>67</td><td>85</td><td>19(+2U)</td><td>ACATTGCCAGGGATTACCATT</td><td>1406.2</td><td>15.1</td><td>1.5</td><td>10.4</td><td>3.3E-25</td><td>3.3E-24</td><td>BT20/MB468</td></tr>
<tr><td>>MI0000264|hsa-mir-7-2&WithFlank&15|+|88611819|88611940@38.59.22;MI0000265|hsa-mir-7-3&WithFlank&19|+|4770664|4770785@37.58.22;MI0000263|hsa-mir-7-1&WithFlank&9|-|83969742|83969863@30.51.22&[MIMAT0000252&hsa-miR-7-5p&offsets|0|-2;m-400&15|+|88611856|88611879&offsets|0|-2];[MIMAT0000252_1&hsa-miR-7-5p&offsets|0|-2;m-402&19|+|4770700|4770723&offsets|0|-2];[MIMAT0000252_2&hsa-miR-7-5p&offsets|0|-2;m-401&9|-|83969811|83969834&offsets|0|-2]|TGGAAGACTAGTGATTTTGTTG</td><td>>MI0000264|hsa-mir-7-2&WithFlank&15|+|88611819|88611940</td><td>38</td><td>59</td><td>22</td><td>TGGAAGACTAGTGATTTTGTTG</td><td>5043.6</td><td>8.2</td><td>0.8</td><td>10.4</td><td>4.1E-25</td><td>3.3E-24</td><td>MB231/MB468</td></tr>
<tr><td>>MI0000456|hsa-mir-140&WithFlank&16|+|69933075|69933186@69.89.21(+1U)&[MIMAT0004597&hsa-miR-140-3p&offsets|+1|+1(+1U);m-45&16|+|69933143|69933164&offsets|0|-1(+1U)]|ACCACAGGGTAGAACCACGGAT</td><td>>MI0000456|hsa-mir-140&WithFlank&16|+|69933075|69933186</td><td>69</td><td>89</td><td>21(+1U)</td><td>ACCACAGGGTAGAACCACGGAT</td><td>894.7</td><td>14.7</td><td>1.4</td><td>10.4</td><td>4.1E-25</td><td>3.4E-24</td><td>MB231/MB468</td></tr>
<tr><td>>MI0000439|hsa-mir-23b&WithFlank&9|+|95085202|95085310@67.85.19&[MIMAT0000418&hsa-miR-23b-3p&offsets|+3|-1;m-42&9|+|95085265|95085285&offsets|+3|+1]|ACATTGCCAGGGATTACCA</td><td>>MI0000439|hsa-mir-23b&WithFlank&9|+|95085202|95085310</td><td>67</td><td>85</td><td>19</td><td>ACATTGCCAGGGATTACCA</td><td>2652.8</td><td>16.0</td><td>1.5</td><td>10.4</td><td>3.3E-25</td><td>3.4E-24</td><td>BT20/MB468</td></tr>
<tr><td>>hg38_21_+_8433172_8446622_RNA45SN1_45S_50ntFlanks@12410.12448.39|TGATCCTTCGATGTCGGCTCTTCCTATCATTGTGAAGCA</td><td>>hg38_21_+_8433172_8446622_RNA45SN1_45S_50ntFlanks</td><td>12410</td><td>12448</td><td>39</td><td>TGATCCTTCGATGTCGGCTCTTCCTATCATTGTGAAGCA</td><td>271.8</td><td>11.0</td><td>1.1</td><td>10.4</td><td>4.2E-25</td><td>3.4E-24</td><td>MB231/MB468</td></tr>
<tr><td>>MI0000441|hsa-mir-30b&WithFlank&8|-|134800514|134800613@23.44.22&[MIMAT0000420&hsa-miR-30b-5p&offsets|0|0;m-76&8|-|134800570|134800591&offsets|0|0]|TGTAAACATCCTACACTCAGCT</td><td>>MI0000441|hsa-mir-30b&WithFlank&8|-|134800514|134800613</td><td>23</td><td>44</td><td>22</td><td>TGTAAACATCCTACACTCAGCT</td><td>18243.3</td><td>9.0</td><td>0.9</td><td>10.3</td><td>4.3E-25</td><td>3.5E-24</td><td>MB231/MB468</td></tr>
<tr><td>>hg38_21_+_8433172_8446622_RNA45SN1_45S_50ntFlanks@4735.4768.34|ACGAAAGTCGGAGGTTCGAAGACGATCAGATACC</td><td>>hg38_21_+_8433172_8446622_RNA45SN1_45S_50ntFlanks</td><td>4735</td><td>4768</td><td>34</td><td>ACGAAAGTCGGAGGTTCGAAGACGATCAGATACC</td><td>752.2</td><td>14.5</td><td>1.4</td><td>10.3</td><td>4.4E-25</td><td>3.5E-24</td><td>MB231/MB468</td></tr>
</table>

<table>
<tr><td>>MI0000093|hsa-mir-92a-1&WithFlank&13|+|91351308|91351397@54.73.20;MI0000094|hsa-mir-92a-2&WithFlank&X|-|134169532|134169618@54.73.20&[MIMAT0000092&hsa-miR-92a-3p&offsets|0|-2;m-9&13|+|91351361|91351382&offsets|0|-2];[MIMAT0000092_1&hsa-miR-92a-3p&offsets|0|-2;m-11&X|-|134169544|134169565&offsets|0|-2]|TATTGCACTTGTCCCGGCCT</td><td>>MI0000093|hsa-mir-92a-1&WithFlank&13|+|91351308|91351397</td><td>54</td><td>73</td><td>20</td><td>TATTGCACTTGTCCCGGCCT</td><td>738.7</td><td>7.8</td><td>0.8</td><td>10.3</td><td>4.4E-25</td><td>3.6E-24</td><td>MB231/MB468</td></tr>
<tr><td>>hg38_21_+_8433172_8446622_RNA45SN1_45S_50ntFlanks@483.506.24|TCGATGTGGTGACGTCGTGCTCTC</td><td>>hg38_21_+_8433172_8446622_RNA45SN1_45S_50ntFlanks</td><td>483</td><td>506</td><td>24</td><td>TCGATGTGGTGACGTCGTGCTCTC</td><td>1474.5</td><td>8.7</td><td>0.8</td><td>10.3</td><td>4.4E-25</td><td>3.6E-24</td><td>MB231/MB468</td></tr>
<tr><td>>MI0006444|hsa-mir-1307&WithFlank&10|-|103394247|103394407@86.107.22(+1A)&[MIMAT0005951&hsa-miR-1307-3p&offsets|0|0(+1A);m-66&10|-|103394300|103394322&offsets|0|-1(+1A)]|ACTCGGCGTGGCGTCGGTCGTGA</td><td>>MI0006444|hsa-mir-1307&WithFlank&10|-|103394247|103394407</td><td>86</td><td>107</td><td>22(+1A)</td><td>ACTCGGCGTGGCGTCGGTCGTGA</td><td>1254.0</td><td>7.1</td><td>0.7</td><td>10.4</td><td>3.6E-25</td><td>3.6E-24</td><td>BT20/MB468</td></tr>
<tr><td>>MI0000108|hsa-mir-103a-2&WithFlank&20|+|3917488|3917577@54.76.23;MI0000109|hsa-mir-103a-1&WithFlank&5|-|168560890|168560979@54.76.23&[MIMAT0000101&hsa-miR-103a-3p&offsets|0|0;m-12&20|+|3917541|3917563&offsets|0|0];[MIMAT0000101_1&hsa-miR-103a-3p&offsets|0|0;m-13&5|-|168560904|168560926&offsets|0|0;m-44220&5|-|168560908|168560926&offsets|0|+4]|AGCAGCATTGTACAGGGCTATGA</td><td>>MI0000108|hsa-mir-103a-2&WithFlank&20|+|3917488|3917577</td><td>54</td><td>76</td><td>23</td><td>AGCAGCATTGTACAGGGCTATGA</td><td>78435.2</td><td>7.3</td><td>0.7</td><td>10.4</td><td>3.6E-25</td><td>3.7E-24</td><td>BT20/MB468</td></tr>
<tr><td>>MI0000272|hsa-mir-182&WithFlank&7|-|129770377|129770498@29.49.21&[MIMAT0000259&hsa-miR-182-5p&offsets|0|-3;m-22&7|-|129770449|129770470&offsets|0|-1]|TTTGGCAATGGTAGAACTCAC</td><td>>MI0000272|hsa-mir-182&WithFlank&7|-|129770377|129770498</td><td>29</td><td>49</td><td>21</td><td>TTTGGCAATGGTAGAACTCAC</td><td>1638.3</td><td>9.0</td><td>0.9</td><td>10.4</td><td>3.7E-25</td><td>3.7E-24</td><td>BT20/MB468</td></tr>
<tr><td>>MI0000650|hsa-mir-200c&WithFlank&12|+|6963693|6963772@50.70.21&[MIMAT0000617&hsa-miR-200c-3p&offsets|0|-2;m-54&12|+|6963742|6963764&offsets|0|-2]|TAATACTGCCGGGTAATGATG</td><td>>MI0000650|hsa-mir-200c&WithFlank&12|+|6963693|6963772</td><td>50</td><td>70</td><td>21</td><td>TAATACTGCCGGGTAATGATG</td><td>7501.7</td><td>9.0</td><td>0.9</td><td>10.4</td><td>3.8E-25</td><td>3.8E-24</td><td>BT20/MB468</td></tr>
<tr><td>>trnaMT_SerGCT_MT_+_12207_12265@14.61.48|AGAACTGCTAACTCATGCCCCCATGTCTAACAACATGGCTTTCTCACC</td><td>>trnaMT_SerGCT_MT_+_12207_12265</td><td>14</td><td>61</td><td>48</td><td>AGAACTGCTAACTCATGCCCCCATGTCTAACAACATGGCTTTCTCACC</td><td>732.7</td><td>14.4</td><td>1.4</td><td>10.3</td><td>4.9E-25</td><td>3.9E-24</td><td>MB231/MB468</td></tr>
<tr><td>>MI0000299|hsa-mir-222&WithFlank&X|-|45747009|45747130@75.97.23(+1U)&[MIMAT0000279&hsa-miR-222-3p&offsets|0|+2(+1U);m-140&X|-|45747033|45747056&offsets|0|-1(+1U)]|AGCTACATCTGGCTACTGGGTCTT</td><td>>MI0000299|hsa-mir-222&WithFlank&X|-|45747009|45747130</td><td>75</td><td>97</td><td>23(+1U)</td><td>AGCTACATCTGGCTACTGGGTCTT</td><td>2726.5</td><td>7.0</td><td>0.7</td><td>10.4</td><td>4.0E-25</td><td>4.0E-24</td><td>BT20/MB468</td></tr>
<tr><td>>trnaMT_ValTAC_MT_+_1602_1670@53.71.19|TAACTTGACCGCTCTGACC</td><td>>trnaMT_ValTAC_MT_+_1602_1670</td><td>53</td><td>71</td><td>19</td><td>TAACTTGACCGCTCTGACC</td><td>810.3</td><td>10.1</td><td>1.0</td><td>10.3</td><td>5.0E-25</td><td>4.1E-24</td><td>MB231/MB468</td></tr>
<tr><td>>hg38_21_+_8433172_8446622_RNA45SN1_45S_50ntFlanks@10389.10407.19|TGAACAGCAGTTGAACATG</td><td>>hg38_21_+_8433172_8446622_RNA45SN1_45S_50ntFlanks</td><td>10389</td><td>10407</td><td>19</td><td>TGAACAGCAGTTGAACATG</td><td>3087.2</td><td>-7.4</td><td>0.7</td><td>-10.3</td><td>4.2E-25</td><td>4.2E-24</td><td>BT20/MB468</td></tr>
<tr><td>>hg38_21_+_8433172_8446622_RNA45SN1_45S_50ntFlanks@6651.6682.32|CGACTCTTAGCGGTGGATCACTCGGCTCGTGC</td><td>>hg38_21_+_8433172_8446622_RNA45SN1_45S_50ntFlanks</td><td>6651</td><td>6682</td><td>32</td><td>CGACTCTTAGCGGTGGATCACTCGGCTCGTGC</td><td>1737.0</td><td>5.5</td><td>0.5</td><td>10.3</td><td>4.2E-25</td><td>4.2E-24</td><td>BT20/MB468</td></tr>
<tr><td>>hg38_MT_+_598_1651_RNR1_12S_50ntFlanks@369.404.36|AATAAAGCTAAAACTCACCTGAGTTGTAAAAAACTC</td><td>>hg38_MT_+_598_1651_RNR1_12S_50ntFlanks</td><td>369</td><td>404</td><td>36</td><td>AATAAAGCTAAAACTCACCTGAGTTGTAAAAAACTC</td><td>714.2</td><td>10.1</td><td>1.0</td><td>10.3</td><td>5.2E-25</td><td>4.2E-24</td><td>MB231/MB468</td></tr>
<tr><td>>hg38_21_+_8433172_8446622_RNA45SN1_45S_50ntFlanks@9566.9597.32|GACGTGCAAATCGGTCGTCCGACCTGGGTATA</td><td>>hg38_21_+_8433172_8446622_RNA45SN1_45S_50ntFlanks</td><td>9566</td><td>9597</td><td>32</td><td>GACGTGCAAATCGGTCGTCCGACCTGGGTATA</td><td>774.0</td><td>-4.7</td><td>0.5</td><td>-10.3</td><td>4.3E-25</td><td>4.3E-24</td><td>BT20/MB468</td></tr>
<tr><td>>hg38_21_+_8433172_8446622_RNA45SN1_45S_50ntFlanks@6651.6673.23|CGACTCTTAGCGGTGGATCACTC</td><td>>hg38_21_+_8433172_8446622_RNA45SN1_45S_50ntFlanks</td><td>6651</td><td>6673</td><td>23</td><td>CGACTCTTAGCGGTGGATCACTC</td><td>1765.7</td><td>5.2</td><td>0.5</td><td>10.3</td><td>4.3E-25</td><td>4.3E-24</td><td>BT20/MB468</td></tr>
<tr><td>>hg38_21_+_8433172_8446622_RNA45SN1_45S_50ntFlanks@475.507.33|CTTCGTGATCGATGTGGTGACGTCGTGCTCTCC</td><td>>hg38_21_+_8433172_8446622_RNA45SN1_45S_50ntFlanks</td><td>475</td><td>507</td><td>33</td><td>CTTCGTGATCGATGTGGTGACGTCGTGCTCTCC</td><td>5790.4</td><td>7.3</td><td>0.7</td><td>10.3</td><td>5.4E-25</td><td>4.4E-24</td><td>MB231/MB468</td></tr>
<tr><td>>hg38_21_+_8433172_8446622_RNA45SN1_45S_50ntFlanks@12995.13038.44|CTCGCTGCGATCTATTGAAAGTCAGCCCTCGACACAAGGGTTTG</td><td>>hg38_21_+_8433172_8446622_RNA45SN1_45S_50ntFlanks</td><td>12995</td><td>13038</td><td>44</td><td>CTCGCTGCGATCTATTGAAAGTCAGCCCTCGACACAAGGGTTTG</td><td>16712.3</td><td>13.6</td><td>1.3</td><td>10.3</td><td>5.5E-25</td><td>4.4E-24</td><td>MB231/MB468</td></tr>
<tr><td>>hg38_21_+_8433172_8446622_RNA45SN1_45S_50ntFlanks@11575.11605.31|ACTTAGAACTGGTGCGGACCAGGGGAATCCG</td><td>>hg38_21_+_8433172_8446622_RNA45SN1_45S_50ntFlanks</td><td>11575</td><td>11605</td><td>31</td><td>ACTTAGAACTGGTGCGGACCAGGGGAATCCG</td><td>965.7</td><td>-6.0</td><td>0.6</td><td>-10.3</td><td>5.6E-25</td><td>4.5E-24</td><td>MB231/MB468</td></tr>
<tr><td>>MI0000078|hsa-mir-22&WithFlank&17|-|1713897|1713993@59.80.22&[MIMAT0000077&hsa-miR-22-3p&offsets|0|0;m-2&17|-|1713914|1713935&offsets|0|0]|AAGCTGCCAGTTGAAGAACTGT</td><td>>MI0000078|hsa-mir-22&WithFlank&17|-|1713897|1713993</td><td>59</td><td>80</td><td>22</td><td>AAGCTGCCAGTTGAAGAACTGT</td><td>22066.3</td><td>8.1</td><td>0.8</td><td>10.3</td><td>5.6E-25</td><td>4.5E-24</td><td>MB231/MB468</td></tr>
<tr><td>>MI0000433|hsa-let-7g&WithFlank&3|-|52268272|52268367@11.31.21(+1C)&[MIMAT0000414&hsa-let-7g-5p&offsets|0|-1(+1C);m-58&3|-|52268336|52268357&offsets|0|-1(+1C)]|TGAGGTAGTAGTTTGTACAGTC</td><td>>MI0000433|hsa-let-7g&WithFlank&3|-|52268272|52268367</td><td>11</td><td>31</td><td>21(+1C)</td><td>TGAGGTAGTAGTTTGTACAGTC</td><td>40675.5</td><td>7.7</td><td>0.7</td><td>10.3</td><td>5.6E-25</td><td>4.5E-24</td><td>MB231/MB468</td></tr>
<tr><td>>hg38_21_+_8433172_8446622_RNA45SN1_45S_50ntFlanks@475.511.37|CTTCGTGATCGATGTGGTGACGTCGTGCTCTCCCGGG</td><td>>hg38_21_+_8433172_8446622_RNA45SN1_45S_50ntFlanks</td><td>475</td><td>511</td><td>37</td><td>CTTCGTGATCGATGTGGTGACGTCGTGCTCTCCCGGG</td><td>3320.6</td><td>6.6</td><td>0.6</td><td>10.3</td><td>4.6E-25</td><td>4.6E-24</td><td>BT20/MB468</td></tr>
<tr><td>>MI0000456|hsa-mir-140&WithFlank&16|+|69933075|69933186@69.89.21&[MIMAT0004597&hsa-miR-140-3p&offsets|+1|+1;m-45&16|+|69933143|69933164&offsets|0|-1]|ACCACAGGGTAGAACCACGGA</td><td>>MI0000456|hsa-mir-140&WithFlank&16|+|69933075|69933186</td><td>69</td><td>89</td><td>21</td><td>ACCACAGGGTAGAACCACGGA</td><td>4204.2</td><td>12.4</td><td>1.2</td><td>10.3</td><td>5.9E-25</td><td>4.7E-24</td><td>MB231/MB468</td></tr>
<tr><td>>trna119_LysCTT_1_-_145395522_145395594@1.33.33,trna11_LysCTT_5_-_180648979_180649051@1.33.33,trna13_LysCTT_6_+_26556774_26556846@1.33.33,trna7_LysCTT_16_+_3225692_3225764@1.33.33,trna9_LysCTT_5_+_180634755_180634827@1.33.33|GCCCGGCTAGCTCAGTCGGTAGAGCATGAGACT</td><td>>trna119_LysCTT_1_-_145395522_145395594</td><td>1</td><td>33</td><td>33</td><td>GCCCGGCTAGCTCAGTCGGTAGAGCATGAGACT</td><td>62476.6</td><td>10.1</td><td>1.0</td><td>10.3</td><td>5.9E-25</td><td>4.7E-24</td><td>MB231/MB468</td></tr>
<tr><td>>am_trnaMT_MetCAT_MT_+_4402_4469@1.47.47|AGTAAGGTCAGCTAAATAAGCTATCGGGCCCATACCCCGAAAATGTT</td><td>>am_trnaMT_MetCAT_MT_+_4402_4469</td><td>1</td><td>47</td><td>47</td><td>AGTAAGGTCAGCTAAATAAGCTATCGGGCCCATACCCCGAAAATGTT</td><td>2294.8</td><td>16.1</td><td>1.6</td><td>10.3</td><td>6.2E-25</td><td>4.9E-24</td><td>MB231/MB468</td></tr>
<tr><td>>hg38_21_+_8433172_8446622_RNA45SN1_45S_50ntFlanks@9574.9610.37|AATCGGTCGTCCGACCTGGGTATAGGGGCGAAAGACT</td><td>>hg38_21_+_8433172_8446622_RNA45SN1_45S_50ntFlanks</td><td>9574</td><td>9610</td><td>37</td><td>AATCGGTCGTCCGACCTGGGTATAGGGGCGAAAGACT</td><td>1414.6</td><td>7.5</td><td>0.7</td><td>10.3</td><td>6.2E-25</td><td>5.0E-24</td><td>MB231/MB468</td></tr>
<tr><td>>MI0000263|hsa-mir-7-1&WithFlank&9|-|83969742|83969863@30.54.25&[MIMAT0000252_2&hsa-miR-7-5p&offsets|0|+1;m-401&9|-|83969811|83969834&offsets|0|+1]|TGGAAGACTAGTGATTTTGTTGTTT</td><td>>MI0000263|hsa-mir-7-1&WithFlank&9|-|83969742|83969863</td><td>30</td><td>54</td><td>25</td><td>TGGAAGACTAGTGATTTTGTTGTTT</td><td>7907.9</td><td>8.0</td><td>0.8</td><td>10.3</td><td>6.2E-25</td><td>5.0E-24</td><td>MB231/MB468</td></tr>
<tr><td>>hg38_21_+_8433172_8446622_RNA45SN1_45S_50ntFlanks@12395.12418.24|AGCGACGTCGCTTTTTGATCCTTC</td><td>>hg38_21_+_8433172_8446622_RNA45SN1_45S_50ntFlanks</td><td>12395</td><td>12418</td><td>24</td><td>AGCGACGTCGCTTTTTGATCCTTC</td><td>831.5</td><td>7.7</td><td>0.7</td><td>10.3</td><td>6.6E-25</td><td>5.3E-24</td><td>MB231/MB468</td></tr>
<tr><td>>MI0000816|hsa-mir-335&WithFlank&7|+|130496105|130496210@58.78.21(+1A)&[MIMAT0004703&hsa-miR-335-3p&offsets|0|-1(+1A);m-159&7|+|130496162|130496183&offsets|0|-1(+1A)]|TTTTTCATTATTGCTCCTGACA</td><td>>MI0000816|hsa-mir-335&WithFlank&7|+|130496105|130496210</td><td>58</td><td>78</td><td>21(+1A)</td><td>TTTTTCATTATTGCTCCTGACA</td><td>1447.7</td><td>8.0</td><td>0.8</td><td>10.3</td><td>6.6E-25</td><td>5.3E-24</td><td>MB231/MB468</td></tr>
<tr><td>>hg38_MT_+_1621_3279_RNR2_16S_50ntFlanks@1428.1457.30|ATTAAAGTCCTACGTGATCTGAGTTCAGAC</td><td>>hg38_MT_+_1621_3279_RNR2_16S_50ntFlanks</td><td>1428</td><td>1457</td><td>30</td><td>ATTAAAGTCCTACGTGATCTGAGTTCAGAC</td><td>448.3</td><td>9.2</td><td>0.9</td><td>10.3</td><td>6.6E-25</td><td>5.3E-24</td><td>MB231/MB468</td></tr>
<tr><td>>MI0000469|hsa-mir-125a&WithFlank&19|+|51693248|51693345@21.43.23(+1U)&[MIMAT0000443&hsa-miR-125a-5p&offsets|0|-1(+1U);m-63&19|+|51693268|51693289&offsets|0|+1(+1U)]|TCCCTGAGACCCTTTAACCTGTGT</td><td>>MI0000469|hsa-mir-125a&WithFlank&19|+|51693248|51693345</td><td>21</td><td>43</td><td>23(+1U)</td><td>TCCCTGAGACCCTTTAACCTGTGT</td><td>2142.6</td><td>6.7</td><td>0.6</td><td>10.3</td><td>6.7E-25</td><td>5.3E-24</td><td>MB231/MB468</td></tr>
<tr><td>>MI0006444|hsa-mir-1307&WithFlank&10|-|103394247|103394407@86.109.24&[MIMAT0005951&hsa-miR-1307-3p&offsets|0|+2;m-66&10|-|103394300|103394322&offsets|0|+1]|ACTCGGCGTGGCGTCGGTCGTGGT</td><td>>MI0006444|hsa-mir-1307&WithFlank&10|-|103394247|103394407</td><td>86</td><td>109</td><td>24</td><td>ACTCGGCGTGGCGTCGGTCGTGGT</td><td>1374.3</td><td>8.1</td><td>0.8</td><td>10.3</td><td>6.7E-25</td><td>5.3E-24</td><td>MB231/MB468</td></tr>
<tr><td>>hg38_21_+_8433172_8446622_RNA45SN1_45S_50ntFlanks@3992.4023.32|GGTGACTCTAGATAACCTCGGGCCGATCGCAC</td><td>>hg38_21_+_8433172_8446622_RNA45SN1_45S_50ntFlanks</td><td>3992</td><td>4023</td><td>32</td><td>GGTGACTCTAGATAACCTCGGGCCGATCGCAC</td><td>1060.6</td><td>7.2</td><td>0.7</td><td>10.3</td><td>6.7E-25</td><td>5.3E-24</td><td>MB231/MB468</td></tr>
<tr><td>>MI0000273|hsa-mir-183&WithFlank&7|-|129774899|129775020@34.55.22&[MIMAT0000261&hsa-miR-183-5p&offsets|+1|+1;m-41&7|-|129774966|129774987&offsets|0|0]|ATGGCACTGGTAGAATTCACTG</td><td>>MI0000273|hsa-mir-183&WithFlank&7|-|129774899|129775020</td><td>34</td><td>55</td><td>22</td><td>ATGGCACTGGTAGAATTCACTG</td><td>8237.1</td><td>7.6</td><td>0.7</td><td>10.3</td><td>5.4E-25</td><td>5.3E-24</td><td>BT20/MB468</td></tr>
<tr><td>>am_MI0000786|hsa-mir-378a&WithFlank&5|+|149732819|149732896@50.70.21(+1U)&[MIMAT0000732&hsa-miR-378a-3p&offsets|+1|0(+1U);m-44&5|+|149732867|149732888&offsets|+1|0(+1U)]|CTGGACTTGGAGTCAGAAGGCT</td><td>>am_MI0000786|hsa-mir-378a&WithFlank&5|+|149732819|149732896</td><td>50</td><td>70</td><td>21(+1U)</td><td>CTGGACTTGGAGTCAGAAGGCT</td><td>685.3</td><td>4.7</td><td>0.5</td><td>10.3</td><td>5.5E-25</td><td>5.4E-24</td><td>BT20/MB468</td></tr>
<tr><td>>hg38_1_-_228634819_228635039_RNA5S12_5S_50ntFlanks@146.170.25|TGGGAATACCGGGTGCTGTAGGCTT</td><td>>hg38_1_-_228634819_228635039_RNA5S12_5S_50ntFlanks</td><td>146</td><td>170</td><td>25</td><td>TGGGAATACCGGGTGCTGTAGGCTT</td><td>2711.6</td><td>7.8</td><td>0.8</td><td>10.3</td><td>6.8E-25</td><td>5.4E-24</td><td>MB231/MB468</td></tr>
<tr><td>>MI0000272|hsa-mir-182&WithFlank&7|-|129770377|129770498@29.51.23&[MIMAT0000259&hsa-miR-182-5p&offsets|0|-1;m-22&7|-|129770449|129770470&offsets|0|+1]|TTTGGCAATGGTAGAACTCACAC</td><td>>MI0000272|hsa-mir-182&WithFlank&7|-|129770377|129770498</td><td>29</td><td>51</td><td>23</td><td>TTTGGCAATGGTAGAACTCACAC</td><td>7798.6</td><td>8.7</td><td>0.8</td><td>10.3</td><td>5.5E-25</td><td>5.4E-24</td><td>BT20/MB468</td></tr>
<tr><td>>hg38_21_+_8433172_8446622_RNA45SN1_45S_50ntFlanks@12073.12095.23|CCGAGGGGCTCTCGCTTCTGGCG</td><td>>hg38_21_+_8433172_8446622_RNA45SN1_45S_50ntFlanks</td><td>12073</td><td>12095</td><td>23</td><td>CCGAGGGGCTCTCGCTTCTGGCG</td><td>1860.6</td><td>-7.6</td><td>0.7</td><td>-10.3</td><td>5.6E-25</td><td>5.5E-24</td><td>BT20/MB468</td></tr>
<tr><td>>hg38_21_+_8433172_8446622_RNA45SN1_45S_50ntFlanks@476.520.45|TTCGTGATCGATGTGGTGACGTCGTGCTCTCCCGGGCCGGGTCCG</td><td>>hg38_21_+_8433172_8446622_RNA45SN1_45S_50ntFlanks</td><td>476</td><td>520</td><td>45</td><td>TTCGTGATCGATGTGGTGACGTCGTGCTCTCCCGGGCCGGGTCCG</td><td>2846.8</td><td>11.8</td><td>1.1</td><td>10.3</td><td>7.0E-25</td><td>5.5E-24</td><td>MB231/MB468</td></tr>
<tr><td>>MI0003591|hsa-mir-584&WithFlank&5|-|149062307|149062415@22.40.19&[MIMAT0003249&hsa-miR-584-5p&offsets|0|-3;m-167&5|-|149062374|149062394&offsets|0|-2]|TTATGGTTTGCCTGGGACT</td><td>>MI0003591|hsa-mir-584&WithFlank&5|-|149062307|149062415</td><td>22</td><td>40</td><td>19</td><td>TTATGGTTTGCCTGGGACT</td><td>2537.8</td><td>10.6</td><td>1.0</td><td>10.3</td><td>7.0E-25</td><td>5.6E-24</td><td>MB231/MB468</td></tr>
</table>

<table>
<tr><th></th><th></th><th></th><th></th><th></th><th></th><th></th><th></th><th></th><th></th><th></th><th></th><th></th></tr>
<tr><td>>trnaMT_GluTTC_MT_-_14674_14742@34.72.39,trnalookalike8_GluTTC_5_-_93905172_93905240@34.72.39|ATATCATTGGTCGTGGTTGTAGTCCGTGCGAGAATACCA</td><td>>trnaMT_GluTTC_MT_-_14674_14742</td><td>34</td><td>72</td><td>39</td><td>ATATCATTGGTCGTGGTTGTAGTCCGTGCGAGAATACCA</td><td>4182.5</td><td>8.4</td><td>0.8</td><td>10.3</td><td>5.8E-25</td><td>5.7E-24</td><td>BT20/MB468</td></tr>
<tr><td>>hg38_21_+_8433172_8446622_RNA45SN1_45S_50ntFlanks@10408.10430.23|GGTCAGTCGGTCCTGAGAGATGG</td><td>>hg38_21_+_8433172_8446622_RNA45SN1_45S_50ntFlanks</td><td>10408</td><td>10430</td><td>23</td><td>GGTCAGTCGGTCCTGAGAGATGG</td><td>851.9</td><td>-6.8</td><td>0.7</td><td>-10.3</td><td>7.2E-25</td><td>5.7E-24</td><td>MB231/MB468</td></tr>
<tr><td>>MI0000077|hsa-mir-21&WithFlank&17|+|59841260|59841343@14.36.23(+1A)&[MIMAT0000076&hsa-miR-21-5p&offsets|0|+1(+1A);m-1&17|+|59841273|59841294&offsets|0|+1(+1A)]|TAGCTTATCAGACTGATGTTGACA</td><td>>MI0000077|hsa-mir-21&WithFlank&17|+|59841260|59841343</td><td>14</td><td>36</td><td>23(+1A)</td><td>TAGCTTATCAGACTGATGTTGACA</td><td>393148.2</td><td>7.5</td><td>0.7</td><td>10.3</td><td>7.2E-25</td><td>5.7E-24</td><td>MB231/MB468</td></tr>
<tr><td>>MI0000816|hsa-mir-335&WithFlank&7|+|130496105|130496210@58.79.22&[MIMAT0004703&hsa-miR-335-3p&offsets|0|0;m-159&7|+|130496162|130496183&offsets|0|0]|TTTTTCATTATTGCTCCTGACC</td><td>>MI0000816|hsa-mir-335&WithFlank&7|+|130496105|130496210</td><td>58</td><td>79</td><td>22</td><td>TTTTTCATTATTGCTCCTGACC</td><td>9739.6</td><td>8.1</td><td>0.8</td><td>10.3</td><td>7.6E-25</td><td>6.0E-24</td><td>MB231/MB468</td></tr>
<tr><td>>trna20_GluTTC_2_-_131094701_131094772@1.32.32,trna5_GluTTC_13_-_41634874_41634945@1.32.32|TCCCATATGGTCTAGCGGTTAGGATTCCTGGT</td><td>>trna20_GluTTC_2_-_131094701_131094772</td><td>1</td><td>32</td><td>32</td><td>TCCCATATGGTCTAGCGGTTAGGATTCCTGGT</td><td>847.3</td><td>14.6</td><td>1.4</td><td>10.3</td><td>7.7E-25</td><td>6.1E-24</td><td>MB231/MB468</td></tr>
<tr><td>>hg38_21_+_8433172_8446622_RNA45SN1_45S_50ntFlanks@5420.5443.24|CCGATTGGATGGTTTAGTGAGGCC</td><td>>hg38_21_+_8433172_8446622_RNA45SN1_45S_50ntFlanks</td><td>5420</td><td>5443</td><td>24</td><td>CCGATTGGATGGTTTAGTGAGGCC</td><td>922.6</td><td>-7.5</td><td>0.7</td><td>-10.3</td><td>6.3E-25</td><td>6.2E-24</td><td>BT20/MB468</td></tr>
<tr><td>>MI0000809|hsa-mir-151a&WithFlank&8|-|140732558|140732659@51.73.23(+1U)&[MIMAT0000757&hsa-miR-151a-3p&offsets|-2|0(+1U);m-30&8|-|140732587|140732607&offsets|-2|0(+1U)]|TACTAGACTGAAGCTCCTTGAGGT</td><td>>MI0000809|hsa-mir-151a&WithFlank&8|-|140732558|140732659</td><td>51</td><td>73</td><td>23(+1U)</td><td>TACTAGACTGAAGCTCCTTGAGGT</td><td>573.3</td><td>7.0</td><td>0.7</td><td>10.3</td><td>6.8E-25</td><td>6.7E-24</td><td>BT20/MB468</td></tr>
<tr><td>>hg38_21_+_8433172_8446622_RNA45SN1_45S_50ntFlanks@9504.9526.23|GTGAACTATGCCTGGGCAGGGCG</td><td>>hg38_21_+_8433172_8446622_RNA45SN1_45S_50ntFlanks</td><td>9504</td><td>9526</td><td>23</td><td>GTGAACTATGCCTGGGCAGGGCG</td><td>3665.5</td><td>-13.4</td><td>1.3</td><td>-10.3</td><td>8.5E-25</td><td>6.7E-24</td><td>MB231/MB468</td></tr>
<tr><td>>MI0000087|hsa-mir-29a&WithFlank&7|-|130876741|130876816@48.70.23&[MIMAT0000086&hsa-miR-29a-3p&offsets|0|+1;m-26&7|-|130876749|130876769&offsets|0|+2]|TAGCACCATCTGAAATCGGTTAT</td><td>>MI0000087|hsa-mir-29a&WithFlank&7|-|130876741|130876816</td><td>48</td><td>70</td><td>23</td><td>TAGCACCATCTGAAATCGGTTAT</td><td>626.9</td><td>12.1</td><td>1.2</td><td>10.3</td><td>8.6E-25</td><td>6.7E-24</td><td>MB231/MB468</td></tr>
<tr><td>>trnaMT_MetCAT_MT_+_4402_4469@38.71.34|CGAAAATGTTGGTTATACCCTTCCCGTACTACCA</td><td>>trnaMT_MetCAT_MT_+_4402_4469</td><td>38</td><td>71</td><td>34</td><td>CGAAAATGTTGGTTATACCCTTCCCGTACTACCA</td><td>592.8</td><td>11.3</td><td>1.1</td><td>10.3</td><td>8.6E-25</td><td>6.7E-24</td><td>MB231/MB468</td></tr>
<tr><td>>am_MI0003639|hsa-mir-625&WithFlank&14|+|65471096|65471192@58.78.21(+1U)&[MIMAT0004808&hsa-miR-625-3p&offsets|0|-1(+1U);m-137&14|+|65471153|65471174&offsets|0|-1(+1U)]|GACTATAGAACTTTCCCCCTCT</td><td>>am_MI0003639|hsa-mir-625&WithFlank&14|+|65471096|65471192</td><td>58</td><td>78</td><td>21(+1U)</td><td>GACTATAGAACTTTCCCCCTCT</td><td>610.3</td><td>12.1</td><td>1.2</td><td>10.3</td><td>8.7E-25</td><td>6.8E-24</td><td>MB231/MB468</td></tr>
<tr><td>>MI0000456|hsa-mir-140&WithFlank&16|+|69933075|69933186@69.88.20&[MIMAT0004597&hsa-miR-140-3p&offsets|+1|0;m-45&16|+|69933143|69933164&offsets|0|-2]|ACCACAGGGTAGAACCACGG</td><td>>MI0000456|hsa-mir-140&WithFlank&16|+|69933075|69933186</td><td>69</td><td>88</td><td>20</td><td>ACCACAGGGTAGAACCACGG</td><td>924.1</td><td>12.7</td><td>1.2</td><td>10.3</td><td>8.7E-25</td><td>6.8E-24</td><td>MB231/MB468</td></tr>
<tr><td>>am_MI0003581|hsa-mir-574&WithFlank&4|+|38868026|38868133@31.53.23&[MIMAT0004795&hsa-miR-574-5p&offsets|0|0;m-213&4|+|38868056|38868078&offsets|0|0]|TGAGTGTGTGTGTGTGAGTGTGT</td><td>>am_MI0003581|hsa-mir-574&WithFlank&4|+|38868026|38868133</td><td>31</td><td>53</td><td>23</td><td>TGAGTGTGTGTGTGTGAGTGTGT</td><td>1420.8</td><td>8.0</td><td>0.8</td><td>10.3</td><td>7.0E-25</td><td>6.8E-24</td><td>BT20/MB468</td></tr>
<tr><td>>hg38_21_+_8433172_8446622_RNA45SN1_45S_50ntFlanks@12416.12448.33|TTCGATGTCGGCTCTTCCTATCATTGTGAAGCA</td><td>>hg38_21_+_8433172_8446622_RNA45SN1_45S_50ntFlanks</td><td>12416</td><td>12448</td><td>33</td><td>TTCGATGTCGGCTCTTCCTATCATTGTGAAGCA</td><td>6289.1</td><td>9.2</td><td>0.9</td><td>10.3</td><td>8.8E-25</td><td>6.9E-24</td><td>MB231/MB468</td></tr>
<tr><td>>hg38_21_+_8433172_8446622_RNA45SN1_45S_50ntFlanks@7973.8009.37|GACGCGACCTCAGATCAGACGTGGCGACCCGCTGAAT</td><td>>hg38_21_+_8433172_8446622_RNA45SN1_45S_50ntFlanks</td><td>7973</td><td>8009</td><td>37</td><td>GACGCGACCTCAGATCAGACGTGGCGACCCGCTGAAT</td><td>1113.5</td><td>6.2</td><td>0.6</td><td>10.3</td><td>7.2E-25</td><td>7.0E-24</td><td>BT20/MB468</td></tr>
<tr><td>>hg38_1_-_228634819_228635039_RNA5S12_5S_50ntFlanks@139.170.32|GACCGCCTGGGAATACCGGGTGCTGTAGGCTT</td><td>>hg38_1_-_228634819_228635039_RNA5S12_5S_50ntFlanks</td><td>139</td><td>170</td><td>32</td><td>GACCGCCTGGGAATACCGGGTGCTGTAGGCTT</td><td>2958.5</td><td>8.0</td><td>0.8</td><td>10.3</td><td>9.1E-25</td><td>7.0E-24</td><td>MB231/MB468</td></tr>
<tr><td>>trna116_GluCTC_1_-_145399233_145399304@1.32.32,trna59_GluCTC_1_+_249168447_249168518@1.32.32,trna71_GluCTC_1_-_161439189_161439260@1.32.32,trna74_GluCTC_1_-_161431809_161431880@1.32.32,trna77_GluCTC_1_-_161424398_161424469@1.32.32,trna77_GluCTC_6_+_28949976_28950047@1.32.32,trna80_GluCTC_1_-_161417018_161417089@1.32.32,trna87_GluCTC_6_-_126101393_126101464@1.32.32|TCCCTGGTGGTCTAGTGGTTAGGATTCGGCGC</td><td>>trna116_GluCTC_1_-_145399233_145399304</td><td>1</td><td>32</td><td>32</td><td>TCCCTGGTGGTCTAGTGGTTAGGATTCGGCGC</td><td>6442.7</td><td>8.3</td><td>0.8</td><td>10.3</td><td>9.1E-25</td><td>7.0E-24</td><td>MB231/MB468</td></tr>
<tr><td>>trna119_LysCTT_1_-_145395522_145395594@4.34.31,trna11_LysCTT_5_-_180648979_180649051@4.34.31,trna13_LysCTT_6_+_26556774_26556846@4.34.31,trna7_LysCTT_16_+_3225692_3225764@4.34.31,trna9_LysCTT_5_+_180634755_180634827@4.34.31|CGGCTAGCTCAGTCGGTAGAGCATGAGACTC</td><td>>trna119_LysCTT_1_-_145395522_145395594</td><td>4</td><td>34</td><td>31</td><td>CGGCTAGCTCAGTCGGTAGAGCATGAGACTC</td><td>843.1</td><td>14.6</td><td>1.4</td><td>10.3</td><td>9.0E-25</td><td>7.0E-24</td><td>MB231/MB468</td></tr>
<tr><td>>hg38_1_-_228634819_228635039_RNA5S12_5S_50ntFlanks@147.171.25|GGGAATACCGGGTGCTGTAGGCTTT</td><td>>hg38_1_-_228634819_228635039_RNA5S12_5S_50ntFlanks</td><td>147</td><td>171</td><td>25</td><td>GGGAATACCGGGTGCTGTAGGCTTT</td><td>1569.2</td><td>5.3</td><td>0.5</td><td>10.3</td><td>7.4E-25</td><td>7.3E-24</td><td>BT20/MB468</td></tr>
<tr><td>>MI0000809|hsa-mir-151a&WithFlank&8|-|140732558|140732659@53.75.23&[MIMAT0000757&hsa-miR-151a-3p&offsets|0|+2;m-30&8|-|140732587|140732607&offsets|0|+2]|CTAGACTGAAGCTCCTTGAGGAC</td><td>>MI0000809|hsa-mir-151a&WithFlank&8|-|140732558|140732659</td><td>53</td><td>75</td><td>23</td><td>CTAGACTGAAGCTCCTTGAGGAC</td><td>625.0</td><td>12.1</td><td>1.2</td><td>10.3</td><td>9.6E-25</td><td>7.4E-24</td><td>MB231/MB468</td></tr>
<tr><td>>trna11_GluTTC_15_-_26327381_26327452@1.36.36,trna3_GluTTC_13_-_45492062_45492133@1.36.36|TCCCACATGGTCTAGCGGTTAGGATTCCTGGTTTTC</td><td>>trna11_GluTTC_15_-_26327381_26327452</td><td>1</td><td>36</td><td>36</td><td>TCCCACATGGTCTAGCGGTTAGGATTCCTGGTTTTC</td><td>1653.7</td><td>15.6</td><td>1.5</td><td>10.3</td><td>9.7E-25</td><td>7.5E-24</td><td>MB231/MB468</td></tr>
<tr><td>>trna152_ValCAC_6_-_27248049_27248121@1.32.32|GCTTCTGTAGTGTAGTGGTTATCACGTTCGCC</td><td>>trna152_ValCAC_6_-_27248049_27248121</td><td>1</td><td>32</td><td>32</td><td>GCTTCTGTAGTGTAGTGGTTATCACGTTCGCC</td><td>9640.4</td><td>12.2</td><td>1.2</td><td>10.3</td><td>9.7E-25</td><td>7.5E-24</td><td>MB231/MB468</td></tr>
<tr><td>>hg38_1_-_228634819_228635039_RNA5S12_5S_50ntFlanks@147.171.25|GGGAATACCGGGTGCTGTAGGCTTT</td><td>>hg38_1_-_228634819_228635039_RNA5S12_5S_50ntFlanks</td><td>147</td><td>171</td><td>25</td><td>GGGAATACCGGGTGCTGTAGGCTTT</td><td>2939.9</td><td>6.5</td><td>0.6</td><td>10.3</td><td>9.8E-25</td><td>7.5E-24</td><td>MB231/MB468</td></tr>
<tr><td>>hg38_21_+_8433172_8446622_RNA45SN1_45S_50ntFlanks@11326.11345.20|CTCGTCGCGCGCGCGTCCGC</td><td>>hg38_21_+_8433172_8446622_RNA45SN1_45S_50ntFlanks</td><td>11326</td><td>11345</td><td>20</td><td>CTCGTCGCGCGCGCGTCCGC</td><td>1830.6</td><td>15.7</td><td>1.5</td><td>10.3</td><td>9.8E-25</td><td>7.6E-24</td><td>MB231/MB468</td></tr>
<tr><td>>MI0000734|hsa-mir-106b&WithFlank&7|-|100093987|100094080@58.78.21(+1A)&[MIMAT0004672&hsa-miR-106b-3p&offsets|0|-1(+1A);m-87&7|-|100094003|100094023&offsets|0|0(+1A)]|CCGCACTGTGGGTACTTGCTGA</td><td>>MI0000734|hsa-mir-106b&WithFlank&7|-|100093987|100094080</td><td>58</td><td>78</td><td>21(+1A)</td><td>CCGCACTGTGGGTACTTGCTGA</td><td>1001.0</td><td>7.9</td><td>0.8</td><td>10.3</td><td>1.0E-24</td><td>7.7E-24</td><td>MB231/MB468</td></tr>
<tr><td>>MI0000078|hsa-mir-22&WithFlank&17|-|1713897|1713993@59.80.22(+1A)&[MIMAT0000077&hsa-miR-22-3p&offsets|0|0(+1A);m-2&17|-|1713914|1713935&offsets|0|0(+1A)]|AAGCTGCCAGTTGAAGAACTGTA</td><td>>MI0000078|hsa-mir-22&WithFlank&17|-|1713897|1713993</td><td>59</td><td>80</td><td>22(+1A)</td><td>AAGCTGCCAGTTGAAGAACTGTA</td><td>650.5</td><td>7.7</td><td>0.7</td><td>10.3</td><td>1.0E-24</td><td>7.9E-24</td><td>MB231/MB468</td></tr>
<tr><td>>hg38_21_+_8433172_8446622_RNA45SN1_45S_50ntFlanks@7978.8017.40|GACCTCAGATCAGACGTGGCGACCCGCTGAATTTAAGCAT</td><td>>hg38_21_+_8433172_8446622_RNA45SN1_45S_50ntFlanks</td><td>7978</td><td>8017</td><td>40</td><td>GACCTCAGATCAGACGTGGCGACCCGCTGAATTTAAGCAT</td><td>505.8</td><td>13.9</td><td>1.4</td><td>10.3</td><td>1.0E-24</td><td>8.0E-24</td><td>MB231/MB468</td></tr>
<tr><td>>hg38_MT_+_1621_3279_RNR2_16S_50ntFlanks@51.71.21|GCTAAACCTAGCCCCAAACCC</td><td>>hg38_MT_+_1621_3279_RNR2_16S_50ntFlanks</td><td>51</td><td>71</td><td>21</td><td>GCTAAACCTAGCCCCAAACCC</td><td>1515.8</td><td>8.3</td><td>0.8</td><td>10.3</td><td>1.0E-24</td><td>8.0E-24</td><td>MB231/MB468</td></tr>
<tr><td>>hg38_21_+_8433172_8446622_RNA45SN1_45S_50ntFlanks@10680.10711.32|GTTCCGGCGGCGTCCGGTGAGCTCTCGCTGGC</td><td>>hg38_21_+_8433172_8446622_RNA45SN1_45S_50ntFlanks</td><td>10680</td><td>10711</td><td>32</td><td>GTTCCGGCGGCGTCCGGTGAGCTCTCGCTGGC</td><td>6593.9</td><td>-6.4</td><td>0.6</td><td>-10.3</td><td>1.1E-24</td><td>8.1E-24</td><td>MB231/MB468</td></tr>
<tr><td>>MI0003560|hsa-mir-92b&WithFlank&1|+|155195171|155195278@67.88.22(+3U)&[MIMAT0003218&hsa-miR-92b-3p&offsets|0|0(+3U);m-115&1|+|155195237|155195258&offsets|0|0(+3U)]|TATTGCACTCGTCCCGGCCTCCTTT</td><td>>MI0003560|hsa-mir-92b&WithFlank&1|+|155195171|155195278</td><td>67</td><td>88</td><td>22(+3U)</td><td>TATTGCACTCGTCCCGGCCTCCTTT</td><td>752.3</td><td>12.4</td><td>1.2</td><td>10.3</td><td>1.1E-24</td><td>8.2E-24</td><td>MB231/MB468</td></tr>
<tr><td>>hg38_21_+_8433172_8446622_RNA45SN1_45S_50ntFlanks@10351.10376.26|CGAGAACTTTGAAGGCCGAAGTGGAG</td><td>>hg38_21_+_8433172_8446622_RNA45SN1_45S_50ntFlanks</td><td>10351</td><td>10376</td><td>26</td><td>CGAGAACTTTGAAGGCCGAAGTGGAG</td><td>1196.9</td><td>-5.8</td><td>0.6</td><td>-10.3</td><td>1.1E-24</td><td>8.4E-24</td><td>MB231/MB468</td></tr>
<tr><td>>MI0000542|hsa-mir-320a&WithFlank&8|-|22244960|22245043@48.69.22(+3U)&[MIMAT0000510&hsa-miR-320a-3p&offsets|0|0(+3U);m-60&8|-|22244975|22244996&offsets|0|0(+3U)]|AAAAGCTGGGTTGAGAGGGCGATTT</td><td>>MI0000542|hsa-mir-320a&WithFlank&8|-|22244960|22245043</td><td>48</td><td>69</td><td>22(+3U)</td><td>AAAAGCTGGGTTGAGAGGGCGATTT</td><td>639.3</td><td>14.0</td><td>1.4</td><td>10.3</td><td>8.8E-25</td><td>8.5E-24</td><td>BT20/MB468</td></tr>
<tr><td>>am_MI0000809|hsa-mir-151a&WithFlank&8|-|140732558|140732659@53.71.19(+1A)&[MIMAT0000757&hsa-miR-151a-3p&offsets|0|-2(+1A);m-30&8|-|140732587|140732607&offsets|0|-2(+1A)]|CTAGACTGAAGCTCCTTGAA</td><td>>am_MI0000809|hsa-mir-151a&WithFlank&8|-|140732558|140732659</td><td>53</td><td>71</td><td>19(+1A)</td><td>CTAGACTGAAGCTCCTTGAA</td><td>1874.9</td><td>13.5</td><td>1.3</td><td>10.3</td><td>8.9E-25</td><td>8.6E-24</td><td>BT20/MB468</td></tr>
<tr><td>>hg38_21_+_8433172_8446622_RNA45SN1_45S_50ntFlanks@10841.10864.24|CGTAACTTCGGGATAAGGATTGGC</td><td>>hg38_21_+_8433172_8446622_RNA45SN1_45S_50ntFlanks</td><td>10841</td><td>10864</td><td>24</td><td>CGTAACTTCGGGATAAGGATTGGC</td><td>2889.0</td><td>-7.2</td><td>0.7</td><td>-10.3</td><td>8.9E-25</td><td>8.6E-24</td><td>BT20/MB468</td></tr>
<tr><td>>hg38_21_+_8433172_8446622_RNA45SN1_45S_50ntFlanks@4679.4720.42|GGCGCAAGACGGACCAGAGCGAAAGCATTTGCCAAGAATGTT</td><td>>hg38_21_+_8433172_8446622_RNA45SN1_45S_50ntFlanks</td><td>4679</td><td>4720</td><td>42</td><td>GGCGCAAGACGGACCAGAGCGAAAGCATTTGCCAAGAATGTT</td><td>296.0</td><td>13.1</td><td>1.3</td><td>10.3</td><td>1.1E-24</td><td>8.7E-24</td><td>MB231/MB468</td></tr>
<tr><td>>MI0000067|hsa-let-7f-1&WithFlank&9|+|94176341|94176439@13.34.22(+1C);MI0000068|hsa-let-7f-2&WithFlank&X|-|53557186|53557280@14.35.22(+1C)&[MIMAT0000067&hsa-let-7f-5p&offsets|0|0(+1C);m-20&9|+|94176353|94176374&offsets|0|0(+1C)];[MIMAT0000067_1&hsa-let-7f-5p&offsets|0|0(+1C);m-19&X|-|53557246|53557267&offsets|0|0(+1C)]|TGAGGTAGTAGATTGTATAGTTC</td><td>>MI0000067|hsa-let-7f-1&WithFlank&9|+|94176341|94176439</td><td>13</td><td>34</td><td>22(+1C)</td><td>TGAGGTAGTAGATTGTATAGTTC</td><td>1020.4</td><td>7.1</td><td>0.7</td><td>10.3</td><td>1.2E-24</td><td>8.9E-24</td><td>MB231/MB468</td></tr>
</table>

<table>
<tr><th></th><th></th><th></th><th></th><th></th><th></th><th></th><th></th><th></th><th></th><th></th><th></th><th></th></tr>
<tr><td>>MI0005531|hsa-mir-450b&WithFlank&X|-|134540179|134540268@17.37.21&[MIMAT0004909&hsa-miR-450b-5p&offsets|0|-1;m-266&X|-|134540232|134540252&offsets|0|0]|TTTTGCAATATGTTCCTGAAT</td><td>>MI0005531|hsa-mir-450b&WithFlank&X|-|134540179|134540268</td><td>17</td><td>37</td><td>21</td><td>TTTTGCAATATGTTCCTGAAT</td><td>866.7</td><td>8.6</td><td>0.8</td><td>10.2</td><td>1.2E-24</td><td>9.1E-24</td><td>MB231/MB468</td></tr>
<tr><td>>MI0000650|hsa-mir-200c&WithFlank&12|+|6963693|6963772@50.70.21&[MIMAT0000617&hsa-miR-200c-3p&offsets|0|-2;m-54&12|+|6963742|6963764&offsets|0|-2]|TAATACTGCCGGGTAATGATG</td><td>>MI0000650|hsa-mir-200c&WithFlank&12|+|6963693|6963772</td><td>50</td><td>70</td><td>21</td><td>TAATACTGCCGGGTAATGATG</td><td>3041.9</td><td>13.1</td><td>1.2</td><td>10.6</td><td>4.9E-26</td><td>9.2E-24</td><td>BT20/MB231</td></tr>
<tr><td>>hg38_21_+_8433172_8446622_RNA45SN1_45S_50ntFlanks@10782.10801.20|GGTGAACAGCCTCTGGCATG</td><td>>hg38_21_+_8433172_8446622_RNA45SN1_45S_50ntFlanks</td><td>10782</td><td>10801</td><td>20</td><td>GGTGAACAGCCTCTGGCATG</td><td>520.7</td><td>-5.2</td><td>0.5</td><td>-10.3</td><td>9.6E-25</td><td>9.2E-24</td><td>BT20/MB468</td></tr>
<tr><td>>MI0000440|hsa-mir-27b&WithFlank&9|+|95085439|95085547@67.86.20&[MIMAT0000419&hsa-miR-27b-3p&offsets|0|-1;m-39&9|+|95085505|95085525&offsets|0|-1]|TTCACAGTGGCTAAGTTCTG</td><td>>MI0000440|hsa-mir-27b&WithFlank&9|+|95085439|95085547</td><td>67</td><td>86</td><td>20</td><td>TTCACAGTGGCTAAGTTCTG</td><td>22980.3</td><td>8.1</td><td>0.8</td><td>10.2</td><td>1.2E-24</td><td>9.3E-24</td><td>MB231/MB468</td></tr>
<tr><td>>hg38_21_+_8433172_8446622_RNA45SN1_45S_50ntFlanks@10410.10429.20|TCAGTCGGTCCTGAGAGATG</td><td>>hg38_21_+_8433172_8446622_RNA45SN1_45S_50ntFlanks</td><td>10410</td><td>10429</td><td>20</td><td>TCAGTCGGTCCTGAGAGATG</td><td>5277.5</td><td>-5.3</td><td>0.5</td><td>-10.2</td><td>1.2E-24</td><td>9.3E-24</td><td>MB231/MB468</td></tr>
<tr><td>>hg38_21_+_8433172_8446622_RNA45SN1_45S_50ntFlanks@12419.12440.22|GATGTCGGCTCTTCCTATCATT</td><td>>hg38_21_+_8433172_8446622_RNA45SN1_45S_50ntFlanks</td><td>12419</td><td>12440</td><td>22</td><td>GATGTCGGCTCTTCCTATCATT</td><td>1508.7</td><td>6.4</td><td>0.6</td><td>10.2</td><td>1.2E-24</td><td>9.4E-24</td><td>MB231/MB468</td></tr>
<tr><td>>hg38_21_+_8433172_8446622_RNA45SN1_45S_50ntFlanks@6651.6672.22|CGACTCTTAGCGGTGGATCACT</td><td>>hg38_21_+_8433172_8446622_RNA45SN1_45S_50ntFlanks</td><td>6651</td><td>6672</td><td>22</td><td>CGACTCTTAGCGGTGGATCACT</td><td>5851.9</td><td>6.4</td><td>0.6</td><td>10.3</td><td>1.0E-24</td><td>9.7E-24</td><td>BT20/MB468</td></tr>
<tr><td>>hg38_MT_+_598_1651_RNR1_12S_50ntFlanks@428.468.41|AAAGTGGCTTTAACATATCTGAACACACAATAGCTAAGACC</td><td>>hg38_MT_+_598_1651_RNR1_12S_50ntFlanks</td><td>428</td><td>468</td><td>41</td><td>AAAGTGGCTTTAACATATCTGAACACACAATAGCTAAGACC</td><td>567.2</td><td>13.8</td><td>1.3</td><td>10.3</td><td>1.0E-24</td><td>9.7E-24</td><td>BT20/MB468</td></tr>
<tr><td>>hg38_21_+_8433172_8446622_RNA45SN1_45S_50ntFlanks@11572.11590.19|CCGACTTAGAACTGGTGCG</td><td>>hg38_21_+_8433172_8446622_RNA45SN1_45S_50ntFlanks</td><td>11572</td><td>11590</td><td>19</td><td>CCGACTTAGAACTGGTGCG</td><td>2217.5</td><td>-4.6</td><td>0.4</td><td>-10.3</td><td>1.0E-24</td><td>9.8E-24</td><td>BT20/MB468</td></tr>
<tr><td>>MI0000061|hsa-let-7a-2&WithFlank&11|-|122146516|122146599@11.32.22;MI0000062|hsa-let-7a-3&WithFlank&22|+|46112743|46112828@10.31.22;MI0000060|hsa-let-7a-1&WithFlank&9|+|94175951|94176042@12.33.22&[MIMAT0000062&hsa-let-7a-5p&offsets|0|0;m-6&11|-|122146568|122146589&offsets|0|0];[MIMAT0000062_1&hsa-let-7a-5p&offsets|0|0;m-5&22|+|46112752|46112773&offsets|0|0];[MIMAT0000062_2&hsa-let-7a-5p&offsets|0|0;m-4&9|+|94175962|94175983&offsets|0|0]|TGAGGTAGTAGGTTGTATAGTT</td><td>>MI0000061|hsa-let-7a-2&WithFlank&11|-|122146516|122146599</td><td>11</td><td>32</td><td>22</td><td>TGAGGTAGTAGGTTGTATAGTT</td><td>329214.0</td><td>8.2</td><td>0.8</td><td>10.2</td><td>1.3E-24</td><td>9.8E-24</td><td>MB231/MB468</td></tr>
<tr><td>>hg38_21_+_8433172_8446622_RNA45SN1_45S_50ntFlanks@6760.6806.47|TTGCGGCCCCGGGTTCCTCCCGGGGCTACGCCTGTCTGAGCGTCGCT</td><td>>hg38_21_+_8433172_8446622_RNA45SN1_45S_50ntFlanks</td><td>6760</td><td>6806</td><td>47</td><td>TTGCGGCCCCGGGTTCCTCCCGGGGCTACGCCTGTCTGAGCGTCGCT</td><td>1080.9</td><td>12.5</td><td>1.2</td><td>10.2</td><td>1.3E-24</td><td>9.9E-24</td><td>MB231/MB468</td></tr>
<tr><td>>trna11_GluTTC_15_-_26327381_26327452@1.35.35,trna3_GluTTC_13_-_45492062_45492133@1.35.35|TCCCACATGGTCTAGCGGTTAGGATTCCTGGTTTT</td><td>>trna11_GluTTC_15_-_26327381_26327452</td><td>1</td><td>35</td><td>35</td><td>TCCCACATGGTCTAGCGGTTAGGATTCCTGGTTTT</td><td>408.6</td><td>11.3</td><td>1.1</td><td>10.3</td><td>1.0E-24</td><td>9.9E-24</td><td>BT20/MB468</td></tr>
<tr><td>>MI0000061|hsa-let-7a-2&WithFlank&11|-|122146516|122146599@11.29.19;MI0000062|hsa-let-7a-3&WithFlank&22|+|46112743|46112828@10.28.19;MI0000060|hsa-let-7a-1&WithFlank&9|+|94175951|94176042@12.30.19&[MIMAT0000062&hsa-let-7a-5p&offsets|0|-3;m-6&11|-|122146568|122146589&offsets|0|-3];[MIMAT0000062_1&hsa-let-7a-5p&offsets|0|-3;m-5&22|+|46112752|46112773&offsets|0|-3];[MIMAT0000062_2&hsa-let-7a-5p&offsets|0|-3;m-4&9|+|94175962|94175983&offsets|0|-3]|TGAGGTAGTAGGTTGTATA</td><td>>MI0000061|hsa-let-7a-2&WithFlank&11|-|122146516|122146599</td><td>11</td><td>29</td><td>19</td><td>TGAGGTAGTAGGTTGTATA</td><td>1470.4</td><td>7.7</td><td>0.8</td><td>10.3</td><td>1.0E-24</td><td>1.0E-23</td><td>BT20/MB468</td></tr>
<tr><td>>hg38_21_+_8433172_8446622_RNA45SN1_45S_50ntFlanks@475.498.24|CTTCGTGATCGATGTGGTGACGTC</td><td>>hg38_21_+_8433172_8446622_RNA45SN1_45S_50ntFlanks</td><td>475</td><td>498</td><td>24</td><td>CTTCGTGATCGATGTGGTGACGTC</td><td>23238.2</td><td>7.9</td><td>0.8</td><td>10.2</td><td>1.3E-24</td><td>1.0E-23</td><td>MB231/MB468</td></tr>
<tr><td>>hg38_21_+_8433172_8446622_RNA45SN1_45S_50ntFlanks@6721.6758.38|AATTGCAGGACACATTGATCATCGACACTTCGAACGCA</td><td>>hg38_21_+_8433172_8446622_RNA45SN1_45S_50ntFlanks</td><td>6721</td><td>6758</td><td>38</td><td>AATTGCAGGACACATTGATCATCGACACTTCGAACGCA</td><td>1115.2</td><td>-5.0</td><td>0.5</td><td>-10.3</td><td>1.0E-24</td><td>1.0E-23</td><td>BT20/MB468</td></tr>
<tr><td>>trna11_GluTTC_15_-_26327381_26327452@1.34.34,trna3_GluTTC_13_-_45492062_45492133@1.34.34|TCCCACATGGTCTAGCGGTTAGGATTCCTGGTTT</td><td>>trna11_GluTTC_15_-_26327381_26327452</td><td>1</td><td>34</td><td>34</td><td>TCCCACATGGTCTAGCGGTTAGGATTCCTGGTTT</td><td>945.5</td><td>14.8</td><td>1.4</td><td>10.2</td><td>1.4E-24</td><td>1.0E-23</td><td>MB231/MB468</td></tr>
<tr><td>>hg38_21_+_8433172_8446622_RNA45SN1_45S_50ntFlanks@6741.6770.30|ATCGACACTTCGAACGCACTTGCGGCCCCG</td><td>>hg38_21_+_8433172_8446622_RNA45SN1_45S_50ntFlanks</td><td>6741</td><td>6770</td><td>30</td><td>ATCGACACTTCGAACGCACTTGCGGCCCCG</td><td>1618.1</td><td>-7.7</td><td>0.8</td><td>-10.3</td><td>1.1E-24</td><td>1.0E-23</td><td>BT20/MB468</td></tr>
<tr><td>>hg38_21_+_8433172_8446622_RNA45SN1_45S_50ntFlanks@476.501.26|TTCGTGATCGATGTGGTGACGTCGTG</td><td>>hg38_21_+_8433172_8446622_RNA45SN1_45S_50ntFlanks</td><td>476</td><td>501</td><td>26</td><td>TTCGTGATCGATGTGGTGACGTCGTG</td><td>4484.4</td><td>7.4</td><td>0.7</td><td>10.2</td><td>1.4E-24</td><td>1.0E-23</td><td>MB231/MB468</td></tr>
<tr><td>>am_MI0003639|hsa-mir-625&WithFlank&14|+|65471096|65471192@58.80.23&[MIMAT0004808&hsa-miR-625-3p&offsets|0|+1;m-137&14|+|65471153|65471174&offsets|0|+1]|GACTATAGAACTTTCCCCCTCAT</td><td>>am_MI0003639|hsa-mir-625&WithFlank&14|+|65471096|65471192</td><td>58</td><td>80</td><td>23</td><td>GACTATAGAACTTTCCCCCTCAT</td><td>1411.2</td><td>12.5</td><td>1.2</td><td>10.2</td><td>1.4E-24</td><td>1.1E-23</td><td>MB231/MB468</td></tr>
<tr><td>>MI0000803|hsa-mir-330&WithFlank&19|-|45638988|45639093@64.86.23(+1U)&[MIMAT0000751&hsa-miR-330-3p&offsets|+1|+1(+1U);m-347&19|-|45639009|45639030&offsets|0|+1(+1U)]|CAAAGCACACGGCCTGCAGAGAGT</td><td>>MI0000803|hsa-mir-330&WithFlank&19|-|45638988|45639093</td><td>64</td><td>86</td><td>23(+1U)</td><td>CAAAGCACACGGCCTGCAGAGAGT</td><td>801.7</td><td>10.6</td><td>1.0</td><td>10.2</td><td>1.4E-24</td><td>1.1E-23</td><td>MB231/MB468</td></tr>
<tr><td>>MI0000078|hsa-mir-22&WithFlank&17|-|1713897|1713993@59.79.21(+1C)&[MIMAT0000077&hsa-miR-22-3p&offsets|0|-1(+1C);m-2&17|-|1713914|1713935&offsets|0|-1(+1C)]|AAGCTGCCAGTTGAAGAACTGC</td><td>>MI0000078|hsa-mir-22&WithFlank&17|-|1713897|1713993</td><td>59</td><td>79</td><td>21(+1C)</td><td>AAGCTGCCAGTTGAAGAACTGC</td><td>3916.1</td><td>8.4</td><td>0.8</td><td>10.3</td><td>1.1E-24</td><td>1.1E-23</td><td>BT20/MB468</td></tr>
<tr><td>>MI0000088|hsa-mir-30a&WithFlank&6|-|71403545|71403627@13.35.23(+1U)&[MIMAT0000087&hsa-miR-30a-5p&offsets|+1|+2(+1U);m-15&6|-|71403595|71403616&offsets|+1|+2(+1U)]|GTAAACATCCTCGACTGGAAGCTT</td><td>>MI0000088|hsa-mir-30a&WithFlank&6|-|71403545|71403627</td><td>13</td><td>35</td><td>23(+1U)</td><td>GTAAACATCCTCGACTGGAAGCTT</td><td>1135.8</td><td>12.0</td><td>1.2</td><td>10.2</td><td>1.4E-24</td><td>1.1E-23</td><td>MB231/MB468</td></tr>
<tr><td>>MI0000542|hsa-mir-320a&WithFlank&8|-|22244960|22245043@49.69.21(+1U)&[MIMAT0000510&hsa-miR-320a-3p&offsets|+1|0(+1U);m-60&8|-|22244975|22244996&offsets|+1|0(+1U)]|AAAGCTGGGTTGAGAGGGCGAT</td><td>>MI0000542|hsa-mir-320a&WithFlank&8|-|22244960|22245043</td><td>49</td><td>69</td><td>21(+1U)</td><td>AAAGCTGGGTTGAGAGGGCGAT</td><td>1335.2</td><td>10.2</td><td>1.0</td><td>10.3</td><td>1.2E-24</td><td>1.1E-23</td><td>BT20/MB468</td></tr>
<tr><td>>hg38_MT_+_1621_3279_RNR2_16S_50ntFlanks@51.88.38|GCTAAACCTAGCCCCAAACCCACTCCACCTTACTACCA</td><td>>hg38_MT_+_1621_3279_RNR2_16S_50ntFlanks</td><td>51</td><td>88</td><td>38</td><td>GCTAAACCTAGCCCCAAACCCACTCCACCTTACTACCA</td><td>3430.6</td><td>10.5</td><td>1.0</td><td>10.2</td><td>1.5E-24</td><td>1.1E-23</td><td>MB231/MB468</td></tr>
<tr><td>>MI0033423|hsa-mir-10399&WithFlank&7|-|139044094|139044163@43.63.21(+1U)&[MIMAT0041630&hsa-miR-10399-3p&offsets|-1|-1(+1U);TJU_CMC_MD2.ID03061.3p-miR&offsets|+3|+2(+1U);m-2657&7|-|139044103|139044124&offsets|+3|+2(+1U)]|TCTCTCGGACAAGCTGTAGGTT</td><td>>MI0033423|hsa-mir-10399&WithFlank&7|-|139044094|139044163</td><td>43</td><td>63</td><td>21(+1U)</td><td>TCTCTCGGACAAGCTGTAGGTT</td><td>796.5</td><td>12.1</td><td>1.2</td><td>10.2</td><td>1.5E-24</td><td>1.1E-23</td><td>MB231/MB468</td></tr>
<tr><td>>trnaMT_ValTAC_MT_+_1602_1670@42.72.31|AGATTTCAACTTAACTTGACCGCTCTGACCA</td><td>>trnaMT_ValTAC_MT_+_1602_1670</td><td>42</td><td>72</td><td>31</td><td>AGATTTCAACTTAACTTGACCGCTCTGACCA</td><td>1849.7</td><td>7.0</td><td>0.7</td><td>10.2</td><td>1.2E-24</td><td>1.1E-23</td><td>BT20/MB468</td></tr>
<tr><td>>hg38_21_+_8433172_8446622_RNA45SN1_45S_50ntFlanks@12974.12991.18|CGTACGTAGCAGAGCAGC</td><td>>hg38_21_+_8433172_8446622_RNA45SN1_45S_50ntFlanks</td><td>12974</td><td>12991</td><td>18</td><td>CGTACGTAGCAGAGCAGC</td><td>4315.6</td><td>-4.9</td><td>0.5</td><td>-10.2</td><td>1.6E-24</td><td>1.2E-23</td><td>MB231/MB468</td></tr>
<tr><td>>MI0000077|hsa-mir-21&WithFlank&17|+|59841260|59841343@13.35.23&[MIMAT0000076&hsa-miR-21-5p&offsets|-1|0;m-1&17|+|59841273|59841294&offsets|-1|0]|GTAGCTTATCAGACTGATGTTGA</td><td>>MI0000077|hsa-mir-21&WithFlank&17|+|59841260|59841343</td><td>13</td><td>35</td><td>23</td><td>GTAGCTTATCAGACTGATGTTGA</td><td>1659.9</td><td>7.1</td><td>0.7</td><td>10.2</td><td>1.6E-24</td><td>1.2E-23</td><td>MB231/MB468</td></tr>
<tr><td>>hg38_21_+_8433172_8446622_RNA45SN1_45S_50ntFlanks@11783.11804.22|CATGAATGGATGAACGAGATTC</td><td>>hg38_21_+_8433172_8446622_RNA45SN1_45S_50ntFlanks</td><td>11783</td><td>11804</td><td>22</td><td>CATGAATGGATGAACGAGATTC</td><td>20548.8</td><td>-5.7</td><td>0.6</td><td>-10.2</td><td>1.6E-24</td><td>1.2E-23</td><td>MB231/MB468</td></tr>
<tr><td>>hg38_21_+_8433172_8446622_RNA45SN1_45S_50ntFlanks@11317.11344.28|ACCCCACGTCTCGTCGCGCGCGCGTCCG</td><td>>hg38_21_+_8433172_8446622_RNA45SN1_45S_50ntFlanks</td><td>11317</td><td>11344</td><td>28</td><td>ACCCCACGTCTCGTCGCGCGCGCGTCCG</td><td>2819.0</td><td>-8.9</td><td>0.9</td><td>-10.2</td><td>1.6E-24</td><td>1.2E-23</td><td>MB231/MB468</td></tr>
<tr><td>>MI0000082|hsa-mir-25&WithFlank&7|-|100093554|100093649@58.78.21(+1G)&[MIMAT0000081&hsa-miR-25-3p&offsets|0|-1(+1G);m-23&7|-|100093571|100093592&offsets|0|-1(+1G)]|CATTGCACTTGTCTCGGTCTGG</td><td>>MI0000082|hsa-mir-25&WithFlank&7|-|100093554|100093649</td><td>58</td><td>78</td><td>21(+1G)</td><td>CATTGCACTTGTCTCGGTCTGG</td><td>5422.2</td><td>7.4</td><td>0.7</td><td>10.2</td><td>1.3E-24</td><td>1.2E-23</td><td>BT20/MB468</td></tr>
<tr><td>>hg38_21_+_8433172_8446622_RNA45SN1_45S_50ntFlanks@5472.5511.40|ACGGCCCTGGCGGAGCGCTGAGAAGACGGTCGAACTTGAC</td><td>>hg38_21_+_8433172_8446622_RNA45SN1_45S_50ntFlanks</td><td>5472</td><td>5511</td><td>40</td><td>ACGGCCCTGGCGGAGCGCTGAGAAGACGGTCGAACTTGAC</td><td>754.2</td><td>12.4</td><td>1.2</td><td>10.2</td><td>1.7E-24</td><td>1.2E-23</td><td>MB231/MB468</td></tr>
<tr><td>>MI0000077|hsa-mir-21&WithFlank&17|+|59841260|59841343@15.36.22(+1A)&[MIMAT0000076&hsa-miR-21-5p&offsets|+1|+1(+1A);m-1&17|+|59841273|59841294&offsets|+1|+1(+1A)]|AGCTTATCAGACTGATGTTGACA</td><td>>MI0000077|hsa-mir-21&WithFlank&17|+|59841260|59841343</td><td>15</td><td>36</td><td>22(+1A)</td><td>AGCTTATCAGACTGATGTTGACA</td><td>578.1</td><td>6.9</td><td>0.7</td><td>10.2</td><td>1.7E-24</td><td>1.3E-23</td><td>MB231/MB468</td></tr>
<tr><td>>MI0003188|hsa-mir-503&WithFlank&X|-|134546322|134546404@12.31.20(+1A)&[MIMAT0002874&hsa-miR-503-5p&offsets|0|-3(+1A);m-297&X|-|134546371|134546393&offsets|0|-3(+1A)]|TAGCAGCGGGAACAGTTCTGA</td><td>>MI0003188|hsa-mir-503&WithFlank&X|-|134546322|134546404</td><td>12</td><td>31</td><td>20(+1A)</td><td>TAGCAGCGGGAACAGTTCTGA</td><td>576.5</td><td>13.8</td><td>1.3</td><td>10.2</td><td>1.3E-24</td><td>1.3E-23</td><td>BT20/MB468</td></tr>
<tr><td>>trnaMT_PheGAA_MT_+_577_647@35.62.28|GAAAATGTTTAGACGGGCTCACATCACC</td><td>>trnaMT_PheGAA_MT_+_577_647</td><td>35</td><td>62</td><td>28</td><td>GAAAATGTTTAGACGGGCTCACATCACC</td><td>716.9</td><td>10.6</td><td>1.0</td><td>10.2</td><td>1.4E-24</td><td>1.3E-23</td><td>BT20/MB468</td></tr>
<tr><td>>hg38_1_-_228634819_228635039_RNA5S12_5S_50ntFlanks@129.171.43|TTGGATGGGAGACCGCCTGGGAATACCGGGTGCTGTAGGCTTT</td><td>>hg38_1_-_228634819_228635039_RNA5S12_5S_50ntFlanks</td><td>129</td><td>171</td><td>43</td><td>TTGGATGGGAGACCGCCTGGGAATACCGGGTGCTGTAGGCTTT</td><td>2871.1</td><td>13.1</td><td>1.3</td><td>10.2</td><td>1.8E-24</td><td>1.3E-23</td><td>MB231/MB468</td></tr>
<tr><td>>MI0000077|hsa-mir-21&WithFlank&17|+|59841260|59841343@14.33.20&[MIMAT0000076&hsa-miR-21-5p&offsets|0|-2;m-1&17|+|59841273|59841294&offsets|0|-2]|TAGCTTATCAGACTGATGTT</td><td>>MI0000077|hsa-mir-21&WithFlank&17|+|59841260|59841343</td><td>14</td><td>33</td><td>20</td><td>TAGCTTATCAGACTGATGTT</td><td>5215.8</td><td>7.7</td><td>0.8</td><td>10.2</td><td>1.4E-24</td><td>1.3E-23</td><td>BT20/MB468</td></tr>
<tr><td>>MI0000746|hsa-mir-99b&WithFlank&19|+|51692606|51692687@13.34.22&[MIMAT0000689&hsa-miR-99b-5p&offsets|0|0;m-10&19|+|51692618|51692639&offsets|0|0]|CACCCGTAGAACCGACCTTGCG</td><td>>MI0000746|hsa-mir-99b&WithFlank&19|+|51692606|51692687</td><td>13</td><td>34</td><td>22</td><td>CACCCGTAGAACCGACCTTGCG</td><td>70763.1</td><td>7.3</td><td>0.7</td><td>10.2</td><td>1.8E-24</td><td>1.4E-23</td><td>MB231/MB468</td></tr>
</table>

<table>
<tr><td>>MI0000283|hsa-mir-203a&WithFlank&14|+|104117399|104117520@72.90.19&[MIMAT0000264&hsa-miR-203a-3p&offsets|+1|-2;m-79&14|+|104117469|104117490&offsets|+1|-2]|TGAAATGTTTAGGACCACT</td><td>>MI0000283|hsa-mir-203a&WithFlank&14|+|104117399|104117520</td><td>72</td><td>90</td><td>19</td><td>TGAAATGTTTAGGACCACT</td><td>1340.8</td><td>8.4</td><td>0.8</td><td>10.2</td><td>1.4E-24</td><td>1.4E-23</td><td>BT20/MB468</td></tr>
<tr><td>>hg38_21_+_8433172_8446622_RNA45SN1_45S_50ntFlanks@7979.8006.28|ACCTCAGATCAGACGTGGCGACCCGCTG</td><td>>hg38_21_+_8433172_8446622_RNA45SN1_45S_50ntFlanks</td><td>7979</td><td>8006</td><td>28</td><td>ACCTCAGATCAGACGTGGCGACCCGCTG</td><td>4449.3</td><td>-6.4</td><td>0.6</td><td>-10.2</td><td>1.5E-24</td><td>1.4E-23</td><td>BT20/MB468</td></tr>
<tr><td>>MI0000809|hsa-mir-151a&WithFlank&8|-|140732558|140732659@53.75.23&[MIMAT0000757&hsa-miR-151a-3p&offsets|0|+2;m-30&8|-|140732587|140732607&offsets|0|+2]|CTAGACTGAAGCTCCTTGAGGAC</td><td>>MI0000809|hsa-mir-151a&WithFlank&8|-|140732558|140732659</td><td>53</td><td>75</td><td>23</td><td>CTAGACTGAAGCTCCTTGAGGAC</td><td>534.9</td><td>11.7</td><td>1.1</td><td>10.2</td><td>1.5E-24</td><td>1.4E-23</td><td>BT20/MB468</td></tr>
<tr><td>>MI0003776|hsa-mir-320b-1&WithFlank&1|+|116671740|116671823@48.67.20;MI0003839|hsa-mir-320b-2&WithFlank&1|-|224257034|224257116@47.66.20;MI0000542|hsa-mir-320a&WithFlank&8|-|22244960|22245043@48.67.20&[MIMAT0005792&hsa-miR-320b&offsets|0|-2;m-183&1|+|116671787|116671806&offsets|0|0];[MIMAT0005792_1&hsa-miR-320b&offsets|0|-2;m-183&1|-|224257051|224257070&offsets|0|0];[MIMAT0000510&hsa-miR-320a-3p&offsets|0|-2;m-60&8|-|22244975|22244996&offsets|0|-2]|AAAAGCTGGGTTGAGAGGGC</td><td>>MI0003776|hsa-mir-320b-1&WithFlank&1|+|116671740|116671823</td><td>48</td><td>67</td><td>20</td><td>AAAAGCTGGGTTGAGAGGGC</td><td>621.4</td><td>7.0</td><td>0.7</td><td>10.2</td><td>1.9E-24</td><td>1.4E-23</td><td>MB231/MB468</td></tr>
<tr><td>>MI0000440|hsa-mir-27b&WithFlank&9|+|95085439|95085547@67.85.19(+1A)&[MIMAT0000419&hsa-miR-27b-3p&offsets|0|-2(+1A);m-39&9|+|95085505|95085525&offsets|0|-2(+1A)]|TTCACAGTGGCTAAGTTCTA</td><td>>MI0000440|hsa-mir-27b&WithFlank&9|+|95085439|95085547</td><td>67</td><td>85</td><td>19(+1A)</td><td>TTCACAGTGGCTAAGTTCTA</td><td>6511.1</td><td>9.9</td><td>1.0</td><td>10.2</td><td>1.5E-24</td><td>1.4E-23</td><td>BT20/MB468</td></tr>
<tr><td>>MI0000103|hsa-mir-101-1&WithFlank&1|-|65058428|65058514@53.73.21(+1U);MI0000739|hsa-mir-101-2&WithFlank&9|+|4850291|4850381@55.75.21(+1U)&[MIMAT0000099&hsa-miR-101-3p&offsets|0|0(+1U);m-17&1|-|65058443|65058462&offsets|0|+1(+1U)];[MIMAT0000099_1&hsa-miR-101-3p&offsets|0|0(+1U);m-16&9|+|4850345|4850364&offsets|0|+1(+1U)]|TACAGTACTGTGATAACTGAAT</td><td>>MI0000103|hsa-mir-101-1&WithFlank&1|-|65058428|65058514</td><td>53</td><td>73</td><td>21(+1U)</td><td>TACAGTACTGTGATAACTGAAT</td><td>3009.8</td><td>9.3</td><td>0.9</td><td>10.2</td><td>1.9E-24</td><td>1.4E-23</td><td>MB231/MB468</td></tr>
<tr><td>>hg38_21_+_8433172_8446622_RNA45SN1_45S_50ntFlanks@6787.6806.20|ACGCCTGTCTGAGCGTCGCT</td><td>>hg38_21_+_8433172_8446622_RNA45SN1_45S_50ntFlanks</td><td>6787</td><td>6806</td><td>20</td><td>ACGCCTGTCTGAGCGTCGCT</td><td>6114.3</td><td>7.0</td><td>0.7</td><td>10.2</td><td>2.0E-24</td><td>1.4E-23</td><td>MB231/MB468</td></tr>
<tr><td>>trnaMT_PheGAA_MT_+_577_647@36.56.21|AAAATGTTTAGACGGGCTCAC</td><td>>trnaMT_PheGAA_MT_+_577_647</td><td>36</td><td>56</td><td>21</td><td>AAAATGTTTAGACGGGCTCAC</td><td>647.0</td><td>12.2</td><td>1.2</td><td>10.2</td><td>2.0E-24</td><td>1.5E-23</td><td>MB231/MB468</td></tr>
<tr><td>>MI0015866|hsa-mir-4326&WithFlank&20|+|63286802|63286872@17.40.24&[MIMAT0016888&hsa-miR-4326&offsets|0|+4;m-339&20|+|63286818|63286841&offsets|0|0]|TGTTCCTCTGTCTCCCAGACTCTG</td><td>>MI0015866|hsa-mir-4326&WithFlank&20|+|63286802|63286872</td><td>17</td><td>40</td><td>24</td><td>TGTTCCTCTGTCTCCCAGACTCTG</td><td>977.1</td><td>14.6</td><td>1.4</td><td>10.2</td><td>1.6E-24</td><td>1.5E-23</td><td>BT20/MB468</td></tr>
<tr><td>>hg38_21_+_8433172_8446622_RNA45SN1_45S_50ntFlanks@12408.12448.41|TTTGATCCTTCGATGTCGGCTCTTCCTATCATTGTGAAGCA</td><td>>hg38_21_+_8433172_8446622_RNA45SN1_45S_50ntFlanks</td><td>12408</td><td>12448</td><td>41</td><td>TTTGATCCTTCGATGTCGGCTCTTCCTATCATTGTGAAGCA</td><td>293.9</td><td>11.1</td><td>1.1</td><td>10.2</td><td>2.0E-24</td><td>1.5E-23</td><td>MB231/MB468</td></tr>
<tr><td>>hg38_21_+_8433172_8446622_RNA45SN1_45S_50ntFlanks@485.516.32|GATGTGGTGACGTCGTGCTCTCCCGGGCCGGG</td><td>>hg38_21_+_8433172_8446622_RNA45SN1_45S_50ntFlanks</td><td>485</td><td>516</td><td>32</td><td>GATGTGGTGACGTCGTGCTCTCCCGGGCCGGG</td><td>1235.0</td><td>14.9</td><td>1.5</td><td>10.2</td><td>1.6E-24</td><td>1.5E-23</td><td>BT20/MB468</td></tr>
<tr><td>>hg38_21_+_8433172_8446622_RNA45SN1_45S_50ntFlanks@12405.12429.25|CTTTTTGATCCTTCGATGTCGGCTC</td><td>>hg38_21_+_8433172_8446622_RNA45SN1_45S_50ntFlanks</td><td>12405</td><td>12429</td><td>25</td><td>CTTTTTGATCCTTCGATGTCGGCTC</td><td>2468.0</td><td>-5.0</td><td>0.5</td><td>-10.2</td><td>2.1E-24</td><td>1.5E-23</td><td>MB231/MB468</td></tr>
<tr><td>>hg38_21_+_8433172_8446622_RNA45SN1_45S_50ntFlanks@3990.4023.34|TTGGTGACTCTAGATAACCTCGGGCCGATCGCAC</td><td>>hg38_21_+_8433172_8446622_RNA45SN1_45S_50ntFlanks</td><td>3990</td><td>4023</td><td>34</td><td>TTGGTGACTCTAGATAACCTCGGGCCGATCGCAC</td><td>700.1</td><td>5.0</td><td>0.5</td><td>10.2</td><td>1.7E-24</td><td>1.5E-23</td><td>BT20/MB468</td></tr>
<tr><td>>MI0000477|hsa-mir-146a&WithFlank&5|+|160485346|160485456@27.46.20(+1C)&[MIMAT0000449&hsa-miR-146a-5p&offsets|0|-2(+1C);m-129&5|+|160485372|160485393&offsets|0|-2(+1C)]|TGAGAACTGAATTCCATGGGC</td><td>>MI0000477|hsa-mir-146a&WithFlank&5|+|160485346|160485456</td><td>27</td><td>46</td><td>20(+1C)</td><td>TGAGAACTGAATTCCATGGGC</td><td>781.1</td><td>14.5</td><td>1.4</td><td>10.2</td><td>2.1E-24</td><td>1.6E-23</td><td>MB231/MB468</td></tr>
<tr><td>>MI0003820|hsa-mir-454&WithFlank&17|-|59137752|59137878@70.94.25&[MIMAT0003885&hsa-miR-454-3p&offsets|0|+2;m-272&17|-|59137787|59137809&offsets|0|+2]|TAGTGCAATATTGCTTATAGGGTTT</td><td>>MI0003820|hsa-mir-454&WithFlank&17|-|59137752|59137878</td><td>70</td><td>94</td><td>25</td><td>TAGTGCAATATTGCTTATAGGGTTT</td><td>2428.2</td><td>8.5</td><td>0.8</td><td>10.2</td><td>2.2E-24</td><td>1.6E-23</td><td>MB231/MB468</td></tr>
<tr><td>>trnaMT_MetCAT_MT_+_4402_4469@34.71.38|ACCCCGAAAATGTTGGTTATACCCTTCCCGTACTACCA</td><td>>trnaMT_MetCAT_MT_+_4402_4469</td><td>34</td><td>71</td><td>38</td><td>ACCCCGAAAATGTTGGTTATACCCTTCCCGTACTACCA</td><td>1007.6</td><td>14.9</td><td>1.5</td><td>10.2</td><td>2.2E-24</td><td>1.6E-23</td><td>MB231/MB468</td></tr>
<tr><td>>MI0000456|hsa-mir-140&WithFlank&16|+|69933075|69933186@69.90.22(+1U)&[MIMAT0004597&hsa-miR-140-3p&offsets|+1|+2(+1U);m-45&16|+|69933143|69933164&offsets|0|0(+1U)]|ACCACAGGGTAGAACCACGGACT</td><td>>MI0000456|hsa-mir-140&WithFlank&16|+|69933075|69933186</td><td>69</td><td>90</td><td>22(+1U)</td><td>ACCACAGGGTAGAACCACGGACT</td><td>3902.8</td><td>12.4</td><td>1.2</td><td>10.2</td><td>2.2E-24</td><td>1.6E-23</td><td>MB231/MB468</td></tr>
<tr><td>>MI0000465|hsa-mir-191&WithFlank&3|-|49020612|49020715@22.44.23(+1C)&[MIMAT0000440&hsa-miR-191-5p&offsets|0|0(+1C);m-48&3|-|49020672|49020694&offsets|0|0(+1C)]|CAACGGAATCCCAAAAGCAGCTGC</td><td>>MI0000465|hsa-mir-191&WithFlank&3|-|49020612|49020715</td><td>22</td><td>44</td><td>23(+1C)</td><td>CAACGGAATCCCAAAAGCAGCTGC</td><td>1471.1</td><td>8.0</td><td>0.8</td><td>10.2</td><td>2.2E-24</td><td>1.6E-23</td><td>MB231/MB468</td></tr>
<tr><td>>hg38_21_+_8433172_8446622_RNA45SN1_45S_50ntFlanks@7980.8006.27|CCTCAGATCAGACGTGGCGACCCGCTG</td><td>>hg38_21_+_8433172_8446622_RNA45SN1_45S_50ntFlanks</td><td>7980</td><td>8006</td><td>27</td><td>CCTCAGATCAGACGTGGCGACCCGCTG</td><td>3119.9</td><td>-7.2</td><td>0.7</td><td>-10.2</td><td>2.2E-24</td><td>1.6E-23</td><td>MB231/MB468</td></tr>
<tr><td>>hg38_21_+_8433172_8446622_RNA45SN1_45S_50ntFlanks@11767.11794.28|TCTAATTAGTGACGCGCATGAATGGATG</td><td>>hg38_21_+_8433172_8446622_RNA45SN1_45S_50ntFlanks</td><td>11767</td><td>11794</td><td>28</td><td>TCTAATTAGTGACGCGCATGAATGGATG</td><td>1183.6</td><td>-4.5</td><td>0.4</td><td>-10.2</td><td>1.7E-24</td><td>1.6E-23</td><td>BT20/MB468</td></tr>
<tr><td>>hg38_21_+_8433172_8446622_RNA45SN1_45S_50ntFlanks@476.522.47|TTCGTGATCGATGTGGTGACGTCGTGCTCTCCCGGGCCGGGTCCGAG</td><td>>hg38_21_+_8433172_8446622_RNA45SN1_45S_50ntFlanks</td><td>476</td><td>522</td><td>47</td><td>TTCGTGATCGATGTGGTGACGTCGTGCTCTCCCGGGCCGGGTCCGAG</td><td>1326.4</td><td>12.5</td><td>1.2</td><td>10.2</td><td>2.3E-24</td><td>1.7E-23</td><td>MB231/MB468</td></tr>
<tr><td>>hg38_MT_+_1621_3279_RNR2_16S_50ntFlanks@51.71.21|GCTAAACCTAGCCCCAAACCC</td><td>>hg38_MT_+_1621_3279_RNR2_16S_50ntFlanks</td><td>51</td><td>71</td><td>21</td><td>GCTAAACCTAGCCCCAAACCC</td><td>1825.7</td><td>8.3</td><td>0.8</td><td>10.2</td><td>1.8E-24</td><td>1.7E-23</td><td>BT20/MB468</td></tr>
<tr><td>>hg38_21_+_8433172_8446622_RNA45SN1_45S_50ntFlanks@475.512.38|CTTCGTGATCGATGTGGTGACGTCGTGCTCTCCCGGGC</td><td>>hg38_21_+_8433172_8446622_RNA45SN1_45S_50ntFlanks</td><td>475</td><td>512</td><td>38</td><td>CTTCGTGATCGATGTGGTGACGTCGTGCTCTCCCGGGC</td><td>843.1</td><td>7.3</td><td>0.7</td><td>10.2</td><td>1.9E-24</td><td>1.7E-23</td><td>BT20/MB468</td></tr>
<tr><td>>hg38_21_+_8433172_8446622_RNA45SN1_45S_50ntFlanks@485.520.36|GATGTGGTGACGTCGTGCTCTCCCGGGCCGGGTCCG</td><td>>hg38_21_+_8433172_8446622_RNA45SN1_45S_50ntFlanks</td><td>485</td><td>520</td><td>36</td><td>GATGTGGTGACGTCGTGCTCTCCCGGGCCGGGTCCG</td><td>1391.3</td><td>15.1</td><td>1.5</td><td>10.2</td><td>1.9E-24</td><td>1.7E-23</td><td>BT20/MB468</td></tr>
<tr><td>>MI0000440|hsa-mir-27b&WithFlank&9|+|95085439|95085547@67.86.20(+1A)&[MIMAT0000419&hsa-miR-27b-3p&offsets|0|-1(+1A);m-39&9|+|95085505|95085525&offsets|0|-1(+1A)]|TTCACAGTGGCTAAGTTCTGA</td><td>>MI0000440|hsa-mir-27b&WithFlank&9|+|95085439|95085547</td><td>67</td><td>86</td><td>20(+1A)</td><td>TTCACAGTGGCTAAGTTCTGA</td><td>24874.1</td><td>9.3</td><td>0.9</td><td>10.2</td><td>1.9E-24</td><td>1.8E-23</td><td>BT20/MB468</td></tr>
<tr><td>>MI0001733|hsa-mir-452&WithFlank&X|-|151959622|151959718@59.80.22&[MIMAT0001636&hsa-miR-452-3p&offsets|-5|-5;m-457&X|-|151959639|151959660&offsets|0|0]|TCAGTCTCATCTGCAAAGAAGT</td><td>>MI0001733|hsa-mir-452&WithFlank&X|-|151959622|151959718</td><td>59</td><td>80</td><td>22</td><td>TCAGTCTCATCTGCAAAGAAGT</td><td>979.6</td><td>14.6</td><td>1.4</td><td>10.2</td><td>1.9E-24</td><td>1.8E-23</td><td>BT20/MB468</td></tr>
<tr><td>>MI0005756|hsa-mir-934&WithFlank&X|+|136550872|136550966@21.43.23&[MIMAT0004977&hsa-miR-934&offsets|0|+1;m-943&X|+|136550892|136550913&offsets|0|+1]|TGTCTACTACTGGAGACACTGGT</td><td>>MI0005756|hsa-mir-934&WithFlank&X|+|136550872|136550966</td><td>21</td><td>43</td><td>23</td><td>TGTCTACTACTGGAGACACTGGT</td><td>2354.1</td><td>14.7</td><td>1.4</td><td>10.5</td><td>1.0E-25</td><td>1.8E-23</td><td>BT20/MB231</td></tr>
<tr><td>>MI0000542|hsa-mir-320a&WithFlank&8|-|22244960|22245043@48.71.24(+1U)&[MIMAT0000510&hsa-miR-320a-3p&offsets|0|+2(+1U);m-60&8|-|22244975|22244996&offsets|0|+2(+1U)]|AAAAGCTGGGTTGAGAGGGCGAAAT</td><td>>MI0000542|hsa-mir-320a&WithFlank&8|-|22244960|22245043</td><td>48</td><td>71</td><td>24(+1U)</td><td>AAAAGCTGGGTTGAGAGGGCGAAAT</td><td>630.6</td><td>8.1</td><td>0.8</td><td>10.2</td><td>2.5E-24</td><td>1.8E-23</td><td>MB231/MB468</td></tr>
<tr><td>>hg38_21_+_8433172_8446622_RNA45SN1_45S_50ntFlanks@9738.9758.21|GAAACGATCTCAACCTATTCT</td><td>>hg38_21_+_8433172_8446622_RNA45SN1_45S_50ntFlanks</td><td>9738</td><td>9758</td><td>21</td><td>GAAACGATCTCAACCTATTCT</td><td>2591.3</td><td>-6.5</td><td>0.6</td><td>-10.2</td><td>2.1E-24</td><td>1.9E-23</td><td>BT20/MB468</td></tr>
<tr><td>>hg38_21_+_8433172_8446622_RNA45SN1_45S_50ntFlanks@6648.6672.25|GTACGACTCTTAGCGGTGGATCACT</td><td>>hg38_21_+_8433172_8446622_RNA45SN1_45S_50ntFlanks</td><td>6648</td><td>6672</td><td>25</td><td>GTACGACTCTTAGCGGTGGATCACT</td><td>658.5</td><td>6.5</td><td>0.6</td><td>10.2</td><td>2.1E-24</td><td>1.9E-23</td><td>BT20/MB468</td></tr>
<tr><td>>hg38_21_+_8433172_8446622_RNA45SN1_45S_50ntFlanks@6698.6720.23|AGCTAGCTGCGAGAATTAATGTG</td><td>>hg38_21_+_8433172_8446622_RNA45SN1_45S_50ntFlanks</td><td>6698</td><td>6720</td><td>23</td><td>AGCTAGCTGCGAGAATTAATGTG</td><td>12582.6</td><td>-5.2</td><td>0.5</td><td>-10.2</td><td>2.1E-24</td><td>1.9E-23</td><td>BT20/MB468</td></tr>
<tr><td>>trna11_GluTTC_15_-_26327381_26327452@1.32.32,trna3_GluTTC_13_-_45492062_45492133@1.32.32|TCCCACATGGTCTAGCGGTTAGGATTCCTGGT</td><td>>trna11_GluTTC_15_-_26327381_26327452</td><td>1</td><td>32</td><td>32</td><td>TCCCACATGGTCTAGCGGTTAGGATTCCTGGT</td><td>1820.5</td><td>13.2</td><td>1.3</td><td>10.2</td><td>2.7E-24</td><td>1.9E-23</td><td>MB231/MB468</td></tr>
<tr><td>>hg38_21_+_8433172_8446622_RNA45SN1_45S_50ntFlanks@5472.5509.38|ACGGCCCTGGCGGAGCGCTGAGAAGACGGTCGAACTTG</td><td>>hg38_21_+_8433172_8446622_RNA45SN1_45S_50ntFlanks</td><td>5472</td><td>5509</td><td>38</td><td>ACGGCCCTGGCGGAGCGCTGAGAAGACGGTCGAACTTG</td><td>1717.3</td><td>7.2</td><td>0.7</td><td>10.2</td><td>2.7E-24</td><td>2.0E-23</td><td>MB231/MB468</td></tr>
<tr><td>>hg38_21_+_8433172_8446622_RNA45SN1_45S_50ntFlanks@4733.4761.29|GAACGAAAGTCGGAGGTTCGAAGACGATC</td><td>>hg38_21_+_8433172_8446622_RNA45SN1_45S_50ntFlanks</td><td>4733</td><td>4761</td><td>29</td><td>GAACGAAAGTCGGAGGTTCGAAGACGATC</td><td>1095.5</td><td>7.0</td><td>0.7</td><td>10.2</td><td>2.7E-24</td><td>2.0E-23</td><td>MB231/MB468</td></tr>
<tr><td>>hg38_21_+_8433172_8446622_RNA45SN1_45S_50ntFlanks@474.506.33|TCTTCGTGATCGATGTGGTGACGTCGTGCTCTC</td><td>>hg38_21_+_8433172_8446622_RNA45SN1_45S_50ntFlanks</td><td>474</td><td>506</td><td>33</td><td>TCTTCGTGATCGATGTGGTGACGTCGTGCTCTC</td><td>1391.4</td><td>8.3</td><td>0.8</td><td>10.2</td><td>2.7E-24</td><td>2.0E-23</td><td>MB231/MB468</td></tr>
<tr><td>>hg38_21_+_8433172_8446622_RNA45SN1_45S_50ntFlanks@476.511.36|TTCGTGATCGATGTGGTGACGTCGTGCTCTCCCGGG</td><td>>hg38_21_+_8433172_8446622_RNA45SN1_45S_50ntFlanks</td><td>476</td><td>511</td><td>36</td><td>TTCGTGATCGATGTGGTGACGTCGTGCTCTCCCGGG</td><td>1314.3</td><td>6.4</td><td>0.6</td><td>10.2</td><td>2.3E-24</td><td>2.1E-23</td><td>BT20/MB468</td></tr>
<tr><td>>MI0000542|hsa-mir-320a&WithFlank&8|-|22244960|22245043@48.68.21(+2U)&[MIMAT0000510&hsa-miR-320a-3p&offsets|0|-1(+2U);m-60&8|-|22244975|22244996&offsets|0|-1(+2U)]|AAAAGCTGGGTTGAGAGGGCGTT</td><td>>MI0000542|hsa-mir-320a&WithFlank&8|-|22244960|22245043</td><td>48</td><td>68</td><td>21(+2U)</td><td>AAAAGCTGGGTTGAGAGGGCGTT</td><td>1673.5</td><td>8.7</td><td>0.9</td><td>10.2</td><td>2.9E-24</td><td>2.1E-23</td><td>MB231/MB468</td></tr>
<tr><td>>hg38_21_+_8433172_8446622_RNA45SN1_45S_50ntFlanks@483.532.50|TCGATGTGGTGACGTCGTGCTCTCCCGGGCCGGGTCCGAGCCGCGACGGG</td><td>>hg38_21_+_8433172_8446622_RNA45SN1_45S_50ntFlanks</td><td>483</td><td>532</td><td>50</td><td>TCGATGTGGTGACGTCGTGCTCTCCCGGGCCGGGTCCGAGCCGCGACGGG</td><td>6923.6</td><td>12.3</td><td>1.2</td><td>10.2</td><td>2.9E-24</td><td>2.1E-23</td><td>MB231/MB468</td></tr>
<tr><td>>hg38_21_+_8433172_8446622_RNA45SN1_45S_50ntFlanks@12395.12415.21|AGCGACGTCGCTTTTTGATCC</td><td>>hg38_21_+_8433172_8446622_RNA45SN1_45S_50ntFlanks</td><td>12395</td><td>12415</td><td>21</td><td>AGCGACGTCGCTTTTTGATCC</td><td>346.8</td><td>8.9</td><td>0.9</td><td>10.2</td><td>2.9E-24</td><td>2.1E-23</td><td>MB231/MB468</td></tr>
</table>

<table>
<tr><th></th><th></th><th></th><th></th><th></th><th></th><th></th><th></th><th></th><th></th><th></th><th></th><th></th></tr>
<tr><td>>MI0000283|hsa-mir-203a&WithFlank&14|+|104117399|104117520@71.92.22&[MIMAT0000264&hsa-miR-203a-3p&offsets|0|0;m-79&14|+|104117469|104117490&offsets|0|0]|GTGAAATGTTTAGGACCACTAG</td><td>>MI0000283|hsa-mir-203a&WithFlank&14|+|104117399|104117520</td><td>71</td><td>92</td><td>22</td><td>GTGAAATGTTTAGGACCACTAG</td><td>2334.1</td><td>14.7</td><td>1.4</td><td>10.5</td><td>1.3E-25</td><td>2.1E-23</td><td>BT20/MB231</td></tr>
<tr><td>>MI0000462|hsa-mir-152&WithFlank&17|-|48037155|48037253@60.80.21(+1U)&[MIMAT0000438&hsa-miR-152-3p&offsets|0|0(+1U);m-97&17|-|48037174|48037194&offsets|0|0(+1U)]|TCAGTGCATGACAGAACTTGGT</td><td>>MI0000462|hsa-mir-152&WithFlank&17|-|48037155|48037253</td><td>60</td><td>80</td><td>21(+1U)</td><td>TCAGTGCATGACAGAACTTGGT</td><td>758.4</td><td>14.2</td><td>1.4</td><td>10.2</td><td>2.3E-24</td><td>2.1E-23</td><td>BT20/MB468</td></tr>
<tr><td>>hg38_21_+_8433172_8446622_RNA45SN1_45S_50ntFlanks@7981.8006.26|CTCAGATCAGACGTGGCGACCCGCTG</td><td>>hg38_21_+_8433172_8446622_RNA45SN1_45S_50ntFlanks</td><td>7981</td><td>8006</td><td>26</td><td>CTCAGATCAGACGTGGCGACCCGCTG</td><td>2564.4</td><td>-7.5</td><td>0.7</td><td>-10.2</td><td>3.0E-24</td><td>2.2E-23</td><td>MB231/MB468</td></tr>
<tr><td>>hg38_21_+_8433172_8446622_RNA45SN1_45S_50ntFlanks@4045.4063.19|CCATTCGAACGTCTGCCCT</td><td>>hg38_21_+_8433172_8446622_RNA45SN1_45S_50ntFlanks</td><td>4045</td><td>4063</td><td>19</td><td>CCATTCGAACGTCTGCCCT</td><td>952.5</td><td>-6.3</td><td>0.6</td><td>-10.2</td><td>2.4E-24</td><td>2.2E-23</td><td>BT20/MB468</td></tr>
<tr><td>>hg38_21_+_8433172_8446622_RNA45SN1_45S_50ntFlanks@7982.8006.25|TCAGATCAGACGTGGCGACCCGCTG</td><td>>hg38_21_+_8433172_8446622_RNA45SN1_45S_50ntFlanks</td><td>7982</td><td>8006</td><td>25</td><td>TCAGATCAGACGTGGCGACCCGCTG</td><td>2023.1</td><td>-6.5</td><td>0.6</td><td>-10.2</td><td>2.4E-24</td><td>2.2E-23</td><td>BT20/MB468</td></tr>
<tr><td>>MI0000542|hsa-mir-320a&WithFlank&8|-|22244960|22245043@48.71.24&[MIMAT0000510&hsa-miR-320a-3p&offsets|0|+2;m-60&8|-|22244975|22244996&offsets|0|+2]|AAAAGCTGGGTTGAGAGGGCGAAA</td><td>>MI0000542|hsa-mir-320a&WithFlank&8|-|22244960|22245043</td><td>48</td><td>71</td><td>24</td><td>AAAAGCTGGGTTGAGAGGGCGAAA</td><td>1315.5</td><td>7.8</td><td>0.8</td><td>10.2</td><td>3.1E-24</td><td>2.2E-23</td><td>MB231/MB468</td></tr>
<tr><td>>MI0000750|hsa-mir-26a-2&WithFlank&12|-|57824603|57824698@20.41.22(+1U);MI0000083|hsa-mir-26a-1&WithFlank&3|+|37969398|37969486@16.37.22(+1U)&[MIMAT0000082&hsa-miR-26a-5p&offsets|0|0(+1U);m-33&12|-|57824658|57824679&offsets|0|0(+1U)];[MIMAT0000082_1&hsa-miR-26a-5p&offsets|0|0(+1U);m-34&3|+|37969413|37969434&offsets|0|0(+1U)]|TTCAAGTAATCCAGGATAGGCTT</td><td>>MI0000750|hsa-mir-26a-2&WithFlank&12|-|57824603|57824698</td><td>20</td><td>41</td><td>22(+1U)</td><td>TTCAAGTAATCCAGGATAGGCTT</td><td>20650.8</td><td>8.4</td><td>0.8</td><td>10.2</td><td>3.2E-24</td><td>2.3E-23</td><td>MB231/MB468</td></tr>
<tr><td>>MI0000736|hsa-mir-30c-1&WithFlank&1|+|40757278|40757378@23.45.23;MI0000254|hsa-mir-30c-2&WithFlank&6|-|71376954|71377037@13.35.23&[MIMAT0000244&hsa-miR-30c-5p&offsets|0|0;m-57&1|+|40757300|40757323&offsets|0|-1];[MIMAT0000244_1&hsa-miR-30c-5p&offsets|0|0;m-56&6|-|71377002|71377025&offsets|0|-1]|TGTAAACATCCTACACTCTCAGC</td><td>>MI0000736|hsa-mir-30c-1&WithFlank&1|+|40757278|40757378</td><td>23</td><td>45</td><td>23</td><td>TGTAAACATCCTACACTCTCAGC</td><td>93241.2</td><td>7.5</td><td>0.7</td><td>10.2</td><td>3.2E-24</td><td>2.3E-23</td><td>MB231/MB468</td></tr>
<tr><td>>MI0001445|hsa-mir-423&WithFlank&17|+|30117073|30117178@58.78.21&[MIMAT0001340&hsa-miR-423-3p&offsets|-1|-3;m-100&17|+|30117131|30117153&offsets|-1|-3]|AAGCTCGGTCTGAGGCCCCTC</td><td>>MI0001445|hsa-mir-423&WithFlank&17|+|30117073|30117178</td><td>58</td><td>78</td><td>21</td><td>AAGCTCGGTCTGAGGCCCCTC</td><td>1360.5</td><td>7.3</td><td>0.7</td><td>10.2</td><td>2.6E-24</td><td>2.4E-23</td><td>BT20/MB468</td></tr>
<tr><td>>MI0000093|hsa-mir-92a-1&WithFlank&13|+|91351308|91351397@54.74.21(+1C);MI0000094|hsa-mir-92a-2&WithFlank&X|-|134169532|134169618@54.74.21(+1C)&[MIMAT0000092&hsa-miR-92a-3p&offsets|0|-1(+1C);m-9&13|+|91351361|91351382&offsets|0|-1(+1C)];[MIMAT0000092_1&hsa-miR-92a-3p&offsets|0|-1(+1C);m-11&X|-|134169544|134169565&offsets|0|-1(+1C)]|TATTGCACTTGTCCCGGCCTGC</td><td>>MI0000093|hsa-mir-92a-1&WithFlank&13|+|91351308|91351397</td><td>54</td><td>74</td><td>21(+1C)</td><td>TATTGCACTTGTCCCGGCCTGC</td><td>24735.0</td><td>7.9</td><td>0.8</td><td>10.1</td><td>3.4E-24</td><td>2.4E-23</td><td>MB231/MB468</td></tr>
<tr><td>>MI0000077|hsa-mir-21&WithFlank&17|+|59841260|59841343@13.37.25&[MIMAT0000076&hsa-miR-21-5p&offsets|-1|+2;m-1&17|+|59841273|59841294&offsets|-1|+2]|GTAGCTTATCAGACTGATGTTGACT</td><td>>MI0000077|hsa-mir-21&WithFlank&17|+|59841260|59841343</td><td>13</td><td>37</td><td>25</td><td>GTAGCTTATCAGACTGATGTTGACT</td><td>1387.5</td><td>10.1</td><td>1.0</td><td>10.1</td><td>3.5E-24</td><td>2.5E-23</td><td>MB231/MB468</td></tr>
<tr><td>>trna111_HisGTG_1_-_147774845_147774916@-1T.22.23,trna118_HisGTG_1_-_145396881_145396952@-1T.22.23,trna16_HisGTG_1_+_146544773_146544844@-1T.22.23,trna1_HisGTG_15_+_45493349_45493420@-1T.22.23,trna21_HisGTG_1_+_147753471_147753542@-1T.22.23,trna33_HisGTG_6_+_27125906_27125977@-1T.22.23,trna7_HisGTG_9_-_14433938_14434009@-1T.22.23,trna8_HisGTG_15_-_45492611_45492682@-1T.22.23,trna9_HisGTG_15_-_45490804_45490875@-1T.22.23|TGCCGTGATCGTATAGTGGTTAG</td><td>>trna111_HisGTG_1_-_147774845_147774916</td><td>-1T</td><td>22</td><td>23</td><td>TGCCGTGATCGTATAGTGGTTAG</td><td>820.5</td><td>12.1</td><td>1.2</td><td>10.1</td><td>3.6E-24</td><td>2.6E-23</td><td>MB231/MB468</td></tr>
<tr><td>>hg38_1_-_228634819_228635039_RNA5S12_5S_50ntFlanks@145.171.27|CTGGGAATACCGGGTGCTGTAGGCTTT</td><td>>hg38_1_-_228634819_228635039_RNA5S12_5S_50ntFlanks</td><td>145</td><td>171</td><td>27</td><td>CTGGGAATACCGGGTGCTGTAGGCTTT</td><td>3226.7</td><td>8.2</td><td>0.8</td><td>10.1</td><td>3.6E-24</td><td>2.6E-23</td><td>MB231/MB468</td></tr>
<tr><td>>MI0000433|hsa-let-7g&WithFlank&3|-|52268272|52268367@11.32.22&[MIMAT0000414&hsa-let-7g-5p&offsets|0|0;m-58&3|-|52268336|52268357&offsets|0|0]|TGAGGTAGTAGTTTGTACAGTT</td><td>>MI0000433|hsa-let-7g&WithFlank&3|-|52268272|52268367</td><td>11</td><td>32</td><td>22</td><td>TGAGGTAGTAGTTTGTACAGTT</td><td>183905.0</td><td>6.4</td><td>0.6</td><td>10.2</td><td>2.9E-24</td><td>2.6E-23</td><td>BT20/MB468</td></tr>
<tr><td>>MI0000650|hsa-mir-200c&WithFlank&12|+|6963693|6963772@50.72.23(+1C)&[MIMAT0000617&hsa-miR-200c-3p&offsets|0|0(+1C);m-54&12|+|6963742|6963764&offsets|0|0(+1C)]|TAATACTGCCGGGTAATGATGGAC</td><td>>MI0000650|hsa-mir-200c&WithFlank&12|+|6963693|6963772</td><td>50</td><td>72</td><td>23(+1C)</td><td>TAATACTGCCGGGTAATGATGGAC</td><td>1494.9</td><td>11.2</td><td>1.1</td><td>10.4</td><td>1.7E-25</td><td>2.6E-23</td><td>BT20/MB231</td></tr>
<tr><td>>hg38_21_+_8433172_8446622_RNA45SN1_45S_50ntFlanks@6738.6770.33|ATCATCGACACTTCGAACGCACTTGCGGCCCCG</td><td>>hg38_21_+_8433172_8446622_RNA45SN1_45S_50ntFlanks</td><td>6738</td><td>6770</td><td>33</td><td>ATCATCGACACTTCGAACGCACTTGCGGCCCCG</td><td>2813.5</td><td>-13.5</td><td>1.3</td><td>-10.2</td><td>2.9E-24</td><td>2.7E-23</td><td>BT20/MB468</td></tr>
<tr><td>>hg38_21_+_8433172_8446622_RNA45SN1_45S_50ntFlanks@12159.12180.22|TTTGACTGGGGCGGTACACCTG</td><td>>hg38_21_+_8433172_8446622_RNA45SN1_45S_50ntFlanks</td><td>12159</td><td>12180</td><td>22</td><td>TTTGACTGGGGCGGTACACCTG</td><td>2435.3</td><td>-9.8</td><td>1.0</td><td>-10.2</td><td>3.0E-24</td><td>2.7E-23</td><td>BT20/MB468</td></tr>
<tr><td>>hg38_21_+_8433172_8446622_RNA45SN1_45S_50ntFlanks@12635.12657.23|AAGCTACCATCTGTGGGATTATG</td><td>>hg38_21_+_8433172_8446622_RNA45SN1_45S_50ntFlanks</td><td>12635</td><td>12657</td><td>23</td><td>AAGCTACCATCTGTGGGATTATG</td><td>688.6</td><td>-5.9</td><td>0.6</td><td>-10.2</td><td>3.0E-24</td><td>2.7E-23</td><td>BT20/MB468</td></tr>
<tr><td>>MI0000066|hsa-let-7e&WithFlank&19|+|51692780|51692870@14.34.21&[MIMAT0000066&hsa-let-7e-5p&offsets|0|-1;m-50&19|+|51692793|51692814&offsets|0|-1]|TGAGGTAGGAGGTTGTATAGT</td><td>>MI0000066|hsa-let-7e&WithFlank&19|+|51692780|51692870</td><td>14</td><td>34</td><td>21</td><td>TGAGGTAGGAGGTTGTATAGT</td><td>4676.7</td><td>7.4</td><td>0.7</td><td>10.2</td><td>3.0E-24</td><td>2.8E-23</td><td>BT20/MB468</td></tr>
<tr><td>>am_trna128_GlyGCC_6_-_27870686_27870756@3.34.32,trna18_GlyGCC_16_+_70822597_70822667@3.34.32,trna19_GlyGCC_16_+_70823410_70823480@3.34.32,trna19_GlyGCC_2_-_157257659_157257729@3.34.32,trna24_GlyGCC_16_-_70812942_70813012@3.34.32,trna25_GlyGCC_16_-_70812114_70812184@3.34.32,trna5_GlyGCC_17_+_8029064_8029134@3.34.32,trna68_GlyGCC_1_-_161493637_161493707@3.34.32|ATTGGTGGTTCAGTGGTAGAATTCTCGCCTGC</td><td>>am_trna128_GlyGCC_6_-_27870686_27870756</td><td>3</td><td>34</td><td>32</td><td>ATTGGTGGTTCAGTGGTAGAATTCTCGCCTGC</td><td>998.6</td><td>12.4</td><td>1.2</td><td>10.1</td><td>3.9E-24</td><td>2.8E-23</td><td>MB231/MB468</td></tr>
<tr><td>>trnaMT_PheGAA_MT_+_577_647@36.60.25|AAAATGTTTAGACGGGCTCACATCA</td><td>>trnaMT_PheGAA_MT_+_577_647</td><td>36</td><td>60</td><td>25</td><td>AAAATGTTTAGACGGGCTCACATCA</td><td>339.2</td><td>13.1</td><td>1.3</td><td>10.2</td><td>3.1E-24</td><td>2.8E-23</td><td>BT20/MB468</td></tr>
<tr><td>>MI0000077|hsa-mir-21&WithFlank&17|+|59841260|59841343@14.33.20(+1U)&[MIMAT0000076&hsa-miR-21-5p&offsets|0|-2(+1U);m-1&17|+|59841273|59841294&offsets|0|-2(+1U)]|TAGCTTATCAGACTGATGTTT</td><td>>MI0000077|hsa-mir-21&WithFlank&17|+|59841260|59841343</td><td>14</td><td>33</td><td>20(+1U)</td><td>TAGCTTATCAGACTGATGTTT</td><td>1271.4</td><td>8.8</td><td>0.9</td><td>10.2</td><td>3.1E-24</td><td>2.8E-23</td><td>BT20/MB468</td></tr>
<tr><td>>MI0000434|hsa-let-7i&WithFlank&12|+|62603680|62603775@12.33.22&[MIMAT0000415&hsa-let-7i-5p&offsets|0|0;m-49&12|+|62603691|62603712&offsets|0|0]|TGAGGTAGTAGTTTGTGCTGTT</td><td>>MI0000434|hsa-let-7i&WithFlank&12|+|62603680|62603775</td><td>12</td><td>33</td><td>22</td><td>TGAGGTAGTAGTTTGTGCTGTT</td><td>728595.2</td><td>8.3</td><td>0.8</td><td>10.1</td><td>3.9E-24</td><td>2.8E-23</td><td>MB231/MB468</td></tr>
<tr><td>>hg38_21_+_8433172_8446622_RNA45SN1_45S_50ntFlanks@475.513.39|CTTCGTGATCGATGTGGTGACGTCGTGCTCTCCCGGGCC</td><td>>hg38_21_+_8433172_8446622_RNA45SN1_45S_50ntFlanks</td><td>475</td><td>513</td><td>39</td><td>CTTCGTGATCGATGTGGTGACGTCGTGCTCTCCCGGGCC</td><td>793.6</td><td>7.6</td><td>0.8</td><td>10.2</td><td>3.1E-24</td><td>2.8E-23</td><td>BT20/MB468</td></tr>
<tr><td>>hg38_21_+_8433172_8446622_RNA45SN1_45S_50ntFlanks@475.524.50|CTTCGTGATCGATGTGGTGACGTCGTGCTCTCCCGGGCCGGGTCCGAGCC</td><td>>hg38_21_+_8433172_8446622_RNA45SN1_45S_50ntFlanks</td><td>475</td><td>524</td><td>50</td><td>CTTCGTGATCGATGTGGTGACGTCGTGCTCTCCCGGGCCGGGTCCGAGCC</td><td>2530.6</td><td>13.7</td><td>1.4</td><td>10.1</td><td>4.0E-24</td><td>2.8E-23</td><td>MB231/MB468</td></tr>
<tr><td>>MI0000077|hsa-mir-21&WithFlank&17|+|59841260|59841343@16.36.21&[MIMAT0000076&hsa-miR-21-5p&offsets|+2|+1;m-1&17|+|59841273|59841294&offsets|+2|+1]|GCTTATCAGACTGATGTTGAC</td><td>>MI0000077|hsa-mir-21&WithFlank&17|+|59841260|59841343</td><td>16</td><td>36</td><td>21</td><td>GCTTATCAGACTGATGTTGAC</td><td>706.1</td><td>12.3</td><td>1.2</td><td>10.1</td><td>4.0E-24</td><td>2.8E-23</td><td>MB231/MB468</td></tr>
<tr><td>>hg38_21_+_8433172_8446622_RNA45SN1_45S_50ntFlanks@4727.4765.39|AATCAAGAACGAAAGTCGGAGGTTCGAAGACGATCAGAT</td><td>>hg38_21_+_8433172_8446622_RNA45SN1_45S_50ntFlanks</td><td>4727</td><td>4765</td><td>39</td><td>AATCAAGAACGAAAGTCGGAGGTTCGAAGACGATCAGAT</td><td>469.4</td><td>6.5</td><td>0.6</td><td>10.2</td><td>3.3E-24</td><td>3.0E-23</td><td>BT20/MB468</td></tr>
<tr><td>>MI0000089|hsa-mir-31&WithFlank&9|-|21512109|21512191@14.36.23&[MIMAT0000089&hsa-miR-31-5p&offsets|0|+2;m-360&9|-|21512157|21512178&offsets|0|+1]|AGGCAAGATGCTGGCATAGCTGT</td><td>>MI0000089|hsa-mir-31&WithFlank&9|-|21512109|21512191</td><td>14</td><td>36</td><td>23</td><td>AGGCAAGATGCTGGCATAGCTGT</td><td>2051.0</td><td>14.5</td><td>1.4</td><td>10.4</td><td>2.0E-25</td><td>3.1E-23</td><td>BT20/MB231</td></tr>
<tr><td>>hg38_21_+_8433172_8446622_RNA45SN1_45S_50ntFlanks@6651.6693.43|CGACTCTTAGCGGTGGATCACTCGGCTCGTGCGTCGATGAAGA</td><td>>hg38_21_+_8433172_8446622_RNA45SN1_45S_50ntFlanks</td><td>6651</td><td>6693</td><td>43</td><td>CGACTCTTAGCGGTGGATCACTCGGCTCGTGCGTCGATGAAGA</td><td>5915.6</td><td>6.8</td><td>0.7</td><td>10.1</td><td>4.3E-24</td><td>3.1E-23</td><td>MB231/MB468</td></tr>
<tr><td>>MI0000061|hsa-let-7a-2&WithFlank&11|-|122146516|122146599@11.30.20(+1G);MI0000062|hsa-let-7a-3&WithFlank&22|+|46112743|46112828@10.29.20(+1G);MI0000060|hsa-let-7a-1&WithFlank&9|+|94175951|94176042@12.31.20(+1G)&[MIMAT0000062&hsa-let-7a-5p&offsets|0|-2(+1G);m-6&11|-|122146568|122146589&offsets|0|-2(+1G)];[MIMAT0000062_1&hsa-let-7a-5p&offsets|0|-2(+1G);m-5&22|+|46112752|46112773&offsets|0|-2(+1G)];[MIMAT0000062_2&hsa-let-7a-5p&offsets|0|-2(+1G);m-4&9|+|94175962|94175983&offsets|0|-2(+1G)]|TGAGGTAGTAGGTTGTATAGG</td><td>>MI0000061|hsa-let-7a-2&WithFlank&11|-|122146516|122146599</td><td>11</td><td>30</td><td>20(+1G)</td><td>TGAGGTAGTAGGTTGTATAGG</td><td>1142.3</td><td>8.3</td><td>0.8</td><td>10.1</td><td>4.4E-24</td><td>3.1E-23</td><td>MB231/MB468</td></tr>
<tr><td>>MI0000736|hsa-mir-30c-1&WithFlank&1|+|40757278|40757378@23.46.24(+1U);MI0000254|hsa-mir-30c-2&WithFlank&6|-|71376954|71377037@13.36.24(+1U)&[MIMAT0000244&hsa-miR-30c-5p&offsets|0|+1(+1U);m-57&1|+|40757300|40757323&offsets|0|0(+1U)];[MIMAT0000244_1&hsa-miR-30c-5p&offsets|0|+1(+1U);m-56&6|-|71377002|71377025&offsets|0|0(+1U)]|TGTAAACATCCTACACTCTCAGCTT</td><td>>MI0000736|hsa-mir-30c-1&WithFlank&1|+|40757278|40757378</td><td>23</td><td>46</td><td>24(+1U)</td><td>TGTAAACATCCTACACTCTCAGCTT</td><td>3994.7</td><td>7.7</td><td>0.8</td><td>10.1</td><td>4.7E-24</td><td>3.4E-23</td><td>MB231/MB468</td></tr>
<tr><td>>MI0000811|hsa-mir-148b&WithFlank&12|+|54337210|54337320@69.91.23&[MIMAT0000759&hsa-miR-148b-3p&offsets|0|+1;m-77&12|+|54337278|54337299&offsets|0|+1]|TCAGTGCATCACAGAACTTTGTC</td><td>>MI0000811|hsa-mir-148b&WithFlank&12|+|54337210|54337320</td><td>69</td><td>91</td><td>23</td><td>TCAGTGCATCACAGAACTTTGTC</td><td>3451.3</td><td>8.5</td><td>0.8</td><td>10.1</td><td>4.7E-24</td><td>3.4E-23</td><td>MB231/MB468</td></tr>
</table>

<table>
<tr><td>>MI0000746|hsa-mir-99b&WithFlank&19|+|51692606|51692687@13.34.22(+1A)&[MIMAT0000689&hsa-miR-99b-5p&offsets|0|0(+1A);m-10&19|+|51692618|51692639&offsets|0|0(+1A)]|CACCCGTAGAACCGACCTTGCGA</td><td>>MI0000746|hsa-mir-99b&WithFlank&19|+|51692606|51692687</td><td>13</td><td>34</td><td>22(+1A)</td><td>CACCCGTAGAACCGACCTTGCGA</td><td>3738.7</td><td>7.8</td><td>0.8</td><td>10.1</td><td>3.8E-24</td><td>3.4E-23</td><td>BT20/MB468</td></tr>
<tr><td>>MI0000811|hsa-mir-148b&WithFlank&12|+|54337210|54337320@69.90.22(+1U)&[MIMAT0000759&hsa-miR-148b-3p&offsets|0|0(+1U);m-77&12|+|54337278|54337299&offsets|0|0(+1U)]|TCAGTGCATCACAGAACTTTGTT</td><td>>MI0000811|hsa-mir-148b&WithFlank&12|+|54337210|54337320</td><td>69</td><td>90</td><td>22(+1U)</td><td>TCAGTGCATCACAGAACTTTGTT</td><td>17981.7</td><td>7.4</td><td>0.7</td><td>10.1</td><td>3.8E-24</td><td>3.4E-23</td><td>BT20/MB468</td></tr>
<tr><td>>MI0000456|hsa-mir-140&WithFlank&16|+|69933075|69933186@69.90.22&[MIMAT0004597&hsa-miR-140-3p&offsets|+1|+2;m-45&16|+|69933143|69933164&offsets|0|0]|ACCACAGGGTAGAACCACGGAC</td><td>>MI0000456|hsa-mir-140&WithFlank&16|+|69933075|69933186</td><td>69</td><td>90</td><td>22</td><td>ACCACAGGGTAGAACCACGGAC</td><td>4696.4</td><td>11.2</td><td>1.1</td><td>10.1</td><td>5.0E-24</td><td>3.5E-23</td><td>MB231/MB468</td></tr>
<tr><td>>MI0000264|hsa-mir-7-2&WithFlank&15|+|88611819|88611940@38.60.23(+1C);MI0000265|hsa-mir-7-3&WithFlank&19|+|4770664|4770785@37.59.23(+1C);MI0000263|hsa-mir-7-1&WithFlank&9|-|83969742|83969863@30.52.23(+1C)&[MIMAT0000252&hsa-miR-7-5p&offsets|0|-1(+1C);m-400&15|+|88611856|88611879&offsets|0|-1(+1C)];[MIMAT0000252_1&hsa-miR-7-5p&offsets|0|-1(+1C);m-402&19|+|4770700|4770723&offsets|0|-1(+1C)];[MIMAT0000252_2&hsa-miR-7-5p&offsets|0|-1(+1C);m-401&9|-|83969811|83969834&offsets|0|-1(+1C)]|TGGAAGACTAGTGATTTTGTTGTC</td><td>>MI0000264|hsa-mir-7-2&WithFlank&15|+|88611819|88611940</td><td>38</td><td>60</td><td>23(+1C)</td><td>TGGAAGACTAGTGATTTTGTTGTC</td><td>26058.2</td><td>7.5</td><td>0.7</td><td>10.1</td><td>5.1E-24</td><td>3.6E-23</td><td>MB231/MB468</td></tr>
<tr><td>>MI0000760|hsa-mir-361&WithFlank&X|-|85903630|85903713@52.75.24&[MIMAT0004682&hsa-miR-361-3p&offsets|+1|+2;m-124&X|-|85903642|85903663&offsets|+1|+3]|CCCCCAGGTGTGATTCTGATTTGC</td><td>>MI0000760|hsa-mir-361&WithFlank&X|-|85903630|85903713</td><td>52</td><td>75</td><td>24</td><td>CCCCCAGGTGTGATTCTGATTTGC</td><td>1023.6</td><td>7.8</td><td>0.8</td><td>10.1</td><td>5.1E-24</td><td>3.6E-23</td><td>MB231/MB468</td></tr>
<tr><td>>hg38_21_+_8433172_8446622_RNA45SN1_45S_50ntFlanks@3840.3879.40|CTCCTACTTGGATAACTGTGGTAATTCTAGAGCTAATACA</td><td>>hg38_21_+_8433172_8446622_RNA45SN1_45S_50ntFlanks</td><td>3840</td><td>3879</td><td>40</td><td>CTCCTACTTGGATAACTGTGGTAATTCTAGAGCTAATACA</td><td>592.5</td><td>13.9</td><td>1.4</td><td>10.1</td><td>4.0E-24</td><td>3.6E-23</td><td>BT20/MB468</td></tr>
<tr><td>>hg38_1_-_228634819_228635039_RNA5S12_5S_50ntFlanks@135.169.35|GGGAGACCGCCTGGGAATACCGGGTGCTGTAGGCT</td><td>>hg38_1_-_228634819_228635039_RNA5S12_5S_50ntFlanks</td><td>135</td><td>169</td><td>35</td><td>GGGAGACCGCCTGGGAATACCGGGTGCTGTAGGCT</td><td>2039.5</td><td>6.4</td><td>0.6</td><td>10.1</td><td>4.1E-24</td><td>3.7E-23</td><td>BT20/MB468</td></tr>
<tr><td>>MI0000082|hsa-mir-25&WithFlank&7|-|100093554|100093649@58.78.21(+1G)&[MIMAT0000081&hsa-miR-25-3p&offsets|0|-1(+1G);m-23&7|-|100093571|100093592&offsets|0|-1(+1G)]|CATTGCACTTGTCTCGGTCTGG</td><td>>MI0000082|hsa-mir-25&WithFlank&7|-|100093554|100093649</td><td>58</td><td>78</td><td>21(+1G)</td><td>CATTGCACTTGTCTCGGTCTGG</td><td>4457.3</td><td>7.4</td><td>0.7</td><td>10.1</td><td>5.2E-24</td><td>3.7E-23</td><td>MB231/MB468</td></tr>
<tr><td>>MI0003591|hsa-mir-584&WithFlank&5|-|149062307|149062415@22.41.20(+1U)&[MIMAT0003249&hsa-miR-584-5p&offsets|0|-2(+1U);m-167&5|-|149062374|149062394&offsets|0|-1(+1U)]|TTATGGTTTGCCTGGGACTGT</td><td>>MI0003591|hsa-mir-584&WithFlank&5|-|149062307|149062415</td><td>22</td><td>41</td><td>20(+1U)</td><td>TTATGGTTTGCCTGGGACTGT</td><td>3196.1</td><td>11.0</td><td>1.1</td><td>10.1</td><td>5.2E-24</td><td>3.7E-23</td><td>MB231/MB468</td></tr>
<tr><td>>trna11_GluTTC_15_-_26327381_26327452@1.34.34,trna3_GluTTC_13_-_45492062_45492133@1.34.34|TCCCACATGGTCTAGCGGTTAGGATTCCTGGTTT</td><td>>trna11_GluTTC_15_-_26327381_26327452</td><td>1</td><td>34</td><td>34</td><td>TCCCACATGGTCTAGCGGTTAGGATTCCTGGTTT</td><td>824.3</td><td>14.3</td><td>1.4</td><td>10.1</td><td>4.1E-24</td><td>3.7E-23</td><td>BT20/MB468</td></tr>
<tr><td>>hg38_21_+_8433172_8446622_RNA45SN1_45S_50ntFlanks@7977.8018.42|CGACCTCAGATCAGACGTGGCGACCCGCTGAATTTAAGCATA</td><td>>hg38_21_+_8433172_8446622_RNA45SN1_45S_50ntFlanks</td><td>7977</td><td>8018</td><td>42</td><td>CGACCTCAGATCAGACGTGGCGACCCGCTGAATTTAAGCATA</td><td>378.0</td><td>13.5</td><td>1.3</td><td>10.1</td><td>5.3E-24</td><td>3.7E-23</td><td>MB231/MB468</td></tr>
<tr><td>>MI0000465|hsa-mir-191&WithFlank&3|-|49020612|49020715@23.44.22&[MIMAT0000440&hsa-miR-191-5p&offsets|+1|0;m-48&3|-|49020672|49020694&offsets|+1|0]|AACGGAATCCCAAAAGCAGCTG</td><td>>MI0000465|hsa-mir-191&WithFlank&3|-|49020612|49020715</td><td>23</td><td>44</td><td>22</td><td>AACGGAATCCCAAAAGCAGCTG</td><td>592.4</td><td>7.2</td><td>0.7</td><td>10.1</td><td>5.3E-24</td><td>3.7E-23</td><td>MB231/MB468</td></tr>
<tr><td>>MI0000440|hsa-mir-27b&WithFlank&9|+|95085439|95085547@67.87.21&[MIMAT0000419&hsa-miR-27b-3p&offsets|0|0;m-39&9|+|95085505|95085525&offsets|0|0]|TTCACAGTGGCTAAGTTCTGC</td><td>>MI0000440|hsa-mir-27b&WithFlank&9|+|95085439|95085547</td><td>67</td><td>87</td><td>21</td><td>TTCACAGTGGCTAAGTTCTGC</td><td>171346.9</td><td>7.5</td><td>0.7</td><td>10.1</td><td>4.1E-24</td><td>3.7E-23</td><td>BT20/MB468</td></tr>
<tr><td>>hg38_21_+_8433172_8446622_RNA45SN1_45S_50ntFlanks@4731.4766.36|AAGAACGAAAGTCGGAGGTTCGAAGACGATCAGATA</td><td>>hg38_21_+_8433172_8446622_RNA45SN1_45S_50ntFlanks</td><td>4731</td><td>4766</td><td>36</td><td>AAGAACGAAAGTCGGAGGTTCGAAGACGATCAGATA</td><td>380.1</td><td>5.8</td><td>0.6</td><td>10.1</td><td>4.2E-24</td><td>3.7E-23</td><td>BT20/MB468</td></tr>
<tr><td>>hg38_21_+_8433172_8446622_RNA45SN1_45S_50ntFlanks@12615.12654.40|GCTGAGGAGCCAATGGGGCGAAGCTACCATCTGTGGGATT</td><td>>hg38_21_+_8433172_8446622_RNA45SN1_45S_50ntFlanks</td><td>12615</td><td>12654</td><td>40</td><td>GCTGAGGAGCCAATGGGGCGAAGCTACCATCTGTGGGATT</td><td>2340.3</td><td>-7.0</td><td>0.7</td><td>-10.1</td><td>4.2E-24</td><td>3.7E-23</td><td>BT20/MB468</td></tr>
<tr><td>>MI0000088|hsa-mir-30a&WithFlank&6|-|71403545|71403627@54.73.20(+2U)&[MIMAT0000088&hsa-miR-30a-3p&offsets|+1|-1(+2U);m-38&6|-|71403554|71403575&offsets|+1|-1(+2U)]|TTTCAGTCGGATGTTTGCAGTT</td><td>>MI0000088|hsa-mir-30a&WithFlank&6|-|71403545|71403627</td><td>54</td><td>73</td><td>20(+2U)</td><td>TTTCAGTCGGATGTTTGCAGTT</td><td>524.1</td><td>7.1</td><td>0.7</td><td>10.1</td><td>4.2E-24</td><td>3.8E-23</td><td>BT20/MB468</td></tr>
<tr><td>>hg38_21_+_8433172_8446622_RNA45SN1_45S_50ntFlanks@476.519.44|TTCGTGATCGATGTGGTGACGTCGTGCTCTCCCGGGCCGGGTCC</td><td>>hg38_21_+_8433172_8446622_RNA45SN1_45S_50ntFlanks</td><td>476</td><td>519</td><td>44</td><td>TTCGTGATCGATGTGGTGACGTCGTGCTCTCCCGGGCCGGGTCC</td><td>1160.2</td><td>12.6</td><td>1.2</td><td>10.1</td><td>5.6E-24</td><td>3.9E-23</td><td>MB231/MB468</td></tr>
<tr><td>>hg38_21_+_8433172_8446622_RNA45SN1_45S_50ntFlanks@3705.3743.39|TACCTGGTTGATCCTGCCAGTAGCATATGCTTGTCTCAA</td><td>>hg38_21_+_8433172_8446622_RNA45SN1_45S_50ntFlanks</td><td>3705</td><td>3743</td><td>39</td><td>TACCTGGTTGATCCTGCCAGTAGCATATGCTTGTCTCAA</td><td>1954.6</td><td>8.1</td><td>0.8</td><td>10.1</td><td>5.6E-24</td><td>3.9E-23</td><td>MB231/MB468</td></tr>
<tr><td>>MI0000063|hsa-let-7b&WithFlank&22|+|46113680|46113774@12.32.21(+1G)&[MIMAT0000063&hsa-let-7b-5p&offsets|0|-1(+1G);m-14&22|+|46113691|46113712&offsets|0|-1(+1G)]|TGAGGTAGTAGGTTGTGTGGTG</td><td>>MI0000063|hsa-let-7b&WithFlank&22|+|46113680|46113774</td><td>12</td><td>32</td><td>21(+1G)</td><td>TGAGGTAGTAGGTTGTGTGGTG</td><td>6373.2</td><td>8.9</td><td>0.9</td><td>10.1</td><td>4.4E-24</td><td>4.0E-23</td><td>BT20/MB468</td></tr>
<tr><td>>MI0000255|hsa-mir-30d&WithFlank&8|-|134804870|134804951@12.32.21&[MIMAT0000245&hsa-miR-30d-5p&offsets|0|-1;m-21&8|-|134804919|134804940&offsets|0|-1]|TGTAAACATCCCCGACTGGAA</td><td>>MI0000255|hsa-mir-30d&WithFlank&8|-|134804870|134804951</td><td>12</td><td>32</td><td>21</td><td>TGTAAACATCCCCGACTGGAA</td><td>855.4</td><td>7.2</td><td>0.7</td><td>10.1</td><td>5.9E-24</td><td>4.1E-23</td><td>MB231/MB468</td></tr>
<tr><td>>MI0000077|hsa-mir-21&WithFlank&17|+|59841260|59841343@14.36.23(+1G)&[MIMAT0000076&hsa-miR-21-5p&offsets|0|+1(+1G);m-1&17|+|59841273|59841294&offsets|0|+1(+1G)]|TAGCTTATCAGACTGATGTTGACG</td><td>>MI0000077|hsa-mir-21&WithFlank&17|+|59841260|59841343</td><td>14</td><td>36</td><td>23(+1G)</td><td>TAGCTTATCAGACTGATGTTGACG</td><td>75588.1</td><td>8.0</td><td>0.8</td><td>10.1</td><td>5.9E-24</td><td>4.2E-23</td><td>MB231/MB468</td></tr>
<tr><td>>hg38_MT_+_1621_3279_RNR2_16S_50ntFlanks@51.90.40|GCTAAACCTAGCCCCAAACCCACTCCACCTTACTACCAGA</td><td>>hg38_MT_+_1621_3279_RNR2_16S_50ntFlanks</td><td>51</td><td>90</td><td>40</td><td>GCTAAACCTAGCCCCAAACCCACTCCACCTTACTACCAGA</td><td>1739.7</td><td>12.6</td><td>1.2</td><td>10.1</td><td>6.0E-24</td><td>4.2E-23</td><td>MB231/MB468</td></tr>
<tr><td>>MI0000285|hsa-mir-205&WithFlank&1|+|209432127|209432248@40.62.23&[MIMAT0000266&hsa-miR-205-5p&offsets|0|+1;m-351&1|+|209432166|209432187&offsets|0|+1]|TCCTTCATTCCACCGGAGTCTGT</td><td>>MI0000285|hsa-mir-205&WithFlank&1|+|209432127|209432248</td><td>40</td><td>62</td><td>23</td><td>TCCTTCATTCCACCGGAGTCTGT</td><td>89973.2</td><td>6.7</td><td>0.7</td><td>10.1</td><td>4.7E-24</td><td>4.2E-23</td><td>BT20/MB468</td></tr>
<tr><td>>MI0001445|hsa-mir-423&WithFlank&17|+|30117073|30117178@58.78.21&[MIMAT0001340&hsa-miR-423-3p&offsets|-1|-3;m-100&17|+|30117131|30117153&offsets|-1|-3]|AAGCTCGGTCTGAGGCCCCTC</td><td>>MI0001445|hsa-mir-423&WithFlank&17|+|30117073|30117178</td><td>58</td><td>78</td><td>21</td><td>AAGCTCGGTCTGAGGCCCCTC</td><td>2015.2</td><td>8.2</td><td>0.8</td><td>10.1</td><td>6.1E-24</td><td>4.3E-23</td><td>MB231/MB468</td></tr>
<tr><td>>trnaMT_GluTTC_MT_-_14674_14742@40.58.19,trnalookalike8_GluTTC_5_-_93905172_93905240@40.58.19|TTGGTCGTGGTTGTAGTCC</td><td>>trnaMT_GluTTC_MT_-_14674_14742</td><td>40</td><td>58</td><td>19</td><td>TTGGTCGTGGTTGTAGTCC</td><td>1898.1</td><td>10.3</td><td>1.0</td><td>10.1</td><td>6.2E-24</td><td>4.3E-23</td><td>MB231/MB468</td></tr>
<tr><td>>hg38_21_+_8433172_8446622_RNA45SN1_45S_50ntFlanks@12996.13016.21|TCGCTGCGATCTATTGAAAGT</td><td>>hg38_21_+_8433172_8446622_RNA45SN1_45S_50ntFlanks</td><td>12996</td><td>13016</td><td>21</td><td>TCGCTGCGATCTATTGAAAGT</td><td>508.8</td><td>6.0</td><td>0.6</td><td>10.1</td><td>4.9E-24</td><td>4.4E-23</td><td>BT20/MB468</td></tr>
<tr><td>>hg38_MT_+_598_1651_RNR1_12S_50ntFlanks@425.452.28|ACGAAAGTGGCTTTAACATATCTGAACA</td><td>>hg38_MT_+_598_1651_RNR1_12S_50ntFlanks</td><td>425</td><td>452</td><td>28</td><td>ACGAAAGTGGCTTTAACATATCTGAACA</td><td>422.8</td><td>9.6</td><td>1.0</td><td>10.1</td><td>5.0E-24</td><td>4.4E-23</td><td>BT20/MB468</td></tr>
<tr><td>>hg38_21_+_8433172_8446622_RNA45SN1_45S_50ntFlanks@13002.13029.28|CGATCTATTGAAAGTCAGCCCTCGACAC</td><td>>hg38_21_+_8433172_8446622_RNA45SN1_45S_50ntFlanks</td><td>13002</td><td>13029</td><td>28</td><td>CGATCTATTGAAAGTCAGCCCTCGACAC</td><td>1784.8</td><td>-12.8</td><td>1.3</td><td>-10.1</td><td>5.0E-24</td><td>4.4E-23</td><td>BT20/MB468</td></tr>
<tr><td>>MI0000270|hsa-mir-181b-1&WithFlank&1|-|198858867|198858988@42.63.22;MI0000683|hsa-mir-181b-2&WithFlank&9|+|124693704|124693804@22.43.22&[MIMAT0000257&hsa-miR-181b-5p&offsets|0|-1;m-73&1|-|198858925|198858947&offsets|0|-1];[MIMAT0000257_1&hsa-miR-181b-5p&offsets|0|-1;m-72&9|+|124693725|124693747&offsets|0|-1]|AACATTCATTGCTGTCGGTGGG</td><td>>MI0000270|hsa-mir-181b-1&WithFlank&1|-|198858867|198858988</td><td>42</td><td>63</td><td>22</td><td>AACATTCATTGCTGTCGGTGGG</td><td>1217.4</td><td>7.5</td><td>0.7</td><td>10.1</td><td>6.4E-24</td><td>4.5E-23</td><td>MB231/MB468</td></tr>
<tr><td>>hg38_21_+_8433172_8446622_RNA45SN1_45S_50ntFlanks@12636.12661.26|AGCTACCATCTGTGGGATTATGACTG</td><td>>hg38_21_+_8433172_8446622_RNA45SN1_45S_50ntFlanks</td><td>12636</td><td>12661</td><td>26</td><td>AGCTACCATCTGTGGGATTATGACTG</td><td>930.3</td><td>-5.0</td><td>0.5</td><td>-10.1</td><td>5.1E-24</td><td>4.5E-23</td><td>BT20/MB468</td></tr>
<tr><td>>MI0000081|hsa-mir-24-2&WithFlank&19|-|13836281|13836365@56.74.19(+1U);MI0000080|hsa-mir-24-1&WithFlank&9|+|95086015|95086094@50.68.19(+1U)&[MIMAT0000080&hsa-miR-24-3p&offsets|0|-3(+1U);m-36&19|-|13836288|13836310&offsets|0|-4(+1U)];[MIMAT0000080_1&hsa-miR-24-3p&offsets|0|-3(+1U);m-35&9|+|95086064|95086086&offsets|0|-4(+1U)]|TGGCTCAGTTCAGCAGGAAT</td><td>>MI0000081|hsa-mir-24-2&WithFlank&19|-|13836281|13836365</td><td>56</td><td>74</td><td>19(+1U)</td><td>TGGCTCAGTTCAGCAGGAAT</td><td>3831.8</td><td>7.3</td><td>0.7</td><td>10.1</td><td>5.1E-24</td><td>4.5E-23</td><td>BT20/MB468</td></tr>
<tr><td>>hg38_21_+_8433172_8446622_RNA45SN1_45S_50ntFlanks@6778.6806.29|CCCGGGGCTACGCCTGTCTGAGCGTCGCT</td><td>>hg38_21_+_8433172_8446622_RNA45SN1_45S_50ntFlanks</td><td>6778</td><td>6806</td><td>29</td><td>CCCGGGGCTACGCCTGTCTGAGCGTCGCT</td><td>2787.6</td><td>6.2</td><td>0.6</td><td>10.1</td><td>6.5E-24</td><td>4.5E-23</td><td>MB231/MB468</td></tr>
<tr><td>>trnaMT_SerGCT_MT_+_12207_12265@1.18.18|GAGAAAGCTCACAAGAAC</td><td>>trnaMT_SerGCT_MT_+_12207_12265</td><td>1</td><td>18</td><td>18</td><td>GAGAAAGCTCACAAGAAC</td><td>20552.5</td><td>10.1</td><td>1.0</td><td>10.1</td><td>6.6E-24</td><td>4.6E-23</td><td>MB231/MB468</td></tr>
<tr><td>>MI0000736|hsa-mir-30c-1&WithFlank&1|+|40757278|40757378@23.45.23(+1A);MI0000254|hsa-mir-30c-2&WithFlank&6|-|71376954|71377037@13.35.23(+1A)&[MIMAT0000244&hsa-miR-30c-5p&offsets|0|0(+1A);m-57&1|+|40757300|40757323&offsets|0|-1(+1A)];[MIMAT0000244_1&hsa-miR-30c-5p&offsets|0|0(+1A);m-56&6|-|71377002|71377025&offsets|0|-1(+1A)]|TGTAAACATCCTACACTCTCAGCA</td><td>>MI0000736|hsa-mir-30c-1&WithFlank&1|+|40757278|40757378</td><td>23</td><td>45</td><td>23(+1A)</td><td>TGTAAACATCCTACACTCTCAGCA</td><td>7162.3</td><td>6.9</td><td>0.7</td><td>10.1</td><td>6.6E-24</td><td>4.6E-23</td><td>MB231/MB468</td></tr>
<tr><td>>hg38_21_+_8433172_8446622_RNA45SN1_45S_50ntFlanks@4732.4768.37|AGAACGAAAGTCGGAGGTTCGAAGACGATCAGATACC</td><td>>hg38_21_+_8433172_8446622_RNA45SN1_45S_50ntFlanks</td><td>4732</td><td>4768</td><td>37</td><td>AGAACGAAAGTCGGAGGTTCGAAGACGATCAGATACC</td><td>313.3</td><td>9.9</td><td>1.0</td><td>10.1</td><td>5.2E-24</td><td>4.6E-23</td><td>BT20/MB468</td></tr>
<tr><td>>trnaMT_SerGCT_MT_+_12207_12265@1.25.25|GAGAAAGCTCACAAGAACTGCTAAC</td><td>>trnaMT_SerGCT_MT_+_12207_12265</td><td>1</td><td>25</td><td>25</td><td>GAGAAAGCTCACAAGAACTGCTAAC</td><td>16750.9</td><td>7.8</td><td>0.8</td><td>10.1</td><td>6.7E-24</td><td>4.7E-23</td><td>MB231/MB468</td></tr>
</table>

<table>
<tr><td>>MI0000266|hsa-mir-10a&WithFlank&17|-|48579832|48579953@28.49.22(+1U)&[MIMAT0000253&hsa-miR-10a-5p&offsets|0|-1(+1U);m-18&17|-|48579905|48579926&offsets|0|0(+1U)]|TACCCTGTAGATCCGAATTTGTT</td><td>>MI0000266|hsa-mir-10a&WithFlank&17|-|48579832|48579953</td><td>28</td><td>49</td><td>22(+1U)</td><td>TACCCTGTAGATCCGAATTTGTT</td><td>824.1</td><td>14.3</td><td>1.4</td><td>10.1</td><td>5.3E-24</td><td>4.7E-23</td><td>BT20/MB468</td></tr>
<tr><td>>hg38_21_+_8433172_8446622_RNA45SN1_45S_50ntFlanks@9477.9497.21|AGCGCACGTGTTAGGACCCGA</td><td>>hg38_21_+_8433172_8446622_RNA45SN1_45S_50ntFlanks</td><td>9477</td><td>9497</td><td>21</td><td>AGCGCACGTGTTAGGACCCGA</td><td>1034.5</td><td>12.4</td><td>1.2</td><td>10.1</td><td>6.9E-24</td><td>4.8E-23</td><td>MB231/MB468</td></tr>
<tr><td>>hg38_21_+_8433172_8446622_RNA45SN1_45S_50ntFlanks@6780.6802.23|CGGGGCTACGCCTGTCTGAGCGT</td><td>>hg38_21_+_8433172_8446622_RNA45SN1_45S_50ntFlanks</td><td>6780</td><td>6802</td><td>23</td><td>CGGGGCTACGCCTGTCTGAGCGT</td><td>877.8</td><td>6.3</td><td>0.6</td><td>10.1</td><td>6.9E-24</td><td>4.8E-23</td><td>MB231/MB468</td></tr>
<tr><td>>hg38_21_+_8433172_8446622_RNA45SN1_45S_50ntFlanks@12159.12179.21|TTTGACTGGGGCGGTACACCT</td><td>>hg38_21_+_8433172_8446622_RNA45SN1_45S_50ntFlanks</td><td>12159</td><td>12179</td><td>21</td><td>TTTGACTGGGGCGGTACACCT</td><td>2875.7</td><td>-9.9</td><td>1.0</td><td>-10.1</td><td>5.5E-24</td><td>4.9E-23</td><td>BT20/MB468</td></tr>
<tr><td>>MI0000067|hsa-let-7f-1&WithFlank&9|+|94176341|94176439@13.33.21(+1C);MI0000068|hsa-let-7f-2&WithFlank&X|-|53557186|53557280@14.34.21(+1C)&[MIMAT0000067&hsa-let-7f-5p&offsets|0|-1(+1C);m-20&9|+|94176353|94176374&offsets|0|-1(+1C)];[MIMAT0000067_1&hsa-let-7f-5p&offsets|0|-1(+1C);m-19&X|-|53557246|53557267&offsets|0|-1(+1C)]|TGAGGTAGTAGATTGTATAGTC</td><td>>MI0000067|hsa-let-7f-1&WithFlank&9|+|94176341|94176439</td><td>13</td><td>33</td><td>21(+1C)</td><td>TGAGGTAGTAGATTGTATAGTC</td><td>44955.7</td><td>7.2</td><td>0.7</td><td>10.1</td><td>7.0E-24</td><td>4.9E-23</td><td>MB231/MB468</td></tr>
<tr><td>>MI0000433|hsa-let-7g&WithFlank&3|-|52268272|52268367@11.30.20(+1C)&[MIMAT0000414&hsa-let-7g-5p&offsets|0|-2(+1C);m-58&3|-|52268336|52268357&offsets|0|-2(+1C)]|TGAGGTAGTAGTTTGTACAGC</td><td>>MI0000433|hsa-let-7g&WithFlank&3|-|52268272|52268367</td><td>11</td><td>30</td><td>20(+1C)</td><td>TGAGGTAGTAGTTTGTACAGC</td><td>1769.2</td><td>8.0</td><td>0.8</td><td>10.1</td><td>7.1E-24</td><td>4.9E-23</td><td>MB231/MB468</td></tr>
<tr><td>>hg38_21_+_8433172_8446622_RNA45SN1_45S_50ntFlanks@11569.11601.33|CAGCCGACTTAGAACTGGTGCGGACCAGGGGAA</td><td>>hg38_21_+_8433172_8446622_RNA45SN1_45S_50ntFlanks</td><td>11569</td><td>11601</td><td>33</td><td>CAGCCGACTTAGAACTGGTGCGGACCAGGGGAA</td><td>2397.7</td><td>-5.8</td><td>0.6</td><td>-10.1</td><td>5.7E-24</td><td>5.0E-23</td><td>BT20/MB468</td></tr>
<tr><td>>MI0000093|hsa-mir-92a-1&WithFlank&13|+|91351308|91351397@54.75.22;MI0000094|hsa-mir-92a-2&WithFlank&X|-|134169532|134169618@54.75.22&[MIMAT0000092&hsa-miR-92a-3p&offsets|0|0;m-9&13|+|91351361|91351382&offsets|0|0];[MIMAT0000092_1&hsa-miR-92a-3p&offsets|0|0;m-11&X|-|134169544|134169565&offsets|0|0]|TATTGCACTTGTCCCGGCCTGT</td><td>>MI0000093|hsa-mir-92a-1&WithFlank&13|+|91351308|91351397</td><td>54</td><td>75</td><td>22</td><td>TATTGCACTTGTCCCGGCCTGT</td><td>197868.0</td><td>8.0</td><td>0.8</td><td>10.1</td><td>7.3E-24</td><td>5.0E-23</td><td>MB231/MB468</td></tr>
<tr><td>>hg38_21_+_8433172_8446622_RNA45SN1_45S_50ntFlanks@8380.8400.21|CAAGAGGGCGTGAAACCGTTA</td><td>>hg38_21_+_8433172_8446622_RNA45SN1_45S_50ntFlanks</td><td>8380</td><td>8400</td><td>21</td><td>CAAGAGGGCGTGAAACCGTTA</td><td>2733.6</td><td>-13.4</td><td>1.3</td><td>-10.1</td><td>5.8E-24</td><td>5.1E-23</td><td>BT20/MB468</td></tr>
<tr><td>>hg38_21_+_8433172_8446622_RNA45SN1_45S_50ntFlanks@6723.6744.22|TTGCAGGACACATTGATCATCG</td><td>>hg38_21_+_8433172_8446622_RNA45SN1_45S_50ntFlanks</td><td>6723</td><td>6744</td><td>22</td><td>TTGCAGGACACATTGATCATCG</td><td>1231.6</td><td>-5.6</td><td>0.6</td><td>-10.1</td><td>5.9E-24</td><td>5.1E-23</td><td>BT20/MB468</td></tr>
<tr><td>>trna10_LysCTT_16_+_3241501_3241573@1.36.36,trna13_LysCTT_14_-_58706613_58706685@1.36.36,trna2_LysCTT_15_+_79152904_79152976@1.36.36|GCCCGGCTAGCTCAGTCGGTAGAGCATGGGACTCTT</td><td>>trna10_LysCTT_16_+_3241501_3241573</td><td>1</td><td>36</td><td>36</td><td>GCCCGGCTAGCTCAGTCGGTAGAGCATGGGACTCTT</td><td>18285.3</td><td>10.9</td><td>1.1</td><td>10.1</td><td>5.9E-24</td><td>5.1E-23</td><td>BT20/MB468</td></tr>
<tr><td>>MI0000082|hsa-mir-25&WithFlank&7|-|100093554|100093649@58.77.20(+1A)&[MIMAT0000081&hsa-miR-25-3p&offsets|0|-2(+1A);m-23&7|-|100093571|100093592&offsets|0|-2(+1A)]|CATTGCACTTGTCTCGGTCTA</td><td>>MI0000082|hsa-mir-25&WithFlank&7|-|100093554|100093649</td><td>58</td><td>77</td><td>20(+1A)</td><td>CATTGCACTTGTCTCGGTCTA</td><td>1971.9</td><td>7.8</td><td>0.8</td><td>10.1</td><td>5.9E-24</td><td>5.2E-23</td><td>BT20/MB468</td></tr>
<tr><td>>hg38_21_+_8433172_8446622_RNA45SN1_45S_50ntFlanks@10405.10424.20|ATGGGTCAGTCGGTCCTGAG</td><td>>hg38_21_+_8433172_8446622_RNA45SN1_45S_50ntFlanks</td><td>10405</td><td>10424</td><td>20</td><td>ATGGGTCAGTCGGTCCTGAG</td><td>1111.6</td><td>6.3</td><td>0.6</td><td>10.1</td><td>6.0E-24</td><td>5.2E-23</td><td>BT20/MB468</td></tr>
<tr><td>>trnaMT_ThrTGT_MT_+_15888_15953@37.69.33|ACCGGAGATGAAAACCTTTTTCCAAGGACACCA</td><td>>trnaMT_ThrTGT_MT_+_15888_15953</td><td>37</td><td>69</td><td>33</td><td>ACCGGAGATGAAAACCTTTTTCCAAGGACACCA</td><td>585.7</td><td>9.5</td><td>0.9</td><td>10.1</td><td>7.7E-24</td><td>5.3E-23</td><td>MB231/MB468</td></tr>
<tr><td>>MI0000087|hsa-mir-29a&WithFlank&7|-|130876741|130876816@48.69.22(+1A)&[MIMAT0000086&hsa-miR-29a-3p&offsets|0|0(+1A);m-26&7|-|130876749|130876769&offsets|0|+1(+1A)]|TAGCACCATCTGAAATCGGTTAA</td><td>>MI0000087|hsa-mir-29a&WithFlank&7|-|130876741|130876816</td><td>48</td><td>69</td><td>22(+1A)</td><td>TAGCACCATCTGAAATCGGTTAA</td><td>507.8</td><td>8.9</td><td>0.9</td><td>10.1</td><td>7.8E-24</td><td>5.3E-23</td><td>MB231/MB468</td></tr>
<tr><td>>am_MI0000786|hsa-mir-378a&WithFlank&5|+|149732819|149732896@50.69.20(+1U)&[MIMAT0000732&hsa-miR-378a-3p&offsets|+1|-1(+1U);m-44&5|+|149732867|149732888&offsets|+1|-1(+1U)]|CTGGACTTGGAGTCAGAAGGT</td><td>>am_MI0000786|hsa-mir-378a&WithFlank&5|+|149732819|149732896</td><td>50</td><td>69</td><td>20(+1U)</td><td>CTGGACTTGGAGTCAGAAGGT</td><td>837.1</td><td>6.0</td><td>0.6</td><td>10.1</td><td>6.2E-24</td><td>5.4E-23</td><td>BT20/MB468</td></tr>
<tr><td>>MI0000440|hsa-mir-27b&WithFlank&9|+|95085439|95085547@66.87.22&[MIMAT0000419&hsa-miR-27b-3p&offsets|-1|0;m-39&9|+|95085505|95085525&offsets|-1|0]|GTTCACAGTGGCTAAGTTCTGC</td><td>>MI0000440|hsa-mir-27b&WithFlank&9|+|95085439|95085547</td><td>66</td><td>87</td><td>22</td><td>GTTCACAGTGGCTAAGTTCTGC</td><td>1058.2</td><td>7.5</td><td>0.7</td><td>10.1</td><td>6.3E-24</td><td>5.4E-23</td><td>BT20/MB468</td></tr>
<tr><td>>hg38_1_-_228634819_228635039_RNA5S12_5S_50ntFlanks@135.170.36|GGGAGACCGCCTGGGAATACCGGGTGCTGTAGGCTT</td><td>>hg38_1_-_228634819_228635039_RNA5S12_5S_50ntFlanks</td><td>135</td><td>170</td><td>36</td><td>GGGAGACCGCCTGGGAATACCGGGTGCTGTAGGCTT</td><td>48400.8</td><td>7.7</td><td>0.8</td><td>10.1</td><td>7.9E-24</td><td>5.4E-23</td><td>MB231/MB468</td></tr>
<tr><td>>MI0003560|hsa-mir-92b&WithFlank&1|+|155195171|155195278@67.86.20(+1U)&[MIMAT0003218&hsa-miR-92b-3p&offsets|0|-2(+1U);m-115&1|+|155195237|155195258&offsets|0|-2(+1U)]|TATTGCACTCGTCCCGGCCTT</td><td>>MI0003560|hsa-mir-92b&WithFlank&1|+|155195171|155195278</td><td>67</td><td>86</td><td>20(+1U)</td><td>TATTGCACTCGTCCCGGCCTT</td><td>815.5</td><td>11.8</td><td>1.2</td><td>10.1</td><td>8.0E-24</td><td>5.5E-23</td><td>MB231/MB468</td></tr>
<tr><td>>hg38_21_+_8433172_8446622_RNA45SN1_45S_50ntFlanks@4729.4767.39|TCAAGAACGAAAGTCGGAGGTTCGAAGACGATCAGATAC</td><td>>hg38_21_+_8433172_8446622_RNA45SN1_45S_50ntFlanks</td><td>4729</td><td>4767</td><td>39</td><td>TCAAGAACGAAAGTCGGAGGTTCGAAGACGATCAGATAC</td><td>310.9</td><td>13.2</td><td>1.3</td><td>10.1</td><td>8.1E-24</td><td>5.6E-23</td><td>MB231/MB468</td></tr>
<tr><td>>MI0000077|hsa-mir-21&WithFlank&17|+|59841260|59841343@14.35.22&[MIMAT0000076&hsa-miR-21-5p&offsets|0|0;m-1&17|+|59841273|59841294&offsets|0|0]|TAGCTTATCAGACTGATGTTGA</td><td>>MI0000077|hsa-mir-21&WithFlank&17|+|59841260|59841343</td><td>14</td><td>35</td><td>22</td><td>TAGCTTATCAGACTGATGTTGA</td><td>274365.7</td><td>7.4</td><td>0.7</td><td>10.1</td><td>8.1E-24</td><td>5.6E-23</td><td>MB231/MB468</td></tr>
<tr><td>>trnaMT_PheGAA_MT_+_577_647@38.62.25|AATGTTTAGACGGGCTCACATCACC</td><td>>trnaMT_PheGAA_MT_+_577_647</td><td>38</td><td>62</td><td>25</td><td>AATGTTTAGACGGGCTCACATCACC</td><td>487.3</td><td>9.5</td><td>0.9</td><td>10.1</td><td>8.1E-24</td><td>5.6E-23</td><td>MB231/MB468</td></tr>
<tr><td>>trna111_HisGTG_1_-_147774845_147774916@-1A.33.34,trna118_HisGTG_1_-_145396881_145396952@-1A.33.34,trna16_HisGTG_1_+_146544773_146544844@-1A.33.34,trna1_HisGTG_15_+_45493349_45493420@-1A.33.34,trna21_HisGTG_1_+_147753471_147753542@-1A.33.34,trna33_HisGTG_6_+_27125906_27125977@-1A.33.34,trna7_HisGTG_9_-_14433938_14434009@-1A.33.34,trna8_HisGTG_15_-_45492611_45492682@-1A.33.34,trna9_HisGTG_15_-_45490804_45490875@-1A.33.34|AGCCGTGATCGTATAGTGGTTAGTACTCTGCGTT</td><td>>trna111_HisGTG_1_-_147774845_147774916</td><td>-1A</td><td>33</td><td>34</td><td>AGCCGTGATCGTATAGTGGTTAGTACTCTGCGTT</td><td>1338.6</td><td>15.3</td><td>1.5</td><td>10.1</td><td>8.2E-24</td><td>5.6E-23</td><td>MB231/MB468</td></tr>
<tr><td>>MI0006444|hsa-mir-1307&WithFlank&10|-|103394247|103394407@86.107.22(+1U)&[MIMAT0005951&hsa-miR-1307-3p&offsets|0|0(+1U);m-66&10|-|103394300|103394322&offsets|0|-1(+1U)]|ACTCGGCGTGGCGTCGGTCGTGT</td><td>>MI0006444|hsa-mir-1307&WithFlank&10|-|103394247|103394407</td><td>86</td><td>107</td><td>22(+1U)</td><td>ACTCGGCGTGGCGTCGGTCGTGT</td><td>1455.2</td><td>7.0</td><td>0.7</td><td>10.1</td><td>6.5E-24</td><td>5.6E-23</td><td>BT20/MB468</td></tr>
<tr><td>>hg38_1_-_228634819_228635039_RNA5S12_5S_50ntFlanks@141.170.30|CCGCCTGGGAATACCGGGTGCTGTAGGCTT</td><td>>hg38_1_-_228634819_228635039_RNA5S12_5S_50ntFlanks</td><td>141</td><td>170</td><td>30</td><td>CCGCCTGGGAATACCGGGTGCTGTAGGCTT</td><td>5314.0</td><td>6.5</td><td>0.6</td><td>10.1</td><td>8.3E-24</td><td>5.7E-23</td><td>MB231/MB468</td></tr>
<tr><td>>MI0000462|hsa-mir-152&WithFlank&17|-|48037155|48037253@60.81.22(+1A)&[MIMAT0000438&hsa-miR-152-3p&offsets|0|+1(+1A);m-97&17|-|48037174|48037194&offsets|0|+1(+1A)]|TCAGTGCATGACAGAACTTGGGA</td><td>>MI0000462|hsa-mir-152&WithFlank&17|-|48037155|48037253</td><td>60</td><td>81</td><td>22(+1A)</td><td>TCAGTGCATGACAGAACTTGGGA</td><td>637.4</td><td>14.0</td><td>1.4</td><td>10.1</td><td>6.6E-24</td><td>5.7E-23</td><td>BT20/MB468</td></tr>
<tr><td>>MI0000087|hsa-mir-29a&WithFlank&7|-|130876741|130876816@47.70.24&[MIMAT0000086&hsa-miR-29a-3p&offsets|-1|+1;m-26&7|-|130876749|130876769&offsets|-1|+2]|CTAGCACCATCTGAAATCGGTTAT</td><td>>MI0000087|hsa-mir-29a&WithFlank&7|-|130876741|130876816</td><td>47</td><td>70</td><td>24</td><td>CTAGCACCATCTGAAATCGGTTAT</td><td>654.0</td><td>11.8</td><td>1.2</td><td>10.1</td><td>8.5E-24</td><td>5.8E-23</td><td>MB231/MB468</td></tr>
<tr><td>>am_MI0003581|hsa-mir-574&WithFlank&4|+|38868026|38868133@31.55.25&[MIMAT0004795&hsa-miR-574-5p&offsets|0|+2;m-213&4|+|38868056|38868078&offsets|0|+2]|TGAGTGTGTGTGTGTGAGTGTGTGT</td><td>>am_MI0003581|hsa-mir-574&WithFlank&4|+|38868026|38868133</td><td>31</td><td>55</td><td>25</td><td>TGAGTGTGTGTGTGTGAGTGTGTGT</td><td>432.7</td><td>7.0</td><td>0.7</td><td>10.1</td><td>6.7E-24</td><td>5.8E-23</td><td>BT20/MB468</td></tr>
<tr><td>>MI0000093|hsa-mir-92a-1&WithFlank&13|+|91351308|91351397@54.76.23(+1U)&[MIMAT0000092&hsa-miR-92a-3p&offsets|0|+1(+1U);m-9&13|+|91351361|91351382&offsets|0|+1(+1U)]|TATTGCACTTGTCCCGGCCTGTTT</td><td>>MI0000093|hsa-mir-92a-1&WithFlank&13|+|91351308|91351397</td><td>54</td><td>76</td><td>23(+1U)</td><td>TATTGCACTTGTCCCGGCCTGTTT</td><td>1575.1</td><td>7.4</td><td>0.7</td><td>10.1</td><td>8.5E-24</td><td>5.8E-23</td><td>MB231/MB468</td></tr>
<tr><td>>hg38_1_-_228634819_228635039_RNA5S12_5S_50ntFlanks@134.170.37|TGGGAGACCGCCTGGGAATACCGGGTGCTGTAGGCTT</td><td>>hg38_1_-_228634819_228635039_RNA5S12_5S_50ntFlanks</td><td>134</td><td>170</td><td>37</td><td>TGGGAGACCGCCTGGGAATACCGGGTGCTGTAGGCTT</td><td>39925.7</td><td>6.0</td><td>0.6</td><td>10.1</td><td>7.0E-24</td><td>6.0E-23</td><td>BT20/MB468</td></tr>
<tr><td>>MI0001733|hsa-mir-452&WithFlank&X|-|151959622|151959718@20.41.22(+1A)&[MIMAT0001635&hsa-miR-452-5p&offsets|0|0(+1A);m-188&X|-|151959677|151959699&offsets|0|-1(+1A)]|AACTGTTTGCAGAGGAAACTGAA</td><td>>MI0001733|hsa-mir-452&WithFlank&X|-|151959622|151959718</td><td>20</td><td>41</td><td>22(+1A)</td><td>AACTGTTTGCAGAGGAAACTGAA</td><td>556.4</td><td>13.8</td><td>1.4</td><td>10.1</td><td>7.0E-24</td><td>6.0E-23</td><td>BT20/MB468</td></tr>
<tr><td>>trnaMT_GluTTC_MT_-_14674_14742@37.71.35,trnalookalike8_GluTTC_5_-_93905172_93905240@37.71.35|TCATTGGTCGTGGTTGTAGTCCGTGCGAGAATACC</td><td>>trnaMT_GluTTC_MT_-_14674_14742</td><td>37</td><td>71</td><td>35</td><td>TCATTGGTCGTGGTTGTAGTCCGTGCGAGAATACC</td><td>1188.9</td><td>11.3</td><td>1.1</td><td>10.1</td><td>8.9E-24</td><td>6.1E-23</td><td>MB231/MB468</td></tr>
</table>

| | | | | | | | | | | | | |
|---|---|---|---|---|---|---|---|---|---|---|---|---|
| >hg38_21_+_8433172_8446622_RNA45SN1_45S_50ntFlanks@12405.12432.28\|CTTTTTGATCCTTCGATGTCGGCTCTTC | >hg38_21_+_8433172_8446622_RNA45SN1_45S_50ntFlanks | 12405 | 12432 | 28 | CTTTTTGATCCTTCGATGTCGGCTCTTC | 5963.6 | -8.4 | 0.8 | -10.1 | 7.1E-24 | 6.1E-23 | BT20/MB468 |
| >MI0000760\|hsa-mir-361&WithFlank&X\|-\|85903630\|85903713@51.73.23&[MIMAT0004682&hsa-miR-361-3p&offsets\|0\|0;m-124&X\|-\|85903642\|85903663&offsets\|0\|+1]\|TCCCCCAGGTGTGATTCTGATTT | >MI0000760\|hsa-mir-361&WithFlank&X\|-\|85903630\|85903713 | 51 | 73 | 23 | TCCCCCAGGTGTGATTCTGATTT | 956.2 | 7.6 | 0.8 | 10.1 | 9.1E-24 | 6.2E-23 | MB231/MB468 |
| >MI0000466\|hsa-mir-9-1&WithFlank&1\|-\|156420335\|156420435@22.43.22;MI0000468\|hsa-mir-9-3&WithFlank&15\|+\|89368011\|89368112@22.43.22;MI0000467\|hsa-mir-9-2&WithFlank&5\|-\|88666847\|88666945@22.43.22&[MIMAT0000441&hsa-miR-9-5p&offsets\|0\|-1;m-104&1\|-\|156420392\|156420414&offsets\|0\|-1];[MIMAT0000441_1&hsa-miR-9-5p&offsets\|0\|-1;m-105&15\|+\|89368032\|89368054&offsets\|0\|-1];[MIMAT0000441_2&hsa-miR-9-5p&offsets\|0\|-1;m-104&5\|-\|88666902\|88666924&offsets\|0\|-1]\|TCTTTGGTTATCTAGCTGTATG | >MI0000466\|hsa-mir-9-1&WithFlank&1\|-\|156420335\|156420435 | 22 | 43 | 22 | TCTTTGGTTATCTAGCTGTATG | 1533.0 | 8.3 | 0.8 | 10.0 | 9.4E-24 | 6.4E-23 | MB231/MB468 |
| >MI0000108\|hsa-mir-103a-2&WithFlank&20\|+\|3917488\|3917577@54.76.23(+1C);MI0000109\|hsa-mir-103a-1&WithFlank&5\|-\|168560890\|168560979@54.76.23(+1C)&[MIMAT0000101&hsa-miR-103a-3p&offsets\|0\|0(+1C);m-12&20\|+\|3917541\|3917563&offsets\|0\|0(+1C)];[MIMAT0000101_1&hsa-miR-103a-3p&offsets\|0\|0(+1C);m-13&5\|-\|168560904\|168560926&offsets\|0\|0(+1C);m-44220&5\|-\|168560908\|168560926&offsets\|0\|+4(+1C)]\|AGCAGCATTGTACAGGGCTATGAC | >MI0000108\|hsa-mir-103a-2&WithFlank&20\|+\|3917488\|3917577 | 54 | 76 | 23(+1C) | AGCAGCATTGTACAGGGCTATGAC | 902.4 | 7.7 | 0.8 | 10.1 | 7.4E-24 | 6.4E-23 | BT20/MB468 |
| >MI0000750\|hsa-mir-26a-2&WithFlank&12\|-\|57824603\|57824698@20.42.23;MI0000083\|hsa-mir-26a-1&WithFlank&3\|+\|37969398\|37969486@16.38.23&[MIMAT0000082&hsa-miR-26a-5p&offsets\|0\|+1;m-33&12\|-\|57824658\|57824679&offsets\|0\|+1];[MIMAT0000082_1&hsa-miR-26a-5p&offsets\|0\|+1;m-34&3\|+\|37969413\|37969434&offsets\|0\|+1]\|TTCAAGTAATCCAGGATAGGCTG | >MI0000750\|hsa-mir-26a-2&WithFlank&12\|-\|57824603\|57824698 | 20 | 42 | 23 | TTCAAGTAATCCAGGATAGGCTG | 1385.7 | 7.5 | 0.7 | 10.0 | 9.6E-24 | 6.5E-23 | MB231/MB468 |
| >hg38_21_+_8433172_8446622_RNA45SN1_45S_50ntFlanks@6671.6698.28\|CTCGGCTCGTGCGTCGATGAAGAACGCA | >hg38_21_+_8433172_8446622_RNA45SN1_45S_50ntFlanks | 6671 | 6698 | 28 | CTCGGCTCGTGCGTCGATGAAGAACGCA | 810.3 | -5.5 | 0.5 | -10.1 | 7.6E-24 | 6.5E-23 | BT20/MB468 |
| >trna111_HisGTG_1_-_147774845_147774916@-1A.26.27,trna118_HisGTG_1_-_145396881_145396952@-1A.26.27,trna16_HisGTG_1_+_146544773_146544844@-1A.26.27,trna1_HisGTG_15_+_45493349_45493420@-1A.26.27,trna21_HisGTG_1_+_147753471_147753542@-1A.26.27,trna33_HisGTG_6_+_27125906_27125977@-1A.26.27,trna7_HisGTG_9_-_14433938_14434009@-1A.26.27,trna8_HisGTG_15_-_45492611_45492682@-1A.26.27,trna9_HisGTG_15_-_45490804_45490875@-1A.26.27\|AGCCGTGATCGTATAGTGGTTAGTACT | >trna111_HisGTG_1_-_147774845_147774916 | -1A | 26 | 27 | AGCCGTGATCGTATAGTGGTTAGTACT | 2322.9 | 9.2 | 0.9 | 10.0 | 9.7E-24 | 6.5E-23 | MB231/MB468 |
| >hg38_21_+_8433172_8446622_RNA45SN1_45S_50ntFlanks@4045.4062.18\|CCATTCGAACGTCTGCCC | >hg38_21_+_8433172_8446622_RNA45SN1_45S_50ntFlanks | 4045 | 4062 | 18 | CCATTCGAACGTCTGCCC | 294.0 | -4.1 | 0.4 | -10.1 | 7.9E-24 | 6.7E-23 | BT20/MB468 |
| >hg38_21_+_8433172_8446622_RNA45SN1_45S_50ntFlanks@12420.12458.39\|ATGTCGGCTCTTCCTATCATTGTGAAGCAGAATTCACCA | >hg38_21_+_8433172_8446622_RNA45SN1_45S_50ntFlanks | 12420 | 12458 | 39 | ATGTCGGCTCTTCCTATCATTGTGAAGCAGAATTCACCA | 823.6 | 5.4 | 0.5 | 10.0 | 9.9E-24 | 6.7E-23 | MB231/MB468 |
| >hg38_21_+_8433172_8446622_RNA45SN1_45S_50ntFlanks@6788.6806.19\|CGCCTGTCTGAGCGTCGCT | >hg38_21_+_8433172_8446622_RNA45SN1_45S_50ntFlanks | 6788 | 6806 | 19 | CGCCTGTCTGAGCGTCGCT | 11745.0 | 6.7 | 0.7 | 10.0 | 1.0E-23 | 6.8E-23 | MB231/MB468 |
| >hg38_21_+_8433172_8446622_RNA45SN1_45S_50ntFlanks@12997.13014.18\|CGCTGCGATCTATTGAAA | >hg38_21_+_8433172_8446622_RNA45SN1_45S_50ntFlanks | 12997 | 13014 | 18 | CGCTGCGATCTATTGAAA | 9640.0 | 6.0 | 0.6 | 10.0 | 1.0E-23 | 6.9E-23 | MB231/MB468 |
| >hg38_21_+_8433172_8446622_RNA45SN1_45S_50ntFlanks@476.507.32\|TTCGTGATCGATGTGGTGACGTCGTGCTCTCC | >hg38_21_+_8433172_8446622_RNA45SN1_45S_50ntFlanks | 476 | 507 | 32 | TTCGTGATCGATGTGGTGACGTCGTGCTCTCC | 1939.3 | 6.2 | 0.6 | 10.1 | 8.1E-24 | 7.0E-23 | BT20/MB468 |
| >trnaMT_SerGCT_MT_+_12207_12265@15.62.48\|GAACTGCTAACTCATGCCCCCATGTCTAACAACATGGCTTTCTCACCA | >trnaMT_SerGCT_MT_+_12207_12265 | 15 | 62 | 48 | GAACTGCTAACTCATGCCCCCATGTCTAACAACATGGCTTTCTCACCA | 1100.6 | 12.5 | 1.2 | 10.0 | 1.0E-23 | 7.1E-23 | MB231/MB468 |
| >hg38_21_+_8433172_8446622_RNA45SN1_45S_50ntFlanks@7977.8010.34\|CGACCTCAGATCAGACGTGGCGACCCGCTGAATT | >hg38_21_+_8433172_8446622_RNA45SN1_45S_50ntFlanks | 7977 | 8010 | 34 | CGACCTCAGATCAGACGTGGCGACCCGCTGAATT | 764.6 | 6.8 | 0.7 | 10.0 | 1.1E-23 | 7.1E-23 | MB231/MB468 |
| >MI0000301\|hsa-mir-224&WithFlank&X\|-\|151958572\|151958664@14.34.21(+1G)&[MIMAT0000281&hsa-miR-224-5p&offsets\|+1\|-3(+1G);m-238&X\|-\|151958630\|151958651&offsets\|0\|-1(+1G)]\|CAAGTCACTAGTGGTTCCGTTG | >MI0000301\|hsa-mir-224&WithFlank&X\|-\|151958572\|151958664 | 14 | 34 | 21(+1G) | CAAGTCACTAGTGGTTCCGTTG | 882.3 | 14.4 | 1.4 | 10.1 | 8.3E-24 | 7.1E-23 | BT20/MB468 |
| >hg38_21_+_8433172_8446622_RNA45SN1_45S_50ntFlanks@475.508.34\|CTTCGTGATCGATGTGGTGACGTCGTGCTCTCCC | >hg38_21_+_8433172_8446622_RNA45SN1_45S_50ntFlanks | 475 | 508 | 34 | CTTCGTGATCGATGTGGTGACGTCGTGCTCTCCC | 1819.4 | 7.3 | 0.7 | 10.0 | 1.1E-23 | 7.1E-23 | MB231/MB468 |
| >hg38_21_+_8433172_8446622_RNA45SN1_45S_50ntFlanks@6719.6740.22\|TGAATTGCAGGACACATTGATC | >hg38_21_+_8433172_8446622_RNA45SN1_45S_50ntFlanks | 6719 | 6740 | 22 | TGAATTGCAGGACACATTGATC | 3711.5 | -4.9 | 0.5 | -10.0 | 1.1E-23 | 7.3E-23 | MB231/MB468 |
| >hg38_21_+_8433172_8446622_RNA45SN1_45S_50ntFlanks@475.503.29\|CTTCGTGATCGATGTGGTGACGTCGTGCT | >hg38_21_+_8433172_8446622_RNA45SN1_45S_50ntFlanks | 475 | 503 | 29 | CTTCGTGATCGATGTGGTGACGTCGTGCT | 7549.6 | 7.0 | 0.7 | 10.0 | 1.1E-23 | 7.3E-23 | MB231/MB468 |
| >hg38_21_+_8433172_8446622_RNA45SN1_45S_50ntFlanks@11661.11681.21\|TGATTTCTGCCCAGTGCTCTG | >hg38_21_+_8433172_8446622_RNA45SN1_45S_50ntFlanks | 11661 | 11681 | 21 | TGATTTCTGCCCAGTGCTCTG | 3104.7 | -5.5 | 0.6 | -10.0 | 1.1E-23 | 7.4E-23 | MB231/MB468 |
| >hg38_21_+_8433172_8446622_RNA45SN1_45S_50ntFlanks@12994.13039.46\|CCTCGCTGCGATCTATTGAAAGTCAGCCCTCGACACAAGGGTTTGT | >hg38_21_+_8433172_8446622_RNA45SN1_45S_50ntFlanks | 12994 | 13039 | 46 | CCTCGCTGCGATCTATTGAAAGTCAGCCCTCGACACAAGGGTTTGT | 162254.7 | 11.3 | 1.1 | 10.0 | 1.1E-23 | 7.5E-23 | MB231/MB468 |
| >MI0003581\|hsa-mir-574&WithFlank&4\|+\|38868026\|38868133@67.88.22&[MIMAT0003239&hsa-miR-574-3p&offsets\|0\|0;m-149&4\|+\|38868092\|38868113&offsets\|0\|0]\|CACGCTCATGCACACACCCACA | >MI0003581\|hsa-mir-574&WithFlank&4\|+\|38868026\|38868133 | 67 | 88 | 22 | CACGCTCATGCACACACCCACA | 2627.7 | 7.7 | 0.8 | 10.0 | 1.1E-23 | 7.5E-23 | MB231/MB468 |
| >hg38_21_+_8433172_8446622_RNA45SN1_45S_50ntFlanks@7977.8019.43\|CGACCTCAGATCAGACGTGGCGACCCGCTGAATTTAAGCATAT | >hg38_21_+_8433172_8446622_RNA45SN1_45S_50ntFlanks | 7977 | 8019 | 43 | CGACCTCAGATCAGACGTGGCGACCCGCTGAATTTAAGCATAT | 622.6 | 14.2 | 1.4 | 10.0 | 1.1E-23 | 7.6E-23 | MB231/MB468 |
| >trna152_ValCAC_6_-_27248049_27248121@1.36.36\|GCTTCTGTAGTGTAGTGGTTATCACGTTCGCCTCAC | >trna152_ValCAC_6_-_27248049_27248121 | 1 | 36 | 36 | GCTTCTGTAGTGTAGTGGTTATCACGTTCGCCTCAC | 1453.9 | 12.9 | 1.3 | 10.0 | 1.2E-23 | 7.7E-23 | MB231/MB468 |
| >MI0000266\|hsa-mir-10a&WithFlank&17\|-\|48579832\|48579953@29.50.22(+1C)&[MIMAT0000253&hsa-miR-10a-5p&offsets\|+1\|0(+1C);m-18&17\|-\|48579905\|48579926&offsets\|+1\|+1(+1C)]\|ACCCTGTAGATCCGAATTTGTGC | >MI0000266\|hsa-mir-10a&WithFlank&17\|-\|48579832\|48579953 | 29 | 50 | 22(+1C) | ACCCTGTAGATCCGAATTTGTGC | 393.6 | 13.3 | 1.3 | 10.1 | 9.1E-24 | 7.8E-23 | BT20/MB468 |
| >trna119_LysCTT_1_-_145395522_145395594@3.34.32,trna11_LysCTT_5_-_180648979_180649051@3.34.32,trna13_LysCTT_6_+_26556774_26556846@3.34.32,trna7_LysCTT_16_+_3225692_3225764@3.34.32,trna9_LysCTT_5_+_180634755_180634827@3.34.32\|CCGGCTAGCTCAGTCGGTAGAGCATGAGACTC | >trna119_LysCTT_1_-_145395522_145395594 | 3 | 34 | 32 | CCGGCTAGCTCAGTCGGTAGAGCATGAGACTC | 693.7 | 12.3 | 1.2 | 10.0 | 1.2E-23 | 7.9E-23 | MB231/MB468 |
| >hg38_21_+_8433172_8446622_RNA45SN1_45S_50ntFlanks@485.517.33\|GATGTGGTGACGTCGTGCTCTCCCGGGCCGGGT | >hg38_21_+_8433172_8446622_RNA45SN1_45S_50ntFlanks | 485 | 517 | 33 | GATGTGGTGACGTCGTGCTCTCCCGGGCCGGGT | 1143.4 | 14.8 | 1.5 | 10.0 | 9.3E-24 | 7.9E-23 | BT20/MB468 |
| >hg38_21_+_8433172_8446622_RNA45SN1_45S_50ntFlanks@476.510.35\|TTCGTGATCGATGTGGTGACGTCGTGCTCTCCCGG | >hg38_21_+_8433172_8446622_RNA45SN1_45S_50ntFlanks | 476 | 510 | 35 | TTCGTGATCGATGTGGTGACGTCGTGCTCTCCCGG | 1758.3 | 6.0 | 0.6 | 10.0 | 9.6E-24 | 8.1E-23 | BT20/MB468 |
| >hg38_21_+_8433172_8446622_RNA45SN1_45S_50ntFlanks@12635.12668.34\|AAGCTACCATCTGTGGGATTATGACTGAACGCCT | >hg38_21_+_8433172_8446622_RNA45SN1_45S_50ntFlanks | 12635 | 12668 | 34 | AAGCTACCATCTGTGGGATTATGACTGAACGCCT | 276.0 | 6.0 | 0.6 | 10.0 | 1.2E-23 | 8.1E-23 | MB231/MB468 |
| >hg38_MT_+_598_1651_RNR1_12S_50ntFlanks@425.456.32\|ACGAAAGTGGCTTTAACATATCTGAACACACA | >hg38_MT_+_598_1651_RNR1_12S_50ntFlanks | 425 | 456 | 32 | ACGAAAGTGGCTTTAACATATCTGAACACACA | 631.6 | 13.9 | 1.4 | 10.0 | 9.6E-24 | 8.2E-23 | BT20/MB468 |
| >MI0003591\|hsa-mir-584&WithFlank&5\|-\|149062307\|149062415@22.41.20(+1U)&[MIMAT0003249&hsa-miR-584-5p&offsets\|0\|-2(+1U);m-167&5\|-\|149062374\|149062394&offsets\|0\|-1(+1U)]\|TTATGGTTTGCCTGGGACTGT | >MI0003591\|hsa-mir-584&WithFlank&5\|-\|149062307\|149062415 | 22 | 41 | 20(+1U) | TTATGGTTTGCCTGGGACTGT | 1859.4 | 10.0 | 1.0 | 10.0 | 9.8E-24 | 8.3E-23 | BT20/MB468 |
| >hg38_1_-_228634819_228635039_RNA5S12_5S_50ntFlanks@138.170.33\|AGACCGCCTGGGAATACCGGGTGCTGTAGGCTT | >hg38_1_-_228634819_228635039_RNA5S12_5S_50ntFlanks | 138 | 170 | 33 | AGACCGCCTGGGAATACCGGGTGCTGTAGGCTT | 2969.2 | 8.2 | 0.8 | 10.0 | 1.3E-23 | 8.5E-23 | MB231/MB468 |
| >MI0000811\|hsa-mir-148b&WithFlank&12\|+\|54337210\|54337320@69.89.21&[MIMAT0000759&hsa-miR-148b-3p&offsets\|0\|-1;m-77&12\|+\|54337278\|54337299&offsets\|0\|-1]\|TCAGTGCATCACAGAACTTTG | >MI0000811\|hsa-mir-148b&WithFlank&12\|+\|54337210\|54337320 | 69 | 89 | 21 | TCAGTGCATCACAGAACTTTG | 1835.0 | 8.4 | 0.8 | 10.0 | 1.3E-23 | 8.5E-23 | MB231/MB468 |
| >MI0000465\|hsa-mir-191&WithFlank&3\|-\|49020612\|49020715@22.44.23(+1A)&[MIMAT0000440&hsa-miR-191-5p&offsets\|0\|0(+1A);m-48&3\|-\|49020672\|49020694&offsets\|0\|0(+1A)]\|CAACGGAATCCCAAAAGCAGCTGA | >MI0000465\|hsa-mir-191&WithFlank&3\|-\|49020612\|49020715 | 22 | 44 | 23(+1A) | CAACGGAATCCCAAAAGCAGCTGA | 828.8 | 6.7 | 0.7 | 10.0 | 1.0E-23 | 8.8E-23 | BT20/MB468 |
| >hg38_21_+_8433172_8446622_RNA45SN1_45S_50ntFlanks@12008.12032.25\|CCGCCGGTGAAATACCACTACTCTG | >hg38_21_+_8433172_8446622_RNA45SN1_45S_50ntFlanks | 12008 | 12032 | 25 | CCGCCGGTGAAATACCACTACTCTG | 5928.3 | -9.0 | 0.9 | -10.0 | 1.4E-23 | 9.0E-23 | MB231/MB468 |
| >hg38_21_+_8433172_8446622_RNA45SN1_45S_50ntFlanks@12405.12426.22\|CTTTTTGATCCTTCGATGTCGG | >hg38_21_+_8433172_8446622_RNA45SN1_45S_50ntFlanks | 12405 | 12426 | 22 | CTTTTTGATCCTTCGATGTCGG | 2765.4 | -13.0 | 1.3 | -10.0 | 1.4E-23 | 9.1E-23 | MB231/MB468 |
| >trna13_ValCAC_19_-_4724647_4724719@1.34.34\|GTTTCCGTAGTGTAGCGGTTATCACATTCGCCTC | >trna13_ValCAC_19_-_4724647_4724719 | 1 | 34 | 34 | GTTTCCGTAGTGTAGCGGTTATCACATTCGCCTC | 1771.0 | 15.7 | 1.6 | 10.0 | 1.4E-23 | 9.5E-23 | MB231/MB468 |
| >MI0000077\|hsa-mir-21&WithFlank&17\|+\|59841260\|59841343@14.36.23(+1A)&[MIMAT0000076&hsa-miR-21-5p&offsets\|0\|+1(+1A);m-1&17\|+\|59841273\|59841294&offsets\|0\|+1(+1A)]\|TAGCTTATCAGACTGATGTTGACA | >MI0000077\|hsa-mir-21&WithFlank&17\|+\|59841260\|59841343 | 14 | 36 | 23(+1A) | TAGCTTATCAGACTGATGTTGACA | 448403.6 | 7.4 | 0.7 | 10.0 | 1.1E-23 | 9.6E-23 | BT20/MB468 |

<table>
<tr><td>>MI0000466|hsa-mir-9-1&WithFlank&1|-|156420335|156420435@22.42.21;MI0000468|hsa-mir-9-3&WithFlank&15|+|89368011|89368112@22.42.21;MI0000467|hsa-mir-9-2&WithFlank&5|-|88666847|88666945@22.42.21&[MIMAT0000441&hsa-miR-9-5p&offsets|0|-2;m-104&1|-|156420392|156420414&offsets|0|-2];[MIMAT0000441_1&hsa-miR-9-5p&offsets|0|-2;m-105&15|+|89368032|89368054&offsets|0|-2];[MIMAT0000441_2&hsa-miR-9-5p&offsets|0|-2;m-104&5|-|88666902|88666924&offsets|0|-2]|TCTTTGGTTATCTAGCTGTAT</td><td>>MI0000466|hsa-mir-9-1&WithFlank&1|-|156420335|156420435</td><td>22</td><td>42</td><td>21</td><td>TCTTTGGTTATCTAGCTGTAT</td><td>581.0</td><td>13.8</td><td>1.4</td><td>10.0</td><td>1.2E-23</td><td>9.8E-23</td><td>BT20/MB468</td></tr>
<tr><td>>hg38_21_+_8433172_8446622_RNA45SN1_45S_50ntFlanks@475.502.28|CTTCGTGATCGATGTGGTGACGTCGTGC</td><td>>hg38_21_+_8433172_8446622_RNA45SN1_45S_50ntFlanks</td><td>475</td><td>502</td><td>28</td><td>CTTCGTGATCGATGTGGTGACGTCGTGC</td><td>1852.5</td><td>6.8</td><td>0.7</td><td>10.0</td><td>1.5E-23</td><td>9.9E-23</td><td>MB231/MB468</td></tr>
<tr><td>>hg38_21_+_8433172_8446622_RNA45SN1_45S_50ntFlanks@6741.6771.31|ATCGACACTTCGAACGCACTTGCGGCCCCGG</td><td>>hg38_21_+_8433172_8446622_RNA45SN1_45S_50ntFlanks</td><td>6741</td><td>6771</td><td>31</td><td>ATCGACACTTCGAACGCACTTGCGGCCCCGG</td><td>1137.7</td><td>-6.0</td><td>0.6</td><td>-10.0</td><td>1.5E-23</td><td>9.9E-23</td><td>MB231/MB468</td></tr>
<tr><td>>MI0000061|hsa-let-7a-2&WithFlank&11|-|122146516|122146599@11.32.22(+1A);MI0000062|hsa-let-7a-3&WithFlank&22|+|46112743|46112828@10.31.22(+1A);MI0000060|hsa-let-7a-1&WithFlank&9|+|94175951|94176042@12.33.22(+1A)&[MIMAT0000062&hsa-let-7a-5p&offsets|0|0(+1A);m-6&11|-|122146568|122146589&offsets|0|0(+1A)];[MIMAT0000062_1&hsa-let-7a-5p&offsets|0|0(+1A);m-5&22|+|46112752|46112773&offsets|0|0(+1A)];[MIMAT0000062_2&hsa-let-7a-5p&offsets|0|0(+1A);m-4&9|+|94175962|94175983&offsets|0|0(+1A)]|TGAGGTAGTAGGTTGTATAGTTA</td><td>>MI0000061|hsa-let-7a-2&WithFlank&11|-|122146516|122146599</td><td>11</td><td>32</td><td>22(+1A)</td><td>TGAGGTAGTAGGTTGTATAGTTA</td><td>3606.5</td><td>7.4</td><td>0.7</td><td>10.0</td><td>1.5E-23</td><td>9.9E-23</td><td>MB231/MB468</td></tr>
<tr><td>>trnaMT_ThrTGT_MT_+_15888_15953@31.69.39|TTGTAAACCGGAGATGAAAACCTTTTTCCAAGGACACCA</td><td>>trnaMT_ThrTGT_MT_+_15888_15953</td><td>31</td><td>69</td><td>39</td><td>TTGTAAACCGGAGATGAAAACCTTTTTCCAAGGACACCA</td><td>1419.1</td><td>15.1</td><td>1.5</td><td>10.0</td><td>1.2E-23</td><td>9.9E-23</td><td>BT20/MB468</td></tr>
<tr><td>>MI0003560|hsa-mir-92b&WithFlank&1|+|155195171|155195278@67.88.22(+2U)&[MIMAT0003218&hsa-miR-92b-3p&offsets|0|0(+2U);m-115&1|+|155195237|155195258&offsets|0|0(+2U)]|TATTGCACTCGTCCCGGCCTCCTT</td><td>>MI0003560|hsa-mir-92b&WithFlank&1|+|155195171|155195278</td><td>67</td><td>88</td><td>22(+2U)</td><td>TATTGCACTCGTCCCGGCCTCCTT</td><td>527.7</td><td>13.7</td><td>1.4</td><td>10.0</td><td>1.2E-23</td><td>9.9E-23</td><td>BT20/MB468</td></tr>
<tr><td>>hg38_21_+_8433172_8446622_RNA45SN1_45S_50ntFlanks@476.513.38|TTCGTGATCGATGTGGTGACGTCGTGCTCTCCCGGGCC</td><td>>hg38_21_+_8433172_8446622_RNA45SN1_45S_50ntFlanks</td><td>476</td><td>513</td><td>38</td><td>TTCGTGATCGATGTGGTGACGTCGTGCTCTCCCGGGCC</td><td>1200.9</td><td>9.9</td><td>1.0</td><td>10.0</td><td>1.6E-23</td><td>1.0E-22</td><td>MB231/MB468</td></tr>
<tr><td>>hg38_21_+_8433172_8446622_RNA45SN1_45S_50ntFlanks@4725.4767.43|TTAATCAAGAACGAAAGTCGGAGGTTCGAAGACGATCAGATAC</td><td>>hg38_21_+_8433172_8446622_RNA45SN1_45S_50ntFlanks</td><td>4725</td><td>4767</td><td>43</td><td>TTAATCAAGAACGAAAGTCGGAGGTTCGAAGACGATCAGATAC</td><td>265.3</td><td>10.9</td><td>1.1</td><td>10.0</td><td>1.6E-23</td><td>1.0E-22</td><td>MB231/MB468</td></tr>
<tr><td>>hg38_21_+_8433172_8446622_RNA45SN1_45S_50ntFlanks@11641.11669.29|CGGCGGGTGTTGACGCGATGTGATTTCTG</td><td>>hg38_21_+_8433172_8446622_RNA45SN1_45S_50ntFlanks</td><td>11641</td><td>11669</td><td>29</td><td>CGGCGGGTGTTGACGCGATGTGATTTCTG</td><td>1668.0</td><td>-9.4</td><td>0.9</td><td>-10.0</td><td>1.2E-23</td><td>1.0E-22</td><td>BT20/MB468</td></tr>
<tr><td>>MI0000746|hsa-mir-99b&WithFlank&19|+|51692606|51692687@51.72.22&[MIMAT0004678&hsa-miR-99b-3p&offsets|0|0;m-151&19|+|51692656|51692678&offsets|0|-1]|CAAGCTCGTGTCTGTGGGTCCG</td><td>>MI0000746|hsa-mir-99b&WithFlank&19|+|51692606|51692687</td><td>51</td><td>72</td><td>22</td><td>CAAGCTCGTGTCTGTGGGTCCG</td><td>4197.2</td><td>6.8</td><td>0.7</td><td>10.0</td><td>1.2E-23</td><td>1.0E-22</td><td>BT20/MB468</td></tr>
<tr><td>>MI0000433|hsa-let-7g&WithFlank&3|-|52268272|52268367@11.30.20(+1C)&[MIMAT0000414&hsa-let-7g-5p&offsets|0|-2(+1C);m-58&3|-|52268336|52268357&offsets|0|-2(+1C)]|TGAGGTAGTAGTTTGTACAGC</td><td>>MI0000433|hsa-let-7g&WithFlank&3|-|52268272|52268367</td><td>11</td><td>30</td><td>20(+1C)</td><td>TGAGGTAGTAGTTTGTACAGC</td><td>1237.7</td><td>7.2</td><td>0.7</td><td>10.0</td><td>1.2E-23</td><td>1.0E-22</td><td>BT20/MB468</td></tr>
<tr><td>>hg38_21_+_8433172_8446622_RNA45SN1_45S_50ntFlanks@10841.10862.22|CGTAACTTCGGGATAAGGATTG</td><td>>hg38_21_+_8433172_8446622_RNA45SN1_45S_50ntFlanks</td><td>10841</td><td>10862</td><td>22</td><td>CGTAACTTCGGGATAAGGATTG</td><td>3780.8</td><td>-7.4</td><td>0.7</td><td>-10.0</td><td>1.3E-23</td><td>1.0E-22</td><td>BT20/MB468</td></tr>
<tr><td>>MI0000066|hsa-let-7e&WithFlank&19|+|51692780|51692870@14.35.22&[MIMAT0000066&hsa-let-7e-5p&offsets|0|0;m-50&19|+|51692793|51692814&offsets|0|0]|TGAGGTAGGAGGTTGTATAGTT</td><td>>MI0000066|hsa-let-7e&WithFlank&19|+|51692780|51692870</td><td>14</td><td>35</td><td>22</td><td>TGAGGTAGGAGGTTGTATAGTT</td><td>31323.6</td><td>7.2</td><td>0.7</td><td>10.0</td><td>1.3E-23</td><td>1.0E-22</td><td>BT20/MB468</td></tr>
<tr><td>>MI0000114|hsa-mir-107&WithFlank&10|-|89592741|89592833@56.76.21;MI0000108|hsa-mir-103a-2&WithFlank&20|+|3917488|3917577@54.74.21;MI0000109|hsa-mir-103a-1&WithFlank&5|-|168560890|168560979@54.74.21&[MIMAT0000104&hsa-miR-107&offsets|0|-2;m-78&10|-|89592758|89592778&offsets|0|0];[MIMAT0000101&hsa-miR-103a-3p&offsets|0|-2;m-12&20|+|3917541|3917563&offsets|0|-2];[MIMAT0000101_1&hsa-miR-103a-3p&offsets|0|-2;m-13&5|-|168560904|168560926&offsets|0|-2;m-44220&5|-|168560908|168560926&offsets|0|+2]|AGCAGCATTGTACAGGGCTAT</td><td>>MI0000114|hsa-mir-107&WithFlank&10|-|89592741|89592833</td><td>56</td><td>76</td><td>21</td><td>AGCAGCATTGTACAGGGCTAT</td><td>2609.6</td><td>8.0</td><td>0.8</td><td>10.0</td><td>1.6E-23</td><td>1.1E-22</td><td>MB231/MB468</td></tr>
<tr><td>>hg38_MT_+_1621_3279_RNR2_16S_50ntFlanks@51.72.22|GCTAAACCTAGCCCCAAACCCA</td><td>>hg38_MT_+_1621_3279_RNR2_16S_50ntFlanks</td><td>51</td><td>72</td><td>22</td><td>GCTAAACCTAGCCCCAAACCCA</td><td>2163.8</td><td>8.5</td><td>0.9</td><td>10.0</td><td>1.3E-23</td><td>1.1E-22</td><td>BT20/MB468</td></tr>
<tr><td>>MI0000239|hsa-mir-197&WithFlank&1|+|109598887|109598973@54.74.21(+1U)&[MIMAT0000227&hsa-miR-197-3p&offsets|0|-1(+1U);m-99&1|+|109598940|109598961&offsets|0|-1(+1U)]|TTCACCACCTTCTCCACCCAGT</td><td>>MI0000239|hsa-mir-197&WithFlank&1|+|109598887|109598973</td><td>54</td><td>74</td><td>21(+1U)</td><td>TTCACCACCTTCTCCACCCAGT</td><td>715.1</td><td>7.1</td><td>0.7</td><td>10.0</td><td>1.6E-23</td><td>1.1E-22</td><td>MB231/MB468</td></tr>
<tr><td>>hg38_21_+_8433172_8446622_RNA45SN1_45S_50ntFlanks@12395.12414.20|AGCGACGTCGCTTTTTGATC</td><td>>hg38_21_+_8433172_8446622_RNA45SN1_45S_50ntFlanks</td><td>12395</td><td>12414</td><td>20</td><td>AGCGACGTCGCTTTTTGATC</td><td>362.6</td><td>3.8</td><td>0.4</td><td>10.0</td><td>1.3E-23</td><td>1.1E-22</td><td>BT20/MB468</td></tr>
<tr><td>>hg38_21_+_8433172_8446622_RNA45SN1_45S_50ntFlanks@6774.6806.33|TCCTCCCGGGGCTACGCCTGTCTGAGCGTCGCT</td><td>>hg38_21_+_8433172_8446622_RNA45SN1_45S_50ntFlanks</td><td>6774</td><td>6806</td><td>33</td><td>TCCTCCCGGGGCTACGCCTGTCTGAGCGTCGCT</td><td>2965.3</td><td>7.2</td><td>0.7</td><td>10.0</td><td>1.6E-23</td><td>1.1E-22</td><td>MB231/MB468</td></tr>
<tr><td>>MI0000760|hsa-mir-361&WithFlank&X|-|85903630|85903713@51.73.23&[MIMAT0004682&hsa-miR-361-3p&offsets|0|0;m-124&X|-|85903642|85903663&offsets|0|+1]|TCCCCCAGGTGTGATTCTGATTT</td><td>>MI0000760|hsa-mir-361&WithFlank&X|-|85903630|85903713</td><td>51</td><td>73</td><td>23</td><td>TCCCCCAGGTGTGATTCTGATTT</td><td>932.3</td><td>7.3</td><td>0.7</td><td>10.0</td><td>1.3E-23</td><td>1.1E-22</td><td>BT20/MB468</td></tr>
<tr><td>>hg38_21_+_8433172_8446622_RNA45SN1_45S_50ntFlanks@4823.4859.37|ATGACCCGCCGGGCAGCTTCCGGGAAACCAAAGTCTT</td><td>>hg38_21_+_8433172_8446622_RNA45SN1_45S_50ntFlanks</td><td>4823</td><td>4859</td><td>37</td><td>ATGACCCGCCGGGCAGCTTCCGGGAAACCAAAGTCTT</td><td>390.9</td><td>11.5</td><td>1.1</td><td>10.0</td><td>1.7E-23</td><td>1.1E-22</td><td>MB231/MB468</td></tr>
<tr><td>>hg38_21_+_8433172_8446622_RNA45SN1_45S_50ntFlanks@485.522.38|GATGTGGTGACGTCGTGCTCTCCCGGGCCGGGTCCGAG</td><td>>hg38_21_+_8433172_8446622_RNA45SN1_45S_50ntFlanks</td><td>485</td><td>522</td><td>38</td><td>GATGTGGTGACGTCGTGCTCTCCCGGGCCGGGTCCGAG</td><td>1439.5</td><td>15.1</td><td>1.5</td><td>10.0</td><td>1.3E-23</td><td>1.1E-22</td><td>BT20/MB468</td></tr>
<tr><td>>MI0000272|hsa-mir-182&WithFlank&7|-|129770377|129770498@29.53.25(+1A)&[MIMAT0000259&hsa-miR-182-5p&offsets|0|+1(+1A);m-22&7|-|129770449|129770470&offsets|0|+3(+1A)]|TTTGGCAATGGTAGAACTCACACTGA</td><td>>MI0000272|hsa-mir-182&WithFlank&7|-|129770377|129770498</td><td>29</td><td>53</td><td>25(+1A)</td><td>TTTGGCAATGGTAGAACTCACACTGA</td><td>4236.4</td><td>9.3</td><td>0.9</td><td>10.0</td><td>1.3E-23</td><td>1.1E-22</td><td>BT20/MB468</td></tr>
<tr><td>>MI0000066|hsa-let-7e&WithFlank&19|+|51692780|51692870@14.34.21&[MIMAT0000066&hsa-let-7e-5p&offsets|0|-1;m-50&19|+|51692793|51692814&offsets|0|-1]|TGAGGTAGGAGGTTGTATAGT</td><td>>MI0000066|hsa-let-7e&WithFlank&19|+|51692780|51692870</td><td>14</td><td>34</td><td>21</td><td>TGAGGTAGGAGGTTGTATAGT</td><td>4178.0</td><td>7.5</td><td>0.8</td><td>10.0</td><td>1.7E-23</td><td>1.1E-22</td><td>MB231/MB468</td></tr>
<tr><td>>MI0000750|hsa-mir-26a-2&WithFlank&12|-|57824603|57824698@20.41.22(+2U);MI0000083|hsa-mir-26a-1&WithFlank&3|+|37969398|37969486@16.37.22(+2U)&[MIMAT0000082&hsa-miR-26a-5p&offsets|0|0(+2U);m-33&12|-|57824658|57824679&offsets|0|0(+2U)];[MIMAT0000082_1&hsa-miR-26a-5p&offsets|0|0(+2U);m-34&3|+|37969413|37969434&offsets|0|0(+2U)]|TTCAAGTAATCCAGGATAGGCTTT</td><td>>MI0000750|hsa-mir-26a-2&WithFlank&12|-|57824603|57824698</td><td>20</td><td>41</td><td>22(+2U)</td><td>TTCAAGTAATCCAGGATAGGCTTT</td><td>3131.0</td><td>6.5</td><td>0.7</td><td>10.0</td><td>1.4E-23</td><td>1.1E-22</td><td>BT20/MB468</td></tr>
<tr><td>>MI0000273|hsa-mir-183&WithFlank&7|-|129774899|129775020@33.54.22(+1A)&[MIMAT0000261&hsa-miR-183-5p&offsets|0|0(+1A);m-41&7|-|129774966|129774987&offsets|-1|-1(+1A)]|TATGGCACTGGTAGAATTCACTA</td><td>>MI0000273|hsa-mir-183&WithFlank&7|-|129774899|129775020</td><td>33</td><td>54</td><td>22(+1A)</td><td>TATGGCACTGGTAGAATTCACTA</td><td>2482.1</td><td>8.6</td><td>0.9</td><td>10.0</td><td>1.4E-23</td><td>1.2E-22</td><td>BT20/MB468</td></tr>
<tr><td>>MI0000086|hsa-mir-28&WithFlank&3|+|188688775|188688872@62.82.21&[MIMAT0004502&hsa-miR-28-3p&offsets|+2|+1;m-28&3|+|188688834|188688855&offsets|+2|+1]|CTAGATTGTGAGCTCCTGGAG</td><td>>MI0000086|hsa-mir-28&WithFlank&3|+|188688775|188688872</td><td>62</td><td>82</td><td>21</td><td>CTAGATTGTGAGCTCCTGGAG</td><td>583.1</td><td>13.8</td><td>1.4</td><td>10.0</td><td>1.4E-23</td><td>1.2E-22</td><td>BT20/MB468</td></tr>
<tr><td>>hg38_MT_+_598_1651_RNR1_12S_50ntFlanks@52.87.36|ATAGGTTTGGTCCTAGCCTTTCTATTAGCTCTTAGT</td><td>>hg38_MT_+_598_1651_RNR1_12S_50ntFlanks</td><td>52</td><td>87</td><td>36</td><td>ATAGGTTTGGTCCTAGCCTTTCTATTAGCTCTTAGT</td><td>1387.5</td><td>7.0</td><td>0.7</td><td>10.0</td><td>1.4E-23</td><td>1.2E-22</td><td>BT20/MB468</td></tr>
<tr><td>>MI0000077|hsa-mir-21&WithFlank&17|+|59841260|59841343@14.33.20(+1A)&[MIMAT0000076&hsa-miR-21-5p&offsets|0|-2(+1A);m-1&17|+|59841273|59841294&offsets|0|-2(+1A)]|TAGCTTATCAGACTGATGTTA</td><td>>MI0000077|hsa-mir-21&WithFlank&17|+|59841260|59841343</td><td>14</td><td>33</td><td>20(+1A)</td><td>TAGCTTATCAGACTGATGTTA</td><td>3432.7</td><td>7.5</td><td>0.8</td><td>10.0</td><td>1.8E-23</td><td>1.2E-22</td><td>MB231/MB468</td></tr>
<tr><td>>hg38_21_+_8433172_8446622_RNA45SN1_45S_50ntFlanks@475.508.34|CTTCGTGATCGATGTGGTGACGTCGTGCTCTCCC</td><td>>hg38_21_+_8433172_8446622_RNA45SN1_45S_50ntFlanks</td><td>475</td><td>508</td><td>34</td><td>CTTCGTGATCGATGTGGTGACGTCGTGCTCTCCC</td><td>1407.4</td><td>6.7</td><td>0.7</td><td>10.0</td><td>1.5E-23</td><td>1.2E-22</td><td>BT20/MB468</td></tr>
<tr><td>>MI0000088|hsa-mir-30a&WithFlank&6|-|71403545|71403627@53.74.22(+1A)&[MIMAT0000088&hsa-miR-30a-3p&offsets|0|0(+1A);m-38&6|-|71403554|71403575&offsets|0|0(+1A)]|CTTTCAGTCGGATGTTTGCAGCA</td><td>>MI0000088|hsa-mir-30a&WithFlank&6|-|71403545|71403627</td><td>53</td><td>74</td><td>22(+1A)</td><td>CTTTCAGTCGGATGTTTGCAGCA</td><td>893.5</td><td>8.0</td><td>0.8</td><td>10.0</td><td>1.9E-23</td><td>1.2E-22</td><td>MB231/MB468</td></tr>
<tr><td>>trnaMT_SerGCT_MT_+_12207_12265@1.21.21|GAGAAAGCTCACAAGAACTGC</td><td>>trnaMT_SerGCT_MT_+_12207_12265</td><td>1</td><td>21</td><td>21</td><td>GAGAAAGCTCACAAGAACTGC</td><td>2916.9</td><td>7.8</td><td>0.8</td><td>10.0</td><td>1.9E-23</td><td>1.2E-22</td><td>MB231/MB468</td></tr>
<tr><td>>MI0001446|hsa-mir-424&WithFlank&X|-|134546608|134546717@54.75.22&[MIMAT0004749&hsa-miR-424-3p&offsets|0|+1;m-305&X|-|134546644|134546664&offsets|0|+1]|CAAAACGTGAGGCGCTGCTATA</td><td>>MI0001446|hsa-mir-424&WithFlank&X|-|134546608|134546717</td><td>54</td><td>75</td><td>22</td><td>CAAAACGTGAGGCGCTGCTATA</td><td>952.3</td><td>11.2</td><td>1.1</td><td>10.0</td><td>1.9E-23</td><td>1.2E-22</td><td>MB231/MB468</td></tr>
<tr><td>>MI0000065|hsa-let-7d&WithFlank&9|+|94178828|94178926@14.36.23&[MIMAT0000065&hsa-let-7d-5p&offsets|0|+1;m-90&9|+|94178841|94178862&offsets|0|+1]|AGAGGTAGTAGGTTGCATAGTTT</td><td>>MI0000065|hsa-let-7d&WithFlank&9|+|94178828|94178926</td><td>14</td><td>36</td><td>23</td><td>AGAGGTAGTAGGTTGCATAGTTT</td><td>695.5</td><td>7.4</td><td>0.7</td><td>10.0</td><td>1.9E-23</td><td>1.2E-22</td><td>MB231/MB468</td></tr>
<tr><td>>MI0000746|hsa-mir-99b&WithFlank&19|+|51692606|51692687@51.73.23&[MIMAT0004678&hsa-miR-99b-3p&offsets|0|+1;m-151&19|+|51692656|51692678&offsets|0|0]|CAAGCTCGTGTCTGTGGGTCCGT</td><td>>MI0000746|hsa-mir-99b&WithFlank&19|+|51692606|51692687</td><td>51</td><td>73</td><td>23</td><td>CAAGCTCGTGTCTGTGGGTCCGT</td><td>2039.4</td><td>7.7</td><td>0.8</td><td>10.0</td><td>2.0E-23</td><td>1.3E-22</td><td>MB231/MB468</td></tr>
<tr><td>>trnaMT_PheGAA_MT_+_577_647@44.74.31|TAGACGGGCTCACATCACCCCATAAACACCA</td><td>>trnaMT_PheGAA_MT_+_577_647</td><td>44</td><td>74</td><td>31</td><td>TAGACGGGCTCACATCACCCCATAAACACCA</td><td>1740.9</td><td>6.7</td><td>0.7</td><td>10.0</td><td>2.0E-23</td><td>1.3E-22</td><td>MB231/MB468</td></tr>
</table>

<table>
<tr><td></td><td></td><td></td><td></td><td></td><td></td><td></td><td></td><td></td><td></td><td></td><td></td><td></td></tr>
<tr><td>>hg38_21_+_8433172_8446622_RNA45SN1_45S_50ntFlanks@4283.4317.35|CTTTAACGAGGATCCATTGGAGGGCAAGTCTGGTG</td><td>>hg38_21_+_8433172_8446622_RNA45SN1_45S_50ntFlanks</td><td>4283</td><td>4317</td><td>35</td><td>CTTTAACGAGGATCCATTGGAGGGCAAGTCTGGTG</td><td>748.5</td><td>-7.3</td><td>0.7</td><td>-10.0</td><td>1.6E-23</td><td>1.3E-22</td><td>BT20/MB468</td></tr>
<tr><td>>MI0000471|hsa-mir-126&WithFlank&9|+|136670596|136670692@57.78.22&[MIMAT0000445&hsa-miR-126-3p&offsets|-1|-1;m-40&9|+|136670653|136670674&offsets|-1|-1]|CTCGTACCGTGAGTAATAATGC</td><td>>MI0000471|hsa-mir-126&WithFlank&9|+|136670596|136670692</td><td>57</td><td>78</td><td>22</td><td>CTCGTACCGTGAGTAATAATGC</td><td>467.7</td><td>13.5</td><td>1.4</td><td>10.0</td><td>1.6E-23</td><td>1.3E-22</td><td>BT20/MB468</td></tr>
<tr><td>>hg38_MT_+_598_1651_RNR1_12S_50ntFlanks@443.480.38|TATCTGAACACACAATAGCTAAGACCCAAACTGGGATT</td><td>>hg38_MT_+_598_1651_RNR1_12S_50ntFlanks</td><td>443</td><td>480</td><td>38</td><td>TATCTGAACACACAATAGCTAAGACCCAAACTGGGATT</td><td>851.7</td><td>12.2</td><td>1.2</td><td>10.0</td><td>2.0E-23</td><td>1.3E-22</td><td>MB231/MB468</td></tr>
<tr><td>>MI0003591|hsa-mir-584&WithFlank&5|-|149062307|149062415@22.41.20&[MIMAT0003249&hsa-miR-584-5p&offsets|0|-2;m-167&5|-|149062374|149062394&offsets|0|-1]|TTATGGTTTGCCTGGGACTG</td><td>>MI0003591|hsa-mir-584&WithFlank&5|-|149062307|149062415</td><td>22</td><td>41</td><td>20</td><td>TTATGGTTTGCCTGGGACTG</td><td>1278.6</td><td>9.9</td><td>1.0</td><td>10.0</td><td>2.1E-23</td><td>1.3E-22</td><td>MB231/MB468</td></tr>
<tr><td>>MI0001445|hsa-mir-423&WithFlank&17|+|30117073|30117178@59.79.21&[MIMAT0001340&hsa-miR-423-3p&offsets|0|-2;m-100&17|+|30117131|30117153&offsets|0|-2]|AGCTCGGTCTGAGGCCCCTCA</td><td>>MI0001445|hsa-mir-423&WithFlank&17|+|30117073|30117178</td><td>59</td><td>79</td><td>21</td><td>AGCTCGGTCTGAGGCCCCTCA</td><td>1464.3</td><td>7.3</td><td>0.7</td><td>10.0</td><td>1.6E-23</td><td>1.3E-22</td><td>BT20/MB468</td></tr>
<tr><td>>hg38_21_+_8433172_8446622_RNA45SN1_45S_50ntFlanks@12935.12956.22|TAAACCATTCGTAGACGACCTG</td><td>>hg38_21_+_8433172_8446622_RNA45SN1_45S_50ntFlanks</td><td>12935</td><td>12956</td><td>22</td><td>TAAACCATTCGTAGACGACCTG</td><td>1448.9</td><td>-5.9</td><td>0.6</td><td>-10.0</td><td>2.1E-23</td><td>1.3E-22</td><td>MB231/MB468</td></tr>
<tr><td>>hg38_1_-_228634819_228635039_RNA5S12_5S_50ntFlanks@153.171.19|ACCGGGTGCTGTAGGCTTT</td><td>>hg38_1_-_228634819_228635039_RNA5S12_5S_50ntFlanks</td><td>153</td><td>171</td><td>19</td><td>ACCGGGTGCTGTAGGCTTT</td><td>23116.1</td><td>8.1</td><td>0.8</td><td>10.0</td><td>2.1E-23</td><td>1.4E-22</td><td>MB231/MB468</td></tr>
<tr><td>>MI0000272|hsa-mir-182&WithFlank&7|-|129770377|129770498@29.54.26(+1U)&[MIMAT0000259&hsa-miR-182-5p&offsets|0|+2(+1U);m-22&7|-|129770449|129770470&offsets|0|+4(+1U)]|TTTGGCAATGGTAGAACTCACACTGGT</td><td>>MI0000272|hsa-mir-182&WithFlank&7|-|129770377|129770498</td><td>29</td><td>54</td><td>26(+1U)</td><td>TTTGGCAATGGTAGAACTCACACTGGT</td><td>1626.3</td><td>7.8</td><td>0.8</td><td>10.0</td><td>1.7E-23</td><td>1.4E-22</td><td>BT20/MB468</td></tr>
<tr><td>>MI0000456|hsa-mir-140&WithFlank&16|+|69933075|69933186@68.90.23(+1U)&[MIMAT0004597&hsa-miR-140-3p&offsets|0|+2(+1U);m-45&16|+|69933143|69933164&offsets|-1|0(+1U)]|TACCACAGGGTAGAACCACGGACT</td><td>>MI0000456|hsa-mir-140&WithFlank&16|+|69933075|69933186</td><td>68</td><td>90</td><td>23(+1U)</td><td>TACCACAGGGTAGAACCACGGACT</td><td>396.2</td><td>13.3</td><td>1.3</td><td>10.0</td><td>1.7E-23</td><td>1.4E-22</td><td>BT20/MB468</td></tr>
<tr><td>>trna152_ValCAC_6_-_27248049_27248121@1.34.34|GCTTCTGTAGTGTAGTGGTTATCACGTTCGCCTC</td><td>>trna152_ValCAC_6_-_27248049_27248121</td><td>1</td><td>34</td><td>34</td><td>GCTTCTGTAGTGTAGTGGTTATCACGTTCGCCTC</td><td>14024.1</td><td>12.6</td><td>1.3</td><td>10.0</td><td>2.1E-23</td><td>1.4E-22</td><td>MB231/MB468</td></tr>
<tr><td>>hg38_21_+_8433172_8446622_RNA45SN1_45S_50ntFlanks@12456.12474.19|CCAAGCGTTGGATTGTTCA</td><td>>hg38_21_+_8433172_8446622_RNA45SN1_45S_50ntFlanks</td><td>12456</td><td>12474</td><td>19</td><td>CCAAGCGTTGGATTGTTCA</td><td>4509.4</td><td>-6.5</td><td>0.7</td><td>-10.0</td><td>2.2E-23</td><td>1.4E-22</td><td>MB231/MB468</td></tr>
<tr><td>>MI0000103|hsa-mir-101-1&WithFlank&1|-|65058428|65058514@53.74.22(+1U);MI0000739|hsa-mir-101-2&WithFlank&9|+|4850291|4850381@55.76.22(+1U)&[MIMAT0000099&hsa-miR-101-3p&offsets|0|+1(+1U);m-17&1|-|65058443|65058462&offsets|0|+2(+1U)];[MIMAT0000099_1&hsa-miR-101-3p&offsets|0|+1(+1U);m-16&9|+|4850345|4850364&offsets|0|+2(+1U)]|TACAGTACTGTGATAACTGAAGT</td><td>>MI0000103|hsa-mir-101-1&WithFlank&1|-|65058428|65058514</td><td>53</td><td>74</td><td>22(+1U)</td><td>TACAGTACTGTGATAACTGAAGT</td><td>2397.0</td><td>8.3</td><td>0.8</td><td>10.0</td><td>2.2E-23</td><td>1.4E-22</td><td>MB231/MB468</td></tr>
<tr><td>>hg38_21_+_8433172_8446622_RNA45SN1_45S_50ntFlanks@11882.11904.23|CTGTTGAGCTTGACTCTAGTCTG</td><td>>hg38_21_+_8433172_8446622_RNA45SN1_45S_50ntFlanks</td><td>11882</td><td>11904</td><td>23</td><td>CTGTTGAGCTTGACTCTAGTCTG</td><td>1056.4</td><td>-6.0</td><td>0.6</td><td>-10.0</td><td>1.8E-23</td><td>1.5E-22</td><td>BT20/MB468</td></tr>
<tr><td>>MI0000063|hsa-let-7b&WithFlank&22|+|46113680|46113774@12.34.23&[MIMAT0000063&hsa-let-7b-5p&offsets|0|+1;m-14&22|+|46113691|46113712&offsets|0|+1]|TGAGGTAGTAGGTTGTGTGGTTT</td><td>>MI0000063|hsa-let-7b&WithFlank&22|+|46113680|46113774</td><td>12</td><td>34</td><td>23</td><td>TGAGGTAGTAGGTTGTGTGGTTT</td><td>5896.4</td><td>7.1</td><td>0.7</td><td>10.0</td><td>2.3E-23</td><td>1.5E-22</td><td>MB231/MB468</td></tr>
<tr><td>>hg38_21_+_8433172_8446622_RNA45SN1_45S_50ntFlanks@12281.12321.41|AATACAGACCGTGAAAGCGGGGCCTCACGATCCTTCTGACC</td><td>>hg38_21_+_8433172_8446622_RNA45SN1_45S_50ntFlanks</td><td>12281</td><td>12321</td><td>41</td><td>AATACAGACCGTGAAAGCGGGGCCTCACGATCCTTCTGACC</td><td>275.0</td><td>11.0</td><td>1.1</td><td>10.0</td><td>2.3E-23</td><td>1.5E-22</td><td>MB231/MB468</td></tr>
<tr><td>>trna111_HisGTG_1_-_147774845_147774916@-1T.26.27,trna118_HisGTG_1_-_145396881_145396952@-1T.26.27,trna16_HisGTG_1_+_146544773_146544844@-1T.26.27,trna1_HisGTG_15_+_45493349_45493420@-1T.26.27,trna21_HisGTG_1_+_147753471_147753542@-1T.26.27,trna33_HisGTG_6_+_27125906_27125977@-1T.26.27,trna7_HisGTG_9_-_14433938_14434009@-1T.26.27,trna8_HisGTG_15_-_45492611_45492682@-1T.26.27,trna9_HisGTG_15_-_45490804_45490875@-1T.26.27|TGCCGTGATCGTATAGTGGTTAGTACT</td><td>>trna111_HisGTG_1_-_147774845_147774916</td><td>-1T</td><td>26</td><td>27</td><td>TGCCGTGATCGTATAGTGGTTAGTACT</td><td>18758.3</td><td>8.8</td><td>0.9</td><td>10.0</td><td>2.3E-23</td><td>1.5E-22</td><td>MB231/MB468</td></tr>
<tr><td>>hg38_MT_+_598_1651_RNR1_12S_50ntFlanks@790.809.20|ACGATAGCCCTTATGAAACT</td><td>>hg38_MT_+_598_1651_RNR1_12S_50ntFlanks</td><td>790</td><td>809</td><td>20</td><td>ACGATAGCCCTTATGAAACT</td><td>636.2</td><td>11.0</td><td>1.1</td><td>10.0</td><td>2.3E-23</td><td>1.5E-22</td><td>MB231/MB468</td></tr>
<tr><td>>MI0000108|hsa-mir-103a-2&WithFlank&20|+|3917488|3917577@54.75.22(+1G);MI0000109|hsa-mir-103a-1&WithFlank&5|-|168560890|168560979@54.75.22(+1G)&[MIMAT0000101&hsa-miR-103a-3p&offsets|0|-1(+1G);m-12&20|+|3917541|3917563&offsets|0|-1(+1G)];[MIMAT0000101_1&hsa-miR-103a-3p&offsets|0|-1(+1G);m-13&5|-|168560904|168560926&offsets|0|-1(+1G);m-44220&5|-|168560908|168560926&offsets|0|+3(+1G)]|AGCAGCATTGTACAGGGCTATGG</td><td>>MI0000108|hsa-mir-103a-2&WithFlank&20|+|3917488|3917577</td><td>54</td><td>75</td><td>22(+1G)</td><td>AGCAGCATTGTACAGGGCTATGG</td><td>3527.0</td><td>7.6</td><td>0.8</td><td>10.0</td><td>2.3E-23</td><td>1.5E-22</td><td>MB231/MB468</td></tr>
<tr><td>>trna119_LysCTT_1_-_145395522_145395594@2.33.32,trna11_LysCTT_5_-_180648979_180649051@2.33.32,trna13_LysCTT_6_+_26556774_26556846@2.33.32,trna7_LysCTT_16_+_3225692_3225764@2.33.32,trna9_LysCTT_5_+_180634755_180634827@2.33.32|CCCGGCTAGCTCAGTCGGTAGAGCATGAGACT</td><td>>trna119_LysCTT_1_-_145395522_145395594</td><td>2</td><td>33</td><td>32</td><td>CCCGGCTAGCTCAGTCGGTAGAGCATGAGACT</td><td>1498.0</td><td>15.5</td><td>1.6</td><td>10.0</td><td>2.3E-23</td><td>1.5E-22</td><td>MB231/MB468</td></tr>
<tr><td>>MI0000446|hsa-mir-125b-1&WithFlank&11|-|122099751|122099850@21.39.19(+1C);MI0000470|hsa-mir-125b-2&WithFlank&21|+|16590231|16590331@23.41.19(+1C)&[MIMAT0000423&hsa-miR-125b-5p&offsets|0|-3(+1C);m-67&11|-|122099809|122099830&offsets|0|-3(+1C)];[MIMAT0000423_1&hsa-miR-125b-5p&offsets|0|-3(+1C);m-68&21|+|16590253|16590274&offsets|0|-3(+1C)]|TCCCTGAGACCCTAACTTGC</td><td>>MI0000446|hsa-mir-125b-1&WithFlank&11|-|122099751|122099850</td><td>21</td><td>39</td><td>19(+1C)</td><td>TCCCTGAGACCCTAACTTGC</td><td>557.4</td><td>11.2</td><td>1.1</td><td>10.0</td><td>2.4E-23</td><td>1.5E-22</td><td>MB231/MB468</td></tr>
<tr><td>>trnaMT_MetCAT_MT_+_4402_4469@36.71.36|CCCGAAAATGTTGGTTATACCCTTCCCGTACTACCA</td><td>>trnaMT_MetCAT_MT_+_4402_4469</td><td>36</td><td>71</td><td>36</td><td>CCCGAAAATGTTGGTTATACCCTTCCCGTACTACCA</td><td>906.6</td><td>12.7</td><td>1.3</td><td>10.0</td><td>2.4E-23</td><td>1.5E-22</td><td>MB231/MB468</td></tr>
<tr><td>>trna20_GluTTC_2_-_131094701_131094772@1.36.36,trna5_GluTTC_13_-_41634874_41634945@1.36.36|TCCCATATGGTCTAGCGGTTAGGATTCCTGGTTTTC</td><td>>trna20_GluTTC_2_-_131094701_131094772</td><td>1</td><td>36</td><td>36</td><td>TCCCATATGGTCTAGCGGTTAGGATTCCTGGTTTTC</td><td>1235.1</td><td>15.2</td><td>1.5</td><td>10.0</td><td>2.4E-23</td><td>1.5E-22</td><td>MB231/MB468</td></tr>
<tr><td>>hg38_21_+_8433172_8446622_RNA45SN1_45S_50ntFlanks@8356.8379.24|GAAAAGAACTTTGAAGAGAGAGTT</td><td>>hg38_21_+_8433172_8446622_RNA45SN1_45S_50ntFlanks</td><td>8356</td><td>8379</td><td>24</td><td>GAAAAGAACTTTGAAGAGAGAGTT</td><td>21418.3</td><td>-7.2</td><td>0.7</td><td>-10.0</td><td>1.9E-23</td><td>1.6E-22</td><td>BT20/MB468</td></tr>
<tr><td>>hg38_21_+_8433172_8446622_RNA45SN1_45S_50ntFlanks@11576.11593.18|CTTAGAACTGGTGCGGAC</td><td>>hg38_21_+_8433172_8446622_RNA45SN1_45S_50ntFlanks</td><td>11576</td><td>11593</td><td>18</td><td>CTTAGAACTGGTGCGGAC</td><td>1021.0</td><td>-6.1</td><td>0.6</td><td>-10.0</td><td>2.0E-23</td><td>1.6E-22</td><td>BT20/MB468</td></tr>
<tr><td>>hg38_21_+_8433172_8446622_RNA45SN1_45S_50ntFlanks@9570.9597.28|TGCAAATCGGTCGTCCGACCTGGGTATA</td><td>>hg38_21_+_8433172_8446622_RNA45SN1_45S_50ntFlanks</td><td>9570</td><td>9597</td><td>28</td><td>TGCAAATCGGTCGTCCGACCTGGGTATA</td><td>2747.5</td><td>-6.9</td><td>0.7</td><td>-10.0</td><td>2.0E-23</td><td>1.6E-22</td><td>BT20/MB468</td></tr>
<tr><td>>hg38_21_+_8433172_8446622_RNA45SN1_45S_50ntFlanks@9568.9597.30|CGTGCAAATCGGTCGTCCGACCTGGGTATA</td><td>>hg38_21_+_8433172_8446622_RNA45SN1_45S_50ntFlanks</td><td>9568</td><td>9597</td><td>30</td><td>CGTGCAAATCGGTCGTCCGACCTGGGTATA</td><td>3570.8</td><td>-8.9</td><td>0.9</td><td>-10.0</td><td>2.0E-23</td><td>1.6E-22</td><td>BT20/MB468</td></tr>
<tr><td>>am_MI0000809|hsa-mir-151a&WithFlank&8|-|140732558|140732659@53.73.21(+2U)&[MIMAT0000757&hsa-miR-151a-3p&offsets|0|0(+2U);m-30&8|-|140732587|140732607&offsets|0|0(+2U)]|CTAGACTGAAGCTCCTTGAGGTT</td><td>>am_MI0000809|hsa-mir-151a&WithFlank&8|-|140732558|140732659</td><td>53</td><td>73</td><td>21(+2U)</td><td>CTAGACTGAAGCTCCTTGAGGTT</td><td>3401.2</td><td>10.4</td><td>1.0</td><td>9.9</td><td>2.6E-23</td><td>1.6E-22</td><td>MB231/MB468</td></tr>
<tr><td>>trna2_LeuTAA_4_-_156384978_156385052@1.18.18|GTTAAGATGGCAGAGCCT</td><td>>trna2_LeuTAA_4_-_156384978_156385052</td><td>1</td><td>18</td><td>18</td><td>GTTAAGATGGCAGAGCCT</td><td>335.9</td><td>10.3</td><td>1.0</td><td>10.0</td><td>2.1E-23</td><td>1.7E-22</td><td>BT20/MB468</td></tr>
<tr><td>>hg38_21_+_8433172_8446622_RNA45SN1_45S_50ntFlanks@4759.4782.24|ATCAGATACCGTCGTAGTTCCGAC</td><td>>hg38_21_+_8433172_8446622_RNA45SN1_45S_50ntFlanks</td><td>4759</td><td>4782</td><td>24</td><td>ATCAGATACCGTCGTAGTTCCGAC</td><td>1731.7</td><td>-12.8</td><td>1.3</td><td>-10.0</td><td>2.1E-23</td><td>1.7E-22</td><td>BT20/MB468</td></tr>
<tr><td>>hg38_MT_+_1621_3279_RNR2_16S_50ntFlanks@644.676.33|AATTGGACCAATCTATCACCCTATAGAAGAACT</td><td>>hg38_MT_+_1621_3279_RNR2_16S_50ntFlanks</td><td>644</td><td>676</td><td>33</td><td>AATTGGACCAATCTATCACCCTATAGAAGAACT</td><td>473.6</td><td>13.5</td><td>1.4</td><td>10.0</td><td>2.1E-23</td><td>1.7E-22</td><td>BT20/MB468</td></tr>
<tr><td>>MI0000093|hsa-mir-92a-1&WithFlank&13|+|91351308|91351397@54.74.21(+1C);MI0000094|hsa-mir-92a-2&WithFlank&X|-|134169532|134169618@54.74.21(+1C)&[MIMAT0000092&hsa-miR-92a-3p&offsets|0|-1(+1C);m-9&13|+|91351361|91351382&offsets|0|-1(+1C)];[MIMAT0000092_1&hsa-miR-92a-3p&offsets|0|-1(+1C);m-11&X|-|134169544|134169565&offsets|0|-1(+1C)]|TATTGCACTTGTCCCGGCCTGC</td><td>>MI0000093|hsa-mir-92a-1&WithFlank&13|+|91351308|91351397</td><td>54</td><td>74</td><td>21(+1C)</td><td>TATTGCACTTGTCCCGGCCTGC</td><td>10688.7</td><td>6.4</td><td>0.6</td><td>10.0</td><td>2.1E-23</td><td>1.7E-22</td><td>BT20/MB468</td></tr>
<tr><td>>MI0000078|hsa-mir-22&WithFlank&17|-|1713897|1713993@59.79.21(+1G)&[MIMAT0000077&hsa-miR-22-3p&offsets|0|-1(+1G);m-2&17|-|1713914|1713935&offsets|0|-1(+1G)]|AAGCTGCCAGTTGAAGAACTGG</td><td>>MI0000078|hsa-mir-22&WithFlank&17|-|1713897|1713993</td><td>59</td><td>79</td><td>21(+1G)</td><td>AAGCTGCCAGTTGAAGAACTGG</td><td>868.0</td><td>8.0</td><td>0.8</td><td>9.9</td><td>2.7E-23</td><td>1.7E-22</td><td>MB231/MB468</td></tr>
<tr><td>>trnaMT_PheGAA_MT_+_577_647@13.34.22|TACCTCCTCAAAGCAATACACT</td><td>>trnaMT_PheGAA_MT_+_577_647</td><td>13</td><td>34</td><td>22</td><td>TACCTCCTCAAAGCAATACACT</td><td>339.3</td><td>9.1</td><td>0.9</td><td>10.0</td><td>2.2E-23</td><td>1.8E-22</td><td>BT20/MB468</td></tr>
<tr><td>>MI0000067|hsa-let-7f-1&WithFlank&9|+|94176341|94176439@13.33.21(+1A);MI0000068|hsa-let-7f-2&WithFlank&X|-|53557186|53557280@14.34.21(+1A)&[MIMAT0000067&hsa-let-7f-5p&offsets|0|-1(+1A);m-20&9|+|94176353|94176374&offsets|0|-1(+1A)];[MIMAT0000067_1&hsa-let-7f-5p&offsets|0|-1(+1A);m-19&X|-|53557246|53557267&offsets|0|-1(+1A)]|TGAGGTAGTAGATTGTATAGTA</td><td>>MI0000067|hsa-let-7f-1&WithFlank&9|+|94176341|94176439</td><td>13</td><td>33</td><td>21(+1A)</td><td>TGAGGTAGTAGATTGTATAGTA</td><td>17955.0</td><td>8.3</td><td>0.8</td><td>10.0</td><td>2.2E-23</td><td>1.8E-22</td><td>BT20/MB468</td></tr>
<tr><td>>MI0000063|hsa-let-7b&WithFlank&22|+|46113680|46113774@12.34.23(+1U)&[MIMAT0000063&hsa-let-7b-5p&offsets|0|+1(+1U);m-14&22|+|46113691|46113712&offsets|0|+1(+1U)]|TGAGGTAGTAGGTTGTGTGGTTTT</td><td>>MI0000063|hsa-let-7b&WithFlank&22|+|46113680|46113774</td><td>12</td><td>34</td><td>23(+1U)</td><td>TGAGGTAGTAGGTTGTGTGGTTTT</td><td>760.4</td><td>7.2</td><td>0.7</td><td>9.9</td><td>2.8E-23</td><td>1.8E-22</td><td>MB231/MB468</td></tr>
</table>

| | | | | | | | | | | | | |
|---|---|---|---|---|---|---|---|---|---|---|---|---|
| >trna14_LysTTT_11_-_59327808_59327880@1.33.33,trna2_LysTTT_17_+_8022473_8022545@1.33.33,trna54_LysTTT_1_+_204475655_204475727@1.33.33,trna5_LysTTT_11_+_59323902_59323974@1.33.33,trna62_LysTTT_1_-_204476158_204476230@1.33.33,trna76_LysTTT_6_+_28918806_28918878@1.33.33\|GCCCGGATAGCTCAGTCGGTAGAGCATCAGACT | >trna14_LysTTT_11_-_59327808_59327880 | 1 | 33 | 33 | GCCCGGATAGCTCAGTCGGTAGAGCATCAGACT | 39262.1 | 10.1 | 1.0 | 10.0 | 2.4E-23 | 1.9E-22 | BT20/MB468 |
| >MI0001445\|hsa-mir-423&WithFlank&17\|+\|30117073\|30117178@59.81.23(+1A)&[MIMAT0001340&hsa-miR-423-3p&offsets\|0\|0(+1A);m-100&17\|+\|30117131\|30117153&offsets\|0\|0(+1A)]\|AGCTCGGTCTGAGGCCCCTCAGTA | >MI0001445\|hsa-mir-423&WithFlank&17\|+\|30117073\|30117178 | 59 | 81 | 23(+1A) | AGCTCGGTCTGAGGCCCCTCAGTA | 2667.0 | 7.5 | 0.8 | 9.9 | 3.0E-23 | 1.9E-22 | MB231/MB468 |
| >hg38_21_+_8433172_8446622_RNA45SN1_45S_50ntFlanks@8357.8377.21\|AAAAGAACTTTGAAGAGAGAG | >hg38_21_+_8433172_8446622_RNA45SN1_45S_50ntFlanks | 8357 | 8377 | 21 | AAAAGAACTTTGAAGAGAGAG | 4713.4 | -5.6 | 0.6 | -10.0 | 2.4E-23 | 1.9E-22 | BT20/MB468 |
| >hg38_21_+_8433172_8446622_RNA45SN1_45S_50ntFlanks@12939.12956.18\|CCATTCGTAGACGACCTG | >hg38_21_+_8433172_8446622_RNA45SN1_45S_50ntFlanks | 12939 | 12956 | 18 | CCATTCGTAGACGACCTG | 978.0 | -6.3 | 0.6 | -10.0 | 2.4E-23 | 1.9E-22 | BT20/MB468 |
| >MI0000086\|hsa-mir-28&WithFlank&3\|+\|188688775\|188688872@20.41.22&[MIMAT0000085&hsa-miR-28-5p&offsets\|0\|0;m-133&3\|+\|188688794\|188688815&offsets\|0\|0]\|AAGGAGCTCACAGTCTATTGAG | >MI0000086\|hsa-mir-28&WithFlank&3\|+\|188688775\|188688872 | 20 | 41 | 22 | AAGGAGCTCACAGTCTATTGAG | 1677.0 | 8.6 | 0.9 | 10.0 | 2.5E-23 | 2.0E-22 | BT20/MB468 |
| >hg38_21_+_8433172_8446622_RNA45SN1_45S_50ntFlanks@11668.11693.26\|TGCCCAGTGCTCTGAATGTCAAAGTG | >hg38_21_+_8433172_8446622_RNA45SN1_45S_50ntFlanks | 11668 | 11693 | 26 | TGCCCAGTGCTCTGAATGTCAAAGTG | 271.1 | 5.1 | 0.5 | 9.9 | 2.5E-23 | 2.0E-22 | BT20/MB468 |
| >hg38_21_+_8433172_8446622_RNA45SN1_45S_50ntFlanks@10681.10710.30\|TTCCGGCGGCGTCCGGTGAGCTCTCGCTGG | >hg38_21_+_8433172_8446622_RNA45SN1_45S_50ntFlanks | 10681 | 10710 | 30 | TTCCGGCGGCGTCCGGTGAGCTCTCGCTGG | 1861.5 | -6.9 | 0.7 | -9.9 | 3.3E-23 | 2.1E-22 | MB231/MB468 |
| >MI0001448\|hsa-mir-425&WithFlank&3\|-\|49020142\|49020240@21.44.24&[MIMAT0003393&hsa-miR-425-5p&offsets\|+1\|+2;m-95&3\|-\|49020200\|49020221&offsets\|+1\|+3]\|ATGACACGATCACTCCCGTTGAGT | >MI0001448\|hsa-mir-425&WithFlank&3\|-\|49020142\|49020240 | 21 | 44 | 24 | ATGACACGATCACTCCCGTTGAGT | 971.9 | 7.6 | 0.8 | 9.9 | 3.4E-23 | 2.2E-22 | MB231/MB468 |
| >hg38_21_+_8433172_8446622_RNA45SN1_45S_50ntFlanks@12416.12449.34\|TTCGATGTCGGCTCTTCCTATCATTGTGAAGCAG | >hg38_21_+_8433172_8446622_RNA45SN1_45S_50ntFlanks | 12416 | 12449 | 34 | TTCGATGTCGGCTCTTCCTATCATTGTGAAGCAG | 380.4 | 4.6 | 0.5 | 9.9 | 2.8E-23 | 2.2E-22 | BT20/MB468 |
| >trnaMT_PheGAA_MT_+_577_647@37.56.20\|AAATGTTTAGACGGGCTCAC | >trnaMT_PheGAA_MT_+_577_647 | 37 | 56 | 20 | AAATGTTTAGACGGGCTCAC | 512.0 | 10.9 | 1.1 | 9.9 | 2.8E-23 | 2.2E-22 | BT20/MB468 |
| >MI0000811\|hsa-mir-148b&WithFlank&12\|+\|54337210\|54337320@69.90.22(+1U)&[MIMAT0000759&hsa-miR-148b-3p&offsets\|0\|0(+1U);m-77&12\|+\|54337278\|54337299&offsets\|0\|0(+1U)]\|TCAGTGCATCACAGAACTTTGTT | >MI0000811\|hsa-mir-148b&WithFlank&12\|+\|54337210\|54337320 | 69 | 90 | 22(+1U) | TCAGTGCATCACAGAACTTTGTT | 31276.0 | 8.5 | 0.9 | 9.9 | 3.6E-23 | 2.3E-22 | MB231/MB468 |
| >hg38_21_+_8433172_8446622_RNA45SN1_45S_50ntFlanks@11669.11688.20\|GCCCAGTGCTCTGAATGTCA | >hg38_21_+_8433172_8446622_RNA45SN1_45S_50ntFlanks | 11669 | 11688 | 20 | GCCCAGTGCTCTGAATGTCA | 797.8 | 6.9 | 0.7 | 9.9 | 3.7E-23 | 2.3E-22 | MB231/MB468 |
| >hg38_21_+_8433172_8446622_RNA45SN1_45S_50ntFlanks@10351.10383.33\|CGAGAACTTTGAAGGCCGAAGTGGAGAAGGGTT | >hg38_21_+_8433172_8446622_RNA45SN1_45S_50ntFlanks | 10351 | 10383 | 33 | CGAGAACTTTGAAGGCCGAAGTGGAGAAGGGTT | 10964.4 | -5.3 | 0.5 | -9.9 | 3.7E-23 | 2.4E-22 | MB231/MB468 |
| >MI0000102\|hsa-mir-100&WithFlank&11\|-\|122152223\|122152314@19.40.22(+1A)&[MIMAT0000098&hsa-miR-100-5p&offsets\|0\|0(+1A);m-27&11\|-\|122152276\|122152296&offsets\|0\|+1(+1A)]\|AACCCGTAGATCCGAACTTGTGA | >MI0000102\|hsa-mir-100&WithFlank&11\|-\|122152223\|122152314 | 19 | 40 | 22(+1A) | AACCCGTAGATCCGAACTTGTGA | 116907.6 | 14.4 | 1.5 | 9.9 | 3.8E-23 | 2.4E-22 | MB231/MB468 |
| >hg38_21_+_8433172_8446622_RNA45SN1_45S_50ntFlanks@12420.12459.40\|ATGTCGGCTCTTCCTATCATTGTGAAGCAGAATTCACCAA | >hg38_21_+_8433172_8446622_RNA45SN1_45S_50ntFlanks | 12420 | 12459 | 40 | ATGTCGGCTCTTCCTATCATTGTGAAGCAGAATTCACCAA | 588.2 | 8.3 | 0.8 | 9.9 | 3.8E-23 | 2.4E-22 | MB231/MB468 |
| >MI0003137\|hsa-mir-193b&WithFlank&16\|+\|14303961\|14304055@57.77.21(+1C)&[MIMAT0002819&hsa-miR-193b-3p&offsets\|0\|-1(+1C);m-166&16\|+\|14304017\|14304038&offsets\|0\|-1(+1C)]\|AACTGGCCCTCAAAGTCCCGCC | >MI0003137\|hsa-mir-193b&WithFlank&16\|+\|14303961\|14304055 | 57 | 77 | 21(+1C) | AACTGGCCCTCAAAGTCCCGCC | 2281.5 | 15.8 | 1.6 | 9.9 | 3.0E-23 | 2.4E-22 | BT20/MB468 |
| >MI0000086\|hsa-mir-28&WithFlank&3\|+\|188688775\|188688872@61.82.22(+1U)&[MIMAT0004502&hsa-miR-28-3p&offsets\|+1\|+1(+1U);m-28&3\|+\|188688834\|188688855&offsets\|+1\|+1(+1U)]\|ACTAGATTGTGAGCTCCTGGAGT | >MI0000086\|hsa-mir-28&WithFlank&3\|+\|188688775\|188688872 | 61 | 82 | 22(+1U) | ACTAGATTGTGAGCTCCTGGAGT | 811.0 | 12.5 | 1.3 | 9.9 | 3.9E-23 | 2.4E-22 | MB231/MB468 |
| >MI0000746\|hsa-mir-99b&WithFlank&19\|+\|51692606\|51692687@13.32.20&[MIMAT0000689&hsa-miR-99b-5p&offsets\|0\|-2;m-10&19\|+\|51692618\|51692639&offsets\|0\|-2]\|CACCCGTAGAACCGACCTTG | >MI0000746\|hsa-mir-99b&WithFlank&19\|+\|51692606\|51692687 | 13 | 32 | 20 | CACCCGTAGAACCGACCTTG | 1333.9 | 8.0 | 0.8 | 9.9 | 3.9E-23 | 2.5E-22 | MB231/MB468 |
| >MI0000816\|hsa-mir-335&WithFlank&7\|+\|130496105\|130496210@57.79.23&[MIMAT0004703&hsa-miR-335-3p&offsets\|-1\|0;m-159&7\|+\|130496162\|130496183&offsets\|-1\|0]\|GTTTTTCATTATTGCTCCTGACC | >MI0000816\|hsa-mir-335&WithFlank&7\|+\|130496105\|130496210 | 57 | 79 | 23 | GTTTTTCATTATTGCTCCTGACC | 778.6 | 7.0 | 0.7 | 9.9 | 3.9E-23 | 2.5E-22 | MB231/MB468 |
| >hg38_21_+_8433172_8446622_RNA45SN1_45S_50ntFlanks@10819.10840.22\|AGGGAAGTCGGCAAGCCGGATC | >hg38_21_+_8433172_8446622_RNA45SN1_45S_50ntFlanks | 10819 | 10840 | 22 | AGGGAAGTCGGCAAGCCGGATC | 3312.5 | -7.7 | 0.8 | -9.9 | 3.2E-23 | 2.6E-22 | BT20/MB468 |
| >MI0000433\|hsa-let-7g&WithFlank&3\|-\|52268272\|52268367@11.31.21(+1G)&[MIMAT0000414&hsa-let-7g-5p&offsets\|0\|-1(+1G);m-58&3\|-\|52268336\|52268357&offsets\|0\|-1(+1G)]\|TGAGGTAGTAGTTTGTACAGTG | >MI0000433\|hsa-let-7g&WithFlank&3\|-\|52268272\|52268367 | 11 | 31 | 21(+1G) | TGAGGTAGTAGTTTGTACAGTG | 8028.5 | 7.1 | 0.7 | 9.9 | 3.3E-23 | 2.6E-22 | BT20/MB468 |
| >hg38_21_+_8433172_8446622_RNA45SN1_45S_50ntFlanks@10411.10431.21\|CAGTCGGTCCTGAGAGATGGG | >hg38_21_+_8433172_8446622_RNA45SN1_45S_50ntFlanks | 10411 | 10431 | 21 | CAGTCGGTCCTGAGAGATGGG | 2595.2 | -7.0 | 0.7 | -9.9 | 4.1E-23 | 2.6E-22 | MB231/MB468 |
| >MI0000269\|hsa-mir-181a-2&WithFlank&9\|+\|124692436\|124692557@83.103.21(+1U)&[MIMAT0004558&hsa-miR-181a-2-3p&offsets\|0\|-1(+1U);m-71&9\|+\|124692518\|124692539&offsets\|0\|-1(+1U)]\|ACCACTGACCGTTGACTGTACT | >MI0000269\|hsa-mir-181a-2&WithFlank&9\|+\|124692436\|124692557 | 83 | 103 | 21(+1U) | ACCACTGACCGTTGACTGTACT | 1393.4 | 10.1 | 1.0 | 9.9 | 4.1E-23 | 2.6E-22 | MB231/MB468 |
| >MI0000264\|hsa-mir-7-2&WithFlank&15\|+\|88611819\|88611940@38.60.23;MI0000265\|hsa-mir-7-3&WithFlank&19\|+\|4770664\|4770785@37.59.23;MI0000263\|hsa-mir-7-1&WithFlank&9\|-\|83969742\|83969863@30.52.23&[MIMAT0000252&hsa-miR-7-5p&offsets\|0\|-1;m-400&15\|+\|88611856\|88611879&offsets\|0\|-1];[MIMAT0000252_1&hsa-miR-7-5p&offsets\|0\|-1;m-402&19\|+\|4770700\|4770723&offsets\|0\|-1];[MIMAT0000252_2&hsa-miR-7-5p&offsets\|0\|-1;m-401&9\|-\|83969811\|83969834&offsets\|0\|-1]\|TGGAAGACTAGTGATTTTGTTGT | >MI0000264\|hsa-mir-7-2&WithFlank&15\|+\|88611819\|88611940 | 38 | 60 | 23 | TGGAAGACTAGTGATTTTGTTGT | 33487.7 | 8.0 | 0.8 | 9.9 | 4.2E-23 | 2.7E-22 | MB231/MB468 |
| >hg38_MT_+_1621_3279_RNR2_16S_50ntFlanks@930.958.29\|AGTGACACATGTTTAACGGCCGCGGTACC | >hg38_MT_+_1621_3279_RNR2_16S_50ntFlanks | 930 | 958 | 29 | AGTGACACATGTTTAACGGCCGCGGTACC | 848.2 | 12.6 | 1.3 | 9.9 | 4.3E-23 | 2.7E-22 | MB231/MB468 |
| >hg38_21_+_8433172_8446622_RNA45SN1_45S_50ntFlanks@6721.6739.19\|AATTGCAGGACACATTGAT | >hg38_21_+_8433172_8446622_RNA45SN1_45S_50ntFlanks | 6721 | 6739 | 19 | AATTGCAGGACACATTGAT | 2268.5 | -5.9 | 0.6 | -9.9 | 4.4E-23 | 2.7E-22 | MB231/MB468 |
| >MI0000542\|hsa-mir-320a&WithFlank&8\|-\|22244960\|22245043@50.70.21(+1U)&[MIMAT0000510&hsa-miR-320a-3p&offsets\|+2\|+1(+1U);m-60&8\|-\|22244975\|22244996&offsets\|+2\|+1(+1U)]\|AAGCTGGGTTGAGAGGGCGAAT | >MI0000542\|hsa-mir-320a&WithFlank&8\|-\|22244960\|22245043 | 50 | 70 | 21(+1U) | AAGCTGGGTTGAGAGGGCGAAT | 418.9 | 13.4 | 1.3 | 9.9 | 3.5E-23 | 2.8E-22 | BT20/MB468 |
| >MI0000089\|hsa-mir-31&WithFlank&9\|-\|21512109\|21512191@14.35.22&[MIMAT0000089&hsa-miR-31-5p&offsets\|0\|+1;m-360&9\|-\|21512157\|21512178&offsets\|0\|0]\|AGGCAAGATGCTGGCATAGCTG | >MI0000089\|hsa-mir-31&WithFlank&9\|-\|21512109\|21512191 | 14 | 35 | 22 | AGGCAAGATGCTGGCATAGCTG | 2788.2 | 8.2 | 0.8 | 9.9 | 3.5E-23 | 2.8E-22 | BT20/MB468 |
| >hg38_21_+_8433172_8446622_RNA45SN1_45S_50ntFlanks@6652.6684.33\|GACTCTTAGCGGTGGATCACTCGGCTCGTGCGT | >hg38_21_+_8433172_8446622_RNA45SN1_45S_50ntFlanks | 6652 | 6684 | 33 | GACTCTTAGCGGTGGATCACTCGGCTCGTGCGT | 1741.5 | 6.5 | 0.7 | 9.9 | 4.4E-23 | 2.8E-22 | MB231/MB468 |
| >MI0000082\|hsa-mir-25&WithFlank&7\|-\|100093554\|100093649@58.79.22(+1A)&[MIMAT0000081&hsa-miR-25-3p&offsets\|0\|0(+1A);m-23&7\|-\|100093571\|100093592&offsets\|0\|0(+1A)]\|CATTGCACTTGTCTCGGTCTGAA | >MI0000082\|hsa-mir-25&WithFlank&7\|-\|100093554\|100093649 | 58 | 79 | 22(+1A) | CATTGCACTTGTCTCGGTCTGAA | 2251.0 | 7.1 | 0.7 | 9.9 | 4.4E-23 | 2.8E-22 | MB231/MB468 |
| >hg38_21_+_8433172_8446622_RNA45SN1_45S_50ntFlanks@9545.9566.22\|GTGGAGGTCCGTAGCGGTCCTG | >hg38_21_+_8433172_8446622_RNA45SN1_45S_50ntFlanks | 9545 | 9566 | 22 | GTGGAGGTCCGTAGCGGTCCTG | 1325.9 | -9.1 | 0.9 | -9.9 | 3.5E-23 | 2.8E-22 | BT20/MB468 |
| >MI0000068\|hsa-let-7f-2&WithFlank&X\|-\|53557186\|53557280@14.36.23&[MIMAT0000067_1&hsa-let-7f-5p&offsets\|0\|+1;m-19&X\|-\|53557246\|53557267&offsets\|0\|+1]\|TGAGGTAGTAGATTGTATAGTTT | >MI0000068\|hsa-let-7f-2&WithFlank&X\|-\|53557186\|53557280 | 14 | 36 | 23 | TGAGGTAGTAGATTGTATAGTTT | 10335.5 | 7.1 | 0.7 | 9.9 | 4.6E-23 | 2.9E-22 | MB231/MB468 |
| >hg38_21_+_8433172_8446622_RNA45SN1_45S_50ntFlanks@9565.9597.33\|TGACGTGCAAATCGGTCGTCCGACCTGGGTATA | >hg38_21_+_8433172_8446622_RNA45SN1_45S_50ntFlanks | 9565 | 9597 | 33 | TGACGTGCAAATCGGTCGTCCGACCTGGGTATA | 1771.8 | -5.4 | 0.5 | -9.9 | 3.7E-23 | 2.9E-22 | BT20/MB468 |
| >hg38_21_+_8433172_8446622_RNA45SN1_45S_50ntFlanks@475.501.27\|CTTCGTGATCGATGTGGTGACGTCGTG | >hg38_21_+_8433172_8446622_RNA45SN1_45S_50ntFlanks | 475 | 501 | 27 | CTTCGTGATCGATGTGGTGACGTCGTG | 1414.4 | 6.5 | 0.7 | 9.9 | 4.7E-23 | 2.9E-22 | MB231/MB468 |
| >MI0000265\|hsa-mir-7-3&WithFlank&19\|+\|4770664\|4770785@37.61.25&[MIMAT0000252_1&hsa-miR-7-5p&offsets\|0\|+1;m-402&19\|+\|4770700\|4770723&offsets\|0\|+1]\|TGGAAGACTAGTGATTTTGTTGTTC | >MI0000265\|hsa-mir-7-3&WithFlank&19\|+\|4770664\|4770785 | 37 | 61 | 25 | TGGAAGACTAGTGATTTTGTTGTTC | 743.0 | 7.7 | 0.8 | 9.9 | 4.8E-23 | 3.0E-22 | MB231/MB468 |
| >MI0000746\|hsa-mir-99b&WithFlank&19\|+\|51692606\|51692687@13.34.22(+1A)&[MIMAT0000689&hsa-miR-99b-5p&offsets\|0\|0(+1A);m-10&19\|+\|51692618\|51692639&offsets\|0\|0(+1A)]\|CACCCGTAGAACCGACCTTGCGA | >MI0000746\|hsa-mir-99b&WithFlank&19\|+\|51692606\|51692687 | 13 | 34 | 22(+1A) | CACCCGTAGAACCGACCTTGCGA | 2062.9 | 7.2 | 0.7 | 9.9 | 4.8E-23 | 3.0E-22 | MB231/MB468 |
| >hg38_21_+_8433172_8446622_RNA45SN1_45S_50ntFlanks@4002.4024.23\|GATAACCTCGGGCCGATCGCACG | >hg38_21_+_8433172_8446622_RNA45SN1_45S_50ntFlanks | 4002 | 4024 | 23 | GATAACCTCGGGCCGATCGCACG | 910.6 | -6.3 | 0.6 | -9.9 | 3.9E-23 | 3.1E-22 | BT20/MB468 |
| >hg38_21_+_8433172_8446622_RNA45SN1_45S_50ntFlanks@10771.10805.35\|CAGGTCTCCAAGGTGAACAGCCTCTGGCATGTTGG | >hg38_21_+_8433172_8446622_RNA45SN1_45S_50ntFlanks | 10771 | 10805 | 35 | CAGGTCTCCAAGGTGAACAGCCTCTGGCATGTTGG | 684.3 | -5.3 | 0.5 | -9.9 | 5.0E-23 | 3.1E-22 | MB231/MB468 |

<table>
<tr><td>>hg38_MT_+_1621_3279_RNR2_16S_50ntFlanks@1349.1388.40|ACAATAGGGTTTACGACCTCGATGTTGGATCAGGACATCC</td><td>>hg38_MT_+_1621_3279_RNR2_16S_50ntFlanks</td><td>1349</td><td>1388</td><td>40</td><td>ACAATAGGGTTTACGACCTCGATGTTGGATCAGGACATCC</td><td>695.5</td><td>14.1</td><td>1.4</td><td>9.9</td><td>4.0E-23</td><td>3.1E-22</td><td>BT20/MB468</td></tr>
<tr><td>>hg38_21_+_8433172_8446622_RNA45SN1_45S_50ntFlanks@6648.6667.20|GTACGACTCTTAGCGGTGGA</td><td>>hg38_21_+_8433172_8446622_RNA45SN1_45S_50ntFlanks</td><td>6648</td><td>6667</td><td>20</td><td>GTACGACTCTTAGCGGTGGA</td><td>505.4</td><td>6.1</td><td>0.6</td><td>9.9</td><td>4.0E-23</td><td>3.1E-22</td><td>BT20/MB468</td></tr>
<tr><td>>hg38_MT_+_598_1651_RNR1_12S_50ntFlanks@52.83.32|ATAGGTTTGGTCCTAGCCTTTCTATTAGCTCT</td><td>>hg38_MT_+_598_1651_RNR1_12S_50ntFlanks</td><td>52</td><td>83</td><td>32</td><td>ATAGGTTTGGTCCTAGCCTTTCTATTAGCTCT</td><td>4063.6</td><td>7.4</td><td>0.7</td><td>9.9</td><td>5.1E-23</td><td>3.2E-22</td><td>MB231/MB468</td></tr>
<tr><td>>hg38_21_+_8433172_8446622_RNA45SN1_45S_50ntFlanks@5527.5550.24|AAAGTCGTAACAAGGTTTCCGTAG</td><td>>hg38_21_+_8433172_8446622_RNA45SN1_45S_50ntFlanks</td><td>5527</td><td>5550</td><td>24</td><td>AAAGTCGTAACAAGGTTTCCGTAG</td><td>694.1</td><td>-6.7</td><td>0.7</td><td>-9.9</td><td>5.1E-23</td><td>3.2E-22</td><td>MB231/MB468</td></tr>
<tr><td>>hg38_21_+_8433172_8446622_RNA45SN1_45S_50ntFlanks@4732.4767.36|AGAACGAAAGTCGGAGGTTCGAAGACGATCAGATAC</td><td>>hg38_21_+_8433172_8446622_RNA45SN1_45S_50ntFlanks</td><td>4732</td><td>4767</td><td>36</td><td>AGAACGAAAGTCGGAGGTTCGAAGACGATCAGATAC</td><td>409.0</td><td>9.2</td><td>0.9</td><td>9.9</td><td>4.1E-23</td><td>3.2E-22</td><td>BT20/MB468</td></tr>
<tr><td>>MI0000734|hsa-mir-106b&WithFlank&7|-|100093987|100094080@58.79.22&[MIMAT0004672&hsa-miR-106b-3p&offsets|0|0;m-87&7|-|100094003|100094023&offsets|0|+1]|CCGCACTGTGGGTACTTGCTGC</td><td>>MI0000734|hsa-mir-106b&WithFlank&7|-|100093987|100094080</td><td>58</td><td>79</td><td>22</td><td>CCGCACTGTGGGTACTTGCTGC</td><td>15562.7</td><td>7.7</td><td>0.8</td><td>9.9</td><td>5.2E-23</td><td>3.2E-22</td><td>MB231/MB468</td></tr>
<tr><td>>MI0000077|hsa-mir-21&WithFlank&17|+|59841260|59841343@14.33.20(+1A)&[MIMAT0000076&hsa-miR-21-5p&offsets|0|-2(+1A);m-1&17|+|59841273|59841294&offsets|0|-2(+1A)]|TAGCTTATCAGACTGATGTTA</td><td>>MI0000077|hsa-mir-21&WithFlank&17|+|59841260|59841343</td><td>14</td><td>33</td><td>20(+1A)</td><td>TAGCTTATCAGACTGATGTTA</td><td>10307.7</td><td>8.8</td><td>0.9</td><td>9.9</td><td>4.3E-23</td><td>3.4E-22</td><td>BT20/MB468</td></tr>
<tr><td>>MI0000760|hsa-mir-361&WithFlank&X|-|85903630|85903713@51.74.24(+1U)&[MIMAT0004682&hsa-miR-361-3p&offsets|0|+1(+1U);m-124&X|-|85903642|85903663&offsets|0|+2(+1U)]|TCCCCCAGGTGTGATTCTGATTTGT</td><td>>MI0000760|hsa-mir-361&WithFlank&X|-|85903630|85903713</td><td>51</td><td>74</td><td>24(+1U)</td><td>TCCCCCAGGTGTGATTCTGATTTGT</td><td>566.8</td><td>11.2</td><td>1.1</td><td>9.9</td><td>5.6E-23</td><td>3.5E-22</td><td>MB231/MB468</td></tr>
<tr><td>>hg38_21_+_8433172_8446622_RNA45SN1_45S_50ntFlanks@475.507.33|CTTCGTGATCGATGTGGTGACGTCGTGCTCTCC</td><td>>hg38_21_+_8433172_8446622_RNA45SN1_45S_50ntFlanks</td><td>475</td><td>507</td><td>33</td><td>CTTCGTGATCGATGTGGTGACGTCGTGCTCTCC</td><td>4081.0</td><td>6.5</td><td>0.7</td><td>9.9</td><td>4.5E-23</td><td>3.5E-22</td><td>BT20/MB468</td></tr>
<tr><td>>hg38_21_+_8433172_8446622_RNA45SN1_45S_50ntFlanks@5472.5490.19|ACGGCCCTGGCGGAGCGCT</td><td>>hg38_21_+_8433172_8446622_RNA45SN1_45S_50ntFlanks</td><td>5472</td><td>5490</td><td>19</td><td>ACGGCCCTGGCGGAGCGCT</td><td>503.7</td><td>6.6</td><td>0.7</td><td>9.9</td><td>4.5E-23</td><td>3.5E-22</td><td>BT20/MB468</td></tr>
<tr><td>>MI0000809|hsa-mir-151a&WithFlank&8|-|140732558|140732659@53.75.23(+1U)&[MIMAT0000757&hsa-miR-151a-3p&offsets|0|+2(+1U);m-30&8|-|140732587|140732607&offsets|0|+2(+1U)]|CTAGACTGAAGCTCCTTGAGGACT</td><td>>MI0000809|hsa-mir-151a&WithFlank&8|-|140732558|140732659</td><td>53</td><td>75</td><td>23(+1U)</td><td>CTAGACTGAAGCTCCTTGAGGACT</td><td>595.1</td><td>13.9</td><td>1.4</td><td>9.9</td><td>4.5E-23</td><td>3.5E-22</td><td>BT20/MB468</td></tr>
<tr><td>>hg38_21_+_8433172_8446622_RNA45SN1_45S_50ntFlanks@12405.12443.39|CTTTTTGATCCTTCGATGTCGGCTCTTCCTATCATTGTG</td><td>>hg38_21_+_8433172_8446622_RNA45SN1_45S_50ntFlanks</td><td>12405</td><td>12443</td><td>39</td><td>CTTTTTGATCCTTCGATGTCGGCTCTTCCTATCATTGTG</td><td>2063.1</td><td>-5.9</td><td>0.6</td><td>-9.9</td><td>4.6E-23</td><td>3.6E-22</td><td>BT20/MB468</td></tr>
<tr><td>>MI0000086|hsa-mir-28&WithFlank&3|+|188688775|188688872@60.80.21(+1U)&[MIMAT0004502&hsa-miR-28-3p&offsets|0|-1(+1U);m-28&3|+|188688834|188688855&offsets|0|-1(+1U)]|CACTAGATTGTGAGCTCCTGGT</td><td>>MI0000086|hsa-mir-28&WithFlank&3|+|188688775|188688872</td><td>60</td><td>80</td><td>21(+1U)</td><td>CACTAGATTGTGAGCTCCTGGT</td><td>1124.8</td><td>12.6</td><td>1.3</td><td>9.9</td><td>5.8E-23</td><td>3.6E-22</td><td>MB231/MB468</td></tr>
<tr><td>>MI0000087|hsa-mir-29a&WithFlank&7|-|130876741|130876816@47.67.21(+1A)&[MIMAT0000086&hsa-miR-29a-3p&offsets|-1|-2(+1A);m-26&7|-|130876749|130876769&offsets|-1|-1(+1A)]|CTAGCACCATCTGAAATCGGTA</td><td>>MI0000087|hsa-mir-29a&WithFlank&7|-|130876741|130876816</td><td>47</td><td>67</td><td>21(+1A)</td><td>CTAGCACCATCTGAAATCGGTA</td><td>918.7</td><td>10.6</td><td>1.1</td><td>9.9</td><td>5.8E-23</td><td>3.6E-22</td><td>MB231/MB468</td></tr>
<tr><td>>MI0003591|hsa-mir-584&WithFlank&5|-|149062307|149062415@22.43.22(+1U)&[MIMAT0003249&hsa-miR-584-5p&offsets|0|0(+1U);m-167&5|-|149062374|149062394&offsets|0|+1(+1U)]|TTATGGTTTGCCTGGGACTGAGT</td><td>>MI0003591|hsa-mir-584&WithFlank&5|-|149062307|149062415</td><td>22</td><td>43</td><td>22(+1U)</td><td>TTATGGTTTGCCTGGGACTGAGT</td><td>894.0</td><td>12.2</td><td>1.2</td><td>9.9</td><td>5.9E-23</td><td>3.6E-22</td><td>MB231/MB468</td></tr>
<tr><td>>hg38_21_+_8433172_8446622_RNA45SN1_45S_50ntFlanks@10391.10409.19|AACAGCAGTTGAACATGGG</td><td>>hg38_21_+_8433172_8446622_RNA45SN1_45S_50ntFlanks</td><td>10391</td><td>10409</td><td>19</td><td>AACAGCAGTTGAACATGGG</td><td>2349.3</td><td>-13.2</td><td>1.3</td><td>-9.9</td><td>4.6E-23</td><td>3.6E-22</td><td>BT20/MB468</td></tr>
<tr><td>>MI0000264|hsa-mir-7-2&WithFlank&15|+|88611819|88611940@38.59.22(+1C);MI0000265|hsa-mir-7-3&WithFlank&19|+|4770664|4770785@37.58.22(+1C);MI0000263|hsa-mir-7-1&WithFlank&9|-|83969742|83969863@30.51.22(+1C)&[MIMAT0000252&hsa-miR-7-5p&offsets|0|-2(+1C);m-400&15|+|88611856|88611879&offsets|0|-2(+1C)];[MIMAT0000252_1&hsa-miR-7-5p&offsets|0|-2(+1C);m-402&19|+|4770700|4770723&offsets|0|-2(+1C)];[MIMAT0000252_2&hsa-miR-7-5p&offsets|0|-2(+1C);m-401&9|-|83969811|83969834&offsets|0|-2(+1C)]|TGGAAGACTAGTGATTTTGTTGC</td><td>>MI0000264|hsa-mir-7-2&WithFlank&15|+|88611819|88611940</td><td>38</td><td>59</td><td>22(+1C)</td><td>TGGAAGACTAGTGATTTTGTTGC</td><td>3987.4</td><td>8.1</td><td>0.8</td><td>9.9</td><td>6.0E-23</td><td>3.7E-22</td><td>MB231/MB468</td></tr>
<tr><td>>hg38_21_+_8433172_8446622_RNA45SN1_45S_50ntFlanks@7974.8013.40|ACGCGACCTCAGATCAGACGTGGCGACCCGCTGAATTTAA</td><td>>hg38_21_+_8433172_8446622_RNA45SN1_45S_50ntFlanks</td><td>7974</td><td>8013</td><td>40</td><td>ACGCGACCTCAGATCAGACGTGGCGACCCGCTGAATTTAA</td><td>2182.3</td><td>7.9</td><td>0.8</td><td>9.9</td><td>6.1E-23</td><td>3.8E-22</td><td>MB231/MB468</td></tr>
<tr><td>>MI0000067|hsa-let-7f-1&WithFlank&9|+|94176341|94176439@13.32.20;MI0000068|hsa-let-7f-2&WithFlank&X|-|53557186|53557280@14.33.20&[MIMAT0000067&hsa-let-7f-5p&offsets|0|-2;m-20&9|+|94176353|94176374&offsets|0|-2];[MIMAT0000067_1&hsa-let-7f-5p&offsets|0|-2;m-19&X|-|53557246|53557267&offsets|0|-2]|TGAGGTAGTAGATTGTATAG</td><td>>MI0000067|hsa-let-7f-1&WithFlank&9|+|94176341|94176439</td><td>13</td><td>32</td><td>20</td><td>TGAGGTAGTAGATTGTATAG</td><td>502.0</td><td>6.9</td><td>0.7</td><td>9.9</td><td>4.9E-23</td><td>3.8E-22</td><td>BT20/MB468</td></tr>
<tr><td>>MI0000441|hsa-mir-30b&WithFlank&8|-|134800514|134800613@23.44.22&[MIMAT0000420&hsa-miR-30b-5p&offsets|0|0;m-76&8|-|134800570|134800591&offsets|0|0]|TGTAAACATCCTACACTCAGCT</td><td>>MI0000441|hsa-mir-30b&WithFlank&8|-|134800514|134800613</td><td>23</td><td>44</td><td>22</td><td>TGTAAACATCCTACACTCAGCT</td><td>58183.3</td><td>10.4</td><td>1.1</td><td>9.9</td><td>4.9E-23</td><td>3.8E-22</td><td>BT20/MB468</td></tr>
<tr><td>>MI0000093|hsa-mir-92a-1&WithFlank&13|+|91351308|91351397@54.74.21(+1A);MI0000094|hsa-mir-92a-2&WithFlank&X|-|134169532|134169618@54.74.21(+1A)&[MIMAT0000092&hsa-miR-92a-3p&offsets|0|-1(+1A);m-9&13|+|91351361|91351382&offsets|0|-1(+1A)];[MIMAT0000092_1&hsa-miR-92a-3p&offsets|0|-1(+1A);m-11&X|-|134169544|134169565&offsets|0|-1(+1A)]|TATTGCACTTGTCCCGGCCTGA</td><td>>MI0000093|hsa-mir-92a-1&WithFlank&13|+|91351308|91351397</td><td>54</td><td>74</td><td>21(+1A)</td><td>TATTGCACTTGTCCCGGCCTGA</td><td>5147.8</td><td>7.4</td><td>0.7</td><td>9.9</td><td>5.1E-23</td><td>3.9E-22</td><td>BT20/MB468</td></tr>
<tr><td>>MI0000077|hsa-mir-21&WithFlank&17|+|59841260|59841343@14.35.22&[MIMAT0000076&hsa-miR-21-5p&offsets|0|0;m-1&17|+|59841273|59841294&offsets|0|0]|TAGCTTATCAGACTGATGTTGA</td><td>>MI0000077|hsa-mir-21&WithFlank&17|+|59841260|59841343</td><td>14</td><td>35</td><td>22</td><td>TAGCTTATCAGACTGATGTTGA</td><td>608718.5</td><td>8.2</td><td>0.8</td><td>9.9</td><td>5.1E-23</td><td>4.0E-22</td><td>BT20/MB468</td></tr>
<tr><td>>hg38_21_+_8433172_8446622_RNA45SN1_45S_50ntFlanks@6732.6773.42|ACATTGATCATCGACACTTCGAACGCACTTGCGGCCCCGGGT</td><td>>hg38_21_+_8433172_8446622_RNA45SN1_45S_50ntFlanks</td><td>6732</td><td>6773</td><td>42</td><td>ACATTGATCATCGACACTTCGAACGCACTTGCGGCCCCGGGT</td><td>2086.5</td><td>10.2</td><td>1.0</td><td>9.9</td><td>6.7E-23</td><td>4.1E-22</td><td>MB231/MB468</td></tr>
<tr><td>>MI0000440|hsa-mir-27b&WithFlank&9|+|95085439|95085547@67.86.20&[MIMAT0000419&hsa-miR-27b-3p&offsets|0|-1;m-39&9|+|95085505|95085525&offsets|0|-1]|TTCACAGTGGCTAAGTTCTG</td><td>>MI0000440|hsa-mir-27b&WithFlank&9|+|95085439|95085547</td><td>67</td><td>86</td><td>20</td><td>TTCACAGTGGCTAAGTTCTG</td><td>29322.5</td><td>8.2</td><td>0.8</td><td>9.9</td><td>5.4E-23</td><td>4.2E-22</td><td>BT20/MB468</td></tr>
<tr><td>>hg38_21_+_8433172_8446622_RNA45SN1_45S_50ntFlanks@11704.11728.25|ATGAAGCGCGGGTAAACGGCGGGAG</td><td>>hg38_21_+_8433172_8446622_RNA45SN1_45S_50ntFlanks</td><td>11704</td><td>11728</td><td>25</td><td>ATGAAGCGCGGGTAAACGGCGGGAG</td><td>657.0</td><td>-6.3</td><td>0.6</td><td>-9.9</td><td>6.8E-23</td><td>4.2E-22</td><td>MB231/MB468</td></tr>
<tr><td>>MI0000746|hsa-mir-99b&WithFlank&19|+|51692606|51692687@13.32.20(+1G)&[MIMAT0000689&hsa-miR-99b-5p&offsets|0|-2(+1G);m-10&19|+|51692618|51692639&offsets|0|-2(+1G)]|CACCCGTAGAACCGACCTTGG</td><td>>MI0000746|hsa-mir-99b&WithFlank&19|+|51692606|51692687</td><td>13</td><td>32</td><td>20(+1G)</td><td>CACCCGTAGAACCGACCTTGG</td><td>2787.0</td><td>16.1</td><td>1.6</td><td>9.9</td><td>5.5E-23</td><td>4.2E-22</td><td>BT20/MB468</td></tr>
<tr><td>>MI0000088|hsa-mir-30a&WithFlank&6|-|71403545|71403627@13.34.22(+1C)&[MIMAT0000087&hsa-miR-30a-5p&offsets|+1|+1(+1C);m-15&6|-|71403595|71403616&offsets|+1|+1(+1C)]|GTAAACATCCTCGACTGGAAGCC</td><td>>MI0000088|hsa-mir-30a&WithFlank&6|-|71403545|71403627</td><td>13</td><td>34</td><td>22(+1C)</td><td>GTAAACATCCTCGACTGGAAGCC</td><td>961.1</td><td>9.8</td><td>1.0</td><td>9.8</td><td>6.9E-23</td><td>4.2E-22</td><td>MB231/MB468</td></tr>
<tr><td>>MI0006444|hsa-mir-1307&WithFlank&10|-|103394247|103394407@87.107.21(+1A)&[MIMAT0005951&hsa-miR-1307-3p&offsets|+1|0(+1A);m-66&10|-|103394300|103394322&offsets|+1|-1(+1A)]|CTCGGCGTGGCGTCGGTCGTGA</td><td>>MI0006444|hsa-mir-1307&WithFlank&10|-|103394247|103394407</td><td>87</td><td>107</td><td>21(+1A)</td><td>CTCGGCGTGGCGTCGGTCGTGA</td><td>647.1</td><td>7.6</td><td>0.8</td><td>9.8</td><td>6.9E-23</td><td>4.3E-22</td><td>MB231/MB468</td></tr>
<tr><td>>hg38_21_+_8433172_8446622_RNA45SN1_45S_50ntFlanks@476.501.26|TTCGTGATCGATGTGGTGACGTCGTG</td><td>>hg38_21_+_8433172_8446622_RNA45SN1_45S_50ntFlanks</td><td>476</td><td>501</td><td>26</td><td>TTCGTGATCGATGTGGTGACGTCGTG</td><td>2144.6</td><td>6.1</td><td>0.6</td><td>9.9</td><td>5.5E-23</td><td>4.3E-22</td><td>BT20/MB468</td></tr>
<tr><td>>hg38_21_+_8433172_8446622_RNA45SN1_45S_50ntFlanks@11572.11591.20|CCGACTTAGAACTGGTGCGG</td><td>>hg38_21_+_8433172_8446622_RNA45SN1_45S_50ntFlanks</td><td>11572</td><td>11591</td><td>20</td><td>CCGACTTAGAACTGGTGCGG</td><td>6454.4</td><td>-6.9</td><td>0.7</td><td>-9.8</td><td>7.0E-23</td><td>4.3E-22</td><td>MB231/MB468</td></tr>
<tr><td>>MI0000299|hsa-mir-222&WithFlank&X|-|45747009|45747130@75.98.24(+1A)&[MIMAT0000279&hsa-miR-222-3p&offsets|0|+3(+1A);m-140&X|-|45747033|45747056&offsets|0|0(+1A)]|AGCTACATCTGGCTACTGGGTCTCA</td><td>>MI0000299|hsa-mir-222&WithFlank&X|-|45747009|45747130</td><td>75</td><td>98</td><td>24(+1A)</td><td>AGCTACATCTGGCTACTGGGTCTCA</td><td>3357.0</td><td>6.8</td><td>0.7</td><td>9.9</td><td>5.7E-23</td><td>4.4E-22</td><td>BT20/MB468</td></tr>
<tr><td>>MI0000266|hsa-mir-10a&WithFlank&17|-|48579832|48579953@28.48.21(+1C)&[MIMAT0000253&hsa-miR-10a-5p&offsets|0|-2(+1C);m-18&17|-|48579905|48579926&offsets|0|-1(+1C)]|TACCCTGTAGATCCGAATTTGC</td><td>>MI0000266|hsa-mir-10a&WithFlank&17|-|48579832|48579953</td><td>28</td><td>48</td><td>21(+1C)</td><td>TACCCTGTAGATCCGAATTTGC</td><td>3246.8</td><td>11.9</td><td>1.2</td><td>9.8</td><td>7.3E-23</td><td>4.5E-22</td><td>MB231/MB468</td></tr>
<tr><td>>hg38_21_+_8433172_8446622_RNA45SN1_45S_50ntFlanks@11729.11747.19|TAACTATGACTCTCTTAAG</td><td>>hg38_21_+_8433172_8446622_RNA45SN1_45S_50ntFlanks</td><td>11729</td><td>11747</td><td>19</td><td>TAACTATGACTCTCTTAAG</td><td>3375.5</td><td>-5.7</td><td>0.6</td><td>-9.9</td><td>5.8E-23</td><td>4.5E-22</td><td>BT20/MB468</td></tr>
<tr><td>>MI0000078|hsa-mir-22&WithFlank&17|-|1713897|1713993@59.80.22&[MIMAT0000077&hsa-miR-22-3p&offsets|0|0;m-2&17|-|1713914|1713935&offsets|0|0]|AAGCTGCCAGTTGAAGAACTGT</td><td>>MI0000078|hsa-mir-22&WithFlank&17|-|1713897|1713993</td><td>59</td><td>80</td><td>22</td><td>AAGCTGCCAGTTGAAGAACTGT</td><td>29100.5</td><td>8.2</td><td>0.8</td><td>9.9</td><td>5.9E-23</td><td>4.5E-22</td><td>BT20/MB468</td></tr>
<tr><td>>hg38_21_+_8433172_8446622_RNA45SN1_45S_50ntFlanks@10389.10410.22|TGAACAGCAGTTGAACATGGGT</td><td>>hg38_21_+_8433172_8446622_RNA45SN1_45S_50ntFlanks</td><td>10389</td><td>10410</td><td>22</td><td>TGAACAGCAGTTGAACATGGGT</td><td>2045.1</td><td>-6.3</td><td>0.6</td><td>-9.8</td><td>7.4E-23</td><td>4.5E-22</td><td>MB231/MB468</td></tr>
<tr><td>>hg38_MT_+_1621_3279_RNR2_16S_50ntFlanks@722.740.19|TGCGTCAGATTAAAACACT</td><td>>hg38_MT_+_1621_3279_RNR2_16S_50ntFlanks</td><td>722</td><td>740</td><td>19</td><td>TGCGTCAGATTAAAACACT</td><td>398.5</td><td>13.3</td><td>1.3</td><td>9.9</td><td>6.0E-23</td><td>4.6E-22</td><td>BT20/MB468</td></tr>
<tr><td>>hg38_21_+_8433172_8446622_RNA45SN1_45S_50ntFlanks@12458.12497.40|AAGCGTTGGATTGTTCACCCACTAATAGGGAACGTGAGCT</td><td>>hg38_21_+_8433172_8446622_RNA45SN1_45S_50ntFlanks</td><td>12458</td><td>12497</td><td>40</td><td>AAGCGTTGGATTGTTCACCCACTAATAGGGAACGTGAGCT</td><td>414.1</td><td>8.1</td><td>0.8</td><td>9.8</td><td>7.6E-23</td><td>4.6E-22</td><td>MB231/MB468</td></tr>
<tr><td>>hg38_21_+_8433172_8446622_RNA45SN1_45S_50ntFlanks@11729.11749.21|TAACTATGACTCTCTTAAGGT</td><td>>hg38_21_+_8433172_8446622_RNA45SN1_45S_50ntFlanks</td><td>11729</td><td>11749</td><td>21</td><td>TAACTATGACTCTCTTAAGGT</td><td>5323.3</td><td>-4.5</td><td>0.5</td><td>-9.8</td><td>7.6E-23</td><td>4.6E-22</td><td>MB231/MB468</td></tr>
</table>

<table>
<tr><td>>trnaMT_SerGCT_MT_+_12207_12265@1.19.19|GAGAAAGCTCACAAGAACT</td><td>>trnaMT_SerGCT_MT_+_12207_12265</td><td>1</td><td>19</td><td>19</td><td>GAGAAAGCTCACAAGAACT</td><td>15321.9</td><td>8.0</td><td>0.8</td><td>9.9</td><td>6.1E-23</td><td>4.7E-22</td><td>BT20/MB468</td></tr>
<tr><td>>hg38_1_-_228634819_228635039_RNA5S12_5S_50ntFlanks@134.173.40|TGGGAGACCGCCTGGGAATACCGGGTGCTGTAGGCTTTTT</td><td>>hg38_1_-_228634819_228635039_RNA5S12_5S_50ntFlanks</td><td>134</td><td>173</td><td>40</td><td>TGGGAGACCGCCTGGGAATACCGGGTGCTGTAGGCTTTTT</td><td>876.9</td><td>14.7</td><td>1.5</td><td>9.8</td><td>7.7E-23</td><td>4.7E-22</td><td>MB231/MB468</td></tr>
<tr><td>>MI0000434|hsa-let-7i&WithFlank&12|+|62603680|62603775@12.31.20(+1G)&[MIMAT0000415&hsa-let-7i-5p&offsets|0|-2(+1G);m-49&12|+|62603691|62603712&offsets|0|-2(+1G)]|TGAGGTAGTAGTTTGTGCTGG</td><td>>MI0000434|hsa-let-7i&WithFlank&12|+|62603680|62603775</td><td>12</td><td>31</td><td>20(+1G)</td><td>TGAGGTAGTAGTTTGTGCTGG</td><td>1385.4</td><td>8.4</td><td>0.9</td><td>9.8</td><td>7.8E-23</td><td>4.8E-22</td><td>MB231/MB468</td></tr>
<tr><td>>hg38_MT_+_1621_3279_RNR2_16S_50ntFlanks@1362.1388.27|CGACCTCGATGTTGGATCAGGACATCC</td><td>>hg38_MT_+_1621_3279_RNR2_16S_50ntFlanks</td><td>1362</td><td>1388</td><td>27</td><td>CGACCTCGATGTTGGATCAGGACATCC</td><td>959.5</td><td>11.8</td><td>1.2</td><td>9.8</td><td>8.1E-23</td><td>4.9E-22</td><td>MB231/MB468</td></tr>
<tr><td>>MI0000255|hsa-mir-30d&WithFlank&8|-|134804870|134804951@12.35.24(+2U)&[MIMAT0000245&hsa-miR-30d-5p&offsets|0|+2(+2U);m-21&8|-|134804919|134804940&offsets|0|+2(+2U)]|TGTAAACATCCCCGACTGGAAGCTTT</td><td>>MI0000255|hsa-mir-30d&WithFlank&8|-|134804870|134804951</td><td>12</td><td>35</td><td>24(+2U)</td><td>TGTAAACATCCCCGACTGGAAGCTTT</td><td>678.1</td><td>7.6</td><td>0.8</td><td>9.9</td><td>6.4E-23</td><td>4.9E-22</td><td>BT20/MB468</td></tr>
<tr><td>>MI0000065|hsa-let-7d&WithFlank&9|+|94178828|94178926@14.34.21(+1C)&[MIMAT0000065&hsa-let-7d-5p&offsets|0|-1(+1C);m-90&9|+|94178841|94178862&offsets|0|-1(+1C)]|AGAGGTAGTAGGTTGCATAGTC</td><td>>MI0000065|hsa-let-7d&WithFlank&9|+|94178828|94178926</td><td>14</td><td>34</td><td>21(+1C)</td><td>AGAGGTAGTAGGTTGCATAGTC</td><td>3502.6</td><td>8.1</td><td>0.8</td><td>9.9</td><td>6.5E-23</td><td>5.0E-22</td><td>BT20/MB468</td></tr>
<tr><td>>hg38_21_+_8433172_8446622_RNA45SN1_45S_50ntFlanks@12617.12653.37|TGAGGAGCCAATGGGGCGAAGCTACCATCTGTGGGAT</td><td>>hg38_21_+_8433172_8446622_RNA45SN1_45S_50ntFlanks</td><td>12617</td><td>12653</td><td>37</td><td>TGAGGAGCCAATGGGGCGAAGCTACCATCTGTGGGAT</td><td>302.2</td><td>-5.7</td><td>0.6</td><td>-9.9</td><td>6.8E-23</td><td>5.2E-22</td><td>BT20/MB468</td></tr>
<tr><td>>hg38_21_+_8433172_8446622_RNA45SN1_45S_50ntFlanks@10786.10823.38|AACAGCCTCTGGCATGTTGGAACAATGTAGGTAAGGGA</td><td>>hg38_21_+_8433172_8446622_RNA45SN1_45S_50ntFlanks</td><td>10786</td><td>10823</td><td>38</td><td>AACAGCCTCTGGCATGTTGGAACAATGTAGGTAAGGGA</td><td>356.3</td><td>-4.7</td><td>0.5</td><td>-9.9</td><td>6.8E-23</td><td>5.2E-22</td><td>BT20/MB468</td></tr>
<tr><td>>MI0000469|hsa-mir-125a&WithFlank&19|+|51693248|51693345@21.44.24&[MIMAT0000443&hsa-miR-125a-5p&offsets|0|0;m-63&19|+|51693268|51693289&offsets|0|+2]|TCCCTGAGACCCTTTAACCTGTGA</td><td>>MI0000469|hsa-mir-125a&WithFlank&19|+|51693248|51693345</td><td>21</td><td>44</td><td>24</td><td>TCCCTGAGACCCTTTAACCTGTGA</td><td>6115.0</td><td>6.2</td><td>0.6</td><td>9.8</td><td>8.6E-23</td><td>5.2E-22</td><td>MB231/MB468</td></tr>
<tr><td>>MI0000077|hsa-mir-21&WithFlank&17|+|59841260|59841343@14.36.23&[MIMAT0000076&hsa-miR-21-5p&offsets|0|+1;m-1&17|+|59841273|59841294&offsets|0|+1]|TAGCTTATCAGACTGATGTTGAC</td><td>>MI0000077|hsa-mir-21&WithFlank&17|+|59841260|59841343</td><td>14</td><td>36</td><td>23</td><td>TAGCTTATCAGACTGATGTTGAC</td><td>4016485.6</td><td>8.6</td><td>0.9</td><td>9.8</td><td>8.7E-23</td><td>5.3E-22</td><td>MB231/MB468</td></tr>
<tr><td>>MI0000061|hsa-let-7a-2&WithFlank&11|-|122146516|122146599@11.32.22;MI0000062|hsa-let-7a-3&WithFlank&22|+|46112743|46112828@10.31.22;MI0000060|hsa-let-7a-1&WithFlank&9|+|94175951|94176042@12.33.22&[MIMAT0000062&hsa-let-7a-5p&offsets|0|0;m-6&11|-|122146568|122146589&offsets|0|0];[MIMAT0000062_1&hsa-let-7a-5p&offsets|0|0;m-5&22|+|46112752|46112773&offsets|0|0];[MIMAT0000062_2&hsa-let-7a-5p&offsets|0|0;m-4&9|+|94175962|94175983&offsets|0|0]|TGAGGTAGTAGGTTGTATAGTT</td><td>>MI0000061|hsa-let-7a-2&WithFlank&11|-|122146516|122146599</td><td>11</td><td>32</td><td>22</td><td>TGAGGTAGTAGGTTGTATAGTT</td><td>191471.0</td><td>7.1</td><td>0.7</td><td>9.8</td><td>7.0E-23</td><td>5.4E-22</td><td>BT20/MB468</td></tr>
<tr><td>>MI0000081|hsa-mir-24-2&WithFlank&19|-|13836281|13836365@56.74.19(+1U);MI0000080|hsa-mir-24-1&WithFlank&9|+|95086015|95086094@50.68.19(+1U)&[MIMAT0000080&hsa-miR-24-3p&offsets|0|-3(+1U);m-36&19|-|13836288|13836310&offsets|0|-4(+1U)];[MIMAT0000080_1&hsa-miR-24-3p&offsets|0|-3(+1U);m-35&9|+|95086064|95086086&offsets|0|-4(+1U)]|TGGCTCAGTTCAGCAGGAAT</td><td>>MI0000081|hsa-mir-24-2&WithFlank&19|-|13836281|13836365</td><td>56</td><td>74</td><td>19(+1U)</td><td>TGGCTCAGTTCAGCAGGAAT</td><td>2062.6</td><td>6.7</td><td>0.7</td><td>9.8</td><td>8.9E-23</td><td>5.4E-22</td><td>MB231/MB468</td></tr>
<tr><td>>hg38_21_+_8433172_8446622_RNA45SN1_45S_50ntFlanks@475.501.27|CTTCGTGATCGATGTGGTGACGTCGTG</td><td>>hg38_21_+_8433172_8446622_RNA45SN1_45S_50ntFlanks</td><td>475</td><td>501</td><td>27</td><td>CTTCGTGATCGATGTGGTGACGTCGTG</td><td>1535.9</td><td>6.3</td><td>0.6</td><td>9.8</td><td>7.1E-23</td><td>5.4E-22</td><td>BT20/MB468</td></tr>
<tr><td>>hg38_21_+_8433172_8446622_RNA45SN1_45S_50ntFlanks@10357.10383.27|CTTTGAAGGCCGAAGTGGAGAAGGGTT</td><td>>hg38_21_+_8433172_8446622_RNA45SN1_45S_50ntFlanks</td><td>10357</td><td>10383</td><td>27</td><td>CTTTGAAGGCCGAAGTGGAGAAGGGTT</td><td>305.6</td><td>-4.0</td><td>0.4</td><td>-9.8</td><td>7.2E-23</td><td>5.5E-22</td><td>BT20/MB468</td></tr>
<tr><td>>MI0000263|hsa-mir-7-1&WithFlank&9|-|83969742|83969863@30.55.26&[MIMAT0000252_2&hsa-miR-7-5p&offsets|0|+2;m-401&9|-|83969811|83969834&offsets|0|+2]|TGGAAGACTAGTGATTTTGTTGTTTT</td><td>>MI0000263|hsa-mir-7-1&WithFlank&9|-|83969742|83969863</td><td>30</td><td>55</td><td>26</td><td>TGGAAGACTAGTGATTTTGTTGTTTT</td><td>1087.4</td><td>7.9</td><td>0.8</td><td>9.8</td><td>9.2E-23</td><td>5.6E-22</td><td>MB231/MB468</td></tr>
<tr><td>>hg38_21_+_8433172_8446622_RNA45SN1_45S_50ntFlanks@12405.12437.33|CTTTTTGATCCTTCGATGTCGGCTCTTCCTATC</td><td>>hg38_21_+_8433172_8446622_RNA45SN1_45S_50ntFlanks</td><td>12405</td><td>12437</td><td>33</td><td>CTTTTTGATCCTTCGATGTCGGCTCTTCCTATC</td><td>1798.4</td><td>-5.1</td><td>0.5</td><td>-9.8</td><td>7.4E-23</td><td>5.6E-22</td><td>BT20/MB468</td></tr>
<tr><td>>MI0000272|hsa-mir-182&WithFlank&7|-|129770377|129770498@29.50.22&[MIMAT0000259&hsa-miR-182-5p&offsets|0|-2;m-22&7|-|129770449|129770470&offsets|0|0]|TTTGGCAATGGTAGAACTCACA</td><td>>MI0000272|hsa-mir-182&WithFlank&7|-|129770377|129770498</td><td>29</td><td>50</td><td>22</td><td>TTTGGCAATGGTAGAACTCACA</td><td>13892.4</td><td>8.7</td><td>0.9</td><td>9.8</td><td>7.4E-23</td><td>5.6E-22</td><td>BT20/MB468</td></tr>
<tr><td>>MI0000811|hsa-mir-148b&WithFlank&12|+|54337210|54337320@69.90.22(+2U)&[MIMAT0000759&hsa-miR-148b-3p&offsets|0|0(+2U);m-77&12|+|54337278|54337299&offsets|0|0(+2U)]|TCAGTGCATCACAGAACTTTGTTT</td><td>>MI0000811|hsa-mir-148b&WithFlank&12|+|54337210|54337320</td><td>69</td><td>90</td><td>22(+2U)</td><td>TCAGTGCATCACAGAACTTTGTTT</td><td>1987.3</td><td>8.3</td><td>0.8</td><td>9.8</td><td>9.7E-23</td><td>5.9E-22</td><td>MB231/MB468</td></tr>
<tr><td>>hg38_21_+_8433172_8446622_RNA45SN1_45S_50ntFlanks@474.505.32|TCTTCGTGATCGATGTGGTGACGTCGTGCTCT</td><td>>hg38_21_+_8433172_8446622_RNA45SN1_45S_50ntFlanks</td><td>474</td><td>505</td><td>32</td><td>TCTTCGTGATCGATGTGGTGACGTCGTGCTCT</td><td>1110.6</td><td>7.2</td><td>0.7</td><td>9.8</td><td>7.8E-23</td><td>5.9E-22</td><td>BT20/MB468</td></tr>
<tr><td>>MI0003834|hsa-mir-769&WithFlank&19|+|46018926|46019055@36.57.22&[MIMAT0003886&hsa-miR-769-5p&offsets|0|0;m-216&19|+|46018961|46018982&offsets|0|0]|TGAGACCTCTGGGTTCTGAGCT</td><td>>MI0003834|hsa-mir-769&WithFlank&19|+|46018926|46019055</td><td>36</td><td>57</td><td>22</td><td>TGAGACCTCTGGGTTCTGAGCT</td><td>4839.3</td><td>8.0</td><td>0.8</td><td>9.8</td><td>9.8E-23</td><td>5.9E-22</td><td>MB231/MB468</td></tr>
<tr><td>>hg38_21_+_8433172_8446622_RNA45SN1_45S_50ntFlanks@3845.3866.22|ACTTGGATAACTGTGGTAATTC</td><td>>hg38_21_+_8433172_8446622_RNA45SN1_45S_50ntFlanks</td><td>3845</td><td>3866</td><td>22</td><td>ACTTGGATAACTGTGGTAATTC</td><td>287.2</td><td>5.8</td><td>0.6</td><td>9.8</td><td>9.9E-23</td><td>6.0E-22</td><td>MB231/MB468</td></tr>
<tr><td>>hg38_21_+_8433172_8446622_RNA45SN1_45S_50ntFlanks@6651.6694.44|CGACTCTTAGCGGTGGATCACTCGGCTCGTGCGTCGATGAAGAA</td><td>>hg38_21_+_8433172_8446622_RNA45SN1_45S_50ntFlanks</td><td>6651</td><td>6694</td><td>44</td><td>CGACTCTTAGCGGTGGATCACTCGGCTCGTGCGTCGATGAAGAA</td><td>3871.4</td><td>6.9</td><td>0.7</td><td>9.8</td><td>1.0E-22</td><td>6.1E-22</td><td>MB231/MB468</td></tr>
<tr><td>>MI0000456|hsa-mir-140&WithFlank&16|+|69933075|69933186@68.88.21&[MIMAT0004597&hsa-miR-140-3p&offsets|0|0;m-45&16|+|69933143|69933164&offsets|-1|-2]|TACCACAGGGTAGAACCACGG</td><td>>MI0000456|hsa-mir-140&WithFlank&16|+|69933075|69933186</td><td>68</td><td>88</td><td>21</td><td>TACCACAGGGTAGAACCACGG</td><td>665.9</td><td>11.5</td><td>1.2</td><td>9.8</td><td>1.0E-22</td><td>6.2E-22</td><td>MB231/MB468</td></tr>
<tr><td>>hg38_21_+_8433172_8446622_RNA45SN1_45S_50ntFlanks@6651.6684.34|CGACTCTTAGCGGTGGATCACTCGGCTCGTGCGT</td><td>>hg38_21_+_8433172_8446622_RNA45SN1_45S_50ntFlanks</td><td>6651</td><td>6684</td><td>34</td><td>CGACTCTTAGCGGTGGATCACTCGGCTCGTGCGT</td><td>3014.9</td><td>4.8</td><td>0.5</td><td>9.8</td><td>8.2E-23</td><td>6.2E-22</td><td>BT20/MB468</td></tr>
<tr><td>>hg38_21_+_8433172_8446622_RNA45SN1_45S_50ntFlanks@6777.6806.30|TCCCGGGGCTACGCCTGTCTGAGCGTCGCT</td><td>>hg38_21_+_8433172_8446622_RNA45SN1_45S_50ntFlanks</td><td>6777</td><td>6806</td><td>30</td><td>TCCCGGGGCTACGCCTGTCTGAGCGTCGCT</td><td>1027.4</td><td>6.0</td><td>0.6</td><td>9.8</td><td>1.0E-22</td><td>6.2E-22</td><td>MB231/MB468</td></tr>
<tr><td>>trnaMT_SerGCT_MT_+_12207_12265@1.25.25|GAGAAAGCTCACAAGAACTGCTAAC</td><td>>trnaMT_SerGCT_MT_+_12207_12265</td><td>1</td><td>25</td><td>25</td><td>GAGAAAGCTCACAAGAACTGCTAAC</td><td>22425.2</td><td>8.0</td><td>0.8</td><td>9.8</td><td>8.2E-23</td><td>6.2E-22</td><td>BT20/MB468</td></tr>
<tr><td>>hg38_21_+_8433172_8446622_RNA45SN1_45S_50ntFlanks@11705.11728.24|TGAAGCGCGGGTAAACGGCGGGAG</td><td>>hg38_21_+_8433172_8446622_RNA45SN1_45S_50ntFlanks</td><td>11705</td><td>11728</td><td>24</td><td>TGAAGCGCGGGTAAACGGCGGGAG</td><td>442.5</td><td>-5.7</td><td>0.6</td><td>-9.8</td><td>1.0E-22</td><td>6.2E-22</td><td>MB231/MB468</td></tr>
<tr><td>>hg38_21_+_8433172_8446622_RNA45SN1_45S_50ntFlanks@12936.12956.21|AAACCATTCGTAGACGACCTG</td><td>>hg38_21_+_8433172_8446622_RNA45SN1_45S_50ntFlanks</td><td>12936</td><td>12956</td><td>21</td><td>AAACCATTCGTAGACGACCTG</td><td>4222.1</td><td>-5.7</td><td>0.6</td><td>-9.8</td><td>1.0E-22</td><td>6.3E-22</td><td>MB231/MB468</td></tr>
<tr><td>>hg38_21_+_8433172_8446622_RNA45SN1_45S_50ntFlanks@10681.10706.26|TTCCGGCGGCGTCCGGTGAGCTCTCG</td><td>>hg38_21_+_8433172_8446622_RNA45SN1_45S_50ntFlanks</td><td>10681</td><td>10706</td><td>26</td><td>TTCCGGCGGCGTCCGGTGAGCTCTCG</td><td>1526.2</td><td>-6.3</td><td>0.6</td><td>-9.8</td><td>1.1E-22</td><td>6.4E-22</td><td>MB231/MB468</td></tr>
<tr><td>>hg38_21_+_8433172_8446622_RNA45SN1_45S_50ntFlanks@474.504.31|TCTTCGTGATCGATGTGGTGACGTCGTGCTC</td><td>>hg38_21_+_8433172_8446622_RNA45SN1_45S_50ntFlanks</td><td>474</td><td>504</td><td>31</td><td>TCTTCGTGATCGATGTGGTGACGTCGTGCTC</td><td>556.1</td><td>7.1</td><td>0.7</td><td>9.8</td><td>8.7E-23</td><td>6.6E-22</td><td>BT20/MB468</td></tr>
<tr><td>>MI0000266|hsa-mir-10a&WithFlank&17|-|48579832|48579953@28.48.21(+1C)&[MIMAT0000253&hsa-miR-10a-5p&offsets|0|-2(+1C);m-18&17|-|48579905|48579926&offsets|0|-1(+1C)]|TACCCTGTAGATCCGAATTTGC</td><td>>MI0000266|hsa-mir-10a&WithFlank&17|-|48579832|48579953</td><td>28</td><td>48</td><td>21(+1C)</td><td>TACCCTGTAGATCCGAATTTGC</td><td>5913.6</td><td>12.4</td><td>1.3</td><td>9.8</td><td>8.8E-23</td><td>6.6E-22</td><td>BT20/MB468</td></tr>
<tr><td>>hg38_21_+_8433172_8446622_RNA45SN1_45S_50ntFlanks@12081.12110.30|CTCTCGCTTCTGGCGCCAAGCGCCCGGCCG</td><td>>hg38_21_+_8433172_8446622_RNA45SN1_45S_50ntFlanks</td><td>12081</td><td>12110</td><td>30</td><td>CTCTCGCTTCTGGCGCCAAGCGCCCGGCCG</td><td>634.3</td><td>-6.3</td><td>0.6</td><td>-9.8</td><td>1.1E-22</td><td>6.7E-22</td><td>MB231/MB468</td></tr>
<tr><td>>MI0000441|hsa-mir-30b&WithFlank&8|-|134800514|134800613@23.43.21(+1C)&[MIMAT0000420&hsa-miR-30b-5p&offsets|0|-1(+1C);m-76&8|-|134800570|134800591&offsets|0|-1(+1C)]|TGTAAACATCCTACACTCAGCC</td><td>>MI0000441|hsa-mir-30b&WithFlank&8|-|134800514|134800613</td><td>23</td><td>43</td><td>21(+1C)</td><td>TGTAAACATCCTACACTCAGCC</td><td>2512.2</td><td>8.8</td><td>0.9</td><td>9.8</td><td>1.1E-22</td><td>6.7E-22</td><td>MB231/MB468</td></tr>
<tr><td>>hg38_21_+_8433172_8446622_RNA45SN1_45S_50ntFlanks@10808.10826.19|CAATGTAGGTAAGGGAAGT</td><td>>hg38_21_+_8433172_8446622_RNA45SN1_45S_50ntFlanks</td><td>10808</td><td>10826</td><td>19</td><td>CAATGTAGGTAAGGGAAGT</td><td>808.9</td><td>-5.9</td><td>0.6</td><td>-9.8</td><td>8.9E-23</td><td>6.7E-22</td><td>BT20/MB468</td></tr>
<tr><td>>hg38_21_+_8433172_8446622_RNA45SN1_45S_50ntFlanks@9534.9565.32|AGGAAACTCTGGTGGAGGTCCGTAGCGGTCCT</td><td>>hg38_21_+_8433172_8446622_RNA45SN1_45S_50ntFlanks</td><td>9534</td><td>9565</td><td>32</td><td>AGGAAACTCTGGTGGAGGTCCGTAGCGGTCCT</td><td>1003.1</td><td>-5.1</td><td>0.5</td><td>-9.8</td><td>9.1E-23</td><td>6.8E-22</td><td>BT20/MB468</td></tr>
<tr><td>>trnaMT_SerGCT_MT_+_12207_12265@1.19.19|GAGAAAGCTCACAAGAACT</td><td>>trnaMT_SerGCT_MT_+_12207_12265</td><td>1</td><td>19</td><td>19</td><td>GAGAAAGCTCACAAGAACT</td><td>12384.3</td><td>7.9</td><td>0.8</td><td>9.8</td><td>1.1E-22</td><td>6.9E-22</td><td>MB231/MB468</td></tr>
<tr><td>>MI0000067|hsa-let-7f-1&WithFlank&9|+|94176341|94176439@13.35.23&[MIMAT0000067&hsa-let-7f-5p&offsets|0|+1;m-20&9|+|94176353|94176374&offsets|0|+1]|TGAGGTAGTAGATTGTATAGTTG</td><td>>MI0000067|hsa-let-7f-1&WithFlank&9|+|94176341|94176439</td><td>13</td><td>35</td><td>23</td><td>TGAGGTAGTAGATTGTATAGTTG</td><td>925.0</td><td>7.2</td><td>0.7</td><td>9.8</td><td>9.2E-23</td><td>6.9E-22</td><td>BT20/MB468</td></tr>
<tr><td>>hg38_21_+_8433172_8446622_RNA45SN1_45S_50ntFlanks@11323.11344.22|CGTCTCGTCGCGCGCGCGTCCG</td><td>>hg38_21_+_8433172_8446622_RNA45SN1_45S_50ntFlanks</td><td>11323</td><td>11344</td><td>22</td><td>CGTCTCGTCGCGCGCGCGTCCG</td><td>5683.1</td><td>-8.8</td><td>0.9</td><td>-9.8</td><td>1.2E-22</td><td>7.0E-22</td><td>MB231/MB468</td></tr>
<tr><td>>trna10_LysCTT_16_+_3241501_3241573@3.35.33,trna13_LysCTT_14_-_58706613_58706685@3.35.33,trna2_LysCTT_15_+_79152904_79152976@3.35.33|CCGGCTAGCTCAGTCGGTAGAGCATGGGACTCT</td><td>>trna10_LysCTT_16_+_3241501_3241573</td><td>3</td><td>35</td><td>33</td><td>CCGGCTAGCTCAGTCGGTAGAGCATGGGACTCT</td><td>1305.6</td><td>11.1</td><td>1.1</td><td>9.8</td><td>1.2E-22</td><td>7.0E-22</td><td>MB231/MB468</td></tr>
<tr><td>>MI0000077|hsa-mir-21&WithFlank&17|+|59841260|59841343@14.34.21(+1U)&[MIMAT0000076&hsa-miR-21-5p&offsets|0|-1(+1U);m-1&17|+|59841273|59841294&offsets|0|-1(+1U)]|TAGCTTATCAGACTGATGTTGT</td><td>>MI0000077|hsa-mir-21&WithFlank&17|+|59841260|59841343</td><td>14</td><td>34</td><td>21(+1U)</td><td>TAGCTTATCAGACTGATGTTGT</td><td>39267.7</td><td>10.4</td><td>1.1</td><td>9.8</td><td>9.4E-23</td><td>7.0E-22</td><td>BT20/MB468</td></tr>
</table>

| | | | | | | | | | | | | |
|---|---|---|---|---|---|---|---|---|---|---|---|---|
| >hg38_21_+_8433172_8446622_RNA45SN1_45S_50ntFlanks@3875.3899.25\|ATACATGCCGACGGGCGCTGACCCC | >hg38_21_+_8433172_8446622_RNA45SN1_45S_50ntFlanks | 3875 | 3899 | 25 | ATACATGCCGACGGGCGCTGACCCC | 577.6 | -4.7 | 0.5 | -9.8 | 9.6E-23 | 7.2E-22 | BT20/MB468 |
| >hg38_21_+_8433172_8446622_RNA45SN1_45S_50ntFlanks@4726.4758.33\|TAATCAAGAACGAAAGTCGGAGGTTCGAAGACG | >hg38_21_+_8433172_8446622_RNA45SN1_45S_50ntFlanks | 4726 | 4758 | 33 | TAATCAAGAACGAAAGTCGGAGGTTCGAAGACG | 1608.9 | -5.3 | 0.5 | -9.8 | 9.6E-23 | 7.2E-22 | BT20/MB468 |
| >MI0000301\|hsa-mir-224&WithFlank&X\|-\|151958572\|151958664@14.38.25&[MIMAT0000281&hsa-miR-224-5p&offsets\|+1\|+1;m-238&X\|-\|151958630\|151958651&offsets\|0\|+3]\|CAAGTCACTAGTGGTTCCGTTTAGT | >MI0000301\|hsa-mir-224&WithFlank&X\|-\|151958572\|151958664 | 14 | 38 | 25 | CAAGTCACTAGTGGTTCCGTTTAGT | 1046.1 | 12.6 | 1.3 | 9.8 | 9.6E-23 | 7.2E-22 | BT20/MB468 |
| >MI0015866\|hsa-mir-4326&WithFlank&20\|+\|63286802\|63286872@17.39.23&[MIMAT0016888&hsa-miR-4326&offsets\|0\|+3;m-339&20\|+\|63286818\|63286841&offsets\|0\|-1]\|TGTTCCTCTGTCTCCCAGACTCT | >MI0015866\|hsa-mir-4326&WithFlank&20\|+\|63286802\|63286872 | 17 | 39 | 23 | TGTTCCTCTGTCTCCCAGACTCT | 646.1 | 14.0 | 1.4 | 9.8 | 9.8E-23 | 7.3E-22 | BT20/MB468 |
| >trnaMT_PheGAA_MT_+_577_647@36.57.22\|AAAATGTTTAGACGGGCTCACA | >trnaMT_PheGAA_MT_+_577_647 | 36 | 57 | 22 | AAAATGTTTAGACGGGCTCACA | 3898.5 | 11.9 | 1.2 | 9.8 | 9.8E-23 | 7.3E-22 | BT20/MB468 |
| >hg38_21_+_8433172_8446622_RNA45SN1_45S_50ntFlanks@10866.10891.26\|CTAAGGGCTGGGTCGGTCGGGCTGGG | >hg38_21_+_8433172_8446622_RNA45SN1_45S_50ntFlanks | 10866 | 10891 | 26 | CTAAGGGCTGGGTCGGTCGGGCTGGG | 332.5 | -5.4 | 0.6 | -9.8 | 1.2E-22 | 7.5E-22 | MB231/MB468 |
| >hg38_21_+_8433172_8446622_RNA45SN1_45S_50ntFlanks@8356.8379.24\|GAAAAGAACTTTGAAGAGAGAGTT | >hg38_21_+_8433172_8446622_RNA45SN1_45S_50ntFlanks | 8356 | 8379 | 24 | GAAAAGAACTTTGAAGAGAGAGTT | 17579.4 | -6.3 | 0.6 | -9.8 | 1.3E-22 | 7.5E-22 | MB231/MB468 |
| >hg38_21_+_8433172_8446622_RNA45SN1_45S_50ntFlanks@485.525.41\|GATGTGGTGACGTCGTGCTCTCCCGGGCCGGGTCCGAGCCG | >hg38_21_+_8433172_8446622_RNA45SN1_45S_50ntFlanks | 485 | 525 | 41 | GATGTGGTGACGTCGTGCTCTCCCGGGCCGGGTCCGAGCCG | 1263.9 | 15.0 | 1.5 | 9.8 | 1.0E-22 | 7.5E-22 | BT20/MB468 |
| >MI0000746\|hsa-mir-99b&WithFlank&19\|+\|51692606\|51692687@51.71.21&[MIMAT0004678&hsa-miR-99b-3p&offsets\|0\|-1;m-151&19\|+\|51692656\|51692678&offsets\|0\|-2]\|CAAGCTCGTGTCTGTGGGTCC | >MI0000746\|hsa-mir-99b&WithFlank&19\|+\|51692606\|51692687 | 51 | 71 | 21 | CAAGCTCGTGTCTGTGGGTCC | 1459.0 | 8.1 | 0.8 | 9.8 | 1.0E-22 | 7.6E-22 | BT20/MB468 |
| >MI0000478\|hsa-mir-149&WithFlank&2\|+\|240455995\|240456095@21.42.22(+1A)&[MIMAT0000450&hsa-miR-149-5p&offsets\|0\|-1(+1A);m-195&2\|+\|240456015\|240456037&offsets\|0\|-1(+1A)]\|TCTGGCTCCGTGTCTTCACTCCA | >MI0000478\|hsa-mir-149&WithFlank&2\|+\|240455995\|240456095 | 21 | 42 | 22(+1A) | TCTGGCTCCGTGTCTTCACTCCA | 1388.7 | 15.1 | 1.5 | 9.8 | 1.0E-22 | 7.7E-22 | BT20/MB468 |
| >MI0000088\|hsa-mir-30a&WithFlank&6\|-\|71403545\|71403627@54.75.22&[MIMAT0000088&hsa-miR-30a-3p&offsets\|+1\|+1;m-38&6\|-\|71403554\|71403575&offsets\|+1\|+1]\|TTTCAGTCGGATGTTTGCAGCT | >MI0000088\|hsa-mir-30a&WithFlank&6\|-\|71403545\|71403627 | 54 | 75 | 22 | TTTCAGTCGGATGTTTGCAGCT | 3207.5 | 10.8 | 1.1 | 9.8 | 1.3E-22 | 7.8E-22 | MB231/MB468 |
| >MI0000087\|hsa-mir-29a&WithFlank&7\|-\|130876741\|130876816@47.67.21&[MIMAT0000086&hsa-miR-29a-3p&offsets\|-1\|-2;m-26&7\|-\|130876749\|130876769&offsets\|-1\|-1]\|CTAGCACCATCTGAAATCGGT | >MI0000087\|hsa-mir-29a&WithFlank&7\|-\|130876741\|130876816 | 47 | 67 | 21 | CTAGCACCATCTGAAATCGGT | 577.9 | 13.8 | 1.4 | 9.8 | 1.1E-22 | 7.8E-22 | BT20/MB468 |
| >MI0000542\|hsa-mir-320a&WithFlank&8\|-\|22244960\|22245043@49.70.22&[MIMAT0000510&hsa-miR-320a-3p&offsets\|+1\|+1;m-60&8\|-\|22244975\|22244996&offsets\|+1\|+1]\|AAAGCTGGGTTGAGAGGGCGAA | >MI0000542\|hsa-mir-320a&WithFlank&8\|-\|22244960\|22245043 | 49 | 70 | 22 | AAAGCTGGGTTGAGAGGGCGAA | 1298.4 | 8.5 | 0.9 | 9.8 | 1.1E-22 | 7.8E-22 | BT20/MB468 |
| >MI0000456\|hsa-mir-140&WithFlank&16\|+\|69933075\|69933186@69.89.21(+1U)&[MIMAT0004597&hsa-miR-140-3p&offsets\|+1\|+1(+1U);m-45&16\|+\|69933143\|69933164&offsets\|0\|-1(+1U)]\|ACCACAGGGTAGAACCACGGAT | >MI0000456\|hsa-mir-140&WithFlank&16\|+\|69933075\|69933186 | 69 | 89 | 21(+1U) | ACCACAGGGTAGAACCACGGAT | 1040.7 | 14.7 | 1.5 | 9.8 | 1.1E-22 | 7.9E-22 | BT20/MB468 |
| >MI0000063\|hsa-let-7b&WithFlank&22\|+\|46113680\|46113774@12.33.22(+1G)&[MIMAT0000063&hsa-let-7b-5p&offsets\|0\|0(+1G);m-14&22\|+\|46113691\|46113712&offsets\|0\|0(+1G)]\|TGAGGTAGTAGGTTGTGTGGTTG | >MI0000063\|hsa-let-7b&WithFlank&22\|+\|46113680\|46113774 | 12 | 33 | 22(+1G) | TGAGGTAGTAGGTTGTGTGGTTG | 1189.9 | 9.8 | 1.0 | 9.8 | 1.1E-22 | 7.9E-22 | BT20/MB468 |
| >hg38_MT_+_598_1651_RNR1_12S_50ntFlanks@428.469.42\|AAAGTGGCTTTAACATATCTGAACACACAATAGCTAAGACCC | >hg38_MT_+_598_1651_RNR1_12S_50ntFlanks | 428 | 469 | 42 | AAAGTGGCTTTAACATATCTGAACACACAATAGCTAAGACCC | 538.4 | 13.7 | 1.4 | 9.8 | 1.1E-22 | 7.9E-22 | BT20/MB468 |
| >hg38_1_-_228634819_228635039_RNA5S12_5S_50ntFlanks@134.169.36\|TGGGAGACCGCCTGGGAATACCGGGTGCTGTAGGCT | >hg38_1_-_228634819_228635039_RNA5S12_5S_50ntFlanks | 134 | 169 | 36 | TGGGAGACCGCCTGGGAATACCGGGTGCTGTAGGCT | 8954.0 | 6.8 | 0.7 | 9.8 | 1.1E-22 | 8.0E-22 | BT20/MB468 |
| >MI0000750\|hsa-mir-26a-2&WithFlank&12\|-\|57824603\|57824698@20.40.21(+1C);MI0000083\|hsa-mir-26a-1&WithFlank&3\|+\|37969398\|37969486@16.36.21(+1C)&[MIMAT0000082&hsa-miR-26a-5p&offsets\|0\|-1(+1C);m-33&12\|-\|57824658\|57824679&offsets\|0\|-1(+1C)];[MIMAT0000082_1&hsa-miR-26a-5p&offsets\|0\|-1(+1C);m-34&3\|+\|37969413\|37969434&offsets\|0\|-1(+1C)]\|TTCAAGTAATCCAGGATAGGCC | >MI0000750\|hsa-mir-26a-2&WithFlank&12\|-\|57824603\|57824698 | 20 | 40 | 21(+1C) | TTCAAGTAATCCAGGATAGGCC | 33319.2 | 7.1 | 0.7 | 9.8 | 1.3E-22 | 8.0E-22 | MB231/MB468 |
| >MI0000433\|hsa-let-7g&WithFlank&3\|-\|52268272\|52268367@10.32.23&[MIMAT0000414&hsa-let-7g-5p&offsets\|-1\|0;m-58&3\|-\|52268336\|52268357&offsets\|-1\|0]\|CTGAGGTAGTAGTTTGTACAGTT | >MI0000433\|hsa-let-7g&WithFlank&3\|-\|52268272\|52268367 | 10 | 32 | 23 | CTGAGGTAGTAGTTTGTACAGTT | 954.6 | 6.4 | 0.6 | 9.8 | 1.1E-22 | 8.1E-22 | BT20/MB468 |
| >MI0000750\|hsa-mir-26a-2&WithFlank&12\|-\|57824603\|57824698@20.39.20(+1U);MI0000083\|hsa-mir-26a-1&WithFlank&3\|+\|37969398\|37969486@16.35.20(+1U)&[MIMAT0000082&hsa-miR-26a-5p&offsets\|0\|-2(+1U);m-33&12\|-\|57824658\|57824679&offsets\|0\|-2(+1U)];[MIMAT0000082_1&hsa-miR-26a-5p&offsets\|0\|-2(+1U);m-34&3\|+\|37969413\|37969434&offsets\|0\|-2(+1U)]\|TTCAAGTAATCCAGGATAGGT | >MI0000750\|hsa-mir-26a-2&WithFlank&12\|-\|57824603\|57824698 | 20 | 39 | 20(+1U) | TTCAAGTAATCCAGGATAGGT | 952.4 | 7.8 | 0.8 | 9.8 | 1.4E-22 | 8.2E-22 | MB231/MB468 |
| >hg38_21_+_8433172_8446622_RNA45SN1_45S_50ntFlanks@12996.13040.45\|TCGCTGCGATCTATTGAAAGTCAGCCCTCGACACAAGGGTTTGTC | >hg38_21_+_8433172_8446622_RNA45SN1_45S_50ntFlanks | 12996 | 13040 | 45 | TCGCTGCGATCTATTGAAAGTCAGCCCTCGACACAAGGGTTTGTC | 2856.9 | 11.8 | 1.2 | 9.8 | 1.4E-22 | 8.2E-22 | MB231/MB468 |
| >am_trna128_GlyGCC_6_-_27870686_27870756@1.33.33,trna18_GlyGCC_16_+_70822597_70822667@1.33.33,trna19_GlyGCC_16_+_70823410_70823480@1.33.33,trna19_GlyGCC_2_-_157257659_157257729@1.33.33,trna24_GlyGCC_16_-_70812942_70813012@1.33.33,trna25_GlyGCC_16_-_70812114_70812184@1.33.33,trna5_GlyGCC_17_+_8029064_8029134@1.33.33,trna68_GlyGCC_1_-_161493637_161493707@1.33.33\|GCATTGGTGGTTCAGTGGTAGAATTCTCGCCTG | >am_trna128_GlyGCC_6_-_27870686_27870756 | 1 | 33 | 33 | GCATTGGTGGTTCAGTGGTAGAATTCTCGCCTG | 4169.7 | 6.2 | 0.6 | 9.8 | 1.1E-22 | 8.3E-22 | BT20/MB468 |
| >MI0000067\|hsa-let-7f-1&WithFlank&9\|+\|94176341\|94176439@13.33.21;MI0000068\|hsa-let-7f-2&WithFlank&X\|-\|53557186\|53557280@14.34.21&[MIMAT0000067&hsa-let-7f-5p&offsets\|0\|-1;m-20&9\|+\|94176353\|94176374&offsets\|0\|-1];[MIMAT0000067_1&hsa-let-7f-5p&offsets\|0\|-1;m-19&X\|-\|53557246\|53557267&offsets\|0\|-1]\|TGAGGTAGTAGATTGTATAGT | >MI0000067\|hsa-let-7f-1&WithFlank&9\|+\|94176341\|94176439 | 13 | 33 | 21 | TGAGGTAGTAGATTGTATAGT | 26043.5 | 6.6 | 0.7 | 9.8 | 1.1E-22 | 8.3E-22 | BT20/MB468 |
| >MI0000086\|hsa-mir-28&WithFlank&3\|+\|188688775\|188688872@60.81.22(+2A)&[MIMAT0004502&hsa-miR-28-3p&offsets\|0\|0(+2A);m-28&3\|+\|188688834\|188688855&offsets\|0\|0(+2A)]\|CACTAGATTGTGAGCTCCTGGAAA | >MI0000086\|hsa-mir-28&WithFlank&3\|+\|188688775\|188688872 | 60 | 81 | 22(+2A) | CACTAGATTGTGAGCTCCTGGAAA | 681.8 | 14.1 | 1.4 | 9.8 | 1.1E-22 | 8.3E-22 | BT20/MB468 |
| >hg38_21_+_8433172_8446622_RNA45SN1_45S_50ntFlanks@10690.10708.19\|CGTCCGGTGAGCTCTCGCT | >hg38_21_+_8433172_8446622_RNA45SN1_45S_50ntFlanks | 10690 | 10708 | 19 | CGTCCGGTGAGCTCTCGCT | 4040.1 | -6.4 | 0.7 | -9.8 | 1.4E-22 | 8.4E-22 | MB231/MB468 |
| >hg38_21_+_8433172_8446622_RNA45SN1_45S_50ntFlanks@12415.12448.34\|CTTCGATGTCGGCTCTTCCTATCATTGTGAAGCA | >hg38_21_+_8433172_8446622_RNA45SN1_45S_50ntFlanks | 12415 | 12448 | 34 | CTTCGATGTCGGCTCTTCCTATCATTGTGAAGCA | 2243.0 | 7.1 | 0.7 | 9.8 | 1.4E-22 | 8.5E-22 | MB231/MB468 |
| >trnaMT_GluTTC_MT_-_14674_14742@38.66.29,trnalookalike8_GluTTC_5_-_93905172_93905240@38.66.29\|CATTGGTCGTGGTTGTAGTCCGTGCGAGA | >trnaMT_GluTTC_MT_-_14674_14742 | 38 | 66 | 29 | CATTGGTCGTGGTTGTAGTCCGTGCGAGA | 343.3 | 7.5 | 0.8 | 9.8 | 1.2E-22 | 8.5E-22 | BT20/MB468 |
| >hg38_21_+_8433172_8446622_RNA45SN1_45S_50ntFlanks@12991.13011.21\|CTCCCTCGCTGCGATCTATTG | >hg38_21_+_8433172_8446622_RNA45SN1_45S_50ntFlanks | 12991 | 13011 | 21 | CTCCCTCGCTGCGATCTATTG | 1271.9 | -7.5 | 0.8 | -9.8 | 1.2E-22 | 8.6E-22 | BT20/MB468 |
| >MI0000750\|hsa-mir-26a-2&WithFlank&12\|-\|57824603\|57824698@20.39.20;MI0000083\|hsa-mir-26a-1&WithFlank&3\|+\|37969398\|37969486@16.35.20&[MIMAT0000082&hsa-miR-26a-5p&offsets\|0\|-2;m-33&12\|-\|57824658\|57824679&offsets\|0\|-2];[MIMAT0000082_1&hsa-miR-26a-5p&offsets\|0\|-2;m-34&3\|+\|37969413\|37969434&offsets\|0\|-2]\|TTCAAGTAATCCAGGATAGG | >MI0000750\|hsa-mir-26a-2&WithFlank&12\|-\|57824603\|57824698 | 20 | 39 | 20 | TTCAAGTAATCCAGGATAGG | 2525.3 | 8.0 | 0.8 | 9.8 | 1.4E-22 | 8.6E-22 | MB231/MB468 |
| >hg38_1_-_228634819_228635039_RNA5S12_5S_50ntFlanks@139.169.31\|GACCGCCTGGGAATACCGGGTGCTGTAGGCT | >hg38_1_-_228634819_228635039_RNA5S12_5S_50ntFlanks | 139 | 169 | 31 | GACCGCCTGGGAATACCGGGTGCTGTAGGCT | 1415.5 | 6.5 | 0.7 | 9.8 | 1.2E-22 | 8.7E-22 | BT20/MB468 |
| >hg38_21_+_8433172_8446622_RNA45SN1_45S_50ntFlanks@6648.6665.18\|GTACGACTCTTAGCGGTG | >hg38_21_+_8433172_8446622_RNA45SN1_45S_50ntFlanks | 6648 | 6665 | 18 | GTACGACTCTTAGCGGTG | 1287.6 | 5.3 | 0.5 | 9.8 | 1.2E-22 | 8.9E-22 | BT20/MB468 |
| >MI0000439\|hsa-mir-23b&WithFlank&9\|+\|95085202\|95085310@67.87.21&[MIMAT0000418&hsa-miR-23b-3p&offsets\|+3\|+1;m-42&9\|+\|95085265\|95085285&offsets\|+3\|+3]\|ACATTGCCAGGGATTACCACG | >MI0000439\|hsa-mir-23b&WithFlank&9\|+\|95085202\|95085310 | 67 | 87 | 21 | ACATTGCCAGGGATTACCACG | 1929.9 | 15.6 | 1.6 | 9.8 | 1.2E-22 | 9.0E-22 | BT20/MB468 |
| >hg38_21_+_8433172_8446622_RNA45SN1_45S_50ntFlanks@11580.11605.26\|GAACTGGTGCGGACCAGGGGAATCCG | >hg38_21_+_8433172_8446622_RNA45SN1_45S_50ntFlanks | 11580 | 11605 | 26 | GAACTGGTGCGGACCAGGGGAATCCG | 1105.7 | -5.9 | 0.6 | -9.8 | 1.2E-22 | 9.0E-22 | BT20/MB468 |
| >hg38_21_+_8433172_8446622_RNA45SN1_45S_50ntFlanks@12416.12440.25\|TTCGATGTCGGCTCTTCCTATCATT | >hg38_21_+_8433172_8446622_RNA45SN1_45S_50ntFlanks | 12416 | 12440 | 25 | TTCGATGTCGGCTCTTCCTATCATT | 2929.4 | 7.6 | 0.8 | 9.8 | 1.5E-22 | 9.1E-22 | MB231/MB468 |
| >hg38_21_+_8433172_8446622_RNA45SN1_45S_50ntFlanks@8380.8397.18\|CAAGAGGGCGTGAAACCG | >hg38_21_+_8433172_8446622_RNA45SN1_45S_50ntFlanks | 8380 | 8397 | 18 | CAAGAGGGCGTGAAACCG | 1190.9 | -7.3 | 0.7 | -9.8 | 1.3E-22 | 9.2E-22 | BT20/MB468 |
| >hg38_21_+_8433172_8446622_RNA45SN1_45S_50ntFlanks@9567.9592.26\|ACGTGCAAATCGGTCGTCCGACCTGG | >hg38_21_+_8433172_8446622_RNA45SN1_45S_50ntFlanks | 9567 | 9592 | 26 | ACGTGCAAATCGGTCGTCCGACCTGG | 744.7 | -6.6 | 0.7 | -9.8 | 1.3E-22 | 9.2E-22 | BT20/MB468 |

<table>
<tr><th></th><th></th><th></th><th></th><th></th><th></th><th></th><th></th><th></th><th></th><th></th><th></th><th></th></tr>
<tr><td>>MI0000762|hsa-mir-362&WithFlank&X|+|50008958|50009034@11.34.24&[MIMAT0000705&hsa-miR-362-5p&offsets|0|0;m-267&X|+|50008968|50008991&offsets|0|0]|AATCCTTGGAACCTAGGTGTGAGT</td><td>>MI0000762|hsa-mir-362&WithFlank&X|+|50008958|50009034</td><td>11</td><td>34</td><td>24</td><td>AATCCTTGGAACCTAGGTGTGAGT</td><td>871.2</td><td>6.7</td><td>0.7</td><td>9.8</td><td>1.3E-22</td><td>9.4E-22</td><td>BT20/MB468</td></tr>
<tr><td>>hg38_21_+_8433172_8446622_RNA45SN1_45S_50ntFlanks@5472.5501.30|ACGGCCCTGGCGGAGCGCTGAGAAGACGGT</td><td>>hg38_21_+_8433172_8446622_RNA45SN1_45S_50ntFlanks</td><td>5472</td><td>5501</td><td>30</td><td>ACGGCCCTGGCGGAGCGCTGAGAAGACGGT</td><td>1350.3</td><td>6.7</td><td>0.7</td><td>9.8</td><td>1.6E-22</td><td>9.4E-22</td><td>MB231/MB468</td></tr>
<tr><td>>hg38_21_+_8433172_8446622_RNA45SN1_45S_50ntFlanks@11323.11343.21|CGTCTCGTCGCGCGCGCGTCC</td><td>>hg38_21_+_8433172_8446622_RNA45SN1_45S_50ntFlanks</td><td>11323</td><td>11343</td><td>21</td><td>CGTCTCGTCGCGCGCGCGTCC</td><td>2751.6</td><td>-6.5</td><td>0.7</td><td>-9.8</td><td>1.6E-22</td><td>9.5E-22</td><td>MB231/MB468</td></tr>
<tr><td>>MI0000736|hsa-mir-30c-1&WithFlank&1|+|40757278|40757378@23.46.24(+1A);MI0000254|hsa-mir-30c-2&WithFlank&6|-|71376954|71377037@13.36.24(+1A)&[MIMAT0000244&hsa-miR-30c-5p&offsets|0|+1(+1A);m-57&1|+|40757300|40757323&offsets|0|0(+1A)];[MIMAT0000244_1&hsa-miR-30c-5p&offsets|0|+1(+1A);m-56&6|-|71377002|71377025&offsets|0|0(+1A)]|TGTAAACATCCTACACTCTCAGCTA</td><td>>MI0000736|hsa-mir-30c-1&WithFlank&1|+|40757278|40757378</td><td>23</td><td>46</td><td>24(+1A)</td><td>TGTAAACATCCTACACTCTCAGCTA</td><td>1240.0</td><td>6.7</td><td>0.7</td><td>9.8</td><td>1.6E-22</td><td>9.5E-22</td><td>MB231/MB468</td></tr>
<tr><td>>hg38_21_+_8433172_8446622_RNA45SN1_45S_50ntFlanks@12414.12448.35|CCTTCGATGTCGGCTCTTCCTATCATTGTGAAGCA</td><td>>hg38_21_+_8433172_8446622_RNA45SN1_45S_50ntFlanks</td><td>12414</td><td>12448</td><td>35</td><td>CCTTCGATGTCGGCTCTTCCTATCATTGTGAAGCA</td><td>668.6</td><td>9.6</td><td>1.0</td><td>9.8</td><td>1.6E-22</td><td>9.5E-22</td><td>MB231/MB468</td></tr>
<tr><td>>trnaMT_PheGAA_MT_+_577_647@35.59.25|GAAAATGTTTAGACGGGCTCACATC</td><td>>trnaMT_PheGAA_MT_+_577_647</td><td>35</td><td>59</td><td>25</td><td>GAAAATGTTTAGACGGGCTCACATC</td><td>854.0</td><td>10.0</td><td>1.0</td><td>9.8</td><td>1.4E-22</td><td>9.8E-22</td><td>BT20/MB468</td></tr>
<tr><td>>hg38_MT_+_1621_3279_RNR2_16S_50ntFlanks@493.517.25|GCTCTTTGGACACTAGGAAAAAACC</td><td>>hg38_MT_+_1621_3279_RNR2_16S_50ntFlanks</td><td>493</td><td>517</td><td>25</td><td>GCTCTTTGGACACTAGGAAAAAACC</td><td>1823.8</td><td>11.0</td><td>1.1</td><td>9.8</td><td>1.7E-22</td><td>9.9E-22</td><td>MB231/MB468</td></tr>
<tr><td>>hg38_21_+_8433172_8446622_RNA45SN1_45S_50ntFlanks@12415.12447.33|CTTCGATGTCGGCTCTTCCTATCATTGTGAAGC</td><td>>hg38_21_+_8433172_8446622_RNA45SN1_45S_50ntFlanks</td><td>12415</td><td>12447</td><td>33</td><td>CTTCGATGTCGGCTCTTCCTATCATTGTGAAGC</td><td>950.9</td><td>6.4</td><td>0.7</td><td>9.8</td><td>1.7E-22</td><td>1.0E-21</td><td>MB231/MB468</td></tr>
<tr><td>>MI0000746|hsa-mir-99b&WithFlank&19|+|51692606|51692687@13.31.19(+1A)&[MIMAT0000689&hsa-miR-99b-5p&offsets|0|-3(+1A);m-10&19|+|51692618|51692639&offsets|0|-3(+1A)]|CACCCGTAGAACCGACCTTA</td><td>>MI0000746|hsa-mir-99b&WithFlank&19|+|51692606|51692687</td><td>13</td><td>31</td><td>19(+1A)</td><td>CACCCGTAGAACCGACCTTA</td><td>644.7</td><td>14.0</td><td>1.4</td><td>9.8</td><td>1.4E-22</td><td>1.0E-21</td><td>BT20/MB468</td></tr>
<tr><td>>MI0000734|hsa-mir-106b&WithFlank&7|-|100093987|100094080@58.80.23&[MIMAT0004672&hsa-miR-106b-3p&offsets|0|+1;m-87&7|-|100094003|100094023&offsets|0|+2]|CCGCACTGTGGGTACTTGCTGCT</td><td>>MI0000734|hsa-mir-106b&WithFlank&7|-|100093987|100094080</td><td>58</td><td>80</td><td>23</td><td>CCGCACTGTGGGTACTTGCTGCT</td><td>662.8</td><td>7.4</td><td>0.8</td><td>9.8</td><td>1.8E-22</td><td>1.0E-21</td><td>MB231/MB468</td></tr>
<tr><td>>hg38_21_+_8433172_8446622_RNA45SN1_45S_50ntFlanks@4722.4758.37|TCATTAATCAAGAACGAAAGTCGGAGGTTCGAAGACG</td><td>>hg38_21_+_8433172_8446622_RNA45SN1_45S_50ntFlanks</td><td>4722</td><td>4758</td><td>37</td><td>TCATTAATCAAGAACGAAAGTCGGAGGTTCGAAGACG</td><td>2807.9</td><td>-6.5</td><td>0.7</td><td>-9.8</td><td>1.5E-22</td><td>1.1E-21</td><td>BT20/MB468</td></tr>
<tr><td>>hg38_21_+_8433172_8446622_RNA45SN1_45S_50ntFlanks@475.522.48|CTTCGTGATCGATGTGGTGACGTCGTGCTCTCCCGGGCCGGGTCCGAG</td><td>>hg38_21_+_8433172_8446622_RNA45SN1_45S_50ntFlanks</td><td>475</td><td>522</td><td>48</td><td>CTTCGTGATCGATGTGGTGACGTCGTGCTCTCCCGGGCCGGGTCCGAG</td><td>1922.6</td><td>11.5</td><td>1.2</td><td>9.8</td><td>1.8E-22</td><td>1.1E-21</td><td>MB231/MB468</td></tr>
<tr><td>>MI0000433|hsa-let-7g&WithFlank&3|-|52268272|52268367@11.32.22&[MIMAT0000414&hsa-let-7g-5p&offsets|0|0;m-58&3|-|52268336|52268357&offsets|0|0]|TGAGGTAGTAGTTTGTACAGTT</td><td>>MI0000433|hsa-let-7g&WithFlank&3|-|52268272|52268367</td><td>11</td><td>32</td><td>22</td><td>TGAGGTAGTAGTTTGTACAGTT</td><td>352727.8</td><td>7.6</td><td>0.8</td><td>9.8</td><td>1.8E-22</td><td>1.1E-21</td><td>MB231/MB468</td></tr>
<tr><td>>hg38_21_+_8433172_8446622_RNA45SN1_45S_50ntFlanks@9534.9566.33|AGGAAACTCTGGTGGAGGTCCGTAGCGGTCCTG</td><td>>hg38_21_+_8433172_8446622_RNA45SN1_45S_50ntFlanks</td><td>9534</td><td>9566</td><td>33</td><td>AGGAAACTCTGGTGGAGGTCCGTAGCGGTCCTG</td><td>1550.5</td><td>-7.1</td><td>0.7</td><td>-9.8</td><td>1.5E-22</td><td>1.1E-21</td><td>BT20/MB468</td></tr>
<tr><td>>hg38_MT_+_1621_3279_RNR2_16S_50ntFlanks@51.95.45|GCTAAACCTAGCCCCAAACCCACTCCACCTTACTACCAGACAACC</td><td>>hg38_MT_+_1621_3279_RNR2_16S_50ntFlanks</td><td>51</td><td>95</td><td>45</td><td>GCTAAACCTAGCCCCAAACCCACTCCACCTTACTACCAGACAACC</td><td>5284.9</td><td>12.7</td><td>1.3</td><td>9.7</td><td>1.9E-22</td><td>1.1E-21</td><td>MB231/MB468</td></tr>
<tr><td>>MI0000266|hsa-mir-10a&WithFlank&17|-|48579832|48579953@29.50.22&[MIMAT0000253&hsa-miR-10a-5p&offsets|+1|0;m-18&17|-|48579905|48579926&offsets|+1|+1]|ACCCTGTAGATCCGAATTTGTG</td><td>>MI0000266|hsa-mir-10a&WithFlank&17|-|48579832|48579953</td><td>29</td><td>50</td><td>22</td><td>ACCCTGTAGATCCGAATTTGTG</td><td>19407.0</td><td>13.3</td><td>1.4</td><td>9.8</td><td>1.6E-22</td><td>1.2E-21</td><td>BT20/MB468</td></tr>
<tr><td>>hg38_21_+_8433172_8446622_RNA45SN1_45S_50ntFlanks@11768.11790.23|CTAATTAGTGACGCGCATGAATG</td><td>>hg38_21_+_8433172_8446622_RNA45SN1_45S_50ntFlanks</td><td>11768</td><td>11790</td><td>23</td><td>CTAATTAGTGACGCGCATGAATG</td><td>976.4</td><td>-5.5</td><td>0.6</td><td>-9.7</td><td>2.0E-22</td><td>1.2E-21</td><td>MB231/MB468</td></tr>
<tr><td>>hg38_21_+_8433172_8446622_RNA45SN1_45S_50ntFlanks@11567.11584.18|AGCAGCCGACTTAGAACT</td><td>>hg38_21_+_8433172_8446622_RNA45SN1_45S_50ntFlanks</td><td>11567</td><td>11584</td><td>18</td><td>AGCAGCCGACTTAGAACT</td><td>3717.1</td><td>6.1</td><td>0.6</td><td>9.8</td><td>1.6E-22</td><td>1.2E-21</td><td>BT20/MB468</td></tr>
<tr><td>>trna119_LysCTT_1_-_145395522_145395594@1.36.36,trna11_LysCTT_5_-_180648979_180649051@1.36.36,trna13_LysCTT_6_+_26556774_26556846@1.36.36,trna7_LysCTT_16_+_3225692_3225764@1.36.36,trna9_LysCTT_5_+_180634755_180634827@1.36.36|GCCCGGCTAGCTCAGTCGGTAGAGCATGAGACTCTT</td><td>>trna119_LysCTT_1_-_145395522_145395594</td><td>1</td><td>36</td><td>36</td><td>GCCCGGCTAGCTCAGTCGGTAGAGCATGAGACTCTT</td><td>4489.5</td><td>8.8</td><td>0.9</td><td>9.8</td><td>1.7E-22</td><td>1.2E-21</td><td>BT20/MB468</td></tr>
<tr><td>>MI0000088|hsa-mir-30a&WithFlank&6|-|71403545|71403627@13.35.23&[MIMAT0000087&hsa-miR-30a-5p&offsets|+1|+2;m-15&6|-|71403595|71403616&offsets|+1|+2]|GTAAACATCCTCGACTGGAAGCT</td><td>>MI0000088|hsa-mir-30a&WithFlank&6|-|71403545|71403627</td><td>13</td><td>35</td><td>23</td><td>GTAAACATCCTCGACTGGAAGCT</td><td>5668.1</td><td>8.4</td><td>0.9</td><td>9.8</td><td>1.7E-22</td><td>1.2E-21</td><td>BT20/MB468</td></tr>
<tr><td>>MI0000433|hsa-let-7g&WithFlank&3|-|52268272|52268367@11.31.21(+1G)&[MIMAT0000414&hsa-let-7g-5p&offsets|0|-1(+1G);m-58&3|-|52268336|52268357&offsets|0|-1(+1G)]|TGAGGTAGTAGTTTGTACAGTG</td><td>>MI0000433|hsa-let-7g&WithFlank&3|-|52268272|52268367</td><td>11</td><td>31</td><td>21(+1G)</td><td>TGAGGTAGTAGTTTGTACAGTG</td><td>9104.6</td><td>7.5</td><td>0.8</td><td>9.7</td><td>2.1E-22</td><td>1.2E-21</td><td>MB231/MB468</td></tr>
<tr><td>>hg38_21_+_8433172_8446622_RNA45SN1_45S_50ntFlanks@4735.4765.31|ACGAAAGTCGGAGGTTCGAAGACGATCAGAT</td><td>>hg38_21_+_8433172_8446622_RNA45SN1_45S_50ntFlanks</td><td>4735</td><td>4765</td><td>31</td><td>ACGAAAGTCGGAGGTTCGAAGACGATCAGAT</td><td>556.7</td><td>9.6</td><td>1.0</td><td>9.7</td><td>2.1E-22</td><td>1.2E-21</td><td>MB231/MB468</td></tr>
<tr><td>>MI0000088|hsa-mir-30a&WithFlank&6|-|71403545|71403627@12.33.22(+1U)&[MIMAT0000087&hsa-miR-30a-5p&offsets|0|0(+1U);m-15&6|-|71403595|71403616&offsets|0|0(+1U)]|TGTAAACATCCTCGACTGGAAGT</td><td>>MI0000088|hsa-mir-30a&WithFlank&6|-|71403545|71403627</td><td>12</td><td>33</td><td>22(+1U)</td><td>TGTAAACATCCTCGACTGGAAGT</td><td>56345.7</td><td>9.5</td><td>1.0</td><td>9.8</td><td>1.8E-22</td><td>1.3E-21</td><td>BT20/MB468</td></tr>
<tr><td>>hg38_21_+_8433172_8446622_RNA45SN1_45S_50ntFlanks@9534.9571.38|AGGAAACTCTGGTGGAGGTCCGTAGCGGTCCTGACGTG</td><td>>hg38_21_+_8433172_8446622_RNA45SN1_45S_50ntFlanks</td><td>9534</td><td>9571</td><td>38</td><td>AGGAAACTCTGGTGGAGGTCCGTAGCGGTCCTGACGTG</td><td>7202.2</td><td>-5.6</td><td>0.6</td><td>-9.8</td><td>1.8E-22</td><td>1.3E-21</td><td>BT20/MB468</td></tr>
<tr><td>>MI0000266|hsa-mir-10a&WithFlank&17|-|48579832|48579953@28.50.23&[MIMAT0000253&hsa-miR-10a-5p&offsets|0|0;m-18&17|-|48579905|48579926&offsets|0|+1]|TACCCTGTAGATCCGAATTTGTG</td><td>>MI0000266|hsa-mir-10a&WithFlank&17|-|48579832|48579953</td><td>28</td><td>50</td><td>23</td><td>TACCCTGTAGATCCGAATTTGTG</td><td>19380.3</td><td>13.1</td><td>1.3</td><td>9.7</td><td>2.2E-22</td><td>1.3E-21</td><td>MB231/MB468</td></tr>
<tr><td>>trna10_LysCTT_16_+_3241501_3241573@4.35.32,trna13_LysCTT_14_-_58706613_58706685@4.35.32,trna2_LysCTT_15_+_79152904_79152976@4.35.32|CGGCTAGCTCAGTCGGTAGAGCATGGGACTCT</td><td>>trna10_LysCTT_16_+_3241501_3241573</td><td>4</td><td>35</td><td>32</td><td>CGGCTAGCTCAGTCGGTAGAGCATGGGACTCT</td><td>1521.7</td><td>11.7</td><td>1.2</td><td>9.7</td><td>2.2E-22</td><td>1.3E-21</td><td>MB231/MB468</td></tr>
<tr><td>>hg38_21_+_8433172_8446622_RNA45SN1_45S_50ntFlanks@476.502.27|TTCGTGATCGATGTGGTGACGTCGTGC</td><td>>hg38_21_+_8433172_8446622_RNA45SN1_45S_50ntFlanks</td><td>476</td><td>502</td><td>27</td><td>TTCGTGATCGATGTGGTGACGTCGTGC</td><td>1439.9</td><td>6.3</td><td>0.7</td><td>9.8</td><td>1.8E-22</td><td>1.3E-21</td><td>BT20/MB468</td></tr>
<tr><td>>hg38_21_+_8433172_8446622_RNA45SN1_45S_50ntFlanks@6786.6807.22|TACGCCTGTCTGAGCGTCGCTT</td><td>>hg38_21_+_8433172_8446622_RNA45SN1_45S_50ntFlanks</td><td>6786</td><td>6807</td><td>22</td><td>TACGCCTGTCTGAGCGTCGCTT</td><td>521.9</td><td>6.6</td><td>0.7</td><td>9.7</td><td>1.8E-22</td><td>1.3E-21</td><td>BT20/MB468</td></tr>
<tr><td>>hg38_21_+_8433172_8446622_RNA45SN1_45S_50ntFlanks@8356.8378.23|GAAAAGAACTTTGAAGAGAGAGT</td><td>>hg38_21_+_8433172_8446622_RNA45SN1_45S_50ntFlanks</td><td>8356</td><td>8378</td><td>23</td><td>GAAAAGAACTTTGAAGAGAGAGT</td><td>4697.8</td><td>-5.2</td><td>0.5</td><td>-9.7</td><td>1.9E-22</td><td>1.3E-21</td><td>BT20/MB468</td></tr>
<tr><td>>hg38_MT_+_598_1651_RNR1_12S_50ntFlanks@368.404.37|CAATAAAGCTAAAACTCACCTGAGTTGTAAAAAACTC</td><td>>hg38_MT_+_598_1651_RNR1_12S_50ntFlanks</td><td>368</td><td>404</td><td>37</td><td>CAATAAAGCTAAAACTCACCTGAGTTGTAAAAAACTC</td><td>762.7</td><td>11.4</td><td>1.2</td><td>9.7</td><td>1.9E-22</td><td>1.3E-21</td><td>BT20/MB468</td></tr>
<tr><td>>hg38_21_+_8433172_8446622_RNA45SN1_45S_50ntFlanks@6651.6678.28|CGACTCTTAGCGGTGGATCACTCGGCTC</td><td>>hg38_21_+_8433172_8446622_RNA45SN1_45S_50ntFlanks</td><td>6651</td><td>6678</td><td>28</td><td>CGACTCTTAGCGGTGGATCACTCGGCTC</td><td>7583.8</td><td>4.0</td><td>0.4</td><td>9.7</td><td>1.9E-22</td><td>1.3E-21</td><td>BT20/MB468</td></tr>
<tr><td>>trnaMT_ValTAC_MT_+_1602_1670@48.70.23|CAACTTAACTTGACCGCTCTGAC</td><td>>trnaMT_ValTAC_MT_+_1602_1670</td><td>48</td><td>70</td><td>23</td><td>CAACTTAACTTGACCGCTCTGAC</td><td>469.0</td><td>11.1</td><td>1.1</td><td>9.7</td><td>1.9E-22</td><td>1.4E-21</td><td>BT20/MB468</td></tr>
<tr><td>>hg38_21_+_8433172_8446622_RNA45SN1_45S_50ntFlanks@10351.10374.24|CGAGAACTTTGAAGGCCGAAGTGG</td><td>>hg38_21_+_8433172_8446622_RNA45SN1_45S_50ntFlanks</td><td>10351</td><td>10374</td><td>24</td><td>CGAGAACTTTGAAGGCCGAAGTGG</td><td>1922.2</td><td>-6.5</td><td>0.7</td><td>-9.7</td><td>2.0E-22</td><td>1.4E-21</td><td>BT20/MB468</td></tr>
<tr><td>>MI0000077|hsa-mir-21&WithFlank&17|+|59841260|59841343@14.35.22(+2U)&[MIMAT0000076&hsa-miR-21-5p&offsets|0|0(+2U);m-1&17|+|59841273|59841294&offsets|0|0(+2U)]|TAGCTTATCAGACTGATGTTGATT</td><td>>MI0000077|hsa-mir-21&WithFlank&17|+|59841260|59841343</td><td>14</td><td>35</td><td>22(+2U)</td><td>TAGCTTATCAGACTGATGTTGATT</td><td>5511.6</td><td>10.0</td><td>1.0</td><td>9.7</td><td>2.0E-22</td><td>1.4E-21</td><td>BT20/MB468</td></tr>
<tr><td>>MI0000465|hsa-mir-191&WithFlank&3|-|49020612|49020715@22.41.20(+1U)&[MIMAT0000440&hsa-miR-191-5p&offsets|0|-3(+1U);m-48&3|-|49020672|49020694&offsets|0|-3(+1U)]|CAACGGAATCCCAAAAGCAGT</td><td>>MI0000465|hsa-mir-191&WithFlank&3|-|49020612|49020715</td><td>22</td><td>41</td><td>20(+1U)</td><td>CAACGGAATCCCAAAAGCAGT</td><td>412.8</td><td>13.3</td><td>1.4</td><td>9.7</td><td>2.0E-22</td><td>1.4E-21</td><td>BT20/MB468</td></tr>
<tr><td>>MI0000255|hsa-mir-30d&WithFlank&8|-|134804870|134804951@12.35.24(+1C)&[MIMAT0000245&hsa-miR-30d-5p&offsets|0|+2(+1C);m-21&8|-|134804919|134804940&offsets|0|+2(+1C)]|TGTAAACATCCCCGACTGGAAGCTC</td><td>>MI0000255|hsa-mir-30d&WithFlank&8|-|134804870|134804951</td><td>12</td><td>35</td><td>24(+1C)</td><td>TGTAAACATCCCCGACTGGAAGCTC</td><td>1406.4</td><td>9.9</td><td>1.0</td><td>9.7</td><td>2.4E-22</td><td>1.4E-21</td><td>MB231/MB468</td></tr>
<tr><td>>am_trnaMT_HisGTG_MT_+_12138_12206@26.44.19|AGATTGTGAATCTGACAAC</td><td>>am_trnaMT_HisGTG_MT_+_12138_12206</td><td>26</td><td>44</td><td>19</td><td>AGATTGTGAATCTGACAAC</td><td>490.3</td><td>7.2</td><td>0.7</td><td>9.7</td><td>2.5E-22</td><td>1.4E-21</td><td>MB231/MB468</td></tr>
<tr><td>>MI0000471|hsa-mir-126&WithFlank&9|+|136670596|136670692@21.41.21&[MIMAT0000444&hsa-miR-126-5p&offsets|0|0;m-114&9|+|136670616|136670636&offsets|0|0]|CATTATTACTTTTGGTACGCG</td><td>>MI0000471|hsa-mir-126&WithFlank&9|+|136670596|136670692</td><td>21</td><td>41</td><td>21</td><td>CATTATTACTTTTGGTACGCG</td><td>965.9</td><td>14.6</td><td>1.5</td><td>9.7</td><td>2.0E-22</td><td>1.4E-21</td><td>BT20/MB468</td></tr>
<tr><td>>hg38_1_-_228634819_228635039_RNA5S12_5S_50ntFlanks@131.171.41|GGATGGGAGACCGCCTGGGAATACCGGGTGCTGTAGGCTTT</td><td>>hg38_1_-_228634819_228635039_RNA5S12_5S_50ntFlanks</td><td>131</td><td>171</td><td>41</td><td>GGATGGGAGACCGCCTGGGAATACCGGGTGCTGTAGGCTTT</td><td>1953.6</td><td>12.3</td><td>1.3</td><td>9.7</td><td>2.5E-22</td><td>1.5E-21</td><td>MB231/MB468</td></tr>
<tr><td>>hg38_21_+_8433172_8446622_RNA45SN1_45S_50ntFlanks@12435.12455.21|ATCATTGTGAAGCAGAATTCA</td><td>>hg38_21_+_8433172_8446622_RNA45SN1_45S_50ntFlanks</td><td>12435</td><td>12455</td><td>21</td><td>ATCATTGTGAAGCAGAATTCA</td><td>1963.2</td><td>-12.9</td><td>1.3</td><td>-9.7</td><td>2.1E-22</td><td>1.5E-21</td><td>BT20/MB468</td></tr>
<tr><td>>hg38_21_+_8433172_8446622_RNA45SN1_45S_50ntFlanks@9575.9596.22|ATCGGTCGTCCGACCTGGGTAT</td><td>>hg38_21_+_8433172_8446622_RNA45SN1_45S_50ntFlanks</td><td>9575</td><td>9596</td><td>22</td><td>ATCGGTCGTCCGACCTGGGTAT</td><td>1930.1</td><td>-12.9</td><td>1.3</td><td>-9.7</td><td>2.1E-22</td><td>1.5E-21</td><td>BT20/MB468</td></tr>
<tr><td>>hg38_21_+_8433172_8446622_RNA45SN1_45S_50ntFlanks@10345.10382.38|TTCAAACGAGAACTTTGAAGGCCGAAGTGGAGAAGGGT</td><td>>hg38_21_+_8433172_8446622_RNA45SN1_45S_50ntFlanks</td><td>10345</td><td>10382</td><td>38</td><td>TTCAAACGAGAACTTTGAAGGCCGAAGTGGAGAAGGGT</td><td>270.4</td><td>8.8</td><td>0.9</td><td>9.7</td><td>2.6E-22</td><td>1.5E-21</td><td>MB231/MB468</td></tr>
</table>

<table>
<tr><td>>hg38_21_+_8433172_8446622_RNA45SN1_45S_50ntFlanks@5446.5466.21|CGGATCGGCCCCGCCGGGGTC</td><td>>hg38_21_+_8433172_8446622_RNA45SN1_45S_50ntFlanks</td><td>5446</td><td>5466</td><td>21</td><td>CGGATCGGCCCCGCCGGGGTC</td><td>3078.4</td><td>-8.1</td><td>0.8</td><td>-9.7</td><td>2.6E-22</td><td>1.5E-21</td><td>MB231/MB468</td></tr>
<tr><td>>MI0000071|hsa-mir-17&WithFlank&13|+|91350599|91350694@20.41.22(+1A)&[MIMAT0000070&hsa-miR-17-5p&offsets|0|-1(+1A);m-88&13|+|91350618|91350640&offsets|0|-1(+1A)]|CAAAGTGCTTACAGTGCAGGTAA</td><td>>MI0000071|hsa-mir-17&WithFlank&13|+|91350599|91350694</td><td>20</td><td>41</td><td>22(+1A)</td><td>CAAAGTGCTTACAGTGCAGGTAA</td><td>2766.0</td><td>6.5</td><td>0.7</td><td>9.7</td><td>2.6E-22</td><td>1.5E-21</td><td>MB231/MB468</td></tr>
<tr><td>>MI0003591|hsa-mir-584&WithFlank&5|-|149062307|149062415@22.42.21(+2U)&[MIMAT0003249&hsa-miR-584-5p&offsets|0|-1(+2U);m-167&5|-|149062374|149062394&offsets|0|0(+2U)]|TTATGGTTTGCCTGGGACTGATT</td><td>>MI0003591|hsa-mir-584&WithFlank&5|-|149062307|149062415</td><td>22</td><td>42</td><td>21(+2U)</td><td>TTATGGTTTGCCTGGGACTGATT</td><td>720.4</td><td>11.9</td><td>1.2</td><td>9.7</td><td>2.6E-22</td><td>1.5E-21</td><td>MB231/MB468</td></tr>
<tr><td>>MI0000093|hsa-mir-92a-1&WithFlank&13|+|91351308|91351397@54.75.22;MI0000094|hsa-mir-92a-2&WithFlank&X|-|134169532|134169618@54.75.22&[MIMAT0000092&hsa-miR-92a-3p&offsets|0|0;m-9&13|+|91351361|91351382&offsets|0|0];[MIMAT0000092_1&hsa-miR-92a-3p&offsets|0|0;m-11&X|-|134169544|134169565&offsets|0|0]|TATTGCACTTGTCCCGGCCTGT</td><td>>MI0000093|hsa-mir-92a-1&WithFlank&13|+|91351308|91351397</td><td>54</td><td>75</td><td>22</td><td>TATTGCACTTGTCCCGGCCTGT</td><td>73217.1</td><td>6.2</td><td>0.6</td><td>9.7</td><td>2.2E-22</td><td>1.5E-21</td><td>BT20/MB468</td></tr>
<tr><td>>hg38_21_+_8433172_8446622_RNA45SN1_45S_50ntFlanks@9545.9574.30|GTGGAGGTCCGTAGCGGTCCTGACGTGCAA</td><td>>hg38_21_+_8433172_8446622_RNA45SN1_45S_50ntFlanks</td><td>9545</td><td>9574</td><td>30</td><td>GTGGAGGTCCGTAGCGGTCCTGACGTGCAA</td><td>615.1</td><td>-6.1</td><td>0.6</td><td>-9.7</td><td>2.2E-22</td><td>1.5E-21</td><td>BT20/MB468</td></tr>
<tr><td>>MI0000750|hsa-mir-26a-2&WithFlank&12|-|57824603|57824698@20.41.22(+1C);MI0000083|hsa-mir-26a-1&WithFlank&3|+|37969398|37969486@16.37.22(+1C)&[MIMAT0000082&hsa-miR-26a-5p&offsets|0|0(+1C);m-33&12|-|57824658|57824679&offsets|0|0(+1C)];[MIMAT0000082_1&hsa-miR-26a-5p&offsets|0|0(+1C);m-34&3|+|37969413|37969434&offsets|0|0(+1C)]|TTCAAGTAATCCAGGATAGGCTC</td><td>>MI0000750|hsa-mir-26a-2&WithFlank&12|-|57824603|57824698</td><td>20</td><td>41</td><td>22(+1C)</td><td>TTCAAGTAATCCAGGATAGGCTC</td><td>2352.2</td><td>8.2</td><td>0.8</td><td>9.7</td><td>2.7E-22</td><td>1.5E-21</td><td>MB231/MB468</td></tr>
<tr><td>>hg38_21_+_8433172_8446622_RNA45SN1_45S_50ntFlanks@10402.10420.19|AACATGGGTCAGTCGGTCC</td><td>>hg38_21_+_8433172_8446622_RNA45SN1_45S_50ntFlanks</td><td>10402</td><td>10420</td><td>19</td><td>AACATGGGTCAGTCGGTCC</td><td>1347.1</td><td>-6.8</td><td>0.7</td><td>-9.7</td><td>2.2E-22</td><td>1.5E-21</td><td>BT20/MB468</td></tr>
<tr><td>>MI0000077|hsa-mir-21&WithFlank&17|+|59841260|59841343@13.36.24&[MIMAT0000076&hsa-miR-21-5p&offsets|-1|+1;m-1&17|+|59841273|59841294&offsets|-1|+1]|GTAGCTTATCAGACTGATGTTGAC</td><td>>MI0000077|hsa-mir-21&WithFlank&17|+|59841260|59841343</td><td>13</td><td>36</td><td>24</td><td>GTAGCTTATCAGACTGATGTTGAC</td><td>961.4</td><td>7.0</td><td>0.7</td><td>9.7</td><td>2.7E-22</td><td>1.6E-21</td><td>MB231/MB468</td></tr>
<tr><td>>MI0000286|hsa-mir-210&WithFlank&11|-|568083|568204@72.93.22&[MIMAT0000267&hsa-miR-210-3p&offsets|0|0;m-102&11|-|568112|568133&offsets|0|0]|CTGTGCGTGTGACAGCGGCTGA</td><td>>MI0000286|hsa-mir-210&WithFlank&11|-|568083|568204</td><td>72</td><td>93</td><td>22</td><td>CTGTGCGTGTGACAGCGGCTGA</td><td>811.6</td><td>7.3</td><td>0.8</td><td>9.7</td><td>2.7E-22</td><td>1.6E-21</td><td>MB231/MB468</td></tr>
<tr><td>>hg38_21_+_8433172_8446622_RNA45SN1_45S_50ntFlanks@8381.8402.22|AAGAGGGCGTGAAACCGTTAAG</td><td>>hg38_21_+_8433172_8446622_RNA45SN1_45S_50ntFlanks</td><td>8381</td><td>8402</td><td>22</td><td>AAGAGGGCGTGAAACCGTTAAG</td><td>1578.7</td><td>-7.6</td><td>0.8</td><td>-9.7</td><td>2.2E-22</td><td>1.6E-21</td><td>BT20/MB468</td></tr>
<tr><td>>MI0000088|hsa-mir-30a&WithFlank&6|-|71403545|71403627@53.74.22&[MIMAT0000088&hsa-miR-30a-3p&offsets|0|0;m-38&6|-|71403554|71403575&offsets|0|0]|CTTTCAGTCGGATGTTTGCAGC</td><td>>MI0000088|hsa-mir-30a&WithFlank&6|-|71403545|71403627</td><td>53</td><td>74</td><td>22</td><td>CTTTCAGTCGGATGTTTGCAGC</td><td>7841.7</td><td>6.4</td><td>0.7</td><td>9.7</td><td>2.3E-22</td><td>1.6E-21</td><td>BT20/MB468</td></tr>
<tr><td>>hg38_1_-_228634819_228635039_RNA5S12_5S_50ntFlanks@145.170.26|CTGGGAATACCGGGTGCTGTAGGCTT</td><td>>hg38_1_-_228634819_228635039_RNA5S12_5S_50ntFlanks</td><td>145</td><td>170</td><td>26</td><td>CTGGGAATACCGGGTGCTGTAGGCTT</td><td>2539.2</td><td>7.8</td><td>0.8</td><td>9.7</td><td>2.8E-22</td><td>1.6E-21</td><td>MB231/MB468</td></tr>
<tr><td>>hg38_21_+_8433172_8446622_RNA45SN1_45S_50ntFlanks@12971.12990.20|TTTCGTACGTAGCAGAGCAG</td><td>>hg38_21_+_8433172_8446622_RNA45SN1_45S_50ntFlanks</td><td>12971</td><td>12990</td><td>20</td><td>TTTCGTACGTAGCAGAGCAG</td><td>433.5</td><td>-5.8</td><td>0.6</td><td>-9.7</td><td>2.8E-22</td><td>1.6E-21</td><td>MB231/MB468</td></tr>
<tr><td>>hg38_21_+_8433172_8446622_RNA45SN1_45S_50ntFlanks@10402.10419.18|AACATGGGTCAGTCGGTC</td><td>>hg38_21_+_8433172_8446622_RNA45SN1_45S_50ntFlanks</td><td>10402</td><td>10419</td><td>18</td><td>AACATGGGTCAGTCGGTC</td><td>625.2</td><td>-6.3</td><td>0.7</td><td>-9.7</td><td>2.4E-22</td><td>1.7E-21</td><td>BT20/MB468</td></tr>
<tr><td>>MI0000079|hsa-mir-23a&WithFlank&19|-|13836581|13836665@54.74.21&[MIMAT0000078&hsa-miR-23a-3p&offsets|+3|+3;m-31&19|-|13836595|13836615&offsets|+3|+3]|ACATTGCCAGGGATTTCCAAC</td><td>>MI0000079|hsa-mir-23a&WithFlank&19|-|13836581|13836665</td><td>54</td><td>74</td><td>21</td><td>ACATTGCCAGGGATTTCCAAC</td><td>461.0</td><td>13.5</td><td>1.4</td><td>9.7</td><td>2.4E-22</td><td>1.7E-21</td><td>BT20/MB468</td></tr>
<tr><td>>hg38_21_+_8433172_8446622_RNA45SN1_45S_50ntFlanks@6650.6668.19|ACGACTCTTAGCGGTGGAT</td><td>>hg38_21_+_8433172_8446622_RNA45SN1_45S_50ntFlanks</td><td>6650</td><td>6668</td><td>19</td><td>ACGACTCTTAGCGGTGGAT</td><td>1607.8</td><td>4.7</td><td>0.5</td><td>9.7</td><td>2.4E-22</td><td>1.7E-21</td><td>BT20/MB468</td></tr>
<tr><td>>hg38_21_+_8433172_8446622_RNA45SN1_45S_50ntFlanks@12405.12433.29|CTTTTTGATCCTTCGATGTCGGCTCTTCC</td><td>>hg38_21_+_8433172_8446622_RNA45SN1_45S_50ntFlanks</td><td>12405</td><td>12433</td><td>29</td><td>CTTTTTGATCCTTCGATGTCGGCTCTTCC</td><td>1960.7</td><td>-4.7</td><td>0.5</td><td>-9.7</td><td>2.4E-22</td><td>1.7E-21</td><td>BT20/MB468</td></tr>
<tr><td>>MI0000093|hsa-mir-92a-1&WithFlank&13|+|91351308|91351397@54.75.22(+1A);MI0000094|hsa-mir-92a-2&WithFlank&X|-|134169532|134169618@54.75.22(+1A)&[MIMAT0000092&hsa-miR-92a-3p&offsets|0|0(+1A);m-9&13|+|91351361|91351382&offsets|0|0(+1A)];[MIMAT0000092_1&hsa-miR-92a-3p&offsets|0|0(+1A);m-11&X|-|134169544|134169565&offsets|0|0(+1A)]|TATTGCACTTGTCCCGGCCTGTA</td><td>>MI0000093|hsa-mir-92a-1&WithFlank&13|+|91351308|91351397</td><td>54</td><td>75</td><td>22(+1A)</td><td>TATTGCACTTGTCCCGGCCTGTA</td><td>25633.7</td><td>6.8</td><td>0.7</td><td>9.7</td><td>3.1E-22</td><td>1.8E-21</td><td>MB231/MB468</td></tr>
<tr><td>>hg38_21_+_8433172_8446622_RNA45SN1_45S_50ntFlanks@4723.4758.36|CATTAATCAAGAACGAAAGTCGGAGGTTCGAAGACG</td><td>>hg38_21_+_8433172_8446622_RNA45SN1_45S_50ntFlanks</td><td>4723</td><td>4758</td><td>36</td><td>CATTAATCAAGAACGAAAGTCGGAGGTTCGAAGACG</td><td>2829.6</td><td>-5.4</td><td>0.6</td><td>-9.7</td><td>2.6E-22</td><td>1.8E-21</td><td>BT20/MB468</td></tr>
<tr><td>>hg38_21_+_8433172_8446622_RNA45SN1_45S_50ntFlanks@6661.6683.23|CGGTGGATCACTCGGCTCGTGCG</td><td>>hg38_21_+_8433172_8446622_RNA45SN1_45S_50ntFlanks</td><td>6661</td><td>6683</td><td>23</td><td>CGGTGGATCACTCGGCTCGTGCG</td><td>4267.5</td><td>7.2</td><td>0.7</td><td>9.7</td><td>3.2E-22</td><td>1.8E-21</td><td>MB231/MB468</td></tr>
<tr><td>>MI0003599|hsa-mir-589&WithFlank&7|-|5495813|5495923@30.52.23&[MIMAT0004799&hsa-miR-589-5p&offsets|0|+1;m-162&7|-|5495874|5495894&offsets|0|+2]|TGAGAACCACGTCTGCTCTGAGC</td><td>>MI0003599|hsa-mir-589&WithFlank&7|-|5495813|5495923</td><td>30</td><td>52</td><td>23</td><td>TGAGAACCACGTCTGCTCTGAGC</td><td>1920.8</td><td>11.6</td><td>1.2</td><td>9.7</td><td>3.2E-22</td><td>1.8E-21</td><td>MB231/MB468</td></tr>
<tr><td>>hg38_21_+_8433172_8446622_RNA45SN1_45S_50ntFlanks@7983.8008.26|CAGATCAGACGTGGCGACCCGCTGAA</td><td>>hg38_21_+_8433172_8446622_RNA45SN1_45S_50ntFlanks</td><td>7983</td><td>8008</td><td>26</td><td>CAGATCAGACGTGGCGACCCGCTGAA</td><td>1579.6</td><td>-5.7</td><td>0.6</td><td>-9.7</td><td>2.6E-22</td><td>1.8E-21</td><td>BT20/MB468</td></tr>
<tr><td>>hg38_21_+_8433172_8446622_RNA45SN1_45S_50ntFlanks@10845.10864.20|ACTTCGGGATAAGGATTGGC</td><td>>hg38_21_+_8433172_8446622_RNA45SN1_45S_50ntFlanks</td><td>10845</td><td>10864</td><td>20</td><td>ACTTCGGGATAAGGATTGGC</td><td>464.2</td><td>-5.9</td><td>0.6</td><td>-9.7</td><td>2.6E-22</td><td>1.8E-21</td><td>BT20/MB468</td></tr>
<tr><td>>MI0000087|hsa-mir-29a&WithFlank&7|-|130876741|130876816@47.67.21(+1G)&[MIMAT0000086&hsa-miR-29a-3p&offsets|-1|-2(+1G);m-26&7|-|130876749|130876769&offsets|-1|-1(+1G)]|CTAGCACCATCTGAAATCGGTG</td><td>>MI0000087|hsa-mir-29a&WithFlank&7|-|130876741|130876816</td><td>47</td><td>67</td><td>21(+1G)</td><td>CTAGCACCATCTGAAATCGGTG</td><td>889.5</td><td>9.6</td><td>1.0</td><td>9.7</td><td>3.2E-22</td><td>1.8E-21</td><td>MB231/MB468</td></tr>
<tr><td>>trna119_LysCTT_1_-_145395522_145395594@1.34.34,trna11_LysCTT_5_-_180648979_180649051@1.34.34,trna13_LysCTT_6_+_26556774_26556846@1.34.34,trna7_LysCTT_16_+_3225692_3225764@1.34.34,trna9_LysCTT_5_+_180634755_180634827@1.34.34|GCCCGGCTAGCTCAGTCGGTAGAGCATGAGACTC</td><td>>trna119_LysCTT_1_-_145395522_145395594</td><td>1</td><td>34</td><td>34</td><td>GCCCGGCTAGCTCAGTCGGTAGAGCATGAGACTC</td><td>3689.1</td><td>7.8</td><td>0.8</td><td>9.7</td><td>2.7E-22</td><td>1.9E-21</td><td>BT20/MB468</td></tr>
<tr><td>>trnaMT_GluTTC_MT_-_14674_14742@39.72.34,trnalookalike8_GluTTC_5_-_93905172_93905240@39.72.34|ATTGGTCGTGGTTGTAGTCCGTGCGAGAATACCA</td><td>>trnaMT_GluTTC_MT_-_14674_14742</td><td>39</td><td>72</td><td>34</td><td>ATTGGTCGTGGTTGTAGTCCGTGCGAGAATACCA</td><td>18494.8</td><td>8.2</td><td>0.8</td><td>9.7</td><td>2.7E-22</td><td>1.9E-21</td><td>BT20/MB468</td></tr>
<tr><td>>MI0000076|hsa-mir-20a&WithFlank&13|+|91351059|91351141@14.35.22(+1U)&[MIMAT0000075&hsa-miR-20a-5p&offsets|0|-1(+1U);m-96&13|+|91351072|91351094&offsets|0|-1(+1U)]|TAAAGTGCTTATAGTGCAGGTAT</td><td>>MI0000076|hsa-mir-20a&WithFlank&13|+|91351059|91351141</td><td>14</td><td>35</td><td>22(+1U)</td><td>TAAAGTGCTTATAGTGCAGGTAT</td><td>618.6</td><td>7.3</td><td>0.8</td><td>9.7</td><td>3.2E-22</td><td>1.9E-21</td><td>MB231/MB468</td></tr>
<tr><td>>MI0000433|hsa-let-7g&WithFlank&3|-|52268272|52268367@11.28.18&[MIMAT0000414&hsa-let-7g-5p&offsets|0|-4;m-58&3|-|52268336|52268357&offsets|0|-4]|TGAGGTAGTAGTTTGTAC</td><td>>MI0000433|hsa-let-7g&WithFlank&3|-|52268272|52268367</td><td>11</td><td>28</td><td>18</td><td>TGAGGTAGTAGTTTGTAC</td><td>10082.8</td><td>8.3</td><td>0.9</td><td>9.7</td><td>3.3E-22</td><td>1.9E-21</td><td>MB231/MB468</td></tr>
<tr><td>>MI0003820|hsa-mir-454&WithFlank&17|-|59137752|59137878@70.93.24&[MIMAT0003885&hsa-miR-454-3p&offsets|0|+1;m-272&17|-|59137787|59137809&offsets|0|+1]|TAGTGCAATATTGCTTATAGGGTT</td><td>>MI0003820|hsa-mir-454&WithFlank&17|-|59137752|59137878</td><td>70</td><td>93</td><td>24</td><td>TAGTGCAATATTGCTTATAGGGTT</td><td>1739.3</td><td>7.8</td><td>0.8</td><td>9.7</td><td>3.3E-22</td><td>1.9E-21</td><td>MB231/MB468</td></tr>
<tr><td>>MI0000108|hsa-mir-103a-2&WithFlank&20|+|3917488|3917577@54.76.23(+1U);MI0000109|hsa-mir-103a-1&WithFlank&5|-|168560890|168560979@54.76.23(+1U)&[MIMAT0000101&hsa-miR-103a-3p&offsets|0|0(+1U);m-12&20|+|3917541|3917563&offsets|0|0(+1U)];[MIMAT0000101_1&hsa-miR-103a-3p&offsets|0|0(+1U);m-13&5|-|168560904|168560926&offsets|0|0(+1U);m-44220&5|-|168560908|168560926&offsets|0|+4(+1U)]|AGCAGCATTGTACAGGGCTATGAT</td><td>>MI0000108|hsa-mir-103a-2&WithFlank&20|+|3917488|3917577</td><td>54</td><td>76</td><td>23(+1U)</td><td>AGCAGCATTGTACAGGGCTATGAT</td><td>7032.8</td><td>7.3</td><td>0.8</td><td>9.7</td><td>2.7E-22</td><td>1.9E-21</td><td>BT20/MB468</td></tr>
<tr><td>>hg38_21_+_8433172_8446622_RNA45SN1_45S_50ntFlanks@12453.12476.24|TCACCAAGCGTTGGATTGTTCACC</td><td>>hg38_21_+_8433172_8446622_RNA45SN1_45S_50ntFlanks</td><td>12453</td><td>12476</td><td>24</td><td>TCACCAAGCGTTGGATTGTTCACC</td><td>838.0</td><td>5.5</td><td>0.6</td><td>9.7</td><td>3.3E-22</td><td>1.9E-21</td><td>MB231/MB468</td></tr>
<tr><td>>hg38_21_+_8433172_8446622_RNA45SN1_45S_50ntFlanks@6741.6770.30|ATCGACACTTCGAACGCACTTGCGGCCCCG</td><td>>hg38_21_+_8433172_8446622_RNA45SN1_45S_50ntFlanks</td><td>6741</td><td>6770</td><td>30</td><td>ATCGACACTTCGAACGCACTTGCGGCCCCG</td><td>1338.4</td><td>-6.0</td><td>0.6</td><td>-9.7</td><td>3.4E-22</td><td>1.9E-21</td><td>MB231/MB468</td></tr>
<tr><td>>hg38_21_+_8433172_8446622_RNA45SN1_45S_50ntFlanks@9575.9597.23|ATCGGTCGTCCGACCTGGGTATA</td><td>>hg38_21_+_8433172_8446622_RNA45SN1_45S_50ntFlanks</td><td>9575</td><td>9597</td><td>23</td><td>ATCGGTCGTCCGACCTGGGTATA</td><td>2258.8</td><td>-13.1</td><td>1.4</td><td>-9.7</td><td>2.8E-22</td><td>1.9E-21</td><td>BT20/MB468</td></tr>
<tr><td>>MI0000065|hsa-let-7d&WithFlank&9|+|94178828|94178926@68.88.21&[MIMAT0004484&hsa-let-7d-3p&offsets|0|-1;m-98&9|+|94178895|94178916&offsets|0|-1]|CTATACGACCTGCTGCCTTTC</td><td>>MI0000065|hsa-let-7d&WithFlank&9|+|94178828|94178926</td><td>68</td><td>88</td><td>21</td><td>CTATACGACCTGCTGCCTTTC</td><td>952.5</td><td>7.5</td><td>0.8</td><td>9.7</td><td>3.5E-22</td><td>2.0E-21</td><td>MB231/MB468</td></tr>
<tr><td>>MI0003760|hsa-mir-671&WithFlank&7|+|151238415|151238544@35.58.24&[MIMAT0003880&hsa-miR-671-5p&offsets|0|+1;m-316&7|+|151238449|151238472&offsets|0|0]|AGGAAGCCCTGGAGGGGCTGGAGG</td><td>>MI0003760|hsa-mir-671&WithFlank&7|+|151238415|151238544</td><td>35</td><td>58</td><td>24</td><td>AGGAAGCCCTGGAGGGGCTGGAGG</td><td>647.1</td><td>7.5</td><td>0.8</td><td>9.7</td><td>3.5E-22</td><td>2.0E-21</td><td>MB231/MB468</td></tr>
<tr><td>>MI0000283|hsa-mir-203a&WithFlank&14|+|104117399|104117520@72.92.21&[MIMAT0000264&hsa-miR-203a-3p&offsets|+1|0;m-79&14|+|104117469|104117490&offsets|+1|0]|TGAAATGTTTAGGACCACTAG</td><td>>MI0000283|hsa-mir-203a&WithFlank&14|+|104117399|104117520</td><td>72</td><td>92</td><td>21</td><td>TGAAATGTTTAGGACCACTAG</td><td>2141.4</td><td>12.6</td><td>1.3</td><td>10.0</td><td>1.4E-23</td><td>2.1E-21</td><td>BT20/MB231</td></tr>
<tr><td>>hg38_21_+_8433172_8446622_RNA45SN1_45S_50ntFlanks@10410.10432.23|TCAGTCGGTCCTGAGAGATGGGC</td><td>>hg38_21_+_8433172_8446622_RNA45SN1_45S_50ntFlanks</td><td>10410</td><td>10432</td><td>23</td><td>TCAGTCGGTCCTGAGAGATGGGC</td><td>666.7</td><td>-4.9</td><td>0.5</td><td>-9.7</td><td>3.1E-22</td><td>2.1E-21</td><td>BT20/MB468</td></tr>
<tr><td>>hg38_21_+_8433172_8446622_RNA45SN1_45S_50ntFlanks@11586.11603.18|GTGCGGACCAGGGGAATC</td><td>>hg38_21_+_8433172_8446622_RNA45SN1_45S_50ntFlanks</td><td>11586</td><td>11603</td><td>18</td><td>GTGCGGACCAGGGGAATC</td><td>1000.0</td><td>-5.7</td><td>0.6</td><td>-9.7</td><td>3.8E-22</td><td>2.1E-21</td><td>MB231/MB468</td></tr>
</table>

<table>
<tr><th></th><th></th><th></th><th></th><th></th><th></th><th></th><th></th><th></th><th></th><th></th><th></th><th></th></tr>
<tr><td>>hg38_21_+_8433172_8446622_RNA45SN1_45S_50ntFlanks@10458.10478.21|ATGGCCTCCGTTGCCCTCGGC</td><td>>hg38_21_+_8433172_8446622_RNA45SN1_45S_50ntFlanks</td><td>10458</td><td>10478</td><td>21</td><td>ATGGCCTCCGTTGCCCTCGGC</td><td>2073.6</td><td>-5.5</td><td>0.6</td><td>-9.7</td><td>3.9E-22</td><td>2.2E-21</td><td>MB231/MB468</td></tr>
<tr><td>>MI0006444|hsa-mir-1307&WithFlank&10|-|103394247|103394407@86.108.23&[MIMAT0005951&hsa-miR-1307-3p&offsets|0|+1;m-66&10|-|103394300|103394322&offsets|0|0]|ACTCGGCGTGGCGTCGGTCGTGG</td><td>>MI0006444|hsa-mir-1307&WithFlank&10|-|103394247|103394407</td><td>86</td><td>108</td><td>23</td><td>ACTCGGCGTGGCGTCGGTCGTGG</td><td>4426.1</td><td>6.2</td><td>0.6</td><td>9.7</td><td>3.2E-22</td><td>2.2E-21</td><td>BT20/MB468</td></tr>
<tr><td>>hg38_21_+_8433172_8446622_RNA45SN1_45S_50ntFlanks@476.508.33|TTCGTGATCGATGTGGTGACGTCGTGCTCTCCC</td><td>>hg38_21_+_8433172_8446622_RNA45SN1_45S_50ntFlanks</td><td>476</td><td>508</td><td>33</td><td>TTCGTGATCGATGTGGTGACGTCGTGCTCTCCC</td><td>601.6</td><td>6.5</td><td>0.7</td><td>9.7</td><td>3.2E-22</td><td>2.2E-21</td><td>BT20/MB468</td></tr>
<tr><td>>MI0000089|hsa-mir-31&WithFlank&9|-|21512109|21512191@14.35.22(+1A)&[MIMAT0000089&hsa-miR-31-5p&offsets|0|+1(+1A);m-360&9|-|21512157|21512178&offsets|0|0(+1A)]|AGGCAAGATGCTGGCATAGCTGA</td><td>>MI0000089|hsa-mir-31&WithFlank&9|-|21512109|21512191</td><td>14</td><td>35</td><td>22(+1A)</td><td>AGGCAAGATGCTGGCATAGCTGA</td><td>946.1</td><td>7.7</td><td>0.8</td><td>9.7</td><td>3.3E-22</td><td>2.3E-21</td><td>BT20/MB468</td></tr>
<tr><td>>hg38_MT_+_598_1651_RNR1_12S_50ntFlanks@905.935.31|AGTATACTTCAAAGGACATTTAACTAAAACC</td><td>>hg38_MT_+_598_1651_RNR1_12S_50ntFlanks</td><td>905</td><td>935</td><td>31</td><td>AGTATACTTCAAAGGACATTTAACTAAAACC</td><td>446.2</td><td>11.4</td><td>1.2</td><td>9.7</td><td>3.3E-22</td><td>2.3E-21</td><td>BT20/MB468</td></tr>
<tr><td>>hg38_21_+_8433172_8446622_RNA45SN1_45S_50ntFlanks@10351.10384.34|CGAGAACTTTGAAGGCCGAAGTGGAGAAGGGTTC</td><td>>hg38_21_+_8433172_8446622_RNA45SN1_45S_50ntFlanks</td><td>10351</td><td>10384</td><td>34</td><td>CGAGAACTTTGAAGGCCGAAGTGGAGAAGGGTTC</td><td>2248.7</td><td>-6.1</td><td>0.6</td><td>-9.7</td><td>3.3E-22</td><td>2.3E-21</td><td>BT20/MB468</td></tr>
<tr><td>>hg38_21_+_8433172_8446622_RNA45SN1_45S_50ntFlanks@475.510.36|CTTCGTGATCGATGTGGTGACGTCGTGCTCTCCCGG</td><td>>hg38_21_+_8433172_8446622_RNA45SN1_45S_50ntFlanks</td><td>475</td><td>510</td><td>36</td><td>CTTCGTGATCGATGTGGTGACGTCGTGCTCTCCCGG</td><td>5051.9</td><td>6.0</td><td>0.6</td><td>9.7</td><td>3.3E-22</td><td>2.3E-21</td><td>BT20/MB468</td></tr>
<tr><td>>MI0003628|hsa-mir-615&WithFlank&12|+|54033944|54034051@67.86.20&[MIMAT0003283&hsa-miR-615-3p&offsets|0|-2;m-495&12|+|54034010|54034030&offsets|0|-1]|TCCGAGCCTGGGTCTCCCTC</td><td>>MI0003628|hsa-mir-615&WithFlank&12|+|54033944|54034051</td><td>67</td><td>86</td><td>20</td><td>TCCGAGCCTGGGTCTCCCTC</td><td>905.8</td><td>7.1</td><td>0.7</td><td>9.7</td><td>3.4E-22</td><td>2.3E-21</td><td>BT20/MB468</td></tr>
<tr><td>>hg38_21_+_8433172_8446622_RNA45SN1_45S_50ntFlanks@475.499.25|CTTCGTGATCGATGTGGTGACGTCG</td><td>>hg38_21_+_8433172_8446622_RNA45SN1_45S_50ntFlanks</td><td>475</td><td>499</td><td>25</td><td>CTTCGTGATCGATGTGGTGACGTCG</td><td>1370.6</td><td>7.4</td><td>0.8</td><td>9.7</td><td>4.1E-22</td><td>2.3E-21</td><td>MB231/MB468</td></tr>
<tr><td>>MI0003591|hsa-mir-584&WithFlank&5|-|149062307|149062415@22.40.19&[MIMAT0003249&hsa-miR-584-5p&offsets|0|-3;m-167&5|-|149062374|149062394&offsets|0|-2]|TTATGGTTTGCCTGGGACT</td><td>>MI0003591|hsa-mir-584&WithFlank&5|-|149062307|149062415</td><td>22</td><td>40</td><td>19</td><td>TTATGGTTTGCCTGGGACT</td><td>1574.0</td><td>9.7</td><td>1.0</td><td>9.7</td><td>3.4E-22</td><td>2.3E-21</td><td>BT20/MB468</td></tr>
<tr><td>>MI0000683|hsa-mir-181b-2&WithFlank&9|+|124693704|124693804@22.46.25&[MIMAT0000257_1&hsa-miR-181b-5p&offsets|0|+2;m-72&9|+|124693725|124693747&offsets|0|+2]|AACATTCATTGCTGTCGGTGGGTTT</td><td>>MI0000683|hsa-mir-181b-2&WithFlank&9|+|124693704|124693804</td><td>22</td><td>46</td><td>25</td><td>AACATTCATTGCTGTCGGTGGGTTT</td><td>2736.8</td><td>7.8</td><td>0.8</td><td>9.7</td><td>4.1E-22</td><td>2.3E-21</td><td>MB231/MB468</td></tr>
<tr><td>>hg38_21_+_8433172_8446622_RNA45SN1_45S_50ntFlanks@476.509.34|TTCGTGATCGATGTGGTGACGTCGTGCTCTCCCG</td><td>>hg38_21_+_8433172_8446622_RNA45SN1_45S_50ntFlanks</td><td>476</td><td>509</td><td>34</td><td>TTCGTGATCGATGTGGTGACGTCGTGCTCTCCCG</td><td>2141.0</td><td>6.3</td><td>0.6</td><td>9.7</td><td>3.5E-22</td><td>2.4E-21</td><td>BT20/MB468</td></tr>
<tr><td>>MI0000067|hsa-let-7f-1&WithFlank&9|+|94176341|94176439@13.34.22;MI0000068|hsa-let-7f-2&WithFlank&X|-|53557186|53557280@14.35.22&[MIMAT0000067&hsa-let-7f-5p&offsets|0|0;m-20&9|+|94176353|94176374&offsets|0|0];[MIMAT0000067_1&hsa-let-7f-5p&offsets|0|0;m-19&X|-|53557246|53557267&offsets|0|0]|TGAGGTAGTAGATTGTATAGTT</td><td>>MI0000067|hsa-let-7f-1&WithFlank&9|+|94176341|94176439</td><td>13</td><td>34</td><td>22</td><td>TGAGGTAGTAGATTGTATAGTT</td><td>329962.1</td><td>6.5</td><td>0.7</td><td>9.7</td><td>3.5E-22</td><td>2.4E-21</td><td>BT20/MB468</td></tr>
<tr><td>>hg38_21_+_8433172_8446622_RNA45SN1_45S_50ntFlanks@11540.11581.42|CGCGGCGCCTCGCCTCGGCCGGCGCCTAGCAGCCGACTTAGA</td><td>>hg38_21_+_8433172_8446622_RNA45SN1_45S_50ntFlanks</td><td>11540</td><td>11581</td><td>42</td><td>CGCGGCGCCTCGCCTCGGCCGGCGCCTAGCAGCCGACTTAGA</td><td>463.2</td><td>5.5</td><td>0.6</td><td>9.7</td><td>4.3E-22</td><td>2.4E-21</td><td>MB231/MB468</td></tr>
<tr><td>>hg38_21_+_8433172_8446622_RNA45SN1_45S_50ntFlanks@3705.3742.38|TACCTGGTTGATCCTGCCAGTAGCATATGCTTGTCTCA</td><td>>hg38_21_+_8433172_8446622_RNA45SN1_45S_50ntFlanks</td><td>3705</td><td>3742</td><td>38</td><td>TACCTGGTTGATCCTGCCAGTAGCATATGCTTGTCTCA</td><td>915.6</td><td>5.5</td><td>0.6</td><td>9.7</td><td>4.4E-22</td><td>2.5E-21</td><td>MB231/MB468</td></tr>
<tr><td>>hg38_21_+_8433172_8446622_RNA45SN1_45S_50ntFlanks@475.509.35|CTTCGTGATCGATGTGGTGACGTCGTGCTCTCCCG</td><td>>hg38_21_+_8433172_8446622_RNA45SN1_45S_50ntFlanks</td><td>475</td><td>509</td><td>35</td><td>CTTCGTGATCGATGTGGTGACGTCGTGCTCTCCCG</td><td>4287.4</td><td>6.5</td><td>0.7</td><td>9.7</td><td>3.7E-22</td><td>2.5E-21</td><td>BT20/MB468</td></tr>
<tr><td>>hg38_21_+_8433172_8446622_RNA45SN1_45S_50ntFlanks@12416.12442.27|TTCGATGTCGGCTCTTCCTATCATTGT</td><td>>hg38_21_+_8433172_8446622_RNA45SN1_45S_50ntFlanks</td><td>12416</td><td>12442</td><td>27</td><td>TTCGATGTCGGCTCTTCCTATCATTGT</td><td>1943.8</td><td>7.6</td><td>0.8</td><td>9.7</td><td>4.5E-22</td><td>2.5E-21</td><td>MB231/MB468</td></tr>
<tr><td>>hg38_21_+_8433172_8446622_RNA45SN1_45S_50ntFlanks@483.506.24|TCGATGTGGTGACGTCGTGCTCTC</td><td>>hg38_21_+_8433172_8446622_RNA45SN1_45S_50ntFlanks</td><td>483</td><td>506</td><td>24</td><td>TCGATGTGGTGACGTCGTGCTCTC</td><td>745.5</td><td>7.4</td><td>0.8</td><td>9.7</td><td>3.8E-22</td><td>2.6E-21</td><td>BT20/MB468</td></tr>
<tr><td>>MI0000065|hsa-let-7d&WithFlank&9|+|94178828|94178926@14.34.21&[MIMAT0000065&hsa-let-7d-5p&offsets|0|-1;m-90&9|+|94178841|94178862&offsets|0|-1]|AGAGGTAGTAGGTTGCATAGT</td><td>>MI0000065|hsa-let-7d&WithFlank&9|+|94178828|94178926</td><td>14</td><td>34</td><td>21</td><td>AGAGGTAGTAGGTTGCATAGT</td><td>3385.6</td><td>7.8</td><td>0.8</td><td>9.7</td><td>4.6E-22</td><td>2.6E-21</td><td>MB231/MB468</td></tr>
<tr><td>>trnaMT_LeuTAA_MT_+_3230_3304@1.30.30|GTTAAGATGGCAGAGCCCGGTAATCGCATA</td><td>>trnaMT_LeuTAA_MT_+_3230_3304</td><td>1</td><td>30</td><td>30</td><td>GTTAAGATGGCAGAGCCCGGTAATCGCATA</td><td>1046.2</td><td>7.4</td><td>0.8</td><td>9.7</td><td>3.8E-22</td><td>2.6E-21</td><td>BT20/MB468</td></tr>
<tr><td>>hg38_21_+_8433172_8446622_RNA45SN1_45S_50ntFlanks@476.521.46|TTCGTGATCGATGTGGTGACGTCGTGCTCTCCCGGGCCGGGTCCGA</td><td>>hg38_21_+_8433172_8446622_RNA45SN1_45S_50ntFlanks</td><td>476</td><td>521</td><td>46</td><td>TTCGTGATCGATGTGGTGACGTCGTGCTCTCCCGGGCCGGGTCCGA</td><td>2540.6</td><td>11.4</td><td>1.2</td><td>9.7</td><td>4.6E-22</td><td>2.6E-21</td><td>MB231/MB468</td></tr>
<tr><td>>trnaMT_SerGCT_MT_+_12207_12265@27.62.36|CATGCCCCCATGTCTAACAACATGGCTTTCTCACCA</td><td>>trnaMT_SerGCT_MT_+_12207_12265</td><td>27</td><td>62</td><td>36</td><td>CATGCCCCCATGTCTAACAACATGGCTTTCTCACCA</td><td>552.3</td><td>9.4</td><td>1.0</td><td>9.7</td><td>4.7E-22</td><td>2.6E-21</td><td>MB231/MB468</td></tr>
<tr><td>>hg38_21_+_8433172_8446622_RNA45SN1_45S_50ntFlanks@12419.12448.30|GATGTCGGCTCTTCCTATCATTGTGAAGCA</td><td>>hg38_21_+_8433172_8446622_RNA45SN1_45S_50ntFlanks</td><td>12419</td><td>12448</td><td>30</td><td>GATGTCGGCTCTTCCTATCATTGTGAAGCA</td><td>4557.2</td><td>8.4</td><td>0.9</td><td>9.7</td><td>4.7E-22</td><td>2.6E-21</td><td>MB231/MB468</td></tr>
<tr><td>>hg38_21_+_8433172_8446622_RNA45SN1_45S_50ntFlanks@7974.7997.24|ACGCGACCTCAGATCAGACGTGGC</td><td>>hg38_21_+_8433172_8446622_RNA45SN1_45S_50ntFlanks</td><td>7974</td><td>7997</td><td>24</td><td>ACGCGACCTCAGATCAGACGTGGC</td><td>1781.9</td><td>6.1</td><td>0.6</td><td>9.7</td><td>3.9E-22</td><td>2.7E-21</td><td>BT20/MB468</td></tr>
<tr><td>>MI0000077|hsa-mir-21&WithFlank&17|+|59841260|59841343@13.35.23&[MIMAT0000076&hsa-miR-21-5p&offsets|-1|0;m-1&17|+|59841273|59841294&offsets|-1|0]|GTAGCTTATCAGACTGATGTTGA</td><td>>MI0000077|hsa-mir-21&WithFlank&17|+|59841260|59841343</td><td>13</td><td>35</td><td>23</td><td>GTAGCTTATCAGACTGATGTTGA</td><td>2044.5</td><td>7.1</td><td>0.7</td><td>9.7</td><td>3.9E-22</td><td>2.7E-21</td><td>BT20/MB468</td></tr>
<tr><td>>hg38_21_+_8433172_8446622_RNA45SN1_45S_50ntFlanks@7985.8006.22|GATCAGACGTGGCGACCCGCTG</td><td>>hg38_21_+_8433172_8446622_RNA45SN1_45S_50ntFlanks</td><td>7985</td><td>8006</td><td>22</td><td>GATCAGACGTGGCGACCCGCTG</td><td>1725.6</td><td>-12.7</td><td>1.3</td><td>-9.7</td><td>4.0E-22</td><td>2.7E-21</td><td>BT20/MB468</td></tr>
<tr><td>>MI0000440|hsa-mir-27b&WithFlank&9|+|95085439|95085547@66.87.22&[MIMAT0000419&hsa-miR-27b-3p&offsets|-1|0;m-39&9|+|95085505|95085525&offsets|-1|0]|GTTCACAGTGGCTAAGTTCTGC</td><td>>MI0000440|hsa-mir-27b&WithFlank&9|+|95085439|95085547</td><td>66</td><td>87</td><td>22</td><td>GTTCACAGTGGCTAAGTTCTGC</td><td>1090.3</td><td>7.9</td><td>0.8</td><td>9.6</td><td>4.9E-22</td><td>2.8E-21</td><td>MB231/MB468</td></tr>
<tr><td>>am_trnaMT_MetCAT_MT_+_4402_4469@1.21.21|AGTAAGGTCAGCTAAATAAGC</td><td>>am_trnaMT_MetCAT_MT_+_4402_4469</td><td>1</td><td>21</td><td>21</td><td>AGTAAGGTCAGCTAAATAAGC</td><td>1693.1</td><td>7.4</td><td>0.8</td><td>9.6</td><td>4.9E-22</td><td>2.8E-21</td><td>MB231/MB468</td></tr>
<tr><td>>MI0000279|hsa-mir-196a-2&WithFlank&12|+|53991732|53991853@31.52.22;MI0000238|hsa-mir-196a-1&WithFlank&17|-|48632484|48632565@13.34.22&[MIMAT0000226&hsa-miR-196a-5p&offsets|0|0;m-261&12|+|53991762|53991783&offsets|0|0];[MIMAT0000226_1&hsa-miR-196a-5p&offsets|0|0;m-268&17|-|48632532|48632553&offsets|0|0]|TAGGTAGTTTCATGTTGTTGGG</td><td>>MI0000279|hsa-mir-196a-2&WithFlank&12|+|53991732|53991853</td><td>31</td><td>52</td><td>22</td><td>TAGGTAGTTTCATGTTGTTGGG</td><td>8150.8</td><td>8.0</td><td>0.8</td><td>9.7</td><td>4.1E-22</td><td>2.8E-21</td><td>BT20/MB468</td></tr>
<tr><td>>hg38_21_+_8433172_8446622_RNA45SN1_45S_50ntFlanks@3985.4002.18|CGGCTTTGGTGACTCTAG</td><td>>hg38_21_+_8433172_8446622_RNA45SN1_45S_50ntFlanks</td><td>3985</td><td>4002</td><td>18</td><td>CGGCTTTGGTGACTCTAG</td><td>2655.5</td><td>-8.5</td><td>0.9</td><td>-9.7</td><td>4.1E-22</td><td>2.8E-21</td><td>BT20/MB468</td></tr>
<tr><td>>MI0000734|hsa-mir-106b&WithFlank&7|-|100093987|100094080@56.76.21(+1C)&[MIMAT0004672&hsa-miR-106b-3p&offsets|-2|-3(+1C);m-87&7|-|100094003|100094023&offsets|-2|-2(+1C)]|TACCGCACTGTGGGTACTTGCC</td><td>>MI0000734|hsa-mir-106b&WithFlank&7|-|100093987|100094080</td><td>56</td><td>76</td><td>21(+1C)</td><td>TACCGCACTGTGGGTACTTGCC</td><td>457.4</td><td>11.5</td><td>1.2</td><td>9.7</td><td>4.2E-22</td><td>2.8E-21</td><td>BT20/MB468</td></tr>
<tr><td>>MI0000434|hsa-let-7i&WithFlank&12|+|62603680|62603775@12.33.22(+1A)&[MIMAT0000415&hsa-let-7i-5p&offsets|0|0(+1A);m-49&12|+|62603691|62603712&offsets|0|0(+1A)]|TGAGGTAGTAGTTTGTGCTGTTA</td><td>>MI0000434|hsa-let-7i&WithFlank&12|+|62603680|62603775</td><td>12</td><td>33</td><td>22(+1A)</td><td>TGAGGTAGTAGTTTGTGCTGTTA</td><td>20230.0</td><td>7.3</td><td>0.8</td><td>9.6</td><td>5.1E-22</td><td>2.9E-21</td><td>MB231/MB468</td></tr>
<tr><td>>MI0000088|hsa-mir-30a&WithFlank&6|-|71403545|71403627@54.74.21(+1A)&[MIMAT0000088&hsa-miR-30a-3p&offsets|+1|0(+1A);m-38&6|-|71403554|71403575&offsets|+1|0(+1A)]|TTTCAGTCGGATGTTTGCAGCA</td><td>>MI0000088|hsa-mir-30a&WithFlank&6|-|71403545|71403627</td><td>54</td><td>74</td><td>21(+1A)</td><td>TTTCAGTCGGATGTTTGCAGCA</td><td>775.7</td><td>10.9</td><td>1.1</td><td>9.6</td><td>5.2E-22</td><td>2.9E-21</td><td>MB231/MB468</td></tr>
<tr><td>>hg38_21_+_8433172_8446622_RNA45SN1_45S_50ntFlanks@5472.5512.41|ACGGCCCTGGCGGAGCGCTGAGAAGACGGTCGAACTTGACT</td><td>>hg38_21_+_8433172_8446622_RNA45SN1_45S_50ntFlanks</td><td>5472</td><td>5512</td><td>41</td><td>ACGGCCCTGGCGGAGCGCTGAGAAGACGGTCGAACTTGACT</td><td>4201.8</td><td>10.2</td><td>1.1</td><td>9.6</td><td>5.4E-22</td><td>3.1E-21</td><td>MB231/MB468</td></tr>
<tr><td>>MI0000076|hsa-mir-20a&WithFlank&13|+|91351059|91351141@14.35.22(+1A)&[MIMAT0000075&hsa-miR-20a-5p&offsets|0|-1(+1A);m-96&13|+|91351072|91351094&offsets|0|-1(+1A)]|TAAAGTGCTTATAGTGCAGGTAA</td><td>>MI0000076|hsa-mir-20a&WithFlank&13|+|91351059|91351141</td><td>14</td><td>35</td><td>22(+1A)</td><td>TAAAGTGCTTATAGTGCAGGTAA</td><td>6891.1</td><td>7.1</td><td>0.7</td><td>9.6</td><td>5.4E-22</td><td>3.1E-21</td><td>MB231/MB468</td></tr>
<tr><td>>MI0000482|hsa-mir-185&WithFlank&22|+|20033133|20033226@21.42.22(+1A)&[MIMAT0000455&hsa-miR-185-5p&offsets|0|0(+1A);m-145&22|+|20033153|20033174&offsets|0|0(+1A)]|TGGAGAGAAAGGCAGTTCCTGAA</td><td>>MI0000482|hsa-mir-185&WithFlank&22|+|20033133|20033226</td><td>21</td><td>42</td><td>22(+1A)</td><td>TGGAGAGAAAGGCAGTTCCTGAA</td><td>621.3</td><td>7.0</td><td>0.7</td><td>9.7</td><td>4.5E-22</td><td>3.1E-21</td><td>BT20/MB468</td></tr>
<tr><td>>trnaMT_SerGCT_MT_+_12207_12265@1.18.18|GAGAAAGCTCACAAGAAC</td><td>>trnaMT_SerGCT_MT_+_12207_12265</td><td>1</td><td>18</td><td>18</td><td>GAGAAAGCTCACAAGAAC</td><td>96934.0</td><td>12.2</td><td>1.3</td><td>9.7</td><td>4.5E-22</td><td>3.1E-21</td><td>BT20/MB468</td></tr>
<tr><td>>MI0000114|hsa-mir-107&WithFlank&10|-|89592741|89592833@56.76.21(+1A);MI0000108|hsa-mir-103a-2&WithFlank&20|+|3917488|3917577@54.74.21(+1A);MI0000109|hsa-mir-103a-1&WithFlank&5|-|168560890|168560979@54.74.21(+1A)&[MIMAT0000104&hsa-miR-107&offsets|0|-2(+1A);m-78&10|-|89592758|89592778&offsets|0|0(+1A)];[MIMAT0000101&hsa-miR-103a-3p&offsets|0|-2(+1A);m-12&20|+|3917541|3917563&offsets|0|-2(+1A)];[MIMAT0000101_1&hsa-miR-103a-3p&offsets|0|-2(+1A);m-13&5|-|168560904|168560926&offsets|0|-2(+1A);m-44220&5|-|168560908|168560926&offsets|0|+2(+1A)]|AGCAGCATTGTACAGGGCTATA</td><td>>MI0000114|hsa-mir-107&WithFlank&10|-|89592741|89592833</td><td>56</td><td>76</td><td>21(+1A)</td><td>AGCAGCATTGTACAGGGCTATA</td><td>3436.3</td><td>8.0</td><td>0.8</td><td>9.7</td><td>4.6E-22</td><td>3.1E-21</td><td>BT20/MB468</td></tr>
<tr><td>>MI0000764|hsa-mir-363&WithFlank&X|-|134169372|134169458@56.77.22&[MIMAT0000707&hsa-miR-363-3p&offsets|0|0;m-178&X|-|134169381|134169403&offsets|0|-1]|AATTGCACGGTATCCATCTGTA</td><td>>MI0000764|hsa-mir-363&WithFlank&X|-|134169372|134169458</td><td>56</td><td>77</td><td>22</td><td>AATTGCACGGTATCCATCTGTA</td><td>1761.1</td><td>14.3</td><td>1.4</td><td>10.0</td><td>2.3E-23</td><td>3.2E-21</td><td>BT20/MB231</td></tr>
<tr><td>>hg38_21_+_8433172_8446622_RNA45SN1_45S_50ntFlanks@7974.8009.36|ACGCGACCTCAGATCAGACGTGGCGACCCGCTGAAT</td><td>>hg38_21_+_8433172_8446622_RNA45SN1_45S_50ntFlanks</td><td>7974</td><td>8009</td><td>36</td><td>ACGCGACCTCAGATCAGACGTGGCGACCCGCTGAAT</td><td>10772.7</td><td>6.7</td><td>0.7</td><td>9.6</td><td>5.7E-22</td><td>3.2E-21</td><td>MB231/MB468</td></tr>
</table>

| | | | | | | | | | | | | |
|---|---|---|---|---|---|---|---|---|---|---|---|---|
| >MI0000234\|hsa-mir-192&WithFlank&11\|-\|64891131\|64891252@30.51.22(+1U)&[MIMAT0000222&hsa-miR-192-5p&offsets\|0\|+1(+1U);m-59&11\|-\|64891203\|64891223&offsets\|0\|+1(+1U)]\|CTGACCTATGAATTGACAGCCAT | >MI0000234\|hsa-mir-192&WithFlank&11\|-\|64891131\|64891252 | 30 | 51 | 22(+1U) | CTGACCTATGAATTGACAGCCAT | 3479.5 | 7.2 | 0.7 | 9.6 | 5.7E-22 | 3.2E-21 | MB231/MB468 |
| >MI0000469\|hsa-mir-125a&WithFlank&19\|+\|51693248\|51693345@21.42.22(+1A)&[MIMAT0000443&hsa-miR-125a-5p&offsets\|0\|-2(+1A);m-63&19\|+\|51693268\|51693289&offsets\|0\|0(+1A)]\|TCCCTGAGACCCTTTAACCTGTA | >MI0000469\|hsa-mir-125a&WithFlank&19\|+\|51693248\|51693345 | 21 | 42 | 22(+1A) | TCCCTGAGACCCTTTAACCTGTA | 2572.5 | 6.5 | 0.7 | 9.6 | 5.9E-22 | 3.3E-21 | MB231/MB468 |
| >hg38_21_+_8433172_8446622_RNA45SN1_45S_50ntFlanks@12973.12992.20\|TCGTACGTAGCAGAGCAGCT | >hg38_21_+_8433172_8446622_RNA45SN1_45S_50ntFlanks | 12973 | 12992 | 20 | TCGTACGTAGCAGAGCAGCT | 1208.5 | -6.2 | 0.6 | -9.7 | 4.9E-22 | 3.3E-21 | BT20/MB468 |
| >hg38_21_+_8433172_8446622_RNA45SN1_45S_50ntFlanks@475.514.40\|CTTCGTGATCGATGTGGTGACGTCGTGCTCTCCCGGGCCG | >hg38_21_+_8433172_8446622_RNA45SN1_45S_50ntFlanks | 475 | 514 | 40 | CTTCGTGATCGATGTGGTGACGTCGTGCTCTCCCGGGCCG | 2625.5 | 7.3 | 0.8 | 9.6 | 4.9E-22 | 3.3E-21 | BT20/MB468 |
| >MI0000434\|hsa-let-7i&WithFlank&12\|+\|62603680\|62603775@12.33.22(+1C)&[MIMAT0000415&hsa-let-7i-5p&offsets\|0\|0(+1C);m-49&12\|+\|62603691\|62603712&offsets\|0\|0(+1C)]\|TGAGGTAGTAGTTTGTGCTGTTC | >MI0000434\|hsa-let-7i&WithFlank&12\|+\|62603680\|62603775 | 12 | 33 | 22(+1C) | TGAGGTAGTAGTTTGTGCTGTTC | 1021.7 | 6.8 | 0.7 | 9.6 | 5.0E-22 | 3.4E-21 | BT20/MB468 |
| >hg38_21_+_8433172_8446622_RNA45SN1_45S_50ntFlanks@6721.6746.26\|AATTGCAGGACACATTGATCATCGAC | >hg38_21_+_8433172_8446622_RNA45SN1_45S_50ntFlanks | 6721 | 6746 | 26 | AATTGCAGGACACATTGATCATCGAC | 1866.9 | -5.2 | 0.5 | -9.6 | 6.1E-22 | 3.4E-21 | MB231/MB468 |
| >MI0000816\|hsa-mir-335&WithFlank&7\|+\|130496105\|130496210@58.78.21&[MIMAT0004703&hsa-miR-335-3p&offsets\|0\|-1;m-159&7\|+\|130496162\|130496183&offsets\|0\|-1]\|TTTTTCATTATTGCTCCTGAC | >MI0000816\|hsa-mir-335&WithFlank&7\|+\|130496105\|130496210 | 58 | 78 | 21 | TTTTTCATTATTGCTCCTGAC | 1875.8 | 8.4 | 0.9 | 9.6 | 6.1E-22 | 3.4E-21 | MB231/MB468 |
| >MI0000285\|hsa-mir-205&WithFlank&1\|+\|209432127\|209432248@40.60.21(+1A)&[MIMAT0000266&hsa-miR-205-5p&offsets\|0\|-1(+1A);m-351&1\|+\|209432166\|209432187&offsets\|0\|-1(+1A)]\|TCCTTCATTCCACCGGAGTCTA | >MI0000285\|hsa-mir-205&WithFlank&1\|+\|209432127\|209432248 | 40 | 60 | 21(+1A) | TCCTTCATTCCACCGGAGTCTA | 13054.7 | 7.3 | 0.8 | 9.6 | 5.2E-22 | 3.5E-21 | BT20/MB468 |
| >MI0000440\|hsa-mir-27b&WithFlank&9\|+\|95085439\|95085547@67.87.21&[MIMAT0000419&hsa-miR-27b-3p&offsets\|0\|0;m-39&9\|+\|95085505\|95085525&offsets\|0\|0]\|TTCACAGTGGCTAAGTTCTGC | >MI0000440\|hsa-mir-27b&WithFlank&9\|+\|95085439\|95085547 | 67 | 87 | 21 | TTCACAGTGGCTAAGTTCTGC | 201189.8 | 8.0 | 0.8 | 9.6 | 6.2E-22 | 3.5E-21 | MB231/MB468 |
| >MI0000065\|hsa-let-7d&WithFlank&9\|+\|94178828\|94178926@68.88.21(+1C)&[MIMAT0004484&hsa-let-7d-3p&offsets\|0\|-1(+1C);m-98&9\|+\|94178895\|94178916&offsets\|0\|-1(+1C)]\|CTATACGACCTGCTGCCTTTCC | >MI0000065\|hsa-let-7d&WithFlank&9\|+\|94178828\|94178926 | 68 | 88 | 21(+1C) | CTATACGACCTGCTGCCTTTCC | 915.2 | 8.0 | 0.8 | 9.6 | 6.3E-22 | 3.5E-21 | MB231/MB468 |
| >MI0000077\|hsa-mir-21&WithFlank&17\|+\|59841260\|59841343@14.35.22(+1G)&[MIMAT0000076&hsa-miR-21-5p&offsets\|0\|0(+1G);m-1&17\|+\|59841273\|59841294&offsets\|0\|0(+1G)]\|TAGCTTATCAGACTGATGTTGAG | >MI0000077\|hsa-mir-21&WithFlank&17\|+\|59841260\|59841343 | 14 | 35 | 22(+1G) | TAGCTTATCAGACTGATGTTGAG | 26133.8 | 9.6 | 1.0 | 9.6 | 5.3E-22 | 3.5E-21 | BT20/MB468 |
| >MI0000063\|hsa-let-7b&WithFlank&22\|+\|46113680\|46113774@12.32.21(+1A)&[MIMAT0000063&hsa-let-7b-5p&offsets\|0\|-1(+1A);m-14&22\|+\|46113691\|46113712&offsets\|0\|-1(+1A)]\|TGAGGTAGTAGGTTGTGTGGTA | >MI0000063\|hsa-let-7b&WithFlank&22\|+\|46113680\|46113774 | 12 | 32 | 21(+1A) | TGAGGTAGTAGGTTGTGTGGTA | 1141.6 | 6.8 | 0.7 | 9.6 | 6.3E-22 | 3.5E-21 | MB231/MB468 |
| >MI0000433\|hsa-let-7g&WithFlank&3\|-\|52268272\|52268367@11.31.21&[MIMAT0000414&hsa-let-7g-5p&offsets\|0\|-1;m-58&3\|-\|52268336\|52268357&offsets\|0\|-1]\|TGAGGTAGTAGTTTGTACAGT | >MI0000433\|hsa-let-7g&WithFlank&3\|-\|52268272\|52268367 | 11 | 31 | 21 | TGAGGTAGTAGTTTGTACAGT | 10063.8 | 6.8 | 0.7 | 9.6 | 5.3E-22 | 3.5E-21 | BT20/MB468 |
| >hg38_21_+_8433172_8446622_RNA45SN1_45S_50ntFlanks@11316.11344.29\|CACCCCACGTCTCGTCGCGCGCGCGTCCG | >hg38_21_+_8433172_8446622_RNA45SN1_45S_50ntFlanks | 11316 | 11344 | 29 | CACCCCACGTCTCGTCGCGCGCGCGTCCG | 5252.3 | -8.1 | 0.8 | -9.6 | 5.3E-22 | 3.6E-21 | BT20/MB468 |
| >hg38_21_+_8433172_8446622_RNA45SN1_45S_50ntFlanks@11318.11344.27\|CCCCACGTCTCGTCGCGCGCGCGTCCG | >hg38_21_+_8433172_8446622_RNA45SN1_45S_50ntFlanks | 11318 | 11344 | 27 | CCCCACGTCTCGTCGCGCGCGCGTCCG | 2278.4 | -12.7 | 1.3 | -9.6 | 6.5E-22 | 3.6E-21 | MB231/MB468 |
| >hg38_21_+_8433172_8446622_RNA45SN1_45S_50ntFlanks@12476.12500.25\|CCACTAATAGGGAACGTGAGCTGGG | >hg38_21_+_8433172_8446622_RNA45SN1_45S_50ntFlanks | 12476 | 12500 | 25 | CCACTAATAGGGAACGTGAGCTGGG | 580.6 | -6.3 | 0.7 | -9.6 | 5.5E-22 | 3.6E-21 | BT20/MB468 |
| >MI0000456\|hsa-mir-140&WithFlank&16\|+\|69933075\|69933186@69.90.22&[MIMAT0004597&hsa-miR-140-3p&offsets\|+1\|+2;m-45&16\|+\|69933143\|69933164&offsets\|0\|0]\|ACCACAGGGTAGAACCACGGAC | >MI0000456\|hsa-mir-140&WithFlank&16\|+\|69933075\|69933186 | 69 | 90 | 22 | ACCACAGGGTAGAACCACGGAC | 3755.5 | 10.7 | 1.1 | 9.6 | 5.5E-22 | 3.6E-21 | BT20/MB468 |
| >MI0000434\|hsa-let-7i&WithFlank&12\|+\|62603680\|62603775@12.32.21(+1C)&[MIMAT0000415&hsa-let-7i-5p&offsets\|0\|-1(+1C);m-49&12\|+\|62603691\|62603712&offsets\|0\|-1(+1C)]\|TGAGGTAGTAGTTTGTGCTGTC | >MI0000434\|hsa-let-7i&WithFlank&12\|+\|62603680\|62603775 | 12 | 32 | 21(+1C) | TGAGGTAGTAGTTTGTGCTGTC | 26600.4 | 6.5 | 0.7 | 9.6 | 5.6E-22 | 3.7E-21 | BT20/MB468 |
| >MI0000108\|hsa-mir-103a-2&WithFlank&20\|+\|3917488\|3917577@54.77.24;MI0000109\|hsa-mir-103a-1&WithFlank&5\|-\|168560890\|168560979@54.77.24&[MIMAT0000101&hsa-miR-103a-3p&offsets\|0\|+1;m-12&20\|+\|3917541\|3917563&offsets\|0\|+1];[MIMAT0000101_1&hsa-miR-103a-3p&offsets\|0\|+1;m-13&5\|-\|168560904\|168560926&offsets\|0\|+1;m-44220&5\|-\|168560908\|168560926&offsets\|0\|+5]\|AGCAGCATTGTACAGGGCTATGAA | >MI0000108\|hsa-mir-103a-2&WithFlank&20\|+\|3917488\|3917577 | 54 | 77 | 24 | AGCAGCATTGTACAGGGCTATGAA | 3896.3 | 8.0 | 0.8 | 9.6 | 5.6E-22 | 3.7E-21 | BT20/MB468 |
| >hg38_21_+_8433172_8446622_RNA45SN1_45S_50ntFlanks@12942.12965.24\|TTCGTAGACGACCTGCTTCTGGGT | >hg38_21_+_8433172_8446622_RNA45SN1_45S_50ntFlanks | 12942 | 12965 | 24 | TTCGTAGACGACCTGCTTCTGGGT | 392.3 | -5.0 | 0.5 | -9.6 | 5.6E-22 | 3.7E-21 | BT20/MB468 |
| >hg38_21_+_8433172_8446622_RNA45SN1_45S_50ntFlanks@4241.4258.18\|CTCTTTCGAGGCCCTGTA | >hg38_21_+_8433172_8446622_RNA45SN1_45S_50ntFlanks | 4241 | 4258 | 18 | CTCTTTCGAGGCCCTGTA | 392.3 | -5.5 | 0.6 | -9.6 | 6.7E-22 | 3.7E-21 | MB231/MB468 |
| >MI0000108\|hsa-mir-103a-2&WithFlank&20\|+\|3917488\|3917577@54.75.22;MI0000109\|hsa-mir-103a-1&WithFlank&5\|-\|168560890\|168560979@54.75.22&[MIMAT0000101&hsa-miR-103a-3p&offsets\|0\|-1;m-12&20\|+\|3917541\|3917563&offsets\|0\|-1];[MIMAT0000101_1&hsa-miR-103a-3p&offsets\|0\|-1;m-13&5\|-\|168560904\|168560926&offsets\|0\|-1;m-44220&5\|-\|168560908\|168560926&offsets\|0\|+3]\|AGCAGCATTGTACAGGGCTATG | >MI0000108\|hsa-mir-103a-2&WithFlank&20\|+\|3917488\|3917577 | 54 | 75 | 22 | AGCAGCATTGTACAGGGCTATG | 17191.6 | 6.8 | 0.7 | 9.6 | 5.7E-22 | 3.8E-21 | BT20/MB468 |
| >MI0000736\|hsa-mir-30c-1&WithFlank&1\|+\|40757278\|40757378@23.48.26;MI0000254\|hsa-mir-30c-2&WithFlank&6\|-\|71376954\|71377037@13.38.26&[MIMAT0000244&hsa-miR-30c-5p&offsets\|0\|+3;m-57&1\|+\|40757300\|40757323&offsets\|0\|+2];[MIMAT0000244_1&hsa-miR-30c-5p&offsets\|0\|+3;m-56&6\|-\|71377002\|71377025&offsets\|0\|+2]\|TGTAAACATCCTACACTCTCAGCTGT | >MI0000736\|hsa-mir-30c-1&WithFlank&1\|+\|40757278\|40757378 | 23 | 48 | 26 | TGTAAACATCCTACACTCTCAGCTGT | 951.0 | 8.0 | 0.8 | 9.6 | 6.9E-22 | 3.8E-21 | MB231/MB468 |
| >MI0000061\|hsa-let-7a-2&WithFlank&11\|-\|122146516\|122146599@11.28.18;MI0000064\|hsa-let-7c&WithFlank&21\|+\|16539822\|16539917@17.34.18;MI0000062\|hsa-let-7a-3&WithFlank&22\|+\|46112743\|46112828@10.27.18;MI0000060\|hsa-let-7a-1&WithFlank&9\|+\|94175951\|94176042@12.29.18&[MIMAT0000062&hsa-let-7a-5p&offsets\|0\|-4;m-6&11\|-\|122146568\|122146589&offsets\|0\|-4];[MIMAT0000064&hsa-let-7c-5p&offsets\|0\|-4;m-37&21\|+\|16539838\|16539859&offsets\|0\|-4];[MIMAT0000062_1&hsa-let-7a-5p&offsets\|0\|-4;m-5&22\|+\|46112752\|46112773&offsets\|0\|-4];[MIMAT0000062_2&hsa-let-7a-5p&offsets\|0\|-4;m-4&9\|+\|94175962\|94175983&offsets\|0\|-4]\|TGAGGTAGTAGGTTGTAT | >MI0000061\|hsa-let-7a-2&WithFlank&11\|-\|122146516\|122146599 | 11 | 28 | 18 | TGAGGTAGTAGGTTGTAT | 8102.8 | 8.3 | 0.9 | 9.6 | 6.9E-22 | 3.8E-21 | MB231/MB468 |
| >MI0000094\|hsa-mir-92a-2&WithFlank&X\|-\|134169532\|134169618@54.76.23&[MIMAT0000092_1&hsa-miR-92a-3p&offsets\|0\|+1;m-11&X\|-\|134169544\|134169565&offsets\|0\|+1]\|TATTGCACTTGTCCCGGCCTGTG | >MI0000094\|hsa-mir-92a-2&WithFlank&X\|-\|134169532\|134169618 | 54 | 76 | 23 | TATTGCACTTGTCCCGGCCTGTG | 1512.4 | 7.0 | 0.7 | 9.6 | 6.9E-22 | 3.8E-21 | MB231/MB468 |
| >am_trna128_GlyGCC_6_-_27870686_27870756@3.35.33,trna18_GlyGCC_16_+_70822597_70822667@3.35.33,trna19_GlyGCC_16_+_70823410_70823480@3.35.33,trna19_GlyGCC_2_-_157257659_157257729@3.35.33,trna24_GlyGCC_16_-_70812942_70813012@3.35.33,trna25_GlyGCC_16_-_70812114_70812184@3.35.33,trna5_GlyGCC_17_+_8029064_8029134@3.35.33,trna68_GlyGCC_1_-_161493637_161493707@3.35.33\|ATTGGTGGTTCAGTGGTAGAATTCTCGCCTGCC | >am_trna128_GlyGCC_6_-_27870686_27870756 | 3 | 35 | 33 | ATTGGTGGTTCAGTGGTAGAATTCTCGCCTGCC | 2161.4 | 11.0 | 1.1 | 9.6 | 6.9E-22 | 3.8E-21 | MB231/MB468 |
| >hg38_21_+_8433172_8446622_RNA45SN1_45S_50ntFlanks@10841.10864.24\|CGTAACTTCGGGATAAGGATTGGC | >hg38_21_+_8433172_8446622_RNA45SN1_45S_50ntFlanks | 10841 | 10864 | 24 | CGTAACTTCGGGATAAGGATTGGC | 2412.2 | -5.9 | 0.6 | -9.6 | 7.0E-22 | 3.9E-21 | MB231/MB468 |
| >hg38_21_+_8433172_8446622_RNA45SN1_45S_50ntFlanks@4734.4761.28\|AACGAAAGTCGGAGGTTCGAAGACGATC | >hg38_21_+_8433172_8446622_RNA45SN1_45S_50ntFlanks | 4734 | 4761 | 28 | AACGAAAGTCGGAGGTTCGAAGACGATC | 661.7 | 5.4 | 0.6 | 9.6 | 7.0E-22 | 3.9E-21 | MB231/MB468 |
| >hg38_21_+_8433172_8446622_RNA45SN1_45S_50ntFlanks@12996.13015.20\|TCGCTGCGATCTATTGAAAG | >hg38_21_+_8433172_8446622_RNA45SN1_45S_50ntFlanks | 12996 | 13015 | 20 | TCGCTGCGATCTATTGAAAG | 1317.7 | 4.9 | 0.5 | 9.6 | 7.0E-22 | 3.9E-21 | MB231/MB468 |
| >hg38_21_+_8433172_8446622_RNA45SN1_45S_50ntFlanks@4003.4025.23\|ATAACCTCGGGCCGATCGCACGC | >hg38_21_+_8433172_8446622_RNA45SN1_45S_50ntFlanks | 4003 | 4025 | 23 | ATAACCTCGGGCCGATCGCACGC | 498.2 | -6.1 | 0.6 | -9.6 | 5.9E-22 | 3.9E-21 | BT20/MB468 |
| >am_MI0000809\|hsa-mir-151a&WithFlank&8\|-\|140732558\|140732659@53.72.20(+1C)&[MIMAT0000757&hsa-miR-151a-3p&offsets\|0\|-1(+1C);m-30&8\|-\|140732587\|140732607&offsets\|0\|-1(+1C)]\|CTAGACTGAAGCTCCTTGAGC | >am_MI0000809\|hsa-mir-151a&WithFlank&8\|-\|140732558\|140732659 | 53 | 72 | 20(+1C) | CTAGACTGAAGCTCCTTGAGC | 2445.6 | 9.7 | 1.0 | 9.6 | 5.9E-22 | 3.9E-21 | BT20/MB468 |

<table>
<tr><td>>hg38_21_+_8433172_8446622_RNA45SN1_45S_50ntFlanks@12405.12429.25|CTTTTTGATCCTTCGATGTCGGCTC</td><td>>hg38_21_+_8433172_8446622_RNA45SN1_45S_50ntFlanks</td><td>12405</td><td>12429</td><td>25</td><td>CTTTTTGATCCTTCGATGTCGGCTC</td><td>2919.6</td><td>-7.1</td><td>0.7</td><td>-9.6</td><td>5.9E-22</td><td>3.9E-21</td><td>BT20/MB468</td></tr>
<tr><td>>hg38_21_+_8433172_8446622_RNA45SN1_45S_50ntFlanks@6775.6806.32|CCTCCCGGGGCTACGCCTGTCTGAGCGTCGCT</td><td>>hg38_21_+_8433172_8446622_RNA45SN1_45S_50ntFlanks</td><td>6775</td><td>6806</td><td>32</td><td>CCTCCCGGGGCTACGCCTGTCTGAGCGTCGCT</td><td>869.4</td><td>5.1</td><td>0.5</td><td>9.6</td><td>7.1E-22</td><td>3.9E-21</td><td>MB231/MB468</td></tr>
<tr><td>>hg38_21_+_8433172_8446622_RNA45SN1_45S_50ntFlanks@11768.11792.25|CTAATTAGTGACGCGCATGAATGGA</td><td>>hg38_21_+_8433172_8446622_RNA45SN1_45S_50ntFlanks</td><td>11768</td><td>11792</td><td>25</td><td>CTAATTAGTGACGCGCATGAATGGA</td><td>1542.2</td><td>-6.1</td><td>0.6</td><td>-9.6</td><td>7.1E-22</td><td>3.9E-21</td><td>MB231/MB468</td></tr>
<tr><td>>MI0000746|hsa-mir-99b&WithFlank&19|+|51692606|51692687@51.71.21(+1A)&[MIMAT0004678&hsa-miR-99b-3p&offsets|0|-1(+1A);m-151&19|+|51692656|51692678&offsets|0|-2(+1A)]|CAAGCTCGTGTCTGTGGGTCCA</td><td>>MI0000746|hsa-mir-99b&WithFlank&19|+|51692606|51692687</td><td>51</td><td>71</td><td>21(+1A)</td><td>CAAGCTCGTGTCTGTGGGTCCA</td><td>1658.3</td><td>8.1</td><td>0.8</td><td>9.6</td><td>6.0E-22</td><td>4.0E-21</td><td>BT20/MB468</td></tr>
<tr><td>>hg38_21_+_8433172_8446622_RNA45SN1_45S_50ntFlanks@6718.6739.22|GTGAATTGCAGGACACATTGAT</td><td>>hg38_21_+_8433172_8446622_RNA45SN1_45S_50ntFlanks</td><td>6718</td><td>6739</td><td>22</td><td>GTGAATTGCAGGACACATTGAT</td><td>722.8</td><td>6.9</td><td>0.7</td><td>9.6</td><td>7.3E-22</td><td>4.0E-21</td><td>MB231/MB468</td></tr>
<tr><td>>MI0000264|hsa-mir-7-2&WithFlank&15|+|88611819|88611940@38.60.23(+1G);MI0000265|hsa-mir-7-3&WithFlank&19|+|4770664|4770785@37.59.23(+1G);MI0000263|hsa-mir-7-1&WithFlank&9|-|83969742|83969863@30.52.23(+1G)&[MIMAT0000252&hsa-miR-7-5p&offsets|0|-1(+1G);m-400&15|+|88611856|88611879&offsets|0|-1(+1G)];[MIMAT0000252_1&hsa-miR-7-5p&offsets|0|-1(+1G);m-402&19|+|4770700|4770723&offsets|0|-1(+1G)];[MIMAT0000252_2&hsa-miR-7-5p&offsets|0|-1(+1G);m-401&9|-|83969811|83969834&offsets|0|-1(+1G)]|TGGAAGACTAGTGATTTTGTTGTG</td><td>>MI0000264|hsa-mir-7-2&WithFlank&15|+|88611819|88611940</td><td>38</td><td>60</td><td>23(+1G)</td><td>TGGAAGACTAGTGATTTTGTTGTG</td><td>7554.5</td><td>7.5</td><td>0.8</td><td>9.6</td><td>7.4E-22</td><td>4.1E-21</td><td>MB231/MB468</td></tr>
<tr><td>>MI0000440|hsa-mir-27b&WithFlank&9|+|95085439|95085547@67.87.21(+1C)&[MIMAT0000419&hsa-miR-27b-3p&offsets|0|0(+1C);m-39&9|+|95085505|95085525&offsets|0|0(+1C)]|TTCACAGTGGCTAAGTTCTGCC</td><td>>MI0000440|hsa-mir-27b&WithFlank&9|+|95085439|95085547</td><td>67</td><td>87</td><td>21(+1C)</td><td>TTCACAGTGGCTAAGTTCTGCC</td><td>629.6</td><td>7.5</td><td>0.8</td><td>9.6</td><td>7.4E-22</td><td>4.1E-21</td><td>MB231/MB468</td></tr>
<tr><td>>hg38_1_-_228634819_228635039_RNA5S12_5S_50ntFlanks@129.169.41|TTGGATGGGAGACCGCCTGGGAATACCGGGTGCTGTAGGCT</td><td>>hg38_1_-_228634819_228635039_RNA5S12_5S_50ntFlanks</td><td>129</td><td>169</td><td>41</td><td>TTGGATGGGAGACCGCCTGGGAATACCGGGTGCTGTAGGCT</td><td>660.3</td><td>14.0</td><td>1.5</td><td>9.6</td><td>6.4E-22</td><td>4.2E-21</td><td>BT20/MB468</td></tr>
<tr><td>>MI0000077|hsa-mir-21&WithFlank&17|+|59841260|59841343@15.37.23&[MIMAT0000076&hsa-miR-21-5p&offsets|+1|+2;m-1&17|+|59841273|59841294&offsets|+1|+2]|AGCTTATCAGACTGATGTTGACT</td><td>>MI0000077|hsa-mir-21&WithFlank&17|+|59841260|59841343</td><td>15</td><td>37</td><td>23</td><td>AGCTTATCAGACTGATGTTGACT</td><td>625.6</td><td>7.0</td><td>0.7</td><td>9.6</td><td>6.4E-22</td><td>4.2E-21</td><td>BT20/MB468</td></tr>
<tr><td>>hg38_21_+_8433172_8446622_RNA45SN1_45S_50ntFlanks@3990.4023.34|TTGGTGACTCTAGATAACCTCGGGCCGATCGCAC</td><td>>hg38_21_+_8433172_8446622_RNA45SN1_45S_50ntFlanks</td><td>3990</td><td>4023</td><td>34</td><td>TTGGTGACTCTAGATAACCTCGGGCCGATCGCAC</td><td>1107.0</td><td>6.0</td><td>0.6</td><td>9.6</td><td>7.7E-22</td><td>4.3E-21</td><td>MB231/MB468</td></tr>
<tr><td>>MI0000471|hsa-mir-126&WithFlank&9|+|136670596|136670692@58.79.22&[MIMAT0000445&hsa-miR-126-3p&offsets|0|0;m-40&9|+|136670653|136670674&offsets|0|0]|TCGTACCGTGAGTAATAATGCG</td><td>>MI0000471|hsa-mir-126&WithFlank&9|+|136670596|136670692</td><td>58</td><td>79</td><td>22</td><td>TCGTACCGTGAGTAATAATGCG</td><td>9611.6</td><td>9.3</td><td>1.0</td><td>9.6</td><td>7.8E-22</td><td>4.3E-21</td><td>MB231/MB468</td></tr>
<tr><td>>MI0000273|hsa-mir-183&WithFlank&7|-|129774899|129775020@34.56.23&[MIMAT0000261&hsa-miR-183-5p&offsets|+1|+2;m-41&7|-|129774966|129774987&offsets|0|+1]|ATGGCACTGGTAGAATTCACTGT</td><td>>MI0000273|hsa-mir-183&WithFlank&7|-|129774899|129775020</td><td>34</td><td>56</td><td>23</td><td>ATGGCACTGGTAGAATTCACTGT</td><td>2420.3</td><td>7.1</td><td>0.7</td><td>9.6</td><td>6.6E-22</td><td>4.3E-21</td><td>BT20/MB468</td></tr>
<tr><td>>trna152_ValCAC_6_-_27248049_27248121@1.33.33|GCTTCTGTAGTGTAGTGGTTATCACGTTCGCCT</td><td>>trna152_ValCAC_6_-_27248049_27248121</td><td>1</td><td>33</td><td>33</td><td>GCTTCTGTAGTGTAGTGGTTATCACGTTCGCCT</td><td>6917.7</td><td>13.2</td><td>1.4</td><td>9.6</td><td>7.9E-22</td><td>4.3E-21</td><td>MB231/MB468</td></tr>
<tr><td>>hg38_21_+_8433172_8446622_RNA45SN1_45S_50ntFlanks@12654.12675.22|TATGACTGAACGCCTCTAAGTC</td><td>>hg38_21_+_8433172_8446622_RNA45SN1_45S_50ntFlanks</td><td>12654</td><td>12675</td><td>22</td><td>TATGACTGAACGCCTCTAAGTC</td><td>2866.2</td><td>-6.9</td><td>0.7</td><td>-9.6</td><td>8.0E-22</td><td>4.4E-21</td><td>MB231/MB468</td></tr>
<tr><td>>MI0000433|hsa-let-7g&WithFlank&3|-|52268272|52268367@10.32.23&[MIMAT0000414&hsa-let-7g-5p&offsets|-1|0;m-58&3|-|52268336|52268357&offsets|-1|0]|CTGAGGTAGTAGTTTGTACAGTT</td><td>>MI0000433|hsa-let-7g&WithFlank&3|-|52268272|52268367</td><td>10</td><td>32</td><td>23</td><td>CTGAGGTAGTAGTTTGTACAGTT</td><td>1480.8</td><td>7.3</td><td>0.8</td><td>9.6</td><td>8.0E-22</td><td>4.4E-21</td><td>MB231/MB468</td></tr>
<tr><td>>am_trnaMT_ProTGG_MT_-_15956_16023@53.71.19|AAGACTTTTTCTCTGACCA</td><td>>am_trnaMT_ProTGG_MT_-_15956_16023</td><td>53</td><td>71</td><td>19</td><td>AAGACTTTTTCTCTGACCA</td><td>11479.5</td><td>5.3</td><td>0.6</td><td>9.6</td><td>8.0E-22</td><td>4.4E-21</td><td>MB231/MB468</td></tr>
<tr><td>>am_MI0000786|hsa-mir-378a&WithFlank&5|+|149732819|149732896@49.70.22(+1G)&[MIMAT0000732&hsa-miR-378a-3p&offsets|0|0(+1G);m-44&5|+|149732867|149732888&offsets|0|0(+1G)]|ACTGGACTTGGAGTCAGAAGGCG</td><td>>am_MI0000786|hsa-mir-378a&WithFlank&5|+|149732819|149732896</td><td>49</td><td>70</td><td>22(+1G)</td><td>ACTGGACTTGGAGTCAGAAGGCG</td><td>570.6</td><td>4.7</td><td>0.5</td><td>9.6</td><td>6.9E-22</td><td>4.5E-21</td><td>BT20/MB468</td></tr>
<tr><td>>MI0000811|hsa-mir-148b&WithFlank&12|+|54337210|54337320@69.90.22&[MIMAT0000759&hsa-miR-148b-3p&offsets|0|0;m-77&12|+|54337278|54337299&offsets|0|0]|TCAGTGCATCACAGAACTTTGT</td><td>>MI0000811|hsa-mir-148b&WithFlank&12|+|54337210|54337320</td><td>69</td><td>90</td><td>22</td><td>TCAGTGCATCACAGAACTTTGT</td><td>32976.4</td><td>8.2</td><td>0.9</td><td>9.6</td><td>8.3E-22</td><td>4.5E-21</td><td>MB231/MB468</td></tr>
<tr><td>>hg38_21_+_8433172_8446622_RNA45SN1_45S_50ntFlanks@11564.11591.28|CCTAGCAGCCGACTTAGAACTGGTGCGG</td><td>>hg38_21_+_8433172_8446622_RNA45SN1_45S_50ntFlanks</td><td>11564</td><td>11591</td><td>28</td><td>CCTAGCAGCCGACTTAGAACTGGTGCGG</td><td>1556.9</td><td>-6.7</td><td>0.7</td><td>-9.6</td><td>7.0E-22</td><td>4.6E-21</td><td>BT20/MB468</td></tr>
<tr><td>>trnaMT_PheGAA_MT_+_577_647@35.54.20|GAAAATGTTTAGACGGGCTC</td><td>>trnaMT_PheGAA_MT_+_577_647</td><td>35</td><td>54</td><td>20</td><td>GAAAATGTTTAGACGGGCTC</td><td>919.7</td><td>9.9</td><td>1.0</td><td>9.6</td><td>7.3E-22</td><td>4.8E-21</td><td>BT20/MB468</td></tr>
<tr><td>>hg38_21_+_8433172_8446622_RNA45SN1_45S_50ntFlanks@10402.10431.30|AACATGGGTCAGTCGGTCCTGAGAGATGGG</td><td>>hg38_21_+_8433172_8446622_RNA45SN1_45S_50ntFlanks</td><td>10402</td><td>10431</td><td>30</td><td>AACATGGGTCAGTCGGTCCTGAGAGATGGG</td><td>621.9</td><td>-6.3</td><td>0.7</td><td>-9.6</td><td>7.4E-22</td><td>4.9E-21</td><td>BT20/MB468</td></tr>
<tr><td>>MI0003560|hsa-mir-92b&WithFlank&1|+|155195171|155195278@67.88.22(+1U)&[MIMAT0003218&hsa-miR-92b-3p&offsets|0|0(+1U);m-115&1|+|155195237|155195258&offsets|0|0(+1U)]|TATTGCACTCGTCCCGGCCTCCT</td><td>>MI0003560|hsa-mir-92b&WithFlank&1|+|155195171|155195278</td><td>67</td><td>88</td><td>22(+1U)</td><td>TATTGCACTCGTCCCGGCCTCCT</td><td>1284.6</td><td>9.6</td><td>1.0</td><td>9.6</td><td>9.0E-22</td><td>4.9E-21</td><td>MB231/MB468</td></tr>
<tr><td>>hg38_21_+_8433172_8446622_RNA45SN1_45S_50ntFlanks@476.503.28|TTCGTGATCGATGTGGTGACGTCGTGCT</td><td>>hg38_21_+_8433172_8446622_RNA45SN1_45S_50ntFlanks</td><td>476</td><td>503</td><td>28</td><td>TTCGTGATCGATGTGGTGACGTCGTGCT</td><td>4601.5</td><td>6.7</td><td>0.7</td><td>9.6</td><td>7.6E-22</td><td>4.9E-21</td><td>BT20/MB468</td></tr>
<tr><td>>hg38_21_+_8433172_8446622_RNA45SN1_45S_50ntFlanks@10392.10409.18|ACAGCAGTTGAACATGGG</td><td>>hg38_21_+_8433172_8446622_RNA45SN1_45S_50ntFlanks</td><td>10392</td><td>10409</td><td>18</td><td>ACAGCAGTTGAACATGGG</td><td>2630.6</td><td>-10.3</td><td>1.1</td><td>-9.6</td><td>7.7E-22</td><td>5.0E-21</td><td>BT20/MB468</td></tr>
<tr><td>>hg38_21_+_8433172_8446622_RNA45SN1_45S_50ntFlanks@5473.5512.40|CGGCCCTGGCGGAGCGCTGAGAAGACGGTCGAACTTGACT</td><td>>hg38_21_+_8433172_8446622_RNA45SN1_45S_50ntFlanks</td><td>5473</td><td>5512</td><td>40</td><td>CGGCCCTGGCGGAGCGCTGAGAAGACGGTCGAACTTGACT</td><td>572.4</td><td>9.3</td><td>1.0</td><td>9.6</td><td>9.3E-22</td><td>5.1E-21</td><td>MB231/MB468</td></tr>
<tr><td>>MI0000462|hsa-mir-152&WithFlank&17|-|48037155|48037253@60.80.21(+1A)&[MIMAT0000438&hsa-miR-152-3p&offsets|0|0(+1A);m-97&17|-|48037174|48037194&offsets|0|0(+1A)]|TCAGTGCATGACAGAACTTGGA</td><td>>MI0000462|hsa-mir-152&WithFlank&17|-|48037155|48037253</td><td>60</td><td>80</td><td>21(+1A)</td><td>TCAGTGCATGACAGAACTTGGA</td><td>666.4</td><td>14.0</td><td>1.5</td><td>9.6</td><td>7.8E-22</td><td>5.1E-21</td><td>BT20/MB468</td></tr>
<tr><td>>hg38_21_+_8433172_8446622_RNA45SN1_45S_50ntFlanks@4732.4767.36|AGAACGAAAGTCGGAGGTTCGAAGACGATCAGATAC</td><td>>hg38_21_+_8433172_8446622_RNA45SN1_45S_50ntFlanks</td><td>4732</td><td>4767</td><td>36</td><td>AGAACGAAAGTCGGAGGTTCGAAGACGATCAGATAC</td><td>1213.5</td><td>11.0</td><td>1.1</td><td>9.6</td><td>9.3E-22</td><td>5.1E-21</td><td>MB231/MB468</td></tr>
<tr><td>>hg38_21_+_8433172_8446622_RNA45SN1_45S_50ntFlanks@12413.12448.36|TCCTTCGATGTCGGCTCTTCCTATCATTGTGAAGCA</td><td>>hg38_21_+_8433172_8446622_RNA45SN1_45S_50ntFlanks</td><td>12413</td><td>12448</td><td>36</td><td>TCCTTCGATGTCGGCTCTTCCTATCATTGTGAAGCA</td><td>292.5</td><td>8.6</td><td>0.9</td><td>9.6</td><td>9.3E-22</td><td>5.1E-21</td><td>MB231/MB468</td></tr>
<tr><td>>MI0003591|hsa-mir-584&WithFlank&5|-|149062307|149062415@22.41.20&[MIMAT0003249&hsa-miR-584-5p&offsets|0|-2;m-167&5|-|149062374|149062394&offsets|0|-1]|TTATGGTTTGCCTGGGACTG</td><td>>MI0003591|hsa-mir-584&WithFlank&5|-|149062307|149062415</td><td>22</td><td>41</td><td>20</td><td>TTATGGTTTGCCTGGGACTG</td><td>851.0</td><td>9.1</td><td>0.9</td><td>9.6</td><td>7.8E-22</td><td>5.1E-21</td><td>BT20/MB468</td></tr>
<tr><td>>trnaMT_GluTTC_MT_-_14674_14742@40.66.27,trnalookalike8_GluTTC_5_-_93905172_93905240@40.66.27|TTGGTCGTGGTTGTAGTCCGTGCGAGA</td><td>>trnaMT_GluTTC_MT_-_14674_14742</td><td>40</td><td>66</td><td>27</td><td>TTGGTCGTGGTTGTAGTCCGTGCGAGA</td><td>711.8</td><td>7.9</td><td>0.8</td><td>9.6</td><td>7.9E-22</td><td>5.1E-21</td><td>BT20/MB468</td></tr>
<tr><td>>MI0000289|hsa-mir-181a-1&WithFlank&1|-|198859038|198859159@30.54.25;MI0000269|hsa-mir-181a-2&WithFlank&9|+|124692436|124692557@45.69.25&[MIMAT0000256&hsa-miR-181a-5p&offsets|0|+2;m-32&1|-|198859108|198859130&offsets|0|+2];[MIMAT0000256_1&hsa-miR-181a-5p&offsets|0|+2;m-32&9|+|124692480|124692502&offsets|0|+2]|AACATTCAACGCTGTCGGTGAGTTT</td><td>>MI0000289|hsa-mir-181a-1&WithFlank&1|-|198859038|198859159</td><td>30</td><td>54</td><td>25</td><td>AACATTCAACGCTGTCGGTGAGTTT</td><td>5563.8</td><td>8.0</td><td>0.8</td><td>9.6</td><td>9.5E-22</td><td>5.2E-21</td><td>MB231/MB468</td></tr>
<tr><td>>trnaMT_HisGTG_MT_+_12138_12206@32.70.39|TGAATCTGACAACAGAGGCTTACGACCCCTTATTTACCC</td><td>>trnaMT_HisGTG_MT_+_12138_12206</td><td>32</td><td>70</td><td>39</td><td>TGAATCTGACAACAGAGGCTTACGACCCCTTATTTACCC</td><td>8072.8</td><td>9.1</td><td>1.0</td><td>9.6</td><td>9.6E-22</td><td>5.2E-21</td><td>MB231/MB468</td></tr>
<tr><td>>trnaMT_LeuTAA_MT_+_3230_3304@1.37.37|GTTAAGATGGCAGAGCCCGGTAATCGCATAAAACTTA</td><td>>trnaMT_LeuTAA_MT_+_3230_3304</td><td>1</td><td>37</td><td>37</td><td>GTTAAGATGGCAGAGCCCGGTAATCGCATAAAACTTA</td><td>852.1</td><td>12.6</td><td>1.3</td><td>9.6</td><td>9.7E-22</td><td>5.3E-21</td><td>MB231/MB468</td></tr>
<tr><td>>MI0000441|hsa-mir-30b&WithFlank&8|-|134800514|134800613@23.43.21(+1C)&[MIMAT0000420&hsa-miR-30b-5p&offsets|0|-1(+1C);m-76&8|-|134800570|134800591&offsets|0|-1(+1C)]|TGTAAACATCCTACACTCAGCC</td><td>>MI0000441|hsa-mir-30b&WithFlank&8|-|134800514|134800613</td><td>23</td><td>43</td><td>21(+1C)</td><td>TGTAAACATCCTACACTCAGCC</td><td>9001.9</td><td>10.4</td><td>1.1</td><td>9.6</td><td>8.3E-22</td><td>5.4E-21</td><td>BT20/MB468</td></tr>
<tr><td>>MI0000299|hsa-mir-222&WithFlank&X|-|45747009|45747130@75.98.24(+1C)&[MIMAT0000279&hsa-miR-222-3p&offsets|0|+3(+1C);m-140&X|-|45747033|45747056&offsets|0|0(+1C)]|AGCTACATCTGGCTACTGGGTCTCC</td><td>>MI0000299|hsa-mir-222&WithFlank&X|-|45747009|45747130</td><td>75</td><td>98</td><td>24(+1C)</td><td>AGCTACATCTGGCTACTGGGTCTCC</td><td>3813.4</td><td>6.3</td><td>0.7</td><td>9.6</td><td>8.3E-22</td><td>5.4E-21</td><td>BT20/MB468</td></tr>
<tr><td>>am_MI0014244|hsa-mir-548x&WithFlank&21|-|18686084|18686170@52.73.22&[MIMAT0015081_1&hsa-miR-548x-3p&offsets|0|+2;m-1121&21|-|18686099|18686119&offsets|0|+1]|TAAAAACTGCAATTACTTTCAC</td><td>>am_MI0014244|hsa-mir-548x&WithFlank&21|-|18686084|18686170</td><td>52</td><td>73</td><td>22</td><td>TAAAAACTGCAATTACTTTCAC</td><td>696.8</td><td>14.1</td><td>1.5</td><td>9.6</td><td>8.4E-22</td><td>5.5E-21</td><td>BT20/MB468</td></tr>
<tr><td>>hg38_21_+_8433172_8446622_RNA45SN1_45S_50ntFlanks@5491.5512.22|GAGAAGACGGTCGAACTTGACT</td><td>>hg38_21_+_8433172_8446622_RNA45SN1_45S_50ntFlanks</td><td>5491</td><td>5512</td><td>22</td><td>GAGAAGACGGTCGAACTTGACT</td><td>787.1</td><td>8.6</td><td>0.9</td><td>9.6</td><td>1.0E-21</td><td>5.5E-21</td><td>MB231/MB468</td></tr>
<tr><td>>hg38_21_+_8433172_8446622_RNA45SN1_45S_50ntFlanks@6652.6693.42|GACTCTTAGCGGTGGATCACTCGGCTCGTGCGTCGATGAAGA</td><td>>hg38_21_+_8433172_8446622_RNA45SN1_45S_50ntFlanks</td><td>6652</td><td>6693</td><td>42</td><td>GACTCTTAGCGGTGGATCACTCGGCTCGTGCGTCGATGAAGA</td><td>1406.2</td><td>6.5</td><td>0.7</td><td>9.6</td><td>1.0E-21</td><td>5.5E-21</td><td>MB231/MB468</td></tr>
</table>

<table>
<tr><td>>trna111_HisGTG_1_-_147774845_147774916@-1G.34.35,trna118_HisGTG_1_-_145396881_145396952@-1G.34.35,trna16_HisGTG_1_+_146544773_146544844@-1G.34.35,trna1_HisGTG_15_+_45493349_45493420@-1G.34.35,trna21_HisGTG_1_+_147753471_147753542@-1G.34.35,trna33_HisGTG_6_+_27125906_27125977@-1G.34.35,trna7_HisGTG_9_-_14433938_14434009@-1G.34.35,trna8_HisGTG_15_-_45492611_45492682@-1G.34.35,trna9_HisGTG_15_-_45490804_45490875@-1G.34.35|GGCCGTGATCGTATAGTGGTTAGTACTCTGCGTTG</td><td>>trna111_HisGTG_1_-_147774845_147774916</td><td>-1G</td><td>34</td><td>35</td><td>GGCCGTGATCGTATAGTGGTTAGTACTCTGCGTTG</td><td>45689.2</td><td>9.4</td><td>1.0</td><td>9.6</td><td>1.0E-21</td><td>5.5E-21</td><td>MB231/MB468</td></tr>
<tr><td>>hg38_21_+_8433172_8446622_RNA45SN1_45S_50ntFlanks@9534.9566.33|AGGAAACTCTGGTGGAGGTCCGTAGCGGTCCTG</td><td>>hg38_21_+_8433172_8446622_RNA45SN1_45S_50ntFlanks</td><td>9534</td><td>9566</td><td>33</td><td>AGGAAACTCTGGTGGAGGTCCGTAGCGGTCCTG</td><td>1281.9</td><td>-6.0</td><td>0.6</td><td>-9.6</td><td>1.0E-21</td><td>5.6E-21</td><td>MB231/MB468</td></tr>
<tr><td>>hg38_MT_+_598_1651_RNR1_12S_50ntFlanks@52.87.36|ATAGGTTTGGTCCTAGCCTTTCTATTAGCTCTTAGT</td><td>>hg38_MT_+_598_1651_RNR1_12S_50ntFlanks</td><td>52</td><td>87</td><td>36</td><td>ATAGGTTTGGTCCTAGCCTTTCTATTAGCTCTTAGT</td><td>727.8</td><td>6.4</td><td>0.7</td><td>9.6</td><td>1.1E-21</td><td>5.9E-21</td><td>MB231/MB468</td></tr>
<tr><td>>MI0000087|hsa-mir-29a&WithFlank&7|-|130876741|130876816@48.67.20(+1G)&[MIMAT0000086&hsa-miR-29a-3p&offsets|0|-2(+1G);m-26&7|-|130876749|130876769&offsets|0|-1(+1G)]|TAGCACCATCTGAAATCGGTG</td><td>>MI0000087|hsa-mir-29a&WithFlank&7|-|130876741|130876816</td><td>48</td><td>67</td><td>20(+1G)</td><td>TAGCACCATCTGAAATCGGTG</td><td>987.4</td><td>9.9</td><td>1.0</td><td>9.6</td><td>1.1E-21</td><td>6.2E-21</td><td>MB231/MB468</td></tr>
<tr><td>>hg38_21_+_8433172_8446622_RNA45SN1_45S_50ntFlanks@12934.12959.26|CTAAACCATTCGTAGACGACCTGCTT</td><td>>hg38_21_+_8433172_8446622_RNA45SN1_45S_50ntFlanks</td><td>12934</td><td>12959</td><td>26</td><td>CTAAACCATTCGTAGACGACCTGCTT</td><td>567.4</td><td>-4.9</td><td>0.5</td><td>-9.6</td><td>9.6E-22</td><td>6.2E-21</td><td>BT20/MB468</td></tr>
<tr><td>>MI0000264|hsa-mir-7-2&WithFlank&15|+|88611819|88611940@38.60.23(+1C);MI0000265|hsa-mir-7-3&WithFlank&19|+|4770664|4770785@37.59.23(+1C);MI0000263|hsa-mir-7-1&WithFlank&9|-|83969742|83969863@30.52.23(+1C)&[MIMAT0000252&hsa-miR-7-5p&offsets|0|-1(+1C);m-400&15|+|88611856|88611879&offsets|0|-1(+1C)];[MIMAT0000252_1&hsa-miR-7-5p&offsets|0|-1(+1C);m-402&19|+|4770700|4770723&offsets|0|-1(+1C)];[MIMAT0000252_2&hsa-miR-7-5p&offsets|0|-1(+1C);m-401&9|-|83969811|83969834&offsets|0|-1(+1C)]|TGGAAGACTAGTGATTTTGTTGTC</td><td>>MI0000264|hsa-mir-7-2&WithFlank&15|+|88611819|88611940</td><td>38</td><td>60</td><td>23(+1C)</td><td>TGGAAGACTAGTGATTTTGTTGTC</td><td>44553.6</td><td>8.0</td><td>0.8</td><td>9.6</td><td>9.7E-22</td><td>6.3E-21</td><td>BT20/MB468</td></tr>
<tr><td>>hg38_21_+_8433172_8446622_RNA45SN1_45S_50ntFlanks@12623.12654.32|GCCAATGGGGCGAAGCTACCATCTGTGGGATT</td><td>>hg38_21_+_8433172_8446622_RNA45SN1_45S_50ntFlanks</td><td>12623</td><td>12654</td><td>32</td><td>GCCAATGGGGCGAAGCTACCATCTGTGGGATT</td><td>207.4</td><td>-5.1</td><td>0.5</td><td>-9.6</td><td>9.9E-22</td><td>6.4E-21</td><td>BT20/MB468</td></tr>
<tr><td>>hg38_21_+_8433172_8446622_RNA45SN1_45S_50ntFlanks@7977.8016.40|CGACCTCAGATCAGACGTGGCGACCCGCTGAATTTAAGCA</td><td>>hg38_21_+_8433172_8446622_RNA45SN1_45S_50ntFlanks</td><td>7977</td><td>8016</td><td>40</td><td>CGACCTCAGATCAGACGTGGCGACCCGCTGAATTTAAGCA</td><td>2055.2</td><td>10.4</td><td>1.1</td><td>9.6</td><td>1.2E-21</td><td>6.5E-21</td><td>MB231/MB468</td></tr>
<tr><td>>hg38_21_+_8433172_8446622_RNA45SN1_45S_50ntFlanks@4588.4614.27|CTATTTTGTTGGTTTTCGGAACTGAGG</td><td>>hg38_21_+_8433172_8446622_RNA45SN1_45S_50ntFlanks</td><td>4588</td><td>4614</td><td>27</td><td>CTATTTTGTTGGTTTTCGGAACTGAGG</td><td>382.2</td><td>-5.2</td><td>0.5</td><td>-9.6</td><td>1.2E-21</td><td>6.6E-21</td><td>MB231/MB468</td></tr>
<tr><td>>hg38_21_+_8433172_8446622_RNA45SN1_45S_50ntFlanks@12405.12436.32|CTTTTTGATCCTTCGATGTCGGCTCTTCCTAT</td><td>>hg38_21_+_8433172_8446622_RNA45SN1_45S_50ntFlanks</td><td>12405</td><td>12436</td><td>32</td><td>CTTTTTGATCCTTCGATGTCGGCTCTTCCTAT</td><td>5521.6</td><td>-9.9</td><td>1.0</td><td>-9.6</td><td>1.0E-21</td><td>6.6E-21</td><td>BT20/MB468</td></tr>
<tr><td>>hg38_21_+_8433172_8446622_RNA45SN1_45S_50ntFlanks@12004.12033.30|CGGGCCGCCGGTGAAATACCACTACTCTGA</td><td>>hg38_21_+_8433172_8446622_RNA45SN1_45S_50ntFlanks</td><td>12004</td><td>12033</td><td>30</td><td>CGGGCCGCCGGTGAAATACCACTACTCTGA</td><td>3568.6</td><td>9.6</td><td>1.0</td><td>9.6</td><td>1.0E-21</td><td>6.7E-21</td><td>BT20/MB468</td></tr>
<tr><td>>MI0000061|hsa-let-7a-2&WithFlank&11|-|122146516|122146599@11.31.21;MI0000062|hsa-let-7a-3&WithFlank&22|+|46112743|46112828@10.30.21;MI0000060|hsa-let-7a-1&WithFlank&9|+|94175951|94176042@12.32.21&[MIMAT0000062&hsa-let-7a-5p&offsets|0|-1;m-6&11|-|122146568|122146589&offsets|0|-1];[MIMAT0000062_1&hsa-let-7a-5p&offsets|0|-1;m-5&22|+|46112752|46112773&offsets|0|-1];[MIMAT0000062_2&hsa-let-7a-5p&offsets|0|-1;m-4&9|+|94175962|94175983&offsets|0|-1]|TGAGGTAGTAGGTTGTATAGT</td><td>>MI0000061|hsa-let-7a-2&WithFlank&11|-|122146516|122146599</td><td>11</td><td>31</td><td>21</td><td>TGAGGTAGTAGGTTGTATAGT</td><td>18239.9</td><td>7.3</td><td>0.8</td><td>9.6</td><td>1.1E-21</td><td>6.8E-21</td><td>BT20/MB468</td></tr>
<tr><td>>MI0000093|hsa-mir-92a-1&WithFlank&13|+|91351308|91351397@54.74.21(+1G);MI0000094|hsa-mir-92a-2&WithFlank&X|-|134169532|134169618@54.74.21(+1G)&[MIMAT0000092&hsa-miR-92a-3p&offsets|0|-1(+1G);m-9&13|+|91351361|91351382&offsets|0|-1(+1G)];[MIMAT0000092_1&hsa-miR-92a-3p&offsets|0|-1(+1G);m-11&X|-|134169544|134169565&offsets|0|-1(+1G)]|TATTGCACTTGTCCCGGCCTGG</td><td>>MI0000093|hsa-mir-92a-1&WithFlank&13|+|91351308|91351397</td><td>54</td><td>74</td><td>21(+1G)</td><td>TATTGCACTTGTCCCGGCCTGG</td><td>8569.3</td><td>7.6</td><td>0.8</td><td>9.6</td><td>1.3E-21</td><td>6.9E-21</td><td>MB231/MB468</td></tr>
<tr><td>>MI0000264|hsa-mir-7-2&WithFlank&15|+|88611819|88611940@38.60.23(+1A);MI0000265|hsa-mir-7-3&WithFlank&19|+|4770664|4770785@37.59.23(+1A);MI0000263|hsa-mir-7-1&WithFlank&9|-|83969742|83969863@30.52.23(+1A)&[MIMAT0000252&hsa-miR-7-5p&offsets|0|-1(+1A);m-400&15|+|88611856|88611879&offsets|0|-1(+1A)];[MIMAT0000252_1&hsa-miR-7-5p&offsets|0|-1(+1A);m-402&19|+|4770700|4770723&offsets|0|-1(+1A)];[MIMAT0000252_2&hsa-miR-7-5p&offsets|0|-1(+1A);m-401&9|-|83969811|83969834&offsets|0|-1(+1A)]|TGGAAGACTAGTGATTTTGTTGTA</td><td>>MI0000264|hsa-mir-7-2&WithFlank&15|+|88611819|88611940</td><td>38</td><td>60</td><td>23(+1A)</td><td>TGGAAGACTAGTGATTTTGTTGTA</td><td>26176.8</td><td>9.7</td><td>1.0</td><td>9.6</td><td>1.1E-21</td><td>7.0E-21</td><td>BT20/MB468</td></tr>
<tr><td>>MI0000272|hsa-mir-182&WithFlank&7|-|129770377|129770498@29.51.23(+1G)&[MIMAT0000259&hsa-miR-182-5p&offsets|0|-1(+1G);m-22&7|-|129770449|129770470&offsets|0|+1(+1G)]|TTTGGCAATGGTAGAACTCACACG</td><td>>MI0000272|hsa-mir-182&WithFlank&7|-|129770377|129770498</td><td>29</td><td>51</td><td>23(+1G)</td><td>TTTGGCAATGGTAGAACTCACACG</td><td>17901.6</td><td>9.4</td><td>1.0</td><td>9.6</td><td>1.1E-21</td><td>7.0E-21</td><td>BT20/MB468</td></tr>
<tr><td>>am_MI0000809|hsa-mir-151a&WithFlank&8|-|140732558|140732659@53.70.18(+3G)&[MIMAT0000757&hsa-miR-151a-3p&offsets|0|-3(+3G);m-30&8|-|140732587|140732607&offsets|0|-3(+3G)]|CTAGACTGAAGCTCCTTGGGG</td><td>>am_MI0000809|hsa-mir-151a&WithFlank&8|-|140732558|140732659</td><td>53</td><td>70</td><td>18(+3G)</td><td>CTAGACTGAAGCTCCTTGGGG</td><td>778.0</td><td>8.7</td><td>0.9</td><td>9.6</td><td>1.1E-21</td><td>7.0E-21</td><td>BT20/MB468</td></tr>
<tr><td>>MI0000077|hsa-mir-21&WithFlank&17|+|59841260|59841343@14.38.25&[MIMAT0000076&hsa-miR-21-5p&offsets|0|+3;m-1&17|+|59841273|59841294&offsets|0|+3]|TAGCTTATCAGACTGATGTTGACTG</td><td>>MI0000077|hsa-mir-21&WithFlank&17|+|59841260|59841343</td><td>14</td><td>38</td><td>25</td><td>TAGCTTATCAGACTGATGTTGACTG</td><td>572.9</td><td>7.5</td><td>0.8</td><td>9.6</td><td>1.1E-21</td><td>7.1E-21</td><td>BT20/MB468</td></tr>
<tr><td>>trnaMT_PheGAA_MT_+_577_647@38.59.22|AATGTTTAGACGGGCTCACATC</td><td>>trnaMT_PheGAA_MT_+_577_647</td><td>38</td><td>59</td><td>22</td><td>AATGTTTAGACGGGCTCACATC</td><td>464.9</td><td>9.2</td><td>1.0</td><td>9.6</td><td>1.1E-21</td><td>7.1E-21</td><td>BT20/MB468</td></tr>
<tr><td>>MI0000061|hsa-let-7a-2&WithFlank&11|-|122146516|122146599@11.32.22(+1C);MI0000062|hsa-let-7a-3&WithFlank&22|+|46112743|46112828@10.31.22(+1C);MI0000060|hsa-let-7a-1&WithFlank&9|+|94175951|94176042@12.33.22(+1C)&[MIMAT0000062&hsa-let-7a-5p&offsets|0|0(+1C);m-6&11|-|122146568|122146589&offsets|0|0(+1C)];[MIMAT0000062_1&hsa-let-7a-5p&offsets|0|0(+1C);m-5&22|+|46112752|46112773&offsets|0|0(+1C)];[MIMAT0000062_2&hsa-let-7a-5p&offsets|0|0(+1C);m-4&9|+|94175962|94175983&offsets|0|0(+1C)]|TGAGGTAGTAGGTTGTATAGTTC</td><td>>MI0000061|hsa-let-7a-2&WithFlank&11|-|122146516|122146599</td><td>11</td><td>32</td><td>22(+1C)</td><td>TGAGGTAGTAGGTTGTATAGTTC</td><td>578.7</td><td>7.3</td><td>0.8</td><td>9.6</td><td>1.1E-21</td><td>7.2E-21</td><td>BT20/MB468</td></tr>
<tr><td>>MI0000465|hsa-mir-191&WithFlank&3|-|49020612|49020715@22.43.22(+1U)&[MIMAT0000440&hsa-miR-191-5p&offsets|0|-1(+1U);m-48&3|-|49020672|49020694&offsets|0|-1(+1U)]|CAACGGAATCCCAAAAGCAGCTT</td><td>>MI0000465|hsa-mir-191&WithFlank&3|-|49020612|49020715</td><td>22</td><td>43</td><td>22(+1U)</td><td>CAACGGAATCCCAAAAGCAGCTT</td><td>784.1</td><td>7.2</td><td>0.8</td><td>9.5</td><td>1.3E-21</td><td>7.2E-21</td><td>MB231/MB468</td></tr>
<tr><td>>hg38_MT_+_598_1651_RNR1_12S_50ntFlanks@52.88.37|ATAGGTTTGGTCCTAGCCTTTCTATTAGCTCTTAGTA</td><td>>hg38_MT_+_598_1651_RNR1_12S_50ntFlanks</td><td>52</td><td>88</td><td>37</td><td>ATAGGTTTGGTCCTAGCCTTTCTATTAGCTCTTAGTA</td><td>2658.5</td><td>7.5</td><td>0.8</td><td>9.5</td><td>1.3E-21</td><td>7.2E-21</td><td>MB231/MB468</td></tr>
<tr><td>>hg38_21_+_8433172_8446622_RNA45SN1_45S_50ntFlanks@10683.10706.24|CCGGCGGCGTCCGGTGAGCTCTCG</td><td>>hg38_21_+_8433172_8446622_RNA45SN1_45S_50ntFlanks</td><td>10683</td><td>10706</td><td>24</td><td>CCGGCGGCGTCCGGTGAGCTCTCG</td><td>4714.8</td><td>-6.4</td><td>0.7</td><td>-9.5</td><td>1.4E-21</td><td>7.3E-21</td><td>MB231/MB468</td></tr>
<tr><td>>MI0000273|hsa-mir-183&WithFlank&7|-|129774899|129775020@33.53.21(+1G)&[MIMAT0000261&hsa-miR-183-5p&offsets|0|-1(+1G);m-41&7|-|129774966|129774987&offsets|-1|-2(+1G)]|TATGGCACTGGTAGAATTCACG</td><td>>MI0000273|hsa-mir-183&WithFlank&7|-|129774899|129775020</td><td>33</td><td>53</td><td>21(+1G)</td><td>TATGGCACTGGTAGAATTCACG</td><td>3290.2</td><td>8.4</td><td>0.9</td><td>9.6</td><td>1.1E-21</td><td>7.3E-21</td><td>BT20/MB468</td></tr>
<tr><td>>hg38_1_-_228634819_228635039_RNA5S12_5S_50ntFlanks@152.169.18|TACCGGGTGCTGTAGGCT</td><td>>hg38_1_-_228634819_228635039_RNA5S12_5S_50ntFlanks</td><td>152</td><td>169</td><td>18</td><td>TACCGGGTGCTGTAGGCT</td><td>708.3</td><td>5.8</td><td>0.6</td><td>9.6</td><td>1.1E-21</td><td>7.3E-21</td><td>BT20/MB468</td></tr>
<tr><td>>hg38_21_+_8433172_8446622_RNA45SN1_45S_50ntFlanks@10458.10477.20|ATGGCCTCCGTTGCCCTCGG</td><td>>hg38_21_+_8433172_8446622_RNA45SN1_45S_50ntFlanks</td><td>10458</td><td>10477</td><td>20</td><td>ATGGCCTCCGTTGCCCTCGG</td><td>2239.9</td><td>-9.8</td><td>1.0</td><td>-9.6</td><td>1.1E-21</td><td>7.3E-21</td><td>BT20/MB468</td></tr>
<tr><td>>hg38_21_+_8433172_8446622_RNA45SN1_45S_50ntFlanks@12419.12457.39|GATGTCGGCTCTTCCTATCATTGTGAAGCAGAATTCACC</td><td>>hg38_21_+_8433172_8446622_RNA45SN1_45S_50ntFlanks</td><td>12419</td><td>12457</td><td>39</td><td>GATGTCGGCTCTTCCTATCATTGTGAAGCAGAATTCACC</td><td>1953.2</td><td>10.6</td><td>1.1</td><td>9.5</td><td>1.4E-21</td><td>7.4E-21</td><td>MB231/MB468</td></tr>
<tr><td>>MI0000434|hsa-let-7i&WithFlank&12|+|62603680|62603775@12.32.21(+1A)&[MIMAT0000415&hsa-let-7i-5p&offsets|0|-1(+1A);m-49&12|+|62603691|62603712&offsets|0|-1(+1A)]|TGAGGTAGTAGTTTGTGCTGTA</td><td>>MI0000434|hsa-let-7i&WithFlank&12|+|62603680|62603775</td><td>12</td><td>32</td><td>21(+1A)</td><td>TGAGGTAGTAGTTTGTGCTGTA</td><td>15624.4</td><td>7.8</td><td>0.8</td><td>9.6</td><td>1.2E-21</td><td>7.4E-21</td><td>BT20/MB468</td></tr>
<tr><td>>hg38_21_+_8433172_8446622_RNA45SN1_45S_50ntFlanks@475.503.29|CTTCGTGATCGATGTGGTGACGTCGTGCT</td><td>>hg38_21_+_8433172_8446622_RNA45SN1_45S_50ntFlanks</td><td>475</td><td>503</td><td>29</td><td>CTTCGTGATCGATGTGGTGACGTCGTGCT</td><td>8350.4</td><td>6.8</td><td>0.7</td><td>9.6</td><td>1.2E-21</td><td>7.4E-21</td><td>BT20/MB468</td></tr>
<tr><td>>MI0000285|hsa-mir-205&WithFlank&1|+|209432127|209432248@40.62.23(+1A)&[MIMAT0000266&hsa-miR-205-5p&offsets|0|+1(+1A);m-351&1|+|209432166|209432187&offsets|0|+1(+1A)]|TCCTTCATTCCACCGGAGTCTGTA</td><td>>MI0000285|hsa-mir-205&WithFlank&1|+|209432127|209432248</td><td>40</td><td>62</td><td>23(+1A)</td><td>TCCTTCATTCCACCGGAGTCTGTA</td><td>1809.4</td><td>14.3</td><td>1.5</td><td>9.9</td><td>5.7E-23</td><td>7.5E-21</td><td>BT20/MB231</td></tr>
<tr><td>>hg38_21_+_8433172_8446622_RNA45SN1_45S_50ntFlanks@11873.11890.18|AAGAAGACCCTGTTGAGC</td><td>>hg38_21_+_8433172_8446622_RNA45SN1_45S_50ntFlanks</td><td>11873</td><td>11890</td><td>18</td><td>AAGAAGACCCTGTTGAGC</td><td>968.0</td><td>-5.3</td><td>0.6</td><td>-9.6</td><td>1.2E-21</td><td>7.5E-21</td><td>BT20/MB468</td></tr>
</table>

<table>
<tr><td>>MI0006444|hsa-mir-1307&WithFlank&10|-|103394247|103394407@86.109.24&[MIMAT0005951&hsa-miR-1307-3p&offsets|0|+2;m-66&10|-|103394300|103394322&offsets|0|+1]|ACTCGGCGTGGCGTCGGTCGTGGT</td><td>>MI0006444|hsa-mir-1307&WithFlank&10|-|103394247|103394407</td><td>86</td><td>109</td><td>24</td><td>ACTCGGCGTGGCGTCGGTCGTGGT</td><td>558.2</td><td>6.5</td><td>0.7</td><td>9.6</td><td>1.2E-21</td><td>7.7E-21</td><td>BT20/MB468</td></tr>
<tr><td>>MI0000466|hsa-mir-9-1&WithFlank&1|-|156420335|156420435@22.42.21(+1A);MI0000468|hsa-mir-9-3&WithFlank&15|+|89368011|89368112@22.42.21(+1A);MI0000467|hsa-mir-9-2&WithFlank&5|-|88666847|88666945@22.42.21(+1A)&[MIMAT0000441&hsa-miR-9-5p&offsets|0|-2(+1A);m-104&1|-|156420392|156420414&offsets|0|-2(+1A)];[MIMAT0000441_1&hsa-miR-9-5p&offsets|0|-2(+1A);m-105&15|+|89368032|89368054&offsets|0|-2(+1A)];[MIMAT0000441_2&hsa-miR-9-5p&offsets|0|-2(+1A);m-104&5|-|88666902|88666924&offsets|0|-2(+1A)]|TCTTTGGTTATCTAGCTGTATA</td><td>>MI0000466|hsa-mir-9-1&WithFlank&1|-|156420335|156420435</td><td>22</td><td>42</td><td>21(+1A)</td><td>TCTTTGGTTATCTAGCTGTATA</td><td>1362.4</td><td>15.1</td><td>1.6</td><td>9.6</td><td>1.3E-21</td><td>8.0E-21</td><td>BT20/MB468</td></tr>
<tr><td>>hg38_21_+_8433172_8446622_RNA45SN1_45S_50ntFlanks@6648.6668.21|GTACGACTCTTAGCGGTGGAT</td><td>>hg38_21_+_8433172_8446622_RNA45SN1_45S_50ntFlanks</td><td>6648</td><td>6668</td><td>21</td><td>GTACGACTCTTAGCGGTGGAT</td><td>1789.1</td><td>4.8</td><td>0.5</td><td>9.6</td><td>1.3E-21</td><td>8.0E-21</td><td>BT20/MB468</td></tr>
<tr><td>>hg38_21_+_8433172_8446622_RNA45SN1_45S_50ntFlanks@5237.5255.19|ACTGACTGGCTCAGCGTGT</td><td>>hg38_21_+_8433172_8446622_RNA45SN1_45S_50ntFlanks</td><td>5237</td><td>5255</td><td>19</td><td>ACTGACTGGCTCAGCGTGT</td><td>384.6</td><td>7.8</td><td>0.8</td><td>9.6</td><td>1.3E-21</td><td>8.1E-21</td><td>BT20/MB468</td></tr>
<tr><td>>MI0000078|hsa-mir-22&WithFlank&17|-|1713897|1713993@59.77.19&[MIMAT0000077&hsa-miR-22-3p&offsets|0|-3;m-2&17|-|1713914|1713935&offsets|0|-3]|AAGCTGCCAGTTGAAGAAC</td><td>>MI0000078|hsa-mir-22&WithFlank&17|-|1713897|1713993</td><td>59</td><td>77</td><td>19</td><td>AAGCTGCCAGTTGAAGAAC</td><td>423.9</td><td>13.4</td><td>1.4</td><td>9.6</td><td>1.3E-21</td><td>8.1E-21</td><td>BT20/MB468</td></tr>
<tr><td>>hg38_21_+_8433172_8446622_RNA45SN1_45S_50ntFlanks@7974.8002.29|ACGCGACCTCAGATCAGACGTGGCGACCC</td><td>>hg38_21_+_8433172_8446622_RNA45SN1_45S_50ntFlanks</td><td>7974</td><td>8002</td><td>29</td><td>ACGCGACCTCAGATCAGACGTGGCGACCC</td><td>428.6</td><td>6.0</td><td>0.6</td><td>9.6</td><td>1.3E-21</td><td>8.1E-21</td><td>BT20/MB468</td></tr>
<tr><td>>MI0000465|hsa-mir-191&WithFlank&3|-|49020612|49020715@22.42.21(+1C)&[MIMAT0000440&hsa-miR-191-5p&offsets|0|-2(+1C);m-48&3|-|49020672|49020694&offsets|0|-2(+1C)]|CAACGGAATCCCAAAAGCAGCC</td><td>>MI0000465|hsa-mir-191&WithFlank&3|-|49020612|49020715</td><td>22</td><td>42</td><td>21(+1C)</td><td>CAACGGAATCCCAAAAGCAGCC</td><td>2767.2</td><td>7.1</td><td>0.7</td><td>9.6</td><td>1.3E-21</td><td>8.2E-21</td><td>BT20/MB468</td></tr>
<tr><td>>MI0000811|hsa-mir-148b&WithFlank&12|+|54337210|54337320@69.89.21(+1C)&[MIMAT0000759&hsa-miR-148b-3p&offsets|0|-1(+1C);m-77&12|+|54337278|54337299&offsets|0|-1(+1C)]|TCAGTGCATCACAGAACTTTGC</td><td>>MI0000811|hsa-mir-148b&WithFlank&12|+|54337210|54337320</td><td>69</td><td>89</td><td>21(+1C)</td><td>TCAGTGCATCACAGAACTTTGC</td><td>4161.5</td><td>8.2</td><td>0.9</td><td>9.6</td><td>1.3E-21</td><td>8.2E-21</td><td>BT20/MB468</td></tr>
<tr><td>>hg38_21_+_8433172_8446622_RNA45SN1_45S_50ntFlanks@4727.4761.35|AATCAAGAACGAAAGTCGGAGGTTCGAAGACGATC</td><td>>hg38_21_+_8433172_8446622_RNA45SN1_45S_50ntFlanks</td><td>4727</td><td>4761</td><td>35</td><td>AATCAAGAACGAAAGTCGGAGGTTCGAAGACGATC</td><td>882.9</td><td>6.4</td><td>0.7</td><td>9.5</td><td>1.5E-21</td><td>8.2E-21</td><td>MB231/MB468</td></tr>
<tr><td>>MI0000289|hsa-mir-181a-1&WithFlank&1|-|198859038|198859159@30.53.24;MI0000269|hsa-mir-181a-2&WithFlank&9|+|124692436|124692557@45.68.24&[MIMAT0000256&hsa-miR-181a-5p&offsets|0|+1;m-32&1|-|198859108|198859130&offsets|0|+1];[MIMAT0000256_1&hsa-miR-181a-5p&offsets|0|+1;m-32&9|+|124692480|124692502&offsets|0|+1]|AACATTCAACGCTGTCGGTGAGTT</td><td>>MI0000289|hsa-mir-181a-1&WithFlank&1|-|198859038|198859159</td><td>30</td><td>53</td><td>24</td><td>AACATTCAACGCTGTCGGTGAGTT</td><td>5485.7</td><td>8.0</td><td>0.8</td><td>9.5</td><td>1.6E-21</td><td>8.5E-21</td><td>MB231/MB468</td></tr>
<tr><td>>trnaMT_ProTGG_MT_-_15956_16023@1.32.32|CAGAGAATAGTTTAAATTAGAATCTTAGCTTT</td><td>>trnaMT_ProTGG_MT_-_15956_16023</td><td>1</td><td>32</td><td>32</td><td>CAGAGAATAGTTTAAATTAGAATCTTAGCTTT</td><td>752.2</td><td>12.0</td><td>1.3</td><td>9.5</td><td>1.6E-21</td><td>8.5E-21</td><td>MB231/MB468</td></tr>
<tr><td>>hg38_21_+_8433172_8446622_RNA45SN1_45S_50ntFlanks@4759.4778.20|ATCAGATACCGTCGTAGTTC</td><td>>hg38_21_+_8433172_8446622_RNA45SN1_45S_50ntFlanks</td><td>4759</td><td>4778</td><td>20</td><td>ATCAGATACCGTCGTAGTTC</td><td>1157.9</td><td>-12.2</td><td>1.3</td><td>-9.5</td><td>1.4E-21</td><td>8.6E-21</td><td>BT20/MB468</td></tr>
<tr><td>>hg38_21_+_8433172_8446622_RNA45SN1_45S_50ntFlanks@11669.11693.25|GCCCAGTGCTCTGAATGTCAAAGTG</td><td>>hg38_21_+_8433172_8446622_RNA45SN1_45S_50ntFlanks</td><td>11669</td><td>11693</td><td>25</td><td>GCCCAGTGCTCTGAATGTCAAAGTG</td><td>262.3</td><td>5.1</td><td>0.5</td><td>9.5</td><td>1.4E-21</td><td>8.7E-21</td><td>BT20/MB468</td></tr>
<tr><td>>MI0000811|hsa-mir-148b&WithFlank&12|+|54337210|54337320@69.90.22&[MIMAT0000759&hsa-miR-148b-3p&offsets|0|0;m-77&12|+|54337278|54337299&offsets|0|0]|TCAGTGCATCACAGAACTTTGT</td><td>>MI0000811|hsa-mir-148b&WithFlank&12|+|54337210|54337320</td><td>69</td><td>90</td><td>22</td><td>TCAGTGCATCACAGAACTTTGT</td><td>32378.0</td><td>7.9</td><td>0.8</td><td>9.5</td><td>1.4E-21</td><td>8.8E-21</td><td>BT20/MB468</td></tr>
<tr><td>>MI0000466|hsa-mir-9-1&WithFlank&1|-|156420335|156420435@22.44.23;MI0000468|hsa-mir-9-3&WithFlank&15|+|89368011|89368112@22.44.23;MI0000467|hsa-mir-9-2&WithFlank&5|-|88666847|88666945@22.44.23&[MIMAT0000441&hsa-miR-9-5p&offsets|0|0;m-104&1|-|156420392|156420414&offsets|0|0];[MIMAT0000441_1&hsa-miR-9-5p&offsets|0|0;m-105&15|+|89368032|89368054&offsets|0|0];[MIMAT0000441_2&hsa-miR-9-5p&offsets|0|0;m-104&5|-|88666902|88666924&offsets|0|0]|TCTTTGGTTATCTAGCTGTATGA</td><td>>MI0000466|hsa-mir-9-1&WithFlank&1|-|156420335|156420435</td><td>22</td><td>44</td><td>23</td><td>TCTTTGGTTATCTAGCTGTATGA</td><td>1149.7</td><td>7.3</td><td>0.8</td><td>9.5</td><td>1.7E-21</td><td>9.0E-21</td><td>MB231/MB468</td></tr>
<tr><td>>MI0000077|hsa-mir-21&WithFlank&17|+|59841260|59841343@14.36.23(+1G)&[MIMAT0000076&hsa-miR-21-5p&offsets|0|+1(+1G);m-1&17|+|59841273|59841294&offsets|0|+1(+1G)]|TAGCTTATCAGACTGATGTTGACG</td><td>>MI0000077|hsa-mir-21&WithFlank&17|+|59841260|59841343</td><td>14</td><td>36</td><td>23(+1G)</td><td>TAGCTTATCAGACTGATGTTGACG</td><td>96852.9</td><td>8.1</td><td>0.9</td><td>9.5</td><td>1.4E-21</td><td>9.1E-21</td><td>BT20/MB468</td></tr>
<tr><td>>hg38_21_+_8433172_8446622_RNA45SN1_45S_50ntFlanks@10392.10409.18|ACAGCAGTTGAACATGGG</td><td>>hg38_21_+_8433172_8446622_RNA45SN1_45S_50ntFlanks</td><td>10392</td><td>10409</td><td>18</td><td>ACAGCAGTTGAACATGGG</td><td>2147.3</td><td>-7.2</td><td>0.8</td><td>-9.5</td><td>1.7E-21</td><td>9.2E-21</td><td>MB231/MB468</td></tr>
<tr><td>>MI0000093|hsa-mir-92a-1&WithFlank&13|+|91351308|91351397@54.75.22(+1A);MI0000094|hsa-mir-92a-2&WithFlank&X|-|134169532|134169618@54.75.22(+1A)&[MIMAT0000092&hsa-miR-92a-3p&offsets|0|0(+1A);m-9&13|+|91351361|91351382&offsets|0|0(+1A)];[MIMAT0000092_1&hsa-miR-92a-3p&offsets|0|0(+1A);m-11&X|-|134169544|134169565&offsets|0|0(+1A)]|TATTGCACTTGTCCCGGCCTGTA</td><td>>MI0000093|hsa-mir-92a-1&WithFlank&13|+|91351308|91351397</td><td>54</td><td>75</td><td>22(+1A)</td><td>TATTGCACTTGTCCCGGCCTGTA</td><td>15940.4</td><td>5.9</td><td>0.6</td><td>9.5</td><td>1.5E-21</td><td>9.2E-21</td><td>BT20/MB468</td></tr>
<tr><td>>hg38_1_-_228634819_228635039_RNA5S12_5S_50ntFlanks@142.170.29|CGCCTGGGAATACCGGGTGCTGTAGGCTT</td><td>>hg38_1_-_228634819_228635039_RNA5S12_5S_50ntFlanks</td><td>142</td><td>170</td><td>29</td><td>CGCCTGGGAATACCGGGTGCTGTAGGCTT</td><td>2583.1</td><td>6.5</td><td>0.7</td><td>9.5</td><td>1.7E-21</td><td>9.3E-21</td><td>MB231/MB468</td></tr>
<tr><td>>MI0000088|hsa-mir-30a&WithFlank&6|-|71403545|71403627@53.73.21(+1A)&[MIMAT0000088&hsa-miR-30a-3p&offsets|0|-1(+1A);m-38&6|-|71403554|71403575&offsets|0|-1(+1A)]|CTTTCAGTCGGATGTTTGCAGA</td><td>>MI0000088|hsa-mir-30a&WithFlank&6|-|71403545|71403627</td><td>53</td><td>73</td><td>21(+1A)</td><td>CTTTCAGTCGGATGTTTGCAGA</td><td>1093.4</td><td>8.2</td><td>0.9</td><td>9.5</td><td>1.5E-21</td><td>9.4E-21</td><td>BT20/MB468</td></tr>
<tr><td>>hg38_21_+_8433172_8446622_RNA45SN1_45S_50ntFlanks@4733.4767.35|GAACGAAAGTCGGAGGTTCGAAGACGATCAGATAC</td><td>>hg38_21_+_8433172_8446622_RNA45SN1_45S_50ntFlanks</td><td>4733</td><td>4767</td><td>35</td><td>GAACGAAAGTCGGAGGTTCGAAGACGATCAGATAC</td><td>1379.3</td><td>11.3</td><td>1.2</td><td>9.5</td><td>1.8E-21</td><td>9.5E-21</td><td>MB231/MB468</td></tr>
<tr><td>>MI0000264|hsa-mir-7-2&WithFlank&15|+|88611819|88611940@38.61.24;MI0000265|hsa-mir-7-3&WithFlank&19|+|4770664|4770785@37.60.24;MI0000263|hsa-mir-7-1&WithFlank&9|-|83969742|83969863@30.53.24&[MIMAT0000252&hsa-miR-7-5p&offsets|0|0;m-400&15|+|88611856|88611879&offsets|0|0];[MIMAT0000252_1&hsa-miR-7-5p&offsets|0|0;m-402&19|+|4770700|4770723&offsets|0|0];[MIMAT0000252_2&hsa-miR-7-5p&offsets|0|0;m-401&9|-|83969811|83969834&offsets|0|0]|TGGAAGACTAGTGATTTTGTTGTT</td><td>>MI0000264|hsa-mir-7-2&WithFlank&15|+|88611819|88611940</td><td>38</td><td>61</td><td>24</td><td>TGGAAGACTAGTGATTTTGTTGTT</td><td>230442.5</td><td>7.4</td><td>0.8</td><td>9.5</td><td>1.8E-21</td><td>9.5E-21</td><td>MB231/MB468</td></tr>
<tr><td>>hg38_1_-_228634819_228635039_RNA5S12_5S_50ntFlanks@152.170.19|TACCGGGTGCTGTAGGCTT</td><td>>hg38_1_-_228634819_228635039_RNA5S12_5S_50ntFlanks</td><td>152</td><td>170</td><td>19</td><td>TACCGGGTGCTGTAGGCTT</td><td>1967.2</td><td>5.6</td><td>0.6</td><td>9.5</td><td>1.5E-21</td><td>9.5E-21</td><td>BT20/MB468</td></tr>
<tr><td>>MI0000063|hsa-let-7b&WithFlank&22|+|46113680|46113774@12.32.21(+1C)&[MIMAT0000063&hsa-let-7b-5p&offsets|0|-1(+1C);m-14&22|+|46113691|46113712&offsets|0|-1(+1C)]|TGAGGTAGTAGGTTGTGTGGTC</td><td>>MI0000063|hsa-let-7b&WithFlank&22|+|46113680|46113774</td><td>12</td><td>32</td><td>21(+1C)</td><td>TGAGGTAGTAGGTTGTGTGGTC</td><td>3466.3</td><td>6.8</td><td>0.7</td><td>9.5</td><td>1.8E-21</td><td>9.6E-21</td><td>MB231/MB468</td></tr>
<tr><td>>MI0000456|hsa-mir-140&WithFlank&16|+|69933075|69933186@69.91.23&[MIMAT0004597&hsa-miR-140-3p&offsets|+1|+3;m-45&16|+|69933143|69933164&offsets|0|+1]|ACCACAGGGTAGAACCACGGACA</td><td>>MI0000456|hsa-mir-140&WithFlank&16|+|69933075|69933186</td><td>69</td><td>91</td><td>23</td><td>ACCACAGGGTAGAACCACGGACA</td><td>1766.5</td><td>10.2</td><td>1.1</td><td>9.5</td><td>1.9E-21</td><td>9.9E-21</td><td>MB231/MB468</td></tr>
<tr><td>>MI0000066|hsa-let-7e&WithFlank&19|+|51692780|51692870@14.34.21(+1C)&[MIMAT0000066&hsa-let-7e-5p&offsets|0|-1(+1C);m-50&19|+|51692793|51692814&offsets|0|-1(+1C)]|TGAGGTAGGAGGTTGTATAGTC</td><td>>MI0000066|hsa-let-7e&WithFlank&19|+|51692780|51692870</td><td>14</td><td>34</td><td>21(+1C)</td><td>TGAGGTAGGAGGTTGTATAGTC</td><td>2642.9</td><td>7.2</td><td>0.8</td><td>9.5</td><td>1.9E-21</td><td>9.9E-21</td><td>MB231/MB468</td></tr>
<tr><td>>hg38_21_+_8433172_8446622_RNA45SN1_45S_50ntFlanks@12405.12434.30|CTTTTTGATCCTTCGATGTCGGCTCTTCCT</td><td>>hg38_21_+_8433172_8446622_RNA45SN1_45S_50ntFlanks</td><td>12405</td><td>12434</td><td>30</td><td>CTTTTTGATCCTTCGATGTCGGCTCTTCCT</td><td>6559.5</td><td>-6.2</td><td>0.7</td><td>-9.5</td><td>1.6E-21</td><td>9.9E-21</td><td>BT20/MB468</td></tr>
<tr><td>>hg38_21_+_8433172_8446622_RNA45SN1_45S_50ntFlanks@7986.8006.21|ATCAGACGTGGCGACCCGCTG</td><td>>hg38_21_+_8433172_8446622_RNA45SN1_45S_50ntFlanks</td><td>7986</td><td>8006</td><td>21</td><td>ATCAGACGTGGCGACCCGCTG</td><td>2603.6</td><td>-9.2</td><td>1.0</td><td>-9.5</td><td>1.9E-21</td><td>1.0E-20</td><td>MB231/MB468</td></tr>
<tr><td>>MI0001448|hsa-mir-425&WithFlank&3|-|49020142|49020240@20.43.24(+1C)&[MIMAT0003393&hsa-miR-425-5p&offsets|0|+1(+1C);m-95&3|-|49020200|49020221&offsets|0|+2(+1C)]|AATGACACGATCACTCCCGTTGAGC</td><td>>MI0001448|hsa-mir-425&WithFlank&3|-|49020142|49020240</td><td>20</td><td>43</td><td>24(+1C)</td><td>AATGACACGATCACTCCCGTTGAGC</td><td>1529.8</td><td>7.8</td><td>0.8</td><td>9.5</td><td>1.6E-21</td><td>1.0E-20</td><td>BT20/MB468</td></tr>
<tr><td>>MI0000283|hsa-mir-203a&WithFlank&14|+|104117399|104117520@72.91.20(+1A)&[MIMAT0000264&hsa-miR-203a-3p&offsets|+1|-1(+1A);m-79&14|+|104117469|104117490&offsets|+1|-1(+1A)]|TGAAATGTTTAGGACCACTAA</td><td>>MI0000283|hsa-mir-203a&WithFlank&14|+|104117399|104117520</td><td>72</td><td>91</td><td>20(+1A)</td><td>TGAAATGTTTAGGACCACTAA</td><td>1150.2</td><td>14.8</td><td>1.6</td><td>9.5</td><td>1.6E-21</td><td>1.0E-20</td><td>BT20/MB468</td></tr>
<tr><td>>hg38_21_+_8433172_8446622_RNA45SN1_45S_50ntFlanks@11705.11728.24|TGAAGCGCGGGTAAACGGCGGGAG</td><td>>hg38_21_+_8433172_8446622_RNA45SN1_45S_50ntFlanks</td><td>11705</td><td>11728</td><td>24</td><td>TGAAGCGCGGGTAAACGGCGGGAG</td><td>526.7</td><td>-7.9</td><td>0.8</td><td>-9.5</td><td>1.6E-21</td><td>1.0E-20</td><td>BT20/MB468</td></tr>
<tr><td>>MI0000071|hsa-mir-17&WithFlank&13|+|91350599|91350694@20.43.24&[MIMAT0000070&hsa-miR-17-5p&offsets|0|+1;m-88&13|+|91350618|91350640&offsets|0|+1]|CAAAGTGCTTACAGTGCAGGTAGT</td><td>>MI0000071|hsa-mir-17&WithFlank&13|+|91350599|91350694</td><td>20</td><td>43</td><td>24</td><td>CAAAGTGCTTACAGTGCAGGTAGT</td><td>10799.0</td><td>7.5</td><td>0.8</td><td>9.5</td><td>1.9E-21</td><td>1.0E-20</td><td>MB231/MB468</td></tr>
<tr><td>>MI0000270|hsa-mir-181b-1&WithFlank&1|-|198858867|198858988@42.63.22(+1C);MI0000683|hsa-mir-181b-2&WithFlank&9|+|124693704|124693804@22.43.22(+1C)&[MIMAT0000257&hsa-miR-181b-5p&offsets|0|-1(+1C);m-73&1|-|198858925|198858947&offsets|0|-1(+1C)];[MIMAT0000257_1&hsa-miR-181b-5p&offsets|0|-1(+1C);m-72&9|+|124693725|124693747&offsets|0|-1(+1C)]|AACATTCATTGCTGTCGGTGGGC</td><td>>MI0000270|hsa-mir-181b-1&WithFlank&1|-|198858867|198858988</td><td>42</td><td>63</td><td>22(+1C)</td><td>AACATTCATTGCTGTCGGTGGGC</td><td>1166.8</td><td>7.7</td><td>0.8</td><td>9.5</td><td>1.9E-21</td><td>1.0E-20</td><td>MB231/MB468</td></tr>
</table>

<table>
<tr><td>>am_MI0000809|hsa-mir-151a&WithFlank&8|-|140732558|140732659@53.73.21(+3U)&[MIMAT0000757&hsa-miR-151a-3p&offsets|0|0(+3U);m-30&8|-|140732587|140732607&offsets|0|0(+3U)]|CTAGACTGAAGCTCCTTGAGGTTT</td><td>>am_MI0000809|hsa-mir-151a&WithFlank&8|-|140732558|140732659</td><td>53</td><td>73</td><td>21(+3U)</td><td>CTAGACTGAAGCTCCTTGAGGTTT</td><td>1016.9</td><td>9.6</td><td>1.0</td><td>9.5</td><td>1.7E-21</td><td>1.0E-20</td><td>BT20/MB468</td></tr>
<tr><td>>MI0000066|hsa-let-7e&WithFlank&19|+|51692780|51692870@14.35.22(+1A)&[MIMAT0000066&hsa-let-7e-5p&offsets|0|0(+1A);m-50&19|+|51692793|51692814&offsets|0|0(+1A)]|TGAGGTAGGAGGTTGTATAGTTA</td><td>>MI0000066|hsa-let-7e&WithFlank&19|+|51692780|51692870</td><td>14</td><td>35</td><td>22(+1A)</td><td>TGAGGTAGGAGGTTGTATAGTTA</td><td>1152.5</td><td>7.2</td><td>0.8</td><td>9.5</td><td>1.7E-21</td><td>1.0E-20</td><td>BT20/MB468</td></tr>
<tr><td>>MI0000077|hsa-mir-21&WithFlank&17|+|59841260|59841343@14.33.20&[MIMAT0000076&hsa-miR-21-5p&offsets|0|-2;m-1&17|+|59841273|59841294&offsets|0|-2]|TAGCTTATCAGACTGATGTT</td><td>>MI0000077|hsa-mir-21&WithFlank&17|+|59841260|59841343</td><td>14</td><td>33</td><td>20</td><td>TAGCTTATCAGACTGATGTT</td><td>1865.6</td><td>6.5</td><td>0.7</td><td>9.5</td><td>2.0E-21</td><td>1.1E-20</td><td>MB231/MB468</td></tr>
<tr><td>>hg38_21_+_8433172_8446622_RNA45SN1_45S_50ntFlanks@4760.4782.23|TCAGATACCGTCGTAGTTCCGAC</td><td>>hg38_21_+_8433172_8446622_RNA45SN1_45S_50ntFlanks</td><td>4760</td><td>4782</td><td>23</td><td>TCAGATACCGTCGTAGTTCCGAC</td><td>911.7</td><td>-8.4</td><td>0.9</td><td>-9.5</td><td>1.7E-21</td><td>1.1E-20</td><td>BT20/MB468</td></tr>
<tr><td>>MI0000650|hsa-mir-200c&WithFlank&12|+|6963693|6963772@50.70.21(+1U)&[MIMAT0000617&hsa-miR-200c-3p&offsets|0|-2(+1U);m-54&12|+|6963742|6963764&offsets|0|-2(+1U)]|TAATACTGCCGGGTAATGATGT</td><td>>MI0000650|hsa-mir-200c&WithFlank&12|+|6963693|6963772</td><td>50</td><td>70</td><td>21(+1U)</td><td>TAATACTGCCGGGTAATGATGT</td><td>3001.6</td><td>9.8</td><td>1.0</td><td>9.5</td><td>1.8E-21</td><td>1.1E-20</td><td>BT20/MB468</td></tr>
<tr><td>>trnaMT_GluTTC_MT_-_14674_14742@34.71.38,trnalookalike8_GluTTC_5_-_93905172_93905240@34.71.38|ATATCATTGGTCGTGGTTGTAGTCCGTGCGAGAATACC</td><td>>trnaMT_GluTTC_MT_-_14674_14742</td><td>34</td><td>71</td><td>38</td><td>ATATCATTGGTCGTGGTTGTAGTCCGTGCGAGAATACC</td><td>1611.0</td><td>10.5</td><td>1.1</td><td>9.5</td><td>2.1E-21</td><td>1.1E-20</td><td>MB231/MB468</td></tr>
<tr><td>>MI0000434|hsa-let-7i&WithFlank&12|+|62603680|62603775@12.31.20(+1A)&[MIMAT0000415&hsa-let-7i-5p&offsets|0|-2(+1A);m-49&12|+|62603691|62603712&offsets|0|-2(+1A)]|TGAGGTAGTAGTTTGTGCTGA</td><td>>MI0000434|hsa-let-7i&WithFlank&12|+|62603680|62603775</td><td>12</td><td>31</td><td>20(+1A)</td><td>TGAGGTAGTAGTTTGTGCTGA</td><td>764.6</td><td>9.3</td><td>1.0</td><td>9.5</td><td>2.1E-21</td><td>1.1E-20</td><td>MB231/MB468</td></tr>
<tr><td>>MI0000086|hsa-mir-28&WithFlank&3|+|188688775|188688872@62.82.21(+1U)&[MIMAT0004502&hsa-miR-28-3p&offsets|+2|+1(+1U);m-28&3|+|188688834|188688855&offsets|+2|+1(+1U)]|CTAGATTGTGAGCTCCTGGAGT</td><td>>MI0000086|hsa-mir-28&WithFlank&3|+|188688775|188688872</td><td>62</td><td>82</td><td>21(+1U)</td><td>CTAGATTGTGAGCTCCTGGAGT</td><td>768.7</td><td>10.6</td><td>1.1</td><td>9.5</td><td>2.1E-21</td><td>1.1E-20</td><td>MB231/MB468</td></tr>
<tr><td>>MI0000439|hsa-mir-23b&WithFlank&9|+|95085202|95085310@65.86.22(+1U)&[MIMAT0000418&hsa-miR-23b-3p&offsets|+1|0(+1U);m-42&9|+|95085265|95085285&offsets|+1|+2(+1U)]|TCACATTGCCAGGGATTACCACT</td><td>>MI0000439|hsa-mir-23b&WithFlank&9|+|95085202|95085310</td><td>65</td><td>86</td><td>22(+1U)</td><td>TCACATTGCCAGGGATTACCACT</td><td>1212.8</td><td>8.1</td><td>0.9</td><td>9.5</td><td>2.1E-21</td><td>1.1E-20</td><td>MB231/MB468</td></tr>
<tr><td>>MI0001445|hsa-mir-423&WithFlank&17|+|30117073|30117178@59.78.20&[MIMAT0001340&hsa-miR-423-3p&offsets|0|-3;m-100&17|+|30117131|30117153&offsets|0|-3]|AGCTCGGTCTGAGGCCCCTC</td><td>>MI0001445|hsa-mir-423&WithFlank&17|+|30117073|30117178</td><td>59</td><td>78</td><td>20</td><td>AGCTCGGTCTGAGGCCCCTC</td><td>4801.0</td><td>7.3</td><td>0.8</td><td>9.5</td><td>1.8E-21</td><td>1.1E-20</td><td>BT20/MB468</td></tr>
<tr><td>>MI0000066|hsa-let-7e&WithFlank&19|+|51692780|51692870@14.36.23&[MIMAT0000066&hsa-let-7e-5p&offsets|0|+1;m-50&19|+|51692793|51692814&offsets|0|+1]|TGAGGTAGGAGGTTGTATAGTTG</td><td>>MI0000066|hsa-let-7e&WithFlank&19|+|51692780|51692870</td><td>14</td><td>36</td><td>23</td><td>TGAGGTAGGAGGTTGTATAGTTG</td><td>1128.2</td><td>7.1</td><td>0.7</td><td>9.5</td><td>1.8E-21</td><td>1.1E-20</td><td>BT20/MB468</td></tr>
<tr><td>>MI0000266|hsa-mir-10a&WithFlank&17|-|48579832|48579953@29.49.21(+1U)&[MIMAT0000253&hsa-miR-10a-5p&offsets|+1|-1(+1U);m-18&17|-|48579905|48579926&offsets|+1|0(+1U)]|ACCCTGTAGATCCGAATTTGTT</td><td>>MI0000266|hsa-mir-10a&WithFlank&17|-|48579832|48579953</td><td>29</td><td>49</td><td>21(+1U)</td><td>ACCCTGTAGATCCGAATTTGTT</td><td>508.7</td><td>13.6</td><td>1.4</td><td>9.5</td><td>1.8E-21</td><td>1.1E-20</td><td>BT20/MB468</td></tr>
<tr><td>>MI0000279|hsa-mir-196a-2&WithFlank&12|+|53991732|53991853@31.52.22(+1U);MI0000238|hsa-mir-196a-1&WithFlank&17|-|48632484|48632565@13.34.22(+1U)&[MIMAT0000226&hsa-miR-196a-5p&offsets|0|0(+1U);m-261&12|+|53991762|53991783&offsets|0|0(+1U)];[MIMAT0000226_1&hsa-miR-196a-5p&offsets|0|0(+1U);m-268&17|-|48632532|48632553&offsets|0|0(+1U)]|TAGGTAGTTTCATGTTGTTGGGT</td><td>>MI0000279|hsa-mir-196a-2&WithFlank&12|+|53991732|53991853</td><td>31</td><td>52</td><td>22(+1U)</td><td>TAGGTAGTTTCATGTTGTTGGGT</td><td>1250.5</td><td>7.9</td><td>0.8</td><td>9.5</td><td>1.9E-21</td><td>1.1E-20</td><td>BT20/MB468</td></tr>
<tr><td>>MI0000440|hsa-mir-27b&WithFlank&9|+|95085439|95085547@67.88.22(+2U)&[MIMAT0000419&hsa-miR-27b-3p&offsets|0|+1(+2U);m-39&9|+|95085505|95085525&offsets|0|+1(+2U)]|TTCACAGTGGCTAAGTTCTGCATT</td><td>>MI0000440|hsa-mir-27b&WithFlank&9|+|95085439|95085547</td><td>67</td><td>88</td><td>22(+2U)</td><td>TTCACAGTGGCTAAGTTCTGCATT</td><td>710.7</td><td>7.2</td><td>0.8</td><td>9.5</td><td>2.2E-21</td><td>1.1E-20</td><td>MB231/MB468</td></tr>
<tr><td>>MI0000650|hsa-mir-200c&WithFlank&12|+|6963693|6963772@50.72.23(+2A)&[MIMAT0000617&hsa-miR-200c-3p&offsets|0|0(+2A);m-54&12|+|6963742|6963764&offsets|0|0(+2A)]|TAATACTGCCGGGTAATGATGGAAA</td><td>>MI0000650|hsa-mir-200c&WithFlank&12|+|6963693|6963772</td><td>50</td><td>72</td><td>23(+2A)</td><td>TAATACTGCCGGGTAATGATGGAAA</td><td>2300.4</td><td>8.3</td><td>0.9</td><td>9.5</td><td>1.9E-21</td><td>1.2E-20</td><td>BT20/MB468</td></tr>
<tr><td>>MI0000482|hsa-mir-185&WithFlank&22|+|20033133|20033226@21.43.23&[MIMAT0000455&hsa-miR-185-5p&offsets|0|+1;m-145&22|+|20033153|20033174&offsets|0|+1]|TGGAGAGAAAGGCAGTTCCTGAT</td><td>>MI0000482|hsa-mir-185&WithFlank&22|+|20033133|20033226</td><td>21</td><td>43</td><td>23</td><td>TGGAGAGAAAGGCAGTTCCTGAT</td><td>815.0</td><td>6.7</td><td>0.7</td><td>9.5</td><td>1.9E-21</td><td>1.2E-20</td><td>BT20/MB468</td></tr>
<tr><td>>trnaMT_GluTTC_MT_-_14674_14742@40.58.19,trnalookalike8_GluTTC_5_-_93905172_93905240@40.58.19|TTGGTCGTGGTTGTAGTCC</td><td>>trnaMT_GluTTC_MT_-_14674_14742</td><td>40</td><td>58</td><td>19</td><td>TTGGTCGTGGTTGTAGTCC</td><td>1171.7</td><td>9.3</td><td>1.0</td><td>9.5</td><td>1.9E-21</td><td>1.2E-20</td><td>BT20/MB468</td></tr>
<tr><td>>hg38_21_+_8433172_8446622_RNA45SN1_45S_50ntFlanks@6652.6694.43|GACTCTTAGCGGTGGATCACTCGGCTCGTGCGTCGATGAAGAA</td><td>>hg38_21_+_8433172_8446622_RNA45SN1_45S_50ntFlanks</td><td>6652</td><td>6694</td><td>43</td><td>GACTCTTAGCGGTGGATCACTCGGCTCGTGCGTCGATGAAGAA</td><td>1043.4</td><td>6.9</td><td>0.7</td><td>9.5</td><td>2.2E-21</td><td>1.2E-20</td><td>MB231/MB468</td></tr>
<tr><td>>MI0000079|hsa-mir-23a&WithFlank&19|-|13836581|13836665@54.72.19(+2U)&[MIMAT0000078&hsa-miR-23a-3p&offsets|+3|+1(+2U);m-31&19|-|13836595|13836615&offsets|+3|+1(+2U)]|ACATTGCCAGGGATTTCCATT</td><td>>MI0000079|hsa-mir-23a&WithFlank&19|-|13836581|13836665</td><td>54</td><td>72</td><td>19(+2U)</td><td>ACATTGCCAGGGATTTCCATT</td><td>624.9</td><td>13.9</td><td>1.5</td><td>9.5</td><td>1.9E-21</td><td>1.2E-20</td><td>BT20/MB468</td></tr>
<tr><td>>MI0000283|hsa-mir-203a&WithFlank&14|+|104117399|104117520@72.92.21(+1U)&[MIMAT0000264&hsa-miR-203a-3p&offsets|+1|0(+1U);m-79&14|+|104117469|104117490&offsets|+1|0(+1U)]|TGAAATGTTTAGGACCACTAGT</td><td>>MI0000283|hsa-mir-203a&WithFlank&14|+|104117399|104117520</td><td>72</td><td>92</td><td>21(+1U)</td><td>TGAAATGTTTAGGACCACTAGT</td><td>1257.7</td><td>13.8</td><td>1.4</td><td>9.8</td><td>9.5E-23</td><td>1.2E-20</td><td>BT20/MB231</td></tr>
<tr><td>>hg38_21_+_8433172_8446622_RNA45SN1_45S_50ntFlanks@476.516.41|TTCGTGATCGATGTGGTGACGTCGTGCTCTCCCGGGCCGGG</td><td>>hg38_21_+_8433172_8446622_RNA45SN1_45S_50ntFlanks</td><td>476</td><td>516</td><td>41</td><td>TTCGTGATCGATGTGGTGACGTCGTGCTCTCCCGGGCCGGG</td><td>2117.7</td><td>7.5</td><td>0.8</td><td>9.5</td><td>2.0E-21</td><td>1.2E-20</td><td>BT20/MB468</td></tr>
<tr><td>>hg38_21_+_8433172_8446622_RNA45SN1_45S_50ntFlanks@11641.11660.20|CGGCGGGTGTTGACGCGATG</td><td>>hg38_21_+_8433172_8446622_RNA45SN1_45S_50ntFlanks</td><td>11641</td><td>11660</td><td>20</td><td>CGGCGGGTGTTGACGCGATG</td><td>10524.1</td><td>-10.7</td><td>1.1</td><td>-9.5</td><td>2.0E-21</td><td>1.2E-20</td><td>BT20/MB468</td></tr>
<tr><td>>hg38_21_+_8433172_8446622_RNA45SN1_45S_50ntFlanks@476.499.24|TTCGTGATCGATGTGGTGACGTCG</td><td>>hg38_21_+_8433172_8446622_RNA45SN1_45S_50ntFlanks</td><td>476</td><td>499</td><td>24</td><td>TTCGTGATCGATGTGGTGACGTCG</td><td>6916.5</td><td>7.3</td><td>0.8</td><td>9.5</td><td>2.3E-21</td><td>1.2E-20</td><td>MB231/MB468</td></tr>
<tr><td>>MI0000070|hsa-mir-16-1&WithFlank&13|-|50048967|50049067@20.41.22(+1C);MI0000115|hsa-mir-16-2&WithFlank&3|+|160404739|160404831@16.37.22(+1C)&[MIMAT0000069&hsa-miR-16-5p&offsets|0|0(+1C);m-62&13|-|50049027|50049048&offsets|0|0(+1C)];[MIMAT0000069_1&hsa-miR-16-5p&offsets|0|0(+1C);m-61&3|+|160404754|160404775&offsets|0|0(+1C)]|TAGCAGCACGTAAATATTGGCGC</td><td>>MI0000070|hsa-mir-16-1&WithFlank&13|-|50048967|50049067</td><td>20</td><td>41</td><td>22(+1C)</td><td>TAGCAGCACGTAAATATTGGCGC</td><td>710.7</td><td>7.6</td><td>0.8</td><td>9.5</td><td>2.4E-21</td><td>1.2E-20</td><td>MB231/MB468</td></tr>
<tr><td>>hg38_21_+_8433172_8446622_RNA45SN1_45S_50ntFlanks@12163.12181.19|ACTGGGGCGGTACACCTGT</td><td>>hg38_21_+_8433172_8446622_RNA45SN1_45S_50ntFlanks</td><td>12163</td><td>12181</td><td>19</td><td>ACTGGGGCGGTACACCTGT</td><td>1394.4</td><td>-7.2</td><td>0.8</td><td>-9.5</td><td>2.4E-21</td><td>1.3E-20</td><td>MB231/MB468</td></tr>
<tr><td>>MI0000542|hsa-mir-320a&WithFlank&8|-|22244960|22245043@48.68.21&[MIMAT0000510&hsa-miR-320a-3p&offsets|0|-1;m-60&8|-|22244975|22244996&offsets|0|-1]|AAAAGCTGGGTTGAGAGGGCG</td><td>>MI0000542|hsa-mir-320a&WithFlank&8|-|22244960|22245043</td><td>48</td><td>68</td><td>21</td><td>AAAAGCTGGGTTGAGAGGGCG</td><td>2355.0</td><td>7.4</td><td>0.8</td><td>9.5</td><td>2.4E-21</td><td>1.3E-20</td><td>MB231/MB468</td></tr>
<tr><td>>MI0000095|hsa-mir-93&WithFlank&7|-|100093762|100093853@17.38.22(+1A)&[MIMAT0000093&hsa-miR-93-5p&offsets|0|-1(+1A);m-29&7|-|100093815|100093837&offsets|0|-1(+1A)]|CAAAGTGCTGTTCGTGCAGGTAA</td><td>>MI0000095|hsa-mir-93&WithFlank&7|-|100093762|100093853</td><td>17</td><td>38</td><td>22(+1A)</td><td>CAAAGTGCTGTTCGTGCAGGTAA</td><td>2681.6</td><td>6.8</td><td>0.7</td><td>9.5</td><td>2.4E-21</td><td>1.3E-20</td><td>MB231/MB468</td></tr>
<tr><td>>hg38_21_+_8433172_8446622_RNA45SN1_45S_50ntFlanks@8176.8204.29|CTTCTGATCGAGGCCCAGCCCGTGGACGG</td><td>>hg38_21_+_8433172_8446622_RNA45SN1_45S_50ntFlanks</td><td>8176</td><td>8204</td><td>29</td><td>CTTCTGATCGAGGCCCAGCCCGTGGACGG</td><td>799.1</td><td>-3.7</td><td>0.4</td><td>-9.5</td><td>2.1E-21</td><td>1.3E-20</td><td>BT20/MB468</td></tr>
<tr><td>>hg38_21_+_8433172_8446622_RNA45SN1_45S_50ntFlanks@3705.3725.21|TACCTGGTTGATCCTGCCAGT</td><td>>hg38_21_+_8433172_8446622_RNA45SN1_45S_50ntFlanks</td><td>3705</td><td>3725</td><td>21</td><td>TACCTGGTTGATCCTGCCAGT</td><td>23522.4</td><td>9.5</td><td>1.0</td><td>9.5</td><td>2.2E-21</td><td>1.3E-20</td><td>BT20/MB468</td></tr>
<tr><td>>MI0005531|hsa-mir-450b&WithFlank&X|-|134540179|134540268@17.37.21&[MIMAT0004909&hsa-miR-450b-5p&offsets|0|-1;m-266&X|-|134540232|134540252&offsets|0|0]|TTTTGCAATATGTTCCTGAAT</td><td>>MI0005531|hsa-mir-450b&WithFlank&X|-|134540179|134540268</td><td>17</td><td>37</td><td>21</td><td>TTTTGCAATATGTTCCTGAAT</td><td>3989.5</td><td>10.5</td><td>1.1</td><td>9.5</td><td>2.2E-21</td><td>1.4E-20</td><td>BT20/MB468</td></tr>
<tr><td>>hg38_21_+_8433172_8446622_RNA45SN1_45S_50ntFlanks@6647.6696.50|CGTACGACTCTTAGCGGTGGATCACTCGGCTCGTGCGTCGATGAAGAACG</td><td>>hg38_21_+_8433172_8446622_RNA45SN1_45S_50ntFlanks</td><td>6647</td><td>6696</td><td>50</td><td>CGTACGACTCTTAGCGGTGGATCACTCGGCTCGTGCGTCGATGAAGAACG</td><td>1506.1</td><td>8.3</td><td>0.9</td><td>9.5</td><td>2.6E-21</td><td>1.4E-20</td><td>MB231/MB468</td></tr>
<tr><td>>MI0001448|hsa-mir-425&WithFlank&3|-|49020142|49020240@20.43.24&[MIMAT0003393&hsa-miR-425-5p&offsets|0|+1;m-95&3|-|49020200|49020221&offsets|0|+2]|AATGACACGATCACTCCCGTTGAG</td><td>>MI0001448|hsa-mir-425&WithFlank&3|-|49020142|49020240</td><td>20</td><td>43</td><td>24</td><td>AATGACACGATCACTCCCGTTGAG</td><td>1561.0</td><td>6.3</td><td>0.7</td><td>9.5</td><td>2.3E-21</td><td>1.4E-20</td><td>BT20/MB468</td></tr>
<tr><td>>MI0000088|hsa-mir-30a&WithFlank&6|-|71403545|71403627@53.75.23&[MIMAT0000088&hsa-miR-30a-3p&offsets|0|+1;m-38&6|-|71403554|71403575&offsets|0|+1]|CTTTCAGTCGGATGTTTGCAGCT</td><td>>MI0000088|hsa-mir-30a&WithFlank&6|-|71403545|71403627</td><td>53</td><td>75</td><td>23</td><td>CTTTCAGTCGGATGTTTGCAGCT</td><td>921.1</td><td>11.9</td><td>1.3</td><td>9.5</td><td>2.7E-21</td><td>1.4E-20</td><td>MB231/MB468</td></tr>
<tr><td>>hg38_21_+_8433172_8446622_RNA45SN1_45S_50ntFlanks@12996.13013.18|TCGCTGCGATCTATTGAA</td><td>>hg38_21_+_8433172_8446622_RNA45SN1_45S_50ntFlanks</td><td>12996</td><td>13013</td><td>18</td><td>TCGCTGCGATCTATTGAA</td><td>5772.9</td><td>4.9</td><td>0.5</td><td>9.5</td><td>2.7E-21</td><td>1.4E-20</td><td>MB231/MB468</td></tr>
<tr><td>>MI0003560|hsa-mir-92b&WithFlank&1|+|155195171|155195278@67.87.21&[MIMAT0003218&hsa-miR-92b-3p&offsets|0|-1;m-115&1|+|155195237|155195258&offsets|0|-1]|TATTGCACTCGTCCCGGCCTC</td><td>>MI0003560|hsa-mir-92b&WithFlank&1|+|155195171|155195278</td><td>67</td><td>87</td><td>21</td><td>TATTGCACTCGTCCCGGCCTC</td><td>2123.6</td><td>9.6</td><td>1.0</td><td>9.5</td><td>2.7E-21</td><td>1.4E-20</td><td>MB231/MB468</td></tr>
<tr><td>>hg38_21_+_8433172_8446622_RNA45SN1_45S_50ntFlanks@12957.12982.26|CTTCTGGGTCGGGGTTTCGTACGTAG</td><td>>hg38_21_+_8433172_8446622_RNA45SN1_45S_50ntFlanks</td><td>12957</td><td>12982</td><td>26</td><td>CTTCTGGGTCGGGGTTTCGTACGTAG</td><td>1212.1</td><td>-8.5</td><td>0.9</td><td>-9.5</td><td>2.3E-21</td><td>1.4E-20</td><td>BT20/MB468</td></tr>
<tr><td>>trnaMT_PheGAA_MT_+_577_647@36.54.19|AAAATGTTTAGACGGGCTC</td><td>>trnaMT_PheGAA_MT_+_577_647</td><td>36</td><td>54</td><td>19</td><td>AAAATGTTTAGACGGGCTC</td><td>1166.8</td><td>11.2</td><td>1.2</td><td>9.5</td><td>2.7E-21</td><td>1.4E-20</td><td>MB231/MB468</td></tr>
</table>

<table>
<tr><th></th><th></th><th></th><th></th><th></th><th></th><th></th><th></th><th></th><th></th><th></th><th></th><th></th></tr>
<tr><td>>hg38_21_+_8433172_8446622_RNA45SN1_45S_50ntFlanks@11668.11703.36|TGCCCAGTGCTCTGAATGTCAAAGTGAAGAAATTCA</td><td>>hg38_21_+_8433172_8446622_RNA45SN1_45S_50ntFlanks</td><td>11668</td><td>11703</td><td>36</td><td>TGCCCAGTGCTCTGAATGTCAAAGTGAAGAAATTCA</td><td>1174.8</td><td>6.4</td><td>0.7</td><td>9.5</td><td>2.8E-21</td><td>1.5E-20</td><td>MB231/MB468</td></tr>
<tr><td>>hg38_21_+_8433172_8446622_RNA45SN1_45S_50ntFlanks@12632.12654.23|GCGAAGCTACCATCTGTGGGATT</td><td>>hg38_21_+_8433172_8446622_RNA45SN1_45S_50ntFlanks</td><td>12632</td><td>12654</td><td>23</td><td>GCGAAGCTACCATCTGTGGGATT</td><td>1640.6</td><td>-5.6</td><td>0.6</td><td>-9.5</td><td>2.8E-21</td><td>1.5E-20</td><td>MB231/MB468</td></tr>
<tr><td>>hg38_21_+_8433172_8446622_RNA45SN1_45S_50ntFlanks@4733.4767.35|GAACGAAAGTCGGAGGTTCGAAGACGATCAGATAC</td><td>>hg38_21_+_8433172_8446622_RNA45SN1_45S_50ntFlanks</td><td>4733</td><td>4767</td><td>35</td><td>GAACGAAAGTCGGAGGTTCGAAGACGATCAGATAC</td><td>295.8</td><td>8.9</td><td>0.9</td><td>9.5</td><td>2.4E-21</td><td>1.5E-20</td><td>BT20/MB468</td></tr>
<tr><td>>hg38_21_+_8433172_8446622_RNA45SN1_45S_50ntFlanks@12407.12426.20|TTTTGATCCTTCGATGTCGG</td><td>>hg38_21_+_8433172_8446622_RNA45SN1_45S_50ntFlanks</td><td>12407</td><td>12426</td><td>20</td><td>TTTTGATCCTTCGATGTCGG</td><td>346.9</td><td>-5.9</td><td>0.6</td><td>-9.5</td><td>2.4E-21</td><td>1.5E-20</td><td>BT20/MB468</td></tr>
<tr><td>>hg38_21_+_8433172_8446622_RNA45SN1_45S_50ntFlanks@7975.8009.35|CGCGACCTCAGATCAGACGTGGCGACCCGCTGAAT</td><td>>hg38_21_+_8433172_8446622_RNA45SN1_45S_50ntFlanks</td><td>7975</td><td>8009</td><td>35</td><td>CGCGACCTCAGATCAGACGTGGCGACCCGCTGAAT</td><td>70880.7</td><td>6.9</td><td>0.7</td><td>9.5</td><td>3.0E-21</td><td>1.5E-20</td><td>MB231/MB468</td></tr>
<tr><td>>MI0000469|hsa-mir-125a&WithFlank&19|+|51693248|51693345@21.42.22&[MIMAT0000443&hsa-miR-125a-5p&offsets|0|-2;m-63&19|+|51693268|51693289&offsets|0|0]|TCCCTGAGACCCTTTAACCTGT</td><td>>MI0000469|hsa-mir-125a&WithFlank&19|+|51693248|51693345</td><td>21</td><td>42</td><td>22</td><td>TCCCTGAGACCCTTTAACCTGT</td><td>10972.4</td><td>7.3</td><td>0.8</td><td>9.5</td><td>2.6E-21</td><td>1.6E-20</td><td>BT20/MB468</td></tr>
<tr><td>>hg38_21_+_8433172_8446622_RNA45SN1_45S_50ntFlanks@11874.11906.33|AGAAGACCCTGTTGAGCTTGACTCTAGTCTGGC</td><td>>hg38_21_+_8433172_8446622_RNA45SN1_45S_50ntFlanks</td><td>11874</td><td>11906</td><td>33</td><td>AGAAGACCCTGTTGAGCTTGACTCTAGTCTGGC</td><td>1063.9</td><td>-6.4</td><td>0.7</td><td>-9.5</td><td>3.0E-21</td><td>1.6E-20</td><td>MB231/MB468</td></tr>
<tr><td>>hg38_21_+_8433172_8446622_RNA45SN1_45S_50ntFlanks@12937.12956.20|AACCATTCGTAGACGACCTG</td><td>>hg38_21_+_8433172_8446622_RNA45SN1_45S_50ntFlanks</td><td>12937</td><td>12956</td><td>20</td><td>AACCATTCGTAGACGACCTG</td><td>3189.4</td><td>-5.4</td><td>0.6</td><td>-9.5</td><td>3.0E-21</td><td>1.6E-20</td><td>MB231/MB468</td></tr>
<tr><td>>MI0000650|hsa-mir-200c&WithFlank&12|+|6963693|6963772@50.69.20&[MIMAT0000617&hsa-miR-200c-3p&offsets|0|-3;m-54&12|+|6963742|6963764&offsets|0|-3]|TAATACTGCCGGGTAATGAT</td><td>>MI0000650|hsa-mir-200c&WithFlank&12|+|6963693|6963772</td><td>50</td><td>69</td><td>20</td><td>TAATACTGCCGGGTAATGAT</td><td>1501.8</td><td>14.1</td><td>1.4</td><td>9.8</td><td>1.3E-22</td><td>1.6E-20</td><td>BT20/MB231</td></tr>
<tr><td>>hg38_21_+_8433172_8446622_RNA45SN1_45S_50ntFlanks@4760.4780.21|TCAGATACCGTCGTAGTTCCG</td><td>>hg38_21_+_8433172_8446622_RNA45SN1_45S_50ntFlanks</td><td>4760</td><td>4780</td><td>21</td><td>TCAGATACCGTCGTAGTTCCG</td><td>3539.2</td><td>-8.5</td><td>0.9</td><td>-9.5</td><td>2.7E-21</td><td>1.6E-20</td><td>BT20/MB468</td></tr>
<tr><td>>hg38_1_-_228634819_228635039_RNA5S12_5S_50ntFlanks@130.169.40|TGGATGGGAGACCGCCTGGGAATACCGGGTGCTGTAGGCT</td><td>>hg38_1_-_228634819_228635039_RNA5S12_5S_50ntFlanks</td><td>130</td><td>169</td><td>40</td><td>TGGATGGGAGACCGCCTGGGAATACCGGGTGCTGTAGGCT</td><td>696.2</td><td>11.6</td><td>1.2</td><td>9.5</td><td>2.7E-21</td><td>1.7E-20</td><td>BT20/MB468</td></tr>
<tr><td>>MI0000067|hsa-let-7f-1&WithFlank&9|+|94176341|94176439@13.34.22;MI0000068|hsa-let-7f-2&WithFlank&X|-|53557186|53557280@14.35.22&[MIMAT0000067&hsa-let-7f-5p&offsets|0|0;m-20&9|+|94176353|94176374&offsets|0|0];[MIMAT0000067_1&hsa-let-7f-5p&offsets|0|0;m-19&X|-|53557246|53557267&offsets|0|0]|TGAGGTAGTAGATTGTATAGTT</td><td>>MI0000067|hsa-let-7f-1&WithFlank&9|+|94176341|94176439</td><td>13</td><td>34</td><td>22</td><td>TGAGGTAGTAGATTGTATAGTT</td><td>415971.0</td><td>7.2</td><td>0.8</td><td>9.5</td><td>3.3E-21</td><td>1.7E-20</td><td>MB231/MB468</td></tr>
<tr><td>>MI0000065|hsa-let-7d&WithFlank&9|+|94178828|94178926@14.34.21(+1C)&[MIMAT0000065&hsa-let-7d-5p&offsets|0|-1(+1C);m-90&9|+|94178841|94178862&offsets|0|-1(+1C)]|AGAGGTAGTAGGTTGCATAGTC</td><td>>MI0000065|hsa-let-7d&WithFlank&9|+|94178828|94178926</td><td>14</td><td>34</td><td>21(+1C)</td><td>AGAGGTAGTAGGTTGCATAGTC</td><td>1745.3</td><td>7.4</td><td>0.8</td><td>9.5</td><td>3.3E-21</td><td>1.7E-20</td><td>MB231/MB468</td></tr>
<tr><td>>hg38_21_+_8433172_8446622_RNA45SN1_45S_50ntFlanks@8195.8213.19|CCGTGGACGGTGTGAGGCC</td><td>>hg38_21_+_8433172_8446622_RNA45SN1_45S_50ntFlanks</td><td>8195</td><td>8213</td><td>19</td><td>CCGTGGACGGTGTGAGGCC</td><td>700.4</td><td>4.5</td><td>0.5</td><td>9.4</td><td>3.4E-21</td><td>1.8E-20</td><td>MB231/MB468</td></tr>
<tr><td>>MI0003591|hsa-mir-584&WithFlank&5|-|149062307|149062415@22.42.21(+1A)&[MIMAT0003249&hsa-miR-584-5p&offsets|0|-1(+1A);m-167&5|-|149062374|149062394&offsets|0|0(+1A)]|TTATGGTTTGCCTGGGACTGAA</td><td>>MI0003591|hsa-mir-584&WithFlank&5|-|149062307|149062415</td><td>22</td><td>42</td><td>21(+1A)</td><td>TTATGGTTTGCCTGGGACTGAA</td><td>656.5</td><td>11.5</td><td>1.2</td><td>9.5</td><td>2.9E-21</td><td>1.8E-20</td><td>BT20/MB468</td></tr>
<tr><td>>hg38_21_+_8433172_8446622_RNA45SN1_45S_50ntFlanks@10680.10712.33|GTTCCGGCGGCGTCCGGTGAGCTCTCGCTGGCC</td><td>>hg38_21_+_8433172_8446622_RNA45SN1_45S_50ntFlanks</td><td>10680</td><td>10712</td><td>33</td><td>GTTCCGGCGGCGTCCGGTGAGCTCTCGCTGGCC</td><td>3060.6</td><td>-5.6</td><td>0.6</td><td>-9.5</td><td>3.0E-21</td><td>1.8E-20</td><td>BT20/MB468</td></tr>
<tr><td>>hg38_MT_+_598_1651_RNR1_12S_50ntFlanks@916.935.20|AAGGACATTTAACTAAAACC</td><td>>hg38_MT_+_598_1651_RNR1_12S_50ntFlanks</td><td>916</td><td>935</td><td>20</td><td>AAGGACATTTAACTAAAACC</td><td>1178.4</td><td>11.7</td><td>1.2</td><td>9.4</td><td>3.5E-21</td><td>1.8E-20</td><td>MB231/MB468</td></tr>
<tr><td>>MI0000066|hsa-let-7e&WithFlank&19|+|51692780|51692870@14.35.22&[MIMAT0000066&hsa-let-7e-5p&offsets|0|0;m-50&19|+|51692793|51692814&offsets|0|0]|TGAGGTAGGAGGTTGTATAGTT</td><td>>MI0000066|hsa-let-7e&WithFlank&19|+|51692780|51692870</td><td>14</td><td>35</td><td>22</td><td>TGAGGTAGGAGGTTGTATAGTT</td><td>23521.9</td><td>7.0</td><td>0.7</td><td>9.4</td><td>3.5E-21</td><td>1.8E-20</td><td>MB231/MB468</td></tr>
<tr><td>>hg38_21_+_8433172_8446622_RNA45SN1_45S_50ntFlanks@4046.4063.18|CATTCGAACGTCTGCCCT</td><td>>hg38_21_+_8433172_8446622_RNA45SN1_45S_50ntFlanks</td><td>4046</td><td>4063</td><td>18</td><td>CATTCGAACGTCTGCCCT</td><td>1586.6</td><td>-5.1</td><td>0.5</td><td>-9.4</td><td>3.5E-21</td><td>1.8E-20</td><td>MB231/MB468</td></tr>
<tr><td>>hg38_21_+_8433172_8446622_RNA45SN1_45S_50ntFlanks@6651.6682.32|CGACTCTTAGCGGTGGATCACTCGGCTCGTGC</td><td>>hg38_21_+_8433172_8446622_RNA45SN1_45S_50ntFlanks</td><td>6651</td><td>6682</td><td>32</td><td>CGACTCTTAGCGGTGGATCACTCGGCTCGTGC</td><td>1935.3</td><td>5.9</td><td>0.6</td><td>9.4</td><td>3.6E-21</td><td>1.9E-20</td><td>MB231/MB468</td></tr>
<tr><td>>hg38_21_+_8433172_8446622_RNA45SN1_45S_50ntFlanks@7974.8011.38|ACGCGACCTCAGATCAGACGTGGCGACCCGCTGAATTT</td><td>>hg38_21_+_8433172_8446622_RNA45SN1_45S_50ntFlanks</td><td>7974</td><td>8011</td><td>38</td><td>ACGCGACCTCAGATCAGACGTGGCGACCCGCTGAATTT</td><td>12316.8</td><td>6.8</td><td>0.7</td><td>9.5</td><td>3.1E-21</td><td>1.9E-20</td><td>BT20/MB468</td></tr>
<tr><td>>hg38_21_+_8433172_8446622_RNA45SN1_45S_50ntFlanks@12635.12653.19|AAGCTACCATCTGTGGGAT</td><td>>hg38_21_+_8433172_8446622_RNA45SN1_45S_50ntFlanks</td><td>12635</td><td>12653</td><td>19</td><td>AAGCTACCATCTGTGGGAT</td><td>3752.9</td><td>-13.9</td><td>1.5</td><td>-9.5</td><td>3.1E-21</td><td>1.9E-20</td><td>BT20/MB468</td></tr>
<tr><td>>hg38_MT_+_598_1651_RNR1_12S_50ntFlanks@428.456.29|AAAGTGGCTTTAACATATCTGAACACACA</td><td>>hg38_MT_+_598_1651_RNR1_12S_50ntFlanks</td><td>428</td><td>456</td><td>29</td><td>AAAGTGGCTTTAACATATCTGAACACACA</td><td>1906.2</td><td>11.6</td><td>1.2</td><td>9.4</td><td>3.7E-21</td><td>1.9E-20</td><td>MB231/MB468</td></tr>
<tr><td>>MI0000746|hsa-mir-99b&WithFlank&19|+|51692606|51692687@13.33.21(+1U)&[MIMAT0000689&hsa-miR-99b-5p&offsets|0|-1(+1U);m-10&19|+|51692618|51692639&offsets|0|-1(+1U)]|CACCCGTAGAACCGACCTTGCT</td><td>>MI0000746|hsa-mir-99b&WithFlank&19|+|51692606|51692687</td><td>13</td><td>33</td><td>21(+1U)</td><td>CACCCGTAGAACCGACCTTGCT</td><td>5127.8</td><td>9.2</td><td>1.0</td><td>9.5</td><td>3.2E-21</td><td>1.9E-20</td><td>BT20/MB468</td></tr>
<tr><td>>am_MI0000809|hsa-mir-151a&WithFlank&8|-|140732558|140732659@17.38.22&[MIMAT0004697&hsa-miR-151a-5p&offsets|0|+1;m-75&8|-|140732623|140732643&offsets|0|+1]|TCGAGGAGCTCACAGTCTAGTA</td><td>>am_MI0000809|hsa-mir-151a&WithFlank&8|-|140732558|140732659</td><td>17</td><td>38</td><td>22</td><td>TCGAGGAGCTCACAGTCTAGTA</td><td>4419.0</td><td>9.6</td><td>1.0</td><td>9.5</td><td>3.2E-21</td><td>1.9E-20</td><td>BT20/MB468</td></tr>
<tr><td>>hg38_1_-_228634819_228635039_RNA5S12_5S_50ntFlanks@139.169.31|GACCGCCTGGGAATACCGGGTGCTGTAGGCT</td><td>>hg38_1_-_228634819_228635039_RNA5S12_5S_50ntFlanks</td><td>139</td><td>169</td><td>31</td><td>GACCGCCTGGGAATACCGGGTGCTGTAGGCT</td><td>1641.2</td><td>7.0</td><td>0.7</td><td>9.4</td><td>3.8E-21</td><td>2.0E-20</td><td>MB231/MB468</td></tr>
<tr><td>>MI0000074|hsa-mir-19b-1&WithFlank&13|+|91351186|91351284@60.81.22;MI0000075|hsa-mir-19b-2&WithFlank&X|-|134169665|134169772@68.89.22&[MIMAT0000074&hsa-miR-19b-3p&offsets|0|-1;m-122&13|+|91351245|91351267&offsets|0|-1];[MIMAT0000074_1&hsa-miR-19b-3p&offsets|0|-1;m-121&X|-|134169683|134169705&offsets|0|-1]|TGTGCAAATCCATGCAAAACTG</td><td>>MI0000074|hsa-mir-19b-1&WithFlank&13|+|91351186|91351284</td><td>60</td><td>81</td><td>22</td><td>TGTGCAAATCCATGCAAAACTG</td><td>1039.1</td><td>6.8</td><td>0.7</td><td>9.4</td><td>3.8E-21</td><td>2.0E-20</td><td>MB231/MB468</td></tr>
<tr><td>>MI0000077|hsa-mir-21&WithFlank&17|+|59841260|59841343@14.34.21(+1G)&[MIMAT0000076&hsa-miR-21-5p&offsets|0|-1(+1G);m-1&17|+|59841273|59841294&offsets|0|-1(+1G)]|TAGCTTATCAGACTGATGTTGG</td><td>>MI0000077|hsa-mir-21&WithFlank&17|+|59841260|59841343</td><td>14</td><td>34</td><td>21(+1G)</td><td>TAGCTTATCAGACTGATGTTGG</td><td>12615.3</td><td>7.4</td><td>0.8</td><td>9.4</td><td>3.8E-21</td><td>2.0E-20</td><td>MB231/MB468</td></tr>
<tr><td>>MI0000270|hsa-mir-181b-1&WithFlank&1|-|198858867|198858988@42.61.20;MI0000683|hsa-mir-181b-2&WithFlank&9|+|124693704|124693804@22.41.20&[MIMAT0000257&hsa-miR-181b-5p&offsets|0|-3;m-73&1|-|198858925|198858947&offsets|0|-3];[MIMAT0000257_1&hsa-miR-181b-5p&offsets|0|-3;m-72&9|+|124693725|124693747&offsets|0|-3]|AACATTCATTGCTGTCGGTG</td><td>>MI0000270|hsa-mir-181b-1&WithFlank&1|-|198858867|198858988</td><td>42</td><td>61</td><td>20</td><td>AACATTCATTGCTGTCGGTG</td><td>736.2</td><td>6.8</td><td>0.7</td><td>9.4</td><td>3.8E-21</td><td>2.0E-20</td><td>MB231/MB468</td></tr>
<tr><td>>MI0000447|hsa-mir-128-1&WithFlank&2|+|135665391|135665484@56.75.20(+1C);MI0000727|hsa-mir-128-2&WithFlank&3|+|35744470|35744565@58.77.20(+1C)&[MIMAT0000424&hsa-miR-128-3p&offsets|0|-1(+1C);m-108&2|+|135665446|135665466&offsets|0|-1(+1C)];[MIMAT0000424_1&hsa-miR-128-3p&offsets|0|-1(+1C);m-117&3|+|35744527|35744547&offsets|0|-1(+1C)]|TCACAGTGAACCGGTCTCTTC</td><td>>MI0000447|hsa-mir-128-1&WithFlank&2|+|135665391|135665484</td><td>56</td><td>75</td><td>20(+1C)</td><td>TCACAGTGAACCGGTCTCTTC</td><td>3183.2</td><td>6.1</td><td>0.6</td><td>9.5</td><td>3.3E-21</td><td>2.0E-20</td><td>BT20/MB468</td></tr>
<tr><td>>hg38_21_+_8433172_8446622_RNA45SN1_45S_50ntFlanks@11781.11798.18|CGCATGAATGGATGAACG</td><td>>hg38_21_+_8433172_8446622_RNA45SN1_45S_50ntFlanks</td><td>11781</td><td>11798</td><td>18</td><td>CGCATGAATGGATGAACG</td><td>761.2</td><td>-5.5</td><td>0.6</td><td>-9.5</td><td>3.4E-21</td><td>2.0E-20</td><td>BT20/MB468</td></tr>
<tr><td>>MI0000811|hsa-mir-148b&WithFlank&12|+|54337210|54337320@69.89.21(+1C)&[MIMAT0000759&hsa-miR-148b-3p&offsets|0|-1(+1C);m-77&12|+|54337278|54337299&offsets|0|-1(+1C)]|TCAGTGCATCACAGAACTTTGC</td><td>>MI0000811|hsa-mir-148b&WithFlank&12|+|54337210|54337320</td><td>69</td><td>89</td><td>21(+1C)</td><td>TCAGTGCATCACAGAACTTTGC</td><td>3632.3</td><td>8.3</td><td>0.9</td><td>9.4</td><td>4.0E-21</td><td>2.1E-20</td><td>MB231/MB468</td></tr>
<tr><td>>hg38_21_+_8433172_8446622_RNA45SN1_45S_50ntFlanks@12420.12455.36|ATGTCGGCTCTTCCTATCATTGTGAAGCAGAATTCA</td><td>>hg38_21_+_8433172_8446622_RNA45SN1_45S_50ntFlanks</td><td>12420</td><td>12455</td><td>36</td><td>ATGTCGGCTCTTCCTATCATTGTGAAGCAGAATTCA</td><td>1836.9</td><td>-5.0</td><td>0.5</td><td>-9.4</td><td>3.5E-21</td><td>2.1E-20</td><td>BT20/MB468</td></tr>
<tr><td>>MI0000446|hsa-mir-125b-1&WithFlank&11|-|122099751|122099850@21.41.21(+1G);MI0000470|hsa-mir-125b-2&WithFlank&21|+|16590231|16590331@23.43.21(+1G)&[MIMAT0000423&hsa-miR-125b-5p&offsets|0|-1(+1G);m-67&11|-|122099809|122099830&offsets|0|-1(+1G)];[MIMAT0000423_1&hsa-miR-125b-5p&offsets|0|-1(+1G);m-68&21|+|16590253|16590274&offsets|0|-1(+1G)]|TCCCTGAGACCCTAACTTGTGG</td><td>>MI0000446|hsa-mir-125b-1&WithFlank&11|-|122099751|122099850</td><td>21</td><td>41</td><td>21(+1G)</td><td>TCCCTGAGACCCTAACTTGTGG</td><td>1893.1</td><td>10.0</td><td>1.1</td><td>9.4</td><td>4.1E-21</td><td>2.1E-20</td><td>MB231/MB468</td></tr>
<tr><td>>MI0000811|hsa-mir-148b&WithFlank&12|+|54337210|54337320@69.90.22(+1A)&[MIMAT0000759&hsa-miR-148b-3p&offsets|0|0(+1A);m-77&12|+|54337278|54337299&offsets|0|0(+1A)]|TCAGTGCATCACAGAACTTTGTA</td><td>>MI0000811|hsa-mir-148b&WithFlank&12|+|54337210|54337320</td><td>69</td><td>90</td><td>22(+1A)</td><td>TCAGTGCATCACAGAACTTTGTA</td><td>1163.0</td><td>7.2</td><td>0.8</td><td>9.4</td><td>4.2E-21</td><td>2.2E-20</td><td>MB231/MB468</td></tr>
<tr><td>>MI0000289|hsa-mir-181a-1&WithFlank&1|-|198859038|198859159@70.90.21(+1U)&[MIMAT0000270&hsa-miR-181a-3p&offsets|0|-1(+1U);m-125&1|-|198859069|198859090&offsets|0|-1(+1U)]|ACCATCGACCGTTGATTGTACT</td><td>>MI0000289|hsa-mir-181a-1&WithFlank&1|-|198859038|198859159</td><td>70</td><td>90</td><td>21(+1U)</td><td>ACCATCGACCGTTGATTGTACT</td><td>1901.1</td><td>10.0</td><td>1.1</td><td>9.4</td><td>4.2E-21</td><td>2.2E-20</td><td>MB231/MB468</td></tr>
<tr><td>>MI0000234|hsa-mir-192&WithFlank&11|-|64891131|64891252@30.50.21&[MIMAT0000222&hsa-miR-192-5p&offsets|0|0;m-59&11|-|64891203|64891223&offsets|0|0]|CTGACCTATGAATTGACAGCC</td><td>>MI0000234|hsa-mir-192&WithFlank&11|-|64891131|64891252</td><td>30</td><td>50</td><td>21</td><td>CTGACCTATGAATTGACAGCC</td><td>2847.7</td><td>6.0</td><td>0.6</td><td>9.4</td><td>3.6E-21</td><td>2.2E-20</td><td>BT20/MB468</td></tr>
<tr><td>>hg38_21_+_8433172_8446622_RNA45SN1_45S_50ntFlanks@9574.9596.23|AATCGGTCGTCCGACCTGGGTAT</td><td>>hg38_21_+_8433172_8446622_RNA45SN1_45S_50ntFlanks</td><td>9574</td><td>9596</td><td>23</td><td>AATCGGTCGTCCGACCTGGGTAT</td><td>4954.8</td><td>-6.1</td><td>0.6</td><td>-9.4</td><td>3.6E-21</td><td>2.2E-20</td><td>BT20/MB468</td></tr>
</table>

<table>
<tr><td>>hg38_21_+_8433172_8446622_RNA45SN1_45S_50ntFlanks@11729.11753.25|TAACTATGACTCTCTTAAGGTAGCC</td><td>>hg38_21_+_8433172_8446622_RNA45SN1_45S_50ntFlanks</td><td>11729</td><td>11753</td><td>25</td><td>TAACTATGACTCTCTTAAGGTAGCC</td><td>2523.2</td><td>-13.3</td><td>1.4</td><td>-9.4</td><td>3.6E-21</td><td>2.2E-20</td><td>BT20/MB468</td></tr>
<tr><td>>MI0001448|hsa-mir-425&WithFlank&3|-|49020142|49020240@20.44.25&[MIMAT0003393&hsa-miR-425-5p&offsets|0|+2;m-95&3|-|49020200|49020221&offsets|0|+3]|AATGACACGATCACTCCCGTTGAGT</td><td>>MI0001448|hsa-mir-425&WithFlank&3|-|49020142|49020240</td><td>20</td><td>44</td><td>25</td><td>AATGACACGATCACTCCCGTTGAGT</td><td>11719.0</td><td>7.7</td><td>0.8</td><td>9.4</td><td>3.7E-21</td><td>2.2E-20</td><td>BT20/MB468</td></tr>
<tr><td>>MI0000456|hsa-mir-140&WithFlank&16|+|69933075|69933186@69.91.23&[MIMAT0004597&hsa-miR-140-3p&offsets|+1|+3;m-45&16|+|69933143|69933164&offsets|0|+1]|ACCACAGGGTAGAACCACGGACA</td><td>>MI0000456|hsa-mir-140&WithFlank&16|+|69933075|69933186</td><td>69</td><td>91</td><td>23</td><td>ACCACAGGGTAGAACCACGGACA</td><td>2823.7</td><td>10.6</td><td>1.1</td><td>9.4</td><td>3.7E-21</td><td>2.2E-20</td><td>BT20/MB468</td></tr>
<tr><td>>MI0000465|hsa-mir-191&WithFlank&3|-|49020612|49020715@22.43.22(+1A)&[MIMAT0000440&hsa-miR-191-5p&offsets|0|-1(+1A);m-48&3|-|49020672|49020694&offsets|0|-1(+1A)]|CAACGGAATCCCAAAAGCAGCTA</td><td>>MI0000465|hsa-mir-191&WithFlank&3|-|49020612|49020715</td><td>22</td><td>43</td><td>22(+1A)</td><td>CAACGGAATCCCAAAAGCAGCTA</td><td>3907.4</td><td>7.4</td><td>0.8</td><td>9.4</td><td>3.8E-21</td><td>2.3E-20</td><td>BT20/MB468</td></tr>
<tr><td>>trnaMT_PheGAA_MT_+_577_647@36.57.22|AAAATGTTTAGACGGGCTCACA</td><td>>trnaMT_PheGAA_MT_+_577_647</td><td>36</td><td>57</td><td>22</td><td>AAAATGTTTAGACGGGCTCACA</td><td>624.1</td><td>9.5</td><td>1.0</td><td>9.4</td><td>4.5E-21</td><td>2.3E-20</td><td>MB231/MB468</td></tr>
<tr><td>>MI0000433|hsa-let-7g&WithFlank&3|-|52268272|52268367@11.32.22(+1A)&[MIMAT0000414&hsa-let-7g-5p&offsets|0|0(+1A);m-58&3|-|52268336|52268357&offsets|0|0(+1A)]|TGAGGTAGTAGTTTGTACAGTTA</td><td>>MI0000433|hsa-let-7g&WithFlank&3|-|52268272|52268367</td><td>11</td><td>32</td><td>22(+1A)</td><td>TGAGGTAGTAGTTTGTACAGTTA</td><td>4125.1</td><td>6.7</td><td>0.7</td><td>9.4</td><td>4.5E-21</td><td>2.3E-20</td><td>MB231/MB468</td></tr>
<tr><td>>hg38_MT_+_598_1651_RNR1_12S_50ntFlanks@905.929.25|AGTATACTTCAAAGGACATTTAACT</td><td>>hg38_MT_+_598_1651_RNR1_12S_50ntFlanks</td><td>905</td><td>929</td><td>25</td><td>AGTATACTTCAAAGGACATTTAACT</td><td>2901.8</td><td>11.2</td><td>1.2</td><td>9.4</td><td>4.6E-21</td><td>2.4E-20</td><td>MB231/MB468</td></tr>
<tr><td>>hg38_21_+_8433172_8446622_RNA45SN1_45S_50ntFlanks@6778.6806.29|CCCGGGGCTACGCCTGTCTGAGCGTCGCT</td><td>>hg38_21_+_8433172_8446622_RNA45SN1_45S_50ntFlanks</td><td>6778</td><td>6806</td><td>29</td><td>CCCGGGGCTACGCCTGTCTGAGCGTCGCT</td><td>1623.7</td><td>5.1</td><td>0.5</td><td>9.4</td><td>4.0E-21</td><td>2.4E-20</td><td>BT20/MB468</td></tr>
<tr><td>>hg38_21_+_8433172_8446622_RNA45SN1_45S_50ntFlanks@10794.10818.25|CTGGCATGTTGGAACAATGTAGGTA</td><td>>hg38_21_+_8433172_8446622_RNA45SN1_45S_50ntFlanks</td><td>10794</td><td>10818</td><td>25</td><td>CTGGCATGTTGGAACAATGTAGGTA</td><td>2842.9</td><td>-7.1</td><td>0.8</td><td>-9.4</td><td>4.8E-21</td><td>2.4E-20</td><td>MB231/MB468</td></tr>
<tr><td>>hg38_21_+_8433172_8446622_RNA45SN1_45S_50ntFlanks@12159.12182.24|TTTGACTGGGGCGGTACACCTGTC</td><td>>hg38_21_+_8433172_8446622_RNA45SN1_45S_50ntFlanks</td><td>12159</td><td>12182</td><td>24</td><td>TTTGACTGGGGCGGTACACCTGTC</td><td>1313.6</td><td>-12.4</td><td>1.3</td><td>-9.4</td><td>4.2E-21</td><td>2.5E-20</td><td>BT20/MB468</td></tr>
<tr><td>>hg38_21_+_8433172_8446622_RNA45SN1_45S_50ntFlanks@12324.12344.21|TTGGGTTTTAAGCAGGAGGTG</td><td>>hg38_21_+_8433172_8446622_RNA45SN1_45S_50ntFlanks</td><td>12324</td><td>12344</td><td>21</td><td>TTGGGTTTTAAGCAGGAGGTG</td><td>1065.7</td><td>-12.1</td><td>1.3</td><td>-9.4</td><td>4.2E-21</td><td>2.5E-20</td><td>BT20/MB468</td></tr>
<tr><td>>hg38_21_+_8433172_8446622_RNA45SN1_45S_50ntFlanks@7984.8008.25|AGATCAGACGTGGCGACCCGCTGAA</td><td>>hg38_21_+_8433172_8446622_RNA45SN1_45S_50ntFlanks</td><td>7984</td><td>8008</td><td>25</td><td>AGATCAGACGTGGCGACCCGCTGAA</td><td>978.5</td><td>-5.2</td><td>0.6</td><td>-9.4</td><td>4.2E-21</td><td>2.5E-20</td><td>BT20/MB468</td></tr>
<tr><td>>hg38_21_+_8433172_8446622_RNA45SN1_45S_50ntFlanks@12615.12653.39|GCTGAGGAGCCAATGGGGCGAAGCTACCATCTGTGGGAT</td><td>>hg38_21_+_8433172_8446622_RNA45SN1_45S_50ntFlanks</td><td>12615</td><td>12653</td><td>39</td><td>GCTGAGGAGCCAATGGGGCGAAGCTACCATCTGTGGGAT</td><td>700.5</td><td>-5.2</td><td>0.5</td><td>-9.4</td><td>4.9E-21</td><td>2.5E-20</td><td>MB231/MB468</td></tr>
<tr><td>>MI0003628|hsa-mir-615&WithFlank&12|+|54033944|54034051@67.88.22&[MIMAT0003283&hsa-miR-615-3p&offsets|0|0;m-495&12|+|54034010|54034030&offsets|0|+1]|TCCGAGCCTGGGTCTCCCTCTT</td><td>>MI0003628|hsa-mir-615&WithFlank&12|+|54033944|54034051</td><td>67</td><td>88</td><td>22</td><td>TCCGAGCCTGGGTCTCCCTCTT</td><td>2494.0</td><td>6.6</td><td>0.7</td><td>9.4</td><td>4.2E-21</td><td>2.5E-20</td><td>BT20/MB468</td></tr>
<tr><td>>hg38_21_+_8433172_8446622_RNA45SN1_45S_50ntFlanks@475.502.28|CTTCGTGATCGATGTGGTGACGTCGTGC</td><td>>hg38_21_+_8433172_8446622_RNA45SN1_45S_50ntFlanks</td><td>475</td><td>502</td><td>28</td><td>CTTCGTGATCGATGTGGTGACGTCGTGC</td><td>1738.0</td><td>6.5</td><td>0.7</td><td>9.4</td><td>4.3E-21</td><td>2.5E-20</td><td>BT20/MB468</td></tr>
<tr><td>>hg38_1_-_228634819_228635039_RNA5S12_5S_50ntFlanks@152.171.20|TACCGGGTGCTGTAGGCTTT</td><td>>hg38_1_-_228634819_228635039_RNA5S12_5S_50ntFlanks</td><td>152</td><td>171</td><td>20</td><td>TACCGGGTGCTGTAGGCTTT</td><td>3837.6</td><td>6.9</td><td>0.7</td><td>9.4</td><td>5.1E-21</td><td>2.6E-20</td><td>MB231/MB468</td></tr>
<tr><td>>hg38_21_+_8433172_8446622_RNA45SN1_45S_50ntFlanks@7977.8011.35|CGACCTCAGATCAGACGTGGCGACCCGCTGAATTT</td><td>>hg38_21_+_8433172_8446622_RNA45SN1_45S_50ntFlanks</td><td>7977</td><td>8011</td><td>35</td><td>CGACCTCAGATCAGACGTGGCGACCCGCTGAATTT</td><td>2384.0</td><td>7.0</td><td>0.7</td><td>9.4</td><td>5.1E-21</td><td>2.6E-20</td><td>MB231/MB468</td></tr>
<tr><td>>hg38_21_+_8433172_8446622_RNA45SN1_45S_50ntFlanks@11564.11582.19|CCTAGCAGCCGACTTAGAA</td><td>>hg38_21_+_8433172_8446622_RNA45SN1_45S_50ntFlanks</td><td>11564</td><td>11582</td><td>19</td><td>CCTAGCAGCCGACTTAGAA</td><td>1055.4</td><td>-5.0</td><td>0.5</td><td>-9.4</td><td>5.1E-21</td><td>2.6E-20</td><td>MB231/MB468</td></tr>
<tr><td>>hg38_21_+_8433172_8446622_RNA45SN1_45S_50ntFlanks@12419.12458.40|GATGTCGGCTCTTCCTATCATTGTGAAGCAGAATTCACCA</td><td>>hg38_21_+_8433172_8446622_RNA45SN1_45S_50ntFlanks</td><td>12419</td><td>12458</td><td>40</td><td>GATGTCGGCTCTTCCTATCATTGTGAAGCAGAATTCACCA</td><td>836.0</td><td>10.8</td><td>1.1</td><td>9.4</td><td>5.1E-21</td><td>2.6E-20</td><td>MB231/MB468</td></tr>
<tr><td>>MI0000081|hsa-mir-24-2&WithFlank&19|-|13836281|13836365@56.77.22(+1U);MI0000080|hsa-mir-24-1&WithFlank&9|+|95086015|95086094@50.71.22(+1U)&[MIMAT0000080&hsa-miR-24-3p&offsets|0|0(+1U);m-36&19|-|13836288|13836310&offsets|0|-1(+1U)];[MIMAT0000080_1&hsa-miR-24-3p&offsets|0|0(+1U);m-35&9|+|95086064|95086086&offsets|0|-1(+1U)]|TGGCTCAGTTCAGCAGGAACAGT</td><td>>MI0000081|hsa-mir-24-2&WithFlank&19|-|13836281|13836365</td><td>56</td><td>77</td><td>22(+1U)</td><td>TGGCTCAGTTCAGCAGGAACAGT</td><td>21403.3</td><td>7.2</td><td>0.8</td><td>9.4</td><td>5.2E-21</td><td>2.6E-20</td><td>MB231/MB468</td></tr>
<tr><td>>MI0000067|hsa-let-7f-1&WithFlank&9|+|94176341|94176439@13.33.21(+1G);MI0000068|hsa-let-7f-2&WithFlank&X|-|53557186|53557280@14.34.21(+1G)&[MIMAT0000067&hsa-let-7f-5p&offsets|0|-1(+1G);m-20&9|+|94176353|94176374&offsets|0|-1(+1G)];[MIMAT0000067_1&hsa-let-7f-5p&offsets|0|-1(+1G);m-19&X|-|53557246|53557267&offsets|0|-1(+1G)]|TGAGGTAGTAGATTGTATAGTG</td><td>>MI0000067|hsa-let-7f-1&WithFlank&9|+|94176341|94176439</td><td>13</td><td>33</td><td>21(+1G)</td><td>TGAGGTAGTAGATTGTATAGTG</td><td>8958.1</td><td>7.1</td><td>0.8</td><td>9.4</td><td>5.2E-21</td><td>2.6E-20</td><td>MB231/MB468</td></tr>
<tr><td>>hg38_21_+_8433172_8446622_RNA45SN1_45S_50ntFlanks@10786.10822.37|AACAGCCTCTGGCATGTTGGAACAATGTAGGTAAGGG</td><td>>hg38_21_+_8433172_8446622_RNA45SN1_45S_50ntFlanks</td><td>10786</td><td>10822</td><td>37</td><td>AACAGCCTCTGGCATGTTGGAACAATGTAGGTAAGGG</td><td>523.3</td><td>-5.7</td><td>0.6</td><td>-9.4</td><td>4.4E-21</td><td>2.6E-20</td><td>BT20/MB468</td></tr>
<tr><td>>hg38_21_+_8433172_8446622_RNA45SN1_45S_50ntFlanks@9568.9595.28|CGTGCAAATCGGTCGTCCGACCTGGGTA</td><td>>hg38_21_+_8433172_8446622_RNA45SN1_45S_50ntFlanks</td><td>9568</td><td>9595</td><td>28</td><td>CGTGCAAATCGGTCGTCCGACCTGGGTA</td><td>1616.7</td><td>-6.8</td><td>0.7</td><td>-9.4</td><td>4.4E-21</td><td>2.6E-20</td><td>BT20/MB468</td></tr>
<tr><td>>hg38_21_+_8433172_8446622_RNA45SN1_45S_50ntFlanks@9572.9597.26|CAAATCGGTCGTCCGACCTGGGTATA</td><td>>hg38_21_+_8433172_8446622_RNA45SN1_45S_50ntFlanks</td><td>9572</td><td>9597</td><td>26</td><td>CAAATCGGTCGTCCGACCTGGGTATA</td><td>7787.7</td><td>-9.2</td><td>1.0</td><td>-9.4</td><td>4.5E-21</td><td>2.7E-20</td><td>BT20/MB468</td></tr>
<tr><td>>MI0000062|hsa-let-7a-3&WithFlank&22|+|46112743|46112828@58.78.21(+1U);MI0000060|hsa-let-7a-1&WithFlank&9|+|94175951|94176042@63.83.21(+1U)&[MIMAT0004481&hsa-let-7a-3p&offsets|0|0(+1U);m-180&22|+|46112800|46112821&offsets|0|-1(+1U)];[MIMAT0004481_1&hsa-let-7a-3p&offsets|0|0(+1U);m-181&9|+|94176013|94176034&offsets|0|-1(+1U)]|CTATACAATCTACTGTCTTTCT</td><td>>MI0000062|hsa-let-7a-3&WithFlank&22|+|46112743|46112828</td><td>58</td><td>78</td><td>21(+1U)</td><td>CTATACAATCTACTGTCTTTCT</td><td>3887.1</td><td>7.1</td><td>0.8</td><td>9.4</td><td>5.3E-21</td><td>2.7E-20</td><td>MB231/MB468</td></tr>
<tr><td>>MI0005756|hsa-mir-934&WithFlank&X|+|136550872|136550966@21.42.22(+1A)&[MIMAT0004977&hsa-miR-934&offsets|0|0(+1A);m-943&X|+|136550892|136550913&offsets|0|0(+1A)]|TGTCTACTACTGGAGACACTGGA</td><td>>MI0005756|hsa-mir-934&WithFlank&X|+|136550872|136550966</td><td>21</td><td>42</td><td>22(+1A)</td><td>TGTCTACTACTGGAGACACTGGA</td><td>18346.4</td><td>10.1</td><td>1.1</td><td>9.4</td><td>4.6E-21</td><td>2.7E-20</td><td>BT20/MB468</td></tr>
<tr><td>>MI0000078|hsa-mir-22&WithFlank&17|-|1713897|1713993@59.78.20&[MIMAT0000077&hsa-miR-22-3p&offsets|0|-2;m-2&17|-|1713914|1713935&offsets|0|-2]|AAGCTGCCAGTTGAAGAACT</td><td>>MI0000078|hsa-mir-22&WithFlank&17|-|1713897|1713993</td><td>59</td><td>78</td><td>20</td><td>AAGCTGCCAGTTGAAGAACT</td><td>1374.9</td><td>9.1</td><td>1.0</td><td>9.4</td><td>5.4E-21</td><td>2.8E-20</td><td>MB231/MB468</td></tr>
<tr><td>>MI0000746|hsa-mir-99b&WithFlank&19|+|51692606|51692687@51.72.22&[MIMAT0004678&hsa-miR-99b-3p&offsets|0|0;m-151&19|+|51692656|51692678&offsets|0|-1]|CAAGCTCGTGTCTGTGGGTCCG</td><td>>MI0000746|hsa-mir-99b&WithFlank&19|+|51692606|51692687</td><td>51</td><td>72</td><td>22</td><td>CAAGCTCGTGTCTGTGGGTCCG</td><td>1869.8</td><td>5.9</td><td>0.6</td><td>9.4</td><td>5.6E-21</td><td>2.8E-20</td><td>MB231/MB468</td></tr>
<tr><td>>trnaMT_GluTTC_MT_-_14674_14742@33.72.40,trnalookalike8_GluTTC_5_-_93905172_93905240@33.72.40|CATATCATTGGTCGTGGTTGTAGTCCGTGCGAGAATACCA</td><td>>trnaMT_GluTTC_MT_-_14674_14742</td><td>33</td><td>72</td><td>40</td><td>CATATCATTGGTCGTGGTTGTAGTCCGTGCGAGAATACCA</td><td>1711.6</td><td>8.7</td><td>0.9</td><td>9.4</td><td>5.6E-21</td><td>2.8E-20</td><td>MB231/MB468</td></tr>
<tr><td>>MI0000086|hsa-mir-28&WithFlank&3|+|188688775|188688872@60.80.21(+1G)&[MIMAT0004502&hsa-miR-28-3p&offsets|0|-1(+1G);m-28&3|+|188688834|188688855&offsets|0|-1(+1G)]|CACTAGATTGTGAGCTCCTGGG</td><td>>MI0000086|hsa-mir-28&WithFlank&3|+|188688775|188688872</td><td>60</td><td>80</td><td>21(+1G)</td><td>CACTAGATTGTGAGCTCCTGGG</td><td>809.9</td><td>12.1</td><td>1.3</td><td>9.4</td><td>5.6E-21</td><td>2.9E-20</td><td>MB231/MB468</td></tr>
<tr><td>>MI0000750|hsa-mir-26a-2&WithFlank&12|-|57824603|57824698@20.40.21(+1A);MI0000083|hsa-mir-26a-1&WithFlank&3|+|37969398|37969486@16.36.21(+1A)&[MIMAT0000082&hsa-miR-26a-5p&offsets|0|-1(+1A);m-33&12|-|57824658|57824679&offsets|0|-1(+1A)];[MIMAT0000082_1&hsa-miR-26a-5p&offsets|0|-1(+1A);m-34&3|+|37969413|37969434&offsets|0|-1(+1A)]|TTCAAGTAATCCAGGATAGGCA</td><td>>MI0000750|hsa-mir-26a-2&WithFlank&12|-|57824603|57824698</td><td>20</td><td>40</td><td>21(+1A)</td><td>TTCAAGTAATCCAGGATAGGCA</td><td>15779.6</td><td>6.7</td><td>0.7</td><td>9.4</td><td>4.9E-21</td><td>2.9E-20</td><td>BT20/MB468</td></tr>
<tr><td>>hg38_21_+_8433172_8446622_RNA45SN1_45S_50ntFlanks@3782.3801.20|CAGTGAAACTGCGAATGGCT</td><td>>hg38_21_+_8433172_8446622_RNA45SN1_45S_50ntFlanks</td><td>3782</td><td>3801</td><td>20</td><td>CAGTGAAACTGCGAATGGCT</td><td>1221.6</td><td>-4.9</td><td>0.5</td><td>-9.4</td><td>5.0E-21</td><td>2.9E-20</td><td>BT20/MB468</td></tr>
<tr><td>>MI0000086|hsa-mir-28&WithFlank&3|+|188688775|188688872@62.82.21(+1U)&[MIMAT0004502&hsa-miR-28-3p&offsets|+2|+1(+1U);m-28&3|+|188688834|188688855&offsets|+2|+1(+1U)]|CTAGATTGTGAGCTCCTGGAGT</td><td>>MI0000086|hsa-mir-28&WithFlank&3|+|188688775|188688872</td><td>62</td><td>82</td><td>21(+1U)</td><td>CTAGATTGTGAGCTCCTGGAGT</td><td>929.8</td><td>10.6</td><td>1.1</td><td>9.4</td><td>5.0E-21</td><td>2.9E-20</td><td>BT20/MB468</td></tr>
<tr><td>>hg38_21_+_8433172_8446622_RNA45SN1_45S_50ntFlanks@12405.12425.21|CTTTTTGATCCTTCGATGTCG</td><td>>hg38_21_+_8433172_8446622_RNA45SN1_45S_50ntFlanks</td><td>12405</td><td>12425</td><td>21</td><td>CTTTTTGATCCTTCGATGTCG</td><td>933.4</td><td>-5.8</td><td>0.6</td><td>-9.4</td><td>5.8E-21</td><td>3.0E-20</td><td>MB231/MB468</td></tr>
<tr><td>>MI0000266|hsa-mir-10a&WithFlank&17|-|48579832|48579953@28.50.23&[MIMAT0000253&hsa-miR-10a-5p&offsets|0|0;m-18&17|-|48579905|48579926&offsets|0|+1]|TACCCTGTAGATCCGAATTTGTG</td><td>>MI0000266|hsa-mir-10a&WithFlank&17|-|48579832|48579953</td><td>28</td><td>50</td><td>23</td><td>TACCCTGTAGATCCGAATTTGTG</td><td>30225.0</td><td>13.4</td><td>1.4</td><td>9.4</td><td>5.1E-21</td><td>3.0E-20</td><td>BT20/MB468</td></tr>
<tr><td>>MI0000465|hsa-mir-191&WithFlank&3|-|49020612|49020715@22.43.22(+1A)&[MIMAT0000440&hsa-miR-191-5p&offsets|0|-1(+1A);m-48&3|-|49020672|49020694&offsets|0|-1(+1A)]|CAACGGAATCCCAAAAGCAGCTA</td><td>>MI0000465|hsa-mir-191&WithFlank&3|-|49020612|49020715</td><td>22</td><td>43</td><td>22(+1A)</td><td>CAACGGAATCCCAAAAGCAGCTA</td><td>2346.9</td><td>7.0</td><td>0.7</td><td>9.4</td><td>6.0E-21</td><td>3.0E-20</td><td>MB231/MB468</td></tr>
<tr><td>>MI0003667|hsa-mir-652&WithFlank&X|+|110055323|110055432@67.87.21(+1U)&[MIMAT0003322&hsa-miR-652-3p&offsets|0|0(+1U);m-148&X|+|110055389|110055410&offsets|0|-1(+1U)]|AATGGCGCCACTAGGGTTGTGT</td><td>>MI0003667|hsa-mir-652&WithFlank&X|+|110055323|110055432</td><td>67</td><td>87</td><td>21(+1U)</td><td>AATGGCGCCACTAGGGTTGTGT</td><td>2153.9</td><td>8.5</td><td>0.9</td><td>9.4</td><td>5.1E-21</td><td>3.0E-20</td><td>BT20/MB468</td></tr>
<tr><td>>MI0000469|hsa-mir-125a&WithFlank&19|+|51693248|51693345@21.42.22&[MIMAT0000443&hsa-miR-125a-5p&offsets|0|-2;m-63&19|+|51693268|51693289&offsets|0|0]|TCCCTGAGACCCTTTAACCTGT</td><td>>MI0000469|hsa-mir-125a&WithFlank&19|+|51693248|51693345</td><td>21</td><td>42</td><td>22</td><td>TCCCTGAGACCCTTTAACCTGT</td><td>4610.3</td><td>6.3</td><td>0.7</td><td>9.4</td><td>6.1E-21</td><td>3.1E-20</td><td>MB231/MB468</td></tr>
</table>

| | | | | | | | | | | | | |
|---|---|---|---|---|---|---|---|---|---|---|---|---|
| >MI0000093\|hsa-mir-92a-1&WithFlank&13\|+\|91351308\|91351397@54.75.22(+1C);MI0000094\|hsa-mir-92a-2&WithFlank&X\|-\|134169532\|134169618@54.75.22(+1C)&[MIMAT0000092&hsa-miR-92a-3p&offsets\|0\|0(+1C);m-9&13\|+\|91351361\|91351382&offsets\|0\|0(+1C)];[MIMAT0000092_1&hsa-miR-92a-3p&offsets\|0\|0(+1C);m-11&X\|-\|134169544\|134169565&offsets\|0\|0(+1C)]\|TATTGCACTTGTCCCGGCCTGTC | >MI0000093\|hsa-mir-92a-1&WithFlank&13\|+\|91351308\|91351397 | 54 | 75 | 22(+1C) | TATTGCACTTGTCCCGGCCTGTC | 661.7 | 5.7 | 0.6 | 9.4 | 5.4E-21 | 3.2E-20 | BT20/MB468 |
| >hg38_1_-_228634819_228635039_RNA5S12_5S_50ntFlanks@147.170.24\|GGGAATACCGGGTGCTGTAGGCTT | >hg38_1_-_228634819_228635039_RNA5S12_5S_50ntFlanks | 147 | 170 | 24 | GGGAATACCGGGTGCTGTAGGCTT | 3173.0 | 6.1 | 0.7 | 9.4 | 6.4E-21 | 3.2E-20 | MB231/MB468 |
| >hg38_21_+_8433172_8446622_RNA45SN1_45S_50ntFlanks@10391.10420.30\|AACAGCAGTTGAACATGGGTCAGTCGGTCC | >hg38_21_+_8433172_8446622_RNA45SN1_45S_50ntFlanks | 10391 | 10420 | 30 | AACAGCAGTTGAACATGGGTCAGTCGGTCC | 316.3 | -4.6 | 0.5 | -9.4 | 6.5E-21 | 3.3E-20 | MB231/MB468 |
| >MI0000440\|hsa-mir-27b&WithFlank&9\|+\|95085439\|95085547@67.85.19&[MIMAT0000419&hsa-miR-27b-3p&offsets\|0\|-2;m-39&9\|+\|95085505\|95085525&offsets\|0\|-2]\|TTCACAGTGGCTAAGTTCT | >MI0000440\|hsa-mir-27b&WithFlank&9\|+\|95085439\|95085547 | 67 | 85 | 19 | TTCACAGTGGCTAAGTTCT | 5061.2 | 7.4 | 0.8 | 9.4 | 6.6E-21 | 3.3E-20 | MB231/MB468 |
| >hg38_21_+_8433172_8446622_RNA45SN1_45S_50ntFlanks@10845.10864.20\|ACTTCGGGATAAGGATTGGC | >hg38_21_+_8433172_8446622_RNA45SN1_45S_50ntFlanks | 10845 | 10864 | 20 | ACTTCGGGATAAGGATTGGC | 390.5 | -5.1 | 0.5 | -9.4 | 6.6E-21 | 3.3E-20 | MB231/MB468 |
| >MI0000067\|hsa-let-7f-1&WithFlank&9\|+\|94176341\|94176439@69.89.21(+1U)&[MIMAT0004486&hsa-let-7f-1-3p&offsets\|0\|-1(+1U);m-275&9\|+\|94176409\|94176430&offsets\|0\|-1(+1U)]\|CTATACAATCTATTGCCTTCCT | >MI0000067\|hsa-let-7f-1&WithFlank&9\|+\|94176341\|94176439 | 69 | 89 | 21(+1U) | CTATACAATCTATTGCCTTCCT | 692.0 | 7.3 | 0.8 | 9.4 | 5.6E-21 | 3.3E-20 | BT20/MB468 |
| >MI0000066\|hsa-let-7e&WithFlank&19\|+\|51692780\|51692870@14.34.21(+1A)&[MIMAT0000066&hsa-let-7e-5p&offsets\|0\|-1(+1A);m-50&19\|+\|51692793\|51692814&offsets\|0\|-1(+1A)]\|TGAGGTAGGAGGTTGTATAGTA | >MI0000066\|hsa-let-7e&WithFlank&19\|+\|51692780\|51692870 | 14 | 34 | 21(+1A) | TGAGGTAGGAGGTTGTATAGTA | 1876.1 | 8.6 | 0.9 | 9.4 | 5.7E-21 | 3.4E-20 | BT20/MB468 |
| >MI0000270\|hsa-mir-181b-1&WithFlank&1\|-\|198858867\|198858988@42.65.24;MI0000683\|hsa-mir-181b-2&WithFlank&9\|+\|124693704\|124693804@22.45.24&[MIMAT0000257&hsa-miR-181b-5p&offsets\|0\|+1;m-73&1\|-\|198858925\|198858947&offsets\|0\|+1];[MIMAT0000257_1&hsa-miR-181b-5p&offsets\|0\|+1;m-72&9\|+\|124693725\|124693747&offsets\|0\|+1]\|AACATTCATTGCTGTCGGTGGGTT | >MI0000270\|hsa-mir-181b-1&WithFlank&1\|-\|198858867\|198858988 | 42 | 65 | 24 | AACATTCATTGCTGTCGGTGGGTT | 5726.0 | 7.2 | 0.8 | 9.4 | 6.7E-21 | 3.4E-20 | MB231/MB468 |
| >hg38_21_+_8433172_8446622_RNA45SN1_45S_50ntFlanks@10362.10383.22\|AAGGCCGAAGTGGAGAAGGGTT | >hg38_21_+_8433172_8446622_RNA45SN1_45S_50ntFlanks | 10362 | 10383 | 22 | AAGGCCGAAGTGGAGAAGGGTT | 1588.0 | -4.8 | 0.5 | -9.4 | 6.9E-21 | 3.5E-20 | MB231/MB468 |
| >hg38_21_+_8433172_8446622_RNA45SN1_45S_50ntFlanks@13007.13039.33\|TATTGAAAGTCAGCCCTCGACACAAGGGTTTGT | >hg38_21_+_8433172_8446622_RNA45SN1_45S_50ntFlanks | 13007 | 13039 | 33 | TATTGAAAGTCAGCCCTCGACACAAGGGTTTGT | 3344.3 | 5.8 | 0.6 | 9.4 | 7.0E-21 | 3.5E-20 | MB231/MB468 |
| >hg38_21_+_8433172_8446622_RNA45SN1_45S_50ntFlanks@11641.11670.30\|CGGCGGGTGTTGACGCGATGTGATTTCTGC | >hg38_21_+_8433172_8446622_RNA45SN1_45S_50ntFlanks | 11641 | 11670 | 30 | CGGCGGGTGTTGACGCGATGTGATTTCTGC | 2691.3 | -10.2 | 1.1 | -9.4 | 6.0E-21 | 3.5E-20 | BT20/MB468 |
| >hg38_21_+_8433172_8446622_RNA45SN1_45S_50ntFlanks@12436.12455.20\|TCATTGTGAAGCAGAATTCA | >hg38_21_+_8433172_8446622_RNA45SN1_45S_50ntFlanks | 12436 | 12455 | 20 | TCATTGTGAAGCAGAATTCA | 4611.6 | -10.0 | 1.1 | -9.4 | 6.0E-21 | 3.5E-20 | BT20/MB468 |
| >MI0000279\|hsa-mir-196a-2&WithFlank&12\|+\|53991732\|53991853@31.51.21(+1A);MI0000238\|hsa-mir-196a-1&WithFlank&17\|-\|48632484\|48632565@13.33.21(+1A)&[MIMAT0000226&hsa-miR-196a-5p&offsets\|0\|-1(+1A);m-261&12\|+\|53991762\|53991783&offsets\|0\|-1(+1A)];[MIMAT0000226_1&hsa-miR-196a-5p&offsets\|0\|-1(+1A);m-268&17\|-\|48632532\|48632553&offsets\|0\|-1(+1A)]\|TAGGTAGTTTCATGTTGTTGGA | >MI0000279\|hsa-mir-196a-2&WithFlank&12\|+\|53991732\|53991853 | 31 | 51 | 21(+1A) | TAGGTAGTTTCATGTTGTTGGA | 2182.7 | 8.9 | 0.9 | 9.4 | 6.0E-21 | 3.5E-20 | BT20/MB468 |
| >MI0000542\|hsa-mir-320a&WithFlank&8\|-\|22244960\|22245043@49.68.20(+2U)&[MIMAT0000510&hsa-miR-320a-3p&offsets\|+1\|-1(+2U);m-60&8\|-\|22244975\|22244996&offsets\|+1\|-1(+2U)]\|AAAGCTGGGTTGAGAGGGCGTT | >MI0000542\|hsa-mir-320a&WithFlank&8\|-\|22244960\|22245043 | 49 | 68 | 20(+2U) | AAAGCTGGGTTGAGAGGGCGTT | 501.8 | 11.6 | 1.2 | 9.4 | 6.0E-21 | 3.5E-20 | BT20/MB468 |
| >MI0000478\|hsa-mir-149&WithFlank&2\|+\|240455995\|240456095@21.43.23&[MIMAT0000450&hsa-miR-149-5p&offsets\|0\|0;m-195&2\|+\|240456015\|240456037&offsets\|0\|0]\|TCTGGCTCCGTGTCTTCACTCCC | >MI0000478\|hsa-mir-149&WithFlank&2\|+\|240455995\|240456095 | 21 | 43 | 23 | TCTGGCTCCGTGTCTTCACTCCC | 3592.8 | 11.8 | 1.3 | 9.4 | 6.1E-21 | 3.6E-20 | BT20/MB468 |
| >MI0000077\|hsa-mir-21&WithFlank&17\|+\|59841260\|59841343@14.32.19&[MIMAT0000076&hsa-miR-21-5p&offsets\|0\|-3;m-1&17\|+\|59841273\|59841294&offsets\|0\|-3]\|TAGCTTATCAGACTGATGT | >MI0000077\|hsa-mir-21&WithFlank&17\|+\|59841260\|59841343 | 14 | 32 | 19 | TAGCTTATCAGACTGATGT | 4041.1 | 7.7 | 0.8 | 9.4 | 6.1E-21 | 3.6E-20 | BT20/MB468 |
| >MI0000767\|hsa-mir-365a&WithFlank&16\|+\|14309279\|14309377@62.83.22;MI0000769\|hsa-mir-365b&WithFlank&17\|+\|31575405\|31575527@74.95.22&[MIMAT0000710&hsa-miR-365a-3p&offsets\|0\|0;m-176&16\|+\|14309340\|14309361&offsets\|0\|0];[MIMAT0022834&hsa-miR-365b-3p&offsets\|0\|0;m-175&17\|+\|31575478\|31575499&offsets\|0\|0]\|TAATGCCCCTAAAAATCCTTAT | >MI0000767\|hsa-mir-365a&WithFlank&16\|+\|14309279\|14309377 | 62 | 83 | 22 | TAATGCCCCTAAAAATCCTTAT | 7206.8 | 9.1 | 1.0 | 9.4 | 6.2E-21 | 3.6E-20 | BT20/MB468 |
| >MI0000279\|hsa-mir-196a-2&WithFlank&12\|+\|53991732\|53991853@31.53.23&[MIMAT0000226&hsa-miR-196a-5p&offsets\|0\|+1;m-261&12\|+\|53991762\|53991783&offsets\|0\|+1]\|TAGGTAGTTTCATGTTGTTGGGA | >MI0000279\|hsa-mir-196a-2&WithFlank&12\|+\|53991732\|53991853 | 31 | 53 | 23 | TAGGTAGTTTCATGTTGTTGGGA | 1423.1 | 14.0 | 1.4 | 9.7 | 3.1E-22 | 3.6E-20 | BT20/MB231 |
| >MI0005566\|hsa-mir-374b&WithFlank&X\|-\|74218541\|74218624@17.38.22&[MIMAT0004955&hsa-miR-374b-5p&offsets\|0\|0;m-153&X\|-\|74218587\|74218608&offsets\|0\|0]\|ATATAATACAACCTGCTAAGTG | >MI0005566\|hsa-mir-374b&WithFlank&X\|-\|74218541\|74218624 | 17 | 38 | 22 | ATATAATACAACCTGCTAAGTG | 3443.3 | 7.5 | 0.8 | 9.4 | 7.4E-21 | 3.7E-20 | MB231/MB468 |
| >MI0005756\|hsa-mir-934&WithFlank&X\|+\|136550872\|136550966@21.40.20(+1A)&[MIMAT0004977&hsa-miR-934&offsets\|0\|-2(+1A);m-943&X\|+\|136550892\|136550913&offsets\|0\|-2(+1A)]\|TGTCTACTACTGGAGACACTA | >MI0005756\|hsa-mir-934&WithFlank&X\|+\|136550872\|136550966 | 21 | 40 | 20(+1A) | TGTCTACTACTGGAGACACTA | 3673.1 | 10.0 | 1.1 | 9.4 | 6.4E-21 | 3.7E-20 | BT20/MB468 |
| >trnaMT_GluTTC_MT_-_14674_14742@44.72.29,trnalookalike8_GluTTC_5_-_93905172_93905240@44.72.29\|TCGTGGTTGTAGTCCGTGCGAGAATACCA | >trnaMT_GluTTC_MT_-_14674_14742 | 44 | 72 | 29 | TCGTGGTTGTAGTCCGTGCGAGAATACCA | 2047.3 | 7.2 | 0.8 | 9.4 | 7.6E-21 | 3.8E-20 | MB231/MB468 |
| >MI0000061\|hsa-let-7a-2&WithFlank&11\|-\|122146516\|122146599@11.31.21(+1G);MI0000062\|hsa-let-7a-3&WithFlank&22\|+\|46112743\|46112828@10.30.21(+1G);MI0000060\|hsa-let-7a-1&WithFlank&9\|+\|94175951\|94176042@12.32.21(+1G)&[MIMAT0000062&hsa-let-7a-5p&offsets\|0\|-1(+1G);m-6&11\|-\|122146568\|122146589&offsets\|0\|-1(+1G)];[MIMAT0000062_1&hsa-let-7a-5p&offsets\|0\|-1(+1G);m-5&22\|+\|46112752\|46112773&offsets\|0\|-1(+1G)];[MIMAT0000062_2&hsa-let-7a-5p&offsets\|0\|-1(+1G);m-4&9\|+\|94175962\|94175983&offsets\|0\|-1(+1G)]\|TGAGGTAGTAGGTTGTATAGTG | >MI0000061\|hsa-let-7a-2&WithFlank&11\|-\|122146516\|122146599 | 11 | 31 | 21(+1G) | TGAGGTAGTAGGTTGTATAGTG | 9448.5 | 8.4 | 0.9 | 9.4 | 6.7E-21 | 3.9E-20 | BT20/MB468 |
| >am_MI0000786\|hsa-mir-378a&WithFlank&5\|+\|149732819\|149732896@49.69.21(+1A)&[MIMAT0000732&hsa-miR-378a-3p&offsets\|0\|-1(+1A);m-44&5\|+\|149732867\|149732888&offsets\|0\|-1(+1A)]\|ACTGGACTTGGAGTCAGAAGGA | >am_MI0000786\|hsa-mir-378a&WithFlank&5\|+\|149732819\|149732896 | 49 | 69 | 21(+1A) | ACTGGACTTGGAGTCAGAAGGA | 3118.4 | 6.1 | 0.6 | 9.4 | 6.7E-21 | 3.9E-20 | BT20/MB468 |
| >hg38_1_-_228634819_228635039_RNA5S12_5S_50ntFlanks@149.170.22\|GAATACCGGGTGCTGTAGGCTT | >hg38_1_-_228634819_228635039_RNA5S12_5S_50ntFlanks | 149 | 170 | 22 | GAATACCGGGTGCTGTAGGCTT | 1953.1 | 5.7 | 0.6 | 9.4 | 6.7E-21 | 3.9E-20 | BT20/MB468 |
| >hg38_21_+_8433172_8446622_RNA45SN1_45S_50ntFlanks@11783.11805.23\|CATGAATGGATGAACGAGATTCC | >hg38_21_+_8433172_8446622_RNA45SN1_45S_50ntFlanks | 11783 | 11805 | 23 | CATGAATGGATGAACGAGATTCC | 2377.1 | -5.6 | 0.6 | -9.4 | 6.9E-21 | 4.0E-20 | BT20/MB468 |
| >hg38_21_+_8433172_8446622_RNA45SN1_45S_50ntFlanks@6661.6700.40\|CGGTGGATCACTCGGCTCGTGCGTCGATGAAGAACGCAGC | >hg38_21_+_8433172_8446622_RNA45SN1_45S_50ntFlanks | 6661 | 6700 | 40 | CGGTGGATCACTCGGCTCGTGCGTCGATGAAGAACGCAGC | 1021.9 | 9.0 | 1.0 | 9.4 | 6.9E-21 | 4.0E-20 | BT20/MB468 |
| >MI0003560\|hsa-mir-92b&WithFlank&1\|+\|155195171\|155195278@67.87.21(+1G)&[MIMAT0003218&hsa-miR-92b-3p&offsets\|0\|-1(+1G);m-115&1\|+\|155195237\|155195258&offsets\|0\|-1(+1G)]\|TATTGCACTCGTCCCGGCCTCG | >MI0003560\|hsa-mir-92b&WithFlank&1\|+\|155195171\|155195278 | 67 | 87 | 21(+1G) | TATTGCACTCGTCCCGGCCTCG | 763.5 | 14.2 | 1.5 | 9.4 | 7.0E-21 | 4.1E-20 | BT20/MB468 |
| >MI0000082\|hsa-mir-25&WithFlank&7\|-\|100093554\|100093649@58.79.22(+1U)&[MIMAT0000081&hsa-miR-25-3p&offsets\|0\|0(+1U);m-23&7\|-\|100093571\|100093592&offsets\|0\|0(+1U)]\|CATTGCACTTGTCTCGGTCTGAT | >MI0000082\|hsa-mir-25&WithFlank&7\|-\|100093554\|100093649 | 58 | 79 | 22(+1U) | CATTGCACTTGTCTCGGTCTGAT | 1796.7 | 6.6 | 0.7 | 9.4 | 7.1E-21 | 4.1E-20 | BT20/MB468 |
| >hg38_21_+_8433172_8446622_RNA45SN1_45S_50ntFlanks@11540.11584.45\|CGCGGCGCCTCGCCTCGGCCGGCGCCTAGCAGCCGACTTAGAACT | >hg38_21_+_8433172_8446622_RNA45SN1_45S_50ntFlanks | 11540 | 11584 | 45 | CGCGGCGCCTCGCCTCGGCCGGCGCCTAGCAGCCGACTTAGAACT | 3674.5 | 11.0 | 1.2 | 9.4 | 8.3E-21 | 4.2E-20 | MB231/MB468 |
| >hg38_21_+_8433172_8446622_RNA45SN1_45S_50ntFlanks@12419.12449.31\|GATGTCGGCTCTTCCTATCATTGTGAAGCAG | >hg38_21_+_8433172_8446622_RNA45SN1_45S_50ntFlanks | 12419 | 12449 | 31 | GATGTCGGCTCTTCCTATCATTGTGAAGCAG | 2501.8 | 6.5 | 0.7 | 9.4 | 8.6E-21 | 4.3E-20 | MB231/MB468 |
| >hg38_21_+_8433172_8446622_RNA45SN1_45S_50ntFlanks@12936.12955.20\|AAACCATTCGTAGACGACCT | >hg38_21_+_8433172_8446622_RNA45SN1_45S_50ntFlanks | 12936 | 12955 | 20 | AAACCATTCGTAGACGACCT | 312.8 | -4.9 | 0.5 | -9.4 | 7.5E-21 | 4.4E-20 | BT20/MB468 |
| >hg38_21_+_8433172_8446622_RNA45SN1_45S_50ntFlanks@11235.11254.20\|CTTCCCGTGGATCGCCCCAG | >hg38_21_+_8433172_8446622_RNA45SN1_45S_50ntFlanks | 11235 | 11254 | 20 | CTTCCCGTGGATCGCCCCAG | 2239.6 | -9.9 | 1.1 | -9.3 | 8.8E-21 | 4.4E-20 | MB231/MB468 |
| >trnaMT_HisGTG_MT_+_12138_12206@32.71.40\|TGAATCTGACAACAGAGGCTTACGACCCCTTATTTACCCC | >trnaMT_HisGTG_MT_+_12138_12206 | 32 | 71 | 40 | TGAATCTGACAACAGAGGCTTACGACCCCTTATTTACCCC | 13097.3 | 8.5 | 0.9 | 9.3 | 8.8E-21 | 4.4E-20 | MB231/MB468 |
| >hg38_21_+_8433172_8446622_RNA45SN1_45S_50ntFlanks@10411.10432.22\|CAGTCGGTCCTGAGAGATGGGC | >hg38_21_+_8433172_8446622_RNA45SN1_45S_50ntFlanks | 10411 | 10432 | 22 | CAGTCGGTCCTGAGAGATGGGC | 1003.8 | -5.3 | 0.6 | -9.4 | 7.7E-21 | 4.5E-20 | BT20/MB468 |

<table>
<tr><td>>hg38_21_+_8433172_8446622_RNA45SN1_45S_50ntFlanks@12996.13036.41|TCGCTGCGATCTATTGAAAGTCAGCCCTCGACACAAGGGTT</td><td>>hg38_21_+_8433172_8446622_RNA45SN1_45S_50ntFlanks</td><td>12996</td><td>13036</td><td>41</td><td>TCGCTGCGATCTATTGAAAGTCAGCCCTCGACACAAGGGTT</td><td>4332.7</td><td>10.0</td><td>1.1</td><td>9.3</td><td>9.1E-21</td><td>4.5E-20</td><td>MB231/MB468</td></tr>
<tr><td>>hg38_21_+_8433172_8446622_RNA45SN1_45S_50ntFlanks@11235.11254.20|CTTCCCGTGGATCGCCCCAG</td><td>>hg38_21_+_8433172_8446622_RNA45SN1_45S_50ntFlanks</td><td>11235</td><td>11254</td><td>20</td><td>CTTCCCGTGGATCGCCCCAG</td><td>2760.5</td><td>-10.0</td><td>1.1</td><td>-9.4</td><td>7.9E-21</td><td>4.6E-20</td><td>BT20/MB468</td></tr>
<tr><td>>hg38_21_+_8433172_8446622_RNA45SN1_45S_50ntFlanks@11667.11687.21|CTGCCCAGTGCTCTGAATGTC</td><td>>hg38_21_+_8433172_8446622_RNA45SN1_45S_50ntFlanks</td><td>11667</td><td>11687</td><td>21</td><td>CTGCCCAGTGCTCTGAATGTC</td><td>289.1</td><td>4.5</td><td>0.5</td><td>9.4</td><td>8.0E-21</td><td>4.6E-20</td><td>BT20/MB468</td></tr>
<tr><td>>trna111_HisGTG_1_-_147774845_147774916@1.35.35,trna118_HisGTG_1_-_145396881_145396952@1.35.35,trna16_HisGTG_1_+_146544773_146544844@1.35.35,trna1_HisGTG_15_+_45493349_45493420@1.35.35,trna21_HisGTG_1_+_147753471_147753542@1.35.35,trna33_HisGTG_6_+_27125906_27125977@1.35.35,trna7_HisGTG_9_-_14433938_14434009@1.35.35,trna8_HisGTG_15_-_45492611_45492682@1.35.35,trna9_HisGTG_15_-_45490804_45490875@1.35.35|GCCGTGATCGTATAGTGGTTAGTACTCTGCGTTGT</td><td>>trna111_HisGTG_1_-_147774845_147774916</td><td>1</td><td>35</td><td>35</td><td>GCCGTGATCGTATAGTGGTTAGTACTCTGCGTTGT</td><td>1441.9</td><td>13.1</td><td>1.4</td><td>9.4</td><td>8.1E-21</td><td>4.7E-20</td><td>BT20/MB468</td></tr>
<tr><td>>MI0000087|hsa-mir-29a&WithFlank&7|-|130876741|130876816@48.67.20(+1A)&[MIMAT0000086&hsa-miR-29a-3p&offsets|0|-2(+1A);m-26&7|-|130876749|130876769&offsets|0|-1(+1A)]|TAGCACCATCTGAAATCGGTA</td><td>>MI0000087|hsa-mir-29a&WithFlank&7|-|130876741|130876816</td><td>48</td><td>67</td><td>20(+1A)</td><td>TAGCACCATCTGAAATCGGTA</td><td>913.4</td><td>10.4</td><td>1.1</td><td>9.3</td><td>9.4E-21</td><td>4.7E-20</td><td>MB231/MB468</td></tr>
<tr><td>>trnaMT_PheGAA_MT_+_577_647@35.54.20|GAAAATGTTTAGACGGGCTC</td><td>>trnaMT_PheGAA_MT_+_577_647</td><td>35</td><td>54</td><td>20</td><td>GAAAATGTTTAGACGGGCTC</td><td>2313.5</td><td>11.4</td><td>1.2</td><td>9.3</td><td>9.4E-21</td><td>4.7E-20</td><td>MB231/MB468</td></tr>
<tr><td>>MI0003820|hsa-mir-454&WithFlank&17|-|59137752|59137878@70.92.23&[MIMAT0003885&hsa-miR-454-3p&offsets|0|0;m-272&17|-|59137787|59137809&offsets|0|0]|TAGTGCAATATTGCTTATAGGGT</td><td>>MI0003820|hsa-mir-454&WithFlank&17|-|59137752|59137878</td><td>70</td><td>92</td><td>23</td><td>TAGTGCAATATTGCTTATAGGGT</td><td>631.1</td><td>7.1</td><td>0.8</td><td>9.3</td><td>9.4E-21</td><td>4.7E-20</td><td>MB231/MB468</td></tr>
<tr><td>>MI0000264|hsa-mir-7-2&WithFlank&15|+|88611819|88611940@38.59.22(+1A);MI0000265|hsa-mir-7-3&WithFlank&19|+|4770664|4770785@37.58.22(+1A);MI0000263|hsa-mir-7-1&WithFlank&9|-|83969742|83969863@30.51.22(+1A)&[MIMAT0000252&hsa-miR-7-5p&offsets|0|-2(+1A);m-400&15|+|88611856|88611879&offsets|0|-2(+1A)];[MIMAT0000252_1&hsa-miR-7-5p&offsets|0|-2(+1A);m-402&19|+|4770700|4770723&offsets|0|-2(+1A)];[MIMAT0000252_2&hsa-miR-7-5p&offsets|0|-2(+1A);m-401&9|-|83969811|83969834&offsets|0|-2(+1A)]|TGGAAGACTAGTGATTTTGTTGA</td><td>>MI0000264|hsa-mir-7-2&WithFlank&15|+|88611819|88611940</td><td>38</td><td>59</td><td>22(+1A)</td><td>TGGAAGACTAGTGATTTTGTTGA</td><td>4551.0</td><td>10.2</td><td>1.1</td><td>9.4</td><td>8.1E-21</td><td>4.7E-20</td><td>BT20/MB468</td></tr>
<tr><td>>MI0000079|hsa-mir-23a&WithFlank&19|-|13836581|13836665@54.71.18(+1G)&[MIMAT0000078&hsa-miR-23a-3p&offsets|+3|0(+1G);m-31&19|-|13836595|13836615&offsets|+3|0(+1G)]|ACATTGCCAGGGATTTCCG</td><td>>MI0000079|hsa-mir-23a&WithFlank&19|-|13836581|13836665</td><td>54</td><td>71</td><td>18(+1G)</td><td>ACATTGCCAGGGATTTCCG</td><td>2177.1</td><td>15.7</td><td>1.7</td><td>9.4</td><td>8.2E-21</td><td>4.7E-20</td><td>BT20/MB468</td></tr>
<tr><td>>hg38_21_+_8433172_8446622_RNA45SN1_45S_50ntFlanks@7975.8013.39|CGCGACCTCAGATCAGACGTGGCGACCCGCTGAATTTAA</td><td>>hg38_21_+_8433172_8446622_RNA45SN1_45S_50ntFlanks</td><td>7975</td><td>8013</td><td>39</td><td>CGCGACCTCAGATCAGACGTGGCGACCCGCTGAATTTAA</td><td>12363.7</td><td>7.8</td><td>0.8</td><td>9.3</td><td>9.5E-21</td><td>4.7E-20</td><td>MB231/MB468</td></tr>
<tr><td>>hg38_21_+_8433172_8446622_RNA45SN1_45S_50ntFlanks@12418.12436.19|CGATGTCGGCTCTTCCTAT</td><td>>hg38_21_+_8433172_8446622_RNA45SN1_45S_50ntFlanks</td><td>12418</td><td>12436</td><td>19</td><td>CGATGTCGGCTCTTCCTAT</td><td>10586.4</td><td>-5.0</td><td>0.5</td><td>-9.3</td><td>9.7E-21</td><td>4.8E-20</td><td>MB231/MB468</td></tr>
<tr><td>>am_trnaMT_ValTAC_MT_+_1602_1670@51.68.18|CTTAACTTGACCGCTCTG</td><td>>am_trnaMT_ValTAC_MT_+_1602_1670</td><td>51</td><td>68</td><td>18</td><td>CTTAACTTGACCGCTCTG</td><td>11567.6</td><td>-8.8</td><td>0.9</td><td>-9.3</td><td>9.9E-21</td><td>4.9E-20</td><td>MB231/MB468</td></tr>
<tr><td>>hg38_21_+_8433172_8446622_RNA45SN1_45S_50ntFlanks@6649.6683.35|TACGACTCTTAGCGGTGGATCACTCGGCTCGTGCG</td><td>>hg38_21_+_8433172_8446622_RNA45SN1_45S_50ntFlanks</td><td>6649</td><td>6683</td><td>35</td><td>TACGACTCTTAGCGGTGGATCACTCGGCTCGTGCG</td><td>3082.0</td><td>6.4</td><td>0.7</td><td>9.3</td><td>1.0E-20</td><td>4.9E-20</td><td>MB231/MB468</td></tr>
<tr><td>>MI0000749|hsa-mir-30e&WithFlank&1|+|40754349|40754452@65.85.21(+2U)&[MIMAT0000693&hsa-miR-30e-3p&offsets|0|-1(+2U);m-25&1|+|40754413|40754434&offsets|0|-1(+2U)]|CTTTCAGTCGGATGTTTACAGTT</td><td>>MI0000749|hsa-mir-30e&WithFlank&1|+|40754349|40754452</td><td>65</td><td>85</td><td>21(+2U)</td><td>CTTTCAGTCGGATGTTTACAGTT</td><td>2740.5</td><td>7.3</td><td>0.8</td><td>9.3</td><td>1.0E-20</td><td>4.9E-20</td><td>MB231/MB468</td></tr>
<tr><td>>MI0000063|hsa-let-7b&WithFlank&22|+|46113680|46113774@12.32.21&[MIMAT0000063&hsa-let-7b-5p&offsets|0|-1;m-14&22|+|46113691|46113712&offsets|0|-1]|TGAGGTAGTAGGTTGTGTGGT</td><td>>MI0000063|hsa-let-7b&WithFlank&22|+|46113680|46113774</td><td>12</td><td>32</td><td>21</td><td>TGAGGTAGTAGGTTGTGTGGT</td><td>3753.8</td><td>6.7</td><td>0.7</td><td>9.3</td><td>1.0E-20</td><td>5.1E-20</td><td>MB231/MB468</td></tr>
<tr><td>>hg38_21_+_8433172_8446622_RNA45SN1_45S_50ntFlanks@12159.12180.22|TTTGACTGGGGCGGTACACCTG</td><td>>hg38_21_+_8433172_8446622_RNA45SN1_45S_50ntFlanks</td><td>12159</td><td>12180</td><td>22</td><td>TTTGACTGGGGCGGTACACCTG</td><td>2013.0</td><td>-12.6</td><td>1.3</td><td>-9.3</td><td>1.0E-20</td><td>5.2E-20</td><td>MB231/MB468</td></tr>
<tr><td>>MI0000070|hsa-mir-16-1&WithFlank&13|-|50048967|50049067@20.39.20(+1U);MI0000115|hsa-mir-16-2&WithFlank&3|+|160404739|160404831@16.35.20(+1U)&[MIMAT0000069&hsa-miR-16-5p&offsets|0|-2(+1U);m-62&13|-|50049027|50049048&offsets|0|-2(+1U)];[MIMAT0000069_1&hsa-miR-16-5p&offsets|0|-2(+1U);m-61&3|+|160404754|160404775&offsets|0|-2(+1U)]|TAGCAGCACGTAAATATTGGT</td><td>>MI0000070|hsa-mir-16-1&WithFlank&13|-|50048967|50049067</td><td>20</td><td>39</td><td>20(+1U)</td><td>TAGCAGCACGTAAATATTGGT</td><td>678.8</td><td>6.7</td><td>0.7</td><td>9.3</td><td>1.1E-20</td><td>5.3E-20</td><td>MB231/MB468</td></tr>
<tr><td>>hg38_21_+_8433172_8446622_RNA45SN1_45S_50ntFlanks@7990.8011.22|GACGTGGCGACCCGCTGAATTT</td><td>>hg38_21_+_8433172_8446622_RNA45SN1_45S_50ntFlanks</td><td>7990</td><td>8011</td><td>22</td><td>GACGTGGCGACCCGCTGAATTT</td><td>3829.7</td><td>6.6</td><td>0.7</td><td>9.3</td><td>1.1E-20</td><td>5.3E-20</td><td>MB231/MB468</td></tr>
<tr><td>>MI0003137|hsa-mir-193b&WithFlank&16|+|14303961|14304055@57.77.21(+1A)&[MIMAT0002819&hsa-miR-193b-3p&offsets|0|-1(+1A);m-166&16|+|14304017|14304038&offsets|0|-1(+1A)]|AACTGGCCCTCAAAGTCCCGCA</td><td>>MI0003137|hsa-mir-193b&WithFlank&16|+|14303961|14304055</td><td>57</td><td>77</td><td>21(+1A)</td><td>AACTGGCCCTCAAAGTCCCGCA</td><td>2795.9</td><td>16.1</td><td>1.7</td><td>9.3</td><td>9.3E-21</td><td>5.4E-20</td><td>BT20/MB468</td></tr>
<tr><td>>MI0000088|hsa-mir-30a&WithFlank&6|-|71403545|71403627@53.73.21&[MIMAT0000088&hsa-miR-30a-3p&offsets|0|-1;m-38&6|-|71403554|71403575&offsets|0|-1]|CTTTCAGTCGGATGTTTGCAG</td><td>>MI0000088|hsa-mir-30a&WithFlank&6|-|71403545|71403627</td><td>53</td><td>73</td><td>21</td><td>CTTTCAGTCGGATGTTTGCAG</td><td>983.0</td><td>6.9</td><td>0.7</td><td>9.3</td><td>9.5E-21</td><td>5.4E-20</td><td>BT20/MB468</td></tr>
<tr><td>>hg38_21_+_8433172_8446622_RNA45SN1_45S_50ntFlanks@12456.12489.34|CCAAGCGTTGGATTGTTCACCCACTAATAGGGAA</td><td>>hg38_21_+_8433172_8446622_RNA45SN1_45S_50ntFlanks</td><td>12456</td><td>12489</td><td>34</td><td>CCAAGCGTTGGATTGTTCACCCACTAATAGGGAA</td><td>895.0</td><td>-6.6</td><td>0.7</td><td>-9.3</td><td>1.1E-20</td><td>5.5E-20</td><td>MB231/MB468</td></tr>
<tr><td>>trna10_AspGTC_12_-_125424193_125424264@36.74.39,trna12_AspGTC_12_-_125411891_125411962@36.74.39,trna144_AspGTC_6_-_27551236_27551307@36.74.39,trna38_AspGTC_17_-_8125556_8125627@36.74.39,trna45_AspGTC_6_+_27447453_27447524@36.74.39,trna48_AspGTC_6_+_27471523_27471594@36.74.39,trna4_AspGTC_12_+_96429799_96429870@36.74.39,trna69_AspGTC_1_-_161440205_161440276@36.74.39,trna72_AspGTC_1_-_161432824_161432895@36.74.39,trna75_AspGTC_1_-_161425414_161425485@36.74.39,trna78_AspGTC_1_-_161418033_161418104@36.74.39,trna81_AspGTC_1_-_161410615_161410686@36.74.39|CACGCGGGAGACCGGGGTTCGATTCCCCGACGGGGAGCC</td><td>>trna10_AspGTC_12_-_125424193_125424264</td><td>36</td><td>74</td><td>39</td><td>CACGCGGGAGACCGGGGTTCGATTCCCCGACGGGGAGCC</td><td>1840.4</td><td>13.3</td><td>1.4</td><td>9.3</td><td>1.1E-20</td><td>5.5E-20</td><td>MB231/MB468</td></tr>
<tr><td>>hg38_21_+_8433172_8446622_RNA45SN1_45S_50ntFlanks@10383.10424.42|TCCATGTGAACAGCAGTTGAACATGGGTCAGTCGGTCCTGAG</td><td>>hg38_21_+_8433172_8446622_RNA45SN1_45S_50ntFlanks</td><td>10383</td><td>10424</td><td>42</td><td>TCCATGTGAACAGCAGTTGAACATGGGTCAGTCGGTCCTGAG</td><td>2668.8</td><td>10.8</td><td>1.2</td><td>9.3</td><td>1.1E-20</td><td>5.6E-20</td><td>MB231/MB468</td></tr>
<tr><td>>MI0000107|hsa-mir-29b-2&WithFlank&1|-|207802437|207802529@58.80.23;MI0000105|hsa-mir-29b-1&WithFlank&7|-|130877453|130877545@57.79.23&[MIMAT0000100&hsa-miR-29b-3p&offsets|0|0;m-81&1|-|207802450|207802472&offsets|0|0];[MIMAT0000100_1&hsa-miR-29b-3p&offsets|0|0;m-82&7|-|130877467|130877489&offsets|0|0]|TAGCACCATTTGAAATCAGTGTT</td><td>>MI0000107|hsa-mir-29b-2&WithFlank&1|-|207802437|207802529</td><td>58</td><td>80</td><td>23</td><td>TAGCACCATTTGAAATCAGTGTT</td><td>2607.1</td><td>9.9</td><td>1.1</td><td>9.3</td><td>1.2E-20</td><td>5.7E-20</td><td>MB231/MB468</td></tr>
<tr><td>>MI0006273|hsa-mir-1180&WithFlank&17|-|19344500|19344580@47.68.22&[MIMAT0005825&hsa-miR-1180-3p&offsets|0|0;m-214&17|-|19344513|19344534&offsets|0|0]|TTTCCGGCTCGCGTGGGTGTGT</td><td>>MI0006273|hsa-mir-1180&WithFlank&17|-|19344500|19344580</td><td>47</td><td>68</td><td>22</td><td>TTTCCGGCTCGCGTGGGTGTGT</td><td>1390.4</td><td>6.5</td><td>0.7</td><td>9.3</td><td>1.2E-20</td><td>5.8E-20</td><td>MB231/MB468</td></tr>
<tr><td>>MI0000456|hsa-mir-140&WithFlank&16|+|69933075|69933186@29.50.22&[MIMAT0000431&hsa-miR-140-5p&offsets|0|0;m-173&16|+|69933103|69933124&offsets|0|0]|CAGTGGTTTTACCCTATGGTAG</td><td>>MI0000456|hsa-mir-140&WithFlank&16|+|69933075|69933186</td><td>29</td><td>50</td><td>22</td><td>CAGTGGTTTTACCCTATGGTAG</td><td>950.0</td><td>7.5</td><td>0.8</td><td>9.3</td><td>1.0E-20</td><td>5.8E-20</td><td>BT20/MB468</td></tr>
<tr><td>>hg38_21_+_8433172_8446622_RNA45SN1_45S_50ntFlanks@6650.6678.29|ACGACTCTTAGCGGTGGATCACTCGGCTC</td><td>>hg38_21_+_8433172_8446622_RNA45SN1_45S_50ntFlanks</td><td>6650</td><td>6678</td><td>29</td><td>ACGACTCTTAGCGGTGGATCACTCGGCTC</td><td>766.6</td><td>4.2</td><td>0.5</td><td>9.3</td><td>1.0E-20</td><td>5.9E-20</td><td>BT20/MB468</td></tr>
<tr><td>>hg38_21_+_8433172_8446622_RNA45SN1_45S_50ntFlanks@11572.11603.32|CCGACTTAGAACTGGTGCGGACCAGGGGAATC</td><td>>hg38_21_+_8433172_8446622_RNA45SN1_45S_50ntFlanks</td><td>11572</td><td>11603</td><td>32</td><td>CCGACTTAGAACTGGTGCGGACCAGGGGAATC</td><td>3732.5</td><td>-5.0</td><td>0.5</td><td>-9.3</td><td>1.2E-20</td><td>5.9E-20</td><td>MB231/MB468</td></tr>
<tr><td>>trnaMT_MetCAT_MT_+_4402_4469@52.70.19|ATACCCTTCCCGTACTACC</td><td>>trnaMT_MetCAT_MT_+_4402_4469</td><td>52</td><td>70</td><td>19</td><td>ATACCCTTCCCGTACTACC</td><td>1977.2</td><td>13.8</td><td>1.5</td><td>9.3</td><td>1.2E-20</td><td>5.9E-20</td><td>MB231/MB468</td></tr>
<tr><td>>MI0000088|hsa-mir-30a&WithFlank&6|-|71403545|71403627@12.33.22(+1A)&[MIMAT0000087&hsa-miR-30a-5p&offsets|0|0(+1A);m-15&6|-|71403595|71403616&offsets|0|0(+1A)]|TGTAAACATCCTCGACTGGAAGA</td><td>>MI0000088|hsa-mir-30a&WithFlank&6|-|71403545|71403627</td><td>12</td><td>33</td><td>22(+1A)</td><td>TGTAAACATCCTCGACTGGAAGA</td><td>18377.2</td><td>9.7</td><td>1.0</td><td>9.3</td><td>1.0E-20</td><td>5.9E-20</td><td>BT20/MB468</td></tr>
<tr><td>>hg38_21_+_8433172_8446622_RNA45SN1_45S_50ntFlanks@11217.11246.30|CTGGACGCGAGCCGGGCCCTTCCCGTGGAT</td><td>>hg38_21_+_8433172_8446622_RNA45SN1_45S_50ntFlanks</td><td>11217</td><td>11246</td><td>30</td><td>CTGGACGCGAGCCGGGCCCTTCCCGTGGAT</td><td>446.3</td><td>7.8</td><td>0.8</td><td>9.3</td><td>1.2E-20</td><td>6.0E-20</td><td>MB231/MB468</td></tr>
<tr><td>>MI0000736|hsa-mir-30c-1&WithFlank&1|+|40757278|40757378@23.42.20;MI0000254|hsa-mir-30c-2&WithFlank&6|-|71376954|71377037@13.32.20&[MIMAT0000244&hsa-miR-30c-5p&offsets|0|-3;m-57&1|+|40757300|40757323&offsets|0|-4];[MIMAT0000244_1&hsa-miR-30c-5p&offsets|0|-3;m-56&6|-|71377002|71377025&offsets|0|-4]|TGTAAACATCCTACACTCTC</td><td>>MI0000736|hsa-mir-30c-1&WithFlank&1|+|40757278|40757378</td><td>23</td><td>42</td><td>20</td><td>TGTAAACATCCTACACTCTC</td><td>3924.2</td><td>7.1</td><td>0.8</td><td>9.3</td><td>1.2E-20</td><td>6.0E-20</td><td>MB231/MB468</td></tr>
<tr><td>>MI0000285|hsa-mir-205&WithFlank&1|+|209432127|209432248@40.59.20&[MIMAT0000266&hsa-miR-205-5p&offsets|0|-2;m-351&1|+|209432166|209432187&offsets|0|-2]|TCCTTCATTCCACCGGAGTC</td><td>>MI0000285|hsa-mir-205&WithFlank&1|+|209432127|209432248</td><td>40</td><td>59</td><td>20</td><td>TCCTTCATTCCACCGGAGTC</td><td>907.5</td><td>6.9</td><td>0.7</td><td>9.3</td><td>1.1E-20</td><td>6.0E-20</td><td>BT20/MB468</td></tr>
</table>

<table>
<tr><td>>MI0000478|hsa-mir-149&WithFlank&2|+|240455995|240456095@21.41.21(+1A)&[MIMAT0000450&hsa-miR-149-5p&offsets|0|-2(+1A);m-195&2|+|240456015|240456037&offsets|0|-2(+1A)]|TCTGGCTCCGTGTCTTCACTCA</td><td>>MI0000478|hsa-mir-149&WithFlank&2|+|240455995|240456095</td><td>21</td><td>41</td><td>21(+1A)</td><td>TCTGGCTCCGTGTCTTCACTCA</td><td>946.6</td><td>14.5</td><td>1.6</td><td>9.3</td><td>1.1E-20</td><td>6.0E-20</td><td>BT20/MB468</td></tr>
<tr><td>>trna162_MetCAT_6_-_26758550_26758622@34.55.22,trna164_MetCAT_6_-_26735574_26735646@34.55.22,trna20_MetCAT_16_+_71460396_71460468@34.55.22,trna21_MetCAT_6_+_26701712_26701784@34.55.22,trna27_MetCAT_6_+_26766444_26766516@34.55.22,trna75_MetCAT_6_+_28912352_28912424@34.55.22,trna97_MetCAT_6_-_28921042_28921114@34.55.22|CATAATCTGAAGGTCCTGAGTT</td><td>>trna162_MetCAT_6_-_26758550_26758622</td><td>34</td><td>55</td><td>22</td><td>CATAATCTGAAGGTCCTGAGTT</td><td>391.2</td><td>6.9</td><td>0.7</td><td>9.3</td><td>1.1E-20</td><td>6.1E-20</td><td>BT20/MB468</td></tr>
<tr><td>>hg38_21_+_8433172_8446622_RNA45SN1_45S_50ntFlanks@5472.5509.38|ACGGCCCTGGCGGAGCGCTGAGAAGACGGTCGAACTTG</td><td>>hg38_21_+_8433172_8446622_RNA45SN1_45S_50ntFlanks</td><td>5472</td><td>5509</td><td>38</td><td>ACGGCCCTGGCGGAGCGCTGAGAAGACGGTCGAACTTG</td><td>765.8</td><td>5.8</td><td>0.6</td><td>9.3</td><td>1.1E-20</td><td>6.2E-20</td><td>BT20/MB468</td></tr>
<tr><td>>MI0000298|hsa-mir-221&WithFlank&X|-|45746151|45746272@31.55.25(+1C)&[MIMAT0004568&hsa-miR-221-5p&offsets|0|+3(+1C);m-405&X|-|45746220|45746242&offsets|0|+2(+1C)]|ACCTGGCATACAATGTAGATTTCTGC</td><td>>MI0000298|hsa-mir-221&WithFlank&X|-|45746151|45746272</td><td>31</td><td>55</td><td>25(+1C)</td><td>ACCTGGCATACAATGTAGATTTCTGC</td><td>1113.1</td><td>9.7</td><td>1.0</td><td>9.3</td><td>1.3E-20</td><td>6.3E-20</td><td>MB231/MB468</td></tr>
<tr><td>>hg38_1_-_228634819_228635039_RNA5S12_5S_50ntFlanks@134.171.38|TGGGAGACCGCCTGGGAATACCGGGTGCTGTAGGCTTT</td><td>>hg38_1_-_228634819_228635039_RNA5S12_5S_50ntFlanks</td><td>134</td><td>171</td><td>38</td><td>TGGGAGACCGCCTGGGAATACCGGGTGCTGTAGGCTTT</td><td>14041.3</td><td>6.5</td><td>0.7</td><td>9.3</td><td>1.1E-20</td><td>6.3E-20</td><td>BT20/MB468</td></tr>
<tr><td>>hg38_21_+_8433172_8446622_RNA45SN1_45S_50ntFlanks@6661.6701.41|CGGTGGATCACTCGGCTCGTGCGTCGATGAAGAACGCAGCT</td><td>>hg38_21_+_8433172_8446622_RNA45SN1_45S_50ntFlanks</td><td>6661</td><td>6701</td><td>41</td><td>CGGTGGATCACTCGGCTCGTGCGTCGATGAAGAACGCAGCT</td><td>750.2</td><td>6.9</td><td>0.7</td><td>9.3</td><td>1.3E-20</td><td>6.5E-20</td><td>MB231/MB468</td></tr>
<tr><td>>MI0016887|hsa-mir-4521&WithFlank&17|+|8186939|8187010@12.35.24&[MIMAT0019058&hsa-miR-4521&offsets|+2|+4;m-765&17|+|8186948|8186969&offsets|+2|+4]|TAAGGAAGTCCTGTGCTCAGTTTT</td><td>>MI0016887|hsa-mir-4521&WithFlank&17|+|8186939|8187010</td><td>12</td><td>35</td><td>24</td><td>TAAGGAAGTCCTGTGCTCAGTTTT</td><td>475.8</td><td>13.5</td><td>1.5</td><td>9.3</td><td>1.1E-20</td><td>6.5E-20</td><td>BT20/MB468</td></tr>
<tr><td>>hg38_21_+_8433172_8446622_RNA45SN1_45S_50ntFlanks@10806.10828.23|AACAATGTAGGTAAGGGAAGTCG</td><td>>hg38_21_+_8433172_8446622_RNA45SN1_45S_50ntFlanks</td><td>10806</td><td>10828</td><td>23</td><td>AACAATGTAGGTAAGGGAAGTCG</td><td>218.8</td><td>-5.1</td><td>0.5</td><td>-9.3</td><td>1.1E-20</td><td>6.5E-20</td><td>BT20/MB468</td></tr>
<tr><td>>hg38_21_+_8433172_8446622_RNA45SN1_45S_50ntFlanks@12415.12457.43|CTTCGATGTCGGCTCTTCCTATCATTGTGAAGCAGAATTCACC</td><td>>hg38_21_+_8433172_8446622_RNA45SN1_45S_50ntFlanks</td><td>12415</td><td>12457</td><td>43</td><td>CTTCGATGTCGGCTCTTCCTATCATTGTGAAGCAGAATTCACC</td><td>527.6</td><td>9.3</td><td>1.0</td><td>9.3</td><td>1.3E-20</td><td>6.6E-20</td><td>MB231/MB468</td></tr>
<tr><td>>MI0005566|hsa-mir-374b&WithFlank&X|-|74218541|74218624@17.39.23&[MIMAT0004955&hsa-miR-374b-5p&offsets|0|+1;m-153&X|-|74218587|74218608&offsets|0|+1]|ATATAATACAACCTGCTAAGTGT</td><td>>MI0005566|hsa-mir-374b&WithFlank&X|-|74218541|74218624</td><td>17</td><td>39</td><td>23</td><td>ATATAATACAACCTGCTAAGTGT</td><td>1086.7</td><td>7.9</td><td>0.8</td><td>9.3</td><td>1.3E-20</td><td>6.6E-20</td><td>MB231/MB468</td></tr>
<tr><td>>hg38_21_+_8433172_8446622_RNA45SN1_45S_50ntFlanks@4759.4783.25|ATCAGATACCGTCGTAGTTCCGACC</td><td>>hg38_21_+_8433172_8446622_RNA45SN1_45S_50ntFlanks</td><td>4759</td><td>4783</td><td>25</td><td>ATCAGATACCGTCGTAGTTCCGACC</td><td>961.1</td><td>-11.9</td><td>1.3</td><td>-9.3</td><td>1.2E-20</td><td>6.6E-20</td><td>BT20/MB468</td></tr>
<tr><td>>hg38_21_+_8433172_8446622_RNA45SN1_45S_50ntFlanks@6780.6806.27|CGGGGCTACGCCTGTCTGAGCGTCGCT</td><td>>hg38_21_+_8433172_8446622_RNA45SN1_45S_50ntFlanks</td><td>6780</td><td>6806</td><td>27</td><td>CGGGGCTACGCCTGTCTGAGCGTCGCT</td><td>780.5</td><td>5.7</td><td>0.6</td><td>9.3</td><td>1.4E-20</td><td>6.6E-20</td><td>MB231/MB468</td></tr>
<tr><td>>MI0000736|hsa-mir-30c-1&WithFlank&1|+|40757278|40757378@23.45.23(+1C);MI0000254|hsa-mir-30c-2&WithFlank&6|-|71376954|71377037@13.35.23(+1C)&[MIMAT0000244&hsa-miR-30c-5p&offsets|0|0(+1C);m-57&1|+|40757300|40757323&offsets|0|-1(+1C)];[MIMAT0000244_1&hsa-miR-30c-5p&offsets|0|0(+1C);m-56&6|-|71377002|71377025&offsets|0|-1(+1C)]|TGTAAACATCCTACACTCTCAGCC</td><td>>MI0000736|hsa-mir-30c-1&WithFlank&1|+|40757278|40757378</td><td>23</td><td>45</td><td>23(+1C)</td><td>TGTAAACATCCTACACTCTCAGCC</td><td>9808.5</td><td>6.8</td><td>0.7</td><td>9.3</td><td>1.4E-20</td><td>6.6E-20</td><td>MB231/MB468</td></tr>
<tr><td>>hg38_21_+_8433172_8446622_RNA45SN1_45S_50ntFlanks@7976.8011.36|GCGACCTCAGATCAGACGTGGCGACCCGCTGAATTT</td><td>>hg38_21_+_8433172_8446622_RNA45SN1_45S_50ntFlanks</td><td>7976</td><td>8011</td><td>36</td><td>GCGACCTCAGATCAGACGTGGCGACCCGCTGAATTT</td><td>1141.9</td><td>9.2</td><td>1.0</td><td>9.3</td><td>1.4E-20</td><td>6.6E-20</td><td>MB231/MB468</td></tr>
<tr><td>>hg38_21_+_8433172_8446622_RNA45SN1_45S_50ntFlanks@12480.12497.18|TAATAGGGAACGTGAGCT</td><td>>hg38_21_+_8433172_8446622_RNA45SN1_45S_50ntFlanks</td><td>12480</td><td>12497</td><td>18</td><td>TAATAGGGAACGTGAGCT</td><td>802.6</td><td>3.6</td><td>0.4</td><td>9.3</td><td>1.2E-20</td><td>6.6E-20</td><td>BT20/MB468</td></tr>
<tr><td>>MI0000077|hsa-mir-21&WithFlank&17|+|59841260|59841343@52.72.21&[MIMAT0004494&hsa-miR-21-3p&offsets|0|0;m-52&17|+|59841311|59841332&offsets|0|-1]|CAACACCAGTCGATGGGCTGT</td><td>>MI0000077|hsa-mir-21&WithFlank&17|+|59841260|59841343</td><td>52</td><td>72</td><td>21</td><td>CAACACCAGTCGATGGGCTGT</td><td>1950.7</td><td>10.1</td><td>1.1</td><td>9.3</td><td>1.4E-20</td><td>6.7E-20</td><td>MB231/MB468</td></tr>
<tr><td>>trnaMT_ValTAC_MT_+_1602_1670@42.71.30|AGATTTCAACTTAACTTGACCGCTCTGACC</td><td>>trnaMT_ValTAC_MT_+_1602_1670</td><td>42</td><td>71</td><td>30</td><td>AGATTTCAACTTAACTTGACCGCTCTGACC</td><td>1293.1</td><td>9.0</td><td>1.0</td><td>9.3</td><td>1.4E-20</td><td>6.9E-20</td><td>MB231/MB468</td></tr>
<tr><td>>trnaMT_PheGAA_MT_+_577_647@38.57.20|AATGTTTAGACGGGCTCACA</td><td>>trnaMT_PheGAA_MT_+_577_647</td><td>38</td><td>57</td><td>20</td><td>AATGTTTAGACGGGCTCACA</td><td>1726.2</td><td>11.0</td><td>1.2</td><td>9.3</td><td>1.2E-20</td><td>7.0E-20</td><td>BT20/MB468</td></tr>
<tr><td>>hg38_21_+_8433172_8446622_RNA45SN1_45S_50ntFlanks@11572.11605.34|CCGACTTAGAACTGGTGCGGACCAGGGGAATCCG</td><td>>hg38_21_+_8433172_8446622_RNA45SN1_45S_50ntFlanks</td><td>11572</td><td>11605</td><td>34</td><td>CCGACTTAGAACTGGTGCGGACCAGGGGAATCCG</td><td>1228.1</td><td>-5.1</td><td>0.6</td><td>-9.3</td><td>1.4E-20</td><td>7.0E-20</td><td>MB231/MB468</td></tr>
<tr><td>>MI0000074|hsa-mir-19b-1&WithFlank&13|+|91351186|91351284@60.81.22(+1U);MI0000075|hsa-mir-19b-2&WithFlank&X|-|134169665|134169772@68.89.22(+1U)&[MIMAT0000074&hsa-miR-19b-3p&offsets|0|-1(+1U);m-122&13|+|91351245|91351267&offsets|0|-1(+1U)];[MIMAT0000074_1&hsa-miR-19b-3p&offsets|0|-1(+1U);m-121&X|-|134169683|134169705&offsets|0|-1(+1U)]|TGTGCAAATCCATGCAAAACTGT</td><td>>MI0000074|hsa-mir-19b-1&WithFlank&13|+|91351186|91351284</td><td>60</td><td>81</td><td>22(+1U)</td><td>TGTGCAAATCCATGCAAAACTGT</td><td>694.7</td><td>6.9</td><td>0.7</td><td>9.3</td><td>1.5E-20</td><td>7.1E-20</td><td>MB231/MB468</td></tr>
<tr><td>>MI0000767|hsa-mir-365a&WithFlank&16|+|14309279|14309377@62.81.20;MI0000769|hsa-mir-365b&WithFlank&17|+|31575405|31575527@74.93.20&[MIMAT0000710&hsa-miR-365a-3p&offsets|0|-2;m-176&16|+|14309340|14309361&offsets|0|-2];[MIMAT0022834&hsa-miR-365b-3p&offsets|0|-2;m-175&17|+|31575478|31575499&offsets|0|-2]|TAATGCCCCTAAAAATCCTT</td><td>>MI0000767|hsa-mir-365a&WithFlank&16|+|14309279|14309377</td><td>62</td><td>81</td><td>20</td><td>TAATGCCCCTAAAAATCCTT</td><td>797.0</td><td>14.3</td><td>1.5</td><td>9.3</td><td>1.2E-20</td><td>7.1E-20</td><td>BT20/MB468</td></tr>
<tr><td>>hg38_21_+_8433172_8446622_RNA45SN1_45S_50ntFlanks@6647.6669.23|CGTACGACTCTTAGCGGTGGATC</td><td>>hg38_21_+_8433172_8446622_RNA45SN1_45S_50ntFlanks</td><td>6647</td><td>6669</td><td>23</td><td>CGTACGACTCTTAGCGGTGGATC</td><td>5507.3</td><td>3.8</td><td>0.4</td><td>9.3</td><td>1.3E-20</td><td>7.2E-20</td><td>BT20/MB468</td></tr>
<tr><td>>MI0000087|hsa-mir-29a&WithFlank&7|-|130876741|130876816@47.67.21(+1A)&[MIMAT0000086&hsa-miR-29a-3p&offsets|-1|-2(+1A);m-26&7|-|130876749|130876769&offsets|-1|-1(+1A)]|CTAGCACCATCTGAAATCGGTA</td><td>>MI0000087|hsa-mir-29a&WithFlank&7|-|130876741|130876816</td><td>47</td><td>67</td><td>21(+1A)</td><td>CTAGCACCATCTGAAATCGGTA</td><td>1308.7</td><td>10.9</td><td>1.2</td><td>9.3</td><td>1.3E-20</td><td>7.3E-20</td><td>BT20/MB468</td></tr>
<tr><td>>MI0000261|hsa-mir-139&WithFlank&11|-|72615057|72615136@13.35.23&[MIMAT0000250&hsa-miR-139-5p&offsets|0|0;m-186&11|-|72615102|72615124&offsets|0|0]|TCTACAGTGCACGTGTCTCCAGT</td><td>>MI0000261|hsa-mir-139&WithFlank&11|-|72615057|72615136</td><td>13</td><td>35</td><td>23</td><td>TCTACAGTGCACGTGTCTCCAGT</td><td>784.4</td><td>11.8</td><td>1.3</td><td>9.3</td><td>1.3E-20</td><td>7.3E-20</td><td>BT20/MB468</td></tr>
<tr><td>>MI0000746|hsa-mir-99b&WithFlank&19|+|51692606|51692687@51.71.21&[MIMAT0004678&hsa-miR-99b-3p&offsets|0|-1;m-151&19|+|51692656|51692678&offsets|0|-2]|CAAGCTCGTGTCTGTGGGTCC</td><td>>MI0000746|hsa-mir-99b&WithFlank&19|+|51692606|51692687</td><td>51</td><td>71</td><td>21</td><td>CAAGCTCGTGTCTGTGGGTCC</td><td>680.3</td><td>7.3</td><td>0.8</td><td>9.3</td><td>1.5E-20</td><td>7.3E-20</td><td>MB231/MB468</td></tr>
<tr><td>>MI0005559|hsa-mir-744&WithFlank&17|+|12081893|12082002@17.36.20(+1U)&[MIMAT0004945&hsa-miR-744-5p&offsets|0|-2(+1U);m-172&17|+|12081909|12081929&offsets|0|-1(+1U)]|TGCGGGGCTAGGGCTAACAGT</td><td>>MI0005559|hsa-mir-744&WithFlank&17|+|12081893|12082002</td><td>17</td><td>36</td><td>20(+1U)</td><td>TGCGGGGCTAGGGCTAACAGT</td><td>789.1</td><td>8.9</td><td>1.0</td><td>9.3</td><td>1.3E-20</td><td>7.4E-20</td><td>BT20/MB468</td></tr>
<tr><td>>hg38_21_+_8433172_8446622_RNA45SN1_45S_50ntFlanks@11785.11803.19|TGAATGGATGAACGAGATT</td><td>>hg38_21_+_8433172_8446622_RNA45SN1_45S_50ntFlanks</td><td>11785</td><td>11803</td><td>19</td><td>TGAATGGATGAACGAGATT</td><td>3473.5</td><td>-6.0</td><td>0.6</td><td>-9.3</td><td>1.3E-20</td><td>7.5E-20</td><td>BT20/MB468</td></tr>
<tr><td>>hg38_21_+_8433172_8446622_RNA45SN1_45S_50ntFlanks@13004.13027.24|ATCTATTGAAAGTCAGCCCTCGAC</td><td>>hg38_21_+_8433172_8446622_RNA45SN1_45S_50ntFlanks</td><td>13004</td><td>13027</td><td>24</td><td>ATCTATTGAAAGTCAGCCCTCGAC</td><td>1831.2</td><td>-12.8</td><td>1.4</td><td>-9.3</td><td>1.3E-20</td><td>7.5E-20</td><td>BT20/MB468</td></tr>
<tr><td>>MI0000076|hsa-mir-20a&WithFlank&13|+|91351059|91351141@14.37.24&[MIMAT0000075&hsa-miR-20a-5p&offsets|0|+1;m-96&13|+|91351072|91351094&offsets|0|+1]|TAAAGTGCTTATAGTGCAGGTAGT</td><td>>MI0000076|hsa-mir-20a&WithFlank&13|+|91351059|91351141</td><td>14</td><td>37</td><td>24</td><td>TAAAGTGCTTATAGTGCAGGTAGT</td><td>6895.2</td><td>7.3</td><td>0.8</td><td>9.3</td><td>1.6E-20</td><td>7.6E-20</td><td>MB231/MB468</td></tr>
<tr><td>>MI0000107|hsa-mir-29b-2&WithFlank&1|-|207802437|207802529@58.79.22(+1C);MI0000105|hsa-mir-29b-1&WithFlank&7|-|130877453|130877545@57.78.22(+1C)&[MIMAT0000100&hsa-miR-29b-3p&offsets|0|-1(+1C);m-81&1|-|207802450|207802472&offsets|0|-1(+1C)];[MIMAT0000100_1&hsa-miR-29b-3p&offsets|0|-1(+1C);m-82&7|-|130877467|130877489&offsets|0|-1(+1C)]|TAGCACCATTTGAAATCAGTGTC</td><td>>MI0000107|hsa-mir-29b-2&WithFlank&1|-|207802437|207802529</td><td>58</td><td>79</td><td>22(+1C)</td><td>TAGCACCATTTGAAATCAGTGTC</td><td>826.3</td><td>14.3</td><td>1.5</td><td>9.3</td><td>1.3E-20</td><td>7.6E-20</td><td>BT20/MB468</td></tr>
<tr><td>>trnaMT_ValTAC_MT_+_1602_1670@48.72.25|CAACTTAACTTGACCGCTCTGACCA</td><td>>trnaMT_ValTAC_MT_+_1602_1670</td><td>48</td><td>72</td><td>25</td><td>CAACTTAACTTGACCGCTCTGACCA</td><td>1776.4</td><td>6.8</td><td>0.7</td><td>9.3</td><td>1.6E-20</td><td>7.6E-20</td><td>MB231/MB468</td></tr>
<tr><td>>hg38_21_+_8433172_8446622_RNA45SN1_45S_50ntFlanks@11784.11803.20|ATGAATGGATGAACGAGATT</td><td>>hg38_21_+_8433172_8446622_RNA45SN1_45S_50ntFlanks</td><td>11784</td><td>11803</td><td>20</td><td>ATGAATGGATGAACGAGATT</td><td>2552.8</td><td>-5.4</td><td>0.6</td><td>-9.3</td><td>1.4E-20</td><td>7.7E-20</td><td>BT20/MB468</td></tr>
<tr><td>>trnaMT_GluTTC_MT_-_14674_14742@46.72.27,trnalookalike8_GluTTC_5_-_93905172_93905240@46.72.27|GTGGTTGTAGTCCGTGCGAGAATACCA</td><td>>trnaMT_GluTTC_MT_-_14674_14742</td><td>46</td><td>72</td><td>27</td><td>GTGGTTGTAGTCCGTGCGAGAATACCA</td><td>1798.8</td><td>10.4</td><td>1.1</td><td>9.3</td><td>1.6E-20</td><td>7.7E-20</td><td>MB231/MB468</td></tr>
<tr><td>>MI0000102|hsa-mir-100&WithFlank&11|-|122152223|122152314@19.40.22&[MIMAT0000098&hsa-miR-100-5p&offsets|0|0;m-27&11|-|122152276|122152296&offsets|0|+1]|AACCCGTAGATCCGAACTTGTG</td><td>>MI0000102|hsa-mir-100&WithFlank&11|-|122152223|122152314</td><td>19</td><td>40</td><td>22</td><td>AACCCGTAGATCCGAACTTGTG</td><td>3020239.3</td><td>15.3</td><td>1.7</td><td>9.3</td><td>1.6E-20</td><td>7.7E-20</td><td>MB231/MB468</td></tr>
<tr><td>>MI0003560|hsa-mir-92b&WithFlank&1|+|155195171|155195278@67.86.20&[MIMAT0003218&hsa-miR-92b-3p&offsets|0|-2;m-115&1|+|155195237|155195258&offsets|0|-2]|TATTGCACTCGTCCCGGCCT</td><td>>MI0003560|hsa-mir-92b&WithFlank&1|+|155195171|155195278</td><td>67</td><td>86</td><td>20</td><td>TATTGCACTCGTCCCGGCCT</td><td>1021.7</td><td>12.2</td><td>1.3</td><td>9.3</td><td>1.4E-20</td><td>7.7E-20</td><td>BT20/MB468</td></tr>
<tr><td>>MI0000063|hsa-let-7b&WithFlank&22|+|46113680|46113774@12.33.22&[MIMAT0000063&hsa-let-7b-5p&offsets|0|0;m-14&22|+|46113691|46113712&offsets|0|0]|TGAGGTAGTAGGTTGTGTGGTT</td><td>>MI0000063|hsa-let-7b&WithFlank&22|+|46113680|46113774</td><td>12</td><td>33</td><td>22</td><td>TGAGGTAGTAGGTTGTGTGGTT</td><td>28700.0</td><td>6.8</td><td>0.7</td><td>9.3</td><td>1.6E-20</td><td>7.8E-20</td><td>MB231/MB468</td></tr>
<tr><td>>hg38_21_+_8433172_8446622_RNA45SN1_45S_50ntFlanks@10382.10424.43|TTCCATGTGAACAGCAGTTGAACATGGGTCAGTCGGTCCTGAG</td><td>>hg38_21_+_8433172_8446622_RNA45SN1_45S_50ntFlanks</td><td>10382</td><td>10424</td><td>43</td><td>TTCCATGTGAACAGCAGTTGAACATGGGTCAGTCGGTCCTGAG</td><td>1012.2</td><td>11.8</td><td>1.3</td><td>9.3</td><td>1.6E-20</td><td>7.8E-20</td><td>MB231/MB468</td></tr>
<tr><td>>am_trnaMT_ValTAC_MT_+_1602_1670@28.57.30|AACTTACACTTAGGAGATTTCAACTTAACT</td><td>>am_trnaMT_ValTAC_MT_+_1602_1670</td><td>28</td><td>57</td><td>30</td><td>AACTTACACTTAGGAGATTTCAACTTAACT</td><td>460.6</td><td>11.5</td><td>1.2</td><td>9.3</td><td>1.4E-20</td><td>7.9E-20</td><td>BT20/MB468</td></tr>
</table>

| | | | | | | | | | | | | |
|---|---|---|---|---|---|---|---|---|---|---|---|---|
| >MI0000438\|hsa-mir-15b&WithFlank&3\|+\|160404582\|160404691@26.47.22&[MIMAT0000417&hsa-miR-15b-5p&offsets\|0\|0;m-110&3\|+\|160404607\|160404628&offsets\|0\|0]\|TAGCAGCACATCATGGTTTACA | >MI0000438\|hsa-mir-15b&WithFlank&3\|+\|160404582\|160404691 | 26 | 47 | 22 | TAGCAGCACATCATGGTTTACA | 4877.2 | 7.1 | 0.8 | 9.3 | 1.7E-20 | 8.0E-20 | MB231/MB468 |
| >hg38_21_+_8433172_8446622_RNA45SN1_45S_50ntFlanks@6734.6774.41\|ATTGATCATCGACACTTCGAACGCACTTGCGGCCCCGGGTT | >hg38_21_+_8433172_8446622_RNA45SN1_45S_50ntFlanks | 6734 | 6774 | 41 | ATTGATCATCGACACTTCGAACGCACTTGCGGCCCCGGGTT | 2087.6 | 11.2 | 1.2 | 9.3 | 1.7E-20 | 8.2E-20 | MB231/MB468 |
| >MI0005559\|hsa-mir-744&WithFlank&17\|+\|12081893\|12082002@17.37.21&[MIMAT0004945&hsa-miR-744-5p&offsets\|0\|-1;m-172&17\|+\|12081909\|12081929&offsets\|0\|0]\|TGCGGGGCTAGGGCTAACAGC | >MI0005559\|hsa-mir-744&WithFlank&17\|+\|12081893\|12082002 | 17 | 37 | 21 | TGCGGGGCTAGGGCTAACAGC | 2623.6 | 7.4 | 0.8 | 9.3 | 1.7E-20 | 8.3E-20 | MB231/MB468 |
| >hg38_21_+_8433172_8446622_RNA45SN1_45S_50ntFlanks@6740.6770.31\|CATCGACACTTCGAACGCACTTGCGGCCCCG | >hg38_21_+_8433172_8446622_RNA45SN1_45S_50ntFlanks | 6740 | 6770 | 31 | CATCGACACTTCGAACGCACTTGCGGCCCCG | 1205.3 | -6.3 | 0.7 | -9.3 | 1.7E-20 | 8.4E-20 | MB231/MB468 |
| >MI0000070\|hsa-mir-16-1&WithFlank&13\|-\|50048967\|50049067@20.40.21(+1A);MI0000115\|hsa-mir-16-2&WithFlank&3\|+\|160404739\|160404831@16.36.21(+1A)&[MIMAT0000069&hsa-miR-16-5p&offsets\|0\|-1(+1A);m-62&13\|-\|50049027\|50049048&offsets\|0\|-1(+1A)];[MIMAT0000069_1&hsa-miR-16-5p&offsets\|0\|-1(+1A);m-61&3\|+\|160404754\|160404775&offsets\|0\|-1(+1A)]\|TAGCAGCACGTAAATATTGGCA | >MI0000070\|hsa-mir-16-1&WithFlank&13\|-\|50048967\|50049067 | 20 | 40 | 21(+1A) | TAGCAGCACGTAAATATTGGCA | 1318.3 | 7.3 | 0.8 | 9.3 | 1.8E-20 | 8.6E-20 | MB231/MB468 |
| >MI0000088\|hsa-mir-30a&WithFlank&6\|-\|71403545\|71403627@54.73.20(+1A)&[MIMAT0000088&hsa-miR-30a-3p&offsets\|+1\|-1(+1A);m-38&6\|-\|71403554\|71403575&offsets\|+1\|-1(+1A)]\|TTTCAGTCGGATGTTTGCAGA | >MI0000088\|hsa-mir-30a&WithFlank&6\|-\|71403545\|71403627 | 54 | 73 | 20(+1A) | TTTCAGTCGGATGTTTGCAGA | 471.7 | 13.5 | 1.5 | 9.3 | 1.5E-20 | 8.6E-20 | BT20/MB468 |
| >MI0000103\|hsa-mir-101-1&WithFlank&1\|-\|65058428\|65058514@53.74.22;MI0000739\|hsa-mir-101-2&WithFlank&9\|+\|4850291\|4850381@55.76.22&[MIMAT0000099&hsa-miR-101-3p&offsets\|0\|+1;m-17&1\|-\|65058443\|65058462&offsets\|0\|+2];[MIMAT0000099_1&hsa-miR-101-3p&offsets\|0\|+1;m-16&9\|+\|4850345\|4850364&offsets\|0\|+2]\|TACAGTACTGTGATAACTGAAG | >MI0000103\|hsa-mir-101-1&WithFlank&1\|-\|65058428\|65058514 | 53 | 74 | 22 | TACAGTACTGTGATAACTGAAG | 7089.6 | 7.7 | 0.8 | 9.3 | 1.9E-20 | 9.0E-20 | MB231/MB468 |
| >MI0000746\|hsa-mir-99b&WithFlank&19\|+\|51692606\|51692687@13.33.21(+1A)&[MIMAT0000689&hsa-miR-99b-5p&offsets\|0\|-1(+1A);m-10&19\|+\|51692618\|51692639&offsets\|0\|-1(+1A)]\|CACCCGTAGAACCGACCTTGCA | >MI0000746\|hsa-mir-99b&WithFlank&19\|+\|51692606\|51692687 | 13 | 33 | 21(+1A) | CACCCGTAGAACCGACCTTGCA | 29990.4 | 8.5 | 0.9 | 9.3 | 1.6E-20 | 9.0E-20 | BT20/MB468 |
| >hg38_21_+_8433172_8446622_RNA45SN1_45S_50ntFlanks@11873.11900.28\|AAGAAGACCCTGTTGAGCTTGACTCTAG | >hg38_21_+_8433172_8446622_RNA45SN1_45S_50ntFlanks | 11873 | 11900 | 28 | AAGAAGACCCTGTTGAGCTTGACTCTAG | 642.8 | -5.6 | 0.6 | -9.3 | 1.9E-20 | 9.0E-20 | MB231/MB468 |
| >hg38_21_+_8433172_8446622_RNA45SN1_45S_50ntFlanks@12006.12033.28\|GGCCGCCGGTGAAATACCACTACTCTGA | >hg38_21_+_8433172_8446622_RNA45SN1_45S_50ntFlanks | 12006 | 12033 | 28 | GGCCGCCGGTGAAATACCACTACTCTGA | 516.4 | 7.1 | 0.8 | 9.3 | 1.6E-20 | 9.0E-20 | BT20/MB468 |
| >MI0000086\|hsa-mir-28&WithFlank&3\|+\|188688775\|188688872@61.82.22(+1U)&[MIMAT0004502&hsa-miR-28-3p&offsets\|+1\|+1(+1U);m-28&3\|+\|188688834\|188688855&offsets\|+1\|+1(+1U)]\|ACTAGATTGTGAGCTCCTGGAGT | >MI0000086\|hsa-mir-28&WithFlank&3\|+\|188688775\|188688872 | 61 | 82 | 22(+1U) | ACTAGATTGTGAGCTCCTGGAGT | 768.6 | 12.2 | 1.3 | 9.3 | 1.6E-20 | 9.0E-20 | BT20/MB468 |
| >hg38_21_+_8433172_8446622_RNA45SN1_45S_50ntFlanks@11569.11601.33\|CAGCCGACTTAGAACTGGTGCGGACCAGGGGAA | >hg38_21_+_8433172_8446622_RNA45SN1_45S_50ntFlanks | 11569 | 11601 | 33 | CAGCCGACTTAGAACTGGTGCGGACCAGGGGAA | 1984.8 | -5.3 | 0.6 | -9.3 | 1.9E-20 | 9.1E-20 | MB231/MB468 |
| >hg38_21_+_8433172_8446622_RNA45SN1_45S_50ntFlanks@10402.10429.28\|AACATGGGTCAGTCGGTCCTGAGAGATG | >hg38_21_+_8433172_8446622_RNA45SN1_45S_50ntFlanks | 10402 | 10429 | 28 | AACATGGGTCAGTCGGTCCTGAGAGATG | 3285.5 | -4.4 | 0.5 | -9.3 | 1.6E-20 | 9.1E-20 | BT20/MB468 |
| >hg38_21_+_8433172_8446622_RNA45SN1_45S_50ntFlanks@10352.10371.20\|GAGAACTTTGAAGGCCGAAG | >hg38_21_+_8433172_8446622_RNA45SN1_45S_50ntFlanks | 10352 | 10371 | 20 | GAGAACTTTGAAGGCCGAAG | 695.0 | 6.7 | 0.7 | 9.3 | 1.9E-20 | 9.1E-20 | MB231/MB468 |
| >trnaMT_ValTAC_MT_+_1602_1670@48.72.25\|CAACTTAACTTGACCGCTCTGACCA | >trnaMT_ValTAC_MT_+_1602_1670 | 48 | 72 | 25 | CAACTTAACTTGACCGCTCTGACCA | 3936.5 | 7.7 | 0.8 | 9.3 | 1.7E-20 | 9.4E-20 | BT20/MB468 |
| >hg38_21_+_8433172_8446622_RNA45SN1_45S_50ntFlanks@12971.12989.19\|TTTCGTACGTAGCAGAGCA | >hg38_21_+_8433172_8446622_RNA45SN1_45S_50ntFlanks | 12971 | 12989 | 19 | TTTCGTACGTAGCAGAGCA | 256.3 | -5.2 | 0.6 | -9.3 | 1.7E-20 | 9.5E-20 | BT20/MB468 |
| >hg38_21_+_8433172_8446622_RNA45SN1_45S_50ntFlanks@485.503.19\|GATGTGGTGACGTCGTGCT | >hg38_21_+_8433172_8446622_RNA45SN1_45S_50ntFlanks | 485 | 503 | 19 | GATGTGGTGACGTCGTGCT | 479.5 | 13.6 | 1.5 | 9.3 | 1.7E-20 | 9.5E-20 | BT20/MB468 |
| >MI0000269\|hsa-mir-181a-2&WithFlank&9\|+\|124692436\|124692557@83.104.22&[MIMAT0004558&hsa-miR-181a-2-3p&offsets\|0\|0;m-71&9\|+\|124692518\|124692539&offsets\|0\|0]\|ACCACTGACCGTTGACTGTACC | >MI0000269\|hsa-mir-181a-2&WithFlank&9\|+\|124692436\|124692557 | 83 | 104 | 22 | ACCACTGACCGTTGACTGTACC | 1305.1 | 7.2 | 0.8 | 9.3 | 1.7E-20 | 9.6E-20 | BT20/MB468 |
| >trnaMT_PheGAA_MT_+_577_647@47.74.28\|ACGGGCTCACATCACCCCATAAACACCA | >trnaMT_PheGAA_MT_+_577_647 | 47 | 74 | 28 | ACGGGCTCACATCACCCCATAAACACCA | 442.4 | 9.4 | 1.0 | 9.3 | 1.7E-20 | 9.6E-20 | BT20/MB468 |
| >hg38_21_+_8433172_8446622_RNA45SN1_45S_50ntFlanks@8380.8402.23\|CAAGAGGGCGTGAAACCGTTAAG | >hg38_21_+_8433172_8446622_RNA45SN1_45S_50ntFlanks | 8380 | 8402 | 23 | CAAGAGGGCGTGAAACCGTTAAG | 1749.3 | -6.2 | 0.7 | -9.3 | 2.0E-20 | 9.7E-20 | MB231/MB468 |
| >hg38_21_+_8433172_8446622_RNA45SN1_45S_50ntFlanks@12619.12654.36\|AGGAGCCAATGGGGCGAAGCTACCATCTGTGGGATT | >hg38_21_+_8433172_8446622_RNA45SN1_45S_50ntFlanks | 12619 | 12654 | 36 | AGGAGCCAATGGGGCGAAGCTACCATCTGTGGGATT | 1000.2 | -5.3 | 0.6 | -9.3 | 1.7E-20 | 9.7E-20 | BT20/MB468 |
| >MI0000065\|hsa-let-7d&WithFlank&9\|+\|94178828\|94178926@14.35.22&[MIMAT0000065&hsa-let-7d-5p&offsets\|0\|0;m-90&9\|+\|94178841\|94178862&offsets\|0\|0]\|AGAGGTAGTAGGTTGCATAGTT | >MI0000065\|hsa-let-7d&WithFlank&9\|+\|94178828\|94178926 | 14 | 35 | 22 | AGAGGTAGTAGGTTGCATAGTT | 16055.5 | 7.3 | 0.8 | 9.3 | 2.1E-20 | 9.9E-20 | MB231/MB468 |
| >hg38_21_+_8433172_8446622_RNA45SN1_45S_50ntFlanks@10400.10424.25\|TGAACATGGGTCAGTCGGTCCTGAG | >hg38_21_+_8433172_8446622_RNA45SN1_45S_50ntFlanks | 10400 | 10424 | 25 | TGAACATGGGTCAGTCGGTCCTGAG | 902.6 | 7.0 | 0.7 | 9.3 | 1.8E-20 | 9.9E-20 | BT20/MB468 |
| >hg38_21_+_8433172_8446622_RNA45SN1_45S_50ntFlanks@7975.8015.41\|CGCGACCTCAGATCAGACGTGGCGACCCGCTGAATTTAAGC | >hg38_21_+_8433172_8446622_RNA45SN1_45S_50ntFlanks | 7975 | 8015 | 41 | CGCGACCTCAGATCAGACGTGGCGACCCGCTGAATTTAAGC | 15550.8 | 9.0 | 1.0 | 9.3 | 1.8E-20 | 9.9E-20 | BT20/MB468 |
| >MI0003591\|hsa-mir-584&WithFlank&5\|-\|149062307\|149062415@22.41.20(+1G)&[MIMAT0003249&hsa-miR-584-5p&offsets\|0\|-2(+1G);m-167&5\|-\|149062374\|149062394&offsets\|0\|-1(+1G)]\|TTATGGTTTGCCTGGGACTGG | >MI0003591\|hsa-mir-584&WithFlank&5\|-\|149062307\|149062415 | 22 | 41 | 20(+1G) | TTATGGTTTGCCTGGGACTGG | 591.4 | 13.9 | 1.5 | 9.3 | 1.9E-20 | 1.0E-19 | BT20/MB468 |
| >MI0003628\|hsa-mir-615&WithFlank&12\|+\|54033944\|54034051@67.87.21&[MIMAT0003283&hsa-miR-615-3p&offsets\|0\|-1;m-495&12\|+\|54034010\|54034030&offsets\|0\|0]\|TCCGAGCCTGGGTCTCCCTCT | >MI0003628\|hsa-mir-615&WithFlank&12\|+\|54033944\|54034051 | 67 | 87 | 21 | TCCGAGCCTGGGTCTCCCTCT | 3740.2 | 6.7 | 0.7 | 9.3 | 1.9E-20 | 1.0E-19 | BT20/MB468 |
| >hg38_21_+_8433172_8446622_RNA45SN1_45S_50ntFlanks@11707.11728.22\|AAGCGCGGGTAAACGGCGGGAG | >hg38_21_+_8433172_8446622_RNA45SN1_45S_50ntFlanks | 11707 | 11728 | 22 | AAGCGCGGGTAAACGGCGGGAG | 228.3 | -5.4 | 0.6 | -9.3 | 1.9E-20 | 1.0E-19 | BT20/MB468 |
| >MI0000285\|hsa-mir-205&WithFlank&1\|+\|209432127\|209432248@40.60.21(+1C)&[MIMAT0000266&hsa-miR-205-5p&offsets\|0\|-1(+1C);m-351&1\|+\|209432166\|209432187&offsets\|0\|-1(+1C)]\|TCCTTCATTCCACCGGAGTCTC | >MI0000285\|hsa-mir-205&WithFlank&1\|+\|209432127\|209432248 | 40 | 60 | 21(+1C) | TCCTTCATTCCACCGGAGTCTC | 484.7 | 6.6 | 0.7 | 9.3 | 1.9E-20 | 1.0E-19 | BT20/MB468 |
| >am_trnaMT_PheGAA_MT_+_577_647@40.58.19\|TGTTTAGACGGGCTCACAT | >am_trnaMT_PheGAA_MT_+_577_647 | 40 | 58 | 19 | TGTTTAGACGGGCTCACAT | 587.6 | 9.6 | 1.0 | 9.3 | 2.2E-20 | 1.1E-19 | MB231/MB468 |
| >hg38_1_-_228634819_228635039_RNA5S12_5S_50ntFlanks@127.170.44\|ACTTGGATGGGAGACCGCCTGGGAATACCGGGTGCTGTAGGCTT | >hg38_1_-_228634819_228635039_RNA5S12_5S_50ntFlanks | 127 | 170 | 44 | ACTTGGATGGGAGACCGCCTGGGAATACCGGGTGCTGTAGGCTT | 2034.0 | 11.7 | 1.3 | 9.3 | 2.2E-20 | 1.1E-19 | MB231/MB468 |
| >MI0000285\|hsa-mir-205&WithFlank&1\|+\|209432127\|209432248@40.59.20(+1A)&[MIMAT0000266&hsa-miR-205-5p&offsets\|0\|-2(+1A);m-351&1\|+\|209432166\|209432187&offsets\|0\|-2(+1A)]\|TCCTTCATTCCACCGGAGTCA | >MI0000285\|hsa-mir-205&WithFlank&1\|+\|209432127\|209432248 | 40 | 59 | 20(+1A) | TCCTTCATTCCACCGGAGTCA | 3470.3 | 8.6 | 0.9 | 9.3 | 2.0E-20 | 1.1E-19 | BT20/MB468 |
| >hg38_21_+_8433172_8446622_RNA45SN1_45S_50ntFlanks@10683.10712.30\|CCGGCGGCGTCCGGTGAGCTCTCGCTGGCC | >hg38_21_+_8433172_8446622_RNA45SN1_45S_50ntFlanks | 10683 | 10712 | 30 | CCGGCGGCGTCCGGTGAGCTCTCGCTGGCC | 1333.7 | -4.9 | 0.5 | -9.3 | 2.0E-20 | 1.1E-19 | BT20/MB468 |
| >MI0000478\|hsa-mir-149&WithFlank&2\|+\|240455995\|240456095@21.40.20&[MIMAT0000450&hsa-miR-149-5p&offsets\|0\|-3;m-195&2\|+\|240456015\|240456037&offsets\|0\|-3]\|TCTGGCTCCGTGTCTTCACT | >MI0000478\|hsa-mir-149&WithFlank&2\|+\|240455995\|240456095 | 21 | 40 | 20 | TCTGGCTCCGTGTCTTCACT | 1284.0 | 15.0 | 1.6 | 9.3 | 2.0E-20 | 1.1E-19 | BT20/MB468 |
| >MI0000298\|hsa-mir-221&WithFlank&X\|-\|45746151\|45746272@31.52.22&[MIMAT0004568&hsa-miR-221-5p&offsets\|0\|0;m-405&X\|-\|45746220\|45746242&offsets\|0\|-1]\|ACCTGGCATACAATGTAGATTT | >MI0000298\|hsa-mir-221&WithFlank&X\|-\|45746151\|45746272 | 31 | 52 | 22 | ACCTGGCATACAATGTAGATTT | 726.4 | 9.4 | 1.0 | 9.2 | 2.3E-20 | 1.1E-19 | MB231/MB468 |
| >MI0000108\|hsa-mir-103a-2&WithFlank&20\|+\|3917488\|3917577@54.75.22(+1U);MI0000109\|hsa-mir-103a-1&WithFlank&5\|-\|168560890\|168560979@54.75.22(+1U)&[MIMAT0000101&hsa-miR-103a-3p&offsets\|0\|-1(+1U);m-12&20\|+\|3917541\|3917563&offsets\|0\|-1(+1U)];[MIMAT0000101_1&hsa-miR-103a-3p&offsets\|0\|-1(+1U);m-13&5\|-\|168560904\|168560926&offsets\|0\|-1(+1U);m-44220&5\|-\|168560908\|168560926&offsets\|0\|+3(+1U)]\|AGCAGCATTGTACAGGGCTATGT | >MI0000108\|hsa-mir-103a-2&WithFlank&20\|+\|3917488\|3917577 | 54 | 75 | 22(+1U) | AGCAGCATTGTACAGGGCTATGT | 5637.2 | 8.0 | 0.9 | 9.3 | 2.0E-20 | 1.1E-19 | BT20/MB468 |
| >hg38_21_+_8433172_8446622_RNA45SN1_45S_50ntFlanks@3849.3866.18\|GGATAACTGTGGTAATTC | >hg38_21_+_8433172_8446622_RNA45SN1_45S_50ntFlanks | 3849 | 3866 | 18 | GGATAACTGTGGTAATTC | 383.0 | 6.2 | 0.7 | 9.2 | 2.4E-20 | 1.1E-19 | MB231/MB468 |
| >MI0000082\|hsa-mir-25&WithFlank&7\|-\|100093554\|100093649@58.79.22(+2A)&[MIMAT0000081&hsa-miR-25-3p&offsets\|0\|0(+2A);m-23&7\|-\|100093571\|100093592&offsets\|0\|0(+2A)]\|CATTGCACTTGTCTCGGTCTGAAA | >MI0000082\|hsa-mir-25&WithFlank&7\|-\|100093554\|100093649 | 58 | 79 | 22(+2A) | CATTGCACTTGTCTCGGTCTGAAA | 905.4 | 6.5 | 0.7 | 9.2 | 2.4E-20 | 1.1E-19 | MB231/MB468 |
| >hg38_1_-_228634819_228635039_RNA5S12_5S_50ntFlanks@138.169.32\|AGACCGCCTGGGAATACCGGGTGCTGTAGGCT | >hg38_1_-_228634819_228635039_RNA5S12_5S_50ntFlanks | 138 | 169 | 32 | AGACCGCCTGGGAATACCGGGTGCTGTAGGCT | 1134.4 | 7.2 | 0.8 | 9.2 | 2.4E-20 | 1.1E-19 | MB231/MB468 |
| >MI0000273\|hsa-mir-183&WithFlank&7\|-\|129774899\|129775020@33.56.24&[MIMAT0000261&hsa-miR-183-5p&offsets\|0\|+2;m-41&7\|-\|129774966\|129774987&offsets\|-1\|+1]\|TATGGCACTGGTAGAATTCACTGT | >MI0000273\|hsa-mir-183&WithFlank&7\|-\|129774899\|129775020 | 33 | 56 | 24 | TATGGCACTGGTAGAATTCACTGT | 1774.2 | 6.7 | 0.7 | 9.3 | 2.1E-20 | 1.2E-19 | BT20/MB468 |

| | | | | | | | | | | | | |
|---|---|---|---|---|---|---|---|---|---|---|---|---|
| >hg38_21_+_8433172_8446622_RNA45SN1_45S_50ntFlanks@6690.6720.31\|AAGAACGCAGCTAGCTGCGAGAATTAATGTG | >hg38_21_+_8433172_8446622_RNA45SN1_45S_50ntFlanks | 6690 | 6720 | 31 | AAGAACGCAGCTAGCTGCGAGAATTAATGTG | 1242.9 | -12.3 | 1.3 | -9.3 | 2.1E-20 | 1.2E-19 | BT20/MB468 |
| >hg38_21_+_8433172_8446622_RNA45SN1_45S_50ntFlanks@12405.12446.42\|CTTTTTGATCCTTCGATGTCGGCTCTTCCTATCATTGTGAAG | >hg38_21_+_8433172_8446622_RNA45SN1_45S_50ntFlanks | 12405 | 12446 | 42 | CTTTTTGATCCTTCGATGTCGGCTCTTCCTATCATTGTGAAG | 398.5 | -6.1 | 0.7 | -9.3 | 2.1E-20 | 1.2E-19 | BT20/MB468 |
| >MI0000082\|hsa-mir-25&WithFlank&7\|-\|100093554\|100093649@58.77.20&[MIMAT0000081&hsa-miR-25-3p&offsets\|0\|-2;m-23&7\|-\|100093571\|100093592&offsets\|0\|-2]\|CATTGCACTTGTCTCGGTCT | >MI0000082\|hsa-mir-25&WithFlank&7\|-\|100093554\|100093649 | 58 | 77 | 20 | CATTGCACTTGTCTCGGTCT | 856.8 | 7.4 | 0.8 | 9.2 | 2.5E-20 | 1.2E-19 | MB231/MB468 |
| >MI0000088\|hsa-mir-30a&WithFlank&6\|-\|71403545\|71403627@54.73.20&[MIMAT0000088&hsa-miR-30a-3p&offsets\|+1\|-1;m-38&6\|-\|71403554\|71403575&offsets\|+1\|-1]\|TTTCAGTCGGATGTTTGCAG | >MI0000088\|hsa-mir-30a&WithFlank&6\|-\|71403545\|71403627 | 54 | 73 | 20 | TTTCAGTCGGATGTTTGCAG | 956.8 | 10.6 | 1.1 | 9.2 | 2.5E-20 | 1.2E-19 | MB231/MB468 |
| >MI0000803\|hsa-mir-330&WithFlank&19\|-\|45638988\|45639093@63.85.23&[MIMAT0000751&hsa-miR-330-3p&offsets\|0\|0;m-347&19\|-\|45639009\|45639030&offsets\|-1\|0]\|GCAAAGCACACGGCCTGCAGAGA | >MI0000803\|hsa-mir-330&WithFlank&19\|-\|45638988\|45639093 | 63 | 85 | 23 | GCAAAGCACACGGCCTGCAGAGA | 429.6 | 6.7 | 0.7 | 9.3 | 2.2E-20 | 1.2E-19 | BT20/MB468 |
| >hg38_MT_+_598_1651_RNR1_12S_50ntFlanks@369.404.36\|AATAAAGCTAAAACTCACCTGAGTTGTAAAAAACTC | >hg38_MT_+_598_1651_RNR1_12S_50ntFlanks | 369 | 404 | 36 | AATAAAGCTAAAACTCACCTGAGTTGTAAAAAACTC | 968.2 | 10.3 | 1.1 | 9.2 | 2.3E-20 | 1.2E-19 | BT20/MB468 |
| >hg38_21_+_8433172_8446622_RNA45SN1_45S_50ntFlanks@10421.10456.36\|TGAGAGATGGGCGAGCGCCGTTCCGAAGGGACGGGC | >hg38_21_+_8433172_8446622_RNA45SN1_45S_50ntFlanks | 10421 | 10456 | 36 | TGAGAGATGGGCGAGCGCCGTTCCGAAGGGACGGGC | 410.3 | -5.0 | 0.5 | -9.2 | 2.3E-20 | 1.2E-19 | BT20/MB468 |
| >hg38_21_+_8433172_8446622_RNA45SN1_45S_50ntFlanks@11561.11585.25\|GCGCCTAGCAGCCGACTTAGAACTG | >hg38_21_+_8433172_8446622_RNA45SN1_45S_50ntFlanks | 11561 | 11585 | 25 | GCGCCTAGCAGCCGACTTAGAACTG | 1421.0 | -6.7 | 0.7 | -9.2 | 2.3E-20 | 1.2E-19 | BT20/MB468 |
| >MI0000077\|hsa-mir-21&WithFlank&17\|+\|59841260\|59841343@15.35.21(+1U)&[MIMAT0000076&hsa-miR-21-5p&offsets\|+1\|0(+1U);m-1&17\|+\|59841273\|59841294&offsets\|+1\|0(+1U)]\|AGCTTATCAGACTGATGTTGAT | >MI0000077\|hsa-mir-21&WithFlank&17\|+\|59841260\|59841343 | 15 | 35 | 21(+1U) | AGCTTATCAGACTGATGTTGAT | 910.9 | 9.6 | 1.0 | 9.2 | 2.3E-20 | 1.2E-19 | BT20/MB468 |
| >MI0000088\|hsa-mir-30a&WithFlank&6\|-\|71403545\|71403627@12.34.23&[MIMAT0000087&hsa-miR-30a-5p&offsets\|0\|+1;m-15&6\|-\|71403595\|71403616&offsets\|0\|+1]\|TGTAAACATCCTCGACTGGAAGC | >MI0000088\|hsa-mir-30a&WithFlank&6\|-\|71403545\|71403627 | 12 | 34 | 23 | TGTAAACATCCTCGACTGGAAGC | 114410.4 | 8.1 | 0.9 | 9.2 | 2.3E-20 | 1.2E-19 | BT20/MB468 |
| >MI0000272\|hsa-mir-182&WithFlank&7\|-\|129770377\|129770498@29.54.26(+1U)&[MIMAT0000259&hsa-miR-182-5p&offsets\|0\|+2(+1U);m-22&7\|-\|129770449\|129770470&offsets\|0\|+4(+1U)]\|TTTGGCAATGGTAGAACTCACACTGGT | >MI0000272\|hsa-mir-182&WithFlank&7\|-\|129770377\|129770498 | 29 | 54 | 26(+1U) | TTTGGCAATGGTAGAACTCACACTGGT | 813.6 | 7.1 | 0.8 | 9.2 | 2.6E-20 | 1.3E-19 | MB231/MB468 |
| >hg38_MT_+_598_1651_RNR1_12S_50ntFlanks@52.83.32\|ATAGGTTTGGTCCTAGCCTTTCTATTAGCTCT | >hg38_MT_+_598_1651_RNR1_12S_50ntFlanks | 52 | 83 | 32 | ATAGGTTTGGTCCTAGCCTTTCTATTAGCTCT | 2945.0 | 6.7 | 0.7 | 9.2 | 2.4E-20 | 1.3E-19 | BT20/MB468 |
| >hg38_21_+_8433172_8446622_RNA45SN1_45S_50ntFlanks@10687.10711.25\|CGGCGTCCGGTGAGCTCTCGCTGGC | >hg38_21_+_8433172_8446622_RNA45SN1_45S_50ntFlanks | 10687 | 10711 | 25 | CGGCGTCCGGTGAGCTCTCGCTGGC | 2718.1 | -5.9 | 0.6 | -9.2 | 2.4E-20 | 1.3E-19 | BT20/MB468 |
| >MI0000089\|hsa-mir-31&WithFlank&9\|-\|21512109\|21512191@14.35.22&[MIMAT0000089&hsa-miR-31-5p&offsets\|0\|+1;m-360&9\|-\|21512157\|21512178&offsets\|0\|0]\|AGGCAAGATGCTGGCATAGCTG | >MI0000089\|hsa-mir-31&WithFlank&9\|-\|21512109\|21512191 | 14 | 35 | 22 | AGGCAAGATGCTGGCATAGCTG | 1112.0 | 13.6 | 1.4 | 9.6 | 1.2E-21 | 1.3E-19 | BT20/MB231 |
| >hg38_21_+_8433172_8446622_RNA45SN1_45S_50ntFlanks@6740.6770.31\|CATCGACACTTCGAACGCACTTGCGGCCCCG | >hg38_21_+_8433172_8446622_RNA45SN1_45S_50ntFlanks | 6740 | 6770 | 31 | CATCGACACTTCGAACGCACTTGCGGCCCCG | 1446.5 | -8.7 | 0.9 | -9.2 | 2.5E-20 | 1.3E-19 | BT20/MB468 |
| >hg38_21_+_8433172_8446622_RNA45SN1_45S_50ntFlanks@10405.10424.20\|ATGGGTCAGTCGGTCCTGAG | >hg38_21_+_8433172_8446622_RNA45SN1_45S_50ntFlanks | 10405 | 10424 | 20 | ATGGGTCAGTCGGTCCTGAG | 932.8 | 6.3 | 0.7 | 9.2 | 2.9E-20 | 1.4E-19 | MB231/MB468 |
| >MI0000078\|hsa-mir-22&WithFlank&17\|-\|1713897\|1713993@59.81.23&[MIMAT0000077&hsa-miR-22-3p&offsets\|0\|+1;m-2&17\|-\|1713914\|1713935&offsets\|0\|+1]\|AAGCTGCCAGTTGAAGAACTGTT | >MI0000078\|hsa-mir-22&WithFlank&17\|-\|1713897\|1713993 | 59 | 81 | 23 | AAGCTGCCAGTTGAAGAACTGTT | 898.5 | 10.6 | 1.1 | 9.2 | 2.9E-20 | 1.4E-19 | MB231/MB468 |
| >trna111_HisGTG_1_-_147774845_147774916@-1T.29.30,trna118_HisGTG_1_-_145396881_145396952@-1T.29.30,trna16_HisGTG_1_+_146544773_146544844@-1T.29.30,trna1_HisGTG_15_+_45493349_45493420@-1T.29.30,trna21_HisGTG_1_+_147753471_147753542@-1T.29.30,trna33_HisGTG_6_+_27125906_27125977@-1T.29.30,trna7_HisGTG_9_-_14433938_14434009@-1T.29.30,trna8_HisGTG_15_-_45492611_45492682@-1T.29.30,trna9_HisGTG_15_-_45490804_45490875@-1T.29.30\|TGCCGTGATCGTATAGTGGTTAGTACTCTG | >trna111_HisGTG_1_-_147774845_147774916 | -1T | 29 | 30 | TGCCGTGATCGTATAGTGGTTAGTACTCTG | 2586.8 | -7.0 | 0.8 | -9.2 | 2.5E-20 | 1.4E-19 | BT20/MB468 |
| >MI0000108\|hsa-mir-103a-2&WithFlank&20\|+\|3917488\|3917577@54.75.22(+1G);MI0000109\|hsa-mir-103a-1&WithFlank&5\|-\|168560890\|168560979@54.75.22(+1G)&[MIMAT0000101&hsa-miR-103a-3p&offsets\|0\|-1(+1G);m-12&20\|+\|3917541\|3917563&offsets\|0\|-1(+1G)];[MIMAT0000101_1&hsa-miR-103a-3p&offsets\|0\|-1(+1G);m-13&5\|-\|168560904\|168560926&offsets\|0\|-1(+1G);m-44220&5\|-\|168560908\|168560926&offsets\|0\|+3(+1G)]\|AGCAGCATTGTACAGGGCTATGG | >MI0000108\|hsa-mir-103a-2&WithFlank&20\|+\|3917488\|3917577 | 54 | 75 | 22(+1G) | AGCAGCATTGTACAGGGCTATGG | 6662.1 | 8.3 | 0.9 | 9.2 | 2.5E-20 | 1.4E-19 | BT20/MB468 |
| >hg38_21_+_8433172_8446622_RNA45SN1_45S_50ntFlanks@11566.11584.19\|TAGCAGCCGACTTAGAACT | >hg38_21_+_8433172_8446622_RNA45SN1_45S_50ntFlanks | 11566 | 11584 | 19 | TAGCAGCCGACTTAGAACT | 920.0 | 6.6 | 0.7 | 9.2 | 2.6E-20 | 1.4E-19 | BT20/MB468 |
| >MI0000750\|hsa-mir-26a-2&WithFlank&12\|-\|57824603\|57824698@20.41.22;MI0000083\|hsa-mir-26a-1&WithFlank&3\|+\|37969398\|37969486@16.37.22&[MIMAT0000082&hsa-miR-26a-5p&offsets\|0\|0;m-33&12\|-\|57824658\|57824679&offsets\|0\|0];[MIMAT0000082_1&hsa-miR-26a-5p&offsets\|0\|0;m-34&3\|+\|37969413\|37969434&offsets\|0\|0]\|TTCAAGTAATCCAGGATAGGCT | >MI0000750\|hsa-mir-26a-2&WithFlank&12\|-\|57824603\|57824698 | 20 | 41 | 22 | TTCAAGTAATCCAGGATAGGCT | 262118.1 | 7.1 | 0.8 | 9.2 | 2.9E-20 | 1.4E-19 | MB231/MB468 |
| >MI0000088\|hsa-mir-30a&WithFlank&6\|-\|71403545\|71403627@53.72.20&[MIMAT0000088&hsa-miR-30a-3p&offsets\|0\|-2;m-38&6\|-\|71403554\|71403575&offsets\|0\|-2]\|CTTTCAGTCGGATGTTTGCA | >MI0000088\|hsa-mir-30a&WithFlank&6\|-\|71403545\|71403627 | 53 | 72 | 20 | CTTTCAGTCGGATGTTTGCA | 646.1 | 9.0 | 1.0 | 9.2 | 3.0E-20 | 1.4E-19 | MB231/MB468 |
| >hg38_1_-_228634819_228635039_RNA5S12_5S_50ntFlanks@137.171.35\|GAGACCGCCTGGGAATACCGGGTGCTGTAGGCTTT | >hg38_1_-_228634819_228635039_RNA5S12_5S_50ntFlanks | 137 | 171 | 35 | GAGACCGCCTGGGAATACCGGGTGCTGTAGGCTTT | 2456.9 | 8.2 | 0.9 | 9.2 | 2.6E-20 | 1.4E-19 | BT20/MB468 |
| >MI0000809\|hsa-mir-151a&WithFlank&8\|-\|140732558\|140732659@53.74.22(+2U)&[MIMAT0000757&hsa-miR-151a-3p&offsets\|0\|+1(+2U);m-30&8\|-\|140732587\|140732607&offsets\|0\|+1(+2U)]\|CTAGACTGAAGCTCCTTGAGGATT | >MI0000809\|hsa-mir-151a&WithFlank&8\|-\|140732558\|140732659 | 53 | 74 | 22(+2U) | CTAGACTGAAGCTCCTTGAGGATT | 989.7 | 9.0 | 1.0 | 9.2 | 2.6E-20 | 1.4E-19 | BT20/MB468 |
| >trnaMT_SerGCT_MT_+_12207_12265@28.62.35\|ATGCCCCCATGTCTAACAACATGGCTTTCTCACCA | >trnaMT_SerGCT_MT_+_12207_12265 | 28 | 62 | 35 | ATGCCCCCATGTCTAACAACATGGCTTTCTCACCA | 549.7 | 11.7 | 1.3 | 9.2 | 2.6E-20 | 1.4E-19 | BT20/MB468 |
| >hg38_21_+_8433172_8446622_RNA45SN1_45S_50ntFlanks@3775.3792.18\|GCCGGTACAGTGAAACTG | >hg38_21_+_8433172_8446622_RNA45SN1_45S_50ntFlanks | 3775 | 3792 | 18 | GCCGGTACAGTGAAACTG | 2090.7 | -7.1 | 0.8 | -9.2 | 2.6E-20 | 1.4E-19 | BT20/MB468 |
| >MI0000279\|hsa-mir-196a-2&WithFlank&12\|+\|53991732\|53991853@31.53.23&[MIMAT0000226&hsa-miR-196a-5p&offsets\|0\|+1;m-261&12\|+\|53991762\|53991783&offsets\|0\|+1]\|TAGGTAGTTTCATGTTGTTGGGA | >MI0000279\|hsa-mir-196a-2&WithFlank&12\|+\|53991732\|53991853 | 31 | 53 | 23 | TAGGTAGTTTCATGTTGTTGGGA | 3781.7 | 8.3 | 0.9 | 9.2 | 2.7E-20 | 1.4E-19 | BT20/MB468 |
| >hg38_21_+_8433172_8446622_RNA45SN1_45S_50ntFlanks@6732.6760.29\|ACATTGATCATCGACACTTCGAACGCACT | >hg38_21_+_8433172_8446622_RNA45SN1_45S_50ntFlanks | 6732 | 6760 | 29 | ACATTGATCATCGACACTTCGAACGCACT | 378.4 | 3.6 | 0.4 | 9.2 | 3.1E-20 | 1.5E-19 | MB231/MB468 |
| >hg38_21_+_8433172_8446622_RNA45SN1_45S_50ntFlanks@11540.11583.44\|CGCGGCGCCTCGCCTCGGCCGGCGCCTAGCAGCCGACTTAGAAC | >hg38_21_+_8433172_8446622_RNA45SN1_45S_50ntFlanks | 11540 | 11583 | 44 | CGCGGCGCCTCGCCTCGGCCGGCGCCTAGCAGCCGACTTAGAAC | 1563.5 | 10.3 | 1.1 | 9.2 | 3.1E-20 | 1.5E-19 | MB231/MB468 |
| >trna111_HisGTG_1_-_147774845_147774916@-1T.33.34,trna118_HisGTG_1_-_145396881_145396952@-1T.33.34,trna16_HisGTG_1_+_146544773_146544844@-1T.33.34,trna1_HisGTG_15_+_45493349_45493420@-1T.33.34,trna21_HisGTG_1_+_147753471_147753542@-1T.33.34,trna33_HisGTG_6_+_27125906_27125977@-1T.33.34,trna7_HisGTG_9_-_14433938_14434009@-1T.33.34,trna8_HisGTG_15_-_45492611_45492682@-1T.33.34,trna9_HisGTG_15_-_45490804_45490875@-1T.33.34\|TGCCGTGATCGTATAGTGGTTAGTACTCTGCGTT | >trna111_HisGTG_1_-_147774845_147774916 | -1T | 33 | 34 | TGCCGTGATCGTATAGTGGTTAGTACTCTGCGTT | 8160.6 | 11.5 | 1.2 | 9.2 | 3.1E-20 | 1.5E-19 | MB231/MB468 |
| >hg38_21_+_8433172_8446622_RNA45SN1_45S_50ntFlanks@10808.10840.33\|CAATGTAGGTAAGGGAAGTCGGCAAGCCGGATC | >hg38_21_+_8433172_8446622_RNA45SN1_45S_50ntFlanks | 10808 | 10840 | 33 | CAATGTAGGTAAGGGAAGTCGGCAAGCCGGATC | 624.3 | -5.9 | 0.6 | -9.2 | 2.8E-20 | 1.5E-19 | BT20/MB468 |
| >hg38_21_+_8433172_8446622_RNA45SN1_45S_50ntFlanks@3705.3736.32\|TACCTGGTTGATCCTGCCAGTAGCATATGCTT | >hg38_21_+_8433172_8446622_RNA45SN1_45S_50ntFlanks | 3705 | 3736 | 32 | TACCTGGTTGATCCTGCCAGTAGCATATGCTT | 4016.3 | 8.3 | 0.9 | 9.2 | 2.8E-20 | 1.5E-19 | BT20/MB468 |
| >trnaMT_SerGCT_MT_+_12207_12265@1.23.23\|GAGAAAGCTCACAAGAACTGCTA | >trnaMT_SerGCT_MT_+_12207_12265 | 1 | 23 | 23 | GAGAAAGCTCACAAGAACTGCTA | 614.7 | 5.4 | 0.6 | 9.2 | 3.2E-20 | 1.5E-19 | MB231/MB468 |
| >MI0000086\|hsa-mir-28&WithFlank&3\|+\|188688775\|188688872@60.82.23(+1U)&[MIMAT0004502&hsa-miR-28-3p&offsets\|0\|+1(+1U);m-28&3\|+\|188688834\|188688855&offsets\|0\|+1(+1U)]\|CACTAGATTGTGAGCTCCTGGAGT | >MI0000086\|hsa-mir-28&WithFlank&3\|+\|188688775\|188688872 | 60 | 82 | 23(+1U) | CACTAGATTGTGAGCTCCTGGAGT | 796.8 | 10.6 | 1.2 | 9.2 | 3.3E-20 | 1.5E-19 | MB231/MB468 |
| >MI0000077\|hsa-mir-21&WithFlank&17\|+\|59841260\|59841343@14.34.21(+1G)&[MIMAT0000076&hsa-miR-21-5p&offsets\|0\|-1(+1G);m-1&17\|+\|59841273\|59841294&offsets\|0\|-1(+1G)]\|TAGCTTATCAGACTGATGTTGG | >MI0000077\|hsa-mir-21&WithFlank&17\|+\|59841260\|59841343 | 14 | 34 | 21(+1G) | TAGCTTATCAGACTGATGTTGG | 56666.4 | 9.4 | 1.0 | 9.2 | 2.9E-20 | 1.5E-19 | BT20/MB468 |
| >MI0000270\|hsa-mir-181b-1&WithFlank&1\|-\|198858867\|198858988@42.64.23;MI0000683\|hsa-mir-181b-2&WithFlank&9\|+\|124693704\|124693804@22.44.23&[MIMAT0000257&hsa-miR-181b-5p&offsets\|0\|0;m-73&1\|-\|198858925\|198858947&offsets\|0\|0];[MIMAT0000257_1&hsa-miR-181b-5p&offsets\|0\|0;m-72&9\|+\|124693725\|124693747&offsets\|0\|0]\|AACATTCATTGCTGTCGGTGGGT | >MI0000270\|hsa-mir-181b-1&WithFlank&1\|-\|198858867\|198858988 | 42 | 64 | 23 | AACATTCATTGCTGTCGGTGGGT | 9366.6 | 7.4 | 0.8 | 9.2 | 3.3E-20 | 1.5E-19 | MB231/MB468 |

| | | | | | | | | | | | | |
|---|---|---|---|---|---|---|---|---|---|---|---|---|
| >MI0000542\|hsa-mir-320a&WithFlank&8\|-\|22244960\|22245043@48.72.25&[MIMAT0000510&hsa-miR-320a-3p&offsets\|0\|+3;m-60&8\|-\|22244975\|22244996&offsets\|0\|+3]\|AAAAGCTGGGTTGAGAGGGCGAAAA | >MI0000542\|hsa-mir-320a&WithFlank&8\|-\|22244960\|22245043 | 48 | 72 | 25 | AAAAGCTGGGTTGAGAGGGCGAAAA | 605.1 | 13.9 | 1.5 | 9.2 | 2.9E-20 | 1.6E-19 | BT20/MB468 |
| >MI0000081\|hsa-mir-24-2&WithFlank&19\|-\|13836281\|13836365@56.77.22(+1C);MI0000080\|hsa-mir-24-1&WithFlank&9\|+\|95086015\|95086094@50.71.22(+1C)&[MIMAT0000080&hsa-miR-24-3p&offsets\|0\|0(+1C);m-36&19\|-\|13836288\|13836310&offsets\|0\|-1(+1C)];[MIMAT0000080_1&hsa-miR-24-3p&offsets\|0\|0(+1C);m-35&9\|+\|95086064\|95086086&offsets\|0\|-1(+1C)]\|TGGCTCAGTTCAGCAGGAACAGC | >MI0000081\|hsa-mir-24-2&WithFlank&19\|-\|13836281\|13836365 | 56 | 77 | 22(+1C) | TGGCTCAGTTCAGCAGGAACAGC | 2399.9 | 7.1 | 0.8 | 9.2 | 3.4E-20 | 1.6E-19 | MB231/MB468 |
| >hg38_21_+_8433172_8446622_RNA45SN1_45S_50ntFlanks@8191.8215.25\|CAGCCCGTGGACGGTGTGAGGCCGG | >hg38_21_+_8433172_8446622_RNA45SN1_45S_50ntFlanks | 8191 | 8215 | 25 | CAGCCCGTGGACGGTGTGAGGCCGG | 747.3 | -5.6 | 0.6 | -9.2 | 3.4E-20 | 1.6E-19 | MB231/MB468 |
| >am_trna10_ValCAC_5_-_180649395_180649467@1.34.34,trna18_ValCAC_5_-_180529253_180529325@1.34.34,trna2_ValCAC_5_+_180524070_180524142@1.34.34,trna6_ValCAC_5_+_180600650_180600722@1.34.34,trna85_ValCAC_1_-_161369490_161369562@1.34.34,trna90_ValCAC_1_-_149684088_149684161@1.34.34,trna98_ValCAC_1_-_149298555_149298627@1.34.34,trna9_ValCAC_6_+_26538282_26538354@1.34.34\|GTTTCCGTAGTGTAGTGGTTATCACGTTCGCCTC | >am_trna10_ValCAC_5_-_180649395_180649467 | 1 | 34 | 34 | GTTTCCGTAGTGTAGTGGTTATCACGTTCGCCTC | 4991.6 | 12.9 | 1.4 | 9.2 | 3.5E-20 | 1.6E-19 | MB231/MB468 |
| >hg38_21_+_8433172_8446622_RNA45SN1_45S_50ntFlanks@5445.5466.22\|TCGGATCGGCCCCGCCGGGGTC | >hg38_21_+_8433172_8446622_RNA45SN1_45S_50ntFlanks | 5445 | 5466 | 22 | TCGGATCGGCCCCGCCGGGGTC | 1815.0 | -5.2 | 0.6 | -9.2 | 3.5E-20 | 1.6E-19 | MB231/MB468 |
| >hg38_21_+_8433172_8446622_RNA45SN1_45S_50ntFlanks@7975.8004.30\|CGCGACCTCAGATCAGACGTGGCGACCCGC | >hg38_21_+_8433172_8446622_RNA45SN1_45S_50ntFlanks | 7975 | 8004 | 30 | CGCGACCTCAGATCAGACGTGGCGACCCGC | 7090.3 | 5.4 | 0.6 | 9.2 | 3.1E-20 | 1.7E-19 | BT20/MB468 |
| >MI0000440\|hsa-mir-27b&WithFlank&9\|+\|95085439\|95085547@67.88.22&[MIMAT0000419&hsa-miR-27b-3p&offsets\|0\|+1;m-39&9\|+\|95085505\|95085525&offsets\|0\|+1]\|TTCACAGTGGCTAAGTTCTGCA | >MI0000440\|hsa-mir-27b&WithFlank&9\|+\|95085439\|95085547 | 67 | 88 | 22 | TTCACAGTGGCTAAGTTCTGCA | 22569.7 | 7.4 | 0.8 | 9.2 | 3.6E-20 | 1.7E-19 | MB231/MB468 |
| >hg38_21_+_8433172_8446622_RNA45SN1_45S_50ntFlanks@12419.12439.21\|GATGTCGGCTCTTCCTATCAT | >hg38_21_+_8433172_8446622_RNA45SN1_45S_50ntFlanks | 12419 | 12439 | 21 | GATGTCGGCTCTTCCTATCAT | 927.8 | 4.7 | 0.5 | 9.2 | 3.1E-20 | 1.7E-19 | BT20/MB468 |
| >hg38_21_+_8433172_8446622_RNA45SN1_45S_50ntFlanks@9605.9622.18\|AAGACTAATCGAACCATC | >hg38_21_+_8433172_8446622_RNA45SN1_45S_50ntFlanks | 9605 | 9622 | 18 | AAGACTAATCGAACCATC | 491.4 | -3.9 | 0.4 | -9.2 | 3.2E-20 | 1.7E-19 | BT20/MB468 |
| >MI0000286\|hsa-mir-210&WithFlank&11\|-\|568083\|568204@72.93.22&[MIMAT0000267&hsa-miR-210-3p&offsets\|0\|0;m-102&11\|-\|568112\|568133&offsets\|0\|0]\|CTGTGCGTGTGACAGCGGCTGA | >MI0000286\|hsa-mir-210&WithFlank&11\|-\|568083\|568204 | 72 | 93 | 22 | CTGTGCGTGTGACAGCGGCTGA | 19125.4 | 11.6 | 1.3 | 9.2 | 3.2E-20 | 1.7E-19 | BT20/MB468 |
| >MI0000272\|hsa-mir-182&WithFlank&7\|-\|129770377\|129770498@29.52.24(+1A)&[MIMAT0000259&hsa-miR-182-5p&offsets\|0\|0(+1A);m-22&7\|-\|129770449\|129770470&offsets\|0\|+2(+1A)]\|TTTGGCAATGGTAGAACTCACACTA | >MI0000272\|hsa-mir-182&WithFlank&7\|-\|129770377\|129770498 | 29 | 52 | 24(+1A) | TTTGGCAATGGTAGAACTCACACTA | 766.1 | 6.5 | 0.7 | 9.2 | 3.7E-20 | 1.8E-19 | MB231/MB468 |
| >hg38_21_+_8433172_8446622_RNA45SN1_45S_50ntFlanks@8170.8187.18\|CAAGTCCTTCTGATCGAG | >hg38_21_+_8433172_8446622_RNA45SN1_45S_50ntFlanks | 8170 | 8187 | 18 | CAAGTCCTTCTGATCGAG | 814.2 | -6.0 | 0.7 | -9.2 | 3.3E-20 | 1.8E-19 | BT20/MB468 |
| >MI0000767\|hsa-mir-365a&WithFlank&16\|+\|14309279\|14309377@62.84.23;MI0000769\|hsa-mir-365b&WithFlank&17\|+\|31575405\|31575527@74.96.23&[MIMAT0000710&hsa-miR-365a-3p&offsets\|0\|+1;m-176&16\|+\|14309340\|14309361&offsets\|0\|+1];[MIMAT0022834&hsa-miR-365b-3p&offsets\|0\|+1;m-175&17\|+\|31575478\|31575499&offsets\|0\|+1]\|TAATGCCCCTAAAAATCCTTATT | >MI0000767\|hsa-mir-365a&WithFlank&16\|+\|14309279\|14309377 | 62 | 84 | 23 | TAATGCCCCTAAAAATCCTTATT | 2539.8 | 9.0 | 1.0 | 9.2 | 3.3E-20 | 1.8E-19 | BT20/MB468 |
| >MI0000466\|hsa-mir-9-1&WithFlank&1\|-\|156420335\|156420435@22.43.22(+1U);MI0000468\|hsa-mir-9-3&WithFlank&15\|+\|89368011\|89368112@22.43.22(+1U);MI0000467\|hsa-mir-9-2&WithFlank&5\|-\|88666847\|88666945@22.43.22(+1U)&[MIMAT0000441&hsa-miR-9-5p&offsets\|0\|-1(+1U);m-104&1\|-\|156420392\|156420414&offsets\|0\|-1(+1U)];[MIMAT0000441_1&hsa-miR-9-5p&offsets\|0\|-1(+1U);m-105&15\|+\|89368032\|89368054&offsets\|0\|-1(+1U)];[MIMAT0000441_2&hsa-miR-9-5p&offsets\|0\|-1(+1U);m-104&5\|-\|88666902\|88666924&offsets\|0\|-1(+1U)]\|TCTTTGGTTATCTAGCTGTATGT | >MI0000466\|hsa-mir-9-1&WithFlank&1\|-\|156420335\|156420435 | 22 | 43 | 22(+1U) | TCTTTGGTTATCTAGCTGTATGT | 809.5 | 14.3 | 1.6 | 9.2 | 3.4E-20 | 1.8E-19 | BT20/MB468 |
| >MI0000750\|hsa-mir-26a-2&WithFlank&12\|-\|57824603\|57824698@20.39.20(+2U);MI0000083\|hsa-mir-26a-1&WithFlank&3\|+\|37969398\|37969486@16.35.20(+2U)&[MIMAT0000082&hsa-miR-26a-5p&offsets\|0\|-2(+2U);m-33&12\|-\|57824658\|57824679&offsets\|0\|-2(+2U)];[MIMAT0000082_1&hsa-miR-26a-5p&offsets\|0\|-2(+2U);m-34&3\|+\|37969413\|37969434&offsets\|0\|-2(+2U)]\|TTCAAGTAATCCAGGATAGGTT | >MI0000750\|hsa-mir-26a-2&WithFlank&12\|-\|57824603\|57824698 | 20 | 39 | 20(+2U) | TTCAAGTAATCCAGGATAGGTT | 1357.0 | 7.4 | 0.8 | 9.2 | 3.9E-20 | 1.8E-19 | MB231/MB468 |
| >MI0000270\|hsa-mir-181b-1&WithFlank&1\|-\|198858867\|198858988@42.63.22(+1A);MI0000683\|hsa-mir-181b-2&WithFlank&9\|+\|124693704\|124693804@22.43.22(+1A)&[MIMAT0000257&hsa-miR-181b-5p&offsets\|0\|-1(+1A);m-73&1\|-\|198858925\|198858947&offsets\|0\|-1(+1A)];[MIMAT0000257_1&hsa-miR-181b-5p&offsets\|0\|-1(+1A);m-72&9\|+\|124693725\|124693747&offsets\|0\|-1(+1A)]\|AACATTCATTGCTGTCGGTGGGA | >MI0000270\|hsa-mir-181b-1&WithFlank&1\|-\|198858867\|198858988 | 42 | 63 | 22(+1A) | AACATTCATTGCTGTCGGTGGGA | 864.1 | 7.2 | 0.8 | 9.2 | 3.9E-20 | 1.8E-19 | MB231/MB468 |
| >hg38_21_+_8433172_8446622_RNA45SN1_45S_50ntFlanks@10794.10818.25\|CTGGCATGTTGGAACAATGTAGGTA | >hg38_21_+_8433172_8446622_RNA45SN1_45S_50ntFlanks | 10794 | 10818 | 25 | CTGGCATGTTGGAACAATGTAGGTA | 3521.0 | -7.0 | 0.8 | -9.2 | 3.5E-20 | 1.9E-19 | BT20/MB468 |
| >hg38_21_+_8433172_8446622_RNA45SN1_45S_50ntFlanks@10786.10818.33\|AACAGCCTCTGGCATGTTGGAACAATGTAGGTA | >hg38_21_+_8433172_8446622_RNA45SN1_45S_50ntFlanks | 10786 | 10818 | 33 | AACAGCCTCTGGCATGTTGGAACAATGTAGGTA | 6753.7 | -6.5 | 0.7 | -9.2 | 3.5E-20 | 1.9E-19 | BT20/MB468 |
| >am_trnaMT_PheGAA_MT_+_577_647@40.59.20\|TGTTTAGACGGGCTCACATC | >am_trnaMT_PheGAA_MT_+_577_647 | 40 | 59 | 20 | TGTTTAGACGGGCTCACATC | 472.2 | 9.0 | 1.0 | 9.2 | 3.6E-20 | 1.9E-19 | BT20/MB468 |
| >trnaMT_PheGAA_MT_+_577_647@43.74.32\|TTAGACGGGCTCACATCACCCCATAAACACCA | >trnaMT_PheGAA_MT_+_577_647 | 43 | 74 | 32 | TTAGACGGGCTCACATCACCCCATAAACACCA | 1107.7 | 10.1 | 1.1 | 9.2 | 4.1E-20 | 1.9E-19 | MB231/MB468 |
| >hg38_21_+_8433172_8446622_RNA45SN1_45S_50ntFlanks@6719.6736.18\|TGAATTGCAGGACACATT | >hg38_21_+_8433172_8446622_RNA45SN1_45S_50ntFlanks | 6719 | 6736 | 18 | TGAATTGCAGGACACATT | 4107.6 | -5.4 | 0.6 | -9.2 | 4.2E-20 | 1.9E-19 | MB231/MB468 |
| >MI0001445\|hsa-mir-423&WithFlank&17\|+\|30117073\|30117178@59.80.22&[MIMAT0001340&hsa-miR-423-3p&offsets\|0\|-1;m-100&17\|+\|30117131\|30117153&offsets\|0\|-1]\|AGCTCGGTCTGAGGCCCCTCAG | >MI0001445\|hsa-mir-423&WithFlank&17\|+\|30117073\|30117178 | 59 | 80 | 22 | AGCTCGGTCTGAGGCCCCTCAG | 1631.2 | 6.4 | 0.7 | 9.2 | 3.7E-20 | 2.0E-19 | BT20/MB468 |
| >MI0000289\|hsa-mir-181a-1&WithFlank&1\|-\|198859038\|198859159@30.49.20;MI0000269\|hsa-mir-181a-2&WithFlank&9\|+\|124692436\|124692557@45.64.20&[MIMAT0000256&hsa-miR-181a-5p&offsets\|0\|-3;m-32&1\|-\|198859108\|198859130&offsets\|0\|-3];[MIMAT0000256_1&hsa-miR-181a-5p&offsets\|0\|-3;m-32&9\|+\|124692480\|124692502&offsets\|0\|-3]\|AACATTCAACGCTGTCGGTG | >MI0000289\|hsa-mir-181a-1&WithFlank&1\|-\|198859038\|198859159 | 30 | 49 | 20 | AACATTCAACGCTGTCGGTG | 1687.7 | 7.2 | 0.8 | 9.2 | 4.2E-20 | 2.0E-19 | MB231/MB468 |
| >MI0000285\|hsa-mir-205&WithFlank&1\|+\|209432127\|209432248@40.60.21&[MIMAT0000266&hsa-miR-205-5p&offsets\|0\|-1;m-351&1\|+\|209432166\|209432187&offsets\|0\|-1]\|TCCTTCATTCCACCGGAGTCT | >MI0000285\|hsa-mir-205&WithFlank&1\|+\|209432127\|209432248 | 40 | 60 | 21 | TCCTTCATTCCACCGGAGTCT | 17703.7 | 7.3 | 0.8 | 9.2 | 3.7E-20 | 2.0E-19 | BT20/MB468 |
| >hg38_21_+_8433172_8446622_RNA45SN1_45S_50ntFlanks@11561.11592.32\|GCGCCTAGCAGCCGACTTAGAACTGGTGCGGA | >hg38_21_+_8433172_8446622_RNA45SN1_45S_50ntFlanks | 11561 | 11592 | 32 | GCGCCTAGCAGCCGACTTAGAACTGGTGCGGA | 1084.2 | -6.2 | 0.7 | -9.2 | 4.3E-20 | 2.0E-19 | MB231/MB468 |
| >hg38_21_+_8433172_8446622_RNA45SN1_45S_50ntFlanks@12932.12956.25\|CGCTAAACCATTCGTAGACGACCTG | >hg38_21_+_8433172_8446622_RNA45SN1_45S_50ntFlanks | 12932 | 12956 | 25 | CGCTAAACCATTCGTAGACGACCTG | 498.8 | -3.5 | 0.4 | -9.2 | 3.8E-20 | 2.0E-19 | BT20/MB468 |
| >hg38_1_-_228634819_228635039_RNA5S12_5S_50ntFlanks@152.171.20\|TACCGGGTGCTGTAGGCTTT | >hg38_1_-_228634819_228635039_RNA5S12_5S_50ntFlanks | 152 | 171 | 20 | TACCGGGTGCTGTAGGCTTT | 1523.4 | 5.3 | 0.6 | 9.2 | 3.8E-20 | 2.0E-19 | BT20/MB468 |
| >hg38_21_+_8433172_8446622_RNA45SN1_45S_50ntFlanks@10687.10706.20\|CGGCGTCCGGTGAGCTCTCG | >hg38_21_+_8433172_8446622_RNA45SN1_45S_50ntFlanks | 10687 | 10706 | 20 | CGGCGTCCGGTGAGCTCTCG | 3238.6 | -6.3 | 0.7 | -9.2 | 3.9E-20 | 2.1E-19 | BT20/MB468 |
| >hg38_21_+_8433172_8446622_RNA45SN1_45S_50ntFlanks@10408.10430.23\|GGTCAGTCGGTCCTGAGAGATGG | >hg38_21_+_8433172_8446622_RNA45SN1_45S_50ntFlanks | 10408 | 10430 | 23 | GGTCAGTCGGTCCTGAGAGATGG | 1024.5 | -12.0 | 1.3 | -9.2 | 3.9E-20 | 2.1E-19 | BT20/MB468 |
| >hg38_21_+_8433172_8446622_RNA45SN1_45S_50ntFlanks@10786.10826.41\|AACAGCCTCTGGCATGTTGGAACAATGTAGGTAAGGGAAGT | >hg38_21_+_8433172_8446622_RNA45SN1_45S_50ntFlanks | 10786 | 10826 | 41 | AACAGCCTCTGGCATGTTGGAACAATGTAGGTAAGGGAAGT | 411.9 | -4.6 | 0.5 | -9.2 | 3.9E-20 | 2.1E-19 | BT20/MB468 |
| >MI0000084\|hsa-mir-26b&WithFlank&2\|+\|218402640\|218402728@18.39.22(+1U)&[MIMAT0000083&hsa-miR-26b-5p&offsets\|0\|+1(+1U);m-53&2\|+\|218402657\|218402678&offsets\|0\|0(+1U)]\|TTCAAGTAATTCAGGATAGGTTT | >MI0000084\|hsa-mir-26b&WithFlank&2\|+\|218402640\|218402728 | 18 | 39 | 22(+1U) | TTCAAGTAATTCAGGATAGGTTT | 1149.4 | 6.9 | 0.8 | 9.2 | 4.5E-20 | 2.1E-19 | MB231/MB468 |
| >MI0000067\|hsa-let-7f-1&WithFlank&9\|+\|94176341\|94176439@13.32.20(+1G);MI0000068\|hsa-let-7f-2&WithFlank&X\|-\|53557186\|53557280@14.33.20(+1G)&[MIMAT0000067&hsa-let-7f-5p&offsets\|0\|-2(+1G);m-20&9\|+\|94176353\|94176374&offsets\|0\|-2(+1G)];[MIMAT0000067_1&hsa-let-7f-5p&offsets\|0\|-2(+1G);m-19&X\|-\|53557246\|53557267&offsets\|0\|-2(+1G)]\|TGAGGTAGTAGATTGTATAGG | >MI0000067\|hsa-let-7f-1&WithFlank&9\|+\|94176341\|94176439 | 13 | 32 | 20(+1G) | TGAGGTAGTAGATTGTATAGG | 1390.6 | 7.3 | 0.8 | 9.2 | 4.5E-20 | 2.1E-19 | MB231/MB468 |
| >hg38_21_+_8433172_8446622_RNA45SN1_45S_50ntFlanks@11661.11679.19\|TGATTTCTGCCCAGTGCTC | >hg38_21_+_8433172_8446622_RNA45SN1_45S_50ntFlanks | 11661 | 11679 | 19 | TGATTTCTGCCCAGTGCTC | 1591.9 | -7.5 | 0.8 | -9.2 | 4.6E-20 | 2.1E-19 | MB231/MB468 |
| >hg38_21_+_8433172_8446622_RNA45SN1_45S_50ntFlanks@10349.10368.20\|AACGAGAACTTTGAAGGCCG | >hg38_21_+_8433172_8446622_RNA45SN1_45S_50ntFlanks | 10349 | 10368 | 20 | AACGAGAACTTTGAAGGCCG | 3486.4 | -3.8 | 0.4 | -9.2 | 4.0E-20 | 2.1E-19 | BT20/MB468 |
| >hg38_21_+_8433172_8446622_RNA45SN1_45S_50ntFlanks@12539.12569.31\|ATGATGTGTTGTTGCCATGGTAATCCTGCTC | >hg38_21_+_8433172_8446622_RNA45SN1_45S_50ntFlanks | 12539 | 12569 | 31 | ATGATGTGTTGTTGCCATGGTAATCCTGCTC | 1329.8 | -7.2 | 0.8 | -9.2 | 4.1E-20 | 2.1E-19 | BT20/MB468 |

<table>
<tr><td>>MI0000088|hsa-mir-30a&WithFlank&6|-|71403545|71403627@12.34.23(+1C)&[MIMAT0000087&hsa-miR-30a-5p&offsets|0|+1(+1C);m-15&6|-|71403595|71403616&offsets|0|+1(+1C)]|TGTAAACATCCTCGACTGGAAGCC</td><td>>MI0000088|hsa-mir-30a&WithFlank&6|-|71403545|71403627</td><td>12</td><td>34</td><td>23(+1C)</td><td>TGTAAACATCCTCGACTGGAAGCC</td><td>47267.8</td><td>8.0</td><td>0.9</td><td>9.2</td><td>4.2E-20</td><td>2.2E-19</td><td>BT20/MB468</td></tr>
<tr><td>>hg38_21_+_8433172_8446622_RNA45SN1_45S_50ntFlanks@11661.11682.22|TGATTTCTGCCCAGTGCTCTGA</td><td>>hg38_21_+_8433172_8446622_RNA45SN1_45S_50ntFlanks</td><td>11661</td><td>11682</td><td>22</td><td>TGATTTCTGCCCAGTGCTCTGA</td><td>465.8</td><td>-5.2</td><td>0.6</td><td>-9.2</td><td>4.7E-20</td><td>2.2E-19</td><td>MB231/MB468</td></tr>
<tr><td>>MI0000750|hsa-mir-26a-2&WithFlank&12|-|57824603|57824698@20.40.21(+1C);MI0000083|hsa-mir-26a-1&WithFlank&3|+|37969398|37969486@16.36.21(+1C)&[MIMAT0000082&hsa-miR-26a-5p&offsets|0|-1(+1C);m-33&12|-|57824658|57824679&offsets|0|-1(+1C)];[MIMAT0000082_1&hsa-miR-26a-5p&offsets|0|-1(+1C);m-34&3|+|37969413|37969434&offsets|0|-1(+1C)]|TTCAAGTAATCCAGGATAGGCC</td><td>>MI0000750|hsa-mir-26a-2&WithFlank&12|-|57824603|57824698</td><td>20</td><td>40</td><td>21(+1C)</td><td>TTCAAGTAATCCAGGATAGGCC</td><td>16151.1</td><td>5.8</td><td>0.6</td><td>9.2</td><td>4.2E-20</td><td>2.2E-19</td><td>BT20/MB468</td></tr>
<tr><td>>hg38_21_+_8433172_8446622_RNA45SN1_45S_50ntFlanks@8380.8402.23|CAAGAGGGCGTGAAACCGTTAAG</td><td>>hg38_21_+_8433172_8446622_RNA45SN1_45S_50ntFlanks</td><td>8380</td><td>8402</td><td>23</td><td>CAAGAGGGCGTGAAACCGTTAAG</td><td>2113.4</td><td>-9.6</td><td>1.0</td><td>-9.2</td><td>4.2E-20</td><td>2.2E-19</td><td>BT20/MB468</td></tr>
<tr><td>>hg38_21_+_8433172_8446622_RNA45SN1_45S_50ntFlanks@9564.9595.32|CTGACGTGCAAATCGGTCGTCCGACCTGGGTA</td><td>>hg38_21_+_8433172_8446622_RNA45SN1_45S_50ntFlanks</td><td>9564</td><td>9595</td><td>32</td><td>CTGACGTGCAAATCGGTCGTCCGACCTGGGTA</td><td>1246.6</td><td>-5.8</td><td>0.6</td><td>-9.2</td><td>4.2E-20</td><td>2.2E-19</td><td>BT20/MB468</td></tr>
<tr><td>>MI0000650|hsa-mir-200c&WithFlank&12|+|6963693|6963772@50.73.24&[MIMAT0000617&hsa-miR-200c-3p&offsets|0|+1;m-54&12|+|6963742|6963764&offsets|0|+1]|TAATACTGCCGGGTAATGATGGAG</td><td>>MI0000650|hsa-mir-200c&WithFlank&12|+|6963693|6963772</td><td>50</td><td>73</td><td>24</td><td>TAATACTGCCGGGTAATGATGGAG</td><td>1194.3</td><td>10.9</td><td>1.2</td><td>9.5</td><td>2.1E-21</td><td>2.2E-19</td><td>BT20/MB231</td></tr>
<tr><td>>hg38_21_+_8433172_8446622_RNA45SN1_45S_50ntFlanks@6646.6674.29|TCGTACGACTCTTAGCGGTGGATCACTCG</td><td>>hg38_21_+_8433172_8446622_RNA45SN1_45S_50ntFlanks</td><td>6646</td><td>6674</td><td>29</td><td>TCGTACGACTCTTAGCGGTGGATCACTCG</td><td>1557.0</td><td>4.8</td><td>0.5</td><td>9.2</td><td>4.3E-20</td><td>2.2E-19</td><td>BT20/MB468</td></tr>
<tr><td>>hg38_1_-_228634819_228635039_RNA5S12_5S_50ntFlanks@138.169.32|AGACCGCCTGGGAATACCGGGTGCTGTAGGCT</td><td>>hg38_1_-_228634819_228635039_RNA5S12_5S_50ntFlanks</td><td>138</td><td>169</td><td>32</td><td>AGACCGCCTGGGAATACCGGGTGCTGTAGGCT</td><td>774.3</td><td>6.4</td><td>0.7</td><td>9.2</td><td>4.3E-20</td><td>2.3E-19</td><td>BT20/MB468</td></tr>
<tr><td>>hg38_21_+_8433172_8446622_RNA45SN1_45S_50ntFlanks@10786.10823.38|AACAGCCTCTGGCATGTTGGAACAATGTAGGTAAGGGA</td><td>>hg38_21_+_8433172_8446622_RNA45SN1_45S_50ntFlanks</td><td>10786</td><td>10823</td><td>38</td><td>AACAGCCTCTGGCATGTTGGAACAATGTAGGTAAGGGA</td><td>294.0</td><td>-4.9</td><td>0.5</td><td>-9.2</td><td>4.9E-20</td><td>2.3E-19</td><td>MB231/MB468</td></tr>
<tr><td>>MI0000065|hsa-let-7d&WithFlank&9|+|94178828|94178926@14.35.22&[MIMAT0000065&hsa-let-7d-5p&offsets|0|0;m-90&9|+|94178841|94178862&offsets|0|0]|AGAGGTAGTAGGTTGCATAGTT</td><td>>MI0000065|hsa-let-7d&WithFlank&9|+|94178828|94178926</td><td>14</td><td>35</td><td>22</td><td>AGAGGTAGTAGGTTGCATAGTT</td><td>25401.2</td><td>7.7</td><td>0.8</td><td>9.2</td><td>4.4E-20</td><td>2.3E-19</td><td>BT20/MB468</td></tr>
<tr><td>>hg38_21_+_8433172_8446622_RNA45SN1_45S_50ntFlanks@8381.8400.20|AAGAGGGCGTGAAACCGTTA</td><td>>hg38_21_+_8433172_8446622_RNA45SN1_45S_50ntFlanks</td><td>8381</td><td>8400</td><td>20</td><td>AAGAGGGCGTGAAACCGTTA</td><td>891.3</td><td>-6.7</td><td>0.7</td><td>-9.2</td><td>5.0E-20</td><td>2.3E-19</td><td>MB231/MB468</td></tr>
<tr><td>>MI0000239|hsa-mir-197&WithFlank&1|+|109598887|109598973@54.76.23&[MIMAT0000227&hsa-miR-197-3p&offsets|0|+1;m-99&1|+|109598940|109598961&offsets|0|+1]|TTCACCACCTTCTCCACCCAGCA</td><td>>MI0000239|hsa-mir-197&WithFlank&1|+|109598887|109598973</td><td>54</td><td>76</td><td>23</td><td>TTCACCACCTTCTCCACCCAGCA</td><td>629.8</td><td>13.9</td><td>1.5</td><td>9.2</td><td>4.5E-20</td><td>2.3E-19</td><td>BT20/MB468</td></tr>
<tr><td>>hg38_21_+_8433172_8446622_RNA45SN1_45S_50ntFlanks@12570.12587.18|AGTACGAGAGGAACCGCA</td><td>>hg38_21_+_8433172_8446622_RNA45SN1_45S_50ntFlanks</td><td>12570</td><td>12587</td><td>18</td><td>AGTACGAGAGGAACCGCA</td><td>961.0</td><td>-11.9</td><td>1.3</td><td>-9.2</td><td>4.7E-20</td><td>2.5E-19</td><td>BT20/MB468</td></tr>
<tr><td>>MI0001446|hsa-mir-424&WithFlank&X|-|134546608|134546717@54.75.22&[MIMAT0004749&hsa-miR-424-3p&offsets|0|+1;m-305&X|-|134546644|134546664&offsets|0|+1]|CAAAACGTGAGGCGCTGCTATA</td><td>>MI0001446|hsa-mir-424&WithFlank&X|-|134546608|134546717</td><td>54</td><td>75</td><td>22</td><td>CAAAACGTGAGGCGCTGCTATA</td><td>686.2</td><td>10.5</td><td>1.1</td><td>9.2</td><td>4.8E-20</td><td>2.5E-19</td><td>BT20/MB468</td></tr>
<tr><td>>hg38_21_+_8433172_8446622_RNA45SN1_45S_50ntFlanks@12415.12448.34|CTTCGATGTCGGCTCTTCCTATCATTGTGAAGCA</td><td>>hg38_21_+_8433172_8446622_RNA45SN1_45S_50ntFlanks</td><td>12415</td><td>12448</td><td>34</td><td>CTTCGATGTCGGCTCTTCCTATCATTGTGAAGCA</td><td>626.2</td><td>5.0</td><td>0.5</td><td>9.2</td><td>4.8E-20</td><td>2.5E-19</td><td>BT20/MB468</td></tr>
<tr><td>>hg38_21_+_8433172_8446622_RNA45SN1_45S_50ntFlanks@10737.10755.19|TAAATCTCGCGCCGGGCCG</td><td>>hg38_21_+_8433172_8446622_RNA45SN1_45S_50ntFlanks</td><td>10737</td><td>10755</td><td>19</td><td>TAAATCTCGCGCCGGGCCG</td><td>626.4</td><td>-6.7</td><td>0.7</td><td>-9.2</td><td>4.8E-20</td><td>2.5E-19</td><td>BT20/MB468</td></tr>
<tr><td>>MI0000098|hsa-mir-96&WithFlank&7|-|129774686|129774775@15.36.22(+1C)&[MIMAT0000095&hsa-miR-96-5p&offsets|0|-1(+1C);m-248&7|-|129774739|129774761&offsets|0|-1(+1C)]|TTTGGCACTAGCACATTTTTGCC</td><td>>MI0000098|hsa-mir-96&WithFlank&7|-|129774686|129774775</td><td>15</td><td>36</td><td>22(+1C)</td><td>TTTGGCACTAGCACATTTTTGCC</td><td>5158.6</td><td>8.1</td><td>0.9</td><td>9.2</td><td>4.9E-20</td><td>2.6E-19</td><td>BT20/MB468</td></tr>
<tr><td>>MI0000093|hsa-mir-92a-1&WithFlank&13|+|91351308|91351397@55.75.21;MI0000094|hsa-mir-92a-2&WithFlank&X|-|134169532|134169618@55.75.21&[MIMAT0000092&hsa-miR-92a-3p&offsets|+1|0;m-9&13|+|91351361|91351382&offsets|+1|0];[MIMAT0000092_1&hsa-miR-92a-3p&offsets|+1|0;m-11&X|-|134169544|134169565&offsets|+1|0]|ATTGCACTTGTCCCGGCCTGT</td><td>>MI0000093|hsa-mir-92a-1&WithFlank&13|+|91351308|91351397</td><td>55</td><td>75</td><td>21</td><td>ATTGCACTTGTCCCGGCCTGT</td><td>2448.5</td><td>7.3</td><td>0.8</td><td>9.2</td><td>5.5E-20</td><td>2.6E-19</td><td>MB231/MB468</td></tr>
<tr><td>>hg38_21_+_8433172_8446622_RNA45SN1_45S_50ntFlanks@12456.12483.28|CCAAGCGTTGGATTGTTCACCCACTAAT</td><td>>hg38_21_+_8433172_8446622_RNA45SN1_45S_50ntFlanks</td><td>12456</td><td>12483</td><td>28</td><td>CCAAGCGTTGGATTGTTCACCCACTAAT</td><td>1439.0</td><td>-4.7</td><td>0.5</td><td>-9.2</td><td>5.5E-20</td><td>2.6E-19</td><td>MB231/MB468</td></tr>
<tr><td>>hg38_21_+_8433172_8446622_RNA45SN1_45S_50ntFlanks@9714.9737.24|CGAATGATTAGAGGTCTTGGGGCC</td><td>>hg38_21_+_8433172_8446622_RNA45SN1_45S_50ntFlanks</td><td>9714</td><td>9737</td><td>24</td><td>CGAATGATTAGAGGTCTTGGGGCC</td><td>471.4</td><td>-7.5</td><td>0.8</td><td>-9.2</td><td>5.0E-20</td><td>2.6E-19</td><td>BT20/MB468</td></tr>
<tr><td>>MI0000078|hsa-mir-22&WithFlank&17|-|1713897|1713993@59.80.22(+1A)&[MIMAT0000077&hsa-miR-22-3p&offsets|0|0(+1A);m-2&17|-|1713914|1713935&offsets|0|0(+1A)]|AAGCTGCCAGTTGAAGAACTGTA</td><td>>MI0000078|hsa-mir-22&WithFlank&17|-|1713897|1713993</td><td>59</td><td>80</td><td>22(+1A)</td><td>AAGCTGCCAGTTGAAGAACTGTA</td><td>1063.1</td><td>8.1</td><td>0.9</td><td>9.2</td><td>5.2E-20</td><td>2.7E-19</td><td>BT20/MB468</td></tr>
<tr><td>>MI0000465|hsa-mir-191&WithFlank&3|-|49020612|49020715@22.42.21(+1A)&[MIMAT0000440&hsa-miR-191-5p&offsets|0|-2(+1A);m-48&3|-|49020672|49020694&offsets|0|-2(+1A)]|CAACGGAATCCCAAAAGCAGCA</td><td>>MI0000465|hsa-mir-191&WithFlank&3|-|49020612|49020715</td><td>22</td><td>42</td><td>21(+1A)</td><td>CAACGGAATCCCAAAAGCAGCA</td><td>2831.6</td><td>8.2</td><td>0.9</td><td>9.2</td><td>5.2E-20</td><td>2.7E-19</td><td>BT20/MB468</td></tr>
<tr><td>>hg38_21_+_8433172_8446622_RNA45SN1_45S_50ntFlanks@10402.10420.19|AACATGGGTCAGTCGGTCC</td><td>>hg38_21_+_8433172_8446622_RNA45SN1_45S_50ntFlanks</td><td>10402</td><td>10420</td><td>19</td><td>AACATGGGTCAGTCGGTCC</td><td>1144.2</td><td>-5.5</td><td>0.6</td><td>-9.1</td><td>5.9E-20</td><td>2.7E-19</td><td>MB231/MB468</td></tr>
<tr><td>>hg38_21_+_8433172_8446622_RNA45SN1_45S_50ntFlanks@12627.12659.33|ATGGGGCGAAGCTACCATCTGTGGGATTATGAC</td><td>>hg38_21_+_8433172_8446622_RNA45SN1_45S_50ntFlanks</td><td>12627</td><td>12659</td><td>33</td><td>ATGGGGCGAAGCTACCATCTGTGGGATTATGAC</td><td>898.6</td><td>-11.8</td><td>1.3</td><td>-9.2</td><td>5.3E-20</td><td>2.8E-19</td><td>BT20/MB468</td></tr>
<tr><td>>MI0000272|hsa-mir-182&WithFlank&7|-|129770377|129770498@29.53.25(+1A)&[MIMAT0000259&hsa-miR-182-5p&offsets|0|+1(+1A);m-22&7|-|129770449|129770470&offsets|0|+3(+1A)]|TTTGGCAATGGTAGAACTCACACTGA</td><td>>MI0000272|hsa-mir-182&WithFlank&7|-|129770377|129770498</td><td>29</td><td>53</td><td>25(+1A)</td><td>TTTGGCAATGGTAGAACTCACACTGA</td><td>578.5</td><td>6.7</td><td>0.7</td><td>9.1</td><td>6.0E-20</td><td>2.8E-19</td><td>MB231/MB468</td></tr>
<tr><td>>MI0000285|hsa-mir-205&WithFlank&1|+|209432127|209432248@40.62.23(+1U)&[MIMAT0000266&hsa-miR-205-5p&offsets|0|+1(+1U);m-351&1|+|209432166|209432187&offsets|0|+1(+1U)]|TCCTTCATTCCACCGGAGTCTGTT</td><td>>MI0000285|hsa-mir-205&WithFlank&1|+|209432127|209432248</td><td>40</td><td>62</td><td>23(+1U)</td><td>TCCTTCATTCCACCGGAGTCTGTT</td><td>1121.8</td><td>13.6</td><td>1.4</td><td>9.5</td><td>2.7E-21</td><td>2.8E-19</td><td>BT20/MB231</td></tr>
<tr><td>>trna119_LysCTT_1_-_145395522_145395594@1.29.29,trna11_LysCTT_5_-_180648979_180649051@1.29.29,trna13_LysCTT_6_+_26556774_26556846@1.29.29,trna32_LysCTT_16_-_3207406_3207478@1.29.29,trna7_LysCTT_16_+_3225692_3225764@1.29.29,trna9_LysCTT_5_+_180634755_180634827@1.29.29|GCCCGGCTAGCTCAGTCGGTAGAGCATGA</td><td>>trna119_LysCTT_1_-_145395522_145395594</td><td>1</td><td>29</td><td>29</td><td>GCCCGGCTAGCTCAGTCGGTAGAGCATGA</td><td>1800.9</td><td>-6.0</td><td>0.7</td><td>-9.2</td><td>5.6E-20</td><td>2.9E-19</td><td>BT20/MB468</td></tr>
<tr><td>>trnaMT_ValTAC_MT_+_1602_1670@53.71.19|TAACTTGACCGCTCTGACC</td><td>>trnaMT_ValTAC_MT_+_1602_1670</td><td>53</td><td>71</td><td>19</td><td>TAACTTGACCGCTCTGACC</td><td>748.1</td><td>9.8</td><td>1.1</td><td>9.2</td><td>5.7E-20</td><td>2.9E-19</td><td>BT20/MB468</td></tr>
<tr><td>>hg38_21_+_8433172_8446622_RNA45SN1_45S_50ntFlanks@476.515.40|TTCGTGATCGATGTGGTGACGTCGTGCTCTCCCGGGCCGG</td><td>>hg38_21_+_8433172_8446622_RNA45SN1_45S_50ntFlanks</td><td>476</td><td>515</td><td>40</td><td>TTCGTGATCGATGTGGTGACGTCGTGCTCTCCCGGGCCGG</td><td>1939.9</td><td>11.0</td><td>1.2</td><td>9.1</td><td>6.3E-20</td><td>2.9E-19</td><td>MB231/MB468</td></tr>
<tr><td>>MI0000285|hsa-mir-205&WithFlank&1|+|209432127|209432248@40.59.20(+1C)&[MIMAT0000266&hsa-miR-205-5p&offsets|0|-2(+1C);m-351&1|+|209432166|209432187&offsets|0|-2(+1C)]|TCCTTCATTCCACCGGAGTCC</td><td>>MI0000285|hsa-mir-205&WithFlank&1|+|209432127|209432248</td><td>40</td><td>59</td><td>20(+1C)</td><td>TCCTTCATTCCACCGGAGTCC</td><td>1082.1</td><td>13.6</td><td>1.4</td><td>9.5</td><td>2.9E-21</td><td>3.0E-19</td><td>BT20/MB231</td></tr>
<tr><td>>hg38_21_+_8433172_8446622_RNA45SN1_45S_50ntFlanks@4728.4758.31|ATCAAGAACGAAAGTCGGAGGTTCGAAGACG</td><td>>hg38_21_+_8433172_8446622_RNA45SN1_45S_50ntFlanks</td><td>4728</td><td>4758</td><td>31</td><td>ATCAAGAACGAAAGTCGGAGGTTCGAAGACG</td><td>1199.1</td><td>-12.2</td><td>1.3</td><td>-9.1</td><td>5.8E-20</td><td>3.0E-19</td><td>BT20/MB468</td></tr>
<tr><td>>am_MI0000786|hsa-mir-378a&WithFlank&5|+|149732819|149732896@49.70.22(+1U)&[MIMAT0000732&hsa-miR-378a-3p&offsets|0|0(+1U);m-44&5|+|149732867|149732888&offsets|0|0(+1U)]|ACTGGACTTGGAGTCAGAAGGCT</td><td>>am_MI0000786|hsa-mir-378a&WithFlank&5|+|149732819|149732896</td><td>49</td><td>70</td><td>22(+1U)</td><td>ACTGGACTTGGAGTCAGAAGGCT</td><td>1709.2</td><td>4.3</td><td>0.5</td><td>9.1</td><td>5.9E-20</td><td>3.0E-19</td><td>BT20/MB468</td></tr>
<tr><td>>trna116_GluCTC_1_-_145399233_145399304@32.70.39,trna71_GluCTC_1_-_161439189_161439260@32.70.39,trna74_GluCTC_1_-_161431809_161431880@32.70.39,trna77_GluCTC_1_-_161424398_161424469@32.70.39,trna77_GluCTC_6_+_28949976_28950047@32.70.39,trna80_GluCTC_1_-_161417018_161417089@32.70.39,trna87_GluCTC_6_-_126101393_126101464@32.70.39|CTCTCACCGCCGCGGCCCGGGTTCGATTCCCGGTCAGGG</td><td>>trna116_GluCTC_1_-_145399233_145399304</td><td>32</td><td>70</td><td>39</td><td>CTCTCACCGCCGCGGCCCGGGTTCGATTCCCGGTCAGGG</td><td>1030.8</td><td>-12.0</td><td>1.3</td><td>-9.1</td><td>5.9E-20</td><td>3.1E-19</td><td>BT20/MB468</td></tr>
<tr><td>>hg38_21_+_8433172_8446622_RNA45SN1_45S_50ntFlanks@10361.10383.23|GAAGGCCGAAGTGGAGAAGGGTT</td><td>>hg38_21_+_8433172_8446622_RNA45SN1_45S_50ntFlanks</td><td>10361</td><td>10383</td><td>23</td><td>GAAGGCCGAAGTGGAGAAGGGTT</td><td>307.6</td><td>-4.9</td><td>0.5</td><td>-9.1</td><td>5.9E-20</td><td>3.1E-19</td><td>BT20/MB468</td></tr>
<tr><td>>trna119_LysCTT_1_-_145395522_145395594@4.36.33,trna11_LysCTT_5_-_180648979_180649051@4.36.33,trna13_LysCTT_6_+_26556774_26556846@4.36.33,trna7_LysCTT_16_+_3225692_3225764@4.36.33,trna9_LysCTT_5_+_180634755_180634827@4.36.33|CGGCTAGCTCAGTCGGTAGAGCATGAGACTCTT</td><td>>trna119_LysCTT_1_-_145395522_145395594</td><td>4</td><td>36</td><td>33</td><td>CGGCTAGCTCAGTCGGTAGAGCATGAGACTCTT</td><td>2673.1</td><td>11.6</td><td>1.3</td><td>9.1</td><td>6.7E-20</td><td>3.1E-19</td><td>MB231/MB468</td></tr>
<tr><td>>hg38_21_+_8433172_8446622_RNA45SN1_45S_50ntFlanks@13002.13039.38|CGATCTATTGAAAGTCAGCCCTCGACACAAGGGTTTGT</td><td>>hg38_21_+_8433172_8446622_RNA45SN1_45S_50ntFlanks</td><td>13002</td><td>13039</td><td>38</td><td>CGATCTATTGAAAGTCAGCCCTCGACACAAGGGTTTGT</td><td>1180.2</td><td>-12.2</td><td>1.3</td><td>-9.1</td><td>6.0E-20</td><td>3.1E-19</td><td>BT20/MB468</td></tr>
</table>

<table>
<tr><td>>MI0001445|hsa-mir-423&WithFlank&17|+|30117073|30117178@58.80.23&[MIMAT0001340&hsa-miR-423-3p&offsets|-1|-1;m-100&17|+|30117131|30117153&offsets|-1|-1]|AAGCTCGGTCTGAGGCCCCTCAG</td><td>>MI0001445|hsa-mir-423&WithFlank&17|+|30117073|30117178</td><td>58</td><td>80</td><td>23</td><td>AAGCTCGGTCTGAGGCCCCTCAG</td><td>796.6</td><td>8.8</td><td>1.0</td><td>9.1</td><td>6.7E-20</td><td>3.1E-19</td><td>MB231/MB468</td></tr>
<tr><td>>MI0000234|hsa-mir-192&WithFlank&11|-|64891131|64891252@30.49.20(+1U)&[MIMAT0000222&hsa-miR-192-5p&offsets|0|-1(+1U);m-59&11|-|64891203|64891223&offsets|0|-1(+1U)]|CTGACCTATGAATTGACAGCT</td><td>>MI0000234|hsa-mir-192&WithFlank&11|-|64891131|64891252</td><td>30</td><td>49</td><td>20(+1U)</td><td>CTGACCTATGAATTGACAGCT</td><td>1520.0</td><td>7.2</td><td>0.8</td><td>9.1</td><td>6.8E-20</td><td>3.1E-19</td><td>MB231/MB468</td></tr>
<tr><td>>trnaMT_PheGAA_MT_+_577_647@38.56.19|AATGTTTAGACGGGCTCAC</td><td>>trnaMT_PheGAA_MT_+_577_647</td><td>38</td><td>56</td><td>19</td><td>AATGTTTAGACGGGCTCAC</td><td>866.1</td><td>9.9</td><td>1.1</td><td>9.1</td><td>6.1E-20</td><td>3.2E-19</td><td>BT20/MB468</td></tr>
<tr><td>>MI0000812|hsa-mir-331&WithFlank&12|+|95308414|95308519@67.87.21(+1U)&[MIMAT0000760&hsa-miR-331-3p&offsets|0|0(+1U);m-192&12|+|95308480|95308501&offsets|0|-1(+1U)]|GCCCCTGGGCCTATCCTAGAAT</td><td>>MI0000812|hsa-mir-331&WithFlank&12|+|95308414|95308519</td><td>67</td><td>87</td><td>21(+1U)</td><td>GCCCCTGGGCCTATCCTAGAAT</td><td>1625.1</td><td>7.4</td><td>0.8</td><td>9.1</td><td>6.1E-20</td><td>3.2E-19</td><td>BT20/MB468</td></tr>
<tr><td>>hg38_21_+_8433172_8446622_RNA45SN1_45S_50ntFlanks@5472.5515.44|ACGGCCCTGGCGGAGCGCTGAGAAGACGGTCGAACTTGACTATC</td><td>>hg38_21_+_8433172_8446622_RNA45SN1_45S_50ntFlanks</td><td>5472</td><td>5515</td><td>44</td><td>ACGGCCCTGGCGGAGCGCTGAGAAGACGGTCGAACTTGACTATC</td><td>764.4</td><td>10.1</td><td>1.1</td><td>9.1</td><td>6.9E-20</td><td>3.2E-19</td><td>MB231/MB468</td></tr>
<tr><td>>MI0000465|hsa-mir-191&WithFlank&3|-|49020612|49020715@22.42.21(+1A)&[MIMAT0000440&hsa-miR-191-5p&offsets|0|-2(+1A);m-48&3|-|49020672|49020694&offsets|0|-2(+1A)]|CAACGGAATCCCAAAAGCAGCA</td><td>>MI0000465|hsa-mir-191&WithFlank&3|-|49020612|49020715</td><td>22</td><td>42</td><td>21(+1A)</td><td>CAACGGAATCCCAAAAGCAGCA</td><td>1167.4</td><td>7.3</td><td>0.8</td><td>9.1</td><td>6.9E-20</td><td>3.2E-19</td><td>MB231/MB468</td></tr>
<tr><td>>MI0000457|hsa-mir-141&WithFlank&12|+|6964091|6964197@65.85.21&[MIMAT0000432&hsa-miR-141-3p&offsets|0|-1;m-134&12|+|6964155|6964175&offsets|0|0]|TAACACTGTCTGGTAAAGATG</td><td>>MI0000457|hsa-mir-141&WithFlank&12|+|6964091|6964197</td><td>65</td><td>85</td><td>21</td><td>TAACACTGTCTGGTAAAGATG</td><td>1338.5</td><td>13.9</td><td>1.5</td><td>9.5</td><td>3.3E-21</td><td>3.2E-19</td><td>BT20/MB231</td></tr>
<tr><td>>MI0000433|hsa-let-7g&WithFlank&3|-|52268272|52268367@11.32.22(+1C)&[MIMAT0000414&hsa-let-7g-5p&offsets|0|0(+1C);m-58&3|-|52268336|52268357&offsets|0|0(+1C)]|TGAGGTAGTAGTTTGTACAGTTC</td><td>>MI0000433|hsa-let-7g&WithFlank&3|-|52268272|52268367</td><td>11</td><td>32</td><td>22(+1C)</td><td>TGAGGTAGTAGTTTGTACAGTTC</td><td>616.3</td><td>8.6</td><td>0.9</td><td>9.1</td><td>7.0E-20</td><td>3.2E-19</td><td>MB231/MB468</td></tr>
<tr><td>>MI0000440|hsa-mir-27b&WithFlank&9|+|95085439|95085547@67.87.21(+1U)&[MIMAT0000419&hsa-miR-27b-3p&offsets|0|0(+1U);m-39&9|+|95085505|95085525&offsets|0|0(+1U)]|TTCACAGTGGCTAAGTTCTGCT</td><td>>MI0000440|hsa-mir-27b&WithFlank&9|+|95085439|95085547</td><td>67</td><td>87</td><td>21(+1U)</td><td>TTCACAGTGGCTAAGTTCTGCT</td><td>4557.0</td><td>7.3</td><td>0.8</td><td>9.1</td><td>7.1E-20</td><td>3.3E-19</td><td>MB231/MB468</td></tr>
<tr><td>>MI0005756|hsa-mir-934&WithFlank&X|+|136550872|136550966@21.42.22&[MIMAT0004977&hsa-miR-934&offsets|0|0;m-943&X|+|136550892|136550913&offsets|0|0]|TGTCTACTACTGGAGACACTGG</td><td>>MI0005756|hsa-mir-934&WithFlank&X|+|136550872|136550966</td><td>21</td><td>42</td><td>22</td><td>TGTCTACTACTGGAGACACTGG</td><td>30632.7</td><td>10.1</td><td>1.1</td><td>9.1</td><td>6.4E-20</td><td>3.3E-19</td><td>BT20/MB468</td></tr>
<tr><td>>hg38_21_+_8433172_8446622_RNA45SN1_45S_50ntFlanks@12617.12653.37|TGAGGAGCCAATGGGGCGAAGCTACCATCTGTGGGAT</td><td>>hg38_21_+_8433172_8446622_RNA45SN1_45S_50ntFlanks</td><td>12617</td><td>12653</td><td>37</td><td>TGAGGAGCCAATGGGGCGAAGCTACCATCTGTGGGAT</td><td>256.2</td><td>-4.6</td><td>0.5</td><td>-9.1</td><td>7.2E-20</td><td>3.3E-19</td><td>MB231/MB468</td></tr>
<tr><td>>hg38_21_+_8433172_8446622_RNA45SN1_45S_50ntFlanks@8380.8405.26|CAAGAGGGCGTGAAACCGTTAAGAGG</td><td>>hg38_21_+_8433172_8446622_RNA45SN1_45S_50ntFlanks</td><td>8380</td><td>8405</td><td>26</td><td>CAAGAGGGCGTGAAACCGTTAAGAGG</td><td>831.2</td><td>-6.9</td><td>0.8</td><td>-9.1</td><td>6.4E-20</td><td>3.3E-19</td><td>BT20/MB468</td></tr>
<tr><td>>hg38_21_+_8433172_8446622_RNA45SN1_45S_50ntFlanks@9545.9566.22|GTGGAGGTCCGTAGCGGTCCTG</td><td>>hg38_21_+_8433172_8446622_RNA45SN1_45S_50ntFlanks</td><td>9545</td><td>9566</td><td>22</td><td>GTGGAGGTCCGTAGCGGTCCTG</td><td>1101.3</td><td>-6.4</td><td>0.7</td><td>-9.1</td><td>7.3E-20</td><td>3.4E-19</td><td>MB231/MB468</td></tr>
<tr><td>>MI0001445|hsa-mir-423&WithFlank&17|+|30117073|30117178@59.81.23(+2U)&[MIMAT0001340&hsa-miR-423-3p&offsets|0|0(+2U);m-100&17|+|30117131|30117153&offsets|0|0(+2U)]|AGCTCGGTCTGAGGCCCCTCAGTTT</td><td>>MI0001445|hsa-mir-423&WithFlank&17|+|30117073|30117178</td><td>59</td><td>81</td><td>23(+2U)</td><td>AGCTCGGTCTGAGGCCCCTCAGTTT</td><td>725.9</td><td>10.3</td><td>1.1</td><td>9.1</td><td>7.3E-20</td><td>3.4E-19</td><td>MB231/MB468</td></tr>
<tr><td>>hg38_21_+_8433172_8446622_RNA45SN1_45S_50ntFlanks@10408.10435.28|GGTCAGTCGGTCCTGAGAGATGGGCGAG</td><td>>hg38_21_+_8433172_8446622_RNA45SN1_45S_50ntFlanks</td><td>10408</td><td>10435</td><td>28</td><td>GGTCAGTCGGTCCTGAGAGATGGGCGAG</td><td>1360.2</td><td>-12.4</td><td>1.4</td><td>-9.1</td><td>6.6E-20</td><td>3.4E-19</td><td>BT20/MB468</td></tr>
<tr><td>>hg38_21_+_8433172_8446622_RNA45SN1_45S_50ntFlanks@11729.11753.25|TAACTATGACTCTCTTAAGGTAGCC</td><td>>hg38_21_+_8433172_8446622_RNA45SN1_45S_50ntFlanks</td><td>11729</td><td>11753</td><td>25</td><td>TAACTATGACTCTCTTAAGGTAGCC</td><td>2060.2</td><td>-6.6</td><td>0.7</td><td>-9.1</td><td>7.6E-20</td><td>3.5E-19</td><td>MB231/MB468</td></tr>
<tr><td>>MI0000285|hsa-mir-205&WithFlank&1|+|209432127|209432248@40.60.21(+1U)&[MIMAT0000266&hsa-miR-205-5p&offsets|0|-1(+1U);m-351&1|+|209432166|209432187&offsets|0|-1(+1U)]|TCCTTCATTCCACCGGAGTCTT</td><td>>MI0000285|hsa-mir-205&WithFlank&1|+|209432127|209432248</td><td>40</td><td>60</td><td>21(+1U)</td><td>TCCTTCATTCCACCGGAGTCTT</td><td>977.4</td><td>13.4</td><td>1.4</td><td>9.4</td><td>3.8E-21</td><td>3.6E-19</td><td>BT20/MB231</td></tr>
<tr><td>>am_trna10_ValCAC_5_-_180649395_180649467@1.32.32,trna12_ValAAC_5_-_180645270_180645342@1.32.32,trna132_ValAAC_6_-_27721179_27721251@1.32.32,trna136_ValAAC_6_-_27648885_27648957@1.32.32,trna139_ValAAC_6_-_27618707_27618779@1.32.32,trna18_ValCAC_5_-_180529253_180529325@1.32.32,trna2_ValAAC_3_+_169490018_169490090@1.32.32,trna2_ValCAC_5_+_180524070_180524142@1.32.32,trna4_ValAAC_5_+_180591154_180591226@1.32.32,trna5_ValAAC_5_+_180596610_180596682@1.32.32,trna6_ValCAC_5_+_180600650_180600722@1.32.32,trna85_ValCAC_1_-_161369490_161369562@1.32.32,trna90_ValCAC_1_-_149684088_149684161@1.32.32,trna98_ValCAC_1_-_149298555_149298627@1.32.32,trna9_ValCAC_6_+_26538282_26538354@1.32.32|GTTTCCGTAGTGTAGTGGTTATCACGTTCGCC</td><td>>am_trna10_ValCAC_5_-_180649395_180649467</td><td>1</td><td>32</td><td>32</td><td>GTTTCCGTAGTGTAGTGGTTATCACGTTCGCC</td><td>3244.8</td><td>7.3</td><td>0.8</td><td>9.1</td><td>6.9E-20</td><td>3.6E-19</td><td>BT20/MB468</td></tr>
<tr><td>>hg38_21_+_8433172_8446622_RNA45SN1_45S_50ntFlanks@11561.11592.32|GCGCCTAGCAGCCGACTTAGAACTGGTGCGGA</td><td>>hg38_21_+_8433172_8446622_RNA45SN1_45S_50ntFlanks</td><td>11561</td><td>11592</td><td>32</td><td>GCGCCTAGCAGCCGACTTAGAACTGGTGCGGA</td><td>1324.1</td><td>-6.4</td><td>0.7</td><td>-9.1</td><td>7.0E-20</td><td>3.6E-19</td><td>BT20/MB468</td></tr>
<tr><td>>hg38_21_+_8433172_8446622_RNA45SN1_45S_50ntFlanks@6746.6770.25|CACTTCGAACGCACTTGCGGCCCCG</td><td>>hg38_21_+_8433172_8446622_RNA45SN1_45S_50ntFlanks</td><td>6746</td><td>6770</td><td>25</td><td>CACTTCGAACGCACTTGCGGCCCCG</td><td>966.2</td><td>-11.9</td><td>1.3</td><td>-9.1</td><td>7.0E-20</td><td>3.6E-19</td><td>BT20/MB468</td></tr>
<tr><td>>MI0003599|hsa-mir-589&WithFlank&7|-|5495813|5495923@30.50.21&[MIMAT0004799&hsa-miR-589-5p&offsets|0|-1;m-162&7|-|5495874|5495894&offsets|0|0]|TGAGAACCACGTCTGCTCTGA</td><td>>MI0003599|hsa-mir-589&WithFlank&7|-|5495813|5495923</td><td>30</td><td>50</td><td>21</td><td>TGAGAACCACGTCTGCTCTGA</td><td>1182.5</td><td>10.6</td><td>1.2</td><td>9.1</td><td>7.9E-20</td><td>3.6E-19</td><td>MB231/MB468</td></tr>
<tr><td>>hg38_21_+_8433172_8446622_RNA45SN1_45S_50ntFlanks@10408.10433.26|GGTCAGTCGGTCCTGAGAGATGGGCG</td><td>>hg38_21_+_8433172_8446622_RNA45SN1_45S_50ntFlanks</td><td>10408</td><td>10433</td><td>26</td><td>GGTCAGTCGGTCCTGAGAGATGGGCG</td><td>983.3</td><td>-11.9</td><td>1.3</td><td>-9.1</td><td>7.1E-20</td><td>3.7E-19</td><td>BT20/MB468</td></tr>
<tr><td>>trnaMT_GluTTC_MT_-_14674_14742@40.66.27,trnalookalike8_GluTTC_5_-_93905172_93905240@40.66.27|TTGGTCGTGGTTGTAGTCCGTGCGAGA</td><td>>trnaMT_GluTTC_MT_-_14674_14742</td><td>40</td><td>66</td><td>27</td><td>TTGGTCGTGGTTGTAGTCCGTGCGAGA</td><td>796.5</td><td>8.3</td><td>0.9</td><td>9.1</td><td>8.0E-20</td><td>3.7E-19</td><td>MB231/MB468</td></tr>
<tr><td>>MI0003560|hsa-mir-92b&WithFlank&1|+|155195171|155195278@67.88.22(+1A)&[MIMAT0003218&hsa-miR-92b-3p&offsets|0|0(+1A);m-115&1|+|155195237|155195258&offsets|0|0(+1A)]|TATTGCACTCGTCCCGGCCTCCA</td><td>>MI0003560|hsa-mir-92b&WithFlank&1|+|155195171|155195278</td><td>67</td><td>88</td><td>22(+1A)</td><td>TATTGCACTCGTCCCGGCCTCCA</td><td>907.8</td><td>12.0</td><td>1.3</td><td>9.1</td><td>7.2E-20</td><td>3.7E-19</td><td>BT20/MB468</td></tr>
<tr><td>>am_MI0003772|hsa-mir-151b&WithFlank&14|-|100109413|100109520@66.85.20(+1C);MI0000809|hsa-mir-151a&WithFlank&8|-|140732558|140732659@17.36.20(+1C)&[MIMAT0010214&hsa-miR-151b&offsets|0|+2(+1C);m-179&14|-|100109436|100109455&offsets|0|0(+1C)];[MIMAT0004697&hsa-miR-151a-5p&offsets|0|-1(+1C);m-75&8|-|140732623|140732643&offsets|0|-1(+1C)]|TCGAGGAGCTCACAGTCTAGC</td><td>>am_MI0003772|hsa-mir-151b&WithFlank&14|-|100109413|100109520</td><td>66</td><td>85</td><td>20(+1C)</td><td>TCGAGGAGCTCACAGTCTAGC</td><td>1143.2</td><td>12.8</td><td>1.4</td><td>9.1</td><td>7.3E-20</td><td>3.7E-19</td><td>BT20/MB468</td></tr>
<tr><td>>hg38_21_+_8433172_8446622_RNA45SN1_45S_50ntFlanks@12456.12489.34|CCAAGCGTTGGATTGTTCACCCACTAATAGGGAA</td><td>>hg38_21_+_8433172_8446622_RNA45SN1_45S_50ntFlanks</td><td>12456</td><td>12489</td><td>34</td><td>CCAAGCGTTGGATTGTTCACCCACTAATAGGGAA</td><td>1060.0</td><td>-12.0</td><td>1.3</td><td>-9.1</td><td>7.3E-20</td><td>3.7E-19</td><td>BT20/MB468</td></tr>
<tr><td>>MI0000750|hsa-mir-26a-2&WithFlank&12|-|57824603|57824698@20.40.21(+1G);MI0000083|hsa-mir-26a-1&WithFlank&3|+|37969398|37969486@16.36.21(+1G)&[MIMAT0000082&hsa-miR-26a-5p&offsets|0|-1(+1G);m-33&12|-|57824658|57824679&offsets|0|-1(+1G)];[MIMAT0000082_1&hsa-miR-26a-5p&offsets|0|-1(+1G);m-34&3|+|37969413|37969434&offsets|0|-1(+1G)]|TTCAAGTAATCCAGGATAGGCG</td><td>>MI0000750|hsa-mir-26a-2&WithFlank&12|-|57824603|57824698</td><td>20</td><td>40</td><td>21(+1G)</td><td>TTCAAGTAATCCAGGATAGGCG</td><td>16628.9</td><td>7.0</td><td>0.8</td><td>9.1</td><td>8.1E-20</td><td>3.7E-19</td><td>MB231/MB468</td></tr>
<tr><td>>hg38_21_+_8433172_8446622_RNA45SN1_45S_50ntFlanks@4735.4767.33|ACGAAAGTCGGAGGTTCGAAGACGATCAGATAC</td><td>>hg38_21_+_8433172_8446622_RNA45SN1_45S_50ntFlanks</td><td>4735</td><td>4767</td><td>33</td><td>ACGAAAGTCGGAGGTTCGAAGACGATCAGATAC</td><td>956.5</td><td>12.3</td><td>1.4</td><td>9.1</td><td>8.2E-20</td><td>3.7E-19</td><td>MB231/MB468</td></tr>
<tr><td>>hg38_21_+_8433172_8446622_RNA45SN1_45S_50ntFlanks@10350.10378.29|ACGAGAACTTTGAAGGCCGAAGTGGAGAA</td><td>>hg38_21_+_8433172_8446622_RNA45SN1_45S_50ntFlanks</td><td>10350</td><td>10378</td><td>29</td><td>ACGAGAACTTTGAAGGCCGAAGTGGAGAA</td><td>373.6</td><td>-4.2</td><td>0.5</td><td>-9.1</td><td>7.3E-20</td><td>3.8E-19</td><td>BT20/MB468</td></tr>
<tr><td>>hg38_21_+_8433172_8446622_RNA45SN1_45S_50ntFlanks@11729.11767.39|TAACTATGACTCTCTTAAGGTAGCCAAATGCCTCGTCAT</td><td>>hg38_21_+_8433172_8446622_RNA45SN1_45S_50ntFlanks</td><td>11729</td><td>11767</td><td>39</td><td>TAACTATGACTCTCTTAAGGTAGCCAAATGCCTCGTCAT</td><td>903.2</td><td>-8.7</td><td>1.0</td><td>-9.1</td><td>7.5E-20</td><td>3.8E-19</td><td>BT20/MB468</td></tr>
<tr><td>>MI0000086|hsa-mir-28&WithFlank&3|+|188688775|188688872@60.80.21(+1U)&[MIMAT0004502&hsa-miR-28-3p&offsets|0|-1(+1U);m-28&3|+|188688834|188688855&offsets|0|-1(+1U)]|CACTAGATTGTGAGCTCCTGGT</td><td>>MI0000086|hsa-mir-28&WithFlank&3|+|188688775|188688872</td><td>60</td><td>80</td><td>21(+1U)</td><td>CACTAGATTGTGAGCTCCTGGT</td><td>1796.9</td><td>13.0</td><td>1.4</td><td>9.1</td><td>7.5E-20</td><td>3.8E-19</td><td>BT20/MB468</td></tr>
<tr><td>>hg38_21_+_8433172_8446622_RNA45SN1_45S_50ntFlanks@9573.9596.24|AAATCGGTCGTCCGACCTGGGTAT</td><td>>hg38_21_+_8433172_8446622_RNA45SN1_45S_50ntFlanks</td><td>9573</td><td>9596</td><td>24</td><td>AAATCGGTCGTCCGACCTGGGTAT</td><td>2807.4</td><td>-6.2</td><td>0.7</td><td>-9.1</td><td>7.5E-20</td><td>3.8E-19</td><td>BT20/MB468</td></tr>
<tr><td>>hg38_21_+_8433172_8446622_RNA45SN1_45S_50ntFlanks@10351.10374.24|CGAGAACTTTGAAGGCCGAAGTGG</td><td>>hg38_21_+_8433172_8446622_RNA45SN1_45S_50ntFlanks</td><td>10351</td><td>10374</td><td>24</td><td>CGAGAACTTTGAAGGCCGAAGTGG</td><td>1638.8</td><td>-4.6</td><td>0.5</td><td>-9.1</td><td>8.4E-20</td><td>3.9E-19</td><td>MB231/MB468</td></tr>
<tr><td>>hg38_21_+_8433172_8446622_RNA45SN1_45S_50ntFlanks@10402.10430.29|AACATGGGTCAGTCGGTCCTGAGAGATGG</td><td>>hg38_21_+_8433172_8446622_RNA45SN1_45S_50ntFlanks</td><td>10402</td><td>10430</td><td>29</td><td>AACATGGGTCAGTCGGTCCTGAGAGATGG</td><td>1548.7</td><td>-5.8</td><td>0.6</td><td>-9.1</td><td>7.6E-20</td><td>3.9E-19</td><td>BT20/MB468</td></tr>
</table>

<table>
<tr><td>>hg38_21_+_8433172_8446622_RNA45SN1_45S_50ntFlanks@10687.10710.24|CGGCGTCCGGTGAGCTCTCGCTGG</td><td>>hg38_21_+_8433172_8446622_RNA45SN1_45S_50ntFlanks</td><td>10687</td><td>10710</td><td>24</td><td>CGGCGTCCGGTGAGCTCTCGCTGG</td><td>4612.2</td><td>-7.6</td><td>0.8</td><td>-9.1</td><td>7.6E-20</td><td>3.9E-19</td><td>BT20/MB468</td></tr>
<tr><td>>MI0000076|hsa-mir-20a&WithFlank&13|+|91351059|91351141@14.36.23(+1A)&[MIMAT0000075&hsa-miR-20a-5p&offsets|0|0(+1A);m-96&13|+|91351072|91351094&offsets|0|0(+1A)]|TAAAGTGCTTATAGTGCAGGTAGA</td><td>>MI0000076|hsa-mir-20a&WithFlank&13|+|91351059|91351141</td><td>14</td><td>36</td><td>23(+1A)</td><td>TAAAGTGCTTATAGTGCAGGTAGA</td><td>15621.9</td><td>7.1</td><td>0.8</td><td>9.1</td><td>8.6E-20</td><td>4.0E-19</td><td>MB231/MB468</td></tr>
<tr><td>>MI0000298|hsa-mir-221&WithFlank&X|-|45746151|45746272@71.90.20(+1C)&[MIMAT0000278&hsa-miR-221-3p&offsets|0|-3(+1C);m-91&X|-|45746181|45746202&offsets|0|-2(+1C)]|AGCTACATTGTCTGCTGGGTC</td><td>>MI0000298|hsa-mir-221&WithFlank&X|-|45746151|45746272</td><td>71</td><td>90</td><td>20(+1C)</td><td>AGCTACATTGTCTGCTGGGTC</td><td>653.2</td><td>8.9</td><td>1.0</td><td>9.1</td><td>8.9E-20</td><td>4.1E-19</td><td>MB231/MB468</td></tr>
<tr><td>>hg38_21_+_8433172_8446622_RNA45SN1_45S_50ntFlanks@12628.12653.26|TGGGGCGAAGCTACCATCTGTGGGAT</td><td>>hg38_21_+_8433172_8446622_RNA45SN1_45S_50ntFlanks</td><td>12628</td><td>12653</td><td>26</td><td>TGGGGCGAAGCTACCATCTGTGGGAT</td><td>1212.8</td><td>-12.2</td><td>1.3</td><td>-9.1</td><td>8.0E-20</td><td>4.1E-19</td><td>BT20/MB468</td></tr>
<tr><td>>am_trnaMT_HisGTG_MT_+_12138_12206@18.38.21|AAAACATCAGATTGTGAATCT</td><td>>am_trnaMT_HisGTG_MT_+_12138_12206</td><td>18</td><td>38</td><td>21</td><td>AAAACATCAGATTGTGAATCT</td><td>655.5</td><td>9.8</td><td>1.1</td><td>9.1</td><td>8.0E-20</td><td>4.1E-19</td><td>BT20/MB468</td></tr>
<tr><td>>MI0000087|hsa-mir-29a&WithFlank&7|-|130876741|130876816@48.68.21(+1U)&[MIMAT0000086&hsa-miR-29a-3p&offsets|0|-1(+1U);m-26&7|-|130876749|130876769&offsets|0|0(+1U)]|TAGCACCATCTGAAATCGGTTT</td><td>>MI0000087|hsa-mir-29a&WithFlank&7|-|130876741|130876816</td><td>48</td><td>68</td><td>21(+1U)</td><td>TAGCACCATCTGAAATCGGTTT</td><td>1367.1</td><td>13.0</td><td>1.4</td><td>9.1</td><td>8.0E-20</td><td>4.1E-19</td><td>BT20/MB468</td></tr>
<tr><td>>MI0000066|hsa-let-7e&WithFlank&19|+|51692780|51692870@14.35.22(+1U)&[MIMAT0000066&hsa-let-7e-5p&offsets|0|0(+1U);m-50&19|+|51692793|51692814&offsets|0|0(+1U)]|TGAGGTAGGAGGTTGTATAGTTT</td><td>>MI0000066|hsa-let-7e&WithFlank&19|+|51692780|51692870</td><td>14</td><td>35</td><td>22(+1U)</td><td>TGAGGTAGGAGGTTGTATAGTTT</td><td>1980.8</td><td>6.9</td><td>0.8</td><td>9.1</td><td>8.9E-20</td><td>4.1E-19</td><td>MB231/MB468</td></tr>
<tr><td>>hg38_21_+_8433172_8446622_RNA45SN1_45S_50ntFlanks@10683.10709.27|CCGGCGGCGTCCGGTGAGCTCTCGCTG</td><td>>hg38_21_+_8433172_8446622_RNA45SN1_45S_50ntFlanks</td><td>10683</td><td>10709</td><td>27</td><td>CCGGCGGCGTCCGGTGAGCTCTCGCTG</td><td>1208.3</td><td>-5.4</td><td>0.6</td><td>-9.1</td><td>9.0E-20</td><td>4.1E-19</td><td>MB231/MB468</td></tr>
<tr><td>>MI0000285|hsa-mir-205&WithFlank&1|+|209432127|209432248@40.59.20(+1C)&[MIMAT0000266&hsa-miR-205-5p&offsets|0|-2(+1C);m-351&1|+|209432166|209432187&offsets|0|-2(+1C)]|TCCTTCATTCCACCGGAGTCC</td><td>>MI0000285|hsa-mir-205&WithFlank&1|+|209432127|209432248</td><td>40</td><td>59</td><td>20(+1C)</td><td>TCCTTCATTCCACCGGAGTCC</td><td>2757.2</td><td>7.4</td><td>0.8</td><td>9.1</td><td>8.2E-20</td><td>4.2E-19</td><td>BT20/MB468</td></tr>
<tr><td>>trnaMT_LeuTAA_MT_+_3230_3304@19.42.24|GGTAATCGCATAAAACTTAAAACT</td><td>>trnaMT_LeuTAA_MT_+_3230_3304</td><td>19</td><td>42</td><td>24</td><td>GGTAATCGCATAAAACTTAAAACT</td><td>447.7</td><td>11.0</td><td>1.2</td><td>9.1</td><td>8.2E-20</td><td>4.2E-19</td><td>BT20/MB468</td></tr>
<tr><td>>hg38_21_+_8433172_8446622_RNA45SN1_45S_50ntFlanks@475.517.43|CTTCGTGATCGATGTGGTGACGTCGTGCTCTCCCGGGCCGGGT</td><td>>hg38_21_+_8433172_8446622_RNA45SN1_45S_50ntFlanks</td><td>475</td><td>517</td><td>43</td><td>CTTCGTGATCGATGTGGTGACGTCGTGCTCTCCCGGGCCGGGT</td><td>2404.7</td><td>8.5</td><td>0.9</td><td>9.1</td><td>8.2E-20</td><td>4.2E-19</td><td>BT20/MB468</td></tr>
<tr><td>>MI0000065|hsa-let-7d&WithFlank&9|+|94178828|94178926@14.34.21&[MIMAT0000065&hsa-let-7d-5p&offsets|0|-1;m-90&9|+|94178841|94178862&offsets|0|-1]|AGAGGTAGTAGGTTGCATAGT</td><td>>MI0000065|hsa-let-7d&WithFlank&9|+|94178828|94178926</td><td>14</td><td>34</td><td>21</td><td>AGAGGTAGTAGGTTGCATAGT</td><td>3968.8</td><td>7.7</td><td>0.8</td><td>9.1</td><td>8.2E-20</td><td>4.2E-19</td><td>BT20/MB468</td></tr>
<tr><td>>hg38_21_+_8433172_8446622_RNA45SN1_45S_50ntFlanks@11577.11605.29|TTAGAACTGGTGCGGACCAGGGGAATCCG</td><td>>hg38_21_+_8433172_8446622_RNA45SN1_45S_50ntFlanks</td><td>11577</td><td>11605</td><td>29</td><td>TTAGAACTGGTGCGGACCAGGGGAATCCG</td><td>411.5</td><td>-5.6</td><td>0.6</td><td>-9.1</td><td>8.3E-20</td><td>4.2E-19</td><td>BT20/MB468</td></tr>
<tr><td>>hg38_21_+_8433172_8446622_RNA45SN1_45S_50ntFlanks@12624.12661.38|CCAATGGGGCGAAGCTACCATCTGTGGGATTATGACTG</td><td>>hg38_21_+_8433172_8446622_RNA45SN1_45S_50ntFlanks</td><td>12624</td><td>12661</td><td>38</td><td>CCAATGGGGCGAAGCTACCATCTGTGGGATTATGACTG</td><td>773.9</td><td>-4.9</td><td>0.5</td><td>-9.1</td><td>8.6E-20</td><td>4.3E-19</td><td>BT20/MB468</td></tr>
<tr><td>>MI0000285|hsa-mir-205&WithFlank&1|+|209432127|209432248@40.61.22(+2A)&[MIMAT0000266&hsa-miR-205-5p&offsets|0|0(+2A);m-351&1|+|209432166|209432187&offsets|0|0(+2A)]|TCCTTCATTCCACCGGAGTCTGAA</td><td>>MI0000285|hsa-mir-205&WithFlank&1|+|209432127|209432248</td><td>40</td><td>61</td><td>22(+2A)</td><td>TCCTTCATTCCACCGGAGTCTGAA</td><td>1169.5</td><td>13.7</td><td>1.5</td><td>9.4</td><td>4.8E-21</td><td>4.4E-19</td><td>BT20/MB231</td></tr>
<tr><td>>hg38_21_+_8433172_8446622_RNA45SN1_45S_50ntFlanks@6719.6747.29|TGAATTGCAGGACACATTGATCATCGACA</td><td>>hg38_21_+_8433172_8446622_RNA45SN1_45S_50ntFlanks</td><td>6719</td><td>6747</td><td>29</td><td>TGAATTGCAGGACACATTGATCATCGACA</td><td>999.3</td><td>-4.2</td><td>0.5</td><td>-9.1</td><td>8.6E-20</td><td>4.4E-19</td><td>BT20/MB468</td></tr>
<tr><td>>hg38_21_+_8433172_8446622_RNA45SN1_45S_50ntFlanks@3705.3740.36|TACCTGGTTGATCCTGCCAGTAGCATATGCTTGTCT</td><td>>hg38_21_+_8433172_8446622_RNA45SN1_45S_50ntFlanks</td><td>3705</td><td>3740</td><td>36</td><td>TACCTGGTTGATCCTGCCAGTAGCATATGCTTGTCT</td><td>545.3</td><td>11.7</td><td>1.3</td><td>9.1</td><td>8.6E-20</td><td>4.4E-19</td><td>BT20/MB468</td></tr>
<tr><td>>MI0000746|hsa-mir-99b&WithFlank&19|+|51692606|51692687@51.73.23(+1U)&[MIMAT0004678&hsa-miR-99b-3p&offsets|0|+1(+1U);m-151&19|+|51692656|51692678&offsets|0|0(+1U)]|CAAGCTCGTGTCTGTGGGTCCGTT</td><td>>MI0000746|hsa-mir-99b&WithFlank&19|+|51692606|51692687</td><td>51</td><td>73</td><td>23(+1U)</td><td>CAAGCTCGTGTCTGTGGGTCCGTT</td><td>728.5</td><td>8.9</td><td>1.0</td><td>9.1</td><td>8.8E-20</td><td>4.4E-19</td><td>BT20/MB468</td></tr>
<tr><td>>hg38_21_+_8433172_8446622_RNA45SN1_45S_50ntFlanks@12157.12180.24|AGTTTGACTGGGGCGGTACACCTG</td><td>>hg38_21_+_8433172_8446622_RNA45SN1_45S_50ntFlanks</td><td>12157</td><td>12180</td><td>24</td><td>AGTTTGACTGGGGCGGTACACCTG</td><td>1030.9</td><td>-12.0</td><td>1.3</td><td>-9.1</td><td>8.9E-20</td><td>4.5E-19</td><td>BT20/MB468</td></tr>
<tr><td>>hg38_21_+_8433172_8446622_RNA45SN1_45S_50ntFlanks@10387.10410.24|TGTGAACAGCAGTTGAACATGGGT</td><td>>hg38_21_+_8433172_8446622_RNA45SN1_45S_50ntFlanks</td><td>10387</td><td>10410</td><td>24</td><td>TGTGAACAGCAGTTGAACATGGGT</td><td>300.6</td><td>-4.1</td><td>0.4</td><td>-9.1</td><td>9.2E-20</td><td>4.6E-19</td><td>BT20/MB468</td></tr>
<tr><td>>hg38_21_+_8433172_8446622_RNA45SN1_45S_50ntFlanks@10411.10428.18|CAGTCGGTCCTGAGAGAT</td><td>>hg38_21_+_8433172_8446622_RNA45SN1_45S_50ntFlanks</td><td>10411</td><td>10428</td><td>18</td><td>CAGTCGGTCCTGAGAGAT</td><td>2159.1</td><td>-5.6</td><td>0.6</td><td>-9.1</td><td>1.0E-19</td><td>4.6E-19</td><td>MB231/MB468</td></tr>
<tr><td>>hg38_1_-_228634819_228635039_RNA5S12_5S_50ntFlanks@144.170.27|CCTGGGAATACCGGGTGCTGTAGGCTT</td><td>>hg38_1_-_228634819_228635039_RNA5S12_5S_50ntFlanks</td><td>144</td><td>170</td><td>27</td><td>CCTGGGAATACCGGGTGCTGTAGGCTT</td><td>5512.6</td><td>6.7</td><td>0.7</td><td>9.1</td><td>9.2E-20</td><td>4.6E-19</td><td>BT20/MB468</td></tr>
<tr><td>>hg38_21_+_8433172_8446622_RNA45SN1_45S_50ntFlanks@7990.8013.24|GACGTGGCGACCCGCTGAATTTAA</td><td>>hg38_21_+_8433172_8446622_RNA45SN1_45S_50ntFlanks</td><td>7990</td><td>8013</td><td>24</td><td>GACGTGGCGACCCGCTGAATTTAA</td><td>426.3</td><td>5.8</td><td>0.6</td><td>9.1</td><td>1.0E-19</td><td>4.7E-19</td><td>MB231/MB468</td></tr>
<tr><td>>MI0000070|hsa-mir-16-1&WithFlank&13|-|50048967|50049067@20.39.20;MI0000115|hsa-mir-16-2&WithFlank&3|+|160404739|160404831@16.35.20&[MIMAT0000069&hsa-miR-16-5p&offsets|0|-2;m-62&13|-|50049027|50049048&offsets|0|-2];[MIMAT0000069_1&hsa-miR-16-5p&offsets|0|-2;m-61&3|+|160404754|160404775&offsets|0|-2]|TAGCAGCACGTAAATATTGG</td><td>>MI0000070|hsa-mir-16-1&WithFlank&13|-|50048967|50049067</td><td>20</td><td>39</td><td>20</td><td>TAGCAGCACGTAAATATTGG</td><td>697.8</td><td>7.2</td><td>0.8</td><td>9.1</td><td>1.1E-19</td><td>4.8E-19</td><td>MB231/MB468</td></tr>
<tr><td>>hg38_21_+_8433172_8446622_RNA45SN1_45S_50ntFlanks@4047.4065.19|ATTCGAACGTCTGCCCTAT</td><td>>hg38_21_+_8433172_8446622_RNA45SN1_45S_50ntFlanks</td><td>4047</td><td>4065</td><td>19</td><td>ATTCGAACGTCTGCCCTAT</td><td>263.4</td><td>-3.9</td><td>0.4</td><td>-9.1</td><td>9.5E-20</td><td>4.8E-19</td><td>BT20/MB468</td></tr>
<tr><td>>hg38_21_+_8433172_8446622_RNA45SN1_45S_50ntFlanks@11874.11893.20|AGAAGACCCTGTTGAGCTTG</td><td>>hg38_21_+_8433172_8446622_RNA45SN1_45S_50ntFlanks</td><td>11874</td><td>11893</td><td>20</td><td>AGAAGACCCTGTTGAGCTTG</td><td>4695.4</td><td>-6.9</td><td>0.8</td><td>-9.1</td><td>9.6E-20</td><td>4.8E-19</td><td>BT20/MB468</td></tr>
<tr><td>>trnaMT_SerGCT_MT_+_12207_12265@23.62.40|AACTCATGCCCCCATGTCTAACAACATGGCTTTCTCACCA</td><td>>trnaMT_SerGCT_MT_+_12207_12265</td><td>23</td><td>62</td><td>40</td><td>AACTCATGCCCCCATGTCTAACAACATGGCTTTCTCACCA</td><td>435.0</td><td>11.4</td><td>1.3</td><td>9.1</td><td>9.6E-20</td><td>4.8E-19</td><td>BT20/MB468</td></tr>
<tr><td>>hg38_21_+_8433172_8446622_RNA45SN1_45S_50ntFlanks@6652.6701.50|GACTCTTAGCGGTGGATCACTCGGCTCGTGCGTCGATGAAGAACGCAGCT</td><td>>hg38_21_+_8433172_8446622_RNA45SN1_45S_50ntFlanks</td><td>6652</td><td>6701</td><td>50</td><td>GACTCTTAGCGGTGGATCACTCGGCTCGTGCGTCGATGAAGAACGCAGCT</td><td>1653.0</td><td>13.1</td><td>1.4</td><td>9.1</td><td>1.1E-19</td><td>4.9E-19</td><td>MB231/MB468</td></tr>
<tr><td>>hg38_21_+_8433172_8446622_RNA45SN1_45S_50ntFlanks@9562.9596.35|TCCTGACGTGCAAATCGGTCGTCCGACCTGGGTAT</td><td>>hg38_21_+_8433172_8446622_RNA45SN1_45S_50ntFlanks</td><td>9562</td><td>9596</td><td>35</td><td>TCCTGACGTGCAAATCGGTCGTCCGACCTGGGTAT</td><td>878.1</td><td>-11.8</td><td>1.3</td><td>-9.1</td><td>9.9E-20</td><td>4.9E-19</td><td>BT20/MB468</td></tr>
<tr><td>>MI0000456|hsa-mir-140&WithFlank&16|+|69933075|69933186@69.90.22(+1G)&[MIMAT0004597&hsa-miR-140-3p&offsets|+1|+2(+1G);m-45&16|+|69933143|69933164&offsets|0|0(+1G)]|ACCACAGGGTAGAACCACGGACG</td><td>>MI0000456|hsa-mir-140&WithFlank&16|+|69933075|69933186</td><td>69</td><td>90</td><td>22(+1G)</td><td>ACCACAGGGTAGAACCACGGACG</td><td>578.1</td><td>13.8</td><td>1.5</td><td>9.1</td><td>1.0E-19</td><td>5.1E-19</td><td>BT20/MB468</td></tr>
<tr><td>>MI0000298|hsa-mir-221&WithFlank&X|-|45746151|45746272@71.92.22(+2U)&[MIMAT0000278&hsa-miR-221-3p&offsets|0|-1(+2U);m-91&X|-|45746181|45746202&offsets|0|0(+2U)]|AGCTACATTGTCTGCTGGGTTTTT</td><td>>MI0000298|hsa-mir-221&WithFlank&X|-|45746151|45746272</td><td>71</td><td>92</td><td>22(+2U)</td><td>AGCTACATTGTCTGCTGGGTTTTT</td><td>1766.1</td><td>9.2</td><td>1.0</td><td>9.1</td><td>1.1E-19</td><td>5.1E-19</td><td>MB231/MB468</td></tr>
<tr><td>>hg38_21_+_8433172_8446622_RNA45SN1_45S_50ntFlanks@11575.11601.27|ACTTAGAACTGGTGCGGACCAGGGGAA</td><td>>hg38_21_+_8433172_8446622_RNA45SN1_45S_50ntFlanks</td><td>11575</td><td>11601</td><td>27</td><td>ACTTAGAACTGGTGCGGACCAGGGGAA</td><td>953.8</td><td>-6.3</td><td>0.7</td><td>-9.1</td><td>1.1E-19</td><td>5.2E-19</td><td>MB231/MB468</td></tr>
<tr><td>>MI0000078|hsa-mir-22&WithFlank&17|-|1713897|1713993@21.41.21&[MIMAT0004495&hsa-miR-22-5p&offsets|0|-1;m-217&17|-|1713953|1713973&offsets|0|0]|AGTTCTTCAGTGGCAAGCTTT</td><td>>MI0000078|hsa-mir-22&WithFlank&17|-|1713897|1713993</td><td>21</td><td>41</td><td>21</td><td>AGTTCTTCAGTGGCAAGCTTT</td><td>1608.6</td><td>8.1</td><td>0.9</td><td>9.1</td><td>1.2E-19</td><td>5.2E-19</td><td>MB231/MB468</td></tr>
<tr><td>>hg38_21_+_8433172_8446622_RNA45SN1_45S_50ntFlanks@12635.12661.27|AAGCTACCATCTGTGGGATTATGACTG</td><td>>hg38_21_+_8433172_8446622_RNA45SN1_45S_50ntFlanks</td><td>12635</td><td>12661</td><td>27</td><td>AAGCTACCATCTGTGGGATTATGACTG</td><td>3034.2</td><td>-5.2</td><td>0.6</td><td>-9.1</td><td>1.1E-19</td><td>5.3E-19</td><td>BT20/MB468</td></tr>
<tr><td>>hg38_21_+_8433172_8446622_RNA45SN1_45S_50ntFlanks@9575.9595.21|ATCGGTCGTCCGACCTGGGTA</td><td>>hg38_21_+_8433172_8446622_RNA45SN1_45S_50ntFlanks</td><td>9575</td><td>9595</td><td>21</td><td>ATCGGTCGTCCGACCTGGGTA</td><td>729.5</td><td>-11.5</td><td>1.3</td><td>-9.1</td><td>1.1E-19</td><td>5.4E-19</td><td>BT20/MB468</td></tr>
<tr><td>>MI0002468|hsa-mir-484&WithFlank&16|+|15643288|15643378@14.35.22&[MIMAT0002174&hsa-miR-484&offsets|0|0;m-127&16|+|15643301|15643321&offsets|0|+1]|TCAGGCTCAGTCCCCTCCCGAT</td><td>>MI0002468|hsa-mir-484&WithFlank&16|+|15643288|15643378</td><td>14</td><td>35</td><td>22</td><td>TCAGGCTCAGTCCCCTCCCGAT</td><td>3887.2</td><td>6.6</td><td>0.7</td><td>9.1</td><td>1.2E-19</td><td>5.4E-19</td><td>MB231/MB468</td></tr>
<tr><td>>MI0000465|hsa-mir-191&WithFlank&3|-|49020612|49020715@22.43.22(+1U)&[MIMAT0000440&hsa-miR-191-5p&offsets|0|-1(+1U);m-48&3|-|49020672|49020694&offsets|0|-1(+1U)]|CAACGGAATCCCAAAAGCAGCTT</td><td>>MI0000465|hsa-mir-191&WithFlank&3|-|49020612|49020715</td><td>22</td><td>43</td><td>22(+1U)</td><td>CAACGGAATCCCAAAAGCAGCTT</td><td>771.9</td><td>6.8</td><td>0.8</td><td>9.1</td><td>1.1E-19</td><td>5.4E-19</td><td>BT20/MB468</td></tr>
<tr><td>>MI0000746|hsa-mir-99b&WithFlank&19|+|51692606|51692687@13.31.19&[MIMAT0000689&hsa-miR-99b-5p&offsets|0|-3;m-10&19|+|51692618|51692639&offsets|0|-3]|CACCCGTAGAACCGACCTT</td><td>>MI0000746|hsa-mir-99b&WithFlank&19|+|51692606|51692687</td><td>13</td><td>31</td><td>19</td><td>CACCCGTAGAACCGACCTT</td><td>435.0</td><td>11.4</td><td>1.3</td><td>9.1</td><td>1.1E-19</td><td>5.4E-19</td><td>BT20/MB468</td></tr>
<tr><td>>hg38_21_+_8433172_8446622_RNA45SN1_45S_50ntFlanks@12416.12433.18|TTCGATGTCGGCTCTTCC</td><td>>hg38_21_+_8433172_8446622_RNA45SN1_45S_50ntFlanks</td><td>12416</td><td>12433</td><td>18</td><td>TTCGATGTCGGCTCTTCC</td><td>3527.6</td><td>6.1</td><td>0.7</td><td>9.1</td><td>1.2E-19</td><td>5.6E-19</td><td>MB231/MB468</td></tr>
<tr><td>>MI0001445|hsa-mir-423&WithFlank&17|+|30117073|30117178@58.80.23(+1C)&[MIMAT0001340&hsa-miR-423-3p&offsets|-1|-1(+1C);m-100&17|+|30117131|30117153&offsets|-1|-1(+1C)]|AAGCTCGGTCTGAGGCCCCTCAGC</td><td>>MI0001445|hsa-mir-423&WithFlank&17|+|30117073|30117178</td><td>58</td><td>80</td><td>23(+1C)</td><td>AAGCTCGGTCTGAGGCCCCTCAGC</td><td>724.7</td><td>8.3</td><td>0.9</td><td>9.1</td><td>1.1E-19</td><td>5.6E-19</td><td>BT20/MB468</td></tr>
<tr><td>>hg38_21_+_8433172_8446622_RNA45SN1_45S_50ntFlanks@12626.12667.42|AATGGGGCGAAGCTACCATCTGTGGGATTATGACTGAACGCC</td><td>>hg38_21_+_8433172_8446622_RNA45SN1_45S_50ntFlanks</td><td>12626</td><td>12667</td><td>42</td><td>AATGGGGCGAAGCTACCATCTGTGGGATTATGACTGAACGCC</td><td>1821.5</td><td>10.5</td><td>1.2</td><td>9.1</td><td>1.3E-19</td><td>5.7E-19</td><td>MB231/MB468</td></tr>
<tr><td>>MI0000746|hsa-mir-99b&WithFlank&19|+|51692606|51692687@13.34.22&[MIMAT0000689&hsa-miR-99b-5p&offsets|0|0;m-10&19|+|51692618|51692639&offsets|0|0]|CACCCGTAGAACCGACCTTGCG</td><td>>MI0000746|hsa-mir-99b&WithFlank&19|+|51692606|51692687</td><td>13</td><td>34</td><td>22</td><td>CACCCGTAGAACCGACCTTGCG</td><td>83550.0</td><td>7.2</td><td>0.8</td><td>9.1</td><td>1.1E-19</td><td>5.7E-19</td><td>BT20/MB468</td></tr>
</table>

<table>
<tr><td>>MI0000272|hsa-mir-182&WithFlank&7|-|129770377|129770498@29.54.26&[MIMAT0000259&hsa-miR-182-5p&offsets|0|+2;m-22&7|-|129770449|129770470&offsets|0|+4]|TTTGGCAATGGTAGAACTCACACTGG</td><td>>MI0000272|hsa-mir-182&WithFlank&7|-|129770377|129770498</td><td>29</td><td>54</td><td>26</td><td>TTTGGCAATGGTAGAACTCACACTGG</td><td>3038.6</td><td>6.5</td><td>0.7</td><td>9.1</td><td>1.3E-19</td><td>5.7E-19</td><td>MB231/MB468</td></tr>
<tr><td>>MI0000234|hsa-mir-192&WithFlank&11|-|64891131|64891252@31.53.23&[MIMAT0000222&hsa-miR-192-5p&offsets|+1|+3;m-59&11|-|64891203|64891223&offsets|+1|+3]|TGACCTATGAATTGACAGCCAGT</td><td>>MI0000234|hsa-mir-192&WithFlank&11|-|64891131|64891252</td><td>31</td><td>53</td><td>23</td><td>TGACCTATGAATTGACAGCCAGT</td><td>1723.2</td><td>6.7</td><td>0.7</td><td>9.1</td><td>1.3E-19</td><td>5.7E-19</td><td>MB231/MB468</td></tr>
<tr><td>>hg38_21_+_8433172_8446622_RNA45SN1_45S_50ntFlanks@3705.3742.38|TACCTGGTTGATCCTGCCAGTAGCATATGCTTGTCTCA</td><td>>hg38_21_+_8433172_8446622_RNA45SN1_45S_50ntFlanks</td><td>3705</td><td>3742</td><td>38</td><td>TACCTGGTTGATCCTGCCAGTAGCATATGCTTGTCTCA</td><td>4527.5</td><td>7.5</td><td>0.8</td><td>9.1</td><td>1.2E-19</td><td>5.8E-19</td><td>BT20/MB468</td></tr>
<tr><td>>MI0000093|hsa-mir-92a-1&WithFlank&13|+|91351308|91351397@54.75.22(+2A);MI0000094|hsa-mir-92a-2&WithFlank&X|-|134169532|134169618@54.75.22(+2A)&[MIMAT0000092&hsa-miR-92a-3p&offsets|0|0(+2A);m-9&13|+|91351361|91351382&offsets|0|0(+2A)];[MIMAT0000092_1&hsa-miR-92a-3p&offsets|0|0(+2A);m-11&X|-|134169544|134169565&offsets|0|0(+2A)]|TATTGCACTTGTCCCGGCCTGTAA</td><td>>MI0000093|hsa-mir-92a-1&WithFlank&13|+|91351308|91351397</td><td>54</td><td>75</td><td>22(+2A)</td><td>TATTGCACTTGTCCCGGCCTGTAA</td><td>1940.5</td><td>6.4</td><td>0.7</td><td>9.1</td><td>1.3E-19</td><td>5.8E-19</td><td>MB231/MB468</td></tr>
<tr><td>>hg38_21_+_8433172_8446622_RNA45SN1_45S_50ntFlanks@9574.9610.37|AATCGGTCGTCCGACCTGGGTATAGGGGCGAAAGACT</td><td>>hg38_21_+_8433172_8446622_RNA45SN1_45S_50ntFlanks</td><td>9574</td><td>9610</td><td>37</td><td>AATCGGTCGTCCGACCTGGGTATAGGGGCGAAAGACT</td><td>600.5</td><td>6.0</td><td>0.7</td><td>9.1</td><td>1.2E-19</td><td>5.9E-19</td><td>BT20/MB468</td></tr>
<tr><td>>MI0000746|hsa-mir-99b&WithFlank&19|+|51692606|51692687@51.70.20&[MIMAT0004678&hsa-miR-99b-3p&offsets|0|-2;m-151&19|+|51692656|51692678&offsets|0|-3]|CAAGCTCGTGTCTGTGGGTC</td><td>>MI0000746|hsa-mir-99b&WithFlank&19|+|51692606|51692687</td><td>51</td><td>70</td><td>20</td><td>CAAGCTCGTGTCTGTGGGTC</td><td>1103.3</td><td>8.3</td><td>0.9</td><td>9.1</td><td>1.2E-19</td><td>5.9E-19</td><td>BT20/MB468</td></tr>
<tr><td>>MI0000264|hsa-mir-7-2&WithFlank&15|+|88611819|88611940@38.61.24;MI0000265|hsa-mir-7-3&WithFlank&19|+|4770664|4770785@37.60.24;MI0000263|hsa-mir-7-1&WithFlank&9|-|83969742|83969863@30.53.24&[MIMAT0000252&hsa-miR-7-5p&offsets|0|0;m-400&15|+|88611856|88611879&offsets|0|0];[MIMAT0000252_1&hsa-miR-7-5p&offsets|0|0;m-402&19|+|4770700|4770723&offsets|0|0];[MIMAT0000252_2&hsa-miR-7-5p&offsets|0|0;m-401&9|-|83969811|83969834&offsets|0|0]|TGGAAGACTAGTGATTTTGTTGTT</td><td>>MI0000264|hsa-mir-7-2&WithFlank&15|+|88611819|88611940</td><td>38</td><td>61</td><td>24</td><td>TGGAAGACTAGTGATTTTGTTGTT</td><td>314428.6</td><td>7.6</td><td>0.8</td><td>9.1</td><td>1.2E-19</td><td>5.9E-19</td><td>BT20/MB468</td></tr>
<tr><td>>MI0000750|hsa-mir-26a-2&WithFlank&12|-|57824603|57824698@20.41.22(+1C);MI0000083|hsa-mir-26a-1&WithFlank&3|+|37969398|37969486@16.37.22(+1C)&[MIMAT0000082&hsa-miR-26a-5p&offsets|0|0(+1C);m-33&12|-|57824658|57824679&offsets|0|0(+1C)];[MIMAT0000082_1&hsa-miR-26a-5p&offsets|0|0(+1C);m-34&3|+|37969413|37969434&offsets|0|0(+1C)]|TTCAAGTAATCCAGGATAGGCTC</td><td>>MI0000750|hsa-mir-26a-2&WithFlank&12|-|57824603|57824698</td><td>20</td><td>41</td><td>22(+1C)</td><td>TTCAAGTAATCCAGGATAGGCTC</td><td>1156.4</td><td>6.9</td><td>0.8</td><td>9.1</td><td>1.2E-19</td><td>5.9E-19</td><td>BT20/MB468</td></tr>
<tr><td>>hg38_21_+_8433172_8446622_RNA45SN1_45S_50ntFlanks@11675.11707.33|TGCTCTGAATGTCAAAGTGAAGAAATTCAATGA</td><td>>hg38_21_+_8433172_8446622_RNA45SN1_45S_50ntFlanks</td><td>11675</td><td>11707</td><td>33</td><td>TGCTCTGAATGTCAAAGTGAAGAAATTCAATGA</td><td>766.4</td><td>6.7</td><td>0.7</td><td>9.1</td><td>1.3E-19</td><td>5.9E-19</td><td>MB231/MB468</td></tr>
<tr><td>>hg38_21_+_8433172_8446622_RNA45SN1_45S_50ntFlanks@12436.12454.19|TCATTGTGAAGCAGAATTC</td><td>>hg38_21_+_8433172_8446622_RNA45SN1_45S_50ntFlanks</td><td>12436</td><td>12454</td><td>19</td><td>TCATTGTGAAGCAGAATTC</td><td>1190.1</td><td>-8.1</td><td>0.9</td><td>-9.1</td><td>1.2E-19</td><td>5.9E-19</td><td>BT20/MB468</td></tr>
<tr><td>>hg38_21_+_8433172_8446622_RNA45SN1_45S_50ntFlanks@10391.10420.30|AACAGCAGTTGAACATGGGTCAGTCGGTCC</td><td>>hg38_21_+_8433172_8446622_RNA45SN1_45S_50ntFlanks</td><td>10391</td><td>10420</td><td>30</td><td>AACAGCAGTTGAACATGGGTCAGTCGGTCC</td><td>367.4</td><td>-7.3</td><td>0.8</td><td>-9.1</td><td>1.2E-19</td><td>6.0E-19</td><td>BT20/MB468</td></tr>
<tr><td>>hg38_21_+_8433172_8446622_RNA45SN1_45S_50ntFlanks@12419.12442.24|GATGTCGGCTCTTCCTATCATTGT</td><td>>hg38_21_+_8433172_8446622_RNA45SN1_45S_50ntFlanks</td><td>12419</td><td>12442</td><td>24</td><td>GATGTCGGCTCTTCCTATCATTGT</td><td>1872.6</td><td>7.3</td><td>0.8</td><td>9.1</td><td>1.3E-19</td><td>6.0E-19</td><td>MB231/MB468</td></tr>
<tr><td>>hg38_21_+_8433172_8446622_RNA45SN1_45S_50ntFlanks@11572.11600.29|CCGACTTAGAACTGGTGCGGACCAGGGGA</td><td>>hg38_21_+_8433172_8446622_RNA45SN1_45S_50ntFlanks</td><td>11572</td><td>11600</td><td>29</td><td>CCGACTTAGAACTGGTGCGGACCAGGGGA</td><td>2410.9</td><td>-4.6</td><td>0.5</td><td>-9.1</td><td>1.2E-19</td><td>6.0E-19</td><td>BT20/MB468</td></tr>
<tr><td>>hg38_21_+_8433172_8446622_RNA45SN1_45S_50ntFlanks@6721.6741.21|AATTGCAGGACACATTGATCA</td><td>>hg38_21_+_8433172_8446622_RNA45SN1_45S_50ntFlanks</td><td>6721</td><td>6741</td><td>21</td><td>AATTGCAGGACACATTGATCA</td><td>646.2</td><td>-6.5</td><td>0.7</td><td>-9.1</td><td>1.2E-19</td><td>6.1E-19</td><td>BT20/MB468</td></tr>
<tr><td>>hg38_21_+_8433172_8446622_RNA45SN1_45S_50ntFlanks@6740.6771.32|CATCGACACTTCGAACGCACTTGCGGCCCCGG</td><td>>hg38_21_+_8433172_8446622_RNA45SN1_45S_50ntFlanks</td><td>6740</td><td>6771</td><td>32</td><td>CATCGACACTTCGAACGCACTTGCGGCCCCGG</td><td>1175.2</td><td>-6.3</td><td>0.7</td><td>-9.1</td><td>1.4E-19</td><td>6.1E-19</td><td>MB231/MB468</td></tr>
<tr><td>>MI0005756|hsa-mir-934&WithFlank&X|+|136550872|136550966@21.41.21&[MIMAT0004977&hsa-miR-934&offsets|0|-1;m-943&X|+|136550892|136550913&offsets|0|-1]|TGTCTACTACTGGAGACACTG</td><td>>MI0005756|hsa-mir-934&WithFlank&X|+|136550872|136550966</td><td>21</td><td>41</td><td>21</td><td>TGTCTACTACTGGAGACACTG</td><td>14424.2</td><td>8.5</td><td>0.9</td><td>9.1</td><td>1.2E-19</td><td>6.2E-19</td><td>BT20/MB468</td></tr>
<tr><td>>MI0005756|hsa-mir-934&WithFlank&X|+|136550872|136550966@21.42.22(+1C)&[MIMAT0004977&hsa-miR-934&offsets|0|0(+1C);m-943&X|+|136550892|136550913&offsets|0|0(+1C)]|TGTCTACTACTGGAGACACTGGC</td><td>>MI0005756|hsa-mir-934&WithFlank&X|+|136550872|136550966</td><td>21</td><td>42</td><td>22(+1C)</td><td>TGTCTACTACTGGAGACACTGGC</td><td>742.7</td><td>7.2</td><td>0.8</td><td>9.1</td><td>1.2E-19</td><td>6.2E-19</td><td>BT20/MB468</td></tr>
<tr><td>>hg38_21_+_8433172_8446622_RNA45SN1_45S_50ntFlanks@6653.6683.31|ACTCTTAGCGGTGGATCACTCGGCTCGTGCG</td><td>>hg38_21_+_8433172_8446622_RNA45SN1_45S_50ntFlanks</td><td>6653</td><td>6683</td><td>31</td><td>ACTCTTAGCGGTGGATCACTCGGCTCGTGCG</td><td>2063.1</td><td>7.0</td><td>0.8</td><td>9.1</td><td>1.4E-19</td><td>6.2E-19</td><td>MB231/MB468</td></tr>
<tr><td>>MI0000299|hsa-mir-222&WithFlank&X|-|45747009|45747130@75.97.23(+3U)&[MIMAT0000279&hsa-miR-222-3p&offsets|0|+2(+3U);m-140&X|-|45747033|45747056&offsets|0|-1(+3U)]|AGCTACATCTGGCTACTGGGTCTTTT</td><td>>MI0000299|hsa-mir-222&WithFlank&X|-|45747009|45747130</td><td>75</td><td>97</td><td>23(+3U)</td><td>AGCTACATCTGGCTACTGGGTCTTTT</td><td>992.8</td><td>11.2</td><td>1.2</td><td>9.1</td><td>1.4E-19</td><td>6.2E-19</td><td>MB231/MB468</td></tr>
<tr><td>>MI0000062|hsa-let-7a-3&WithFlank&22|+|46112743|46112828@58.79.22;MI0000060|hsa-let-7a-1&WithFlank&9|+|94175951|94176042@63.84.22&[MIMAT0004481&hsa-let-7a-3p&offsets|0|+1;m-180&22|+|46112800|46112821&offsets|0|0];[MIMAT0004481_1&hsa-let-7a-3p&offsets|0|+1;m-181&9|+|94176013|94176034&offsets|0|0]|CTATACAATCTACTGTCTTTCC</td><td>>MI0000062|hsa-let-7a-3&WithFlank&22|+|46112743|46112828</td><td>58</td><td>79</td><td>22</td><td>CTATACAATCTACTGTCTTTCC</td><td>647.2</td><td>6.7</td><td>0.7</td><td>9.1</td><td>1.4E-19</td><td>6.3E-19</td><td>MB231/MB468</td></tr>
<tr><td>>MI0000811|hsa-mir-148b&WithFlank&12|+|54337210|54337320@69.90.22(+1G)&[MIMAT0000759&hsa-miR-148b-3p&offsets|0|0(+1G);m-77&12|+|54337278|54337299&offsets|0|0(+1G)]|TCAGTGCATCACAGAACTTTGTG</td><td>>MI0000811|hsa-mir-148b&WithFlank&12|+|54337210|54337320</td><td>69</td><td>90</td><td>22(+1G)</td><td>TCAGTGCATCACAGAACTTTGTG</td><td>939.5</td><td>8.0</td><td>0.9</td><td>9.1</td><td>1.4E-19</td><td>6.3E-19</td><td>MB231/MB468</td></tr>
<tr><td>>MI0000272|hsa-mir-182&WithFlank&7|-|129770377|129770498@29.53.25(+1U)&[MIMAT0000259&hsa-miR-182-5p&offsets|0|+1(+1U);m-22&7|-|129770449|129770470&offsets|0|+3(+1U)]|TTTGGCAATGGTAGAACTCACACTGT</td><td>>MI0000272|hsa-mir-182&WithFlank&7|-|129770377|129770498</td><td>29</td><td>53</td><td>25(+1U)</td><td>TTTGGCAATGGTAGAACTCACACTGT</td><td>3393.9</td><td>7.5</td><td>0.8</td><td>9.1</td><td>1.4E-19</td><td>6.3E-19</td><td>MB231/MB468</td></tr>
<tr><td>>MI0000811|hsa-mir-148b&WithFlank&12|+|54337210|54337320@69.90.22(+1A)&[MIMAT0000759&hsa-miR-148b-3p&offsets|0|0(+1A);m-77&12|+|54337278|54337299&offsets|0|0(+1A)]|TCAGTGCATCACAGAACTTTGTA</td><td>>MI0000811|hsa-mir-148b&WithFlank&12|+|54337210|54337320</td><td>69</td><td>90</td><td>22(+1A)</td><td>TCAGTGCATCACAGAACTTTGTA</td><td>2006.8</td><td>7.7</td><td>0.8</td><td>9.1</td><td>1.3E-19</td><td>6.4E-19</td><td>BT20/MB468</td></tr>
<tr><td>>am_MI0000809|hsa-mir-151a&WithFlank&8|-|140732558|140732659@53.73.21(+2U)&[MIMAT0000757&hsa-miR-151a-3p&offsets|0|0(+2U);m-30&8|-|140732587|140732607&offsets|0|0(+2U)]|CTAGACTGAAGCTCCTTGAGGTT</td><td>>am_MI0000809|hsa-mir-151a&WithFlank&8|-|140732558|140732659</td><td>53</td><td>73</td><td>21(+2U)</td><td>CTAGACTGAAGCTCCTTGAGGTT</td><td>2263.6</td><td>9.6</td><td>1.1</td><td>9.1</td><td>1.3E-19</td><td>6.6E-19</td><td>BT20/MB468</td></tr>
<tr><td>>MI0000264|hsa-mir-7-2&WithFlank&15|+|88611819|88611940@38.59.22(+1C);MI0000265|hsa-mir-7-3&WithFlank&19|+|4770664|4770785@37.58.22(+1C);MI0000263|hsa-mir-7-1&WithFlank&9|-|83969742|83969863@30.51.22(+1C)&[MIMAT0000252&hsa-miR-7-5p&offsets|0|-2(+1C);m-400&15|+|88611856|88611879&offsets|0|-2(+1C)];[MIMAT0000252_1&hsa-miR-7-5p&offsets|0|-2(+1C);m-402&19|+|4770700|4770723&offsets|0|-2(+1C)];[MIMAT0000252_2&hsa-miR-7-5p&offsets|0|-2(+1C);m-401&9|-|83969811|83969834&offsets|0|-2(+1C)]|TGGAAGACTAGTGATTTTGTTGC</td><td>>MI0000264|hsa-mir-7-2&WithFlank&15|+|88611819|88611940</td><td>38</td><td>59</td><td>22(+1C)</td><td>TGGAAGACTAGTGATTTTGTTGC</td><td>7555.0</td><td>8.8</td><td>1.0</td><td>9.1</td><td>1.4E-19</td><td>6.9E-19</td><td>BT20/MB468</td></tr>
<tr><td>>hg38_21_+_8433172_8446622_RNA45SN1_45S_50ntFlanks@6748.6770.23|CTTCGAACGCACTTGCGGCCCCG</td><td>>hg38_21_+_8433172_8446622_RNA45SN1_45S_50ntFlanks</td><td>6748</td><td>6770</td><td>23</td><td>CTTCGAACGCACTTGCGGCCCCG</td><td>276.4</td><td>-5.6</td><td>0.6</td><td>-9.1</td><td>1.4E-19</td><td>6.9E-19</td><td>BT20/MB468</td></tr>
<tr><td>>hg38_21_+_8433172_8446622_RNA45SN1_45S_50ntFlanks@3882.3899.18|CCGACGGGCGCTGACCCC</td><td>>hg38_21_+_8433172_8446622_RNA45SN1_45S_50ntFlanks</td><td>3882</td><td>3899</td><td>18</td><td>CCGACGGGCGCTGACCCC</td><td>1830.3</td><td>-6.7</td><td>0.7</td><td>-9.1</td><td>1.4E-19</td><td>6.9E-19</td><td>BT20/MB468</td></tr>
<tr><td>>hg38_21_+_8433172_8446622_RNA45SN1_45S_50ntFlanks@10819.10840.22|AGGGAAGTCGGCAAGCCGGATC</td><td>>hg38_21_+_8433172_8446622_RNA45SN1_45S_50ntFlanks</td><td>10819</td><td>10840</td><td>22</td><td>AGGGAAGTCGGCAAGCCGGATC</td><td>2764.7</td><td>-5.0</td><td>0.6</td><td>-9.0</td><td>1.5E-19</td><td>7.0E-19</td><td>MB231/MB468</td></tr>
<tr><td>>trnaMT_SerGCT_MT_+_12207_12265@29.62.34|TGCCCCCATGTCTAACAACATGGCTTTCTCACCA</td><td>>trnaMT_SerGCT_MT_+_12207_12265</td><td>29</td><td>62</td><td>34</td><td>TGCCCCCATGTCTAACAACATGGCTTTCTCACCA</td><td>435.7</td><td>11.4</td><td>1.3</td><td>9.1</td><td>1.4E-19</td><td>7.0E-19</td><td>BT20/MB468</td></tr>
<tr><td>>hg38_MT_+_1621_3279_RNR2_16S_50ntFlanks@1427.1458.32|GATTAAAGTCCTACGTGATCTGAGTTCAGACC</td><td>>hg38_MT_+_1621_3279_RNR2_16S_50ntFlanks</td><td>1427</td><td>1458</td><td>32</td><td>GATTAAAGTCCTACGTGATCTGAGTTCAGACC</td><td>920.5</td><td>9.8</td><td>1.1</td><td>9.0</td><td>1.6E-19</td><td>7.0E-19</td><td>MB231/MB468</td></tr>
<tr><td>>hg38_21_+_8433172_8446622_RNA45SN1_45S_50ntFlanks@3985.4002.18|CGGCTTTGGTGACTCTAG</td><td>>hg38_21_+_8433172_8446622_RNA45SN1_45S_50ntFlanks</td><td>3985</td><td>4002</td><td>18</td><td>CGGCTTTGGTGACTCTAG</td><td>2243.8</td><td>-10.8</td><td>1.2</td><td>-9.0</td><td>1.6E-19</td><td>7.1E-19</td><td>MB231/MB468</td></tr>
<tr><td>>am_trnaMT_PheGAA_MT_+_577_647@41.59.19|GTTTAGACGGGCTCACATC</td><td>>am_trnaMT_PheGAA_MT_+_577_647</td><td>41</td><td>59</td><td>19</td><td>GTTTAGACGGGCTCACATC</td><td>671.5</td><td>9.3</td><td>1.0</td><td>9.0</td><td>1.6E-19</td><td>7.2E-19</td><td>MB231/MB468</td></tr>
<tr><td>>MI0005756|hsa-mir-934&WithFlank&X|+|136550872|136550966@21.43.23&[MIMAT0004977&hsa-miR-934&offsets|0|+1;m-943&X|+|136550892|136550913&offsets|0|+1]|TGTCTACTACTGGAGACACTGGT</td><td>>MI0005756|hsa-mir-934&WithFlank&X|+|136550872|136550966</td><td>21</td><td>43</td><td>23</td><td>TGTCTACTACTGGAGACACTGGT</td><td>5849.0</td><td>6.9</td><td>0.8</td><td>9.0</td><td>1.5E-19</td><td>7.2E-19</td><td>BT20/MB468</td></tr>
<tr><td>>hg38_21_+_8433172_8446622_RNA45SN1_45S_50ntFlanks@10458.10477.20|ATGGCCTCCGTTGCCCTCGG</td><td>>hg38_21_+_8433172_8446622_RNA45SN1_45S_50ntFlanks</td><td>10458</td><td>10477</td><td>20</td><td>ATGGCCTCCGTTGCCCTCGG</td><td>1825.9</td><td>-12.4</td><td>1.4</td><td>-9.0</td><td>1.6E-19</td><td>7.2E-19</td><td>MB231/MB468</td></tr>
<tr><td>>MI0000650|hsa-mir-200c&WithFlank&12|+|6963693|6963772@49.72.24&[MIMAT0000617&hsa-miR-200c-3p&offsets|-1|0;m-54&12|+|6963742|6963764&offsets|-1|0]|CTAATACTGCCGGGTAATGATGGA</td><td>>MI0000650|hsa-mir-200c&WithFlank&12|+|6963693|6963772</td><td>49</td><td>72</td><td>24</td><td>CTAATACTGCCGGGTAATGATGGA</td><td>1159.0</td><td>7.3</td><td>0.8</td><td>9.0</td><td>1.5E-19</td><td>7.2E-19</td><td>BT20/MB468</td></tr>
</table>

<table>
<tr><th></th><th></th><th></th><th></th><th></th><th></th><th></th><th></th><th></th><th></th><th></th><th></th><th></th></tr>
<tr><td>>hg38_21_+_8433172_8446622_RNA45SN1_45S_50ntFlanks@4734.4767.34|AACGAAAGTCGGAGGTTCGAAGACGATCAGATAC</td><td>>hg38_21_+_8433172_8446622_RNA45SN1_45S_50ntFlanks</td><td>4734</td><td>4767</td><td>34</td><td>AACGAAAGTCGGAGGTTCGAAGACGATCAGATAC</td><td>589.0</td><td>9.9</td><td>1.1</td><td>9.0</td><td>1.6E-19</td><td>7.3E-19</td><td>MB231/MB468</td></tr>
<tr><td>>hg38_21_+_8433172_8446622_RNA45SN1_45S_50ntFlanks@10411.10433.23|CAGTCGGTCCTGAGAGATGGGCG</td><td>>hg38_21_+_8433172_8446622_RNA45SN1_45S_50ntFlanks</td><td>10411</td><td>10433</td><td>23</td><td>CAGTCGGTCCTGAGAGATGGGCG</td><td>1262.2</td><td>-8.8</td><td>1.0</td><td>-9.0</td><td>1.5E-19</td><td>7.3E-19</td><td>BT20/MB468</td></tr>
<tr><td>>MI0000825|hsa-mir-345&WithFlank&14|+|100307853|100307962@24.45.22&[MIMAT0000772&hsa-miR-345-5p&offsets|0|0;m-193&14|+|100307876|100307896&offsets|0|+1]|GCTGACTCCTAGTCCAGGGCTC</td><td>>MI0000825|hsa-mir-345&WithFlank&14|+|100307853|100307962</td><td>24</td><td>45</td><td>22</td><td>GCTGACTCCTAGTCCAGGGCTC</td><td>913.6</td><td>7.4</td><td>0.8</td><td>9.0</td><td>1.6E-19</td><td>7.3E-19</td><td>MB231/MB468</td></tr>
<tr><td>>hg38_21_+_8433172_8446622_RNA45SN1_45S_50ntFlanks@9604.9621.18|AAAGACTAATCGAACCAT</td><td>>hg38_21_+_8433172_8446622_RNA45SN1_45S_50ntFlanks</td><td>9604</td><td>9621</td><td>18</td><td>AAAGACTAATCGAACCAT</td><td>297.8</td><td>-4.1</td><td>0.5</td><td>-9.0</td><td>1.5E-19</td><td>7.4E-19</td><td>BT20/MB468</td></tr>
<tr><td>>hg38_21_+_8433172_8446622_RNA45SN1_45S_50ntFlanks@9738.9759.22|GAAACGATCTCAACCTATTCTC</td><td>>hg38_21_+_8433172_8446622_RNA45SN1_45S_50ntFlanks</td><td>9738</td><td>9759</td><td>22</td><td>GAAACGATCTCAACCTATTCTC</td><td>503.0</td><td>-5.4</td><td>0.6</td><td>-9.0</td><td>1.5E-19</td><td>7.4E-19</td><td>BT20/MB468</td></tr>
<tr><td>>MI0000098|hsa-mir-96&WithFlank&7|-|129774686|129774775@15.36.22(+1A)&[MIMAT0000095&hsa-miR-96-5p&offsets|0|-1(+1A);m-248&7|-|129774739|129774761&offsets|0|-1(+1A)]|TTTGGCACTAGCACATTTTTGCA</td><td>>MI0000098|hsa-mir-96&WithFlank&7|-|129774686|129774775</td><td>15</td><td>36</td><td>22(+1A)</td><td>TTTGGCACTAGCACATTTTTGCA</td><td>6547.8</td><td>9.2</td><td>1.0</td><td>9.0</td><td>1.6E-19</td><td>7.7E-19</td><td>BT20/MB468</td></tr>
<tr><td>>trnaMT_GluTTC_MT_-_14674_14742@41.72.32,trnalookalike8_GluTTC_5_-_93905172_93905240@41.72.32|TGGTCGTGGTTGTAGTCCGTGCGAGAATACCA</td><td>>trnaMT_GluTTC_MT_-_14674_14742</td><td>41</td><td>72</td><td>32</td><td>TGGTCGTGGTTGTAGTCCGTGCGAGAATACCA</td><td>3195.1</td><td>5.7</td><td>0.6</td><td>9.0</td><td>1.6E-19</td><td>7.7E-19</td><td>BT20/MB468</td></tr>
<tr><td>>hg38_21_+_8433172_8446622_RNA45SN1_45S_50ntFlanks@12939.12956.18|CCATTCGTAGACGACCTG</td><td>>hg38_21_+_8433172_8446622_RNA45SN1_45S_50ntFlanks</td><td>12939</td><td>12956</td><td>18</td><td>CCATTCGTAGACGACCTG</td><td>839.7</td><td>-4.5</td><td>0.5</td><td>-9.0</td><td>1.7E-19</td><td>7.7E-19</td><td>MB231/MB468</td></tr>
<tr><td>>MI0000746|hsa-mir-99b&WithFlank&19|+|51692606|51692687@51.73.23(+1U)&[MIMAT0004678&hsa-miR-99b-3p&offsets|0|+1(+1U);m-151&19|+|51692656|51692678&offsets|0|0(+1U)]|CAAGCTCGTGTCTGTGGGTCCGTT</td><td>>MI0000746|hsa-mir-99b&WithFlank&19|+|51692606|51692687</td><td>51</td><td>73</td><td>23(+1U)</td><td>CAAGCTCGTGTCTGTGGGTCCGTT</td><td>679.3</td><td>9.0</td><td>1.0</td><td>9.0</td><td>1.7E-19</td><td>7.7E-19</td><td>MB231/MB468</td></tr>
<tr><td>>MI0000440|hsa-mir-27b&WithFlank&9|+|95085439|95085547@67.88.22(+1U)&[MIMAT0000419&hsa-miR-27b-3p&offsets|0|+1(+1U);m-39&9|+|95085505|95085525&offsets|0|+1(+1U)]|TTCACAGTGGCTAAGTTCTGCAT</td><td>>MI0000440|hsa-mir-27b&WithFlank&9|+|95085439|95085547</td><td>67</td><td>88</td><td>22(+1U)</td><td>TTCACAGTGGCTAAGTTCTGCAT</td><td>1399.3</td><td>7.8</td><td>0.9</td><td>9.0</td><td>1.7E-19</td><td>7.8E-19</td><td>MB231/MB468</td></tr>
<tr><td>>MI0000750|hsa-mir-26a-2&WithFlank&12|-|57824603|57824698@20.41.22(+1A);MI0000083|hsa-mir-26a-1&WithFlank&3|+|37969398|37969486@16.37.22(+1A)&[MIMAT0000082&hsa-miR-26a-5p&offsets|0|0(+1A);m-33&12|-|57824658|57824679&offsets|0|0(+1A)];[MIMAT0000082_1&hsa-miR-26a-5p&offsets|0|0(+1A);m-34&3|+|37969413|37969434&offsets|0|0(+1A)]|TTCAAGTAATCCAGGATAGGCTA</td><td>>MI0000750|hsa-mir-26a-2&WithFlank&12|-|57824603|57824698</td><td>20</td><td>41</td><td>22(+1A)</td><td>TTCAAGTAATCCAGGATAGGCTA</td><td>3408.4</td><td>7.0</td><td>0.8</td><td>9.0</td><td>1.7E-19</td><td>7.8E-19</td><td>MB231/MB468</td></tr>
<tr><td>>MI0006444|hsa-mir-1307&WithFlank&10|-|103394247|103394407@87.109.23&[MIMAT0005951&hsa-miR-1307-3p&offsets|+1|+2;m-66&10|-|103394300|103394322&offsets|+1|+1]|CTCGGCGTGGCGTCGGTCGTGGT</td><td>>MI0006444|hsa-mir-1307&WithFlank&10|-|103394247|103394407</td><td>87</td><td>109</td><td>23</td><td>CTCGGCGTGGCGTCGGTCGTGGT</td><td>771.1</td><td>8.8</td><td>1.0</td><td>9.0</td><td>1.8E-19</td><td>7.8E-19</td><td>MB231/MB468</td></tr>
<tr><td>>hg38_MT_+_598_1651_RNR1_12S_50ntFlanks@367.404.38|CCAATAAAGCTAAAACTCACCTGAGTTGTAAAAAACTC</td><td>>hg38_MT_+_598_1651_RNR1_12S_50ntFlanks</td><td>367</td><td>404</td><td>38</td><td>CCAATAAAGCTAAAACTCACCTGAGTTGTAAAAAACTC</td><td>736.0</td><td>10.1</td><td>1.1</td><td>9.0</td><td>1.8E-19</td><td>7.8E-19</td><td>MB231/MB468</td></tr>
<tr><td>>MI0000456|hsa-mir-140&WithFlank&16|+|69933075|69933186@69.88.20&[MIMAT0004597&hsa-miR-140-3p&offsets|+1|0;m-45&16|+|69933143|69933164&offsets|0|-2]|ACCACAGGGTAGAACCACGG</td><td>>MI0000456|hsa-mir-140&WithFlank&16|+|69933075|69933186</td><td>69</td><td>88</td><td>20</td><td>ACCACAGGGTAGAACCACGG</td><td>752.0</td><td>12.2</td><td>1.3</td><td>9.0</td><td>1.6E-19</td><td>7.8E-19</td><td>BT20/MB468</td></tr>
<tr><td>>MI0000078|hsa-mir-22&WithFlank&17|-|1713897|1713993@59.79.21(+1G)&[MIMAT0000077&hsa-miR-22-3p&offsets|0|-1(+1G);m-2&17|-|1713914|1713935&offsets|0|-1(+1G)]|AAGCTGCCAGTTGAAGAACTGG</td><td>>MI0000078|hsa-mir-22&WithFlank&17|-|1713897|1713993</td><td>59</td><td>79</td><td>21(+1G)</td><td>AAGCTGCCAGTTGAAGAACTGG</td><td>2186.9</td><td>9.1</td><td>1.0</td><td>9.0</td><td>1.6E-19</td><td>7.8E-19</td><td>BT20/MB468</td></tr>
<tr><td>>hg38_1_-_228634819_228635039_RNA5S12_5S_50ntFlanks@84.103.20|CTCGTCTGATCTCGGAAGCT</td><td>>hg38_1_-_228634819_228635039_RNA5S12_5S_50ntFlanks</td><td>84</td><td>103</td><td>20</td><td>CTCGTCTGATCTCGGAAGCT</td><td>1029.2</td><td>-5.7</td><td>0.6</td><td>-9.0</td><td>1.6E-19</td><td>8.0E-19</td><td>BT20/MB468</td></tr>
<tr><td>>hg38_21_+_8433172_8446622_RNA45SN1_45S_50ntFlanks@7975.8001.27|CGCGACCTCAGATCAGACGTGGCGACC</td><td>>hg38_21_+_8433172_8446622_RNA45SN1_45S_50ntFlanks</td><td>7975</td><td>8001</td><td>27</td><td>CGCGACCTCAGATCAGACGTGGCGACC</td><td>1673.9</td><td>7.2</td><td>0.8</td><td>9.0</td><td>1.7E-19</td><td>8.0E-19</td><td>BT20/MB468</td></tr>
<tr><td>>MI0000736|hsa-mir-30c-1&WithFlank&1|+|40757278|40757378@23.44.22;MI0000254|hsa-mir-30c-2&WithFlank&6|-|71376954|71377037@13.34.22&[MIMAT0000244&hsa-miR-30c-5p&offsets|0|-1;m-57&1|+|40757300|40757323&offsets|0|-2];[MIMAT0000244_1&hsa-miR-30c-5p&offsets|0|-1;m-56&6|-|71377002|71377025&offsets|0|-2]|TGTAAACATCCTACACTCTCAG</td><td>>MI0000736|hsa-mir-30c-1&WithFlank&1|+|40757278|40757378</td><td>23</td><td>44</td><td>22</td><td>TGTAAACATCCTACACTCTCAG</td><td>601.0</td><td>6.9</td><td>0.8</td><td>9.0</td><td>1.8E-19</td><td>8.1E-19</td><td>MB231/MB468</td></tr>
<tr><td>>hg38_21_+_8433172_8446622_RNA45SN1_45S_50ntFlanks@12365.12382.18|ATAACTGGCTTGTGGCGG</td><td>>hg38_21_+_8433172_8446622_RNA45SN1_45S_50ntFlanks</td><td>12365</td><td>12382</td><td>18</td><td>ATAACTGGCTTGTGGCGG</td><td>799.2</td><td>-11.6</td><td>1.3</td><td>-9.0</td><td>1.7E-19</td><td>8.1E-19</td><td>BT20/MB468</td></tr>
<tr><td>>hg38_21_+_8433172_8446622_RNA45SN1_45S_50ntFlanks@11669.11702.34|GCCCAGTGCTCTGAATGTCAAAGTGAAGAAATTC</td><td>>hg38_21_+_8433172_8446622_RNA45SN1_45S_50ntFlanks</td><td>11669</td><td>11702</td><td>34</td><td>GCCCAGTGCTCTGAATGTCAAAGTGAAGAAATTC</td><td>2294.2</td><td>6.7</td><td>0.7</td><td>9.0</td><td>1.8E-19</td><td>8.2E-19</td><td>MB231/MB468</td></tr>
<tr><td>>hg38_21_+_8433172_8446622_RNA45SN1_45S_50ntFlanks@6746.6771.26|CACTTCGAACGCACTTGCGGCCCCGG</td><td>>hg38_21_+_8433172_8446622_RNA45SN1_45S_50ntFlanks</td><td>6746</td><td>6771</td><td>26</td><td>CACTTCGAACGCACTTGCGGCCCCGG</td><td>682.7</td><td>-11.4</td><td>1.3</td><td>-9.0</td><td>1.7E-19</td><td>8.2E-19</td><td>BT20/MB468</td></tr>
<tr><td>>trnaMT_PheGAA_MT_+_577_647@37.57.21|AAATGTTTAGACGGGCTCACA</td><td>>trnaMT_PheGAA_MT_+_577_647</td><td>37</td><td>57</td><td>21</td><td>AAATGTTTAGACGGGCTCACA</td><td>496.3</td><td>10.6</td><td>1.2</td><td>9.0</td><td>1.7E-19</td><td>8.2E-19</td><td>BT20/MB468</td></tr>
<tr><td>>MI0000095|hsa-mir-93&WithFlank&7|-|100093762|100093853@17.39.23(+1A)&[MIMAT0000093&hsa-miR-93-5p&offsets|0|0(+1A);m-29&7|-|100093815|100093837&offsets|0|0(+1A)]|CAAAGTGCTGTTCGTGCAGGTAGA</td><td>>MI0000095|hsa-mir-93&WithFlank&7|-|100093762|100093853</td><td>17</td><td>39</td><td>23(+1A)</td><td>CAAAGTGCTGTTCGTGCAGGTAGA</td><td>7163.5</td><td>7.0</td><td>0.8</td><td>9.0</td><td>1.9E-19</td><td>8.2E-19</td><td>MB231/MB468</td></tr>
<tr><td>>hg38_21_+_8433172_8446622_RNA45SN1_45S_50ntFlanks@8196.8215.20|CGTGGACGGTGTGAGGCCGG</td><td>>hg38_21_+_8433172_8446622_RNA45SN1_45S_50ntFlanks</td><td>8196</td><td>8215</td><td>20</td><td>CGTGGACGGTGTGAGGCCGG</td><td>1270.6</td><td>-5.1</td><td>0.6</td><td>-9.0</td><td>1.7E-19</td><td>8.3E-19</td><td>BT20/MB468</td></tr>
<tr><td>>MI0000255|hsa-mir-30d&WithFlank&8|-|134804870|134804951@12.35.24(+1A)&[MIMAT0000245&hsa-miR-30d-5p&offsets|0|+2(+1A);m-21&8|-|134804919|134804940&offsets|0|+2(+1A)]|TGTAAACATCCCCGACTGGAAGCTA</td><td>>MI0000255|hsa-mir-30d&WithFlank&8|-|134804870|134804951</td><td>12</td><td>35</td><td>24(+1A)</td><td>TGTAAACATCCCCGACTGGAAGCTA</td><td>1152.5</td><td>9.1</td><td>1.0</td><td>9.0</td><td>1.9E-19</td><td>8.4E-19</td><td>MB231/MB468</td></tr>
<tr><td>>hg38_MT_+_1621_3279_RNR2_16S_50ntFlanks@51.92.42|GCTAAACCTAGCCCCAAACCCACTCCACCTTACTACCAGACA</td><td>>hg38_MT_+_1621_3279_RNR2_16S_50ntFlanks</td><td>51</td><td>92</td><td>42</td><td>GCTAAACCTAGCCCCAAACCCACTCCACCTTACTACCAGACA</td><td>683.9</td><td>14.1</td><td>1.6</td><td>9.0</td><td>1.7E-19</td><td>8.4E-19</td><td>BT20/MB468</td></tr>
<tr><td>>MI0005756|hsa-mir-934&WithFlank&X|+|136550872|136550966@21.42.22(+2A)&[MIMAT0004977&hsa-miR-934&offsets|0|0(+2A);m-943&X|+|136550892|136550913&offsets|0|0(+2A)]|TGTCTACTACTGGAGACACTGGAA</td><td>>MI0005756|hsa-mir-934&WithFlank&X|+|136550872|136550966</td><td>21</td><td>42</td><td>22(+2A)</td><td>TGTCTACTACTGGAGACACTGGAA</td><td>4317.6</td><td>9.7</td><td>1.1</td><td>9.0</td><td>1.7E-19</td><td>8.4E-19</td><td>BT20/MB468</td></tr>
<tr><td>>hg38_21_+_8433172_8446622_RNA45SN1_45S_50ntFlanks@12437.12473.37|CATTGTGAAGCAGAATTCACCAAGCGTTGGATTGTTC</td><td>>hg38_21_+_8433172_8446622_RNA45SN1_45S_50ntFlanks</td><td>12437</td><td>12473</td><td>37</td><td>CATTGTGAAGCAGAATTCACCAAGCGTTGGATTGTTC</td><td>1411.3</td><td>-5.0</td><td>0.6</td><td>-9.0</td><td>1.7E-19</td><td>8.5E-19</td><td>BT20/MB468</td></tr>
<tr><td>>MI0000727|hsa-mir-128-2&WithFlank&3|+|35744470|35744565@58.79.22&[MIMAT0000424_1&hsa-miR-128-3p&offsets|0|+1;m-117&3|+|35744527|35744547&offsets|0|+1]|TCACAGTGAACCGGTCTCTTTC</td><td>>MI0000727|hsa-mir-128-2&WithFlank&3|+|35744470|35744565</td><td>58</td><td>79</td><td>22</td><td>TCACAGTGAACCGGTCTCTTTC</td><td>634.9</td><td>5.6</td><td>0.6</td><td>9.0</td><td>1.7E-19</td><td>8.5E-19</td><td>BT20/MB468</td></tr>
<tr><td>>MI0000077|hsa-mir-21&WithFlank&17|+|59841260|59841343@14.34.21&[MIMAT0000076&hsa-miR-21-5p&offsets|0|-1;m-1&17|+|59841273|59841294&offsets|0|-1]|TAGCTTATCAGACTGATGTTG</td><td>>MI0000077|hsa-mir-21&WithFlank&17|+|59841260|59841343</td><td>14</td><td>34</td><td>21</td><td>TAGCTTATCAGACTGATGTTG</td><td>48761.6</td><td>7.3</td><td>0.8</td><td>9.0</td><td>1.8E-19</td><td>8.5E-19</td><td>BT20/MB468</td></tr>
<tr><td>>MI0000471|hsa-mir-126&WithFlank&9|+|136670596|136670692@59.78.20&[MIMAT0000445&hsa-miR-126-3p&offsets|+1|-1;m-40&9|+|136670653|136670674&offsets|+1|-1]|CGTACCGTGAGTAATAATGC</td><td>>MI0000471|hsa-mir-126&WithFlank&9|+|136670596|136670692</td><td>59</td><td>78</td><td>20</td><td>CGTACCGTGAGTAATAATGC</td><td>1300.8</td><td>10.7</td><td>1.2</td><td>9.0</td><td>1.9E-19</td><td>8.5E-19</td><td>MB231/MB468</td></tr>
<tr><td>>am_MI0000809|hsa-mir-151a&WithFlank&8|-|140732558|140732659@53.71.19&[MIMAT0000757&hsa-miR-151a-3p&offsets|0|-2;m-30&8|-|140732587|140732607&offsets|0|-2]|CTAGACTGAAGCTCCTTGA</td><td>>am_MI0000809|hsa-mir-151a&WithFlank&8|-|140732558|140732659</td><td>53</td><td>71</td><td>19</td><td>CTAGACTGAAGCTCCTTGA</td><td>1481.7</td><td>9.0</td><td>1.0</td><td>9.0</td><td>1.9E-19</td><td>8.6E-19</td><td>MB231/MB468</td></tr>
<tr><td>>hg38_1_-_228634819_228635039_RNA5S12_5S_50ntFlanks@137.170.34|GAGACCGCCTGGGAATACCGGGTGCTGTAGGCTT</td><td>>hg38_1_-_228634819_228635039_RNA5S12_5S_50ntFlanks</td><td>137</td><td>170</td><td>34</td><td>GAGACCGCCTGGGAATACCGGGTGCTGTAGGCTT</td><td>1409.2</td><td>9.3</td><td>1.0</td><td>9.0</td><td>2.0E-19</td><td>8.7E-19</td><td>MB231/MB468</td></tr>
<tr><td>>hg38_1_-_228634819_228635039_RNA5S12_5S_50ntFlanks@145.170.26|CTGGGAATACCGGGTGCTGTAGGCTT</td><td>>hg38_1_-_228634819_228635039_RNA5S12_5S_50ntFlanks</td><td>145</td><td>170</td><td>26</td><td>CTGGGAATACCGGGTGCTGTAGGCTT</td><td>1409.0</td><td>6.7</td><td>0.7</td><td>9.0</td><td>1.8E-19</td><td>8.9E-19</td><td>BT20/MB468</td></tr>
<tr><td>>MI0001445|hsa-mir-423&WithFlank&17|+|30117073|30117178@23.44.22&[MIMAT0004748&hsa-miR-423-5p&offsets|0|-1;m-141&17|+|30117095|30117117&offsets|0|-1]|TGAGGGGCAGAGAGCGAGACTT</td><td>>MI0001445|hsa-mir-423&WithFlank&17|+|30117073|30117178</td><td>23</td><td>44</td><td>22</td><td>TGAGGGGCAGAGAGCGAGACTT</td><td>1671.0</td><td>9.7</td><td>1.1</td><td>9.0</td><td>2.1E-19</td><td>9.3E-19</td><td>MB231/MB468</td></tr>
<tr><td>>MI0000270|hsa-mir-181b-1&WithFlank&1|-|198858867|198858988@42.64.23(+1C);MI0000683|hsa-mir-181b-2&WithFlank&9|+|124693704|124693804@22.44.23(+1C)&[MIMAT0000257&hsa-miR-181b-5p&offsets|0|0(+1C);m-73&1|-|198858925|198858947&offsets|0|0(+1C)];[MIMAT0000257_1&hsa-miR-181b-5p&offsets|0|0(+1C);m-72&9|+|124693725|124693747&offsets|0|0(+1C)]|AACATTCATTGCTGTCGGTGGGTC</td><td>>MI0000270|hsa-mir-181b-1&WithFlank&1|-|198858867|198858988</td><td>42</td><td>64</td><td>23(+1C)</td><td>AACATTCATTGCTGTCGGTGGGTC</td><td>703.3</td><td>7.3</td><td>0.8</td><td>9.0</td><td>2.1E-19</td><td>9.3E-19</td><td>MB231/MB468</td></tr>
<tr><td>>trnaMT_SerGCT_MT_+_12207_12265@1.38.38|GAGAAAGCTCACAAGAACTGCTAACTCATGCCCCCATG</td><td>>trnaMT_SerGCT_MT_+_12207_12265</td><td>1</td><td>38</td><td>38</td><td>GAGAAAGCTCACAAGAACTGCTAACTCATGCCCCCATG</td><td>994.7</td><td>5.4</td><td>0.6</td><td>9.0</td><td>2.1E-19</td><td>9.5E-19</td><td>MB231/MB468</td></tr>
<tr><td>>hg38_21_+_8433172_8446622_RNA45SN1_45S_50ntFlanks@11873.11905.33|AAGAAGACCCTGTTGAGCTTGACTCTAGTCTGG</td><td>>hg38_21_+_8433172_8446622_RNA45SN1_45S_50ntFlanks</td><td>11873</td><td>11905</td><td>33</td><td>AAGAAGACCCTGTTGAGCTTGACTCTAGTCTGG</td><td>484.9</td><td>-7.7</td><td>0.9</td><td>-9.0</td><td>2.1E-19</td><td>9.5E-19</td><td>MB231/MB468</td></tr>
<tr><td>>hg38_21_+_8433172_8446622_RNA45SN1_45S_50ntFlanks@11999.12031.33|CCCTGCGGGCCGCCGGTGAAATACCACTACTCT</td><td>>hg38_21_+_8433172_8446622_RNA45SN1_45S_50ntFlanks</td><td>11999</td><td>12031</td><td>33</td><td>CCCTGCGGGCCGCCGGTGAAATACCACTACTCT</td><td>653.3</td><td>-11.3</td><td>1.3</td><td>-9.0</td><td>2.0E-19</td><td>9.5E-19</td><td>BT20/MB468</td></tr>
</table>

<table>
<tr><td>>am_trnaMT_PheGAA_MT_+_577_647@40.62.23|TGTTTAGACGGGCTCACATCACC</td><td>>am_trnaMT_PheGAA_MT_+_577_647</td><td>40</td><td>62</td><td>23</td><td>TGTTTAGACGGGCTCACATCACC</td><td>621.8</td><td>9.1</td><td>1.0</td><td>9.0</td><td>2.2E-19</td><td>9.6E-19</td><td>MB231/MB468</td></tr>
<tr><td>>hg38_21_+_8433172_8446622_RNA45SN1_45S_50ntFlanks@3988.4023.36|CTTTGGTGACTCTAGATAACCTCGGGCCGATCGCAC</td><td>>hg38_21_+_8433172_8446622_RNA45SN1_45S_50ntFlanks</td><td>3988</td><td>4023</td><td>36</td><td>CTTTGGTGACTCTAGATAACCTCGGGCCGATCGCAC</td><td>1277.1</td><td>6.7</td><td>0.7</td><td>9.0</td><td>2.2E-19</td><td>9.8E-19</td><td>MB231/MB468</td></tr>
<tr><td>>MI0000456|hsa-mir-140&WithFlank&16|+|69933075|69933186@68.91.24&[MIMAT0004597&hsa-miR-140-3p&offsets|0|+3;m-45&16|+|69933143|69933164&offsets|-1|+1]|TACCACAGGGTAGAACCACGGACA</td><td>>MI0000456|hsa-mir-140&WithFlank&16|+|69933075|69933186</td><td>68</td><td>91</td><td>24</td><td>TACCACAGGGTAGAACCACGGACA</td><td>819.1</td><td>10.5</td><td>1.2</td><td>9.0</td><td>2.2E-19</td><td>9.8E-19</td><td>MB231/MB468</td></tr>
<tr><td>>MI0000447|hsa-mir-128-1&WithFlank&2|+|135665391|135665484@56.75.20(+1A);MI0000727|hsa-mir-128-2&WithFlank&3|+|35744470|35744565@58.77.20(+1A)&[MIMAT0000424&hsa-miR-128-3p&offsets|0|-1(+1A);m-108&2|+|135665446|135665466&offsets|0|-1(+1A)];[MIMAT0000424_1&hsa-miR-128-3p&offsets|0|-1(+1A);m-117&3|+|35744527|35744547&offsets|0|-1(+1A)]|TCACAGTGAACCGGTCTCTTA</td><td>>MI0000447|hsa-mir-128-1&WithFlank&2|+|135665391|135665484</td><td>56</td><td>75</td><td>20(+1A)</td><td>TCACAGTGAACCGGTCTCTTA</td><td>1261.5</td><td>7.0</td><td>0.8</td><td>9.0</td><td>2.1E-19</td><td>1.0E-18</td><td>BT20/MB468</td></tr>
<tr><td>>MI0000462|hsa-mir-152&WithFlank&17|-|48037155|48037253@60.79.20(+1A)&[MIMAT0000438&hsa-miR-152-3p&offsets|0|-1(+1A);m-97&17|-|48037174|48037194&offsets|0|-1(+1A)]|TCAGTGCATGACAGAACTTGA</td><td>>MI0000462|hsa-mir-152&WithFlank&17|-|48037155|48037253</td><td>60</td><td>79</td><td>20(+1A)</td><td>TCAGTGCATGACAGAACTTGA</td><td>1133.2</td><td>14.8</td><td>1.6</td><td>9.0</td><td>2.1E-19</td><td>1.0E-18</td><td>BT20/MB468</td></tr>
<tr><td>>am_MI0000809|hsa-mir-151a&WithFlank&8|-|140732558|140732659@53.72.20(+1C)&[MIMAT0000757&hsa-miR-151a-3p&offsets|0|-1(+1C);m-30&8|-|140732587|140732607&offsets|0|-1(+1C)]|CTAGACTGAAGCTCCTTGAGC</td><td>>am_MI0000809|hsa-mir-151a&WithFlank&8|-|140732558|140732659</td><td>53</td><td>72</td><td>20(+1C)</td><td>CTAGACTGAAGCTCCTTGAGC</td><td>2020.8</td><td>9.6</td><td>1.1</td><td>9.0</td><td>2.3E-19</td><td>1.0E-18</td><td>MB231/MB468</td></tr>
<tr><td>>hg38_21_+_8433172_8446622_RNA45SN1_45S_50ntFlanks@6721.6738.18|AATTGCAGGACACATTGA</td><td>>hg38_21_+_8433172_8446622_RNA45SN1_45S_50ntFlanks</td><td>6721</td><td>6738</td><td>18</td><td>AATTGCAGGACACATTGA</td><td>358.7</td><td>-5.9</td><td>0.7</td><td>-9.0</td><td>2.4E-19</td><td>1.0E-18</td><td>MB231/MB468</td></tr>
<tr><td>>MI0000440|hsa-mir-27b&WithFlank&9|+|95085439|95085547@67.86.20(+1G)&[MIMAT0000419&hsa-miR-27b-3p&offsets|0|-1(+1G);m-39&9|+|95085505|95085525&offsets|0|-1(+1G)]|TTCACAGTGGCTAAGTTCTGG</td><td>>MI0000440|hsa-mir-27b&WithFlank&9|+|95085439|95085547</td><td>67</td><td>86</td><td>20(+1G)</td><td>TTCACAGTGGCTAAGTTCTGG</td><td>1256.6</td><td>8.0</td><td>0.9</td><td>9.0</td><td>2.4E-19</td><td>1.0E-18</td><td>MB231/MB468</td></tr>
<tr><td>>MI0000465|hsa-mir-191&WithFlank&3|-|49020612|49020715@22.43.22&[MIMAT0000440&hsa-miR-191-5p&offsets|0|-1;m-48&3|-|49020672|49020694&offsets|0|-1]|CAACGGAATCCCAAAAGCAGCT</td><td>>MI0000465|hsa-mir-191&WithFlank&3|-|49020612|49020715</td><td>22</td><td>43</td><td>22</td><td>CAACGGAATCCCAAAAGCAGCT</td><td>18599.6</td><td>6.9</td><td>0.8</td><td>9.0</td><td>2.2E-19</td><td>1.0E-18</td><td>BT20/MB468</td></tr>
<tr><td>>MI0001448|hsa-mir-425&WithFlank&3|-|49020142|49020240@20.43.24(+1G)&[MIMAT0003393&hsa-miR-425-5p&offsets|0|+1(+1G);m-95&3|-|49020200|49020221&offsets|0|+2(+1G)]|AATGACACGATCACTCCCGTTGAGG</td><td>>MI0001448|hsa-mir-425&WithFlank&3|-|49020142|49020240</td><td>20</td><td>43</td><td>24(+1G)</td><td>AATGACACGATCACTCCCGTTGAGG</td><td>909.7</td><td>9.6</td><td>1.1</td><td>9.0</td><td>2.4E-19</td><td>1.0E-18</td><td>MB231/MB468</td></tr>
<tr><td>>hg38_1_-_228634819_228635039_RNA5S12_5S_50ntFlanks@153.170.18|ACCGGGTGCTGTAGGCTT</td><td>>hg38_1_-_228634819_228635039_RNA5S12_5S_50ntFlanks</td><td>153</td><td>170</td><td>18</td><td>ACCGGGTGCTGTAGGCTT</td><td>24012.4</td><td>7.8</td><td>0.9</td><td>9.0</td><td>2.4E-19</td><td>1.1E-18</td><td>MB231/MB468</td></tr>
<tr><td>>hg38_21_+_8433172_8446622_RNA45SN1_45S_50ntFlanks@5527.5550.24|AAAGTCGTAACAAGGTTTCCGTAG</td><td>>hg38_21_+_8433172_8446622_RNA45SN1_45S_50ntFlanks</td><td>5527</td><td>5550</td><td>24</td><td>AAAGTCGTAACAAGGTTTCCGTAG</td><td>835.9</td><td>-11.7</td><td>1.3</td><td>-9.0</td><td>2.2E-19</td><td>1.1E-18</td><td>BT20/MB468</td></tr>
<tr><td>>MI0000279|hsa-mir-196a-2&WithFlank&12|+|53991732|53991853@31.51.21;MI0000238|hsa-mir-196a-1&WithFlank&17|-|48632484|48632565@13.33.21&[MIMAT0000226&hsa-miR-196a-5p&offsets|0|-1;m-261&12|+|53991762|53991783&offsets|0|-1];[MIMAT0000226_1&hsa-miR-196a-5p&offsets|0|-1;m-268&17|-|48632532|48632553&offsets|0|-1]|TAGGTAGTTTCATGTTGTTGG</td><td>>MI0000279|hsa-mir-196a-2&WithFlank&12|+|53991732|53991853</td><td>31</td><td>51</td><td>21</td><td>TAGGTAGTTTCATGTTGTTGG</td><td>927.1</td><td>13.4</td><td>1.4</td><td>9.3</td><td>1.2E-20</td><td>1.1E-18</td><td>BT20/MB231</td></tr>
<tr><td>>MI0000070|hsa-mir-16-1&WithFlank&13|-|50048967|50049067@20.42.23;MI0000115|hsa-mir-16-2&WithFlank&3|+|160404739|160404831@16.38.23&[MIMAT0000069&hsa-miR-16-5p&offsets|0|+1;m-62&13|-|50049027|50049048&offsets|0|+1];[MIMAT0000069_1&hsa-miR-16-5p&offsets|0|+1;m-61&3|+|160404754|160404775&offsets|0|+1]|TAGCAGCACGTAAATATTGGCGT</td><td>>MI0000070|hsa-mir-16-1&WithFlank&13|-|50048967|50049067</td><td>20</td><td>42</td><td>23</td><td>TAGCAGCACGTAAATATTGGCGT</td><td>5898.8</td><td>7.2</td><td>0.8</td><td>9.0</td><td>2.4E-19</td><td>1.1E-18</td><td>MB231/MB468</td></tr>
<tr><td>>hg38_21_+_8433172_8446622_RNA45SN1_45S_50ntFlanks@3993.4023.31|GTGACTCTAGATAACCTCGGGCCGATCGCAC</td><td>>hg38_21_+_8433172_8446622_RNA45SN1_45S_50ntFlanks</td><td>3993</td><td>4023</td><td>31</td><td>GTGACTCTAGATAACCTCGGGCCGATCGCAC</td><td>765.4</td><td>4.5</td><td>0.5</td><td>9.0</td><td>2.2E-19</td><td>1.1E-18</td><td>BT20/MB468</td></tr>
<tr><td>>trnaMT_GluTTC_MT_-_14674_14742@32.72.41,trnalookalike8_GluTTC_5_-_93905172_93905240@32.72.41|TCATATCATTGGTCGTGGTTGTAGTCCGTGCGAGAATACCA</td><td>>trnaMT_GluTTC_MT_-_14674_14742</td><td>32</td><td>72</td><td>41</td><td>TCATATCATTGGTCGTGGTTGTAGTCCGTGCGAGAATACCA</td><td>919.1</td><td>10.4</td><td>1.2</td><td>9.0</td><td>2.2E-19</td><td>1.1E-18</td><td>BT20/MB468</td></tr>
<tr><td>>MI0000088|hsa-mir-30a&WithFlank&6|-|71403545|71403627@12.35.24&[MIMAT0000087&hsa-miR-30a-5p&offsets|0|+2;m-15&6|-|71403595|71403616&offsets|0|+2]|TGTAAACATCCTCGACTGGAAGCT</td><td>>MI0000088|hsa-mir-30a&WithFlank&6|-|71403545|71403627</td><td>12</td><td>35</td><td>24</td><td>TGTAAACATCCTCGACTGGAAGCT</td><td>298276.3</td><td>7.8</td><td>0.9</td><td>9.0</td><td>2.2E-19</td><td>1.1E-18</td><td>BT20/MB468</td></tr>
<tr><td>>hg38_21_+_8433172_8446622_RNA45SN1_45S_50ntFlanks@485.527.43|GATGTGGTGACGTCGTGCTCTCCCGGGCCGGGTCCGAGCCGCG</td><td>>hg38_21_+_8433172_8446622_RNA45SN1_45S_50ntFlanks</td><td>485</td><td>527</td><td>43</td><td>GATGTGGTGACGTCGTGCTCTCCCGGGCCGGGTCCGAGCCGCG</td><td>876.7</td><td>14.4</td><td>1.6</td><td>9.0</td><td>2.2E-19</td><td>1.1E-18</td><td>BT20/MB468</td></tr>
<tr><td>>am_trna117_GlyTCC_1_-_145397864_145397935@1.30.30,trna45_GlyTCC_1_+_161500903_161500974@1.30.30,trna70_GlyTCC_1_-_161439547_161439618@1.30.30,trna73_GlyTCC_1_-_161432166_161432237@1.30.30,trna76_GlyTCC_1_-_161424756_161424827@1.30.30,trna79_GlyTCC_1_-_161417375_161417446@1.30.30|GCGTTGGTGGTATAGTGGTGAGCATAGCTG</td><td>>am_trna117_GlyTCC_1_-_145397864_145397935</td><td>1</td><td>30</td><td>30</td><td>GCGTTGGTGGTATAGTGGTGAGCATAGCTG</td><td>1309.9</td><td>-5.8</td><td>0.6</td><td>-9.0</td><td>2.3E-19</td><td>1.1E-18</td><td>BT20/MB468</td></tr>
<tr><td>>hg38_21_+_8433172_8446622_RNA45SN1_45S_50ntFlanks@6650.6677.28|ACGACTCTTAGCGGTGGATCACTCGGCT</td><td>>hg38_21_+_8433172_8446622_RNA45SN1_45S_50ntFlanks</td><td>6650</td><td>6677</td><td>28</td><td>ACGACTCTTAGCGGTGGATCACTCGGCT</td><td>641.3</td><td>3.8</td><td>0.4</td><td>9.0</td><td>2.3E-19</td><td>1.1E-18</td><td>BT20/MB468</td></tr>
<tr><td>>MI0000434|hsa-let-7i&WithFlank&12|+|62603680|62603775@12.33.22&[MIMAT0000415&hsa-let-7i-5p&offsets|0|0;m-49&12|+|62603691|62603712&offsets|0|0]|TGAGGTAGTAGTTTGTGCTGTT</td><td>>MI0000434|hsa-let-7i&WithFlank&12|+|62603680|62603775</td><td>12</td><td>33</td><td>22</td><td>TGAGGTAGTAGTTTGTGCTGTT</td><td>194232.5</td><td>6.1</td><td>0.7</td><td>9.0</td><td>2.3E-19</td><td>1.1E-18</td><td>BT20/MB468</td></tr>
<tr><td>>hg38_21_+_8433172_8446622_RNA45SN1_45S_50ntFlanks@12633.12652.20|CGAAGCTACCATCTGTGGGA</td><td>>hg38_21_+_8433172_8446622_RNA45SN1_45S_50ntFlanks</td><td>12633</td><td>12652</td><td>20</td><td>CGAAGCTACCATCTGTGGGA</td><td>780.4</td><td>-11.6</td><td>1.3</td><td>-9.0</td><td>2.4E-19</td><td>1.1E-18</td><td>BT20/MB468</td></tr>
<tr><td>>hg38_21_+_8433172_8446622_RNA45SN1_45S_50ntFlanks@6772.6799.28|GTTCCTCCCGGGGCTACGCCTGTCTGAG</td><td>>hg38_21_+_8433172_8446622_RNA45SN1_45S_50ntFlanks</td><td>6772</td><td>6799</td><td>28</td><td>GTTCCTCCCGGGGCTACGCCTGTCTGAG</td><td>862.8</td><td>-11.7</td><td>1.3</td><td>-9.0</td><td>2.4E-19</td><td>1.1E-18</td><td>BT20/MB468</td></tr>
<tr><td>>hg38_21_+_8433172_8446622_RNA45SN1_45S_50ntFlanks@10391.10419.29|AACAGCAGTTGAACATGGGTCAGTCGGTC</td><td>>hg38_21_+_8433172_8446622_RNA45SN1_45S_50ntFlanks</td><td>10391</td><td>10419</td><td>29</td><td>AACAGCAGTTGAACATGGGTCAGTCGGTC</td><td>283.4</td><td>-4.6</td><td>0.5</td><td>-9.0</td><td>2.6E-19</td><td>1.1E-18</td><td>MB231/MB468</td></tr>
<tr><td>>trna10_LysCTT_16_+_3241501_3241573@4.33.30,trna13_LysCTT_14_-_58706613_58706685@4.33.30,trna2_LysCTT_15_+_79152904_79152976@4.33.30|CGGCTAGCTCAGTCGGTAGAGCATGGGACT</td><td>>trna10_LysCTT_16_+_3241501_3241573</td><td>4</td><td>33</td><td>30</td><td>CGGCTAGCTCAGTCGGTAGAGCATGGGACT</td><td>2389.6</td><td>11.5</td><td>1.3</td><td>9.0</td><td>2.6E-19</td><td>1.1E-18</td><td>MB231/MB468</td></tr>
<tr><td>>hg38_21_+_8433172_8446622_RNA45SN1_45S_50ntFlanks@10388.10410.23|GTGAACAGCAGTTGAACATGGGT</td><td>>hg38_21_+_8433172_8446622_RNA45SN1_45S_50ntFlanks</td><td>10388</td><td>10410</td><td>23</td><td>GTGAACAGCAGTTGAACATGGGT</td><td>398.9</td><td>-5.4</td><td>0.6</td><td>-9.0</td><td>2.6E-19</td><td>1.1E-18</td><td>MB231/MB468</td></tr>
<tr><td>>MI0000289|hsa-mir-181a-1&WithFlank&1|-|198859038|198859159@30.51.22(+1A);MI0000269|hsa-mir-181a-2&WithFlank&9|+|124692436|124692557@45.66.22(+1A)&[MIMAT0000256&hsa-miR-181a-5p&offsets|0|-1(+1A);m-32&1|-|198859108|198859130&offsets|0|-1(+1A)];[MIMAT0000256_1&hsa-miR-181a-5p&offsets|0|-1(+1A);m-32&9|+|124692480|124692502&offsets|0|-1(+1A)]|AACATTCAACGCTGTCGGTGAGA</td><td>>MI0000289|hsa-mir-181a-1&WithFlank&1|-|198859038|198859159</td><td>30</td><td>51</td><td>22(+1A)</td><td>AACATTCAACGCTGTCGGTGAGA</td><td>1111.8</td><td>6.8</td><td>0.8</td><td>9.0</td><td>2.6E-19</td><td>1.1E-18</td><td>MB231/MB468</td></tr>
<tr><td>>hg38_21_+_8433172_8446622_RNA45SN1_45S_50ntFlanks@12417.12447.31|TCGATGTCGGCTCTTCCTATCATTGTGAAGC</td><td>>hg38_21_+_8433172_8446622_RNA45SN1_45S_50ntFlanks</td><td>12417</td><td>12447</td><td>31</td><td>TCGATGTCGGCTCTTCCTATCATTGTGAAGC</td><td>647.0</td><td>5.5</td><td>0.6</td><td>9.0</td><td>2.6E-19</td><td>1.1E-18</td><td>MB231/MB468</td></tr>
<tr><td>>hg38_21_+_8433172_8446622_RNA45SN1_45S_50ntFlanks@6648.6694.47|GTACGACTCTTAGCGGTGGATCACTCGGCTCGTGCGTCGATGAAGAA</td><td>>hg38_21_+_8433172_8446622_RNA45SN1_45S_50ntFlanks</td><td>6648</td><td>6694</td><td>47</td><td>GTACGACTCTTAGCGGTGGATCACTCGGCTCGTGCGTCGATGAAGAA</td><td>412.6</td><td>8.5</td><td>0.9</td><td>9.0</td><td>2.6E-19</td><td>1.1E-18</td><td>MB231/MB468</td></tr>
<tr><td>>MI0000061|hsa-let-7a-2&WithFlank&11|-|122146516|122146599@11.33.23;MI0000062|hsa-let-7a-3&WithFlank&22|+|46112743|46112828@10.32.23;MI0000060|hsa-let-7a-1&WithFlank&9|+|94175951|94176042@12.34.23&[MIMAT0000062&hsa-let-7a-5p&offsets|0|+1;m-6&11|-|122146568|122146589&offsets|0|+1];[MIMAT0000062_1&hsa-let-7a-5p&offsets|0|+1;m-5&22|+|46112752|46112773&offsets|0|+1];[MIMAT0000062_2&hsa-let-7a-5p&offsets|0|+1;m-4&9|+|94175962|94175983&offsets|0|+1]|TGAGGTAGTAGGTTGTATAGTTT</td><td>>MI0000061|hsa-let-7a-2&WithFlank&11|-|122146516|122146599</td><td>11</td><td>33</td><td>23</td><td>TGAGGTAGTAGGTTGTATAGTTT</td><td>4264.3</td><td>6.9</td><td>0.8</td><td>9.0</td><td>2.4E-19</td><td>1.2E-18</td><td>BT20/MB468</td></tr>
<tr><td>>hg38_21_+_8433172_8446622_RNA45SN1_45S_50ntFlanks@476.517.42|TTCGTGATCGATGTGGTGACGTCGTGCTCTCCCGGGCCGGGT</td><td>>hg38_21_+_8433172_8446622_RNA45SN1_45S_50ntFlanks</td><td>476</td><td>517</td><td>42</td><td>TTCGTGATCGATGTGGTGACGTCGTGCTCTCCCGGGCCGGGT</td><td>1316.1</td><td>7.9</td><td>0.9</td><td>9.0</td><td>2.5E-19</td><td>1.2E-18</td><td>BT20/MB468</td></tr>
<tr><td>>MI0000067|hsa-let-7f-1&WithFlank&9|+|94176341|94176439@13.30.18;MI0000068|hsa-let-7f-2&WithFlank&X|-|53557186|53557280@14.31.18&[MIMAT0000067&hsa-let-7f-5p&offsets|0|-4;m-20&9|+|94176353|94176374&offsets|0|-4];[MIMAT0000067_1&hsa-let-7f-5p&offsets|0|-4;m-19&X|-|53557246|53557267&offsets|0|-4]|TGAGGTAGTAGATTGTAT</td><td>>MI0000067|hsa-let-7f-1&WithFlank&9|+|94176341|94176439</td><td>13</td><td>30</td><td>18</td><td>TGAGGTAGTAGATTGTAT</td><td>11569.2</td><td>7.5</td><td>0.8</td><td>9.0</td><td>2.7E-19</td><td>1.2E-18</td><td>MB231/MB468</td></tr>
<tr><td>>am_trnaMT_LysTTT_MT_+_8295_8364@29.52.24,trnalookalike7_LysTTT_1_+_568844_568913@29.52.24|TTTAAGTTAAAGATTAAGAGAACC</td><td>>am_trnaMT_LysTTT_MT_+_8295_8364</td><td>29</td><td>52</td><td>24</td><td>TTTAAGTTAAAGATTAAGAGAACC</td><td>818.6</td><td>9.7</td><td>1.1</td><td>9.0</td><td>2.5E-19</td><td>1.2E-18</td><td>BT20/MB468</td></tr>
<tr><td>>hg38_21_+_8433172_8446622_RNA45SN1_45S_50ntFlanks@10350.10381.32|ACGAGAACTTTGAAGGCCGAAGTGGAGAAGGG</td><td>>hg38_21_+_8433172_8446622_RNA45SN1_45S_50ntFlanks</td><td>10350</td><td>10381</td><td>32</td><td>ACGAGAACTTTGAAGGCCGAAGTGGAGAAGGG</td><td>190.5</td><td>-3.9</td><td>0.4</td><td>-9.0</td><td>2.5E-19</td><td>1.2E-18</td><td>BT20/MB468</td></tr>
</table>

| | | | | | | | | | | | | |
|---|---|---|---|---|---|---|---|---|---|---|---|---|
| >MI0000234\|hsa-mir-192&WithFlank&11\|-\|64891131\|64891252@30.49.20(+1A)&[MIMAT0000222&hsa-miR-192-5p&offsets\|0\|-1(+1A);m-59&11\|-\|64891203\|64891223&offsets\|0\|-1(+1A)]\|CTGACCTATGAATTGACAGCA | >MI0000234\|hsa-mir-192&WithFlank&11\|-\|64891131\|64891252 | 30 | 49 | 20(+1A) | CTGACCTATGAATTGACAGCA | 882.3 | 7.4 | 0.8 | 9.0 | 2.5E-19 | 1.2E-18 | BT20/MB468 |
| >am_trnaMT_PheGAA_MT_+_577_647@40.58.19\|TGTTTAGACGGGCTCACAT | >am_trnaMT_PheGAA_MT_+_577_647 | 40 | 58 | 19 | TGTTTAGACGGGCTCACAT | 407.3 | 8.9 | 1.0 | 9.0 | 2.5E-19 | 1.2E-18 | BT20/MB468 |
| >trnaMT_PheGAA_MT_+_577_647@38.62.25\|AATGTTTAGACGGGCTCACATCACC | >trnaMT_PheGAA_MT_+_577_647 | 38 | 62 | 25 | AATGTTTAGACGGGCTCACATCACC | 448.9 | 9.1 | 1.0 | 9.0 | 2.6E-19 | 1.2E-18 | BT20/MB468 |
| >MI0000061\|hsa-let-7a-2&WithFlank&11\|-\|122146516\|122146599@11.30.20(+1G);MI0000062\|hsa-let-7a-3&WithFlank&22\|+\|46112743\|46112828@10.29.20(+1G);MI0000060\|hsa-let-7a-1&WithFlank&9\|+\|94175951\|94176042@12.31.20(+1G)&[MIMAT0000062&hsa-let-7a-5p&offsets\|0\|-2(+1G);m-6&11\|-\|122146568\|122146589&offsets\|0\|-2(+1G)];[MIMAT0000062_1&hsa-let-7a-5p&offsets\|0\|-2(+1G);m-5&22\|+\|46112752\|46112773&offsets\|0\|-2(+1G)];[MIMAT0000062_2&hsa-let-7a-5p&offsets\|0\|-2(+1G);m-4&9\|+\|94175962\|94175983&offsets\|0\|-2(+1G)]\|TGAGGTAGTAGGTTGTATAGG | >MI0000061\|hsa-let-7a-2&WithFlank&11\|-\|122146516\|122146599 | 11 | 30 | 20(+1G) | TGAGGTAGTAGGTTGTATAGG | 1251.9 | 8.2 | 0.9 | 9.0 | 2.6E-19 | 1.2E-18 | BT20/MB468 |
| >MI0000743\|hsa-mir-34c&WithFlank&11\|+\|111513433\|111513521@19.41.23&[MIMAT0000686&hsa-miR-34c-5p&offsets\|0\|0;m-336&11\|+\|111513451\|111513473&offsets\|0\|0]\|AGGCAGTGTAGTTAGCTGATTGC | >MI0000743\|hsa-mir-34c&WithFlank&11\|+\|111513433\|111513521 | 19 | 41 | 23 | AGGCAGTGTAGTTAGCTGATTGC | 901.7 | -13.2 | 1.4 | -9.3 | 1.5E-20 | 1.3E-18 | BT20/MB231 |
| >hg38_21_+_8433172_8446622_RNA45SN1_45S_50ntFlanks@6743.6771.29\|CGACACTTCGAACGCACTTGCGGCCCCGG | >hg38_21_+_8433172_8446622_RNA45SN1_45S_50ntFlanks | 6743 | 6771 | 29 | CGACACTTCGAACGCACTTGCGGCCCCGG | 1582.7 | -7.1 | 0.8 | -9.0 | 2.7E-19 | 1.3E-18 | BT20/MB468 |
| >MI0000465\|hsa-mir-191&WithFlank&3\|-\|49020612\|49020715@22.44.23(+1C)&[MIMAT0000440&hsa-miR-191-5p&offsets\|0\|0(+1C);m-48&3\|-\|49020672\|49020694&offsets\|0\|0(+1C)]\|CAACGGAATCCCAAAAGCAGCTGC | >MI0000465\|hsa-mir-191&WithFlank&3\|-\|49020612\|49020715 | 22 | 44 | 23(+1C) | CAACGGAATCCCAAAAGCAGCTGC | 590.6 | 6.4 | 0.7 | 9.0 | 2.7E-19 | 1.3E-18 | BT20/MB468 |
| >hg38_21_+_8433172_8446622_RNA45SN1_45S_50ntFlanks@6648.6678.31\|GTACGACTCTTAGCGGTGGATCACTCGGCTC | >hg38_21_+_8433172_8446622_RNA45SN1_45S_50ntFlanks | 6648 | 6678 | 31 | GTACGACTCTTAGCGGTGGATCACTCGGCTC | 935.0 | 4.8 | 0.5 | 9.0 | 2.7E-19 | 1.3E-18 | BT20/MB468 |
| >MI0000760\|hsa-mir-361&WithFlank&X\|-\|85903630\|85903713@51.74.24&[MIMAT0004682&hsa-miR-361-3p&offsets\|0\|+1;m-124&X\|-\|85903642\|85903663&offsets\|0\|+2]\|TCCCCCAGGTGTGATTCTGATTTG | >MI0000760\|hsa-mir-361&WithFlank&X\|-\|85903630\|85903713 | 51 | 74 | 24 | TCCCCCAGGTGTGATTCTGATTTG | 590.3 | 6.5 | 0.7 | 9.0 | 2.7E-19 | 1.3E-18 | BT20/MB468 |
| >hg38_21_+_8433172_8446622_RNA45SN1_45S_50ntFlanks@11781.11804.24\|CGCATGAATGGATGAACGAGATTC | >hg38_21_+_8433172_8446622_RNA45SN1_45S_50ntFlanks | 11781 | 11804 | 24 | CGCATGAATGGATGAACGAGATTC | 2431.2 | -6.5 | 0.7 | -9.0 | 3.0E-19 | 1.3E-18 | MB231/MB468 |
| >hg38_21_+_8433172_8446622_RNA45SN1_45S_50ntFlanks@6719.6746.28\|TGAATTGCAGGACACATTGATCATCGAC | >hg38_21_+_8433172_8446622_RNA45SN1_45S_50ntFlanks | 6719 | 6746 | 28 | TGAATTGCAGGACACATTGATCATCGAC | 804.1 | -4.5 | 0.5 | -9.0 | 3.0E-19 | 1.3E-18 | MB231/MB468 |
| >hg38_21_+_8433172_8446622_RNA45SN1_45S_50ntFlanks@6719.6744.26\|TGAATTGCAGGACACATTGATCATCG | >hg38_21_+_8433172_8446622_RNA45SN1_45S_50ntFlanks | 6719 | 6744 | 26 | TGAATTGCAGGACACATTGATCATCG | 7856.9 | -5.5 | 0.6 | -9.0 | 3.0E-19 | 1.3E-18 | MB231/MB468 |
| >hg38_21_+_8433172_8446622_RNA45SN1_45S_50ntFlanks@10346.10368.23\|TCAAACGAGAACTTTGAAGGCCG | >hg38_21_+_8433172_8446622_RNA45SN1_45S_50ntFlanks | 10346 | 10368 | 23 | TCAAACGAGAACTTTGAAGGCCG | 1962.0 | -4.0 | 0.4 | -9.0 | 2.8E-19 | 1.3E-18 | BT20/MB468 |
| >hg38_21_+_8433172_8446622_RNA45SN1_45S_50ntFlanks@8381.8400.20\|AAGAGGGCGTGAAACCGTTA | >hg38_21_+_8433172_8446622_RNA45SN1_45S_50ntFlanks | 8381 | 8400 | 20 | AAGAGGGCGTGAAACCGTTA | 1080.7 | -8.9 | 1.0 | -9.0 | 2.8E-19 | 1.3E-18 | BT20/MB468 |
| >MI0000070\|hsa-mir-16-1&WithFlank&13\|-\|50048967\|50049067@20.41.22;MI0000115\|hsa-mir-16-2&WithFlank&3\|+\|160404739\|160404831@16.37.22&[MIMAT0000069&hsa-miR-16-5p&offsets\|0\|0;m-62&13\|-\|50049027\|50049048&offsets\|0\|0];[MIMAT0000069_1&hsa-miR-16-5p&offsets\|0\|0;m-61&3\|+\|160404754\|160404775&offsets\|0\|0]\|TAGCAGCACGTAAATATTGGCG | >MI0000070\|hsa-mir-16-1&WithFlank&13\|-\|50048967\|50049067 | 20 | 41 | 22 | TAGCAGCACGTAAATATTGGCG | 6830.1 | 7.1 | 0.8 | 9.0 | 3.0E-19 | 1.3E-18 | MB231/MB468 |
| >trnaMT_PheGAA_MT_+_577_647@35.56.22\|GAAAATGTTTAGACGGGCTCAC | >trnaMT_PheGAA_MT_+_577_647 | 35 | 56 | 22 | GAAAATGTTTAGACGGGCTCAC | 1387.1 | 11.4 | 1.3 | 9.0 | 2.8E-19 | 1.3E-18 | BT20/MB468 |
| >hg38_21_+_8433172_8446622_RNA45SN1_45S_50ntFlanks@6747.6770.24\|ACTTCGAACGCACTTGCGGCCCCG | >hg38_21_+_8433172_8446622_RNA45SN1_45S_50ntFlanks | 6747 | 6770 | 24 | ACTTCGAACGCACTTGCGGCCCCG | 388.7 | -5.6 | 0.6 | -9.0 | 3.1E-19 | 1.3E-18 | MB231/MB468 |
| >hg38_21_+_8433172_8446622_RNA45SN1_45S_50ntFlanks@6652.6676.25\|GACTCTTAGCGGTGGATCACTCGGC | >hg38_21_+_8433172_8446622_RNA45SN1_45S_50ntFlanks | 6652 | 6676 | 25 | GACTCTTAGCGGTGGATCACTCGGC | 1145.2 | 6.9 | 0.8 | 9.0 | 3.1E-19 | 1.3E-18 | MB231/MB468 |
| >MI0000067\|hsa-let-7f-1&WithFlank&9\|+\|94176341\|94176439@13.34.22(+1A);MI0000068\|hsa-let-7f-2&WithFlank&X\|-\|53557186\|53557280@14.35.22(+1A)&[MIMAT0000067&hsa-let-7f-5p&offsets\|0\|0(+1A);m-20&9\|+\|94176353\|94176374&offsets\|0\|0(+1A)];[MIMAT0000067_1&hsa-let-7f-5p&offsets\|0\|0(+1A);m-19&X\|-\|53557246\|53557267&offsets\|0\|0(+1A)]\|TGAGGTAGTAGATTGTATAGTTA | >MI0000067\|hsa-let-7f-1&WithFlank&9\|+\|94176341\|94176439 | 13 | 34 | 22(+1A) | TGAGGTAGTAGATTGTATAGTTA | 4050.6 | 6.2 | 0.7 | 9.0 | 3.1E-19 | 1.3E-18 | MB231/MB468 |
| >MI0001445\|hsa-mir-423&WithFlank&17\|+\|30117073\|30117178@58.81.24(+1U)&[MIMAT0001340&hsa-miR-423-3p&offsets\|-1\|0(+1U);m-100&17\|+\|30117131\|30117153&offsets\|-1\|0(+1U)]\|AAGCTCGGTCTGAGGCCCCTCAGTT | >MI0001445\|hsa-mir-423&WithFlank&17\|+\|30117073\|30117178 | 58 | 81 | 24(+1U) | AAGCTCGGTCTGAGGCCCCTCAGTT | 1241.2 | 9.6 | 1.1 | 9.0 | 3.1E-19 | 1.3E-18 | MB231/MB468 |
| >hg38_21_+_8433172_8446622_RNA45SN1_45S_50ntFlanks@11641.11665.25\|CGGCGGGTGTTGACGCGATGTGATT | >hg38_21_+_8433172_8446622_RNA45SN1_45S_50ntFlanks | 11641 | 11665 | 25 | CGGCGGGTGTTGACGCGATGTGATT | 3695.2 | -7.6 | 0.9 | -9.0 | 2.9E-19 | 1.4E-18 | BT20/MB468 |
| >hg38_21_+_8433172_8446622_RNA45SN1_45S_50ntFlanks@11323.11342.20\|CGTCTCGTCGCGCGCGCGTC | >hg38_21_+_8433172_8446622_RNA45SN1_45S_50ntFlanks | 11323 | 11342 | 20 | CGTCTCGTCGCGCGCGCGTC | 3059.1 | -8.1 | 0.9 | -9.0 | 3.2E-19 | 1.4E-18 | MB231/MB468 |
| >hg38_21_+_8433172_8446622_RNA45SN1_45S_50ntFlanks@11781.11806.26\|CGCATGAATGGATGAACGAGATTCCC | >hg38_21_+_8433172_8446622_RNA45SN1_45S_50ntFlanks | 11781 | 11806 | 26 | CGCATGAATGGATGAACGAGATTCCC | 677.2 | -11.4 | 1.3 | -9.0 | 2.9E-19 | 1.4E-18 | BT20/MB468 |
| >MI0000087\|hsa-mir-29a&WithFlank&7\|-\|130876741\|130876816@47.67.21(+1G)&[MIMAT0000086&hsa-miR-29a-3p&offsets\|-1\|-2(+1G);m-26&7\|-\|130876749\|130876769&offsets\|-1\|-1(+1G)]\|CTAGCACCATCTGAAATCGGTG | >MI0000087\|hsa-mir-29a&WithFlank&7\|-\|130876741\|130876816 | 47 | 67 | 21(+1G) | CTAGCACCATCTGAAATCGGTG | 1060.4 | 9.6 | 1.1 | 9.0 | 2.9E-19 | 1.4E-18 | BT20/MB468 |
| >MI0000434\|hsa-let-7i&WithFlank&12\|+\|62603680\|62603775@12.34.23&[MIMAT0000415&hsa-let-7i-5p&offsets\|0\|+1;m-49&12\|+\|62603691\|62603712&offsets\|0\|+1]\|TGAGGTAGTAGTTTGTGCTGTTG | >MI0000434\|hsa-let-7i&WithFlank&12\|+\|62603680\|62603775 | 12 | 34 | 23 | TGAGGTAGTAGTTTGTGCTGTTG | 1010.6 | 7.0 | 0.8 | 9.0 | 2.9E-19 | 1.4E-18 | BT20/MB468 |
| >hg38_21_+_8433172_8446622_RNA45SN1_45S_50ntFlanks@6651.6683.33\|CGACTCTTAGCGGTGGATCACTCGGCTCGTGCG | >hg38_21_+_8433172_8446622_RNA45SN1_45S_50ntFlanks | 6651 | 6683 | 33 | CGACTCTTAGCGGTGGATCACTCGGCTCGTGCG | 16977.9 | 6.2 | 0.7 | 9.0 | 3.2E-19 | 1.4E-18 | MB231/MB468 |
| >am_MI0000809\|hsa-mir-151a&WithFlank&8\|-\|140732558\|140732659@52.72.21(+1A)&[MIMAT0000757&hsa-miR-151a-3p&offsets\|-1\|-1(+1A);m-30&8\|-\|140732587\|140732607&offsets\|-1\|-1(+1A)]\|ACTAGACTGAAGCTCCTTGAGA | >am_MI0000809\|hsa-mir-151a&WithFlank&8\|-\|140732558\|140732659 | 52 | 72 | 21(+1A) | ACTAGACTGAAGCTCCTTGAGA | 511.1 | 10.6 | 1.2 | 9.0 | 3.0E-19 | 1.4E-18 | BT20/MB468 |
| >MI0000077\|hsa-mir-21&WithFlank&17\|+\|59841260\|59841343@14.34.21&[MIMAT0000076&hsa-miR-21-5p&offsets\|0\|-1;m-1&17\|+\|59841273\|59841294&offsets\|0\|-1]\|TAGCTTATCAGACTGATGTTG | >MI0000077\|hsa-mir-21&WithFlank&17\|+\|59841260\|59841343 | 14 | 34 | 21 | TAGCTTATCAGACTGATGTTG | 33299.2 | 7.0 | 0.8 | 9.0 | 3.3E-19 | 1.4E-18 | MB231/MB468 |
| >MI0000061\|hsa-let-7a-2&WithFlank&11\|-\|122146516\|122146599@11.32.22(+1A);MI0000062\|hsa-let-7a-3&WithFlank&22\|+\|46112743\|46112828@10.31.22(+1A);MI0000060\|hsa-let-7a-1&WithFlank&9\|+\|94175951\|94176042@12.33.22(+1A)&[MIMAT0000062&hsa-let-7a-5p&offsets\|0\|0(+1A);m-6&11\|-\|122146568\|122146589&offsets\|0\|0(+1A)];[MIMAT0000062_1&hsa-let-7a-5p&offsets\|0\|0(+1A);m-5&22\|+\|46112752\|46112773&offsets\|0\|0(+1A)];[MIMAT0000062_2&hsa-let-7a-5p&offsets\|0\|0(+1A);m-4&9\|+\|94175962\|94175983&offsets\|0\|0(+1A)]\|TGAGGTAGTAGGTTGTATAGTTA | >MI0000061\|hsa-let-7a-2&WithFlank&11\|-\|122146516\|122146599 | 11 | 32 | 22(+1A) | TGAGGTAGTAGGTTGTATAGTTA | 3890.9 | 7.2 | 0.8 | 9.0 | 3.0E-19 | 1.4E-18 | BT20/MB468 |
| >hg38_21_+_8433172_8446622_RNA45SN1_45S_50ntFlanks@9572.9595.24\|CAAATCGGTCGTCCGACCTGGGTA | >hg38_21_+_8433172_8446622_RNA45SN1_45S_50ntFlanks | 9572 | 9595 | 24 | CAAATCGGTCGTCCGACCTGGGTA | 2689.9 | -5.9 | 0.7 | -9.0 | 3.3E-19 | 1.4E-18 | MB231/MB468 |
| >MI0005566\|hsa-mir-374b&WithFlank&X\|-\|74218541\|74218624@17.37.21&[MIMAT0004955&hsa-miR-374b-5p&offsets\|0\|-1;m-153&X\|-\|74218587\|74218608&offsets\|0\|-1]\|ATATAATACAACCTGCTAAGT | >MI0005566\|hsa-mir-374b&WithFlank&X\|-\|74218541\|74218624 | 17 | 37 | 21 | ATATAATACAACCTGCTAAGT | 2676.9 | 7.2 | 0.8 | 9.0 | 3.3E-19 | 1.4E-18 | MB231/MB468 |
| >MI0003834\|hsa-mir-769&WithFlank&19\|+\|46018926\|46019055@36.57.22&[MIMAT0003886&hsa-miR-769-5p&offsets\|0\|0;m-216&19\|+\|46018961\|46018982&offsets\|0\|0]\|TGAGACCTCTGGGTTCTGAGCT | >MI0003834\|hsa-mir-769&WithFlank&19\|+\|46018926\|46019055 | 36 | 57 | 22 | TGAGACCTCTGGGTTCTGAGCT | 7121.6 | 8.3 | 0.9 | 9.0 | 3.1E-19 | 1.4E-18 | BT20/MB468 |
| >trna10_MetCAT_8_-_124169470_124169542@31.51.21,trna22_MetCAT_16_-_87417628_87417700@31.51.21\|TCTCATAATCTGAAGGTCGTG | >trna10_MetCAT_8_-_124169470_124169542 | 31 | 51 | 21 | TCTCATAATCTGAAGGTCGTG | 978.0 | -11.9 | 1.3 | -9.0 | 3.1E-19 | 1.5E-18 | BT20/MB468 |
| >MI0000272\|hsa-mir-182&WithFlank&7\|-\|129770377\|129770498@29.52.24(+1U)&[MIMAT0000259&hsa-miR-182-5p&offsets\|0\|0(+1U);m-22&7\|-\|129770449\|129770470&offsets\|0\|+2(+1U)]\|TTTGGCAATGGTAGAACTCACACTT | >MI0000272\|hsa-mir-182&WithFlank&7\|-\|129770377\|129770498 | 29 | 52 | 24(+1U) | TTTGGCAATGGTAGAACTCACACTT | 1133.0 | 9.6 | 1.1 | 9.0 | 3.1E-19 | 1.5E-18 | BT20/MB468 |
| >hg38_21_+_8433172_8446622_RNA45SN1_45S_50ntFlanks@11641.11669.29\|CGGCGGGTGTTGACGCGATGTGATTTCTG | >hg38_21_+_8433172_8446622_RNA45SN1_45S_50ntFlanks | 11641 | 11669 | 29 | CGGCGGGTGTTGACGCGATGTGATTTCTG | 1381.9 | -8.1 | 0.9 | -9.0 | 3.4E-19 | 1.5E-18 | MB231/MB468 |
| >hg38_21_+_8433172_8446622_RNA45SN1_45S_50ntFlanks@11894.11912.19\|ACTCTAGTCTGGCACGGTG | >hg38_21_+_8433172_8446622_RNA45SN1_45S_50ntFlanks | 11894 | 11912 | 19 | ACTCTAGTCTGGCACGGTG | 1273.6 | -8.5 | 1.0 | -9.0 | 3.2E-19 | 1.5E-18 | BT20/MB468 |
| >trna15_ValAAC_5_-_180615416_180615488@1.32.32\|GTTTCCGTAGTGTAGTGGTCATCACGTTCGCC | >trna15_ValAAC_5_-_180615416_180615488 | 1 | 32 | 32 | GTTTCCGTAGTGTAGTGGTCATCACGTTCGCC | 1685.9 | 10.2 | 1.1 | 9.0 | 3.5E-19 | 1.5E-18 | MB231/MB468 |
| >hg38_21_+_8433172_8446622_RNA45SN1_45S_50ntFlanks@6700.6719.20\|CTAGCTGCGAGAATTAATGT | >hg38_21_+_8433172_8446622_RNA45SN1_45S_50ntFlanks | 6700 | 6719 | 20 | CTAGCTGCGAGAATTAATGT | 414.4 | -5.7 | 0.6 | -9.0 | 3.3E-19 | 1.5E-18 | BT20/MB468 |

<table>
<tr><th></th><th></th><th></th><th></th><th></th><th></th><th></th><th></th><th></th><th></th><th></th><th></th><th></th></tr>
<tr><td>>MI0000088|hsa-mir-30a&WithFlank&6|-|71403545|71403627@12.32.21(+1A)&[MIMAT0000087&hsa-miR-30a-5p&offsets|0|-1(+1A);m-15&6|-|71403595|71403616&offsets|0|-1(+1A)]|TGTAAACATCCTCGACTGGAAA</td><td>>MI0000088|hsa-mir-30a&WithFlank&6|-|71403545|71403627</td><td>12</td><td>32</td><td>21(+1A)</td><td>TGTAAACATCCTCGACTGGAAA</td><td>5706.1</td><td>10.2</td><td>1.1</td><td>9.0</td><td>3.3E-19</td><td>1.5E-18</td><td>BT20/MB468</td></tr>
<tr><td>>MI0000071|hsa-mir-17&WithFlank&13|+|91350599|91350694@20.42.23(+1C)&[MIMAT0000070&hsa-miR-17-5p&offsets|0|0(+1C);m-88&13|+|91350618|91350640&offsets|0|0(+1C)]|CAAAGTGCTTACAGTGCAGGTAGC</td><td>>MI0000071|hsa-mir-17&WithFlank&13|+|91350599|91350694</td><td>20</td><td>42</td><td>23(+1C)</td><td>CAAAGTGCTTACAGTGCAGGTAGC</td><td>1268.9</td><td>7.3</td><td>0.8</td><td>8.9</td><td>3.6E-19</td><td>1.5E-18</td><td>MB231/MB468</td></tr>
<tr><td>>MI0000736|hsa-mir-30c-1&WithFlank&1|+|40757278|40757378@23.46.24;MI0000254|hsa-mir-30c-2&WithFlank&6|-|71376954|71377037@13.36.24&[MIMAT0000244&hsa-miR-30c-5p&offsets|0|+1;m-57&1|+|40757300|40757323&offsets|0|0];[MIMAT0000244_1&hsa-miR-30c-5p&offsets|0|+1;m-56&6|-|71377002|71377025&offsets|0|0]|TGTAAACATCCTACACTCTCAGCT</td><td>>MI0000736|hsa-mir-30c-1&WithFlank&1|+|40757278|40757378</td><td>23</td><td>46</td><td>24</td><td>TGTAAACATCCTACACTCTCAGCT</td><td>72170.4</td><td>6.7</td><td>0.7</td><td>8.9</td><td>3.6E-19</td><td>1.5E-18</td><td>MB231/MB468</td></tr>
<tr><td>>am_MI0015825|hsa-mir-378c&WithFlank&10|-|130962582|130962674@17.37.21&[MIMAT0016847&hsa-miR-378c&offsets|0|-4;m-130&10|-|130962639|130962658&offsets|0|+1]|ACTGGACTTGGAGTCAGAAGA</td><td>>am_MI0015825|hsa-mir-378c&WithFlank&10|-|130962582|130962674</td><td>17</td><td>37</td><td>21</td><td>ACTGGACTTGGAGTCAGAAGA</td><td>1201.5</td><td>6.8</td><td>0.8</td><td>9.0</td><td>3.4E-19</td><td>1.6E-18</td><td>BT20/MB468</td></tr>
<tr><td>>MI0000749|hsa-mir-30e&WithFlank&1|+|40754349|40754452@65.85.21(+1U)&[MIMAT0000693&hsa-miR-30e-3p&offsets|0|-1(+1U);m-25&1|+|40754413|40754434&offsets|0|-1(+1U)]|CTTTCAGTCGGATGTTTACAGT</td><td>>MI0000749|hsa-mir-30e&WithFlank&1|+|40754349|40754452</td><td>65</td><td>85</td><td>21(+1U)</td><td>CTTTCAGTCGGATGTTTACAGT</td><td>8531.0</td><td>6.8</td><td>0.8</td><td>8.9</td><td>3.6E-19</td><td>1.6E-18</td><td>MB231/MB468</td></tr>
<tr><td>>MI0000286|hsa-mir-210&WithFlank&11|-|568083|568204@72.93.22(+1C)&[MIMAT0000267&hsa-miR-210-3p&offsets|0|0(+1C);m-102&11|-|568112|568133&offsets|0|0(+1C)]|CTGTGCGTGTGACAGCGGCTGAC</td><td>>MI0000286|hsa-mir-210&WithFlank&11|-|568083|568204</td><td>72</td><td>93</td><td>22(+1C)</td><td>CTGTGCGTGTGACAGCGGCTGAC</td><td>1579.7</td><td>15.3</td><td>1.7</td><td>9.0</td><td>3.5E-19</td><td>1.6E-18</td><td>BT20/MB468</td></tr>
<tr><td>>hg38_21_+_8433172_8446622_RNA45SN1_45S_50ntFlanks@8381.8399.19|AAGAGGGCGTGAAACCGTT</td><td>>hg38_21_+_8433172_8446622_RNA45SN1_45S_50ntFlanks</td><td>8381</td><td>8399</td><td>19</td><td>AAGAGGGCGTGAAACCGTT</td><td>285.1</td><td>-5.9</td><td>0.7</td><td>-9.0</td><td>3.5E-19</td><td>1.6E-18</td><td>BT20/MB468</td></tr>
<tr><td>>hg38_21_+_8433172_8446622_RNA45SN1_45S_50ntFlanks@6651.6687.37|CGACTCTTAGCGGTGGATCACTCGGCTCGTGCGTCGA</td><td>>hg38_21_+_8433172_8446622_RNA45SN1_45S_50ntFlanks</td><td>6651</td><td>6687</td><td>37</td><td>CGACTCTTAGCGGTGGATCACTCGGCTCGTGCGTCGA</td><td>3127.7</td><td>6.6</td><td>0.7</td><td>8.9</td><td>3.8E-19</td><td>1.6E-18</td><td>MB231/MB468</td></tr>
<tr><td>>hg38_21_+_8433172_8446622_RNA45SN1_45S_50ntFlanks@4000.4024.25|TAGATAACCTCGGGCCGATCGCACG</td><td>>hg38_21_+_8433172_8446622_RNA45SN1_45S_50ntFlanks</td><td>4000</td><td>4024</td><td>25</td><td>TAGATAACCTCGGGCCGATCGCACG</td><td>280.2</td><td>-4.1</td><td>0.5</td><td>-8.9</td><td>3.6E-19</td><td>1.7E-18</td><td>BT20/MB468</td></tr>
<tr><td>>MI0000272|hsa-mir-182&WithFlank&7|-|129770377|129770498@29.50.22(+1U)&[MIMAT0000259&hsa-miR-182-5p&offsets|0|-2(+1U);m-22&7|-|129770449|129770470&offsets|0|0(+1U)]|TTTGGCAATGGTAGAACTCACAT</td><td>>MI0000272|hsa-mir-182&WithFlank&7|-|129770377|129770498</td><td>29</td><td>50</td><td>22(+1U)</td><td>TTTGGCAATGGTAGAACTCACAT</td><td>2133.4</td><td>10.1</td><td>1.1</td><td>8.9</td><td>3.6E-19</td><td>1.7E-18</td><td>BT20/MB468</td></tr>
<tr><td>>hg38_21_+_8433172_8446622_RNA45SN1_45S_50ntFlanks@5125.5145.21|AGCGGTCGGCGTCCCCCAACT</td><td>>hg38_21_+_8433172_8446622_RNA45SN1_45S_50ntFlanks</td><td>5125</td><td>5145</td><td>21</td><td>AGCGGTCGGCGTCCCCCAACT</td><td>480.5</td><td>13.6</td><td>1.5</td><td>8.9</td><td>3.8E-19</td><td>1.7E-18</td><td>BT20/MB468</td></tr>
<tr><td>>am_trnaMT_ValTAC_MT_+_1602_1670@28.52.25|AACTTACACTTAGGAGATTTCAACT</td><td>>am_trnaMT_ValTAC_MT_+_1602_1670</td><td>28</td><td>52</td><td>25</td><td>AACTTACACTTAGGAGATTTCAACT</td><td>502.1</td><td>9.3</td><td>1.0</td><td>8.9</td><td>3.8E-19</td><td>1.8E-18</td><td>BT20/MB468</td></tr>
<tr><td>>hg38_21_+_8433172_8446622_RNA45SN1_45S_50ntFlanks@12617.12654.38|TGAGGAGCCAATGGGGCGAAGCTACCATCTGTGGGATT</td><td>>hg38_21_+_8433172_8446622_RNA45SN1_45S_50ntFlanks</td><td>12617</td><td>12654</td><td>38</td><td>TGAGGAGCCAATGGGGCGAAGCTACCATCTGTGGGATT</td><td>615.2</td><td>-5.5</td><td>0.6</td><td>-8.9</td><td>3.9E-19</td><td>1.8E-18</td><td>BT20/MB468</td></tr>
<tr><td>>am_trnaMT_PheGAA_MT_+_577_647@42.59.18|TTTAGACGGGCTCACATC</td><td>>am_trnaMT_PheGAA_MT_+_577_647</td><td>42</td><td>59</td><td>18</td><td>TTTAGACGGGCTCACATC</td><td>333.5</td><td>8.6</td><td>1.0</td><td>8.9</td><td>3.9E-19</td><td>1.8E-18</td><td>BT20/MB468</td></tr>
<tr><td>>hg38_21_+_8433172_8446622_RNA45SN1_45S_50ntFlanks@5421.5443.23|CGATTGGATGGTTTAGTGAGGCC</td><td>>hg38_21_+_8433172_8446622_RNA45SN1_45S_50ntFlanks</td><td>5421</td><td>5443</td><td>23</td><td>CGATTGGATGGTTTAGTGAGGCC</td><td>397.0</td><td>-7.4</td><td>0.8</td><td>-8.9</td><td>3.9E-19</td><td>1.8E-18</td><td>BT20/MB468</td></tr>
<tr><td>>hg38_21_+_8433172_8446622_RNA45SN1_45S_50ntFlanks@4557.4587.31|CAGCTAGGAATAATGGAATAGGACCGCGGTT</td><td>>hg38_21_+_8433172_8446622_RNA45SN1_45S_50ntFlanks</td><td>4557</td><td>4587</td><td>31</td><td>CAGCTAGGAATAATGGAATAGGACCGCGGTT</td><td>228.0</td><td>-5.5</td><td>0.6</td><td>-8.9</td><td>3.9E-19</td><td>1.8E-18</td><td>BT20/MB468</td></tr>
<tr><td>>hg38_1_-_228634819_228635039_RNA5S12_5S_50ntFlanks@147.169.23|GGGAATACCGGGTGCTGTAGGCT</td><td>>hg38_1_-_228634819_228635039_RNA5S12_5S_50ntFlanks</td><td>147</td><td>169</td><td>23</td><td>GGGAATACCGGGTGCTGTAGGCT</td><td>658.7</td><td>5.7</td><td>0.6</td><td>8.9</td><td>4.3E-19</td><td>1.8E-18</td><td>MB231/MB468</td></tr>
<tr><td>>hg38_21_+_8433172_8446622_RNA45SN1_45S_50ntFlanks@7974.8015.42|ACGCGACCTCAGATCAGACGTGGCGACCCGCTGAATTTAAGC</td><td>>hg38_21_+_8433172_8446622_RNA45SN1_45S_50ntFlanks</td><td>7974</td><td>8015</td><td>42</td><td>ACGCGACCTCAGATCAGACGTGGCGACCCGCTGAATTTAAGC</td><td>1467.0</td><td>8.7</td><td>1.0</td><td>8.9</td><td>4.3E-19</td><td>1.8E-18</td><td>MB231/MB468</td></tr>
<tr><td>>hg38_21_+_8433172_8446622_RNA45SN1_45S_50ntFlanks@11729.11750.22|TAACTATGACTCTCTTAAGGTA</td><td>>hg38_21_+_8433172_8446622_RNA45SN1_45S_50ntFlanks</td><td>11729</td><td>11750</td><td>22</td><td>TAACTATGACTCTCTTAAGGTA</td><td>1299.7</td><td>-5.0</td><td>0.6</td><td>-8.9</td><td>4.0E-19</td><td>1.8E-18</td><td>BT20/MB468</td></tr>
<tr><td>>MI0000812|hsa-mir-331&WithFlank&12|+|95308414|95308519@67.87.21(+1U)&[MIMAT0000760&hsa-miR-331-3p&offsets|0|0(+1U);m-192&12|+|95308480|95308501&offsets|0|-1(+1U)]|GCCCCTGGGCCTATCCTAGAAT</td><td>>MI0000812|hsa-mir-331&WithFlank&12|+|95308414|95308519</td><td>67</td><td>87</td><td>21(+1U)</td><td>GCCCCTGGGCCTATCCTAGAAT</td><td>1644.0</td><td>7.7</td><td>0.9</td><td>8.9</td><td>4.4E-19</td><td>1.9E-18</td><td>MB231/MB468</td></tr>
<tr><td>>hg38_21_+_8433172_8446622_RNA45SN1_45S_50ntFlanks@6648.6677.30|GTACGACTCTTAGCGGTGGATCACTCGGCT</td><td>>hg38_21_+_8433172_8446622_RNA45SN1_45S_50ntFlanks</td><td>6648</td><td>6677</td><td>30</td><td>GTACGACTCTTAGCGGTGGATCACTCGGCT</td><td>582.9</td><td>4.5</td><td>0.5</td><td>8.9</td><td>4.1E-19</td><td>1.9E-18</td><td>BT20/MB468</td></tr>
<tr><td>>MI0000076|hsa-mir-20a&WithFlank&13|+|91351059|91351141@14.36.23(+1C)&[MIMAT0000075&hsa-miR-20a-5p&offsets|0|0(+1C);m-96&13|+|91351072|91351094&offsets|0|0(+1C)]|TAAAGTGCTTATAGTGCAGGTAGC</td><td>>MI0000076|hsa-mir-20a&WithFlank&13|+|91351059|91351141</td><td>14</td><td>36</td><td>23(+1C)</td><td>TAAAGTGCTTATAGTGCAGGTAGC</td><td>801.2</td><td>7.4</td><td>0.8</td><td>8.9</td><td>4.5E-19</td><td>1.9E-18</td><td>MB231/MB468</td></tr>
<tr><td>>hg38_21_+_8433172_8446622_RNA45SN1_45S_50ntFlanks@12635.12653.19|AAGCTACCATCTGTGGGAT</td><td>>hg38_21_+_8433172_8446622_RNA45SN1_45S_50ntFlanks</td><td>12635</td><td>12653</td><td>19</td><td>AAGCTACCATCTGTGGGAT</td><td>3040.3</td><td>-9.9</td><td>1.1</td><td>-8.9</td><td>4.5E-19</td><td>1.9E-18</td><td>MB231/MB468</td></tr>
<tr><td>>hg38_21_+_8433172_8446622_RNA45SN1_45S_50ntFlanks@12619.12653.35|AGGAGCCAATGGGGCGAAGCTACCATCTGTGGGAT</td><td>>hg38_21_+_8433172_8446622_RNA45SN1_45S_50ntFlanks</td><td>12619</td><td>12653</td><td>35</td><td>AGGAGCCAATGGGGCGAAGCTACCATCTGTGGGAT</td><td>407.7</td><td>-6.6</td><td>0.7</td><td>-8.9</td><td>4.5E-19</td><td>1.9E-18</td><td>MB231/MB468</td></tr>
<tr><td>>MI0001448|hsa-mir-425&WithFlank&3|-|49020142|49020240@20.41.22&[MIMAT0003393&hsa-miR-425-5p&offsets|0|-1;m-95&3|-|49020200|49020221&offsets|0|0]|AATGACACGATCACTCCCGTTG</td><td>>MI0001448|hsa-mir-425&WithFlank&3|-|49020142|49020240</td><td>20</td><td>41</td><td>22</td><td>AATGACACGATCACTCCCGTTG</td><td>682.3</td><td>7.7</td><td>0.9</td><td>8.9</td><td>4.2E-19</td><td>1.9E-18</td><td>BT20/MB468</td></tr>
<tr><td>>hg38_21_+_8433172_8446622_RNA45SN1_45S_50ntFlanks@12163.12181.19|ACTGGGGCGGTACACCTGT</td><td>>hg38_21_+_8433172_8446622_RNA45SN1_45S_50ntFlanks</td><td>12163</td><td>12181</td><td>19</td><td>ACTGGGGCGGTACACCTGT</td><td>1694.3</td><td>-9.5</td><td>1.1</td><td>-8.9</td><td>4.3E-19</td><td>2.0E-18</td><td>BT20/MB468</td></tr>
<tr><td>>MI0000077|hsa-mir-21&WithFlank&17|+|59841260|59841343@52.72.21&[MIMAT0004494&hsa-miR-21-3p&offsets|0|0;m-52&17|+|59841311|59841332&offsets|0|-1]|CAACACCAGTCGATGGGCTGT</td><td>>MI0000077|hsa-mir-21&WithFlank&17|+|59841260|59841343</td><td>52</td><td>72</td><td>21</td><td>CAACACCAGTCGATGGGCTGT</td><td>4100.2</td><td>10.9</td><td>1.2</td><td>8.9</td><td>4.4E-19</td><td>2.0E-18</td><td>BT20/MB468</td></tr>
<tr><td>>trna152_ValCAC_6_-_27248049_27248121@1.33.33|GCTTCTGTAGTGTAGTGGTTATCACGTTCGCCT</td><td>>trna152_ValCAC_6_-_27248049_27248121</td><td>1</td><td>33</td><td>33</td><td>GCTTCTGTAGTGTAGTGGTTATCACGTTCGCCT</td><td>374.5</td><td>8.8</td><td>1.0</td><td>8.9</td><td>4.4E-19</td><td>2.0E-18</td><td>BT20/MB468</td></tr>
<tr><td>>am_MI0000809|hsa-mir-151a&WithFlank&8|-|140732558|140732659@53.70.18(+3G)&[MIMAT0000757&hsa-miR-151a-3p&offsets|0|-3(+3G);m-30&8|-|140732587|140732607&offsets|0|-3(+3G)]|CTAGACTGAAGCTCCTTGGGG</td><td>>am_MI0000809|hsa-mir-151a&WithFlank&8|-|140732558|140732659</td><td>53</td><td>70</td><td>18(+3G)</td><td>CTAGACTGAAGCTCCTTGGGG</td><td>743.2</td><td>8.9</td><td>1.0</td><td>8.9</td><td>4.7E-19</td><td>2.0E-18</td><td>MB231/MB468</td></tr>
<tr><td>>hg38_21_+_8433172_8446622_RNA45SN1_45S_50ntFlanks@11540.11582.43|CGCGGCGCCTCGCCTCGGCCGGCGCCTAGCAGCCGACTTAGAA</td><td>>hg38_21_+_8433172_8446622_RNA45SN1_45S_50ntFlanks</td><td>11540</td><td>11582</td><td>43</td><td>CGCGGCGCCTCGCCTCGGCCGGCGCCTAGCAGCCGACTTAGAA</td><td>1005.3</td><td>5.8</td><td>0.7</td><td>8.9</td><td>4.7E-19</td><td>2.0E-18</td><td>MB231/MB468</td></tr>
<tr><td>>hg38_21_+_8433172_8446622_RNA45SN1_45S_50ntFlanks@11675.11709.35|TGCTCTGAATGTCAAAGTGAAGAAATTCAATGAAG</td><td>>hg38_21_+_8433172_8446622_RNA45SN1_45S_50ntFlanks</td><td>11675</td><td>11709</td><td>35</td><td>TGCTCTGAATGTCAAAGTGAAGAAATTCAATGAAG</td><td>692.7</td><td>6.2</td><td>0.7</td><td>8.9</td><td>4.7E-19</td><td>2.0E-18</td><td>MB231/MB468</td></tr>
<tr><td>>MI0000071|hsa-mir-17&WithFlank&13|+|91350599|91350694@20.42.23(+1A)&[MIMAT0000070&hsa-miR-17-5p&offsets|0|0(+1A);m-88&13|+|91350618|91350640&offsets|0|0(+1A)]|CAAAGTGCTTACAGTGCAGGTAGA</td><td>>MI0000071|hsa-mir-17&WithFlank&13|+|91350599|91350694</td><td>20</td><td>42</td><td>23(+1A)</td><td>CAAAGTGCTTACAGTGCAGGTAGA</td><td>16588.7</td><td>6.8</td><td>0.8</td><td>8.9</td><td>4.7E-19</td><td>2.0E-18</td><td>MB231/MB468</td></tr>
<tr><td>>trna119_LysCTT_1_-_145395522_145395594@4.35.32,trna11_LysCTT_5_-_180648979_180649051@4.35.32,trna13_LysCTT_6_+_26556774_26556846@4.35.32,trna7_LysCTT_16_+_3225692_3225764@4.35.32,trna9_LysCTT_5_+_180634755_180634827@4.35.32|CGGCTAGCTCAGTCGGTAGAGCATGAGACTCT</td><td>>trna119_LysCTT_1_-_145395522_145395594</td><td>4</td><td>35</td><td>32</td><td>CGGCTAGCTCAGTCGGTAGAGCATGAGACTCT</td><td>4280.6</td><td>11.4</td><td>1.3</td><td>8.9</td><td>4.7E-19</td><td>2.0E-18</td><td>MB231/MB468</td></tr>
<tr><td>>hg38_21_+_8433172_8446622_RNA45SN1_45S_50ntFlanks@11564.11591.28|CCTAGCAGCCGACTTAGAACTGGTGCGG</td><td>>hg38_21_+_8433172_8446622_RNA45SN1_45S_50ntFlanks</td><td>11564</td><td>11591</td><td>28</td><td>CCTAGCAGCCGACTTAGAACTGGTGCGG</td><td>1281.6</td><td>-6.4</td><td>0.7</td><td>-8.9</td><td>4.8E-19</td><td>2.0E-18</td><td>MB231/MB468</td></tr>
<tr><td>>hg38_21_+_8433172_8446622_RNA45SN1_45S_50ntFlanks@9574.9597.24|AATCGGTCGTCCGACCTGGGTATA</td><td>>hg38_21_+_8433172_8446622_RNA45SN1_45S_50ntFlanks</td><td>9574</td><td>9597</td><td>24</td><td>AATCGGTCGTCCGACCTGGGTATA</td><td>5593.2</td><td>-5.9</td><td>0.7</td><td>-8.9</td><td>4.4E-19</td><td>2.0E-18</td><td>BT20/MB468</td></tr>
<tr><td>>hg38_21_+_8433172_8446622_RNA45SN1_45S_50ntFlanks@12419.12440.22|GATGTCGGCTCTTCCTATCATT</td><td>>hg38_21_+_8433172_8446622_RNA45SN1_45S_50ntFlanks</td><td>12419</td><td>12440</td><td>22</td><td>GATGTCGGCTCTTCCTATCATT</td><td>368.9</td><td>4.0</td><td>0.5</td><td>8.9</td><td>4.5E-19</td><td>2.1E-18</td><td>BT20/MB468</td></tr>
<tr><td>>MI0000285|hsa-mir-205&WithFlank&1|+|209432127|209432248@40.59.20(+1A)&[MIMAT0000266&hsa-miR-205-5p&offsets|0|-2(+1A);m-351&1|+|209432166|209432187&offsets|0|-2(+1A)]|TCCTTCATTCCACCGGAGTCA</td><td>>MI0000285|hsa-mir-205&WithFlank&1|+|209432127|209432248</td><td>40</td><td>59</td><td>20(+1A)</td><td>TCCTTCATTCCACCGGAGTCA</td><td>1228.4</td><td>13.8</td><td>1.5</td><td>9.2</td><td>2.5E-20</td><td>2.1E-18</td><td>BT20/MB231</td></tr>
<tr><td>>hg38_21_+_8433172_8446622_RNA45SN1_45S_50ntFlanks@6671.6697.27|CTCGGCTCGTGCGTCGATGAAGAACGC</td><td>>hg38_21_+_8433172_8446622_RNA45SN1_45S_50ntFlanks</td><td>6671</td><td>6697</td><td>27</td><td>CTCGGCTCGTGCGTCGATGAAGAACGC</td><td>433.9</td><td>-4.0</td><td>0.5</td><td>-8.9</td><td>4.5E-19</td><td>2.1E-18</td><td>BT20/MB468</td></tr>
<tr><td>>MI0000070|hsa-mir-16-1&WithFlank&13|-|50048967|50049067@20.40.21(+1A);MI0000115|hsa-mir-16-2&WithFlank&3|+|160404739|160404831@16.36.21(+1A)&[MIMAT0000069&hsa-miR-16-5p&offsets|0|-1(+1A);m-62&13|-|50049027|50049048&offsets|0|-1(+1A)];[MIMAT0000069_1&hsa-miR-16-5p&offsets|0|-1(+1A);m-61&3|+|160404754|160404775&offsets|0|-1(+1A)]|TAGCAGCACGTAAATATTGGCA</td><td>>MI0000070|hsa-mir-16-1&WithFlank&13|-|50048967|50049067</td><td>20</td><td>40</td><td>21(+1A)</td><td>TAGCAGCACGTAAATATTGGCA</td><td>14765.0</td><td>10.5</td><td>1.2</td><td>8.9</td><td>4.5E-19</td><td>2.1E-18</td><td>BT20/MB468</td></tr>
</table>

| | | | | | | | | | | | | |
|---|---|---|---|---|---|---|---|---|---|---|---|---|
| >trna14_LysTTT_11_-_59327808_59327880@1.35.35,trna2_LysTTT_17_+_8022473_8022545@1.35.35,trna54_LysTTT_1_+_204475655_204475727@1.35.35,trna5_LysTTT_11_+_59323902_59323974@1.35.35,trna62_LysTTT_1_-_204476158_204476230@1.35.35,trna76_LysTTT_6_+_28918806_28918878@1.35.35\|GCCCGGATAGCTCAGTCGGTAGAGCATCAGACTTT | >trna14_LysTTT_11_-_59327808_59327880 | 1 | 35 | 35 | GCCCGGATAGCTCAGTCGGTAGAGCATCAGACTTT | 3031.0 | 14.2 | 1.6 | 8.9 | 4.5E-19 | 2.1E-18 | BT20/MB468 |
| >hg38_21_+_8433172_8446622_RNA45SN1_45S_50ntFlanks@12957.12982.26\|CTTCTGGGTCGGGGTTTCGTACGTAG | >hg38_21_+_8433172_8446622_RNA45SN1_45S_50ntFlanks | 12957 | 12982 | 26 | CTTCTGGGTCGGGGTTTCGTACGTAG | 1012.5 | -11.6 | 1.3 | -8.9 | 5.0E-19 | 2.1E-18 | MB231/MB468 |
| >MI0000746\|hsa-mir-99b&WithFlank&19\|+\|51692606\|51692687@13.33.21(+1C)&[MIMAT0000689&hsa-miR-99b-5p&offsets\|0\|-1(+1C);m-10&19\|+\|51692618\|51692639&offsets\|0\|-1(+1C)]\|CACCCGTAGAACCGACCTTGCC | >MI0000746\|hsa-mir-99b&WithFlank&19\|+\|51692606\|51692687 | 13 | 33 | 21(+1C) | CACCCGTAGAACCGACCTTGCC | 1504.9 | 8.4 | 0.9 | 8.9 | 4.7E-19 | 2.1E-18 | BT20/MB468 |
| >hg38_21_+_8433172_8446622_RNA45SN1_45S_50ntFlanks@12449.12480.32\|GAATTCACCAAGCGTTGGATTGTTCACCCACT | >hg38_21_+_8433172_8446622_RNA45SN1_45S_50ntFlanks | 12449 | 12480 | 32 | GAATTCACCAAGCGTTGGATTGTTCACCCACT | 744.9 | 9.7 | 1.1 | 8.9 | 5.0E-19 | 2.1E-18 | MB231/MB468 |
| >MI0000750\|hsa-mir-26a-2&WithFlank&12\|-\|57824603\|57824698@20.41.22(+1U);MI0000083\|hsa-mir-26a-1&WithFlank&3\|+\|37969398\|37969486@16.37.22(+1U)&[MIMAT0000082&hsa-miR-26a-5p&offsets\|0\|0(+1U);m-33&12\|-\|57824658\|57824679&offsets\|0\|0(+1U)];[MIMAT0000082_1&hsa-miR-26a-5p&offsets\|0\|0(+1U);m-34&3\|+\|37969413\|37969434&offsets\|0\|0(+1U)]\|TTCAAGTAATCCAGGATAGGCTT | >MI0000750\|hsa-mir-26a-2&WithFlank&12\|-\|57824603\|57824698 | 20 | 41 | 22(+1U) | TTCAAGTAATCCAGGATAGGCTT | 8084.0 | 6.7 | 0.8 | 8.9 | 4.7E-19 | 2.2E-18 | BT20/MB468 |
| >hg38_21_+_8433172_8446622_RNA45SN1_45S_50ntFlanks@8191.8212.22\|CAGCCCGTGGACGGTGTGAGGC | >hg38_21_+_8433172_8446622_RNA45SN1_45S_50ntFlanks | 8191 | 8212 | 22 | CAGCCCGTGGACGGTGTGAGGC | 579.4 | -4.7 | 0.5 | -8.9 | 4.7E-19 | 2.2E-18 | BT20/MB468 |
| >hg38_21_+_8433172_8446622_RNA45SN1_45S_50ntFlanks@6649.6692.44\|TACGACTCTTAGCGGTGGATCACTCGGCTCGTGCGTCGATGAAG | >hg38_21_+_8433172_8446622_RNA45SN1_45S_50ntFlanks | 6649 | 6692 | 44 | TACGACTCTTAGCGGTGGATCACTCGGCTCGTGCGTCGATGAAG | 444.3 | 4.8 | 0.5 | 8.9 | 5.1E-19 | 2.2E-18 | MB231/MB468 |
| >hg38_21_+_8433172_8446622_RNA45SN1_45S_50ntFlanks@12157.12182.26\|AGTTTGACTGGGGCGGTACACCTGTC | >hg38_21_+_8433172_8446622_RNA45SN1_45S_50ntFlanks | 12157 | 12182 | 26 | AGTTTGACTGGGGCGGTACACCTGTC | 480.8 | -5.5 | 0.6 | -8.9 | 5.1E-19 | 2.2E-18 | MB231/MB468 |
| >MI0000298\|hsa-mir-221&WithFlank&X\|-\|45746151\|45746272@71.93.23(+1C)&[MIMAT0000278&hsa-miR-221-3p&offsets\|0\|0(+1C);m-91&X\|-\|45746181\|45746202&offsets\|0\|+1(+1C)]\|AGCTACATTGTCTGCTGGGTTTCC | >MI0000298\|hsa-mir-221&WithFlank&X\|-\|45746151\|45746272 | 71 | 93 | 23(+1C) | AGCTACATTGTCTGCTGGGTTTCC | 939.5 | 5.5 | 0.6 | 8.9 | 4.8E-19 | 2.2E-18 | BT20/MB468 |
| >hg38_21_+_8433172_8446622_RNA45SN1_45S_50ntFlanks@9534.9561.28\|AGGAAACTCTGGTGGAGGTCCGTAGCGG | >hg38_21_+_8433172_8446622_RNA45SN1_45S_50ntFlanks | 9534 | 9561 | 28 | AGGAAACTCTGGTGGAGGTCCGTAGCGG | 1386.0 | -6.3 | 0.7 | -8.9 | 5.2E-19 | 2.2E-18 | MB231/MB468 |
| >MI0000066\|hsa-let-7e&WithFlank&19\|+\|51692780\|51692870@14.36.23&[MIMAT0000066&hsa-let-7e-5p&offsets\|0\|+1;m-50&19\|+\|51692793\|51692814&offsets\|0\|+1]\|TGAGGTAGGAGGTTGTATAGTTG | >MI0000066\|hsa-let-7e&WithFlank&19\|+\|51692780\|51692870 | 14 | 36 | 23 | TGAGGTAGGAGGTTGTATAGTTG | 694.6 | 6.6 | 0.7 | 8.9 | 5.3E-19 | 2.2E-18 | MB231/MB468 |
| >hg38_MT_+_1621_3279_RNR2_16S_50ntFlanks@51.87.37\|GCTAAACCTAGCCCCAAACCCACTCCACCTTACTACC | >hg38_MT_+_1621_3279_RNR2_16S_50ntFlanks | 51 | 87 | 37 | GCTAAACCTAGCCCCAAACCCACTCCACCTTACTACC | 1441.2 | 9.4 | 1.1 | 8.9 | 5.0E-19 | 2.3E-18 | BT20/MB468 |
| >trnaMT_PheGAA_MT_+_577_647@36.62.27\|AAAATGTTTAGACGGGCTCACATCACC | >trnaMT_PheGAA_MT_+_577_647 | 36 | 62 | 27 | AAAATGTTTAGACGGGCTCACATCACC | 585.8 | 10.8 | 1.2 | 8.9 | 5.0E-19 | 2.3E-18 | BT20/MB468 |
| >hg38_21_+_8433172_8446622_RNA45SN1_45S_50ntFlanks@12436.12455.20\|TCATTGTGAAGCAGAATTCA | >hg38_21_+_8433172_8446622_RNA45SN1_45S_50ntFlanks | 12436 | 12455 | 20 | TCATTGTGAAGCAGAATTCA | 3793.0 | -7.0 | 0.8 | -8.9 | 5.4E-19 | 2.3E-18 | MB231/MB468 |
| >MI0000234\|hsa-mir-192&WithFlank&11\|-\|64891131\|64891252@31.51.21(+1U)&[MIMAT0000222&hsa-miR-192-5p&offsets\|+1\|+1(+1U);m-59&11\|-\|64891203\|64891223&offsets\|+1\|+1(+1U)]\|TGACCTATGAATTGACAGCCAT | >MI0000234\|hsa-mir-192&WithFlank&11\|-\|64891131\|64891252 | 31 | 51 | 21(+1U) | TGACCTATGAATTGACAGCCAT | 799.5 | 6.4 | 0.7 | 8.9 | 5.4E-19 | 2.3E-18 | MB231/MB468 |
| >MI0000440\|hsa-mir-27b&WithFlank&9\|+\|95085439\|95085547@67.87.21(+1G)&[MIMAT0000419&hsa-miR-27b-3p&offsets\|0\|0(+1G);m-39&9\|+\|95085505\|95085525&offsets\|0\|0(+1G)]\|TTCACAGTGGCTAAGTTCTGCG | >MI0000440\|hsa-mir-27b&WithFlank&9\|+\|95085439\|95085547 | 67 | 87 | 21(+1G) | TTCACAGTGGCTAAGTTCTGCG | 1777.6 | 7.3 | 0.8 | 8.9 | 5.5E-19 | 2.3E-18 | MB231/MB468 |
| >MI0000255\|hsa-mir-30d&WithFlank&8\|-\|134804870\|134804951@12.35.24(+1C)&[MIMAT0000245&hsa-miR-30d-5p&offsets\|0\|+2(+1C);m-21&8\|-\|134804919\|134804940&offsets\|0\|+2(+1C)]\|TGTAAACATCCCCGACTGGAAGCTC | >MI0000255\|hsa-mir-30d&WithFlank&8\|-\|134804870\|134804951 | 12 | 35 | 24(+1C) | TGTAAACATCCCCGACTGGAAGCTC | 847.7 | 8.9 | 1.0 | 8.9 | 5.1E-19 | 2.3E-18 | BT20/MB468 |
| >hg38_21_+_8433172_8446622_RNA45SN1_45S_50ntFlanks@6645.6669.25\|CTCGTACGACTCTTAGCGGTGGATC | >hg38_21_+_8433172_8446622_RNA45SN1_45S_50ntFlanks | 6645 | 6669 | 25 | CTCGTACGACTCTTAGCGGTGGATC | 5536.5 | 3.9 | 0.4 | 8.9 | 5.3E-19 | 2.4E-18 | BT20/MB468 |
| >MI0000456\|hsa-mir-140&WithFlank&16\|+\|69933075\|69933186@69.89.21&[MIMAT0004597&hsa-miR-140-3p&offsets\|+1\|+1;m-45&16\|+\|69933143\|69933164&offsets\|0\|-1]\|ACCACAGGGTAGAACCACGGA | >MI0000456\|hsa-mir-140&WithFlank&16\|+\|69933075\|69933186 | 69 | 89 | 21 | ACCACAGGGTAGAACCACGGA | 4549.2 | 12.2 | 1.4 | 8.9 | 5.3E-19 | 2.4E-18 | BT20/MB468 |
| >hg38_21_+_8433172_8446622_RNA45SN1_45S_50ntFlanks@12570.12587.18\|AGTACGAGAGGAACCGCA | >hg38_21_+_8433172_8446622_RNA45SN1_45S_50ntFlanks | 12570 | 12587 | 18 | AGTACGAGAGGAACCGCA | 817.0 | -5.8 | 0.7 | -8.9 | 5.7E-19 | 2.4E-18 | MB231/MB468 |
| >hg38_21_+_8433172_8446622_RNA45SN1_45S_50ntFlanks@4759.4779.21\|ATCAGATACCGTCGTAGTTCC | >hg38_21_+_8433172_8446622_RNA45SN1_45S_50ntFlanks | 4759 | 4779 | 21 | ATCAGATACCGTCGTAGTTCC | 593.5 | -11.2 | 1.3 | -8.9 | 5.4E-19 | 2.4E-18 | BT20/MB468 |
| >MI0000760\|hsa-mir-361&WithFlank&X\|-\|85903630\|85903713@12.33.22&[MIMAT0000703&hsa-miR-361-5p&offsets\|0\|0;m-89&X\|-\|85903681\|85903702&offsets\|0\|0]\|TTATCAGAATCTCCAGGGGTAC | >MI0000760\|hsa-mir-361&WithFlank&X\|-\|85903630\|85903713 | 12 | 33 | 22 | TTATCAGAATCTCCAGGGGTAC | 819.4 | 6.6 | 0.7 | 8.9 | 5.8E-19 | 2.5E-18 | MB231/MB468 |
| >hg38_21_+_8433172_8446622_RNA45SN1_45S_50ntFlanks@5420.5445.26\|CCGATTGGATGGTTTAGTGAGGCCCT | >hg38_21_+_8433172_8446622_RNA45SN1_45S_50ntFlanks | 5420 | 5445 | 26 | CCGATTGGATGGTTTAGTGAGGCCCT | 542.1 | -7.7 | 0.9 | -8.9 | 5.5E-19 | 2.5E-18 | BT20/MB468 |
| >hg38_21_+_8433172_8446622_RNA45SN1_45S_50ntFlanks@10400.10424.25\|TGAACATGGGTCAGTCGGTCCTGAG | >hg38_21_+_8433172_8446622_RNA45SN1_45S_50ntFlanks | 10400 | 10424 | 25 | TGAACATGGGTCAGTCGGTCCTGAG | 909.1 | 7.2 | 0.8 | 8.9 | 5.9E-19 | 2.5E-18 | MB231/MB468 |
| >hg38_MT_+_598_1651_RNR1_12S_50ntFlanks@608.633.26\|TGTTCTGTAATCGATAAACCCCGATC | >hg38_MT_+_598_1651_RNR1_12S_50ntFlanks | 608 | 633 | 26 | TGTTCTGTAATCGATAAACCCCGATC | 1140.9 | 10.3 | 1.2 | 8.9 | 6.0E-19 | 2.5E-18 | MB231/MB468 |
| >hg38_21_+_8433172_8446622_RNA45SN1_45S_50ntFlanks@6741.6772.32\|ATCGACACTTCGAACGCACTTGCGGCCCCGGG | >hg38_21_+_8433172_8446622_RNA45SN1_45S_50ntFlanks | 6741 | 6772 | 32 | ATCGACACTTCGAACGCACTTGCGGCCCCGGG | 352.8 | -6.1 | 0.7 | -8.9 | 5.6E-19 | 2.6E-18 | BT20/MB468 |
| >hg38_1_-_228634819_228635039_RNA5S12_5S_50ntFlanks@134.172.39\|TGGGAGACCGCCTGGGAATACCGGGTGCTGTAGGCTTTT | >hg38_1_-_228634819_228635039_RNA5S12_5S_50ntFlanks | 134 | 172 | 39 | TGGGAGACCGCCTGGGAATACCGGGTGCTGTAGGCTTTT | 3987.5 | 11.9 | 1.3 | 8.9 | 6.3E-19 | 2.6E-18 | MB231/MB468 |
| >MI0000283\|hsa-mir-203a&WithFlank&14\|+\|104117399\|104117520@72.93.22&[MIMAT0000264&hsa-miR-203a-3p&offsets\|+1\|+1;m-79&14\|+\|104117469\|104117490&offsets\|+1\|+1]\|TGAAATGTTTAGGACCACTAGA | >MI0000283\|hsa-mir-203a&WithFlank&14\|+\|104117399\|104117520 | 72 | 93 | 22 | TGAAATGTTTAGGACCACTAGA | 1045.6 | 13.5 | 1.5 | 9.2 | 3.3E-20 | 2.7E-18 | BT20/MB231 |
| >MI0000750\|hsa-mir-26a-2&WithFlank&12\|-\|57824603\|57824698@20.42.23;MI0000083\|hsa-mir-26a-1&WithFlank&3\|+\|37969398\|37969486@16.38.23&[MIMAT0000082&hsa-miR-26a-5p&offsets\|0\|+1;m-33&12\|-\|57824658\|57824679&offsets\|0\|+1];[MIMAT0000082_1&hsa-miR-26a-5p&offsets\|0\|+1;m-34&3\|+\|37969413\|37969434&offsets\|0\|+1]\|TTCAAGTAATCCAGGATAGGCTG | >MI0000750\|hsa-mir-26a-2&WithFlank&12\|-\|57824603\|57824698 | 20 | 42 | 23 | TTCAAGTAATCCAGGATAGGCTG | 1130.3 | 6.9 | 0.8 | 8.9 | 5.9E-19 | 2.7E-18 | BT20/MB468 |
| >MI0000066\|hsa-let-7e&WithFlank&19\|+\|51692780\|51692870@14.35.22(+1U)&[MIMAT0000066&hsa-let-7e-5p&offsets\|0\|0(+1U);m-50&19\|+\|51692793\|51692814&offsets\|0\|0(+1U)]\|TGAGGTAGGAGGTTGTATAGTTT | >MI0000066\|hsa-let-7e&WithFlank&19\|+\|51692780\|51692870 | 14 | 35 | 22(+1U) | TGAGGTAGGAGGTTGTATAGTTT | 1692.2 | 6.4 | 0.7 | 8.9 | 6.0E-19 | 2.7E-18 | BT20/MB468 |
| >hg38_21_+_8433172_8446622_RNA45SN1_45S_50ntFlanks@4823.4851.29\|ATGACCCGCCGGGCAGCTTCCGGGAAACC | >hg38_21_+_8433172_8446622_RNA45SN1_45S_50ntFlanks | 4823 | 4851 | 29 | ATGACCCGCCGGGCAGCTTCCGGGAAACC | 597.6 | 9.6 | 1.1 | 8.9 | 6.5E-19 | 2.8E-18 | MB231/MB468 |
| >hg38_21_+_8433172_8446622_RNA45SN1_45S_50ntFlanks@12416.12453.38\|TTCGATGTCGGCTCTTCCTATCATTGTGAAGCAGAATT | >hg38_21_+_8433172_8446622_RNA45SN1_45S_50ntFlanks | 12416 | 12453 | 38 | TTCGATGTCGGCTCTTCCTATCATTGTGAAGCAGAATT | 1120.2 | 9.9 | 1.1 | 8.9 | 6.6E-19 | 2.8E-18 | MB231/MB468 |
| >hg38_21_+_8433172_8446622_RNA45SN1_45S_50ntFlanks@11572.11602.31\|CCGACTTAGAACTGGTGCGGACCAGGGGAAT | >hg38_21_+_8433172_8446622_RNA45SN1_45S_50ntFlanks | 11572 | 11602 | 31 | CCGACTTAGAACTGGTGCGGACCAGGGGAAT | 3681.3 | -5.1 | 0.6 | -8.9 | 6.2E-19 | 2.8E-18 | BT20/MB468 |
| >am_MI0000809\|hsa-mir-151a&WithFlank&8\|-\|140732558\|140732659@17.38.22&[MIMAT0004697&hsa-miR-151a-5p&offsets\|0\|+1;m-75&8\|-\|140732623\|140732643&offsets\|0\|+1]\|TCGAGGAGCTCACAGTCTAGTA | >am_MI0000809\|hsa-mir-151a&WithFlank&8\|-\|140732558\|140732659 | 17 | 38 | 22 | TCGAGGAGCTCACAGTCTAGTA | 795.1 | 7.4 | 0.8 | 8.9 | 6.7E-19 | 2.8E-18 | MB231/MB468 |
| >hg38_21_+_8433172_8446622_RNA45SN1_45S_50ntFlanks@12457.12487.31\|CAAGCGTTGGATTGTTCACCCACTAATAGGG | >hg38_21_+_8433172_8446622_RNA45SN1_45S_50ntFlanks | 12457 | 12487 | 31 | CAAGCGTTGGATTGTTCACCCACTAATAGGG | 1605.6 | -5.8 | 0.6 | -8.9 | 6.8E-19 | 2.9E-18 | MB231/MB468 |
| >hg38_MT_+_1621_3279_RNR2_16S_50ntFlanks@1370.1389.20\|ATGTTGGATCAGGACATCCC | >hg38_MT_+_1621_3279_RNR2_16S_50ntFlanks | 1370 | 1389 | 20 | ATGTTGGATCAGGACATCCC | 488.8 | 13.6 | 1.5 | 8.9 | 6.4E-19 | 2.9E-18 | BT20/MB468 |
| >hg38_21_+_8433172_8446622_RNA45SN1_45S_50ntFlanks@12417.12448.32\|TCGATGTCGGCTCTTCCTATCATTGTGAAGCA | >hg38_21_+_8433172_8446622_RNA45SN1_45S_50ntFlanks | 12417 | 12448 | 32 | TCGATGTCGGCTCTTCCTATCATTGTGAAGCA | 1570.7 | 6.3 | 0.7 | 8.9 | 7.0E-19 | 2.9E-18 | MB231/MB468 |
| >MI0000095\|hsa-mir-93&WithFlank&7\|-\|100093762\|100093853@17.39.23(+1A)&[MIMAT0000093&hsa-miR-93-5p&offsets\|0\|0(+1A);m-29&7\|-\|100093815\|100093837&offsets\|0\|0(+1A)]\|CAAAGTGCTGTTCGTGCAGGTAGA | >MI0000095\|hsa-mir-93&WithFlank&7\|-\|100093762\|100093853 | 17 | 39 | 23(+1A) | CAAAGTGCTGTTCGTGCAGGTAGA | 4987.7 | 6.2 | 0.7 | 8.9 | 6.5E-19 | 2.9E-18 | BT20/MB468 |
| >MI0000465\|hsa-mir-191&WithFlank&3\|-\|49020612\|49020715@22.42.21(+1C)&[MIMAT0000440&hsa-miR-191-5p&offsets\|0\|-2(+1C);m-48&3\|-\|49020672\|49020694&offsets\|0\|-2(+1C)]\|CAACGGAATCCCAAAAGCAGCC | >MI0000465\|hsa-mir-191&WithFlank&3\|-\|49020612\|49020715 | 22 | 42 | 21(+1C) | CAACGGAATCCCAAAAGCAGCC | 1928.5 | 6.9 | 0.8 | 8.9 | 7.0E-19 | 2.9E-18 | MB231/MB468 |

| | | | | | | | | | | | | |
|---|---|---|---|---|---|---|---|---|---|---|---|---|
| >hg38_21_+_8433172_8446622_RNA45SN1_45S_50ntFlanks@11768.11791.24\|CTAATTAGTGACGCGCATGAATGG | >hg38_21_+_8433172_8446622_RNA45SN1_45S_50ntFlanks | 11768 | 11791 | 24 | CTAATTAGTGACGCGCATGAATGG | 609.6 | -11.2 | 1.3 | -8.9 | 6.5E-19 | 3.0E-18 | BT20/MB468 |
| >MI0000440\|hsa-mir-27b&WithFlank&9\|+\|95085439\|95085547@67.88.22&[MIMAT0000419&hsa-miR-27b-3p&offsets\|0\|+1;m-39&9\|+\|95085505\|95085525&offsets\|0\|+1]\|TTCACAGTGGCTAAGTTCTGCA | >MI0000440\|hsa-mir-27b&WithFlank&9\|+\|95085439\|95085547 | 67 | 88 | 22 | TTCACAGTGGCTAAGTTCTGCA | 21335.7 | 7.0 | 0.8 | 8.9 | 6.7E-19 | 3.0E-18 | BT20/MB468 |
| >MI0000264\|hsa-mir-7-2&WithFlank&15\|+\|88611819\|88611940@38.60.23;MI0000265\|hsa-mir-7-3&WithFlank&19\|+\|4770664\|4770785@37.59.23;MI0000263\|hsa-mir-7-1&WithFlank&9\|-\|83969742\|83969863@30.52.23&[MIMAT0000252&hsa-miR-7-5p&offsets\|0\|-1;m-400&15\|+\|88611856\|88611879&offsets\|0\|-1];[MIMAT0000252_1&hsa-miR-7-5p&offsets\|0\|-1;m-402&19\|+\|4770700\|4770723&offsets\|0\|-1];[MIMAT0000252_2&hsa-miR-7-5p&offsets\|0\|-1;m-401&9\|-\|83969811\|83969834&offsets\|0\|-1]\|TGGAAGACTAGTGATTTTGTTGT | >MI0000264\|hsa-mir-7-2&WithFlank&15\|+\|88611819\|88611940 | 38 | 60 | 23 | TGGAAGACTAGTGATTTTGTTGT | 55826.6 | 8.4 | 0.9 | 8.9 | 6.8E-19 | 3.1E-18 | BT20/MB468 |
| >hg38_21_+_8433172_8446622_RNA45SN1_45S_50ntFlanks@6777.6802.26\|TCCCGGGGCTACGCCTGTCTGAGCGT | >hg38_21_+_8433172_8446622_RNA45SN1_45S_50ntFlanks | 6777 | 6802 | 26 | TCCCGGGGCTACGCCTGTCTGAGCGT | 567.8 | 6.1 | 0.7 | 8.9 | 7.3E-19 | 3.1E-18 | MB231/MB468 |
| >MI0000279\|hsa-mir-196a-2&WithFlank&12\|+\|53991732\|53991853@31.51.21;MI0000238\|hsa-mir-196a-1&WithFlank&17\|-\|48632484\|48632565@13.33.21&[MIMAT0000226&hsa-miR-196a-5p&offsets\|0\|-1;m-261&12\|+\|53991762\|53991783&offsets\|0\|-1];[MIMAT0000226_1&hsa-miR-196a-5p&offsets\|0\|-1;m-268&17\|-\|48632532\|48632553&offsets\|0\|-1]\|TAGGTAGTTTCATGTTGTTGG | >MI0000279\|hsa-mir-196a-2&WithFlank&12\|+\|53991732\|53991853 | 31 | 51 | 21 | TAGGTAGTTTCATGTTGTTGG | 2369.5 | 7.3 | 0.8 | 8.9 | 6.9E-19 | 3.1E-18 | BT20/MB468 |
| >hg38_21_+_8433172_8446622_RNA45SN1_45S_50ntFlanks@475.523.49\|CTTCGTGATCGATGTGGTGACGTCGTGCTCTCCCGGGCCGGGTCCGAGC | >hg38_21_+_8433172_8446622_RNA45SN1_45S_50ntFlanks | 475 | 523 | 49 | CTTCGTGATCGATGTGGTGACGTCGTGCTCTCCCGGGCCGGGTCCGAGC | 2186.1 | 11.2 | 1.3 | 8.9 | 7.4E-19 | 3.1E-18 | MB231/MB468 |
| >hg38_21_+_8433172_8446622_RNA45SN1_45S_50ntFlanks@12163.12180.18\|ACTGGGGCGGTACACCTG | >hg38_21_+_8433172_8446622_RNA45SN1_45S_50ntFlanks | 12163 | 12180 | 18 | ACTGGGGCGGTACACCTG | 2432.2 | -9.4 | 1.1 | -8.9 | 6.9E-19 | 3.1E-18 | BT20/MB468 |
| >MI0000456\|hsa-mir-140&WithFlank&16\|+\|69933075\|69933186@68.89.22&[MIMAT0004597&hsa-miR-140-3p&offsets\|0\|+1;m-45&16\|+\|69933143\|69933164&offsets\|-1\|-1]\|TACCACAGGGTAGAACCACGGA | >MI0000456\|hsa-mir-140&WithFlank&16\|+\|69933075\|69933186 | 68 | 89 | 22 | TACCACAGGGTAGAACCACGGA | 2404.7 | 11.4 | 1.3 | 8.9 | 7.5E-19 | 3.1E-18 | MB231/MB468 |
| >MI0000469\|hsa-mir-125a&WithFlank&19\|+\|51693248\|51693345@21.43.23&[MIMAT0000443&hsa-miR-125a-5p&offsets\|0\|-1;m-63&19\|+\|51693268\|51693289&offsets\|0\|+1]\|TCCCTGAGACCCTTTAACCTGTG | >MI0000469\|hsa-mir-125a&WithFlank&19\|+\|51693248\|51693345 | 21 | 43 | 23 | TCCCTGAGACCCTTTAACCTGTG | 50336.0 | 6.9 | 0.8 | 8.9 | 7.3E-19 | 3.3E-18 | BT20/MB468 |
| >trnaMT_PheGAA_MT_+_577_647@1.20.20\|GTTTATGTAGCTTACCTCCT | >trnaMT_PheGAA_MT_+_577_647 | 1 | 20 | 20 | GTTTATGTAGCTTACCTCCT | 1119.3 | 8.8 | 1.0 | 8.9 | 7.9E-19 | 3.3E-18 | MB231/MB468 |
| >MI0006444\|hsa-mir-1307&WithFlank&10\|-\|103394247\|103394407@87.108.22&[MIMAT0005951&hsa-miR-1307-3p&offsets\|+1\|+1;m-66&10\|-\|103394300\|103394322&offsets\|+1\|0]\|CTCGGCGTGGCGTCGGTCGTGG | >MI0006444\|hsa-mir-1307&WithFlank&10\|-\|103394247\|103394407 | 87 | 108 | 22 | CTCGGCGTGGCGTCGGTCGTGG | 1581.4 | 6.0 | 0.7 | 8.9 | 7.4E-19 | 3.3E-18 | BT20/MB468 |
| >trnaMT_SerGCT_MT_+_12207_12265@1.21.21\|GAGAAAGCTCACAAGAACTGC | >trnaMT_SerGCT_MT_+_12207_12265 | 1 | 21 | 21 | GAGAAAGCTCACAAGAACTGC | 2006.3 | 7.0 | 0.8 | 8.9 | 7.5E-19 | 3.4E-18 | BT20/MB468 |
| >MI0000255\|hsa-mir-30d&WithFlank&8\|-\|134804870\|134804951@12.35.24(+3U)&[MIMAT0000245&hsa-miR-30d-5p&offsets\|0\|+2(+3U);m-21&8\|-\|134804919\|134804940&offsets\|0\|+2(+3U)]\|TGTAAACATCCCCGACTGGAAGCTTTT | >MI0000255\|hsa-mir-30d&WithFlank&8\|-\|134804870\|134804951 | 12 | 35 | 24(+3U) | TGTAAACATCCCCGACTGGAAGCTTTT | 1599.3 | 10.8 | 1.2 | 8.9 | 8.2E-19 | 3.5E-18 | MB231/MB468 |
| >hg38_21_+_8433172_8446622_RNA45SN1_45S_50ntFlanks@12457.12475.19\|CAAGCGTTGGATTGTTCAC | >hg38_21_+_8433172_8446622_RNA45SN1_45S_50ntFlanks | 12457 | 12475 | 19 | CAAGCGTTGGATTGTTCAC | 1121.2 | -7.3 | 0.8 | -8.9 | 7.7E-19 | 3.5E-18 | BT20/MB468 |
| >hg38_21_+_8433172_8446622_RNA45SN1_45S_50ntFlanks@10735.10755.21\|TGTAAATCTCGCGCCGGGCCG | >hg38_21_+_8433172_8446622_RNA45SN1_45S_50ntFlanks | 10735 | 10755 | 21 | TGTAAATCTCGCGCCGGGCCG | 611.3 | -6.8 | 0.8 | -8.9 | 7.7E-19 | 3.5E-18 | BT20/MB468 |
| >MI0000542\|hsa-mir-320a&WithFlank&8\|-\|22244960\|22245043@48.68.21(+1G)&[MIMAT0000510&hsa-miR-320a-3p&offsets\|0\|-1(+1G);m-60&8\|-\|22244975\|22244996&offsets\|0\|-1(+1G)]\|AAAAGCTGGGTTGAGAGGGCGG | >MI0000542\|hsa-mir-320a&WithFlank&8\|-\|22244960\|22245043 | 48 | 68 | 21(+1G) | AAAAGCTGGGTTGAGAGGGCGG | 3486.5 | 10.6 | 1.2 | 8.9 | 7.8E-19 | 3.5E-18 | BT20/MB468 |
| >hg38_21_+_8433172_8446622_RNA45SN1_45S_50ntFlanks@10684.10709.26\|CGGCGGCGTCCGGTGAGCTCTCGCTG | >hg38_21_+_8433172_8446622_RNA45SN1_45S_50ntFlanks | 10684 | 10709 | 26 | CGGCGGCGTCCGGTGAGCTCTCGCTG | 1869.9 | -4.2 | 0.5 | -8.9 | 8.4E-19 | 3.5E-18 | MB231/MB468 |
| >hg38_21_+_8433172_8446622_RNA45SN1_45S_50ntFlanks@12002.12032.31\|TGCGGGCCGCCGGTGAAATACCACTACTCTG | >hg38_21_+_8433172_8446622_RNA45SN1_45S_50ntFlanks | 12002 | 12032 | 31 | TGCGGGCCGCCGGTGAAATACCACTACTCTG | 2670.1 | -7.2 | 0.8 | -8.9 | 7.9E-19 | 3.5E-18 | BT20/MB468 |
| >trna10_AspGTC_12_-_125424193_125424264@1.34.34,trna12_AspGTC_12_-_125411891_125411962@1.34.34,trna38_AspGTC_17_-_8125556_8125627@1.34.34,trna45_AspGTC_6_+_27447453_27447524@1.34.34,trna48_AspGTC_6_+_27471523_27471594@1.34.34,trna4_AspGTC_12_+_96429799_96429870@1.34.34,trna69_AspGTC_1_-_161440205_161440276@1.34.34,trna72_AspGTC_1_-_161432824_161432895@1.34.34,trna75_AspGTC_1_-_161425414_161425485@1.34.34,trna78_AspGTC_1_-_161418033_161418104@1.34.34,trna81_AspGTC_1_-_161410615_161410686@1.34.34\|TCCTCGTTAGTATAGTGGTGAGTATCCCCGCCTG | >trna10_AspGTC_12_-_125424193_125424264 | 1 | 34 | 34 | TCCTCGTTAGTATAGTGGTGAGTATCCCCGCCTG | 2037.1 | 8.4 | 0.9 | 8.9 | 8.5E-19 | 3.5E-18 | MB231/MB468 |
| >hg38_1_-_228634819_228635039_RNA5S12_5S_50ntFlanks@139.170.32\|GACCGCCTGGGAATACCGGGTGCTGTAGGCTT | >hg38_1_-_228634819_228635039_RNA5S12_5S_50ntFlanks | 139 | 170 | 32 | GACCGCCTGGGAATACCGGGTGCTGTAGGCTT | 1976.7 | 7.1 | 0.8 | 8.9 | 8.0E-19 | 3.6E-18 | BT20/MB468 |
| >hg38_21_+_8433172_8446622_RNA45SN1_45S_50ntFlanks@5491.5516.26\|GAGAAGACGGTCGAACTTGACTATCT | >hg38_21_+_8433172_8446622_RNA45SN1_45S_50ntFlanks | 5491 | 5516 | 26 | GAGAAGACGGTCGAACTTGACTATCT | 13052.1 | 10.5 | 1.2 | 8.9 | 8.0E-19 | 3.6E-18 | BT20/MB468 |
| >MI0000264\|hsa-mir-7-2&WithFlank&15\|+\|88611819\|88611940@38.61.24(+1A);MI0000265\|hsa-mir-7-3&WithFlank&19\|+\|4770664\|4770785@37.60.24(+1A);MI0000263\|hsa-mir-7-1&WithFlank&9\|-\|83969742\|83969863@30.53.24(+1A)&[MIMAT0000252&hsa-miR-7-5p&offsets\|0\|0(+1A);m-400&15\|+\|88611856\|88611879&offsets\|0\|0(+1A)];[MIMAT0000252_1&hsa-miR-7-5p&offsets\|0\|0(+1A);m-402&19\|+\|4770700\|4770723&offsets\|0\|0(+1A)];[MIMAT0000252_2&hsa-miR-7-5p&offsets\|0\|0(+1A);m-401&9\|-\|83969811\|83969834&offsets\|0\|0(+1A)]\|TGGAAGACTAGTGATTTTGTTGTTA | >MI0000264\|hsa-mir-7-2&WithFlank&15\|+\|88611819\|88611940 | 38 | 61 | 24(+1A) | TGGAAGACTAGTGATTTTGTTGTTA | 3467.9 | 7.9 | 0.9 | 8.9 | 8.0E-19 | 3.6E-18 | BT20/MB468 |
| >hg38_21_+_8433172_8446622_RNA45SN1_45S_50ntFlanks@7973.8011.39\|GACGCGACCTCAGATCAGACGTGGCGACCCGCTGAATTT | >hg38_21_+_8433172_8446622_RNA45SN1_45S_50ntFlanks | 7973 | 8011 | 39 | GACGCGACCTCAGATCAGACGTGGCGACCCGCTGAATTT | 1676.0 | 7.2 | 0.8 | 8.9 | 8.2E-19 | 3.7E-18 | BT20/MB468 |
| >MI0001652\|hsa-mir-450a-1&WithFlank&X\|-\|134540335\|134540437@24.43.20;MI0003187\|hsa-mir-450a-2&WithFlank&X\|-\|134540502\|134540613@29.48.20&[MIMAT0001545&hsa-miR-450a-5p&offsets\|0\|-2;m-288&X\|-\|134540393\|134540414&offsets\|0\|-2];[MIMAT0001545_1&hsa-miR-450a-5p&offsets\|0\|-2;m-287&X\|-\|134540564\|134540585&offsets\|0\|-2]\|TTTTGCGATGTGTTCCTAAT | >MI0001652\|hsa-mir-450a-1&WithFlank&X\|-\|134540335\|134540437 | 24 | 43 | 20 | TTTTGCGATGTGTTCCTAAT | 911.1 | 12.0 | 1.4 | 8.9 | 8.2E-19 | 3.7E-18 | BT20/MB468 |
| >MI0003205\|hsa-mir-532&WithFlank&X\|+\|50003142\|50003244@26.47.22&[MIMAT0002888&hsa-miR-532-5p&offsets\|0\|0;m-65&X\|+\|50003167\|50003188&offsets\|0\|0]\|CATGCCTTGAGTGTAGGACCGT | >MI0003205\|hsa-mir-532&WithFlank&X\|+\|50003142\|50003244 | 26 | 47 | 22 | CATGCCTTGAGTGTAGGACCGT | 8484.8 | 6.3 | 0.7 | 8.8 | 8.9E-19 | 3.7E-18 | MB231/MB468 |
| >MI0000471\|hsa-mir-126&WithFlank&9\|+\|136670596\|136670692@58.78.21(+1U)&[MIMAT0000445&hsa-miR-126-3p&offsets\|0\|-1(+1U);m-40&9\|+\|136670653\|136670674&offsets\|0\|-1(+1U)]\|TCGTACCGTGAGTAATAATGCT | >MI0000471\|hsa-mir-126&WithFlank&9\|+\|136670596\|136670692 | 58 | 78 | 21(+1U) | TCGTACCGTGAGTAATAATGCT | 448.3 | 11.4 | 1.3 | 8.9 | 8.3E-19 | 3.7E-18 | BT20/MB468 |
| >hg38_21_+_8433172_8446622_RNA45SN1_45S_50ntFlanks@11885.11905.21\|TTGAGCTTGACTCTAGTCTGG | >hg38_21_+_8433172_8446622_RNA45SN1_45S_50ntFlanks | 11885 | 11905 | 21 | TTGAGCTTGACTCTAGTCTGG | 888.8 | -8.6 | 1.0 | -8.9 | 8.3E-19 | 3.7E-18 | BT20/MB468 |
| >MI0000746\|hsa-mir-99b&WithFlank&19\|+\|51692606\|51692687@13.34.22(+1C)&[MIMAT0000689&hsa-miR-99b-5p&offsets\|0\|0(+1C);m-10&19\|+\|51692618\|51692639&offsets\|0\|0(+1C)]\|CACCCGTAGAACCGACCTTGCGC | >MI0000746\|hsa-mir-99b&WithFlank&19\|+\|51692606\|51692687 | 13 | 34 | 22(+1C) | CACCCGTAGAACCGACCTTGCGC | 670.7 | 8.9 | 1.0 | 8.8 | 9.0E-19 | 3.8E-18 | MB231/MB468 |
| >hg38_21_+_8433172_8446622_RNA45SN1_45S_50ntFlanks@10687.10710.24\|CGGCGTCCGGTGAGCTCTCGCTGG | >hg38_21_+_8433172_8446622_RNA45SN1_45S_50ntFlanks | 10687 | 10710 | 24 | CGGCGTCCGGTGAGCTCTCGCTGG | 3868.8 | -7.0 | 0.8 | -8.8 | 9.0E-19 | 3.8E-18 | MB231/MB468 |
| >hg38_21_+_8433172_8446622_RNA45SN1_45S_50ntFlanks@11316.11342.27\|CACCCCACGTCTCGTCGCGCGCGCGTC | >hg38_21_+_8433172_8446622_RNA45SN1_45S_50ntFlanks | 11316 | 11342 | 27 | CACCCCACGTCTCGTCGCGCGCGCGTC | 1481.4 | -12.1 | 1.4 | -8.8 | 9.1E-19 | 3.8E-18 | MB231/MB468 |
| >MI0000077\|hsa-mir-21&WithFlank&17\|+\|59841260\|59841343@52.73.22&[MIMAT0004494&hsa-miR-21-3p&offsets\|0\|+1;m-52&17\|+\|59841311\|59841332&offsets\|0\|0]\|CAACACCAGTCGATGGGCTGTC | >MI0000077\|hsa-mir-21&WithFlank&17\|+\|59841260\|59841343 | 52 | 73 | 22 | CAACACCAGTCGATGGGCTGTC | 962.1 | 7.5 | 0.9 | 8.8 | 9.2E-19 | 3.8E-18 | MB231/MB468 |
| >am_trnaMT_TyrGTA_MT_-_5826_5891@1.33.33\|GGTAAAATGGCTGAGTGAAGCATTGGACTGTAA | >am_trnaMT_TyrGTA_MT_-_5826_5891 | 1 | 33 | 33 | GGTAAAATGGCTGAGTGAAGCATTGGACTGTAA | 852.3 | 10.7 | 1.2 | 8.8 | 9.5E-19 | 4.0E-18 | MB231/MB468 |
| >MI0000465\|hsa-mir-191&WithFlank&3\|-\|49020612\|49020715@22.43.22&[MIMAT0000440&hsa-miR-191-5p&offsets\|0\|-1;m-48&3\|-\|49020672\|49020694&offsets\|0\|-1]\|CAACGGAATCCCAAAAGCAGCT | >MI0000465\|hsa-mir-191&WithFlank&3\|-\|49020612\|49020715 | 22 | 43 | 22 | CAACGGAATCCCAAAAGCAGCT | 14679.1 | 6.8 | 0.8 | 8.8 | 9.6E-19 | 4.0E-18 | MB231/MB468 |
| >hg38_21_+_8433172_8446622_RNA45SN1_45S_50ntFlanks@6719.6752.34\|TGAATTGCAGGACACATTGATCATCGACACTTCG | >hg38_21_+_8433172_8446622_RNA45SN1_45S_50ntFlanks | 6719 | 6752 | 34 | TGAATTGCAGGACACATTGATCATCGACACTTCG | 221.5 | -5.0 | 0.6 | -8.8 | 9.0E-19 | 4.0E-18 | BT20/MB468 |

<table>
<tr><th></th><th></th><th></th><th></th><th></th><th></th><th></th><th></th><th></th><th></th><th></th><th></th><th></th></tr>
<tr><td>>MI0000767|hsa-mir-365a&WithFlank&16|+|14309279|14309377@62.82.21;MI0000769|hsa-mir-365b&WithFlank&17|+|31575405|31575527@74.94.21&[MIMAT0000710&hsa-miR-365a-3p&offsets|0|-1;m-176&16|+|14309340|14309361&offsets|0|-1];[MIMAT0022834&hsa-miR-365b-3p&offsets|0|-1;m-175&17|+|31575478|31575499&offsets|0|-1]|TAATGCCCCTAAAAATCCTTA</td><td>>MI0000767|hsa-mir-365a&WithFlank&16|+|14309279|14309377</td><td>62</td><td>82</td><td>21</td><td>TAATGCCCCTAAAAATCCTTA</td><td>1682.6</td><td>13.3</td><td>1.5</td><td>8.8</td><td>9.2E-19</td><td>4.1E-18</td><td>BT20/MB468</td></tr>
<tr><td>>hg38_21_+_8433172_8446622_RNA45SN1_45S_50ntFlanks@7975.8018.44|CGCGACCTCAGATCAGACGTGGCGACCCGCTGAATTTAAGCATA</td><td>>hg38_21_+_8433172_8446622_RNA45SN1_45S_50ntFlanks</td><td>7975</td><td>8018</td><td>44</td><td>CGCGACCTCAGATCAGACGTGGCGACCCGCTGAATTTAAGCATA</td><td>3564.2</td><td>8.5</td><td>1.0</td><td>8.8</td><td>9.9E-19</td><td>4.1E-18</td><td>MB231/MB468</td></tr>
<tr><td>>hg38_21_+_8433172_8446622_RNA45SN1_45S_50ntFlanks@6644.6665.22|CCTCGTACGACTCTTAGCGGTG</td><td>>hg38_21_+_8433172_8446622_RNA45SN1_45S_50ntFlanks</td><td>6644</td><td>6665</td><td>22</td><td>CCTCGTACGACTCTTAGCGGTG</td><td>634.2</td><td>5.4</td><td>0.6</td><td>8.8</td><td>9.3E-19</td><td>4.1E-18</td><td>BT20/MB468</td></tr>
<tr><td>>trnaMT_ValTAC_MT_+_1602_1670@50.70.21|ACTTAACTTGACCGCTCTGAC</td><td>>trnaMT_ValTAC_MT_+_1602_1670</td><td>50</td><td>70</td><td>21</td><td>ACTTAACTTGACCGCTCTGAC</td><td>433.6</td><td>9.2</td><td>1.0</td><td>8.8</td><td>9.3E-19</td><td>4.1E-18</td><td>BT20/MB468</td></tr>
<tr><td>>hg38_21_+_8433172_8446622_RNA45SN1_45S_50ntFlanks@5473.5516.44|CGGCCCTGGCGGAGCGCTGAGAAGACGGTCGAACTTGACTATCT</td><td>>hg38_21_+_8433172_8446622_RNA45SN1_45S_50ntFlanks</td><td>5473</td><td>5516</td><td>44</td><td>CGGCCCTGGCGGAGCGCTGAGAAGACGGTCGAACTTGACTATCT</td><td>3800.3</td><td>11.3</td><td>1.3</td><td>8.8</td><td>1.0E-18</td><td>4.2E-18</td><td>MB231/MB468</td></tr>
<tr><td>>hg38_21_+_8433172_8446622_RNA45SN1_45S_50ntFlanks@6719.6745.27|TGAATTGCAGGACACATTGATCATCGA</td><td>>hg38_21_+_8433172_8446622_RNA45SN1_45S_50ntFlanks</td><td>6719</td><td>6745</td><td>27</td><td>TGAATTGCAGGACACATTGATCATCGA</td><td>542.8</td><td>-11.1</td><td>1.3</td><td>-8.8</td><td>9.4E-19</td><td>4.2E-18</td><td>BT20/MB468</td></tr>
<tr><td>>hg38_21_+_8433172_8446622_RNA45SN1_45S_50ntFlanks@9567.9597.31|ACGTGCAAATCGGTCGTCCGACCTGGGTATA</td><td>>hg38_21_+_8433172_8446622_RNA45SN1_45S_50ntFlanks</td><td>9567</td><td>9597</td><td>31</td><td>ACGTGCAAATCGGTCGTCCGACCTGGGTATA</td><td>3424.7</td><td>-9.2</td><td>1.0</td><td>-8.8</td><td>9.4E-19</td><td>4.2E-18</td><td>BT20/MB468</td></tr>
<tr><td>>hg38_21_+_8433172_8446622_RNA45SN1_45S_50ntFlanks@4449.4471.23|CCTCTCGGCGCCCCCTCGATGCT</td><td>>hg38_21_+_8433172_8446622_RNA45SN1_45S_50ntFlanks</td><td>4449</td><td>4471</td><td>23</td><td>CCTCTCGGCGCCCCCTCGATGCT</td><td>435.6</td><td>-7.6</td><td>0.9</td><td>-8.8</td><td>9.5E-19</td><td>4.2E-18</td><td>BT20/MB468</td></tr>
<tr><td>>MI0000085|hsa-mir-27a&WithFlank&19|-|13836434|13836523@57.75.19(+1U)&[MIMAT0000084&hsa-miR-27a-3p&offsets|0|-2(+1U);m-43&19|-|13836447|13836467&offsets|0|-2(+1U)]|TTCACAGTGGCTAAGTTCCT</td><td>>MI0000085|hsa-mir-27a&WithFlank&19|-|13836434|13836523</td><td>57</td><td>75</td><td>19(+1U)</td><td>TTCACAGTGGCTAAGTTCCT</td><td>956.2</td><td>6.9</td><td>0.8</td><td>8.8</td><td>1.0E-18</td><td>4.3E-18</td><td>MB231/MB468</td></tr>
<tr><td>>MI0000264|hsa-mir-7-2&WithFlank&15|+|88611819|88611940@38.57.20(+1C);MI0000265|hsa-mir-7-3&WithFlank&19|+|4770664|4770785@37.56.20(+1C);MI0000263|hsa-mir-7-1&WithFlank&9|-|83969742|83969863@30.49.20(+1C)&[MIMAT0000252&hsa-miR-7-5p&offsets|0|-4(+1C);m-400&15|+|88611856|88611879&offsets|0|-4(+1C)];[MIMAT0000252_1&hsa-miR-7-5p&offsets|0|-4(+1C);m-402&19|+|4770700|4770723&offsets|0|-4(+1C)];[MIMAT0000252_2&hsa-miR-7-5p&offsets|0|-4(+1C);m-401&9|-|83969811|83969834&offsets|0|-4(+1C)]|TGGAAGACTAGTGATTTTGTC</td><td>>MI0000264|hsa-mir-7-2&WithFlank&15|+|88611819|88611940</td><td>38</td><td>57</td><td>20(+1C)</td><td>TGGAAGACTAGTGATTTTGTC</td><td>756.0</td><td>10.1</td><td>1.1</td><td>8.8</td><td>1.0E-18</td><td>4.3E-18</td><td>MB231/MB468</td></tr>
<tr><td>>hg38_MT_+_598_1651_RNR1_12S_50ntFlanks@904.929.26|AAGTATACTTCAAAGGACATTTAACT</td><td>>hg38_MT_+_598_1651_RNR1_12S_50ntFlanks</td><td>904</td><td>929</td><td>26</td><td>AAGTATACTTCAAAGGACATTTAACT</td><td>479.2</td><td>11.1</td><td>1.3</td><td>8.8</td><td>9.8E-19</td><td>4.3E-18</td><td>BT20/MB468</td></tr>
<tr><td>>hg38_21_+_8433172_8446622_RNA45SN1_45S_50ntFlanks@11768.11804.37|CTAATTAGTGACGCGCATGAATGGATGAACGAGATTC</td><td>>hg38_21_+_8433172_8446622_RNA45SN1_45S_50ntFlanks</td><td>11768</td><td>11804</td><td>37</td><td>CTAATTAGTGACGCGCATGAATGGATGAACGAGATTC</td><td>437.6</td><td>-5.2</td><td>0.6</td><td>-8.8</td><td>1.1E-18</td><td>4.4E-18</td><td>MB231/MB468</td></tr>
<tr><td>>trnaMT_PheGAA_MT_+_577_647@38.56.19|AATGTTTAGACGGGCTCAC</td><td>>trnaMT_PheGAA_MT_+_577_647</td><td>38</td><td>56</td><td>19</td><td>AATGTTTAGACGGGCTCAC</td><td>746.0</td><td>9.9</td><td>1.1</td><td>8.8</td><td>1.1E-18</td><td>4.4E-18</td><td>MB231/MB468</td></tr>
<tr><td>>hg38_21_+_8433172_8446622_RNA45SN1_45S_50ntFlanks@6745.6771.27|ACACTTCGAACGCACTTGCGGCCCCGG</td><td>>hg38_21_+_8433172_8446622_RNA45SN1_45S_50ntFlanks</td><td>6745</td><td>6771</td><td>27</td><td>ACACTTCGAACGCACTTGCGGCCCCGG</td><td>3075.8</td><td>-9.0</td><td>1.0</td><td>-8.8</td><td>1.0E-18</td><td>4.4E-18</td><td>BT20/MB468</td></tr>
<tr><td>>MI0000086|hsa-mir-28&WithFlank&3|+|188688775|188688872@60.80.21(+1G)&[MIMAT0004502&hsa-miR-28-3p&offsets|0|-1(+1G);m-28&3|+|188688834|188688855&offsets|0|-1(+1G)]|CACTAGATTGTGAGCTCCTGGG</td><td>>MI0000086|hsa-mir-28&WithFlank&3|+|188688775|188688872</td><td>60</td><td>80</td><td>21(+1G)</td><td>CACTAGATTGTGAGCTCCTGGG</td><td>2138.1</td><td>13.2</td><td>1.5</td><td>8.8</td><td>1.0E-18</td><td>4.5E-18</td><td>BT20/MB468</td></tr>
<tr><td>>hg38_21_+_8433172_8446622_RNA45SN1_45S_50ntFlanks@10840.10868.29|CCGTAACTTCGGGATAAGGATTGGCTCTA</td><td>>hg38_21_+_8433172_8446622_RNA45SN1_45S_50ntFlanks</td><td>10840</td><td>10868</td><td>29</td><td>CCGTAACTTCGGGATAAGGATTGGCTCTA</td><td>537.3</td><td>-4.9</td><td>0.6</td><td>-8.8</td><td>1.0E-18</td><td>4.5E-18</td><td>BT20/MB468</td></tr>
<tr><td>>hg38_21_+_8433172_8446622_RNA45SN1_45S_50ntFlanks@10375.10407.33|AGAAGGGTTCCATGTGAACAGCAGTTGAACATG</td><td>>hg38_21_+_8433172_8446622_RNA45SN1_45S_50ntFlanks</td><td>10375</td><td>10407</td><td>33</td><td>AGAAGGGTTCCATGTGAACAGCAGTTGAACATG</td><td>286.4</td><td>-5.7</td><td>0.6</td><td>-8.8</td><td>1.1E-18</td><td>4.5E-18</td><td>MB231/MB468</td></tr>
<tr><td>>trna129_MetCAT_6_-_27870271_27870342@31.51.21,trna142_MetCAT_6_-_27560600_27560671@31.51.21,trna150_MetCAT_6_-_27300764_27300835@31.51.21,trna169_MetCAT_6_-_26330529_26330600@31.51.21,trna171_MetCAT_6_-_26313352_26313423@31.51.21,trna20_MetCAT_17_-_80452597_80452668@31.51.21,trna2_MetCAT_6_+_26286754_26286825@31.51.21,trna32_MetCAT_1_+_153643726_153643797@31.51.21,trna61_MetCAT_6_+_27745664_27745735@31.51.21|CCCATAACCCAGAGGTCGATG</td><td>>trna129_MetCAT_6_-_27870271_27870342</td><td>31</td><td>51</td><td>21</td><td>CCCATAACCCAGAGGTCGATG</td><td>1305.0</td><td>-12.3</td><td>1.4</td><td>-8.8</td><td>1.0E-18</td><td>4.5E-18</td><td>BT20/MB468</td></tr>
<tr><td>>hg38_21_+_8433172_8446622_RNA45SN1_45S_50ntFlanks@11270.11295.26|CGGCCGCCCCCGGGGAGCCCGGCGGG</td><td>>hg38_21_+_8433172_8446622_RNA45SN1_45S_50ntFlanks</td><td>11270</td><td>11295</td><td>26</td><td>CGGCCGCCCCCGGGGAGCCCGGCGGG</td><td>680.3</td><td>-11.4</td><td>1.3</td><td>-8.8</td><td>1.0E-18</td><td>4.6E-18</td><td>BT20/MB468</td></tr>
<tr><td>>hg38_21_+_8433172_8446622_RNA45SN1_45S_50ntFlanks@10683.10711.29|CCGGCGGCGTCCGGTGAGCTCTCGCTGGC</td><td>>hg38_21_+_8433172_8446622_RNA45SN1_45S_50ntFlanks</td><td>10683</td><td>10711</td><td>29</td><td>CCGGCGGCGTCCGGTGAGCTCTCGCTGGC</td><td>1166.9</td><td>-6.0</td><td>0.7</td><td>-8.8</td><td>1.0E-18</td><td>4.6E-18</td><td>BT20/MB468</td></tr>
<tr><td>>MI0000095|hsa-mir-93&WithFlank&7|-|100093762|100093853@17.39.23(+1G)&[MIMAT0000093&hsa-miR-93-5p&offsets|0|0(+1G);m-29&7|-|100093815|100093837&offsets|0|0(+1G)]|CAAAGTGCTGTTCGTGCAGGTAGG</td><td>>MI0000095|hsa-mir-93&WithFlank&7|-|100093762|100093853</td><td>17</td><td>39</td><td>23(+1G)</td><td>CAAAGTGCTGTTCGTGCAGGTAGG</td><td>835.4</td><td>6.9</td><td>0.8</td><td>8.8</td><td>1.1E-18</td><td>4.6E-18</td><td>MB231/MB468</td></tr>
<tr><td>>MI0000283|hsa-mir-203a&WithFlank&14|+|104117399|104117520@72.90.19&[MIMAT0000264&hsa-miR-203a-3p&offsets|+1|-2;m-79&14|+|104117469|104117490&offsets|+1|-2]|TGAAATGTTTAGGACCACT</td><td>>MI0000283|hsa-mir-203a&WithFlank&14|+|104117399|104117520</td><td>72</td><td>90</td><td>19</td><td>TGAAATGTTTAGGACCACT</td><td>591.4</td><td>12.7</td><td>1.4</td><td>9.1</td><td>5.8E-20</td><td>4.6E-18</td><td>BT20/MB231</td></tr>
<tr><td>>hg38_21_+_8433172_8446622_RNA45SN1_45S_50ntFlanks@4760.4782.23|TCAGATACCGTCGTAGTTCCGAC</td><td>>hg38_21_+_8433172_8446622_RNA45SN1_45S_50ntFlanks</td><td>4760</td><td>4782</td><td>23</td><td>TCAGATACCGTCGTAGTTCCGAC</td><td>772.0</td><td>-6.3</td><td>0.7</td><td>-8.8</td><td>1.1E-18</td><td>4.7E-18</td><td>MB231/MB468</td></tr>
<tr><td>>trna111_HisGTG_1_-_147774845_147774916@-1G.33.34,trna118_HisGTG_1_-_145396881_145396952@-1G.33.34,trna16_HisGTG_1_+_146544773_146544844@-1G.33.34,trna1_HisGTG_15_+_45493349_45493420@-1G.33.34,trna21_HisGTG_1_+_147753471_147753542@-1G.33.34,trna33_HisGTG_6_+_27125906_27125977@-1G.33.34,trna7_HisGTG_9_-_14433938_14434009@-1G.33.34,trna8_HisGTG_15_-_45492611_45492682@-1G.33.34,trna9_HisGTG_15_-_45490804_45490875@-1G.33.34|GGCCGTGATCGTATAGTGGTTAGTACTCTGCGTT</td><td>>trna111_HisGTG_1_-_147774845_147774916</td><td>-1G</td><td>33</td><td>34</td><td>GGCCGTGATCGTATAGTGGTTAGTACTCTGCGTT</td><td>5586.5</td><td>9.0</td><td>1.0</td><td>8.8</td><td>1.1E-18</td><td>4.7E-18</td><td>BT20/MB468</td></tr>
<tr><td>>MI0000650|hsa-mir-200c&WithFlank&12|+|6963693|6963772@50.72.23(+2A)&[MIMAT0000617&hsa-miR-200c-3p&offsets|0|0(+2A);m-54&12|+|6963742|6963764&offsets|0|0(+2A)]|TAATACTGCCGGGTAATGATGGAAA</td><td>>MI0000650|hsa-mir-200c&WithFlank&12|+|6963693|6963772</td><td>50</td><td>72</td><td>23(+2A)</td><td>TAATACTGCCGGGTAATGATGGAAA</td><td>914.8</td><td>13.3</td><td>1.5</td><td>9.1</td><td>6.1E-20</td><td>4.7E-18</td><td>BT20/MB231</td></tr>
<tr><td>>MI0000433|hsa-let-7g&WithFlank&3|-|52268272|52268367@11.28.18&[MIMAT0000414&hsa-let-7g-5p&offsets|0|-4;m-58&3|-|52268336|52268357&offsets|0|-4]|TGAGGTAGTAGTTTGTAC</td><td>>MI0000433|hsa-let-7g&WithFlank&3|-|52268272|52268367</td><td>11</td><td>28</td><td>18</td><td>TGAGGTAGTAGTTTGTAC</td><td>4384.7</td><td>6.7</td><td>0.8</td><td>8.8</td><td>1.1E-18</td><td>4.8E-18</td><td>BT20/MB468</td></tr>
<tr><td>>MI0000115|hsa-mir-16-2&WithFlank&3|+|160404739|160404831@58.79.22&[MIMAT0004518&hsa-miR-16-2-3p&offsets|-1|-1;m-252&3|+|160404796|160404816&offsets|0|+1]|ACCAATATTACTGTGCTGCTTT</td><td>>MI0000115|hsa-mir-16-2&WithFlank&3|+|160404739|160404831</td><td>58</td><td>79</td><td>22</td><td>ACCAATATTACTGTGCTGCTTT</td><td>2273.0</td><td>6.6</td><td>0.8</td><td>8.8</td><td>1.2E-18</td><td>5.0E-18</td><td>MB231/MB468</td></tr>
<tr><td>>hg38_21_+_8433172_8446622_RNA45SN1_45S_50ntFlanks@11666.11702.37|TCTGCCCAGTGCTCTGAATGTCAAAGTGAAGAAATTC</td><td>>hg38_21_+_8433172_8446622_RNA45SN1_45S_50ntFlanks</td><td>11666</td><td>11702</td><td>37</td><td>TCTGCCCAGTGCTCTGAATGTCAAAGTGAAGAAATTC</td><td>981.6</td><td>5.5</td><td>0.6</td><td>8.8</td><td>1.2E-18</td><td>5.0E-18</td><td>MB231/MB468</td></tr>
<tr><td>>MI0000465|hsa-mir-191&WithFlank&3|-|49020612|49020715@22.44.23&[MIMAT0000440&hsa-miR-191-5p&offsets|0|0;m-48&3|-|49020672|49020694&offsets|0|0]|CAACGGAATCCCAAAAGCAGCTG</td><td>>MI0000465|hsa-mir-191&WithFlank&3|-|49020612|49020715</td><td>22</td><td>44</td><td>23</td><td>CAACGGAATCCCAAAAGCAGCTG</td><td>22597.8</td><td>6.6</td><td>0.8</td><td>8.8</td><td>1.2E-18</td><td>5.0E-18</td><td>MB231/MB468</td></tr>
<tr><td>>MI0000750|hsa-mir-26a-2&WithFlank&12|-|57824603|57824698@20.40.21;MI0000083|hsa-mir-26a-1&WithFlank&3|+|37969398|37969486@16.36.21&[MIMAT0000082&hsa-miR-26a-5p&offsets|0|-1;m-33&12|-|57824658|57824679&offsets|0|-1];[MIMAT0000082_1&hsa-miR-26a-5p&offsets|0|-1;m-34&3|+|37969413|37969434&offsets|0|-1]|TTCAAGTAATCCAGGATAGGC</td><td>>MI0000750|hsa-mir-26a-2&WithFlank&12|-|57824603|57824698</td><td>20</td><td>40</td><td>21</td><td>TTCAAGTAATCCAGGATAGGC</td><td>3632.4</td><td>7.3</td><td>0.8</td><td>8.8</td><td>1.3E-18</td><td>5.4E-18</td><td>MB231/MB468</td></tr>
<tr><td>>hg38_21_+_8433172_8446622_RNA45SN1_45S_50ntFlanks@4730.4767.38|CAAGAACGAAAGTCGGAGGTTCGAAGACGATCAGATAC</td><td>>hg38_21_+_8433172_8446622_RNA45SN1_45S_50ntFlanks</td><td>4730</td><td>4767</td><td>38</td><td>CAAGAACGAAAGTCGGAGGTTCGAAGACGATCAGATAC</td><td>786.8</td><td>10.1</td><td>1.2</td><td>8.8</td><td>1.3E-18</td><td>5.4E-18</td><td>MB231/MB468</td></tr>
<tr><td>>hg38_21_+_8433172_8446622_RNA45SN1_45S_50ntFlanks@5484.5514.31|GAGCGCTGAGAAGACGGTCGAACTTGACTAT</td><td>>hg38_21_+_8433172_8446622_RNA45SN1_45S_50ntFlanks</td><td>5484</td><td>5514</td><td>31</td><td>GAGCGCTGAGAAGACGGTCGAACTTGACTAT</td><td>1489.7</td><td>-5.2</td><td>0.6</td><td>-8.8</td><td>1.3E-18</td><td>5.4E-18</td><td>MB231/MB468</td></tr>
<tr><td>>hg38_1_-_228634819_228635039_RNA5S12_5S_50ntFlanks@149.171.23|GAATACCGGGTGCTGTAGGCTTT</td><td>>hg38_1_-_228634819_228635039_RNA5S12_5S_50ntFlanks</td><td>149</td><td>171</td><td>23</td><td>GAATACCGGGTGCTGTAGGCTTT</td><td>3520.9</td><td>6.8</td><td>0.8</td><td>8.8</td><td>1.4E-18</td><td>5.6E-18</td><td>MB231/MB468</td></tr>
<tr><td>>MI0000095|hsa-mir-93&WithFlank&7|-|100093762|100093853@17.40.24&[MIMAT0000093&hsa-miR-93-5p&offsets|0|+1;m-29&7|-|100093815|100093837&offsets|0|+1]|CAAAGTGCTGTTCGTGCAGGTAGT</td><td>>MI0000095|hsa-mir-93&WithFlank&7|-|100093762|100093853</td><td>17</td><td>40</td><td>24</td><td>CAAAGTGCTGTTCGTGCAGGTAGT</td><td>12220.0</td><td>7.3</td><td>0.8</td><td>8.8</td><td>1.4E-18</td><td>5.6E-18</td><td>MB231/MB468</td></tr>
<tr><td>>hg38_21_+_8433172_8446622_RNA45SN1_45S_50ntFlanks@8177.8195.19|TTCTGATCGAGGCCCAGCC</td><td>>hg38_21_+_8433172_8446622_RNA45SN1_45S_50ntFlanks</td><td>8177</td><td>8195</td><td>19</td><td>TTCTGATCGAGGCCCAGCC</td><td>317.8</td><td>3.9</td><td>0.4</td><td>8.8</td><td>1.3E-18</td><td>5.7E-18</td><td>BT20/MB468</td></tr>
</table>

<table>
<tr><td>>hg38_21_+_8433172_8446622_RNA45SN1_45S_50ntFlanks@7991.8008.18|ACGTGGCGACCCGCTGAA</td><td>>hg38_21_+_8433172_8446622_RNA45SN1_45S_50ntFlanks</td><td>7991</td><td>8008</td><td>18</td><td>ACGTGGCGACCCGCTGAA</td><td>6076.2</td><td>-5.1</td><td>0.6</td><td>-8.8</td><td>1.3E-18</td><td>5.7E-18</td><td>BT20/MB468</td></tr>
<tr><td>>hg38_21_+_8433172_8446622_RNA45SN1_45S_50ntFlanks@11894.11912.19|ACTCTAGTCTGGCACGGTG</td><td>>hg38_21_+_8433172_8446622_RNA45SN1_45S_50ntFlanks</td><td>11894</td><td>11912</td><td>19</td><td>ACTCTAGTCTGGCACGGTG</td><td>1050.5</td><td>-11.6</td><td>1.3</td><td>-8.8</td><td>1.4E-18</td><td>5.8E-18</td><td>MB231/MB468</td></tr>
<tr><td>>MI0000088|hsa-mir-30a&WithFlank&6|-|71403545|71403627@53.73.21(+1G)&[MIMAT0000088&hsa-miR-30a-3p&offsets|0|-1(+1G);m-38&6|-|71403554|71403575&offsets|0|-1(+1G)]|CTTTCAGTCGGATGTTTGCAGG</td><td>>MI0000088|hsa-mir-30a&WithFlank&6|-|71403545|71403627</td><td>53</td><td>73</td><td>21(+1G)</td><td>CTTTCAGTCGGATGTTTGCAGG</td><td>804.3</td><td>10.1</td><td>1.1</td><td>8.8</td><td>1.4E-18</td><td>5.8E-18</td><td>MB231/MB468</td></tr>
<tr><td>>hg38_21_+_8433172_8446622_RNA45SN1_45S_50ntFlanks@11578.11605.28|TAGAACTGGTGCGGACCAGGGGAATCCG</td><td>>hg38_21_+_8433172_8446622_RNA45SN1_45S_50ntFlanks</td><td>11578</td><td>11605</td><td>28</td><td>TAGAACTGGTGCGGACCAGGGGAATCCG</td><td>344.3</td><td>-4.6</td><td>0.5</td><td>-8.8</td><td>1.3E-18</td><td>5.8E-18</td><td>BT20/MB468</td></tr>
<tr><td>>hg38_21_+_8433172_8446622_RNA45SN1_45S_50ntFlanks@12159.12181.23|TTTGACTGGGGCGGTACACCTGT</td><td>>hg38_21_+_8433172_8446622_RNA45SN1_45S_50ntFlanks</td><td>12159</td><td>12181</td><td>23</td><td>TTTGACTGGGGCGGTACACCTGT</td><td>1399.3</td><td>-7.7</td><td>0.9</td><td>-8.8</td><td>1.3E-18</td><td>5.8E-18</td><td>BT20/MB468</td></tr>
<tr><td>>MI0000082|hsa-mir-25&WithFlank&7|-|100093554|100093649@58.79.22(+2A)&[MIMAT0000081&hsa-miR-25-3p&offsets|0|0(+2A);m-23&7|-|100093571|100093592&offsets|0|0(+2A)]|CATTGCACTTGTCTCGGTCTGAAA</td><td>>MI0000082|hsa-mir-25&WithFlank&7|-|100093554|100093649</td><td>58</td><td>79</td><td>22(+2A)</td><td>CATTGCACTTGTCTCGGTCTGAAA</td><td>1043.2</td><td>6.5</td><td>0.7</td><td>8.8</td><td>1.3E-18</td><td>5.9E-18</td><td>BT20/MB468</td></tr>
<tr><td>>hg38_21_+_8433172_8446622_RNA45SN1_45S_50ntFlanks@5477.5516.40|CCTGGCGGAGCGCTGAGAAGACGGTCGAACTTGACTATCT</td><td>>hg38_21_+_8433172_8446622_RNA45SN1_45S_50ntFlanks</td><td>5477</td><td>5516</td><td>40</td><td>CCTGGCGGAGCGCTGAGAAGACGGTCGAACTTGACTATCT</td><td>259.6</td><td>12.7</td><td>1.4</td><td>8.8</td><td>1.3E-18</td><td>5.9E-18</td><td>BT20/MB468</td></tr>
<tr><td>>hg38_21_+_8433172_8446622_RNA45SN1_45S_50ntFlanks@10351.10378.28|CGAGAACTTTGAAGGCCGAAGTGGAGAA</td><td>>hg38_21_+_8433172_8446622_RNA45SN1_45S_50ntFlanks</td><td>10351</td><td>10378</td><td>28</td><td>CGAGAACTTTGAAGGCCGAAGTGGAGAA</td><td>900.0</td><td>-5.4</td><td>0.6</td><td>-8.8</td><td>1.3E-18</td><td>5.9E-18</td><td>BT20/MB468</td></tr>
<tr><td>>hg38_MT_+_1621_3279_RNR2_16S_50ntFlanks@460.492.33|TTGTAAATTTAACTGTTAGTCCAAAGAGGAACA</td><td>>hg38_MT_+_1621_3279_RNR2_16S_50ntFlanks</td><td>460</td><td>492</td><td>33</td><td>TTGTAAATTTAACTGTTAGTCCAAAGAGGAACA</td><td>1917.2</td><td>10.5</td><td>1.2</td><td>8.8</td><td>1.5E-18</td><td>6.1E-18</td><td>MB231/MB468</td></tr>
<tr><td>>hg38_21_+_8433172_8446622_RNA45SN1_45S_50ntFlanks@12456.12483.28|CCAAGCGTTGGATTGTTCACCCACTAAT</td><td>>hg38_21_+_8433172_8446622_RNA45SN1_45S_50ntFlanks</td><td>12456</td><td>12483</td><td>28</td><td>CCAAGCGTTGGATTGTTCACCCACTAAT</td><td>1697.6</td><td>-6.1</td><td>0.7</td><td>-8.8</td><td>1.4E-18</td><td>6.1E-18</td><td>BT20/MB468</td></tr>
<tr><td>>MI0000469|hsa-mir-125a&WithFlank&19|+|51693248|51693345@21.44.24&[MIMAT0000443&hsa-miR-125a-5p&offsets|0|0;m-63&19|+|51693268|51693289&offsets|0|+2]|TCCCTGAGACCCTTTAACCTGTGA</td><td>>MI0000469|hsa-mir-125a&WithFlank&19|+|51693248|51693345</td><td>21</td><td>44</td><td>24</td><td>TCCCTGAGACCCTTTAACCTGTGA</td><td>9023.0</td><td>6.5</td><td>0.7</td><td>8.8</td><td>1.4E-18</td><td>6.2E-18</td><td>BT20/MB468</td></tr>
<tr><td>>MI0000082|hsa-mir-25&WithFlank&7|-|100093554|100093649@58.78.21(+1C)&[MIMAT0000081&hsa-miR-25-3p&offsets|0|-1(+1C);m-23&7|-|100093571|100093592&offsets|0|-1(+1C)]|CATTGCACTTGTCTCGGTCTGC</td><td>>MI0000082|hsa-mir-25&WithFlank&7|-|100093554|100093649</td><td>58</td><td>78</td><td>21(+1C)</td><td>CATTGCACTTGTCTCGGTCTGC</td><td>935.4</td><td>8.7</td><td>1.0</td><td>8.8</td><td>1.4E-18</td><td>6.3E-18</td><td>BT20/MB468</td></tr>
<tr><td>>hg38_21_+_8433172_8446622_RNA45SN1_45S_50ntFlanks@3984.4002.19|GCGGCTTTGGTGACTCTAG</td><td>>hg38_21_+_8433172_8446622_RNA45SN1_45S_50ntFlanks</td><td>3984</td><td>4002</td><td>19</td><td>GCGGCTTTGGTGACTCTAG</td><td>627.8</td><td>-11.3</td><td>1.3</td><td>-8.8</td><td>1.4E-18</td><td>6.3E-18</td><td>BT20/MB468</td></tr>
<tr><td>>hg38_21_+_8433172_8446622_RNA45SN1_45S_50ntFlanks@11572.11601.30|CCGACTTAGAACTGGTGCGGACCAGGGGAA</td><td>>hg38_21_+_8433172_8446622_RNA45SN1_45S_50ntFlanks</td><td>11572</td><td>11601</td><td>30</td><td>CCGACTTAGAACTGGTGCGGACCAGGGGAA</td><td>5667.9</td><td>-5.7</td><td>0.6</td><td>-8.8</td><td>1.5E-18</td><td>6.3E-18</td><td>MB231/MB468</td></tr>
<tr><td>>hg38_21_+_8433172_8446622_RNA45SN1_45S_50ntFlanks@12997.13015.19|CGCTGCGATCTATTGAAAG</td><td>>hg38_21_+_8433172_8446622_RNA45SN1_45S_50ntFlanks</td><td>12997</td><td>13015</td><td>19</td><td>CGCTGCGATCTATTGAAAG</td><td>2058.1</td><td>4.8</td><td>0.5</td><td>8.8</td><td>1.5E-18</td><td>6.3E-18</td><td>MB231/MB468</td></tr>
<tr><td>>MI0003560|hsa-mir-92b&WithFlank&1|+|155195171|155195278@67.86.20(+1U)&[MIMAT0003218&hsa-miR-92b-3p&offsets|0|-2(+1U);m-115&1|+|155195237|155195258&offsets|0|-2(+1U)]|TATTGCACTCGTCCCGGCCTT</td><td>>MI0003560|hsa-mir-92b&WithFlank&1|+|155195171|155195278</td><td>67</td><td>86</td><td>20(+1U)</td><td>TATTGCACTCGTCCCGGCCTT</td><td>820.1</td><td>11.5</td><td>1.3</td><td>8.8</td><td>1.5E-18</td><td>6.5E-18</td><td>BT20/MB468</td></tr>
<tr><td>>hg38_21_+_8433172_8446622_RNA45SN1_45S_50ntFlanks@12995.13036.42|CTCGCTGCGATCTATTGAAAGTCAGCCCTCGACACAAGGGTT</td><td>>hg38_21_+_8433172_8446622_RNA45SN1_45S_50ntFlanks</td><td>12995</td><td>13036</td><td>42</td><td>CTCGCTGCGATCTATTGAAAGTCAGCCCTCGACACAAGGGTT</td><td>3051.9</td><td>10.0</td><td>1.1</td><td>8.8</td><td>1.5E-18</td><td>6.5E-18</td><td>BT20/MB468</td></tr>
<tr><td>>hg38_21_+_8433172_8446622_RNA45SN1_45S_50ntFlanks@9575.9603.29|ATCGGTCGTCCGACCTGGGTATAGGGGCG</td><td>>hg38_21_+_8433172_8446622_RNA45SN1_45S_50ntFlanks</td><td>9575</td><td>9603</td><td>29</td><td>ATCGGTCGTCCGACCTGGGTATAGGGGCG</td><td>507.9</td><td>-11.0</td><td>1.2</td><td>-8.8</td><td>1.5E-18</td><td>6.6E-18</td><td>BT20/MB468</td></tr>
<tr><td>>hg38_21_+_8433172_8446622_RNA45SN1_45S_50ntFlanks@10679.10711.33|GGTTCCGGCGGCGTCCGGTGAGCTCTCGCTGGC</td><td>>hg38_21_+_8433172_8446622_RNA45SN1_45S_50ntFlanks</td><td>10679</td><td>10711</td><td>33</td><td>GGTTCCGGCGGCGTCCGGTGAGCTCTCGCTGGC</td><td>5973.1</td><td>-8.0</td><td>0.9</td><td>-8.8</td><td>1.5E-18</td><td>6.6E-18</td><td>BT20/MB468</td></tr>
<tr><td>>hg38_21_+_8433172_8446622_RNA45SN1_45S_50ntFlanks@10466.10497.32|CGTTGCCCTCGGCCGATCGAAAGGGAGTCGGG</td><td>>hg38_21_+_8433172_8446622_RNA45SN1_45S_50ntFlanks</td><td>10466</td><td>10497</td><td>32</td><td>CGTTGCCCTCGGCCGATCGAAAGGGAGTCGGG</td><td>542.5</td><td>-11.1</td><td>1.3</td><td>-8.8</td><td>1.5E-18</td><td>6.7E-18</td><td>BT20/MB468</td></tr>
<tr><td>>hg38_21_+_8433172_8446622_RNA45SN1_45S_50ntFlanks@5494.5516.23|AAGACGGTCGAACTTGACTATCT</td><td>>hg38_21_+_8433172_8446622_RNA45SN1_45S_50ntFlanks</td><td>5494</td><td>5516</td><td>23</td><td>AAGACGGTCGAACTTGACTATCT</td><td>1178.6</td><td>7.0</td><td>0.8</td><td>8.8</td><td>1.6E-18</td><td>6.7E-18</td><td>MB231/MB468</td></tr>
<tr><td>>hg38_21_+_8433172_8446622_RNA45SN1_45S_50ntFlanks@11161.11178.18|CACGGTCCCCCGCGAGGG</td><td>>hg38_21_+_8433172_8446622_RNA45SN1_45S_50ntFlanks</td><td>11161</td><td>11178</td><td>18</td><td>CACGGTCCCCCGCGAGGG</td><td>1126.2</td><td>-11.7</td><td>1.3</td><td>-8.8</td><td>1.6E-18</td><td>6.7E-18</td><td>MB231/MB468</td></tr>
<tr><td>>MI0001445|hsa-mir-423&WithFlank&17|+|30117073|30117178@59.77.19(+1U)&[MIMAT0001340&hsa-miR-423-3p&offsets|0|-4(+1U);m-100&17|+|30117131|30117153&offsets|0|-4(+1U)]|AGCTCGGTCTGAGGCCCCTT</td><td>>MI0001445|hsa-mir-423&WithFlank&17|+|30117073|30117178</td><td>59</td><td>77</td><td>19(+1U)</td><td>AGCTCGGTCTGAGGCCCCTT</td><td>1155.5</td><td>9.4</td><td>1.1</td><td>8.8</td><td>1.7E-18</td><td>6.8E-18</td><td>MB231/MB468</td></tr>
<tr><td>>hg38_21_+_8433172_8446622_RNA45SN1_45S_50ntFlanks@12073.12090.18|CCGAGGGGCTCTCGCTTC</td><td>>hg38_21_+_8433172_8446622_RNA45SN1_45S_50ntFlanks</td><td>12073</td><td>12090</td><td>18</td><td>CCGAGGGGCTCTCGCTTC</td><td>368.6</td><td>-4.5</td><td>0.5</td><td>-8.8</td><td>1.7E-18</td><td>6.8E-18</td><td>MB231/MB468</td></tr>
<tr><td>>MI0000434|hsa-let-7i&WithFlank&12|+|62603680|62603775@12.31.20(+2C)&[MIMAT0000415&hsa-let-7i-5p&offsets|0|-2(+2C);m-49&12|+|62603691|62603712&offsets|0|-2(+2C)]|TGAGGTAGTAGTTTGTGCTGCC</td><td>>MI0000434|hsa-let-7i&WithFlank&12|+|62603680|62603775</td><td>12</td><td>31</td><td>20(+2C)</td><td>TGAGGTAGTAGTTTGTGCTGCC</td><td>963.1</td><td>9.7</td><td>1.1</td><td>8.8</td><td>1.7E-18</td><td>6.9E-18</td><td>MB231/MB468</td></tr>
<tr><td>>hg38_21_+_8433172_8446622_RNA45SN1_45S_50ntFlanks@10391.10409.19|AACAGCAGTTGAACATGGG</td><td>>hg38_21_+_8433172_8446622_RNA45SN1_45S_50ntFlanks</td><td>10391</td><td>10409</td><td>19</td><td>AACAGCAGTTGAACATGGG</td><td>1930.4</td><td>-10.5</td><td>1.2</td><td>-8.8</td><td>1.7E-18</td><td>6.9E-18</td><td>MB231/MB468</td></tr>
<tr><td>>hg38_21_+_8433172_8446622_RNA45SN1_45S_50ntFlanks@10410.10433.24|TCAGTCGGTCCTGAGAGATGGGCG</td><td>>hg38_21_+_8433172_8446622_RNA45SN1_45S_50ntFlanks</td><td>10410</td><td>10433</td><td>24</td><td>TCAGTCGGTCCTGAGAGATGGGCG</td><td>852.4</td><td>-6.3</td><td>0.7</td><td>-8.8</td><td>1.7E-18</td><td>7.0E-18</td><td>MB231/MB468</td></tr>
<tr><td>>hg38_21_+_8433172_8446622_RNA45SN1_45S_50ntFlanks@11316.11336.21|CACCCCACGTCTCGTCGCGCG</td><td>>hg38_21_+_8433172_8446622_RNA45SN1_45S_50ntFlanks</td><td>11316</td><td>11336</td><td>21</td><td>CACCCCACGTCTCGTCGCGCG</td><td>898.6</td><td>-8.9</td><td>1.0</td><td>-8.8</td><td>1.8E-18</td><td>7.2E-18</td><td>MB231/MB468</td></tr>
<tr><td>>hg38_21_+_8433172_8446622_RNA45SN1_45S_50ntFlanks@11585.11616.32|GGTGCGGACCAGGGGAATCCGACTGTTTAATT</td><td>>hg38_21_+_8433172_8446622_RNA45SN1_45S_50ntFlanks</td><td>11585</td><td>11616</td><td>32</td><td>GGTGCGGACCAGGGGAATCCGACTGTTTAATT</td><td>372.9</td><td>8.5</td><td>1.0</td><td>8.8</td><td>1.8E-18</td><td>7.3E-18</td><td>MB231/MB468</td></tr>
<tr><td>>hg38_MT_+_1621_3279_RNR2_16S_50ntFlanks@1363.1388.26|GACCTCGATGTTGGATCAGGACATCC</td><td>>hg38_MT_+_1621_3279_RNR2_16S_50ntFlanks</td><td>1363</td><td>1388</td><td>26</td><td>GACCTCGATGTTGGATCAGGACATCC</td><td>870.6</td><td>10.0</td><td>1.1</td><td>8.8</td><td>1.8E-18</td><td>7.3E-18</td><td>MB231/MB468</td></tr>
<tr><td>>MI0000077|hsa-mir-21&WithFlank&17|+|59841260|59841343@14.35.22(+2A)&[MIMAT0000076&hsa-miR-21-5p&offsets|0|0(+2A);m-1&17|+|59841273|59841294&offsets|0|0(+2A)]|TAGCTTATCAGACTGATGTTGAAA</td><td>>MI0000077|hsa-mir-21&WithFlank&17|+|59841260|59841343</td><td>14</td><td>35</td><td>22(+2A)</td><td>TAGCTTATCAGACTGATGTTGAAA</td><td>2230.4</td><td>6.7</td><td>0.8</td><td>8.8</td><td>1.8E-18</td><td>7.3E-18</td><td>MB231/MB468</td></tr>
<tr><td>>hg38_21_+_8433172_8446622_RNA45SN1_45S_50ntFlanks@12567.12587.21|CTCAGTACGAGAGGAACCGCA</td><td>>hg38_21_+_8433172_8446622_RNA45SN1_45S_50ntFlanks</td><td>12567</td><td>12587</td><td>21</td><td>CTCAGTACGAGAGGAACCGCA</td><td>400.9</td><td>-7.4</td><td>0.8</td><td>-8.8</td><td>1.8E-18</td><td>7.3E-18</td><td>MB231/MB468</td></tr>
<tr><td>>hg38_21_+_8433172_8446622_RNA45SN1_45S_50ntFlanks@11669.11692.24|GCCCAGTGCTCTGAATGTCAAAGT</td><td>>hg38_21_+_8433172_8446622_RNA45SN1_45S_50ntFlanks</td><td>11669</td><td>11692</td><td>24</td><td>GCCCAGTGCTCTGAATGTCAAAGT</td><td>487.9</td><td>7.5</td><td>0.9</td><td>8.8</td><td>1.8E-18</td><td>7.3E-18</td><td>MB231/MB468</td></tr>
<tr><td>>hg38_21_+_8433172_8446622_RNA45SN1_45S_50ntFlanks@5472.5489.18|ACGGCCCTGGCGGAGCGC</td><td>>hg38_21_+_8433172_8446622_RNA45SN1_45S_50ntFlanks</td><td>5472</td><td>5489</td><td>18</td><td>ACGGCCCTGGCGGAGCGC</td><td>1760.6</td><td>10.5</td><td>1.2</td><td>8.8</td><td>1.8E-18</td><td>7.3E-18</td><td>MB231/MB468</td></tr>
<tr><td>>hg38_21_+_8433172_8446622_RNA45SN1_45S_50ntFlanks@12405.12425.21|CTTTTTGATCCTTCGATGTCG</td><td>>hg38_21_+_8433172_8446622_RNA45SN1_45S_50ntFlanks</td><td>12405</td><td>12425</td><td>21</td><td>CTTTTTGATCCTTCGATGTCG</td><td>1125.4</td><td>-8.7</td><td>1.0</td><td>-8.8</td><td>1.7E-18</td><td>7.4E-18</td><td>BT20/MB468</td></tr>
<tr><td>>hg38_21_+_8433172_8446622_RNA45SN1_45S_50ntFlanks@10845.10863.19|ACTTCGGGATAAGGATTGG</td><td>>hg38_21_+_8433172_8446622_RNA45SN1_45S_50ntFlanks</td><td>10845</td><td>10863</td><td>19</td><td>ACTTCGGGATAAGGATTGG</td><td>392.0</td><td>-6.5</td><td>0.7</td><td>-8.8</td><td>1.7E-18</td><td>7.5E-18</td><td>BT20/MB468</td></tr>
<tr><td>>hg38_1_-_228634819_228635039_RNA5S12_5S_50ntFlanks@153.171.19|ACCGGGTGCTGTAGGCTTT</td><td>>hg38_1_-_228634819_228635039_RNA5S12_5S_50ntFlanks</td><td>153</td><td>171</td><td>19</td><td>ACCGGGTGCTGTAGGCTTT</td><td>21635.8</td><td>7.7</td><td>0.9</td><td>8.8</td><td>1.7E-18</td><td>7.5E-18</td><td>BT20/MB468</td></tr>
<tr><td>>MI0000264|hsa-mir-7-2&WithFlank&15|+|88611819|88611940@38.59.22(+1G);MI0000265|hsa-mir-7-3&WithFlank&19|+|4770664|4770785@37.58.22(+1G);MI0000263|hsa-mir-7-1&WithFlank&9|-|83969742|83969863@30.51.22(+1G)&[MIMAT0000252&hsa-miR-7-5p&offsets|0|-2(+1G);m-400&15|+|88611856|88611879&offsets|0|-2(+1G)];[MIMAT0000252_1&hsa-miR-7-5p&offsets|0|-2(+1G);m-402&19|+|4770700|4770723&offsets|0|-2(+1G)];[MIMAT0000252_2&hsa-miR-7-5p&offsets|0|-2(+1G);m-401&9|-|83969811|83969834&offsets|0|-2(+1G)]|TGGAAGACTAGTGATTTTGTTGG</td><td>>MI0000264|hsa-mir-7-2&WithFlank&15|+|88611819|88611940</td><td>38</td><td>59</td><td>22(+1G)</td><td>TGGAAGACTAGTGATTTTGTTGG</td><td>1581.6</td><td>7.9</td><td>0.9</td><td>8.8</td><td>1.9E-18</td><td>7.5E-18</td><td>MB231/MB468</td></tr>
<tr><td>>MI0003205|hsa-mir-532&WithFlank&X|+|50003142|50003244@26.46.21(+1C)&[MIMAT0002888&hsa-miR-532-5p&offsets|0|-1(+1C);m-65&X|+|50003167|50003188&offsets|0|-1(+1C)]|CATGCCTTGAGTGTAGGACCGC</td><td>>MI0003205|hsa-mir-532&WithFlank&X|+|50003142|50003244</td><td>26</td><td>46</td><td>21(+1C)</td><td>CATGCCTTGAGTGTAGGACCGC</td><td>968.0</td><td>6.3</td><td>0.7</td><td>8.8</td><td>1.9E-18</td><td>7.7E-18</td><td>MB231/MB468</td></tr>
<tr><td>>hg38_21_+_8433172_8446622_RNA45SN1_45S_50ntFlanks@6742.6771.30|TCGACACTTCGAACGCACTTGCGGCCCCGG</td><td>>hg38_21_+_8433172_8446622_RNA45SN1_45S_50ntFlanks</td><td>6742</td><td>6771</td><td>30</td><td>TCGACACTTCGAACGCACTTGCGGCCCCGG</td><td>314.7</td><td>-5.6</td><td>0.6</td><td>-8.8</td><td>1.8E-18</td><td>7.8E-18</td><td>BT20/MB468</td></tr>
<tr><td>>hg38_21_+_8433172_8446622_RNA45SN1_45S_50ntFlanks@11729.11752.24|TAACTATGACTCTCTTAAGGTAGC</td><td>>hg38_21_+_8433172_8446622_RNA45SN1_45S_50ntFlanks</td><td>11729</td><td>11752</td><td>24</td><td>TAACTATGACTCTCTTAAGGTAGC</td><td>2019.8</td><td>-6.2</td><td>0.7</td><td>-8.8</td><td>1.9E-18</td><td>8.2E-18</td><td>BT20/MB468</td></tr>
<tr><td>>hg38_21_+_8433172_8446622_RNA45SN1_45S_50ntFlanks@10866.10890.25|CTAAGGGCTGGGTCGGTCGGGCTGG</td><td>>hg38_21_+_8433172_8446622_RNA45SN1_45S_50ntFlanks</td><td>10866</td><td>10890</td><td>25</td><td>CTAAGGGCTGGGTCGGTCGGGCTGG</td><td>266.4</td><td>-4.6</td><td>0.5</td><td>-8.8</td><td>2.0E-18</td><td>8.3E-18</td><td>MB231/MB468</td></tr>
<tr><td>>hg38_21_+_8433172_8446622_RNA45SN1_45S_50ntFlanks@7978.8010.33|GACCTCAGATCAGACGTGGCGACCCGCTGAATT</td><td>>hg38_21_+_8433172_8446622_RNA45SN1_45S_50ntFlanks</td><td>7978</td><td>8010</td><td>33</td><td>GACCTCAGATCAGACGTGGCGACCCGCTGAATT</td><td>406.4</td><td>8.4</td><td>1.0</td><td>8.8</td><td>2.1E-18</td><td>8.4E-18</td><td>MB231/MB468</td></tr>
</table>

<table>
<tr><td></td><td></td><td></td><td></td><td></td><td></td><td></td><td></td><td></td><td></td><td></td><td></td><td></td></tr>
<tr><td>>MI0000081|hsa-mir-24-2&WithFlank&19|-|13836281|13836365@56.76.21(+1A);MI0000080|hsa-mir-24-1&WithFlank&9|+|95086015|95086094@50.70.21(+1A)&[MIMAT0000080&hsa-miR-24-3p&offsets|0|-1(+1A);m-36&19|-|13836288|13836310&offsets|0|-2(+1A)];[MIMAT0000080_1&hsa-miR-24-3p&offsets|0|-1(+1A);m-35&9|+|95086064|95086086&offsets|0|-2(+1A)]|TGGCTCAGTTCAGCAGGAACAA</td><td>>MI0000081|hsa-mir-24-2&WithFlank&19|-|13836281|13836365</td><td>56</td><td>76</td><td>21(+1A)</td><td>TGGCTCAGTTCAGCAGGAACAA</td><td>2932.9</td><td>6.2</td><td>0.7</td><td>8.8</td><td>2.1E-18</td><td>8.4E-18</td><td>MB231/MB468</td></tr>
<tr><td>>hg38_1_-_228634819_228635039_RNA5S12_5S_50ntFlanks@153.170.18|ACCGGGTGCTGTAGGCTT</td><td>>hg38_1_-_228634819_228635039_RNA5S12_5S_50ntFlanks</td><td>153</td><td>170</td><td>18</td><td>ACCGGGTGCTGTAGGCTT</td><td>34926.9</td><td>8.0</td><td>0.9</td><td>8.8</td><td>1.9E-18</td><td>8.4E-18</td><td>BT20/MB468</td></tr>
<tr><td>>MI0000087|hsa-mir-29a&WithFlank&7|-|130876741|130876816@48.67.20&[MIMAT0000086&hsa-miR-29a-3p&offsets|0|-2;m-26&7|-|130876749|130876769&offsets|0|-1]|TAGCACCATCTGAAATCGGT</td><td>>MI0000087|hsa-mir-29a&WithFlank&7|-|130876741|130876816</td><td>48</td><td>67</td><td>20</td><td>TAGCACCATCTGAAATCGGT</td><td>1329.5</td><td>10.4</td><td>1.2</td><td>8.8</td><td>2.1E-18</td><td>8.4E-18</td><td>MB231/MB468</td></tr>
<tr><td>>hg38_21_+_8433172_8446622_RNA45SN1_45S_50ntFlanks@10362.10388.27|AAGGCCGAAGTGGAGAAGGGTTCCATG</td><td>>hg38_21_+_8433172_8446622_RNA45SN1_45S_50ntFlanks</td><td>10362</td><td>10388</td><td>27</td><td>AAGGCCGAAGTGGAGAAGGGTTCCATG</td><td>2961.2</td><td>-13.5</td><td>1.5</td><td>-8.8</td><td>2.0E-18</td><td>8.8E-18</td><td>BT20/MB468</td></tr>
<tr><td>>MI0000071|hsa-mir-17&WithFlank&13|+|91350599|91350694@20.40.21&[MIMAT0000070&hsa-miR-17-5p&offsets|0|-2;m-88&13|+|91350618|91350640&offsets|0|-2]|CAAAGTGCTTACAGTGCAGGT</td><td>>MI0000071|hsa-mir-17&WithFlank&13|+|91350599|91350694</td><td>20</td><td>40</td><td>21</td><td>CAAAGTGCTTACAGTGCAGGT</td><td>3035.5</td><td>6.6</td><td>0.8</td><td>8.7</td><td>2.2E-18</td><td>8.9E-18</td><td>MB231/MB468</td></tr>
<tr><td>>hg38_21_+_8433172_8446622_RNA45SN1_45S_50ntFlanks@11233.11254.22|CCCTTCCCGTGGATCGCCCCAG</td><td>>hg38_21_+_8433172_8446622_RNA45SN1_45S_50ntFlanks</td><td>11233</td><td>11254</td><td>22</td><td>CCCTTCCCGTGGATCGCCCCAG</td><td>971.1</td><td>-11.9</td><td>1.4</td><td>-8.8</td><td>2.1E-18</td><td>8.9E-18</td><td>BT20/MB468</td></tr>
<tr><td>>hg38_21_+_8433172_8446622_RNA45SN1_45S_50ntFlanks@9533.9553.21|GAGGAAACTCTGGTGGAGGTC</td><td>>hg38_21_+_8433172_8446622_RNA45SN1_45S_50ntFlanks</td><td>9533</td><td>9553</td><td>21</td><td>GAGGAAACTCTGGTGGAGGTC</td><td>6287.9</td><td>-5.6</td><td>0.6</td><td>-8.8</td><td>2.1E-18</td><td>9.1E-18</td><td>BT20/MB468</td></tr>
<tr><td>>hg38_21_+_8433172_8446622_RNA45SN1_45S_50ntFlanks@12631.12654.24|GGCGAAGCTACCATCTGTGGGATT</td><td>>hg38_21_+_8433172_8446622_RNA45SN1_45S_50ntFlanks</td><td>12631</td><td>12654</td><td>24</td><td>GGCGAAGCTACCATCTGTGGGATT</td><td>388.2</td><td>-4.8</td><td>0.6</td><td>-8.7</td><td>2.3E-18</td><td>9.2E-18</td><td>MB231/MB468</td></tr>
<tr><td>>MI0000542|hsa-mir-320a&WithFlank&8|-|22244960|22245043@49.71.23&[MIMAT0000510&hsa-miR-320a-3p&offsets|+1|+2;m-60&8|-|22244975|22244996&offsets|+1|+2]|AAAGCTGGGTTGAGAGGGCGAAA</td><td>>MI0000542|hsa-mir-320a&WithFlank&8|-|22244960|22245043</td><td>49</td><td>71</td><td>23</td><td>AAAGCTGGGTTGAGAGGGCGAAA</td><td>691.8</td><td>11.6</td><td>1.3</td><td>8.7</td><td>2.2E-18</td><td>9.3E-18</td><td>BT20/MB468</td></tr>
<tr><td>>hg38_21_+_8433172_8446622_RNA45SN1_45S_50ntFlanks@5476.5516.41|CCCTGGCGGAGCGCTGAGAAGACGGTCGAACTTGACTATCT</td><td>>hg38_21_+_8433172_8446622_RNA45SN1_45S_50ntFlanks</td><td>5476</td><td>5516</td><td>41</td><td>CCCTGGCGGAGCGCTGAGAAGACGGTCGAACTTGACTATCT</td><td>259.1</td><td>12.7</td><td>1.4</td><td>8.7</td><td>2.2E-18</td><td>9.4E-18</td><td>BT20/MB468</td></tr>
<tr><td>>hg38_21_+_8433172_8446622_RNA45SN1_45S_50ntFlanks@6652.6700.49|GACTCTTAGCGGTGGATCACTCGGCTCGTGCGTCGATGAAGAACGCAGC</td><td>>hg38_21_+_8433172_8446622_RNA45SN1_45S_50ntFlanks</td><td>6652</td><td>6700</td><td>49</td><td>GACTCTTAGCGGTGGATCACTCGGCTCGTGCGTCGATGAAGAACGCAGC</td><td>2298.6</td><td>11.4</td><td>1.3</td><td>8.7</td><td>2.3E-18</td><td>9.4E-18</td><td>MB231/MB468</td></tr>
<tr><td>>MI0000077|hsa-mir-21&WithFlank&17|+|59841260|59841343@14.39.26(+1U)&[MIMAT0000076&hsa-miR-21-5p&offsets|0|+4(+1U);m-1&17|+|59841273|59841294&offsets|0|+4(+1U)]|TAGCTTATCAGACTGATGTTGACTGTT</td><td>>MI0000077|hsa-mir-21&WithFlank&17|+|59841260|59841343</td><td>14</td><td>39</td><td>26(+1U)</td><td>TAGCTTATCAGACTGATGTTGACTGTT</td><td>821.6</td><td>6.6</td><td>0.8</td><td>8.7</td><td>2.3E-18</td><td>9.4E-18</td><td>MB231/MB468</td></tr>
<tr><td>>MI0000095|hsa-mir-93&WithFlank&7|-|100093762|100093853@17.39.23(+1C)&[MIMAT0000093&hsa-miR-93-5p&offsets|0|0(+1C);m-29&7|-|100093815|100093837&offsets|0|0(+1C)]|CAAAGTGCTGTTCGTGCAGGTAGC</td><td>>MI0000095|hsa-mir-93&WithFlank&7|-|100093762|100093853</td><td>17</td><td>39</td><td>23(+1C)</td><td>CAAAGTGCTGTTCGTGCAGGTAGC</td><td>1449.4</td><td>7.4</td><td>0.8</td><td>8.7</td><td>2.4E-18</td><td>9.5E-18</td><td>MB231/MB468</td></tr>
<tr><td>>MI0000264|hsa-mir-7-2&WithFlank&15|+|88611819|88611940@38.58.21;MI0000265|hsa-mir-7-3&WithFlank&19|+|4770664|4770785@37.57.21;MI0000263|hsa-mir-7-1&WithFlank&9|-|83969742|83969863@30.50.21&[MIMAT0000252&hsa-miR-7-5p&offsets|0|-3;m-400&15|+|88611856|88611879&offsets|0|-3];[MIMAT0000252_1&hsa-miR-7-5p&offsets|0|-3;m-402&19|+|4770700|4770723&offsets|0|-3];[MIMAT0000252_2&hsa-miR-7-5p&offsets|0|-3;m-401&9|-|83969811|83969834&offsets|0|-3]|TGGAAGACTAGTGATTTTGTT</td><td>>MI0000264|hsa-mir-7-2&WithFlank&15|+|88611819|88611940</td><td>38</td><td>58</td><td>21</td><td>TGGAAGACTAGTGATTTTGTT</td><td>5027.2</td><td>8.0</td><td>0.9</td><td>8.7</td><td>2.2E-18</td><td>9.6E-18</td><td>BT20/MB468</td></tr>
<tr><td>>hg38_21_+_8433172_8446622_RNA45SN1_45S_50ntFlanks@12157.12179.23|AGTTTGACTGGGGCGGTACACCT</td><td>>hg38_21_+_8433172_8446622_RNA45SN1_45S_50ntFlanks</td><td>12157</td><td>12179</td><td>23</td><td>AGTTTGACTGGGGCGGTACACCT</td><td>507.0</td><td>-6.2</td><td>0.7</td><td>-8.7</td><td>2.4E-18</td><td>9.7E-18</td><td>MB231/MB468</td></tr>
<tr><td>>hg38_21_+_8433172_8446622_RNA45SN1_45S_50ntFlanks@11576.11605.30|CTTAGAACTGGTGCGGACCAGGGGAATCCG</td><td>>hg38_21_+_8433172_8446622_RNA45SN1_45S_50ntFlanks</td><td>11576</td><td>11605</td><td>30</td><td>CTTAGAACTGGTGCGGACCAGGGGAATCCG</td><td>887.7</td><td>-11.8</td><td>1.3</td><td>-8.7</td><td>2.3E-18</td><td>9.7E-18</td><td>BT20/MB468</td></tr>
<tr><td>>MI0000067|hsa-let-7f-1&WithFlank&9|+|94176341|94176439@13.33.21(+1G);MI0000068|hsa-let-7f-2&WithFlank&X|-|53557186|53557280@14.34.21(+1G)&[MIMAT0000067&hsa-let-7f-5p&offsets|0|-1(+1G);m-20&9|+|94176353|94176374&offsets|0|-1(+1G)];[MIMAT0000067_1&hsa-let-7f-5p&offsets|0|-1(+1G);m-19&X|-|53557246|53557267&offsets|0|-1(+1G)]|TGAGGTAGTAGATTGTATAGTG</td><td>>MI0000067|hsa-let-7f-1&WithFlank&9|+|94176341|94176439</td><td>13</td><td>33</td><td>21(+1G)</td><td>TGAGGTAGTAGATTGTATAGTG</td><td>16081.9</td><td>7.7</td><td>0.9</td><td>8.7</td><td>2.3E-18</td><td>9.9E-18</td><td>BT20/MB468</td></tr>
<tr><td>>MI0000438|hsa-mir-15b&WithFlank&3|+|160404582|160404691@26.46.21&[MIMAT0000417&hsa-miR-15b-5p&offsets|0|-1;m-110&3|+|160404607|160404628&offsets|0|-1]|TAGCAGCACATCATGGTTTAC</td><td>>MI0000438|hsa-mir-15b&WithFlank&3|+|160404582|160404691</td><td>26</td><td>46</td><td>21</td><td>TAGCAGCACATCATGGTTTAC</td><td>912.2</td><td>7.0</td><td>0.8</td><td>8.7</td><td>2.5E-18</td><td>9.9E-18</td><td>MB231/MB468</td></tr>
<tr><td>>MI0000070|hsa-mir-16-1&WithFlank&13|-|50048967|50049067@20.41.22(+1C);MI0000115|hsa-mir-16-2&WithFlank&3|+|160404739|160404831@16.37.22(+1C)&[MIMAT0000069&hsa-miR-16-5p&offsets|0|0(+1C);m-62&13|-|50049027|50049048&offsets|0|0(+1C)];[MIMAT0000069_1&hsa-miR-16-5p&offsets|0|0(+1C);m-61&3|+|160404754|160404775&offsets|0|0(+1C)]|TAGCAGCACGTAAATATTGGCGC</td><td>>MI0000070|hsa-mir-16-1&WithFlank&13|-|50048967|50049067</td><td>20</td><td>41</td><td>22(+1C)</td><td>TAGCAGCACGTAAATATTGGCGC</td><td>2665.6</td><td>9.2</td><td>1.1</td><td>8.7</td><td>2.3E-18</td><td>1.0E-17</td><td>BT20/MB468</td></tr>
<tr><td>>hg38_21_+_8433172_8446622_RNA45SN1_45S_50ntFlanks@12933.12956.24|GCTAAACCATTCGTAGACGACCTG</td><td>>hg38_21_+_8433172_8446622_RNA45SN1_45S_50ntFlanks</td><td>12933</td><td>12956</td><td>24</td><td>GCTAAACCATTCGTAGACGACCTG</td><td>757.8</td><td>-5.8</td><td>0.7</td><td>-8.7</td><td>2.3E-18</td><td>1.0E-17</td><td>BT20/MB468</td></tr>
<tr><td>>hg38_21_+_8433172_8446622_RNA45SN1_45S_50ntFlanks@12615.12653.39|GCTGAGGAGCCAATGGGGCGAAGCTACCATCTGTGGGAT</td><td>>hg38_21_+_8433172_8446622_RNA45SN1_45S_50ntFlanks</td><td>12615</td><td>12653</td><td>39</td><td>GCTGAGGAGCCAATGGGGCGAAGCTACCATCTGTGGGAT</td><td>824.6</td><td>-7.9</td><td>0.9</td><td>-8.7</td><td>2.4E-18</td><td>1.0E-17</td><td>BT20/MB468</td></tr>
<tr><td>>hg38_21_+_8433172_8446622_RNA45SN1_45S_50ntFlanks@12463.12486.24|TTGGATTGTTCACCCACTAATAGG</td><td>>hg38_21_+_8433172_8446622_RNA45SN1_45S_50ntFlanks</td><td>12463</td><td>12486</td><td>24</td><td>TTGGATTGTTCACCCACTAATAGG</td><td>428.3</td><td>3.9</td><td>0.5</td><td>8.7</td><td>2.5E-18</td><td>1.0E-17</td><td>MB231/MB468</td></tr>
<tr><td>>hg38_21_+_8433172_8446622_RNA45SN1_45S_50ntFlanks@6652.6683.32|GACTCTTAGCGGTGGATCACTCGGCTCGTGCG</td><td>>hg38_21_+_8433172_8446622_RNA45SN1_45S_50ntFlanks</td><td>6652</td><td>6683</td><td>32</td><td>GACTCTTAGCGGTGGATCACTCGGCTCGTGCG</td><td>9234.9</td><td>6.9</td><td>0.8</td><td>8.7</td><td>2.6E-18</td><td>1.0E-17</td><td>MB231/MB468</td></tr>
<tr><td>>hg38_21_+_8433172_8446622_RNA45SN1_45S_50ntFlanks@9567.9594.28|ACGTGCAAATCGGTCGTCCGACCTGGGT</td><td>>hg38_21_+_8433172_8446622_RNA45SN1_45S_50ntFlanks</td><td>9567</td><td>9594</td><td>28</td><td>ACGTGCAAATCGGTCGTCCGACCTGGGT</td><td>317.4</td><td>-4.4</td><td>0.5</td><td>-8.7</td><td>2.4E-18</td><td>1.0E-17</td><td>BT20/MB468</td></tr>
<tr><td>>hg38_21_+_8433172_8446622_RNA45SN1_45S_50ntFlanks@6772.6803.32|GTTCCTCCCGGGGCTACGCCTGTCTGAGCGTC</td><td>>hg38_21_+_8433172_8446622_RNA45SN1_45S_50ntFlanks</td><td>6772</td><td>6803</td><td>32</td><td>GTTCCTCCCGGGGCTACGCCTGTCTGAGCGTC</td><td>374.5</td><td>-5.4</td><td>0.6</td><td>-8.7</td><td>2.4E-18</td><td>1.0E-17</td><td>BT20/MB468</td></tr>
<tr><td>>MI0000086|hsa-mir-28&WithFlank&3|+|188688775|188688872@60.79.20(+1A)&[MIMAT0004502&hsa-miR-28-3p&offsets|0|-2(+1A);m-28&3|+|188688834|188688855&offsets|0|-2(+1A)]|CACTAGATTGTGAGCTCCTGA</td><td>>MI0000086|hsa-mir-28&WithFlank&3|+|188688775|188688872</td><td>60</td><td>79</td><td>20(+1A)</td><td>CACTAGATTGTGAGCTCCTGA</td><td>1003.4</td><td>12.6</td><td>1.4</td><td>8.7</td><td>2.4E-18</td><td>1.0E-17</td><td>BT20/MB468</td></tr>
<tr><td>>MI0000750|hsa-mir-26a-2&WithFlank&12|-|57824603|57824698@20.39.20(+1U);MI0000083|hsa-mir-26a-1&WithFlank&3|+|37969398|37969486@16.35.20(+1U)&[MIMAT0000082&hsa-miR-26a-5p&offsets|0|-2(+1U);m-33&12|-|57824658|57824679&offsets|0|-2(+1U)];[MIMAT0000082_1&hsa-miR-26a-5p&offsets|0|-2(+1U);m-34&3|+|37969413|37969434&offsets|0|-2(+1U)]|TTCAAGTAATCCAGGATAGGT</td><td>>MI0000750|hsa-mir-26a-2&WithFlank&12|-|57824603|57824698</td><td>20</td><td>39</td><td>20(+1U)</td><td>TTCAAGTAATCCAGGATAGGT</td><td>1231.6</td><td>7.9</td><td>0.9</td><td>8.7</td><td>2.4E-18</td><td>1.0E-17</td><td>BT20/MB468</td></tr>
<tr><td>>hg38_21_+_8433172_8446622_RNA45SN1_45S_50ntFlanks@7982.8006.25|TCAGATCAGACGTGGCGACCCGCTG</td><td>>hg38_21_+_8433172_8446622_RNA45SN1_45S_50ntFlanks</td><td>7982</td><td>8006</td><td>25</td><td>TCAGATCAGACGTGGCGACCCGCTG</td><td>1707.8</td><td>-6.2</td><td>0.7</td><td>-8.7</td><td>2.6E-18</td><td>1.1E-17</td><td>MB231/MB468</td></tr>
<tr><td>>hg38_21_+_8433172_8446622_RNA45SN1_45S_50ntFlanks@12420.12446.27|ATGTCGGCTCTTCCTATCATTGTGAAG</td><td>>hg38_21_+_8433172_8446622_RNA45SN1_45S_50ntFlanks</td><td>12420</td><td>12446</td><td>27</td><td>ATGTCGGCTCTTCCTATCATTGTGAAG</td><td>3971.7</td><td>-5.0</td><td>0.6</td><td>-8.7</td><td>2.5E-18</td><td>1.1E-17</td><td>BT20/MB468</td></tr>
<tr><td>>hg38_21_+_8433172_8446622_RNA45SN1_45S_50ntFlanks@10408.10429.22|GGTCAGTCGGTCCTGAGAGATG</td><td>>hg38_21_+_8433172_8446622_RNA45SN1_45S_50ntFlanks</td><td>10408</td><td>10429</td><td>22</td><td>GGTCAGTCGGTCCTGAGAGATG</td><td>1996.8</td><td>-5.2</td><td>0.6</td><td>-8.7</td><td>2.7E-18</td><td>1.1E-17</td><td>MB231/MB468</td></tr>
<tr><td>>hg38_21_+_8433172_8446622_RNA45SN1_45S_50ntFlanks@7986.8003.18|ATCAGACGTGGCGACCCG</td><td>>hg38_21_+_8433172_8446622_RNA45SN1_45S_50ntFlanks</td><td>7986</td><td>8003</td><td>18</td><td>ATCAGACGTGGCGACCCG</td><td>871.3</td><td>-11.8</td><td>1.3</td><td>-8.7</td><td>2.6E-18</td><td>1.1E-17</td><td>BT20/MB468</td></tr>
<tr><td>>hg38_21_+_8433172_8446622_RNA45SN1_45S_50ntFlanks@12324.12344.21|TTGGGTTTTAAGCAGGAGGTG</td><td>>hg38_21_+_8433172_8446622_RNA45SN1_45S_50ntFlanks</td><td>12324</td><td>12344</td><td>21</td><td>TTGGGTTTTAAGCAGGAGGTG</td><td>892.1</td><td>-11.4</td><td>1.3</td><td>-8.7</td><td>2.7E-18</td><td>1.1E-17</td><td>MB231/MB468</td></tr>
<tr><td>>hg38_21_+_8433172_8446622_RNA45SN1_45S_50ntFlanks@7979.8006.28|ACCTCAGATCAGACGTGGCGACCCGCTG</td><td>>hg38_21_+_8433172_8446622_RNA45SN1_45S_50ntFlanks</td><td>7979</td><td>8006</td><td>28</td><td>ACCTCAGATCAGACGTGGCGACCCGCTG</td><td>3714.7</td><td>-6.1</td><td>0.7</td><td>-8.7</td><td>2.8E-18</td><td>1.1E-17</td><td>MB231/MB468</td></tr>
<tr><td>>hg38_21_+_8433172_8446622_RNA45SN1_45S_50ntFlanks@485.526.42|GATGTGGTGACGTCGTGCTCTCCCGGGCCGGGTCCGAGCCGC</td><td>>hg38_21_+_8433172_8446622_RNA45SN1_45S_50ntFlanks</td><td>485</td><td>526</td><td>42</td><td>GATGTGGTGACGTCGTGCTCTCCCGGGCCGGGTCCGAGCCGC</td><td>453.9</td><td>13.5</td><td>1.5</td><td>8.7</td><td>2.7E-18</td><td>1.1E-17</td><td>BT20/MB468</td></tr>
<tr><td>>MI0000811|hsa-mir-148b&WithFlank&12|+|54337210|54337320@69.89.21(+1G)&[MIMAT0000759&hsa-miR-148b-3p&offsets|0|-1(+1G);m-77&12|+|54337278|54337299&offsets|0|-1(+1G)]|TCAGTGCATCACAGAACTTTGG</td><td>>MI0000811|hsa-mir-148b&WithFlank&12|+|54337210|54337320</td><td>69</td><td>89</td><td>21(+1G)</td><td>TCAGTGCATCACAGAACTTTGG</td><td>1489.5</td><td>8.3</td><td>0.9</td><td>8.7</td><td>2.8E-18</td><td>1.1E-17</td><td>MB231/MB468</td></tr>
<tr><td>>hg38_21_+_8433172_8446622_RNA45SN1_45S_50ntFlanks@10408.10433.26|GGTCAGTCGGTCCTGAGAGATGGGCG</td><td>>hg38_21_+_8433172_8446622_RNA45SN1_45S_50ntFlanks</td><td>10408</td><td>10433</td><td>26</td><td>GGTCAGTCGGTCCTGAGAGATGGGCG</td><td>822.9</td><td>-5.5</td><td>0.6</td><td>-8.7</td><td>2.9E-18</td><td>1.1E-17</td><td>MB231/MB468</td></tr>
</table>

<table>
<tr><td>>hg38_21_+_8433172_8446622_RNA45SN1_45S_50ntFlanks@6746.6770.25|CACTTCGAACGCACTTGCGGCCCCCG</td><td>>hg38_21_+_8433172_8446622_RNA45SN1_45S_50ntFlanks</td><td>6746</td><td>6770</td><td>25</td><td>CACTTCGAACGCACTTGCGGCCCCG</td><td>800.8</td><td>-7.9</td><td>0.9</td><td>-8.7</td><td>2.9E-18</td><td>1.2E-17</td><td>MB231/MB468</td></tr>
<tr><td>>MI0000750|hsa-mir-26a-2&WithFlank&12|-|57824603|57824698@20.43.24;MI0000083|hsa-mir-26a-1&WithFlank&3|+|37969398|37969486@16.39.24&[MIMAT0000082&hsa-miR-26a-5p&offsets|0|+2;m-33&12|-|57824658|57824679&offsets|0|+2];[MIMAT0000082_1&hsa-miR-26a-5p&offsets|0|+2;m-34&3|+|37969413|37969434&offsets|0|+2]|TTCAAGTAATCCAGGATAGGCTGT</td><td>>MI0000750|hsa-mir-26a-2&WithFlank&12|-|57824603|57824698</td><td>20</td><td>43</td><td>24</td><td>TTCAAGTAATCCAGGATAGGCTGT</td><td>1246.3</td><td>9.3</td><td>1.1</td><td>8.7</td><td>2.9E-18</td><td>1.2E-17</td><td>MB231/MB468</td></tr>
<tr><td>>hg38_21_+_8433172_8446622_RNA45SN1_45S_50ntFlanks@11199.11225.27|CCGGCGGCGGCGGCGACTCTGGACGCG</td><td>>hg38_21_+_8433172_8446622_RNA45SN1_45S_50ntFlanks</td><td>11199</td><td>11225</td><td>27</td><td>CCGGCGGCGGCGGCGACTCTGGACGCG</td><td>859.3</td><td>-11.3</td><td>1.3</td><td>-8.7</td><td>2.9E-18</td><td>1.2E-17</td><td>MB231/MB468</td></tr>
<tr><td>>MI0000440|hsa-mir-27b&WithFlank&9|+|95085439|95085547@67.87.21(+2U)&[MIMAT0000419&hsa-miR-27b-3p&offsets|0|0(+2U);m-39&9|+|95085505|95085525&offsets|0|0(+2U)]|TTCACAGTGGCTAAGTTCTGCTT</td><td>>MI0000440|hsa-mir-27b&WithFlank&9|+|95085439|95085547</td><td>67</td><td>87</td><td>21(+2U)</td><td>TTCACAGTGGCTAAGTTCTGCTT</td><td>1816.6</td><td>7.5</td><td>0.9</td><td>8.7</td><td>2.9E-18</td><td>1.2E-17</td><td>MB231/MB468</td></tr>
<tr><td>>MI0006444|hsa-mir-1307&WithFlank&10|-|103394247|103394407@86.107.22&[MIMAT0005951&hsa-miR-1307-3p&offsets|0|0;m-66&10|-|103394300|103394322&offsets|0|-1]|ACTCGGCGTGGCGTCGGTCGTG</td><td>>MI0006444|hsa-mir-1307&WithFlank&10|-|103394247|103394407</td><td>86</td><td>107</td><td>22</td><td>ACTCGGCGTGGCGTCGGTCGTG</td><td>890.0</td><td>5.7</td><td>0.7</td><td>8.7</td><td>2.7E-18</td><td>1.2E-17</td><td>BT20/MB468</td></tr>
<tr><td>>am_trna10_ValCAC_5_-_180649395_180649467@1.29.29,trna12_ValAAC_5_-_180645270_180645342@1.29.29,trna132_ValAAC_6_-_27721179_27721251@1.29.29,trna136_ValAAC_6_-_27648885_27648957@1.29.29,trna139_ValAAC_6_-_27618707_27618779@1.29.29,trna18_ValCAC_5_-_180529253_180529325@1.29.29,trna2_ValAAC_3_+_169490018_169490090@1.29.29,trna2_ValCAC_5_+_180524070_180524142@1.29.29,trna4_ValAAC_5_+_180591154_180591226@1.29.29,trna5_ValAAC_5_+_180596610_180596682@1.29.29,trna6_ValCAC_5_+_180600650_180600722@1.29.29,trna85_ValCAC_1_-_161369490_161369562@1.29.29,trna90_ValCAC_1_-_149684088_149684161@1.29.29,trna98_ValCAC_1_-_149298555_149298627@1.29.29,trna9_ValCAC_6_+_26538282_26538354@1.29.29|GTTTCCGTAGTGTAGTGGTTATCACGTTC</td><td>>am_trna10_ValCAC_5_-_180649395_180649467</td><td>1</td><td>29</td><td>29</td><td>GTTTCCGTAGTGTAGTGGTTATCACGTTC</td><td>16930.6</td><td>-6.9</td><td>0.8</td><td>-8.7</td><td>2.8E-18</td><td>1.2E-17</td><td>BT20/MB468</td></tr>
<tr><td>>MI0000440|hsa-mir-27b&WithFlank&9|+|95085439|95085547@67.86.20(+1G)&[MIMAT0000419&hsa-miR-27b-3p&offsets|0|-1(+1G);m-39&9|+|95085505|95085525&offsets|0|-1(+1G)]|TTCACAGTGGCTAAGTTCTGG</td><td>>MI0000440|hsa-mir-27b&WithFlank&9|+|95085439|95085547</td><td>67</td><td>86</td><td>20(+1G)</td><td>TTCACAGTGGCTAAGTTCTGG</td><td>4603.2</td><td>9.6</td><td>1.1</td><td>8.7</td><td>2.8E-18</td><td>1.2E-17</td><td>BT20/MB468</td></tr>
<tr><td>>hg38_21_+_8433172_8446622_RNA45SN1_45S_50ntFlanks@11316.11338.23|CACCCCACGTCTCGTCGCGCGCG</td><td>>hg38_21_+_8433172_8446622_RNA45SN1_45S_50ntFlanks</td><td>11316</td><td>11338</td><td>23</td><td>CACCCCACGTCTCGTCGCGCGCG</td><td>1020.1</td><td>-11.6</td><td>1.3</td><td>-8.7</td><td>3.0E-18</td><td>1.2E-17</td><td>MB231/MB468</td></tr>
<tr><td>>MI0000077|hsa-mir-21&WithFlank&17|+|59841260|59841343@14.32.19&[MIMAT0000076&hsa-miR-21-5p&offsets|0|-3;m-1&17|+|59841273|59841294&offsets|0|-3]|TAGCTTATCAGACTGATGT</td><td>>MI0000077|hsa-mir-21&WithFlank&17|+|59841260|59841343</td><td>14</td><td>32</td><td>19</td><td>TAGCTTATCAGACTGATGT</td><td>2007.3</td><td>7.0</td><td>0.8</td><td>8.7</td><td>3.0E-18</td><td>1.2E-17</td><td>MB231/MB468</td></tr>
<tr><td>>hg38_21_+_8433172_8446622_RNA45SN1_45S_50ntFlanks@4367.4389.23|CAGTTAAAAAGCTCGTAGTTGGA</td><td>>hg38_21_+_8433172_8446622_RNA45SN1_45S_50ntFlanks</td><td>4367</td><td>4389</td><td>23</td><td>CAGTTAAAAAGCTCGTAGTTGGA</td><td>1309.9</td><td>-6.2</td><td>0.7</td><td>-8.7</td><td>3.0E-18</td><td>1.2E-17</td><td>MB231/MB468</td></tr>
<tr><td>>MI0000289|hsa-mir-181a-1&WithFlank&1|-|198859038|198859159@30.52.23;MI0000269|hsa-mir-181a-2&WithFlank&9|+|124692436|124692557@45.67.23&[MIMAT0000256&hsa-miR-181a-5p&offsets|0|0;m-32&1|-|198859108|198859130&offsets|0|0];[MIMAT0000256_1&hsa-miR-181a-5p&offsets|0|0;m-32&9|+|124692480|124692502&offsets|0|0]|AACATTCAACGCTGTCGGTGAGT</td><td>>MI0000289|hsa-mir-181a-1&WithFlank&1|-|198859038|198859159</td><td>30</td><td>52</td><td>23</td><td>AACATTCAACGCTGTCGGTGAGT</td><td>15901.1</td><td>7.1</td><td>0.8</td><td>8.7</td><td>3.1E-18</td><td>1.2E-17</td><td>MB231/MB468</td></tr>
<tr><td>>hg38_21_+_8433172_8446622_RNA45SN1_45S_50ntFlanks@3841.3879.39|TCCTACTTGGATAACTGTGGTAATTCTAGAGCTAATACA</td><td>>hg38_21_+_8433172_8446622_RNA45SN1_45S_50ntFlanks</td><td>3841</td><td>3879</td><td>39</td><td>TCCTACTTGGATAACTGTGGTAATTCTAGAGCTAATACA</td><td>471.9</td><td>11.5</td><td>1.3</td><td>8.7</td><td>2.9E-18</td><td>1.2E-17</td><td>BT20/MB468</td></tr>
<tr><td>>hg38_21_+_8433172_8446622_RNA45SN1_45S_50ntFlanks@12157.12182.26|AGTTTGACTGGGGCGGTACACCTGTC</td><td>>hg38_21_+_8433172_8446622_RNA45SN1_45S_50ntFlanks</td><td>12157</td><td>12182</td><td>26</td><td>AGTTTGACTGGGGCGGTACACCTGTC</td><td>569.0</td><td>-7.7</td><td>0.9</td><td>-8.7</td><td>3.0E-18</td><td>1.3E-17</td><td>BT20/MB468</td></tr>
<tr><td>>MI0000465|hsa-mir-191&WithFlank&3|-|49020612|49020715@22.45.24&[MIMAT0000440&hsa-miR-191-5p&offsets|0|+1;m-48&3|-|49020672|49020694&offsets|0|+1]|CAACGGAATCCCAAAAGCAGCTGT</td><td>>MI0000465|hsa-mir-191&WithFlank&3|-|49020612|49020715</td><td>22</td><td>45</td><td>24</td><td>CAACGGAATCCCAAAAGCAGCTGT</td><td>4329.1</td><td>6.3</td><td>0.7</td><td>8.7</td><td>3.0E-18</td><td>1.3E-17</td><td>BT20/MB468</td></tr>
<tr><td>>hg38_21_+_8433172_8446622_RNA45SN1_45S_50ntFlanks@11561.11591.31|GCGCCTAGCAGCCGACTTAGAACTGGTGCGG</td><td>>hg38_21_+_8433172_8446622_RNA45SN1_45S_50ntFlanks</td><td>11561</td><td>11591</td><td>31</td><td>GCGCCTAGCAGCCGACTTAGAACTGGTGCGG</td><td>951.1</td><td>-6.6</td><td>0.8</td><td>-8.7</td><td>3.2E-18</td><td>1.3E-17</td><td>MB231/MB468</td></tr>
<tr><td>>hg38_21_+_8433172_8446622_RNA45SN1_45S_50ntFlanks@10458.10478.21|ATGGCCTCCGTTGCCCTCGGC</td><td>>hg38_21_+_8433172_8446622_RNA45SN1_45S_50ntFlanks</td><td>10458</td><td>10478</td><td>21</td><td>ATGGCCTCCGTTGCCCTCGGC</td><td>2496.6</td><td>-7.1</td><td>0.8</td><td>-8.7</td><td>3.0E-18</td><td>1.3E-17</td><td>BT20/MB468</td></tr>
<tr><td>>MI0003776|hsa-mir-320b-1&WithFlank&1|+|116671740|116671823@48.66.19;MI0003839|hsa-mir-320b-2&WithFlank&1|-|224257034|224257116@47.65.19;MI0003778|hsa-mir-320c-1&WithFlank&18|+|21683512|21683595@48.66.19;MI0008191|hsa-mir-320c-2&WithFlank&18|+|24321669|24321752@48.66.19;MI0000542|hsa-mir-320a&WithFlank&8|-|22244960|22245043@48.66.19&[MIMAT0005792&hsa-miR-320b&offsets|0|-3;m-183&1|+|116671787|116671806&offsets|0|-1];[MIMAT0005792_1&hsa-miR-320b&offsets|0|-3;m-183&1|-|224257051|224257070&offsets|0|-1];[MIMAT0005793&hsa-miR-320c&offsets|0|-1;m-222&18|+|21683559|21683578&offsets|0|-1];[MIMAT0005793_1&hsa-miR-320c&offsets|0|-1;m-219&18|+|24321716|24321735&offsets|0|-1];[MIMAT0000510&hsa-miR-320a-3p&offsets|0|-3;m-60&8|-|22244975|22244996&offsets|0|-3]|AAAAGCTGGGTTGAGAGGG</td><td>>MI0003776|hsa-mir-320b-1&WithFlank&1|+|116671740|116671823</td><td>48</td><td>66</td><td>19</td><td>AAAAGCTGGGTTGAGAGGG</td><td>653.3</td><td>10.9</td><td>1.3</td><td>8.7</td><td>3.0E-18</td><td>1.3E-17</td><td>BT20/MB468</td></tr>
<tr><td>>hg38_1_-_228634819_228635039_RNA5S12_5S_50ntFlanks@144.169.26|CCTGGGAATACCGGGTGCTGTAGGCT</td><td>>hg38_1_-_228634819_228635039_RNA5S12_5S_50ntFlanks</td><td>144</td><td>169</td><td>26</td><td>CCTGGGAATACCGGGTGCTGTAGGCT</td><td>761.6</td><td>8.3</td><td>1.0</td><td>8.7</td><td>3.3E-18</td><td>1.3E-17</td><td>MB231/MB468</td></tr>
<tr><td>>hg38_21_+_8433172_8446622_RNA45SN1_45S_50ntFlanks@11321.11342.22|CACGTCTCGTCGCGCGCGCGTC</td><td>>hg38_21_+_8433172_8446622_RNA45SN1_45S_50ntFlanks</td><td>11321</td><td>11342</td><td>22</td><td>CACGTCTCGTCGCGCGCGCGTC</td><td>730.9</td><td>-6.1</td><td>0.7</td><td>-8.7</td><td>3.4E-18</td><td>1.3E-17</td><td>MB231/MB468</td></tr>
<tr><td>>hg38_21_+_8433172_8446622_RNA45SN1_45S_50ntFlanks@11563.11584.22|GCCTAGCAGCCGACTTAGAACT</td><td>>hg38_21_+_8433172_8446622_RNA45SN1_45S_50ntFlanks</td><td>11563</td><td>11584</td><td>22</td><td>GCCTAGCAGCCGACTTAGAACT</td><td>629.4</td><td>9.1</td><td>1.1</td><td>8.7</td><td>3.4E-18</td><td>1.4E-17</td><td>MB231/MB468</td></tr>
<tr><td>>hg38_21_+_8433172_8446622_RNA45SN1_45S_50ntFlanks@6652.6684.33|GACTCTTAGCGGTGGATCACTCGGCTCGTGCGT</td><td>>hg38_21_+_8433172_8446622_RNA45SN1_45S_50ntFlanks</td><td>6652</td><td>6684</td><td>33</td><td>GACTCTTAGCGGTGGATCACTCGGCTCGTGCGT</td><td>532.3</td><td>4.5</td><td>0.5</td><td>8.7</td><td>3.2E-18</td><td>1.4E-17</td><td>BT20/MB468</td></tr>
<tr><td>>MI0000065|hsa-let-7d&WithFlank&9|+|94178828|94178926@68.88.21&[MIMAT0004484&hsa-let-7d-3p&offsets|0|-1;m-98&9|+|94178895|94178916&offsets|0|-1]|CTATACGACCTGCTGCCTTTC</td><td>>MI0000065|hsa-let-7d&WithFlank&9|+|94178828|94178926</td><td>68</td><td>88</td><td>21</td><td>CTATACGACCTGCTGCCTTTC</td><td>662.4</td><td>6.7</td><td>0.8</td><td>8.7</td><td>3.2E-18</td><td>1.4E-17</td><td>BT20/MB468</td></tr>
<tr><td>>MI0000067|hsa-let-7f-1&WithFlank&9|+|94176341|94176439@13.30.18;MI0000068|hsa-let-7f-2&WithFlank&X|-|53557186|53557280@14.31.18&[MIMAT0000067&hsa-let-7f-5p&offsets|0|-4;m-20&9|+|94176353|94176374&offsets|0|-4];[MIMAT0000067_1&hsa-let-7f-5p&offsets|0|-4;m-19&X|-|53557246|53557267&offsets|0|-4]|TGAGGTAGTAGATTGTAT</td><td>>MI0000067|hsa-let-7f-1&WithFlank&9|+|94176341|94176439</td><td>13</td><td>30</td><td>18</td><td>TGAGGTAGTAGATTGTAT</td><td>7397.4</td><td>6.6</td><td>0.8</td><td>8.7</td><td>3.2E-18</td><td>1.4E-17</td><td>BT20/MB468</td></tr>
<tr><td>>hg38_21_+_8433172_8446622_RNA45SN1_45S_50ntFlanks@10806.10826.21|AACAATGTAGGTAAGGGAAGT</td><td>>hg38_21_+_8433172_8446622_RNA45SN1_45S_50ntFlanks</td><td>10806</td><td>10826</td><td>21</td><td>AACAATGTAGGTAAGGGAAGT</td><td>280.0</td><td>-4.7</td><td>0.5</td><td>-8.7</td><td>3.4E-18</td><td>1.4E-17</td><td>MB231/MB468</td></tr>
<tr><td>>hg38_21_+_8433172_8446622_RNA45SN1_45S_50ntFlanks@9568.9595.28|CGTGCAAATCGGTCGTCCGACCTGGGTA</td><td>>hg38_21_+_8433172_8446622_RNA45SN1_45S_50ntFlanks</td><td>9568</td><td>9595</td><td>28</td><td>CGTGCAAATCGGTCGTCCGACCTGGGTA</td><td>1393.4</td><td>-5.5</td><td>0.6</td><td>-8.7</td><td>3.5E-18</td><td>1.4E-17</td><td>MB231/MB468</td></tr>
<tr><td>>hg38_21_+_8433172_8446622_RNA45SN1_45S_50ntFlanks@6650.6683.34|ACGACTCTTAGCGGTGGATCACTCGGCTCGTGCG</td><td>>hg38_21_+_8433172_8446622_RNA45SN1_45S_50ntFlanks</td><td>6650</td><td>6683</td><td>34</td><td>ACGACTCTTAGCGGTGGATCACTCGGCTCGTGCG</td><td>3624.4</td><td>6.3</td><td>0.7</td><td>8.7</td><td>3.5E-18</td><td>1.4E-17</td><td>MB231/MB468</td></tr>
<tr><td>>hg38_21_+_8433172_8446622_RNA45SN1_45S_50ntFlanks@6733.6770.38|CATTGATCATCGACACTTCGAACGCACTTGCGGCCCCG</td><td>>hg38_21_+_8433172_8446622_RNA45SN1_45S_50ntFlanks</td><td>6733</td><td>6770</td><td>38</td><td>CATTGATCATCGACACTTCGAACGCACTTGCGGCCCCG</td><td>947.5</td><td>-4.2</td><td>0.5</td><td>-8.7</td><td>3.4E-18</td><td>1.4E-17</td><td>BT20/MB468</td></tr>
<tr><td>>hg38_21_+_8433172_8446622_RNA45SN1_45S_50ntFlanks@9569.9596.28|GTGCAAATCGGTCGTCCGACCTGGGTAT</td><td>>hg38_21_+_8433172_8446622_RNA45SN1_45S_50ntFlanks</td><td>9569</td><td>9596</td><td>28</td><td>GTGCAAATCGGTCGTCCGACCTGGGTAT</td><td>449.0</td><td>-6.1</td><td>0.7</td><td>-8.7</td><td>3.4E-18</td><td>1.4E-17</td><td>BT20/MB468</td></tr>
<tr><td>>MI0000088|hsa-mir-30a&WithFlank&6|-|71403545|71403627@12.34.23(+1A)&[MIMAT0000087&hsa-miR-30a-5p&offsets|0|+1(+1A);m-15&6|-|71403595|71403616&offsets|0|+1(+1A)]|TGTAAACATCCTCGACTGGAAGCA</td><td>>MI0000088|hsa-mir-30a&WithFlank&6|-|71403545|71403627</td><td>12</td><td>34</td><td>23(+1A)</td><td>TGTAAACATCCTCGACTGGAAGCA</td><td>57090.4</td><td>8.8</td><td>1.0</td><td>8.7</td><td>3.4E-18</td><td>1.4E-17</td><td>BT20/MB468</td></tr>
<tr><td>>hg38_21_+_8433172_8446622_RNA45SN1_45S_50ntFlanks@3705.3728.24|TACCTGGTTGATCCTGCCAGTAGC</td><td>>hg38_21_+_8433172_8446622_RNA45SN1_45S_50ntFlanks</td><td>3705</td><td>3728</td><td>24</td><td>TACCTGGTTGATCCTGCCAGTAGC</td><td>5515.0</td><td>8.5</td><td>1.0</td><td>8.7</td><td>3.4E-18</td><td>1.5E-17</td><td>BT20/MB468</td></tr>
<tr><td>>MI0000239|hsa-mir-197&WithFlank&1|+|109598887|109598973@54.75.22&[MIMAT0000227&hsa-miR-197-3p&offsets|0|0;m-99&1|+|109598940|109598961&offsets|0|0]|TTCACCACCTTCTCCACCCAGC</td><td>>MI0000239|hsa-mir-197&WithFlank&1|+|109598887|109598973</td><td>54</td><td>75</td><td>22</td><td>TTCACCACCTTCTCCACCCAGC</td><td>2956.0</td><td>6.7</td><td>0.8</td><td>8.7</td><td>3.7E-18</td><td>1.5E-17</td><td>MB231/MB468</td></tr>
<tr><td>>MI0000098|hsa-mir-96&WithFlank&7|-|129774686|129774775@15.37.23&[MIMAT0000095&hsa-miR-96-5p&offsets|0|0;m-248&7|-|129774739|129774761&offsets|0|0]|TTTGGCACTAGCACATTTTTGCT</td><td>>MI0000098|hsa-mir-96&WithFlank&7|-|129774686|129774775</td><td>15</td><td>37</td><td>23</td><td>TTTGGCACTAGCACATTTTTGCT</td><td>32647.7</td><td>7.8</td><td>0.9</td><td>8.7</td><td>3.5E-18</td><td>1.5E-17</td><td>BT20/MB468</td></tr>
</table>

<table>
<tr><th></th><th></th><th></th><th></th><th></th><th></th><th></th><th></th><th></th><th></th><th></th><th></th><th></th></tr>
<tr><td>>trna116_GluCTC_1_-_145399233_145399304@3.33.31,trna59_GluCTC_1_+_249168447_249168518@3.33.31,trna71_GluCTC_1_-_161439189_161439260@3.33.31,trna74_GluCTC_1_-_161431809_161431880@3.33.31,trna77_GluCTC_1_-_161424398_161424469@3.33.31,trna77_GluCTC_6_+_28949976_28950047@3.33.31,trna80_GluCTC_1_-_161417018_161417089@3.33.31,trna87_GluCTC_6_-_126101393_126101464@3.33.31|CCTGGTGGTCTAGTGGTTAGGATTCGGCGCT</td><td>>trna116_GluCTC_1_-_145399233_145399304</td><td>3</td><td>33</td><td>31</td><td>CCTGGTGGTCTAGTGGTTAGGATTCGGCGCT</td><td>682.4</td><td>9.7</td><td>1.1</td><td>8.7</td><td>3.7E-18</td><td>1.5E-17</td><td>MB231/MB468</td></tr>
<tr><td>>hg38_21_+_8433172_8446622_RNA45SN1_45S_50ntFlanks@6648.6693.46|GTACGACTCTTAGCGGTGGATCACTCGGCTCGTGCGTCGATGAAGA</td><td>>hg38_21_+_8433172_8446622_RNA45SN1_45S_50ntFlanks</td><td>6648</td><td>6693</td><td>46</td><td>GTACGACTCTTAGCGGTGGATCACTCGGCTCGTGCGTCGATGAAGA</td><td>605.6</td><td>7.6</td><td>0.9</td><td>8.7</td><td>3.7E-18</td><td>1.5E-17</td><td>MB231/MB468</td></tr>
<tr><td>>MI0000081|hsa-mir-24-2&WithFlank&19|-|13836281|13836365@18.41.24&[MIMAT0004497&hsa-miR-24-2-5p&offsets|-1|+1;m-226&19|-|13836326|13836348&offsets|0|+1]|GTGCCTACTGAGCTGAAACACAGT</td><td>>MI0000081|hsa-mir-24-2&WithFlank&19|-|13836281|13836365</td><td>18</td><td>41</td><td>24</td><td>GTGCCTACTGAGCTGAAACACAGT</td><td>1143.4</td><td>6.2</td><td>0.7</td><td>8.7</td><td>3.8E-18</td><td>1.5E-17</td><td>MB231/MB468</td></tr>
<tr><td>>hg38_21_+_8433172_8446622_RNA45SN1_45S_50ntFlanks@11576.11603.28|CTTAGAACTGGTGCGGACCAGGGGAATC</td><td>>hg38_21_+_8433172_8446622_RNA45SN1_45S_50ntFlanks</td><td>11576</td><td>11603</td><td>28</td><td>CTTAGAACTGGTGCGGACCAGGGGAATC</td><td>3020.9</td><td>-6.6</td><td>0.8</td><td>-8.7</td><td>3.6E-18</td><td>1.5E-17</td><td>BT20/MB468</td></tr>
<tr><td>>hg38_21_+_8433172_8446622_RNA45SN1_45S_50ntFlanks@12635.12659.25|AAGCTACCATCTGTGGGATTATGAC</td><td>>hg38_21_+_8433172_8446622_RNA45SN1_45S_50ntFlanks</td><td>12635</td><td>12659</td><td>25</td><td>AAGCTACCATCTGTGGGATTATGAC</td><td>1301.0</td><td>-4.9</td><td>0.6</td><td>-8.7</td><td>3.8E-18</td><td>1.5E-17</td><td>MB231/MB468</td></tr>
<tr><td>>hg38_21_+_8433172_8446622_RNA45SN1_45S_50ntFlanks@11566.11583.18|TAGCAGCCGACTTAGAAC</td><td>>hg38_21_+_8433172_8446622_RNA45SN1_45S_50ntFlanks</td><td>11566</td><td>11583</td><td>18</td><td>TAGCAGCCGACTTAGAAC</td><td>699.1</td><td>7.9</td><td>0.9</td><td>8.7</td><td>3.8E-18</td><td>1.5E-17</td><td>MB231/MB468</td></tr>
<tr><td>>hg38_21_+_8433172_8446622_RNA45SN1_45S_50ntFlanks@9562.9597.36|TCCTGACGTGCAAATCGGTCGTCCGACCTGGGTATA</td><td>>hg38_21_+_8433172_8446622_RNA45SN1_45S_50ntFlanks</td><td>9562</td><td>9597</td><td>36</td><td>TCCTGACGTGCAAATCGGTCGTCCGACCTGGGTATA</td><td>399.1</td><td>-5.1</td><td>0.6</td><td>-8.7</td><td>3.9E-18</td><td>1.5E-17</td><td>MB231/MB468</td></tr>
<tr><td>>MI0000076|hsa-mir-20a&WithFlank&13|+|91351059|91351141@14.36.23&[MIMAT0000075&hsa-miR-20a-5p&offsets|0|0;m-96&13|+|91351072|91351094&offsets|0|0]|TAAAGTGCTTATAGTGCAGGTAG</td><td>>MI0000076|hsa-mir-20a&WithFlank&13|+|91351059|91351141</td><td>14</td><td>36</td><td>23</td><td>TAAAGTGCTTATAGTGCAGGTAG</td><td>78110.8</td><td>6.8</td><td>0.8</td><td>8.7</td><td>3.9E-18</td><td>1.5E-17</td><td>MB231/MB468</td></tr>
<tr><td>>MI0000816|hsa-mir-335&WithFlank&7|+|130496105|130496210@58.78.21(+1U)&[MIMAT0004703&hsa-miR-335-3p&offsets|0|-1(+1U);m-159&7|+|130496162|130496183&offsets|0|-1(+1U)]|TTTTTCATTATTGCTCCTGACT</td><td>>MI0000816|hsa-mir-335&WithFlank&7|+|130496105|130496210</td><td>58</td><td>78</td><td>21(+1U)</td><td>TTTTTCATTATTGCTCCTGACT</td><td>2004.5</td><td>7.4</td><td>0.8</td><td>8.7</td><td>3.7E-18</td><td>1.5E-17</td><td>BT20/MB468</td></tr>
<tr><td>>am_trnaMT_ProTGG_MT_-_15956_16023@52.70.19|AAAGACTTTTTCTCTGACC</td><td>>am_trnaMT_ProTGG_MT_-_15956_16023</td><td>52</td><td>70</td><td>19</td><td>AAAGACTTTTTCTCTGACC</td><td>5250.4</td><td>11.2</td><td>1.3</td><td>8.7</td><td>3.9E-18</td><td>1.6E-17</td><td>MB231/MB468</td></tr>
<tr><td>>hg38_21_+_8433172_8446622_RNA45SN1_45S_50ntFlanks@8342.8378.37|CGTAAGGGAAAGTTGAAAAGAACTTTGAAGAGAGAGT</td><td>>hg38_21_+_8433172_8446622_RNA45SN1_45S_50ntFlanks</td><td>8342</td><td>8378</td><td>37</td><td>CGTAAGGGAAAGTTGAAAAGAACTTTGAAGAGAGAGT</td><td>1080.3</td><td>10.0</td><td>1.2</td><td>8.7</td><td>3.9E-18</td><td>1.6E-17</td><td>MB231/MB468</td></tr>
<tr><td>>trnaMT_SerGCT_MT_+_12207_12265@1.23.23|GAGAAAGCTCACAAGAACTGCTA</td><td>>trnaMT_SerGCT_MT_+_12207_12265</td><td>1</td><td>23</td><td>23</td><td>GAGAAAGCTCACAAGAACTGCTA</td><td>966.3</td><td>5.8</td><td>0.7</td><td>8.7</td><td>3.7E-18</td><td>1.6E-17</td><td>BT20/MB468</td></tr>
<tr><td>>hg38_21_+_8433172_8446622_RNA45SN1_45S_50ntFlanks@12626.12669.44|AATGGGGCGAAGCTACCATCTGTGGGATTATGACTGAACGCCTC</td><td>>hg38_21_+_8433172_8446622_RNA45SN1_45S_50ntFlanks</td><td>12626</td><td>12669</td><td>44</td><td>AATGGGGCGAAGCTACCATCTGTGGGATTATGACTGAACGCCTC</td><td>331.0</td><td>7.6</td><td>0.9</td><td>8.7</td><td>4.0E-18</td><td>1.6E-17</td><td>MB231/MB468</td></tr>
<tr><td>>MI0000434|hsa-let-7i&WithFlank&12|+|62603680|62603775@12.32.21&[MIMAT0000415&hsa-let-7i-5p&offsets|0|-1;m-49&12|+|62603691|62603712&offsets|0|-1]|TGAGGTAGTAGTTTGTGCTGT</td><td>>MI0000434|hsa-let-7i&WithFlank&12|+|62603680|62603775</td><td>12</td><td>32</td><td>21</td><td>TGAGGTAGTAGTTTGTGCTGT</td><td>9856.6</td><td>6.8</td><td>0.8</td><td>8.7</td><td>3.7E-18</td><td>1.6E-17</td><td>BT20/MB468</td></tr>
<tr><td>>hg38_21_+_8433172_8446622_RNA45SN1_45S_50ntFlanks@6647.6683.37|CGTACGACTCTTAGCGGTGGATCACTCGGCTCGTGCG</td><td>>hg38_21_+_8433172_8446622_RNA45SN1_45S_50ntFlanks</td><td>6647</td><td>6683</td><td>37</td><td>CGTACGACTCTTAGCGGTGGATCACTCGGCTCGTGCG</td><td>863.8</td><td>10.0</td><td>1.2</td><td>8.7</td><td>4.0E-18</td><td>1.6E-17</td><td>MB231/MB468</td></tr>
<tr><td>>hg38_21_+_8433172_8446622_RNA45SN1_45S_50ntFlanks@9570.9595.26|TGCAAATCGGTCGTCCGACCTGGGTA</td><td>>hg38_21_+_8433172_8446622_RNA45SN1_45S_50ntFlanks</td><td>9570</td><td>9595</td><td>26</td><td>TGCAAATCGGTCGTCCGACCTGGGTA</td><td>1746.6</td><td>-4.9</td><td>0.6</td><td>-8.7</td><td>4.0E-18</td><td>1.6E-17</td><td>MB231/MB468</td></tr>
<tr><td>>hg38_21_+_8433172_8446622_RNA45SN1_45S_50ntFlanks@11668.11686.19|TGCCCAGTGCTCTGAATGT</td><td>>hg38_21_+_8433172_8446622_RNA45SN1_45S_50ntFlanks</td><td>11668</td><td>11686</td><td>19</td><td>TGCCCAGTGCTCTGAATGT</td><td>523.3</td><td>8.9</td><td>1.0</td><td>8.7</td><td>4.1E-18</td><td>1.6E-17</td><td>MB231/MB468</td></tr>
<tr><td>>MI0000077|hsa-mir-21&WithFlank&17|+|59841260|59841343@14.37.24(+1U)&[MIMAT0000076&hsa-miR-21-5p&offsets|0|+2(+1U);m-1&17|+|59841273|59841294&offsets|0|+2(+1U)]|TAGCTTATCAGACTGATGTTGACTT</td><td>>MI0000077|hsa-mir-21&WithFlank&17|+|59841260|59841343</td><td>14</td><td>37</td><td>24(+1U)</td><td>TAGCTTATCAGACTGATGTTGACTT</td><td>830.6</td><td>10.1</td><td>1.2</td><td>8.7</td><td>4.1E-18</td><td>1.6E-17</td><td>MB231/MB468</td></tr>
<tr><td>>hg38_21_+_8433172_8446622_RNA45SN1_45S_50ntFlanks@8357.8379.23|AAAAGAACTTTGAAGAGAGAGTT</td><td>>hg38_21_+_8433172_8446622_RNA45SN1_45S_50ntFlanks</td><td>8357</td><td>8379</td><td>23</td><td>AAAAGAACTTTGAAGAGAGAGTT</td><td>64051.6</td><td>-7.6</td><td>0.9</td><td>-8.7</td><td>3.9E-18</td><td>1.6E-17</td><td>BT20/MB468</td></tr>
<tr><td>>hg38_21_+_8433172_8446622_RNA45SN1_45S_50ntFlanks@3875.3898.24|ATACATGCCGACGGGCGCTGACCC</td><td>>hg38_21_+_8433172_8446622_RNA45SN1_45S_50ntFlanks</td><td>3875</td><td>3898</td><td>24</td><td>ATACATGCCGACGGGCGCTGACCC</td><td>1666.6</td><td>-4.7</td><td>0.5</td><td>-8.7</td><td>4.2E-18</td><td>1.7E-17</td><td>MB231/MB468</td></tr>
<tr><td>>hg38_21_+_8433172_8446622_RNA45SN1_45S_50ntFlanks@9534.9562.29|AGGAAACTCTGGTGGAGGTCCGTAGCGGT</td><td>>hg38_21_+_8433172_8446622_RNA45SN1_45S_50ntFlanks</td><td>9534</td><td>9562</td><td>29</td><td>AGGAAACTCTGGTGGAGGTCCGTAGCGGT</td><td>1148.4</td><td>-5.1</td><td>0.6</td><td>-8.7</td><td>4.0E-18</td><td>1.7E-17</td><td>BT20/MB468</td></tr>
<tr><td>>hg38_1_-_228634819_228635039_RNA5S12_5S_50ntFlanks@136.171.36|GGAGACCGCCTGGGAATACCGGGTGCTGTAGGCTTT</td><td>>hg38_1_-_228634819_228635039_RNA5S12_5S_50ntFlanks</td><td>136</td><td>171</td><td>36</td><td>GGAGACCGCCTGGGAATACCGGGTGCTGTAGGCTTT</td><td>8246.8</td><td>6.6</td><td>0.8</td><td>8.7</td><td>4.3E-18</td><td>1.7E-17</td><td>MB231/MB468</td></tr>
<tr><td>>hg38_21_+_8433172_8446622_RNA45SN1_45S_50ntFlanks@10771.10806.36|CAGGTCTCCAAGGTGAACAGCCTCTGGCATGTTGGA</td><td>>hg38_21_+_8433172_8446622_RNA45SN1_45S_50ntFlanks</td><td>10771</td><td>10806</td><td>36</td><td>CAGGTCTCCAAGGTGAACAGCCTCTGGCATGTTGGA</td><td>525.2</td><td>-11.0</td><td>1.3</td><td>-8.7</td><td>4.0E-18</td><td>1.7E-17</td><td>BT20/MB468</td></tr>
<tr><td>>MI0003667|hsa-mir-652&WithFlank&X|+|110055323|110055432@67.87.21&[MIMAT0003322&hsa-miR-652-3p&offsets|0|0;m-148&X|+|110055389|110055410&offsets|0|-1]|AATGGCGCCACTAGGGTTGTG</td><td>>MI0003667|hsa-mir-652&WithFlank&X|+|110055323|110055432</td><td>67</td><td>87</td><td>21</td><td>AATGGCGCCACTAGGGTTGTG</td><td>604.8</td><td>7.5</td><td>0.9</td><td>8.7</td><td>4.0E-18</td><td>1.7E-17</td><td>BT20/MB468</td></tr>
<tr><td>>hg38_21_+_8433172_8446622_RNA45SN1_45S_50ntFlanks@4731.4766.36|AAGAACGAAAGTCGGAGGTTCGAAGACGATCAGATA</td><td>>hg38_21_+_8433172_8446622_RNA45SN1_45S_50ntFlanks</td><td>4731</td><td>4766</td><td>36</td><td>AAGAACGAAAGTCGGAGGTTCGAAGACGATCAGATA</td><td>700.3</td><td>6.9</td><td>0.8</td><td>8.7</td><td>4.3E-18</td><td>1.7E-17</td><td>MB231/MB468</td></tr>
<tr><td>>hg38_21_+_8433172_8446622_RNA45SN1_45S_50ntFlanks@11670.11699.30|CCCAGTGCTCTGAATGTCAAAGTGAAGAAA</td><td>>hg38_21_+_8433172_8446622_RNA45SN1_45S_50ntFlanks</td><td>11670</td><td>11699</td><td>30</td><td>CCCAGTGCTCTGAATGTCAAAGTGAAGAAA</td><td>208.3</td><td>-4.9</td><td>0.6</td><td>-8.7</td><td>4.0E-18</td><td>1.7E-17</td><td>BT20/MB468</td></tr>
<tr><td>>trnaMT_ProTGG_MT_-_15956_16023@40.58.19|AATGGTGGAGTTAAAGACT</td><td>>trnaMT_ProTGG_MT_-_15956_16023</td><td>40</td><td>58</td><td>19</td><td>AATGGTGGAGTTAAAGACT</td><td>1473.0</td><td>10.3</td><td>1.2</td><td>8.7</td><td>4.3E-18</td><td>1.7E-17</td><td>MB231/MB468</td></tr>
<tr><td>>hg38_21_+_8433172_8446622_RNA45SN1_45S_50ntFlanks@12539.12573.35|ATGATGTGTTGTTGCCATGGTAATCCTGCTCAGTA</td><td>>hg38_21_+_8433172_8446622_RNA45SN1_45S_50ntFlanks</td><td>12539</td><td>12573</td><td>35</td><td>ATGATGTGTTGTTGCCATGGTAATCCTGCTCAGTA</td><td>166.3</td><td>-4.9</td><td>0.6</td><td>-8.7</td><td>4.1E-18</td><td>1.7E-17</td><td>BT20/MB468</td></tr>
<tr><td>>MI0000266|hsa-mir-10a&WithFlank&17|-|48579832|48579953@28.50.23(+1A)&[MIMAT0000253&hsa-miR-10a-5p&offsets|0|0(+1A);m-18&17|-|48579905|48579926&offsets|0|+1(+1A)]|TACCCTGTAGATCCGAATTTGTGA</td><td>>MI0000266|hsa-mir-10a&WithFlank&17|-|48579832|48579953</td><td>28</td><td>50</td><td>23(+1A)</td><td>TACCCTGTAGATCCGAATTTGTGA</td><td>2989.3</td><td>12.4</td><td>1.4</td><td>8.7</td><td>4.1E-18</td><td>1.7E-17</td><td>BT20/MB468</td></tr>
<tr><td>>trnaMT_GluTTC_MT_-_14674_14742@39.71.33,trnalookalike8_GluTTC_5_-_93905172_93905240@39.71.33|ATTGGTCGTGGTTGTAGTCCGTGCGAGAATACC</td><td>>trnaMT_GluTTC_MT_-_14674_14742</td><td>39</td><td>71</td><td>33</td><td>ATTGGTCGTGGTTGTAGTCCGTGCGAGAATACC</td><td>4837.1</td><td>11.0</td><td>1.3</td><td>8.7</td><td>4.4E-18</td><td>1.7E-17</td><td>MB231/MB468</td></tr>
<tr><td>>hg38_21_+_8433172_8446622_RNA45SN1_45S_50ntFlanks@6646.6671.26|TCGTACGACTCTTAGCGGTGGATCAC</td><td>>hg38_21_+_8433172_8446622_RNA45SN1_45S_50ntFlanks</td><td>6646</td><td>6671</td><td>26</td><td>TCGTACGACTCTTAGCGGTGGATCAC</td><td>612.6</td><td>4.6</td><td>0.5</td><td>8.7</td><td>4.2E-18</td><td>1.7E-17</td><td>BT20/MB468</td></tr>
<tr><td>>hg38_21_+_8433172_8446622_RNA45SN1_45S_50ntFlanks@12417.12450.34|TCGATGTCGGCTCTTCCTATCATTGTGAAGCAGA</td><td>>hg38_21_+_8433172_8446622_RNA45SN1_45S_50ntFlanks</td><td>12417</td><td>12450</td><td>34</td><td>TCGATGTCGGCTCTTCCTATCATTGTGAAGCAGA</td><td>335.3</td><td>4.0</td><td>0.5</td><td>8.7</td><td>4.5E-18</td><td>1.8E-17</td><td>MB231/MB468</td></tr>
<tr><td>>hg38_21_+_8433172_8446622_RNA45SN1_45S_50ntFlanks@11569.11603.35|CAGCCGACTTAGAACTGGTGCGGACCAGGGGAATC</td><td>>hg38_21_+_8433172_8446622_RNA45SN1_45S_50ntFlanks</td><td>11569</td><td>11603</td><td>35</td><td>CAGCCGACTTAGAACTGGTGCGGACCAGGGGAATC</td><td>1580.1</td><td>-5.0</td><td>0.6</td><td>-8.7</td><td>4.5E-18</td><td>1.8E-17</td><td>MB231/MB468</td></tr>
<tr><td>>MI0000085|hsa-mir-27a&WithFlank&19|-|13836434|13836523@57.74.18;MI0000440|hsa-mir-27b&WithFlank&9|+|95085439|95085547@67.84.18&[MIMAT0000084&hsa-miR-27a-3p&offsets|0|-3;m-43&19|-|13836447|13836467&offsets|0|-3];[MIMAT0000419&hsa-miR-27b-3p&offsets|0|-3;m-39&9|+|95085505|95085525&offsets|0|-3]|TTCACAGTGGCTAAGTTC</td><td>>MI0000085|hsa-mir-27a&WithFlank&19|-|13836434|13836523</td><td>57</td><td>74</td><td>18</td><td>TTCACAGTGGCTAAGTTC</td><td>1613.7</td><td>7.4</td><td>0.8</td><td>8.7</td><td>4.5E-18</td><td>1.8E-17</td><td>MB231/MB468</td></tr>
<tr><td>>hg38_21_+_8433172_8446622_RNA45SN1_45S_50ntFlanks@12163.12182.20|ACTGGGGCGGTACACCTGTC</td><td>>hg38_21_+_8433172_8446622_RNA45SN1_45S_50ntFlanks</td><td>12163</td><td>12182</td><td>20</td><td>ACTGGGGCGGTACACCTGTC</td><td>2760.0</td><td>-5.4</td><td>0.6</td><td>-8.7</td><td>4.6E-18</td><td>1.8E-17</td><td>MB231/MB468</td></tr>
<tr><td>>MI0000081|hsa-mir-24-2&WithFlank&19|-|13836281|13836365@56.78.23;MI0000080|hsa-mir-24-1&WithFlank&9|+|95086015|95086094@50.72.23&[MIMAT0000080&hsa-miR-24-3p&offsets|0|+1;m-36&19|-|13836288|13836310&offsets|0|0];[MIMAT0000080_1&hsa-miR-24-3p&offsets|0|+1;m-35&9|+|95086064|95086086&offsets|0|0]|TGGCTCAGTTCAGCAGGAACAGG</td><td>>MI0000081|hsa-mir-24-2&WithFlank&19|-|13836281|13836365</td><td>56</td><td>78</td><td>23</td><td>TGGCTCAGTTCAGCAGGAACAGG</td><td>970.7</td><td>7.0</td><td>0.8</td><td>8.7</td><td>4.7E-18</td><td>1.8E-17</td><td>MB231/MB468</td></tr>
<tr><td>>MI0000073|hsa-mir-19a&WithFlank&13|+|91350885|91350978@55.78.24&[MIMAT0000073&hsa-miR-19a-3p&offsets|0|+1;m-201&13|+|91350939|91350961&offsets|0|+1]|TGTGCAAATCTATGCAAAACTGAT</td><td>>MI0000073|hsa-mir-19a&WithFlank&13|+|91350885|91350978</td><td>55</td><td>78</td><td>24</td><td>TGTGCAAATCTATGCAAAACTGAT</td><td>799.1</td><td>6.4</td><td>0.7</td><td>8.7</td><td>4.8E-18</td><td>1.9E-17</td><td>MB231/MB468</td></tr>
<tr><td>>trnaMT_MetCAT_MT_+_4402_4469@52.71.20|ATACCCTTCCCGTACTACCA</td><td>>trnaMT_MetCAT_MT_+_4402_4469</td><td>52</td><td>71</td><td>20</td><td>ATACCCTTCCCGTACTACCA</td><td>2667.8</td><td>11.4</td><td>1.3</td><td>8.7</td><td>4.9E-18</td><td>1.9E-17</td><td>MB231/MB468</td></tr>
<tr><td>>hg38_21_+_8433172_8446622_RNA45SN1_45S_50ntFlanks@6719.6749.31|TGAATTGCAGGACACATTGATCATCGACACT</td><td>>hg38_21_+_8433172_8446622_RNA45SN1_45S_50ntFlanks</td><td>6719</td><td>6749</td><td>31</td><td>TGAATTGCAGGACACATTGATCATCGACACT</td><td>306.7</td><td>-3.8</td><td>0.4</td><td>-8.7</td><td>4.6E-18</td><td>1.9E-17</td><td>BT20/MB468</td></tr>
<tr><td>>MI0000088|hsa-mir-30a&WithFlank&6|-|71403545|71403627@12.31.20&[MIMAT0000087&hsa-miR-30a-5p&offsets|0|-2;m-15&6|-|71403595|71403616&offsets|0|-2]|TGTAAACATCCTCGACTGGA</td><td>>MI0000088|hsa-mir-30a&WithFlank&6|-|71403545|71403627</td><td>12</td><td>31</td><td>20</td><td>TGTAAACATCCTCGACTGGA</td><td>4546.0</td><td>7.0</td><td>0.8</td><td>8.7</td><td>4.7E-18</td><td>2.0E-17</td><td>BT20/MB468</td></tr>
<tr><td>>hg38_21_+_8433172_8446622_RNA45SN1_45S_50ntFlanks@12936.12953.18|AAACCATTCGTAGACGAC</td><td>>hg38_21_+_8433172_8446622_RNA45SN1_45S_50ntFlanks</td><td>12936</td><td>12953</td><td>18</td><td>AAACCATTCGTAGACGAC</td><td>632.0</td><td>-5.4</td><td>0.6</td><td>-8.7</td><td>4.7E-18</td><td>2.0E-17</td><td>BT20/MB468</td></tr>
<tr><td>>hg38_21_+_8433172_8446622_RNA45SN1_45S_50ntFlanks@12003.12034.32|GCGGGCCGCCGGTGAAATACCACTACTCTGAT</td><td>>hg38_21_+_8433172_8446622_RNA45SN1_45S_50ntFlanks</td><td>12003</td><td>12034</td><td>32</td><td>GCGGGCCGCCGGTGAAATACCACTACTCTGAT</td><td>2275.3</td><td>7.7</td><td>0.9</td><td>8.7</td><td>5.1E-18</td><td>2.0E-17</td><td>MB231/MB468</td></tr>
</table>

<table>
<tr><td>>hg38_21_+_8433172_8446622_RNA45SN1_45S_50ntFlanks@12627.12659.33|ATGGGGCGAAGCTACCATCTGTGGGATTATGAC</td><td>>hg38_21_+_8433172_8446622_RNA45SN1_45S_50ntFlanks</td><td>12627</td><td>12659</td><td>33</td><td>ATGGGGCGAAGCTACCATCTGTGGGATTATGAC</td><td>764.1</td><td>-4.7</td><td>0.5</td><td>-8.7</td><td>5.1E-18</td><td>2.0E-17</td><td>MB231/MB468</td></tr>
<tr><td>>hg38_21_+_8433172_8446622_RNA45SN1_45S_50ntFlanks@7986.8005.20|ATCAGACGTGGCGACCCGCT</td><td>>hg38_21_+_8433172_8446622_RNA45SN1_45S_50ntFlanks</td><td>7986</td><td>8005</td><td>20</td><td>ATCAGACGTGGCGACCCGCT</td><td>658.6</td><td>-11.4</td><td>1.3</td><td>-8.7</td><td>4.7E-18</td><td>2.0E-17</td><td>BT20/MB468</td></tr>
<tr><td>>hg38_21_+_8433172_8446622_RNA45SN1_45S_50ntFlanks@10843.10865.23|TAACTTCGGGATAAGGATTGGCT</td><td>>hg38_21_+_8433172_8446622_RNA45SN1_45S_50ntFlanks</td><td>10843</td><td>10865</td><td>23</td><td>TAACTTCGGGATAAGGATTGGCT</td><td>887.8</td><td>-6.9</td><td>0.8</td><td>-8.7</td><td>4.7E-18</td><td>2.0E-17</td><td>BT20/MB468</td></tr>
<tr><td>>MI0000076|hsa-mir-20a&WithFlank&13|+|91351059|91351141@14.34.21&[MIMAT0000075&hsa-miR-20a-5p&offsets|0|-2;m-96&13|+|91351072|91351094&offsets|0|-2]|TAAAGTGCTTATAGTGCAGGT</td><td>>MI0000076|hsa-mir-20a&WithFlank&13|+|91351059|91351141</td><td>14</td><td>34</td><td>21</td><td>TAAAGTGCTTATAGTGCAGGT</td><td>4652.0</td><td>7.1</td><td>0.8</td><td>8.7</td><td>5.1E-18</td><td>2.0E-17</td><td>MB231/MB468</td></tr>
<tr><td>>hg38_21_+_8433172_8446622_RNA45SN1_45S_50ntFlanks@10402.10435.34|AACATGGGTCAGTCGGTCCTGAGAGATGGGCGAG</td><td>>hg38_21_+_8433172_8446622_RNA45SN1_45S_50ntFlanks</td><td>10402</td><td>10435</td><td>34</td><td>AACATGGGTCAGTCGGTCCTGAGAGATGGGCGAG</td><td>399.3</td><td>-6.7</td><td>0.8</td><td>-8.7</td><td>5.1E-18</td><td>2.0E-17</td><td>MB231/MB468</td></tr>
<tr><td>>hg38_21_+_8433172_8446622_RNA45SN1_45S_50ntFlanks@11785.11804.20|TGAATGGATGAACGAGATTC</td><td>>hg38_21_+_8433172_8446622_RNA45SN1_45S_50ntFlanks</td><td>11785</td><td>11804</td><td>20</td><td>TGAATGGATGAACGAGATTC</td><td>8258.9</td><td>-5.1</td><td>0.6</td><td>-8.7</td><td>5.1E-18</td><td>2.0E-17</td><td>MB231/MB468</td></tr>
<tr><td>>MI0000114|hsa-mir-107&WithFlank&10|-|89592741|89592833@56.76.21;MI0000108|hsa-mir-103a-2&WithFlank&20|+|3917488|3917577@54.74.21;MI0000109|hsa-mir-103a-1&WithFlank&5|-|168560890|168560979@54.74.21&[MIMAT0000104&hsa-miR-107&offsets|0|-2;m-78&10|-|89592758|89592778&offsets|0|0];[MIMAT0000101&hsa-miR-103a-3p&offsets|0|-2;m-12&20|+|3917541|3917563&offsets|0|-2];[MIMAT0000101_1&hsa-miR-103a-3p&offsets|0|-2;m-13&5|-|168560904|168560926&offsets|0|-2;m-44220&5|-|168560908|168560926&offsets|0|+2]|AGCAGCATTGTACAGGGCTAT</td><td>>MI0000114|hsa-mir-107&WithFlank&10|-|89592741|89592833</td><td>56</td><td>76</td><td>21</td><td>AGCAGCATTGTACAGGGCTAT</td><td>2370.8</td><td>7.6</td><td>0.9</td><td>8.7</td><td>4.9E-18</td><td>2.0E-17</td><td>BT20/MB468</td></tr>
<tr><td>>MI0000264|hsa-mir-7-2&WithFlank&15|+|88611819|88611940@38.60.23(+1G);MI0000265|hsa-mir-7-3&WithFlank&19|+|4770664|4770785@37.59.23(+1G);MI0000263|hsa-mir-7-1&WithFlank&9|-|83969742|83969863@30.52.23(+1G)&[MIMAT0000252&hsa-miR-7-5p&offsets|0|-1(+1G);m-400&15|+|88611856|88611879&offsets|0|-1(+1G)];[MIMAT0000252_1&hsa-miR-7-5p&offsets|0|-1(+1G);m-402&19|+|4770700|4770723&offsets|0|-1(+1G)];[MIMAT0000252_2&hsa-miR-7-5p&offsets|0|-1(+1G);m-401&9|-|83969811|83969834&offsets|0|-1(+1G)]|TGGAAGACTAGTGATTTTGTTGTG</td><td>>MI0000264|hsa-mir-7-2&WithFlank&15|+|88611819|88611940</td><td>38</td><td>60</td><td>23(+1G)</td><td>TGGAAGACTAGTGATTTTGTTGTG</td><td>25002.7</td><td>9.0</td><td>1.0</td><td>8.7</td><td>5.0E-18</td><td>2.1E-17</td><td>BT20/MB468</td></tr>
<tr><td>>MI0000079|hsa-mir-23a&WithFlank&19|-|13836581|13836665@52.74.23&[MIMAT0000078&hsa-miR-23a-3p&offsets|+1|+3;m-31&19|-|13836595|13836615&offsets|+1|+3]|TCACATTGCCAGGGATTTCCAAC</td><td>>MI0000079|hsa-mir-23a&WithFlank&19|-|13836581|13836665</td><td>52</td><td>74</td><td>23</td><td>TCACATTGCCAGGGATTTCCAAC</td><td>837.8</td><td>7.1</td><td>0.8</td><td>8.6</td><td>5.3E-18</td><td>2.1E-17</td><td>MB231/MB468</td></tr>
<tr><td>>MI0000066|hsa-let-7e&WithFlank&19|+|51692780|51692870@14.34.21(+1G)&[MIMAT0000066&hsa-let-7e-5p&offsets|0|-1(+1G);m-50&19|+|51692793|51692814&offsets|0|-1(+1G)]|TGAGGTAGGAGGTTGTATAGTG</td><td>>MI0000066|hsa-let-7e&WithFlank&19|+|51692780|51692870</td><td>14</td><td>34</td><td>21(+1G)</td><td>TGAGGTAGGAGGTTGTATAGTG</td><td>1620.2</td><td>8.4</td><td>1.0</td><td>8.7</td><td>5.0E-18</td><td>2.1E-17</td><td>BT20/MB468</td></tr>
<tr><td>>hg38_21_+_8433172_8446622_RNA45SN1_45S_50ntFlanks@475.516.42|CTTCGTGATCGATGTGGTGACGTCGTGCTCTCCCGGGCCGGG</td><td>>hg38_21_+_8433172_8446622_RNA45SN1_45S_50ntFlanks</td><td>475</td><td>516</td><td>42</td><td>CTTCGTGATCGATGTGGTGACGTCGTGCTCTCCCGGGCCGGG</td><td>4430.8</td><td>7.8</td><td>0.9</td><td>8.7</td><td>5.0E-18</td><td>2.1E-17</td><td>BT20/MB468</td></tr>
<tr><td>>MI0000114|hsa-mir-107&WithFlank&10|-|89592741|89592833@56.76.21(+1U);MI0000108|hsa-mir-103a-2&WithFlank&20|+|3917488|3917577@54.74.21(+1U);MI0000109|hsa-mir-103a-1&WithFlank&5|-|168560890|168560979@54.74.21(+1U)&[MIMAT0000104&hsa-miR-107&offsets|0|-2(+1U);m-78&10|-|89592758|89592778&offsets|0|0(+1U)];[MIMAT0000101&hsa-miR-103a-3p&offsets|0|-2(+1U);m-12&20|+|3917541|3917563&offsets|0|-2(+1U)];[MIMAT0000101_1&hsa-miR-103a-3p&offsets|0|-2(+1U);m-13&5|-|168560904|168560926&offsets|0|-2(+1U);m-44220&5|-|168560908|168560926&offsets|0|+2(+1U)]|AGCAGCATTGTACAGGGCTATT</td><td>>MI0000114|hsa-mir-107&WithFlank&10|-|89592741|89592833</td><td>56</td><td>76</td><td>21(+1U)</td><td>AGCAGCATTGTACAGGGCTATT</td><td>771.2</td><td>7.7</td><td>0.9</td><td>8.7</td><td>5.0E-18</td><td>2.1E-17</td><td>BT20/MB468</td></tr>
<tr><td>>MI0000434|hsa-let-7i&WithFlank&12|+|62603680|62603775@12.31.20(+1C)&[MIMAT0000415&hsa-let-7i-5p&offsets|0|-2(+1C);m-49&12|+|62603691|62603712&offsets|0|-2(+1C)]|TGAGGTAGTAGTTTGTGCTGC</td><td>>MI0000434|hsa-let-7i&WithFlank&12|+|62603680|62603775</td><td>12</td><td>31</td><td>20(+1C)</td><td>TGAGGTAGTAGTTTGTGCTGC</td><td>1229.8</td><td>7.1</td><td>0.8</td><td>8.7</td><td>5.0E-18</td><td>2.1E-17</td><td>BT20/MB468</td></tr>
<tr><td>>MI0000298|hsa-mir-221&WithFlank&X|-|45746151|45746272@31.53.23(+1C)&[MIMAT0004568&hsa-miR-221-5p&offsets|0|+1(+1C);m-405&X|-|45746220|45746242&offsets|0|0(+1C)]|ACCTGGCATACAATGTAGATTTCC</td><td>>MI0000298|hsa-mir-221&WithFlank&X|-|45746151|45746272</td><td>31</td><td>53</td><td>23(+1C)</td><td>ACCTGGCATACAATGTAGATTTCC</td><td>1042.2</td><td>10.1</td><td>1.2</td><td>8.6</td><td>5.5E-18</td><td>2.1E-17</td><td>MB231/MB468</td></tr>
<tr><td>>hg38_21_+_8433172_8446622_RNA45SN1_45S_50ntFlanks@6646.6685.40|TCGTACGACTCTTAGCGGTGGATCACTCGGCTCGTGCGTC</td><td>>hg38_21_+_8433172_8446622_RNA45SN1_45S_50ntFlanks</td><td>6646</td><td>6685</td><td>40</td><td>TCGTACGACTCTTAGCGGTGGATCACTCGGCTCGTGCGTC</td><td>14704.6</td><td>8.2</td><td>0.9</td><td>8.6</td><td>5.2E-18</td><td>2.1E-17</td><td>BT20/MB468</td></tr>
<tr><td>>hg38_21_+_8433172_8446622_RNA45SN1_45S_50ntFlanks@4006.4024.19|ACCTCGGGCCGATCGCACG</td><td>>hg38_21_+_8433172_8446622_RNA45SN1_45S_50ntFlanks</td><td>4006</td><td>4024</td><td>19</td><td>ACCTCGGGCCGATCGCACG</td><td>599.4</td><td>-5.7</td><td>0.7</td><td>-8.6</td><td>5.2E-18</td><td>2.2E-17</td><td>BT20/MB468</td></tr>
<tr><td>>hg38_21_+_8433172_8446622_RNA45SN1_45S_50ntFlanks@12459.12487.29|AGCGTTGGATTGTTCACCCACTAATAGGG</td><td>>hg38_21_+_8433172_8446622_RNA45SN1_45S_50ntFlanks</td><td>12459</td><td>12487</td><td>29</td><td>AGCGTTGGATTGTTCACCCACTAATAGGG</td><td>708.9</td><td>-11.5</td><td>1.3</td><td>-8.6</td><td>5.2E-18</td><td>2.2E-17</td><td>BT20/MB468</td></tr>
<tr><td>>MI0005756|hsa-mir-934&WithFlank&X|+|136550872|136550966@21.42.22(+2A)&[MIMAT0004977&hsa-miR-934&offsets|0|0(+2A);m-943&X|+|136550892|136550913&offsets|0|0(+2A)]|TGTCTACTACTGGAGACACTGGAA</td><td>>MI0005756|hsa-mir-934&WithFlank&X|+|136550872|136550966</td><td>21</td><td>42</td><td>22(+2A)</td><td>TGTCTACTACTGGAGACACTGGAA</td><td>1431.0</td><td>14.0</td><td>1.6</td><td>9.0</td><td>2.9E-19</td><td>2.2E-17</td><td>BT20/MB231</td></tr>
<tr><td>>MI0000063|hsa-let-7b&WithFlank&22|+|46113680|46113774@12.33.22(+1A)&[MIMAT0000063&hsa-let-7b-5p&offsets|0|0(+1A);m-14&22|+|46113691|46113712&offsets|0|0(+1A)]|TGAGGTAGTAGGTTGTGTGGTTA</td><td>>MI0000063|hsa-let-7b&WithFlank&22|+|46113680|46113774</td><td>12</td><td>33</td><td>22(+1A)</td><td>TGAGGTAGTAGGTTGTGTGGTTA</td><td>1643.7</td><td>6.3</td><td>0.7</td><td>8.6</td><td>5.6E-18</td><td>2.2E-17</td><td>MB231/MB468</td></tr>
<tr><td>>hg38_21_+_8433172_8446622_RNA45SN1_45S_50ntFlanks@12157.12181.25|AGTTTGACTGGGGCGGTACACCTGT</td><td>>hg38_21_+_8433172_8446622_RNA45SN1_45S_50ntFlanks</td><td>12157</td><td>12181</td><td>25</td><td>AGTTTGACTGGGGCGGTACACCTGT</td><td>584.1</td><td>-11.2</td><td>1.3</td><td>-8.6</td><td>5.3E-18</td><td>2.2E-17</td><td>BT20/MB468</td></tr>
<tr><td>>hg38_21_+_8433172_8446622_RNA45SN1_45S_50ntFlanks@12971.12991.21|TTTCGTACGTAGCAGAGCAGC</td><td>>hg38_21_+_8433172_8446622_RNA45SN1_45S_50ntFlanks</td><td>12971</td><td>12991</td><td>21</td><td>TTTCGTACGTAGCAGAGCAGC</td><td>407.2</td><td>-7.1</td><td>0.8</td><td>-8.6</td><td>5.4E-18</td><td>2.2E-17</td><td>BT20/MB468</td></tr>
<tr><td>>hg38_21_+_8433172_8446622_RNA45SN1_45S_50ntFlanks@6734.6773.40|ATTGATCATCGACACTTCGAACGCACTTGCGGCCCCGGGT</td><td>>hg38_21_+_8433172_8446622_RNA45SN1_45S_50ntFlanks</td><td>6734</td><td>6773</td><td>40</td><td>ATTGATCATCGACACTTCGAACGCACTTGCGGCCCCGGGT</td><td>2646.4</td><td>10.8</td><td>1.3</td><td>8.6</td><td>5.7E-18</td><td>2.2E-17</td><td>MB231/MB468</td></tr>
<tr><td>>hg38_21_+_8433172_8446622_RNA45SN1_45S_50ntFlanks@8114.8143.30|GCGCGGGACATGTGGCGTACGGAAGACCCG</td><td>>hg38_21_+_8433172_8446622_RNA45SN1_45S_50ntFlanks</td><td>8114</td><td>8143</td><td>30</td><td>GCGCGGGACATGTGGCGTACGGAAGACCCG</td><td>806.0</td><td>-11.6</td><td>1.3</td><td>-8.6</td><td>5.4E-18</td><td>2.2E-17</td><td>BT20/MB468</td></tr>
<tr><td>>hg38_21_+_8433172_8446622_RNA45SN1_45S_50ntFlanks@10408.10435.28|GGTCAGTCGGTCCTGAGAGATGGGCGAG</td><td>>hg38_21_+_8433172_8446622_RNA45SN1_45S_50ntFlanks</td><td>10408</td><td>10435</td><td>28</td><td>GGTCAGTCGGTCCTGAGAGATGGGCGAG</td><td>1110.6</td><td>-11.7</td><td>1.4</td><td>-8.6</td><td>5.8E-18</td><td>2.3E-17</td><td>MB231/MB468</td></tr>
<tr><td>>hg38_MT_+_1621_3279_RNR2_16S_50ntFlanks@1216.1262.47|CAACCTCCGAGCAGTACATGCTAAGACTTCACCAGTCAAAGCGAACT</td><td>>hg38_MT_+_1621_3279_RNR2_16S_50ntFlanks</td><td>1216</td><td>1262</td><td>47</td><td>CAACCTCCGAGCAGTACATGCTAAGACTTCACCAGTCAAAGCGAACT</td><td>540.7</td><td>11.7</td><td>1.4</td><td>8.6</td><td>5.7E-18</td><td>2.3E-17</td><td>BT20/MB468</td></tr>
<tr><td>>MI0000438|hsa-mir-15b&WithFlank&3|+|160404582|160404691@64.84.21&[MIMAT0004586&hsa-miR-15b-3p&offsets|0|-1;m-204&3|+|160404645|160404665&offsets|0|0]|CGAATCATTATTTGCTGCTCT</td><td>>MI0000438|hsa-mir-15b&WithFlank&3|+|160404582|160404691</td><td>64</td><td>84</td><td>21</td><td>CGAATCATTATTTGCTGCTCT</td><td>892.2</td><td>6.7</td><td>0.8</td><td>8.6</td><td>6.1E-18</td><td>2.3E-17</td><td>MB231/MB468</td></tr>
<tr><td>>hg38_21_+_8433172_8446622_RNA45SN1_45S_50ntFlanks@12071.12095.25|CCCCGAGGGGCTCTCGCTTCTGGCG</td><td>>hg38_21_+_8433172_8446622_RNA45SN1_45S_50ntFlanks</td><td>12071</td><td>12095</td><td>25</td><td>CCCCGAGGGGCTCTCGCTTCTGGCG</td><td>1320.8</td><td>-6.7</td><td>0.8</td><td>-8.6</td><td>5.7E-18</td><td>2.3E-17</td><td>BT20/MB468</td></tr>
<tr><td>>MI0000114|hsa-mir-107&WithFlank&10|-|89592741|89592833@56.77.22&[MIMAT0000104&hsa-miR-107&offsets|0|-1;m-78&10|-|89592758|89592778&offsets|0|+1]|AGCAGCATTGTACAGGGCTATC</td><td>>MI0000114|hsa-mir-107&WithFlank&10|-|89592741|89592833</td><td>56</td><td>77</td><td>22</td><td>AGCAGCATTGTACAGGGCTATC</td><td>2320.9</td><td>6.9</td><td>0.8</td><td>8.6</td><td>6.1E-18</td><td>2.4E-17</td><td>MB231/MB468</td></tr>
<tr><td>>hg38_21_+_8433172_8446622_RNA45SN1_45S_50ntFlanks@12654.12674.21|TATGACTGAACGCCTCTAAGT</td><td>>hg38_21_+_8433172_8446622_RNA45SN1_45S_50ntFlanks</td><td>12654</td><td>12674</td><td>21</td><td>TATGACTGAACGCCTCTAAGT</td><td>897.7</td><td>-5.4</td><td>0.6</td><td>-8.6</td><td>6.1E-18</td><td>2.4E-17</td><td>MB231/MB468</td></tr>
<tr><td>>hg38_21_+_8433172_8446622_RNA45SN1_45S_50ntFlanks@7975.8017.43|CGCGACCTCAGATCAGACGTGGCGACCCGCTGAATTTAAGCAT</td><td>>hg38_21_+_8433172_8446622_RNA45SN1_45S_50ntFlanks</td><td>7975</td><td>8017</td><td>43</td><td>CGCGACCTCAGATCAGACGTGGCGACCCGCTGAATTTAAGCAT</td><td>4702.5</td><td>7.4</td><td>0.9</td><td>8.6</td><td>6.2E-18</td><td>2.4E-17</td><td>MB231/MB468</td></tr>
<tr><td>>hg38_21_+_8433172_8446622_RNA45SN1_45S_50ntFlanks@4003.4042.40|ATAACCTCGGGCCGATCGCACGCCCCCCGTGGCGGCGACG</td><td>>hg38_21_+_8433172_8446622_RNA45SN1_45S_50ntFlanks</td><td>4003</td><td>4042</td><td>40</td><td>ATAACCTCGGGCCGATCGCACGCCCCCCGTGGCGGCGACG</td><td>435.3</td><td>-6.3</td><td>0.7</td><td>-8.6</td><td>5.7E-18</td><td>2.4E-17</td><td>BT20/MB468</td></tr>
<tr><td>>trnaMT_ValTAC_MT_+_1602_1670@49.72.24|AACTTAACTTGACCGCTCTGACCA</td><td>>trnaMT_ValTAC_MT_+_1602_1670</td><td>49</td><td>72</td><td>24</td><td>AACTTAACTTGACCGCTCTGACCA</td><td>2519.3</td><td>6.4</td><td>0.7</td><td>8.6</td><td>5.8E-18</td><td>2.4E-17</td><td>BT20/MB468</td></tr>
<tr><td>>MI0000070|hsa-mir-16-1&WithFlank&13|-|50048967|50049067@20.39.20;MI0000115|hsa-mir-16-2&WithFlank&3|+|160404739|160404831@16.35.20&[MIMAT0000069&hsa-miR-16-5p&offsets|0|-2;m-62&13|-|50049027|50049048&offsets|0|-2];[MIMAT0000069_1&hsa-miR-16-5p&offsets|0|-2;m-61&3|+|160404754|160404775&offsets|0|-2]|TAGCAGCACGTAAATATTGG</td><td>>MI0000070|hsa-mir-16-1&WithFlank&13|-|50048967|50049067</td><td>20</td><td>39</td><td>20</td><td>TAGCAGCACGTAAATATTGG</td><td>981.0</td><td>7.5</td><td>0.9</td><td>8.6</td><td>5.8E-18</td><td>2.4E-17</td><td>BT20/MB468</td></tr>
<tr><td>>MI0000439|hsa-mir-23b&WithFlank&9|+|95085202|95085310@65.86.22(+1U)&[MIMAT0000418&hsa-miR-23b-3p&offsets|+1|0(+1U);m-42&9|+|95085265|95085285&offsets|+1|+2(+1U)]|TCACATTGCCAGGGATTACCACT</td><td>>MI0000439|hsa-mir-23b&WithFlank&9|+|95085202|95085310</td><td>65</td><td>86</td><td>22(+1U)</td><td>TCACATTGCCAGGGATTACCACT</td><td>833.8</td><td>7.3</td><td>0.8</td><td>8.6</td><td>5.8E-18</td><td>2.4E-17</td><td>BT20/MB468</td></tr>
<tr><td>>hg38_21_+_8433172_8446622_RNA45SN1_45S_50ntFlanks@7990.8017.28|GACGTGGCGACCCGCTGAATTTAAGCAT</td><td>>hg38_21_+_8433172_8446622_RNA45SN1_45S_50ntFlanks</td><td>7990</td><td>8017</td><td>28</td><td>GACGTGGCGACCCGCTGAATTTAAGCAT</td><td>677.0</td><td>6.3</td><td>0.7</td><td>8.6</td><td>6.3E-18</td><td>2.4E-17</td><td>MB231/MB468</td></tr>
<tr><td>>hg38_21_+_8433172_8446622_RNA45SN1_45S_50ntFlanks@7990.8010.21|GACGTGGCGACCCGCTGAATT</td><td>>hg38_21_+_8433172_8446622_RNA45SN1_45S_50ntFlanks</td><td>7990</td><td>8010</td><td>21</td><td>GACGTGGCGACCCGCTGAATT</td><td>1688.7</td><td>6.4</td><td>0.7</td><td>8.6</td><td>6.3E-18</td><td>2.4E-17</td><td>MB231/MB468</td></tr>
</table>

| | | | | | | | | | | | | |
|---|---|---|---|---|---|---|---|---|---|---|---|---|
| >hg38_21_+_8433172_8446622_RNA45SN1_45S_50ntFlanks@10843.10868.26\|TAACTTCGGGATAAGGATTGGCTCTA | >hg38_21_+_8433172_8446622_RNA45SN1_45S_50ntFlanks | 10843 | 10868 | 26 | TAACTTCGGGATAAGGATTGGCTCTA | 805.8 | -7.0 | 0.8 | -8.6 | 5.9E-18 | 2.4E-17 | BT20/MB468 |
| >am_MI0000786\|hsa-mir-378a&WithFlank&5\|+\|149732819\|149732896@49.69.21&[MIMAT0000732&hsa-miR-378a-3p&offsets\|0\|-1;m-44&5\|+\|149732867\|149732888&offsets\|0\|-1]\|ACTGGACTTGGAGTCAGAAGG | >am_MI0000786\|hsa-mir-378a&WithFlank&5\|+\|149732819\|149732896 | 49 | 69 | 21 | ACTGGACTTGGAGTCAGAAGG | 5489.7 | 5.3 | 0.6 | 8.6 | 5.9E-18 | 2.4E-17 | BT20/MB468 |
| >hg38_21_+_8433172_8446622_RNA45SN1_45S_50ntFlanks@10683.10706.24\|CCGGCGGCGTCCGGTGAGCTCTCG | >hg38_21_+_8433172_8446622_RNA45SN1_45S_50ntFlanks | 10683 | 10706 | 24 | CCGGCGGCGTCCGGTGAGCTCTCG | 5742.1 | -6.0 | 0.7 | -8.6 | 5.9E-18 | 2.4E-17 | BT20/MB468 |
| >hg38_21_+_8433172_8446622_RNA45SN1_45S_50ntFlanks@475.521.47\|CTTCGTGATCGATGTGGTGACGTCGTGCTCTCCCGGGCCGGGTCCGA | >hg38_21_+_8433172_8446622_RNA45SN1_45S_50ntFlanks | 475 | 521 | 47 | CTTCGTGATCGATGTGGTGACGTCGTGCTCTCCCGGGCCGGGTCCGA | 1528.0 | 8.8 | 1.0 | 8.6 | 6.0E-18 | 2.5E-17 | BT20/MB468 |
| >MI0000234\|hsa-mir-192&WithFlank&11\|-\|64891131\|64891252@30.51.22(+1U)&[MIMAT0000222&hsa-miR-192-5p&offsets\|0\|+1(+1U);m-59&11\|-\|64891203\|64891223&offsets\|0\|+1(+1U)]\|CTGACCTATGAATTGACAGCCAT | >MI0000234\|hsa-mir-192&WithFlank&11\|-\|64891131\|64891252 | 30 | 51 | 22(+1U) | CTGACCTATGAATTGACAGCCAT | 1018.7 | 5.1 | 0.6 | 8.6 | 6.0E-18 | 2.5E-17 | BT20/MB468 |
| >MI0000433\|hsa-let-7g&WithFlank&3\|-\|52268272\|52268367@11.32.22(+1A)&[MIMAT0000414&hsa-let-7g-5p&offsets\|0\|0(+1A);m-58&3\|-\|52268336\|52268357&offsets\|0\|0(+1A)]\|TGAGGTAGTAGTTTGTACAGTTA | >MI0000433\|hsa-let-7g&WithFlank&3\|-\|52268272\|52268367 | 11 | 32 | 22(+1A) | TGAGGTAGTAGTTTGTACAGTTA | 3443.8 | 6.1 | 0.7 | 8.6 | 6.0E-18 | 2.5E-17 | BT20/MB468 |
| >hg38_21_+_8433172_8446622_RNA45SN1_45S_50ntFlanks@12324.12348.25\|TTGGGTTTTAAGCAGGAGGTGTCAG | >hg38_21_+_8433172_8446622_RNA45SN1_45S_50ntFlanks | 12324 | 12348 | 25 | TTGGGTTTTAAGCAGGAGGTGTCAG | 492.4 | -10.9 | 1.3 | -8.6 | 6.0E-18 | 2.5E-17 | BT20/MB468 |
| >MI0000103\|hsa-mir-101-1&WithFlank&1\|-\|65058428\|65058514@53.74.22;MI0000739\|hsa-mir-101-2&WithFlank&9\|+\|4850291\|4850381@55.76.22&[MIMAT0000099&hsa-miR-101-3p&offsets\|0\|+1;m-17&1\|-\|65058443\|65058462&offsets\|0\|+2];[MIMAT0000099_1&hsa-miR-101-3p&offsets\|0\|+1;m-16&9\|+\|4850345\|4850364&offsets\|0\|+2]\|TACAGTACTGTGATAACTGAAG | >MI0000103\|hsa-mir-101-1&WithFlank&1\|-\|65058428\|65058514 | 53 | 74 | 22 | TACAGTACTGTGATAACTGAAG | 33856.7 | 9.6 | 1.1 | 8.6 | 6.1E-18 | 2.5E-17 | BT20/MB468 |
| >hg38_21_+_8433172_8446622_RNA45SN1_45S_50ntFlanks@9568.9592.25\|CGTGCAAATCGGTCGTCCGACCTGG | >hg38_21_+_8433172_8446622_RNA45SN1_45S_50ntFlanks | 9568 | 9592 | 25 | CGTGCAAATCGGTCGTCCGACCTGG | 603.0 | -11.2 | 1.3 | -8.6 | 6.2E-18 | 2.5E-17 | BT20/MB468 |
| >hg38_21_+_8433172_8446622_RNA45SN1_45S_50ntFlanks@483.507.25\|TCGATGTGGTGACGTCGTGCTCTCC | >hg38_21_+_8433172_8446622_RNA45SN1_45S_50ntFlanks | 483 | 507 | 25 | TCGATGTGGTGACGTCGTGCTCTCC | 844.2 | 9.0 | 1.0 | 8.6 | 6.6E-18 | 2.5E-17 | MB231/MB468 |
| >hg38_21_+_8433172_8446622_RNA45SN1_45S_50ntFlanks@8175.8216.42\|CCTTCTGATCGAGGCCCAGCCCGTGGACGGTGTGAGGCCGGT | >hg38_21_+_8433172_8446622_RNA45SN1_45S_50ntFlanks | 8175 | 8216 | 42 | CCTTCTGATCGAGGCCCAGCCCGTGGACGGTGTGAGGCCGGT | 341.3 | 9.8 | 1.1 | 8.6 | 6.3E-18 | 2.6E-17 | BT20/MB468 |
| >hg38_21_+_8433172_8446622_RNA45SN1_45S_50ntFlanks@6652.6682.31\|GACTCTTAGCGGTGGATCACTCGGCTCGTGC | >hg38_21_+_8433172_8446622_RNA45SN1_45S_50ntFlanks | 6652 | 6682 | 31 | GACTCTTAGCGGTGGATCACTCGGCTCGTGC | 351.7 | 4.9 | 0.6 | 8.6 | 6.3E-18 | 2.6E-17 | BT20/MB468 |
| >hg38_MT_+_1621_3279_RNR2_16S_50ntFlanks@459.492.34\|CTTGTAAATTTAACTGTTAGTCCAAAGAGGAACA | >hg38_MT_+_1621_3279_RNR2_16S_50ntFlanks | 459 | 492 | 34 | CTTGTAAATTTAACTGTTAGTCCAAAGAGGAACA | 2674.0 | 10.4 | 1.2 | 8.6 | 6.7E-18 | 2.6E-17 | MB231/MB468 |
| >hg38_21_+_8433172_8446622_RNA45SN1_45S_50ntFlanks@7984.8008.25\|AGATCAGACGTGGCGACCCGCTGAA | >hg38_21_+_8433172_8446622_RNA45SN1_45S_50ntFlanks | 7984 | 8008 | 25 | AGATCAGACGTGGCGACCCGCTGAA | 822.7 | -5.3 | 0.6 | -8.6 | 6.7E-18 | 2.6E-17 | MB231/MB468 |
| >hg38_21_+_8433172_8446622_RNA45SN1_45S_50ntFlanks@12971.12990.20\|TTTCGTACGTAGCAGAGCAG | >hg38_21_+_8433172_8446622_RNA45SN1_45S_50ntFlanks | 12971 | 12990 | 20 | TTTCGTACGTAGCAGAGCAG | 514.3 | -11.0 | 1.3 | -8.6 | 6.4E-18 | 2.6E-17 | BT20/MB468 |
| >hg38_21_+_8433172_8446622_RNA45SN1_45S_50ntFlanks@12632.12653.22\|GCGAAGCTACCATCTGTGGGAT | >hg38_21_+_8433172_8446622_RNA45SN1_45S_50ntFlanks | 12632 | 12653 | 22 | GCGAAGCTACCATCTGTGGGAT | 963.0 | -11.9 | 1.4 | -8.6 | 6.5E-18 | 2.6E-17 | BT20/MB468 |
| >MI0000749\|hsa-mir-30e&WithFlank&1\|+\|40754349\|40754452@65.85.21(+1U)&[MIMAT0000693&hsa-miR-30e-3p&offsets\|0\|-1(+1U);m-25&1\|+\|40754413\|40754434&offsets\|0\|-1(+1U)]\|CTTTCAGTCGGATGTTTACAGT | >MI0000749\|hsa-mir-30e&WithFlank&1\|+\|40754349\|40754452 | 65 | 85 | 21(+1U) | CTTTCAGTCGGATGTTTACAGT | 6987.0 | 6.3 | 0.7 | 8.6 | 6.6E-18 | 2.7E-17 | BT20/MB468 |
| >MI0000441\|hsa-mir-30b&WithFlank&8\|-\|134800514\|134800613@23.43.21&[MIMAT0000420&hsa-miR-30b-5p&offsets\|0\|-1;m-76&8\|-\|134800570\|134800591&offsets\|0\|-1]\|TGTAAACATCCTACACTCAGC | >MI0000441\|hsa-mir-30b&WithFlank&8\|-\|134800514\|134800613 | 23 | 43 | 21 | TGTAAACATCCTACACTCAGC | 650.3 | 8.9 | 1.0 | 8.6 | 7.1E-18 | 2.7E-17 | MB231/MB468 |
| >MI0000809\|hsa-mir-151a&WithFlank&8\|-\|140732558\|140732659@53.74.22(+1G)&[MIMAT0000757&hsa-miR-151a-3p&offsets\|0\|+1(+1G);m-30&8\|-\|140732587\|140732607&offsets\|0\|+1(+1G)]\|CTAGACTGAAGCTCCTTGAGGAG | >MI0000809\|hsa-mir-151a&WithFlank&8\|-\|140732558\|140732659 | 53 | 74 | 22(+1G) | CTAGACTGAAGCTCCTTGAGGAG | 812.0 | 9.1 | 1.1 | 8.6 | 6.7E-18 | 2.7E-17 | BT20/MB468 |
| >hg38_21_+_8433172_8446622_RNA45SN1_45S_50ntFlanks@7976.8009.34\|GCGACCTCAGATCAGACGTGGCGACCCGCTGAAT | >hg38_21_+_8433172_8446622_RNA45SN1_45S_50ntFlanks | 7976 | 8009 | 34 | GCGACCTCAGATCAGACGTGGCGACCCGCTGAAT | 730.1 | 8.2 | 1.0 | 8.6 | 7.2E-18 | 2.8E-17 | MB231/MB468 |
| >hg38_21_+_8433172_8446622_RNA45SN1_45S_50ntFlanks@7977.8017.41\|CGACCTCAGATCAGACGTGGCGACCCGCTGAATTTAAGCAT | >hg38_21_+_8433172_8446622_RNA45SN1_45S_50ntFlanks | 7977 | 8017 | 41 | CGACCTCAGATCAGACGTGGCGACCCGCTGAATTTAAGCAT | 794.1 | 9.1 | 1.1 | 8.6 | 7.2E-18 | 2.8E-17 | MB231/MB468 |
| >MI0003560\|hsa-mir-92b&WithFlank&1\|+\|155195171\|155195278@67.87.21&[MIMAT0003218&hsa-miR-92b-3p&offsets\|0\|-1;m-115&1\|+\|155195237\|155195258&offsets\|0\|-1]\|TATTGCACTCGTCCCGGCCTC | >MI0003560\|hsa-mir-92b&WithFlank&1\|+\|155195171\|155195278 | 67 | 87 | 21 | TATTGCACTCGTCCCGGCCTC | 2061.3 | 9.3 | 1.1 | 8.6 | 6.9E-18 | 2.8E-17 | BT20/MB468 |
| >hg38_21_+_8433172_8446622_RNA45SN1_45S_50ntFlanks@3899.3919.21\|CCTTCGCGGGGGGGATGCGTG | >hg38_21_+_8433172_8446622_RNA45SN1_45S_50ntFlanks | 3899 | 3919 | 21 | CCTTCGCGGGGGGGATGCGTG | 436.0 | -6.9 | 0.8 | -8.6 | 7.5E-18 | 2.9E-17 | MB231/MB468 |
| >hg38_21_+_8433172_8446622_RNA45SN1_45S_50ntFlanks@10402.10431.30\|AACATGGGTCAGTCGGTCCTGAGAGATGGG | >hg38_21_+_8433172_8446622_RNA45SN1_45S_50ntFlanks | 10402 | 10431 | 30 | AACATGGGTCAGTCGGTCCTGAGAGATGGG | 539.5 | -3.9 | 0.5 | -8.6 | 7.5E-18 | 2.9E-17 | MB231/MB468 |
| >hg38_21_+_8433172_8446622_RNA45SN1_45S_50ntFlanks@3985.4005.21\|CGGCTTTGGTGACTCTAGATA | >hg38_21_+_8433172_8446622_RNA45SN1_45S_50ntFlanks | 3985 | 4005 | 21 | CGGCTTTGGTGACTCTAGATA | 439.2 | -5.4 | 0.6 | -8.6 | 7.1E-18 | 2.9E-17 | BT20/MB468 |
| >MI0001445\|hsa-mir-423&WithFlank&17\|+\|30117073\|30117178@58.80.23(+1C)&[MIMAT0001340&hsa-miR-423-3p&offsets\|-1\|-1(+1C);m-100&17\|+\|30117131\|30117153&offsets\|-1\|-1(+1C)]\|AAGCTCGGTCTGAGGCCCCTCAGC | >MI0001445\|hsa-mir-423&WithFlank&17\|+\|30117073\|30117178 | 58 | 80 | 23(+1C) | AAGCTCGGTCTGAGGCCCCTCAGC | 1075.9 | 9.1 | 1.1 | 8.6 | 7.6E-18 | 2.9E-17 | MB231/MB468 |
| >MI0000471\|hsa-mir-126&WithFlank&9\|+\|136670596\|136670692@58.79.22&[MIMAT0000445&hsa-miR-126-3p&offsets\|0\|0;m-40&9\|+\|136670653\|136670674&offsets\|0\|0]\|TCGTACCGTGAGTAATAATGCG | >MI0000471\|hsa-mir-126&WithFlank&9\|+\|136670596\|136670692 | 58 | 79 | 22 | TCGTACCGTGAGTAATAATGCG | 9596.8 | 9.0 | 1.0 | 8.6 | 7.1E-18 | 2.9E-17 | BT20/MB468 |
| >am_trnaMT_ValTAC_MT_+_1602_1670@28.47.20\|AACTTACACTTAGGAGATTT | >am_trnaMT_ValTAC_MT_+_1602_1670 | 28 | 47 | 20 | AACTTACACTTAGGAGATTT | 830.4 | 12.3 | 1.4 | 8.6 | 7.3E-18 | 3.0E-17 | BT20/MB468 |
| >MI0000088\|hsa-mir-30a&WithFlank&6\|-\|71403545\|71403627@12.33.22&[MIMAT0000087&hsa-miR-30a-5p&offsets\|0\|0;m-15&6\|-\|71403595\|71403616&offsets\|0\|0]\|TGTAAACATCCTCGACTGGAAG | >MI0000088\|hsa-mir-30a&WithFlank&6\|-\|71403545\|71403627 | 12 | 33 | 22 | TGTAAACATCCTCGACTGGAAG | 29392.8 | 8.6 | 1.0 | 8.6 | 7.3E-18 | 3.0E-17 | BT20/MB468 |
| >hg38_21_+_8433172_8446622_RNA45SN1_45S_50ntFlanks@12628.12653.26\|TGGGGCGAAGCTACCATCTGTGGGAT | >hg38_21_+_8433172_8446622_RNA45SN1_45S_50ntFlanks | 12628 | 12653 | 26 | TGGGGCGAAGCTACCATCTGTGGGAT | 993.5 | -11.5 | 1.3 | -8.6 | 7.9E-18 | 3.0E-17 | MB231/MB468 |
| >hg38_21_+_8433172_8446622_RNA45SN1_45S_50ntFlanks@11666.11687.22\|TCTGCCCAGTGCTCTGAATGTC | >hg38_21_+_8433172_8446622_RNA45SN1_45S_50ntFlanks | 11666 | 11687 | 22 | TCTGCCCAGTGCTCTGAATGTC | 1383.8 | 5.3 | 0.6 | 8.6 | 8.0E-18 | 3.1E-17 | MB231/MB468 |
| >hg38_21_+_8433172_8446622_RNA45SN1_45S_50ntFlanks@12405.12422.18\|CTTTTTGATCCTTCGATG | >hg38_21_+_8433172_8446622_RNA45SN1_45S_50ntFlanks | 12405 | 12422 | 18 | CTTTTTGATCCTTCGATG | 873.9 | -11.8 | 1.4 | -8.6 | 7.6E-18 | 3.1E-17 | BT20/MB468 |
| >MI0000299\|hsa-mir-222&WithFlank&X\|-\|45747009\|45747130@75.99.25(+1U)&[MIMAT0000279&hsa-miR-222-3p&offsets\|0\|+4(+1U);m-140&X\|-\|45747033\|45747056&offsets\|0\|+1(+1U)]\|AGCTACATCTGGCTACTGGGTCTCTT | >MI0000299\|hsa-mir-222&WithFlank&X\|-\|45747009\|45747130 | 75 | 99 | 25(+1U) | AGCTACATCTGGCTACTGGGTCTCTT | 1950.0 | 6.5 | 0.8 | 8.6 | 7.6E-18 | 3.1E-17 | BT20/MB468 |
| >hg38_21_+_8433172_8446622_RNA45SN1_45S_50ntFlanks@12957.12979.23\|CTTCTGGGTCGGGGTTTCGTACG | >hg38_21_+_8433172_8446622_RNA45SN1_45S_50ntFlanks | 12957 | 12979 | 23 | CTTCTGGGTCGGGGTTTCGTACG | 338.2 | -5.2 | 0.6 | -8.6 | 8.1E-18 | 3.1E-17 | MB231/MB468 |
| >MI0001446\|hsa-mir-424&WithFlank&X\|-\|134546608\|134546717@17.37.21&[MIMAT0001341&hsa-miR-424-5p&offsets\|0\|-1;m-152&X\|-\|134546681\|134546701&offsets\|0\|0]\|CAGCAGCAATTCATGTTTTGA | >MI0001446\|hsa-mir-424&WithFlank&X\|-\|134546608\|134546717 | 17 | 37 | 21 | CAGCAGCAATTCATGTTTTGA | 3399.4 | 14.3 | 1.7 | 8.6 | 8.0E-18 | 3.2E-17 | BT20/MB468 |
| >hg38_21_+_8433172_8446622_RNA45SN1_45S_50ntFlanks@11784.11804.21\|ATGAATGGATGAACGAGATTC | >hg38_21_+_8433172_8446622_RNA45SN1_45S_50ntFlanks | 11784 | 11804 | 21 | ATGAATGGATGAACGAGATTC | 5007.6 | -4.5 | 0.5 | -8.6 | 8.7E-18 | 3.3E-17 | MB231/MB468 |
| >hg38_21_+_8433172_8446622_RNA45SN1_45S_50ntFlanks@4760.4778.19\|TCAGATACCGTCGTAGTTC | >hg38_21_+_8433172_8446622_RNA45SN1_45S_50ntFlanks | 4760 | 4778 | 19 | TCAGATACCGTCGTAGTTC | 504.0 | -11.0 | 1.3 | -8.6 | 8.4E-18 | 3.4E-17 | BT20/MB468 |
| >hg38_21_+_8433172_8446622_RNA45SN1_45S_50ntFlanks@10411.10432.22\|CAGTCGGTCCTGAGAGATGGGC | >hg38_21_+_8433172_8446622_RNA45SN1_45S_50ntFlanks | 10411 | 10432 | 22 | CAGTCGGTCCTGAGAGATGGGC | 836.7 | -5.5 | 0.6 | -8.6 | 9.0E-18 | 3.5E-17 | MB231/MB468 |
| >hg38_21_+_8433172_8446622_RNA45SN1_45S_50ntFlanks@11315.11344.30\|CCACCCCACGTCTCGTCGCGCGCGCGTCCG | >hg38_21_+_8433172_8446622_RNA45SN1_45S_50ntFlanks | 11315 | 11344 | 30 | CCACCCCACGTCTCGTCGCGCGCGCGTCCG | 936.8 | -11.4 | 1.3 | -8.6 | 9.0E-18 | 3.5E-17 | MB231/MB468 |
| >hg38_MT_+_1621_3279_RNR2_16S_50ntFlanks@493.518.26\|GCTCTTTGGACACTAGGAAAAAACCT | >hg38_MT_+_1621_3279_RNR2_16S_50ntFlanks | 493 | 518 | 26 | GCTCTTTGGACACTAGGAAAAAACCT | 2724.4 | 10.4 | 1.2 | 8.6 | 9.1E-18 | 3.5E-17 | MB231/MB468 |
| >trna152_ValCAC_6_-_27248049_27248121@1.29.29\|GCTTCTGTAGTGTAGTGGTTATCACGTTC | >trna152_ValCAC_6_-_27248049_27248121 | 1 | 29 | 29 | GCTTCTGTAGTGTAGTGGTTATCACGTTC | 9702.3 | -6.5 | 0.8 | -8.6 | 8.5E-18 | 3.5E-17 | BT20/MB468 |
| >hg38_1_-_228634819_228635039_RNA5S12_5S_50ntFlanks@149.169.21\|GAATACCGGGTGCTGTAGGCT | >hg38_1_-_228634819_228635039_RNA5S12_5S_50ntFlanks | 149 | 169 | 21 | GAATACCGGGTGCTGTAGGCT | 800.3 | 6.1 | 0.7 | 8.6 | 8.6E-18 | 3.5E-17 | BT20/MB468 |
| >hg38_21_+_8433172_8446622_RNA45SN1_45S_50ntFlanks@4731.4768.38\|AAGAACGAAAGTCGGAGGTTCGAAGACGATCAGATACC | >hg38_21_+_8433172_8446622_RNA45SN1_45S_50ntFlanks | 4731 | 4768 | 38 | AAGAACGAAAGTCGGAGGTTCGAAGACGATCAGATACC | 501.6 | 10.1 | 1.2 | 8.6 | 8.6E-18 | 3.5E-17 | BT20/MB468 |
| >hg38_21_+_8433172_8446622_RNA45SN1_45S_50ntFlanks@4555.4587.33\|CGCAGCTAGGAATAATGGAATAGGACCGCGGTT | >hg38_21_+_8433172_8446622_RNA45SN1_45S_50ntFlanks | 4555 | 4587 | 33 | CGCAGCTAGGAATAATGGAATAGGACCGCGGTT | 140.5 | -4.7 | 0.5 | -8.6 | 8.6E-18 | 3.5E-17 | BT20/MB468 |

<table>
<tr><td>>MI0000650|hsa-mir-200c&WithFlank&12|+|6963693|6963772@50.69.20(+1A)&[MIMAT0000617&hsa-miR-200c-3p&offsets|0|-3(+1A);m-54&12|+|6963742|6963764&offsets|0|-3(+1A)]|TAATACTGCCGGGTAATGATA</td><td>>MI0000650|hsa-mir-200c&WithFlank&12|+|6963693|6963772</td><td>50</td><td>69</td><td>20(+1A)</td><td>TAATACTGCCGGGTAATGATA</td><td>1636.0</td><td>11.1</td><td>1.3</td><td>8.6</td><td>8.6E-18</td><td>3.5E-17</td><td>BT20/MB468</td></tr>
<tr><td>>hg38_21_+_8433172_8446622_RNA45SN1_45S_50ntFlanks@11781.11806.26|CGCATGAATGGATGAACGAGATTCCC</td><td>>hg38_21_+_8433172_8446622_RNA45SN1_45S_50ntFlanks</td><td>11781</td><td>11806</td><td>26</td><td>CGCATGAATGGATGAACGAGATTCCC</td><td>567.8</td><td>-6.9</td><td>0.8</td><td>-8.6</td><td>9.2E-18</td><td>3.5E-17</td><td>MB231/MB468</td></tr>
<tr><td>>hg38_21_+_8433172_8446622_RNA45SN1_45S_50ntFlanks@11641.11662.22|CGGCGGGTGTTGACGCGATGTG</td><td>>hg38_21_+_8433172_8446622_RNA45SN1_45S_50ntFlanks</td><td>11641</td><td>11662</td><td>22</td><td>CGGCGGGTGTTGACGCGATGTG</td><td>1400.9</td><td>-5.4</td><td>0.6</td><td>-8.6</td><td>9.4E-18</td><td>3.6E-17</td><td>MB231/MB468</td></tr>
<tr><td>>trna16_TyrGTA_14_-_21131351_21131444@59.76.18,trna17_TyrGTA_14_-_21128117_21128210@59.76.18,trna19_TyrGTA_14_-_21121258_21121351@59.76.18,trna2_TyrGTA_2_+_27273650_27273738@59.76.18,trna4_TyrGTA_8_+_67025602_67025694@59.76.18,trna5_TyrGTA_14_+_21151432_21151520@59.76.18,trna5_TyrGTA_8_+_67026223_67026311@59.76.18|TTCCGGCTCGAAGGACCA</td><td>>trna16_TyrGTA_14_-_21131351_21131444</td><td>59</td><td>76</td><td>18</td><td>TTCCGGCTCGAAGGACCA</td><td>4013.9</td><td>7.3</td><td>0.9</td><td>8.6</td><td>8.9E-18</td><td>3.6E-17</td><td>BT20/MB468</td></tr>
<tr><td>>hg38_1_-_228634819_228635039_RNA5S12_5S_50ntFlanks@133.171.39|ATGGGAGACCGCCTGGGAATACCGGGTGCTGTAGGCTTT</td><td>>hg38_1_-_228634819_228635039_RNA5S12_5S_50ntFlanks</td><td>133</td><td>171</td><td>39</td><td>ATGGGAGACCGCCTGGGAATACCGGGTGCTGTAGGCTTT</td><td>5148.0</td><td>6.0</td><td>0.7</td><td>8.6</td><td>8.9E-18</td><td>3.6E-17</td><td>BT20/MB468</td></tr>
<tr><td>>MI0000469|hsa-mir-125a&WithFlank&19|+|51693248|51693345@21.43.23(+1U)&[MIMAT0000443&hsa-miR-125a-5p&offsets|0|-1(+1U);m-63&19|+|51693268|51693289&offsets|0|+1(+1U)]|TCCCTGAGACCCTTTAACCTGTGT</td><td>>MI0000469|hsa-mir-125a&WithFlank&19|+|51693248|51693345</td><td>21</td><td>43</td><td>23(+1U)</td><td>TCCCTGAGACCCTTTAACCTGTGT</td><td>2522.9</td><td>6.6</td><td>0.8</td><td>8.6</td><td>9.1E-18</td><td>3.7E-17</td><td>BT20/MB468</td></tr>
<tr><td>>hg38_21_+_8433172_8446622_RNA45SN1_45S_50ntFlanks@4003.4024.22|ATAACCTCGGGCCGATCGCACG</td><td>>hg38_21_+_8433172_8446622_RNA45SN1_45S_50ntFlanks</td><td>4003</td><td>4024</td><td>22</td><td>ATAACCTCGGGCCGATCGCACG</td><td>1486.4</td><td>-5.0</td><td>0.6</td><td>-8.6</td><td>9.7E-18</td><td>3.7E-17</td><td>MB231/MB468</td></tr>
<tr><td>>hg38_21_+_8433172_8446622_RNA45SN1_45S_50ntFlanks@4731.4767.37|AAGAACGAAAGTCGGAGGTTCGAAGACGATCAGATAC</td><td>>hg38_21_+_8433172_8446622_RNA45SN1_45S_50ntFlanks</td><td>4731</td><td>4767</td><td>37</td><td>AAGAACGAAAGTCGGAGGTTCGAAGACGATCAGATAC</td><td>1599.5</td><td>10.4</td><td>1.2</td><td>8.6</td><td>9.8E-18</td><td>3.7E-17</td><td>MB231/MB468</td></tr>
<tr><td>>hg38_21_+_8433172_8446622_RNA45SN1_45S_50ntFlanks@6733.6773.41|CATTGATCATCGACACTTCGAACGCACTTGCGGCCCCGGGT</td><td>>hg38_21_+_8433172_8446622_RNA45SN1_45S_50ntFlanks</td><td>6733</td><td>6773</td><td>41</td><td>CATTGATCATCGACACTTCGAACGCACTTGCGGCCCCGGGT</td><td>1086.5</td><td>9.0</td><td>1.0</td><td>8.6</td><td>9.8E-18</td><td>3.7E-17</td><td>MB231/MB468</td></tr>
<tr><td>>MI0000483|hsa-mir-186&WithFlank&1|-|71067625|71067722@21.41.21(+1A)&[MIMAT0000456&hsa-miR-186-5p&offsets|0|-1(+1A);m-84&1|-|71067681|71067702&offsets|0|-1(+1A)]|CAAAGAATTCTCCTTTTGGGCA</td><td>>MI0000483|hsa-mir-186&WithFlank&1|-|71067625|71067722</td><td>21</td><td>41</td><td>21(+1A)</td><td>CAAAGAATTCTCCTTTTGGGCA</td><td>1287.6</td><td>8.2</td><td>1.0</td><td>8.6</td><td>9.3E-18</td><td>3.7E-17</td><td>BT20/MB468</td></tr>
<tr><td>>MI0000750|hsa-mir-26a-2&WithFlank&12|-|57824603|57824698@20.41.22(+3U);MI0000083|hsa-mir-26a-1&WithFlank&3|+|37969398|37969486@16.37.22(+3U)&[MIMAT0000082&hsa-miR-26a-5p&offsets|0|0(+3U);m-33&12|-|57824658|57824679&offsets|0|0(+3U)];[MIMAT0000082_1&hsa-miR-26a-5p&offsets|0|0(+3U);m-34&3|+|37969413|37969434&offsets|0|0(+3U)]|TTCAAGTAATCCAGGATAGGCTTTT</td><td>>MI0000750|hsa-mir-26a-2&WithFlank&12|-|57824603|57824698</td><td>20</td><td>41</td><td>22(+3U)</td><td>TTCAAGTAATCCAGGATAGGCTTTT</td><td>747.6</td><td>9.7</td><td>1.1</td><td>8.6</td><td>9.9E-18</td><td>3.8E-17</td><td>MB231/MB468</td></tr>
<tr><td>>hg38_21_+_8433172_8446622_RNA45SN1_45S_50ntFlanks@9570.9596.27|TGCAAATCGGTCGTCCGACCTGGGTAT</td><td>>hg38_21_+_8433172_8446622_RNA45SN1_45S_50ntFlanks</td><td>9570</td><td>9596</td><td>27</td><td>TGCAAATCGGTCGTCCGACCTGGGTAT</td><td>2306.8</td><td>-6.7</td><td>0.8</td><td>-8.6</td><td>1.0E-17</td><td>3.8E-17</td><td>MB231/MB468</td></tr>
<tr><td>>hg38_21_+_8433172_8446622_RNA45SN1_45S_50ntFlanks@13022.13039.18|CTCGACACAAGGGTTTGT</td><td>>hg38_21_+_8433172_8446622_RNA45SN1_45S_50ntFlanks</td><td>13022</td><td>13039</td><td>18</td><td>CTCGACACAAGGGTTTGT</td><td>1938.8</td><td>4.3</td><td>0.5</td><td>8.6</td><td>1.0E-17</td><td>3.9E-17</td><td>MB231/MB468</td></tr>
<tr><td>>MI0000736|hsa-mir-30c-1&WithFlank&1|+|40757278|40757378@23.45.23(+1G);MI0000254|hsa-mir-30c-2&WithFlank&6|-|71376954|71377037@13.35.23(+1G)&[MIMAT0000244&hsa-miR-30c-5p&offsets|0|0(+1G);m-57&1|+|40757300|40757323&offsets|0|-1(+1G)];[MIMAT0000244_1&hsa-miR-30c-5p&offsets|0|0(+1G);m-56&6|-|71377002|71377025&offsets|0|-1(+1G)]|TGTAAACATCCTACACTCTCAGCG</td><td>>MI0000736|hsa-mir-30c-1&WithFlank&1|+|40757278|40757378</td><td>23</td><td>45</td><td>23(+1G)</td><td>TGTAAACATCCTACACTCTCAGCG</td><td>5150.3</td><td>6.6</td><td>0.8</td><td>8.6</td><td>1.0E-17</td><td>3.9E-17</td><td>MB231/MB468</td></tr>
<tr><td>>hg38_21_+_8433172_8446622_RNA45SN1_45S_50ntFlanks@11569.11600.32|CAGCCGACTTAGAACTGGTGCGGACCAGGGGA</td><td>>hg38_21_+_8433172_8446622_RNA45SN1_45S_50ntFlanks</td><td>11569</td><td>11600</td><td>32</td><td>CAGCCGACTTAGAACTGGTGCGGACCAGGGGA</td><td>1111.9</td><td>-4.4</td><td>0.5</td><td>-8.6</td><td>9.6E-18</td><td>3.9E-17</td><td>BT20/MB468</td></tr>
<tr><td>>trnaMT_GluTTC_MT_-_14674_14742@38.61.24,trnalookalike8_GluTTC_5_-_93905172_93905240@38.61.24|CATTGGTCGTGGTTGTAGTCCGTG</td><td>>trnaMT_GluTTC_MT_-_14674_14742</td><td>38</td><td>61</td><td>24</td><td>CATTGGTCGTGGTTGTAGTCCGTG</td><td>361.6</td><td>8.6</td><td>1.0</td><td>8.6</td><td>9.6E-18</td><td>3.9E-17</td><td>BT20/MB468</td></tr>
<tr><td>>hg38_21_+_8433172_8446622_RNA45SN1_45S_50ntFlanks@9739.9757.19|AAACGATCTCAACCTATTC</td><td>>hg38_21_+_8433172_8446622_RNA45SN1_45S_50ntFlanks</td><td>9739</td><td>9757</td><td>19</td><td>AAACGATCTCAACCTATTC</td><td>723.5</td><td>-8.0</td><td>0.9</td><td>-8.6</td><td>9.7E-18</td><td>3.9E-17</td><td>BT20/MB468</td></tr>
<tr><td>>hg38_21_+_8433172_8446622_RNA45SN1_45S_50ntFlanks@10690.10710.21|CGTCCGGTGAGCTCTCGCTGG</td><td>>hg38_21_+_8433172_8446622_RNA45SN1_45S_50ntFlanks</td><td>10690</td><td>10710</td><td>21</td><td>CGTCCGGTGAGCTCTCGCTGG</td><td>652.1</td><td>-7.9</td><td>0.9</td><td>-8.6</td><td>9.9E-18</td><td>4.0E-17</td><td>BT20/MB468</td></tr>
<tr><td>>hg38_21_+_8433172_8446622_RNA45SN1_45S_50ntFlanks@12641.12659.19|CCATCTGTGGGATTATGAC</td><td>>hg38_21_+_8433172_8446622_RNA45SN1_45S_50ntFlanks</td><td>12641</td><td>12659</td><td>19</td><td>CCATCTGTGGGATTATGAC</td><td>676.5</td><td>-6.1</td><td>0.7</td><td>-8.6</td><td>9.9E-18</td><td>4.0E-17</td><td>BT20/MB468</td></tr>
<tr><td>>MI0000805|hsa-mir-342&WithFlank&14|+|100109649|100109759@67.90.24&[MIMAT0000753&hsa-miR-342-3p&offsets|0|+1;m-128&14|+|100109715|100109737&offsets|0|+1]|TCTCACACAGAAATCGCACCCGTC</td><td>>MI0000805|hsa-mir-342&WithFlank&14|+|100109649|100109759</td><td>67</td><td>90</td><td>24</td><td>TCTCACACAGAAATCGCACCCGTC</td><td>699.0</td><td>6.5</td><td>0.8</td><td>8.6</td><td>1.1E-17</td><td>4.0E-17</td><td>MB231/MB468</td></tr>
<tr><td>>hg38_21_+_8433172_8446622_RNA45SN1_45S_50ntFlanks@10821.10840.20|GGAAGTCGGCAAGCCGGATC</td><td>>hg38_21_+_8433172_8446622_RNA45SN1_45S_50ntFlanks</td><td>10821</td><td>10840</td><td>20</td><td>GGAAGTCGGCAAGCCGGATC</td><td>2336.2</td><td>-5.5</td><td>0.6</td><td>-8.6</td><td>1.0E-17</td><td>4.1E-17</td><td>BT20/MB468</td></tr>
<tr><td>>hg38_21_+_8433172_8446622_RNA45SN1_45S_50ntFlanks@12449.12476.28|GAATTCACCAAGCGTTGGATTGTTCACC</td><td>>hg38_21_+_8433172_8446622_RNA45SN1_45S_50ntFlanks</td><td>12449</td><td>12476</td><td>28</td><td>GAATTCACCAAGCGTTGGATTGTTCACC</td><td>1129.8</td><td>8.9</td><td>1.0</td><td>8.6</td><td>1.1E-17</td><td>4.1E-17</td><td>MB231/MB468</td></tr>
<tr><td>>MI0000070|hsa-mir-16-1&WithFlank&13|-|50048967|50049067@20.41.22;MI0000115|hsa-mir-16-2&WithFlank&3|+|160404739|160404831@16.37.22&[MIMAT0000069&hsa-miR-16-5p&offsets|0|0;m-62&13|-|50049027|50049048&offsets|0|0];[MIMAT0000069_1&hsa-miR-16-5p&offsets|0|0;m-61&3|+|160404754|160404775&offsets|0|0]|TAGCAGCACGTAAATATTGGCG</td><td>>MI0000070|hsa-mir-16-1&WithFlank&13|-|50048967|50049067</td><td>20</td><td>41</td><td>22</td><td>TAGCAGCACGTAAATATTGGCG</td><td>33941.3</td><td>9.2</td><td>1.1</td><td>8.6</td><td>1.0E-17</td><td>4.1E-17</td><td>BT20/MB468</td></tr>
<tr><td>>am_MI0015825|hsa-mir-378c&WithFlank&10|-|130962582|130962674@17.37.21&[MIMAT0016847&hsa-miR-378c&offsets|0|-4;m-130&10|-|130962639|130962658&offsets|0|+1]|ACTGGACTTGGAGTCAGAAGA</td><td>>am_MI0015825|hsa-mir-378c&WithFlank&10|-|130962582|130962674</td><td>17</td><td>37</td><td>21</td><td>ACTGGACTTGGAGTCAGAAGA</td><td>664.7</td><td>6.2</td><td>0.7</td><td>8.6</td><td>1.1E-17</td><td>4.1E-17</td><td>MB231/MB468</td></tr>
<tr><td>>hg38_21_+_8433172_8446622_RNA45SN1_45S_50ntFlanks@9604.9622.19|AAAGACTAATCGAACCATC</td><td>>hg38_21_+_8433172_8446622_RNA45SN1_45S_50ntFlanks</td><td>9604</td><td>9622</td><td>19</td><td>AAAGACTAATCGAACCATC</td><td>939.3</td><td>-4.1</td><td>0.5</td><td>-8.6</td><td>1.0E-17</td><td>4.1E-17</td><td>BT20/MB468</td></tr>
<tr><td>>hg38_21_+_8433172_8446622_RNA45SN1_45S_50ntFlanks@12654.12674.21|TATGACTGAACGCCTCTAAGT</td><td>>hg38_21_+_8433172_8446622_RNA45SN1_45S_50ntFlanks</td><td>12654</td><td>12674</td><td>21</td><td>TATGACTGAACGCCTCTAAGT</td><td>1044.9</td><td>-6.3</td><td>0.7</td><td>-8.6</td><td>1.0E-17</td><td>4.1E-17</td><td>BT20/MB468</td></tr>
<tr><td>>MI0000811|hsa-mir-148b&WithFlank&12|+|54337210|54337320@69.90.22(+2U)&[MIMAT0000759&hsa-miR-148b-3p&offsets|0|0(+2U);m-77&12|+|54337278|54337299&offsets|0|0(+2U)]|TCAGTGCATCACAGAACTTTGTTT</td><td>>MI0000811|hsa-mir-148b&WithFlank&12|+|54337210|54337320</td><td>69</td><td>90</td><td>22(+2U)</td><td>TCAGTGCATCACAGAACTTTGTTT</td><td>1025.4</td><td>7.0</td><td>0.8</td><td>8.6</td><td>1.0E-17</td><td>4.1E-17</td><td>BT20/MB468</td></tr>
<tr><td>>hg38_21_+_8433172_8446622_RNA45SN1_45S_50ntFlanks@12622.12654.33|AGCCAATGGGGCGAAGCTACCATCTGTGGGATT</td><td>>hg38_21_+_8433172_8446622_RNA45SN1_45S_50ntFlanks</td><td>12622</td><td>12654</td><td>33</td><td>AGCCAATGGGGCGAAGCTACCATCTGTGGGATT</td><td>251.1</td><td>-4.3</td><td>0.5</td><td>-8.6</td><td>1.1E-17</td><td>4.1E-17</td><td>MB231/MB468</td></tr>
<tr><td>>MI0000456|hsa-mir-140&WithFlank&16|+|69933075|69933186@68.88.21&[MIMAT0004597&hsa-miR-140-3p&offsets|0|0;m-45&16|+|69933143|69933164&offsets|-1|-2]|TACCACAGGGTAGAACCACGG</td><td>>MI0000456|hsa-mir-140&WithFlank&16|+|69933075|69933186</td><td>68</td><td>88</td><td>21</td><td>TACCACAGGGTAGAACCACGG</td><td>660.2</td><td>11.2</td><td>1.3</td><td>8.6</td><td>1.0E-17</td><td>4.1E-17</td><td>BT20/MB468</td></tr>
<tr><td>>hg38_MT_+_598_1651_RNR1_12S_50ntFlanks@52.88.37|ATAGGTTTGGTCCTAGCCTTTCTATTAGCTCTTAGTA</td><td>>hg38_MT_+_598_1651_RNR1_12S_50ntFlanks</td><td>52</td><td>88</td><td>37</td><td>ATAGGTTTGGTCCTAGCCTTTCTATTAGCTCTTAGTA</td><td>2894.2</td><td>7.4</td><td>0.9</td><td>8.6</td><td>1.1E-17</td><td>4.2E-17</td><td>BT20/MB468</td></tr>
<tr><td>>MI0000107|hsa-mir-29b-2&WithFlank&1|-|207802437|207802529@58.80.23;MI0000105|hsa-mir-29b-1&WithFlank&7|-|130877453|130877545@57.79.23&[MIMAT0000100&hsa-miR-29b-3p&offsets|0|0;m-81&1|-|207802450|207802472&offsets|0|0];[MIMAT0000100_1&hsa-miR-29b-3p&offsets|0|0;m-82&7|-|130877467|130877489&offsets|0|0]|TAGCACCATTTGAAATCAGTGTT</td><td>>MI0000107|hsa-mir-29b-2&WithFlank&1|-|207802437|207802529</td><td>58</td><td>80</td><td>23</td><td>TAGCACCATTTGAAATCAGTGTT</td><td>6061.5</td><td>10.8</td><td>1.3</td><td>8.6</td><td>1.1E-17</td><td>4.3E-17</td><td>BT20/MB468</td></tr>
<tr><td>>MI0000299|hsa-mir-222&WithFlank&X|-|45747009|45747130@75.96.22(+1A)&[MIMAT0000279&hsa-miR-222-3p&offsets|0|+1(+1A);m-140&X|-|45747033|45747056&offsets|0|-2(+1A)]|AGCTACATCTGGCTACTGGGTCA</td><td>>MI0000299|hsa-mir-222&WithFlank&X|-|45747009|45747130</td><td>75</td><td>96</td><td>22(+1A)</td><td>AGCTACATCTGGCTACTGGGTCA</td><td>1194.5</td><td>7.6</td><td>0.9</td><td>8.6</td><td>1.1E-17</td><td>4.3E-17</td><td>BT20/MB468</td></tr>
<tr><td>>hg38_21_+_8433172_8446622_RNA45SN1_45S_50ntFlanks@12366.12383.18|TAACTGGCTTGTGGCGGC</td><td>>hg38_21_+_8433172_8446622_RNA45SN1_45S_50ntFlanks</td><td>12366</td><td>12383</td><td>18</td><td>TAACTGGCTTGTGGCGGC</td><td>1182.4</td><td>-7.7</td><td>0.9</td><td>-8.6</td><td>1.1E-17</td><td>4.3E-17</td><td>MB231/MB468</td></tr>
<tr><td>>MI0000270|hsa-mir-181b-1&WithFlank&1|-|198858867|198858988@42.62.21;MI0000683|hsa-mir-181b-2&WithFlank&9|+|124693704|124693804@22.42.21&[MIMAT0000257&hsa-miR-181b-5p&offsets|0|-2;m-73&1|-|198858925|198858947&offsets|0|-2];[MIMAT0000257_1&hsa-miR-181b-5p&offsets|0|-2;m-72&9|+|124693725|124693747&offsets|0|-2]|AACATTCATTGCTGTCGGTGG</td><td>>MI0000270|hsa-mir-181b-1&WithFlank&1|-|198858867|198858988</td><td>42</td><td>62</td><td>21</td><td>AACATTCATTGCTGTCGGTGG</td><td>1381.3</td><td>6.9</td><td>0.8</td><td>8.6</td><td>1.1E-17</td><td>4.4E-17</td><td>MB231/MB468</td></tr>
<tr><td>>hg38_21_+_8433172_8446622_RNA45SN1_45S_50ntFlanks@485.532.48|GATGTGGTGACGTCGTGCTCTCCCGGGCCGGGTCCGAGCCGCGACGGG</td><td>>hg38_21_+_8433172_8446622_RNA45SN1_45S_50ntFlanks</td><td>485</td><td>532</td><td>48</td><td>GATGTGGTGACGTCGTGCTCTCCCGGGCCGGGTCCGAGCCGCGACGGG</td><td>36982.9</td><td>30.0</td><td>3.5</td><td>8.6</td><td>1.2E-17</td><td>4.4E-17</td><td>MB231/MB468</td></tr>
<tr><td>>hg38_21_+_8433172_8446622_RNA45SN1_45S_50ntFlanks@6732.6774.43|ACATTGATCATCGACACTTCGAACGCACTTGCGGCCCCGGGTT</td><td>>hg38_21_+_8433172_8446622_RNA45SN1_45S_50ntFlanks</td><td>6732</td><td>6774</td><td>43</td><td>ACATTGATCATCGACACTTCGAACGCACTTGCGGCCCCGGGTT</td><td>1393.4</td><td>10.5</td><td>1.2</td><td>8.6</td><td>1.2E-17</td><td>4.4E-17</td><td>MB231/MB468</td></tr>
</table>

<table>
<tr><th></th><th></th><th></th><th></th><th></th><th></th><th></th><th></th><th></th><th></th><th></th><th></th><th></th></tr>
<tr><td>>MI0000542|hsa-mir-320a&WithFlank&8|-|22244960|22245043@48.69.22(+1G)&[MIMAT0000510&hsa-miR-320a-3p&offsets|0|0(+1G);m-60&8|-|22244975|22244996&offsets|0|0(+1G)]|AAAAGCTGGGTTGAGAGGGCGAG</td><td>>MI0000542|hsa-mir-320a&WithFlank&8|-|22244960|22245043</td><td>48</td><td>69</td><td>22(+1G)</td><td>AAAAGCTGGGTTGAGAGGGCGAG</td><td>2936.7</td><td>11.5</td><td>1.3</td><td>8.6</td><td>1.1E-17</td><td>4.4E-17</td><td>BT20/MB468</td></tr>
<tr><td>>MI0000272|hsa-mir-182&WithFlank&7|-|129770377|129770498@29.51.23(+1A)&[MIMAT0000259&hsa-miR-182-5p&offsets|0|-1(+1A);m-22&7|-|129770449|129770470&offsets|0|+1(+1A)]|TTTGGCAATGGTAGAACTCACACA</td><td>>MI0000272|hsa-mir-182&WithFlank&7|-|129770377|129770498</td><td>29</td><td>51</td><td>23(+1A)</td><td>TTTGGCAATGGTAGAACTCACACA</td><td>1302.3</td><td>6.2</td><td>0.7</td><td>8.6</td><td>1.2E-17</td><td>4.5E-17</td><td>MB231/MB468</td></tr>
<tr><td>>hg38_21_+_8433172_8446622_RNA45SN1_45S_50ntFlanks@9716.9737.22|AATGATTAGAGGTCTTGGGGCC</td><td>>hg38_21_+_8433172_8446622_RNA45SN1_45S_50ntFlanks</td><td>9716</td><td>9737</td><td>22</td><td>AATGATTAGAGGTCTTGGGGCC</td><td>259.3</td><td>-3.1</td><td>0.4</td><td>-8.6</td><td>1.1E-17</td><td>4.5E-17</td><td>BT20/MB468</td></tr>
<tr><td>>hg38_21_+_8433172_8446622_RNA45SN1_45S_50ntFlanks@8176.8197.22|CTTCTGATCGAGGCCCAGCCCG</td><td>>hg38_21_+_8433172_8446622_RNA45SN1_45S_50ntFlanks</td><td>8176</td><td>8197</td><td>22</td><td>CTTCTGATCGAGGCCCAGCCCG</td><td>510.0</td><td>-3.9</td><td>0.5</td><td>-8.6</td><td>1.1E-17</td><td>4.5E-17</td><td>BT20/MB468</td></tr>
<tr><td>>MI0000477|hsa-mir-146a&WithFlank&5|+|160485346|160485456@27.48.22&[MIMAT0000449&hsa-miR-146a-5p&offsets|0|0;m-129&5|+|160485372|160485393&offsets|0|0]|TGAGAACTGAATTCCATGGGTT</td><td>>MI0000477|hsa-mir-146a&WithFlank&5|+|160485346|160485456</td><td>27</td><td>48</td><td>22</td><td>TGAGAACTGAATTCCATGGGTT</td><td>15401.6</td><td>-8.4</td><td>0.9</td><td>-8.9</td><td>6.4E-19</td><td>4.6E-17</td><td>BT20/MB231</td></tr>
<tr><td>>MI0000650|hsa-mir-200c&WithFlank&12|+|6963693|6963772@50.71.22(+1C)&[MIMAT0000617&hsa-miR-200c-3p&offsets|0|-1(+1C);m-54&12|+|6963742|6963764&offsets|0|-1(+1C)]|TAATACTGCCGGGTAATGATGGC</td><td>>MI0000650|hsa-mir-200c&WithFlank&12|+|6963693|6963772</td><td>50</td><td>71</td><td>22(+1C)</td><td>TAATACTGCCGGGTAATGATGGC</td><td>746.6</td><td>13.0</td><td>1.5</td><td>8.9</td><td>6.2E-19</td><td>4.6E-17</td><td>BT20/MB231</td></tr>
<tr><td>>MI0000286|hsa-mir-210&WithFlank&11|-|568083|568204@72.93.22(+3A)&[MIMAT0000267&hsa-miR-210-3p&offsets|0|0(+3A);m-102&11|-|568112|568133&offsets|0|0(+3A)]|CTGTGCGTGTGACAGCGGCTGAAAA</td><td>>MI0000286|hsa-mir-210&WithFlank&11|-|568083|568204</td><td>72</td><td>93</td><td>22(+3A)</td><td>CTGTGCGTGTGACAGCGGCTGAAAA</td><td>749.1</td><td>11.7</td><td>1.4</td><td>8.6</td><td>1.1E-17</td><td>4.6E-17</td><td>BT20/MB468</td></tr>
<tr><td>>MI0000440|hsa-mir-27b&WithFlank&9|+|95085439|95085547@67.88.22(+1U)&[MIMAT0000419&hsa-miR-27b-3p&offsets|0|+1(+1U);m-39&9|+|95085505|95085525&offsets|0|+1(+1U)]|TTCACAGTGGCTAAGTTCTGCAT</td><td>>MI0000440|hsa-mir-27b&WithFlank&9|+|95085439|95085547</td><td>67</td><td>88</td><td>22(+1U)</td><td>TTCACAGTGGCTAAGTTCTGCAT</td><td>1173.2</td><td>7.2</td><td>0.8</td><td>8.6</td><td>1.1E-17</td><td>4.6E-17</td><td>BT20/MB468</td></tr>
<tr><td>>MI0000067|hsa-let-7f-1&WithFlank&9|+|94176341|94176439@13.34.22(+1C);MI0000068|hsa-let-7f-2&WithFlank&X|-|53557186|53557280@14.35.22(+1C)&[MIMAT0000067&hsa-let-7f-5p&offsets|0|0(+1C);m-20&9|+|94176353|94176374&offsets|0|0(+1C)];[MIMAT0000067_1&hsa-let-7f-5p&offsets|0|0(+1C);m-19&X|-|53557246|53557267&offsets|0|0(+1C)]|TGAGGTAGTAGATTGTATAGTTC</td><td>>MI0000067|hsa-let-7f-1&WithFlank&9|+|94176341|94176439</td><td>13</td><td>34</td><td>22(+1C)</td><td>TGAGGTAGTAGATTGTATAGTTC</td><td>620.2</td><td>6.1</td><td>0.7</td><td>8.6</td><td>1.2E-17</td><td>4.6E-17</td><td>BT20/MB468</td></tr>
<tr><td>>am_trnaMT_TyrGTA_MT_-_5826_5891@1.31.31|GGTAAAATGGCTGAGTGAAGCATTGGACTGT</td><td>>am_trnaMT_TyrGTA_MT_-_5826_5891</td><td>1</td><td>31</td><td>31</td><td>GGTAAAATGGCTGAGTGAAGCATTGGACTGT</td><td>587.6</td><td>10.0</td><td>1.2</td><td>8.5</td><td>1.2E-17</td><td>4.7E-17</td><td>MB231/MB468</td></tr>
<tr><td>>trnaMT_GluTTC_MT_-_14674_14742@39.58.20,trnalookalike8_GluTTC_5_-_93905172_93905240@39.58.20|ATTGGTCGTGGTTGTAGTCC</td><td>>trnaMT_GluTTC_MT_-_14674_14742</td><td>39</td><td>58</td><td>20</td><td>ATTGGTCGTGGTTGTAGTCC</td><td>1387.2</td><td>10.2</td><td>1.2</td><td>8.5</td><td>1.3E-17</td><td>4.8E-17</td><td>MB231/MB468</td></tr>
<tr><td>>MI0000471|hsa-mir-126&WithFlank&9|+|136670596|136670692@57.79.23&[MIMAT0000445&hsa-miR-126-3p&offsets|-1|0;m-40&9|+|136670653|136670674&offsets|-1|0]|CTCGTACCGTGAGTAATAATGCG</td><td>>MI0000471|hsa-mir-126&WithFlank&9|+|136670596|136670692</td><td>57</td><td>79</td><td>23</td><td>CTCGTACCGTGAGTAATAATGCG</td><td>1537.1</td><td>10.5</td><td>1.2</td><td>8.5</td><td>1.3E-17</td><td>4.8E-17</td><td>MB231/MB468</td></tr>
<tr><td>>trnaMT_ValTAC_MT_+_1602_1670@55.72.18|ACTTGACCGCTCTGACCA</td><td>>trnaMT_ValTAC_MT_+_1602_1670</td><td>55</td><td>72</td><td>18</td><td>ACTTGACCGCTCTGACCA</td><td>4727.2</td><td>6.3</td><td>0.7</td><td>8.6</td><td>1.2E-17</td><td>4.8E-17</td><td>BT20/MB468</td></tr>
<tr><td>>am_trnaMT_LysTTT_MT_+_8295_8364@34.52.19,trnalookalike7_LysTTT_1_+_568844_568913@34.52.19|GTTAAAGATTAAGAGAACC</td><td>>am_trnaMT_LysTTT_MT_+_8295_8364</td><td>34</td><td>52</td><td>19</td><td>GTTAAAGATTAAGAGAACC</td><td>853.7</td><td>9.7</td><td>1.1</td><td>8.6</td><td>1.2E-17</td><td>4.9E-17</td><td>BT20/MB468</td></tr>
<tr><td>>trna16_TyrGTA_14_-_21131351_21131444@59.76.18,trna17_TyrGTA_14_-_21128117_21128210@59.76.18,trna19_TyrGTA_14_-_21121258_21121351@59.76.18,trna2_TyrGTA_2_+_27273650_27273738@59.76.18,trna4_TyrGTA_8_+_67025602_67025694@59.76.18,trna5_TyrGTA_14_+_21151432_21151520@59.76.18,trna5_TyrGTA_8_+_67026223_67026311@59.76.18|TTCCGGCTCGAAGGACCA</td><td>>trna16_TyrGTA_14_-_21131351_21131444</td><td>59</td><td>76</td><td>18</td><td>TTCCGGCTCGAAGGACCA</td><td>2205.6</td><td>6.7</td><td>0.8</td><td>8.5</td><td>1.3E-17</td><td>4.9E-17</td><td>MB231/MB468</td></tr>
<tr><td>>hg38_21_+_8433172_8446622_RNA45SN1_45S_50ntFlanks@12627.12667.41|ATGGGGCGAAGCTACCATCTGTGGGATTATGACTGAACGCC</td><td>>hg38_21_+_8433172_8446622_RNA45SN1_45S_50ntFlanks</td><td>12627</td><td>12667</td><td>41</td><td>ATGGGGCGAAGCTACCATCTGTGGGATTATGACTGAACGCC</td><td>733.1</td><td>8.4</td><td>1.0</td><td>8.5</td><td>1.3E-17</td><td>4.9E-17</td><td>MB231/MB468</td></tr>
<tr><td>>trnaMT_ThrTGT_MT_+_15888_15953@37.69.33|ACCGGAGATGAAAACCTTTTTCCAAGGACACCA</td><td>>trnaMT_ThrTGT_MT_+_15888_15953</td><td>37</td><td>69</td><td>33</td><td>ACCGGAGATGAAAACCTTTTTCCAAGGACACCA</td><td>612.6</td><td>9.4</td><td>1.1</td><td>8.5</td><td>1.2E-17</td><td>4.9E-17</td><td>BT20/MB468</td></tr>
<tr><td>>hg38_21_+_8433172_8446622_RNA45SN1_45S_50ntFlanks@11561.11586.26|GCGCCTAGCAGCCGACTTAGAACTGG</td><td>>hg38_21_+_8433172_8446622_RNA45SN1_45S_50ntFlanks</td><td>11561</td><td>11586</td><td>26</td><td>GCGCCTAGCAGCCGACTTAGAACTGG</td><td>520.1</td><td>-5.5</td><td>0.6</td><td>-8.5</td><td>1.3E-17</td><td>5.0E-17</td><td>BT20/MB468</td></tr>
<tr><td>>hg38_21_+_8433172_8446622_RNA45SN1_45S_50ntFlanks@9307.9330.24|AAGGAGTCTAACACGTGCGCGAGT</td><td>>hg38_21_+_8433172_8446622_RNA45SN1_45S_50ntFlanks</td><td>9307</td><td>9330</td><td>24</td><td>AAGGAGTCTAACACGTGCGCGAGT</td><td>873.6</td><td>7.8</td><td>0.9</td><td>8.5</td><td>1.3E-17</td><td>5.1E-17</td><td>MB231/MB468</td></tr>
<tr><td>>MI0000811|hsa-mir-148b&WithFlank&12|+|54337210|54337320@69.89.21(+1G)&[MIMAT0000759&hsa-miR-148b-3p&offsets|0|-1(+1G);m-77&12|+|54337278|54337299&offsets|0|-1(+1G)]|TCAGTGCATCACAGAACTTTGG</td><td>>MI0000811|hsa-mir-148b&WithFlank&12|+|54337210|54337320</td><td>69</td><td>89</td><td>21(+1G)</td><td>TCAGTGCATCACAGAACTTTGG</td><td>3083.0</td><td>9.0</td><td>1.1</td><td>8.5</td><td>1.3E-17</td><td>5.1E-17</td><td>BT20/MB468</td></tr>
<tr><td>>hg38_21_+_8433172_8446622_RNA45SN1_45S_50ntFlanks@7974.8016.43|ACGCGACCTCAGATCAGACGTGGCGACCCGCTGAATTTAAGCA</td><td>>hg38_21_+_8433172_8446622_RNA45SN1_45S_50ntFlanks</td><td>7974</td><td>8016</td><td>43</td><td>ACGCGACCTCAGATCAGACGTGGCGACCCGCTGAATTTAAGCA</td><td>2864.7</td><td>8.9</td><td>1.0</td><td>8.5</td><td>1.4E-17</td><td>5.1E-17</td><td>MB231/MB468</td></tr>
<tr><td>>hg38_21_+_8433172_8446622_RNA45SN1_45S_50ntFlanks@12415.12438.24|CTTCGATGTCGGCTCTTCCTATCA</td><td>>hg38_21_+_8433172_8446622_RNA45SN1_45S_50ntFlanks</td><td>12415</td><td>12438</td><td>24</td><td>CTTCGATGTCGGCTCTTCCTATCA</td><td>667.9</td><td>4.2</td><td>0.5</td><td>8.5</td><td>1.4E-17</td><td>5.2E-17</td><td>MB231/MB468</td></tr>
<tr><td>>MI0000298|hsa-mir-221&WithFlank&X|-|45746151|45746272@71.92.22(+1U)&[MIMAT0000278&hsa-miR-221-3p&offsets|0|-1(+1U);m-91&X|-|45746181|45746202&offsets|0|0(+1U)]|AGCTACATTGTCTGCTGGGTTTT</td><td>>MI0000298|hsa-mir-221&WithFlank&X|-|45746151|45746272</td><td>71</td><td>92</td><td>22(+1U)</td><td>AGCTACATTGTCTGCTGGGTTTT</td><td>2581.3</td><td>5.9</td><td>0.7</td><td>8.5</td><td>1.3E-17</td><td>5.2E-17</td><td>BT20/MB468</td></tr>
<tr><td>>MI0000483|hsa-mir-186&WithFlank&1|-|71067625|71067722@21.41.21(+1C)&[MIMAT0000456&hsa-miR-186-5p&offsets|0|-1(+1C);m-84&1|-|71067681|71067702&offsets|0|-1(+1C)]|CAAAGAATTCTCCTTTTGGGCC</td><td>>MI0000483|hsa-mir-186&WithFlank&1|-|71067625|71067722</td><td>21</td><td>41</td><td>21(+1C)</td><td>CAAAGAATTCTCCTTTTGGGCC</td><td>803.4</td><td>7.0</td><td>0.8</td><td>8.5</td><td>1.3E-17</td><td>5.2E-17</td><td>BT20/MB468</td></tr>
<tr><td>>hg38_21_+_8433172_8446622_RNA45SN1_45S_50ntFlanks@12567.12587.21|CTCAGTACGAGAGGAACCGCA</td><td>>hg38_21_+_8433172_8446622_RNA45SN1_45S_50ntFlanks</td><td>12567</td><td>12587</td><td>21</td><td>CTCAGTACGAGAGGAACCGCA</td><td>481.9</td><td>-10.9</td><td>1.3</td><td>-8.5</td><td>1.3E-17</td><td>5.3E-17</td><td>BT20/MB468</td></tr>
<tr><td>>hg38_21_+_8433172_8446622_RNA45SN1_45S_50ntFlanks@7980.8006.27|CCTCAGATCAGACGTGGCGACCCGCTG</td><td>>hg38_21_+_8433172_8446622_RNA45SN1_45S_50ntFlanks</td><td>7980</td><td>8006</td><td>27</td><td>CCTCAGATCAGACGTGGCGACCCGCTG</td><td>3722.6</td><td>-7.4</td><td>0.9</td><td>-8.5</td><td>1.3E-17</td><td>5.3E-17</td><td>BT20/MB468</td></tr>
<tr><td>>MI0000440|hsa-mir-27b&WithFlank&9|+|95085439|95085547@67.87.21(+1G)&[MIMAT0000419&hsa-miR-27b-3p&offsets|0|0(+1G);m-39&9|+|95085505|95085525&offsets|0|0(+1G)]|TTCACAGTGGCTAAGTTCTGCG</td><td>>MI0000440|hsa-mir-27b&WithFlank&9|+|95085439|95085547</td><td>67</td><td>87</td><td>21(+1G)</td><td>TTCACAGTGGCTAAGTTCTGCG</td><td>3297.0</td><td>8.0</td><td>0.9</td><td>8.5</td><td>1.3E-17</td><td>5.3E-17</td><td>BT20/MB468</td></tr>
<tr><td>>am_trna10_LysCTT_16_+_3241501_3241573@31.51.21,trna119_LysCTT_1_-_145395522_145395594@31.51.21,trna11_LysCTT_5_-_180648979_180649051@31.51.21,trna13_LysCTT_16_+_3246153_3246225@31.51.21,trna13_LysCTT_6_+_26556774_26556846@31.51.21,trna30_LysCTT_16_-_3230555_3230627@31.51.21,trna5_LysCTT_16_+_3214939_3215011@31.51.21,trna7_LysCTT_16_+_3225692_3225764@31.51.21,trna8_LysCTT_1_+_39970195_39970267@31.51.21,trna9_LysCTT_5_+_180634755_180634827@31.51.21|ACTCTTAATCTCAGGGTCGTG</td><td>>am_trna10_LysCTT_16_+_3241501_3241573</td><td>31</td><td>51</td><td>21</td><td>ACTCTTAATCTCAGGGTCGTG</td><td>597.4</td><td>-11.2</td><td>1.3</td><td>-8.5</td><td>1.3E-17</td><td>5.3E-17</td><td>BT20/MB468</td></tr>
<tr><td>>hg38_21_+_8433172_8446622_RNA45SN1_45S_50ntFlanks@6646.6669.24|TCGTACGACTCTTAGCGGTGGATC</td><td>>hg38_21_+_8433172_8446622_RNA45SN1_45S_50ntFlanks</td><td>6646</td><td>6669</td><td>24</td><td>TCGTACGACTCTTAGCGGTGGATC</td><td>37278.7</td><td>4.1</td><td>0.5</td><td>8.5</td><td>1.3E-17</td><td>5.3E-17</td><td>BT20/MB468</td></tr>
<tr><td>>MI0000438|hsa-mir-15b&WithFlank&3|+|160404582|160404691@26.47.22&[MIMAT0000417&hsa-miR-15b-5p&offsets|0|0;m-110&3|+|160404607|160404628&offsets|0|0]|TAGCAGCACATCATGGTTTACA</td><td>>MI0000438|hsa-mir-15b&WithFlank&3|+|160404582|160404691</td><td>26</td><td>47</td><td>22</td><td>TAGCAGCACATCATGGTTTACA</td><td>8626.5</td><td>7.7</td><td>0.9</td><td>8.5</td><td>1.4E-17</td><td>5.3E-17</td><td>BT20/MB468</td></tr>
<tr><td>>MI0001445|hsa-mir-423&WithFlank&17|+|30117073|30117178@23.44.22(+1C)&[MIMAT0004748&hsa-miR-423-5p&offsets|0|-1(+1C);m-141&17|+|30117095|30117117&offsets|0|-1(+1C)]|TGAGGGGCAGAGAGCGAGACTTC</td><td>>MI0001445|hsa-mir-423&WithFlank&17|+|30117073|30117178</td><td>23</td><td>44</td><td>22(+1C)</td><td>TGAGGGGCAGAGAGCGAGACTTC</td><td>951.7</td><td>8.7</td><td>1.0</td><td>8.5</td><td>1.4E-17</td><td>5.3E-17</td><td>BT20/MB468</td></tr>
<tr><td>>hg38_21_+_8433172_8446622_RNA45SN1_45S_50ntFlanks@6777.6806.30|TCCCGGGGCTACGCCTGTCTGAGCGTCGCT</td><td>>hg38_21_+_8433172_8446622_RNA45SN1_45S_50ntFlanks</td><td>6777</td><td>6806</td><td>30</td><td>TCCCGGGGCTACGCCTGTCTGAGCGTCGCT</td><td>643.6</td><td>5.0</td><td>0.6</td><td>8.5</td><td>1.4E-17</td><td>5.4E-17</td><td>BT20/MB468</td></tr>
<tr><td>>MI0000456|hsa-mir-140&WithFlank&16|+|69933075|69933186@68.90.23&[MIMAT0004597&hsa-miR-140-3p&offsets|0|+2;m-45&16|+|69933143|69933164&offsets|-1|0]|TACCACAGGGTAGAACCACGGAC</td><td>>MI0000456|hsa-mir-140&WithFlank&16|+|69933075|69933186</td><td>68</td><td>90</td><td>23</td><td>TACCACAGGGTAGAACCACGGAC</td><td>2149.0</td><td>10.9</td><td>1.3</td><td>8.5</td><td>1.4E-17</td><td>5.4E-17</td><td>MB231/MB468</td></tr>
<tr><td>>hg38_21_+_8433172_8446622_RNA45SN1_45S_50ntFlanks@12157.12180.24|AGTTTGACTGGGGCGGTACACCTG</td><td>>hg38_21_+_8433172_8446622_RNA45SN1_45S_50ntFlanks</td><td>12157</td><td>12180</td><td>24</td><td>AGTTTGACTGGGGCGGTACACCTG</td><td>857.2</td><td>-11.3</td><td>1.3</td><td>-8.5</td><td>1.4E-17</td><td>5.4E-17</td><td>MB231/MB468</td></tr>
<tr><td>>hg38_21_+_8433172_8446622_RNA45SN1_45S_50ntFlanks@10807.10826.20|ACAATGTAGGTAAGGGAAGT</td><td>>hg38_21_+_8433172_8446622_RNA45SN1_45S_50ntFlanks</td><td>10807</td><td>10826</td><td>20</td><td>ACAATGTAGGTAAGGGAAGT</td><td>205.1</td><td>-5.3</td><td>0.6</td><td>-8.5</td><td>1.4E-17</td><td>5.4E-17</td><td>BT20/MB468</td></tr>
</table>

| | | | | | | | | | | | | |
|---|---|---|---|---|---|---|---|---|---|---|---|---|
| >MI0000447\|hsa-mir-128-1&WithFlank&2\|+\|135665391\|135665484@56.76.21;MI0000727\|hsa-mir-128-2&WithFlank&3\|+\|35744470\|35744565@58.78.21&[MIMAT0000424&hsa-miR-128-3p&offsets\|0\|0;m-108&2\|+\|135665446\|135665466&offsets\|0\|0];[MIMAT0000424_1&hsa-miR-128-3p&offsets\|0\|0;m-117&3\|+\|35744527\|35744547&offsets\|0\|0]\|TCACAGTGAACCGGTCTCTTT | >MI0000447\|hsa-mir-128-1&WithFlank&2\|+\|135665391\|135665484 | 56 | 76 | 21 | TCACAGTGAACCGGTCTCTTT | 26171.1 | 5.7 | 0.7 | 8.5 | 1.4E-17 | 5.4E-17 | BT20/MB468 |
| >MI0001448\|hsa-mir-425&WithFlank&3\|-\|49020142\|49020240@21.44.24&[MIMAT0003393&hsa-miR-425-5p&offsets\|+1\|+2;m-95&3\|-\|49020200\|49020221&offsets\|+1\|+3]\|ATGACACGATCACTCCCGTTGAGT | >MI0001448\|hsa-mir-425&WithFlank&3\|-\|49020142\|49020240 | 21 | 44 | 24 | ATGACACGATCACTCCCGTTGAGT | 1403.3 | 7.9 | 0.9 | 8.5 | 1.4E-17 | 5.5E-17 | BT20/MB468 |
| >hg38_21_+_8433172_8446622_RNA45SN1_45S_50ntFlanks@12366.12383.18\|TAACTGGCTTGTGGCGGC | >hg38_21_+_8433172_8446622_RNA45SN1_45S_50ntFlanks | 12366 | 12383 | 18 | TAACTGGCTTGTGGCGGC | 1444.1 | -8.0 | 0.9 | -8.5 | 1.4E-17 | 5.6E-17 | BT20/MB468 |
| >hg38_21_+_8433172_8446622_RNA45SN1_45S_50ntFlanks@12435.12455.21\|ATCATTGTGAAGCAGAATTCA | >hg38_21_+_8433172_8446622_RNA45SN1_45S_50ntFlanks | 12435 | 12455 | 21 | ATCATTGTGAAGCAGAATTCA | 1617.5 | -8.4 | 1.0 | -8.5 | 1.5E-17 | 5.6E-17 | MB231/MB468 |
| >MI0001445\|hsa-mir-423&WithFlank&17\|+\|30117073\|30117178@23.44.22&[MIMAT0004748&hsa-miR-423-5p&offsets\|0\|-1;m-141&17\|+\|30117095\|30117117&offsets\|0\|-1]\|TGAGGGGCAGAGAGCGAGACTT | >MI0001445\|hsa-mir-423&WithFlank&17\|+\|30117073\|30117178 | 23 | 44 | 22 | TGAGGGGCAGAGAGCGAGACTT | 928.2 | 8.6 | 1.0 | 8.5 | 1.4E-17 | 5.6E-17 | BT20/MB468 |
| >hg38_MT_+_1621_3279_RNR2_16S_50ntFlanks@644.663.20\|AATTGGACCAATCTATCACC | >hg38_MT_+_1621_3279_RNR2_16S_50ntFlanks | 644 | 663 | 20 | AATTGGACCAATCTATCACC | 376.1 | 9.8 | 1.2 | 8.5 | 1.4E-17 | 5.6E-17 | BT20/MB468 |
| >MI0000103\|hsa-mir-101-1&WithFlank&1\|-\|65058428\|65058514@52.73.22;MI0000739\|hsa-mir-101-2&WithFlank&9\|+\|4850291\|4850381@54.75.22&[MIMAT0000099&hsa-miR-101-3p&offsets\|-1\|0;m-17&1\|-\|65058443\|65058462&offsets\|-1\|+1];[MIMAT0000099_1&hsa-miR-101-3p&offsets\|-1\|0;m-16&9\|+\|4850345\|4850364&offsets\|-1\|+1]\|GTACAGTACTGTGATAACTGAA | >MI0000103\|hsa-mir-101-1&WithFlank&1\|-\|65058428\|65058514 | 52 | 73 | 22 | GTACAGTACTGTGATAACTGAA | 5884.6 | 6.5 | 0.8 | 8.5 | 1.5E-17 | 5.7E-17 | MB231/MB468 |
| >hg38_21_+_8433172_8446622_RNA45SN1_45S_50ntFlanks@11874.11903.30\|AGAAGACCCTGTTGAGCTTGACTCTAGTCT | >hg38_21_+_8433172_8446622_RNA45SN1_45S_50ntFlanks | 11874 | 11903 | 30 | AGAAGACCCTGTTGAGCTTGACTCTAGTCT | 219.7 | -4.7 | 0.6 | -8.5 | 1.5E-17 | 5.8E-17 | MB231/MB468 |
| >MI0000650\|hsa-mir-200c&WithFlank&12\|+\|6963693\|6963772@49.71.23&[MIMAT0000617&hsa-miR-200c-3p&offsets\|-1\|-1;m-54&12\|+\|6963742\|6963764&offsets\|-1\|-1]\|CTAATACTGCCGGGTAATGATGG | >MI0000650\|hsa-mir-200c&WithFlank&12\|+\|6963693\|6963772 | 49 | 71 | 23 | CTAATACTGCCGGGTAATGATGG | 525.5 | 12.5 | 1.4 | 8.9 | 8.4E-19 | 5.9E-17 | BT20/MB231 |
| >MI0000108\|hsa-mir-103a-2&WithFlank&20\|+\|3917488\|3917577@56.76.21;MI0000109\|hsa-mir-103a-1&WithFlank&5\|-\|168560890\|168560979@56.76.21&[MIMAT0000101&hsa-miR-103a-3p&offsets\|+2\|0;m-12&20\|+\|3917541\|3917563&offsets\|+2\|0];[MIMAT0000101_1&hsa-miR-103a-3p&offsets\|+2\|0;m-13&5\|-\|168560904\|168560926&offsets\|+2\|0;m-44220&5\|-\|168560908\|168560926&offsets\|+2\|+4]\|CAGCATTGTACAGGGCTATGA | >MI0000108\|hsa-mir-103a-2&WithFlank&20\|+\|3917488\|3917577 | 56 | 76 | 21 | CAGCATTGTACAGGGCTATGA | 1042.5 | 6.8 | 0.8 | 8.5 | 1.6E-17 | 6.0E-17 | MB231/MB468 |
| >MI0000299\|hsa-mir-222&WithFlank&X\|-\|45747009\|45747130@75.98.24&[MIMAT0000279&hsa-miR-222-3p&offsets\|0\|+3;m-140&X\|-\|45747033\|45747056&offsets\|0\|0]\|AGCTACATCTGGCTACTGGGTCTC | >MI0000299\|hsa-mir-222&WithFlank&X\|-\|45747009\|45747130 | 75 | 98 | 24 | AGCTACATCTGGCTACTGGGTCTC | 9663.1 | 6.2 | 0.7 | 8.5 | 1.5E-17 | 6.0E-17 | BT20/MB468 |
| >hg38_21_+_8433172_8446622_RNA45SN1_45S_50ntFlanks@11769.11786.18\|TAATTAGTGACGCGCATG | >hg38_21_+_8433172_8446622_RNA45SN1_45S_50ntFlanks | 11769 | 11786 | 18 | TAATTAGTGACGCGCATG | 355.4 | -5.8 | 0.7 | -8.5 | 1.6E-17 | 6.2E-17 | BT20/MB468 |
| >MI0000272\|hsa-mir-182&WithFlank&7\|-\|129770377\|129770498@29.53.25&[MIMAT0000259&hsa-miR-182-5p&offsets\|0\|+1;m-22&7\|-\|129770449\|129770470&offsets\|0\|+3]\|TTTGGCAATGGTAGAACTCACACTG | >MI0000272\|hsa-mir-182&WithFlank&7\|-\|129770377\|129770498 | 29 | 53 | 25 | TTTGGCAATGGTAGAACTCACACTG | 6760.1 | 6.3 | 0.7 | 8.5 | 1.7E-17 | 6.2E-17 | MB231/MB468 |
| >hg38_21_+_8433172_8446622_RNA45SN1_45S_50ntFlanks@12405.12424.20\|CTTTTTGATCCTTCGATGTC | >hg38_21_+_8433172_8446622_RNA45SN1_45S_50ntFlanks | 12405 | 12424 | 20 | CTTTTTGATCCTTCGATGTC | 1135.7 | -7.1 | 0.8 | -8.5 | 1.7E-17 | 6.2E-17 | MB231/MB468 |
| >hg38_21_+_8433172_8446622_RNA45SN1_45S_50ntFlanks@12419.12452.34\|GATGTCGGCTCTTCCTATCATTGTGAAGCAGAAT | >hg38_21_+_8433172_8446622_RNA45SN1_45S_50ntFlanks | 12419 | 12452 | 34 | GATGTCGGCTCTTCCTATCATTGTGAAGCAGAAT | 2180.1 | 9.0 | 1.1 | 8.5 | 1.7E-17 | 6.2E-17 | MB231/MB468 |
| >am_trnaMT_MetCAT_MT_+_4402_4469@1.21.21\|AGTAAGGTCAGCTAAATAAGC | >am_trnaMT_MetCAT_MT_+_4402_4469 | 1 | 21 | 21 | AGTAAGGTCAGCTAAATAAGC | 1201.5 | 6.7 | 0.8 | 8.5 | 1.6E-17 | 6.4E-17 | BT20/MB468 |
| >trna116_GluCTC_1_-_145399233_145399304@32.70.39,trna71_GluCTC_1_-_161439189_161439260@32.70.39,trna74_GluCTC_1_-_161431809_161431880@32.70.39,trna77_GluCTC_1_-_161424398_161424469@32.70.39,trna77_GluCTC_6_+_28949976_28950047@32.70.39,trna80_GluCTC_1_-_161417018_161417089@32.70.39,trna87_GluCTC_6_-_126101393_126101464@32.70.39\|CTCTCACCGCCGCGGCCCGGGTTCGATTCCCGGTCAGGG | >trna116_GluCTC_1_-_145399233_145399304 | 32 | 70 | 39 | CTCTCACCGCCGCGGCCCGGGTTCGATTCCCGGTCAGGG | 869.2 | -11.3 | 1.3 | -8.5 | 1.7E-17 | 6.4E-17 | MB231/MB468 |
| >MI0000457\|hsa-mir-141&WithFlank&12\|+\|6964091\|6964197@23.44.22&[MIMAT0004598&hsa-miR-141-5p&offsets\|0\|0;m-221&12\|+\|6964113\|6964133&offsets\|0\|+1]\|CATCTTCCAGTACAGTGTTGGA | >MI0000457\|hsa-mir-141&WithFlank&12\|+\|6964091\|6964197 | 23 | 44 | 22 | CATCTTCCAGTACAGTGTTGGA | 1403.4 | 8.3 | 1.0 | 8.5 | 1.6E-17 | 6.4E-17 | BT20/MB468 |
| >MI0005756\|hsa-mir-934&WithFlank&X\|+\|136550872\|136550966@21.40.20(+1A)&[MIMAT0004977&hsa-miR-934&offsets\|0\|-2(+1A);m-943&X\|+\|136550892\|136550913&offsets\|0\|-2(+1A)]\|TGTCTACTACTGGAGACACTA | >MI0005756\|hsa-mir-934&WithFlank&X\|+\|136550872\|136550966 | 21 | 40 | 20(+1A) | TGTCTACTACTGGAGACACTA | 1227.2 | 13.8 | 1.6 | 8.8 | 9.6E-19 | 6.6E-17 | BT20/MB231 |
| >MI0000234\|hsa-mir-192&WithFlank&11\|-\|64891131\|64891252@30.50.21&[MIMAT0000222&hsa-miR-192-5p&offsets\|0\|0;m-59&11\|-\|64891203\|64891223&offsets\|0\|0]\|CTGACCTATGAATTGACAGCC | >MI0000234\|hsa-mir-192&WithFlank&11\|-\|64891131\|64891252 | 30 | 50 | 21 | CTGACCTATGAATTGACAGCC | 3800.4 | 6.7 | 0.8 | 8.5 | 1.8E-17 | 6.6E-17 | MB231/MB468 |
| >MI0000279\|hsa-mir-196a-2&WithFlank&12\|+\|53991732\|53991853@31.51.21(+1A);MI0000238\|hsa-mir-196a-1&WithFlank&17\|-\|48632484\|48632565@13.33.21(+1A)&[MIMAT0000226&hsa-miR-196a-5p&offsets\|0\|-1(+1A);m-261&12\|+\|53991762\|53991783&offsets\|0\|-1(+1A)];[MIMAT0000226_1&hsa-miR-196a-5p&offsets\|0\|-1(+1A);m-268&17\|-\|48632532\|48632553&offsets\|0\|-1(+1A)]\|TAGGTAGTTTCATGTTGTTGGA | >MI0000279\|hsa-mir-196a-2&WithFlank&12\|+\|53991732\|53991853 | 31 | 51 | 21(+1A) | TAGGTAGTTTCATGTTGTTGGA | 779.2 | 13.1 | 1.5 | 8.8 | 1.0E-18 | 6.7E-17 | BT20/MB231 |
| >hg38_21_+_8433172_8446622_RNA45SN1_45S_50ntFlanks@6748.6770.23\|CTTCGAACGCACTTGCGGCCCCG | >hg38_21_+_8433172_8446622_RNA45SN1_45S_50ntFlanks | 6748 | 6770 | 23 | CTTCGAACGCACTTGCGGCCCCG | 233.9 | -4.5 | 0.5 | -8.5 | 1.8E-17 | 6.8E-17 | MB231/MB468 |
| >MI0000748\|hsa-mir-130b&WithFlank&22\|+\|21653298\|21653391@19.40.22&[MIMAT0004680&hsa-miR-130b-5p&offsets\|0\|+1;m-234&22\|+\|21653316\|21653337&offsets\|0\|0]\|ACTCTTTCCCTGTTGCACTACT | >MI0000748\|hsa-mir-130b&WithFlank&22\|+\|21653298\|21653391 | 19 | 40 | 22 | ACTCTTTCCCTGTTGCACTACT | 1265.1 | 7.1 | 0.8 | 8.5 | 1.7E-17 | 6.8E-17 | BT20/MB468 |
| >MI0000065\|hsa-let-7d&WithFlank&9\|+\|94178828\|94178926@14.36.23&[MIMAT0000065&hsa-let-7d-5p&offsets\|0\|+1;m-90&9\|+\|94178841\|94178862&offsets\|0\|+1]\|AGAGGTAGTAGGTTGCATAGTTT | >MI0000065\|hsa-let-7d&WithFlank&9\|+\|94178828\|94178926 | 14 | 36 | 23 | AGAGGTAGTAGGTTGCATAGTTT | 840.1 | 7.4 | 0.9 | 8.5 | 1.7E-17 | 6.8E-17 | BT20/MB468 |
| >hg38_21_+_8433172_8446622_RNA45SN1_45S_50ntFlanks@4763.4780.18\|GATACCGTCGTAGTTCCG | >hg38_21_+_8433172_8446622_RNA45SN1_45S_50ntFlanks | 4763 | 4780 | 18 | GATACCGTCGTAGTTCCG | 756.3 | -6.5 | 0.8 | -8.5 | 1.8E-17 | 6.9E-17 | BT20/MB468 |
| >MI0000811\|hsa-mir-148b&WithFlank&12\|+\|54337210\|54337320@69.90.22(+1G)&[MIMAT0000759&hsa-miR-148b-3p&offsets\|0\|0(+1G);m-77&12\|+\|54337278\|54337299&offsets\|0\|0(+1G)]\|TCAGTGCATCACAGAACTTTGTG | >MI0000811\|hsa-mir-148b&WithFlank&12\|+\|54337210\|54337320 | 69 | 90 | 22(+1G) | TCAGTGCATCACAGAACTTTGTG | 1336.4 | 8.2 | 1.0 | 8.5 | 1.8E-17 | 6.9E-17 | BT20/MB468 |
| >hg38_21_+_8433172_8446622_RNA45SN1_45S_50ntFlanks@3705.3728.24\|TACCTGGTTGATCCTGCCAGTAGC | >hg38_21_+_8433172_8446622_RNA45SN1_45S_50ntFlanks | 3705 | 3728 | 24 | TACCTGGTTGATCCTGCCAGTAGC | 769.4 | 5.9 | 0.7 | 8.5 | 1.8E-17 | 6.9E-17 | MB231/MB468 |
| >hg38_MT_+_598_1651_RNR1_12S_50ntFlanks@719.744.26\|TAAAGACGTTAGGTCAAGGTGTAGCC | >hg38_MT_+_598_1651_RNR1_12S_50ntFlanks | 719 | 744 | 26 | TAAAGACGTTAGGTCAAGGTGTAGCC | 1700.0 | 7.4 | 0.9 | 8.5 | 1.8E-17 | 6.9E-17 | MB231/MB468 |
| >hg38_1_-_228634819_228635039_RNA5S12_5S_50ntFlanks@131.170.40\|GGATGGGAGACCGCCTGGGAATACCGGGTGCTGTAGGCTT | >hg38_1_-_228634819_228635039_RNA5S12_5S_50ntFlanks | 131 | 170 | 40 | GGATGGGAGACCGCCTGGGAATACCGGGTGCTGTAGGCTT | 1932.0 | 10.6 | 1.2 | 8.5 | 1.9E-17 | 7.0E-17 | MB231/MB468 |
| >MI0002468\|hsa-mir-484&WithFlank&16\|+\|15643288\|15643378@14.35.22(+1U)&[MIMAT0002174&hsa-miR-484&offsets\|0\|0(+1U);m-127&16\|+\|15643301\|15643321&offsets\|0\|+1(+1U)]\|TCAGGCTCAGTCCCCTCCCGATT | >MI0002468\|hsa-mir-484&WithFlank&16\|+\|15643288\|15643378 | 14 | 35 | 22(+1U) | TCAGGCTCAGTCCCCTCCCGATT | 1799.4 | 6.7 | 0.8 | 8.5 | 1.9E-17 | 7.1E-17 | MB231/MB468 |
| >hg38_21_+_8433172_8446622_RNA45SN1_45S_50ntFlanks@12996.13017.22\|TCGCTGCGATCTATTGAAAGTC | >hg38_21_+_8433172_8446622_RNA45SN1_45S_50ntFlanks | 12996 | 13017 | 22 | TCGCTGCGATCTATTGAAAGTC | 834.7 | 6.2 | 0.7 | 8.5 | 1.8E-17 | 7.1E-17 | BT20/MB468 |
| >MI0000082\|hsa-mir-25&WithFlank&7\|-\|100093554\|100093649@58.77.20&[MIMAT0000081&hsa-miR-25-3p&offsets\|0\|-2;m-23&7\|-\|100093571\|100093592&offsets\|0\|-2]\|CATTGCACTTGTCTCGGTCT | >MI0000082\|hsa-mir-25&WithFlank&7\|-\|100093554\|100093649 | 58 | 77 | 20 | CATTGCACTTGTCTCGGTCT | 936.7 | 7.2 | 0.8 | 8.5 | 1.8E-17 | 7.2E-17 | BT20/MB468 |
| >trna10_MetCAT_8_-_124169470_124169542@31.51.21,trna22_MetCAT_16_-_87417628_87417700@31.51.21\|TCTCATAATCTGAAGGTCGTG | >trna10_MetCAT_8_-_124169470_124169542 | 31 | 51 | 21 | TCTCATAATCTGAAGGTCGTG | 805.3 | -8.0 | 0.9 | -8.5 | 1.9E-17 | 7.2E-17 | MB231/MB468 |
| >MI0000440\|hsa-mir-27b&WithFlank&9\|+\|95085439\|95085547@67.88.22(+1A)&[MIMAT0000419&hsa-miR-27b-3p&offsets\|0\|+1(+1A);m-39&9\|+\|95085505\|95085525&offsets\|0\|+1(+1A)]\|TTCACAGTGGCTAAGTTCTGCAA | >MI0000440\|hsa-mir-27b&WithFlank&9\|+\|95085439\|95085547 | 67 | 88 | 22(+1A) | TTCACAGTGGCTAAGTTCTGCAA | 1130.2 | 6.6 | 0.8 | 8.5 | 2.0E-17 | 7.3E-17 | MB231/MB468 |
| >MI0000085\|hsa-mir-27a&WithFlank&19\|-\|13836434\|13836523@57.75.19&[MIMAT0000084&hsa-miR-27a-3p&offsets\|0\|-2;m-43&19\|-\|13836447\|13836467&offsets\|0\|-2]\|TTCACAGTGGCTAAGTTCC | >MI0000085\|hsa-mir-27a&WithFlank&19\|-\|13836434\|13836523 | 57 | 75 | 19 | TTCACAGTGGCTAAGTTCC | 1813.4 | 6.6 | 0.8 | 8.5 | 2.0E-17 | 7.3E-17 | MB231/MB468 |
| >MI0000434\|hsa-let-7i&WithFlank&12\|+\|62603680\|62603775@12.33.22(+1U)&[MIMAT0000415&hsa-let-7i-5p&offsets\|0\|0(+1U);m-49&12\|+\|62603691\|62603712&offsets\|0\|0(+1U)]\|TGAGGTAGTAGTTTGTGCTGTTT | >MI0000434\|hsa-let-7i&WithFlank&12\|+\|62603680\|62603775 | 12 | 33 | 22(+1U) | TGAGGTAGTAGTTTGTGCTGTTT | 7388.9 | 6.2 | 0.7 | 8.5 | 1.9E-17 | 7.4E-17 | BT20/MB468 |
| >am_MI0003581\|hsa-mir-574&WithFlank&4\|+\|38868026\|38868133@31.54.24&[MIMAT0004795&hsa-miR-574-5p&offsets\|0\|+1;m-213&4\|+\|38868056\|38868078&offsets\|0\|+1]\|TGAGTGTGTGTGTGTGAGTGTGTG | >am_MI0003581\|hsa-mir-574&WithFlank&4\|+\|38868026\|38868133 | 31 | 54 | 24 | TGAGTGTGTGTGTGTGAGTGTGTG | 687.8 | 7.0 | 0.8 | 8.5 | 1.9E-17 | 7.4E-17 | BT20/MB468 |
| >hg38_21_+_8433172_8446622_RNA45SN1_45S_50ntFlanks@10351.10390.40\|CGAGAACTTTGAAGGCCGAAGTGGAGAAGGGTTCCATGTG | >hg38_21_+_8433172_8446622_RNA45SN1_45S_50ntFlanks | 10351 | 10390 | 40 | CGAGAACTTTGAAGGCCGAAGTGGAGAAGGGTTCCATGTG | 4777.2 | -6.0 | 0.7 | -8.5 | 1.9E-17 | 7.4E-17 | BT20/MB468 |

<table>
<tr><th></th><th></th><th></th><th></th><th></th><th></th><th></th><th></th><th></th><th></th><th></th><th></th><th></th></tr>
<tr><td>>MI0000108|hsa-mir-103a-2&WithFlank&20|+|3917488|3917577@54.75.22(+2U);MI0000109|hsa-mir-103a-1&WithFlank&5|-|168560890|168560979@54.75.22(+2U)&[MIMAT0000101&hsa-miR-103a-3p&offsets|0|-1(+2U);m-12&20|+|3917541|3917563&offsets|0|-1(+2U)];[MIMAT0000101_1&hsa-miR-103a-3p&offsets|0|-1(+2U);m-13&5|-|168560904|168560926&offsets|0|-1(+2U);m-44220&5|-|168560908|168560926&offsets|0|+3(+2U)]|AGCAGCATTGTACAGGGCTATGTT</td><td>>MI0000108|hsa-mir-103a-2&WithFlank&20|+|3917488|3917577</td><td>54</td><td>75</td><td>22(+2U)</td><td>AGCAGCATTGTACAGGGCTATGTT</td><td>808.2</td><td>8.5</td><td>1.0</td><td>8.5</td><td>2.0E-17</td><td>7.4E-17</td><td>MB231/MB468</td></tr>
<tr><td>>hg38_MT_+_1621_3279_RNR2_16S_50ntFlanks@1438.1457.20|TACGTGATCTGAGTTCAGAC</td><td>>hg38_MT_+_1621_3279_RNR2_16S_50ntFlanks</td><td>1438</td><td>1457</td><td>20</td><td>TACGTGATCTGAGTTCAGAC</td><td>1298.8</td><td>10.0</td><td>1.2</td><td>8.5</td><td>2.0E-17</td><td>7.4E-17</td><td>MB231/MB468</td></tr>
<tr><td>>MI0000095|hsa-mir-93&WithFlank&7|-|100093762|100093853@17.38.22&[MIMAT0000093&hsa-miR-93-5p&offsets|0|-1;m-29&7|-|100093815|100093837&offsets|0|-1]|CAAAGTGCTGTTCGTGCAGGTA</td><td>>MI0000095|hsa-mir-93&WithFlank&7|-|100093762|100093853</td><td>17</td><td>38</td><td>22</td><td>CAAAGTGCTGTTCGTGCAGGTA</td><td>2038.7</td><td>7.1</td><td>0.8</td><td>8.5</td><td>2.0E-17</td><td>7.5E-17</td><td>MB231/MB468</td></tr>
<tr><td>>trnaMT_HisGTG_MT_+_12138_12206@32.69.38|TGAATCTGACAACAGAGGCTTACGACCCCTTATTTACC</td><td>>trnaMT_HisGTG_MT_+_12138_12206</td><td>32</td><td>69</td><td>38</td><td>TGAATCTGACAACAGAGGCTTACGACCCCTTATTTACC</td><td>2208.8</td><td>9.4</td><td>1.1</td><td>8.5</td><td>2.0E-17</td><td>7.5E-17</td><td>MB231/MB468</td></tr>
<tr><td>>MI0000469|hsa-mir-125a&WithFlank&19|+|51693248|51693345@21.43.23&[MIMAT0000443&hsa-miR-125a-5p&offsets|0|-1;m-63&19|+|51693268|51693289&offsets|0|+1]|TCCCTGAGACCCTTTAACCTGTG</td><td>>MI0000469|hsa-mir-125a&WithFlank&19|+|51693248|51693345</td><td>21</td><td>43</td><td>23</td><td>TCCCTGAGACCCTTTAACCTGTG</td><td>26036.5</td><td>6.2</td><td>0.7</td><td>8.5</td><td>2.1E-17</td><td>7.7E-17</td><td>MB231/MB468</td></tr>
<tr><td>>hg38_21_+_8433172_8446622_RNA45SN1_45S_50ntFlanks@12449.12473.25|GAATTCACCAAGCGTTGGATTGTTC</td><td>>hg38_21_+_8433172_8446622_RNA45SN1_45S_50ntFlanks</td><td>12449</td><td>12473</td><td>25</td><td>GAATTCACCAAGCGTTGGATTGTTC</td><td>285.5</td><td>7.0</td><td>0.8</td><td>8.5</td><td>2.1E-17</td><td>7.8E-17</td><td>MB231/MB468</td></tr>
<tr><td>>hg38_21_+_8433172_8446622_RNA45SN1_45S_50ntFlanks@4130.4162.33|ATTCCGGAGAGGGAGCCTGAGAAACGGCTACCA</td><td>>hg38_21_+_8433172_8446622_RNA45SN1_45S_50ntFlanks</td><td>4130</td><td>4162</td><td>33</td><td>ATTCCGGAGAGGGAGCCTGAGAAACGGCTACCA</td><td>404.1</td><td>-7.3</td><td>0.9</td><td>-8.5</td><td>2.0E-17</td><td>7.8E-17</td><td>BT20/MB468</td></tr>
<tr><td>>hg38_21_+_8433172_8446622_RNA45SN1_45S_50ntFlanks@12637.12654.18|GCTACCATCTGTGGGATT</td><td>>hg38_21_+_8433172_8446622_RNA45SN1_45S_50ntFlanks</td><td>12637</td><td>12654</td><td>18</td><td>GCTACCATCTGTGGGATT</td><td>1100.8</td><td>-6.4</td><td>0.8</td><td>-8.5</td><td>2.0E-17</td><td>7.8E-17</td><td>BT20/MB468</td></tr>
<tr><td>>hg38_21_+_8433172_8446622_RNA45SN1_45S_50ntFlanks@4069.4095.27|CTTTCGATGGTAGTCGCCGTGCCTACC</td><td>>hg38_21_+_8433172_8446622_RNA45SN1_45S_50ntFlanks</td><td>4069</td><td>4095</td><td>27</td><td>CTTTCGATGGTAGTCGCCGTGCCTACC</td><td>184.3</td><td>-4.8</td><td>0.6</td><td>-8.5</td><td>2.0E-17</td><td>7.8E-17</td><td>BT20/MB468</td></tr>
<tr><td>>MI0000089|hsa-mir-31&WithFlank&9|-|21512109|21512191@14.34.21&[MIMAT0000089&hsa-miR-31-5p&offsets|0|0;m-360&9|-|21512157|21512178&offsets|0|-1]|AGGCAAGATGCTGGCATAGCT</td><td>>MI0000089|hsa-mir-31&WithFlank&9|-|21512109|21512191</td><td>14</td><td>34</td><td>21</td><td>AGGCAAGATGCTGGCATAGCT</td><td>671.1</td><td>12.9</td><td>1.5</td><td>8.8</td><td>1.2E-18</td><td>7.9E-17</td><td>BT20/MB231</td></tr>
<tr><td>>MI0000298|hsa-mir-221&WithFlank&X|-|45746151|45746272@31.54.24&[MIMAT0004568&hsa-miR-221-5p&offsets|0|+2;m-405&X|-|45746220|45746242&offsets|0|+1]|ACCTGGCATACAATGTAGATTTCT</td><td>>MI0000298|hsa-mir-221&WithFlank&X|-|45746151|45746272</td><td>31</td><td>54</td><td>24</td><td>ACCTGGCATACAATGTAGATTTCT</td><td>1128.2</td><td>5.8</td><td>0.7</td><td>8.5</td><td>2.0E-17</td><td>7.9E-17</td><td>BT20/MB468</td></tr>
<tr><td>>hg38_21_+_8433172_8446622_RNA45SN1_45S_50ntFlanks@12625.12654.30|CAATGGGGCGAAGCTACCATCTGTGGGATT</td><td>>hg38_21_+_8433172_8446622_RNA45SN1_45S_50ntFlanks</td><td>12625</td><td>12654</td><td>30</td><td>CAATGGGGCGAAGCTACCATCTGTGGGATT</td><td>3398.2</td><td>-4.5</td><td>0.5</td><td>-8.5</td><td>2.1E-17</td><td>7.9E-17</td><td>MB231/MB468</td></tr>
<tr><td>>hg38_21_+_8433172_8446622_RNA45SN1_45S_50ntFlanks@4823.4844.22|ATGACCCGCCGGGCAGCTTCCG</td><td>>hg38_21_+_8433172_8446622_RNA45SN1_45S_50ntFlanks</td><td>4823</td><td>4844</td><td>22</td><td>ATGACCCGCCGGGCAGCTTCCG</td><td>314.4</td><td>5.6</td><td>0.7</td><td>8.5</td><td>2.2E-17</td><td>8.0E-17</td><td>MB231/MB468</td></tr>
<tr><td>>hg38_21_+_8433172_8446622_RNA45SN1_45S_50ntFlanks@12405.12422.18|CTTTTTGATCCTTCGATG</td><td>>hg38_21_+_8433172_8446622_RNA45SN1_45S_50ntFlanks</td><td>12405</td><td>12422</td><td>18</td><td>CTTTTTGATCCTTCGATG</td><td>713.6</td><td>-8.6</td><td>1.0</td><td>-8.5</td><td>2.2E-17</td><td>8.1E-17</td><td>MB231/MB468</td></tr>
<tr><td>>hg38_21_+_8433172_8446622_RNA45SN1_45S_50ntFlanks@11873.11890.18|AAGAAGACCCTGTTGAGC</td><td>>hg38_21_+_8433172_8446622_RNA45SN1_45S_50ntFlanks</td><td>11873</td><td>11890</td><td>18</td><td>AAGAAGACCCTGTTGAGC</td><td>814.2</td><td>-4.9</td><td>0.6</td><td>-8.5</td><td>2.2E-17</td><td>8.1E-17</td><td>MB231/MB468</td></tr>
<tr><td>>hg38_21_+_8433172_8446622_RNA45SN1_45S_50ntFlanks@12416.12452.37|TTCGATGTCGGCTCTTCCTATCATTGTGAAGCAGAAT</td><td>>hg38_21_+_8433172_8446622_RNA45SN1_45S_50ntFlanks</td><td>12416</td><td>12452</td><td>37</td><td>TTCGATGTCGGCTCTTCCTATCATTGTGAAGCAGAAT</td><td>1333.4</td><td>9.7</td><td>1.1</td><td>8.5</td><td>2.2E-17</td><td>8.1E-17</td><td>MB231/MB468</td></tr>
<tr><td>>hg38_21_+_8433172_8446622_RNA45SN1_45S_50ntFlanks@6721.6742.22|AATTGCAGGACACATTGATCAT</td><td>>hg38_21_+_8433172_8446622_RNA45SN1_45S_50ntFlanks</td><td>6721</td><td>6742</td><td>22</td><td>AATTGCAGGACACATTGATCAT</td><td>707.1</td><td>-7.0</td><td>0.8</td><td>-8.5</td><td>2.1E-17</td><td>8.2E-17</td><td>BT20/MB468</td></tr>
<tr><td>>hg38_21_+_8433172_8446622_RNA45SN1_45S_50ntFlanks@10806.10823.18|AACAATGTAGGTAAGGGA</td><td>>hg38_21_+_8433172_8446622_RNA45SN1_45S_50ntFlanks</td><td>10806</td><td>10823</td><td>18</td><td>AACAATGTAGGTAAGGGA</td><td>363.6</td><td>-6.2</td><td>0.7</td><td>-8.5</td><td>2.2E-17</td><td>8.2E-17</td><td>MB231/MB468</td></tr>
<tr><td>>hg38_21_+_8433172_8446622_RNA45SN1_45S_50ntFlanks@6650.6667.18|ACGACTCTTAGCGGTGGA</td><td>>hg38_21_+_8433172_8446622_RNA45SN1_45S_50ntFlanks</td><td>6650</td><td>6667</td><td>18</td><td>ACGACTCTTAGCGGTGGA</td><td>397.1</td><td>5.4</td><td>0.6</td><td>8.5</td><td>2.1E-17</td><td>8.2E-17</td><td>BT20/MB468</td></tr>
<tr><td>>hg38_21_+_8433172_8446622_RNA45SN1_45S_50ntFlanks@12994.13038.45|CCTCGCTGCGATCTATTGAAAGTCAGCCCTCGACACAAGGGTTTG</td><td>>hg38_21_+_8433172_8446622_RNA45SN1_45S_50ntFlanks</td><td>12994</td><td>13038</td><td>45</td><td>CCTCGCTGCGATCTATTGAAAGTCAGCCCTCGACACAAGGGTTTG</td><td>1604.3</td><td>11.1</td><td>1.3</td><td>8.5</td><td>2.2E-17</td><td>8.3E-17</td><td>MB231/MB468</td></tr>
<tr><td>>hg38_21_+_8433172_8446622_RNA45SN1_45S_50ntFlanks@12631.12654.24|GGCGAAGCTACCATCTGTGGGATT</td><td>>hg38_21_+_8433172_8446622_RNA45SN1_45S_50ntFlanks</td><td>12631</td><td>12654</td><td>24</td><td>GGCGAAGCTACCATCTGTGGGATT</td><td>454.5</td><td>-10.8</td><td>1.3</td><td>-8.5</td><td>2.1E-17</td><td>8.3E-17</td><td>BT20/MB468</td></tr>
<tr><td>>hg38_21_+_8433172_8446622_RNA45SN1_45S_50ntFlanks@12418.12452.35|CGATGTCGGCTCTTCCTATCATTGTGAAGCAGAAT</td><td>>hg38_21_+_8433172_8446622_RNA45SN1_45S_50ntFlanks</td><td>12418</td><td>12452</td><td>35</td><td>CGATGTCGGCTCTTCCTATCATTGTGAAGCAGAAT</td><td>571.7</td><td>5.2</td><td>0.6</td><td>8.5</td><td>2.3E-17</td><td>8.4E-17</td><td>MB231/MB468</td></tr>
<tr><td>>hg38_21_+_8433172_8446622_RNA45SN1_45S_50ntFlanks@4443.4471.29|CCCTTGCCTCTCGGCGCCCCCTCGATGCT</td><td>>hg38_21_+_8433172_8446622_RNA45SN1_45S_50ntFlanks</td><td>4443</td><td>4471</td><td>29</td><td>CCCTTGCCTCTCGGCGCCCCCTCGATGCT</td><td>736.0</td><td>-11.5</td><td>1.4</td><td>-8.5</td><td>2.2E-17</td><td>8.4E-17</td><td>BT20/MB468</td></tr>
<tr><td>>MI0000289|hsa-mir-181a-1&WithFlank&1|-|198859038|198859159@30.51.22(+1C);MI0000269|hsa-mir-181a-2&WithFlank&9|+|124692436|124692557@45.66.22(+1C)&[MIMAT0000256&hsa-miR-181a-5p&offsets|0|-1(+1C);m-32&1|-|198859108|198859130&offsets|0|-1(+1C)];[MIMAT0000256_1&hsa-miR-181a-5p&offsets|0|-1(+1C);m-32&9|+|124692480|124692502&offsets|0|-1(+1C)]|AACATTCAACGCTGTCGGTGAGC</td><td>>MI0000289|hsa-mir-181a-1&WithFlank&1|-|198859038|198859159</td><td>30</td><td>51</td><td>22(+1C)</td><td>AACATTCAACGCTGTCGGTGAGC</td><td>1797.1</td><td>7.0</td><td>0.8</td><td>8.5</td><td>2.3E-17</td><td>8.5E-17</td><td>MB231/MB468</td></tr>
<tr><td>>hg38_21_+_8433172_8446622_RNA45SN1_45S_50ntFlanks@4733.4766.34|GAACGAAAGTCGGAGGTTCGAAGACGATCAGATA</td><td>>hg38_21_+_8433172_8446622_RNA45SN1_45S_50ntFlanks</td><td>4733</td><td>4766</td><td>34</td><td>GAACGAAAGTCGGAGGTTCGAAGACGATCAGATA</td><td>407.7</td><td>8.6</td><td>1.0</td><td>8.5</td><td>2.3E-17</td><td>8.5E-17</td><td>MB231/MB468</td></tr>
<tr><td>>hg38_1_-_228634819_228635039_RNA5S12_5S_50ntFlanks@135.172.38|GGGAGACCGCCTGGGAATACCGGGTGCTGTAGGCTTTT</td><td>>hg38_1_-_228634819_228635039_RNA5S12_5S_50ntFlanks</td><td>135</td><td>172</td><td>38</td><td>GGGAGACCGCCTGGGAATACCGGGTGCTGTAGGCTTTT</td><td>1003.3</td><td>8.6</td><td>1.0</td><td>8.5</td><td>2.3E-17</td><td>8.5E-17</td><td>MB231/MB468</td></tr>
<tr><td>>hg38_21_+_8433172_8446622_RNA45SN1_45S_50ntFlanks@12412.12432.21|ATCCTTCGATGTCGGCTCTTC</td><td>>hg38_21_+_8433172_8446622_RNA45SN1_45S_50ntFlanks</td><td>12412</td><td>12432</td><td>21</td><td>ATCCTTCGATGTCGGCTCTTC</td><td>593.4</td><td>-11.2</td><td>1.3</td><td>-8.5</td><td>2.2E-17</td><td>8.6E-17</td><td>BT20/MB468</td></tr>
<tr><td>>hg38_21_+_8433172_8446622_RNA45SN1_45S_50ntFlanks@6773.6799.27|TTCCTCCCGGGGCTACGCCTGTCTGAG</td><td>>hg38_21_+_8433172_8446622_RNA45SN1_45S_50ntFlanks</td><td>6773</td><td>6799</td><td>27</td><td>TTCCTCCCGGGGCTACGCCTGTCTGAG</td><td>355.8</td><td>-7.0</td><td>0.8</td><td>-8.5</td><td>2.4E-17</td><td>8.6E-17</td><td>MB231/MB468</td></tr>
<tr><td>>MI0000279|hsa-mir-196a-2&WithFlank&12|+|53991732|53991853@31.52.22(+1U);MI0000238|hsa-mir-196a-1&WithFlank&17|-|48632484|48632565@13.34.22(+1U)&[MIMAT0000226&hsa-miR-196a-5p&offsets|0|0(+1U);m-261&12|+|53991762|53991783&offsets|0|0(+1U)];[MIMAT0000226_1&hsa-miR-196a-5p&offsets|0|0(+1U);m-268&17|-|48632532|48632553&offsets|0|0(+1U)]|TAGGTAGTTTCATGTTGTTGGGT</td><td>>MI0000279|hsa-mir-196a-2&WithFlank&12|+|53991732|53991853</td><td>31</td><td>52</td><td>22(+1U)</td><td>TAGGTAGTTTCATGTTGTTGGGT</td><td>488.3</td><td>12.4</td><td>1.4</td><td>8.8</td><td>1.3E-18</td><td>8.6E-17</td><td>BT20/MB231</td></tr>
<tr><td>>hg38_21_+_8433172_8446622_RNA45SN1_45S_50ntFlanks@9545.9564.20|GTGGAGGTCCGTAGCGGTCC</td><td>>hg38_21_+_8433172_8446622_RNA45SN1_45S_50ntFlanks</td><td>9545</td><td>9564</td><td>20</td><td>GTGGAGGTCCGTAGCGGTCC</td><td>563.6</td><td>-7.8</td><td>0.9</td><td>-8.5</td><td>2.3E-17</td><td>8.8E-17</td><td>BT20/MB468</td></tr>
<tr><td>>hg38_21_+_8433172_8446622_RNA45SN1_45S_50ntFlanks@12616.12653.38|CTGAGGAGCCAATGGGGCGAAGCTACCATCTGTGGGAT</td><td>>hg38_21_+_8433172_8446622_RNA45SN1_45S_50ntFlanks</td><td>12616</td><td>12653</td><td>38</td><td>CTGAGGAGCCAATGGGGCGAAGCTACCATCTGTGGGAT</td><td>226.6</td><td>-6.6</td><td>0.8</td><td>-8.5</td><td>2.3E-17</td><td>8.8E-17</td><td>BT20/MB468</td></tr>
<tr><td>>hg38_21_+_8433172_8446622_RNA45SN1_45S_50ntFlanks@12133.12150.18|CTCCGGGGACAGTGCCAG</td><td>>hg38_21_+_8433172_8446622_RNA45SN1_45S_50ntFlanks</td><td>12133</td><td>12150</td><td>18</td><td>CTCCGGGGACAGTGCCAG</td><td>1141.1</td><td>-12.1</td><td>1.4</td><td>-8.5</td><td>2.3E-17</td><td>8.9E-17</td><td>BT20/MB468</td></tr>
<tr><td>>hg38_21_+_8433172_8446622_RNA45SN1_45S_50ntFlanks@13004.13028.25|ATCTATTGAAAGTCAGCCCTCGACA</td><td>>hg38_21_+_8433172_8446622_RNA45SN1_45S_50ntFlanks</td><td>13004</td><td>13028</td><td>25</td><td>ATCTATTGAAAGTCAGCCCTCGACA</td><td>828.8</td><td>-11.7</td><td>1.4</td><td>-8.5</td><td>2.3E-17</td><td>8.9E-17</td><td>BT20/MB468</td></tr>
<tr><td>>hg38_21_+_8433172_8446622_RNA45SN1_45S_50ntFlanks@11208.11231.24|GCGGCGACTCTGGACGCGAGCCGG</td><td>>hg38_21_+_8433172_8446622_RNA45SN1_45S_50ntFlanks</td><td>11208</td><td>11231</td><td>24</td><td>GCGGCGACTCTGGACGCGAGCCGG</td><td>748.9</td><td>-5.2</td><td>0.6</td><td>-8.5</td><td>2.4E-17</td><td>9.0E-17</td><td>MB231/MB468</td></tr>
<tr><td>>hg38_21_+_8433172_8446622_RNA45SN1_45S_50ntFlanks@6786.6806.21|TACGCCTGTCTGAGCGTCGCT</td><td>>hg38_21_+_8433172_8446622_RNA45SN1_45S_50ntFlanks</td><td>6786</td><td>6806</td><td>21</td><td>TACGCCTGTCTGAGCGTCGCT</td><td>4626.5</td><td>7.4</td><td>0.9</td><td>8.5</td><td>2.4E-17</td><td>9.3E-17</td><td>BT20/MB468</td></tr>
<tr><td>>hg38_21_+_8433172_8446622_RNA45SN1_45S_50ntFlanks@12997.13039.43|CGCTGCGATCTATTGAAAGTCAGCCCTCGACACAAGGGTTTGT</td><td>>hg38_21_+_8433172_8446622_RNA45SN1_45S_50ntFlanks</td><td>12997</td><td>13039</td><td>43</td><td>CGCTGCGATCTATTGAAAGTCAGCCCTCGACACAAGGGTTTGT</td><td>68395.8</td><td>7.8</td><td>0.9</td><td>8.5</td><td>2.4E-17</td><td>9.4E-17</td><td>BT20/MB468</td></tr>
<tr><td>>hg38_21_+_8433172_8446622_RNA45SN1_45S_50ntFlanks@10402.10433.32|AACATGGGTCAGTCGGTCCTGAGAGATGGGCG</td><td>>hg38_21_+_8433172_8446622_RNA45SN1_45S_50ntFlanks</td><td>10402</td><td>10433</td><td>32</td><td>AACATGGGTCAGTCGGTCCTGAGAGATGGGCG</td><td>320.9</td><td>-5.3</td><td>0.6</td><td>-8.5</td><td>2.4E-17</td><td>9.4E-17</td><td>BT20/MB468</td></tr>
<tr><td>>hg38_21_+_8433172_8446622_RNA45SN1_45S_50ntFlanks@11641.11670.30|CGGCGGGTGTTGACGCGATGTGATTTCTGC</td><td>>hg38_21_+_8433172_8446622_RNA45SN1_45S_50ntFlanks</td><td>11641</td><td>11670</td><td>30</td><td>CGGCGGGTGTTGACGCGATGTGATTTCTGC</td><td>2191.2</td><td>-8.2</td><td>1.0</td><td>-8.5</td><td>2.6E-17</td><td>9.4E-17</td><td>MB231/MB468</td></tr>
<tr><td>>hg38_21_+_8433172_8446622_RNA45SN1_45S_50ntFlanks@11661.11679.19|TGATTTCTGCCCAGTGCTC</td><td>>hg38_21_+_8433172_8446622_RNA45SN1_45S_50ntFlanks</td><td>11661</td><td>11679</td><td>19</td><td>TGATTTCTGCCCAGTGCTC</td><td>1959.1</td><td>-9.3</td><td>1.1</td><td>-8.5</td><td>2.5E-17</td><td>9.4E-17</td><td>BT20/MB468</td></tr>
<tr><td>>hg38_MT_+_598_1651_RNR1_12S_50ntFlanks@428.456.29|AAAGTGGCTTTAACATATCTGAACACACA</td><td>>hg38_MT_+_598_1651_RNR1_12S_50ntFlanks</td><td>428</td><td>456</td><td>29</td><td>AAAGTGGCTTTAACATATCTGAACACACA</td><td>1470.7</td><td>11.1</td><td>1.3</td><td>8.5</td><td>2.5E-17</td><td>9.6E-17</td><td>BT20/MB468</td></tr>
<tr><td>>hg38_21_+_8433172_8446622_RNA45SN1_45S_50ntFlanks@6742.6771.30|TCGACACTTCGAACGCACTTGCGGCCCCGG</td><td>>hg38_21_+_8433172_8446622_RNA45SN1_45S_50ntFlanks</td><td>6742</td><td>6771</td><td>30</td><td>TCGACACTTCGAACGCACTTGCGGCCCCGG</td><td>260.8</td><td>-5.2</td><td>0.6</td><td>-8.5</td><td>2.6E-17</td><td>9.7E-17</td><td>MB231/MB468</td></tr>
<tr><td>>hg38_21_+_8433172_8446622_RNA45SN1_45S_50ntFlanks@8552.8569.18|CTCCCCGGAGGGGGCGGG</td><td>>hg38_21_+_8433172_8446622_RNA45SN1_45S_50ntFlanks</td><td>8552</td><td>8569</td><td>18</td><td>CTCCCCGGAGGGGGCGGG</td><td>544.7</td><td>-11.1</td><td>1.3</td><td>-8.5</td><td>2.5E-17</td><td>9.7E-17</td><td>BT20/MB468</td></tr>
<tr><td>>hg38_21_+_8433172_8446622_RNA45SN1_45S_50ntFlanks@10806.10823.18|AACAATGTAGGTAAGGGA</td><td>>hg38_21_+_8433172_8446622_RNA45SN1_45S_50ntFlanks</td><td>10806</td><td>10823</td><td>18</td><td>AACAATGTAGGTAAGGGA</td><td>432.4</td><td>-10.7</td><td>1.3</td><td>-8.5</td><td>2.6E-17</td><td>9.9E-17</td><td>BT20/MB468</td></tr>
<tr><td>>hg38_21_+_8433172_8446622_RNA45SN1_45S_50ntFlanks@11886.11904.19|TGAGCTTGACTCTAGTCTG</td><td>>hg38_21_+_8433172_8446622_RNA45SN1_45S_50ntFlanks</td><td>11886</td><td>11904</td><td>19</td><td>TGAGCTTGACTCTAGTCTG</td><td>654.9</td><td>-8.4</td><td>1.0</td><td>-8.5</td><td>2.7E-17</td><td>9.9E-17</td><td>MB231/MB468</td></tr>
<tr><td>>hg38_21_+_8433172_8446622_RNA45SN1_45S_50ntFlanks@4005.4024.20|AACCTCGGGCCGATCGCACG</td><td>>hg38_21_+_8433172_8446622_RNA45SN1_45S_50ntFlanks</td><td>4005</td><td>4024</td><td>20</td><td>AACCTCGGGCCGATCGCACG</td><td>545.0</td><td>-4.0</td><td>0.5</td><td>-8.5</td><td>2.6E-17</td><td>1.0E-16</td><td>BT20/MB468</td></tr>
<tr><td>>MI0000065|hsa-let-7d&WithFlank&9|+|94178828|94178926@68.88.21(+1A)&[MIMAT0004484&hsa-let-7d-3p&offsets|0|-1(+1A);m-98&9|+|94178895|94178916&offsets|0|-1(+1A)]|CTATACGACCTGCTGCCTTTCA</td><td>>MI0000065|hsa-let-7d&WithFlank&9|+|94178828|94178926</td><td>68</td><td>88</td><td>21(+1A)</td><td>CTATACGACCTGCTGCCTTTCA</td><td>865.4</td><td>9.3</td><td>1.1</td><td>8.5</td><td>2.6E-17</td><td>1.0E-16</td><td>BT20/MB468</td></tr>
<tr><td>>MI0000542|hsa-mir-320a&WithFlank&8|-|22244960|22245043@49.69.21&[MIMAT0000510&hsa-miR-320a-3p&offsets|+1|0;m-60&8|-|22244975|22244996&offsets|+1|0]|AAAGCTGGGTTGAGAGGGCGA</td><td>>MI0000542|hsa-mir-320a&WithFlank&8|-|22244960|22245043</td><td>49</td><td>69</td><td>21</td><td>AAAGCTGGGTTGAGAGGGCGA</td><td>1936.2</td><td>9.2</td><td>1.1</td><td>8.5</td><td>2.6E-17</td><td>1.0E-16</td><td>BT20/MB468</td></tr>
<tr><td>>MI0000440|hsa-mir-27b&WithFlank&9|+|95085439|95085547@67.88.22(+1A)&[MIMAT0000419&hsa-miR-27b-3p&offsets|0|+1(+1A);m-39&9|+|95085505|95085525&offsets|0|+1(+1A)]|TTCACAGTGGCTAAGTTCTGCAA</td><td>>MI0000440|hsa-mir-27b&WithFlank&9|+|95085439|95085547</td><td>67</td><td>88</td><td>22(+1A)</td><td>TTCACAGTGGCTAAGTTCTGCAA</td><td>1602.1</td><td>6.8</td><td>0.8</td><td>8.5</td><td>2.6E-17</td><td>1.0E-16</td><td>BT20/MB468</td></tr>
</table>

<table>
<tr><td>>MI0000077|hsa-mir-21&WithFlank&17|+|59841260|59841343@14.35.22(+2A)&[MIMAT0000076&hsa-miR-21-5p&offsets|0|0(+2A);m-1&17|+|59841273|59841294&offsets|0|0(+2A)]|TAGCTTATCAGACTGATGTTGAAA</td><td>>MI0000077|hsa-mir-21&WithFlank&17|+|59841260|59841343</td><td>14</td><td>35</td><td>22(+2A)</td><td>TAGCTTATCAGACTGATGTTGAAA</td><td>4274.1</td><td>7.4</td><td>0.9</td><td>8.5</td><td>2.7E-17</td><td>1.0E-16</td><td>BT20/MB468</td></tr>
<tr><td>>hg38_21_+_8433172_8446622_RNA45SN1_45S_50ntFlanks@9553.9570.18|CCGTAGCGGTCCTGACGT</td><td>>hg38_21_+_8433172_8446622_RNA45SN1_45S_50ntFlanks</td><td>9553</td><td>9570</td><td>18</td><td>CCGTAGCGGTCCTGACGT</td><td>591.0</td><td>3.8</td><td>0.5</td><td>8.5</td><td>2.9E-17</td><td>1.1E-16</td><td>MB231/MB468</td></tr>
<tr><td>>hg38_21_+_8433172_8446622_RNA45SN1_45S_50ntFlanks@6722.6745.24|ATTGCAGGACACATTGATCATCGA</td><td>>hg38_21_+_8433172_8446622_RNA45SN1_45S_50ntFlanks</td><td>6722</td><td>6745</td><td>24</td><td>ATTGCAGGACACATTGATCATCGA</td><td>196.4</td><td>-4.9</td><td>0.6</td><td>-8.5</td><td>2.9E-17</td><td>1.1E-16</td><td>BT20/MB468</td></tr>
<tr><td>>MI0000441|hsa-mir-30b&WithFlank&8|-|134800514|134800613@23.43.21(+1A)&[MIMAT0000420&hsa-miR-30b-5p&offsets|0|-1(+1A);m-76&8|-|134800570|134800591&offsets|0|-1(+1A)]|TGTAAACATCCTACACTCAGCA</td><td>>MI0000441|hsa-mir-30b&WithFlank&8|-|134800514|134800613</td><td>23</td><td>43</td><td>21(+1A)</td><td>TGTAAACATCCTACACTCAGCA</td><td>12190.3</td><td>12.2</td><td>1.4</td><td>8.5</td><td>2.9E-17</td><td>1.1E-16</td><td>BT20/MB468</td></tr>
<tr><td>>MI0000283|hsa-mir-203a&WithFlank&14|+|104117399|104117520@71.90.20&[MIMAT0000264&hsa-miR-203a-3p&offsets|0|-2;m-79&14|+|104117469|104117490&offsets|0|-2]|GTGAAATGTTTAGGACCACT</td><td>>MI0000283|hsa-mir-203a&WithFlank&14|+|104117399|104117520</td><td>71</td><td>90</td><td>20</td><td>GTGAAATGTTTAGGACCACT</td><td>693.0</td><td>11.0</td><td>1.3</td><td>8.8</td><td>1.8E-18</td><td>1.1E-16</td><td>BT20/MB231</td></tr>
<tr><td>>hg38_21_+_8433172_8446622_RNA45SN1_45S_50ntFlanks@5484.5514.31|GAGCGCTGAGAAGACGGTCGAACTTGACTAT</td><td>>hg38_21_+_8433172_8446622_RNA45SN1_45S_50ntFlanks</td><td>5484</td><td>5514</td><td>31</td><td>GAGCGCTGAGAAGACGGTCGAACTTGACTAT</td><td>1758.2</td><td>-5.8</td><td>0.7</td><td>-8.4</td><td>2.9E-17</td><td>1.1E-16</td><td>BT20/MB468</td></tr>
<tr><td>>hg38_21_+_8433172_8446622_RNA45SN1_45S_50ntFlanks@3705.3722.18|TACCTGGTTGATCCTGCC</td><td>>hg38_21_+_8433172_8446622_RNA45SN1_45S_50ntFlanks</td><td>3705</td><td>3722</td><td>18</td><td>TACCTGGTTGATCCTGCC</td><td>12099.3</td><td>4.9</td><td>0.6</td><td>8.4</td><td>2.9E-17</td><td>1.1E-16</td><td>BT20/MB468</td></tr>
<tr><td>>MI0000465|hsa-mir-191&WithFlank&3|-|49020612|49020715@22.42.21(+1G)&[MIMAT0000440&hsa-miR-191-5p&offsets|0|-2(+1G);m-48&3|-|49020672|49020694&offsets|0|-2(+1G)]|CAACGGAATCCCAAAAGCAGCG</td><td>>MI0000465|hsa-mir-191&WithFlank&3|-|49020612|49020715</td><td>22</td><td>42</td><td>21(+1G)</td><td>CAACGGAATCCCAAAAGCAGCG</td><td>889.8</td><td>7.0</td><td>0.8</td><td>8.4</td><td>3.1E-17</td><td>1.1E-16</td><td>MB231/MB468</td></tr>
<tr><td>>trnaMT_ProTGG_MT_-_15956_16023@51.71.21|TAAAGACTTTTTCTCTGACCA</td><td>>trnaMT_ProTGG_MT_-_15956_16023</td><td>51</td><td>71</td><td>21</td><td>TAAAGACTTTTTCTCTGACCA</td><td>1302.3</td><td>6.8</td><td>0.8</td><td>8.4</td><td>3.0E-17</td><td>1.1E-16</td><td>BT20/MB468</td></tr>
<tr><td>>MI0000802|hsa-mir-340&WithFlank&5|-|180015297|180015403@22.42.21&[MIMAT0004692&hsa-miR-340-5p&offsets|0|-1;m-220&5|-|180015361|180015382&offsets|0|-1]|TTATAAAGCAATGAGACTGAT</td><td>>MI0000802|hsa-mir-340&WithFlank&5|-|180015297|180015403</td><td>22</td><td>42</td><td>21</td><td>TTATAAAGCAATGAGACTGAT</td><td>571.2</td><td>6.0</td><td>0.7</td><td>8.4</td><td>3.1E-17</td><td>1.1E-16</td><td>MB231/MB468</td></tr>
<tr><td>>hg38_21_+_8433172_8446622_RNA45SN1_45S_50ntFlanks@4728.4751.24|ATCAAGAACGAAAGTCGGAGGTTC</td><td>>hg38_21_+_8433172_8446622_RNA45SN1_45S_50ntFlanks</td><td>4728</td><td>4751</td><td>24</td><td>ATCAAGAACGAAAGTCGGAGGTTC</td><td>423.3</td><td>-10.7</td><td>1.3</td><td>-8.4</td><td>3.0E-17</td><td>1.1E-16</td><td>BT20/MB468</td></tr>
<tr><td>>MI0000085|hsa-mir-27a&WithFlank&19|-|13836434|13836523@57.75.19(+1A)&[MIMAT0000084&hsa-miR-27a-3p&offsets|0|-2(+1A);m-43&19|-|13836447|13836467&offsets|0|-2(+1A)]|TTCACAGTGGCTAAGTTCCA</td><td>>MI0000085|hsa-mir-27a&WithFlank&19|-|13836434|13836523</td><td>57</td><td>75</td><td>19(+1A)</td><td>TTCACAGTGGCTAAGTTCCA</td><td>6591.1</td><td>6.6</td><td>0.8</td><td>8.4</td><td>3.1E-17</td><td>1.1E-16</td><td>MB231/MB468</td></tr>
<tr><td>>hg38_21_+_8433172_8446622_RNA45SN1_45S_50ntFlanks@4759.4786.28|ATCAGATACCGTCGTAGTTCCGACCATA</td><td>>hg38_21_+_8433172_8446622_RNA45SN1_45S_50ntFlanks</td><td>4759</td><td>4786</td><td>28</td><td>ATCAGATACCGTCGTAGTTCCGACCATA</td><td>618.5</td><td>-11.3</td><td>1.3</td><td>-8.4</td><td>3.0E-17</td><td>1.2E-16</td><td>BT20/MB468</td></tr>
<tr><td>>hg38_21_+_8433172_8446622_RNA45SN1_45S_50ntFlanks@3874.3899.26|AATACATGCCGACGGGCGCTGACCCC</td><td>>hg38_21_+_8433172_8446622_RNA45SN1_45S_50ntFlanks</td><td>3874</td><td>3899</td><td>26</td><td>AATACATGCCGACGGGCGCTGACCCC</td><td>508.6</td><td>-4.5</td><td>0.5</td><td>-8.4</td><td>3.1E-17</td><td>1.2E-16</td><td>BT20/MB468</td></tr>
<tr><td>>hg38_21_+_8433172_8446622_RNA45SN1_45S_50ntFlanks@11235.11253.19|CTTCCCGTGGATCGCCCCA</td><td>>hg38_21_+_8433172_8446622_RNA45SN1_45S_50ntFlanks</td><td>11235</td><td>11253</td><td>19</td><td>CTTCCCGTGGATCGCCCCA</td><td>867.8</td><td>-11.3</td><td>1.3</td><td>-8.4</td><td>3.2E-17</td><td>1.2E-16</td><td>MB231/MB468</td></tr>
<tr><td>>MI0000456|hsa-mir-140&WithFlank&16|+|69933075|69933186@69.90.22(+1U)&[MIMAT0004597&hsa-miR-140-3p&offsets|+1|+2(+1U);m-45&16|+|69933143|69933164&offsets|0|0(+1U)]|ACCACAGGGTAGAACCACGGACT</td><td>>MI0000456|hsa-mir-140&WithFlank&16|+|69933075|69933186</td><td>69</td><td>90</td><td>22(+1U)</td><td>ACCACAGGGTAGAACCACGGACT</td><td>1672.5</td><td>10.9</td><td>1.3</td><td>8.4</td><td>3.1E-17</td><td>1.2E-16</td><td>BT20/MB468</td></tr>
<tr><td>>MI0000462|hsa-mir-152&WithFlank&17|-|48037155|48037253@60.81.22&[MIMAT0000438&hsa-miR-152-3p&offsets|0|+1;m-97&17|-|48037174|48037194&offsets|0|+1]|TCAGTGCATGACAGAACTTGGG</td><td>>MI0000462|hsa-mir-152&WithFlank&17|-|48037155|48037253</td><td>60</td><td>81</td><td>22</td><td>TCAGTGCATGACAGAACTTGGG</td><td>1752.8</td><td>11.8</td><td>1.4</td><td>8.4</td><td>3.1E-17</td><td>1.2E-16</td><td>BT20/MB468</td></tr>
<tr><td>>hg38_21_+_8433172_8446622_RNA45SN1_45S_50ntFlanks@10423.10453.31|AGAGATGGGCGAGCGCCGTTCCGAAGGGACG</td><td>>hg38_21_+_8433172_8446622_RNA45SN1_45S_50ntFlanks</td><td>10423</td><td>10453</td><td>31</td><td>AGAGATGGGCGAGCGCCGTTCCGAAGGGACG</td><td>172.4</td><td>-3.6</td><td>0.4</td><td>-8.4</td><td>3.2E-17</td><td>1.2E-16</td><td>BT20/MB468</td></tr>
<tr><td>>trnaMT_GluTTC_MT_-_14674_14742@41.71.31,trnalookalike8_GluTTC_5_-_93905172_93905240@41.71.31|TGGTCGTGGTTGTAGTCCGTGCGAGAATACC</td><td>>trnaMT_GluTTC_MT_-_14674_14742</td><td>41</td><td>71</td><td>31</td><td>TGGTCGTGGTTGTAGTCCGTGCGAGAATACC</td><td>960.5</td><td>9.6</td><td>1.1</td><td>8.4</td><td>3.3E-17</td><td>1.2E-16</td><td>MB231/MB468</td></tr>
<tr><td>>hg38_21_+_8433172_8446622_RNA45SN1_45S_50ntFlanks@12934.12952.19|CTAAACCATTCGTAGACGA</td><td>>hg38_21_+_8433172_8446622_RNA45SN1_45S_50ntFlanks</td><td>12934</td><td>12952</td><td>19</td><td>CTAAACCATTCGTAGACGA</td><td>511.7</td><td>-11.0</td><td>1.3</td><td>-8.4</td><td>3.2E-17</td><td>1.2E-16</td><td>BT20/MB468</td></tr>
<tr><td>>MI0000065|hsa-let-7d&WithFlank&9|+|94178828|94178926@14.35.22(+1A)&[MIMAT0000065&hsa-let-7d-5p&offsets|0|0(+1A);m-90&9|+|94178841|94178862&offsets|0|0(+1A)]|AGAGGTAGTAGGTTGCATAGTTA</td><td>>MI0000065|hsa-let-7d&WithFlank&9|+|94178828|94178926</td><td>14</td><td>35</td><td>22(+1A)</td><td>AGAGGTAGTAGGTTGCATAGTTA</td><td>711.1</td><td>7.5</td><td>0.9</td><td>8.4</td><td>3.2E-17</td><td>1.2E-16</td><td>BT20/MB468</td></tr>
<tr><td>>hg38_21_+_8433172_8446622_RNA45SN1_45S_50ntFlanks@10806.10826.21|AACAATGTAGGTAAGGGAAGT</td><td>>hg38_21_+_8433172_8446622_RNA45SN1_45S_50ntFlanks</td><td>10806</td><td>10826</td><td>21</td><td>AACAATGTAGGTAAGGGAAGT</td><td>333.2</td><td>-5.2</td><td>0.6</td><td>-8.4</td><td>3.2E-17</td><td>1.2E-16</td><td>BT20/MB468</td></tr>
<tr><td>>MI0005756|hsa-mir-934&WithFlank&X|+|136550872|136550966@21.41.21(+1U)&[MIMAT0004977&hsa-miR-934&offsets|0|-1(+1U);m-943&X|+|136550892|136550913&offsets|0|-1(+1U)]|TGTCTACTACTGGAGACACTGT</td><td>>MI0005756|hsa-mir-934&WithFlank&X|+|136550872|136550966</td><td>21</td><td>41</td><td>21(+1U)</td><td>TGTCTACTACTGGAGACACTGT</td><td>2917.6</td><td>9.2</td><td>1.1</td><td>8.4</td><td>3.3E-17</td><td>1.2E-16</td><td>BT20/MB468</td></tr>
<tr><td>>hg38_21_+_8433172_8446622_RNA45SN1_45S_50ntFlanks@8170.8197.28|CAAGTCCTTCTGATCGAGGCCCAGCCCG</td><td>>hg38_21_+_8433172_8446622_RNA45SN1_45S_50ntFlanks</td><td>8170</td><td>8197</td><td>28</td><td>CAAGTCCTTCTGATCGAGGCCCAGCCCG</td><td>662.1</td><td>-6.0</td><td>0.7</td><td>-8.4</td><td>3.4E-17</td><td>1.2E-16</td><td>MB231/MB468</td></tr>
<tr><td>>hg38_21_+_8433172_8446622_RNA45SN1_45S_50ntFlanks@10680.10706.27|GTTCCGGCGGCGTCCGGTGAGCTCTCG</td><td>>hg38_21_+_8433172_8446622_RNA45SN1_45S_50ntFlanks</td><td>10680</td><td>10706</td><td>27</td><td>GTTCCGGCGGCGTCCGGTGAGCTCTCG</td><td>6878.1</td><td>-6.2</td><td>0.7</td><td>-8.4</td><td>3.4E-17</td><td>1.3E-16</td><td>MB231/MB468</td></tr>
<tr><td>>hg38_21_+_8433172_8446622_RNA45SN1_45S_50ntFlanks@11727.11748.22|AGTAACTATGACTCTCTTAAGG</td><td>>hg38_21_+_8433172_8446622_RNA45SN1_45S_50ntFlanks</td><td>11727</td><td>11748</td><td>22</td><td>AGTAACTATGACTCTCTTAAGG</td><td>450.3</td><td>4.1</td><td>0.5</td><td>8.4</td><td>3.5E-17</td><td>1.3E-16</td><td>MB231/MB468</td></tr>
<tr><td>>hg38_21_+_8433172_8446622_RNA45SN1_45S_50ntFlanks@10768.10785.18|CAGCAGGTCTCCAAGGTG</td><td>>hg38_21_+_8433172_8446622_RNA45SN1_45S_50ntFlanks</td><td>10768</td><td>10785</td><td>18</td><td>CAGCAGGTCTCCAAGGTG</td><td>701.6</td><td>-5.5</td><td>0.7</td><td>-8.4</td><td>3.4E-17</td><td>1.3E-16</td><td>BT20/MB468</td></tr>
<tr><td>>MI0000074|hsa-mir-19b-1&WithFlank&13|+|91351186|91351284@60.82.23;MI0000075|hsa-mir-19b-2&WithFlank&X|-|134169665|134169772@68.90.23&[MIMAT0000074&hsa-miR-19b-3p&offsets|0|0;m-122&13|+|91351245|91351267&offsets|0|0];[MIMAT0000074_1&hsa-miR-19b-3p&offsets|0|0;m-121&X|-|134169683|134169705&offsets|0|0]|TGTGCAAATCCATGCAAAACTGA</td><td>>MI0000074|hsa-mir-19b-1&WithFlank&13|+|91351186|91351284</td><td>60</td><td>82</td><td>23</td><td>TGTGCAAATCCATGCAAAACTGA</td><td>2406.1</td><td>7.1</td><td>0.8</td><td>8.4</td><td>3.5E-17</td><td>1.3E-16</td><td>MB231/MB468</td></tr>
<tr><td>>MI0000084|hsa-mir-26b&WithFlank&2|+|218402640|218402728@18.39.22&[MIMAT0000083&hsa-miR-26b-5p&offsets|0|+1;m-53&2|+|218402657|218402678&offsets|0|0]|TTCAAGTAATTCAGGATAGGTT</td><td>>MI0000084|hsa-mir-26b&WithFlank&2|+|218402640|218402728</td><td>18</td><td>39</td><td>22</td><td>TTCAAGTAATTCAGGATAGGTT</td><td>18868.7</td><td>6.5</td><td>0.8</td><td>8.4</td><td>3.6E-17</td><td>1.3E-16</td><td>MB231/MB468</td></tr>
<tr><td>>hg38_21_+_8433172_8446622_RNA45SN1_45S_50ntFlanks@485.529.45|GATGTGGTGACGTCGTGCTCTCCCGGGCCGGGTCCGAGCCGCGAC</td><td>>hg38_21_+_8433172_8446622_RNA45SN1_45S_50ntFlanks</td><td>485</td><td>529</td><td>45</td><td>GATGTGGTGACGTCGTGCTCTCCCGGGCCGGGTCCGAGCCGCGAC</td><td>472.1</td><td>13.5</td><td>1.6</td><td>8.4</td><td>3.4E-17</td><td>1.3E-16</td><td>BT20/MB468</td></tr>
<tr><td>>hg38_21_+_8433172_8446622_RNA45SN1_45S_50ntFlanks@12489.12509.21|ACGTGAGCTGGGTTTAGACCG</td><td>>hg38_21_+_8433172_8446622_RNA45SN1_45S_50ntFlanks</td><td>12489</td><td>12509</td><td>21</td><td>ACGTGAGCTGGGTTTAGACCG</td><td>756.1</td><td>-5.0</td><td>0.6</td><td>-8.4</td><td>3.6E-17</td><td>1.3E-16</td><td>MB231/MB468</td></tr>
<tr><td>>MI0000067|hsa-let-7f-1&WithFlank&9|+|94176341|94176439@13.32.20(+1G);MI0000068|hsa-let-7f-2&WithFlank&X|-|53557186|53557280@14.33.20(+1G)&[MIMAT0000067&hsa-let-7f-5p&offsets|0|-2(+1G);m-20&9|+|94176353|94176374&offsets|0|-2(+1G)];[MIMAT0000067_1&hsa-let-7f-5p&offsets|0|-2(+1G);m-19&X|-|53557246|53557267&offsets|0|-2(+1G)]|TGAGGTAGTAGATTGTATAGG</td><td>>MI0000067|hsa-let-7f-1&WithFlank&9|+|94176341|94176439</td><td>13</td><td>32</td><td>20(+1G)</td><td>TGAGGTAGTAGATTGTATAGG</td><td>1687.5</td><td>7.3</td><td>0.9</td><td>8.4</td><td>3.5E-17</td><td>1.3E-16</td><td>BT20/MB468</td></tr>
<tr><td>>hg38_21_+_8433172_8446622_RNA45SN1_45S_50ntFlanks@12488.12521.34|AACGTGAGCTGGGTTTAGACCGTCGTGAGACAGG</td><td>>hg38_21_+_8433172_8446622_RNA45SN1_45S_50ntFlanks</td><td>12488</td><td>12521</td><td>34</td><td>AACGTGAGCTGGGTTTAGACCGTCGTGAGACAGG</td><td>601.4</td><td>-11.2</td><td>1.3</td><td>-8.4</td><td>3.5E-17</td><td>1.3E-16</td><td>BT20/MB468</td></tr>
<tr><td>>hg38_21_+_8433172_8446622_RNA45SN1_45S_50ntFlanks@6651.6687.37|CGACTCTTAGCGGTGGATCACTCGGCTCGTGCGTCGA</td><td>>hg38_21_+_8433172_8446622_RNA45SN1_45S_50ntFlanks</td><td>6651</td><td>6687</td><td>37</td><td>CGACTCTTAGCGGTGGATCACTCGGCTCGTGCGTCGA</td><td>1997.9</td><td>5.6</td><td>0.7</td><td>8.4</td><td>3.6E-17</td><td>1.4E-16</td><td>BT20/MB468</td></tr>
<tr><td>>hg38_21_+_8433172_8446622_RNA45SN1_45S_50ntFlanks@12458.12487.30|AAGCGTTGGATTGTTCACCCACTAATAGGG</td><td>>hg38_21_+_8433172_8446622_RNA45SN1_45S_50ntFlanks</td><td>12458</td><td>12487</td><td>30</td><td>AAGCGTTGGATTGTTCACCCACTAATAGGG</td><td>618.0</td><td>-6.0</td><td>0.7</td><td>-8.4</td><td>3.6E-17</td><td>1.4E-16</td><td>BT20/MB468</td></tr>
<tr><td>>hg38_21_+_8433172_8446622_RNA45SN1_45S_50ntFlanks@11885.11905.21|TTGAGCTTGACTCTAGTCTGG</td><td>>hg38_21_+_8433172_8446622_RNA45SN1_45S_50ntFlanks</td><td>11885</td><td>11905</td><td>21</td><td>TTGAGCTTGACTCTAGTCTGG</td><td>729.2</td><td>-11.1</td><td>1.3</td><td>-8.4</td><td>3.8E-17</td><td>1.4E-16</td><td>MB231/MB468</td></tr>
<tr><td>>hg38_21_+_8433172_8446622_RNA45SN1_45S_50ntFlanks@12619.12653.35|AGGAGCCAATGGGGCGAAGCTACCATCTGTGGGAT</td><td>>hg38_21_+_8433172_8446622_RNA45SN1_45S_50ntFlanks</td><td>12619</td><td>12653</td><td>35</td><td>AGGAGCCAATGGGGCGAAGCTACCATCTGTGGGAT</td><td>485.1</td><td>-7.3</td><td>0.9</td><td>-8.4</td><td>3.7E-17</td><td>1.4E-16</td><td>BT20/MB468</td></tr>
<tr><td>>hg38_21_+_8433172_8446622_RNA45SN1_45S_50ntFlanks@11561.11588.28|GCGCCTAGCAGCCGACTTAGAACTGGTG</td><td>>hg38_21_+_8433172_8446622_RNA45SN1_45S_50ntFlanks</td><td>11561</td><td>11588</td><td>28</td><td>GCGCCTAGCAGCCGACTTAGAACTGGTG</td><td>459.0</td><td>-4.1</td><td>0.5</td><td>-8.4</td><td>3.8E-17</td><td>1.4E-16</td><td>MB231/MB468</td></tr>
<tr><td>>MI0000736|hsa-mir-30c-1&WithFlank&1|+|40757278|40757378@23.44.22(+1U);MI0000254|hsa-mir-30c-2&WithFlank&6|-|71376954|71377037@13.34.22(+1U)&[MIMAT0000244&hsa-miR-30c-5p&offsets|0|-1(+1U);m-57&1|+|40757300|40757323&offsets|0|-2(+1U)];[MIMAT0000244_1&hsa-miR-30c-5p&offsets|0|-1(+1U);m-56&6|-|71377002|71377025&offsets|0|-2(+1U)]|TGTAAACATCCTACACTCTCAGT</td><td>>MI0000736|hsa-mir-30c-1&WithFlank&1|+|40757278|40757378</td><td>23</td><td>44</td><td>22(+1U)</td><td>TGTAAACATCCTACACTCTCAGT</td><td>11761.4</td><td>6.6</td><td>0.8</td><td>8.4</td><td>3.7E-17</td><td>1.4E-16</td><td>BT20/MB468</td></tr>
<tr><td>>MI0000298|hsa-mir-221&WithFlank&X|-|45746151|45746272@71.91.21(+1A)&[MIMAT0000278&hsa-miR-221-3p&offsets|0|-2(+1A);m-91&X|-|45746181|45746202&offsets|0|-1(+1A)]|AGCTACATTGTCTGCTGGGTTA</td><td>>MI0000298|hsa-mir-221&WithFlank&X|-|45746151|45746272</td><td>71</td><td>91</td><td>21(+1A)</td><td>AGCTACATTGTCTGCTGGGTTA</td><td>890.7</td><td>7.3</td><td>0.9</td><td>8.4</td><td>3.8E-17</td><td>1.4E-16</td><td>BT20/MB468</td></tr>
</table>

<table>
<tr><td>>MI0001445|hsa-mir-423&WithFlank&17|+|30117073|30117178@58.81.24(+1U)&[MIMAT0001340&hsa-miR-423-3p&offsets|-1|0(+1U);m-100&17|+|30117131|30117153&offsets|-1|0(+1U)]|AAGCTCGGTCTGAGGCCCCTCAGTT</td><td>>MI0001445|hsa-mir-423&WithFlank&17|+|30117073|30117178</td><td>58</td><td>81</td><td>24(+1U)</td><td>AAGCTCGGTCTGAGGCCCCTCAGTT</td><td>465.4</td><td>7.9</td><td>0.9</td><td>8.4</td><td>3.8E-17</td><td>1.4E-16</td><td>BT20/MB468</td></tr>
<tr><td>>hg38_21_+_8433172_8446622_RNA45SN1_45S_50ntFlanks@12072.12095.24|CCCGAGGGGCTCTCGCTTCTGGCG</td><td>>hg38_21_+_8433172_8446622_RNA45SN1_45S_50ntFlanks</td><td>12072</td><td>12095</td><td>24</td><td>CCCGAGGGGCTCTCGCTTCTGGCG</td><td>981.8</td><td>-6.6</td><td>0.8</td><td>-8.4</td><td>3.8E-17</td><td>1.4E-16</td><td>BT20/MB468</td></tr>
<tr><td>>hg38_21_+_8433172_8446622_RNA45SN1_45S_50ntFlanks@5489.5512.24|CTGAGAAGACGGTCGAACTTGACT</td><td>>hg38_21_+_8433172_8446622_RNA45SN1_45S_50ntFlanks</td><td>5489</td><td>5512</td><td>24</td><td>CTGAGAAGACGGTCGAACTTGACT</td><td>796.9</td><td>6.2</td><td>0.7</td><td>8.4</td><td>4.0E-17</td><td>1.4E-16</td><td>MB231/MB468</td></tr>
<tr><td>>hg38_21_+_8433172_8446622_RNA45SN1_45S_50ntFlanks@4731.4761.31|AAGAACGAAAGTCGGAGGTTCGAAGACGATC</td><td>>hg38_21_+_8433172_8446622_RNA45SN1_45S_50ntFlanks</td><td>4731</td><td>4761</td><td>31</td><td>AAGAACGAAAGTCGGAGGTTCGAAGACGATC</td><td>751.4</td><td>3.7</td><td>0.4</td><td>8.4</td><td>3.9E-17</td><td>1.5E-16</td><td>BT20/MB468</td></tr>
<tr><td>>hg38_21_+_8433172_8446622_RNA45SN1_45S_50ntFlanks@12000.12032.33|CCTGCGGGCCGCCGGTGAAATACCACTACTCTG</td><td>>hg38_21_+_8433172_8446622_RNA45SN1_45S_50ntFlanks</td><td>12000</td><td>12032</td><td>33</td><td>CCTGCGGGCCGCCGGTGAAATACCACTACTCTG</td><td>235.5</td><td>-6.6</td><td>0.8</td><td>-8.4</td><td>3.9E-17</td><td>1.5E-16</td><td>BT20/MB468</td></tr>
<tr><td>>MI0000255|hsa-mir-30d&WithFlank&8|-|134804870|134804951@12.33.22(+2U)&[MIMAT0000245&hsa-miR-30d-5p&offsets|0|0(+2U);m-21&8|-|134804919|134804940&offsets|0|0(+2U)]|TGTAAACATCCCCGACTGGAAGTT</td><td>>MI0000255|hsa-mir-30d&WithFlank&8|-|134804870|134804951</td><td>12</td><td>33</td><td>22(+2U)</td><td>TGTAAACATCCCCGACTGGAAGTT</td><td>735.6</td><td>10.0</td><td>1.2</td><td>8.4</td><td>4.0E-17</td><td>1.5E-16</td><td>MB231/MB468</td></tr>
<tr><td>>trna129_MetCAT_6_-_27870271_27870342@31.51.21,trna142_MetCAT_6_-_27560600_27560671@31.51.21,trna150_MetCAT_6_-_27300764_27300835@31.51.21,trna169_MetCAT_6_-_26330529_26330600@31.51.21,trna171_MetCAT_6_-_26313352_26313423@31.51.21,trna20_MetCAT_17_-_80452597_80452668@31.51.21,trna2_MetCAT_6_+_26286754_26286825@31.51.21,trna32_MetCAT_1_+_153643726_153643797@31.51.21,trna61_MetCAT_6_+_27745664_27745735@31.51.21|CCCATAACCCAGAGGTCGATG</td><td>>trna129_MetCAT_6_-_27870271_27870342</td><td>31</td><td>51</td><td>21</td><td>CCCATAACCCAGAGGTCGATG</td><td>1059.8</td><td>-11.6</td><td>1.4</td><td>-8.4</td><td>4.0E-17</td><td>1.5E-16</td><td>MB231/MB468</td></tr>
<tr><td>>hg38_21_+_8433172_8446622_RNA45SN1_45S_50ntFlanks@8168.8187.20|CCCAAGTCCTTCTGATCGAG</td><td>>hg38_21_+_8433172_8446622_RNA45SN1_45S_50ntFlanks</td><td>8168</td><td>8187</td><td>20</td><td>CCCAAGTCCTTCTGATCGAG</td><td>492.9</td><td>-6.8</td><td>0.8</td><td>-8.4</td><td>3.9E-17</td><td>1.5E-16</td><td>BT20/MB468</td></tr>
<tr><td>>hg38_21_+_8433172_8446622_RNA45SN1_45S_50ntFlanks@9562.9596.35|TCCTGACGTGCAAATCGGTCGTCCGACCTGGGTAT</td><td>>hg38_21_+_8433172_8446622_RNA45SN1_45S_50ntFlanks</td><td>9562</td><td>9596</td><td>35</td><td>TCCTGACGTGCAAATCGGTCGTCCGACCTGGGTAT</td><td>737.7</td><td>-5.8</td><td>0.7</td><td>-8.4</td><td>4.1E-17</td><td>1.5E-16</td><td>MB231/MB468</td></tr>
<tr><td>>hg38_21_+_8433172_8446622_RNA45SN1_45S_50ntFlanks@7984.8006.23|AGATCAGACGTGGCGACCCGCTG</td><td>>hg38_21_+_8433172_8446622_RNA45SN1_45S_50ntFlanks</td><td>7984</td><td>8006</td><td>23</td><td>AGATCAGACGTGGCGACCCGCTG</td><td>2663.0</td><td>-7.9</td><td>0.9</td><td>-8.4</td><td>4.0E-17</td><td>1.5E-16</td><td>BT20/MB468</td></tr>
<tr><td>>MI0000465|hsa-mir-191&WithFlank&3|-|49020612|49020715@22.44.23&[MIMAT0000440&hsa-miR-191-5p&offsets|0|0;m-48&3|-|49020672|49020694&offsets|0|0]|CAACGGAATCCCAAAAGCAGCTG</td><td>>MI0000465|hsa-mir-191&WithFlank&3|-|49020612|49020715</td><td>22</td><td>44</td><td>23</td><td>CAACGGAATCCCAAAAGCAGCTG</td><td>20718.8</td><td>6.2</td><td>0.7</td><td>8.4</td><td>4.0E-17</td><td>1.5E-16</td><td>BT20/MB468</td></tr>
<tr><td>>MI0000285|hsa-mir-205&WithFlank&1|+|209432127|209432248@40.61.22(+2A)&[MIMAT0000266&hsa-miR-205-5p&offsets|0|0(+2A);m-351&1|+|209432166|209432187&offsets|0|0(+2A)]|TCCTTCATTCCACCGGAGTCTGAA</td><td>>MI0000285|hsa-mir-205&WithFlank&1|+|209432127|209432248</td><td>40</td><td>61</td><td>22(+2A)</td><td>TCCTTCATTCCACCGGAGTCTGAA</td><td>2669.2</td><td>6.3</td><td>0.7</td><td>8.4</td><td>4.1E-17</td><td>1.5E-16</td><td>BT20/MB468</td></tr>
<tr><td>>hg38_21_+_8433172_8446622_RNA45SN1_45S_50ntFlanks@12636.12654.19|AGCTACCATCTGTGGGATT</td><td>>hg38_21_+_8433172_8446622_RNA45SN1_45S_50ntFlanks</td><td>12636</td><td>12654</td><td>19</td><td>AGCTACCATCTGTGGGATT</td><td>2128.2</td><td>-6.3</td><td>0.7</td><td>-8.4</td><td>4.2E-17</td><td>1.5E-16</td><td>MB231/MB468</td></tr>
<tr><td>>hg38_21_+_8433172_8446622_RNA45SN1_45S_50ntFlanks@7977.8009.33|CGACCTCAGATCAGACGTGGCGACCCGCTGAAT</td><td>>hg38_21_+_8433172_8446622_RNA45SN1_45S_50ntFlanks</td><td>7977</td><td>8009</td><td>33</td><td>CGACCTCAGATCAGACGTGGCGACCCGCTGAAT</td><td>451.9</td><td>4.7</td><td>0.6</td><td>8.4</td><td>4.3E-17</td><td>1.5E-16</td><td>MB231/MB468</td></tr>
<tr><td>>MI0003820|hsa-mir-454&WithFlank&17|-|59137752|59137878@70.93.24&[MIMAT0003885&hsa-miR-454-3p&offsets|0|+1;m-272&17|-|59137787|59137809&offsets|0|+1]|TAGTGCAATATTGCTTATAGGGTT</td><td>>MI0003820|hsa-mir-454&WithFlank&17|-|59137752|59137878</td><td>70</td><td>93</td><td>24</td><td>TAGTGCAATATTGCTTATAGGGTT</td><td>2144.5</td><td>7.8</td><td>0.9</td><td>8.4</td><td>4.1E-17</td><td>1.6E-16</td><td>BT20/MB468</td></tr>
<tr><td>>hg38_21_+_8433172_8446622_RNA45SN1_45S_50ntFlanks@12157.12179.23|AGTTTGACTGGGGCGGTACACCT</td><td>>hg38_21_+_8433172_8446622_RNA45SN1_45S_50ntFlanks</td><td>12157</td><td>12179</td><td>23</td><td>AGTTTGACTGGGGCGGTACACCT</td><td>604.1</td><td>-11.2</td><td>1.3</td><td>-8.4</td><td>4.2E-17</td><td>1.6E-16</td><td>BT20/MB468</td></tr>
<tr><td>>trnaMT_ValTAC_MT_+_1602_1670@53.72.20|TAACTTGACCGCTCTGACCA</td><td>>trnaMT_ValTAC_MT_+_1602_1670</td><td>53</td><td>72</td><td>20</td><td>TAACTTGACCGCTCTGACCA</td><td>3618.0</td><td>10.7</td><td>1.3</td><td>8.4</td><td>4.2E-17</td><td>1.6E-16</td><td>BT20/MB468</td></tr>
<tr><td>>hg38_21_+_8433172_8446622_RNA45SN1_45S_50ntFlanks@4588.4614.27|CTATTTTGTTGGTTTTCGGAACTGAGG</td><td>>hg38_21_+_8433172_8446622_RNA45SN1_45S_50ntFlanks</td><td>4588</td><td>4614</td><td>27</td><td>CTATTTTGTTGGTTTTCGGAACTGAGG</td><td>459.8</td><td>-4.9</td><td>0.6</td><td>-8.4</td><td>4.2E-17</td><td>1.6E-16</td><td>BT20/MB468</td></tr>
<tr><td>>am_MI0000786|hsa-mir-378a&WithFlank&5|+|149732819|149732896@49.70.22(+1A)&[MIMAT0000732&hsa-miR-378a-3p&offsets|0|0(+1A);m-44&5|+|149732867|149732888&offsets|0|0(+1A)]|ACTGGACTTGGAGTCAGAAGGCA</td><td>>am_MI0000786|hsa-mir-378a&WithFlank&5|+|149732819|149732896</td><td>49</td><td>70</td><td>22(+1A)</td><td>ACTGGACTTGGAGTCAGAAGGCA</td><td>2773.2</td><td>4.9</td><td>0.6</td><td>8.4</td><td>4.2E-17</td><td>1.6E-16</td><td>BT20/MB468</td></tr>
<tr><td>>am_trnaMT_PheGAA_MT_+_577_647@41.59.19|GTTTAGACGGGCTCACATC</td><td>>am_trnaMT_PheGAA_MT_+_577_647</td><td>41</td><td>59</td><td>19</td><td>GTTTAGACGGGCTCACATC</td><td>907.3</td><td>9.5</td><td>1.1</td><td>8.4</td><td>4.3E-17</td><td>1.6E-16</td><td>BT20/MB468</td></tr>
<tr><td>>hg38_21_+_8433172_8446622_RNA45SN1_45S_50ntFlanks@5490.5516.27|TGAGAAGACGGTCGAACTTGACTATCT</td><td>>hg38_21_+_8433172_8446622_RNA45SN1_45S_50ntFlanks</td><td>5490</td><td>5516</td><td>27</td><td>TGAGAAGACGGTCGAACTTGACTATCT</td><td>1919.1</td><td>10.4</td><td>1.2</td><td>8.4</td><td>4.4E-17</td><td>1.6E-16</td><td>MB231/MB468</td></tr>
<tr><td>>hg38_21_+_8433172_8446622_RNA45SN1_45S_50ntFlanks@12476.12500.25|CCACTAATAGGGAACGTGAGCTGGG</td><td>>hg38_21_+_8433172_8446622_RNA45SN1_45S_50ntFlanks</td><td>12476</td><td>12500</td><td>25</td><td>CCACTAATAGGGAACGTGAGCTGGG</td><td>495.4</td><td>-4.7</td><td>0.6</td><td>-8.4</td><td>4.4E-17</td><td>1.6E-16</td><td>MB231/MB468</td></tr>
<tr><td>>hg38_21_+_8433172_8446622_RNA45SN1_45S_50ntFlanks@11890.11907.18|CTTGACTCTAGTCTGGCA</td><td>>hg38_21_+_8433172_8446622_RNA45SN1_45S_50ntFlanks</td><td>11890</td><td>11907</td><td>18</td><td>CTTGACTCTAGTCTGGCA</td><td>671.6</td><td>-5.3</td><td>0.6</td><td>-8.4</td><td>4.3E-17</td><td>1.6E-16</td><td>BT20/MB468</td></tr>
<tr><td>>hg38_21_+_8433172_8446622_RNA45SN1_45S_50ntFlanks@6746.6771.26|CACTTCGAACGCACTTGCGGCCCCGG</td><td>>hg38_21_+_8433172_8446622_RNA45SN1_45S_50ntFlanks</td><td>6746</td><td>6771</td><td>26</td><td>CACTTCGAACGCACTTGCGGCCCCGG</td><td>574.1</td><td>-10.7</td><td>1.3</td><td>-8.4</td><td>4.4E-17</td><td>1.6E-16</td><td>MB231/MB468</td></tr>
<tr><td>>MI0001445|hsa-mir-423&WithFlank&17|+|30117073|30117178@23.44.22(+1C)&[MIMAT0004748&hsa-miR-423-5p&offsets|0|-1(+1C);m-141&17|+|30117095|30117117&offsets|0|-1(+1C)]|TGAGGGGCAGAGAGCGAGACTTC</td><td>>MI0001445|hsa-mir-423&WithFlank&17|+|30117073|30117178</td><td>23</td><td>44</td><td>22(+1C)</td><td>TGAGGGGCAGAGAGCGAGACTTC</td><td>946.2</td><td>8.9</td><td>1.1</td><td>8.4</td><td>4.5E-17</td><td>1.6E-16</td><td>MB231/MB468</td></tr>
<tr><td>>hg38_21_+_8433172_8446622_RNA45SN1_45S_50ntFlanks@11999.12031.33|CCCTGCGGGCCGCCGGTGAAATACCACTACTCT</td><td>>hg38_21_+_8433172_8446622_RNA45SN1_45S_50ntFlanks</td><td>11999</td><td>12031</td><td>33</td><td>CCCTGCGGGCCGCCGGTGAAATACCACTACTCT</td><td>547.3</td><td>-10.7</td><td>1.3</td><td>-8.4</td><td>4.5E-17</td><td>1.6E-16</td><td>MB231/MB468</td></tr>
<tr><td>>MI0003560|hsa-mir-92b&WithFlank&1|+|155195171|155195278@67.88.22(+1U)&[MIMAT0003218&hsa-miR-92b-3p&offsets|0|0(+1U);m-115&1|+|155195237|155195258&offsets|0|0(+1U)]|TATTGCACTCGTCCCGGCCTCCT</td><td>>MI0003560|hsa-mir-92b&WithFlank&1|+|155195171|155195278</td><td>67</td><td>88</td><td>22(+1U)</td><td>TATTGCACTCGTCCCGGCCTCCT</td><td>1312.6</td><td>9.3</td><td>1.1</td><td>8.4</td><td>4.3E-17</td><td>1.6E-16</td><td>BT20/MB468</td></tr>
<tr><td>>hg38_1_-_228634819_228635039_RNA5S12_5S_50ntFlanks@51.84.34|GTCTACGGCCATACCACCCTGAACGCGCCCGATC</td><td>>hg38_1_-_228634819_228635039_RNA5S12_5S_50ntFlanks</td><td>51</td><td>84</td><td>34</td><td>GTCTACGGCCATACCACCCTGAACGCGCCCGATC</td><td>5377.6</td><td>6.3</td><td>0.7</td><td>8.4</td><td>4.4E-17</td><td>1.6E-16</td><td>BT20/MB468</td></tr>
<tr><td>>hg38_21_+_8433172_8446622_RNA45SN1_45S_50ntFlanks@12626.12653.28|AATGGGGCGAAGCTACCATCTGTGGGAT</td><td>>hg38_21_+_8433172_8446622_RNA45SN1_45S_50ntFlanks</td><td>12626</td><td>12653</td><td>28</td><td>AATGGGGCGAAGCTACCATCTGTGGGAT</td><td>4055.6</td><td>-9.2</td><td>1.1</td><td>-8.4</td><td>4.4E-17</td><td>1.6E-16</td><td>BT20/MB468</td></tr>
<tr><td>>hg38_21_+_8433172_8446622_RNA45SN1_45S_50ntFlanks@11613.11640.28|AATTAAAACAAAGCATCGCGAAGGCCCG</td><td>>hg38_21_+_8433172_8446622_RNA45SN1_45S_50ntFlanks</td><td>11613</td><td>11640</td><td>28</td><td>AATTAAAACAAAGCATCGCGAAGGCCCG</td><td>477.5</td><td>-8.0</td><td>0.9</td><td>-8.4</td><td>4.6E-17</td><td>1.6E-16</td><td>MB231/MB468</td></tr>
<tr><td>>hg38_21_+_8433172_8446622_RNA45SN1_45S_50ntFlanks@11594.11611.18|CAGGGGAATCCGACTGTT</td><td>>hg38_21_+_8433172_8446622_RNA45SN1_45S_50ntFlanks</td><td>11594</td><td>11611</td><td>18</td><td>CAGGGGAATCCGACTGTT</td><td>1764.4</td><td>-3.9</td><td>0.5</td><td>-8.4</td><td>4.4E-17</td><td>1.6E-16</td><td>BT20/MB468</td></tr>
<tr><td>>hg38_21_+_8433172_8446622_RNA45SN1_45S_50ntFlanks@12625.12653.29|CAATGGGGCGAAGCTACCATCTGTGGGAT</td><td>>hg38_21_+_8433172_8446622_RNA45SN1_45S_50ntFlanks</td><td>12625</td><td>12653</td><td>29</td><td>CAATGGGGCGAAGCTACCATCTGTGGGAT</td><td>1210.0</td><td>-5.8</td><td>0.7</td><td>-8.4</td><td>4.6E-17</td><td>1.7E-16</td><td>MB231/MB468</td></tr>
<tr><td>>hg38_21_+_8433172_8446622_RNA45SN1_45S_50ntFlanks@12997.13017.21|CGCTGCGATCTATTGAAAGTC</td><td>>hg38_21_+_8433172_8446622_RNA45SN1_45S_50ntFlanks</td><td>12997</td><td>13017</td><td>21</td><td>CGCTGCGATCTATTGAAAGTC</td><td>1356.0</td><td>6.0</td><td>0.7</td><td>8.4</td><td>4.5E-17</td><td>1.7E-16</td><td>BT20/MB468</td></tr>
<tr><td>>hg38_21_+_8433172_8446622_RNA45SN1_45S_50ntFlanks@4045.4063.19|CCATTCGAACGTCTGCCCT</td><td>>hg38_21_+_8433172_8446622_RNA45SN1_45S_50ntFlanks</td><td>4045</td><td>4063</td><td>19</td><td>CCATTCGAACGTCTGCCCT</td><td>809.8</td><td>-4.8</td><td>0.6</td><td>-8.4</td><td>4.7E-17</td><td>1.7E-16</td><td>MB231/MB468</td></tr>
<tr><td>>MI0000103|hsa-mir-101-1&WithFlank&1|-|65058428|65058514@53.73.21(+1A);MI0000739|hsa-mir-101-2&WithFlank&9|+|4850291|4850381@55.75.21(+1A)&[MIMAT0000099&hsa-miR-101-3p&offsets|0|0(+1A);m-17&1|-|65058443|65058462&offsets|0|+1(+1A)];[MIMAT0000099_1&hsa-miR-101-3p&offsets|0|0(+1A);m-16&9|+|4850345|4850364&offsets|0|+1(+1A)]|TACAGTACTGTGATAACTGAAA</td><td>>MI0000103|hsa-mir-101-1&WithFlank&1|-|65058428|65058514</td><td>53</td><td>73</td><td>21(+1A)</td><td>TACAGTACTGTGATAACTGAAA</td><td>1071.5</td><td>7.0</td><td>0.8</td><td>8.4</td><td>4.7E-17</td><td>1.7E-16</td><td>MB231/MB468</td></tr>
<tr><td>>hg38_21_+_8433172_8446622_RNA45SN1_45S_50ntFlanks@10771.10797.27|CAGGTCTCCAAGGTGAACAGCCTCTGG</td><td>>hg38_21_+_8433172_8446622_RNA45SN1_45S_50ntFlanks</td><td>10771</td><td>10797</td><td>27</td><td>CAGGTCTCCAAGGTGAACAGCCTCTGG</td><td>538.9</td><td>-11.1</td><td>1.3</td><td>-8.4</td><td>4.6E-17</td><td>1.7E-16</td><td>BT20/MB468</td></tr>
<tr><td>>hg38_1_-_228634819_228635039_RNA5S12_5S_50ntFlanks@149.171.23|GAATACCGGGTGCTGTAGGCTTT</td><td>>hg38_1_-_228634819_228635039_RNA5S12_5S_50ntFlanks</td><td>149</td><td>171</td><td>23</td><td>GAATACCGGGTGCTGTAGGCTTT</td><td>1673.7</td><td>5.5</td><td>0.6</td><td>8.4</td><td>4.6E-17</td><td>1.7E-16</td><td>BT20/MB468</td></tr>
<tr><td>>hg38_21_+_8433172_8446622_RNA45SN1_45S_50ntFlanks@12033.12053.21|ATCGTTTTTTCACTGACCCGG</td><td>>hg38_21_+_8433172_8446622_RNA45SN1_45S_50ntFlanks</td><td>12033</td><td>12053</td><td>21</td><td>ATCGTTTTTTCACTGACCCGG</td><td>571.2</td><td>-11.2</td><td>1.3</td><td>-8.4</td><td>4.6E-17</td><td>1.7E-16</td><td>BT20/MB468</td></tr>
<tr><td>>MI0000433|hsa-let-7g&WithFlank&3|-|52268272|52268367@11.33.23&[MIMAT0000414&hsa-let-7g-5p&offsets|0|+1;m-58&3|-|52268336|52268357&offsets|0|+1]|TGAGGTAGTAGTTTGTACAGTTT</td><td>>MI0000433|hsa-let-7g&WithFlank&3|-|52268272|52268367</td><td>11</td><td>33</td><td>23</td><td>TGAGGTAGTAGTTTGTACAGTTT</td><td>2430.7</td><td>6.3</td><td>0.7</td><td>8.4</td><td>4.7E-17</td><td>1.7E-16</td><td>BT20/MB468</td></tr>
</table>

<table>
<tr><td>>am_trna134_GluTTC_1_-_16861774_16861845@1.29.29,trna5_GluTTC_1_+_17199078_17199149@1.29.29,trna84_GluTTC_1_-_161391883_161391954@1.29.29,trna94_GluTTC_1_-_149664355_149664427@1.29.29|TCCCTGGTGGTCTAGTGGCTAGGATTCGG</td><td>>am_trna134_GluTTC_1_-_16861774_16861845</td><td>1</td><td>29</td><td>29</td><td>TCCCTGGTGGTCTAGTGGCTAGGATTCGG</td><td>812.3</td><td>-7.2</td><td>0.9</td><td>-8.4</td><td>4.8E-17</td><td>1.7E-16</td><td>MB231/MB468</td></tr>
<tr><td>>hg38_21_+_8433172_8446622_RNA45SN1_45S_50ntFlanks@6738.6772.35|ATCATCGACACTTCGAACGCACTTGCGGCCCCGGG</td><td>>hg38_21_+_8433172_8446622_RNA45SN1_45S_50ntFlanks</td><td>6738</td><td>6772</td><td>35</td><td>ATCATCGACACTTCGAACGCACTTGCGGCCCCGGG</td><td>538.9</td><td>-11.1</td><td>1.3</td><td>-8.4</td><td>4.7E-17</td><td>1.8E-16</td><td>BT20/MB468</td></tr>
<tr><td>>hg38_21_+_8433172_8446622_RNA45SN1_45S_50ntFlanks@10362.10384.23|AAGGCCGAAGTGGAGAAGGGTTC</td><td>>hg38_21_+_8433172_8446622_RNA45SN1_45S_50ntFlanks</td><td>10362</td><td>10384</td><td>23</td><td>AAGGCCGAAGTGGAGAAGGGTTC</td><td>609.0</td><td>-7.8</td><td>0.9</td><td>-8.4</td><td>4.8E-17</td><td>1.8E-16</td><td>BT20/MB468</td></tr>
<tr><td>>hg38_MT_+_1621_3279_RNR2_16S_50ntFlanks@1439.1457.19|ACGTGATCTGAGTTCAGAC</td><td>>hg38_MT_+_1621_3279_RNR2_16S_50ntFlanks</td><td>1439</td><td>1457</td><td>19</td><td>ACGTGATCTGAGTTCAGAC</td><td>461.5</td><td>8.7</td><td>1.0</td><td>8.4</td><td>5.0E-17</td><td>1.8E-16</td><td>MB231/MB468</td></tr>
<tr><td>>hg38_21_+_8433172_8446622_RNA45SN1_45S_50ntFlanks@10349.10377.29|AACGAGAACTTTGAAGGCCGAAGTGGAGA</td><td>>hg38_21_+_8433172_8446622_RNA45SN1_45S_50ntFlanks</td><td>10349</td><td>10377</td><td>29</td><td>AACGAGAACTTTGAAGGCCGAAGTGGAGA</td><td>945.8</td><td>-3.0</td><td>0.4</td><td>-8.4</td><td>4.8E-17</td><td>1.8E-16</td><td>BT20/MB468</td></tr>
<tr><td>>MI0000081|hsa-mir-24-2&WithFlank&19|-|13836281|13836365@56.75.20;MI0000080|hsa-mir-24-1&WithFlank&9|+|95086015|95086094@50.69.20&[MIMAT0000080&hsa-miR-24-3p&offsets|0|-2;m-36&19|-|13836288|13836310&offsets|0|-3];[MIMAT0000080_1&hsa-miR-24-3p&offsets|0|-2;m-35&9|+|95086064|95086086&offsets|0|-3]|TGGCTCAGTTCAGCAGGAAC</td><td>>MI0000081|hsa-mir-24-2&WithFlank&19|-|13836281|13836365</td><td>56</td><td>75</td><td>20</td><td>TGGCTCAGTTCAGCAGGAAC</td><td>31016.9</td><td>6.4</td><td>0.8</td><td>8.4</td><td>5.0E-17</td><td>1.8E-16</td><td>MB231/MB468</td></tr>
<tr><td>>trnaMT_ValTAC_MT_+_1602_1670@54.71.18|AACTTGACCGCTCTGACC</td><td>>trnaMT_ValTAC_MT_+_1602_1670</td><td>54</td><td>71</td><td>18</td><td>AACTTGACCGCTCTGACC</td><td>1402.6</td><td>10.1</td><td>1.2</td><td>8.4</td><td>5.1E-17</td><td>1.8E-16</td><td>MB231/MB468</td></tr>
<tr><td>>am_MI0006370|hsa-mir-1303&WithFlank&5|+|154685770|154685867@57.79.23&[MIMAT0005891&hsa-miR-1303&offsets|-1|0;m-516&5|+|154685826|154685848&offsets|0|0]|TTTTAGAGACGGGGTCTTGCTCT</td><td>>am_MI0006370|hsa-mir-1303&WithFlank&5|+|154685770|154685867</td><td>57</td><td>79</td><td>23</td><td>TTTTAGAGACGGGGTCTTGCTCT</td><td>453.3</td><td>5.8</td><td>0.7</td><td>8.4</td><td>4.9E-17</td><td>1.8E-16</td><td>BT20/MB468</td></tr>
<tr><td>>MI0000103|hsa-mir-101-1&WithFlank&1|-|65058428|65058514@53.74.22(+1U);MI0000739|hsa-mir-101-2&WithFlank&9|+|4850291|4850381@55.76.22(+1U)&[MIMAT0000099&hsa-miR-101-3p&offsets|0|+1(+1U);m-17&1|-|65058443|65058462&offsets|0|+2(+1U)];[MIMAT0000099_1&hsa-miR-101-3p&offsets|0|+1(+1U);m-16&9|+|4850345|4850364&offsets|0|+2(+1U)]|TACAGTACTGTGATAACTGAAGT</td><td>>MI0000103|hsa-mir-101-1&WithFlank&1|-|65058428|65058514</td><td>53</td><td>74</td><td>22(+1U)</td><td>TACAGTACTGTGATAACTGAAGT</td><td>3149.3</td><td>8.4</td><td>1.0</td><td>8.4</td><td>5.0E-17</td><td>1.8E-16</td><td>BT20/MB468</td></tr>
<tr><td>>hg38_21_+_8433172_8446622_RNA45SN1_45S_50ntFlanks@6743.6771.29|CGACACTTCGAACGCACTTGCGGCCCCGG</td><td>>hg38_21_+_8433172_8446622_RNA45SN1_45S_50ntFlanks</td><td>6743</td><td>6771</td><td>29</td><td>CGACACTTCGAACGCACTTGCGGCCCCGG</td><td>1308.1</td><td>-5.9</td><td>0.7</td><td>-8.4</td><td>5.1E-17</td><td>1.8E-16</td><td>MB231/MB468</td></tr>
<tr><td>>hg38_21_+_8433172_8446622_RNA45SN1_45S_50ntFlanks@13004.13025.22|ATCTATTGAAAGTCAGCCCTCG</td><td>>hg38_21_+_8433172_8446622_RNA45SN1_45S_50ntFlanks</td><td>13004</td><td>13025</td><td>22</td><td>ATCTATTGAAAGTCAGCCCTCG</td><td>696.5</td><td>-11.4</td><td>1.4</td><td>-8.4</td><td>5.0E-17</td><td>1.8E-16</td><td>BT20/MB468</td></tr>
<tr><td>>MI0000471|hsa-mir-126&WithFlank&9|+|136670596|136670692@58.77.20(+1U)&[MIMAT0000445&hsa-miR-126-3p&offsets|0|-2(+1U);m-40&9|+|136670653|136670674&offsets|0|-2(+1U)]|TCGTACCGTGAGTAATAATGT</td><td>>MI0000471|hsa-mir-126&WithFlank&9|+|136670596|136670692</td><td>58</td><td>77</td><td>20(+1U)</td><td>TCGTACCGTGAGTAATAATGT</td><td>2019.4</td><td>10.5</td><td>1.3</td><td>8.4</td><td>5.2E-17</td><td>1.9E-16</td><td>MB231/MB468</td></tr>
<tr><td>>hg38_21_+_8433172_8446622_RNA45SN1_45S_50ntFlanks@10808.10825.18|CAATGTAGGTAAGGGAAG</td><td>>hg38_21_+_8433172_8446622_RNA45SN1_45S_50ntFlanks</td><td>10808</td><td>10825</td><td>18</td><td>CAATGTAGGTAAGGGAAG</td><td>847.7</td><td>-11.7</td><td>1.4</td><td>-8.4</td><td>5.1E-17</td><td>1.9E-16</td><td>BT20/MB468</td></tr>
<tr><td>>hg38_21_+_8433172_8446622_RNA45SN1_45S_50ntFlanks@3773.3792.20|CGGCCGGTACAGTGAAACTG</td><td>>hg38_21_+_8433172_8446622_RNA45SN1_45S_50ntFlanks</td><td>3773</td><td>3792</td><td>20</td><td>CGGCCGGTACAGTGAAACTG</td><td>228.4</td><td>-5.4</td><td>0.6</td><td>-8.4</td><td>5.1E-17</td><td>1.9E-16</td><td>BT20/MB468</td></tr>
<tr><td>>trnaMT_SerGCT_MT_+_12207_12265@14.62.49|AGAACTGCTAACTCATGCCCCCATGTCTAACAACATGGCTTTCTCACCA</td><td>>trnaMT_SerGCT_MT_+_12207_12265</td><td>14</td><td>62</td><td>49</td><td>AGAACTGCTAACTCATGCCCCCATGTCTAACAACATGGCTTTCTCACCA</td><td>1120.5</td><td>14.8</td><td>1.8</td><td>8.4</td><td>5.1E-17</td><td>1.9E-16</td><td>BT20/MB468</td></tr>
<tr><td>>MI0000070|hsa-mir-16-1&WithFlank&13|-|50048967|50049067@20.39.20(+1U);MI0000115|hsa-mir-16-2&WithFlank&3|+|160404739|160404831@16.35.20(+1U)&[MIMAT0000069&hsa-miR-16-5p&offsets|0|-2(+1U);m-62&13|-|50049027|50049048&offsets|0|-2(+1U)];[MIMAT0000069_1&hsa-miR-16-5p&offsets|0|-2(+1U);m-61&3|+|160404754|160404775&offsets|0|-2(+1U)]|TAGCAGCACGTAAATATTGGT</td><td>>MI0000070|hsa-mir-16-1&WithFlank&13|-|50048967|50049067</td><td>20</td><td>39</td><td>20(+1U)</td><td>TAGCAGCACGTAAATATTGGT</td><td>2597.7</td><td>8.3</td><td>1.0</td><td>8.4</td><td>5.1E-17</td><td>1.9E-16</td><td>BT20/MB468</td></tr>
<tr><td>>hg38_21_+_8433172_8446622_RNA45SN1_45S_50ntFlanks@12454.12477.24|CACCAAGCGTTGGATTGTTCACCC</td><td>>hg38_21_+_8433172_8446622_RNA45SN1_45S_50ntFlanks</td><td>12454</td><td>12477</td><td>24</td><td>CACCAAGCGTTGGATTGTTCACCC</td><td>365.6</td><td>3.8</td><td>0.5</td><td>8.4</td><td>5.3E-17</td><td>1.9E-16</td><td>MB231/MB468</td></tr>
<tr><td>>hg38_21_+_8433172_8446622_RNA45SN1_45S_50ntFlanks@12637.12659.23|GCTACCATCTGTGGGATTATGAC</td><td>>hg38_21_+_8433172_8446622_RNA45SN1_45S_50ntFlanks</td><td>12637</td><td>12659</td><td>23</td><td>GCTACCATCTGTGGGATTATGAC</td><td>331.1</td><td>-4.6</td><td>0.5</td><td>-8.4</td><td>5.3E-17</td><td>1.9E-16</td><td>BT20/MB468</td></tr>
<tr><td>>am_MI0000786|hsa-mir-378a&WithFlank&5|+|149732819|149732896@50.70.21&[MIMAT0000732&hsa-miR-378a-3p&offsets|+1|0;m-44&5|+|149732867|149732888&offsets|+1|0]|CTGGACTTGGAGTCAGAAGGC</td><td>>am_MI0000786|hsa-mir-378a&WithFlank&5|+|149732819|149732896</td><td>50</td><td>70</td><td>21</td><td>CTGGACTTGGAGTCAGAAGGC</td><td>2096.7</td><td>5.0</td><td>0.6</td><td>8.4</td><td>5.3E-17</td><td>1.9E-16</td><td>BT20/MB468</td></tr>
<tr><td>>hg38_21_+_8433172_8446622_RNA45SN1_45S_50ntFlanks@12934.12972.39|CTAAACCATTCGTAGACGACCTGCTTCTGGGTCGGGGTT</td><td>>hg38_21_+_8433172_8446622_RNA45SN1_45S_50ntFlanks</td><td>12934</td><td>12972</td><td>39</td><td>CTAAACCATTCGTAGACGACCTGCTTCTGGGTCGGGGTT</td><td>1000.6</td><td>-5.8</td><td>0.7</td><td>-8.4</td><td>5.4E-17</td><td>2.0E-16</td><td>MB231/MB468</td></tr>
<tr><td>>am_MI0016746|hsa-mir-548o-2&WithFlank&20|+|38516557|38516638@51.71.21(+1U);MI0006402|hsa-mir-548o&WithFlank&7|-|102405736|102405861@93.113.21(+1U)&[MIMAT0005919&hsa-miR-548o-3p&offsets|0|-1(+1U);m-431&20|+|38516607|38516628&offsets|0|-1(+1U)];[MIMAT0005919_1&hsa-miR-548o-3p&offsets|0|-1(+1U);m-430&7|-|102405748|102405769&offsets|0|-1(+1U)]|CCAAAACTGCAGTTACTTTTGT</td><td>>am_MI0016746|hsa-mir-548o-2&WithFlank&20|+|38516557|38516638</td><td>51</td><td>71</td><td>21(+1U)</td><td>CCAAAACTGCAGTTACTTTTGT</td><td>512.5</td><td>10.5</td><td>1.2</td><td>8.4</td><td>5.3E-17</td><td>2.0E-16</td><td>BT20/MB468</td></tr>
<tr><td>>MI0000108|hsa-mir-103a-2&WithFlank&20|+|3917488|3917577@54.77.24(+1U);MI0000109|hsa-mir-103a-1&WithFlank&5|-|168560890|168560979@54.77.24(+1U)&[MIMAT0000101&hsa-miR-103a-3p&offsets|0|+1(+1U);m-12&20|+|3917541|3917563&offsets|0|+1(+1U)];[MIMAT0000101_1&hsa-miR-103a-3p&offsets|0|+1(+1U);m-13&5|-|168560904|168560926&offsets|0|+1(+1U);m-44220&5|-|168560908|168560926&offsets|0|+5(+1U)]|AGCAGCATTGTACAGGGCTATGAAT</td><td>>MI0000108|hsa-mir-103a-2&WithFlank&20|+|3917488|3917577</td><td>54</td><td>77</td><td>24(+1U)</td><td>AGCAGCATTGTACAGGGCTATGAAT</td><td>834.0</td><td>7.2</td><td>0.9</td><td>8.4</td><td>5.4E-17</td><td>2.0E-16</td><td>BT20/MB468</td></tr>
<tr><td>>am_trnaMT_PheGAA_MT_+_577_647@43.62.20|TTAGACGGGCTCACATCACC</td><td>>am_trnaMT_PheGAA_MT_+_577_647</td><td>43</td><td>62</td><td>20</td><td>TTAGACGGGCTCACATCACC</td><td>429.4</td><td>8.3</td><td>1.0</td><td>8.4</td><td>5.6E-17</td><td>2.0E-16</td><td>MB231/MB468</td></tr>
<tr><td>>MI0000283|hsa-mir-203a&WithFlank&14|+|104117399|104117520@71.92.22&[MIMAT0000264&hsa-miR-203a-3p&offsets|0|0;m-79&14|+|104117469|104117490&offsets|0|0]|GTGAAATGTTTAGGACCACTAG</td><td>>MI0000283|hsa-mir-203a&WithFlank&14|+|104117399|104117520</td><td>71</td><td>92</td><td>22</td><td>GTGAAATGTTTAGGACCACTAG</td><td>5737.0</td><td>8.3</td><td>1.0</td><td>8.4</td><td>5.5E-17</td><td>2.0E-16</td><td>BT20/MB468</td></tr>
<tr><td>>hg38_21_+_8433172_8446622_RNA45SN1_45S_50ntFlanks@11613.11640.28|AATTAAAACAAAGCATCGCGAAGGCCCG</td><td>>hg38_21_+_8433172_8446622_RNA45SN1_45S_50ntFlanks</td><td>11613</td><td>11640</td><td>28</td><td>AATTAAAACAAAGCATCGCGAAGGCCCG</td><td>578.8</td><td>-11.2</td><td>1.3</td><td>-8.4</td><td>5.5E-17</td><td>2.0E-16</td><td>BT20/MB468</td></tr>
<tr><td>>MI0000084|hsa-mir-26b&WithFlank&2|+|218402640|218402728@18.38.21(+1A)&[MIMAT0000083&hsa-miR-26b-5p&offsets|0|0(+1A);m-53&2|+|218402657|218402678&offsets|0|-1(+1A)]|TTCAAGTAATTCAGGATAGGTA</td><td>>MI0000084|hsa-mir-26b&WithFlank&2|+|218402640|218402728</td><td>18</td><td>38</td><td>21(+1A)</td><td>TTCAAGTAATTCAGGATAGGTA</td><td>597.4</td><td>6.6</td><td>0.8</td><td>8.4</td><td>5.7E-17</td><td>2.0E-16</td><td>MB231/MB468</td></tr>
<tr><td>>MI0000434|hsa-let-7i&WithFlank&12|+|62603680|62603775@12.33.22(+2U)&[MIMAT0000415&hsa-let-7i-5p&offsets|0|0(+2U);m-49&12|+|62603691|62603712&offsets|0|0(+2U)]|TGAGGTAGTAGTTTGTGCTGTTTT</td><td>>MI0000434|hsa-let-7i&WithFlank&12|+|62603680|62603775</td><td>12</td><td>33</td><td>22(+2U)</td><td>TGAGGTAGTAGTTTGTGCTGTTTT</td><td>1097.5</td><td>10.0</td><td>1.2</td><td>8.4</td><td>5.7E-17</td><td>2.0E-16</td><td>MB231/MB468</td></tr>
<tr><td>>hg38_21_+_8433172_8446622_RNA45SN1_45S_50ntFlanks@11718.11742.25|AACGGCGGGAGTAACTATGACTCTC</td><td>>hg38_21_+_8433172_8446622_RNA45SN1_45S_50ntFlanks</td><td>11718</td><td>11742</td><td>25</td><td>AACGGCGGGAGTAACTATGACTCTC</td><td>1323.9</td><td>-5.6</td><td>0.7</td><td>-8.4</td><td>5.6E-17</td><td>2.1E-16</td><td>BT20/MB468</td></tr>
<tr><td>>hg38_21_+_8433172_8446622_RNA45SN1_45S_50ntFlanks@4728.4752.25|ATCAAGAACGAAAGTCGGAGGTTCG</td><td>>hg38_21_+_8433172_8446622_RNA45SN1_45S_50ntFlanks</td><td>4728</td><td>4752</td><td>25</td><td>ATCAAGAACGAAAGTCGGAGGTTCG</td><td>362.7</td><td>-10.5</td><td>1.3</td><td>-8.4</td><td>5.6E-17</td><td>2.1E-16</td><td>BT20/MB468</td></tr>
<tr><td>>MI0000285|hsa-mir-205&WithFlank&1|+|209432127|209432248@40.59.20(+1G)&[MIMAT0000266&hsa-miR-205-5p&offsets|0|-2(+1G);m-351&1|+|209432166|209432187&offsets|0|-2(+1G)]|TCCTTCATTCCACCGGAGTCG</td><td>>MI0000285|hsa-mir-205&WithFlank&1|+|209432127|209432248</td><td>40</td><td>59</td><td>20(+1G)</td><td>TCCTTCATTCCACCGGAGTCG</td><td>2384.3</td><td>8.4</td><td>1.0</td><td>8.4</td><td>5.7E-17</td><td>2.1E-16</td><td>BT20/MB468</td></tr>
<tr><td>>hg38_21_+_8433172_8446622_RNA45SN1_45S_50ntFlanks@3875.3898.24|ATACATGCCGACGGGCGCTGACCC</td><td>>hg38_21_+_8433172_8446622_RNA45SN1_45S_50ntFlanks</td><td>3875</td><td>3898</td><td>24</td><td>ATACATGCCGACGGGCGCTGACCC</td><td>1968.9</td><td>-6.4</td><td>0.8</td><td>-8.4</td><td>5.7E-17</td><td>2.1E-16</td><td>BT20/MB468</td></tr>
<tr><td>>MI0000542|hsa-mir-320a&WithFlank&8|-|22244960|22245043@48.69.22(+1C)&[MIMAT0000510&hsa-miR-320a-3p&offsets|0|0(+1C);m-60&8|-|22244975|22244996&offsets|0|0(+1C)]|AAAAGCTGGGTTGAGAGGGCGAC</td><td>>MI0000542|hsa-mir-320a&WithFlank&8|-|22244960|22245043</td><td>48</td><td>69</td><td>22(+1C)</td><td>AAAAGCTGGGTTGAGAGGGCGAC</td><td>2841.8</td><td>11.0</td><td>1.3</td><td>8.4</td><td>5.7E-17</td><td>2.1E-16</td><td>BT20/MB468</td></tr>
<tr><td>>hg38_21_+_8433172_8446622_RNA45SN1_45S_50ntFlanks@6651.6676.26|CGACTCTTAGCGGTGGATCACTCGGC</td><td>>hg38_21_+_8433172_8446622_RNA45SN1_45S_50ntFlanks</td><td>6651</td><td>6676</td><td>26</td><td>CGACTCTTAGCGGTGGATCACTCGGC</td><td>3955.1</td><td>5.9</td><td>0.7</td><td>8.4</td><td>5.9E-17</td><td>2.1E-16</td><td>MB231/MB468</td></tr>
<tr><td>>MI0000441|hsa-mir-30b&WithFlank&8|-|134800514|134800613@23.43.21(+1A)&[MIMAT0000420&hsa-miR-30b-5p&offsets|0|-1(+1A);m-76&8|-|134800570|134800591&offsets|0|-1(+1A)]|TGTAAACATCCTACACTCAGCA</td><td>>MI0000441|hsa-mir-30b&WithFlank&8|-|134800514|134800613</td><td>23</td><td>43</td><td>21(+1A)</td><td>TGTAAACATCCTACACTCAGCA</td><td>1660.9</td><td>9.6</td><td>1.2</td><td>8.4</td><td>5.9E-17</td><td>2.1E-16</td><td>MB231/MB468</td></tr>
<tr><td>>hg38_21_+_8433172_8446622_RNA45SN1_45S_50ntFlanks@11575.11600.26|ACTTAGAACTGGTGCGGACCAGGGGA</td><td>>hg38_21_+_8433172_8446622_RNA45SN1_45S_50ntFlanks</td><td>11575</td><td>11600</td><td>26</td><td>ACTTAGAACTGGTGCGGACCAGGGGA</td><td>642.6</td><td>-5.0</td><td>0.6</td><td>-8.4</td><td>5.8E-17</td><td>2.1E-16</td><td>BT20/MB468</td></tr>
</table>

<table>
<tr><td>>hg38_21_+_8433172_8446622_RNA45SN1_45S_50ntFlanks@9738.9755.18|GAAACGATCTCAACCTAT</td><td>>hg38_21_+_8433172_8446622_RNA45SN1_45S_50ntFlanks</td><td>9738</td><td>9755</td><td>18</td><td>GAAACGATCTCAACCTAT</td><td>657.2</td><td>-11.4</td><td>1.4</td><td>-8.4</td><td>5.9E-17</td><td>2.2E-16</td><td>BT20/MB468</td></tr>
<tr><td>>hg38_21_+_8433172_8446622_RNA45SN1_45S_50ntFlanks@10820.10840.21|GGGAAGTCGGCAAGCCGGATC</td><td>>hg38_21_+_8433172_8446622_RNA45SN1_45S_50ntFlanks</td><td>10820</td><td>10840</td><td>21</td><td>GGGAAGTCGGCAAGCCGGATC</td><td>1016.4</td><td>-5.1</td><td>0.6</td><td>-8.4</td><td>6.0E-17</td><td>2.2E-16</td><td>BT20/MB468</td></tr>
<tr><td>>hg38_21_+_8433172_8446622_RNA45SN1_45S_50ntFlanks@11588.11611.24|GCGGACCAGGGGAATCCGACTGTT</td><td>>hg38_21_+_8433172_8446622_RNA45SN1_45S_50ntFlanks</td><td>11588</td><td>11611</td><td>24</td><td>GCGGACCAGGGGAATCCGACTGTT</td><td>704.6</td><td>4.1</td><td>0.5</td><td>8.4</td><td>6.1E-17</td><td>2.2E-16</td><td>MB231/MB468</td></tr>
<tr><td>>hg38_21_+_8433172_8446622_RNA45SN1_45S_50ntFlanks@9504.9525.22|GTGAACTATGCCTGGGCAGGGC</td><td>>hg38_21_+_8433172_8446622_RNA45SN1_45S_50ntFlanks</td><td>9504</td><td>9525</td><td>22</td><td>GTGAACTATGCCTGGGCAGGGC</td><td>797.2</td><td>-7.0</td><td>0.8</td><td>-8.4</td><td>6.0E-17</td><td>2.2E-16</td><td>BT20/MB468</td></tr>
<tr><td>>MI0000283|hsa-mir-203a&WithFlank&14|+|104117399|104117520@71.93.23&[MIMAT0000264&hsa-miR-203a-3p&offsets|0|+1;m-79&14|+|104117469|104117490&offsets|0|+1]|GTGAAATGTTTAGGACCACTAGA</td><td>>MI0000283|hsa-mir-203a&WithFlank&14|+|104117399|104117520</td><td>71</td><td>93</td><td>23</td><td>GTGAAATGTTTAGGACCACTAGA</td><td>513.4</td><td>12.5</td><td>1.4</td><td>8.7</td><td>3.6E-18</td><td>2.2E-16</td><td>BT20/MB231</td></tr>
<tr><td>>hg38_21_+_8433172_8446622_RNA45SN1_45S_50ntFlanks@11564.11586.23|CCTAGCAGCCGACTTAGAACTGG</td><td>>hg38_21_+_8433172_8446622_RNA45SN1_45S_50ntFlanks</td><td>11564</td><td>11586</td><td>23</td><td>CCTAGCAGCCGACTTAGAACTGG</td><td>585.4</td><td>-3.6</td><td>0.4</td><td>-8.4</td><td>6.1E-17</td><td>2.2E-16</td><td>BT20/MB468</td></tr>
<tr><td>>hg38_21_+_8433172_8446622_RNA45SN1_45S_50ntFlanks@12458.12496.39|AAGCGTTGGATTGTTCACCCACTAATAGGGAACGTGAGC</td><td>>hg38_21_+_8433172_8446622_RNA45SN1_45S_50ntFlanks</td><td>12458</td><td>12496</td><td>39</td><td>AAGCGTTGGATTGTTCACCCACTAATAGGGAACGTGAGC</td><td>666.4</td><td>7.8</td><td>0.9</td><td>8.4</td><td>6.2E-17</td><td>2.2E-16</td><td>MB231/MB468</td></tr>
<tr><td>>MI0000465|hsa-mir-191&WithFlank&3|-|49020612|49020715@22.42.21(+1G)&[MIMAT0000440&hsa-miR-191-5p&offsets|0|-2(+1G);m-48&3|-|49020672|49020694&offsets|0|-2(+1G)]|CAACGGAATCCCAAAAGCAGCG</td><td>>MI0000465|hsa-mir-191&WithFlank&3|-|49020612|49020715</td><td>22</td><td>42</td><td>21(+1G)</td><td>CAACGGAATCCCAAAAGCAGCG</td><td>2029.2</td><td>7.9</td><td>0.9</td><td>8.4</td><td>6.1E-17</td><td>2.2E-16</td><td>BT20/MB468</td></tr>
<tr><td>>MI0000650|hsa-mir-200c&WithFlank&12|+|6963693|6963772@50.69.20(+1A)&[MIMAT0000617&hsa-miR-200c-3p&offsets|0|-3(+1A);m-54&12|+|6963742|6963764&offsets|0|-3(+1A)]|TAATACTGCCGGGTAATGATA</td><td>>MI0000650|hsa-mir-200c&WithFlank&12|+|6963693|6963772</td><td>50</td><td>69</td><td>20(+1A)</td><td>TAATACTGCCGGGTAATGATA</td><td>594.2</td><td>12.7</td><td>1.5</td><td>8.7</td><td>3.7E-18</td><td>2.3E-16</td><td>BT20/MB231</td></tr>
<tr><td>>hg38_21_+_8433172_8446622_RNA45SN1_45S_50ntFlanks@8177.8195.19|TTCTGATCGAGGCCCAGCC</td><td>>hg38_21_+_8433172_8446622_RNA45SN1_45S_50ntFlanks</td><td>8177</td><td>8195</td><td>19</td><td>TTCTGATCGAGGCCCAGCC</td><td>787.1</td><td>5.5</td><td>0.7</td><td>8.4</td><td>6.4E-17</td><td>2.3E-16</td><td>MB231/MB468</td></tr>
<tr><td>>hg38_21_+_8433172_8446622_RNA45SN1_45S_50ntFlanks@6773.6799.27|TTCCTCCCGGGGCTACGCCTGTCTGAG</td><td>>hg38_21_+_8433172_8446622_RNA45SN1_45S_50ntFlanks</td><td>6773</td><td>6799</td><td>27</td><td>TTCCTCCCGGGGCTACGCCTGTCTGAG</td><td>427.3</td><td>-10.7</td><td>1.3</td><td>-8.4</td><td>6.2E-17</td><td>2.3E-16</td><td>BT20/MB468</td></tr>
<tr><td>>hg38_21_+_8433172_8446622_RNA45SN1_45S_50ntFlanks@5472.5507.36|ACGGCCCTGGCGGAGCGCTGAGAAGACGGTCGAACT</td><td>>hg38_21_+_8433172_8446622_RNA45SN1_45S_50ntFlanks</td><td>5472</td><td>5507</td><td>36</td><td>ACGGCCCTGGCGGAGCGCTGAGAAGACGGTCGAACT</td><td>635.5</td><td>8.4</td><td>1.0</td><td>8.4</td><td>6.4E-17</td><td>2.3E-16</td><td>MB231/MB468</td></tr>
<tr><td>>hg38_21_+_8433172_8446622_RNA45SN1_45S_50ntFlanks@11886.11905.20|TGAGCTTGACTCTAGTCTGG</td><td>>hg38_21_+_8433172_8446622_RNA45SN1_45S_50ntFlanks</td><td>11886</td><td>11905</td><td>20</td><td>TGAGCTTGACTCTAGTCTGG</td><td>449.8</td><td>-10.8</td><td>1.3</td><td>-8.4</td><td>6.5E-17</td><td>2.4E-16</td><td>BT20/MB468</td></tr>
<tr><td>>MI0000289|hsa-mir-181a-1&WithFlank&1|-|198859038|198859159@30.51.22(+1G);MI0000269|hsa-mir-181a-2&WithFlank&9|+|124692436|124692557@45.66.22(+1G)&[MIMAT0000256&hsa-miR-181a-5p&offsets|0|-1(+1G);m-32&1|-|198859108|198859130&offsets|0|-1(+1G)];[MIMAT0000256_1&hsa-miR-181a-5p&offsets|0|-1(+1G);m-32&9|+|124692480|124692502&offsets|0|-1(+1G)]|AACATTCAACGCTGTCGGTGAGG</td><td>>MI0000289|hsa-mir-181a-1&WithFlank&1|-|198859038|198859159</td><td>30</td><td>51</td><td>22(+1G)</td><td>AACATTCAACGCTGTCGGTGAGG</td><td>820.4</td><td>7.1</td><td>0.9</td><td>8.4</td><td>6.7E-17</td><td>2.4E-16</td><td>MB231/MB468</td></tr>
<tr><td>>hg38_21_+_8433172_8446622_RNA45SN1_45S_50ntFlanks@4003.4026.24|ATAACCTCGGGCCGATCGCACGCC</td><td>>hg38_21_+_8433172_8446622_RNA45SN1_45S_50ntFlanks</td><td>4003</td><td>4026</td><td>24</td><td>ATAACCTCGGGCCGATCGCACGCC</td><td>389.5</td><td>-10.6</td><td>1.3</td><td>-8.4</td><td>6.6E-17</td><td>2.4E-16</td><td>BT20/MB468</td></tr>
<tr><td>>MI0005756|hsa-mir-934&WithFlank&X|+|136550872|136550966@21.42.22(+1G)&[MIMAT0004977&hsa-miR-934&offsets|0|0(+1G);m-943&X|+|136550892|136550913&offsets|0|0(+1G)]|TGTCTACTACTGGAGACACTGGG</td><td>>MI0005756|hsa-mir-934&WithFlank&X|+|136550872|136550966</td><td>21</td><td>42</td><td>22(+1G)</td><td>TGTCTACTACTGGAGACACTGGG</td><td>2162.8</td><td>9.4</td><td>1.1</td><td>8.4</td><td>6.6E-17</td><td>2.4E-16</td><td>BT20/MB468</td></tr>
<tr><td>>hg38_21_+_8433172_8446622_RNA45SN1_45S_50ntFlanks@7986.8012.27|ATCAGACGTGGCGACCCGCTGAATTTA</td><td>>hg38_21_+_8433172_8446622_RNA45SN1_45S_50ntFlanks</td><td>7986</td><td>8012</td><td>27</td><td>ATCAGACGTGGCGACCCGCTGAATTTA</td><td>528.3</td><td>-11.0</td><td>1.3</td><td>-8.4</td><td>6.7E-17</td><td>2.5E-16</td><td>BT20/MB468</td></tr>
<tr><td>>MI0003776|hsa-mir-320b-1&WithFlank&1|+|116671740|116671823@48.66.19(+1A);MI0003839|hsa-mir-320b-2&WithFlank&1|-|224257034|224257116@47.65.19(+1A);MI0003778|hsa-mir-320c-1&WithFlank&18|+|21683512|21683595@48.66.19(+1A);MI0008191|hsa-mir-320c-2&WithFlank&18|+|24321669|24321752@48.66.19(+1A);MI0000542|hsa-mir-320a&WithFlank&8|-|22244960|22245043@48.66.19(+1A)&[MIMAT0005792&hsa-miR-320b&offsets|0|-3(+1A);m-183&1|+|116671787|116671806&offsets|0|-1(+1A)];[MIMAT0005792_1&hsa-miR-320b&offsets|0|-3(+1A);m-183&1|-|224257051|224257070&offsets|0|-1(+1A)];[MIMAT0005793&hsa-miR-320c&offsets|0|-1(+1A);m-222&18|+|21683559|21683578&offsets|0|-1(+1A)];[MIMAT0005793_1&hsa-miR-320c&offsets|0|-1(+1A);m-219&18|+|24321716|24321735&offsets|0|-1(+1A)];[MIMAT0000510&hsa-miR-320a-3p&offsets|0|-3(+1A);m-60&8|-|22244975|22244996&offsets|0|-3(+1A)]|AAAAGCTGGGTTGAGAGGGA</td><td>>MI0003776|hsa-mir-320b-1&WithFlank&1|+|116671740|116671823</td><td>48</td><td>66</td><td>19(+1A)</td><td>AAAAGCTGGGTTGAGAGGGA</td><td>720.0</td><td>11.1</td><td>1.3</td><td>8.4</td><td>6.8E-17</td><td>2.5E-16</td><td>BT20/MB468</td></tr>
<tr><td>>MI0000070|hsa-mir-16-1&WithFlank&13|-|50048967|50049067@20.40.21;MI0000115|hsa-mir-16-2&WithFlank&3|+|160404739|160404831@16.36.21&[MIMAT0000069&hsa-miR-16-5p&offsets|0|-1;m-62&13|-|50049027|50049048&offsets|0|-1];[MIMAT0000069_1&hsa-miR-16-5p&offsets|0|-1;m-61&3|+|160404754|160404775&offsets|0|-1]|TAGCAGCACGTAAATATTGGC</td><td>>MI0000070|hsa-mir-16-1&WithFlank&13|-|50048967|50049067</td><td>20</td><td>40</td><td>21</td><td>TAGCAGCACGTAAATATTGGC</td><td>2513.9</td><td>6.3</td><td>0.8</td><td>8.3</td><td>6.9E-17</td><td>2.5E-16</td><td>MB231/MB468</td></tr>
<tr><td>>trnaMT_PheGAA_MT_+_577_647@39.59.21|ATGTTTAGACGGGCTCACATC</td><td>>trnaMT_PheGAA_MT_+_577_647</td><td>39</td><td>59</td><td>21</td><td>ATGTTTAGACGGGCTCACATC</td><td>643.4</td><td>8.8</td><td>1.1</td><td>8.3</td><td>6.9E-17</td><td>2.5E-16</td><td>MB231/MB468</td></tr>
<tr><td>>MI0000094|hsa-mir-92a-2&WithFlank&X|-|134169532|134169618@54.76.23&[MIMAT0000092_1&hsa-miR-92a-3p&offsets|0|+1;m-11&X|-|134169544|134169565&offsets|0|+1]|TATTGCACTTGTCCCGGCCTGTG</td><td>>MI0000094|hsa-mir-92a-2&WithFlank&X|-|134169532|134169618</td><td>54</td><td>76</td><td>23</td><td>TATTGCACTTGTCCCGGCCTGTG</td><td>1190.3</td><td>6.4</td><td>0.8</td><td>8.4</td><td>6.8E-17</td><td>2.5E-16</td><td>BT20/MB468</td></tr>
<tr><td>>hg38_21_+_8433172_8446622_RNA45SN1_45S_50ntFlanks@12785.12805.21|CCCCGCGCGCGCGGGAGGGCG</td><td>>hg38_21_+_8433172_8446622_RNA45SN1_45S_50ntFlanks</td><td>12785</td><td>12805</td><td>21</td><td>CCCCGCGCGCGCGGGAGGGCG</td><td>535.4</td><td>-10.6</td><td>1.3</td><td>-8.3</td><td>7.0E-17</td><td>2.5E-16</td><td>MB231/MB468</td></tr>
<tr><td>>MI0000082|hsa-mir-25&WithFlank&7|-|100093554|100093649@58.78.21(+1C)&[MIMAT0000081&hsa-miR-25-3p&offsets|0|-1(+1C);m-23&7|-|100093571|100093592&offsets|0|-1(+1C)]|CATTGCACTTGTCTCGGTCTGC</td><td>>MI0000082|hsa-mir-25&WithFlank&7|-|100093554|100093649</td><td>58</td><td>78</td><td>21(+1C)</td><td>CATTGCACTTGTCTCGGTCTGC</td><td>932.8</td><td>8.9</td><td>1.1</td><td>8.3</td><td>7.0E-17</td><td>2.5E-16</td><td>MB231/MB468</td></tr>
<tr><td>>MI0000285|hsa-mir-205&WithFlank&1|+|209432127|209432248@40.61.22&[MIMAT0000266&hsa-miR-205-5p&offsets|0|0;m-351&1|+|209432166|209432187&offsets|0|0]|TCCTTCATTCCACCGGAGTCTG</td><td>>MI0000285|hsa-mir-205&WithFlank&1|+|209432127|209432248</td><td>40</td><td>61</td><td>22</td><td>TCCTTCATTCCACCGGAGTCTG</td><td>61363.6</td><td>6.1</td><td>0.7</td><td>8.3</td><td>6.8E-17</td><td>2.5E-16</td><td>BT20/MB468</td></tr>
<tr><td>>MI0000071|hsa-mir-17&WithFlank&13|+|91350599|91350694@20.42.23&[MIMAT0000070&hsa-miR-17-5p&offsets|0|0;m-88&13|+|91350618|91350640&offsets|0|0]|CAAAGTGCTTACAGTGCAGGTAG</td><td>>MI0000071|hsa-mir-17&WithFlank&13|+|91350599|91350694</td><td>20</td><td>42</td><td>23</td><td>CAAAGTGCTTACAGTGCAGGTAG</td><td>31160.2</td><td>6.1</td><td>0.7</td><td>8.3</td><td>7.0E-17</td><td>2.5E-16</td><td>MB231/MB468</td></tr>
<tr><td>>hg38_21_+_8433172_8446622_RNA45SN1_45S_50ntFlanks@12942.12972.31|TTCGTAGACGACCTGCTTCTGGGTCGGGGTT</td><td>>hg38_21_+_8433172_8446622_RNA45SN1_45S_50ntFlanks</td><td>12942</td><td>12972</td><td>31</td><td>TTCGTAGACGACCTGCTTCTGGGTCGGGGTT</td><td>418.7</td><td>-7.4</td><td>0.9</td><td>-8.3</td><td>6.9E-17</td><td>2.5E-16</td><td>BT20/MB468</td></tr>
<tr><td>>hg38_1_-_228634819_228635039_RNA5S12_5S_50ntFlanks@152.170.19|TACCGGGTGCTGTAGGCTT</td><td>>hg38_1_-_228634819_228635039_RNA5S12_5S_50ntFlanks</td><td>152</td><td>170</td><td>19</td><td>TACCGGGTGCTGTAGGCTT</td><td>3315.9</td><td>6.6</td><td>0.8</td><td>8.3</td><td>7.0E-17</td><td>2.5E-16</td><td>MB231/MB468</td></tr>
<tr><td>>hg38_21_+_8433172_8446622_RNA45SN1_45S_50ntFlanks@12624.12653.30|CCAATGGGGCGAAGCTACCATCTGTGGGAT</td><td>>hg38_21_+_8433172_8446622_RNA45SN1_45S_50ntFlanks</td><td>12624</td><td>12653</td><td>30</td><td>CCAATGGGGCGAAGCTACCATCTGTGGGAT</td><td>461.1</td><td>-10.8</td><td>1.3</td><td>-8.3</td><td>6.9E-17</td><td>2.5E-16</td><td>BT20/MB468</td></tr>
<tr><td>>hg38_21_+_8433172_8446622_RNA45SN1_45S_50ntFlanks@6772.6799.28|GTTCCTCCCGGGGCTACGCCTGTCTGAG</td><td>>hg38_21_+_8433172_8446622_RNA45SN1_45S_50ntFlanks</td><td>6772</td><td>6799</td><td>28</td><td>GTTCCTCCCGGGGCTACGCCTGTCTGAG</td><td>723.0</td><td>-7.6</td><td>0.9</td><td>-8.3</td><td>7.1E-17</td><td>2.5E-16</td><td>MB231/MB468</td></tr>
<tr><td>>hg38_21_+_8433172_8446622_RNA45SN1_45S_50ntFlanks@7974.8004.31|ACGCGACCTCAGATCAGACGTGGCGACCCGC</td><td>>hg38_21_+_8433172_8446622_RNA45SN1_45S_50ntFlanks</td><td>7974</td><td>8004</td><td>31</td><td>ACGCGACCTCAGATCAGACGTGGCGACCCGC</td><td>877.8</td><td>5.1</td><td>0.6</td><td>8.3</td><td>7.0E-17</td><td>2.5E-16</td><td>BT20/MB468</td></tr>
<tr><td>>MI0000433|hsa-let-7g&WithFlank&3|-|52268272|52268367@11.30.20(+1G)&[MIMAT0000414&hsa-let-7g-5p&offsets|0|-2(+1G);m-58&3|-|52268336|52268357&offsets|0|-2(+1G)]|TGAGGTAGTAGTTTGTACAGG</td><td>>MI0000433|hsa-let-7g&WithFlank&3|-|52268272|52268367</td><td>11</td><td>30</td><td>20(+1G)</td><td>TGAGGTAGTAGTTTGTACAGG</td><td>692.0</td><td>8.7</td><td>1.0</td><td>8.3</td><td>7.1E-17</td><td>2.5E-16</td><td>MB231/MB468</td></tr>
<tr><td>>hg38_21_+_8433172_8446622_RNA45SN1_45S_50ntFlanks@11768.11791.24|CTAATTAGTGACGCGCATGAATGG</td><td>>hg38_21_+_8433172_8446622_RNA45SN1_45S_50ntFlanks</td><td>11768</td><td>11791</td><td>24</td><td>CTAATTAGTGACGCGCATGAATGG</td><td>506.8</td><td>-10.6</td><td>1.3</td><td>-8.3</td><td>7.2E-17</td><td>2.5E-16</td><td>MB231/MB468</td></tr>
<tr><td>>hg38_21_+_8433172_8446622_RNA45SN1_45S_50ntFlanks@10679.10710.32|GGTTCCGGCGGCGTCCGGTGAGCTCTCGCTGG</td><td>>hg38_21_+_8433172_8446622_RNA45SN1_45S_50ntFlanks</td><td>10679</td><td>10710</td><td>32</td><td>GGTTCCGGCGGCGTCCGGTGAGCTCTCGCTGG</td><td>33416.2</td><td>-7.8</td><td>0.9</td><td>-8.3</td><td>7.0E-17</td><td>2.5E-16</td><td>BT20/MB468</td></tr>
<tr><td>>hg38_21_+_8433172_8446622_RNA45SN1_45S_50ntFlanks@12357.12376.20|CCACAGGGATAACTGGCTTG</td><td>>hg38_21_+_8433172_8446622_RNA45SN1_45S_50ntFlanks</td><td>12357</td><td>12376</td><td>20</td><td>CCACAGGGATAACTGGCTTG</td><td>827.7</td><td>-11.3</td><td>1.4</td><td>-8.3</td><td>7.2E-17</td><td>2.6E-16</td><td>MB231/MB468</td></tr>
<tr><td>>MI0000264|hsa-mir-7-2&WithFlank&15|+|88611819|88611940@38.59.22;MI0000265|hsa-mir-7-3&WithFlank&19|+|4770664|4770785@37.58.22;MI0000263|hsa-mir-7-1&WithFlank&9|-|83969742|83969863@30.51.22&[MIMAT0000252&hsa-miR-7-5p&offsets|0|-2;m-400&15|+|88611856|88611879&offsets|0|-2];[MIMAT0000252_1&hsa-miR-7-5p&offsets|0|-2;m-402&19|+|4770700|4770723&offsets|0|-2];[MIMAT0000252_2&hsa-miR-7-5p&offsets|0|-2;m-401&9|-|83969811|83969834&offsets|0|-2]|TGGAAGACTAGTGATTTTGTTG</td><td>>MI0000264|hsa-mir-7-2&WithFlank&15|+|88611819|88611940</td><td>38</td><td>59</td><td>22</td><td>TGGAAGACTAGTGATTTTGTTG</td><td>5379.4</td><td>8.0</td><td>1.0</td><td>8.3</td><td>7.1E-17</td><td>2.6E-16</td><td>BT20/MB468</td></tr>
</table>

| | | | | | | | | | | | | |
|---|---|---|---|---|---|---|---|---|---|---|---|---|
| >hg38_21_+_8433172_8446622_RNA45SN1_45S_50ntFlanks@9575.9596.22\|ATCGGTCGTCCGACCTGGGTAT | >hg38_21_+_8433172_8446622_RNA45SN1_45S_50ntFlanks | 9575 | 9596 | 22 | ATCGGTCGTCCGACCTGGGTAT | 1592.8 | -8.2 | 1.0 | -8.3 | 7.2E-17 | 2.6E-16 | MB231/MB468 |
| >hg38_21_+_8433172_8446622_RNA45SN1_45S_50ntFlanks@8105.8143.39\|CGCGGCGGGGCGCGGGACATGTGGCGTACGGAAGACCCG | >hg38_21_+_8433172_8446622_RNA45SN1_45S_50ntFlanks | 8105 | 8143 | 39 | CGCGGCGGGGCGCGGGACATGTGGCGTACGGAAGACCCG | 2102.7 | -7.0 | 0.8 | -8.3 | 7.3E-17 | 2.6E-16 | MB231/MB468 |
| >MI0000447\|hsa-mir-128-1&WithFlank&2\|+\|135665391\|135665484@56.76.21(+1A);MI0000727\|hsa-mir-128-2&WithFlank&3\|+\|35744470\|35744565@58.78.21(+1A)&[MIMAT0000424&hsa-miR-128-3p&offsets\|0\|0(+1A);m-108&2\|+\|135665446\|135665466&offsets\|0\|0(+1A)];[MIMAT0000424_1&hsa-miR-128-3p&offsets\|0\|0(+1A);m-117&3\|+\|35744527\|35744547&offsets\|0\|0(+1A)]\|TCACAGTGAACCGGTCTCTTTA | >MI0000447\|hsa-mir-128-1&WithFlank&2\|+\|135665391\|135665484 | 56 | 76 | 21(+1A) | TCACAGTGAACCGGTCTCTTTA | 4096.0 | 5.5 | 0.7 | 8.3 | 7.2E-17 | 2.6E-16 | BT20/MB468 |
| >hg38_21_+_8433172_8446622_RNA45SN1_45S_50ntFlanks@11573.11601.29\|CGACTTAGAACTGGTGCGGACCAGGGGAA | >hg38_21_+_8433172_8446622_RNA45SN1_45S_50ntFlanks | 11573 | 11601 | 29 | CGACTTAGAACTGGTGCGGACCAGGGGAA | 671.4 | -4.6 | 0.5 | -8.3 | 7.2E-17 | 2.6E-16 | BT20/MB468 |
| >hg38_21_+_8433172_8446622_RNA45SN1_45S_50ntFlanks@10690.10708.19\|CGTCCGGTGAGCTCTCGCT | >hg38_21_+_8433172_8446622_RNA45SN1_45S_50ntFlanks | 10690 | 10708 | 19 | CGTCCGGTGAGCTCTCGCT | 4867.5 | -6.4 | 0.8 | -8.3 | 7.3E-17 | 2.6E-16 | BT20/MB468 |
| >MI0000103\|hsa-mir-101-1&WithFlank&1\|-\|65058428\|65058514@53.72.20;MI0000739\|hsa-mir-101-2&WithFlank&9\|+\|4850291\|4850381@55.74.20&[MIMAT0000099&hsa-miR-101-3p&offsets\|0\|-1;m-17&1\|-\|65058443\|65058462&offsets\|0\|0];[MIMAT0000099_1&hsa-miR-101-3p&offsets\|0\|-1;m-16&9\|+\|4850345\|4850364&offsets\|0\|0]\|TACAGTACTGTGATAACTGA | >MI0000103\|hsa-mir-101-1&WithFlank&1\|-\|65058428\|65058514 | 53 | 72 | 20 | TACAGTACTGTGATAACTGA | 3698.2 | 6.6 | 0.8 | 8.3 | 7.6E-17 | 2.7E-16 | MB231/MB468 |
| >hg38_21_+_8433172_8446622_RNA45SN1_45S_50ntFlanks@11564.11603.40\|CCTAGCAGCCGACTTAGAACTGGTGCGGACCAGGGGAATC | >hg38_21_+_8433172_8446622_RNA45SN1_45S_50ntFlanks | 11564 | 11603 | 40 | CCTAGCAGCCGACTTAGAACTGGTGCGGACCAGGGGAATC | 723.3 | -5.6 | 0.7 | -8.3 | 7.5E-17 | 2.7E-16 | BT20/MB468 |
| >MI0001445\|hsa-mir-423&WithFlank&17\|+\|30117073\|30117178@59.77.19(+1U)&[MIMAT0001340&hsa-miR-423-3p&offsets\|0\|-4(+1U);m-100&17\|+\|30117131\|30117153&offsets\|0\|-4(+1U)]\|AGCTCGGTCTGAGGCCCCTT | >MI0001445\|hsa-mir-423&WithFlank&17\|+\|30117073\|30117178 | 59 | 77 | 19(+1U) | AGCTCGGTCTGAGGCCCCTT | 820.8 | 8.6 | 1.0 | 8.3 | 7.6E-17 | 2.7E-16 | BT20/MB468 |
| >hg38_21_+_8433172_8446622_RNA45SN1_45S_50ntFlanks@12423.12457.35\|TCGGCTCTTCCTATCATTGTGAAGCAGAATTCACC | >hg38_21_+_8433172_8446622_RNA45SN1_45S_50ntFlanks | 12423 | 12457 | 35 | TCGGCTCTTCCTATCATTGTGAAGCAGAATTCACC | 549.5 | 7.5 | 0.9 | 8.3 | 7.7E-17 | 2.7E-16 | MB231/MB468 |
| >MI0000063\|hsa-let-7b&WithFlank&22\|+\|46113680\|46113774@12.32.21(+1G)&[MIMAT0000063&hsa-let-7b-5p&offsets\|0\|-1(+1G);m-14&22\|+\|46113691\|46113712&offsets\|0\|-1(+1G)]\|TGAGGTAGTAGGTTGTGTGGTG | >MI0000063\|hsa-let-7b&WithFlank&22\|+\|46113680\|46113774 | 12 | 32 | 21(+1G) | TGAGGTAGTAGGTTGTGTGGTG | 962.8 | 6.5 | 0.8 | 8.3 | 7.8E-17 | 2.7E-16 | MB231/MB468 |
| >hg38_21_+_8433172_8446622_RNA45SN1_45S_50ntFlanks@11576.11603.28\|CTTAGAACTGGTGCGGACCAGGGGAATC | >hg38_21_+_8433172_8446622_RNA45SN1_45S_50ntFlanks | 11576 | 11603 | 28 | CTTAGAACTGGTGCGGACCAGGGGAATC | 2499.3 | -5.6 | 0.7 | -8.3 | 7.8E-17 | 2.7E-16 | MB231/MB468 |
| >hg38_21_+_8433172_8446622_RNA45SN1_45S_50ntFlanks@10393.10419.27\|CAGCAGTTGAACATGGGTCAGTCGGTC | >hg38_21_+_8433172_8446622_RNA45SN1_45S_50ntFlanks | 10393 | 10419 | 27 | CAGCAGTTGAACATGGGTCAGTCGGTC | 254.9 | -4.5 | 0.5 | -8.3 | 7.7E-17 | 2.8E-16 | BT20/MB468 |
| >hg38_21_+_8433172_8446622_RNA45SN1_45S_50ntFlanks@10479.10497.19\|CGATCGAAAGGGAGTCGGG | >hg38_21_+_8433172_8446622_RNA45SN1_45S_50ntFlanks | 10479 | 10497 | 19 | CGATCGAAAGGGAGTCGGG | 423.0 | -10.7 | 1.3 | -8.3 | 7.7E-17 | 2.8E-16 | BT20/MB468 |
| >hg38_21_+_8433172_8446622_RNA45SN1_45S_50ntFlanks@11233.11254.22\|CCCTTCCCGTGGATCGCCCCAG | >hg38_21_+_8433172_8446622_RNA45SN1_45S_50ntFlanks | 11233 | 11254 | 22 | CCCTTCCCGTGGATCGCCCCAG | 793.3 | -11.2 | 1.3 | -8.3 | 7.9E-17 | 2.8E-16 | MB231/MB468 |
| >MI0000071\|hsa-mir-17&WithFlank&13\|+\|91350599\|91350694@20.42.23(+1G)&[MIMAT0000070&hsa-miR-17-5p&offsets\|0\|0(+1G);m-88&13\|+\|91350618\|91350640&offsets\|0\|0(+1G)]\|CAAAGTGCTTACAGTGCAGGTAGG | >MI0000071\|hsa-mir-17&WithFlank&13\|+\|91350599\|91350694 | 20 | 42 | 23(+1G) | CAAAGTGCTTACAGTGCAGGTAGG | 1090.2 | 6.9 | 0.8 | 8.3 | 8.0E-17 | 2.8E-16 | MB231/MB468 |
| >hg38_21_+_8433172_8446622_RNA45SN1_45S_50ntFlanks@11729.11767.39\|TAACTATGACTCTCTTAAGGTAGCCAAATGCCTCGTCAT | >hg38_21_+_8433172_8446622_RNA45SN1_45S_50ntFlanks | 11729 | 11767 | 39 | TAACTATGACTCTCTTAAGGTAGCCAAATGCCTCGTCAT | 762.5 | -4.9 | 0.6 | -8.3 | 8.1E-17 | 2.9E-16 | MB231/MB468 |
| >MI0000447\|hsa-mir-128-1&WithFlank&2\|+\|135665391\|135665484@56.77.22&[MIMAT0000424&hsa-miR-128-3p&offsets\|0\|+1;m-108&2\|+\|135665446\|135665466&offsets\|0\|+1]\|TCACAGTGAACCGGTCTCTTTT | >MI0000447\|hsa-mir-128-1&WithFlank&2\|+\|135665391\|135665484 | 56 | 77 | 22 | TCACAGTGAACCGGTCTCTTTT | 6410.3 | 5.3 | 0.6 | 8.3 | 7.9E-17 | 2.9E-16 | BT20/MB468 |
| >MI0000289\|hsa-mir-181a-1&WithFlank&1\|-\|198859038\|198859159@70.90.21(+1U)&[MIMAT0000270&hsa-miR-181a-3p&offsets\|0\|-1(+1U);m-125&1\|-\|198859069\|198859090&offsets\|0\|-1(+1U)]\|ACCATCGACCGTTGATTGTACT | >MI0000289\|hsa-mir-181a-1&WithFlank&1\|-\|198859038\|198859159 | 70 | 90 | 21(+1U) | ACCATCGACCGTTGATTGTACT | 545.1 | -12.4 | 1.4 | -8.7 | 4.9E-18 | 2.9E-16 | BT20/MB231 |
| >MI0000650\|hsa-mir-200c&WithFlank&12\|+\|6963693\|6963772@49.72.24&[MIMAT0000617&hsa-miR-200c-3p&offsets\|-1\|0;m-54&12\|+\|6963742\|6963764&offsets\|-1\|0]\|CTAATACTGCCGGGTAATGATGGA | >MI0000650\|hsa-mir-200c&WithFlank&12\|+\|6963693\|6963772 | 49 | 72 | 24 | CTAATACTGCCGGGTAATGATGGA | 453.2 | 12.3 | 1.4 | 8.7 | 5.0E-18 | 2.9E-16 | BT20/MB231 |
| >hg38_21_+_8433172_8446622_RNA45SN1_45S_50ntFlanks@11874.11903.30\|AGAAGACCCTGTTGAGCTTGACTCTAGTCT | >hg38_21_+_8433172_8446622_RNA45SN1_45S_50ntFlanks | 11874 | 11903 | 30 | AGAAGACCCTGTTGAGCTTGACTCTAGTCT | 266.0 | -4.9 | 0.6 | -8.3 | 8.0E-17 | 2.9E-16 | BT20/MB468 |
| >hg38_21_+_8433172_8446622_RNA45SN1_45S_50ntFlanks@11581.11605.25\|AACTGGTGCGGACCAGGGGAATCCG | >hg38_21_+_8433172_8446622_RNA45SN1_45S_50ntFlanks | 11581 | 11605 | 25 | AACTGGTGCGGACCAGGGGAATCCG | 1511.3 | -4.6 | 0.6 | -8.3 | 8.3E-17 | 2.9E-16 | MB231/MB468 |
| >hg38_21_+_8433172_8446622_RNA45SN1_45S_50ntFlanks@6745.6769.25\|ACACTTCGAACGCACTTGCGGCCCC | >hg38_21_+_8433172_8446622_RNA45SN1_45S_50ntFlanks | 6745 | 6769 | 25 | ACACTTCGAACGCACTTGCGGCCCC | 181.0 | -4.8 | 0.6 | -8.3 | 8.2E-17 | 2.9E-16 | BT20/MB468 |
| >hg38_21_+_8433172_8446622_RNA45SN1_45S_50ntFlanks@12539.12566.28\|ATGATGTGTTGTTGCCATGGTAATCCTG | >hg38_21_+_8433172_8446622_RNA45SN1_45S_50ntFlanks | 12539 | 12566 | 28 | ATGATGTGTTGTTGCCATGGTAATCCTG | 844.6 | -8.0 | 1.0 | -8.3 | 8.4E-17 | 3.0E-16 | MB231/MB468 |
| >hg38_21_+_8433172_8446622_RNA45SN1_45S_50ntFlanks@12366.12384.19\|TAACTGGCTTGTGGCGGCC | >hg38_21_+_8433172_8446622_RNA45SN1_45S_50ntFlanks | 12366 | 12384 | 19 | TAACTGGCTTGTGGCGGCC | 430.8 | -10.7 | 1.3 | -8.3 | 8.3E-17 | 3.0E-16 | BT20/MB468 |
| >MI0000088\|hsa-mir-30a&WithFlank&6\|-\|71403545\|71403627@13.34.22(+1C)&[MIMAT0000087&hsa-miR-30a-5p&offsets\|+1\|+1(+1C);m-15&6\|-\|71403595\|71403616&offsets\|+1\|+1(+1C)]\|GTAAACATCCTCGACTGGAAGCC | >MI0000088\|hsa-mir-30a&WithFlank&6\|-\|71403545\|71403627 | 13 | 34 | 22(+1C) | GTAAACATCCTCGACTGGAAGCC | 946.3 | 9.6 | 1.2 | 8.3 | 8.6E-17 | 3.1E-16 | BT20/MB468 |
| >am_trnaMT_ArgTCG_MT_+_10405_10469@28.48.21\|ATTTCGACTCATTAAATTATG | >am_trnaMT_ArgTCG_MT_+_10405_10469 | 28 | 48 | 21 | ATTTCGACTCATTAAATTATG | 593.0 | -5.1 | 0.6 | -8.3 | 8.8E-17 | 3.1E-16 | MB231/MB468 |
| >hg38_21_+_8433172_8446622_RNA45SN1_45S_50ntFlanks@6787.6807.21\|ACGCCTGTCTGAGCGTCGCTT | >hg38_21_+_8433172_8446622_RNA45SN1_45S_50ntFlanks | 6787 | 6807 | 21 | ACGCCTGTCTGAGCGTCGCTT | 712.1 | 6.3 | 0.8 | 8.3 | 8.8E-17 | 3.1E-16 | MB231/MB468 |
| >hg38_21_+_8433172_8446622_RNA45SN1_45S_50ntFlanks@12044.12070.27\|ACTGACCCGGTGAGGCGGGGGGGCGAG | >hg38_21_+_8433172_8446622_RNA45SN1_45S_50ntFlanks | 12044 | 12070 | 27 | ACTGACCCGGTGAGGCGGGGGGGCGAG | 362.9 | -10.5 | 1.3 | -8.3 | 8.7E-17 | 3.1E-16 | BT20/MB468 |
| >hg38_21_+_8433172_8446622_RNA45SN1_45S_50ntFlanks@9562.9597.36\|TCCTGACGTGCAAATCGGTCGTCCGACCTGGGTATA | >hg38_21_+_8433172_8446622_RNA45SN1_45S_50ntFlanks | 9562 | 9597 | 36 | TCCTGACGTGCAAATCGGTCGTCCGACCTGGGTATA | 476.2 | -7.5 | 0.9 | -8.3 | 8.9E-17 | 3.2E-16 | BT20/MB468 |
| >hg38_21_+_8433172_8446622_RNA45SN1_45S_50ntFlanks@5444.5466.23\|CTCGGATCGGCCCCGCCGGGGTC | >hg38_21_+_8433172_8446622_RNA45SN1_45S_50ntFlanks | 5444 | 5466 | 23 | CTCGGATCGGCCCCGCCGGGGTC | 6963.0 | -6.2 | 0.7 | -8.3 | 9.2E-17 | 3.2E-16 | MB231/MB468 |
| >hg38_1_-_228634819_228635039_RNA5S12_5S_50ntFlanks@133.170.38\|ATGGGAGACCGCCTGGGAATACCGGGTGCTGTAGGCTT | >hg38_1_-_228634819_228635039_RNA5S12_5S_50ntFlanks | 133 | 170 | 38 | ATGGGAGACCGCCTGGGAATACCGGGTGCTGTAGGCTT | 10932.2 | 5.1 | 0.6 | 8.3 | 9.0E-17 | 3.2E-16 | BT20/MB468 |
| >am_trna128_GlyGCC_6_-_27870686_27870756@1.35.35,trna18_GlyGCC_16_+_70822597_70822667@1.35.35,trna19_GlyGCC_16_+_70823410_70823480@1.35.35,trna19_GlyGCC_2_-_157257659_157257729@1.35.35,trna24_GlyGCC_16_-_70812942_70813012@1.35.35,trna25_GlyGCC_16_-_70812114_70812184@1.35.35,trna5_GlyGCC_17_+_8029064_8029134@1.35.35,trna68_GlyGCC_1_-_161493637_161493707@1.35.35\|GCATTGGTGGTTCAGTGGTAGAATTCTCGCCTGCC | >am_trna128_GlyGCC_6_-_27870686_27870756 | 1 | 35 | 35 | GCATTGGTGGTTCAGTGGTAGAATTCTCGCCTGCC | 7177.8 | 8.5 | 1.0 | 8.3 | 9.0E-17 | 3.2E-16 | BT20/MB468 |
| >MI0000457\|hsa-mir-141&WithFlank&12\|+\|6964091\|6964197@23.44.22&[MIMAT0004598&hsa-miR-141-5p&offsets\|0\|0;m-221&12\|+\|6964113\|6964133&offsets\|0\|+1]\|CATCTTCCAGTACAGTGTTGGA | >MI0000457\|hsa-mir-141&WithFlank&12\|+\|6964091\|6964197 | 23 | 44 | 22 | CATCTTCCAGTACAGTGTTGGA | 512.6 | 12.5 | 1.4 | 8.6 | 5.7E-18 | 3.3E-16 | BT20/MB231 |
| >MI0000650\|hsa-mir-200c&WithFlank&12\|+\|6963693\|6963772@50.70.21(+1U)&[MIMAT0000617&hsa-miR-200c-3p&offsets\|0\|-2(+1U);m-54&12\|+\|6963742\|6963764&offsets\|0\|-2(+1U)]\|TAATACTGCCGGGTAATGATGT | >MI0000650\|hsa-mir-200c&WithFlank&12\|+\|6963693\|6963772 | 50 | 70 | 21(+1U) | TAATACTGCCGGGTAATGATGT | 1061.6 | 11.6 | 1.3 | 8.6 | 5.8E-18 | 3.3E-16 | BT20/MB231 |
| >MI0000089\|hsa-mir-31&WithFlank&9\|-\|21512109\|21512191@14.35.22(+1A)&[MIMAT0000089&hsa-miR-31-5p&offsets\|0\|+1(+1A);m-360&9\|-\|21512157\|21512178&offsets\|0\|0(+1A)]\|AGGCAAGATGCTGGCATAGCTGA | >MI0000089\|hsa-mir-31&WithFlank&9\|-\|21512109\|21512191 | 14 | 35 | 22(+1A) | AGGCAAGATGCTGGCATAGCTGA | 391.3 | 12.1 | 1.4 | 8.6 | 6.0E-18 | 3.3E-16 | BT20/MB231 |
| >hg38_21_+_8433172_8446622_RNA45SN1_45S_50ntFlanks@12159.12182.24\|TTTGACTGGGGCGGTACACCTGTC | >hg38_21_+_8433172_8446622_RNA45SN1_45S_50ntFlanks | 12159 | 12182 | 24 | TTTGACTGGGGCGGTACACCTGTC | 1094.2 | -7.2 | 0.9 | -8.3 | 9.4E-17 | 3.3E-16 | MB231/MB468 |
| >hg38_21_+_8433172_8446622_RNA45SN1_45S_50ntFlanks@3705.3745.41\|TACCTGGTTGATCCTGCCAGTAGCATATGCTTGTCTCAAAG | >hg38_21_+_8433172_8446622_RNA45SN1_45S_50ntFlanks | 3705 | 3745 | 41 | TACCTGGTTGATCCTGCCAGTAGCATATGCTTGTCTCAAAG | 1491.0 | 7.6 | 0.9 | 8.3 | 9.3E-17 | 3.3E-16 | BT20/MB468 |
| >MI0000734\|hsa-mir-106b&WithFlank&7\|-\|100093987\|100094080@58.77.20(+1A)&[MIMAT0004672&hsa-miR-106b-3p&offsets\|0\|-2(+1A);m-87&7\|-\|100094003\|100094023&offsets\|0\|-1(+1A)]\|CCGCACTGTGGGTACTTGCTA | >MI0000734\|hsa-mir-106b&WithFlank&7\|-\|100093987\|100094080 | 58 | 77 | 20(+1A) | CCGCACTGTGGGTACTTGCTA | 637.6 | 9.5 | 1.1 | 8.3 | 9.5E-17 | 3.3E-16 | MB231/MB468 |

<table>
<tr><td>>MI0000088|hsa-mir-30a&WithFlank&6|-|71403545|71403627@12.33.22(+2U)&[MIMAT0000087&hsa-miR-30a-5p&offsets|0|0(+2U);m-15&6|-|71403595|71403616&offsets|0|0(+2U)]|TGTAAACATCCTCGACTGGAAGTT</td><td>>MI0000088|hsa-mir-30a&WithFlank&6|-|71403545|71403627</td><td>12</td><td>33</td><td>22(+2U)</td><td>TGTAAACATCCTCGACTGGAAGTT</td><td>1858.4</td><td>9.4</td><td>1.1</td><td>8.3</td><td>9.6E-17</td><td>3.4E-16</td><td>MB231/MB468</td></tr>
<tr><td>>MI0000465|hsa-mir-191&WithFlank&3|-|49020612|49020715@22.42.21&[MIMAT0000440&hsa-miR-191-5p&offsets|0|-2;m-48&3|-|49020672|49020694&offsets|0|-2]|CAACGGAATCCCAAAAGCAGC</td><td>>MI0000465|hsa-mir-191&WithFlank&3|-|49020612|49020715</td><td>22</td><td>42</td><td>21</td><td>CAACGGAATCCCAAAAGCAGC</td><td>945.2</td><td>6.6</td><td>0.8</td><td>8.3</td><td>9.7E-17</td><td>3.4E-16</td><td>MB231/MB468</td></tr>
<tr><td>>hg38_21_+_8433172_8446622_RNA45SN1_45S_50ntFlanks@9504.9525.22|GTGAACTATGCCTGGGCAGGGC</td><td>>hg38_21_+_8433172_8446622_RNA45SN1_45S_50ntFlanks</td><td>9504</td><td>9525</td><td>22</td><td>GTGAACTATGCCTGGGCAGGGC</td><td>673.4</td><td>-5.3</td><td>0.6</td><td>-8.3</td><td>9.7E-17</td><td>3.4E-16</td><td>MB231/MB468</td></tr>
<tr><td>>hg38_21_+_8433172_8446622_RNA45SN1_45S_50ntFlanks@3899.3919.21|CCTTCGCGGGGGGGATGCGTG</td><td>>hg38_21_+_8433172_8446622_RNA45SN1_45S_50ntFlanks</td><td>3899</td><td>3919</td><td>21</td><td>CCTTCGCGGGGGGGATGCGTG</td><td>528.2</td><td>-5.4</td><td>0.7</td><td>-8.3</td><td>9.6E-17</td><td>3.4E-16</td><td>BT20/MB468</td></tr>
<tr><td>>hg38_21_+_8433172_8446622_RNA45SN1_45S_50ntFlanks@12615.12654.40|GCTGAGGAGCCAATGGGGCGAAGCTACCATCTGTGGGATT</td><td>>hg38_21_+_8433172_8446622_RNA45SN1_45S_50ntFlanks</td><td>12615</td><td>12654</td><td>40</td><td>GCTGAGGAGCCAATGGGGCGAAGCTACCATCTGTGGGATT</td><td>2026.8</td><td>-4.5</td><td>0.5</td><td>-8.3</td><td>9.8E-17</td><td>3.4E-16</td><td>MB231/MB468</td></tr>
<tr><td>>MI0000264|hsa-mir-7-2&WithFlank&15|+|88611819|88611940@38.59.22(+1G);MI0000265|hsa-mir-7-3&WithFlank&19|+|4770664|4770785@37.58.22(+1G);MI0000263|hsa-mir-7-1&WithFlank&9|-|83969742|83969863@30.51.22(+1G)&[MIMAT0000252&hsa-miR-7-5p&offsets|0|-2(+1G);m-400&15|+|88611856|88611879&offsets|0|-2(+1G)];[MIMAT0000252_1&hsa-miR-7-5p&offsets|0|-2(+1G);m-402&19|+|4770700|4770723&offsets|0|-2(+1G)];[MIMAT0000252_2&hsa-miR-7-5p&offsets|0|-2(+1G);m-401&9|-|83969811|83969834&offsets|0|-2(+1G)]|TGGAAGACTAGTGATTTTGTTGG</td><td>>MI0000264|hsa-mir-7-2&WithFlank&15|+|88611819|88611940</td><td>38</td><td>59</td><td>22(+1G)</td><td>TGGAAGACTAGTGATTTTGTTGG</td><td>5796.3</td><td>9.5</td><td>1.1</td><td>8.3</td><td>9.7E-17</td><td>3.5E-16</td><td>BT20/MB468</td></tr>
<tr><td>>hg38_1_-_228634819_228635039_RNA5S12_5S_50ntFlanks@132.171.40|GATGGGAGACCGCCTGGGAATACCGGGTGCTGTAGGCTTT</td><td>>hg38_1_-_228634819_228635039_RNA5S12_5S_50ntFlanks</td><td>132</td><td>171</td><td>40</td><td>GATGGGAGACCGCCTGGGAATACCGGGTGCTGTAGGCTTT</td><td>13692.8</td><td>5.8</td><td>0.7</td><td>8.3</td><td>9.9E-17</td><td>3.5E-16</td><td>MB231/MB468</td></tr>
<tr><td>>MI0000750|hsa-mir-26a-2&WithFlank&12|-|57824603|57824698@20.41.22;MI0000083|hsa-mir-26a-1&WithFlank&3|+|37969398|37969486@16.37.22&[MIMAT0000082&hsa-miR-26a-5p&offsets|0|0;m-33&12|-|57824658|57824679&offsets|0|0];[MIMAT0000082_1&hsa-miR-26a-5p&offsets|0|0;m-34&3|+|37969413|37969434&offsets|0|0]|TTCAAGTAATCCAGGATAGGCT</td><td>>MI0000750|hsa-mir-26a-2&WithFlank&12|-|57824603|57824698</td><td>20</td><td>41</td><td>22</td><td>TTCAAGTAATCCAGGATAGGCT</td><td>112468.0</td><td>5.6</td><td>0.7</td><td>8.3</td><td>9.7E-17</td><td>3.5E-16</td><td>BT20/MB468</td></tr>
<tr><td>>hg38_21_+_8433172_8446622_RNA45SN1_45S_50ntFlanks@12676.12703.28|AGAATCCCGCCCAGGCGGAACGATACGG</td><td>>hg38_21_+_8433172_8446622_RNA45SN1_45S_50ntFlanks</td><td>12676</td><td>12703</td><td>28</td><td>AGAATCCCGCCCAGGCGGAACGATACGG</td><td>421.2</td><td>-10.7</td><td>1.3</td><td>-8.3</td><td>1.0E-16</td><td>3.6E-16</td><td>BT20/MB468</td></tr>
<tr><td>>trna10_LysCTT_16_+_3241501_3241573@1.34.34,trna13_LysCTT_14_-_58706613_58706685@1.34.34,trna2_LysCTT_15_+_79152904_79152976@1.34.34|GCCCGGCTAGCTCAGTCGGTAGAGCATGGGACTC</td><td>>trna10_LysCTT_16_+_3241501_3241573</td><td>1</td><td>34</td><td>34</td><td>GCCCGGCTAGCTCAGTCGGTAGAGCATGGGACTC</td><td>9627.8</td><td>10.9</td><td>1.3</td><td>8.3</td><td>1.0E-16</td><td>3.6E-16</td><td>MB231/MB468</td></tr>
<tr><td>>hg38_21_+_8433172_8446622_RNA45SN1_45S_50ntFlanks@4733.4761.29|GAACGAAAGTCGGAGGTTCGAAGACGATC</td><td>>hg38_21_+_8433172_8446622_RNA45SN1_45S_50ntFlanks</td><td>4733</td><td>4761</td><td>29</td><td>GAACGAAAGTCGGAGGTTCGAAGACGATC</td><td>320.6</td><td>4.9</td><td>0.6</td><td>8.3</td><td>1.0E-16</td><td>3.6E-16</td><td>BT20/MB468</td></tr>
<tr><td>>hg38_21_+_8433172_8446622_RNA45SN1_45S_50ntFlanks@12991.13016.26|CTCCCTCGCTGCGATCTATTGAAAGT</td><td>>hg38_21_+_8433172_8446622_RNA45SN1_45S_50ntFlanks</td><td>12991</td><td>13016</td><td>26</td><td>CTCCCTCGCTGCGATCTATTGAAAGT</td><td>407.8</td><td>-3.1</td><td>0.4</td><td>-8.3</td><td>1.0E-16</td><td>3.6E-16</td><td>BT20/MB468</td></tr>
<tr><td>>hg38_21_+_8433172_8446622_RNA45SN1_45S_50ntFlanks@6719.6745.27|TGAATTGCAGGACACATTGATCATCGA</td><td>>hg38_21_+_8433172_8446622_RNA45SN1_45S_50ntFlanks</td><td>6719</td><td>6745</td><td>27</td><td>TGAATTGCAGGACACATTGATCATCGA</td><td>458.9</td><td>-6.1</td><td>0.7</td><td>-8.3</td><td>1.0E-16</td><td>3.7E-16</td><td>MB231/MB468</td></tr>
<tr><td>>hg38_21_+_8433172_8446622_RNA45SN1_45S_50ntFlanks@10684.10711.28|CGGCGGCGTCCGGTGAGCTCTCGCTGGC</td><td>>hg38_21_+_8433172_8446622_RNA45SN1_45S_50ntFlanks</td><td>10684</td><td>10711</td><td>28</td><td>CGGCGGCGTCCGGTGAGCTCTCGCTGGC</td><td>2708.7</td><td>-5.4</td><td>0.6</td><td>-8.3</td><td>1.0E-16</td><td>3.7E-16</td><td>BT20/MB468</td></tr>
<tr><td>>hg38_MT_+_598_1651_RNR1_12S_50ntFlanks@608.633.26|TGTTCTGTAATCGATAAACCCCGATC</td><td>>hg38_MT_+_598_1651_RNR1_12S_50ntFlanks</td><td>608</td><td>633</td><td>26</td><td>TGTTCTGTAATCGATAAACCCCGATC</td><td>336.2</td><td>8.4</td><td>1.0</td><td>8.3</td><td>1.0E-16</td><td>3.7E-16</td><td>BT20/MB468</td></tr>
<tr><td>>hg38_21_+_8433172_8446622_RNA45SN1_45S_50ntFlanks@11768.11786.19|CTAATTAGTGACGCGCATG</td><td>>hg38_21_+_8433172_8446622_RNA45SN1_45S_50ntFlanks</td><td>11768</td><td>11786</td><td>19</td><td>CTAATTAGTGACGCGCATG</td><td>736.6</td><td>-6.5</td><td>0.8</td><td>-8.3</td><td>1.0E-16</td><td>3.7E-16</td><td>BT20/MB468</td></tr>
<tr><td>>MI0003820|hsa-mir-454&WithFlank&17|-|59137752|59137878@70.92.23&[MIMAT0003885&hsa-miR-454-3p&offsets|0|0;m-272&17|-|59137787|59137809&offsets|0|0]|TAGTGCAATATTGCTTATAGGGT</td><td>>MI0003820|hsa-mir-454&WithFlank&17|-|59137752|59137878</td><td>70</td><td>92</td><td>23</td><td>TAGTGCAATATTGCTTATAGGGT</td><td>1851.7</td><td>8.4</td><td>1.0</td><td>8.3</td><td>1.0E-16</td><td>3.7E-16</td><td>BT20/MB468</td></tr>
<tr><td>>hg38_1_-_228634819_228635039_RNA5S12_5S_50ntFlanks@140.171.32|ACCGCCTGGGAATACCGGGTGCTGTAGGCTTT</td><td>>hg38_1_-_228634819_228635039_RNA5S12_5S_50ntFlanks</td><td>140</td><td>171</td><td>32</td><td>ACCGCCTGGGAATACCGGGTGCTGTAGGCTTT</td><td>14941.3</td><td>6.3</td><td>0.8</td><td>8.3</td><td>1.1E-16</td><td>3.7E-16</td><td>MB231/MB468</td></tr>
<tr><td>>hg38_21_+_8433172_8446622_RNA45SN1_45S_50ntFlanks@11704.11728.25|ATGAAGCGCGGGTAAACGGCGGGAG</td><td>>hg38_21_+_8433172_8446622_RNA45SN1_45S_50ntFlanks</td><td>11704</td><td>11728</td><td>25</td><td>ATGAAGCGCGGGTAAACGGCGGGAG</td><td>780.6</td><td>-7.6</td><td>0.9</td><td>-8.3</td><td>1.1E-16</td><td>3.8E-16</td><td>BT20/MB468</td></tr>
<tr><td>>hg38_21_+_8433172_8446622_RNA45SN1_45S_50ntFlanks@11569.11605.37|CAGCCGACTTAGAACTGGTGCGGACCAGGGGAATCCG</td><td>>hg38_21_+_8433172_8446622_RNA45SN1_45S_50ntFlanks</td><td>11569</td><td>11605</td><td>37</td><td>CAGCCGACTTAGAACTGGTGCGGACCAGGGGAATCCG</td><td>245.4</td><td>-5.0</td><td>0.6</td><td>-8.3</td><td>1.1E-16</td><td>3.8E-16</td><td>MB231/MB468</td></tr>
<tr><td>>hg38_21_+_8433172_8446622_RNA45SN1_45S_50ntFlanks@11270.11295.26|CGGCCGCCCCCGGGGAGCCCGGCGGG</td><td>>hg38_21_+_8433172_8446622_RNA45SN1_45S_50ntFlanks</td><td>11270</td><td>11295</td><td>26</td><td>CGGCCGCCCCCGGGGAGCCCGGCGGG</td><td>570.2</td><td>-10.7</td><td>1.3</td><td>-8.3</td><td>1.1E-16</td><td>3.9E-16</td><td>MB231/MB468</td></tr>
<tr><td>>hg38_21_+_8433172_8446622_RNA45SN1_45S_50ntFlanks@12418.12441.24|CGATGTCGGCTCTTCCTATCATTG</td><td>>hg38_21_+_8433172_8446622_RNA45SN1_45S_50ntFlanks</td><td>12418</td><td>12441</td><td>24</td><td>CGATGTCGGCTCTTCCTATCATTG</td><td>866.1</td><td>-4.8</td><td>0.6</td><td>-8.3</td><td>1.1E-16</td><td>3.9E-16</td><td>BT20/MB468</td></tr>
<tr><td>>MI0000434|hsa-let-7i&WithFlank&12|+|62603680|62603775@12.33.22(+1A)&[MIMAT0000415&hsa-let-7i-5p&offsets|0|0(+1A);m-49&12|+|62603691|62603712&offsets|0|0(+1A)]|TGAGGTAGTAGTTTGTGCTGTTA</td><td>>MI0000434|hsa-let-7i&WithFlank&12|+|62603680|62603775</td><td>12</td><td>33</td><td>22(+1A)</td><td>TGAGGTAGTAGTTTGTGCTGTTA</td><td>11542.9</td><td>6.2</td><td>0.7</td><td>8.3</td><td>1.1E-16</td><td>3.9E-16</td><td>BT20/MB468</td></tr>
<tr><td>>MI0000298|hsa-mir-221&WithFlank&X|-|45746151|45746272@71.93.23(+1U)&[MIMAT0000278&hsa-miR-221-3p&offsets|0|0(+1U);m-91&X|-|45746181|45746202&offsets|0|+1(+1U)]|AGCTACATTGTCTGCTGGGTTTCT</td><td>>MI0000298|hsa-mir-221&WithFlank&X|-|45746151|45746272</td><td>71</td><td>93</td><td>23(+1U)</td><td>AGCTACATTGTCTGCTGGGTTTCT</td><td>6174.7</td><td>5.3</td><td>0.6</td><td>8.3</td><td>1.1E-16</td><td>3.9E-16</td><td>BT20/MB468</td></tr>
<tr><td>>MI0000440|hsa-mir-27b&WithFlank&9|+|95085439|95085547@67.85.19&[MIMAT0000419&hsa-miR-27b-3p&offsets|0|-2;m-39&9|+|95085505|95085525&offsets|0|-2]|TTCACAGTGGCTAAGTTCT</td><td>>MI0000440|hsa-mir-27b&WithFlank&9|+|95085439|95085547</td><td>67</td><td>85</td><td>19</td><td>TTCACAGTGGCTAAGTTCT</td><td>4484.2</td><td>6.9</td><td>0.8</td><td>8.3</td><td>1.1E-16</td><td>3.9E-16</td><td>BT20/MB468</td></tr>
<tr><td>>hg38_21_+_8433172_8446622_RNA45SN1_45S_50ntFlanks@12636.12654.19|AGCTACCATCTGTGGGATT</td><td>>hg38_21_+_8433172_8446622_RNA45SN1_45S_50ntFlanks</td><td>12636</td><td>12654</td><td>19</td><td>AGCTACCATCTGTGGGATT</td><td>2585.6</td><td>-9.3</td><td>1.1</td><td>-8.3</td><td>1.1E-16</td><td>3.9E-16</td><td>BT20/MB468</td></tr>
<tr><td>>am_MI0000786|hsa-mir-378a&WithFlank&5|+|149732819|149732896@49.70.22(+1U)&[MIMAT0000732&hsa-miR-378a-3p&offsets|0|0(+1U);m-44&5|+|149732867|149732888&offsets|0|0(+1U)]|ACTGGACTTGGAGTCAGAAGGCT</td><td>>am_MI0000786|hsa-mir-378a&WithFlank&5|+|149732819|149732896</td><td>49</td><td>70</td><td>22(+1U)</td><td>ACTGGACTTGGAGTCAGAAGGCT</td><td>3790.6</td><td>5.8</td><td>0.7</td><td>8.3</td><td>1.1E-16</td><td>4.0E-16</td><td>MB231/MB468</td></tr>
<tr><td>>MI0000095|hsa-mir-93&WithFlank&7|-|100093762|100093853@17.37.21&[MIMAT0000093&hsa-miR-93-5p&offsets|0|-2;m-29&7|-|100093815|100093837&offsets|0|-2]|CAAAGTGCTGTTCGTGCAGGT</td><td>>MI0000095|hsa-mir-93&WithFlank&7|-|100093762|100093853</td><td>17</td><td>37</td><td>21</td><td>CAAAGTGCTGTTCGTGCAGGT</td><td>3445.8</td><td>7.1</td><td>0.9</td><td>8.3</td><td>1.2E-16</td><td>4.0E-16</td><td>MB231/MB468</td></tr>
<tr><td>>MI0000076|hsa-mir-20a&WithFlank&13|+|91351059|91351141@14.35.22&[MIMAT0000075&hsa-miR-20a-5p&offsets|0|-1;m-96&13|+|91351072|91351094&offsets|0|-1]|TAAAGTGCTTATAGTGCAGGTA</td><td>>MI0000076|hsa-mir-20a&WithFlank&13|+|91351059|91351141</td><td>14</td><td>35</td><td>22</td><td>TAAAGTGCTTATAGTGCAGGTA</td><td>1503.3</td><td>6.7</td><td>0.8</td><td>8.3</td><td>1.2E-16</td><td>4.0E-16</td><td>MB231/MB468</td></tr>
<tr><td>>hg38_21_+_8433172_8446622_RNA45SN1_45S_50ntFlanks@12416.12440.25|TTCGATGTCGGCTCTTCCTATCATT</td><td>>hg38_21_+_8433172_8446622_RNA45SN1_45S_50ntFlanks</td><td>12416</td><td>12440</td><td>25</td><td>TTCGATGTCGGCTCTTCCTATCATT</td><td>682.5</td><td>5.2</td><td>0.6</td><td>8.3</td><td>1.1E-16</td><td>4.0E-16</td><td>BT20/MB468</td></tr>
<tr><td>>trnaMT_PheGAA_MT_+_577_647@1.18.18|GTTTATGTAGCTTACCTC</td><td>>trnaMT_PheGAA_MT_+_577_647</td><td>1</td><td>18</td><td>18</td><td>GTTTATGTAGCTTACCTC</td><td>1160.4</td><td>6.7</td><td>0.8</td><td>8.3</td><td>1.1E-16</td><td>4.1E-16</td><td>BT20/MB468</td></tr>
<tr><td>>hg38_21_+_8433172_8446622_RNA45SN1_45S_50ntFlanks@8357.8387.31|AAAAGAACTTTGAAGAGAGAGTTCAAGAGGG</td><td>>hg38_21_+_8433172_8446622_RNA45SN1_45S_50ntFlanks</td><td>8357</td><td>8387</td><td>31</td><td>AAAAGAACTTTGAAGAGAGAGTTCAAGAGGG</td><td>158.5</td><td>-3.7</td><td>0.4</td><td>-8.3</td><td>1.2E-16</td><td>4.1E-16</td><td>BT20/MB468</td></tr>
<tr><td>>hg38_21_+_8433172_8446622_RNA45SN1_45S_50ntFlanks@13002.13023.22|CGATCTATTGAAAGTCAGCCCT</td><td>>hg38_21_+_8433172_8446622_RNA45SN1_45S_50ntFlanks</td><td>13002</td><td>13023</td><td>22</td><td>CGATCTATTGAAAGTCAGCCCT</td><td>497.9</td><td>-11.0</td><td>1.3</td><td>-8.3</td><td>1.2E-16</td><td>4.2E-16</td><td>BT20/MB468</td></tr>
<tr><td>>hg38_21_+_8433172_8446622_RNA45SN1_45S_50ntFlanks@6732.6773.42|ACATTGATCATCGACACTTCGAACGCACTTGCGGCCCCGGGT</td><td>>hg38_21_+_8433172_8446622_RNA45SN1_45S_50ntFlanks</td><td>6732</td><td>6773</td><td>42</td><td>ACATTGATCATCGACACTTCGAACGCACTTGCGGCCCCGGGT</td><td>1075.5</td><td>9.0</td><td>1.1</td><td>8.3</td><td>1.2E-16</td><td>4.2E-16</td><td>BT20/MB468</td></tr>
<tr><td>>hg38_21_+_8433172_8446622_RNA45SN1_45S_50ntFlanks@12420.12460.41|ATGTCGGCTCTTCCTATCATTGTGAAGCAGAATTCACCAAG</td><td>>hg38_21_+_8433172_8446622_RNA45SN1_45S_50ntFlanks</td><td>12420</td><td>12460</td><td>41</td><td>ATGTCGGCTCTTCCTATCATTGTGAAGCAGAATTCACCAAG</td><td>586.7</td><td>4.9</td><td>0.6</td><td>8.3</td><td>1.2E-16</td><td>4.2E-16</td><td>MB231/MB468</td></tr>
<tr><td>>hg38_21_+_8433172_8446622_RNA45SN1_45S_50ntFlanks@6652.6687.36|GACTCTTAGCGGTGGATCACTCGGCTCGTGCGTCGA</td><td>>hg38_21_+_8433172_8446622_RNA45SN1_45S_50ntFlanks</td><td>6652</td><td>6687</td><td>36</td><td>GACTCTTAGCGGTGGATCACTCGGCTCGTGCGTCGA</td><td>2889.8</td><td>6.3</td><td>0.8</td><td>8.3</td><td>1.2E-16</td><td>4.2E-16</td><td>MB231/MB468</td></tr>
<tr><td>>hg38_21_+_8433172_8446622_RNA45SN1_45S_50ntFlanks@10574.10608.35|AAGCCGGCGGGAGCCCCGGGGAGAGTTCTCTTTTC</td><td>>hg38_21_+_8433172_8446622_RNA45SN1_45S_50ntFlanks</td><td>10574</td><td>10608</td><td>35</td><td>AAGCCGGCGGGAGCCCCGGGGAGAGTTCTCTTTTC</td><td>486.4</td><td>-6.4</td><td>0.8</td><td>-8.3</td><td>1.2E-16</td><td>4.2E-16</td><td>MB231/MB468</td></tr>
<tr><td>>hg38_21_+_8433172_8446622_RNA45SN1_45S_50ntFlanks@12044.12068.25|ACTGACCCGGTGAGGCGGGGGGGCG</td><td>>hg38_21_+_8433172_8446622_RNA45SN1_45S_50ntFlanks</td><td>12044</td><td>12068</td><td>25</td><td>ACTGACCCGGTGAGGCGGGGGGGCG</td><td>260.3</td><td>-6.5</td><td>0.8</td><td>-8.3</td><td>1.2E-16</td><td>4.2E-16</td><td>BT20/MB468</td></tr>
<tr><td>>hg38_21_+_8433172_8446622_RNA45SN1_45S_50ntFlanks@10391.10419.29|AACAGCAGTTGAACATGGGTCAGTCGGTC</td><td>>hg38_21_+_8433172_8446622_RNA45SN1_45S_50ntFlanks</td><td>10391</td><td>10419</td><td>29</td><td>AACAGCAGTTGAACATGGGTCAGTCGGTC</td><td>326.5</td><td>-10.3</td><td>1.2</td><td>-8.3</td><td>1.2E-16</td><td>4.2E-16</td><td>BT20/MB468</td></tr>
<tr><td>>am_trnaMT_ProTGG_MT_-_15956_16023@53.71.19|AAGACTTTTTCTCTGACCA</td><td>>am_trnaMT_ProTGG_MT_-_15956_16023</td><td>53</td><td>71</td><td>19</td><td>AAGACTTTTTCTCTGACCA</td><td>11276.7</td><td>5.1</td><td>0.6</td><td>8.3</td><td>1.2E-16</td><td>4.2E-16</td><td>BT20/MB468</td></tr>
<tr><td>>trnaMT_SerGCT_MT_+_12207_12265@1.26.26|GAGAAAGCTCACAAGAACTGCTAACT</td><td>>trnaMT_SerGCT_MT_+_12207_12265</td><td>1</td><td>26</td><td>26</td><td>GAGAAAGCTCACAAGAACTGCTAACT</td><td>10831.0</td><td>6.5</td><td>0.8</td><td>8.3</td><td>1.2E-16</td><td>4.2E-16</td><td>MB231/MB468</td></tr>
<tr><td>>hg38_21_+_8433172_8446622_RNA45SN1_45S_50ntFlanks@8472.8492.21|CGGCCCGGCGGATCTTTCCCG</td><td>>hg38_21_+_8433172_8446622_RNA45SN1_45S_50ntFlanks</td><td>8472</td><td>8492</td><td>21</td><td>CGGCCCGGCGGATCTTTCCCG</td><td>574.0</td><td>-11.2</td><td>1.3</td><td>-8.3</td><td>1.2E-16</td><td>4.3E-16</td><td>BT20/MB468</td></tr>
</table>

<table>
<tr><td>>hg38_21_+_8433172_8446622_RNA45SN1_45S_50ntFlanks@7986.8008.23|ATCAGACGTGGCGACCCGCTGAA</td><td>>hg38_21_+_8433172_8446622_RNA45SN1_45S_50ntFlanks</td><td>7986</td><td>8008</td><td>23</td><td>ATCAGACGTGGCGACCCGCTGAA</td><td>437.8</td><td>-10.8</td><td>1.3</td><td>-8.3</td><td>1.2E-16</td><td>4.3E-16</td><td>BT20/MB468</td></tr>
<tr><td>>hg38_21_+_8433172_8446622_RNA45SN1_45S_50ntFlanks@3984.4002.19|GCGGCTTTGGTGACTCTAG</td><td>>hg38_21_+_8433172_8446622_RNA45SN1_45S_50ntFlanks</td><td>3984</td><td>4002</td><td>19</td><td>GCGGCTTTGGTGACTCTAG</td><td>523.4</td><td>-10.6</td><td>1.3</td><td>-8.3</td><td>1.3E-16</td><td>4.4E-16</td><td>MB231/MB468</td></tr>
<tr><td>>MI0000095|hsa-mir-93&WithFlank&7|-|100093762|100093853@17.39.23&[MIMAT0000093&hsa-miR-93-5p&offsets|0|0;m-29&7|-|100093815|100093837&offsets|0|0]|CAAAGTGCTGTTCGTGCAGGTAG</td><td>>MI0000095|hsa-mir-93&WithFlank&7|-|100093762|100093853</td><td>17</td><td>39</td><td>23</td><td>CAAAGTGCTGTTCGTGCAGGTAG</td><td>29841.4</td><td>6.4</td><td>0.8</td><td>8.3</td><td>1.3E-16</td><td>4.4E-16</td><td>MB231/MB468</td></tr>
<tr><td>>hg38_21_+_8433172_8446622_RNA45SN1_45S_50ntFlanks@3802.3834.33|CATTAAATCAGTTATGGTTCCTTTGGTCGCTCG</td><td>>hg38_21_+_8433172_8446622_RNA45SN1_45S_50ntFlanks</td><td>3802</td><td>3834</td><td>33</td><td>CATTAAATCAGTTATGGTTCCTTTGGTCGCTCG</td><td>231.5</td><td>-4.5</td><td>0.5</td><td>-8.3</td><td>1.2E-16</td><td>4.4E-16</td><td>BT20/MB468</td></tr>
<tr><td>>MI0000447|hsa-mir-128-1&WithFlank&2|+|135665391|135665484@56.75.20(+1G);MI0000727|hsa-mir-128-2&WithFlank&3|+|35744470|35744565@58.77.20(+1G)&[MIMAT0000424&hsa-miR-128-3p&offsets|0|-1(+1G);m-108&2|+|135665446|135665466&offsets|0|-1(+1G)];[MIMAT0000424_1&hsa-miR-128-3p&offsets|0|-1(+1G);m-117&3|+|35744527|35744547&offsets|0|-1(+1G)]|TCACAGTGAACCGGTCTCTTG</td><td>>MI0000447|hsa-mir-128-1&WithFlank&2|+|135665391|135665484</td><td>56</td><td>75</td><td>20(+1G)</td><td>TCACAGTGAACCGGTCTCTTG</td><td>837.8</td><td>7.1</td><td>0.9</td><td>8.3</td><td>1.2E-16</td><td>4.4E-16</td><td>BT20/MB468</td></tr>
<tr><td>>hg38_21_+_8433172_8446622_RNA45SN1_45S_50ntFlanks@11564.11581.18|CCTAGCAGCCGACTTAGA</td><td>>hg38_21_+_8433172_8446622_RNA45SN1_45S_50ntFlanks</td><td>11564</td><td>11581</td><td>18</td><td>CCTAGCAGCCGACTTAGA</td><td>800.4</td><td>-3.8</td><td>0.5</td><td>-8.3</td><td>1.3E-16</td><td>4.4E-16</td><td>MB231/MB468</td></tr>
<tr><td>>MI0000542|hsa-mir-320a&WithFlank&8|-|22244960|22245043@49.69.21(+2U)&[MIMAT0000510&hsa-miR-320a-3p&offsets|+1|0(+2U);m-60&8|-|22244975|22244996&offsets|+1|0(+2U)]|AAAGCTGGGTTGAGAGGGCGATT</td><td>>MI0000542|hsa-mir-320a&WithFlank&8|-|22244960|22245043</td><td>49</td><td>69</td><td>21(+2U)</td><td>AAAGCTGGGTTGAGAGGGCGATT</td><td>535.6</td><td>10.6</td><td>1.3</td><td>8.3</td><td>1.3E-16</td><td>4.5E-16</td><td>BT20/MB468</td></tr>
<tr><td>>hg38_21_+_8433172_8446622_RNA45SN1_45S_50ntFlanks@10690.10710.21|CGTCCGGTGAGCTCTCGCTGG</td><td>>hg38_21_+_8433172_8446622_RNA45SN1_45S_50ntFlanks</td><td>10690</td><td>10710</td><td>21</td><td>CGTCCGGTGAGCTCTCGCTGG</td><td>536.6</td><td>-10.6</td><td>1.3</td><td>-8.3</td><td>1.3E-16</td><td>4.5E-16</td><td>MB231/MB468</td></tr>
<tr><td>>hg38_21_+_8433172_8446622_RNA45SN1_45S_50ntFlanks@5472.5513.42|ACGGCCCTGGCGGAGCGCTGAGAAGACGGTCGAACTTGACTA</td><td>>hg38_21_+_8433172_8446622_RNA45SN1_45S_50ntFlanks</td><td>5472</td><td>5513</td><td>42</td><td>ACGGCCCTGGCGGAGCGCTGAGAAGACGGTCGAACTTGACTA</td><td>782.0</td><td>8.7</td><td>1.0</td><td>8.3</td><td>1.3E-16</td><td>4.5E-16</td><td>MB231/MB468</td></tr>
<tr><td>>hg38_21_+_8433172_8446622_RNA45SN1_45S_50ntFlanks@11873.11893.21|AAGAAGACCCTGTTGAGCTTG</td><td>>hg38_21_+_8433172_8446622_RNA45SN1_45S_50ntFlanks</td><td>11873</td><td>11893</td><td>21</td><td>AAGAAGACCCTGTTGAGCTTG</td><td>4456.3</td><td>-5.9</td><td>0.7</td><td>-8.3</td><td>1.3E-16</td><td>4.5E-16</td><td>BT20/MB468</td></tr>
<tr><td>>MI0000483|hsa-mir-186&WithFlank&1|-|71067625|71067722@21.40.20&[MIMAT0000456&hsa-miR-186-5p&offsets|0|-2;m-84&1|-|71067681|71067702&offsets|0|-2]|CAAAGAATTCTCCTTTTGGG</td><td>>MI0000483|hsa-mir-186&WithFlank&1|-|71067625|71067722</td><td>21</td><td>40</td><td>20</td><td>CAAAGAATTCTCCTTTTGGG</td><td>737.7</td><td>6.0</td><td>0.7</td><td>8.3</td><td>1.3E-16</td><td>4.5E-16</td><td>MB231/MB468</td></tr>
<tr><td>>hg38_21_+_8433172_8446622_RNA45SN1_45S_50ntFlanks@12151.12182.32|GTGGGGAGTTTGACTGGGGCGGTACACCTGTC</td><td>>hg38_21_+_8433172_8446622_RNA45SN1_45S_50ntFlanks</td><td>12151</td><td>12182</td><td>32</td><td>GTGGGGAGTTTGACTGGGGCGGTACACCTGTC</td><td>371.5</td><td>-10.5</td><td>1.3</td><td>-8.3</td><td>1.3E-16</td><td>4.6E-16</td><td>BT20/MB468</td></tr>
<tr><td>>MI0000298|hsa-mir-221&WithFlank&X|-|45746151|45746272@31.53.23&[MIMAT0004568&hsa-miR-221-5p&offsets|0|+1;m-405&X|-|45746220|45746242&offsets|0|0]|ACCTGGCATACAATGTAGATTTC</td><td>>MI0000298|hsa-mir-221&WithFlank&X|-|45746151|45746272</td><td>31</td><td>53</td><td>23</td><td>ACCTGGCATACAATGTAGATTTC</td><td>542.8</td><td>5.5</td><td>0.7</td><td>8.3</td><td>1.3E-16</td><td>4.7E-16</td><td>BT20/MB468</td></tr>
<tr><td>>MI0000447|hsa-mir-128-1&WithFlank&2|+|135665391|135665484@56.74.19;MI0000727|hsa-mir-128-2&WithFlank&3|+|35744470|35744565@58.76.19&[MIMAT0000424&hsa-miR-128-3p&offsets|0|-2;m-108&2|+|135665446|135665466&offsets|0|-2];[MIMAT0000424_1&hsa-miR-128-3p&offsets|0|-2;m-117&3|+|35744527|35744547&offsets|0|-2]|TCACAGTGAACCGGTCTCT</td><td>>MI0000447|hsa-mir-128-1&WithFlank&2|+|135665391|135665484</td><td>56</td><td>74</td><td>19</td><td>TCACAGTGAACCGGTCTCT</td><td>449.1</td><td>6.7</td><td>0.8</td><td>8.3</td><td>1.3E-16</td><td>4.7E-16</td><td>BT20/MB468</td></tr>
<tr><td>>hg38_21_+_8433172_8446622_RNA45SN1_45S_50ntFlanks@12476.12499.24|CCACTAATAGGGAACGTGAGCTGG</td><td>>hg38_21_+_8433172_8446622_RNA45SN1_45S_50ntFlanks</td><td>12476</td><td>12499</td><td>24</td><td>CCACTAATAGGGAACGTGAGCTGG</td><td>368.1</td><td>-10.5</td><td>1.3</td><td>-8.3</td><td>1.3E-16</td><td>4.7E-16</td><td>BT20/MB468</td></tr>
<tr><td>>hg38_1_-_228634819_228635039_RNA5S12_5S_50ntFlanks@84.101.18|CTCGTCTGATCTCGGAAG</td><td>>hg38_1_-_228634819_228635039_RNA5S12_5S_50ntFlanks</td><td>84</td><td>101</td><td>18</td><td>CTCGTCTGATCTCGGAAG</td><td>1513.5</td><td>-5.4</td><td>0.7</td><td>-8.3</td><td>1.4E-16</td><td>4.7E-16</td><td>MB231/MB468</td></tr>
<tr><td>>MI0000483|hsa-mir-186&WithFlank&1|-|71067625|71067722@21.41.21&[MIMAT0000456&hsa-miR-186-5p&offsets|0|-1;m-84&1|-|71067681|71067702&offsets|0|-1]|CAAAGAATTCTCCTTTTGGGC</td><td>>MI0000483|hsa-mir-186&WithFlank&1|-|71067625|71067722</td><td>21</td><td>41</td><td>21</td><td>CAAAGAATTCTCCTTTTGGGC</td><td>885.7</td><td>6.6</td><td>0.8</td><td>8.3</td><td>1.4E-16</td><td>4.8E-16</td><td>BT20/MB468</td></tr>
<tr><td>>MI0000066|hsa-let-7e&WithFlank&19|+|51692780|51692870@14.33.20(+1C)&[MIMAT0000066&hsa-let-7e-5p&offsets|0|-2(+1C);m-50&19|+|51692793|51692814&offsets|0|-2(+1C)]|TGAGGTAGGAGGTTGTATAGC</td><td>>MI0000066|hsa-let-7e&WithFlank&19|+|51692780|51692870</td><td>14</td><td>33</td><td>20(+1C)</td><td>TGAGGTAGGAGGTTGTATAGC</td><td>598.1</td><td>8.7</td><td>1.0</td><td>8.3</td><td>1.4E-16</td><td>4.9E-16</td><td>BT20/MB468</td></tr>
<tr><td>>hg38_21_+_8433172_8446622_RNA45SN1_45S_50ntFlanks@12407.12426.20|TTTTGATCCTTCGATGTCGG</td><td>>hg38_21_+_8433172_8446622_RNA45SN1_45S_50ntFlanks</td><td>12407</td><td>12426</td><td>20</td><td>TTTTGATCCTTCGATGTCGG</td><td>290.8</td><td>-6.3</td><td>0.8</td><td>-8.3</td><td>1.4E-16</td><td>4.9E-16</td><td>MB231/MB468</td></tr>
<tr><td>>hg38_21_+_8433172_8446622_RNA45SN1_45S_50ntFlanks@485.508.24|GATGTGGTGACGTCGTGCTCTCCC</td><td>>hg38_21_+_8433172_8446622_RNA45SN1_45S_50ntFlanks</td><td>485</td><td>508</td><td>24</td><td>GATGTGGTGACGTCGTGCTCTCCC</td><td>4227.5</td><td>30.0</td><td>3.6</td><td>8.3</td><td>1.4E-16</td><td>4.9E-16</td><td>MB231/MB468</td></tr>
<tr><td>>hg38_21_+_8433172_8446622_RNA45SN1_45S_50ntFlanks@485.509.25|GATGTGGTGACGTCGTGCTCTCCCG</td><td>>hg38_21_+_8433172_8446622_RNA45SN1_45S_50ntFlanks</td><td>485</td><td>509</td><td>25</td><td>GATGTGGTGACGTCGTGCTCTCCCG</td><td>5213.0</td><td>30.0</td><td>3.6</td><td>8.3</td><td>1.4E-16</td><td>4.9E-16</td><td>MB231/MB468</td></tr>
<tr><td>>hg38_21_+_8433172_8446622_RNA45SN1_45S_50ntFlanks@485.515.31|GATGTGGTGACGTCGTGCTCTCCCGGGCCGG</td><td>>hg38_21_+_8433172_8446622_RNA45SN1_45S_50ntFlanks</td><td>485</td><td>515</td><td>31</td><td>GATGTGGTGACGTCGTGCTCTCCCGGGCCGG</td><td>4366.4</td><td>30.0</td><td>3.6</td><td>8.3</td><td>1.4E-16</td><td>4.9E-16</td><td>MB231/MB468</td></tr>
<tr><td>>hg38_21_+_8433172_8446622_RNA45SN1_45S_50ntFlanks@485.521.37|GATGTGGTGACGTCGTGCTCTCCCGGGCCGGGTCCGA</td><td>>hg38_21_+_8433172_8446622_RNA45SN1_45S_50ntFlanks</td><td>485</td><td>521</td><td>37</td><td>GATGTGGTGACGTCGTGCTCTCCCGGGCCGGGTCCGA</td><td>3433.7</td><td>30.0</td><td>3.6</td><td>8.3</td><td>1.4E-16</td><td>4.9E-16</td><td>MB231/MB468</td></tr>
<tr><td>>hg38_21_+_8433172_8446622_RNA45SN1_45S_50ntFlanks@485.525.41|GATGTGGTGACGTCGTGCTCTCCCGGGCCGGGTCCGAGCCG</td><td>>hg38_21_+_8433172_8446622_RNA45SN1_45S_50ntFlanks</td><td>485</td><td>525</td><td>41</td><td>GATGTGGTGACGTCGTGCTCTCCCGGGCCGGGTCCGAGCCG</td><td>4092.9</td><td>30.0</td><td>3.6</td><td>8.3</td><td>1.4E-16</td><td>4.9E-16</td><td>MB231/MB468</td></tr>
<tr><td>>hg38_21_+_8433172_8446622_RNA45SN1_45S_50ntFlanks@485.527.43|GATGTGGTGACGTCGTGCTCTCCCGGGCCGGGTCCGAGCCGCG</td><td>>hg38_21_+_8433172_8446622_RNA45SN1_45S_50ntFlanks</td><td>485</td><td>527</td><td>43</td><td>GATGTGGTGACGTCGTGCTCTCCCGGGCCGGGTCCGAGCCGCG</td><td>3221.4</td><td>30.0</td><td>3.6</td><td>8.3</td><td>1.4E-16</td><td>4.9E-16</td><td>MB231/MB468</td></tr>
<tr><td>>hg38_21_+_8433172_8446622_RNA45SN1_45S_50ntFlanks@485.528.44|GATGTGGTGACGTCGTGCTCTCCCGGGCCGGGTCCGAGCCGCGA</td><td>>hg38_21_+_8433172_8446622_RNA45SN1_45S_50ntFlanks</td><td>485</td><td>528</td><td>44</td><td>GATGTGGTGACGTCGTGCTCTCCCGGGCCGGGTCCGAGCCGCGA</td><td>2082.3</td><td>30.0</td><td>3.6</td><td>8.3</td><td>1.4E-16</td><td>4.9E-16</td><td>MB231/MB468</td></tr>
<tr><td>>hg38_21_+_8433172_8446622_RNA45SN1_45S_50ntFlanks@485.530.46|GATGTGGTGACGTCGTGCTCTCCCGGGCCGGGTCCGAGCCGCGACG</td><td>>hg38_21_+_8433172_8446622_RNA45SN1_45S_50ntFlanks</td><td>485</td><td>530</td><td>46</td><td>GATGTGGTGACGTCGTGCTCTCCCGGGCCGGGTCCGAGCCGCGACG</td><td>3192.1</td><td>30.0</td><td>3.6</td><td>8.3</td><td>1.4E-16</td><td>4.9E-16</td><td>MB231/MB468</td></tr>
<tr><td>>hg38_21_+_8433172_8446622_RNA45SN1_45S_50ntFlanks@485.531.47|GATGTGGTGACGTCGTGCTCTCCCGGGCCGGGTCCGAGCCGCGACGG</td><td>>hg38_21_+_8433172_8446622_RNA45SN1_45S_50ntFlanks</td><td>485</td><td>531</td><td>47</td><td>GATGTGGTGACGTCGTGCTCTCCCGGGCCGGGTCCGAGCCGCGACGG</td><td>4101.3</td><td>30.0</td><td>3.6</td><td>8.3</td><td>1.4E-16</td><td>4.9E-16</td><td>MB231/MB468</td></tr>
<tr><td>>MI0000079|hsa-mir-23a&WithFlank&19|-|13836581|13836665@54.71.18&[MIMAT0000078&hsa-miR-23a-3p&offsets|+3|0;m-31&19|-|13836595|13836615&offsets|+3|0]|ACATTGCCAGGGATTTCC</td><td>>MI0000079|hsa-mir-23a&WithFlank&19|-|13836581|13836665</td><td>54</td><td>71</td><td>18</td><td>ACATTGCCAGGGATTTCC</td><td>19437.9</td><td>30.0</td><td>3.6</td><td>8.3</td><td>1.4E-16</td><td>4.9E-16</td><td>MB231/MB468</td></tr>
<tr><td>>MI0000079|hsa-mir-23a&WithFlank&19|-|13836581|13836665@54.72.19(+1U)&[MIMAT0000078&hsa-miR-23a-3p&offsets|+3|+1(+1U);m-31&19|-|13836595|13836615&offsets|+3|+1(+1U)]|ACATTGCCAGGGATTTCCAT</td><td>>MI0000079|hsa-mir-23a&WithFlank&19|-|13836581|13836665</td><td>54</td><td>72</td><td>19(+1U)</td><td>ACATTGCCAGGGATTTCCAT</td><td>3723.0</td><td>30.0</td><td>3.6</td><td>8.3</td><td>1.4E-16</td><td>4.9E-16</td><td>MB231/MB468</td></tr>
<tr><td>>MI0000079|hsa-mir-23a&WithFlank&19|-|13836581|13836665@54.72.19(+2U)&[MIMAT0000078&hsa-miR-23a-3p&offsets|+3|+1(+2U);m-31&19|-|13836595|13836615&offsets|+3|+1(+2U)]|ACATTGCCAGGGATTTCCATT</td><td>>MI0000079|hsa-mir-23a&WithFlank&19|-|13836581|13836665</td><td>54</td><td>72</td><td>19(+2U)</td><td>ACATTGCCAGGGATTTCCATT</td><td>3840.9</td><td>30.0</td><td>3.6</td><td>8.3</td><td>1.4E-16</td><td>4.9E-16</td><td>MB231/MB468</td></tr>
<tr><td>>MI0000079|hsa-mir-23a&WithFlank&19|-|13836581|13836665@54.72.19&[MIMAT0000078&hsa-miR-23a-3p&offsets|+3|+1;m-31&19|-|13836595|13836615&offsets|+3|+1]|ACATTGCCAGGGATTTCCA</td><td>>MI0000079|hsa-mir-23a&WithFlank&19|-|13836581|13836665</td><td>54</td><td>72</td><td>19</td><td>ACATTGCCAGGGATTTCCA</td><td>22491.1</td><td>30.0</td><td>3.6</td><td>8.3</td><td>1.4E-16</td><td>4.9E-16</td><td>MB231/MB468</td></tr>
<tr><td>>MI0000079|hsa-mir-23a&WithFlank&19|-|13836581|13836665@54.73.20(+1U)&[MIMAT0000078&hsa-miR-23a-3p&offsets|+3|+2(+1U);m-31&19|-|13836595|13836615&offsets|+3|+2(+1U)]|ACATTGCCAGGGATTTCCAAT</td><td>>MI0000079|hsa-mir-23a&WithFlank&19|-|13836581|13836665</td><td>54</td><td>73</td><td>20(+1U)</td><td>ACATTGCCAGGGATTTCCAAT</td><td>17374.0</td><td>30.0</td><td>3.6</td><td>8.3</td><td>1.4E-16</td><td>4.9E-16</td><td>MB231/MB468</td></tr>
<tr><td>>MI0000079|hsa-mir-23a&WithFlank&19|-|13836581|13836665@54.73.20&[MIMAT0000078&hsa-miR-23a-3p&offsets|+3|+2;m-31&19|-|13836595|13836615&offsets|+3|+2]|ACATTGCCAGGGATTTCCAA</td><td>>MI0000079|hsa-mir-23a&WithFlank&19|-|13836581|13836665</td><td>54</td><td>73</td><td>20</td><td>ACATTGCCAGGGATTTCCAA</td><td>13045.2</td><td>30.0</td><td>3.6</td><td>8.3</td><td>1.4E-16</td><td>4.9E-16</td><td>MB231/MB468</td></tr>
<tr><td>>MI0000081|hsa-mir-24-2&WithFlank&19|-|13836281|13836365@56.77.22(+1A);MI0000080|hsa-mir-24-1&WithFlank&9|+|95086015|95086094@50.71.22(+1A)&[MIMAT0000080&hsa-miR-24-3p&offsets|0|0(+1A);m-36&19|-|13836288|13836310&offsets|0|-1(+1A)];[MIMAT0000080_1&hsa-miR-24-3p&offsets|0|0(+1A);m-35&9|+|95086064|95086086&offsets|0|-1(+1A)]|TGGCTCAGTTCAGCAGGAACAGA</td><td>>MI0000081|hsa-mir-24-2&WithFlank&19|-|13836281|13836365</td><td>56</td><td>77</td><td>22(+1A)</td><td>TGGCTCAGTTCAGCAGGAACAGA</td><td>875.5</td><td>5.8</td><td>0.7</td><td>8.3</td><td>1.4E-16</td><td>4.9E-16</td><td>MB231/MB468</td></tr>
<tr><td>>MI0000439|hsa-mir-23b&WithFlank&9|+|95085202|95085310@67.84.18&[MIMAT0000418&hsa-miR-23b-3p&offsets|+3|-2;m-42&9|+|95085265|95085285&offsets|+3|0]|ACATTGCCAGGGATTACC</td><td>>MI0000439|hsa-mir-23b&WithFlank&9|+|95085202|95085310</td><td>67</td><td>84</td><td>18</td><td>ACATTGCCAGGGATTACC</td><td>2371.9</td><td>30.0</td><td>3.6</td><td>8.3</td><td>1.4E-16</td><td>4.9E-16</td><td>MB231/MB468</td></tr>
<tr><td>>MI0000439|hsa-mir-23b&WithFlank&9|+|95085202|95085310@67.85.19(+2U)&[MIMAT0000418&hsa-miR-23b-3p&offsets|+3|-1(+2U);m-42&9|+|95085265|95085285&offsets|+3|+1(+2U)]|ACATTGCCAGGGATTACCATT</td><td>>MI0000439|hsa-mir-23b&WithFlank&9|+|95085202|95085310</td><td>67</td><td>85</td><td>19(+2U)</td><td>ACATTGCCAGGGATTACCATT</td><td>2643.9</td><td>30.0</td><td>3.6</td><td>8.3</td><td>1.4E-16</td><td>4.9E-16</td><td>MB231/MB468</td></tr>
<tr><td>>MI0000439|hsa-mir-23b&WithFlank&9|+|95085202|95085310@67.86.20(+1U)&[MIMAT0000418&hsa-miR-23b-3p&offsets|+3|0(+1U);m-42&9|+|95085265|95085285&offsets|+3|+2(+1U)]|ACATTGCCAGGGATTACCACT</td><td>>MI0000439|hsa-mir-23b&WithFlank&9|+|95085202|95085310</td><td>67</td><td>86</td><td>20(+1U)</td><td>ACATTGCCAGGGATTACCACT</td><td>18479.5</td><td>30.0</td><td>3.6</td><td>8.3</td><td>1.4E-16</td><td>4.9E-16</td><td>MB231/MB468</td></tr>
<tr><td>>MI0000439|hsa-mir-23b&WithFlank&9|+|95085202|95085310@67.86.20&[MIMAT0000418&hsa-miR-23b-3p&offsets|+3|0;m-42&9|+|95085265|95085285&offsets|+3|+2]|ACATTGCCAGGGATTACCAC</td><td>>MI0000439|hsa-mir-23b&WithFlank&9|+|95085202|95085310</td><td>67</td><td>86</td><td>20</td><td>ACATTGCCAGGGATTACCAC</td><td>5929.0</td><td>30.0</td><td>3.6</td><td>8.3</td><td>1.4E-16</td><td>4.9E-16</td><td>MB231/MB468</td></tr>
</table>

<table>
<tr><td>>MI0000750|hsa-mir-26a-2&WithFlank&12|-|57824603|57824698@23.41.19;MI0000083|hsa-mir-26a-1&WithFlank&3|+|37969398|37969486@19.37.19&[MIMAT0000082&hsa-miR-26a-5p&offsets|+3|0;m-33&12|-|57824658|57824679&offsets|+3|0];[MIMAT0000082_1&hsa-miR-26a-5p&offsets|+3|0;m-34&3|+|37969413|37969434&offsets|+3|0]|AAGTAATCCAGGATAGGCT</td><td>>MI0000750|hsa-mir-26a-2&WithFlank&12|-|57824603|57824698</td><td>23</td><td>41</td><td>19</td><td>AAGTAATCCAGGATAGGCT</td><td>2586.7</td><td>30.0</td><td>3.6</td><td>8.3</td><td>1.4E-16</td><td>4.9E-16</td><td>MB231/MB468</td></tr>
<tr><td>>hg38_21_+_8433172_8446622_RNA45SN1_45S_50ntFlanks@11564.11601.38|CCTAGCAGCCGACTTAGAACTGGTGCGGACCAGGGGAA</td><td>>hg38_21_+_8433172_8446622_RNA45SN1_45S_50ntFlanks</td><td>11564</td><td>11601</td><td>38</td><td>CCTAGCAGCCGACTTAGAACTGGTGCGGACCAGGGGAA</td><td>1179.0</td><td>-5.4</td><td>0.7</td><td>-8.3</td><td>1.4E-16</td><td>4.9E-16</td><td>BT20/MB468</td></tr>
<tr><td>>hg38_21_+_8433172_8446622_RNA45SN1_45S_50ntFlanks@4734.4761.28|AACGAAAGTCGGAGGTTCGAAGACGATC</td><td>>hg38_21_+_8433172_8446622_RNA45SN1_45S_50ntFlanks</td><td>4734</td><td>4761</td><td>28</td><td>AACGAAAGTCGGAGGTTCGAAGACGATC</td><td>333.1</td><td>4.1</td><td>0.5</td><td>8.3</td><td>1.4E-16</td><td>4.9E-16</td><td>BT20/MB468</td></tr>
<tr><td>>MI0000079|hsa-mir-23a&WithFlank&19|-|13836581|13836665@54.73.20(+2U)&[MIMAT0000078&hsa-miR-23a-3p&offsets|+3|+2(+2U);m-31&19|-|13836595|13836615&offsets|+3|+2(+2U)]|ACATTGCCAGGGATTTCCAATT</td><td>>MI0000079|hsa-mir-23a&WithFlank&19|-|13836581|13836665</td><td>54</td><td>73</td><td>20(+2U)</td><td>ACATTGCCAGGGATTTCCAATT</td><td>2104.9</td><td>30.0</td><td>3.6</td><td>8.3</td><td>1.5E-16</td><td>5.0E-16</td><td>MB231/MB468</td></tr>
<tr><td>>hg38_21_+_8433172_8446622_RNA45SN1_45S_50ntFlanks@10845.10868.24|ACTTCGGGATAAGGATTGGCTCTA</td><td>>hg38_21_+_8433172_8446622_RNA45SN1_45S_50ntFlanks</td><td>10845</td><td>10868</td><td>24</td><td>ACTTCGGGATAAGGATTGGCTCTA</td><td>916.5</td><td>-6.1</td><td>0.7</td><td>-8.3</td><td>1.5E-16</td><td>5.0E-16</td><td>MB231/MB468</td></tr>
<tr><td>>hg38_21_+_8433172_8446622_RNA45SN1_45S_50ntFlanks@9564.9596.33|CTGACGTGCAAATCGGTCGTCCGACCTGGGTAT</td><td>>hg38_21_+_8433172_8446622_RNA45SN1_45S_50ntFlanks</td><td>9564</td><td>9596</td><td>33</td><td>CTGACGTGCAAATCGGTCGTCCGACCTGGGTAT</td><td>11176.1</td><td>-5.3</td><td>0.6</td><td>-8.3</td><td>1.5E-16</td><td>5.1E-16</td><td>MB231/MB468</td></tr>
<tr><td>>hg38_21_+_8433172_8446622_RNA45SN1_45S_50ntFlanks@12072.12090.19|CCCGAGGGGCTCTCGCTTC</td><td>>hg38_21_+_8433172_8446622_RNA45SN1_45S_50ntFlanks</td><td>12072</td><td>12090</td><td>19</td><td>CCCGAGGGGCTCTCGCTTC</td><td>185.5</td><td>-5.0</td><td>0.6</td><td>-8.3</td><td>1.4E-16</td><td>5.1E-16</td><td>BT20/MB468</td></tr>
<tr><td>>hg38_MT_+_598_1651_RNR1_12S_50ntFlanks@905.929.25|AGTATACTTCAAAGGACATTTAACT</td><td>>hg38_MT_+_598_1651_RNR1_12S_50ntFlanks</td><td>905</td><td>929</td><td>25</td><td>AGTATACTTCAAAGGACATTTAACT</td><td>1659.9</td><td>10.3</td><td>1.2</td><td>8.3</td><td>1.5E-16</td><td>5.1E-16</td><td>BT20/MB468</td></tr>
<tr><td>>hg38_21_+_8433172_8446622_RNA45SN1_45S_50ntFlanks@11212.11232.21|CGACTCTGGACGCGAGCCGGG</td><td>>hg38_21_+_8433172_8446622_RNA45SN1_45S_50ntFlanks</td><td>11212</td><td>11232</td><td>21</td><td>CGACTCTGGACGCGAGCCGGG</td><td>1110.9</td><td>-9.7</td><td>1.2</td><td>-8.3</td><td>1.5E-16</td><td>5.2E-16</td><td>MB231/MB468</td></tr>
<tr><td>>hg38_21_+_8433172_8446622_RNA45SN1_45S_50ntFlanks@12934.12966.33|CTAAACCATTCGTAGACGACCTGCTTCTGGGTC</td><td>>hg38_21_+_8433172_8446622_RNA45SN1_45S_50ntFlanks</td><td>12934</td><td>12966</td><td>33</td><td>CTAAACCATTCGTAGACGACCTGCTTCTGGGTC</td><td>269.1</td><td>-4.7</td><td>0.6</td><td>-8.3</td><td>1.5E-16</td><td>5.2E-16</td><td>MB231/MB468</td></tr>
<tr><td>>hg38_21_+_8433172_8446622_RNA45SN1_45S_50ntFlanks@6661.6685.25|CGGTGGATCACTCGGCTCGTGCGTC</td><td>>hg38_21_+_8433172_8446622_RNA45SN1_45S_50ntFlanks</td><td>6661</td><td>6685</td><td>25</td><td>CGGTGGATCACTCGGCTCGTGCGTC</td><td>728.5</td><td>4.8</td><td>0.6</td><td>8.3</td><td>1.5E-16</td><td>5.2E-16</td><td>BT20/MB468</td></tr>
<tr><td>>hg38_21_+_8433172_8446622_RNA45SN1_45S_50ntFlanks@11707.11728.22|AAGCGCGGGTAAACGGCGGGAG</td><td>>hg38_21_+_8433172_8446622_RNA45SN1_45S_50ntFlanks</td><td>11707</td><td>11728</td><td>22</td><td>AAGCGCGGGTAAACGGCGGGAG</td><td>194.4</td><td>-4.2</td><td>0.5</td><td>-8.3</td><td>1.5E-16</td><td>5.2E-16</td><td>MB231/MB468</td></tr>
<tr><td>>trna10_ValCAC_5_-_180649395_180649467@59.76.18,trna12_ValAAC_5_-_180645270_180645342@59.76.18,trna132_ValAAC_6_-_27721179_27721251@59.76.18,trna136_ValAAC_6_-_27648885_27648957@59.76.18,trna15_ValAAC_5_-_180615416_180615488@59.76.18,trna18_ValCAC_5_-_180529253_180529325@59.76.18,trna2_ValAAC_3_+_169490018_169490090@59.76.18,trna2_ValCAC_5_+_180524070_180524142@59.76.18,trna4_ValAAC_5_+_180591154_180591226@59.76.18,trna5_ValAAC_5_+_180596610_180596682@59.76.18,trna6_ValCAC_5_+_180600650_180600722@59.76.18,trna85_ValCAC_1_-_161369490_161369562@59.76.18,trna90_ValCAC_1_-_149684088_149684161@60.77.18,trna9_ValCAC_6_+_26538282_26538354@59.76.18|AACCGGGCGGAAACACCA</td><td>>trna10_ValCAC_5_-_180649395_180649467</td><td>59</td><td>76</td><td>18</td><td>AACCGGGCGGAAACACCA</td><td>1222.9</td><td>5.9</td><td>0.7</td><td>8.3</td><td>1.5E-16</td><td>5.2E-16</td><td>BT20/MB468</td></tr>
<tr><td>>MI0000093|hsa-mir-92a-1&WithFlank&13|+|91351308|91351397@54.74.21(+1G);MI0000094|hsa-mir-92a-2&WithFlank&X|-|134169532|134169618@54.74.21(+1G)&[MIMAT0000092&hsa-miR-92a-3p&offsets|0|-1(+1G);m-9&13|+|91351361|91351382&offsets|0|-1(+1G)];[MIMAT0000092_1&hsa-miR-92a-3p&offsets|0|-1(+1G);m-11&X|-|134169544|134169565&offsets|0|-1(+1G)]|TATTGCACTTGTCCCGGCCTGG</td><td>>MI0000093|hsa-mir-92a-1&WithFlank&13|+|91351308|91351397</td><td>54</td><td>74</td><td>21(+1G)</td><td>TATTGCACTTGTCCCGGCCTGG</td><td>5570.0</td><td>6.7</td><td>0.8</td><td>8.3</td><td>1.5E-16</td><td>5.2E-16</td><td>BT20/MB468</td></tr>
<tr><td>>MI0000061|hsa-let-7a-2&WithFlank&11|-|122146516|122146599@11.28.18;MI0000064|hsa-let-7c&WithFlank&21|+|16539822|16539917@17.34.18;MI0000062|hsa-let-7a-3&WithFlank&22|+|46112743|46112828@10.27.18;MI0000060|hsa-let-7a-1&WithFlank&9|+|94175951|94176042@12.29.18&[MIMAT0000062&hsa-let-7a-5p&offsets|0|-4;m-6&11|-|122146568|122146589&offsets|0|-4];[MIMAT0000064&hsa-let-7c-5p&offsets|0|-4;m-37&21|+|16539838|16539859&offsets|0|-4];[MIMAT0000062_1&hsa-let-7a-5p&offsets|0|-4;m-5&22|+|46112752|46112773&offsets|0|-4];[MIMAT0000062_2&hsa-let-7a-5p&offsets|0|-4;m-4&9|+|94175962|94175983&offsets|0|-4]|TGAGGTAGTAGGTTGTAT</td><td>>MI0000061|hsa-let-7a-2&WithFlank&11|-|122146516|122146599</td><td>11</td><td>28</td><td>18</td><td>TGAGGTAGTAGGTTGTAT</td><td>3700.4</td><td>6.8</td><td>0.8</td><td>8.3</td><td>1.5E-16</td><td>5.3E-16</td><td>BT20/MB468</td></tr>
<tr><td>>MI0000081|hsa-mir-24-2&WithFlank&19|-|13836281|13836365@56.77.22(+1C);MI0000080|hsa-mir-24-1&WithFlank&9|+|95086015|95086094@50.71.22(+1C)&[MIMAT0000080&hsa-miR-24-3p&offsets|0|0(+1C);m-36&19|-|13836288|13836310&offsets|0|-1(+1C)];[MIMAT0000080_1&hsa-miR-24-3p&offsets|0|0(+1C);m-35&9|+|95086064|95086086&offsets|0|-1(+1C)]|TGGCTCAGTTCAGCAGGAACAGC</td><td>>MI0000081|hsa-mir-24-2&WithFlank&19|-|13836281|13836365</td><td>56</td><td>77</td><td>22(+1C)</td><td>TGGCTCAGTTCAGCAGGAACAGC</td><td>3203.9</td><td>7.2</td><td>0.9</td><td>8.3</td><td>1.5E-16</td><td>5.3E-16</td><td>BT20/MB468</td></tr>
<tr><td>>hg38_21_+_8433172_8446622_RNA45SN1_45S_50ntFlanks@12971.12991.21|TTTCGTACGTAGCAGAGCAGC</td><td>>hg38_21_+_8433172_8446622_RNA45SN1_45S_50ntFlanks</td><td>12971</td><td>12991</td><td>21</td><td>TTTCGTACGTAGCAGAGCAGC</td><td>351.8</td><td>-4.2</td><td>0.5</td><td>-8.3</td><td>1.5E-16</td><td>5.3E-16</td><td>MB231/MB468</td></tr>
<tr><td>>hg38_21_+_8433172_8446622_RNA45SN1_45S_50ntFlanks@11569.11605.37|CAGCCGACTTAGAACTGGTGCGGACCAGGGGAATCCG</td><td>>hg38_21_+_8433172_8446622_RNA45SN1_45S_50ntFlanks</td><td>11569</td><td>11605</td><td>37</td><td>CAGCCGACTTAGAACTGGTGCGGACCAGGGGAATCCG</td><td>297.7</td><td>-5.2</td><td>0.6</td><td>-8.3</td><td>1.5E-16</td><td>5.3E-16</td><td>BT20/MB468</td></tr>
<tr><td>>hg38_21_+_8433172_8446622_RNA45SN1_45S_50ntFlanks@10727.10755.29|GGAGAGGGTGTAAATCTCGCGCCGGGCCG</td><td>>hg38_21_+_8433172_8446622_RNA45SN1_45S_50ntFlanks</td><td>10727</td><td>10755</td><td>29</td><td>GGAGAGGGTGTAAATCTCGCGCCGGGCCG</td><td>563.0</td><td>-11.1</td><td>1.3</td><td>-8.3</td><td>1.6E-16</td><td>5.4E-16</td><td>BT20/MB468</td></tr>
<tr><td>>hg38_21_+_8433172_8446622_RNA45SN1_45S_50ntFlanks@11576.11605.30|CTTAGAACTGGTGCGGACCAGGGGAATCCG</td><td>>hg38_21_+_8433172_8446622_RNA45SN1_45S_50ntFlanks</td><td>11576</td><td>11605</td><td>30</td><td>CTTAGAACTGGTGCGGACCAGGGGAATCCG</td><td>740.2</td><td>-5.7</td><td>0.7</td><td>-8.2</td><td>1.6E-16</td><td>5.4E-16</td><td>MB231/MB468</td></tr>
<tr><td>>hg38_21_+_8433172_8446622_RNA45SN1_45S_50ntFlanks@485.522.38|GATGTGGTGACGTCGTGCTCTCCCGGGCCGGGTCCGAG</td><td>>hg38_21_+_8433172_8446622_RNA45SN1_45S_50ntFlanks</td><td>485</td><td>522</td><td>38</td><td>GATGTGGTGACGTCGTGCTCTCCCGGGCCGGGTCCGAG</td><td>2324.8</td><td>30.0</td><td>3.6</td><td>8.2</td><td>1.6E-16</td><td>5.4E-16</td><td>MB231/MB468</td></tr>
<tr><td>>hg38_21_+_8433172_8446622_RNA45SN1_45S_50ntFlanks@11318.11342.25|CCCCACGTCTCGTCGCGCGCGCGTC</td><td>>hg38_21_+_8433172_8446622_RNA45SN1_45S_50ntFlanks</td><td>11318</td><td>11342</td><td>25</td><td>CCCCACGTCTCGTCGCGCGCGCGTC</td><td>534.4</td><td>-10.6</td><td>1.3</td><td>-8.2</td><td>1.6E-16</td><td>5.4E-16</td><td>MB231/MB468</td></tr>
<tr><td>>MI0000440|hsa-mir-27b&WithFlank&9|+|95085439|95085547@67.87.21(+1C)&[MIMAT0000419&hsa-miR-27b-3p&offsets|0|0(+1C);m-39&9|+|95085505|95085525&offsets|0|0(+1C)]|TTCACAGTGGCTAAGTTCTGCC</td><td>>MI0000440|hsa-mir-27b&WithFlank&9|+|95085439|95085547</td><td>67</td><td>87</td><td>21(+1C)</td><td>TTCACAGTGGCTAAGTTCTGCC</td><td>906.5</td><td>7.8</td><td>0.9</td><td>8.3</td><td>1.6E-16</td><td>5.5E-16</td><td>BT20/MB468</td></tr>
<tr><td>>hg38_21_+_8433172_8446622_RNA45SN1_45S_50ntFlanks@8357.8375.19|AAAAGAACTTTGAAGAGAG</td><td>>hg38_21_+_8433172_8446622_RNA45SN1_45S_50ntFlanks</td><td>8357</td><td>8375</td><td>19</td><td>AAAAGAACTTTGAAGAGAG</td><td>5052.6</td><td>-5.4</td><td>0.7</td><td>-8.3</td><td>1.6E-16</td><td>5.5E-16</td><td>BT20/MB468</td></tr>
<tr><td>>hg38_21_+_8433172_8446622_RNA45SN1_45S_50ntFlanks@4731.4761.31|AAGAACGAAAGTCGGAGGTTCGAAGACGATC</td><td>>hg38_21_+_8433172_8446622_RNA45SN1_45S_50ntFlanks</td><td>4731</td><td>4761</td><td>31</td><td>AAGAACGAAAGTCGGAGGTTCGAAGACGATC</td><td>2460.0</td><td>5.7</td><td>0.7</td><td>8.2</td><td>1.6E-16</td><td>5.5E-16</td><td>MB231/MB468</td></tr>
<tr><td>>am_trna106_HisGTG_1_-_149155828_149155899@7.29.23,trna111_HisGTG_1_-_147774845_147774916@7.29.23,trna118_HisGTG_1_-_145396881_145396952@7.29.23,trna16_HisGTG_1_+_146544773_146544844@7.29.23,trna1_HisGTG_15_+_45493349_45493420@7.29.23,trna21_HisGTG_1_+_147753471_147753542@7.29.23,trna33_HisGTG_6_+_27125906_27125977@7.29.23,trna7_HisGTG_9_-_14433938_14434009@7.29.23,trna8_HisGTG_15_-_45492611_45492682@7.29.23,trna9_HisGTG_15_-_45490804_45490875@7.29.23|ATCGTATAGTGGTTAGTACTCTG</td><td>>am_trna106_HisGTG_1_-_149155828_149155899</td><td>7</td><td>29</td><td>23</td><td>ATCGTATAGTGGTTAGTACTCTG</td><td>419.0</td><td>-10.7</td><td>1.3</td><td>-8.2</td><td>1.6E-16</td><td>5.6E-16</td><td>BT20/MB468</td></tr>
<tr><td>>MI0005756|hsa-mir-934&WithFlank&X|+|136550872|136550966@21.41.21(+1U)&[MIMAT0004977&hsa-miR-934&offsets|0|-1(+1U);m-943&X|+|136550892|136550913&offsets|0|-1(+1U)]|TGTCTACTACTGGAGACACTGT</td><td>>MI0005756|hsa-mir-934&WithFlank&X|+|136550872|136550966</td><td>21</td><td>41</td><td>21(+1U)</td><td>TGTCTACTACTGGAGACACTGT</td><td>955.5</td><td>13.4</td><td>1.6</td><td>8.6</td><td>1.0E-17</td><td>5.6E-16</td><td>BT20/MB231</td></tr>
<tr><td>>hg38_21_+_8433172_8446622_RNA45SN1_45S_50ntFlanks@6721.6738.18|AATTGCAGGACACATTGA</td><td>>hg38_21_+_8433172_8446622_RNA45SN1_45S_50ntFlanks</td><td>6721</td><td>6738</td><td>18</td><td>AATTGCAGGACACATTGA</td><td>431.4</td><td>-10.7</td><td>1.3</td><td>-8.2</td><td>1.7E-16</td><td>5.8E-16</td><td>BT20/MB468</td></tr>
<tr><td>>hg38_21_+_8433172_8446622_RNA45SN1_45S_50ntFlanks@485.529.45|GATGTGGTGACGTCGTGCTCTCCCGGGCCGGGTCCGAGCCGCGAC</td><td>>hg38_21_+_8433172_8446622_RNA45SN1_45S_50ntFlanks</td><td>485</td><td>529</td><td>45</td><td>GATGTGGTGACGTCGTGCTCTCCCGGGCCGGGTCCGAGCCGCGAC</td><td>3775.4</td><td>29.9</td><td>3.6</td><td>8.2</td><td>1.7E-16</td><td>5.8E-16</td><td>MB231/MB468</td></tr>
<tr><td>>hg38_21_+_8433172_8446622_RNA45SN1_45S_50ntFlanks@11769.11794.26|TAATTAGTGACGCGCATGAATGGATG</td><td>>hg38_21_+_8433172_8446622_RNA45SN1_45S_50ntFlanks</td><td>11769</td><td>11794</td><td>26</td><td>TAATTAGTGACGCGCATGAATGGATG</td><td>982.9</td><td>-3.4</td><td>0.4</td><td>-8.2</td><td>1.7E-16</td><td>5.8E-16</td><td>BT20/MB468</td></tr>
<tr><td>>hg38_21_+_8433172_8446622_RNA45SN1_45S_50ntFlanks@6778.6807.30|CCCGGGGCTACGCCTGTCTGAGCGTCGCTT</td><td>>hg38_21_+_8433172_8446622_RNA45SN1_45S_50ntFlanks</td><td>6778</td><td>6807</td><td>30</td><td>CCCGGGGCTACGCCTGTCTGAGCGTCGCTT</td><td>485.5</td><td>5.3</td><td>0.6</td><td>8.2</td><td>1.7E-16</td><td>5.8E-16</td><td>MB231/MB468</td></tr>
<tr><td>>hg38_21_+_8433172_8446622_RNA45SN1_45S_50ntFlanks@10389.10409.21|TGAACAGCAGTTGAACATGGG</td><td>>hg38_21_+_8433172_8446622_RNA45SN1_45S_50ntFlanks</td><td>10389</td><td>10409</td><td>21</td><td>TGAACAGCAGTTGAACATGGG</td><td>750.0</td><td>-8.1</td><td>1.0</td><td>-8.2</td><td>1.7E-16</td><td>5.9E-16</td><td>BT20/MB468</td></tr>
<tr><td>>hg38_21_+_8433172_8446622_RNA45SN1_45S_50ntFlanks@12416.12438.23|TTCGATGTCGGCTCTTCCTATCA</td><td>>hg38_21_+_8433172_8446622_RNA45SN1_45S_50ntFlanks</td><td>12416</td><td>12438</td><td>23</td><td>TTCGATGTCGGCTCTTCCTATCA</td><td>686.4</td><td>5.6</td><td>0.7</td><td>8.2</td><td>1.7E-16</td><td>5.9E-16</td><td>BT20/MB468</td></tr>
</table>

<table>
<tr><td>>hg38_21_+_8433172_8446622_RNA45SN1_45S_50ntFlanks@10690.10709.20|CGTCCGGTGAGCTCTCGCTG</td><td>>hg38_21_+_8433172_8446622_RNA45SN1_45S_50ntFlanks</td><td>10690</td><td>10709</td><td>20</td><td>CGTCCGGTGAGCTCTCGCTG</td><td>1264.9</td><td>-5.5</td><td>0.7</td><td>-8.2</td><td>1.7E-16</td><td>5.9E-16</td><td>BT20/MB468</td></tr>
<tr><td>>hg38_21_+_8433172_8446622_RNA45SN1_45S_50ntFlanks@51.71.21|GCTGACACGCTGTCCTCTGGC</td><td>>hg38_21_+_8433172_8446622_RNA45SN1_45S_50ntFlanks</td><td>51</td><td>71</td><td>21</td><td>GCTGACACGCTGTCCTCTGGC</td><td>645.2</td><td>3.7</td><td>0.4</td><td>8.2</td><td>1.8E-16</td><td>5.9E-16</td><td>MB231/MB468</td></tr>
<tr><td>>trna116_GluCTC_1_-_145399233_145399304@1.39.39,trna59_GluCTC_1_+_249168447_249168518@1.39.39,trna71_GluCTC_1_-_161439189_161439260@1.39.39,trna74_GluCTC_1_-_161431809_161431880@1.39.39,trna77_GluCTC_1_-_161424398_161424469@1.39.39,trna77_GluCTC_6_+_28949976_28950047@1.39.39,trna80_GluCTC_1_-_161417018_161417089@1.39.39,trna87_GluCTC_6_-_126101393_126101464@1.39.39|TCCCTGGTGGTCTAGTGGTTAGGATTCGGCGCTCTCACC</td><td>>trna116_GluCTC_1_-_145399233_145399304</td><td>1</td><td>39</td><td>39</td><td>TCCCTGGTGGTCTAGTGGTTAGGATTCGGCGCTCTCACC</td><td>2477.8</td><td>6.8</td><td>0.8</td><td>8.2</td><td>1.8E-16</td><td>6.1E-16</td><td>BT20/MB468</td></tr>
<tr><td>>MI0000084|hsa-mir-26b&WithFlank&2|+|218402640|218402728@18.38.21(+1C)&[MIMAT0000083&hsa-miR-26b-5p&offsets|0|0(+1C);m-53&2|+|218402657|218402678&offsets|0|-1(+1C)]|TTCAAGTAATTCAGGATAGGTC</td><td>>MI0000084|hsa-mir-26b&WithFlank&2|+|218402640|218402728</td><td>18</td><td>38</td><td>21(+1C)</td><td>TTCAAGTAATTCAGGATAGGTC</td><td>1977.6</td><td>6.3</td><td>0.8</td><td>8.2</td><td>1.8E-16</td><td>6.1E-16</td><td>MB231/MB468</td></tr>
<tr><td>>am_trnaMT_LysTTT_MT_+_8295_8364@30.52.23,trnalookalike7_LysTTT_1_+_568844_568913@30.52.23|TTAAGTTAAAGATTAAGAGAACC</td><td>>am_trnaMT_LysTTT_MT_+_8295_8364</td><td>30</td><td>52</td><td>23</td><td>TTAAGTTAAAGATTAAGAGAACC</td><td>818.8</td><td>9.3</td><td>1.1</td><td>8.2</td><td>1.8E-16</td><td>6.1E-16</td><td>BT20/MB468</td></tr>
<tr><td>>am_MI0000809|hsa-mir-151a&WithFlank&8|-|140732558|140732659@17.37.21&[MIMAT0004697&hsa-miR-151a-5p&offsets|0|0;m-75&8|-|140732623|140732643&offsets|0|0]|TCGAGGAGCTCACAGTCTAGT</td><td>>am_MI0000809|hsa-mir-151a&WithFlank&8|-|140732558|140732659</td><td>17</td><td>37</td><td>21</td><td>TCGAGGAGCTCACAGTCTAGT</td><td>9317.6</td><td>10.4</td><td>1.3</td><td>8.2</td><td>1.8E-16</td><td>6.1E-16</td><td>BT20/MB468</td></tr>
<tr><td>>hg38_21_+_8433172_8446622_RNA45SN1_45S_50ntFlanks@6721.6744.24|AATTGCAGGACACATTGATCATCG</td><td>>hg38_21_+_8433172_8446622_RNA45SN1_45S_50ntFlanks</td><td>6721</td><td>6744</td><td>24</td><td>AATTGCAGGACACATTGATCATCG</td><td>16817.7</td><td>-5.7</td><td>0.7</td><td>-8.2</td><td>1.8E-16</td><td>6.2E-16</td><td>MB231/MB468</td></tr>
<tr><td>>hg38_21_+_8433172_8446622_RNA45SN1_45S_50ntFlanks@11027.11051.25|CTCTCTCTCTCTCTCCCCCGCTCCC</td><td>>hg38_21_+_8433172_8446622_RNA45SN1_45S_50ntFlanks</td><td>11027</td><td>11051</td><td>25</td><td>CTCTCTCTCTCTCTCCCCCGCTCCC</td><td>454.9</td><td>-10.4</td><td>1.3</td><td>-8.2</td><td>1.9E-16</td><td>6.3E-16</td><td>MB231/MB468</td></tr>
<tr><td>>MI0000103|hsa-mir-101-1&WithFlank&1|-|65058428|65058514@53.73.21(+1U);MI0000739|hsa-mir-101-2&WithFlank&9|+|4850291|4850381@55.75.21(+1U)&[MIMAT0000099&hsa-miR-101-3p&offsets|0|0(+1U);m-17&1|-|65058443|65058462&offsets|0|+1(+1U)];[MIMAT0000099_1&hsa-miR-101-3p&offsets|0|0(+1U);m-16&9|+|4850345|4850364&offsets|0|+1(+1U)]|TACAGTACTGTGATAACTGAAT</td><td>>MI0000103|hsa-mir-101-1&WithFlank&1|-|65058428|65058514</td><td>53</td><td>73</td><td>21(+1U)</td><td>TACAGTACTGTGATAACTGAAT</td><td>3146.2</td><td>9.0</td><td>1.1</td><td>8.2</td><td>1.8E-16</td><td>6.4E-16</td><td>BT20/MB468</td></tr>
<tr><td>>hg38_21_+_8433172_8446622_RNA45SN1_45S_50ntFlanks@11575.11600.26|ACTTAGAACTGGTGCGGACCAGGGGA</td><td>>hg38_21_+_8433172_8446622_RNA45SN1_45S_50ntFlanks</td><td>11575</td><td>11600</td><td>26</td><td>ACTTAGAACTGGTGCGGACCAGGGGA</td><td>529.4</td><td>-4.8</td><td>0.6</td><td>-8.2</td><td>1.9E-16</td><td>6.4E-16</td><td>MB231/MB468</td></tr>
<tr><td>>MI0000750|hsa-mir-26a-2&WithFlank&12|-|57824603|57824698@20.40.21(+1G);MI0000083|hsa-mir-26a-1&WithFlank&3|+|37969398|37969486@16.36.21(+1G)&[MIMAT0000082&hsa-miR-26a-5p&offsets|0|-1(+1G);m-33&12|-|57824658|57824679&offsets|0|-1(+1G)];[MIMAT0000082_1&hsa-miR-26a-5p&offsets|0|-1(+1G);m-34&3|+|37969413|37969434&offsets|0|-1(+1G)]|TTCAAGTAATCCAGGATAGGCG</td><td>>MI0000750|hsa-mir-26a-2&WithFlank&12|-|57824603|57824698</td><td>20</td><td>40</td><td>21(+1G)</td><td>TTCAAGTAATCCAGGATAGGCG</td><td>10303.7</td><td>6.0</td><td>0.7</td><td>8.2</td><td>1.9E-16</td><td>6.5E-16</td><td>BT20/MB468</td></tr>
<tr><td>>hg38_21_+_8433172_8446622_RNA45SN1_45S_50ntFlanks@12627.12657.31|ATGGGGCGAAGCTACCATCTGTGGGATTATG</td><td>>hg38_21_+_8433172_8446622_RNA45SN1_45S_50ntFlanks</td><td>12627</td><td>12657</td><td>31</td><td>ATGGGGCGAAGCTACCATCTGTGGGATTATG</td><td>506.4</td><td>-7.3</td><td>0.9</td><td>-8.2</td><td>1.9E-16</td><td>6.5E-16</td><td>BT20/MB468</td></tr>
<tr><td>>MI0000542|hsa-mir-320a&WithFlank&8|-|22244960|22245043@48.68.21(+1C)&[MIMAT0000510&hsa-miR-320a-3p&offsets|0|-1(+1C);m-60&8|-|22244975|22244996&offsets|0|-1(+1C)]|AAAAGCTGGGTTGAGAGGGCGC</td><td>>MI0000542|hsa-mir-320a&WithFlank&8|-|22244960|22245043</td><td>48</td><td>68</td><td>21(+1C)</td><td>AAAAGCTGGGTTGAGAGGGCGC</td><td>1433.2</td><td>10.4</td><td>1.3</td><td>8.2</td><td>1.9E-16</td><td>6.5E-16</td><td>BT20/MB468</td></tr>
<tr><td>>hg38_21_+_8433172_8446622_RNA45SN1_45S_50ntFlanks@12347.12365.19|AGAAAAGTTACCACAGGGA</td><td>>hg38_21_+_8433172_8446622_RNA45SN1_45S_50ntFlanks</td><td>12347</td><td>12365</td><td>19</td><td>AGAAAAGTTACCACAGGGA</td><td>353.3</td><td>-10.5</td><td>1.3</td><td>-8.2</td><td>1.9E-16</td><td>6.6E-16</td><td>BT20/MB468</td></tr>
<tr><td>>hg38_21_+_8433172_8446622_RNA45SN1_45S_50ntFlanks@7978.8011.34|GACCTCAGATCAGACGTGGCGACCCGCTGAATTT</td><td>>hg38_21_+_8433172_8446622_RNA45SN1_45S_50ntFlanks</td><td>7978</td><td>8011</td><td>34</td><td>GACCTCAGATCAGACGTGGCGACCCGCTGAATTT</td><td>1142.2</td><td>7.9</td><td>1.0</td><td>8.2</td><td>2.0E-16</td><td>6.6E-16</td><td>MB231/MB468</td></tr>
<tr><td>>MI0000072|hsa-mir-18a&WithFlank&13|+|91350745|91350827@12.34.23&[MIMAT0000072&hsa-miR-18a-5p&offsets|0|0;m-262&13|+|91350756|91350778&offsets|0|0]|TAAGGTGCATCTAGTGCAGATAG</td><td>>MI0000072|hsa-mir-18a&WithFlank&13|+|91350745|91350827</td><td>12</td><td>34</td><td>23</td><td>TAAGGTGCATCTAGTGCAGATAG</td><td>1192.8</td><td>6.3</td><td>0.8</td><td>8.2</td><td>2.0E-16</td><td>6.6E-16</td><td>MB231/MB468</td></tr>
<tr><td>>MI0000085|hsa-mir-27a&WithFlank&19|-|13836434|13836523@16.36.21&[MIMAT0004501&hsa-miR-27a-5p&offsets|0|-1;m-270&19|-|13836487|13836508&offsets|0|-1]|AGGGCTTAGCTGCTTGTGAGC</td><td>>MI0000085|hsa-mir-27a&WithFlank&19|-|13836434|13836523</td><td>16</td><td>36</td><td>21</td><td>AGGGCTTAGCTGCTTGTGAGC</td><td>941.4</td><td>7.8</td><td>0.9</td><td>8.2</td><td>2.0E-16</td><td>6.6E-16</td><td>MB231/MB468</td></tr>
<tr><td>>hg38_21_+_8433172_8446622_RNA45SN1_45S_50ntFlanks@485.526.42|GATGTGGTGACGTCGTGCTCTCCCGGGCCGGGTCCGAGCCGC</td><td>>hg38_21_+_8433172_8446622_RNA45SN1_45S_50ntFlanks</td><td>485</td><td>526</td><td>42</td><td>GATGTGGTGACGTCGTGCTCTCCCGGGCCGGGTCCGAGCCGC</td><td>1854.5</td><td>29.9</td><td>3.6</td><td>8.2</td><td>2.0E-16</td><td>6.7E-16</td><td>MB231/MB468</td></tr>
<tr><td>>hg38_21_+_8433172_8446622_RNA45SN1_45S_50ntFlanks@6747.6771.25|ACTTCGAACGCACTTGCGGCCCCGG</td><td>>hg38_21_+_8433172_8446622_RNA45SN1_45S_50ntFlanks</td><td>6747</td><td>6771</td><td>25</td><td>ACTTCGAACGCACTTGCGGCCCCGG</td><td>245.7</td><td>-5.7</td><td>0.7</td><td>-8.2</td><td>2.0E-16</td><td>6.7E-16</td><td>MB231/MB468</td></tr>
<tr><td>>MI0000061|hsa-let-7a-2&WithFlank&11|-|122146516|122146599@11.32.22(+1G);MI0000062|hsa-let-7a-3&WithFlank&22|+|46112743|46112828@10.31.22(+1G);MI0000060|hsa-let-7a-1&WithFlank&9|+|94175951|94176042@12.33.22(+1G)&[MIMAT0000062&hsa-let-7a-5p&offsets|0|0(+1G);m-6&11|-|122146568|122146589&offsets|0|0(+1G)];[MIMAT0000062_1&hsa-let-7a-5p&offsets|0|0(+1G);m-5&22|+|46112752|46112773&offsets|0|0(+1G)];[MIMAT0000062_2&hsa-let-7a-5p&offsets|0|0(+1G);m-4&9|+|94175962|94175983&offsets|0|0(+1G)]|TGAGGTAGTAGGTTGTATAGTTG</td><td>>MI0000061|hsa-let-7a-2&WithFlank&11|-|122146516|122146599</td><td>11</td><td>32</td><td>22(+1G)</td><td>TGAGGTAGTAGGTTGTATAGTTG</td><td>590.7</td><td>8.6</td><td>1.0</td><td>8.2</td><td>1.9E-16</td><td>6.7E-16</td><td>BT20/MB468</td></tr>
<tr><td>>hg38_21_+_8433172_8446622_RNA45SN1_45S_50ntFlanks@12940.12966.27|CATTCGTAGACGACCTGCTTCTGGGTC</td><td>>hg38_21_+_8433172_8446622_RNA45SN1_45S_50ntFlanks</td><td>12940</td><td>12966</td><td>27</td><td>CATTCGTAGACGACCTGCTTCTGGGTC</td><td>573.8</td><td>-7.7</td><td>0.9</td><td>-8.2</td><td>2.0E-16</td><td>6.7E-16</td><td>BT20/MB468</td></tr>
<tr><td>>hg38_21_+_8433172_8446622_RNA45SN1_45S_50ntFlanks@10786.10818.33|AACAGCCTCTGGCATGTTGGAACAATGTAGGTA</td><td>>hg38_21_+_8433172_8446622_RNA45SN1_45S_50ntFlanks</td><td>10786</td><td>10818</td><td>33</td><td>AACAGCCTCTGGCATGTTGGAACAATGTAGGTA</td><td>5598.2</td><td>-5.3</td><td>0.6</td><td>-8.2</td><td>2.0E-16</td><td>6.7E-16</td><td>MB231/MB468</td></tr>
<tr><td>>MI0000285|hsa-mir-205&WithFlank&1|+|209432127|209432248@40.61.22(+1A)&[MIMAT0000266&hsa-miR-205-5p&offsets|0|0(+1A);m-351&1|+|209432166|209432187&offsets|0|0(+1A)]|TCCTTCATTCCACCGGAGTCTGA</td><td>>MI0000285|hsa-mir-205&WithFlank&1|+|209432127|209432248</td><td>40</td><td>61</td><td>22(+1A)</td><td>TCCTTCATTCCACCGGAGTCTGA</td><td>41853.0</td><td>14.6</td><td>1.7</td><td>8.5</td><td>1.3E-17</td><td>6.8E-16</td><td>BT20/MB231</td></tr>
<tr><td>>trnaMT_PheGAA_MT_+_577_647@43.74.32|TTAGACGGGCTCACATCACCCCATAAACACCA</td><td>>trnaMT_PheGAA_MT_+_577_647</td><td>43</td><td>74</td><td>32</td><td>TTAGACGGGCTCACATCACCCCATAAACACCA</td><td>842.9</td><td>9.5</td><td>1.2</td><td>8.2</td><td>2.0E-16</td><td>6.9E-16</td><td>BT20/MB468</td></tr>
<tr><td>>hg38_21_+_8433172_8446622_RNA45SN1_45S_50ntFlanks@7978.8016.39|GACCTCAGATCAGACGTGGCGACCCGCTGAATTTAAGCA</td><td>>hg38_21_+_8433172_8446622_RNA45SN1_45S_50ntFlanks</td><td>7978</td><td>8016</td><td>39</td><td>GACCTCAGATCAGACGTGGCGACCCGCTGAATTTAAGCA</td><td>725.0</td><td>9.9</td><td>1.2</td><td>8.2</td><td>2.0E-16</td><td>6.9E-16</td><td>MB231/MB468</td></tr>
<tr><td>>hg38_21_+_8433172_8446622_RNA45SN1_45S_50ntFlanks@12416.12447.32|TTCGATGTCGGCTCTTCCTATCATTGTGAAGC</td><td>>hg38_21_+_8433172_8446622_RNA45SN1_45S_50ntFlanks</td><td>12416</td><td>12447</td><td>32</td><td>TTCGATGTCGGCTCTTCCTATCATTGTGAAGC</td><td>2411.6</td><td>9.1</td><td>1.1</td><td>8.2</td><td>2.1E-16</td><td>6.9E-16</td><td>MB231/MB468</td></tr>
<tr><td>>hg38_21_+_8433172_8446622_RNA45SN1_45S_50ntFlanks@5492.5516.25|AGAAGACGGTCGAACTTGACTATCT</td><td>>hg38_21_+_8433172_8446622_RNA45SN1_45S_50ntFlanks</td><td>5492</td><td>5516</td><td>25</td><td>AGAAGACGGTCGAACTTGACTATCT</td><td>3437.4</td><td>6.1</td><td>0.7</td><td>8.2</td><td>2.1E-16</td><td>7.0E-16</td><td>MB231/MB468</td></tr>
<tr><td>>hg38_21_+_8433172_8446622_RNA45SN1_45S_50ntFlanks@10375.10407.33|AGAAGGGTTCCATGTGAACAGCAGTTGAACATG</td><td>>hg38_21_+_8433172_8446622_RNA45SN1_45S_50ntFlanks</td><td>10375</td><td>10407</td><td>33</td><td>AGAAGGGTTCCATGTGAACAGCAGTTGAACATG</td><td>335.2</td><td>-10.4</td><td>1.3</td><td>-8.2</td><td>2.0E-16</td><td>7.0E-16</td><td>BT20/MB468</td></tr>
<tr><td>>MI0000439|hsa-mir-23b&WithFlank&9|+|95085202|95085310@67.86.20(+1C)&[MIMAT0000418&hsa-miR-23b-3p&offsets|+3|0(+1C);m-42&9|+|95085265|95085285&offsets|+3|+2(+1C)]|ACATTGCCAGGGATTACCACC</td><td>>MI0000439|hsa-mir-23b&WithFlank&9|+|95085202|95085310</td><td>67</td><td>86</td><td>20(+1C)</td><td>ACATTGCCAGGGATTACCACC</td><td>2461.6</td><td>29.8</td><td>3.6</td><td>8.2</td><td>2.1E-16</td><td>7.1E-16</td><td>MB231/MB468</td></tr>
<tr><td>>hg38_21_+_8433172_8446622_RNA45SN1_45S_50ntFlanks@8176.8203.28|CTTCTGATCGAGGCCCAGCCCGTGGACG</td><td>>hg38_21_+_8433172_8446622_RNA45SN1_45S_50ntFlanks</td><td>8176</td><td>8203</td><td>28</td><td>CTTCTGATCGAGGCCCAGCCCGTGGACG</td><td>2021.2</td><td>-3.4</td><td>0.4</td><td>-8.2</td><td>2.1E-16</td><td>7.2E-16</td><td>BT20/MB468</td></tr>
<tr><td>>MI0000103|hsa-mir-101-1&WithFlank&1|-|65058428|65058514@53.73.21;MI0000739|hsa-mir-101-2&WithFlank&9|+|4850291|4850381@55.75.21&[MIMAT0000099&hsa-miR-101-3p&offsets|0|0;m-17&1|-|65058443|65058462&offsets|0|+1];[MIMAT0000099_1&hsa-miR-101-3p&offsets|0|0;m-16&9|+|4850345|4850364&offsets|0|+1]|TACAGTACTGTGATAACTGAA</td><td>>MI0000103|hsa-mir-101-1&WithFlank&1|-|65058428|65058514</td><td>53</td><td>73</td><td>21</td><td>TACAGTACTGTGATAACTGAA</td><td>2295.9</td><td>6.7</td><td>0.8</td><td>8.2</td><td>2.1E-16</td><td>7.2E-16</td><td>MB231/MB468</td></tr>
<tr><td>>MI0000081|hsa-mir-24-2&WithFlank&19|-|13836281|13836365@56.75.20;MI0000080|hsa-mir-24-1&WithFlank&9|+|95086015|95086094@50.69.20&[MIMAT0000080&hsa-miR-24-3p&offsets|0|-2;m-36&19|-|13836288|13836310&offsets|0|-3];[MIMAT0000080_1&hsa-miR-24-3p&offsets|0|-2;m-35&9|+|95086064|95086086&offsets|0|-3]|TGGCTCAGTTCAGCAGGAAC</td><td>>MI0000081|hsa-mir-24-2&WithFlank&19|-|13836281|13836365</td><td>56</td><td>75</td><td>20</td><td>TGGCTCAGTTCAGCAGGAAC</td><td>27017.6</td><td>5.9</td><td>0.7</td><td>8.2</td><td>2.1E-16</td><td>7.2E-16</td><td>BT20/MB468</td></tr>
<tr><td>>hg38_21_+_8433172_8446622_RNA45SN1_45S_50ntFlanks@3838.3860.23|CTCTCCTACTTGGATAACTGTGG</td><td>>hg38_21_+_8433172_8446622_RNA45SN1_45S_50ntFlanks</td><td>3838</td><td>3860</td><td>23</td><td>CTCTCCTACTTGGATAACTGTGG</td><td>592.7</td><td>-3.8</td><td>0.5</td><td>-8.2</td><td>2.1E-16</td><td>7.2E-16</td><td>BT20/MB468</td></tr>
<tr><td>>hg38_21_+_8433172_8446622_RNA45SN1_45S_50ntFlanks@11534.11553.20|GTTCCGCGCGGCGCCTCGCC</td><td>>hg38_21_+_8433172_8446622_RNA45SN1_45S_50ntFlanks</td><td>11534</td><td>11553</td><td>20</td><td>GTTCCGCGCGGCGCCTCGCC</td><td>643.7</td><td>-11.3</td><td>1.4</td><td>-8.2</td><td>2.1E-16</td><td>7.3E-16</td><td>BT20/MB468</td></tr>
</table>

| | | | | | | | | | | | | |
|---|---|---|---|---|---|---|---|---|---|---|---|---|
| >MI0000095\|hsa-mir-93&WithFlank&7\|-\|100093762\|100093853@17.38.22(+1A)&[MIMAT0000093&hsa-miR-93-5p&offsets\|0\|-1(+1A);m-29&7\|-\|100093815\|100093837&offsets\|0\|-1(+1A)]\|CAAAGTGCTGTTCGTGCAGGTAA | >MI0000095\|hsa-mir-93&WithFlank&7\|-\|100093762\|100093853 | 17 | 38 | 22(+1A) | CAAAGTGCTGTTCGTGCAGGTAA | 4475.0 | 7.2 | 0.9 | 8.2 | 2.1E-16 | 7.3E-16 | BT20/MB468 |
| >hg38_21_+_8433172_8446622_RNA45SN1_45S_50ntFlanks@6747.6771.25\|ACTTCGAACGCACTTGCGGCCCCGG | >hg38_21_+_8433172_8446622_RNA45SN1_45S_50ntFlanks | 6747 | 6771 | 25 | ACTTCGAACGCACTTGCGGCCCCGG | 286.6 | -10.2 | 1.2 | -8.2 | 2.2E-16 | 7.4E-16 | BT20/MB468 |
| >hg38_21_+_8433172_8446622_RNA45SN1_45S_50ntFlanks@9563.9596.34\|CCTGACGTGCAAATCGGTCGTCCGACCTGGGTAT | >hg38_21_+_8433172_8446622_RNA45SN1_45S_50ntFlanks | 9563 | 9596 | 34 | CCTGACGTGCAAATCGGTCGTCCGACCTGGGTAT | 986.9 | -7.9 | 1.0 | -8.2 | 2.2E-16 | 7.4E-16 | BT20/MB468 |
| >hg38_21_+_8433172_8446622_RNA45SN1_45S_50ntFlanks@485.520.36\|GATGTGGTGACGTCGTGCTCTCCCGGGCCGGGTCCG | >hg38_21_+_8433172_8446622_RNA45SN1_45S_50ntFlanks | 485 | 520 | 36 | GATGTGGTGACGTCGTGCTCTCCCGGGCCGGGTCCG | 3583.9 | 29.8 | 3.6 | 8.2 | 2.2E-16 | 7.4E-16 | MB231/MB468 |
| >hg38_21_+_8433172_8446622_RNA45SN1_45S_50ntFlanks@10839.10865.27\|TCCGTAACTTCGGGATAAGGATTGGCT | >hg38_21_+_8433172_8446622_RNA45SN1_45S_50ntFlanks | 10839 | 10865 | 27 | TCCGTAACTTCGGGATAAGGATTGGCT | 573.3 | -8.0 | 1.0 | -8.2 | 2.2E-16 | 7.4E-16 | BT20/MB468 |
| >hg38_21_+_8433172_8446622_RNA45SN1_45S_50ntFlanks@6649.6668.20\|TACGACTCTTAGCGGTGGAT | >hg38_21_+_8433172_8446622_RNA45SN1_45S_50ntFlanks | 6649 | 6668 | 20 | TACGACTCTTAGCGGTGGAT | 861.7 | 4.0 | 0.5 | 8.2 | 2.2E-16 | 7.5E-16 | BT20/MB468 |
| >hg38_21_+_8433172_8446622_RNA45SN1_45S_50ntFlanks@4759.4778.20\|ATCAGATACCGTCGTAGTTC | >hg38_21_+_8433172_8446622_RNA45SN1_45S_50ntFlanks | 4759 | 4778 | 20 | ATCAGATACCGTCGTAGTTC | 970.1 | -7.6 | 0.9 | -8.2 | 2.2E-16 | 7.5E-16 | MB231/MB468 |
| >hg38_21_+_8433172_8446622_RNA45SN1_45S_50ntFlanks@13007.13025.19\|TATTGAAAGTCAGCCCTCG | >hg38_21_+_8433172_8446622_RNA45SN1_45S_50ntFlanks | 13007 | 13025 | 19 | TATTGAAAGTCAGCCCTCG | 722.1 | -6.9 | 0.8 | -8.2 | 2.2E-16 | 7.5E-16 | BT20/MB468 |
| >hg38_21_+_8433172_8446622_RNA45SN1_45S_50ntFlanks@10349.10374.26\|AACGAGAACTTTGAAGGCCGAAGTGG | >hg38_21_+_8433172_8446622_RNA45SN1_45S_50ntFlanks | 10349 | 10374 | 26 | AACGAGAACTTTGAAGGCCGAAGTGG | 618.9 | -3.4 | 0.4 | -8.2 | 2.2E-16 | 7.6E-16 | BT20/MB468 |
| >hg38_21_+_8433172_8446622_RNA45SN1_45S_50ntFlanks@11576.11601.26\|CTTAGAACTGGTGCGGACCAGGGGAA | >hg38_21_+_8433172_8446622_RNA45SN1_45S_50ntFlanks | 11576 | 11601 | 26 | CTTAGAACTGGTGCGGACCAGGGGAA | 994.6 | -8.5 | 1.0 | -8.2 | 2.3E-16 | 7.6E-16 | MB231/MB468 |
| >MI0000650\|hsa-mir-200c&WithFlank&12\|+\|6963693\|6963772@49.70.22&[MIMAT0000617&hsa-miR-200c-3p&offsets\|-1\|-2;m-54&12\|+\|6963742\|6963764&offsets\|-1\|-2]\|CTAATACTGCCGGGTAATGATG | >MI0000650\|hsa-mir-200c&WithFlank&12\|+\|6963693\|6963772 | 49 | 70 | 22 | CTAATACTGCCGGGTAATGATG | 961.7 | 11.5 | 1.4 | 8.2 | 2.3E-16 | 7.8E-16 | BT20/MB468 |
| >hg38_21_+_8433172_8446622_RNA45SN1_45S_50ntFlanks@6721.6753.33\|AATTGCAGGACACATTGATCATCGACACTTCGA | >hg38_21_+_8433172_8446622_RNA45SN1_45S_50ntFlanks | 6721 | 6753 | 33 | AATTGCAGGACACATTGATCATCGACACTTCGA | 120.9 | -4.7 | 0.6 | -8.2 | 2.3E-16 | 7.8E-16 | BT20/MB468 |
| >MI0000285\|hsa-mir-205&WithFlank&1\|+\|209432127\|209432248@40.59.20(+1G)&[MIMAT0000266&hsa-miR-205-5p&offsets\|0\|-2(+1G);m-351&1\|+\|209432166\|209432187&offsets\|0\|-2(+1G)]\|TCCTTCATTCCACCGGAGTCG | >MI0000285\|hsa-mir-205&WithFlank&1\|+\|209432127\|209432248 | 40 | 59 | 20(+1G) | TCCTTCATTCCACCGGAGTCG | 858.6 | 13.2 | 1.6 | 8.5 | 1.5E-17 | 7.8E-16 | BT20/MB231 |
| >MI0000079\|hsa-mir-23a&WithFlank&19\|-\|13836581\|13836665@54.74.21&[MIMAT0000078&hsa-miR-23a-3p&offsets\|+3\|+3;m-31&19\|-\|13836595\|13836615&offsets\|+3\|+3]\|ACATTGCCAGGGATTTCCAAC | >MI0000079\|hsa-mir-23a&WithFlank&19\|-\|13836581\|13836665 | 54 | 74 | 21 | ACATTGCCAGGGATTTCCAAC | 1750.3 | 29.8 | 3.6 | 8.2 | 2.4E-16 | 7.9E-16 | MB231/MB468 |
| >MI0000085\|hsa-mir-27a&WithFlank&19\|-\|13836434\|13836523@57.76.20(+3U)&[MIMAT0000084&hsa-miR-27a-3p&offsets\|0\|-1(+3U);m-43&19\|-\|13836447\|13836467&offsets\|0\|-1(+3U)]\|TTCACAGTGGCTAAGTTCCGTTT | >MI0000085\|hsa-mir-27a&WithFlank&19\|-\|13836434\|13836523 | 57 | 76 | 20(+3U) | TTCACAGTGGCTAAGTTCCGTTT | 3492.0 | 6.8 | 0.8 | 8.2 | 2.4E-16 | 8.0E-16 | MB231/MB468 |
| >hg38_MT_+_598_1651_RNR1_12S_50ntFlanks@916.935.20\|AAGGACATTTAACTAAAACC | >hg38_MT_+_598_1651_RNR1_12S_50ntFlanks | 916 | 935 | 20 | AAGGACATTTAACTAAAACC | 822.6 | 11.0 | 1.3 | 8.2 | 2.3E-16 | 8.0E-16 | BT20/MB468 |
| >hg38_21_+_8433172_8446622_RNA45SN1_45S_50ntFlanks@8114.8143.30\|GCGCGGGACATGTGGCGTACGGAAGACCCG | >hg38_21_+_8433172_8446622_RNA45SN1_45S_50ntFlanks | 8114 | 8143 | 30 | GCGCGGGACATGTGGCGTACGGAAGACCCG | 670.3 | -11.0 | 1.3 | -8.2 | 2.4E-16 | 8.0E-16 | MB231/MB468 |
| >hg38_21_+_8433172_8446622_RNA45SN1_45S_50ntFlanks@6737.6770.34\|GATCATCGACACTTCGAACGCACTTGCGGCCCCG | >hg38_21_+_8433172_8446622_RNA45SN1_45S_50ntFlanks | 6737 | 6770 | 34 | GATCATCGACACTTCGAACGCACTTGCGGCCCCG | 453.9 | -10.8 | 1.3 | -8.2 | 2.3E-16 | 8.0E-16 | BT20/MB468 |
| >hg38_21_+_8433172_8446622_RNA45SN1_45S_50ntFlanks@4773.4793.21\|TAGTTCCGACCATAAACGATG | >hg38_21_+_8433172_8446622_RNA45SN1_45S_50ntFlanks | 4773 | 4793 | 21 | TAGTTCCGACCATAAACGATG | 1573.2 | -5.8 | 0.7 | -8.2 | 2.4E-16 | 8.0E-16 | BT20/MB468 |
| >hg38_21_+_8433172_8446622_RNA45SN1_45S_50ntFlanks@4657.4689.33\|CTAGAGGTGAAATTCTTGGACCGGCGCAAGACG | >hg38_21_+_8433172_8446622_RNA45SN1_45S_50ntFlanks | 4657 | 4689 | 33 | CTAGAGGTGAAATTCTTGGACCGGCGCAAGACG | 1966.3 | -6.8 | 0.8 | -8.2 | 2.4E-16 | 8.1E-16 | MB231/MB468 |
| >hg38_21_+_8433172_8446622_RNA45SN1_45S_50ntFlanks@10866.10891.26\|CTAAGGGCTGGGTCGGTCGGGCTGGG | >hg38_21_+_8433172_8446622_RNA45SN1_45S_50ntFlanks | 10866 | 10891 | 26 | CTAAGGGCTGGGTCGGTCGGGCTGGG | 397.4 | -5.4 | 0.7 | -8.2 | 2.4E-16 | 8.1E-16 | BT20/MB468 |
| >hg38_21_+_8433172_8446622_RNA45SN1_45S_50ntFlanks@10808.10825.18\|CAATGTAGGTAAGGGAAG | >hg38_21_+_8433172_8446622_RNA45SN1_45S_50ntFlanks | 10808 | 10825 | 18 | CAATGTAGGTAAGGGAAG | 689.3 | -8.5 | 1.0 | -8.2 | 2.4E-16 | 8.1E-16 | MB231/MB468 |
| >MI0000285\|hsa-mir-205&WithFlank&1\|+\|209432127\|209432248@40.62.23(+1U)&[MIMAT0000266&hsa-miR-205-5p&offsets\|0\|+1(+1U);m-351&1\|+\|209432166\|209432187&offsets\|0\|+1(+1U)]\|TCCTTCATTCCACCGGAGTCTGTT | >MI0000285\|hsa-mir-205&WithFlank&1\|+\|209432127\|209432248 | 40 | 62 | 23(+1U) | TCCTTCATTCCACCGGAGTCTGTT | 2464.6 | 6.1 | 0.7 | 8.2 | 2.4E-16 | 8.2E-16 | BT20/MB468 |
| >MI0000439\|hsa-mir-23b&WithFlank&9\|+\|95085202\|95085310@67.86.20(+1A)&[MIMAT0000418&hsa-miR-23b-3p&offsets\|+3\|0(+1A);m-42&9\|+\|95085265\|95085285&offsets\|+3\|+2(+1A)]\|ACATTGCCAGGGATTACCACA | >MI0000439\|hsa-mir-23b&WithFlank&9\|+\|95085202\|95085310 | 67 | 86 | 20(+1A) | ACATTGCCAGGGATTACCACA | 1725.3 | 29.8 | 3.6 | 8.2 | 2.5E-16 | 8.3E-16 | MB231/MB468 |
| >MI0000298\|hsa-mir-221&WithFlank&X\|-\|45746151\|45746272@71.93.23(+1G)&[MIMAT0000278&hsa-miR-221-3p&offsets\|0\|0(+1G);m-91&X\|-\|45746181\|45746202&offsets\|0\|+1(+1G)]\|AGCTACATTGTCTGCTGGGTTTCG | >MI0000298\|hsa-mir-221&WithFlank&X\|-\|45746151\|45746272 | 71 | 93 | 23(+1G) | AGCTACATTGTCTGCTGGGTTTCG | 695.0 | 5.5 | 0.7 | 8.2 | 2.5E-16 | 8.4E-16 | BT20/MB468 |
| >MI0001448\|hsa-mir-425&WithFlank&3\|-\|49020142\|49020240@20.42.23&[MIMAT0003393&hsa-miR-425-5p&offsets\|0\|0;m-95&3\|-\|49020200\|49020221&offsets\|0\|+1]\|AATGACACGATCACTCCCGTTGA | >MI0001448\|hsa-mir-425&WithFlank&3\|-\|49020142\|49020240 | 20 | 42 | 23 | AATGACACGATCACTCCCGTTGA | 718.1 | 7.3 | 0.9 | 8.2 | 2.5E-16 | 8.4E-16 | BT20/MB468 |
| >hg38_21_+_8433172_8446622_RNA45SN1_45S_50ntFlanks@12126.12150.25\|CGACCCGCTCCGGGGACAGTGCCAG | >hg38_21_+_8433172_8446622_RNA45SN1_45S_50ntFlanks | 12126 | 12150 | 25 | CGACCCGCTCCGGGGACAGTGCCAG | 423.2 | -10.7 | 1.3 | -8.2 | 2.5E-16 | 8.4E-16 | BT20/MB468 |
| >MI0000093\|hsa-mir-92a-1&WithFlank&13\|+\|91351308\|91351397@55.75.21;MI0000094\|hsa-mir-92a-2&WithFlank&X\|-\|134169532\|134169618@55.75.21&[MIMAT0000092&hsa-miR-92a-3p&offsets\|+1\|0;m-9&13\|+\|91351361\|91351382&offsets\|+1\|0];[MIMAT0000092_1&hsa-miR-92a-3p&offsets\|+1\|0;m-11&X\|-\|134169544\|134169565&offsets\|+1\|0]\|ATTGCACTTGTCCCGGCCTGT | >MI0000093\|hsa-mir-92a-1&WithFlank&13\|+\|91351308\|91351397 | 55 | 75 | 21 | ATTGCACTTGTCCCGGCCTGT | 791.8 | 5.3 | 0.7 | 8.2 | 2.5E-16 | 8.5E-16 | BT20/MB468 |
| >hg38_21_+_8433172_8446622_RNA45SN1_45S_50ntFlanks@11612.11640.29\|TAATTAAAACAAAGCATCGCGAAGGCCCG | >hg38_21_+_8433172_8446622_RNA45SN1_45S_50ntFlanks | 11612 | 11640 | 29 | TAATTAAAACAAAGCATCGCGAAGGCCCG | 457.0 | -10.8 | 1.3 | -8.2 | 2.5E-16 | 8.5E-16 | BT20/MB468 |
| >hg38_21_+_8433172_8446622_RNA45SN1_45S_50ntFlanks@11564.11601.38\|CCTAGCAGCCGACTTAGAACTGGTGCGGACCAGGGGAA | >hg38_21_+_8433172_8446622_RNA45SN1_45S_50ntFlanks | 11564 | 11601 | 38 | CCTAGCAGCCGACTTAGAACTGGTGCGGACCAGGGGAA | 963.9 | -5.4 | 0.7 | -8.2 | 2.5E-16 | 8.5E-16 | MB231/MB468 |
| >hg38_21_+_8433172_8446622_RNA45SN1_45S_50ntFlanks@3984.4001.18\|GCGGCTTTGGTGACTCTA | >hg38_21_+_8433172_8446622_RNA45SN1_45S_50ntFlanks | 3984 | 4001 | 18 | GCGGCTTTGGTGACTCTA | 798.5 | -6.1 | 0.7 | -8.2 | 2.5E-16 | 8.5E-16 | BT20/MB468 |
| >hg38_21_+_8433172_8446622_RNA45SN1_45S_50ntFlanks@3705.3739.35\|TACCTGGTTGATCCTGCCAGTAGCATATGCTTGTC | >hg38_21_+_8433172_8446622_RNA45SN1_45S_50ntFlanks | 3705 | 3739 | 35 | TACCTGGTTGATCCTGCCAGTAGCATATGCTTGTC | 2861.7 | 6.4 | 0.8 | 8.2 | 2.6E-16 | 8.7E-16 | BT20/MB468 |
| >MI0000298\|hsa-mir-221&WithFlank&X\|-\|45746151\|45746272@71.91.21(+2C)&[MIMAT0000278&hsa-miR-221-3p&offsets\|0\|-2(+2C);m-91&X\|-\|45746181\|45746202&offsets\|0\|-1(+2C)]\|AGCTACATTGTCTGCTGGGTTCC | >MI0000298\|hsa-mir-221&WithFlank&X\|-\|45746151\|45746272 | 71 | 91 | 21(+2C) | AGCTACATTGTCTGCTGGGTTCC | 639.2 | 8.3 | 1.0 | 8.2 | 2.6E-16 | 8.8E-16 | MB231/MB468 |
| >hg38_21_+_8433172_8446622_RNA45SN1_45S_50ntFlanks@11768.11786.19\|CTAATTAGTGACGCGCATG | >hg38_21_+_8433172_8446622_RNA45SN1_45S_50ntFlanks | 11768 | 11786 | 19 | CTAATTAGTGACGCGCATG | 601.8 | -7.4 | 0.9 | -8.2 | 2.6E-16 | 8.8E-16 | MB231/MB468 |
| >hg38_21_+_8433172_8446622_RNA45SN1_45S_50ntFlanks@12126.12146.21\|CGACCCGCTCCGGGGACAGTG | >hg38_21_+_8433172_8446622_RNA45SN1_45S_50ntFlanks | 12126 | 12146 | 21 | CGACCCGCTCCGGGGACAGTG | 357.0 | -10.5 | 1.3 | -8.2 | 2.6E-16 | 8.8E-16 | BT20/MB468 |
| >hg38_21_+_8433172_8446622_RNA45SN1_45S_50ntFlanks@10786.10825.40\|AACAGCCTCTGGCATGTTGGAACAATGTAGGTAAGGGAAG | >hg38_21_+_8433172_8446622_RNA45SN1_45S_50ntFlanks | 10786 | 10825 | 40 | AACAGCCTCTGGCATGTTGGAACAATGTAGGTAAGGGAAG | 372.4 | -5.9 | 0.7 | -8.2 | 2.6E-16 | 8.8E-16 | BT20/MB468 |
| >hg38_21_+_8433172_8446622_RNA45SN1_45S_50ntFlanks@12416.12437.22\|TTCGATGTCGGCTCTTCCTATC | >hg38_21_+_8433172_8446622_RNA45SN1_45S_50ntFlanks | 12416 | 12437 | 22 | TTCGATGTCGGCTCTTCCTATC | 4593.0 | 5.1 | 0.6 | 8.2 | 2.7E-16 | 8.9E-16 | MB231/MB468 |
| >hg38_21_+_8433172_8446622_RNA45SN1_45S_50ntFlanks@10409.10430.22\|GTCAGTCGGTCCTGAGAGATGG | >hg38_21_+_8433172_8446622_RNA45SN1_45S_50ntFlanks | 10409 | 10430 | 22 | GTCAGTCGGTCCTGAGAGATGG | 386.9 | -3.9 | 0.5 | -8.2 | 2.7E-16 | 9.0E-16 | MB231/MB468 |
| >hg38_21_+_8433172_8446622_RNA45SN1_45S_50ntFlanks@12432.12455.24\|CCTATCATTGTGAAGCAGAATTCA | >hg38_21_+_8433172_8446622_RNA45SN1_45S_50ntFlanks | 12432 | 12455 | 24 | CCTATCATTGTGAAGCAGAATTCA | 219.2 | -6.0 | 0.7 | -8.2 | 2.7E-16 | 9.0E-16 | MB231/MB468 |
| >hg38_21_+_8433172_8446622_RNA45SN1_45S_50ntFlanks@9567.9592.26\|ACGTGCAAATCGGTCGTCCGACCTGG | >hg38_21_+_8433172_8446622_RNA45SN1_45S_50ntFlanks | 9567 | 9592 | 26 | ACGTGCAAATCGGTCGTCCGACCTGG | 628.8 | -5.5 | 0.7 | -8.2 | 2.7E-16 | 9.1E-16 | MB231/MB468 |
| >hg38_21_+_8433172_8446622_RNA45SN1_45S_50ntFlanks@12450.12480.31\|AATTCACCAAGCGTTGGATTGTTCACCCACT | >hg38_21_+_8433172_8446622_RNA45SN1_45S_50ntFlanks | 12450 | 12480 | 31 | AATTCACCAAGCGTTGGATTGTTCACCCACT | 463.9 | 7.5 | 0.9 | 8.2 | 2.8E-16 | 9.3E-16 | MB231/MB468 |
| >MI0000477\|hsa-mir-146a&WithFlank&5\|+\|160485346\|160485456@27.48.22(+1U)&[MIMAT0000449&hsa-miR-146a-5p&offsets\|0\|0(+1U);m-129&5\|+\|160485372\|160485393&offsets\|0\|0(+1U)]\|TGAGAACTGAATTCCATGGGTTT | >MI0000477\|hsa-mir-146a&WithFlank&5\|+\|160485346\|160485456 | 27 | 48 | 22(+1U) | TGAGAACTGAATTCCATGGGTTT | 632.2 | -12.7 | 1.5 | -8.5 | 1.8E-17 | 9.3E-16 | BT20/MB231 |
| >hg38_21_+_8433172_8446622_RNA45SN1_45S_50ntFlanks@12635.12667.33\|AAGCTACCATCTGTGGGATTATGACTGAACGCC | >hg38_21_+_8433172_8446622_RNA45SN1_45S_50ntFlanks | 12635 | 12667 | 33 | AAGCTACCATCTGTGGGATTATGACTGAACGCC | 538.2 | 7.7 | 0.9 | 8.2 | 2.8E-16 | 9.3E-16 | MB231/MB468 |
| >hg38_21_+_8433172_8446622_RNA45SN1_45S_50ntFlanks@11874.11905.32\|AGAAGACCCTGTTGAGCTTGACTCTAGTCTGG | >hg38_21_+_8433172_8446622_RNA45SN1_45S_50ntFlanks | 11874 | 11905 | 32 | AGAAGACCCTGTTGAGCTTGACTCTAGTCTGG | 428.1 | -10.3 | 1.3 | -8.2 | 2.8E-16 | 9.4E-16 | MB231/MB468 |

<table>
<tr><td>>MI0000447|hsa-mir-128-1&WithFlank&2|+|135665391|135665484@56.75.20;MI0000727|hsa-mir-128-2&WithFlank&3|+|35744470|35744565@58.77.20&[MIMAT0000424&hsa-miR-128-3p&offsets|0|-1;m-108&2|+|135665446|135665466&offsets|0|-1];[MIMAT0000424_1&hsa-miR-128-3p&offsets|0|-1;m-117&3|+|35744527|35744547&offsets|0|-1]|TCACAGTGAACCGGTCTCTT</td><td>>MI0000447|hsa-mir-128-1&WithFlank&2|+|135665391|135665484</td><td>56</td><td>75</td><td>20</td><td>TCACAGTGAACCGGTCTCTT</td><td>2561.5</td><td>6.1</td><td>0.7</td><td>8.2</td><td>2.9E-16</td><td>9.5E-16</td><td>MB231/MB468</td></tr>
<tr><td>>hg38_21_+_8433172_8446622_RNA45SN1_45S_50ntFlanks@485.524.40|GATGTGGTGACGTCGTGCTCTCCCGGGCCGGGTCCGAGCC</td><td>>hg38_21_+_8433172_8446622_RNA45SN1_45S_50ntFlanks</td><td>485</td><td>524</td><td>40</td><td>GATGTGGTGACGTCGTGCTCTCCCGGGCCGGGTCCGAGCC</td><td>1833.8</td><td>29.7</td><td>3.6</td><td>8.2</td><td>2.9E-16</td><td>9.6E-16</td><td>MB231/MB468</td></tr>
<tr><td>>hg38_21_+_8433172_8446622_RNA45SN1_45S_50ntFlanks@11885.11904.20|TTGAGCTTGACTCTAGTCTG</td><td>>hg38_21_+_8433172_8446622_RNA45SN1_45S_50ntFlanks</td><td>11885</td><td>11904</td><td>20</td><td>TTGAGCTTGACTCTAGTCTG</td><td>1278.4</td><td>-8.1</td><td>1.0</td><td>-8.2</td><td>2.9E-16</td><td>9.6E-16</td><td>MB231/MB468</td></tr>
<tr><td>>MI0000081|hsa-mir-24-2&WithFlank&19|-|13836281|13836365@56.76.21(+1A);MI0000080|hsa-mir-24-1&WithFlank&9|+|95086015|95086094@50.70.21(+1A)&[MIMAT0000080&hsa-miR-24-3p&offsets|0|-1(+1A);m-36&19|-|13836288|13836310&offsets|0|-2(+1A)];[MIMAT0000080_1&hsa-miR-24-3p&offsets|0|-1(+1A);m-35&9|+|95086064|95086086&offsets|0|-2(+1A)]|TGGCTCAGTTCAGCAGGAACAA</td><td>>MI0000081|hsa-mir-24-2&WithFlank&19|-|13836281|13836365</td><td>56</td><td>76</td><td>21(+1A)</td><td>TGGCTCAGTTCAGCAGGAACAA</td><td>4217.5</td><td>6.4</td><td>0.8</td><td>8.2</td><td>2.8E-16</td><td>9.6E-16</td><td>BT20/MB468</td></tr>
<tr><td>>hg38_21_+_8433172_8446622_RNA45SN1_45S_50ntFlanks@6778.6802.25|CCCGGGGCTACGCCTGTCTGAGCGT</td><td>>hg38_21_+_8433172_8446622_RNA45SN1_45S_50ntFlanks</td><td>6778</td><td>6802</td><td>25</td><td>CCCGGGGCTACGCCTGTCTGAGCGT</td><td>3112.8</td><td>7.0</td><td>0.9</td><td>8.2</td><td>2.9E-16</td><td>9.6E-16</td><td>MB231/MB468</td></tr>
<tr><td>>hg38_21_+_8433172_8446622_RNA45SN1_45S_50ntFlanks@12456.12477.22|CCAAGCGTTGGATTGTTCACCC</td><td>>hg38_21_+_8433172_8446622_RNA45SN1_45S_50ntFlanks</td><td>12456</td><td>12477</td><td>22</td><td>CCAAGCGTTGGATTGTTCACCC</td><td>6317.2</td><td>-4.4</td><td>0.5</td><td>-8.2</td><td>2.9E-16</td><td>9.6E-16</td><td>MB231/MB468</td></tr>
<tr><td>>hg38_21_+_8433172_8446622_RNA45SN1_45S_50ntFlanks@7989.8016.28|AGACGTGGCGACCCGCTGAATTTAAGCA</td><td>>hg38_21_+_8433172_8446622_RNA45SN1_45S_50ntFlanks</td><td>7989</td><td>8016</td><td>28</td><td>AGACGTGGCGACCCGCTGAATTTAAGCA</td><td>1263.6</td><td>5.2</td><td>0.6</td><td>8.2</td><td>2.9E-16</td><td>9.7E-16</td><td>MB231/MB468</td></tr>
<tr><td>>hg38_21_+_8433172_8446622_RNA45SN1_45S_50ntFlanks@11874.11907.34|AGAAGACCCTGTTGAGCTTGACTCTAGTCTGGCA</td><td>>hg38_21_+_8433172_8446622_RNA45SN1_45S_50ntFlanks</td><td>11874</td><td>11907</td><td>34</td><td>AGAAGACCCTGTTGAGCTTGACTCTAGTCTGGCA</td><td>285.1</td><td>-10.1</td><td>1.2</td><td>-8.2</td><td>2.9E-16</td><td>9.9E-16</td><td>BT20/MB468</td></tr>
<tr><td>>MI0003776|hsa-mir-320b-1&WithFlank&1|+|116671740|116671823@48.68.21;MI0003839|hsa-mir-320b-2&WithFlank&1|-|224257034|224257116@47.67.21&[MIMAT0005792&hsa-miR-320b&offsets|0|-1;m-183&1|+|116671787|116671806&offsets|0|+1];[MIMAT0005792_1&hsa-miR-320b&offsets|0|-1;m-183&1|-|224257051|224257070&offsets|0|+1]|AAAAGCTGGGTTGAGAGGGCA</td><td>>MI0003776|hsa-mir-320b-1&WithFlank&1|+|116671740|116671823</td><td>48</td><td>68</td><td>21</td><td>AAAAGCTGGGTTGAGAGGGCA</td><td>3312.2</td><td>11.2</td><td>1.4</td><td>8.2</td><td>3.0E-16</td><td>1.0E-15</td><td>BT20/MB468</td></tr>
<tr><td>>hg38_21_+_8433172_8446622_RNA45SN1_45S_50ntFlanks@475.497.23|CTTCGTGATCGATGTGGTGACGT</td><td>>hg38_21_+_8433172_8446622_RNA45SN1_45S_50ntFlanks</td><td>475</td><td>497</td><td>23</td><td>CTTCGTGATCGATGTGGTGACGT</td><td>13015.1</td><td>6.6</td><td>0.8</td><td>8.2</td><td>3.0E-16</td><td>1.0E-15</td><td>BT20/MB468</td></tr>
<tr><td>>hg38_21_+_8433172_8446622_RNA45SN1_45S_50ntFlanks@11665.11702.38|TTCTGCCCAGTGCTCTGAATGTCAAAGTGAAGAAATTC</td><td>>hg38_21_+_8433172_8446622_RNA45SN1_45S_50ntFlanks</td><td>11665</td><td>11702</td><td>38</td><td>TTCTGCCCAGTGCTCTGAATGTCAAAGTGAAGAAATTC</td><td>624.6</td><td>7.7</td><td>0.9</td><td>8.2</td><td>3.1E-16</td><td>1.0E-15</td><td>MB231/MB468</td></tr>
<tr><td>>hg38_21_+_8433172_8446622_RNA45SN1_45S_50ntFlanks@12991.13019.29|CTCCCTCGCTGCGATCTATTGAAAGTCAG</td><td>>hg38_21_+_8433172_8446622_RNA45SN1_45S_50ntFlanks</td><td>12991</td><td>13019</td><td>29</td><td>CTCCCTCGCTGCGATCTATTGAAAGTCAG</td><td>450.8</td><td>-10.8</td><td>1.3</td><td>-8.2</td><td>3.0E-16</td><td>1.0E-15</td><td>BT20/MB468</td></tr>
<tr><td>>MI0000434|hsa-let-7i&WithFlank&12|+|62603680|62603775@12.32.21(+1G)&[MIMAT0000415&hsa-let-7i-5p&offsets|0|-1(+1G);m-49&12|+|62603691|62603712&offsets|0|-1(+1G)]|TGAGGTAGTAGTTTGTGCTGTG</td><td>>MI0000434|hsa-let-7i&WithFlank&12|+|62603680|62603775</td><td>12</td><td>32</td><td>21(+1G)</td><td>TGAGGTAGTAGTTTGTGCTGTG</td><td>13515.8</td><td>7.2</td><td>0.9</td><td>8.2</td><td>3.1E-16</td><td>1.0E-15</td><td>BT20/MB468</td></tr>
<tr><td>>MI0000088|hsa-mir-30a&WithFlank&6|-|71403545|71403627@12.32.21&[MIMAT0000087&hsa-miR-30a-5p&offsets|0|-1;m-15&6|-|71403595|71403616&offsets|0|-1]|TGTAAACATCCTCGACTGGAA</td><td>>MI0000088|hsa-mir-30a&WithFlank&6|-|71403545|71403627</td><td>12</td><td>32</td><td>21</td><td>TGTAAACATCCTCGACTGGAA</td><td>4845.3</td><td>7.9</td><td>1.0</td><td>8.2</td><td>3.1E-16</td><td>1.0E-15</td><td>BT20/MB468</td></tr>
<tr><td>>hg38_21_+_8433172_8446622_RNA45SN1_45S_50ntFlanks@8380.8403.24|CAAGAGGGCGTGAAACCGTTAAGA</td><td>>hg38_21_+_8433172_8446622_RNA45SN1_45S_50ntFlanks</td><td>8380</td><td>8403</td><td>24</td><td>CAAGAGGGCGTGAAACCGTTAAGA</td><td>580.2</td><td>-5.1</td><td>0.6</td><td>-8.2</td><td>3.1E-16</td><td>1.0E-15</td><td>BT20/MB468</td></tr>
<tr><td>>am_MI0000786|hsa-mir-378a&WithFlank&5|+|149732819|149732896@49.69.21(+1U)&[MIMAT0000732&hsa-miR-378a-3p&offsets|0|-1(+1U);m-44&5|+|149732867|149732888&offsets|0|-1(+1U)]|ACTGGACTTGGAGTCAGAAGGT</td><td>>am_MI0000786|hsa-mir-378a&WithFlank&5|+|149732819|149732896</td><td>49</td><td>69</td><td>21(+1U)</td><td>ACTGGACTTGGAGTCAGAAGGT</td><td>7541.9</td><td>6.0</td><td>0.7</td><td>8.2</td><td>3.1E-16</td><td>1.0E-15</td><td>MB231/MB468</td></tr>
<tr><td>>hg38_21_+_8433172_8446622_RNA45SN1_45S_50ntFlanks@11710.11728.19|CGCGGGTAAACGGCGGGAG</td><td>>hg38_21_+_8433172_8446622_RNA45SN1_45S_50ntFlanks</td><td>11710</td><td>11728</td><td>19</td><td>CGCGGGTAAACGGCGGGAG</td><td>260.6</td><td>-6.5</td><td>0.8</td><td>-8.2</td><td>3.2E-16</td><td>1.0E-15</td><td>MB231/MB468</td></tr>
<tr><td>>trnaMT_ValTAC_MT_+_1602_1670@29.72.44|ACTTACACTTAGGAGATTTCAACTTAACTTGACCGCTCTGACCA</td><td>>trnaMT_ValTAC_MT_+_1602_1670</td><td>29</td><td>72</td><td>44</td><td>ACTTACACTTAGGAGATTTCAACTTAACTTGACCGCTCTGACCA</td><td>522.7</td><td>9.0</td><td>1.1</td><td>8.2</td><td>3.2E-16</td><td>1.1E-15</td><td>MB231/MB468</td></tr>
<tr><td>>hg38_21_+_8433172_8446622_RNA45SN1_45S_50ntFlanks@9554.9574.21|CGTAGCGGTCCTGACGTGCAA</td><td>>hg38_21_+_8433172_8446622_RNA45SN1_45S_50ntFlanks</td><td>9554</td><td>9574</td><td>21</td><td>CGTAGCGGTCCTGACGTGCAA</td><td>2189.7</td><td>-3.7</td><td>0.4</td><td>-8.2</td><td>3.1E-16</td><td>1.1E-15</td><td>BT20/MB468</td></tr>
<tr><td>>MI0000285|hsa-mir-205&WithFlank&1|+|209432127|209432248@40.59.20&[MIMAT0000266&hsa-miR-205-5p&offsets|0|-2;m-351&1|+|209432166|209432187&offsets|0|-2]|TCCTTCATTCCACCGGAGTC</td><td>>MI0000285|hsa-mir-205&WithFlank&1|+|209432127|209432248</td><td>40</td><td>59</td><td>20</td><td>TCCTTCATTCCACCGGAGTC</td><td>414.9</td><td>12.2</td><td>1.4</td><td>8.5</td><td>2.1E-17</td><td>1.1E-15</td><td>BT20/MB231</td></tr>
<tr><td>>hg38_21_+_8433172_8446622_RNA45SN1_45S_50ntFlanks@12626.12659.34|AATGGGGCGAAGCTACCATCTGTGGGATTATGAC</td><td>>hg38_21_+_8433172_8446622_RNA45SN1_45S_50ntFlanks</td><td>12626</td><td>12659</td><td>34</td><td>AATGGGGCGAAGCTACCATCTGTGGGATTATGAC</td><td>484.4</td><td>-5.5</td><td>0.7</td><td>-8.2</td><td>3.2E-16</td><td>1.1E-15</td><td>BT20/MB468</td></tr>
<tr><td>>hg38_21_+_8433172_8446622_RNA45SN1_45S_50ntFlanks@10786.10815.30|AACAGCCTCTGGCATGTTGGAACAATGTAG</td><td>>hg38_21_+_8433172_8446622_RNA45SN1_45S_50ntFlanks</td><td>10786</td><td>10815</td><td>30</td><td>AACAGCCTCTGGCATGTTGGAACAATGTAG</td><td>217.4</td><td>-4.2</td><td>0.5</td><td>-8.2</td><td>3.2E-16</td><td>1.1E-15</td><td>BT20/MB468</td></tr>
<tr><td>>hg38_1_-_228634819_228635039_RNA5S12_5S_50ntFlanks@138.170.33|AGACCGCCTGGGAATACCGGGTGCTGTAGGCTT</td><td>>hg38_1_-_228634819_228635039_RNA5S12_5S_50ntFlanks</td><td>138</td><td>170</td><td>33</td><td>AGACCGCCTGGGAATACCGGGTGCTGTAGGCTT</td><td>1732.9</td><td>7.2</td><td>0.9</td><td>8.2</td><td>3.2E-16</td><td>1.1E-15</td><td>BT20/MB468</td></tr>
<tr><td>>am_trnaMT_ArgTCG_MT_+_10405_10469@28.48.21|ATTTCGACTCATTAAATTATG</td><td>>am_trnaMT_ArgTCG_MT_+_10405_10469</td><td>28</td><td>48</td><td>21</td><td>ATTTCGACTCATTAAATTATG</td><td>698.4</td><td>-6.2</td><td>0.8</td><td>-8.2</td><td>3.2E-16</td><td>1.1E-15</td><td>BT20/MB468</td></tr>
<tr><td>>MI0000093|hsa-mir-92a-1&WithFlank&13|+|91351308|91351397@54.76.23&[MIMAT0000092&hsa-miR-92a-3p&offsets|0|+1;m-9&13|+|91351361|91351382&offsets|0|+1]|TATTGCACTTGTCCCGGCCTGTT</td><td>>MI0000093|hsa-mir-92a-1&WithFlank&13|+|91351308|91351397</td><td>54</td><td>76</td><td>23</td><td>TATTGCACTTGTCCCGGCCTGTT</td><td>4699.5</td><td>5.4</td><td>0.7</td><td>8.2</td><td>3.2E-16</td><td>1.1E-15</td><td>BT20/MB468</td></tr>
<tr><td>>MI0000067|hsa-let-7f-1&WithFlank&9|+|94176341|94176439@13.34.22(+1A);MI0000068|hsa-let-7f-2&WithFlank&X|-|53557186|53557280@14.35.22(+1A)&[MIMAT0000067&hsa-let-7f-5p&offsets|0|0(+1A);m-20&9|+|94176353|94176374&offsets|0|0(+1A)];[MIMAT0000067_1&hsa-let-7f-5p&offsets|0|0(+1A);m-19&X|-|53557246|53557267&offsets|0|0(+1A)]|TGAGGTAGTAGATTGTATAGTTA</td><td>>MI0000067|hsa-let-7f-1&WithFlank&9|+|94176341|94176439</td><td>13</td><td>34</td><td>22(+1A)</td><td>TGAGGTAGTAGATTGTATAGTTA</td><td>6396.4</td><td>6.6</td><td>0.8</td><td>8.2</td><td>3.2E-16</td><td>1.1E-15</td><td>BT20/MB468</td></tr>
<tr><td>>trnaMT_ValTAC_MT_+_1602_1670@52.72.21|TTAACTTGACCGCTCTGACCA</td><td>>trnaMT_ValTAC_MT_+_1602_1670</td><td>52</td><td>72</td><td>21</td><td>TTAACTTGACCGCTCTGACCA</td><td>611.5</td><td>8.7</td><td>1.1</td><td>8.2</td><td>3.3E-16</td><td>1.1E-15</td><td>MB231/MB468</td></tr>
<tr><td>>hg38_21_+_8433172_8446622_RNA45SN1_45S_50ntFlanks@4003.4020.18|ATAACCTCGGGCCGATCG</td><td>>hg38_21_+_8433172_8446622_RNA45SN1_45S_50ntFlanks</td><td>4003</td><td>4020</td><td>18</td><td>ATAACCTCGGGCCGATCG</td><td>425.2</td><td>-4.9</td><td>0.6</td><td>-8.2</td><td>3.3E-16</td><td>1.1E-15</td><td>MB231/MB468</td></tr>
<tr><td>>hg38_1_-_228634819_228635039_RNA5S12_5S_50ntFlanks@145.171.27|CTGGGAATACCGGGTGCTGTAGGCTTT</td><td>>hg38_1_-_228634819_228635039_RNA5S12_5S_50ntFlanks</td><td>145</td><td>171</td><td>27</td><td>CTGGGAATACCGGGTGCTGTAGGCTTT</td><td>1071.7</td><td>6.3</td><td>0.8</td><td>8.2</td><td>3.2E-16</td><td>1.1E-15</td><td>BT20/MB468</td></tr>
<tr><td>>MI0016005|hsa-mir-3615&WithFlank&17|+|74748607|74748705@57.78.22&[MIMAT0017994&hsa-miR-3615&offsets|0|+1;m-271&17|+|74748663|74748684&offsets|0|0]|TCTCTCGGCTCCTCGCGGCTCG</td><td>>MI0016005|hsa-mir-3615&WithFlank&17|+|74748607|74748705</td><td>57</td><td>78</td><td>22</td><td>TCTCTCGGCTCCTCGCGGCTCG</td><td>1395.8</td><td>6.2</td><td>0.8</td><td>8.2</td><td>3.3E-16</td><td>1.1E-15</td><td>MB231/MB468</td></tr>
<tr><td>>hg38_21_+_8433172_8446622_RNA45SN1_45S_50ntFlanks@12441.12473.33|GTGAAGCAGAATTCACCAAGCGTTGGATTGTTC</td><td>>hg38_21_+_8433172_8446622_RNA45SN1_45S_50ntFlanks</td><td>12441</td><td>12473</td><td>33</td><td>GTGAAGCAGAATTCACCAAGCGTTGGATTGTTC</td><td>312.3</td><td>4.0</td><td>0.5</td><td>8.2</td><td>3.4E-16</td><td>1.1E-15</td><td>MB231/MB468</td></tr>
<tr><td>>hg38_1_-_228634819_228635039_RNA5S12_5S_50ntFlanks@51.79.29|GTCTACGGCCATACCACCCTGAACGCGCC</td><td>>hg38_1_-_228634819_228635039_RNA5S12_5S_50ntFlanks</td><td>51</td><td>79</td><td>29</td><td>GTCTACGGCCATACCACCCTGAACGCGCC</td><td>596.1</td><td>5.3</td><td>0.7</td><td>8.2</td><td>3.3E-16</td><td>1.1E-15</td><td>BT20/MB468</td></tr>
<tr><td>>MI0000434|hsa-let-7i&WithFlank&12|+|62603680|62603775@12.31.20&[MIMAT0000415&hsa-let-7i-5p&offsets|0|-2;m-49&12|+|62603691|62603712&offsets|0|-2]|TGAGGTAGTAGTTTGTGCTG</td><td>>MI0000434|hsa-let-7i&WithFlank&12|+|62603680|62603775</td><td>12</td><td>31</td><td>20</td><td>TGAGGTAGTAGTTTGTGCTG</td><td>672.2</td><td>6.6</td><td>0.8</td><td>8.2</td><td>3.4E-16</td><td>1.1E-15</td><td>BT20/MB468</td></tr>
<tr><td>>hg38_21_+_8433172_8446622_RNA45SN1_45S_50ntFlanks@11874.11909.36|AGAAGACCCTGTTGAGCTTGACTCTAGTCTGGCACG</td><td>>hg38_21_+_8433172_8446622_RNA45SN1_45S_50ntFlanks</td><td>11874</td><td>11909</td><td>36</td><td>AGAAGACCCTGTTGAGCTTGACTCTAGTCTGGCACG</td><td>206.4</td><td>-6.3</td><td>0.8</td><td>-8.2</td><td>3.4E-16</td><td>1.1E-15</td><td>BT20/MB468</td></tr>
<tr><td>>MI0000081|hsa-mir-24-2&WithFlank&19|-|13836281|13836365@56.77.22(+1U);MI0000080|hsa-mir-24-1&WithFlank&9|+|95086015|95086094@50.71.22(+1U)&[MIMAT0000080&hsa-miR-24-3p&offsets|0|0(+1U);m-36&19|-|13836288|13836310&offsets|0|-1(+1U)];[MIMAT0000080_1&hsa-miR-24-3p&offsets|0|0(+1U);m-35&9|+|95086064|95086086&offsets|0|-1(+1U)]|TGGCTCAGTTCAGCAGGAACAGT</td><td>>MI0000081|hsa-mir-24-2&WithFlank&19|-|13836281|13836365</td><td>56</td><td>77</td><td>22(+1U)</td><td>TGGCTCAGTTCAGCAGGAACAGT</td><td>25353.7</td><td>7.1</td><td>0.9</td><td>8.2</td><td>3.4E-16</td><td>1.1E-15</td><td>BT20/MB468</td></tr>
<tr><td>>am_trnaMT_PheGAA_MT_+_577_647@41.60.20|GTTTAGACGGGCTCACATCA</td><td>>am_trnaMT_PheGAA_MT_+_577_647</td><td>41</td><td>60</td><td>20</td><td>GTTTAGACGGGCTCACATCA</td><td>306.8</td><td>9.4</td><td>1.1</td><td>8.2</td><td>3.5E-16</td><td>1.2E-15</td><td>BT20/MB468</td></tr>
<tr><td>>MI0000298|hsa-mir-221&WithFlank&X|-|45746151|45746272@71.91.21(+1C)&[MIMAT0000278&hsa-miR-221-3p&offsets|0|-2(+1C);m-91&X|-|45746181|45746202&offsets|0|-1(+1C)]|AGCTACATTGTCTGCTGGGTTC</td><td>>MI0000298|hsa-mir-221&WithFlank&X|-|45746151|45746272</td><td>71</td><td>91</td><td>21(+1C)</td><td>AGCTACATTGTCTGCTGGGTTC</td><td>1894.2</td><td>6.5</td><td>0.8</td><td>8.2</td><td>3.5E-16</td><td>1.2E-15</td><td>BT20/MB468</td></tr>
<tr><td>>hg38_21_+_8433172_8446622_RNA45SN1_45S_50ntFlanks@12151.12180.30|GTGGGGAGTTTGACTGGGGCGGTACACCTG</td><td>>hg38_21_+_8433172_8446622_RNA45SN1_45S_50ntFlanks</td><td>12151</td><td>12180</td><td>30</td><td>GTGGGGAGTTTGACTGGGGCGGTACACCTG</td><td>417.2</td><td>-10.7</td><td>1.3</td><td>-8.2</td><td>3.5E-16</td><td>1.2E-15</td><td>BT20/MB468</td></tr>
<tr><td>>hg38_21_+_8433172_8446622_RNA45SN1_45S_50ntFlanks@12941.12966.26|ATTCGTAGACGACCTGCTTCTGGGTC</td><td>>hg38_21_+_8433172_8446622_RNA45SN1_45S_50ntFlanks</td><td>12941</td><td>12966</td><td>26</td><td>ATTCGTAGACGACCTGCTTCTGGGTC</td><td>658.7</td><td>-11.4</td><td>1.4</td><td>-8.2</td><td>3.5E-16</td><td>1.2E-15</td><td>BT20/MB468</td></tr>
</table>

| | | | | | | | | | | | | |
|---|---|---|---|---|---|---|---|---|---|---|---|---|
| >MI0000079\|hsa-mir-23a&WithFlank&19\|-\|13836581\|13836665@54.71.18(+1G)&[MIMAT0000078&hsa-miR-23a-3p&offsets\|+3\|0(+1G);m-31&19\|-\|13836595\|13836615&offsets\|+3\|0(+1G)]\|ACATTGCCAGGGATTTCCG | >MI0000079\|hsa-mir-23a&WithFlank&19\|-\|13836581\|13836665 | 54 | 71 | 18(+1G) | ACATTGCCAGGGATTTCCG | 1532.8 | 29.6 | 3.6 | 8.2 | 3.6E-16 | 1.2E-15 | MB231/MB468 |
| >hg38_21_+_8433172_8446622_RNA45SN1_45S_50ntFlanks@12073.12090.18\|CCGAGGGGCTCTCGCTTC | >hg38_21_+_8433172_8446622_RNA45SN1_45S_50ntFlanks | 12073 | 12090 | 18 | CCGAGGGGCTCTCGCTTC | 438.8 | -5.3 | 0.6 | -8.2 | 3.6E-16 | 1.2E-15 | BT20/MB468 |
| >MI0000285\|hsa-mir-205&WithFlank&1\|+\|209432127\|209432248@40.62.23(+1A)&[MIMAT0000266&hsa-miR-205-5p&offsets\|0\|+1(+1A);m-351&1\|+\|209432166\|209432187&offsets\|0\|+1(+1A)]\|TCCTTCATTCCACCGGAGTCTGTA | >MI0000285\|hsa-mir-205&WithFlank&1\|+\|209432127\|209432248 | 40 | 62 | 23(+1A) | TCCTTCATTCCACCGGAGTCTGTA | 4032.5 | 6.1 | 0.8 | 8.2 | 3.6E-16 | 1.2E-15 | BT20/MB468 |
| >MI0000469\|hsa-mir-125a&WithFlank&19\|+\|51693248\|51693345@21.43.23(+1G)&[MIMAT0000443&hsa-miR-125a-5p&offsets\|0\|-1(+1G);m-63&19\|+\|51693268\|51693289&offsets\|0\|+1(+1G)]\|TCCCTGAGACCCTTTAACCTGTGG | >MI0000469\|hsa-mir-125a&WithFlank&19\|+\|51693248\|51693345 | 21 | 43 | 23(+1G) | TCCCTGAGACCCTTTAACCTGTGG | 689.8 | 7.1 | 0.9 | 8.2 | 3.6E-16 | 1.2E-15 | BT20/MB468 |
| >MI0000070\|hsa-mir-16-1&WithFlank&13\|-\|50048967\|50049067@20.42.23;MI0000115\|hsa-mir-16-2&WithFlank&3\|+\|160404739\|160404831@16.38.23&[MIMAT0000069&hsa-miR-16-5p&offsets\|0\|+1;m-62&13\|-\|50049027\|50049048&offsets\|0\|+1];[MIMAT0000069_1&hsa-miR-16-5p&offsets\|0\|+1;m-61&3\|+\|160404754\|160404775&offsets\|0\|+1]\|TAGCAGCACGTAAATATTGGCGT | >MI0000070\|hsa-mir-16-1&WithFlank&13\|-\|50048967\|50049067 | 20 | 42 | 23 | TAGCAGCACGTAAATATTGGCGT | 19909.5 | 8.6 | 1.1 | 8.1 | 3.6E-16 | 1.2E-15 | BT20/MB468 |
| >MI0000447\|hsa-mir-128-1&WithFlank&2\|+\|135665391\|135665484@56.77.22&[MIMAT0000424&hsa-miR-128-3p&offsets\|0\|+1;m-108&2\|+\|135665446\|135665466&offsets\|0\|+1]\|TCACAGTGAACCGGTCTCTTTT | >MI0000447\|hsa-mir-128-1&WithFlank&2\|+\|135665391\|135665484 | 56 | 77 | 22 | TCACAGTGAACCGGTCTCTTTT | 6863.3 | 5.7 | 0.7 | 8.1 | 3.7E-16 | 1.2E-15 | MB231/MB468 |
| >hg38_21_+_8433172_8446622_RNA45SN1_45S_50ntFlanks@11918.11950.33\|ACATGAGAGGTGTAGAATAAGTGGGAGGCCCCC | >hg38_21_+_8433172_8446622_RNA45SN1_45S_50ntFlanks | 11918 | 11950 | 33 | ACATGAGAGGTGTAGAATAAGTGGGAGGCCCCC | 305.9 | -5.9 | 0.7 | -8.1 | 3.7E-16 | 1.2E-15 | BT20/MB468 |
| >MI0000234\|hsa-mir-192&WithFlank&11\|-\|64891131\|64891252@30.51.22&[MIMAT0000222&hsa-miR-192-5p&offsets\|0\|+1;m-59&11\|-\|64891203\|64891223&offsets\|0\|+1]\|CTGACCTATGAATTGACAGCCA | >MI0000234\|hsa-mir-192&WithFlank&11\|-\|64891131\|64891252 | 30 | 51 | 22 | CTGACCTATGAATTGACAGCCA | 804.1 | 6.6 | 0.8 | 8.1 | 3.8E-16 | 1.2E-15 | MB231/MB468 |
| >hg38_21_+_8433172_8446622_RNA45SN1_45S_50ntFlanks@6744.6770.27\|GACACTTCGAACGCACTTGCGGCCCCG | >hg38_21_+_8433172_8446622_RNA45SN1_45S_50ntFlanks | 6744 | 6770 | 27 | GACACTTCGAACGCACTTGCGGCCCCG | 1361.2 | -6.6 | 0.8 | -8.1 | 3.8E-16 | 1.3E-15 | BT20/MB468 |
| >am_trna10_LysCTT_16_+_3241501_3241573@31.51.21,trna119_LysCTT_1_-_145395522_145395594@31.51.21,trna11_LysCTT_5_-_180648979_180649051@31.51.21,trna13_LysCTT_16_+_3246153_3246225@31.51.21,trna13_LysCTT_6_+_26556774_26556846@31.51.21,trna30_LysCTT_16_-_3230555_3230627@31.51.21,trna5_LysCTT_16_+_3214939_3215011@31.51.21,trna7_LysCTT_16_+_3225692_3225764@31.51.21,trna8_LysCTT_1_+_39970195_39970267@31.51.21,trna9_LysCTT_5_+_180634755_180634827@31.51.21\|ACTCTTAATCTCAGGGTCGTG | >am_trna10_LysCTT_16_+_3241501_3241573 | 31 | 51 | 21 | ACTCTTAATCTCAGGGTCGTG | 491.2 | -10.5 | 1.3 | -8.1 | 3.8E-16 | 1.3E-15 | MB231/MB468 |
| >MI0000465\|hsa-mir-191&WithFlank&3\|-\|49020612\|49020715@22.42.21&[MIMAT0000440&hsa-miR-191-5p&offsets\|0\|-2;m-48&3\|-\|49020672\|49020694&offsets\|0\|-2]\|CAACGGAATCCCAAAAGCAGC | >MI0000465\|hsa-mir-191&WithFlank&3\|-\|49020612\|49020715 | 22 | 42 | 21 | CAACGGAATCCCAAAAGCAGC | 1155.2 | 6.6 | 0.8 | 8.1 | 3.8E-16 | 1.3E-15 | BT20/MB468 |
| >MI0000483\|hsa-mir-186&WithFlank&1\|-\|71067625\|71067722@21.42.22&[MIMAT0000456&hsa-miR-186-5p&offsets\|0\|0;m-84&1\|-\|71067681\|71067702&offsets\|0\|0]\|CAAAGAATTCTCCTTTTGGGCT | >MI0000483\|hsa-mir-186&WithFlank&1\|-\|71067625\|71067722 | 21 | 42 | 22 | CAAAGAATTCTCCTTTTGGGCT | 5533.9 | 6.7 | 0.8 | 8.1 | 3.8E-16 | 1.3E-15 | BT20/MB468 |
| >hg38_21_+_8433172_8446622_RNA45SN1_45S_50ntFlanks@12636.12653.18\|AGCTACCATCTGTGGGAT | >hg38_21_+_8433172_8446622_RNA45SN1_45S_50ntFlanks | 12636 | 12653 | 18 | AGCTACCATCTGTGGGAT | 1212.5 | -12.2 | 1.5 | -8.1 | 3.9E-16 | 1.3E-15 | BT20/MB468 |
| >trna10_AspGTC_12_-_125424193_125424264@36.75.40,trna12_AspGTC_12_-_125411891_125411962@36.75.40,trna144_AspGTC_6_-_27551236_27551307@36.75.40,trna38_AspGTC_17_-_8125556_8125627@36.75.40,trna45_AspGTC_6_+_27447453_27447524@36.75.40,trna48_AspGTC_6_+_27471523_27471594@36.75.40,trna4_AspGTC_12_+_96429799_96429870@36.75.40,trna69_AspGTC_1_-_161440205_161440276@36.75.40,trna72_AspGTC_1_-_161432824_161432895@36.75.40,trna75_AspGTC_1_-_161425414_161425485@36.75.40,trna78_AspGTC_1_-_161418033_161418104@36.75.40,trna81_AspGTC_1_-_161410615_161410686@36.75.40\|CACGCGGGAGACCGGGGTTCGATTCCCCGACGGGGAGCCA | >trna10_AspGTC_12_-_125424193_125424264 | 36 | 75 | 40 | CACGCGGGAGACCGGGGTTCGATTCCCCGACGGGGAGCCA | 2319.9 | 9.8 | 1.2 | 8.1 | 3.9E-16 | 1.3E-15 | MB231/MB468 |
| >MI0000103\|hsa-mir-101-1&WithFlank&1\|-\|65058428\|65058514@53.73.21(+1A);MI0000739\|hsa-mir-101-2&WithFlank&9\|+\|4850291\|4850381@55.75.21(+1A)&[MIMAT0000099&hsa-miR-101-3p&offsets\|0\|0(+1A);m-17&1\|-\|65058443\|65058462&offsets\|0\|+1(+1A)];[MIMAT0000099_1&hsa-miR-101-3p&offsets\|0\|0(+1A);m-16&9\|+\|4850345\|4850364&offsets\|0\|+1(+1A)]\|TACAGTACTGTGATAACTGAAA | >MI0000103\|hsa-mir-101-1&WithFlank&1\|-\|65058428\|65058514 | 53 | 73 | 21(+1A) | TACAGTACTGTGATAACTGAAA | 10803.0 | 10.0 | 1.2 | 8.1 | 3.9E-16 | 1.3E-15 | BT20/MB468 |
| >hg38_21_+_8433172_8446622_RNA45SN1_45S_50ntFlanks@11718.11740.23\|AACGGCGGGAGTAACTATGACTC | >hg38_21_+_8433172_8446622_RNA45SN1_45S_50ntFlanks | 11718 | 11740 | 23 | AACGGCGGGAGTAACTATGACTC | 1673.5 | -7.5 | 0.9 | -8.1 | 3.9E-16 | 1.3E-15 | BT20/MB468 |
| >am_trnaMT_TyrGTA_MT_-_5826_5891@1.32.32\|GGTAAAATGGCTGAGTGAAGCATTGGACTGTA | >am_trnaMT_TyrGTA_MT_-_5826_5891 | 1 | 32 | 32 | GGTAAAATGGCTGAGTGAAGCATTGGACTGTA | 5163.0 | 10.2 | 1.3 | 8.1 | 4.0E-16 | 1.3E-15 | MB231/MB468 |
| >trnaMT_HisGTG_MT_+_12138_12206@32.63.32\|TGAATCTGACAACAGAGGCTTACGACCCCTTA | >trnaMT_HisGTG_MT_+_12138_12206 | 32 | 63 | 32 | TGAATCTGACAACAGAGGCTTACGACCCCTTA | 2264.3 | 7.5 | 0.9 | 8.1 | 4.0E-16 | 1.3E-15 | MB231/MB468 |
| >hg38_21_+_8433172_8446622_RNA45SN1_45S_50ntFlanks@5489.5512.24\|CTGAGAAGACGGTCGAACTTGACT | >hg38_21_+_8433172_8446622_RNA45SN1_45S_50ntFlanks | 5489 | 5512 | 24 | CTGAGAAGACGGTCGAACTTGACT | 460.0 | 5.2 | 0.6 | 8.1 | 4.0E-16 | 1.3E-15 | BT20/MB468 |
| >hg38_21_+_8433172_8446622_RNA45SN1_45S_50ntFlanks@485.519.35\|GATGTGGTGACGTCGTGCTCTCCCGGGCCGGGTCC | >hg38_21_+_8433172_8446622_RNA45SN1_45S_50ntFlanks | 485 | 519 | 35 | GATGTGGTGACGTCGTGCTCTCCCGGGCCGGGTCC | 1481.8 | 29.5 | 3.6 | 8.1 | 4.0E-16 | 1.3E-15 | MB231/MB468 |
| >hg38_21_+_8433172_8446622_RNA45SN1_45S_50ntFlanks@11733.11751.19\|TATGACTCTCTTAAGGTAG | >hg38_21_+_8433172_8446622_RNA45SN1_45S_50ntFlanks | 11733 | 11751 | 19 | TATGACTCTCTTAAGGTAG | 306.3 | -10.3 | 1.3 | -8.1 | 4.0E-16 | 1.3E-15 | BT20/MB468 |
| >trnaMT_ProTGG_MT_-_15956_16023@51.70.20\|TAAAGACTTTTTCTCTGACC | >trnaMT_ProTGG_MT_-_15956_16023 | 51 | 70 | 20 | TAAAGACTTTTTCTCTGACC | 747.3 | 9.2 | 1.1 | 8.1 | 4.0E-16 | 1.3E-15 | MB231/MB468 |
| >MI0000067\|hsa-let-7f-1&WithFlank&9\|+\|94176341\|94176439@13.32.20(+1A);MI0000068\|hsa-let-7f-2&WithFlank&X\|-\|53557186\|53557280@14.33.20(+1A)&[MIMAT0000067&hsa-let-7f-5p&offsets\|0\|-2(+1A);m-20&9\|+\|94176353\|94176374&offsets\|0\|-2(+1A)];[MIMAT0000067_1&hsa-let-7f-5p&offsets\|0\|-2(+1A);m-19&X\|-\|53557246\|53557267&offsets\|0\|-2(+1A)]\|TGAGGTAGTAGATTGTATAGA | >MI0000067\|hsa-let-7f-1&WithFlank&9\|+\|94176341\|94176439 | 13 | 32 | 20(+1A) | TGAGGTAGTAGATTGTATAGA | 566.2 | 7.8 | 1.0 | 8.1 | 4.1E-16 | 1.3E-15 | MB231/MB468 |
| >hg38_21_+_8433172_8446622_RNA45SN1_45S_50ntFlanks@12324.12348.25\|TTGGGTTTTAAGCAGGAGGTGTCAG | >hg38_21_+_8433172_8446622_RNA45SN1_45S_50ntFlanks | 12324 | 12348 | 25 | TTGGGTTTTAAGCAGGAGGTGTCAG | 409.1 | -10.3 | 1.3 | -8.1 | 4.1E-16 | 1.3E-15 | MB231/MB468 |
| >hg38_21_+_8433172_8446622_RNA45SN1_45S_50ntFlanks@13004.13028.25\|ATCTATTGAAAGTCAGCCCTCGACA | >hg38_21_+_8433172_8446622_RNA45SN1_45S_50ntFlanks | 13004 | 13028 | 25 | ATCTATTGAAAGTCAGCCCTCGACA | 674.5 | -11.0 | 1.3 | -8.1 | 4.1E-16 | 1.4E-15 | MB231/MB468 |
| >hg38_21_+_8433172_8446622_RNA45SN1_45S_50ntFlanks@4445.4471.27\|CTTGCCTCTCGGCGCCCCCTCGATGCT | >hg38_21_+_8433172_8446622_RNA45SN1_45S_50ntFlanks | 4445 | 4471 | 27 | CTTGCCTCTCGGCGCCCCCTCGATGCT | 262.3 | -10.0 | 1.2 | -8.1 | 4.1E-16 | 1.4E-15 | BT20/MB468 |
| >hg38_21_+_8433172_8446622_RNA45SN1_45S_50ntFlanks@4443.4471.29\|CCCTTGCCTCTCGGCGCCCCCTCGATGCT | >hg38_21_+_8433172_8446622_RNA45SN1_45S_50ntFlanks | 4443 | 4471 | 29 | CCCTTGCCTCTCGGCGCCCCCTCGATGCT | 600.9 | -10.8 | 1.3 | -8.1 | 4.2E-16 | 1.4E-15 | MB231/MB468 |
| >hg38_21_+_8433172_8446622_RNA45SN1_45S_50ntFlanks@12944.12964.21\|CGTAGACGACCTGCTTCTGGG | >hg38_21_+_8433172_8446622_RNA45SN1_45S_50ntFlanks | 12944 | 12964 | 21 | CGTAGACGACCTGCTTCTGGG | 304.7 | -5.5 | 0.7 | -8.1 | 4.2E-16 | 1.4E-15 | BT20/MB468 |
| >MI0000750\|hsa-mir-26a-2&WithFlank&12\|-\|57824603\|57824698@20.41.22(+1A);MI0000083\|hsa-mir-26a-1&WithFlank&3\|+\|37969398\|37969486@16.37.22(+1A)&[MIMAT0000082&hsa-miR-26a-5p&offsets\|0\|0(+1A);m-33&12\|-\|57824658\|57824679&offsets\|0\|0(+1A)];[MIMAT0000082_1&hsa-miR-26a-5p&offsets\|0\|0(+1A);m-34&3\|+\|37969413\|37969434&offsets\|0\|0(+1A)]\|TTCAAGTAATCCAGGATAGGCTA | >MI0000750\|hsa-mir-26a-2&WithFlank&12\|-\|57824603\|57824698 | 20 | 41 | 22(+1A) | TTCAAGTAATCCAGGATAGGCTA | 2985.5 | 6.5 | 0.8 | 8.1 | 4.2E-16 | 1.4E-15 | BT20/MB468 |
| >MI0000298\|hsa-mir-221&WithFlank&X\|-\|45746151\|45746272@71.92.22&[MIMAT0000278&hsa-miR-221-3p&offsets\|0\|-1;m-91&X\|-\|45746181\|45746202&offsets\|0\|0]\|AGCTACATTGTCTGCTGGGTTT | >MI0000298\|hsa-mir-221&WithFlank&X\|-\|45746151\|45746272 | 71 | 92 | 22 | AGCTACATTGTCTGCTGGGTTT | 13041.8 | 6.0 | 0.7 | 8.1 | 4.2E-16 | 1.4E-15 | BT20/MB468 |
| >hg38_21_+_8433172_8446622_RNA45SN1_45S_50ntFlanks@11733.11751.19\|TATGACTCTCTTAAGGTAG | >hg38_21_+_8433172_8446622_RNA45SN1_45S_50ntFlanks | 11733 | 11751 | 19 | TATGACTCTCTTAAGGTAG | 261.7 | -4.3 | 0.5 | -8.1 | 4.3E-16 | 1.4E-15 | MB231/MB468 |
| >hg38_1_-_228634819_228635039_RNA5S12_5S_50ntFlanks@138.171.34\|AGACCGCCTGGGAATACCGGGTGCTGTAGGCTTT | >hg38_1_-_228634819_228635039_RNA5S12_5S_50ntFlanks | 138 | 171 | 34 | AGACCGCCTGGGAATACCGGGTGCTGTAGGCTTT | 2168.3 | 7.8 | 1.0 | 8.1 | 4.3E-16 | 1.4E-15 | BT20/MB468 |
| >hg38_21_+_8433172_8446622_RNA45SN1_45S_50ntFlanks@4588.4616.29\|CTATTTTGTTGGTTTTCGGAACTGAGGCC | >hg38_21_+_8433172_8446622_RNA45SN1_45S_50ntFlanks | 4588 | 4616 | 29 | CTATTTTGTTGGTTTTCGGAACTGAGGCC | 963.4 | -5.7 | 0.7 | -8.1 | 4.3E-16 | 1.4E-15 | BT20/MB468 |
| >hg38_21_+_8433172_8446622_RNA45SN1_45S_50ntFlanks@9567.9586.20\|ACGTGCAAATCGGTCGTCCG | >hg38_21_+_8433172_8446622_RNA45SN1_45S_50ntFlanks | 9567 | 9586 | 20 | ACGTGCAAATCGGTCGTCCG | 406.8 | -6.0 | 0.7 | -8.1 | 4.3E-16 | 1.4E-15 | BT20/MB468 |

<table>
<tr><td>>MI0000447|hsa-mir-128-1&WithFlank&2|+|135665391|135665484@56.78.23&[MIMAT0000424&hsa-miR-128-3p&offsets|0|+2;m-108&2|+|135665446|135665466&offsets|0|+2]|TCACAGTGAACCGGTCTCTTTTT</td><td>>MI0000447|hsa-mir-128-1&WithFlank&2|+|135665391|135665484</td><td>56</td><td>78</td><td>23</td><td>TCACAGTGAACCGGTCTCTTTTT</td><td>851.8</td><td>5.7</td><td>0.7</td><td>8.1</td><td>4.4E-16</td><td>1.4E-15</td><td>MB231/MB468</td></tr>
<tr><td>>hg38_21_+_8433172_8446622_RNA45SN1_45S_50ntFlanks@9565.9596.32|TGACGTGCAAATCGGTCGTCCGACCTGGGTAT</td><td>>hg38_21_+_8433172_8446622_RNA45SN1_45S_50ntFlanks</td><td>9565</td><td>9596</td><td>32</td><td>TGACGTGCAAATCGGTCGTCCGACCTGGGTAT</td><td>4817.8</td><td>-7.2</td><td>0.9</td><td>-8.1</td><td>4.4E-16</td><td>1.5E-15</td><td>BT20/MB468</td></tr>
<tr><td>>MI0000298|hsa-mir-221&WithFlank&X|-|45746151|45746272@71.90.20&[MIMAT0000278&hsa-miR-221-3p&offsets|0|-3;m-91&X|-|45746181|45746202&offsets|0|-2]|AGCTACATTGTCTGCTGGGT</td><td>>MI0000298|hsa-mir-221&WithFlank&X|-|45746151|45746272</td><td>71</td><td>90</td><td>20</td><td>AGCTACATTGTCTGCTGGGT</td><td>473.9</td><td>6.0</td><td>0.7</td><td>8.1</td><td>4.5E-16</td><td>1.5E-15</td><td>BT20/MB468</td></tr>
<tr><td>>hg38_21_+_8433172_8446622_RNA45SN1_45S_50ntFlanks@10391.10408.18|AACAGCAGTTGAACATGG</td><td>>hg38_21_+_8433172_8446622_RNA45SN1_45S_50ntFlanks</td><td>10391</td><td>10408</td><td>18</td><td>AACAGCAGTTGAACATGG</td><td>585.2</td><td>-4.7</td><td>0.6</td><td>-8.1</td><td>4.5E-16</td><td>1.5E-15</td><td>MB231/MB468</td></tr>
<tr><td>>am_MI0000786|hsa-mir-378a&WithFlank&5|+|149732819|149732896@49.70.22&[MIMAT0000732&hsa-miR-378a-3p&offsets|0|0;m-44&5|+|149732867|149732888&offsets|0|0]|ACTGGACTTGGAGTCAGAAGGC</td><td>>am_MI0000786|hsa-mir-378a&WithFlank&5|+|149732819|149732896</td><td>49</td><td>70</td><td>22</td><td>ACTGGACTTGGAGTCAGAAGGC</td><td>21541.4</td><td>4.9</td><td>0.6</td><td>8.1</td><td>4.5E-16</td><td>1.5E-15</td><td>BT20/MB468</td></tr>
<tr><td>>MI0005531|hsa-mir-450b&WithFlank&X|-|134540179|134540268@17.38.22&[MIMAT0004909&hsa-miR-450b-5p&offsets|0|0;m-266&X|-|134540232|134540252&offsets|0|+1]|TTTTGCAATATGTTCCTGAATA</td><td>>MI0005531|hsa-mir-450b&WithFlank&X|-|134540179|134540268</td><td>17</td><td>38</td><td>22</td><td>TTTTGCAATATGTTCCTGAATA</td><td>1642.8</td><td>10.3</td><td>1.3</td><td>8.1</td><td>4.5E-16</td><td>1.5E-15</td><td>BT20/MB468</td></tr>
<tr><td>>am_trna2_GlyGCC_21_-_18827107_18827177@57.74.18,trna35_GlyGCC_1_+_161413094_161413164@57.74.18,trna37_GlyGCC_1_+_161420467_161420537@57.74.18,trna39_GlyGCC_1_+_161427898_161427968@57.74.18,trna41_GlyGCC_1_+_161435258_161435328@57.74.18|TTCCCGGCCCATGCACCA</td><td>>am_trna2_GlyGCC_21_-_18827107_18827177</td><td>57</td><td>74</td><td>18</td><td>TTCCCGGCCCATGCACCA</td><td>467.5</td><td>10.4</td><td>1.2</td><td>8.4</td><td>3.1E-17</td><td>1.5E-15</td><td>BT20/MB231</td></tr>
<tr><td>>hg38_21_+_8433172_8446622_RNA45SN1_45S_50ntFlanks@6747.6770.24|ACTTCGAACGCACTTGCGGCCCCG</td><td>>hg38_21_+_8433172_8446622_RNA45SN1_45S_50ntFlanks</td><td>6747</td><td>6770</td><td>24</td><td>ACTTCGAACGCACTTGCGGCCCCG</td><td>465.5</td><td>-7.2</td><td>0.9</td><td>-8.1</td><td>4.6E-16</td><td>1.5E-15</td><td>BT20/MB468</td></tr>
<tr><td>>hg38_21_+_8433172_8446622_RNA45SN1_45S_50ntFlanks@10767.10785.19|GCAGCAGGTCTCCAAGGTG</td><td>>hg38_21_+_8433172_8446622_RNA45SN1_45S_50ntFlanks</td><td>10767</td><td>10785</td><td>19</td><td>GCAGCAGGTCTCCAAGGTG</td><td>204.1</td><td>-3.7</td><td>0.5</td><td>-8.1</td><td>4.6E-16</td><td>1.5E-15</td><td>BT20/MB468</td></tr>
<tr><td>>hg38_1_-_228634819_228635039_RNA5S12_5S_50ntFlanks@84.103.20|CTCGTCTGATCTCGGAAGCT</td><td>>hg38_1_-_228634819_228635039_RNA5S12_5S_50ntFlanks</td><td>84</td><td>103</td><td>20</td><td>CTCGTCTGATCTCGGAAGCT</td><td>857.9</td><td>-5.7</td><td>0.7</td><td>-8.1</td><td>4.7E-16</td><td>1.5E-15</td><td>MB231/MB468</td></tr>
<tr><td>>hg38_21_+_8433172_8446622_RNA45SN1_45S_50ntFlanks@10771.10805.35|CAGGTCTCCAAGGTGAACAGCCTCTGGCATGTTGG</td><td>>hg38_21_+_8433172_8446622_RNA45SN1_45S_50ntFlanks</td><td>10771</td><td>10805</td><td>35</td><td>CAGGTCTCCAAGGTGAACAGCCTCTGGCATGTTGG</td><td>805.3</td><td>-7.7</td><td>1.0</td><td>-8.1</td><td>4.7E-16</td><td>1.5E-15</td><td>BT20/MB468</td></tr>
<tr><td>>MI0000108|hsa-mir-103a-2&WithFlank&20|+|3917488|3917577@54.76.23(+2U);MI0000109|hsa-mir-103a-1&WithFlank&5|-|168560890|168560979@54.76.23(+2U)&[MIMAT0000101&hsa-miR-103a-3p&offsets|0|0(+2U);m-12&20|+|3917541|3917563&offsets|0|0(+2U)];[MIMAT0000101_1&hsa-miR-103a-3p&offsets|0|0(+2U);m-13&5|-|168560904|168560926&offsets|0|0(+2U);m-44220&5|-|168560908|168560926&offsets|0|+4(+2U)]|AGCAGCATTGTACAGGGCTATGATT</td><td>>MI0000108|hsa-mir-103a-2&WithFlank&20|+|3917488|3917577</td><td>54</td><td>76</td><td>23(+2U)</td><td>AGCAGCATTGTACAGGGCTATGATT</td><td>865.8</td><td>5.9</td><td>0.7</td><td>8.1</td><td>4.7E-16</td><td>1.5E-15</td><td>BT20/MB468</td></tr>
<tr><td>>MI0000076|hsa-mir-20a&WithFlank&13|+|91351059|91351141@14.36.23(+1G)&[MIMAT0000075&hsa-miR-20a-5p&offsets|0|0(+1G);m-96&13|+|91351072|91351094&offsets|0|0(+1G)]|TAAAGTGCTTATAGTGCAGGTAGG</td><td>>MI0000076|hsa-mir-20a&WithFlank&13|+|91351059|91351141</td><td>14</td><td>36</td><td>23(+1G)</td><td>TAAAGTGCTTATAGTGCAGGTAGG</td><td>998.3</td><td>7.0</td><td>0.9</td><td>8.1</td><td>4.7E-16</td><td>1.5E-15</td><td>MB231/MB468</td></tr>
<tr><td>>hg38_21_+_8433172_8446622_RNA45SN1_45S_50ntFlanks@4722.4752.31|TCATTAATCAAGAACGAAAGTCGGAGGTTCG</td><td>>hg38_21_+_8433172_8446622_RNA45SN1_45S_50ntFlanks</td><td>4722</td><td>4752</td><td>31</td><td>TCATTAATCAAGAACGAAAGTCGGAGGTTCG</td><td>531.4</td><td>-11.0</td><td>1.4</td><td>-8.1</td><td>4.8E-16</td><td>1.6E-15</td><td>BT20/MB468</td></tr>
<tr><td>>hg38_21_+_8433172_8446622_RNA45SN1_45S_50ntFlanks@8154.8181.28|CCGCTCGTGGGGGGCCCAAGTCCTTCTG</td><td>>hg38_21_+_8433172_8446622_RNA45SN1_45S_50ntFlanks</td><td>8154</td><td>8181</td><td>28</td><td>CCGCTCGTGGGGGGCCCAAGTCCTTCTG</td><td>1041.8</td><td>-5.7</td><td>0.7</td><td>-8.1</td><td>4.8E-16</td><td>1.6E-15</td><td>BT20/MB468</td></tr>
<tr><td>>MI0000088|hsa-mir-30a&WithFlank&6|-|71403545|71403627@54.75.22&[MIMAT0000088&hsa-miR-30a-3p&offsets|+1|+1;m-38&6|-|71403554|71403575&offsets|+1|+1]|TTTCAGTCGGATGTTTGCAGCT</td><td>>MI0000088|hsa-mir-30a&WithFlank&6|-|71403545|71403627</td><td>54</td><td>75</td><td>22</td><td>TTTCAGTCGGATGTTTGCAGCT</td><td>570.8</td><td>8.0</td><td>1.0</td><td>8.1</td><td>4.8E-16</td><td>1.6E-15</td><td>BT20/MB468</td></tr>
<tr><td>>hg38_21_+_8433172_8446622_RNA45SN1_45S_50ntFlanks@7975.8016.42|CGCGACCTCAGATCAGACGTGGCGACCCGCTGAATTTAAGCA</td><td>>hg38_21_+_8433172_8446622_RNA45SN1_45S_50ntFlanks</td><td>7975</td><td>8016</td><td>42</td><td>CGCGACCTCAGATCAGACGTGGCGACCCGCTGAATTTAAGCA</td><td>7581.5</td><td>6.9</td><td>0.8</td><td>8.1</td><td>4.8E-16</td><td>1.6E-15</td><td>BT20/MB468</td></tr>
<tr><td>>hg38_21_+_8433172_8446622_RNA45SN1_45S_50ntFlanks@3993.4023.31|GTGACTCTAGATAACCTCGGGCCGATCGCAC</td><td>>hg38_21_+_8433172_8446622_RNA45SN1_45S_50ntFlanks</td><td>3993</td><td>4023</td><td>31</td><td>GTGACTCTAGATAACCTCGGGCCGATCGCAC</td><td>1827.7</td><td>6.1</td><td>0.7</td><td>8.1</td><td>4.9E-16</td><td>1.6E-15</td><td>MB231/MB468</td></tr>
<tr><td>>MI0000063|hsa-let-7b&WithFlank&22|+|46113680|46113774@12.31.20(+1A)&[MIMAT0000063&hsa-let-7b-5p&offsets|0|-2(+1A);m-14&22|+|46113691|46113712&offsets|0|-2(+1A)]|TGAGGTAGTAGGTTGTGTGGA</td><td>>MI0000063|hsa-let-7b&WithFlank&22|+|46113680|46113774</td><td>12</td><td>31</td><td>20(+1A)</td><td>TGAGGTAGTAGGTTGTGTGGA</td><td>823.3</td><td>11.4</td><td>1.4</td><td>8.1</td><td>4.9E-16</td><td>1.6E-15</td><td>BT20/MB468</td></tr>
<tr><td>>MI0000439|hsa-mir-23b&WithFlank&9|+|95085202|95085310@67.85.19(+1U)&[MIMAT0000418&hsa-miR-23b-3p&offsets|+3|-1(+1U);m-42&9|+|95085265|95085285&offsets|+3|+1(+1U)]|ACATTGCCAGGGATTACCAT</td><td>>MI0000439|hsa-mir-23b&WithFlank&9|+|95085202|95085310</td><td>67</td><td>85</td><td>19(+1U)</td><td>ACATTGCCAGGGATTACCAT</td><td>1492.5</td><td>29.5</td><td>3.6</td><td>8.1</td><td>4.9E-16</td><td>1.6E-15</td><td>MB231/MB468</td></tr>
<tr><td>>hg38_21_+_8433172_8446622_RNA45SN1_45S_50ntFlanks@12638.12661.24|CTACCATCTGTGGGATTATGACTG</td><td>>hg38_21_+_8433172_8446622_RNA45SN1_45S_50ntFlanks</td><td>12638</td><td>12661</td><td>24</td><td>CTACCATCTGTGGGATTATGACTG</td><td>1019.9</td><td>-4.8</td><td>0.6</td><td>-8.1</td><td>4.9E-16</td><td>1.6E-15</td><td>BT20/MB468</td></tr>
<tr><td>>hg38_21_+_8433172_8446622_RNA45SN1_45S_50ntFlanks@10574.10608.35|AAGCCGGCGGGAGCCCCGGGGAGAGTTCTCTTTTC</td><td>>hg38_21_+_8433172_8446622_RNA45SN1_45S_50ntFlanks</td><td>10574</td><td>10608</td><td>35</td><td>AAGCCGGCGGGAGCCCCGGGGAGAGTTCTCTTTTC</td><td>592.4</td><td>-11.2</td><td>1.4</td><td>-8.1</td><td>5.0E-16</td><td>1.6E-15</td><td>BT20/MB468</td></tr>
<tr><td>>hg38_21_+_8433172_8446622_RNA45SN1_45S_50ntFlanks@6743.6770.28|CGACACTTCGAACGCACTTGCGGCCCCG</td><td>>hg38_21_+_8433172_8446622_RNA45SN1_45S_50ntFlanks</td><td>6743</td><td>6770</td><td>28</td><td>CGACACTTCGAACGCACTTGCGGCCCCG</td><td>1207.2</td><td>-5.7</td><td>0.7</td><td>-8.1</td><td>5.0E-16</td><td>1.6E-15</td><td>MB231/MB468</td></tr>
<tr><td>>MI0000811|hsa-mir-148b&WithFlank&12|+|54337210|54337320@69.89.21(+1A)&[MIMAT0000759&hsa-miR-148b-3p&offsets|0|-1(+1A);m-77&12|+|54337278|54337299&offsets|0|-1(+1A)]|TCAGTGCATCACAGAACTTTGA</td><td>>MI0000811|hsa-mir-148b&WithFlank&12|+|54337210|54337320</td><td>69</td><td>89</td><td>21(+1A)</td><td>TCAGTGCATCACAGAACTTTGA</td><td>685.6</td><td>8.5</td><td>1.0</td><td>8.1</td><td>5.0E-16</td><td>1.6E-15</td><td>MB231/MB468</td></tr>
<tr><td>>hg38_21_+_8433172_8446622_RNA45SN1_45S_50ntFlanks@10845.10868.24|ACTTCGGGATAAGGATTGGCTCTA</td><td>>hg38_21_+_8433172_8446622_RNA45SN1_45S_50ntFlanks</td><td>10845</td><td>10868</td><td>24</td><td>ACTTCGGGATAAGGATTGGCTCTA</td><td>1120.8</td><td>-6.7</td><td>0.8</td><td>-8.1</td><td>5.0E-16</td><td>1.6E-15</td><td>BT20/MB468</td></tr>
<tr><td>>MI0000440|hsa-mir-27b&WithFlank&9|+|95085439|95085547@67.85.19(+2U)&[MIMAT0000419&hsa-miR-27b-3p&offsets|0|-2(+2U);m-39&9|+|95085505|95085525&offsets|0|-2(+2U)]|TTCACAGTGGCTAAGTTCTTT</td><td>>MI0000440|hsa-mir-27b&WithFlank&9|+|95085439|95085547</td><td>67</td><td>85</td><td>19(+2U)</td><td>TTCACAGTGGCTAAGTTCTTT</td><td>560.0</td><td>6.4</td><td>0.8</td><td>8.1</td><td>5.0E-16</td><td>1.6E-15</td><td>BT20/MB468</td></tr>
<tr><td>>hg38_21_+_8433172_8446622_RNA45SN1_45S_50ntFlanks@10388.10424.37|GTGAACAGCAGTTGAACATGGGTCAGTCGGTCCTGAG</td><td>>hg38_21_+_8433172_8446622_RNA45SN1_45S_50ntFlanks</td><td>10388</td><td>10424</td><td>37</td><td>GTGAACAGCAGTTGAACATGGGTCAGTCGGTCCTGAG</td><td>984.5</td><td>5.6</td><td>0.7</td><td>8.1</td><td>5.1E-16</td><td>1.6E-15</td><td>MB231/MB468</td></tr>
<tr><td>>hg38_21_+_8433172_8446622_RNA45SN1_45S_50ntFlanks@12480.12497.18|TAATAGGGAACGTGAGCT</td><td>>hg38_21_+_8433172_8446622_RNA45SN1_45S_50ntFlanks</td><td>12480</td><td>12497</td><td>18</td><td>TAATAGGGAACGTGAGCT</td><td>1535.0</td><td>4.9</td><td>0.6</td><td>8.1</td><td>5.1E-16</td><td>1.7E-15</td><td>MB231/MB468</td></tr>
<tr><td>>am_trna128_GlyGCC_6_-_27870686_27870756@4.31.28,trna133_GlyCCC_1_-_16872434_16872504@4.31.28,trna18_GlyGCC_16_+_70822597_70822667@4.31.28,trna19_GlyGCC_16_+_70823410_70823480@4.31.28,trna19_GlyGCC_2_-_157257659_157257729@4.31.28,trna24_GlyGCC_16_-_70812942_70813012@4.31.28,trna25_GlyGCC_16_-_70812114_70812184@4.31.28,trna4_GlyCCC_1_+_17188416_17188486@4.31.28,trna5_GlyGCC_17_+_8029064_8029134@4.31.28,trna68_GlyGCC_1_-_161493637_161493707@4.31.28|TTGGTGGTTCAGTGGTAGAATTCTCGCC</td><td>>am_trna128_GlyGCC_6_-_27870686_27870756</td><td>4</td><td>31</td><td>28</td><td>TTGGTGGTTCAGTGGTAGAATTCTCGCC</td><td>656.3</td><td>9.5</td><td>1.2</td><td>8.1</td><td>5.1E-16</td><td>1.7E-15</td><td>MB231/MB468</td></tr>
<tr><td>>hg38_21_+_8433172_8446622_RNA45SN1_45S_50ntFlanks@6663.6681.19|GTGGATCACTCGGCTCGTG</td><td>>hg38_21_+_8433172_8446622_RNA45SN1_45S_50ntFlanks</td><td>6663</td><td>6681</td><td>19</td><td>GTGGATCACTCGGCTCGTG</td><td>1134.3</td><td>-5.6</td><td>0.7</td><td>-8.1</td><td>5.1E-16</td><td>1.7E-15</td><td>MB231/MB468</td></tr>
<tr><td>>MI0000103|hsa-mir-101-1&WithFlank&1|-|65058428|65058514@52.72.21;MI0000739|hsa-mir-101-2&WithFlank&9|+|4850291|4850381@54.74.21&[MIMAT0000099&hsa-miR-101-3p&offsets|-1|-1;m-17&1|-|65058443|65058462&offsets|-1|0];[MIMAT0000099_1&hsa-miR-101-3p&offsets|-1|-1;m-16&9|+|4850345|4850364&offsets|-1|0]|GTACAGTACTGTGATAACTGA</td><td>>MI0000103|hsa-mir-101-1&WithFlank&1|-|65058428|65058514</td><td>52</td><td>72</td><td>21</td><td>GTACAGTACTGTGATAACTGA</td><td>2262.4</td><td>6.5</td><td>0.8</td><td>8.1</td><td>5.2E-16</td><td>1.7E-15</td><td>MB231/MB468</td></tr>
<tr><td>>MI0000264|hsa-mir-7-2&WithFlank&15|+|88611819|88611940@38.57.20;MI0000265|hsa-mir-7-3&WithFlank&19|+|4770664|4770785@37.56.20;MI0000263|hsa-mir-7-1&WithFlank&9|-|83969742|83969863@30.49.20&[MIMAT0000252&hsa-miR-7-5p&offsets|0|-4;m-400&15|+|88611856|88611879&offsets|0|-4];[MIMAT0000252_1&hsa-miR-7-5p&offsets|0|-4;m-402&19|+|4770700|4770723&offsets|0|-4];[MIMAT0000252_2&hsa-miR-7-5p&offsets|0|-4;m-401&9|-|83969811|83969834&offsets|0|-4]|TGGAAGACTAGTGATTTTGT</td><td>>MI0000264|hsa-mir-7-2&WithFlank&15|+|88611819|88611940</td><td>38</td><td>57</td><td>20</td><td>TGGAAGACTAGTGATTTTGT</td><td>662.9</td><td>8.7</td><td>1.1</td><td>8.1</td><td>5.2E-16</td><td>1.7E-15</td><td>MB231/MB468</td></tr>
<tr><td>>hg38_21_+_8433172_8446622_RNA45SN1_45S_50ntFlanks@6708.6731.24|GAGAATTAATGTGAATTGCAGGAC</td><td>>hg38_21_+_8433172_8446622_RNA45SN1_45S_50ntFlanks</td><td>6708</td><td>6731</td><td>24</td><td>GAGAATTAATGTGAATTGCAGGAC</td><td>554.9</td><td>7.5</td><td>0.9</td><td>8.1</td><td>5.2E-16</td><td>1.7E-15</td><td>MB231/MB468</td></tr>
<tr><td>>hg38_21_+_8433172_8446622_RNA45SN1_45S_50ntFlanks@12432.12455.24|CCTATCATTGTGAAGCAGAATTCA</td><td>>hg38_21_+_8433172_8446622_RNA45SN1_45S_50ntFlanks</td><td>12432</td><td>12455</td><td>24</td><td>CCTATCATTGTGAAGCAGAATTCA</td><td>260.4</td><td>-10.0</td><td>1.2</td><td>-8.1</td><td>5.3E-16</td><td>1.7E-15</td><td>BT20/MB468</td></tr>
<tr><td>>MI0000734|hsa-mir-106b&WithFlank&7|-|100093987|100094080@58.76.19&[MIMAT0004672&hsa-miR-106b-3p&offsets|0|-3;m-87&7|-|100094003|100094023&offsets|0|-2]|CCGCACTGTGGGTACTTGC</td><td>>MI0000734|hsa-mir-106b&WithFlank&7|-|100093987|100094080</td><td>58</td><td>76</td><td>19</td><td>CCGCACTGTGGGTACTTGC</td><td>592.3</td><td>9.2</td><td>1.1</td><td>8.1</td><td>5.3E-16</td><td>1.7E-15</td><td>MB231/MB468</td></tr>
</table>

<table>
<tr><td>>MI0000650|hsa-mir-200c&WithFlank&12|+|6963693|6963772@50.70.21(+2A)&[MIMAT0000617&hsa-miR-200c-3p&offsets|0|-2(+2A);m-54&12|+|6963742|6963764&offsets|0|-2(+2A)]|TAATACTGCCGGGTAATGATGAA</td><td>>MI0000650|hsa-mir-200c&WithFlank&12|+|6963693|6963772</td><td>50</td><td>70</td><td>21(+2A)</td><td>TAATACTGCCGGGTAATGATGAA</td><td>1341.3</td><td>12.6</td><td>1.5</td><td>8.1</td><td>5.3E-16</td><td>1.7E-15</td><td>BT20/MB468</td></tr>
<tr><td>>hg38_21_+_8433172_8446622_RNA45SN1_45S_50ntFlanks@11575.11594.20|ACTTAGAACTGGTGCGGACC</td><td>>hg38_21_+_8433172_8446622_RNA45SN1_45S_50ntFlanks</td><td>11575</td><td>11594</td><td>20</td><td>ACTTAGAACTGGTGCGGACC</td><td>1045.3</td><td>-3.6</td><td>0.4</td><td>-8.1</td><td>5.3E-16</td><td>1.7E-15</td><td>BT20/MB468</td></tr>
<tr><td>>trnaMT_ProTGG_MT_-_15956_16023@1.32.32|CAGAGAATAGTTTAAATTAGAATCTTAGCTTT</td><td>>trnaMT_ProTGG_MT_-_15956_16023</td><td>1</td><td>32</td><td>32</td><td>CAGAGAATAGTTTAAATTAGAATCTTAGCTTT</td><td>1175.3</td><td>12.4</td><td>1.5</td><td>8.1</td><td>5.3E-16</td><td>1.7E-15</td><td>BT20/MB468</td></tr>
<tr><td>>hg38_21_+_8433172_8446622_RNA45SN1_45S_50ntFlanks@11766.11791.26|ATCTAATTAGTGACGCGCATGAATGG</td><td>>hg38_21_+_8433172_8446622_RNA45SN1_45S_50ntFlanks</td><td>11766</td><td>11791</td><td>26</td><td>ATCTAATTAGTGACGCGCATGAATGG</td><td>288.2</td><td>-10.2</td><td>1.3</td><td>-8.1</td><td>5.3E-16</td><td>1.7E-15</td><td>BT20/MB468</td></tr>
<tr><td>>hg38_21_+_8433172_8446622_RNA45SN1_45S_50ntFlanks@6721.6757.37|AATTGCAGGACACATTGATCATCGACACTTCGAACGC</td><td>>hg38_21_+_8433172_8446622_RNA45SN1_45S_50ntFlanks</td><td>6721</td><td>6757</td><td>37</td><td>AATTGCAGGACACATTGATCATCGACACTTCGAACGC</td><td>162.3</td><td>-4.7</td><td>0.6</td><td>-8.1</td><td>5.4E-16</td><td>1.8E-15</td><td>BT20/MB468</td></tr>
<tr><td>>hg38_21_+_8433172_8446622_RNA45SN1_45S_50ntFlanks@6788.6807.20|CGCCTGTCTGAGCGTCGCTT</td><td>>hg38_21_+_8433172_8446622_RNA45SN1_45S_50ntFlanks</td><td>6788</td><td>6807</td><td>20</td><td>CGCCTGTCTGAGCGTCGCTT</td><td>1946.3</td><td>6.0</td><td>0.7</td><td>8.1</td><td>5.4E-16</td><td>1.8E-15</td><td>MB231/MB468</td></tr>
<tr><td>>MI0000081|hsa-mir-24-2&WithFlank&19|-|13836281|13836365@56.74.19(+1A);MI0000080|hsa-mir-24-1&WithFlank&9|+|95086015|95086094@50.68.19(+1A)&[MIMAT0000080&hsa-miR-24-3p&offsets|0|-3(+1A);m-36&19|-|13836288|13836310&offsets|0|-4(+1A)];[MIMAT0000080_1&hsa-miR-24-3p&offsets|0|-3(+1A);m-35&9|+|95086064|95086086&offsets|0|-4(+1A)]|TGGCTCAGTTCAGCAGGAAA</td><td>>MI0000081|hsa-mir-24-2&WithFlank&19|-|13836281|13836365</td><td>56</td><td>74</td><td>19(+1A)</td><td>TGGCTCAGTTCAGCAGGAAA</td><td>982.5</td><td>7.7</td><td>0.9</td><td>8.1</td><td>5.5E-16</td><td>1.8E-15</td><td>BT20/MB468</td></tr>
<tr><td>>hg38_21_+_8433172_8446622_RNA45SN1_45S_50ntFlanks@5420.5445.26|CCGATTGGATGGTTTAGTGAGGCCCT</td><td>>hg38_21_+_8433172_8446622_RNA45SN1_45S_50ntFlanks</td><td>5420</td><td>5445</td><td>26</td><td>CCGATTGGATGGTTTAGTGAGGCCCT</td><td>458.1</td><td>-10.4</td><td>1.3</td><td>-8.1</td><td>5.5E-16</td><td>1.8E-15</td><td>MB231/MB468</td></tr>
<tr><td>>hg38_21_+_8433172_8446622_RNA45SN1_45S_50ntFlanks@11210.11232.23|GGCGACTCTGGACGCGAGCCGGG</td><td>>hg38_21_+_8433172_8446622_RNA45SN1_45S_50ntFlanks</td><td>11210</td><td>11232</td><td>23</td><td>GGCGACTCTGGACGCGAGCCGGG</td><td>650.2</td><td>-8.4</td><td>1.0</td><td>-8.1</td><td>5.6E-16</td><td>1.8E-15</td><td>MB231/MB468</td></tr>
<tr><td>>hg38_21_+_8433172_8446622_RNA45SN1_45S_50ntFlanks@12357.12375.19|CCACAGGGATAACTGGCTT</td><td>>hg38_21_+_8433172_8446622_RNA45SN1_45S_50ntFlanks</td><td>12357</td><td>12375</td><td>19</td><td>CCACAGGGATAACTGGCTT</td><td>415.0</td><td>-10.3</td><td>1.3</td><td>-8.1</td><td>5.6E-16</td><td>1.8E-15</td><td>MB231/MB468</td></tr>
<tr><td>>trnaMT_ValTAC_MT_+_1602_1670@53.72.20|TAACTTGACCGCTCTGACCA</td><td>>trnaMT_ValTAC_MT_+_1602_1670</td><td>53</td><td>72</td><td>20</td><td>TAACTTGACCGCTCTGACCA</td><td>1321.1</td><td>9.4</td><td>1.2</td><td>8.1</td><td>5.6E-16</td><td>1.8E-15</td><td>MB231/MB468</td></tr>
<tr><td>>hg38_21_+_8433172_8446622_RNA45SN1_45S_50ntFlanks@10362.10390.29|AAGGCCGAAGTGGAGAAGGGTTCCATGTG</td><td>>hg38_21_+_8433172_8446622_RNA45SN1_45S_50ntFlanks</td><td>10362</td><td>10390</td><td>29</td><td>AAGGCCGAAGTGGAGAAGGGTTCCATGTG</td><td>1588.6</td><td>-8.9</td><td>1.1</td><td>-8.1</td><td>5.6E-16</td><td>1.8E-15</td><td>BT20/MB468</td></tr>
<tr><td>>hg38_MT_+_1621_3279_RNR2_16S_50ntFlanks@1438.1458.21|TACGTGATCTGAGTTCAGACC</td><td>>hg38_MT_+_1621_3279_RNR2_16S_50ntFlanks</td><td>1438</td><td>1458</td><td>21</td><td>TACGTGATCTGAGTTCAGACC</td><td>748.3</td><td>9.3</td><td>1.2</td><td>8.1</td><td>5.7E-16</td><td>1.8E-15</td><td>MB231/MB468</td></tr>
<tr><td>>hg38_21_+_8433172_8446622_RNA45SN1_45S_50ntFlanks@10408.10432.25|GGTCAGTCGGTCCTGAGAGATGGGC</td><td>>hg38_21_+_8433172_8446622_RNA45SN1_45S_50ntFlanks</td><td>10408</td><td>10432</td><td>25</td><td>GGTCAGTCGGTCCTGAGAGATGGGC</td><td>351.2</td><td>-10.4</td><td>1.3</td><td>-8.1</td><td>5.7E-16</td><td>1.8E-15</td><td>BT20/MB468</td></tr>
<tr><td>>hg38_21_+_8433172_8446622_RNA45SN1_45S_50ntFlanks@11208.11227.20|GCGGCGACTCTGGACGCGAG</td><td>>hg38_21_+_8433172_8446622_RNA45SN1_45S_50ntFlanks</td><td>11208</td><td>11227</td><td>20</td><td>GCGGCGACTCTGGACGCGAG</td><td>930.1</td><td>-8.0</td><td>1.0</td><td>-8.1</td><td>5.8E-16</td><td>1.9E-15</td><td>MB231/MB468</td></tr>
<tr><td>>hg38_MT_+_1621_3279_RNR2_16S_50ntFlanks@51.84.34|GCTAAACCTAGCCCCAAACCCACTCCACCTTACT</td><td>>hg38_MT_+_1621_3279_RNR2_16S_50ntFlanks</td><td>51</td><td>84</td><td>34</td><td>GCTAAACCTAGCCCCAAACCCACTCCACCTTACT</td><td>1345.1</td><td>8.7</td><td>1.1</td><td>8.1</td><td>5.8E-16</td><td>1.9E-15</td><td>MB231/MB468</td></tr>
<tr><td>>MI0003599|hsa-mir-589&WithFlank&7|-|5495813|5495923@30.50.21&[MIMAT0004799&hsa-miR-589-5p&offsets|0|-1;m-162&7|-|5495874|5495894&offsets|0|0]|TGAGAACCACGTCTGCTCTGA</td><td>>MI0003599|hsa-mir-589&WithFlank&7|-|5495813|5495923</td><td>30</td><td>50</td><td>21</td><td>TGAGAACCACGTCTGCTCTGA</td><td>806.1</td><td>9.9</td><td>1.2</td><td>8.1</td><td>5.8E-16</td><td>1.9E-15</td><td>BT20/MB468</td></tr>
<tr><td>>hg38_21_+_8433172_8446622_RNA45SN1_45S_50ntFlanks@10869.10901.33|AGGGCTGGGTCGGTCGGGCTGGGGCGCGAAGCG</td><td>>hg38_21_+_8433172_8446622_RNA45SN1_45S_50ntFlanks</td><td>10869</td><td>10901</td><td>33</td><td>AGGGCTGGGTCGGTCGGGCTGGGGCGCGAAGCG</td><td>201.5</td><td>-6.4</td><td>0.8</td><td>-8.1</td><td>5.9E-16</td><td>1.9E-15</td><td>MB231/MB468</td></tr>
<tr><td>>MI0000483|hsa-mir-186&WithFlank&1|-|71067625|71067722@21.42.22&[MIMAT0000456&hsa-miR-186-5p&offsets|0|0;m-84&1|-|71067681|71067702&offsets|0|0]|CAAAGAATTCTCCTTTTGGGCT</td><td>>MI0000483|hsa-mir-186&WithFlank&1|-|71067625|71067722</td><td>21</td><td>42</td><td>22</td><td>CAAAGAATTCTCCTTTTGGGCT</td><td>2578.4</td><td>5.9</td><td>0.7</td><td>8.1</td><td>5.9E-16</td><td>1.9E-15</td><td>MB231/MB468</td></tr>
<tr><td>>hg38_21_+_8433172_8446622_RNA45SN1_45S_50ntFlanks@12151.12182.32|GTGGGGAGTTTGACTGGGGCGGTACACCTGTC</td><td>>hg38_21_+_8433172_8446622_RNA45SN1_45S_50ntFlanks</td><td>12151</td><td>12182</td><td>32</td><td>GTGGGGAGTTTGACTGGGGCGGTACACCTGTC</td><td>311.3</td><td>-6.4</td><td>0.8</td><td>-8.1</td><td>6.0E-16</td><td>1.9E-15</td><td>MB231/MB468</td></tr>
<tr><td>>MI0000736|hsa-mir-30c-1&WithFlank&1|+|40757278|40757378@23.44.22(+1A);MI0000254|hsa-mir-30c-2&WithFlank&6|-|71376954|71377037@13.34.22(+1A)&[MIMAT0000244&hsa-miR-30c-5p&offsets|0|-1(+1A);m-57&1|+|40757300|40757323&offsets|0|-2(+1A)];[MIMAT0000244_1&hsa-miR-30c-5p&offsets|0|-1(+1A);m-56&6|-|71377002|71377025&offsets|0|-2(+1A)]|TGTAAACATCCTACACTCTCAGA</td><td>>MI0000736|hsa-mir-30c-1&WithFlank&1|+|40757278|40757378</td><td>23</td><td>44</td><td>22(+1A)</td><td>TGTAAACATCCTACACTCTCAGA</td><td>3456.6</td><td>7.0</td><td>0.9</td><td>8.1</td><td>6.0E-16</td><td>1.9E-15</td><td>BT20/MB468</td></tr>
<tr><td>>hg38_1_-_228634819_228635039_RNA5S12_5S_50ntFlanks@133.170.38|ATGGGAGACCGCCTGGGAATACCGGGTGCTGTAGGCTT</td><td>>hg38_1_-_228634819_228635039_RNA5S12_5S_50ntFlanks</td><td>133</td><td>170</td><td>38</td><td>ATGGGAGACCGCCTGGGAATACCGGGTGCTGTAGGCTT</td><td>14279.5</td><td>5.7</td><td>0.7</td><td>8.1</td><td>6.0E-16</td><td>1.9E-15</td><td>MB231/MB468</td></tr>
<tr><td>>hg38_21_+_8433172_8446622_RNA45SN1_45S_50ntFlanks@4759.4781.23|ATCAGATACCGTCGTAGTTCCGA</td><td>>hg38_21_+_8433172_8446622_RNA45SN1_45S_50ntFlanks</td><td>4759</td><td>4781</td><td>23</td><td>ATCAGATACCGTCGTAGTTCCGA</td><td>253.7</td><td>-10.0</td><td>1.2</td><td>-8.1</td><td>6.0E-16</td><td>2.0E-15</td><td>BT20/MB468</td></tr>
<tr><td>>am_MI0000786|hsa-mir-378a&WithFlank&5|+|149732819|149732896@50.70.21(+1U)&[MIMAT0000732&hsa-miR-378a-3p&offsets|+1|0(+1U);m-44&5|+|149732867|149732888&offsets|+1|0(+1U)]|CTGGACTTGGAGTCAGAAGGCT</td><td>>am_MI0000786|hsa-mir-378a&WithFlank&5|+|149732819|149732896</td><td>50</td><td>70</td><td>21(+1U)</td><td>CTGGACTTGGAGTCAGAAGGCT</td><td>1206.7</td><td>5.8</td><td>0.7</td><td>8.1</td><td>6.1E-16</td><td>2.0E-15</td><td>MB231/MB468</td></tr>
<tr><td>>hg38_21_+_8433172_8446622_RNA45SN1_45S_50ntFlanks@11675.11702.28|TGCTCTGAATGTCAAAGTGAAGAAATTC</td><td>>hg38_21_+_8433172_8446622_RNA45SN1_45S_50ntFlanks</td><td>11675</td><td>11702</td><td>28</td><td>TGCTCTGAATGTCAAAGTGAAGAAATTC</td><td>2262.7</td><td>4.9</td><td>0.6</td><td>8.1</td><td>6.1E-16</td><td>2.0E-15</td><td>MB231/MB468</td></tr>
<tr><td>>trnaMT_LeuTAA_MT_+_3230_3304@18.42.25|CGGTAATCGCATAAAACTTAAAACT</td><td>>trnaMT_LeuTAA_MT_+_3230_3304</td><td>18</td><td>42</td><td>25</td><td>CGGTAATCGCATAAAACTTAAAACT</td><td>905.2</td><td>11.4</td><td>1.4</td><td>8.1</td><td>6.1E-16</td><td>2.0E-15</td><td>BT20/MB468</td></tr>
<tr><td>>hg38_21_+_8433172_8446622_RNA45SN1_45S_50ntFlanks@12627.12652.26|ATGGGGCGAAGCTACCATCTGTGGGA</td><td>>hg38_21_+_8433172_8446622_RNA45SN1_45S_50ntFlanks</td><td>12627</td><td>12652</td><td>26</td><td>ATGGGGCGAAGCTACCATCTGTGGGA</td><td>311.6</td><td>-10.3</td><td>1.3</td><td>-8.1</td><td>6.1E-16</td><td>2.0E-15</td><td>BT20/MB468</td></tr>
<tr><td>>hg38_21_+_8433172_8446622_RNA45SN1_45S_50ntFlanks@12632.12657.26|GCGAAGCTACCATCTGTGGGATTATG</td><td>>hg38_21_+_8433172_8446622_RNA45SN1_45S_50ntFlanks</td><td>12632</td><td>12657</td><td>26</td><td>GCGAAGCTACCATCTGTGGGATTATG</td><td>112.1</td><td>-4.3</td><td>0.5</td><td>-8.1</td><td>6.1E-16</td><td>2.0E-15</td><td>BT20/MB468</td></tr>
<tr><td>>hg38_21_+_8433172_8446622_RNA45SN1_45S_50ntFlanks@10839.10865.27|TCCGTAACTTCGGGATAAGGATTGGCT</td><td>>hg38_21_+_8433172_8446622_RNA45SN1_45S_50ntFlanks</td><td>10839</td><td>10865</td><td>27</td><td>TCCGTAACTTCGGGATAAGGATTGGCT</td><td>482.2</td><td>-4.9</td><td>0.6</td><td>-8.1</td><td>6.2E-16</td><td>2.0E-15</td><td>MB231/MB468</td></tr>
<tr><td>>hg38_21_+_8433172_8446622_RNA45SN1_45S_50ntFlanks@11161.11179.19|CACGGTCCCCCGCGAGGGG</td><td>>hg38_21_+_8433172_8446622_RNA45SN1_45S_50ntFlanks</td><td>11161</td><td>11179</td><td>19</td><td>CACGGTCCCCCGCGAGGGG</td><td>613.9</td><td>-10.8</td><td>1.3</td><td>-8.1</td><td>6.2E-16</td><td>2.0E-15</td><td>MB231/MB468</td></tr>
<tr><td>>MI0000084|hsa-mir-26b&WithFlank&2|+|218402640|218402728@18.38.21(+1C)&[MIMAT0000083&hsa-miR-26b-5p&offsets|0|0(+1C);m-53&2|+|218402657|218402678&offsets|0|-1(+1C)]|TTCAAGTAATTCAGGATAGGTC</td><td>>MI0000084|hsa-mir-26b&WithFlank&2|+|218402640|218402728</td><td>18</td><td>38</td><td>21(+1C)</td><td>TTCAAGTAATTCAGGATAGGTC</td><td>4816.0</td><td>7.3</td><td>0.9</td><td>8.1</td><td>6.2E-16</td><td>2.0E-15</td><td>BT20/MB468</td></tr>
<tr><td>>hg38_21_+_8433172_8446622_RNA45SN1_45S_50ntFlanks@10392.10420.29|ACAGCAGTTGAACATGGGTCAGTCGGTCC</td><td>>hg38_21_+_8433172_8446622_RNA45SN1_45S_50ntFlanks</td><td>10392</td><td>10420</td><td>29</td><td>ACAGCAGTTGAACATGGGTCAGTCGGTCC</td><td>316.0</td><td>-4.3</td><td>0.5</td><td>-8.1</td><td>6.2E-16</td><td>2.0E-15</td><td>MB231/MB468</td></tr>
<tr><td>>am_MI0015825|hsa-mir-378c&WithFlank&10|-|130962582|130962674@17.36.20;MI0000786|hsa-mir-378a&WithFlank&5|+|149732819|149732896@49.68.20&[MIMAT0016847&hsa-miR-378c&offsets|0|-5;m-130&10|-|130962639|130962658&offsets|0|0];[MIMAT0000732&hsa-miR-378a-3p&offsets|0|-2;m-44&5|+|149732867|149732888&offsets|0|-2]|ACTGGACTTGGAGTCAGAAG</td><td>>am_MI0015825|hsa-mir-378c&WithFlank&10|-|130962582|130962674</td><td>17</td><td>36</td><td>20</td><td>ACTGGACTTGGAGTCAGAAG</td><td>830.4</td><td>5.9</td><td>0.7</td><td>8.1</td><td>6.2E-16</td><td>2.0E-15</td><td>MB231/MB468</td></tr>
<tr><td>>hg38_1_-_228634819_228635039_RNA5S12_5S_50ntFlanks@151.171.21|ATACCGGGTGCTGTAGGCTTT</td><td>>hg38_1_-_228634819_228635039_RNA5S12_5S_50ntFlanks</td><td>151</td><td>171</td><td>21</td><td>ATACCGGGTGCTGTAGGCTTT</td><td>3121.0</td><td>6.1</td><td>0.8</td><td>8.1</td><td>6.3E-16</td><td>2.0E-15</td><td>MB231/MB468</td></tr>
<tr><td>>MI0000456|hsa-mir-140&WithFlank&16|+|69933075|69933186@68.91.24&[MIMAT0004597&hsa-miR-140-3p&offsets|0|+3;m-45&16|+|69933143|69933164&offsets|-1|+1]|TACCACAGGGTAGAACCACGGACA</td><td>>MI0000456|hsa-mir-140&WithFlank&16|+|69933075|69933186</td><td>68</td><td>91</td><td>24</td><td>TACCACAGGGTAGAACCACGGACA</td><td>864.3</td><td>10.2</td><td>1.3</td><td>8.1</td><td>6.3E-16</td><td>2.0E-15</td><td>BT20/MB468</td></tr>
<tr><td>>hg38_21_+_8433172_8446622_RNA45SN1_45S_50ntFlanks@9504.9527.24|GTGAACTATGCCTGGGCAGGGCGA</td><td>>hg38_21_+_8433172_8446622_RNA45SN1_45S_50ntFlanks</td><td>9504</td><td>9527</td><td>24</td><td>GTGAACTATGCCTGGGCAGGGCGA</td><td>325.4</td><td>-10.3</td><td>1.3</td><td>-8.1</td><td>6.3E-16</td><td>2.1E-15</td><td>BT20/MB468</td></tr>
<tr><td>>MI0000749|hsa-mir-30e&WithFlank&1|+|40754349|40754452@65.85.21&[MIMAT0000693&hsa-miR-30e-3p&offsets|0|-1;m-25&1|+|40754413|40754434&offsets|0|-1]|CTTTCAGTCGGATGTTTACAG</td><td>>MI0000749|hsa-mir-30e&WithFlank&1|+|40754349|40754452</td><td>65</td><td>85</td><td>21</td><td>CTTTCAGTCGGATGTTTACAG</td><td>944.6</td><td>6.4</td><td>0.8</td><td>8.1</td><td>6.4E-16</td><td>2.1E-15</td><td>MB231/MB468</td></tr>
<tr><td>>hg38_21_+_8433172_8446622_RNA45SN1_45S_50ntFlanks@12458.12475.18|AAGCGTTGGATTGTTCAC</td><td>>hg38_21_+_8433172_8446622_RNA45SN1_45S_50ntFlanks</td><td>12458</td><td>12475</td><td>18</td><td>AAGCGTTGGATTGTTCAC</td><td>459.2</td><td>-4.6</td><td>0.6</td><td>-8.1</td><td>6.4E-16</td><td>2.1E-15</td><td>BT20/MB468</td></tr>
<tr><td>>hg38_21_+_8433172_8446622_RNA45SN1_45S_50ntFlanks@11587.11616.30|TGCGGACCAGGGGAATCCGACTGTTTAATT</td><td>>hg38_21_+_8433172_8446622_RNA45SN1_45S_50ntFlanks</td><td>11587</td><td>11616</td><td>30</td><td>TGCGGACCAGGGGAATCCGACTGTTTAATT</td><td>387.8</td><td>5.4</td><td>0.7</td><td>8.1</td><td>6.5E-16</td><td>2.1E-15</td><td>MB231/MB468</td></tr>
<tr><td>>hg38_21_+_8433172_8446622_RNA45SN1_45S_50ntFlanks@3994.4023.30|TGACTCTAGATAACCTCGGGCCGATCGCAC</td><td>>hg38_21_+_8433172_8446622_RNA45SN1_45S_50ntFlanks</td><td>3994</td><td>4023</td><td>30</td><td>TGACTCTAGATAACCTCGGGCCGATCGCAC</td><td>385.3</td><td>6.9</td><td>0.9</td><td>8.1</td><td>6.5E-16</td><td>2.1E-15</td><td>MB231/MB468</td></tr>
<tr><td>>hg38_21_+_8433172_8446622_RNA45SN1_45S_50ntFlanks@485.528.44|GATGTGGTGACGTCGTGCTCTCCCGGGCCGGGTCCGAGCCGCGA</td><td>>hg38_21_+_8433172_8446622_RNA45SN1_45S_50ntFlanks</td><td>485</td><td>528</td><td>44</td><td>GATGTGGTGACGTCGTGCTCTCCCGGGCCGGGTCCGAGCCGCGA</td><td>543.0</td><td>13.7</td><td>1.7</td><td>8.1</td><td>6.4E-16</td><td>2.1E-15</td><td>BT20/MB468</td></tr>
<tr><td>>hg38_21_+_8433172_8446622_RNA45SN1_45S_50ntFlanks@4003.4020.18|ATAACCTCGGGCCGATCG</td><td>>hg38_21_+_8433172_8446622_RNA45SN1_45S_50ntFlanks</td><td>4003</td><td>4020</td><td>18</td><td>ATAACCTCGGGCCGATCG</td><td>499.3</td><td>-5.7</td><td>0.7</td><td>-8.1</td><td>6.5E-16</td><td>2.1E-15</td><td>BT20/MB468</td></tr>
<tr><td>>MI0000283|hsa-mir-203a&WithFlank&14|+|104117399|104117520@71.91.21(+1A)&[MIMAT0000264&hsa-miR-203a-3p&offsets|0|-1(+1A);m-79&14|+|104117469|104117490&offsets|0|-1(+1A)]|GTGAAATGTTTAGGACCACTAA</td><td>>MI0000283|hsa-mir-203a&WithFlank&14|+|104117399|104117520</td><td>71</td><td>91</td><td>21(+1A)</td><td>GTGAAATGTTTAGGACCACTAA</td><td>442.7</td><td>12.3</td><td>1.5</td><td>8.4</td><td>4.4E-17</td><td>2.1E-15</td><td>BT20/MB231</td></tr>
</table>

| | | | | | | | | | | | | |
|---|---|---|---|---|---|---|---|---|---|---|---|---|
| >hg38_21_+_8433172_8446622_RNA45SN1_45S_50ntFlanks@8380.8401.22\|CAAGAGGGCGTGAAACCGTTAA | >hg38_21_+_8433172_8446622_RNA45SN1_45S_50ntFlanks | 8380 | 8401 | 22 | CAAGAGGGCGTGAAACCGTTAA | 510.8 | -11.0 | 1.4 | -8.1 | 6.6E-16 | 2.1E-15 | BT20/MB468 |
| >hg38_21_+_8433172_8446622_RNA45SN1_45S_50ntFlanks@12451.12477.27\|ATTCACCAAGCGTTGGATTGTTCACCC | >hg38_21_+_8433172_8446622_RNA45SN1_45S_50ntFlanks | 12451 | 12477 | 27 | ATTCACCAAGCGTTGGATTGTTCACCC | 509.0 | 7.3 | 0.9 | 8.1 | 6.7E-16 | 2.1E-15 | MB231/MB468 |
| >hg38_21_+_8433172_8446622_RNA45SN1_45S_50ntFlanks@4763.4780.18\|GATACCGTCGTAGTTCCG | >hg38_21_+_8433172_8446622_RNA45SN1_45S_50ntFlanks | 4763 | 4780 | 18 | GATACCGTCGTAGTTCCG | 633.0 | -4.7 | 0.6 | -8.1 | 6.7E-16 | 2.1E-15 | MB231/MB468 |
| >hg38_21_+_8433172_8446622_RNA45SN1_45S_50ntFlanks@12995.13039.45\|CTCGCTGCGATCTATTGAAAGTCAGCCCTCGACACAAGGGTTTGT | >hg38_21_+_8433172_8446622_RNA45SN1_45S_50ntFlanks | 12995 | 13039 | 45 | CTCGCTGCGATCTATTGAAAGTCAGCCCTCGACACAAGGGTTTGT | 114935.2 | 8.4 | 1.0 | 8.1 | 6.6E-16 | 2.1E-15 | BT20/MB468 |
| >hg38_21_+_8433172_8446622_RNA45SN1_45S_50ntFlanks@10783.10818.36\|GTGAACAGCCTCTGGCATGTTGGAACAATGTAGGTA | >hg38_21_+_8433172_8446622_RNA45SN1_45S_50ntFlanks | 10783 | 10818 | 36 | GTGAACAGCCTCTGGCATGTTGGAACAATGTAGGTA | 1187.1 | -5.9 | 0.7 | -8.1 | 6.6E-16 | 2.1E-15 | BT20/MB468 |
| >MI0000299\|hsa-mir-222&WithFlank&X\|-\|45747009\|45747130@75.99.25(+1A)&[MIMAT0000279&hsa-miR-222-3p&offsets\|0\|+4(+1A);m-140&X\|-\|45747033\|45747056&offsets\|0\|+1(+1A)]\|AGCTACATCTGGCTACTGGGTCTCTA | >MI0000299\|hsa-mir-222&WithFlank&X\|-\|45747009\|45747130 | 75 | 99 | 25(+1A) | AGCTACATCTGGCTACTGGGTCTCTA | 1419.5 | 9.9 | 1.2 | 8.1 | 6.7E-16 | 2.1E-15 | MB231/MB468 |
| >hg38_21_+_8433172_8446622_RNA45SN1_45S_50ntFlanks@8552.8569.18\|CTCCCCGGAGGGGGCGGG | >hg38_21_+_8433172_8446622_RNA45SN1_45S_50ntFlanks | 8552 | 8569 | 18 | CTCCCCGGAGGGGGCGGG | 448.4 | -10.4 | 1.3 | -8.1 | 6.8E-16 | 2.2E-15 | MB231/MB468 |
| >trna20_GluTTC_2_-_131094701_131094772@1.33.33,trna5_GluTTC_13_-_41634874_41634945@1.33.33\|TCCCATATGGTCTAGCGGTTAGGATTCCTGGTT | >trna20_GluTTC_2_-_131094701_131094772 | 1 | 33 | 33 | TCCCATATGGTCTAGCGGTTAGGATTCCTGGTT | 933.2 | 10.7 | 1.3 | 8.1 | 6.8E-16 | 2.2E-15 | BT20/MB468 |
| >hg38_21_+_8433172_8446622_RNA45SN1_45S_50ntFlanks@12934.12955.22\|CTAAACCATTCGTAGACGACCT | >hg38_21_+_8433172_8446622_RNA45SN1_45S_50ntFlanks | 12934 | 12955 | 22 | CTAAACCATTCGTAGACGACCT | 297.1 | -4.0 | 0.5 | -8.1 | 7.0E-16 | 2.2E-15 | MB231/MB468 |
| >MI0005559\|hsa-mir-744&WithFlank&17\|+\|12081893\|12082002@17.38.22&[MIMAT0004945&hsa-miR-744-5p&offsets\|0\|0;m-172&17\|+\|12081909\|12081929&offsets\|0\|+1]\|TGCGGGGCTAGGGCTAACAGCA | >MI0005559\|hsa-mir-744&WithFlank&17\|+\|12081893\|12082002 | 17 | 38 | 22 | TGCGGGGCTAGGGCTAACAGCA | 1785.8 | 6.7 | 0.8 | 8.1 | 6.9E-16 | 2.2E-15 | BT20/MB468 |
| >hg38_21_+_8433172_8446622_RNA45SN1_45S_50ntFlanks@485.518.34\|GATGTGGTGACGTCGTGCTCTCCCGGGCCGGGTC | >hg38_21_+_8433172_8446622_RNA45SN1_45S_50ntFlanks | 485 | 518 | 34 | GATGTGGTGACGTCGTGCTCTCCCGGGCCGGGTC | 1242.2 | 29.3 | 3.6 | 8.1 | 7.0E-16 | 2.2E-15 | MB231/MB468 |
| >hg38_21_+_8433172_8446622_RNA45SN1_45S_50ntFlanks@10572.10591.20\|AGAAGCCGGCGGGAGCCCCG | >hg38_21_+_8433172_8446622_RNA45SN1_45S_50ntFlanks | 10572 | 10591 | 20 | AGAAGCCGGCGGGAGCCCCG | 771.8 | -11.6 | 1.4 | -8.1 | 7.0E-16 | 2.3E-15 | BT20/MB468 |
| >hg38_21_+_8433172_8446622_RNA45SN1_45S_50ntFlanks@6719.6746.28\|TGAATTGCAGGACACATTGATCATCGAC | >hg38_21_+_8433172_8446622_RNA45SN1_45S_50ntFlanks | 6719 | 6746 | 28 | TGAATTGCAGGACACATTGATCATCGAC | 944.6 | -6.1 | 0.8 | -8.1 | 7.0E-16 | 2.3E-15 | BT20/MB468 |
| >MI0000650\|hsa-mir-200c&WithFlank&12\|+\|6963693\|6963772@50.71.22(+1C)&[MIMAT0000617&hsa-miR-200c-3p&offsets\|0\|-1(+1C);m-54&12\|+\|6963742\|6963764&offsets\|0\|-1(+1C)]\|TAATACTGCCGGGTAATGATGGC | >MI0000650\|hsa-mir-200c&WithFlank&12\|+\|6963693\|6963772 | 50 | 71 | 22(+1C) | TAATACTGCCGGGTAATGATGGC | 2039.7 | 9.7 | 1.2 | 8.1 | 7.0E-16 | 2.3E-15 | BT20/MB468 |
| >hg38_21_+_8433172_8446622_RNA45SN1_45S_50ntFlanks@9568.9592.25\|CGTGCAAATCGGTCGTCCGACCTGG | >hg38_21_+_8433172_8446622_RNA45SN1_45S_50ntFlanks | 9568 | 9592 | 25 | CGTGCAAATCGGTCGTCCGACCTGG | 509.8 | -6.9 | 0.9 | -8.1 | 7.1E-16 | 2.3E-15 | MB231/MB468 |
| >hg38_21_+_8433172_8446622_RNA45SN1_45S_50ntFlanks@4283.4317.35\|CTTTAACGAGGATCCATTGGAGGGCAAGTCTGGTG | >hg38_21_+_8433172_8446622_RNA45SN1_45S_50ntFlanks | 4283 | 4317 | 35 | CTTTAACGAGGATCCATTGGAGGGCAAGTCTGGTG | 647.6 | -5.3 | 0.7 | -8.1 | 7.1E-16 | 2.3E-15 | MB231/MB468 |
| >am_trnaMT_HisGTG_MT_+_12138_12206@26.44.19\|AGATTGTGAATCTGACAAC | >am_trnaMT_HisGTG_MT_+_12138_12206 | 26 | 44 | 19 | AGATTGTGAATCTGACAAC | 1296.9 | 8.3 | 1.0 | 8.1 | 7.1E-16 | 2.3E-15 | BT20/MB468 |
| >hg38_21_+_8433172_8446622_RNA45SN1_45S_50ntFlanks@11027.11050.24\|CTCTCTCTCTCTCTCCCCCGCTCC | >hg38_21_+_8433172_8446622_RNA45SN1_45S_50ntFlanks | 11027 | 11050 | 24 | CTCTCTCTCTCTCTCCCCCGCTCC | 438.9 | -10.4 | 1.3 | -8.1 | 7.2E-16 | 2.3E-15 | MB231/MB468 |
| >hg38_21_+_8433172_8446622_RNA45SN1_45S_50ntFlanks@11890.11912.23\|CTTGACTCTAGTCTGGCACGGTG | >hg38_21_+_8433172_8446622_RNA45SN1_45S_50ntFlanks | 11890 | 11912 | 23 | CTTGACTCTAGTCTGGCACGGTG | 624.0 | -5.1 | 0.6 | -8.1 | 7.2E-16 | 2.3E-15 | BT20/MB468 |
| >hg38_21_+_8433172_8446622_RNA45SN1_45S_50ntFlanks@8182.8203.22\|ATCGAGGCCCAGCCCGTGGACG | >hg38_21_+_8433172_8446622_RNA45SN1_45S_50ntFlanks | 8182 | 8203 | 22 | ATCGAGGCCCAGCCCGTGGACG | 247.1 | -9.9 | 1.2 | -8.1 | 7.2E-16 | 2.3E-15 | BT20/MB468 |
| >MI0000283\|hsa-mir-203a&WithFlank&14\|+\|104117399\|104117520@72.93.22&[MIMAT0000264&hsa-miR-203a-3p&offsets\|+1\|+1;m-79&14\|+\|104117469\|104117490&offsets\|+1\|+1]\|TGAAATGTTTAGGACCACTAGA | >MI0000283\|hsa-mir-203a&WithFlank&14\|+\|104117399\|104117520 | 72 | 93 | 22 | TGAAATGTTTAGGACCACTAGA | 2783.3 | 10.4 | 1.3 | 8.1 | 7.2E-16 | 2.3E-15 | BT20/MB468 |
| >hg38_21_+_8433172_8446622_RNA45SN1_45S_50ntFlanks@9572.9594.23\|CAAATCGGTCGTCCGACCTGGGT | >hg38_21_+_8433172_8446622_RNA45SN1_45S_50ntFlanks | 9572 | 9594 | 23 | CAAATCGGTCGTCCGACCTGGGT | 522.8 | -4.8 | 0.6 | -8.1 | 7.3E-16 | 2.3E-15 | MB231/MB468 |
| >hg38_21_+_8433172_8446622_RNA45SN1_45S_50ntFlanks@8198.8215.18\|TGGACGGTGTGAGGCCGG | >hg38_21_+_8433172_8446622_RNA45SN1_45S_50ntFlanks | 8198 | 8215 | 18 | TGGACGGTGTGAGGCCGG | 2234.8 | -6.8 | 0.8 | -8.1 | 7.3E-16 | 2.3E-15 | BT20/MB468 |
| >MI0000077\|hsa-mir-21&WithFlank&17\|+\|59841260\|59841343@13.37.25&[MIMAT0000076&hsa-miR-21-5p&offsets\|-1\|+2;m-1&17\|+\|59841273\|59841294&offsets\|-1\|+2]\|GTAGCTTATCAGACTGATGTTGACT | >MI0000077\|hsa-mir-21&WithFlank&17\|+\|59841260\|59841343 | 13 | 37 | 25 | GTAGCTTATCAGACTGATGTTGACT | 500.3 | 8.4 | 1.0 | 8.1 | 7.3E-16 | 2.4E-15 | BT20/MB468 |
| >hg38_21_+_8433172_8446622_RNA45SN1_45S_50ntFlanks@485.517.33\|GATGTGGTGACGTCGTGCTCTCCCGGGCCGGGT | >hg38_21_+_8433172_8446622_RNA45SN1_45S_50ntFlanks | 485 | 517 | 33 | GATGTGGTGACGTCGTGCTCTCCCGGGCCGGGT | 1869.4 | 29.3 | 3.6 | 8.1 | 7.4E-16 | 2.4E-15 | MB231/MB468 |
| >hg38_21_+_8433172_8446622_RNA45SN1_45S_50ntFlanks@13004.13025.22\|ATCTATTGAAAGTCAGCCCTCG | >hg38_21_+_8433172_8446622_RNA45SN1_45S_50ntFlanks | 13004 | 13025 | 22 | ATCTATTGAAAGTCAGCCCTCG | 568.2 | -10.7 | 1.3 | -8.1 | 7.6E-16 | 2.4E-15 | MB231/MB468 |
| >MI0000650\|hsa-mir-200c&WithFlank&12\|+\|6963693\|6963772@49.70.22&[MIMAT0000617&hsa-miR-200c-3p&offsets\|-1\|-2;m-54&12\|+\|6963742\|6963764&offsets\|-1\|-2]\|CTAATACTGCCGGGTAATGATG | >MI0000650\|hsa-mir-200c&WithFlank&12\|+\|6963693\|6963772 | 49 | 70 | 22 | CTAATACTGCCGGGTAATGATG | 362.8 | 12.0 | 1.4 | 8.4 | 5.1E-17 | 2.4E-15 | BT20/MB231 |
| >hg38_21_+_8433172_8446622_RNA45SN1_45S_50ntFlanks@8170.8197.28\|CAAGTCCTTCTGATCGAGGCCCAGCCCG | >hg38_21_+_8433172_8446622_RNA45SN1_45S_50ntFlanks | 8170 | 8197 | 28 | CAAGTCCTTCTGATCGAGGCCCAGCCCG | 793.6 | -7.7 | 1.0 | -8.1 | 7.6E-16 | 2.5E-15 | BT20/MB468 |
| >hg38_21_+_8433172_8446622_RNA45SN1_45S_50ntFlanks@9540.9571.32\|CTCTGGTGGAGGTCCGTAGCGGTCCTGACGTG | >hg38_21_+_8433172_8446622_RNA45SN1_45S_50ntFlanks | 9540 | 9571 | 32 | CTCTGGTGGAGGTCCGTAGCGGTCCTGACGTG | 380.8 | -3.2 | 0.4 | -8.1 | 7.7E-16 | 2.5E-15 | BT20/MB468 |
| >hg38_21_+_8433172_8446622_RNA45SN1_45S_50ntFlanks@12416.12454.39\|TTCGATGTCGGCTCTTCCTATCATTGTGAAGCAGAATTC | >hg38_21_+_8433172_8446622_RNA45SN1_45S_50ntFlanks | 12416 | 12454 | 39 | TTCGATGTCGGCTCTTCCTATCATTGTGAAGCAGAATTC | 920.3 | 7.9 | 1.0 | 8.1 | 7.8E-16 | 2.5E-15 | MB231/MB468 |
| >trna152_ValCAC_6_-_27248049_27248121@1.32.32\|GCTTCTGTAGTGTAGTGGTTATCACGTTCGCC | >trna152_ValCAC_6_-_27248049_27248121 | 1 | 32 | 32 | GCTTCTGTAGTGTAGTGGTTATCACGTTCGCC | 601.3 | 8.0 | 1.0 | 8.1 | 7.8E-16 | 2.5E-15 | BT20/MB468 |
| >hg38_21_+_8433172_8446622_RNA45SN1_45S_50ntFlanks@6651.6691.41\|CGACTCTTAGCGGTGGATCACTCGGCTCGTGCGTCGATGAA | >hg38_21_+_8433172_8446622_RNA45SN1_45S_50ntFlanks | 6651 | 6691 | 41 | CGACTCTTAGCGGTGGATCACTCGGCTCGTGCGTCGATGAA | 579.1 | 5.4 | 0.7 | 8.1 | 7.8E-16 | 2.5E-15 | BT20/MB468 |
| >trnaMT_PheGAA_MT_+_577_647@39.60.22\|ATGTTTAGACGGGCTCACATCA | >trnaMT_PheGAA_MT_+_577_647 | 39 | 60 | 22 | ATGTTTAGACGGGCTCACATCA | 248.6 | 9.1 | 1.1 | 8.1 | 7.9E-16 | 2.5E-15 | BT20/MB468 |
| >hg38_21_+_8433172_8446622_RNA45SN1_45S_50ntFlanks@12439.12472.34\|TTGTGAAGCAGAATTCACCAAGCGTTGGATTGTT | >hg38_21_+_8433172_8446622_RNA45SN1_45S_50ntFlanks | 12439 | 12472 | 34 | TTGTGAAGCAGAATTCACCAAGCGTTGGATTGTT | 446.3 | 7.4 | 0.9 | 8.1 | 7.9E-16 | 2.5E-15 | MB231/MB468 |
| >hg38_21_+_8433172_8446622_RNA45SN1_45S_50ntFlanks@9567.9597.31\|ACGTGCAAATCGGTCGTCCGACCTGGGTATA | >hg38_21_+_8433172_8446622_RNA45SN1_45S_50ntFlanks | 9567 | 9597 | 31 | ACGTGCAAATCGGTCGTCCGACCTGGGTATA | 2896.5 | -6.2 | 0.8 | -8.1 | 8.0E-16 | 2.5E-15 | MB231/MB468 |
| >hg38_21_+_8433172_8446622_RNA45SN1_45S_50ntFlanks@8381.8405.25\|AAGAGGGCGTGAAACCGTTAAGAGG | >hg38_21_+_8433172_8446622_RNA45SN1_45S_50ntFlanks | 8381 | 8405 | 25 | AAGAGGGCGTGAAACCGTTAAGAGG | 278.2 | -5.4 | 0.7 | -8.1 | 8.0E-16 | 2.5E-15 | BT20/MB468 |
| >hg38_21_+_8433172_8446622_RNA45SN1_45S_50ntFlanks@9004.9026.23\|TCTCCCCCGCGGGGGCGCGCCGG | >hg38_21_+_8433172_8446622_RNA45SN1_45S_50ntFlanks | 9004 | 9026 | 23 | TCTCCCCCGCGGGGGCGCGCCGG | 621.9 | -10.9 | 1.3 | -8.1 | 8.1E-16 | 2.6E-15 | MB231/MB468 |
| >MI0000089\|hsa-mir-31&WithFlank&9\|-\|21512109\|21512191@14.35.22(+1C)&[MIMAT0000089&hsa-miR-31-5p&offsets\|0\|+1(+1C);m-360&9\|-\|21512157\|21512178&offsets\|0\|0(+1C)]\|AGGCAAGATGCTGGCATAGCTGC | >MI0000089\|hsa-mir-31&WithFlank&9\|-\|21512109\|21512191 | 14 | 35 | 22(+1C) | AGGCAAGATGCTGGCATAGCTGC | 289.0 | 11.7 | 1.4 | 8.4 | 5.5E-17 | 2.6E-15 | BT20/MB231 |
| >hg38_21_+_8433172_8446622_RNA45SN1_45S_50ntFlanks@7977.8010.34\|CGACCTCAGATCAGACGTGGCGACCCGCTGAATT | >hg38_21_+_8433172_8446622_RNA45SN1_45S_50ntFlanks | 7977 | 8010 | 34 | CGACCTCAGATCAGACGTGGCGACCCGCTGAATT | 256.2 | 5.0 | 0.6 | 8.1 | 8.1E-16 | 2.6E-15 | BT20/MB468 |
| >MI0000234\|hsa-mir-192&WithFlank&11\|-\|64891131\|64891252@30.49.20&[MIMAT0000222&hsa-miR-192-5p&offsets\|0\|-1;m-59&11\|-\|64891203\|64891223&offsets\|0\|-1]\|CTGACCTATGAATTGACAGC | >MI0000234\|hsa-mir-192&WithFlank&11\|-\|64891131\|64891252 | 30 | 49 | 20 | CTGACCTATGAATTGACAGC | 596.8 | 6.6 | 0.8 | 8.1 | 8.1E-16 | 2.6E-15 | BT20/MB468 |
| >hg38_21_+_8433172_8446622_RNA45SN1_45S_50ntFlanks@12416.12448.33\|TTCGATGTCGGCTCTTCCTATCATTGTGAAGCA | >hg38_21_+_8433172_8446622_RNA45SN1_45S_50ntFlanks | 12416 | 12448 | 33 | TTCGATGTCGGCTCTTCCTATCATTGTGAAGCA | 1067.8 | 6.4 | 0.8 | 8.1 | 8.2E-16 | 2.6E-15 | BT20/MB468 |
| >MI0000086\|hsa-mir-28&WithFlank&3\|+\|188688775\|188688872@60.82.23(+1U)&[MIMAT0004502&hsa-miR-28-3p&offsets\|0\|+1(+1U);m-28&3\|+\|188688834\|188688855&offsets\|0\|+1(+1U)]\|CACTAGATTGTGAGCTCCTGGAGT | >MI0000086\|hsa-mir-28&WithFlank&3\|+\|188688775\|188688872 | 60 | 82 | 23(+1U) | CACTAGATTGTGAGCTCCTGGAGT | 1021.4 | 10.8 | 1.3 | 8.1 | 8.2E-16 | 2.6E-15 | BT20/MB468 |
| >MI0000746\|hsa-mir-99b&WithFlank&19\|+\|51692606\|51692687@51.71.21(+1U)&[MIMAT0004678&hsa-miR-99b-3p&offsets\|0\|-1(+1U);m-151&19\|+\|51692656\|51692678&offsets\|0\|-2(+1U)]\|CAAGCTCGTGTCTGTGGGTCCT | >MI0000746\|hsa-mir-99b&WithFlank&19\|+\|51692606\|51692687 | 51 | 71 | 21(+1U) | CAAGCTCGTGTCTGTGGGTCCT | 513.7 | 9.5 | 1.2 | 8.1 | 8.2E-16 | 2.6E-15 | BT20/MB468 |
| >MI0025242\|hsa-mir-7706&WithFlank&15\|+\|85380590\|85380668@50.71.22(+1U)&[MIMAT0030021&hsa-miR-7706&offsets\|0\|-2(+1U);m-390&15\|+\|85380639\|85380662&offsets\|0\|-2(+1U)]\|TGAAGCGCCTGTGCTCTGCCGAT | >MI0025242\|hsa-mir-7706&WithFlank&15\|+\|85380590\|85380668 | 50 | 71 | 22(+1U) | TGAAGCGCCTGTGCTCTGCCGAT | 397.6 | 7.7 | 1.0 | 8.1 | 8.3E-16 | 2.6E-15 | BT20/MB468 |
| >hg38_21_+_8433172_8446622_RNA45SN1_45S_50ntFlanks@12635.12652.18\|AAGCTACCATCTGTGGGA | >hg38_21_+_8433172_8446622_RNA45SN1_45S_50ntFlanks | 12635 | 12652 | 18 | AAGCTACCATCTGTGGGA | 373.0 | -10.5 | 1.3 | -8.0 | 8.3E-16 | 2.6E-15 | BT20/MB468 |
| >trnaMT_SerGCT_MT_+_12207_12265@27.62.36\|CATGCCCCCATGTCTAACAACATGGCTTTCTCACCA | >trnaMT_SerGCT_MT_+_12207_12265 | 27 | 62 | 36 | CATGCCCCCATGTCTAACAACATGGCTTTCTCACCA | 659.5 | 9.4 | 1.2 | 8.0 | 8.3E-16 | 2.7E-15 | BT20/MB468 |

| | | | | | | | | | | | | |
|---|---|---|---|---|---|---|---|---|---|---|---|---|
| >hg38_21_+_8433172_8446622_RNA45SN1_45S_50ntFlanks@10362.10390.29\|AAGGCCGAAGTGGAGAAGGGTTCCATGTG | >hg38_21_+_8433172_8446622_RNA45SN1_45S_50ntFlanks | 10362 | 10390 | 29 | AAGGCCGAAGTGGAGAAGGGTTCCATGTG | 1303.0 | -5.9 | 0.7 | -8.0 | 8.4E-16 | 2.7E-15 | MB231/MB468 |
| >hg38_21_+_8433172_8446622_RNA45SN1_45S_50ntFlanks@10922.10946.25\|CTGGACGAGGCGCCGCCGCCCCCCC | >hg38_21_+_8433172_8446622_RNA45SN1_45S_50ntFlanks | 10922 | 10946 | 25 | CTGGACGAGGCGCCGCCGCCCCCCC | 370.8 | -10.1 | 1.3 | -8.0 | 8.4E-16 | 2.7E-15 | MB231/MB468 |
| >hg38_21_+_8433172_8446622_RNA45SN1_45S_50ntFlanks@12008.12043.36\|CCGCCGGTGAAATACCACTACTCTGATCGTTTTTTC | >hg38_21_+_8433172_8446622_RNA45SN1_45S_50ntFlanks | 12008 | 12043 | 36 | CCGCCGGTGAAATACCACTACTCTGATCGTTTTTTC | 1067.4 | -12.1 | 1.5 | -8.0 | 8.4E-16 | 2.7E-15 | BT20/MB468 |
| >hg38_21_+_8433172_8446622_RNA45SN1_45S_50ntFlanks@12678.12703.26\|AATCCCGCCCAGGCGGAACGATACGG | >hg38_21_+_8433172_8446622_RNA45SN1_45S_50ntFlanks | 12678 | 12703 | 26 | AATCCCGCCCAGGCGGAACGATACGG | 296.7 | -10.2 | 1.3 | -8.0 | 8.4E-16 | 2.7E-15 | BT20/MB468 |
| >hg38_21_+_8433172_8446622_RNA45SN1_45S_50ntFlanks@11784.11806.23\|ATGAATGGATGAACGAGATTCCC | >hg38_21_+_8433172_8446622_RNA45SN1_45S_50ntFlanks | 11784 | 11806 | 23 | ATGAATGGATGAACGAGATTCCC | 1028.1 | -5.4 | 0.7 | -8.0 | 8.5E-16 | 2.7E-15 | BT20/MB468 |
| >hg38_21_+_8433172_8446622_RNA45SN1_45S_50ntFlanks@12464.12496.33\|TGGATTGTTCACCCACTAATAGGGAACGTGAGC | >hg38_21_+_8433172_8446622_RNA45SN1_45S_50ntFlanks | 12464 | 12496 | 33 | TGGATTGTTCACCCACTAATAGGGAACGTGAGC | 345.1 | 6.7 | 0.8 | 8.0 | 8.6E-16 | 2.7E-15 | MB231/MB468 |
| >MI0000273\|hsa-mir-183&WithFlank&7\|-\|129774899\|129775020@33.56.24&[MIMAT0000261&hsa-miR-183-5p&offsets\|0\|+2;m-41&7\|-\|129774966\|129774987&offsets\|-1\|+1]\|TATGGCACTGGTAGAATTCACTGT | >MI0000273\|hsa-mir-183&WithFlank&7\|-\|129774899\|129775020 | 33 | 56 | 24 | TATGGCACTGGTAGAATTCACTGT | 1217.7 | 6.5 | 0.8 | 8.0 | 8.7E-16 | 2.7E-15 | MB231/MB468 |
| >hg38_21_+_8433172_8446622_RNA45SN1_45S_50ntFlanks@9569.9595.27\|GTGCAAATCGGTCGTCCGACCTGGGTA | >hg38_21_+_8433172_8446622_RNA45SN1_45S_50ntFlanks | 9569 | 9595 | 27 | GTGCAAATCGGTCGTCCGACCTGGGTA | 273.4 | -2.9 | 0.4 | -8.0 | 8.7E-16 | 2.8E-15 | BT20/MB468 |
| >MI0000078\|hsa-mir-22&WithFlank&17\|-\|1713897\|1713993@59.78.20&[MIMAT0000077&hsa-miR-22-3p&offsets\|0\|-2;m-2&17\|-\|1713914\|1713935&offsets\|0\|-2]\|AAGCTGCCAGTTGAAGAACT | >MI0000078\|hsa-mir-22&WithFlank&17\|-\|1713897\|1713993 | 59 | 78 | 20 | AAGCTGCCAGTTGAAGAACT | 1510.7 | 9.0 | 1.1 | 8.0 | 8.7E-16 | 2.8E-15 | BT20/MB468 |
| >MI0000081\|hsa-mir-24-2&WithFlank&19\|-\|13836281\|13836365@56.77.22;MI0000080\|hsa-mir-24-1&WithFlank&9\|+\|95086015\|95086094@50.71.22&[MIMAT0000080&hsa-miR-24-3p&offsets\|0\|0;m-36&19\|-\|13836288\|13836310&offsets\|0\|-1];[MIMAT0000080_1&hsa-miR-24-3p&offsets\|0\|0;m-35&9\|+\|95086064\|95086086&offsets\|0\|-1]\|TGGCTCAGTTCAGCAGGAACAG | >MI0000081\|hsa-mir-24-2&WithFlank&19\|-\|13836281\|13836365 | 56 | 77 | 22 | TGGCTCAGTTCAGCAGGAACAG | 30556.7 | 5.9 | 0.7 | 8.0 | 8.7E-16 | 2.8E-15 | MB231/MB468 |
| >hg38_21_+_8433172_8446622_RNA45SN1_45S_50ntFlanks@12605.12625.21\|TATGTGCTTGGCTGAGGAGCC | >hg38_21_+_8433172_8446622_RNA45SN1_45S_50ntFlanks | 12605 | 12625 | 21 | TATGTGCTTGGCTGAGGAGCC | 5999.1 | -4.5 | 0.6 | -8.0 | 8.8E-16 | 2.8E-15 | MB231/MB468 |
| >hg38_21_+_8433172_8446622_RNA45SN1_45S_50ntFlanks@12416.12456.41\|TTCGATGTCGGCTCTTCCTATCATTGTGAAGCAGAATTCAC | >hg38_21_+_8433172_8446622_RNA45SN1_45S_50ntFlanks | 12416 | 12456 | 41 | TTCGATGTCGGCTCTTCCTATCATTGTGAAGCAGAATTCAC | 539.8 | 7.5 | 0.9 | 8.0 | 8.9E-16 | 2.8E-15 | MB231/MB468 |
| >hg38_21_+_8433172_8446622_RNA45SN1_45S_50ntFlanks@9504.9524.21\|GTGAACTATGCCTGGGCAGGG | >hg38_21_+_8433172_8446622_RNA45SN1_45S_50ntFlanks | 9504 | 9524 | 21 | GTGAACTATGCCTGGGCAGGG | 348.3 | -10.4 | 1.3 | -8.0 | 8.9E-16 | 2.8E-15 | BT20/MB468 |
| >hg38_21_+_8433172_8446622_RNA45SN1_45S_50ntFlanks@7989.8015.27\|AGACGTGGCGACCCGCTGAATTTAAGC | >hg38_21_+_8433172_8446622_RNA45SN1_45S_50ntFlanks | 7989 | 8015 | 27 | AGACGTGGCGACCCGCTGAATTTAAGC | 595.2 | 6.5 | 0.8 | 8.0 | 8.9E-16 | 2.8E-15 | MB231/MB468 |
| >hg38_21_+_8433172_8446622_RNA45SN1_45S_50ntFlanks@6739.6770.32\|TCATCGACACTTCGAACGCACTTGCGGCCCCG | >hg38_21_+_8433172_8446622_RNA45SN1_45S_50ntFlanks | 6739 | 6770 | 32 | TCATCGACACTTCGAACGCACTTGCGGCCCCG | 391.9 | -10.6 | 1.3 | -8.0 | 8.9E-16 | 2.8E-15 | BT20/MB468 |
| >MI0000088\|hsa-mir-30a&WithFlank&6\|-\|71403545\|71403627@12.34.23(+1G)&[MIMAT0000087&hsa-miR-30a-5p&offsets\|0\|+1(+1G);m-15&6\|-\|71403595\|71403616&offsets\|0\|+1(+1G)]\|TGTAAACATCCTCGACTGGAAGCG | >MI0000088\|hsa-mir-30a&WithFlank&6\|-\|71403545\|71403627 | 12 | 34 | 23(+1G) | TGTAAACATCCTCGACTGGAAGCG | 41502.7 | 8.6 | 1.1 | 8.0 | 9.0E-16 | 2.8E-15 | BT20/MB468 |
| >hg38_21_+_8433172_8446622_RNA45SN1_45S_50ntFlanks@8198.8224.27\|TGGACGGTGTGAGGCCGGTAGCGGCCC | >hg38_21_+_8433172_8446622_RNA45SN1_45S_50ntFlanks | 8198 | 8224 | 27 | TGGACGGTGTGAGGCCGGTAGCGGCCC | 181.6 | -4.1 | 0.5 | -8.0 | 9.1E-16 | 2.9E-15 | BT20/MB468 |
| >trna111_HisGTG_1_-_147774845_147774916@-1T.24.25,trna118_HisGTG_1_-_145396881_145396952@-1T.24.25,trna16_HisGTG_1_+_146544773_146544844@-1T.24.25,trna1_HisGTG_15_+_45493349_45493420@-1T.24.25,trna21_HisGTG_1_+_147753471_147753542@-1T.24.25,trna33_HisGTG_6_+_27125906_27125977@-1T.24.25,trna7_HisGTG_9_-_14433938_14434009@-1T.24.25,trna8_HisGTG_15_-_45492611_45492682@-1T.24.25,trna9_HisGTG_15_-_45490804_45490875@-1T.24.25\|TGCCGTGATCGTATAGTGGTTAGTA | >trna111_HisGTG_1_-_147774845_147774916 | -1T | 24 | 25 | TGCCGTGATCGTATAGTGGTTAGTA | 833.1 | 6.7 | 0.8 | 8.0 | 9.1E-16 | 2.9E-15 | MB231/MB468 |
| >hg38_21_+_8433172_8446622_RNA45SN1_45S_50ntFlanks@9738.9755.18\|GAAACGATCTCAACCTAT | >hg38_21_+_8433172_8446622_RNA45SN1_45S_50ntFlanks | 9738 | 9755 | 18 | GAAACGATCTCAACCTAT | 537.9 | -10.6 | 1.3 | -8.0 | 9.3E-16 | 2.9E-15 | MB231/MB468 |
| >MI0000087\|hsa-mir-29a&WithFlank&7\|-\|130876741\|130876816@48.69.22(+1A)&[MIMAT0000086&hsa-miR-29a-3p&offsets\|0\|0(+1A);m-26&7\|-\|130876749\|130876769&offsets\|0\|+1(+1A)]\|TAGCACCATCTGAAATCGGTTAA | >MI0000087\|hsa-mir-29a&WithFlank&7\|-\|130876741\|130876816 | 48 | 69 | 22(+1A) | TAGCACCATCTGAAATCGGTTAA | 817.7 | 9.3 | 1.2 | 8.0 | 9.3E-16 | 2.9E-15 | BT20/MB468 |
| >MI0000749\|hsa-mir-30e&WithFlank&1\|+\|40754349\|40754452@65.85.21(+2U)&[MIMAT0000693&hsa-miR-30e-3p&offsets\|0\|-1(+2U);m-25&1\|+\|40754413\|40754434&offsets\|0\|-1(+2U)]\|CTTTCAGTCGGATGTTTACAGTT | >MI0000749\|hsa-mir-30e&WithFlank&1\|+\|40754349\|40754452 | 65 | 85 | 21(+2U) | CTTTCAGTCGGATGTTTACAGTT | 933.4 | 5.4 | 0.7 | 8.0 | 9.4E-16 | 3.0E-15 | BT20/MB468 |
| >MI0000255\|hsa-mir-30d&WithFlank&8\|-\|134804870\|134804951@12.35.24(+1A)&[MIMAT0000245&hsa-miR-30d-5p&offsets\|0\|+2(+1A);m-21&8\|-\|134804919\|134804940&offsets\|0\|+2(+1A)]\|TGTAAACATCCCCGACTGGAAGCTA | >MI0000255\|hsa-mir-30d&WithFlank&8\|-\|134804870\|134804951 | 12 | 35 | 24(+1A) | TGTAAACATCCCCGACTGGAAGCTA | 1340.0 | 9.0 | 1.1 | 8.0 | 9.4E-16 | 3.0E-15 | BT20/MB468 |
| >hg38_21_+_8433172_8446622_RNA45SN1_45S_50ntFlanks@12033.12053.21\|ATCGTTTTTTCACTGACCCGG | >hg38_21_+_8433172_8446622_RNA45SN1_45S_50ntFlanks | 12033 | 12053 | 21 | ATCGTTTTTTCACTGACCCGG | 469.5 | -10.5 | 1.3 | -8.0 | 9.5E-16 | 3.0E-15 | MB231/MB468 |
| >hg38_21_+_8433172_8446622_RNA45SN1_45S_50ntFlanks@12412.12443.32\|ATCCTTCGATGTCGGCTCTTCCTATCATTGTG | >hg38_21_+_8433172_8446622_RNA45SN1_45S_50ntFlanks | 12412 | 12443 | 32 | ATCCTTCGATGTCGGCTCTTCCTATCATTGTG | 341.9 | -10.4 | 1.3 | -8.0 | 9.5E-16 | 3.0E-15 | BT20/MB468 |
| >hg38_21_+_8433172_8446622_RNA45SN1_45S_50ntFlanks@9504.9523.20\|GTGAACTATGCCTGGGCAGG | >hg38_21_+_8433172_8446622_RNA45SN1_45S_50ntFlanks | 9504 | 9523 | 20 | GTGAACTATGCCTGGGCAGG | 1283.5 | -7.8 | 1.0 | -8.0 | 9.5E-16 | 3.0E-15 | MB231/MB468 |
| >hg38_21_+_8433172_8446622_RNA45SN1_45S_50ntFlanks@11567.11594.28\|AGCAGCCGACTTAGAACTGGTGCGGACC | >hg38_21_+_8433172_8446622_RNA45SN1_45S_50ntFlanks | 11567 | 11594 | 28 | AGCAGCCGACTTAGAACTGGTGCGGACC | 573.6 | 3.8 | 0.5 | 8.0 | 9.6E-16 | 3.0E-15 | MB231/MB468 |
| >hg38_21_+_8433172_8446622_RNA45SN1_45S_50ntFlanks@10798.10818.21\|CATGTTGGAACAATGTAGGTA | >hg38_21_+_8433172_8446622_RNA45SN1_45S_50ntFlanks | 10798 | 10818 | 21 | CATGTTGGAACAATGTAGGTA | 2768.0 | -5.8 | 0.7 | -8.0 | 9.6E-16 | 3.0E-15 | BT20/MB468 |
| >hg38_21_+_8433172_8446622_RNA45SN1_45S_50ntFlanks@51.74.24\|GCTGACACGCTGTCCTCTGGCGAC | >hg38_21_+_8433172_8446622_RNA45SN1_45S_50ntFlanks | 51 | 74 | 24 | GCTGACACGCTGTCCTCTGGCGAC | 740.3 | 6.5 | 0.8 | 8.0 | 9.8E-16 | 3.1E-15 | MB231/MB468 |
| >hg38_21_+_8433172_8446622_RNA45SN1_45S_50ntFlanks@10866.10884.19\|CTAAGGGCTGGGTCGGTCG | >hg38_21_+_8433172_8446622_RNA45SN1_45S_50ntFlanks | 10866 | 10884 | 19 | CTAAGGGCTGGGTCGGTCG | 393.3 | -10.2 | 1.3 | -8.0 | 9.9E-16 | 3.1E-15 | MB231/MB468 |
| >MI0000439\|hsa-mir-23b&WithFlank&9\|+\|95085202\|95085310@67.85.19&[MIMAT0000418&hsa-miR-23b-3p&offsets\|+3\|-1;m-42&9\|+\|95085265\|95085285&offsets\|+3\|+1]\|ACATTGCCAGGGATTACCA | >MI0000439\|hsa-mir-23b&WithFlank&9\|+\|95085202\|95085310 | 67 | 85 | 19 | ACATTGCCAGGGATTACCA | 1110.0 | 29.1 | 3.6 | 8.0 | 1.0E-15 | 3.2E-15 | MB231/MB468 |
| >hg38_21_+_8433172_8446622_RNA45SN1_45S_50ntFlanks@4760.4783.24\|TCAGATACCGTCGTAGTTCCGACC | >hg38_21_+_8433172_8446622_RNA45SN1_45S_50ntFlanks | 4760 | 4783 | 24 | TCAGATACCGTCGTAGTTCCGACC | 555.1 | -4.4 | 0.5 | -8.0 | 1.0E-15 | 3.2E-15 | BT20/MB468 |
| >hg38_21_+_8433172_8446622_RNA45SN1_45S_50ntFlanks@4472.4494.23\|CTTAGCTGAGTGTCCCGCGGGGC | >hg38_21_+_8433172_8446622_RNA45SN1_45S_50ntFlanks | 4472 | 4494 | 23 | CTTAGCTGAGTGTCCCGCGGGGC | 354.5 | -10.5 | 1.3 | -8.0 | 1.0E-15 | 3.2E-15 | BT20/MB468 |
| >MI0000087\|hsa-mir-29a&WithFlank&7\|-\|130876741\|130876816@48.67.20(+1G)&[MIMAT0000086&hsa-miR-29a-3p&offsets\|0\|-2(+1G);m-26&7\|-\|130876749\|130876769&offsets\|0\|-1(+1G)]\|TAGCACCATCTGAAATCGGTG | >MI0000087\|hsa-mir-29a&WithFlank&7\|-\|130876741\|130876816 | 48 | 67 | 20(+1G) | TAGCACCATCTGAAATCGGTG | 2135.2 | 10.7 | 1.3 | 8.0 | 1.0E-15 | 3.2E-15 | BT20/MB468 |
| >hg38_21_+_8433172_8446622_RNA45SN1_45S_50ntFlanks@475.518.44\|CTTCGTGATCGATGTGGTGACGTCGTGCTCTCCCGGGCCGGGTC | >hg38_21_+_8433172_8446622_RNA45SN1_45S_50ntFlanks | 475 | 518 | 44 | CTTCGTGATCGATGTGGTGACGTCGTGCTCTCCCGGGCCGGGTC | 1190.9 | 7.7 | 1.0 | 8.0 | 1.0E-15 | 3.2E-15 | BT20/MB468 |
| >MI0000286\|hsa-mir-210&WithFlank&11\|-\|568083\|568204@72.94.23&[MIMAT0000267&hsa-miR-210-3p&offsets\|0\|+1;m-102&11\|-\|568112\|568133&offsets\|0\|+1]\|CTGTGCGTGTGACAGCGGCTGAT | >MI0000286\|hsa-mir-210&WithFlank&11\|-\|568083\|568204 | 72 | 94 | 23 | CTGTGCGTGTGACAGCGGCTGAT | 588.7 | 8.9 | 1.1 | 8.0 | 1.0E-15 | 3.2E-15 | MB231/MB468 |
| >MI0000299\|hsa-mir-222&WithFlank&X\|-\|45747009\|45747130@75.96.22(+1G)&[MIMAT0000279&hsa-miR-222-3p&offsets\|0\|+1(+1G);m-140&X\|-\|45747033\|45747056&offsets\|0\|-2(+1G)]\|AGCTACATCTGGCTACTGGGTCG | >MI0000299\|hsa-mir-222&WithFlank&X\|-\|45747009\|45747130 | 75 | 96 | 22(+1G) | AGCTACATCTGGCTACTGGGTCG | 699.2 | 7.1 | 0.9 | 8.0 | 1.0E-15 | 3.2E-15 | BT20/MB468 |
| >MI0001445\|hsa-mir-423&WithFlank&17\|+\|30117073\|30117178@59.82.24&[MIMAT0001340&hsa-miR-423-3p&offsets\|0\|+1;m-100&17\|+\|30117131\|30117153&offsets\|0\|+1]\|AGCTCGGTCTGAGGCCCCTCAGTC | >MI0001445\|hsa-mir-423&WithFlank&17\|+\|30117073\|30117178 | 59 | 82 | 24 | AGCTCGGTCTGAGGCCCCTCAGTC | 890.2 | 9.6 | 1.2 | 8.0 | 1.0E-15 | 3.2E-15 | MB231/MB468 |
| >MI0000482\|hsa-mir-185&WithFlank&22\|+\|20033133\|20033226@21.42.22&[MIMAT0000455&hsa-miR-185-5p&offsets\|0\|0;m-145&22\|+\|20033153\|20033174&offsets\|0\|0]\|TGGAGAGAAAGGCAGTTCCTGA | >MI0000482\|hsa-mir-185&WithFlank&22\|+\|20033133\|20033226 | 21 | 42 | 22 | TGGAGAGAAAGGCAGTTCCTGA | 3158.9 | 5.9 | 0.7 | 8.0 | 1.0E-15 | 3.3E-15 | BT20/MB468 |
| >hg38_21_+_8433172_8446622_RNA45SN1_45S_50ntFlanks@11739.11768.30\|TCTCTTAAGGTAGCCAAATGCCTCGTCATC | >hg38_21_+_8433172_8446622_RNA45SN1_45S_50ntFlanks | 11739 | 11768 | 30 | TCTCTTAAGGTAGCCAAATGCCTCGTCATC | 263.1 | -10.0 | 1.3 | -8.0 | 1.1E-15 | 3.3E-15 | BT20/MB468 |
| >hg38_21_+_8433172_8446622_RNA45SN1_45S_50ntFlanks@11576.11600.25\|CTTAGAACTGGTGCGGACCAGGGGA | >hg38_21_+_8433172_8446622_RNA45SN1_45S_50ntFlanks | 11576 | 11600 | 25 | CTTAGAACTGGTGCGGACCAGGGGA | 493.2 | -5.2 | 0.6 | -8.0 | 1.1E-15 | 3.4E-15 | MB231/MB468 |

<table>
<tr><td>>hg38_21_+_8433172_8446622_RNA45SN1_45S_50ntFlanks@4008.4042.35|CTCGGGCCGATCGCACGCCCCCCGTGGCGGCGACG</td><td>>hg38_21_+_8433172_8446622_RNA45SN1_45S_50ntFlanks</td><td>4008</td><td>4042</td><td>35</td><td>CTCGGGCCGATCGCACGCCCCCCGTGGCGGCGACG</td><td>179.9</td><td>-3.7</td><td>0.5</td><td>-8.0</td><td>1.1E-15</td><td>3.4E-15</td><td>BT20/MB468</td></tr>
<tr><td>>hg38_21_+_8433172_8446622_RNA45SN1_45S_50ntFlanks@10410.10428.19|TCAGTCGGTCCTGAGAGAT</td><td>>hg38_21_+_8433172_8446622_RNA45SN1_45S_50ntFlanks</td><td>10410</td><td>10428</td><td>19</td><td>TCAGTCGGTCCTGAGAGAT</td><td>489.7</td><td>-4.6</td><td>0.6</td><td>-8.0</td><td>1.1E-15</td><td>3.4E-15</td><td>MB231/MB468</td></tr>
<tr><td>>hg38_21_+_8433172_8446622_RNA45SN1_45S_50ntFlanks@475.495.21|CTTCGTGATCGATGTGGTGAC</td><td>>hg38_21_+_8433172_8446622_RNA45SN1_45S_50ntFlanks</td><td>475</td><td>495</td><td>21</td><td>CTTCGTGATCGATGTGGTGAC</td><td>1064.0</td><td>5.6</td><td>0.7</td><td>8.0</td><td>1.1E-15</td><td>3.4E-15</td><td>BT20/MB468</td></tr>
<tr><td>>hg38_21_+_8433172_8446622_RNA45SN1_45S_50ntFlanks@6772.6803.32|GTTCCTCCCGGGGCTACGCCTGTCTGAGCGTC</td><td>>hg38_21_+_8433172_8446622_RNA45SN1_45S_50ntFlanks</td><td>6772</td><td>6803</td><td>32</td><td>GTTCCTCCCGGGGCTACGCCTGTCTGAGCGTC</td><td>318.3</td><td>-4.0</td><td>0.5</td><td>-8.0</td><td>1.1E-15</td><td>3.5E-15</td><td>MB231/MB468</td></tr>
<tr><td>>hg38_21_+_8433172_8446622_RNA45SN1_45S_50ntFlanks@10684.10710.27|CGGCGGCGTCCGGTGAGCTCTCGCTGG</td><td>>hg38_21_+_8433172_8446622_RNA45SN1_45S_50ntFlanks</td><td>10684</td><td>10710</td><td>27</td><td>CGGCGGCGTCCGGTGAGCTCTCGCTGG</td><td>6187.2</td><td>-4.3</td><td>0.5</td><td>-8.0</td><td>1.1E-15</td><td>3.5E-15</td><td>MB231/MB468</td></tr>
<tr><td>>hg38_21_+_8433172_8446622_RNA45SN1_45S_50ntFlanks@4560.4587.28|CTAGGAATAATGGAATAGGACCGCGGTT</td><td>>hg38_21_+_8433172_8446622_RNA45SN1_45S_50ntFlanks</td><td>4560</td><td>4587</td><td>28</td><td>CTAGGAATAATGGAATAGGACCGCGGTT</td><td>276.2</td><td>-10.1</td><td>1.3</td><td>-8.0</td><td>1.1E-15</td><td>3.5E-15</td><td>BT20/MB468</td></tr>
<tr><td>>hg38_21_+_8433172_8446622_RNA45SN1_45S_50ntFlanks@10681.10709.29|TTCCGGCGGCGTCCGGTGAGCTCTCGCTG</td><td>>hg38_21_+_8433172_8446622_RNA45SN1_45S_50ntFlanks</td><td>10681</td><td>10709</td><td>29</td><td>TTCCGGCGGCGTCCGGTGAGCTCTCGCTG</td><td>396.8</td><td>-5.5</td><td>0.7</td><td>-8.0</td><td>1.1E-15</td><td>3.5E-15</td><td>MB231/MB468</td></tr>
<tr><td>>am_trna111_HisGTG_1_-_147774845_147774916@-1G.26.27,trna118_HisGTG_1_-_145396881_145396952@-1G.26.27,trna16_HisGTG_1_+_146544773_146544844@-1G.26.27,trna1_HisGTG_15_+_45493349_45493420@-1G.26.27,trna21_HisGTG_1_+_147753471_147753542@-1G.26.27,trna33_HisGTG_6_+_27125906_27125977@-1G.26.27,trna7_HisGTG_9_-_14433938_14434009@-1G.26.27,trna8_HisGTG_15_-_45492611_45492682@-1G.26.27,trna9_HisGTG_15_-_45490804_45490875@-1G.26.27|GGCCGTGATCGTATAGTGGTTAGTACT</td><td>>am_trna111_HisGTG_1_-_147774845_147774916</td><td>-1G</td><td>26</td><td>27</td><td>GGCCGTGATCGTATAGTGGTTAGTACT</td><td>4913.9</td><td>6.6</td><td>0.8</td><td>8.0</td><td>1.1E-15</td><td>3.5E-15</td><td>MB231/MB468</td></tr>
<tr><td>>MI0000273|hsa-mir-183&WithFlank&7|-|129774899|129775020@34.55.22(+1A)&[MIMAT0000261&hsa-miR-183-5p&offsets|+1|+1(+1A);m-41&7|-|129774966|129774987&offsets|0|0(+1A)]|ATGGCACTGGTAGAATTCACTGA</td><td>>MI0000273|hsa-mir-183&WithFlank&7|-|129774899|129775020</td><td>34</td><td>55</td><td>22(+1A)</td><td>ATGGCACTGGTAGAATTCACTGA</td><td>449.2</td><td>9.3</td><td>1.2</td><td>8.0</td><td>1.1E-15</td><td>3.5E-15</td><td>BT20/MB468</td></tr>
<tr><td>>trna116_GluCTC_1_-_145399233_145399304@35.75.41,trna134_GluTTC_1_-_16861774_16861845@35.75.41,trna71_GluCTC_1_-_161439189_161439260@35.75.41,trna74_GluCTC_1_-_161431809_161431880@35.75.41,trna77_GluCTC_1_-_161424398_161424469@35.75.41,trna77_GluCTC_6_+_28949976_28950047@35.75.41,trna80_GluCTC_1_-_161417018_161417089@35.75.41,trna84_GluTTC_1_-_161391883_161391954@35.75.41,trna87_GluCTC_6_-_126101393_126101464@35.75.41|TCACCGCCGCGGCCCGGGTTCGATTCCCGGTCAGGGAACCA</td><td>>trna116_GluCTC_1_-_145399233_145399304</td><td>35</td><td>75</td><td>41</td><td>TCACCGCCGCGGCCCGGGTTCGATTCCCGGTCAGGGAACCA</td><td>989.0</td><td>7.4</td><td>0.9</td><td>8.0</td><td>1.1E-15</td><td>3.5E-15</td><td>MB231/MB468</td></tr>
<tr><td>>hg38_21_+_8433172_8446622_RNA45SN1_45S_50ntFlanks@8324.8356.33|CCGATAGTCAACAAGTACCGTAAGGGAAAGTTG</td><td>>hg38_21_+_8433172_8446622_RNA45SN1_45S_50ntFlanks</td><td>8324</td><td>8356</td><td>33</td><td>CCGATAGTCAACAAGTACCGTAAGGGAAAGTTG</td><td>245.9</td><td>-9.9</td><td>1.2</td><td>-8.0</td><td>1.1E-15</td><td>3.6E-15</td><td>BT20/MB468</td></tr>
<tr><td>>hg38_21_+_8433172_8446622_RNA45SN1_45S_50ntFlanks@10808.10840.33|CAATGTAGGTAAGGGAAGTCGGCAAGCCGGATC</td><td>>hg38_21_+_8433172_8446622_RNA45SN1_45S_50ntFlanks</td><td>10808</td><td>10840</td><td>33</td><td>CAATGTAGGTAAGGGAAGTCGGCAAGCCGGATC</td><td>522.6</td><td>-4.8</td><td>0.6</td><td>-8.0</td><td>1.1E-15</td><td>3.6E-15</td><td>MB231/MB468</td></tr>
<tr><td>>hg38_21_+_8433172_8446622_RNA45SN1_45S_50ntFlanks@12416.12444.29|TTCGATGTCGGCTCTTCCTATCATTGTGA</td><td>>hg38_21_+_8433172_8446622_RNA45SN1_45S_50ntFlanks</td><td>12416</td><td>12444</td><td>29</td><td>TTCGATGTCGGCTCTTCCTATCATTGTGA</td><td>2446.1</td><td>5.5</td><td>0.7</td><td>8.0</td><td>1.1E-15</td><td>3.6E-15</td><td>MB231/MB468</td></tr>
<tr><td>>hg38_21_+_8433172_8446622_RNA45SN1_45S_50ntFlanks@12934.12965.32|CTAAACCATTCGTAGACGACCTGCTTCTGGGT</td><td>>hg38_21_+_8433172_8446622_RNA45SN1_45S_50ntFlanks</td><td>12934</td><td>12965</td><td>32</td><td>CTAAACCATTCGTAGACGACCTGCTTCTGGGT</td><td>423.4</td><td>-4.8</td><td>0.6</td><td>-8.0</td><td>1.1E-15</td><td>3.6E-15</td><td>BT20/MB468</td></tr>
<tr><td>>hg38_21_+_8433172_8446622_RNA45SN1_45S_50ntFlanks@12133.12150.18|CTCCGGGGACAGTGCCAG</td><td>>hg38_21_+_8433172_8446622_RNA45SN1_45S_50ntFlanks</td><td>12133</td><td>12150</td><td>18</td><td>CTCCGGGGACAGTGCCAG</td><td>927.5</td><td>-8.7</td><td>1.1</td><td>-8.0</td><td>1.1E-15</td><td>3.6E-15</td><td>MB231/MB468</td></tr>
<tr><td>>MI0000650|hsa-mir-200c&WithFlank&12|+|6963693|6963772@50.72.23(+2U)&[MIMAT0000617&hsa-miR-200c-3p&offsets|0|0(+2U);m-54&12|+|6963742|6963764&offsets|0|0(+2U)]|TAATACTGCCGGGTAATGATGGATT</td><td>>MI0000650|hsa-mir-200c&WithFlank&12|+|6963693|6963772</td><td>50</td><td>72</td><td>23(+2U)</td><td>TAATACTGCCGGGTAATGATGGATT</td><td>373.3</td><td>8.5</td><td>1.0</td><td>8.3</td><td>7.9E-17</td><td>3.6E-15</td><td>BT20/MB231</td></tr>
<tr><td>>MI0005756|hsa-mir-934&WithFlank&X|+|136550872|136550966@21.42.22(+1C)&[MIMAT0004977&hsa-miR-934&offsets|0|0(+1C);m-943&X|+|136550892|136550913&offsets|0|0(+1C)]|TGTCTACTACTGGAGACACTGGC</td><td>>MI0005756|hsa-mir-934&WithFlank&X|+|136550872|136550966</td><td>21</td><td>42</td><td>22(+1C)</td><td>TGTCTACTACTGGAGACACTGGC</td><td>303.2</td><td>11.7</td><td>1.4</td><td>8.3</td><td>7.8E-17</td><td>3.6E-15</td><td>BT20/MB231</td></tr>
<tr><td>>MI0000079|hsa-mir-23a&WithFlank&19|-|13836581|13836665@54.73.20(+1A)&[MIMAT0000078&hsa-miR-23a-3p&offsets|+3|+2(+1A);m-31&19|-|13836595|13836615&offsets|+3|+2(+1A)]|ACATTGCCAGGGATTTCCAAA</td><td>>MI0000079|hsa-mir-23a&WithFlank&19|-|13836581|13836665</td><td>54</td><td>73</td><td>20(+1A)</td><td>ACATTGCCAGGGATTTCCAAA</td><td>1052.5</td><td>29.1</td><td>3.6</td><td>8.0</td><td>1.2E-15</td><td>3.6E-15</td><td>MB231/MB468</td></tr>
<tr><td>>hg38_21_+_8433172_8446622_RNA45SN1_45S_50ntFlanks@3771.3792.22|CACGGCCGGTACAGTGAAACTG</td><td>>hg38_21_+_8433172_8446622_RNA45SN1_45S_50ntFlanks</td><td>3771</td><td>3792</td><td>22</td><td>CACGGCCGGTACAGTGAAACTG</td><td>330.4</td><td>-6.9</td><td>0.9</td><td>-8.0</td><td>1.2E-15</td><td>3.6E-15</td><td>BT20/MB468</td></tr>
<tr><td>>hg38_21_+_8433172_8446622_RNA45SN1_45S_50ntFlanks@12456.12481.26|CCAAGCGTTGGATTGTTCACCCACTA</td><td>>hg38_21_+_8433172_8446622_RNA45SN1_45S_50ntFlanks</td><td>12456</td><td>12481</td><td>26</td><td>CCAAGCGTTGGATTGTTCACCCACTA</td><td>986.9</td><td>-5.3</td><td>0.7</td><td>-8.0</td><td>1.2E-15</td><td>3.6E-15</td><td>MB231/MB468</td></tr>
<tr><td>>am_MI0000786|hsa-mir-378a&WithFlank&5|+|149732819|149732896@50.69.20(+1U)&[MIMAT0000732&hsa-miR-378a-3p&offsets|+1|-1(+1U);m-44&5|+|149732867|149732888&offsets|+1|-1(+1U)]|CTGGACTTGGAGTCAGAAGGT</td><td>>am_MI0000786|hsa-mir-378a&WithFlank&5|+|149732819|149732896</td><td>50</td><td>69</td><td>20(+1U)</td><td>CTGGACTTGGAGTCAGAAGGT</td><td>600.0</td><td>5.8</td><td>0.7</td><td>8.0</td><td>1.2E-15</td><td>3.7E-15</td><td>MB231/MB468</td></tr>
<tr><td>>MI0000299|hsa-mir-222&WithFlank&X|-|45747009|45747130@75.96.22&[MIMAT0000279&hsa-miR-222-3p&offsets|0|+1;m-140&X|-|45747033|45747056&offsets|0|-2]|AGCTACATCTGGCTACTGGGTC</td><td>>MI0000299|hsa-mir-222&WithFlank&X|-|45747009|45747130</td><td>75</td><td>96</td><td>22</td><td>AGCTACATCTGGCTACTGGGTC</td><td>1230.3</td><td>6.1</td><td>0.8</td><td>8.0</td><td>1.2E-15</td><td>3.7E-15</td><td>BT20/MB468</td></tr>
<tr><td>>hg38_MT_+_1621_3279_RNR2_16S_50ntFlanks@51.81.31|GCTAAACCTAGCCCCAAACCCACTCCACCTT</td><td>>hg38_MT_+_1621_3279_RNR2_16S_50ntFlanks</td><td>51</td><td>81</td><td>31</td><td>GCTAAACCTAGCCCCAAACCCACTCCACCTT</td><td>620.0</td><td>8.1</td><td>1.0</td><td>8.0</td><td>1.2E-15</td><td>3.7E-15</td><td>BT20/MB468</td></tr>
<tr><td>>hg38_21_+_8433172_8446622_RNA45SN1_45S_50ntFlanks@3843.3860.18|CTACTTGGATAACTGTGG</td><td>>hg38_21_+_8433172_8446622_RNA45SN1_45S_50ntFlanks</td><td>3843</td><td>3860</td><td>18</td><td>CTACTTGGATAACTGTGG</td><td>256.0</td><td>-3.8</td><td>0.5</td><td>-8.0</td><td>1.2E-15</td><td>3.8E-15</td><td>BT20/MB468</td></tr>
<tr><td>>hg38_21_+_8433172_8446622_RNA45SN1_45S_50ntFlanks@12589.12614.26|GTTCAGACATTTGGTGTATGTGCTTG</td><td>>hg38_21_+_8433172_8446622_RNA45SN1_45S_50ntFlanks</td><td>12589</td><td>12614</td><td>26</td><td>GTTCAGACATTTGGTGTATGTGCTTG</td><td>506.4</td><td>-11.0</td><td>1.4</td><td>-8.0</td><td>1.2E-15</td><td>3.8E-15</td><td>BT20/MB468</td></tr>
<tr><td>>hg38_21_+_8433172_8446622_RNA45SN1_45S_50ntFlanks@12001.12031.31|CTGCGGGCCGCCGGTGAAATACCACTACTCT</td><td>>hg38_21_+_8433172_8446622_RNA45SN1_45S_50ntFlanks</td><td>12001</td><td>12031</td><td>31</td><td>CTGCGGGCCGCCGGTGAAATACCACTACTCT</td><td>244.4</td><td>-4.3</td><td>0.5</td><td>-8.0</td><td>1.2E-15</td><td>3.8E-15</td><td>BT20/MB468</td></tr>
<tr><td>>MI0000746|hsa-mir-99b&WithFlank&19|+|51692606|51692687@13.34.22(+1C)&[MIMAT0000689&hsa-miR-99b-5p&offsets|0|0(+1C);m-10&19|+|51692618|51692639&offsets|0|0(+1C)]|CACCCGTAGAACCGACCTTGCGC</td><td>>MI0000746|hsa-mir-99b&WithFlank&19|+|51692606|51692687</td><td>13</td><td>34</td><td>22(+1C)</td><td>CACCCGTAGAACCGACCTTGCGC</td><td>672.8</td><td>8.6</td><td>1.1</td><td>8.0</td><td>1.2E-15</td><td>3.9E-15</td><td>BT20/MB468</td></tr>
<tr><td>>trna111_HisGTG_1_-_147774845_147774916@-1A.29.30,trna118_HisGTG_1_-_145396881_145396952@-1A.29.30,trna16_HisGTG_1_+_146544773_146544844@-1A.29.30,trna1_HisGTG_15_+_45493349_45493420@-1A.29.30,trna21_HisGTG_1_+_147753471_147753542@-1A.29.30,trna33_HisGTG_6_+_27125906_27125977@-1A.29.30,trna7_HisGTG_9_-_14433938_14434009@-1A.29.30,trna8_HisGTG_15_-_45492611_45492682@-1A.29.30,trna9_HisGTG_15_-_45490804_45490875@-1A.29.30|AGCCGTGATCGTATAGTGGTTAGTACTCTG</td><td>>trna111_HisGTG_1_-_147774845_147774916</td><td>-1A</td><td>29</td><td>30</td><td>AGCCGTGATCGTATAGTGGTTAGTACTCTG</td><td>322.4</td><td>-10.3</td><td>1.3</td><td>-8.0</td><td>1.2E-15</td><td>3.9E-15</td><td>BT20/MB468</td></tr>
<tr><td>>MI0000289|hsa-mir-181a-1&WithFlank&1|-|198859038|198859159@30.50.21;MI0000269|hsa-mir-181a-2&WithFlank&9|+|124692436|124692557@45.65.21&[MIMAT0000256&hsa-miR-181a-5p&offsets|0|-2;m-32&1|-|198859108|198859130&offsets|0|-2];[MIMAT0000256_1&hsa-miR-181a-5p&offsets|0|-2;m-32&9|+|124692480|124692502&offsets|0|-2]|AACATTCAACGCTGTCGGTGA</td><td>>MI0000289|hsa-mir-181a-1&WithFlank&1|-|198859038|198859159</td><td>30</td><td>50</td><td>21</td><td>AACATTCAACGCTGTCGGTGA</td><td>1597.6</td><td>6.4</td><td>0.8</td><td>8.0</td><td>1.2E-15</td><td>3.9E-15</td><td>MB231/MB468</td></tr>
<tr><td>>MI0016887|hsa-mir-4521&WithFlank&17|+|8186939|8187010@14.34.21&[MIMAT0019058&hsa-miR-4521&offsets|+4|+3;m-765&17|+|8186948|8186969&offsets|+4|+3]|AGGAAGTCCTGTGCTCAGTTT</td><td>>MI0016887|hsa-mir-4521&WithFlank&17|+|8186939|8187010</td><td>14</td><td>34</td><td>21</td><td>AGGAAGTCCTGTGCTCAGTTT</td><td>565.1</td><td>9.6</td><td>1.2</td><td>8.0</td><td>1.2E-15</td><td>3.9E-15</td><td>BT20/MB468</td></tr>
<tr><td>>hg38_21_+_8433172_8446622_RNA45SN1_45S_50ntFlanks@10408.10431.24|GGTCAGTCGGTCCTGAGAGATGGG</td><td>>hg38_21_+_8433172_8446622_RNA45SN1_45S_50ntFlanks</td><td>10408</td><td>10431</td><td>24</td><td>GGTCAGTCGGTCCTGAGAGATGGG</td><td>371.6</td><td>-10.5</td><td>1.3</td><td>-8.0</td><td>1.3E-15</td><td>3.9E-15</td><td>BT20/MB468</td></tr>
<tr><td>>hg38_21_+_8433172_8446622_RNA45SN1_45S_50ntFlanks@9504.9533.30|GTGAACTATGCCTGGGCAGGGCGAAGCCAG</td><td>>hg38_21_+_8433172_8446622_RNA45SN1_45S_50ntFlanks</td><td>9504</td><td>9533</td><td>30</td><td>GTGAACTATGCCTGGGCAGGGCGAAGCCAG</td><td>739.8</td><td>-11.5</td><td>1.4</td><td>-8.0</td><td>1.3E-15</td><td>3.9E-15</td><td>BT20/MB468</td></tr>
<tr><td>>hg38_1_-_228634819_228635039_RNA5S12_5S_50ntFlanks@152.169.18|TACCGGGTGCTGTAGGCT</td><td>>hg38_1_-_228634819_228635039_RNA5S12_5S_50ntFlanks</td><td>152</td><td>169</td><td>18</td><td>TACCGGGTGCTGTAGGCT</td><td>703.5</td><td>6.0</td><td>0.8</td><td>8.0</td><td>1.3E-15</td><td>3.9E-15</td><td>MB231/MB468</td></tr>
<tr><td>>hg38_21_+_8433172_8446622_RNA45SN1_45S_50ntFlanks@13002.13025.24|CGATCTATTGAAAGTCAGCCCTCG</td><td>>hg38_21_+_8433172_8446622_RNA45SN1_45S_50ntFlanks</td><td>13002</td><td>13025</td><td>24</td><td>CGATCTATTGAAAGTCAGCCCTCG</td><td>297.1</td><td>-10.2</td><td>1.3</td><td>-8.0</td><td>1.3E-15</td><td>4.0E-15</td><td>BT20/MB468</td></tr>
<tr><td>>hg38_21_+_8433172_8446622_RNA45SN1_45S_50ntFlanks@8198.8225.28|TGGACGGTGTGAGGCCGGTAGCGGCCCC</td><td>>hg38_21_+_8433172_8446622_RNA45SN1_45S_50ntFlanks</td><td>8198</td><td>8225</td><td>28</td><td>TGGACGGTGTGAGGCCGGTAGCGGCCCC</td><td>227.1</td><td>-4.5</td><td>0.6</td><td>-8.0</td><td>1.3E-15</td><td>4.1E-15</td><td>BT20/MB468</td></tr>
<tr><td>>hg38_21_+_8433172_8446622_RNA45SN1_45S_50ntFlanks@4240.4258.19|ACTCTTTCGAGGCCCTGTA</td><td>>hg38_21_+_8433172_8446622_RNA45SN1_45S_50ntFlanks</td><td>4240</td><td>4258</td><td>19</td><td>ACTCTTTCGAGGCCCTGTA</td><td>196.0</td><td>-4.7</td><td>0.6</td><td>-8.0</td><td>1.3E-15</td><td>4.1E-15</td><td>MB231/MB468</td></tr>
</table>

| | | | | | | | | | | | | |
|---|---|---|---|---|---|---|---|---|---|---|---|---|
| >hg38_21_+_8433172_8446622_RNA45SN1_45S_50ntFlanks@8191.8214.24\|CAGCCCGTGGACGGTGTGAGGCCG | >hg38_21_+_8433172_8446622_RNA45SN1_45S_50ntFlanks | 8191 | 8214 | 24 | CAGCCCGTGGACGGTGTGAGGCCG | 531.1 | -4.6 | 0.6 | -8.0 | 1.3E-15 | 4.1E-15 | BT20/MB468 |
| >MI0000283\|hsa-mir-203a&WithFlank&14\|+\|104117399\|104117520@71.90.20&[MIMAT0000264&hsa-miR-203a-3p&offsets\|0\|-2;m-79&14\|+\|104117469\|104117490&offsets\|0\|-2]\|GTGAAATGTTTAGGACCACT | >MI0000283\|hsa-mir-203a&WithFlank&14\|+\|104117399\|104117520 | 71 | 90 | 20 | GTGAAATGTTTAGGACCACT | 1682.6 | 7.5 | 0.9 | 8.0 | 1.3E-15 | 4.2E-15 | BT20/MB468 |
| >MI0000764\|hsa-mir-363&WithFlank&X\|-\|134169372\|134169458@56.76.21&[MIMAT0000707&hsa-miR-363-3p&offsets\|0\|-1;m-178&X\|-\|134169381\|134169403&offsets\|0\|-2]\|AATTGCACGGTATCCATCTGT | >MI0000764\|hsa-mir-363&WithFlank&X\|-\|134169372\|134169458 | 56 | 76 | 21 | AATTGCACGGTATCCATCTGT | 564.7 | 6.8 | 0.8 | 8.0 | 1.3E-15 | 4.2E-15 | BT20/MB468 |
| >MI0000078\|hsa-mir-22&WithFlank&17\|-\|1713897\|1713993@21.42.22&[MIMAT0004495&hsa-miR-22-5p&offsets\|0\|0;m-217&17\|-\|1713953\|1713973&offsets\|0\|+1]\|AGTTCTTCAGTGGCAAGCTTTA | >MI0000078\|hsa-mir-22&WithFlank&17\|-\|1713897\|1713993 | 21 | 42 | 22 | AGTTCTTCAGTGGCAAGCTTTA | 856.3 | 9.5 | 1.2 | 8.0 | 1.3E-15 | 4.2E-15 | BT20/MB468 |
| >MI0000746\|hsa-mir-99b&WithFlank&19\|+\|51692606\|51692687@13.32.20(+2U)&[MIMAT0000689&hsa-miR-99b-5p&offsets\|0\|-2(+2U);m-10&19\|+\|51692618\|51692639&offsets\|0\|-2(+2U)]\|CACCCGTAGAACCGACCTTGTT | >MI0000746\|hsa-mir-99b&WithFlank&19\|+\|51692606\|51692687 | 13 | 32 | 20(+2U) | CACCCGTAGAACCGACCTTGTT | 2168.3 | 9.0 | 1.1 | 8.3 | 9.4E-17 | 4.2E-15 | BT20/MB231 |
| >hg38_21_+_8433172_8446622_RNA45SN1_45S_50ntFlanks@10240.10259.20\|CTACGCCGCGACGAGTAGGA | >hg38_21_+_8433172_8446622_RNA45SN1_45S_50ntFlanks | 10240 | 10259 | 20 | CTACGCCGCGACGAGTAGGA | 465.6 | 5.8 | 0.7 | 8.0 | 1.4E-15 | 4.2E-15 | BT20/MB468 |
| >MI0003834\|hsa-mir-769&WithFlank&19\|+\|46018926\|46019055@36.56.21(+1C)&[MIMAT0003886&hsa-miR-769-5p&offsets\|0\|-1(+1C);m-216&19\|+\|46018961\|46018982&offsets\|0\|-1(+1C)]\|TGAGACCTCTGGGTTCTGAGCC | >MI0003834\|hsa-mir-769&WithFlank&19\|+\|46018926\|46019055 | 36 | 56 | 21(+1C) | TGAGACCTCTGGGTTCTGAGCC | 681.0 | 8.5 | 1.1 | 8.0 | 1.4E-15 | 4.4E-15 | MB231/MB468 |
| >hg38_21_+_8433172_8446622_RNA45SN1_45S_50ntFlanks@9545.9563.19\|GTGGAGGTCCGTAGCGGTC | >hg38_21_+_8433172_8446622_RNA45SN1_45S_50ntFlanks | 9545 | 9563 | 19 | GTGGAGGTCCGTAGCGGTC | 2244.8 | -5.5 | 0.7 | -8.0 | 1.4E-15 | 4.5E-15 | MB231/MB468 |
| >hg38_21_+_8433172_8446622_RNA45SN1_45S_50ntFlanks@6656.6681.26\|CTTAGCGGTGGATCACTCGGCTCGTG | >hg38_21_+_8433172_8446622_RNA45SN1_45S_50ntFlanks | 6656 | 6681 | 26 | CTTAGCGGTGGATCACTCGGCTCGTG | 1845.0 | -4.6 | 0.6 | -8.0 | 1.5E-15 | 4.5E-15 | BT20/MB468 |
| >hg38_21_+_8433172_8446622_RNA45SN1_45S_50ntFlanks@12942.12966.25\|TTCGTAGACGACCTGCTTCTGGGTC | >hg38_21_+_8433172_8446622_RNA45SN1_45S_50ntFlanks | 12942 | 12966 | 25 | TTCGTAGACGACCTGCTTCTGGGTC | 625.7 | -11.3 | 1.4 | -8.0 | 1.5E-15 | 4.5E-15 | BT20/MB468 |
| >MI0000750\|hsa-mir-26a-2&WithFlank&12\|-\|57824603\|57824698@20.39.20;MI0000083\|hsa-mir-26a-1&WithFlank&3\|+\|37969398\|37969486@16.35.20&[MIMAT0000082&hsa-miR-26a-5p&offsets\|0\|-2;m-33&12\|-\|57824658\|57824679&offsets\|0\|-2];[MIMAT0000082_1&hsa-miR-26a-5p&offsets\|0\|-2;m-34&3\|+\|37969413\|37969434&offsets\|0\|-2]\|TTCAAGTAATCCAGGATAGG | >MI0000750\|hsa-mir-26a-2&WithFlank&12\|-\|57824603\|57824698 | 20 | 39 | 20 | TTCAAGTAATCCAGGATAGG | 2075.8 | 7.4 | 0.9 | 8.0 | 1.5E-15 | 4.5E-15 | BT20/MB468 |
| >hg38_21_+_8433172_8446622_RNA45SN1_45S_50ntFlanks@8154.8181.28\|CCGCTCGTGGGGGGCCCAAGTCCTTCTG | >hg38_21_+_8433172_8446622_RNA45SN1_45S_50ntFlanks | 8154 | 8181 | 28 | CCGCTCGTGGGGGGCCCAAGTCCTTCTG | 893.0 | -4.3 | 0.5 | -8.0 | 1.5E-15 | 4.5E-15 | MB231/MB468 |
| >hg38_21_+_8433172_8446622_RNA45SN1_45S_50ntFlanks@6651.6700.50\|CGACTCTTAGCGGTGGATCACTCGGCTCGTGCGTCGATGAAGAACGCAGC | >hg38_21_+_8433172_8446622_RNA45SN1_45S_50ntFlanks | 6651 | 6700 | 50 | CGACTCTTAGCGGTGGATCACTCGGCTCGTGCGTCGATGAAGAACGCAGC | 1965.3 | 8.6 | 1.1 | 8.0 | 1.5E-15 | 4.6E-15 | BT20/MB468 |
| >hg38_21_+_8433172_8446622_RNA45SN1_45S_50ntFlanks@7985.8008.24\|GATCAGACGTGGCGACCCGCTGAA | >hg38_21_+_8433172_8446622_RNA45SN1_45S_50ntFlanks | 7985 | 8008 | 24 | GATCAGACGTGGCGACCCGCTGAA | 323.5 | -10.3 | 1.3 | -8.0 | 1.5E-15 | 4.6E-15 | BT20/MB468 |
| >hg38_21_+_8433172_8446622_RNA45SN1_45S_50ntFlanks@9554.9573.20\|CGTAGCGGTCCTGACGTGCA | >hg38_21_+_8433172_8446622_RNA45SN1_45S_50ntFlanks | 9554 | 9573 | 20 | CGTAGCGGTCCTGACGTGCA | 10620.8 | -3.1 | 0.4 | -8.0 | 1.5E-15 | 4.6E-15 | BT20/MB468 |
| >hg38_21_+_8433172_8446622_RNA45SN1_45S_50ntFlanks@3873.3898.26\|TAATACATGCCGACGGGCGCTGACCC | >hg38_21_+_8433172_8446622_RNA45SN1_45S_50ntFlanks | 3873 | 3898 | 26 | TAATACATGCCGACGGGCGCTGACCC | 1219.3 | -6.8 | 0.8 | -8.0 | 1.5E-15 | 4.7E-15 | BT20/MB468 |
| >MI0000433\|hsa-let-7g&WithFlank&3\|-\|52268272\|52268367@11.29.19&[MIMAT0000414&hsa-let-7g-5p&offsets\|0\|-3;m-58&3\|-\|52268336\|52268357&offsets\|0\|-3]\|TGAGGTAGTAGTTTGTACA | >MI0000433\|hsa-let-7g&WithFlank&3\|-\|52268272\|52268367 | 11 | 29 | 19 | TGAGGTAGTAGTTTGTACA | 493.5 | 7.7 | 1.0 | 8.0 | 1.5E-15 | 4.8E-15 | BT20/MB468 |
| >hg38_21_+_8433172_8446622_RNA45SN1_45S_50ntFlanks@12998.13039.42\|GCTGCGATCTATTGAAAGTCAGCCCTCGACACAAGGGTTTGT | >hg38_21_+_8433172_8446622_RNA45SN1_45S_50ntFlanks | 12998 | 13039 | 42 | GCTGCGATCTATTGAAAGTCAGCCCTCGACACAAGGGTTTGT | 13085.0 | 8.5 | 1.1 | 8.0 | 1.5E-15 | 4.8E-15 | BT20/MB468 |
| >trna116_GluCTC_1_-_145399233_145399304@37.74.38,trna134_GluTTC_1_-_16861774_16861845@37.74.38,trna71_GluCTC_1_-_161439189_161439260@37.74.38,trna74_GluCTC_1_-_161431809_161431880@37.74.38,trna77_GluCTC_1_-_161424398_161424469@37.74.38,trna77_GluCTC_6_+_28949976_28950047@37.74.38,trna80_GluCTC_1_-_161417018_161417089@37.74.38,trna84_GluTTC_1_-_161391883_161391954@37.74.38,trna87_GluCTC_6_-_126101393_126101464@37.74.38\|ACCGCCGCGGCCCGGGTTCGATTCCCGGTCAGGGAACC | >trna116_GluCTC_1_-_145399233_145399304 | 37 | 74 | 38 | ACCGCCGCGGCCCGGGTTCGATTCCCGGTCAGGGAACC | 9542.0 | 11.0 | 1.4 | 8.0 | 1.5E-15 | 4.8E-15 | MB231/MB468 |
| >MI0000438\|hsa-mir-15b&WithFlank&3\|+\|160404582\|160404691@26.46.21&[MIMAT0000417&hsa-miR-15b-5p&offsets\|0\|-1;m-110&3\|+\|160404607\|160404628&offsets\|0\|-1]\|TAGCAGCACATCATGGTTTAC | >MI0000438\|hsa-mir-15b&WithFlank&3\|+\|160404582\|160404691 | 26 | 46 | 21 | TAGCAGCACATCATGGTTTAC | 1250.3 | 7.2 | 0.9 | 8.0 | 1.6E-15 | 4.8E-15 | BT20/MB468 |
| >hg38_21_+_8433172_8446622_RNA45SN1_45S_50ntFlanks@12456.12474.19\|CCAAGCGTTGGATTGTTCA | >hg38_21_+_8433172_8446622_RNA45SN1_45S_50ntFlanks | 12456 | 12474 | 19 | CCAAGCGTTGGATTGTTCA | 5441.8 | -7.1 | 0.9 | -8.0 | 1.6E-15 | 4.8E-15 | BT20/MB468 |
| >MI0000815\|hsa-mir-339&WithFlank&7\|-\|1022927\|1023032@21.43.23&[MIMAT0000764&hsa-miR-339-5p&offsets\|0\|0;m-163&7\|-\|1022990\|1023012&offsets\|0\|0]\|TCCCTGTCCTCCAGGAGCTCACG | >MI0000815\|hsa-mir-339&WithFlank&7\|-\|1022927\|1023032 | 21 | 43 | 23 | TCCCTGTCCTCCAGGAGCTCACG | 1419.7 | 6.4 | 0.8 | 8.0 | 1.6E-15 | 4.9E-15 | MB231/MB468 |
| >hg38_21_+_8433172_8446622_RNA45SN1_45S_50ntFlanks@4976.4993.18\|CCGGACACGGACAGGATT | >hg38_21_+_8433172_8446622_RNA45SN1_45S_50ntFlanks | 4976 | 4993 | 18 | CCGGACACGGACAGGATT | 408.0 | 7.8 | 1.0 | 8.0 | 1.6E-15 | 4.9E-15 | BT20/MB468 |
| >MI0000273\|hsa-mir-183&WithFlank&7\|-\|129774899\|129775020@34.53.20(+1G)&[MIMAT0000261&hsa-miR-183-5p&offsets\|+1\|-1(+1G);m-41&7\|-\|129774966\|129774987&offsets\|0\|-2(+1G)]\|ATGGCACTGGTAGAATTCACG | >MI0000273\|hsa-mir-183&WithFlank&7\|-\|129774899\|129775020 | 34 | 53 | 20(+1G) | ATGGCACTGGTAGAATTCACG | 887.1 | 10.2 | 1.3 | 8.0 | 1.6E-15 | 4.9E-15 | BT20/MB468 |
| >trnaMT_ValTAC_MT_+_1602_1670@41.72.32\|GAGATTTCAACTTAACTTGACCGCTCTGACCA | >trnaMT_ValTAC_MT_+_1602_1670 | 41 | 72 | 32 | GAGATTTCAACTTAACTTGACCGCTCTGACCA | 640.7 | 5.8 | 0.7 | 8.0 | 1.6E-15 | 4.9E-15 | MB231/MB468 |
| >hg38_21_+_8433172_8446622_RNA45SN1_45S_50ntFlanks@10922.10939.18\|CTGGACGAGGCGCCGCCG | >hg38_21_+_8433172_8446622_RNA45SN1_45S_50ntFlanks | 10922 | 10939 | 18 | CTGGACGAGGCGCCGCCG | 761.8 | -7.7 | 1.0 | -8.0 | 1.6E-15 | 4.9E-15 | MB231/MB468 |
| >trna10_AspGTC_12_-_125424193_125424264@37.73.37,trna12_AspGTC_12_-_125411891_125411962@37.73.37,trna144_AspGTC_6_-_27551236_27551307@37.73.37,trna38_AspGTC_17_-_8125556_8125627@37.73.37,trna45_AspGTC_6_+_27447453_27447524@37.73.37,trna48_AspGTC_6_+_27471523_27471594@37.73.37,trna4_AspGTC_12_+_96429799_96429870@37.73.37,trna69_AspGTC_1_-_161440205_161440276@37.73.37,trna72_AspGTC_1_-_161432824_161432895@37.73.37,trna75_AspGTC_1_-_161425414_161425485@37.73.37,trna78_AspGTC_1_-_161418033_161418104@37.73.37,trna81_AspGTC_1_-_161410615_161410686@37.73.37\|ACGCGGGAGACCGGGGTTCGATTCCCCGACGGGGAGC | >trna10_AspGTC_12_-_125424193_125424264 | 37 | 73 | 37 | ACGCGGGAGACCGGGGTTCGATTCCCCGACGGGGAGC | 5002.7 | 8.9 | 1.1 | 8.0 | 1.6E-15 | 5.0E-15 | MB231/MB468 |
| >hg38_21_+_8433172_8446622_RNA45SN1_45S_50ntFlanks@10408.10426.19\|GGTCAGTCGGTCCTGAGAG | >hg38_21_+_8433172_8446622_RNA45SN1_45S_50ntFlanks | 10408 | 10426 | 19 | GGTCAGTCGGTCCTGAGAG | 505.2 | -11.0 | 1.4 | -8.0 | 1.6E-15 | 5.0E-15 | BT20/MB468 |
| >hg38_21_+_8433172_8446622_RNA45SN1_45S_50ntFlanks@11821.11844.24\|CTATCCAGCGAAACCACAGCCAAG | >hg38_21_+_8433172_8446622_RNA45SN1_45S_50ntFlanks | 11821 | 11844 | 24 | CTATCCAGCGAAACCACAGCCAAG | 228.7 | -9.8 | 1.2 | -8.0 | 1.7E-15 | 5.1E-15 | BT20/MB468 |
| >hg38_MT_+_1621_3279_RNR2_16S_50ntFlanks@1216.1248.33\|CAACCTCCGAGCAGTACATGCTAAGACTTCACC | >hg38_MT_+_1621_3279_RNR2_16S_50ntFlanks | 1216 | 1248 | 33 | CAACCTCCGAGCAGTACATGCTAAGACTTCACC | 1249.1 | 9.8 | 1.2 | 8.0 | 1.6E-15 | 5.1E-15 | MB231/MB468 |
| >hg38_21_+_8433172_8446622_RNA45SN1_45S_50ntFlanks@10846.10868.23\|CTTCGGGATAAGGATTGGCTCTA | >hg38_21_+_8433172_8446622_RNA45SN1_45S_50ntFlanks | 10846 | 10868 | 23 | CTTCGGGATAAGGATTGGCTCTA | 874.2 | -6.9 | 0.9 | -8.0 | 1.7E-15 | 5.1E-15 | BT20/MB468 |
| >MI0000255\|hsa-mir-30d&WithFlank&8\|-\|134804870\|134804951@12.33.22(+1G)&[MIMAT0000245&hsa-miR-30d-5p&offsets\|0\|0(+1G);m-21&8\|-\|134804919\|134804940&offsets\|0\|0(+1G)]\|TGTAAACATCCCCGACTGGAAGG | >MI0000255\|hsa-mir-30d&WithFlank&8\|-\|134804870\|134804951 | 12 | 33 | 22(+1G) | TGTAAACATCCCCGACTGGAAGG | 1414.9 | 10.6 | 1.3 | 8.0 | 1.7E-15 | 5.1E-15 | BT20/MB468 |
| >hg38_21_+_8433172_8446622_RNA45SN1_45S_50ntFlanks@12459.12477.19\|AGCGTTGGATTGTTCACCC | >hg38_21_+_8433172_8446622_RNA45SN1_45S_50ntFlanks | 12459 | 12477 | 19 | AGCGTTGGATTGTTCACCC | 686.3 | -4.4 | 0.5 | -8.0 | 1.7E-15 | 5.1E-15 | BT20/MB468 |
| >MI0000440\|hsa-mir-27b&WithFlank&9\|+\|95085439\|95085547@67.87.21(+1U)&[MIMAT0000419&hsa-miR-27b-3p&offsets\|0\|0(+1U);m-39&9\|+\|95085505\|95085525&offsets\|0\|0(+1U)]\|TTCACAGTGGCTAAGTTCTGCT | >MI0000440\|hsa-mir-27b&WithFlank&9\|+\|95085439\|95085547 | 67 | 87 | 21(+1U) | TTCACAGTGGCTAAGTTCTGCT | 5767.0 | 7.3 | 0.9 | 8.0 | 1.7E-15 | 5.2E-15 | BT20/MB468 |
| >MI0016887\|hsa-mir-4521&WithFlank&17\|+\|8186939\|8187010@14.35.22&[MIMAT0019058&hsa-miR-4521&offsets\|+4\|+4;m-765&17\|+\|8186948\|8186969&offsets\|+4\|+4]\|AGGAAGTCCTGTGCTCAGTTTT | >MI0016887\|hsa-mir-4521&WithFlank&17\|+\|8186939\|8187010 | 14 | 35 | 22 | AGGAAGTCCTGTGCTCAGTTTT | 545.8 | 10.8 | 1.4 | 8.0 | 1.7E-15 | 5.2E-15 | BT20/MB468 |
| >hg38_21_+_8433172_8446622_RNA45SN1_45S_50ntFlanks@4590.4614.25\|ATTTTGTTGGTTTTCGGAACTGAGG | >hg38_21_+_8433172_8446622_RNA45SN1_45S_50ntFlanks | 4590 | 4614 | 25 | ATTTTGTTGGTTTTCGGAACTGAGG | 141.2 | -3.8 | 0.5 | -8.0 | 1.7E-15 | 5.3E-15 | BT20/MB468 |
| >MI0000289\|hsa-mir-181a-1&WithFlank&1\|-\|198859038\|198859159@30.51.22;MI0000269\|hsa-mir-181a-2&WithFlank&9\|+\|124692436\|124692557@45.66.22&[MIMAT0000256&hsa-miR-181a-5p&offsets\|0\|-1;m-32&1\|-\|198859108\|198859130&offsets\|0\|-1];[MIMAT0000256_1&hsa-miR-181a-5p&offsets\|0\|-1;m-32&9\|+\|124692480\|124692502&offsets\|0\|-1]\|AACATTCAACGCTGTCGGTGAG | >MI0000289\|hsa-mir-181a-1&WithFlank&1\|-\|198859038\|198859159 | 30 | 51 | 22 | AACATTCAACGCTGTCGGTGAG | 1735.4 | 6.7 | 0.8 | 8.0 | 1.7E-15 | 5.3E-15 | MB231/MB468 |
| >hg38_21_+_8433172_8446622_RNA45SN1_45S_50ntFlanks@10771.10797.27\|CAGGTCTCCAAGGTGAACAGCCTCTGG | >hg38_21_+_8433172_8446622_RNA45SN1_45S_50ntFlanks | 10771 | 10797 | 27 | CAGGTCTCCAAGGTGAACAGCCTCTGG | 447.5 | -6.7 | 0.8 | -8.0 | 1.7E-15 | 5.3E-15 | MB231/MB468 |

<table>
<tr><th></th><th></th><th></th><th></th><th></th><th></th><th></th><th></th><th></th><th></th><th></th><th></th><th></th></tr>
<tr><td>>hg38_21_+_8433172_8446622_RNA45SN1_45S_50ntFlanks@10375.10406.32|AGAAGGGTTCCATGTGAACAGCAGTTGAACAT</td><td>>hg38_21_+_8433172_8446622_RNA45SN1_45S_50ntFlanks</td><td>10375</td><td>10406</td><td>32</td><td>AGAAGGGTTCCATGTGAACAGCAGTTGAACAT</td><td>260.3</td><td>-10.0</td><td>1.3</td><td>-8.0</td><td>1.8E-15</td><td>5.4E-15</td><td>BT20/MB468</td></tr>
<tr><td>>hg38_21_+_8433172_8446622_RNA45SN1_45S_50ntFlanks@11571.11590.20|GCCGACTTAGAACTGGTGCG</td><td>>hg38_21_+_8433172_8446622_RNA45SN1_45S_50ntFlanks</td><td>11571</td><td>11590</td><td>20</td><td>GCCGACTTAGAACTGGTGCG</td><td>543.1</td><td>4.6</td><td>0.6</td><td>8.0</td><td>1.7E-15</td><td>5.4E-15</td><td>MB231/MB468</td></tr>
<tr><td>>hg38_21_+_8433172_8446622_RNA45SN1_45S_50ntFlanks@10420.10443.24|CTGAGAGATGGGCGAGCGCCGTTC</td><td>>hg38_21_+_8433172_8446622_RNA45SN1_45S_50ntFlanks</td><td>10420</td><td>10443</td><td>24</td><td>CTGAGAGATGGGCGAGCGCCGTTC</td><td>320.1</td><td>-7.2</td><td>0.9</td><td>-8.0</td><td>1.8E-15</td><td>5.4E-15</td><td>BT20/MB468</td></tr>
<tr><td>>hg38_21_+_8433172_8446622_RNA45SN1_45S_50ntFlanks@12636.12657.22|AGCTACCATCTGTGGGATTATG</td><td>>hg38_21_+_8433172_8446622_RNA45SN1_45S_50ntFlanks</td><td>12636</td><td>12657</td><td>22</td><td>AGCTACCATCTGTGGGATTATG</td><td>266.8</td><td>-10.1</td><td>1.3</td><td>-8.0</td><td>1.8E-15</td><td>5.4E-15</td><td>BT20/MB468</td></tr>
<tr><td>>am_trnaMT_PheGAA_MT_+_577_647@41.58.18|GTTTAGACGGGCTCACAT</td><td>>am_trnaMT_PheGAA_MT_+_577_647</td><td>41</td><td>58</td><td>18</td><td>GTTTAGACGGGCTCACAT</td><td>1228.1</td><td>9.6</td><td>1.2</td><td>8.0</td><td>1.7E-15</td><td>5.4E-15</td><td>MB231/MB468</td></tr>
<tr><td>>MI0000108|hsa-mir-103a-2&WithFlank&20|+|3917488|3917577@54.78.25&[MIMAT0000101&hsa-miR-103a-3p&offsets|0|+2;m-12&20|+|3917541|3917563&offsets|0|+2]|AGCAGCATTGTACAGGGCTATGAAA</td><td>>MI0000108|hsa-mir-103a-2&WithFlank&20|+|3917488|3917577</td><td>54</td><td>78</td><td>25</td><td>AGCAGCATTGTACAGGGCTATGAAA</td><td>762.7</td><td>7.4</td><td>0.9</td><td>8.0</td><td>1.8E-15</td><td>5.5E-15</td><td>BT20/MB468</td></tr>
<tr><td>>hg38_21_+_8433172_8446622_RNA45SN1_45S_50ntFlanks@12627.12652.26|ATGGGGCGAAGCTACCATCTGTGGGA</td><td>>hg38_21_+_8433172_8446622_RNA45SN1_45S_50ntFlanks</td><td>12627</td><td>12652</td><td>26</td><td>ATGGGGCGAAGCTACCATCTGTGGGA</td><td>267.6</td><td>-6.1</td><td>0.8</td><td>-8.0</td><td>1.8E-15</td><td>5.6E-15</td><td>MB231/MB468</td></tr>
<tr><td>>hg38_21_+_8433172_8446622_RNA45SN1_45S_50ntFlanks@10806.10840.35|AACAATGTAGGTAAGGGAAGTCGGCAAGCCGGATC</td><td>>hg38_21_+_8433172_8446622_RNA45SN1_45S_50ntFlanks</td><td>10806</td><td>10840</td><td>35</td><td>AACAATGTAGGTAAGGGAAGTCGGCAAGCCGGATC</td><td>252.3</td><td>-4.3</td><td>0.5</td><td>-8.0</td><td>1.8E-15</td><td>5.6E-15</td><td>BT20/MB468</td></tr>
<tr><td>>hg38_21_+_8433172_8446622_RNA45SN1_45S_50ntFlanks@10572.10608.37|AGAAGCCGGCGGGAGCCCCGGGGAGAGTTCTCTTTTC</td><td>>hg38_21_+_8433172_8446622_RNA45SN1_45S_50ntFlanks</td><td>10572</td><td>10608</td><td>37</td><td>AGAAGCCGGCGGGAGCCCCGGGGAGAGTTCTCTTTTC</td><td>447.5</td><td>-10.8</td><td>1.4</td><td>-8.0</td><td>1.8E-15</td><td>5.6E-15</td><td>BT20/MB468</td></tr>
<tr><td>>hg38_21_+_8433172_8446622_RNA45SN1_45S_50ntFlanks@12624.12653.30|CCAATGGGGCGAAGCTACCATCTGTGGGAT</td><td>>hg38_21_+_8433172_8446622_RNA45SN1_45S_50ntFlanks</td><td>12624</td><td>12653</td><td>30</td><td>CCAATGGGGCGAAGCTACCATCTGTGGGAT</td><td>380.0</td><td>-10.1</td><td>1.3</td><td>-8.0</td><td>1.8E-15</td><td>5.7E-15</td><td>MB231/MB468</td></tr>
<tr><td>>hg38_21_+_8433172_8446622_RNA45SN1_45S_50ntFlanks@10348.10369.22|AAACGAGAACTTTGAAGGCCGA</td><td>>hg38_21_+_8433172_8446622_RNA45SN1_45S_50ntFlanks</td><td>10348</td><td>10369</td><td>22</td><td>AAACGAGAACTTTGAAGGCCGA</td><td>923.1</td><td>3.9</td><td>0.5</td><td>8.0</td><td>1.8E-15</td><td>5.7E-15</td><td>MB231/MB468</td></tr>
<tr><td>>hg38_21_+_8433172_8446622_RNA45SN1_45S_50ntFlanks@10387.10424.38|TGTGAACAGCAGTTGAACATGGGTCAGTCGGTCCTGAG</td><td>>hg38_21_+_8433172_8446622_RNA45SN1_45S_50ntFlanks</td><td>10387</td><td>10424</td><td>38</td><td>TGTGAACAGCAGTTGAACATGGGTCAGTCGGTCCTGAG</td><td>1218.0</td><td>8.6</td><td>1.1</td><td>8.0</td><td>1.8E-15</td><td>5.7E-15</td><td>MB231/MB468</td></tr>
<tr><td>>hg38_21_+_8433172_8446622_RNA45SN1_45S_50ntFlanks@11668.11704.37|TGCCCAGTGCTCTGAATGTCAAAGTGAAGAAATTCAA</td><td>>hg38_21_+_8433172_8446622_RNA45SN1_45S_50ntFlanks</td><td>11668</td><td>11704</td><td>37</td><td>TGCCCAGTGCTCTGAATGTCAAAGTGAAGAAATTCAA</td><td>339.0</td><td>6.8</td><td>0.9</td><td>8.0</td><td>1.8E-15</td><td>5.7E-15</td><td>MB231/MB468</td></tr>
<tr><td>>hg38_21_+_8433172_8446622_RNA45SN1_45S_50ntFlanks@11874.11904.31|AGAAGACCCTGTTGAGCTTGACTCTAGTCTG</td><td>>hg38_21_+_8433172_8446622_RNA45SN1_45S_50ntFlanks</td><td>11874</td><td>11904</td><td>31</td><td>AGAAGACCCTGTTGAGCTTGACTCTAGTCTG</td><td>428.5</td><td>-10.3</td><td>1.3</td><td>-8.0</td><td>1.9E-15</td><td>5.7E-15</td><td>MB231/MB468</td></tr>
<tr><td>>hg38_21_+_8433172_8446622_RNA45SN1_45S_50ntFlanks@12159.12178.20|TTTGACTGGGGCGGTACACC</td><td>>hg38_21_+_8433172_8446622_RNA45SN1_45S_50ntFlanks</td><td>12159</td><td>12178</td><td>20</td><td>TTTGACTGGGGCGGTACACC</td><td>411.4</td><td>-5.6</td><td>0.7</td><td>-7.9</td><td>1.9E-15</td><td>5.7E-15</td><td>BT20/MB468</td></tr>
<tr><td>>MI0000062|hsa-let-7a-3&WithFlank&22|+|46112743|46112828@58.79.22;MI0000060|hsa-let-7a-1&WithFlank&9|+|94175951|94176042@63.84.22&[MIMAT0004481&hsa-let-7a-3p&offsets|0|+1;m-180&22|+|46112800|46112821&offsets|0|0];[MIMAT0004481_1&hsa-let-7a-3p&offsets|0|+1;m-181&9|+|94176013|94176034&offsets|0|0]|CTATACAATCTACTGTCTTTCC</td><td>>MI0000062|hsa-let-7a-3&WithFlank&22|+|46112743|46112828</td><td>58</td><td>79</td><td>22</td><td>CTATACAATCTACTGTCTTTCC</td><td>690.6</td><td>6.5</td><td>0.8</td><td>7.9</td><td>1.9E-15</td><td>5.7E-15</td><td>BT20/MB468</td></tr>
<tr><td>>hg38_21_+_8433172_8446622_RNA45SN1_45S_50ntFlanks@12622.12654.33|AGCCAATGGGGCGAAGCTACCATCTGTGGGATT</td><td>>hg38_21_+_8433172_8446622_RNA45SN1_45S_50ntFlanks</td><td>12622</td><td>12654</td><td>33</td><td>AGCCAATGGGGCGAAGCTACCATCTGTGGGATT</td><td>292.6</td><td>-4.8</td><td>0.6</td><td>-7.9</td><td>1.9E-15</td><td>5.8E-15</td><td>BT20/MB468</td></tr>
<tr><td>>hg38_21_+_8433172_8446622_RNA45SN1_45S_50ntFlanks@12161.12179.19|TGACTGGGGCGGTACACCT</td><td>>hg38_21_+_8433172_8446622_RNA45SN1_45S_50ntFlanks</td><td>12161</td><td>12179</td><td>19</td><td>TGACTGGGGCGGTACACCT</td><td>404.6</td><td>-6.1</td><td>0.8</td><td>-7.9</td><td>1.9E-15</td><td>5.8E-15</td><td>BT20/MB468</td></tr>
<tr><td>>am_trna162_MetCAT_6_-_26758550_26758622@34.54.21,trna164_MetCAT_6_-_26735574_26735646@34.54.21,trna20_MetCAT_16_+_71460396_71460468@34.54.21,trna21_MetCAT_6_+_26701712_26701784@34.54.21,trna27_MetCAT_6_+_26766444_26766516@34.54.21,trna75_MetCAT_6_+_28912352_28912424@34.54.21,trna97_MetCAT_6_-_28921042_28921114@34.54.21|CATAATCTGAAGGTCCTGAGT</td><td>>am_trna162_MetCAT_6_-_26758550_26758622</td><td>34</td><td>54</td><td>21</td><td>CATAATCTGAAGGTCCTGAGT</td><td>2684.0</td><td>5.1</td><td>0.6</td><td>7.9</td><td>1.9E-15</td><td>5.9E-15</td><td>BT20/MB468</td></tr>
<tr><td>>hg38_21_+_8433172_8446622_RNA45SN1_45S_50ntFlanks@11886.11905.20|TGAGCTTGACTCTAGTCTGG</td><td>>hg38_21_+_8433172_8446622_RNA45SN1_45S_50ntFlanks</td><td>11886</td><td>11905</td><td>20</td><td>TGAGCTTGACTCTAGTCTGG</td><td>371.3</td><td>-10.1</td><td>1.3</td><td>-7.9</td><td>1.9E-15</td><td>5.9E-15</td><td>MB231/MB468</td></tr>
<tr><td>>hg38_21_+_8433172_8446622_RNA45SN1_45S_50ntFlanks@6737.6771.35|GATCATCGACACTTCGAACGCACTTGCGGCCCCGG</td><td>>hg38_21_+_8433172_8446622_RNA45SN1_45S_50ntFlanks</td><td>6737</td><td>6771</td><td>35</td><td>GATCATCGACACTTCGAACGCACTTGCGGCCCCGG</td><td>288.6</td><td>-10.2</td><td>1.3</td><td>-7.9</td><td>1.9E-15</td><td>5.9E-15</td><td>BT20/MB468</td></tr>
<tr><td>>hg38_21_+_8433172_8446622_RNA45SN1_45S_50ntFlanks@10349.10383.35|AACGAGAACTTTGAAGGCCGAAGTGGAGAAGGGTT</td><td>>hg38_21_+_8433172_8446622_RNA45SN1_45S_50ntFlanks</td><td>10349</td><td>10383</td><td>35</td><td>AACGAGAACTTTGAAGGCCGAAGTGGAGAAGGGTT</td><td>2607.4</td><td>-3.2</td><td>0.4</td><td>-7.9</td><td>2.0E-15</td><td>6.0E-15</td><td>BT20/MB468</td></tr>
<tr><td>>hg38_21_+_8433172_8446622_RNA45SN1_45S_50ntFlanks@4367.4394.28|CAGTTAAAAAGCTCGTAGTTGGATCTTG</td><td>>hg38_21_+_8433172_8446622_RNA45SN1_45S_50ntFlanks</td><td>4367</td><td>4394</td><td>28</td><td>CAGTTAAAAAGCTCGTAGTTGGATCTTG</td><td>252.7</td><td>-10.0</td><td>1.3</td><td>-7.9</td><td>2.0E-15</td><td>6.1E-15</td><td>BT20/MB468</td></tr>
<tr><td>>hg38_21_+_8433172_8446622_RNA45SN1_45S_50ntFlanks@7976.8010.35|GCGACCTCAGATCAGACGTGGCGACCCGCTGAATT</td><td>>hg38_21_+_8433172_8446622_RNA45SN1_45S_50ntFlanks</td><td>7976</td><td>8010</td><td>35</td><td>GCGACCTCAGATCAGACGTGGCGACCCGCTGAATT</td><td>341.5</td><td>8.7</td><td>1.1</td><td>7.9</td><td>2.0E-15</td><td>6.1E-15</td><td>BT20/MB468</td></tr>
<tr><td>>trnaMT_ValTAC_MT_+_1602_1670@47.72.26|TCAACTTAACTTGACCGCTCTGACCA</td><td>>trnaMT_ValTAC_MT_+_1602_1670</td><td>47</td><td>72</td><td>26</td><td>TCAACTTAACTTGACCGCTCTGACCA</td><td>474.6</td><td>7.4</td><td>0.9</td><td>7.9</td><td>2.0E-15</td><td>6.1E-15</td><td>BT20/MB468</td></tr>
<tr><td>>hg38_21_+_8433172_8446622_RNA45SN1_45S_50ntFlanks@10807.10826.20|ACAATGTAGGTAAGGGAAGT</td><td>>hg38_21_+_8433172_8446622_RNA45SN1_45S_50ntFlanks</td><td>10807</td><td>10826</td><td>20</td><td>ACAATGTAGGTAAGGGAAGT</td><td>172.2</td><td>-4.7</td><td>0.6</td><td>-7.9</td><td>2.0E-15</td><td>6.2E-15</td><td>MB231/MB468</td></tr>
<tr><td>>MI0000108|hsa-mir-103a-2&WithFlank&20|+|3917488|3917577@54.76.23(+1G);MI0000109|hsa-mir-103a-1&WithFlank&5|-|168560890|168560979@54.76.23(+1G)&[MIMAT0000101&hsa-miR-103a-3p&offsets|0|0(+1G);m-12&20|+|3917541|3917563&offsets|0|0(+1G)];[MIMAT0000101_1&hsa-miR-103a-3p&offsets|0|0(+1G);m-13&5|-|168560904|168560926&offsets|0|0(+1G);m-44220&5|-|168560908|168560926&offsets|0|+4(+1G)]|AGCAGCATTGTACAGGGCTATGAG</td><td>>MI0000108|hsa-mir-103a-2&WithFlank&20|+|3917488|3917577</td><td>54</td><td>76</td><td>23(+1G)</td><td>AGCAGCATTGTACAGGGCTATGAG</td><td>774.6</td><td>8.9</td><td>1.1</td><td>7.9</td><td>2.1E-15</td><td>6.3E-15</td><td>BT20/MB468</td></tr>
<tr><td>>hg38_21_+_8433172_8446622_RNA45SN1_45S_50ntFlanks@12415.12449.35|CTTCGATGTCGGCTCTTCCTATCATTGTGAAGCAG</td><td>>hg38_21_+_8433172_8446622_RNA45SN1_45S_50ntFlanks</td><td>12415</td><td>12449</td><td>35</td><td>CTTCGATGTCGGCTCTTCCTATCATTGTGAAGCAG</td><td>1093.8</td><td>4.8</td><td>0.6</td><td>7.9</td><td>2.0E-15</td><td>6.3E-15</td><td>MB231/MB468</td></tr>
<tr><td>>hg38_21_+_8433172_8446622_RNA45SN1_45S_50ntFlanks@10846.10864.19|CTTCGGGATAAGGATTGGC</td><td>>hg38_21_+_8433172_8446622_RNA45SN1_45S_50ntFlanks</td><td>10846</td><td>10864</td><td>19</td><td>CTTCGGGATAAGGATTGGC</td><td>293.5</td><td>-4.4</td><td>0.6</td><td>-7.9</td><td>2.0E-15</td><td>6.3E-15</td><td>MB231/MB468</td></tr>
<tr><td>>hg38_21_+_8433172_8446622_RNA45SN1_45S_50ntFlanks@11235.11269.35|CTTCCCGTGGATCGCCCCAGCTGCGGCGGGCGTCG</td><td>>hg38_21_+_8433172_8446622_RNA45SN1_45S_50ntFlanks</td><td>11235</td><td>11269</td><td>35</td><td>CTTCCCGTGGATCGCCCCAGCTGCGGCGGGCGTCG</td><td>360.0</td><td>-10.1</td><td>1.3</td><td>-7.9</td><td>2.1E-15</td><td>6.3E-15</td><td>MB231/MB468</td></tr>
<tr><td>>hg38_21_+_8433172_8446622_RNA45SN1_45S_50ntFlanks@11571.11594.24|GCCGACTTAGAACTGGTGCGGACC</td><td>>hg38_21_+_8433172_8446622_RNA45SN1_45S_50ntFlanks</td><td>11571</td><td>11594</td><td>24</td><td>GCCGACTTAGAACTGGTGCGGACC</td><td>898.7</td><td>5.3</td><td>0.7</td><td>7.9</td><td>2.1E-15</td><td>6.3E-15</td><td>MB231/MB468</td></tr>
<tr><td>>MI0000273|hsa-mir-183&WithFlank&7|-|129774899|129775020@33.53.21(+1C)&[MIMAT0000261&hsa-miR-183-5p&offsets|0|-1(+1C);m-41&7|-|129774966|129774987&offsets|-1|-2(+1C)]|TATGGCACTGGTAGAATTCACC</td><td>>MI0000273|hsa-mir-183&WithFlank&7|-|129774899|129775020</td><td>33</td><td>53</td><td>21(+1C)</td><td>TATGGCACTGGTAGAATTCACC</td><td>1334.9</td><td>5.9</td><td>0.7</td><td>7.9</td><td>2.1E-15</td><td>6.3E-15</td><td>MB231/MB468</td></tr>
<tr><td>>hg38_1_-_228634819_228635039_RNA5S12_5S_50ntFlanks@133.169.37|ATGGGAGACCGCCTGGGAATACCGGGTGCTGTAGGCT</td><td>>hg38_1_-_228634819_228635039_RNA5S12_5S_50ntFlanks</td><td>133</td><td>169</td><td>37</td><td>ATGGGAGACCGCCTGGGAATACCGGGTGCTGTAGGCT</td><td>3292.2</td><td>5.8</td><td>0.7</td><td>7.9</td><td>2.1E-15</td><td>6.4E-15</td><td>BT20/MB468</td></tr>
<tr><td>>hg38_21_+_8433172_8446622_RNA45SN1_45S_50ntFlanks@10099.10117.19|CGCGCGCGCGTGTGGTGTG</td><td>>hg38_21_+_8433172_8446622_RNA45SN1_45S_50ntFlanks</td><td>10099</td><td>10117</td><td>19</td><td>CGCGCGCGCGTGTGGTGTG</td><td>317.6</td><td>-9.9</td><td>1.2</td><td>-7.9</td><td>2.1E-15</td><td>6.5E-15</td><td>MB231/MB468</td></tr>
<tr><td>>MI0000810|hsa-mir-135b&WithFlank&1|-|205448296|205448404@22.43.22&[MIMAT0000758&hsa-miR-135b-5p&offsets|0|-1;m-321&1|-|205448361|205448383&offsets|0|-1]|TATGGCTTTTCATTCCTATGTG</td><td>>MI0000810|hsa-mir-135b&WithFlank&1|-|205448296|205448404</td><td>22</td><td>43</td><td>22</td><td>TATGGCTTTTCATTCCTATGTG</td><td>564.3</td><td>6.0</td><td>0.8</td><td>7.9</td><td>2.1E-15</td><td>6.5E-15</td><td>BT20/MB468</td></tr>
<tr><td>>hg38_21_+_8433172_8446622_RNA45SN1_45S_50ntFlanks@4003.4025.23|ATAACCTCGGGCCGATCGCACGC</td><td>>hg38_21_+_8433172_8446622_RNA45SN1_45S_50ntFlanks</td><td>4003</td><td>4025</td><td>23</td><td>ATAACCTCGGGCCGATCGCACGC</td><td>420.0</td><td>-6.7</td><td>0.8</td><td>-7.9</td><td>2.1E-15</td><td>6.5E-15</td><td>MB231/MB468</td></tr>
<tr><td>>hg38_21_+_8433172_8446622_RNA45SN1_45S_50ntFlanks@8472.8492.21|CGGCCCGGCGGATCTTTCCCG</td><td>>hg38_21_+_8433172_8446622_RNA45SN1_45S_50ntFlanks</td><td>8472</td><td>8492</td><td>21</td><td>CGGCCCGGCGGATCTTTCCCG</td><td>474.7</td><td>-10.5</td><td>1.3</td><td>-7.9</td><td>2.1E-15</td><td>6.6E-15</td><td>MB231/MB468</td></tr>
<tr><td>>trnaMT_LysTTT_MT_+_8295_8364@1.36.36,trnalookalike7_LysTTT_1_+_568844_568913@1.36.36|CACTGTAAAGCTAACTTAGCATTAACCTTTTAAGTT</td><td>>trnaMT_LysTTT_MT_+_8295_8364</td><td>1</td><td>36</td><td>36</td><td>CACTGTAAAGCTAACTTAGCATTAACCTTTTAAGTT</td><td>693.7</td><td>5.9</td><td>0.7</td><td>7.9</td><td>2.2E-15</td><td>6.6E-15</td><td>BT20/MB468</td></tr>
<tr><td>>MI0003137|hsa-mir-193b&WithFlank&16|+|14303961|14304055@57.77.21(+1G)&[MIMAT0002819&hsa-miR-193b-3p&offsets|0|-1(+1G);m-166&16|+|14304017|14304038&offsets|0|-1(+1G)]|AACTGGCCCTCAAAGTCCCGCG</td><td>>MI0003137|hsa-mir-193b&WithFlank&16|+|14303961|14304055</td><td>57</td><td>77</td><td>21(+1G)</td><td>AACTGGCCCTCAAAGTCCCGCG</td><td>2047.5</td><td>13.6</td><td>1.7</td><td>7.9</td><td>2.2E-15</td><td>6.6E-15</td><td>BT20/MB468</td></tr>
<tr><td>>hg38_21_+_8433172_8446622_RNA45SN1_45S_50ntFlanks@10401.10439.39|GAACATGGGTCAGTCGGTCCTGAGAGATGGGCGAGCGCC</td><td>>hg38_21_+_8433172_8446622_RNA45SN1_45S_50ntFlanks</td><td>10401</td><td>10439</td><td>39</td><td>GAACATGGGTCAGTCGGTCCTGAGAGATGGGCGAGCGCC</td><td>573.1</td><td>8.7</td><td>1.1</td><td>7.9</td><td>2.2E-15</td><td>6.7E-15</td><td>MB231/MB468</td></tr>
<tr><td>>hg38_21_+_8433172_8446622_RNA45SN1_45S_50ntFlanks@6651.6690.40|CGACTCTTAGCGGTGGATCACTCGGCTCGTGCGTCGATGA</td><td>>hg38_21_+_8433172_8446622_RNA45SN1_45S_50ntFlanks</td><td>6651</td><td>6690</td><td>40</td><td>CGACTCTTAGCGGTGGATCACTCGGCTCGTGCGTCGATGA</td><td>3171.9</td><td>4.8</td><td>0.6</td><td>7.9</td><td>2.2E-15</td><td>6.7E-15</td><td>MB231/MB468</td></tr>
</table>

<table>
<tr><td>>hg38_21_+_8433172_8446622_RNA45SN1_45S_50ntFlanks@12081.12110.30|CTCTCGCTTCTGGCGCCAAGCGCCCGGCCG</td><td>>hg38_21_+_8433172_8446622_RNA45SN1_45S_50ntFlanks</td><td>12081</td><td>12110</td><td>30</td><td>CTCTCGCTTCTGGCGCCAAGCGCCCGGCCG</td><td>763.8</td><td>-7.6</td><td>1.0</td><td>-7.9</td><td>2.2E-15</td><td>6.8E-15</td><td>BT20/MB468</td></tr>
<tr><td>>MI0000283|hsa-mir-203a&WithFlank&14|+|104117399|104117520@72.91.20(+1A)&[MIMAT0000264&hsa-miR-203a-3p&offsets|+1|-1(+1A);m-79&14|+|104117469|104117490&offsets|+1|-1(+1A)]|TGAAATGTTTAGGACCACTAA</td><td>>MI0000283|hsa-mir-203a&WithFlank&14|+|104117399|104117520</td><td>72</td><td>91</td><td>20(+1A)</td><td>TGAAATGTTTAGGACCACTAA</td><td>413.5</td><td>12.2</td><td>1.5</td><td>8.3</td><td>1.5E-16</td><td>6.8E-15</td><td>BT20/MB231</td></tr>
<tr><td>>hg38_21_+_8433172_8446622_RNA45SN1_45S_50ntFlanks@12628.12661.34|TGGGGCGAAGCTACCATCTGTGGGATTATGACTG</td><td>>hg38_21_+_8433172_8446622_RNA45SN1_45S_50ntFlanks</td><td>12628</td><td>12661</td><td>34</td><td>TGGGGCGAAGCTACCATCTGTGGGATTATGACTG</td><td>432.6</td><td>-4.1</td><td>0.5</td><td>-7.9</td><td>2.3E-15</td><td>6.9E-15</td><td>MB231/MB468</td></tr>
<tr><td>>hg38_21_+_8433172_8446622_RNA45SN1_45S_50ntFlanks@10687.10705.19|CGGCGTCCGGTGAGCTCTC</td><td>>hg38_21_+_8433172_8446622_RNA45SN1_45S_50ntFlanks</td><td>10687</td><td>10705</td><td>19</td><td>CGGCGTCCGGTGAGCTCTC</td><td>148.5</td><td>-4.3</td><td>0.5</td><td>-7.9</td><td>2.3E-15</td><td>7.0E-15</td><td>BT20/MB468</td></tr>
<tr><td>>MI0000457|hsa-mir-141&WithFlank&12|+|6964091|6964197@65.85.21&[MIMAT0000432&hsa-miR-141-3p&offsets|0|-1;m-134&12|+|6964155|6964175&offsets|0|0]|TAACACTGTCTGGTAAAGATG</td><td>>MI0000457|hsa-mir-141&WithFlank&12|+|6964091|6964197</td><td>65</td><td>85</td><td>21</td><td>TAACACTGTCTGGTAAAGATG</td><td>3725.6</td><td>7.9</td><td>1.0</td><td>7.9</td><td>2.3E-15</td><td>7.0E-15</td><td>BT20/MB468</td></tr>
<tr><td>>MI0000440|hsa-mir-27b&WithFlank&9|+|95085439|95085547@67.87.21(+2U)&[MIMAT0000419&hsa-miR-27b-3p&offsets|0|0(+2U);m-39&9|+|95085505|95085525&offsets|0|0(+2U)]|TTCACAGTGGCTAAGTTCTGCTT</td><td>>MI0000440|hsa-mir-27b&WithFlank&9|+|95085439|95085547</td><td>67</td><td>87</td><td>21(+2U)</td><td>TTCACAGTGGCTAAGTTCTGCTT</td><td>1213.9</td><td>6.6</td><td>0.8</td><td>7.9</td><td>2.3E-15</td><td>7.0E-15</td><td>BT20/MB468</td></tr>
<tr><td>>MI0000088|hsa-mir-30a&WithFlank&6|-|71403545|71403627@12.35.24(+1U)&[MIMAT0000087&hsa-miR-30a-5p&offsets|0|+2(+1U);m-15&6|-|71403595|71403616&offsets|0|+2(+1U)]|TGTAAACATCCTCGACTGGAAGCTT</td><td>>MI0000088|hsa-mir-30a&WithFlank&6|-|71403545|71403627</td><td>12</td><td>35</td><td>24(+1U)</td><td>TGTAAACATCCTCGACTGGAAGCTT</td><td>7108.2</td><td>7.2</td><td>0.9</td><td>7.9</td><td>2.3E-15</td><td>7.0E-15</td><td>BT20/MB468</td></tr>
<tr><td>>trna10_AspGTC_12_-_125424193_125424264@35.74.40,trna12_AspGTC_12_-_125411891_125411962@35.74.40,trna144_AspGTC_6_-_27551236_27551307@35.74.40,trna38_AspGTC_17_-_8125556_8125627@35.74.40,trna45_AspGTC_6_+_27447453_27447524@35.74.40,trna48_AspGTC_6_+_27471523_27471594@35.74.40,trna4_AspGTC_12_+_96429799_96429870@35.74.40,trna69_AspGTC_1_-_161440205_161440276@35.74.40,trna72_AspGTC_1_-_161432824_161432895@35.74.40,trna75_AspGTC_1_-_161425414_161425485@35.74.40,trna78_AspGTC_1_-_161418033_161418104@35.74.40,trna81_AspGTC_1_-_161410615_161410686@35.74.40|TCACGCGGGAGACCGGGGTTCGATTCCCCGACGGGGAGCC</td><td>>trna10_AspGTC_12_-_125424193_125424264</td><td>35</td><td>74</td><td>40</td><td>TCACGCGGGAGACCGGGGTTCGATTCCCCGACGGGGAGCC</td><td>2119.0</td><td>11.1</td><td>1.4</td><td>7.9</td><td>2.3E-15</td><td>7.0E-15</td><td>MB231/MB468</td></tr>
<tr><td>>hg38_21_+_8433172_8446622_RNA45SN1_45S_50ntFlanks@10786.10816.31|AACAGCCTCTGGCATGTTGGAACAATGTAGG</td><td>>hg38_21_+_8433172_8446622_RNA45SN1_45S_50ntFlanks</td><td>10786</td><td>10816</td><td>31</td><td>AACAGCCTCTGGCATGTTGGAACAATGTAGG</td><td>821.8</td><td>-4.0</td><td>0.5</td><td>-7.9</td><td>2.3E-15</td><td>7.1E-15</td><td>MB231/MB468</td></tr>
<tr><td>>hg38_21_+_8433172_8446622_RNA45SN1_45S_50ntFlanks@12412.12436.25|ATCCTTCGATGTCGGCTCTTCCTAT</td><td>>hg38_21_+_8433172_8446622_RNA45SN1_45S_50ntFlanks</td><td>12412</td><td>12436</td><td>25</td><td>ATCCTTCGATGTCGGCTCTTCCTAT</td><td>330.4</td><td>-10.4</td><td>1.3</td><td>-7.9</td><td>2.3E-15</td><td>7.1E-15</td><td>BT20/MB468</td></tr>
<tr><td>>hg38_1_-_228634819_228635039_RNA5S12_5S_50ntFlanks@151.170.20|ATACCGGGTGCTGTAGGCTT</td><td>>hg38_1_-_228634819_228635039_RNA5S12_5S_50ntFlanks</td><td>151</td><td>170</td><td>20</td><td>ATACCGGGTGCTGTAGGCTT</td><td>1931.2</td><td>5.1</td><td>0.6</td><td>7.9</td><td>2.3E-15</td><td>7.1E-15</td><td>BT20/MB468</td></tr>
<tr><td>>hg38_21_+_8433172_8446622_RNA45SN1_45S_50ntFlanks@8104.8143.40|CCGCGGCGGGGCGCGGGACATGTGGCGTACGGAAGACCCG</td><td>>hg38_21_+_8433172_8446622_RNA45SN1_45S_50ntFlanks</td><td>8104</td><td>8143</td><td>40</td><td>CCGCGGCGGGGCGCGGGACATGTGGCGTACGGAAGACCCG</td><td>2549.6</td><td>-7.0</td><td>0.9</td><td>-7.9</td><td>2.3E-15</td><td>7.1E-15</td><td>BT20/MB468</td></tr>
<tr><td>>MI0000103|hsa-mir-101-1&WithFlank&1|-|65058428|65058514@52.73.22(+1A);MI0000739|hsa-mir-101-2&WithFlank&9|+|4850291|4850381@54.75.22(+1A)&[MIMAT0000099&hsa-miR-101-3p&offsets|-1|0(+1A);m-17&1|-|65058443|65058462&offsets|-1|+1(+1A)];[MIMAT0000099_1&hsa-miR-101-3p&offsets|-1|0(+1A);m-16&9|+|4850345|4850364&offsets|-1|+1(+1A)]|GTACAGTACTGTGATAACTGAAA</td><td>>MI0000103|hsa-mir-101-1&WithFlank&1|-|65058428|65058514</td><td>52</td><td>73</td><td>22(+1A)</td><td>GTACAGTACTGTGATAACTGAAA</td><td>1135.1</td><td>6.4</td><td>0.8</td><td>7.9</td><td>2.3E-15</td><td>7.1E-15</td><td>MB231/MB468</td></tr>
<tr><td>>hg38_21_+_8433172_8446622_RNA45SN1_45S_50ntFlanks@3985.4005.21|CGGCTTTGGTGACTCTAGATA</td><td>>hg38_21_+_8433172_8446622_RNA45SN1_45S_50ntFlanks</td><td>3985</td><td>4005</td><td>21</td><td>CGGCTTTGGTGACTCTAGATA</td><td>362.0</td><td>-10.1</td><td>1.3</td><td>-7.9</td><td>2.3E-15</td><td>7.1E-15</td><td>MB231/MB468</td></tr>
<tr><td>>hg38_21_+_8433172_8446622_RNA45SN1_45S_50ntFlanks@6649.6696.48|TACGACTCTTAGCGGTGGATCACTCGGCTCGTGCGTCGATGAAGAACG</td><td>>hg38_21_+_8433172_8446622_RNA45SN1_45S_50ntFlanks</td><td>6649</td><td>6696</td><td>48</td><td>TACGACTCTTAGCGGTGGATCACTCGGCTCGTGCGTCGATGAAGAACG</td><td>1023.0</td><td>7.9</td><td>1.0</td><td>7.9</td><td>2.3E-15</td><td>7.1E-15</td><td>MB231/MB468</td></tr>
<tr><td>>hg38_21_+_8433172_8446622_RNA45SN1_45S_50ntFlanks@11534.11553.20|GTTCCGCGCGGCGCCTCGCC</td><td>>hg38_21_+_8433172_8446622_RNA45SN1_45S_50ntFlanks</td><td>11534</td><td>11553</td><td>20</td><td>GTTCCGCGCGGCGCCTCGCC</td><td>526.2</td><td>-10.6</td><td>1.3</td><td>-7.9</td><td>2.3E-15</td><td>7.2E-15</td><td>MB231/MB468</td></tr>
<tr><td>>hg38_21_+_8433172_8446622_RNA45SN1_45S_50ntFlanks@11641.11668.28|CGGCGGGTGTTGACGCGATGTGATTTCT</td><td>>hg38_21_+_8433172_8446622_RNA45SN1_45S_50ntFlanks</td><td>11641</td><td>11668</td><td>28</td><td>CGGCGGGTGTTGACGCGATGTGATTTCT</td><td>327.9</td><td>-5.5</td><td>0.7</td><td>-7.9</td><td>2.4E-15</td><td>7.3E-15</td><td>BT20/MB468</td></tr>
<tr><td>>MI0000456|hsa-mir-140&WithFlank&16|+|69933075|69933186@68.89.22&[MIMAT0004597&hsa-miR-140-3p&offsets|0|+1;m-45&16|+|69933143|69933164&offsets|-1|-1]|TACCACAGGGTAGAACCACGGA</td><td>>MI0000456|hsa-mir-140&WithFlank&16|+|69933075|69933186</td><td>68</td><td>89</td><td>22</td><td>TACCACAGGGTAGAACCACGGA</td><td>2913.5</td><td>11.3</td><td>1.4</td><td>7.9</td><td>2.4E-15</td><td>7.3E-15</td><td>BT20/MB468</td></tr>
<tr><td>>hg38_21_+_8433172_8446622_RNA45SN1_45S_50ntFlanks@10479.10497.19|CGATCGAAAGGGAGTCGGG</td><td>>hg38_21_+_8433172_8446622_RNA45SN1_45S_50ntFlanks</td><td>10479</td><td>10497</td><td>19</td><td>CGATCGAAAGGGAGTCGGG</td><td>348.9</td><td>-10.0</td><td>1.3</td><td>-7.9</td><td>2.4E-15</td><td>7.3E-15</td><td>MB231/MB468</td></tr>
<tr><td>>hg38_21_+_8433172_8446622_RNA45SN1_45S_50ntFlanks@12616.12653.38|CTGAGGAGCCAATGGGGCGAAGCTACCATCTGTGGGAT</td><td>>hg38_21_+_8433172_8446622_RNA45SN1_45S_50ntFlanks</td><td>12616</td><td>12653</td><td>38</td><td>CTGAGGAGCCAATGGGGCGAAGCTACCATCTGTGGGAT</td><td>198.2</td><td>-3.9</td><td>0.5</td><td>-7.9</td><td>2.4E-15</td><td>7.3E-15</td><td>MB231/MB468</td></tr>
<tr><td>>hg38_21_+_8433172_8446622_RNA45SN1_45S_50ntFlanks@12366.12384.19|TAACTGGCTTGTGGCGGCC</td><td>>hg38_21_+_8433172_8446622_RNA45SN1_45S_50ntFlanks</td><td>12366</td><td>12384</td><td>19</td><td>TAACTGGCTTGTGGCGGCC</td><td>355.3</td><td>-10.0</td><td>1.3</td><td>-7.9</td><td>2.4E-15</td><td>7.4E-15</td><td>MB231/MB468</td></tr>
<tr><td>>trna6_TrpCCA_12_+_98898030_98898101@30.52.23|ACTCCAGATCAGAAGGCTGCGTG</td><td>>trna6_TrpCCA_12_+_98898030_98898101</td><td>30</td><td>52</td><td>23</td><td>ACTCCAGATCAGAAGGCTGCGTG</td><td>261.5</td><td>-10.0</td><td>1.3</td><td>-7.9</td><td>2.4E-15</td><td>7.4E-15</td><td>BT20/MB468</td></tr>
<tr><td>>MI0003591|hsa-mir-584&WithFlank&5|-|149062307|149062415@22.42.21(+1U)&[MIMAT0003249&hsa-miR-584-5p&offsets|0|-1(+1U);m-167&5|-|149062374|149062394&offsets|0|0(+1U)]|TTATGGTTTGCCTGGGACTGAT</td><td>>MI0003591|hsa-mir-584&WithFlank&5|-|149062307|149062415</td><td>22</td><td>42</td><td>21(+1U)</td><td>TTATGGTTTGCCTGGGACTGAT</td><td>803.7</td><td>10.7</td><td>1.3</td><td>7.9</td><td>2.5E-15</td><td>7.4E-15</td><td>BT20/MB468</td></tr>
<tr><td>>MI0005566|hsa-mir-374b&WithFlank&X|-|74218541|74218624@17.37.21&[MIMAT0004955&hsa-miR-374b-5p&offsets|0|-1;m-153&X|-|74218587|74218608&offsets|0|-1]|ATATAATACAACCTGCTAAGT</td><td>>MI0005566|hsa-mir-374b&WithFlank&X|-|74218541|74218624</td><td>17</td><td>37</td><td>21</td><td>ATATAATACAACCTGCTAAGT</td><td>4687.2</td><td>7.8</td><td>1.0</td><td>7.9</td><td>2.5E-15</td><td>7.5E-15</td><td>BT20/MB468</td></tr>
<tr><td>>MI0000299|hsa-mir-222&WithFlank&X|-|45747009|45747130@75.95.21&[MIMAT0000279&hsa-miR-222-3p&offsets|0|0;m-140&X|-|45747033|45747056&offsets|0|-3]|AGCTACATCTGGCTACTGGGT</td><td>>MI0000299|hsa-mir-222&WithFlank&X|-|45747009|45747130</td><td>75</td><td>95</td><td>21</td><td>AGCTACATCTGGCTACTGGGT</td><td>2280.3</td><td>6.3</td><td>0.8</td><td>7.9</td><td>2.5E-15</td><td>7.5E-15</td><td>BT20/MB468</td></tr>
<tr><td>>hg38_21_+_8433172_8446622_RNA45SN1_45S_50ntFlanks@10727.10755.29|GGAGAGGGTGTAAATCTCGCGCCGGGCCG</td><td>>hg38_21_+_8433172_8446622_RNA45SN1_45S_50ntFlanks</td><td>10727</td><td>10755</td><td>29</td><td>GGAGAGGGTGTAAATCTCGCGCCGGGCCG</td><td>464.4</td><td>-10.4</td><td>1.3</td><td>-7.9</td><td>2.5E-15</td><td>7.5E-15</td><td>MB231/MB468</td></tr>
<tr><td>>hg38_21_+_8433172_8446622_RNA45SN1_45S_50ntFlanks@6650.6695.46|ACGACTCTTAGCGGTGGATCACTCGGCTCGTGCGTCGATGAAGAAC</td><td>>hg38_21_+_8433172_8446622_RNA45SN1_45S_50ntFlanks</td><td>6650</td><td>6695</td><td>46</td><td>ACGACTCTTAGCGGTGGATCACTCGGCTCGTGCGTCGATGAAGAAC</td><td>619.1</td><td>9.5</td><td>1.2</td><td>7.9</td><td>2.5E-15</td><td>7.5E-15</td><td>BT20/MB468</td></tr>
<tr><td>>hg38_21_+_8433172_8446622_RNA45SN1_45S_50ntFlanks@6734.6774.41|ATTGATCATCGACACTTCGAACGCACTTGCGGCCCCGGGTT</td><td>>hg38_21_+_8433172_8446622_RNA45SN1_45S_50ntFlanks</td><td>6734</td><td>6774</td><td>41</td><td>ATTGATCATCGACACTTCGAACGCACTTGCGGCCCCGGGTT</td><td>870.2</td><td>9.7</td><td>1.2</td><td>7.9</td><td>2.5E-15</td><td>7.5E-15</td><td>BT20/MB468</td></tr>
<tr><td>>hg38_21_+_8433172_8446622_RNA45SN1_45S_50ntFlanks@10348.10368.21|AAACGAGAACTTTGAAGGCCG</td><td>>hg38_21_+_8433172_8446622_RNA45SN1_45S_50ntFlanks</td><td>10348</td><td>10368</td><td>21</td><td>AAACGAGAACTTTGAAGGCCG</td><td>1558.8</td><td>-3.3</td><td>0.4</td><td>-7.9</td><td>2.5E-15</td><td>7.5E-15</td><td>BT20/MB468</td></tr>
<tr><td>>trnaMT_ValTAC_MT_+_1602_1670@49.72.24|AACTTAACTTGACCGCTCTGACCA</td><td>>trnaMT_ValTAC_MT_+_1602_1670</td><td>49</td><td>72</td><td>24</td><td>AACTTAACTTGACCGCTCTGACCA</td><td>697.6</td><td>4.7</td><td>0.6</td><td>7.9</td><td>2.5E-15</td><td>7.6E-15</td><td>MB231/MB468</td></tr>
<tr><td>>hg38_21_+_8433172_8446622_RNA45SN1_45S_50ntFlanks@12934.12955.22|CTAAACCATTCGTAGACGACCT</td><td>>hg38_21_+_8433172_8446622_RNA45SN1_45S_50ntFlanks</td><td>12934</td><td>12955</td><td>22</td><td>CTAAACCATTCGTAGACGACCT</td><td>342.7</td><td>-6.6</td><td>0.8</td><td>-7.9</td><td>2.6E-15</td><td>7.7E-15</td><td>BT20/MB468</td></tr>
<tr><td>>MI0000089|hsa-mir-31&WithFlank&9|-|21512109|21512191@14.34.21&[MIMAT0000089&hsa-miR-31-5p&offsets|0|0;m-360&9|-|21512157|21512178&offsets|0|-1]|AGGCAAGATGCTGGCATAGCT</td><td>>MI0000089|hsa-mir-31&WithFlank&9|-|21512109|21512191</td><td>14</td><td>34</td><td>21</td><td>AGGCAAGATGCTGGCATAGCT</td><td>1835.7</td><td>9.6</td><td>1.2</td><td>7.9</td><td>2.6E-15</td><td>7.7E-15</td><td>BT20/MB468</td></tr>
<tr><td>>trna116_GluCTC_1_-_145399233_145399304@1.31.31,trna59_GluCTC_1_+_249168447_249168518@1.31.31,trna71_GluCTC_1_-_161439189_161439260@1.31.31,trna74_GluCTC_1_-_161431809_161431880@1.31.31,trna77_GluCTC_1_-_161424398_161424469@1.31.31,trna77_GluCTC_6_+_28949976_28950047@1.31.31,trna80_GluCTC_1_-_161417018_161417089@1.31.31,trna87_GluCTC_6_-_126101393_126101464@1.31.31|TCCCTGGTGGTCTAGTGGTTAGGATTCGGCG</td><td>>trna116_GluCTC_1_-_145399233_145399304</td><td>1</td><td>31</td><td>31</td><td>TCCCTGGTGGTCTAGTGGTTAGGATTCGGCG</td><td>83940.0</td><td>-5.3</td><td>0.7</td><td>-7.9</td><td>2.6E-15</td><td>7.7E-15</td><td>BT20/MB468</td></tr>
<tr><td>>hg38_21_+_8433172_8446622_RNA45SN1_45S_50ntFlanks@6728.6756.29|GGACACATTGATCATCGACACTTCGAACG</td><td>>hg38_21_+_8433172_8446622_RNA45SN1_45S_50ntFlanks</td><td>6728</td><td>6756</td><td>29</td><td>GGACACATTGATCATCGACACTTCGAACG</td><td>446.3</td><td>4.9</td><td>0.6</td><td>7.9</td><td>2.5E-15</td><td>7.7E-15</td><td>MB231/MB468</td></tr>
<tr><td>>MI0000438|hsa-mir-15b&WithFlank&3|+|160404582|160404691@64.85.22&[MIMAT0004586&hsa-miR-15b-3p&offsets|0|0;m-204&3|+|160404645|160404665&offsets|0|+1]|CGAATCATTATTTGCTGCTCTA</td><td>>MI0000438|hsa-mir-15b&WithFlank&3|+|160404582|160404691</td><td>64</td><td>85</td><td>22</td><td>CGAATCATTATTTGCTGCTCTA</td><td>453.2</td><td>6.5</td><td>0.8</td><td>7.9</td><td>2.6E-15</td><td>7.8E-15</td><td>BT20/MB468</td></tr>
<tr><td>>am_MI0000815|hsa-mir-339&WithFlank&7|-|1022927|1023032@21.42.22(+1A)&[MIMAT0000764&hsa-miR-339-5p&offsets|0|-1(+1A);m-163&7|-|1022990|1023012&offsets|0|-1(+1A)]|TCCCTGTCCTCCAGGAGCTCACA</td><td>>am_MI0000815|hsa-mir-339&WithFlank&7|-|1022927|1023032</td><td>21</td><td>42</td><td>22(+1A)</td><td>TCCCTGTCCTCCAGGAGCTCACA</td><td>928.8</td><td>7.7</td><td>1.0</td><td>7.9</td><td>2.6E-15</td><td>7.9E-15</td><td>BT20/MB468</td></tr>
<tr><td>>am_trnaMT_ValTAC_MT_+_1602_1670@48.65.18|CAACTTAACTTGACCGCT</td><td>>am_trnaMT_ValTAC_MT_+_1602_1670</td><td>48</td><td>65</td><td>18</td><td>CAACTTAACTTGACCGCT</td><td>434.9</td><td>8.7</td><td>1.1</td><td>7.9</td><td>2.6E-15</td><td>7.9E-15</td><td>BT20/MB468</td></tr>
<tr><td>>MI0000266|hsa-mir-10a&WithFlank&17|-|48579832|48579953@28.49.22&[MIMAT0000253&hsa-miR-10a-5p&offsets|0|-1;m-18&17|-|48579905|48579926&offsets|0|0]|TACCCTGTAGATCCGAATTTGT</td><td>>MI0000266|hsa-mir-10a&WithFlank&17|-|48579832|48579953</td><td>28</td><td>49</td><td>22</td><td>TACCCTGTAGATCCGAATTTGT</td><td>44384.4</td><td>12.6</td><td>1.6</td><td>7.9</td><td>2.6E-15</td><td>7.9E-15</td><td>BT20/MB468</td></tr>
</table>

<table>
<tr><td>>hg38_21_+_8433172_8446622_RNA45SN1_45S_50ntFlanks@11768.11803.36|CTAATTAGTGACGCGCATGAATGGATGAACGAGATT</td><td>>hg38_21_+_8433172_8446622_RNA45SN1_45S_50ntFlanks</td><td>11768</td><td>11803</td><td>36</td><td>CTAATTAGTGACGCGCATGAATGGATGAACGAGATT</td><td>1495.2</td><td>-5.2</td><td>0.7</td><td>-7.9</td><td>2.6E-15</td><td>7.9E-15</td><td>MB231/MB468</td></tr>
<tr><td>>hg38_21_+_8433172_8446622_RNA45SN1_45S_50ntFlanks@12954.12972.19|CTGCTTCTGGGTCGGGGTT</td><td>>hg38_21_+_8433172_8446622_RNA45SN1_45S_50ntFlanks</td><td>12954</td><td>12972</td><td>19</td><td>CTGCTTCTGGGTCGGGGTT</td><td>454.3</td><td>-7.0</td><td>0.9</td><td>-7.9</td><td>2.6E-15</td><td>8.0E-15</td><td>MB231/MB468</td></tr>
<tr><td>>hg38_21_+_8433172_8446622_RNA45SN1_45S_50ntFlanks@476.497.22|TTCGTGATCGATGTGGTGACGT</td><td>>hg38_21_+_8433172_8446622_RNA45SN1_45S_50ntFlanks</td><td>476</td><td>497</td><td>22</td><td>TTCGTGATCGATGTGGTGACGT</td><td>1148.0</td><td>6.8</td><td>0.9</td><td>7.9</td><td>2.7E-15</td><td>8.0E-15</td><td>BT20/MB468</td></tr>
<tr><td>>hg38_21_+_8433172_8446622_RNA45SN1_45S_50ntFlanks@3705.3744.40|TACCTGGTTGATCCTGCCAGTAGCATATGCTTGTCTCAAA</td><td>>hg38_21_+_8433172_8446622_RNA45SN1_45S_50ntFlanks</td><td>3705</td><td>3744</td><td>40</td><td>TACCTGGTTGATCCTGCCAGTAGCATATGCTTGTCTCAAA</td><td>383.5</td><td>7.0</td><td>0.9</td><td>7.9</td><td>2.6E-15</td><td>8.0E-15</td><td>MB231/MB468</td></tr>
<tr><td>>am_trnaMT_ValTAC_MT_+_1602_1670@29.52.24|ACTTACACTTAGGAGATTTCAACT</td><td>>am_trnaMT_ValTAC_MT_+_1602_1670</td><td>29</td><td>52</td><td>24</td><td>ACTTACACTTAGGAGATTTCAACT</td><td>470.6</td><td>8.7</td><td>1.1</td><td>7.9</td><td>2.7E-15</td><td>8.0E-15</td><td>BT20/MB468</td></tr>
<tr><td>>hg38_21_+_8433172_8446622_RNA45SN1_45S_50ntFlanks@11575.11593.19|ACTTAGAACTGGTGCGGAC</td><td>>hg38_21_+_8433172_8446622_RNA45SN1_45S_50ntFlanks</td><td>11575</td><td>11593</td><td>19</td><td>ACTTAGAACTGGTGCGGAC</td><td>1171.4</td><td>-4.1</td><td>0.5</td><td>-7.9</td><td>2.6E-15</td><td>8.0E-15</td><td>MB231/MB468</td></tr>
<tr><td>>MI0000098|hsa-mir-96&WithFlank&7|-|129774686|129774775@15.36.22(+1G)&[MIMAT0000095&hsa-miR-96-5p&offsets|0|-1(+1G);m-248&7|-|129774739|129774761&offsets|0|-1(+1G)]|TTTGGCACTAGCACATTTTTGCG</td><td>>MI0000098|hsa-mir-96&WithFlank&7|-|129774686|129774775</td><td>15</td><td>36</td><td>22(+1G)</td><td>TTTGGCACTAGCACATTTTTGCG</td><td>4822.4</td><td>8.8</td><td>1.1</td><td>7.9</td><td>2.7E-15</td><td>8.2E-15</td><td>BT20/MB468</td></tr>
<tr><td>>trnaMT_ValTAC_MT_+_1602_1670@42.71.30|AGATTTCAACTTAACTTGACCGCTCTGACC</td><td>>trnaMT_ValTAC_MT_+_1602_1670</td><td>42</td><td>71</td><td>30</td><td>AGATTTCAACTTAACTTGACCGCTCTGACC</td><td>709.1</td><td>7.9</td><td>1.0</td><td>7.9</td><td>2.7E-15</td><td>8.2E-15</td><td>BT20/MB468</td></tr>
<tr><td>>am_trna106_HisGTG_1_-_149155828_149155899@7.28.22,trna111_HisGTG_1_-_147774845_147774916@7.28.22,trna118_HisGTG_1_-_145396881_145396952@7.28.22,trna16_HisGTG_1_+_146544773_146544844@7.28.22,trna1_HisGTG_15_+_45493349_45493420@7.28.22,trna21_HisGTG_1_+_147753471_147753542@7.28.22,trna33_HisGTG_6_+_27125906_27125977@7.28.22,trna7_HisGTG_9_-_14433938_14434009@7.28.22,trna8_HisGTG_15_-_45492611_45492682@7.28.22,trna9_HisGTG_15_-_45490804_45490875@7.28.22|ATCGTATAGTGGTTAGTACTCT</td><td>>am_trna106_HisGTG_1_-_149155828_149155899</td><td>7</td><td>28</td><td>22</td><td>ATCGTATAGTGGTTAGTACTCT</td><td>519.1</td><td>-11.0</td><td>1.4</td><td>-7.9</td><td>2.8E-15</td><td>8.3E-15</td><td>BT20/MB468</td></tr>
<tr><td>>hg38_21_+_8433172_8446622_RNA45SN1_45S_50ntFlanks@6773.6803.31|TTCCTCCCGGGGCTACGCCTGTCTGAGCGTC</td><td>>hg38_21_+_8433172_8446622_RNA45SN1_45S_50ntFlanks</td><td>6773</td><td>6803</td><td>31</td><td>TTCCTCCCGGGGCTACGCCTGTCTGAGCGTC</td><td>1342.5</td><td>-5.3</td><td>0.7</td><td>-7.9</td><td>2.8E-15</td><td>8.3E-15</td><td>BT20/MB468</td></tr>
<tr><td>>MI0000071|hsa-mir-17&WithFlank&13|+|91350599|91350694@20.41.22&[MIMAT0000070&hsa-miR-17-5p&offsets|0|-1;m-88&13|+|91350618|91350640&offsets|0|-1]|CAAAGTGCTTACAGTGCAGGTA</td><td>>MI0000071|hsa-mir-17&WithFlank&13|+|91350599|91350694</td><td>20</td><td>41</td><td>22</td><td>CAAAGTGCTTACAGTGCAGGTA</td><td>2112.1</td><td>6.3</td><td>0.8</td><td>7.9</td><td>2.7E-15</td><td>8.3E-15</td><td>MB231/MB468</td></tr>
<tr><td>>hg38_21_+_8433172_8446622_RNA45SN1_45S_50ntFlanks@10388.10410.23|GTGAACAGCAGTTGAACATGGGT</td><td>>hg38_21_+_8433172_8446622_RNA45SN1_45S_50ntFlanks</td><td>10388</td><td>10410</td><td>23</td><td>GTGAACAGCAGTTGAACATGGGT</td><td>482.0</td><td>-5.8</td><td>0.7</td><td>-7.9</td><td>2.8E-15</td><td>8.4E-15</td><td>BT20/MB468</td></tr>
<tr><td>>hg38_21_+_8433172_8446622_RNA45SN1_45S_50ntFlanks@12627.12657.31|ATGGGGCGAAGCTACCATCTGTGGGATTATG</td><td>>hg38_21_+_8433172_8446622_RNA45SN1_45S_50ntFlanks</td><td>12627</td><td>12657</td><td>31</td><td>ATGGGGCGAAGCTACCATCTGTGGGATTATG</td><td>439.5</td><td>-4.0</td><td>0.5</td><td>-7.9</td><td>2.8E-15</td><td>8.4E-15</td><td>MB231/MB468</td></tr>
<tr><td>>trnaMT_GluTTC_MT_-_14674_14742@39.58.20,trnalookalike8_GluTTC_5_-_93905172_93905240@39.58.20|ATTGGTCGTGGTTGTAGTCC</td><td>>trnaMT_GluTTC_MT_-_14674_14742</td><td>39</td><td>58</td><td>20</td><td>ATTGGTCGTGGTTGTAGTCC</td><td>1082.8</td><td>9.6</td><td>1.2</td><td>7.9</td><td>2.8E-15</td><td>8.4E-15</td><td>BT20/MB468</td></tr>
<tr><td>>hg38_21_+_8433172_8446622_RNA45SN1_45S_50ntFlanks@10410.10433.24|TCAGTCGGTCCTGAGAGATGGGCG</td><td>>hg38_21_+_8433172_8446622_RNA45SN1_45S_50ntFlanks</td><td>10410</td><td>10433</td><td>24</td><td>TCAGTCGGTCCTGAGAGATGGGCG</td><td>1029.5</td><td>-7.1</td><td>0.9</td><td>-7.9</td><td>2.8E-15</td><td>8.5E-15</td><td>BT20/MB468</td></tr>
<tr><td>>hg38_21_+_8433172_8446622_RNA45SN1_45S_50ntFlanks@5421.5443.23|CGATTGGATGGTTTAGTGAGGCC</td><td>>hg38_21_+_8433172_8446622_RNA45SN1_45S_50ntFlanks</td><td>5421</td><td>5443</td><td>23</td><td>CGATTGGATGGTTTAGTGAGGCC</td><td>333.8</td><td>-10.0</td><td>1.3</td><td>-7.9</td><td>2.8E-15</td><td>8.5E-15</td><td>MB231/MB468</td></tr>
<tr><td>>hg38_21_+_8433172_8446622_RNA45SN1_45S_50ntFlanks@12934.12953.20|CTAAACCATTCGTAGACGAC</td><td>>hg38_21_+_8433172_8446622_RNA45SN1_45S_50ntFlanks</td><td>12934</td><td>12953</td><td>20</td><td>CTAAACCATTCGTAGACGAC</td><td>644.9</td><td>-4.6</td><td>0.6</td><td>-7.9</td><td>2.8E-15</td><td>8.5E-15</td><td>MB231/MB468</td></tr>
<tr><td>>hg38_21_+_8433172_8446622_RNA45SN1_45S_50ntFlanks@13002.13020.19|CGATCTATTGAAAGTCAGC</td><td>>hg38_21_+_8433172_8446622_RNA45SN1_45S_50ntFlanks</td><td>13002</td><td>13020</td><td>19</td><td>CGATCTATTGAAAGTCAGC</td><td>243.5</td><td>-9.9</td><td>1.3</td><td>-7.9</td><td>2.9E-15</td><td>8.6E-15</td><td>BT20/MB468</td></tr>
<tr><td>>MI0005756|hsa-mir-934&WithFlank&X|+|136550872|136550966@21.40.20&[MIMAT0004977&hsa-miR-934&offsets|0|-2;m-943&X|+|136550892|136550913&offsets|0|-2]|TGTCTACTACTGGAGACACT</td><td>>MI0005756|hsa-mir-934&WithFlank&X|+|136550872|136550966</td><td>21</td><td>40</td><td>20</td><td>TGTCTACTACTGGAGACACT</td><td>452.2</td><td>12.3</td><td>1.5</td><td>8.2</td><td>2.0E-16</td><td>8.6E-15</td><td>BT20/MB231</td></tr>
<tr><td>>MI0000078|hsa-mir-22&WithFlank&17|-|1713897|1713993@21.41.21&[MIMAT0004495&hsa-miR-22-5p&offsets|0|-1;m-217&17|-|1713953|1713973&offsets|0|0]|AGTTCTTCAGTGGCAAGCTTT</td><td>>MI0000078|hsa-mir-22&WithFlank&17|-|1713897|1713993</td><td>21</td><td>41</td><td>21</td><td>AGTTCTTCAGTGGCAAGCTTT</td><td>1079.4</td><td>7.2</td><td>0.9</td><td>7.9</td><td>2.9E-15</td><td>8.6E-15</td><td>BT20/MB468</td></tr>
<tr><td>>trnaMT_ValTAC_MT_+_1602_1670@50.72.23|ACTTAACTTGACCGCTCTGACCA</td><td>>trnaMT_ValTAC_MT_+_1602_1670</td><td>50</td><td>72</td><td>23</td><td>ACTTAACTTGACCGCTCTGACCA</td><td>4448.9</td><td>5.5</td><td>0.7</td><td>7.9</td><td>2.9E-15</td><td>8.7E-15</td><td>BT20/MB468</td></tr>
<tr><td>>hg38_21_+_8433172_8446622_RNA45SN1_45S_50ntFlanks@4005.4042.38|AACCTCGGGCCGATCGCACGCCCCCCGTGGCGGCGACG</td><td>>hg38_21_+_8433172_8446622_RNA45SN1_45S_50ntFlanks</td><td>4005</td><td>4042</td><td>38</td><td>AACCTCGGGCCGATCGCACGCCCCCCGTGGCGGCGACG</td><td>172.6</td><td>-4.4</td><td>0.6</td><td>-7.9</td><td>2.9E-15</td><td>8.8E-15</td><td>BT20/MB468</td></tr>
<tr><td>>hg38_21_+_8433172_8446622_RNA45SN1_45S_50ntFlanks@10512.10533.22|TCCGGAGTGGCGGAGATGGGCG</td><td>>hg38_21_+_8433172_8446622_RNA45SN1_45S_50ntFlanks</td><td>10512</td><td>10533</td><td>22</td><td>TCCGGAGTGGCGGAGATGGGCG</td><td>793.4</td><td>-8.7</td><td>1.1</td><td>-7.9</td><td>2.9E-15</td><td>8.8E-15</td><td>MB231/MB468</td></tr>
<tr><td>>MI0003820|hsa-mir-454&WithFlank&17|-|59137752|59137878@70.94.25&[MIMAT0003885&hsa-miR-454-3p&offsets|0|+2;m-272&17|-|59137787|59137809&offsets|0|+2]|TAGTGCAATATTGCTTATAGGGTTT</td><td>>MI0003820|hsa-mir-454&WithFlank&17|-|59137752|59137878</td><td>70</td><td>94</td><td>25</td><td>TAGTGCAATATTGCTTATAGGGTTT</td><td>1881.1</td><td>7.8</td><td>1.0</td><td>7.9</td><td>3.0E-15</td><td>8.9E-15</td><td>BT20/MB468</td></tr>
<tr><td>>hg38_21_+_8433172_8446622_RNA45SN1_45S_50ntFlanks@7988.8019.32|CAGACGTGGCGACCCGCTGAATTTAAGCATAT</td><td>>hg38_21_+_8433172_8446622_RNA45SN1_45S_50ntFlanks</td><td>7988</td><td>8019</td><td>32</td><td>CAGACGTGGCGACCCGCTGAATTTAAGCATAT</td><td>366.0</td><td>5.7</td><td>0.7</td><td>7.9</td><td>2.9E-15</td><td>8.9E-15</td><td>MB231/MB468</td></tr>
<tr><td>>hg38_21_+_8433172_8446622_RNA45SN1_45S_50ntFlanks@12991.13015.25|CTCCCTCGCTGCGATCTATTGAAAG</td><td>>hg38_21_+_8433172_8446622_RNA45SN1_45S_50ntFlanks</td><td>12991</td><td>13015</td><td>25</td><td>CTCCCTCGCTGCGATCTATTGAAAG</td><td>2381.8</td><td>-5.2</td><td>0.7</td><td>-7.9</td><td>3.0E-15</td><td>9.1E-15</td><td>MB231/MB468</td></tr>
<tr><td>>hg38_21_+_8433172_8446622_RNA45SN1_45S_50ntFlanks@10362.10384.23|AAGGCCGAAGTGGAGAAGGGTTC</td><td>>hg38_21_+_8433172_8446622_RNA45SN1_45S_50ntFlanks</td><td>10362</td><td>10384</td><td>23</td><td>AAGGCCGAAGTGGAGAAGGGTTC</td><td>520.1</td><td>-4.2</td><td>0.5</td><td>-7.9</td><td>3.0E-15</td><td>9.2E-15</td><td>MB231/MB468</td></tr>
<tr><td>>trna111_HisGTG_1_-_147774845_147774916@-1T.25.26,trna118_HisGTG_1_-_145396881_145396952@-1T.25.26,trna16_HisGTG_1_+_146544773_146544844@-1T.25.26,trna1_HisGTG_15_+_45493349_45493420@-1T.25.26,trna21_HisGTG_1_+_147753471_147753542@-1T.25.26,trna33_HisGTG_6_+_27125906_27125977@-1T.25.26,trna7_HisGTG_9_-_14433938_14434009@-1T.25.26,trna8_HisGTG_15_-_45492611_45492682@-1T.25.26,trna9_HisGTG_15_-_45490804_45490875@-1T.25.26|TGCCGTGATCGTATAGTGGTTAGTAC</td><td>>trna111_HisGTG_1_-_147774845_147774916</td><td>-1T</td><td>25</td><td>26</td><td>TGCCGTGATCGTATAGTGGTTAGTAC</td><td>1506.2</td><td>6.5</td><td>0.8</td><td>7.9</td><td>3.0E-15</td><td>9.2E-15</td><td>MB231/MB468</td></tr>
<tr><td>>hg38_21_+_8433172_8446622_RNA45SN1_45S_50ntFlanks@11534.11560.27|GTTCCGCGCGGCGCCTCGCCTCGGCCG</td><td>>hg38_21_+_8433172_8446622_RNA45SN1_45S_50ntFlanks</td><td>11534</td><td>11560</td><td>27</td><td>GTTCCGCGCGGCGCCTCGCCTCGGCCG</td><td>288.9</td><td>-10.2</td><td>1.3</td><td>-7.9</td><td>3.1E-15</td><td>9.2E-15</td><td>BT20/MB468</td></tr>
<tr><td>>hg38_21_+_8433172_8446622_RNA45SN1_45S_50ntFlanks@10350.10368.19|ACGAGAACTTTGAAGGCCG</td><td>>hg38_21_+_8433172_8446622_RNA45SN1_45S_50ntFlanks</td><td>10350</td><td>10368</td><td>19</td><td>ACGAGAACTTTGAAGGCCG</td><td>7381.0</td><td>-4.6</td><td>0.6</td><td>-7.9</td><td>3.1E-15</td><td>9.4E-15</td><td>MB231/MB468</td></tr>
<tr><td>>MI0003666|hsa-mir-651&WithFlank&X|+|8126959|8127067@22.43.22&[MIMAT0003321&hsa-miR-651-5p&offsets|0|0;m-399&X|+|8126980|8127001&offsets|0|0]|TTTAGGATAAGCTTGACTTTTG</td><td>>MI0003666|hsa-mir-651&WithFlank&X|+|8126959|8127067</td><td>22</td><td>43</td><td>22</td><td>TTTAGGATAAGCTTGACTTTTG</td><td>1229.2</td><td>9.0</td><td>1.1</td><td>7.9</td><td>3.1E-15</td><td>9.4E-15</td><td>MB231/MB468</td></tr>
<tr><td>>hg38_21_+_8433172_8446622_RNA45SN1_45S_50ntFlanks@4733.4765.33|GAACGAAAGTCGGAGGTTCGAAGACGATCAGAT</td><td>>hg38_21_+_8433172_8446622_RNA45SN1_45S_50ntFlanks</td><td>4733</td><td>4765</td><td>33</td><td>GAACGAAAGTCGGAGGTTCGAAGACGATCAGAT</td><td>744.2</td><td>8.8</td><td>1.1</td><td>7.9</td><td>3.1E-15</td><td>9.4E-15</td><td>MB231/MB468</td></tr>
<tr><td>>am_MI0000786|hsa-mir-378a&WithFlank&5|+|149732819|149732896@49.69.21&[MIMAT0000732&hsa-miR-378a-3p&offsets|0|-1;m-44&5|+|149732867|149732888&offsets|0|-1]|ACTGGACTTGGAGTCAGAAGG</td><td>>am_MI0000786|hsa-mir-378a&WithFlank&5|+|149732819|149732896</td><td>49</td><td>69</td><td>21</td><td>ACTGGACTTGGAGTCAGAAGG</td><td>5604.5</td><td>5.7</td><td>0.7</td><td>7.9</td><td>3.1E-15</td><td>9.5E-15</td><td>MB231/MB468</td></tr>
<tr><td>>hg38_21_+_8433172_8446622_RNA45SN1_45S_50ntFlanks@10808.10830.23|CAATGTAGGTAAGGGAAGTCGGC</td><td>>hg38_21_+_8433172_8446622_RNA45SN1_45S_50ntFlanks</td><td>10808</td><td>10830</td><td>23</td><td>CAATGTAGGTAAGGGAAGTCGGC</td><td>198.5</td><td>-4.3</td><td>0.5</td><td>-7.9</td><td>3.2E-15</td><td>9.6E-15</td><td>BT20/MB468</td></tr>
<tr><td>>hg38_21_+_8433172_8446622_RNA45SN1_45S_50ntFlanks@11767.11791.25|TCTAATTAGTGACGCGCATGAATGG</td><td>>hg38_21_+_8433172_8446622_RNA45SN1_45S_50ntFlanks</td><td>11767</td><td>11791</td><td>25</td><td>TCTAATTAGTGACGCGCATGAATGG</td><td>159.1</td><td>-4.2</td><td>0.5</td><td>-7.9</td><td>3.2E-15</td><td>9.8E-15</td><td>MB231/MB468</td></tr>
<tr><td>>hg38_21_+_8433172_8446622_RNA45SN1_45S_50ntFlanks@10458.10497.40|ATGGCCTCCGTTGCCCTCGGCCGATCGAAAGGGAGTCGGG</td><td>>hg38_21_+_8433172_8446622_RNA45SN1_45S_50ntFlanks</td><td>10458</td><td>10497</td><td>40</td><td>ATGGCCTCCGTTGCCCTCGGCCGATCGAAAGGGAGTCGGG</td><td>228.0</td><td>-9.8</td><td>1.2</td><td>-7.9</td><td>3.3E-15</td><td>9.9E-15</td><td>BT20/MB468</td></tr>
<tr><td>>hg38_21_+_8433172_8446622_RNA45SN1_45S_50ntFlanks@11767.11803.37|TCTAATTAGTGACGCGCATGAATGGATGAACGAGATT</td><td>>hg38_21_+_8433172_8446622_RNA45SN1_45S_50ntFlanks</td><td>11767</td><td>11803</td><td>37</td><td>TCTAATTAGTGACGCGCATGAATGGATGAACGAGATT</td><td>461.6</td><td>-3.4</td><td>0.4</td><td>-7.9</td><td>3.3E-15</td><td>1.0E-14</td><td>BT20/MB468</td></tr>
<tr><td>>hg38_21_+_8433172_8446622_RNA45SN1_45S_50ntFlanks@12934.12957.24|CTAAACCATTCGTAGACGACCTGC</td><td>>hg38_21_+_8433172_8446622_RNA45SN1_45S_50ntFlanks</td><td>12934</td><td>12957</td><td>24</td><td>CTAAACCATTCGTAGACGACCTGC</td><td>233.7</td><td>-3.8</td><td>0.5</td><td>-7.9</td><td>3.4E-15</td><td>1.0E-14</td><td>BT20/MB468</td></tr>
<tr><td>>hg38_21_+_8433172_8446622_RNA45SN1_45S_50ntFlanks@4367.4394.28|CAGTTAAAAAGCTCGTAGTTGGATCTTG</td><td>>hg38_21_+_8433172_8446622_RNA45SN1_45S_50ntFlanks</td><td>4367</td><td>4394</td><td>28</td><td>CAGTTAAAAAGCTCGTAGTTGGATCTTG</td><td>212.2</td><td>-6.0</td><td>0.8</td><td>-7.9</td><td>3.3E-15</td><td>1.0E-14</td><td>MB231/MB468</td></tr>
<tr><td>>hg38_21_+_8433172_8446622_RNA45SN1_45S_50ntFlanks@4731.4767.37|AAGAACGAAAGTCGGAGGTTCGAAGACGATCAGATAC</td><td>>hg38_21_+_8433172_8446622_RNA45SN1_45S_50ntFlanks</td><td>4731</td><td>4767</td><td>37</td><td>AAGAACGAAAGTCGGAGGTTCGAAGACGATCAGATAC</td><td>637.3</td><td>8.9</td><td>1.1</td><td>7.9</td><td>3.4E-15</td><td>1.0E-14</td><td>BT20/MB468</td></tr>
<tr><td>>MI0000299|hsa-mir-222&WithFlank&X|-|45747009|45747130@75.97.23&[MIMAT0000279&hsa-miR-222-3p&offsets|0|+2;m-140&X|-|45747033|45747056&offsets|0|-1]|AGCTACATCTGGCTACTGGGTCT</td><td>>MI0000299|hsa-mir-222&WithFlank&X|-|45747009|45747130</td><td>75</td><td>97</td><td>23</td><td>AGCTACATCTGGCTACTGGGTCT</td><td>5706.8</td><td>6.4</td><td>0.8</td><td>7.9</td><td>3.4E-15</td><td>1.0E-14</td><td>BT20/MB468</td></tr>
<tr><td>>MI0016887|hsa-mir-4521&WithFlank&17|+|8186939|8187010@13.35.23&[MIMAT0019058&hsa-miR-4521&offsets|+3|+4;m-765&17|+|8186948|8186969&offsets|+3|+4]|AAGGAAGTCCTGTGCTCAGTTTT</td><td>>MI0016887|hsa-mir-4521&WithFlank&17|+|8186939|8187010</td><td>13</td><td>35</td><td>23</td><td>AAGGAAGTCCTGTGCTCAGTTTT</td><td>905.1</td><td>10.2</td><td>1.3</td><td>7.9</td><td>3.4E-15</td><td>1.0E-14</td><td>BT20/MB468</td></tr>
</table>

<table>
<tr><th></th><th></th><th></th><th></th><th></th><th></th><th></th><th></th><th></th><th></th><th></th><th></th><th></th></tr>
<tr><td>>hg38_21_+_8433172_8446622_RNA45SN1_45S_50ntFlanks@6645.6674.30|CTCGTACGACTCTTAGCGGTGGATCACTCG</td><td>>hg38_21_+_8433172_8446622_RNA45SN1_45S_50ntFlanks</td><td>6645</td><td>6674</td><td>30</td><td>CTCGTACGACTCTTAGCGGTGGATCACTCG</td><td>444.6</td><td>4.0</td><td>0.5</td><td>7.9</td><td>3.4E-15</td><td>1.0E-14</td><td>BT20/MB468</td></tr>
<tr><td>>hg38_21_+_8433172_8446622_RNA45SN1_45S_50ntFlanks@12417.12441.25|TCGATGTCGGCTCTTCCTATCATTG</td><td>>hg38_21_+_8433172_8446622_RNA45SN1_45S_50ntFlanks</td><td>12417</td><td>12441</td><td>25</td><td>TCGATGTCGGCTCTTCCTATCATTG</td><td>812.1</td><td>-3.6</td><td>0.5</td><td>-7.9</td><td>3.5E-15</td><td>1.0E-14</td><td>BT20/MB468</td></tr>
<tr><td>>hg38_21_+_8433172_8446622_RNA45SN1_45S_50ntFlanks@12456.12479.24|CCAAGCGTTGGATTGTTCACCCAC</td><td>>hg38_21_+_8433172_8446622_RNA45SN1_45S_50ntFlanks</td><td>12456</td><td>12479</td><td>24</td><td>CCAAGCGTTGGATTGTTCACCCAC</td><td>661.1</td><td>-5.8</td><td>0.7</td><td>-7.9</td><td>3.5E-15</td><td>1.0E-14</td><td>BT20/MB468</td></tr>
<tr><td>>MI0000782|hsa-mir-374a&WithFlank&X|-|74287280|74287363@48.69.22&[MIMAT0004688&hsa-miR-374a-3p&offsets|0|0;m-74&X|-|74287295|74287316&offsets|0|0]|CTTATCAGATTGTATTGTAATT</td><td>>MI0000782|hsa-mir-374a&WithFlank&X|-|74287280|74287363</td><td>48</td><td>69</td><td>22</td><td>CTTATCAGATTGTATTGTAATT</td><td>1910.7</td><td>6.8</td><td>0.9</td><td>7.9</td><td>3.4E-15</td><td>1.0E-14</td><td>MB231/MB468</td></tr>
<tr><td>>MI0000441|hsa-mir-30b&WithFlank&8|-|134800514|134800613@23.43.21(+1G)&[MIMAT0000420&hsa-miR-30b-5p&offsets|0|-1(+1G);m-76&8|-|134800570|134800591&offsets|0|-1(+1G)]|TGTAAACATCCTACACTCAGCG</td><td>>MI0000441|hsa-mir-30b&WithFlank&8|-|134800514|134800613</td><td>23</td><td>43</td><td>21(+1G)</td><td>TGTAAACATCCTACACTCAGCG</td><td>9150.7</td><td>11.7</td><td>1.5</td><td>7.9</td><td>3.5E-15</td><td>1.0E-14</td><td>BT20/MB468</td></tr>
<tr><td>>hg38_21_+_8433172_8446622_RNA45SN1_45S_50ntFlanks@12636.12653.18|AGCTACCATCTGTGGGAT</td><td>>hg38_21_+_8433172_8446622_RNA45SN1_45S_50ntFlanks</td><td>12636</td><td>12653</td><td>18</td><td>AGCTACCATCTGTGGGAT</td><td>977.8</td><td>-11.5</td><td>1.5</td><td>-7.9</td><td>3.5E-15</td><td>1.0E-14</td><td>MB231/MB468</td></tr>
<tr><td>>hg38_21_+_8433172_8446622_RNA45SN1_45S_50ntFlanks@4003.4026.24|ATAACCTCGGGCCGATCGCACGCC</td><td>>hg38_21_+_8433172_8446622_RNA45SN1_45S_50ntFlanks</td><td>4003</td><td>4026</td><td>24</td><td>ATAACCTCGGGCCGATCGCACGCC</td><td>324.7</td><td>-9.9</td><td>1.3</td><td>-7.9</td><td>3.5E-15</td><td>1.1E-14</td><td>MB231/MB468</td></tr>
<tr><td>>hg38_21_+_8433172_8446622_RNA45SN1_45S_50ntFlanks@9736.9765.30|CCGAAACGATCTCAACCTATTCTCAAACTT</td><td>>hg38_21_+_8433172_8446622_RNA45SN1_45S_50ntFlanks</td><td>9736</td><td>9765</td><td>30</td><td>CCGAAACGATCTCAACCTATTCTCAAACTT</td><td>283.8</td><td>-10.1</td><td>1.3</td><td>-7.9</td><td>3.6E-15</td><td>1.1E-14</td><td>BT20/MB468</td></tr>
<tr><td>>hg38_21_+_8433172_8446622_RNA45SN1_45S_50ntFlanks@12931.12956.26|GCGCTAAACCATTCGTAGACGACCTG</td><td>>hg38_21_+_8433172_8446622_RNA45SN1_45S_50ntFlanks</td><td>12931</td><td>12956</td><td>26</td><td>GCGCTAAACCATTCGTAGACGACCTG</td><td>419.0</td><td>-4.7</td><td>0.6</td><td>-7.9</td><td>3.6E-15</td><td>1.1E-14</td><td>BT20/MB468</td></tr>
<tr><td>>hg38_21_+_8433172_8446622_RNA45SN1_45S_50ntFlanks@11584.11616.33|TGGTGCGGACCAGGGGAATCCGACTGTTTAATT</td><td>>hg38_21_+_8433172_8446622_RNA45SN1_45S_50ntFlanks</td><td>11584</td><td>11616</td><td>33</td><td>TGGTGCGGACCAGGGGAATCCGACTGTTTAATT</td><td>384.6</td><td>7.7</td><td>1.0</td><td>7.9</td><td>3.6E-15</td><td>1.1E-14</td><td>BT20/MB468</td></tr>
<tr><td>>hg38_21_+_8433172_8446622_RNA45SN1_45S_50ntFlanks@10802.10822.21|TTGGAACAATGTAGGTAAGGG</td><td>>hg38_21_+_8433172_8446622_RNA45SN1_45S_50ntFlanks</td><td>10802</td><td>10822</td><td>21</td><td>TTGGAACAATGTAGGTAAGGG</td><td>125.7</td><td>-4.6</td><td>0.6</td><td>-7.9</td><td>3.7E-15</td><td>1.1E-14</td><td>BT20/MB468</td></tr>
<tr><td>>hg38_21_+_8433172_8446622_RNA45SN1_45S_50ntFlanks@10841.10858.18|CGTAACTTCGGGATAAGG</td><td>>hg38_21_+_8433172_8446622_RNA45SN1_45S_50ntFlanks</td><td>10841</td><td>10858</td><td>18</td><td>CGTAACTTCGGGATAAGG</td><td>994.3</td><td>-6.4</td><td>0.8</td><td>-7.9</td><td>3.7E-15</td><td>1.1E-14</td><td>BT20/MB468</td></tr>
<tr><td>>hg38_21_+_8433172_8446622_RNA45SN1_45S_50ntFlanks@11999.12030.32|CCCTGCGGGCCGCCGGTGAAATACCACTACTC</td><td>>hg38_21_+_8433172_8446622_RNA45SN1_45S_50ntFlanks</td><td>11999</td><td>12030</td><td>32</td><td>CCCTGCGGGCCGCCGGTGAAATACCACTACTC</td><td>228.9</td><td>-9.8</td><td>1.3</td><td>-7.9</td><td>3.7E-15</td><td>1.1E-14</td><td>BT20/MB468</td></tr>
<tr><td>>hg38_21_+_8433172_8446622_RNA45SN1_45S_50ntFlanks@6739.6770.32|TCATCGACACTTCGAACGCACTTGCGGCCCCG</td><td>>hg38_21_+_8433172_8446622_RNA45SN1_45S_50ntFlanks</td><td>6739</td><td>6770</td><td>32</td><td>TCATCGACACTTCGAACGCACTTGCGGCCCCG</td><td>323.8</td><td>-6.6</td><td>0.8</td><td>-7.9</td><td>3.6E-15</td><td>1.1E-14</td><td>MB231/MB468</td></tr>
<tr><td>>am_MI0000815|hsa-mir-339&WithFlank&7|-|1022927|1023032@21.42.22&[MIMAT0000764&hsa-miR-339-5p&offsets|0|-1;m-163&7|-|1022990|1023012&offsets|0|-1]|TCCCTGTCCTCCAGGAGCTCAC</td><td>>am_MI0000815|hsa-mir-339&WithFlank&7|-|1022927|1023032</td><td>21</td><td>42</td><td>22</td><td>TCCCTGTCCTCCAGGAGCTCAC</td><td>727.7</td><td>7.2</td><td>0.9</td><td>7.9</td><td>3.7E-15</td><td>1.1E-14</td><td>BT20/MB468</td></tr>
<tr><td>>am_trnaMT_ValTAC_MT_+_1602_1670@28.48.21|AACTTACACTTAGGAGATTTC</td><td>>am_trnaMT_ValTAC_MT_+_1602_1670</td><td>28</td><td>48</td><td>21</td><td>AACTTACACTTAGGAGATTTC</td><td>478.7</td><td>9.0</td><td>1.1</td><td>7.9</td><td>3.8E-15</td><td>1.1E-14</td><td>BT20/MB468</td></tr>
<tr><td>>trna10_AlaCGC_6_+_26553731_26553802@58.75.18,trna119_AlaCGC_6_-_28641613_28641684@58.75.18,trna13_AlaCGC_2_+_157257281_157257352@58.75.18,trna13_AlaTGC_12_-_125406301_125406372@58.75.18,trna66_AlaTGC_6_+_28611222_28611293@58.75.18,trna8_AlaTGC_12_+_125424512_125424583@58.75.18,trna8_AlaTGC_5_+_180633868_180633939@58.75.18|TCCCCGGCATCTCCACCA</td><td>>trna10_AlaCGC_6_+_26553731_26553802</td><td>58</td><td>75</td><td>18</td><td>TCCCCGGCATCTCCACCA</td><td>5513.3</td><td>7.0</td><td>0.9</td><td>7.9</td><td>3.7E-15</td><td>1.1E-14</td><td>MB231/MB468</td></tr>
<tr><td>>hg38_21_+_8433172_8446622_RNA45SN1_45S_50ntFlanks@10392.10420.29|ACAGCAGTTGAACATGGGTCAGTCGGTCC</td><td>>hg38_21_+_8433172_8446622_RNA45SN1_45S_50ntFlanks</td><td>10392</td><td>10420</td><td>29</td><td>ACAGCAGTTGAACATGGGTCAGTCGGTCC</td><td>371.1</td><td>-7.0</td><td>0.9</td><td>-7.9</td><td>3.8E-15</td><td>1.1E-14</td><td>BT20/MB468</td></tr>
<tr><td>>MI0000269|hsa-mir-181a-2&WithFlank&9|+|124692436|124692557@83.103.21(+1U)&[MIMAT0004558&hsa-miR-181a-2-3p&offsets|0|-1(+1U);m-71&9|+|124692518|124692539&offsets|0|-1(+1U)]|ACCACTGACCGTTGACTGTACT</td><td>>MI0000269|hsa-mir-181a-2&WithFlank&9|+|124692436|124692557</td><td>83</td><td>103</td><td>21(+1U)</td><td>ACCACTGACCGTTGACTGTACT</td><td>771.7</td><td>8.9</td><td>1.1</td><td>7.9</td><td>3.8E-15</td><td>1.1E-14</td><td>BT20/MB468</td></tr>
<tr><td>>MI0000749|hsa-mir-30e&WithFlank&1|+|40754349|40754452@65.86.22&[MIMAT0000693&hsa-miR-30e-3p&offsets|0|0;m-25&1|+|40754413|40754434&offsets|0|0]|CTTTCAGTCGGATGTTTACAGC</td><td>>MI0000749|hsa-mir-30e&WithFlank&1|+|40754349|40754452</td><td>65</td><td>86</td><td>22</td><td>CTTTCAGTCGGATGTTTACAGC</td><td>6294.5</td><td>5.9</td><td>0.7</td><td>7.9</td><td>3.7E-15</td><td>1.1E-14</td><td>MB231/MB468</td></tr>
<tr><td>>MI0000273|hsa-mir-183&WithFlank&7|-|129774899|129775020@34.53.20(+1A)&[MIMAT0000261&hsa-miR-183-5p&offsets|+1|-1(+1A);m-41&7|-|129774966|129774987&offsets|0|-2(+1A)]|ATGGCACTGGTAGAATTCACA</td><td>>MI0000273|hsa-mir-183&WithFlank&7|-|129774899|129775020</td><td>34</td><td>53</td><td>20(+1A)</td><td>ATGGCACTGGTAGAATTCACA</td><td>1000.1</td><td>10.2</td><td>1.3</td><td>7.9</td><td>3.8E-15</td><td>1.1E-14</td><td>BT20/MB468</td></tr>
<tr><td>>hg38_21_+_8433172_8446622_RNA45SN1_45S_50ntFlanks@12423.12443.21|TCGGCTCTTCCTATCATTGTG</td><td>>hg38_21_+_8433172_8446622_RNA45SN1_45S_50ntFlanks</td><td>12423</td><td>12443</td><td>21</td><td>TCGGCTCTTCCTATCATTGTG</td><td>309.3</td><td>-3.5</td><td>0.4</td><td>-7.9</td><td>3.8E-15</td><td>1.1E-14</td><td>BT20/MB468</td></tr>
<tr><td>>hg38_21_+_8433172_8446622_RNA45SN1_45S_50ntFlanks@10787.10816.30|ACAGCCTCTGGCATGTTGGAACAATGTAGG</td><td>>hg38_21_+_8433172_8446622_RNA45SN1_45S_50ntFlanks</td><td>10787</td><td>10816</td><td>30</td><td>ACAGCCTCTGGCATGTTGGAACAATGTAGG</td><td>326.4</td><td>-3.3</td><td>0.4</td><td>-7.9</td><td>3.8E-15</td><td>1.1E-14</td><td>BT20/MB468</td></tr>
<tr><td>>trnaMT_HisGTG_MT_+_12138_12206@28.72.45|ATTGTGAATCTGACAACAGAGGCTTACGACCCCTTATTTACCCCA</td><td>>trnaMT_HisGTG_MT_+_12138_12206</td><td>28</td><td>72</td><td>45</td><td>ATTGTGAATCTGACAACAGAGGCTTACGACCCCTTATTTACCCCA</td><td>1147.2</td><td>7.5</td><td>0.9</td><td>7.9</td><td>3.8E-15</td><td>1.1E-14</td><td>MB231/MB468</td></tr>
<tr><td>>trna112_GlnCTG_1_-_147737382_147737453@30.50.21,trna131_GlnCTG_6_-_27759135_27759206@30.50.21,trna146_GlnCTG_6_-_27515531_27515602@30.50.21,trna15_GlnCTG_1_+_145963304_145963375@30.50.21,trna19_GlnCTG_1_+_147505038_147505109@30.50.21,trna1_GlnCTG_6_+_18836402_18836473@30.50.21,trna28_GlnCTG_1_+_149186125_149186196@30.50.21,trna3_GlnCTG_17_+_8023070_8023141@30.50.21,trna49_GlnCTG_6_+_27487308_27487379@30.50.21,trna7_GlnCTG_15_-_66161400_66161471@30.50.21,trna99_GlnCTG_6_-_28909378_28909449@30.50.21|GACTCTGAATCCAGCGATCCG</td><td>>trna112_GlnCTG_1_-_147737382_147737453</td><td>30</td><td>50</td><td>21</td><td>GACTCTGAATCCAGCGATCCG</td><td>390.2</td><td>-5.8</td><td>0.7</td><td>-7.9</td><td>3.9E-15</td><td>1.1E-14</td><td>BT20/MB468</td></tr>
<tr><td>>hg38_21_+_8433172_8446622_RNA45SN1_45S_50ntFlanks@12589.12614.26|GTTCAGACATTTGGTGTATGTGCTTG</td><td>>hg38_21_+_8433172_8446622_RNA45SN1_45S_50ntFlanks</td><td>12589</td><td>12614</td><td>26</td><td>GTTCAGACATTTGGTGTATGTGCTTG</td><td>413.9</td><td>-7.5</td><td>1.0</td><td>-7.9</td><td>3.9E-15</td><td>1.2E-14</td><td>MB231/MB468</td></tr>
<tr><td>>hg38_21_+_8433172_8446622_RNA45SN1_45S_50ntFlanks@12605.12623.19|TATGTGCTTGGCTGAGGAG</td><td>>hg38_21_+_8433172_8446622_RNA45SN1_45S_50ntFlanks</td><td>12605</td><td>12623</td><td>19</td><td>TATGTGCTTGGCTGAGGAG</td><td>299.9</td><td>-4.8</td><td>0.6</td><td>-7.9</td><td>4.0E-15</td><td>1.2E-14</td><td>BT20/MB468</td></tr>
<tr><td>>hg38_21_+_8433172_8446622_RNA45SN1_45S_50ntFlanks@11890.11912.23|CTTGACTCTAGTCTGGCACGGTG</td><td>>hg38_21_+_8433172_8446622_RNA45SN1_45S_50ntFlanks</td><td>11890</td><td>11912</td><td>23</td><td>CTTGACTCTAGTCTGGCACGGTG</td><td>513.9</td><td>-6.3</td><td>0.8</td><td>-7.9</td><td>3.9E-15</td><td>1.2E-14</td><td>MB231/MB468</td></tr>
<tr><td>>hg38_21_+_8433172_8446622_RNA45SN1_45S_50ntFlanks@10351.10381.31|CGAGAACTTTGAAGGCCGAAGTGGAGAAGGG</td><td>>hg38_21_+_8433172_8446622_RNA45SN1_45S_50ntFlanks</td><td>10351</td><td>10381</td><td>31</td><td>CGAGAACTTTGAAGGCCGAAGTGGAGAAGGG</td><td>424.5</td><td>-4.6</td><td>0.6</td><td>-7.9</td><td>4.0E-15</td><td>1.2E-14</td><td>BT20/MB468</td></tr>
<tr><td>>hg38_21_+_8433172_8446622_RNA45SN1_45S_50ntFlanks@12008.12034.27|CCGCCGGTGAAATACCACTACTCTGAT</td><td>>hg38_21_+_8433172_8446622_RNA45SN1_45S_50ntFlanks</td><td>12008</td><td>12034</td><td>27</td><td>CCGCCGGTGAAATACCACTACTCTGAT</td><td>208.1</td><td>-6.2</td><td>0.8</td><td>-7.9</td><td>4.0E-15</td><td>1.2E-14</td><td>BT20/MB468</td></tr>
<tr><td>>hg38_21_+_8433172_8446622_RNA45SN1_45S_50ntFlanks@12996.13036.41|TCGCTGCGATCTATTGAAAGTCAGCCCTCGACACAAGGGTT</td><td>>hg38_21_+_8433172_8446622_RNA45SN1_45S_50ntFlanks</td><td>12996</td><td>13036</td><td>41</td><td>TCGCTGCGATCTATTGAAAGTCAGCCCTCGACACAAGGGTT</td><td>2934.5</td><td>9.2</td><td>1.2</td><td>7.9</td><td>4.0E-15</td><td>1.2E-14</td><td>BT20/MB468</td></tr>
<tr><td>>MI0005559|hsa-mir-744&WithFlank&17|+|12081893|12082002@17.38.22&[MIMAT0004945&hsa-miR-744-5p&offsets|0|0;m-172&17|+|12081909|12081929&offsets|0|+1]|TGCGGGGCTAGGGCTAACAGCA</td><td>>MI0005559|hsa-mir-744&WithFlank&17|+|12081893|12082002</td><td>17</td><td>38</td><td>22</td><td>TGCGGGGCTAGGGCTAACAGCA</td><td>1070.2</td><td>6.2</td><td>0.8</td><td>7.9</td><td>3.9E-15</td><td>1.2E-14</td><td>MB231/MB468</td></tr>
<tr><td>>hg38_21_+_8433172_8446622_RNA45SN1_45S_50ntFlanks@12657.12674.18|GACTGAACGCCTCTAAGT</td><td>>hg38_21_+_8433172_8446622_RNA45SN1_45S_50ntFlanks</td><td>12657</td><td>12674</td><td>18</td><td>GACTGAACGCCTCTAAGT</td><td>442.1</td><td>-3.3</td><td>0.4</td><td>-7.9</td><td>4.0E-15</td><td>1.2E-14</td><td>MB231/MB468</td></tr>
<tr><td>>MI0000736|hsa-mir-30c-1&WithFlank&1|+|40757278|40757378@23.44.22(+1G);MI0000254|hsa-mir-30c-2&WithFlank&6|-|71376954|71377037@13.34.22(+1G)&[MIMAT0000244&hsa-miR-30c-5p&offsets|0|-1(+1G);m-57&1|+|40757300|40757323&offsets|0|-2(+1G)];[MIMAT0000244_1&hsa-miR-30c-5p&offsets|0|-1(+1G);m-56&6|-|71377002|71377025&offsets|0|-2(+1G)]|TGTAAACATCCTACACTCTCAGG</td><td>>MI0000736|hsa-mir-30c-1&WithFlank&1|+|40757278|40757378</td><td>23</td><td>44</td><td>22(+1G)</td><td>TGTAAACATCCTACACTCTCAGG</td><td>651.1</td><td>8.2</td><td>1.0</td><td>7.9</td><td>4.0E-15</td><td>1.2E-14</td><td>MB231/MB468</td></tr>
<tr><td>>hg38_21_+_8433172_8446622_RNA45SN1_45S_50ntFlanks@9554.9584.31|CGTAGCGGTCCTGACGTGCAAATCGGTCGTC</td><td>>hg38_21_+_8433172_8446622_RNA45SN1_45S_50ntFlanks</td><td>9554</td><td>9584</td><td>31</td><td>CGTAGCGGTCCTGACGTGCAAATCGGTCGTC</td><td>1635.0</td><td>-4.7</td><td>0.6</td><td>-7.9</td><td>4.1E-15</td><td>1.2E-14</td><td>BT20/MB468</td></tr>
<tr><td>>hg38_21_+_8433172_8446622_RNA45SN1_45S_50ntFlanks@3782.3801.20|CAGTGAAACTGCGAATGGCT</td><td>>hg38_21_+_8433172_8446622_RNA45SN1_45S_50ntFlanks</td><td>3782</td><td>3801</td><td>20</td><td>CAGTGAAACTGCGAATGGCT</td><td>1032.5</td><td>-4.2</td><td>0.5</td><td>-7.9</td><td>4.0E-15</td><td>1.2E-14</td><td>MB231/MB468</td></tr>
<tr><td>>hg38_21_+_8433172_8446622_RNA45SN1_45S_50ntFlanks@11540.11586.47|CGCGGCGCCTCGCCTCGGCCGGCGCCTAGCAGCCGACTTAGAACTGG</td><td>>hg38_21_+_8433172_8446622_RNA45SN1_45S_50ntFlanks</td><td>11540</td><td>11586</td><td>47</td><td>CGCGGCGCCTCGCCTCGGCCGGCGCCTAGCAGCCGACTTAGAACTGG</td><td>677.9</td><td>7.6</td><td>1.0</td><td>7.9</td><td>4.0E-15</td><td>1.2E-14</td><td>MB231/MB468</td></tr>
<tr><td>>MI0000273|hsa-mir-183&WithFlank&7|-|129774899|129775020@33.54.22&[MIMAT0000261&hsa-miR-183-5p&offsets|0|0;m-41&7|-|129774966|129774987&offsets|-1|-1]|TATGGCACTGGTAGAATTCACT</td><td>>MI0000273|hsa-mir-183&WithFlank&7|-|129774899|129775020</td><td>33</td><td>54</td><td>22</td><td>TATGGCACTGGTAGAATTCACT</td><td>11169.4</td><td>5.9</td><td>0.7</td><td>7.9</td><td>4.0E-15</td><td>1.2E-14</td><td>MB231/MB468</td></tr>
<tr><td>>MI0000100|hsa-mir-98&WithFlank&X|-|53556217|53556347@28.48.21&[MIMAT0000096&hsa-miR-98-5p&offsets|0|-1;m-143&X|-|53556299|53556320&offsets|0|-1]|TGAGGTAGTAAGTTGTATTGT</td><td>>MI0000100|hsa-mir-98&WithFlank&X|-|53556217|53556347</td><td>28</td><td>48</td><td>21</td><td>TGAGGTAGTAAGTTGTATTGT</td><td>1082.0</td><td>6.0</td><td>0.8</td><td>7.9</td><td>4.0E-15</td><td>1.2E-14</td><td>MB231/MB468</td></tr>
<tr><td>>hg38_21_+_8433172_8446622_RNA45SN1_45S_50ntFlanks@5471.5516.46|CACGGCCCTGGCGGAGCGCTGAGAAGACGGTCGAACTTGACTATCT</td><td>>hg38_21_+_8433172_8446622_RNA45SN1_45S_50ntFlanks</td><td>5471</td><td>5516</td><td>46</td><td>CACGGCCCTGGCGGAGCGCTGAGAAGACGGTCGAACTTGACTATCT</td><td>5318.2</td><td>10.6</td><td>1.4</td><td>7.9</td><td>4.0E-15</td><td>1.2E-14</td><td>MB231/MB468</td></tr>
</table>

| | | | | | | | | | | | | |
|---|---|---|---|---|---|---|---|---|---|---|---|---|
| >hg38_21_+_8433172_8446622_RNA45SN1_45S_50ntFlanks@8182.8210.29\|ATCGAGGCCCAGCCCGTGGACGGTGTGAG | >hg38_21_+_8433172_8446622_RNA45SN1_45S_50ntFlanks | 8182 | 8210 | 29 | ATCGAGGCCCAGCCCGTGGACGGTGTGAG | 220.6 | -9.8 | 1.2 | -7.9 | 4.1E-15 | 1.2E-14 | BT20/MB468 |
| >hg38_21_+_8433172_8446622_RNA45SN1_45S_50ntFlanks@10841.10862.22\|CGTAACTTCGGGATAAGGATTG | >hg38_21_+_8433172_8446622_RNA45SN1_45S_50ntFlanks | 10841 | 10862 | 22 | CGTAACTTCGGGATAAGGATTG | 3093.4 | -5.6 | 0.7 | -7.9 | 4.1E-15 | 1.2E-14 | MB231/MB468 |
| >hg38_21_+_8433172_8446622_RNA45SN1_45S_50ntFlanks@10841.10868.28\|CGTAACTTCGGGATAAGGATTGGCTCTA | >hg38_21_+_8433172_8446622_RNA45SN1_45S_50ntFlanks | 10841 | 10868 | 28 | CGTAACTTCGGGATAAGGATTGGCTCTA | 6703.0 | -4.5 | 0.6 | -7.8 | 4.2E-15 | 1.2E-14 | BT20/MB468 |
| >hg38_21_+_8433172_8446622_RNA45SN1_45S_50ntFlanks@10781.10805.25\|AGGTGAACAGCCTCTGGCATGTTGG | >hg38_21_+_8433172_8446622_RNA45SN1_45S_50ntFlanks | 10781 | 10805 | 25 | AGGTGAACAGCCTCTGGCATGTTGG | 300.6 | -10.2 | 1.3 | -7.8 | 4.2E-15 | 1.2E-14 | BT20/MB468 |
| >hg38_21_+_8433172_8446622_RNA45SN1_45S_50ntFlanks@11759.11777.19\|CCTCGTCATCTAATTAGTG | >hg38_21_+_8433172_8446622_RNA45SN1_45S_50ntFlanks | 11759 | 11777 | 19 | CCTCGTCATCTAATTAGTG | 380.1 | -10.6 | 1.3 | -7.8 | 4.3E-15 | 1.3E-14 | BT20/MB468 |
| >MI0003205\|hsa-mir-532&WithFlank&X\|+\|50003142\|50003244@26.46.21(+1G)&[MIMAT0002888&hsa-miR-532-5p&offsets\|0\|-1(+1G);m-65&X\|+\|50003167\|50003188&offsets\|0\|-1(+1G)]\|CATGCCTTGAGTGTAGGACCGG | >MI0003205\|hsa-mir-532&WithFlank&X\|+\|50003142\|50003244 | 26 | 46 | 21(+1G) | CATGCCTTGAGTGTAGGACCGG | 557.4 | 6.2 | 0.8 | 7.8 | 4.3E-15 | 1.3E-14 | BT20/MB468 |
| >hg38_MT_+_1621_3279_RNR2_16S_50ntFlanks@51.84.34\|GCTAAACCTAGCCCCAAACCCACTCCACCTTACT | >hg38_MT_+_1621_3279_RNR2_16S_50ntFlanks | 51 | 84 | 34 | GCTAAACCTAGCCCCAAACCCACTCCACCTTACT | 1669.4 | 8.8 | 1.1 | 7.8 | 4.4E-15 | 1.3E-14 | BT20/MB468 |
| >hg38_21_+_8433172_8446622_RNA45SN1_45S_50ntFlanks@11575.11601.27\|ACTTAGAACTGGTGCGGACCAGGGGAA | >hg38_21_+_8433172_8446622_RNA45SN1_45S_50ntFlanks | 11575 | 11601 | 27 | ACTTAGAACTGGTGCGGACCAGGGGAA | 1171.4 | -5.5 | 0.7 | -7.8 | 4.4E-15 | 1.3E-14 | BT20/MB468 |
| >hg38_21_+_8433172_8446622_RNA45SN1_45S_50ntFlanks@10781.10818.38\|AGGTGAACAGCCTCTGGCATGTTGGAACAATGTAGGTA | >hg38_21_+_8433172_8446622_RNA45SN1_45S_50ntFlanks | 10781 | 10818 | 38 | AGGTGAACAGCCTCTGGCATGTTGGAACAATGTAGGTA | 1702.4 | -5.8 | 0.7 | -7.8 | 4.5E-15 | 1.3E-14 | BT20/MB468 |
| >hg38_21_+_8433172_8446622_RNA45SN1_45S_50ntFlanks@12973.13001.29\|TCGTACGTAGCAGAGCAGCTCCCTCGCTG | >hg38_21_+_8433172_8446622_RNA45SN1_45S_50ntFlanks | 12973 | 13001 | 29 | TCGTACGTAGCAGAGCAGCTCCCTCGCTG | 228.6 | -9.8 | 1.3 | -7.8 | 4.5E-15 | 1.3E-14 | BT20/MB468 |
| >MI0003581\|hsa-mir-574&WithFlank&4\|+\|38868026\|38868133@67.88.22&[MIMAT0003239&hsa-miR-574-3p&offsets\|0\|0;m-149&4\|+\|38868092\|38868113&offsets\|0\|0]\|CACGCTCATGCACACACCCACA | >MI0003581\|hsa-mir-574&WithFlank&4\|+\|38868026\|38868133 | 67 | 88 | 22 | CACGCTCATGCACACACCCACA | 1560.8 | 6.7 | 0.9 | 7.8 | 4.5E-15 | 1.3E-14 | BT20/MB468 |
| >hg38_21_+_8433172_8446622_RNA45SN1_45S_50ntFlanks@10352.10372.21\|GAGAACTTTGAAGGCCGAAGT | >hg38_21_+_8433172_8446622_RNA45SN1_45S_50ntFlanks | 10352 | 10372 | 21 | GAGAACTTTGAAGGCCGAAGT | 791.9 | 7.9 | 1.0 | 7.8 | 4.4E-15 | 1.3E-14 | MB231/MB468 |
| >am_trna106_HisGTG_1_-_149155828_149155899@7.29.23,trna111_HisGTG_1_-_147774845_147774916@7.29.23,trna118_HisGTG_1_-_145396881_145396952@7.29.23,trna16_HisGTG_1_+_146544773_146544844@7.29.23,trna1_HisGTG_15_+_45493349_45493420@7.29.23,trna21_HisGTG_1_+_147753471_147753542@7.29.23,trna33_HisGTG_6_+_27125906_27125977@7.29.23,trna7_HisGTG_9_-_14433938_14434009@7.29.23,trna8_HisGTG_15_-_45492611_45492682@7.29.23,trna9_HisGTG_15_-_45490804_45490875@7.29.23\|ATCGTATAGTGGTTAGTACTCTG | >am_trna106_HisGTG_1_-_149155828_149155899 | 7 | 29 | 23 | ATCGTATAGTGGTTAGTACTCTG | 346.9 | -10.0 | 1.3 | -7.8 | 4.4E-15 | 1.3E-14 | MB231/MB468 |
| >MI0000438\|hsa-mir-15b&WithFlank&3\|+\|160404582\|160404691@64.84.21&[MIMAT0004586&hsa-miR-15b-3p&offsets\|0\|-1;m-204&3\|+\|160404645\|160404665&offsets\|0\|0]\|CGAATCATTATTTGCTGCTCT | >MI0000438\|hsa-mir-15b&WithFlank&3\|+\|160404582\|160404691 | 64 | 84 | 21 | CGAATCATTATTTGCTGCTCT | 959.3 | 6.5 | 0.8 | 7.8 | 4.6E-15 | 1.3E-14 | BT20/MB468 |
| >hg38_21_+_8433172_8446622_RNA45SN1_45S_50ntFlanks@3854.3872.19\|ACTGTGGTAATTCTAGAGC | >hg38_21_+_8433172_8446622_RNA45SN1_45S_50ntFlanks | 3854 | 3872 | 19 | ACTGTGGTAATTCTAGAGC | 653.2 | -4.2 | 0.5 | -7.8 | 4.6E-15 | 1.4E-14 | BT20/MB468 |
| >hg38_21_+_8433172_8446622_RNA45SN1_45S_50ntFlanks@10683.10710.28\|CCGGCGGCGTCCGGTGAGCTCTCGCTGG | >hg38_21_+_8433172_8446622_RNA45SN1_45S_50ntFlanks | 10683 | 10710 | 28 | CCGGCGGCGTCCGGTGAGCTCTCGCTGG | 335.7 | -10.4 | 1.3 | -7.8 | 4.6E-15 | 1.4E-14 | BT20/MB468 |
| >hg38_21_+_8433172_8446622_RNA45SN1_45S_50ntFlanks@12044.12070.27\|ACTGACCCGGTGAGGCGGGGGGGCGAG | >hg38_21_+_8433172_8446622_RNA45SN1_45S_50ntFlanks | 12044 | 12070 | 27 | ACTGACCCGGTGAGGCGGGGGGGCGAG | 301.4 | -9.8 | 1.3 | -7.8 | 4.6E-15 | 1.4E-14 | MB231/MB468 |
| >hg38_21_+_8433172_8446622_RNA45SN1_45S_50ntFlanks@12437.12454.18\|CATTGTGAAGCAGAATTC | >hg38_21_+_8433172_8446622_RNA45SN1_45S_50ntFlanks | 12437 | 12454 | 18 | CATTGTGAAGCAGAATTC | 2191.9 | -4.3 | 0.6 | -7.8 | 4.6E-15 | 1.4E-14 | MB231/MB468 |
| >hg38_21_+_8433172_8446622_RNA45SN1_45S_50ntFlanks@3840.3861.22\|CTCCTACTTGGATAACTGTGGT | >hg38_21_+_8433172_8446622_RNA45SN1_45S_50ntFlanks | 3840 | 3861 | 22 | CTCCTACTTGGATAACTGTGGT | 552.7 | 4.1 | 0.5 | 7.8 | 4.7E-15 | 1.4E-14 | BT20/MB468 |
| >hg38_21_+_8433172_8446622_RNA45SN1_45S_50ntFlanks@10843.10865.23\|TAACTTCGGGATAAGGATTGGCT | >hg38_21_+_8433172_8446622_RNA45SN1_45S_50ntFlanks | 10843 | 10865 | 23 | TAACTTCGGGATAAGGATTGGCT | 738.7 | -4.9 | 0.6 | -7.8 | 4.6E-15 | 1.4E-14 | MB231/MB468 |
| >MI0000477\|hsa-mir-146a&WithFlank&5\|+\|160485346\|160485456@27.50.24&[MIMAT0000449&hsa-miR-146a-5p&offsets\|0\|+2;m-129&5\|+\|160485372\|160485393&offsets\|0\|+2]\|TGAGAACTGAATTCCATGGGTTGT | >MI0000477\|hsa-mir-146a&WithFlank&5\|+\|160485346\|160485456 | 27 | 50 | 24 | TGAGAACTGAATTCCATGGGTTGT | 429.0 | -12.1 | 1.5 | -8.2 | 3.2E-16 | 1.4E-14 | BT20/MB231 |
| >hg38_21_+_8433172_8446622_RNA45SN1_45S_50ntFlanks@9565.9603.39\|TGACGTGCAAATCGGTCGTCCGACCTGGGTATAGGGGCG | >hg38_21_+_8433172_8446622_RNA45SN1_45S_50ntFlanks | 9565 | 9603 | 39 | TGACGTGCAAATCGGTCGTCCGACCTGGGTATAGGGGCG | 284.1 | 4.5 | 0.6 | 7.8 | 4.7E-15 | 1.4E-14 | MB231/MB468 |
| >hg38_21_+_8433172_8446622_RNA45SN1_45S_50ntFlanks@10794.10822.29\|CTGGCATGTTGGAACAATGTAGGTAAGGG | >hg38_21_+_8433172_8446622_RNA45SN1_45S_50ntFlanks | 10794 | 10822 | 29 | CTGGCATGTTGGAACAATGTAGGTAAGGG | 320.4 | -10.3 | 1.3 | -7.8 | 4.8E-15 | 1.4E-14 | BT20/MB468 |
| >hg38_21_+_8433172_8446622_RNA45SN1_45S_50ntFlanks@10687.10709.23\|CGGCGTCCGGTGAGCTCTCGCTG | >hg38_21_+_8433172_8446622_RNA45SN1_45S_50ntFlanks | 10687 | 10709 | 23 | CGGCGTCCGGTGAGCTCTCGCTG | 2429.5 | -5.6 | 0.7 | -7.8 | 4.7E-15 | 1.4E-14 | MB231/MB468 |
| >hg38_21_+_8433172_8446622_RNA45SN1_45S_50ntFlanks@10407.10429.23\|GGGTCAGTCGGTCCTGAGAGATG | >hg38_21_+_8433172_8446622_RNA45SN1_45S_50ntFlanks | 10407 | 10429 | 23 | GGGTCAGTCGGTCCTGAGAGATG | 209.8 | -5.2 | 0.7 | -7.8 | 4.9E-15 | 1.4E-14 | BT20/MB468 |
| >hg38_21_+_8433172_8446622_RNA45SN1_45S_50ntFlanks@11573.11605.33\|CGACTTAGAACTGGTGCGGACCAGGGGAATCCG | >hg38_21_+_8433172_8446622_RNA45SN1_45S_50ntFlanks | 11573 | 11605 | 33 | CGACTTAGAACTGGTGCGGACCAGGGGAATCCG | 212.8 | -3.7 | 0.5 | -7.8 | 4.9E-15 | 1.4E-14 | BT20/MB468 |
| >hg38_21_+_8433172_8446622_RNA45SN1_45S_50ntFlanks@11766.11790.25\|ATCTAATTAGTGACGCGCATGAATG | >hg38_21_+_8433172_8446622_RNA45SN1_45S_50ntFlanks | 11766 | 11790 | 25 | ATCTAATTAGTGACGCGCATGAATG | 211.9 | -9.7 | 1.2 | -7.8 | 4.9E-15 | 1.4E-14 | BT20/MB468 |
| >hg38_21_+_8433172_8446622_RNA45SN1_45S_50ntFlanks@11612.11640.29\|TAATTAAAACAAAGCATCGCGAAGGCCCG | >hg38_21_+_8433172_8446622_RNA45SN1_45S_50ntFlanks | 11612 | 11640 | 29 | TAATTAAAACAAAGCATCGCGAAGGCCCG | 377.7 | -10.1 | 1.3 | -7.8 | 4.9E-15 | 1.5E-14 | MB231/MB468 |
| >hg38_21_+_8433172_8446622_RNA45SN1_45S_50ntFlanks@12126.12150.25\|CGACCCGCTCCGGGGACAGTGCCAG | >hg38_21_+_8433172_8446622_RNA45SN1_45S_50ntFlanks | 12126 | 12150 | 25 | CGACCCGCTCCGGGGACAGTGCCAG | 347.7 | -10.0 | 1.3 | -7.8 | 4.9E-15 | 1.5E-14 | MB231/MB468 |
| >trna20_GluTTC_2_-_131094701_131094772@1.30.30,trna5_GluTTC_13_-_41634874_41634945@1.30.30\|TCCCATATGGTCTAGCGGTTAGGATTCCTG | >trna20_GluTTC_2_-_131094701_131094772 | 1 | 30 | 30 | TCCCATATGGTCTAGCGGTTAGGATTCCTG | 241.4 | -5.0 | 0.6 | -7.8 | 5.0E-15 | 1.5E-14 | BT20/MB468 |
| >hg38_21_+_8433172_8446622_RNA45SN1_45S_50ntFlanks@12412.12434.23\|ATCCTTCGATGTCGGCTCTTCCT | >hg38_21_+_8433172_8446622_RNA45SN1_45S_50ntFlanks | 12412 | 12434 | 23 | ATCCTTCGATGTCGGCTCTTCCT | 334.2 | -10.4 | 1.3 | -7.8 | 5.0E-15 | 1.5E-14 | BT20/MB468 |
| >hg38_1_-_228634819_228635039_RNA5S12_5S_50ntFlanks@139.171.33\|GACCGCCTGGGAATACCGGGTGCTGTAGGCTTT | >hg38_1_-_228634819_228635039_RNA5S12_5S_50ntFlanks | 139 | 171 | 33 | GACCGCCTGGGAATACCGGGTGCTGTAGGCTTT | 2232.2 | 7.8 | 1.0 | 7.8 | 5.0E-15 | 1.5E-14 | BT20/MB468 |
| >hg38_21_+_8433172_8446622_RNA45SN1_45S_50ntFlanks@4588.4616.29\|CTATTTTGTTGGTTTTCGGAACTGAGGCC | >hg38_21_+_8433172_8446622_RNA45SN1_45S_50ntFlanks | 4588 | 4616 | 29 | CTATTTTGTTGGTTTTCGGAACTGAGGCC | 823.5 | -5.1 | 0.6 | -7.8 | 4.9E-15 | 1.5E-14 | MB231/MB468 |
| >hg38_21_+_8433172_8446622_RNA45SN1_45S_50ntFlanks@6745.6772.28\|ACACTTCGAACGCACTTGCGGCCCCGGG | >hg38_21_+_8433172_8446622_RNA45SN1_45S_50ntFlanks | 6745 | 6772 | 28 | ACACTTCGAACGCACTTGCGGCCCCGGG | 328.0 | -9.9 | 1.3 | -7.8 | 4.9E-15 | 1.5E-14 | MB231/MB468 |
| >hg38_21_+_8433172_8446622_RNA45SN1_45S_50ntFlanks@4738.4757.20\|AAAGTCGGAGGTTCGAAGAC | >hg38_21_+_8433172_8446622_RNA45SN1_45S_50ntFlanks | 4738 | 4757 | 20 | AAAGTCGGAGGTTCGAAGAC | 531.1 | 4.4 | 0.6 | 7.8 | 4.9E-15 | 1.5E-14 | MB231/MB468 |
| >hg38_1_-_228634819_228635039_RNA5S12_5S_50ntFlanks@149.170.22\|GAATACCGGGTGCTGTAGGCTT | >hg38_1_-_228634819_228635039_RNA5S12_5S_50ntFlanks | 149 | 170 | 22 | GAATACCGGGTGCTGTAGGCTT | 2488.9 | 6.3 | 0.8 | 7.8 | 5.0E-15 | 1.5E-14 | MB231/MB468 |
| >hg38_21_+_8433172_8446622_RNA45SN1_45S_50ntFlanks@12991.13013.23\|CTCCCTCGCTGCGATCTATTGAA | >hg38_21_+_8433172_8446622_RNA45SN1_45S_50ntFlanks | 12991 | 13013 | 23 | CTCCCTCGCTGCGATCTATTGAA | 5876.7 | -5.6 | 0.7 | -7.8 | 5.0E-15 | 1.5E-14 | MB231/MB468 |
| >hg38_21_+_8433172_8446622_RNA45SN1_45S_50ntFlanks@12416.12450.35\|TTCGATGTCGGCTCTTCCTATCATTGTGAAGCAGA | >hg38_21_+_8433172_8446622_RNA45SN1_45S_50ntFlanks | 12416 | 12450 | 35 | TTCGATGTCGGCTCTTCCTATCATTGTGAAGCAGA | 917.0 | 8.0 | 1.0 | 7.8 | 5.0E-15 | 1.5E-14 | MB231/MB468 |
| >hg38_21_+_8433172_8446622_RNA45SN1_45S_50ntFlanks@10587.10607.21\|CCCCGGGGAGAGTTCTCTTTT | >hg38_21_+_8433172_8446622_RNA45SN1_45S_50ntFlanks | 10587 | 10607 | 21 | CCCCGGGGAGAGTTCTCTTTT | 347.4 | -5.7 | 0.7 | -7.8 | 5.1E-15 | 1.5E-14 | BT20/MB468 |
| >hg38_21_+_8433172_8446622_RNA45SN1_45S_50ntFlanks@12632.12653.22\|GCGAAGCTACCATCTGTGGGAT | >hg38_21_+_8433172_8446622_RNA45SN1_45S_50ntFlanks | 12632 | 12653 | 22 | GCGAAGCTACCATCTGTGGGAT | 793.9 | -7.2 | 0.9 | -7.8 | 5.0E-15 | 1.5E-14 | MB231/MB468 |
| >MI0000749\|hsa-mir-30e&WithFlank&1\|+\|40754349\|40754452@65.85.21(+1A)&[MIMAT0000693&hsa-miR-30e-3p&offsets\|0\|-1(+1A);m-25&1\|+\|40754413\|40754434&offsets\|0\|-1(+1A)]\|CTTTCAGTCGGATGTTTACAGA | >MI0000749\|hsa-mir-30e&WithFlank&1\|+\|40754349\|40754452 | 65 | 85 | 21(+1A) | CTTTCAGTCGGATGTTTACAGA | 724.2 | 6.8 | 0.9 | 7.8 | 5.1E-15 | 1.5E-14 | BT20/MB468 |
| >hg38_21_+_8433172_8446622_RNA45SN1_45S_50ntFlanks@12157.12181.25\|AGTTTGACTGGGGCGGTACACCTGT | >hg38_21_+_8433172_8446622_RNA45SN1_45S_50ntFlanks | 12157 | 12181 | 25 | AGTTTGACTGGGGCGGTACACCTGT | 489.6 | -8.7 | 1.1 | -7.8 | 5.1E-15 | 1.5E-14 | MB231/MB468 |
| >MI0000749\|hsa-mir-30e&WithFlank&1\|+\|40754349\|40754452@23.46.24(+2U)&[MIMAT0000692&hsa-miR-30e-5p&offsets\|0\|+2(+2U);m-24&1\|+\|40754371\|40754394&offsets\|0\|0(+2U)]\|TGTAAACATCCTTGACTGGAAGCTTT | >MI0000749\|hsa-mir-30e&WithFlank&1\|+\|40754349\|40754452 | 23 | 46 | 24(+2U) | TGTAAACATCCTTGACTGGAAGCTTT | 777.6 | 6.0 | 0.8 | 7.8 | 5.1E-15 | 1.5E-14 | MB231/MB468 |
| >hg38_21_+_8433172_8446622_RNA45SN1_45S_50ntFlanks@11621.11640.20\|CAAAGCATCGCGAAGGCCCG | >hg38_21_+_8433172_8446622_RNA45SN1_45S_50ntFlanks | 11621 | 11640 | 20 | CAAAGCATCGCGAAGGCCCG | 349.8 | -10.4 | 1.3 | -7.8 | 5.3E-15 | 1.5E-14 | BT20/MB468 |

| | | | | | | | | | | | | |
|---|---|---|---|---|---|---|---|---|---|---|---|---|
| >hg38_21_+_8433172_8446622_RNA45SN1_45S_50ntFlanks@6738.6772.35\|ATCATCGACACTTCGAACGCACTTGCGGCCCCGGG | >hg38_21_+_8433172_8446622_RNA45SN1_45S_50ntFlanks | 6738 | 6772 | 35 | ATCATCGACACTTCGAACGCACTTGCGGCCCCGGG | 447.1 | -6.8 | 0.9 | -7.8 | 5.1E-15 | 1.5E-14 | MB231/MB468 |
| >hg38_21_+_8433172_8446622_RNA45SN1_45S_50ntFlanks@13002.13023.22\|CGATCTATTGAAAGTCAGCCCT | >hg38_21_+_8433172_8446622_RNA45SN1_45S_50ntFlanks | 13002 | 13023 | 22 | CGATCTATTGAAAGTCAGCCCT | 426.1 | -10.3 | 1.3 | -7.8 | 5.2E-15 | 1.5E-14 | MB231/MB468 |
| >hg38_21_+_8433172_8446622_RNA45SN1_45S_50ntFlanks@476.514.39\|TTCGTGATCGATGTGGTGACGTCGTGCTCTCCCGGGCCG | >hg38_21_+_8433172_8446622_RNA45SN1_45S_50ntFlanks | 476 | 514 | 39 | TTCGTGATCGATGTGGTGACGTCGTGCTCTCCCGGGCCG | 1056.7 | 9.6 | 1.2 | 7.8 | 5.2E-15 | 1.6E-14 | MB231/MB468 |
| >am_MI0000748\|hsa-mir-130b&WithFlank&22\|+\|21653298\|21653391@19.39.21&[MIMAT0004680&hsa-miR-130b-5p&offsets\|0\|0;m-234&22\|+\|21653316\|21653337&offsets\|0\|-1]\|ACTCTTTCCCTGTTGCACTAC | >am_MI0000748\|hsa-mir-130b&WithFlank&22\|+\|21653298\|21653391 | 19 | 39 | 21 | ACTCTTTCCCTGTTGCACTAC | 1297.2 | 8.3 | 1.1 | 7.8 | 5.2E-15 | 1.6E-14 | MB231/MB468 |
| >hg38_21_+_8433172_8446622_RNA45SN1_45S_50ntFlanks@11620.11640.21\|ACAAAGCATCGCGAAGGCCCG | >hg38_21_+_8433172_8446622_RNA45SN1_45S_50ntFlanks | 11620 | 11640 | 21 | ACAAAGCATCGCGAAGGCCCG | 315.1 | -7.2 | 0.9 | -7.8 | 5.4E-15 | 1.6E-14 | BT20/MB468 |
| >hg38_21_+_8433172_8446622_RNA45SN1_45S_50ntFlanks@12640.12659.20\|ACCATCTGTGGGATTATGAC | >hg38_21_+_8433172_8446622_RNA45SN1_45S_50ntFlanks | 12640 | 12659 | 20 | ACCATCTGTGGGATTATGAC | 315.8 | -3.0 | 0.4 | -7.8 | 5.5E-15 | 1.6E-14 | BT20/MB468 |
| >hg38_21_+_8433172_8446622_RNA45SN1_45S_50ntFlanks@10355.10383.29\|AACTTTGAAGGCCGAAGTGGAGAAGGGTT | >hg38_21_+_8433172_8446622_RNA45SN1_45S_50ntFlanks | 10355 | 10383 | 29 | AACTTTGAAGGCCGAAGTGGAGAAGGGTT | 181.0 | -3.1 | 0.4 | -7.8 | 5.5E-15 | 1.6E-14 | BT20/MB468 |
| >trnaMT_SerGCT_MT_+_12207_12265@1.26.26\|GAGAAAGCTCACAAGAACTGCTAACT | >trnaMT_SerGCT_MT_+_12207_12265 | 1 | 26 | 26 | GAGAAAGCTCACAAGAACTGCTAACT | 11568.4 | 6.4 | 0.8 | 7.8 | 5.6E-15 | 1.6E-14 | BT20/MB468 |
| >hg38_21_+_8433172_8446622_RNA45SN1_45S_50ntFlanks@7986.8008.23\|ATCAGACGTGGCGACCCGCTGAA | >hg38_21_+_8433172_8446622_RNA45SN1_45S_50ntFlanks | 7986 | 8008 | 23 | ATCAGACGTGGCGACCCGCTGAA | 370.0 | -10.1 | 1.3 | -7.8 | 5.5E-15 | 1.6E-14 | MB231/MB468 |
| >hg38_21_+_8433172_8446622_RNA45SN1_45S_50ntFlanks@9505.9526.22\|TGAACTATGCCTGGGCAGGGCG | >hg38_21_+_8433172_8446622_RNA45SN1_45S_50ntFlanks | 9505 | 9526 | 22 | TGAACTATGCCTGGGCAGGGCG | 251.3 | -10.0 | 1.3 | -7.8 | 5.7E-15 | 1.7E-14 | BT20/MB468 |
| >MI0000650\|hsa-mir-200c&WithFlank&12\|+\|6963693\|6963772@50.71.22(+2U)&[MIMAT0000617&hsa-miR-200c-3p&offsets\|0\|-1(+2U);m-54&12\|+\|6963742\|6963764&offsets\|0\|-1(+2U)]\|TAATACTGCCGGGTAATGATGGTT | >MI0000650\|hsa-mir-200c&WithFlank&12\|+\|6963693\|6963772 | 50 | 71 | 22(+2U) | TAATACTGCCGGGTAATGATGGTT | 526.1 | 8.7 | 1.1 | 7.8 | 5.7E-15 | 1.7E-14 | BT20/MB468 |
| >hg38_21_+_8433172_8446622_RNA45SN1_45S_50ntFlanks@12476.12493.18\|CCACTAATAGGGAACGTG | >hg38_21_+_8433172_8446622_RNA45SN1_45S_50ntFlanks | 12476 | 12493 | 18 | CCACTAATAGGGAACGTG | 307.8 | -3.9 | 0.5 | -7.8 | 5.6E-15 | 1.7E-14 | MB231/MB468 |
| >MI0000440\|hsa-mir-27b&WithFlank&9\|+\|95085439\|95085547@67.85.19(+1U)&[MIMAT0000419&hsa-miR-27b-3p&offsets\|0\|-2(+1U);m-39&9\|+\|95085505\|95085525&offsets\|0\|-2(+1U)]\|TTCACAGTGGCTAAGTTCTT | >MI0000440\|hsa-mir-27b&WithFlank&9\|+\|95085439\|95085547 | 67 | 85 | 19(+1U) | TTCACAGTGGCTAAGTTCTT | 648.3 | 9.9 | 1.3 | 7.8 | 5.8E-15 | 1.7E-14 | BT20/MB468 |
| >hg38_21_+_8433172_8446622_RNA45SN1_45S_50ntFlanks@11641.11664.24\|CGGCGGGTGTTGACGCGATGTGAT | >hg38_21_+_8433172_8446622_RNA45SN1_45S_50ntFlanks | 11641 | 11664 | 24 | CGGCGGGTGTTGACGCGATGTGAT | 672.6 | -5.3 | 0.7 | -7.8 | 5.6E-15 | 1.7E-14 | MB231/MB468 |
| >hg38_21_+_8433172_8446622_RNA45SN1_45S_50ntFlanks@6733.6773.41\|CATTGATCATCGACACTTCGAACGCACTTGCGGCCCCGGGT | >hg38_21_+_8433172_8446622_RNA45SN1_45S_50ntFlanks | 6733 | 6773 | 41 | CATTGATCATCGACACTTCGAACGCACTTGCGGCCCCGGGT | 587.4 | 7.9 | 1.0 | 7.8 | 5.8E-15 | 1.7E-14 | BT20/MB468 |
| >hg38_21_+_8433172_8446622_RNA45SN1_45S_50ntFlanks@10830.10858.29\|CAAGCCGGATCCGTAACTTCGGGATAAGG | >hg38_21_+_8433172_8446622_RNA45SN1_45S_50ntFlanks | 10830 | 10858 | 29 | CAAGCCGGATCCGTAACTTCGGGATAAGG | 299.5 | -10.2 | 1.3 | -7.8 | 5.9E-15 | 1.7E-14 | BT20/MB468 |
| >hg38_21_+_8433172_8446622_RNA45SN1_45S_50ntFlanks@12418.12435.18\|CGATGTCGGCTCTTCCTA | >hg38_21_+_8433172_8446622_RNA45SN1_45S_50ntFlanks | 12418 | 12435 | 18 | CGATGTCGGCTCTTCCTA | 6893.3 | -4.9 | 0.6 | -7.8 | 5.7E-15 | 1.7E-14 | MB231/MB468 |
| >hg38_21_+_8433172_8446622_RNA45SN1_45S_50ntFlanks@11161.11188.28\|CACGGTCCCCCGCGAGGGGGGCCCGGGC | >hg38_21_+_8433172_8446622_RNA45SN1_45S_50ntFlanks | 11161 | 11188 | 28 | CACGGTCCCCCGCGAGGGGGGCCCGGGC | 246.5 | -9.9 | 1.3 | -7.8 | 5.9E-15 | 1.7E-14 | BT20/MB468 |
| >hg38_21_+_8433172_8446622_RNA45SN1_45S_50ntFlanks@11316.11343.28\|CACCCCACGTCTCGTCGCGCGCGCGTCC | >hg38_21_+_8433172_8446622_RNA45SN1_45S_50ntFlanks | 11316 | 11343 | 28 | CACCCCACGTCTCGTCGCGCGCGCGTCC | 329.1 | -9.9 | 1.3 | -7.8 | 5.8E-15 | 1.7E-14 | MB231/MB468 |
| >MI0000081\|hsa-mir-24-2&WithFlank&19\|-\|13836281\|13836365@56.77.22(+1A);MI0000080\|hsa-mir-24-1&WithFlank&9\|+\|95086015\|95086094@50.71.22(+1A)&[MIMAT0000080&hsa-miR-24-3p&offsets\|0\|0(+1A);m-36&19\|-\|13836288\|13836310&offsets\|0\|-1(+1A)];[MIMAT0000080_1&hsa-miR-24-3p&offsets\|0\|0(+1A);m-35&9\|+\|95086064\|95086086&offsets\|0\|-1(+1A)]\|TGGCTCAGTTCAGCAGGAACAGA | >MI0000081\|hsa-mir-24-2&WithFlank&19\|-\|13836281\|13836365 | 56 | 77 | 22(+1A) | TGGCTCAGTTCAGCAGGAACAGA | 3540.1 | 7.6 | 1.0 | 7.8 | 5.9E-15 | 1.7E-14 | BT20/MB468 |
| >hg38_21_+_8433172_8446622_RNA45SN1_45S_50ntFlanks@475.496.22\|CTTCGTGATCGATGTGGTGACG | >hg38_21_+_8433172_8446622_RNA45SN1_45S_50ntFlanks | 475 | 496 | 22 | CTTCGTGATCGATGTGGTGACG | 5887.1 | 6.0 | 0.8 | 7.8 | 5.9E-15 | 1.7E-14 | BT20/MB468 |
| >MI0000483\|hsa-mir-186&WithFlank&1\|-\|71067625\|71067722@21.44.24&[MIMAT0000456&hsa-miR-186-5p&offsets\|0\|+2;m-84&1\|-\|71067681\|71067702&offsets\|0\|+2]\|CAAAGAATTCTCCTTTTGGGCTTT | >MI0000483\|hsa-mir-186&WithFlank&1\|-\|71067625\|71067722 | 21 | 44 | 24 | CAAAGAATTCTCCTTTTGGGCTTT | 767.7 | 6.5 | 0.8 | 7.8 | 5.8E-15 | 1.7E-14 | MB231/MB468 |
| >trna10_ValCAC_5_-_180649395_180649467@59.76.18,trna12_ValAAC_5_-_180645270_180645342@59.76.18,trna132_ValAAC_6_-_27721179_27721251@59.76.18,trna136_ValAAC_6_-_27648885_27648957@59.76.18,trna15_ValAAC_5_-_180615416_180615488@59.76.18,trna18_ValCAC_5_-_180529253_180529325@59.76.18,trna2_ValAAC_3_+_169490018_169490090@59.76.18,trna2_ValCAC_5_+_180524070_180524142@59.76.18,trna4_ValAAC_5_+_180591154_180591226@59.76.18,trna5_ValAAC_5_+_180596610_180596682@59.76.18,trna6_ValCAC_5_+_180600650_180600722@59.76.18,trna85_ValCAC_1_-_161369490_161369562@59.76.18,trna90_ValCAC_1_-_149684088_149684161@60.77.18,trna9_ValCAC_6_+_26538282_26538354@59.76.18\|AACCGGGCGGAAACACCA | >trna10_ValCAC_5_-_180649395_180649467 | 59 | 76 | 18 | AACCGGGCGGAAACACCA | 854.6 | 5.7 | 0.7 | 7.8 | 5.8E-15 | 1.7E-14 | MB231/MB468 |
| >hg38_21_+_8433172_8446622_RNA45SN1_45S_50ntFlanks@12420.12452.33\|ATGTCGGCTCTTCCTATCATTGTGAAGCAGAAT | >hg38_21_+_8433172_8446622_RNA45SN1_45S_50ntFlanks | 12420 | 12452 | 33 | ATGTCGGCTCTTCCTATCATTGTGAAGCAGAAT | 718.0 | 4.6 | 0.6 | 7.8 | 5.8E-15 | 1.7E-14 | MB231/MB468 |
| >hg38_21_+_8433172_8446622_RNA45SN1_45S_50ntFlanks@10795.10818.24\|TGGCATGTTGGAACAATGTAGGTA | >hg38_21_+_8433172_8446622_RNA45SN1_45S_50ntFlanks | 10795 | 10818 | 24 | TGGCATGTTGGAACAATGTAGGTA | 1025.6 | -5.2 | 0.7 | -7.8 | 6.0E-15 | 1.7E-14 | BT20/MB468 |
| >hg38_21_+_8433172_8446622_RNA45SN1_45S_50ntFlanks@12008.12043.36\|CCGCCGGTGAAATACCACTACTCTGATCGTTTTTTC | >hg38_21_+_8433172_8446622_RNA45SN1_45S_50ntFlanks | 12008 | 12043 | 36 | CCGCCGGTGAAATACCACTACTCTGATCGTTTTTTC | 856.8 | -11.3 | 1.4 | -7.8 | 5.9E-15 | 1.7E-14 | MB231/MB468 |
| >hg38_21_+_8433172_8446622_RNA45SN1_45S_50ntFlanks@10351.10384.34\|CGAGAACTTTGAAGGCCGAAGTGGAGAAGGGTTC | >hg38_21_+_8433172_8446622_RNA45SN1_45S_50ntFlanks | 10351 | 10384 | 34 | CGAGAACTTTGAAGGCCGAAGTGGAGAAGGGTTC | 1910.7 | -5.0 | 0.6 | -7.8 | 5.9E-15 | 1.7E-14 | MB231/MB468 |
| >trnaMT_GlnTTG_MT_-_4329_4400@1.33.33,trnalookalike2_GlnTTG_1_-_564879_564950@1.33.33\|TAGGATGGGGTGTGATAGGTGGCACGGAGAATT | >trnaMT_GlnTTG_MT_-_4329_4400 | 1 | 33 | 33 | TAGGATGGGGTGTGATAGGTGGCACGGAGAATT | 1387.0 | 11.6 | 1.5 | 7.8 | 6.1E-15 | 1.8E-14 | BT20/MB468 |
| >hg38_21_+_8433172_8446622_RNA45SN1_45S_50ntFlanks@11027.11049.23\|CTCTCTCTCTCTCTCCCCCGCTC | >hg38_21_+_8433172_8446622_RNA45SN1_45S_50ntFlanks | 11027 | 11049 | 23 | CTCTCTCTCTCTCTCCCCCGCTC | 293.3 | -9.8 | 1.3 | -7.8 | 6.0E-15 | 1.8E-14 | MB231/MB468 |
| >hg38_21_+_8433172_8446622_RNA45SN1_45S_50ntFlanks@10869.10902.34\|AGGGCTGGGTCGGTCGGGCTGGGGCGCGAAGCGG | >hg38_21_+_8433172_8446622_RNA45SN1_45S_50ntFlanks | 10869 | 10902 | 34 | AGGGCTGGGTCGGTCGGGCTGGGGCGCGAAGCGG | 344.3 | -8.2 | 1.1 | -7.8 | 6.0E-15 | 1.8E-14 | MB231/MB468 |
| >hg38_21_+_8433172_8446622_RNA45SN1_45S_50ntFlanks@12633.12657.25\|CGAAGCTACCATCTGTGGGATTATG | >hg38_21_+_8433172_8446622_RNA45SN1_45S_50ntFlanks | 12633 | 12657 | 25 | CGAAGCTACCATCTGTGGGATTATG | 500.7 | -4.3 | 0.6 | -7.8 | 6.0E-15 | 1.8E-14 | MB231/MB468 |
| >am_MI0003776\|hsa-mir-320b-1&WithFlank&1\|+\|116671740\|116671823@48.65.18;MI0003839\|hsa-mir-320b-2&WithFlank&1\|-\|224257034\|224257116@47.64.18;MI0008190\|hsa-mir-320d-1&WithFlank&13\|-\|40727810\|40727893@48.65.18;MI0003778\|hsa-mir-320c-1&WithFlank&18\|+\|21683512\|21683595@48.65.18;MI0008191\|hsa-mir-320c-2&WithFlank&18\|+\|24321669\|24321752@48.65.18;MI0000542\|hsa-mir-320a&WithFlank&8\|-\|22244960\|22245043@48.65.18;MI0008192\|hsa-mir-320d-2&WithFlank&X\|-\|140926154\|140926237@48.65.18&[MIMAT0005792&hsa-miR-320b&offsets\|0\|-4;m-183&1\|+\|116671787\|116671806&offsets\|0\|-2];[MIMAT0005792_1&hsa-miR-320b&offsets\|0\|-4;m-183&1\|-\|224257051\|224257070&offsets\|0\|-2];[MIMAT0006764&hsa-miR-320d&offsets\|0\|-1;m-246&13\|-\|40727829\|40727846&offsets\|0\|0];[MIMAT0005793&hsa-miR-320c&offsets\|0\|-2;m-222&18\|+\|21683559\|21683578&offsets\|0\|-2];[MIMAT0005793_1&hsa-miR-320c&offsets\|0\|-2;m-219&18\|+\|24321716\|24321735&offsets\|0\|-2];[MIMAT0000510&hsa-miR-320a-3p&offsets\|0\|-4;m-60&8\|-\|22244975\|22244996&offsets\|0\|-4];[MIMAT0006764_1&hsa-miR-320d&offsets\|0\|-1;m-253&X\|-\|140926173\|140926190&offsets\|0\|0]\|AAAAGCTGGGTTGAGAGG | >am_MI0003776\|hsa-mir-320b-1&WithFlank&1\|+\|116671740\|116671823 | 48 | 65 | 18 | AAAAGCTGGGTTGAGAGG | 746.6 | 10.5 | 1.4 | 7.8 | 6.2E-15 | 1.8E-14 | BT20/MB468 |
| >hg38_21_+_8433172_8446622_RNA45SN1_45S_50ntFlanks@10362.10388.27\|AAGGCCGAAGTGGAGAAGGGTTCCATG | >hg38_21_+_8433172_8446622_RNA45SN1_45S_50ntFlanks | 10362 | 10388 | 27 | AAGGCCGAAGTGGAGAAGGGTTCCATG | 2364.7 | -9.3 | 1.2 | -7.8 | 6.1E-15 | 1.8E-14 | MB231/MB468 |
| >am_MI0000786\|hsa-mir-378a&WithFlank&5\|+\|149732819\|149732896@49.69.21(+1A)&[MIMAT0000732&hsa-miR-378a-3p&offsets\|0\|-1(+1A);m-44&5\|+\|149732867\|149732888&offsets\|0\|-1(+1A)]\|ACTGGACTTGGAGTCAGAAGGA | >am_MI0000786\|hsa-mir-378a&WithFlank&5\|+\|149732819\|149732896 | 49 | 69 | 21(+1A) | ACTGGACTTGGAGTCAGAAGGA | 2009.6 | 5.7 | 0.7 | 7.8 | 6.1E-15 | 1.8E-14 | MB231/MB468 |

<table>
<tr><th></th><th></th><th></th><th></th><th></th><th></th><th></th><th></th><th></th><th></th><th></th><th></th><th></th></tr>
<tr><td>>trna111_HisGTG_1_-_147774845_147774916@1.32.32,trna118_HisGTG_1_-_145396881_145396952@1.32.32,trna16_HisGTG_1_+_146544773_146544844@1.32.32,trna1_HisGTG_15_+_45493349_45493420@1.32.32,trna21_HisGTG_1_+_147753471_147753542@1.32.32,trna33_HisGTG_6_+_27125906_27125977@1.32.32,trna7_HisGTG_9_-_14433938_14434009@1.32.32,trna8_HisGTG_15_-_45492611_45492682@1.32.32,trna9_HisGTG_15_-_45490804_45490875@1.32.32|GCCGTGATCGTATAGTGGTTAGTACTCTGCGT</td><td>>trna111_HisGTG_1_-_147774845_147774916</td><td>1</td><td>32</td><td>32</td><td>GCCGTGATCGTATAGTGGTTAGTACTCTGCGT</td><td>1169.9</td><td>8.6</td><td>1.1</td><td>7.8</td><td>6.1E-15</td><td>1.8E-14</td><td>MB231/MB468</td></tr>
<tr><td>>hg38_21_+_8433172_8446622_RNA45SN1_45S_50ntFlanks@11270.11293.24|CGGCCGCCCCCGGGGAGCCCGGCG</td><td>>hg38_21_+_8433172_8446622_RNA45SN1_45S_50ntFlanks</td><td>11270</td><td>11293</td><td>24</td><td>CGGCCGCCCCCGGGGAGCCCGGCG</td><td>229.6</td><td>-9.8</td><td>1.3</td><td>-7.8</td><td>6.3E-15</td><td>1.8E-14</td><td>BT20/MB468</td></tr>
<tr><td>>hg38_21_+_8433172_8446622_RNA45SN1_45S_50ntFlanks@12539.12566.28|ATGATGTGTTGTTGCCATGGTAATCCTG</td><td>>hg38_21_+_8433172_8446622_RNA45SN1_45S_50ntFlanks</td><td>12539</td><td>12566</td><td>28</td><td>ATGATGTGTTGTTGCCATGGTAATCCTG</td><td>1021.2</td><td>-8.0</td><td>1.0</td><td>-7.8</td><td>6.3E-15</td><td>1.8E-14</td><td>BT20/MB468</td></tr>
<tr><td>>MI0000077|hsa-mir-21&WithFlank&17|+|59841260|59841343@14.34.21(+2U)&[MIMAT0000076&hsa-miR-21-5p&offsets|0|-1(+2U);m-1&17|+|59841273|59841294&offsets|0|-1(+2U)]|TAGCTTATCAGACTGATGTTGTT</td><td>>MI0000077|hsa-mir-21&WithFlank&17|+|59841260|59841343</td><td>14</td><td>34</td><td>21(+2U)</td><td>TAGCTTATCAGACTGATGTTGTT</td><td>622.0</td><td>9.2</td><td>1.2</td><td>7.8</td><td>6.2E-15</td><td>1.8E-14</td><td>MB231/MB468</td></tr>
<tr><td>>hg38_21_+_8433172_8446622_RNA45SN1_45S_50ntFlanks@10689.10709.21|GCGTCCGGTGAGCTCTCGCTG</td><td>>hg38_21_+_8433172_8446622_RNA45SN1_45S_50ntFlanks</td><td>10689</td><td>10709</td><td>21</td><td>GCGTCCGGTGAGCTCTCGCTG</td><td>400.7</td><td>-8.4</td><td>1.1</td><td>-7.8</td><td>6.3E-15</td><td>1.8E-14</td><td>MB231/MB468</td></tr>
<tr><td>>hg38_21_+_8433172_8446622_RNA45SN1_45S_50ntFlanks@4472.4497.26|CTTAGCTGAGTGTCCCGCGGGGCCCG</td><td>>hg38_21_+_8433172_8446622_RNA45SN1_45S_50ntFlanks</td><td>4472</td><td>4497</td><td>26</td><td>CTTAGCTGAGTGTCCCGCGGGGCCCG</td><td>216.7</td><td>-9.8</td><td>1.3</td><td>-7.8</td><td>6.5E-15</td><td>1.9E-14</td><td>BT20/MB468</td></tr>
<tr><td>>hg38_21_+_8433172_8446622_RNA45SN1_45S_50ntFlanks@9738.9765.28|GAAACGATCTCAACCTATTCTCAAACTT</td><td>>hg38_21_+_8433172_8446622_RNA45SN1_45S_50ntFlanks</td><td>9738</td><td>9765</td><td>28</td><td>GAAACGATCTCAACCTATTCTCAAACTT</td><td>1102.3</td><td>-5.3</td><td>0.7</td><td>-7.8</td><td>6.5E-15</td><td>1.9E-14</td><td>BT20/MB468</td></tr>
<tr><td>>hg38_21_+_8433172_8446622_RNA45SN1_45S_50ntFlanks@10781.10805.25|AGGTGAACAGCCTCTGGCATGTTGG</td><td>>hg38_21_+_8433172_8446622_RNA45SN1_45S_50ntFlanks</td><td>10781</td><td>10805</td><td>25</td><td>AGGTGAACAGCCTCTGGCATGTTGG</td><td>249.0</td><td>-6.7</td><td>0.9</td><td>-7.8</td><td>6.4E-15</td><td>1.9E-14</td><td>MB231/MB468</td></tr>
<tr><td>>hg38_21_+_8433172_8446622_RNA45SN1_45S_50ntFlanks@4002.4020.19|GATAACCTCGGGCCGATCG</td><td>>hg38_21_+_8433172_8446622_RNA45SN1_45S_50ntFlanks</td><td>4002</td><td>4020</td><td>19</td><td>GATAACCTCGGGCCGATCG</td><td>218.7</td><td>-4.6</td><td>0.6</td><td>-7.8</td><td>6.5E-15</td><td>1.9E-14</td><td>BT20/MB468</td></tr>
<tr><td>>MI0000088|hsa-mir-30a&WithFlank&6|-|71403545|71403627@12.35.24(+1C)&[MIMAT0000087&hsa-miR-30a-5p&offsets|0|+2(+1C);m-15&6|-|71403595|71403616&offsets|0|+2(+1C)]|TGTAAACATCCTCGACTGGAAGCTC</td><td>>MI0000088|hsa-mir-30a&WithFlank&6|-|71403545|71403627</td><td>12</td><td>35</td><td>24(+1C)</td><td>TGTAAACATCCTCGACTGGAAGCTC</td><td>954.2</td><td>7.4</td><td>1.0</td><td>7.8</td><td>6.5E-15</td><td>1.9E-14</td><td>BT20/MB468</td></tr>
<tr><td>>hg38_21_+_8433172_8446622_RNA45SN1_45S_50ntFlanks@10389.10409.21|TGAACAGCAGTTGAACATGGG</td><td>>hg38_21_+_8433172_8446622_RNA45SN1_45S_50ntFlanks</td><td>10389</td><td>10409</td><td>21</td><td>TGAACAGCAGTTGAACATGGG</td><td>615.8</td><td>-8.9</td><td>1.1</td><td>-7.8</td><td>6.4E-15</td><td>1.9E-14</td><td>MB231/MB468</td></tr>
<tr><td>>trnaMT_MetCAT_MT_+_4402_4469@50.71.22|TTATACCCTTCCCGTACTACCA</td><td>>trnaMT_MetCAT_MT_+_4402_4469</td><td>50</td><td>71</td><td>22</td><td>TTATACCCTTCCCGTACTACCA</td><td>626.7</td><td>7.9</td><td>1.0</td><td>7.8</td><td>6.4E-15</td><td>1.9E-14</td><td>MB231/MB468</td></tr>
<tr><td>>am_trna2_GlyGCC_21_-_18827107_18827177@57.74.18,trna35_GlyGCC_1_+_161413094_161413164@57.74.18,trna37_GlyGCC_1_+_161420467_161420537@57.74.18,trna39_GlyGCC_1_+_161427898_161427968@57.74.18,trna41_GlyGCC_1_+_161435258_161435328@57.74.18|TTCCCGGCCCATGCACCA</td><td>>am_trna2_GlyGCC_21_-_18827107_18827177</td><td>57</td><td>74</td><td>18</td><td>TTCCCGGCCCATGCACCA</td><td>1086.9</td><td>10.9</td><td>1.4</td><td>7.8</td><td>6.6E-15</td><td>1.9E-14</td><td>BT20/MB468</td></tr>
<tr><td>>hg38_21_+_8433172_8446622_RNA45SN1_45S_50ntFlanks@10727.10757.31|GGAGAGGGTGTAAATCTCGCGCCGGGCCGTA</td><td>>hg38_21_+_8433172_8446622_RNA45SN1_45S_50ntFlanks</td><td>10727</td><td>10757</td><td>31</td><td>GGAGAGGGTGTAAATCTCGCGCCGGGCCGTA</td><td>216.5</td><td>-9.8</td><td>1.3</td><td>-7.8</td><td>6.7E-15</td><td>1.9E-14</td><td>BT20/MB468</td></tr>
<tr><td>>hg38_21_+_8433172_8446622_RNA45SN1_45S_50ntFlanks@5073.5098.26|AATTCCGATAACGAACGAGACTCTGG</td><td>>hg38_21_+_8433172_8446622_RNA45SN1_45S_50ntFlanks</td><td>5073</td><td>5098</td><td>26</td><td>AATTCCGATAACGAACGAGACTCTGG</td><td>327.5</td><td>-3.6</td><td>0.5</td><td>-7.8</td><td>6.7E-15</td><td>1.9E-14</td><td>BT20/MB468</td></tr>
<tr><td>>hg38_21_+_8433172_8446622_RNA45SN1_45S_50ntFlanks@10404.10430.27|CATGGGTCAGTCGGTCCTGAGAGATGG</td><td>>hg38_21_+_8433172_8446622_RNA45SN1_45S_50ntFlanks</td><td>10404</td><td>10430</td><td>27</td><td>CATGGGTCAGTCGGTCCTGAGAGATGG</td><td>256.4</td><td>-4.2</td><td>0.5</td><td>-7.8</td><td>6.5E-15</td><td>1.9E-14</td><td>MB231/MB468</td></tr>
<tr><td>>MI0000088|hsa-mir-30a&WithFlank&6|-|71403545|71403627@12.33.22(+1G)&[MIMAT0000087&hsa-miR-30a-5p&offsets|0|0(+1G);m-15&6|-|71403595|71403616&offsets|0|0(+1G)]|TGTAAACATCCTCGACTGGAAGG</td><td>>MI0000088|hsa-mir-30a&WithFlank&6|-|71403545|71403627</td><td>12</td><td>33</td><td>22(+1G)</td><td>TGTAAACATCCTCGACTGGAAGG</td><td>755.8</td><td>8.4</td><td>1.1</td><td>7.8</td><td>6.6E-15</td><td>1.9E-14</td><td>MB231/MB468</td></tr>
<tr><td>>hg38_21_+_8433172_8446622_RNA45SN1_45S_50ntFlanks@6645.6668.24|CTCGTACGACTCTTAGCGGTGGAT</td><td>>hg38_21_+_8433172_8446622_RNA45SN1_45S_50ntFlanks</td><td>6645</td><td>6668</td><td>24</td><td>CTCGTACGACTCTTAGCGGTGGAT</td><td>1412.2</td><td>4.7</td><td>0.6</td><td>7.8</td><td>6.7E-15</td><td>2.0E-14</td><td>MB231/MB468</td></tr>
<tr><td>>trnaMT_PheGAA_MT_+_577_647@39.56.18|ATGTTTAGACGGGCTCAC</td><td>>trnaMT_PheGAA_MT_+_577_647</td><td>39</td><td>56</td><td>18</td><td>ATGTTTAGACGGGCTCAC</td><td>5451.1</td><td>9.4</td><td>1.2</td><td>7.8</td><td>6.9E-15</td><td>2.0E-14</td><td>MB231/MB468</td></tr>
<tr><td>>hg38_21_+_8433172_8446622_RNA45SN1_45S_50ntFlanks@6745.6762.18|ACACTTCGAACGCACTTG</td><td>>hg38_21_+_8433172_8446622_RNA45SN1_45S_50ntFlanks</td><td>6745</td><td>6762</td><td>18</td><td>ACACTTCGAACGCACTTG</td><td>271.9</td><td>-6.9</td><td>0.9</td><td>-7.8</td><td>7.1E-15</td><td>2.0E-14</td><td>BT20/MB468</td></tr>
<tr><td>>hg38_21_+_8433172_8446622_RNA45SN1_45S_50ntFlanks@12347.12365.19|AGAAAAGTTACCACAGGGA</td><td>>hg38_21_+_8433172_8446622_RNA45SN1_45S_50ntFlanks</td><td>12347</td><td>12365</td><td>19</td><td>AGAAAAGTTACCACAGGGA</td><td>292.0</td><td>-9.8</td><td>1.3</td><td>-7.8</td><td>7.1E-15</td><td>2.1E-14</td><td>MB231/MB468</td></tr>
<tr><td>>hg38_21_+_8433172_8446622_RNA45SN1_45S_50ntFlanks@12151.12180.30|GTGGGGAGTTTGACTGGGGCGGTACACCTG</td><td>>hg38_21_+_8433172_8446622_RNA45SN1_45S_50ntFlanks</td><td>12151</td><td>12180</td><td>30</td><td>GTGGGGAGTTTGACTGGGGCGGTACACCTG</td><td>344.4</td><td>-10.0</td><td>1.3</td><td>-7.8</td><td>7.1E-15</td><td>2.1E-14</td><td>MB231/MB468</td></tr>
<tr><td>>MI0000289|hsa-mir-181a-1&WithFlank&1|-|198859038|198859159@70.90.21(+1A)&[MIMAT0000270&hsa-miR-181a-3p&offsets|0|-1(+1A);m-125&1|-|198859069|198859090&offsets|0|-1(+1A)]|ACCATCGACCGTTGATTGTACA</td><td>>MI0000289|hsa-mir-181a-1&WithFlank&1|-|198859038|198859159</td><td>70</td><td>90</td><td>21(+1A)</td><td>ACCATCGACCGTTGATTGTACA</td><td>718.5</td><td>9.5</td><td>1.2</td><td>7.8</td><td>7.2E-15</td><td>2.1E-14</td><td>MB231/MB468</td></tr>
<tr><td>>hg38_21_+_8433172_8446622_RNA45SN1_45S_50ntFlanks@6744.6771.28|GACACTTCGAACGCACTTGCGGCCCCGG</td><td>>hg38_21_+_8433172_8446622_RNA45SN1_45S_50ntFlanks</td><td>6744</td><td>6771</td><td>28</td><td>GACACTTCGAACGCACTTGCGGCCCCGG</td><td>888.1</td><td>-6.1</td><td>0.8</td><td>-7.8</td><td>7.4E-15</td><td>2.1E-14</td><td>BT20/MB468</td></tr>
<tr><td>>hg38_21_+_8433172_8446622_RNA45SN1_45S_50ntFlanks@12419.12453.35|GATGTCGGCTCTTCCTATCATTGTGAAGCAGAATT</td><td>>hg38_21_+_8433172_8446622_RNA45SN1_45S_50ntFlanks</td><td>12419</td><td>12453</td><td>35</td><td>GATGTCGGCTCTTCCTATCATTGTGAAGCAGAATT</td><td>1419.7</td><td>8.2</td><td>1.1</td><td>7.8</td><td>7.3E-15</td><td>2.1E-14</td><td>MB231/MB468</td></tr>
<tr><td>>hg38_21_+_8433172_8446622_RNA45SN1_45S_50ntFlanks@11787.11804.18|AATGGATGAACGAGATTC</td><td>>hg38_21_+_8433172_8446622_RNA45SN1_45S_50ntFlanks</td><td>11787</td><td>11804</td><td>18</td><td>AATGGATGAACGAGATTC</td><td>6637.3</td><td>-4.3</td><td>0.6</td><td>-7.8</td><td>7.3E-15</td><td>2.1E-14</td><td>MB231/MB468</td></tr>
<tr><td>>MI0005756|hsa-mir-934&WithFlank&X|+|136550872|136550966@21.41.21(+1A)&[MIMAT0004977&hsa-miR-934&offsets|0|-1(+1A);m-943&X|+|136550892|136550913&offsets|0|-1(+1A)]|TGTCTACTACTGGAGACACTGA</td><td>>MI0005756|hsa-mir-934&WithFlank&X|+|136550872|136550966</td><td>21</td><td>41</td><td>21(+1A)</td><td>TGTCTACTACTGGAGACACTGA</td><td>9348.1</td><td>11.7</td><td>1.5</td><td>7.8</td><td>7.5E-15</td><td>2.1E-14</td><td>BT20/MB468</td></tr>
<tr><td>>MI0000084|hsa-mir-26b&WithFlank&2|+|218402640|218402728@18.39.22&[MIMAT0000083&hsa-miR-26b-5p&offsets|0|+1;m-53&2|+|218402657|218402678&offsets|0|0]|TTCAAGTAATTCAGGATAGGTT</td><td>>MI0000084|hsa-mir-26b&WithFlank&2|+|218402640|218402728</td><td>18</td><td>39</td><td>22</td><td>TTCAAGTAATTCAGGATAGGTT</td><td>35579.5</td><td>7.1</td><td>0.9</td><td>7.8</td><td>7.5E-15</td><td>2.1E-14</td><td>BT20/MB468</td></tr>
<tr><td>>MI0000440|hsa-mir-27b&WithFlank&9|+|95085439|95085547@67.86.20(+2U)&[MIMAT0000419&hsa-miR-27b-3p&offsets|0|-1(+2U);m-39&9|+|95085505|95085525&offsets|0|-1(+2U)]|TTCACAGTGGCTAAGTTCTGTT</td><td>>MI0000440|hsa-mir-27b&WithFlank&9|+|95085439|95085547</td><td>67</td><td>86</td><td>20(+2U)</td><td>TTCACAGTGGCTAAGTTCTGTT</td><td>590.5</td><td>8.6</td><td>1.1</td><td>7.8</td><td>7.5E-15</td><td>2.1E-14</td><td>BT20/MB468</td></tr>
<tr><td>>MI0000239|hsa-mir-197&WithFlank&1|+|109598887|109598973@54.74.21(+1A)&[MIMAT0000227&hsa-miR-197-3p&offsets|0|-1(+1A);m-99&1|+|109598940|109598961&offsets|0|-1(+1A)]|TTCACCACCTTCTCCACCCAGA</td><td>>MI0000239|hsa-mir-197&WithFlank&1|+|109598887|109598973</td><td>54</td><td>74</td><td>21(+1A)</td><td>TTCACCACCTTCTCCACCCAGA</td><td>2548.3</td><td>11.2</td><td>1.4</td><td>7.8</td><td>7.5E-15</td><td>2.2E-14</td><td>BT20/MB468</td></tr>
<tr><td>>MI0000447|hsa-mir-128-1&WithFlank&2|+|135665391|135665484@56.75.20;MI0000727|hsa-mir-128-2&WithFlank&3|+|35744470|35744565@58.77.20&[MIMAT0000424&hsa-miR-128-3p&offsets|0|-1;m-108&2|+|135665446|135665466&offsets|0|-1];[MIMAT0000424_1&hsa-miR-128-3p&offsets|0|-1;m-117&3|+|35744527|35744547&offsets|0|-1]|TCACAGTGAACCGGTCTCTT</td><td>>MI0000447|hsa-mir-128-1&WithFlank&2|+|135665391|135665484</td><td>56</td><td>75</td><td>20</td><td>TCACAGTGAACCGGTCTCTT</td><td>2628.7</td><td>5.8</td><td>0.7</td><td>7.8</td><td>7.5E-15</td><td>2.2E-14</td><td>BT20/MB468</td></tr>
<tr><td>>hg38_21_+_8433172_8446622_RNA45SN1_45S_50ntFlanks@12640.12667.28|ACCATCTGTGGGATTATGACTGAACGCC</td><td>>hg38_21_+_8433172_8446622_RNA45SN1_45S_50ntFlanks</td><td>12640</td><td>12667</td><td>28</td><td>ACCATCTGTGGGATTATGACTGAACGCC</td><td>536.4</td><td>8.7</td><td>1.1</td><td>7.8</td><td>7.4E-15</td><td>2.2E-14</td><td>MB231/MB468</td></tr>
<tr><td>>hg38_21_+_8433172_8446622_RNA45SN1_45S_50ntFlanks@9504.9533.30|GTGAACTATGCCTGGGCAGGGCGAAGCCAG</td><td>>hg38_21_+_8433172_8446622_RNA45SN1_45S_50ntFlanks</td><td>9504</td><td>9533</td><td>30</td><td>GTGAACTATGCCTGGGCAGGGCGAAGCCAG</td><td>597.8</td><td>-10.8</td><td>1.4</td><td>-7.8</td><td>7.4E-15</td><td>2.2E-14</td><td>MB231/MB468</td></tr>
<tr><td>>hg38_21_+_8433172_8446622_RNA45SN1_45S_50ntFlanks@9534.9552.19|AGGAAACTCTGGTGGAGGT</td><td>>hg38_21_+_8433172_8446622_RNA45SN1_45S_50ntFlanks</td><td>9534</td><td>9552</td><td>19</td><td>AGGAAACTCTGGTGGAGGT</td><td>2842.1</td><td>-4.5</td><td>0.6</td><td>-7.8</td><td>7.4E-15</td><td>2.2E-14</td><td>MB231/MB468</td></tr>
<tr><td>>hg38_21_+_8433172_8446622_RNA45SN1_45S_50ntFlanks@12957.12990.34|CTTCTGGGTCGGGGTTTCGTACGTAGCAGAGCAG</td><td>>hg38_21_+_8433172_8446622_RNA45SN1_45S_50ntFlanks</td><td>12957</td><td>12990</td><td>34</td><td>CTTCTGGGTCGGGGTTTCGTACGTAGCAGAGCAG</td><td>306.0</td><td>-6.4</td><td>0.8</td><td>-7.8</td><td>7.6E-15</td><td>2.2E-14</td><td>BT20/MB468</td></tr>
<tr><td>>hg38_21_+_8433172_8446622_RNA45SN1_45S_50ntFlanks@4449.4472.24|CCTCTCGGCGCCCCCTCGATGCTC</td><td>>hg38_21_+_8433172_8446622_RNA45SN1_45S_50ntFlanks</td><td>4449</td><td>4472</td><td>24</td><td>CCTCTCGGCGCCCCCTCGATGCTC</td><td>256.0</td><td>-10.0</td><td>1.3</td><td>-7.8</td><td>7.7E-15</td><td>2.2E-14</td><td>BT20/MB468</td></tr>
<tr><td>>hg38_21_+_8433172_8446622_RNA45SN1_45S_50ntFlanks@12488.12521.34|AACGTGAGCTGGGTTTAGACCGTCGTGAGACAGG</td><td>>hg38_21_+_8433172_8446622_RNA45SN1_45S_50ntFlanks</td><td>12488</td><td>12521</td><td>34</td><td>AACGTGAGCTGGGTTTAGACCGTCGTGAGACAGG</td><td>513.7</td><td>-7.2</td><td>0.9</td><td>-7.8</td><td>7.5E-15</td><td>2.2E-14</td><td>MB231/MB468</td></tr>
<tr><td>>MI0000734|hsa-mir-106b&WithFlank&7|-|100093987|100094080@18.38.21&[MIMAT0000680&hsa-miR-106b-5p&offsets|0|0;m-103&7|-|100094043|100094063&offsets|0|0]|TAAAGTGCTGACAGTGCAGAT</td><td>>MI0000734|hsa-mir-106b&WithFlank&7|-|100093987|100094080</td><td>18</td><td>38</td><td>21</td><td>TAAAGTGCTGACAGTGCAGAT</td><td>2737.3</td><td>6.8</td><td>0.9</td><td>7.8</td><td>7.5E-15</td><td>2.2E-14</td><td>MB231/MB468</td></tr>
<tr><td>>hg38_21_+_8433172_8446622_RNA45SN1_45S_50ntFlanks@8104.8143.40|CCGCGGCGGGGCGCGGGACATGTGGCGTACGGAAGACCCG</td><td>>hg38_21_+_8433172_8446622_RNA45SN1_45S_50ntFlanks</td><td>8104</td><td>8143</td><td>40</td><td>CCGCGGCGGGGCGCGGGACATGTGGCGTACGGAAGACCCG</td><td>2136.1</td><td>-6.1</td><td>0.8</td><td>-7.8</td><td>7.6E-15</td><td>2.2E-14</td><td>MB231/MB468</td></tr>
<tr><td>>hg38_21_+_8433172_8446622_RNA45SN1_45S_50ntFlanks@12991.13019.29|CTCCCTCGCTGCGATCTATTGAAAGTCAG</td><td>>hg38_21_+_8433172_8446622_RNA45SN1_45S_50ntFlanks</td><td>12991</td><td>13019</td><td>29</td><td>CTCCCTCGCTGCGATCTATTGAAAGTCAG</td><td>377.6</td><td>-10.1</td><td>1.3</td><td>-7.8</td><td>7.6E-15</td><td>2.2E-14</td><td>MB231/MB468</td></tr>
</table>

| | | | | | | | | | | | | |
|---|---|---|---|---|---|---|---|---|---|---|---|---|
| >MI0003834\|hsa-mir-769&WithFlank&19\|+\|46018926\|46019055@36.57.22(+1U)&[MIMAT0003886&hsa-miR-769-5p&offsets\|0\|0(+1U);m-216&19\|+\|46018961\|46018982&offsets\|0\|0(+1U)]\|TGAGACCTCTGGGTTCTGAGCTT | >MI0003834\|hsa-mir-769&WithFlank&19\|+\|46018926\|46019055 | 36 | 57 | 22(+1U) | TGAGACCTCTGGGTTCTGAGCTT | 666.8 | 9.2 | 1.2 | 7.8 | 7.8E-15 | 2.2E-14 | BT20/MB468 |
| >MI0000476\|hsa-mir-138-1&WithFlank&3\|+\|44114206\|44114316@29.52.24&[MIMAT0000430_1&hsa-miR-138-5p&offsets\|0\|+1;m-497&3\|+\|44114234\|44114256&offsets\|0\|+1]\|AGCTGGTGTTGTGAATCAGGCCGT | >MI0000476\|hsa-mir-138-1&WithFlank&3\|+\|44114206\|44114316 | 29 | 52 | 24 | AGCTGGTGTTGTGAATCAGGCCGT | 303.0 | -11.6 | 1.4 | -8.1 | 5.2E-16 | 2.2E-14 | BT20/MB231 |
| >hg38_21_+_8433172_8446622_RNA45SN1_45S_50ntFlanks@12453.12477.25\|TCACCAAGCGTTGGATTGTTCACCC | >hg38_21_+_8433172_8446622_RNA45SN1_45S_50ntFlanks | 12453 | 12477 | 25 | TCACCAAGCGTTGGATTGTTCACCC | 500.8 | 7.2 | 0.9 | 7.8 | 7.6E-15 | 2.2E-14 | MB231/MB468 |
| >hg38_21_+_8433172_8446622_RNA45SN1_45S_50ntFlanks@7989.8006.18\|AGACGTGGCGACCCGCTG | >hg38_21_+_8433172_8446622_RNA45SN1_45S_50ntFlanks | 7989 | 8006 | 18 | AGACGTGGCGACCCGCTG | 1844.6 | -4.8 | 0.6 | -7.8 | 7.9E-15 | 2.2E-14 | BT20/MB468 |
| >MI0000469\|hsa-mir-125a&WithFlank&19\|+\|51693248\|51693345@21.42.22(+1U)&[MIMAT0000443&hsa-miR-125a-5p&offsets\|0\|-2(+1U);m-63&19\|+\|51693268\|51693289&offsets\|0\|0(+1U)]\|TCCCTGAGACCCTTTAACCTGTT | >MI0000469\|hsa-mir-125a&WithFlank&19\|+\|51693248\|51693345 | 21 | 42 | 22(+1U) | TCCCTGAGACCCTTTAACCTGTT | 878.8 | 8.4 | 1.1 | 7.8 | 8.0E-15 | 2.3E-14 | BT20/MB468 |
| >hg38_21_+_8433172_8446622_RNA45SN1_45S_50ntFlanks@9504.9522.19\|GTGAACTATGCCTGGGCAG | >hg38_21_+_8433172_8446622_RNA45SN1_45S_50ntFlanks | 9504 | 9522 | 19 | GTGAACTATGCCTGGGCAG | 613.7 | -6.0 | 0.8 | -7.8 | 8.0E-15 | 2.3E-14 | BT20/MB468 |
| >hg38_21_+_8433172_8446622_RNA45SN1_45S_50ntFlanks@9567.9594.28\|ACGTGCAAATCGGTCGTCCGACCTGGGT | >hg38_21_+_8433172_8446622_RNA45SN1_45S_50ntFlanks | 9567 | 9594 | 28 | ACGTGCAAATCGGTCGTCCGACCTGGGT | 269.7 | -4.6 | 0.6 | -7.8 | 7.9E-15 | 2.3E-14 | MB231/MB468 |
| >hg38_21_+_8433172_8446622_RNA45SN1_45S_50ntFlanks@5525.5550.26\|TAAAAGTCGTAACAAGGTTTCCGTAG | >hg38_21_+_8433172_8446622_RNA45SN1_45S_50ntFlanks | 5525 | 5550 | 26 | TAAAAGTCGTAACAAGGTTTCCGTAG | 208.4 | -9.7 | 1.2 | -7.8 | 8.1E-15 | 2.3E-14 | BT20/MB468 |
| >am_trnaMT_PheGAA_MT_+_577_647@40.62.23\|TGTTTAGACGGGCTCACATCACC | >am_trnaMT_PheGAA_MT_+_577_647 | 40 | 62 | 23 | TGTTTAGACGGGCTCACATCACC | 448.9 | 8.5 | 1.1 | 7.8 | 8.1E-15 | 2.3E-14 | BT20/MB468 |
| >MI0000063\|hsa-let-7b&WithFlank&22\|+\|46113680\|46113774@12.33.22(+2A)&[MIMAT0000063&hsa-let-7b-5p&offsets\|0\|0(+2A);m-14&22\|+\|46113691\|46113712&offsets\|0\|0(+2A)]\|TGAGGTAGTAGGTTGTGTGGTTAA | >MI0000063\|hsa-let-7b&WithFlank&22\|+\|46113680\|46113774 | 12 | 33 | 22(+2A) | TGAGGTAGTAGGTTGTGTGGTTAA | 581.5 | 8.3 | 1.1 | 7.8 | 8.1E-15 | 2.3E-14 | BT20/MB468 |
| >hg38_21_+_8433172_8446622_RNA45SN1_45S_50ntFlanks@10735.10755.21\|TGTAAATCTCGCGCCGGGCCG | >hg38_21_+_8433172_8446622_RNA45SN1_45S_50ntFlanks | 10735 | 10755 | 21 | TGTAAATCTCGCGCCGGGCCG | 505.8 | -6.5 | 0.8 | -7.8 | 8.0E-15 | 2.3E-14 | MB231/MB468 |
| >hg38_21_+_8433172_8446622_RNA45SN1_45S_50ntFlanks@4732.4761.30\|AGAACGAAAGTCGGAGGTTCGAAGACGATC | >hg38_21_+_8433172_8446622_RNA45SN1_45S_50ntFlanks | 4732 | 4761 | 30 | AGAACGAAAGTCGGAGGTTCGAAGACGATC | 1054.6 | 5.9 | 0.8 | 7.8 | 8.0E-15 | 2.3E-14 | MB231/MB468 |
| >MI0005566\|hsa-mir-374b&WithFlank&X\|-\|74218541\|74218624@17.38.22&[MIMAT0004955&hsa-miR-374b-5p&offsets\|0\|0;m-153&X\|-\|74218587\|74218608&offsets\|0\|0]\|ATATAATACAACCTGCTAAGTG | >MI0005566\|hsa-mir-374b&WithFlank&X\|-\|74218541\|74218624 | 17 | 38 | 22 | ATATAATACAACCTGCTAAGTG | 4228.4 | 7.5 | 1.0 | 7.8 | 8.2E-15 | 2.3E-14 | BT20/MB468 |
| >hg38_21_+_8433172_8446622_RNA45SN1_45S_50ntFlanks@3854.3871.18\|ACTGTGGTAATTCTAGAG | >hg38_21_+_8433172_8446622_RNA45SN1_45S_50ntFlanks | 3854 | 3871 | 18 | ACTGTGGTAATTCTAGAG | 329.8 | -4.7 | 0.6 | -7.8 | 8.0E-15 | 2.3E-14 | MB231/MB468 |
| >hg38_21_+_8433172_8446622_RNA45SN1_45S_50ntFlanks@4064.4081.18\|ATCAACTTTCGATGGTAG | >hg38_21_+_8433172_8446622_RNA45SN1_45S_50ntFlanks | 4064 | 4081 | 18 | ATCAACTTTCGATGGTAG | 585.6 | -11.2 | 1.4 | -7.8 | 8.3E-15 | 2.4E-14 | BT20/MB468 |
| >hg38_1_-_228634819_228635039_RNA5S12_5S_50ntFlanks@131.169.39\|GGATGGGAGACCGCCTGGGAATACCGGGTGCTGTAGGCT | >hg38_1_-_228634819_228635039_RNA5S12_5S_50ntFlanks | 131 | 169 | 39 | GGATGGGAGACCGCCTGGGAATACCGGGTGCTGTAGGCT | 341.9 | 9.4 | 1.2 | 7.8 | 8.3E-15 | 2.4E-14 | BT20/MB468 |
| >MI0000299\|hsa-mir-222&WithFlank&X\|-\|45747009\|45747130@75.99.25(+2U)&[MIMAT0000279&hsa-miR-222-3p&offsets\|0\|+4(+2U);m-140&X\|-\|45747033\|45747056&offsets\|0\|+1(+2U)]\|AGCTACATCTGGCTACTGGGTCTCTTT | >MI0000299\|hsa-mir-222&WithFlank&X\|-\|45747009\|45747130 | 75 | 99 | 25(+2U) | AGCTACATCTGGCTACTGGGTCTCTTT | 853.5 | 6.3 | 0.8 | 7.8 | 8.3E-15 | 2.4E-14 | BT20/MB468 |
| >trna116_GluCTC_1_-_145399233_145399304@32.71.40,trna71_GluCTC_1_-_161439189_161439260@32.71.40,trna74_GluCTC_1_-_161431809_161431880@32.71.40,trna77_GluCTC_1_-_161424398_161424469@32.71.40,trna77_GluCTC_6_+_28949976_28950047@32.71.40,trna80_GluCTC_1_-_161417018_161417089@32.71.40,trna87_GluCTC_6_-_126101393_126101464@32.71.40\|CTCTCACCGCCGCGGCCCGGGTTCGATTCCCGGTCAGGGA | >trna116_GluCTC_1_-_145399233_145399304 | 32 | 71 | 40 | CTCTCACCGCCGCGGCCCGGGTTCGATTCCCGGTCAGGGA | 296.1 | -10.2 | 1.3 | -7.8 | 8.4E-15 | 2.4E-14 | BT20/MB468 |
| >hg38_21_+_8433172_8446622_RNA45SN1_45S_50ntFlanks@11677.11702.26\|CTCTGAATGTCAAAGTGAAGAAATTC | >hg38_21_+_8433172_8446622_RNA45SN1_45S_50ntFlanks | 11677 | 11702 | 26 | CTCTGAATGTCAAAGTGAAGAAATTC | 486.8 | -4.4 | 0.6 | -7.8 | 8.4E-15 | 2.4E-14 | BT20/MB468 |
| >MI0000441\|hsa-mir-30b&WithFlank&8\|-\|134800514\|134800613@23.43.21(+1G)&[MIMAT0000420&hsa-miR-30b-5p&offsets\|0\|-1(+1G);m-76&8\|-\|134800570\|134800591&offsets\|0\|-1(+1G)]\|TGTAAACATCCTACACTCAGCG | >MI0000441\|hsa-mir-30b&WithFlank&8\|-\|134800514\|134800613 | 23 | 43 | 21(+1G) | TGTAAACATCCTACACTCAGCG | 1425.9 | 9.3 | 1.2 | 7.8 | 8.3E-15 | 2.4E-14 | MB231/MB468 |
| >hg38_21_+_8433172_8446622_RNA45SN1_45S_50ntFlanks@12418.12448.31\|CGATGTCGGCTCTTCCTATCATTGTGAAGCA | >hg38_21_+_8433172_8446622_RNA45SN1_45S_50ntFlanks | 12418 | 12448 | 31 | CGATGTCGGCTCTTCCTATCATTGTGAAGCA | 1188.0 | 5.1 | 0.7 | 7.8 | 8.3E-15 | 2.4E-14 | MB231/MB468 |
| >hg38_1_-_228634819_228635039_RNA5S12_5S_50ntFlanks@84.112.29\|CTCGTCTGATCTCGGAAGCTAAGCAGGGT | >hg38_1_-_228634819_228635039_RNA5S12_5S_50ntFlanks | 84 | 112 | 29 | CTCGTCTGATCTCGGAAGCTAAGCAGGGT | 253.3 | -5.6 | 0.7 | -7.8 | 8.6E-15 | 2.5E-14 | MB231/MB468 |
| >hg38_21_+_8433172_8446622_RNA45SN1_45S_50ntFlanks@12476.12499.24\|CCACTAATAGGGAACGTGAGCTGG | >hg38_21_+_8433172_8446622_RNA45SN1_45S_50ntFlanks | 12476 | 12499 | 24 | CCACTAATAGGGAACGTGAGCTGG | 310.9 | -5.6 | 0.7 | -7.8 | 8.7E-15 | 2.5E-14 | MB231/MB468 |
| >hg38_21_+_8433172_8446622_RNA45SN1_45S_50ntFlanks@12934.12952.19\|CTAAACCATTCGTAGACGA | >hg38_21_+_8433172_8446622_RNA45SN1_45S_50ntFlanks | 12934 | 12952 | 19 | CTAAACCATTCGTAGACGA | 424.9 | -6.5 | 0.8 | -7.8 | 8.7E-15 | 2.5E-14 | MB231/MB468 |
| >hg38_21_+_8433172_8446622_RNA45SN1_45S_50ntFlanks@12633.12661.29\|CGAAGCTACCATCTGTGGGATTATGACTG | >hg38_21_+_8433172_8446622_RNA45SN1_45S_50ntFlanks | 12633 | 12661 | 29 | CGAAGCTACCATCTGTGGGATTATGACTG | 1428.6 | -3.9 | 0.5 | -7.8 | 8.7E-15 | 2.5E-14 | MB231/MB468 |
| >trna116_GluCTC_1_-_145399233_145399304@37.75.39,trna134_GluTTC_1_-_16861774_16861845@37.75.39,trna71_GluCTC_1_-_161439189_161439260@37.75.39,trna74_GluCTC_1_-_161431809_161431880@37.75.39,trna77_GluCTC_1_-_161424398_161424469@37.75.39,trna77_GluCTC_6_+_28949976_28950047@37.75.39,trna80_GluCTC_1_-_161417018_161417089@37.75.39,trna84_GluTTC_1_-_161391883_161391954@37.75.39,trna87_GluCTC_6_-_126101393_126101464@37.75.39\|ACCGCCGCGGCCCGGGTTCGATTCCCGGTCAGGGAACCA | >trna116_GluCTC_1_-_145399233_145399304 | 37 | 75 | 39 | ACCGCCGCGGCCCGGGTTCGATTCCCGGTCAGGGAACCA | 9442.0 | 9.0 | 1.2 | 7.8 | 8.8E-15 | 2.6E-14 | MB231/MB468 |
| >am_trna143_LysTTT_6_-_27559593_27559665@6.24.19,trna14_LysTTT_11_-_59327808_59327880@6.24.19,trna2_LysTTT_17_+_8022473_8022545@6.24.19,trna54_LysTTT_1_+_204475655_204475727@6.24.19,trna5_LysTTT_11_+_59323902_59323974@6.24.19,trna62_LysTTT_1_-_204476158_204476230@6.24.19,trna76_LysTTT_6_+_28918806_28918878@6.24.19\|GATAGCTCAGTCGGTAGAG | >am_trna143_LysTTT_6_-_27559593_27559665 | 6 | 24 | 19 | GATAGCTCAGTCGGTAGAG | 363.0 | -10.5 | 1.4 | -7.8 | 9.0E-15 | 2.6E-14 | BT20/MB468 |
| >hg38_21_+_8433172_8446622_RNA45SN1_45S_50ntFlanks@13006.13028.23\|CTATTGAAAGTCAGCCCTCGACA | >hg38_21_+_8433172_8446622_RNA45SN1_45S_50ntFlanks | 13006 | 13028 | 23 | CTATTGAAAGTCAGCCCTCGACA | 610.4 | -11.2 | 1.5 | -7.8 | 9.1E-15 | 2.6E-14 | BT20/MB468 |
| >hg38_21_+_8433172_8446622_RNA45SN1_45S_50ntFlanks@10846.10864.19\|CTTCGGGATAAGGATTGGC | >hg38_21_+_8433172_8446622_RNA45SN1_45S_50ntFlanks | 10846 | 10864 | 19 | CTTCGGGATAAGGATTGGC | 347.3 | -7.1 | 0.9 | -7.8 | 9.1E-15 | 2.6E-14 | BT20/MB468 |
| >hg38_21_+_8433172_8446622_RNA45SN1_45S_50ntFlanks@12416.12442.27\|TTCGATGTCGGCTCTTCCTATCATTGT | >hg38_21_+_8433172_8446622_RNA45SN1_45S_50ntFlanks | 12416 | 12442 | 27 | TTCGATGTCGGCTCTTCCTATCATTGT | 426.2 | 5.1 | 0.7 | 7.8 | 9.1E-15 | 2.6E-14 | BT20/MB468 |
| >hg38_21_+_8433172_8446622_RNA45SN1_45S_50ntFlanks@5472.5506.35\|ACGGCCCTGGCGGAGCGCTGAGAAGACGGTCGAAC | >hg38_21_+_8433172_8446622_RNA45SN1_45S_50ntFlanks | 5472 | 5506 | 35 | ACGGCCCTGGCGGAGCGCTGAGAAGACGGTCGAAC | 1038.4 | 8.6 | 1.1 | 7.8 | 8.9E-15 | 2.6E-14 | MB231/MB468 |
| >hg38_21_+_8433172_8446622_RNA45SN1_45S_50ntFlanks@9534.9571.38\|AGGAAACTCTGGTGGAGGTCCGTAGCGGTCCTGACGTG | >hg38_21_+_8433172_8446622_RNA45SN1_45S_50ntFlanks | 9534 | 9571 | 38 | AGGAAACTCTGGTGGAGGTCCGTAGCGGTCCTGACGTG | 6289.4 | -3.7 | 0.5 | -7.8 | 8.9E-15 | 2.6E-14 | MB231/MB468 |
| >hg38_21_+_8433172_8446622_RNA45SN1_45S_50ntFlanks@12073.12095.23\|CCGAGGGGCTCTCGCTTCTGGCG | >hg38_21_+_8433172_8446622_RNA45SN1_45S_50ntFlanks | 12073 | 12095 | 23 | CCGAGGGGCTCTCGCTTCTGGCG | 1550.7 | -5.8 | 0.8 | -7.8 | 9.0E-15 | 2.6E-14 | MB231/MB468 |
| >MI0000650\|hsa-mir-200c&WithFlank&12\|+\|6963693\|6963772@50.70.21(+2A)&[MIMAT0000617&hsa-miR-200c-3p&offsets\|0\|-2(+2A);m-54&12\|+\|6963742\|6963764&offsets\|0\|-2(+2A)]\|TAATACTGCCGGGTAATGATGAA | >MI0000650\|hsa-mir-200c&WithFlank&12\|+\|6963693\|6963772 | 50 | 70 | 21(+2A) | TAATACTGCCGGGTAATGATGAA | 448.2 | 12.3 | 1.5 | 8.1 | 6.2E-16 | 2.6E-14 | BT20/MB231 |
| >hg38_21_+_8433172_8446622_RNA45SN1_45S_50ntFlanks@12044.12072.29\|ACTGACCCGGTGAGGCGGGGGGGCGAGCC | >hg38_21_+_8433172_8446622_RNA45SN1_45S_50ntFlanks | 12044 | 12072 | 29 | ACTGACCCGGTGAGGCGGGGGGGCGAGCC | 196.4 | -9.6 | 1.2 | -7.7 | 9.2E-15 | 2.6E-14 | BT20/MB468 |
| >hg38_21_+_8433172_8446622_RNA45SN1_45S_50ntFlanks@6738.6756.19\|ATCATCGACACTTCGAACG | >hg38_21_+_8433172_8446622_RNA45SN1_45S_50ntFlanks | 6738 | 6756 | 19 | ATCATCGACACTTCGAACG | 631.7 | -11.3 | 1.5 | -7.7 | 9.2E-15 | 2.6E-14 | BT20/MB468 |
| >hg38_MT_+_598_1651_RNR1_12S_50ntFlanks@606.633.28\|CCTGTTCTGTAATCGATAAACCCCGATC | >hg38_MT_+_598_1651_RNR1_12S_50ntFlanks | 606 | 633 | 28 | CCTGTTCTGTAATCGATAAACCCCGATC | 899.3 | 9.2 | 1.2 | 7.8 | 9.1E-15 | 2.6E-14 | MB231/MB468 |
| >hg38_21_+_8433172_8446622_RNA45SN1_45S_50ntFlanks@3854.3881.28\|ACTGTGGTAATTCTAGAGCTAATACATG | >hg38_21_+_8433172_8446622_RNA45SN1_45S_50ntFlanks | 3854 | 3881 | 28 | ACTGTGGTAATTCTAGAGCTAATACATG | 1748.3 | -6.0 | 0.8 | -7.7 | 9.3E-15 | 2.6E-14 | BT20/MB468 |
| >hg38_21_+_8433172_8446622_RNA45SN1_45S_50ntFlanks@4128.4162.35\|CGATTCCGGAGAGGGAGCCTGAGAAACGGCTACCA | >hg38_21_+_8433172_8446622_RNA45SN1_45S_50ntFlanks | 4128 | 4162 | 35 | CGATTCCGGAGAGGGAGCCTGAGAAACGGCTACCA | 149.4 | -5.8 | 0.8 | -7.7 | 9.3E-15 | 2.6E-14 | BT20/MB468 |

| | | | | | | | | | | | | |
|---|---|---|---|---|---|---|---|---|---|---|---|---|
| >hg38_21_+_8433172_8446622_RNA45SN1_45S_50ntFlanks@10805.10823.19\|GAACAATGTAGGTAAGGGA | >hg38_21_+_8433172_8446622_RNA45SN1_45S_50ntFlanks | 10805 | 10823 | 19 | GAACAATGTAGGTAAGGGA | 212.6 | -9.7 | 1.3 | -7.7 | 9.4E-15 | 2.6E-14 | BT20/MB468 |
| >hg38_21_+_8433172_8446622_RNA45SN1_45S_50ntFlanks@9545.9562.18\|GTGGAGGTCCGTAGCGGT | >hg38_21_+_8433172_8446622_RNA45SN1_45S_50ntFlanks | 9545 | 9562 | 18 | GTGGAGGTCCGTAGCGGT | 170.2 | -4.2 | 0.5 | -7.7 | 9.4E-15 | 2.7E-14 | BT20/MB468 |
| >MI0003834\|hsa-mir-769&WithFlank&19\|+\|46018926\|46019055@36.56.21(+1A)&[MIMAT0003886&hsa-miR-769-5p&offsets\|0\|-1(+1A);m-216&19\|+\|46018961\|46018982&offsets\|0\|-1(+1A)]\|TGAGACCTCTGGGTTCTGAGCA | >MI0003834\|hsa-mir-769&WithFlank&19\|+\|46018926\|46019055 | 36 | 56 | 21(+1A) | TGAGACCTCTGGGTTCTGAGCA | 1463.0 | 11.2 | 1.4 | 7.7 | 9.4E-15 | 2.7E-14 | BT20/MB468 |
| >hg38_21_+_8433172_8446622_RNA45SN1_45S_50ntFlanks@11767.11791.25\|TCTAATTAGTGACGCGCATGAATGG | >hg38_21_+_8433172_8446622_RNA45SN1_45S_50ntFlanks | 11767 | 11791 | 25 | TCTAATTAGTGACGCGCATGAATGG | 184.9 | -9.5 | 1.2 | -7.7 | 9.4E-15 | 2.7E-14 | BT20/MB468 |
| >hg38_21_+_8433172_8446622_RNA45SN1_45S_50ntFlanks@12960.12982.23\|CTGGGTCGGGGTTTCGTACGTAG | >hg38_21_+_8433172_8446622_RNA45SN1_45S_50ntFlanks | 12960 | 12982 | 23 | CTGGGTCGGGGTTTCGTACGTAG | 138.6 | -4.4 | 0.6 | -7.7 | 9.5E-15 | 2.7E-14 | BT20/MB468 |
| >hg38_21_+_8433172_8446622_RNA45SN1_45S_50ntFlanks@11569.11602.34\|CAGCCGACTTAGAACTGGTGCGGACCAGGGGAAT | >hg38_21_+_8433172_8446622_RNA45SN1_45S_50ntFlanks | 11569 | 11602 | 34 | CAGCCGACTTAGAACTGGTGCGGACCAGGGGAAT | 1451.4 | -4.5 | 0.6 | -7.7 | 9.6E-15 | 2.7E-14 | BT20/MB468 |
| >MI0000272\|hsa-mir-182&WithFlank&7\|-\|129770377\|129770498@29.54.26(+1A)&[MIMAT0000259&hsa-miR-182-5p&offsets\|0\|+2(+1A);m-22&7\|-\|129770449\|129770470&offsets\|0\|+4(+1A)]\|TTTGGCAATGGTAGAACTCACACTGGA | >MI0000272\|hsa-mir-182&WithFlank&7\|-\|129770377\|129770498 | 29 | 54 | 26(+1A) | TTTGGCAATGGTAGAACTCACACTGGA | 501.8 | 9.6 | 1.2 | 7.7 | 9.7E-15 | 2.7E-14 | BT20/MB468 |
| >hg38_21_+_8433172_8446622_RNA45SN1_45S_50ntFlanks@12365.12383.19\|ATAACTGGCTTGTGGCGGC | >hg38_21_+_8433172_8446622_RNA45SN1_45S_50ntFlanks | 12365 | 12383 | 19 | ATAACTGGCTTGTGGCGGC | 213.0 | -9.7 | 1.3 | -7.7 | 9.7E-15 | 2.7E-14 | BT20/MB468 |
| >MI0000478\|hsa-mir-149&WithFlank&2\|+\|240455995\|240456095@21.41.21&[MIMAT0000450&hsa-miR-149-5p&offsets\|0\|-2;m-195&2\|+\|240456015\|240456037&offsets\|0\|-2]\|TCTGGCTCCGTGTCTTCACTC | >MI0000478\|hsa-mir-149&WithFlank&2\|+\|240455995\|240456095 | 21 | 41 | 21 | TCTGGCTCCGTGTCTTCACTC | 1397.4 | 11.5 | 1.5 | 7.7 | 9.8E-15 | 2.8E-14 | BT20/MB468 |
| >hg38_21_+_8433172_8446622_RNA45SN1_45S_50ntFlanks@11769.11791.23\|TAATTAGTGACGCGCATGAATGG | >hg38_21_+_8433172_8446622_RNA45SN1_45S_50ntFlanks | 11769 | 11791 | 23 | TAATTAGTGACGCGCATGAATGG | 791.1 | -3.9 | 0.5 | -7.7 | 9.5E-15 | 2.8E-14 | MB231/MB468 |
| >am_MI0005763\|hsa-mir-941-1&WithFlank&20\|+\|63919443\|63919526@53.74.22(+1G);MI0005764\|hsa-mir-941-2&WithFlank&20\|+\|63919499\|63919582@53.74.22(+1G);MI0005765\|hsa-mir-941-3&WithFlank&20\|+\|63919555\|63919638@53.74.22(+1G);MI0005766\|hsa-mir-941-4&WithFlank&20\|+\|63919750\|63919833@53.74.22(+1G);MI0031520\|hsa-mir-941-5&WithFlank&20\|+\|63919862\|63919945@53.74.22(+1G)&[MIMAT0004984&hsa-miR-941&offsets\|0\|-1(+1G);m-263&20\|+\|63919495\|63919517&offsets\|0\|-1(+1G)];[MIMAT0004984_1&hsa-miR-941&offsets\|0\|-1(+1G);m-263&20\|+\|63919551\|63919573&offsets\|0\|-1(+1G)];[MIMAT0004984_2&hsa-miR-941&offsets\|0\|-1(+1G);m-263&20\|+\|63919607\|63919629&offsets\|0\|-1(+1G)];[MIMAT0004984_3&hsa-miR-941&offsets\|0\|-1(+1G);m-263&20\|+\|63919802\|63919824&offsets\|0\|-1(+1G)];[MIMAT0004984_4&hsa-miR-941&offsets\|0\|-1(+1G);m-263&20\|+\|63919914\|63919936&offsets\|0\|-1(+1G)]\|CACCCGGCTGTGTGCACATGTGG | >am_MI0005763\|hsa-mir-941-1&WithFlank&20\|+\|63919443\|63919526 | 53 | 74 | 22(+1G) | CACCCGGCTGTGTGCACATGTGG | 447.8 | 10.5 | 1.4 | 7.7 | 9.8E-15 | 2.8E-14 | BT20/MB468 |
| >MI0000815\|hsa-mir-339&WithFlank&7\|-\|1022927\|1023032@21.44.24&[MIMAT0000764&hsa-miR-339-5p&offsets\|0\|+1;m-163&7\|-\|1022990\|1023012&offsets\|0\|+1]\|TCCCTGTCCTCCAGGAGCTCACGT | >MI0000815\|hsa-mir-339&WithFlank&7\|-\|1022927\|1023032 | 21 | 44 | 24 | TCCCTGTCCTCCAGGAGCTCACGT | 680.5 | 7.7 | 1.0 | 7.7 | 9.6E-15 | 2.8E-14 | MB231/MB468 |
| >hg38_21_+_8433172_8446622_RNA45SN1_45S_50ntFlanks@3840.3865.26\|CTCCTACTTGGATAACTGTGGTAATT | >hg38_21_+_8433172_8446622_RNA45SN1_45S_50ntFlanks | 3840 | 3865 | 26 | CTCCTACTTGGATAACTGTGGTAATT | 412.6 | 4.4 | 0.6 | 7.7 | 9.9E-15 | 2.8E-14 | BT20/MB468 |
| >hg38_21_+_8433172_8446622_RNA45SN1_45S_50ntFlanks@6670.6699.30\|ACTCGGCTCGTGCGTCGATGAAGAACGCAG | >hg38_21_+_8433172_8446622_RNA45SN1_45S_50ntFlanks | 6670 | 6699 | 30 | ACTCGGCTCGTGCGTCGATGAAGAACGCAG | 438.4 | -3.4 | 0.4 | -7.7 | 1.0E-14 | 2.8E-14 | BT20/MB468 |
| >hg38_21_+_8433172_8446622_RNA45SN1_45S_50ntFlanks@12996.13039.44\|TCGCTGCGATCTATTGAAAGTCAGCCCTCGACACAAGGGTTTGT | >hg38_21_+_8433172_8446622_RNA45SN1_45S_50ntFlanks | 12996 | 13039 | 44 | TCGCTGCGATCTATTGAAAGTCAGCCCTCGACACAAGGGTTTGT | 76411.8 | 7.6 | 1.0 | 7.7 | 1.0E-14 | 2.8E-14 | BT20/MB468 |
| >hg38_1_-_228634819_228635039_RNA5S12_5S_50ntFlanks@136.170.35\|GGAGACCGCCTGGGAATACCGGGTGCTGTAGGCTT | >hg38_1_-_228634819_228635039_RNA5S12_5S_50ntFlanks | 136 | 170 | 35 | GGAGACCGCCTGGGAATACCGGGTGCTGTAGGCTT | 4276.1 | 5.9 | 0.8 | 7.7 | 9.9E-15 | 2.9E-14 | MB231/MB468 |
| >hg38_21_+_8433172_8446622_RNA45SN1_45S_50ntFlanks@10914.10936.23\|CGCCGCGGCTGGACGAGGCGCCG | >hg38_21_+_8433172_8446622_RNA45SN1_45S_50ntFlanks | 10914 | 10936 | 23 | CGCCGCGGCTGGACGAGGCGCCG | 377.9 | -10.1 | 1.3 | -7.7 | 9.9E-15 | 2.9E-14 | MB231/MB468 |
| >hg38_21_+_8433172_8446622_RNA45SN1_45S_50ntFlanks@12934.12966.33\|CTAAACCATTCGTAGACGACCTGCTTCTGGGTC | >hg38_21_+_8433172_8446622_RNA45SN1_45S_50ntFlanks | 12934 | 12966 | 33 | CTAAACCATTCGTAGACGACCTGCTTCTGGGTC | 319.4 | -6.9 | 0.9 | -7.7 | 1.0E-14 | 2.9E-14 | BT20/MB468 |
| >hg38_21_+_8433172_8446622_RNA45SN1_45S_50ntFlanks@12126.12146.21\|CGACCCGCTCCGGGGACAGTG | >hg38_21_+_8433172_8446622_RNA45SN1_45S_50ntFlanks | 12126 | 12146 | 21 | CGACCCGCTCCGGGGACAGTG | 296.9 | -9.8 | 1.3 | -7.7 | 1.0E-14 | 2.9E-14 | MB231/MB468 |
| >hg38_21_+_8433172_8446622_RNA45SN1_45S_50ntFlanks@12078.12095.18\|GGGCTCTCGCTTCTGGCG | >hg38_21_+_8433172_8446622_RNA45SN1_45S_50ntFlanks | 12078 | 12095 | 18 | GGGCTCTCGCTTCTGGCG | 343.7 | -7.3 | 0.9 | -7.7 | 1.0E-14 | 2.9E-14 | BT20/MB468 |
| >hg38_21_+_8433172_8446622_RNA45SN1_45S_50ntFlanks@10350.10377.28\|ACGAGAACTTTGAAGGCCGAAGTGGAGA | >hg38_21_+_8433172_8446622_RNA45SN1_45S_50ntFlanks | 10350 | 10377 | 28 | ACGAGAACTTTGAAGGCCGAAGTGGAGA | 1673.4 | -3.7 | 0.5 | -7.7 | 1.0E-14 | 3.0E-14 | MB231/MB468 |
| >hg38_21_+_8433172_8446622_RNA45SN1_45S_50ntFlanks@11317.11336.20\|ACCCCACGTCTCGTCGCGCG | >hg38_21_+_8433172_8446622_RNA45SN1_45S_50ntFlanks | 11317 | 11336 | 20 | ACCCCACGTCTCGTCGCGCG | 434.7 | -10.3 | 1.3 | -7.7 | 1.0E-14 | 3.0E-14 | MB231/MB468 |
| >hg38_21_+_8433172_8446622_RNA45SN1_45S_50ntFlanks@10683.10711.29\|CCGGCGGCGTCCGGTGAGCTCTCGCTGGC | >hg38_21_+_8433172_8446622_RNA45SN1_45S_50ntFlanks | 10683 | 10711 | 29 | CCGGCGGCGTCCGGTGAGCTCTCGCTGGC | 969.8 | -4.7 | 0.6 | -7.7 | 1.0E-14 | 3.0E-14 | MB231/MB468 |
| >hg38_21_+_8433172_8446622_RNA45SN1_45S_50ntFlanks@12539.12567.29\|ATGATGTGTTGTTGCCATGGTAATCCTGC | >hg38_21_+_8433172_8446622_RNA45SN1_45S_50ntFlanks | 12539 | 12567 | 29 | ATGATGTGTTGTTGCCATGGTAATCCTGC | 507.2 | -5.9 | 0.8 | -7.7 | 1.1E-14 | 3.0E-14 | BT20/MB468 |
| >trnaMT_ValTAC_MT_+_1602_1670@52.72.21\|TTAACTTGACCGCTCTGACCA | >trnaMT_ValTAC_MT_+_1602_1670 | 52 | 72 | 21 | TTAACTTGACCGCTCTGACCA | 1385.8 | 9.7 | 1.3 | 7.7 | 1.1E-14 | 3.1E-14 | BT20/MB468 |
| >hg38_21_+_8433172_8446622_RNA45SN1_45S_50ntFlanks@10786.10820.35\|AACAGCCTCTGGCATGTTGGAACAATGTAGGTAAG | >hg38_21_+_8433172_8446622_RNA45SN1_45S_50ntFlanks | 10786 | 10820 | 35 | AACAGCCTCTGGCATGTTGGAACAATGTAGGTAAG | 297.5 | -3.5 | 0.5 | -7.7 | 1.1E-14 | 3.1E-14 | BT20/MB468 |
| >MI0000077\|hsa-mir-21&WithFlank&17\|+\|59841260\|59841343@14.39.26&[MIMAT0000076&hsa-miR-21-5p&offsets\|0\|+4;m-1&17\|+\|59841273\|59841294&offsets\|0\|+4]\|TAGCTTATCAGACTGATGTTGACTGT | >MI0000077\|hsa-mir-21&WithFlank&17\|+\|59841260\|59841343 | 14 | 39 | 26 | TAGCTTATCAGACTGATGTTGACTGT | 1497.9 | 9.6 | 1.2 | 7.7 | 1.1E-14 | 3.1E-14 | BT20/MB468 |
| >hg38_21_+_8433172_8446622_RNA45SN1_45S_50ntFlanks@12621.12654.34\|GAGCCAATGGGGCGAAGCTACCATCTGTGGGATT | >hg38_21_+_8433172_8446622_RNA45SN1_45S_50ntFlanks | 12621 | 12654 | 34 | GAGCCAATGGGGCGAAGCTACCATCTGTGGGATT | 132.7 | -4.3 | 0.6 | -7.7 | 1.1E-14 | 3.1E-14 | MB231/MB468 |
| >hg38_21_+_8433172_8446622_RNA45SN1_45S_50ntFlanks@12991.13015.25\|CTCCCTCGCTGCGATCTATTGAAAG | >hg38_21_+_8433172_8446622_RNA45SN1_45S_50ntFlanks | 12991 | 13015 | 25 | CTCCCTCGCTGCGATCTATTGAAAG | 2856.6 | -7.3 | 0.9 | -7.7 | 1.1E-14 | 3.1E-14 | BT20/MB468 |
| >hg38_21_+_8433172_8446622_RNA45SN1_45S_50ntFlanks@12463.12496.34\|TTGGATTGTTCACCCACTAATAGGGAACGTGAGC | >hg38_21_+_8433172_8446622_RNA45SN1_45S_50ntFlanks | 12463 | 12496 | 34 | TTGGATTGTTCACCCACTAATAGGGAACGTGAGC | 932.5 | 7.7 | 1.0 | 7.7 | 1.1E-14 | 3.1E-14 | MB231/MB468 |
| >MI0003834\|hsa-mir-769&WithFlank&19\|+\|46018926\|46019055@36.56.21(+1G)&[MIMAT0003886&hsa-miR-769-5p&offsets\|0\|-1(+1G);m-216&19\|+\|46018961\|46018982&offsets\|0\|-1(+1G)]\|TGAGACCTCTGGGTTCTGAGCG | >MI0003834\|hsa-mir-769&WithFlank&19\|+\|46018926\|46019055 | 36 | 56 | 21(+1G) | TGAGACCTCTGGGTTCTGAGCG | 1113.3 | 12.7 | 1.6 | 7.7 | 1.1E-14 | 3.1E-14 | BT20/MB468 |
| >hg38_21_+_8433172_8446622_RNA45SN1_45S_50ntFlanks@10401.10424.24\|GAACATGGGTCAGTCGGTCCTGAG | >hg38_21_+_8433172_8446622_RNA45SN1_45S_50ntFlanks | 10401 | 10424 | 24 | GAACATGGGTCAGTCGGTCCTGAG | 1997.2 | 5.0 | 0.6 | 7.7 | 1.1E-14 | 3.1E-14 | MB231/MB468 |
| >MI0000273\|hsa-mir-183&WithFlank&7\|-\|129774899\|129775020@34.54.21&[MIMAT0000261&hsa-miR-183-5p&offsets\|+1\|0;m-41&7\|-\|129774966\|129774987&offsets\|0\|-1]\|ATGGCACTGGTAGAATTCACT | >MI0000273\|hsa-mir-183&WithFlank&7\|-\|129774899\|129775020 | 34 | 54 | 21 | ATGGCACTGGTAGAATTCACT | 2124.6 | 5.8 | 0.7 | 7.7 | 1.1E-14 | 3.2E-14 | MB231/MB468 |
| >hg38_21_+_8433172_8446622_RNA45SN1_45S_50ntFlanks@4722.4751.30\|TCATTAATCAAGAACGAAAGTCGGAGGTTC | >hg38_21_+_8433172_8446622_RNA45SN1_45S_50ntFlanks | 4722 | 4751 | 30 | TCATTAATCAAGAACGAAAGTCGGAGGTTC | 430.9 | -7.4 | 1.0 | -7.7 | 1.1E-14 | 3.2E-14 | BT20/MB468 |
| >hg38_21_+_8433172_8446622_RNA45SN1_45S_50ntFlanks@11234.11254.21\|CCTTCCCGTGGATCGCCCCAG | >hg38_21_+_8433172_8446622_RNA45SN1_45S_50ntFlanks | 11234 | 11254 | 21 | CCTTCCCGTGGATCGCCCCAG | 284.2 | -10.1 | 1.3 | -7.7 | 1.1E-14 | 3.2E-14 | BT20/MB468 |
| >trnaMT_PheGAA_MT_+_577_647@1.34.34\|GTTTATGTAGCTTACCTCCTCAAAGCAATACACT | >trnaMT_PheGAA_MT_+_577_647 | 1 | 34 | 34 | GTTTATGTAGCTTACCTCCTCAAAGCAATACACT | 1635.5 | 9.8 | 1.3 | 7.7 | 1.1E-14 | 3.2E-14 | MB231/MB468 |
| >hg38_21_+_8433172_8446622_RNA45SN1_45S_50ntFlanks@10344.10373.30\|ATTCAAACGAGAACTTTGAAGGCCGAAGTG | >hg38_21_+_8433172_8446622_RNA45SN1_45S_50ntFlanks | 10344 | 10373 | 30 | ATTCAAACGAGAACTTTGAAGGCCGAAGTG | 482.8 | 7.1 | 0.9 | 7.7 | 1.1E-14 | 3.2E-14 | MB231/MB468 |
| >hg38_21_+_8433172_8446622_RNA45SN1_45S_50ntFlanks@4103.4121.19\|CCACGGGTGACGGGGAATC | >hg38_21_+_8433172_8446622_RNA45SN1_45S_50ntFlanks | 4103 | 4121 | 19 | CCACGGGTGACGGGGAATC | 223.8 | -9.8 | 1.3 | -7.7 | 1.2E-14 | 3.3E-14 | BT20/MB468 |
| >hg38_21_+_8433172_8446622_RNA45SN1_45S_50ntFlanks@10807.10828.22\|ACAATGTAGGTAAGGGAAGTCG | >hg38_21_+_8433172_8446622_RNA45SN1_45S_50ntFlanks | 10807 | 10828 | 22 | ACAATGTAGGTAAGGGAAGTCG | 136.5 | -4.4 | 0.6 | -7.7 | 1.2E-14 | 3.3E-14 | BT20/MB468 |
| >MI0000077\|hsa-mir-21&WithFlank&17\|+\|59841260\|59841343@14.34.21(+1C)&[MIMAT0000076&hsa-miR-21-5p&offsets\|0\|-1(+1C);m-1&17\|+\|59841273\|59841294&offsets\|0\|-1(+1C)]\|TAGCTTATCAGACTGATGTTGC | >MI0000077\|hsa-mir-21&WithFlank&17\|+\|59841260\|59841343 | 14 | 34 | 21(+1C) | TAGCTTATCAGACTGATGTTGC | 753.6 | 8.3 | 1.1 | 7.7 | 1.1E-14 | 3.3E-14 | MB231/MB468 |
| >MI0000283\|hsa-mir-203a&WithFlank&14\|+\|104117399\|104117520@71.92.22(+1U)&[MIMAT0000264&hsa-miR-203a-3p&offsets\|0\|0(+1U);m-79&14\|+\|104117469\|104117490&offsets\|0\|0(+1U)]\|GTGAAATGTTTAGGACCACTAGT | >MI0000283\|hsa-mir-203a&WithFlank&14\|+\|104117399\|104117520 | 71 | 92 | 22(+1U) | GTGAAATGTTTAGGACCACTAGT | 227.5 | 11.3 | 1.4 | 8.1 | 8.0E-16 | 3.3E-14 | BT20/MB231 |

<table>
<tr><td>>am_trna2_GlyGCC_21_-_18827107_18827177@1.27.27,trna35_GlyGCC_1_+_161413094_161413164@1.27.27,trna37_GlyGCC_1_+_161420467_161420537@1.27.27,trna39_GlyGCC_1_+_161427898_161427968@1.27.27,trna41_GlyGCC_1_+_161435258_161435328@1.27.27|GCATGGGTGGTTCAGTGGTAGAATTCT</td><td>>am_trna2_GlyGCC_21_-_18827107_18827177</td><td>1</td><td>27</td><td>27</td><td>GCATGGGTGGTTCAGTGGTAGAATTCT</td><td>1180.6</td><td>7.5</td><td>0.9</td><td>8.1</td><td>8.2E-16</td><td>3.3E-14</td><td>BT20/MB231</td></tr>
<tr><td>>hg38_21_+_8433172_8446622_RNA45SN1_45S_50ntFlanks@12417.12435.19|TCGATGTCGGCTCTTCCTA</td><td>>hg38_21_+_8433172_8446622_RNA45SN1_45S_50ntFlanks</td><td>12417</td><td>12435</td><td>19</td><td>TCGATGTCGGCTCTTCCTA</td><td>2610.6</td><td>-3.9</td><td>0.5</td><td>-7.7</td><td>1.2E-14</td><td>3.4E-14</td><td>MB231/MB468</td></tr>
<tr><td>>hg38_21_+_8433172_8446622_RNA45SN1_45S_50ntFlanks@475.498.24|CTTCGTGATCGATGTGGTGACGTC</td><td>>hg38_21_+_8433172_8446622_RNA45SN1_45S_50ntFlanks</td><td>475</td><td>498</td><td>24</td><td>CTTCGTGATCGATGTGGTGACGTC</td><td>14194.9</td><td>6.9</td><td>0.9</td><td>7.7</td><td>1.2E-14</td><td>3.4E-14</td><td>BT20/MB468</td></tr>
<tr><td>>am_trna162_MetCAT_6_-_26758550_26758622@31.53.23,trna164_MetCAT_6_-_26735574_26735646@31.53.23,trna20_MetCAT_16_+_71460396_71460468@31.53.23,trna21_MetCAT_6_+_26701712_26701784@31.53.23,trna27_MetCAT_6_+_26766444_26766516@31.53.23,trna75_MetCAT_6_+_28912352_28912424@31.53.23,trna97_MetCAT_6_-_28921042_28921114@31.53.23|TCTCATAATCTGAAGGTCCTGAG</td><td>>am_trna162_MetCAT_6_-_26758550_26758622</td><td>31</td><td>53</td><td>23</td><td>TCTCATAATCTGAAGGTCCTGAG</td><td>365.6</td><td>-10.5</td><td>1.4</td><td>-7.7</td><td>1.2E-14</td><td>3.4E-14</td><td>BT20/MB468</td></tr>
<tr><td>>MI0000750|hsa-mir-26a-2&WithFlank&12|-|57824603|57824698@20.40.21;MI0000083|hsa-mir-26a-1&WithFlank&3|+|37969398|37969486@16.36.21&[MIMAT0000082&hsa-miR-26a-5p&offsets|0|-1;m-33&12|-|57824658|57824679&offsets|0|-1];[MIMAT0000082_1&hsa-miR-26a-5p&offsets|0|-1;m-34&3|+|37969413|37969434&offsets|0|-1]|TTCAAGTAATCCAGGATAGGC</td><td>>MI0000750|hsa-mir-26a-2&WithFlank&12|-|57824603|57824698</td><td>20</td><td>40</td><td>21</td><td>TTCAAGTAATCCAGGATAGGC</td><td>2734.3</td><td>6.6</td><td>0.9</td><td>7.7</td><td>1.2E-14</td><td>3.4E-14</td><td>BT20/MB468</td></tr>
<tr><td>>hg38_21_+_8433172_8446622_RNA45SN1_45S_50ntFlanks@12008.12033.26|CCGCCGGTGAAATACCACTACTCTGA</td><td>>hg38_21_+_8433172_8446622_RNA45SN1_45S_50ntFlanks</td><td>12008</td><td>12033</td><td>26</td><td>CCGCCGGTGAAATACCACTACTCTGA</td><td>248.3</td><td>-5.6</td><td>0.7</td><td>-7.7</td><td>1.2E-14</td><td>3.5E-14</td><td>MB231/MB468</td></tr>
<tr><td>>hg38_21_+_8433172_8446622_RNA45SN1_45S_50ntFlanks@3930.3959.30|ATCAAAACCAACCCGGTCAGCCCCTCTCCG</td><td>>hg38_21_+_8433172_8446622_RNA45SN1_45S_50ntFlanks</td><td>3930</td><td>3959</td><td>30</td><td>ATCAAAACCAACCCGGTCAGCCCCTCTCCG</td><td>184.9</td><td>-9.5</td><td>1.2</td><td>-7.7</td><td>1.2E-14</td><td>3.5E-14</td><td>BT20/MB468</td></tr>
<tr><td>>hg38_21_+_8433172_8446622_RNA45SN1_45S_50ntFlanks@11621.11640.20|CAAAGCATCGCGAAGGCCCG</td><td>>hg38_21_+_8433172_8446622_RNA45SN1_45S_50ntFlanks</td><td>11621</td><td>11640</td><td>20</td><td>CAAAGCATCGCGAAGGCCCG</td><td>289.7</td><td>-7.2</td><td>0.9</td><td>-7.7</td><td>1.2E-14</td><td>3.5E-14</td><td>MB231/MB468</td></tr>
<tr><td>>MI0000811|hsa-mir-148b&WithFlank&12|+|54337210|54337320@69.89.21(+1A)&[MIMAT0000759&hsa-miR-148b-3p&offsets|0|-1(+1A);m-77&12|+|54337278|54337299&offsets|0|-1(+1A)]|TCAGTGCATCACAGAACTTTGA</td><td>>MI0000811|hsa-mir-148b&WithFlank&12|+|54337210|54337320</td><td>69</td><td>89</td><td>21(+1A)</td><td>TCAGTGCATCACAGAACTTTGA</td><td>2228.9</td><td>9.9</td><td>1.3</td><td>7.7</td><td>1.3E-14</td><td>3.5E-14</td><td>BT20/MB468</td></tr>
<tr><td>>MI0000088|hsa-mir-30a&WithFlank&6|-|71403545|71403627@53.71.19&[MIMAT0000088&hsa-miR-30a-3p&offsets|0|-3;m-38&6|-|71403554|71403575&offsets|0|-3]|CTTTCAGTCGGATGTTTGC</td><td>>MI0000088|hsa-mir-30a&WithFlank&6|-|71403545|71403627</td><td>53</td><td>71</td><td>19</td><td>CTTTCAGTCGGATGTTTGC</td><td>1146.3</td><td>6.3</td><td>0.8</td><td>7.7</td><td>1.3E-14</td><td>3.5E-14</td><td>BT20/MB468</td></tr>
<tr><td>>trnaMT_ValTAC_MT_+_1602_1670@54.71.18|AACTTGACCGCTCTGACC</td><td>>trnaMT_ValTAC_MT_+_1602_1670</td><td>54</td><td>71</td><td>18</td><td>AACTTGACCGCTCTGACC</td><td>852.5</td><td>9.2</td><td>1.2</td><td>7.7</td><td>1.3E-14</td><td>3.5E-14</td><td>BT20/MB468</td></tr>
<tr><td>>hg38_21_+_8433172_8446622_RNA45SN1_45S_50ntFlanks@8380.8404.25|CAAGAGGGCGTGAAACCGTTAAGAG</td><td>>hg38_21_+_8433172_8446622_RNA45SN1_45S_50ntFlanks</td><td>8380</td><td>8404</td><td>25</td><td>CAAGAGGGCGTGAAACCGTTAAGAG</td><td>382.4</td><td>-10.6</td><td>1.4</td><td>-7.7</td><td>1.3E-14</td><td>3.5E-14</td><td>BT20/MB468</td></tr>
<tr><td>>hg38_21_+_8433172_8446622_RNA45SN1_45S_50ntFlanks@12004.12035.32|CGGGCCGCCGGTGAAATACCACTACTCTGATC</td><td>>hg38_21_+_8433172_8446622_RNA45SN1_45S_50ntFlanks</td><td>12004</td><td>12035</td><td>32</td><td>CGGGCCGCCGGTGAAATACCACTACTCTGATC</td><td>1558.0</td><td>9.1</td><td>1.2</td><td>7.7</td><td>1.3E-14</td><td>3.5E-14</td><td>BT20/MB468</td></tr>
<tr><td>>hg38_1_-_228634819_228635039_RNA5S12_5S_50ntFlanks@130.170.41|TGGATGGGAGACCGCCTGGGAATACCGGGTGCTGTAGGCTT</td><td>>hg38_1_-_228634819_228635039_RNA5S12_5S_50ntFlanks</td><td>130</td><td>170</td><td>41</td><td>TGGATGGGAGACCGCCTGGGAATACCGGGTGCTGTAGGCTT</td><td>1879.4</td><td>11.3</td><td>1.5</td><td>7.7</td><td>1.3E-14</td><td>3.6E-14</td><td>BT20/MB468</td></tr>
<tr><td>>hg38_21_+_8433172_8446622_RNA45SN1_45S_50ntFlanks@12940.12972.33|CATTCGTAGACGACCTGCTTCTGGGTCGGGGTT</td><td>>hg38_21_+_8433172_8446622_RNA45SN1_45S_50ntFlanks</td><td>12940</td><td>12972</td><td>33</td><td>CATTCGTAGACGACCTGCTTCTGGGTCGGGGTT</td><td>219.8</td><td>-4.4</td><td>0.6</td><td>-7.7</td><td>1.2E-14</td><td>3.6E-14</td><td>MB231/MB468</td></tr>
<tr><td>>MI0000061|hsa-let-7a-2&WithFlank&11|-|122146516|122146599@11.30.20;MI0000062|hsa-let-7a-3&WithFlank&22|+|46112743|46112828@10.29.20;MI0000060|hsa-let-7a-1&WithFlank&9|+|94175951|94176042@12.31.20&[MIMAT0000062&hsa-let-7a-5p&offsets|0|-2;m-6&11|-|122146568|122146589&offsets|0|-2];[MIMAT0000062_1&hsa-let-7a-5p&offsets|0|-2;m-5&22|+|46112752|46112773&offsets|0|-2];[MIMAT0000062_2&hsa-let-7a-5p&offsets|0|-2;m-4&9|+|94175962|94175983&offsets|0|-2]|TGAGGTAGTAGGTTGTATAG</td><td>>MI0000061|hsa-let-7a-2&WithFlank&11|-|122146516|122146599</td><td>11</td><td>30</td><td>20</td><td>TGAGGTAGTAGGTTGTATAG</td><td>557.0</td><td>8.8</td><td>1.1</td><td>7.7</td><td>1.3E-14</td><td>3.6E-14</td><td>MB231/MB468</td></tr>
<tr><td>>hg38_21_+_8433172_8446622_RNA45SN1_45S_50ntFlanks@11566.11586.21|TAGCAGCCGACTTAGAACTGG</td><td>>hg38_21_+_8433172_8446622_RNA45SN1_45S_50ntFlanks</td><td>11566</td><td>11586</td><td>21</td><td>TAGCAGCCGACTTAGAACTGG</td><td>1053.2</td><td>5.1</td><td>0.7</td><td>7.7</td><td>1.3E-14</td><td>3.6E-14</td><td>MB231/MB468</td></tr>
<tr><td>>hg38_21_+_8433172_8446622_RNA45SN1_45S_50ntFlanks@12658.12675.18|ACTGAACGCCTCTAAGTC</td><td>>hg38_21_+_8433172_8446622_RNA45SN1_45S_50ntFlanks</td><td>12658</td><td>12675</td><td>18</td><td>ACTGAACGCCTCTAAGTC</td><td>704.5</td><td>-5.6</td><td>0.7</td><td>-7.7</td><td>1.3E-14</td><td>3.6E-14</td><td>BT20/MB468</td></tr>
<tr><td>>hg38_21_+_8433172_8446622_RNA45SN1_45S_50ntFlanks@11316.11334.19|CACCCCACGTCTCGTCGCG</td><td>>hg38_21_+_8433172_8446622_RNA45SN1_45S_50ntFlanks</td><td>11316</td><td>11334</td><td>19</td><td>CACCCCACGTCTCGTCGCG</td><td>765.2</td><td>-5.5</td><td>0.7</td><td>-7.7</td><td>1.3E-14</td><td>3.6E-14</td><td>MB231/MB468</td></tr>
<tr><td>>hg38_21_+_8433172_8446622_RNA45SN1_45S_50ntFlanks@10389.10408.20|TGAACAGCAGTTGAACATGG</td><td>>hg38_21_+_8433172_8446622_RNA45SN1_45S_50ntFlanks</td><td>10389</td><td>10408</td><td>20</td><td>TGAACAGCAGTTGAACATGG</td><td>259.1</td><td>-6.8</td><td>0.9</td><td>-7.7</td><td>1.3E-14</td><td>3.6E-14</td><td>BT20/MB468</td></tr>
<tr><td>>hg38_21_+_8433172_8446622_RNA45SN1_45S_50ntFlanks@12927.12956.30|CCTGGCGCTAAACCATTCGTAGACGACCTG</td><td>>hg38_21_+_8433172_8446622_RNA45SN1_45S_50ntFlanks</td><td>12927</td><td>12956</td><td>30</td><td>CCTGGCGCTAAACCATTCGTAGACGACCTG</td><td>192.1</td><td>-9.6</td><td>1.2</td><td>-7.7</td><td>1.3E-14</td><td>3.6E-14</td><td>BT20/MB468</td></tr>
<tr><td>>MI0000073|hsa-mir-19a&WithFlank&13|+|91350885|91350978@55.76.22(+1U)&[MIMAT0000073&hsa-miR-19a-3p&offsets|0|-1(+1U);m-201&13|+|91350939|91350961&offsets|0|-1(+1U)]|TGTGCAAATCTATGCAAAACTGT</td><td>>MI0000073|hsa-mir-19a&WithFlank&13|+|91350885|91350978</td><td>55</td><td>76</td><td>22(+1U)</td><td>TGTGCAAATCTATGCAAAACTGT</td><td>624.5</td><td>5.6</td><td>0.7</td><td>7.7</td><td>1.3E-14</td><td>3.6E-14</td><td>MB231/MB468</td></tr>
<tr><td>>hg38_21_+_8433172_8446622_RNA45SN1_45S_50ntFlanks@12418.12447.30|CGATGTCGGCTCTTCCTATCATTGTGAAGC</td><td>>hg38_21_+_8433172_8446622_RNA45SN1_45S_50ntFlanks</td><td>12418</td><td>12447</td><td>30</td><td>CGATGTCGGCTCTTCCTATCATTGTGAAGC</td><td>525.6</td><td>4.3</td><td>0.6</td><td>7.7</td><td>1.3E-14</td><td>3.7E-14</td><td>MB231/MB468</td></tr>
<tr><td>>hg38_21_+_8433172_8446622_RNA45SN1_45S_50ntFlanks@11876.11906.31|AAGACCCTGTTGAGCTTGACTCTAGTCTGGC</td><td>>hg38_21_+_8433172_8446622_RNA45SN1_45S_50ntFlanks</td><td>11876</td><td>11906</td><td>31</td><td>AAGACCCTGTTGAGCTTGACTCTAGTCTGGC</td><td>309.7</td><td>-9.9</td><td>1.3</td><td>-7.7</td><td>1.3E-14</td><td>3.7E-14</td><td>MB231/MB468</td></tr>
<tr><td>>MI0000255|hsa-mir-30d&WithFlank&8|-|134804870|134804951@12.33.22(+2U)&[MIMAT0000245&hsa-miR-30d-5p&offsets|0|0(+2U);m-21&8|-|134804919|134804940&offsets|0|0(+2U)]|TGTAAACATCCCCGACTGGAAGTT</td><td>>MI0000255|hsa-mir-30d&WithFlank&8|-|134804870|134804951</td><td>12</td><td>33</td><td>22(+2U)</td><td>TGTAAACATCCCCGACTGGAAGTT</td><td>2269.5</td><td>11.3</td><td>1.5</td><td>7.7</td><td>1.3E-14</td><td>3.7E-14</td><td>BT20/MB468</td></tr>
<tr><td>>MI0000805|hsa-mir-342&WithFlank&14|+|100109649|100109759@67.90.24&[MIMAT0000753&hsa-miR-342-3p&offsets|0|+1;m-128&14|+|100109715|100109737&offsets|0|+1]|TCTCACACAGAAATCGCACCCGTC</td><td>>MI0000805|hsa-mir-342&WithFlank&14|+|100109649|100109759</td><td>67</td><td>90</td><td>24</td><td>TCTCACACAGAAATCGCACCCGTC</td><td>778.6</td><td>6.3</td><td>0.8</td><td>7.7</td><td>1.3E-14</td><td>3.7E-14</td><td>BT20/MB468</td></tr>
<tr><td>>MI0000085|hsa-mir-27a&WithFlank&19|-|13836434|13836523@57.77.21(+1U)&[MIMAT0000084&hsa-miR-27a-3p&offsets|0|0(+1U);m-43&19|-|13836447|13836467&offsets|0|0(+1U)]|TTCACAGTGGCTAAGTTCCGCT</td><td>>MI0000085|hsa-mir-27a&WithFlank&19|-|13836434|13836523</td><td>57</td><td>77</td><td>21(+1U)</td><td>TTCACAGTGGCTAAGTTCCGCT</td><td>980.4</td><td>6.5</td><td>0.8</td><td>7.7</td><td>1.3E-14</td><td>3.7E-14</td><td>MB231/MB468</td></tr>
<tr><td>>am_trnaMT_SerGCT_MT_+_12207_12265@14.32.19|AGAACTGCTAACTCATGCC</td><td>>am_trnaMT_SerGCT_MT_+_12207_12265</td><td>14</td><td>32</td><td>19</td><td>AGAACTGCTAACTCATGCC</td><td>738.3</td><td>8.9</td><td>1.2</td><td>7.7</td><td>1.3E-14</td><td>3.7E-14</td><td>BT20/MB468</td></tr>
<tr><td>>hg38_21_+_8433172_8446622_RNA45SN1_45S_50ntFlanks@10366.10383.18|CCGAAGTGGAGAAGGGTT</td><td>>hg38_21_+_8433172_8446622_RNA45SN1_45S_50ntFlanks</td><td>10366</td><td>10383</td><td>18</td><td>CCGAAGTGGAGAAGGGTT</td><td>100.4</td><td>-4.3</td><td>0.6</td><td>-7.7</td><td>1.4E-14</td><td>3.7E-14</td><td>BT20/MB468</td></tr>
<tr><td>>MI0000750|hsa-mir-26a-2&WithFlank&12|-|57824603|57824698@20.39.20(+2U);MI0000083|hsa-mir-26a-1&WithFlank&3|+|37969398|37969486@16.35.20(+2U)&[MIMAT0000082&hsa-miR-26a-5p&offsets|0|-2(+2U);m-33&12|-|57824658|57824679&offsets|0|-2(+2U)];[MIMAT0000082_1&hsa-miR-26a-5p&offsets|0|-2(+2U);m-34&3|+|37969413|37969434&offsets|0|-2(+2U)]|TTCAAGTAATCCAGGATAGGTT</td><td>>MI0000750|hsa-mir-26a-2&WithFlank&12|-|57824603|57824698</td><td>20</td><td>39</td><td>20(+2U)</td><td>TTCAAGTAATCCAGGATAGGTT</td><td>1086.6</td><td>6.8</td><td>0.9</td><td>7.7</td><td>1.4E-14</td><td>3.8E-14</td><td>BT20/MB468</td></tr>
<tr><td>>am_trnaMT_PheGAA_MT_+_577_647@42.62.21|TTTAGACGGGCTCACATCACC</td><td>>am_trnaMT_PheGAA_MT_+_577_647</td><td>42</td><td>62</td><td>21</td><td>TTTAGACGGGCTCACATCACC</td><td>552.0</td><td>8.3</td><td>1.1</td><td>7.7</td><td>1.3E-14</td><td>3.8E-14</td><td>MB231/MB468</td></tr>
<tr><td>>MI0000078|hsa-mir-22&WithFlank&17|-|1713897|1713993@59.79.21(+1A)&[MIMAT0000077&hsa-miR-22-3p&offsets|0|-1(+1A);m-2&17|-|1713914|1713935&offsets|0|-1(+1A)]|AAGCTGCCAGTTGAAGAACTGA</td><td>>MI0000078|hsa-mir-22&WithFlank&17|-|1713897|1713993</td><td>59</td><td>79</td><td>21(+1A)</td><td>AAGCTGCCAGTTGAAGAACTGA</td><td>1649.0</td><td>9.5</td><td>1.2</td><td>7.7</td><td>1.4E-14</td><td>3.8E-14</td><td>BT20/MB468</td></tr>
<tr><td>>hg38_21_+_8433172_8446622_RNA45SN1_45S_50ntFlanks@11876.11905.30|AAGACCCTGTTGAGCTTGACTCTAGTCTGG</td><td>>hg38_21_+_8433172_8446622_RNA45SN1_45S_50ntFlanks</td><td>11876</td><td>11905</td><td>30</td><td>AAGACCCTGTTGAGCTTGACTCTAGTCTGG</td><td>268.5</td><td>-6.9</td><td>0.9</td><td>-7.7</td><td>1.4E-14</td><td>3.8E-14</td><td>BT20/MB468</td></tr>
<tr><td>>hg38_21_+_8433172_8446622_RNA45SN1_45S_50ntFlanks@10866.10894.29|CTAAGGGCTGGGTCGGTCGGGCTGGGGCG</td><td>>hg38_21_+_8433172_8446622_RNA45SN1_45S_50ntFlanks</td><td>10866</td><td>10894</td><td>29</td><td>CTAAGGGCTGGGTCGGTCGGGCTGGGGCG</td><td>242.1</td><td>-4.6</td><td>0.6</td><td>-7.7</td><td>1.4E-14</td><td>3.8E-14</td><td>BT20/MB468</td></tr>
<tr><td>>hg38_21_+_8433172_8446622_RNA45SN1_45S_50ntFlanks@4472.4493.22|CTTAGCTGAGTGTCCCGCGGGG</td><td>>hg38_21_+_8433172_8446622_RNA45SN1_45S_50ntFlanks</td><td>4472</td><td>4493</td><td>22</td><td>CTTAGCTGAGTGTCCCGCGGGG</td><td>148.6</td><td>-5.8</td><td>0.8</td><td>-7.7</td><td>1.4E-14</td><td>3.8E-14</td><td>BT20/MB468</td></tr>
<tr><td>>am_trna162_MetCAT_6_-_26758550_26758622@31.51.21,trna164_MetCAT_6_-_26735574_26735646@31.51.21,trna20_MetCAT_16_+_71460396_71460468@31.51.21,trna21_MetCAT_6_+_26701712_26701784@31.51.21,trna27_MetCAT_6_+_26766444_26766516@31.51.21,trna75_MetCAT_6_+_28912352_28912424@31.51.21,trna97_MetCAT_6_-_28921042_28921114@31.51.21|TCTCATAATCTGAAGGTCCTG</td><td>>am_trna162_MetCAT_6_-_26758550_26758622</td><td>31</td><td>51</td><td>21</td><td>TCTCATAATCTGAAGGTCCTG</td><td>444.8</td><td>-10.8</td><td>1.4</td><td>-7.7</td><td>1.4E-14</td><td>3.9E-14</td><td>BT20/MB468</td></tr>
<tr><td>>MI0000085|hsa-mir-27a&WithFlank&19|-|13836434|13836523@57.76.20&[MIMAT0000084&hsa-miR-27a-3p&offsets|0|-1;m-43&19|-|13836447|13836467&offsets|0|-1]|TTCACAGTGGCTAAGTTCCG</td><td>>MI0000085|hsa-mir-27a&WithFlank&19|-|13836434|13836523</td><td>57</td><td>76</td><td>20</td><td>TTCACAGTGGCTAAGTTCCG</td><td>39765.7</td><td>6.3</td><td>0.8</td><td>7.7</td><td>1.4E-14</td><td>3.9E-14</td><td>MB231/MB468</td></tr>
</table>

| | | | | | | | | | | | | |
|---|---|---|---|---|---|---|---|---|---|---|---|---|
| >trna119_LysCTT_1_-_145395522_145395594@1.31.31,trna11_LysCTT_5_-_180648979_180649051@1.31.31,trna13_LysCTT_6_+_26556774_26556846@1.31.31,trna32_LysCTT_16_-_3207406_3207478@1.31.31,trna7_LysCTT_16_+_3225692_3225764@1.31.31,trna9_LysCTT_5_+_180634755_180634827@1.31.31\|GCCCGGCTAGCTCAGTCGGTAGAGCATGAGA | >trna119_LysCTT_1_-_145395522_145395594 | 1 | 31 | 31 | GCCCGGCTAGCTCAGTCGGTAGAGCATGAGA | 4890.4 | -5.1 | 0.7 | -7.7 | 1.4E-14 | 3.9E-14 | BT20/MB468 |
| >MI0000471\|hsa-mir-126&WithFlank&9\|+\|136670596\|136670692@59.78.20&[MIMAT0000445&hsa-miR-126-3p&offsets\|+1\|-1;m-40&9\|+\|136670653\|136670674&offsets\|+1\|-1]\|CGTACCGTGAGTAATAATGC | >MI0000471\|hsa-mir-126&WithFlank&9\|+\|136670596\|136670692 | 59 | 78 | 20 | CGTACCGTGAGTAATAATGC | 1146.6 | 10.2 | 1.3 | 7.7 | 1.4E-14 | 4.0E-14 | BT20/MB468 |
| >MI0000767\|hsa-mir-365a&WithFlank&16\|+\|14309279\|14309377@62.81.20(+2U);MI0000769\|hsa-mir-365b&WithFlank&17\|+\|31575405\|31575527@74.93.20(+2U)&[MIMAT0000710&hsa-miR-365a-3p&offsets\|0\|-2(+2U);m-176&16\|+\|14309340\|14309361&offsets\|0\|-2(+2U)];[MIMAT0022834&hsa-miR-365b-3p&offsets\|0\|-2(+2U);m-175&17\|+\|31575478\|31575499&offsets\|0\|-2(+2U)]\|TAATGCCCCTAAAAATCCTTTT | >MI0000767\|hsa-mir-365a&WithFlank&16\|+\|14309279\|14309377 | 62 | 81 | 20(+2U) | TAATGCCCCTAAAAATCCTTTT | 1144.8 | 11.5 | 1.5 | 7.7 | 1.4E-14 | 4.0E-14 | BT20/MB468 |
| >hg38_21_+_8433172_8446622_RNA45SN1_45S_50ntFlanks@10805.10822.18\|GAACAATGTAGGTAAGGG | >hg38_21_+_8433172_8446622_RNA45SN1_45S_50ntFlanks | 10805 | 10822 | 18 | GAACAATGTAGGTAAGGG | 198.0 | -5.7 | 0.7 | -7.7 | 1.4E-14 | 4.0E-14 | MB231/MB468 |
| >hg38_21_+_8433172_8446622_RNA45SN1_45S_50ntFlanks@12162.12180.19\|GACTGGGGCGGTACACCTG | >hg38_21_+_8433172_8446622_RNA45SN1_45S_50ntFlanks | 12162 | 12180 | 19 | GACTGGGGCGGTACACCTG | 374.6 | -7.4 | 1.0 | -7.7 | 1.5E-14 | 4.0E-14 | BT20/MB468 |
| >hg38_21_+_8433172_8446622_RNA45SN1_45S_50ntFlanks@10369.10407.39\|AAGTGGAGAAGGGTTCCATGTGAACAGCAGTTGAACATG | >hg38_21_+_8433172_8446622_RNA45SN1_45S_50ntFlanks | 10369 | 10407 | 39 | AAGTGGAGAAGGGTTCCATGTGAACAGCAGTTGAACATG | 307.3 | -10.3 | 1.3 | -7.7 | 1.5E-14 | 4.0E-14 | BT20/MB468 |
| >hg38_21_+_8433172_8446622_RNA45SN1_45S_50ntFlanks@8472.8491.20\|CGGCCCGGCGGATCTTTCCC | >hg38_21_+_8433172_8446622_RNA45SN1_45S_50ntFlanks | 8472 | 8491 | 20 | CGGCCCGGCGGATCTTTCCC | 330.2 | -10.4 | 1.3 | -7.7 | 1.5E-14 | 4.0E-14 | BT20/MB468 |
| >hg38_21_+_8433172_8446622_RNA45SN1_45S_50ntFlanks@6699.6718.20\|GCTAGCTGCGAGAATTAATG | >hg38_21_+_8433172_8446622_RNA45SN1_45S_50ntFlanks | 6699 | 6718 | 20 | GCTAGCTGCGAGAATTAATG | 9533.8 | -4.8 | 0.6 | -7.7 | 1.5E-14 | 4.1E-14 | MB231/MB468 |
| >hg38_21_+_8433172_8446622_RNA45SN1_45S_50ntFlanks@8174.8202.29\|TCCTTCTGATCGAGGCCCAGCCCGTGGAC | >hg38_21_+_8433172_8446622_RNA45SN1_45S_50ntFlanks | 8174 | 8202 | 29 | TCCTTCTGATCGAGGCCCAGCCCGTGGAC | 249.2 | 6.8 | 0.9 | 7.7 | 1.5E-14 | 4.1E-14 | BT20/MB468 |
| >hg38_21_+_8433172_8446622_RNA45SN1_45S_50ntFlanks@11641.11659.19\|CGGCGGGTGTTGACGCGAT | >hg38_21_+_8433172_8446622_RNA45SN1_45S_50ntFlanks | 11641 | 11659 | 19 | CGGCGGGTGTTGACGCGAT | 248.4 | -9.9 | 1.3 | -7.7 | 1.5E-14 | 4.2E-14 | BT20/MB468 |
| >hg38_21_+_8433172_8446622_RNA45SN1_45S_50ntFlanks@8472.8497.26\|CGGCCCGGCGGATCTTTCCCGCCCCC | >hg38_21_+_8433172_8446622_RNA45SN1_45S_50ntFlanks | 8472 | 8497 | 26 | CGGCCCGGCGGATCTTTCCCGCCCCC | 231.2 | -9.8 | 1.3 | -7.7 | 1.5E-14 | 4.2E-14 | BT20/MB468 |
| >trna12_TrpCCA_17_+_19411494_19411565@30.52.23,trna168_TrpCCA_6_-_26331672_26331743@30.52.23,trna170_TrpCCA_6_-_26319330_26319401@30.52.23,trna1_TrpCCA_7_+_99067307_99067378@30.52.23,trna39_TrpCCA_17_-_8124187_8124258@30.52.23,trna6_TrpCCA_17_+_8089676_8089747@30.52.23\|ACTCCAGATCAGAAGGTTGCGTG | >trna12_TrpCCA_17_+_19411494_19411565 | 30 | 52 | 23 | ACTCCAGATCAGAAGGTTGCGTG | 239.0 | -9.9 | 1.3 | -7.7 | 1.5E-14 | 4.2E-14 | BT20/MB468 |
| >hg38_21_+_8433172_8446622_RNA45SN1_45S_50ntFlanks@10806.10825.20\|AACAATGTAGGTAAGGGAAG | >hg38_21_+_8433172_8446622_RNA45SN1_45S_50ntFlanks | 10806 | 10825 | 20 | AACAATGTAGGTAAGGGAAG | 307.6 | -10.3 | 1.3 | -7.7 | 1.6E-14 | 4.3E-14 | BT20/MB468 |
| >hg38_21_+_8433172_8446622_RNA45SN1_45S_50ntFlanks@10101.10127.27\|CGCGCGCGTGTGGTGTGCGTCGGAGGG | >hg38_21_+_8433172_8446622_RNA45SN1_45S_50ntFlanks | 10101 | 10127 | 27 | CGCGCGCGTGTGGTGTGCGTCGGAGGG | 383.1 | -10.2 | 1.3 | -7.7 | 1.5E-14 | 4.3E-14 | MB231/MB468 |
| >hg38_21_+_8433172_8446622_RNA45SN1_45S_50ntFlanks@10411.10435.25\|CAGTCGGTCCTGAGAGATGGGCGAG | >hg38_21_+_8433172_8446622_RNA45SN1_45S_50ntFlanks | 10411 | 10435 | 25 | CAGTCGGTCCTGAGAGATGGGCGAG | 402.4 | -10.2 | 1.3 | -7.7 | 1.5E-14 | 4.3E-14 | MB231/MB468 |
| >hg38_21_+_8433172_8446622_RNA45SN1_45S_50ntFlanks@11564.11603.40\|CCTAGCAGCCGACTTAGAACTGGTGCGGACCAGGGGAATC | >hg38_21_+_8433172_8446622_RNA45SN1_45S_50ntFlanks | 11564 | 11603 | 40 | CCTAGCAGCCGACTTAGAACTGGTGCGGACCAGGGGAATC | 606.3 | -4.5 | 0.6 | -7.7 | 1.6E-14 | 4.4E-14 | MB231/MB468 |
| >hg38_21_+_8433172_8446622_RNA45SN1_45S_50ntFlanks@10808.10828.21\|CAATGTAGGTAAGGGAAGTCG | >hg38_21_+_8433172_8446622_RNA45SN1_45S_50ntFlanks | 10808 | 10828 | 21 | CAATGTAGGTAAGGGAAGTCG | 411.3 | -5.8 | 0.8 | -7.7 | 1.6E-14 | 4.4E-14 | MB231/MB468 |
| >MI0000234\|hsa-mir-192&WithFlank&11\|-\|64891131\|64891252@30.51.22&[MIMAT0000222&hsa-miR-192-5p&offsets\|0\|+1;m-59&11\|-\|64891203\|64891223&offsets\|0\|+1]\|CTGACCTATGAATTGACAGCCA | >MI0000234\|hsa-mir-192&WithFlank&11\|-\|64891131\|64891252 | 30 | 51 | 22 | CTGACCTATGAATTGACAGCCA | 698.5 | 6.1 | 0.8 | 7.7 | 1.6E-14 | 4.5E-14 | BT20/MB468 |
| >hg38_21_+_8433172_8446622_RNA45SN1_45S_50ntFlanks@10350.10376.27\|ACGAGAACTTTGAAGGCCGAAGTGGAG | >hg38_21_+_8433172_8446622_RNA45SN1_45S_50ntFlanks | 10350 | 10376 | 27 | ACGAGAACTTTGAAGGCCGAAGTGGAG | 987.5 | -4.1 | 0.5 | -7.7 | 1.6E-14 | 4.5E-14 | MB231/MB468 |
| >hg38_21_+_8433172_8446622_RNA45SN1_45S_50ntFlanks@10572.10607.36\|AGAAGCCGGCGGGAGCCCCGGGGAGAGTTCTCTTTT | >hg38_21_+_8433172_8446622_RNA45SN1_45S_50ntFlanks | 10572 | 10607 | 36 | AGAAGCCGGCGGGAGCCCCGGGGAGAGTTCTCTTTT | 232.9 | -9.9 | 1.3 | -7.7 | 1.7E-14 | 4.6E-14 | BT20/MB468 |
| >hg38_21_+_8433172_8446622_RNA45SN1_45S_50ntFlanks@5494.5516.23\|AAGACGGTCGAACTTGACTATCT | >hg38_21_+_8433172_8446622_RNA45SN1_45S_50ntFlanks | 5494 | 5516 | 23 | AAGACGGTCGAACTTGACTATCT | 1172.1 | 6.7 | 0.9 | 7.7 | 1.7E-14 | 4.6E-14 | BT20/MB468 |
| >hg38_21_+_8433172_8446622_RNA45SN1_45S_50ntFlanks@12405.12423.19\|CTTTTTGATCCTTCGATGT | >hg38_21_+_8433172_8446622_RNA45SN1_45S_50ntFlanks | 12405 | 12423 | 19 | CTTTTTGATCCTTCGATGT | 286.1 | -10.2 | 1.3 | -7.7 | 1.7E-14 | 4.7E-14 | BT20/MB468 |
| >hg38_21_+_8433172_8446622_RNA45SN1_45S_50ntFlanks@11874.11908.35\|AGAAGACCCTGTTGAGCTTGACTCTAGTCTGGCAC | >hg38_21_+_8433172_8446622_RNA45SN1_45S_50ntFlanks | 11874 | 11908 | 35 | AGAAGACCCTGTTGAGCTTGACTCTAGTCTGGCAC | 297.1 | -4.8 | 0.6 | -7.7 | 1.6E-14 | 4.7E-14 | MB231/MB468 |
| >MI0000299\|hsa-mir-222&WithFlank&X\|-\|45747009\|45747130@75.96.22(+1C)&[MIMAT0000279&hsa-miR-222-3p&offsets\|0\|+1(+1C);m-140&X\|-\|45747033\|45747056&offsets\|0\|-2(+1C)]\|AGCTACATCTGGCTACTGGGTCC | >MI0000299\|hsa-mir-222&WithFlank&X\|-\|45747009\|45747130 | 75 | 96 | 22(+1C) | AGCTACATCTGGCTACTGGGTCC | 904.9 | 6.5 | 0.9 | 7.7 | 1.7E-14 | 4.7E-14 | BT20/MB468 |
| >MI0016887\|hsa-mir-4521&WithFlank&17\|+\|8186939\|8187010@10.34.25&[MIMAT0019058&hsa-miR-4521&offsets\|0\|+3;m-765&17\|+\|8186948\|8186969&offsets\|0\|+3]\|GCTAAGGAAGTCCTGTGCTCAGTTT | >MI0016887\|hsa-mir-4521&WithFlank&17\|+\|8186939\|8187010 | 10 | 34 | 25 | GCTAAGGAAGTCCTGTGCTCAGTTT | 1059.8 | 5.9 | 0.8 | 7.7 | 1.7E-14 | 4.7E-14 | MB231/MB468 |
| >am_MI0015825\|hsa-mir-378c&WithFlank&10\|-\|130962582\|130962674@17.36.20;MI0000786\|hsa-mir-378a&WithFlank&5\|+\|149732819\|149732896@49.68.20&[MIMAT0016847&hsa-miR-378c&offsets\|0\|-5;m-130&10\|-\|130962639\|130962658&offsets\|0\|0];[MIMAT0000732&hsa-miR-378a-3p&offsets\|0\|-2;m-44&5\|+\|149732867\|149732888&offsets\|0\|-2]\|ACTGGACTTGGAGTCAGAAG | >am_MI0015825\|hsa-mir-378c&WithFlank&10\|-\|130962582\|130962674 | 17 | 36 | 20 | ACTGGACTTGGAGTCAGAAG | 1279.9 | 6.3 | 0.8 | 7.7 | 1.7E-14 | 4.7E-14 | BT20/MB468 |
| >hg38_21_+_8433172_8446622_RNA45SN1_45S_50ntFlanks@10840.10865.26\|CCGTAACTTCGGGATAAGGATTGGCT | >hg38_21_+_8433172_8446622_RNA45SN1_45S_50ntFlanks | 10840 | 10865 | 26 | CCGTAACTTCGGGATAAGGATTGGCT | 1665.1 | -3.8 | 0.5 | -7.7 | 1.7E-14 | 4.7E-14 | MB231/MB468 |
| >hg38_21_+_8433172_8446622_RNA45SN1_45S_50ntFlanks@10572.10608.37\|AGAAGCCGGCGGGAGCCCCGGGGAGAGTTCTCTTTTC | >hg38_21_+_8433172_8446622_RNA45SN1_45S_50ntFlanks | 10572 | 10608 | 37 | AGAAGCCGGCGGGAGCCCCGGGGAGAGTTCTCTTTTC | 365.8 | -10.1 | 1.3 | -7.7 | 1.7E-14 | 4.8E-14 | MB231/MB468 |
| >hg38_21_+_8433172_8446622_RNA45SN1_45S_50ntFlanks@12635.12652.18\|AAGCTACCATCTGTGGGA | >hg38_21_+_8433172_8446622_RNA45SN1_45S_50ntFlanks | 12635 | 12652 | 18 | AAGCTACCATCTGTGGGA | 308.1 | -9.8 | 1.3 | -7.7 | 1.7E-14 | 4.8E-14 | MB231/MB468 |
| >MI0000108\|hsa-mir-103a-2&WithFlank&20\|+\|3917488\|3917577@54.75.22(+1C);MI0000109\|hsa-mir-103a-1&WithFlank&5\|-\|168560890\|168560979@54.75.22(+1C)&[MIMAT0000101&hsa-miR-103a-3p&offsets\|0\|-1(+1C);m-12&20\|+\|3917541\|3917563&offsets\|0\|-1(+1C)];[MIMAT0000101_1&hsa-miR-103a-3p&offsets\|0\|-1(+1C);m-13&5\|-\|168560904\|168560926&offsets\|0\|-1(+1C);m-44220&5\|-\|168560908\|168560926&offsets\|0\|+3(+1C)]\|AGCAGCATTGTACAGGGCTATGC | >MI0000108\|hsa-mir-103a-2&WithFlank&20\|+\|3917488\|3917577 | 54 | 75 | 22(+1C) | AGCAGCATTGTACAGGGCTATGC | 576.3 | 8.4 | 1.1 | 7.7 | 1.7E-14 | 4.8E-14 | BT20/MB468 |
| >hg38_21_+_8433172_8446622_RNA45SN1_45S_50ntFlanks@11661.11678.18\|TGATTTCTGCCCAGTGCT | >hg38_21_+_8433172_8446622_RNA45SN1_45S_50ntFlanks | 11661 | 11678 | 18 | TGATTTCTGCCCAGTGCT | 285.4 | -10.1 | 1.3 | -7.7 | 1.8E-14 | 4.8E-14 | BT20/MB468 |
| >trnaMT_ValTAC_MT_+_1602_1670@46.72.27\|TTCAACTTAACTTGACCGCTCTGACCA | >trnaMT_ValTAC_MT_+_1602_1670 | 46 | 72 | 27 | TTCAACTTAACTTGACCGCTCTGACCA | 602.4 | 10.1 | 1.3 | 7.7 | 1.8E-14 | 4.8E-14 | BT20/MB468 |
| >hg38_21_+_8433172_8446622_RNA45SN1_45S_50ntFlanks@6789.6806.18\|GCCTGTCTGAGCGTCGCT | >hg38_21_+_8433172_8446622_RNA45SN1_45S_50ntFlanks | 6789 | 6806 | 18 | GCCTGTCTGAGCGTCGCT | 1783.7 | 5.8 | 0.8 | 7.7 | 1.8E-14 | 4.8E-14 | BT20/MB468 |
| >hg38_21_+_8433172_8446622_RNA45SN1_45S_50ntFlanks@12499.12518.20\|GGTTTAGACCGTCGTGAGAC | >hg38_21_+_8433172_8446622_RNA45SN1_45S_50ntFlanks | 12499 | 12518 | 20 | GGTTTAGACCGTCGTGAGAC | 268.8 | -10.1 | 1.3 | -7.7 | 1.8E-14 | 4.9E-14 | BT20/MB468 |
| >am_trnaMT_LysTTT_MT_+_8295_8364@27.52.26,trnalookalike7_LysTTT_1_+_568844_568913@27.52.26\|CTTTTAAGTTAAAGATTAAGAGAACC | >am_trnaMT_LysTTT_MT_+_8295_8364 | 27 | 52 | 26 | CTTTTAAGTTAAAGATTAAGAGAACC | 579.4 | 9.1 | 1.2 | 7.7 | 1.8E-14 | 4.9E-14 | BT20/MB468 |
| >hg38_21_+_8433172_8446622_RNA45SN1_45S_50ntFlanks@11027.11053.27\|CTCTCTCTCTCTCTCCCCCGCTCCCCG | >hg38_21_+_8433172_8446622_RNA45SN1_45S_50ntFlanks | 11027 | 11053 | 27 | CTCTCTCTCTCTCTCCCCCGCTCCCCG | 321.6 | -9.9 | 1.3 | -7.7 | 1.7E-14 | 4.9E-14 | MB231/MB468 |
| >hg38_21_+_8433172_8446622_RNA45SN1_45S_50ntFlanks@10678.10712.35\|CGGTTCCGGCGGCGTCCGGTGAGCTCTCGCTGGCC | >hg38_21_+_8433172_8446622_RNA45SN1_45S_50ntFlanks | 10678 | 10712 | 35 | CGGTTCCGGCGGCGTCCGGTGAGCTCTCGCTGGCC | 6819.7 | -6.1 | 0.8 | -7.7 | 1.8E-14 | 4.9E-14 | BT20/MB468 |
| >hg38_MT_+_1621_3279_RNR2_16S_50ntFlanks@1427.1458.32\|GATTAAAGTCCTACGTGATCTGAGTTCAGACC | >hg38_MT_+_1621_3279_RNR2_16S_50ntFlanks | 1427 | 1458 | 32 | GATTAAAGTCCTACGTGATCTGAGTTCAGACC | 297.7 | 7.9 | 1.0 | 7.7 | 1.8E-14 | 4.9E-14 | BT20/MB468 |
| >hg38_21_+_8433172_8446622_RNA45SN1_45S_50ntFlanks@10498.10516.19\|TTCAGATCCCCGAATCCGG | >hg38_21_+_8433172_8446622_RNA45SN1_45S_50ntFlanks | 10498 | 10516 | 19 | TTCAGATCCCCGAATCCGG | 203.0 | -9.7 | 1.3 | -7.7 | 1.8E-14 | 4.9E-14 | BT20/MB468 |
| >hg38_21_+_8433172_8446622_RNA45SN1_45S_50ntFlanks@13008.13025.18\|ATTGAAAGTCAGCCCTCG | >hg38_21_+_8433172_8446622_RNA45SN1_45S_50ntFlanks | 13008 | 13025 | 18 | ATTGAAAGTCAGCCCTCG | 1291.8 | -6.3 | 0.8 | -7.7 | 1.8E-14 | 5.0E-14 | BT20/MB468 |
| >MI0000743\|hsa-mir-34c&WithFlank&11\|+\|111513433\|111513521@19.40.22(+1U)&[MIMAT0000686&hsa-miR-34c-5p&offsets\|0\|-1(+1U);m-336&11\|+\|111513451\|111513473&offsets\|0\|-1(+1U)]\|AGGCAGTGTAGTTAGCTGATTGT | >MI0000743\|hsa-mir-34c&WithFlank&11\|+\|111513433\|111513521 | 19 | 40 | 22(+1U) | AGGCAGTGTAGTTAGCTGATTGT | 230.5 | -11.2 | 1.4 | -8.0 | 1.2E-15 | 5.0E-14 | BT20/MB231 |
| >hg38_21_+_8433172_8446622_RNA45SN1_45S_50ntFlanks@10782.10801.20\|GGTGAACAGCCTCTGGCATG | >hg38_21_+_8433172_8446622_RNA45SN1_45S_50ntFlanks | 10782 | 10801 | 20 | GGTGAACAGCCTCTGGCATG | 457.5 | -3.2 | 0.4 | -7.7 | 1.8E-14 | 5.0E-14 | MB231/MB468 |

| | | | | | | | | | | | | |
|---|---|---|---|---|---|---|---|---|---|---|---|---|
| >MI0000767\|hsa-mir-365a&WithFlank&16\|+\|14309279\|14309377@62.83.22;MI0000769\|hsa-mir-365b&WithFlank&17\|+\|31575405\|31575527@74.95.22&[MIMAT0000710&hsa-miR-365a-3p&offsets\|0\|0;m-176&16\|+\|14309340\|14309361&offsets\|0\|0];[MIMAT0022834&hsa-miR-365b-3p&offsets\|0\|0;m-175&17\|+\|31575478\|31575499&offsets\|0\|0]\|TAATGCCCCTAAAAATCCTTAT | >MI0000767\|hsa-mir-365a&WithFlank&16\|+\|14309279\|14309377 | 62 | 83 | 22 | TAATGCCCCTAAAAATCCTTAT | 573.2 | 5.7 | 0.7 | 7.7 | 1.8E-14 | 5.0E-14 | MB231/MB468 |
| >hg38_1_-_228634819_228635039_RNA5S12_5S_50ntFlanks@144.171.28\|CCTGGGAATACCGGGTGCTGTAGGCTTT | >hg38_1_-_228634819_228635039_RNA5S12_5S_50ntFlanks | 144 | 171 | 28 | CCTGGGAATACCGGGTGCTGTAGGCTTT | 4532.4 | 5.9 | 0.8 | 7.7 | 1.8E-14 | 5.0E-14 | BT20/MB468 |
| >am_trna119_LysCTT_1_-_145395522_145395594@16.33.18,trna11_LysCTT_5_-_180648979_180649051@16.33.18,trna128_LysCTT_1_-_55423542_55423614@16.33.18,trna13_LysCTT_6_+_26556774_26556846@16.33.18,trna6_LysCTT_19_-_52425396_52425468@16.33.18,trna7_LysCTT_16_+_3225692_3225764@16.33.18,trna9_LysCTT_5_+_180634755_180634827@16.33.18\|TCGGTAGAGCATGAGACT | >am_trna119_LysCTT_1_-_145395522_145395594 | 16 | 33 | 18 | TCGGTAGAGCATGAGACT | 1634.9 | 10.0 | 1.3 | 7.7 | 1.8E-14 | 5.1E-14 | MB231/MB468 |
| >hg38_21_+_8433172_8446622_RNA45SN1_45S_50ntFlanks@3873.3898.26\|TAATACATGCCGACGGGCGCTGACCC | >hg38_21_+_8433172_8446622_RNA45SN1_45S_50ntFlanks | 3873 | 3898 | 26 | TAATACATGCCGACGGGCGCTGACCC | 1024.4 | -4.7 | 0.6 | -7.7 | 1.8E-14 | 5.1E-14 | MB231/MB468 |
| >trnaMT_PheGAA_MT_+_577_647@39.56.18\|ATGTTTAGACGGGCTCAC | >trnaMT_PheGAA_MT_+_577_647 | 39 | 56 | 18 | ATGTTTAGACGGGCTCAC | 5166.4 | 9.1 | 1.2 | 7.7 | 1.9E-14 | 5.1E-14 | BT20/MB468 |
| >MI0000272\|hsa-mir-182&WithFlank&7\|-\|129770377\|129770498@29.52.24&[MIMAT0000259&hsa-miR-182-5p&offsets\|0\|0;m-22&7\|-\|129770449\|129770470&offsets\|0\|+2]\|TTTGGCAATGGTAGAACTCACACT | >MI0000272\|hsa-mir-182&WithFlank&7\|-\|129770377\|129770498 | 29 | 52 | 24 | TTTGGCAATGGTAGAACTCACACT | 22661.9 | 5.9 | 0.8 | 7.7 | 1.8E-14 | 5.1E-14 | MB231/MB468 |
| >trna10_AspGTC_12_-_125424193_125424264@32.72.41,trna12_AspGTC_12_-_125411891_125411962@32.72.41,trna144_AspGTC_6_-_27551236_27551307@32.72.41,trna38_AspGTC_17_-_8125556_8125627@32.72.41,trna45_AspGTC_6_+_27447453_27447524@32.72.41,trna48_AspGTC_6_+_27471523_27471594@32.72.41,trna4_AspGTC_12_+_96429799_96429870@32.72.41,trna69_AspGTC_1_-_161440205_161440276@32.72.41,trna72_AspGTC_1_-_161432824_161432895@32.72.41,trna75_AspGTC_1_-_161425414_161425485@32.72.41,trna78_AspGTC_1_-_161418033_161418104@32.72.41,trna81_AspGTC_1_-_161410615_161410686@32.72.41\|CTGTCACGCGGGAGACCGGGGTTCGATTCCCCGACGGGGAG | >trna10_AspGTC_12_-_125424193_125424264 | 32 | 72 | 41 | CTGTCACGCGGGAGACCGGGGTTCGATTCCCCGACGGGGAG | 307.3 | -7.1 | 0.9 | -7.7 | 1.9E-14 | 5.2E-14 | BT20/MB468 |
| >hg38_21_+_8433172_8446622_RNA45SN1_45S_50ntFlanks@8182.8211.30\|ATCGAGGCCCAGCCCGTGGACGGTGTGAGG | >hg38_21_+_8433172_8446622_RNA45SN1_45S_50ntFlanks | 8182 | 8211 | 30 | ATCGAGGCCCAGCCCGTGGACGGTGTGAGG | 193.2 | -9.6 | 1.3 | -7.7 | 1.9E-14 | 5.3E-14 | BT20/MB468 |
| >MI0000087\|hsa-mir-29a&WithFlank&7\|-\|130876741\|130876816@48.67.20(+1A)&[MIMAT0000086&hsa-miR-29a-3p&offsets\|0\|-2(+1A);m-26&7\|-\|130876749\|130876769&offsets\|0\|-1(+1A)]\|TAGCACCATCTGAAATCGGTA | >MI0000087\|hsa-mir-29a&WithFlank&7\|-\|130876741\|130876816 | 48 | 67 | 20(+1A) | TAGCACCATCTGAAATCGGTA | 2368.3 | 11.4 | 1.5 | 7.7 | 2.0E-14 | 5.3E-14 | BT20/MB468 |
| >hg38_21_+_8433172_8446622_RNA45SN1_45S_50ntFlanks@7983.8007.25\|CAGATCAGACGTGGCGACCCGCTGA | >hg38_21_+_8433172_8446622_RNA45SN1_45S_50ntFlanks | 7983 | 8007 | 25 | CAGATCAGACGTGGCGACCCGCTGA | 270.3 | -5.6 | 0.7 | -7.7 | 1.9E-14 | 5.3E-14 | MB231/MB468 |
| >hg38_21_+_8433172_8446622_RNA45SN1_45S_50ntFlanks@10729.10755.27\|AGAGGGTGTAAATCTCGCGCCGGGCCG | >hg38_21_+_8433172_8446622_RNA45SN1_45S_50ntFlanks | 10729 | 10755 | 27 | AGAGGGTGTAAATCTCGCGCCGGGCCG | 221.7 | -9.8 | 1.3 | -7.7 | 2.0E-14 | 5.3E-14 | BT20/MB468 |
| >hg38_21_+_8433172_8446622_RNA45SN1_45S_50ntFlanks@12954.12972.19\|CTGCTTCTGGGTCGGGGTT | >hg38_21_+_8433172_8446622_RNA45SN1_45S_50ntFlanks | 12954 | 12972 | 19 | CTGCTTCTGGGTCGGGGTT | 552.0 | -7.5 | 1.0 | -7.7 | 2.0E-14 | 5.4E-14 | BT20/MB468 |
| >hg38_21_+_8433172_8446622_RNA45SN1_45S_50ntFlanks@6700.6717.18\|CTAGCTGCGAGAATTAAT | >hg38_21_+_8433172_8446622_RNA45SN1_45S_50ntFlanks | 6700 | 6717 | 18 | CTAGCTGCGAGAATTAAT | 171.7 | -9.4 | 1.2 | -7.7 | 2.0E-14 | 5.4E-14 | BT20/MB468 |
| >hg38_21_+_8433172_8446622_RNA45SN1_45S_50ntFlanks@10392.10419.28\|ACAGCAGTTGAACATGGGTCAGTCGGTC | >hg38_21_+_8433172_8446622_RNA45SN1_45S_50ntFlanks | 10392 | 10419 | 28 | ACAGCAGTTGAACATGGGTCAGTCGGTC | 400.8 | -4.0 | 0.5 | -7.7 | 1.9E-14 | 5.4E-14 | MB231/MB468 |
| >hg38_21_+_8433172_8446622_RNA45SN1_45S_50ntFlanks@6645.6667.23\|CTCGTACGACTCTTAGCGGTGGA | >hg38_21_+_8433172_8446622_RNA45SN1_45S_50ntFlanks | 6645 | 6667 | 23 | CTCGTACGACTCTTAGCGGTGGA | 1879.5 | 5.9 | 0.8 | 7.7 | 1.9E-14 | 5.4E-14 | MB231/MB468 |
| >hg38_MT_+_1621_3279_RNR2_16S_50ntFlanks@51.94.44\|GCTAAACCTAGCCCCAAACCCACTCCACCTTACTACCAGACAAC | >hg38_MT_+_1621_3279_RNR2_16S_50ntFlanks | 51 | 94 | 44 | GCTAAACCTAGCCCCAAACCCACTCCACCTTACTACCAGACAAC | 613.2 | 11.9 | 1.5 | 7.7 | 2.0E-14 | 5.4E-14 | BT20/MB468 |
| >hg38_1_-_228634819_228635039_RNA5S12_5S_50ntFlanks@131.171.41\|GGATGGGAGACCGCCTGGGAATACCGGGTGCTGTAGGCTTT | >hg38_1_-_228634819_228635039_RNA5S12_5S_50ntFlanks | 131 | 171 | 41 | GGATGGGAGACCGCCTGGGAATACCGGGTGCTGTAGGCTTT | 805.6 | 10.8 | 1.4 | 7.7 | 2.0E-14 | 5.4E-14 | BT20/MB468 |
| >hg38_21_+_8433172_8446622_RNA45SN1_45S_50ntFlanks@11233.11250.18\|CCCTTCCCGTGGATCGCC | >hg38_21_+_8433172_8446622_RNA45SN1_45S_50ntFlanks | 11233 | 11250 | 18 | CCCTTCCCGTGGATCGCC | 606.6 | -11.2 | 1.5 | -7.7 | 2.0E-14 | 5.5E-14 | BT20/MB468 |
| >hg38_21_+_8433172_8446622_RNA45SN1_45S_50ntFlanks@11768.11804.37\|CTAATTAGTGACGCGCATGAATGGATGAACGAGATTC | >hg38_21_+_8433172_8446622_RNA45SN1_45S_50ntFlanks | 11768 | 11804 | 37 | CTAATTAGTGACGCGCATGAATGGATGAACGAGATTC | 524.4 | -7.4 | 1.0 | -7.6 | 2.0E-14 | 5.5E-14 | BT20/MB468 |
| >hg38_21_+_8433172_8446622_RNA45SN1_45S_50ntFlanks@6652.6677.26\|GACTCTTAGCGGTGGATCACTCGGCT | >hg38_21_+_8433172_8446622_RNA45SN1_45S_50ntFlanks | 6652 | 6677 | 26 | GACTCTTAGCGGTGGATCACTCGGCT | 654.2 | 3.4 | 0.4 | 7.6 | 2.0E-14 | 5.5E-14 | BT20/MB468 |
| >MI0000746\|hsa-mir-99b&WithFlank&19\|+\|51692606\|51692687@51.70.20(+1U)&[MIMAT0004678&hsa-miR-99b-3p&offsets\|0\|-2(+1U);m-151&19\|+\|51692656\|51692678&offsets\|0\|-3(+1U)]\|CAAGCTCGTGTCTGTGGGTCT | >MI0000746\|hsa-mir-99b&WithFlank&19\|+\|51692606\|51692687 | 51 | 70 | 20(+1U) | CAAGCTCGTGTCTGTGGGTCT | 1238.7 | 9.3 | 1.2 | 7.6 | 2.1E-14 | 5.6E-14 | BT20/MB468 |
| >hg38_21_+_8433172_8446622_RNA45SN1_45S_50ntFlanks@9504.9527.24\|GTGAACTATGCCTGGGCAGGGCGA | >hg38_21_+_8433172_8446622_RNA45SN1_45S_50ntFlanks | 9504 | 9527 | 24 | GTGAACTATGCCTGGGCAGGGCGA | 269.4 | -9.6 | 1.3 | -7.6 | 2.1E-14 | 5.8E-14 | MB231/MB468 |
| >hg38_21_+_8433172_8446622_RNA45SN1_45S_50ntFlanks@9504.9524.21\|GTGAACTATGCCTGGGCAGGG | >hg38_21_+_8433172_8446622_RNA45SN1_45S_50ntFlanks | 9504 | 9524 | 21 | GTGAACTATGCCTGGGCAGGG | 287.6 | -9.7 | 1.3 | -7.6 | 2.1E-14 | 5.8E-14 | MB231/MB468 |
| >MI0000298\|hsa-mir-221&WithFlank&X\|-\|45746151\|45746272@71.91.21&[MIMAT0000278&hsa-miR-221-3p&offsets\|0\|-2;m-91&X\|-\|45746181\|45746202&offsets\|0\|-1]\|AGCTACATTGTCTGCTGGGTT | >MI0000298\|hsa-mir-221&WithFlank&X\|-\|45746151\|45746272 | 71 | 91 | 21 | AGCTACATTGTCTGCTGGGTT | 1289.3 | 6.1 | 0.8 | 7.6 | 2.2E-14 | 5.9E-14 | BT20/MB468 |
| >trnaMT_PheGAA_MT_+_577_647@35.53.19\|GAAAATGTTTAGACGGGCT | >trnaMT_PheGAA_MT_+_577_647 | 35 | 53 | 19 | GAAAATGTTTAGACGGGCT | 841.1 | 5.8 | 0.8 | 7.6 | 2.2E-14 | 5.9E-14 | BT20/MB468 |
| >hg38_21_+_8433172_8446622_RNA45SN1_45S_50ntFlanks@12957.12990.34\|CTTCTGGGTCGGGGTTTCGTACGTAGCAGAGCAG | >hg38_21_+_8433172_8446622_RNA45SN1_45S_50ntFlanks | 12957 | 12990 | 34 | CTTCTGGGTCGGGGTTTCGTACGTAGCAGAGCAG | 254.2 | -9.6 | 1.3 | -7.6 | 2.1E-14 | 5.9E-14 | MB231/MB468 |
| >hg38_21_+_8433172_8446622_RNA45SN1_45S_50ntFlanks@12991.13012.22\|CTCCCTCGCTGCGATCTATTGA | >hg38_21_+_8433172_8446622_RNA45SN1_45S_50ntFlanks | 12991 | 13012 | 22 | CTCCCTCGCTGCGATCTATTGA | 1618.4 | -4.2 | 0.6 | -7.6 | 2.1E-14 | 6.0E-14 | MB231/MB468 |
| >am_MI0014244\|hsa-mir-548x&WithFlank&21\|-\|18686084\|18686170@52.73.22&[MIMAT0015081_1&hsa-miR-548x-3p&offsets\|0\|+2;m-1121&21\|-\|18686099\|18686119&offsets\|0\|+1]\|TAAAAACTGCAATTACTTTCAC | >am_MI0014244\|hsa-mir-548x&WithFlank&21\|-\|18686084\|18686170 | 52 | 73 | 22 | TAAAAACTGCAATTACTTTCAC | 264.1 | 11.5 | 1.4 | 8.0 | 1.5E-15 | 6.0E-14 | BT20/MB231 |
| >MI0000089\|hsa-mir-31&WithFlank&9\|-\|21512109\|21512191@14.35.22(+1C)&[MIMAT0000089&hsa-miR-31-5p&offsets\|0\|+1(+1C);m-360&9\|-\|21512157\|21512178&offsets\|0\|0(+1C)]\|AGGCAAGATGCTGGCATAGCTGC | >MI0000089\|hsa-mir-31&WithFlank&9\|-\|21512109\|21512191 | 14 | 35 | 22(+1C) | AGGCAAGATGCTGGCATAGCTGC | 686.4 | 10.4 | 1.4 | 7.6 | 2.3E-14 | 6.2E-14 | BT20/MB468 |
| >hg38_21_+_8433172_8446622_RNA45SN1_45S_50ntFlanks@11918.11951.34\|ACATGAGAGGTGTAGAATAAGTGGGAGGCCCCCG | >hg38_21_+_8433172_8446622_RNA45SN1_45S_50ntFlanks | 11918 | 11951 | 34 | ACATGAGAGGTGTAGAATAAGTGGGAGGCCCCCG | 289.1 | -7.1 | 0.9 | -7.6 | 2.3E-14 | 6.2E-14 | BT20/MB468 |
| >MI0000802\|hsa-mir-340&WithFlank&5\|-\|180015297\|180015403@22.42.21&[MIMAT0004692&hsa-miR-340-5p&offsets\|0\|-1;m-220&5\|-\|180015361\|180015382&offsets\|0\|-1]\|TTATAAAGCAATGAGACTGAT | >MI0000802\|hsa-mir-340&WithFlank&5\|-\|180015297\|180015403 | 22 | 42 | 21 | TTATAAAGCAATGAGACTGAT | 2504.9 | 7.9 | 1.0 | 7.6 | 2.3E-14 | 6.3E-14 | BT20/MB468 |
| >hg38_21_+_8433172_8446622_RNA45SN1_45S_50ntFlanks@9352.9370.19\|CCGTGGCGCAATGAAGGTG | >hg38_21_+_8433172_8446622_RNA45SN1_45S_50ntFlanks | 9352 | 9370 | 19 | CCGTGGCGCAATGAAGGTG | 987.4 | -8.9 | 1.2 | -7.6 | 2.3E-14 | 6.3E-14 | BT20/MB468 |
| >hg38_21_+_8433172_8446622_RNA45SN1_45S_50ntFlanks@12937.12966.30\|AACCATTCGTAGACGACCTGCTTCTGGGTC | >hg38_21_+_8433172_8446622_RNA45SN1_45S_50ntFlanks | 12937 | 12966 | 30 | AACCATTCGTAGACGACCTGCTTCTGGGTC | 225.3 | -6.4 | 0.8 | -7.6 | 2.3E-14 | 6.3E-14 | BT20/MB468 |
| >hg38_21_+_8433172_8446622_RNA45SN1_45S_50ntFlanks@10805.10822.18\|GAACAATGTAGGTAAGGG | >hg38_21_+_8433172_8446622_RNA45SN1_45S_50ntFlanks | 10805 | 10822 | 18 | GAACAATGTAGGTAAGGG | 236.5 | -6.4 | 0.8 | -7.6 | 2.3E-14 | 6.3E-14 | BT20/MB468 |
| >hg38_21_+_8433172_8446622_RNA45SN1_45S_50ntFlanks@6739.6771.33\|TCATCGACACTTCGAACGCACTTGCGGCCCCGG | >hg38_21_+_8433172_8446622_RNA45SN1_45S_50ntFlanks | 6739 | 6771 | 33 | TCATCGACACTTCGAACGCACTTGCGGCCCCGG | 355.3 | -6.7 | 0.9 | -7.6 | 2.4E-14 | 6.4E-14 | BT20/MB468 |
| >trna130_GlnTTG_6_-_27763640_27763711@1.29.29,trna173_GlnTTG_6_-_26311975_26312046@1.29.29,trna174_GlnTTG_6_-_26311424_26311495@1.29.29\|GGCCCCATGGTGTAATGGTTAGCACTCTG | >trna130_GlnTTG_6_-_27763640_27763711 | 1 | 29 | 29 | GGCCCCATGGTGTAATGGTTAGCACTCTG | 555.7 | -5.9 | 0.8 | -7.6 | 2.3E-14 | 6.4E-14 | MB231/MB468 |
| >MI0000103\|hsa-mir-101-1&WithFlank&1\|-\|65058428\|65058514@53.72.20;MI0000739\|hsa-mir-101-2&WithFlank&9\|+\|4850291\|4850381@55.74.20&[MIMAT0000099&hsa-miR-101-3p&offsets\|0\|-1;m-17&1\|-\|65058443\|65058462&offsets\|0\|0];[MIMAT0000099_1&hsa-miR-101-3p&offsets\|0\|-1;m-16&9\|+\|4850345\|4850364&offsets\|0\|0]\|TACAGTACTGTGATAACTGA | >MI0000103\|hsa-mir-101-1&WithFlank&1\|-\|65058428\|65058514 | 53 | 72 | 20 | TACAGTACTGTGATAACTGA | 5514.7 | 6.9 | 0.9 | 7.6 | 2.4E-14 | 6.4E-14 | BT20/MB468 |
| >hg38_21_+_8433172_8446622_RNA45SN1_45S_50ntFlanks@6645.6676.32\|CTCGTACGACTCTTAGCGGTGGATCACTCGGC | >hg38_21_+_8433172_8446622_RNA45SN1_45S_50ntFlanks | 6645 | 6676 | 32 | CTCGTACGACTCTTAGCGGTGGATCACTCGGC | 374.9 | 6.9 | 0.9 | 7.6 | 2.4E-14 | 6.4E-14 | BT20/MB468 |
| >hg38_21_+_8433172_8446622_RNA45SN1_45S_50ntFlanks@9352.9370.19\|CCGTGGCGCAATGAAGGTG | >hg38_21_+_8433172_8446622_RNA45SN1_45S_50ntFlanks | 9352 | 9370 | 19 | CCGTGGCGCAATGAAGGTG | 840.4 | -6.5 | 0.8 | -7.6 | 2.3E-14 | 6.5E-14 | MB231/MB468 |

| | | | | | | | | | | | | |
|---|---|---|---|---|---|---|---|---|---|---|---|---|
| >hg38_21_+_8433172_8446622_RNA45SN1_45S_50ntFlanks@12625.12659.35\|CAATGGGGCGAAGCTACCATCTGTGGGATTATGAC | >hg38_21_+_8433172_8446622_RNA45SN1_45S_50ntFlanks | 12625 | 12659 | 35 | CAATGGGGCGAAGCTACCATCTGTGGGATTATGAC | 124.9 | -4.3 | 0.6 | -7.6 | 2.4E-14 | 6.5E-14 | BT20/MB468 |
| >MI0000434\|hsa-let-7i&WithFlank&12\|+\|62603680\|62603775@12.31.20(+1G)&[MIMAT0000415&hsa-let-7i-5p&offsets\|0\|-2(+1G);m-49&12\|+\|62603691\|62603712&offsets\|0\|-2(+1G)]\|TGAGGTAGTAGTTTGTGCTGG | >MI0000434\|hsa-let-7i&WithFlank&12\|+\|62603680\|62603775 | 12 | 31 | 20(+1G) | TGAGGTAGTAGTTTGTGCTGG | 793.7 | 7.3 | 1.0 | 7.6 | 2.4E-14 | 6.5E-14 | BT20/MB468 |
| >hg38_21_+_8433172_8446622_RNA45SN1_45S_50ntFlanks@12405.12434.30\|CTTTTTGATCCTTCGATGTCGGCTCTTCCT | >hg38_21_+_8433172_8446622_RNA45SN1_45S_50ntFlanks | 12405 | 12434 | 30 | CTTTTTGATCCTTCGATGTCGGCTCTTCCT | 5551.5 | -5.1 | 0.7 | -7.6 | 2.3E-14 | 6.5E-14 | MB231/MB468 |
| >hg38_21_+_8433172_8446622_RNA45SN1_45S_50ntFlanks@9572.9601.30\|CAAATCGGTCGTCCGACCTGGGTATAGGGG | >hg38_21_+_8433172_8446622_RNA45SN1_45S_50ntFlanks | 9572 | 9601 | 30 | CAAATCGGTCGTCCGACCTGGGTATAGGGG | 312.2 | -3.2 | 0.4 | -7.6 | 2.4E-14 | 6.5E-14 | BT20/MB468 |
| >hg38_21_+_8433172_8446622_RNA45SN1_45S_50ntFlanks@11578.11605.28\|TAGAACTGGTGCGGACCAGGGGAATCCG | >hg38_21_+_8433172_8446622_RNA45SN1_45S_50ntFlanks | 11578 | 11605 | 28 | TAGAACTGGTGCGGACCAGGGGAATCCG | 286.8 | -4.3 | 0.6 | -7.6 | 2.3E-14 | 6.6E-14 | MB231/MB468 |
| >am_trna130_GlnTTG_6_-_27763640_27763711@13.30.18,trna146_GlnCTG_6_-_27515531_27515602@13.30.18,trna16_GlnTTG_17_+_47269890_47269961@13.30.18,trna173_GlnTTG_6_-_26311975_26312046@13.30.18,trna174_GlnTTG_6_-_26311424_26311495@13.30.18,trna1_GlnCTG_6_+_18836402_18836473@13.30.18,trna3_GlnCTG_17_+_8023070_8023141@13.30.18,trna42_GlnCTG_6_+_27263212_27263283@13.30.18,trna49_GlnCTG_6_+_27487308_27487379@13.30.18,trna64_GlnTTG_6_+_28557156_28557227@13.30.18,trna7_GlnCTG_15_-_66161400_66161471@13.30.18,trna84_GlnTTG_6_+_145503859_145503930@13.30.18,trna99_GlnCTG_6_-_28909378_28909449@13.30.18\|TAATGGTTAGCACTCTGG | >am_trna130_GlnTTG_6_-_27763640_27763711 | 13 | 30 | 18 | TAATGGTTAGCACTCTGG | 184.9 | -4.3 | 0.6 | -7.6 | 2.4E-14 | 6.6E-14 | BT20/MB468 |
| >hg38_21_+_8433172_8446622_RNA45SN1_45S_50ntFlanks@6690.6720.31\|AAGAACGCAGCTAGCTGCGAGAATTAATGTG | >hg38_21_+_8433172_8446622_RNA45SN1_45S_50ntFlanks | 6690 | 6720 | 31 | AAGAACGCAGCTAGCTGCGAGAATTAATGTG | 1074.4 | -4.4 | 0.6 | -7.6 | 2.4E-14 | 6.6E-14 | MB231/MB468 |
| >hg38_21_+_8433172_8446622_RNA45SN1_45S_50ntFlanks@10384.10401.18\|CCATGTGAACAGCAGTTG | >hg38_21_+_8433172_8446622_RNA45SN1_45S_50ntFlanks | 10384 | 10401 | 18 | CCATGTGAACAGCAGTTG | 291.0 | -10.2 | 1.3 | -7.6 | 2.5E-14 | 6.6E-14 | BT20/MB468 |
| >hg38_21_+_8433172_8446622_RNA45SN1_45S_50ntFlanks@10737.10755.19\|TAAATCTCGCGCCGGGCCG | >hg38_21_+_8433172_8446622_RNA45SN1_45S_50ntFlanks | 10737 | 10755 | 19 | TAAATCTCGCGCCGGGCCG | 518.1 | -8.7 | 1.1 | -7.6 | 2.4E-14 | 6.7E-14 | MB231/MB468 |
| >hg38_21_+_8433172_8446622_RNA45SN1_45S_50ntFlanks@10468.10497.30\|TTGCCCTCGGCCGATCGAAAGGGAGTCGGG | >hg38_21_+_8433172_8446622_RNA45SN1_45S_50ntFlanks | 10468 | 10497 | 30 | TTGCCCTCGGCCGATCGAAAGGGAGTCGGG | 289.2 | -7.0 | 0.9 | -7.6 | 2.5E-14 | 6.7E-14 | BT20/MB468 |
| >hg38_21_+_8433172_8446622_RNA45SN1_45S_50ntFlanks@7990.8018.29\|GACGTGGCGACCCGCTGAATTTAAGCATA | >hg38_21_+_8433172_8446622_RNA45SN1_45S_50ntFlanks | 7990 | 8018 | 29 | GACGTGGCGACCCGCTGAATTTAAGCATA | 390.5 | 6.9 | 0.9 | 7.6 | 2.4E-14 | 6.8E-14 | MB231/MB468 |
| >MI0000273\|hsa-mir-183&WithFlank&7\|-\|129774899\|129775020@34.56.23&[MIMAT0000261&hsa-miR-183-5p&offsets\|+1\|+2;m-41&7\|-\|129774966\|129774987&offsets\|0\|+1]\|ATGGCACTGGTAGAATTCACTGT | >MI0000273\|hsa-mir-183&WithFlank&7\|-\|129774899\|129775020 | 34 | 56 | 23 | ATGGCACTGGTAGAATTCACTGT | 1046.6 | 6.2 | 0.8 | 7.6 | 2.4E-14 | 6.8E-14 | MB231/MB468 |
| >MI0000077\|hsa-mir-21&WithFlank&17\|+\|59841260\|59841343@13.36.24&[MIMAT0000076&hsa-miR-21-5p&offsets\|-1\|+1;m-1&17\|+\|59841273\|59841294&offsets\|-1\|+1]\|GTAGCTTATCAGACTGATGTTGAC | >MI0000077\|hsa-mir-21&WithFlank&17\|+\|59841260\|59841343 | 13 | 36 | 24 | GTAGCTTATCAGACTGATGTTGAC | 510.3 | 5.8 | 0.8 | 7.6 | 2.5E-14 | 6.8E-14 | BT20/MB468 |
| >hg38_21_+_8433172_8446622_RNA45SN1_45S_50ntFlanks@13004.13027.24\|ATCTATTGAAAGTCAGCCCTCGAC | >hg38_21_+_8433172_8446622_RNA45SN1_45S_50ntFlanks | 13004 | 13027 | 24 | ATCTATTGAAAGTCAGCCCTCGAC | 1495.1 | -8.0 | 1.0 | -7.6 | 2.4E-14 | 6.9E-14 | MB231/MB468 |
| >MI0000081\|hsa-mir-24-2&WithFlank&19\|-\|13836281\|13836365@56.78.23;MI0000080\|hsa-mir-24-1&WithFlank&9\|+\|95086015\|95086094@50.72.23&[MIMAT0000080&hsa-miR-24-3p&offsets\|0\|+1;m-36&19\|-\|13836288\|13836310&offsets\|0\|0];[MIMAT0000080_1&hsa-miR-24-3p&offsets\|0\|+1;m-35&9\|+\|95086064\|95086086&offsets\|0\|0]\|TGGCTCAGTTCAGCAGGAACAGG | >MI0000081\|hsa-mir-24-2&WithFlank&19\|-\|13836281\|13836365 | 56 | 78 | 23 | TGGCTCAGTTCAGCAGGAACAGG | 2487.2 | 8.1 | 1.1 | 7.6 | 2.6E-14 | 6.9E-14 | BT20/MB468 |
| >MI0000084\|hsa-mir-26b&WithFlank&2\|+\|218402640\|218402728@18.38.21(+1A)&[MIMAT0000083&hsa-miR-26b-5p&offsets\|0\|0(+1A);m-53&2\|+\|218402657\|218402678&offsets\|0\|-1(+1A)]\|TTCAAGTAATTCAGGATAGGTA | >MI0000084\|hsa-mir-26b&WithFlank&2\|+\|218402640\|218402728 | 18 | 38 | 21(+1A) | TTCAAGTAATTCAGGATAGGTA | 3193.8 | 8.7 | 1.1 | 7.6 | 2.6E-14 | 6.9E-14 | BT20/MB468 |
| >hg38_21_+_8433172_8446622_RNA45SN1_45S_50ntFlanks@8357.8375.19\|AAAAGAACTTTGAAGAGAG | >hg38_21_+_8433172_8446622_RNA45SN1_45S_50ntFlanks | 8357 | 8375 | 19 | AAAAGAACTTTGAAGAGAG | 4282.8 | -4.6 | 0.6 | -7.6 | 2.5E-14 | 6.9E-14 | MB231/MB468 |
| >MI0000447\|hsa-mir-128-1&WithFlank&2\|+\|135665391\|135665484@56.75.20(+1C);MI0000727\|hsa-mir-128-2&WithFlank&3\|+\|35744470\|35744565@58.77.20(+1C)&[MIMAT0000424&hsa-miR-128-3p&offsets\|0\|-1(+1C);m-108&2\|+\|135665446\|135665466&offsets\|0\|-1(+1C)];[MIMAT0000424_1&hsa-miR-128-3p&offsets\|0\|-1(+1C);m-117&3\|+\|35744527\|35744547&offsets\|0\|-1(+1C)]\|TCACAGTGAACCGGTCTCTTC | >MI0000447\|hsa-mir-128-1&WithFlank&2\|+\|135665391\|135665484 | 56 | 75 | 20(+1C) | TCACAGTGAACCGGTCTCTTC | 1529.3 | 5.3 | 0.7 | 7.6 | 2.5E-14 | 7.0E-14 | MB231/MB468 |
| >hg38_21_+_8433172_8446622_RNA45SN1_45S_50ntFlanks@11572.11592.21\|CCGACTTAGAACTGGTGCGGA | >hg38_21_+_8433172_8446622_RNA45SN1_45S_50ntFlanks | 11572 | 11592 | 21 | CCGACTTAGAACTGGTGCGGA | 777.9 | -5.3 | 0.7 | -7.6 | 2.5E-14 | 7.0E-14 | MB231/MB468 |
| >hg38_21_+_8433172_8446622_RNA45SN1_45S_50ntFlanks@12008.12030.23\|CCGCCGGTGAAATACCACTACTC | >hg38_21_+_8433172_8446622_RNA45SN1_45S_50ntFlanks | 12008 | 12030 | 23 | CCGCCGGTGAAATACCACTACTC | 684.7 | -9.0 | 1.2 | -7.6 | 2.5E-14 | 7.1E-14 | MB231/MB468 |
| >hg38_21_+_8433172_8446622_RNA45SN1_45S_50ntFlanks@10571.10591.21\|GAGAAGCCGGCGGGAGCCCCG | >hg38_21_+_8433172_8446622_RNA45SN1_45S_50ntFlanks | 10571 | 10591 | 21 | GAGAAGCCGGCGGGAGCCCCG | 484.1 | -10.9 | 1.4 | -7.6 | 2.7E-14 | 7.1E-14 | BT20/MB468 |
| >hg38_21_+_8433172_8446622_RNA45SN1_45S_50ntFlanks@4760.4778.19\|TCAGATACCGTCGTAGTTC | >hg38_21_+_8433172_8446622_RNA45SN1_45S_50ntFlanks | 4760 | 4778 | 19 | TCAGATACCGTCGTAGTTC | 434.5 | -4.6 | 0.6 | -7.6 | 2.5E-14 | 7.1E-14 | MB231/MB468 |
| >MI0016887\|hsa-mir-4521&WithFlank&17\|+\|8186939\|8187010@13.34.22&[MIMAT0019058&hsa-miR-4521&offsets\|+3\|+3;m-765&17\|+\|8186948\|8186969&offsets\|+3\|+3]\|AAGGAAGTCCTGTGCTCAGTTT | >MI0016887\|hsa-mir-4521&WithFlank&17\|+\|8186939\|8187010 | 13 | 34 | 22 | AAGGAAGTCCTGTGCTCAGTTT | 847.6 | 11.0 | 1.4 | 7.6 | 2.7E-14 | 7.2E-14 | BT20/MB468 |
| >trnaMT_ValTAC_MT_+_1602_1670@55.72.18\|ACTTGACCGCTCTGACCA | >trnaMT_ValTAC_MT_+_1602_1670 | 55 | 72 | 18 | ACTTGACCGCTCTGACCA | 1989.5 | 5.4 | 0.7 | 7.6 | 2.6E-14 | 7.2E-14 | MB231/MB468 |
| >MI0000650\|hsa-mir-200c&WithFlank&12\|+\|6963693\|6963772@50.72.23(+2U)&[MIMAT0000617&hsa-miR-200c-3p&offsets\|0\|0(+2U);m-54&12\|+\|6963742\|6963764&offsets\|0\|0(+2U)]\|TAATACTGCCGGGTAATGATGGATT | >MI0000650\|hsa-mir-200c&WithFlank&12\|+\|6963693\|6963772 | 50 | 72 | 23(+2U) | TAATACTGCCGGGTAATGATGGATT | 851.3 | 7.0 | 0.9 | 7.6 | 2.7E-14 | 7.2E-14 | BT20/MB468 |
| >hg38_21_+_8433172_8446622_RNA45SN1_45S_50ntFlanks@10794.10816.23\|CTGGCATGTTGGAACAATGTAGG | >hg38_21_+_8433172_8446622_RNA45SN1_45S_50ntFlanks | 10794 | 10816 | 23 | CTGGCATGTTGGAACAATGTAGG | 1733.2 | -5.2 | 0.7 | -7.6 | 2.7E-14 | 7.2E-14 | BT20/MB468 |
| >hg38_21_+_8433172_8446622_RNA45SN1_45S_50ntFlanks@4726.4759.34\|TAATCAAGAACGAAAGTCGGAGGTTCGAAGACGA | >hg38_21_+_8433172_8446622_RNA45SN1_45S_50ntFlanks | 4726 | 4759 | 34 | TAATCAAGAACGAAAGTCGGAGGTTCGAAGACGA | 620.0 | -5.3 | 0.7 | -7.6 | 2.7E-14 | 7.2E-14 | BT20/MB468 |
| >hg38_1_-_228634819_228635039_RNA5S12_5S_50ntFlanks@51.85.35\|GTCTACGGCCATACCACCCTGAACGCGCCCGATCT | >hg38_1_-_228634819_228635039_RNA5S12_5S_50ntFlanks | 51 | 85 | 35 | GTCTACGGCCATACCACCCTGAACGCGCCCGATCT | 754.9 | 5.4 | 0.7 | 7.6 | 2.7E-14 | 7.2E-14 | BT20/MB468 |
| >hg38_21_+_8433172_8446622_RNA45SN1_45S_50ntFlanks@4759.4793.35\|ATCAGATACCGTCGTAGTTCCGACCATAAACGATG | >hg38_21_+_8433172_8446622_RNA45SN1_45S_50ntFlanks | 4759 | 4793 | 35 | ATCAGATACCGTCGTAGTTCCGACCATAAACGATG | 175.6 | -9.4 | 1.2 | -7.6 | 2.7E-14 | 7.3E-14 | BT20/MB468 |
| >hg38_21_+_8433172_8446622_RNA45SN1_45S_50ntFlanks@6742.6770.29\|TCGACACTTCGAACGCACTTGCGGCCCCG | >hg38_21_+_8433172_8446622_RNA45SN1_45S_50ntFlanks | 6742 | 6770 | 29 | TCGACACTTCGAACGCACTTGCGGCCCCG | 285.9 | -6.4 | 0.8 | -7.6 | 2.6E-14 | 7.3E-14 | MB231/MB468 |
| >hg38_21_+_8433172_8446622_RNA45SN1_45S_50ntFlanks@6654.6683.30\|CTCTTAGCGGTGGATCACTCGGCTCGTGCG | >hg38_21_+_8433172_8446622_RNA45SN1_45S_50ntFlanks | 6654 | 6683 | 30 | CTCTTAGCGGTGGATCACTCGGCTCGTGCG | 889.8 | 5.7 | 0.8 | 7.6 | 2.6E-14 | 7.3E-14 | MB231/MB468 |
| >hg38_21_+_8433172_8446622_RNA45SN1_45S_50ntFlanks@9736.9754.19\|CCGAAACGATCTCAACCTA | >hg38_21_+_8433172_8446622_RNA45SN1_45S_50ntFlanks | 9736 | 9754 | 19 | CCGAAACGATCTCAACCTA | 405.8 | -6.2 | 0.8 | -7.6 | 2.6E-14 | 7.3E-14 | MB231/MB468 |
| >hg38_21_+_8433172_8446622_RNA45SN1_45S_50ntFlanks@8472.8494.23\|CGGCCCGGCGGATCTTTCCCGCC | >hg38_21_+_8433172_8446622_RNA45SN1_45S_50ntFlanks | 8472 | 8494 | 23 | CGGCCCGGCGGATCTTTCCCGCC | 380.7 | -10.6 | 1.4 | -7.6 | 2.7E-14 | 7.3E-14 | BT20/MB468 |
| >hg38_21_+_8433172_8446622_RNA45SN1_45S_50ntFlanks@476.515.40\|TTCGTGATCGATGTGGTGACGTCGTGCTCTCCCGGGCCGG | >hg38_21_+_8433172_8446622_RNA45SN1_45S_50ntFlanks | 476 | 515 | 40 | TTCGTGATCGATGTGGTGACGTCGTGCTCTCCCGGGCCGG | 426.8 | 8.6 | 1.1 | 7.6 | 2.8E-14 | 7.4E-14 | BT20/MB468 |
| >hg38_21_+_8433172_8446622_RNA45SN1_45S_50ntFlanks@12405.12424.20\|CTTTTTGATCCTTCGATGTC | >hg38_21_+_8433172_8446622_RNA45SN1_45S_50ntFlanks | 12405 | 12424 | 20 | CTTTTTGATCCTTCGATGTC | 1392.2 | -8.4 | 1.1 | -7.6 | 2.8E-14 | 7.4E-14 | BT20/MB468 |
| >hg38_21_+_8433172_8446622_RNA45SN1_45S_50ntFlanks@9568.9594.27\|CGTGCAAATCGGTCGTCCGACCTGGGT | >hg38_21_+_8433172_8446622_RNA45SN1_45S_50ntFlanks | 9568 | 9594 | 27 | CGTGCAAATCGGTCGTCCGACCTGGGT | 344.1 | -4.0 | 0.5 | -7.6 | 2.8E-14 | 7.5E-14 | BT20/MB468 |
| >MI0000265\|hsa-mir-7-3&WithFlank&19\|+\|4770664\|4770785@37.61.25&[MIMAT0000252_1&hsa-miR-7-5p&offsets\|0\|+1;m-402&19\|+\|4770700\|4770723&offsets\|0\|+1]\|TGGAAGACTAGTGATTTTGTTGTTC | >MI0000265\|hsa-mir-7-3&WithFlank&19\|+\|4770664\|4770785 | 37 | 61 | 25 | TGGAAGACTAGTGATTTTGTTGTTC | 485.0 | 6.7 | 0.9 | 7.6 | 2.8E-14 | 7.5E-14 | BT20/MB468 |
| >hg38_21_+_8433172_8446622_RNA45SN1_45S_50ntFlanks@11766.11791.26\|ATCTAATTAGTGACGCGCATGAATGG | >hg38_21_+_8433172_8446622_RNA45SN1_45S_50ntFlanks | 11766 | 11791 | 26 | ATCTAATTAGTGACGCGCATGAATGG | 238.9 | -9.5 | 1.2 | -7.6 | 2.7E-14 | 7.5E-14 | MB231/MB468 |
| >hg38_21_+_8433172_8446622_RNA45SN1_45S_50ntFlanks@8182.8206.25\|ATCGAGGCCCAGCCCGTGGACGGTG | >hg38_21_+_8433172_8446622_RNA45SN1_45S_50ntFlanks | 8182 | 8206 | 25 | ATCGAGGCCCAGCCCGTGGACGGTG | 194.9 | -9.6 | 1.3 | -7.6 | 2.8E-14 | 7.5E-14 | BT20/MB468 |
| >hg38_21_+_8433172_8446622_RNA45SN1_45S_50ntFlanks@12415.12432.18\|CTTCGATGTCGGCTCTTC | >hg38_21_+_8433172_8446622_RNA45SN1_45S_50ntFlanks | 12415 | 12432 | 18 | CTTCGATGTCGGCTCTTC | 2962.1 | -3.5 | 0.5 | -7.6 | 2.8E-14 | 7.6E-14 | BT20/MB468 |
| >hg38_21_+_8433172_8446622_RNA45SN1_45S_50ntFlanks@10781.10807.27\|AGGTGAACAGCCTCTGGCATGTTGGAA | >hg38_21_+_8433172_8446622_RNA45SN1_45S_50ntFlanks | 10781 | 10807 | 27 | AGGTGAACAGCCTCTGGCATGTTGGAA | 333.8 | -10.4 | 1.4 | -7.6 | 2.8E-14 | 7.6E-14 | BT20/MB468 |
| >hg38_21_+_8433172_8446622_RNA45SN1_45S_50ntFlanks@12044.12071.28\|ACTGACCCGGTGAGGCGGGGGGGCGAGC | >hg38_21_+_8433172_8446622_RNA45SN1_45S_50ntFlanks | 12044 | 12071 | 28 | ACTGACCCGGTGAGGCGGGGGGGCGAGC | 170.6 | -9.4 | 1.2 | -7.6 | 2.9E-14 | 7.6E-14 | BT20/MB468 |

<table>
<tr><th></th><th></th><th></th><th></th><th></th><th></th><th></th><th></th><th></th><th></th><th></th><th></th><th></th></tr>
<tr><td>>hg38_21_+_8433172_8446622_RNA45SN1_45S_50ntFlanks@11318.11338.21|CCCCACGTCTCGTCGCGCGCG</td><td>>hg38_21_+_8433172_8446622_RNA45SN1_45S_50ntFlanks</td><td>11318</td><td>11338</td><td>21</td><td>CCCCACGTCTCGTCGCGCGCG</td><td>307.1</td><td>-9.8</td><td>1.3</td><td>-7.6</td><td>2.7E-14</td><td>7.6E-14</td><td>MB231/MB468</td></tr>
<tr><td>>hg38_21_+_8433172_8446622_RNA45SN1_45S_50ntFlanks@12151.12181.31|GTGGGGAGTTTGACTGGGGCGGTACACCTGT</td><td>>hg38_21_+_8433172_8446622_RNA45SN1_45S_50ntFlanks</td><td>12151</td><td>12181</td><td>31</td><td>GTGGGGAGTTTGACTGGGGCGGTACACCTGT</td><td>187.6</td><td>-9.5</td><td>1.3</td><td>-7.6</td><td>2.9E-14</td><td>7.7E-14</td><td>BT20/MB468</td></tr>
<tr><td>>hg38_21_+_8433172_8446622_RNA45SN1_45S_50ntFlanks@4570.4587.18|TGGAATAGGACCGCGGTT</td><td>>hg38_21_+_8433172_8446622_RNA45SN1_45S_50ntFlanks</td><td>4570</td><td>4587</td><td>18</td><td>TGGAATAGGACCGCGGTT</td><td>295.2</td><td>-3.8</td><td>0.5</td><td>-7.6</td><td>2.9E-14</td><td>7.7E-14</td><td>BT20/MB468</td></tr>
<tr><td>>hg38_21_+_8433172_8446622_RNA45SN1_45S_50ntFlanks@11209.11230.22|CGGCGACTCTGGACGCGAGCCG</td><td>>hg38_21_+_8433172_8446622_RNA45SN1_45S_50ntFlanks</td><td>11209</td><td>11230</td><td>22</td><td>CGGCGACTCTGGACGCGAGCCG</td><td>543.3</td><td>-4.5</td><td>0.6</td><td>-7.6</td><td>2.8E-14</td><td>7.7E-14</td><td>MB231/MB468</td></tr>
<tr><td>>trna111_HisGTG_1_-_147774845_147774916@-1T.34.35,trna118_HisGTG_1_-_145396881_145396952@-1T.34.35,trna16_HisGTG_1_+_146544773_146544844@-1T.34.35,trna1_HisGTG_15_+_45493349_45493420@-1T.34.35,trna21_HisGTG_1_+_147753471_147753542@-1T.34.35,trna33_HisGTG_6_+_27125906_27125977@-1T.34.35,trna7_HisGTG_9_-_14433938_14434009@-1T.34.35,trna8_HisGTG_15_-_45492611_45492682@-1T.34.35,trna9_HisGTG_15_-_45490804_45490875@-1T.34.35|TGCCGTGATCGTATAGTGGTTAGTACTCTGCGTTG</td><td>>trna111_HisGTG_1_-_147774845_147774916</td><td>-1T</td><td>34</td><td>35</td><td>TGCCGTGATCGTATAGTGGTTAGTACTCTGCGTTG</td><td>5391.9</td><td>7.5</td><td>1.0</td><td>7.6</td><td>2.8E-14</td><td>7.8E-14</td><td>MB231/MB468</td></tr>
<tr><td>>hg38_21_+_8433172_8446622_RNA45SN1_45S_50ntFlanks@10412.10430.19|AGTCGGTCCTGAGAGATGG</td><td>>hg38_21_+_8433172_8446622_RNA45SN1_45S_50ntFlanks</td><td>10412</td><td>10430</td><td>19</td><td>AGTCGGTCCTGAGAGATGG</td><td>174.0</td><td>-5.7</td><td>0.8</td><td>-7.6</td><td>3.0E-14</td><td>7.9E-14</td><td>BT20/MB468</td></tr>
<tr><td>>hg38_21_+_8433172_8446622_RNA45SN1_45S_50ntFlanks@12991.13027.37|CTCCCTCGCTGCGATCTATTGAAAGTCAGCCCTCGAC</td><td>>hg38_21_+_8433172_8446622_RNA45SN1_45S_50ntFlanks</td><td>12991</td><td>13027</td><td>37</td><td>CTCCCTCGCTGCGATCTATTGAAAGTCAGCCCTCGAC</td><td>292.8</td><td>-10.2</td><td>1.3</td><td>-7.6</td><td>3.0E-14</td><td>7.9E-14</td><td>BT20/MB468</td></tr>
<tr><td>>hg38_21_+_8433172_8446622_RNA45SN1_45S_50ntFlanks@10408.10425.18|GGTCAGTCGGTCCTGAGA</td><td>>hg38_21_+_8433172_8446622_RNA45SN1_45S_50ntFlanks</td><td>10408</td><td>10425</td><td>18</td><td>GGTCAGTCGGTCCTGAGA</td><td>210.9</td><td>-6.6</td><td>0.9</td><td>-7.6</td><td>3.0E-14</td><td>7.9E-14</td><td>BT20/MB468</td></tr>
<tr><td>>hg38_21_+_8433172_8446622_RNA45SN1_45S_50ntFlanks@11580.11605.26|GAACTGGTGCGGACCAGGGGAATCCG</td><td>>hg38_21_+_8433172_8446622_RNA45SN1_45S_50ntFlanks</td><td>11580</td><td>11605</td><td>26</td><td>GAACTGGTGCGGACCAGGGGAATCCG</td><td>922.4</td><td>-4.9</td><td>0.7</td><td>-7.6</td><td>2.8E-14</td><td>7.9E-14</td><td>MB231/MB468</td></tr>
<tr><td>>hg38_21_+_8433172_8446622_RNA45SN1_45S_50ntFlanks@4728.4759.32|ATCAAGAACGAAAGTCGGAGGTTCGAAGACGA</td><td>>hg38_21_+_8433172_8446622_RNA45SN1_45S_50ntFlanks</td><td>4728</td><td>4759</td><td>32</td><td>ATCAAGAACGAAAGTCGGAGGTTCGAAGACGA</td><td>225.5</td><td>-9.8</td><td>1.3</td><td>-7.6</td><td>3.0E-14</td><td>7.9E-14</td><td>BT20/MB468</td></tr>
<tr><td>>hg38_21_+_8433172_8446622_RNA45SN1_45S_50ntFlanks@12630.12654.25|GGGCGAAGCTACCATCTGTGGGATT</td><td>>hg38_21_+_8433172_8446622_RNA45SN1_45S_50ntFlanks</td><td>12630</td><td>12654</td><td>25</td><td>GGGCGAAGCTACCATCTGTGGGATT</td><td>197.8</td><td>-9.6</td><td>1.3</td><td>-7.6</td><td>3.0E-14</td><td>7.9E-14</td><td>BT20/MB468</td></tr>
<tr><td>>hg38_21_+_8433172_8446622_RNA45SN1_45S_50ntFlanks@10425.10456.32|AGATGGGCGAGCGCCGTTCCGAAGGGACGGGC</td><td>>hg38_21_+_8433172_8446622_RNA45SN1_45S_50ntFlanks</td><td>10425</td><td>10456</td><td>32</td><td>AGATGGGCGAGCGCCGTTCCGAAGGGACGGGC</td><td>222.5</td><td>-4.3</td><td>0.6</td><td>-7.6</td><td>3.0E-14</td><td>7.9E-14</td><td>BT20/MB468</td></tr>
<tr><td>>am_trnaMT_GluTTC_MT_-_14674_14742@27.52.26,trnalookalike8_GluTTC_5_-_93905172_93905240@27.52.26|GTTTTTCATATCATTGGTCGTGGTTG</td><td>>am_trnaMT_GluTTC_MT_-_14674_14742</td><td>27</td><td>52</td><td>26</td><td>GTTTTTCATATCATTGGTCGTGGTTG</td><td>295.1</td><td>-10.2</td><td>1.3</td><td>-7.6</td><td>3.0E-14</td><td>7.9E-14</td><td>BT20/MB468</td></tr>
<tr><td>>hg38_21_+_8433172_8446622_RNA45SN1_45S_50ntFlanks@10873.10902.30|CTGGGTCGGTCGGGCTGGGGCGCGAAGCGG</td><td>>hg38_21_+_8433172_8446622_RNA45SN1_45S_50ntFlanks</td><td>10873</td><td>10902</td><td>30</td><td>CTGGGTCGGTCGGGCTGGGGCGCGAAGCGG</td><td>259.2</td><td>-9.6</td><td>1.3</td><td>-7.6</td><td>2.8E-14</td><td>8.0E-14</td><td>MB231/MB468</td></tr>
<tr><td>>MI0000263|hsa-mir-7-1&WithFlank&9|-|83969742|83969863@30.54.25&[MIMAT0000252_2&hsa-miR-7-5p&offsets|0|+1;m-401&9|-|83969811|83969834&offsets|0|+1]|TGGAAGACTAGTGATTTTGTTGTTT</td><td>>MI0000263|hsa-mir-7-1&WithFlank&9|-|83969742|83969863</td><td>30</td><td>54</td><td>25</td><td>TGGAAGACTAGTGATTTTGTTGTTT</td><td>3773.1</td><td>6.6</td><td>0.9</td><td>7.6</td><td>3.0E-14</td><td>8.0E-14</td><td>BT20/MB468</td></tr>
<tr><td>>hg38_1_-_228634819_228635039_RNA5S12_5S_50ntFlanks@51.90.40|GTCTACGGCCATACCACCCTGAACGCGCCCGATCTCGTCT</td><td>>hg38_1_-_228634819_228635039_RNA5S12_5S_50ntFlanks</td><td>51</td><td>90</td><td>40</td><td>GTCTACGGCCATACCACCCTGAACGCGCCCGATCTCGTCT</td><td>519.6</td><td>9.1</td><td>1.2</td><td>7.6</td><td>3.0E-14</td><td>8.1E-14</td><td>BT20/MB468</td></tr>
<tr><td>>hg38_21_+_8433172_8446622_RNA45SN1_45S_50ntFlanks@482.505.24|ATCGATGTGGTGACGTCGTGCTCT</td><td>>hg38_21_+_8433172_8446622_RNA45SN1_45S_50ntFlanks</td><td>482</td><td>505</td><td>24</td><td>ATCGATGTGGTGACGTCGTGCTCT</td><td>202.4</td><td>-9.7</td><td>1.3</td><td>-7.6</td><td>3.1E-14</td><td>8.1E-14</td><td>BT20/MB468</td></tr>
<tr><td>>hg38_21_+_8433172_8446622_RNA45SN1_45S_50ntFlanks@6646.6683.38|TCGTACGACTCTTAGCGGTGGATCACTCGGCTCGTGCG</td><td>>hg38_21_+_8433172_8446622_RNA45SN1_45S_50ntFlanks</td><td>6646</td><td>6683</td><td>38</td><td>TCGTACGACTCTTAGCGGTGGATCACTCGGCTCGTGCG</td><td>2121.4</td><td>7.2</td><td>0.9</td><td>7.6</td><td>3.1E-14</td><td>8.1E-14</td><td>BT20/MB468</td></tr>
<tr><td>>MI0000089|hsa-mir-31&WithFlank&9|-|21512109|21512191@14.34.21(+1A)&[MIMAT0000089&hsa-miR-31-5p&offsets|0|0(+1A);m-360&9|-|21512157|21512178&offsets|0|-1(+1A)]|AGGCAAGATGCTGGCATAGCTA</td><td>>MI0000089|hsa-mir-31&WithFlank&9|-|21512109|21512191</td><td>14</td><td>34</td><td>21(+1A)</td><td>AGGCAAGATGCTGGCATAGCTA</td><td>270.0</td><td>11.6</td><td>1.5</td><td>7.9</td><td>2.1E-15</td><td>8.1E-14</td><td>BT20/MB231</td></tr>
<tr><td>>MI0000077|hsa-mir-21&WithFlank&17|+|59841260|59841343@14.34.21(+2C)&[MIMAT0000076&hsa-miR-21-5p&offsets|0|-1(+2C);m-1&17|+|59841273|59841294&offsets|0|-1(+2C)]|TAGCTTATCAGACTGATGTTGCC</td><td>>MI0000077|hsa-mir-21&WithFlank&17|+|59841260|59841343</td><td>14</td><td>34</td><td>21(+2C)</td><td>TAGCTTATCAGACTGATGTTGCC</td><td>2452.1</td><td>7.1</td><td>0.9</td><td>7.6</td><td>3.1E-14</td><td>8.2E-14</td><td>BT20/MB468</td></tr>
<tr><td>>hg38_21_+_8433172_8446622_RNA45SN1_45S_50ntFlanks@7986.8017.32|ATCAGACGTGGCGACCCGCTGAATTTAAGCAT</td><td>>hg38_21_+_8433172_8446622_RNA45SN1_45S_50ntFlanks</td><td>7986</td><td>8017</td><td>32</td><td>ATCAGACGTGGCGACCCGCTGAATTTAAGCAT</td><td>215.4</td><td>-9.7</td><td>1.3</td><td>-7.6</td><td>3.1E-14</td><td>8.2E-14</td><td>BT20/MB468</td></tr>
<tr><td>>MI0000816|hsa-mir-335&WithFlank&7|+|130496105|130496210@58.78.21(+1A)&[MIMAT0004703&hsa-miR-335-3p&offsets|0|-1(+1A);m-159&7|+|130496162|130496183&offsets|0|-1(+1A)]|TTTTTCATTATTGCTCCTGACA</td><td>>MI0000816|hsa-mir-335&WithFlank&7|+|130496105|130496210</td><td>58</td><td>78</td><td>21(+1A)</td><td>TTTTTCATTATTGCTCCTGACA</td><td>1117.5</td><td>7.3</td><td>1.0</td><td>7.6</td><td>3.1E-14</td><td>8.2E-14</td><td>BT20/MB468</td></tr>
<tr><td>>am_trnaMT_HisGTG_MT_+_12138_12206@19.38.20|AAACATCAGATTGTGAATCT</td><td>>am_trnaMT_HisGTG_MT_+_12138_12206</td><td>19</td><td>38</td><td>20</td><td>AAACATCAGATTGTGAATCT</td><td>542.4</td><td>8.7</td><td>1.2</td><td>7.6</td><td>3.1E-14</td><td>8.2E-14</td><td>BT20/MB468</td></tr>
<tr><td>>hg38_1_-_228634819_228635039_RNA5S12_5S_50ntFlanks@51.91.41|GTCTACGGCCATACCACCCTGAACGCGCCCGATCTCGTCTG</td><td>>hg38_1_-_228634819_228635039_RNA5S12_5S_50ntFlanks</td><td>51</td><td>91</td><td>41</td><td>GTCTACGGCCATACCACCCTGAACGCGCCCGATCTCGTCTG</td><td>1082.3</td><td>8.7</td><td>1.1</td><td>7.6</td><td>3.1E-14</td><td>8.2E-14</td><td>BT20/MB468</td></tr>
<tr><td>>hg38_21_+_8433172_8446622_RNA45SN1_45S_50ntFlanks@12412.12431.20|ATCCTTCGATGTCGGCTCTT</td><td>>hg38_21_+_8433172_8446622_RNA45SN1_45S_50ntFlanks</td><td>12412</td><td>12431</td><td>20</td><td>ATCCTTCGATGTCGGCTCTT</td><td>288.8</td><td>-10.2</td><td>1.3</td><td>-7.6</td><td>3.1E-14</td><td>8.2E-14</td><td>BT20/MB468</td></tr>
<tr><td>>hg38_21_+_8433172_8446622_RNA45SN1_45S_50ntFlanks@9720.9737.18|ATTAGAGGTCTTGGGGCC</td><td>>hg38_21_+_8433172_8446622_RNA45SN1_45S_50ntFlanks</td><td>9720</td><td>9737</td><td>18</td><td>ATTAGAGGTCTTGGGGCC</td><td>213.3</td><td>-9.7</td><td>1.3</td><td>-7.6</td><td>3.1E-14</td><td>8.3E-14</td><td>BT20/MB468</td></tr>
<tr><td>>hg38_21_+_8433172_8446622_RNA45SN1_45S_50ntFlanks@12994.13036.43|CCTCGCTGCGATCTATTGAAAGTCAGCCCTCGACACAAGGGTT</td><td>>hg38_21_+_8433172_8446622_RNA45SN1_45S_50ntFlanks</td><td>12994</td><td>13036</td><td>43</td><td>CCTCGCTGCGATCTATTGAAAGTCAGCCCTCGACACAAGGGTT</td><td>579.6</td><td>8.7</td><td>1.1</td><td>7.6</td><td>3.0E-14</td><td>8.4E-14</td><td>MB231/MB468</td></tr>
<tr><td>>hg38_21_+_8433172_8446622_RNA45SN1_45S_50ntFlanks@13002.13028.27|CGATCTATTGAAAGTCAGCCCTCGACA</td><td>>hg38_21_+_8433172_8446622_RNA45SN1_45S_50ntFlanks</td><td>13002</td><td>13028</td><td>27</td><td>CGATCTATTGAAAGTCAGCCCTCGACA</td><td>2643.0</td><td>-8.0</td><td>1.0</td><td>-7.6</td><td>3.0E-14</td><td>8.4E-14</td><td>MB231/MB468</td></tr>
<tr><td>>hg38_21_+_8433172_8446622_RNA45SN1_45S_50ntFlanks@11199.11227.29|CCGGCGGCGGCGGCGACTCTGGACGCGAG</td><td>>hg38_21_+_8433172_8446622_RNA45SN1_45S_50ntFlanks</td><td>11199</td><td>11227</td><td>29</td><td>CCGGCGGCGGCGGCGACTCTGGACGCGAG</td><td>259.2</td><td>-9.6</td><td>1.3</td><td>-7.6</td><td>3.0E-14</td><td>8.5E-14</td><td>MB231/MB468</td></tr>
<tr><td>>hg38_1_-_228634819_228635039_RNA5S12_5S_50ntFlanks@84.101.18|CTCGTCTGATCTCGGAAG</td><td>>hg38_1_-_228634819_228635039_RNA5S12_5S_50ntFlanks</td><td>84</td><td>101</td><td>18</td><td>CTCGTCTGATCTCGGAAG</td><td>1813.2</td><td>-5.3</td><td>0.7</td><td>-7.6</td><td>3.2E-14</td><td>8.5E-14</td><td>BT20/MB468</td></tr>
<tr><td>>am_trnaMT_ProTGG_MT_-_15956_16023@52.70.19|AAAGACTTTTTCTCTGACC</td><td>>am_trnaMT_ProTGG_MT_-_15956_16023</td><td>52</td><td>70</td><td>19</td><td>AAAGACTTTTTCTCTGACC</td><td>4187.1</td><td>10.7</td><td>1.4</td><td>7.6</td><td>3.2E-14</td><td>8.5E-14</td><td>BT20/MB468</td></tr>
<tr><td>>hg38_21_+_8433172_8446622_RNA45SN1_45S_50ntFlanks@6745.6772.28|ACACTTCGAACGCACTTGCGGCCCCGGG</td><td>>hg38_21_+_8433172_8446622_RNA45SN1_45S_50ntFlanks</td><td>6745</td><td>6772</td><td>28</td><td>ACACTTCGAACGCACTTGCGGCCCCGGG</td><td>398.7</td><td>-6.8</td><td>0.9</td><td>-7.6</td><td>3.2E-14</td><td>8.5E-14</td><td>BT20/MB468</td></tr>
<tr><td>>hg38_21_+_8433172_8446622_RNA45SN1_45S_50ntFlanks@12417.12454.38|TCGATGTCGGCTCTTCCTATCATTGTGAAGCAGAATTC</td><td>>hg38_21_+_8433172_8446622_RNA45SN1_45S_50ntFlanks</td><td>12417</td><td>12454</td><td>38</td><td>TCGATGTCGGCTCTTCCTATCATTGTGAAGCAGAATTC</td><td>394.8</td><td>4.1</td><td>0.5</td><td>7.6</td><td>3.1E-14</td><td>8.6E-14</td><td>MB231/MB468</td></tr>
<tr><td>>MI0003834|hsa-mir-769&WithFlank&19|+|46018926|46019055@36.56.21&[MIMAT0003886&hsa-miR-769-5p&offsets|0|-1;m-216&19|+|46018961|46018982&offsets|0|-1]|TGAGACCTCTGGGTTCTGAGC</td><td>>MI0003834|hsa-mir-769&WithFlank&19|+|46018926|46019055</td><td>36</td><td>56</td><td>21</td><td>TGAGACCTCTGGGTTCTGAGC</td><td>1211.8</td><td>8.5</td><td>1.1</td><td>7.6</td><td>3.1E-14</td><td>8.7E-14</td><td>MB231/MB468</td></tr>
<tr><td>>MI0001519|hsa-mir-20b&WithFlank&X|-|134169803|134169883@12.34.23&[MIMAT0001413&hsa-miR-20b-5p&offsets|0|0;m-241&X|-|134169850|134169872&offsets|0|0]|CAAAGTGCTCATAGTGCAGGTAG</td><td>>MI0001519|hsa-mir-20b&WithFlank&X|-|134169803|134169883</td><td>12</td><td>34</td><td>23</td><td>CAAAGTGCTCATAGTGCAGGTAG</td><td>241.0</td><td>11.4</td><td>1.4</td><td>7.9</td><td>2.3E-15</td><td>8.7E-14</td><td>BT20/MB231</td></tr>
<tr><td>>hg38_21_+_8433172_8446622_RNA45SN1_45S_50ntFlanks@12363.12382.20|GGATAACTGGCTTGTGGCGG</td><td>>hg38_21_+_8433172_8446622_RNA45SN1_45S_50ntFlanks</td><td>12363</td><td>12382</td><td>20</td><td>GGATAACTGGCTTGTGGCGG</td><td>355.1</td><td>-10.5</td><td>1.4</td><td>-7.6</td><td>3.3E-14</td><td>8.8E-14</td><td>BT20/MB468</td></tr>
<tr><td>>MI0000065|hsa-let-7d&WithFlank&9|+|94178828|94178926@68.88.21(+1G)&[MIMAT0004484&hsa-let-7d-3p&offsets|0|-1(+1G);m-98&9|+|94178895|94178916&offsets|0|-1(+1G)]|CTATACGACCTGCTGCCTTTCG</td><td>>MI0000065|hsa-let-7d&WithFlank&9|+|94178828|94178926</td><td>68</td><td>88</td><td>21(+1G)</td><td>CTATACGACCTGCTGCCTTTCG</td><td>572.1</td><td>9.5</td><td>1.2</td><td>7.6</td><td>3.3E-14</td><td>8.8E-14</td><td>BT20/MB468</td></tr>
<tr><td>>hg38_21_+_8433172_8446622_RNA45SN1_45S_50ntFlanks@4657.4682.26|CTAGAGGTGAAATTCTTGGACCGGCG</td><td>>hg38_21_+_8433172_8446622_RNA45SN1_45S_50ntFlanks</td><td>4657</td><td>4682</td><td>26</td><td>CTAGAGGTGAAATTCTTGGACCGGCG</td><td>397.8</td><td>-10.6</td><td>1.4</td><td>-7.6</td><td>3.3E-14</td><td>8.8E-14</td><td>BT20/MB468</td></tr>
<tr><td>>hg38_21_+_8433172_8446622_RNA45SN1_45S_50ntFlanks@4064.4081.18|ATCAACTTTCGATGGTAG</td><td>>hg38_21_+_8433172_8446622_RNA45SN1_45S_50ntFlanks</td><td>4064</td><td>4081</td><td>18</td><td>ATCAACTTTCGATGGTAG</td><td>472.2</td><td>-10.5</td><td>1.4</td><td>-7.6</td><td>3.2E-14</td><td>8.9E-14</td><td>MB231/MB468</td></tr>
<tr><td>>hg38_21_+_8433172_8446622_RNA45SN1_45S_50ntFlanks@11586.11605.20|GTGCGGACCAGGGGAATCCG</td><td>>hg38_21_+_8433172_8446622_RNA45SN1_45S_50ntFlanks</td><td>11586</td><td>11605</td><td>20</td><td>GTGCGGACCAGGGGAATCCG</td><td>249.1</td><td>-10.0</td><td>1.3</td><td>-7.6</td><td>3.4E-14</td><td>8.9E-14</td><td>BT20/MB468</td></tr>
<tr><td>>hg38_21_+_8433172_8446622_RNA45SN1_45S_50ntFlanks@12628.12659.32|TGGGGCGAAGCTACCATCTGTGGGATTATGAC</td><td>>hg38_21_+_8433172_8446622_RNA45SN1_45S_50ntFlanks</td><td>12628</td><td>12659</td><td>32</td><td>TGGGGCGAAGCTACCATCTGTGGGATTATGAC</td><td>223.5</td><td>-9.8</td><td>1.3</td><td>-7.6</td><td>3.4E-14</td><td>8.9E-14</td><td>BT20/MB468</td></tr>
<tr><td>>hg38_21_+_8433172_8446622_RNA45SN1_45S_50ntFlanks@11207.11225.19|GGCGGCGACTCTGGACGCG</td><td>>hg38_21_+_8433172_8446622_RNA45SN1_45S_50ntFlanks</td><td>11207</td><td>11225</td><td>19</td><td>GGCGGCGACTCTGGACGCG</td><td>288.7</td><td>-4.7</td><td>0.6</td><td>-7.6</td><td>3.2E-14</td><td>8.9E-14</td><td>MB231/MB468</td></tr>
<tr><td>>hg38_21_+_8433172_8446622_RNA45SN1_45S_50ntFlanks@10731.10752.22|AGGGTGTAAATCTCGCGCCGGG</td><td>>hg38_21_+_8433172_8446622_RNA45SN1_45S_50ntFlanks</td><td>10731</td><td>10752</td><td>22</td><td>AGGGTGTAAATCTCGCGCCGGG</td><td>205.6</td><td>-9.7</td><td>1.3</td><td>-7.6</td><td>3.4E-14</td><td>9.0E-14</td><td>BT20/MB468</td></tr>
</table>

| | | | | | | | | | | | | |
|---|---|---|---|---|---|---|---|---|---|---|---|---|
| >trna130_GlnTTG_6_-_27763640_27763711@1.29.29,trna173_GlnTTG_6_-_26311975_26312046@1.29.29,trna174_GlnTTG_6_-_26311424_26311495@1.29.29\|GGCCCCATGGTGTAATGGTTAGCACTCTG | >trna130_GlnTTG_6_-_27763640_27763711 | 1 | 29 | 29 | GGCCCCATGGTGTAATGGTTAGCACTCTG | 650.5 | -8.2 | 1.1 | -7.6 | 3.4E-14 | 9.0E-14 | BT20/MB468 |
| >hg38_21_+_8433172_8446622_RNA45SN1_45S_50ntFlanks@11027.11060.34\|CTCTCTCTCTCTCTCCCCCGCTCCCCGTCCTCCC | >hg38_21_+_8433172_8446622_RNA45SN1_45S_50ntFlanks | 11027 | 11060 | 34 | CTCTCTCTCTCTCTCCCCCGCTCCCCGTCCTCCC | 262.6 | -9.6 | 1.3 | -7.6 | 3.2E-14 | 9.0E-14 | MB231/MB468 |
| >MI0000070\|hsa-mir-16-1&WithFlank&13\|-\|50048967\|50049067@20.40.21;MI0000115\|hsa-mir-16-2&WithFlank&3\|+\|160404739\|160404831@16.36.21&[MIMAT0000069&hsa-miR-16-5p&offsets\|0\|-1;m-62&13\|-\|50049027\|50049048&offsets\|0\|-1];[MIMAT0000069_1&hsa-miR-16-5p&offsets\|0\|-1;m-61&3\|+\|160404754\|160404775&offsets\|0\|-1]\|TAGCAGCACGTAAATATTGGC | >MI0000070\|hsa-mir-16-1&WithFlank&13\|-\|50048967\|50049067 | 20 | 40 | 21 | TAGCAGCACGTAAATATTGGC | 5164.6 | 7.1 | 0.9 | 7.6 | 3.5E-14 | 9.1E-14 | BT20/MB468 |
| >MI0000289\|hsa-mir-181a-1&WithFlank&1\|-\|198859038\|198859159@30.53.24(+1C);MI0000269\|hsa-mir-181a-2&WithFlank&9\|+\|124692436\|124692557@45.68.24(+1C)&[MIMAT0000256&hsa-miR-181a-5p&offsets\|0\|+1(+1C);m-32&1\|-\|198859108\|198859130&offsets\|0\|+1(+1C)];[MIMAT0000256_1&hsa-miR-181a-5p&offsets\|0\|+1(+1C);m-32&9\|+\|124692480\|124692502&offsets\|0\|+1(+1C)]\|AACATTCAACGCTGTCGGTGAGTTC | >MI0000289\|hsa-mir-181a-1&WithFlank&1\|-\|198859038\|198859159 | 30 | 53 | 24(+1C) | AACATTCAACGCTGTCGGTGAGTTC | 623.5 | 8.1 | 1.1 | 7.6 | 3.3E-14 | 9.1E-14 | MB231/MB468 |
| >hg38_21_+_8433172_8446622_RNA45SN1_45S_50ntFlanks@6779.6806.28\|CCGGGGCTACGCCTGTCTGAGCGTCGCT | >hg38_21_+_8433172_8446622_RNA45SN1_45S_50ntFlanks | 6779 | 6806 | 28 | CCGGGGCTACGCCTGTCTGAGCGTCGCT | 1252.6 | 4.7 | 0.6 | 7.6 | 3.3E-14 | 9.1E-14 | MB231/MB468 |
| >hg38_21_+_8433172_8446622_RNA45SN1_45S_50ntFlanks@12457.12477.21\|CAAGCGTTGGATTGTTCACCC | >hg38_21_+_8433172_8446622_RNA45SN1_45S_50ntFlanks | 12457 | 12477 | 21 | CAAGCGTTGGATTGTTCACCC | 3036.3 | -4.8 | 0.6 | -7.6 | 3.5E-14 | 9.1E-14 | BT20/MB468 |
| >hg38_21_+_8433172_8446622_RNA45SN1_45S_50ntFlanks@10866.10892.27\|CTAAGGGCTGGGTCGGTCGGGCTGGGG | >hg38_21_+_8433172_8446622_RNA45SN1_45S_50ntFlanks | 10866 | 10892 | 27 | CTAAGGGCTGGGTCGGTCGGGCTGGGG | 312.6 | -8.1 | 1.1 | -7.6 | 3.3E-14 | 9.2E-14 | MB231/MB468 |
| >hg38_21_+_8433172_8446622_RNA45SN1_45S_50ntFlanks@12973.12992.20\|TCGTACGTAGCAGAGCAGCT | >hg38_21_+_8433172_8446622_RNA45SN1_45S_50ntFlanks | 12973 | 12992 | 20 | TCGTACGTAGCAGAGCAGCT | 1072.1 | -3.5 | 0.5 | -7.6 | 3.3E-14 | 9.3E-14 | MB231/MB468 |
| >hg38_21_+_8433172_8446622_RNA45SN1_45S_50ntFlanks@12151.12179.29\|GTGGGGAGTTTGACTGGGGCGGTACACCT | >hg38_21_+_8433172_8446622_RNA45SN1_45S_50ntFlanks | 12151 | 12179 | 29 | GTGGGGAGTTTGACTGGGGCGGTACACCT | 239.0 | -9.9 | 1.3 | -7.6 | 3.6E-14 | 9.4E-14 | BT20/MB468 |
| >MI0000093\|hsa-mir-92a-1&WithFlank&13\|+\|91351308\|91351397@54.75.22(+2A);MI0000094\|hsa-mir-92a-2&WithFlank&X\|-\|134169532\|134169618@54.75.22(+2A)&[MIMAT0000092&hsa-miR-92a-3p&offsets\|0\|0(+2A);m-9&13\|+\|91351361\|91351382&offsets\|0\|0(+2A)];[MIMAT0000092_1&hsa-miR-92a-3p&offsets\|0\|0(+2A);m-11&X\|-\|134169544\|134169565&offsets\|0\|0(+2A)]\|TATTGCACTTGTCCCGGCCTGTAA | >MI0000093\|hsa-mir-92a-1&WithFlank&13\|+\|91351308\|91351397 | 54 | 75 | 22(+2A) | TATTGCACTTGTCCCGGCCTGTAA | 1112.1 | 5.3 | 0.7 | 7.6 | 3.6E-14 | 9.4E-14 | BT20/MB468 |
| >hg38_21_+_8433172_8446622_RNA45SN1_45S_50ntFlanks@10817.10840.24\|TAAGGGAAGTCGGCAAGCCGGATC | >hg38_21_+_8433172_8446622_RNA45SN1_45S_50ntFlanks | 10817 | 10840 | 24 | TAAGGGAAGTCGGCAAGCCGGATC | 176.5 | -5.9 | 0.8 | -7.6 | 3.6E-14 | 9.5E-14 | BT20/MB468 |
| >MI0000068\|hsa-let-7f-2&WithFlank&X\|-\|53557186\|53557280@14.36.23&[MIMAT0000067_1&hsa-let-7f-5p&offsets\|0\|+1;m-19&X\|-\|53557246\|53557267&offsets\|0\|+1]\|TGAGGTAGTAGATTGTATAGTTT | >MI0000068\|hsa-let-7f-2&WithFlank&X\|-\|53557186\|53557280 | 14 | 36 | 23 | TGAGGTAGTAGATTGTATAGTTT | 4673.4 | 5.7 | 0.7 | 7.6 | 3.6E-14 | 9.5E-14 | BT20/MB468 |
| >hg38_21_+_8433172_8446622_RNA45SN1_45S_50ntFlanks@11881.11905.25\|CCTGTTGAGCTTGACTCTAGTCTGG | >hg38_21_+_8433172_8446622_RNA45SN1_45S_50ntFlanks | 11881 | 11905 | 25 | CCTGTTGAGCTTGACTCTAGTCTGG | 168.7 | -9.4 | 1.2 | -7.6 | 3.6E-14 | 9.5E-14 | BT20/MB468 |
| >hg38_21_+_8433172_8446622_RNA45SN1_45S_50ntFlanks@12676.12708.33\|AGAATCCCGCCCAGGCGGAACGATACGGCAGCG | >hg38_21_+_8433172_8446622_RNA45SN1_45S_50ntFlanks | 12676 | 12708 | 33 | AGAATCCCGCCCAGGCGGAACGATACGGCAGCG | 172.2 | -9.4 | 1.2 | -7.6 | 3.6E-14 | 9.5E-14 | BT20/MB468 |
| >trnaMT_PheGAA_MT_+_577_647@56.74.19\|CATCACCCCATAAACACCA | >trnaMT_PheGAA_MT_+_577_647 | 56 | 74 | 19 | CATCACCCCATAAACACCA | 1557.8 | 8.5 | 1.1 | 7.6 | 3.4E-14 | 9.5E-14 | MB231/MB468 |
| >hg38_21_+_8433172_8446622_RNA45SN1_45S_50ntFlanks@3785.3812.28\|TGAAACTGCGAATGGCTCATTAAATCAG | >hg38_21_+_8433172_8446622_RNA45SN1_45S_50ntFlanks | 3785 | 3812 | 28 | TGAAACTGCGAATGGCTCATTAAATCAG | 222.1 | -9.8 | 1.3 | -7.6 | 3.7E-14 | 9.6E-14 | BT20/MB468 |
| >trna112_GlnCTG_1_-_147737382_147737453@30.50.21,trna131_GlnCTG_6_-_27759135_27759206@30.50.21,trna146_GlnCTG_6_-_27515531_27515602@30.50.21,trna15_GlnCTG_1_+_145963304_145963375@30.50.21,trna19_GlnCTG_1_+_147505038_147505109@30.50.21,trna1_GlnCTG_6_+_18836402_18836473@30.50.21,trna28_GlnCTG_1_+_149186125_149186196@30.50.21,trna3_GlnCTG_17_+_8023070_8023141@30.50.21,trna49_GlnCTG_6_+_27487308_27487379@30.50.21,trna7_GlnCTG_15_-_66161400_66161471@30.50.21,trna99_GlnCTG_6_-_28909378_28909449@30.50.21\|GACTCTGAATCCAGCGATCCG | >trna112_GlnCTG_1_-_147737382_147737453 | 30 | 50 | 21 | GACTCTGAATCCAGCGATCCG | 318.3 | -9.9 | 1.3 | -7.6 | 3.5E-14 | 9.6E-14 | MB231/MB468 |
| >hg38_1_-_228634819_228635039_RNA5S12_5S_50ntFlanks@128.169.42\|CTTGGATGGGAGACCGCCTGGGAATACCGGGTGCTGTAGGCT | >hg38_1_-_228634819_228635039_RNA5S12_5S_50ntFlanks | 128 | 169 | 42 | CTTGGATGGGAGACCGCCTGGGAATACCGGGTGCTGTAGGCT | 496.5 | 10.8 | 1.4 | 7.6 | 3.7E-14 | 9.7E-14 | BT20/MB468 |
| >hg38_21_+_8433172_8446622_RNA45SN1_45S_50ntFlanks@11781.11805.25\|CGCATGAATGGATGAACGAGATTCC | >hg38_21_+_8433172_8446622_RNA45SN1_45S_50ntFlanks | 11781 | 11805 | 25 | CGCATGAATGGATGAACGAGATTCC | 476.7 | -6.3 | 0.8 | -7.6 | 3.7E-14 | 9.7E-14 | BT20/MB468 |
| >hg38_21_+_8433172_8446622_RNA45SN1_45S_50ntFlanks@8170.8188.19\|CAAGTCCTTCTGATCGAGG | >hg38_21_+_8433172_8446622_RNA45SN1_45S_50ntFlanks | 8170 | 8188 | 19 | CAAGTCCTTCTGATCGAGG | 821.8 | -11.7 | 1.5 | -7.6 | 3.7E-14 | 9.7E-14 | BT20/MB468 |
| >am_trna134_GluTTC_1_-_16861774_16861845@1.28.28,trna5_GluTTC_1_+_17199078_17199149@1.28.28,trna84_GluTTC_1_-_161391883_161391954@1.28.28,trna94_GluTTC_1_-_149664355_149664427@1.28.28\|TCCCTGGTGGTCTAGTGGCTAGGATTCG | >am_trna134_GluTTC_1_-_16861774_16861845 | 1 | 28 | 28 | TCCCTGGTGGTCTAGTGGCTAGGATTCG | 324.3 | -6.4 | 0.8 | -7.6 | 3.5E-14 | 9.7E-14 | MB231/MB468 |
| >MI0000078\|hsa-mir-22&WithFlank&17\|-\|1713897\|1713993@59.81.23&[MIMAT0000077&hsa-miR-22-3p&offsets\|0\|+1;m-2&17\|-\|1713914\|1713935&offsets\|0\|+1]\|AAGCTGCCAGTTGAAGAACTGTT | >MI0000078\|hsa-mir-22&WithFlank&17\|-\|1713897\|1713993 | 59 | 81 | 23 | AAGCTGCCAGTTGAAGAACTGTT | 690.1 | 9.9 | 1.3 | 7.6 | 3.7E-14 | 9.7E-14 | BT20/MB468 |
| >hg38_21_+_8433172_8446622_RNA45SN1_45S_50ntFlanks@4759.4777.19\|ATCAGATACCGTCGTAGTT | >hg38_21_+_8433172_8446622_RNA45SN1_45S_50ntFlanks | 4759 | 4777 | 19 | ATCAGATACCGTCGTAGTT | 167.3 | -9.4 | 1.2 | -7.6 | 3.7E-14 | 9.8E-14 | BT20/MB468 |
| >hg38_21_+_8433172_8446622_RNA45SN1_45S_50ntFlanks@10687.10712.26\|CGGCGTCCGGTGAGCTCTCGCTGGCC | >hg38_21_+_8433172_8446622_RNA45SN1_45S_50ntFlanks | 10687 | 10712 | 26 | CGGCGTCCGGTGAGCTCTCGCTGGCC | 742.1 | -5.8 | 0.8 | -7.6 | 3.7E-14 | 9.8E-14 | BT20/MB468 |
| >hg38_21_+_8433172_8446622_RNA45SN1_45S_50ntFlanks@6664.6681.18\|TGGATCACTCGGCTCGTG | >hg38_21_+_8433172_8446622_RNA45SN1_45S_50ntFlanks | 6664 | 6681 | 18 | TGGATCACTCGGCTCGTG | 1127.8 | -5.1 | 0.7 | -7.6 | 3.5E-14 | 9.8E-14 | MB231/MB468 |
| >hg38_1_-_228634819_228635039_RNA5S12_5S_50ntFlanks@151.169.19\|ATACCGGGTGCTGTAGGCT | >hg38_1_-_228634819_228635039_RNA5S12_5S_50ntFlanks | 151 | 169 | 19 | ATACCGGGTGCTGTAGGCT | 802.9 | 5.6 | 0.7 | 7.6 | 3.8E-14 | 9.8E-14 | BT20/MB468 |
| >am_trnaMT_LysTTT_MT_+_8295_8364@33.52.20,trnalookalike7_LysTTT_1_+_568844_568913@33.52.20\|AGTTAAAGATTAAGAGAACC | >am_trnaMT_LysTTT_MT_+_8295_8364 | 33 | 52 | 20 | AGTTAAAGATTAAGAGAACC | 495.4 | 8.8 | 1.2 | 7.6 | 3.8E-14 | 9.9E-14 | BT20/MB468 |
| >MI0000084\|hsa-mir-26b&WithFlank&2\|+\|218402640\|218402728@18.38.21(+1G)&[MIMAT0000083&hsa-miR-26b-5p&offsets\|0\|0(+1G);m-53&2\|+\|218402657\|218402678&offsets\|0\|-1(+1G)]\|TTCAAGTAATTCAGGATAGGTG | >MI0000084\|hsa-mir-26b&WithFlank&2\|+\|218402640\|218402728 | 18 | 38 | 21(+1G) | TTCAAGTAATTCAGGATAGGTG | 2788.6 | 8.6 | 1.1 | 7.6 | 3.8E-14 | 9.9E-14 | BT20/MB468 |
| >hg38_21_+_8433172_8446622_RNA45SN1_45S_50ntFlanks@11591.11616.26\|GACCAGGGGAATCCGACTGTTTAATT | >hg38_21_+_8433172_8446622_RNA45SN1_45S_50ntFlanks | 11591 | 11616 | 26 | GACCAGGGGAATCCGACTGTTTAATT | 286.4 | 4.7 | 0.6 | 7.6 | 3.6E-14 | 9.9E-14 | MB231/MB468 |
| >MI0000100\|hsa-mir-98&WithFlank&X\|-\|53556217\|53556347@28.48.21(+1C)&[MIMAT0000096&hsa-miR-98-5p&offsets\|0\|-1(+1C);m-143&X\|-\|53556299\|53556320&offsets\|0\|-1(+1C)]\|TGAGGTAGTAAGTTGTATTGTC | >MI0000100\|hsa-mir-98&WithFlank&X\|-\|53556217\|53556347 | 28 | 48 | 21(+1C) | TGAGGTAGTAAGTTGTATTGTC | 1614.0 | 5.7 | 0.7 | 7.6 | 3.6E-14 | 9.9E-14 | MB231/MB468 |
| >hg38_21_+_8433172_8446622_RNA45SN1_45S_50ntFlanks@11576.11593.18\|CTTAGAACTGGTGCGGAC | >hg38_21_+_8433172_8446622_RNA45SN1_45S_50ntFlanks | 11576 | 11593 | 18 | CTTAGAACTGGTGCGGAC | 861.9 | -4.2 | 0.6 | -7.6 | 3.6E-14 | 1.0E-13 | MB231/MB468 |
| >hg38_21_+_8433172_8446622_RNA45SN1_45S_50ntFlanks@12409.12433.25\|TTGATCCTTCGATGTCGGCTCTTCC | >hg38_21_+_8433172_8446622_RNA45SN1_45S_50ntFlanks | 12409 | 12433 | 25 | TTGATCCTTCGATGTCGGCTCTTCC | 567.4 | 4.9 | 0.6 | 7.6 | 3.6E-14 | 1.0E-13 | MB231/MB468 |
| >MI0000272\|hsa-mir-182&WithFlank&7\|-\|129770377\|129770498@29.51.23(+1C)&[MIMAT0000259&hsa-miR-182-5p&offsets\|0\|-1(+1C);m-22&7\|-\|129770449\|129770470&offsets\|0\|+1(+1C)]\|TTTGGCAATGGTAGAACTCACACC | >MI0000272\|hsa-mir-182&WithFlank&7\|-\|129770377\|129770498 | 29 | 51 | 23(+1C) | TTTGGCAATGGTAGAACTCACACC | 2716.5 | 6.0 | 0.8 | 7.6 | 3.6E-14 | 1.0E-13 | MB231/MB468 |
| >hg38_21_+_8433172_8446622_RNA45SN1_45S_50ntFlanks@5420.5444.25\|CCGATTGGATGGTTTAGTGAGGCCC | >hg38_21_+_8433172_8446622_RNA45SN1_45S_50ntFlanks | 5420 | 5444 | 25 | CCGATTGGATGGTTTAGTGAGGCCC | 189.2 | -9.6 | 1.3 | -7.6 | 3.8E-14 | 1.0E-13 | BT20/MB468 |
| >hg38_21_+_8433172_8446622_RNA45SN1_45S_50ntFlanks@12459.12487.29\|AGCGTTGGATTGTTCACCCACTAATAGGG | >hg38_21_+_8433172_8446622_RNA45SN1_45S_50ntFlanks | 12459 | 12487 | 29 | AGCGTTGGATTGTTCACCCACTAATAGGG | 595.4 | -6.4 | 0.8 | -7.6 | 3.7E-14 | 1.0E-13 | MB231/MB468 |
| >MI0000456\|hsa-mir-140&WithFlank&16\|+\|69933075\|69933186@68.90.23&[MIMAT0004597&hsa-miR-140-3p&offsets\|0\|+2;m-45&16\|+\|69933143\|69933164&offsets\|-1\|0]\|TACCACAGGGTAGAACCACGGAC | >MI0000456\|hsa-mir-140&WithFlank&16\|+\|69933075\|69933186 | 68 | 90 | 23 | TACCACAGGGTAGAACCACGGAC | 1234.2 | 9.8 | 1.3 | 7.6 | 3.9E-14 | 1.0E-13 | BT20/MB468 |
| >hg38_21_+_8433172_8446622_RNA45SN1_45S_50ntFlanks@11876.11905.30\|AAGACCCTGTTGAGCTTGACTCTAGTCTGG | >hg38_21_+_8433172_8446622_RNA45SN1_45S_50ntFlanks | 11876 | 11905 | 30 | AAGACCCTGTTGAGCTTGACTCTAGTCTGG | 223.0 | -6.1 | 0.8 | -7.6 | 3.7E-14 | 1.0E-13 | MB231/MB468 |
| >hg38_21_+_8433172_8446622_RNA45SN1_45S_50ntFlanks@3840.3862.23\|CTCCTACTTGGATAACTGTGGTA | >hg38_21_+_8433172_8446622_RNA45SN1_45S_50ntFlanks | 3840 | 3862 | 23 | CTCCTACTTGGATAACTGTGGTA | 579.7 | 4.1 | 0.5 | 7.6 | 3.9E-14 | 1.0E-13 | BT20/MB468 |

| | | | | | | | | | | | | |
|---|---|---|---|---|---|---|---|---|---|---|---|---|
| >MI0000471\|hsa-mir-126&WithFlank&9\|+\|136670596\|136670692@57.79.23&[MIMAT0000445&hsa-miR-126-3p&offsets\|-1\|0;m-40&9\|+\|136670653\|136670674&offsets\|-1\|0]\|CTCGTACCGTGAGTAATAATGCG | >MI0000471\|hsa-mir-126&WithFlank&9\|+\|136670596\|136670692 | 57 | 79 | 23 | CTCGTACCGTGAGTAATAATGCG | 1521.5 | 10.2 | 1.3 | 7.6 | 4.0E-14 | 1.0E-13 | BT20/MB468 |
| >MI0000471\|hsa-mir-126&WithFlank&9\|+\|136670596\|136670692@58.77.20(+1U)&[MIMAT0000445&hsa-miR-126-3p&offsets\|0\|-2(+1U);m-40&9\|+\|136670653\|136670674&offsets\|0\|-2(+1U)]\|TCGTACCGTGAGTAATAATGT | >MI0000471\|hsa-mir-126&WithFlank&9\|+\|136670596\|136670692 | 58 | 77 | 20(+1U) | TCGTACCGTGAGTAATAATGT | 2289.3 | 10.4 | 1.4 | 7.6 | 4.0E-14 | 1.0E-13 | BT20/MB468 |
| >hg38_21_+_8433172_8446622_RNA45SN1_45S_50ntFlanks@12996.13014.19\|TCGCTGCGATCTATTGAAA | >hg38_21_+_8433172_8446622_RNA45SN1_45S_50ntFlanks | 12996 | 13014 | 19 | TCGCTGCGATCTATTGAAA | 877.5 | 3.6 | 0.5 | 7.6 | 4.0E-14 | 1.0E-13 | BT20/MB468 |
| >trnaMT_PheGAA_MT_+_577_647@1.35.35\|GTTTATGTAGCTTACCTCCTCAAAGCAATACACTG | >trnaMT_PheGAA_MT_+_577_647 | 1 | 35 | 35 | GTTTATGTAGCTTACCTCCTCAAAGCAATACACTG | 848.0 | 6.6 | 0.9 | 7.6 | 3.8E-14 | 1.0E-13 | MB231/MB468 |
| >MI0000815\|hsa-mir-339&WithFlank&7\|-\|1022927\|1023032@21.43.23&[MIMAT0000764&hsa-miR-339-5p&offsets\|0\|0;m-163&7\|-\|1022990\|1023012&offsets\|0\|0]\|TCCCTGTCCTCCAGGAGCTCACG | >MI0000815\|hsa-mir-339&WithFlank&7\|-\|1022927\|1023032 | 21 | 43 | 23 | TCCCTGTCCTCCAGGAGCTCACG | 2078.6 | 6.6 | 0.9 | 7.6 | 4.0E-14 | 1.0E-13 | BT20/MB468 |
| >hg38_21_+_8433172_8446622_RNA45SN1_45S_50ntFlanks@6787.6806.20\|ACGCCTGTCTGAGCGTCGCT | >hg38_21_+_8433172_8446622_RNA45SN1_45S_50ntFlanks | 6787 | 6806 | 20 | ACGCCTGTCTGAGCGTCGCT | 2704.4 | 5.5 | 0.7 | 7.6 | 4.1E-14 | 1.1E-13 | BT20/MB468 |
| >hg38_21_+_8433172_8446622_RNA45SN1_45S_50ntFlanks@10821.10838.18\|GGAAGTCGGCAAGCCGGA | >hg38_21_+_8433172_8446622_RNA45SN1_45S_50ntFlanks | 10821 | 10838 | 18 | GGAAGTCGGCAAGCCGGA | 596.9 | -4.6 | 0.6 | -7.6 | 4.1E-14 | 1.1E-13 | BT20/MB468 |
| >hg38_21_+_8433172_8446622_RNA45SN1_45S_50ntFlanks@10866.10902.37\|CTAAGGGCTGGGTCGGTCGGGCTGGGGCGCGAAGCGG | >hg38_21_+_8433172_8446622_RNA45SN1_45S_50ntFlanks | 10866 | 10902 | 37 | CTAAGGGCTGGGTCGGTCGGGCTGGGGCGCGAAGCGG | 320.6 | -9.9 | 1.3 | -7.6 | 3.9E-14 | 1.1E-13 | MB231/MB468 |
| >hg38_21_+_8433172_8446622_RNA45SN1_45S_50ntFlanks@4761.4780.20\|CAGATACCGTCGTAGTTCCG | >hg38_21_+_8433172_8446622_RNA45SN1_45S_50ntFlanks | 4761 | 4780 | 20 | CAGATACCGTCGTAGTTCCG | 227.9 | -9.8 | 1.3 | -7.6 | 4.1E-14 | 1.1E-13 | BT20/MB468 |
| >MI0000286\|hsa-mir-210&WithFlank&11\|-\|568083\|568204@72.93.22(+1A)&[MIMAT0000267&hsa-miR-210-3p&offsets\|0\|0(+1A);m-102&11\|-\|568112\|568133&offsets\|0\|0(+1A)]\|CTGTGCGTGTGACAGCGGCTGAA | >MI0000286\|hsa-mir-210&WithFlank&11\|-\|568083\|568204 | 72 | 93 | 22(+1A) | CTGTGCGTGTGACAGCGGCTGAA | 13709.7 | 12.8 | 1.7 | 7.6 | 4.1E-14 | 1.1E-13 | BT20/MB468 |
| >hg38_21_+_8433172_8446622_RNA45SN1_45S_50ntFlanks@12456.12482.27\|CCAAGCGTTGGATTGTTCACCCACTAA | >hg38_21_+_8433172_8446622_RNA45SN1_45S_50ntFlanks | 12456 | 12482 | 27 | CCAAGCGTTGGATTGTTCACCCACTAA | 178.1 | -9.5 | 1.3 | -7.6 | 4.1E-14 | 1.1E-13 | BT20/MB468 |
| >hg38_21_+_8433172_8446622_RNA45SN1_45S_50ntFlanks@9504.9521.18\|GTGAACTATGCCTGGGCA | >hg38_21_+_8433172_8446622_RNA45SN1_45S_50ntFlanks | 9504 | 9521 | 18 | GTGAACTATGCCTGGGCA | 1290.1 | -6.2 | 0.8 | -7.6 | 4.1E-14 | 1.1E-13 | BT20/MB468 |
| >MI0000065\|hsa-let-7d&WithFlank&9\|+\|94178828\|94178926@14.34.21(+1G)&[MIMAT0000065&hsa-let-7d-5p&offsets\|0\|-1(+1G);m-90&9\|+\|94178841\|94178862&offsets\|0\|-1(+1G)]\|AGAGGTAGTAGGTTGCATAGTG | >MI0000065\|hsa-let-7d&WithFlank&9\|+\|94178828\|94178926 | 14 | 34 | 21(+1G) | AGAGGTAGTAGGTTGCATAGTG | 1557.4 | 9.7 | 1.3 | 7.6 | 4.1E-14 | 1.1E-13 | BT20/MB468 |
| >hg38_21_+_8433172_8446622_RNA45SN1_45S_50ntFlanks@10786.10822.37\|AACAGCCTCTGGCATGTTGGAACAATGTAGGTAAGGG | >hg38_21_+_8433172_8446622_RNA45SN1_45S_50ntFlanks | 10786 | 10822 | 37 | AACAGCCTCTGGCATGTTGGAACAATGTAGGTAAGGG | 461.7 | -3.4 | 0.4 | -7.6 | 3.9E-14 | 1.1E-13 | MB231/MB468 |
| >MI0000063\|hsa-let-7b&WithFlank&22\|+\|46113680\|46113774@12.31.20(+1C)&[MIMAT0000063&hsa-let-7b-5p&offsets\|0\|-2(+1C);m-14&22\|+\|46113691\|46113712&offsets\|0\|-2(+1C)]\|TGAGGTAGTAGGTTGTGTGGC | >MI0000063\|hsa-let-7b&WithFlank&22\|+\|46113680\|46113774 | 12 | 31 | 20(+1C) | TGAGGTAGTAGGTTGTGTGGC | 1464.4 | 8.4 | 1.1 | 7.6 | 4.1E-14 | 1.1E-13 | BT20/MB468 |
| >hg38_21_+_8433172_8446622_RNA45SN1_45S_50ntFlanks@12936.12966.31\|AAACCATTCGTAGACGACCTGCTTCTGGGTC | >hg38_21_+_8433172_8446622_RNA45SN1_45S_50ntFlanks | 12936 | 12966 | 31 | AAACCATTCGTAGACGACCTGCTTCTGGGTC | 239.2 | -9.9 | 1.3 | -7.6 | 4.2E-14 | 1.1E-13 | BT20/MB468 |
| >hg38_21_+_8433172_8446622_RNA45SN1_45S_50ntFlanks@12488.12520.33\|AACGTGAGCTGGGTTTAGACCGTCGTGAGACAG | >hg38_21_+_8433172_8446622_RNA45SN1_45S_50ntFlanks | 12488 | 12520 | 33 | AACGTGAGCTGGGTTTAGACCGTCGTGAGACAG | 370.7 | -6.7 | 0.9 | -7.6 | 4.2E-14 | 1.1E-13 | BT20/MB468 |
| >hg38_21_+_8433172_8446622_RNA45SN1_45S_50ntFlanks@10425.10453.29\|AGATGGGCGAGCGCCGTTCCGAAGGGACG | >hg38_21_+_8433172_8446622_RNA45SN1_45S_50ntFlanks | 10425 | 10453 | 29 | AGATGGGCGAGCGCCGTTCCGAAGGGACG | 169.2 | -5.7 | 0.8 | -7.6 | 4.2E-14 | 1.1E-13 | BT20/MB468 |
| >hg38_21_+_8433172_8446622_RNA45SN1_45S_50ntFlanks@12456.12473.18\|CCAAGCGTTGGATTGTTC | >hg38_21_+_8433172_8446622_RNA45SN1_45S_50ntFlanks | 12456 | 12473 | 18 | CCAAGCGTTGGATTGTTC | 6690.7 | -6.5 | 0.9 | -7.6 | 4.2E-14 | 1.1E-13 | BT20/MB468 |
| >hg38_21_+_8433172_8446622_RNA45SN1_45S_50ntFlanks@10808.10828.21\|CAATGTAGGTAAGGGAAGTCG | >hg38_21_+_8433172_8446622_RNA45SN1_45S_50ntFlanks | 10808 | 10828 | 21 | CAATGTAGGTAAGGGAAGTCG | 493.6 | -7.0 | 0.9 | -7.6 | 4.2E-14 | 1.1E-13 | BT20/MB468 |
| >hg38_21_+_8433172_8446622_RNA45SN1_45S_50ntFlanks@10843.10868.26\|TAACTTCGGGATAAGGATTGGCTCTA | >hg38_21_+_8433172_8446622_RNA45SN1_45S_50ntFlanks | 10843 | 10868 | 26 | TAACTTCGGGATAAGGATTGGCTCTA | 660.8 | -5.6 | 0.7 | -7.6 | 4.0E-14 | 1.1E-13 | MB231/MB468 |
| >hg38_21_+_8433172_8446622_RNA45SN1_45S_50ntFlanks@10513.10533.21\|CCGGAGTGGCGGAGATGGGCG | >hg38_21_+_8433172_8446622_RNA45SN1_45S_50ntFlanks | 10513 | 10533 | 21 | CCGGAGTGGCGGAGATGGGCG | 764.5 | -6.3 | 0.8 | -7.6 | 4.0E-14 | 1.1E-13 | MB231/MB468 |
| >MI0001445\|hsa-mir-423&WithFlank&17\|+\|30117073\|30117178@59.82.24&[MIMAT0001340&hsa-miR-423-3p&offsets\|0\|+1;m-100&17\|+\|30117131\|30117153&offsets\|0\|+1]\|AGCTCGGTCTGAGGCCCCTCAGTC | >MI0001445\|hsa-mir-423&WithFlank&17\|+\|30117073\|30117178 | 59 | 82 | 24 | AGCTCGGTCTGAGGCCCCTCAGTC | 390.4 | 8.2 | 1.1 | 7.6 | 4.3E-14 | 1.1E-13 | BT20/MB468 |
| >hg38_21_+_8433172_8446622_RNA45SN1_45S_50ntFlanks@5125.5158.34\|AGCGGTCGGCGTCCCCCAACTTCTTAGAGGGACA | >hg38_21_+_8433172_8446622_RNA45SN1_45S_50ntFlanks | 5125 | 5158 | 34 | AGCGGTCGGCGTCCCCCAACTTCTTAGAGGGACA | 502.5 | 7.6 | 1.0 | 7.5 | 4.4E-14 | 1.1E-13 | BT20/MB468 |
| >hg38_21_+_8433172_8446622_RNA45SN1_45S_50ntFlanks@10393.10420.28\|CAGCAGTTGAACATGGGTCAGTCGGTCC | >hg38_21_+_8433172_8446622_RNA45SN1_45S_50ntFlanks | 10393 | 10420 | 28 | CAGCAGTTGAACATGGGTCAGTCGGTCC | 176.1 | -9.5 | 1.3 | -7.5 | 4.4E-14 | 1.1E-13 | BT20/MB468 |
| >hg38_MT_+_1621_3279_RNR2_16S_50ntFlanks@1428.1458.31\|ATTAAAGTCCTACGTGATCTGAGTTCAGACC | >hg38_MT_+_1621_3279_RNR2_16S_50ntFlanks | 1428 | 1458 | 31 | ATTAAAGTCCTACGTGATCTGAGTTCAGACC | 1766.1 | 9.6 | 1.3 | 7.6 | 4.1E-14 | 1.1E-13 | MB231/MB468 |
| >hg38_21_+_8433172_8446622_RNA45SN1_45S_50ntFlanks@12942.12965.24\|TTCGTAGACGACCTGCTTCTGGGT | >hg38_21_+_8433172_8446622_RNA45SN1_45S_50ntFlanks | 12942 | 12965 | 24 | TTCGTAGACGACCTGCTTCTGGGT | 337.3 | -3.5 | 0.5 | -7.6 | 4.1E-14 | 1.1E-13 | MB231/MB468 |
| >hg38_21_+_8433172_8446622_RNA45SN1_45S_50ntFlanks@4472.4494.23\|CTTAGCTGAGTGTCCCGCGGGGC | >hg38_21_+_8433172_8446622_RNA45SN1_45S_50ntFlanks | 4472 | 4494 | 23 | CTTAGCTGAGTGTCCCGCGGGGC | 302.7 | -9.8 | 1.3 | -7.6 | 4.1E-14 | 1.1E-13 | MB231/MB468 |
| >hg38_21_+_8433172_8446622_RNA45SN1_45S_50ntFlanks@11873.11899.27\|AAGAAGACCCTGTTGAGCTTGACTCTA | >hg38_21_+_8433172_8446622_RNA45SN1_45S_50ntFlanks | 11873 | 11899 | 27 | AAGAAGACCCTGTTGAGCTTGACTCTA | 244.6 | -4.0 | 0.5 | -7.6 | 4.1E-14 | 1.1E-13 | MB231/MB468 |
| >hg38_21_+_8433172_8446622_RNA45SN1_45S_50ntFlanks@12635.12657.23\|AAGCTACCATCTGTGGGATTATG | >hg38_21_+_8433172_8446622_RNA45SN1_45S_50ntFlanks | 12635 | 12657 | 23 | AAGCTACCATCTGTGGGATTATG | 589.6 | -4.1 | 0.5 | -7.6 | 4.1E-14 | 1.1E-13 | MB231/MB468 |
| >hg38_21_+_8433172_8446622_RNA45SN1_45S_50ntFlanks@11208.11234.27\|GCGGCGACTCTGGACGCGAGCCGGGCC | >hg38_21_+_8433172_8446622_RNA45SN1_45S_50ntFlanks | 11208 | 11234 | 27 | GCGGCGACTCTGGACGCGAGCCGGGCC | 1895.1 | -7.4 | 1.0 | -7.6 | 4.2E-14 | 1.1E-13 | MB231/MB468 |
| >hg38_21_+_8433172_8446622_RNA45SN1_45S_50ntFlanks@3705.3739.35\|TACCTGGTTGATCCTGCCAGTAGCATATGCTTGTC | >hg38_21_+_8433172_8446622_RNA45SN1_45S_50ntFlanks | 3705 | 3739 | 35 | TACCTGGTTGATCCTGCCAGTAGCATATGCTTGTC | 522.7 | 4.1 | 0.5 | 7.6 | 4.2E-14 | 1.2E-13 | MB231/MB468 |
| >hg38_21_+_8433172_8446622_RNA45SN1_45S_50ntFlanks@10386.10424.39\|ATGTGAACAGCAGTTGAACATGGGTCAGTCGGTCCTGAG | >hg38_21_+_8433172_8446622_RNA45SN1_45S_50ntFlanks | 10386 | 10424 | 39 | ATGTGAACAGCAGTTGAACATGGGTCAGTCGGTCCTGAG | 1125.7 | 8.3 | 1.1 | 7.6 | 4.2E-14 | 1.2E-13 | MB231/MB468 |
| >MI0000273\|hsa-mir-183&WithFlank&7\|-\|129774899\|129775020@33.55.23&[MIMAT0000261&hsa-miR-183-5p&offsets\|0\|+1;m-41&7\|-\|129774966\|129774987&offsets\|-1\|0]\|TATGGCACTGGTAGAATTCACTG | >MI0000273\|hsa-mir-183&WithFlank&7\|-\|129774899\|129775020 | 33 | 55 | 23 | TATGGCACTGGTAGAATTCACTG | 2281.2 | 5.5 | 0.7 | 7.6 | 4.2E-14 | 1.2E-13 | MB231/MB468 |
| >MI0000103\|hsa-mir-101-1&WithFlank&1\|-\|65058428\|65058514@52.72.21;MI0000739\|hsa-mir-101-2&WithFlank&9\|+\|4850291\|4850381@54.74.21&[MIMAT0000099&hsa-miR-101-3p&offsets\|-1\|-1;m-17&1\|-\|65058443\|65058462&offsets\|-1\|0];[MIMAT0000099_1&hsa-miR-101-3p&offsets\|-1\|-1;m-16&9\|+\|4850345\|4850364&offsets\|-1\|0]\|GTACAGTACTGTGATAACTGA | >MI0000103\|hsa-mir-101-1&WithFlank&1\|-\|65058428\|65058514 | 52 | 72 | 21 | GTACAGTACTGTGATAACTGA | 12772.6 | 8.7 | 1.2 | 7.5 | 4.5E-14 | 1.2E-13 | BT20/MB468 |
| >hg38_21_+_8433172_8446622_RNA45SN1_45S_50ntFlanks@11821.11842.22\|CTATCCAGCGAAACCACAGCCA | >hg38_21_+_8433172_8446622_RNA45SN1_45S_50ntFlanks | 11821 | 11842 | 22 | CTATCCAGCGAAACCACAGCCA | 157.1 | -9.3 | 1.2 | -7.5 | 4.5E-14 | 1.2E-13 | BT20/MB468 |
| >hg38_21_+_8433172_8446622_RNA45SN1_45S_50ntFlanks@11580.11606.27\|GAACTGGTGCGGACCAGGGGAATCCGA | >hg38_21_+_8433172_8446622_RNA45SN1_45S_50ntFlanks | 11580 | 11606 | 27 | GAACTGGTGCGGACCAGGGGAATCCGA | 144.8 | -4.5 | 0.6 | -7.5 | 4.5E-14 | 1.2E-13 | BT20/MB468 |
| >MI0000285\|hsa-mir-205&WithFlank&1\|+\|209432127\|209432248@40.60.21(+1C)&[MIMAT0000266&hsa-miR-205-5p&offsets\|0\|-1(+1C);m-351&1\|+\|209432166\|209432187&offsets\|0\|-1(+1C)]\|TCCTTCATTCCACCGGAGTCTC | >MI0000285\|hsa-mir-205&WithFlank&1\|+\|209432127\|209432248 | 40 | 60 | 21(+1C) | TCCTTCATTCCACCGGAGTCTC | 205.1 | 11.2 | 1.4 | 7.9 | 3.1E-15 | 1.2E-13 | BT20/MB231 |
| >hg38_MT_+_1621_3279_RNR2_16S_50ntFlanks@51.88.38\|GCTAAACCTAGCCCCAAACCCACTCCACCTTACTACCA | >hg38_MT_+_1621_3279_RNR2_16S_50ntFlanks | 51 | 88 | 38 | GCTAAACCTAGCCCCAAACCCACTCCACCTTACTACCA | 1089.8 | 8.6 | 1.1 | 7.5 | 4.6E-14 | 1.2E-13 | BT20/MB468 |
| >hg38_21_+_8433172_8446622_RNA45SN1_45S_50ntFlanks@10572.10589.18\|AGAAGCCGGCGGGAGCCC | >hg38_21_+_8433172_8446622_RNA45SN1_45S_50ntFlanks | 10572 | 10589 | 18 | AGAAGCCGGCGGGAGCCC | 160.9 | -6.2 | 0.8 | -7.5 | 4.7E-14 | 1.2E-13 | BT20/MB468 |
| >MI0000285\|hsa-mir-205&WithFlank&1\|+\|209432127\|209432248@40.61.22(+1A)&[MIMAT0000266&hsa-miR-205-5p&offsets\|0\|0(+1A);m-351&1\|+\|209432166\|209432187&offsets\|0\|0(+1A)]\|TCCTTCATTCCACCGGAGTCTGA | >MI0000285\|hsa-mir-205&WithFlank&1\|+\|209432127\|209432248 | 40 | 61 | 22(+1A) | TCCTTCATTCCACCGGAGTCTGA | 1339.3 | -7.8 | 1.0 | -7.5 | 4.4E-14 | 1.2E-13 | MB231/MB468 |
| >hg38_21_+_8433172_8446622_RNA45SN1_45S_50ntFlanks@12624.12659.36\|CCAATGGGGCGAAGCTACCATCTGTGGGATTATGAC | >hg38_21_+_8433172_8446622_RNA45SN1_45S_50ntFlanks | 12624 | 12659 | 36 | CCAATGGGGCGAAGCTACCATCTGTGGGATTATGAC | 197.5 | -9.6 | 1.3 | -7.5 | 4.7E-14 | 1.2E-13 | BT20/MB468 |
| >hg38_21_+_8433172_8446622_RNA45SN1_45S_50ntFlanks@4759.4780.22\|ATCAGATACCGTCGTAGTTCCG | >hg38_21_+_8433172_8446622_RNA45SN1_45S_50ntFlanks | 4759 | 4780 | 22 | ATCAGATACCGTCGTAGTTCCG | 6191.0 | -8.5 | 1.1 | -7.5 | 4.4E-14 | 1.2E-13 | MB231/MB468 |
| >hg38_21_+_8433172_8446622_RNA45SN1_45S_50ntFlanks@5421.5445.25\|CGATTGGATGGTTTAGTGAGGCCCT | >hg38_21_+_8433172_8446622_RNA45SN1_45S_50ntFlanks | 5421 | 5445 | 25 | CGATTGGATGGTTTAGTGAGGCCCT | 224.7 | -9.4 | 1.2 | -7.5 | 4.5E-14 | 1.2E-13 | MB231/MB468 |
| >hg38_21_+_8433172_8446622_RNA45SN1_45S_50ntFlanks@4560.4579.20\|CTAGGAATAATGGAATAGGA | >hg38_21_+_8433172_8446622_RNA45SN1_45S_50ntFlanks | 4560 | 4579 | 20 | CTAGGAATAATGGAATAGGA | 246.4 | -9.9 | 1.3 | -7.5 | 4.8E-14 | 1.2E-13 | BT20/MB468 |

<table>
<tr><th></th><th></th><th></th><th></th><th></th><th></th><th></th><th></th><th></th><th></th><th></th><th></th><th></th></tr>
<tr><td>>MI0000077|hsa-mir-21&WithFlank&17|+|59841260|59841343@14.34.21(+1C)&[MIMAT0000076&hsa-miR-21-5p&offsets|0|-1(+1C);m-1&17|+|59841273|59841294&offsets|0|-1(+1C)]|TAGCTTATCAGACTGATGTTGC</td><td>>MI0000077|hsa-mir-21&WithFlank&17|+|59841260|59841343</td><td>14</td><td>34</td><td>21(+1C)</td><td>TAGCTTATCAGACTGATGTTGC</td><td>3520.9</td><td>10.2</td><td>1.4</td><td>7.5</td><td>4.8E-14</td><td>1.2E-13</td><td>BT20/MB468</td></tr>
<tr><td>>hg38_21_+_8433172_8446622_RNA45SN1_45S_50ntFlanks@5467.5484.18|GGCCCACGGCCCTGGCGG</td><td>>hg38_21_+_8433172_8446622_RNA45SN1_45S_50ntFlanks</td><td>5467</td><td>5484</td><td>18</td><td>GGCCCACGGCCCTGGCGG</td><td>208.1</td><td>-9.7</td><td>1.3</td><td>-7.5</td><td>4.8E-14</td><td>1.2E-13</td><td>BT20/MB468</td></tr>
<tr><td>>hg38_21_+_8433172_8446622_RNA45SN1_45S_50ntFlanks@12433.12455.23|CTATCATTGTGAAGCAGAATTCA</td><td>>hg38_21_+_8433172_8446622_RNA45SN1_45S_50ntFlanks</td><td>12433</td><td>12455</td><td>23</td><td>CTATCATTGTGAAGCAGAATTCA</td><td>186.2</td><td>-9.5</td><td>1.3</td><td>-7.5</td><td>4.8E-14</td><td>1.2E-13</td><td>BT20/MB468</td></tr>
<tr><td>>hg38_21_+_8433172_8446622_RNA45SN1_45S_50ntFlanks@10392.10430.39|ACAGCAGTTGAACATGGGTCAGTCGGTCCTGAGAGATGG</td><td>>hg38_21_+_8433172_8446622_RNA45SN1_45S_50ntFlanks</td><td>10392</td><td>10430</td><td>39</td><td>ACAGCAGTTGAACATGGGTCAGTCGGTCCTGAGAGATGG</td><td>657.3</td><td>-11.4</td><td>1.5</td><td>-7.5</td><td>4.8E-14</td><td>1.2E-13</td><td>BT20/MB468</td></tr>
<tr><td>>hg38_MT_+_1621_3279_RNR2_16S_50ntFlanks@51.80.30|GCTAAACCTAGCCCCAAACCCACTCCACCT</td><td>>hg38_MT_+_1621_3279_RNR2_16S_50ntFlanks</td><td>51</td><td>80</td><td>30</td><td>GCTAAACCTAGCCCCAAACCCACTCCACCT</td><td>467.8</td><td>8.6</td><td>1.1</td><td>7.5</td><td>4.9E-14</td><td>1.2E-13</td><td>BT20/MB468</td></tr>
<tr><td>>hg38_21_+_8433172_8446622_RNA45SN1_45S_50ntFlanks@9534.9554.21|AGGAAACTCTGGTGGAGGTCC</td><td>>hg38_21_+_8433172_8446622_RNA45SN1_45S_50ntFlanks</td><td>9534</td><td>9554</td><td>21</td><td>AGGAAACTCTGGTGGAGGTCC</td><td>340.5</td><td>-10.4</td><td>1.4</td><td>-7.5</td><td>4.9E-14</td><td>1.3E-13</td><td>BT20/MB468</td></tr>
<tr><td>>hg38_1_-_228634819_228635039_RNA5S12_5S_50ntFlanks@51.86.36|GTCTACGGCCATACCACCCTGAACGCGCCCGATCTC</td><td>>hg38_1_-_228634819_228635039_RNA5S12_5S_50ntFlanks</td><td>51</td><td>86</td><td>36</td><td>GTCTACGGCCATACCACCCTGAACGCGCCCGATCTC</td><td>632.3</td><td>8.1</td><td>1.1</td><td>7.5</td><td>4.9E-14</td><td>1.3E-13</td><td>BT20/MB468</td></tr>
<tr><td>>hg38_21_+_8433172_8446622_RNA45SN1_45S_50ntFlanks@12539.12569.31|ATGATGTGTTGTTGCCATGGTAATCCTGCTC</td><td>>hg38_21_+_8433172_8446622_RNA45SN1_45S_50ntFlanks</td><td>12539</td><td>12569</td><td>31</td><td>ATGATGTGTTGTTGCCATGGTAATCCTGCTC</td><td>1119.0</td><td>-7.1</td><td>0.9</td><td>-7.5</td><td>4.6E-14</td><td>1.3E-13</td><td>MB231/MB468</td></tr>
<tr><td>>hg38_21_+_8433172_8446622_RNA45SN1_45S_50ntFlanks@11885.11912.28|TTGAGCTTGACTCTAGTCTGGCACGGTG</td><td>>hg38_21_+_8433172_8446622_RNA45SN1_45S_50ntFlanks</td><td>11885</td><td>11912</td><td>28</td><td>TTGAGCTTGACTCTAGTCTGGCACGGTG</td><td>162.5</td><td>-9.3</td><td>1.2</td><td>-7.5</td><td>4.9E-14</td><td>1.3E-13</td><td>BT20/MB468</td></tr>
<tr><td>>hg38_21_+_8433172_8446622_RNA45SN1_45S_50ntFlanks@10510.10533.24|AATCCGGAGTGGCGGAGATGGGCG</td><td>>hg38_21_+_8433172_8446622_RNA45SN1_45S_50ntFlanks</td><td>10510</td><td>10533</td><td>24</td><td>AATCCGGAGTGGCGGAGATGGGCG</td><td>559.5</td><td>-6.5</td><td>0.9</td><td>-7.5</td><td>4.6E-14</td><td>1.3E-13</td><td>MB231/MB468</td></tr>
<tr><td>>hg38_21_+_8433172_8446622_RNA45SN1_45S_50ntFlanks@12003.12032.30|GCGGGCCGCCGGTGAAATACCACTACTCTG</td><td>>hg38_21_+_8433172_8446622_RNA45SN1_45S_50ntFlanks</td><td>12003</td><td>12032</td><td>30</td><td>GCGGGCCGCCGGTGAAATACCACTACTCTG</td><td>1350.2</td><td>-4.4</td><td>0.6</td><td>-7.5</td><td>5.0E-14</td><td>1.3E-13</td><td>BT20/MB468</td></tr>
<tr><td>>hg38_21_+_8433172_8446622_RNA45SN1_45S_50ntFlanks@12419.12445.27|GATGTCGGCTCTTCCTATCATTGTGAA</td><td>>hg38_21_+_8433172_8446622_RNA45SN1_45S_50ntFlanks</td><td>12419</td><td>12445</td><td>27</td><td>GATGTCGGCTCTTCCTATCATTGTGAA</td><td>449.6</td><td>4.2</td><td>0.6</td><td>7.5</td><td>4.7E-14</td><td>1.3E-13</td><td>MB231/MB468</td></tr>
<tr><td>>hg38_1_-_228634819_228635039_RNA5S12_5S_50ntFlanks@51.87.37|GTCTACGGCCATACCACCCTGAACGCGCCCGATCTCG</td><td>>hg38_1_-_228634819_228635039_RNA5S12_5S_50ntFlanks</td><td>51</td><td>87</td><td>37</td><td>GTCTACGGCCATACCACCCTGAACGCGCCCGATCTCG</td><td>1918.2</td><td>11.7</td><td>1.6</td><td>7.5</td><td>5.0E-14</td><td>1.3E-13</td><td>BT20/MB468</td></tr>
<tr><td>>hg38_21_+_8433172_8446622_RNA45SN1_45S_50ntFlanks@12457.12487.31|CAAGCGTTGGATTGTTCACCCACTAATAGGG</td><td>>hg38_21_+_8433172_8446622_RNA45SN1_45S_50ntFlanks</td><td>12457</td><td>12487</td><td>31</td><td>CAAGCGTTGGATTGTTCACCCACTAATAGGG</td><td>1931.7</td><td>-8.5</td><td>1.1</td><td>-7.5</td><td>5.1E-14</td><td>1.3E-13</td><td>BT20/MB468</td></tr>
<tr><td>>hg38_21_+_8433172_8446622_RNA45SN1_45S_50ntFlanks@13004.13023.20|ATCTATTGAAAGTCAGCCCT</td><td>>hg38_21_+_8433172_8446622_RNA45SN1_45S_50ntFlanks</td><td>13004</td><td>13023</td><td>20</td><td>ATCTATTGAAAGTCAGCCCT</td><td>164.9</td><td>-9.4</td><td>1.2</td><td>-7.5</td><td>5.1E-14</td><td>1.3E-13</td><td>BT20/MB468</td></tr>
<tr><td>>MI0000272|hsa-mir-182&WithFlank&7|-|129770377|129770498@29.49.21(+1G)&[MIMAT0000259&hsa-miR-182-5p&offsets|0|-3(+1G);m-22&7|-|129770449|129770470&offsets|0|-1(+1G)]|TTTGGCAATGGTAGAACTCACG</td><td>>MI0000272|hsa-mir-182&WithFlank&7|-|129770377|129770498</td><td>29</td><td>49</td><td>21(+1G)</td><td>TTTGGCAATGGTAGAACTCACG</td><td>1476.2</td><td>10.9</td><td>1.5</td><td>7.5</td><td>5.1E-14</td><td>1.3E-13</td><td>BT20/MB468</td></tr>
<tr><td>>MI0000081|hsa-mir-24-2&WithFlank&19|-|13836281|13836365@56.76.21;MI0000080|hsa-mir-24-1&WithFlank&9|+|95086015|95086094@50.70.21&[MIMAT0000080&hsa-miR-24-3p&offsets|0|-1;m-36&19|-|13836288|13836310&offsets|0|-2];[MIMAT0000080_1&hsa-miR-24-3p&offsets|0|-1;m-35&9|+|95086064|95086086&offsets|0|-2]|TGGCTCAGTTCAGCAGGAACA</td><td>>MI0000081|hsa-mir-24-2&WithFlank&19|-|13836281|13836365</td><td>56</td><td>76</td><td>21</td><td>TGGCTCAGTTCAGCAGGAACA</td><td>8032.9</td><td>5.4</td><td>0.7</td><td>7.5</td><td>4.8E-14</td><td>1.3E-13</td><td>MB231/MB468</td></tr>
<tr><td>>am_trna2_ArgCCT_16_+_3202901_3202973@14.31.18,trna3_ArgCCT_7_+_139025446_139025518@14.31.18|AATGGATAAGGCATTGGC</td><td>>am_trna2_ArgCCT_16_+_3202901_3202973</td><td>14</td><td>31</td><td>18</td><td>AATGGATAAGGCATTGGC</td><td>276.7</td><td>-10.1</td><td>1.3</td><td>-7.5</td><td>5.1E-14</td><td>1.3E-13</td><td>BT20/MB468</td></tr>
<tr><td>>hg38_21_+_8433172_8446622_RNA45SN1_45S_50ntFlanks@13002.13025.24|CGATCTATTGAAAGTCAGCCCTCG</td><td>>hg38_21_+_8433172_8446622_RNA45SN1_45S_50ntFlanks</td><td>13002</td><td>13025</td><td>24</td><td>CGATCTATTGAAAGTCAGCCCTCG</td><td>247.3</td><td>-9.5</td><td>1.3</td><td>-7.5</td><td>4.8E-14</td><td>1.3E-13</td><td>MB231/MB468</td></tr>
<tr><td>>hg38_21_+_8433172_8446622_RNA45SN1_45S_50ntFlanks@10681.10711.31|TTCCGGCGGCGTCCGGTGAGCTCTCGCTGGC</td><td>>hg38_21_+_8433172_8446622_RNA45SN1_45S_50ntFlanks</td><td>10681</td><td>10711</td><td>31</td><td>TTCCGGCGGCGTCCGGTGAGCTCTCGCTGGC</td><td>967.2</td><td>-6.5</td><td>0.9</td><td>-7.5</td><td>5.2E-14</td><td>1.3E-13</td><td>BT20/MB468</td></tr>
<tr><td>>hg38_21_+_8433172_8446622_RNA45SN1_45S_50ntFlanks@6733.6771.39|CATTGATCATCGACACTTCGAACGCACTTGCGGCCCCGG</td><td>>hg38_21_+_8433172_8446622_RNA45SN1_45S_50ntFlanks</td><td>6733</td><td>6771</td><td>39</td><td>CATTGATCATCGACACTTCGAACGCACTTGCGGCCCCGG</td><td>793.0</td><td>-3.9</td><td>0.5</td><td>-7.5</td><td>5.2E-14</td><td>1.3E-13</td><td>BT20/MB468</td></tr>
<tr><td>>trna111_HisGTG_1_-_147774845_147774916@-1T.33.34,trna118_HisGTG_1_-_145396881_145396952@-1T.33.34,trna16_HisGTG_1_+_146544773_146544844@-1T.33.34,trna1_HisGTG_15_+_45493349_45493420@-1T.33.34,trna21_HisGTG_1_+_147753471_147753542@-1T.33.34,trna33_HisGTG_6_+_27125906_27125977@-1T.33.34,trna7_HisGTG_9_-_14433938_14434009@-1T.33.34,trna8_HisGTG_15_-_45492611_45492682@-1T.33.34,trna9_HisGTG_15_-_45490804_45490875@-1T.33.34|TGCCGTGATCGTATAGTGGTTAGTACTCTGCGTT</td><td>>trna111_HisGTG_1_-_147774845_147774916</td><td>-1T</td><td>33</td><td>34</td><td>TGCCGTGATCGTATAGTGGTTAGTACTCTGCGTT</td><td>1165.8</td><td>8.4</td><td>1.1</td><td>7.5</td><td>5.2E-14</td><td>1.3E-13</td><td>BT20/MB468</td></tr>
<tr><td>>MI0000084|hsa-mir-26b&WithFlank&2|+|218402640|218402728@18.38.21&[MIMAT0000083&hsa-miR-26b-5p&offsets|0|0;m-53&2|+|218402657|218402678&offsets|0|-1]|TTCAAGTAATTCAGGATAGGT</td><td>>MI0000084|hsa-mir-26b&WithFlank&2|+|218402640|218402728</td><td>18</td><td>38</td><td>21</td><td>TTCAAGTAATTCAGGATAGGT</td><td>2215.3</td><td>6.0</td><td>0.8</td><td>7.5</td><td>4.9E-14</td><td>1.3E-13</td><td>MB231/MB468</td></tr>
<tr><td>>MI0000734|hsa-mir-106b&WithFlank&7|-|100093987|100094080@18.39.22&[MIMAT0000680&hsa-miR-106b-5p&offsets|0|+1;m-103&7|-|100094043|100094063&offsets|0|+1]|TAAAGTGCTGACAGTGCAGATA</td><td>>MI0000734|hsa-mir-106b&WithFlank&7|-|100093987|100094080</td><td>18</td><td>39</td><td>22</td><td>TAAAGTGCTGACAGTGCAGATA</td><td>3634.3</td><td>6.1</td><td>0.8</td><td>7.5</td><td>4.9E-14</td><td>1.3E-13</td><td>MB231/MB468</td></tr>
<tr><td>>MI0000085|hsa-mir-27a&WithFlank&19|-|13836434|13836523@57.77.21&[MIMAT0000084&hsa-miR-27a-3p&offsets|0|0;m-43&19|-|13836447|13836467&offsets|0|0]|TTCACAGTGGCTAAGTTCCGC</td><td>>MI0000085|hsa-mir-27a&WithFlank&19|-|13836434|13836523</td><td>57</td><td>77</td><td>21</td><td>TTCACAGTGGCTAAGTTCCGC</td><td>6969.8</td><td>6.0</td><td>0.8</td><td>7.5</td><td>4.9E-14</td><td>1.3E-13</td><td>MB231/MB468</td></tr>
<tr><td>>hg38_MT_+_1621_3279_RNR2_16S_50ntFlanks@1370.1388.19|ATGTTGGATCAGGACATCC</td><td>>hg38_MT_+_1621_3279_RNR2_16S_50ntFlanks</td><td>1370</td><td>1388</td><td>19</td><td>ATGTTGGATCAGGACATCC</td><td>1560.0</td><td>9.8</td><td>1.3</td><td>7.5</td><td>4.9E-14</td><td>1.3E-13</td><td>MB231/MB468</td></tr>
<tr><td>>hg38_21_+_8433172_8446622_RNA45SN1_45S_50ntFlanks@10808.10829.22|CAATGTAGGTAAGGGAAGTCGG</td><td>>hg38_21_+_8433172_8446622_RNA45SN1_45S_50ntFlanks</td><td>10808</td><td>10829</td><td>22</td><td>CAATGTAGGTAAGGGAAGTCGG</td><td>184.5</td><td>-9.5</td><td>1.3</td><td>-7.5</td><td>5.3E-14</td><td>1.4E-13</td><td>BT20/MB468</td></tr>
<tr><td>>hg38_21_+_8433172_8446622_RNA45SN1_45S_50ntFlanks@10587.10607.21|CCCCGGGGAGAGTTCTCTTTT</td><td>>hg38_21_+_8433172_8446622_RNA45SN1_45S_50ntFlanks</td><td>10587</td><td>10607</td><td>21</td><td>CCCCGGGGAGAGTTCTCTTTT</td><td>281.3</td><td>-9.7</td><td>1.3</td><td>-7.5</td><td>5.0E-14</td><td>1.4E-13</td><td>MB231/MB468</td></tr>
<tr><td>>MI0000469|hsa-mir-125a&WithFlank&19|+|51693248|51693345@21.41.21&[MIMAT0000443&hsa-miR-125a-5p&offsets|0|-3;m-63&19|+|51693268|51693289&offsets|0|-1]|TCCCTGAGACCCTTTAACCTG</td><td>>MI0000469|hsa-mir-125a&WithFlank&19|+|51693248|51693345</td><td>21</td><td>41</td><td>21</td><td>TCCCTGAGACCCTTTAACCTG</td><td>1058.2</td><td>8.6</td><td>1.1</td><td>7.5</td><td>5.3E-14</td><td>1.4E-13</td><td>BT20/MB468</td></tr>
<tr><td>>trnaMT_PheGAA_MT_+_577_647@39.59.21|ATGTTTAGACGGGCTCACATC</td><td>>trnaMT_PheGAA_MT_+_577_647</td><td>39</td><td>59</td><td>21</td><td>ATGTTTAGACGGGCTCACATC</td><td>711.6</td><td>8.8</td><td>1.2</td><td>7.5</td><td>5.4E-14</td><td>1.4E-13</td><td>BT20/MB468</td></tr>
<tr><td>>hg38_21_+_8433172_8446622_RNA45SN1_45S_50ntFlanks@12998.13017.20|GCTGCGATCTATTGAAAGTC</td><td>>hg38_21_+_8433172_8446622_RNA45SN1_45S_50ntFlanks</td><td>12998</td><td>13017</td><td>20</td><td>GCTGCGATCTATTGAAAGTC</td><td>580.0</td><td>8.3</td><td>1.1</td><td>7.5</td><td>5.0E-14</td><td>1.4E-13</td><td>MB231/MB468</td></tr>
<tr><td>>hg38_21_+_8433172_8446622_RNA45SN1_45S_50ntFlanks@6773.6793.21|TTCCTCCCGGGGCTACGCCTG</td><td>>hg38_21_+_8433172_8446622_RNA45SN1_45S_50ntFlanks</td><td>6773</td><td>6793</td><td>21</td><td>TTCCTCCCGGGGCTACGCCTG</td><td>291.2</td><td>-9.8</td><td>1.3</td><td>-7.5</td><td>5.1E-14</td><td>1.4E-13</td><td>MB231/MB468</td></tr>
<tr><td>>hg38_21_+_8433172_8446622_RNA45SN1_45S_50ntFlanks@6646.6672.27|TCGTACGACTCTTAGCGGTGGATCACT</td><td>>hg38_21_+_8433172_8446622_RNA45SN1_45S_50ntFlanks</td><td>6646</td><td>6672</td><td>27</td><td>TCGTACGACTCTTAGCGGTGGATCACT</td><td>2505.6</td><td>5.7</td><td>0.8</td><td>7.5</td><td>5.1E-14</td><td>1.4E-13</td><td>MB231/MB468</td></tr>
<tr><td>>hg38_21_+_8433172_8446622_RNA45SN1_45S_50ntFlanks@4472.4496.25|CTTAGCTGAGTGTCCCGCGGGGCCC</td><td>>hg38_21_+_8433172_8446622_RNA45SN1_45S_50ntFlanks</td><td>4472</td><td>4496</td><td>25</td><td>CTTAGCTGAGTGTCCCGCGGGGCCC</td><td>182.3</td><td>-9.5</td><td>1.3</td><td>-7.5</td><td>5.5E-14</td><td>1.4E-13</td><td>BT20/MB468</td></tr>
<tr><td>>hg38_21_+_8433172_8446622_RNA45SN1_45S_50ntFlanks@9564.9594.31|CTGACGTGCAAATCGGTCGTCCGACCTGGGT</td><td>>hg38_21_+_8433172_8446622_RNA45SN1_45S_50ntFlanks</td><td>9564</td><td>9594</td><td>31</td><td>CTGACGTGCAAATCGGTCGTCCGACCTGGGT</td><td>366.6</td><td>-3.9</td><td>0.5</td><td>-7.5</td><td>5.5E-14</td><td>1.4E-13</td><td>BT20/MB468</td></tr>
<tr><td>>hg38_21_+_8433172_8446622_RNA45SN1_45S_50ntFlanks@4723.4759.37|CATTAATCAAGAACGAAAGTCGGAGGTTCGAAGACGA</td><td>>hg38_21_+_8433172_8446622_RNA45SN1_45S_50ntFlanks</td><td>4723</td><td>4759</td><td>37</td><td>CATTAATCAAGAACGAAAGTCGGAGGTTCGAAGACGA</td><td>678.1</td><td>-5.1</td><td>0.7</td><td>-7.5</td><td>5.5E-14</td><td>1.4E-13</td><td>BT20/MB468</td></tr>
<tr><td>>am_trnaMT_ValTAC_MT_+_1602_1670@51.68.18|CTTAACTTGACCGCTCTG</td><td>>am_trnaMT_ValTAC_MT_+_1602_1670</td><td>51</td><td>68</td><td>18</td><td>CTTAACTTGACCGCTCTG</td><td>13303.0</td><td>-7.1</td><td>0.9</td><td>-7.5</td><td>5.6E-14</td><td>1.4E-13</td><td>BT20/MB468</td></tr>
<tr><td>>MI0000286|hsa-mir-210&WithFlank&11|-|568083|568204@72.94.23&[MIMAT0000267&hsa-miR-210-3p&offsets|0|+1;m-102&11|-|568112|568133&offsets|0|+1]|CTGTGCGTGTGACAGCGGCTGAT</td><td>>MI0000286|hsa-mir-210&WithFlank&11|-|568083|568204</td><td>72</td><td>94</td><td>23</td><td>CTGTGCGTGTGACAGCGGCTGAT</td><td>8577.8</td><td>12.5</td><td>1.7</td><td>7.5</td><td>5.6E-14</td><td>1.4E-13</td><td>BT20/MB468</td></tr>
<tr><td>>hg38_21_+_8433172_8446622_RNA45SN1_45S_50ntFlanks@11315.11342.28|CCACCCCACGTCTCGTCGCGCGCGCGTC</td><td>>hg38_21_+_8433172_8446622_RNA45SN1_45S_50ntFlanks</td><td>11315</td><td>11342</td><td>28</td><td>CCACCCCACGTCTCGTCGCGCGCGCGTC</td><td>219.2</td><td>-9.4</td><td>1.2</td><td>-7.5</td><td>5.3E-14</td><td>1.4E-13</td><td>MB231/MB468</td></tr>
<tr><td>>MI0000739|hsa-mir-101-2&WithFlank&9|+|4850291|4850381@55.77.23&[MIMAT0000099_1&hsa-miR-101-3p&offsets|0|+2;m-16&9|+|4850345|4850364&offsets|0|+3]|TACAGTACTGTGATAACTGAAGA</td><td>>MI0000739|hsa-mir-101-2&WithFlank&9|+|4850291|4850381</td><td>55</td><td>77</td><td>23</td><td>TACAGTACTGTGATAACTGAAGA</td><td>2242.2</td><td>8.2</td><td>1.1</td><td>7.5</td><td>5.7E-14</td><td>1.4E-13</td><td>BT20/MB468</td></tr>
<tr><td>>MI0000098|hsa-mir-96&WithFlank&7|-|129774686|129774775@15.36.22(+1C)&[MIMAT0000095&hsa-miR-96-5p&offsets|0|-1(+1C);m-248&7|-|129774739|129774761&offsets|0|-1(+1C)]|TTTGGCACTAGCACATTTTTGCC</td><td>>MI0000098|hsa-mir-96&WithFlank&7|-|129774686|129774775</td><td>15</td><td>36</td><td>22(+1C)</td><td>TTTGGCACTAGCACATTTTTGCC</td><td>643.4</td><td>5.4</td><td>0.7</td><td>7.5</td><td>5.3E-14</td><td>1.4E-13</td><td>MB231/MB468</td></tr>
<tr><td>>hg38_21_+_8433172_8446622_RNA45SN1_45S_50ntFlanks@12416.12445.30|TTCGATGTCGGCTCTTCCTATCATTGTGAA</td><td>>hg38_21_+_8433172_8446622_RNA45SN1_45S_50ntFlanks</td><td>12416</td><td>12445</td><td>30</td><td>TTCGATGTCGGCTCTTCCTATCATTGTGAA</td><td>955.2</td><td>5.0</td><td>0.7</td><td>7.5</td><td>5.3E-14</td><td>1.4E-13</td><td>MB231/MB468</td></tr>
<tr><td>>hg38_21_+_8433172_8446622_RNA45SN1_45S_50ntFlanks@13006.13025.20|CTATTGAAAGTCAGCCCTCG</td><td>>hg38_21_+_8433172_8446622_RNA45SN1_45S_50ntFlanks</td><td>13006</td><td>13025</td><td>20</td><td>CTATTGAAAGTCAGCCCTCG</td><td>614.0</td><td>-6.3</td><td>0.8</td><td>-7.5</td><td>5.3E-14</td><td>1.5E-13</td><td>MB231/MB468</td></tr>
</table>

<table>
<tr><td>>hg38_21_+_8433172_8446622_RNA45SN1_45S_50ntFlanks@11873.11908.36|AAGAAGACCCTGTTGAGCTTGACTCTAGTCTGGCAC</td><td>>hg38_21_+_8433172_8446622_RNA45SN1_45S_50ntFlanks</td><td>11873</td><td>11908</td><td>36</td><td>AAGAAGACCCTGTTGAGCTTGACTCTAGTCTGGCAC</td><td>232.0</td><td>-6.2</td><td>0.8</td><td>-7.5</td><td>5.7E-14</td><td>1.5E-13</td><td>BT20/MB468</td></tr>
<tr><td>>hg38_21_+_8433172_8446622_RNA45SN1_45S_50ntFlanks@6735.6770.36|TTGATCATCGACACTTCGAACGCACTTGCGGCCCCG</td><td>>hg38_21_+_8433172_8446622_RNA45SN1_45S_50ntFlanks</td><td>6735</td><td>6770</td><td>36</td><td>TTGATCATCGACACTTCGAACGCACTTGCGGCCCCG</td><td>232.0</td><td>-4.7</td><td>0.6</td><td>-7.5</td><td>5.7E-14</td><td>1.5E-13</td><td>BT20/MB468</td></tr>
<tr><td>>hg38_21_+_8433172_8446622_RNA45SN1_45S_50ntFlanks@8176.8217.42|CTTCTGATCGAGGCCCAGCCCGTGGACGGTGTGAGGCCGGTA</td><td>>hg38_21_+_8433172_8446622_RNA45SN1_45S_50ntFlanks</td><td>8176</td><td>8217</td><td>42</td><td>CTTCTGATCGAGGCCCAGCCCGTGGACGGTGTGAGGCCGGTA</td><td>425.4</td><td>4.7</td><td>0.6</td><td>7.5</td><td>5.4E-14</td><td>1.5E-13</td><td>MB231/MB468</td></tr>
<tr><td>>hg38_21_+_8433172_8446622_RNA45SN1_45S_50ntFlanks@6734.6773.40|ATTGATCATCGACACTTCGAACGCACTTGCGGCCCCGGGT</td><td>>hg38_21_+_8433172_8446622_RNA45SN1_45S_50ntFlanks</td><td>6734</td><td>6773</td><td>40</td><td>ATTGATCATCGACACTTCGAACGCACTTGCGGCCCCGGGT</td><td>1241.7</td><td>9.5</td><td>1.3</td><td>7.5</td><td>5.8E-14</td><td>1.5E-13</td><td>BT20/MB468</td></tr>
<tr><td>>hg38_21_+_8433172_8446622_RNA45SN1_45S_50ntFlanks@6719.6750.32|TGAATTGCAGGACACATTGATCATCGACACTT</td><td>>hg38_21_+_8433172_8446622_RNA45SN1_45S_50ntFlanks</td><td>6719</td><td>6750</td><td>32</td><td>TGAATTGCAGGACACATTGATCATCGACACTT</td><td>112.9</td><td>-4.1</td><td>0.5</td><td>-7.5</td><td>5.8E-14</td><td>1.5E-13</td><td>BT20/MB468</td></tr>
<tr><td>>hg38_21_+_8433172_8446622_RNA45SN1_45S_50ntFlanks@4129.4162.34|GATTCCGGAGAGGGAGCCTGAGAAACGGCTACCA</td><td>>hg38_21_+_8433172_8446622_RNA45SN1_45S_50ntFlanks</td><td>4129</td><td>4162</td><td>34</td><td>GATTCCGGAGAGGGAGCCTGAGAAACGGCTACCA</td><td>148.0</td><td>-9.2</td><td>1.2</td><td>-7.5</td><td>5.9E-14</td><td>1.5E-13</td><td>BT20/MB468</td></tr>
<tr><td>>hg38_21_+_8433172_8446622_RNA45SN1_45S_50ntFlanks@6661.6687.27|CGGTGGATCACTCGGCTCGTGCGTCGA</td><td>>hg38_21_+_8433172_8446622_RNA45SN1_45S_50ntFlanks</td><td>6661</td><td>6687</td><td>27</td><td>CGGTGGATCACTCGGCTCGTGCGTCGA</td><td>1026.8</td><td>5.5</td><td>0.7</td><td>7.5</td><td>5.5E-14</td><td>1.5E-13</td><td>MB231/MB468</td></tr>
<tr><td>>hg38_21_+_8433172_8446622_RNA45SN1_45S_50ntFlanks@5524.5550.27|GTAAAAGTCGTAACAAGGTTTCCGTAG</td><td>>hg38_21_+_8433172_8446622_RNA45SN1_45S_50ntFlanks</td><td>5524</td><td>5550</td><td>27</td><td>GTAAAAGTCGTAACAAGGTTTCCGTAG</td><td>166.8</td><td>-9.4</td><td>1.2</td><td>-7.5</td><td>5.9E-14</td><td>1.5E-13</td><td>BT20/MB468</td></tr>
<tr><td>>trnaMT_PheGAA_MT_+_577_647@39.62.24|ATGTTTAGACGGGCTCACATCACC</td><td>>trnaMT_PheGAA_MT_+_577_647</td><td>39</td><td>62</td><td>24</td><td>ATGTTTAGACGGGCTCACATCACC</td><td>930.2</td><td>8.9</td><td>1.2</td><td>7.5</td><td>5.5E-14</td><td>1.5E-13</td><td>MB231/MB468</td></tr>
<tr><td>>hg38_21_+_8433172_8446622_RNA45SN1_45S_50ntFlanks@12942.12964.23|TTCGTAGACGACCTGCTTCTGGG</td><td>>hg38_21_+_8433172_8446622_RNA45SN1_45S_50ntFlanks</td><td>12942</td><td>12964</td><td>23</td><td>TTCGTAGACGACCTGCTTCTGGG</td><td>161.9</td><td>-3.9</td><td>0.5</td><td>-7.5</td><td>5.6E-14</td><td>1.5E-13</td><td>MB231/MB468</td></tr>
<tr><td>>am_trna146_GlnCTG_6_-_27515531_27515602@1.29.29,trna1_GlnCTG_6_+_18836402_18836473@1.29.29,trna3_GlnCTG_17_+_8023070_8023141@1.29.29,trna42_GlnCTG_6_+_27263212_27263283@1.29.29,trna49_GlnCTG_6_+_27487308_27487379@1.29.29,trna7_GlnCTG_15_-_66161400_66161471@1.29.29,trna99_GlnCTG_6_-_28909378_28909449@1.29.29|GGTTCCATGGTGTAATGGTTAGCACTCTG</td><td>>am_trna146_GlnCTG_6_-_27515531_27515602</td><td>1</td><td>29</td><td>29</td><td>GGTTCCATGGTGTAATGGTTAGCACTCTG</td><td>8849.0</td><td>-5.4</td><td>0.7</td><td>-7.5</td><td>5.6E-14</td><td>1.5E-13</td><td>MB231/MB468</td></tr>
<tr><td>>MI0000542|hsa-mir-320a&WithFlank&8|-|22244960|22245043@48.69.22(+1C)&[MIMAT0000510&hsa-miR-320a-3p&offsets|0|0(+1C);m-60&8|-|22244975|22244996&offsets|0|0(+1C)]|AAAAGCTGGGTTGAGAGGGCGAC</td><td>>MI0000542|hsa-mir-320a&WithFlank&8|-|22244960|22245043</td><td>48</td><td>69</td><td>22(+1C)</td><td>AAAAGCTGGGTTGAGAGGGCGAC</td><td>912.1</td><td>9.7</td><td>1.3</td><td>7.5</td><td>5.6E-14</td><td>1.5E-13</td><td>MB231/MB468</td></tr>
<tr><td>>MI0000095|hsa-mir-93&WithFlank&7|-|100093762|100093853@17.39.23(+1C)&[MIMAT0000093&hsa-miR-93-5p&offsets|0|0(+1C);m-29&7|-|100093815|100093837&offsets|0|0(+1C)]|CAAAGTGCTGTTCGTGCAGGTAGC</td><td>>MI0000095|hsa-mir-93&WithFlank&7|-|100093762|100093853</td><td>17</td><td>39</td><td>23(+1C)</td><td>CAAAGTGCTGTTCGTGCAGGTAGC</td><td>539.1</td><td>5.7</td><td>0.8</td><td>7.5</td><td>6.0E-14</td><td>1.5E-13</td><td>BT20/MB468</td></tr>
<tr><td>>hg38_21_+_8433172_8446622_RNA45SN1_45S_50ntFlanks@3939.3959.21|AACCCGGTCAGCCCCTCTCCG</td><td>>hg38_21_+_8433172_8446622_RNA45SN1_45S_50ntFlanks</td><td>3939</td><td>3959</td><td>21</td><td>AACCCGGTCAGCCCCTCTCCG</td><td>417.4</td><td>-10.3</td><td>1.4</td><td>-7.5</td><td>5.6E-14</td><td>1.5E-13</td><td>MB231/MB468</td></tr>
<tr><td>>MI0000447|hsa-mir-128-1&WithFlank&2|+|135665391|135665484@56.76.21;MI0000727|hsa-mir-128-2&WithFlank&3|+|35744470|35744565@58.78.21&[MIMAT0000424&hsa-miR-128-3p&offsets|0|0;m-108&2|+|135665446|135665466&offsets|0|0];[MIMAT0000424_1&hsa-miR-128-3p&offsets|0|0;m-117&3|+|35744527|35744547&offsets|0|0]|TCACAGTGAACCGGTCTCTTT</td><td>>MI0000447|hsa-mir-128-1&WithFlank&2|+|135665391|135665484</td><td>56</td><td>76</td><td>21</td><td>TCACAGTGAACCGGTCTCTTT</td><td>15701.2</td><td>5.2</td><td>0.7</td><td>7.5</td><td>5.6E-14</td><td>1.5E-13</td><td>MB231/MB468</td></tr>
<tr><td>>hg38_21_+_8433172_8446622_RNA45SN1_45S_50ntFlanks@3711.3737.27|GTTGATCCTGCCAGTAGCATATGCTTG</td><td>>hg38_21_+_8433172_8446622_RNA45SN1_45S_50ntFlanks</td><td>3711</td><td>3737</td><td>27</td><td>GTTGATCCTGCCAGTAGCATATGCTTG</td><td>501.0</td><td>-5.8</td><td>0.8</td><td>-7.5</td><td>6.0E-14</td><td>1.5E-13</td><td>BT20/MB468</td></tr>
<tr><td>>hg38_21_+_8433172_8446622_RNA45SN1_45S_50ntFlanks@6667.6686.20|ATCACTCGGCTCGTGCGTCG</td><td>>hg38_21_+_8433172_8446622_RNA45SN1_45S_50ntFlanks</td><td>6667</td><td>6686</td><td>20</td><td>ATCACTCGGCTCGTGCGTCG</td><td>203.3</td><td>-9.7</td><td>1.3</td><td>-7.5</td><td>6.1E-14</td><td>1.5E-13</td><td>BT20/MB468</td></tr>
<tr><td>>hg38_21_+_8433172_8446622_RNA45SN1_45S_50ntFlanks@12606.12625.20|ATGTGCTTGGCTGAGGAGCC</td><td>>hg38_21_+_8433172_8446622_RNA45SN1_45S_50ntFlanks</td><td>12606</td><td>12625</td><td>20</td><td>ATGTGCTTGGCTGAGGAGCC</td><td>624.5</td><td>-4.0</td><td>0.5</td><td>-7.5</td><td>6.1E-14</td><td>1.5E-13</td><td>BT20/MB468</td></tr>
<tr><td>>hg38_21_+_8433172_8446622_RNA45SN1_45S_50ntFlanks@11848.11872.25|ACGGGCTTGGCGGAATCAGCGGGGA</td><td>>hg38_21_+_8433172_8446622_RNA45SN1_45S_50ntFlanks</td><td>11848</td><td>11872</td><td>25</td><td>ACGGGCTTGGCGGAATCAGCGGGGA</td><td>130.1</td><td>-4.2</td><td>0.6</td><td>-7.5</td><td>6.1E-14</td><td>1.5E-13</td><td>BT20/MB468</td></tr>
<tr><td>>MI0000088|hsa-mir-30a&WithFlank&6|-|71403545|71403627@12.35.24(+1A)&[MIMAT0000087&hsa-miR-30a-5p&offsets|0|+2(+1A);m-15&6|-|71403595|71403616&offsets|0|+2(+1A)]|TGTAAACATCCTCGACTGGAAGCTA</td><td>>MI0000088|hsa-mir-30a&WithFlank&6|-|71403545|71403627</td><td>12</td><td>35</td><td>24(+1A)</td><td>TGTAAACATCCTCGACTGGAAGCTA</td><td>2108.0</td><td>7.7</td><td>1.0</td><td>7.5</td><td>6.1E-14</td><td>1.6E-13</td><td>BT20/MB468</td></tr>
<tr><td>>hg38_21_+_8433172_8446622_RNA45SN1_45S_50ntFlanks@11999.12029.31|CCCTGCGGGCCGCCGGTGAAATACCACTACT</td><td>>hg38_21_+_8433172_8446622_RNA45SN1_45S_50ntFlanks</td><td>11999</td><td>12029</td><td>31</td><td>CCCTGCGGGCCGCCGGTGAAATACCACTACT</td><td>127.8</td><td>-5.8</td><td>0.8</td><td>-7.5</td><td>6.1E-14</td><td>1.6E-13</td><td>BT20/MB468</td></tr>
<tr><td>>hg38_21_+_8433172_8446622_RNA45SN1_45S_50ntFlanks@11847.11872.26|AACGGGCTTGGCGGAATCAGCGGGGA</td><td>>hg38_21_+_8433172_8446622_RNA45SN1_45S_50ntFlanks</td><td>11847</td><td>11872</td><td>26</td><td>AACGGGCTTGGCGGAATCAGCGGGGA</td><td>199.7</td><td>-5.4</td><td>0.7</td><td>-7.5</td><td>5.8E-14</td><td>1.6E-13</td><td>MB231/MB468</td></tr>
<tr><td>>MI0000471|hsa-mir-126&WithFlank&9|+|136670596|136670692@58.78.21(+1A)&[MIMAT0000445&hsa-miR-126-3p&offsets|0|-1(+1A);m-40&9|+|136670653|136670674&offsets|0|-1(+1A)]|TCGTACCGTGAGTAATAATGCA</td><td>>MI0000471|hsa-mir-126&WithFlank&9|+|136670596|136670692</td><td>58</td><td>78</td><td>21(+1A)</td><td>TCGTACCGTGAGTAATAATGCA</td><td>1660.1</td><td>10.0</td><td>1.3</td><td>7.5</td><td>5.8E-14</td><td>1.6E-13</td><td>MB231/MB468</td></tr>
<tr><td>>MI0000433|hsa-let-7g&WithFlank&3|-|52268272|52268367@11.30.20(+1G)&[MIMAT0000414&hsa-let-7g-5p&offsets|0|-2(+1G);m-58&3|-|52268336|52268357&offsets|0|-2(+1G)]|TGAGGTAGTAGTTTGTACAGG</td><td>>MI0000433|hsa-let-7g&WithFlank&3|-|52268272|52268367</td><td>11</td><td>30</td><td>20(+1G)</td><td>TGAGGTAGTAGTTTGTACAGG</td><td>592.9</td><td>8.2</td><td>1.1</td><td>7.5</td><td>6.2E-14</td><td>1.6E-13</td><td>BT20/MB468</td></tr>
<tr><td>>hg38_21_+_8433172_8446622_RNA45SN1_45S_50ntFlanks@12324.12345.22|TTGGGTTTTAAGCAGGAGGTGT</td><td>>hg38_21_+_8433172_8446622_RNA45SN1_45S_50ntFlanks</td><td>12324</td><td>12345</td><td>22</td><td>TTGGGTTTTAAGCAGGAGGTGT</td><td>274.8</td><td>-10.1</td><td>1.3</td><td>-7.5</td><td>6.2E-14</td><td>1.6E-13</td><td>BT20/MB468</td></tr>
<tr><td>>hg38_21_+_8433172_8446622_RNA45SN1_45S_50ntFlanks@10384.10406.23|CCATGTGAACAGCAGTTGAACAT</td><td>>hg38_21_+_8433172_8446622_RNA45SN1_45S_50ntFlanks</td><td>10384</td><td>10406</td><td>23</td><td>CCATGTGAACAGCAGTTGAACAT</td><td>191.6</td><td>-9.6</td><td>1.3</td><td>-7.5</td><td>6.2E-14</td><td>1.6E-13</td><td>BT20/MB468</td></tr>
<tr><td>>hg38_21_+_8433172_8446622_RNA45SN1_45S_50ntFlanks@10681.10710.30|TTCCGGCGGCGTCCGGTGAGCTCTCGCTGG</td><td>>hg38_21_+_8433172_8446622_RNA45SN1_45S_50ntFlanks</td><td>10681</td><td>10710</td><td>30</td><td>TTCCGGCGGCGTCCGGTGAGCTCTCGCTGG</td><td>2277.1</td><td>-7.2</td><td>1.0</td><td>-7.5</td><td>6.2E-14</td><td>1.6E-13</td><td>BT20/MB468</td></tr>
<tr><td>>hg38_21_+_8433172_8446622_RNA45SN1_45S_50ntFlanks@4283.4314.32|CTTTAACGAGGATCCATTGGAGGGCAAGTCTG</td><td>>hg38_21_+_8433172_8446622_RNA45SN1_45S_50ntFlanks</td><td>4283</td><td>4314</td><td>32</td><td>CTTTAACGAGGATCCATTGGAGGGCAAGTCTG</td><td>162.4</td><td>-9.3</td><td>1.2</td><td>-7.5</td><td>6.2E-14</td><td>1.6E-13</td><td>BT20/MB468</td></tr>
<tr><td>>hg38_21_+_8433172_8446622_RNA45SN1_45S_50ntFlanks@8170.8212.43|CAAGTCCTTCTGATCGAGGCCCAGCCCGTGGACGGTGTGAGGC</td><td>>hg38_21_+_8433172_8446622_RNA45SN1_45S_50ntFlanks</td><td>8170</td><td>8212</td><td>43</td><td>CAAGTCCTTCTGATCGAGGCCCAGCCCGTGGACGGTGTGAGGC</td><td>401.2</td><td>-4.9</td><td>0.7</td><td>-7.5</td><td>6.2E-14</td><td>1.6E-13</td><td>BT20/MB468</td></tr>
<tr><td>>hg38_21_+_8433172_8446622_RNA45SN1_45S_50ntFlanks@10841.10863.23|CGTAACTTCGGGATAAGGATTGG</td><td>>hg38_21_+_8433172_8446622_RNA45SN1_45S_50ntFlanks</td><td>10841</td><td>10863</td><td>23</td><td>CGTAACTTCGGGATAAGGATTGG</td><td>1752.9</td><td>-5.3</td><td>0.7</td><td>-7.5</td><td>5.8E-14</td><td>1.6E-13</td><td>MB231/MB468</td></tr>
<tr><td>>hg38_21_+_8433172_8446622_RNA45SN1_45S_50ntFlanks@6719.6738.20|TGAATTGCAGGACACATTGA</td><td>>hg38_21_+_8433172_8446622_RNA45SN1_45S_50ntFlanks</td><td>6719</td><td>6738</td><td>20</td><td>TGAATTGCAGGACACATTGA</td><td>179.7</td><td>-9.5</td><td>1.3</td><td>-7.5</td><td>6.3E-14</td><td>1.6E-13</td><td>BT20/MB468</td></tr>
<tr><td>>hg38_21_+_8433172_8446622_RNA45SN1_45S_50ntFlanks@10410.10435.26|TCAGTCGGTCCTGAGAGATGGGCGAG</td><td>>hg38_21_+_8433172_8446622_RNA45SN1_45S_50ntFlanks</td><td>10410</td><td>10435</td><td>26</td><td>TCAGTCGGTCCTGAGAGATGGGCGAG</td><td>340.7</td><td>-10.4</td><td>1.4</td><td>-7.5</td><td>6.3E-14</td><td>1.6E-13</td><td>BT20/MB468</td></tr>
<tr><td>>hg38_21_+_8433172_8446622_RNA45SN1_45S_50ntFlanks@12044.12068.25|ACTGACCCGGTGAGGCGGGGGGGCG</td><td>>hg38_21_+_8433172_8446622_RNA45SN1_45S_50ntFlanks</td><td>12044</td><td>12068</td><td>25</td><td>ACTGACCCGGTGAGGCGGGGGGGCG</td><td>214.9</td><td>-9.3</td><td>1.2</td><td>-7.5</td><td>5.9E-14</td><td>1.6E-13</td><td>MB231/MB468</td></tr>
<tr><td>>MI0000103|hsa-mir-101-1&WithFlank&1|-|65058428|65058514@52.73.22(+1A);MI0000739|hsa-mir-101-2&WithFlank&9|+|4850291|4850381@54.75.22(+1A)&[MIMAT0000099&hsa-miR-101-3p&offsets|-1|0(+1A);m-17&1|-|65058443|65058462&offsets|-1|+1(+1A)];[MIMAT0000099_1&hsa-miR-101-3p&offsets|-1|0(+1A);m-16&9|+|4850345|4850364&offsets|-1|+1(+1A)]|GTACAGTACTGTGATAACTGAAA</td><td>>MI0000103|hsa-mir-101-1&WithFlank&1|-|65058428|65058514</td><td>52</td><td>73</td><td>22(+1A)</td><td>GTACAGTACTGTGATAACTGAAA</td><td>6847.4</td><td>8.7</td><td>1.2</td><td>7.5</td><td>6.4E-14</td><td>1.6E-13</td><td>BT20/MB468</td></tr>
<tr><td>>MI0000650|hsa-mir-200c&WithFlank&12|+|6963693|6963772@50.70.21(+1C)&[MIMAT0000617&hsa-miR-200c-3p&offsets|0|-2(+1C);m-54&12|+|6963742|6963764&offsets|0|-2(+1C)]|TAATACTGCCGGGTAATGATGC</td><td>>MI0000650|hsa-mir-200c&WithFlank&12|+|6963693|6963772</td><td>50</td><td>70</td><td>21(+1C)</td><td>TAATACTGCCGGGTAATGATGC</td><td>239.9</td><td>11.4</td><td>1.5</td><td>7.8</td><td>4.3E-15</td><td>1.6E-13</td><td>BT20/MB231</td></tr>
<tr><td>>hg38_21_+_8433172_8446622_RNA45SN1_45S_50ntFlanks@10845.10866.22|ACTTCGGGATAAGGATTGGCTC</td><td>>hg38_21_+_8433172_8446622_RNA45SN1_45S_50ntFlanks</td><td>10845</td><td>10866</td><td>22</td><td>ACTTCGGGATAAGGATTGGCTC</td><td>278.0</td><td>-5.5</td><td>0.7</td><td>-7.5</td><td>6.4E-14</td><td>1.6E-13</td><td>BT20/MB468</td></tr>
<tr><td>>hg38_21_+_8433172_8446622_RNA45SN1_45S_50ntFlanks@11575.11605.31|ACTTAGAACTGGTGCGGACCAGGGGAATCCG</td><td>>hg38_21_+_8433172_8446622_RNA45SN1_45S_50ntFlanks</td><td>11575</td><td>11605</td><td>31</td><td>ACTTAGAACTGGTGCGGACCAGGGGAATCCG</td><td>1154.0</td><td>-7.7</td><td>1.0</td><td>-7.5</td><td>6.6E-14</td><td>1.7E-13</td><td>BT20/MB468</td></tr>
<tr><td>>trnaMT_ValTAC_MT_+_1602_1670@28.72.45|AACTTACACTTAGGAGATTTCAACTTAACTTGACCGCTCTGACCA</td><td>>trnaMT_ValTAC_MT_+_1602_1670</td><td>28</td><td>72</td><td>45</td><td>AACTTACACTTAGGAGATTTCAACTTAACTTGACCGCTCTGACCA</td><td>696.3</td><td>10.0</td><td>1.3</td><td>7.5</td><td>6.6E-14</td><td>1.7E-13</td><td>BT20/MB468</td></tr>
<tr><td>>hg38_21_+_8433172_8446622_RNA45SN1_45S_50ntFlanks@11317.11338.22|ACCCCACGTCTCGTCGCGCGCG</td><td>>hg38_21_+_8433172_8446622_RNA45SN1_45S_50ntFlanks</td><td>11317</td><td>11338</td><td>22</td><td>ACCCCACGTCTCGTCGCGCGCG</td><td>496.9</td><td>-8.6</td><td>1.1</td><td>-7.5</td><td>6.2E-14</td><td>1.7E-13</td><td>MB231/MB468</td></tr>
<tr><td>>hg38_21_+_8433172_8446622_RNA45SN1_45S_50ntFlanks@11233.11250.18|CCCTTCCCGTGGATCGCC</td><td>>hg38_21_+_8433172_8446622_RNA45SN1_45S_50ntFlanks</td><td>11233</td><td>11250</td><td>18</td><td>CCCTTCCCGTGGATCGCC</td><td>488.7</td><td>-10.5</td><td>1.4</td><td>-7.5</td><td>6.2E-14</td><td>1.7E-13</td><td>MB231/MB468</td></tr>
<tr><td>>hg38_21_+_8433172_8446622_RNA45SN1_45S_50ntFlanks@4297.4321.25|CATTGGAGGGCAAGTCTGGTGCCAG</td><td>>hg38_21_+_8433172_8446622_RNA45SN1_45S_50ntFlanks</td><td>4297</td><td>4321</td><td>25</td><td>CATTGGAGGGCAAGTCTGGTGCCAG</td><td>225.6</td><td>-6.0</td><td>0.8</td><td>-7.5</td><td>6.8E-14</td><td>1.7E-13</td><td>BT20/MB468</td></tr>
</table>

<table>
<tr><th></th><th></th><th></th><th></th><th></th><th></th><th></th><th></th><th></th><th></th><th></th><th></th><th></th></tr>
<tr><td>>hg38_21_+_8433172_8446622_RNA45SN1_45S_50ntFlanks@10384.10404.21|CCATGTGAACAGCAGTTGAAC</td><td>>hg38_21_+_8433172_8446622_RNA45SN1_45S_50ntFlanks</td><td>10384</td><td>10404</td><td>21</td><td>CCATGTGAACAGCAGTTGAAC</td><td>203.5</td><td>-4.2</td><td>0.6</td><td>-7.5</td><td>6.8E-14</td><td>1.7E-13</td><td>BT20/MB468</td></tr>
<tr><td>>hg38_21_+_8433172_8446622_RNA45SN1_45S_50ntFlanks@10349.10376.28|AACGAGAACTTTGAAGGCCGAAGTGGAG</td><td>>hg38_21_+_8433172_8446622_RNA45SN1_45S_50ntFlanks</td><td>10349</td><td>10376</td><td>28</td><td>AACGAGAACTTTGAAGGCCGAAGTGGAG</td><td>549.3</td><td>-3.5</td><td>0.5</td><td>-7.5</td><td>6.9E-14</td><td>1.7E-13</td><td>BT20/MB468</td></tr>
<tr><td>>MI0000736|hsa-mir-30c-1&WithFlank&1|+|40757278|40757378@23.45.23(+1A);MI0000254|hsa-mir-30c-2&WithFlank&6|-|71376954|71377037@13.35.23(+1A)&[MIMAT0000244&hsa-miR-30c-5p&offsets|0|0(+1A);m-57&1|+|40757300|40757323&offsets|0|-1(+1A)];[MIMAT0000244_1&hsa-miR-30c-5p&offsets|0|0(+1A);m-56&6|-|71377002|71377025&offsets|0|-1(+1A)]|TGTAAACATCCTACACTCTCAGCA</td><td>>MI0000736|hsa-mir-30c-1&WithFlank&1|+|40757278|40757378</td><td>23</td><td>45</td><td>23(+1A)</td><td>TGTAAACATCCTACACTCTCAGCA</td><td>7710.5</td><td>6.8</td><td>0.9</td><td>7.5</td><td>6.9E-14</td><td>1.7E-13</td><td>BT20/MB468</td></tr>
<tr><td>>MI0000095|hsa-mir-93&WithFlank&7|-|100093762|100093853@17.39.23(+1G)&[MIMAT0000093&hsa-miR-93-5p&offsets|0|0(+1G);m-29&7|-|100093815|100093837&offsets|0|0(+1G)]|CAAAGTGCTGTTCGTGCAGGTAGG</td><td>>MI0000095|hsa-mir-93&WithFlank&7|-|100093762|100093853</td><td>17</td><td>39</td><td>23(+1G)</td><td>CAAAGTGCTGTTCGTGCAGGTAGG</td><td>699.2</td><td>6.3</td><td>0.8</td><td>7.5</td><td>6.9E-14</td><td>1.7E-13</td><td>BT20/MB468</td></tr>
<tr><td>>hg38_21_+_8433172_8446622_RNA45SN1_45S_50ntFlanks@6651.6685.35|CGACTCTTAGCGGTGGATCACTCGGCTCGTGCGTC</td><td>>hg38_21_+_8433172_8446622_RNA45SN1_45S_50ntFlanks</td><td>6651</td><td>6685</td><td>35</td><td>CGACTCTTAGCGGTGGATCACTCGGCTCGTGCGTC</td><td>1723.2</td><td>3.8</td><td>0.5</td><td>7.5</td><td>6.5E-14</td><td>1.8E-13</td><td>MB231/MB468</td></tr>
<tr><td>>hg38_21_+_8433172_8446622_RNA45SN1_45S_50ntFlanks@11534.11551.18|GTTCCGCGCGGCGCCTCG</td><td>>hg38_21_+_8433172_8446622_RNA45SN1_45S_50ntFlanks</td><td>11534</td><td>11551</td><td>18</td><td>GTTCCGCGCGGCGCCTCG</td><td>1013.1</td><td>-6.9</td><td>0.9</td><td>-7.5</td><td>7.0E-14</td><td>1.8E-13</td><td>BT20/MB468</td></tr>
<tr><td>>trna16_GlnTTG_17_+_47269890_47269961@1.29.29,trna64_GlnTTG_6_+_28557156_28557227@1.29.29,trna84_GlnTTG_6_+_145503859_145503930@1.29.29|GGTCCCATGGTGTAATGGTTAGCACTCTG</td><td>>trna16_GlnTTG_17_+_47269890_47269961</td><td>1</td><td>29</td><td>29</td><td>GGTCCCATGGTGTAATGGTTAGCACTCTG</td><td>1703.8</td><td>-6.6</td><td>0.9</td><td>-7.5</td><td>6.6E-14</td><td>1.8E-13</td><td>MB231/MB468</td></tr>
<tr><td>>am_trnaMT_SerGCT_MT_+_12207_12265@14.32.19|AGAACTGCTAACTCATGCC</td><td>>am_trnaMT_SerGCT_MT_+_12207_12265</td><td>14</td><td>32</td><td>19</td><td>AGAACTGCTAACTCATGCC</td><td>885.6</td><td>9.4</td><td>1.2</td><td>7.5</td><td>6.6E-14</td><td>1.8E-13</td><td>MB231/MB468</td></tr>
<tr><td>>trnaMT_GluTTC_MT_-_14674_14742@46.72.27,trnalookalike8_GluTTC_5_-_93905172_93905240@46.72.27|GTGGTTGTAGTCCGTGCGAGAATACCA</td><td>>trnaMT_GluTTC_MT_-_14674_14742</td><td>46</td><td>72</td><td>27</td><td>GTGGTTGTAGTCCGTGCGAGAATACCA</td><td>913.8</td><td>9.3</td><td>1.2</td><td>7.5</td><td>7.1E-14</td><td>1.8E-13</td><td>BT20/MB468</td></tr>
<tr><td>>am_trnaMT_PheGAA_MT_+_577_647@41.58.18|GTTTAGACGGGCTCACAT</td><td>>am_trnaMT_PheGAA_MT_+_577_647</td><td>41</td><td>58</td><td>18</td><td>GTTTAGACGGGCTCACAT</td><td>1021.2</td><td>9.2</td><td>1.2</td><td>7.5</td><td>7.2E-14</td><td>1.8E-13</td><td>BT20/MB468</td></tr>
<tr><td>>hg38_21_+_8433172_8446622_RNA45SN1_45S_50ntFlanks@6771.6799.29|GGTTCCTCCCGGGGCTACGCCTGTCTGAG</td><td>>hg38_21_+_8433172_8446622_RNA45SN1_45S_50ntFlanks</td><td>6771</td><td>6799</td><td>29</td><td>GGTTCCTCCCGGGGCTACGCCTGTCTGAG</td><td>155.2</td><td>-4.7</td><td>0.6</td><td>-7.5</td><td>7.2E-14</td><td>1.8E-13</td><td>BT20/MB468</td></tr>
<tr><td>>hg38_MT_+_598_1651_RNR1_12S_50ntFlanks@916.936.21|AAGGACATTTAACTAAAACCC</td><td>>hg38_MT_+_598_1651_RNR1_12S_50ntFlanks</td><td>916</td><td>936</td><td>21</td><td>AAGGACATTTAACTAAAACCC</td><td>506.4</td><td>9.9</td><td>1.3</td><td>7.5</td><td>7.2E-14</td><td>1.8E-13</td><td>BT20/MB468</td></tr>
<tr><td>>hg38_21_+_8433172_8446622_RNA45SN1_45S_50ntFlanks@6651.6677.27|CGACTCTTAGCGGTGGATCACTCGGCT</td><td>>hg38_21_+_8433172_8446622_RNA45SN1_45S_50ntFlanks</td><td>6651</td><td>6677</td><td>27</td><td>CGACTCTTAGCGGTGGATCACTCGGCT</td><td>6207.4</td><td>3.9</td><td>0.5</td><td>7.5</td><td>7.2E-14</td><td>1.8E-13</td><td>BT20/MB468</td></tr>
<tr><td>>MI0000093|hsa-mir-92a-1&WithFlank&13|+|91351308|91351397@54.74.21;MI0000094|hsa-mir-92a-2&WithFlank&X|-|134169532|134169618@54.74.21&[MIMAT0000092&hsa-miR-92a-3p&offsets|0|-1;m-9&13|+|91351361|91351382&offsets|0|-1];[MIMAT0000092_1&hsa-miR-92a-3p&offsets|0|-1;m-11&X|-|134169544|134169565&offsets|0|-1]|TATTGCACTTGTCCCGGCCTG</td><td>>MI0000093|hsa-mir-92a-1&WithFlank&13|+|91351308|91351397</td><td>54</td><td>74</td><td>21</td><td>TATTGCACTTGTCCCGGCCTG</td><td>1665.3</td><td>6.4</td><td>0.9</td><td>7.5</td><td>7.4E-14</td><td>1.9E-13</td><td>BT20/MB468</td></tr>
<tr><td>>MI0003139|hsa-mir-181d&WithFlank&19|+|13874869|13875017@42.64.23&[MIMAT0002821&hsa-miR-181d-5p&offsets|0|0;m-194&19|+|13874910|13874932&offsets|0|0]|AACATTCATTGTTGTCGGTGGGT</td><td>>MI0003139|hsa-mir-181d&WithFlank&19|+|13874869|13875017</td><td>42</td><td>64</td><td>23</td><td>AACATTCATTGTTGTCGGTGGGT</td><td>548.3</td><td>5.5</td><td>0.7</td><td>7.5</td><td>7.5E-14</td><td>1.9E-13</td><td>BT20/MB468</td></tr>
<tr><td>>hg38_21_+_8433172_8446622_RNA45SN1_45S_50ntFlanks@3705.3741.37|TACCTGGTTGATCCTGCCAGTAGCATATGCTTGTCTC</td><td>>hg38_21_+_8433172_8446622_RNA45SN1_45S_50ntFlanks</td><td>3705</td><td>3741</td><td>37</td><td>TACCTGGTTGATCCTGCCAGTAGCATATGCTTGTCTC</td><td>1225.7</td><td>4.7</td><td>0.6</td><td>7.5</td><td>6.9E-14</td><td>1.9E-13</td><td>MB231/MB468</td></tr>
<tr><td>>hg38_1_-_228634819_228635039_RNA5S12_5S_50ntFlanks@154.171.18|CCGGGTGCTGTAGGCTTT</td><td>>hg38_1_-_228634819_228635039_RNA5S12_5S_50ntFlanks</td><td>154</td><td>171</td><td>18</td><td>CCGGGTGCTGTAGGCTTT</td><td>3944.7</td><td>5.6</td><td>0.8</td><td>7.5</td><td>7.0E-14</td><td>1.9E-13</td><td>MB231/MB468</td></tr>
<tr><td>>hg38_21_+_8433172_8446622_RNA45SN1_45S_50ntFlanks@9572.9592.21|CAAATCGGTCGTCCGACCTGG</td><td>>hg38_21_+_8433172_8446622_RNA45SN1_45S_50ntFlanks</td><td>9572</td><td>9592</td><td>21</td><td>CAAATCGGTCGTCCGACCTGG</td><td>183.9</td><td>-9.5</td><td>1.3</td><td>-7.5</td><td>7.6E-14</td><td>1.9E-13</td><td>BT20/MB468</td></tr>
<tr><td>>hg38_21_+_8433172_8446622_RNA45SN1_45S_50ntFlanks@10794.10822.29|CTGGCATGTTGGAACAATGTAGGTAAGGG</td><td>>hg38_21_+_8433172_8446622_RNA45SN1_45S_50ntFlanks</td><td>10794</td><td>10822</td><td>29</td><td>CTGGCATGTTGGAACAATGTAGGTAAGGG</td><td>262.3</td><td>-9.6</td><td>1.3</td><td>-7.5</td><td>7.1E-14</td><td>1.9E-13</td><td>MB231/MB468</td></tr>
<tr><td>>hg38_21_+_8433172_8446622_RNA45SN1_45S_50ntFlanks@10411.10443.33|CAGTCGGTCCTGAGAGATGGGCGAGCGCCGTTC</td><td>>hg38_21_+_8433172_8446622_RNA45SN1_45S_50ntFlanks</td><td>10411</td><td>10443</td><td>33</td><td>CAGTCGGTCCTGAGAGATGGGCGAGCGCCGTTC</td><td>649.9</td><td>-7.3</td><td>1.0</td><td>-7.5</td><td>7.6E-14</td><td>1.9E-13</td><td>BT20/MB468</td></tr>
<tr><td>>am_trna11_ProAGG_16_+_3241989_3242060@1.29.29,trna12_ProAGG_6_+_26555498_26555569@1.29.29,trna12_ProTGG_11_-_75946869_75946940@1.29.29,trna14_ProTGG_5_-_180615854_180615925@1.29.29,trna22_ProAGG_14_-_21081560_21081631@1.29.29,trna23_ProAGG_14_-_21077495_21077566@1.29.29,trna28_ProTGG_16_-_3234133_3234204@1.29.29,trna29_ProAGG_16_-_3232635_3232706@1.29.29,trna2_ProAGG_7_+_128423504_128423575@1.29.29,trna30_ProCGG_6_+_27059521_27059592@1.29.29,trna37_ProCGG_17_-_8126151_8126222@1.29.29,trna3_ProTGG_16_+_3208923_3208994@1.29.29,trna52_ProCGG_1_+_167683962_167684033@1.29.29,trna65_ProAGG_1_-_167684725_167684796@1.29.29,trna6_ProCGG_16_+_3222049_3222120@1.29.29,trna6_ProTGG_14_+_21152175_21152246@1.29.29,trna8_ProTGG_16_+_3238094_3238165@1.29.29,trna9_ProAGG_11_+_75946557_75946628@1.29.29,trna9_ProAGG_16_+_3239634_3239705@1.29.29|GGCTCGTTGGTCTAGGGGTATGATTCTCG</td><td>>am_trna11_ProAGG_16_+_3241989_3242060</td><td>1</td><td>29</td><td>29</td><td>GGCTCGTTGGTCTAGGGGTATGATTCTCG</td><td>1528.4</td><td>-6.5</td><td>0.9</td><td>-7.5</td><td>7.7E-14</td><td>1.9E-13</td><td>BT20/MB468</td></tr>
<tr><td>>hg38_21_+_8433172_8446622_RNA45SN1_45S_50ntFlanks@10420.10456.37|CTGAGAGATGGGCGAGCGCCGTTCCGAAGGGACGGGC</td><td>>hg38_21_+_8433172_8446622_RNA45SN1_45S_50ntFlanks</td><td>10420</td><td>10456</td><td>37</td><td>CTGAGAGATGGGCGAGCGCCGTTCCGAAGGGACGGGC</td><td>820.4</td><td>-4.8</td><td>0.6</td><td>-7.5</td><td>7.7E-14</td><td>1.9E-13</td><td>BT20/MB468</td></tr>
<tr><td>>hg38_21_+_8433172_8446622_RNA45SN1_45S_50ntFlanks@4759.4781.23|ATCAGATACCGTCGTAGTTCCGA</td><td>>hg38_21_+_8433172_8446622_RNA45SN1_45S_50ntFlanks</td><td>4759</td><td>4781</td><td>23</td><td>ATCAGATACCGTCGTAGTTCCGA</td><td>213.6</td><td>-9.3</td><td>1.2</td><td>-7.5</td><td>7.2E-14</td><td>1.9E-13</td><td>MB231/MB468</td></tr>
<tr><td>>hg38_21_+_8433172_8446622_RNA45SN1_45S_50ntFlanks@10408.10425.18|GGTCAGTCGGTCCTGAGA</td><td>>hg38_21_+_8433172_8446622_RNA45SN1_45S_50ntFlanks</td><td>10408</td><td>10425</td><td>18</td><td>GGTCAGTCGGTCCTGAGA</td><td>176.6</td><td>-4.6</td><td>0.6</td><td>-7.5</td><td>7.2E-14</td><td>1.9E-13</td><td>MB231/MB468</td></tr>
<tr><td>>MI0000441|hsa-mir-30b&WithFlank&8|-|134800514|134800613@23.43.21&[MIMAT0000420&hsa-miR-30b-5p&offsets|0|-1;m-76&8|-|134800570|134800591&offsets|0|-1]|TGTAAACATCCTACACTCAGC</td><td>>MI0000441|hsa-mir-30b&WithFlank&8|-|134800514|134800613</td><td>23</td><td>43</td><td>21</td><td>TGTAAACATCCTACACTCAGC</td><td>1643.8</td><td>9.9</td><td>1.3</td><td>7.5</td><td>7.7E-14</td><td>1.9E-13</td><td>BT20/MB468</td></tr>
<tr><td>>hg38_21_+_8433172_8446622_RNA45SN1_45S_50ntFlanks@3870.3898.29|AGCTAATACATGCCGACGGGCGCTGACCC</td><td>>hg38_21_+_8433172_8446622_RNA45SN1_45S_50ntFlanks</td><td>3870</td><td>3898</td><td>29</td><td>AGCTAATACATGCCGACGGGCGCTGACCC</td><td>219.0</td><td>-9.8</td><td>1.3</td><td>-7.5</td><td>7.8E-14</td><td>1.9E-13</td><td>BT20/MB468</td></tr>
<tr><td>>hg38_21_+_8433172_8446622_RNA45SN1_45S_50ntFlanks@11534.11560.27|GTTCCGCGCGGCGCCTCGCCTCGGCCG</td><td>>hg38_21_+_8433172_8446622_RNA45SN1_45S_50ntFlanks</td><td>11534</td><td>11560</td><td>27</td><td>GTTCCGCGCGGCGCCTCGCCTCGGCCG</td><td>237.5</td><td>-9.5</td><td>1.3</td><td>-7.5</td><td>7.4E-14</td><td>2.0E-13</td><td>MB231/MB468</td></tr>
<tr><td>>MI0000111|hsa-mir-105-1&WithFlank&X|-|152392213|152392305@19.41.23;MI0000112|hsa-mir-105-2&WithFlank&X|-|152394406|152394498@19.41.23&[MIMAT0000102&hsa-miR-105-5p&offsets|0|0;m-735&X|-|152392266|152392287&offsets|0|+1];[MIMAT0000102_1&hsa-miR-105-5p&offsets|0|0;m-735&X|-|152394459|152394480&offsets|0|+1]|TCAAATGCTCAGACTCCTGTGGT</td><td>>MI0000111|hsa-mir-105-1&WithFlank&X|-|152392213|152392305</td><td>19</td><td>41</td><td>23</td><td>TCAAATGCTCAGACTCCTGTGGT</td><td>209.8</td><td>11.2</td><td>1.4</td><td>7.8</td><td>5.3E-15</td><td>2.0E-13</td><td>BT20/MB231</td></tr>
<tr><td>>MI0000062|hsa-let-7a-3&WithFlank&22|+|46112743|46112828@58.78.21(+1U);MI0000060|hsa-let-7a-1&WithFlank&9|+|94175951|94176042@63.83.21(+1U)&[MIMAT0004481&hsa-let-7a-3p&offsets|0|0(+1U);m-180&22|+|46112800|46112821&offsets|0|-1(+1U)];[MIMAT0004481_1&hsa-let-7a-3p&offsets|0|0(+1U);m-181&9|+|94176013|94176034&offsets|0|-1(+1U)]|CTATACAATCTACTGTCTTTCT</td><td>>MI0000062|hsa-let-7a-3&WithFlank&22|+|46112743|46112828</td><td>58</td><td>78</td><td>21(+1U)</td><td>CTATACAATCTACTGTCTTTCT</td><td>2725.2</td><td>6.3</td><td>0.8</td><td>7.5</td><td>8.0E-14</td><td>2.0E-13</td><td>BT20/MB468</td></tr>
<tr><td>>MI0000760|hsa-mir-361&WithFlank&X|-|85903630|85903713@12.33.22&[MIMAT0000703&hsa-miR-361-5p&offsets|0|0;m-89&X|-|85903681|85903702&offsets|0|0]|TTATCAGAATCTCCAGGGGTAC</td><td>>MI0000760|hsa-mir-361&WithFlank&X|-|85903630|85903713</td><td>12</td><td>33</td><td>22</td><td>TTATCAGAATCTCCAGGGGTAC</td><td>628.1</td><td>6.0</td><td>0.8</td><td>7.5</td><td>8.0E-14</td><td>2.0E-13</td><td>BT20/MB468</td></tr>
<tr><td>>MI0000802|hsa-mir-340&WithFlank&5|-|180015297|180015403@22.43.22&[MIMAT0004692&hsa-miR-340-5p&offsets|0|0;m-220&5|-|180015361|180015382&offsets|0|0]|TTATAAAGCAATGAGACTGATT</td><td>>MI0000802|hsa-mir-340&WithFlank&5|-|180015297|180015403</td><td>22</td><td>43</td><td>22</td><td>TTATAAAGCAATGAGACTGATT</td><td>5723.8</td><td>5.7</td><td>0.8</td><td>7.5</td><td>7.5E-14</td><td>2.0E-13</td><td>MB231/MB468</td></tr>
<tr><td>>hg38_21_+_8433172_8446622_RNA45SN1_45S_50ntFlanks@10402.10430.29|AACATGGGTCAGTCGGTCCTGAGAGATGG</td><td>>hg38_21_+_8433172_8446622_RNA45SN1_45S_50ntFlanks</td><td>10402</td><td>10430</td><td>29</td><td>AACATGGGTCAGTCGGTCCTGAGAGATGG</td><td>1318.3</td><td>-4.5</td><td>0.6</td><td>-7.5</td><td>7.6E-14</td><td>2.1E-13</td><td>MB231/MB468</td></tr>
<tr><td>>hg38_21_+_8433172_8446622_RNA45SN1_45S_50ntFlanks@10410.10443.34|TCAGTCGGTCCTGAGAGATGGGCGAGCGCCGTTC</td><td>>hg38_21_+_8433172_8446622_RNA45SN1_45S_50ntFlanks</td><td>10410</td><td>10443</td><td>34</td><td>TCAGTCGGTCCTGAGAGATGGGCGAGCGCCGTTC</td><td>225.7</td><td>-6.3</td><td>0.8</td><td>-7.5</td><td>8.2E-14</td><td>2.1E-13</td><td>BT20/MB468</td></tr>
<tr><td>>hg38_21_+_8433172_8446622_RNA45SN1_45S_50ntFlanks@8380.8405.26|CAAGAGGGCGTGAAACCGTTAAGAGG</td><td>>hg38_21_+_8433172_8446622_RNA45SN1_45S_50ntFlanks</td><td>8380</td><td>8405</td><td>26</td><td>CAAGAGGGCGTGAAACCGTTAAGAGG</td><td>705.0</td><td>-4.8</td><td>0.6</td><td>-7.5</td><td>7.7E-14</td><td>2.1E-13</td><td>MB231/MB468</td></tr>
<tr><td>>am_trna143_LysTTT_6_-_27559593_27559665@6.24.19,trna14_LysTTT_11_-_59327808_59327880@6.24.19,trna2_LysTTT_17_+_8022473_8022545@6.24.19,trna54_LysTTT_1_+_204475655_204475727@6.24.19,trna5_LysTTT_11_+_59323902_59323974@6.24.19,trna62_LysTTT_1_-_204476158_204476230@6.24.19,trna76_LysTTT_6_+_28918806_28918878@6.24.19|GATAGCTCAGTCGGTAGAG</td><td>>am_trna143_LysTTT_6_-_27559593_27559665</td><td>6</td><td>24</td><td>19</td><td>GATAGCTCAGTCGGTAGAG</td><td>295.6</td><td>-9.8</td><td>1.3</td><td>-7.5</td><td>7.7E-14</td><td>2.1E-13</td><td>MB231/MB468</td></tr>
<tr><td>>hg38_21_+_8433172_8446622_RNA45SN1_45S_50ntFlanks@10410.10428.19|TCAGTCGGTCCTGAGAGAT</td><td>>hg38_21_+_8433172_8446622_RNA45SN1_45S_50ntFlanks</td><td>10410</td><td>10428</td><td>19</td><td>TCAGTCGGTCCTGAGAGAT</td><td>579.0</td><td>-4.8</td><td>0.6</td><td>-7.5</td><td>8.3E-14</td><td>2.1E-13</td><td>BT20/MB468</td></tr>
<tr><td>>hg38_MT_+_1621_3279_RNR2_16S_50ntFlanks@1404.1421.18|TATTAAAGGTTCGTTTGT</td><td>>hg38_MT_+_1621_3279_RNR2_16S_50ntFlanks</td><td>1404</td><td>1421</td><td>18</td><td>TATTAAAGGTTCGTTTGT</td><td>1176.0</td><td>5.7</td><td>0.8</td><td>7.5</td><td>7.7E-14</td><td>2.1E-13</td><td>MB231/MB468</td></tr>
<tr><td>>hg38_21_+_8433172_8446622_RNA45SN1_45S_50ntFlanks@11235.11252.18|CTTCCCGTGGATCGCCCC</td><td>>hg38_21_+_8433172_8446622_RNA45SN1_45S_50ntFlanks</td><td>11235</td><td>11252</td><td>18</td><td>CTTCCCGTGGATCGCCCC</td><td>1209.4</td><td>-5.5</td><td>0.7</td><td>-7.5</td><td>8.3E-14</td><td>2.1E-13</td><td>BT20/MB468</td></tr>
</table>

<table>
<tr><td></td><td></td><td></td><td></td><td></td><td></td><td></td><td></td><td></td><td></td><td></td><td></td><td></td></tr>
<tr><td>>hg38_21_+_8433172_8446622_RNA45SN1_45S_50ntFlanks@12488.12518.31|AACGTGAGCTGGGTTTAGACCGTCGTGAGAC</td><td>>hg38_21_+_8433172_8446622_RNA45SN1_45S_50ntFlanks</td><td>12488</td><td>12518</td><td>31</td><td>AACGTGAGCTGGGTTTAGACCGTCGTGAGAC</td><td>301.3</td><td>-4.0</td><td>0.5</td><td>-7.5</td><td>7.8E-14</td><td>2.1E-13</td><td>MB231/MB468</td></tr>
<tr><td>>MI0000298|hsa-mir-221&WithFlank&X|-|45746151|45746272@71.92.22(+1A)&[MIMAT0000278&hsa-miR-221-3p&offsets|0|-1(+1A);m-91&X|-|45746181|45746202&offsets|0|0(+1A)]|AGCTACATTGTCTGCTGGGTTTA</td><td>>MI0000298|hsa-mir-221&WithFlank&X|-|45746151|45746272</td><td>71</td><td>92</td><td>22(+1A)</td><td>AGCTACATTGTCTGCTGGGTTTA</td><td>1012.8</td><td>6.0</td><td>0.8</td><td>7.5</td><td>8.5E-14</td><td>2.1E-13</td><td>BT20/MB468</td></tr>
<tr><td>>hg38_21_+_8433172_8446622_RNA45SN1_45S_50ntFlanks@11849.11872.24|CGGGCTTGGCGGAATCAGCGGGGA</td><td>>hg38_21_+_8433172_8446622_RNA45SN1_45S_50ntFlanks</td><td>11849</td><td>11872</td><td>24</td><td>CGGGCTTGGCGGAATCAGCGGGGA</td><td>210.9</td><td>-9.3</td><td>1.2</td><td>-7.5</td><td>7.9E-14</td><td>2.1E-13</td><td>MB231/MB468</td></tr>
<tr><td>>hg38_21_+_8433172_8446622_RNA45SN1_45S_50ntFlanks@12991.13014.24|CTCCCTCGCTGCGATCTATTGAAA</td><td>>hg38_21_+_8433172_8446622_RNA45SN1_45S_50ntFlanks</td><td>12991</td><td>13014</td><td>24</td><td>CTCCCTCGCTGCGATCTATTGAAA</td><td>1879.1</td><td>-6.4</td><td>0.9</td><td>-7.5</td><td>8.5E-14</td><td>2.1E-13</td><td>BT20/MB468</td></tr>
<tr><td>>hg38_21_+_8433172_8446622_RNA45SN1_45S_50ntFlanks@10420.10457.38|CTGAGAGATGGGCGAGCGCCGTTCCGAAGGGACGGGCG</td><td>>hg38_21_+_8433172_8446622_RNA45SN1_45S_50ntFlanks</td><td>10420</td><td>10457</td><td>38</td><td>CTGAGAGATGGGCGAGCGCCGTTCCGAAGGGACGGGCG</td><td>182.2</td><td>-9.5</td><td>1.3</td><td>-7.5</td><td>8.6E-14</td><td>2.1E-13</td><td>BT20/MB468</td></tr>
<tr><td>>MI0003776|hsa-mir-320b-1&WithFlank&1|+|116671740|116671823@48.67.20(+1U);MI0003839|hsa-mir-320b-2&WithFlank&1|-|224257034|224257116@47.66.20(+1U);MI0000542|hsa-mir-320a&WithFlank&8|-|22244960|22245043@48.67.20(+1U)&[MIMAT0005792&hsa-miR-320b&offsets|0|-2(+1U);m-183&1|+|116671787|116671806&offsets|0|0(+1U)];[MIMAT0005792_1&hsa-miR-320b&offsets|0|-2(+1U);m-183&1|-|224257051|224257070&offsets|0|0(+1U)];[MIMAT0000510&hsa-miR-320a-3p&offsets|0|-2(+1U);m-60&8|-|22244975|22244996&offsets|0|-2(+1U)]|AAAAGCTGGGTTGAGAGGGCT</td><td>>MI0003776|hsa-mir-320b-1&WithFlank&1|+|116671740|116671823</td><td>48</td><td>67</td><td>20(+1U)</td><td>AAAAGCTGGGTTGAGAGGGCT</td><td>1058.3</td><td>9.9</td><td>1.3</td><td>7.5</td><td>8.6E-14</td><td>2.1E-13</td><td>BT20/MB468</td></tr>
<tr><td>>hg38_21_+_8433172_8446622_RNA45SN1_45S_50ntFlanks@4163.4182.20|CATCCAAGGAAGGCAGCAGG</td><td>>hg38_21_+_8433172_8446622_RNA45SN1_45S_50ntFlanks</td><td>4163</td><td>4182</td><td>20</td><td>CATCCAAGGAAGGCAGCAGG</td><td>665.3</td><td>-7.4</td><td>1.0</td><td>-7.5</td><td>8.6E-14</td><td>2.2E-13</td><td>BT20/MB468</td></tr>
<tr><td>>MI0001448|hsa-mir-425&WithFlank&3|-|49020142|49020240@20.43.24(+1G)&[MIMAT0003393&hsa-miR-425-5p&offsets|0|+1(+1G);m-95&3|-|49020200|49020221&offsets|0|+2(+1G)]|AATGACACGATCACTCCCGTTGAGG</td><td>>MI0001448|hsa-mir-425&WithFlank&3|-|49020142|49020240</td><td>20</td><td>43</td><td>24(+1G)</td><td>AATGACACGATCACTCCCGTTGAGG</td><td>1112.2</td><td>9.6</td><td>1.3</td><td>7.5</td><td>8.7E-14</td><td>2.2E-13</td><td>BT20/MB468</td></tr>
<tr><td>>hg38_21_+_8433172_8446622_RNA45SN1_45S_50ntFlanks@10347.10372.26|CAAACGAGAACTTTGAAGGCCGAAGT</td><td>>hg38_21_+_8433172_8446622_RNA45SN1_45S_50ntFlanks</td><td>10347</td><td>10372</td><td>26</td><td>CAAACGAGAACTTTGAAGGCCGAAGT</td><td>302.3</td><td>4.6</td><td>0.6</td><td>7.5</td><td>8.1E-14</td><td>2.2E-13</td><td>MB231/MB468</td></tr>
<tr><td>>am_trnaMT_PheGAA_MT_+_577_647@40.57.18|TGTTTAGACGGGCTCACA</td><td>>am_trnaMT_PheGAA_MT_+_577_647</td><td>40</td><td>57</td><td>18</td><td>TGTTTAGACGGGCTCACA</td><td>1115.0</td><td>9.0</td><td>1.2</td><td>7.5</td><td>8.1E-14</td><td>2.2E-13</td><td>MB231/MB468</td></tr>
<tr><td>>hg38_21_+_8433172_8446622_RNA45SN1_45S_50ntFlanks@10678.10711.34|CGGTTCCGGCGGCGTCCGGTGAGCTCTCGCTGGC</td><td>>hg38_21_+_8433172_8446622_RNA45SN1_45S_50ntFlanks</td><td>10678</td><td>10711</td><td>34</td><td>CGGTTCCGGCGGCGTCCGGTGAGCTCTCGCTGGC</td><td>32570.6</td><td>-6.7</td><td>0.9</td><td>-7.5</td><td>8.8E-14</td><td>2.2E-13</td><td>BT20/MB468</td></tr>
<tr><td>>trna2_GlyGCC_21_-_18827107_18827177@1.31.31,trna35_GlyGCC_1_+_161413094_161413164@1.31.31,trna37_GlyGCC_1_+_161420467_161420537@1.31.31,trna39_GlyGCC_1_+_161427898_161427968@1.31.31,trna41_GlyGCC_1_+_161435258_161435328@1.31.31|GCATGGGTGGTTCAGTGGTAGAATTCTCGCC</td><td>>trna2_GlyGCC_21_-_18827107_18827177</td><td>1</td><td>31</td><td>31</td><td>GCATGGGTGGTTCAGTGGTAGAATTCTCGCC</td><td>30985.9</td><td>6.2</td><td>0.8</td><td>7.8</td><td>6.0E-15</td><td>2.2E-13</td><td>BT20/MB231</td></tr>
<tr><td>>hg38_21_+_8433172_8446622_RNA45SN1_45S_50ntFlanks@13002.13026.25|CGATCTATTGAAAGTCAGCCCTCGA</td><td>>hg38_21_+_8433172_8446622_RNA45SN1_45S_50ntFlanks</td><td>13002</td><td>13026</td><td>25</td><td>CGATCTATTGAAAGTCAGCCCTCGA</td><td>157.4</td><td>-9.3</td><td>1.2</td><td>-7.5</td><td>8.9E-14</td><td>2.2E-13</td><td>BT20/MB468</td></tr>
<tr><td>>MI0002468|hsa-mir-484&WithFlank&16|+|15643288|15643378@14.35.22&[MIMAT0002174&hsa-miR-484&offsets|0|0;m-127&16|+|15643301|15643321&offsets|0|+1]|TCAGGCTCAGTCCCCTCCCGAT</td><td>>MI0002468|hsa-mir-484&WithFlank&16|+|15643288|15643378</td><td>14</td><td>35</td><td>22</td><td>TCAGGCTCAGTCCCCTCCCGAT</td><td>2967.6</td><td>5.9</td><td>0.8</td><td>7.5</td><td>8.9E-14</td><td>2.2E-13</td><td>BT20/MB468</td></tr>
<tr><td>>MI0000764|hsa-mir-363&WithFlank&X|-|134169372|134169458@56.76.21&[MIMAT0000707&hsa-miR-363-3p&offsets|0|-1;m-178&X|-|134169381|134169403&offsets|0|-2]|AATTGCACGGTATCCATCTGT</td><td>>MI0000764|hsa-mir-363&WithFlank&X|-|134169372|134169458</td><td>56</td><td>76</td><td>21</td><td>AATTGCACGGTATCCATCTGT</td><td>214.2</td><td>11.2</td><td>1.4</td><td>7.8</td><td>6.1E-15</td><td>2.2E-13</td><td>BT20/MB231</td></tr>
<tr><td>>hg38_21_+_8433172_8446622_RNA45SN1_45S_50ntFlanks@12407.12435.29|TTTTGATCCTTCGATGTCGGCTCTTCCTA</td><td>>hg38_21_+_8433172_8446622_RNA45SN1_45S_50ntFlanks</td><td>12407</td><td>12435</td><td>29</td><td>TTTTGATCCTTCGATGTCGGCTCTTCCTA</td><td>314.8</td><td>-4.7</td><td>0.6</td><td>-7.5</td><td>9.0E-14</td><td>2.2E-13</td><td>BT20/MB468</td></tr>
<tr><td>>MI0000115|hsa-mir-16-2&WithFlank&3|+|160404739|160404831@58.79.22&[MIMAT0004518&hsa-miR-16-2-3p&offsets|-1|-1;m-252&3|+|160404796|160404816&offsets|0|+1]|ACCAATATTACTGTGCTGCTTT</td><td>>MI0000115|hsa-mir-16-2&WithFlank&3|+|160404739|160404831</td><td>58</td><td>79</td><td>22</td><td>ACCAATATTACTGTGCTGCTTT</td><td>1996.2</td><td>6.1</td><td>0.8</td><td>7.5</td><td>9.0E-14</td><td>2.2E-13</td><td>BT20/MB468</td></tr>
<tr><td>>hg38_21_+_8433172_8446622_RNA45SN1_45S_50ntFlanks@11233.11252.20|CCCTTCCCGTGGATCGCCCC</td><td>>hg38_21_+_8433172_8446622_RNA45SN1_45S_50ntFlanks</td><td>11233</td><td>11252</td><td>20</td><td>CCCTTCCCGTGGATCGCCCC</td><td>256.0</td><td>-10.0</td><td>1.3</td><td>-7.5</td><td>9.1E-14</td><td>2.2E-13</td><td>BT20/MB468</td></tr>
<tr><td>>hg38_21_+_8433172_8446622_RNA45SN1_45S_50ntFlanks@8324.8356.33|CCGATAGTCAACAAGTACCGTAAGGGAAAGTTG</td><td>>hg38_21_+_8433172_8446622_RNA45SN1_45S_50ntFlanks</td><td>8324</td><td>8356</td><td>33</td><td>CCGATAGTCAACAAGTACCGTAAGGGAAAGTTG</td><td>204.9</td><td>-9.3</td><td>1.2</td><td>-7.5</td><td>8.5E-14</td><td>2.3E-13</td><td>MB231/MB468</td></tr>
<tr><td>>hg38_21_+_8433172_8446622_RNA45SN1_45S_50ntFlanks@12419.12438.20|GATGTCGGCTCTTCCTATCA</td><td>>hg38_21_+_8433172_8446622_RNA45SN1_45S_50ntFlanks</td><td>12419</td><td>12438</td><td>20</td><td>GATGTCGGCTCTTCCTATCA</td><td>781.4</td><td>4.7</td><td>0.6</td><td>7.5</td><td>9.2E-14</td><td>2.3E-13</td><td>BT20/MB468</td></tr>
<tr><td>>hg38_21_+_8433172_8446622_RNA45SN1_45S_50ntFlanks@3986.4023.38|GGCTTTGGTGACTCTAGATAACCTCGGGCCGATCGCAC</td><td>>hg38_21_+_8433172_8446622_RNA45SN1_45S_50ntFlanks</td><td>3986</td><td>4023</td><td>38</td><td>GGCTTTGGTGACTCTAGATAACCTCGGGCCGATCGCAC</td><td>811.6</td><td>7.7</td><td>1.0</td><td>7.5</td><td>8.7E-14</td><td>2.3E-13</td><td>MB231/MB468</td></tr>
<tr><td>>MI0000469|hsa-mir-125a&WithFlank&19|+|51693248|51693345@21.41.21(+1G)&[MIMAT0000443&hsa-miR-125a-5p&offsets|0|-3(+1G);m-63&19|+|51693268|51693289&offsets|0|-1(+1G)]|TCCCTGAGACCCTTTAACCTGG</td><td>>MI0000469|hsa-mir-125a&WithFlank&19|+|51693248|51693345</td><td>21</td><td>41</td><td>21(+1G)</td><td>TCCCTGAGACCCTTTAACCTGG</td><td>622.0</td><td>8.3</td><td>1.1</td><td>7.4</td><td>9.4E-14</td><td>2.3E-13</td><td>BT20/MB468</td></tr>
<tr><td>>MI0000089|hsa-mir-31&WithFlank&9|-|21512109|21512191@14.34.21(+1A)&[MIMAT0000089&hsa-miR-31-5p&offsets|0|0(+1A);m-360&9|-|21512157|21512178&offsets|0|-1(+1A)]|AGGCAAGATGCTGGCATAGCTA</td><td>>MI0000089|hsa-mir-31&WithFlank&9|-|21512109|21512191</td><td>14</td><td>34</td><td>21(+1A)</td><td>AGGCAAGATGCTGGCATAGCTA</td><td>742.2</td><td>10.9</td><td>1.5</td><td>7.4</td><td>9.4E-14</td><td>2.3E-13</td><td>BT20/MB468</td></tr>
<tr><td>>hg38_1_-_228634819_228635039_RNA5S12_5S_50ntFlanks@141.169.29|CCGCCTGGGAATACCGGGTGCTGTAGGCT</td><td>>hg38_1_-_228634819_228635039_RNA5S12_5S_50ntFlanks</td><td>141</td><td>169</td><td>29</td><td>CCGCCTGGGAATACCGGGTGCTGTAGGCT</td><td>2419.5</td><td>5.3</td><td>0.7</td><td>7.5</td><td>8.8E-14</td><td>2.4E-13</td><td>MB231/MB468</td></tr>
<tr><td>>hg38_21_+_8433172_8446622_RNA45SN1_45S_50ntFlanks@10574.10607.34|AAGCCGGCGGGAGCCCCGGGGAGAGTTCTCTTTT</td><td>>hg38_21_+_8433172_8446622_RNA45SN1_45S_50ntFlanks</td><td>10574</td><td>10607</td><td>34</td><td>AAGCCGGCGGGAGCCCCGGGGAGAGTTCTCTTTT</td><td>212.2</td><td>-9.7</td><td>1.3</td><td>-7.4</td><td>9.5E-14</td><td>2.4E-13</td><td>BT20/MB468</td></tr>
<tr><td>>hg38_21_+_8433172_8446622_RNA45SN1_45S_50ntFlanks@10781.10806.26|AGGTGAACAGCCTCTGGCATGTTGGA</td><td>>hg38_21_+_8433172_8446622_RNA45SN1_45S_50ntFlanks</td><td>10781</td><td>10806</td><td>26</td><td>AGGTGAACAGCCTCTGGCATGTTGGA</td><td>235.0</td><td>-9.9</td><td>1.3</td><td>-7.4</td><td>9.6E-14</td><td>2.4E-13</td><td>BT20/MB468</td></tr>
<tr><td>>hg38_21_+_8433172_8446622_RNA45SN1_45S_50ntFlanks@10829.10858.30|GCAAGCCGGATCCGTAACTTCGGGATAAGG</td><td>>hg38_21_+_8433172_8446622_RNA45SN1_45S_50ntFlanks</td><td>10829</td><td>10858</td><td>30</td><td>GCAAGCCGGATCCGTAACTTCGGGATAAGG</td><td>202.5</td><td>-4.9</td><td>0.7</td><td>-7.4</td><td>9.6E-14</td><td>2.4E-13</td><td>BT20/MB468</td></tr>
<tr><td>>hg38_21_+_8433172_8446622_RNA45SN1_45S_50ntFlanks@9714.9735.22|CGAATGATTAGAGGTCTTGGGG</td><td>>hg38_21_+_8433172_8446622_RNA45SN1_45S_50ntFlanks</td><td>9714</td><td>9735</td><td>22</td><td>CGAATGATTAGAGGTCTTGGGG</td><td>145.5</td><td>-6.0</td><td>0.8</td><td>-7.4</td><td>9.8E-14</td><td>2.4E-13</td><td>BT20/MB468</td></tr>
<tr><td>>hg38_21_+_8433172_8446622_RNA45SN1_45S_50ntFlanks@5487.5514.28|CGCTGAGAAGACGGTCGAACTTGACTAT</td><td>>hg38_21_+_8433172_8446622_RNA45SN1_45S_50ntFlanks</td><td>5487</td><td>5514</td><td>28</td><td>CGCTGAGAAGACGGTCGAACTTGACTAT</td><td>1606.8</td><td>-4.9</td><td>0.7</td><td>-7.4</td><td>9.9E-14</td><td>2.4E-13</td><td>BT20/MB468</td></tr>
<tr><td>>MI0000471|hsa-mir-126&WithFlank&9|+|136670596|136670692@59.79.21&[MIMAT0000445&hsa-miR-126-3p&offsets|+1|0;m-40&9|+|136670653|136670674&offsets|+1|0]|CGTACCGTGAGTAATAATGCG</td><td>>MI0000471|hsa-mir-126&WithFlank&9|+|136670596|136670692</td><td>59</td><td>79</td><td>21</td><td>CGTACCGTGAGTAATAATGCG</td><td>682.3</td><td>10.4</td><td>1.4</td><td>7.4</td><td>9.9E-14</td><td>2.4E-13</td><td>BT20/MB468</td></tr>
<tr><td>>hg38_21_+_8433172_8446622_RNA45SN1_45S_50ntFlanks@10850.10868.19|GGGATAAGGATTGGCTCTA</td><td>>hg38_21_+_8433172_8446622_RNA45SN1_45S_50ntFlanks</td><td>10850</td><td>10868</td><td>19</td><td>GGGATAAGGATTGGCTCTA</td><td>119.7</td><td>-3.9</td><td>0.5</td><td>-7.4</td><td>9.9E-14</td><td>2.5E-13</td><td>BT20/MB468</td></tr>
<tr><td>>hg38_21_+_8433172_8446622_RNA45SN1_45S_50ntFlanks@6743.6772.30|CGACACTTCGAACGCACTTGCGGCCCCGGG</td><td>>hg38_21_+_8433172_8446622_RNA45SN1_45S_50ntFlanks</td><td>6743</td><td>6772</td><td>30</td><td>CGACACTTCGAACGCACTTGCGGCCCCGGG</td><td>308.3</td><td>-5.9</td><td>0.8</td><td>-7.4</td><td>1.0E-13</td><td>2.5E-13</td><td>BT20/MB468</td></tr>
<tr><td>>hg38_21_+_8433172_8446622_RNA45SN1_45S_50ntFlanks@10843.10862.20|TAACTTCGGGATAAGGATTG</td><td>>hg38_21_+_8433172_8446622_RNA45SN1_45S_50ntFlanks</td><td>10843</td><td>10862</td><td>20</td><td>TAACTTCGGGATAAGGATTG</td><td>420.7</td><td>-4.3</td><td>0.6</td><td>-7.5</td><td>9.3E-14</td><td>2.5E-13</td><td>MB231/MB468</td></tr>
<tr><td>>hg38_21_+_8433172_8446622_RNA45SN1_45S_50ntFlanks@6790.6807.18|CCTGTCTGAGCGTCGCTT</td><td>>hg38_21_+_8433172_8446622_RNA45SN1_45S_50ntFlanks</td><td>6790</td><td>6807</td><td>18</td><td>CCTGTCTGAGCGTCGCTT</td><td>1742.5</td><td>5.7</td><td>0.8</td><td>7.4</td><td>9.4E-14</td><td>2.5E-13</td><td>MB231/MB468</td></tr>
<tr><td>>MI0000074|hsa-mir-19b-1&WithFlank&13|+|91351186|91351284@60.81.22;MI0000075|hsa-mir-19b-2&WithFlank&X|-|134169665|134169772@68.89.22&[MIMAT0000074&hsa-miR-19b-3p&offsets|0|-1;m-122&13|+|91351245|91351267&offsets|0|-1];[MIMAT0000074_1&hsa-miR-19b-3p&offsets|0|-1;m-121&X|-|134169683|134169705&offsets|0|-1]|TGTGCAAATCCATGCAAAACTG</td><td>>MI0000074|hsa-mir-19b-1&WithFlank&13|+|91351186|91351284</td><td>60</td><td>81</td><td>22</td><td>TGTGCAAATCCATGCAAAACTG</td><td>1652.2</td><td>7.2</td><td>1.0</td><td>7.4</td><td>1.0E-13</td><td>2.5E-13</td><td>BT20/MB468</td></tr>
<tr><td>>hg38_21_+_8433172_8446622_RNA45SN1_45S_50ntFlanks@10826.10844.19|TCGGCAAGCCGGATCCGTA</td><td>>hg38_21_+_8433172_8446622_RNA45SN1_45S_50ntFlanks</td><td>10826</td><td>10844</td><td>19</td><td>TCGGCAAGCCGGATCCGTA</td><td>432.3</td><td>-10.7</td><td>1.4</td><td>-7.4</td><td>1.0E-13</td><td>2.5E-13</td><td>BT20/MB468</td></tr>
<tr><td>>hg38_21_+_8433172_8446622_RNA45SN1_45S_50ntFlanks@11766.11803.38|ATCTAATTAGTGACGCGCATGAATGGATGAACGAGATT</td><td>>hg38_21_+_8433172_8446622_RNA45SN1_45S_50ntFlanks</td><td>11766</td><td>11803</td><td>38</td><td>ATCTAATTAGTGACGCGCATGAATGGATGAACGAGATT</td><td>217.3</td><td>-9.8</td><td>1.3</td><td>-7.4</td><td>1.0E-13</td><td>2.5E-13</td><td>BT20/MB468</td></tr>
<tr><td>>hg38_1_-_228634819_228635039_RNA5S12_5S_50ntFlanks@51.84.34|GTCTACGGCCATACCACCCTGAACGCGCCCGATC</td><td>>hg38_1_-_228634819_228635039_RNA5S12_5S_50ntFlanks</td><td>51</td><td>84</td><td>34</td><td>GTCTACGGCCATACCACCCTGAACGCGCCCGATC</td><td>3163.6</td><td>5.8</td><td>0.8</td><td>7.4</td><td>9.4E-14</td><td>2.5E-13</td><td>MB231/MB468</td></tr>
<tr><td>>hg38_21_+_8433172_8446622_RNA45SN1_45S_50ntFlanks@12356.12376.21|ACCACAGGGATAACTGGCTTG</td><td>>hg38_21_+_8433172_8446622_RNA45SN1_45S_50ntFlanks</td><td>12356</td><td>12376</td><td>21</td><td>ACCACAGGGATAACTGGCTTG</td><td>255.8</td><td>-9.6</td><td>1.3</td><td>-7.4</td><td>9.5E-14</td><td>2.5E-13</td><td>MB231/MB468</td></tr>
</table>

<table>
<tr><th></th><th></th><th></th><th></th><th></th><th></th><th></th><th></th><th></th><th></th><th></th><th></th><th></th></tr>
<tr><td>>am_trna162_MetCAT_6_-_26758550_26758622@31.53.23,trna164_MetCAT_6_-_26735574_26735646@31.53.23,trna20_MetCAT_16_+_71460396_71460468@31.53.23,trna21_MetCAT_6_+_26701712_26701784@31.53.23,trna27_MetCAT_6_+_26766444_26766516@31.53.23,trna75_MetCAT_6_+_28912352_28912424@31.53.23,trna97_MetCAT_6_-_28921042_28921114@31.53.23|TCTCATAATCTGAAGGTCCTGAG</td><td>>am_trna162_MetCAT_6_-_26758550_26758622</td><td>31</td><td>53</td><td>23</td><td>TCTCATAATCTGAAGGTCCTGAG</td><td>297.9</td><td>-9.8</td><td>1.3</td><td>-7.4</td><td>9.5E-14</td><td>2.5E-13</td><td>MB231/MB468</td></tr>
<tr><td>>hg38_21_+_8433172_8446622_RNA45SN1_45S_50ntFlanks@12539.12565.27|ATGATGTGTTGTTGCCATGGTAATCCT</td><td>>hg38_21_+_8433172_8446622_RNA45SN1_45S_50ntFlanks</td><td>12539</td><td>12565</td><td>27</td><td>ATGATGTGTTGTTGCCATGGTAATCCT</td><td>209.5</td><td>-9.7</td><td>1.3</td><td>-7.4</td><td>1.0E-13</td><td>2.5E-13</td><td>BT20/MB468</td></tr>
<tr><td>>hg38_21_+_8433172_8446622_RNA45SN1_45S_50ntFlanks@12399.12419.21|ACGTCGCTTTTTGATCCTTCG</td><td>>hg38_21_+_8433172_8446622_RNA45SN1_45S_50ntFlanks</td><td>12399</td><td>12419</td><td>21</td><td>ACGTCGCTTTTTGATCCTTCG</td><td>170.3</td><td>-4.0</td><td>0.5</td><td>-7.4</td><td>1.0E-13</td><td>2.6E-13</td><td>BT20/MB468</td></tr>
<tr><td>>hg38_21_+_8433172_8446622_RNA45SN1_45S_50ntFlanks@12993.13039.47|CCCTCGCTGCGATCTATTGAAAGTCAGCCCTCGACACAAGGGTTTGT</td><td>>hg38_21_+_8433172_8446622_RNA45SN1_45S_50ntFlanks</td><td>12993</td><td>13039</td><td>47</td><td>CCCTCGCTGCGATCTATTGAAAGTCAGCCCTCGACACAAGGGTTTGT</td><td>13475.7</td><td>7.4</td><td>1.0</td><td>7.4</td><td>9.6E-14</td><td>2.6E-13</td><td>MB231/MB468</td></tr>
<tr><td>>hg38_21_+_8433172_8446622_RNA45SN1_45S_50ntFlanks@8182.8204.23|ATCGAGGCCCAGCCCGTGGACGG</td><td>>hg38_21_+_8433172_8446622_RNA45SN1_45S_50ntFlanks</td><td>8182</td><td>8204</td><td>23</td><td>ATCGAGGCCCAGCCCGTGGACGG</td><td>159.8</td><td>-9.3</td><td>1.3</td><td>-7.4</td><td>1.0E-13</td><td>2.6E-13</td><td>BT20/MB468</td></tr>
<tr><td>>hg38_21_+_8433172_8446622_RNA45SN1_45S_50ntFlanks@10408.10434.27|GGTCAGTCGGTCCTGAGAGATGGGCGA</td><td>>hg38_21_+_8433172_8446622_RNA45SN1_45S_50ntFlanks</td><td>10408</td><td>10434</td><td>27</td><td>GGTCAGTCGGTCCTGAGAGATGGGCGA</td><td>152.3</td><td>-9.2</td><td>1.2</td><td>-7.4</td><td>1.0E-13</td><td>2.6E-13</td><td>BT20/MB468</td></tr>
<tr><td>>hg38_21_+_8433172_8446622_RNA45SN1_45S_50ntFlanks@12360.12382.23|CAGGGATAACTGGCTTGTGGCGG</td><td>>hg38_21_+_8433172_8446622_RNA45SN1_45S_50ntFlanks</td><td>12360</td><td>12382</td><td>23</td><td>CAGGGATAACTGGCTTGTGGCGG</td><td>189.9</td><td>-6.4</td><td>0.9</td><td>-7.4</td><td>1.1E-13</td><td>2.6E-13</td><td>BT20/MB468</td></tr>
<tr><td>>hg38_21_+_8433172_8446622_RNA45SN1_45S_50ntFlanks@6719.6741.23|TGAATTGCAGGACACATTGATCA</td><td>>hg38_21_+_8433172_8446622_RNA45SN1_45S_50ntFlanks</td><td>6719</td><td>6741</td><td>23</td><td>TGAATTGCAGGACACATTGATCA</td><td>370.4</td><td>-3.3</td><td>0.4</td><td>-7.4</td><td>9.7E-14</td><td>2.6E-13</td><td>MB231/MB468</td></tr>
<tr><td>>hg38_21_+_8433172_8446622_RNA45SN1_45S_50ntFlanks@7986.8004.19|ATCAGACGTGGCGACCCGC</td><td>>hg38_21_+_8433172_8446622_RNA45SN1_45S_50ntFlanks</td><td>7986</td><td>8004</td><td>19</td><td>ATCAGACGTGGCGACCCGC</td><td>245.7</td><td>-9.9</td><td>1.3</td><td>-7.4</td><td>1.1E-13</td><td>2.6E-13</td><td>BT20/MB468</td></tr>
<tr><td>>hg38_21_+_8433172_8446622_RNA45SN1_45S_50ntFlanks@10392.10419.28|ACAGCAGTTGAACATGGGTCAGTCGGTC</td><td>>hg38_21_+_8433172_8446622_RNA45SN1_45S_50ntFlanks</td><td>10392</td><td>10419</td><td>28</td><td>ACAGCAGTTGAACATGGGTCAGTCGGTC</td><td>465.4</td><td>-6.9</td><td>0.9</td><td>-7.4</td><td>1.1E-13</td><td>2.6E-13</td><td>BT20/MB468</td></tr>
<tr><td>>hg38_21_+_8433172_8446622_RNA45SN1_45S_50ntFlanks@8170.8188.19|CAAGTCCTTCTGATCGAGG</td><td>>hg38_21_+_8433172_8446622_RNA45SN1_45S_50ntFlanks</td><td>8170</td><td>8188</td><td>19</td><td>CAAGTCCTTCTGATCGAGG</td><td>656.4</td><td>-10.9</td><td>1.5</td><td>-7.4</td><td>9.9E-14</td><td>2.6E-13</td><td>MB231/MB468</td></tr>
<tr><td>>hg38_21_+_8433172_8446622_RNA45SN1_45S_50ntFlanks@4995.5024.30|ACAGATTGATAGCTCTTTCTCGATTCCGTG</td><td>>hg38_21_+_8433172_8446622_RNA45SN1_45S_50ntFlanks</td><td>4995</td><td>5024</td><td>30</td><td>ACAGATTGATAGCTCTTTCTCGATTCCGTG</td><td>301.5</td><td>-4.9</td><td>0.7</td><td>-7.4</td><td>1.1E-13</td><td>2.6E-13</td><td>BT20/MB468</td></tr>
<tr><td>>hg38_21_+_8433172_8446622_RNA45SN1_45S_50ntFlanks@8182.8208.27|ATCGAGGCCCAGCCCGTGGACGGTGTG</td><td>>hg38_21_+_8433172_8446622_RNA45SN1_45S_50ntFlanks</td><td>8182</td><td>8208</td><td>27</td><td>ATCGAGGCCCAGCCCGTGGACGGTGTG</td><td>148.8</td><td>-9.2</td><td>1.2</td><td>-7.4</td><td>1.1E-13</td><td>2.6E-13</td><td>BT20/MB468</td></tr>
<tr><td>>hg38_21_+_8433172_8446622_RNA45SN1_45S_50ntFlanks@12396.12415.20|GCGACGTCGCTTTTTGATCC</td><td>>hg38_21_+_8433172_8446622_RNA45SN1_45S_50ntFlanks</td><td>12396</td><td>12415</td><td>20</td><td>GCGACGTCGCTTTTTGATCC</td><td>337.4</td><td>4.1</td><td>0.5</td><td>7.4</td><td>1.0E-13</td><td>2.7E-13</td><td>MB231/MB468</td></tr>
<tr><td>>hg38_21_+_8433172_8446622_RNA45SN1_45S_50ntFlanks@12539.12557.19|ATGATGTGTTGTTGCCATG</td><td>>hg38_21_+_8433172_8446622_RNA45SN1_45S_50ntFlanks</td><td>12539</td><td>12557</td><td>19</td><td>ATGATGTGTTGTTGCCATG</td><td>175.8</td><td>-9.5</td><td>1.3</td><td>-7.4</td><td>1.1E-13</td><td>2.7E-13</td><td>BT20/MB468</td></tr>
<tr><td>>hg38_21_+_8433172_8446622_RNA45SN1_45S_50ntFlanks@6670.6698.29|ACTCGGCTCGTGCGTCGATGAAGAACGCA</td><td>>hg38_21_+_8433172_8446622_RNA45SN1_45S_50ntFlanks</td><td>6670</td><td>6698</td><td>29</td><td>ACTCGGCTCGTGCGTCGATGAAGAACGCA</td><td>234.4</td><td>-3.3</td><td>0.4</td><td>-7.4</td><td>1.1E-13</td><td>2.7E-13</td><td>BT20/MB468</td></tr>
<tr><td>>hg38_21_+_8433172_8446622_RNA45SN1_45S_50ntFlanks@10471.10488.18|CCCTCGGCCGATCGAAAG</td><td>>hg38_21_+_8433172_8446622_RNA45SN1_45S_50ntFlanks</td><td>10471</td><td>10488</td><td>18</td><td>CCCTCGGCCGATCGAAAG</td><td>502.5</td><td>-4.5</td><td>0.6</td><td>-7.4</td><td>1.1E-13</td><td>2.7E-13</td><td>BT20/MB468</td></tr>
<tr><td>>hg38_21_+_8433172_8446622_RNA45SN1_45S_50ntFlanks@11873.11907.35|AAGAAGACCCTGTTGAGCTTGACTCTAGTCTGGCA</td><td>>hg38_21_+_8433172_8446622_RNA45SN1_45S_50ntFlanks</td><td>11873</td><td>11907</td><td>35</td><td>AAGAAGACCCTGTTGAGCTTGACTCTAGTCTGGCA</td><td>212.2</td><td>-9.3</td><td>1.3</td><td>-7.4</td><td>1.0E-13</td><td>2.7E-13</td><td>MB231/MB468</td></tr>
<tr><td>>hg38_21_+_8433172_8446622_RNA45SN1_45S_50ntFlanks@10794.10823.30|CTGGCATGTTGGAACAATGTAGGTAAGGGA</td><td>>hg38_21_+_8433172_8446622_RNA45SN1_45S_50ntFlanks</td><td>10794</td><td>10823</td><td>30</td><td>CTGGCATGTTGGAACAATGTAGGTAAGGGA</td><td>216.7</td><td>-6.4</td><td>0.9</td><td>-7.4</td><td>1.1E-13</td><td>2.7E-13</td><td>BT20/MB468</td></tr>
<tr><td>>hg38_21_+_8433172_8446622_RNA45SN1_45S_50ntFlanks@10916.10933.18|CCGCGGCTGGACGAGGCG</td><td>>hg38_21_+_8433172_8446622_RNA45SN1_45S_50ntFlanks</td><td>10916</td><td>10933</td><td>18</td><td>CCGCGGCTGGACGAGGCG</td><td>213.6</td><td>-5.3</td><td>0.7</td><td>-7.4</td><td>1.0E-13</td><td>2.7E-13</td><td>MB231/MB468</td></tr>
<tr><td>>am_trna111_HisGTG_1_-_147774845_147774916@1.29.29,trna118_HisGTG_1_-_145396881_145396952@1.29.29,trna16_HisGTG_1_+_146544773_146544844@1.29.29,trna1_HisGTG_15_+_45493349_45493420@1.29.29,trna21_HisGTG_1_+_147753471_147753542@1.29.29,trna33_HisGTG_6_+_27125906_27125977@1.29.29,trna7_HisGTG_9_-_14433938_14434009@1.29.29,trna8_HisGTG_15_-_45492611_45492682@1.29.29,trna9_HisGTG_15_-_45490804_45490875@1.29.29|GCCGTGATCGTATAGTGGTTAGTACTCTG</td><td>>am_trna111_HisGTG_1_-_147774845_147774916</td><td>1</td><td>29</td><td>29</td><td>GCCGTGATCGTATAGTGGTTAGTACTCTG</td><td>4354.6</td><td>-5.4</td><td>0.7</td><td>-7.4</td><td>1.1E-13</td><td>2.8E-13</td><td>BT20/MB468</td></tr>
<tr><td>>hg38_21_+_8433172_8446622_RNA45SN1_45S_50ntFlanks@11314.11344.31|CCCACCCCACGTCTCGTCGCGCGCGCGTCCG</td><td>>hg38_21_+_8433172_8446622_RNA45SN1_45S_50ntFlanks</td><td>11314</td><td>11344</td><td>31</td><td>CCCACCCCACGTCTCGTCGCGCGCGCGTCCG</td><td>215.3</td><td>-9.3</td><td>1.3</td><td>-7.4</td><td>1.0E-13</td><td>2.8E-13</td><td>MB231/MB468</td></tr>
<tr><td>>hg38_21_+_8433172_8446622_RNA45SN1_45S_50ntFlanks@4006.4025.20|ACCTCGGGCCGATCGCACGC</td><td>>hg38_21_+_8433172_8446622_RNA45SN1_45S_50ntFlanks</td><td>4006</td><td>4025</td><td>20</td><td>ACCTCGGGCCGATCGCACGC</td><td>266.3</td><td>-6.1</td><td>0.8</td><td>-7.4</td><td>1.1E-13</td><td>2.8E-13</td><td>BT20/MB468</td></tr>
<tr><td>>hg38_21_+_8433172_8446622_RNA45SN1_45S_50ntFlanks@6721.6752.32|AATTGCAGGACACATTGATCATCGACACTTCG</td><td>>hg38_21_+_8433172_8446622_RNA45SN1_45S_50ntFlanks</td><td>6721</td><td>6752</td><td>32</td><td>AATTGCAGGACACATTGATCATCGACACTTCG</td><td>336.8</td><td>-3.0</td><td>0.4</td><td>-7.4</td><td>1.1E-13</td><td>2.8E-13</td><td>BT20/MB468</td></tr>
<tr><td>>hg38_21_+_8433172_8446622_RNA45SN1_45S_50ntFlanks@3787.3812.26|AAACTGCGAATGGCTCATTAAATCAG</td><td>>hg38_21_+_8433172_8446622_RNA45SN1_45S_50ntFlanks</td><td>3787</td><td>3812</td><td>26</td><td>AAACTGCGAATGGCTCATTAAATCAG</td><td>218.0</td><td>-9.8</td><td>1.3</td><td>-7.4</td><td>1.1E-13</td><td>2.8E-13</td><td>BT20/MB468</td></tr>
<tr><td>>hg38_21_+_8433172_8446622_RNA45SN1_45S_50ntFlanks@10805.10826.22|GAACAATGTAGGTAAGGGAAGT</td><td>>hg38_21_+_8433172_8446622_RNA45SN1_45S_50ntFlanks</td><td>10805</td><td>10826</td><td>22</td><td>GAACAATGTAGGTAAGGGAAGT</td><td>166.6</td><td>-9.4</td><td>1.3</td><td>-7.4</td><td>1.1E-13</td><td>2.8E-13</td><td>BT20/MB468</td></tr>
<tr><td>>hg38_21_+_8433172_8446622_RNA45SN1_45S_50ntFlanks@10571.10591.21|GAGAAGCCGGCGGGAGCCCCG</td><td>>hg38_21_+_8433172_8446622_RNA45SN1_45S_50ntFlanks</td><td>10571</td><td>10591</td><td>21</td><td>GAGAAGCCGGCGGGAGCCCCG</td><td>394.1</td><td>-10.2</td><td>1.4</td><td>-7.4</td><td>1.0E-13</td><td>2.8E-13</td><td>MB231/MB468</td></tr>
<tr><td>>MI0000100|hsa-mir-98&WithFlank&X|-|53556217|53556347@28.49.22&[MIMAT0000096&hsa-miR-98-5p&offsets|0|0;m-143&X|-|53556299|53556320&offsets|0|0]|TGAGGTAGTAAGTTGTATTGTT</td><td>>MI0000100|hsa-mir-98&WithFlank&X|-|53556217|53556347</td><td>28</td><td>49</td><td>22</td><td>TGAGGTAGTAAGTTGTATTGTT</td><td>13629.8</td><td>5.5</td><td>0.7</td><td>7.4</td><td>1.1E-13</td><td>2.8E-13</td><td>MB231/MB468</td></tr>
<tr><td>>MI0000103|hsa-mir-101-1&WithFlank&1|-|65058428|65058514@53.72.20(+2U);MI0000739|hsa-mir-101-2&WithFlank&9|+|4850291|4850381@55.74.20(+2U)&[MIMAT0000099&hsa-miR-101-3p&offsets|0|-1(+2U);m-17&1|-|65058443|65058462&offsets|0|0(+2U)];[MIMAT0000099_1&hsa-miR-101-3p&offsets|0|-1(+2U);m-16&9|+|4850345|4850364&offsets|0|0(+2U)]|TACAGTACTGTGATAACTGATT</td><td>>MI0000103|hsa-mir-101-1&WithFlank&1|-|65058428|65058514</td><td>53</td><td>72</td><td>20(+2U)</td><td>TACAGTACTGTGATAACTGATT</td><td>877.1</td><td>9.9</td><td>1.3</td><td>7.4</td><td>1.1E-13</td><td>2.8E-13</td><td>MB231/MB468</td></tr>
<tr><td>>hg38_21_+_8433172_8446622_RNA45SN1_45S_50ntFlanks@12405.12428.24|CTTTTTGATCCTTCGATGTCGGCT</td><td>>hg38_21_+_8433172_8446622_RNA45SN1_45S_50ntFlanks</td><td>12405</td><td>12428</td><td>24</td><td>CTTTTTGATCCTTCGATGTCGGCT</td><td>125.8</td><td>-5.4</td><td>0.7</td><td>-7.4</td><td>1.2E-13</td><td>2.8E-13</td><td>BT20/MB468</td></tr>
<tr><td>>hg38_21_+_8433172_8446622_RNA45SN1_45S_50ntFlanks@7986.8016.31|ATCAGACGTGGCGACCCGCTGAATTTAAGCA</td><td>>hg38_21_+_8433172_8446622_RNA45SN1_45S_50ntFlanks</td><td>7986</td><td>8016</td><td>31</td><td>ATCAGACGTGGCGACCCGCTGAATTTAAGCA</td><td>184.7</td><td>-9.5</td><td>1.3</td><td>-7.4</td><td>1.2E-13</td><td>2.9E-13</td><td>BT20/MB468</td></tr>
<tr><td>>hg38_21_+_8433172_8446622_RNA45SN1_45S_50ntFlanks@12449.12480.32|GAATTCACCAAGCGTTGGATTGTTCACCCACT</td><td>>hg38_21_+_8433172_8446622_RNA45SN1_45S_50ntFlanks</td><td>12449</td><td>12480</td><td>32</td><td>GAATTCACCAAGCGTTGGATTGTTCACCCACT</td><td>633.0</td><td>9.3</td><td>1.3</td><td>7.4</td><td>1.2E-13</td><td>2.9E-13</td><td>BT20/MB468</td></tr>
<tr><td>>trna10_AspGTC_12_-_125424193_125424264@35.73.39,trna12_AspGTC_12_-_125411891_125411962@35.73.39,trna144_AspGTC_6_-_27551236_27551307@35.73.39,trna38_AspGTC_17_-_8125556_8125627@35.73.39,trna45_AspGTC_6_+_27447453_27447524@35.73.39,trna48_AspGTC_6_+_27471523_27471594@35.73.39,trna4_AspGTC_12_+_96429799_96429870@35.73.39,trna69_AspGTC_1_-_161440205_161440276@35.73.39,trna72_AspGTC_1_-_161432824_161432895@35.73.39,trna75_AspGTC_1_-_161425414_161425485@35.73.39,trna78_AspGTC_1_-_161418033_161418104@35.73.39,trna81_AspGTC_1_-_161410615_161410686@35.73.39|TCACGCGGGAGACCGGGGTTCGATTCCCCGACGGGGAGC</td><td>>trna10_AspGTC_12_-_125424193_125424264</td><td>35</td><td>73</td><td>39</td><td>TCACGCGGGAGACCGGGGTTCGATTCCCCGACGGGGAGC</td><td>1229.5</td><td>7.3</td><td>1.0</td><td>7.4</td><td>1.1E-13</td><td>2.9E-13</td><td>MB231/MB468</td></tr>
<tr><td>>MI0000440|hsa-mir-27b&WithFlank&9|+|95085439|95085547@66.86.21(+1U)&[MIMAT0000419&hsa-miR-27b-3p&offsets|-1|-1(+1U);m-39&9|+|95085505|95085525&offsets|-1|-1(+1U)]|GTTCACAGTGGCTAAGTTCTGT</td><td>>MI0000440|hsa-mir-27b&WithFlank&9|+|95085439|95085547</td><td>66</td><td>86</td><td>21(+1U)</td><td>GTTCACAGTGGCTAAGTTCTGT</td><td>402.0</td><td>8.7</td><td>1.2</td><td>7.4</td><td>1.2E-13</td><td>2.9E-13</td><td>BT20/MB468</td></tr>
<tr><td>>hg38_21_+_8433172_8446622_RNA45SN1_45S_50ntFlanks@475.520.46|CTTCGTGATCGATGTGGTGACGTCGTGCTCTCCCGGGCCGGGTCCG</td><td>>hg38_21_+_8433172_8446622_RNA45SN1_45S_50ntFlanks</td><td>475</td><td>520</td><td>46</td><td>CTTCGTGATCGATGTGGTGACGTCGTGCTCTCCCGGGCCGGGTCCG</td><td>771.3</td><td>9.8</td><td>1.3</td><td>7.4</td><td>1.2E-13</td><td>2.9E-13</td><td>BT20/MB468</td></tr>
<tr><td>>MI0000081|hsa-mir-24-2&WithFlank&19|-|13836281|13836365@56.77.22;MI0000080|hsa-mir-24-1&WithFlank&9|+|95086015|95086094@50.71.22&[MIMAT0000080&hsa-miR-24-3p&offsets|0|0;m-36&19|-|13836288|13836310&offsets|0|-1];[MIMAT0000080_1&hsa-miR-24-3p&offsets|0|0;m-35&9|+|95086064|95086086&offsets|0|-1]|TGGCTCAGTTCAGCAGGAACAG</td><td>>MI0000081|hsa-mir-24-2&WithFlank&19|-|13836281|13836365</td><td>56</td><td>77</td><td>22</td><td>TGGCTCAGTTCAGCAGGAACAG</td><td>21079.7</td><td>5.1</td><td>0.7</td><td>7.4</td><td>1.2E-13</td><td>3.0E-13</td><td>BT20/MB468</td></tr>
<tr><td>>hg38_21_+_8433172_8446622_RNA45SN1_45S_50ntFlanks@6647.6676.30|CGTACGACTCTTAGCGGTGGATCACTCGGC</td><td>>hg38_21_+_8433172_8446622_RNA45SN1_45S_50ntFlanks</td><td>6647</td><td>6676</td><td>30</td><td>CGTACGACTCTTAGCGGTGGATCACTCGGC</td><td>616.0</td><td>7.4</td><td>1.0</td><td>7.4</td><td>1.2E-13</td><td>3.0E-13</td><td>BT20/MB468</td></tr>
<tr><td>>hg38_21_+_8433172_8446622_RNA45SN1_45S_50ntFlanks@6719.6742.24|TGAATTGCAGGACACATTGATCAT</td><td>>hg38_21_+_8433172_8446622_RNA45SN1_45S_50ntFlanks</td><td>6719</td><td>6742</td><td>24</td><td>TGAATTGCAGGACACATTGATCAT</td><td>658.8</td><td>-8.2</td><td>1.1</td><td>-7.4</td><td>1.2E-13</td><td>3.0E-13</td><td>BT20/MB468</td></tr>
<tr><td>>hg38_21_+_8433172_8446622_RNA45SN1_45S_50ntFlanks@11739.11758.20|TCTCTTAAGGTAGCCAAATG</td><td>>hg38_21_+_8433172_8446622_RNA45SN1_45S_50ntFlanks</td><td>11739</td><td>11758</td><td>20</td><td>TCTCTTAAGGTAGCCAAATG</td><td>268.1</td><td>-10.1</td><td>1.4</td><td>-7.4</td><td>1.3E-13</td><td>3.1E-13</td><td>BT20/MB468</td></tr>
</table>

| | | | | | | | | | | | | |
|---|---|---|---|---|---|---|---|---|---|---|---|---|
| >hg38_21_+_8433172_8446622_RNA45SN1_45S_50ntFlanks@11318.11343.26\|CCCCACGTCTCGTCGCGCGCGCGTCC | >hg38_21_+_8433172_8446622_RNA45SN1_45S_50ntFlanks | 11318 | 11343 | 26 | CCCCACGTCTCGTCGCGCGCGCGTCC | 246.5 | -9.5 | 1.3 | -7.4 | 1.2E-13 | 3.1E-13 | MB231/MB468 |
| >hg38_21_+_8433172_8446622_RNA45SN1_45S_50ntFlanks@11769.11804.36\|TAATTAGTGACGCGCATGAATGGATGAACGAGATTC | >hg38_21_+_8433172_8446622_RNA45SN1_45S_50ntFlanks | 11769 | 11804 | 36 | TAATTAGTGACGCGCATGAATGGATGAACGAGATTC | 400.0 | -4.8 | 0.7 | -7.4 | 1.3E-13 | 3.1E-13 | BT20/MB468 |
| >hg38_21_+_8433172_8446622_RNA45SN1_45S_50ntFlanks@11575.11599.25\|ACTTAGAACTGGTGCGGACCAGGGG | >hg38_21_+_8433172_8446622_RNA45SN1_45S_50ntFlanks | 11575 | 11599 | 25 | ACTTAGAACTGGTGCGGACCAGGGG | 168.5 | -6.0 | 0.8 | -7.4 | 1.3E-13 | 3.1E-13 | BT20/MB468 |
| >hg38_21_+_8433172_8446622_RNA45SN1_45S_50ntFlanks@9545.9573.29\|GTGGAGGTCCGTAGCGGTCCTGACGTGCA | >hg38_21_+_8433172_8446622_RNA45SN1_45S_50ntFlanks | 9545 | 9573 | 29 | GTGGAGGTCCGTAGCGGTCCTGACGTGCA | 878.4 | -3.4 | 0.5 | -7.4 | 1.3E-13 | 3.1E-13 | BT20/MB468 |
| >am_trnaMT_PheGAA_MT_+_577_647@44.62.19\|TAGACGGGCTCACATCACC | >am_trnaMT_PheGAA_MT_+_577_647 | 44 | 62 | 19 | TAGACGGGCTCACATCACC | 1058.1 | 5.4 | 0.7 | 7.4 | 1.2E-13 | 3.1E-13 | MB231/MB468 |
| >hg38_1_-_228634819_228635039_RNA5S12_5S_50ntFlanks@146.169.24\|TGGGAATACCGGGTGCTGTAGGCT | >hg38_1_-_228634819_228635039_RNA5S12_5S_50ntFlanks | 146 | 169 | 24 | TGGGAATACCGGGTGCTGTAGGCT | 810.6 | 7.7 | 1.0 | 7.4 | 1.3E-13 | 3.1E-13 | BT20/MB468 |
| >hg38_21_+_8433172_8446622_RNA45SN1_45S_50ntFlanks@9534.9555.22\|AGGAAACTCTGGTGGAGGTCCG | >hg38_21_+_8433172_8446622_RNA45SN1_45S_50ntFlanks | 9534 | 9555 | 22 | AGGAAACTCTGGTGGAGGTCCG | 245.4 | -9.9 | 1.3 | -7.4 | 1.3E-13 | 3.1E-13 | BT20/MB468 |
| >hg38_21_+_8433172_8446622_RNA45SN1_45S_50ntFlanks@11769.11792.24\|TAATTAGTGACGCGCATGAATGGA | >hg38_21_+_8433172_8446622_RNA45SN1_45S_50ntFlanks | 11769 | 11792 | 24 | TAATTAGTGACGCGCATGAATGGA | 521.3 | -4.8 | 0.7 | -7.4 | 1.2E-13 | 3.1E-13 | MB231/MB468 |
| >hg38_21_+_8433172_8446622_RNA45SN1_45S_50ntFlanks@10411.10433.23\|CAGTCGGTCCTGAGAGATGGGCG | >hg38_21_+_8433172_8446622_RNA45SN1_45S_50ntFlanks | 10411 | 10433 | 23 | CAGTCGGTCCTGAGAGATGGGCG | 1044.3 | -7.4 | 1.0 | -7.4 | 1.2E-13 | 3.1E-13 | MB231/MB468 |
| >MI0000084\|hsa-mir-26b&WithFlank&2\|+\|218402640\|218402728@18.39.22(+1U)&[MIMAT0000083&hsa-miR-26b-5p&offsets\|0\|+1(+1U);m-53&2\|+\|218402657\|218402678&offsets\|0\|0(+1U)]\|TTCAAGTAATTCAGGATAGGTTT | >MI0000084\|hsa-mir-26b&WithFlank&2\|+\|218402640\|218402728 | 18 | 39 | 22(+1U) | TTCAAGTAATTCAGGATAGGTTT | 1390.0 | 6.9 | 0.9 | 7.4 | 1.3E-13 | 3.2E-13 | BT20/MB468 |
| >MI0003205\|hsa-mir-532&WithFlank&X\|+\|50003142\|50003244@26.46.21(+1A)&[MIMAT0002888&hsa-miR-532-5p&offsets\|0\|-1(+1A);m-65&X\|+\|50003167\|50003188&offsets\|0\|-1(+1A)]\|CATGCCTTGAGTGTAGGACCGA | >MI0003205\|hsa-mir-532&WithFlank&X\|+\|50003142\|50003244 | 26 | 46 | 21(+1A) | CATGCCTTGAGTGTAGGACCGA | 433.5 | 7.6 | 1.0 | 7.4 | 1.3E-13 | 3.2E-13 | BT20/MB468 |
| >am_trna13_LysCTT_14_-_58706613_58706685@31.51.21,trna2_LysCTT_15_+_79152904_79152976@31.51.21,trna4_LysCTT_18_-_43669271_43669343@31.51.21\|ACTCTTAATCCCAGGGTCGTG | >am_trna13_LysCTT_14_-_58706613_58706685 | 31 | 51 | 21 | ACTCTTAATCCCAGGGTCGTG | 256.8 | -9.6 | 1.3 | -7.4 | 1.2E-13 | 3.2E-13 | MB231/MB468 |
| >hg38_21_+_8433172_8446622_RNA45SN1_45S_50ntFlanks@12365.12382.18\|ATAACTGGCTTGTGGCGG | >hg38_21_+_8433172_8446622_RNA45SN1_45S_50ntFlanks | 12365 | 12382 | 18 | ATAACTGGCTTGTGGCGG | 669.6 | -8.1 | 1.1 | -7.4 | 1.2E-13 | 3.2E-13 | MB231/MB468 |
| >trna6_TrpCCA_12_+_98898030_98898101@30.52.23\|ACTCCAGATCAGAAGGCTGCGTG | >trna6_TrpCCA_12_+_98898030_98898101 | 30 | 52 | 23 | ACTCCAGATCAGAAGGCTGCGTG | 218.8 | -9.3 | 1.3 | -7.4 | 1.2E-13 | 3.2E-13 | MB231/MB468 |
| >MI0000085\|hsa-mir-27a&WithFlank&19\|-\|13836434\|13836523@57.76.20(+1A)&[MIMAT0000084&hsa-miR-27a-3p&offsets\|0\|-1(+1A);m-43&19\|-\|13836447\|13836467&offsets\|0\|-1(+1A)]\|TTCACAGTGGCTAAGTTCCGA | >MI0000085\|hsa-mir-27a&WithFlank&19\|-\|13836434\|13836523 | 57 | 76 | 20(+1A) | TTCACAGTGGCTAAGTTCCGA | 8297.8 | 5.9 | 0.8 | 7.4 | 1.2E-13 | 3.3E-13 | MB231/MB468 |
| >hg38_21_+_8433172_8446622_RNA45SN1_45S_50ntFlanks@11576.11601.26\|CTTAGAACTGGTGCGGACCAGGGGAA | >hg38_21_+_8433172_8446622_RNA45SN1_45S_50ntFlanks | 11576 | 11601 | 26 | CTTAGAACTGGTGCGGACCAGGGGAA | 1220.9 | -8.2 | 1.1 | -7.4 | 1.4E-13 | 3.3E-13 | BT20/MB468 |
| >hg38_21_+_8433172_8446622_RNA45SN1_45S_50ntFlanks@6651.6693.43\|CGACTCTTAGCGGTGGATCACTCGGCTCGTGCGTCGATGAAGA | >hg38_21_+_8433172_8446622_RNA45SN1_45S_50ntFlanks | 6651 | 6693 | 43 | CGACTCTTAGCGGTGGATCACTCGGCTCGTGCGTCGATGAAGA | 5178.9 | 6.3 | 0.8 | 7.4 | 1.4E-13 | 3.3E-13 | BT20/MB468 |
| >trnaMT_PheGAA_MT_+_577_647@39.57.19\|ATGTTTAGACGGGCTCACA | >trnaMT_PheGAA_MT_+_577_647 | 39 | 57 | 19 | ATGTTTAGACGGGCTCACA | 3070.6 | 10.1 | 1.4 | 7.4 | 1.4E-13 | 3.4E-13 | BT20/MB468 |
| >hg38_21_+_8433172_8446622_RNA45SN1_45S_50ntFlanks@3840.3868.29\|CTCCTACTTGGATAACTGTGGTAATTCTA | >hg38_21_+_8433172_8446622_RNA45SN1_45S_50ntFlanks | 3840 | 3868 | 29 | CTCCTACTTGGATAACTGTGGTAATTCTA | 634.8 | 3.4 | 0.5 | 7.4 | 1.4E-13 | 3.4E-13 | BT20/MB468 |
| >trna10_AspGTC_12_-_125424193_125424264@1.30.30,trna12_AspGTC_12_-_125411891_125411962@1.30.30,trna38_AspGTC_17_-_8125556_8125627@1.30.30,trna45_AspGTC_6_+_27447453_27447524@1.30.30,trna48_AspGTC_6_+_27471523_27471594@1.30.30,trna4_AspGTC_12_+_96429799_96429870@1.30.30,trna69_AspGTC_1_-_161440205_161440276@1.30.30,trna72_AspGTC_1_-_161432824_161432895@1.30.30,trna75_AspGTC_1_-_161425414_161425485@1.30.30,trna78_AspGTC_1_-_161418033_161418104@1.30.30,trna81_AspGTC_1_-_161410615_161410686@1.30.30\|TCCTCGTTAGTATAGTGGTGAGTATCCCCG | >trna10_AspGTC_12_-_125424193_125424264 | 1 | 30 | 30 | TCCTCGTTAGTATAGTGGTGAGTATCCCCG | 341.8 | -4.3 | 0.6 | -7.4 | 1.3E-13 | 3.4E-13 | MB231/MB468 |
| >hg38_21_+_8433172_8446622_RNA45SN1_45S_50ntFlanks@11661.11678.18\|TGATTTCTGCCCAGTGCT | >hg38_21_+_8433172_8446622_RNA45SN1_45S_50ntFlanks | 11661 | 11678 | 18 | TGATTTCTGCCCAGTGCT | 235.9 | -6.4 | 0.9 | -7.4 | 1.3E-13 | 3.4E-13 | MB231/MB468 |
| >am_trna13_LysCTT_14_-_58706613_58706685@31.51.21,trna2_LysCTT_15_+_79152904_79152976@31.51.21,trna4_LysCTT_18_-_43669271_43669343@31.51.21\|ACTCTTAATCCCAGGGTCGTG | >am_trna13_LysCTT_14_-_58706613_58706685 | 31 | 51 | 21 | ACTCTTAATCCCAGGGTCGTG | 315.8 | -7.1 | 1.0 | -7.4 | 1.4E-13 | 3.4E-13 | BT20/MB468 |
| >hg38_21_+_8433172_8446622_RNA45SN1_45S_50ntFlanks@11739.11768.30\|TCTCTTAAGGTAGCCAAATGCCTCGTCATC | >hg38_21_+_8433172_8446622_RNA45SN1_45S_50ntFlanks | 11739 | 11768 | 30 | TCTCTTAAGGTAGCCAAATGCCTCGTCATC | 228.1 | -4.0 | 0.5 | -7.4 | 1.3E-13 | 3.5E-13 | MB231/MB468 |
| >hg38_21_+_8433172_8446622_RNA45SN1_45S_50ntFlanks@12435.12454.20\|ATCATTGTGAAGCAGAATTC | >hg38_21_+_8433172_8446622_RNA45SN1_45S_50ntFlanks | 12435 | 12454 | 20 | ATCATTGTGAAGCAGAATTC | 425.4 | -10.7 | 1.5 | -7.4 | 1.4E-13 | 3.5E-13 | BT20/MB468 |
| >trna10_AspGTC_12_-_125424193_125424264@32.72.41,trna12_AspGTC_12_-_125411891_125411962@32.72.41,trna144_AspGTC_6_-_27551236_27551307@32.72.41,trna38_AspGTC_17_-_8125556_8125627@32.72.41,trna45_AspGTC_6_+_27447453_27447524@32.72.41,trna48_AspGTC_6_+_27471523_27471594@32.72.41,trna4_AspGTC_12_+_96429799_96429870@32.72.41,trna69_AspGTC_1_-_161440205_161440276@32.72.41,trna72_AspGTC_1_-_161432824_161432895@32.72.41,trna75_AspGTC_1_-_161425414_161425485@32.72.41,trna78_AspGTC_1_-_161418033_161418104@32.72.41,trna81_AspGTC_1_-_161410615_161410686@32.72.41\|CTGTCACGCGGGAGACCGGGGTTCGATTCCCCGACGGGGAG | >trna10_AspGTC_12_-_125424193_125424264 | 32 | 72 | 41 | CTGTCACGCGGGAGACCGGGGTTCGATTCCCCGACGGGGAG | 252.7 | -9.6 | 1.3 | -7.4 | 1.3E-13 | 3.5E-13 | MB231/MB468 |
| >hg38_21_+_8433172_8446622_RNA45SN1_45S_50ntFlanks@13002.13020.19\|CGATCTATTGAAAGTCAGC | >hg38_21_+_8433172_8446622_RNA45SN1_45S_50ntFlanks | 13002 | 13020 | 19 | CGATCTATTGAAAGTCAGC | 201.6 | -9.2 | 1.2 | -7.4 | 1.3E-13 | 3.5E-13 | MB231/MB468 |
| >hg38_21_+_8433172_8446622_RNA45SN1_45S_50ntFlanks@3984.4001.18\|GCGGCTTTGGTGACTCTA | >hg38_21_+_8433172_8446622_RNA45SN1_45S_50ntFlanks | 3984 | 4001 | 18 | GCGGCTTTGGTGACTCTA | 670.5 | -6.9 | 0.9 | -7.4 | 1.3E-13 | 3.5E-13 | MB231/MB468 |
| >MI0000253\|hsa-mir-148a&WithFlank&7\|-\|25949913\|25949992@50.70.21&[MIMAT0000243&hsa-miR-148a-3p&offsets\|0\|-1;m-7&7\|-\|25949922\|25949943&offsets\|0\|-1]\|TCAGTGCACTACAGAACTTTG | >MI0000253\|hsa-mir-148a&WithFlank&7\|-\|25949913\|25949992 | 50 | 70 | 21 | TCAGTGCACTACAGAACTTTG | 2631.1 | 9.4 | 1.3 | 7.4 | 1.4E-13 | 3.5E-13 | BT20/MB468 |
| >hg38_21_+_8433172_8446622_RNA45SN1_45S_50ntFlanks@10574.10596.23\|AAGCCGGCGGGAGCCCCGGGGAG | >hg38_21_+_8433172_8446622_RNA45SN1_45S_50ntFlanks | 10574 | 10596 | 23 | AAGCCGGCGGGAGCCCCGGGGAG | 184.0 | -9.5 | 1.3 | -7.4 | 1.5E-13 | 3.5E-13 | BT20/MB468 |
| >hg38_21_+_8433172_8446622_RNA45SN1_45S_50ntFlanks@6719.6743.25\|TGAATTGCAGGACACATTGATCATC | >hg38_21_+_8433172_8446622_RNA45SN1_45S_50ntFlanks | 6719 | 6743 | 25 | TGAATTGCAGGACACATTGATCATC | 5658.7 | -4.3 | 0.6 | -7.4 | 1.3E-13 | 3.6E-13 | MB231/MB468 |
| >hg38_21_+_8433172_8446622_RNA45SN1_45S_50ntFlanks@5490.5512.23\|TGAGAAGACGGTCGAACTTGACT | >hg38_21_+_8433172_8446622_RNA45SN1_45S_50ntFlanks | 5490 | 5512 | 23 | TGAGAAGACGGTCGAACTTGACT | 1115.0 | 9.3 | 1.3 | 7.4 | 1.4E-13 | 3.6E-13 | MB231/MB468 |
| >trnaMT_LysTTT_MT_+_8295_8364@1.20.20,trnalookalike7_LysTTT_1_+_568844_568913@1.20.20\|CACTGTAAAGCTAACTTAGC | >trnaMT_LysTTT_MT_+_8295_8364 | 1 | 20 | 20 | CACTGTAAAGCTAACTTAGC | 925.2 | 8.9 | 1.2 | 7.4 | 1.5E-13 | 3.6E-13 | BT20/MB468 |
| >hg38_21_+_8433172_8446622_RNA45SN1_45S_50ntFlanks@10681.10711.31\|TTCCGGCGGCGTCCGGTGAGCTCTCGCTGGC | >hg38_21_+_8433172_8446622_RNA45SN1_45S_50ntFlanks | 10681 | 10711 | 31 | TTCCGGCGGCGTCCGGTGAGCTCTCGCTGGC | 795.1 | -5.5 | 0.7 | -7.4 | 1.4E-13 | 3.6E-13 | MB231/MB468 |
| >hg38_21_+_8433172_8446622_RNA45SN1_45S_50ntFlanks@4006.4025.20\|ACCTCGGGCCGATCGCACGC | >hg38_21_+_8433172_8446622_RNA45SN1_45S_50ntFlanks | 4006 | 4025 | 20 | ACCTCGGGCCGATCGCACGC | 237.7 | -3.2 | 0.4 | -7.4 | 1.4E-13 | 3.6E-13 | MB231/MB468 |
| >hg38_21_+_8433172_8446622_RNA45SN1_45S_50ntFlanks@6646.6668.23\|TCGTACGACTCTTAGCGGTGGAT | >hg38_21_+_8433172_8446622_RNA45SN1_45S_50ntFlanks | 6646 | 6668 | 23 | TCGTACGACTCTTAGCGGTGGAT | 5069.3 | 4.1 | 0.6 | 7.4 | 1.4E-13 | 3.6E-13 | MB231/MB468 |
| >hg38_21_+_8433172_8446622_RNA45SN1_45S_50ntFlanks@8357.8378.22\|AAAAGAACTTTGAAGAGAGAGT | >hg38_21_+_8433172_8446622_RNA45SN1_45S_50ntFlanks | 8357 | 8378 | 22 | AAAAGAACTTTGAAGAGAGAGT | 16569.6 | -4.4 | 0.6 | -7.4 | 1.4E-13 | 3.6E-13 | MB231/MB468 |
| >hg38_21_+_8433172_8446622_RNA45SN1_45S_50ntFlanks@11666.11693.28\|TCTGCCCAGTGCTCTGAATGTCAAAGTG | >hg38_21_+_8433172_8446622_RNA45SN1_45S_50ntFlanks | 11666 | 11693 | 28 | TCTGCCCAGTGCTCTGAATGTCAAAGTG | 392.2 | 4.5 | 0.6 | 7.4 | 1.4E-13 | 3.6E-13 | MB231/MB468 |
| >trna101_AlaAGC_6_-_28831462_28831533@58.75.18,trna102_AlaAGC_6_-_28806221_28806292@58.75.18,trna110_AlaTGC_6_-_28757547_28757618@58.75.18,trna113_AlaTGC_6_-_28726141_28726212@58.75.18,trna65_AlaAGC_6_+_28574933_28575004@58.75.18\|TCCCCGGCACCTCCACCA | >trna101_AlaAGC_6_-_28831462_28831533 | 58 | 75 | 18 | TCCCCGGCACCTCCACCA | 4476.4 | 6.6 | 0.9 | 7.4 | 1.4E-13 | 3.7E-13 | MB231/MB468 |

<table>
<tr><td>>hg38_21_+_8433172_8446622_RNA45SN1_45S_50ntFlanks@10822.10840.19|GAAGTCGGCAAGCCGGATC</td><td>>hg38_21_+_8433172_8446622_RNA45SN1_45S_50ntFlanks</td><td>10822</td><td>10840</td><td>19</td><td>GAAGTCGGCAAGCCGGATC</td><td>276.2</td><td>-5.9</td><td>0.8</td><td>-7.4</td><td>1.5E-13</td><td>3.7E-13</td><td>BT20/MB468</td></tr>
<tr><td>>hg38_21_+_8433172_8446622_RNA45SN1_45S_50ntFlanks@9198.9222.25|GTTCTCTCGGGGCCACGCGCGCGTC</td><td>>hg38_21_+_8433172_8446622_RNA45SN1_45S_50ntFlanks</td><td>9198</td><td>9222</td><td>25</td><td>GTTCTCTCGGGGCCACGCGCGCGTC</td><td>299.2</td><td>-9.8</td><td>1.3</td><td>-7.4</td><td>1.4E-13</td><td>3.7E-13</td><td>MB231/MB468</td></tr>
<tr><td>>hg38_21_+_8433172_8446622_RNA45SN1_45S_50ntFlanks@12386.12414.29|AGCGTTCATAGCGACGTCGCTTTTTGATC</td><td>>hg38_21_+_8433172_8446622_RNA45SN1_45S_50ntFlanks</td><td>12386</td><td>12414</td><td>29</td><td>AGCGTTCATAGCGACGTCGCTTTTTGATC</td><td>158.5</td><td>-5.8</td><td>0.8</td><td>-7.4</td><td>1.6E-13</td><td>3.8E-13</td><td>BT20/MB468</td></tr>
<tr><td>>MI0000734|hsa-mir-106b&WithFlank&7|-|100093987|100094080@58.77.20(+1A)&[MIMAT0004672&hsa-miR-106b-3p&offsets|0|-2(+1A);m-87&7|-|100094003|100094023&offsets|0|-1(+1A)]|CCGCACTGTGGGTACTTGCTA</td><td>>MI0000734|hsa-mir-106b&WithFlank&7|-|100093987|100094080</td><td>58</td><td>77</td><td>20(+1A)</td><td>CCGCACTGTGGGTACTTGCTA</td><td>962.1</td><td>9.8</td><td>1.3</td><td>7.4</td><td>1.6E-13</td><td>3.8E-13</td><td>BT20/MB468</td></tr>
<tr><td>>hg38_21_+_8433172_8446622_RNA45SN1_45S_50ntFlanks@12159.12183.25|TTTGACTGGGGCGGTACACCTGTCA</td><td>>hg38_21_+_8433172_8446622_RNA45SN1_45S_50ntFlanks</td><td>12159</td><td>12183</td><td>25</td><td>TTTGACTGGGGCGGTACACCTGTCA</td><td>230.4</td><td>-9.8</td><td>1.3</td><td>-7.4</td><td>1.6E-13</td><td>3.8E-13</td><td>BT20/MB468</td></tr>
<tr><td>>hg38_21_+_8433172_8446622_RNA45SN1_45S_50ntFlanks@11783.11803.21|CATGAATGGATGAACGAGATT</td><td>>hg38_21_+_8433172_8446622_RNA45SN1_45S_50ntFlanks</td><td>11783</td><td>11803</td><td>21</td><td>CATGAATGGATGAACGAGATT</td><td>7216.5</td><td>-4.3</td><td>0.6</td><td>-7.4</td><td>1.5E-13</td><td>3.8E-13</td><td>MB231/MB468</td></tr>
<tr><td>>hg38_21_+_8433172_8446622_RNA45SN1_45S_50ntFlanks@3771.3792.22|CACGGCCGGTACAGTGAAACTG</td><td>>hg38_21_+_8433172_8446622_RNA45SN1_45S_50ntFlanks</td><td>3771</td><td>3792</td><td>22</td><td>CACGGCCGGTACAGTGAAACTG</td><td>292.5</td><td>-3.9</td><td>0.5</td><td>-7.4</td><td>1.5E-13</td><td>3.8E-13</td><td>MB231/MB468</td></tr>
<tr><td>>MI0000487|hsa-mir-193a&WithFlank&17|+|31559990|31560089@27.48.22&[MIMAT0004614&hsa-miR-193a-5p&offsets|0|0;m-123&17|+|31560016|31560037&offsets|0|0]|TGGGTCTTTGCGGGCGAGATGA</td><td>>MI0000487|hsa-mir-193a&WithFlank&17|+|31559990|31560089</td><td>27</td><td>48</td><td>22</td><td>TGGGTCTTTGCGGGCGAGATGA</td><td>677.7</td><td>7.7</td><td>1.0</td><td>7.4</td><td>1.6E-13</td><td>3.8E-13</td><td>BT20/MB468</td></tr>
<tr><td>>hg38_21_+_8433172_8446622_RNA45SN1_45S_50ntFlanks@11572.11599.28|CCGACTTAGAACTGGTGCGGACCAGGGG</td><td>>hg38_21_+_8433172_8446622_RNA45SN1_45S_50ntFlanks</td><td>11572</td><td>11599</td><td>28</td><td>CCGACTTAGAACTGGTGCGGACCAGGGG</td><td>335.7</td><td>-3.1</td><td>0.4</td><td>-7.4</td><td>1.6E-13</td><td>3.9E-13</td><td>BT20/MB468</td></tr>
<tr><td>>hg38_21_+_8433172_8446622_RNA45SN1_45S_50ntFlanks@3876.3899.24|TACATGCCGACGGGCGCTGACCCC</td><td>>hg38_21_+_8433172_8446622_RNA45SN1_45S_50ntFlanks</td><td>3876</td><td>3899</td><td>24</td><td>TACATGCCGACGGGCGCTGACCCC</td><td>197.8</td><td>-2.9</td><td>0.4</td><td>-7.4</td><td>1.6E-13</td><td>3.9E-13</td><td>BT20/MB468</td></tr>
<tr><td>>hg38_21_+_8433172_8446622_RNA45SN1_45S_50ntFlanks@3777.3801.25|CGGTACAGTGAAACTGCGAATGGCT</td><td>>hg38_21_+_8433172_8446622_RNA45SN1_45S_50ntFlanks</td><td>3777</td><td>3801</td><td>25</td><td>CGGTACAGTGAAACTGCGAATGGCT</td><td>462.3</td><td>-4.1</td><td>0.6</td><td>-7.4</td><td>1.7E-13</td><td>4.0E-13</td><td>BT20/MB468</td></tr>
<tr><td>>hg38_21_+_8433172_8446622_RNA45SN1_45S_50ntFlanks@11888.11905.18|AGCTTGACTCTAGTCTGG</td><td>>hg38_21_+_8433172_8446622_RNA45SN1_45S_50ntFlanks</td><td>11888</td><td>11905</td><td>18</td><td>AGCTTGACTCTAGTCTGG</td><td>182.5</td><td>-9.5</td><td>1.3</td><td>-7.4</td><td>1.7E-13</td><td>4.0E-13</td><td>BT20/MB468</td></tr>
<tr><td>>MI0000542|hsa-mir-320a&WithFlank&8|-|22244960|22245043@49.68.20&[MIMAT0000510&hsa-miR-320a-3p&offsets|+1|-1;m-60&8|-|22244975|22244996&offsets|+1|-1]|AAAGCTGGGTTGAGAGGGCG</td><td>>MI0000542|hsa-mir-320a&WithFlank&8|-|22244960|22245043</td><td>49</td><td>68</td><td>20</td><td>AAAGCTGGGTTGAGAGGGCG</td><td>583.2</td><td>9.2</td><td>1.2</td><td>7.4</td><td>1.7E-13</td><td>4.0E-13</td><td>BT20/MB468</td></tr>
<tr><td>>hg38_21_+_8433172_8446622_RNA45SN1_45S_50ntFlanks@3775.3794.20|GCCGGTACAGTGAAACTGCG</td><td>>hg38_21_+_8433172_8446622_RNA45SN1_45S_50ntFlanks</td><td>3775</td><td>3794</td><td>20</td><td>GCCGGTACAGTGAAACTGCG</td><td>187.7</td><td>-3.6</td><td>0.5</td><td>-7.4</td><td>1.7E-13</td><td>4.1E-13</td><td>BT20/MB468</td></tr>
<tr><td>>hg38_21_+_8433172_8446622_RNA45SN1_45S_50ntFlanks@12991.13028.38|CTCCCTCGCTGCGATCTATTGAAAGTCAGCCCTCGACA</td><td>>hg38_21_+_8433172_8446622_RNA45SN1_45S_50ntFlanks</td><td>12991</td><td>13028</td><td>38</td><td>CTCCCTCGCTGCGATCTATTGAAAGTCAGCCCTCGACA</td><td>201.0</td><td>-9.6</td><td>1.3</td><td>-7.4</td><td>1.7E-13</td><td>4.1E-13</td><td>BT20/MB468</td></tr>
<tr><td>>hg38_21_+_8433172_8446622_RNA45SN1_45S_50ntFlanks@11233.11253.21|CCCTTCCCGTGGATCGCCCCA</td><td>>hg38_21_+_8433172_8446622_RNA45SN1_45S_50ntFlanks</td><td>11233</td><td>11253</td><td>21</td><td>CCCTTCCCGTGGATCGCCCCA</td><td>246.3</td><td>-9.9</td><td>1.3</td><td>-7.4</td><td>1.7E-13</td><td>4.1E-13</td><td>BT20/MB468</td></tr>
<tr><td>>hg38_21_+_8433172_8446622_RNA45SN1_45S_50ntFlanks@11792.11810.19|ATGAACGAGATTCCCACTG</td><td>>hg38_21_+_8433172_8446622_RNA45SN1_45S_50ntFlanks</td><td>11792</td><td>11810</td><td>19</td><td>ATGAACGAGATTCCCACTG</td><td>211.0</td><td>-9.7</td><td>1.3</td><td>-7.4</td><td>1.7E-13</td><td>4.1E-13</td><td>BT20/MB468</td></tr>
<tr><td>>hg38_21_+_8433172_8446622_RNA45SN1_45S_50ntFlanks@12991.13008.18|CTCCCTCGCTGCGATCTA</td><td>>hg38_21_+_8433172_8446622_RNA45SN1_45S_50ntFlanks</td><td>12991</td><td>13008</td><td>18</td><td>CTCCCTCGCTGCGATCTA</td><td>307.2</td><td>-10.3</td><td>1.4</td><td>-7.4</td><td>1.7E-13</td><td>4.1E-13</td><td>BT20/MB468</td></tr>
<tr><td>>hg38_21_+_8433172_8446622_RNA45SN1_45S_50ntFlanks@9571.9596.26|GCAAATCGGTCGTCCGACCTGGGTAT</td><td>>hg38_21_+_8433172_8446622_RNA45SN1_45S_50ntFlanks</td><td>9571</td><td>9596</td><td>26</td><td>GCAAATCGGTCGTCCGACCTGGGTAT</td><td>211.8</td><td>-3.7</td><td>0.5</td><td>-7.4</td><td>1.6E-13</td><td>4.1E-13</td><td>MB231/MB468</td></tr>
<tr><td>>hg38_21_+_8433172_8446622_RNA45SN1_45S_50ntFlanks@12456.12482.27|CCAAGCGTTGGATTGTTCACCCACTAA</td><td>>hg38_21_+_8433172_8446622_RNA45SN1_45S_50ntFlanks</td><td>12456</td><td>12482</td><td>27</td><td>CCAAGCGTTGGATTGTTCACCCACTAA</td><td>149.1</td><td>-6.0</td><td>0.8</td><td>-7.4</td><td>1.6E-13</td><td>4.2E-13</td><td>MB231/MB468</td></tr>
<tr><td>>hg38_21_+_8433172_8446622_RNA45SN1_45S_50ntFlanks@8472.8494.23|CGGCCCGGCGGATCTTTCCCGCC</td><td>>hg38_21_+_8433172_8446622_RNA45SN1_45S_50ntFlanks</td><td>8472</td><td>8494</td><td>23</td><td>CGGCCCGGCGGATCTTTCCCGCC</td><td>310.2</td><td>-9.9</td><td>1.3</td><td>-7.4</td><td>1.6E-13</td><td>4.2E-13</td><td>MB231/MB468</td></tr>
<tr><td>>am_MI0015825|hsa-mir-378c&WithFlank&10|-|130962582|130962674@17.34.18;MI0016749|hsa-mir-378d-1&WithFlank&4|-|5923269|5923334@37.54.18;MI0000786|hsa-mir-378a&WithFlank&5|+|149732819|149732896@49.66.18;MI0003840|hsa-mir-378d-2&WithFlank&8|-|93916016|93916125@28.45.18&[MIMAT0016847&hsa-miR-378c&offsets|0|-7;m-130&10|-|130962639|130962658&offsets|0|-2];[MIMAT0018926&hsa-miR-378d&offsets|0|-2;m-228&4|-|5923281|5923298&offsets|0|0];[MIMAT0000732&hsa-miR-378a-3p&offsets|0|-4;m-44&5|+|149732867|149732888&offsets|0|-4];[MIMAT0018926_1&hsa-miR-378d&offsets|0|-2;m-227&8|-|93916081|93916098&offsets|0|0]|ACTGGACTTGGAGTCAGA</td><td>>am_MI0015825|hsa-mir-378c&WithFlank&10|-|130962582|130962674</td><td>17</td><td>34</td><td>18</td><td>ACTGGACTTGGAGTCAGA</td><td>594.1</td><td>5.6</td><td>0.8</td><td>7.4</td><td>1.6E-13</td><td>4.3E-13</td><td>MB231/MB468</td></tr>
<tr><td>>hg38_21_+_8433172_8446622_RNA45SN1_45S_50ntFlanks@11873.11894.22|AAGAAGACCCTGTTGAGCTTGA</td><td>>hg38_21_+_8433172_8446622_RNA45SN1_45S_50ntFlanks</td><td>11873</td><td>11894</td><td>22</td><td>AAGAAGACCCTGTTGAGCTTGA</td><td>146.9</td><td>-3.6</td><td>0.5</td><td>-7.4</td><td>1.8E-13</td><td>4.3E-13</td><td>BT20/MB468</td></tr>
<tr><td>>hg38_21_+_8433172_8446622_RNA45SN1_45S_50ntFlanks@10420.10453.34|CTGAGAGATGGGCGAGCGCCGTTCCGAAGGGACG</td><td>>hg38_21_+_8433172_8446622_RNA45SN1_45S_50ntFlanks</td><td>10420</td><td>10453</td><td>34</td><td>CTGAGAGATGGGCGAGCGCCGTTCCGAAGGGACG</td><td>503.5</td><td>-4.6</td><td>0.6</td><td>-7.4</td><td>1.8E-13</td><td>4.4E-13</td><td>BT20/MB468</td></tr>
<tr><td>>hg38_21_+_8433172_8446622_RNA45SN1_45S_50ntFlanks@8472.8491.20|CGGCCCGGCGGATCTTTCCC</td><td>>hg38_21_+_8433172_8446622_RNA45SN1_45S_50ntFlanks</td><td>8472</td><td>8491</td><td>20</td><td>CGGCCCGGCGGATCTTTCCC</td><td>272.6</td><td>-9.7</td><td>1.3</td><td>-7.4</td><td>1.7E-13</td><td>4.4E-13</td><td>MB231/MB468</td></tr>
<tr><td>>hg38_21_+_8433172_8446622_RNA45SN1_45S_50ntFlanks@9736.9765.30|CCGAAACGATCTCAACCTATTCTCAAACTT</td><td>>hg38_21_+_8433172_8446622_RNA45SN1_45S_50ntFlanks</td><td>9736</td><td>9765</td><td>30</td><td>CCGAAACGATCTCAACCTATTCTCAAACTT</td><td>241.9</td><td>-9.5</td><td>1.3</td><td>-7.4</td><td>1.7E-13</td><td>4.4E-13</td><td>MB231/MB468</td></tr>
<tr><td>>hg38_21_+_8433172_8446622_RNA45SN1_45S_50ntFlanks@5449.5466.18|ATCGGCCCCGCCGGGGTC</td><td>>hg38_21_+_8433172_8446622_RNA45SN1_45S_50ntFlanks</td><td>5449</td><td>5466</td><td>18</td><td>ATCGGCCCCGCCGGGGTC</td><td>134.3</td><td>-9.1</td><td>1.2</td><td>-7.4</td><td>1.8E-13</td><td>4.4E-13</td><td>BT20/MB468</td></tr>
<tr><td>>hg38_21_+_8433172_8446622_RNA45SN1_45S_50ntFlanks@11918.11951.34|ACATGAGAGGTGTAGAATAAGTGGGAGGCCCCCG</td><td>>hg38_21_+_8433172_8446622_RNA45SN1_45S_50ntFlanks</td><td>11918</td><td>11951</td><td>34</td><td>ACATGAGAGGTGTAGAATAAGTGGGAGGCCCCCG</td><td>235.5</td><td>-9.5</td><td>1.3</td><td>-7.4</td><td>1.7E-13</td><td>4.4E-13</td><td>MB231/MB468</td></tr>
<tr><td>>hg38_21_+_8433172_8446622_RNA45SN1_45S_50ntFlanks@10770.10805.36|GCAGGTCTCCAAGGTGAACAGCCTCTGGCATGTTGG</td><td>>hg38_21_+_8433172_8446622_RNA45SN1_45S_50ntFlanks</td><td>10770</td><td>10805</td><td>36</td><td>GCAGGTCTCCAAGGTGAACAGCCTCTGGCATGTTGG</td><td>128.4</td><td>-5.8</td><td>0.8</td><td>-7.4</td><td>1.8E-13</td><td>4.4E-13</td><td>BT20/MB468</td></tr>
<tr><td>>hg38_21_+_8433172_8446622_RNA45SN1_45S_50ntFlanks@12073.12096.24|CCGAGGGGCTCTCGCTTCTGGCGC</td><td>>hg38_21_+_8433172_8446622_RNA45SN1_45S_50ntFlanks</td><td>12073</td><td>12096</td><td>24</td><td>CCGAGGGGCTCTCGCTTCTGGCGC</td><td>843.0</td><td>-5.9</td><td>0.8</td><td>-7.4</td><td>1.9E-13</td><td>4.5E-13</td><td>BT20/MB468</td></tr>
<tr><td>>hg38_21_+_8433172_8446622_RNA45SN1_45S_50ntFlanks@4728.4748.21|ATCAAGAACGAAAGTCGGAGG</td><td>>hg38_21_+_8433172_8446622_RNA45SN1_45S_50ntFlanks</td><td>4728</td><td>4748</td><td>21</td><td>ATCAAGAACGAAAGTCGGAGG</td><td>170.5</td><td>-9.4</td><td>1.3</td><td>-7.4</td><td>1.9E-13</td><td>4.5E-13</td><td>BT20/MB468</td></tr>
<tr><td>>MI0002470|hsa-mir-486-1&WithFlank&8|-|41660435|41660514@10.31.22(+1U);MI0023622|hsa-mir-486-2&WithFlank&8|+|41660438|41660513@7.28.22(+1U)&[MIMAT0002177&hsa-miR-486-5p&offsets|0|0(+1U);m-107&8|-|41660484|41660505&offsets|0|0(+1U)];[MIMAT0002177_1&hsa-miR-486-5p&offsets|0|0(+1U);TJU_CMC_MD2.ID03138.5p-miR&offsets|0|-2(+1U);m-106&8|+|41660444|41660465&offsets|0|0(+1U)]|TCCTGTACTGAGCTGCCCCGAGT</td><td>>MI0002470|hsa-mir-486-1&WithFlank&8|-|41660435|41660514</td><td>10</td><td>31</td><td>22(+1U)</td><td>TCCTGTACTGAGCTGCCCCGAGT</td><td>281.7</td><td>-6.8</td><td>0.9</td><td>-7.7</td><td>1.3E-14</td><td>4.5E-13</td><td>BT20/MB231</td></tr>
<tr><td>>MI0000264|hsa-mir-7-2&WithFlank&15|+|88611819|88611940@38.57.20(+1C);MI0000265|hsa-mir-7-3&WithFlank&19|+|4770664|4770785@37.56.20(+1C);MI0000263|hsa-mir-7-1&WithFlank&9|-|83969742|83969863@30.49.20(+1C)&[MIMAT0000252&hsa-miR-7-5p&offsets|0|-4(+1C);m-400&15|+|88611856|88611879&offsets|0|-4(+1C)];[MIMAT0000252_1&hsa-miR-7-5p&offsets|0|-4(+1C);m-402&19|+|4770700|4770723&offsets|0|-4(+1C)];[MIMAT0000252_2&hsa-miR-7-5p&offsets|0|-4(+1C);m-401&9|-|83969811|83969834&offsets|0|-4(+1C)]|TGGAAGACTAGTGATTTTGTC</td><td>>MI0000264|hsa-mir-7-2&WithFlank&15|+|88611819|88611940</td><td>38</td><td>57</td><td>20(+1C)</td><td>TGGAAGACTAGTGATTTTGTC</td><td>708.7</td><td>9.7</td><td>1.3</td><td>7.4</td><td>1.9E-13</td><td>4.5E-13</td><td>BT20/MB468</td></tr>
<tr><td>>hg38_21_+_8433172_8446622_RNA45SN1_45S_50ntFlanks@10806.10825.20|AACAATGTAGGTAAGGGAAG</td><td>>hg38_21_+_8433172_8446622_RNA45SN1_45S_50ntFlanks</td><td>10806</td><td>10825</td><td>20</td><td>AACAATGTAGGTAAGGGAAG</td><td>251.0</td><td>-9.5</td><td>1.3</td><td>-7.4</td><td>1.7E-13</td><td>4.6E-13</td><td>MB231/MB468</td></tr>
<tr><td>>MI0000283|hsa-mir-203a&WithFlank&14|+|104117399|104117520@72.92.21(+1C)&[MIMAT0000264&hsa-miR-203a-3p&offsets|+1|0(+1C);m-79&14|+|104117469|104117490&offsets|+1|0(+1C)]|TGAAATGTTTAGGACCACTAGC</td><td>>MI0000283|hsa-mir-203a&WithFlank&14|+|104117399|104117520</td><td>72</td><td>92</td><td>21(+1C)</td><td>TGAAATGTTTAGGACCACTAGC</td><td>171.9</td><td>10.9</td><td>1.4</td><td>7.7</td><td>1.3E-14</td><td>4.6E-13</td><td>BT20/MB231</td></tr>
<tr><td>>hg38_21_+_8433172_8446622_RNA45SN1_45S_50ntFlanks@10351.10379.29|CGAGAACTTTGAAGGCCGAAGTGGAGAAG</td><td>>hg38_21_+_8433172_8446622_RNA45SN1_45S_50ntFlanks</td><td>10351</td><td>10379</td><td>29</td><td>CGAGAACTTTGAAGGCCGAAGTGGAGAAG</td><td>581.5</td><td>-3.6</td><td>0.5</td><td>-7.4</td><td>1.9E-13</td><td>4.6E-13</td><td>BT20/MB468</td></tr>
<tr><td>>hg38_21_+_8433172_8446622_RNA45SN1_45S_50ntFlanks@8380.8397.18|CAAGAGGGCGTGAAACCG</td><td>>hg38_21_+_8433172_8446622_RNA45SN1_45S_50ntFlanks</td><td>8380</td><td>8397</td><td>18</td><td>CAAGAGGGCGTGAAACCG</td><td>1003.8</td><td>-5.3</td><td>0.7</td><td>-7.4</td><td>1.7E-13</td><td>4.6E-13</td><td>MB231/MB468</td></tr>
<tr><td>>hg38_21_+_8433172_8446622_RNA45SN1_45S_50ntFlanks@10781.10807.27|AGGTGAACAGCCTCTGGCATGTTGGAA</td><td>>hg38_21_+_8433172_8446622_RNA45SN1_45S_50ntFlanks</td><td>10781</td><td>10807</td><td>27</td><td>AGGTGAACAGCCTCTGGCATGTTGGAA</td><td>273.2</td><td>-6.9</td><td>0.9</td><td>-7.4</td><td>1.8E-13</td><td>4.6E-13</td><td>MB231/MB468</td></tr>
<tr><td>>hg38_21_+_8433172_8446622_RNA45SN1_45S_50ntFlanks@11318.11336.19|CCCCACGTCTCGTCGCGCG</td><td>>hg38_21_+_8433172_8446622_RNA45SN1_45S_50ntFlanks</td><td>11318</td><td>11336</td><td>19</td><td>CCCCACGTCTCGTCGCGCG</td><td>316.2</td><td>-9.9</td><td>1.3</td><td>-7.4</td><td>1.8E-13</td><td>4.6E-13</td><td>MB231/MB468</td></tr>
<tr><td>>hg38_21_+_8433172_8446622_RNA45SN1_45S_50ntFlanks@11752.11780.29|CCAAATGCCTCGTCATCTAATTAGTGACG</td><td>>hg38_21_+_8433172_8446622_RNA45SN1_45S_50ntFlanks</td><td>11752</td><td>11780</td><td>29</td><td>CCAAATGCCTCGTCATCTAATTAGTGACG</td><td>178.6</td><td>-9.5</td><td>1.3</td><td>-7.4</td><td>2.0E-13</td><td>4.7E-13</td><td>BT20/MB468</td></tr>
</table>

<table>
<tr><th></th><th></th><th></th><th></th><th></th><th></th><th></th><th></th><th></th><th></th><th></th><th></th><th></th></tr>
<tr><td>>MI0000289|hsa-mir-181a-1&WithFlank&1|-|198859038|198859159@30.52.23(+1C);MI0000269|hsa-mir-181a-2&WithFlank&9|+|124692436|124692557@45.67.23(+1C)&[MIMAT0000256&hsa-miR-181a-5p&offsets|0|0(+1C);m-32&1|-|198859108|198859130&offsets|0|0(+1C)];[MIMAT0000256_1&hsa-miR-181a-5p&offsets|0|0(+1C);m-32&9|+|124692480|124692502&offsets|0|0(+1C)]|AACATTCAACGCTGTCGGTGAGTC</td><td>>MI0000289|hsa-mir-181a-1&WithFlank&1|-|198859038|198859159</td><td>30</td><td>52</td><td>23(+1C)</td><td>AACATTCAACGCTGTCGGTGAGTC</td><td>672.6</td><td>8.9</td><td>1.2</td><td>7.4</td><td>1.8E-13</td><td>4.7E-13</td><td>MB231/MB468</td></tr>
<tr><td>>am_trna162_MetCAT_6_-_26758550_26758622@34.54.21,trna164_MetCAT_6_-_26735574_26735646@34.54.21,trna20_MetCAT_16_+_71460396_71460468@34.54.21,trna21_MetCAT_6_+_26701712_26701784@34.54.21,trna27_MetCAT_6_+_26766444_26766516@34.54.21,trna75_MetCAT_6_+_28912352_28912424@34.54.21,trna97_MetCAT_6_-_28921042_28921114@34.54.21|CATAATCTGAAGGTCCTGAGT</td><td>>am_trna162_MetCAT_6_-_26758550_26758622</td><td>34</td><td>54</td><td>21</td><td>CATAATCTGAAGGTCCTGAGT</td><td>3787.9</td><td>5.9</td><td>0.8</td><td>7.4</td><td>1.8E-13</td><td>4.7E-13</td><td>MB231/MB468</td></tr>
<tr><td>>hg38_21_+_8433172_8446622_RNA45SN1_45S_50ntFlanks@4445.4471.27|CTTGCCTCTCGGCGCCCCCTCGATGCT</td><td>>hg38_21_+_8433172_8446622_RNA45SN1_45S_50ntFlanks</td><td>4445</td><td>4471</td><td>27</td><td>CTTGCCTCTCGGCGCCCCCTCGATGCT</td><td>223.2</td><td>-7.6</td><td>1.0</td><td>-7.4</td><td>1.8E-13</td><td>4.7E-13</td><td>MB231/MB468</td></tr>
<tr><td>>hg38_21_+_8433172_8446622_RNA45SN1_45S_50ntFlanks@5487.5514.28|CGCTGAGAAGACGGTCGAACTTGACTAT</td><td>>hg38_21_+_8433172_8446622_RNA45SN1_45S_50ntFlanks</td><td>5487</td><td>5514</td><td>28</td><td>CGCTGAGAAGACGGTCGAACTTGACTAT</td><td>1329.6</td><td>-5.8</td><td>0.8</td><td>-7.4</td><td>1.8E-13</td><td>4.7E-13</td><td>MB231/MB468</td></tr>
<tr><td>>hg38_21_+_8433172_8446622_RNA45SN1_45S_50ntFlanks@11569.11592.24|CAGCCGACTTAGAACTGGTGCGGA</td><td>>hg38_21_+_8433172_8446622_RNA45SN1_45S_50ntFlanks</td><td>11569</td><td>11592</td><td>24</td><td>CAGCCGACTTAGAACTGGTGCGGA</td><td>440.0</td><td>-4.9</td><td>0.7</td><td>-7.4</td><td>1.8E-13</td><td>4.7E-13</td><td>MB231/MB468</td></tr>
<tr><td>>hg38_21_+_8433172_8446622_RNA45SN1_45S_50ntFlanks@11234.11254.21|CCTTCCCGTGGATCGCCCCAG</td><td>>hg38_21_+_8433172_8446622_RNA45SN1_45S_50ntFlanks</td><td>11234</td><td>11254</td><td>21</td><td>CCTTCCCGTGGATCGCCCCAG</td><td>232.7</td><td>-9.4</td><td>1.3</td><td>-7.4</td><td>1.8E-13</td><td>4.7E-13</td><td>MB231/MB468</td></tr>
<tr><td>>am_MI0000786|hsa-mir-378a&WithFlank&5|+|149732819|149732896@49.70.22&[MIMAT0000732&hsa-miR-378a-3p&offsets|0|0;m-44&5|+|149732867|149732888&offsets|0|0]|ACTGGACTTGGAGTCAGAAGGC</td><td>>am_MI0000786|hsa-mir-378a&WithFlank&5|+|149732819|149732896</td><td>49</td><td>70</td><td>22</td><td>ACTGGACTTGGAGTCAGAAGGC</td><td>28210.1</td><td>5.6</td><td>0.8</td><td>7.4</td><td>1.8E-13</td><td>4.7E-13</td><td>MB231/MB468</td></tr>
<tr><td>>hg38_21_+_8433172_8446622_RNA45SN1_45S_50ntFlanks@10369.10407.39|AAGTGGAGAAGGGTTCCATGTGAACAGCAGTTGAACATG</td><td>>hg38_21_+_8433172_8446622_RNA45SN1_45S_50ntFlanks</td><td>10369</td><td>10407</td><td>39</td><td>AAGTGGAGAAGGGTTCCATGTGAACAGCAGTTGAACATG</td><td>257.0</td><td>-4.8</td><td>0.7</td><td>-7.4</td><td>1.8E-13</td><td>4.8E-13</td><td>MB231/MB468</td></tr>
<tr><td>>hg38_21_+_8433172_8446622_RNA45SN1_45S_50ntFlanks@12991.13009.19|CTCCCTCGCTGCGATCTAT</td><td>>hg38_21_+_8433172_8446622_RNA45SN1_45S_50ntFlanks</td><td>12991</td><td>13009</td><td>19</td><td>CTCCCTCGCTGCGATCTAT</td><td>264.5</td><td>-10.0</td><td>1.4</td><td>-7.3</td><td>2.0E-13</td><td>4.8E-13</td><td>BT20/MB468</td></tr>
<tr><td>>MI0000095|hsa-mir-93&WithFlank&7|-|100093762|100093853@17.38.22&[MIMAT0000093&hsa-miR-93-5p&offsets|0|-1;m-29&7|-|100093815|100093837&offsets|0|-1]|CAAAGTGCTGTTCGTGCAGGTA</td><td>>MI0000095|hsa-mir-93&WithFlank&7|-|100093762|100093853</td><td>17</td><td>38</td><td>22</td><td>CAAAGTGCTGTTCGTGCAGGTA</td><td>1872.4</td><td>6.7</td><td>0.9</td><td>7.3</td><td>2.0E-13</td><td>4.8E-13</td><td>BT20/MB468</td></tr>
<tr><td>>hg38_21_+_8433172_8446622_RNA45SN1_45S_50ntFlanks@10350.10383.34|ACGAGAACTTTGAAGGCCGAAGTGGAGAAGGGTT</td><td>>hg38_21_+_8433172_8446622_RNA45SN1_45S_50ntFlanks</td><td>10350</td><td>10383</td><td>34</td><td>ACGAGAACTTTGAAGGCCGAAGTGGAGAAGGGTT</td><td>4681.3</td><td>-3.6</td><td>0.5</td><td>-7.4</td><td>1.9E-13</td><td>4.8E-13</td><td>MB231/MB468</td></tr>
<tr><td>>hg38_21_+_8433172_8446622_RNA45SN1_45S_50ntFlanks@9534.9564.31|AGGAAACTCTGGTGGAGGTCCGTAGCGGTCC</td><td>>hg38_21_+_8433172_8446622_RNA45SN1_45S_50ntFlanks</td><td>9534</td><td>9564</td><td>31</td><td>AGGAAACTCTGGTGGAGGTCCGTAGCGGTCC</td><td>956.5</td><td>-4.0</td><td>0.5</td><td>-7.4</td><td>1.9E-13</td><td>4.9E-13</td><td>MB231/MB468</td></tr>
<tr><td>>hg38_21_+_8433172_8446622_RNA45SN1_45S_50ntFlanks@12151.12183.33|GTGGGGAGTTTGACTGGGGCGGTACACCTGTCA</td><td>>hg38_21_+_8433172_8446622_RNA45SN1_45S_50ntFlanks</td><td>12151</td><td>12183</td><td>33</td><td>GTGGGGAGTTTGACTGGGGCGGTACACCTGTCA</td><td>156.7</td><td>-9.3</td><td>1.3</td><td>-7.3</td><td>2.0E-13</td><td>4.9E-13</td><td>BT20/MB468</td></tr>
<tr><td>>hg38_21_+_8433172_8446622_RNA45SN1_45S_50ntFlanks@10845.10862.18|ACTTCGGGATAAGGATTG</td><td>>hg38_21_+_8433172_8446622_RNA45SN1_45S_50ntFlanks</td><td>10845</td><td>10862</td><td>18</td><td>ACTTCGGGATAAGGATTG</td><td>766.4</td><td>-7.8</td><td>1.1</td><td>-7.3</td><td>2.0E-13</td><td>4.9E-13</td><td>BT20/MB468</td></tr>
<tr><td>>hg38_21_+_8433172_8446622_RNA45SN1_45S_50ntFlanks@11561.11603.43|GCGCCTAGCAGCCGACTTAGAACTGGTGCGGACCAGGGGAATC</td><td>>hg38_21_+_8433172_8446622_RNA45SN1_45S_50ntFlanks</td><td>11561</td><td>11603</td><td>43</td><td>GCGCCTAGCAGCCGACTTAGAACTGGTGCGGACCAGGGGAATC</td><td>283.4</td><td>-5.7</td><td>0.8</td><td>-7.4</td><td>1.9E-13</td><td>4.9E-13</td><td>MB231/MB468</td></tr>
<tr><td>>hg38_21_+_8433172_8446622_RNA45SN1_45S_50ntFlanks@12510.12527.18|TCGTGAGACAGGTTAGTT</td><td>>hg38_21_+_8433172_8446622_RNA45SN1_45S_50ntFlanks</td><td>12510</td><td>12527</td><td>18</td><td>TCGTGAGACAGGTTAGTT</td><td>386.5</td><td>-7.5</td><td>1.0</td><td>-7.3</td><td>2.1E-13</td><td>4.9E-13</td><td>BT20/MB468</td></tr>
<tr><td>>MI0000802|hsa-mir-340&WithFlank&5|-|180015297|180015403@22.43.22&[MIMAT0004692&hsa-miR-340-5p&offsets|0|0;m-220&5|-|180015361|180015382&offsets|0|0]|TTATAAAGCAATGAGACTGATT</td><td>>MI0000802|hsa-mir-340&WithFlank&5|-|180015297|180015403</td><td>22</td><td>43</td><td>22</td><td>TTATAAAGCAATGAGACTGATT</td><td>26618.9</td><td>7.6</td><td>1.0</td><td>7.3</td><td>2.1E-13</td><td>5.0E-13</td><td>BT20/MB468</td></tr>
<tr><td>>hg38_21_+_8433172_8446622_RNA45SN1_45S_50ntFlanks@3930.3959.30|ATCAAAACCAACCCGGTCAGCCCCTCTCCG</td><td>>hg38_21_+_8433172_8446622_RNA45SN1_45S_50ntFlanks</td><td>3930</td><td>3959</td><td>30</td><td>ATCAAAACCAACCCGGTCAGCCCCTCTCCG</td><td>156.1</td><td>-5.3</td><td>0.7</td><td>-7.4</td><td>1.9E-13</td><td>5.1E-13</td><td>MB231/MB468</td></tr>
<tr><td>>hg38_21_+_8433172_8446622_RNA45SN1_45S_50ntFlanks@12324.12349.26|TTGGGTTTTAAGCAGGAGGTGTCAGA</td><td>>hg38_21_+_8433172_8446622_RNA45SN1_45S_50ntFlanks</td><td>12324</td><td>12349</td><td>26</td><td>TTGGGTTTTAAGCAGGAGGTGTCAGA</td><td>143.9</td><td>-9.2</td><td>1.2</td><td>-7.3</td><td>2.1E-13</td><td>5.1E-13</td><td>BT20/MB468</td></tr>
<tr><td>>trna116_GluCTC_1_-_145399233_145399304@32.71.40,trna71_GluCTC_1_-_161439189_161439260@32.71.40,trna74_GluCTC_1_-_161431809_161431880@32.71.40,trna77_GluCTC_1_-_161424398_161424469@32.71.40,trna77_GluCTC_6_+_28949976_28950047@32.71.40,trna80_GluCTC_1_-_161417018_161417089@32.71.40,trna87_GluCTC_6_-_126101393_126101464@32.71.40|CTCTCACCGCCGCGGCCCGGGTTCGATTCCCGGTCAGGGA</td><td>>trna116_GluCTC_1_-_145399233_145399304</td><td>32</td><td>71</td><td>40</td><td>CTCTCACCGCCGCGGCCCGGGTTCGATTCCCGGTCAGGGA</td><td>248.7</td><td>-9.5</td><td>1.3</td><td>-7.4</td><td>1.9E-13</td><td>5.1E-13</td><td>MB231/MB468</td></tr>
<tr><td>>hg38_21_+_8433172_8446622_RNA45SN1_45S_50ntFlanks@7993.8016.24|GTGGCGACCCGCTGAATTTAAGCA</td><td>>hg38_21_+_8433172_8446622_RNA45SN1_45S_50ntFlanks</td><td>7993</td><td>8016</td><td>24</td><td>GTGGCGACCCGCTGAATTTAAGCA</td><td>573.6</td><td>5.4</td><td>0.7</td><td>7.4</td><td>2.0E-13</td><td>5.1E-13</td><td>MB231/MB468</td></tr>
<tr><td>>hg38_21_+_8433172_8446622_RNA45SN1_45S_50ntFlanks@11669.11703.35|GCCCAGTGCTCTGAATGTCAAAGTGAAGAAATTCA</td><td>>hg38_21_+_8433172_8446622_RNA45SN1_45S_50ntFlanks</td><td>11669</td><td>11703</td><td>35</td><td>GCCCAGTGCTCTGAATGTCAAAGTGAAGAAATTCA</td><td>775.3</td><td>7.4</td><td>1.0</td><td>7.4</td><td>2.0E-13</td><td>5.1E-13</td><td>MB231/MB468</td></tr>
<tr><td>>MI0000447|hsa-mir-128-1&WithFlank&2|+|135665391|135665484@56.76.21(+2A);MI0000727|hsa-mir-128-2&WithFlank&3|+|35744470|35744565@58.78.21(+2A)&[MIMAT0000424&hsa-miR-128-3p&offsets|0|0(+2A);m-108&2|+|135665446|135665466&offsets|0|0(+2A)];[MIMAT0000424_1&hsa-miR-128-3p&offsets|0|0(+2A);m-117&3|+|35744527|35744547&offsets|0|0(+2A)]|TCACAGTGAACCGGTCTCTTTAA</td><td>>MI0000447|hsa-mir-128-1&WithFlank&2|+|135665391|135665484</td><td>56</td><td>76</td><td>21(+2A)</td><td>TCACAGTGAACCGGTCTCTTTAA</td><td>1746.3</td><td>4.9</td><td>0.7</td><td>7.3</td><td>2.2E-13</td><td>5.2E-13</td><td>BT20/MB468</td></tr>
<tr><td>>hg38_21_+_8433172_8446622_RNA45SN1_45S_50ntFlanks@11593.11613.21|CCAGGGGAATCCGACTGTTTA</td><td>>hg38_21_+_8433172_8446622_RNA45SN1_45S_50ntFlanks</td><td>11593</td><td>11613</td><td>21</td><td>CCAGGGGAATCCGACTGTTTA</td><td>354.1</td><td>-3.7</td><td>0.5</td><td>-7.3</td><td>2.2E-13</td><td>5.2E-13</td><td>BT20/MB468</td></tr>
<tr><td>>MI0000285|hsa-mir-205&WithFlank&1|+|209432127|209432248@40.61.22(+3A)&[MIMAT0000266&hsa-miR-205-5p&offsets|0|0(+3A);m-351&1|+|209432166|209432187&offsets|0|0(+3A)]|TCCTTCATTCCACCGGAGTCTGAAA</td><td>>MI0000285|hsa-mir-205&WithFlank&1|+|209432127|209432248</td><td>40</td><td>61</td><td>22(+3A)</td><td>TCCTTCATTCCACCGGAGTCTGAAA</td><td>206.0</td><td>11.2</td><td>1.5</td><td>7.7</td><td>1.5E-14</td><td>5.2E-13</td><td>BT20/MB231</td></tr>
<tr><td>>MI0000283|hsa-mir-203a&WithFlank&14|+|104117399|104117520@72.92.21(+1C)&[MIMAT0000264&hsa-miR-203a-3p&offsets|+1|0(+1C);m-79&14|+|104117469|104117490&offsets|+1|0(+1C)]|TGAAATGTTTAGGACCACTAGC</td><td>>MI0000283|hsa-mir-203a&WithFlank&14|+|104117399|104117520</td><td>72</td><td>92</td><td>21(+1C)</td><td>TGAAATGTTTAGGACCACTAGC</td><td>389.3</td><td>9.6</td><td>1.3</td><td>7.3</td><td>2.2E-13</td><td>5.2E-13</td><td>BT20/MB468</td></tr>
<tr><td>>hg38_21_+_8433172_8446622_RNA45SN1_45S_50ntFlanks@4443.4472.30|CCCTTGCCTCTCGGCGCCCCCTCGATGCTC</td><td>>hg38_21_+_8433172_8446622_RNA45SN1_45S_50ntFlanks</td><td>4443</td><td>4472</td><td>30</td><td>CCCTTGCCTCTCGGCGCCCCCTCGATGCTC</td><td>312.2</td><td>-10.3</td><td>1.4</td><td>-7.3</td><td>2.2E-13</td><td>5.2E-13</td><td>BT20/MB468</td></tr>
<tr><td>>hg38_21_+_8433172_8446622_RNA45SN1_45S_50ntFlanks@9575.9594.20|ATCGGTCGTCCGACCTGGGT</td><td>>hg38_21_+_8433172_8446622_RNA45SN1_45S_50ntFlanks</td><td>9575</td><td>9594</td><td>20</td><td>ATCGGTCGTCCGACCTGGGT</td><td>146.4</td><td>-9.2</td><td>1.3</td><td>-7.3</td><td>2.2E-13</td><td>5.2E-13</td><td>BT20/MB468</td></tr>
<tr><td>>hg38_21_+_8433172_8446622_RNA45SN1_45S_50ntFlanks@10458.10497.40|ATGGCCTCCGTTGCCCTCGGCCGATCGAAAGGGAGTCGGG</td><td>>hg38_21_+_8433172_8446622_RNA45SN1_45S_50ntFlanks</td><td>10458</td><td>10497</td><td>40</td><td>ATGGCCTCCGTTGCCCTCGGCCGATCGAAAGGGAGTCGGG</td><td>189.6</td><td>-9.1</td><td>1.2</td><td>-7.3</td><td>2.0E-13</td><td>5.3E-13</td><td>MB231/MB468</td></tr>
<tr><td>>hg38_21_+_8433172_8446622_RNA45SN1_45S_50ntFlanks@12161.12180.20|TGACTGGGGCGGTACACCTG</td><td>>hg38_21_+_8433172_8446622_RNA45SN1_45S_50ntFlanks</td><td>12161</td><td>12180</td><td>20</td><td>TGACTGGGGCGGTACACCTG</td><td>307.5</td><td>-6.1</td><td>0.8</td><td>-7.3</td><td>2.0E-13</td><td>5.3E-13</td><td>MB231/MB468</td></tr>
<tr><td>>hg38_21_+_8433172_8446622_RNA45SN1_45S_50ntFlanks@7985.8012.28|GATCAGACGTGGCGACCCGCTGAATTTA</td><td>>hg38_21_+_8433172_8446622_RNA45SN1_45S_50ntFlanks</td><td>7985</td><td>8012</td><td>28</td><td>GATCAGACGTGGCGACCCGCTGAATTTA</td><td>175.1</td><td>-9.4</td><td>1.3</td><td>-7.3</td><td>2.2E-13</td><td>5.3E-13</td><td>BT20/MB468</td></tr>
<tr><td>>hg38_21_+_8433172_8446622_RNA45SN1_45S_50ntFlanks@12942.12973.32|TTCGTAGACGACCTGCTTCTGGGTCGGGGTTT</td><td>>hg38_21_+_8433172_8446622_RNA45SN1_45S_50ntFlanks</td><td>12942</td><td>12973</td><td>32</td><td>TTCGTAGACGACCTGCTTCTGGGTCGGGGTTT</td><td>204.0</td><td>-5.0</td><td>0.7</td><td>-7.3</td><td>2.2E-13</td><td>5.4E-13</td><td>BT20/MB468</td></tr>
<tr><td>>hg38_21_+_8433172_8446622_RNA45SN1_45S_50ntFlanks@9564.9592.29|CTGACGTGCAAATCGGTCGTCCGACCTGG</td><td>>hg38_21_+_8433172_8446622_RNA45SN1_45S_50ntFlanks</td><td>9564</td><td>9592</td><td>29</td><td>CTGACGTGCAAATCGGTCGTCCGACCTGG</td><td>951.5</td><td>-4.0</td><td>0.5</td><td>-7.3</td><td>2.1E-13</td><td>5.4E-13</td><td>MB231/MB468</td></tr>
<tr><td>>hg38_21_+_8433172_8446622_RNA45SN1_45S_50ntFlanks@10512.10533.22|TCCGGAGTGGCGGAGATGGGCG</td><td>>hg38_21_+_8433172_8446622_RNA45SN1_45S_50ntFlanks</td><td>10512</td><td>10533</td><td>22</td><td>TCCGGAGTGGCGGAGATGGGCG</td><td>982.9</td><td>-8.4</td><td>1.1</td><td>-7.3</td><td>2.3E-13</td><td>5.4E-13</td><td>BT20/MB468</td></tr>
<tr><td>>hg38_21_+_8433172_8446622_RNA45SN1_45S_50ntFlanks@6647.6667.21|CGTACGACTCTTAGCGGTGGA</td><td>>hg38_21_+_8433172_8446622_RNA45SN1_45S_50ntFlanks</td><td>6647</td><td>6667</td><td>21</td><td>CGTACGACTCTTAGCGGTGGA</td><td>1105.0</td><td>5.4</td><td>0.7</td><td>7.3</td><td>2.1E-13</td><td>5.4E-13</td><td>MB231/MB468</td></tr>
<tr><td>>hg38_21_+_8433172_8446622_RNA45SN1_45S_50ntFlanks@10423.10456.34|AGAGATGGGCGAGCGCCGTTCCGAAGGGACGGGC</td><td>>hg38_21_+_8433172_8446622_RNA45SN1_45S_50ntFlanks</td><td>10423</td><td>10456</td><td>34</td><td>AGAGATGGGCGAGCGCCGTTCCGAAGGGACGGGC</td><td>258.7</td><td>-2.9</td><td>0.4</td><td>-7.3</td><td>2.3E-13</td><td>5.4E-13</td><td>BT20/MB468</td></tr>
<tr><td>>MI0000802|hsa-mir-340&WithFlank&5|-|180015297|180015403@22.42.21(+1C)&[MIMAT0004692&hsa-miR-340-5p&offsets|0|-1(+1C);m-220&5|-|180015361|180015382&offsets|0|-1(+1C)]|TTATAAAGCAATGAGACTGATC</td><td>>MI0000802|hsa-mir-340&WithFlank&5|-|180015297|180015403</td><td>22</td><td>42</td><td>21(+1C)</td><td>TTATAAAGCAATGAGACTGATC</td><td>2711.6</td><td>8.0</td><td>1.1</td><td>7.3</td><td>2.3E-13</td><td>5.4E-13</td><td>BT20/MB468</td></tr>
<tr><td>>hg38_21_+_8433172_8446622_RNA45SN1_45S_50ntFlanks@12126.12148.23|CGACCCGCTCCGGGGACAGTGCC</td><td>>hg38_21_+_8433172_8446622_RNA45SN1_45S_50ntFlanks</td><td>12126</td><td>12148</td><td>23</td><td>CGACCCGCTCCGGGGACAGTGCC</td><td>162.8</td><td>-9.3</td><td>1.3</td><td>-7.3</td><td>2.3E-13</td><td>5.4E-13</td><td>BT20/MB468</td></tr>
<tr><td>>hg38_21_+_8433172_8446622_RNA45SN1_45S_50ntFlanks@11161.11188.28|CACGGTCCCCCGCGAGGGGGGCCCGGGC</td><td>>hg38_21_+_8433172_8446622_RNA45SN1_45S_50ntFlanks</td><td>11161</td><td>11188</td><td>28</td><td>CACGGTCCCCCGCGAGGGGGGCCCGGGC</td><td>204.2</td><td>-9.2</td><td>1.3</td><td>-7.3</td><td>2.1E-13</td><td>5.5E-13</td><td>MB231/MB468</td></tr>
</table>

<table>
<tr><td>>MI0000253|hsa-mir-148a&WithFlank&7|-|25949913|25949992@50.72.23(+1A)&[MIMAT0000243&hsa-miR-148a-3p&offsets|0|+1(+1A);m-7&7|-|25949922|25949943&offsets|0|+1(+1A)]|TCAGTGCACTACAGAACTTTGTCA</td><td>>MI0000253|hsa-mir-148a&WithFlank&7|-|25949913|25949992</td><td>50</td><td>72</td><td>23(+1A)</td><td>TCAGTGCACTACAGAACTTTGTCA</td><td>811.3</td><td>8.6</td><td>1.2</td><td>7.3</td><td>2.3E-13</td><td>5.5E-13</td><td>BT20/MB468</td></tr>
<tr><td>>am_trna106_PheGAA_6_-_28775610_28775682@31.52.22,trna109_PheGAA_6_-_28758499_28758571@31.52.22,trna11_PheGAA_12_-_125412389_125412461@31.52.22,trna13_PheGAA_11_-_59333853_59333925@31.52.22,trna14_PheGAA_19_-_1383361_1383433@31.52.22,trna15_PheGAA_11_-_59324970_59325042@31.52.22,trna1_PheGAA_13_-_95201904_95201976@31.52.22,trna72_PheGAA_6_+_28732164_28732236@31.52.22,trna96_PheGAA_6_-_28949449_28949521@31.52.22|ACTGAAGATCTAAAGGTCCCTG</td><td>>am_trna106_PheGAA_6_-_28775610_28775682</td><td>31</td><td>52</td><td>22</td><td>ACTGAAGATCTAAAGGTCCCTG</td><td>183.7</td><td>-9.5</td><td>1.3</td><td>-7.3</td><td>2.3E-13</td><td>5.5E-13</td><td>BT20/MB468</td></tr>
<tr><td>>hg38_21_+_8433172_8446622_RNA45SN1_45S_50ntFlanks@9552.9571.20|TCCGTAGCGGTCCTGACGTG</td><td>>hg38_21_+_8433172_8446622_RNA45SN1_45S_50ntFlanks</td><td>9552</td><td>9571</td><td>20</td><td>TCCGTAGCGGTCCTGACGTG</td><td>596.1</td><td>-3.7</td><td>0.5</td><td>-7.3</td><td>2.3E-13</td><td>5.5E-13</td><td>BT20/MB468</td></tr>
<tr><td>>hg38_21_+_8433172_8446622_RNA45SN1_45S_50ntFlanks@9570.9594.25|TGCAAATCGGTCGTCCGACCTGGGT</td><td>>hg38_21_+_8433172_8446622_RNA45SN1_45S_50ntFlanks</td><td>9570</td><td>9594</td><td>25</td><td>TGCAAATCGGTCGTCCGACCTGGGT</td><td>274.7</td><td>-3.9</td><td>0.5</td><td>-7.3</td><td>2.1E-13</td><td>5.5E-13</td><td>MB231/MB468</td></tr>
<tr><td>>hg38_21_+_8433172_8446622_RNA45SN1_45S_50ntFlanks@12363.12382.20|GGATAACTGGCTTGTGGCGG</td><td>>hg38_21_+_8433172_8446622_RNA45SN1_45S_50ntFlanks</td><td>12363</td><td>12382</td><td>20</td><td>GGATAACTGGCTTGTGGCGG</td><td>289.6</td><td>-9.8</td><td>1.3</td><td>-7.3</td><td>2.2E-13</td><td>5.6E-13</td><td>MB231/MB468</td></tr>
<tr><td>>hg38_21_+_8433172_8446622_RNA45SN1_45S_50ntFlanks@4002.4024.23|GATAACCTCGGGCCGATCGCACG</td><td>>hg38_21_+_8433172_8446622_RNA45SN1_45S_50ntFlanks</td><td>4002</td><td>4024</td><td>23</td><td>GATAACCTCGGGCCGATCGCACG</td><td>799.2</td><td>-3.8</td><td>0.5</td><td>-7.3</td><td>2.2E-13</td><td>5.6E-13</td><td>MB231/MB468</td></tr>
<tr><td>>MI0000489|hsa-mir-195&WithFlank&17|-|7017609|7017707@21.42.22&[MIMAT0000461&hsa-miR-195-5p&offsets|0|+1;m-191&17|-|7017666|7017687&offsets|0|0]|TAGCAGCACAGAAATATTGGCA</td><td>>MI0000489|hsa-mir-195&WithFlank&17|-|7017609|7017707</td><td>21</td><td>42</td><td>22</td><td>TAGCAGCACAGAAATATTGGCA</td><td>1205.1</td><td>10.4</td><td>1.4</td><td>7.3</td><td>2.4E-13</td><td>5.7E-13</td><td>BT20/MB468</td></tr>
<tr><td>>trnaMT_PheGAA_MT_+_577_647@56.74.19|CATCACCCCATAAACACCA</td><td>>trnaMT_PheGAA_MT_+_577_647</td><td>56</td><td>74</td><td>19</td><td>CATCACCCCATAAACACCA</td><td>1863.3</td><td>8.5</td><td>1.2</td><td>7.3</td><td>2.4E-13</td><td>5.7E-13</td><td>BT20/MB468</td></tr>
<tr><td>>hg38_21_+_8433172_8446622_RNA45SN1_45S_50ntFlanks@11593.11612.20|CCAGGGGAATCCGACTGTTT</td><td>>hg38_21_+_8433172_8446622_RNA45SN1_45S_50ntFlanks</td><td>11593</td><td>11612</td><td>20</td><td>CCAGGGGAATCCGACTGTTT</td><td>1108.9</td><td>-4.5</td><td>0.6</td><td>-7.3</td><td>2.4E-13</td><td>5.7E-13</td><td>BT20/MB468</td></tr>
<tr><td>>hg38_21_+_8433172_8446622_RNA45SN1_45S_50ntFlanks@10351.10371.21|CGAGAACTTTGAAGGCCGAAG</td><td>>hg38_21_+_8433172_8446622_RNA45SN1_45S_50ntFlanks</td><td>10351</td><td>10371</td><td>21</td><td>CGAGAACTTTGAAGGCCGAAG</td><td>880.5</td><td>-3.8</td><td>0.5</td><td>-7.3</td><td>2.4E-13</td><td>5.7E-13</td><td>BT20/MB468</td></tr>
<tr><td>>hg38_21_+_8433172_8446622_RNA45SN1_45S_50ntFlanks@6652.6678.27|GACTCTTAGCGGTGGATCACTCGGCTC</td><td>>hg38_21_+_8433172_8446622_RNA45SN1_45S_50ntFlanks</td><td>6652</td><td>6678</td><td>27</td><td>GACTCTTAGCGGTGGATCACTCGGCTC</td><td>1326.0</td><td>3.7</td><td>0.5</td><td>7.3</td><td>2.4E-13</td><td>5.7E-13</td><td>BT20/MB468</td></tr>
<tr><td>>hg38_21_+_8433172_8446622_RNA45SN1_45S_50ntFlanks@12638.12659.22|CTACCATCTGTGGGATTATGAC</td><td>>hg38_21_+_8433172_8446622_RNA45SN1_45S_50ntFlanks</td><td>12638</td><td>12659</td><td>22</td><td>CTACCATCTGTGGGATTATGAC</td><td>808.1</td><td>-6.0</td><td>0.8</td><td>-7.3</td><td>2.4E-13</td><td>5.7E-13</td><td>BT20/MB468</td></tr>
<tr><td>>hg38_21_+_8433172_8446622_RNA45SN1_45S_50ntFlanks@13007.13039.33|TATTGAAAGTCAGCCCTCGACACAAGGGTTTGT</td><td>>hg38_21_+_8433172_8446622_RNA45SN1_45S_50ntFlanks</td><td>13007</td><td>13039</td><td>33</td><td>TATTGAAAGTCAGCCCTCGACACAAGGGTTTGT</td><td>2895.7</td><td>5.3</td><td>0.7</td><td>7.3</td><td>2.4E-13</td><td>5.7E-13</td><td>BT20/MB468</td></tr>
<tr><td>>hg38_21_+_8433172_8446622_RNA45SN1_45S_50ntFlanks@11593.11613.21|CCAGGGGAATCCGACTGTTTA</td><td>>hg38_21_+_8433172_8446622_RNA45SN1_45S_50ntFlanks</td><td>11593</td><td>11613</td><td>21</td><td>CCAGGGGAATCCGACTGTTTA</td><td>293.8</td><td>-3.5</td><td>0.5</td><td>-7.3</td><td>2.2E-13</td><td>5.8E-13</td><td>MB231/MB468</td></tr>
<tr><td>>hg38_1_-_228634819_228635039_RNA5S12_5S_50ntFlanks@131.170.40|GGATGGGAGACCGCCTGGGAATACCGGGTGCTGTAGGCTT</td><td>>hg38_1_-_228634819_228635039_RNA5S12_5S_50ntFlanks</td><td>131</td><td>170</td><td>40</td><td>GGATGGGAGACCGCCTGGGAATACCGGGTGCTGTAGGCTT</td><td>886.7</td><td>9.2</td><td>1.3</td><td>7.3</td><td>2.4E-13</td><td>5.8E-13</td><td>BT20/MB468</td></tr>
<tr><td>>hg38_21_+_8433172_8446622_RNA45SN1_45S_50ntFlanks@9545.9567.23|GTGGAGGTCCGTAGCGGTCCTGA</td><td>>hg38_21_+_8433172_8446622_RNA45SN1_45S_50ntFlanks</td><td>9545</td><td>9567</td><td>23</td><td>GTGGAGGTCCGTAGCGGTCCTGA</td><td>224.4</td><td>-4.7</td><td>0.6</td><td>-7.3</td><td>2.2E-13</td><td>5.8E-13</td><td>MB231/MB468</td></tr>
<tr><td>>hg38_21_+_8433172_8446622_RNA45SN1_45S_50ntFlanks@10781.10804.24|AGGTGAACAGCCTCTGGCATGTTG</td><td>>hg38_21_+_8433172_8446622_RNA45SN1_45S_50ntFlanks</td><td>10781</td><td>10804</td><td>24</td><td>AGGTGAACAGCCTCTGGCATGTTG</td><td>276.0</td><td>-10.1</td><td>1.4</td><td>-7.3</td><td>2.5E-13</td><td>5.8E-13</td><td>BT20/MB468</td></tr>
<tr><td>>hg38_21_+_8433172_8446622_RNA45SN1_45S_50ntFlanks@8356.8377.22|GAAAAGAACTTTGAAGAGAGAG</td><td>>hg38_21_+_8433172_8446622_RNA45SN1_45S_50ntFlanks</td><td>8356</td><td>8377</td><td>22</td><td>GAAAAGAACTTTGAAGAGAGAG</td><td>1084.5</td><td>-4.7</td><td>0.6</td><td>-7.3</td><td>2.5E-13</td><td>5.9E-13</td><td>BT20/MB468</td></tr>
<tr><td>>hg38_21_+_8433172_8446622_RNA45SN1_45S_50ntFlanks@10408.10431.24|GGTCAGTCGGTCCTGAGAGATGGG</td><td>>hg38_21_+_8433172_8446622_RNA45SN1_45S_50ntFlanks</td><td>10408</td><td>10431</td><td>24</td><td>GGTCAGTCGGTCCTGAGAGATGGG</td><td>307.4</td><td>-8.0</td><td>1.1</td><td>-7.3</td><td>2.3E-13</td><td>5.9E-13</td><td>MB231/MB468</td></tr>
<tr><td>>hg38_21_+_8433172_8446622_RNA45SN1_45S_50ntFlanks@9505.9526.22|TGAACTATGCCTGGGCAGGGCG</td><td>>hg38_21_+_8433172_8446622_RNA45SN1_45S_50ntFlanks</td><td>9505</td><td>9526</td><td>22</td><td>TGAACTATGCCTGGGCAGGGCG</td><td>209.9</td><td>-9.3</td><td>1.3</td><td>-7.3</td><td>2.3E-13</td><td>6.0E-13</td><td>MB231/MB468</td></tr>
<tr><td>>MI0000816|hsa-mir-335&WithFlank&7|+|130496105|130496210@58.79.22&[MIMAT0004703&hsa-miR-335-3p&offsets|0|0;m-159&7|+|130496162|130496183&offsets|0|0]|TTTTTCATTATTGCTCCTGACC</td><td>>MI0000816|hsa-mir-335&WithFlank&7|+|130496105|130496210</td><td>58</td><td>79</td><td>22</td><td>TTTTTCATTATTGCTCCTGACC</td><td>3624.2</td><td>6.4</td><td>0.9</td><td>7.3</td><td>2.5E-13</td><td>6.0E-13</td><td>BT20/MB468</td></tr>
<tr><td>>hg38_21_+_8433172_8446622_RNA45SN1_45S_50ntFlanks@11999.12030.32|CCCTGCGGGCCGCCGGTGAAATACCACTACTC</td><td>>hg38_21_+_8433172_8446622_RNA45SN1_45S_50ntFlanks</td><td>11999</td><td>12030</td><td>32</td><td>CCCTGCGGGCCGCCGGTGAAATACCACTACTC</td><td>191.2</td><td>-9.2</td><td>1.2</td><td>-7.3</td><td>2.4E-13</td><td>6.1E-13</td><td>MB231/MB468</td></tr>
<tr><td>>hg38_21_+_8433172_8446622_RNA45SN1_45S_50ntFlanks@12415.12435.21|CTTCGATGTCGGCTCTTCCTA</td><td>>hg38_21_+_8433172_8446622_RNA45SN1_45S_50ntFlanks</td><td>12415</td><td>12435</td><td>21</td><td>CTTCGATGTCGGCTCTTCCTA</td><td>1268.4</td><td>-3.8</td><td>0.5</td><td>-7.3</td><td>2.4E-13</td><td>6.1E-13</td><td>MB231/MB468</td></tr>
<tr><td>>hg38_21_+_8433172_8446622_RNA45SN1_45S_50ntFlanks@11786.11803.18|GAATGGATGAACGAGATT</td><td>>hg38_21_+_8433172_8446622_RNA45SN1_45S_50ntFlanks</td><td>11786</td><td>11803</td><td>18</td><td>GAATGGATGAACGAGATT</td><td>1182.6</td><td>-4.5</td><td>0.6</td><td>-7.3</td><td>2.6E-13</td><td>6.2E-13</td><td>BT20/MB468</td></tr>
<tr><td>>hg38_21_+_8433172_8446622_RNA45SN1_45S_50ntFlanks@10794.10816.23|CTGGCATGTTGGAACAATGTAGG</td><td>>hg38_21_+_8433172_8446622_RNA45SN1_45S_50ntFlanks</td><td>10794</td><td>10816</td><td>23</td><td>CTGGCATGTTGGAACAATGTAGG</td><td>1423.4</td><td>-4.9</td><td>0.7</td><td>-7.3</td><td>2.4E-13</td><td>6.2E-13</td><td>MB231/MB468</td></tr>
<tr><td>>hg38_21_+_8433172_8446622_RNA45SN1_45S_50ntFlanks@6650.6692.43|ACGACTCTTAGCGGTGGATCACTCGGCTCGTGCGTCGATGAAG</td><td>>hg38_21_+_8433172_8446622_RNA45SN1_45S_50ntFlanks</td><td>6650</td><td>6692</td><td>43</td><td>ACGACTCTTAGCGGTGGATCACTCGGCTCGTGCGTCGATGAAG</td><td>988.5</td><td>4.7</td><td>0.6</td><td>7.3</td><td>2.4E-13</td><td>6.2E-13</td><td>MB231/MB468</td></tr>
<tr><td>>hg38_21_+_8433172_8446622_RNA45SN1_45S_50ntFlanks@9504.9523.20|GTGAACTATGCCTGGGCAGG</td><td>>hg38_21_+_8433172_8446622_RNA45SN1_45S_50ntFlanks</td><td>9504</td><td>9523</td><td>20</td><td>GTGAACTATGCCTGGGCAGG</td><td>1557.4</td><td>-6.8</td><td>0.9</td><td>-7.3</td><td>2.6E-13</td><td>6.2E-13</td><td>BT20/MB468</td></tr>
<tr><td>>hg38_21_+_8433172_8446622_RNA45SN1_45S_50ntFlanks@12412.12435.24|ATCCTTCGATGTCGGCTCTTCCTA</td><td>>hg38_21_+_8433172_8446622_RNA45SN1_45S_50ntFlanks</td><td>12412</td><td>12435</td><td>24</td><td>ATCCTTCGATGTCGGCTCTTCCTA</td><td>185.3</td><td>-9.5</td><td>1.3</td><td>-7.3</td><td>2.6E-13</td><td>6.2E-13</td><td>BT20/MB468</td></tr>
<tr><td>>hg38_21_+_8433172_8446622_RNA45SN1_45S_50ntFlanks@9572.9598.27|CAAATCGGTCGTCCGACCTGGGTATAG</td><td>>hg38_21_+_8433172_8446622_RNA45SN1_45S_50ntFlanks</td><td>9572</td><td>9598</td><td>27</td><td>CAAATCGGTCGTCCGACCTGGGTATAG</td><td>171.4</td><td>-9.4</td><td>1.3</td><td>-7.3</td><td>2.7E-13</td><td>6.3E-13</td><td>BT20/MB468</td></tr>
<tr><td>>am_MI0000809|hsa-mir-151a&WithFlank&8|-|140732558|140732659@17.37.21&[MIMAT0004697&hsa-miR-151a-5p&offsets|0|0;m-75&8|-|140732623|140732643&offsets|0|0]|TCGAGGAGCTCACAGTCTAGT</td><td>>am_MI0000809|hsa-mir-151a&WithFlank&8|-|140732558|140732659</td><td>17</td><td>37</td><td>21</td><td>TCGAGGAGCTCACAGTCTAGT</td><td>2242.3</td><td>8.6</td><td>1.2</td><td>7.3</td><td>2.4E-13</td><td>6.3E-13</td><td>MB231/MB468</td></tr>
<tr><td>>hg38_21_+_8433172_8446622_RNA45SN1_45S_50ntFlanks@11319.11344.26|CCCACGTCTCGTCGCGCGCGCGTCCG</td><td>>hg38_21_+_8433172_8446622_RNA45SN1_45S_50ntFlanks</td><td>11319</td><td>11344</td><td>26</td><td>CCCACGTCTCGTCGCGCGCGCGTCCG</td><td>873.8</td><td>-6.5</td><td>0.9</td><td>-7.3</td><td>2.4E-13</td><td>6.3E-13</td><td>MB231/MB468</td></tr>
<tr><td>>hg38_21_+_8433172_8446622_RNA45SN1_45S_50ntFlanks@4002.4026.25|GATAACCTCGGGCCGATCGCACGCC</td><td>>hg38_21_+_8433172_8446622_RNA45SN1_45S_50ntFlanks</td><td>4002</td><td>4026</td><td>25</td><td>GATAACCTCGGGCCGATCGCACGCC</td><td>130.1</td><td>-9.0</td><td>1.2</td><td>-7.3</td><td>2.7E-13</td><td>6.3E-13</td><td>BT20/MB468</td></tr>
<tr><td>>MI0000650|hsa-mir-200c&WithFlank&12|+|6963693|6963772@50.71.22(+2U)&[MIMAT0000617&hsa-miR-200c-3p&offsets|0|-1(+2U);m-54&12|+|6963742|6963764&offsets|0|-1(+2U)]|TAATACTGCCGGGTAATGATGGTT</td><td>>MI0000650|hsa-mir-200c&WithFlank&12|+|6963693|6963772</td><td>50</td><td>71</td><td>22(+2U)</td><td>TAATACTGCCGGGTAATGATGGTT</td><td>198.1</td><td>11.1</td><td>1.5</td><td>7.7</td><td>1.9E-14</td><td>6.4E-13</td><td>BT20/MB231</td></tr>
<tr><td>>hg38_21_+_8433172_8446622_RNA45SN1_45S_50ntFlanks@3985.4004.20|CGGCTTTGGTGACTCTAGAT</td><td>>hg38_21_+_8433172_8446622_RNA45SN1_45S_50ntFlanks</td><td>3985</td><td>4004</td><td>20</td><td>CGGCTTTGGTGACTCTAGAT</td><td>141.8</td><td>-9.1</td><td>1.3</td><td>-7.3</td><td>2.7E-13</td><td>6.5E-13</td><td>BT20/MB468</td></tr>
<tr><td>>hg38_21_+_8433172_8446622_RNA45SN1_45S_50ntFlanks@10411.10453.43|CAGTCGGTCCTGAGAGATGGGCGAGCGCCGTTCCGAAGGGACG</td><td>>hg38_21_+_8433172_8446622_RNA45SN1_45S_50ntFlanks</td><td>10411</td><td>10453</td><td>43</td><td>CAGTCGGTCCTGAGAGATGGGCGAGCGCCGTTCCGAAGGGACG</td><td>317.2</td><td>-4.2</td><td>0.6</td><td>-7.3</td><td>2.7E-13</td><td>6.5E-13</td><td>BT20/MB468</td></tr>
<tr><td>>hg38_21_+_8433172_8446622_RNA45SN1_45S_50ntFlanks@6651.6697.47|CGACTCTTAGCGGTGGATCACTCGGCTCGTGCGTCGATGAAGAACGC</td><td>>hg38_21_+_8433172_8446622_RNA45SN1_45S_50ntFlanks</td><td>6651</td><td>6697</td><td>47</td><td>CGACTCTTAGCGGTGGATCACTCGGCTCGTGCGTCGATGAAGAACGC</td><td>8699.6</td><td>5.1</td><td>0.7</td><td>7.3</td><td>2.5E-13</td><td>6.5E-13</td><td>MB231/MB468</td></tr>
<tr><td>>hg38_21_+_8433172_8446622_RNA45SN1_45S_50ntFlanks@4759.4782.24|ATCAGATACCGTCGTAGTTCCGAC</td><td>>hg38_21_+_8433172_8446622_RNA45SN1_45S_50ntFlanks</td><td>4759</td><td>4782</td><td>24</td><td>ATCAGATACCGTCGTAGTTCCGAC</td><td>1442.1</td><td>-7.4</td><td>1.0</td><td>-7.3</td><td>2.5E-13</td><td>6.5E-13</td><td>MB231/MB468</td></tr>
<tr><td>>hg38_21_+_8433172_8446622_RNA45SN1_45S_50ntFlanks@6776.6806.31|CTCCCGGGGCTACGCCTGTCTGAGCGTCGCT</td><td>>hg38_21_+_8433172_8446622_RNA45SN1_45S_50ntFlanks</td><td>6776</td><td>6806</td><td>31</td><td>CTCCCGGGGCTACGCCTGTCTGAGCGTCGCT</td><td>3922.0</td><td>4.2</td><td>0.6</td><td>7.3</td><td>2.5E-13</td><td>6.5E-13</td><td>MB231/MB468</td></tr>
<tr><td>>hg38_21_+_8433172_8446622_RNA45SN1_45S_50ntFlanks@12935.12955.21|TAAACCATTCGTAGACGACCT</td><td>>hg38_21_+_8433172_8446622_RNA45SN1_45S_50ntFlanks</td><td>12935</td><td>12955</td><td>21</td><td>TAAACCATTCGTAGACGACCT</td><td>107.5</td><td>-3.8</td><td>0.5</td><td>-7.3</td><td>2.8E-13</td><td>6.6E-13</td><td>BT20/MB468</td></tr>
<tr><td>>hg38_21_+_8433172_8446622_RNA45SN1_45S_50ntFlanks@11317.11342.26|ACCCCACGTCTCGTCGCGCGCGCGTC</td><td>>hg38_21_+_8433172_8446622_RNA45SN1_45S_50ntFlanks</td><td>11317</td><td>11342</td><td>26</td><td>ACCCCACGTCTCGTCGCGCGCGCGTC</td><td>690.8</td><td>-8.1</td><td>1.1</td><td>-7.3</td><td>2.6E-13</td><td>6.6E-13</td><td>MB231/MB468</td></tr>
<tr><td>>hg38_21_+_8433172_8446622_RNA45SN1_45S_50ntFlanks@11270.11293.24|CGGCCGCCCCCGGGGAGCCCGGCG</td><td>>hg38_21_+_8433172_8446622_RNA45SN1_45S_50ntFlanks</td><td>11270</td><td>11293</td><td>24</td><td>CGGCCGCCCCCGGGGAGCCCGGCG</td><td>189.8</td><td>-9.1</td><td>1.3</td><td>-7.3</td><td>2.6E-13</td><td>6.6E-13</td><td>MB231/MB468</td></tr>
<tr><td>>hg38_21_+_8433172_8446622_RNA45SN1_45S_50ntFlanks@4367.4398.32|CAGTTAAAAAGCTCGTAGTTGGATCTTGGGAG</td><td>>hg38_21_+_8433172_8446622_RNA45SN1_45S_50ntFlanks</td><td>4367</td><td>4398</td><td>32</td><td>CAGTTAAAAAGCTCGTAGTTGGATCTTGGGAG</td><td>353.4</td><td>-7.0</td><td>1.0</td><td>-7.3</td><td>2.8E-13</td><td>6.7E-13</td><td>BT20/MB468</td></tr>
<tr><td>>hg38_21_+_8433172_8446622_RNA45SN1_45S_50ntFlanks@6772.6797.26|GTTCCTCCCGGGGCTACGCCTGTCTG</td><td>>hg38_21_+_8433172_8446622_RNA45SN1_45S_50ntFlanks</td><td>6772</td><td>6797</td><td>26</td><td>GTTCCTCCCGGGGCTACGCCTGTCTG</td><td>169.8</td><td>-5.7</td><td>0.8</td><td>-7.3</td><td>2.6E-13</td><td>6.7E-13</td><td>MB231/MB468</td></tr>
<tr><td>>hg38_21_+_8433172_8446622_RNA45SN1_45S_50ntFlanks@4367.4391.25|CAGTTAAAAAGCTCGTAGTTGGATC</td><td>>hg38_21_+_8433172_8446622_RNA45SN1_45S_50ntFlanks</td><td>4367</td><td>4391</td><td>25</td><td>CAGTTAAAAAGCTCGTAGTTGGATC</td><td>1076.5</td><td>-4.4</td><td>0.6</td><td>-7.3</td><td>2.7E-13</td><td>6.8E-13</td><td>MB231/MB468</td></tr>
</table>

<table>
<tr><th></th><th></th><th></th><th></th><th></th><th></th><th></th><th></th><th></th><th></th><th></th><th></th><th></th></tr>
<tr><td>>MI0000469|hsa-mir-125a&WithFlank&19|+|51693248|51693345@21.41.21(+1A)&[MIMAT0000443&hsa-miR-125a-5p&offsets|0|-3(+1A);m-63&19|+|51693268|51693289&offsets|0|-1(+1A)]|TCCCTGAGACCCTTTAACCTGA</td><td>>MI0000469|hsa-mir-125a&WithFlank&19|+|51693248|51693345</td><td>21</td><td>41</td><td>21(+1A)</td><td>TCCCTGAGACCCTTTAACCTGA</td><td>506.8</td><td>9.3</td><td>1.3</td><td>7.3</td><td>2.9E-13</td><td>6.8E-13</td><td>BT20/MB468</td></tr>
<tr><td>>hg38_21_+_8433172_8446622_RNA45SN1_45S_50ntFlanks@9573.9597.25|AAATCGGTCGTCCGACCTGGGTATA</td><td>>hg38_21_+_8433172_8446622_RNA45SN1_45S_50ntFlanks</td><td>9573</td><td>9597</td><td>25</td><td>AAATCGGTCGTCCGACCTGGGTATA</td><td>2578.1</td><td>-5.0</td><td>0.7</td><td>-7.3</td><td>2.7E-13</td><td>6.8E-13</td><td>MB231/MB468</td></tr>
<tr><td>>hg38_21_+_8433172_8446622_RNA45SN1_45S_50ntFlanks@7989.8010.22|AGACGTGGCGACCCGCTGAATT</td><td>>hg38_21_+_8433172_8446622_RNA45SN1_45S_50ntFlanks</td><td>7989</td><td>8010</td><td>22</td><td>AGACGTGGCGACCCGCTGAATT</td><td>8586.5</td><td>6.8</td><td>0.9</td><td>7.3</td><td>2.7E-13</td><td>6.9E-13</td><td>MB231/MB468</td></tr>
<tr><td>>hg38_21_+_8433172_8446622_RNA45SN1_45S_50ntFlanks@10463.10497.35|CTCCGTTGCCCTCGGCCGATCGAAAGGGAGTCGGG</td><td>>hg38_21_+_8433172_8446622_RNA45SN1_45S_50ntFlanks</td><td>10463</td><td>10497</td><td>35</td><td>CTCCGTTGCCCTCGGCCGATCGAAAGGGAGTCGGG</td><td>130.0</td><td>-9.0</td><td>1.2</td><td>-7.3</td><td>2.9E-13</td><td>6.9E-13</td><td>BT20/MB468</td></tr>
<tr><td>>am_trna11_ProAGG_16_+_3241989_3242060@1.28.28,trna12_ProAGG_6_+_26555498_26555569@1.28.28,trna12_ProTGG_11_-_75946869_75946940@1.28.28,trna14_ProTGG_5_-_180615854_180615925@1.28.28,trna22_ProAGG_14_-_21081560_21081631@1.28.28,trna23_ProAGG_14_-_21077495_21077566@1.28.28,trna28_ProTGG_16_-_3234133_3234204@1.28.28,trna29_ProAGG_16_-_3232635_3232706@1.28.28,trna2_ProAGG_7_+_128423504_128423575@1.28.28,trna30_ProCGG_6_+_27059521_27059592@1.28.28,trna37_ProCGG_17_-_8126151_8126222@1.28.28,trna3_ProTGG_16_+_3208923_3208994@1.28.28,trna52_ProCGG_1_+_167683962_167684033@1.28.28,trna65_ProAGG_1_-_167684725_167684796@1.28.28,trna6_ProCGG_16_+_3222049_3222120@1.28.28,trna6_ProTGG_14_+_21152175_21152246@1.28.28,trna8_ProTGG_16_+_3238094_3238165@1.28.28,trna9_ProAGG_11_+_75946557_75946628@1.28.28,trna9_ProAGG_16_+_3239634_3239705@1.28.28|GGCTCGTTGGTCTAGGGGTATGATTCTC</td><td>>am_trna11_ProAGG_16_+_3241989_3242060</td><td>1</td><td>28</td><td>28</td><td>GGCTCGTTGGTCTAGGGGTATGATTCTC</td><td>509.8</td><td>-6.1</td><td>0.8</td><td>-7.3</td><td>2.7E-13</td><td>7.0E-13</td><td>MB231/MB468</td></tr>
<tr><td>>hg38_21_+_8433172_8446622_RNA45SN1_45S_50ntFlanks@6689.6720.32|GAAGAACGCAGCTAGCTGCGAGAATTAATGTG</td><td>>hg38_21_+_8433172_8446622_RNA45SN1_45S_50ntFlanks</td><td>6689</td><td>6720</td><td>32</td><td>GAAGAACGCAGCTAGCTGCGAGAATTAATGTG</td><td>130.0</td><td>-4.8</td><td>0.7</td><td>-7.3</td><td>3.0E-13</td><td>7.1E-13</td><td>BT20/MB468</td></tr>
<tr><td>>hg38_21_+_8433172_8446622_RNA45SN1_45S_50ntFlanks@9567.9595.29|ACGTGCAAATCGGTCGTCCGACCTGGGTA</td><td>>hg38_21_+_8433172_8446622_RNA45SN1_45S_50ntFlanks</td><td>9567</td><td>9595</td><td>29</td><td>ACGTGCAAATCGGTCGTCCGACCTGGGTA</td><td>1401.5</td><td>-5.0</td><td>0.7</td><td>-7.3</td><td>2.8E-13</td><td>7.1E-13</td><td>MB231/MB468</td></tr>
<tr><td>>trna10_LysCTT_16_+_3241501_3241573@1.34.34,trna13_LysCTT_14_-_58706613_58706685@1.34.34,trna2_LysCTT_15_+_79152904_79152976@1.34.34|GCCCGGCTAGCTCAGTCGGTAGAGCATGGGACTC</td><td>>trna10_LysCTT_16_+_3241501_3241573</td><td>1</td><td>34</td><td>34</td><td>GCCCGGCTAGCTCAGTCGGTAGAGCATGGGACTC</td><td>5797.4</td><td>9.9</td><td>1.4</td><td>7.3</td><td>3.0E-13</td><td>7.1E-13</td><td>BT20/MB468</td></tr>
<tr><td>>MI0000084|hsa-mir-26b&WithFlank&2|+|218402640|218402728@18.38.21&[MIMAT0000083&hsa-miR-26b-5p&offsets|0|0;m-53&2|+|218402657|218402678&offsets|0|-1]|TTCAAGTAATTCAGGATAGGT</td><td>>MI0000084|hsa-mir-26b&WithFlank&2|+|218402640|218402728</td><td>18</td><td>38</td><td>21</td><td>TTCAAGTAATTCAGGATAGGT</td><td>5635.7</td><td>7.1</td><td>1.0</td><td>7.3</td><td>3.0E-13</td><td>7.1E-13</td><td>BT20/MB468</td></tr>
<tr><td>>hg38_21_+_8433172_8446622_RNA45SN1_45S_50ntFlanks@10574.10591.18|AAGCCGGCGGGAGCCCCG</td><td>>hg38_21_+_8433172_8446622_RNA45SN1_45S_50ntFlanks</td><td>10574</td><td>10591</td><td>18</td><td>AAGCCGGCGGGAGCCCCG</td><td>367.2</td><td>-10.5</td><td>1.4</td><td>-7.3</td><td>3.0E-13</td><td>7.1E-13</td><td>BT20/MB468</td></tr>
<tr><td>>hg38_21_+_8433172_8446622_RNA45SN1_45S_50ntFlanks@10388.10424.37|GTGAACAGCAGTTGAACATGGGTCAGTCGGTCCTGAG</td><td>>hg38_21_+_8433172_8446622_RNA45SN1_45S_50ntFlanks</td><td>10388</td><td>10424</td><td>37</td><td>GTGAACAGCAGTTGAACATGGGTCAGTCGGTCCTGAG</td><td>1473.1</td><td>6.0</td><td>0.8</td><td>7.3</td><td>3.0E-13</td><td>7.2E-13</td><td>BT20/MB468</td></tr>
<tr><td>>hg38_21_+_8433172_8446622_RNA45SN1_45S_50ntFlanks@10808.10829.22|CAATGTAGGTAAGGGAAGTCGG</td><td>>hg38_21_+_8433172_8446622_RNA45SN1_45S_50ntFlanks</td><td>10808</td><td>10829</td><td>22</td><td>CAATGTAGGTAAGGGAAGTCGG</td><td>154.2</td><td>-6.3</td><td>0.9</td><td>-7.3</td><td>2.8E-13</td><td>7.2E-13</td><td>MB231/MB468</td></tr>
<tr><td>>hg38_21_+_8433172_8446622_RNA45SN1_45S_50ntFlanks@10384.10401.18|CCATGTGAACAGCAGTTG</td><td>>hg38_21_+_8433172_8446622_RNA45SN1_45S_50ntFlanks</td><td>10384</td><td>10401</td><td>18</td><td>CCATGTGAACAGCAGTTG</td><td>237.4</td><td>-9.5</td><td>1.3</td><td>-7.3</td><td>2.8E-13</td><td>7.2E-13</td><td>MB231/MB468</td></tr>
<tr><td>>hg38_21_+_8433172_8446622_RNA45SN1_45S_50ntFlanks@12405.12427.23|CTTTTTGATCCTTCGATGTCGGC</td><td>>hg38_21_+_8433172_8446622_RNA45SN1_45S_50ntFlanks</td><td>12405</td><td>12427</td><td>23</td><td>CTTTTTGATCCTTCGATGTCGGC</td><td>140.0</td><td>-9.1</td><td>1.3</td><td>-7.3</td><td>3.1E-13</td><td>7.3E-13</td><td>BT20/MB468</td></tr>
<tr><td>>hg38_21_+_8433172_8446622_RNA45SN1_45S_50ntFlanks@11619.11640.22|AACAAAGCATCGCGAAGGCCCG</td><td>>hg38_21_+_8433172_8446622_RNA45SN1_45S_50ntFlanks</td><td>11619</td><td>11640</td><td>22</td><td>AACAAAGCATCGCGAAGGCCCG</td><td>185.4</td><td>-9.5</td><td>1.3</td><td>-7.3</td><td>3.1E-13</td><td>7.3E-13</td><td>BT20/MB468</td></tr>
<tr><td>>hg38_21_+_8433172_8446622_RNA45SN1_45S_50ntFlanks@10731.10755.25|AGGGTGTAAATCTCGCGCCGGGCCG</td><td>>hg38_21_+_8433172_8446622_RNA45SN1_45S_50ntFlanks</td><td>10731</td><td>10755</td><td>25</td><td>AGGGTGTAAATCTCGCGCCGGGCCG</td><td>209.1</td><td>-9.3</td><td>1.3</td><td>-7.3</td><td>2.8E-13</td><td>7.3E-13</td><td>MB231/MB468</td></tr>
<tr><td>>hg38_21_+_8433172_8446622_RNA45SN1_45S_50ntFlanks@12412.12441.30|ATCCTTCGATGTCGGCTCTTCCTATCATTG</td><td>>hg38_21_+_8433172_8446622_RNA45SN1_45S_50ntFlanks</td><td>12412</td><td>12441</td><td>30</td><td>ATCCTTCGATGTCGGCTCTTCCTATCATTG</td><td>149.1</td><td>-9.2</td><td>1.3</td><td>-7.3</td><td>3.1E-13</td><td>7.4E-13</td><td>BT20/MB468</td></tr>
<tr><td>>hg38_21_+_8433172_8446622_RNA45SN1_45S_50ntFlanks@10427.10453.27|ATGGGCGAGCGCCGTTCCGAAGGGACG</td><td>>hg38_21_+_8433172_8446622_RNA45SN1_45S_50ntFlanks</td><td>10427</td><td>10453</td><td>27</td><td>ATGGGCGAGCGCCGTTCCGAAGGGACG</td><td>393.6</td><td>-5.1</td><td>0.7</td><td>-7.3</td><td>3.1E-13</td><td>7.4E-13</td><td>BT20/MB468</td></tr>
<tr><td>>hg38_21_+_8433172_8446622_RNA45SN1_45S_50ntFlanks@7975.8000.26|CGCGACCTCAGATCAGACGTGGCGAC</td><td>>hg38_21_+_8433172_8446622_RNA45SN1_45S_50ntFlanks</td><td>7975</td><td>8000</td><td>26</td><td>CGCGACCTCAGATCAGACGTGGCGAC</td><td>668.7</td><td>7.6</td><td>1.0</td><td>7.3</td><td>3.1E-13</td><td>7.4E-13</td><td>BT20/MB468</td></tr>
<tr><td>>hg38_21_+_8433172_8446622_RNA45SN1_45S_50ntFlanks@12628.12657.30|TGGGGCGAAGCTACCATCTGTGGGATTATG</td><td>>hg38_21_+_8433172_8446622_RNA45SN1_45S_50ntFlanks</td><td>12628</td><td>12657</td><td>30</td><td>TGGGGCGAAGCTACCATCTGTGGGATTATG</td><td>144.7</td><td>-9.2</td><td>1.3</td><td>-7.3</td><td>3.1E-13</td><td>7.4E-13</td><td>BT20/MB468</td></tr>
<tr><td>>hg38_21_+_8433172_8446622_RNA45SN1_45S_50ntFlanks@12568.12587.20|TCAGTACGAGAGGAACCGCA</td><td>>hg38_21_+_8433172_8446622_RNA45SN1_45S_50ntFlanks</td><td>12568</td><td>12587</td><td>20</td><td>TCAGTACGAGAGGAACCGCA</td><td>134.4</td><td>-9.1</td><td>1.2</td><td>-7.3</td><td>3.2E-13</td><td>7.4E-13</td><td>BT20/MB468</td></tr>
<tr><td>>hg38_21_+_8433172_8446622_RNA45SN1_45S_50ntFlanks@482.506.25|ATCGATGTGGTGACGTCGTGCTCTC</td><td>>hg38_21_+_8433172_8446622_RNA45SN1_45S_50ntFlanks</td><td>482</td><td>506</td><td>25</td><td>ATCGATGTGGTGACGTCGTGCTCTC</td><td>147.5</td><td>-9.2</td><td>1.3</td><td>-7.3</td><td>3.2E-13</td><td>7.5E-13</td><td>BT20/MB468</td></tr>
<tr><td>>am_trna2_GlyTCC_19_+_4724082_4724153@1.30.30|GCGTTGGTGGTATAGTGGTTAGCATAGCTG</td><td>>am_trna2_GlyTCC_19_+_4724082_4724153</td><td>1</td><td>30</td><td>30</td><td>GCGTTGGTGGTATAGTGGTTAGCATAGCTG</td><td>222.6</td><td>-9.8</td><td>1.3</td><td>-7.3</td><td>3.2E-13</td><td>7.5E-13</td><td>BT20/MB468</td></tr>
<tr><td>>hg38_21_+_8433172_8446622_RNA45SN1_45S_50ntFlanks@12934.12951.18|CTAAACCATTCGTAGACG</td><td>>hg38_21_+_8433172_8446622_RNA45SN1_45S_50ntFlanks</td><td>12934</td><td>12951</td><td>18</td><td>CTAAACCATTCGTAGACG</td><td>802.6</td><td>-6.6</td><td>0.9</td><td>-7.3</td><td>3.2E-13</td><td>7.5E-13</td><td>BT20/MB468</td></tr>
<tr><td>>hg38_21_+_8433172_8446622_RNA45SN1_45S_50ntFlanks@10571.10596.26|GAGAAGCCGGCGGGAGCCCCGGGGAG</td><td>>hg38_21_+_8433172_8446622_RNA45SN1_45S_50ntFlanks</td><td>10571</td><td>10596</td><td>26</td><td>GAGAAGCCGGCGGGAGCCCCGGGGAG</td><td>151.8</td><td>-9.2</td><td>1.3</td><td>-7.3</td><td>3.2E-13</td><td>7.5E-13</td><td>BT20/MB468</td></tr>
<tr><td>>hg38_21_+_8433172_8446622_RNA45SN1_45S_50ntFlanks@4567.4587.21|TAATGGAATAGGACCGCGGTT</td><td>>hg38_21_+_8433172_8446622_RNA45SN1_45S_50ntFlanks</td><td>4567</td><td>4587</td><td>21</td><td>TAATGGAATAGGACCGCGGTT</td><td>354.3</td><td>-4.0</td><td>0.6</td><td>-7.3</td><td>3.0E-13</td><td>7.6E-13</td><td>MB231/MB468</td></tr>
<tr><td>>hg38_21_+_8433172_8446622_RNA45SN1_45S_50ntFlanks@4722.4752.31|TCATTAATCAAGAACGAAAGTCGGAGGTTCG</td><td>>hg38_21_+_8433172_8446622_RNA45SN1_45S_50ntFlanks</td><td>4722</td><td>4752</td><td>31</td><td>TCATTAATCAAGAACGAAAGTCGGAGGTTCG</td><td>468.9</td><td>-4.9</td><td>0.7</td><td>-7.3</td><td>3.0E-13</td><td>7.6E-13</td><td>MB231/MB468</td></tr>
<tr><td>>trnaMT_PheGAA_MT_+_577_647@55.74.20|ACATCACCCCATAAACACCA</td><td>>trnaMT_PheGAA_MT_+_577_647</td><td>55</td><td>74</td><td>20</td><td>ACATCACCCCATAAACACCA</td><td>793.6</td><td>9.4</td><td>1.3</td><td>7.3</td><td>3.0E-13</td><td>7.8E-13</td><td>MB231/MB468</td></tr>
<tr><td>>MI0000285|hsa-mir-205&WithFlank&1|+|209432127|209432248@40.61.22&[MIMAT0000266&hsa-miR-205-5p&offsets|0|0;m-351&1|+|209432166|209432187&offsets|0|0]|TCCTTCATTCCACCGGAGTCTG</td><td>>MI0000285|hsa-mir-205&WithFlank&1|+|209432127|209432248</td><td>40</td><td>61</td><td>22</td><td>TCCTTCATTCCACCGGAGTCTG</td><td>936.4</td><td>-7.5</td><td>1.0</td><td>-7.3</td><td>3.0E-13</td><td>7.8E-13</td><td>MB231/MB468</td></tr>
<tr><td>>hg38_21_+_8433172_8446622_RNA45SN1_45S_50ntFlanks@12459.12478.20|AGCGTTGGATTGTTCACCCA</td><td>>hg38_21_+_8433172_8446622_RNA45SN1_45S_50ntFlanks</td><td>12459</td><td>12478</td><td>20</td><td>AGCGTTGGATTGTTCACCCA</td><td>607.4</td><td>-3.7</td><td>0.5</td><td>-7.3</td><td>3.1E-13</td><td>7.8E-13</td><td>MB231/MB468</td></tr>
<tr><td>>hg38_21_+_8433172_8446622_RNA45SN1_45S_50ntFlanks@9736.9758.23|CCGAAACGATCTCAACCTATTCT</td><td>>hg38_21_+_8433172_8446622_RNA45SN1_45S_50ntFlanks</td><td>9736</td><td>9758</td><td>23</td><td>CCGAAACGATCTCAACCTATTCT</td><td>127.5</td><td>-9.0</td><td>1.2</td><td>-7.3</td><td>3.4E-13</td><td>7.9E-13</td><td>BT20/MB468</td></tr>
<tr><td>>hg38_21_+_8433172_8446622_RNA45SN1_45S_50ntFlanks@6651.6674.24|CGACTCTTAGCGGTGGATCACTCG</td><td>>hg38_21_+_8433172_8446622_RNA45SN1_45S_50ntFlanks</td><td>6651</td><td>6674</td><td>24</td><td>CGACTCTTAGCGGTGGATCACTCG</td><td>1361.3</td><td>3.9</td><td>0.5</td><td>7.3</td><td>3.4E-13</td><td>8.0E-13</td><td>BT20/MB468</td></tr>
<tr><td>>hg38_21_+_8433172_8446622_RNA45SN1_45S_50ntFlanks@11581.11603.23|AACTGGTGCGGACCAGGGGAATC</td><td>>hg38_21_+_8433172_8446622_RNA45SN1_45S_50ntFlanks</td><td>11581</td><td>11603</td><td>23</td><td>AACTGGTGCGGACCAGGGGAATC</td><td>4022.4</td><td>-4.9</td><td>0.7</td><td>-7.3</td><td>3.4E-13</td><td>8.0E-13</td><td>BT20/MB468</td></tr>
<tr><td>>hg38_21_+_8433172_8446622_RNA45SN1_45S_50ntFlanks@10423.10457.35|AGAGATGGGCGAGCGCCGTTCCGAAGGGACGGGCG</td><td>>hg38_21_+_8433172_8446622_RNA45SN1_45S_50ntFlanks</td><td>10423</td><td>10457</td><td>35</td><td>AGAGATGGGCGAGCGCCGTTCCGAAGGGACGGGCG</td><td>145.4</td><td>-9.2</td><td>1.3</td><td>-7.3</td><td>3.4E-13</td><td>8.0E-13</td><td>BT20/MB468</td></tr>
<tr><td>>MI0000087|hsa-mir-29a&WithFlank&7|-|130876741|130876816@48.67.20&[MIMAT0000086&hsa-miR-29a-3p&offsets|0|-2;m-26&7|-|130876749|130876769&offsets|0|-1]|TAGCACCATCTGAAATCGGT</td><td>>MI0000087|hsa-mir-29a&WithFlank&7|-|130876741|130876816</td><td>48</td><td>67</td><td>20</td><td>TAGCACCATCTGAAATCGGT</td><td>1305.0</td><td>10.1</td><td>1.4</td><td>7.3</td><td>3.5E-13</td><td>8.1E-13</td><td>BT20/MB468</td></tr>
<tr><td>>hg38_21_+_8433172_8446622_RNA45SN1_45S_50ntFlanks@10408.10428.21|GGTCAGTCGGTCCTGAGAGAT</td><td>>hg38_21_+_8433172_8446622_RNA45SN1_45S_50ntFlanks</td><td>10408</td><td>10428</td><td>21</td><td>GGTCAGTCGGTCCTGAGAGAT</td><td>211.9</td><td>-6.1</td><td>0.8</td><td>-7.3</td><td>3.5E-13</td><td>8.1E-13</td><td>BT20/MB468</td></tr>
<tr><td>>hg38_21_+_8433172_8446622_RNA45SN1_45S_50ntFlanks@3705.3744.40|TACCTGGTTGATCCTGCCAGTAGCATATGCTTGTCTCAAA</td><td>>hg38_21_+_8433172_8446622_RNA45SN1_45S_50ntFlanks</td><td>3705</td><td>3744</td><td>40</td><td>TACCTGGTTGATCCTGCCAGTAGCATATGCTTGTCTCAAA</td><td>2240.3</td><td>9.3</td><td>1.3</td><td>7.3</td><td>3.5E-13</td><td>8.2E-13</td><td>BT20/MB468</td></tr>
<tr><td>>MI0000285|hsa-mir-205&WithFlank&1|+|209432127|209432248@40.61.22(+3A)&[MIMAT0000266&hsa-miR-205-5p&offsets|0|0(+3A);m-351&1|+|209432166|209432187&offsets|0|0(+3A)]|TCCTTCATTCCACCGGAGTCTGAAA</td><td>>MI0000285|hsa-mir-205&WithFlank&1|+|209432127|209432248</td><td>40</td><td>61</td><td>22(+3A)</td><td>TCCTTCATTCCACCGGAGTCTGAAA</td><td>509.2</td><td>6.1</td><td>0.8</td><td>7.3</td><td>3.5E-13</td><td>8.3E-13</td><td>BT20/MB468</td></tr>
<tr><td>>MI0000650|hsa-mir-200c&WithFlank&12|+|6963693|6963772@49.71.23&[MIMAT0000617&hsa-miR-200c-3p&offsets|-1|-1;m-54&12|+|6963742|6963764&offsets|-1|-1]|CTAATACTGCCGGGTAATGATGG</td><td>>MI0000650|hsa-mir-200c&WithFlank&12|+|6963693|6963772</td><td>49</td><td>71</td><td>23</td><td>CTAATACTGCCGGGTAATGATGG</td><td>1344.3</td><td>8.7</td><td>1.2</td><td>7.3</td><td>3.6E-13</td><td>8.3E-13</td><td>BT20/MB468</td></tr>
<tr><td>>hg38_21_+_8433172_8446622_RNA45SN1_45S_50ntFlanks@3711.3737.27|GTTGATCCTGCCAGTAGCATATGCTTG</td><td>>hg38_21_+_8433172_8446622_RNA45SN1_45S_50ntFlanks</td><td>3711</td><td>3737</td><td>27</td><td>GTTGATCCTGCCAGTAGCATATGCTTG</td><td>431.2</td><td>-4.3</td><td>0.6</td><td>-7.3</td><td>3.3E-13</td><td>8.3E-13</td><td>MB231/MB468</td></tr>
</table>

| | | | | | | | | | | | | |
|---|---|---|---|---|---|---|---|---|---|---|---|---|
| >hg38_21_+_8433172_8446622_RNA45SN1_45S_50ntFlanks@11199.11225.27\|CCGGCGGCGGCGGCGACTCTGGACGCG | >hg38_21_+_8433172_8446622_RNA45SN1_45S_50ntFlanks | 11199 | 11225 | 27 | CCGGCGGCGGCGGCGACTCTGGACGCG | 1038.9 | -7.5 | 1.0 | -7.3 | 3.6E-13 | 8.5E-13 | BT20/MB468 |
| >am_trna2_LeuTAA_X_-_55207755_55207829@1.18.18,trna4_LeuTAA_20_-_55933428_55933502@1.18.18,trna60_LeuTAA_1_-_238106956_238107030@1.18.18,trna9_LeuTAA_2_+_117781417_117781491@1.18.18,trnaMT_LeuTAA_MT_+_3230_3304@1.18.18\|GTTAAGATGGCAGAGCCC | >am_trna2_LeuTAA_X_-_55207755_55207829 | 1 | 18 | 18 | GTTAAGATGGCAGAGCCC | 559.5 | 9.8 | 1.4 | 7.3 | 3.7E-13 | 8.5E-13 | BT20/MB468 |
| >hg38_21_+_8433172_8446622_RNA45SN1_45S_50ntFlanks@11760.11777.18\|CTCGTCATCTAATTAGTG | >hg38_21_+_8433172_8446622_RNA45SN1_45S_50ntFlanks | 11760 | 11777 | 18 | CTCGTCATCTAATTAGTG | 141.8 | -9.1 | 1.3 | -7.3 | 3.7E-13 | 8.6E-13 | BT20/MB468 |
| >hg38_21_+_8433172_8446622_RNA45SN1_45S_50ntFlanks@12329.12346.18\|TTTTAAGCAGGAGGTGTC | >hg38_21_+_8433172_8446622_RNA45SN1_45S_50ntFlanks | 12329 | 12346 | 18 | TTTTAAGCAGGAGGTGTC | 252.1 | -6.3 | 0.9 | -7.3 | 3.7E-13 | 8.6E-13 | BT20/MB468 |
| >hg38_21_+_8433172_8446622_RNA45SN1_45S_50ntFlanks@12418.12443.26\|CGATGTCGGCTCTTCCTATCATTGTG | >hg38_21_+_8433172_8446622_RNA45SN1_45S_50ntFlanks | 12418 | 12443 | 26 | CGATGTCGGCTCTTCCTATCATTGTG | 3716.2 | -4.5 | 0.6 | -7.3 | 3.7E-13 | 8.6E-13 | BT20/MB468 |
| >hg38_21_+_8433172_8446622_RNA45SN1_45S_50ntFlanks@9507.9526.20\|AACTATGCCTGGGCAGGGCG | >hg38_21_+_8433172_8446622_RNA45SN1_45S_50ntFlanks | 9507 | 9526 | 20 | AACTATGCCTGGGCAGGGCG | 149.8 | -9.2 | 1.3 | -7.3 | 3.7E-13 | 8.6E-13 | BT20/MB468 |
| >hg38_21_+_8433172_8446622_RNA45SN1_45S_50ntFlanks@10398.10424.27\|GTTGAACATGGGTCAGTCGGTCCTGAG | >hg38_21_+_8433172_8446622_RNA45SN1_45S_50ntFlanks | 10398 | 10424 | 27 | GTTGAACATGGGTCAGTCGGTCCTGAG | 1114.9 | 7.8 | 1.1 | 7.3 | 3.5E-13 | 8.9E-13 | MB231/MB468 |
| >MI0003834\|hsa-mir-769&WithFlank&19\|+\|46018926\|46019055@36.56.21(+1C)&[MIMAT0003886&hsa-miR-769-5p&offsets\|0\|-1(+1C);m-216&19\|+\|46018961\|46018982&offsets\|0\|-1(+1C)]\|TGAGACCTCTGGGTTCTGAGCC | >MI0003834\|hsa-mir-769&WithFlank&19\|+\|46018926\|46019055 | 36 | 56 | 21(+1C) | TGAGACCTCTGGGTTCTGAGCC | 1106.5 | 8.9 | 1.2 | 7.3 | 3.8E-13 | 8.9E-13 | BT20/MB468 |
| >hg38_21_+_8433172_8446622_RNA45SN1_45S_50ntFlanks@12973.13001.29\|TCGTACGTAGCAGAGCAGCTCCCTCGCTG | >hg38_21_+_8433172_8446622_RNA45SN1_45S_50ntFlanks | 12973 | 13001 | 29 | TCGTACGTAGCAGAGCAGCTCCCTCGCTG | 193.4 | -9.2 | 1.3 | -7.3 | 3.5E-13 | 9.0E-13 | MB231/MB468 |
| >hg38_21_+_8433172_8446622_RNA45SN1_45S_50ntFlanks@3802.3829.28\|CATTAAATCAGTTATGGTTCCTTTGGTC | >hg38_21_+_8433172_8446622_RNA45SN1_45S_50ntFlanks | 3802 | 3829 | 28 | CATTAAATCAGTTATGGTTCCTTTGGTC | 177.4 | -5.8 | 0.8 | -7.3 | 3.9E-13 | 9.0E-13 | BT20/MB468 |
| >hg38_21_+_8433172_8446622_RNA45SN1_45S_50ntFlanks@12478.12497.20\|ACTAATAGGGAACGTGAGCT | >hg38_21_+_8433172_8446622_RNA45SN1_45S_50ntFlanks | 12478 | 12497 | 20 | ACTAATAGGGAACGTGAGCT | 1361.6 | 2.6 | 0.4 | 7.3 | 3.9E-13 | 9.0E-13 | BT20/MB468 |
| >hg38_21_+_8433172_8446622_RNA45SN1_45S_50ntFlanks@4723.4751.29\|CATTAATCAAGAACGAAAGTCGGAGGTTC | >hg38_21_+_8433172_8446622_RNA45SN1_45S_50ntFlanks | 4723 | 4751 | 29 | CATTAATCAAGAACGAAAGTCGGAGGTTC | 449.6 | -4.8 | 0.7 | -7.3 | 3.9E-13 | 9.1E-13 | BT20/MB468 |
| >MI0000734\|hsa-mir-106b&WithFlank&7\|-\|100093987\|100094080@18.37.20&[MIMAT0000680&hsa-miR-106b-5p&offsets\|0\|-1;m-103&7\|-\|100094043\|100094063&offsets\|0\|-1]\|TAAAGTGCTGACAGTGCAGA | >MI0000734\|hsa-mir-106b&WithFlank&7\|-\|100093987\|100094080 | 18 | 37 | 20 | TAAAGTGCTGACAGTGCAGA | 1421.7 | 8.1 | 1.1 | 7.3 | 3.9E-13 | 9.1E-13 | BT20/MB468 |
| >trnaMT_SerGCT_MT_+_12207_12265@1.20.20\|GAGAAAGCTCACAAGAACTG | >trnaMT_SerGCT_MT_+_12207_12265 | 1 | 20 | 20 | GAGAAAGCTCACAAGAACTG | 1458.5 | 4.0 | 0.5 | 7.3 | 3.9E-13 | 9.1E-13 | BT20/MB468 |
| >hg38_21_+_8433172_8446622_RNA45SN1_45S_50ntFlanks@11577.11605.29\|TTAGAACTGGTGCGGACCAGGGGAATCCG | >hg38_21_+_8433172_8446622_RNA45SN1_45S_50ntFlanks | 11577 | 11605 | 29 | TTAGAACTGGTGCGGACCAGGGGAATCCG | 345.6 | -5.5 | 0.8 | -7.3 | 3.6E-13 | 9.1E-13 | MB231/MB468 |
| >hg38_21_+_8433172_8446622_RNA45SN1_45S_50ntFlanks@12934.12972.39\|CTAAACCATTCGTAGACGACCTGCTTCTGGGTCGGGGTT | >hg38_21_+_8433172_8446622_RNA45SN1_45S_50ntFlanks | 12934 | 12972 | 39 | CTAAACCATTCGTAGACGACCTGCTTCTGGGTCGGGGTT | 1220.9 | -7.3 | 1.0 | -7.3 | 3.9E-13 | 9.2E-13 | BT20/MB468 |
| >hg38_21_+_8433172_8446622_RNA45SN1_45S_50ntFlanks@10410.10435.26\|TCAGTCGGTCCTGAGAGATGGGCGAG | >hg38_21_+_8433172_8446622_RNA45SN1_45S_50ntFlanks | 10410 | 10435 | 26 | TCAGTCGGTCCTGAGAGATGGGCGAG | 276.6 | -9.7 | 1.3 | -7.3 | 3.6E-13 | 9.2E-13 | MB231/MB468 |
| >hg38_21_+_8433172_8446622_RNA45SN1_45S_50ntFlanks@6661.6687.27\|CGGTGGATCACTCGGCTCGTGCGTCGA | >hg38_21_+_8433172_8446622_RNA45SN1_45S_50ntFlanks | 6661 | 6687 | 27 | CGGTGGATCACTCGGCTCGTGCGTCGA | 1733.8 | 6.0 | 0.8 | 7.3 | 4.0E-13 | 9.3E-13 | BT20/MB468 |
| >trna10_LysCTT_16_+_3241501_3241573@1.29.29,trna13_LysCTT_14_-_58706613_58706685@1.29.29,trna2_LysCTT_15_+_79152904_79152976@1.29.29\|GCCCGGCTAGCTCAGTCGGTAGAGCATGG | >trna10_LysCTT_16_+_3241501_3241573 | 1 | 29 | 29 | GCCCGGCTAGCTCAGTCGGTAGAGCATGG | 595.5 | -5.8 | 0.8 | -7.3 | 4.0E-13 | 9.3E-13 | BT20/MB468 |
| >hg38_1_-_228634819_228635039_RNA5S12_5S_50ntFlanks@84.112.29\|CTCGTCTGATCTCGGAAGCTAAGCAGGGT | >hg38_1_-_228634819_228635039_RNA5S12_5S_50ntFlanks | 84 | 112 | 29 | CTCGTCTGATCTCGGAAGCTAAGCAGGGT | 303.9 | -5.8 | 0.8 | -7.3 | 4.0E-13 | 9.3E-13 | BT20/MB468 |
| >hg38_21_+_8433172_8446622_RNA45SN1_45S_50ntFlanks@4472.4492.21\|CTTAGCTGAGTGTCCCGCGGG | >hg38_21_+_8433172_8446622_RNA45SN1_45S_50ntFlanks | 4472 | 4492 | 21 | CTTAGCTGAGTGTCCCGCGGG | 172.2 | -6.1 | 0.8 | -7.3 | 4.1E-13 | 9.5E-13 | BT20/MB468 |
| >MI0000253\|hsa-mir-148a&WithFlank&7\|-\|25949913\|25949992@50.69.20(+2U)&[MIMAT0000243&hsa-miR-148a-3p&offsets\|0\|-2(+2U);m-7&7\|-\|25949922\|25949943&offsets\|0\|-2(+2U)]\|TCAGTGCACTACAGAACTTTTT | >MI0000253\|hsa-mir-148a&WithFlank&7\|-\|25949913\|25949992 | 50 | 69 | 20(+2U) | TCAGTGCACTACAGAACTTTTT | 550.6 | 8.4 | 1.2 | 7.3 | 4.1E-13 | 9.5E-13 | BT20/MB468 |
| >hg38_21_+_8433172_8446622_RNA45SN1_45S_50ntFlanks@12934.12953.20\|CTAAACCATTCGTAGACGAC | >hg38_21_+_8433172_8446622_RNA45SN1_45S_50ntFlanks | 12934 | 12953 | 20 | CTAAACCATTCGTAGACGAC | 767.5 | -7.6 | 1.0 | -7.3 | 4.1E-13 | 9.5E-13 | BT20/MB468 |
| >hg38_21_+_8433172_8446622_RNA45SN1_45S_50ntFlanks@11199.11220.22\|CCGGCGGCGGCGGCGACTCTGG | >hg38_21_+_8433172_8446622_RNA45SN1_45S_50ntFlanks | 11199 | 11220 | 22 | CCGGCGGCGGCGGCGACTCTGG | 193.4 | -9.2 | 1.3 | -7.3 | 3.7E-13 | 9.5E-13 | MB231/MB468 |
| >hg38_21_+_8433172_8446622_RNA45SN1_45S_50ntFlanks@12937.12966.30\|AACCATTCGTAGACGACCTGCTTCTGGGTC | >hg38_21_+_8433172_8446622_RNA45SN1_45S_50ntFlanks | 12937 | 12966 | 30 | AACCATTCGTAGACGACCTGCTTCTGGGTC | 184.7 | -9.1 | 1.3 | -7.3 | 3.7E-13 | 9.6E-13 | MB231/MB468 |
| >hg38_21_+_8433172_8446622_RNA45SN1_45S_50ntFlanks@10107.10127.21\|CGTGTGGTGTGCGTCGGAGGG | >hg38_21_+_8433172_8446622_RNA45SN1_45S_50ntFlanks | 10107 | 10127 | 21 | CGTGTGGTGTGCGTCGGAGGG | 189.6 | -9.1 | 1.3 | -7.3 | 3.8E-13 | 9.6E-13 | MB231/MB468 |
| >MI0000542\|hsa-mir-320a&WithFlank&8\|-\|22244960\|22245043@50.70.21&[MIMAT0000510&hsa-miR-320a-3p&offsets\|+2\|+1;m-60&8\|-\|22244975\|22244996&offsets\|+2\|+1]\|AAGCTGGGTTGAGAGGGCGAA | >MI0000542\|hsa-mir-320a&WithFlank&8\|-\|22244960\|22245043 | 50 | 70 | 21 | AAGCTGGGTTGAGAGGGCGAA | 460.6 | 10.2 | 1.4 | 7.3 | 4.1E-13 | 9.6E-13 | BT20/MB468 |
| >hg38_1_-_228634819_228635039_RNA5S12_5S_50ntFlanks@137.170.34\|GAGACCGCCTGGGAATACCGGGTGCTGTAGGCTT | >hg38_1_-_228634819_228635039_RNA5S12_5S_50ntFlanks | 137 | 170 | 34 | GAGACCGCCTGGGAATACCGGGTGCTGTAGGCTT | 1122.5 | 8.7 | 1.2 | 7.3 | 4.2E-13 | 9.6E-13 | BT20/MB468 |
| >hg38_21_+_8433172_8446622_RNA45SN1_45S_50ntFlanks@11209.11234.26\|CGGCGACTCTGGACGCGAGCCGGGCC | >hg38_21_+_8433172_8446622_RNA45SN1_45S_50ntFlanks | 11209 | 11234 | 26 | CGGCGACTCTGGACGCGAGCCGGGCC | 1429.3 | -5.6 | 0.8 | -7.3 | 3.8E-13 | 9.7E-13 | MB231/MB468 |
| >hg38_21_+_8433172_8446622_RNA45SN1_45S_50ntFlanks@10797.10818.22\|GCATGTTGGAACAATGTAGGTA | >hg38_21_+_8433172_8446622_RNA45SN1_45S_50ntFlanks | 10797 | 10818 | 22 | GCATGTTGGAACAATGTAGGTA | 3394.7 | -5.4 | 0.7 | -7.2 | 4.2E-13 | 9.8E-13 | BT20/MB468 |
| >hg38_21_+_8433172_8446622_RNA45SN1_45S_50ntFlanks@11874.11912.39\|AGAAGACCCTGTTGAGCTTGACTCTAGTCTGGCACGGTG | >hg38_21_+_8433172_8446622_RNA45SN1_45S_50ntFlanks | 11874 | 11912 | 39 | AGAAGACCCTGTTGAGCTTGACTCTAGTCTGGCACGGTG | 128.3 | -9.0 | 1.2 | -7.2 | 4.2E-13 | 9.8E-13 | BT20/MB468 |
| >MI0000102\|hsa-mir-100&WithFlank&11\|-\|122152223\|122152314@19.40.22(+1C)&[MIMAT0000098&hsa-miR-100-5p&offsets\|0\|0(+1C);m-27&11\|-\|122152276\|122152296&offsets\|0\|+1(+1C)]\|AACCCGTAGATCCGAACTTGTGC | >MI0000102\|hsa-mir-100&WithFlank&11\|-\|122152223\|122152314 | 19 | 40 | 22(+1C) | AACCCGTAGATCCGAACTTGTGC | 27731.2 | 16.0 | 2.2 | 7.3 | 3.9E-13 | 9.9E-13 | MB231/MB468 |
| >hg38_21_+_8433172_8446622_RNA45SN1_45S_50ntFlanks@12974.13001.28\|CGTACGTAGCAGAGCAGCTCCCTCGCTG | >hg38_21_+_8433172_8446622_RNA45SN1_45S_50ntFlanks | 12974 | 13001 | 28 | CGTACGTAGCAGAGCAGCTCCCTCGCTG | 126.6 | -9.0 | 1.2 | -7.2 | 4.3E-13 | 9.9E-13 | BT20/MB468 |
| >hg38_21_+_8433172_8446622_RNA45SN1_45S_50ntFlanks@12499.12518.20\|GGTTTAGACCGTCGTGAGAC | >hg38_21_+_8433172_8446622_RNA45SN1_45S_50ntFlanks | 12499 | 12518 | 20 | GGTTTAGACCGTCGTGAGAC | 224.5 | -9.4 | 1.3 | -7.3 | 3.9E-13 | 9.9E-13 | MB231/MB468 |
| >hg38_21_+_8433172_8446622_RNA45SN1_45S_50ntFlanks@3705.3748.44\|TACCTGGTTGATCCTGCCAGTAGCATATGCTTGTCTCAAAGATT | >hg38_21_+_8433172_8446622_RNA45SN1_45S_50ntFlanks | 3705 | 3748 | 44 | TACCTGGTTGATCCTGCCAGTAGCATATGCTTGTCTCAAAGATT | 19227.6 | 11.8 | 1.6 | 7.2 | 4.3E-13 | 1.0E-12 | BT20/MB468 |
| >hg38_21_+_8433172_8446622_RNA45SN1_45S_50ntFlanks@12081.12106.26\|CTCTCGCTTCTGGCGCCAAGCGCCCG | >hg38_21_+_8433172_8446622_RNA45SN1_45S_50ntFlanks | 12081 | 12106 | 26 | CTCTCGCTTCTGGCGCCAAGCGCCCG | 358.9 | -4.6 | 0.6 | -7.3 | 3.9E-13 | 1.0E-12 | MB231/MB468 |
| >hg38_21_+_8433172_8446622_RNA45SN1_45S_50ntFlanks@12071.12095.25\|CCCCGAGGGGCTCTCGCTTCTGGCG | >hg38_21_+_8433172_8446622_RNA45SN1_45S_50ntFlanks | 12071 | 12095 | 25 | CCCCGAGGGGCTCTCGCTTCTGGCG | 1114.3 | -4.4 | 0.6 | -7.3 | 3.9E-13 | 1.0E-12 | MB231/MB468 |
| >hg38_21_+_8433172_8446622_RNA45SN1_45S_50ntFlanks@11641.11659.19\|CGGCGGGTGTTGACGCGAT | >hg38_21_+_8433172_8446622_RNA45SN1_45S_50ntFlanks | 11641 | 11659 | 19 | CGGCGGGTGTTGACGCGAT | 206.0 | -9.3 | 1.3 | -7.3 | 4.0E-13 | 1.0E-12 | MB231/MB468 |
| >hg38_21_+_8433172_8446622_RNA45SN1_45S_50ntFlanks@12946.12964.19\|TAGACGACCTGCTTCTGGG | >hg38_21_+_8433172_8446622_RNA45SN1_45S_50ntFlanks | 12946 | 12964 | 19 | TAGACGACCTGCTTCTGGG | 187.1 | -4.5 | 0.6 | -7.2 | 4.4E-13 | 1.0E-12 | BT20/MB468 |
| >hg38_21_+_8433172_8446622_RNA45SN1_45S_50ntFlanks@12935.12953.19\|TAAACCATTCGTAGACGAC | >hg38_21_+_8433172_8446622_RNA45SN1_45S_50ntFlanks | 12935 | 12953 | 19 | TAAACCATTCGTAGACGAC | 226.7 | -4.8 | 0.7 | -7.2 | 4.5E-13 | 1.0E-12 | BT20/MB468 |
| >MI0000483\|hsa-mir-186&WithFlank&1\|-\|71067625\|71067722@21.41.21(+1G)&[MIMAT0000456&hsa-miR-186-5p&offsets\|0\|-1(+1G);m-84&1\|-\|71067681\|71067702&offsets\|0\|-1(+1G)]\|CAAAGAATTCTCCTTTTGGGCG | >MI0000483\|hsa-mir-186&WithFlank&1\|-\|71067625\|71067722 | 21 | 41 | 21(+1G) | CAAAGAATTCTCCTTTTGGGCG | 821.1 | 7.5 | 1.0 | 7.2 | 4.4E-13 | 1.0E-12 | BT20/MB468 |
| >hg38_21_+_8433172_8446622_RNA45SN1_45S_50ntFlanks@10869.10889.21\|AGGGCTGGGTCGGTCGGGCTG | >hg38_21_+_8433172_8446622_RNA45SN1_45S_50ntFlanks | 10869 | 10889 | 21 | AGGGCTGGGTCGGTCGGGCTG | 260.6 | -6.3 | 0.9 | -7.3 | 4.0E-13 | 1.0E-12 | MB231/MB468 |
| >hg38_21_+_8433172_8446622_RNA45SN1_45S_50ntFlanks@10421.10457.37\|TGAGAGATGGGCGAGCGCCGTTCCGAAGGGACGGGCG | >hg38_21_+_8433172_8446622_RNA45SN1_45S_50ntFlanks | 10421 | 10457 | 37 | TGAGAGATGGGCGAGCGCCGTTCCGAAGGGACGGGCG | 127.6 | -9.0 | 1.2 | -7.2 | 4.5E-13 | 1.0E-12 | BT20/MB468 |
| >hg38_21_+_8433172_8446622_RNA45SN1_45S_50ntFlanks@10771.10796.26\|CAGGTCTCCAAGGTGAACAGCCTCTG | >hg38_21_+_8433172_8446622_RNA45SN1_45S_50ntFlanks | 10771 | 10796 | 26 | CAGGTCTCCAAGGTGAACAGCCTCTG | 659.7 | -4.8 | 0.7 | -7.2 | 4.5E-13 | 1.0E-12 | BT20/MB468 |
| >hg38_21_+_8433172_8446622_RNA45SN1_45S_50ntFlanks@10816.10840.25\|GTAAGGGAAGTCGGCAAGCCGGATC | >hg38_21_+_8433172_8446622_RNA45SN1_45S_50ntFlanks | 10816 | 10840 | 25 | GTAAGGGAAGTCGGCAAGCCGGATC | 158.1 | -4.0 | 0.6 | -7.2 | 4.5E-13 | 1.0E-12 | BT20/MB468 |

<table>
<tr><th></th><th></th><th></th><th></th><th></th><th></th><th></th><th></th><th></th><th></th><th></th><th></th><th></th></tr>
<tr><td>>hg38_21_+_8433172_8446622_RNA45SN1_45S_50ntFlanks@4103.4121.19|CCACGGGTGACGGGGAATC</td><td>>hg38_21_+_8433172_8446622_RNA45SN1_45S_50ntFlanks</td><td>4103</td><td>4121</td><td>19</td><td>CCACGGGTGACGGGGAATC</td><td>189.4</td><td>-5.1</td><td>0.7</td><td>-7.3</td><td>4.1E-13</td><td>1.0E-12</td><td>MB231/MB468</td></tr>
<tr><td>>hg38_21_+_8433172_8446622_RNA45SN1_45S_50ntFlanks@10377.10407.31|AAGGGTTCCATGTGAACAGCAGTTGAACATG</td><td>>hg38_21_+_8433172_8446622_RNA45SN1_45S_50ntFlanks</td><td>10377</td><td>10407</td><td>31</td><td>AAGGGTTCCATGTGAACAGCAGTTGAACATG</td><td>118.2</td><td>-8.9</td><td>1.2</td><td>-7.2</td><td>4.5E-13</td><td>1.0E-12</td><td>BT20/MB468</td></tr>
<tr><td>>hg38_21_+_8433172_8446622_RNA45SN1_45S_50ntFlanks@10409.10433.25|GTCAGTCGGTCCTGAGAGATGGGCG</td><td>>hg38_21_+_8433172_8446622_RNA45SN1_45S_50ntFlanks</td><td>10409</td><td>10433</td><td>25</td><td>GTCAGTCGGTCCTGAGAGATGGGCG</td><td>221.2</td><td>-3.4</td><td>0.5</td><td>-7.2</td><td>4.5E-13</td><td>1.0E-12</td><td>BT20/MB468</td></tr>
<tr><td>>hg38_21_+_8433172_8446622_RNA45SN1_45S_50ntFlanks@9576.9596.21|TCGGTCGTCCGACCTGGGTAT</td><td>>hg38_21_+_8433172_8446622_RNA45SN1_45S_50ntFlanks</td><td>9576</td><td>9596</td><td>21</td><td>TCGGTCGTCCGACCTGGGTAT</td><td>177.2</td><td>-9.5</td><td>1.3</td><td>-7.2</td><td>4.5E-13</td><td>1.0E-12</td><td>BT20/MB468</td></tr>
<tr><td>>hg38_21_+_8433172_8446622_RNA45SN1_45S_50ntFlanks@10436.10453.18|CGCCGTTCCGAAGGGACG</td><td>>hg38_21_+_8433172_8446622_RNA45SN1_45S_50ntFlanks</td><td>10436</td><td>10453</td><td>18</td><td>CGCCGTTCCGAAGGGACG</td><td>358.1</td><td>-10.5</td><td>1.4</td><td>-7.2</td><td>4.6E-13</td><td>1.0E-12</td><td>BT20/MB468</td></tr>
<tr><td>>MI0000749|hsa-mir-30e&WithFlank&1|+|40754349|40754452@65.83.19&[MIMAT0000693&hsa-miR-30e-3p&offsets|0|-3;m-25&1|+|40754413|40754434&offsets|0|-3]|CTTTCAGTCGGATGTTTAC</td><td>>MI0000749|hsa-mir-30e&WithFlank&1|+|40754349|40754452</td><td>65</td><td>83</td><td>19</td><td>CTTTCAGTCGGATGTTTAC</td><td>2109.3</td><td>6.0</td><td>0.8</td><td>7.3</td><td>4.1E-13</td><td>1.1E-12</td><td>MB231/MB468</td></tr>
<tr><td>>hg38_21_+_8433172_8446622_RNA45SN1_45S_50ntFlanks@6775.6806.32|CCTCCCGGGGCTACGCCTGTCTGAGCGTCGCT</td><td>>hg38_21_+_8433172_8446622_RNA45SN1_45S_50ntFlanks</td><td>6775</td><td>6806</td><td>32</td><td>CCTCCCGGGGCTACGCCTGTCTGAGCGTCGCT</td><td>1116.4</td><td>5.2</td><td>0.7</td><td>7.2</td><td>4.6E-13</td><td>1.1E-12</td><td>BT20/MB468</td></tr>
<tr><td>>MI0000103|hsa-mir-101-1&WithFlank&1|-|65058428|65058514@53.73.21;MI0000739|hsa-mir-101-2&WithFlank&9|+|4850291|4850381@55.75.21&[MIMAT0000099&hsa-miR-101-3p&offsets|0|0;m-17&1|-|65058443|65058462&offsets|0|+1];[MIMAT0000099_1&hsa-miR-101-3p&offsets|0|0;m-16&9|+|4850345|4850364&offsets|0|+1]|TACAGTACTGTGATAACTGAA</td><td>>MI0000103|hsa-mir-101-1&WithFlank&1|-|65058428|65058514</td><td>53</td><td>73</td><td>21</td><td>TACAGTACTGTGATAACTGAA</td><td>5968.3</td><td>7.8</td><td>1.1</td><td>7.2</td><td>4.6E-13</td><td>1.1E-12</td><td>BT20/MB468</td></tr>
<tr><td>>hg38_21_+_8433172_8446622_RNA45SN1_45S_50ntFlanks@7983.8008.26|CAGATCAGACGTGGCGACCCGCTGAA</td><td>>hg38_21_+_8433172_8446622_RNA45SN1_45S_50ntFlanks</td><td>7983</td><td>8008</td><td>26</td><td>CAGATCAGACGTGGCGACCCGCTGAA</td><td>1350.4</td><td>-4.9</td><td>0.7</td><td>-7.2</td><td>4.2E-13</td><td>1.1E-12</td><td>MB231/MB468</td></tr>
<tr><td>>hg38_21_+_8433172_8446622_RNA45SN1_45S_50ntFlanks@12419.12444.26|GATGTCGGCTCTTCCTATCATTGTGA</td><td>>hg38_21_+_8433172_8446622_RNA45SN1_45S_50ntFlanks</td><td>12419</td><td>12444</td><td>26</td><td>GATGTCGGCTCTTCCTATCATTGTGA</td><td>1377.9</td><td>4.7</td><td>0.6</td><td>7.2</td><td>4.2E-13</td><td>1.1E-12</td><td>MB231/MB468</td></tr>
<tr><td>>hg38_21_+_8433172_8446622_RNA45SN1_45S_50ntFlanks@8357.8377.21|AAAAGAACTTTGAAGAGAGAG</td><td>>hg38_21_+_8433172_8446622_RNA45SN1_45S_50ntFlanks</td><td>8357</td><td>8377</td><td>21</td><td>AAAAGAACTTTGAAGAGAGAG</td><td>4024.6</td><td>-4.0</td><td>0.6</td><td>-7.2</td><td>4.2E-13</td><td>1.1E-12</td><td>MB231/MB468</td></tr>
<tr><td>>hg38_21_+_8433172_8446622_RNA45SN1_45S_50ntFlanks@12623.12654.32|GCCAATGGGGCGAAGCTACCATCTGTGGGATT</td><td>>hg38_21_+_8433172_8446622_RNA45SN1_45S_50ntFlanks</td><td>12623</td><td>12654</td><td>32</td><td>GCCAATGGGGCGAAGCTACCATCTGTGGGATT</td><td>180.1</td><td>-3.8</td><td>0.5</td><td>-7.2</td><td>4.2E-13</td><td>1.1E-12</td><td>MB231/MB468</td></tr>
<tr><td>>MI0000077|hsa-mir-21&WithFlank&17|+|59841260|59841343@52.73.22&[MIMAT0004494&hsa-miR-21-3p&offsets|0|+1;m-52&17|+|59841311|59841332&offsets|0|0]|CAACACCAGTCGATGGGCTGTC</td><td>>MI0000077|hsa-mir-21&WithFlank&17|+|59841260|59841343</td><td>52</td><td>73</td><td>22</td><td>CAACACCAGTCGATGGGCTGTC</td><td>1652.7</td><td>8.0</td><td>1.1</td><td>7.2</td><td>4.7E-13</td><td>1.1E-12</td><td>BT20/MB468</td></tr>
<tr><td>>hg38_21_+_8433172_8446622_RNA45SN1_45S_50ntFlanks@4560.4587.28|CTAGGAATAATGGAATAGGACCGCGGTT</td><td>>hg38_21_+_8433172_8446622_RNA45SN1_45S_50ntFlanks</td><td>4560</td><td>4587</td><td>28</td><td>CTAGGAATAATGGAATAGGACCGCGGTT</td><td>234.6</td><td>-7.6</td><td>1.1</td><td>-7.2</td><td>4.3E-13</td><td>1.1E-12</td><td>MB231/MB468</td></tr>
<tr><td>>hg38_21_+_8433172_8446622_RNA45SN1_45S_50ntFlanks@4283.4307.25|CTTTAACGAGGATCCATTGGAGGGC</td><td>>hg38_21_+_8433172_8446622_RNA45SN1_45S_50ntFlanks</td><td>4283</td><td>4307</td><td>25</td><td>CTTTAACGAGGATCCATTGGAGGGC</td><td>145.8</td><td>-9.2</td><td>1.3</td><td>-7.2</td><td>4.7E-13</td><td>1.1E-12</td><td>BT20/MB468</td></tr>
<tr><td>>am_trna11_ArgACG_3_-_45730491_45730563@39.56.18,trna138_ArgACG_6_-_27638344_27638416@39.56.18,trna156_ArgACG_6_-_27181623_27181695@39.56.18,trna36_ArgACG_6_+_27182952_27183024@39.56.18|TCAGAAGATTCTAGGTTC</td><td>>am_trna11_ArgACG_3_-_45730491_45730563</td><td>39</td><td>56</td><td>18</td><td>TCAGAAGATTCTAGGTTC</td><td>148.9</td><td>-9.2</td><td>1.3</td><td>-7.2</td><td>4.7E-13</td><td>1.1E-12</td><td>BT20/MB468</td></tr>
<tr><td>>hg38_21_+_8433172_8446622_RNA45SN1_45S_50ntFlanks@12388.12406.19|CGTTCATAGCGACGTCGCT</td><td>>hg38_21_+_8433172_8446622_RNA45SN1_45S_50ntFlanks</td><td>12388</td><td>12406</td><td>19</td><td>CGTTCATAGCGACGTCGCT</td><td>361.1</td><td>3.8</td><td>0.5</td><td>7.2</td><td>4.3E-13</td><td>1.1E-12</td><td>MB231/MB468</td></tr>
<tr><td>>hg38_21_+_8433172_8446622_RNA45SN1_45S_50ntFlanks@12456.12473.18|CCAAGCGTTGGATTGTTC</td><td>>hg38_21_+_8433172_8446622_RNA45SN1_45S_50ntFlanks</td><td>12456</td><td>12473</td><td>18</td><td>CCAAGCGTTGGATTGTTC</td><td>5567.3</td><td>-5.1</td><td>0.7</td><td>-7.2</td><td>4.3E-13</td><td>1.1E-12</td><td>MB231/MB468</td></tr>
<tr><td>>hg38_1_-_228634819_228635039_RNA5S12_5S_50ntFlanks@154.171.18|CCGGGTGCTGTAGGCTTT</td><td>>hg38_1_-_228634819_228635039_RNA5S12_5S_50ntFlanks</td><td>154</td><td>171</td><td>18</td><td>CCGGGTGCTGTAGGCTTT</td><td>2707.7</td><td>4.8</td><td>0.7</td><td>7.2</td><td>4.8E-13</td><td>1.1E-12</td><td>BT20/MB468</td></tr>
<tr><td>>hg38_21_+_8433172_8446622_RNA45SN1_45S_50ntFlanks@12417.12452.36|TCGATGTCGGCTCTTCCTATCATTGTGAAGCAGAAT</td><td>>hg38_21_+_8433172_8446622_RNA45SN1_45S_50ntFlanks</td><td>12417</td><td>12452</td><td>36</td><td>TCGATGTCGGCTCTTCCTATCATTGTGAAGCAGAAT</td><td>545.8</td><td>6.8</td><td>0.9</td><td>7.2</td><td>4.4E-13</td><td>1.1E-12</td><td>MB231/MB468</td></tr>
<tr><td>>hg38_21_+_8433172_8446622_RNA45SN1_45S_50ntFlanks@10841.10875.35|CGTAACTTCGGGATAAGGATTGGCTCTAAGGGCTG</td><td>>hg38_21_+_8433172_8446622_RNA45SN1_45S_50ntFlanks</td><td>10841</td><td>10875</td><td>35</td><td>CGTAACTTCGGGATAAGGATTGGCTCTAAGGGCTG</td><td>285.1</td><td>-5.5</td><td>0.8</td><td>-7.2</td><td>4.8E-13</td><td>1.1E-12</td><td>BT20/MB468</td></tr>
<tr><td>>MI0003628|hsa-mir-615&WithFlank&12|+|54033944|54034051@67.85.19&[MIMAT0003283&hsa-miR-615-3p&offsets|0|-3;m-495&12|+|54034010|54034030&offsets|0|-2]|TCCGAGCCTGGGTCTCCCT</td><td>>MI0003628|hsa-mir-615&WithFlank&12|+|54033944|54034051</td><td>67</td><td>85</td><td>19</td><td>TCCGAGCCTGGGTCTCCCT</td><td>511.9</td><td>7.4</td><td>1.0</td><td>7.2</td><td>4.8E-13</td><td>1.1E-12</td><td>BT20/MB468</td></tr>
<tr><td>>hg38_21_+_8433172_8446622_RNA45SN1_45S_50ntFlanks@4102.4121.20|ACCACGGGTGACGGGGAATC</td><td>>hg38_21_+_8433172_8446622_RNA45SN1_45S_50ntFlanks</td><td>4102</td><td>4121</td><td>20</td><td>ACCACGGGTGACGGGGAATC</td><td>557.5</td><td>-5.9</td><td>0.8</td><td>-7.2</td><td>4.9E-13</td><td>1.1E-12</td><td>BT20/MB468</td></tr>
<tr><td>>MI0000298|hsa-mir-221&WithFlank&X|-|45746151|45746272@31.56.26&[MIMAT0004568&hsa-miR-221-5p&offsets|0|+4;m-405&X|-|45746220|45746242&offsets|0|+3]|ACCTGGCATACAATGTAGATTTCTGT</td><td>>MI0000298|hsa-mir-221&WithFlank&X|-|45746151|45746272</td><td>31</td><td>56</td><td>26</td><td>ACCTGGCATACAATGTAGATTTCTGT</td><td>783.8</td><td>5.0</td><td>0.7</td><td>7.2</td><td>4.9E-13</td><td>1.1E-12</td><td>BT20/MB468</td></tr>
<tr><td>>hg38_21_+_8433172_8446622_RNA45SN1_45S_50ntFlanks@10808.10830.23|CAATGTAGGTAAGGGAAGTCGGC</td><td>>hg38_21_+_8433172_8446622_RNA45SN1_45S_50ntFlanks</td><td>10808</td><td>10830</td><td>23</td><td>CAATGTAGGTAAGGGAAGTCGGC</td><td>165.0</td><td>-5.0</td><td>0.7</td><td>-7.2</td><td>4.4E-13</td><td>1.1E-12</td><td>MB231/MB468</td></tr>
<tr><td>>hg38_21_+_8433172_8446622_RNA45SN1_45S_50ntFlanks@5525.5550.26|TAAAAGTCGTAACAAGGTTTCCGTAG</td><td>>hg38_21_+_8433172_8446622_RNA45SN1_45S_50ntFlanks</td><td>5525</td><td>5550</td><td>26</td><td>TAAAAGTCGTAACAAGGTTTCCGTAG</td><td>179.4</td><td>-4.1</td><td>0.6</td><td>-7.2</td><td>4.4E-13</td><td>1.1E-12</td><td>MB231/MB468</td></tr>
<tr><td>>hg38_21_+_8433172_8446622_RNA45SN1_45S_50ntFlanks@10807.10825.19|ACAATGTAGGTAAGGGAAG</td><td>>hg38_21_+_8433172_8446622_RNA45SN1_45S_50ntFlanks</td><td>10807</td><td>10825</td><td>19</td><td>ACAATGTAGGTAAGGGAAG</td><td>199.7</td><td>-9.6</td><td>1.3</td><td>-7.2</td><td>4.9E-13</td><td>1.1E-12</td><td>BT20/MB468</td></tr>
<tr><td>>hg38_21_+_8433172_8446622_RNA45SN1_45S_50ntFlanks@12405.12423.19|CTTTTTGATCCTTCGATGT</td><td>>hg38_21_+_8433172_8446622_RNA45SN1_45S_50ntFlanks</td><td>12405</td><td>12423</td><td>19</td><td>CTTTTTGATCCTTCGATGT</td><td>238.3</td><td>-5.6</td><td>0.8</td><td>-7.2</td><td>4.5E-13</td><td>1.1E-12</td><td>MB231/MB468</td></tr>
<tr><td>>MI0000272|hsa-mir-182&WithFlank&7|-|129770377|129770498@29.50.22&[MIMAT0000259&hsa-miR-182-5p&offsets|0|-2;m-22&7|-|129770449|129770470&offsets|0|0]|TTTGGCAATGGTAGAACTCACA</td><td>>MI0000272|hsa-mir-182&WithFlank&7|-|129770377|129770498</td><td>29</td><td>50</td><td>22</td><td>TTTGGCAATGGTAGAACTCACA</td><td>1315.5</td><td>5.6</td><td>0.8</td><td>7.2</td><td>4.5E-13</td><td>1.1E-12</td><td>MB231/MB468</td></tr>
<tr><td>>MI0000081|hsa-mir-24-2&WithFlank&19|-|13836281|13836365@56.75.20(+1G);MI0000080|hsa-mir-24-1&WithFlank&9|+|95086015|95086094@50.69.20(+1G)&[MIMAT0000080&hsa-miR-24-3p&offsets|0|-2(+1G);m-36&19|-|13836288|13836310&offsets|0|-3(+1G)];[MIMAT0000080_1&hsa-miR-24-3p&offsets|0|-2(+1G);m-35&9|+|95086064|95086086&offsets|0|-3(+1G)]|TGGCTCAGTTCAGCAGGAACG</td><td>>MI0000081|hsa-mir-24-2&WithFlank&19|-|13836281|13836365</td><td>56</td><td>75</td><td>20(+1G)</td><td>TGGCTCAGTTCAGCAGGAACG</td><td>796.9</td><td>6.3</td><td>0.9</td><td>7.2</td><td>4.9E-13</td><td>1.1E-12</td><td>BT20/MB468</td></tr>
<tr><td>>am_trna106_HisGTG_1_-_149155828_149155899@7.28.22,trna111_HisGTG_1_-_147774845_147774916@7.28.22,trna118_HisGTG_1_-_145396881_145396952@7.28.22,trna16_HisGTG_1_+_146544773_146544844@7.28.22,trna1_HisGTG_15_+_45493349_45493420@7.28.22,trna21_HisGTG_1_+_147753471_147753542@7.28.22,trna33_HisGTG_6_+_27125906_27125977@7.28.22,trna7_HisGTG_9_-_14433938_14434009@7.28.22,trna8_HisGTG_15_-_45492611_45492682@7.28.22,trna9_HisGTG_15_-_45490804_45490875@7.28.22|ATCGTATAGTGGTTAGTACTCT</td><td>>am_trna106_HisGTG_1_-_149155828_149155899</td><td>7</td><td>28</td><td>22</td><td>ATCGTATAGTGGTTAGTACTCT</td><td>426.5</td><td>-6.9</td><td>0.9</td><td>-7.2</td><td>4.5E-13</td><td>1.1E-12</td><td>MB231/MB468</td></tr>
<tr><td>>hg38_21_+_8433172_8446622_RNA45SN1_45S_50ntFlanks@12033.12052.20|ATCGTTTTTTCACTGACCCG</td><td>>hg38_21_+_8433172_8446622_RNA45SN1_45S_50ntFlanks</td><td>12033</td><td>12052</td><td>20</td><td>ATCGTTTTTTCACTGACCCG</td><td>184.7</td><td>-9.5</td><td>1.3</td><td>-7.2</td><td>5.0E-13</td><td>1.1E-12</td><td>BT20/MB468</td></tr>
<tr><td>>MI0000482|hsa-mir-185&WithFlank&22|+|20033133|20033226@21.42.22&[MIMAT0000455&hsa-miR-185-5p&offsets|0|0;m-145&22|+|20033153|20033174&offsets|0|0]|TGGAGAGAAAGGCAGTTCCTGA</td><td>>MI0000482|hsa-mir-185&WithFlank&22|+|20033133|20033226</td><td>21</td><td>42</td><td>22</td><td>TGGAGAGAAAGGCAGTTCCTGA</td><td>1961.4</td><td>5.5</td><td>0.8</td><td>7.2</td><td>4.5E-13</td><td>1.1E-12</td><td>MB231/MB468</td></tr>
<tr><td>>MI0003684|hsa-mir-660&WithFlank&X|+|50013235|50013343@22.44.23&[MIMAT0003338&hsa-miR-660-5p&offsets|0|+1;m-157&X|+|50013256|50013278&offsets|0|0]|TACCCATTGCATATCGGAGTTGT</td><td>>MI0003684|hsa-mir-660&WithFlank&X|+|50013235|50013343</td><td>22</td><td>44</td><td>23</td><td>TACCCATTGCATATCGGAGTTGT</td><td>952.5</td><td>5.7</td><td>0.8</td><td>7.2</td><td>4.5E-13</td><td>1.1E-12</td><td>MB231/MB468</td></tr>
<tr><td>>hg38_21_+_8433172_8446622_RNA45SN1_45S_50ntFlanks@10727.10757.31|GGAGAGGGTGTAAATCTCGCGCCGGGCCGTA</td><td>>hg38_21_+_8433172_8446622_RNA45SN1_45S_50ntFlanks</td><td>10727</td><td>10757</td><td>31</td><td>GGAGAGGGTGTAAATCTCGCGCCGGGCCGTA</td><td>181.7</td><td>-9.1</td><td>1.3</td><td>-7.2</td><td>4.5E-13</td><td>1.1E-12</td><td>MB231/MB468</td></tr>
<tr><td>>hg38_21_+_8433172_8446622_RNA45SN1_45S_50ntFlanks@11759.11777.19|CCTCGTCATCTAATTAGTG</td><td>>hg38_21_+_8433172_8446622_RNA45SN1_45S_50ntFlanks</td><td>11759</td><td>11777</td><td>19</td><td>CCTCGTCATCTAATTAGTG</td><td>312.9</td><td>-6.3</td><td>0.9</td><td>-7.2</td><td>4.5E-13</td><td>1.1E-12</td><td>MB231/MB468</td></tr>
<tr><td>>hg38_21_+_8433172_8446622_RNA45SN1_45S_50ntFlanks@5472.5494.23|ACGGCCCTGGCGGAGCGCTGAGA</td><td>>hg38_21_+_8433172_8446622_RNA45SN1_45S_50ntFlanks</td><td>5472</td><td>5494</td><td>23</td><td>ACGGCCCTGGCGGAGCGCTGAGA</td><td>389.6</td><td>6.0</td><td>0.8</td><td>7.2</td><td>5.0E-13</td><td>1.1E-12</td><td>BT20/MB468</td></tr>
<tr><td>>hg38_21_+_8433172_8446622_RNA45SN1_45S_50ntFlanks@10420.10443.24|CTGAGAGATGGGCGAGCGCCGTTC</td><td>>hg38_21_+_8433172_8446622_RNA45SN1_45S_50ntFlanks</td><td>10420</td><td>10443</td><td>24</td><td>CTGAGAGATGGGCGAGCGCCGTTC</td><td>283.2</td><td>-4.2</td><td>0.6</td><td>-7.2</td><td>4.6E-13</td><td>1.2E-12</td><td>MB231/MB468</td></tr>
<tr><td>>hg38_21_+_8433172_8446622_RNA45SN1_45S_50ntFlanks@10402.10433.32|AACATGGGTCAGTCGGTCCTGAGAGATGGGCG</td><td>>hg38_21_+_8433172_8446622_RNA45SN1_45S_50ntFlanks</td><td>10402</td><td>10433</td><td>32</td><td>AACATGGGTCAGTCGGTCCTGAGAGATGGGCG</td><td>277.8</td><td>-3.5</td><td>0.5</td><td>-7.2</td><td>4.6E-13</td><td>1.2E-12</td><td>MB231/MB468</td></tr>
<tr><td>>hg38_21_+_8433172_8446622_RNA45SN1_45S_50ntFlanks@11564.11588.25|CCTAGCAGCCGACTTAGAACTGGTG</td><td>>hg38_21_+_8433172_8446622_RNA45SN1_45S_50ntFlanks</td><td>11564</td><td>11588</td><td>25</td><td>CCTAGCAGCCGACTTAGAACTGGTG</td><td>1003.3</td><td>-4.5</td><td>0.6</td><td>-7.2</td><td>5.1E-13</td><td>1.2E-12</td><td>BT20/MB468</td></tr>
<tr><td>>hg38_21_+_8433172_8446622_RNA45SN1_45S_50ntFlanks@9574.9596.23|AATCGGTCGTCCGACCTGGGTAT</td><td>>hg38_21_+_8433172_8446622_RNA45SN1_45S_50ntFlanks</td><td>9574</td><td>9596</td><td>23</td><td>AATCGGTCGTCCGACCTGGGTAT</td><td>4188.3</td><td>-4.5</td><td>0.6</td><td>-7.2</td><td>4.6E-13</td><td>1.2E-12</td><td>MB231/MB468</td></tr>
<tr><td>>hg38_21_+_8433172_8446622_RNA45SN1_45S_50ntFlanks@10396.10424.29|CAGTTGAACATGGGTCAGTCGGTCCTGAG</td><td>>hg38_21_+_8433172_8446622_RNA45SN1_45S_50ntFlanks</td><td>10396</td><td>10424</td><td>29</td><td>CAGTTGAACATGGGTCAGTCGGTCCTGAG</td><td>2746.7</td><td>4.8</td><td>0.7</td><td>7.2</td><td>4.6E-13</td><td>1.2E-12</td><td>MB231/MB468</td></tr>
<tr><td>>trna17_ValTAC_11_-_59318102_59318174@1.30.30,trna4_ValTAC_X_-_18693029_18693101@1.30.30|GGTTCCATAGTGTAGTGGTTATCACGTCTG</td><td>>trna17_ValTAC_11_-_59318102_59318174</td><td>1</td><td>30</td><td>30</td><td>GGTTCCATAGTGTAGTGGTTATCACGTCTG</td><td>321.9</td><td>-10.3</td><td>1.4</td><td>-7.2</td><td>5.1E-13</td><td>1.2E-12</td><td>BT20/MB468</td></tr>
<tr><td>>hg38_21_+_8433172_8446622_RNA45SN1_45S_50ntFlanks@10866.10889.24|CTAAGGGCTGGGTCGGTCGGGCTG</td><td>>hg38_21_+_8433172_8446622_RNA45SN1_45S_50ntFlanks</td><td>10866</td><td>10889</td><td>24</td><td>CTAAGGGCTGGGTCGGTCGGGCTG</td><td>181.5</td><td>-9.1</td><td>1.3</td><td>-7.2</td><td>4.7E-13</td><td>1.2E-12</td><td>MB231/MB468</td></tr>
</table>

| | | | | | | | | | | | | |
|---|---|---|---|---|---|---|---|---|---|---|---|---|
| >hg38_21_+_8433172_8446622_RNA45SN1_45S_50ntFlanks@10805.10823.19\|GAACAATGTAGGTAAGGGA | >hg38_21_+_8433172_8446622_RNA45SN1_45S_50ntFlanks | 10805 | 10823 | 19 | GAACAATGTAGGTAAGGGA | 176.6 | -9.0 | 1.2 | -7.2 | 4.7E-13 | 1.2E-12 | MB231/MB468 |
| >am_trnaMT_GluTTC_MT_-_14674_14742@27.52.26,trnalookalike8_GluTTC_5_-_93905172_93905240@27.52.26\|GTTTTTCATATCATTGGTCGTGGTTG | >am_trnaMT_GluTTC_MT_-_14674_14742 | 27 | 52 | 26 | GTTTTTCATATCATTGGTCGTGGTTG | 248.5 | -9.5 | 1.3 | -7.2 | 4.7E-13 | 1.2E-12 | MB231/MB468 |
| >hg38_21_+_8433172_8446622_RNA45SN1_45S_50ntFlanks@11821.11844.24\|CTATCCAGCGAAACCACAGCCAAG | >hg38_21_+_8433172_8446622_RNA45SN1_45S_50ntFlanks | 11821 | 11844 | 24 | CTATCCAGCGAAACCACAGCCAAG | 193.1 | -7.4 | 1.0 | -7.2 | 4.8E-13 | 1.2E-12 | MB231/MB468 |
| >hg38_21_+_8433172_8446622_RNA45SN1_45S_50ntFlanks@12539.12563.25\|ATGATGTGTTGTTGCCATGGTAATC | >hg38_21_+_8433172_8446622_RNA45SN1_45S_50ntFlanks | 12539 | 12563 | 25 | ATGATGTGTTGTTGCCATGGTAATC | 383.7 | -5.6 | 0.8 | -7.2 | 4.8E-13 | 1.2E-12 | MB231/MB468 |
| >trna12_TrpCCA_17_+_19411494_19411565@30.52.23,trna168_TrpCCA_6_-_26331672_26331743@30.52.23,trna170_TrpCCA_6_-_26319330_26319401@30.52.23,trna1_TrpCCA_7_+_99067307_99067378@30.52.23,trna39_TrpCCA_17_-_8124187_8124258@30.52.23,trna6_TrpCCA_17_+_8089676_8089747@30.52.23\|ACTCCAGATCAGAAGGTTGCGTG | >trna12_TrpCCA_17_+_19411494_19411565 | 30 | 52 | 23 | ACTCCAGATCAGAAGGTTGCGTG | 199.3 | -9.2 | 1.3 | -7.2 | 4.9E-13 | 1.2E-12 | MB231/MB468 |
| >MI0000111\|hsa-mir-105-1&WithFlank&X\|-\|152392213\|152392305@21.42.22;MI0000112\|hsa-mir-105-2&WithFlank&X\|-\|152394406\|152394498@21.42.22&[MIMAT0000102&hsa-miR-105-5p&offsets\|+2\|+1;m-735&X\|-\|152392266\|152392287&offsets\|+2\|+2];[MIMAT0000102_1&hsa-miR-105-5p&offsets\|+2\|+1;m-735&X\|-\|152394459\|152394480&offsets\|+2\|+2]\|AAATGCTCAGACTCCTGTGGTG | >MI0000111\|hsa-mir-105-1&WithFlank&X\|-\|152392213\|152392305 | 21 | 42 | 22 | AAATGCTCAGACTCCTGTGGTG | 166.2 | 10.9 | 1.4 | 7.6 | 3.6E-14 | 1.2E-12 | BT20/MB231 |
| >hg38_21_+_8433172_8446622_RNA45SN1_45S_50ntFlanks@13005.13027.23\|TCTATTGAAAGTCAGCCCTCGAC | >hg38_21_+_8433172_8446622_RNA45SN1_45S_50ntFlanks | 13005 | 13027 | 23 | TCTATTGAAAGTCAGCCCTCGAC | 271.1 | -10.1 | 1.4 | -7.2 | 5.4E-13 | 1.2E-12 | BT20/MB468 |
| >hg38_21_+_8433172_8446622_RNA45SN1_45S_50ntFlanks@8472.8497.26\|CGGCCCGGCGGATCTTTCCCGCCCCC | >hg38_21_+_8433172_8446622_RNA45SN1_45S_50ntFlanks | 8472 | 8497 | 26 | CGGCCCGGCGGATCTTTCCCGCCCCC | 191.8 | -9.2 | 1.3 | -7.2 | 4.9E-13 | 1.2E-12 | MB231/MB468 |
| >hg38_21_+_8433172_8446622_RNA45SN1_45S_50ntFlanks@9534.9562.29\|AGGAAACTCTGGTGGAGGTCCGTAGCGGT | >hg38_21_+_8433172_8446622_RNA45SN1_45S_50ntFlanks | 9534 | 9562 | 29 | AGGAAACTCTGGTGGAGGTCCGTAGCGGT | 975.2 | -4.0 | 0.6 | -7.2 | 4.9E-13 | 1.2E-12 | MB231/MB468 |
| >hg38_21_+_8433172_8446622_RNA45SN1_45S_50ntFlanks@10798.10818.21\|CATGTTGGAACAATGTAGGTA | >hg38_21_+_8433172_8446622_RNA45SN1_45S_50ntFlanks | 10798 | 10818 | 21 | CATGTTGGAACAATGTAGGTA | 2274.0 | -4.9 | 0.7 | -7.2 | 5.0E-13 | 1.2E-12 | MB231/MB468 |
| >hg38_21_+_8433172_8446622_RNA45SN1_45S_50ntFlanks@10805.10840.36\|GAACAATGTAGGTAAGGGAAGTCGGCAAGCCGGATC | >hg38_21_+_8433172_8446622_RNA45SN1_45S_50ntFlanks | 10805 | 10840 | 36 | GAACAATGTAGGTAAGGGAAGTCGGCAAGCCGGATC | 115.8 | -3.8 | 0.5 | -7.2 | 5.5E-13 | 1.3E-12 | BT20/MB468 |
| >hg38_21_+_8433172_8446622_RNA45SN1_45S_50ntFlanks@11670.11699.30\|CCCAGTGCTCTGAATGTCAAAGTGAAGAAA | >hg38_21_+_8433172_8446622_RNA45SN1_45S_50ntFlanks | 11670 | 11699 | 30 | CCCAGTGCTCTGAATGTCAAAGTGAAGAAA | 177.8 | -4.2 | 0.6 | -7.2 | 5.0E-13 | 1.3E-12 | MB231/MB468 |
| >hg38_21_+_8433172_8446622_RNA45SN1_45S_50ntFlanks@12324.12355.32\|TTGGGTTTTAAGCAGGAGGTGTCAGAAAAGTT | >hg38_21_+_8433172_8446622_RNA45SN1_45S_50ntFlanks | 12324 | 12355 | 32 | TTGGGTTTTAAGCAGGAGGTGTCAGAAAAGTT | 120.4 | -8.9 | 1.2 | -7.2 | 5.5E-13 | 1.3E-12 | BT20/MB468 |
| >hg38_21_+_8433172_8446622_RNA45SN1_45S_50ntFlanks@7986.8012.27\|ATCAGACGTGGCGACCCGCTGAATTTA | >hg38_21_+_8433172_8446622_RNA45SN1_45S_50ntFlanks | 7986 | 8012 | 27 | ATCAGACGTGGCGACCCGCTGAATTTA | 449.7 | -6.5 | 0.9 | -7.2 | 5.1E-13 | 1.3E-12 | MB231/MB468 |
| >hg38_21_+_8433172_8446622_RNA45SN1_45S_50ntFlanks@12932.12956.25\|CGCTAAACCATTCGTAGACGACCTG | >hg38_21_+_8433172_8446622_RNA45SN1_45S_50ntFlanks | 12932 | 12956 | 25 | CGCTAAACCATTCGTAGACGACCTG | 418.3 | -3.7 | 0.5 | -7.2 | 5.1E-13 | 1.3E-12 | MB231/MB468 |
| >hg38_21_+_8433172_8446622_RNA45SN1_45S_50ntFlanks@4760.4777.18\|TCAGATACCGTCGTAGTT | >hg38_21_+_8433172_8446622_RNA45SN1_45S_50ntFlanks | 4760 | 4777 | 18 | TCAGATACCGTCGTAGTT | 128.7 | -9.0 | 1.2 | -7.2 | 5.6E-13 | 1.3E-12 | BT20/MB468 |
| >trna15_ValAAC_5_-_180615416_180615488@1.29.29\|GTTTCCGTAGTGTAGTGGTCATCACGTTC | >trna15_ValAAC_5_-_180615416_180615488 | 1 | 29 | 29 | GTTTCCGTAGTGTAGTGGTCATCACGTTC | 2014.9 | -8.6 | 1.2 | -7.2 | 5.6E-13 | 1.3E-12 | BT20/MB468 |
| >hg38_21_+_8433172_8446622_RNA45SN1_45S_50ntFlanks@6648.6692.45\|GTACGACTCTTAGCGGTGGATCACTCGGCTCGTGCGTCGATGAAG | >hg38_21_+_8433172_8446622_RNA45SN1_45S_50ntFlanks | 6648 | 6692 | 45 | GTACGACTCTTAGCGGTGGATCACTCGGCTCGTGCGTCGATGAAG | 939.3 | 5.0 | 0.7 | 7.2 | 5.1E-13 | 1.3E-12 | MB231/MB468 |
| >hg38_21_+_8433172_8446622_RNA45SN1_45S_50ntFlanks@4297.4320.24\|CATTGGAGGGCAAGTCTGGTGCCA | >hg38_21_+_8433172_8446622_RNA45SN1_45S_50ntFlanks | 4297 | 4320 | 24 | CATTGGAGGGCAAGTCTGGTGCCA | 154.9 | -9.3 | 1.3 | -7.2 | 5.6E-13 | 1.3E-12 | BT20/MB468 |
| >hg38_21_+_8433172_8446622_RNA45SN1_45S_50ntFlanks@10683.10709.27\|CCGGCGGCGTCCGGTGAGCTCTCGCTG | >hg38_21_+_8433172_8446622_RNA45SN1_45S_50ntFlanks | 10683 | 10709 | 27 | CCGGCGGCGTCCGGTGAGCTCTCGCTG | 1459.0 | -6.2 | 0.9 | -7.2 | 5.7E-13 | 1.3E-12 | BT20/MB468 |
| >hg38_21_+_8433172_8446622_RNA45SN1_45S_50ntFlanks@9180.9197.18\|CCGTCGGGCCCGGGGGAG | >hg38_21_+_8433172_8446622_RNA45SN1_45S_50ntFlanks | 9180 | 9197 | 18 | CCGTCGGGCCCGGGGGAG | 193.0 | -9.6 | 1.3 | -7.2 | 5.7E-13 | 1.3E-12 | BT20/MB468 |
| >MI0005756\|hsa-mir-934&WithFlank&X\|+\|136550872\|136550966@21.42.22(+1A)&[MIMAT0004977&hsa-miR-934&offsets\|0\|0(+1A);m-943&X\|+\|136550892\|136550913&offsets\|0\|0(+1A)]\|TGTCTACTACTGGAGACACTGGA | >MI0005756\|hsa-mir-934&WithFlank&X\|+\|136550872\|136550966 | 21 | 42 | 22(+1A) | TGTCTACTACTGGAGACACTGGA | 6133.5 | 16.1 | 2.1 | 7.6 | 3.8E-14 | 1.3E-12 | BT20/MB231 |
| >hg38_21_+_8433172_8446622_RNA45SN1_45S_50ntFlanks@10838.10865.28\|ATCCGTAACTTCGGGATAAGGATTGGCT | >hg38_21_+_8433172_8446622_RNA45SN1_45S_50ntFlanks | 10838 | 10865 | 28 | ATCCGTAACTTCGGGATAAGGATTGGCT | 175.0 | -9.4 | 1.3 | -7.2 | 5.8E-13 | 1.3E-12 | BT20/MB468 |
| >hg38_21_+_8433172_8446622_RNA45SN1_45S_50ntFlanks@12934.12970.37\|CTAAACCATTCGTAGACGACCTGCTTCTGGGTCGGGG | >hg38_21_+_8433172_8446622_RNA45SN1_45S_50ntFlanks | 12934 | 12970 | 37 | CTAAACCATTCGTAGACGACCTGCTTCTGGGTCGGGG | 232.6 | -9.9 | 1.4 | -7.2 | 5.8E-13 | 1.3E-12 | BT20/MB468 |
| >hg38_21_+_8433172_8446622_RNA45SN1_45S_50ntFlanks@11209.11227.19\|CGGCGACTCTGGACGCGAG | >hg38_21_+_8433172_8446622_RNA45SN1_45S_50ntFlanks | 11209 | 11227 | 19 | CGGCGACTCTGGACGCGAG | 682.6 | -6.6 | 0.9 | -7.2 | 5.3E-13 | 1.3E-12 | MB231/MB468 |
| >hg38_21_+_8433172_8446622_RNA45SN1_45S_50ntFlanks@12457.12475.19\|CAAGCGTTGGATTGTTCAC | >hg38_21_+_8433172_8446622_RNA45SN1_45S_50ntFlanks | 12457 | 12475 | 19 | CAAGCGTTGGATTGTTCAC | 936.3 | -4.7 | 0.7 | -7.2 | 5.3E-13 | 1.3E-12 | MB231/MB468 |
| >hg38_21_+_8433172_8446622_RNA45SN1_45S_50ntFlanks@11215.11234.20\|CTCTGGACGCGAGCCGGGCC | >hg38_21_+_8433172_8446622_RNA45SN1_45S_50ntFlanks | 11215 | 11234 | 20 | CTCTGGACGCGAGCCGGGCC | 127.2 | -9.0 | 1.2 | -7.2 | 5.8E-13 | 1.3E-12 | BT20/MB468 |
| >hg38_21_+_8433172_8446622_RNA45SN1_45S_50ntFlanks@9738.9758.21\|GAAACGATCTCAACCTATTCT | >hg38_21_+_8433172_8446622_RNA45SN1_45S_50ntFlanks | 9738 | 9758 | 21 | GAAACGATCTCAACCTATTCT | 2162.4 | -4.5 | 0.6 | -7.2 | 5.3E-13 | 1.3E-12 | MB231/MB468 |
| >MI0000289\|hsa-mir-181a-1&WithFlank&1\|-\|198859038\|198859159@70.90.21(+1A)&[MIMAT0000270&hsa-miR-181a-3p&offsets\|0\|-1(+1A);m-125&1\|-\|198859069\|198859090&offsets\|0\|-1(+1A)]\|ACCATCGACCGTTGATTGTACA | >MI0000289\|hsa-mir-181a-1&WithFlank&1\|-\|198859038\|198859159 | 70 | 90 | 21(+1A) | ACCATCGACCGTTGATTGTACA | 206.7 | -11.0 | 1.5 | -7.6 | 4.0E-14 | 1.3E-12 | BT20/MB231 |
| >hg38_21_+_8433172_8446622_RNA45SN1_45S_50ntFlanks@12940.12966.27\|CATTCGTAGACGACCTGCTTCTGGGTC | >hg38_21_+_8433172_8446622_RNA45SN1_45S_50ntFlanks | 12940 | 12966 | 27 | CATTCGTAGACGACCTGCTTCTGGGTC | 494.4 | -4.1 | 0.6 | -7.2 | 5.4E-13 | 1.3E-12 | MB231/MB468 |
| >hg38_21_+_8433172_8446622_RNA45SN1_45S_50ntFlanks@6743.6770.28\|CGACACTTCGAACGCACTTGCGGCCCCG | >hg38_21_+_8433172_8446622_RNA45SN1_45S_50ntFlanks | 6743 | 6770 | 28 | CGACACTTCGAACGCACTTGCGGCCCCG | 1467.2 | -8.2 | 1.1 | -7.2 | 5.9E-13 | 1.3E-12 | BT20/MB468 |
| >hg38_1_-_228634819_228635039_RNA5S12_5S_50ntFlanks@51.94.44\|GTCTACGGCCATACCACCCTGAACGCGCCCGATCTCGTCTGATC | >hg38_1_-_228634819_228635039_RNA5S12_5S_50ntFlanks | 51 | 94 | 44 | GTCTACGGCCATACCACCCTGAACGCGCCCGATCTCGTCTGATC | 1274.7 | 12.2 | 1.7 | 7.2 | 5.9E-13 | 1.3E-12 | BT20/MB468 |
| >hg38_21_+_8433172_8446622_RNA45SN1_45S_50ntFlanks@9714.9737.24\|CGAATGATTAGAGGTCTTGGGGCC | >hg38_21_+_8433172_8446622_RNA45SN1_45S_50ntFlanks | 9714 | 9737 | 24 | CGAATGATTAGAGGTCTTGGGGCC | 403.3 | -5.5 | 0.8 | -7.2 | 5.4E-13 | 1.3E-12 | MB231/MB468 |
| >hg38_21_+_8433172_8446622_RNA45SN1_45S_50ntFlanks@12328.12346.19\|GTTTTAAGCAGGAGGTGTC | >hg38_21_+_8433172_8446622_RNA45SN1_45S_50ntFlanks | 12328 | 12346 | 19 | GTTTTAAGCAGGAGGTGTC | 139.5 | -9.1 | 1.3 | -7.2 | 6.0E-13 | 1.4E-12 | BT20/MB468 |
| >hg38_21_+_8433172_8446622_RNA45SN1_45S_50ntFlanks@12633.12656.24\|CGAAGCTACCATCTGTGGGATTAT | >hg38_21_+_8433172_8446622_RNA45SN1_45S_50ntFlanks | 12633 | 12656 | 24 | CGAAGCTACCATCTGTGGGATTAT | 130.8 | -9.0 | 1.3 | -7.2 | 6.0E-13 | 1.4E-12 | BT20/MB468 |
| >hg38_21_+_8433172_8446622_RNA45SN1_45S_50ntFlanks@10513.10533.21\|CCGGAGTGGCGGAGATGGGCG | >hg38_21_+_8433172_8446622_RNA45SN1_45S_50ntFlanks | 10513 | 10533 | 21 | CCGGAGTGGCGGAGATGGGCG | 938.7 | -6.5 | 0.9 | -7.2 | 6.0E-13 | 1.4E-12 | BT20/MB468 |
| >am_trnaMT_ValTAC_MT_+_1602_1670@49.67.19\|AACTTAACTTGACCGCTCT | >am_trnaMT_ValTAC_MT_+_1602_1670 | 49 | 67 | 19 | AACTTAACTTGACCGCTCT | 4878.1 | 7.4 | 1.0 | 7.2 | 6.0E-13 | 1.4E-12 | BT20/MB468 |
| >hg38_21_+_8433172_8446622_RNA45SN1_45S_50ntFlanks@9545.9567.23\|GTGGAGGTCCGTAGCGGTCCTGA | >hg38_21_+_8433172_8446622_RNA45SN1_45S_50ntFlanks | 9545 | 9567 | 23 | GTGGAGGTCCGTAGCGGTCCTGA | 269.1 | -6.6 | 0.9 | -7.2 | 6.0E-13 | 1.4E-12 | BT20/MB468 |
| >hg38_21_+_8433172_8446622_RNA45SN1_45S_50ntFlanks@4472.4497.26\|CTTAGCTGAGTGTCCCGCGGGGCCCG | >hg38_21_+_8433172_8446622_RNA45SN1_45S_50ntFlanks | 4472 | 4497 | 26 | CTTAGCTGAGTGTCCCGCGGGGCCCG | 183.7 | -9.1 | 1.3 | -7.2 | 5.4E-13 | 1.4E-12 | MB231/MB468 |
| >hg38_21_+_8433172_8446622_RNA45SN1_45S_50ntFlanks@8388.8405.18\|CGTGAAACCGTTAAGAGG | >hg38_21_+_8433172_8446622_RNA45SN1_45S_50ntFlanks | 8388 | 8405 | 18 | CGTGAAACCGTTAAGAGG | 155.2 | -9.3 | 1.3 | -7.2 | 6.1E-13 | 1.4E-12 | BT20/MB468 |
| >hg38_21_+_8433172_8446622_RNA45SN1_45S_50ntFlanks@7989.8011.23\|AGACGTGGCGACCCGCTGAATTT | >hg38_21_+_8433172_8446622_RNA45SN1_45S_50ntFlanks | 7989 | 8011 | 23 | AGACGTGGCGACCCGCTGAATTT | 15130.1 | 6.6 | 0.9 | 7.2 | 5.5E-13 | 1.4E-12 | MB231/MB468 |
| >hg38_21_+_8433172_8446622_RNA45SN1_45S_50ntFlanks@8380.8401.22\|CAAGAGGGCGTGAAACCGTTAA | >hg38_21_+_8433172_8446622_RNA45SN1_45S_50ntFlanks | 8380 | 8401 | 22 | CAAGAGGGCGTGAAACCGTTAA | 422.4 | -6.0 | 0.8 | -7.2 | 5.6E-13 | 1.4E-12 | MB231/MB468 |
| >hg38_21_+_8433172_8446622_RNA45SN1_45S_50ntFlanks@11586.11605.20\|GTGCGGACCAGGGGAATCCG | >hg38_21_+_8433172_8446622_RNA45SN1_45S_50ntFlanks | 11586 | 11605 | 20 | GTGCGGACCAGGGGAATCCG | 203.8 | -9.2 | 1.3 | -7.2 | 5.6E-13 | 1.4E-12 | MB231/MB468 |
| >am_MI0000786\|hsa-mir-378a&WithFlank&5\|+\|149732819\|149732896@49.69.21(+1G)&[MIMAT0000732&hsa-miR-378a-3p&offsets\|0\|-1(+1G);m-44&5\|+\|149732867\|149732888&offsets\|0\|-1(+1G)]\|ACTGGACTTGGAGTCAGAAGGG | >am_MI0000786\|hsa-mir-378a&WithFlank&5\|+\|149732819\|149732896 | 49 | 69 | 21(+1G) | ACTGGACTTGGAGTCAGAAGGG | 454.6 | 6.0 | 0.8 | 7.2 | 6.2E-13 | 1.4E-12 | BT20/MB468 |

| | | | | | | | | | | | | |
|---|---|---|---|---|---|---|---|---|---|---|---|---|
| >MI0000088\|hsa-mir-30a&WithFlank&6\|-\|71403545\|71403627@12.35.24(+2U)&[MIMAT0000087&hsa-miR-30a-5p&offsets\|0\|+2(+2U);m-15&6\|-\|71403595\|71403616&offsets\|0\|+2(+2U)]\|TGTAAACATCCTCGACTGGAAGCTTT | >MI0000088\|hsa-mir-30a&WithFlank&6\|-\|71403545\|71403627 | 12 | 35 | 24(+2U) | TGTAAACATCCTCGACTGGAAGCTTT | 792.4 | 6.0 | 0.8 | 7.2 | 6.2E-13 | 1.4E-12 | BT20/MB468 |
| >hg38_21_+_8433172_8446622_RNA45SN1_45S_50ntFlanks@13009.13027.19\|TTGAAAGTCAGCCCTCGAC | >hg38_21_+_8433172_8446622_RNA45SN1_45S_50ntFlanks | 13009 | 13027 | 19 | TTGAAAGTCAGCCCTCGAC | 410.8 | -10.7 | 1.5 | -7.2 | 6.2E-13 | 1.4E-12 | BT20/MB468 |
| >hg38_21_+_8433172_8446622_RNA45SN1_45S_50ntFlanks@10391.10408.18\|AACAGCAGTTGAACATGG | >hg38_21_+_8433172_8446622_RNA45SN1_45S_50ntFlanks | 10391 | 10408 | 18 | AACAGCAGTTGAACATGG | 697.0 | -6.6 | 0.9 | -7.2 | 6.3E-13 | 1.4E-12 | BT20/MB468 |
| >hg38_21_+_8433172_8446622_RNA45SN1_45S_50ntFlanks@11534.11552.19\|GTTCCGCGCGGCGCCTCGC | >hg38_21_+_8433172_8446622_RNA45SN1_45S_50ntFlanks | 11534 | 11552 | 19 | GTTCCGCGCGGCGCCTCGC | 597.2 | -7.6 | 1.1 | -7.2 | 6.3E-13 | 1.4E-12 | BT20/MB468 |
| >hg38_21_+_8433172_8446622_RNA45SN1_45S_50ntFlanks@10866.10901.36\|CTAAGGGCTGGGTCGGTCGGGCTGGGGCGCGAAGCG | >hg38_21_+_8433172_8446622_RNA45SN1_45S_50ntFlanks | 10866 | 10901 | 36 | CTAAGGGCTGGGTCGGTCGGGCTGGGGCGCGAAGCG | 251.4 | -9.5 | 1.3 | -7.2 | 5.7E-13 | 1.4E-12 | MB231/MB468 |
| >hg38_21_+_8433172_8446622_RNA45SN1_45S_50ntFlanks@4264.4282.19\|AATGAGTCCACTTTAAATC | >hg38_21_+_8433172_8446622_RNA45SN1_45S_50ntFlanks | 4264 | 4282 | 19 | AATGAGTCCACTTTAAATC | 139.8 | -9.1 | 1.3 | -7.2 | 6.3E-13 | 1.4E-12 | BT20/MB468 |
| >MI0000100\|hsa-mir-98&WithFlank&X\|-\|53556217\|53556347@28.48.21(+1A)&[MIMAT0000096&hsa-miR-98-5p&offsets\|0\|-1(+1A);m-143&X\|-\|53556299\|53556320&offsets\|0\|-1(+1A)]\|TGAGGTAGTAAGTTGTATTGTA | >MI0000100\|hsa-mir-98&WithFlank&X\|-\|53556217\|53556347 | 28 | 48 | 21(+1A) | TGAGGTAGTAAGTTGTATTGTA | 874.3 | 7.1 | 1.0 | 7.2 | 6.3E-13 | 1.4E-12 | BT20/MB468 |
| >am_MI0003772\|hsa-mir-151b&WithFlank&14\|-\|100109413\|100109520@66.85.20(+1G);MI0000809\|hsa-mir-151a&WithFlank&8\|-\|140732558\|140732659@17.36.20(+1G)&[MIMAT0010214&hsa-miR-151b&offsets\|0\|+2(+1G);m-179&14\|-\|100109436\|100109455&offsets\|0\|0(+1G)];[MIMAT0004697&hsa-miR-151a-5p&offsets\|0\|-1(+1G);m-75&8\|-\|140732623\|140732643&offsets\|0\|-1(+1G)]\|TCGAGGAGCTCACAGTCTAGG | >am_MI0003772\|hsa-mir-151b&WithFlank&14\|-\|100109413\|100109520 | 66 | 85 | 20(+1G) | TCGAGGAGCTCACAGTCTAGG | 877.6 | 11.6 | 1.6 | 7.2 | 6.3E-13 | 1.4E-12 | BT20/MB468 |
| >hg38_21_+_8433172_8446622_RNA45SN1_45S_50ntFlanks@10682.10706.25\|TCCGGCGGCGTCCGGTGAGCTCTCG | >hg38_21_+_8433172_8446622_RNA45SN1_45S_50ntFlanks | 10682 | 10706 | 25 | TCCGGCGGCGTCCGGTGAGCTCTCG | 1048.7 | -4.7 | 0.7 | -7.2 | 5.8E-13 | 1.4E-12 | MB231/MB468 |
| >hg38_21_+_8433172_8446622_RNA45SN1_45S_50ntFlanks@12078.12095.18\|GGGCTCTCGCTTCTGGCG | >hg38_21_+_8433172_8446622_RNA45SN1_45S_50ntFlanks | 12078 | 12095 | 18 | GGGCTCTCGCTTCTGGCG | 288.5 | -4.6 | 0.6 | -7.2 | 5.8E-13 | 1.4E-12 | MB231/MB468 |
| >hg38_21_+_8433172_8446622_RNA45SN1_45S_50ntFlanks@10384.10403.20\|CCATGTGAACAGCAGTTGAA | >hg38_21_+_8433172_8446622_RNA45SN1_45S_50ntFlanks | 10384 | 10403 | 20 | CCATGTGAACAGCAGTTGAA | 242.1 | -9.9 | 1.4 | -7.2 | 6.4E-13 | 1.4E-12 | BT20/MB468 |
| >hg38_21_+_8433172_8446622_RNA45SN1_45S_50ntFlanks@3705.3735.31\|TACCTGGTTGATCCTGCCAGTAGCATATGCT | >hg38_21_+_8433172_8446622_RNA45SN1_45S_50ntFlanks | 3705 | 3735 | 31 | TACCTGGTTGATCCTGCCAGTAGCATATGCT | 1580.3 | 9.0 | 1.3 | 7.2 | 6.4E-13 | 1.5E-12 | BT20/MB468 |
| >MI0005756\|hsa-mir-934&WithFlank&X\|+\|136550872\|136550966@21.43.23(+1U)&[MIMAT0004977&hsa-miR-934&offsets\|0\|+1(+1U);m-943&X\|+\|136550892\|136550913&offsets\|0\|+1(+1U)]\|TGTCTACTACTGGAGACACTGGTT | >MI0005756\|hsa-mir-934&WithFlank&X\|+\|136550872\|136550966 | 21 | 43 | 23(+1U) | TGTCTACTACTGGAGACACTGGTT | 196.5 | 11.1 | 1.5 | 7.5 | 4.4E-14 | 1.5E-12 | BT20/MB231 |
| >hg38_21_+_8433172_8446622_RNA45SN1_45S_50ntFlanks@11620.11640.21\|ACAAAGCATCGCGAAGGCCCG | >hg38_21_+_8433172_8446622_RNA45SN1_45S_50ntFlanks | 11620 | 11640 | 21 | ACAAAGCATCGCGAAGGCCCG | 265.4 | -5.5 | 0.8 | -7.2 | 5.9E-13 | 1.5E-12 | MB231/MB468 |
| >hg38_21_+_8433172_8446622_RNA45SN1_45S_50ntFlanks@10866.10902.37\|CTAAGGGCTGGGTCGGTCGGGCTGGGGCGCGAAGCGG | >hg38_21_+_8433172_8446622_RNA45SN1_45S_50ntFlanks | 10866 | 10902 | 37 | CTAAGGGCTGGGTCGGTCGGGCTGGGGCGCGAAGCGG | 377.1 | -6.8 | 0.9 | -7.2 | 6.5E-13 | 1.5E-12 | BT20/MB468 |
| >hg38_21_+_8433172_8446622_RNA45SN1_45S_50ntFlanks@10361.10390.30\|GAAGGCCGAAGTGGAGAAGGGTTCCATGTG | >hg38_21_+_8433172_8446622_RNA45SN1_45S_50ntFlanks | 10361 | 10390 | 30 | GAAGGCCGAAGTGGAGAAGGGTTCCATGTG | 257.9 | -10.0 | 1.4 | -7.2 | 6.6E-13 | 1.5E-12 | BT20/MB468 |
| >hg38_21_+_8433172_8446622_RNA45SN1_45S_50ntFlanks@10826.10858.33\|TCGGCAAGCCGGATCCGTAACTTCGGGATAAGG | >hg38_21_+_8433172_8446622_RNA45SN1_45S_50ntFlanks | 10826 | 10858 | 33 | TCGGCAAGCCGGATCCGTAACTTCGGGATAAGG | 406.3 | -10.7 | 1.5 | -7.2 | 6.6E-13 | 1.5E-12 | BT20/MB468 |
| >hg38_21_+_8433172_8446622_RNA45SN1_45S_50ntFlanks@4560.4577.18\|CTAGGAATAATGGAATAG | >hg38_21_+_8433172_8446622_RNA45SN1_45S_50ntFlanks | 4560 | 4577 | 18 | CTAGGAATAATGGAATAG | 165.6 | -9.4 | 1.3 | -7.2 | 6.6E-13 | 1.5E-12 | BT20/MB468 |
| >hg38_21_+_8433172_8446622_RNA45SN1_45S_50ntFlanks@10729.10755.27\|AGAGGGTGTAAATCTCGCGCCGGGCCG | >hg38_21_+_8433172_8446622_RNA45SN1_45S_50ntFlanks | 10729 | 10755 | 27 | AGAGGGTGTAAATCTCGCGCCGGGCCG | 183.0 | -9.1 | 1.3 | -7.2 | 6.0E-13 | 1.5E-12 | MB231/MB468 |
| >hg38_21_+_8433172_8446622_RNA45SN1_45S_50ntFlanks@4006.4026.21\|ACCTCGGGCCGATCGCACGCC | >hg38_21_+_8433172_8446622_RNA45SN1_45S_50ntFlanks | 4006 | 4026 | 21 | ACCTCGGGCCGATCGCACGCC | 150.0 | -9.2 | 1.3 | -7.2 | 6.7E-13 | 1.5E-12 | BT20/MB468 |
| >hg38_21_+_8433172_8446622_RNA45SN1_45S_50ntFlanks@12044.12072.29\|ACTGACCCGGTGAGGCGGGGGGGCGAGCC | >hg38_21_+_8433172_8446622_RNA45SN1_45S_50ntFlanks | 12044 | 12072 | 29 | ACTGACCCGGTGAGGCGGGGGGGCGAGCC | 163.3 | -8.9 | 1.2 | -7.2 | 6.1E-13 | 1.5E-12 | MB231/MB468 |
| >hg38_21_+_8433172_8446622_RNA45SN1_45S_50ntFlanks@10572.10591.20\|AGAAGCCGGCGGGAGCCCCG | >hg38_21_+_8433172_8446622_RNA45SN1_45S_50ntFlanks | 10572 | 10591 | 20 | AGAAGCCGGCGGGAGCCCCG | 631.1 | -8.9 | 1.2 | -7.2 | 6.1E-13 | 1.5E-12 | MB231/MB468 |
| >hg38_21_+_8433172_8446622_RNA45SN1_45S_50ntFlanks@11576.11594.19\|CTTAGAACTGGTGCGGACC | >hg38_21_+_8433172_8446622_RNA45SN1_45S_50ntFlanks | 11576 | 11594 | 19 | CTTAGAACTGGTGCGGACC | 810.3 | -3.8 | 0.5 | -7.2 | 6.8E-13 | 1.5E-12 | BT20/MB468 |
| >hg38_21_+_8433172_8446622_RNA45SN1_45S_50ntFlanks@12621.12654.34\|GAGCCAATGGGGCGAAGCTACCATCTGTGGGATT | >hg38_21_+_8433172_8446622_RNA45SN1_45S_50ntFlanks | 12621 | 12654 | 34 | GAGCCAATGGGGCGAAGCTACCATCTGTGGGATT | 154.4 | -5.4 | 0.8 | -7.2 | 6.8E-13 | 1.5E-12 | BT20/MB468 |
| >hg38_21_+_8433172_8446622_RNA45SN1_45S_50ntFlanks@8170.8195.26\|CAAGTCCTTCTGATCGAGGCCCAGCC | >hg38_21_+_8433172_8446622_RNA45SN1_45S_50ntFlanks | 8170 | 8195 | 26 | CAAGTCCTTCTGATCGAGGCCCAGCC | 844.3 | -5.8 | 0.8 | -7.2 | 6.8E-13 | 1.5E-12 | BT20/MB468 |
| >am_trnaMT_LysTTT_MT_+_8295_8364@28.52.25,trnalookalike7_LysTTT_1_+_568844_568913@28.52.25\|TTTTAAGTTAAAGATTAAGAGAACC | >am_trnaMT_LysTTT_MT_+_8295_8364 | 28 | 52 | 25 | TTTTAAGTTAAAGATTAAGAGAACC | 929.5 | 9.0 | 1.3 | 7.2 | 6.9E-13 | 1.6E-12 | BT20/MB468 |
| >trnaMT_HisGTG_MT_+_12138_12206@32.72.41\|TGAATCTGACAACAGAGGCTTACGACCCCTTATTTACCCCA | >trnaMT_HisGTG_MT_+_12138_12206 | 32 | 72 | 41 | TGAATCTGACAACAGAGGCTTACGACCCCTTATTTACCCCA | 5778.5 | 9.1 | 1.3 | 7.2 | 6.4E-13 | 1.6E-12 | MB231/MB468 |
| >hg38_21_+_8433172_8446622_RNA45SN1_45S_50ntFlanks@6773.6797.25\|TTCCTCCCGGGGCTACGCCTGTCTG | >hg38_21_+_8433172_8446622_RNA45SN1_45S_50ntFlanks | 6773 | 6797 | 25 | TTCCTCCCGGGGCTACGCCTGTCTG | 223.3 | -6.4 | 0.9 | -7.2 | 7.1E-13 | 1.6E-12 | BT20/MB468 |
| >hg38_21_+_8433172_8446622_RNA45SN1_45S_50ntFlanks@9574.9598.25\|AATCGGTCGTCCGACCTGGGTATAG | >hg38_21_+_8433172_8446622_RNA45SN1_45S_50ntFlanks | 9574 | 9598 | 25 | AATCGGTCGTCCGACCTGGGTATAG | 144.1 | -3.9 | 0.5 | -7.2 | 7.1E-13 | 1.6E-12 | BT20/MB468 |
| >hg38_21_+_8433172_8446622_RNA45SN1_45S_50ntFlanks@12974.12992.19\|CGTACGTAGCAGAGCAGCT | >hg38_21_+_8433172_8446622_RNA45SN1_45S_50ntFlanks | 12974 | 12992 | 19 | CGTACGTAGCAGAGCAGCT | 449.7 | -4.8 | 0.7 | -7.2 | 7.1E-13 | 1.6E-12 | BT20/MB468 |
| >hg38_21_+_8433172_8446622_RNA45SN1_45S_50ntFlanks@11315.11338.24\|CCACCCCACGTCTCGTCGCGCGCG | >hg38_21_+_8433172_8446622_RNA45SN1_45S_50ntFlanks | 11315 | 11338 | 24 | CCACCCCACGTCTCGTCGCGCGCG | 231.3 | -9.4 | 1.3 | -7.2 | 6.4E-13 | 1.6E-12 | MB231/MB468 |
| >hg38_21_+_8433172_8446622_RNA45SN1_45S_50ntFlanks@5485.5514.30\|AGCGCTGAGAAGACGGTCGAACTTGACTAT | >hg38_21_+_8433172_8446622_RNA45SN1_45S_50ntFlanks | 5485 | 5514 | 30 | AGCGCTGAGAAGACGGTCGAACTTGACTAT | 1555.9 | -5.2 | 0.7 | -7.2 | 6.4E-13 | 1.6E-12 | MB231/MB468 |
| >hg38_21_+_8433172_8446622_RNA45SN1_45S_50ntFlanks@4163.4184.22\|CATCCAAGGAAGGCAGCAGGCG | >hg38_21_+_8433172_8446622_RNA45SN1_45S_50ntFlanks | 4163 | 4184 | 22 | CATCCAAGGAAGGCAGCAGGCG | 446.1 | -6.9 | 1.0 | -7.2 | 7.1E-13 | 1.6E-12 | BT20/MB468 |
| >hg38_21_+_8433172_8446622_RNA45SN1_45S_50ntFlanks@10597.10617.21\|AGTTCTCTTTTCTTTGTGAAG | >hg38_21_+_8433172_8446622_RNA45SN1_45S_50ntFlanks | 10597 | 10617 | 21 | AGTTCTCTTTTCTTTGTGAAG | 142.3 | -9.1 | 1.3 | -7.2 | 7.2E-13 | 1.6E-12 | BT20/MB468 |
| >hg38_21_+_8433172_8446622_RNA45SN1_45S_50ntFlanks@12081.12108.28\|CTCTCGCTTCTGGCGCCAAGCGCCCGGC | >hg38_21_+_8433172_8446622_RNA45SN1_45S_50ntFlanks | 12081 | 12108 | 28 | CTCTCGCTTCTGGCGCCAAGCGCCCGGC | 383.7 | -4.6 | 0.6 | -7.2 | 6.5E-13 | 1.6E-12 | MB231/MB468 |
| >hg38_21_+_8433172_8446622_RNA45SN1_45S_50ntFlanks@12655.12673.19\|ATGACTGAACGCCTCTAAG | >hg38_21_+_8433172_8446622_RNA45SN1_45S_50ntFlanks | 12655 | 12673 | 19 | ATGACTGAACGCCTCTAAG | 400.8 | -6.9 | 1.0 | -7.2 | 7.2E-13 | 1.6E-12 | BT20/MB468 |
| >hg38_21_+_8433172_8446622_RNA45SN1_45S_50ntFlanks@11567.11587.21\|AGCAGCCGACTTAGAACTGGT | >hg38_21_+_8433172_8446622_RNA45SN1_45S_50ntFlanks | 11567 | 11587 | 21 | AGCAGCCGACTTAGAACTGGT | 936.8 | 3.7 | 0.5 | 7.2 | 6.5E-13 | 1.6E-12 | MB231/MB468 |
| >hg38_21_+_8433172_8446622_RNA45SN1_45S_50ntFlanks@10805.10826.22\|GAACAATGTAGGTAAGGGAAGT | >hg38_21_+_8433172_8446622_RNA45SN1_45S_50ntFlanks | 10805 | 10826 | 22 | GAACAATGTAGGTAAGGGAAGT | 138.6 | -6.2 | 0.9 | -7.2 | 6.6E-13 | 1.6E-12 | MB231/MB468 |
| >hg38_21_+_8433172_8446622_RNA45SN1_45S_50ntFlanks@11561.11582.22\|GCGCCTAGCAGCCGACTTAGAA | >hg38_21_+_8433172_8446622_RNA45SN1_45S_50ntFlanks | 11561 | 11582 | 22 | GCGCCTAGCAGCCGACTTAGAA | 243.2 | -9.9 | 1.4 | -7.2 | 7.3E-13 | 1.6E-12 | BT20/MB468 |
| >hg38_1_-_228634819_228635039_RNA5S12_5S_50ntFlanks@133.169.37\|ATGGGAGACCGCCTGGGAATACCGGGTGCTGTAGGCT | >hg38_1_-_228634819_228635039_RNA5S12_5S_50ntFlanks | 133 | 169 | 37 | ATGGGAGACCGCCTGGGAATACCGGGTGCTGTAGGCT | 1995.3 | 5.3 | 0.7 | 7.2 | 6.7E-13 | 1.7E-12 | MB231/MB468 |
| >hg38_21_+_8433172_8446622_RNA45SN1_45S_50ntFlanks@9166.9197.32\|CCGGGCGGGTCGCGCCGTCGGGCCCGGGGGAG | >hg38_21_+_8433172_8446622_RNA45SN1_45S_50ntFlanks | 9166 | 9197 | 32 | CCGGGCGGGTCGCGCCGTCGGGCCCGGGGGAG | 230.3 | -9.4 | 1.3 | -7.2 | 6.8E-13 | 1.7E-12 | MB231/MB468 |
| >trna2_GlyGCC_21_-_18827107_18827177@1.29.29,trna35_GlyGCC_1_+_161413094_161413164@1.29.29,trna37_GlyGCC_1_+_161420467_161420537@1.29.29,trna39_GlyGCC_1_+_161427898_161427968@1.29.29,trna41_GlyGCC_1_+_161435258_161435328@1.29.29\|GCATGGGTGGTTCAGTGGTAGAATTCTCG | >trna2_GlyGCC_21_-_18827107_18827177 | 1 | 29 | 29 | GCATGGGTGGTTCAGTGGTAGAATTCTCG | 9041.0 | -7.5 | 1.0 | -7.2 | 6.8E-13 | 1.7E-12 | MB231/MB468 |
| >MI0006273\|hsa-mir-1180&WithFlank&17\|-\|19344500\|19344580@47.67.21(+1C)&[MIMAT0005825&hsa-miR-1180-3p&offsets\|0\|-1(+1C);m-214&17\|-\|19344513\|19344534&offsets\|0\|-1(+1C)]\|TTTCCGGCTCGCGTGGGTGTGC | >MI0006273\|hsa-mir-1180&WithFlank&17\|-\|19344500\|19344580 | 47 | 67 | 21(+1C) | TTTCCGGCTCGCGTGGGTGTGC | 473.5 | 8.6 | 1.2 | 7.2 | 7.5E-13 | 1.7E-12 | BT20/MB468 |
| >hg38_21_+_8433172_8446622_RNA45SN1_45S_50ntFlanks@11573.11593.21\|CGACTTAGAACTGGTGCGGAC | >hg38_21_+_8433172_8446622_RNA45SN1_45S_50ntFlanks | 11573 | 11593 | 21 | CGACTTAGAACTGGTGCGGAC | 291.1 | -3.1 | 0.4 | -7.2 | 7.5E-13 | 1.7E-12 | BT20/MB468 |
| >hg38_21_+_8433172_8446622_RNA45SN1_45S_50ntFlanks@12456.12481.26\|CCAAGCGTTGGATTGTTCACCCACTA | >hg38_21_+_8433172_8446622_RNA45SN1_45S_50ntFlanks | 12456 | 12481 | 26 | CCAAGCGTTGGATTGTTCACCCACTA | 1185.2 | -7.9 | 1.1 | -7.2 | 7.5E-13 | 1.7E-12 | BT20/MB468 |

<table>
<tr><td>>hg38_21_+_8433172_8446622_RNA45SN1_45S_50ntFlanks@4564.4587.24|GAATAATGGAATAGGACCGCGGTT</td><td>>hg38_21_+_8433172_8446622_RNA45SN1_45S_50ntFlanks</td><td>4564</td><td>4587</td><td>24</td><td>GAATAATGGAATAGGACCGCGGTT</td><td>148.0</td><td>-9.2</td><td>1.3</td><td>-7.2</td><td>7.5E-13</td><td>1.7E-12</td><td>BT20/MB468</td></tr>
<tr><td>>hg38_21_+_8433172_8446622_RNA45SN1_45S_50ntFlanks@11614.11640.27|ATTAAAACAAAGCATCGCGAAGGCCCG</td><td>>hg38_21_+_8433172_8446622_RNA45SN1_45S_50ntFlanks</td><td>11614</td><td>11640</td><td>27</td><td>ATTAAAACAAAGCATCGCGAAGGCCCG</td><td>174.1</td><td>-9.4</td><td>1.3</td><td>-7.2</td><td>7.5E-13</td><td>1.7E-12</td><td>BT20/MB468</td></tr>
<tr><td>>hg38_21_+_8433172_8446622_RNA45SN1_45S_50ntFlanks@11786.11804.19|GAATGGATGAACGAGATTC</td><td>>hg38_21_+_8433172_8446622_RNA45SN1_45S_50ntFlanks</td><td>11786</td><td>11804</td><td>19</td><td>GAATGGATGAACGAGATTC</td><td>1481.9</td><td>-6.5</td><td>0.9</td><td>-7.2</td><td>7.5E-13</td><td>1.7E-12</td><td>BT20/MB468</td></tr>
<tr><td>>hg38_21_+_8433172_8446622_RNA45SN1_45S_50ntFlanks@5524.5550.27|GTAAAAGTCGTAACAAGGTTTCCGTAG</td><td>>hg38_21_+_8433172_8446622_RNA45SN1_45S_50ntFlanks</td><td>5524</td><td>5550</td><td>27</td><td>GTAAAAGTCGTAACAAGGTTTCCGTAG</td><td>143.8</td><td>-3.9</td><td>0.5</td><td>-7.2</td><td>6.8E-13</td><td>1.7E-12</td><td>MB231/MB468</td></tr>
<tr><td>>hg38_21_+_8433172_8446622_RNA45SN1_45S_50ntFlanks@11115.11133.19|CGGCAGGGGCCGGCGGCGG</td><td>>hg38_21_+_8433172_8446622_RNA45SN1_45S_50ntFlanks</td><td>11115</td><td>11133</td><td>19</td><td>CGGCAGGGGCCGGCGGCGG</td><td>124.3</td><td>-9.0</td><td>1.2</td><td>-7.2</td><td>7.6E-13</td><td>1.7E-12</td><td>BT20/MB468</td></tr>
<tr><td>>hg38_21_+_8433172_8446622_RNA45SN1_45S_50ntFlanks@3775.3792.18|GCCGGTACAGTGAAACTG</td><td>>hg38_21_+_8433172_8446622_RNA45SN1_45S_50ntFlanks</td><td>3775</td><td>3792</td><td>18</td><td>GCCGGTACAGTGAAACTG</td><td>1810.6</td><td>-5.1</td><td>0.7</td><td>-7.2</td><td>6.9E-13</td><td>1.7E-12</td><td>MB231/MB468</td></tr>
<tr><td>>hg38_21_+_8433172_8446622_RNA45SN1_45S_50ntFlanks@10735.10752.18|TGTAAATCTCGCGCCGGG</td><td>>hg38_21_+_8433172_8446622_RNA45SN1_45S_50ntFlanks</td><td>10735</td><td>10752</td><td>18</td><td>TGTAAATCTCGCGCCGGG</td><td>165.3</td><td>-9.4</td><td>1.3</td><td>-7.2</td><td>7.6E-13</td><td>1.7E-12</td><td>BT20/MB468</td></tr>
<tr><td>>hg38_21_+_8433172_8446622_RNA45SN1_45S_50ntFlanks@6650.6668.19|ACGACTCTTAGCGGTGGAT</td><td>>hg38_21_+_8433172_8446622_RNA45SN1_45S_50ntFlanks</td><td>6650</td><td>6668</td><td>19</td><td>ACGACTCTTAGCGGTGGAT</td><td>678.3</td><td>3.6</td><td>0.5</td><td>7.2</td><td>6.9E-13</td><td>1.7E-12</td><td>MB231/MB468</td></tr>
<tr><td>>hg38_21_+_8433172_8446622_RNA45SN1_45S_50ntFlanks@12539.12563.25|ATGATGTGTTGTTGCCATGGTAATC</td><td>>hg38_21_+_8433172_8446622_RNA45SN1_45S_50ntFlanks</td><td>12539</td><td>12563</td><td>25</td><td>ATGATGTGTTGTTGCCATGGTAATC</td><td>453.6</td><td>-6.7</td><td>0.9</td><td>-7.2</td><td>7.6E-13</td><td>1.7E-12</td><td>BT20/MB468</td></tr>
<tr><td>>hg38_21_+_8433172_8446622_RNA45SN1_45S_50ntFlanks@11535.11552.18|TTCCGCGCGGCGCCTCGC</td><td>>hg38_21_+_8433172_8446622_RNA45SN1_45S_50ntFlanks</td><td>11535</td><td>11552</td><td>18</td><td>TTCCGCGCGGCGCCTCGC</td><td>147.4</td><td>-9.2</td><td>1.3</td><td>-7.2</td><td>7.6E-13</td><td>1.7E-12</td><td>BT20/MB468</td></tr>
<tr><td>>hg38_21_+_8433172_8446622_RNA45SN1_45S_50ntFlanks@4738.4765.28|AAAGTCGGAGGTTCGAAGACGATCAGAT</td><td>>hg38_21_+_8433172_8446622_RNA45SN1_45S_50ntFlanks</td><td>4738</td><td>4765</td><td>28</td><td>AAAGTCGGAGGTTCGAAGACGATCAGAT</td><td>669.8</td><td>6.9</td><td>1.0</td><td>7.2</td><td>7.0E-13</td><td>1.7E-12</td><td>MB231/MB468</td></tr>
<tr><td>>hg38_21_+_8433172_8446622_RNA45SN1_45S_50ntFlanks@7986.8011.26|ATCAGACGTGGCGACCCGCTGAATTT</td><td>>hg38_21_+_8433172_8446622_RNA45SN1_45S_50ntFlanks</td><td>7986</td><td>8011</td><td>26</td><td>ATCAGACGTGGCGACCCGCTGAATTT</td><td>155.2</td><td>-9.3</td><td>1.3</td><td>-7.2</td><td>7.7E-13</td><td>1.7E-12</td><td>BT20/MB468</td></tr>
<tr><td>>MI0000079|hsa-mir-23a&WithFlank&19|-|13836581|13836665@51.72.22&[MIMAT0000078&hsa-miR-23a-3p&offsets|0|+1;m-31&19|-|13836595|13836615&offsets|0|+1]|ATCACATTGCCAGGGATTTCCA</td><td>>MI0000079|hsa-mir-23a&WithFlank&19|-|13836581|13836665</td><td>51</td><td>72</td><td>22</td><td>ATCACATTGCCAGGGATTTCCA</td><td>183.6</td><td>-9.5</td><td>1.3</td><td>-7.2</td><td>7.8E-13</td><td>1.7E-12</td><td>BT20/MB468</td></tr>
<tr><td>>hg38_21_+_8433172_8446622_RNA45SN1_45S_50ntFlanks@9004.9026.23|TCTCCCCCGCGGGGGCGCGCCGG</td><td>>hg38_21_+_8433172_8446622_RNA45SN1_45S_50ntFlanks</td><td>9004</td><td>9026</td><td>23</td><td>TCTCCCCCGCGGGGGCGCGCCGG</td><td>753.7</td><td>-7.6</td><td>1.1</td><td>-7.2</td><td>7.8E-13</td><td>1.7E-12</td><td>BT20/MB468</td></tr>
<tr><td>>MI0000736|hsa-mir-30c-1&WithFlank&1|+|40757278|40757378@23.45.23(+1C);MI0000254|hsa-mir-30c-2&WithFlank&6|-|71376954|71377037@13.35.23(+1C)&[MIMAT0000244&hsa-miR-30c-5p&offsets|0|0(+1C);m-57&1|+|40757300|40757323&offsets|0|-1(+1C)];[MIMAT0000244_1&hsa-miR-30c-5p&offsets|0|0(+1C);m-56&6|-|71377002|71377025&offsets|0|-1(+1C)]|TGTAAACATCCTACACTCTCAGCC</td><td>>MI0000736|hsa-mir-30c-1&WithFlank&1|+|40757278|40757378</td><td>23</td><td>45</td><td>23(+1C)</td><td>TGTAAACATCCTACACTCTCAGCC</td><td>6143.2</td><td>5.8</td><td>0.8</td><td>7.2</td><td>7.8E-13</td><td>1.7E-12</td><td>BT20/MB468</td></tr>
<tr><td>>MI0000086|hsa-mir-28&WithFlank&3|+|188688775|188688872@60.81.22(+1U)&[MIMAT0004502&hsa-miR-28-3p&offsets|0|0(+1U);m-28&3|+|188688834|188688855&offsets|0|0(+1U)]|CACTAGATTGTGAGCTCCTGGAT</td><td>>MI0000086|hsa-mir-28&WithFlank&3|+|188688775|188688872</td><td>60</td><td>81</td><td>22(+1U)</td><td>CACTAGATTGTGAGCTCCTGGAT</td><td>1296.9</td><td>10.8</td><td>1.5</td><td>7.2</td><td>7.8E-13</td><td>1.7E-12</td><td>BT20/MB468</td></tr>
<tr><td>>hg38_21_+_8433172_8446622_RNA45SN1_45S_50ntFlanks@12936.12966.31|AAACCATTCGTAGACGACCTGCTTCTGGGTC</td><td>>hg38_21_+_8433172_8446622_RNA45SN1_45S_50ntFlanks</td><td>12936</td><td>12966</td><td>31</td><td>AAACCATTCGTAGACGACCTGCTTCTGGGTC</td><td>204.1</td><td>-4.4</td><td>0.6</td><td>-7.2</td><td>7.1E-13</td><td>1.8E-12</td><td>MB231/MB468</td></tr>
<tr><td>>hg38_21_+_8433172_8446622_RNA45SN1_45S_50ntFlanks@5488.5516.29|GCTGAGAAGACGGTCGAACTTGACTATCT</td><td>>hg38_21_+_8433172_8446622_RNA45SN1_45S_50ntFlanks</td><td>5488</td><td>5516</td><td>29</td><td>GCTGAGAAGACGGTCGAACTTGACTATCT</td><td>1138.0</td><td>7.4</td><td>1.0</td><td>7.2</td><td>7.8E-13</td><td>1.8E-12</td><td>BT20/MB468</td></tr>
<tr><td>>hg38_21_+_8433172_8446622_RNA45SN1_45S_50ntFlanks@12151.12179.29|GTGGGGAGTTTGACTGGGGCGGTACACCT</td><td>>hg38_21_+_8433172_8446622_RNA45SN1_45S_50ntFlanks</td><td>12151</td><td>12179</td><td>29</td><td>GTGGGGAGTTTGACTGGGGCGGTACACCT</td><td>196.9</td><td>-9.2</td><td>1.3</td><td>-7.2</td><td>7.2E-13</td><td>1.8E-12</td><td>MB231/MB468</td></tr>
<tr><td>>hg38_21_+_8433172_8446622_RNA45SN1_45S_50ntFlanks@7974.7997.24|ACGCGACCTCAGATCAGACGTGGC</td><td>>hg38_21_+_8433172_8446622_RNA45SN1_45S_50ntFlanks</td><td>7974</td><td>7997</td><td>24</td><td>ACGCGACCTCAGATCAGACGTGGC</td><td>876.1</td><td>5.3</td><td>0.7</td><td>7.2</td><td>7.2E-13</td><td>1.8E-12</td><td>MB231/MB468</td></tr>
<tr><td>>MI0002468|hsa-mir-484&WithFlank&16|+|15643288|15643378@14.35.22(+1U)&[MIMAT0002174&hsa-miR-484&offsets|0|0(+1U);m-127&16|+|15643301|15643321&offsets|0|+1(+1U)]|TCAGGCTCAGTCCCCTCCCGATT</td><td>>MI0002468|hsa-mir-484&WithFlank&16|+|15643288|15643378</td><td>14</td><td>35</td><td>22(+1U)</td><td>TCAGGCTCAGTCCCCTCCCGATT</td><td>1146.7</td><td>5.8</td><td>0.8</td><td>7.2</td><td>8.0E-13</td><td>1.8E-12</td><td>BT20/MB468</td></tr>
<tr><td>>hg38_21_+_8433172_8446622_RNA45SN1_45S_50ntFlanks@6649.6668.20|TACGACTCTTAGCGGTGGAT</td><td>>hg38_21_+_8433172_8446622_RNA45SN1_45S_50ntFlanks</td><td>6649</td><td>6668</td><td>20</td><td>TACGACTCTTAGCGGTGGAT</td><td>496.1</td><td>3.4</td><td>0.5</td><td>7.2</td><td>7.3E-13</td><td>1.8E-12</td><td>MB231/MB468</td></tr>
<tr><td>>hg38_21_+_8433172_8446622_RNA45SN1_45S_50ntFlanks@11783.11800.18|CATGAATGGATGAACGAG</td><td>>hg38_21_+_8433172_8446622_RNA45SN1_45S_50ntFlanks</td><td>11783</td><td>11800</td><td>18</td><td>CATGAATGGATGAACGAG</td><td>406.2</td><td>-6.9</td><td>1.0</td><td>-7.2</td><td>8.2E-13</td><td>1.8E-12</td><td>BT20/MB468</td></tr>
<tr><td>>hg38_21_+_8433172_8446622_RNA45SN1_45S_50ntFlanks@12477.12498.22|CACTAATAGGGAACGTGAGCTG</td><td>>hg38_21_+_8433172_8446622_RNA45SN1_45S_50ntFlanks</td><td>12477</td><td>12498</td><td>22</td><td>CACTAATAGGGAACGTGAGCTG</td><td>1086.1</td><td>-3.1</td><td>0.4</td><td>-7.2</td><td>8.2E-13</td><td>1.8E-12</td><td>BT20/MB468</td></tr>
<tr><td>>hg38_21_+_8433172_8446622_RNA45SN1_45S_50ntFlanks@5492.5516.25|AGAAGACGGTCGAACTTGACTATCT</td><td>>hg38_21_+_8433172_8446622_RNA45SN1_45S_50ntFlanks</td><td>5492</td><td>5516</td><td>25</td><td>AGAAGACGGTCGAACTTGACTATCT</td><td>15609.0</td><td>8.1</td><td>1.1</td><td>7.2</td><td>8.2E-13</td><td>1.8E-12</td><td>BT20/MB468</td></tr>
<tr><td>>hg38_21_+_8433172_8446622_RNA45SN1_45S_50ntFlanks@3782.3812.31|CAGTGAAACTGCGAATGGCTCATTAAATCAG</td><td>>hg38_21_+_8433172_8446622_RNA45SN1_45S_50ntFlanks</td><td>3782</td><td>3812</td><td>31</td><td>CAGTGAAACTGCGAATGGCTCATTAAATCAG</td><td>134.8</td><td>-9.1</td><td>1.3</td><td>-7.2</td><td>8.2E-13</td><td>1.8E-12</td><td>BT20/MB468</td></tr>
<tr><td>>hg38_21_+_8433172_8446622_RNA45SN1_45S_50ntFlanks@6647.6668.22|CGTACGACTCTTAGCGGTGGAT</td><td>>hg38_21_+_8433172_8446622_RNA45SN1_45S_50ntFlanks</td><td>6647</td><td>6668</td><td>22</td><td>CGTACGACTCTTAGCGGTGGAT</td><td>1188.5</td><td>4.2</td><td>0.6</td><td>7.2</td><td>7.4E-13</td><td>1.8E-12</td><td>MB231/MB468</td></tr>
<tr><td>>hg38_21_+_8433172_8446622_RNA45SN1_45S_50ntFlanks@8177.8202.26|TTCTGATCGAGGCCCAGCCCGTGGAC</td><td>>hg38_21_+_8433172_8446622_RNA45SN1_45S_50ntFlanks</td><td>8177</td><td>8202</td><td>26</td><td>TTCTGATCGAGGCCCAGCCCGTGGAC</td><td>359.6</td><td>6.5</td><td>0.9</td><td>7.2</td><td>7.6E-13</td><td>1.9E-12</td><td>MB231/MB468</td></tr>
<tr><td>>hg38_21_+_8433172_8446622_RNA45SN1_45S_50ntFlanks@11316.11340.25|CACCCCACGTCTCGTCGCGCGCGCG</td><td>>hg38_21_+_8433172_8446622_RNA45SN1_45S_50ntFlanks</td><td>11316</td><td>11340</td><td>25</td><td>CACCCCACGTCTCGTCGCGCGCGCG</td><td>196.4</td><td>-9.2</td><td>1.3</td><td>-7.2</td><td>7.6E-13</td><td>1.9E-12</td><td>MB231/MB468</td></tr>
<tr><td>>hg38_21_+_8433172_8446622_RNA45SN1_45S_50ntFlanks@10107.10125.19|CGTGTGGTGTGCGTCGGAG</td><td>>hg38_21_+_8433172_8446622_RNA45SN1_45S_50ntFlanks</td><td>10107</td><td>10125</td><td>19</td><td>CGTGTGGTGTGCGTCGGAG</td><td>194.5</td><td>-9.2</td><td>1.3</td><td>-7.2</td><td>7.6E-13</td><td>1.9E-12</td><td>MB231/MB468</td></tr>
<tr><td>>MI0000103|hsa-mir-101-1&WithFlank&1|-|65058428|65058514@52.73.22;MI0000739|hsa-mir-101-2&WithFlank&9|+|4850291|4850381@54.75.22&[MIMAT0000099&hsa-miR-101-3p&offsets|-1|0;m-17&1|-|65058443|65058462&offsets|-1|+1];[MIMAT0000099_1&hsa-miR-101-3p&offsets|-1|0;m-16&9|+|4850345|4850364&offsets|-1|+1]|GTACAGTACTGTGATAACTGAA</td><td>>MI0000103|hsa-mir-101-1&WithFlank&1|-|65058428|65058514</td><td>52</td><td>73</td><td>22</td><td>GTACAGTACTGTGATAACTGAA</td><td>17618.8</td><td>7.8</td><td>1.1</td><td>7.2</td><td>8.5E-13</td><td>1.9E-12</td><td>BT20/MB468</td></tr>
<tr><td>>hg38_21_+_8433172_8446622_RNA45SN1_45S_50ntFlanks@9546.9572.27|TGGAGGTCCGTAGCGGTCCTGACGTGC</td><td>>hg38_21_+_8433172_8446622_RNA45SN1_45S_50ntFlanks</td><td>9546</td><td>9572</td><td>27</td><td>TGGAGGTCCGTAGCGGTCCTGACGTGC</td><td>319.0</td><td>3.5</td><td>0.5</td><td>7.2</td><td>7.7E-13</td><td>1.9E-12</td><td>MB231/MB468</td></tr>
<tr><td>>hg38_21_+_8433172_8446622_RNA45SN1_45S_50ntFlanks@4007.4042.36|CCTCGGGCCGATCGCACGCCCCCCGTGGCGGCGACG</td><td>>hg38_21_+_8433172_8446622_RNA45SN1_45S_50ntFlanks</td><td>4007</td><td>4042</td><td>36</td><td>CCTCGGGCCGATCGCACGCCCCCCGTGGCGGCGACG</td><td>154.5</td><td>-2.5</td><td>0.4</td><td>-7.2</td><td>8.5E-13</td><td>1.9E-12</td><td>BT20/MB468</td></tr>
<tr><td>>MI0000782|hsa-mir-374a&WithFlank&X|-|74287280|74287363@18.39.22&[MIMAT0000727&hsa-miR-374a-5p&offsets|0|0;m-184&X|-|74287325|74287346&offsets|0|0]|TTATAATACAACCTGATAAGTG</td><td>>MI0000782|hsa-mir-374a&WithFlank&X|-|74287280|74287363</td><td>18</td><td>39</td><td>22</td><td>TTATAATACAACCTGATAAGTG</td><td>2116.0</td><td>5.7</td><td>0.8</td><td>7.2</td><td>7.8E-13</td><td>1.9E-12</td><td>MB231/MB468</td></tr>
<tr><td>>hg38_21_+_8433172_8446622_RNA45SN1_45S_50ntFlanks@6646.6664.19|TCGTACGACTCTTAGCGGT</td><td>>hg38_21_+_8433172_8446622_RNA45SN1_45S_50ntFlanks</td><td>6646</td><td>6664</td><td>19</td><td>TCGTACGACTCTTAGCGGT</td><td>15642.5</td><td>4.9</td><td>0.7</td><td>7.2</td><td>7.8E-13</td><td>1.9E-12</td><td>MB231/MB468</td></tr>
<tr><td>>hg38_21_+_8433172_8446622_RNA45SN1_45S_50ntFlanks@6744.6771.28|GACACTTCGAACGCACTTGCGGCCCCGG</td><td>>hg38_21_+_8433172_8446622_RNA45SN1_45S_50ntFlanks</td><td>6744</td><td>6771</td><td>28</td><td>GACACTTCGAACGCACTTGCGGCCCCGG</td><td>745.8</td><td>-4.5</td><td>0.6</td><td>-7.2</td><td>7.8E-13</td><td>1.9E-12</td><td>MB231/MB468</td></tr>
<tr><td>>hg38_21_+_8433172_8446622_RNA45SN1_45S_50ntFlanks@9576.9597.22|TCGGTCGTCCGACCTGGGTATA</td><td>>hg38_21_+_8433172_8446622_RNA45SN1_45S_50ntFlanks</td><td>9576</td><td>9597</td><td>22</td><td>TCGGTCGTCCGACCTGGGTATA</td><td>190.4</td><td>-9.6</td><td>1.3</td><td>-7.2</td><td>8.7E-13</td><td>1.9E-12</td><td>BT20/MB468</td></tr>
<tr><td>>hg38_21_+_8433172_8446622_RNA45SN1_45S_50ntFlanks@11884.11904.21|GTTGAGCTTGACTCTAGTCTG</td><td>>hg38_21_+_8433172_8446622_RNA45SN1_45S_50ntFlanks</td><td>11884</td><td>11904</td><td>21</td><td>GTTGAGCTTGACTCTAGTCTG</td><td>171.2</td><td>-9.4</td><td>1.3</td><td>-7.1</td><td>8.7E-13</td><td>1.9E-12</td><td>BT20/MB468</td></tr>
<tr><td>>am_trna117_GlyTCC_1_-_145397864_145397935@1.29.29,trna45_GlyTCC_1_+_161500903_161500974@1.29.29,trna70_GlyTCC_1_-_161439547_161439618@1.29.29,trna73_GlyTCC_1_-_161432166_161432237@1.29.29,trna76_GlyTCC_1_-_161424756_161424827@1.29.29,trna79_GlyTCC_1_-_161417375_161417446@1.29.29|GCGTTGGTGGTATAGTGGTGAGCATAGCT</td><td>>am_trna117_GlyTCC_1_-_145397864_145397935</td><td>1</td><td>29</td><td>29</td><td>GCGTTGGTGGTATAGTGGTGAGCATAGCT</td><td>1089.8</td><td>-4.7</td><td>0.7</td><td>-7.1</td><td>8.7E-13</td><td>1.9E-12</td><td>BT20/MB468</td></tr>
<tr><td>>hg38_21_+_8433172_8446622_RNA45SN1_45S_50ntFlanks@10427.10457.31|ATGGGCGAGCGCCGTTCCGAAGGGACGGGCG</td><td>>hg38_21_+_8433172_8446622_RNA45SN1_45S_50ntFlanks</td><td>10427</td><td>10457</td><td>31</td><td>ATGGGCGAGCGCCGTTCCGAAGGGACGGGCG</td><td>168.3</td><td>-9.4</td><td>1.3</td><td>-7.1</td><td>8.8E-13</td><td>2.0E-12</td><td>BT20/MB468</td></tr>
<tr><td>>hg38_21_+_8433172_8446622_RNA45SN1_45S_50ntFlanks@10678.10710.33|CGGTTCCGGCGGCGTCCGGTGAGCTCTCGCTGG</td><td>>hg38_21_+_8433172_8446622_RNA45SN1_45S_50ntFlanks</td><td>10678</td><td>10710</td><td>33</td><td>CGGTTCCGGCGGCGTCCGGTGAGCTCTCGCTGG</td><td>168089.3</td><td>-7.0</td><td>1.0</td><td>-7.1</td><td>8.9E-13</td><td>2.0E-12</td><td>BT20/MB468</td></tr>
<tr><td>>hg38_21_+_8433172_8446622_RNA45SN1_45S_50ntFlanks@11561.11603.43|GCGCCTAGCAGCCGACTTAGAACTGGTGCGGACCAGGGGAATC</td><td>>hg38_21_+_8433172_8446622_RNA45SN1_45S_50ntFlanks</td><td>11561</td><td>11603</td><td>43</td><td>GCGCCTAGCAGCCGACTTAGAACTGGTGCGGACCAGGGGAATC</td><td>352.5</td><td>-5.3</td><td>0.7</td><td>-7.1</td><td>8.9E-13</td><td>2.0E-12</td><td>BT20/MB468</td></tr>
<tr><td>>MI0000102|hsa-mir-100&WithFlank&11|-|122152223|122152314@19.40.22(+1U)&[MIMAT0000098&hsa-miR-100-5p&offsets|0|0(+1U);m-27&11|-|122152276|122152296&offsets|0|+1(+1U)]|AACCCGTAGATCCGAACTTGTGT</td><td>>MI0000102|hsa-mir-100&WithFlank&11|-|122152223|122152314</td><td>19</td><td>40</td><td>22(+1U)</td><td>AACCCGTAGATCCGAACTTGTGT</td><td>63208.6</td><td>-6.7</td><td>0.9</td><td>-7.5</td><td>6.1E-14</td><td>2.0E-12</td><td>BT20/MB231</td></tr>
<tr><td>>hg38_21_+_8433172_8446622_RNA45SN1_45S_50ntFlanks@10498.10516.19|TTCAGATCCCCGAATCCGG</td><td>>hg38_21_+_8433172_8446622_RNA45SN1_45S_50ntFlanks</td><td>10498</td><td>10516</td><td>19</td><td>TTCAGATCCCCGAATCCGG</td><td>168.4</td><td>-9.0</td><td>1.3</td><td>-7.2</td><td>8.1E-13</td><td>2.0E-12</td><td>MB231/MB468</td></tr>
</table>

<table>
<tr><td></td><td></td><td></td><td></td><td></td><td></td><td></td><td></td><td></td><td></td><td></td><td></td><td></td></tr>
<tr><td>>hg38_21_+_8433172_8446622_RNA45SN1_45S_50ntFlanks@11212.11230.19|CGACTCTGGACGCGAGCCG</td><td>>hg38_21_+_8433172_8446622_RNA45SN1_45S_50ntFlanks</td><td>11212</td><td>11230</td><td>19</td><td>CGACTCTGGACGCGAGCCG</td><td>158.3</td><td>-4.9</td><td>0.7</td><td>-7.1</td><td>9.2E-13</td><td>2.0E-12</td><td>BT20/MB468</td></tr>
<tr><td>>hg38_21_+_8433172_8446622_RNA45SN1_45S_50ntFlanks@2959.2979.21|CGACGTGTGGCGTGGGTCGAC</td><td>>hg38_21_+_8433172_8446622_RNA45SN1_45S_50ntFlanks</td><td>2959</td><td>2979</td><td>21</td><td>CGACGTGTGGCGTGGGTCGAC</td><td>135.7</td><td>-9.1</td><td>1.3</td><td>-7.1</td><td>9.2E-13</td><td>2.0E-12</td><td>BT20/MB468</td></tr>
<tr><td>>hg38_21_+_8433172_8446622_RNA45SN1_45S_50ntFlanks@12073.12110.38|CCGAGGGGCTCTCGCTTCTGGCGCCAAGCGCCCGGCCG</td><td>>hg38_21_+_8433172_8446622_RNA45SN1_45S_50ntFlanks</td><td>12073</td><td>12110</td><td>38</td><td>CCGAGGGGCTCTCGCTTCTGGCGCCAAGCGCCCGGCCG</td><td>184.5</td><td>-5.5</td><td>0.8</td><td>-7.2</td><td>8.3E-13</td><td>2.0E-12</td><td>MB231/MB468</td></tr>
<tr><td>>hg38_21_+_8433172_8446622_RNA45SN1_45S_50ntFlanks@10458.10495.38|ATGGCCTCCGTTGCCCTCGGCCGATCGAAAGGGAGTCG</td><td>>hg38_21_+_8433172_8446622_RNA45SN1_45S_50ntFlanks</td><td>10458</td><td>10495</td><td>38</td><td>ATGGCCTCCGTTGCCCTCGGCCGATCGAAAGGGAGTCG</td><td>243.2</td><td>-9.9</td><td>1.4</td><td>-7.1</td><td>9.2E-13</td><td>2.0E-12</td><td>BT20/MB468</td></tr>
<tr><td>>MI0005756|hsa-mir-934&WithFlank&X|+|136550872|136550966@21.42.22&[MIMAT0004977&hsa-miR-934&offsets|0|0;m-943&X|+|136550892|136550913&offsets|0|0]|TGTCTACTACTGGAGACACTGG</td><td>>MI0005756|hsa-mir-934&WithFlank&X|+|136550872|136550966</td><td>21</td><td>42</td><td>22</td><td>TGTCTACTACTGGAGACACTGG</td><td>9997.0</td><td>16.8</td><td>2.2</td><td>7.5</td><td>6.3E-14</td><td>2.0E-12</td><td>BT20/MB231</td></tr>
<tr><td>>hg38_21_+_8433172_8446622_RNA45SN1_45S_50ntFlanks@8381.8401.21|AAGAGGGCGTGAAACCGTTAA</td><td>>hg38_21_+_8433172_8446622_RNA45SN1_45S_50ntFlanks</td><td>8381</td><td>8401</td><td>21</td><td>AAGAGGGCGTGAAACCGTTAA</td><td>169.3</td><td>-9.4</td><td>1.3</td><td>-7.1</td><td>9.3E-13</td><td>2.1E-12</td><td>BT20/MB468</td></tr>
<tr><td>>hg38_21_+_8433172_8446622_RNA45SN1_45S_50ntFlanks@11209.11231.23|CGGCGACTCTGGACGCGAGCCGG</td><td>>hg38_21_+_8433172_8446622_RNA45SN1_45S_50ntFlanks</td><td>11209</td><td>11231</td><td>23</td><td>CGGCGACTCTGGACGCGAGCCGG</td><td>692.1</td><td>-4.4</td><td>0.6</td><td>-7.2</td><td>8.4E-13</td><td>2.1E-12</td><td>MB231/MB468</td></tr>
<tr><td>>hg38_21_+_8433172_8446622_RNA45SN1_45S_50ntFlanks@9534.9560.27|AGGAAACTCTGGTGGAGGTCCGTAGCG</td><td>>hg38_21_+_8433172_8446622_RNA45SN1_45S_50ntFlanks</td><td>9534</td><td>9560</td><td>27</td><td>AGGAAACTCTGGTGGAGGTCCGTAGCG</td><td>406.1</td><td>-3.7</td><td>0.5</td><td>-7.1</td><td>9.4E-13</td><td>2.1E-12</td><td>BT20/MB468</td></tr>
<tr><td>>hg38_21_+_8433172_8446622_RNA45SN1_45S_50ntFlanks@12927.12956.30|CCTGGCGCTAAACCATTCGTAGACGACCTG</td><td>>hg38_21_+_8433172_8446622_RNA45SN1_45S_50ntFlanks</td><td>12927</td><td>12956</td><td>30</td><td>CCTGGCGCTAAACCATTCGTAGACGACCTG</td><td>160.0</td><td>-8.9</td><td>1.2</td><td>-7.2</td><td>8.5E-13</td><td>2.1E-12</td><td>MB231/MB468</td></tr>
<tr><td>>MI0000095|hsa-mir-93&WithFlank&7|-|100093762|100093853@17.37.21&[MIMAT0000093&hsa-miR-93-5p&offsets|0|-2;m-29&7|-|100093815|100093837&offsets|0|-2]|CAAAGTGCTGTTCGTGCAGGT</td><td>>MI0000095|hsa-mir-93&WithFlank&7|-|100093762|100093853</td><td>17</td><td>37</td><td>21</td><td>CAAAGTGCTGTTCGTGCAGGT</td><td>2164.8</td><td>6.2</td><td>0.9</td><td>7.1</td><td>9.4E-13</td><td>2.1E-12</td><td>BT20/MB468</td></tr>
<tr><td>>hg38_21_+_8433172_8446622_RNA45SN1_45S_50ntFlanks@6646.6676.31|TCGTACGACTCTTAGCGGTGGATCACTCGGC</td><td>>hg38_21_+_8433172_8446622_RNA45SN1_45S_50ntFlanks</td><td>6646</td><td>6676</td><td>31</td><td>TCGTACGACTCTTAGCGGTGGATCACTCGGC</td><td>912.6</td><td>5.3</td><td>0.7</td><td>7.2</td><td>8.5E-13</td><td>2.1E-12</td><td>MB231/MB468</td></tr>
<tr><td>>trna129_MetCAT_6_-_27870271_27870342@32.51.20,trna142_MetCAT_6_-_27560600_27560671@32.51.20,trna150_MetCAT_6_-_27300764_27300835@32.51.20,trna169_MetCAT_6_-_26330529_26330600@32.51.20,trna171_MetCAT_6_-_26313352_26313423@32.51.20,trna20_MetCAT_17_-_80452597_80452668@32.51.20,trna2_MetCAT_6_+_26286754_26286825@32.51.20,trna32_MetCAT_1_+_153643726_153643797@32.51.20,trna61_MetCAT_6_+_27745664_27745735@32.51.20|CCATAACCCAGAGGTCGATG</td><td>>trna129_MetCAT_6_-_27870271_27870342</td><td>32</td><td>51</td><td>20</td><td>CCATAACCCAGAGGTCGATG</td><td>246.9</td><td>-9.9</td><td>1.4</td><td>-7.1</td><td>9.4E-13</td><td>2.1E-12</td><td>BT20/MB468</td></tr>
<tr><td>>hg38_21_+_8433172_8446622_RNA45SN1_45S_50ntFlanks@11669.11692.24|GCCCAGTGCTCTGAATGTCAAAGT</td><td>>hg38_21_+_8433172_8446622_RNA45SN1_45S_50ntFlanks</td><td>11669</td><td>11692</td><td>24</td><td>GCCCAGTGCTCTGAATGTCAAAGT</td><td>280.1</td><td>6.4</td><td>0.9</td><td>7.1</td><td>9.5E-13</td><td>2.1E-12</td><td>BT20/MB468</td></tr>
<tr><td>>hg38_21_+_8433172_8446622_RNA45SN1_45S_50ntFlanks@4723.4752.30|CATTAATCAAGAACGAAAGTCGGAGGTTCG</td><td>>hg38_21_+_8433172_8446622_RNA45SN1_45S_50ntFlanks</td><td>4723</td><td>4752</td><td>30</td><td>CATTAATCAAGAACGAAAGTCGGAGGTTCG</td><td>557.3</td><td>-6.4</td><td>0.9</td><td>-7.1</td><td>9.5E-13</td><td>2.1E-12</td><td>BT20/MB468</td></tr>
<tr><td>>hg38_21_+_8433172_8446622_RNA45SN1_45S_50ntFlanks@4853.4883.31|AAGTCTTTGGGTTCCGGGGGGAGTATGGTTG</td><td>>hg38_21_+_8433172_8446622_RNA45SN1_45S_50ntFlanks</td><td>4853</td><td>4883</td><td>31</td><td>AAGTCTTTGGGTTCCGGGGGGAGTATGGTTG</td><td>124.6</td><td>-9.0</td><td>1.3</td><td>-7.1</td><td>9.5E-13</td><td>2.1E-12</td><td>BT20/MB468</td></tr>
<tr><td>>hg38_21_+_8433172_8446622_RNA45SN1_45S_50ntFlanks@10869.10892.24|AGGGCTGGGTCGGTCGGGCTGGGG</td><td>>hg38_21_+_8433172_8446622_RNA45SN1_45S_50ntFlanks</td><td>10869</td><td>10892</td><td>24</td><td>AGGGCTGGGTCGGTCGGGCTGGGG</td><td>274.3</td><td>-9.7</td><td>1.4</td><td>-7.2</td><td>8.6E-13</td><td>2.1E-12</td><td>MB231/MB468</td></tr>
<tr><td>>hg38_21_+_8433172_8446622_RNA45SN1_45S_50ntFlanks@6722.6740.19|ATTGCAGGACACATTGATC</td><td>>hg38_21_+_8433172_8446622_RNA45SN1_45S_50ntFlanks</td><td>6722</td><td>6740</td><td>19</td><td>ATTGCAGGACACATTGATC</td><td>1184.5</td><td>-4.5</td><td>0.6</td><td>-7.1</td><td>9.7E-13</td><td>2.1E-12</td><td>BT20/MB468</td></tr>
<tr><td>>hg38_21_+_8433172_8446622_RNA45SN1_45S_50ntFlanks@10795.10818.24|TGGCATGTTGGAACAATGTAGGTA</td><td>>hg38_21_+_8433172_8446622_RNA45SN1_45S_50ntFlanks</td><td>10795</td><td>10818</td><td>24</td><td>TGGCATGTTGGAACAATGTAGGTA</td><td>846.7</td><td>-4.5</td><td>0.6</td><td>-7.1</td><td>8.7E-13</td><td>2.1E-12</td><td>MB231/MB468</td></tr>
<tr><td>>trna2_GlyGCC_21_-_18827107_18827177@1.28.28,trna35_GlyGCC_1_+_161413094_161413164@1.28.28,trna37_GlyGCC_1_+_161420467_161420537@1.28.28,trna39_GlyGCC_1_+_161427898_161427968@1.28.28,trna41_GlyGCC_1_+_161435258_161435328@1.28.28|GCATGGGTGGTTCAGTGGTAGAATTCTC</td><td>>trna2_GlyGCC_21_-_18827107_18827177</td><td>1</td><td>28</td><td>28</td><td>GCATGGGTGGTTCAGTGGTAGAATTCTC</td><td>8602.6</td><td>-7.5</td><td>1.0</td><td>-7.1</td><td>8.7E-13</td><td>2.1E-12</td><td>MB231/MB468</td></tr>
<tr><td>>MI0000750|hsa-mir-26a-2&WithFlank&12|-|57824603|57824698@20.43.24;MI0000083|hsa-mir-26a-1&WithFlank&3|+|37969398|37969486@16.39.24&[MIMAT0000082&hsa-miR-26a-5p&offsets|0|+2;m-33&12|-|57824658|57824679&offsets|0|+2];[MIMAT0000082_1&hsa-miR-26a-5p&offsets|0|+2;m-34&3|+|37969413|37969434&offsets|0|+2]|TTCAAGTAATCCAGGATAGGCTGT</td><td>>MI0000750|hsa-mir-26a-2&WithFlank&12|-|57824603|57824698</td><td>20</td><td>43</td><td>24</td><td>TTCAAGTAATCCAGGATAGGCTGT</td><td>457.5</td><td>7.6</td><td>1.1</td><td>7.1</td><td>9.7E-13</td><td>2.2E-12</td><td>BT20/MB468</td></tr>
<tr><td>>MI0000095|hsa-mir-93&WithFlank&7|-|100093762|100093853@17.40.24&[MIMAT0000093&hsa-miR-93-5p&offsets|0|+1;m-29&7|-|100093815|100093837&offsets|0|+1]|CAAAGTGCTGTTCGTGCAGGTAGT</td><td>>MI0000095|hsa-mir-93&WithFlank&7|-|100093762|100093853</td><td>17</td><td>40</td><td>24</td><td>CAAAGTGCTGTTCGTGCAGGTAGT</td><td>4188.6</td><td>5.4</td><td>0.8</td><td>7.1</td><td>9.8E-13</td><td>2.2E-12</td><td>BT20/MB468</td></tr>
<tr><td>>hg38_21_+_8433172_8446622_RNA45SN1_45S_50ntFlanks@3899.3920.22|CCTTCGCGGGGGGGATGCGTGC</td><td>>hg38_21_+_8433172_8446622_RNA45SN1_45S_50ntFlanks</td><td>3899</td><td>3920</td><td>22</td><td>CCTTCGCGGGGGGGATGCGTGC</td><td>156.3</td><td>-8.9</td><td>1.2</td><td>-7.1</td><td>8.8E-13</td><td>2.2E-12</td><td>MB231/MB468</td></tr>
<tr><td>>hg38_21_+_8433172_8446622_RNA45SN1_45S_50ntFlanks@11577.11603.27|TTAGAACTGGTGCGGACCAGGGGAATC</td><td>>hg38_21_+_8433172_8446622_RNA45SN1_45S_50ntFlanks</td><td>11577</td><td>11603</td><td>27</td><td>TTAGAACTGGTGCGGACCAGGGGAATC</td><td>810.2</td><td>-4.4</td><td>0.6</td><td>-7.1</td><td>8.9E-13</td><td>2.2E-12</td><td>MB231/MB468</td></tr>
<tr><td>>hg38_1_-_228634819_228635039_RNA5S12_5S_50ntFlanks@129.170.42|TTGGATGGGAGACCGCCTGGGAATACCGGGTGCTGTAGGCTT</td><td>>hg38_1_-_228634819_228635039_RNA5S12_5S_50ntFlanks</td><td>129</td><td>170</td><td>42</td><td>TTGGATGGGAGACCGCCTGGGAATACCGGGTGCTGTAGGCTT</td><td>1763.7</td><td>11.2</td><td>1.6</td><td>7.1</td><td>9.9E-13</td><td>2.2E-12</td><td>BT20/MB468</td></tr>
<tr><td>>hg38_21_+_8433172_8446622_RNA45SN1_45S_50ntFlanks@11235.11255.21|CTTCCCGTGGATCGCCCCAGC</td><td>>hg38_21_+_8433172_8446622_RNA45SN1_45S_50ntFlanks</td><td>11235</td><td>11255</td><td>21</td><td>CTTCCCGTGGATCGCCCCAGC</td><td>253.6</td><td>-4.2</td><td>0.6</td><td>-7.1</td><td>9.9E-13</td><td>2.2E-12</td><td>BT20/MB468</td></tr>
<tr><td>>MI0000734|hsa-mir-106b&WithFlank&7|-|100093987|100094080@18.38.21&[MIMAT0000680&hsa-miR-106b-5p&offsets|0|0;m-103&7|-|100094043|100094063&offsets|0|0]|TAAAGTGCTGACAGTGCAGAT</td><td>>MI0000734|hsa-mir-106b&WithFlank&7|-|100093987|100094080</td><td>18</td><td>38</td><td>21</td><td>TAAAGTGCTGACAGTGCAGAT</td><td>8491.7</td><td>8.1</td><td>1.1</td><td>7.1</td><td>9.9E-13</td><td>2.2E-12</td><td>BT20/MB468</td></tr>
<tr><td>>hg38_21_+_8433172_8446622_RNA45SN1_45S_50ntFlanks@4724.4752.29|ATTAATCAAGAACGAAAGTCGGAGGTTCG</td><td>>hg38_21_+_8433172_8446622_RNA45SN1_45S_50ntFlanks</td><td>4724</td><td>4752</td><td>29</td><td>ATTAATCAAGAACGAAAGTCGGAGGTTCG</td><td>523.6</td><td>-6.3</td><td>0.9</td><td>-7.1</td><td>1.0E-12</td><td>2.2E-12</td><td>BT20/MB468</td></tr>
<tr><td>>hg38_21_+_8433172_8446622_RNA45SN1_45S_50ntFlanks@10766.10797.32|CGCAGCAGGTCTCCAAGGTGAACAGCCTCTGG</td><td>>hg38_21_+_8433172_8446622_RNA45SN1_45S_50ntFlanks</td><td>10766</td><td>10797</td><td>32</td><td>CGCAGCAGGTCTCCAAGGTGAACAGCCTCTGG</td><td>113.5</td><td>-8.8</td><td>1.2</td><td>-7.1</td><td>1.0E-12</td><td>2.2E-12</td><td>BT20/MB468</td></tr>
<tr><td>>hg38_21_+_8433172_8446622_RNA45SN1_45S_50ntFlanks@10680.10712.33|GTTCCGGCGGCGTCCGGTGAGCTCTCGCTGGCC</td><td>>hg38_21_+_8433172_8446622_RNA45SN1_45S_50ntFlanks</td><td>10680</td><td>10712</td><td>33</td><td>GTTCCGGCGGCGTCCGGTGAGCTCTCGCTGGCC</td><td>2614.2</td><td>-3.9</td><td>0.5</td><td>-7.1</td><td>9.1E-13</td><td>2.2E-12</td><td>MB231/MB468</td></tr>
<tr><td>>hg38_21_+_8433172_8446622_RNA45SN1_45S_50ntFlanks@12488.12505.18|AACGTGAGCTGGGTTTAG</td><td>>hg38_21_+_8433172_8446622_RNA45SN1_45S_50ntFlanks</td><td>12488</td><td>12505</td><td>18</td><td>AACGTGAGCTGGGTTTAG</td><td>202.7</td><td>-4.6</td><td>0.6</td><td>-7.1</td><td>1.0E-12</td><td>2.3E-12</td><td>BT20/MB468</td></tr>
<tr><td>>hg38_21_+_8433172_8446622_RNA45SN1_45S_50ntFlanks@10801.10818.18|GTTGGAACAATGTAGGTA</td><td>>hg38_21_+_8433172_8446622_RNA45SN1_45S_50ntFlanks</td><td>10801</td><td>10818</td><td>18</td><td>GTTGGAACAATGTAGGTA</td><td>357.8</td><td>-10.5</td><td>1.5</td><td>-7.1</td><td>1.0E-12</td><td>2.3E-12</td><td>BT20/MB468</td></tr>
<tr><td>>hg38_21_+_8433172_8446622_RNA45SN1_45S_50ntFlanks@11641.11661.21|CGGCGGGTGTTGACGCGATGT</td><td>>hg38_21_+_8433172_8446622_RNA45SN1_45S_50ntFlanks</td><td>11641</td><td>11661</td><td>21</td><td>CGGCGGGTGTTGACGCGATGT</td><td>137.0</td><td>-9.1</td><td>1.3</td><td>-7.1</td><td>1.0E-12</td><td>2.3E-12</td><td>BT20/MB468</td></tr>
<tr><td>>hg38_21_+_8433172_8446622_RNA45SN1_45S_50ntFlanks@6661.6683.23|CGGTGGATCACTCGGCTCGTGCG</td><td>>hg38_21_+_8433172_8446622_RNA45SN1_45S_50ntFlanks</td><td>6661</td><td>6683</td><td>23</td><td>CGGTGGATCACTCGGCTCGTGCG</td><td>1947.7</td><td>5.8</td><td>0.8</td><td>7.1</td><td>1.0E-12</td><td>2.3E-12</td><td>BT20/MB468</td></tr>
<tr><td>>hg38_21_+_8433172_8446622_RNA45SN1_45S_50ntFlanks@3785.3812.28|TGAAACTGCGAATGGCTCATTAAATCAG</td><td>>hg38_21_+_8433172_8446622_RNA45SN1_45S_50ntFlanks</td><td>3785</td><td>3812</td><td>28</td><td>TGAAACTGCGAATGGCTCATTAAATCAG</td><td>183.4</td><td>-9.1</td><td>1.3</td><td>-7.1</td><td>9.3E-13</td><td>2.3E-12</td><td>MB231/MB468</td></tr>
<tr><td>>hg38_MT_+_598_1651_RNR1_12S_50ntFlanks@428.452.25|AAAGTGGCTTTAACATATCTGAACA</td><td>>hg38_MT_+_598_1651_RNR1_12S_50ntFlanks</td><td>428</td><td>452</td><td>25</td><td>AAAGTGGCTTTAACATATCTGAACA</td><td>612.5</td><td>8.7</td><td>1.2</td><td>7.1</td><td>1.0E-12</td><td>2.3E-12</td><td>BT20/MB468</td></tr>
<tr><td>>hg38_21_+_8433172_8446622_RNA45SN1_45S_50ntFlanks@11593.11612.20|CCAGGGGAATCCGACTGTTT</td><td>>hg38_21_+_8433172_8446622_RNA45SN1_45S_50ntFlanks</td><td>11593</td><td>11612</td><td>20</td><td>CCAGGGGAATCCGACTGTTT</td><td>911.6</td><td>-4.2</td><td>0.6</td><td>-7.1</td><td>9.4E-13</td><td>2.3E-12</td><td>MB231/MB468</td></tr>
<tr><td>>hg38_21_+_8433172_8446622_RNA45SN1_45S_50ntFlanks@12324.12356.33|TTGGGTTTTAAGCAGGAGGTGTCAGAAAAGTTA</td><td>>hg38_21_+_8433172_8446622_RNA45SN1_45S_50ntFlanks</td><td>12324</td><td>12356</td><td>33</td><td>TTGGGTTTTAAGCAGGAGGTGTCAGAAAAGTTA</td><td>106.3</td><td>-8.7</td><td>1.2</td><td>-7.1</td><td>1.0E-12</td><td>2.3E-12</td><td>BT20/MB468</td></tr>
<tr><td>>hg38_21_+_8433172_8446622_RNA45SN1_45S_50ntFlanks@11618.11640.23|AAACAAAGCATCGCGAAGGCCCG</td><td>>hg38_21_+_8433172_8446622_RNA45SN1_45S_50ntFlanks</td><td>11618</td><td>11640</td><td>23</td><td>AAACAAAGCATCGCGAAGGCCCG</td><td>150.4</td><td>-9.2</td><td>1.3</td><td>-7.1</td><td>1.0E-12</td><td>2.3E-12</td><td>BT20/MB468</td></tr>
<tr><td>>hg38_21_+_8433172_8446622_RNA45SN1_45S_50ntFlanks@9551.9569.19|GTCCGTAGCGGTCCTGACG</td><td>>hg38_21_+_8433172_8446622_RNA45SN1_45S_50ntFlanks</td><td>9551</td><td>9569</td><td>19</td><td>GTCCGTAGCGGTCCTGACG</td><td>563.0</td><td>4.6</td><td>0.6</td><td>7.1</td><td>9.4E-13</td><td>2.3E-12</td><td>MB231/MB468</td></tr>
<tr><td>>am_MI0000786|hsa-mir-378a&WithFlank&5|+|149732819|149732896@50.70.21&[MIMAT0000732&hsa-miR-378a-3p&offsets|+1|0;m-44&5|+|149732867|149732888&offsets|+1|0]|CTGGACTTGGAGTCAGAAGGC</td><td>>am_MI0000786|hsa-mir-378a&WithFlank&5|+|149732819|149732896</td><td>50</td><td>70</td><td>21</td><td>CTGGACTTGGAGTCAGAAGGC</td><td>2370.2</td><td>5.5</td><td>0.8</td><td>7.1</td><td>9.5E-13</td><td>2.3E-12</td><td>MB231/MB468</td></tr>
<tr><td>>trnaMT_GluTTC_MT_-_14674_14742@39.66.28,trnalookalike8_GluTTC_5_-_93905172_93905240@39.66.28|ATTGGTCGTGGTTGTAGTCCGTGCGAGA</td><td>>trnaMT_GluTTC_MT_-_14674_14742</td><td>39</td><td>66</td><td>28</td><td>ATTGGTCGTGGTTGTAGTCCGTGCGAGA</td><td>567.6</td><td>8.3</td><td>1.2</td><td>7.1</td><td>1.1E-12</td><td>2.3E-12</td><td>BT20/MB468</td></tr>
<tr><td>>hg38_21_+_8433172_8446622_RNA45SN1_45S_50ntFlanks@8552.8577.26|CTCCCCGGAGGGGGCGGGCTCCGGCG</td><td>>hg38_21_+_8433172_8446622_RNA45SN1_45S_50ntFlanks</td><td>8552</td><td>8577</td><td>26</td><td>CTCCCCGGAGGGGGCGGGCTCCGGCG</td><td>159.8</td><td>-8.9</td><td>1.2</td><td>-7.1</td><td>9.6E-13</td><td>2.4E-12</td><td>MB231/MB468</td></tr>
</table>

<table>
<tr><td>>hg38_21_+_8433172_8446622_RNA45SN1_45S_50ntFlanks@6773.6797.25|TTCCTCCCGGGGCTACGCCTGTCTG</td><td>>hg38_21_+_8433172_8446622_RNA45SN1_45S_50ntFlanks</td><td>6773</td><td>6797</td><td>25</td><td>TTCCTCCCGGGGCTACGCCTGTCTG</td><td>183.2</td><td>-9.1</td><td>1.3</td><td>-7.1</td><td>9.7E-13</td><td>2.4E-12</td><td>MB231/MB468</td></tr>
<tr><td>>hg38_21_+_8433172_8446622_RNA45SN1_45S_50ntFlanks@6743.6772.30|CGACACTTCGAACGCACTTGCGGCCCCGGG</td><td>>hg38_21_+_8433172_8446622_RNA45SN1_45S_50ntFlanks</td><td>6743</td><td>6772</td><td>30</td><td>CGACACTTCGAACGCACTTGCGGCCCCGGG</td><td>253.8</td><td>-5.0</td><td>0.7</td><td>-7.1</td><td>9.8E-13</td><td>2.4E-12</td><td>MB231/MB468</td></tr>
<tr><td>>hg38_21_+_8433172_8446622_RNA45SN1_45S_50ntFlanks@8171.8197.27|AAGTCCTTCTGATCGAGGCCCAGCCCG</td><td>>hg38_21_+_8433172_8446622_RNA45SN1_45S_50ntFlanks</td><td>8171</td><td>8197</td><td>27</td><td>AAGTCCTTCTGATCGAGGCCCAGCCCG</td><td>190.6</td><td>-4.5</td><td>0.6</td><td>-7.1</td><td>1.1E-12</td><td>2.4E-12</td><td>BT20/MB468</td></tr>
<tr><td>>hg38_21_+_8433172_8446622_RNA45SN1_45S_50ntFlanks@10794.10823.30|CTGGCATGTTGGAACAATGTAGGTAAGGGA</td><td>>hg38_21_+_8433172_8446622_RNA45SN1_45S_50ntFlanks</td><td>10794</td><td>10823</td><td>30</td><td>CTGGCATGTTGGAACAATGTAGGTAAGGGA</td><td>176.7</td><td>-9.0</td><td>1.3</td><td>-7.1</td><td>9.8E-13</td><td>2.4E-12</td><td>MB231/MB468</td></tr>
<tr><td>>hg38_21_+_8433172_8446622_RNA45SN1_45S_50ntFlanks@9720.9737.18|ATTAGAGGTCTTGGGGCC</td><td>>hg38_21_+_8433172_8446622_RNA45SN1_45S_50ntFlanks</td><td>9720</td><td>9737</td><td>18</td><td>ATTAGAGGTCTTGGGGCC</td><td>176.6</td><td>-9.0</td><td>1.3</td><td>-7.1</td><td>9.8E-13</td><td>2.4E-12</td><td>MB231/MB468</td></tr>
<tr><td>>MI0000746|hsa-mir-99b&WithFlank&19|+|51692606|51692687@51.72.22(+1C)&[MIMAT0004678&hsa-miR-99b-3p&offsets|0|0(+1C);m-151&19|+|51692656|51692678&offsets|0|-1(+1C)]|CAAGCTCGTGTCTGTGGGTCCGC</td><td>>MI0000746|hsa-mir-99b&WithFlank&19|+|51692606|51692687</td><td>51</td><td>72</td><td>22(+1C)</td><td>CAAGCTCGTGTCTGTGGGTCCGC</td><td>424.3</td><td>9.4</td><td>1.3</td><td>7.1</td><td>1.1E-12</td><td>2.4E-12</td><td>BT20/MB468</td></tr>
<tr><td>>hg38_21_+_8433172_8446622_RNA45SN1_45S_50ntFlanks@12417.12438.22|TCGATGTCGGCTCTTCCTATCA</td><td>>hg38_21_+_8433172_8446622_RNA45SN1_45S_50ntFlanks</td><td>12417</td><td>12438</td><td>22</td><td>TCGATGTCGGCTCTTCCTATCA</td><td>1241.3</td><td>4.0</td><td>0.6</td><td>7.1</td><td>9.8E-13</td><td>2.4E-12</td><td>MB231/MB468</td></tr>
<tr><td>>hg38_21_+_8433172_8446622_RNA45SN1_45S_50ntFlanks@10414.10431.18|TCGGTCCTGAGAGATGGG</td><td>>hg38_21_+_8433172_8446622_RNA45SN1_45S_50ntFlanks</td><td>10414</td><td>10431</td><td>18</td><td>TCGGTCCTGAGAGATGGG</td><td>371.0</td><td>-5.6</td><td>0.8</td><td>-7.1</td><td>1.1E-12</td><td>2.4E-12</td><td>BT20/MB468</td></tr>
<tr><td>>MI0000102|hsa-mir-100&WithFlank&11|-|122152223|122152314@20.39.20&[MIMAT0000098&hsa-miR-100-5p&offsets|+1|-1;m-27&11|-|122152276|122152296&offsets|+1|0]|ACCCGTAGATCCGAACTTGT</td><td>>MI0000102|hsa-mir-100&WithFlank&11|-|122152223|122152314</td><td>20</td><td>39</td><td>20</td><td>ACCCGTAGATCCGAACTTGT</td><td>296.2</td><td>-7.0</td><td>0.9</td><td>-7.5</td><td>7.5E-14</td><td>2.4E-12</td><td>BT20/MB231</td></tr>
<tr><td>>hg38_21_+_8433172_8446622_RNA45SN1_45S_50ntFlanks@10574.10591.18|AAGCCGGCGGGAGCCCCG</td><td>>hg38_21_+_8433172_8446622_RNA45SN1_45S_50ntFlanks</td><td>10574</td><td>10591</td><td>18</td><td>AAGCCGGCGGGAGCCCCG</td><td>297.8</td><td>-9.8</td><td>1.4</td><td>-7.1</td><td>9.9E-13</td><td>2.4E-12</td><td>MB231/MB468</td></tr>
<tr><td>>trnaMT_ValTAC_MT_+_1602_1670@50.72.23|ACTTAACTTGACCGCTCTGACCA</td><td>>trnaMT_ValTAC_MT_+_1602_1670</td><td>50</td><td>72</td><td>23</td><td>ACTTAACTTGACCGCTCTGACCA</td><td>1426.1</td><td>4.0</td><td>0.6</td><td>7.1</td><td>9.9E-13</td><td>2.4E-12</td><td>MB231/MB468</td></tr>
<tr><td>>hg38_21_+_8433172_8446622_RNA45SN1_45S_50ntFlanks@9565.9592.28|TGACGTGCAAATCGGTCGTCCGACCTGG</td><td>>hg38_21_+_8433172_8446622_RNA45SN1_45S_50ntFlanks</td><td>9565</td><td>9592</td><td>28</td><td>TGACGTGCAAATCGGTCGTCCGACCTGG</td><td>480.0</td><td>-3.6</td><td>0.5</td><td>-7.1</td><td>1.1E-12</td><td>2.4E-12</td><td>BT20/MB468</td></tr>
<tr><td>>hg38_21_+_8433172_8446622_RNA45SN1_45S_50ntFlanks@11739.11767.29|TCTCTTAAGGTAGCCAAATGCCTCGTCAT</td><td>>hg38_21_+_8433172_8446622_RNA45SN1_45S_50ntFlanks</td><td>11739</td><td>11767</td><td>29</td><td>TCTCTTAAGGTAGCCAAATGCCTCGTCAT</td><td>156.6</td><td>-9.3</td><td>1.3</td><td>-7.1</td><td>1.1E-12</td><td>2.4E-12</td><td>BT20/MB468</td></tr>
<tr><td>>hg38_21_+_8433172_8446622_RNA45SN1_45S_50ntFlanks@3866.3898.33|CTAGAGCTAATACATGCCGACGGGCGCTGACCC</td><td>>hg38_21_+_8433172_8446622_RNA45SN1_45S_50ntFlanks</td><td>3866</td><td>3898</td><td>33</td><td>CTAGAGCTAATACATGCCGACGGGCGCTGACCC</td><td>166.1</td><td>-9.4</td><td>1.3</td><td>-7.1</td><td>1.1E-12</td><td>2.4E-12</td><td>BT20/MB468</td></tr>
<tr><td>>hg38_21_+_8433172_8446622_RNA45SN1_45S_50ntFlanks@12606.12626.21|ATGTGCTTGGCTGAGGAGCCA</td><td>>hg38_21_+_8433172_8446622_RNA45SN1_45S_50ntFlanks</td><td>12606</td><td>12626</td><td>21</td><td>ATGTGCTTGGCTGAGGAGCCA</td><td>315.6</td><td>-4.0</td><td>0.6</td><td>-7.1</td><td>1.0E-12</td><td>2.5E-12</td><td>MB231/MB468</td></tr>
<tr><td>>MI0000088|hsa-mir-30a&WithFlank&6|-|71403545|71403627@12.33.22(+1G)&[MIMAT0000087&hsa-miR-30a-5p&offsets|0|0(+1G);m-15&6|-|71403595|71403616&offsets|0|0(+1G)]|TGTAAACATCCTCGACTGGAAGG</td><td>>MI0000088|hsa-mir-30a&WithFlank&6|-|71403545|71403627</td><td>12</td><td>33</td><td>22(+1G)</td><td>TGTAAACATCCTCGACTGGAAGG</td><td>3762.0</td><td>10.4</td><td>1.5</td><td>7.1</td><td>1.1E-12</td><td>2.5E-12</td><td>BT20/MB468</td></tr>
<tr><td>>hg38_21_+_8433172_8446622_RNA45SN1_45S_50ntFlanks@11233.11252.20|CCCTTCCCGTGGATCGCCCC</td><td>>hg38_21_+_8433172_8446622_RNA45SN1_45S_50ntFlanks</td><td>11233</td><td>11252</td><td>20</td><td>CCCTTCCCGTGGATCGCCCC</td><td>208.8</td><td>-9.3</td><td>1.3</td><td>-7.1</td><td>1.0E-12</td><td>2.5E-12</td><td>MB231/MB468</td></tr>
<tr><td>>hg38_21_+_8433172_8446622_RNA45SN1_45S_50ntFlanks@12163.12180.18|ACTGGGGCGGTACACCTG</td><td>>hg38_21_+_8433172_8446622_RNA45SN1_45S_50ntFlanks</td><td>12163</td><td>12180</td><td>18</td><td>ACTGGGGCGGTACACCTG</td><td>2000.3</td><td>-7.2</td><td>1.0</td><td>-7.1</td><td>1.0E-12</td><td>2.5E-12</td><td>MB231/MB468</td></tr>
<tr><td>>hg38_21_+_8433172_8446622_RNA45SN1_45S_50ntFlanks@10361.10388.28|GAAGGCCGAAGTGGAGAAGGGTTCCATG</td><td>>hg38_21_+_8433172_8446622_RNA45SN1_45S_50ntFlanks</td><td>10361</td><td>10388</td><td>28</td><td>GAAGGCCGAAGTGGAGAAGGGTTCCATG</td><td>480.6</td><td>-10.9</td><td>1.5</td><td>-7.1</td><td>1.1E-12</td><td>2.5E-12</td><td>BT20/MB468</td></tr>
<tr><td>>hg38_21_+_8433172_8446622_RNA45SN1_45S_50ntFlanks@12638.12659.22|CTACCATCTGTGGGATTATGAC</td><td>>hg38_21_+_8433172_8446622_RNA45SN1_45S_50ntFlanks</td><td>12638</td><td>12659</td><td>22</td><td>CTACCATCTGTGGGATTATGAC</td><td>670.3</td><td>-5.2</td><td>0.7</td><td>-7.1</td><td>1.0E-12</td><td>2.5E-12</td><td>MB231/MB468</td></tr>
<tr><td>>hg38_21_+_8433172_8446622_RNA45SN1_45S_50ntFlanks@10768.10797.30|CAGCAGGTCTCCAAGGTGAACAGCCTCTGG</td><td>>hg38_21_+_8433172_8446622_RNA45SN1_45S_50ntFlanks</td><td>10768</td><td>10797</td><td>30</td><td>CAGCAGGTCTCCAAGGTGAACAGCCTCTGG</td><td>139.6</td><td>-5.6</td><td>0.8</td><td>-7.1</td><td>1.0E-12</td><td>2.5E-12</td><td>MB231/MB468</td></tr>
<tr><td>>am_trnaMT_PheGAA_MT_+_577_647@40.57.18|TGTTTAGACGGGCTCACA</td><td>>am_trnaMT_PheGAA_MT_+_577_647</td><td>40</td><td>57</td><td>18</td><td>TGTTTAGACGGGCTCACA</td><td>1458.5</td><td>9.2</td><td>1.3</td><td>7.1</td><td>1.2E-12</td><td>2.5E-12</td><td>BT20/MB468</td></tr>
<tr><td>>hg38_21_+_8433172_8446622_RNA45SN1_45S_50ntFlanks@10357.10390.34|CTTTGAAGGCCGAAGTGGAGAAGGGTTCCATGTG</td><td>>hg38_21_+_8433172_8446622_RNA45SN1_45S_50ntFlanks</td><td>10357</td><td>10390</td><td>34</td><td>CTTTGAAGGCCGAAGTGGAGAAGGGTTCCATGTG</td><td>220.1</td><td>-5.0</td><td>0.7</td><td>-7.1</td><td>1.2E-12</td><td>2.5E-12</td><td>BT20/MB468</td></tr>
<tr><td>>hg38_1_-_228634819_228635039_RNA5S12_5S_50ntFlanks@151.170.20|ATACCGGGTGCTGTAGGCTT</td><td>>hg38_1_-_228634819_228635039_RNA5S12_5S_50ntFlanks</td><td>151</td><td>170</td><td>20</td><td>ATACCGGGTGCTGTAGGCTT</td><td>2457.6</td><td>5.7</td><td>0.8</td><td>7.1</td><td>1.1E-12</td><td>2.6E-12</td><td>MB231/MB468</td></tr>
<tr><td>>hg38_21_+_8433172_8446622_RNA45SN1_45S_50ntFlanks@10683.10710.28|CCGGCGGCGTCCGGTGAGCTCTCGCTGG</td><td>>hg38_21_+_8433172_8446622_RNA45SN1_45S_50ntFlanks</td><td>10683</td><td>10710</td><td>28</td><td>CCGGCGGCGTCCGGTGAGCTCTCGCTGG</td><td>277.2</td><td>-5.9</td><td>0.8</td><td>-7.1</td><td>1.1E-12</td><td>2.6E-12</td><td>MB231/MB468</td></tr>
<tr><td>>hg38_21_+_8433172_8446622_RNA45SN1_45S_50ntFlanks@12405.12430.26|CTTTTTGATCCTTCGATGTCGGCTCT</td><td>>hg38_21_+_8433172_8446622_RNA45SN1_45S_50ntFlanks</td><td>12405</td><td>12430</td><td>26</td><td>CTTTTTGATCCTTCGATGTCGGCTCT</td><td>762.2</td><td>-4.5</td><td>0.6</td><td>-7.1</td><td>1.2E-12</td><td>2.6E-12</td><td>BT20/MB468</td></tr>
<tr><td>>hg38_21_+_8433172_8446622_RNA45SN1_45S_50ntFlanks@4724.4759.36|ATTAATCAAGAACGAAAGTCGGAGGTTCGAAGACGA</td><td>>hg38_21_+_8433172_8446622_RNA45SN1_45S_50ntFlanks</td><td>4724</td><td>4759</td><td>36</td><td>ATTAATCAAGAACGAAAGTCGGAGGTTCGAAGACGA</td><td>1233.1</td><td>-5.7</td><td>0.8</td><td>-7.1</td><td>1.2E-12</td><td>2.6E-12</td><td>BT20/MB468</td></tr>
<tr><td>>hg38_21_+_8433172_8446622_RNA45SN1_45S_50ntFlanks@4738.4767.30|AAAGTCGGAGGTTCGAAGACGATCAGATAC</td><td>>hg38_21_+_8433172_8446622_RNA45SN1_45S_50ntFlanks</td><td>4738</td><td>4767</td><td>30</td><td>AAAGTCGGAGGTTCGAAGACGATCAGATAC</td><td>1376.5</td><td>9.4</td><td>1.3</td><td>7.1</td><td>1.1E-12</td><td>2.6E-12</td><td>MB231/MB468</td></tr>
<tr><td>>hg38_21_+_8433172_8446622_RNA45SN1_45S_50ntFlanks@11601.11619.19|ATCCGACTGTTTAATTAAA</td><td>>hg38_21_+_8433172_8446622_RNA45SN1_45S_50ntFlanks</td><td>11601</td><td>11619</td><td>19</td><td>ATCCGACTGTTTAATTAAA</td><td>122.5</td><td>-8.9</td><td>1.3</td><td>-7.1</td><td>1.2E-12</td><td>2.6E-12</td><td>BT20/MB468</td></tr>
<tr><td>>hg38_21_+_8433172_8446622_RNA45SN1_45S_50ntFlanks@12934.12969.36|CTAAACCATTCGTAGACGACCTGCTTCTGGGTCGGG</td><td>>hg38_21_+_8433172_8446622_RNA45SN1_45S_50ntFlanks</td><td>12934</td><td>12969</td><td>36</td><td>CTAAACCATTCGTAGACGACCTGCTTCTGGGTCGGG</td><td>195.2</td><td>-9.6</td><td>1.4</td><td>-7.1</td><td>1.2E-12</td><td>2.6E-12</td><td>BT20/MB468</td></tr>
<tr><td>>hg38_21_+_8433172_8446622_RNA45SN1_45S_50ntFlanks@11783.11802.20|CATGAATGGATGAACGAGAT</td><td>>hg38_21_+_8433172_8446622_RNA45SN1_45S_50ntFlanks</td><td>11783</td><td>11802</td><td>20</td><td>CATGAATGGATGAACGAGAT</td><td>291.1</td><td>-3.0</td><td>0.4</td><td>-7.1</td><td>1.2E-12</td><td>2.6E-12</td><td>BT20/MB468</td></tr>
<tr><td>>hg38_21_+_8433172_8446622_RNA45SN1_45S_50ntFlanks@6788.6806.19|CGCCTGTCTGAGCGTCGCT</td><td>>hg38_21_+_8433172_8446622_RNA45SN1_45S_50ntFlanks</td><td>6788</td><td>6806</td><td>19</td><td>CGCCTGTCTGAGCGTCGCT</td><td>5132.6</td><td>5.3</td><td>0.7</td><td>7.1</td><td>1.2E-12</td><td>2.6E-12</td><td>BT20/MB468</td></tr>
<tr><td>>hg38_21_+_8433172_8446622_RNA45SN1_45S_50ntFlanks@4443.4472.30|CCCTTGCCTCTCGGCGCCCCCTCGATGCTC</td><td>>hg38_21_+_8433172_8446622_RNA45SN1_45S_50ntFlanks</td><td>4443</td><td>4472</td><td>30</td><td>CCCTTGCCTCTCGGCGCCCCCTCGATGCTC</td><td>252.8</td><td>-9.6</td><td>1.3</td><td>-7.1</td><td>1.1E-12</td><td>2.6E-12</td><td>MB231/MB468</td></tr>
<tr><td>>hg38_21_+_8433172_8446622_RNA45SN1_45S_50ntFlanks@11199.11223.25|CCGGCGGCGGCGGCGACTCTGGACG</td><td>>hg38_21_+_8433172_8446622_RNA45SN1_45S_50ntFlanks</td><td>11199</td><td>11223</td><td>25</td><td>CCGGCGGCGGCGGCGACTCTGGACG</td><td>149.5</td><td>-8.8</td><td>1.2</td><td>-7.1</td><td>1.1E-12</td><td>2.7E-12</td><td>MB231/MB468</td></tr>
<tr><td>>hg38_21_+_8433172_8446622_RNA45SN1_45S_50ntFlanks@4560.4578.19|CTAGGAATAATGGAATAGG</td><td>>hg38_21_+_8433172_8446622_RNA45SN1_45S_50ntFlanks</td><td>4560</td><td>4578</td><td>19</td><td>CTAGGAATAATGGAATAGG</td><td>139.4</td><td>-9.1</td><td>1.3</td><td>-7.1</td><td>1.2E-12</td><td>2.7E-12</td><td>BT20/MB468</td></tr>
<tr><td>>hg38_1_-_228634819_228635039_RNA5S12_5S_50ntFlanks@127.170.44|ACTTGGATGGGAGACCGCCTGGGAATACCGGGTGCTGTAGGCTT</td><td>>hg38_1_-_228634819_228635039_RNA5S12_5S_50ntFlanks</td><td>127</td><td>170</td><td>44</td><td>ACTTGGATGGGAGACCGCCTGGGAATACCGGGTGCTGTAGGCTT</td><td>916.7</td><td>10.3</td><td>1.4</td><td>7.1</td><td>1.2E-12</td><td>2.7E-12</td><td>BT20/MB468</td></tr>
<tr><td>>hg38_21_+_8433172_8446622_RNA45SN1_45S_50ntFlanks@11592.11611.20|ACCAGGGGAATCCGACTGTT</td><td>>hg38_21_+_8433172_8446622_RNA45SN1_45S_50ntFlanks</td><td>11592</td><td>11611</td><td>20</td><td>ACCAGGGGAATCCGACTGTT</td><td>411.1</td><td>-4.7</td><td>0.7</td><td>-7.1</td><td>1.1E-12</td><td>2.7E-12</td><td>MB231/MB468</td></tr>
<tr><td>>hg38_21_+_8433172_8446622_RNA45SN1_45S_50ntFlanks@11575.11606.32|ACTTAGAACTGGTGCGGACCAGGGGAATCCGA</td><td>>hg38_21_+_8433172_8446622_RNA45SN1_45S_50ntFlanks</td><td>11575</td><td>11606</td><td>32</td><td>ACTTAGAACTGGTGCGGACCAGGGGAATCCGA</td><td>170.9</td><td>-3.9</td><td>0.6</td><td>-7.1</td><td>1.2E-12</td><td>2.7E-12</td><td>BT20/MB468</td></tr>
<tr><td>>MI0000268|hsa-mir-34a&WithFlank&1|-|9151662|9151783@28.50.23&[MIMAT0000255&hsa-miR-34a-5p&offsets|0|+1;m-119&1|-|9151735|9151756&offsets|0|+1]|TGGCAGTGTCTTAGCTGGTTGTT</td><td>>MI0000268|hsa-mir-34a&WithFlank&1|-|9151662|9151783</td><td>28</td><td>50</td><td>23</td><td>TGGCAGTGTCTTAGCTGGTTGTT</td><td>1175.0</td><td>7.2</td><td>1.0</td><td>7.1</td><td>1.2E-12</td><td>2.7E-12</td><td>BT20/MB468</td></tr>
<tr><td>>hg38_21_+_8433172_8446622_RNA45SN1_45S_50ntFlanks@3731.3748.18|ATGCTTGTCTCAAAGATT</td><td>>hg38_21_+_8433172_8446622_RNA45SN1_45S_50ntFlanks</td><td>3731</td><td>3748</td><td>18</td><td>ATGCTTGTCTCAAAGATT</td><td>503.9</td><td>9.1</td><td>1.3</td><td>7.1</td><td>1.3E-12</td><td>2.7E-12</td><td>BT20/MB468</td></tr>
<tr><td>>hg38_21_+_8433172_8446622_RNA45SN1_45S_50ntFlanks@6738.6758.21|ATCATCGACACTTCGAACGCA</td><td>>hg38_21_+_8433172_8446622_RNA45SN1_45S_50ntFlanks</td><td>6738</td><td>6758</td><td>21</td><td>ATCATCGACACTTCGAACGCA</td><td>185.1</td><td>-9.5</td><td>1.3</td><td>-7.1</td><td>1.3E-12</td><td>2.7E-12</td><td>BT20/MB468</td></tr>
<tr><td>>MI0003780|hsa-mir-1296&WithFlank&10|-|63372951|63373054@22.43.22&[MIMAT0005794&hsa-miR-1296-5p&offsets|0|0;m-304&10|-|63373012|63373033&offsets|0|0]|TTAGGGCCCTGGCTCCATCTCC</td><td>>MI0003780|hsa-mir-1296&WithFlank&10|-|63372951|63373054</td><td>22</td><td>43</td><td>22</td><td>TTAGGGCCCTGGCTCCATCTCC</td><td>1042.1</td><td>8.5</td><td>1.2</td><td>7.1</td><td>1.3E-12</td><td>2.8E-12</td><td>BT20/MB468</td></tr>
<tr><td>>hg38_21_+_8433172_8446622_RNA45SN1_45S_50ntFlanks@10689.10710.22|GCGTCCGGTGAGCTCTCGCTGG</td><td>>hg38_21_+_8433172_8446622_RNA45SN1_45S_50ntFlanks</td><td>10689</td><td>10710</td><td>22</td><td>GCGTCCGGTGAGCTCTCGCTGG</td><td>118.0</td><td>-8.9</td><td>1.3</td><td>-7.1</td><td>1.3E-12</td><td>2.8E-12</td><td>BT20/MB468</td></tr>
<tr><td>>hg38_21_+_8433172_8446622_RNA45SN1_45S_50ntFlanks@10731.10752.22|AGGGTGTAAATCTCGCGCCGGG</td><td>>hg38_21_+_8433172_8446622_RNA45SN1_45S_50ntFlanks</td><td>10731</td><td>10752</td><td>22</td><td>AGGGTGTAAATCTCGCGCCGGG</td><td>170.3</td><td>-9.0</td><td>1.3</td><td>-7.1</td><td>1.2E-12</td><td>2.8E-12</td><td>MB231/MB468</td></tr>
<tr><td>>hg38_21_+_8433172_8446622_RNA45SN1_45S_50ntFlanks@10461.10497.37|GCCTCCGTTGCCCTCGGCCGATCGAAAGGGAGTCGGG</td><td>>hg38_21_+_8433172_8446622_RNA45SN1_45S_50ntFlanks</td><td>10461</td><td>10497</td><td>37</td><td>GCCTCCGTTGCCCTCGGCCGATCGAAAGGGAGTCGGG</td><td>125.7</td><td>-9.0</td><td>1.3</td><td>-7.1</td><td>1.3E-12</td><td>2.8E-12</td><td>BT20/MB468</td></tr>
<tr><td>>hg38_21_+_8433172_8446622_RNA45SN1_45S_50ntFlanks@6772.6793.22|GTTCCTCCCGGGGCTACGCCTG</td><td>>hg38_21_+_8433172_8446622_RNA45SN1_45S_50ntFlanks</td><td>6772</td><td>6793</td><td>22</td><td>GTTCCTCCCGGGGCTACGCCTG</td><td>161.0</td><td>-9.3</td><td>1.3</td><td>-7.1</td><td>1.3E-12</td><td>2.8E-12</td><td>BT20/MB468</td></tr>
<tr><td>>MI0000285|hsa-mir-205&WithFlank&1|+|209432127|209432248@40.61.22(+1A)&[MIMAT0000266&hsa-miR-205-5p&offsets|0|0(+1A);m-351&1|+|209432166|209432187&offsets|0|0(+1A)]|TCCTTCATTCCACCGGAGTCTGA</td><td>>MI0000285|hsa-mir-205&WithFlank&1|+|209432127|209432248</td><td>40</td><td>61</td><td>22(+1A)</td><td>TCCTTCATTCCACCGGAGTCTGA</td><td>86324.2</td><td>6.1</td><td>0.9</td><td>7.1</td><td>1.3E-12</td><td>2.8E-12</td><td>BT20/MB468</td></tr>
<tr><td>>trnaMT_PheGAA_MT_+_577_647@39.57.19|ATGTTTAGACGGGCTCACA</td><td>>trnaMT_PheGAA_MT_+_577_647</td><td>39</td><td>57</td><td>19</td><td>ATGTTTAGACGGGCTCACA</td><td>1145.8</td><td>8.8</td><td>1.2</td><td>7.1</td><td>1.2E-12</td><td>2.8E-12</td><td>MB231/MB468</td></tr>
<tr><td>>hg38_21_+_8433172_8446622_RNA45SN1_45S_50ntFlanks@12643.12661.19|ATCTGTGGGATTATGACTG</td><td>>hg38_21_+_8433172_8446622_RNA45SN1_45S_50ntFlanks</td><td>12643</td><td>12661</td><td>19</td><td>ATCTGTGGGATTATGACTG</td><td>150.0</td><td>-9.2</td><td>1.3</td><td>-7.1</td><td>1.3E-12</td><td>2.8E-12</td><td>BT20/MB468</td></tr>
<tr><td>>hg38_21_+_8433172_8446622_RNA45SN1_45S_50ntFlanks@4043.4063.21|ACCCATTCGAACGTCTGCCCT</td><td>>hg38_21_+_8433172_8446622_RNA45SN1_45S_50ntFlanks</td><td>4043</td><td>4063</td><td>21</td><td>ACCCATTCGAACGTCTGCCCT</td><td>107.0</td><td>-8.7</td><td>1.2</td><td>-7.1</td><td>1.3E-12</td><td>2.8E-12</td><td>BT20/MB468</td></tr>
<tr><td>>hg38_21_+_8433172_8446622_RNA45SN1_45S_50ntFlanks@4724.4752.29|ATTAATCAAGAACGAAAGTCGGAGGTTCG</td><td>>hg38_21_+_8433172_8446622_RNA45SN1_45S_50ntFlanks</td><td>4724</td><td>4752</td><td>29</td><td>ATTAATCAAGAACGAAAGTCGGAGGTTCG</td><td>451.2</td><td>-4.8</td><td>0.7</td><td>-7.1</td><td>1.2E-12</td><td>2.9E-12</td><td>MB231/MB468</td></tr>
</table>

<table>
<tr><td>>hg38_21_+_8433172_8446622_RNA45SN1_45S_50ntFlanks@8168.8188.21|CCCAAGTCCTTCTGATCGAGG</td><td>>hg38_21_+_8433172_8446622_RNA45SN1_45S_50ntFlanks</td><td>8168</td><td>8188</td><td>21</td><td>CCCAAGTCCTTCTGATCGAGG</td><td>452.3</td><td>-10.8</td><td>1.5</td><td>-7.1</td><td>1.3E-12</td><td>2.9E-12</td><td>BT20/MB468</td></tr>
<tr><td>>hg38_21_+_8433172_8446622_RNA45SN1_45S_50ntFlanks@10116.10136.21|TGCGTCGGAGGGCGGCGGCGG</td><td>>hg38_21_+_8433172_8446622_RNA45SN1_45S_50ntFlanks</td><td>10116</td><td>10136</td><td>21</td><td>TGCGTCGGAGGGCGGCGGCGG</td><td>148.3</td><td>-8.8</td><td>1.2</td><td>-7.1</td><td>1.2E-12</td><td>2.9E-12</td><td>MB231/MB468</td></tr>
<tr><td>>trnaMT_GluTTC_MT_-_14674_14742@39.63.25,trnalookalike8_GluTTC_5_-_93905172_93905240@39.63.25|ATTGGTCGTGGTTGTAGTCCGTGCG</td><td>>trnaMT_GluTTC_MT_-_14674_14742</td><td>39</td><td>63</td><td>25</td><td>ATTGGTCGTGGTTGTAGTCCGTGCG</td><td>542.9</td><td>8.3</td><td>1.2</td><td>7.1</td><td>1.3E-12</td><td>2.9E-12</td><td>BT20/MB468</td></tr>
<tr><td>>trnaMT_GluTTC_MT_-_14674_14742@33.72.40,trnalookalike8_GluTTC_5_-_93905172_93905240@33.72.40|CATATCATTGGTCGTGGTTGTAGTCCGTGCGAGAATACCA</td><td>>trnaMT_GluTTC_MT_-_14674_14742</td><td>33</td><td>72</td><td>40</td><td>CATATCATTGGTCGTGGTTGTAGTCCGTGCGAGAATACCA</td><td>813.8</td><td>7.4</td><td>1.0</td><td>7.1</td><td>1.3E-12</td><td>2.9E-12</td><td>BT20/MB468</td></tr>
<tr><td>>MI0000255|hsa-mir-30d&WithFlank&8|-|134804870|134804951@12.36.25&[MIMAT0000245&hsa-miR-30d-5p&offsets|0|+3;m-21&8|-|134804919|134804940&offsets|0|+3]|TGTAAACATCCCCGACTGGAAGCTG</td><td>>MI0000255|hsa-mir-30d&WithFlank&8|-|134804870|134804951</td><td>12</td><td>36</td><td>25</td><td>TGTAAACATCCCCGACTGGAAGCTG</td><td>599.9</td><td>9.3</td><td>1.3</td><td>7.1</td><td>1.4E-12</td><td>2.9E-12</td><td>BT20/MB468</td></tr>
<tr><td>>hg38_MT_+_1621_3279_RNR2_16S_50ntFlanks@1438.1457.20|TACGTGATCTGAGTTCAGAC</td><td>>hg38_MT_+_1621_3279_RNR2_16S_50ntFlanks</td><td>1438</td><td>1457</td><td>20</td><td>TACGTGATCTGAGTTCAGAC</td><td>417.4</td><td>8.2</td><td>1.2</td><td>7.1</td><td>1.4E-12</td><td>2.9E-12</td><td>BT20/MB468</td></tr>
<tr><td>>MI0000084|hsa-mir-26b&WithFlank&2|+|218402640|218402728@18.39.22(+1A)&[MIMAT0000083&hsa-miR-26b-5p&offsets|0|+1(+1A);m-53&2|+|218402657|218402678&offsets|0|0(+1A)]|TTCAAGTAATTCAGGATAGGTTA</td><td>>MI0000084|hsa-mir-26b&WithFlank&2|+|218402640|218402728</td><td>18</td><td>39</td><td>22(+1A)</td><td>TTCAAGTAATTCAGGATAGGTTA</td><td>1050.3</td><td>7.4</td><td>1.0</td><td>7.1</td><td>1.4E-12</td><td>2.9E-12</td><td>BT20/MB468</td></tr>
<tr><td>>hg38_21_+_8433172_8446622_RNA45SN1_45S_50ntFlanks@9736.9757.22|CCGAAACGATCTCAACCTATTC</td><td>>hg38_21_+_8433172_8446622_RNA45SN1_45S_50ntFlanks</td><td>9736</td><td>9757</td><td>22</td><td>CCGAAACGATCTCAACCTATTC</td><td>113.8</td><td>-8.8</td><td>1.2</td><td>-7.1</td><td>1.4E-12</td><td>3.0E-12</td><td>BT20/MB468</td></tr>
<tr><td>>hg38_21_+_8433172_8446622_RNA45SN1_45S_50ntFlanks@9534.9559.26|AGGAAACTCTGGTGGAGGTCCGTAGC</td><td>>hg38_21_+_8433172_8446622_RNA45SN1_45S_50ntFlanks</td><td>9534</td><td>9559</td><td>26</td><td>AGGAAACTCTGGTGGAGGTCCGTAGC</td><td>113.1</td><td>-3.7</td><td>0.5</td><td>-7.1</td><td>1.4E-12</td><td>3.0E-12</td><td>BT20/MB468</td></tr>
<tr><td>>hg38_21_+_8433172_8446622_RNA45SN1_45S_50ntFlanks@5467.5484.18|GGCCCACGGCCCTGGCGG</td><td>>hg38_21_+_8433172_8446622_RNA45SN1_45S_50ntFlanks</td><td>5467</td><td>5484</td><td>18</td><td>GGCCCACGGCCCTGGCGG</td><td>171.0</td><td>-9.0</td><td>1.3</td><td>-7.1</td><td>1.2E-12</td><td>3.0E-12</td><td>MB231/MB468</td></tr>
<tr><td>>hg38_21_+_8433172_8446622_RNA45SN1_45S_50ntFlanks@4002.4042.41|GATAACCTCGGGCCGATCGCACGCCCCCCGTGGCGGCGACG</td><td>>hg38_21_+_8433172_8446622_RNA45SN1_45S_50ntFlanks</td><td>4002</td><td>4042</td><td>41</td><td>GATAACCTCGGGCCGATCGCACGCCCCCCGTGGCGGCGACG</td><td>138.7</td><td>-9.1</td><td>1.3</td><td>-7.1</td><td>1.4E-12</td><td>3.0E-12</td><td>BT20/MB468</td></tr>
<tr><td>>hg38_21_+_8433172_8446622_RNA45SN1_45S_50ntFlanks@12957.12987.31|CTTCTGGGTCGGGGTTTCGTACGTAGCAGAG</td><td>>hg38_21_+_8433172_8446622_RNA45SN1_45S_50ntFlanks</td><td>12957</td><td>12987</td><td>31</td><td>CTTCTGGGTCGGGGTTTCGTACGTAGCAGAG</td><td>219.0</td><td>-3.9</td><td>0.6</td><td>-7.1</td><td>1.2E-12</td><td>3.0E-12</td><td>MB231/MB468</td></tr>
<tr><td>>hg38_21_+_8433172_8446622_RNA45SN1_45S_50ntFlanks@8179.8211.33|CTGATCGAGGCCCAGCCCGTGGACGGTGTGAGG</td><td>>hg38_21_+_8433172_8446622_RNA45SN1_45S_50ntFlanks</td><td>8179</td><td>8211</td><td>33</td><td>CTGATCGAGGCCCAGCCCGTGGACGGTGTGAGG</td><td>211.3</td><td>-9.3</td><td>1.3</td><td>-7.1</td><td>1.3E-12</td><td>3.0E-12</td><td>MB231/MB468</td></tr>
<tr><td>>hg38_21_+_8433172_8446622_RNA45SN1_45S_50ntFlanks@11568.11611.44|GCAGCCGACTTAGAACTGGTGCGGACCAGGGGAATCCGACTGTT</td><td>>hg38_21_+_8433172_8446622_RNA45SN1_45S_50ntFlanks</td><td>11568</td><td>11611</td><td>44</td><td>GCAGCCGACTTAGAACTGGTGCGGACCAGGGGAATCCGACTGTT</td><td>3138.1</td><td>6.4</td><td>0.9</td><td>7.1</td><td>1.4E-12</td><td>3.0E-12</td><td>BT20/MB468</td></tr>
<tr><td>>hg38_21_+_8433172_8446622_RNA45SN1_45S_50ntFlanks@11792.11810.19|ATGAACGAGATTCCCACTG</td><td>>hg38_21_+_8433172_8446622_RNA45SN1_45S_50ntFlanks</td><td>11792</td><td>11810</td><td>19</td><td>ATGAACGAGATTCCCACTG</td><td>176.2</td><td>-4.8</td><td>0.7</td><td>-7.1</td><td>1.3E-12</td><td>3.1E-12</td><td>MB231/MB468</td></tr>
<tr><td>>hg38_21_+_8433172_8446622_RNA45SN1_45S_50ntFlanks@4240.4258.19|ACTCTTTCGAGGCCCTGTA</td><td>>hg38_21_+_8433172_8446622_RNA45SN1_45S_50ntFlanks</td><td>4240</td><td>4258</td><td>19</td><td>ACTCTTTCGAGGCCCTGTA</td><td>233.0</td><td>-6.1</td><td>0.9</td><td>-7.1</td><td>1.4E-12</td><td>3.1E-12</td><td>BT20/MB468</td></tr>
<tr><td>>hg38_21_+_8433172_8446622_RNA45SN1_45S_50ntFlanks@12008.12039.32|CCGCCGGTGAAATACCACTACTCTGATCGTTT</td><td>>hg38_21_+_8433172_8446622_RNA45SN1_45S_50ntFlanks</td><td>12008</td><td>12039</td><td>32</td><td>CCGCCGGTGAAATACCACTACTCTGATCGTTT</td><td>124.4</td><td>-3.7</td><td>0.5</td><td>-7.1</td><td>1.4E-12</td><td>3.1E-12</td><td>BT20/MB468</td></tr>
<tr><td>>hg38_21_+_8433172_8446622_RNA45SN1_45S_50ntFlanks@3802.3834.33|CATTAAATCAGTTATGGTTCCTTTGGTCGCTCG</td><td>>hg38_21_+_8433172_8446622_RNA45SN1_45S_50ntFlanks</td><td>3802</td><td>3834</td><td>33</td><td>CATTAAATCAGTTATGGTTCCTTTGGTCGCTCG</td><td>203.4</td><td>-3.4</td><td>0.5</td><td>-7.1</td><td>1.3E-12</td><td>3.1E-12</td><td>MB231/MB468</td></tr>
<tr><td>>hg38_21_+_8433172_8446622_RNA45SN1_45S_50ntFlanks@12991.13008.18|CTCCCTCGCTGCGATCTA</td><td>>hg38_21_+_8433172_8446622_RNA45SN1_45S_50ntFlanks</td><td>12991</td><td>13008</td><td>18</td><td>CTCCCTCGCTGCGATCTA</td><td>249.9</td><td>-7.0</td><td>1.0</td><td>-7.1</td><td>1.3E-12</td><td>3.1E-12</td><td>MB231/MB468</td></tr>
<tr><td>>hg38_21_+_8433172_8446622_RNA45SN1_45S_50ntFlanks@11845.11872.28|GGAACGGGCTTGGCGGAATCAGCGGGGA</td><td>>hg38_21_+_8433172_8446622_RNA45SN1_45S_50ntFlanks</td><td>11845</td><td>11872</td><td>28</td><td>GGAACGGGCTTGGCGGAATCAGCGGGGA</td><td>199.9</td><td>-7.4</td><td>1.0</td><td>-7.1</td><td>1.3E-12</td><td>3.1E-12</td><td>MB231/MB468</td></tr>
<tr><td>>hg38_21_+_8433172_8446622_RNA45SN1_45S_50ntFlanks@11845.11872.28|GGAACGGGCTTGGCGGAATCAGCGGGGA</td><td>>hg38_21_+_8433172_8446622_RNA45SN1_45S_50ntFlanks</td><td>11845</td><td>11872</td><td>28</td><td>GGAACGGGCTTGGCGGAATCAGCGGGGA</td><td>239.7</td><td>-6.0</td><td>0.8</td><td>-7.1</td><td>1.4E-12</td><td>3.1E-12</td><td>BT20/MB468</td></tr>
<tr><td>>hg38_21_+_8433172_8446622_RNA45SN1_45S_50ntFlanks@12000.12032.33|CCTGCGGGCCGCCGGTGAAATACCACTACTCTG</td><td>>hg38_21_+_8433172_8446622_RNA45SN1_45S_50ntFlanks</td><td>12000</td><td>12032</td><td>33</td><td>CCTGCGGGCCGCCGGTGAAATACCACTACTCTG</td><td>200.5</td><td>-7.4</td><td>1.0</td><td>-7.1</td><td>1.3E-12</td><td>3.1E-12</td><td>MB231/MB468</td></tr>
<tr><td>>hg38_21_+_8433172_8446622_RNA45SN1_45S_50ntFlanks@10101.10125.25|CGCGCGCGTGTGGTGTGCGTCGGAG</td><td>>hg38_21_+_8433172_8446622_RNA45SN1_45S_50ntFlanks</td><td>10101</td><td>10125</td><td>25</td><td>CGCGCGCGTGTGGTGTGCGTCGGAG</td><td>261.5</td><td>-9.6</td><td>1.4</td><td>-7.1</td><td>1.3E-12</td><td>3.1E-12</td><td>MB231/MB468</td></tr>
<tr><td>>hg38_21_+_8433172_8446622_RNA45SN1_45S_50ntFlanks@11212.11231.20|CGACTCTGGACGCGAGCCGG</td><td>>hg38_21_+_8433172_8446622_RNA45SN1_45S_50ntFlanks</td><td>11212</td><td>11231</td><td>20</td><td>CGACTCTGGACGCGAGCCGG</td><td>161.1</td><td>-5.8</td><td>0.8</td><td>-7.1</td><td>1.3E-12</td><td>3.1E-12</td><td>MB231/MB468</td></tr>
<tr><td>>MI0000749|hsa-mir-30e&WithFlank&1|+|40754349|40754452@23.46.24(+1U)&[MIMAT0000692&hsa-miR-30e-5p&offsets|0|+2(+1U);m-24&1|+|40754371|40754394&offsets|0|0(+1U)]|TGTAAACATCCTTGACTGGAAGCTT</td><td>>MI0000749|hsa-mir-30e&WithFlank&1|+|40754349|40754452</td><td>23</td><td>46</td><td>24(+1U)</td><td>TGTAAACATCCTTGACTGGAAGCTT</td><td>1442.2</td><td>5.6</td><td>0.8</td><td>7.1</td><td>1.3E-12</td><td>3.1E-12</td><td>MB231/MB468</td></tr>
<tr><td>>hg38_21_+_8433172_8446622_RNA45SN1_45S_50ntFlanks@10878.10901.24|TCGGTCGGGCTGGGGCGCGAAGCG</td><td>>hg38_21_+_8433172_8446622_RNA45SN1_45S_50ntFlanks</td><td>10878</td><td>10901</td><td>24</td><td>TCGGTCGGGCTGGGGCGCGAAGCG</td><td>141.2</td><td>-6.2</td><td>0.9</td><td>-7.1</td><td>1.3E-12</td><td>3.2E-12</td><td>MB231/MB468</td></tr>
<tr><td>>hg38_21_+_8433172_8446622_RNA45SN1_45S_50ntFlanks@13006.13025.20|CTATTGAAAGTCAGCCCTCG</td><td>>hg38_21_+_8433172_8446622_RNA45SN1_45S_50ntFlanks</td><td>13006</td><td>13025</td><td>20</td><td>CTATTGAAAGTCAGCCCTCG</td><td>757.3</td><td>-8.2</td><td>1.2</td><td>-7.1</td><td>1.5E-12</td><td>3.2E-12</td><td>BT20/MB468</td></tr>
<tr><td>>hg38_21_+_8433172_8446622_RNA45SN1_45S_50ntFlanks@7986.8014.29|ATCAGACGTGGCGACCCGCTGAATTTAAG</td><td>>hg38_21_+_8433172_8446622_RNA45SN1_45S_50ntFlanks</td><td>7986</td><td>8014</td><td>29</td><td>ATCAGACGTGGCGACCCGCTGAATTTAAG</td><td>129.2</td><td>-9.0</td><td>1.3</td><td>-7.1</td><td>1.5E-12</td><td>3.2E-12</td><td>BT20/MB468</td></tr>
<tr><td>>MI0000065|hsa-let-7d&WithFlank&9|+|94178828|94178926@14.34.21(+1A)&[MIMAT0000065&hsa-let-7d-5p&offsets|0|-1(+1A);m-90&9|+|94178841|94178862&offsets|0|-1(+1A)]|AGAGGTAGTAGGTTGCATAGTA</td><td>>MI0000065|hsa-let-7d&WithFlank&9|+|94178828|94178926</td><td>14</td><td>34</td><td>21(+1A)</td><td>AGAGGTAGTAGGTTGCATAGTA</td><td>1618.5</td><td>10.4</td><td>1.5</td><td>7.1</td><td>1.5E-12</td><td>3.2E-12</td><td>BT20/MB468</td></tr>
<tr><td>>hg38_1_-_228634819_228635039_RNA5S12_5S_50ntFlanks@51.92.42|GTCTACGGCCATACCACCCTGAACGCGCCCGATCTCGTCTGA</td><td>>hg38_1_-_228634819_228635039_RNA5S12_5S_50ntFlanks</td><td>51</td><td>92</td><td>42</td><td>GTCTACGGCCATACCACCCTGAACGCGCCCGATCTCGTCTGA</td><td>488.1</td><td>10.5</td><td>1.5</td><td>7.1</td><td>1.5E-12</td><td>3.2E-12</td><td>BT20/MB468</td></tr>
<tr><td>>hg38_1_-_228634819_228635039_RNA5S12_5S_50ntFlanks@129.171.43|TTGGATGGGAGACCGCCTGGGAATACCGGGTGCTGTAGGCTTT</td><td>>hg38_1_-_228634819_228635039_RNA5S12_5S_50ntFlanks</td><td>129</td><td>171</td><td>43</td><td>TTGGATGGGAGACCGCCTGGGAATACCGGGTGCTGTAGGCTTT</td><td>1024.8</td><td>11.4</td><td>1.6</td><td>7.1</td><td>1.5E-12</td><td>3.2E-12</td><td>BT20/MB468</td></tr>
<tr><td>>am_trna10_LysCTT_16_+_3241501_3241573@32.51.20,trna119_LysCTT_1_-_145395522_145395594@32.51.20,trna11_LysCTT_5_-_180648979_180649051@32.51.20,trna13_LysCTT_16_+_3246153_3246225@32.51.20,trna13_LysCTT_6_+_26556774_26556846@32.51.20,trna30_LysCTT_16_-_3230555_3230627@32.51.20,trna5_LysCTT_16_+_3214939_3215011@32.51.20,trna7_LysCTT_16_+_3225692_3225764@32.51.20,trna8_LysCTT_1_+_39970195_39970267@32.51.20,trna9_LysCTT_5_+_180634755_180634827@32.51.20|CTCTTAATCTCAGGGTCGTG</td><td>>am_trna10_LysCTT_16_+_3241501_3241573</td><td>32</td><td>51</td><td>20</td><td>CTCTTAATCTCAGGGTCGTG</td><td>170.6</td><td>-9.4</td><td>1.3</td><td>-7.1</td><td>1.5E-12</td><td>3.2E-12</td><td>BT20/MB468</td></tr>
<tr><td>>hg38_21_+_8433172_8446622_RNA45SN1_45S_50ntFlanks@12934.12961.28|CTAAACCATTCGTAGACGACCTGCTTCT</td><td>>hg38_21_+_8433172_8446622_RNA45SN1_45S_50ntFlanks</td><td>12934</td><td>12961</td><td>28</td><td>CTAAACCATTCGTAGACGACCTGCTTCT</td><td>427.5</td><td>-5.2</td><td>0.7</td><td>-7.1</td><td>1.5E-12</td><td>3.2E-12</td><td>BT20/MB468</td></tr>
<tr><td>>hg38_21_+_8433172_8446622_RNA45SN1_45S_50ntFlanks@4759.4776.18|ATCAGATACCGTCGTAGT</td><td>>hg38_21_+_8433172_8446622_RNA45SN1_45S_50ntFlanks</td><td>4759</td><td>4776</td><td>18</td><td>ATCAGATACCGTCGTAGT</td><td>170.8</td><td>-9.4</td><td>1.3</td><td>-7.1</td><td>1.5E-12</td><td>3.3E-12</td><td>BT20/MB468</td></tr>
<tr><td>>hg38_21_+_8433172_8446622_RNA45SN1_45S_50ntFlanks@11602.11620.19|TCCGACTGTTTAATTAAAA</td><td>>hg38_21_+_8433172_8446622_RNA45SN1_45S_50ntFlanks</td><td>11602</td><td>11620</td><td>19</td><td>TCCGACTGTTTAATTAAAA</td><td>599.9</td><td>-4.2</td><td>0.6</td><td>-7.1</td><td>1.5E-12</td><td>3.3E-12</td><td>BT20/MB468</td></tr>
<tr><td>>hg38_21_+_8433172_8446622_RNA45SN1_45S_50ntFlanks@10436.10453.18|CGCCGTTCCGAAGGGACG</td><td>>hg38_21_+_8433172_8446622_RNA45SN1_45S_50ntFlanks</td><td>10436</td><td>10453</td><td>18</td><td>CGCCGTTCCGAAGGGACG</td><td>288.7</td><td>-9.7</td><td>1.4</td><td>-7.1</td><td>1.4E-12</td><td>3.3E-12</td><td>MB231/MB468</td></tr>
<tr><td>>hg38_21_+_8433172_8446622_RNA45SN1_45S_50ntFlanks@9738.9759.22|GAAACGATCTCAACCTATTCTC</td><td>>hg38_21_+_8433172_8446622_RNA45SN1_45S_50ntFlanks</td><td>9738</td><td>9759</td><td>22</td><td>GAAACGATCTCAACCTATTCTC</td><td>443.9</td><td>-3.2</td><td>0.4</td><td>-7.1</td><td>1.4E-12</td><td>3.3E-12</td><td>MB231/MB468</td></tr>
<tr><td>>hg38_21_+_8433172_8446622_RNA45SN1_45S_50ntFlanks@11452.11472.21|CTCCTCGGCGCGCGGCGGCGG</td><td>>hg38_21_+_8433172_8446622_RNA45SN1_45S_50ntFlanks</td><td>11452</td><td>11472</td><td>21</td><td>CTCCTCGGCGCGCGGCGGCGG</td><td>169.5</td><td>-9.0</td><td>1.3</td><td>-7.1</td><td>1.4E-12</td><td>3.3E-12</td><td>MB231/MB468</td></tr>
<tr><td>>hg38_21_+_8433172_8446622_RNA45SN1_45S_50ntFlanks@12151.12181.31|GTGGGGAGTTTGACTGGGGCGGTACACCTGT</td><td>>hg38_21_+_8433172_8446622_RNA45SN1_45S_50ntFlanks</td><td>12151</td><td>12181</td><td>31</td><td>GTGGGGAGTTTGACTGGGGCGGTACACCTGT</td><td>155.6</td><td>-8.9</td><td>1.2</td><td>-7.1</td><td>1.4E-12</td><td>3.3E-12</td><td>MB231/MB468</td></tr>
<tr><td>>hg38_21_+_8433172_8446622_RNA45SN1_45S_50ntFlanks@11886.11903.18|TGAGCTTGACTCTAGTCT</td><td>>hg38_21_+_8433172_8446622_RNA45SN1_45S_50ntFlanks</td><td>11886</td><td>11903</td><td>18</td><td>TGAGCTTGACTCTAGTCT</td><td>213.1</td><td>-6.3</td><td>0.9</td><td>-7.1</td><td>1.5E-12</td><td>3.3E-12</td><td>BT20/MB468</td></tr>
<tr><td>>hg38_21_+_8433172_8446622_RNA45SN1_45S_50ntFlanks@8379.8402.24|TCAAGAGGGCGTGAAACCGTTAAG</td><td>>hg38_21_+_8433172_8446622_RNA45SN1_45S_50ntFlanks</td><td>8379</td><td>8402</td><td>24</td><td>TCAAGAGGGCGTGAAACCGTTAAG</td><td>199.2</td><td>-9.6</td><td>1.4</td><td>-7.1</td><td>1.6E-12</td><td>3.4E-12</td><td>BT20/MB468</td></tr>
<tr><td>>hg38_21_+_8433172_8446622_RNA45SN1_45S_50ntFlanks@10781.10806.26|AGGTGAACAGCCTCTGGCATGTTGGA</td><td>>hg38_21_+_8433172_8446622_RNA45SN1_45S_50ntFlanks</td><td>10781</td><td>10806</td><td>26</td><td>AGGTGAACAGCCTCTGGCATGTTGGA</td><td>192.6</td><td>-9.2</td><td>1.3</td><td>-7.1</td><td>1.4E-12</td><td>3.4E-12</td><td>MB231/MB468</td></tr>
<tr><td>>hg38_21_+_8433172_8446622_RNA45SN1_45S_50ntFlanks@10772.10805.34|AGGTCTCCAAGGTGAACAGCCTCTGGCATGTTGG</td><td>>hg38_21_+_8433172_8446622_RNA45SN1_45S_50ntFlanks</td><td>10772</td><td>10805</td><td>34</td><td>AGGTCTCCAAGGTGAACAGCCTCTGGCATGTTGG</td><td>103.4</td><td>-3.7</td><td>0.5</td><td>-7.1</td><td>1.6E-12</td><td>3.5E-12</td><td>BT20/MB468</td></tr>
<tr><td>>hg38_21_+_8433172_8446622_RNA45SN1_45S_50ntFlanks@7975.7997.23|CGCGACCTCAGATCAGACGTGGC</td><td>>hg38_21_+_8433172_8446622_RNA45SN1_45S_50ntFlanks</td><td>7975</td><td>7997</td><td>23</td><td>CGCGACCTCAGATCAGACGTGGC</td><td>6025.8</td><td>5.5</td><td>0.8</td><td>7.1</td><td>1.4E-12</td><td>3.5E-12</td><td>MB231/MB468</td></tr>
<tr><td>>hg38_21_+_8433172_8446622_RNA45SN1_45S_50ntFlanks@13008.13039.32|ATTGAAAGTCAGCCCTCGACACAAGGGTTTGT</td><td>>hg38_21_+_8433172_8446622_RNA45SN1_45S_50ntFlanks</td><td>13008</td><td>13039</td><td>32</td><td>ATTGAAAGTCAGCCCTCGACACAAGGGTTTGT</td><td>3762.5</td><td>4.8</td><td>0.7</td><td>7.1</td><td>1.6E-12</td><td>3.5E-12</td><td>BT20/MB468</td></tr>
</table>

<table>
<tr><td>>hg38_21_+_8433172_8446622_RNA45SN1_45S_50ntFlanks@11918.11950.33|ACATGAGAGGTGTAGAATAAGTGGGAGGCCCCC</td><td>>hg38_21_+_8433172_8446622_RNA45SN1_45S_50ntFlanks</td><td>11918</td><td>11950</td><td>33</td><td>ACATGAGAGGTGTAGAATAAGTGGGAGGCCCCC</td><td>250.1</td><td>-7.7</td><td>1.1</td><td>-7.1</td><td>1.5E-12</td><td>3.5E-12</td><td>MB231/MB468</td></tr>
<tr><td>>hg38_21_+_8433172_8446622_RNA45SN1_45S_50ntFlanks@10679.10705.27|GGTTCCGGCGGCGTCCGGTGAGCTCTC</td><td>>hg38_21_+_8433172_8446622_RNA45SN1_45S_50ntFlanks</td><td>10679</td><td>10705</td><td>27</td><td>GGTTCCGGCGGCGTCCGGTGAGCTCTC</td><td>224.8</td><td>-7.6</td><td>1.1</td><td>-7.1</td><td>1.5E-12</td><td>3.5E-12</td><td>MB231/MB468</td></tr>
<tr><td>>hg38_21_+_8433172_8446622_RNA45SN1_45S_50ntFlanks@9534.9555.22|AGGAAACTCTGGTGGAGGTCCG</td><td>>hg38_21_+_8433172_8446622_RNA45SN1_45S_50ntFlanks</td><td>9534</td><td>9555</td><td>22</td><td>AGGAAACTCTGGTGGAGGTCCG</td><td>200.1</td><td>-9.2</td><td>1.3</td><td>-7.1</td><td>1.5E-12</td><td>3.5E-12</td><td>MB231/MB468</td></tr>
<tr><td>>hg38_21_+_8433172_8446622_RNA45SN1_45S_50ntFlanks@12637.12654.18|GCTACCATCTGTGGGATT</td><td>>hg38_21_+_8433172_8446622_RNA45SN1_45S_50ntFlanks</td><td>12637</td><td>12654</td><td>18</td><td>GCTACCATCTGTGGGATT</td><td>899.4</td><td>-5.8</td><td>0.8</td><td>-7.1</td><td>1.5E-12</td><td>3.5E-12</td><td>MB231/MB468</td></tr>
<tr><td>>hg38_21_+_8433172_8446622_RNA45SN1_45S_50ntFlanks@10387.10424.38|TGTGAACAGCAGTTGAACATGGGTCAGTCGGTCCTGAG</td><td>>hg38_21_+_8433172_8446622_RNA45SN1_45S_50ntFlanks</td><td>10387</td><td>10424</td><td>38</td><td>TGTGAACAGCAGTTGAACATGGGTCAGTCGGTCCTGAG</td><td>881.0</td><td>7.9</td><td>1.1</td><td>7.1</td><td>1.7E-12</td><td>3.6E-12</td><td>BT20/MB468</td></tr>
<tr><td>>hg38_21_+_8433172_8446622_RNA45SN1_45S_50ntFlanks@6667.6699.33|ATCACTCGGCTCGTGCGTCGATGAAGAACGCAG</td><td>>hg38_21_+_8433172_8446622_RNA45SN1_45S_50ntFlanks</td><td>6667</td><td>6699</td><td>33</td><td>ATCACTCGGCTCGTGCGTCGATGAAGAACGCAG</td><td>122.2</td><td>-8.9</td><td>1.3</td><td>-7.1</td><td>1.7E-12</td><td>3.6E-12</td><td>BT20/MB468</td></tr>
<tr><td>>hg38_21_+_8433172_8446622_RNA45SN1_45S_50ntFlanks@12994.13013.20|CCTCGCTGCGATCTATTGAA</td><td>>hg38_21_+_8433172_8446622_RNA45SN1_45S_50ntFlanks</td><td>12994</td><td>13013</td><td>20</td><td>CCTCGCTGCGATCTATTGAA</td><td>702.3</td><td>-4.8</td><td>0.7</td><td>-7.1</td><td>1.7E-12</td><td>3.6E-12</td><td>BT20/MB468</td></tr>
<tr><td>>MI0000095|hsa-mir-93&WithFlank&7|-|100093762|100093853@17.39.23&[MIMAT0000093&hsa-miR-93-5p&offsets|0|0;m-29&7|-|100093815|100093837&offsets|0|0]|CAAAGTGCTGTTCGTGCAGGTAG</td><td>>MI0000095|hsa-mir-93&WithFlank&7|-|100093762|100093853</td><td>17</td><td>39</td><td>23</td><td>CAAAGTGCTGTTCGTGCAGGTAG</td><td>22074.5</td><td>5.7</td><td>0.8</td><td>7.1</td><td>1.7E-12</td><td>3.6E-12</td><td>BT20/MB468</td></tr>
<tr><td>>hg38_21_+_8433172_8446622_RNA45SN1_45S_50ntFlanks@9573.9602.30|AAATCGGTCGTCCGACCTGGGTATAGGGGC</td><td>>hg38_21_+_8433172_8446622_RNA45SN1_45S_50ntFlanks</td><td>9573</td><td>9602</td><td>30</td><td>AAATCGGTCGTCCGACCTGGGTATAGGGGC</td><td>384.9</td><td>3.5</td><td>0.5</td><td>7.1</td><td>1.5E-12</td><td>3.6E-12</td><td>MB231/MB468</td></tr>
<tr><td>>MI0000086|hsa-mir-28&WithFlank&3|+|188688775|188688872@20.40.21&[MIMAT0000085&hsa-miR-28-5p&offsets|0|-1;m-133&3|+|188688794|188688815&offsets|0|-1]|AAGGAGCTCACAGTCTATTGA</td><td>>MI0000086|hsa-mir-28&WithFlank&3|+|188688775|188688872</td><td>20</td><td>40</td><td>21</td><td>AAGGAGCTCACAGTCTATTGA</td><td>801.1</td><td>9.3</td><td>1.3</td><td>7.1</td><td>1.7E-12</td><td>3.6E-12</td><td>BT20/MB468</td></tr>
<tr><td>>hg38_21_+_8433172_8446622_RNA45SN1_45S_50ntFlanks@12161.12179.19|TGACTGGGGCGGTACACCT</td><td>>hg38_21_+_8433172_8446622_RNA45SN1_45S_50ntFlanks</td><td>12161</td><td>12179</td><td>19</td><td>TGACTGGGGCGGTACACCT</td><td>333.5</td><td>-6.3</td><td>0.9</td><td>-7.1</td><td>1.5E-12</td><td>3.6E-12</td><td>MB231/MB468</td></tr>
<tr><td>>hg38_21_+_8433172_8446622_RNA45SN1_45S_50ntFlanks@11874.11907.34|AGAAGACCCTGTTGAGCTTGACTCTAGTCTGGCA</td><td>>hg38_21_+_8433172_8446622_RNA45SN1_45S_50ntFlanks</td><td>11874</td><td>11907</td><td>34</td><td>AGAAGACCCTGTTGAGCTTGACTCTAGTCTGGCA</td><td>241.9</td><td>-6.9</td><td>1.0</td><td>-7.1</td><td>1.5E-12</td><td>3.6E-12</td><td>MB231/MB468</td></tr>
<tr><td>>hg38_21_+_8433172_8446622_RNA45SN1_45S_50ntFlanks@7973.8012.40|GACGCGACCTCAGATCAGACGTGGCGACCCGCTGAATTTA</td><td>>hg38_21_+_8433172_8446622_RNA45SN1_45S_50ntFlanks</td><td>7973</td><td>8012</td><td>40</td><td>GACGCGACCTCAGATCAGACGTGGCGACCCGCTGAATTTA</td><td>814.5</td><td>5.2</td><td>0.7</td><td>7.1</td><td>1.5E-12</td><td>3.6E-12</td><td>MB231/MB468</td></tr>
<tr><td>>hg38_21_+_8433172_8446622_RNA45SN1_45S_50ntFlanks@12159.12183.25|TTTGACTGGGGCGGTACACCTGTCA</td><td>>hg38_21_+_8433172_8446622_RNA45SN1_45S_50ntFlanks</td><td>12159</td><td>12183</td><td>25</td><td>TTTGACTGGGGCGGTACACCTGTCA</td><td>192.2</td><td>-4.9</td><td>0.7</td><td>-7.1</td><td>1.5E-12</td><td>3.6E-12</td><td>MB231/MB468</td></tr>
<tr><td>>MI0000085|hsa-mir-27a&WithFlank&19|-|13836434|13836523@57.76.20(+1U)&[MIMAT0000084&hsa-miR-27a-3p&offsets|0|-1(+1U);m-43&19|-|13836447|13836467&offsets|0|-1(+1U)]|TTCACAGTGGCTAAGTTCCGT</td><td>>MI0000085|hsa-mir-27a&WithFlank&19|-|13836434|13836523</td><td>57</td><td>76</td><td>20(+1U)</td><td>TTCACAGTGGCTAAGTTCCGT</td><td>18057.6</td><td>5.9</td><td>0.8</td><td>7.1</td><td>1.5E-12</td><td>3.6E-12</td><td>MB231/MB468</td></tr>
<tr><td>>hg38_21_+_8433172_8446622_RNA45SN1_45S_50ntFlanks@13004.13029.26|ATCTATTGAAAGTCAGCCCTCGACAC</td><td>>hg38_21_+_8433172_8446622_RNA45SN1_45S_50ntFlanks</td><td>13004</td><td>13029</td><td>26</td><td>ATCTATTGAAAGTCAGCCCTCGACAC</td><td>153.7</td><td>-9.3</td><td>1.3</td><td>-7.1</td><td>1.7E-12</td><td>3.6E-12</td><td>BT20/MB468</td></tr>
<tr><td>>hg38_21_+_8433172_8446622_RNA45SN1_45S_50ntFlanks@4005.4026.22|AACCTCGGGCCGATCGCACGCC</td><td>>hg38_21_+_8433172_8446622_RNA45SN1_45S_50ntFlanks</td><td>4005</td><td>4026</td><td>22</td><td>AACCTCGGGCCGATCGCACGCC</td><td>135.5</td><td>-9.1</td><td>1.3</td><td>-7.1</td><td>1.7E-12</td><td>3.6E-12</td><td>BT20/MB468</td></tr>
<tr><td>>hg38_21_+_8433172_8446622_RNA45SN1_45S_50ntFlanks@6737.6771.35|GATCATCGACACTTCGAACGCACTTGCGGCCCCGG</td><td>>hg38_21_+_8433172_8446622_RNA45SN1_45S_50ntFlanks</td><td>6737</td><td>6771</td><td>35</td><td>GATCATCGACACTTCGAACGCACTTGCGGCCCCGG</td><td>241.7</td><td>-5.6</td><td>0.8</td><td>-7.1</td><td>1.5E-12</td><td>3.6E-12</td><td>MB231/MB468</td></tr>
<tr><td>>MI0000070|hsa-mir-16-1&WithFlank&13|-|50048967|50049067@20.40.21(+1U);MI0000115|hsa-mir-16-2&WithFlank&3|+|160404739|160404831@16.36.21(+1U)&[MIMAT0000069&hsa-miR-16-5p&offsets|0|-1(+1U);m-62&13|-|50049027|50049048&offsets|0|-1(+1U)];[MIMAT0000069_1&hsa-miR-16-5p&offsets|0|-1(+1U);m-61&3|+|160404754|160404775&offsets|0|-1(+1U)]|TAGCAGCACGTAAATATTGGCT</td><td>>MI0000070|hsa-mir-16-1&WithFlank&13|-|50048967|50049067</td><td>20</td><td>40</td><td>21(+1U)</td><td>TAGCAGCACGTAAATATTGGCT</td><td>3428.1</td><td>11.8</td><td>1.7</td><td>7.1</td><td>1.7E-12</td><td>3.7E-12</td><td>BT20/MB468</td></tr>
<tr><td>>hg38_21_+_8433172_8446622_RNA45SN1_45S_50ntFlanks@8381.8403.23|AAGAGGGCGTGAAACCGTTAAGA</td><td>>hg38_21_+_8433172_8446622_RNA45SN1_45S_50ntFlanks</td><td>8381</td><td>8403</td><td>23</td><td>AAGAGGGCGTGAAACCGTTAAGA</td><td>245.2</td><td>-4.6</td><td>0.7</td><td>-7.1</td><td>1.7E-12</td><td>3.7E-12</td><td>BT20/MB468</td></tr>
<tr><td>>hg38_21_+_8433172_8446622_RNA45SN1_45S_50ntFlanks@4449.4471.23|CCTCTCGGCGCCCCCTCGATGCT</td><td>>hg38_21_+_8433172_8446622_RNA45SN1_45S_50ntFlanks</td><td>4449</td><td>4471</td><td>23</td><td>CCTCTCGGCGCCCCCTCGATGCT</td><td>370.1</td><td>-6.0</td><td>0.8</td><td>-7.1</td><td>1.5E-12</td><td>3.7E-12</td><td>MB231/MB468</td></tr>
<tr><td>>hg38_21_+_8433172_8446622_RNA45SN1_45S_50ntFlanks@11235.11269.35|CTTCCCGTGGATCGCCCCAGCTGCGGCGGGCGTCG</td><td>>hg38_21_+_8433172_8446622_RNA45SN1_45S_50ntFlanks</td><td>11235</td><td>11269</td><td>35</td><td>CTTCCCGTGGATCGCCCCAGCTGCGGCGGGCGTCG</td><td>437.8</td><td>-6.7</td><td>0.9</td><td>-7.1</td><td>1.7E-12</td><td>3.7E-12</td><td>BT20/MB468</td></tr>
<tr><td>>hg38_MT_+_598_1651_RNR1_12S_50ntFlanks@367.404.38|CCAATAAAGCTAAAACTCACCTGAGTTGTAAAAAACTC</td><td>>hg38_MT_+_598_1651_RNR1_12S_50ntFlanks</td><td>367</td><td>404</td><td>38</td><td>CCAATAAAGCTAAAACTCACCTGAGTTGTAAAAAACTC</td><td>459.9</td><td>9.1</td><td>1.3</td><td>7.1</td><td>1.7E-12</td><td>3.7E-12</td><td>BT20/MB468</td></tr>
<tr><td>>hg38_21_+_8433172_8446622_RNA45SN1_45S_50ntFlanks@8381.8402.22|AAGAGGGCGTGAAACCGTTAAG</td><td>>hg38_21_+_8433172_8446622_RNA45SN1_45S_50ntFlanks</td><td>8381</td><td>8402</td><td>22</td><td>AAGAGGGCGTGAAACCGTTAAG</td><td>1344.5</td><td>-4.2</td><td>0.6</td><td>-7.1</td><td>1.6E-12</td><td>3.7E-12</td><td>MB231/MB468</td></tr>
<tr><td>>hg38_21_+_8433172_8446622_RNA45SN1_45S_50ntFlanks@11601.11620.20|ATCCGACTGTTTAATTAAAA</td><td>>hg38_21_+_8433172_8446622_RNA45SN1_45S_50ntFlanks</td><td>11601</td><td>11620</td><td>20</td><td>ATCCGACTGTTTAATTAAAA</td><td>112.5</td><td>-8.8</td><td>1.2</td><td>-7.1</td><td>1.8E-12</td><td>3.8E-12</td><td>BT20/MB468</td></tr>
<tr><td>>hg38_21_+_8433172_8446622_RNA45SN1_45S_50ntFlanks@10826.10844.19|TCGGCAAGCCGGATCCGTA</td><td>>hg38_21_+_8433172_8446622_RNA45SN1_45S_50ntFlanks</td><td>10826</td><td>10844</td><td>19</td><td>TCGGCAAGCCGGATCCGTA</td><td>349.4</td><td>-7.2</td><td>1.0</td><td>-7.1</td><td>1.6E-12</td><td>3.8E-12</td><td>MB231/MB468</td></tr>
<tr><td>>MI0000749|hsa-mir-30e&WithFlank&1|+|40754349|40754452@65.85.21&[MIMAT0000693&hsa-miR-30e-3p&offsets|0|-1;m-25&1|+|40754413|40754434&offsets|0|-1]|CTTTCAGTCGGATGTTTACAG</td><td>>MI0000749|hsa-mir-30e&WithFlank&1|+|40754349|40754452</td><td>65</td><td>85</td><td>21</td><td>CTTTCAGTCGGATGTTTACAG</td><td>789.5</td><td>5.9</td><td>0.8</td><td>7.0</td><td>1.8E-12</td><td>3.9E-12</td><td>BT20/MB468</td></tr>
<tr><td>>hg38_21_+_8433172_8446622_RNA45SN1_45S_50ntFlanks@12991.13009.19|CTCCCTCGCTGCGATCTAT</td><td>>hg38_21_+_8433172_8446622_RNA45SN1_45S_50ntFlanks</td><td>12991</td><td>13009</td><td>19</td><td>CTCCCTCGCTGCGATCTAT</td><td>214.9</td><td>-9.3</td><td>1.3</td><td>-7.1</td><td>1.6E-12</td><td>3.9E-12</td><td>MB231/MB468</td></tr>
<tr><td>>hg38_1_-_228634819_228635039_RNA5S12_5S_50ntFlanks@144.169.26|CCTGGGAATACCGGGTGCTGTAGGCT</td><td>>hg38_1_-_228634819_228635039_RNA5S12_5S_50ntFlanks</td><td>144</td><td>169</td><td>26</td><td>CCTGGGAATACCGGGTGCTGTAGGCT</td><td>873.6</td><td>8.2</td><td>1.2</td><td>7.0</td><td>1.8E-12</td><td>3.9E-12</td><td>BT20/MB468</td></tr>
<tr><td>>hg38_21_+_8433172_8446622_RNA45SN1_45S_50ntFlanks@12324.12343.20|TTGGGTTTTAAGCAGGAGGT</td><td>>hg38_21_+_8433172_8446622_RNA45SN1_45S_50ntFlanks</td><td>12324</td><td>12343</td><td>20</td><td>TTGGGTTTTAAGCAGGAGGT</td><td>102.5</td><td>-8.7</td><td>1.2</td><td>-7.0</td><td>1.8E-12</td><td>3.9E-12</td><td>BT20/MB468</td></tr>
<tr><td>>hg38_21_+_8433172_8446622_RNA45SN1_45S_50ntFlanks@11161.11179.19|CACGGTCCCCCGCGAGGGG</td><td>>hg38_21_+_8433172_8446622_RNA45SN1_45S_50ntFlanks</td><td>11161</td><td>11179</td><td>19</td><td>CACGGTCCCCCGCGAGGGG</td><td>746.3</td><td>-7.5</td><td>1.1</td><td>-7.0</td><td>1.8E-12</td><td>3.9E-12</td><td>BT20/MB468</td></tr>
<tr><td>>am_trnaMT_ValTAC_MT_+_1602_1670@38.62.25|TAGGAGATTTCAACTTAACTTGACC</td><td>>am_trnaMT_ValTAC_MT_+_1602_1670</td><td>38</td><td>62</td><td>25</td><td>TAGGAGATTTCAACTTAACTTGACC</td><td>691.4</td><td>8.9</td><td>1.3</td><td>7.0</td><td>1.8E-12</td><td>3.9E-12</td><td>BT20/MB468</td></tr>
<tr><td>>hg38_21_+_8433172_8446622_RNA45SN1_45S_50ntFlanks@8168.8197.30|CCCAAGTCCTTCTGATCGAGGCCCAGCCCG</td><td>>hg38_21_+_8433172_8446622_RNA45SN1_45S_50ntFlanks</td><td>8168</td><td>8197</td><td>30</td><td>CCCAAGTCCTTCTGATCGAGGCCCAGCCCG</td><td>160.0</td><td>-9.3</td><td>1.3</td><td>-7.0</td><td>1.8E-12</td><td>3.9E-12</td><td>BT20/MB468</td></tr>
<tr><td>>hg38_21_+_8433172_8446622_RNA45SN1_45S_50ntFlanks@10402.10426.25|AACATGGGTCAGTCGGTCCTGAGAG</td><td>>hg38_21_+_8433172_8446622_RNA45SN1_45S_50ntFlanks</td><td>10402</td><td>10426</td><td>25</td><td>AACATGGGTCAGTCGGTCCTGAGAG</td><td>451.5</td><td>-4.8</td><td>0.7</td><td>-7.0</td><td>1.8E-12</td><td>3.9E-12</td><td>BT20/MB468</td></tr>
<tr><td>>hg38_21_+_8433172_8446622_RNA45SN1_45S_50ntFlanks@12954.12982.29|CTGCTTCTGGGTCGGGGTTTCGTACGTAG</td><td>>hg38_21_+_8433172_8446622_RNA45SN1_45S_50ntFlanks</td><td>12954</td><td>12982</td><td>29</td><td>CTGCTTCTGGGTCGGGGTTTCGTACGTAG</td><td>118.7</td><td>-8.9</td><td>1.3</td><td>-7.0</td><td>1.8E-12</td><td>4.0E-12</td><td>BT20/MB468</td></tr>
<tr><td>>MI0000736|hsa-mir-30c-1&WithFlank&1|+|40757278|40757378@23.45.23;MI0000254|hsa-mir-30c-2&WithFlank&6|-|71376954|71377037@13.35.23&[MIMAT0000244&hsa-miR-30c-5p&offsets|0|0;m-57&1|+|40757300|40757323&offsets|0|-1];[MIMAT0000244_1&hsa-miR-30c-5p&offsets|0|0;m-56&6|-|71377002|71377025&offsets|0|-1]|TGTAAACATCCTACACTCTCAGC</td><td>>MI0000736|hsa-mir-30c-1&WithFlank&1|+|40757278|40757378</td><td>23</td><td>45</td><td>23</td><td>TGTAAACATCCTACACTCTCAGC</td><td>29241.7</td><td>5.5</td><td>0.8</td><td>7.0</td><td>1.9E-12</td><td>4.0E-12</td><td>BT20/MB468</td></tr>
<tr><td>>hg38_1_-_228634819_228635039_RNA5S12_5S_50ntFlanks@140.170.31|ACCGCCTGGGAATACCGGGTGCTGTAGGCTT</td><td>>hg38_1_-_228634819_228635039_RNA5S12_5S_50ntFlanks</td><td>140</td><td>170</td><td>31</td><td>ACCGCCTGGGAATACCGGGTGCTGTAGGCTT</td><td>39159.0</td><td>5.3</td><td>0.8</td><td>7.1</td><td>1.7E-12</td><td>4.0E-12</td><td>MB231/MB468</td></tr>
<tr><td>>hg38_21_+_8433172_8446622_RNA45SN1_45S_50ntFlanks@6721.6741.21|AATTGCAGGACACATTGATCA</td><td>>hg38_21_+_8433172_8446622_RNA45SN1_45S_50ntFlanks</td><td>6721</td><td>6741</td><td>21</td><td>AATTGCAGGACACATTGATCA</td><td>549.7</td><td>-4.0</td><td>0.6</td><td>-7.1</td><td>1.7E-12</td><td>4.0E-12</td><td>MB231/MB468</td></tr>
<tr><td>>hg38_21_+_8433172_8446622_RNA45SN1_45S_50ntFlanks@12419.12448.30|GATGTCGGCTCTTCCTATCATTGTGAAGCA</td><td>>hg38_21_+_8433172_8446622_RNA45SN1_45S_50ntFlanks</td><td>12419</td><td>12448</td><td>30</td><td>GATGTCGGCTCTTCCTATCATTGTGAAGCA</td><td>761.2</td><td>5.5</td><td>0.8</td><td>7.0</td><td>1.9E-12</td><td>4.0E-12</td><td>BT20/MB468</td></tr>
<tr><td>>hg38_21_+_8433172_8446622_RNA45SN1_45S_50ntFlanks@6742.6770.29|TCGACACTTCGAACGCACTTGCGGCCCCG</td><td>>hg38_21_+_8433172_8446622_RNA45SN1_45S_50ntFlanks</td><td>6742</td><td>6770</td><td>29</td><td>TCGACACTTCGAACGCACTTGCGGCCCCG</td><td>347.4</td><td>-6.8</td><td>1.0</td><td>-7.0</td><td>1.9E-12</td><td>4.0E-12</td><td>BT20/MB468</td></tr>
<tr><td>>hg38_21_+_8433172_8446622_RNA45SN1_45S_50ntFlanks@12488.12525.38|AACGTGAGCTGGGTTTAGACCGTCGTGAGACAGGTTAG</td><td>>hg38_21_+_8433172_8446622_RNA45SN1_45S_50ntFlanks</td><td>12488</td><td>12525</td><td>38</td><td>AACGTGAGCTGGGTTTAGACCGTCGTGAGACAGGTTAG</td><td>135.1</td><td>-9.1</td><td>1.3</td><td>-7.0</td><td>1.9E-12</td><td>4.0E-12</td><td>BT20/MB468</td></tr>
<tr><td>>hg38_21_+_8433172_8446622_RNA45SN1_45S_50ntFlanks@10395.10424.30|GCAGTTGAACATGGGTCAGTCGGTCCTGAG</td><td>>hg38_21_+_8433172_8446622_RNA45SN1_45S_50ntFlanks</td><td>10395</td><td>10424</td><td>30</td><td>GCAGTTGAACATGGGTCAGTCGGTCCTGAG</td><td>1108.3</td><td>5.1</td><td>0.7</td><td>7.1</td><td>1.7E-12</td><td>4.0E-12</td><td>MB231/MB468</td></tr>
<tr><td>>hg38_21_+_8433172_8446622_RNA45SN1_45S_50ntFlanks@10761.10785.25|ATATCCGCAGCAGGTCTCCAAGGTG</td><td>>hg38_21_+_8433172_8446622_RNA45SN1_45S_50ntFlanks</td><td>10761</td><td>10785</td><td>25</td><td>ATATCCGCAGCAGGTCTCCAAGGTG</td><td>113.8</td><td>-8.8</td><td>1.3</td><td>-7.0</td><td>1.9E-12</td><td>4.0E-12</td><td>BT20/MB468</td></tr>
<tr><td>>trnaMT_HisGTG_MT_+_12138_12206@32.67.36|TGAATCTGACAACAGAGGCTTACGACCCCTTATTTA</td><td>>trnaMT_HisGTG_MT_+_12138_12206</td><td>32</td><td>67</td><td>36</td><td>TGAATCTGACAACAGAGGCTTACGACCCCTTATTTA</td><td>868.0</td><td>8.1</td><td>1.1</td><td>7.1</td><td>1.7E-12</td><td>4.0E-12</td><td>MB231/MB468</td></tr>
</table>

<table>
<tr><td>>hg38_21_+_8433172_8446622_RNA45SN1_45S_50ntFlanks@13002.13030.29|CGATCTATTGAAAGTCAGCCCTCGACACA</td><td>>hg38_21_+_8433172_8446622_RNA45SN1_45S_50ntFlanks</td><td>13002</td><td>13030</td><td>29</td><td>CGATCTATTGAAAGTCAGCCCTCGACACA</td><td>127.0</td><td>-9.0</td><td>1.3</td><td>-7.0</td><td>1.9E-12</td><td>4.1E-12</td><td>BT20/MB468</td></tr>
<tr><td>>hg38_21_+_8433172_8446622_RNA45SN1_45S_50ntFlanks@9575.9605.31|ATCGGTCGTCCGACCTGGGTATAGGGGCGAA</td><td>>hg38_21_+_8433172_8446622_RNA45SN1_45S_50ntFlanks</td><td>9575</td><td>9605</td><td>31</td><td>ATCGGTCGTCCGACCTGGGTATAGGGGCGAA</td><td>158.3</td><td>-9.3</td><td>1.3</td><td>-7.0</td><td>1.9E-12</td><td>4.1E-12</td><td>BT20/MB468</td></tr>
<tr><td>>hg38_21_+_8433172_8446622_RNA45SN1_45S_50ntFlanks@10727.10752.26|GGAGAGGGTGTAAATCTCGCGCCGGG</td><td>>hg38_21_+_8433172_8446622_RNA45SN1_45S_50ntFlanks</td><td>10727</td><td>10752</td><td>26</td><td>GGAGAGGGTGTAAATCTCGCGCCGGG</td><td>131.3</td><td>-9.0</td><td>1.3</td><td>-7.0</td><td>1.9E-12</td><td>4.1E-12</td><td>BT20/MB468</td></tr>
<tr><td>>hg38_1_-_228634819_228635039_RNA5S12_5S_50ntFlanks@142.169.28|CGCCTGGGAATACCGGGTGCTGTAGGCT</td><td>>hg38_1_-_228634819_228635039_RNA5S12_5S_50ntFlanks</td><td>142</td><td>169</td><td>28</td><td>CGCCTGGGAATACCGGGTGCTGTAGGCT</td><td>508.0</td><td>5.3</td><td>0.8</td><td>7.0</td><td>1.9E-12</td><td>4.1E-12</td><td>BT20/MB468</td></tr>
<tr><td>>hg38_21_+_8433172_8446622_RNA45SN1_45S_50ntFlanks@10788.10816.29|CAGCCTCTGGCATGTTGGAACAATGTAGG</td><td>>hg38_21_+_8433172_8446622_RNA45SN1_45S_50ntFlanks</td><td>10788</td><td>10816</td><td>29</td><td>CAGCCTCTGGCATGTTGGAACAATGTAGG</td><td>484.9</td><td>-3.5</td><td>0.5</td><td>-7.0</td><td>1.9E-12</td><td>4.1E-12</td><td>BT20/MB468</td></tr>
<tr><td>>hg38_21_+_8433172_8446622_RNA45SN1_45S_50ntFlanks@11702.11728.27|CAATGAAGCGCGGGTAAACGGCGGGAG</td><td>>hg38_21_+_8433172_8446622_RNA45SN1_45S_50ntFlanks</td><td>11702</td><td>11728</td><td>27</td><td>CAATGAAGCGCGGGTAAACGGCGGGAG</td><td>104.6</td><td>-8.7</td><td>1.2</td><td>-7.0</td><td>1.9E-12</td><td>4.1E-12</td><td>BT20/MB468</td></tr>
<tr><td>>hg38_21_+_8433172_8446622_RNA45SN1_45S_50ntFlanks@6722.6745.24|ATTGCAGGACACATTGATCATCGA</td><td>>hg38_21_+_8433172_8446622_RNA45SN1_45S_50ntFlanks</td><td>6722</td><td>6745</td><td>24</td><td>ATTGCAGGACACATTGATCATCGA</td><td>171.1</td><td>-3.4</td><td>0.5</td><td>-7.1</td><td>1.7E-12</td><td>4.1E-12</td><td>MB231/MB468</td></tr>
<tr><td>>hg38_21_+_8433172_8446622_RNA45SN1_45S_50ntFlanks@12934.12951.18|CTAAACCATTCGTAGACG</td><td>>hg38_21_+_8433172_8446622_RNA45SN1_45S_50ntFlanks</td><td>12934</td><td>12951</td><td>18</td><td>CTAAACCATTCGTAGACG</td><td>691.1</td><td>-3.8</td><td>0.5</td><td>-7.1</td><td>1.7E-12</td><td>4.1E-12</td><td>MB231/MB468</td></tr>
<tr><td>>hg38_21_+_8433172_8446622_RNA45SN1_45S_50ntFlanks@10781.10802.22|AGGTGAACAGCCTCTGGCATGT</td><td>>hg38_21_+_8433172_8446622_RNA45SN1_45S_50ntFlanks</td><td>10781</td><td>10802</td><td>22</td><td>AGGTGAACAGCCTCTGGCATGT</td><td>233.4</td><td>-3.5</td><td>0.5</td><td>-7.0</td><td>1.9E-12</td><td>4.1E-12</td><td>BT20/MB468</td></tr>
<tr><td>>hg38_21_+_8433172_8446622_RNA45SN1_45S_50ntFlanks@10782.10818.37|GGTGAACAGCCTCTGGCATGTTGGAACAATGTAGGTA</td><td>>hg38_21_+_8433172_8446622_RNA45SN1_45S_50ntFlanks</td><td>10782</td><td>10818</td><td>37</td><td>GGTGAACAGCCTCTGGCATGTTGGAACAATGTAGGTA</td><td>286.0</td><td>-4.6</td><td>0.7</td><td>-7.1</td><td>1.7E-12</td><td>4.2E-12</td><td>MB231/MB468</td></tr>
<tr><td>>hg38_21_+_8433172_8446622_RNA45SN1_45S_50ntFlanks@10738.10756.19|AAATCTCGCGCCGGGCCGT</td><td>>hg38_21_+_8433172_8446622_RNA45SN1_45S_50ntFlanks</td><td>10738</td><td>10756</td><td>19</td><td>AAATCTCGCGCCGGGCCGT</td><td>1174.1</td><td>6.3</td><td>0.9</td><td>7.1</td><td>1.8E-12</td><td>4.2E-12</td><td>MB231/MB468</td></tr>
<tr><td>>trna116_GluCTC_1_-_145399233_145399304@36.75.40,trna134_GluTTC_1_-_16861774_16861845@36.75.40,trna71_GluCTC_1_-_161439189_161439260@36.75.40,trna74_GluCTC_1_-_161431809_161431880@36.75.40,trna77_GluCTC_1_-_161424398_161424469@36.75.40,trna77_GluCTC_6_+_28949976_28950047@36.75.40,trna80_GluCTC_1_-_161417018_161417089@36.75.40,trna84_GluTTC_1_-_161391883_161391954@36.75.40,trna87_GluCTC_6_-_126101393_126101464@36.75.40|CACCGCCGCGGCCCGGGTTCGATTCCCGGTCAGGGAACCA</td><td>>trna116_GluCTC_1_-_145399233_145399304</td><td>36</td><td>75</td><td>40</td><td>CACCGCCGCGGCCCGGGTTCGATTCCCGGTCAGGGAACCA</td><td>1193.7</td><td>6.8</td><td>1.0</td><td>7.1</td><td>1.8E-12</td><td>4.2E-12</td><td>MB231/MB468</td></tr>
<tr><td>>hg38_21_+_8433172_8446622_RNA45SN1_45S_50ntFlanks@7989.8017.29|AGACGTGGCGACCCGCTGAATTTAAGCAT</td><td>>hg38_21_+_8433172_8446622_RNA45SN1_45S_50ntFlanks</td><td>7989</td><td>8017</td><td>29</td><td>AGACGTGGCGACCCGCTGAATTTAAGCAT</td><td>931.2</td><td>4.7</td><td>0.7</td><td>7.1</td><td>1.8E-12</td><td>4.2E-12</td><td>MB231/MB468</td></tr>
<tr><td>>hg38_21_+_8433172_8446622_RNA45SN1_45S_50ntFlanks@10770.10797.28|GCAGGTCTCCAAGGTGAACAGCCTCTGG</td><td>>hg38_21_+_8433172_8446622_RNA45SN1_45S_50ntFlanks</td><td>10770</td><td>10797</td><td>28</td><td>GCAGGTCTCCAAGGTGAACAGCCTCTGG</td><td>106.0</td><td>-8.7</td><td>1.2</td><td>-7.0</td><td>2.0E-12</td><td>4.2E-12</td><td>BT20/MB468</td></tr>
<tr><td>>hg38_21_+_8433172_8446622_RNA45SN1_45S_50ntFlanks@11051.11075.25|CCGTCCTCCCCCCTCCCCGGGGGAG</td><td>>hg38_21_+_8433172_8446622_RNA45SN1_45S_50ntFlanks</td><td>11051</td><td>11075</td><td>25</td><td>CCGTCCTCCCCCCTCCCCGGGGGAG</td><td>147.9</td><td>-9.2</td><td>1.3</td><td>-7.0</td><td>2.0E-12</td><td>4.3E-12</td><td>BT20/MB468</td></tr>
<tr><td>>hg38_21_+_8433172_8446622_RNA45SN1_45S_50ntFlanks@10378.10407.30|AGGGTTCCATGTGAACAGCAGTTGAACATG</td><td>>hg38_21_+_8433172_8446622_RNA45SN1_45S_50ntFlanks</td><td>10378</td><td>10407</td><td>30</td><td>AGGGTTCCATGTGAACAGCAGTTGAACATG</td><td>128.9</td><td>-9.0</td><td>1.3</td><td>-7.0</td><td>2.0E-12</td><td>4.3E-12</td><td>BT20/MB468</td></tr>
<tr><td>>hg38_21_+_8433172_8446622_RNA45SN1_45S_50ntFlanks@11321.11338.18|CACGTCTCGTCGCGCGCG</td><td>>hg38_21_+_8433172_8446622_RNA45SN1_45S_50ntFlanks</td><td>11321</td><td>11338</td><td>18</td><td>CACGTCTCGTCGCGCGCG</td><td>399.5</td><td>-7.4</td><td>1.1</td><td>-7.0</td><td>1.8E-12</td><td>4.3E-12</td><td>MB231/MB468</td></tr>
<tr><td>>hg38_21_+_8433172_8446622_RNA45SN1_45S_50ntFlanks@11233.11253.21|CCCTTCCCGTGGATCGCCCCA</td><td>>hg38_21_+_8433172_8446622_RNA45SN1_45S_50ntFlanks</td><td>11233</td><td>11253</td><td>21</td><td>CCCTTCCCGTGGATCGCCCCA</td><td>200.8</td><td>-9.2</td><td>1.3</td><td>-7.0</td><td>1.8E-12</td><td>4.3E-12</td><td>MB231/MB468</td></tr>
<tr><td>>trna10_AlaCGC_6_+_26553731_26553802@58.75.18,trna119_AlaCGC_6_-_28641613_28641684@58.75.18,trna13_AlaCGC_2_+_157257281_157257352@58.75.18,trna13_AlaTGC_12_-_125406301_125406372@58.75.18,trna66_AlaTGC_6_+_28611222_28611293@58.75.18,trna8_AlaTGC_12_+_125424512_125424583@58.75.18,trna8_AlaTGC_5_+_180633868_180633939@58.75.18|TCCCCGGCATCTCCACCA</td><td>>trna10_AlaCGC_6_+_26553731_26553802</td><td>58</td><td>75</td><td>18</td><td>TCCCCGGCATCTCCACCA</td><td>3949.3</td><td>6.2</td><td>0.9</td><td>7.0</td><td>2.0E-12</td><td>4.3E-12</td><td>BT20/MB468</td></tr>
<tr><td>>hg38_21_+_8433172_8446622_RNA45SN1_45S_50ntFlanks@4444.4471.28|CCTTGCCTCTCGGCGCCCCCTCGATGCT</td><td>>hg38_21_+_8433172_8446622_RNA45SN1_45S_50ntFlanks</td><td>4444</td><td>4471</td><td>28</td><td>CCTTGCCTCTCGGCGCCCCCTCGATGCT</td><td>162.6</td><td>-9.3</td><td>1.3</td><td>-7.0</td><td>2.0E-12</td><td>4.3E-12</td><td>BT20/MB468</td></tr>
<tr><td>>am_trna128_GlyGCC_6_-_27870686_27870756@1.36.36,trna18_GlyGCC_16_+_70822597_70822667@1.36.36,trna19_GlyGCC_16_+_70823410_70823480@1.36.36,trna19_GlyGCC_2_-_157257659_157257729@1.36.36,trna24_GlyGCC_16_-_70812942_70813012@1.36.36,trna25_GlyGCC_16_-_70812114_70812184@1.36.36,trna5_GlyGCC_17_+_8029064_8029134@1.36.36,trna68_GlyGCC_1_-_161493637_161493707@1.36.36|GCATTGGTGGTTCAGTGGTAGAATTCTCGCCTGCCA</td><td>>am_trna128_GlyGCC_6_-_27870686_27870756</td><td>1</td><td>36</td><td>36</td><td>GCATTGGTGGTTCAGTGGTAGAATTCTCGCCTGCCA</td><td>453.1</td><td>5.4</td><td>0.8</td><td>7.0</td><td>2.0E-12</td><td>4.3E-12</td><td>BT20/MB468</td></tr>
<tr><td>>hg38_21_+_8433172_8446622_RNA45SN1_45S_50ntFlanks@9504.9529.26|GTGAACTATGCCTGGGCAGGGCGAAG</td><td>>hg38_21_+_8433172_8446622_RNA45SN1_45S_50ntFlanks</td><td>9504</td><td>9529</td><td>26</td><td>GTGAACTATGCCTGGGCAGGGCGAAG</td><td>191.1</td><td>-9.6</td><td>1.4</td><td>-7.0</td><td>2.0E-12</td><td>4.3E-12</td><td>BT20/MB468</td></tr>
<tr><td>>am_trnaMT_PheGAA_MT_+_577_647@41.62.22|GTTTAGACGGGCTCACATCACC</td><td>>am_trnaMT_PheGAA_MT_+_577_647</td><td>41</td><td>62</td><td>22</td><td>GTTTAGACGGGCTCACATCACC</td><td>1167.1</td><td>8.7</td><td>1.2</td><td>7.0</td><td>1.8E-12</td><td>4.3E-12</td><td>MB231/MB468</td></tr>
<tr><td>>hg38_21_+_8433172_8446622_RNA45SN1_45S_50ntFlanks@10574.10607.34|AAGCCGGCGGGAGCCCCGGGGAGAGTTCTCTTTT</td><td>>hg38_21_+_8433172_8446622_RNA45SN1_45S_50ntFlanks</td><td>10574</td><td>10607</td><td>34</td><td>AAGCCGGCGGGAGCCCCGGGGAGAGTTCTCTTTT</td><td>173.9</td><td>-9.0</td><td>1.3</td><td>-7.0</td><td>1.8E-12</td><td>4.4E-12</td><td>MB231/MB468</td></tr>
<tr><td>>hg38_21_+_8433172_8446622_RNA45SN1_45S_50ntFlanks@10375.10406.32|AGAAGGGTTCCATGTGAACAGCAGTTGAACAT</td><td>>hg38_21_+_8433172_8446622_RNA45SN1_45S_50ntFlanks</td><td>10375</td><td>10406</td><td>32</td><td>AGAAGGGTTCCATGTGAACAGCAGTTGAACAT</td><td>226.6</td><td>-4.1</td><td>0.6</td><td>-7.0</td><td>1.8E-12</td><td>4.4E-12</td><td>MB231/MB468</td></tr>
<tr><td>>trna14_LysTTT_11_-_59327808_59327880@1.32.32,trna2_LysTTT_17_+_8022473_8022545@1.32.32,trna54_LysTTT_1_+_204475655_204475727@1.32.32,trna5_LysTTT_11_+_59323902_59323974@1.32.32,trna62_LysTTT_1_-_204476158_204476230@1.32.32,trna76_LysTTT_6_+_28918806_28918878@1.32.32|GCCCGGATAGCTCAGTCGGTAGAGCATCAGAC</td><td>>trna14_LysTTT_11_-_59327808_59327880</td><td>1</td><td>32</td><td>32</td><td>GCCCGGATAGCTCAGTCGGTAGAGCATCAGAC</td><td>714.6</td><td>5.1</td><td>0.7</td><td>7.0</td><td>1.9E-12</td><td>4.4E-12</td><td>MB231/MB468</td></tr>
<tr><td>>hg38_21_+_8433172_8446622_RNA45SN1_45S_50ntFlanks@11273.11293.21|CCGCCCCCGGGGAGCCCGGCG</td><td>>hg38_21_+_8433172_8446622_RNA45SN1_45S_50ntFlanks</td><td>11273</td><td>11293</td><td>21</td><td>CCGCCCCCGGGGAGCCCGGCG</td><td>122.2</td><td>-8.9</td><td>1.3</td><td>-7.0</td><td>2.1E-12</td><td>4.4E-12</td><td>BT20/MB468</td></tr>
<tr><td>>hg38_21_+_8433172_8446622_RNA45SN1_45S_50ntFlanks@11767.11804.38|TCTAATTAGTGACGCGCATGAATGGATGAACGAGATTC</td><td>>hg38_21_+_8433172_8446622_RNA45SN1_45S_50ntFlanks</td><td>11767</td><td>11804</td><td>38</td><td>TCTAATTAGTGACGCGCATGAATGGATGAACGAGATTC</td><td>99.8</td><td>-4.0</td><td>0.6</td><td>-7.0</td><td>2.1E-12</td><td>4.4E-12</td><td>BT20/MB468</td></tr>
<tr><td>>am_trnaMT_ValTAC_MT_+_1602_1670@38.57.20|TAGGAGATTTCAACTTAACT</td><td>>am_trnaMT_ValTAC_MT_+_1602_1670</td><td>38</td><td>57</td><td>20</td><td>TAGGAGATTTCAACTTAACT</td><td>626.9</td><td>8.4</td><td>1.2</td><td>7.0</td><td>2.1E-12</td><td>4.5E-12</td><td>BT20/MB468</td></tr>
<tr><td>>hg38_21_+_8433172_8446622_RNA45SN1_45S_50ntFlanks@8194.8225.32|CCCGTGGACGGTGTGAGGCCGGTAGCGGCCCC</td><td>>hg38_21_+_8433172_8446622_RNA45SN1_45S_50ntFlanks</td><td>8194</td><td>8225</td><td>32</td><td>CCCGTGGACGGTGTGAGGCCGGTAGCGGCCCC</td><td>124.1</td><td>-8.9</td><td>1.3</td><td>-7.0</td><td>2.1E-12</td><td>4.5E-12</td><td>BT20/MB468</td></tr>
<tr><td>>hg38_21_+_8433172_8446622_RNA45SN1_45S_50ntFlanks@9534.9554.21|AGGAAACTCTGGTGGAGGTCC</td><td>>hg38_21_+_8433172_8446622_RNA45SN1_45S_50ntFlanks</td><td>9534</td><td>9554</td><td>21</td><td>AGGAAACTCTGGTGGAGGTCC</td><td>278.0</td><td>-6.4</td><td>0.9</td><td>-7.0</td><td>1.9E-12</td><td>4.5E-12</td><td>MB231/MB468</td></tr>
<tr><td>>hg38_21_+_8433172_8446622_RNA45SN1_45S_50ntFlanks@10869.10902.34|AGGGCTGGGTCGGTCGGGCTGGGGCGCGAAGCGG</td><td>>hg38_21_+_8433172_8446622_RNA45SN1_45S_50ntFlanks</td><td>10869</td><td>10902</td><td>34</td><td>AGGGCTGGGTCGGTCGGGCTGGGGCGCGAAGCGG</td><td>414.3</td><td>-6.5</td><td>0.9</td><td>-7.0</td><td>2.1E-12</td><td>4.5E-12</td><td>BT20/MB468</td></tr>
<tr><td>>MI0000734|hsa-mir-106b&WithFlank&7|-|100093987|100094080@18.39.22&[MIMAT0000680&hsa-miR-106b-5p&offsets|0|+1;m-103&7|-|100094043|100094063&offsets|0|+1]|TAAAGTGCTGACAGTGCAGATA</td><td>>MI0000734|hsa-mir-106b&WithFlank&7|-|100093987|100094080</td><td>18</td><td>39</td><td>22</td><td>TAAAGTGCTGACAGTGCAGATA</td><td>15721.0</td><td>8.0</td><td>1.1</td><td>7.0</td><td>2.1E-12</td><td>4.5E-12</td><td>BT20/MB468</td></tr>
<tr><td>>hg38_21_+_8433172_8446622_RNA45SN1_45S_50ntFlanks@12957.12980.24|CTTCTGGGTCGGGGTTTCGTACGT</td><td>>hg38_21_+_8433172_8446622_RNA45SN1_45S_50ntFlanks</td><td>12957</td><td>12980</td><td>24</td><td>CTTCTGGGTCGGGGTTTCGTACGT</td><td>107.1</td><td>-8.7</td><td>1.2</td><td>-7.0</td><td>2.1E-12</td><td>4.5E-12</td><td>BT20/MB468</td></tr>
<tr><td>>hg38_21_+_8433172_8446622_RNA45SN1_45S_50ntFlanks@6722.6746.25|ATTGCAGGACACATTGATCATCGAC</td><td>>hg38_21_+_8433172_8446622_RNA45SN1_45S_50ntFlanks</td><td>6722</td><td>6746</td><td>25</td><td>ATTGCAGGACACATTGATCATCGAC</td><td>250.8</td><td>-3.6</td><td>0.5</td><td>-7.0</td><td>2.1E-12</td><td>4.6E-12</td><td>BT20/MB468</td></tr>
<tr><td>>hg38_21_+_8433172_8446622_RNA45SN1_45S_50ntFlanks@11891.11912.22|TTGACTCTAGTCTGGCACGGTG</td><td>>hg38_21_+_8433172_8446622_RNA45SN1_45S_50ntFlanks</td><td>11891</td><td>11912</td><td>22</td><td>TTGACTCTAGTCTGGCACGGTG</td><td>675.9</td><td>-5.1</td><td>0.7</td><td>-7.0</td><td>1.9E-12</td><td>4.6E-12</td><td>MB231/MB468</td></tr>
<tr><td>>hg38_21_+_8433172_8446622_RNA45SN1_45S_50ntFlanks@12874.12891.18|GCTGGAAAGGCGGCCGCC</td><td>>hg38_21_+_8433172_8446622_RNA45SN1_45S_50ntFlanks</td><td>12874</td><td>12891</td><td>18</td><td>GCTGGAAAGGCGGCCGCC</td><td>186.2</td><td>-9.5</td><td>1.4</td><td>-7.0</td><td>2.2E-12</td><td>4.6E-12</td><td>BT20/MB468</td></tr>
<tr><td>>am_trna117_GlyTCC_1_-_145397864_145397935@1.29.29,trna45_GlyTCC_1_+_161500903_161500974@1.29.29,trna70_GlyTCC_1_-_161439547_161439618@1.29.29,trna73_GlyTCC_1_-_161432166_161432237@1.29.29,trna76_GlyTCC_1_-_161424756_161424827@1.29.29,trna79_GlyTCC_1_-_161417375_161417446@1.29.29|GCGTTGGTGGTATAGTGGTGAGCATAGCT</td><td>>am_trna117_GlyTCC_1_-_145397864_145397935</td><td>1</td><td>29</td><td>29</td><td>GCGTTGGTGGTATAGTGGTGAGCATAGCT</td><td>904.1</td><td>-5.5</td><td>0.8</td><td>-7.0</td><td>1.9E-12</td><td>4.6E-12</td><td>MB231/MB468</td></tr>
</table>

<table>
<tr><td>>am_trna128_GlyGCC_6_-_27870686_27870756@1.28.28,trna133_GlyCCC_1_-_16872434_16872504@1.28.28,trna18_GlyGCC_16_+_70822597_70822667@1.28.28,trna19_GlyGCC_16_+_70823410_70823480@1.28.28,trna19_GlyGCC_2_-_157257659_157257729@1.28.28,trna24_GlyGCC_16_-_70812942_70813012@1.28.28,trna25_GlyGCC_16_-_70812114_70812184@1.28.28,trna4_GlyCCC_1_+_17188416_17188486@1.28.28,trna5_GlyGCC_17_+_8029064_8029134@1.28.28,trna68_GlyGCC_1_-_161493637_161493707@1.28.28|GCATTGGTGGTTCAGTGGTAGAATTCTC</td><td>>am_trna128_GlyGCC_6_-_27870686_27870756</td><td>1</td><td>28</td><td>28</td><td>GCATTGGTGGTTCAGTGGTAGAATTCTC</td><td>35234.6</td><td>-4.7</td><td>0.7</td><td>-7.0</td><td>2.2E-12</td><td>4.6E-12</td><td>BT20/MB468</td></tr>
<tr><td>>hg38_21_+_8433172_8446622_RNA45SN1_45S_50ntFlanks@11564.11600.37|CCTAGCAGCCGACTTAGAACTGGTGCGGACCAGGGGA</td><td>>hg38_21_+_8433172_8446622_RNA45SN1_45S_50ntFlanks</td><td>11564</td><td>11600</td><td>37</td><td>CCTAGCAGCCGACTTAGAACTGGTGCGGACCAGGGGA</td><td>487.9</td><td>-3.7</td><td>0.5</td><td>-7.0</td><td>2.2E-12</td><td>4.7E-12</td><td>BT20/MB468</td></tr>
<tr><td>>hg38_21_+_8433172_8446622_RNA45SN1_45S_50ntFlanks@7991.8019.29|ACGTGGCGACCCGCTGAATTTAAGCATAT</td><td>>hg38_21_+_8433172_8446622_RNA45SN1_45S_50ntFlanks</td><td>7991</td><td>8019</td><td>29</td><td>ACGTGGCGACCCGCTGAATTTAAGCATAT</td><td>489.0</td><td>6.1</td><td>0.9</td><td>7.0</td><td>2.0E-12</td><td>4.7E-12</td><td>MB231/MB468</td></tr>
<tr><td>>hg38_21_+_8433172_8446622_RNA45SN1_45S_50ntFlanks@10787.10818.32|ACAGCCTCTGGCATGTTGGAACAATGTAGGTA</td><td>>hg38_21_+_8433172_8446622_RNA45SN1_45S_50ntFlanks</td><td>10787</td><td>10818</td><td>32</td><td>ACAGCCTCTGGCATGTTGGAACAATGTAGGTA</td><td>1504.6</td><td>-4.5</td><td>0.6</td><td>-7.0</td><td>2.0E-12</td><td>4.7E-12</td><td>MB231/MB468</td></tr>
<tr><td>>hg38_21_+_8433172_8446622_RNA45SN1_45S_50ntFlanks@10805.10828.24|GAACAATGTAGGTAAGGGAAGTCG</td><td>>hg38_21_+_8433172_8446622_RNA45SN1_45S_50ntFlanks</td><td>10805</td><td>10828</td><td>24</td><td>GAACAATGTAGGTAAGGGAAGTCG</td><td>117.2</td><td>-5.5</td><td>0.8</td><td>-7.0</td><td>2.2E-12</td><td>4.7E-12</td><td>BT20/MB468</td></tr>
<tr><td>>hg38_21_+_8433172_8446622_RNA45SN1_45S_50ntFlanks@11766.11794.29|ATCTAATTAGTGACGCGCATGAATGGATG</td><td>>hg38_21_+_8433172_8446622_RNA45SN1_45S_50ntFlanks</td><td>11766</td><td>11794</td><td>29</td><td>ATCTAATTAGTGACGCGCATGAATGGATG</td><td>123.3</td><td>-8.9</td><td>1.3</td><td>-7.0</td><td>2.2E-12</td><td>4.8E-12</td><td>BT20/MB468</td></tr>
<tr><td>>hg38_21_+_8433172_8446622_RNA45SN1_45S_50ntFlanks@8168.8188.21|CCCAAGTCCTTCTGATCGAGG</td><td>>hg38_21_+_8433172_8446622_RNA45SN1_45S_50ntFlanks</td><td>8168</td><td>8188</td><td>21</td><td>CCCAAGTCCTTCTGATCGAGG</td><td>360.9</td><td>-10.1</td><td>1.4</td><td>-7.0</td><td>2.0E-12</td><td>4.8E-12</td><td>MB231/MB468</td></tr>
<tr><td>>hg38_21_+_8433172_8446622_RNA45SN1_45S_50ntFlanks@10412.10430.19|AGTCGGTCCTGAGAGATGG</td><td>>hg38_21_+_8433172_8446622_RNA45SN1_45S_50ntFlanks</td><td>10412</td><td>10430</td><td>19</td><td>AGTCGGTCCTGAGAGATGG</td><td>144.3</td><td>-8.7</td><td>1.2</td><td>-7.0</td><td>2.0E-12</td><td>4.8E-12</td><td>MB231/MB468</td></tr>
<tr><td>>hg38_21_+_8433172_8446622_RNA45SN1_45S_50ntFlanks@3985.4026.42|CGGCTTTGGTGACTCTAGATAACCTCGGGCCGATCGCACGCC</td><td>>hg38_21_+_8433172_8446622_RNA45SN1_45S_50ntFlanks</td><td>3985</td><td>4026</td><td>42</td><td>CGGCTTTGGTGACTCTAGATAACCTCGGGCCGATCGCACGCC</td><td>639.4</td><td>-11.3</td><td>1.6</td><td>-7.0</td><td>2.3E-12</td><td>4.8E-12</td><td>BT20/MB468</td></tr>
<tr><td>>hg38_21_+_8433172_8446622_RNA45SN1_45S_50ntFlanks@12433.12455.23|CTATCATTGTGAAGCAGAATTCA</td><td>>hg38_21_+_8433172_8446622_RNA45SN1_45S_50ntFlanks</td><td>12433</td><td>12455</td><td>23</td><td>CTATCATTGTGAAGCAGAATTCA</td><td>154.6</td><td>-8.8</td><td>1.3</td><td>-7.0</td><td>2.0E-12</td><td>4.8E-12</td><td>MB231/MB468</td></tr>
<tr><td>>hg38_21_+_8433172_8446622_RNA45SN1_45S_50ntFlanks@12636.12661.26|AGCTACCATCTGTGGGATTATGACTG</td><td>>hg38_21_+_8433172_8446622_RNA45SN1_45S_50ntFlanks</td><td>12636</td><td>12661</td><td>26</td><td>AGCTACCATCTGTGGGATTATGACTG</td><td>804.8</td><td>-3.5</td><td>0.5</td><td>-7.0</td><td>2.0E-12</td><td>4.8E-12</td><td>MB231/MB468</td></tr>
<tr><td>>hg38_21_+_8433172_8446622_RNA45SN1_45S_50ntFlanks@10482.10512.31|TCGAAAGGGAGTCGGGTTCAGATCCCCGAAT</td><td>>hg38_21_+_8433172_8446622_RNA45SN1_45S_50ntFlanks</td><td>10482</td><td>10512</td><td>31</td><td>TCGAAAGGGAGTCGGGTTCAGATCCCCGAAT</td><td>162.2</td><td>-9.3</td><td>1.3</td><td>-7.0</td><td>2.3E-12</td><td>4.8E-12</td><td>BT20/MB468</td></tr>
<tr><td>>hg38_1_-_228634819_228635039_RNA5S12_5S_50ntFlanks@136.170.35|GGAGACCGCCTGGGAATACCGGGTGCTGTAGGCTT</td><td>>hg38_1_-_228634819_228635039_RNA5S12_5S_50ntFlanks</td><td>136</td><td>170</td><td>35</td><td>GGAGACCGCCTGGGAATACCGGGTGCTGTAGGCTT</td><td>2925.4</td><td>5.1</td><td>0.7</td><td>7.0</td><td>2.3E-12</td><td>4.8E-12</td><td>BT20/MB468</td></tr>
<tr><td>>hg38_21_+_8433172_8446622_RNA45SN1_45S_50ntFlanks@10384.10406.23|CCATGTGAACAGCAGTTGAACAT</td><td>>hg38_21_+_8433172_8446622_RNA45SN1_45S_50ntFlanks</td><td>10384</td><td>10406</td><td>23</td><td>CCATGTGAACAGCAGTTGAACAT</td><td>158.5</td><td>-8.9</td><td>1.3</td><td>-7.0</td><td>2.1E-12</td><td>4.9E-12</td><td>MB231/MB468</td></tr>
<tr><td>>hg38_21_+_8433172_8446622_RNA45SN1_45S_50ntFlanks@12003.12033.31|GCGGGCCGCCGGTGAAATACCACTACTCTGA</td><td>>hg38_21_+_8433172_8446622_RNA45SN1_45S_50ntFlanks</td><td>12003</td><td>12033</td><td>31</td><td>GCGGGCCGCCGGTGAAATACCACTACTCTGA</td><td>1679.2</td><td>6.1</td><td>0.9</td><td>7.0</td><td>2.1E-12</td><td>4.9E-12</td><td>MB231/MB468</td></tr>
<tr><td>>hg38_21_+_8433172_8446622_RNA45SN1_45S_50ntFlanks@10348.10372.25|AAACGAGAACTTTGAAGGCCGAAGT</td><td>>hg38_21_+_8433172_8446622_RNA45SN1_45S_50ntFlanks</td><td>10348</td><td>10372</td><td>25</td><td>AAACGAGAACTTTGAAGGCCGAAGT</td><td>2128.1</td><td>4.8</td><td>0.7</td><td>7.0</td><td>2.1E-12</td><td>4.9E-12</td><td>MB231/MB468</td></tr>
<tr><td>>hg38_21_+_8433172_8446622_RNA45SN1_45S_50ntFlanks@10807.10840.34|ACAATGTAGGTAAGGGAAGTCGGCAAGCCGGATC</td><td>>hg38_21_+_8433172_8446622_RNA45SN1_45S_50ntFlanks</td><td>10807</td><td>10840</td><td>34</td><td>ACAATGTAGGTAAGGGAAGTCGGCAAGCCGGATC</td><td>146.4</td><td>-5.3</td><td>0.8</td><td>-7.0</td><td>2.4E-12</td><td>5.0E-12</td><td>BT20/MB468</td></tr>
<tr><td>>hg38_21_+_8433172_8446622_RNA45SN1_45S_50ntFlanks@10384.10405.22|CCATGTGAACAGCAGTTGAACA</td><td>>hg38_21_+_8433172_8446622_RNA45SN1_45S_50ntFlanks</td><td>10384</td><td>10405</td><td>22</td><td>CCATGTGAACAGCAGTTGAACA</td><td>509.6</td><td>-3.2</td><td>0.5</td><td>-7.0</td><td>2.1E-12</td><td>5.0E-12</td><td>MB231/MB468</td></tr>
<tr><td>>hg38_21_+_8433172_8446622_RNA45SN1_45S_50ntFlanks@11579.11603.25|AGAACTGGTGCGGACCAGGGGAATC</td><td>>hg38_21_+_8433172_8446622_RNA45SN1_45S_50ntFlanks</td><td>11579</td><td>11603</td><td>25</td><td>AGAACTGGTGCGGACCAGGGGAATC</td><td>2241.5</td><td>-4.3</td><td>0.6</td><td>-7.0</td><td>2.1E-12</td><td>5.1E-12</td><td>MB231/MB468</td></tr>
<tr><td>>hg38_21_+_8433172_8446622_RNA45SN1_45S_50ntFlanks@8115.8143.29|CGCGGGACATGTGGCGTACGGAAGACCCG</td><td>>hg38_21_+_8433172_8446622_RNA45SN1_45S_50ntFlanks</td><td>8115</td><td>8143</td><td>29</td><td>CGCGGGACATGTGGCGTACGGAAGACCCG</td><td>1235.7</td><td>-6.9</td><td>1.0</td><td>-7.0</td><td>2.2E-12</td><td>5.1E-12</td><td>MB231/MB468</td></tr>
<tr><td>>trnaMT_ProTGG_MT_-_15956_16023@40.58.19|AATGGTGGAGTTAAAGACT</td><td>>trnaMT_ProTGG_MT_-_15956_16023</td><td>40</td><td>58</td><td>19</td><td>AATGGTGGAGTTAAAGACT</td><td>571.3</td><td>8.8</td><td>1.3</td><td>7.0</td><td>2.4E-12</td><td>5.1E-12</td><td>BT20/MB468</td></tr>
<tr><td>>hg38_21_+_8433172_8446622_RNA45SN1_45S_50ntFlanks@10404.10435.32|CATGGGTCAGTCGGTCCTGAGAGATGGGCGAG</td><td>>hg38_21_+_8433172_8446622_RNA45SN1_45S_50ntFlanks</td><td>10404</td><td>10435</td><td>32</td><td>CATGGGTCAGTCGGTCCTGAGAGATGGGCGAG</td><td>111.2</td><td>-8.8</td><td>1.3</td><td>-7.0</td><td>2.4E-12</td><td>5.1E-12</td><td>BT20/MB468</td></tr>
<tr><td>>hg38_21_+_8433172_8446622_RNA45SN1_45S_50ntFlanks@8380.8404.25|CAAGAGGGCGTGAAACCGTTAAGAG</td><td>>hg38_21_+_8433172_8446622_RNA45SN1_45S_50ntFlanks</td><td>8380</td><td>8404</td><td>25</td><td>CAAGAGGGCGTGAAACCGTTAAGAG</td><td>318.0</td><td>-5.9</td><td>0.8</td><td>-7.0</td><td>2.2E-12</td><td>5.2E-12</td><td>MB231/MB468</td></tr>
<tr><td>>hg38_21_+_8433172_8446622_RNA45SN1_45S_50ntFlanks@12632.12659.28|GCGAAGCTACCATCTGTGGGATTATGAC</td><td>>hg38_21_+_8433172_8446622_RNA45SN1_45S_50ntFlanks</td><td>12632</td><td>12659</td><td>28</td><td>GCGAAGCTACCATCTGTGGGATTATGAC</td><td>246.2</td><td>-4.4</td><td>0.6</td><td>-7.0</td><td>2.4E-12</td><td>5.2E-12</td><td>BT20/MB468</td></tr>
<tr><td>>hg38_21_+_8433172_8446622_RNA45SN1_45S_50ntFlanks@11873.11899.27|AAGAAGACCCTGTTGAGCTTGACTCTA</td><td>>hg38_21_+_8433172_8446622_RNA45SN1_45S_50ntFlanks</td><td>11873</td><td>11899</td><td>27</td><td>AAGAAGACCCTGTTGAGCTTGACTCTA</td><td>300.5</td><td>-3.3</td><td>0.5</td><td>-7.0</td><td>2.5E-12</td><td>5.2E-12</td><td>BT20/MB468</td></tr>
<tr><td>>am_trna129_MetCAT_6_-_27870271_27870342@34.51.18,trna142_MetCAT_6_-_27560600_27560671@34.51.18,trna150_MetCAT_6_-_27300764_27300835@34.51.18,trna169_MetCAT_6_-_26330529_26330600@34.51.18,trna171_MetCAT_6_-_26313352_26313423@34.51.18,trna20_MetCAT_17_-_80452597_80452668@34.51.18,trna2_MetCAT_6_+_26286754_26286825@34.51.18,trna32_MetCAT_1_+_153643726_153643797@34.51.18,trna61_MetCAT_6_+_27745664_27745735@34.51.18|ATAACCCAGAGGTCGATG</td><td>>am_trna129_MetCAT_6_-_27870271_27870342</td><td>34</td><td>51</td><td>18</td><td>ATAACCCAGAGGTCGATG</td><td>257.6</td><td>-10.0</td><td>1.4</td><td>-7.0</td><td>2.5E-12</td><td>5.2E-12</td><td>BT20/MB468</td></tr>
<tr><td>>hg38_21_+_8433172_8446622_RNA45SN1_45S_50ntFlanks@12588.12610.23|GGTTCAGACATTTGGTGTATGTG</td><td>>hg38_21_+_8433172_8446622_RNA45SN1_45S_50ntFlanks</td><td>12588</td><td>12610</td><td>23</td><td>GGTTCAGACATTTGGTGTATGTG</td><td>124.8</td><td>-9.0</td><td>1.3</td><td>-7.0</td><td>2.5E-12</td><td>5.3E-12</td><td>BT20/MB468</td></tr>
<tr><td>>hg38_21_+_8433172_8446622_RNA45SN1_45S_50ntFlanks@12488.12505.18|AACGTGAGCTGGGTTTAG</td><td>>hg38_21_+_8433172_8446622_RNA45SN1_45S_50ntFlanks</td><td>12488</td><td>12505</td><td>18</td><td>AACGTGAGCTGGGTTTAG</td><td>169.5</td><td>-5.0</td><td>0.7</td><td>-7.0</td><td>2.2E-12</td><td>5.3E-12</td><td>MB231/MB468</td></tr>
<tr><td>>hg38_21_+_8433172_8446622_RNA45SN1_45S_50ntFlanks@7983.8003.21|CAGATCAGACGTGGCGACCCG</td><td>>hg38_21_+_8433172_8446622_RNA45SN1_45S_50ntFlanks</td><td>7983</td><td>8003</td><td>21</td><td>CAGATCAGACGTGGCGACCCG</td><td>222.7</td><td>-5.5</td><td>0.8</td><td>-7.0</td><td>2.2E-12</td><td>5.3E-12</td><td>MB231/MB468</td></tr>
<tr><td>>hg38_21_+_8433172_8446622_RNA45SN1_45S_50ntFlanks@12933.12956.24|GCTAAACCATTCGTAGACGACCTG</td><td>>hg38_21_+_8433172_8446622_RNA45SN1_45S_50ntFlanks</td><td>12933</td><td>12956</td><td>24</td><td>GCTAAACCATTCGTAGACGACCTG</td><td>653.3</td><td>-3.8</td><td>0.5</td><td>-7.0</td><td>2.2E-12</td><td>5.3E-12</td><td>MB231/MB468</td></tr>
<tr><td>>hg38_21_+_8433172_8446622_RNA45SN1_45S_50ntFlanks@3705.3741.37|TACCTGGTTGATCCTGCCAGTAGCATATGCTTGTCTC</td><td>>hg38_21_+_8433172_8446622_RNA45SN1_45S_50ntFlanks</td><td>3705</td><td>3741</td><td>37</td><td>TACCTGGTTGATCCTGCCAGTAGCATATGCTTGTCTC</td><td>6069.0</td><td>6.8</td><td>1.0</td><td>7.0</td><td>2.5E-12</td><td>5.3E-12</td><td>BT20/MB468</td></tr>
<tr><td>>hg38_21_+_8433172_8446622_RNA45SN1_45S_50ntFlanks@6667.6686.20|ATCACTCGGCTCGTGCGTCG</td><td>>hg38_21_+_8433172_8446622_RNA45SN1_45S_50ntFlanks</td><td>6667</td><td>6686</td><td>20</td><td>ATCACTCGGCTCGTGCGTCG</td><td>171.2</td><td>-9.0</td><td>1.3</td><td>-7.0</td><td>2.3E-12</td><td>5.3E-12</td><td>MB231/MB468</td></tr>
<tr><td>>MI0000264|hsa-mir-7-2&WithFlank&15|+|88611819|88611940@38.57.20;MI0000265|hsa-mir-7-3&WithFlank&19|+|4770664|4770785@37.56.20;MI0000263|hsa-mir-7-1&WithFlank&9|-|83969742|83969863@30.49.20&[MIMAT0000252&hsa-miR-7-5p&offsets|0|-4;m-400&15|+|88611856|88611879&offsets|0|-4];[MIMAT0000252_1&hsa-miR-7-5p&offsets|0|-4;m-402&19|+|4770700|4770723&offsets|0|-4];[MIMAT0000252_2&hsa-miR-7-5p&offsets|0|-4;m-401&9|-|83969811|83969834&offsets|0|-4]|TGGAAGACTAGTGATTTTGT</td><td>>MI0000264|hsa-mir-7-2&WithFlank&15|+|88611819|88611940</td><td>38</td><td>57</td><td>20</td><td>TGGAAGACTAGTGATTTTGT</td><td>748.8</td><td>8.5</td><td>1.2</td><td>7.0</td><td>2.5E-12</td><td>5.3E-12</td><td>BT20/MB468</td></tr>
<tr><td>>hg38_21_+_8433172_8446622_RNA45SN1_45S_50ntFlanks@11561.11585.25|GCGCCTAGCAGCCGACTTAGAACTG</td><td>>hg38_21_+_8433172_8446622_RNA45SN1_45S_50ntFlanks</td><td>11561</td><td>11585</td><td>25</td><td>GCGCCTAGCAGCCGACTTAGAACTG</td><td>1204.3</td><td>-4.3</td><td>0.6</td><td>-7.0</td><td>2.3E-12</td><td>5.4E-12</td><td>MB231/MB468</td></tr>
<tr><td>>hg38_21_+_8433172_8446622_RNA45SN1_45S_50ntFlanks@3593.3610.18|CCCGTCCGTCCGTCCGTC</td><td>>hg38_21_+_8433172_8446622_RNA45SN1_45S_50ntFlanks</td><td>3593</td><td>3610</td><td>18</td><td>CCCGTCCGTCCGTCCGTC</td><td>135.0</td><td>-9.1</td><td>1.3</td><td>-7.0</td><td>2.6E-12</td><td>5.4E-12</td><td>BT20/MB468</td></tr>
<tr><td>>hg38_21_+_8433172_8446622_RNA45SN1_45S_50ntFlanks@8737.8754.18|CCCGTCCGTCCGTCCGTC</td><td>>hg38_21_+_8433172_8446622_RNA45SN1_45S_50ntFlanks</td><td>8737</td><td>8754</td><td>18</td><td>CCCGTCCGTCCGTCCGTC</td><td>135.0</td><td>-9.1</td><td>1.3</td><td>-7.0</td><td>2.6E-12</td><td>5.4E-12</td><td>BT20/MB468</td></tr>
<tr><td>>hg38_21_+_8433172_8446622_RNA45SN1_45S_50ntFlanks@6721.6750.30|AATTGCAGGACACATTGATCATCGACACTT</td><td>>hg38_21_+_8433172_8446622_RNA45SN1_45S_50ntFlanks</td><td>6721</td><td>6750</td><td>30</td><td>AATTGCAGGACACATTGATCATCGACACTT</td><td>311.7</td><td>-4.2</td><td>0.6</td><td>-7.0</td><td>2.6E-12</td><td>5.4E-12</td><td>BT20/MB468</td></tr>
<tr><td>>hg38_21_+_8433172_8446622_RNA45SN1_45S_50ntFlanks@10571.10608.38|GAGAAGCCGGCGGGAGCCCCGGGGAGAGTTCTCTTTTC</td><td>>hg38_21_+_8433172_8446622_RNA45SN1_45S_50ntFlanks</td><td>10571</td><td>10608</td><td>38</td><td>GAGAAGCCGGCGGGAGCCCCGGGGAGAGTTCTCTTTTC</td><td>140.7</td><td>-9.1</td><td>1.3</td><td>-7.0</td><td>2.6E-12</td><td>5.4E-12</td><td>BT20/MB468</td></tr>
<tr><td>>hg38_21_+_8433172_8446622_RNA45SN1_45S_50ntFlanks@6732.6774.43|ACATTGATCATCGACACTTCGAACGCACTTGCGGCCCCGGGTT</td><td>>hg38_21_+_8433172_8446622_RNA45SN1_45S_50ntFlanks</td><td>6732</td><td>6774</td><td>43</td><td>ACATTGATCATCGACACTTCGAACGCACTTGCGGCCCCGGGTT</td><td>701.5</td><td>9.2</td><td>1.3</td><td>7.0</td><td>2.6E-12</td><td>5.4E-12</td><td>BT20/MB468</td></tr>
<tr><td>>hg38_21_+_8433172_8446622_RNA45SN1_45S_50ntFlanks@7986.8018.33|ATCAGACGTGGCGACCCGCTGAATTTAAGCATA</td><td>>hg38_21_+_8433172_8446622_RNA45SN1_45S_50ntFlanks</td><td>7986</td><td>8018</td><td>33</td><td>ATCAGACGTGGCGACCCGCTGAATTTAAGCATA</td><td>114.8</td><td>-8.8</td><td>1.3</td><td>-7.0</td><td>2.6E-12</td><td>5.4E-12</td><td>BT20/MB468</td></tr>
</table>

<table>
<tr><th></th><th></th><th></th><th></th><th></th><th></th><th></th><th></th><th></th><th></th><th></th><th></th><th></th></tr>
<tr><td>>hg38_21_+_8433172_8446622_RNA45SN1_45S_50ntFlanks@4128.4162.35|CGATTCCGGAGAGGGAGCCTGAGAAACGGCTACCA</td><td>>hg38_21_+_8433172_8446622_RNA45SN1_45S_50ntFlanks</td><td>4128</td><td>4162</td><td>35</td><td>CGATTCCGGAGAGGGAGCCTGAGAAACGGCTACCA</td><td>126.1</td><td>-6.0</td><td>0.9</td><td>-7.0</td><td>2.3E-12</td><td>5.4E-12</td><td>MB231/MB468</td></tr>
<tr><td>>trna116_GluCTC_1_-_145399233_145399304@32.73.42,trna71_GluCTC_1_-_161439189_161439260@32.73.42,trna74_GluCTC_1_-_161431809_161431880@32.73.42,trna77_GluCTC_1_-_161424398_161424469@32.73.42,trna77_GluCTC_6_+_28949976_28950047@32.73.42,trna80_GluCTC_1_-_161417018_161417089@32.73.42,trna87_GluCTC_6_-_126101393_126101464@32.73.42|CTCTCACCGCCGCGGCCCGGGTTCGATTCCCGGTCAGGGAAC</td><td>>trna116_GluCTC_1_-_145399233_145399304</td><td>32</td><td>73</td><td>42</td><td>CTCTCACCGCCGCGGCCCGGGTTCGATTCCCGGTCAGGGAAC</td><td>180.4</td><td>-9.5</td><td>1.4</td><td>-7.0</td><td>2.6E-12</td><td>5.5E-12</td><td>BT20/MB468</td></tr>
<tr><td>>hg38_21_+_8433172_8446622_RNA45SN1_45S_50ntFlanks@10786.10821.36|AACAGCCTCTGGCATGTTGGAACAATGTAGGTAAGG</td><td>>hg38_21_+_8433172_8446622_RNA45SN1_45S_50ntFlanks</td><td>10786</td><td>10821</td><td>36</td><td>AACAGCCTCTGGCATGTTGGAACAATGTAGGTAAGG</td><td>319.0</td><td>-2.7</td><td>0.4</td><td>-7.0</td><td>2.6E-12</td><td>5.5E-12</td><td>BT20/MB468</td></tr>
<tr><td>>hg38_21_+_8433172_8446622_RNA45SN1_45S_50ntFlanks@12676.12708.33|AGAATCCCGCCCAGGCGGAACGATACGGCAGCG</td><td>>hg38_21_+_8433172_8446622_RNA45SN1_45S_50ntFlanks</td><td>12676</td><td>12708</td><td>33</td><td>AGAATCCCGCCCAGGCGGAACGATACGGCAGCG</td><td>143.6</td><td>-8.7</td><td>1.2</td><td>-7.0</td><td>2.3E-12</td><td>5.5E-12</td><td>MB231/MB468</td></tr>
<tr><td>>hg38_21_+_8433172_8446622_RNA45SN1_45S_50ntFlanks@10832.10862.31|AGCCGGATCCGTAACTTCGGGATAAGGATTG</td><td>>hg38_21_+_8433172_8446622_RNA45SN1_45S_50ntFlanks</td><td>10832</td><td>10862</td><td>31</td><td>AGCCGGATCCGTAACTTCGGGATAAGGATTG</td><td>139.0</td><td>-9.1</td><td>1.3</td><td>-7.0</td><td>2.6E-12</td><td>5.5E-12</td><td>BT20/MB468</td></tr>
<tr><td>>MI0000098|hsa-mir-96&WithFlank&7|-|129774686|129774775@15.37.23&[MIMAT0000095&hsa-miR-96-5p&offsets|0|0;m-248&7|-|129774739|129774761&offsets|0|0]|TTTGGCACTAGCACATTTTTGCT</td><td>>MI0000098|hsa-mir-96&WithFlank&7|-|129774686|129774775</td><td>15</td><td>37</td><td>23</td><td>TTTGGCACTAGCACATTTTTGCT</td><td>4688.2</td><td>5.2</td><td>0.7</td><td>7.0</td><td>2.4E-12</td><td>5.5E-12</td><td>MB231/MB468</td></tr>
<tr><td>>hg38_21_+_8433172_8446622_RNA45SN1_45S_50ntFlanks@11535.11553.19|TTCCGCGCGGCGCCTCGCC</td><td>>hg38_21_+_8433172_8446622_RNA45SN1_45S_50ntFlanks</td><td>11535</td><td>11553</td><td>19</td><td>TTCCGCGCGGCGCCTCGCC</td><td>120.8</td><td>-8.9</td><td>1.3</td><td>-7.0</td><td>2.6E-12</td><td>5.5E-12</td><td>BT20/MB468</td></tr>
<tr><td>>hg38_21_+_8433172_8446622_RNA45SN1_45S_50ntFlanks@11847.11872.26|AACGGGCTTGGCGGAATCAGCGGGGA</td><td>>hg38_21_+_8433172_8446622_RNA45SN1_45S_50ntFlanks</td><td>11847</td><td>11872</td><td>26</td><td>AACGGGCTTGGCGGAATCAGCGGGGA</td><td>251.0</td><td>-3.3</td><td>0.5</td><td>-7.0</td><td>2.6E-12</td><td>5.5E-12</td><td>BT20/MB468</td></tr>
<tr><td>>hg38_21_+_8433172_8446622_RNA45SN1_45S_50ntFlanks@12804.12821.18|CGCGTGCCCCGCCGCGCG</td><td>>hg38_21_+_8433172_8446622_RNA45SN1_45S_50ntFlanks</td><td>12804</td><td>12821</td><td>18</td><td>CGCGTGCCCCGCCGCGCG</td><td>97.3</td><td>-8.6</td><td>1.2</td><td>-7.0</td><td>2.7E-12</td><td>5.6E-12</td><td>BT20/MB468</td></tr>
<tr><td>>hg38_21_+_8433172_8446622_RNA45SN1_45S_50ntFlanks@12942.12972.31|TTCGTAGACGACCTGCTTCTGGGTCGGGGTT</td><td>>hg38_21_+_8433172_8446622_RNA45SN1_45S_50ntFlanks</td><td>12942</td><td>12972</td><td>31</td><td>TTCGTAGACGACCTGCTTCTGGGTCGGGGTT</td><td>352.6</td><td>-5.4</td><td>0.8</td><td>-7.0</td><td>2.4E-12</td><td>5.6E-12</td><td>MB231/MB468</td></tr>
<tr><td>>hg38_21_+_8433172_8446622_RNA45SN1_45S_50ntFlanks@8154.8178.25|CCGCTCGTGGGGGGCCCAAGTCCTT</td><td>>hg38_21_+_8433172_8446622_RNA45SN1_45S_50ntFlanks</td><td>8154</td><td>8178</td><td>25</td><td>CCGCTCGTGGGGGGCCCAAGTCCTT</td><td>129.2</td><td>-9.0</td><td>1.3</td><td>-7.0</td><td>2.7E-12</td><td>5.6E-12</td><td>BT20/MB468</td></tr>
<tr><td>>MI0002470|hsa-mir-486-1&WithFlank&8|-|41660435|41660514@10.31.22;MI0023622|hsa-mir-486-2&WithFlank&8|+|41660438|41660513@7.28.22&[MIMAT0002177&hsa-miR-486-5p&offsets|0|0;m-107&8|-|41660484|41660505&offsets|0|0];[MIMAT0002177_1&hsa-miR-486-5p&offsets|0|0;TJU_CMC_MD2.ID03138.5p-miR&offsets|0|-2;m-106&8|+|41660444|41660465&offsets|0|0]|TCCTGTACTGAGCTGCCCCGAG</td><td>>MI0002470|hsa-mir-486-1&WithFlank&8|-|41660435|41660514</td><td>10</td><td>31</td><td>22</td><td>TCCTGTACTGAGCTGCCCCGAG</td><td>197.0</td><td>-6.7</td><td>0.9</td><td>-7.4</td><td>1.8E-13</td><td>5.6E-12</td><td>BT20/MB231</td></tr>
<tr><td>>hg38_21_+_8433172_8446622_RNA45SN1_45S_50ntFlanks@11741.11758.18|TCTTAAGGTAGCCAAATG</td><td>>hg38_21_+_8433172_8446622_RNA45SN1_45S_50ntFlanks</td><td>11741</td><td>11758</td><td>18</td><td>TCTTAAGGTAGCCAAATG</td><td>284.6</td><td>-10.1</td><td>1.5</td><td>-7.0</td><td>2.7E-12</td><td>5.6E-12</td><td>BT20/MB468</td></tr>
<tr><td>>hg38_21_+_8433172_8446622_RNA45SN1_45S_50ntFlanks@4752.4782.31|GAAGACGATCAGATACCGTCGTAGTTCCGAC</td><td>>hg38_21_+_8433172_8446622_RNA45SN1_45S_50ntFlanks</td><td>4752</td><td>4782</td><td>31</td><td>GAAGACGATCAGATACCGTCGTAGTTCCGAC</td><td>109.9</td><td>-8.8</td><td>1.3</td><td>-7.0</td><td>2.7E-12</td><td>5.6E-12</td><td>BT20/MB468</td></tr>
<tr><td>>hg38_21_+_8433172_8446622_RNA45SN1_45S_50ntFlanks@12928.12956.29|CTGGCGCTAAACCATTCGTAGACGACCTG</td><td>>hg38_21_+_8433172_8446622_RNA45SN1_45S_50ntFlanks</td><td>12928</td><td>12956</td><td>29</td><td>CTGGCGCTAAACCATTCGTAGACGACCTG</td><td>102.6</td><td>-8.7</td><td>1.2</td><td>-7.0</td><td>2.7E-12</td><td>5.6E-12</td><td>BT20/MB468</td></tr>
<tr><td>>hg38_21_+_8433172_8446622_RNA45SN1_45S_50ntFlanks@3986.4023.38|GGCTTTGGTGACTCTAGATAACCTCGGGCCGATCGCAC</td><td>>hg38_21_+_8433172_8446622_RNA45SN1_45S_50ntFlanks</td><td>3986</td><td>4023</td><td>38</td><td>GGCTTTGGTGACTCTAGATAACCTCGGGCCGATCGCAC</td><td>624.6</td><td>7.0</td><td>1.0</td><td>7.0</td><td>2.7E-12</td><td>5.7E-12</td><td>BT20/MB468</td></tr>
<tr><td>>hg38_21_+_8433172_8446622_RNA45SN1_45S_50ntFlanks@10351.10388.38|CGAGAACTTTGAAGGCCGAAGTGGAGAAGGGTTCCATG</td><td>>hg38_21_+_8433172_8446622_RNA45SN1_45S_50ntFlanks</td><td>10351</td><td>10388</td><td>38</td><td>CGAGAACTTTGAAGGCCGAAGTGGAGAAGGGTTCCATG</td><td>5026.6</td><td>-6.0</td><td>0.9</td><td>-7.0</td><td>2.4E-12</td><td>5.7E-12</td><td>MB231/MB468</td></tr>
<tr><td>>hg38_21_+_8433172_8446622_RNA45SN1_45S_50ntFlanks@3873.3899.27|TAATACATGCCGACGGGCGCTGACCCC</td><td>>hg38_21_+_8433172_8446622_RNA45SN1_45S_50ntFlanks</td><td>3873</td><td>3899</td><td>27</td><td>TAATACATGCCGACGGGCGCTGACCCC</td><td>280.3</td><td>-3.2</td><td>0.5</td><td>-7.0</td><td>2.4E-12</td><td>5.7E-12</td><td>MB231/MB468</td></tr>
<tr><td>>hg38_21_+_8433172_8446622_RNA45SN1_45S_50ntFlanks@10808.10827.20|CAATGTAGGTAAGGGAAGTC</td><td>>hg38_21_+_8433172_8446622_RNA45SN1_45S_50ntFlanks</td><td>10808</td><td>10827</td><td>20</td><td>CAATGTAGGTAAGGGAAGTC</td><td>268.5</td><td>-6.4</td><td>0.9</td><td>-7.0</td><td>2.8E-12</td><td>5.8E-12</td><td>BT20/MB468</td></tr>
<tr><td>>hg38_21_+_8433172_8446622_RNA45SN1_45S_50ntFlanks@12539.12565.27|ATGATGTGTTGTTGCCATGGTAATCCT</td><td>>hg38_21_+_8433172_8446622_RNA45SN1_45S_50ntFlanks</td><td>12539</td><td>12565</td><td>27</td><td>ATGATGTGTTGTTGCCATGGTAATCCT</td><td>174.5</td><td>-9.0</td><td>1.3</td><td>-7.0</td><td>2.5E-12</td><td>5.9E-12</td><td>MB231/MB468</td></tr>
<tr><td>>hg38_21_+_8433172_8446622_RNA45SN1_45S_50ntFlanks@7987.8006.20|TCAGACGTGGCGACCCGCTG</td><td>>hg38_21_+_8433172_8446622_RNA45SN1_45S_50ntFlanks</td><td>7987</td><td>8006</td><td>20</td><td>TCAGACGTGGCGACCCGCTG</td><td>129.3</td><td>-9.0</td><td>1.3</td><td>-7.0</td><td>2.8E-12</td><td>5.9E-12</td><td>BT20/MB468</td></tr>
<tr><td>>hg38_21_+_8433172_8446622_RNA45SN1_45S_50ntFlanks@13006.13027.22|CTATTGAAAGTCAGCCCTCGAC</td><td>>hg38_21_+_8433172_8446622_RNA45SN1_45S_50ntFlanks</td><td>13006</td><td>13027</td><td>22</td><td>CTATTGAAAGTCAGCCCTCGAC</td><td>877.6</td><td>-8.4</td><td>1.2</td><td>-7.0</td><td>2.8E-12</td><td>5.9E-12</td><td>BT20/MB468</td></tr>
<tr><td>>hg38_1_-_228634819_228635039_RNA5S12_5S_50ntFlanks@151.171.21|ATACCGGGTGCTGTAGGCTTT</td><td>>hg38_1_-_228634819_228635039_RNA5S12_5S_50ntFlanks</td><td>151</td><td>171</td><td>21</td><td>ATACCGGGTGCTGTAGGCTTT</td><td>1516.9</td><td>4.8</td><td>0.7</td><td>7.0</td><td>2.8E-12</td><td>5.9E-12</td><td>BT20/MB468</td></tr>
<tr><td>>hg38_21_+_8433172_8446622_RNA45SN1_45S_50ntFlanks@5417.5443.27|CTACCGATTGGATGGTTTAGTGAGGCC</td><td>>hg38_21_+_8433172_8446622_RNA45SN1_45S_50ntFlanks</td><td>5417</td><td>5443</td><td>27</td><td>CTACCGATTGGATGGTTTAGTGAGGCC</td><td>125.4</td><td>-9.0</td><td>1.3</td><td>-7.0</td><td>2.8E-12</td><td>5.9E-12</td><td>BT20/MB468</td></tr>
<tr><td>>hg38_21_+_8433172_8446622_RNA45SN1_45S_50ntFlanks@12044.12071.28|ACTGACCCGGTGAGGCGGGGGGGCGAGC</td><td>>hg38_21_+_8433172_8446622_RNA45SN1_45S_50ntFlanks</td><td>12044</td><td>12071</td><td>28</td><td>ACTGACCCGGTGAGGCGGGGGGGCGAGC</td><td>143.4</td><td>-8.7</td><td>1.2</td><td>-7.0</td><td>2.5E-12</td><td>6.0E-12</td><td>MB231/MB468</td></tr>
<tr><td>>hg38_21_+_8433172_8446622_RNA45SN1_45S_50ntFlanks@4160.4182.23|CCACATCCAAGGAAGGCAGCAGG</td><td>>hg38_21_+_8433172_8446622_RNA45SN1_45S_50ntFlanks</td><td>4160</td><td>4182</td><td>23</td><td>CCACATCCAAGGAAGGCAGCAGG</td><td>160.6</td><td>-6.2</td><td>0.9</td><td>-7.0</td><td>2.9E-12</td><td>6.0E-12</td><td>BT20/MB468</td></tr>
<tr><td>>hg38_21_+_8433172_8446622_RNA45SN1_45S_50ntFlanks@12126.12151.26|CGACCCGCTCCGGGGACAGTGCCAGG</td><td>>hg38_21_+_8433172_8446622_RNA45SN1_45S_50ntFlanks</td><td>12126</td><td>12151</td><td>26</td><td>CGACCCGCTCCGGGGACAGTGCCAGG</td><td>119.5</td><td>-8.9</td><td>1.3</td><td>-7.0</td><td>2.9E-12</td><td>6.0E-12</td><td>BT20/MB468</td></tr>
<tr><td>>hg38_21_+_8433172_8446622_RNA45SN1_45S_50ntFlanks@7988.8006.19|CAGACGTGGCGACCCGCTG</td><td>>hg38_21_+_8433172_8446622_RNA45SN1_45S_50ntFlanks</td><td>7988</td><td>8006</td><td>19</td><td>CAGACGTGGCGACCCGCTG</td><td>149.2</td><td>-9.2</td><td>1.3</td><td>-7.0</td><td>2.9E-12</td><td>6.0E-12</td><td>BT20/MB468</td></tr>
<tr><td>>hg38_21_+_8433172_8446622_RNA45SN1_45S_50ntFlanks@5472.5501.30|ACGGCCCTGGCGGAGCGCTGAGAAGACGGT</td><td>>hg38_21_+_8433172_8446622_RNA45SN1_45S_50ntFlanks</td><td>5472</td><td>5501</td><td>30</td><td>ACGGCCCTGGCGGAGCGCTGAGAAGACGGT</td><td>1877.7</td><td>6.9</td><td>1.0</td><td>7.0</td><td>2.9E-12</td><td>6.1E-12</td><td>BT20/MB468</td></tr>
<tr><td>>hg38_21_+_8433172_8446622_RNA45SN1_45S_50ntFlanks@12633.12652.20|CGAAGCTACCATCTGTGGGA</td><td>>hg38_21_+_8433172_8446622_RNA45SN1_45S_50ntFlanks</td><td>12633</td><td>12652</td><td>20</td><td>CGAAGCTACCATCTGTGGGA</td><td>663.7</td><td>-6.1</td><td>0.9</td><td>-7.0</td><td>2.6E-12</td><td>6.2E-12</td><td>MB231/MB468</td></tr>
<tr><td>>hg38_21_+_8433172_8446622_RNA45SN1_45S_50ntFlanks@12439.12473.35|TTGTGAAGCAGAATTCACCAAGCGTTGGATTGTTC</td><td>>hg38_21_+_8433172_8446622_RNA45SN1_45S_50ntFlanks</td><td>12439</td><td>12473</td><td>35</td><td>TTGTGAAGCAGAATTCACCAAGCGTTGGATTGTTC</td><td>1011.5</td><td>4.8</td><td>0.7</td><td>7.0</td><td>2.6E-12</td><td>6.2E-12</td><td>MB231/MB468</td></tr>
<tr><td>>hg38_21_+_8433172_8446622_RNA45SN1_45S_50ntFlanks@12081.12107.27|CTCTCGCTTCTGGCGCCAAGCGCCCGG</td><td>>hg38_21_+_8433172_8446622_RNA45SN1_45S_50ntFlanks</td><td>12081</td><td>12107</td><td>27</td><td>CTCTCGCTTCTGGCGCCAAGCGCCCGG</td><td>350.9</td><td>-4.3</td><td>0.6</td><td>-7.0</td><td>2.7E-12</td><td>6.2E-12</td><td>MB231/MB468</td></tr>
<tr><td>>hg38_21_+_8433172_8446622_RNA45SN1_45S_50ntFlanks@12634.12653.20|GAAGCTACCATCTGTGGGAT</td><td>>hg38_21_+_8433172_8446622_RNA45SN1_45S_50ntFlanks</td><td>12634</td><td>12653</td><td>20</td><td>GAAGCTACCATCTGTGGGAT</td><td>211.7</td><td>-9.7</td><td>1.4</td><td>-7.0</td><td>3.0E-12</td><td>6.3E-12</td><td>BT20/MB468</td></tr>
<tr><td>>hg38_21_+_8433172_8446622_RNA45SN1_45S_50ntFlanks@10914.10946.33|CGCCGCGGCTGGACGAGGCGCCGCCGCCCCCCC</td><td>>hg38_21_+_8433172_8446622_RNA45SN1_45S_50ntFlanks</td><td>10914</td><td>10946</td><td>33</td><td>CGCCGCGGCTGGACGAGGCGCCGCCGCCCCCCC</td><td>113.1</td><td>-8.8</td><td>1.3</td><td>-7.0</td><td>3.0E-12</td><td>6.3E-12</td><td>BT20/MB468</td></tr>
<tr><td>>hg38_21_+_8433172_8446622_RNA45SN1_45S_50ntFlanks@10678.10695.18|CGGTTCCGGCGGCGTCCG</td><td>>hg38_21_+_8433172_8446622_RNA45SN1_45S_50ntFlanks</td><td>10678</td><td>10695</td><td>18</td><td>CGGTTCCGGCGGCGTCCG</td><td>102.8</td><td>-8.7</td><td>1.2</td><td>-7.0</td><td>3.0E-12</td><td>6.3E-12</td><td>BT20/MB468</td></tr>
<tr><td>>MI0000782|hsa-mir-374a&WithFlank&X|-|74287280|74287363@18.38.21&[MIMAT0000727&hsa-miR-374a-5p&offsets|0|-1;m-184&X|-|74287325|74287346&offsets|0|-1]|TTATAATACAACCTGATAAGT</td><td>>MI0000782|hsa-mir-374a&WithFlank&X|-|74287280|74287363</td><td>18</td><td>38</td><td>21</td><td>TTATAATACAACCTGATAAGT</td><td>783.0</td><td>5.2</td><td>0.7</td><td>7.0</td><td>2.7E-12</td><td>6.4E-12</td><td>MB231/MB468</td></tr>
<tr><td>>am_trnaMT_PheGAA_MT_+_577_647@44.62.19|TAGACGGGCTCACATCACC</td><td>>am_trnaMT_PheGAA_MT_+_577_647</td><td>44</td><td>62</td><td>19</td><td>TAGACGGGCTCACATCACC</td><td>1149.4</td><td>5.4</td><td>0.8</td><td>7.0</td><td>3.1E-12</td><td>6.5E-12</td><td>BT20/MB468</td></tr>
<tr><td>>MI0000102|hsa-mir-100&WithFlank&11|-|122152223|122152314@19.40.22&[MIMAT0000098&hsa-miR-100-5p&offsets|0|0;m-27&11|-|122152276|122152296&offsets|0|+1]|AACCCGTAGATCCGAACTTGTG</td><td>>MI0000102|hsa-mir-100&WithFlank&11|-|122152223|122152314</td><td>19</td><td>40</td><td>22</td><td>AACCCGTAGATCCGAACTTGTG</td><td>863377.7</td><td>-6.0</td><td>0.8</td><td>-7.3</td><td>2.1E-13</td><td>6.5E-12</td><td>BT20/MB231</td></tr>
<tr><td>>MI0000749|hsa-mir-30e&WithFlank&1|+|40754349|40754452@65.86.22&[MIMAT0000693&hsa-miR-30e-3p&offsets|0|0;m-25&1|+|40754413|40754434&offsets|0|0]|CTTTCAGTCGGATGTTTACAGC</td><td>>MI0000749|hsa-mir-30e&WithFlank&1|+|40754349|40754452</td><td>65</td><td>86</td><td>22</td><td>CTTTCAGTCGGATGTTTACAGC</td><td>3347.7</td><td>4.6</td><td>0.7</td><td>7.0</td><td>3.1E-12</td><td>6.5E-12</td><td>BT20/MB468</td></tr>
<tr><td>>MI0000270|hsa-mir-181b-1&WithFlank&1|-|198858867|198858988@42.63.22(+1A);MI0000683|hsa-mir-181b-2&WithFlank&9|+|124693704|124693804@22.43.22(+1A)&[MIMAT0000257&hsa-miR-181b-5p&offsets|0|-1(+1A);m-73&1|-|198858925|198858947&offsets|0|-1(+1A)];[MIMAT0000257_1&hsa-miR-181b-5p&offsets|0|-1(+1A);m-72&9|+|124693725|124693747&offsets|0|-1(+1A)]|AACATTCATTGCTGTCGGTGGGA</td><td>>MI0000270|hsa-mir-181b-1&WithFlank&1|-|198858867|198858988</td><td>42</td><td>63</td><td>22(+1A)</td><td>AACATTCATTGCTGTCGGTGGGA</td><td>419.1</td><td>5.9</td><td>0.8</td><td>7.0</td><td>3.1E-12</td><td>6.5E-12</td><td>BT20/MB468</td></tr>
<tr><td>>hg38_21_+_8433172_8446622_RNA45SN1_45S_50ntFlanks@12938.12955.18|ACCATTCGTAGACGACCT</td><td>>hg38_21_+_8433172_8446622_RNA45SN1_45S_50ntFlanks</td><td>12938</td><td>12955</td><td>18</td><td>ACCATTCGTAGACGACCT</td><td>97.7</td><td>-3.6</td><td>0.5</td><td>-7.0</td><td>3.1E-12</td><td>6.5E-12</td><td>BT20/MB468</td></tr>
</table>

| | | | | | | | | | | | | |
|---|---|---|---|---|---|---|---|---|---|---|---|---|
| >trna10_AspGTC_12_-_125424193_125424264@36.73.38,trna12_AspGTC_12_-_125411891_125411962@36.73.38,trna144_AspGTC_6_-_27551236_27551307@36.73.38,trna38_AspGTC_17_-_8125556_8125627@36.73.38,trna45_AspGTC_6_+_27447453_27447524@36.73.38,trna48_AspGTC_6_+_27471523_27471594@36.73.38,trna4_AspGTC_12_+_96429799_96429870@36.73.38,trna69_AspGTC_1_-_161440205_161440276@36.73.38,trna72_AspGTC_1_-_161432824_161432895@36.73.38,trna75_AspGTC_1_-_161425414_161425485@36.73.38,trna78_AspGTC_1_-_161418033_161418104@36.73.38,trna81_AspGTC_1_-_161410615_161410686@36.73.38\|CACGCGGGAGACCGGGGTTCGATTCCCCGACGGGGAGC | >trna10_AspGTC_12_-_125424193_125424264 | 36 | 73 | 38 | CACGCGGGAGACCGGGGTTCGATTCCCCGACGGGGAGC | 1235.2 | 8.7 | 1.2 | 7.0 | 2.8E-12 | 6.5E-12 | MB231/MB468 |
| >MI0000086\|hsa-mir-28&WithFlank&3\|+\|188688775\|188688872@20.41.22(+1A)&[MIMAT0000085&hsa-miR-28-5p&offsets\|0\|0(+1A);m-133&3\|+\|188688794\|188688815&offsets\|0\|0(+1A)]\|AAGGAGCTCACAGTCTATTGAGA | >MI0000086\|hsa-mir-28&WithFlank&3\|+\|188688775\|188688872 | 20 | 41 | 22(+1A) | AAGGAGCTCACAGTCTATTGAGA | 1038.3 | 9.7 | 1.4 | 7.0 | 3.1E-12 | 6.5E-12 | BT20/MB468 |
| >MI0000077\|hsa-mir-21&WithFlank&17\|+\|59841260\|59841343@15.35.21&[MIMAT0000076&hsa-miR-21-5p&offsets\|+1\|0;m-1&17\|+\|59841273\|59841294&offsets\|+1\|0]\|AGCTTATCAGACTGATGTTGA | >MI0000077\|hsa-mir-21&WithFlank&17\|+\|59841260\|59841343 | 15 | 35 | 21 | AGCTTATCAGACTGATGTTGA | 526.4 | 10.0 | 1.4 | 7.0 | 3.2E-12 | 6.6E-12 | BT20/MB468 |
| >hg38_21_+_8433172_8446622_RNA45SN1_45S_50ntFlanks@6651.6692.42\|CGACTCTTAGCGGTGGATCACTCGGCTCGTGCGTCGATGAAG | >hg38_21_+_8433172_8446622_RNA45SN1_45S_50ntFlanks | 6651 | 6692 | 42 | CGACTCTTAGCGGTGGATCACTCGGCTCGTGCGTCGATGAAG | 9746.1 | 4.2 | 0.6 | 7.0 | 2.8E-12 | 6.6E-12 | MB231/MB468 |
| >hg38_21_+_8433172_8446622_RNA45SN1_45S_50ntFlanks@11729.11766.38\|TAACTATGACTCTCTTAAGGTAGCCAAATGCCTCGTCA | >hg38_21_+_8433172_8446622_RNA45SN1_45S_50ntFlanks | 11729 | 11766 | 38 | TAACTATGACTCTCTTAAGGTAGCCAAATGCCTCGTCA | 150.0 | -9.2 | 1.3 | -7.0 | 3.2E-12 | 6.6E-12 | BT20/MB468 |
| >hg38_21_+_8433172_8446622_RNA45SN1_45S_50ntFlanks@11315.11332.18\|CCACCCCACGTCTCGTCG | >hg38_21_+_8433172_8446622_RNA45SN1_45S_50ntFlanks | 11315 | 11332 | 18 | CCACCCCACGTCTCGTCG | 286.3 | -9.7 | 1.4 | -7.0 | 2.8E-12 | 6.6E-12 | MB231/MB468 |
| >MI0003666\|hsa-mir-651&WithFlank&X\|+\|8126959\|8127067@22.43.22&[MIMAT0003321&hsa-miR-651-5p&offsets\|0\|0;m-399&X\|+\|8126980\|8127001&offsets\|0\|0]\|TTTAGGATAAGCTTGACTTTTG | >MI0003666\|hsa-mir-651&WithFlank&X\|+\|8126959\|8127067 | 22 | 43 | 22 | TTTAGGATAAGCTTGACTTTTG | 1434.8 | 8.9 | 1.3 | 7.0 | 3.2E-12 | 6.7E-12 | BT20/MB468 |
| >hg38_21_+_8433172_8446622_RNA45SN1_45S_50ntFlanks@10408.10426.19\|GGTCAGTCGGTCCTGAGAG | >hg38_21_+_8433172_8446622_RNA45SN1_45S_50ntFlanks | 10408 | 10426 | 19 | GGTCAGTCGGTCCTGAGAG | 413.7 | -6.4 | 0.9 | -7.0 | 2.8E-12 | 6.7E-12 | MB231/MB468 |
| >hg38_21_+_8433172_8446622_RNA45SN1_45S_50ntFlanks@4044.4063.20\|CCCATTCGAACGTCTGCCCT | >hg38_21_+_8433172_8446622_RNA45SN1_45S_50ntFlanks | 4044 | 4063 | 20 | CCCATTCGAACGTCTGCCCT | 99.4 | -8.6 | 1.2 | -7.0 | 3.2E-12 | 6.8E-12 | BT20/MB468 |
| >hg38_21_+_8433172_8446622_RNA45SN1_45S_50ntFlanks@12329.12346.18\|TTTTAAGCAGGAGGTGTC | >hg38_21_+_8433172_8446622_RNA45SN1_45S_50ntFlanks | 12329 | 12346 | 18 | TTTTAAGCAGGAGGTGTC | 213.3 | -4.3 | 0.6 | -7.0 | 2.9E-12 | 6.8E-12 | MB231/MB468 |
| >hg38_21_+_8433172_8446622_RNA45SN1_45S_50ntFlanks@8196.8215.20\|CGTGGACGGTGTGAGGCCGG | >hg38_21_+_8433172_8446622_RNA45SN1_45S_50ntFlanks | 8196 | 8215 | 20 | CGTGGACGGTGTGAGGCCGG | 1081.5 | -3.9 | 0.6 | -7.0 | 2.9E-12 | 6.8E-12 | MB231/MB468 |
| >hg38_21_+_8433172_8446622_RNA45SN1_45S_50ntFlanks@9166.9197.32\|CCGGGCGGGTCGCGCCGTCGGGCCCGGGGGAG | >hg38_21_+_8433172_8446622_RNA45SN1_45S_50ntFlanks | 9166 | 9197 | 32 | CCGGGCGGGTCGCGCCGTCGGGCCCGGGGGAG | 283.7 | -6.8 | 1.0 | -7.0 | 3.3E-12 | 6.8E-12 | BT20/MB468 |
| >hg38_21_+_8433172_8446622_RNA45SN1_45S_50ntFlanks@6652.6687.36\|GACTCTTAGCGGTGGATCACTCGGCTCGTGCGTCGA | >hg38_21_+_8433172_8446622_RNA45SN1_45S_50ntFlanks | 6652 | 6687 | 36 | GACTCTTAGCGGTGGATCACTCGGCTCGTGCGTCGA | 1548.3 | 5.1 | 0.7 | 7.0 | 3.3E-12 | 6.8E-12 | BT20/MB468 |
| >hg38_21_+_8433172_8446622_RNA45SN1_45S_50ntFlanks@4002.4025.24\|GATAACCTCGGGCCGATCGCACGC | >hg38_21_+_8433172_8446622_RNA45SN1_45S_50ntFlanks | 4002 | 4025 | 24 | GATAACCTCGGGCCGATCGCACGC | 192.7 | -6.0 | 0.9 | -7.0 | 3.3E-12 | 6.8E-12 | BT20/MB468 |
| >hg38_21_+_8433172_8446622_RNA45SN1_45S_50ntFlanks@8737.8758.22\|CCCGTCCGTCCGTCCGTCCGTC | >hg38_21_+_8433172_8446622_RNA45SN1_45S_50ntFlanks | 8737 | 8758 | 22 | CCCGTCCGTCCGTCCGTCCGTC | 147.7 | -9.2 | 1.3 | -7.0 | 3.3E-12 | 6.8E-12 | BT20/MB468 |
| >hg38_21_+_8433172_8446622_RNA45SN1_45S_50ntFlanks@9736.9755.20\|CCGAAACGATCTCAACCTAT | >hg38_21_+_8433172_8446622_RNA45SN1_45S_50ntFlanks | 9736 | 9755 | 20 | CCGAAACGATCTCAACCTAT | 195.5 | -4.8 | 0.7 | -7.0 | 2.9E-12 | 6.8E-12 | MB231/MB468 |
| >hg38_21_+_8433172_8446622_RNA45SN1_45S_50ntFlanks@7989.8018.30\|AGACGTGGCGACCCGCTGAATTTAAGCATA | >hg38_21_+_8433172_8446622_RNA45SN1_45S_50ntFlanks | 7989 | 8018 | 30 | AGACGTGGCGACCCGCTGAATTTAAGCATA | 370.3 | 4.2 | 0.6 | 7.0 | 2.9E-12 | 6.9E-12 | MB231/MB468 |
| >hg38_21_+_8433172_8446622_RNA45SN1_45S_50ntFlanks@10384.10405.22\|CCATGTGAACAGCAGTTGAACA | >hg38_21_+_8433172_8446622_RNA45SN1_45S_50ntFlanks | 10384 | 10405 | 22 | CCATGTGAACAGCAGTTGAACA | 579.3 | -4.7 | 0.7 | -7.0 | 3.3E-12 | 6.9E-12 | BT20/MB468 |
| >hg38_21_+_8433172_8446622_RNA45SN1_45S_50ntFlanks@12638.12657.20\|CTACCATCTGTGGGATTATG | >hg38_21_+_8433172_8446622_RNA45SN1_45S_50ntFlanks | 12638 | 12657 | 20 | CTACCATCTGTGGGATTATG | 313.9 | -7.1 | 1.0 | -7.0 | 3.3E-12 | 6.9E-12 | BT20/MB468 |
| >hg38_21_+_8433172_8446622_RNA45SN1_45S_50ntFlanks@11881.11905.25\|CCTGTTGAGCTTGACTCTAGTCTGG | >hg38_21_+_8433172_8446622_RNA45SN1_45S_50ntFlanks | 11881 | 11905 | 25 | CCTGTTGAGCTTGACTCTAGTCTGG | 141.7 | -8.7 | 1.2 | -7.0 | 2.9E-12 | 6.9E-12 | MB231/MB468 |
| >hg38_21_+_8433172_8446622_RNA45SN1_45S_50ntFlanks@12358.12376.19\|CACAGGGATAACTGGCTTG | >hg38_21_+_8433172_8446622_RNA45SN1_45S_50ntFlanks | 12358 | 12376 | 19 | CACAGGGATAACTGGCTTG | 143.5 | -9.2 | 1.3 | -7.0 | 3.3E-12 | 6.9E-12 | BT20/MB468 |
| >MI0000108\|hsa-mir-103a-2&WithFlank&20\|+\|3917488\|3917577@56.76.21;MI0000109\|hsa-mir-103a-1&WithFlank&5\|-\|168560890\|168560979@56.76.21&[MIMAT0000101&hsa-miR-103a-3p&offsets\|+2\|0;m-12&20\|+\|3917541\|3917563&offsets\|+2\|0];[MIMAT0000101_1&hsa-miR-103a-3p&offsets\|+2\|0;m-13&5\|-\|168560904\|168560926&offsets\|+2\|0;m-44220&5\|-\|168560908\|168560926&offsets\|+2\|+4]\|CAGCATTGTACAGGGCTATGA | >MI0000108\|hsa-mir-103a-2&WithFlank&20\|+\|3917488\|3917577 | 56 | 76 | 21 | CAGCATTGTACAGGGCTATGA | 839.9 | 6.2 | 0.9 | 7.0 | 3.3E-12 | 6.9E-12 | BT20/MB468 |
| >MI0003834\|hsa-mir-769&WithFlank&19\|+\|46018926\|46019055@36.56.21&[MIMAT0003886&hsa-miR-769-5p&offsets\|0\|-1;m-216&19\|+\|46018961\|46018982&offsets\|0\|-1]\|TGAGACCTCTGGGTTCTGAGC | >MI0003834\|hsa-mir-769&WithFlank&19\|+\|46018926\|46019055 | 36 | 56 | 21 | TGAGACCTCTGGGTTCTGAGC | 1749.4 | 8.8 | 1.3 | 7.0 | 3.4E-12 | 7.1E-12 | BT20/MB468 |
| >hg38_21_+_8433172_8446622_RNA45SN1_45S_50ntFlanks@6790.6807.18\|CCTGTCTGAGCGTCGCTT | >hg38_21_+_8433172_8446622_RNA45SN1_45S_50ntFlanks | 6790 | 6807 | 18 | CCTGTCTGAGCGTCGCTT | 1097.5 | 4.8 | 0.7 | 7.0 | 3.4E-12 | 7.1E-12 | BT20/MB468 |
| >hg38_21_+_8433172_8446622_RNA45SN1_45S_50ntFlanks@4322.4339.18\|CAGCCGCGGTAATTCCAG | >hg38_21_+_8433172_8446622_RNA45SN1_45S_50ntFlanks | 4322 | 4339 | 18 | CAGCCGCGGTAATTCCAG | 124.7 | -9.0 | 1.3 | -7.0 | 3.4E-12 | 7.1E-12 | BT20/MB468 |
| >hg38_21_+_8433172_8446622_RNA45SN1_45S_50ntFlanks@10808.10826.19\|CAATGTAGGTAAGGGAAGT | >hg38_21_+_8433172_8446622_RNA45SN1_45S_50ntFlanks | 10808 | 10826 | 19 | CAATGTAGGTAAGGGAAGT | 663.4 | -6.2 | 0.9 | -7.0 | 3.1E-12 | 7.2E-12 | MB231/MB468 |
| >hg38_21_+_8433172_8446622_RNA45SN1_45S_50ntFlanks@3874.3898.25\|AATACATGCCGACGGGCGCTGACCC | >hg38_21_+_8433172_8446622_RNA45SN1_45S_50ntFlanks | 3874 | 3898 | 25 | AATACATGCCGACGGGCGCTGACCC | 1747.0 | -4.4 | 0.6 | -7.0 | 3.1E-12 | 7.2E-12 | MB231/MB468 |
| >MI0000471\|hsa-mir-126&WithFlank&9\|+\|136670596\|136670692@58.78.21(+1A)&[MIMAT0000445&hsa-miR-126-3p&offsets\|0\|-1(+1A);m-40&9\|+\|136670653\|136670674&offsets\|0\|-1(+1A)]\|TCGTACCGTGAGTAATAATGCA | >MI0000471\|hsa-mir-126&WithFlank&9\|+\|136670596\|136670692 | 58 | 78 | 21(+1A) | TCGTACCGTGAGTAATAATGCA | 2821.3 | 10.4 | 1.5 | 7.0 | 3.5E-12 | 7.2E-12 | BT20/MB468 |
| >hg38_21_+_8433172_8446622_RNA45SN1_45S_50ntFlanks@12636.12659.24\|AGCTACCATCTGTGGGATTATGAC | >hg38_21_+_8433172_8446622_RNA45SN1_45S_50ntFlanks | 12636 | 12659 | 24 | AGCTACCATCTGTGGGATTATGAC | 569.9 | -4.4 | 0.6 | -7.0 | 3.1E-12 | 7.2E-12 | MB231/MB468 |
| >hg38_21_+_8433172_8446622_RNA45SN1_45S_50ntFlanks@12574.12604.31\|CGAGAGGAACCGCAGGTTCAGACATTTGGTG | >hg38_21_+_8433172_8446622_RNA45SN1_45S_50ntFlanks | 12574 | 12604 | 31 | CGAGAGGAACCGCAGGTTCAGACATTTGGTG | 105.1 | -8.7 | 1.3 | -7.0 | 3.5E-12 | 7.2E-12 | BT20/MB468 |
| >hg38_21_+_8433172_8446622_RNA45SN1_45S_50ntFlanks@10349.10372.24\|AACGAGAACTTTGAAGGCCGAAGT | >hg38_21_+_8433172_8446622_RNA45SN1_45S_50ntFlanks | 10349 | 10372 | 24 | AACGAGAACTTTGAAGGCCGAAGT | 3011.9 | 5.1 | 0.7 | 7.0 | 3.1E-12 | 7.3E-12 | MB231/MB468 |
| >hg38_21_+_8433172_8446622_RNA45SN1_45S_50ntFlanks@12412.12437.26\|ATCCTTCGATGTCGGCTCTTCCTATC | >hg38_21_+_8433172_8446622_RNA45SN1_45S_50ntFlanks | 12412 | 12437 | 26 | ATCCTTCGATGTCGGCTCTTCCTATC | 126.4 | -9.0 | 1.3 | -7.0 | 3.5E-12 | 7.3E-12 | BT20/MB468 |
| >hg38_21_+_8433172_8446622_RNA45SN1_45S_50ntFlanks@7984.8007.24\|AGATCAGACGTGGCGACCCGCTGA | >hg38_21_+_8433172_8446622_RNA45SN1_45S_50ntFlanks | 7984 | 8007 | 24 | AGATCAGACGTGGCGACCCGCTGA | 179.7 | -4.7 | 0.7 | -7.0 | 3.6E-12 | 7.4E-12 | BT20/MB468 |
| >hg38_21_+_8433172_8446622_RNA45SN1_45S_50ntFlanks@6738.6769.32\|ATCATCGACACTTCGAACGCACTTGCGGCCCC | >hg38_21_+_8433172_8446622_RNA45SN1_45S_50ntFlanks | 6738 | 6769 | 32 | ATCATCGACACTTCGAACGCACTTGCGGCCCC | 116.0 | -8.9 | 1.3 | -7.0 | 3.6E-12 | 7.4E-12 | BT20/MB468 |
| >hg38_21_+_8433172_8446622_RNA45SN1_45S_50ntFlanks@10479.10512.34\|CGATCGAAAGGGAGTCGGGTTCAGATCCCCGAAT | >hg38_21_+_8433172_8446622_RNA45SN1_45S_50ntFlanks | 10479 | 10512 | 34 | CGATCGAAAGGGAGTCGGGTTCAGATCCCCGAAT | 138.4 | -9.1 | 1.3 | -7.0 | 3.6E-12 | 7.4E-12 | BT20/MB468 |
| >hg38_21_+_8433172_8446622_RNA45SN1_45S_50ntFlanks@12419.12437.19\|GATGTCGGCTCTTCCTATC | >hg38_21_+_8433172_8446622_RNA45SN1_45S_50ntFlanks | 12419 | 12437 | 19 | GATGTCGGCTCTTCCTATC | 8391.0 | 4.3 | 0.6 | 7.0 | 3.2E-12 | 7.4E-12 | MB231/MB468 |
| >MI0000102\|hsa-mir-100&WithFlank&11\|-\|122152223\|122152314@22.40.19&[MIMAT0000098&hsa-miR-100-5p&offsets\|+3\|0;m-27&11\|-\|122152276\|122152296&offsets\|+3\|+1]\|CCGTAGATCCGAACTTGTG | >MI0000102\|hsa-mir-100&WithFlank&11\|-\|122152223\|122152314 | 22 | 40 | 19 | CCGTAGATCCGAACTTGTG | 3210.7 | 16.6 | 2.4 | 7.0 | 3.2E-12 | 7.4E-12 | MB231/MB468 |
| >hg38_21_+_8433172_8446622_RNA45SN1_45S_50ntFlanks@10351.10388.38\|CGAGAACTTTGAAGGCCGAAGTGGAGAAGGGTTCCATG | >hg38_21_+_8433172_8446622_RNA45SN1_45S_50ntFlanks | 10351 | 10388 | 38 | CGAGAACTTTGAAGGCCGAAGTGGAGAAGGGTTCCATG | 6289.4 | -6.7 | 1.0 | -7.0 | 3.6E-12 | 7.4E-12 | BT20/MB468 |
| >hg38_21_+_8433172_8446622_RNA45SN1_45S_50ntFlanks@12499.12521.23\|GGTTTAGACCGTCGTGAGACAGG | >hg38_21_+_8433172_8446622_RNA45SN1_45S_50ntFlanks | 12499 | 12521 | 23 | GGTTTAGACCGTCGTGAGACAGG | 105.8 | -8.7 | 1.3 | -7.0 | 3.6E-12 | 7.4E-12 | BT20/MB468 |
| >hg38_21_+_8433172_8446622_RNA45SN1_45S_50ntFlanks@11885.11912.28\|TTGAGCTTGACTCTAGTCTGGCACGGTG | >hg38_21_+_8433172_8446622_RNA45SN1_45S_50ntFlanks | 11885 | 11912 | 28 | TTGAGCTTGACTCTAGTCTGGCACGGTG | 135.1 | -8.7 | 1.2 | -7.0 | 3.2E-12 | 7.4E-12 | MB231/MB468 |
| >MI0002468\|hsa-mir-484&WithFlank&16\|+\|15643288\|15643378@14.36.23&[MIMAT0002174&hsa-miR-484&offsets\|0\|+1;m-127&16\|+\|15643301\|15643321&offsets\|0\|+2]\|TCAGGCTCAGTCCCCTCCCGATA | >MI0002468\|hsa-mir-484&WithFlank&16\|+\|15643288\|15643378 | 14 | 36 | 23 | TCAGGCTCAGTCCCCTCCCGATA | 552.6 | 5.8 | 0.8 | 7.0 | 3.6E-12 | 7.4E-12 | BT20/MB468 |
| >hg38_21_+_8433172_8446622_RNA45SN1_45S_50ntFlanks@4367.4391.25\|CAGTTAAAAAGCTCGTAGTTGGATC | >hg38_21_+_8433172_8446622_RNA45SN1_45S_50ntFlanks | 4367 | 4391 | 25 | CAGTTAAAAAGCTCGTAGTTGGATC | 1257.9 | -5.1 | 0.7 | -7.0 | 3.6E-12 | 7.4E-12 | BT20/MB468 |

| | | | | | | | | | | | | |
|---|---|---|---|---|---|---|---|---|---|---|---|---|
| >hg38_21_+_8433172_8446622_RNA45SN1_45S_50ntFlanks@12970.12990.21\|GTTTCGTACGTAGCAGAGCAG | >hg38_21_+_8433172_8446622_RNA45SN1_45S_50ntFlanks | 12970 | 12990 | 21 | GTTTCGTACGTAGCAGAGCAG | 155.2 | -9.3 | 1.3 | -6.9 | 3.7E-12 | 7.6E-12 | BT20/MB468 |
| >MI0000085\|hsa-mir-27a&WithFlank&19\|-\|13836434\|13836523@57.76.20(+2U)&[MIMAT0000084&hsa-miR-27a-3p&offsets\|0\|-1(+2U);m-43&19\|-\|13836447\|13836467&offsets\|0\|-1(+2U)]\|TTCACAGTGGCTAAGTTCCGTT | >MI0000085\|hsa-mir-27a&WithFlank&19\|-\|13836434\|13836523 | 57 | 76 | 20(+2U) | TTCACAGTGGCTAAGTTCCGTT | 7157.8 | 5.6 | 0.8 | 7.0 | 3.3E-12 | 7.6E-12 | MB231/MB468 |
| >hg38_21_+_8433172_8446622_RNA45SN1_45S_50ntFlanks@9567.9588.22\|ACGTGCAAATCGGTCGTCCGAC | >hg38_21_+_8433172_8446622_RNA45SN1_45S_50ntFlanks | 9567 | 9588 | 22 | ACGTGCAAATCGGTCGTCCGAC | 134.6 | -9.1 | 1.3 | -6.9 | 3.7E-12 | 7.7E-12 | BT20/MB468 |
| >hg38_21_+_8433172_8446622_RNA45SN1_45S_50ntFlanks@10805.10825.21\|GAACAATGTAGGTAAGGGAAG | >hg38_21_+_8433172_8446622_RNA45SN1_45S_50ntFlanks | 10805 | 10825 | 21 | GAACAATGTAGGTAAGGGAAG | 153.4 | -9.3 | 1.3 | -6.9 | 3.7E-12 | 7.7E-12 | BT20/MB468 |
| >hg38_21_+_8433172_8446622_RNA45SN1_45S_50ntFlanks@12159.12181.23\|TTTGACTGGGGCGGTACACCTGT | >hg38_21_+_8433172_8446622_RNA45SN1_45S_50ntFlanks | 12159 | 12181 | 23 | TTTGACTGGGGCGGTACACCTGT | 1161.7 | -8.0 | 1.2 | -7.0 | 3.3E-12 | 7.7E-12 | MB231/MB468 |
| >hg38_21_+_8433172_8446622_RNA45SN1_45S_50ntFlanks@4046.4063.18\|CATTCGAACGTCTGCCCT | >hg38_21_+_8433172_8446622_RNA45SN1_45S_50ntFlanks | 4046 | 4063 | 18 | CATTCGAACGTCTGCCCT | 1919.9 | -4.4 | 0.6 | -6.9 | 3.8E-12 | 7.7E-12 | BT20/MB468 |
| >hg38_21_+_8433172_8446622_RNA45SN1_45S_50ntFlanks@13004.13023.20\|ATCTATTGAAAGTCAGCCCT | >hg38_21_+_8433172_8446622_RNA45SN1_45S_50ntFlanks | 13004 | 13023 | 20 | ATCTATTGAAAGTCAGCCCT | 137.5 | -8.7 | 1.2 | -7.0 | 3.3E-12 | 7.8E-12 | MB231/MB468 |
| >hg38_21_+_8433172_8446622_RNA45SN1_45S_50ntFlanks@12160.12180.21\|TTGACTGGGGCGGTACACCTG | >hg38_21_+_8433172_8446622_RNA45SN1_45S_50ntFlanks | 12160 | 12180 | 21 | TTGACTGGGGCGGTACACCTG | 123.6 | -8.9 | 1.3 | -6.9 | 3.8E-12 | 7.8E-12 | BT20/MB468 |
| >hg38_21_+_8433172_8446622_RNA45SN1_45S_50ntFlanks@9305.9338.34\|CCAAGGAGTCTAACACGTGCGCGAGTCGGGGGCT | >hg38_21_+_8433172_8446622_RNA45SN1_45S_50ntFlanks | 9305 | 9338 | 34 | CCAAGGAGTCTAACACGTGCGCGAGTCGGGGGCT | 301.6 | -5.3 | 0.8 | -6.9 | 3.8E-12 | 7.8E-12 | BT20/MB468 |
| >hg38_21_+_8433172_8446622_RNA45SN1_45S_50ntFlanks@10844.10865.22\|AACTTCGGGATAAGGATTGGCT | >hg38_21_+_8433172_8446622_RNA45SN1_45S_50ntFlanks | 10844 | 10865 | 22 | AACTTCGGGATAAGGATTGGCT | 289.0 | -4.6 | 0.7 | -7.0 | 3.4E-12 | 7.9E-12 | MB231/MB468 |
| >hg38_21_+_8433172_8446622_RNA45SN1_45S_50ntFlanks@5152.5185.34\|AGGGACAAGTGGCGTTCAGCCACCCGAGATTGAG | >hg38_21_+_8433172_8446622_RNA45SN1_45S_50ntFlanks | 5152 | 5185 | 34 | AGGGACAAGTGGCGTTCAGCCACCCGAGATTGAG | 658.3 | -11.4 | 1.6 | -6.9 | 3.8E-12 | 7.9E-12 | BT20/MB468 |
| >MI0000272\|hsa-mir-182&WithFlank&7\|-\|129770377\|129770498@29.51.23&[MIMAT0000259&hsa-miR-182-5p&offsets\|0\|-1;m-22&7\|-\|129770449\|129770470&offsets\|0\|+1]\|TTTGGCAATGGTAGAACTCACAC | >MI0000272\|hsa-mir-182&WithFlank&7\|-\|129770377\|129770498 | 29 | 51 | 23 | TTTGGCAATGGTAGAACTCACAC | 785.9 | 5.6 | 0.8 | 7.0 | 3.4E-12 | 7.9E-12 | MB231/MB468 |
| >hg38_21_+_8433172_8446622_RNA45SN1_45S_50ntFlanks@11452.11478.27\|CTCCTCGGCGCGCGGCGGCGGCGGCGG | >hg38_21_+_8433172_8446622_RNA45SN1_45S_50ntFlanks | 11452 | 11478 | 27 | CTCCTCGGCGCGCGGCGGCGGCGGCGG | 151.6 | -8.8 | 1.3 | -7.0 | 3.4E-12 | 7.9E-12 | MB231/MB468 |
| >hg38_21_+_8433172_8446622_RNA45SN1_45S_50ntFlanks@12157.12183.27\|AGTTTGACTGGGGCGGTACACCTGTCA | >hg38_21_+_8433172_8446622_RNA45SN1_45S_50ntFlanks | 12157 | 12183 | 27 | AGTTTGACTGGGGCGGTACACCTGTCA | 104.1 | -8.7 | 1.3 | -6.9 | 3.9E-12 | 8.1E-12 | BT20/MB468 |
| >hg38_21_+_8433172_8446622_RNA45SN1_45S_50ntFlanks@10845.10863.19\|ACTTCGGGATAAGGATTGG | >hg38_21_+_8433172_8446622_RNA45SN1_45S_50ntFlanks | 10845 | 10863 | 19 | ACTTCGGGATAAGGATTGG | 332.6 | -4.4 | 0.6 | -7.0 | 3.5E-12 | 8.1E-12 | MB231/MB468 |
| >hg38_21_+_8433172_8446622_RNA45SN1_45S_50ntFlanks@10873.10894.22\|CTGGGTCGGTCGGGCTGGGGCG | >hg38_21_+_8433172_8446622_RNA45SN1_45S_50ntFlanks | 10873 | 10894 | 22 | CTGGGTCGGTCGGGCTGGGGCG | 143.7 | -5.4 | 0.8 | -7.0 | 3.5E-12 | 8.1E-12 | MB231/MB468 |
| >hg38_21_+_8433172_8446622_RNA45SN1_45S_50ntFlanks@6652.6690.39\|GACTCTTAGCGGTGGATCACTCGGCTCGTGCGTCGATGA | >hg38_21_+_8433172_8446622_RNA45SN1_45S_50ntFlanks | 6652 | 6690 | 39 | GACTCTTAGCGGTGGATCACTCGGCTCGTGCGTCGATGA | 927.4 | 5.0 | 0.7 | 7.0 | 3.5E-12 | 8.1E-12 | MB231/MB468 |
| >hg38_21_+_8433172_8446622_RNA45SN1_45S_50ntFlanks@10386.10410.25\|ATGTGAACAGCAGTTGAACATGGGT | >hg38_21_+_8433172_8446622_RNA45SN1_45S_50ntFlanks | 10386 | 10410 | 25 | ATGTGAACAGCAGTTGAACATGGGT | 236.5 | -4.0 | 0.6 | -6.9 | 4.0E-12 | 8.1E-12 | BT20/MB468 |
| >hg38_21_+_8433172_8446622_RNA45SN1_45S_50ntFlanks@10412.10429.18\|AGTCGGTCCTGAGAGATG | >hg38_21_+_8433172_8446622_RNA45SN1_45S_50ntFlanks | 10412 | 10429 | 18 | AGTCGGTCCTGAGAGATG | 166.4 | -3.1 | 0.4 | -7.0 | 3.5E-12 | 8.2E-12 | MB231/MB468 |
| >trna116_GluCTC_1_-_145399233_145399304@39.74.36,trna134_GluTTC_1_-_16861774_16861845@39.74.36,trna71_GluCTC_1_-_161439189_161439260@39.74.36,trna74_GluCTC_1_-_161431809_161431880@39.74.36,trna77_GluCTC_1_-_161424398_161424469@39.74.36,trna77_GluCTC_6_+_28949976_28950047@39.74.36,trna80_GluCTC_1_-_161417018_161417089@39.74.36,trna84_GluTTC_1_-_161391883_161391954@39.74.36,trna87_GluCTC_6_-_126101393_126101464@39.74.36\|CGCCGCGGCCCGGGTTCGATTCCCGGTCAGGGAACC | >trna116_GluCTC_1_-_145399233_145399304 | 39 | 74 | 36 | CGCCGCGGCCCGGGTTCGATTCCCGGTCAGGGAACC | 869.7 | 9.2 | 1.3 | 7.0 | 3.5E-12 | 8.2E-12 | MB231/MB468 |
| >hg38_21_+_8433172_8446622_RNA45SN1_45S_50ntFlanks@10343.10373.31\|TATTCAAACGAGAACTTTGAAGGCCGAAGTG | >hg38_21_+_8433172_8446622_RNA45SN1_45S_50ntFlanks | 10343 | 10373 | 31 | TATTCAAACGAGAACTTTGAAGGCCGAAGTG | 354.9 | 6.4 | 0.9 | 7.0 | 3.6E-12 | 8.3E-12 | MB231/MB468 |
| >hg38_21_+_8433172_8446622_RNA45SN1_45S_50ntFlanks@3874.3898.25\|AATACATGCCGACGGGCGCTGACCC | >hg38_21_+_8433172_8446622_RNA45SN1_45S_50ntFlanks | 3874 | 3898 | 25 | AATACATGCCGACGGGCGCTGACCC | 2086.5 | -6.2 | 0.9 | -6.9 | 4.1E-12 | 8.4E-12 | BT20/MB468 |
| >hg38_21_+_8433172_8446622_RNA45SN1_45S_50ntFlanks@12415.12439.25\|CTTCGATGTCGGCTCTTCCTATCAT | >hg38_21_+_8433172_8446622_RNA45SN1_45S_50ntFlanks | 12415 | 12439 | 25 | CTTCGATGTCGGCTCTTCCTATCAT | 416.0 | 3.4 | 0.5 | 7.0 | 3.6E-12 | 8.4E-12 | MB231/MB468 |
| >hg38_21_+_8433172_8446622_RNA45SN1_45S_50ntFlanks@6646.6695.50\|TCGTACGACTCTTAGCGGTGGATCACTCGGCTCGTGCGTCGATGAAGAAC | >hg38_21_+_8433172_8446622_RNA45SN1_45S_50ntFlanks | 6646 | 6695 | 50 | TCGTACGACTCTTAGCGGTGGATCACTCGGCTCGTGCGTCGATGAAGAAC | 2587.4 | 10.0 | 1.4 | 6.9 | 4.1E-12 | 8.4E-12 | BT20/MB468 |
| >hg38_21_+_8433172_8446622_RNA45SN1_45S_50ntFlanks@9566.9596.31\|GACGTGCAAATCGGTCGTCCGACCTGGGTAT | >hg38_21_+_8433172_8446622_RNA45SN1_45S_50ntFlanks | 9566 | 9596 | 31 | GACGTGCAAATCGGTCGTCCGACCTGGGTAT | 1904.8 | -5.8 | 0.8 | -6.9 | 4.1E-12 | 8.4E-12 | BT20/MB468 |
| >hg38_21_+_8433172_8446622_RNA45SN1_45S_50ntFlanks@10866.10884.19\|CTAAGGGCTGGGTCGGTCG | >hg38_21_+_8433172_8446622_RNA45SN1_45S_50ntFlanks | 10866 | 10884 | 19 | CTAAGGGCTGGGTCGGTCG | 479.5 | -6.7 | 1.0 | -6.9 | 4.1E-12 | 8.4E-12 | BT20/MB468 |
| >hg38_21_+_8433172_8446622_RNA45SN1_45S_50ntFlanks@7975.8013.39\|CGCGACCTCAGATCAGACGTGGCGACCCGCTGAATTTAA | >hg38_21_+_8433172_8446622_RNA45SN1_45S_50ntFlanks | 7975 | 8013 | 39 | CGCGACCTCAGATCAGACGTGGCGACCCGCTGAATTTAA | 4634.5 | 6.1 | 0.9 | 6.9 | 4.1E-12 | 8.4E-12 | BT20/MB468 |
| >hg38_21_+_8433172_8446622_RNA45SN1_45S_50ntFlanks@10827.10844.18\|CGGCAAGCCGGATCCGTA | >hg38_21_+_8433172_8446622_RNA45SN1_45S_50ntFlanks | 10827 | 10844 | 18 | CGGCAAGCCGGATCCGTA | 135.9 | -9.1 | 1.3 | -6.9 | 4.1E-12 | 8.4E-12 | BT20/MB468 |
| >hg38_21_+_8433172_8446622_RNA45SN1_45S_50ntFlanks@5473.5501.29\|CGGCCCTGGCGGAGCGCTGAGAAGACGGT | >hg38_21_+_8433172_8446622_RNA45SN1_45S_50ntFlanks | 5473 | 5501 | 29 | CGGCCCTGGCGGAGCGCTGAGAAGACGGT | 704.6 | 7.0 | 1.0 | 6.9 | 4.1E-12 | 8.4E-12 | BT20/MB468 |
| >hg38_21_+_8433172_8446622_RNA45SN1_45S_50ntFlanks@11821.11842.22\|CTATCCAGCGAAACCACAGCCA | >hg38_21_+_8433172_8446622_RNA45SN1_45S_50ntFlanks | 11821 | 11842 | 22 | CTATCCAGCGAAACCACAGCCA | 130.9 | -8.6 | 1.2 | -7.0 | 3.6E-12 | 8.4E-12 | MB231/MB468 |
| >hg38_21_+_8433172_8446622_RNA45SN1_45S_50ntFlanks@12476.12493.18\|CCACTAATAGGGAACGTG | >hg38_21_+_8433172_8446622_RNA45SN1_45S_50ntFlanks | 12476 | 12493 | 18 | CCACTAATAGGGAACGTG | 353.3 | -6.4 | 0.9 | -6.9 | 4.1E-12 | 8.5E-12 | BT20/MB468 |
| >hg38_21_+_8433172_8446622_RNA45SN1_45S_50ntFlanks@5446.5467.22\|CGGATCGGCCCCGCCGGGGTCG | >hg38_21_+_8433172_8446622_RNA45SN1_45S_50ntFlanks | 5446 | 5467 | 22 | CGGATCGGCCCCGCCGGGGTCG | 130.1 | -9.0 | 1.3 | -6.9 | 4.2E-12 | 8.6E-12 | BT20/MB468 |
| >trna23_ArgCCG_17_-_66016013_66016085@14.31.18\|AATGGATAAGGCATCAGC | >trna23_ArgCCG_17_-_66016013_66016085 | 14 | 31 | 18 | AATGGATAAGGCATCAGC | 167.1 | -9.4 | 1.4 | -6.9 | 4.3E-12 | 8.8E-12 | BT20/MB468 |
| >hg38_21_+_8433172_8446622_RNA45SN1_45S_50ntFlanks@10574.10596.23\|AAGCCGGCGGGAGCCCCGGGGAG | >hg38_21_+_8433172_8446622_RNA45SN1_45S_50ntFlanks | 10574 | 10596 | 23 | AAGCCGGCGGGAGCCCCGGGGAG | 151.2 | -8.8 | 1.3 | -6.9 | 3.8E-12 | 8.8E-12 | MB231/MB468 |
| >hg38_21_+_8433172_8446622_RNA45SN1_45S_50ntFlanks@10841.10858.18\|CGTAACTTCGGGATAAGG | >hg38_21_+_8433172_8446622_RNA45SN1_45S_50ntFlanks | 10841 | 10858 | 18 | CGTAACTTCGGGATAAGG | 805.5 | -6.0 | 0.9 | -6.9 | 3.8E-12 | 8.8E-12 | MB231/MB468 |
| >hg38_21_+_8433172_8446622_RNA45SN1_45S_50ntFlanks@6737.6756.20\|GATCATCGACACTTCGAACG | >hg38_21_+_8433172_8446622_RNA45SN1_45S_50ntFlanks | 6737 | 6756 | 20 | GATCATCGACACTTCGAACG | 124.3 | -8.9 | 1.3 | -6.9 | 4.3E-12 | 8.8E-12 | BT20/MB468 |
| >MI0000273\|hsa-mir-183&WithFlank&7\|-\|129774899\|129775020@34.55.22&[MIMAT0000261&hsa-miR-183-5p&offsets\|+1\|+1;m-41&7\|-\|129774966\|129774987&offsets\|0\|0]\|ATGGCACTGGTAGAATTCACTG | >MI0000273\|hsa-mir-183&WithFlank&7\|-\|129774899\|129775020 | 34 | 55 | 22 | ATGGCACTGGTAGAATTCACTG | 1572.7 | 5.4 | 0.8 | 6.9 | 3.8E-12 | 8.9E-12 | MB231/MB468 |
| >hg38_21_+_8433172_8446622_RNA45SN1_45S_50ntFlanks@8472.8490.19\|CGGCCCGGCGGATCTTTCC | >hg38_21_+_8433172_8446622_RNA45SN1_45S_50ntFlanks | 8472 | 8490 | 19 | CGGCCCGGCGGATCTTTCC | 138.5 | -9.1 | 1.3 | -6.9 | 4.4E-12 | 8.9E-12 | BT20/MB468 |
| >hg38_21_+_8433172_8446622_RNA45SN1_45S_50ntFlanks@12324.12342.19\|TTGGGTTTTAAGCAGGAGG | >hg38_21_+_8433172_8446622_RNA45SN1_45S_50ntFlanks | 12324 | 12342 | 19 | TTGGGTTTTAAGCAGGAGG | 110.8 | -8.8 | 1.3 | -6.9 | 4.4E-12 | 9.0E-12 | BT20/MB468 |
| >MI0000067\|hsa-let-7f-1&WithFlank&9\|+\|94176341\|94176439@13.32.20(+1A);MI0000068\|hsa-let-7f-2&WithFlank&X\|-\|53557186\|53557280@14.33.20(+1A)&[MIMAT0000067&hsa-let-7f-5p&offsets\|0\|-2(+1A);m-20&9\|+\|94176353\|94176374&offsets\|0\|-2(+1A)];[MIMAT0000067_1&hsa-let-7f-5p&offsets\|0\|-2(+1A);m-19&X\|-\|53557246\|53557267&offsets\|0\|-2(+1A)]\|TGAGGTAGTAGATTGTATAGA | >MI0000067\|hsa-let-7f-1&WithFlank&9\|+\|94176341\|94176439 | 13 | 32 | 20(+1A) | TGAGGTAGTAGATTGTATAGA | 1008.1 | 8.4 | 1.2 | 6.9 | 4.4E-12 | 9.0E-12 | BT20/MB468 |
| >MI0000746\|hsa-mir-99b&WithFlank&19\|+\|51692606\|51692687@14.34.21&[MIMAT0000689&hsa-miR-99b-5p&offsets\|+1\|0;m-10&19\|+\|51692618\|51692639&offsets\|+1\|0]\|ACCCGTAGAACCGACCTTGCG | >MI0000746\|hsa-mir-99b&WithFlank&19\|+\|51692606\|51692687 | 14 | 34 | 21 | ACCCGTAGAACCGACCTTGCG | 723.1 | 7.7 | 1.1 | 6.9 | 4.4E-12 | 9.1E-12 | BT20/MB468 |
| >hg38_21_+_8433172_8446622_RNA45SN1_45S_50ntFlanks@6646.6685.40\|TCGTACGACTCTTAGCGGTGGATCACTCGGCTCGTGCGTC | >hg38_21_+_8433172_8446622_RNA45SN1_45S_50ntFlanks | 6646 | 6685 | 40 | TCGTACGACTCTTAGCGGTGGATCACTCGGCTCGTGCGTC | 1584.0 | 5.2 | 0.7 | 6.9 | 3.9E-12 | 9.1E-12 | MB231/MB468 |
| >hg38_21_+_8433172_8446622_RNA45SN1_45S_50ntFlanks@12324.12345.22\|TTGGGTTTTAAGCAGGAGGTGT | >hg38_21_+_8433172_8446622_RNA45SN1_45S_50ntFlanks | 12324 | 12345 | 22 | TTGGGTTTTAAGCAGGAGGTGT | 240.6 | -5.1 | 0.7 | -6.9 | 3.9E-12 | 9.1E-12 | MB231/MB468 |
| >hg38_21_+_8433172_8446622_RNA45SN1_45S_50ntFlanks@7981.8008.28\|CTCAGATCAGACGTGGCGACCCGCTGAA | >hg38_21_+_8433172_8446622_RNA45SN1_45S_50ntFlanks | 7981 | 8008 | 28 | CTCAGATCAGACGTGGCGACCCGCTGAA | 541.9 | -4.0 | 0.6 | -6.9 | 4.5E-12 | 9.1E-12 | BT20/MB468 |
| >hg38_21_+_8433172_8446622_RNA45SN1_45S_50ntFlanks@12992.13011.20\|TCCCTCGCTGCGATCTATTG | >hg38_21_+_8433172_8446622_RNA45SN1_45S_50ntFlanks | 12992 | 13011 | 20 | TCCCTCGCTGCGATCTATTG | 216.9 | -6.5 | 0.9 | -6.9 | 4.5E-12 | 9.2E-12 | BT20/MB468 |

| | | | | | | | | | | | | |
|---|---|---|---|---|---|---|---|---|---|---|---|---|
| >hg38_21_+_8433172_8446622_RNA45SN1_45S_50ntFlanks@12360.12382.23\|CAGGGATAACTGGCTTGTGGCGG | >hg38_21_+_8433172_8446622_RNA45SN1_45S_50ntFlanks | 12360 | 12382 | 23 | CAGGGATAACTGGCTTGTGGCGG | 158.8 | -8.9 | 1.3 | -6.9 | 4.0E-12 | 9.2E-12 | MB231/MB468 |
| >hg38_21_+_8433172_8446622_RNA45SN1_45S_50ntFlanks@11675.11706.32\|TGCTCTGAATGTCAAAGTGAAGAAATTCAATG | >hg38_21_+_8433172_8446622_RNA45SN1_45S_50ntFlanks | 11675 | 11706 | 32 | TGCTCTGAATGTCAAAGTGAAGAAATTCAATG | 1160.6 | 5.4 | 0.8 | 6.9 | 4.0E-12 | 9.2E-12 | MB231/MB468 |
| >MI0000488\|hsa-mir-194-1&WithFlank&1\|-\|220118151\|220118247@21.42.22;MI0000732\|hsa-mir-194-2&WithFlank&11\|-\|64891349\|64891445@21.42.22&[MIMAT0000460&hsa-miR-194-5p&offsets\|0\|0;m-131&1\|-\|220118206\|220118227&offsets\|0\|0];[MIMAT0000460_1&hsa-miR-194-5p&offsets\|0\|0;m-126&11\|-\|64891404\|64891425&offsets\|0\|0]\|TGTAACAGCAACTCCATGTGGA | >MI0000488\|hsa-mir-194-1&WithFlank&1\|-\|220118151\|220118247 | 21 | 42 | 22 | TGTAACAGCAACTCCATGTGGA | 601.0 | 4.9 | 0.7 | 6.9 | 4.0E-12 | 9.2E-12 | MB231/MB468 |
| >hg38_21_+_8433172_8446622_RNA45SN1_45S_50ntFlanks@12159.12178.20\|TTTGACTGGGGCGGTACACC | >hg38_21_+_8433172_8446622_RNA45SN1_45S_50ntFlanks | 12159 | 12178 | 20 | TTTGACTGGGGCGGTACACC | 358.3 | -4.1 | 0.6 | -6.9 | 4.0E-12 | 9.3E-12 | MB231/MB468 |
| >trna10_LysCTT_16_+_3241501_3241573@1.28.28,trna119_LysCTT_1_-_145395522_145395594@1.28.28,trna11_LysCTT_5_-_180648979_180649051@1.28.28,trna13_LysCTT_14_-_58706613_58706685@1.28.28,trna13_LysCTT_6_+_26556774_26556846@1.28.28,trna2_LysCTT_15_+_79152904_79152976@1.28.28,trna32_LysCTT_16_-_3207406_3207478@1.28.28,trna7_LysCTT_16_+_3225692_3225764@1.28.28,trna9_LysCTT_5_+_180634755_180634827@1.28.28\|GCCCGGCTAGCTCAGTCGGTAGAGCATG | >trna10_LysCTT_16_+_3241501_3241573 | 1 | 28 | 28 | GCCCGGCTAGCTCAGTCGGTAGAGCATG | 3329.8 | -5.0 | 0.7 | -6.9 | 4.5E-12 | 9.3E-12 | BT20/MB468 |
| >hg38_21_+_8433172_8446622_RNA45SN1_45S_50ntFlanks@3776.3801.26\|CCGGTACAGTGAAACTGCGAATGGCT | >hg38_21_+_8433172_8446622_RNA45SN1_45S_50ntFlanks | 3776 | 3801 | 26 | CCGGTACAGTGAAACTGCGAATGGCT | 351.0 | -3.8 | 0.5 | -6.9 | 4.6E-12 | 9.3E-12 | BT20/MB468 |
| >hg38_21_+_8433172_8446622_RNA45SN1_45S_50ntFlanks@12035.12053.19\|CGTTTTTTCACTGACCCGG | >hg38_21_+_8433172_8446622_RNA45SN1_45S_50ntFlanks | 12035 | 12053 | 19 | CGTTTTTTCACTGACCCGG | 127.0 | -9.0 | 1.3 | -6.9 | 4.6E-12 | 9.3E-12 | BT20/MB468 |
| >hg38_21_+_8433172_8446622_RNA45SN1_45S_50ntFlanks@11552.11574.23\|CCTCGGCCGGCGCCTAGCAGCCG | >hg38_21_+_8433172_8446622_RNA45SN1_45S_50ntFlanks | 11552 | 11574 | 23 | CCTCGGCCGGCGCCTAGCAGCCG | 98.1 | -5.4 | 0.8 | -6.9 | 4.6E-12 | 9.4E-12 | BT20/MB468 |
| >hg38_21_+_8433172_8446622_RNA45SN1_45S_50ntFlanks@4738.4761.24\|AAAGTCGGAGGTTCGAAGACGATC | >hg38_21_+_8433172_8446622_RNA45SN1_45S_50ntFlanks | 4738 | 4761 | 24 | AAAGTCGGAGGTTCGAAGACGATC | 1249.6 | 4.8 | 0.7 | 6.9 | 4.1E-12 | 9.4E-12 | MB231/MB468 |
| >hg38_21_+_8433172_8446622_RNA45SN1_45S_50ntFlanks@4760.4793.34\|TCAGATACCGTCGTAGTTCCGACCATAAACGATG | >hg38_21_+_8433172_8446622_RNA45SN1_45S_50ntFlanks | 4760 | 4793 | 34 | TCAGATACCGTCGTAGTTCCGACCATAAACGATG | 133.7 | -3.5 | 0.5 | -6.9 | 4.6E-12 | 9.4E-12 | BT20/MB468 |
| >hg38_21_+_8433172_8446622_RNA45SN1_45S_50ntFlanks@4069.4098.30\|CTTTCGATGGTAGTCGCCGTGCCTACCATG | >hg38_21_+_8433172_8446622_RNA45SN1_45S_50ntFlanks | 4069 | 4098 | 30 | CTTTCGATGGTAGTCGCCGTGCCTACCATG | 148.2 | -9.2 | 1.3 | -6.9 | 4.6E-12 | 9.4E-12 | BT20/MB468 |
| >hg38_21_+_8433172_8446622_RNA45SN1_45S_50ntFlanks@10481.10498.18\|ATCGAAAGGGAGTCGGGT | >hg38_21_+_8433172_8446622_RNA45SN1_45S_50ntFlanks | 10481 | 10498 | 18 | ATCGAAAGGGAGTCGGGT | 117.9 | -8.9 | 1.3 | -6.9 | 4.6E-12 | 9.4E-12 | BT20/MB468 |
| >hg38_21_+_8433172_8446622_RNA45SN1_45S_50ntFlanks@12654.12671.18\|TATGACTGAACGCCTCTA | >hg38_21_+_8433172_8446622_RNA45SN1_45S_50ntFlanks | 12654 | 12671 | 18 | TATGACTGAACGCCTCTA | 157.1 | -4.0 | 0.6 | -6.9 | 4.6E-12 | 9.4E-12 | BT20/MB468 |
| >hg38_21_+_8433172_8446622_RNA45SN1_45S_50ntFlanks@11752.11777.26\|CCAAATGCCTCGTCATCTAATTAGTG | >hg38_21_+_8433172_8446622_RNA45SN1_45S_50ntFlanks | 11752 | 11777 | 26 | CCAAATGCCTCGTCATCTAATTAGTG | 91.5 | -8.5 | 1.2 | -6.9 | 4.7E-12 | 9.6E-12 | BT20/MB468 |
| >hg38_21_+_8433172_8446622_RNA45SN1_45S_50ntFlanks@10781.10803.23\|AGGTGAACAGCCTCTGGCATGTT | >hg38_21_+_8433172_8446622_RNA45SN1_45S_50ntFlanks | 10781 | 10803 | 23 | AGGTGAACAGCCTCTGGCATGTT | 283.2 | -4.4 | 0.6 | -6.9 | 4.7E-12 | 9.6E-12 | BT20/MB468 |
| >hg38_21_+_8433172_8446622_RNA45SN1_45S_50ntFlanks@6645.6662.18\|CTCGTACGACTCTTAGCG | >hg38_21_+_8433172_8446622_RNA45SN1_45S_50ntFlanks | 6645 | 6662 | 18 | CTCGTACGACTCTTAGCG | 4644.7 | 3.5 | 0.5 | 6.9 | 4.7E-12 | 9.6E-12 | BT20/MB468 |
| >hg38_21_+_8433172_8446622_RNA45SN1_45S_50ntFlanks@12461.12487.27\|CGTTGGATTGTTCACCCACTAATAGGG | >hg38_21_+_8433172_8446622_RNA45SN1_45S_50ntFlanks | 12461 | 12487 | 27 | CGTTGGATTGTTCACCCACTAATAGGG | 174.4 | -9.0 | 1.3 | -6.9 | 4.2E-12 | 9.6E-12 | MB231/MB468 |
| >hg38_21_+_8433172_8446622_RNA45SN1_45S_50ntFlanks@12634.12654.21\|GAAGCTACCATCTGTGGGATT | >hg38_21_+_8433172_8446622_RNA45SN1_45S_50ntFlanks | 12634 | 12654 | 21 | GAAGCTACCATCTGTGGGATT | 404.2 | -5.4 | 0.8 | -6.9 | 4.7E-12 | 9.7E-12 | BT20/MB468 |
| >hg38_21_+_8433172_8446622_RNA45SN1_45S_50ntFlanks@11564.11581.18\|CCTAGCAGCCGACTTAGA | >hg38_21_+_8433172_8446622_RNA45SN1_45S_50ntFlanks | 11564 | 11581 | 18 | CCTAGCAGCCGACTTAGA | 927.6 | -5.9 | 0.9 | -6.9 | 4.8E-12 | 9.8E-12 | BT20/MB468 |
| >trnaMT_GluTTC_MT_-_14674_14742@40.57.18,trnalookalike8_GluTTC_5_-_93905172_93905240@40.57.18\|TTGGTCGTGGTTGTAGTC | >trnaMT_GluTTC_MT_-_14674_14742 | 40 | 57 | 18 | TTGGTCGTGGTTGTAGTC | 538.7 | 8.5 | 1.2 | 6.9 | 4.8E-12 | 9.8E-12 | BT20/MB468 |
| >hg38_21_+_8433172_8446622_RNA45SN1_45S_50ntFlanks@8342.8379.38\|CGTAAGGGAAAGTTGAAAAGAACTTTGAAGAGAGAGTT | >hg38_21_+_8433172_8446622_RNA45SN1_45S_50ntFlanks | 8342 | 8379 | 38 | CGTAAGGGAAAGTTGAAAAGAACTTTGAAGAGAGAGTT | 875.1 | 8.6 | 1.2 | 6.9 | 4.3E-12 | 9.9E-12 | MB231/MB468 |
| >hg38_21_+_8433172_8446622_RNA45SN1_45S_50ntFlanks@3705.3748.44\|TACCTGGTTGATCCTGCCAGTAGCATATGCTTGTCTCAAAGATT | >hg38_21_+_8433172_8446622_RNA45SN1_45S_50ntFlanks | 3705 | 3748 | 44 | TACCTGGTTGATCCTGCCAGTAGCATATGCTTGTCTCAAAGATT | 1027.5 | 7.8 | 1.1 | 6.9 | 4.3E-12 | 9.9E-12 | MB231/MB468 |
| >hg38_21_+_8433172_8446622_RNA45SN1_45S_50ntFlanks@11580.11603.24\|GAACTGGTGCGGACCAGGGGAATC | >hg38_21_+_8433172_8446622_RNA45SN1_45S_50ntFlanks | 11580 | 11603 | 24 | GAACTGGTGCGGACCAGGGGAATC | 2994.5 | -4.4 | 0.6 | -6.9 | 4.9E-12 | 1.0E-11 | BT20/MB468 |
| >hg38_21_+_8433172_8446622_RNA45SN1_45S_50ntFlanks@3705.3725.21\|TACCTGGTTGATCCTGCCAGT | >hg38_21_+_8433172_8446622_RNA45SN1_45S_50ntFlanks | 3705 | 3725 | 21 | TACCTGGTTGATCCTGCCAGT | 1312.0 | 5.6 | 0.8 | 6.9 | 4.3E-12 | 1.0E-11 | MB231/MB468 |
| >hg38_21_+_8433172_8446622_RNA45SN1_45S_50ntFlanks@12539.12573.35\|ATGATGTGTTGTTGCCATGGTAATCCTGCTCAGTA | >hg38_21_+_8433172_8446622_RNA45SN1_45S_50ntFlanks | 12539 | 12573 | 35 | ATGATGTGTTGTTGCCATGGTAATCCTGCTCAGTA | 137.3 | -8.7 | 1.3 | -6.9 | 4.3E-12 | 1.0E-11 | MB231/MB468 |
| >hg38_21_+_8433172_8446622_RNA45SN1_45S_50ntFlanks@4160.4184.25\|CCACATCCAAGGAAGGCAGCAGGCG | >hg38_21_+_8433172_8446622_RNA45SN1_45S_50ntFlanks | 4160 | 4184 | 25 | CCACATCCAAGGAAGGCAGCAGGCG | 113.9 | -8.8 | 1.3 | -6.9 | 4.9E-12 | 1.0E-11 | BT20/MB468 |
| >hg38_21_+_8433172_8446622_RNA45SN1_45S_50ntFlanks@4399.4423.25\|CGGGCGGGCGGTCCGCCGCGAGGCG | >hg38_21_+_8433172_8446622_RNA45SN1_45S_50ntFlanks | 4399 | 4423 | 25 | CGGGCGGGCGGTCCGCCGCGAGGCG | 108.8 | -8.8 | 1.3 | -6.9 | 4.9E-12 | 1.0E-11 | BT20/MB468 |
| >hg38_21_+_8433172_8446622_RNA45SN1_45S_50ntFlanks@10808.10827.20\|CAATGTAGGTAAGGGAAGTC | >hg38_21_+_8433172_8446622_RNA45SN1_45S_50ntFlanks | 10808 | 10827 | 20 | CAATGTAGGTAAGGGAAGTC | 225.1 | -4.3 | 0.6 | -6.9 | 4.3E-12 | 1.0E-11 | MB231/MB468 |
| >hg38_21_+_8433172_8446622_RNA45SN1_45S_50ntFlanks@10728.10755.28\|GAGAGGGTGTAAATCTCGCGCCGGGCCG | >hg38_21_+_8433172_8446622_RNA45SN1_45S_50ntFlanks | 10728 | 10755 | 28 | GAGAGGGTGTAAATCTCGCGCCGGGCCG | 131.8 | -9.0 | 1.3 | -6.9 | 4.9E-12 | 1.0E-11 | BT20/MB468 |
| >hg38_21_+_8433172_8446622_RNA45SN1_45S_50ntFlanks@6647.6664.18\|CGTACGACTCTTAGCGGT | >hg38_21_+_8433172_8446622_RNA45SN1_45S_50ntFlanks | 6647 | 6664 | 18 | CGTACGACTCTTAGCGGT | 2420.8 | 5.1 | 0.7 | 6.9 | 4.4E-12 | 1.0E-11 | MB231/MB468 |
| >hg38_21_+_8433172_8446622_RNA45SN1_45S_50ntFlanks@11888.11905.18\|AGCTTGACTCTAGTCTGG | >hg38_21_+_8433172_8446622_RNA45SN1_45S_50ntFlanks | 11888 | 11905 | 18 | AGCTTGACTCTAGTCTGG | 150.3 | -8.8 | 1.3 | -6.9 | 4.4E-12 | 1.0E-11 | MB231/MB468 |
| >hg38_21_+_8433172_8446622_RNA45SN1_45S_50ntFlanks@4445.4472.28\|CTTGCCTCTCGGCGCCCCCTCGATGCTC | >hg38_21_+_8433172_8446622_RNA45SN1_45S_50ntFlanks | 4445 | 4472 | 28 | CTTGCCTCTCGGCGCCCCCTCGATGCTC | 120.2 | -8.9 | 1.3 | -6.9 | 5.0E-12 | 1.0E-11 | BT20/MB468 |
| >hg38_21_+_8433172_8446622_RNA45SN1_45S_50ntFlanks@7979.7996.18\|ACCTCAGATCAGACGTGG | >hg38_21_+_8433172_8446622_RNA45SN1_45S_50ntFlanks | 7979 | 7996 | 18 | ACCTCAGATCAGACGTGG | 102.4 | -8.7 | 1.3 | -6.9 | 5.0E-12 | 1.0E-11 | BT20/MB468 |
| >hg38_21_+_8433172_8446622_RNA45SN1_45S_50ntFlanks@10116.10135.20\|TGCGTCGGAGGGCGGCGGCG | >hg38_21_+_8433172_8446622_RNA45SN1_45S_50ntFlanks | 10116 | 10135 | 20 | TGCGTCGGAGGGCGGCGGCG | 130.3 | -8.6 | 1.2 | -6.9 | 4.4E-12 | 1.0E-11 | MB231/MB468 |
| >hg38_21_+_8433172_8446622_RNA45SN1_45S_50ntFlanks@10420.10457.38\|CTGAGAGATGGGCGAGCGCCGTTCCGAAGGGACGGGCG | >hg38_21_+_8433172_8446622_RNA45SN1_45S_50ntFlanks | 10420 | 10457 | 38 | CTGAGAGATGGGCGAGCGCCGTTCCGAAGGGACGGGCG | 154.5 | -8.8 | 1.3 | -6.9 | 4.4E-12 | 1.0E-11 | MB231/MB468 |
| >trna10_AspGTC_12_-_125424193_125424264@39.75.37,trna12_AspGTC_12_-_125411891_125411962@39.75.37,trna144_AspGTC_6_-_27551236_27551307@39.75.37,trna38_AspGTC_17_-_8125556_8125627@39.75.37,trna45_AspGTC_6_+_27447453_27447524@39.75.37,trna48_AspGTC_6_+_27471523_27471594@39.75.37,trna4_AspGTC_12_+_96429799_96429870@39.75.37,trna69_AspGTC_1_-_161440205_161440276@39.75.37,trna72_AspGTC_1_-_161432824_161432895@39.75.37,trna75_AspGTC_1_-_161425414_161425485@39.75.37,trna78_AspGTC_1_-_161418033_161418104@39.75.37,trna81_AspGTC_1_-_161410615_161410686@39.75.37\|GCGGGAGACCGGGGTTCGATTCCCCGACGGGGAGCCA | >trna10_AspGTC_12_-_125424193_125424264 | 39 | 75 | 37 | GCGGGAGACCGGGGTTCGATTCCCCGACGGGGAGCCA | 1267.6 | 6.0 | 0.9 | 6.9 | 4.4E-12 | 1.0E-11 | MB231/MB468 |
| >hg38_21_+_8433172_8446622_RNA45SN1_45S_50ntFlanks@11882.11900.19\|CTGTTGAGCTTGACTCTAG | >hg38_21_+_8433172_8446622_RNA45SN1_45S_50ntFlanks | 11882 | 11900 | 19 | CTGTTGAGCTTGACTCTAG | 258.8 | -10.0 | 1.4 | -6.9 | 5.0E-12 | 1.0E-11 | BT20/MB468 |
| >hg38_21_+_8433172_8446622_RNA45SN1_45S_50ntFlanks@10806.10829.24\|AACAATGTAGGTAAGGGAAGTCGG | >hg38_21_+_8433172_8446622_RNA45SN1_45S_50ntFlanks | 10806 | 10829 | 24 | AACAATGTAGGTAAGGGAAGTCGG | 100.3 | -5.3 | 0.8 | -6.9 | 5.0E-12 | 1.0E-11 | BT20/MB468 |
| >hg38_21_+_8433172_8446622_RNA45SN1_45S_50ntFlanks@9572.9603.32\|CAAATCGGTCGTCCGACCTGGGTATAGGGGCG | >hg38_21_+_8433172_8446622_RNA45SN1_45S_50ntFlanks | 9572 | 9603 | 32 | CAAATCGGTCGTCCGACCTGGGTATAGGGGCG | 1633.7 | -4.0 | 0.6 | -6.9 | 5.0E-12 | 1.0E-11 | BT20/MB468 |
| >hg38_21_+_8433172_8446622_RNA45SN1_45S_50ntFlanks@10351.10382.32\|CGAGAACTTTGAAGGCCGAAGTGGAGAAGGGT | >hg38_21_+_8433172_8446622_RNA45SN1_45S_50ntFlanks | 10351 | 10382 | 32 | CGAGAACTTTGAAGGCCGAAGTGGAGAAGGGT | 863.0 | -3.1 | 0.4 | -6.9 | 5.1E-12 | 1.0E-11 | BT20/MB468 |
| >hg38_21_+_8433172_8446622_RNA45SN1_45S_50ntFlanks@10914.10936.23\|CGCCGCGGCTGGACGAGGCGCCG | >hg38_21_+_8433172_8446622_RNA45SN1_45S_50ntFlanks | 10914 | 10936 | 23 | CGCCGCGGCTGGACGAGGCGCCG | 465.1 | -7.0 | 1.0 | -6.9 | 5.2E-12 | 1.0E-11 | BT20/MB468 |
| >hg38_21_+_8433172_8446622_RNA45SN1_45S_50ntFlanks@9568.9594.27\|CGTGCAAATCGGTCGTCCGACCTGGGT | >hg38_21_+_8433172_8446622_RNA45SN1_45S_50ntFlanks | 9568 | 9594 | 27 | CGTGCAAATCGGTCGTCCGACCTGGGT | 291.7 | -4.2 | 0.6 | -6.9 | 4.6E-12 | 1.1E-11 | MB231/MB468 |
| >hg38_21_+_8433172_8446622_RNA45SN1_45S_50ntFlanks@5493.5516.24\|GAAGACGGTCGAACTTGACTATCT | >hg38_21_+_8433172_8446622_RNA45SN1_45S_50ntFlanks | 5493 | 5516 | 24 | GAAGACGGTCGAACTTGACTATCT | 676.0 | 8.5 | 1.2 | 6.9 | 4.6E-12 | 1.1E-11 | MB231/MB468 |

<table>
<tr><td>>trna111_HisGTG_1_-_147774845_147774916@1.32.32,trna118_HisGTG_1_-_145396881_145396952@1.32.32,trna16_HisGTG_1_+_146544773_146544844@1.32.32,trna1_HisGTG_15_+_45493349_45493420@1.32.32,trna21_HisGTG_1_+_147753471_147753542@1.32.32,trna33_HisGTG_6_+_27125906_27125977@1.32.32,trna7_HisGTG_9_-_14433938_14434009@1.32.32,trna8_HisGTG_15_-_45492611_45492682@1.32.32,trna9_HisGTG_15_-_45490804_45490875@1.32.32|GCCGTGATCGTATAGTGGTTAGTACTCTGCGT</td><td>>trna111_HisGTG_1_-_147774845_147774916</td><td>1</td><td>32</td><td>32</td><td>GCCGTGATCGTATAGTGGTTAGTACTCTGCGT</td><td>513.2</td><td>7.1</td><td>1.0</td><td>6.9</td><td>5.2E-12</td><td>1.1E-11</td><td>BT20/MB468</td></tr>
<tr><td>>hg38_21_+_8433172_8446622_RNA45SN1_45S_50ntFlanks@12324.12347.24|TTGGGTTTTAAGCAGGAGGTGTCA</td><td>>hg38_21_+_8433172_8446622_RNA45SN1_45S_50ntFlanks</td><td>12324</td><td>12347</td><td>24</td><td>TTGGGTTTTAAGCAGGAGGTGTCA</td><td>95.0</td><td>-8.6</td><td>1.2</td><td>-6.9</td><td>5.2E-12</td><td>1.1E-11</td><td>BT20/MB468</td></tr>
<tr><td>>hg38_21_+_8433172_8446622_RNA45SN1_45S_50ntFlanks@12457.12488.32|CAAGCGTTGGATTGTTCACCCACTAATAGGGA</td><td>>hg38_21_+_8433172_8446622_RNA45SN1_45S_50ntFlanks</td><td>12457</td><td>12488</td><td>32</td><td>CAAGCGTTGGATTGTTCACCCACTAATAGGGA</td><td>537.4</td><td>-4.6</td><td>0.7</td><td>-6.9</td><td>4.6E-12</td><td>1.1E-11</td><td>MB231/MB468</td></tr>
<tr><td>>hg38_21_+_8433172_8446622_RNA45SN1_45S_50ntFlanks@12459.12484.26|AGCGTTGGATTGTTCACCCACTAATA</td><td>>hg38_21_+_8433172_8446622_RNA45SN1_45S_50ntFlanks</td><td>12459</td><td>12484</td><td>26</td><td>AGCGTTGGATTGTTCACCCACTAATA</td><td>177.1</td><td>-9.5</td><td>1.4</td><td>-6.9</td><td>5.3E-12</td><td>1.1E-11</td><td>BT20/MB468</td></tr>
<tr><td>>hg38_21_+_8433172_8446622_RNA45SN1_45S_50ntFlanks@12162.12179.18|GACTGGGGCGGTACACCT</td><td>>hg38_21_+_8433172_8446622_RNA45SN1_45S_50ntFlanks</td><td>12162</td><td>12179</td><td>18</td><td>GACTGGGGCGGTACACCT</td><td>205.1</td><td>-9.7</td><td>1.4</td><td>-6.9</td><td>5.3E-12</td><td>1.1E-11</td><td>BT20/MB468</td></tr>
<tr><td>>MI0000065|hsa-let-7d&WithFlank&9|+|94178828|94178926@14.33.20(+1C)&[MIMAT0000065&hsa-let-7d-5p&offsets|0|-2(+1C);m-90&9|+|94178841|94178862&offsets|0|-2(+1C)]|AGAGGTAGTAGGTTGCATAGC</td><td>>MI0000065|hsa-let-7d&WithFlank&9|+|94178828|94178926</td><td>14</td><td>33</td><td>20(+1C)</td><td>AGAGGTAGTAGGTTGCATAGC</td><td>540.1</td><td>8.9</td><td>1.3</td><td>6.9</td><td>5.3E-12</td><td>1.1E-11</td><td>BT20/MB468</td></tr>
<tr><td>>am_trna162_MetCAT_6_-_26758550_26758622@31.52.22,trna164_MetCAT_6_-_26735574_26735646@31.52.22,trna20_MetCAT_16_+_71460396_71460468@31.52.22,trna21_MetCAT_6_+_26701712_26701784@31.52.22,trna27_MetCAT_6_+_26766444_26766516@31.52.22,trna75_MetCAT_6_+_28912352_28912424@31.52.22,trna97_MetCAT_6_-_28921042_28921114@31.52.22|TCTCATAATCTGAAGGTCCTGA</td><td>>am_trna162_MetCAT_6_-_26758550_26758622</td><td>31</td><td>52</td><td>22</td><td>TCTCATAATCTGAAGGTCCTGA</td><td>118.4</td><td>-8.9</td><td>1.3</td><td>-6.9</td><td>5.3E-12</td><td>1.1E-11</td><td>BT20/MB468</td></tr>
<tr><td>>hg38_21_+_8433172_8446622_RNA45SN1_45S_50ntFlanks@12073.12096.24|CCGAGGGGCTCTCGCTTCTGGCGC</td><td>>hg38_21_+_8433172_8446622_RNA45SN1_45S_50ntFlanks</td><td>12073</td><td>12096</td><td>24</td><td>CCGAGGGGCTCTCGCTTCTGGCGC</td><td>714.1</td><td>-4.0</td><td>0.6</td><td>-6.9</td><td>4.7E-12</td><td>1.1E-11</td><td>MB231/MB468</td></tr>
<tr><td>>MI0000746|hsa-mir-99b&WithFlank&19|+|51692606|51692687@51.69.19&[MIMAT0004678&hsa-miR-99b-3p&offsets|0|-3;m-151&19|+|51692656|51692678&offsets|0|-4]|CAAGCTCGTGTCTGTGGGT</td><td>>MI0000746|hsa-mir-99b&WithFlank&19|+|51692606|51692687</td><td>51</td><td>69</td><td>19</td><td>CAAGCTCGTGTCTGTGGGT</td><td>1001.5</td><td>8.9</td><td>1.3</td><td>6.9</td><td>5.3E-12</td><td>1.1E-11</td><td>BT20/MB468</td></tr>
<tr><td>>hg38_21_+_8433172_8446622_RNA45SN1_45S_50ntFlanks@10781.10801.21|AGGTGAACAGCCTCTGGCATG</td><td>>hg38_21_+_8433172_8446622_RNA45SN1_45S_50ntFlanks</td><td>10781</td><td>10801</td><td>21</td><td>AGGTGAACAGCCTCTGGCATG</td><td>585.5</td><td>-3.9</td><td>0.6</td><td>-6.9</td><td>4.7E-12</td><td>1.1E-11</td><td>MB231/MB468</td></tr>
<tr><td>>hg38_21_+_8433172_8446622_RNA45SN1_45S_50ntFlanks@8115.8143.29|CGCGGGACATGTGGCGTACGGAAGACCCG</td><td>>hg38_21_+_8433172_8446622_RNA45SN1_45S_50ntFlanks</td><td>8115</td><td>8143</td><td>29</td><td>CGCGGGACATGTGGCGTACGGAAGACCCG</td><td>1508.8</td><td>-5.0</td><td>0.7</td><td>-6.9</td><td>5.4E-12</td><td>1.1E-11</td><td>BT20/MB468</td></tr>
<tr><td>>hg38_21_+_8433172_8446622_RNA45SN1_45S_50ntFlanks@4160.4178.19|CCACATCCAAGGAAGGCAG</td><td>>hg38_21_+_8433172_8446622_RNA45SN1_45S_50ntFlanks</td><td>4160</td><td>4178</td><td>19</td><td>CCACATCCAAGGAAGGCAG</td><td>99.3</td><td>-8.6</td><td>1.3</td><td>-6.9</td><td>5.4E-12</td><td>1.1E-11</td><td>BT20/MB468</td></tr>
<tr><td>>hg38_21_+_8433172_8446622_RNA45SN1_45S_50ntFlanks@6664.6681.18|TGGATCACTCGGCTCGTG</td><td>>hg38_21_+_8433172_8446622_RNA45SN1_45S_50ntFlanks</td><td>6664</td><td>6681</td><td>18</td><td>TGGATCACTCGGCTCGTG</td><td>1345.3</td><td>-4.7</td><td>0.7</td><td>-6.9</td><td>5.4E-12</td><td>1.1E-11</td><td>BT20/MB468</td></tr>
<tr><td>>hg38_21_+_8433172_8446622_RNA45SN1_45S_50ntFlanks@13002.13026.25|CGATCTATTGAAAGTCAGCCCTCGA</td><td>>hg38_21_+_8433172_8446622_RNA45SN1_45S_50ntFlanks</td><td>13002</td><td>13026</td><td>25</td><td>CGATCTATTGAAAGTCAGCCCTCGA</td><td>130.4</td><td>-8.6</td><td>1.2</td><td>-6.9</td><td>4.8E-12</td><td>1.1E-11</td><td>MB231/MB468</td></tr>
<tr><td>>hg38_21_+_8433172_8446622_RNA45SN1_45S_50ntFlanks@12934.12970.37|CTAAACCATTCGTAGACGACCTGCTTCTGGGTCGGGG</td><td>>hg38_21_+_8433172_8446622_RNA45SN1_45S_50ntFlanks</td><td>12934</td><td>12970</td><td>37</td><td>CTAAACCATTCGTAGACGACCTGCTTCTGGGTCGGGG</td><td>188.5</td><td>-9.1</td><td>1.3</td><td>-6.9</td><td>4.8E-12</td><td>1.1E-11</td><td>MB231/MB468</td></tr>
<tr><td>>hg38_21_+_8433172_8446622_RNA45SN1_45S_50ntFlanks@5490.5516.27|TGAGAAGACGGTCGAACTTGACTATCT</td><td>>hg38_21_+_8433172_8446622_RNA45SN1_45S_50ntFlanks</td><td>5490</td><td>5516</td><td>27</td><td>TGAGAAGACGGTCGAACTTGACTATCT</td><td>2635.0</td><td>10.7</td><td>1.5</td><td>6.9</td><td>5.5E-12</td><td>1.1E-11</td><td>BT20/MB468</td></tr>
<tr><td>>MI0000085|hsa-mir-27a&WithFlank&19|-|13836434|13836523@57.75.19&[MIMAT0000084&hsa-miR-27a-3p&offsets|0|-2;m-43&19|-|13836447|13836467&offsets|0|-2]|TTCACAGTGGCTAAGTTCC</td><td>>MI0000085|hsa-mir-27a&WithFlank&19|-|13836434|13836523</td><td>57</td><td>75</td><td>19</td><td>TTCACAGTGGCTAAGTTCC</td><td>1241.4</td><td>5.8</td><td>0.8</td><td>6.9</td><td>5.5E-12</td><td>1.1E-11</td><td>BT20/MB468</td></tr>
<tr><td>>hg38_21_+_8433172_8446622_RNA45SN1_45S_50ntFlanks@12154.12182.29|GGGAGTTTGACTGGGGCGGTACACCTGTC</td><td>>hg38_21_+_8433172_8446622_RNA45SN1_45S_50ntFlanks</td><td>12154</td><td>12182</td><td>29</td><td>GGGAGTTTGACTGGGGCGGTACACCTGTC</td><td>105.9</td><td>-8.7</td><td>1.3</td><td>-6.9</td><td>5.6E-12</td><td>1.1E-11</td><td>BT20/MB468</td></tr>
<tr><td>>hg38_21_+_8433172_8446622_RNA45SN1_45S_50ntFlanks@11894.11920.27|ACTCTAGTCTGGCACGGTGAAGAGACA</td><td>>hg38_21_+_8433172_8446622_RNA45SN1_45S_50ntFlanks</td><td>11894</td><td>11920</td><td>27</td><td>ACTCTAGTCTGGCACGGTGAAGAGACA</td><td>143.5</td><td>-9.2</td><td>1.3</td><td>-6.9</td><td>5.6E-12</td><td>1.1E-11</td><td>BT20/MB468</td></tr>
<tr><td>>hg38_21_+_8433172_8446622_RNA45SN1_45S_50ntFlanks@11873.11912.40|AAGAAGACCCTGTTGAGCTTGACTCTAGTCTGGCACGGTG</td><td>>hg38_21_+_8433172_8446622_RNA45SN1_45S_50ntFlanks</td><td>11873</td><td>11912</td><td>40</td><td>AAGAAGACCCTGTTGAGCTTGACTCTAGTCTGGCACGGTG</td><td>111.7</td><td>-8.8</td><td>1.3</td><td>-6.9</td><td>5.6E-12</td><td>1.1E-11</td><td>BT20/MB468</td></tr>
<tr><td>>hg38_MT_+_598_1651_RNR1_12S_50ntFlanks@52.77.26|ATAGGTTTGGTCCTAGCCTTTCTATT</td><td>>hg38_MT_+_598_1651_RNR1_12S_50ntFlanks</td><td>52</td><td>77</td><td>26</td><td>ATAGGTTTGGTCCTAGCCTTTCTATT</td><td>653.8</td><td>7.5</td><td>1.1</td><td>6.9</td><td>4.9E-12</td><td>1.1E-11</td><td>MB231/MB468</td></tr>
<tr><td>>hg38_21_+_8433172_8446622_RNA45SN1_45S_50ntFlanks@11661.11682.22|TGATTTCTGCCCAGTGCTCTGA</td><td>>hg38_21_+_8433172_8446622_RNA45SN1_45S_50ntFlanks</td><td>11661</td><td>11682</td><td>22</td><td>TGATTTCTGCCCAGTGCTCTGA</td><td>566.1</td><td>-4.5</td><td>0.7</td><td>-6.9</td><td>5.6E-12</td><td>1.1E-11</td><td>BT20/MB468</td></tr>
<tr><td>>hg38_21_+_8433172_8446622_RNA45SN1_45S_50ntFlanks@5472.5512.41|ACGGCCCTGGCGGAGCGCTGAGAAGACGGTCGAACTTGACT</td><td>>hg38_21_+_8433172_8446622_RNA45SN1_45S_50ntFlanks</td><td>5472</td><td>5512</td><td>41</td><td>ACGGCCCTGGCGGAGCGCTGAGAAGACGGTCGAACTTGACT</td><td>735.2</td><td>7.4</td><td>1.1</td><td>6.9</td><td>5.7E-12</td><td>1.1E-11</td><td>BT20/MB468</td></tr>
<tr><td>>hg38_MT_+_598_1651_RNR1_12S_50ntFlanks@428.455.28|AAAGTGGCTTTAACATATCTGAACACAC</td><td>>hg38_MT_+_598_1651_RNR1_12S_50ntFlanks</td><td>428</td><td>455</td><td>28</td><td>AAAGTGGCTTTAACATATCTGAACACAC</td><td>572.5</td><td>9.5</td><td>1.4</td><td>6.9</td><td>5.7E-12</td><td>1.1E-11</td><td>BT20/MB468</td></tr>
<tr><td>>hg38_21_+_8433172_8446622_RNA45SN1_45S_50ntFlanks@4069.4095.27|CTTTCGATGGTAGTCGCCGTGCCTACC</td><td>>hg38_21_+_8433172_8446622_RNA45SN1_45S_50ntFlanks</td><td>4069</td><td>4095</td><td>27</td><td>CTTTCGATGGTAGTCGCCGTGCCTACC</td><td>162.3</td><td>-3.3</td><td>0.5</td><td>-6.9</td><td>5.0E-12</td><td>1.2E-11</td><td>MB231/MB468</td></tr>
<tr><td>>hg38_21_+_8433172_8446622_RNA45SN1_45S_50ntFlanks@12944.12972.29|CGTAGACGACCTGCTTCTGGGTCGGGGTT</td><td>>hg38_21_+_8433172_8446622_RNA45SN1_45S_50ntFlanks</td><td>12944</td><td>12972</td><td>29</td><td>CGTAGACGACCTGCTTCTGGGTCGGGGTT</td><td>149.8</td><td>-4.4</td><td>0.6</td><td>-6.9</td><td>5.1E-12</td><td>1.2E-11</td><td>MB231/MB468</td></tr>
<tr><td>>MI0000483|hsa-mir-186&WithFlank&1|-|71067625|71067722@21.43.23&[MIMAT0000456&hsa-miR-186-5p&offsets|0|+1;m-84&1|-|71067681|71067702&offsets|0|+1]|CAAAGAATTCTCCTTTTGGGCTT</td><td>>MI0000483|hsa-mir-186&WithFlank&1|-|71067625|71067722</td><td>21</td><td>43</td><td>23</td><td>CAAAGAATTCTCCTTTTGGGCTT</td><td>4042.8</td><td>7.0</td><td>1.0</td><td>6.9</td><td>5.8E-12</td><td>1.2E-11</td><td>BT20/MB468</td></tr>
<tr><td>>hg38_21_+_8433172_8446622_RNA45SN1_45S_50ntFlanks@7990.8011.22|GACGTGGCGACCCGCTGAATTT</td><td>>hg38_21_+_8433172_8446622_RNA45SN1_45S_50ntFlanks</td><td>7990</td><td>8011</td><td>22</td><td>GACGTGGCGACCCGCTGAATTT</td><td>1099.3</td><td>4.5</td><td>0.7</td><td>6.9</td><td>5.8E-12</td><td>1.2E-11</td><td>BT20/MB468</td></tr>
<tr><td>>hg38_21_+_8433172_8446622_RNA45SN1_45S_50ntFlanks@11575.11599.25|ACTTAGAACTGGTGCGGACCAGGGG</td><td>>hg38_21_+_8433172_8446622_RNA45SN1_45S_50ntFlanks</td><td>11575</td><td>11599</td><td>25</td><td>ACTTAGAACTGGTGCGGACCAGGGG</td><td>138.6</td><td>-8.7</td><td>1.3</td><td>-6.9</td><td>5.1E-12</td><td>1.2E-11</td><td>MB231/MB468</td></tr>
<tr><td>>hg38_21_+_8433172_8446622_RNA45SN1_45S_50ntFlanks@6651.6668.18|CGACTCTTAGCGGTGGAT</td><td>>hg38_21_+_8433172_8446622_RNA45SN1_45S_50ntFlanks</td><td>6651</td><td>6668</td><td>18</td><td>CGACTCTTAGCGGTGGAT</td><td>6324.6</td><td>3.7</td><td>0.5</td><td>6.9</td><td>5.1E-12</td><td>1.2E-11</td><td>MB231/MB468</td></tr>
<tr><td>>hg38_21_+_8433172_8446622_RNA45SN1_45S_50ntFlanks@11641.11663.23|CGGCGGGTGTTGACGCGATGTGA</td><td>>hg38_21_+_8433172_8446622_RNA45SN1_45S_50ntFlanks</td><td>11641</td><td>11663</td><td>23</td><td>CGGCGGGTGTTGACGCGATGTGA</td><td>330.8</td><td>-8.0</td><td>1.2</td><td>-6.9</td><td>5.2E-12</td><td>1.2E-11</td><td>MB231/MB468</td></tr>
<tr><td>>hg38_21_+_8433172_8446622_RNA45SN1_45S_50ntFlanks@11874.11910.37|AGAAGACCCTGTTGAGCTTGACTCTAGTCTGGCACGG</td><td>>hg38_21_+_8433172_8446622_RNA45SN1_45S_50ntFlanks</td><td>11874</td><td>11910</td><td>37</td><td>AGAAGACCCTGTTGAGCTTGACTCTAGTCTGGCACGG</td><td>149.6</td><td>-8.8</td><td>1.3</td><td>-6.9</td><td>5.2E-12</td><td>1.2E-11</td><td>MB231/MB468</td></tr>
<tr><td>>hg38_21_+_8433172_8446622_RNA45SN1_45S_50ntFlanks@10403.10420.18|ACATGGGTCAGTCGGTCC</td><td>>hg38_21_+_8433172_8446622_RNA45SN1_45S_50ntFlanks</td><td>10403</td><td>10420</td><td>18</td><td>ACATGGGTCAGTCGGTCC</td><td>131.1</td><td>-4.0</td><td>0.6</td><td>-6.9</td><td>5.9E-12</td><td>1.2E-11</td><td>BT20/MB468</td></tr>
<tr><td>>hg38_21_+_8433172_8446622_RNA45SN1_45S_50ntFlanks@9567.9584.18|ACGTGCAAATCGGTCGTC</td><td>>hg38_21_+_8433172_8446622_RNA45SN1_45S_50ntFlanks</td><td>9567</td><td>9584</td><td>18</td><td>ACGTGCAAATCGGTCGTC</td><td>355.9</td><td>-4.8</td><td>0.7</td><td>-6.9</td><td>5.9E-12</td><td>1.2E-11</td><td>BT20/MB468</td></tr>
<tr><td>>hg38_21_+_8433172_8446622_RNA45SN1_45S_50ntFlanks@12685.12703.19|CCCAGGCGGAACGATACGG</td><td>>hg38_21_+_8433172_8446622_RNA45SN1_45S_50ntFlanks</td><td>12685</td><td>12703</td><td>19</td><td>CCCAGGCGGAACGATACGG</td><td>113.8</td><td>-8.8</td><td>1.3</td><td>-6.9</td><td>5.9E-12</td><td>1.2E-11</td><td>BT20/MB468</td></tr>
<tr><td>>MI0000081|hsa-mir-24-2&WithFlank&19|-|13836281|13836365@56.76.21;MI0000080|hsa-mir-24-1&WithFlank&9|+|95086015|95086094@50.70.21&[MIMAT0000080&hsa-miR-24-3p&offsets|0|-1;m-36&19|-|13836288|13836310&offsets|0|-2];[MIMAT0000080_1&hsa-miR-24-3p&offsets|0|-1;m-35&9|+|95086064|95086086&offsets|0|-2]|TGGCTCAGTTCAGCAGGAACA</td><td>>MI0000081|hsa-mir-24-2&WithFlank&19|-|13836281|13836365</td><td>56</td><td>76</td><td>21</td><td>TGGCTCAGTTCAGCAGGAACA</td><td>8853.9</td><td>5.2</td><td>0.8</td><td>6.9</td><td>6.0E-12</td><td>1.2E-11</td><td>BT20/MB468</td></tr>
<tr><td>>MI0000085|hsa-mir-27a&WithFlank&19|-|13836434|13836523@57.76.20(+1G)&[MIMAT0000084&hsa-miR-27a-3p&offsets|0|-1(+1G);m-43&19|-|13836447|13836467&offsets|0|-1(+1G)]|TTCACAGTGGCTAAGTTCCGG</td><td>>MI0000085|hsa-mir-27a&WithFlank&19|-|13836434|13836523</td><td>57</td><td>76</td><td>20(+1G)</td><td>TTCACAGTGGCTAAGTTCCGG</td><td>1059.0</td><td>5.6</td><td>0.8</td><td>6.9</td><td>5.2E-12</td><td>1.2E-11</td><td>MB231/MB468</td></tr>
<tr><td>>hg38_21_+_8433172_8446622_RNA45SN1_45S_50ntFlanks@10845.10862.18|ACTTCGGGATAAGGATTG</td><td>>hg38_21_+_8433172_8446622_RNA45SN1_45S_50ntFlanks</td><td>10845</td><td>10862</td><td>18</td><td>ACTTCGGGATAAGGATTG</td><td>641.4</td><td>-4.5</td><td>0.7</td><td>-6.9</td><td>5.3E-12</td><td>1.2E-11</td><td>MB231/MB468</td></tr>
<tr><td>>hg38_21_+_8433172_8446622_RNA45SN1_45S_50ntFlanks@11752.11780.29|CCAAATGCCTCGTCATCTAATTAGTGACG</td><td>>hg38_21_+_8433172_8446622_RNA45SN1_45S_50ntFlanks</td><td>11752</td><td>11780</td><td>29</td><td>CCAAATGCCTCGTCATCTAATTAGTGACG</td><td>147.3</td><td>-8.8</td><td>1.3</td><td>-6.9</td><td>5.3E-12</td><td>1.2E-11</td><td>MB231/MB468</td></tr>
<tr><td>>hg38_21_+_8433172_8446622_RNA45SN1_45S_50ntFlanks@12128.12146.19|ACCCGCTCCGGGGACAGTG</td><td>>hg38_21_+_8433172_8446622_RNA45SN1_45S_50ntFlanks</td><td>12128</td><td>12146</td><td>19</td><td>ACCCGCTCCGGGGACAGTG</td><td>90.8</td><td>-8.5</td><td>1.2</td><td>-6.9</td><td>6.0E-12</td><td>1.2E-11</td><td>BT20/MB468</td></tr>
<tr><td>>hg38_21_+_8433172_8446622_RNA45SN1_45S_50ntFlanks@11874.11909.36|AGAAGACCCTGTTGAGCTTGACTCTAGTCTGGCACG</td><td>>hg38_21_+_8433172_8446622_RNA45SN1_45S_50ntFlanks</td><td>11874</td><td>11909</td><td>36</td><td>AGAAGACCCTGTTGAGCTTGACTCTAGTCTGGCACG</td><td>175.8</td><td>-7.2</td><td>1.0</td><td>-6.9</td><td>5.4E-12</td><td>1.2E-11</td><td>MB231/MB468</td></tr>
<tr><td>>hg38_21_+_8433172_8446622_RNA45SN1_45S_50ntFlanks@13005.13028.24|TCTATTGAAAGTCAGCCCTCGACA</td><td>>hg38_21_+_8433172_8446622_RNA45SN1_45S_50ntFlanks</td><td>13005</td><td>13028</td><td>24</td><td>TCTATTGAAAGTCAGCCCTCGACA</td><td>169.5</td><td>-9.4</td><td>1.4</td><td>-6.9</td><td>6.2E-12</td><td>1.2E-11</td><td>BT20/MB468</td></tr>
<tr><td>>hg38_21_+_8433172_8446622_RNA45SN1_45S_50ntFlanks@12161.12181.21|TGACTGGGGCGGTACACCTGT</td><td>>hg38_21_+_8433172_8446622_RNA45SN1_45S_50ntFlanks</td><td>12161</td><td>12181</td><td>21</td><td>TGACTGGGGCGGTACACCTGT</td><td>149.0</td><td>-8.8</td><td>1.3</td><td>-6.9</td><td>5.4E-12</td><td>1.2E-11</td><td>MB231/MB468</td></tr>
<tr><td>>hg38_21_+_8433172_8446622_RNA45SN1_45S_50ntFlanks@12045.12070.26|CTGACCCGGTGAGGCGGGGGGGCGAG</td><td>>hg38_21_+_8433172_8446622_RNA45SN1_45S_50ntFlanks</td><td>12045</td><td>12070</td><td>26</td><td>CTGACCCGGTGAGGCGGGGGGGCGAG</td><td>104.2</td><td>-8.7</td><td>1.3</td><td>-6.9</td><td>6.2E-12</td><td>1.2E-11</td><td>BT20/MB468</td></tr>
<tr><td>>hg38_21_+_8433172_8446622_RNA45SN1_45S_50ntFlanks@9540.9571.32|CTCTGGTGGAGGTCCGTAGCGGTCCTGACGTG</td><td>>hg38_21_+_8433172_8446622_RNA45SN1_45S_50ntFlanks</td><td>9540</td><td>9571</td><td>32</td><td>CTCTGGTGGAGGTCCGTAGCGGTCCTGACGTG</td><td>326.4</td><td>-2.9</td><td>0.4</td><td>-6.9</td><td>5.5E-12</td><td>1.2E-11</td><td>MB231/MB468</td></tr>
</table>

| | | | | | | | | | | | | |
|---|---|---|---|---|---|---|---|---|---|---|---|---|
| >hg38_21_+_8433172_8446622_RNA45SN1_45S_50ntFlanks@10357.10376.20\|CTTTGAAGGCCGAAGTGGAG | >hg38_21_+_8433172_8446622_RNA45SN1_45S_50ntFlanks | 10357 | 10376 | 20 | CTTTGAAGGCCGAAGTGGAG | 105.9 | -4.2 | 0.6 | -6.9 | 6.2E-12 | 1.2E-11 | BT20/MB468 |
| >hg38_21_+_8433172_8446622_RNA45SN1_45S_50ntFlanks@10869.10901.33\|AGGGCTGGGTCGGTCGGGCTGGGGCGCGAAGCG | >hg38_21_+_8433172_8446622_RNA45SN1_45S_50ntFlanks | 10869 | 10901 | 33 | AGGGCTGGGTCGGTCGGGCTGGGGCGCGAAGCG | 242.7 | -5.8 | 0.8 | -6.9 | 6.3E-12 | 1.3E-11 | BT20/MB468 |
| >hg38_21_+_8433172_8446622_RNA45SN1_45S_50ntFlanks@13002.13027.26\|CGATCTATTGAAAGTCAGCCCTCGAC | >hg38_21_+_8433172_8446622_RNA45SN1_45S_50ntFlanks | 13002 | 13027 | 26 | CGATCTATTGAAAGTCAGCCCTCGAC | 5178.7 | -8.0 | 1.2 | -6.9 | 5.5E-12 | 1.3E-11 | MB231/MB468 |
| >hg38_21_+_8433172_8446622_RNA45SN1_45S_50ntFlanks@10101.10126.26\|CGCGCGCGTGTGGTGTGCGTCGGAGG | >hg38_21_+_8433172_8446622_RNA45SN1_45S_50ntFlanks | 10101 | 10126 | 26 | CGCGCGCGTGTGGTGTGCGTCGGAGG | 262.6 | -7.7 | 1.1 | -6.9 | 5.6E-12 | 1.3E-11 | MB231/MB468 |
| >hg38_21_+_8433172_8446622_RNA45SN1_45S_50ntFlanks@8179.8211.33\|CTGATCGAGGCCCAGCCCGTGGACGGTGTGAGG | >hg38_21_+_8433172_8446622_RNA45SN1_45S_50ntFlanks | 8179 | 8211 | 33 | CTGATCGAGGCCCAGCCCGTGGACGGTGTGAGG | 261.4 | -6.7 | 1.0 | -6.9 | 6.4E-12 | 1.3E-11 | BT20/MB468 |
| >hg38_21_+_8433172_8446622_RNA45SN1_45S_50ntFlanks@11323.11340.18\|CGTCTCGTCGCGCGCGCG | >hg38_21_+_8433172_8446622_RNA45SN1_45S_50ntFlanks | 11323 | 11340 | 18 | CGTCTCGTCGCGCGCGCG | 340.2 | -5.8 | 0.8 | -6.9 | 5.6E-12 | 1.3E-11 | MB231/MB468 |
| >hg38_21_+_8433172_8446622_RNA45SN1_45S_50ntFlanks@5423.5443.21\|ATTGGATGGTTTAGTGAGGCC | >hg38_21_+_8433172_8446622_RNA45SN1_45S_50ntFlanks | 5423 | 5443 | 21 | ATTGGATGGTTTAGTGAGGCC | 112.4 | -5.6 | 0.8 | -6.9 | 6.4E-12 | 1.3E-11 | BT20/MB468 |
| >MI0000816\|hsa-mir-335&WithFlank&7\|+\|130496105\|130496210@22.43.22&[MIMAT0000765&hsa-miR-335-5p&offsets\|0\|-1;m-249&7\|+\|130496126\|130496147&offsets\|0\|0]\|TCAAGAGCAATAACGAAAAATG | >MI0000816\|hsa-mir-335&WithFlank&7\|+\|130496105\|130496210 | 22 | 43 | 22 | TCAAGAGCAATAACGAAAAATG | 501.8 | 6.5 | 0.9 | 6.9 | 6.4E-12 | 1.3E-11 | BT20/MB468 |
| >hg38_21_+_8433172_8446622_RNA45SN1_45S_50ntFlanks@13005.13025.21\|TCTATTGAAAGTCAGCCCTCG | >hg38_21_+_8433172_8446622_RNA45SN1_45S_50ntFlanks | 13005 | 13025 | 21 | TCTATTGAAAGTCAGCCCTCG | 162.0 | -9.3 | 1.4 | -6.9 | 6.5E-12 | 1.3E-11 | BT20/MB468 |
| >hg38_21_+_8433172_8446622_RNA45SN1_45S_50ntFlanks@12937.12955.19\|AACCATTCGTAGACGACCT | >hg38_21_+_8433172_8446622_RNA45SN1_45S_50ntFlanks | 12937 | 12955 | 19 | AACCATTCGTAGACGACCT | 216.0 | -3.0 | 0.4 | -6.9 | 5.8E-12 | 1.3E-11 | MB231/MB468 |
| >hg38_21_+_8433172_8446622_RNA45SN1_45S_50ntFlanks@12367.12384.18\|AACTGGCTTGTGGCGGCC | >hg38_21_+_8433172_8446622_RNA45SN1_45S_50ntFlanks | 12367 | 12384 | 18 | AACTGGCTTGTGGCGGCC | 162.7 | -3.5 | 0.5 | -6.9 | 6.6E-12 | 1.3E-11 | BT20/MB468 |
| >hg38_21_+_8433172_8446622_RNA45SN1_45S_50ntFlanks@4473.4494.22\|TTAGCTGAGTGTCCCGCGGGGC | >hg38_21_+_8433172_8446622_RNA45SN1_45S_50ntFlanks | 4473 | 4494 | 22 | TTAGCTGAGTGTCCCGCGGGGC | 93.0 | -8.5 | 1.2 | -6.9 | 6.6E-12 | 1.3E-11 | BT20/MB468 |
| >hg38_21_+_8433172_8446622_RNA45SN1_45S_50ntFlanks@12539.12557.19\|ATGATGTGTTGTTGCCATG | >hg38_21_+_8433172_8446622_RNA45SN1_45S_50ntFlanks | 12539 | 12557 | 19 | ATGATGTGTTGTTGCCATG | 149.4 | -8.8 | 1.3 | -6.9 | 5.9E-12 | 1.3E-11 | MB231/MB468 |
| >hg38_21_+_8433172_8446622_RNA45SN1_45S_50ntFlanks@7985.8005.21\|GATCAGACGTGGCGACCCGCT | >hg38_21_+_8433172_8446622_RNA45SN1_45S_50ntFlanks | 7985 | 8005 | 21 | GATCAGACGTGGCGACCCGCT | 136.4 | -9.1 | 1.3 | -6.9 | 6.7E-12 | 1.3E-11 | BT20/MB468 |
| >trna129_MetCAT_6_-_27870271_27870342@32.51.20,trna142_MetCAT_6_-_27560600_27560671@32.51.20,trna150_MetCAT_6_-_27300764_27300835@32.51.20,trna169_MetCAT_6_-_26330529_26330600@32.51.20,trna171_MetCAT_6_-_26313352_26313423@32.51.20,trna20_MetCAT_17_-_80452597_80452668@32.51.20,trna2_MetCAT_6_+_26286754_26286825@32.51.20,trna32_MetCAT_1_+_153643726_153643797@32.51.20,trna61_MetCAT_6_+_27745664_27745735@32.51.20\|CCATAACCCAGAGGTCGATG | >trna129_MetCAT_6_-_27870271_27870342 | 32 | 51 | 20 | CCATAACCCAGAGGTCGATG | 199.6 | -9.2 | 1.3 | -6.9 | 5.9E-12 | 1.3E-11 | MB231/MB468 |
| >hg38_21_+_8433172_8446622_RNA45SN1_45S_50ntFlanks@7992.8017.26\|CGTGGCGACCCGCTGAATTTAAGCAT | >hg38_21_+_8433172_8446622_RNA45SN1_45S_50ntFlanks | 7992 | 8017 | 26 | CGTGGCGACCCGCTGAATTTAAGCAT | 312.0 | 3.8 | 0.6 | 6.9 | 5.9E-12 | 1.3E-11 | MB231/MB468 |
| >hg38_21_+_8433172_8446622_RNA45SN1_45S_50ntFlanks@10916.10936.21\|CCGCGGCTGGACGAGGCGCCG | >hg38_21_+_8433172_8446622_RNA45SN1_45S_50ntFlanks | 10916 | 10936 | 21 | CCGCGGCTGGACGAGGCGCCG | 185.5 | -9.1 | 1.3 | -6.9 | 5.9E-12 | 1.3E-11 | MB231/MB468 |
| >hg38_21_+_8433172_8446622_RNA45SN1_45S_50ntFlanks@12628.12659.32\|TGGGGCGAAGCTACCATCTGTGGGATTATGAC | >hg38_21_+_8433172_8446622_RNA45SN1_45S_50ntFlanks | 12628 | 12659 | 32 | TGGGGCGAAGCTACCATCTGTGGGATTATGAC | 185.9 | -5.5 | 0.8 | -6.9 | 5.9E-12 | 1.3E-11 | MB231/MB468 |
| >hg38_21_+_8433172_8446622_RNA45SN1_45S_50ntFlanks@3991.4023.33\|TGGTGACTCTAGATAACCTCGGGCCGATCGCAC | >hg38_21_+_8433172_8446622_RNA45SN1_45S_50ntFlanks | 3991 | 4023 | 33 | TGGTGACTCTAGATAACCTCGGGCCGATCGCAC | 503.4 | 4.2 | 0.6 | 6.9 | 5.9E-12 | 1.3E-11 | MB231/MB468 |
| >hg38_21_+_8433172_8446622_RNA45SN1_45S_50ntFlanks@3771.3799.29\|CACGGCCGGTACAGTGAAACTGCGAATGG | >hg38_21_+_8433172_8446622_RNA45SN1_45S_50ntFlanks | 3771 | 3799 | 29 | CACGGCCGGTACAGTGAAACTGCGAATGG | 87.4 | -3.9 | 0.6 | -6.9 | 6.7E-12 | 1.3E-11 | BT20/MB468 |
| >am_trnaMT_MetCAT_MT_+_4402_4469@1.18.18\|AGTAAGGTCAGCTAAATA | >am_trnaMT_MetCAT_MT_+_4402_4469 | 1 | 18 | 18 | AGTAAGGTCAGCTAAATA | 815.5 | 9.1 | 1.3 | 6.9 | 6.8E-12 | 1.4E-11 | BT20/MB468 |
| >hg38_21_+_8433172_8446622_RNA45SN1_45S_50ntFlanks@6723.6745.23\|TTGCAGGACACATTGATCATCGA | >hg38_21_+_8433172_8446622_RNA45SN1_45S_50ntFlanks | 6723 | 6745 | 23 | TTGCAGGACACATTGATCATCGA | 107.0 | -8.7 | 1.3 | -6.9 | 6.8E-12 | 1.4E-11 | BT20/MB468 |
| >hg38_21_+_8433172_8446622_RNA45SN1_45S_50ntFlanks@12892.12933.42\|CCCTCGCCCGTCACGCACCGCACGTTCGTGGGGAACCTGGCG | >hg38_21_+_8433172_8446622_RNA45SN1_45S_50ntFlanks | 12892 | 12933 | 42 | CCCTCGCCCGTCACGCACCGCACGTTCGTGGGGAACCTGGCG | 148.0 | -9.2 | 1.3 | -6.9 | 6.8E-12 | 1.4E-11 | BT20/MB468 |
| >hg38_21_+_8433172_8446622_RNA45SN1_45S_50ntFlanks@10516.10533.18\|GAGTGGCGGAGATGGGCG | >hg38_21_+_8433172_8446622_RNA45SN1_45S_50ntFlanks | 10516 | 10533 | 18 | GAGTGGCGGAGATGGGCG | 253.1 | -7.7 | 1.1 | -6.9 | 6.0E-12 | 1.4E-11 | MB231/MB468 |
| >hg38_21_+_8433172_8446622_RNA45SN1_45S_50ntFlanks@11998.12032.35\|GCCCTGCGGGCCGCCGGTGAAATACCACTACTCTG | >hg38_21_+_8433172_8446622_RNA45SN1_45S_50ntFlanks | 11998 | 12032 | 35 | GCCCTGCGGGCCGCCGGTGAAATACCACTACTCTG | 100.5 | -8.6 | 1.3 | -6.9 | 6.9E-12 | 1.4E-11 | BT20/MB468 |
| >hg38_21_+_8433172_8446622_RNA45SN1_45S_50ntFlanks@12624.12652.29\|CCAATGGGGCGAAGCTACCATCTGTGGGA | >hg38_21_+_8433172_8446622_RNA45SN1_45S_50ntFlanks | 12624 | 12652 | 29 | CCAATGGGGCGAAGCTACCATCTGTGGGA | 110.7 | -8.8 | 1.3 | -6.9 | 6.9E-12 | 1.4E-11 | BT20/MB468 |
| >hg38_21_+_8433172_8446622_RNA45SN1_45S_50ntFlanks@10731.10755.25\|AGGGTGTAAATCTCGCGCCGGGCCG | >hg38_21_+_8433172_8446622_RNA45SN1_45S_50ntFlanks | 10731 | 10755 | 25 | AGGGTGTAAATCTCGCGCCGGGCCG | 255.7 | -6.2 | 0.9 | -6.9 | 6.9E-12 | 1.4E-11 | BT20/MB468 |
| >hg38_21_+_8433172_8446622_RNA45SN1_45S_50ntFlanks@10408.10434.27\|GGTCAGTCGGTCCTGAGAGATGGGCGA | >hg38_21_+_8433172_8446622_RNA45SN1_45S_50ntFlanks | 10408 | 10434 | 27 | GGTCAGTCGGTCCTGAGAGATGGGCGA | 126.5 | -8.6 | 1.2 | -6.9 | 6.1E-12 | 1.4E-11 | MB231/MB468 |
| >hg38_21_+_8433172_8446622_RNA45SN1_45S_50ntFlanks@10128.10147.20\|CGGCGGCGGCGGCGGCGGCG | >hg38_21_+_8433172_8446622_RNA45SN1_45S_50ntFlanks | 10128 | 10147 | 20 | CGGCGGCGGCGGCGGCGGCG | 117.2 | -5.1 | 0.8 | -6.9 | 7.0E-12 | 1.4E-11 | BT20/MB468 |
| >hg38_21_+_8433172_8446622_RNA45SN1_45S_50ntFlanks@11795.11820.26\|AACGAGATTCCCACTGTCCCTACCTA | >hg38_21_+_8433172_8446622_RNA45SN1_45S_50ntFlanks | 11795 | 11820 | 26 | AACGAGATTCCCACTGTCCCTACCTA | 101.1 | -8.7 | 1.3 | -6.9 | 7.0E-12 | 1.4E-11 | BT20/MB468 |
| >hg38_21_+_8433172_8446622_RNA45SN1_45S_50ntFlanks@8845.8864.20\|CGGCGGCGGCGGCGGCGGCG | >hg38_21_+_8433172_8446622_RNA45SN1_45S_50ntFlanks | 8845 | 8864 | 20 | CGGCGGCGGCGGCGGCGGCG | 117.2 | -5.1 | 0.8 | -6.9 | 7.0E-12 | 1.4E-11 | BT20/MB468 |
| >hg38_21_+_8433172_8446622_RNA45SN1_45S_50ntFlanks@12417.12455.39\|TCGATGTCGGCTCTTCCTATCATTGTGAAGCAGAATTCA | >hg38_21_+_8433172_8446622_RNA45SN1_45S_50ntFlanks | 12417 | 12455 | 39 | TCGATGTCGGCTCTTCCTATCATTGTGAAGCAGAATTCA | 407.5 | -3.6 | 0.5 | -6.9 | 7.0E-12 | 1.4E-11 | BT20/MB468 |
| >hg38_21_+_8433172_8446622_RNA45SN1_45S_50ntFlanks@10512.10534.23\|TCCGGAGTGGCGGAGATGGGCGC | >hg38_21_+_8433172_8446622_RNA45SN1_45S_50ntFlanks | 10512 | 10534 | 23 | TCCGGAGTGGCGGAGATGGGCGC | 151.1 | -9.2 | 1.3 | -6.9 | 7.0E-12 | 1.4E-11 | BT20/MB468 |
| >hg38_21_+_8433172_8446622_RNA45SN1_45S_50ntFlanks@12628.12652.25\|TGGGGCGAAGCTACCATCTGTGGGA | >hg38_21_+_8433172_8446622_RNA45SN1_45S_50ntFlanks | 12628 | 12652 | 25 | TGGGGCGAAGCTACCATCTGTGGGA | 87.0 | -8.4 | 1.2 | -6.9 | 7.0E-12 | 1.4E-11 | BT20/MB468 |
| >hg38_21_+_8433172_8446622_RNA45SN1_45S_50ntFlanks@6722.6743.22\|ATTGCAGGACACATTGATCATC | >hg38_21_+_8433172_8446622_RNA45SN1_45S_50ntFlanks | 6722 | 6743 | 22 | ATTGCAGGACACATTGATCATC | 1072.8 | -3.9 | 0.6 | -6.9 | 7.0E-12 | 1.4E-11 | BT20/MB468 |
| >hg38_21_+_8433172_8446622_RNA45SN1_45S_50ntFlanks@6666.6689.24\|GATCACTCGGCTCGTGCGTCGATG | >hg38_21_+_8433172_8446622_RNA45SN1_45S_50ntFlanks | 6666 | 6689 | 24 | GATCACTCGGCTCGTGCGTCGATG | 90.5 | -8.5 | 1.2 | -6.9 | 7.0E-12 | 1.4E-11 | BT20/MB468 |
| >hg38_21_+_8433172_8446622_RNA45SN1_45S_50ntFlanks@12628.12657.30\|TGGGGCGAAGCTACCATCTGTGGGATTATG | >hg38_21_+_8433172_8446622_RNA45SN1_45S_50ntFlanks | 12628 | 12657 | 30 | TGGGGCGAAGCTACCATCTGTGGGATTATG | 121.9 | -6.0 | 0.9 | -6.9 | 6.2E-12 | 1.4E-11 | MB231/MB468 |
| >am_trnaMT_ValTAC_MT_+_1602_1670@28.46.19\|AACTTACACTTAGGAGATT | >am_trnaMT_ValTAC_MT_+_1602_1670 | 28 | 46 | 19 | AACTTACACTTAGGAGATT | 440.4 | 8.6 | 1.2 | 6.9 | 7.1E-12 | 1.4E-11 | BT20/MB468 |
| >hg38_21_+_8433172_8446622_RNA45SN1_45S_50ntFlanks@11212.11231.20\|CGACTCTGGACGCGAGCCGG | >hg38_21_+_8433172_8446622_RNA45SN1_45S_50ntFlanks | 11212 | 11231 | 20 | CGACTCTGGACGCGAGCCGG | 195.6 | -5.9 | 0.9 | -6.9 | 7.1E-12 | 1.4E-11 | BT20/MB468 |
| >hg38_21_+_8433172_8446622_RNA45SN1_45S_50ntFlanks@10788.10815.28\|CAGCCTCTGGCATGTTGGAACAATGTAG | >hg38_21_+_8433172_8446622_RNA45SN1_45S_50ntFlanks | 10788 | 10815 | 28 | CAGCCTCTGGCATGTTGGAACAATGTAG | 285.4 | -3.4 | 0.5 | -6.9 | 7.1E-12 | 1.4E-11 | BT20/MB468 |
| >MI0000077\|hsa-mir-21&WithFlank&17\|+\|59841260\|59841343@14.34.21(+2U)&[MIMAT0000076&hsa-miR-21-5p&offsets\|0\|-1(+2U);m-1&17\|+\|59841273\|59841294&offsets\|0\|-1(+2U)]\|TAGCTTATCAGACTGATGTTGTT | >MI0000077\|hsa-mir-21&WithFlank&17\|+\|59841260\|59841343 | 14 | 34 | 21(+2U) | TAGCTTATCAGACTGATGTTGTT | 1339.2 | 10.0 | 1.5 | 6.9 | 7.2E-12 | 1.4E-11 | BT20/MB468 |
| >hg38_21_+_8433172_8446622_RNA45SN1_45S_50ntFlanks@10461.10480.20\|GCCTCCGTTGCCCTCGGCCG | >hg38_21_+_8433172_8446622_RNA45SN1_45S_50ntFlanks | 10461 | 10480 | 20 | GCCTCCGTTGCCCTCGGCCG | 84.1 | -5.2 | 0.8 | -6.9 | 7.2E-12 | 1.4E-11 | BT20/MB468 |
| >hg38_21_+_8433172_8446622_RNA45SN1_45S_50ntFlanks@12360.12384.25\|CAGGGATAACTGGCTTGTGGCGGCC | >hg38_21_+_8433172_8446622_RNA45SN1_45S_50ntFlanks | 12360 | 12384 | 25 | CAGGGATAACTGGCTTGTGGCGGCC | 139.9 | -4.1 | 0.6 | -6.9 | 7.2E-12 | 1.4E-11 | BT20/MB468 |
| >hg38_21_+_8433172_8446622_RNA45SN1_45S_50ntFlanks@11534.11555.22\|GTTCCGCGCGGCGCCTCGCCTC | >hg38_21_+_8433172_8446622_RNA45SN1_45S_50ntFlanks | 11534 | 11555 | 22 | GTTCCGCGCGGCGCCTCGCCTC | 142.9 | -6.0 | 0.9 | -6.9 | 7.2E-12 | 1.4E-11 | BT20/MB468 |
| >hg38_21_+_8433172_8446622_RNA45SN1_45S_50ntFlanks@12511.12538.28\|CGTGAGACAGGTTAGTTTTACCCTACTG | >hg38_21_+_8433172_8446622_RNA45SN1_45S_50ntFlanks | 12511 | 12538 | 28 | CGTGAGACAGGTTAGTTTTACCCTACTG | 103.8 | -8.7 | 1.3 | -6.9 | 7.2E-12 | 1.4E-11 | BT20/MB468 |

| | | | | | | | | | | | |
|---|---|---|---|---|---|---|---|---|---|---|---|
| >hg38_21_+_8433172_8446622_RNA45SN1_45S_50ntFlanks@11199.11223.25\|CCGGCGGCGGCGGCGACTCTGGACG | >hg38_21_+_8433172_8446622_RNA45SN1_45S_50ntFlanks | 11199 | 11223 | 25 | CCGGCGGCGGCGGCGACTCTGGACG | 181.4 | -5.4 | 0.8 | -6.9 | 7.2E-12 | 1.4E-11 | BT20/MB468 |
| >hg38_21_+_8433172_8446622_RNA45SN1_45S_50ntFlanks@12008.12031.24\|CCGCCGGTGAAATACCACTACTCT | >hg38_21_+_8433172_8446622_RNA45SN1_45S_50ntFlanks | 12008 | 12031 | 24 | CCGCCGGTGAAATACCACTACTCT | 1188.6 | -4.9 | 0.7 | -6.9 | 6.4E-12 | 1.5E-11 | MB231/MB468 |
| >am_trna106_PheGAA_6_-_28775610_28775682@31.52.22,trna109_PheGAA_6_-_28758499_28758571@31.52.22,trna11_PheGAA_12_-_125412389_125412461@31.52.22,trna13_PheGAA_11_-_59333853_59333925@31.52.22,trna14_PheGAA_19_-_1383361_1383433@31.52.22,trna15_PheGAA_11_-_59324970_59325042@31.52.22,trna1_PheGAA_13_-_95201904_95201976@31.52.22,trna72_PheGAA_6_+_28732164_28732236@31.52.22,trna96_PheGAA_6_-_28949449_28949521@31.52.22\|ACTGAAGATCTAAAGGTCCCTG | >am_trna106_PheGAA_6_-_28775610_28775682 | 31 | 52 | 22 | ACTGAAGATCTAAAGGTCCCTG | 153.3 | -8.8 | 1.3 | -6.9 | 6.4E-12 | 1.5E-11 | MB231/MB468 |
| >hg38_21_+_8433172_8446622_RNA45SN1_45S_50ntFlanks@10572.10589.18\|AGAAGCCGGCGGGAGCCC | >hg38_21_+_8433172_8446622_RNA45SN1_45S_50ntFlanks | 10572 | 10589 | 18 | AGAAGCCGGCGGGAGCCC | 131.9 | -8.6 | 1.3 | -6.9 | 6.5E-12 | 1.5E-11 | MB231/MB468 |
| >hg38_21_+_8433172_8446622_RNA45SN1_45S_50ntFlanks@4047.4067.21\|ATTCGAACGTCTGCCCTATCA | >hg38_21_+_8433172_8446622_RNA45SN1_45S_50ntFlanks | 4047 | 4067 | 21 | ATTCGAACGTCTGCCCTATCA | 196.9 | -2.9 | 0.4 | -6.8 | 7.4E-12 | 1.5E-11 | BT20/MB468 |
| >hg38_21_+_8433172_8446622_RNA45SN1_45S_50ntFlanks@11115.11132.18\|CGGCAGGGGCCGGCGGCG | >hg38_21_+_8433172_8446622_RNA45SN1_45S_50ntFlanks | 11115 | 11132 | 18 | CGGCAGGGGCCGGCGGCG | 104.8 | -8.7 | 1.3 | -6.8 | 7.4E-12 | 1.5E-11 | BT20/MB468 |
| >hg38_21_+_8433172_8446622_RNA45SN1_45S_50ntFlanks@11577.11603.27\|TTAGAACTGGTGCGGACCAGGGGAATC | >hg38_21_+_8433172_8446622_RNA45SN1_45S_50ntFlanks | 11577 | 11603 | 27 | TTAGAACTGGTGCGGACCAGGGGAATC | 957.2 | -5.0 | 0.7 | -6.8 | 7.4E-12 | 1.5E-11 | BT20/MB468 |
| >hg38_21_+_8433172_8446622_RNA45SN1_45S_50ntFlanks@11885.11904.20\|TTGAGCTTGACTCTAGTCTG | >hg38_21_+_8433172_8446622_RNA45SN1_45S_50ntFlanks | 11885 | 11904 | 20 | TTGAGCTTGACTCTAGTCTG | 1566.0 | -8.6 | 1.3 | -6.8 | 7.5E-12 | 1.5E-11 | BT20/MB468 |
| >trna10_LysCTT_16_+_3241501_3241573@1.33.33,trna13_LysCTT_14_-_58706613_58706685@1.33.33,trna2_LysCTT_15_+_79152904_79152976@1.33.33\|GCCCGGCTAGCTCAGTCGGTAGAGCATGGGACT | >trna10_LysCTT_16_+_3241501_3241573 | 1 | 33 | 33 | GCCCGGCTAGCTCAGTCGGTAGAGCATGGGACT | 1625.5 | 9.0 | 1.3 | 6.9 | 6.5E-12 | 1.5E-11 | MB231/MB468 |
| >trna11_GluTTC_15_-_26327381_26327452@1.30.30,trna3_GluTTC_13_-_45492062_45492133@1.30.30\|TCCCACATGGTCTAGCGGTTAGGATTCCTG | >trna11_GluTTC_15_-_26327381_26327452 | 1 | 30 | 30 | TCCCACATGGTCTAGCGGTTAGGATTCCTG | 1542.0 | -4.2 | 0.6 | -6.8 | 7.5E-12 | 1.5E-11 | BT20/MB468 |
| >hg38_21_+_8433172_8446622_RNA45SN1_45S_50ntFlanks@13004.13039.36\|ATCTATTGAAAGTCAGCCCTCGACACAAGGGTTTGT | >hg38_21_+_8433172_8446622_RNA45SN1_45S_50ntFlanks | 13004 | 13039 | 36 | ATCTATTGAAAGTCAGCCCTCGACACAAGGGTTTGT | 124.2 | -8.9 | 1.3 | -6.8 | 7.6E-12 | 1.5E-11 | BT20/MB468 |
| >hg38_21_+_8433172_8446622_RNA45SN1_45S_50ntFlanks@12033.12055.23\|ATCGTTTTTTCACTGACCCGGTG | >hg38_21_+_8433172_8446622_RNA45SN1_45S_50ntFlanks | 12033 | 12055 | 23 | ATCGTTTTTTCACTGACCCGGTG | 106.6 | -8.7 | 1.3 | -6.8 | 7.6E-12 | 1.5E-11 | BT20/MB468 |
| >hg38_21_+_8433172_8446622_RNA45SN1_45S_50ntFlanks@12473.12496.24\|CACCCACTAATAGGGAACGTGAGC | >hg38_21_+_8433172_8446622_RNA45SN1_45S_50ntFlanks | 12473 | 12496 | 24 | CACCCACTAATAGGGAACGTGAGC | 932.6 | 4.0 | 0.6 | 6.9 | 6.6E-12 | 1.5E-11 | MB231/MB468 |
| >hg38_21_+_8433172_8446622_RNA45SN1_45S_50ntFlanks@11873.11897.25\|AAGAAGACCCTGTTGAGCTTGACTC | >hg38_21_+_8433172_8446622_RNA45SN1_45S_50ntFlanks | 11873 | 11897 | 25 | AAGAAGACCCTGTTGAGCTTGACTC | 523.8 | -4.1 | 0.6 | -6.9 | 6.7E-12 | 1.5E-11 | MB231/MB468 |
| >hg38_21_+_8433172_8446622_RNA45SN1_45S_50ntFlanks@10186.10207.22\|TCCTCCCCTCCTCCCGCCCACG | >hg38_21_+_8433172_8446622_RNA45SN1_45S_50ntFlanks | 10186 | 10207 | 22 | TCCTCCCCTCCTCCCGCCCACG | 136.1 | -8.7 | 1.3 | -6.9 | 6.8E-12 | 1.5E-11 | MB231/MB468 |
| >hg38_21_+_8433172_8446622_RNA45SN1_45S_50ntFlanks@12995.13012.18\|CTCGCTGCGATCTATTGA | >hg38_21_+_8433172_8446622_RNA45SN1_45S_50ntFlanks | 12995 | 13012 | 18 | CTCGCTGCGATCTATTGA | 883.9 | 3.3 | 0.5 | 6.9 | 6.9E-12 | 1.6E-11 | MB231/MB468 |
| >hg38_21_+_8433172_8446622_RNA45SN1_45S_50ntFlanks@6728.6752.25\|GGACACATTGATCATCGACACTTCG | >hg38_21_+_8433172_8446622_RNA45SN1_45S_50ntFlanks | 6728 | 6752 | 25 | GGACACATTGATCATCGACACTTCG | 302.9 | 4.6 | 0.7 | 6.9 | 7.0E-12 | 1.6E-11 | MB231/MB468 |
| >hg38_21_+_8433172_8446622_RNA45SN1_45S_50ntFlanks@12488.12512.25\|AACGTGAGCTGGGTTTAGACCGTCG | >hg38_21_+_8433172_8446622_RNA45SN1_45S_50ntFlanks | 12488 | 12512 | 25 | AACGTGAGCTGGGTTTAGACCGTCG | 182.7 | -5.2 | 0.8 | -6.9 | 7.0E-12 | 1.6E-11 | MB231/MB468 |
| >hg38_21_+_8433172_8446622_RNA45SN1_45S_50ntFlanks@12588.12616.29\|GGTTCAGACATTTGGTGTATGTGCTTGGC | >hg38_21_+_8433172_8446622_RNA45SN1_45S_50ntFlanks | 12588 | 12616 | 29 | GGTTCAGACATTTGGTGTATGTGCTTGGC | 103.5 | -8.7 | 1.3 | -6.8 | 8.3E-12 | 1.6E-11 | BT20/MB468 |
| >hg38_21_+_8433172_8446622_RNA45SN1_45S_50ntFlanks@4047.4081.35\|ATTCGAACGTCTGCCCTATCAACTTTCGATGGTAG | >hg38_21_+_8433172_8446622_RNA45SN1_45S_50ntFlanks | 4047 | 4081 | 35 | ATTCGAACGTCTGCCCTATCAACTTTCGATGGTAG | 93.7 | -8.5 | 1.3 | -6.8 | 8.3E-12 | 1.6E-11 | BT20/MB468 |
| >hg38_21_+_8433172_8446622_RNA45SN1_45S_50ntFlanks@10922.10947.26\|CTGGACGAGGCGCCGCCGCCCCCCCC | >hg38_21_+_8433172_8446622_RNA45SN1_45S_50ntFlanks | 10922 | 10947 | 26 | CTGGACGAGGCGCCGCCGCCCCCCCC | 93.5 | -3.8 | 0.6 | -6.8 | 8.3E-12 | 1.6E-11 | BT20/MB468 |
| >hg38_21_+_8433172_8446622_RNA45SN1_45S_50ntFlanks@4006.4042.37\|ACCTCGGGCCGATCGCACGCCCCCCGTGGCGGCGACG | >hg38_21_+_8433172_8446622_RNA45SN1_45S_50ntFlanks | 4006 | 4042 | 37 | ACCTCGGGCCGATCGCACGCCCCCCGTGGCGGCGACG | 211.1 | -3.7 | 0.5 | -6.8 | 8.4E-12 | 1.7E-11 | BT20/MB468 |
| >hg38_21_+_8433172_8446622_RNA45SN1_45S_50ntFlanks@10866.10901.36\|CTAAGGGCTGGGTCGGTCGGGCTGGGGCGCGAAGCG | >hg38_21_+_8433172_8446622_RNA45SN1_45S_50ntFlanks | 10866 | 10901 | 36 | CTAAGGGCTGGGTCGGTCGGGCTGGGGCGCGAAGCG | 294.4 | -6.6 | 1.0 | -6.8 | 8.4E-12 | 1.7E-11 | BT20/MB468 |
| >hg38_21_+_8433172_8446622_RNA45SN1_45S_50ntFlanks@12126.12148.23\|CGACCCGCTCCGGGGACAGTGCC | >hg38_21_+_8433172_8446622_RNA45SN1_45S_50ntFlanks | 12126 | 12148 | 23 | CGACCCGCTCCGGGGACAGTGCC | 134.1 | -8.6 | 1.3 | -6.9 | 7.3E-12 | 1.7E-11 | MB231/MB468 |
| >hg38_21_+_8433172_8446622_RNA45SN1_45S_50ntFlanks@5489.5516.28\|CTGAGAAGACGGTCGAACTTGACTATCT | >hg38_21_+_8433172_8446622_RNA45SN1_45S_50ntFlanks | 5489 | 5516 | 28 | CTGAGAAGACGGTCGAACTTGACTATCT | 3560.0 | 5.0 | 0.7 | 6.9 | 7.3E-12 | 1.7E-11 | MB231/MB468 |
| >hg38_21_+_8433172_8446622_RNA45SN1_45S_50ntFlanks@11321.11344.24\|CACGTCTCGTCGCGCGCGCGTCCG | >hg38_21_+_8433172_8446622_RNA45SN1_45S_50ntFlanks | 11321 | 11344 | 24 | CACGTCTCGTCGCGCGCGCGTCCG | 2868.6 | -7.1 | 1.0 | -6.9 | 7.4E-12 | 1.7E-11 | MB231/MB468 |
| >hg38_21_+_8433172_8446622_RNA45SN1_45S_50ntFlanks@11873.11904.32\|AAGAAGACCCTGTTGAGCTTGACTCTAGTCTG | >hg38_21_+_8433172_8446622_RNA45SN1_45S_50ntFlanks | 11873 | 11904 | 32 | AAGAAGACCCTGTTGAGCTTGACTCTAGTCTG | 531.1 | -7.7 | 1.1 | -6.9 | 7.4E-12 | 1.7E-11 | MB231/MB468 |
| >hg38_21_+_8433172_8446622_RNA45SN1_45S_50ntFlanks@10410.10431.22\|TCAGTCGGTCCTGAGAGATGGG | >hg38_21_+_8433172_8446622_RNA45SN1_45S_50ntFlanks | 10410 | 10431 | 22 | TCAGTCGGTCCTGAGAGATGGG | 1303.7 | -5.2 | 0.8 | -6.8 | 7.4E-12 | 1.7E-11 | MB231/MB468 |
| >MI0000650\|hsa-mir-200c&WithFlank&12\|+\|6963693\|6963772@50.70.21(+1C)&[MIMAT0000617&hsa-miR-200c-3p&offsets\|0\|-2(+1C);m-54&12\|+\|6963742\|6963764&offsets\|0\|-2(+1C)]\|TAATACTGCCGGGTAATGATGC | >MI0000650\|hsa-mir-200c&WithFlank&12\|+\|6963693\|6963772 | 50 | 70 | 21(+1C) | TAATACTGCCGGGTAATGATGC | 649.5 | 10.3 | 1.5 | 6.8 | 8.5E-12 | 1.7E-11 | BT20/MB468 |
| >hg38_21_+_8433172_8446622_RNA45SN1_45S_50ntFlanks@12638.12657.20\|CTACCATCTGTGGGATTATG | >hg38_21_+_8433172_8446622_RNA45SN1_45S_50ntFlanks | 12638 | 12657 | 20 | CTACCATCTGTGGGATTATG | 258.9 | -6.2 | 0.9 | -6.8 | 7.4E-12 | 1.7E-11 | MB231/MB468 |
| >hg38_21_+_8433172_8446622_RNA45SN1_45S_50ntFlanks@12938.12966.29\|ACCATTCGTAGACGACCTGCTTCTGGGTC | >hg38_21_+_8433172_8446622_RNA45SN1_45S_50ntFlanks | 12938 | 12966 | 29 | ACCATTCGTAGACGACCTGCTTCTGGGTC | 119.3 | -8.9 | 1.3 | -6.8 | 8.6E-12 | 1.7E-11 | BT20/MB468 |
| >hg38_1_-_228634819_228635039_RNA5S12_5S_50ntFlanks@145.169.25\|CTGGGAATACCGGGTGCTGTAGGCT | >hg38_1_-_228634819_228635039_RNA5S12_5S_50ntFlanks | 145 | 169 | 25 | CTGGGAATACCGGGTGCTGTAGGCT | 565.1 | 7.3 | 1.1 | 6.8 | 8.6E-12 | 1.7E-11 | BT20/MB468 |
| >hg38_21_+_8433172_8446622_RNA45SN1_45S_50ntFlanks@12412.12433.22\|ATCCTTCGATGTCGGCTCTTCC | >hg38_21_+_8433172_8446622_RNA45SN1_45S_50ntFlanks | 12412 | 12433 | 22 | ATCCTTCGATGTCGGCTCTTCC | 120.0 | -8.9 | 1.3 | -6.8 | 8.7E-12 | 1.7E-11 | BT20/MB468 |
| >hg38_21_+_8433172_8446622_RNA45SN1_45S_50ntFlanks@3861.3881.21\|TAATTCTAGAGCTAATACATG | >hg38_21_+_8433172_8446622_RNA45SN1_45S_50ntFlanks | 3861 | 3881 | 21 | TAATTCTAGAGCTAATACATG | 197.4 | -9.6 | 1.4 | -6.8 | 8.7E-12 | 1.7E-11 | BT20/MB468 |
| >hg38_21_+_8433172_8446622_RNA45SN1_45S_50ntFlanks@12450.12480.31\|AATTCACCAAGCGTTGGATTGTTCACCCACT | >hg38_21_+_8433172_8446622_RNA45SN1_45S_50ntFlanks | 12450 | 12480 | 31 | AATTCACCAAGCGTTGGATTGTTCACCCACT | 422.3 | 7.1 | 1.0 | 6.8 | 8.7E-12 | 1.7E-11 | BT20/MB468 |
| >hg38_21_+_8433172_8446622_RNA45SN1_45S_50ntFlanks@12991.13028.38\|CTCCCTCGCTGCGATCTATTGAAAGTCAGCCCTCGACA | >hg38_21_+_8433172_8446622_RNA45SN1_45S_50ntFlanks | 12991 | 13028 | 38 | CTCCCTCGCTGCGATCTATTGAAAGTCAGCCCTCGACA | 166.4 | -5.7 | 0.8 | -6.8 | 7.6E-12 | 1.7E-11 | MB231/MB468 |
| >am_trnaMT_TyrGTA_MT_-_5826_5891@1.32.32\|GGTAAAATGGCTGAGTGAAGCATTGGACTGTA | >am_trnaMT_TyrGTA_MT_-_5826_5891 | 1 | 32 | 32 | GGTAAAATGGCTGAGTGAAGCATTGGACTGTA | 1835.0 | 8.4 | 1.2 | 6.8 | 8.8E-12 | 1.7E-11 | BT20/MB468 |
| >hg38_21_+_8433172_8446622_RNA45SN1_45S_50ntFlanks@12152.12180.29\|TGGGGAGTTTGACTGGGGCGGTACACCTG | >hg38_21_+_8433172_8446622_RNA45SN1_45S_50ntFlanks | 12152 | 12180 | 29 | TGGGGAGTTTGACTGGGGCGGTACACCTG | 104.8 | -8.7 | 1.3 | -6.8 | 8.8E-12 | 1.7E-11 | BT20/MB468 |
| >hg38_21_+_8433172_8446622_RNA45SN1_45S_50ntFlanks@11873.11906.34\|AAGAAGACCCTGTTGAGCTTGACTCTAGTCTGGC | >hg38_21_+_8433172_8446622_RNA45SN1_45S_50ntFlanks | 11873 | 11906 | 34 | AAGAAGACCCTGTTGAGCTTGACTCTAGTCTGGC | 1075.1 | -6.6 | 1.0 | -6.8 | 8.8E-12 | 1.7E-11 | BT20/MB468 |
| >MI0000488\|hsa-mir-194-1&WithFlank&1\|-\|220118151\|220118247@21.42.22;MI0000732\|hsa-mir-194-2&WithFlank&11\|-\|64891349\|64891445@21.42.22&[MIMAT0000460&hsa-miR-194-5p&offsets\|0\|0;m-131&1\|-\|220118206\|220118227&offsets\|0\|0];[MIMAT0000460_1&hsa-miR-194-5p&offsets\|0\|0;m-126&11\|-\|64891404\|64891425&offsets\|0\|0]\|TGTAACAGCAACTCCATGTGGA | >MI0000488\|hsa-mir-194-1&WithFlank&1\|-\|220118151\|220118247 | 21 | 42 | 22 | TGTAACAGCAACTCCATGTGGA | 1589.0 | 6.1 | 0.9 | 6.8 | 8.8E-12 | 1.7E-11 | BT20/MB468 |
| >hg38_21_+_8433172_8446622_RNA45SN1_45S_50ntFlanks@12786.12805.20\|CCCGCGCGCGCGGGAGGGCG | >hg38_21_+_8433172_8446622_RNA45SN1_45S_50ntFlanks | 12786 | 12805 | 20 | CCCGCGCGCGCGGGAGGGCG | 84.8 | -8.4 | 1.2 | -6.8 | 8.8E-12 | 1.7E-11 | BT20/MB468 |
| >hg38_21_+_8433172_8446622_RNA45SN1_45S_50ntFlanks@12083.12110.28\|CTCGCTTCTGGCGCCAAGCGCCCGGCCG | >hg38_21_+_8433172_8446622_RNA45SN1_45S_50ntFlanks | 12083 | 12110 | 28 | CTCGCTTCTGGCGCCAAGCGCCCGGCCG | 111.5 | -8.8 | 1.3 | -6.8 | 8.9E-12 | 1.7E-11 | BT20/MB468 |

| | | | | | | | | | | | | |
|---|---|---|---|---|---|---|---|---|---|---|---|---|
| >trna10_AspGTC_12_-_125424193_125424264@32.70.39,trna12_AspGTC_12_-_125411891_125411962@32.70.39,trna144_AspGTC_6_-_27551236_27551307@32.70.39,trna38_AspGTC_17_-_8125556_8125627@32.70.39,trna45_AspGTC_6_+_27447453_27447524@32.70.39,trna48_AspGTC_6_+_27471523_27471594@32.70.39,trna4_AspGTC_12_+_96429799_96429870@32.70.39,trna69_AspGTC_1_-_161440205_161440276@32.70.39,trna72_AspGTC_1_-_161432824_161432895@32.70.39,trna75_AspGTC_1_-_161425414_161425485@32.70.39,trna78_AspGTC_1_-_161418033_161418104@32.70.39,trna81_AspGTC_1_-_161410615_161410686@32.70.39\|CTGTCACGCGGGAGACCGGGGTTCGATTCCCCGACGGGG | >trna10_AspGTC_12_-_125424193_125424264 | 32 | 70 | 39 | CTGTCACGCGGGAGACCGGGGTTCGATTCCCCGACGGGG | 103.7 | -8.7 | 1.3 | -6.8 | 8.9E-12 | 1.8E-11 | BT20/MB468 |
| >hg38_21_+_8433172_8446622_RNA45SN1_45S_50ntFlanks@4588.4615.28\|CTATTTTGTTGGTTTTCGGAACTGAGGC | >hg38_21_+_8433172_8446622_RNA45SN1_45S_50ntFlanks | 4588 | 4615 | 28 | CTATTTTGTTGGTTTTCGGAACTGAGGC | 152.9 | -6.0 | 0.9 | -6.8 | 8.9E-12 | 1.8E-11 | BT20/MB468 |
| >hg38_21_+_8433172_8446622_RNA45SN1_45S_50ntFlanks@11190.11207.18\|CCCGGGGGGCCGGCGGCG | >hg38_21_+_8433172_8446622_RNA45SN1_45S_50ntFlanks | 11190 | 11207 | 18 | CCCGGGGGGCCGGCGGCG | 140.4 | -8.7 | 1.3 | -6.8 | 7.9E-12 | 1.8E-11 | MB231/MB468 |
| >hg38_21_+_8433172_8446622_RNA45SN1_45S_50ntFlanks@12154.12180.27\|GGGAGTTTGACTGGGGCGGTACACCTG | >hg38_21_+_8433172_8446622_RNA45SN1_45S_50ntFlanks | 12154 | 12180 | 27 | GGGAGTTTGACTGGGGCGGTACACCTG | 102.7 | -8.7 | 1.3 | -6.8 | 9.0E-12 | 1.8E-11 | BT20/MB468 |
| >hg38_21_+_8433172_8446622_RNA45SN1_45S_50ntFlanks@12347.12366.20\|AGAAAAGTTACCACAGGGAT | >hg38_21_+_8433172_8446622_RNA45SN1_45S_50ntFlanks | 12347 | 12366 | 20 | AGAAAAGTTACCACAGGGAT | 96.6 | -8.6 | 1.3 | -6.8 | 9.1E-12 | 1.8E-11 | BT20/MB468 |
| >hg38_21_+_8433172_8446622_RNA45SN1_45S_50ntFlanks@3900.3919.20\|CTTCGCGGGGGGGATGCGTG | >hg38_21_+_8433172_8446622_RNA45SN1_45S_50ntFlanks | 3900 | 3919 | 20 | CTTCGCGGGGGGGATGCGTG | 124.1 | -5.4 | 0.8 | -6.8 | 8.1E-12 | 1.8E-11 | MB231/MB468 |
| >hg38_21_+_8433172_8446622_RNA45SN1_45S_50ntFlanks@11876.11893.18\|AAGACCCTGTTGAGCTTG | >hg38_21_+_8433172_8446622_RNA45SN1_45S_50ntFlanks | 11876 | 11893 | 18 | AAGACCCTGTTGAGCTTG | 1872.4 | -4.9 | 0.7 | -6.8 | 8.1E-12 | 1.8E-11 | MB231/MB468 |
| >hg38_21_+_8433172_8446622_RNA45SN1_45S_50ntFlanks@7974.8015.42\|ACGCGACCTCAGATCAGACGTGGCGACCCGCTGAATTTAAGC | >hg38_21_+_8433172_8446622_RNA45SN1_45S_50ntFlanks | 7974 | 8015 | 42 | ACGCGACCTCAGATCAGACGTGGCGACCCGCTGAATTTAAGC | 1742.7 | 8.6 | 1.3 | 6.8 | 9.4E-12 | 1.9E-11 | BT20/MB468 |
| >hg38_21_+_8433172_8446622_RNA45SN1_45S_50ntFlanks@10678.10696.19\|CGGTTCCGGCGGCGTCCGG | >hg38_21_+_8433172_8446622_RNA45SN1_45S_50ntFlanks | 10678 | 10696 | 19 | CGGTTCCGGCGGCGTCCGG | 82.5 | -8.4 | 1.2 | -6.8 | 9.5E-12 | 1.9E-11 | BT20/MB468 |
| >hg38_21_+_8433172_8446622_RNA45SN1_45S_50ntFlanks@11718.11738.21\|AACGGCGGGAGTAACTATGAC | >hg38_21_+_8433172_8446622_RNA45SN1_45S_50ntFlanks | 11718 | 11738 | 21 | AACGGCGGGAGTAACTATGAC | 738.9 | -8.4 | 1.2 | -6.8 | 9.5E-12 | 1.9E-11 | BT20/MB468 |
| >hg38_21_+_8433172_8446622_RNA45SN1_45S_50ntFlanks@9739.9757.19\|AAACGATCTCAACCTATTC | >hg38_21_+_8433172_8446622_RNA45SN1_45S_50ntFlanks | 9739 | 9757 | 19 | AAACGATCTCAACCTATTC | 630.3 | -3.8 | 0.6 | -6.8 | 8.3E-12 | 1.9E-11 | MB231/MB468 |
| >hg38_21_+_8433172_8446622_RNA45SN1_45S_50ntFlanks@3769.3786.18\|CGCACGGCCGGTACAGTG | >hg38_21_+_8433172_8446622_RNA45SN1_45S_50ntFlanks | 3769 | 3786 | 18 | CGCACGGCCGGTACAGTG | 151.6 | -4.1 | 0.6 | -6.8 | 9.5E-12 | 1.9E-11 | BT20/MB468 |
| >hg38_21_+_8433172_8446622_RNA45SN1_45S_50ntFlanks@12999.13039.41\|CTGCGATCTATTGAAAGTCAGCCCTCGACACAAGGGTTTGT | >hg38_21_+_8433172_8446622_RNA45SN1_45S_50ntFlanks | 12999 | 13039 | 41 | CTGCGATCTATTGAAAGTCAGCCCTCGACACAAGGGTTTGT | 3192.1 | 5.1 | 0.7 | 6.8 | 9.6E-12 | 1.9E-11 | BT20/MB468 |
| >hg38_1_-_228634819_228635039_RNA5S12_5S_50ntFlanks@141.170.30\|CCGCCTGGGAATACCGGGTGCTGTAGGCTT | >hg38_1_-_228634819_228635039_RNA5S12_5S_50ntFlanks | 141 | 170 | 30 | CCGCCTGGGAATACCGGGTGCTGTAGGCTT | 5054.1 | 6.2 | 0.9 | 6.8 | 9.6E-12 | 1.9E-11 | BT20/MB468 |
| >hg38_21_+_8433172_8446622_RNA45SN1_45S_50ntFlanks@10511.10533.23\|ATCCGGAGTGGCGGAGATGGGCG | >hg38_21_+_8433172_8446622_RNA45SN1_45S_50ntFlanks | 10511 | 10533 | 23 | ATCCGGAGTGGCGGAGATGGGCG | 136.0 | -9.1 | 1.3 | -6.8 | 9.6E-12 | 1.9E-11 | BT20/MB468 |
| >MI0000102\|hsa-mir-100&WithFlank&11\|-\|122152223\|122152314@19.40.22(+1C)&[MIMAT0000098&hsa-miR-100-5p&offsets\|0\|0(+1C);m-27&11\|-\|122152276\|122152296&offsets\|0\|+1(+1C)]\|AACCCGTAGATCCGAACTTGTGC | >MI0000102\|hsa-mir-100&WithFlank&11\|-\|122152223\|122152314 | 19 | 40 | 22(+1C) | AACCCGTAGATCCGAACTTGTGC | 7436.0 | -6.5 | 0.9 | -7.2 | 6.1E-13 | 1.9E-11 | BT20/MB231 |
| >hg38_21_+_8433172_8446622_RNA45SN1_45S_50ntFlanks@10408.10428.21\|GGTCAGTCGGTCCTGAGAGAT | >hg38_21_+_8433172_8446622_RNA45SN1_45S_50ntFlanks | 10408 | 10428 | 21 | GGTCAGTCGGTCCTGAGAGAT | 186.8 | -3.2 | 0.5 | -6.8 | 8.4E-12 | 1.9E-11 | MB231/MB468 |
| >hg38_21_+_8433172_8446622_RNA45SN1_45S_50ntFlanks@9505.9523.19\|TGAACTATGCCTGGGCAGG | >hg38_21_+_8433172_8446622_RNA45SN1_45S_50ntFlanks | 9505 | 9523 | 19 | TGAACTATGCCTGGGCAGG | 123.6 | -5.2 | 0.8 | -6.8 | 9.6E-12 | 1.9E-11 | BT20/MB468 |
| >hg38_21_+_8433172_8446622_RNA45SN1_45S_50ntFlanks@6645.6664.20\|CTCGTACGACTCTTAGCGGT | >hg38_21_+_8433172_8446622_RNA45SN1_45S_50ntFlanks | 6645 | 6664 | 20 | CTCGTACGACTCTTAGCGGT | 2240.9 | 5.1 | 0.8 | 6.8 | 8.4E-12 | 1.9E-11 | MB231/MB468 |
| >hg38_21_+_8433172_8446622_RNA45SN1_45S_50ntFlanks@12033.12052.20\|ATCGTTTTTTCACTGACCCG | >hg38_21_+_8433172_8446622_RNA45SN1_45S_50ntFlanks | 12033 | 12052 | 20 | ATCGTTTTTTCACTGACCCG | 151.0 | -8.8 | 1.3 | -6.8 | 8.4E-12 | 1.9E-11 | MB231/MB468 |
| >hg38_21_+_8433172_8446622_RNA45SN1_45S_50ntFlanks@10912.10932.21\|CGCGCCGCGGCTGGACGAGGC | >hg38_21_+_8433172_8446622_RNA45SN1_45S_50ntFlanks | 10912 | 10932 | 21 | CGCGCCGCGGCTGGACGAGGC | 238.6 | -9.5 | 1.4 | -6.8 | 8.5E-12 | 1.9E-11 | MB231/MB468 |
| >MI0000285\|hsa-mir-205&WithFlank&1\|+\|209432127\|209432248@40.62.23&[MIMAT0000266&hsa-miR-205-5p&offsets\|0\|+1;m-351&1\|+\|209432166\|209432187&offsets\|0\|+1]\|TCCTTCATTCCACCGGAGTCTGT | >MI0000285\|hsa-mir-205&WithFlank&1\|+\|209432127\|209432248 | 40 | 62 | 23 | TCCTTCATTCCACCGGAGTCTGT | 928.7 | -7.1 | 1.0 | -6.8 | 8.5E-12 | 1.9E-11 | MB231/MB468 |
| >hg38_21_+_8433172_8446622_RNA45SN1_45S_50ntFlanks@9534.9569.36\|AGGAAACTCTGGTGGAGGTCCGTAGCGGTCCTGACG | >hg38_21_+_8433172_8446622_RNA45SN1_45S_50ntFlanks | 9534 | 9569 | 36 | AGGAAACTCTGGTGGAGGTCCGTAGCGGTCCTGACG | 1937.1 | -5.0 | 0.7 | -6.8 | 9.8E-12 | 1.9E-11 | BT20/MB468 |
| >hg38_21_+_8433172_8446622_RNA45SN1_45S_50ntFlanks@11517.11539.23\|CCGCCCCCGGGGCCGCGGTTCCG | >hg38_21_+_8433172_8446622_RNA45SN1_45S_50ntFlanks | 11517 | 11539 | 23 | CCGCCCCCGGGGCCGCGGTTCCG | 95.5 | -8.6 | 1.3 | -6.8 | 1.0E-11 | 2.0E-11 | BT20/MB468 |
| >hg38_21_+_8433172_8446622_RNA45SN1_45S_50ntFlanks@12633.12651.19\|CGAAGCTACCATCTGTGGG | >hg38_21_+_8433172_8446622_RNA45SN1_45S_50ntFlanks | 12633 | 12651 | 19 | CGAAGCTACCATCTGTGGG | 700.6 | -5.3 | 0.8 | -6.8 | 1.0E-11 | 2.0E-11 | BT20/MB468 |
| >hg38_21_+_8433172_8446622_RNA45SN1_45S_50ntFlanks@12637.12659.23\|GCTACCATCTGTGGGATTATGAC | >hg38_21_+_8433172_8446622_RNA45SN1_45S_50ntFlanks | 12637 | 12659 | 23 | GCTACCATCTGTGGGATTATGAC | 286.4 | -3.3 | 0.5 | -6.8 | 8.7E-12 | 2.0E-11 | MB231/MB468 |
| >hg38_21_+_8433172_8446622_RNA45SN1_45S_50ntFlanks@6698.6717.20\|AGCTAGCTGCGAGAATTAAT | >hg38_21_+_8433172_8446622_RNA45SN1_45S_50ntFlanks | 6698 | 6717 | 20 | AGCTAGCTGCGAGAATTAAT | 169.2 | -4.4 | 0.6 | -6.8 | 1.0E-11 | 2.0E-11 | BT20/MB468 |
| >hg38_21_+_8433172_8446622_RNA45SN1_45S_50ntFlanks@10687.10712.26\|CGGCGTCCGGTGAGCTCTCGCTGGCC | >hg38_21_+_8433172_8446622_RNA45SN1_45S_50ntFlanks | 10687 | 10712 | 26 | CGGCGTCCGGTGAGCTCTCGCTGGCC | 643.2 | -4.3 | 0.6 | -6.8 | 8.8E-12 | 2.0E-11 | MB231/MB468 |
| >hg38_21_+_8433172_8446622_RNA45SN1_45S_50ntFlanks@4367.4389.23\|CAGTTAAAAAGCTCGTAGTTGGA | >hg38_21_+_8433172_8446622_RNA45SN1_45S_50ntFlanks | 4367 | 4389 | 23 | CAGTTAAAAAGCTCGTAGTTGGA | 1543.9 | -8.3 | 1.2 | -6.8 | 1.0E-11 | 2.0E-11 | BT20/MB468 |
| >hg38_21_+_8433172_8446622_RNA45SN1_45S_50ntFlanks@9573.9597.25\|AAATCGGTCGTCCGACCTGGGTATA | >hg38_21_+_8433172_8446622_RNA45SN1_45S_50ntFlanks | 9573 | 9597 | 25 | AAATCGGTCGTCCGACCTGGGTATA | 3084.0 | -6.2 | 0.9 | -6.8 | 1.0E-11 | 2.0E-11 | BT20/MB468 |
| >hg38_21_+_8433172_8446622_RNA45SN1_45S_50ntFlanks@12588.12614.27\|GGTTCAGACATTTGGTGTATGTGCTTG | >hg38_21_+_8433172_8446622_RNA45SN1_45S_50ntFlanks | 12588 | 12614 | 27 | GGTTCAGACATTTGGTGTATGTGCTTG | 138.3 | -9.1 | 1.3 | -6.8 | 1.0E-11 | 2.0E-11 | BT20/MB468 |
| >hg38_21_+_8433172_8446622_RNA45SN1_45S_50ntFlanks@9570.9592.23\|TGCAAATCGGTCGTCCGACCTGG | >hg38_21_+_8433172_8446622_RNA45SN1_45S_50ntFlanks | 9570 | 9592 | 23 | TGCAAATCGGTCGTCCGACCTGG | 121.1 | -5.5 | 0.8 | -6.8 | 1.0E-11 | 2.0E-11 | BT20/MB468 |
| >hg38_1_-_228634819_228635039_RNA5S12_5S_50ntFlanks@51.82.32\|GTCTACGGCCATACCACCCTGAACGCGCCCGA | >hg38_1_-_228634819_228635039_RNA5S12_5S_50ntFlanks | 51 | 82 | 32 | GTCTACGGCCATACCACCCTGAACGCGCCCGA | 826.3 | 4.9 | 0.7 | 6.8 | 1.0E-11 | 2.0E-11 | BT20/MB468 |
| >hg38_21_+_8433172_8446622_RNA45SN1_45S_50ntFlanks@12160.12179.20\|TTGACTGGGGCGGTACACCT | >hg38_21_+_8433172_8446622_RNA45SN1_45S_50ntFlanks | 12160 | 12179 | 20 | TTGACTGGGGCGGTACACCT | 147.2 | -6.1 | 0.9 | -6.8 | 1.0E-11 | 2.0E-11 | BT20/MB468 |
| >hg38_21_+_8433172_8446622_RNA45SN1_45S_50ntFlanks@11737.11768.32\|ACTCTCTTAAGGTAGCCAAATGCCTCGTCATC | >hg38_21_+_8433172_8446622_RNA45SN1_45S_50ntFlanks | 11737 | 11768 | 32 | ACTCTCTTAAGGTAGCCAAATGCCTCGTCATC | 98.2 | -8.6 | 1.3 | -6.8 | 1.0E-11 | 2.0E-11 | BT20/MB468 |
| >hg38_21_+_8433172_8446622_RNA45SN1_45S_50ntFlanks@476.495.20\|TTCGTGATCGATGTGGTGAC | >hg38_21_+_8433172_8446622_RNA45SN1_45S_50ntFlanks | 476 | 495 | 20 | TTCGTGATCGATGTGGTGAC | 839.4 | 5.4 | 0.8 | 6.8 | 1.0E-11 | 2.0E-11 | BT20/MB468 |
| >hg38_21_+_8433172_8446622_RNA45SN1_45S_50ntFlanks@8182.8223.42\|ATCGAGGCCCAGCCCGTGGACGGTGTGAGGCCGGTAGCGGCC | >hg38_21_+_8433172_8446622_RNA45SN1_45S_50ntFlanks | 8182 | 8223 | 42 | ATCGAGGCCCAGCCCGTGGACGGTGTGAGGCCGGTAGCGGCC | 110.2 | -8.8 | 1.3 | -6.8 | 1.0E-11 | 2.0E-11 | BT20/MB468 |
| >hg38_21_+_8433172_8446622_RNA45SN1_45S_50ntFlanks@8113.8143.31\|GGCGCGGGACATGTGGCGTACGGAAGACCCG | >hg38_21_+_8433172_8446622_RNA45SN1_45S_50ntFlanks | 8113 | 8143 | 31 | GGCGCGGGACATGTGGCGTACGGAAGACCCG | 138.0 | -9.1 | 1.3 | -6.8 | 1.1E-11 | 2.1E-11 | BT20/MB468 |
| >hg38_21_+_8433172_8446622_RNA45SN1_45S_50ntFlanks@9556.9582.27\|TAGCGGTCCTGACGTGCAAATCGGTCG | >hg38_21_+_8433172_8446622_RNA45SN1_45S_50ntFlanks | 9556 | 9582 | 27 | TAGCGGTCCTGACGTGCAAATCGGTCG | 477.2 | -4.0 | 0.6 | -6.8 | 9.2E-12 | 2.1E-11 | MB231/MB468 |
| >hg38_21_+_8433172_8446622_RNA45SN1_45S_50ntFlanks@11843.11872.30\|AGGGAACGGGCTTGGCGGAATCAGCGGGGA | >hg38_21_+_8433172_8446622_RNA45SN1_45S_50ntFlanks | 11843 | 11872 | 30 | AGGGAACGGGCTTGGCGGAATCAGCGGGGA | 113.8 | -5.9 | 0.9 | -6.8 | 9.2E-12 | 2.1E-11 | MB231/MB468 |
| >hg38_21_+_8433172_8446622_RNA45SN1_45S_50ntFlanks@9736.9755.20\|CCGAAACGATCTCAACCTAT | >hg38_21_+_8433172_8446622_RNA45SN1_45S_50ntFlanks | 9736 | 9755 | 20 | CCGAAACGATCTCAACCTAT | 229.0 | -5.8 | 0.8 | -6.8 | 1.1E-11 | 2.1E-11 | BT20/MB468 |
| >hg38_21_+_8433172_8446622_RNA45SN1_45S_50ntFlanks@8325.8356.32\|CGATAGTCAACAAGTACCGTAAGGGAAAGTTG | >hg38_21_+_8433172_8446622_RNA45SN1_45S_50ntFlanks | 8325 | 8356 | 32 | CGATAGTCAACAAGTACCGTAAGGGAAAGTTG | 99.3 | -8.6 | 1.3 | -6.8 | 1.1E-11 | 2.1E-11 | BT20/MB468 |

<table>
<tr><td>>trna111_HisGTG_1_-_147774845_147774916@-1A.34.35,trna118_HisGTG_1_-_145396881_145396952@-1A.34.35,trna16_HisGTG_1_+_146544773_146544844@-1A.34.35,trna1_HisGTG_15_+_45493349_45493420@-1A.34.35,trna21_HisGTG_1_+_147753471_147753542@-1A.34.35,trna33_HisGTG_6_+_27125906_27125977@-1A.34.35,trna7_HisGTG_9_-_14433938_14434009@-1A.34.35,trna8_HisGTG_15_-_45492611_45492682@-1A.34.35,trna9_HisGTG_15_-_45490804_45490875@-1A.34.35|AGCCGTGATCGTATAGTGGTTAGTACTCTGCGTTG</td><td>>trna111_HisGTG_1_-_147774845_147774916</td><td>-1A</td><td>34</td><td>35</td><td>AGCCGTGATCGTATAGTGGTTAGTACTCTGCGTTG</td><td>1131.9</td><td>8.1</td><td>1.2</td><td>6.8</td><td>9.3E-12</td><td>2.1E-11</td><td>MB231/MB468</td></tr>
<tr><td>>hg38_21_+_8433172_8446622_RNA45SN1_45S_50ntFlanks@10425.10457.33|AGATGGGCGAGCGCCGTTCCGAAGGGACGGGCG</td><td>>hg38_21_+_8433172_8446622_RNA45SN1_45S_50ntFlanks</td><td>10425</td><td>10457</td><td>33</td><td>AGATGGGCGAGCGCCGTTCCGAAGGGACGGGCG</td><td>110.2</td><td>-8.8</td><td>1.3</td><td>-6.8</td><td>1.1E-11</td><td>2.1E-11</td><td>BT20/MB468</td></tr>
<tr><td>>trnaMT_SerGCT_MT_+_12207_12265@1.20.20|GAGAAAGCTCACAAGAACTG</td><td>>trnaMT_SerGCT_MT_+_12207_12265</td><td>1</td><td>20</td><td>20</td><td>GAGAAAGCTCACAAGAACTG</td><td>1302.9</td><td>4.1</td><td>0.6</td><td>6.8</td><td>9.4E-12</td><td>2.1E-11</td><td>MB231/MB468</td></tr>
<tr><td>>hg38_21_+_8433172_8446622_RNA45SN1_45S_50ntFlanks@9503.9526.24|GGTGAACTATGCCTGGGCAGGGCG</td><td>>hg38_21_+_8433172_8446622_RNA45SN1_45S_50ntFlanks</td><td>9503</td><td>9526</td><td>24</td><td>GGTGAACTATGCCTGGGCAGGGCG</td><td>98.5</td><td>-8.6</td><td>1.3</td><td>-6.8</td><td>1.1E-11</td><td>2.1E-11</td><td>BT20/MB468</td></tr>
<tr><td>>MI0005756|hsa-mir-934&WithFlank&X|+|136550872|136550966@21.40.20&[MIMAT0004977&hsa-miR-934&offsets|0|-2;m-943&X|+|136550892|136550913&offsets|0|-2]|TGTCTACTACTGGAGACACT</td><td>>MI0005756|hsa-mir-934&WithFlank&X|+|136550872|136550966</td><td>21</td><td>40</td><td>20</td><td>TGTCTACTACTGGAGACACT</td><td>1324.2</td><td>8.7</td><td>1.3</td><td>6.8</td><td>1.1E-11</td><td>2.1E-11</td><td>BT20/MB468</td></tr>
<tr><td>>hg38_21_+_8433172_8446622_RNA45SN1_45S_50ntFlanks@12490.12520.31|CGTGAGCTGGGTTTAGACCGTCGTGAGACAG</td><td>>hg38_21_+_8433172_8446622_RNA45SN1_45S_50ntFlanks</td><td>12490</td><td>12520</td><td>31</td><td>CGTGAGCTGGGTTTAGACCGTCGTGAGACAG</td><td>116.6</td><td>-5.9</td><td>0.9</td><td>-6.8</td><td>9.7E-12</td><td>2.2E-11</td><td>MB231/MB468</td></tr>
<tr><td>>hg38_21_+_8433172_8446622_RNA45SN1_45S_50ntFlanks@5417.5445.29|CTACCGATTGGATGGTTTAGTGAGGCCCT</td><td>>hg38_21_+_8433172_8446622_RNA45SN1_45S_50ntFlanks</td><td>5417</td><td>5445</td><td>29</td><td>CTACCGATTGGATGGTTTAGTGAGGCCCT</td><td>151.9</td><td>-8.8</td><td>1.3</td><td>-6.8</td><td>9.7E-12</td><td>2.2E-11</td><td>MB231/MB468</td></tr>
<tr><td>>hg38_21_+_8433172_8446622_RNA45SN1_45S_50ntFlanks@12605.12624.20|TATGTGCTTGGCTGAGGAGC</td><td>>hg38_21_+_8433172_8446622_RNA45SN1_45S_50ntFlanks</td><td>12605</td><td>12624</td><td>20</td><td>TATGTGCTTGGCTGAGGAGC</td><td>1683.1</td><td>-3.7</td><td>0.5</td><td>-6.8</td><td>1.1E-11</td><td>2.2E-11</td><td>BT20/MB468</td></tr>
<tr><td>>hg38_21_+_8433172_8446622_RNA45SN1_45S_50ntFlanks@10845.10866.22|ACTTCGGGATAAGGATTGGCTC</td><td>>hg38_21_+_8433172_8446622_RNA45SN1_45S_50ntFlanks</td><td>10845</td><td>10866</td><td>22</td><td>ACTTCGGGATAAGGATTGGCTC</td><td>235.0</td><td>-3.9</td><td>0.6</td><td>-6.8</td><td>9.7E-12</td><td>2.2E-11</td><td>MB231/MB468</td></tr>
<tr><td>>hg38_21_+_8433172_8446622_RNA45SN1_45S_50ntFlanks@4588.4608.21|CTATTTTGTTGGTTTTCGGAA</td><td>>hg38_21_+_8433172_8446622_RNA45SN1_45S_50ntFlanks</td><td>4588</td><td>4608</td><td>21</td><td>CTATTTTGTTGGTTTTCGGAA</td><td>185.1</td><td>-9.5</td><td>1.4</td><td>-6.8</td><td>1.1E-11</td><td>2.2E-11</td><td>BT20/MB468</td></tr>
<tr><td>>hg38_21_+_8433172_8446622_RNA45SN1_45S_50ntFlanks@4734.4765.32|AACGAAAGTCGGAGGTTCGAAGACGATCAGAT</td><td>>hg38_21_+_8433172_8446622_RNA45SN1_45S_50ntFlanks</td><td>4734</td><td>4765</td><td>32</td><td>AACGAAAGTCGGAGGTTCGAAGACGATCAGAT</td><td>366.9</td><td>7.9</td><td>1.2</td><td>6.8</td><td>1.1E-11</td><td>2.2E-11</td><td>BT20/MB468</td></tr>
<tr><td>>hg38_21_+_8433172_8446622_RNA45SN1_45S_50ntFlanks@9569.9597.29|GTGCAAATCGGTCGTCCGACCTGGGTATA</td><td>>hg38_21_+_8433172_8446622_RNA45SN1_45S_50ntFlanks</td><td>9569</td><td>9597</td><td>29</td><td>GTGCAAATCGGTCGTCCGACCTGGGTATA</td><td>863.3</td><td>-4.3</td><td>0.6</td><td>-6.8</td><td>1.1E-11</td><td>2.2E-11</td><td>BT20/MB468</td></tr>
<tr><td>>hg38_21_+_8433172_8446622_RNA45SN1_45S_50ntFlanks@11876.11906.31|AAGACCCTGTTGAGCTTGACTCTAGTCTGGC</td><td>>hg38_21_+_8433172_8446622_RNA45SN1_45S_50ntFlanks</td><td>11876</td><td>11906</td><td>31</td><td>AAGACCCTGTTGAGCTTGACTCTAGTCTGGC</td><td>382.2</td><td>-6.0</td><td>0.9</td><td>-6.8</td><td>1.1E-11</td><td>2.2E-11</td><td>BT20/MB468</td></tr>
<tr><td>>trna101_AlaAGC_6_-_28831462_28831533@58.75.18,trna102_AlaAGC_6_-_28806221_28806292@58.75.18,trna110_AlaTGC_6_-_28757547_28757618@58.75.18,trna113_AlaTGC_6_-_28726141_28726212@58.75.18,trna65_AlaAGC_6_+_28574933_28575004@58.75.18|TCCCCGGCACCTCCACCA</td><td>>trna101_AlaAGC_6_-_28831462_28831533</td><td>58</td><td>75</td><td>18</td><td>TCCCCGGCACCTCCACCA</td><td>3362.7</td><td>5.9</td><td>0.9</td><td>6.8</td><td>1.1E-11</td><td>2.2E-11</td><td>BT20/MB468</td></tr>
<tr><td>>trna10_AspGTC_12_-_125424193_125424264@37.75.39,trna12_AspGTC_12_-_125411891_125411962@37.75.39,trna144_AspGTC_6_-_27551236_27551307@37.75.39,trna38_AspGTC_17_-_8125556_8125627@37.75.39,trna45_AspGTC_6_+_27447453_27447524@37.75.39,trna48_AspGTC_6_+_27471523_27471594@37.75.39,trna4_AspGTC_12_+_96429799_96429870@37.75.39,trna69_AspGTC_1_-_161440205_161440276@37.75.39,trna72_AspGTC_1_-_161432824_161432895@37.75.39,trna75_AspGTC_1_-_161425414_161425485@37.75.39,trna78_AspGTC_1_-_161418033_161418104@37.75.39,trna81_AspGTC_1_-_161410615_161410686@37.75.39|ACGCGGGAGACCGGGGTTCGATTCCCCGACGGGGAGCCA</td><td>>trna10_AspGTC_12_-_125424193_125424264</td><td>37</td><td>75</td><td>39</td><td>ACGCGGGAGACCGGGGTTCGATTCCCCGACGGGGAGCCA</td><td>3065.3</td><td>9.2</td><td>1.4</td><td>6.8</td><td>9.9E-12</td><td>2.2E-11</td><td>MB231/MB468</td></tr>
<tr><td>>hg38_21_+_8433172_8446622_RNA45SN1_45S_50ntFlanks@6646.6667.22|TCGTACGACTCTTAGCGGTGGA</td><td>>hg38_21_+_8433172_8446622_RNA45SN1_45S_50ntFlanks</td><td>6646</td><td>6667</td><td>22</td><td>TCGTACGACTCTTAGCGGTGGA</td><td>4588.7</td><td>5.1</td><td>0.7</td><td>6.8</td><td>1.0E-11</td><td>2.2E-11</td><td>MB231/MB468</td></tr>
<tr><td>>MI0000272|hsa-mir-182&WithFlank&7|-|129770377|129770498@29.51.23(+1G)&[MIMAT0000259&hsa-miR-182-5p&offsets|0|-1(+1G);m-22&7|-|129770449|129770470&offsets|0|+1(+1G)]|TTTGGCAATGGTAGAACTCACACG</td><td>>MI0000272|hsa-mir-182&WithFlank&7|-|129770377|129770498</td><td>29</td><td>51</td><td>23(+1G)</td><td>TTTGGCAATGGTAGAACTCACACG</td><td>1178.8</td><td>5.7</td><td>0.8</td><td>6.8</td><td>1.0E-11</td><td>2.2E-11</td><td>MB231/MB468</td></tr>
<tr><td>>hg38_21_+_8433172_8446622_RNA45SN1_45S_50ntFlanks@7986.8007.22|ATCAGACGTGGCGACCCGCTGA</td><td>>hg38_21_+_8433172_8446622_RNA45SN1_45S_50ntFlanks</td><td>7986</td><td>8007</td><td>22</td><td>ATCAGACGTGGCGACCCGCTGA</td><td>105.8</td><td>-8.7</td><td>1.3</td><td>-6.8</td><td>1.2E-11</td><td>2.2E-11</td><td>BT20/MB468</td></tr>
<tr><td>>hg38_21_+_8433172_8446622_RNA45SN1_45S_50ntFlanks@12935.12952.18|TAAACCATTCGTAGACGA</td><td>>hg38_21_+_8433172_8446622_RNA45SN1_45S_50ntFlanks</td><td>12935</td><td>12952</td><td>18</td><td>TAAACCATTCGTAGACGA</td><td>133.6</td><td>-3.6</td><td>0.5</td><td>-6.8</td><td>1.0E-11</td><td>2.3E-11</td><td>MB231/MB468</td></tr>
<tr><td>>hg38_21_+_8433172_8446622_RNA45SN1_45S_50ntFlanks@9568.9586.19|CGTGCAAATCGGTCGTCCG</td><td>>hg38_21_+_8433172_8446622_RNA45SN1_45S_50ntFlanks</td><td>9568</td><td>9586</td><td>19</td><td>CGTGCAAATCGGTCGTCCG</td><td>202.1</td><td>-4.0</td><td>0.6</td><td>-6.8</td><td>1.0E-11</td><td>2.3E-11</td><td>MB231/MB468</td></tr>
<tr><td>>hg38_21_+_8433172_8446622_RNA45SN1_45S_50ntFlanks@11766.11786.21|ATCTAATTAGTGACGCGCATG</td><td>>hg38_21_+_8433172_8446622_RNA45SN1_45S_50ntFlanks</td><td>11766</td><td>11786</td><td>21</td><td>ATCTAATTAGTGACGCGCATG</td><td>111.3</td><td>-8.8</td><td>1.3</td><td>-6.8</td><td>1.2E-11</td><td>2.3E-11</td><td>BT20/MB468</td></tr>
<tr><td>>hg38_21_+_8433172_8446622_RNA45SN1_45S_50ntFlanks@3738.3756.19|TCTCAAAGATTAAGCCATG</td><td>>hg38_21_+_8433172_8446622_RNA45SN1_45S_50ntFlanks</td><td>3738</td><td>3756</td><td>19</td><td>TCTCAAAGATTAAGCCATG</td><td>99.0</td><td>-8.6</td><td>1.3</td><td>-6.8</td><td>1.2E-11</td><td>2.3E-11</td><td>BT20/MB468</td></tr>
<tr><td>>hg38_21_+_8433172_8446622_RNA45SN1_45S_50ntFlanks@4588.4617.30|CTATTTTGTTGGTTTTCGGAACTGAGGCCA</td><td>>hg38_21_+_8433172_8446622_RNA45SN1_45S_50ntFlanks</td><td>4588</td><td>4617</td><td>30</td><td>CTATTTTGTTGGTTTTCGGAACTGAGGCCA</td><td>85.9</td><td>-8.4</td><td>1.2</td><td>-6.8</td><td>1.2E-11</td><td>2.3E-11</td><td>BT20/MB468</td></tr>
<tr><td>>hg38_21_+_8433172_8446622_RNA45SN1_45S_50ntFlanks@11666.11688.23|TCTGCCCAGTGCTCTGAATGTCA</td><td>>hg38_21_+_8433172_8446622_RNA45SN1_45S_50ntFlanks</td><td>11666</td><td>11688</td><td>23</td><td>TCTGCCCAGTGCTCTGAATGTCA</td><td>408.2</td><td>4.4</td><td>0.7</td><td>6.8</td><td>1.0E-11</td><td>2.3E-11</td><td>MB231/MB468</td></tr>
<tr><td>>hg38_21_+_8433172_8446622_RNA45SN1_45S_50ntFlanks@4472.4493.22|CTTAGCTGAGTGTCCCGCGGGG</td><td>>hg38_21_+_8433172_8446622_RNA45SN1_45S_50ntFlanks</td><td>4472</td><td>4493</td><td>22</td><td>CTTAGCTGAGTGTCCCGCGGGG</td><td>124.0</td><td>-8.5</td><td>1.3</td><td>-6.8</td><td>1.0E-11</td><td>2.3E-11</td><td>MB231/MB468</td></tr>
<tr><td>>hg38_21_+_8433172_8446622_RNA45SN1_45S_50ntFlanks@10689.10706.18|GCGTCCGGTGAGCTCTCG</td><td>>hg38_21_+_8433172_8446622_RNA45SN1_45S_50ntFlanks</td><td>10689</td><td>10706</td><td>18</td><td>GCGTCCGGTGAGCTCTCG</td><td>1278.2</td><td>-7.8</td><td>1.1</td><td>-6.8</td><td>1.2E-11</td><td>2.3E-11</td><td>BT20/MB468</td></tr>
<tr><td>>hg38_21_+_8433172_8446622_RNA45SN1_45S_50ntFlanks@10843.10862.20|TAACTTCGGGATAAGGATTG</td><td>>hg38_21_+_8433172_8446622_RNA45SN1_45S_50ntFlanks</td><td>10843</td><td>10862</td><td>20</td><td>TAACTTCGGGATAAGGATTG</td><td>499.6</td><td>-6.2</td><td>0.9</td><td>-6.8</td><td>1.2E-11</td><td>2.3E-11</td><td>BT20/MB468</td></tr>
<tr><td>>trna111_HisGTG_1_-_147774845_147774916@1.34.34,trna118_HisGTG_1_-_145396881_145396952@1.34.34,trna16_HisGTG_1_+_146544773_146544844@1.34.34,trna1_HisGTG_15_+_45493349_45493420@1.34.34,trna21_HisGTG_1_+_147753471_147753542@1.34.34,trna33_HisGTG_6_+_27125906_27125977@1.34.34,trna7_HisGTG_9_-_14433938_14434009@1.34.34,trna8_HisGTG_15_-_45492611_45492682@1.34.34,trna9_HisGTG_15_-_45490804_45490875@1.34.34|GCCGTGATCGTATAGTGGTTAGTACTCTGCGTTG</td><td>>trna111_HisGTG_1_-_147774845_147774916</td><td>1</td><td>34</td><td>34</td><td>GCCGTGATCGTATAGTGGTTAGTACTCTGCGTTG</td><td>6754.9</td><td>6.6</td><td>1.0</td><td>6.8</td><td>1.0E-11</td><td>2.3E-11</td><td>MB231/MB468</td></tr>
<tr><td>>hg38_21_+_8433172_8446622_RNA45SN1_45S_50ntFlanks@10783.10818.36|GTGAACAGCCTCTGGCATGTTGGAACAATGTAGGTA</td><td>>hg38_21_+_8433172_8446622_RNA45SN1_45S_50ntFlanks</td><td>10783</td><td>10818</td><td>36</td><td>GTGAACAGCCTCTGGCATGTTGGAACAATGTAGGTA</td><td>987.1</td><td>-4.4</td><td>0.6</td><td>-6.8</td><td>1.0E-11</td><td>2.3E-11</td><td>MB231/MB468</td></tr>
<tr><td>>hg38_21_+_8433172_8446622_RNA45SN1_45S_50ntFlanks@4002.4020.19|GATAACCTCGGGCCGATCG</td><td>>hg38_21_+_8433172_8446622_RNA45SN1_45S_50ntFlanks</td><td>4002</td><td>4020</td><td>19</td><td>GATAACCTCGGGCCGATCG</td><td>189.9</td><td>-3.5</td><td>0.5</td><td>-6.8</td><td>1.0E-11</td><td>2.3E-11</td><td>MB231/MB468</td></tr>
<tr><td>>hg38_21_+_8433172_8446622_RNA45SN1_45S_50ntFlanks@12440.12473.34|TGTGAAGCAGAATTCACCAAGCGTTGGATTGTTC</td><td>>hg38_21_+_8433172_8446622_RNA45SN1_45S_50ntFlanks</td><td>12440</td><td>12473</td><td>34</td><td>TGTGAAGCAGAATTCACCAAGCGTTGGATTGTTC</td><td>454.0</td><td>6.4</td><td>0.9</td><td>6.8</td><td>1.1E-11</td><td>2.4E-11</td><td>MB231/MB468</td></tr>
<tr><td>>hg38_21_+_8433172_8446622_RNA45SN1_45S_50ntFlanks@6651.6672.22|CGACTCTTAGCGGTGGATCACT</td><td>>hg38_21_+_8433172_8446622_RNA45SN1_45S_50ntFlanks</td><td>6651</td><td>6672</td><td>22</td><td>CGACTCTTAGCGGTGGATCACT</td><td>1982.8</td><td>5.1</td><td>0.7</td><td>6.8</td><td>1.1E-11</td><td>2.4E-11</td><td>MB231/MB468</td></tr>
<tr><td>>hg38_21_+_8433172_8446622_RNA45SN1_45S_50ntFlanks@9539.9563.25|ACTCTGGTGGAGGTCCGTAGCGGTC</td><td>>hg38_21_+_8433172_8446622_RNA45SN1_45S_50ntFlanks</td><td>9539</td><td>9563</td><td>25</td><td>ACTCTGGTGGAGGTCCGTAGCGGTC</td><td>148.3</td><td>-9.2</td><td>1.4</td><td>-6.8</td><td>1.2E-11</td><td>2.4E-11</td><td>BT20/MB468</td></tr>
<tr><td>>hg38_21_+_8433172_8446622_RNA45SN1_45S_50ntFlanks@12941.12960.20|ATTCGTAGACGACCTGCTTC</td><td>>hg38_21_+_8433172_8446622_RNA45SN1_45S_50ntFlanks</td><td>12941</td><td>12960</td><td>20</td><td>ATTCGTAGACGACCTGCTTC</td><td>796.7</td><td>3.9</td><td>0.6</td><td>6.8</td><td>1.1E-11</td><td>2.4E-11</td><td>MB231/MB468</td></tr>
<tr><td>>hg38_21_+_8433172_8446622_RNA45SN1_45S_50ntFlanks@11874.11900.27|AGAAGACCCTGTTGAGCTTGACTCTAG</td><td>>hg38_21_+_8433172_8446622_RNA45SN1_45S_50ntFlanks</td><td>11874</td><td>11900</td><td>27</td><td>AGAAGACCCTGTTGAGCTTGACTCTAG</td><td>573.5</td><td>-5.7</td><td>0.8</td><td>-6.8</td><td>1.1E-11</td><td>2.4E-11</td><td>MB231/MB468</td></tr>
<tr><td>>hg38_21_+_8433172_8446622_RNA45SN1_45S_50ntFlanks@11581.11616.36|AACTGGTGCGGACCAGGGGAATCCGACTGTTTAATT</td><td>>hg38_21_+_8433172_8446622_RNA45SN1_45S_50ntFlanks</td><td>11581</td><td>11616</td><td>36</td><td>AACTGGTGCGGACCAGGGGAATCCGACTGTTTAATT</td><td>579.9</td><td>4.4</td><td>0.6</td><td>6.8</td><td>1.1E-11</td><td>2.4E-11</td><td>MB231/MB468</td></tr>
<tr><td>>trnaMT_GluTTC_MT_-_14674_14742@40.63.24,trnalookalike8_GluTTC_5_-_93905172_93905240@40.63.24|TTGGTCGTGGTTGTAGTCCGTGCG</td><td>>trnaMT_GluTTC_MT_-_14674_14742</td><td>40</td><td>63</td><td>24</td><td>TTGGTCGTGGTTGTAGTCCGTGCG</td><td>473.6</td><td>8.0</td><td>1.2</td><td>6.8</td><td>1.3E-11</td><td>2.4E-11</td><td>BT20/MB468</td></tr>
<tr><td>>hg38_21_+_8433172_8446622_RNA45SN1_45S_50ntFlanks@12457.12488.32|CAAGCGTTGGATTGTTCACCCACTAATAGGGA</td><td>>hg38_21_+_8433172_8446622_RNA45SN1_45S_50ntFlanks</td><td>12457</td><td>12488</td><td>32</td><td>CAAGCGTTGGATTGTTCACCCACTAATAGGGA</td><td>644.2</td><td>-6.7</td><td>1.0</td><td>-6.8</td><td>1.3E-11</td><td>2.5E-11</td><td>BT20/MB468</td></tr>
<tr><td>>hg38_21_+_8433172_8446622_RNA45SN1_45S_50ntFlanks@12654.12673.20|TATGACTGAACGCCTCTAAG</td><td>>hg38_21_+_8433172_8446622_RNA45SN1_45S_50ntFlanks</td><td>12654</td><td>12673</td><td>20</td><td>TATGACTGAACGCCTCTAAG</td><td>105.3</td><td>-8.7</td><td>1.3</td><td>-6.8</td><td>1.3E-11</td><td>2.5E-11</td><td>BT20/MB468</td></tr>
</table>

| | | | | | | | | | | | | |
|---|---|---|---|---|---|---|---|---|---|---|---|---|
| >hg38_21_+_8433172_8446622_RNA45SN1_45S_50ntFlanks@5526.5550.25\|AAAAGTCGTAACAAGGTTTCCGTAG | >hg38_21_+_8433172_8446622_RNA45SN1_45S_50ntFlanks | 5526 | 5550 | 25 | AAAAGTCGTAACAAGGTTTCCGTAG | 158.7 | -4.8 | 0.7 | -6.8 | 1.3E-11 | 2.5E-11 | BT20/MB468 |
| >hg38_21_+_8433172_8446622_RNA45SN1_45S_50ntFlanks@7983.8004.22\|CAGATCAGACGTGGCGACCCGC | >hg38_21_+_8433172_8446622_RNA45SN1_45S_50ntFlanks | 7983 | 8004 | 22 | CAGATCAGACGTGGCGACCCGC | 409.6 | -5.1 | 0.8 | -6.8 | 1.1E-11 | 2.5E-11 | MB231/MB468 |
| >trnaMT_GluTTC_MT_-_14674_14742@44.72.29,trnalookalike8_GluTTC_5_-_93905172_93905240@44.72.29\|TCGTGGTTGTAGTCCGTGCGAGAATACCA | >trnaMT_GluTTC_MT_-_14674_14742 | 44 | 72 | 29 | TCGTGGTTGTAGTCCGTGCGAGAATACCA | 452.0 | 4.7 | 0.7 | 6.8 | 1.3E-11 | 2.5E-11 | BT20/MB468 |
| >hg38_21_+_8433172_8446622_RNA45SN1_45S_50ntFlanks@12944.12964.21\|CGTAGACGACCTGCTTCTGGG | >hg38_21_+_8433172_8446622_RNA45SN1_45S_50ntFlanks | 12944 | 12964 | 21 | CGTAGACGACCTGCTTCTGGG | 262.1 | -3.9 | 0.6 | -6.8 | 1.1E-11 | 2.5E-11 | MB231/MB468 |
| >hg38_21_+_8433172_8446622_RNA45SN1_45S_50ntFlanks@3775.3799.25\|GCCGGTACAGTGAAACTGCGAATGG | >hg38_21_+_8433172_8446622_RNA45SN1_45S_50ntFlanks | 3775 | 3799 | 25 | GCCGGTACAGTGAAACTGCGAATGG | 96.7 | -8.6 | 1.3 | -6.8 | 1.3E-11 | 2.5E-11 | BT20/MB468 |
| >hg38_21_+_8433172_8446622_RNA45SN1_45S_50ntFlanks@10409.10431.23\|GTCAGTCGGTCCTGAGAGATGGG | >hg38_21_+_8433172_8446622_RNA45SN1_45S_50ntFlanks | 10409 | 10431 | 23 | GTCAGTCGGTCCTGAGAGATGGG | 157.9 | -2.9 | 0.4 | -6.8 | 1.1E-11 | 2.5E-11 | MB231/MB468 |
| >hg38_21_+_8433172_8446622_RNA45SN1_45S_50ntFlanks@6751.6770.20\|CGAACGCACTTGCGGCCCCG | >hg38_21_+_8433172_8446622_RNA45SN1_45S_50ntFlanks | 6751 | 6770 | 20 | CGAACGCACTTGCGGCCCCG | 87.7 | -8.4 | 1.2 | -6.8 | 1.3E-11 | 2.5E-11 | BT20/MB468 |
| >hg38_21_+_8433172_8446622_RNA45SN1_45S_50ntFlanks@3876.3898.23\|TACATGCCGACGGGCGCTGACCC | >hg38_21_+_8433172_8446622_RNA45SN1_45S_50ntFlanks | 3876 | 3898 | 23 | TACATGCCGACGGGCGCTGACCC | 752.0 | -4.3 | 0.6 | -6.8 | 1.3E-11 | 2.5E-11 | BT20/MB468 |
| >hg38_21_+_8433172_8446622_RNA45SN1_45S_50ntFlanks@10680.10710.31\|GTTCCGGCGGCGTCCGGTGAGCTCTCGCTGG | >hg38_21_+_8433172_8446622_RNA45SN1_45S_50ntFlanks | 10680 | 10710 | 31 | GTTCCGGCGGCGTCCGGTGAGCTCTCGCTGG | 18722.2 | -6.6 | 1.0 | -6.8 | 1.3E-11 | 2.5E-11 | BT20/MB468 |
| >hg38_21_+_8433172_8446622_RNA45SN1_45S_50ntFlanks@12406.12436.31\|TTTTTGATCCTTCGATGTCGGCTCTTCCTAT | >hg38_21_+_8433172_8446622_RNA45SN1_45S_50ntFlanks | 12406 | 12436 | 31 | TTTTTGATCCTTCGATGTCGGCTCTTCCTAT | 217.9 | -4.4 | 0.7 | -6.8 | 1.3E-11 | 2.5E-11 | BT20/MB468 |
| >hg38_21_+_8433172_8446622_RNA45SN1_45S_50ntFlanks@12412.12446.35\|ATCCTTCGATGTCGGCTCTTCCTATCATTGTGAAG | >hg38_21_+_8433172_8446622_RNA45SN1_45S_50ntFlanks | 12412 | 12446 | 35 | ATCCTTCGATGTCGGCTCTTCCTATCATTGTGAAG | 88.4 | -8.5 | 1.2 | -6.8 | 1.3E-11 | 2.5E-11 | BT20/MB468 |
| >hg38_21_+_8433172_8446622_RNA45SN1_45S_50ntFlanks@10421.10443.23\|TGAGAGATGGGCGAGCGCCGTTC | >hg38_21_+_8433172_8446622_RNA45SN1_45S_50ntFlanks | 10421 | 10443 | 23 | TGAGAGATGGGCGAGCGCCGTTC | 93.9 | -8.5 | 1.3 | -6.8 | 1.3E-11 | 2.6E-11 | BT20/MB468 |
| >hg38_21_+_8433172_8446622_RNA45SN1_45S_50ntFlanks@11739.11758.20\|TCTCTTAAGGTAGCCAAATG | >hg38_21_+_8433172_8446622_RNA45SN1_45S_50ntFlanks | 11739 | 11758 | 20 | TCTCTTAAGGTAGCCAAATG | 226.7 | -4.3 | 0.6 | -6.8 | 1.2E-11 | 2.6E-11 | MB231/MB468 |
| >hg38_21_+_8433172_8446622_RNA45SN1_45S_50ntFlanks@4657.4688.32\|CTAGAGGTGAAATTCTTGGACCGGCGCAAGAC | >hg38_21_+_8433172_8446622_RNA45SN1_45S_50ntFlanks | 4657 | 4688 | 32 | CTAGAGGTGAAATTCTTGGACCGGCGCAAGAC | 186.9 | -9.5 | 1.4 | -6.8 | 1.3E-11 | 2.6E-11 | BT20/MB468 |
| >hg38_21_+_8433172_8446622_RNA45SN1_45S_50ntFlanks@10402.10435.34\|AACATGGGTCAGTCGGTCCTGAGAGATGGGCGAG | >hg38_21_+_8433172_8446622_RNA45SN1_45S_50ntFlanks | 10402 | 10435 | 34 | AACATGGGTCAGTCGGTCCTGAGAGATGGGCGAG | 481.5 | -6.6 | 1.0 | -6.8 | 1.3E-11 | 2.6E-11 | BT20/MB468 |
| >hg38_21_+_8433172_8446622_RNA45SN1_45S_50ntFlanks@11452.11472.21\|CTCCTCGGCGCGCGGCGGCGG | >hg38_21_+_8433172_8446622_RNA45SN1_45S_50ntFlanks | 11452 | 11472 | 21 | CTCCTCGGCGCGCGGCGGCGG | 208.2 | -5.9 | 0.9 | -6.8 | 1.4E-11 | 2.6E-11 | BT20/MB468 |
| >hg38_21_+_8433172_8446622_RNA45SN1_45S_50ntFlanks@11873.11900.28\|AAGAAGACCCTGTTGAGCTTGACTCTAG | >hg38_21_+_8433172_8446622_RNA45SN1_45S_50ntFlanks | 11873 | 11900 | 28 | AAGAAGACCCTGTTGAGCTTGACTCTAG | 778.2 | -6.2 | 0.9 | -6.8 | 1.4E-11 | 2.6E-11 | BT20/MB468 |
| >hg38_21_+_8433172_8446622_RNA45SN1_45S_50ntFlanks@11769.11803.35\|TAATTAGTGACGCGCATGAATGGATGAACGAGATT | >hg38_21_+_8433172_8446622_RNA45SN1_45S_50ntFlanks | 11769 | 11803 | 35 | TAATTAGTGACGCGCATGAATGGATGAACGAGATT | 623.7 | -2.8 | 0.4 | -6.8 | 1.4E-11 | 2.6E-11 | BT20/MB468 |
| >MI0000273\|hsa-mir-183&WithFlank&7\|-\|129774899\|129775020@33.53.21&[MIMAT0000261&hsa-miR-183-5p&offsets\|0\|-1;m-41&7\|-\|129774966\|129774987&offsets\|-1\|-2]\|TATGGCACTGGTAGAATTCAC | >MI0000273\|hsa-mir-183&WithFlank&7\|-\|129774899\|129775020 | 33 | 53 | 21 | TATGGCACTGGTAGAATTCAC | 503.6 | 8.5 | 1.3 | 6.8 | 1.4E-11 | 2.6E-11 | BT20/MB468 |
| >trna10_AspGTC_12_-_125424193_125424264@39.74.36,trna12_AspGTC_12_-_125411891_125411962@39.74.36,trna144_AspGTC_6_-_27551236_27551307@39.74.36,trna38_AspGTC_17_-_8125556_8125627@39.74.36,trna45_AspGTC_6_+_27447453_27447524@39.74.36,trna48_AspGTC_6_+_27471523_27471594@39.74.36,trna4_AspGTC_12_+_96429799_96429870@39.74.36,trna69_AspGTC_1_-_161440205_161440276@39.74.36,trna72_AspGTC_1_-_161432824_161432895@39.74.36,trna75_AspGTC_1_-_161425414_161425485@39.74.36,trna78_AspGTC_1_-_161418033_161418104@39.74.36,trna81_AspGTC_1_-_161410615_161410686@39.74.36\|GCGGGAGACCGGGGTTCGATTCCCCGACGGGGAGCC | >trna10_AspGTC_12_-_125424193_125424264 | 39 | 74 | 36 | GCGGGAGACCGGGGTTCGATTCCCCGACGGGGAGCC | 838.9 | 8.0 | 1.2 | 6.8 | 1.2E-11 | 2.6E-11 | MB231/MB468 |
| >hg38_21_+_8433172_8446622_RNA45SN1_45S_50ntFlanks@10571.10596.26\|GAGAAGCCGGCGGGAGCCCCGGGGAG | >hg38_21_+_8433172_8446622_RNA45SN1_45S_50ntFlanks | 10571 | 10596 | 26 | GAGAAGCCGGCGGGAGCCCCGGGGAG | 125.4 | -8.5 | 1.3 | -6.8 | 1.2E-11 | 2.7E-11 | MB231/MB468 |
| >hg38_21_+_8433172_8446622_RNA45SN1_45S_50ntFlanks@12347.12364.18\|AGAAAAGTTACCACAGGG | >hg38_21_+_8433172_8446622_RNA45SN1_45S_50ntFlanks | 12347 | 12364 | 18 | AGAAAAGTTACCACAGGG | 80.0 | -8.3 | 1.2 | -6.8 | 1.4E-11 | 2.7E-11 | BT20/MB468 |
| >trna9_ArgTCT_1_+_94313129_94313213@1.29.29\|GGCTCCGTGGCGCAATGGATAGCGCATTG | >trna9_ArgTCT_1_+_94313129_94313213 | 1 | 29 | 29 | GGCTCCGTGGCGCAATGGATAGCGCATTG | 129.0 | -9.0 | 1.3 | -6.8 | 1.4E-11 | 2.7E-11 | BT20/MB468 |
| >MI0001448\|hsa-mir-425&WithFlank&3\|-\|49020142\|49020240@20.43.24(+1A)&[MIMAT0003393&hsa-miR-425-5p&offsets\|0\|+1(+1A);m-95&3\|-\|49020200\|49020221&offsets\|0\|+2(+1A)]\|AATGACACGATCACTCCCGTTGAGA | >MI0001448\|hsa-mir-425&WithFlank&3\|-\|49020142\|49020240 | 20 | 43 | 24(+1A) | AATGACACGATCACTCCCGTTGAGA | 766.0 | 8.4 | 1.2 | 6.8 | 1.4E-11 | 2.7E-11 | BT20/MB468 |
| >hg38_21_+_8433172_8446622_RNA45SN1_45S_50ntFlanks@10843.10863.21\|TAACTTCGGGATAAGGATTGG | >hg38_21_+_8433172_8446622_RNA45SN1_45S_50ntFlanks | 10843 | 10863 | 21 | TAACTTCGGGATAAGGATTGG | 187.1 | -6.0 | 0.9 | -6.8 | 1.4E-11 | 2.7E-11 | BT20/MB468 |
| >hg38_21_+_8433172_8446622_RNA45SN1_45S_50ntFlanks@10781.10808.28\|AGGTGAACAGCCTCTGGCATGTTGGAAC | >hg38_21_+_8433172_8446622_RNA45SN1_45S_50ntFlanks | 10781 | 10808 | 28 | AGGTGAACAGCCTCTGGCATGTTGGAAC | 117.1 | -8.9 | 1.3 | -6.8 | 1.4E-11 | 2.7E-11 | BT20/MB468 |
| >hg38_21_+_8433172_8446622_RNA45SN1_45S_50ntFlanks@10829.10852.24\|GCAAGCCGGATCCGTAACTTCGGG | >hg38_21_+_8433172_8446622_RNA45SN1_45S_50ntFlanks | 10829 | 10852 | 24 | GCAAGCCGGATCCGTAACTTCGGG | 327.0 | -3.8 | 0.6 | -6.8 | 1.4E-11 | 2.7E-11 | BT20/MB468 |
| >hg38_21_+_8433172_8446622_RNA45SN1_45S_50ntFlanks@11535.11552.18\|TTCCGCGCGGCGCCTCGC | >hg38_21_+_8433172_8446622_RNA45SN1_45S_50ntFlanks | 11535 | 11552 | 18 | TTCCGCGCGGCGCCTCGC | 124.9 | -4.3 | 0.6 | -6.8 | 1.2E-11 | 2.8E-11 | MB231/MB468 |
| >hg38_21_+_8433172_8446622_RNA45SN1_45S_50ntFlanks@12417.12442.26\|TCGATGTCGGCTCTTCCTATCATTGT | >hg38_21_+_8433172_8446622_RNA45SN1_45S_50ntFlanks | 12417 | 12442 | 26 | TCGATGTCGGCTCTTCCTATCATTGT | 620.9 | 4.0 | 0.6 | 6.8 | 1.2E-11 | 2.8E-11 | MB231/MB468 |
| >hg38_21_+_8433172_8446622_RNA45SN1_45S_50ntFlanks@7985.8007.23\|GATCAGACGTGGCGACCCGCTGA | >hg38_21_+_8433172_8446622_RNA45SN1_45S_50ntFlanks | 7985 | 8007 | 23 | GATCAGACGTGGCGACCCGCTGA | 101.3 | -8.7 | 1.3 | -6.8 | 1.4E-11 | 2.8E-11 | BT20/MB468 |
| >am_trnaMT_PheGAA_MT_+_577_647@43.62.20\|TTAGACGGGCTCACATCACC | >am_trnaMT_PheGAA_MT_+_577_647 | 43 | 62 | 20 | TTAGACGGGCTCACATCACC | 296.4 | 7.6 | 1.1 | 6.8 | 1.4E-11 | 2.8E-11 | BT20/MB468 |
| >hg38_21_+_8433172_8446622_RNA45SN1_45S_50ntFlanks@4555.4587.33\|CGCAGCTAGGAATAATGGAATAGGACCGCGGTT | >hg38_21_+_8433172_8446622_RNA45SN1_45S_50ntFlanks | 4555 | 4587 | 33 | CGCAGCTAGGAATAATGGAATAGGACCGCGGTT | 114.7 | -8.4 | 1.2 | -6.8 | 1.3E-11 | 2.8E-11 | MB231/MB468 |
| >hg38_21_+_8433172_8446622_RNA45SN1_45S_50ntFlanks@4738.4766.29\|AAAGTCGGAGGTTCGAAGACGATCAGATA | >hg38_21_+_8433172_8446622_RNA45SN1_45S_50ntFlanks | 4738 | 4766 | 29 | AAAGTCGGAGGTTCGAAGACGATCAGATA | 644.2 | 5.8 | 0.9 | 6.8 | 1.3E-11 | 2.8E-11 | MB231/MB468 |
| >hg38_21_+_8433172_8446622_RNA45SN1_45S_50ntFlanks@12457.12478.22\|CAAGCGTTGGATTGTTCACCCA | >hg38_21_+_8433172_8446622_RNA45SN1_45S_50ntFlanks | 12457 | 12478 | 22 | CAAGCGTTGGATTGTTCACCCA | 3008.9 | -4.3 | 0.6 | -6.8 | 1.3E-11 | 2.8E-11 | MB231/MB468 |
| >hg38_21_+_8433172_8446622_RNA45SN1_45S_50ntFlanks@8194.8215.22\|CCCGTGGACGGTGTGAGGCCGG | >hg38_21_+_8433172_8446622_RNA45SN1_45S_50ntFlanks | 8194 | 8215 | 22 | CCCGTGGACGGTGTGAGGCCGG | 1190.7 | -5.4 | 0.8 | -6.8 | 1.3E-11 | 2.8E-11 | MB231/MB468 |
| >hg38_21_+_8433172_8446622_RNA45SN1_45S_50ntFlanks@12539.12558.20\|ATGATGTGTTGTTGCCATGG | >hg38_21_+_8433172_8446622_RNA45SN1_45S_50ntFlanks | 12539 | 12558 | 20 | ATGATGTGTTGTTGCCATGG | 92.4 | -8.5 | 1.3 | -6.8 | 1.5E-11 | 2.8E-11 | BT20/MB468 |
| >hg38_21_+_8433172_8446622_RNA45SN1_45S_50ntFlanks@10478.10512.35\|CCGATCGAAAGGGAGTCGGGTTCAGATCCCCGAAT | >hg38_21_+_8433172_8446622_RNA45SN1_45S_50ntFlanks | 10478 | 10512 | 35 | CCGATCGAAAGGGAGTCGGGTTCAGATCCCCGAAT | 97.7 | -8.6 | 1.3 | -6.7 | 1.5E-11 | 2.9E-11 | BT20/MB468 |
| >hg38_21_+_8433172_8446622_RNA45SN1_45S_50ntFlanks@12971.12988.18\|TTTCGTACGTAGCAGAGC | >hg38_21_+_8433172_8446622_RNA45SN1_45S_50ntFlanks | 12971 | 12988 | 18 | TTTCGTACGTAGCAGAGC | 114.7 | -3.0 | 0.5 | -6.7 | 1.5E-11 | 2.9E-11 | BT20/MB468 |
| >hg38_21_+_8433172_8446622_RNA45SN1_45S_50ntFlanks@12935.12966.32\|TAAACCATTCGTAGACGACCTGCTTCTGGGTC | >hg38_21_+_8433172_8446622_RNA45SN1_45S_50ntFlanks | 12935 | 12966 | 32 | TAAACCATTCGTAGACGACCTGCTTCTGGGTC | 92.6 | -8.5 | 1.3 | -6.7 | 1.5E-11 | 2.9E-11 | BT20/MB468 |
| >hg38_21_+_8433172_8446622_RNA45SN1_45S_50ntFlanks@6666.6686.21\|GATCACTCGGCTCGTGCGTCG | >hg38_21_+_8433172_8446622_RNA45SN1_45S_50ntFlanks | 6666 | 6686 | 21 | GATCACTCGGCTCGTGCGTCG | 104.0 | -8.7 | 1.3 | -6.7 | 1.5E-11 | 2.9E-11 | BT20/MB468 |
| >hg38_21_+_8433172_8446622_RNA45SN1_45S_50ntFlanks@9507.9526.20\|AACTATGCCTGGGCAGGGCG | >hg38_21_+_8433172_8446622_RNA45SN1_45S_50ntFlanks | 9507 | 9526 | 20 | AACTATGCCTGGGCAGGGCG | 123.4 | -8.5 | 1.3 | -6.8 | 1.3E-11 | 2.9E-11 | MB231/MB468 |
| >hg38_21_+_8433172_8446622_RNA45SN1_45S_50ntFlanks@12324.12349.26\|TTGGGTTTTAAGCAGGAGGTGTCAGA | >hg38_21_+_8433172_8446622_RNA45SN1_45S_50ntFlanks | 12324 | 12349 | 26 | TTGGGTTTTAAGCAGGAGGTGTCAGA | 120.4 | -8.5 | 1.3 | -6.8 | 1.3E-11 | 2.9E-11 | MB231/MB468 |
| >trna14_LysTTT_11_-_59327808_59327880@1.36.36,trna2_LysTTT_17_+_8022473_8022545@1.36.36,trna54_LysTTT_1_+_204475655_204475727@1.36.36,trna5_LysTTT_11_+_59323902_59323974@1.36.36,trna62_LysTTT_1_-_204476158_204476230@1.36.36,trna76_LysTTT_6_+_28918806_28918878@1.36.36\|GCCCGGATAGCTCAGTCGGTAGAGCATCAGACTTTT | >trna14_LysTTT_11_-_59327808_59327880 | 1 | 36 | 36 | GCCCGGATAGCTCAGTCGGTAGAGCATCAGACTTTT | 3618.0 | 10.7 | 1.6 | 6.7 | 1.5E-11 | 2.9E-11 | BT20/MB468 |

| | | | | | | | | | | | | |
|---|---|---|---|---|---|---|---|---|---|---|---|---|
| >hg38_21_+_8433172_8446622_RNA45SN1_45S_50ntFlanks@11255.11281.27\|CTGCGGCGGGCGTCGCGGCCGCCCCCG | >hg38_21_+_8433172_8446622_RNA45SN1_45S_50ntFlanks | 11255 | 11281 | 27 | CTGCGGCGGGCGTCGCGGCCGCCCCCG | 111.6 | -8.8 | 1.3 | -6.7 | 1.5E-11 | 3.0E-11 | BT20/MB468 |
| >hg38_21_+_8433172_8446622_RNA45SN1_45S_50ntFlanks@12936.12955.20\|AAACCATTCGTAGACGACCT | >hg38_21_+_8433172_8446622_RNA45SN1_45S_50ntFlanks | 12936 | 12955 | 20 | AAACCATTCGTAGACGACCT | 282.1 | -3.0 | 0.4 | -6.8 | 1.3E-11 | 3.0E-11 | MB231/MB468 |
| >trna10_AspGTC_12_-_125424193_125424264@31.72.42,trna12_AspGTC_12_-_125411891_125411962@31.72.42,trna38_AspGTC_17_-_8125556_8125627@31.72.42,trna45_AspGTC_6_+_27447453_27447524@31.72.42,trna48_AspGTC_6_+_27471523_27471594@31.72.42,trna4_AspGTC_12_+_96429799_96429870@31.72.42,trna69_AspGTC_1_-_161440205_161440276@31.72.42,trna72_AspGTC_1_-_161432824_161432895@31.72.42,trna75_AspGTC_1_-_161425414_161425485@31.72.42,trna78_AspGTC_1_-_161418033_161418104@31.72.42,trna81_AspGTC_1_-_161410615_161410686@31.72.42\|CCTGTCACGCGGGAGACCGGGGTTCGATTCCCCGACGGGGAG | >trna10_AspGTC_12_-_125424193_125424264 | 31 | 72 | 42 | CCTGTCACGCGGGAGACCGGGGTTCGATTCCCCGACGGGGAG | 198.3 | -9.2 | 1.4 | -6.8 | 1.3E-11 | 3.0E-11 | MB231/MB468 |
| >hg38_21_+_8433172_8446622_RNA45SN1_45S_50ntFlanks@11769.11790.22\|TAATTAGTGACGCGCATGAATG | >hg38_21_+_8433172_8446622_RNA45SN1_45S_50ntFlanks | 11769 | 11790 | 22 | TAATTAGTGACGCGCATGAATG | 531.2 | -3.6 | 0.5 | -6.7 | 1.5E-11 | 3.0E-11 | BT20/MB468 |
| >hg38_21_+_8433172_8446622_RNA45SN1_45S_50ntFlanks@4726.4752.27\|TAATCAAGAACGAAAGTCGGAGGTTCG | >hg38_21_+_8433172_8446622_RNA45SN1_45S_50ntFlanks | 4726 | 4752 | 27 | TAATCAAGAACGAAAGTCGGAGGTTCG | 227.0 | -5.9 | 0.9 | -6.7 | 1.5E-11 | 3.0E-11 | BT20/MB468 |
| >hg38_21_+_8433172_8446622_RNA45SN1_45S_50ntFlanks@11104.11123.20\|AAGGGTCGGGGCGGCAGGGG | >hg38_21_+_8433172_8446622_RNA45SN1_45S_50ntFlanks | 11104 | 11123 | 20 | AAGGGTCGGGGCGGCAGGGG | 131.9 | -8.6 | 1.3 | -6.8 | 1.4E-11 | 3.0E-11 | MB231/MB468 |
| >hg38_21_+_8433172_8446622_RNA45SN1_45S_50ntFlanks@4722.4748.27\|TCATTAATCAAGAACGAAAGTCGGAGG | >hg38_21_+_8433172_8446622_RNA45SN1_45S_50ntFlanks | 4722 | 4748 | 27 | TCATTAATCAAGAACGAAAGTCGGAGG | 369.7 | -4.7 | 0.7 | -6.7 | 1.6E-11 | 3.0E-11 | BT20/MB468 |
| >hg38_21_+_8433172_8446622_RNA45SN1_45S_50ntFlanks@12626.12657.32\|AATGGGGCGAAGCTACCATCTGTGGGATTATG | >hg38_21_+_8433172_8446622_RNA45SN1_45S_50ntFlanks | 12626 | 12657 | 32 | AATGGGGCGAAGCTACCATCTGTGGGATTATG | 349.0 | -4.9 | 0.7 | -6.7 | 1.6E-11 | 3.1E-11 | BT20/MB468 |
| >hg38_21_+_8433172_8446622_RNA45SN1_45S_50ntFlanks@10571.10589.19\|GAGAAGCCGGCGGGAGCCC | >hg38_21_+_8433172_8446622_RNA45SN1_45S_50ntFlanks | 10571 | 10589 | 19 | GAGAAGCCGGCGGGAGCCC | 106.3 | -8.7 | 1.3 | -6.7 | 1.6E-11 | 3.1E-11 | BT20/MB468 |
| >hg38_21_+_8433172_8446622_RNA45SN1_45S_50ntFlanks@12631.12653.23\|GGCGAAGCTACCATCTGTGGGAT | >hg38_21_+_8433172_8446622_RNA45SN1_45S_50ntFlanks | 12631 | 12653 | 23 | GGCGAAGCTACCATCTGTGGGAT | 137.0 | -9.1 | 1.3 | -6.7 | 1.6E-11 | 3.1E-11 | BT20/MB468 |
| >hg38_21_+_8433172_8446622_RNA45SN1_45S_50ntFlanks@11537.11560.24\|CCGCGCGGCGCCTCGCCTCGGCCG | >hg38_21_+_8433172_8446622_RNA45SN1_45S_50ntFlanks | 11537 | 11560 | 24 | CCGCGCGGCGCCTCGCCTCGGCCG | 94.6 | -8.6 | 1.3 | -6.7 | 1.6E-11 | 3.1E-11 | BT20/MB468 |
| >MI0000088\|hsa-mir-30a&WithFlank&6\|-\|71403545\|71403627@12.33.22(+2U)&[MIMAT0000087&hsa-miR-30a-5p&offsets\|0\|0(+2U);m-15&6\|-\|71403595\|71403616&offsets\|0\|0(+2U)]\|TGTAAACATCCTCGACTGGAAGTT | >MI0000088\|hsa-mir-30a&WithFlank&6\|-\|71403545\|71403627 | 12 | 33 | 22(+2U) | TGTAAACATCCTCGACTGGAAGTT | 3605.8 | 10.1 | 1.5 | 6.7 | 1.6E-11 | 3.1E-11 | BT20/MB468 |
| >hg38_21_+_8433172_8446622_RNA45SN1_45S_50ntFlanks@12991.13010.20\|CTCCCTCGCTGCGATCTATT | >hg38_21_+_8433172_8446622_RNA45SN1_45S_50ntFlanks | 12991 | 13010 | 20 | CTCCCTCGCTGCGATCTATT | 415.6 | -5.4 | 0.8 | -6.7 | 1.6E-11 | 3.1E-11 | BT20/MB468 |
| >MI0000736\|hsa-mir-30c-1&WithFlank&1\|+\|40757278\|40757378@23.46.24;MI0000254\|hsa-mir-30c-2&WithFlank&6\|-\|71376954\|71377037@13.36.24&[MIMAT0000244&hsa-miR-30c-5p&offsets\|0\|+1;m-57&1\|+\|40757300\|40757323&offsets\|0\|0];[MIMAT0000244_1&hsa-miR-30c-5p&offsets\|0\|+1;m-56&6\|-\|71377002\|71377025&offsets\|0\|0]\|TGTAAACATCCTACACTCTCAGCT | >MI0000736\|hsa-mir-30c-1&WithFlank&1\|+\|40757278\|40757378 | 23 | 46 | 24 | TGTAAACATCCTACACTCTCAGCT | 40090.0 | 5.5 | 0.8 | 6.7 | 1.6E-11 | 3.1E-11 | BT20/MB468 |
| >hg38_21_+_8433172_8446622_RNA45SN1_45S_50ntFlanks@10444.10462.19\|CGAAGGGACGGGCGATGGC | >hg38_21_+_8433172_8446622_RNA45SN1_45S_50ntFlanks | 10444 | 10462 | 19 | CGAAGGGACGGGCGATGGC | 156.8 | -3.9 | 0.6 | -6.7 | 1.6E-11 | 3.1E-11 | BT20/MB468 |
| >MI0000483\|hsa-mir-186&WithFlank&1\|-\|71067625\|71067722@21.43.23&[MIMAT0000456&hsa-miR-186-5p&offsets\|0\|+1;m-84&1\|-\|71067681\|71067702&offsets\|0\|+1]\|CAAAGAATTCTCCTTTTGGGCTT | >MI0000483\|hsa-mir-186&WithFlank&1\|-\|71067625\|71067722 | 21 | 43 | 23 | CAAAGAATTCTCCTTTTGGGCTT | 1156.5 | 5.5 | 0.8 | 6.8 | 1.4E-11 | 3.1E-11 | MB231/MB468 |
| >hg38_21_+_8433172_8446622_RNA45SN1_45S_50ntFlanks@11741.11759.19\|TCTTAAGGTAGCCAAATGC | >hg38_21_+_8433172_8446622_RNA45SN1_45S_50ntFlanks | 11741 | 11759 | 19 | TCTTAAGGTAGCCAAATGC | 176.2 | -9.5 | 1.4 | -6.7 | 1.6E-11 | 3.2E-11 | BT20/MB468 |
| >MI0003591\|hsa-mir-584&WithFlank&5\|-\|149062307\|149062415@22.43.22&[MIMAT0003249&hsa-miR-584-5p&offsets\|0\|0;m-167&5\|-\|149062374\|149062394&offsets\|0\|+1]\|TTATGGTTTGCCTGGGACTGAG | >MI0003591\|hsa-mir-584&WithFlank&5\|-\|149062307\|149062415 | 22 | 43 | 22 | TTATGGTTTGCCTGGGACTGAG | 503.1 | 9.6 | 1.4 | 6.7 | 1.6E-11 | 3.2E-11 | BT20/MB468 |
| >MI0000085\|hsa-mir-27a&WithFlank&19\|-\|13836434\|13836523@57.75.19(+1U)&[MIMAT0000084&hsa-miR-27a-3p&offsets\|0\|-2(+1U);m-43&19\|-\|13836447\|13836467&offsets\|0\|-2(+1U)]\|TTCACAGTGGCTAAGTTCCT | >MI0000085\|hsa-mir-27a&WithFlank&19\|-\|13836434\|13836523 | 57 | 75 | 19(+1U) | TTCACAGTGGCTAAGTTCCT | 683.7 | 6.1 | 0.9 | 6.7 | 1.7E-11 | 3.2E-11 | BT20/MB468 |
| >hg38_21_+_8433172_8446622_RNA45SN1_45S_50ntFlanks@8196.8224.29\|CGTGGACGGTGTGAGGCCGGTAGCGGCCC | >hg38_21_+_8433172_8446622_RNA45SN1_45S_50ntFlanks | 8196 | 8224 | 29 | CGTGGACGGTGTGAGGCCGGTAGCGGCCC | 476.2 | -3.4 | 0.5 | -6.7 | 1.7E-11 | 3.2E-11 | BT20/MB468 |
| >hg38_21_+_8433172_8446622_RNA45SN1_45S_50ntFlanks@11853.11872.20\|CTTGGCGGAATCAGCGGGGA | >hg38_21_+_8433172_8446622_RNA45SN1_45S_50ntFlanks | 11853 | 11872 | 20 | CTTGGCGGAATCAGCGGGGA | 123.8 | -8.9 | 1.3 | -6.7 | 1.7E-11 | 3.2E-11 | BT20/MB468 |
| >MI0003190\|hsa-mir-505&WithFlank&X\|-\|139924142\|139924237@57.80.24&[MIMAT0002876&hsa-miR-505-3p&offsets\|0\|+2;m-196&X\|-\|139924160\|139924181&offsets\|0\|+2]\|CGTCAACACTTGCTGGTTTCCTCT | >MI0003190\|hsa-mir-505&WithFlank&X\|-\|139924142\|139924237 | 57 | 80 | 24 | CGTCAACACTTGCTGGTTTCCTCT | 447.0 | 8.6 | 1.3 | 6.7 | 1.7E-11 | 3.2E-11 | BT20/MB468 |
| >hg38_1_-_228634819_228635039_RNA5S12_5S_50ntFlanks@136.171.36\|GGAGACCGCCTGGGAATACCGGGTGCTGTAGGCTTT | >hg38_1_-_228634819_228635039_RNA5S12_5S_50ntFlanks | 136 | 171 | 36 | GGAGACCGCCTGGGAATACCGGGTGCTGTAGGCTTT | 2711.9 | 4.8 | 0.7 | 6.7 | 1.7E-11 | 3.2E-11 | BT20/MB468 |
| >hg38_21_+_8433172_8446622_RNA45SN1_45S_50ntFlanks@11641.11663.23\|CGGCGGGTGTTGACGCGATGTGA | >hg38_21_+_8433172_8446622_RNA45SN1_45S_50ntFlanks | 11641 | 11663 | 23 | CGGCGGGTGTTGACGCGATGTGA | 399.5 | -7.0 | 1.0 | -6.7 | 1.7E-11 | 3.2E-11 | BT20/MB468 |
| >hg38_21_+_8433172_8446622_RNA45SN1_45S_50ntFlanks@12008.12030.23\|CCGCCGGTGAAATACCACTACTC | >hg38_21_+_8433172_8446622_RNA45SN1_45S_50ntFlanks | 12008 | 12030 | 23 | CCGCCGGTGAAATACCACTACTC | 840.9 | -6.7 | 1.0 | -6.7 | 1.7E-11 | 3.3E-11 | BT20/MB468 |
| >hg38_21_+_8433172_8446622_RNA45SN1_45S_50ntFlanks@6771.6799.29\|GGTTCCTCCCGGGGCTACGCCTGTCTGAG | >hg38_21_+_8433172_8446622_RNA45SN1_45S_50ntFlanks | 6771 | 6799 | 29 | GGTTCCTCCCGGGGCTACGCCTGTCTGAG | 125.7 | -8.6 | 1.3 | -6.8 | 1.5E-11 | 3.3E-11 | MB231/MB468 |
| >hg38_21_+_8433172_8446622_RNA45SN1_45S_50ntFlanks@5449.5466.18\|ATCGGCCCCGCCGGGGTC | >hg38_21_+_8433172_8446622_RNA45SN1_45S_50ntFlanks | 5449 | 5466 | 18 | ATCGGCCCCGCCGGGGTC | 112.1 | -8.4 | 1.2 | -6.8 | 1.5E-11 | 3.3E-11 | MB231/MB468 |
| >MI0000085\|hsa-mir-27a&WithFlank&19\|-\|13836434\|13836523@57.74.18(+1A);MI0000440\|hsa-mir-27b&WithFlank&9\|+\|95085439\|95085547@67.84.18(+1A)&[MIMAT0000084&hsa-miR-27a-3p&offsets\|0\|-3(+1A);m-43&19\|-\|13836447\|13836467&offsets\|0\|-3(+1A)];[MIMAT0000419&hsa-miR-27b-3p&offsets\|0\|-3(+1A);m-39&9\|+\|95085505\|95085525&offsets\|0\|-3(+1A)]\|TTCACAGTGGCTAAGTTCA | >MI0000085\|hsa-mir-27a&WithFlank&19\|-\|13836434\|13836523 | 57 | 74 | 18(+1A) | TTCACAGTGGCTAAGTTCA | 683.8 | 8.4 | 1.2 | 6.7 | 1.7E-11 | 3.3E-11 | BT20/MB468 |
| >hg38_21_+_8433172_8446622_RNA45SN1_45S_50ntFlanks@12151.12183.33\|GTGGGGAGTTTGACTGGGGCGGTACACCTGTCA | >hg38_21_+_8433172_8446622_RNA45SN1_45S_50ntFlanks | 12151 | 12183 | 33 | GTGGGGAGTTTGACTGGGGCGGTACACCTGTCA | 134.7 | -3.8 | 0.6 | -6.7 | 1.5E-11 | 3.3E-11 | MB231/MB468 |
| >hg38_21_+_8433172_8446622_RNA45SN1_45S_50ntFlanks@10794.10826.33\|CTGGCATGTTGGAACAATGTAGGTAAGGGAAGT | >hg38_21_+_8433172_8446622_RNA45SN1_45S_50ntFlanks | 10794 | 10826 | 33 | CTGGCATGTTGGAACAATGTAGGTAAGGGAAGT | 210.6 | -5.5 | 0.8 | -6.7 | 1.5E-11 | 3.3E-11 | MB231/MB468 |
| >MI0000234\|hsa-mir-192&WithFlank&11\|-\|64891131\|64891252@31.53.23&[MIMAT0000222&hsa-miR-192-5p&offsets\|+1\|+3;m-59&11\|-\|64891203\|64891223&offsets\|+1\|+3]\|TGACCTATGAATTGACAGCCAGT | >MI0000234\|hsa-mir-192&WithFlank&11\|-\|64891131\|64891252 | 31 | 53 | 23 | TGACCTATGAATTGACAGCCAGT | 597.1 | 4.8 | 0.7 | 6.7 | 1.7E-11 | 3.3E-11 | BT20/MB468 |
| >hg38_21_+_8433172_8446622_RNA45SN1_45S_50ntFlanks@12457.12476.20\|CAAGCGTTGGATTGTTCACC | >hg38_21_+_8433172_8446622_RNA45SN1_45S_50ntFlanks | 12457 | 12476 | 20 | CAAGCGTTGGATTGTTCACC | 5615.1 | -4.3 | 0.6 | -6.7 | 1.7E-11 | 3.3E-11 | BT20/MB468 |
| >hg38_21_+_8433172_8446622_RNA45SN1_45S_50ntFlanks@8475.8492.18\|CCCGGCGGATCTTTCCCG | >hg38_21_+_8433172_8446622_RNA45SN1_45S_50ntFlanks | 8475 | 8492 | 18 | CCCGGCGGATCTTTCCCG | 121.3 | -8.9 | 1.3 | -6.7 | 1.7E-11 | 3.3E-11 | BT20/MB468 |
| >hg38_21_+_8433172_8446622_RNA45SN1_45S_50ntFlanks@6753.6770.18\|AACGCACTTGCGGCCCCG | >hg38_21_+_8433172_8446622_RNA45SN1_45S_50ntFlanks | 6753 | 6770 | 18 | AACGCACTTGCGGCCCCG | 89.7 | -8.5 | 1.3 | -6.7 | 1.7E-11 | 3.3E-11 | BT20/MB468 |
| >hg38_21_+_8433172_8446622_RNA45SN1_45S_50ntFlanks@12072.12096.25\|CCCGAGGGGCTCTCGCTTCTGGCGC | >hg38_21_+_8433172_8446622_RNA45SN1_45S_50ntFlanks | 12072 | 12096 | 25 | CCCGAGGGGCTCTCGCTTCTGGCGC | 292.2 | -4.7 | 0.7 | -6.7 | 1.7E-11 | 3.3E-11 | BT20/MB468 |
| >hg38_21_+_8433172_8446622_RNA45SN1_45S_50ntFlanks@12957.12979.23\|CTTCTGGGTCGGGGTTTCGTACG | >hg38_21_+_8433172_8446622_RNA45SN1_45S_50ntFlanks | 12957 | 12979 | 23 | CTTCTGGGTCGGGGTTTCGTACG | 403.6 | -6.3 | 0.9 | -6.7 | 1.8E-11 | 3.4E-11 | BT20/MB468 |
| >hg38_21_+_8433172_8446622_RNA45SN1_45S_50ntFlanks@12456.12490.35\|CCAAGCGTTGGATTGTTCACCCACTAATAGGGAAC | >hg38_21_+_8433172_8446622_RNA45SN1_45S_50ntFlanks | 12456 | 12490 | 35 | CCAAGCGTTGGATTGTTCACCCACTAATAGGGAAC | 541.0 | -4.1 | 0.6 | -6.7 | 1.8E-11 | 3.4E-11 | BT20/MB468 |
| >hg38_21_+_8433172_8446622_RNA45SN1_45S_50ntFlanks@12405.12427.23\|CTTTTTGATCCTTCGATGTCGGC | >hg38_21_+_8433172_8446622_RNA45SN1_45S_50ntFlanks | 12405 | 12427 | 23 | CTTTTTGATCCTTCGATGTCGGC | 116.1 | -8.4 | 1.3 | -6.7 | 1.6E-11 | 3.4E-11 | MB231/MB468 |
| >hg38_1_-_228634819_228635039_RNA5S12_5S_50ntFlanks@130.171.42\|TGGATGGGAGACCGCCTGGGAATACCGGGTGCTGTAGGCTTT | >hg38_1_-_228634819_228635039_RNA5S12_5S_50ntFlanks | 130 | 171 | 42 | TGGATGGGAGACCGCCTGGGAATACCGGGTGCTGTAGGCTTT | 1034.5 | 10.6 | 1.6 | 6.7 | 1.8E-11 | 3.4E-11 | BT20/MB468 |
| >hg38_21_+_8433172_8446622_RNA45SN1_45S_50ntFlanks@11583.11605.23\|CTGGTGCGGACCAGGGGAATCCG | >hg38_21_+_8433172_8446622_RNA45SN1_45S_50ntFlanks | 11583 | 11605 | 23 | CTGGTGCGGACCAGGGGAATCCG | 570.1 | -4.3 | 0.6 | -6.7 | 1.6E-11 | 3.4E-11 | MB231/MB468 |
| >hg38_21_+_8433172_8446622_RNA45SN1_45S_50ntFlanks@12489.12521.33\|ACGTGAGCTGGGTTTAGACCGTCGTGAGACAGG | >hg38_21_+_8433172_8446622_RNA45SN1_45S_50ntFlanks | 12489 | 12521 | 33 | ACGTGAGCTGGGTTTAGACCGTCGTGAGACAGG | 132.3 | -5.8 | 0.9 | -6.7 | 1.8E-11 | 3.5E-11 | BT20/MB468 |

<table>
<tr><td>>hg38_21_+_8433172_8446622_RNA45SN1_45S_50ntFlanks@9504.9521.18|GTGAACTATGCCTGGGCA</td><td>>hg38_21_+_8433172_8446622_RNA45SN1_45S_50ntFlanks</td><td>9504</td><td>9521</td><td>18</td><td>GTGAACTATGCCTGGGCA</td><td>1093.3</td><td>-4.0</td><td>0.6</td><td>-6.7</td><td>1.6E-11</td><td>3.5E-11</td><td>MB231/MB468</td></tr>
<tr><td>>hg38_21_+_8433172_8446622_RNA45SN1_45S_50ntFlanks@11161.11189.29|CACGGTCCCCCGCGAGGGGGGCCCGGGCA</td><td>>hg38_21_+_8433172_8446622_RNA45SN1_45S_50ntFlanks</td><td>11161</td><td>11189</td><td>29</td><td>CACGGTCCCCCGCGAGGGGGGCCCGGGCA</td><td>110.7</td><td>-8.8</td><td>1.3</td><td>-6.7</td><td>1.8E-11</td><td>3.5E-11</td><td>BT20/MB468</td></tr>
<tr><td>>am_trna11_ArgACG_3_-_45730491_45730563@39.56.18,trna138_ArgACG_6_-_27638344_27638416@39.56.18,trna156_ArgACG_6_-_27181623_27181695@39.56.18,trna36_ArgACG_6_+_27182952_27183024@39.56.18|TCAGAAGATTCTAGGTTC</td><td>>am_trna11_ArgACG_3_-_45730491_45730563</td><td>39</td><td>56</td><td>18</td><td>TCAGAAGATTCTAGGTTC</td><td>122.7</td><td>-8.5</td><td>1.3</td><td>-6.7</td><td>1.6E-11</td><td>3.5E-11</td><td>MB231/MB468</td></tr>
<tr><td>>hg38_21_+_8433172_8446622_RNA45SN1_45S_50ntFlanks@13003.13027.25|GATCTATTGAAAGTCAGCCCTCGAC</td><td>>hg38_21_+_8433172_8446622_RNA45SN1_45S_50ntFlanks</td><td>13003</td><td>13027</td><td>25</td><td>GATCTATTGAAAGTCAGCCCTCGAC</td><td>115.9</td><td>-8.9</td><td>1.3</td><td>-6.7</td><td>1.8E-11</td><td>3.5E-11</td><td>BT20/MB468</td></tr>
<tr><td>>hg38_21_+_8433172_8446622_RNA45SN1_45S_50ntFlanks@9533.9563.31|GAGGAAACTCTGGTGGAGGTCCGTAGCGGTC</td><td>>hg38_21_+_8433172_8446622_RNA45SN1_45S_50ntFlanks</td><td>9533</td><td>9563</td><td>31</td><td>GAGGAAACTCTGGTGGAGGTCCGTAGCGGTC</td><td>1070.0</td><td>-4.6</td><td>0.7</td><td>-6.7</td><td>1.8E-11</td><td>3.5E-11</td><td>BT20/MB468</td></tr>
<tr><td>>MI0000446|hsa-mir-125b-1&WithFlank&11|-|122099751|122099850@21.43.23(+1U)&[MIMAT0000423&hsa-miR-125b-5p&offsets|0|+1(+1U);m-67&11|-|122099809|122099830&offsets|0|+1(+1U)]|TCCCTGAGACCCTAACTTGTGATT</td><td>>MI0000446|hsa-mir-125b-1&WithFlank&11|-|122099751|122099850</td><td>21</td><td>43</td><td>23(+1U)</td><td>TCCCTGAGACCCTAACTTGTGATT</td><td>329.7</td><td>-7.4</td><td>1.0</td><td>-7.1</td><td>1.1E-12</td><td>3.5E-11</td><td>BT20/MB231</td></tr>
<tr><td>>MI0000239|hsa-mir-197&WithFlank&1|+|109598887|109598973@54.74.21&[MIMAT0000227&hsa-miR-197-3p&offsets|0|-1;m-99&1|+|109598940|109598961&offsets|0|-1]|TTCACCACCTTCTCCACCCAG</td><td>>MI0000239|hsa-mir-197&WithFlank&1|+|109598887|109598973</td><td>54</td><td>74</td><td>21</td><td>TTCACCACCTTCTCCACCCAG</td><td>718.4</td><td>10.3</td><td>1.5</td><td>6.7</td><td>1.9E-11</td><td>3.5E-11</td><td>BT20/MB468</td></tr>
<tr><td>>hg38_21_+_8433172_8446622_RNA45SN1_45S_50ntFlanks@11641.11679.39|CGGCGGGTGTTGACGCGATGTGATTTCTGCCCAGTGCTC</td><td>>hg38_21_+_8433172_8446622_RNA45SN1_45S_50ntFlanks</td><td>11641</td><td>11679</td><td>39</td><td>CGGCGGGTGTTGACGCGATGTGATTTCTGCCCAGTGCTC</td><td>160.0</td><td>-9.3</td><td>1.4</td><td>-6.7</td><td>1.9E-11</td><td>3.5E-11</td><td>BT20/MB468</td></tr>
<tr><td>>hg38_21_+_8433172_8446622_RNA45SN1_45S_50ntFlanks@10735.10752.18|TGTAAATCTCGCGCCGGG</td><td>>hg38_21_+_8433172_8446622_RNA45SN1_45S_50ntFlanks</td><td>10735</td><td>10752</td><td>18</td><td>TGTAAATCTCGCGCCGGG</td><td>135.9</td><td>-8.7</td><td>1.3</td><td>-6.7</td><td>1.6E-11</td><td>3.5E-11</td><td>MB231/MB468</td></tr>
<tr><td>>hg38_21_+_8433172_8446622_RNA45SN1_45S_50ntFlanks@10393.10420.28|CAGCAGTTGAACATGGGTCAGTCGGTCC</td><td>>hg38_21_+_8433172_8446622_RNA45SN1_45S_50ntFlanks</td><td>10393</td><td>10420</td><td>28</td><td>CAGCAGTTGAACATGGGTCAGTCGGTCC</td><td>157.7</td><td>-3.4</td><td>0.5</td><td>-6.7</td><td>1.6E-11</td><td>3.6E-11</td><td>MB231/MB468</td></tr>
<tr><td>>hg38_21_+_8433172_8446622_RNA45SN1_45S_50ntFlanks@11577.11601.25|TTAGAACTGGTGCGGACCAGGGGAA</td><td>>hg38_21_+_8433172_8446622_RNA45SN1_45S_50ntFlanks</td><td>11577</td><td>11601</td><td>25</td><td>TTAGAACTGGTGCGGACCAGGGGAA</td><td>237.9</td><td>-5.3</td><td>0.8</td><td>-6.7</td><td>1.6E-11</td><td>3.6E-11</td><td>MB231/MB468</td></tr>
<tr><td>>MI0000816|hsa-mir-335&WithFlank&7|+|130496105|130496210@58.78.21&[MIMAT0004703&hsa-miR-335-3p&offsets|0|-1;m-159&7|+|130496162|130496183&offsets|0|-1]|TTTTTCATTATTGCTCCTGAC</td><td>>MI0000816|hsa-mir-335&WithFlank&7|+|130496105|130496210</td><td>58</td><td>78</td><td>21</td><td>TTTTTCATTATTGCTCCTGAC</td><td>636.6</td><td>6.5</td><td>1.0</td><td>6.7</td><td>1.9E-11</td><td>3.6E-11</td><td>BT20/MB468</td></tr>
<tr><td>>hg38_21_+_8433172_8446622_RNA45SN1_45S_50ntFlanks@7990.8016.27|GACGTGGCGACCCGCTGAATTTAAGCA</td><td>>hg38_21_+_8433172_8446622_RNA45SN1_45S_50ntFlanks</td><td>7990</td><td>8016</td><td>27</td><td>GACGTGGCGACCCGCTGAATTTAAGCA</td><td>1171.4</td><td>7.3</td><td>1.1</td><td>6.7</td><td>1.6E-11</td><td>3.6E-11</td><td>MB231/MB468</td></tr>
<tr><td>>hg38_21_+_8433172_8446622_RNA45SN1_45S_50ntFlanks@10427.10457.31|ATGGGCGAGCGCCGTTCCGAAGGGACGGGCG</td><td>>hg38_21_+_8433172_8446622_RNA45SN1_45S_50ntFlanks</td><td>10427</td><td>10457</td><td>31</td><td>ATGGGCGAGCGCCGTTCCGAAGGGACGGGCG</td><td>138.8</td><td>-6.2</td><td>0.9</td><td>-6.7</td><td>1.6E-11</td><td>3.6E-11</td><td>MB231/MB468</td></tr>
<tr><td>>hg38_21_+_8433172_8446622_RNA45SN1_45S_50ntFlanks@10572.10596.25|AGAAGCCGGCGGGAGCCCCGGGGAG</td><td>>hg38_21_+_8433172_8446622_RNA45SN1_45S_50ntFlanks</td><td>10572</td><td>10596</td><td>25</td><td>AGAAGCCGGCGGGAGCCCCGGGGAG</td><td>143.6</td><td>-5.5</td><td>0.8</td><td>-6.7</td><td>1.9E-11</td><td>3.6E-11</td><td>BT20/MB468</td></tr>
<tr><td>>hg38_21_+_8433172_8446622_RNA45SN1_45S_50ntFlanks@10690.10711.22|CGTCCGGTGAGCTCTCGCTGGC</td><td>>hg38_21_+_8433172_8446622_RNA45SN1_45S_50ntFlanks</td><td>10690</td><td>10711</td><td>22</td><td>CGTCCGGTGAGCTCTCGCTGGC</td><td>103.5</td><td>-8.7</td><td>1.3</td><td>-6.7</td><td>1.9E-11</td><td>3.7E-11</td><td>BT20/MB468</td></tr>
<tr><td>>hg38_21_+_8433172_8446622_RNA45SN1_45S_50ntFlanks@11876.11904.29|AAGACCCTGTTGAGCTTGACTCTAGTCTG</td><td>>hg38_21_+_8433172_8446622_RNA45SN1_45S_50ntFlanks</td><td>11876</td><td>11904</td><td>29</td><td>AAGACCCTGTTGAGCTTGACTCTAGTCTG</td><td>152.0</td><td>-9.2</td><td>1.4</td><td>-6.7</td><td>1.9E-11</td><td>3.7E-11</td><td>BT20/MB468</td></tr>
<tr><td>>hg38_21_+_8433172_8446622_RNA45SN1_45S_50ntFlanks@12456.12479.24|CCAAGCGTTGGATTGTTCACCCAC</td><td>>hg38_21_+_8433172_8446622_RNA45SN1_45S_50ntFlanks</td><td>12456</td><td>12479</td><td>24</td><td>CCAAGCGTTGGATTGTTCACCCAC</td><td>550.5</td><td>-4.6</td><td>0.7</td><td>-6.7</td><td>1.7E-11</td><td>3.7E-11</td><td>MB231/MB468</td></tr>
<tr><td>>hg38_21_+_8433172_8446622_RNA45SN1_45S_50ntFlanks@11579.11606.28|AGAACTGGTGCGGACCAGGGGAATCCGA</td><td>>hg38_21_+_8433172_8446622_RNA45SN1_45S_50ntFlanks</td><td>11579</td><td>11606</td><td>28</td><td>AGAACTGGTGCGGACCAGGGGAATCCGA</td><td>151.4</td><td>-3.5</td><td>0.5</td><td>-6.7</td><td>1.9E-11</td><td>3.7E-11</td><td>BT20/MB468</td></tr>
<tr><td>>hg38_21_+_8433172_8446622_RNA45SN1_45S_50ntFlanks@10794.10826.33|CTGGCATGTTGGAACAATGTAGGTAAGGGAAGT</td><td>>hg38_21_+_8433172_8446622_RNA45SN1_45S_50ntFlanks</td><td>10794</td><td>10826</td><td>33</td><td>CTGGCATGTTGGAACAATGTAGGTAAGGGAAGT</td><td>267.6</td><td>-3.8</td><td>0.6</td><td>-6.7</td><td>1.9E-11</td><td>3.7E-11</td><td>BT20/MB468</td></tr>
<tr><td>>hg38_21_+_8433172_8446622_RNA45SN1_45S_50ntFlanks@4554.4572.19|CCGCAGCTAGGAATAATGG</td><td>>hg38_21_+_8433172_8446622_RNA45SN1_45S_50ntFlanks</td><td>4554</td><td>4572</td><td>19</td><td>CCGCAGCTAGGAATAATGG</td><td>106.7</td><td>-8.7</td><td>1.3</td><td>-6.7</td><td>1.9E-11</td><td>3.7E-11</td><td>BT20/MB468</td></tr>
<tr><td>>trna112_GlnCTG_1_-_147737382_147737453@31.50.20,trna131_GlnCTG_6_-_27759135_27759206@31.50.20,trna146_GlnCTG_6_-_27515531_27515602@31.50.20,trna15_GlnCTG_1_+_145963304_145963375@31.50.20,trna19_GlnCTG_1_+_147505038_147505109@31.50.20,trna1_GlnCTG_6_+_18836402_18836473@31.50.20,trna28_GlnCTG_1_+_149186125_149186196@31.50.20,trna3_GlnCTG_17_+_8023070_8023141@31.50.20,trna49_GlnCTG_6_+_27487308_27487379@31.50.20,trna7_GlnCTG_15_-_66161400_66161471@31.50.20,trna99_GlnCTG_6_-_28909378_28909449@31.50.20|ACTCTGAATCCAGCGATCCG</td><td>>trna112_GlnCTG_1_-_147737382_147737453</td><td>31</td><td>50</td><td>20</td><td>ACTCTGAATCCAGCGATCCG</td><td>159.3</td><td>-8.9</td><td>1.3</td><td>-6.7</td><td>1.7E-11</td><td>3.7E-11</td><td>MB231/MB468</td></tr>
<tr><td>>hg38_21_+_8433172_8446622_RNA45SN1_45S_50ntFlanks@12499.12517.19|GGTTTAGACCGTCGTGAGA</td><td>>hg38_21_+_8433172_8446622_RNA45SN1_45S_50ntFlanks</td><td>12499</td><td>12517</td><td>19</td><td>GGTTTAGACCGTCGTGAGA</td><td>93.3</td><td>-8.5</td><td>1.3</td><td>-6.7</td><td>2.0E-11</td><td>3.8E-11</td><td>BT20/MB468</td></tr>
<tr><td>>hg38_21_+_8433172_8446622_RNA45SN1_45S_50ntFlanks@9562.9592.31|TCCTGACGTGCAAATCGGTCGTCCGACCTGG</td><td>>hg38_21_+_8433172_8446622_RNA45SN1_45S_50ntFlanks</td><td>9562</td><td>9592</td><td>31</td><td>TCCTGACGTGCAAATCGGTCGTCCGACCTGG</td><td>126.1</td><td>-5.4</td><td>0.8</td><td>-6.7</td><td>2.0E-11</td><td>3.8E-11</td><td>BT20/MB468</td></tr>
<tr><td>>hg38_21_+_8433172_8446622_RNA45SN1_45S_50ntFlanks@11579.11603.25|AGAACTGGTGCGGACCAGGGGAATC</td><td>>hg38_21_+_8433172_8446622_RNA45SN1_45S_50ntFlanks</td><td>11579</td><td>11603</td><td>25</td><td>AGAACTGGTGCGGACCAGGGGAATC</td><td>2677.7</td><td>-4.5</td><td>0.7</td><td>-6.7</td><td>2.0E-11</td><td>3.8E-11</td><td>BT20/MB468</td></tr>
<tr><td>>MI0000746|hsa-mir-99b&WithFlank&19|+|51692606|51692687@13.35.23&[MIMAT0000689&hsa-miR-99b-5p&offsets|0|+1;m-10&19|+|51692618|51692639&offsets|0|+1]|CACCCGTAGAACCGACCTTGCGG</td><td>>MI0000746|hsa-mir-99b&WithFlank&19|+|51692606|51692687</td><td>13</td><td>35</td><td>23</td><td>CACCCGTAGAACCGACCTTGCGG</td><td>723.7</td><td>8.6</td><td>1.3</td><td>6.7</td><td>2.0E-11</td><td>3.8E-11</td><td>BT20/MB468</td></tr>
<tr><td>>hg38_21_+_8433172_8446622_RNA45SN1_45S_50ntFlanks@11728.11753.26|GTAACTATGACTCTCTTAAGGTAGCC</td><td>>hg38_21_+_8433172_8446622_RNA45SN1_45S_50ntFlanks</td><td>11728</td><td>11753</td><td>26</td><td>GTAACTATGACTCTCTTAAGGTAGCC</td><td>196.4</td><td>-9.6</td><td>1.4</td><td>-6.7</td><td>2.0E-11</td><td>3.8E-11</td><td>BT20/MB468</td></tr>
<tr><td>>hg38_21_+_8433172_8446622_RNA45SN1_45S_50ntFlanks@10351.10381.31|CGAGAACTTTGAAGGCCGAAGTGGAGAAGGG</td><td>>hg38_21_+_8433172_8446622_RNA45SN1_45S_50ntFlanks</td><td>10351</td><td>10381</td><td>31</td><td>CGAGAACTTTGAAGGCCGAAGTGGAGAAGGG</td><td>372.4</td><td>-3.1</td><td>0.5</td><td>-6.7</td><td>1.7E-11</td><td>3.8E-11</td><td>MB231/MB468</td></tr>
<tr><td>>hg38_21_+_8433172_8446622_RNA45SN1_45S_50ntFlanks@12407.12435.29|TTTTGATCCTTCGATGTCGGCTCTTCCTA</td><td>>hg38_21_+_8433172_8446622_RNA45SN1_45S_50ntFlanks</td><td>12407</td><td>12435</td><td>29</td><td>TTTTGATCCTTCGATGTCGGCTCTTCCTA</td><td>277.7</td><td>-3.2</td><td>0.5</td><td>-6.7</td><td>1.7E-11</td><td>3.8E-11</td><td>MB231/MB468</td></tr>
<tr><td>>hg38_21_+_8433172_8446622_RNA45SN1_45S_50ntFlanks@8381.8401.21|AAGAGGGCGTGAAACCGTTAA</td><td>>hg38_21_+_8433172_8446622_RNA45SN1_45S_50ntFlanks</td><td>8381</td><td>8401</td><td>21</td><td>AAGAGGGCGTGAAACCGTTAA</td><td>139.4</td><td>-6.2</td><td>0.9</td><td>-6.7</td><td>1.7E-11</td><td>3.8E-11</td><td>MB231/MB468</td></tr>
<tr><td>>am_MI0000786|hsa-mir-378a&WithFlank&5|+|149732819|149732896@49.70.22(+1A)&[MIMAT0000732&hsa-miR-378a-3p&offsets|0|0(+1A);m-44&5|+|149732867|149732888&offsets|0|0(+1A)]|ACTGGACTTGGAGTCAGAAGGCA</td><td>>am_MI0000786|hsa-mir-378a&WithFlank&5|+|149732819|149732896</td><td>49</td><td>70</td><td>22(+1A)</td><td>ACTGGACTTGGAGTCAGAAGGCA</td><td>2172.9</td><td>4.8</td><td>0.7</td><td>6.7</td><td>1.8E-11</td><td>3.9E-11</td><td>MB231/MB468</td></tr>
<tr><td>>MI0016887|hsa-mir-4521&WithFlank&17|+|8186939|8187010@10.34.25&[MIMAT0019058&hsa-miR-4521&offsets|0|+3;m-765&17|+|8186948|8186969&offsets|0|+3]|GCTAAGGAAGTCCTGTGCTCAGTTT</td><td>>MI0016887|hsa-mir-4521&WithFlank&17|+|8186939|8187010</td><td>10</td><td>34</td><td>25</td><td>GCTAAGGAAGTCCTGTGCTCAGTTT</td><td>2939.4</td><td>7.1</td><td>1.1</td><td>6.7</td><td>2.0E-11</td><td>3.9E-11</td><td>BT20/MB468</td></tr>
<tr><td>>hg38_21_+_8433172_8446622_RNA45SN1_45S_50ntFlanks@11703.11727.25|AATGAAGCGCGGGTAAACGGCGGGA</td><td>>hg38_21_+_8433172_8446622_RNA45SN1_45S_50ntFlanks</td><td>11703</td><td>11727</td><td>25</td><td>AATGAAGCGCGGGTAAACGGCGGGA</td><td>164.4</td><td>-3.6</td><td>0.5</td><td>-6.7</td><td>2.1E-11</td><td>3.9E-11</td><td>BT20/MB468</td></tr>
<tr><td>>MI0005756|hsa-mir-934&WithFlank&X|+|136550872|136550966@21.43.23(+1U)&[MIMAT0004977&hsa-miR-934&offsets|0|+1(+1U);m-943&X|+|136550892|136550913&offsets|0|+1(+1U)]|TGTCTACTACTGGAGACACTGGTT</td><td>>MI0005756|hsa-mir-934&WithFlank&X|+|136550872|136550966</td><td>21</td><td>43</td><td>23(+1U)</td><td>TGTCTACTACTGGAGACACTGGTT</td><td>542.4</td><td>7.9</td><td>1.2</td><td>6.7</td><td>2.1E-11</td><td>3.9E-11</td><td>BT20/MB468</td></tr>
<tr><td>>am_trnaMT_GlyTCC_MT_+_9991_10058@28.48.21|ACTTCCAATTAACTAGTTTTG</td><td>>am_trnaMT_GlyTCC_MT_+_9991_10058</td><td>28</td><td>48</td><td>21</td><td>ACTTCCAATTAACTAGTTTTG</td><td>115.9</td><td>-8.9</td><td>1.3</td><td>-6.7</td><td>2.1E-11</td><td>3.9E-11</td><td>BT20/MB468</td></tr>
<tr><td>>hg38_21_+_8433172_8446622_RNA45SN1_45S_50ntFlanks@10878.10902.25|TCGGTCGGGCTGGGGCGCGAAGCGG</td><td>>hg38_21_+_8433172_8446622_RNA45SN1_45S_50ntFlanks</td><td>10878</td><td>10902</td><td>25</td><td>TCGGTCGGGCTGGGGCGCGAAGCGG</td><td>162.5</td><td>-7.1</td><td>1.1</td><td>-6.7</td><td>1.8E-11</td><td>3.9E-11</td><td>MB231/MB468</td></tr>
<tr><td>>hg38_21_+_8433172_8446622_RNA45SN1_45S_50ntFlanks@11873.11894.22|AAGAAGACCCTGTTGAGCTTGA</td><td>>hg38_21_+_8433172_8446622_RNA45SN1_45S_50ntFlanks</td><td>11873</td><td>11894</td><td>22</td><td>AAGAAGACCCTGTTGAGCTTGA</td><td>114.1</td><td>-8.4</td><td>1.3</td><td>-6.7</td><td>1.8E-11</td><td>3.9E-11</td><td>MB231/MB468</td></tr>
<tr><td>>hg38_21_+_8433172_8446622_RNA45SN1_45S_50ntFlanks@10684.10706.23|CGGCGGCGTCCGGTGAGCTCTCG</td><td>>hg38_21_+_8433172_8446622_RNA45SN1_45S_50ntFlanks</td><td>10684</td><td>10706</td><td>23</td><td>CGGCGGCGTCCGGTGAGCTCTCG</td><td>4823.8</td><td>-6.7</td><td>1.0</td><td>-6.7</td><td>2.1E-11</td><td>3.9E-11</td><td>BT20/MB468</td></tr>
<tr><td>>hg38_21_+_8433172_8446622_RNA45SN1_45S_50ntFlanks@6721.6770.50|AATTGCAGGACACATTGATCATCGACACTTCGAACGCACTTGCGGCCCCG</td><td>>hg38_21_+_8433172_8446622_RNA45SN1_45S_50ntFlanks</td><td>6721</td><td>6770</td><td>50</td><td>AATTGCAGGACACATTGATCATCGACACTTCGAACGCACTTGCGGCCCCG</td><td>204.0</td><td>-9.7</td><td>1.4</td><td>-6.7</td><td>2.1E-11</td><td>3.9E-11</td><td>BT20/MB468</td></tr>
<tr><td>>hg38_1_-_228634819_228635039_RNA5S12_5S_50ntFlanks@126.170.45|TACTTGGATGGGAGACCGCCTGGGAATACCGGGTGCTGTAGGCTT</td><td>>hg38_1_-_228634819_228635039_RNA5S12_5S_50ntFlanks</td><td>126</td><td>170</td><td>45</td><td>TACTTGGATGGGAGACCGCCTGGGAATACCGGGTGCTGTAGGCTT</td><td>1821.0</td><td>8.7</td><td>1.3</td><td>6.7</td><td>1.8E-11</td><td>4.0E-11</td><td>MB231/MB468</td></tr>
<tr><td>>hg38_21_+_8433172_8446622_RNA45SN1_45S_50ntFlanks@8170.8187.18|CAAGTCCTTCTGATCGAG</td><td>>hg38_21_+_8433172_8446622_RNA45SN1_45S_50ntFlanks</td><td>8170</td><td>8187</td><td>18</td><td>CAAGTCCTTCTGATCGAG</td><td>685.9</td><td>-4.1</td><td>0.6</td><td>-6.7</td><td>1.8E-11</td><td>4.0E-11</td><td>MB231/MB468</td></tr>
<tr><td>>hg38_21_+_8433172_8446622_RNA45SN1_45S_50ntFlanks@11886.11903.18|TGAGCTTGACTCTAGTCT</td><td>>hg38_21_+_8433172_8446622_RNA45SN1_45S_50ntFlanks</td><td>11886</td><td>11903</td><td>18</td><td>TGAGCTTGACTCTAGTCT</td><td>176.6</td><td>-7.2</td><td>1.1</td><td>-6.7</td><td>1.8E-11</td><td>4.0E-11</td><td>MB231/MB468</td></tr>
</table>

<table>
<tr><td>>hg38_21_+_8433172_8446622_RNA45SN1_45S_50ntFlanks@10689.10711.23|GCGTCCGGTGAGCTCTCGCTGGC</td><td>>hg38_21_+_8433172_8446622_RNA45SN1_45S_50ntFlanks</td><td>10689</td><td>10711</td><td>23</td><td>GCGTCCGGTGAGCTCTCGCTGGC</td><td>104.5</td><td>-8.7</td><td>1.3</td><td>-6.7</td><td>2.1E-11</td><td>4.0E-11</td><td>BT20/MB468</td></tr>
<tr><td>>hg38_21_+_8433172_8446622_RNA45SN1_45S_50ntFlanks@12434.12455.22|TATCATTGTGAAGCAGAATTCA</td><td>>hg38_21_+_8433172_8446622_RNA45SN1_45S_50ntFlanks</td><td>12434</td><td>12455</td><td>22</td><td>TATCATTGTGAAGCAGAATTCA</td><td>228.6</td><td>-3.2</td><td>0.5</td><td>-6.7</td><td>1.8E-11</td><td>4.0E-11</td><td>MB231/MB468</td></tr>
<tr><td>>hg38_21_+_8433172_8446622_RNA45SN1_45S_50ntFlanks@13008.13027.20|ATTGAAAGTCAGCCCTCGAC</td><td>>hg38_21_+_8433172_8446622_RNA45SN1_45S_50ntFlanks</td><td>13008</td><td>13027</td><td>20</td><td>ATTGAAAGTCAGCCCTCGAC</td><td>688.1</td><td>-8.3</td><td>1.2</td><td>-6.7</td><td>2.2E-11</td><td>4.1E-11</td><td>BT20/MB468</td></tr>
<tr><td>>hg38_21_+_8433172_8446622_RNA45SN1_45S_50ntFlanks@12450.12476.27|AATTCACCAAGCGTTGGATTGTTCACC</td><td>>hg38_21_+_8433172_8446622_RNA45SN1_45S_50ntFlanks</td><td>12450</td><td>12476</td><td>27</td><td>AATTCACCAAGCGTTGGATTGTTCACC</td><td>493.7</td><td>6.9</td><td>1.0</td><td>6.7</td><td>1.9E-11</td><td>4.1E-11</td><td>MB231/MB468</td></tr>
<tr><td>>hg38_21_+_8433172_8446622_RNA45SN1_45S_50ntFlanks@10389.10406.18|TGAACAGCAGTTGAACAT</td><td>>hg38_21_+_8433172_8446622_RNA45SN1_45S_50ntFlanks</td><td>10389</td><td>10406</td><td>18</td><td>TGAACAGCAGTTGAACAT</td><td>114.7</td><td>-8.8</td><td>1.3</td><td>-6.7</td><td>2.2E-11</td><td>4.1E-11</td><td>BT20/MB468</td></tr>
<tr><td>>hg38_21_+_8433172_8446622_RNA45SN1_45S_50ntFlanks@12939.12966.28|CCATTCGTAGACGACCTGCTTCTGGGTC</td><td>>hg38_21_+_8433172_8446622_RNA45SN1_45S_50ntFlanks</td><td>12939</td><td>12966</td><td>28</td><td>CCATTCGTAGACGACCTGCTTCTGGGTC</td><td>122.7</td><td>-5.7</td><td>0.9</td><td>-6.7</td><td>2.2E-11</td><td>4.1E-11</td><td>BT20/MB468</td></tr>
<tr><td>>hg38_21_+_8433172_8446622_RNA45SN1_45S_50ntFlanks@10821.10838.18|GGAAGTCGGCAAGCCGGA</td><td>>hg38_21_+_8433172_8446622_RNA45SN1_45S_50ntFlanks</td><td>10821</td><td>10838</td><td>18</td><td>GGAAGTCGGCAAGCCGGA</td><td>498.4</td><td>-3.8</td><td>0.6</td><td>-6.7</td><td>1.9E-11</td><td>4.1E-11</td><td>MB231/MB468</td></tr>
<tr><td>>hg38_21_+_8433172_8446622_RNA45SN1_45S_50ntFlanks@11782.11804.23|GCATGAATGGATGAACGAGATTC</td><td>>hg38_21_+_8433172_8446622_RNA45SN1_45S_50ntFlanks</td><td>11782</td><td>11804</td><td>23</td><td>GCATGAATGGATGAACGAGATTC</td><td>175.4</td><td>-3.0</td><td>0.5</td><td>-6.7</td><td>2.2E-11</td><td>4.1E-11</td><td>BT20/MB468</td></tr>
<tr><td>>hg38_21_+_8433172_8446622_RNA45SN1_45S_50ntFlanks@11882.11900.19|CTGTTGAGCTTGACTCTAG</td><td>>hg38_21_+_8433172_8446622_RNA45SN1_45S_50ntFlanks</td><td>11882</td><td>11900</td><td>19</td><td>CTGTTGAGCTTGACTCTAG</td><td>210.6</td><td>-9.3</td><td>1.4</td><td>-6.7</td><td>1.9E-11</td><td>4.2E-11</td><td>MB231/MB468</td></tr>
<tr><td>>hg38_21_+_8433172_8446622_RNA45SN1_45S_50ntFlanks@11874.11899.26|AGAAGACCCTGTTGAGCTTGACTCTA</td><td>>hg38_21_+_8433172_8446622_RNA45SN1_45S_50ntFlanks</td><td>11874</td><td>11899</td><td>26</td><td>AGAAGACCCTGTTGAGCTTGACTCTA</td><td>293.5</td><td>-5.1</td><td>0.8</td><td>-6.7</td><td>1.9E-11</td><td>4.2E-11</td><td>MB231/MB468</td></tr>
<tr><td>>hg38_21_+_8433172_8446622_RNA45SN1_45S_50ntFlanks@7980.8008.29|CCTCAGATCAGACGTGGCGACCCGCTGAA</td><td>>hg38_21_+_8433172_8446622_RNA45SN1_45S_50ntFlanks</td><td>7980</td><td>8008</td><td>29</td><td>CCTCAGATCAGACGTGGCGACCCGCTGAA</td><td>686.6</td><td>-5.4</td><td>0.8</td><td>-6.7</td><td>2.2E-11</td><td>4.2E-11</td><td>BT20/MB468</td></tr>
<tr><td>>hg38_21_+_8433172_8446622_RNA45SN1_45S_50ntFlanks@5515.5533.19|CTAGAGGAAGTAAAAGTCG</td><td>>hg38_21_+_8433172_8446622_RNA45SN1_45S_50ntFlanks</td><td>5515</td><td>5533</td><td>19</td><td>CTAGAGGAAGTAAAAGTCG</td><td>103.0</td><td>-8.7</td><td>1.3</td><td>-6.7</td><td>2.2E-11</td><td>4.2E-11</td><td>BT20/MB468</td></tr>
<tr><td>>hg38_21_+_8433172_8446622_RNA45SN1_45S_50ntFlanks@11212.11232.21|CGACTCTGGACGCGAGCCGGG</td><td>>hg38_21_+_8433172_8446622_RNA45SN1_45S_50ntFlanks</td><td>11212</td><td>11232</td><td>21</td><td>CGACTCTGGACGCGAGCCGGG</td><td>1349.2</td><td>-6.6</td><td>1.0</td><td>-6.7</td><td>2.2E-11</td><td>4.2E-11</td><td>BT20/MB468</td></tr>
<tr><td>>hg38_21_+_8433172_8446622_RNA45SN1_45S_50ntFlanks@12474.12496.23|ACCCACTAATAGGGAACGTGAGC</td><td>>hg38_21_+_8433172_8446622_RNA45SN1_45S_50ntFlanks</td><td>12474</td><td>12496</td><td>23</td><td>ACCCACTAATAGGGAACGTGAGC</td><td>2870.3</td><td>4.0</td><td>0.6</td><td>6.7</td><td>1.9E-11</td><td>4.3E-11</td><td>MB231/MB468</td></tr>
<tr><td>>MI0000079|hsa-mir-23a&WithFlank&19|-|13836581|13836665@51.71.21&[MIMAT0000078&hsa-miR-23a-3p&offsets|0|0;m-31&19|-|13836595|13836615&offsets|0|0]|ATCACATTGCCAGGGATTTCC</td><td>>MI0000079|hsa-mir-23a&WithFlank&19|-|13836581|13836665</td><td>51</td><td>71</td><td>21</td><td>ATCACATTGCCAGGGATTTCC</td><td>127.5</td><td>-9.0</td><td>1.3</td><td>-6.7</td><td>2.3E-11</td><td>4.3E-11</td><td>BT20/MB468</td></tr>
<tr><td>>hg38_21_+_8433172_8446622_RNA45SN1_45S_50ntFlanks@11569.11602.34|CAGCCGACTTAGAACTGGTGCGGACCAGGGGAAT</td><td>>hg38_21_+_8433172_8446622_RNA45SN1_45S_50ntFlanks</td><td>11569</td><td>11602</td><td>34</td><td>CAGCCGACTTAGAACTGGTGCGGACCAGGGGAAT</td><td>1226.7</td><td>-3.7</td><td>0.6</td><td>-6.7</td><td>2.0E-11</td><td>4.3E-11</td><td>MB231/MB468</td></tr>
<tr><td>>MI0000085|hsa-mir-27a&WithFlank&19|-|13836434|13836523@57.74.18;MI0000440|hsa-mir-27b&WithFlank&9|+|95085439|95085547@67.84.18&[MIMAT0000084&hsa-miR-27a-3p&offsets|0|-3;m-43&19|-|13836447|13836467&offsets|0|-3];[MIMAT0000419&hsa-miR-27b-3p&offsets|0|-3;m-39&9|+|95085505|95085525&offsets|0|-3]|TTCACAGTGGCTAAGTTC</td><td>>MI0000085|hsa-mir-27a&WithFlank&19|-|13836434|13836523</td><td>57</td><td>74</td><td>18</td><td>TTCACAGTGGCTAAGTTC</td><td>696.4</td><td>5.8</td><td>0.9</td><td>6.7</td><td>2.3E-11</td><td>4.3E-11</td><td>BT20/MB468</td></tr>
<tr><td>>hg38_21_+_8433172_8446622_RNA45SN1_45S_50ntFlanks@9454.9503.50|CCGCGCCGGGGAGGTGGAGCACGAGCGCACGTGTTAGGACCCGAAAGATG</td><td>>hg38_21_+_8433172_8446622_RNA45SN1_45S_50ntFlanks</td><td>9454</td><td>9503</td><td>50</td><td>CCGCGCCGGGGAGGTGGAGCACGAGCGCACGTGTTAGGACCCGAAAGATG</td><td>192.6</td><td>-9.6</td><td>1.4</td><td>-6.7</td><td>2.3E-11</td><td>4.3E-11</td><td>BT20/MB468</td></tr>
<tr><td>>trna2_GlyGCC_21_-_18827107_18827177@1.29.29,trna35_GlyGCC_1_+_161413094_161413164@1.29.29,trna37_GlyGCC_1_+_161420467_161420537@1.29.29,trna39_GlyGCC_1_+_161427898_161427968@1.29.29,trna41_GlyGCC_1_+_161435258_161435328@1.29.29|GCATGGGTGGTTCAGTGGTAGAATTCTCG</td><td>>trna2_GlyGCC_21_-_18827107_18827177</td><td>1</td><td>29</td><td>29</td><td>GCATGGGTGGTTCAGTGGTAGAATTCTCG</td><td>11277.0</td><td>-4.0</td><td>0.6</td><td>-6.7</td><td>2.3E-11</td><td>4.3E-11</td><td>BT20/MB468</td></tr>
<tr><td>>hg38_21_+_8433172_8446622_RNA45SN1_45S_50ntFlanks@4370.4398.29|TTAAAAAGCTCGTAGTTGGATCTTGGGAG</td><td>>hg38_21_+_8433172_8446622_RNA45SN1_45S_50ntFlanks</td><td>4370</td><td>4398</td><td>29</td><td>TTAAAAAGCTCGTAGTTGGATCTTGGGAG</td><td>125.6</td><td>-8.5</td><td>1.3</td><td>-6.7</td><td>2.0E-11</td><td>4.3E-11</td><td>MB231/MB468</td></tr>
<tr><td>>hg38_21_+_8433172_8446622_RNA45SN1_45S_50ntFlanks@9736.9754.19|CCGAAACGATCTCAACCTA</td><td>>hg38_21_+_8433172_8446622_RNA45SN1_45S_50ntFlanks</td><td>9736</td><td>9754</td><td>19</td><td>CCGAAACGATCTCAACCTA</td><td>472.6</td><td>-6.7</td><td>1.0</td><td>-6.7</td><td>2.3E-11</td><td>4.4E-11</td><td>BT20/MB468</td></tr>
<tr><td>>hg38_21_+_8433172_8446622_RNA45SN1_45S_50ntFlanks@9504.9529.26|GTGAACTATGCCTGGGCAGGGCGAAG</td><td>>hg38_21_+_8433172_8446622_RNA45SN1_45S_50ntFlanks</td><td>9504</td><td>9529</td><td>26</td><td>GTGAACTATGCCTGGGCAGGGCGAAG</td><td>155.1</td><td>-8.9</td><td>1.3</td><td>-6.7</td><td>2.0E-11</td><td>4.4E-11</td><td>MB231/MB468</td></tr>
<tr><td>>MI0000077|hsa-mir-21&WithFlank&17|+|59841260|59841343@14.32.19(+1C)&[MIMAT0000076&hsa-miR-21-5p&offsets|0|-3(+1C);m-1&17|+|59841273|59841294&offsets|0|-3(+1C)]|TAGCTTATCAGACTGATGTC</td><td>>MI0000077|hsa-mir-21&WithFlank&17|+|59841260|59841343</td><td>14</td><td>32</td><td>19(+1C)</td><td>TAGCTTATCAGACTGATGTC</td><td>698.0</td><td>8.9</td><td>1.3</td><td>6.7</td><td>2.4E-11</td><td>4.5E-11</td><td>BT20/MB468</td></tr>
<tr><td>>hg38_21_+_8433172_8446622_RNA45SN1_45S_50ntFlanks@9714.9735.22|CGAATGATTAGAGGTCTTGGGG</td><td>>hg38_21_+_8433172_8446622_RNA45SN1_45S_50ntFlanks</td><td>9714</td><td>9735</td><td>22</td><td>CGAATGATTAGAGGTCTTGGGG</td><td>122.2</td><td>-8.5</td><td>1.3</td><td>-6.7</td><td>2.1E-11</td><td>4.5E-11</td><td>MB231/MB468</td></tr>
<tr><td>>MI0000102|hsa-mir-100&WithFlank&11|-|122152223|122152314@19.40.22(+2U)&[MIMAT0000098&hsa-miR-100-5p&offsets|0|0(+2U);m-27&11|-|122152276|122152296&offsets|0|+1(+2U)]|AACCCGTAGATCCGAACTTGTGTT</td><td>>MI0000102|hsa-mir-100&WithFlank&11|-|122152223|122152314</td><td>19</td><td>40</td><td>22(+2U)</td><td>AACCCGTAGATCCGAACTTGTGTT</td><td>644.1</td><td>-6.6</td><td>0.9</td><td>-7.1</td><td>1.5E-12</td><td>4.5E-11</td><td>BT20/MB231</td></tr>
<tr><td>>hg38_21_+_8433172_8446622_RNA45SN1_45S_50ntFlanks@12634.12653.20|GAAGCTACCATCTGTGGGAT</td><td>>hg38_21_+_8433172_8446622_RNA45SN1_45S_50ntFlanks</td><td>12634</td><td>12653</td><td>20</td><td>GAAGCTACCATCTGTGGGAT</td><td>171.9</td><td>-9.0</td><td>1.3</td><td>-6.7</td><td>2.1E-11</td><td>4.6E-11</td><td>MB231/MB468</td></tr>
<tr><td>>MI0000782|hsa-mir-374a&WithFlank&X|-|74287280|74287363@48.69.22&[MIMAT0004688&hsa-miR-374a-3p&offsets|0|0;m-74&X|-|74287295|74287316&offsets|0|0]|CTTATCAGATTGTATTGTAATT</td><td>>MI0000782|hsa-mir-374a&WithFlank&X|-|74287280|74287363</td><td>48</td><td>69</td><td>22</td><td>CTTATCAGATTGTATTGTAATT</td><td>6710.0</td><td>8.3</td><td>1.2</td><td>6.7</td><td>2.4E-11</td><td>4.6E-11</td><td>BT20/MB468</td></tr>
<tr><td>>hg38_21_+_8433172_8446622_RNA45SN1_45S_50ntFlanks@13006.13027.22|CTATTGAAAGTCAGCCCTCGAC</td><td>>hg38_21_+_8433172_8446622_RNA45SN1_45S_50ntFlanks</td><td>13006</td><td>13027</td><td>22</td><td>CTATTGAAAGTCAGCCCTCGAC</td><td>707.7</td><td>-6.8</td><td>1.0</td><td>-6.7</td><td>2.1E-11</td><td>4.6E-11</td><td>MB231/MB468</td></tr>
<tr><td>>hg38_21_+_8433172_8446622_RNA45SN1_45S_50ntFlanks@12407.12440.34|TTTTGATCCTTCGATGTCGGCTCTTCCTATCATT</td><td>>hg38_21_+_8433172_8446622_RNA45SN1_45S_50ntFlanks</td><td>12407</td><td>12440</td><td>34</td><td>TTTTGATCCTTCGATGTCGGCTCTTCCTATCATT</td><td>338.0</td><td>4.3</td><td>0.6</td><td>6.7</td><td>2.1E-11</td><td>4.6E-11</td><td>MB231/MB468</td></tr>
<tr><td>>hg38_21_+_8433172_8446622_RNA45SN1_45S_50ntFlanks@4761.4780.20|CAGATACCGTCGTAGTTCCG</td><td>>hg38_21_+_8433172_8446622_RNA45SN1_45S_50ntFlanks</td><td>4761</td><td>4780</td><td>20</td><td>CAGATACCGTCGTAGTTCCG</td><td>189.7</td><td>-5.4</td><td>0.8</td><td>-6.7</td><td>2.1E-11</td><td>4.6E-11</td><td>MB231/MB468</td></tr>
<tr><td>>hg38_21_+_8433172_8446622_RNA45SN1_45S_50ntFlanks@10905.10936.32|CTGGGCGCGCGCCGCGGCTGGACGAGGCGCCG</td><td>>hg38_21_+_8433172_8446622_RNA45SN1_45S_50ntFlanks</td><td>10905</td><td>10936</td><td>32</td><td>CTGGGCGCGCGCCGCGGCTGGACGAGGCGCCG</td><td>151.2</td><td>-8.8</td><td>1.3</td><td>-6.7</td><td>2.1E-11</td><td>4.7E-11</td><td>MB231/MB468</td></tr>
<tr><td>>hg38_21_+_8433172_8446622_RNA45SN1_45S_50ntFlanks@12569.12587.19|CAGTACGAGAGGAACCGCA</td><td>>hg38_21_+_8433172_8446622_RNA45SN1_45S_50ntFlanks</td><td>12569</td><td>12587</td><td>19</td><td>CAGTACGAGAGGAACCGCA</td><td>146.1</td><td>-3.2</td><td>0.5</td><td>-6.7</td><td>2.5E-11</td><td>4.7E-11</td><td>BT20/MB468</td></tr>
<tr><td>>hg38_21_+_8433172_8446622_RNA45SN1_45S_50ntFlanks@6667.6689.23|ATCACTCGGCTCGTGCGTCGATG</td><td>>hg38_21_+_8433172_8446622_RNA45SN1_45S_50ntFlanks</td><td>6667</td><td>6689</td><td>23</td><td>ATCACTCGGCTCGTGCGTCGATG</td><td>88.0</td><td>-8.5</td><td>1.3</td><td>-6.7</td><td>2.5E-11</td><td>4.7E-11</td><td>BT20/MB468</td></tr>
<tr><td>>hg38_21_+_8433172_8446622_RNA45SN1_45S_50ntFlanks@11729.11754.26|TAACTATGACTCTCTTAAGGTAGCCA</td><td>>hg38_21_+_8433172_8446622_RNA45SN1_45S_50ntFlanks</td><td>11729</td><td>11754</td><td>26</td><td>TAACTATGACTCTCTTAAGGTAGCCA</td><td>262.1</td><td>-10.0</td><td>1.5</td><td>-6.7</td><td>2.5E-11</td><td>4.7E-11</td><td>BT20/MB468</td></tr>
<tr><td>>hg38_21_+_8433172_8446622_RNA45SN1_45S_50ntFlanks@10786.10828.43|AACAGCCTCTGGCATGTTGGAACAATGTAGGTAAGGGAAGTCG</td><td>>hg38_21_+_8433172_8446622_RNA45SN1_45S_50ntFlanks</td><td>10786</td><td>10828</td><td>43</td><td>AACAGCCTCTGGCATGTTGGAACAATGTAGGTAAGGGAAGTCG</td><td>101.1</td><td>-8.7</td><td>1.3</td><td>-6.7</td><td>2.5E-11</td><td>4.8E-11</td><td>BT20/MB468</td></tr>
<tr><td>>hg38_21_+_8433172_8446622_RNA45SN1_45S_50ntFlanks@4367.4398.32|CAGTTAAAAAGCTCGTAGTTGGATCTTGGGAG</td><td>>hg38_21_+_8433172_8446622_RNA45SN1_45S_50ntFlanks</td><td>4367</td><td>4398</td><td>32</td><td>CAGTTAAAAAGCTCGTAGTTGGATCTTGGGAG</td><td>291.1</td><td>-7.0</td><td>1.0</td><td>-6.7</td><td>2.2E-11</td><td>4.8E-11</td><td>MB231/MB468</td></tr>
<tr><td>>hg38_21_+_8433172_8446622_RNA45SN1_45S_50ntFlanks@8170.8196.27|CAAGTCCTTCTGATCGAGGCCCAGCCC</td><td>>hg38_21_+_8433172_8446622_RNA45SN1_45S_50ntFlanks</td><td>8170</td><td>8196</td><td>27</td><td>CAAGTCCTTCTGATCGAGGCCCAGCCC</td><td>91.7</td><td>-3.6</td><td>0.5</td><td>-6.7</td><td>2.5E-11</td><td>4.8E-11</td><td>BT20/MB468</td></tr>
<tr><td>>hg38_21_+_8433172_8446622_RNA45SN1_45S_50ntFlanks@10423.10457.35|AGAGATGGGCGAGCGCCGTTCCGAAGGGACGGGCG</td><td>>hg38_21_+_8433172_8446622_RNA45SN1_45S_50ntFlanks</td><td>10423</td><td>10457</td><td>35</td><td>AGAGATGGGCGAGCGCCGTTCCGAAGGGACGGGCG</td><td>122.3</td><td>-5.2</td><td>0.8</td><td>-6.7</td><td>2.2E-11</td><td>4.8E-11</td><td>MB231/MB468</td></tr>
<tr><td>>hg38_21_+_8433172_8446622_RNA45SN1_45S_50ntFlanks@11570.11601.32|AGCCGACTTAGAACTGGTGCGGACCAGGGGAA</td><td>>hg38_21_+_8433172_8446622_RNA45SN1_45S_50ntFlanks</td><td>11570</td><td>11601</td><td>32</td><td>AGCCGACTTAGAACTGGTGCGGACCAGGGGAA</td><td>571.8</td><td>-3.5</td><td>0.5</td><td>-6.7</td><td>2.6E-11</td><td>4.9E-11</td><td>BT20/MB468</td></tr>
<tr><td>>hg38_21_+_8433172_8446622_RNA45SN1_45S_50ntFlanks@5420.5444.25|CCGATTGGATGGTTTAGTGAGGCCC</td><td>>hg38_21_+_8433172_8446622_RNA45SN1_45S_50ntFlanks</td><td>5420</td><td>5444</td><td>25</td><td>CCGATTGGATGGTTTAGTGAGGCCC</td><td>161.7</td><td>-7.1</td><td>1.1</td><td>-6.7</td><td>2.2E-11</td><td>4.9E-11</td><td>MB231/MB468</td></tr>
<tr><td>>hg38_21_+_8433172_8446622_RNA45SN1_45S_50ntFlanks@12420.12448.29|ATGTCGGCTCTTCCTATCATTGTGAAGCA</td><td>>hg38_21_+_8433172_8446622_RNA45SN1_45S_50ntFlanks</td><td>12420</td><td>12448</td><td>29</td><td>ATGTCGGCTCTTCCTATCATTGTGAAGCA</td><td>2342.0</td><td>4.5</td><td>0.7</td><td>6.7</td><td>2.2E-11</td><td>4.9E-11</td><td>MB231/MB468</td></tr>
<tr><td>>am_trna10_LysCTT_16_+_3241501_3241573@32.51.20,trna119_LysCTT_1_-_145395522_145395594@32.51.20,trna11_LysCTT_5_-_180648979_180649051@32.51.20,trna13_LysCTT_16_+_3246153_3246225@32.51.20,trna13_LysCTT_6_+_26556774_26556846@32.51.20,trna30_LysCTT_16_-_3230555_3230627@32.51.20,trna5_LysCTT_16_+_3214939_3215011@32.51.20,trna7_LysCTT_16_+_3225692_3225764@32.51.20,trna8_LysCTT_1_+_39970195_39970267@32.51.20,trna9_LysCTT_5_+_180634755_180634827@32.51.20|CTCTTAATCTCAGGGTCGTG</td><td>>am_trna10_LysCTT_16_+_3241501_3241573</td><td>32</td><td>51</td><td>20</td><td>CTCTTAATCTCAGGGTCGTG</td><td>139.3</td><td>-8.7</td><td>1.3</td><td>-6.7</td><td>2.3E-11</td><td>4.9E-11</td><td>MB231/MB468</td></tr>
<tr><td>>hg38_21_+_8433172_8446622_RNA45SN1_45S_50ntFlanks@12399.12433.35|ACGTCGCTTTTTGATCCTTCGATGTCGGCTCTTCC</td><td>>hg38_21_+_8433172_8446622_RNA45SN1_45S_50ntFlanks</td><td>12399</td><td>12433</td><td>35</td><td>ACGTCGCTTTTTGATCCTTCGATGTCGGCTCTTCC</td><td>545.0</td><td>5.4</td><td>0.8</td><td>6.7</td><td>2.3E-11</td><td>4.9E-11</td><td>MB231/MB468</td></tr>
<tr><td>>hg38_21_+_8433172_8446622_RNA45SN1_45S_50ntFlanks@12874.12891.18|GCTGGAAAGGCGGCCGCC</td><td>>hg38_21_+_8433172_8446622_RNA45SN1_45S_50ntFlanks</td><td>12874</td><td>12891</td><td>18</td><td>GCTGGAAAGGCGGCCGCC</td><td>151.6</td><td>-8.8</td><td>1.3</td><td>-6.7</td><td>2.3E-11</td><td>5.0E-11</td><td>MB231/MB468</td></tr>
</table>

<table>
<tr><td>>hg38_21_+_8433172_8446622_RNA45SN1_45S_50ntFlanks@4657.4686.30|CTAGAGGTGAAATTCTTGGACCGGCGCAAG</td><td>>hg38_21_+_8433172_8446622_RNA45SN1_45S_50ntFlanks</td><td>4657</td><td>4686</td><td>30</td><td>CTAGAGGTGAAATTCTTGGACCGGCGCAAG</td><td>130.0</td><td>-9.0</td><td>1.4</td><td>-6.7</td><td>2.7E-11</td><td>5.0E-11</td><td>BT20/MB468</td></tr>
<tr><td>>hg38_21_+_8433172_8446622_RNA45SN1_45S_50ntFlanks@11076.11103.28|CGCCGCGTGGGGGCGGCGGCGGGGGGAG</td><td>>hg38_21_+_8433172_8446622_RNA45SN1_45S_50ntFlanks</td><td>11076</td><td>11103</td><td>28</td><td>CGCCGCGTGGGGGCGGCGGCGGGGGGAG</td><td>122.1</td><td>-8.9</td><td>1.3</td><td>-6.7</td><td>2.7E-11</td><td>5.0E-11</td><td>BT20/MB468</td></tr>
<tr><td>>hg38_21_+_8433172_8446622_RNA45SN1_45S_50ntFlanks@9554.9576.23|CGTAGCGGTCCTGACGTGCAAAT</td><td>>hg38_21_+_8433172_8446622_RNA45SN1_45S_50ntFlanks</td><td>9554</td><td>9576</td><td>23</td><td>CGTAGCGGTCCTGACGTGCAAAT</td><td>2350.6</td><td>4.3</td><td>0.6</td><td>6.7</td><td>2.3E-11</td><td>5.0E-11</td><td>MB231/MB468</td></tr>
<tr><td>>hg38_21_+_8433172_8446622_RNA45SN1_45S_50ntFlanks@10440.10460.21|GTTCCGAAGGGACGGGCGATG</td><td>>hg38_21_+_8433172_8446622_RNA45SN1_45S_50ntFlanks</td><td>10440</td><td>10460</td><td>21</td><td>GTTCCGAAGGGACGGGCGATG</td><td>336.1</td><td>4.1</td><td>0.6</td><td>6.7</td><td>2.3E-11</td><td>5.0E-11</td><td>MB231/MB468</td></tr>
<tr><td>>MI0000085|hsa-mir-27a&WithFlank&19|-|13836434|13836523@16.37.22&[MIMAT0004501&hsa-miR-27a-5p&offsets|0|0;m-270&19|-|13836487|13836508&offsets|0|0]|AGGGCTTAGCTGCTTGTGAGCA</td><td>>MI0000085|hsa-mir-27a&WithFlank&19|-|13836434|13836523</td><td>16</td><td>37</td><td>22</td><td>AGGGCTTAGCTGCTTGTGAGCA</td><td>1119.8</td><td>6.7</td><td>1.0</td><td>6.7</td><td>2.3E-11</td><td>5.0E-11</td><td>MB231/MB468</td></tr>
<tr><td>>hg38_21_+_8433172_8446622_RNA45SN1_45S_50ntFlanks@8157.8178.22|CTCGTGGGGGGCCCAAGTCCTT</td><td>>hg38_21_+_8433172_8446622_RNA45SN1_45S_50ntFlanks</td><td>8157</td><td>8178</td><td>22</td><td>CTCGTGGGGGGCCCAAGTCCTT</td><td>161.5</td><td>-8.9</td><td>1.3</td><td>-6.7</td><td>2.3E-11</td><td>5.1E-11</td><td>MB231/MB468</td></tr>
<tr><td>>hg38_21_+_8433172_8446622_RNA45SN1_45S_50ntFlanks@13009.13028.20|TTGAAAGTCAGCCCTCGACA</td><td>>hg38_21_+_8433172_8446622_RNA45SN1_45S_50ntFlanks</td><td>13009</td><td>13028</td><td>20</td><td>TTGAAAGTCAGCCCTCGACA</td><td>187.9</td><td>-9.5</td><td>1.4</td><td>-6.7</td><td>2.7E-11</td><td>5.1E-11</td><td>BT20/MB468</td></tr>
<tr><td>>trnaMT_LeuTAA_MT_+_3230_3304@19.44.26|GGTAATCGCATAAAACTTAAAACTTT</td><td>>trnaMT_LeuTAA_MT_+_3230_3304</td><td>19</td><td>44</td><td>26</td><td>GGTAATCGCATAAAACTTAAAACTTT</td><td>299.6</td><td>4.7</td><td>0.7</td><td>6.7</td><td>2.7E-11</td><td>5.1E-11</td><td>BT20/MB468</td></tr>
<tr><td>>hg38_21_+_8433172_8446622_RNA45SN1_45S_50ntFlanks@11594.11613.20|CAGGGGAATCCGACTGTTTA</td><td>>hg38_21_+_8433172_8446622_RNA45SN1_45S_50ntFlanks</td><td>11594</td><td>11613</td><td>20</td><td>CAGGGGAATCCGACTGTTTA</td><td>235.0</td><td>-2.8</td><td>0.4</td><td>-6.7</td><td>2.7E-11</td><td>5.1E-11</td><td>BT20/MB468</td></tr>
<tr><td>>hg38_21_+_8433172_8446622_RNA45SN1_45S_50ntFlanks@3777.3799.23|CGGTACAGTGAAACTGCGAATGG</td><td>>hg38_21_+_8433172_8446622_RNA45SN1_45S_50ntFlanks</td><td>3777</td><td>3799</td><td>23</td><td>CGGTACAGTGAAACTGCGAATGG</td><td>76.7</td><td>-3.9</td><td>0.6</td><td>-6.7</td><td>2.7E-11</td><td>5.1E-11</td><td>BT20/MB468</td></tr>
<tr><td>>hg38_21_+_8433172_8446622_RNA45SN1_45S_50ntFlanks@9574.9602.29|AATCGGTCGTCCGACCTGGGTATAGGGGC</td><td>>hg38_21_+_8433172_8446622_RNA45SN1_45S_50ntFlanks</td><td>9574</td><td>9602</td><td>29</td><td>AATCGGTCGTCCGACCTGGGTATAGGGGC</td><td>1228.6</td><td>4.3</td><td>0.6</td><td>6.7</td><td>2.4E-11</td><td>5.1E-11</td><td>MB231/MB468</td></tr>
<tr><td>>hg38_21_+_8433172_8446622_RNA45SN1_45S_50ntFlanks@11739.11767.29|TCTCTTAAGGTAGCCAAATGCCTCGTCAT</td><td>>hg38_21_+_8433172_8446622_RNA45SN1_45S_50ntFlanks</td><td>11739</td><td>11767</td><td>29</td><td>TCTCTTAAGGTAGCCAAATGCCTCGTCAT</td><td>128.3</td><td>-8.6</td><td>1.3</td><td>-6.7</td><td>2.4E-11</td><td>5.1E-11</td><td>MB231/MB468</td></tr>
<tr><td>>hg38_21_+_8433172_8446622_RNA45SN1_45S_50ntFlanks@10597.10617.21|AGTTCTCTTTTCTTTGTGAAG</td><td>>hg38_21_+_8433172_8446622_RNA45SN1_45S_50ntFlanks</td><td>10597</td><td>10617</td><td>21</td><td>AGTTCTCTTTTCTTTGTGAAG</td><td>117.3</td><td>-8.5</td><td>1.3</td><td>-6.7</td><td>2.4E-11</td><td>5.2E-11</td><td>MB231/MB468</td></tr>
<tr><td>>hg38_21_+_8433172_8446622_RNA45SN1_45S_50ntFlanks@6647.6680.34|CGTACGACTCTTAGCGGTGGATCACTCGGCTCGT</td><td>>hg38_21_+_8433172_8446622_RNA45SN1_45S_50ntFlanks</td><td>6647</td><td>6680</td><td>34</td><td>CGTACGACTCTTAGCGGTGGATCACTCGGCTCGT</td><td>932.9</td><td>3.0</td><td>0.4</td><td>6.7</td><td>2.8E-11</td><td>5.2E-11</td><td>BT20/MB468</td></tr>
<tr><td>>hg38_21_+_8433172_8446622_RNA45SN1_45S_50ntFlanks@12162.12181.20|GACTGGGGCGGTACACCTGT</td><td>>hg38_21_+_8433172_8446622_RNA45SN1_45S_50ntFlanks</td><td>12162</td><td>12181</td><td>20</td><td>GACTGGGGCGGTACACCTGT</td><td>167.9</td><td>-6.2</td><td>0.9</td><td>-6.7</td><td>2.8E-11</td><td>5.2E-11</td><td>BT20/MB468</td></tr>
<tr><td>>MI0003129|hsa-mir-146b&WithFlank&10|+|102436506|102436590@15.38.24&[MIMAT0002809&hsa-miR-146b-5p&offsets|0|+1;m-70&10|+|102436520|102436543&offsets|0|0]|TGAGAACTGAATTCCATAGGCTGT</td><td>>MI0003129|hsa-mir-146b&WithFlank&10|+|102436506|102436590</td><td>15</td><td>38</td><td>24</td><td>TGAGAACTGAATTCCATAGGCTGT</td><td>1063.9</td><td>5.9</td><td>0.9</td><td>6.7</td><td>2.4E-11</td><td>5.2E-11</td><td>MB231/MB468</td></tr>
<tr><td>>hg38_21_+_8433172_8446622_RNA45SN1_45S_50ntFlanks@7977.8011.35|CGACCTCAGATCAGACGTGGCGACCCGCTGAATTT</td><td>>hg38_21_+_8433172_8446622_RNA45SN1_45S_50ntFlanks</td><td>7977</td><td>8011</td><td>35</td><td>CGACCTCAGATCAGACGTGGCGACCCGCTGAATTT</td><td>503.9</td><td>4.5</td><td>0.7</td><td>6.7</td><td>2.8E-11</td><td>5.2E-11</td><td>BT20/MB468</td></tr>
<tr><td>>hg38_21_+_8433172_8446622_RNA45SN1_45S_50ntFlanks@12510.12538.29|TCGTGAGACAGGTTAGTTTTACCCTACTG</td><td>>hg38_21_+_8433172_8446622_RNA45SN1_45S_50ntFlanks</td><td>12510</td><td>12538</td><td>29</td><td>TCGTGAGACAGGTTAGTTTTACCCTACTG</td><td>112.9</td><td>-8.8</td><td>1.3</td><td>-6.7</td><td>2.8E-11</td><td>5.3E-11</td><td>BT20/MB468</td></tr>
<tr><td>>trna116_GluCTC_1_-_145399233_145399304@32.72.41,trna71_GluCTC_1_-_161439189_161439260@32.72.41,trna74_GluCTC_1_-_161431809_161431880@32.72.41,trna77_GluCTC_1_-_161424398_161424469@32.72.41,trna77_GluCTC_6_+_28949976_28950047@32.72.41,trna80_GluCTC_1_-_161417018_161417089@32.72.41,trna87_GluCTC_6_-_126101393_126101464@32.72.41|CTCTCACCGCCGCGGCCCGGGTTCGATTCCCGGTCAGGGAA</td><td>>trna116_GluCTC_1_-_145399233_145399304</td><td>32</td><td>72</td><td>41</td><td>CTCTCACCGCCGCGGCCCGGGTTCGATTCCCGGTCAGGGAA</td><td>95.3</td><td>-8.6</td><td>1.3</td><td>-6.7</td><td>2.8E-11</td><td>5.3E-11</td><td>BT20/MB468</td></tr>
<tr><td>>hg38_21_+_8433172_8446622_RNA45SN1_45S_50ntFlanks@12406.12431.26|TTTTTGATCCTTCGATGTCGGCTCTT</td><td>>hg38_21_+_8433172_8446622_RNA45SN1_45S_50ntFlanks</td><td>12406</td><td>12431</td><td>26</td><td>TTTTTGATCCTTCGATGTCGGCTCTT</td><td>271.5</td><td>-4.2</td><td>0.6</td><td>-6.7</td><td>2.8E-11</td><td>5.3E-11</td><td>BT20/MB468</td></tr>
<tr><td>>hg38_21_+_8433172_8446622_RNA45SN1_45S_50ntFlanks@10781.10801.21|AGGTGAACAGCCTCTGGCATG</td><td>>hg38_21_+_8433172_8446622_RNA45SN1_45S_50ntFlanks</td><td>10781</td><td>10801</td><td>21</td><td>AGGTGAACAGCCTCTGGCATG</td><td>700.1</td><td>-4.4</td><td>0.7</td><td>-6.7</td><td>2.8E-11</td><td>5.3E-11</td><td>BT20/MB468</td></tr>
<tr><td>>hg38_21_+_8433172_8446622_RNA45SN1_45S_50ntFlanks@9534.9561.28|AGGAAACTCTGGTGGAGGTCCGTAGCGG</td><td>>hg38_21_+_8433172_8446622_RNA45SN1_45S_50ntFlanks</td><td>9534</td><td>9561</td><td>28</td><td>AGGAAACTCTGGTGGAGGTCCGTAGCGG</td><td>1692.9</td><td>-7.8</td><td>1.2</td><td>-6.7</td><td>2.9E-11</td><td>5.4E-11</td><td>BT20/MB468</td></tr>
<tr><td>>hg38_21_+_8433172_8446622_RNA45SN1_45S_50ntFlanks@10388.10409.22|GTGAACAGCAGTTGAACATGGG</td><td>>hg38_21_+_8433172_8446622_RNA45SN1_45S_50ntFlanks</td><td>10388</td><td>10409</td><td>22</td><td>GTGAACAGCAGTTGAACATGGG</td><td>114.3</td><td>-8.8</td><td>1.3</td><td>-6.7</td><td>2.9E-11</td><td>5.4E-11</td><td>BT20/MB468</td></tr>
<tr><td>>hg38_21_+_8433172_8446622_RNA45SN1_45S_50ntFlanks@12158.12180.23|GTTTGACTGGGGCGGTACACCTG</td><td>>hg38_21_+_8433172_8446622_RNA45SN1_45S_50ntFlanks</td><td>12158</td><td>12180</td><td>23</td><td>GTTTGACTGGGGCGGTACACCTG</td><td>107.6</td><td>-8.7</td><td>1.3</td><td>-6.7</td><td>2.9E-11</td><td>5.4E-11</td><td>BT20/MB468</td></tr>
<tr><td>>hg38_21_+_8433172_8446622_RNA45SN1_45S_50ntFlanks@11561.11591.31|GCGCCTAGCAGCCGACTTAGAACTGGTGCGG</td><td>>hg38_21_+_8433172_8446622_RNA45SN1_45S_50ntFlanks</td><td>11561</td><td>11591</td><td>31</td><td>GCGCCTAGCAGCCGACTTAGAACTGGTGCGG</td><td>1170.4</td><td>-7.1</td><td>1.1</td><td>-6.7</td><td>2.9E-11</td><td>5.4E-11</td><td>BT20/MB468</td></tr>
<tr><td>>hg38_21_+_8433172_8446622_RNA45SN1_45S_50ntFlanks@10830.10858.29|CAAGCCGGATCCGTAACTTCGGGATAAGG</td><td>>hg38_21_+_8433172_8446622_RNA45SN1_45S_50ntFlanks</td><td>10830</td><td>10858</td><td>29</td><td>CAAGCCGGATCCGTAACTTCGGGATAAGG</td><td>252.4</td><td>-5.1</td><td>0.8</td><td>-6.7</td><td>2.5E-11</td><td>5.4E-11</td><td>MB231/MB468</td></tr>
<tr><td>>hg38_21_+_8433172_8446622_RNA45SN1_45S_50ntFlanks@12002.12032.31|TGCGGGCCGCCGGTGAAATACCACTACTCTG</td><td>>hg38_21_+_8433172_8446622_RNA45SN1_45S_50ntFlanks</td><td>12002</td><td>12032</td><td>31</td><td>TGCGGGCCGCCGGTGAAATACCACTACTCTG</td><td>2263.5</td><td>-4.9</td><td>0.7</td><td>-6.7</td><td>2.5E-11</td><td>5.5E-11</td><td>MB231/MB468</td></tr>
<tr><td>>MI0000077|hsa-mir-21&WithFlank&17|+|59841260|59841343@13.34.22&[MIMAT0000076&hsa-miR-21-5p&offsets|-1|-1;m-1&17|+|59841273|59841294&offsets|-1|-1]|GTAGCTTATCAGACTGATGTTG</td><td>>MI0000077|hsa-mir-21&WithFlank&17|+|59841260|59841343</td><td>13</td><td>34</td><td>22</td><td>GTAGCTTATCAGACTGATGTTG</td><td>417.0</td><td>5.9</td><td>0.9</td><td>6.6</td><td>3.0E-11</td><td>5.5E-11</td><td>BT20/MB468</td></tr>
<tr><td>>hg38_21_+_8433172_8446622_RNA45SN1_45S_50ntFlanks@3872.3898.27|CTAATACATGCCGACGGGCGCTGACCC</td><td>>hg38_21_+_8433172_8446622_RNA45SN1_45S_50ntFlanks</td><td>3872</td><td>3898</td><td>27</td><td>CTAATACATGCCGACGGGCGCTGACCC</td><td>486.9</td><td>-5.4</td><td>0.8</td><td>-6.7</td><td>2.6E-11</td><td>5.5E-11</td><td>MB231/MB468</td></tr>
<tr><td>>hg38_21_+_8433172_8446622_RNA45SN1_45S_50ntFlanks@10869.10886.18|AGGGCTGGGTCGGTCGGG</td><td>>hg38_21_+_8433172_8446622_RNA45SN1_45S_50ntFlanks</td><td>10869</td><td>10886</td><td>18</td><td>AGGGCTGGGTCGGTCGGG</td><td>176.2</td><td>-9.0</td><td>1.4</td><td>-6.7</td><td>2.6E-11</td><td>5.6E-11</td><td>MB231/MB468</td></tr>
<tr><td>>hg38_21_+_8433172_8446622_RNA45SN1_45S_50ntFlanks@11881.11900.20|CCTGTTGAGCTTGACTCTAG</td><td>>hg38_21_+_8433172_8446622_RNA45SN1_45S_50ntFlanks</td><td>11881</td><td>11900</td><td>20</td><td>CCTGTTGAGCTTGACTCTAG</td><td>129.6</td><td>-8.6</td><td>1.3</td><td>-6.7</td><td>2.6E-11</td><td>5.6E-11</td><td>MB231/MB468</td></tr>
<tr><td>>hg38_21_+_8433172_8446622_RNA45SN1_45S_50ntFlanks@10463.10483.21|CTCCGTTGCCCTCGGCCGATC</td><td>>hg38_21_+_8433172_8446622_RNA45SN1_45S_50ntFlanks</td><td>10463</td><td>10483</td><td>21</td><td>CTCCGTTGCCCTCGGCCGATC</td><td>143.6</td><td>-4.5</td><td>0.7</td><td>-6.6</td><td>3.0E-11</td><td>5.6E-11</td><td>BT20/MB468</td></tr>
<tr><td>>hg38_21_+_8433172_8446622_RNA45SN1_45S_50ntFlanks@6700.6718.19|CTAGCTGCGAGAATTAATG</td><td>>hg38_21_+_8433172_8446622_RNA45SN1_45S_50ntFlanks</td><td>6700</td><td>6718</td><td>19</td><td>CTAGCTGCGAGAATTAATG</td><td>5570.5</td><td>-4.3</td><td>0.6</td><td>-6.7</td><td>2.6E-11</td><td>5.7E-11</td><td>MB231/MB468</td></tr>
<tr><td>>hg38_21_+_8433172_8446622_RNA45SN1_45S_50ntFlanks@10463.10497.35|CTCCGTTGCCCTCGGCCGATCGAAAGGGAGTCGGG</td><td>>hg38_21_+_8433172_8446622_RNA45SN1_45S_50ntFlanks</td><td>10463</td><td>10497</td><td>35</td><td>CTCCGTTGCCCTCGGCCGATCGAAAGGGAGTCGGG</td><td>109.5</td><td>-8.4</td><td>1.3</td><td>-6.7</td><td>2.6E-11</td><td>5.7E-11</td><td>MB231/MB468</td></tr>
<tr><td>>hg38_21_+_8433172_8446622_RNA45SN1_45S_50ntFlanks@11210.11233.24|GGCGACTCTGGACGCGAGCCGGGC</td><td>>hg38_21_+_8433172_8446622_RNA45SN1_45S_50ntFlanks</td><td>11210</td><td>11233</td><td>24</td><td>GGCGACTCTGGACGCGAGCCGGGC</td><td>214.2</td><td>-4.2</td><td>0.6</td><td>-6.7</td><td>2.6E-11</td><td>5.7E-11</td><td>MB231/MB468</td></tr>
<tr><td>>hg38_21_+_8433172_8446622_RNA45SN1_45S_50ntFlanks@12627.12651.25|ATGGGGCGAAGCTACCATCTGTGGG</td><td>>hg38_21_+_8433172_8446622_RNA45SN1_45S_50ntFlanks</td><td>12627</td><td>12651</td><td>25</td><td>ATGGGGCGAAGCTACCATCTGTGGG</td><td>75.5</td><td>-8.2</td><td>1.2</td><td>-6.6</td><td>3.1E-11</td><td>5.7E-11</td><td>BT20/MB468</td></tr>
<tr><td>>MI0000264|hsa-mir-7-2&WithFlank&15|+|88611819|88611940@38.58.21(+1A);MI0000265|hsa-mir-7-3&WithFlank&19|+|4770664|4770785@37.57.21(+1A);MI0000263|hsa-mir-7-1&WithFlank&9|-|83969742|83969863@30.50.21(+1A)&[MIMAT0000252&hsa-miR-7-5p&offsets|0|-3(+1A);m-400&15|+|88611856|88611879&offsets|0|-3(+1A)];[MIMAT0000252_1&hsa-miR-7-5p&offsets|0|-3(+1A);m-402&19|+|4770700|4770723&offsets|0|-3(+1A)];[MIMAT0000252_2&hsa-miR-7-5p&offsets|0|-3(+1A);m-401&9|-|83969811|83969834&offsets|0|-3(+1A)]|TGGAAGACTAGTGATTTTGTTA</td><td>>MI0000264|hsa-mir-7-2&WithFlank&15|+|88611819|88611940</td><td>38</td><td>58</td><td>21(+1A)</td><td>TGGAAGACTAGTGATTTTGTTA</td><td>1448.4</td><td>10.2</td><td>1.5</td><td>6.6</td><td>3.1E-11</td><td>5.7E-11</td><td>BT20/MB468</td></tr>
<tr><td>>hg38_21_+_8433172_8446622_RNA45SN1_45S_50ntFlanks@11570.11601.32|AGCCGACTTAGAACTGGTGCGGACCAGGGGAA</td><td>>hg38_21_+_8433172_8446622_RNA45SN1_45S_50ntFlanks</td><td>11570</td><td>11601</td><td>32</td><td>AGCCGACTTAGAACTGGTGCGGACCAGGGGAA</td><td>476.5</td><td>-3.4</td><td>0.5</td><td>-6.7</td><td>2.6E-11</td><td>5.7E-11</td><td>MB231/MB468</td></tr>
<tr><td>>hg38_21_+_8433172_8446622_RNA45SN1_45S_50ntFlanks@4160.4182.23|CCACATCCAAGGAAGGCAGCAGG</td><td>>hg38_21_+_8433172_8446622_RNA45SN1_45S_50ntFlanks</td><td>4160</td><td>4182</td><td>23</td><td>CCACATCCAAGGAAGGCAGCAGG</td><td>136.3</td><td>-8.7</td><td>1.3</td><td>-6.7</td><td>2.6E-11</td><td>5.7E-11</td><td>MB231/MB468</td></tr>
<tr><td>>hg38_MT_+_1621_3279_RNR2_16S_50ntFlanks@460.492.33|TTGTAAATTTAACTGTTAGTCCAAAGAGGAACA</td><td>>hg38_MT_+_1621_3279_RNR2_16S_50ntFlanks</td><td>460</td><td>492</td><td>33</td><td>TTGTAAATTTAACTGTTAGTCCAAAGAGGAACA</td><td>925.2</td><td>9.3</td><td>1.4</td><td>6.6</td><td>3.1E-11</td><td>5.8E-11</td><td>BT20/MB468</td></tr>
<tr><td>>hg38_21_+_8433172_8446622_RNA45SN1_45S_50ntFlanks@11874.11912.39|AGAAGACCCTGTTGAGCTTGACTCTAGTCTGGCACGGTG</td><td>>hg38_21_+_8433172_8446622_RNA45SN1_45S_50ntFlanks</td><td>11874</td><td>11912</td><td>39</td><td>AGAAGACCCTGTTGAGCTTGACTCTAGTCTGGCACGGTG</td><td>106.8</td><td>-8.3</td><td>1.2</td><td>-6.7</td><td>2.7E-11</td><td>5.8E-11</td><td>MB231/MB468</td></tr>
<tr><td>>hg38_21_+_8433172_8446622_RNA45SN1_45S_50ntFlanks@11641.11666.26|CGGCGGGTGTTGACGCGATGTGATTT</td><td>>hg38_21_+_8433172_8446622_RNA45SN1_45S_50ntFlanks</td><td>11641</td><td>11666</td><td>26</td><td>CGGCGGGTGTTGACGCGATGTGATTT</td><td>127.6</td><td>-9.0</td><td>1.4</td><td>-6.6</td><td>3.1E-11</td><td>5.8E-11</td><td>BT20/MB468</td></tr>
</table>

| | | | | | | | | | | | | |
|---|---|---|---|---|---|---|---|---|---|---|---|---|
| >hg38_21_+_8433172_8446622_RNA45SN1_45S_50ntFlanks@3787.3812.26\|AAACTGCGAATGGCTCATTAAATCAG | >hg38_21_+_8433172_8446622_RNA45SN1_45S_50ntFlanks | 3787 | 3812 | 26 | AAACTGCGAATGGCTCATTAAATCAG | 185.3 | -4.3 | 0.6 | -6.7 | 2.7E-11 | 5.8E-11 | MB231/MB468 |
| >hg38_21_+_8433172_8446622_RNA45SN1_45S_50ntFlanks@8423.8442.20\|CGCAGTCCGCCCGGAGGATT | >hg38_21_+_8433172_8446622_RNA45SN1_45S_50ntFlanks | 8423 | 8442 | 20 | CGCAGTCCGCCCGGAGGATT | 193.6 | -3.7 | 0.6 | -6.6 | 3.1E-11 | 5.8E-11 | BT20/MB468 |
| >hg38_21_+_8433172_8446622_RNA45SN1_45S_50ntFlanks@10882.10902.21\|TCGGGCTGGGGCGCGAAGCGG | >hg38_21_+_8433172_8446622_RNA45SN1_45S_50ntFlanks | 10882 | 10902 | 21 | TCGGGCTGGGGCGCGAAGCGG | 141.0 | -4.8 | 0.7 | -6.7 | 2.7E-11 | 5.9E-11 | MB231/MB468 |
| >hg38_21_+_8433172_8446622_RNA45SN1_45S_50ntFlanks@11999.12033.35\|CCCTGCGGGCCGCCGGTGAAATACCACTACTCTGA | >hg38_21_+_8433172_8446622_RNA45SN1_45S_50ntFlanks | 11999 | 12033 | 35 | CCCTGCGGGCCGCCGGTGAAATACCACTACTCTGA | 113.1 | -8.4 | 1.3 | -6.7 | 2.7E-11 | 5.9E-11 | MB231/MB468 |
| >hg38_21_+_8433172_8446622_RNA45SN1_45S_50ntFlanks@7990.8019.30\|GACGTGGCGACCCGCTGAATTTAAGCATAT | >hg38_21_+_8433172_8446622_RNA45SN1_45S_50ntFlanks | 7990 | 8019 | 30 | GACGTGGCGACCCGCTGAATTTAAGCATAT | 1076.5 | 8.9 | 1.3 | 6.7 | 2.7E-11 | 5.9E-11 | MB231/MB468 |
| >hg38_21_+_8433172_8446622_RNA45SN1_45S_50ntFlanks@476.499.24\|TTCGTGATCGATGTGGTGACGTCG | >hg38_21_+_8433172_8446622_RNA45SN1_45S_50ntFlanks | 476 | 499 | 24 | TTCGTGATCGATGTGGTGACGTCG | 2974.0 | 5.8 | 0.9 | 6.6 | 3.2E-11 | 5.9E-11 | BT20/MB468 |
| >hg38_21_+_8433172_8446622_RNA45SN1_45S_50ntFlanks@12354.12376.23\|TTACCACAGGGATAACTGGCTTG | >hg38_21_+_8433172_8446622_RNA45SN1_45S_50ntFlanks | 12354 | 12376 | 23 | TTACCACAGGGATAACTGGCTTG | 127.8 | -9.0 | 1.4 | -6.6 | 3.2E-11 | 5.9E-11 | BT20/MB468 |
| >hg38_21_+_8433172_8446622_RNA45SN1_45S_50ntFlanks@11562.11592.31\|CGCCTAGCAGCCGACTTAGAACTGGTGCGGA | >hg38_21_+_8433172_8446622_RNA45SN1_45S_50ntFlanks | 11562 | 11592 | 31 | CGCCTAGCAGCCGACTTAGAACTGGTGCGGA | 771.8 | -4.2 | 0.6 | -6.7 | 2.8E-11 | 6.0E-11 | MB231/MB468 |
| >hg38_MT_+_1621_3279_RNR2_16S_50ntFlanks@51.95.45\|GCTAAACCTAGCCCCAAACCCACTCCACCTTACTACCAGACAACC | >hg38_MT_+_1621_3279_RNR2_16S_50ntFlanks | 51 | 95 | 45 | GCTAAACCTAGCCCCAAACCCACTCCACCTTACTACCAGACAACC | 1349.0 | 10.4 | 1.6 | 6.6 | 3.3E-11 | 6.0E-11 | BT20/MB468 |
| >hg38_21_+_8433172_8446622_RNA45SN1_45S_50ntFlanks@6772.6793.22\|GTTCCTCCCGGGGCTACGCCTG | >hg38_21_+_8433172_8446622_RNA45SN1_45S_50ntFlanks | 6772 | 6793 | 22 | GTTCCTCCCGGGGCTACGCCTG | 133.9 | -8.6 | 1.3 | -6.7 | 2.8E-11 | 6.1E-11 | MB231/MB468 |
| >hg38_21_+_8433172_8446622_RNA45SN1_45S_50ntFlanks@10411.10443.33\|CAGTCGGTCCTGAGAGATGGGCGAGCGCCGTTC | >hg38_21_+_8433172_8446622_RNA45SN1_45S_50ntFlanks | 10411 | 10443 | 33 | CAGTCGGTCCTGAGAGATGGGCGAGCGCCGTTC | 564.2 | -4.6 | 0.7 | -6.7 | 2.8E-11 | 6.1E-11 | MB231/MB468 |
| >hg38_21_+_8433172_8446622_RNA45SN1_45S_50ntFlanks@10794.10825.32\|CTGGCATGTTGGAACAATGTAGGTAAGGGAAG | >hg38_21_+_8433172_8446622_RNA45SN1_45S_50ntFlanks | 10794 | 10825 | 32 | CTGGCATGTTGGAACAATGTAGGTAAGGGAAG | 329.3 | -4.3 | 0.7 | -6.6 | 3.3E-11 | 6.1E-11 | BT20/MB468 |
| >hg38_21_+_8433172_8446622_RNA45SN1_45S_50ntFlanks@4069.4089.21\|CTTTCGATGGTAGTCGCCGTG | >hg38_21_+_8433172_8446622_RNA45SN1_45S_50ntFlanks | 4069 | 4089 | 21 | CTTTCGATGGTAGTCGCCGTG | 172.2 | -4.4 | 0.7 | -6.6 | 3.3E-11 | 6.1E-11 | BT20/MB468 |
| >hg38_21_+_8433172_8446622_RNA45SN1_45S_50ntFlanks@7985.8004.20\|GATCAGACGTGGCGACCCGC | >hg38_21_+_8433172_8446622_RNA45SN1_45S_50ntFlanks | 7985 | 8004 | 20 | GATCAGACGTGGCGACCCGC | 136.1 | -9.1 | 1.4 | -6.6 | 3.3E-11 | 6.2E-11 | BT20/MB468 |
| >hg38_21_+_8433172_8446622_RNA45SN1_45S_50ntFlanks@11641.11664.24\|CGGCGGGTGTTGACGCGATGTGAT | >hg38_21_+_8433172_8446622_RNA45SN1_45S_50ntFlanks | 11641 | 11664 | 24 | CGGCGGGTGTTGACGCGATGTGAT | 812.4 | -6.0 | 0.9 | -6.6 | 3.3E-11 | 6.2E-11 | BT20/MB468 |
| >hg38_21_+_8433172_8446622_RNA45SN1_45S_50ntFlanks@9027.9064.38\|CGTCTCCTCGTGGGGGGGCCGGGCCACCCCTCCCACGG | >hg38_21_+_8433172_8446622_RNA45SN1_45S_50ntFlanks | 9027 | 9064 | 38 | CGTCTCCTCGTGGGGGGGCCGGGCCACCCCTCCCACGG | 131.7 | -8.6 | 1.3 | -6.7 | 2.9E-11 | 6.2E-11 | MB231/MB468 |
| >hg38_21_+_8433172_8446622_RNA45SN1_45S_50ntFlanks@12654.12671.18\|TATGACTGAACGCCTCTA | >hg38_21_+_8433172_8446622_RNA45SN1_45S_50ntFlanks | 12654 | 12671 | 18 | TATGACTGAACGCCTCTA | 125.9 | -8.6 | 1.3 | -6.7 | 2.9E-11 | 6.2E-11 | MB231/MB468 |
| >hg38_21_+_8433172_8446622_RNA45SN1_45S_50ntFlanks@12490.12521.32\|CGTGAGCTGGGTTTAGACCGTCGTGAGACAGG | >hg38_21_+_8433172_8446622_RNA45SN1_45S_50ntFlanks | 12490 | 12521 | 32 | CGTGAGCTGGGTTTAGACCGTCGTGAGACAGG | 121.7 | -8.5 | 1.3 | -6.7 | 2.9E-11 | 6.2E-11 | MB231/MB468 |
| >hg38_21_+_8433172_8446622_RNA45SN1_45S_50ntFlanks@6650.6676.27\|ACGACTCTTAGCGGTGGATCACTCGGC | >hg38_21_+_8433172_8446622_RNA45SN1_45S_50ntFlanks | 6650 | 6676 | 27 | ACGACTCTTAGCGGTGGATCACTCGGC | 572.2 | 6.9 | 1.0 | 6.6 | 3.3E-11 | 6.2E-11 | BT20/MB468 |
| >hg38_21_+_8433172_8446622_RNA45SN1_45S_50ntFlanks@12406.12435.30\|TTTTTGATCCTTCGATGTCGGCTCTTCCTA | >hg38_21_+_8433172_8446622_RNA45SN1_45S_50ntFlanks | 12406 | 12435 | 30 | TTTTTGATCCTTCGATGTCGGCTCTTCCTA | 106.4 | -5.3 | 0.8 | -6.6 | 3.4E-11 | 6.3E-11 | BT20/MB468 |
| >hg38_21_+_8433172_8446622_RNA45SN1_45S_50ntFlanks@12974.13001.28\|CGTACGTAGCAGAGCAGCTCCCTCGCTG | >hg38_21_+_8433172_8446622_RNA45SN1_45S_50ntFlanks | 12974 | 13001 | 28 | CGTACGTAGCAGAGCAGCTCCCTCGCTG | 105.5 | -8.3 | 1.2 | -6.7 | 2.9E-11 | 6.3E-11 | MB231/MB468 |
| >hg38_21_+_8433172_8446622_RNA45SN1_45S_50ntFlanks@12156.12180.25\|GAGTTTGACTGGGGCGGTACACCTG | >hg38_21_+_8433172_8446622_RNA45SN1_45S_50ntFlanks | 12156 | 12180 | 25 | GAGTTTGACTGGGGCGGTACACCTG | 99.9 | -8.6 | 1.3 | -6.6 | 3.4E-11 | 6.3E-11 | BT20/MB468 |
| >hg38_21_+_8433172_8446622_RNA45SN1_45S_50ntFlanks@10377.10407.31\|AAGGGTTCCATGTGAACAGCAGTTGAACATG | >hg38_21_+_8433172_8446622_RNA45SN1_45S_50ntFlanks | 10377 | 10407 | 31 | AAGGGTTCCATGTGAACAGCAGTTGAACATG | 100.6 | -5.1 | 0.8 | -6.6 | 2.9E-11 | 6.3E-11 | MB231/MB468 |
| >hg38_21_+_8433172_8446622_RNA45SN1_45S_50ntFlanks@6738.6760.23\|ATCATCGACACTTCGAACGCACT | >hg38_21_+_8433172_8446622_RNA45SN1_45S_50ntFlanks | 6738 | 6760 | 23 | ATCATCGACACTTCGAACGCACT | 115.2 | -8.8 | 1.3 | -6.6 | 3.4E-11 | 6.4E-11 | BT20/MB468 |
| >hg38_21_+_8433172_8446622_RNA45SN1_45S_50ntFlanks@12957.12981.25\|CTTCTGGGTCGGGGTTTCGTACGTA | >hg38_21_+_8433172_8446622_RNA45SN1_45S_50ntFlanks | 12957 | 12981 | 25 | CTTCTGGGTCGGGGTTTCGTACGTA | 109.2 | -5.8 | 0.9 | -6.6 | 3.0E-11 | 6.4E-11 | MB231/MB468 |
| >trna12_ArgCCT_16_+_3243918_3243990@34.51.18,trna18_ArgCCT_17_+_73030001_73030073@34.51.18,trna21_ArgCCT_17_-_73030526_73030598@34.51.18,trna2_ArgCCT_16_+_3202901_3202973@34.51.18,trna3_ArgCCT_7_+_139025446_139025518@34.51.18\|CCTAAGCCAGGGATTGTG | >trna12_ArgCCT_16_+_3243918_3243990 | 34 | 51 | 18 | CCTAAGCCAGGGATTGTG | 105.1 | -8.7 | 1.3 | -6.6 | 3.5E-11 | 6.4E-11 | BT20/MB468 |
| >hg38_21_+_8433172_8446622_RNA45SN1_45S_50ntFlanks@6653.6698.46\|ACTCTTAGCGGTGGATCACTCGGCTCGTGCGTCGATGAAGAACGCA | >hg38_21_+_8433172_8446622_RNA45SN1_45S_50ntFlanks | 6653 | 6698 | 46 | ACTCTTAGCGGTGGATCACTCGGCTCGTGCGTCGATGAAGAACGCA | 616.8 | 4.5 | 0.7 | 6.6 | 3.0E-11 | 6.4E-11 | MB231/MB468 |
| >hg38_1_-_228634819_228635039_RNA5S12_5S_50ntFlanks@150.171.22\|AATACCGGGTGCTGTAGGCTTT | >hg38_1_-_228634819_228635039_RNA5S12_5S_50ntFlanks | 150 | 171 | 22 | AATACCGGGTGCTGTAGGCTTT | 4896.7 | 5.4 | 0.8 | 6.6 | 3.0E-11 | 6.4E-11 | MB231/MB468 |
| >hg38_21_+_8433172_8446622_RNA45SN1_45S_50ntFlanks@11320.11342.23\|CCACGTCTCGTCGCGCGCGCGTC | >hg38_21_+_8433172_8446622_RNA45SN1_45S_50ntFlanks | 11320 | 11342 | 23 | CCACGTCTCGTCGCGCGCGCGTC | 514.3 | -6.1 | 0.9 | -6.6 | 3.0E-11 | 6.5E-11 | MB231/MB468 |
| >trna111_HisGTG_1_-_147774845_147774916@1.34.34,trna118_HisGTG_1_-_145396881_145396952@1.34.34,trna16_HisGTG_1_+_146544773_146544844@1.34.34,trna1_HisGTG_15_+_45493349_45493420@1.34.34,trna21_HisGTG_1_+_147753471_147753542@1.34.34,trna33_HisGTG_6_+_27125906_27125977@1.34.34,trna7_HisGTG_9_-_14433938_14434009@1.34.34,trna8_HisGTG_15_-_45492611_45492682@1.34.34,trna9_HisGTG_15_-_45490804_45490875@1.34.34\|GCCGTGATCGTATAGTGGTTAGTACTCTGCGTTG | >trna111_HisGTG_1_-_147774845_147774916 | 1 | 34 | 34 | GCCGTGATCGTATAGTGGTTAGTACTCTGCGTTG | 3178.7 | 5.2 | 0.8 | 6.6 | 3.5E-11 | 6.5E-11 | BT20/MB468 |
| >hg38_21_+_8433172_8446622_RNA45SN1_45S_50ntFlanks@11694.11717.24\|AAGAAATTCAATGAAGCGCGGGTA | >hg38_21_+_8433172_8446622_RNA45SN1_45S_50ntFlanks | 11694 | 11717 | 24 | AAGAAATTCAATGAAGCGCGGGTA | 269.9 | -4.0 | 0.6 | -6.6 | 3.5E-11 | 6.6E-11 | BT20/MB468 |
| >hg38_21_+_8433172_8446622_RNA45SN1_45S_50ntFlanks@8168.8197.30\|CCCAAGTCCTTCTGATCGAGGCCCAGCCCG | >hg38_21_+_8433172_8446622_RNA45SN1_45S_50ntFlanks | 8168 | 8197 | 30 | CCCAAGTCCTTCTGATCGAGGCCCAGCCCG | 130.6 | -8.6 | 1.3 | -6.6 | 3.0E-11 | 6.6E-11 | MB231/MB468 |
| >hg38_21_+_8433172_8446622_RNA45SN1_45S_50ntFlanks@3853.3881.29\|AACTGTGGTAATTCTAGAGCTAATACATG | >hg38_21_+_8433172_8446622_RNA45SN1_45S_50ntFlanks | 3853 | 3881 | 29 | AACTGTGGTAATTCTAGAGCTAATACATG | 554.5 | -4.0 | 0.6 | -6.6 | 3.5E-11 | 6.6E-11 | BT20/MB468 |
| >hg38_21_+_8433172_8446622_RNA45SN1_45S_50ntFlanks@10680.10711.32\|GTTCCGGCGGCGTCCGGTGAGCTCTCGCTGGC | >hg38_21_+_8433172_8446622_RNA45SN1_45S_50ntFlanks | 10680 | 10711 | 32 | GTTCCGGCGGCGTCCGGTGAGCTCTCGCTGGC | 8038.0 | -7.2 | 1.1 | -6.6 | 3.6E-11 | 6.6E-11 | BT20/MB468 |
| >hg38_21_+_8433172_8446622_RNA45SN1_45S_50ntFlanks@3992.4024.33\|GGTGACTCTAGATAACCTCGGGCCGATCGCACG | >hg38_21_+_8433172_8446622_RNA45SN1_45S_50ntFlanks | 3992 | 4024 | 33 | GGTGACTCTAGATAACCTCGGGCCGATCGCACG | 512.1 | 4.3 | 0.6 | 6.6 | 3.1E-11 | 6.7E-11 | MB231/MB468 |
| >hg38_21_+_8433172_8446622_RNA45SN1_45S_50ntFlanks@4817.4835.19\|ATTCCCATGACCCGCCGGG | >hg38_21_+_8433172_8446622_RNA45SN1_45S_50ntFlanks | 4817 | 4835 | 19 | ATTCCCATGACCCGCCGGG | 76.5 | -8.3 | 1.2 | -6.6 | 3.6E-11 | 6.7E-11 | BT20/MB468 |
| >hg38_21_+_8433172_8446622_RNA45SN1_45S_50ntFlanks@6690.6718.29\|AAGAACGCAGCTAGCTGCGAGAATTAATG | >hg38_21_+_8433172_8446622_RNA45SN1_45S_50ntFlanks | 6690 | 6718 | 29 | AAGAACGCAGCTAGCTGCGAGAATTAATG | 113.6 | -8.8 | 1.3 | -6.6 | 3.6E-11 | 6.7E-11 | BT20/MB468 |
| >hg38_21_+_8433172_8446622_RNA45SN1_45S_50ntFlanks@9554.9595.42\|CGTAGCGGTCCTGACGTGCAAATCGGTCGTCCGACCTGGGTA | >hg38_21_+_8433172_8446622_RNA45SN1_45S_50ntFlanks | 9554 | 9595 | 42 | CGTAGCGGTCCTGACGTGCAAATCGGTCGTCCGACCTGGGTA | 86.0 | -8.4 | 1.3 | -6.6 | 3.6E-11 | 6.7E-11 | BT20/MB468 |
| >hg38_21_+_8433172_8446622_RNA45SN1_45S_50ntFlanks@4444.4471.28\|CCTTGCCTCTCGGCGCCCCCTCGATGCT | >hg38_21_+_8433172_8446622_RNA45SN1_45S_50ntFlanks | 4444 | 4471 | 28 | CCTTGCCTCTCGGCGCCCCCTCGATGCT | 132.6 | -8.6 | 1.3 | -6.6 | 3.1E-11 | 6.7E-11 | MB231/MB468 |
| >hg38_21_+_8433172_8446622_RNA45SN1_45S_50ntFlanks@8191.8213.23\|CAGCCCGTGGACGGTGTGAGGCC | >hg38_21_+_8433172_8446622_RNA45SN1_45S_50ntFlanks | 8191 | 8213 | 23 | CAGCCCGTGGACGGTGTGAGGCC | 551.0 | -3.2 | 0.5 | -6.6 | 3.7E-11 | 6.9E-11 | BT20/MB468 |
| >hg38_21_+_8433172_8446622_RNA45SN1_45S_50ntFlanks@13004.13029.26\|ATCTATTGAAAGTCAGCCCTCGACAC | >hg38_21_+_8433172_8446622_RNA45SN1_45S_50ntFlanks | 13004 | 13029 | 26 | ATCTATTGAAAGTCAGCCCTCGACAC | 125.5 | -8.5 | 1.3 | -6.6 | 3.2E-11 | 6.9E-11 | MB231/MB468 |
| >trna59_GluCTC_1_+_249168447_249168518@32.70.39\|CTCTCACCGCCGCGGCCCGGGTTCGATTCCCGGTCAGGA | >trna59_GluCTC_1_+_249168447_249168518 | 32 | 70 | 39 | CTCTCACCGCCGCGGCCCGGGTTCGATTCCCGGTCAGGA | 87.0 | -8.4 | 1.3 | -6.6 | 3.7E-11 | 6.9E-11 | BT20/MB468 |
| >hg38_21_+_8433172_8446622_RNA45SN1_45S_50ntFlanks@11641.11661.21\|CGGCGGGTGTTGACGCGATGT | >hg38_21_+_8433172_8446622_RNA45SN1_45S_50ntFlanks | 11641 | 11661 | 21 | CGGCGGGTGTTGACGCGATGT | 114.9 | -5.9 | 0.9 | -6.6 | 3.2E-11 | 6.9E-11 | MB231/MB468 |
| >hg38_21_+_8433172_8446622_RNA45SN1_45S_50ntFlanks@7988.8007.20\|CAGACGTGGCGACCCGCTGA | >hg38_21_+_8433172_8446622_RNA45SN1_45S_50ntFlanks | 7988 | 8007 | 20 | CAGACGTGGCGACCCGCTGA | 507.5 | -3.8 | 0.6 | -6.6 | 3.8E-11 | 6.9E-11 | BT20/MB468 |
| >hg38_21_+_8433172_8446622_RNA45SN1_45S_50ntFlanks@11215.11234.20\|CTCTGGACGCGAGCCGGGCC | >hg38_21_+_8433172_8446622_RNA45SN1_45S_50ntFlanks | 11215 | 11234 | 20 | CTCTGGACGCGAGCCGGGCC | 105.7 | -8.3 | 1.3 | -6.6 | 3.3E-11 | 7.0E-11 | MB231/MB468 |
| >hg38_21_+_8433172_8446622_RNA45SN1_45S_50ntFlanks@5420.5438.19\|CCGATTGGATGGTTTAGTG | >hg38_21_+_8433172_8446622_RNA45SN1_45S_50ntFlanks | 5420 | 5438 | 19 | CCGATTGGATGGTTTAGTG | 123.4 | -8.9 | 1.4 | -6.6 | 3.8E-11 | 7.1E-11 | BT20/MB468 |

<table>
<tr><td>>hg38_21_+_8433172_8446622_RNA45SN1_45S_50ntFlanks@12490.12521.32|CGTGAGCTGGGTTTAGACCGTCGTGAGACAGG</td><td>>hg38_21_+_8433172_8446622_RNA45SN1_45S_50ntFlanks</td><td>12490</td><td>12521</td><td>32</td><td>CGTGAGCTGGGTTTAGACCGTCGTGAGACAGG</td><td>145.9</td><td>-5.5</td><td>0.8</td><td>-6.6</td><td>3.8E-11</td><td>7.1E-11</td><td>BT20/MB468</td></tr>
<tr><td>>hg38_21_+_8433172_8446622_RNA45SN1_45S_50ntFlanks@12996.13038.43|TCGCTGCGATCTATTGAAAGTCAGCCCTCGACACAAGGGTTTG</td><td>>hg38_21_+_8433172_8446622_RNA45SN1_45S_50ntFlanks</td><td>12996</td><td>13038</td><td>43</td><td>TCGCTGCGATCTATTGAAAGTCAGCCCTCGACACAAGGGTTTG</td><td>1304.3</td><td>9.9</td><td>1.5</td><td>6.6</td><td>3.8E-11</td><td>7.1E-11</td><td>BT20/MB468</td></tr>
<tr><td>>hg38_21_+_8433172_8446622_RNA45SN1_45S_50ntFlanks@4264.4282.19|AATGAGTCCACTTTAAATC</td><td>>hg38_21_+_8433172_8446622_RNA45SN1_45S_50ntFlanks</td><td>4264</td><td>4282</td><td>19</td><td>AATGAGTCCACTTTAAATC</td><td>118.2</td><td>-8.5</td><td>1.3</td><td>-6.6</td><td>3.3E-11</td><td>7.1E-11</td><td>MB231/MB468</td></tr>
<tr><td>>trna10_LysCTT_16_+_3241501_3241573@1.30.30,trna13_LysCTT_14_-_58706613_58706685@1.30.30,trna2_LysCTT_15_+_79152904_79152976@1.30.30|GCCCGGCTAGCTCAGTCGGTAGAGCATGGG</td><td>>trna10_LysCTT_16_+_3241501_3241573</td><td>1</td><td>30</td><td>30</td><td>GCCCGGCTAGCTCAGTCGGTAGAGCATGGG</td><td>139.1</td><td>-8.7</td><td>1.3</td><td>-6.6</td><td>3.3E-11</td><td>7.1E-11</td><td>MB231/MB468</td></tr>
<tr><td>>MI0016887|hsa-mir-4521&WithFlank&17|+|8186939|8187010@10.35.26&[MIMAT0019058&hsa-miR-4521&offsets|0|+4;m-765&17|+|8186948|8186969&offsets|0|+4]|GCTAAGGAAGTCCTGTGCTCAGTTTT</td><td>>MI0016887|hsa-mir-4521&WithFlank&17|+|8186939|8187010</td><td>10</td><td>35</td><td>26</td><td>GCTAAGGAAGTCCTGTGCTCAGTTTT</td><td>1776.0</td><td>5.4</td><td>0.8</td><td>6.6</td><td>3.3E-11</td><td>7.1E-11</td><td>MB231/MB468</td></tr>
<tr><td>>hg38_21_+_8433172_8446622_RNA45SN1_45S_50ntFlanks@13005.13027.23|TCTATTGAAAGTCAGCCCTCGAC</td><td>>hg38_21_+_8433172_8446622_RNA45SN1_45S_50ntFlanks</td><td>13005</td><td>13027</td><td>23</td><td>TCTATTGAAAGTCAGCCCTCGAC</td><td>222.7</td><td>-6.0</td><td>0.9</td><td>-6.6</td><td>3.3E-11</td><td>7.1E-11</td><td>MB231/MB468</td></tr>
<tr><td>>hg38_21_+_8433172_8446622_RNA45SN1_45S_50ntFlanks@9027.9059.33|CGTCTCCTCGTGGGGGGGCCGGGCCACCCCTCC</td><td>>hg38_21_+_8433172_8446622_RNA45SN1_45S_50ntFlanks</td><td>9027</td><td>9059</td><td>33</td><td>CGTCTCCTCGTGGGGGGGCCGGGCCACCCCTCC</td><td>97.4</td><td>-8.6</td><td>1.3</td><td>-6.6</td><td>3.9E-11</td><td>7.1E-11</td><td>BT20/MB468</td></tr>
<tr><td>>MI0000102|hsa-mir-100&WithFlank&11|-|122152223|122152314@19.39.21&[MIMAT0000098&hsa-miR-100-5p&offsets|0|-1;m-27&11|-|122152276|122152296&offsets|0|0]|AACCCGTAGATCCGAACTTGT</td><td>>MI0000102|hsa-mir-100&WithFlank&11|-|122152223|122152314</td><td>19</td><td>39</td><td>21</td><td>AACCCGTAGATCCGAACTTGT</td><td>173046.6</td><td>-6.1</td><td>0.9</td><td>-7.0</td><td>2.4E-12</td><td>7.1E-11</td><td>BT20/MB231</td></tr>
<tr><td>>hg38_21_+_8433172_8446622_RNA45SN1_45S_50ntFlanks@11641.11668.28|CGGCGGGTGTTGACGCGATGTGATTTCT</td><td>>hg38_21_+_8433172_8446622_RNA45SN1_45S_50ntFlanks</td><td>11641</td><td>11668</td><td>28</td><td>CGGCGGGTGTTGACGCGATGTGATTTCT</td><td>283.4</td><td>-3.5</td><td>0.5</td><td>-6.6</td><td>3.3E-11</td><td>7.1E-11</td><td>MB231/MB468</td></tr>
<tr><td>>hg38_21_+_8433172_8446622_RNA45SN1_45S_50ntFlanks@11768.11798.31|CTAATTAGTGACGCGCATGAATGGATGAACG</td><td>>hg38_21_+_8433172_8446622_RNA45SN1_45S_50ntFlanks</td><td>11768</td><td>11798</td><td>31</td><td>CTAATTAGTGACGCGCATGAATGGATGAACG</td><td>94.6</td><td>-8.6</td><td>1.3</td><td>-6.6</td><td>3.9E-11</td><td>7.2E-11</td><td>BT20/MB468</td></tr>
<tr><td>>hg38_21_+_8433172_8446622_RNA45SN1_45S_50ntFlanks@10410.10453.44|TCAGTCGGTCCTGAGAGATGGGCGAGCGCCGTTCCGAAGGGACG</td><td>>hg38_21_+_8433172_8446622_RNA45SN1_45S_50ntFlanks</td><td>10410</td><td>10453</td><td>44</td><td>TCAGTCGGTCCTGAGAGATGGGCGAGCGCCGTTCCGAAGGGACG</td><td>112.1</td><td>-4.5</td><td>0.7</td><td>-6.6</td><td>3.9E-11</td><td>7.2E-11</td><td>BT20/MB468</td></tr>
<tr><td>>hg38_MT_+_1621_3279_RNR2_16S_50ntFlanks@1370.1388.19|ATGTTGGATCAGGACATCC</td><td>>hg38_MT_+_1621_3279_RNR2_16S_50ntFlanks</td><td>1370</td><td>1388</td><td>19</td><td>ATGTTGGATCAGGACATCC</td><td>945.7</td><td>8.9</td><td>1.3</td><td>6.6</td><td>3.9E-11</td><td>7.3E-11</td><td>BT20/MB468</td></tr>
<tr><td>>MI0000749|hsa-mir-30e&WithFlank&1|+|40754349|40754452@23.45.23(+1A)&[MIMAT0000692&hsa-miR-30e-5p&offsets|0|+1(+1A);m-24&1|+|40754371|40754394&offsets|0|-1(+1A)]|TGTAAACATCCTTGACTGGAAGCA</td><td>>MI0000749|hsa-mir-30e&WithFlank&1|+|40754349|40754452</td><td>23</td><td>45</td><td>23(+1A)</td><td>TGTAAACATCCTTGACTGGAAGCA</td><td>1312.3</td><td>4.9</td><td>0.7</td><td>6.6</td><td>3.4E-11</td><td>7.3E-11</td><td>MB231/MB468</td></tr>
<tr><td>>hg38_21_+_8433172_8446622_RNA45SN1_45S_50ntFlanks@12488.12518.31|AACGTGAGCTGGGTTTAGACCGTCGTGAGAC</td><td>>hg38_21_+_8433172_8446622_RNA45SN1_45S_50ntFlanks</td><td>12488</td><td>12518</td><td>31</td><td>AACGTGAGCTGGGTTTAGACCGTCGTGAGAC</td><td>344.0</td><td>-6.1</td><td>0.9</td><td>-6.6</td><td>4.0E-11</td><td>7.3E-11</td><td>BT20/MB468</td></tr>
<tr><td>>hg38_21_+_8433172_8446622_RNA45SN1_45S_50ntFlanks@11999.12029.31|CCCTGCGGGCCGCCGGTGAAATACCACTACT</td><td>>hg38_21_+_8433172_8446622_RNA45SN1_45S_50ntFlanks</td><td>11999</td><td>12029</td><td>31</td><td>CCCTGCGGGCCGCCGGTGAAATACCACTACT</td><td>105.8</td><td>-8.3</td><td>1.3</td><td>-6.6</td><td>3.4E-11</td><td>7.4E-11</td><td>MB231/MB468</td></tr>
<tr><td>>hg38_21_+_8433172_8446622_RNA45SN1_45S_50ntFlanks@4443.4470.28|CCCTTGCCTCTCGGCGCCCCCTCGATGC</td><td>>hg38_21_+_8433172_8446622_RNA45SN1_45S_50ntFlanks</td><td>4443</td><td>4470</td><td>28</td><td>CCCTTGCCTCTCGGCGCCCCCTCGATGC</td><td>101.6</td><td>-8.7</td><td>1.3</td><td>-6.6</td><td>4.0E-11</td><td>7.4E-11</td><td>BT20/MB468</td></tr>
<tr><td>>hg38_21_+_8433172_8446622_RNA45SN1_45S_50ntFlanks@12991.13012.22|CTCCCTCGCTGCGATCTATTGA</td><td>>hg38_21_+_8433172_8446622_RNA45SN1_45S_50ntFlanks</td><td>12991</td><td>13012</td><td>22</td><td>CTCCCTCGCTGCGATCTATTGA</td><td>1903.1</td><td>-6.2</td><td>0.9</td><td>-6.6</td><td>4.0E-11</td><td>7.4E-11</td><td>BT20/MB468</td></tr>
<tr><td>>trna111_HisGTG_1_-_147774845_147774916@2.30.29,trna118_HisGTG_1_-_145396881_145396952@2.30.29,trna16_HisGTG_1_+_146544773_146544844@2.30.29,trna1_HisGTG_15_+_45493349_45493420@2.30.29,trna21_HisGTG_1_+_147753471_147753542@2.30.29,trna33_HisGTG_6_+_27125906_27125977@2.30.29,trna7_HisGTG_9_-_14433938_14434009@2.30.29,trna8_HisGTG_15_-_45492611_45492682@2.30.29,trna9_HisGTG_15_-_45490804_45490875@2.30.29|CCGTGATCGTATAGTGGTTAGTACTCTGC</td><td>>trna111_HisGTG_1_-_147774845_147774916</td><td>2</td><td>30</td><td>29</td><td>CCGTGATCGTATAGTGGTTAGTACTCTGC</td><td>601.7</td><td>5.2</td><td>0.8</td><td>6.6</td><td>4.0E-11</td><td>7.5E-11</td><td>BT20/MB468</td></tr>
<tr><td>>hg38_21_+_8433172_8446622_RNA45SN1_45S_50ntFlanks@11564.11600.37|CCTAGCAGCCGACTTAGAACTGGTGCGGACCAGGGGA</td><td>>hg38_21_+_8433172_8446622_RNA45SN1_45S_50ntFlanks</td><td>11564</td><td>11600</td><td>37</td><td>CCTAGCAGCCGACTTAGAACTGGTGCGGACCAGGGGA</td><td>402.3</td><td>-3.7</td><td>0.6</td><td>-6.6</td><td>3.5E-11</td><td>7.5E-11</td><td>MB231/MB468</td></tr>
<tr><td>>MI0000285|hsa-mir-205&WithFlank&1|+|209432127|209432248@40.60.21&[MIMAT0000266&hsa-miR-205-5p&offsets|0|-1;m-351&1|+|209432166|209432187&offsets|0|-1]|TCCTTCATTCCACCGGAGTCT</td><td>>MI0000285|hsa-mir-205&WithFlank&1|+|209432127|209432248</td><td>40</td><td>60</td><td>21</td><td>TCCTTCATTCCACCGGAGTCT</td><td>124.3</td><td>-8.5</td><td>1.3</td><td>-6.6</td><td>3.5E-11</td><td>7.5E-11</td><td>MB231/MB468</td></tr>
<tr><td>>hg38_21_+_8433172_8446622_RNA45SN1_45S_50ntFlanks@7983.8005.23|CAGATCAGACGTGGCGACCCGCT</td><td>>hg38_21_+_8433172_8446622_RNA45SN1_45S_50ntFlanks</td><td>7983</td><td>8005</td><td>23</td><td>CAGATCAGACGTGGCGACCCGCT</td><td>362.3</td><td>-4.0</td><td>0.6</td><td>-6.6</td><td>3.5E-11</td><td>7.5E-11</td><td>MB231/MB468</td></tr>
<tr><td>>hg38_21_+_8433172_8446622_RNA45SN1_45S_50ntFlanks@11785.11806.22|TGAATGGATGAACGAGATTCCC</td><td>>hg38_21_+_8433172_8446622_RNA45SN1_45S_50ntFlanks</td><td>11785</td><td>11806</td><td>22</td><td>TGAATGGATGAACGAGATTCCC</td><td>1064.6</td><td>-8.1</td><td>1.2</td><td>-6.6</td><td>4.1E-11</td><td>7.5E-11</td><td>BT20/MB468</td></tr>
<tr><td>>hg38_21_+_8433172_8446622_RNA45SN1_45S_50ntFlanks@10873.10894.22|CTGGGTCGGTCGGGCTGGGGCG</td><td>>hg38_21_+_8433172_8446622_RNA45SN1_45S_50ntFlanks</td><td>10873</td><td>10894</td><td>22</td><td>CTGGGTCGGTCGGGCTGGGGCG</td><td>178.0</td><td>-4.1</td><td>0.6</td><td>-6.6</td><td>4.1E-11</td><td>7.5E-11</td><td>BT20/MB468</td></tr>
<tr><td>>trna116_GluCTC_1_-_145399233_145399304@32.73.42,trna71_GluCTC_1_-_161439189_161439260@32.73.42,trna74_GluCTC_1_-_161431809_161431880@32.73.42,trna77_GluCTC_1_-_161424398_161424469@32.73.42,trna77_GluCTC_6_+_28949976_28950047@32.73.42,trna80_GluCTC_1_-_161417018_161417089@32.73.42,trna87_GluCTC_6_-_126101393_126101464@32.73.42|CTCTCACCGCCGCGGCCCGGGTTCGATTCCCGGTCAGGGAAC</td><td>>trna116_GluCTC_1_-_145399233_145399304</td><td>32</td><td>73</td><td>42</td><td>CTCTCACCGCCGCGGCCCGGGTTCGATTCCCGGTCAGGGAAC</td><td>151.4</td><td>-8.8</td><td>1.3</td><td>-6.6</td><td>3.6E-11</td><td>7.7E-11</td><td>MB231/MB468</td></tr>
<tr><td>>hg38_21_+_8433172_8446622_RNA45SN1_45S_50ntFlanks@12162.12179.18|GACTGGGGCGGTACACCT</td><td>>hg38_21_+_8433172_8446622_RNA45SN1_45S_50ntFlanks</td><td>12162</td><td>12179</td><td>18</td><td>GACTGGGGCGGTACACCT</td><td>167.8</td><td>-9.0</td><td>1.4</td><td>-6.6</td><td>3.6E-11</td><td>7.7E-11</td><td>MB231/MB468</td></tr>
<tr><td>>trna10_AspGTC_12_-_125424193_125424264@35.75.41,trna12_AspGTC_12_-_125411891_125411962@35.75.41,trna144_AspGTC_6_-_27551236_27551307@35.75.41,trna38_AspGTC_17_-_8125556_8125627@35.75.41,trna45_AspGTC_6_+_27447453_27447524@35.75.41,trna48_AspGTC_6_+_27471523_27471594@35.75.41,trna4_AspGTC_12_+_96429799_96429870@35.75.41,trna69_AspGTC_1_-_161440205_161440276@35.75.41,trna72_AspGTC_1_-_161432824_161432895@35.75.41,trna75_AspGTC_1_-_161425414_161425485@35.75.41,trna78_AspGTC_1_-_161418033_161418104@35.75.41,trna81_AspGTC_1_-_161410615_161410686@35.75.41|TCACGCGGGAGACCGGGGTTCGATTCCCCGACGGGGAGCCA</td><td>>trna10_AspGTC_12_-_125424193_125424264</td><td>35</td><td>75</td><td>41</td><td>TCACGCGGGAGACCGGGGTTCGATTCCCCGACGGGGAGCCA</td><td>5254.0</td><td>9.1</td><td>1.4</td><td>6.6</td><td>3.6E-11</td><td>7.7E-11</td><td>MB231/MB468</td></tr>
<tr><td>>hg38_21_+_8433172_8446622_RNA45SN1_45S_50ntFlanks@3780.3801.22|TACAGTGAAACTGCGAATGGCT</td><td>>hg38_21_+_8433172_8446622_RNA45SN1_45S_50ntFlanks</td><td>3780</td><td>3801</td><td>22</td><td>TACAGTGAAACTGCGAATGGCT</td><td>208.3</td><td>-3.6</td><td>0.5</td><td>-6.6</td><td>4.2E-11</td><td>7.8E-11</td><td>BT20/MB468</td></tr>
<tr><td>>hg38_21_+_8433172_8446622_RNA45SN1_45S_50ntFlanks@11617.11640.24|AAAACAAAGCATCGCGAAGGCCCG</td><td>>hg38_21_+_8433172_8446622_RNA45SN1_45S_50ntFlanks</td><td>11617</td><td>11640</td><td>24</td><td>AAAACAAAGCATCGCGAAGGCCCG</td><td>95.5</td><td>-8.6</td><td>1.3</td><td>-6.6</td><td>4.3E-11</td><td>8.0E-11</td><td>BT20/MB468</td></tr>
<tr><td>>am_trna11_ProAGG_16_+_3241989_3242060@1.29.29,trna12_ProAGG_6_+_26555498_26555569@1.29.29,trna12_ProTGG_11_-_75946869_75946940@1.29.29,trna14_ProTGG_5_-_180615854_180615925@1.29.29,trna22_ProAGG_14_-_21081560_21081631@1.29.29,trna23_ProAGG_14_-_21077495_21077566@1.29.29,trna28_ProTGG_16_-_3234133_3234204@1.29.29,trna29_ProAGG_16_-_3232635_3232706@1.29.29,trna2_ProAGG_7_+_128423504_128423575@1.29.29,trna30_ProCGG_6_+_27059521_27059592@1.29.29,trna37_ProCGG_17_-_8126151_8126222@1.29.29,trna3_ProTGG_16_+_3208923_3208994@1.29.29,trna52_ProCGG_1_+_167683962_167684033@1.29.29,trna65_ProAGG_1_-_167684725_167684796@1.29.29,trna6_ProCGG_16_+_3222049_3222120@1.29.29,trna6_ProTGG_14_+_21152175_21152246@1.29.29,trna8_ProTGG_16_+_3238094_3238165@1.29.29,trna9_ProAGG_11_+_75946557_75946628@1.29.29,trna9_ProAGG_16_+_3239634_3239705@1.29.29|GGCTCGTTGGTCTAGGGGTATGATTCTCG</td><td>>am_trna11_ProAGG_16_+_3241989_3242060</td><td>1</td><td>29</td><td>29</td><td>GGCTCGTTGGTCTAGGGGTATGATTCTCG</td><td>1281.8</td><td>-6.3</td><td>0.9</td><td>-6.6</td><td>3.7E-11</td><td>8.0E-11</td><td>MB231/MB468</td></tr>
<tr><td>>hg38_21_+_8433172_8446622_RNA45SN1_45S_50ntFlanks@6661.6696.36|CGGTGGATCACTCGGCTCGTGCGTCGATGAAGAACG</td><td>>hg38_21_+_8433172_8446622_RNA45SN1_45S_50ntFlanks</td><td>6661</td><td>6696</td><td>36</td><td>CGGTGGATCACTCGGCTCGTGCGTCGATGAAGAACG</td><td>777.3</td><td>3.3</td><td>0.5</td><td>6.6</td><td>3.7E-11</td><td>8.0E-11</td><td>MB231/MB468</td></tr>
<tr><td>>hg38_21_+_8433172_8446622_RNA45SN1_45S_50ntFlanks@12973.13000.28|TCGTACGTAGCAGAGCAGCTCCCTCGCT</td><td>>hg38_21_+_8433172_8446622_RNA45SN1_45S_50ntFlanks</td><td>12973</td><td>13000</td><td>28</td><td>TCGTACGTAGCAGAGCAGCTCCCTCGCT</td><td>101.1</td><td>-3.4</td><td>0.5</td><td>-6.6</td><td>4.3E-11</td><td>8.0E-11</td><td>BT20/MB468</td></tr>
<tr><td>>hg38_21_+_8433172_8446622_RNA45SN1_45S_50ntFlanks@12386.12414.29|AGCGTTCATAGCGACGTCGCTTTTTGATC</td><td>>hg38_21_+_8433172_8446622_RNA45SN1_45S_50ntFlanks</td><td>12386</td><td>12414</td><td>29</td><td>AGCGTTCATAGCGACGTCGCTTTTTGATC</td><td>137.2</td><td>-3.8</td><td>0.6</td><td>-6.6</td><td>3.8E-11</td><td>8.0E-11</td><td>MB231/MB468</td></tr>
<tr><td>>hg38_21_+_8433172_8446622_RNA45SN1_45S_50ntFlanks@12642.12659.18|CATCTGTGGGATTATGAC</td><td>>hg38_21_+_8433172_8446622_RNA45SN1_45S_50ntFlanks</td><td>12642</td><td>12659</td><td>18</td><td>CATCTGTGGGATTATGAC</td><td>194.3</td><td>-4.7</td><td>0.7</td><td>-6.6</td><td>4.4E-11</td><td>8.0E-11</td><td>BT20/MB468</td></tr>
<tr><td>>hg38_21_+_8433172_8446622_RNA45SN1_45S_50ntFlanks@12539.12560.22|ATGATGTGTTGTTGCCATGGTA</td><td>>hg38_21_+_8433172_8446622_RNA45SN1_45S_50ntFlanks</td><td>12539</td><td>12560</td><td>22</td><td>ATGATGTGTTGTTGCCATGGTA</td><td>114.6</td><td>-8.4</td><td>1.3</td><td>-6.6</td><td>3.8E-11</td><td>8.1E-11</td><td>MB231/MB468</td></tr>
</table>

<table>
<tr><th></th><th></th><th></th><th></th><th></th><th></th><th></th><th></th><th></th><th></th><th></th><th></th><th></th></tr>
<tr><td>>hg38_21_+_8433172_8446622_RNA45SN1_45S_50ntFlanks@9027.9065.39|CGTCTCCTCGTGGGGGGGCCGGGCCACCCCTCCCACGGC</td><td>>hg38_21_+_8433172_8446622_RNA45SN1_45S_50ntFlanks</td><td>9027</td><td>9065</td><td>39</td><td>CGTCTCCTCGTGGGGGGGCCGGGCCACCCCTCCCACGGC</td><td>111.1</td><td>-8.8</td><td>1.3</td><td>-6.6</td><td>4.4E-11</td><td>8.1E-11</td><td>BT20/MB468</td></tr>
<tr><td>>hg38_21_+_8433172_8446622_RNA45SN1_45S_50ntFlanks@10756.10785.30|TACCCATATCCGCAGCAGGTCTCCAAGGTG</td><td>>hg38_21_+_8433172_8446622_RNA45SN1_45S_50ntFlanks</td><td>10756</td><td>10785</td><td>30</td><td>TACCCATATCCGCAGCAGGTCTCCAAGGTG</td><td>84.6</td><td>-8.4</td><td>1.3</td><td>-6.6</td><td>4.4E-11</td><td>8.1E-11</td><td>BT20/MB468</td></tr>
<tr><td>>hg38_21_+_8433172_8446622_RNA45SN1_45S_50ntFlanks@10407.10429.23|GGGTCAGTCGGTCCTGAGAGATG</td><td>>hg38_21_+_8433172_8446622_RNA45SN1_45S_50ntFlanks</td><td>10407</td><td>10429</td><td>23</td><td>GGGTCAGTCGGTCCTGAGAGATG</td><td>180.5</td><td>-3.4</td><td>0.5</td><td>-6.6</td><td>3.8E-11</td><td>8.2E-11</td><td>MB231/MB468</td></tr>
<tr><td>>hg38_21_+_8433172_8446622_RNA45SN1_45S_50ntFlanks@4567.4587.21|TAATGGAATAGGACCGCGGTT</td><td>>hg38_21_+_8433172_8446622_RNA45SN1_45S_50ntFlanks</td><td>4567</td><td>4587</td><td>21</td><td>TAATGGAATAGGACCGCGGTT</td><td>414.6</td><td>-4.6</td><td>0.7</td><td>-6.6</td><td>4.5E-11</td><td>8.2E-11</td><td>BT20/MB468</td></tr>
<tr><td>>hg38_21_+_8433172_8446622_RNA45SN1_45S_50ntFlanks@11534.11555.22|GTTCCGCGCGGCGCCTCGCCTC</td><td>>hg38_21_+_8433172_8446622_RNA45SN1_45S_50ntFlanks</td><td>11534</td><td>11555</td><td>22</td><td>GTTCCGCGCGGCGCCTCGCCTC</td><td>116.2</td><td>-8.4</td><td>1.3</td><td>-6.6</td><td>3.9E-11</td><td>8.2E-11</td><td>MB231/MB468</td></tr>
<tr><td>>am_trnaMT_PheGAA_MT_+_577_647@42.62.21|TTTAGACGGGCTCACATCACC</td><td>>am_trnaMT_PheGAA_MT_+_577_647</td><td>42</td><td>62</td><td>21</td><td>TTTAGACGGGCTCACATCACC</td><td>533.5</td><td>8.1</td><td>1.2</td><td>6.6</td><td>4.5E-11</td><td>8.2E-11</td><td>BT20/MB468</td></tr>
<tr><td>>hg38_21_+_8433172_8446622_RNA45SN1_45S_50ntFlanks@3769.3792.24|CGCACGGCCGGTACAGTGAAACTG</td><td>>hg38_21_+_8433172_8446622_RNA45SN1_45S_50ntFlanks</td><td>3769</td><td>3792</td><td>24</td><td>CGCACGGCCGGTACAGTGAAACTG</td><td>194.0</td><td>-4.4</td><td>0.7</td><td>-6.6</td><td>4.5E-11</td><td>8.3E-11</td><td>BT20/MB468</td></tr>
<tr><td>>hg38_21_+_8433172_8446622_RNA45SN1_45S_50ntFlanks@11887.11904.18|GAGCTTGACTCTAGTCTG</td><td>>hg38_21_+_8433172_8446622_RNA45SN1_45S_50ntFlanks</td><td>11887</td><td>11904</td><td>18</td><td>GAGCTTGACTCTAGTCTG</td><td>101.1</td><td>-8.7</td><td>1.3</td><td>-6.6</td><td>4.5E-11</td><td>8.3E-11</td><td>BT20/MB468</td></tr>
<tr><td>>hg38_21_+_8433172_8446622_RNA45SN1_45S_50ntFlanks@12709.12742.34|CCGCGGAGCCTCGGTTGGCCTCGGATAGCCGGTC</td><td>>hg38_21_+_8433172_8446622_RNA45SN1_45S_50ntFlanks</td><td>12709</td><td>12742</td><td>34</td><td>CCGCGGAGCCTCGGTTGGCCTCGGATAGCCGGTC</td><td>96.5</td><td>-8.6</td><td>1.3</td><td>-6.6</td><td>4.5E-11</td><td>8.3E-11</td><td>BT20/MB468</td></tr>
<tr><td>>hg38_21_+_8433172_8446622_RNA45SN1_45S_50ntFlanks@12457.12486.30|CAAGCGTTGGATTGTTCACCCACTAATAGG</td><td>>hg38_21_+_8433172_8446622_RNA45SN1_45S_50ntFlanks</td><td>12457</td><td>12486</td><td>30</td><td>CAAGCGTTGGATTGTTCACCCACTAATAGG</td><td>492.9</td><td>-4.1</td><td>0.6</td><td>-6.6</td><td>4.6E-11</td><td>8.3E-11</td><td>BT20/MB468</td></tr>
<tr><td>>MI0000114|hsa-mir-107&WithFlank&10|-|89592741|89592833@56.77.22&[MIMAT0000104&hsa-miR-107&offsets|0|-1;m-78&10|-|89592758|89592778&offsets|0|+1]|AGCAGCATTGTACAGGGCTATC</td><td>>MI0000114|hsa-mir-107&WithFlank&10|-|89592741|89592833</td><td>56</td><td>77</td><td>22</td><td>AGCAGCATTGTACAGGGCTATC</td><td>1181.6</td><td>5.6</td><td>0.8</td><td>6.6</td><td>4.6E-11</td><td>8.4E-11</td><td>BT20/MB468</td></tr>
<tr><td>>hg38_21_+_8433172_8446622_RNA45SN1_45S_50ntFlanks@12416.12433.18|TTCGATGTCGGCTCTTCC</td><td>>hg38_21_+_8433172_8446622_RNA45SN1_45S_50ntFlanks</td><td>12416</td><td>12433</td><td>18</td><td>TTCGATGTCGGCTCTTCC</td><td>881.7</td><td>3.7</td><td>0.6</td><td>6.6</td><td>4.6E-11</td><td>8.4E-11</td><td>BT20/MB468</td></tr>
<tr><td>>hg38_21_+_8433172_8446622_RNA45SN1_45S_50ntFlanks@12712.12751.40|CGGAGCCTCGGTTGGCCTCGGATAGCCGGTCCCCCGCCTG</td><td>>hg38_21_+_8433172_8446622_RNA45SN1_45S_50ntFlanks</td><td>12712</td><td>12751</td><td>40</td><td>CGGAGCCTCGGTTGGCCTCGGATAGCCGGTCCCCCGCCTG</td><td>110.2</td><td>-8.8</td><td>1.3</td><td>-6.6</td><td>4.6E-11</td><td>8.4E-11</td><td>BT20/MB468</td></tr>
<tr><td>>hg38_21_+_8433172_8446622_RNA45SN1_45S_50ntFlanks@7984.8012.29|AGATCAGACGTGGCGACCCGCTGAATTTA</td><td>>hg38_21_+_8433172_8446622_RNA45SN1_45S_50ntFlanks</td><td>7984</td><td>8012</td><td>29</td><td>AGATCAGACGTGGCGACCCGCTGAATTTA</td><td>141.1</td><td>-4.2</td><td>0.6</td><td>-6.6</td><td>4.6E-11</td><td>8.5E-11</td><td>BT20/MB468</td></tr>
<tr><td>>trna10_AspGTC_12_-_125424193_125424264@1.30.30,trna12_AspGTC_12_-_125411891_125411962@1.30.30,trna38_AspGTC_17_-_8125556_8125627@1.30.30,trna45_AspGTC_6_+_27447453_27447524@1.30.30,trna48_AspGTC_6_+_27471523_27471594@1.30.30,trna4_AspGTC_12_+_96429799_96429870@1.30.30,trna69_AspGTC_1_-_161440205_161440276@1.30.30,trna72_AspGTC_1_-_161432824_161432895@1.30.30,trna75_AspGTC_1_-_161425414_161425485@1.30.30,trna78_AspGTC_1_-_161418033_161418104@1.30.30,trna81_AspGTC_1_-_161410615_161410686@1.30.30|TCCTCGTTAGTATAGTGGTGAGTATCCCCG</td><td>>trna10_AspGTC_12_-_125424193_125424264</td><td>1</td><td>30</td><td>30</td><td>TCCTCGTTAGTATAGTGGTGAGTATCCCCG</td><td>394.3</td><td>-6.4</td><td>1.0</td><td>-6.6</td><td>4.6E-11</td><td>8.5E-11</td><td>BT20/MB468</td></tr>
<tr><td>>MI0000071|hsa-mir-17&WithFlank&13|+|91350599|91350694@20.41.22(+1A)&[MIMAT0000070&hsa-miR-17-5p&offsets|0|-1(+1A);m-88&13|+|91350618|91350640&offsets|0|-1(+1A)]|CAAAGTGCTTACAGTGCAGGTAA</td><td>>MI0000071|hsa-mir-17&WithFlank&13|+|91350599|91350694</td><td>20</td><td>41</td><td>22(+1A)</td><td>CAAAGTGCTTACAGTGCAGGTAA</td><td>3074.9</td><td>6.3</td><td>1.0</td><td>6.6</td><td>4.7E-11</td><td>8.6E-11</td><td>BT20/MB468</td></tr>
<tr><td>>hg38_21_+_8433172_8446622_RNA45SN1_45S_50ntFlanks@12490.12509.20|CGTGAGCTGGGTTTAGACCG</td><td>>hg38_21_+_8433172_8446622_RNA45SN1_45S_50ntFlanks</td><td>12490</td><td>12509</td><td>20</td><td>CGTGAGCTGGGTTTAGACCG</td><td>102.5</td><td>-8.3</td><td>1.3</td><td>-6.6</td><td>4.0E-11</td><td>8.6E-11</td><td>MB231/MB468</td></tr>
<tr><td>>MI0000063|hsa-let-7b&WithFlank&22|+|46113680|46113774@12.31.20(+1G)&[MIMAT0000063&hsa-let-7b-5p&offsets|0|-2(+1G);m-14&22|+|46113691|46113712&offsets|0|-2(+1G)]|TGAGGTAGTAGGTTGTGTGGG</td><td>>MI0000063|hsa-let-7b&WithFlank&22|+|46113680|46113774</td><td>12</td><td>31</td><td>20(+1G)</td><td>TGAGGTAGTAGGTTGTGTGGG</td><td>910.6</td><td>9.4</td><td>1.4</td><td>6.6</td><td>4.7E-11</td><td>8.6E-11</td><td>BT20/MB468</td></tr>
<tr><td>>hg38_21_+_8433172_8446622_RNA45SN1_45S_50ntFlanks@12997.13038.42|CGCTGCGATCTATTGAAAGTCAGCCCTCGACACAAGGGTTTG</td><td>>hg38_21_+_8433172_8446622_RNA45SN1_45S_50ntFlanks</td><td>12997</td><td>13038</td><td>42</td><td>CGCTGCGATCTATTGAAAGTCAGCCCTCGACACAAGGGTTTG</td><td>1079.6</td><td>9.4</td><td>1.4</td><td>6.6</td><td>4.8E-11</td><td>8.7E-11</td><td>BT20/MB468</td></tr>
<tr><td>>hg38_21_+_8433172_8446622_RNA45SN1_45S_50ntFlanks@12479.12509.31|CTAATAGGGAACGTGAGCTGGGTTTAGACCG</td><td>>hg38_21_+_8433172_8446622_RNA45SN1_45S_50ntFlanks</td><td>12479</td><td>12509</td><td>31</td><td>CTAATAGGGAACGTGAGCTGGGTTTAGACCG</td><td>147.5</td><td>-4.8</td><td>0.7</td><td>-6.6</td><td>4.8E-11</td><td>8.8E-11</td><td>BT20/MB468</td></tr>
<tr><td>>hg38_21_+_8433172_8446622_RNA45SN1_45S_50ntFlanks@11572.11602.31|CCGACTTAGAACTGGTGCGGACCAGGGGAAT</td><td>>hg38_21_+_8433172_8446622_RNA45SN1_45S_50ntFlanks</td><td>11572</td><td>11602</td><td>31</td><td>CCGACTTAGAACTGGTGCGGACCAGGGGAAT</td><td>3118.5</td><td>-3.9</td><td>0.6</td><td>-6.6</td><td>4.2E-11</td><td>8.9E-11</td><td>MB231/MB468</td></tr>
<tr><td>>hg38_21_+_8433172_8446622_RNA45SN1_45S_50ntFlanks@8472.8489.18|CGGCCCGGCGGATCTTTC</td><td>>hg38_21_+_8433172_8446622_RNA45SN1_45S_50ntFlanks</td><td>8472</td><td>8489</td><td>18</td><td>CGGCCCGGCGGATCTTTC</td><td>95.5</td><td>-8.6</td><td>1.3</td><td>-6.6</td><td>4.9E-11</td><td>8.9E-11</td><td>BT20/MB468</td></tr>
<tr><td>>hg38_21_+_8433172_8446622_RNA45SN1_45S_50ntFlanks@12624.12659.36|CCAATGGGGCGAAGCTACCATCTGTGGGATTATGAC</td><td>>hg38_21_+_8433172_8446622_RNA45SN1_45S_50ntFlanks</td><td>12624</td><td>12659</td><td>36</td><td>CCAATGGGGCGAAGCTACCATCTGTGGGATTATGAC</td><td>176.4</td><td>-3.4</td><td>0.5</td><td>-6.6</td><td>4.2E-11</td><td>9.0E-11</td><td>MB231/MB468</td></tr>
<tr><td>>MI0000253|hsa-mir-148a&WithFlank&7|-|25949913|25949992@50.71.22(+1A)&[MIMAT0000243&hsa-miR-148a-3p&offsets|0|0(+1A);m-7&7|-|25949922|25949943&offsets|0|0(+1A)]|TCAGTGCACTACAGAACTTTGTA</td><td>>MI0000253|hsa-mir-148a&WithFlank&7|-|25949913|25949992</td><td>50</td><td>71</td><td>22(+1A)</td><td>TCAGTGCACTACAGAACTTTGTA</td><td>920.9</td><td>5.9</td><td>0.9</td><td>7.0</td><td>3.0E-12</td><td>9.0E-11</td><td>BT20/MB231</td></tr>
<tr><td>>hg38_21_+_8433172_8446622_RNA45SN1_45S_50ntFlanks@4472.4496.25|CTTAGCTGAGTGTCCCGCGGGGCCC</td><td>>hg38_21_+_8433172_8446622_RNA45SN1_45S_50ntFlanks</td><td>4472</td><td>4496</td><td>25</td><td>CTTAGCTGAGTGTCCCGCGGGGCCC</td><td>157.0</td><td>-7.0</td><td>1.1</td><td>-6.6</td><td>4.2E-11</td><td>9.0E-11</td><td>MB231/MB468</td></tr>
<tr><td>>hg38_21_+_8433172_8446622_RNA45SN1_45S_50ntFlanks@7983.8007.25|CAGATCAGACGTGGCGACCCGCTGA</td><td>>hg38_21_+_8433172_8446622_RNA45SN1_45S_50ntFlanks</td><td>7983</td><td>8007</td><td>25</td><td>CAGATCAGACGTGGCGACCCGCTGA</td><td>322.3</td><td>-4.8</td><td>0.7</td><td>-6.6</td><td>5.0E-11</td><td>9.1E-11</td><td>BT20/MB468</td></tr>
<tr><td>>hg38_21_+_8433172_8446622_RNA45SN1_45S_50ntFlanks@3782.3802.21|CAGTGAAACTGCGAATGGCTC</td><td>>hg38_21_+_8433172_8446622_RNA45SN1_45S_50ntFlanks</td><td>3782</td><td>3802</td><td>21</td><td>CAGTGAAACTGCGAATGGCTC</td><td>160.1</td><td>-3.6</td><td>0.5</td><td>-6.6</td><td>5.0E-11</td><td>9.1E-11</td><td>BT20/MB468</td></tr>
<tr><td>>hg38_21_+_8433172_8446622_RNA45SN1_45S_50ntFlanks@11552.11590.39|CCTCGGCCGGCGCCTAGCAGCCGACTTAGAACTGGTGCG</td><td>>hg38_21_+_8433172_8446622_RNA45SN1_45S_50ntFlanks</td><td>11552</td><td>11590</td><td>39</td><td>CCTCGGCCGGCGCCTAGCAGCCGACTTAGAACTGGTGCG</td><td>394.4</td><td>3.7</td><td>0.6</td><td>6.6</td><td>4.4E-11</td><td>9.3E-11</td><td>MB231/MB468</td></tr>
<tr><td>>hg38_21_+_8433172_8446622_RNA45SN1_45S_50ntFlanks@4738.4768.31|AAAGTCGGAGGTTCGAAGACGATCAGATACC</td><td>>hg38_21_+_8433172_8446622_RNA45SN1_45S_50ntFlanks</td><td>4738</td><td>4768</td><td>31</td><td>AAAGTCGGAGGTTCGAAGACGATCAGATACC</td><td>1366.7</td><td>9.1</td><td>1.4</td><td>6.6</td><td>4.4E-11</td><td>9.3E-11</td><td>MB231/MB468</td></tr>
<tr><td>>trna116_GluCTC_1_-_145399233_145399304@32.69.38,trna59_GluCTC_1_+_249168447_249168518@32.69.38,trna71_GluCTC_1_-_161439189_161439260@32.69.38,trna74_GluCTC_1_-_161431809_161431880@32.69.38,trna77_GluCTC_1_-_161424398_161424469@32.69.38,trna77_GluCTC_6_+_28949976_28950047@32.69.38,trna80_GluCTC_1_-_161417018_161417089@32.69.38,trna87_GluCTC_6_-_126101393_126101464@32.69.38|CTCTCACCGCCGCGGCCCGGGTTCGATTCCCGGTCAGG</td><td>>trna116_GluCTC_1_-_145399233_145399304</td><td>32</td><td>69</td><td>38</td><td>CTCTCACCGCCGCGGCCCGGGTTCGATTCCCGGTCAGG</td><td>121.5</td><td>-8.5</td><td>1.3</td><td>-6.6</td><td>4.4E-11</td><td>9.3E-11</td><td>MB231/MB468</td></tr>
<tr><td>>hg38_21_+_8433172_8446622_RNA45SN1_45S_50ntFlanks@9532.9553.22|AGAGGAAACTCTGGTGGAGGTC</td><td>>hg38_21_+_8433172_8446622_RNA45SN1_45S_50ntFlanks</td><td>9532</td><td>9553</td><td>22</td><td>AGAGGAAACTCTGGTGGAGGTC</td><td>284.6</td><td>6.4</td><td>1.0</td><td>6.6</td><td>4.4E-11</td><td>9.3E-11</td><td>MB231/MB468</td></tr>
<tr><td>>hg38_21_+_8433172_8446622_RNA45SN1_45S_50ntFlanks@5417.5445.29|CTACCGATTGGATGGTTTAGTGAGGCCCT</td><td>>hg38_21_+_8433172_8446622_RNA45SN1_45S_50ntFlanks</td><td>5417</td><td>5445</td><td>29</td><td>CTACCGATTGGATGGTTTAGTGAGGCCCT</td><td>180.3</td><td>-5.9</td><td>0.9</td><td>-6.6</td><td>5.1E-11</td><td>9.4E-11</td><td>BT20/MB468</td></tr>
<tr><td>>hg38_21_+_8433172_8446622_RNA45SN1_45S_50ntFlanks@11571.11612.42|GCCGACTTAGAACTGGTGCGGACCAGGGGAATCCGACTGTTT</td><td>>hg38_21_+_8433172_8446622_RNA45SN1_45S_50ntFlanks</td><td>11571</td><td>11612</td><td>42</td><td>GCCGACTTAGAACTGGTGCGGACCAGGGGAATCCGACTGTTT</td><td>349.5</td><td>5.4</td><td>0.8</td><td>6.6</td><td>4.5E-11</td><td>9.5E-11</td><td>MB231/MB468</td></tr>
<tr><td>>hg38_21_+_8433172_8446622_RNA45SN1_45S_50ntFlanks@7984.8007.24|AGATCAGACGTGGCGACCCGCTGA</td><td>>hg38_21_+_8433172_8446622_RNA45SN1_45S_50ntFlanks</td><td>7984</td><td>8007</td><td>24</td><td>AGATCAGACGTGGCGACCCGCTGA</td><td>151.8</td><td>-5.5</td><td>0.8</td><td>-6.6</td><td>4.5E-11</td><td>9.5E-11</td><td>MB231/MB468</td></tr>
<tr><td>>hg38_21_+_8433172_8446622_RNA45SN1_45S_50ntFlanks@11881.11903.23|CCTGTTGAGCTTGACTCTAGTCT</td><td>>hg38_21_+_8433172_8446622_RNA45SN1_45S_50ntFlanks</td><td>11881</td><td>11903</td><td>23</td><td>CCTGTTGAGCTTGACTCTAGTCT</td><td>297.7</td><td>-3.0</td><td>0.5</td><td>-6.6</td><td>4.5E-11</td><td>9.5E-11</td><td>MB231/MB468</td></tr>
<tr><td>>trnaMT_GluTTC_MT_-_14674_14742@39.71.33,trnalookalike8_GluTTC_5_-_93905172_93905240@39.71.33|ATTGGTCGTGGTTGTAGTCCGTGCGAGAATACC</td><td>>trnaMT_GluTTC_MT_-_14674_14742</td><td>39</td><td>71</td><td>33</td><td>ATTGGTCGTGGTTGTAGTCCGTGCGAGAATACC</td><td>1219.4</td><td>8.8</td><td>1.3</td><td>6.6</td><td>5.2E-11</td><td>9.6E-11</td><td>BT20/MB468</td></tr>
<tr><td>>hg38_21_+_8433172_8446622_RNA45SN1_45S_50ntFlanks@4760.4786.27|TCAGATACCGTCGTAGTTCCGACCATA</td><td>>hg38_21_+_8433172_8446622_RNA45SN1_45S_50ntFlanks</td><td>4760</td><td>4786</td><td>27</td><td>TCAGATACCGTCGTAGTTCCGACCATA</td><td>361.2</td><td>-4.8</td><td>0.7</td><td>-6.6</td><td>5.3E-11</td><td>9.6E-11</td><td>BT20/MB468</td></tr>
<tr><td>>hg38_21_+_8433172_8446622_RNA45SN1_45S_50ntFlanks@7989.8019.31|AGACGTGGCGACCCGCTGAATTTAAGCATAT</td><td>>hg38_21_+_8433172_8446622_RNA45SN1_45S_50ntFlanks</td><td>7989</td><td>8019</td><td>31</td><td>AGACGTGGCGACCCGCTGAATTTAAGCATAT</td><td>868.7</td><td>8.4</td><td>1.3</td><td>6.6</td><td>4.6E-11</td><td>9.7E-11</td><td>MB231/MB468</td></tr>
<tr><td>>hg38_21_+_8433172_8446622_RNA45SN1_45S_50ntFlanks@11561.11601.41|GCGCCTAGCAGCCGACTTAGAACTGGTGCGGACCAGGGGAA</td><td>>hg38_21_+_8433172_8446622_RNA45SN1_45S_50ntFlanks</td><td>11561</td><td>11601</td><td>41</td><td>GCGCCTAGCAGCCGACTTAGAACTGGTGCGGACCAGGGGAA</td><td>343.4</td><td>-5.9</td><td>0.9</td><td>-6.6</td><td>4.6E-11</td><td>9.7E-11</td><td>MB231/MB468</td></tr>
<tr><td>>hg38_21_+_8433172_8446622_RNA45SN1_45S_50ntFlanks@11115.11133.19|CGGCAGGGGCCGGCGGCGG</td><td>>hg38_21_+_8433172_8446622_RNA45SN1_45S_50ntFlanks</td><td>11115</td><td>11133</td><td>19</td><td>CGGCAGGGGCCGGCGGCGG</td><td>103.8</td><td>-8.3</td><td>1.3</td><td>-6.6</td><td>4.6E-11</td><td>9.7E-11</td><td>MB231/MB468</td></tr>
<tr><td>>MI0000114|hsa-mir-107&WithFlank&10|-|89592741|89592833@56.78.23&[MIMAT0000104&hsa-miR-107&offsets|0|0;m-78&10|-|89592758|89592778&offsets|0|+2]|AGCAGCATTGTACAGGGCTATCA</td><td>>MI0000114|hsa-mir-107&WithFlank&10|-|89592741|89592833</td><td>56</td><td>78</td><td>23</td><td>AGCAGCATTGTACAGGGCTATCA</td><td>904.6</td><td>5.2</td><td>0.8</td><td>6.6</td><td>4.6E-11</td><td>9.7E-11</td><td>MB231/MB468</td></tr>
</table>

<table>
<tr><th></th><th></th><th></th><th></th><th></th><th></th><th></th><th></th><th></th><th></th><th></th><th></th><th></th></tr>
<tr><td>>hg38_21_+_8433172_8446622_RNA45SN1_45S_50ntFlanks@10408.10427.20|GGTCAGTCGGTCCTGAGAGA</td><td>>hg38_21_+_8433172_8446622_RNA45SN1_45S_50ntFlanks</td><td>10408</td><td>10427</td><td>20</td><td>GGTCAGTCGGTCCTGAGAGA</td><td>112.7</td><td>-8.8</td><td>1.3</td><td>-6.6</td><td>5.3E-11</td><td>9.7E-11</td><td>BT20/MB468</td></tr>
<tr><td>>hg38_21_+_8433172_8446622_RNA45SN1_45S_50ntFlanks@8845.8867.23|CGGCGGCGGCGGCGGCGGCGGGA</td><td>>hg38_21_+_8433172_8446622_RNA45SN1_45S_50ntFlanks</td><td>8845</td><td>8867</td><td>23</td><td>CGGCGGCGGCGGCGGCGGCGGGA</td><td>86.6</td><td>-8.4</td><td>1.3</td><td>-6.6</td><td>5.4E-11</td><td>9.8E-11</td><td>BT20/MB468</td></tr>
<tr><td>>hg38_21_+_8433172_8446622_RNA45SN1_45S_50ntFlanks@12875.12892.18|CTGGAAAGGCGGCCGCCC</td><td>>hg38_21_+_8433172_8446622_RNA45SN1_45S_50ntFlanks</td><td>12875</td><td>12892</td><td>18</td><td>CTGGAAAGGCGGCCGCCC</td><td>98.8</td><td>-8.6</td><td>1.3</td><td>-6.6</td><td>5.4E-11</td><td>9.8E-11</td><td>BT20/MB468</td></tr>
<tr><td>>trnaMT_PheGAA_MT_+_577_647@42.74.33|TTTAGACGGGCTCACATCACCCCATAAACACCA</td><td>>trnaMT_PheGAA_MT_+_577_647</td><td>42</td><td>74</td><td>33</td><td>TTTAGACGGGCTCACATCACCCCATAAACACCA</td><td>864.7</td><td>10.0</td><td>1.5</td><td>6.6</td><td>5.4E-11</td><td>9.8E-11</td><td>BT20/MB468</td></tr>
<tr><td>>hg38_21_+_8433172_8446622_RNA45SN1_45S_50ntFlanks@10587.10608.22|CCCCGGGGAGAGTTCTCTTTTC</td><td>>hg38_21_+_8433172_8446622_RNA45SN1_45S_50ntFlanks</td><td>10587</td><td>10608</td><td>22</td><td>CCCCGGGGAGAGTTCTCTTTTC</td><td>145.3</td><td>-6.0</td><td>0.9</td><td>-6.6</td><td>5.4E-11</td><td>9.9E-11</td><td>BT20/MB468</td></tr>
<tr><td>>hg38_21_+_8433172_8446622_RNA45SN1_45S_50ntFlanks@10398.10424.27|GTTGAACATGGGTCAGTCGGTCCTGAG</td><td>>hg38_21_+_8433172_8446622_RNA45SN1_45S_50ntFlanks</td><td>10398</td><td>10424</td><td>27</td><td>GTTGAACATGGGTCAGTCGGTCCTGAG</td><td>1055.4</td><td>7.5</td><td>1.1</td><td>6.6</td><td>5.5E-11</td><td>1.0E-10</td><td>BT20/MB468</td></tr>
<tr><td>>trnaMT_LysTTT_MT_+_8295_8364@1.26.26,trnalookalike7_LysTTT_1_+_568844_568913@1.26.26|CACTGTAAAGCTAACTTAGCATTAAC</td><td>>trnaMT_LysTTT_MT_+_8295_8364</td><td>1</td><td>26</td><td>26</td><td>CACTGTAAAGCTAACTTAGCATTAAC</td><td>1107.1</td><td>4.5</td><td>0.7</td><td>6.6</td><td>4.7E-11</td><td>1.0E-10</td><td>MB231/MB468</td></tr>
<tr><td>>hg38_21_+_8433172_8446622_RNA45SN1_45S_50ntFlanks@10392.10435.44|ACAGCAGTTGAACATGGGTCAGTCGGTCCTGAGAGATGGGCGAG</td><td>>hg38_21_+_8433172_8446622_RNA45SN1_45S_50ntFlanks</td><td>10392</td><td>10435</td><td>44</td><td>ACAGCAGTTGAACATGGGTCAGTCGGTCCTGAGAGATGGGCGAG</td><td>424.3</td><td>-10.7</td><td>1.6</td><td>-6.6</td><td>5.5E-11</td><td>1.0E-10</td><td>BT20/MB468</td></tr>
<tr><td>>hg38_21_+_8433172_8446622_RNA45SN1_45S_50ntFlanks@10838.10868.31|ATCCGTAACTTCGGGATAAGGATTGGCTCTA</td><td>>hg38_21_+_8433172_8446622_RNA45SN1_45S_50ntFlanks</td><td>10838</td><td>10868</td><td>31</td><td>ATCCGTAACTTCGGGATAAGGATTGGCTCTA</td><td>132.0</td><td>-9.0</td><td>1.4</td><td>-6.6</td><td>5.5E-11</td><td>1.0E-10</td><td>BT20/MB468</td></tr>
<tr><td>>hg38_21_+_8433172_8446622_RNA45SN1_45S_50ntFlanks@476.521.46|TTCGTGATCGATGTGGTGACGTCGTGCTCTCCCGGGCCGGGTCCGA</td><td>>hg38_21_+_8433172_8446622_RNA45SN1_45S_50ntFlanks</td><td>476</td><td>521</td><td>46</td><td>TTCGTGATCGATGTGGTGACGTCGTGCTCTCCCGGGCCGGGTCCGA</td><td>756.7</td><td>9.4</td><td>1.4</td><td>6.6</td><td>5.5E-11</td><td>1.0E-10</td><td>BT20/MB468</td></tr>
<tr><td>>hg38_21_+_8433172_8446622_RNA45SN1_45S_50ntFlanks@10587.10608.22|CCCCGGGGAGAGTTCTCTTTTC</td><td>>hg38_21_+_8433172_8446622_RNA45SN1_45S_50ntFlanks</td><td>10587</td><td>10608</td><td>22</td><td>CCCCGGGGAGAGTTCTCTTTTC</td><td>117.7</td><td>-8.5</td><td>1.3</td><td>-6.6</td><td>4.7E-11</td><td>1.0E-10</td><td>MB231/MB468</td></tr>
<tr><td>>hg38_21_+_8433172_8446622_RNA45SN1_45S_50ntFlanks@7986.8015.30|ATCAGACGTGGCGACCCGCTGAATTTAAGC</td><td>>hg38_21_+_8433172_8446622_RNA45SN1_45S_50ntFlanks</td><td>7986</td><td>8015</td><td>30</td><td>ATCAGACGTGGCGACCCGCTGAATTTAAGC</td><td>84.2</td><td>-8.4</td><td>1.3</td><td>-6.6</td><td>5.6E-11</td><td>1.0E-10</td><td>BT20/MB468</td></tr>
<tr><td>>hg38_21_+_8433172_8446622_RNA45SN1_45S_50ntFlanks@10806.10827.22|AACAATGTAGGTAAGGGAAGTC</td><td>>hg38_21_+_8433172_8446622_RNA45SN1_45S_50ntFlanks</td><td>10806</td><td>10827</td><td>22</td><td>AACAATGTAGGTAAGGGAAGTC</td><td>116.5</td><td>-4.1</td><td>0.6</td><td>-6.6</td><td>5.6E-11</td><td>1.0E-10</td><td>BT20/MB468</td></tr>
<tr><td>>hg38_21_+_8433172_8446622_RNA45SN1_45S_50ntFlanks@6726.6744.19|CAGGACACATTGATCATCG</td><td>>hg38_21_+_8433172_8446622_RNA45SN1_45S_50ntFlanks</td><td>6726</td><td>6744</td><td>19</td><td>CAGGACACATTGATCATCG</td><td>294.7</td><td>-7.0</td><td>1.1</td><td>-6.6</td><td>5.7E-11</td><td>1.0E-10</td><td>BT20/MB468</td></tr>
<tr><td>>trnaMT_SerGCT_MT_+_12207_12265@13.62.50|AAGAACTGCTAACTCATGCCCCCATGTCTAACAACATGGCTTTCTCACCA</td><td>>trnaMT_SerGCT_MT_+_12207_12265</td><td>13</td><td>62</td><td>50</td><td>AAGAACTGCTAACTCATGCCCCCATGTCTAACAACATGGCTTTCTCACCA</td><td>2119.1</td><td>16.0</td><td>2.4</td><td>6.6</td><td>4.9E-11</td><td>1.0E-10</td><td>MB231/MB468</td></tr>
<tr><td>>hg38_21_+_8433172_8446622_RNA45SN1_45S_50ntFlanks@11569.11591.23|CAGCCGACTTAGAACTGGTGCGG</td><td>>hg38_21_+_8433172_8446622_RNA45SN1_45S_50ntFlanks</td><td>11569</td><td>11591</td><td>23</td><td>CAGCCGACTTAGAACTGGTGCGG</td><td>180.7</td><td>-5.8</td><td>0.9</td><td>-6.6</td><td>4.9E-11</td><td>1.0E-10</td><td>MB231/MB468</td></tr>
<tr><td>>hg38_21_+_8433172_8446622_RNA45SN1_45S_50ntFlanks@12931.12956.26|GCGCTAAACCATTCGTAGACGACCTG</td><td>>hg38_21_+_8433172_8446622_RNA45SN1_45S_50ntFlanks</td><td>12931</td><td>12956</td><td>26</td><td>GCGCTAAACCATTCGTAGACGACCTG</td><td>359.0</td><td>-3.9</td><td>0.6</td><td>-6.6</td><td>5.0E-11</td><td>1.1E-10</td><td>MB231/MB468</td></tr>
<tr><td>>hg38_21_+_8433172_8446622_RNA45SN1_45S_50ntFlanks@6709.6734.26|AGAATTAATGTGAATTGCAGGACACA</td><td>>hg38_21_+_8433172_8446622_RNA45SN1_45S_50ntFlanks</td><td>6709</td><td>6734</td><td>26</td><td>AGAATTAATGTGAATTGCAGGACACA</td><td>90.1</td><td>-8.5</td><td>1.3</td><td>-6.5</td><td>5.8E-11</td><td>1.1E-10</td><td>BT20/MB468</td></tr>
<tr><td>>trna2_ArgCCT_16_+_3202901_3202973@13.30.18,trna3_ArgCCT_7_+_139025446_139025518@13.30.18|TAATGGATAAGGCATTGG</td><td>>trna2_ArgCCT_16_+_3202901_3202973</td><td>13</td><td>30</td><td>18</td><td>TAATGGATAAGGCATTGG</td><td>120.7</td><td>-8.9</td><td>1.4</td><td>-6.5</td><td>5.9E-11</td><td>1.1E-10</td><td>BT20/MB468</td></tr>
<tr><td>>hg38_21_+_8433172_8446622_RNA45SN1_45S_50ntFlanks@12408.12433.26|TTTGATCCTTCGATGTCGGCTCTTCC</td><td>>hg38_21_+_8433172_8446622_RNA45SN1_45S_50ntFlanks</td><td>12408</td><td>12433</td><td>26</td><td>TTTGATCCTTCGATGTCGGCTCTTCC</td><td>473.8</td><td>4.0</td><td>0.6</td><td>6.6</td><td>5.0E-11</td><td>1.1E-10</td><td>MB231/MB468</td></tr>
<tr><td>>hg38_21_+_8433172_8446622_RNA45SN1_45S_50ntFlanks@12712.12742.31|CGGAGCCTCGGTTGGCCTCGGATAGCCGGTC</td><td>>hg38_21_+_8433172_8446622_RNA45SN1_45S_50ntFlanks</td><td>12712</td><td>12742</td><td>31</td><td>CGGAGCCTCGGTTGGCCTCGGATAGCCGGTC</td><td>91.1</td><td>-8.5</td><td>1.3</td><td>-6.5</td><td>5.9E-11</td><td>1.1E-10</td><td>BT20/MB468</td></tr>
<tr><td>>hg38_21_+_8433172_8446622_RNA45SN1_45S_50ntFlanks@11614.11640.27|ATTAAAACAAAGCATCGCGAAGGCCCG</td><td>>hg38_21_+_8433172_8446622_RNA45SN1_45S_50ntFlanks</td><td>11614</td><td>11640</td><td>27</td><td>ATTAAAACAAAGCATCGCGAAGGCCCG</td><td>144.7</td><td>-5.7</td><td>0.9</td><td>-6.6</td><td>5.1E-11</td><td>1.1E-10</td><td>MB231/MB468</td></tr>
<tr><td>>hg38_21_+_8433172_8446622_RNA45SN1_45S_50ntFlanks@10781.10804.24|AGGTGAACAGCCTCTGGCATGTTG</td><td>>hg38_21_+_8433172_8446622_RNA45SN1_45S_50ntFlanks</td><td>10781</td><td>10804</td><td>24</td><td>AGGTGAACAGCCTCTGGCATGTTG</td><td>226.4</td><td>-6.0</td><td>0.9</td><td>-6.6</td><td>5.2E-11</td><td>1.1E-10</td><td>MB231/MB468</td></tr>
<tr><td>>MI0000736|hsa-mir-30c-1&WithFlank&1|+|40757278|40757378@23.45.23(+1G);MI0000254|hsa-mir-30c-2&WithFlank&6|-|71376954|71377037@13.35.23(+1G)&[MIMAT0000244&hsa-miR-30c-5p&offsets|0|0(+1G);m-57&1|+|40757300|40757323&offsets|0|-1(+1G)];[MIMAT0000244_1&hsa-miR-30c-5p&offsets|0|0(+1G);m-56&6|-|71377002|71377025&offsets|0|-1(+1G)]|TGTAAACATCCTACACTCTCAGCG</td><td>>MI0000736|hsa-mir-30c-1&WithFlank&1|+|40757278|40757378</td><td>23</td><td>45</td><td>23(+1G)</td><td>TGTAAACATCCTACACTCTCAGCG</td><td>5040.0</td><td>6.3</td><td>1.0</td><td>6.5</td><td>6.0E-11</td><td>1.1E-10</td><td>BT20/MB468</td></tr>
<tr><td>>hg38_21_+_8433172_8446622_RNA45SN1_45S_50ntFlanks@10393.10430.38|CAGCAGTTGAACATGGGTCAGTCGGTCCTGAGAGATGG</td><td>>hg38_21_+_8433172_8446622_RNA45SN1_45S_50ntFlanks</td><td>10393</td><td>10430</td><td>38</td><td>CAGCAGTTGAACATGGGTCAGTCGGTCCTGAGAGATGG</td><td>392.2</td><td>-5.6</td><td>0.9</td><td>-6.5</td><td>6.0E-11</td><td>1.1E-10</td><td>BT20/MB468</td></tr>
<tr><td>>hg38_21_+_8433172_8446622_RNA45SN1_45S_50ntFlanks@12324.12355.32|TTGGGTTTTAAGCAGGAGGTGTCAGAAAAGTT</td><td>>hg38_21_+_8433172_8446622_RNA45SN1_45S_50ntFlanks</td><td>12324</td><td>12355</td><td>32</td><td>TTGGGTTTTAAGCAGGAGGTGTCAGAAAAGTT</td><td>101.6</td><td>-8.2</td><td>1.3</td><td>-6.6</td><td>5.2E-11</td><td>1.1E-10</td><td>MB231/MB468</td></tr>
<tr><td>>hg38_21_+_8433172_8446622_RNA45SN1_45S_50ntFlanks@10404.10429.26|CATGGGTCAGTCGGTCCTGAGAGATG</td><td>>hg38_21_+_8433172_8446622_RNA45SN1_45S_50ntFlanks</td><td>10404</td><td>10429</td><td>26</td><td>CATGGGTCAGTCGGTCCTGAGAGATG</td><td>1036.2</td><td>-3.0</td><td>0.5</td><td>-6.5</td><td>6.1E-11</td><td>1.1E-10</td><td>BT20/MB468</td></tr>
<tr><td>>hg38_21_+_8433172_8446622_RNA45SN1_45S_50ntFlanks@7975.8004.30|CGCGACCTCAGATCAGACGTGGCGACCCGC</td><td>>hg38_21_+_8433172_8446622_RNA45SN1_45S_50ntFlanks</td><td>7975</td><td>8004</td><td>30</td><td>CGCGACCTCAGATCAGACGTGGCGACCCGC</td><td>3459.5</td><td>4.6</td><td>0.7</td><td>6.6</td><td>5.3E-11</td><td>1.1E-10</td><td>MB231/MB468</td></tr>
<tr><td>>hg38_21_+_8433172_8446622_RNA45SN1_45S_50ntFlanks@10761.10780.20|ATATCCGCAGCAGGTCTCCA</td><td>>hg38_21_+_8433172_8446622_RNA45SN1_45S_50ntFlanks</td><td>10761</td><td>10780</td><td>20</td><td>ATATCCGCAGCAGGTCTCCA</td><td>115.9</td><td>-8.8</td><td>1.4</td><td>-6.5</td><td>6.2E-11</td><td>1.1E-10</td><td>BT20/MB468</td></tr>
<tr><td>>MI0000063|hsa-let-7b&WithFlank&22|+|46113680|46113774@66.86.21(+1U)&[MIMAT0004482&hsa-let-7b-3p&offsets|0|-1(+1U);m-171&22|+|46113745|46113766&offsets|0|-1(+1U)]|CTATACAACCTACTGCCTTCCT</td><td>>MI0000063|hsa-let-7b&WithFlank&22|+|46113680|46113774</td><td>66</td><td>86</td><td>21(+1U)</td><td>CTATACAACCTACTGCCTTCCT</td><td>563.0</td><td>7.0</td><td>1.1</td><td>6.5</td><td>6.2E-11</td><td>1.1E-10</td><td>BT20/MB468</td></tr>
<tr><td>>hg38_21_+_8433172_8446622_RNA45SN1_45S_50ntFlanks@4557.4587.31|CAGCTAGGAATAATGGAATAGGACCGCGGTT</td><td>>hg38_21_+_8433172_8446622_RNA45SN1_45S_50ntFlanks</td><td>4557</td><td>4587</td><td>31</td><td>CAGCTAGGAATAATGGAATAGGACCGCGGTT</td><td>194.7</td><td>-5.0</td><td>0.8</td><td>-6.6</td><td>5.4E-11</td><td>1.1E-10</td><td>MB231/MB468</td></tr>
<tr><td>>hg38_21_+_8433172_8446622_RNA45SN1_45S_50ntFlanks@4555.4572.18|CGCAGCTAGGAATAATGG</td><td>>hg38_21_+_8433172_8446622_RNA45SN1_45S_50ntFlanks</td><td>4555</td><td>4572</td><td>18</td><td>CGCAGCTAGGAATAATGG</td><td>405.0</td><td>-4.6</td><td>0.7</td><td>-6.5</td><td>6.3E-11</td><td>1.2E-10</td><td>BT20/MB468</td></tr>
<tr><td>>hg38_21_+_8433172_8446622_RNA45SN1_45S_50ntFlanks@10461.10497.37|GCCTCCGTTGCCCTCGGCCGATCGAAAGGGAGTCGGG</td><td>>hg38_21_+_8433172_8446622_RNA45SN1_45S_50ntFlanks</td><td>10461</td><td>10497</td><td>37</td><td>GCCTCCGTTGCCCTCGGCCGATCGAAAGGGAGTCGGG</td><td>104.2</td><td>-8.3</td><td>1.3</td><td>-6.6</td><td>5.5E-11</td><td>1.2E-10</td><td>MB231/MB468</td></tr>
<tr><td>>hg38_21_+_8433172_8446622_RNA45SN1_45S_50ntFlanks@4783.4800.18|CATAAACGATGCCGACCG</td><td>>hg38_21_+_8433172_8446622_RNA45SN1_45S_50ntFlanks</td><td>4783</td><td>4800</td><td>18</td><td>CATAAACGATGCCGACCG</td><td>81.5</td><td>-8.3</td><td>1.3</td><td>-6.5</td><td>6.4E-11</td><td>1.2E-10</td><td>BT20/MB468</td></tr>
<tr><td>>hg38_21_+_8433172_8446622_RNA45SN1_45S_50ntFlanks@11051.11075.25|CCGTCCTCCCCCCTCCCCGGGGGAG</td><td>>hg38_21_+_8433172_8446622_RNA45SN1_45S_50ntFlanks</td><td>11051</td><td>11075</td><td>25</td><td>CCGTCCTCCCCCCTCCCCGGGGGAG</td><td>124.8</td><td>-8.5</td><td>1.3</td><td>-6.6</td><td>5.5E-11</td><td>1.2E-10</td><td>MB231/MB468</td></tr>
<tr><td>>hg38_21_+_8433172_8446622_RNA45SN1_45S_50ntFlanks@10805.10825.21|GAACAATGTAGGTAAGGGAAG</td><td>>hg38_21_+_8433172_8446622_RNA45SN1_45S_50ntFlanks</td><td>10805</td><td>10825</td><td>21</td><td>GAACAATGTAGGTAAGGGAAG</td><td>125.0</td><td>-8.5</td><td>1.3</td><td>-6.6</td><td>5.6E-11</td><td>1.2E-10</td><td>MB231/MB468</td></tr>
<tr><td>>hg38_21_+_8433172_8446622_RNA45SN1_45S_50ntFlanks@4283.4307.25|CTTTAACGAGGATCCATTGGAGGGC</td><td>>hg38_21_+_8433172_8446622_RNA45SN1_45S_50ntFlanks</td><td>4283</td><td>4307</td><td>25</td><td>CTTTAACGAGGATCCATTGGAGGGC</td><td>126.6</td><td>-5.2</td><td>0.8</td><td>-6.6</td><td>5.6E-11</td><td>1.2E-10</td><td>MB231/MB468</td></tr>
<tr><td>>hg38_21_+_8433172_8446622_RNA45SN1_45S_50ntFlanks@9027.9058.32|CGTCTCCTCGTGGGGGGGCCGGGCCACCCCTC</td><td>>hg38_21_+_8433172_8446622_RNA45SN1_45S_50ntFlanks</td><td>9027</td><td>9058</td><td>32</td><td>CGTCTCCTCGTGGGGGGGCCGGGCCACCCCTC</td><td>92.1</td><td>-8.5</td><td>1.3</td><td>-6.5</td><td>6.5E-11</td><td>1.2E-10</td><td>BT20/MB468</td></tr>
<tr><td>>hg38_21_+_8433172_8446622_RNA45SN1_45S_50ntFlanks@5423.5445.23|ATTGGATGGTTTAGTGAGGCCCT</td><td>>hg38_21_+_8433172_8446622_RNA45SN1_45S_50ntFlanks</td><td>5423</td><td>5445</td><td>23</td><td>ATTGGATGGTTTAGTGAGGCCCT</td><td>113.2</td><td>-8.4</td><td>1.3</td><td>-6.6</td><td>5.6E-11</td><td>1.2E-10</td><td>MB231/MB468</td></tr>
<tr><td>>hg38_21_+_8433172_8446622_RNA45SN1_45S_50ntFlanks@10922.10941.20|CTGGACGAGGCGCCGCCGCC</td><td>>hg38_21_+_8433172_8446622_RNA45SN1_45S_50ntFlanks</td><td>10922</td><td>10941</td><td>20</td><td>CTGGACGAGGCGCCGCCGCC</td><td>263.6</td><td>-6.9</td><td>1.0</td><td>-6.6</td><td>5.6E-11</td><td>1.2E-10</td><td>MB231/MB468</td></tr>
<tr><td>>hg38_21_+_8433172_8446622_RNA45SN1_45S_50ntFlanks@4069.4094.26|CTTTCGATGGTAGTCGCCGTGCCTAC</td><td>>hg38_21_+_8433172_8446622_RNA45SN1_45S_50ntFlanks</td><td>4069</td><td>4094</td><td>26</td><td>CTTTCGATGGTAGTCGCCGTGCCTAC</td><td>108.7</td><td>-3.6</td><td>0.5</td><td>-6.5</td><td>6.5E-11</td><td>1.2E-10</td><td>BT20/MB468</td></tr>
<tr><td>>trna10_GlyTCC_17_+_8124866_8124937@1.30.30|GCGTTGGTGGTATAGTGGTAAGCATAGCTG</td><td>>trna10_GlyTCC_17_+_8124866_8124937</td><td>1</td><td>30</td><td>30</td><td>GCGTTGGTGGTATAGTGGTAAGCATAGCTG</td><td>186.1</td><td>-7.2</td><td>1.1</td><td>-6.6</td><td>5.6E-11</td><td>1.2E-10</td><td>MB231/MB468</td></tr>
<tr><td>>hg38_21_+_8433172_8446622_RNA45SN1_45S_50ntFlanks@10389.10407.19|TGAACAGCAGTTGAACATG</td><td>>hg38_21_+_8433172_8446622_RNA45SN1_45S_50ntFlanks</td><td>10389</td><td>10407</td><td>19</td><td>TGAACAGCAGTTGAACATG</td><td>2555.6</td><td>-5.0</td><td>0.8</td><td>-6.6</td><td>5.7E-11</td><td>1.2E-10</td><td>MB231/MB468</td></tr>
<tr><td>>hg38_21_+_8433172_8446622_RNA45SN1_45S_50ntFlanks@12415.12439.25|CTTCGATGTCGGCTCTTCCTATCAT</td><td>>hg38_21_+_8433172_8446622_RNA45SN1_45S_50ntFlanks</td><td>12415</td><td>12439</td><td>25</td><td>CTTCGATGTCGGCTCTTCCTATCAT</td><td>878.1</td><td>4.3</td><td>0.7</td><td>6.5</td><td>6.6E-11</td><td>1.2E-10</td><td>BT20/MB468</td></tr>
<tr><td>>hg38_21_+_8433172_8446622_RNA45SN1_45S_50ntFlanks@10128.10147.20|CGGCGGCGGCGGCGGCGGCG</td><td>>hg38_21_+_8433172_8446622_RNA45SN1_45S_50ntFlanks</td><td>10128</td><td>10147</td><td>20</td><td>CGGCGGCGGCGGCGGCGGCG</td><td>96.1</td><td>-8.2</td><td>1.2</td><td>-6.6</td><td>5.7E-11</td><td>1.2E-10</td><td>MB231/MB468</td></tr>
<tr><td>>hg38_21_+_8433172_8446622_RNA45SN1_45S_50ntFlanks@8845.8864.20|CGGCGGCGGCGGCGGCGGCG</td><td>>hg38_21_+_8433172_8446622_RNA45SN1_45S_50ntFlanks</td><td>8845</td><td>8864</td><td>20</td><td>CGGCGGCGGCGGCGGCGGCG</td><td>96.1</td><td>-8.2</td><td>1.2</td><td>-6.6</td><td>5.7E-11</td><td>1.2E-10</td><td>MB231/MB468</td></tr>
<tr><td>>hg38_21_+_8433172_8446622_RNA45SN1_45S_50ntFlanks@8194.8211.18|CCCGTGGACGGTGTGAGG</td><td>>hg38_21_+_8433172_8446622_RNA45SN1_45S_50ntFlanks</td><td>8194</td><td>8211</td><td>18</td><td>CCCGTGGACGGTGTGAGG</td><td>694.8</td><td>-5.3</td><td>0.8</td><td>-6.5</td><td>6.6E-11</td><td>1.2E-10</td><td>BT20/MB468</td></tr>
</table>

<table>
<tr><td></td><td></td><td></td><td></td><td></td><td></td><td></td><td></td><td></td><td></td><td></td><td></td><td></td></tr>
<tr><td>>hg38_21_+_8433172_8446622_RNA45SN1_45S_50ntFlanks@8737.8758.22|CCCGTCCGTCCGTCCGTCCGTC</td><td>>hg38_21_+_8433172_8446622_RNA45SN1_45S_50ntFlanks</td><td>8737</td><td>8758</td><td>22</td><td>CCCGTCCGTCCGTCCGTCCGTC</td><td>120.6</td><td>-8.5</td><td>1.3</td><td>-6.6</td><td>5.8E-11</td><td>1.2E-10</td><td>MB231/MB468</td></tr>
<tr><td>>hg38_21_+_8433172_8446622_RNA45SN1_45S_50ntFlanks@8105.8143.39|CGCGGCGGGGCGCGGGACATGTGGCGTACGGAAGACCCG</td><td>>hg38_21_+_8433172_8446622_RNA45SN1_45S_50ntFlanks</td><td>8105</td><td>8143</td><td>39</td><td>CGCGGCGGGGCGCGGGACATGTGGCGTACGGAAGACCCG</td><td>2494.0</td><td>-6.7</td><td>1.0</td><td>-6.5</td><td>6.7E-11</td><td>1.2E-10</td><td>BT20/MB468</td></tr>
<tr><td>>hg38_21_+_8433172_8446622_RNA45SN1_45S_50ntFlanks@11587.11605.19|TGCGGACCAGGGGAATCCG</td><td>>hg38_21_+_8433172_8446622_RNA45SN1_45S_50ntFlanks</td><td>11587</td><td>11605</td><td>19</td><td>TGCGGACCAGGGGAATCCG</td><td>143.4</td><td>-5.6</td><td>0.9</td><td>-6.5</td><td>6.7E-11</td><td>1.2E-10</td><td>BT20/MB468</td></tr>
<tr><td>>hg38_21_+_8433172_8446622_RNA45SN1_45S_50ntFlanks@6731.6758.28|CACATTGATCATCGACACTTCGAACGCA</td><td>>hg38_21_+_8433172_8446622_RNA45SN1_45S_50ntFlanks</td><td>6731</td><td>6758</td><td>28</td><td>CACATTGATCATCGACACTTCGAACGCA</td><td>208.1</td><td>-3.0</td><td>0.5</td><td>-6.5</td><td>6.8E-11</td><td>1.2E-10</td><td>BT20/MB468</td></tr>
<tr><td>>hg38_21_+_8433172_8446622_RNA45SN1_45S_50ntFlanks@12510.12527.18|TCGTGAGACAGGTTAGTT</td><td>>hg38_21_+_8433172_8446622_RNA45SN1_45S_50ntFlanks</td><td>12510</td><td>12527</td><td>18</td><td>TCGTGAGACAGGTTAGTT</td><td>317.5</td><td>-5.9</td><td>0.9</td><td>-6.5</td><td>5.9E-11</td><td>1.2E-10</td><td>MB231/MB468</td></tr>
<tr><td>>hg38_21_+_8433172_8446622_RNA45SN1_45S_50ntFlanks@11568.11611.44|GCAGCCGACTTAGAACTGGTGCGGACCAGGGGAATCCGACTGTT</td><td>>hg38_21_+_8433172_8446622_RNA45SN1_45S_50ntFlanks</td><td>11568</td><td>11611</td><td>44</td><td>GCAGCCGACTTAGAACTGGTGCGGACCAGGGGAATCCGACTGTT</td><td>961.8</td><td>5.0</td><td>0.8</td><td>6.5</td><td>5.9E-11</td><td>1.2E-10</td><td>MB231/MB468</td></tr>
<tr><td>>hg38_21_+_8433172_8446622_RNA45SN1_45S_50ntFlanks@10794.10815.22|CTGGCATGTTGGAACAATGTAG</td><td>>hg38_21_+_8433172_8446622_RNA45SN1_45S_50ntFlanks</td><td>10794</td><td>10815</td><td>22</td><td>CTGGCATGTTGGAACAATGTAG</td><td>468.7</td><td>-5.2</td><td>0.8</td><td>-6.5</td><td>6.9E-11</td><td>1.2E-10</td><td>BT20/MB468</td></tr>
<tr><td>>hg38_21_+_8433172_8446622_RNA45SN1_45S_50ntFlanks@3931.3959.29|TCAAAACCAACCCGGTCAGCCCCTCTCCG</td><td>>hg38_21_+_8433172_8446622_RNA45SN1_45S_50ntFlanks</td><td>3931</td><td>3959</td><td>29</td><td>TCAAAACCAACCCGGTCAGCCCCTCTCCG</td><td>98.2</td><td>-8.2</td><td>1.3</td><td>-6.5</td><td>5.9E-11</td><td>1.2E-10</td><td>MB231/MB468</td></tr>
<tr><td>>hg38_21_+_8433172_8446622_RNA45SN1_45S_50ntFlanks@10571.10608.38|GAGAAGCCGGCGGGAGCCCCGGGGAGAGTTCTCTTTTC</td><td>>hg38_21_+_8433172_8446622_RNA45SN1_45S_50ntFlanks</td><td>10571</td><td>10608</td><td>38</td><td>GAGAAGCCGGCGGGAGCCCCGGGGAGAGTTCTCTTTTC</td><td>115.8</td><td>-8.4</td><td>1.3</td><td>-6.5</td><td>5.9E-11</td><td>1.2E-10</td><td>MB231/MB468</td></tr>
<tr><td>>hg38_21_+_8433172_8446622_RNA45SN1_45S_50ntFlanks@10866.10886.21|CTAAGGGCTGGGTCGGTCGGG</td><td>>hg38_21_+_8433172_8446622_RNA45SN1_45S_50ntFlanks</td><td>10866</td><td>10886</td><td>21</td><td>CTAAGGGCTGGGTCGGTCGGG</td><td>143.5</td><td>-6.9</td><td>1.1</td><td>-6.5</td><td>5.9E-11</td><td>1.2E-10</td><td>MB231/MB468</td></tr>
<tr><td>>hg38_21_+_8433172_8446622_RNA45SN1_45S_50ntFlanks@12004.12034.31|CGGGCCGCCGGTGAAATACCACTACTCTGAT</td><td>>hg38_21_+_8433172_8446622_RNA45SN1_45S_50ntFlanks</td><td>12004</td><td>12034</td><td>31</td><td>CGGGCCGCCGGTGAAATACCACTACTCTGAT</td><td>736.2</td><td>7.9</td><td>1.2</td><td>6.5</td><td>6.9E-11</td><td>1.2E-10</td><td>BT20/MB468</td></tr>
<tr><td>>hg38_21_+_8433172_8446622_RNA45SN1_45S_50ntFlanks@8380.8399.20|CAAGAGGGCGTGAAACCGTT</td><td>>hg38_21_+_8433172_8446622_RNA45SN1_45S_50ntFlanks</td><td>8380</td><td>8399</td><td>20</td><td>CAAGAGGGCGTGAAACCGTT</td><td>605.3</td><td>-5.7</td><td>0.9</td><td>-6.5</td><td>5.9E-11</td><td>1.2E-10</td><td>MB231/MB468</td></tr>
<tr><td>>hg38_21_+_8433172_8446622_RNA45SN1_45S_50ntFlanks@12417.12449.33|TCGATGTCGGCTCTTCCTATCATTGTGAAGCAG</td><td>>hg38_21_+_8433172_8446622_RNA45SN1_45S_50ntFlanks</td><td>12417</td><td>12449</td><td>33</td><td>TCGATGTCGGCTCTTCCTATCATTGTGAAGCAG</td><td>440.2</td><td>3.6</td><td>0.6</td><td>6.5</td><td>6.0E-11</td><td>1.3E-10</td><td>MB231/MB468</td></tr>
<tr><td>>hg38_21_+_8433172_8446622_RNA45SN1_45S_50ntFlanks@12954.12982.29|CTGCTTCTGGGTCGGGGTTTCGTACGTAG</td><td>>hg38_21_+_8433172_8446622_RNA45SN1_45S_50ntFlanks</td><td>12954</td><td>12982</td><td>29</td><td>CTGCTTCTGGGTCGGGGTTTCGTACGTAG</td><td>101.7</td><td>-4.1</td><td>0.6</td><td>-6.5</td><td>6.0E-11</td><td>1.3E-10</td><td>MB231/MB468</td></tr>
<tr><td>>hg38_21_+_8433172_8446622_RNA45SN1_45S_50ntFlanks@11729.11754.26|TAACTATGACTCTCTTAAGGTAGCCA</td><td>>hg38_21_+_8433172_8446622_RNA45SN1_45S_50ntFlanks</td><td>11729</td><td>11754</td><td>26</td><td>TAACTATGACTCTCTTAAGGTAGCCA</td><td>209.5</td><td>-9.3</td><td>1.4</td><td>-6.5</td><td>6.0E-11</td><td>1.3E-10</td><td>MB231/MB468</td></tr>
<tr><td>>hg38_21_+_8433172_8446622_RNA45SN1_45S_50ntFlanks@10383.10424.42|TCCATGTGAACAGCAGTTGAACATGGGTCAGTCGGTCCTGAG</td><td>>hg38_21_+_8433172_8446622_RNA45SN1_45S_50ntFlanks</td><td>10383</td><td>10424</td><td>42</td><td>TCCATGTGAACAGCAGTTGAACATGGGTCAGTCGGTCCTGAG</td><td>739.2</td><td>8.7</td><td>1.3</td><td>6.5</td><td>6.9E-11</td><td>1.3E-10</td><td>BT20/MB468</td></tr>
<tr><td>>hg38_21_+_8433172_8446622_RNA45SN1_45S_50ntFlanks@4724.4751.28|ATTAATCAAGAACGAAAGTCGGAGGTTC</td><td>>hg38_21_+_8433172_8446622_RNA45SN1_45S_50ntFlanks</td><td>4724</td><td>4751</td><td>28</td><td>ATTAATCAAGAACGAAAGTCGGAGGTTC</td><td>505.7</td><td>-4.2</td><td>0.6</td><td>-6.5</td><td>7.0E-11</td><td>1.3E-10</td><td>BT20/MB468</td></tr>
<tr><td>>hg38_21_+_8433172_8446622_RNA45SN1_45S_50ntFlanks@13005.13028.24|TCTATTGAAAGTCAGCCCTCGACA</td><td>>hg38_21_+_8433172_8446622_RNA45SN1_45S_50ntFlanks</td><td>13005</td><td>13028</td><td>24</td><td>TCTATTGAAAGTCAGCCCTCGACA</td><td>137.3</td><td>-8.7</td><td>1.3</td><td>-6.5</td><td>6.0E-11</td><td>1.3E-10</td><td>MB231/MB468</td></tr>
<tr><td>>hg38_21_+_8433172_8446622_RNA45SN1_45S_50ntFlanks@7992.8016.25|CGTGGCGACCCGCTGAATTTAAGCA</td><td>>hg38_21_+_8433172_8446622_RNA45SN1_45S_50ntFlanks</td><td>7992</td><td>8016</td><td>25</td><td>CGTGGCGACCCGCTGAATTTAAGCA</td><td>1222.8</td><td>4.8</td><td>0.7</td><td>6.5</td><td>6.1E-11</td><td>1.3E-10</td><td>MB231/MB468</td></tr>
<tr><td>>hg38_21_+_8433172_8446622_RNA45SN1_45S_50ntFlanks@9605.9622.18|AAGACTAATCGAACCATC</td><td>>hg38_21_+_8433172_8446622_RNA45SN1_45S_50ntFlanks</td><td>9605</td><td>9622</td><td>18</td><td>AAGACTAATCGAACCATC</td><td>427.2</td><td>-3.2</td><td>0.5</td><td>-6.5</td><td>6.1E-11</td><td>1.3E-10</td><td>MB231/MB468</td></tr>
<tr><td>>hg38_21_+_8433172_8446622_RNA45SN1_45S_50ntFlanks@12415.12442.28|CTTCGATGTCGGCTCTTCCTATCATTGT</td><td>>hg38_21_+_8433172_8446622_RNA45SN1_45S_50ntFlanks</td><td>12415</td><td>12442</td><td>28</td><td>CTTCGATGTCGGCTCTTCCTATCATTGT</td><td>597.0</td><td>4.2</td><td>0.6</td><td>6.5</td><td>6.1E-11</td><td>1.3E-10</td><td>MB231/MB468</td></tr>
<tr><td>>MI0000100|hsa-mir-98&WithFlank&X|-|53556217|53556347@28.48.21(+1C)&[MIMAT0000096&hsa-miR-98-5p&offsets|0|-1(+1C);m-143&X|-|53556299|53556320&offsets|0|-1(+1C)]|TGAGGTAGTAAGTTGTATTGTC</td><td>>MI0000100|hsa-mir-98&WithFlank&X|-|53556217|53556347</td><td>28</td><td>48</td><td>21(+1C)</td><td>TGAGGTAGTAAGTTGTATTGTC</td><td>1520.5</td><td>5.3</td><td>0.8</td><td>6.5</td><td>7.1E-11</td><td>1.3E-10</td><td>BT20/MB468</td></tr>
<tr><td>>hg38_21_+_8433172_8446622_RNA45SN1_45S_50ntFlanks@8194.8214.21|CCCGTGGACGGTGTGAGGCCG</td><td>>hg38_21_+_8433172_8446622_RNA45SN1_45S_50ntFlanks</td><td>8194</td><td>8214</td><td>21</td><td>CCCGTGGACGGTGTGAGGCCG</td><td>548.2</td><td>-3.4</td><td>0.5</td><td>-6.5</td><td>6.2E-11</td><td>1.3E-10</td><td>MB231/MB468</td></tr>
<tr><td>>hg38_21_+_8433172_8446622_RNA45SN1_45S_50ntFlanks@6671.6698.28|CTCGGCTCGTGCGTCGATGAAGAACGCA</td><td>>hg38_21_+_8433172_8446622_RNA45SN1_45S_50ntFlanks</td><td>6671</td><td>6698</td><td>28</td><td>CTCGGCTCGTGCGTCGATGAAGAACGCA</td><td>706.3</td><td>-3.4</td><td>0.5</td><td>-6.5</td><td>6.2E-11</td><td>1.3E-10</td><td>MB231/MB468</td></tr>
<tr><td>>hg38_21_+_8433172_8446622_RNA45SN1_45S_50ntFlanks@4064.4101.38|ATCAACTTTCGATGGTAGTCGCCGTGCCTACCATGGTG</td><td>>hg38_21_+_8433172_8446622_RNA45SN1_45S_50ntFlanks</td><td>4064</td><td>4101</td><td>38</td><td>ATCAACTTTCGATGGTAGTCGCCGTGCCTACCATGGTG</td><td>100.1</td><td>-8.6</td><td>1.3</td><td>-6.5</td><td>7.2E-11</td><td>1.3E-10</td><td>BT20/MB468</td></tr>
<tr><td>>hg38_21_+_8433172_8446622_RNA45SN1_45S_50ntFlanks@10727.10752.26|GGAGAGGGTGTAAATCTCGCGCCGGG</td><td>>hg38_21_+_8433172_8446622_RNA45SN1_45S_50ntFlanks</td><td>10727</td><td>10752</td><td>26</td><td>GGAGAGGGTGTAAATCTCGCGCCGGG</td><td>108.8</td><td>-8.3</td><td>1.3</td><td>-6.5</td><td>6.2E-11</td><td>1.3E-10</td><td>MB231/MB468</td></tr>
<tr><td>>hg38_21_+_8433172_8446622_RNA45SN1_45S_50ntFlanks@10873.10901.29|CTGGGTCGGTCGGGCTGGGGCGCGAAGCG</td><td>>hg38_21_+_8433172_8446622_RNA45SN1_45S_50ntFlanks</td><td>10873</td><td>10901</td><td>29</td><td>CTGGGTCGGTCGGGCTGGGGCGCGAAGCG</td><td>98.0</td><td>-8.2</td><td>1.3</td><td>-6.5</td><td>6.3E-11</td><td>1.3E-10</td><td>MB231/MB468</td></tr>
<tr><td>>hg38_MT_+_1621_3279_RNR2_16S_50ntFlanks@51.73.23|GCTAAACCTAGCCCCAAACCCAC</td><td>>hg38_MT_+_1621_3279_RNR2_16S_50ntFlanks</td><td>51</td><td>73</td><td>23</td><td>GCTAAACCTAGCCCCAAACCCAC</td><td>873.4</td><td>7.6</td><td>1.2</td><td>6.5</td><td>7.3E-11</td><td>1.3E-10</td><td>BT20/MB468</td></tr>
<tr><td>>hg38_1_-_228634819_228635039_RNA5S12_5S_50ntFlanks@150.169.20|AATACCGGGTGCTGTAGGCT</td><td>>hg38_1_-_228634819_228635039_RNA5S12_5S_50ntFlanks</td><td>150</td><td>169</td><td>20</td><td>AATACCGGGTGCTGTAGGCT</td><td>965.6</td><td>4.8</td><td>0.7</td><td>6.5</td><td>7.3E-11</td><td>1.3E-10</td><td>BT20/MB468</td></tr>
<tr><td>>hg38_21_+_8433172_8446622_RNA45SN1_45S_50ntFlanks@3899.3920.22|CCTTCGCGGGGGGGATGCGTGC</td><td>>hg38_21_+_8433172_8446622_RNA45SN1_45S_50ntFlanks</td><td>3899</td><td>3920</td><td>22</td><td>CCTTCGCGGGGGGGATGCGTGC</td><td>190.9</td><td>-5.4</td><td>0.8</td><td>-6.5</td><td>7.3E-11</td><td>1.3E-10</td><td>BT20/MB468</td></tr>
<tr><td>>hg38_21_+_8433172_8446622_RNA45SN1_45S_50ntFlanks@10572.10607.36|AGAAGCCGGCGGGAGCCCCGGGGAGAGTTCTCTTTT</td><td>>hg38_21_+_8433172_8446622_RNA45SN1_45S_50ntFlanks</td><td>10572</td><td>10607</td><td>36</td><td>AGAAGCCGGCGGGAGCCCCGGGGAGAGTTCTCTTTT</td><td>195.4</td><td>-6.5</td><td>1.0</td><td>-6.5</td><td>6.3E-11</td><td>1.3E-10</td><td>MB231/MB468</td></tr>
<tr><td>>hg38_21_+_8433172_8446622_RNA45SN1_45S_50ntFlanks@10498.10521.24|TTCAGATCCCCGAATCCGGAGTGG</td><td>>hg38_21_+_8433172_8446622_RNA45SN1_45S_50ntFlanks</td><td>10498</td><td>10521</td><td>24</td><td>TTCAGATCCCCGAATCCGGAGTGG</td><td>257.7</td><td>-10.0</td><td>1.5</td><td>-6.5</td><td>7.4E-11</td><td>1.3E-10</td><td>BT20/MB468</td></tr>
<tr><td>>hg38_21_+_8433172_8446622_RNA45SN1_45S_50ntFlanks@12072.12089.18|CCCGAGGGGCTCTCGCTT</td><td>>hg38_21_+_8433172_8446622_RNA45SN1_45S_50ntFlanks</td><td>12072</td><td>12089</td><td>18</td><td>CCCGAGGGGCTCTCGCTT</td><td>113.7</td><td>-5.1</td><td>0.8</td><td>-6.5</td><td>7.5E-11</td><td>1.3E-10</td><td>BT20/MB468</td></tr>
<tr><td>>hg38_21_+_8433172_8446622_RNA45SN1_45S_50ntFlanks@3593.3610.18|CCCGTCCGTCCGTCCGTC</td><td>>hg38_21_+_8433172_8446622_RNA45SN1_45S_50ntFlanks</td><td>3593</td><td>3610</td><td>18</td><td>CCCGTCCGTCCGTCCGTC</td><td>111.0</td><td>-8.4</td><td>1.3</td><td>-6.5</td><td>6.5E-11</td><td>1.3E-10</td><td>MB231/MB468</td></tr>
<tr><td>>hg38_21_+_8433172_8446622_RNA45SN1_45S_50ntFlanks@8737.8754.18|CCCGTCCGTCCGTCCGTC</td><td>>hg38_21_+_8433172_8446622_RNA45SN1_45S_50ntFlanks</td><td>8737</td><td>8754</td><td>18</td><td>CCCGTCCGTCCGTCCGTC</td><td>111.0</td><td>-8.4</td><td>1.3</td><td>-6.5</td><td>6.5E-11</td><td>1.3E-10</td><td>MB231/MB468</td></tr>
<tr><td>>hg38_21_+_8433172_8446622_RNA45SN1_45S_50ntFlanks@12634.12654.21|GAAGCTACCATCTGTGGGATT</td><td>>hg38_21_+_8433172_8446622_RNA45SN1_45S_50ntFlanks</td><td>12634</td><td>12654</td><td>21</td><td>GAAGCTACCATCTGTGGGATT</td><td>335.4</td><td>-4.5</td><td>0.7</td><td>-6.5</td><td>6.5E-11</td><td>1.4E-10</td><td>MB231/MB468</td></tr>
<tr><td>>hg38_21_+_8433172_8446622_RNA45SN1_45S_50ntFlanks@10689.10710.22|GCGTCCGGTGAGCTCTCGCTGG</td><td>>hg38_21_+_8433172_8446622_RNA45SN1_45S_50ntFlanks</td><td>10689</td><td>10710</td><td>22</td><td>GCGTCCGGTGAGCTCTCGCTGG</td><td>97.8</td><td>-8.2</td><td>1.3</td><td>-6.5</td><td>6.5E-11</td><td>1.4E-10</td><td>MB231/MB468</td></tr>
<tr><td>>hg38_21_+_8433172_8446622_RNA45SN1_45S_50ntFlanks@8107.8143.37|CGGCGGGGCGCGGGACATGTGGCGTACGGAAGACCCG</td><td>>hg38_21_+_8433172_8446622_RNA45SN1_45S_50ntFlanks</td><td>8107</td><td>8143</td><td>37</td><td>CGGCGGGGCGCGGGACATGTGGCGTACGGAAGACCCG</td><td>142.7</td><td>-8.7</td><td>1.3</td><td>-6.5</td><td>6.5E-11</td><td>1.4E-10</td><td>MB231/MB468</td></tr>
<tr><td>>hg38_21_+_8433172_8446622_RNA45SN1_45S_50ntFlanks@12407.12436.30|TTTTGATCCTTCGATGTCGGCTCTTCCTAT</td><td>>hg38_21_+_8433172_8446622_RNA45SN1_45S_50ntFlanks</td><td>12407</td><td>12436</td><td>30</td><td>TTTTGATCCTTCGATGTCGGCTCTTCCTAT</td><td>901.1</td><td>-5.1</td><td>0.8</td><td>-6.5</td><td>7.5E-11</td><td>1.4E-10</td><td>BT20/MB468</td></tr>
<tr><td>>hg38_21_+_8433172_8446622_RNA45SN1_45S_50ntFlanks@12001.12030.30|CTGCGGGCCGCCGGTGAAATACCACTACTC</td><td>>hg38_21_+_8433172_8446622_RNA45SN1_45S_50ntFlanks</td><td>12001</td><td>12030</td><td>30</td><td>CTGCGGGCCGCCGGTGAAATACCACTACTC</td><td>105.2</td><td>-5.3</td><td>0.8</td><td>-6.5</td><td>7.5E-11</td><td>1.4E-10</td><td>BT20/MB468</td></tr>
<tr><td>>hg38_21_+_8433172_8446622_RNA45SN1_45S_50ntFlanks@6738.6771.34|ATCATCGACACTTCGAACGCACTTGCGGCCCCGG</td><td>>hg38_21_+_8433172_8446622_RNA45SN1_45S_50ntFlanks</td><td>6738</td><td>6771</td><td>34</td><td>ATCATCGACACTTCGAACGCACTTGCGGCCCCGG</td><td>2153.6</td><td>-7.2</td><td>1.1</td><td>-6.5</td><td>6.5E-11</td><td>1.4E-10</td><td>MB231/MB468</td></tr>
<tr><td>>hg38_21_+_8433172_8446622_RNA45SN1_45S_50ntFlanks@4853.4883.31|AAGTCTTTGGGTTCCGGGGGGAGTATGGTTG</td><td>>hg38_21_+_8433172_8446622_RNA45SN1_45S_50ntFlanks</td><td>4853</td><td>4883</td><td>31</td><td>AAGTCTTTGGGTTCCGGGGGGAGTATGGTTG</td><td>105.7</td><td>-8.3</td><td>1.3</td><td>-6.5</td><td>6.5E-11</td><td>1.4E-10</td><td>MB231/MB468</td></tr>
<tr><td>>hg38_21_+_8433172_8446622_RNA45SN1_45S_50ntFlanks@9562.9595.34|TCCTGACGTGCAAATCGGTCGTCCGACCTGGGTA</td><td>>hg38_21_+_8433172_8446622_RNA45SN1_45S_50ntFlanks</td><td>9562</td><td>9595</td><td>34</td><td>TCCTGACGTGCAAATCGGTCGTCCGACCTGGGTA</td><td>175.1</td><td>-5.5</td><td>0.8</td><td>-6.5</td><td>7.6E-11</td><td>1.4E-10</td><td>BT20/MB468</td></tr>
<tr><td>>hg38_21_+_8433172_8446622_RNA45SN1_45S_50ntFlanks@11783.11810.28|CATGAATGGATGAACGAGATTCCCACTG</td><td>>hg38_21_+_8433172_8446622_RNA45SN1_45S_50ntFlanks</td><td>11783</td><td>11810</td><td>28</td><td>CATGAATGGATGAACGAGATTCCCACTG</td><td>118.5</td><td>-8.9</td><td>1.4</td><td>-6.5</td><td>7.6E-11</td><td>1.4E-10</td><td>BT20/MB468</td></tr>
<tr><td>>hg38_21_+_8433172_8446622_RNA45SN1_45S_50ntFlanks@475.522.48|CTTCGTGATCGATGTGGTGACGTCGTGCTCTCCCGGGCCGGGTCCGAG</td><td>>hg38_21_+_8433172_8446622_RNA45SN1_45S_50ntFlanks</td><td>475</td><td>522</td><td>48</td><td>CTTCGTGATCGATGTGGTGACGTCGTGCTCTCCCGGGCCGGGTCCGAG</td><td>615.1</td><td>9.5</td><td>1.5</td><td>6.5</td><td>7.7E-11</td><td>1.4E-10</td><td>BT20/MB468</td></tr>
<tr><td>>hg38_21_+_8433172_8446622_RNA45SN1_45S_50ntFlanks@12638.12661.24|CTACCATCTGTGGGATTATGACTG</td><td>>hg38_21_+_8433172_8446622_RNA45SN1_45S_50ntFlanks</td><td>12638</td><td>12661</td><td>24</td><td>CTACCATCTGTGGGATTATGACTG</td><td>884.5</td><td>-3.5</td><td>0.5</td><td>-6.5</td><td>6.7E-11</td><td>1.4E-10</td><td>MB231/MB468</td></tr>
<tr><td>>hg38_21_+_8433172_8446622_RNA45SN1_45S_50ntFlanks@11209.11233.25|CGGCGACTCTGGACGCGAGCCGGGC</td><td>>hg38_21_+_8433172_8446622_RNA45SN1_45S_50ntFlanks</td><td>11209</td><td>11233</td><td>25</td><td>CGGCGACTCTGGACGCGAGCCGGGC</td><td>819.9</td><td>-4.2</td><td>0.6</td><td>-6.5</td><td>6.7E-11</td><td>1.4E-10</td><td>MB231/MB468</td></tr>
<tr><td>>MI0000294|hsa-mir-218-1&WithFlank&4|+|20528269|20528390@31.52.22(+1U);MI0000295|hsa-mir-218-2&WithFlank&5|-|168768140|168768261@31.52.22(+1U)&[MIMAT0000275&hsa-miR-218-5p&offsets|0|+1(+1U);m-185&4|+|20528299|20528319&offsets|0|+1(+1U)];[MIMAT0000275_1&hsa-miR-218-5p&offsets|0|+1(+1U);m-187&5|-|168768211|168768231&offsets|0|+1(+1U)]|TTGTGCTTGATCTAACCATGTGT</td><td>>MI0000294|hsa-mir-218-1&WithFlank&4|+|20528269|20528390</td><td>31</td><td>52</td><td>22(+1U)</td><td>TTGTGCTTGATCTAACCATGTGT</td><td>215.0</td><td>-7.4</td><td>1.1</td><td>-6.9</td><td>4.8E-12</td><td>1.4E-10</td><td>BT20/MB231</td></tr>
</table>

| | | | | | | | | | | | | |
|---|---|---|---|---|---|---|---|---|---|---|---|---|
| >hg38_21_+_8433172_8446622_RNA45SN1_45S_50ntFlanks@6651.6694.44\|CGACTCTTAGCGGTGGATCACTCGGCTCGTGCGTCGATGAAGAA | >hg38_21_+_8433172_8446622_RNA45SN1_45S_50ntFlanks | 6651 | 6694 | 44 | CGACTCTTAGCGGTGGATCACTCGGCTCGTGCGTCGATGAAGAA | 2135.4 | 5.7 | 0.9 | 6.5 | 7.8E-11 | 1.4E-10 | BT20/MB468 |
| >MI0000081\|hsa-mir-24-2&WithFlank&19\|-\|13836281\|13836365@56.73.18;MI0000080\|hsa-mir-24-1&WithFlank&9\|+\|95086015\|95086094@50.67.18&[MIMAT0000080&hsa-miR-24-3p&offsets\|0\|-4;m-36&19\|-\|13836288\|13836310&offsets\|0\|-5];[MIMAT0000080_1&hsa-miR-24-3p&offsets\|0\|-4;m-35&9\|+\|95086064\|95086086&offsets\|0\|-5]\|TGGCTCAGTTCAGCAGGA | >MI0000081\|hsa-mir-24-2&WithFlank&19\|-\|13836281\|13836365 | 56 | 73 | 18 | TGGCTCAGTTCAGCAGGA | 407.8 | 8.5 | 1.3 | 6.5 | 7.9E-11 | 1.4E-10 | BT20/MB468 |
| >hg38_21_+_8433172_8446622_RNA45SN1_45S_50ntFlanks@10409.10435.27\|GTCAGTCGGTCCTGAGAGATGGGCGAG | >hg38_21_+_8433172_8446622_RNA45SN1_45S_50ntFlanks | 10409 | 10435 | 27 | GTCAGTCGGTCCTGAGAGATGGGCGAG | 154.5 | -5.5 | 0.8 | -6.5 | 6.8E-11 | 1.4E-10 | MB231/MB468 |
| >hg38_21_+_8433172_8446622_RNA45SN1_45S_50ntFlanks@482.507.26\|ATCGATGTGGTGACGTCGTGCTCTCC | >hg38_21_+_8433172_8446622_RNA45SN1_45S_50ntFlanks | 482 | 507 | 26 | ATCGATGTGGTGACGTCGTGCTCTCC | 73.2 | -8.2 | 1.3 | -6.5 | 8.0E-11 | 1.4E-10 | BT20/MB468 |
| >hg38_21_+_8433172_8446622_RNA45SN1_45S_50ntFlanks@11641.11667.27\|CGGCGGGTGTTGACGCGATGTGATTTC | >hg38_21_+_8433172_8446622_RNA45SN1_45S_50ntFlanks | 11641 | 11667 | 27 | CGGCGGGTGTTGACGCGATGTGATTTC | 459.9 | -5.4 | 0.8 | -6.5 | 8.0E-11 | 1.4E-10 | BT20/MB468 |
| >hg38_21_+_8433172_8446622_RNA45SN1_45S_50ntFlanks@10684.10712.29\|CGGCGGCGTCCGGTGAGCTCTCGCTGGCC | >hg38_21_+_8433172_8446622_RNA45SN1_45S_50ntFlanks | 10684 | 10712 | 29 | CGGCGGCGTCCGGTGAGCTCTCGCTGGCC | 309.3 | -4.1 | 0.6 | -6.5 | 8.0E-11 | 1.4E-10 | BT20/MB468 |
| >hg38_21_+_8433172_8446622_RNA45SN1_45S_50ntFlanks@6738.6770.33\|ATCATCGACACTTCGAACGCACTTGCGGCCCCG | >hg38_21_+_8433172_8446622_RNA45SN1_45S_50ntFlanks | 6738 | 6770 | 33 | ATCATCGACACTTCGAACGCACTTGCGGCCCCG | 2324.5 | -7.3 | 1.1 | -6.5 | 6.9E-11 | 1.4E-10 | MB231/MB468 |
| >hg38_21_+_8433172_8446622_RNA45SN1_45S_50ntFlanks@10784.10818.35\|TGAACAGCCTCTGGCATGTTGGAACAATGTAGGTA | >hg38_21_+_8433172_8446622_RNA45SN1_45S_50ntFlanks | 10784 | 10818 | 35 | TGAACAGCCTCTGGCATGTTGGAACAATGTAGGTA | 438.0 | -3.7 | 0.6 | -6.5 | 8.1E-11 | 1.5E-10 | BT20/MB468 |
| >hg38_21_+_8433172_8446622_RNA45SN1_45S_50ntFlanks@12071.12096.26\|CCCCGAGGGGCTCTCGCTTCTGGCGC | >hg38_21_+_8433172_8446622_RNA45SN1_45S_50ntFlanks | 12071 | 12096 | 26 | CCCCGAGGGGCTCTCGCTTCTGGCGC | 424.3 | -5.3 | 0.8 | -6.5 | 8.1E-11 | 1.5E-10 | BT20/MB468 |
| >hg38_21_+_8433172_8446622_RNA45SN1_45S_50ntFlanks@10508.10533.26\|CGAATCCGGAGTGGCGGAGATGGGCG | >hg38_21_+_8433172_8446622_RNA45SN1_45S_50ntFlanks | 10508 | 10533 | 26 | CGAATCCGGAGTGGCGGAGATGGGCG | 115.4 | -8.4 | 1.3 | -6.5 | 7.0E-11 | 1.5E-10 | MB231/MB468 |
| >hg38_21_+_8433172_8446622_RNA45SN1_45S_50ntFlanks@12416.12446.31\|TTCGATGTCGGCTCTTCCTATCATTGTGAAG | >hg38_21_+_8433172_8446622_RNA45SN1_45S_50ntFlanks | 12416 | 12446 | 31 | TTCGATGTCGGCTCTTCCTATCATTGTGAAG | 2442.1 | 5.1 | 0.8 | 6.5 | 7.0E-11 | 1.5E-10 | MB231/MB468 |
| >hg38_21_+_8433172_8446622_RNA45SN1_45S_50ntFlanks@8115.8140.26\|CGCGGGACATGTGGCGTACGGAAGAC | >hg38_21_+_8433172_8446622_RNA45SN1_45S_50ntFlanks | 8115 | 8140 | 26 | CGCGGGACATGTGGCGTACGGAAGAC | 188.2 | -7.3 | 1.1 | -6.5 | 7.0E-11 | 1.5E-10 | MB231/MB468 |
| >hg38_21_+_8433172_8446622_RNA45SN1_45S_50ntFlanks@9504.9531.28\|GTGAACTATGCCTGGGCAGGGCGAAGCC | >hg38_21_+_8433172_8446622_RNA45SN1_45S_50ntFlanks | 9504 | 9531 | 28 | GTGAACTATGCCTGGGCAGGGCGAAGCC | 112.5 | -8.8 | 1.4 | -6.5 | 8.2E-11 | 1.5E-10 | BT20/MB468 |
| >hg38_21_+_8433172_8446622_RNA45SN1_45S_50ntFlanks@12473.12497.25\|CACCCACTAATAGGGAACGTGAGCT | >hg38_21_+_8433172_8446622_RNA45SN1_45S_50ntFlanks | 12473 | 12497 | 25 | CACCCACTAATAGGGAACGTGAGCT | 761.9 | 4.3 | 0.7 | 6.5 | 7.1E-11 | 1.5E-10 | MB231/MB468 |
| >hg38_21_+_8433172_8446622_RNA45SN1_45S_50ntFlanks@10845.10867.23\|ACTTCGGGATAAGGATTGGCTCT | >hg38_21_+_8433172_8446622_RNA45SN1_45S_50ntFlanks | 10845 | 10867 | 23 | ACTTCGGGATAAGGATTGGCTCT | 378.6 | -4.0 | 0.6 | -6.5 | 8.2E-11 | 1.5E-10 | BT20/MB468 |
| >hg38_21_+_8433172_8446622_RNA45SN1_45S_50ntFlanks@9556.9582.27\|TAGCGGTCCTGACGTGCAAATCGGTCG | >hg38_21_+_8433172_8446622_RNA45SN1_45S_50ntFlanks | 9556 | 9582 | 27 | TAGCGGTCCTGACGTGCAAATCGGTCG | 561.9 | -7.0 | 1.1 | -6.5 | 8.2E-11 | 1.5E-10 | BT20/MB468 |
| >hg38_21_+_8433172_8446622_RNA45SN1_45S_50ntFlanks@13010.13027.18\|TGAAAGTCAGCCCTCGAC | >hg38_21_+_8433172_8446622_RNA45SN1_45S_50ntFlanks | 13010 | 13027 | 18 | TGAAAGTCAGCCCTCGAC | 162.3 | -9.3 | 1.4 | -6.5 | 8.2E-11 | 1.5E-10 | BT20/MB468 |
| >hg38_21_+_8433172_8446622_RNA45SN1_45S_50ntFlanks@12633.12659.27\|CGAAGCTACCATCTGTGGGATTATGAC | >hg38_21_+_8433172_8446622_RNA45SN1_45S_50ntFlanks | 12633 | 12659 | 27 | CGAAGCTACCATCTGTGGGATTATGAC | 826.6 | -4.2 | 0.6 | -6.5 | 7.1E-11 | 1.5E-10 | MB231/MB468 |
| >hg38_21_+_8433172_8446622_RNA45SN1_45S_50ntFlanks@4069.4098.30\|CTTTCGATGGTAGTCGCCGTGCCTACCATG | >hg38_21_+_8433172_8446622_RNA45SN1_45S_50ntFlanks | 4069 | 4098 | 30 | CTTTCGATGGTAGTCGCCGTGCCTACCATG | 120.7 | -8.5 | 1.3 | -6.5 | 7.1E-11 | 1.5E-10 | MB231/MB468 |
| >trna116_GluCTC_1_-_145399233_145399304@32.75.44,trna71_GluCTC_1_-_161439189_161439260@32.75.44,trna74_GluCTC_1_-_161431809_161431880@32.75.44,trna77_GluCTC_1_-_161424398_161424469@32.75.44,trna77_GluCTC_6_+_28949976_28950047@32.75.44,trna80_GluCTC_1_-_161417018_161417089@32.75.44,trna87_GluCTC_6_-_126101393_126101464@32.75.44\|CTCTCACCGCCGCGGCCCGGGTTCGATTCCCGGTCAGGGAACCA | >trna116_GluCTC_1_-_145399233_145399304 | 32 | 75 | 44 | CTCTCACCGCCGCGGCCCGGGTTCGATTCCCGGTCAGGGAACCA | 124.9 | -9.0 | 1.4 | -6.5 | 8.3E-11 | 1.5E-10 | BT20/MB468 |
| >hg38_21_+_8433172_8446622_RNA45SN1_45S_50ntFlanks@9739.9759.21\|AAACGATCTCAACCTATTCTC | >hg38_21_+_8433172_8446622_RNA45SN1_45S_50ntFlanks | 9739 | 9759 | 21 | AAACGATCTCAACCTATTCTC | 420.4 | -3.6 | 0.5 | -6.5 | 8.3E-11 | 1.5E-10 | BT20/MB468 |
| >MI0000253\|hsa-mir-148a&WithFlank&7\|-\|25949913\|25949992@50.70.21(+1A)&[MIMAT0000243&hsa-miR-148a-3p&offsets\|0\|-1(+1A);m-7&7\|-\|25949922\|25949943&offsets\|0\|-1(+1A)]\|TCAGTGCACTACAGAACTTTGA | >MI0000253\|hsa-mir-148a&WithFlank&7\|-\|25949913\|25949992 | 50 | 70 | 21(+1A) | TCAGTGCACTACAGAACTTTGA | 13307.8 | 5.3 | 0.8 | 6.9 | 5.1E-12 | 1.5E-10 | BT20/MB231 |
| >hg38_21_+_8433172_8446622_RNA45SN1_45S_50ntFlanks@6648.6669.22\|GTACGACTCTTAGCGGTGGATC | >hg38_21_+_8433172_8446622_RNA45SN1_45S_50ntFlanks | 6648 | 6669 | 22 | GTACGACTCTTAGCGGTGGATC | 1939.7 | 3.4 | 0.5 | 6.5 | 8.4E-11 | 1.5E-10 | BT20/MB468 |
| >hg38_21_+_8433172_8446622_RNA45SN1_45S_50ntFlanks@12444.12467.24\|AAGCAGAATTCACCAAGCGTTGGA | >hg38_21_+_8433172_8446622_RNA45SN1_45S_50ntFlanks | 12444 | 12467 | 24 | AAGCAGAATTCACCAAGCGTTGGA | 156.9 | -9.3 | 1.4 | -6.5 | 8.4E-11 | 1.5E-10 | BT20/MB468 |
| >hg38_21_+_8433172_8446622_RNA45SN1_45S_50ntFlanks@10794.10815.22\|CTGGCATGTTGGAACAATGTAG | >hg38_21_+_8433172_8446622_RNA45SN1_45S_50ntFlanks | 10794 | 10815 | 22 | CTGGCATGTTGGAACAATGTAG | 375.6 | -5.6 | 0.9 | -6.5 | 7.3E-11 | 1.5E-10 | MB231/MB468 |
| >hg38_21_+_8433172_8446622_RNA45SN1_45S_50ntFlanks@10392.10418.27\|ACAGCAGTTGAACATGGGTCAGTCGGT | >hg38_21_+_8433172_8446622_RNA45SN1_45S_50ntFlanks | 10392 | 10418 | 27 | ACAGCAGTTGAACATGGGTCAGTCGGT | 150.2 | -2.9 | 0.5 | -6.5 | 8.5E-11 | 1.5E-10 | BT20/MB468 |
| >hg38_21_+_8433172_8446622_RNA45SN1_45S_50ntFlanks@11769.11792.24\|TAATTAGTGACGCGCATGAATGGA | >hg38_21_+_8433172_8446622_RNA45SN1_45S_50ntFlanks | 11769 | 11792 | 24 | TAATTAGTGACGCGCATGAATGGA | 616.9 | -6.7 | 1.0 | -6.5 | 8.5E-11 | 1.5E-10 | BT20/MB468 |
| >hg38_21_+_8433172_8446622_RNA45SN1_45S_50ntFlanks@11534.11552.19\|GTTCCGCGCGGCGCCTCGC | >hg38_21_+_8433172_8446622_RNA45SN1_45S_50ntFlanks | 11534 | 11552 | 19 | GTTCCGCGCGGCGCCTCGC | 489.7 | -6.3 | 1.0 | -6.5 | 7.4E-11 | 1.5E-10 | MB231/MB468 |
| >hg38_21_+_8433172_8446622_RNA45SN1_45S_50ntFlanks@10357.10377.21\|CTTTGAAGGCCGAAGTGGAGA | >hg38_21_+_8433172_8446622_RNA45SN1_45S_50ntFlanks | 10357 | 10377 | 21 | CTTTGAAGGCCGAAGTGGAGA | 222.7 | -3.8 | 0.6 | -6.5 | 8.6E-11 | 1.5E-10 | BT20/MB468 |
| >hg38_21_+_8433172_8446622_RNA45SN1_45S_50ntFlanks@7978.8019.42\|GACCTCAGATCAGACGTGGCGACCCGCTGAATTTAAGCATAT | >hg38_21_+_8433172_8446622_RNA45SN1_45S_50ntFlanks | 7978 | 8019 | 42 | GACCTCAGATCAGACGTGGCGACCCGCTGAATTTAAGCATAT | 335.0 | 13.3 | 2.0 | 6.5 | 7.5E-11 | 1.6E-10 | MB231/MB468 |
| >hg38_21_+_8433172_8446622_RNA45SN1_45S_50ntFlanks@8154.8178.25\|CCGCTCGTGGGGGGCCCAAGTCCTT | >hg38_21_+_8433172_8446622_RNA45SN1_45S_50ntFlanks | 8154 | 8178 | 25 | CCGCTCGTGGGGGGCCCAAGTCCTT | 106.4 | -8.3 | 1.3 | -6.5 | 7.5E-11 | 1.6E-10 | MB231/MB468 |
| >hg38_21_+_8433172_8446622_RNA45SN1_45S_50ntFlanks@12478.12497.20\|ACTAATAGGGAACGTGAGCT | >hg38_21_+_8433172_8446622_RNA45SN1_45S_50ntFlanks | 12478 | 12497 | 20 | ACTAATAGGGAACGTGAGCT | 2031.6 | 3.6 | 0.6 | 6.5 | 7.6E-11 | 1.6E-10 | MB231/MB468 |
| >hg38_21_+_8433172_8446622_RNA45SN1_45S_50ntFlanks@11881.11904.24\|CCTGTTGAGCTTGACTCTAGTCTG | >hg38_21_+_8433172_8446622_RNA45SN1_45S_50ntFlanks | 11881 | 11904 | 24 | CCTGTTGAGCTTGACTCTAGTCTG | 1568.9 | -6.5 | 1.0 | -6.5 | 8.9E-11 | 1.6E-10 | BT20/MB468 |
| >hg38_21_+_8433172_8446622_RNA45SN1_45S_50ntFlanks@11273.11293.21\|CCGCCCCCGGGGAGCCCGGCG | >hg38_21_+_8433172_8446622_RNA45SN1_45S_50ntFlanks | 11273 | 11293 | 21 | CCGCCCCCGGGGAGCCCGGCG | 100.9 | -8.2 | 1.3 | -6.5 | 7.7E-11 | 1.6E-10 | MB231/MB468 |
| >hg38_21_+_8433172_8446622_RNA45SN1_45S_50ntFlanks@12957.12981.25\|CTTCTGGGTCGGGGTTTCGTACGTA | >hg38_21_+_8433172_8446622_RNA45SN1_45S_50ntFlanks | 12957 | 12981 | 25 | CTTCTGGGTCGGGGTTTCGTACGTA | 130.5 | -5.2 | 0.8 | -6.5 | 9.0E-11 | 1.6E-10 | BT20/MB468 |
| >hg38_21_+_8433172_8446622_RNA45SN1_45S_50ntFlanks@10679.10712.34\|GGTTCCGGCGGCGTCCGGTGAGCTCTCGCTGGCC | >hg38_21_+_8433172_8446622_RNA45SN1_45S_50ntFlanks | 10679 | 10712 | 34 | GGTTCCGGCGGCGTCCGGTGAGCTCTCGCTGGCC | 238.9 | -5.6 | 0.9 | -6.5 | 7.8E-11 | 1.6E-10 | MB231/MB468 |
| >hg38_21_+_8433172_8446622_RNA45SN1_45S_50ntFlanks@7974.8014.41\|ACGCGACCTCAGATCAGACGTGGCGACCCGCTGAATTTAAG | >hg38_21_+_8433172_8446622_RNA45SN1_45S_50ntFlanks | 7974 | 8014 | 41 | ACGCGACCTCAGATCAGACGTGGCGACCCGCTGAATTTAAG | 909.0 | 5.2 | 0.8 | 6.5 | 7.8E-11 | 1.6E-10 | MB231/MB468 |
| >hg38_21_+_8433172_8446622_RNA45SN1_45S_50ntFlanks@11766.11794.29\|ATCTAATTAGTGACGCGCATGAATGGATG | >hg38_21_+_8433172_8446622_RNA45SN1_45S_50ntFlanks | 11766 | 11794 | 29 | ATCTAATTAGTGACGCGCATGAATGGATG | 101.7 | -8.2 | 1.3 | -6.5 | 7.8E-11 | 1.6E-10 | MB231/MB468 |
| >hg38_21_+_8433172_8446622_RNA45SN1_45S_50ntFlanks@12588.12610.23\|GGTTCAGACATTTGGTGTATGTG | >hg38_21_+_8433172_8446622_RNA45SN1_45S_50ntFlanks | 12588 | 12610 | 23 | GGTTCAGACATTTGGTGTATGTG | 102.6 | -8.3 | 1.3 | -6.5 | 7.8E-11 | 1.6E-10 | MB231/MB468 |
| >hg38_21_+_8433172_8446622_RNA45SN1_45S_50ntFlanks@12476.12509.34\|CCACTAATAGGGAACGTGAGCTGGGTTTAGACCG | >hg38_21_+_8433172_8446622_RNA45SN1_45S_50ntFlanks | 12476 | 12509 | 34 | CCACTAATAGGGAACGTGAGCTGGGTTTAGACCG | 108.8 | -8.8 | 1.4 | -6.5 | 9.1E-11 | 1.6E-10 | BT20/MB468 |
| >MI0000283\|hsa-mir-203a&WithFlank&14\|+\|104117399\|104117520@71.93.23&[MIMAT0000264&hsa-miR-203a-3p&offsets\|0\|+1;m-79&14\|+\|104117469\|104117490&offsets\|0\|+1]\|GTGAAATGTTTAGGACCACTAGA | >MI0000283\|hsa-mir-203a&WithFlank&14\|+\|104117399\|104117520 | 71 | 93 | 23 | GTGAAATGTTTAGGACCACTAGA | 1345.4 | 9.4 | 1.4 | 6.5 | 9.1E-11 | 1.6E-10 | BT20/MB468 |
| >hg38_21_+_8433172_8446622_RNA45SN1_45S_50ntFlanks@4003.4042.40\|ATAACCTCGGGCCGATCGCACGCCCCCCGTGGCGGCGACG | >hg38_21_+_8433172_8446622_RNA45SN1_45S_50ntFlanks | 4003 | 4042 | 40 | ATAACCTCGGGCCGATCGCACGCCCCCCGTGGCGGCGACG | 385.2 | -4.2 | 0.6 | -6.5 | 7.9E-11 | 1.6E-10 | MB231/MB468 |
| >hg38_21_+_8433172_8446622_RNA45SN1_45S_50ntFlanks@4723.4752.30\|CATTAATCAAGAACGAAAGTCGGAGGTTCG | >hg38_21_+_8433172_8446622_RNA45SN1_45S_50ntFlanks | 4723 | 4752 | 30 | CATTAATCAAGAACGAAAGTCGGAGGTTCG | 482.8 | -4.6 | 0.7 | -6.5 | 7.9E-11 | 1.6E-10 | MB231/MB468 |

<table>
<tr><th></th><th></th><th></th><th></th><th></th><th></th><th></th><th></th><th></th><th></th><th></th><th></th><th></th></tr>
<tr><td>>hg38_21_+_8433172_8446622_RNA45SN1_45S_50ntFlanks@7975.8014.40|CGCGACCTCAGATCAGACGTGGCGACCCGCTGAATTTAAG</td><td>>hg38_21_+_8433172_8446622_RNA45SN1_45S_50ntFlanks</td><td>7975</td><td>8014</td><td>40</td><td>CGCGACCTCAGATCAGACGTGGCGACCCGCTGAATTTAAG</td><td>7755.8</td><td>5.2</td><td>0.8</td><td>6.5</td><td>8.1E-11</td><td>1.7E-10</td><td>MB231/MB468</td></tr>
<tr><td>>MI0000076|hsa-mir-20a&WithFlank&13|+|91351059|91351141@14.35.22(+1A)&[MIMAT0000075&hsa-miR-20a-5p&offsets|0|-1(+1A);m-96&13|+|91351072|91351094&offsets|0|-1(+1A)]|TAAAGTGCTTATAGTGCAGGTAA</td><td>>MI0000076|hsa-mir-20a&WithFlank&13|+|91351059|91351141</td><td>14</td><td>35</td><td>22(+1A)</td><td>TAAAGTGCTTATAGTGCAGGTAA</td><td>6937.0</td><td>6.8</td><td>1.1</td><td>6.5</td><td>9.4E-11</td><td>1.7E-10</td><td>BT20/MB468</td></tr>
<tr><td>>hg38_21_+_8433172_8446622_RNA45SN1_45S_50ntFlanks@12992.13013.22|TCCCTCGCTGCGATCTATTGAA</td><td>>hg38_21_+_8433172_8446622_RNA45SN1_45S_50ntFlanks</td><td>12992</td><td>13013</td><td>22</td><td>TCCCTCGCTGCGATCTATTGAA</td><td>886.5</td><td>-4.6</td><td>0.7</td><td>-6.5</td><td>9.4E-11</td><td>1.7E-10</td><td>BT20/MB468</td></tr>
<tr><td>>hg38_21_+_8433172_8446622_RNA45SN1_45S_50ntFlanks@12459.12486.28|AGCGTTGGATTGTTCACCCACTAATAGG</td><td>>hg38_21_+_8433172_8446622_RNA45SN1_45S_50ntFlanks</td><td>12459</td><td>12486</td><td>28</td><td>AGCGTTGGATTGTTCACCCACTAATAGG</td><td>137.4</td><td>-5.6</td><td>0.9</td><td>-6.5</td><td>9.4E-11</td><td>1.7E-10</td><td>BT20/MB468</td></tr>
<tr><td>>hg38_21_+_8433172_8446622_RNA45SN1_45S_50ntFlanks@12874.12892.19|GCTGGAAAGGCGGCCGCCC</td><td>>hg38_21_+_8433172_8446622_RNA45SN1_45S_50ntFlanks</td><td>12874</td><td>12892</td><td>19</td><td>GCTGGAAAGGCGGCCGCCC</td><td>88.1</td><td>-8.5</td><td>1.3</td><td>-6.5</td><td>9.4E-11</td><td>1.7E-10</td><td>BT20/MB468</td></tr>
<tr><td>>hg38_21_+_8433172_8446622_RNA45SN1_45S_50ntFlanks@11759.11780.22|CCTCGTCATCTAATTAGTGACG</td><td>>hg38_21_+_8433172_8446622_RNA45SN1_45S_50ntFlanks</td><td>11759</td><td>11780</td><td>22</td><td>CCTCGTCATCTAATTAGTGACG</td><td>203.0</td><td>-9.7</td><td>1.5</td><td>-6.5</td><td>9.5E-11</td><td>1.7E-10</td><td>BT20/MB468</td></tr>
<tr><td>>hg38_21_+_8433172_8446622_RNA45SN1_45S_50ntFlanks@4390.4408.19|TCTTGGGAGCGGGCGGGCG</td><td>>hg38_21_+_8433172_8446622_RNA45SN1_45S_50ntFlanks</td><td>4390</td><td>4408</td><td>19</td><td>TCTTGGGAGCGGGCGGGCG</td><td>89.8</td><td>-8.5</td><td>1.3</td><td>-6.5</td><td>9.5E-11</td><td>1.7E-10</td><td>BT20/MB468</td></tr>
<tr><td>>hg38_21_+_8433172_8446622_RNA45SN1_45S_50ntFlanks@10805.10828.24|GAACAATGTAGGTAAGGGAAGTCG</td><td>>hg38_21_+_8433172_8446622_RNA45SN1_45S_50ntFlanks</td><td>10805</td><td>10828</td><td>24</td><td>GAACAATGTAGGTAAGGGAAGTCG</td><td>97.5</td><td>-5.3</td><td>0.8</td><td>-6.5</td><td>8.2E-11</td><td>1.7E-10</td><td>MB231/MB468</td></tr>
<tr><td>>hg38_21_+_8433172_8446622_RNA45SN1_45S_50ntFlanks@10420.10455.36|CTGAGAGATGGGCGAGCGCCGTTCCGAAGGGACGGG</td><td>>hg38_21_+_8433172_8446622_RNA45SN1_45S_50ntFlanks</td><td>10420</td><td>10455</td><td>36</td><td>CTGAGAGATGGGCGAGCGCCGTTCCGAAGGGACGGG</td><td>98.0</td><td>-8.6</td><td>1.3</td><td>-6.5</td><td>9.5E-11</td><td>1.7E-10</td><td>BT20/MB468</td></tr>
<tr><td>>hg38_21_+_8433172_8446622_RNA45SN1_45S_50ntFlanks@8190.8212.23|CCAGCCCGTGGACGGTGTGAGGC</td><td>>hg38_21_+_8433172_8446622_RNA45SN1_45S_50ntFlanks</td><td>8190</td><td>8212</td><td>23</td><td>CCAGCCCGTGGACGGTGTGAGGC</td><td>512.5</td><td>-4.8</td><td>0.7</td><td>-6.5</td><td>9.5E-11</td><td>1.7E-10</td><td>BT20/MB468</td></tr>
<tr><td>>MI0000090|hsa-mir-32&WithFlank&9|-|109046223|109046304@12.33.22&[MIMAT0000090&hsa-miR-32-5p&offsets|0|0;m-208&9|-|109046273|109046293&offsets|0|+1]|TATTGCACATTACTAAGTTGCA</td><td>>MI0000090|hsa-mir-32&WithFlank&9|-|109046223|109046304</td><td>12</td><td>33</td><td>22</td><td>TATTGCACATTACTAAGTTGCA</td><td>2277.4</td><td>5.9</td><td>0.9</td><td>6.5</td><td>8.2E-11</td><td>1.7E-10</td><td>MB231/MB468</td></tr>
<tr><td>>hg38_21_+_8433172_8446622_RNA45SN1_45S_50ntFlanks@13006.13028.23|CTATTGAAAGTCAGCCCTCGACA</td><td>>hg38_21_+_8433172_8446622_RNA45SN1_45S_50ntFlanks</td><td>13006</td><td>13028</td><td>23</td><td>CTATTGAAAGTCAGCCCTCGACA</td><td>495.0</td><td>-6.6</td><td>1.0</td><td>-6.5</td><td>8.2E-11</td><td>1.7E-10</td><td>MB231/MB468</td></tr>
<tr><td>>MI0000285|hsa-mir-205&WithFlank&1|+|209432127|209432248@40.61.22(+1C)&[MIMAT0000266&hsa-miR-205-5p&offsets|0|0(+1C);m-351&1|+|209432166|209432187&offsets|0|0(+1C)]|TCCTTCATTCCACCGGAGTCTGC</td><td>>MI0000285|hsa-mir-205&WithFlank&1|+|209432127|209432248</td><td>40</td><td>61</td><td>22(+1C)</td><td>TCCTTCATTCCACCGGAGTCTGC</td><td>116.8</td><td>-8.4</td><td>1.3</td><td>-6.5</td><td>8.3E-11</td><td>1.7E-10</td><td>MB231/MB468</td></tr>
<tr><td>>hg38_21_+_8433172_8446622_RNA45SN1_45S_50ntFlanks@12446.12476.31|GCAGAATTCACCAAGCGTTGGATTGTTCACC</td><td>>hg38_21_+_8433172_8446622_RNA45SN1_45S_50ntFlanks</td><td>12446</td><td>12476</td><td>31</td><td>GCAGAATTCACCAAGCGTTGGATTGTTCACC</td><td>817.9</td><td>5.0</td><td>0.8</td><td>6.5</td><td>8.3E-11</td><td>1.7E-10</td><td>MB231/MB468</td></tr>
<tr><td>>hg38_1_-_228634819_228635039_RNA5S12_5S_50ntFlanks@132.171.40|GATGGGAGACCGCCTGGGAATACCGGGTGCTGTAGGCTTT</td><td>>hg38_1_-_228634819_228635039_RNA5S12_5S_50ntFlanks</td><td>132</td><td>171</td><td>40</td><td>GATGGGAGACCGCCTGGGAATACCGGGTGCTGTAGGCTTT</td><td>6610.3</td><td>4.5</td><td>0.7</td><td>6.5</td><td>9.6E-11</td><td>1.7E-10</td><td>BT20/MB468</td></tr>
<tr><td>>hg38_21_+_8433172_8446622_RNA45SN1_45S_50ntFlanks@8168.8187.20|CCCAAGTCCTTCTGATCGAG</td><td>>hg38_21_+_8433172_8446622_RNA45SN1_45S_50ntFlanks</td><td>8168</td><td>8187</td><td>20</td><td>CCCAAGTCCTTCTGATCGAG</td><td>417.8</td><td>-3.9</td><td>0.6</td><td>-6.5</td><td>8.3E-11</td><td>1.7E-10</td><td>MB231/MB468</td></tr>
<tr><td>>hg38_21_+_8433172_8446622_RNA45SN1_45S_50ntFlanks@9570.9592.23|TGCAAATCGGTCGTCCGACCTGG</td><td>>hg38_21_+_8433172_8446622_RNA45SN1_45S_50ntFlanks</td><td>9570</td><td>9592</td><td>23</td><td>TGCAAATCGGTCGTCCGACCTGG</td><td>98.8</td><td>-8.2</td><td>1.3</td><td>-6.5</td><td>8.3E-11</td><td>1.7E-10</td><td>MB231/MB468</td></tr>
<tr><td>>hg38_21_+_8433172_8446622_RNA45SN1_45S_50ntFlanks@7986.8009.24|ATCAGACGTGGCGACCCGCTGAAT</td><td>>hg38_21_+_8433172_8446622_RNA45SN1_45S_50ntFlanks</td><td>7986</td><td>8009</td><td>24</td><td>ATCAGACGTGGCGACCCGCTGAAT</td><td>78.8</td><td>-8.3</td><td>1.3</td><td>-6.5</td><td>9.7E-11</td><td>1.7E-10</td><td>BT20/MB468</td></tr>
<tr><td>>hg38_21_+_8433172_8446622_RNA45SN1_45S_50ntFlanks@7975.8020.46|CGCGACCTCAGATCAGACGTGGCGACCCGCTGAATTTAAGCATATT</td><td>>hg38_21_+_8433172_8446622_RNA45SN1_45S_50ntFlanks</td><td>7975</td><td>8020</td><td>46</td><td>CGCGACCTCAGATCAGACGTGGCGACCCGCTGAATTTAAGCATATT</td><td>1279.1</td><td>15.2</td><td>2.3</td><td>6.5</td><td>8.4E-11</td><td>1.7E-10</td><td>MB231/MB468</td></tr>
<tr><td>>MI0000111|hsa-mir-105-1&WithFlank&X|-|152392213|152392305@21.42.22;MI0000112|hsa-mir-105-2&WithFlank&X|-|152394406|152394498@21.42.22&[MIMAT0000102&hsa-miR-105-5p&offsets|+2|+1;m-735&X|-|152392266|152392287&offsets|+2|+2];[MIMAT0000102_1&hsa-miR-105-5p&offsets|+2|+1;m-735&X|-|152394459|152394480&offsets|+2|+2]|AAATGCTCAGACTCCTGTGGTG</td><td>>MI0000111|hsa-mir-105-1&WithFlank&X|-|152392213|152392305</td><td>21</td><td>42</td><td>22</td><td>AAATGCTCAGACTCCTGTGGTG</td><td>417.3</td><td>5.3</td><td>0.8</td><td>6.5</td><td>9.8E-11</td><td>1.7E-10</td><td>BT20/MB468</td></tr>
<tr><td>>hg38_21_+_8433172_8446622_RNA45SN1_45S_50ntFlanks@11718.11740.23|AACGGCGGGAGTAACTATGACTC</td><td>>hg38_21_+_8433172_8446622_RNA45SN1_45S_50ntFlanks</td><td>11718</td><td>11740</td><td>23</td><td>AACGGCGGGAGTAACTATGACTC</td><td>1379.3</td><td>-4.9</td><td>0.7</td><td>-6.5</td><td>8.5E-11</td><td>1.7E-10</td><td>MB231/MB468</td></tr>
<tr><td>>hg38_21_+_8433172_8446622_RNA45SN1_45S_50ntFlanks@12941.12961.21|ATTCGTAGACGACCTGCTTCT</td><td>>hg38_21_+_8433172_8446622_RNA45SN1_45S_50ntFlanks</td><td>12941</td><td>12961</td><td>21</td><td>ATTCGTAGACGACCTGCTTCT</td><td>672.4</td><td>3.6</td><td>0.6</td><td>6.5</td><td>8.5E-11</td><td>1.7E-10</td><td>MB231/MB468</td></tr>
<tr><td>>trna17_ValTAC_11_-_59318102_59318174@1.34.34,trna4_ValTAC_X_-_18693029_18693101@1.34.34|GGTTCCATAGTGTAGTGGTTATCACGTCTGCTTT</td><td>>trna17_ValTAC_11_-_59318102_59318174</td><td>1</td><td>34</td><td>34</td><td>GGTTCCATAGTGTAGTGGTTATCACGTCTGCTTT</td><td>1034.1</td><td>14.9</td><td>2.3</td><td>6.5</td><td>8.5E-11</td><td>1.8E-10</td><td>MB231/MB468</td></tr>
<tr><td>>hg38_21_+_8433172_8446622_RNA45SN1_45S_50ntFlanks@4163.4184.22|CATCCAAGGAAGGCAGCAGGCG</td><td>>hg38_21_+_8433172_8446622_RNA45SN1_45S_50ntFlanks</td><td>4163</td><td>4184</td><td>22</td><td>CATCCAAGGAAGGCAGCAGGCG</td><td>383.7</td><td>-5.9</td><td>0.9</td><td>-6.5</td><td>8.5E-11</td><td>1.8E-10</td><td>MB231/MB468</td></tr>
<tr><td>>hg38_21_+_8433172_8446622_RNA45SN1_45S_50ntFlanks@10403.10424.22|ACATGGGTCAGTCGGTCCTGAG</td><td>>hg38_21_+_8433172_8446622_RNA45SN1_45S_50ntFlanks</td><td>10403</td><td>10424</td><td>22</td><td>ACATGGGTCAGTCGGTCCTGAG</td><td>958.9</td><td>4.2</td><td>0.7</td><td>6.5</td><td>1.0E-10</td><td>1.8E-10</td><td>BT20/MB468</td></tr>
<tr><td>>hg38_21_+_8433172_8446622_RNA45SN1_45S_50ntFlanks@12991.13027.37|CTCCCTCGCTGCGATCTATTGAAAGTCAGCCCTCGAC</td><td>>hg38_21_+_8433172_8446622_RNA45SN1_45S_50ntFlanks</td><td>12991</td><td>13027</td><td>37</td><td>CTCCCTCGCTGCGATCTATTGAAAGTCAGCCCTCGAC</td><td>244.7</td><td>-4.7</td><td>0.7</td><td>-6.5</td><td>8.6E-11</td><td>1.8E-10</td><td>MB231/MB468</td></tr>
<tr><td>>hg38_21_+_8433172_8446622_RNA45SN1_45S_50ntFlanks@10682.10706.25|TCCGGCGGCGTCCGGTGAGCTCTCG</td><td>>hg38_21_+_8433172_8446622_RNA45SN1_45S_50ntFlanks</td><td>10682</td><td>10706</td><td>25</td><td>TCCGGCGGCGTCCGGTGAGCTCTCG</td><td>1282.0</td><td>-4.6</td><td>0.7</td><td>-6.5</td><td>1.0E-10</td><td>1.8E-10</td><td>BT20/MB468</td></tr>
<tr><td>>hg38_21_+_8433172_8446622_RNA45SN1_45S_50ntFlanks@12489.12521.33|ACGTGAGCTGGGTTTAGACCGTCGTGAGACAGG</td><td>>hg38_21_+_8433172_8446622_RNA45SN1_45S_50ntFlanks</td><td>12489</td><td>12521</td><td>33</td><td>ACGTGAGCTGGGTTTAGACCGTCGTGAGACAGG</td><td>111.1</td><td>-8.4</td><td>1.3</td><td>-6.5</td><td>8.7E-11</td><td>1.8E-10</td><td>MB231/MB468</td></tr>
<tr><td>>hg38_21_+_8433172_8446622_RNA45SN1_45S_50ntFlanks@12630.12654.25|GGGCGAAGCTACCATCTGTGGGATT</td><td>>hg38_21_+_8433172_8446622_RNA45SN1_45S_50ntFlanks</td><td>12630</td><td>12654</td><td>25</td><td>GGGCGAAGCTACCATCTGTGGGATT</td><td>169.3</td><td>-4.6</td><td>0.7</td><td>-6.5</td><td>8.7E-11</td><td>1.8E-10</td><td>MB231/MB468</td></tr>
<tr><td>>hg38_21_+_8433172_8446622_RNA45SN1_45S_50ntFlanks@9033.9059.27|CTCGTGGGGGGGCCGGGCCACCCCTCC</td><td>>hg38_21_+_8433172_8446622_RNA45SN1_45S_50ntFlanks</td><td>9033</td><td>9059</td><td>27</td><td>CTCGTGGGGGGGCCGGGCCACCCCTCC</td><td>137.7</td><td>-8.7</td><td>1.3</td><td>-6.5</td><td>8.7E-11</td><td>1.8E-10</td><td>MB231/MB468</td></tr>
<tr><td>>hg38_21_+_8433172_8446622_RNA45SN1_45S_50ntFlanks@12477.12497.21|CACTAATAGGGAACGTGAGCT</td><td>>hg38_21_+_8433172_8446622_RNA45SN1_45S_50ntFlanks</td><td>12477</td><td>12497</td><td>21</td><td>CACTAATAGGGAACGTGAGCT</td><td>2582.4</td><td>3.6</td><td>0.6</td><td>6.5</td><td>8.8E-11</td><td>1.8E-10</td><td>MB231/MB468</td></tr>
<tr><td>>hg38_21_+_8433172_8446622_RNA45SN1_45S_50ntFlanks@9352.9377.26|CCGTGGCGCAATGAAGGTGAAGGCCG</td><td>>hg38_21_+_8433172_8446622_RNA45SN1_45S_50ntFlanks</td><td>9352</td><td>9377</td><td>26</td><td>CCGTGGCGCAATGAAGGTGAAGGCCG</td><td>89.8</td><td>-8.5</td><td>1.3</td><td>-6.5</td><td>1.0E-10</td><td>1.8E-10</td><td>BT20/MB468</td></tr>
<tr><td>>hg38_21_+_8433172_8446622_RNA45SN1_45S_50ntFlanks@10787.10818.32|ACAGCCTCTGGCATGTTGGAACAATGTAGGTA</td><td>>hg38_21_+_8433172_8446622_RNA45SN1_45S_50ntFlanks</td><td>10787</td><td>10818</td><td>32</td><td>ACAGCCTCTGGCATGTTGGAACAATGTAGGTA</td><td>1834.1</td><td>-4.8</td><td>0.7</td><td>-6.5</td><td>1.0E-10</td><td>1.8E-10</td><td>BT20/MB468</td></tr>
<tr><td>>hg38_21_+_8433172_8446622_RNA45SN1_45S_50ntFlanks@4764.4781.18|ATACCGTCGTAGTTCCGA</td><td>>hg38_21_+_8433172_8446622_RNA45SN1_45S_50ntFlanks</td><td>4764</td><td>4781</td><td>18</td><td>ATACCGTCGTAGTTCCGA</td><td>113.7</td><td>-3.5</td><td>0.5</td><td>-6.5</td><td>1.0E-10</td><td>1.8E-10</td><td>BT20/MB468</td></tr>
<tr><td>>hg38_21_+_8433172_8446622_RNA45SN1_45S_50ntFlanks@12419.12442.24|GATGTCGGCTCTTCCTATCATTGT</td><td>>hg38_21_+_8433172_8446622_RNA45SN1_45S_50ntFlanks</td><td>12419</td><td>12442</td><td>24</td><td>GATGTCGGCTCTTCCTATCATTGT</td><td>386.2</td><td>4.8</td><td>0.7</td><td>6.5</td><td>1.0E-10</td><td>1.8E-10</td><td>BT20/MB468</td></tr>
<tr><td>>hg38_MT_+_598_1651_RNR1_12S_50ntFlanks@416.462.47|AAATAGACTACGAAAGTGGCTTTAACATATCTGAACACACAATAGCT</td><td>>hg38_MT_+_598_1651_RNR1_12S_50ntFlanks</td><td>416</td><td>462</td><td>47</td><td>AAATAGACTACGAAAGTGGCTTTAACATATCTGAACACACAATAGCT</td><td>732.0</td><td>14.4</td><td>2.2</td><td>6.5</td><td>9.0E-11</td><td>1.8E-10</td><td>MB231/MB468</td></tr>
<tr><td>>hg38_21_+_8433172_8446622_RNA45SN1_45S_50ntFlanks@12637.12661.25|GCTACCATCTGTGGGATTATGACTG</td><td>>hg38_21_+_8433172_8446622_RNA45SN1_45S_50ntFlanks</td><td>12637</td><td>12661</td><td>25</td><td>GCTACCATCTGTGGGATTATGACTG</td><td>432.8</td><td>-3.6</td><td>0.6</td><td>-6.5</td><td>1.0E-10</td><td>1.9E-10</td><td>BT20/MB468</td></tr>
<tr><td>>hg38_MT_+_1621_3279_RNR2_16S_50ntFlanks@459.492.34|CTTGTAAATTTAACTGTTAGTCCAAAGAGGAACA</td><td>>hg38_MT_+_1621_3279_RNR2_16S_50ntFlanks</td><td>459</td><td>492</td><td>34</td><td>CTTGTAAATTTAACTGTTAGTCCAAAGAGGAACA</td><td>942.7</td><td>8.7</td><td>1.4</td><td>6.5</td><td>1.0E-10</td><td>1.9E-10</td><td>BT20/MB468</td></tr>
<tr><td>>hg38_21_+_8433172_8446622_RNA45SN1_45S_50ntFlanks@10512.10534.23|TCCGGAGTGGCGGAGATGGGCGC</td><td>>hg38_21_+_8433172_8446622_RNA45SN1_45S_50ntFlanks</td><td>10512</td><td>10534</td><td>23</td><td>TCCGGAGTGGCGGAGATGGGCGC</td><td>123.0</td><td>-8.5</td><td>1.3</td><td>-6.5</td><td>9.1E-11</td><td>1.9E-10</td><td>MB231/MB468</td></tr>
<tr><td>>hg38_21_+_8433172_8446622_RNA45SN1_45S_50ntFlanks@9553.9573.21|CCGTAGCGGTCCTGACGTGCA</td><td>>hg38_21_+_8433172_8446622_RNA45SN1_45S_50ntFlanks</td><td>9553</td><td>9573</td><td>21</td><td>CCGTAGCGGTCCTGACGTGCA</td><td>474.4</td><td>3.3</td><td>0.5</td><td>6.5</td><td>9.1E-11</td><td>1.9E-10</td><td>MB231/MB468</td></tr>
<tr><td>>hg38_21_+_8433172_8446622_RNA45SN1_45S_50ntFlanks@11535.11553.19|TTCCGCGCGGCGCCTCGCC</td><td>>hg38_21_+_8433172_8446622_RNA45SN1_45S_50ntFlanks</td><td>11535</td><td>11553</td><td>19</td><td>TTCCGCGCGGCGCCTCGCC</td><td>99.6</td><td>-8.2</td><td>1.3</td><td>-6.5</td><td>9.1E-11</td><td>1.9E-10</td><td>MB231/MB468</td></tr>
<tr><td>>hg38_21_+_8433172_8446622_RNA45SN1_45S_50ntFlanks@7980.8003.24|CCTCAGATCAGACGTGGCGACCCG</td><td>>hg38_21_+_8433172_8446622_RNA45SN1_45S_50ntFlanks</td><td>7980</td><td>8003</td><td>24</td><td>CCTCAGATCAGACGTGGCGACCCG</td><td>221.3</td><td>-3.9</td><td>0.6</td><td>-6.5</td><td>1.1E-10</td><td>1.9E-10</td><td>BT20/MB468</td></tr>
<tr><td>>hg38_21_+_8433172_8446622_RNA45SN1_45S_50ntFlanks@7984.8005.22|AGATCAGACGTGGCGACCCGCT</td><td>>hg38_21_+_8433172_8446622_RNA45SN1_45S_50ntFlanks</td><td>7984</td><td>8005</td><td>22</td><td>AGATCAGACGTGGCGACCCGCT</td><td>176.5</td><td>-4.7</td><td>0.7</td><td>-6.5</td><td>9.2E-11</td><td>1.9E-10</td><td>MB231/MB468</td></tr>
<tr><td>>hg38_21_+_8433172_8446622_RNA45SN1_45S_50ntFlanks@5421.5445.25|CGATTGGATGGTTTAGTGAGGCCCT</td><td>>hg38_21_+_8433172_8446622_RNA45SN1_45S_50ntFlanks</td><td>5421</td><td>5445</td><td>25</td><td>CGATTGGATGGTTTAGTGAGGCCCT</td><td>270.9</td><td>-5.8</td><td>0.9</td><td>-6.5</td><td>1.1E-10</td><td>1.9E-10</td><td>BT20/MB468</td></tr>
<tr><td>>hg38_21_+_8433172_8446622_RNA45SN1_45S_50ntFlanks@8106.8143.38|GCGGCGGGGCGCGGGACATGTGGCGTACGGAAGACCCG</td><td>>hg38_21_+_8433172_8446622_RNA45SN1_45S_50ntFlanks</td><td>8106</td><td>8143</td><td>38</td><td>GCGGCGGGGCGCGGGACATGTGGCGTACGGAAGACCCG</td><td>110.2</td><td>-8.8</td><td>1.4</td><td>-6.5</td><td>1.1E-10</td><td>1.9E-10</td><td>BT20/MB468</td></tr>
</table>

<table>
<tr><td></td><td></td><td></td><td></td><td></td><td></td><td></td><td></td><td></td><td></td><td></td><td></td><td></td></tr>
<tr><td>>hg38_21_+_8433172_8446622_RNA45SN1_45S_50ntFlanks@4498.4517.20|AAGCGTTTACTTTGAAAAAA</td><td>>hg38_21_+_8433172_8446622_RNA45SN1_45S_50ntFlanks</td><td>4498</td><td>4517</td><td>20</td><td>AAGCGTTTACTTTGAAAAAA</td><td>291.2</td><td>-4.0</td><td>0.6</td><td>-6.5</td><td>9.3E-11</td><td>1.9E-10</td><td>MB231/MB468</td></tr>
<tr><td>>hg38_21_+_8433172_8446622_RNA45SN1_45S_50ntFlanks@12513.12538.26|TGAGACAGGTTAGTTTTACCCTACTG</td><td>>hg38_21_+_8433172_8446622_RNA45SN1_45S_50ntFlanks</td><td>12513</td><td>12538</td><td>26</td><td>TGAGACAGGTTAGTTTTACCCTACTG</td><td>124.8</td><td>-9.0</td><td>1.4</td><td>-6.5</td><td>1.1E-10</td><td>1.9E-10</td><td>BT20/MB468</td></tr>
<tr><td>>hg38_21_+_8433172_8446622_RNA45SN1_45S_50ntFlanks@9465.9503.39|AGGTGGAGCACGAGCGCACGTGTTAGGACCCGAAAGATG</td><td>>hg38_21_+_8433172_8446622_RNA45SN1_45S_50ntFlanks</td><td>9465</td><td>9503</td><td>39</td><td>AGGTGGAGCACGAGCGCACGTGTTAGGACCCGAAAGATG</td><td>77.8</td><td>-8.3</td><td>1.3</td><td>-6.5</td><td>1.1E-10</td><td>1.9E-10</td><td>BT20/MB468</td></tr>
<tr><td>>hg38_21_+_8433172_8446622_RNA45SN1_45S_50ntFlanks@3793.3810.18|CGAATGGCTCATTAAATC</td><td>>hg38_21_+_8433172_8446622_RNA45SN1_45S_50ntFlanks</td><td>3793</td><td>3810</td><td>18</td><td>CGAATGGCTCATTAAATC</td><td>150.4</td><td>-5.3</td><td>0.8</td><td>-6.5</td><td>1.1E-10</td><td>1.9E-10</td><td>BT20/MB468</td></tr>
<tr><td>>hg38_21_+_8433172_8446622_RNA45SN1_45S_50ntFlanks@11783.11807.25|CATGAATGGATGAACGAGATTCCCA</td><td>>hg38_21_+_8433172_8446622_RNA45SN1_45S_50ntFlanks</td><td>11783</td><td>11807</td><td>25</td><td>CATGAATGGATGAACGAGATTCCCA</td><td>274.4</td><td>-6.5</td><td>1.0</td><td>-6.5</td><td>1.1E-10</td><td>1.9E-10</td><td>BT20/MB468</td></tr>
<tr><td>>hg38_21_+_8433172_8446622_RNA45SN1_45S_50ntFlanks@11567.11586.20|AGCAGCCGACTTAGAACTGG</td><td>>hg38_21_+_8433172_8446622_RNA45SN1_45S_50ntFlanks</td><td>11567</td><td>11586</td><td>20</td><td>AGCAGCCGACTTAGAACTGG</td><td>3459.2</td><td>3.7</td><td>0.6</td><td>6.5</td><td>9.4E-11</td><td>1.9E-10</td><td>MB231/MB468</td></tr>
<tr><td>>hg38_21_+_8433172_8446622_RNA45SN1_45S_50ntFlanks@12417.12443.27|TCGATGTCGGCTCTTCCTATCATTGTG</td><td>>hg38_21_+_8433172_8446622_RNA45SN1_45S_50ntFlanks</td><td>12417</td><td>12443</td><td>27</td><td>TCGATGTCGGCTCTTCCTATCATTGTG</td><td>2098.8</td><td>-3.1</td><td>0.5</td><td>-6.5</td><td>1.1E-10</td><td>1.9E-10</td><td>BT20/MB468</td></tr>
<tr><td>>hg38_21_+_8433172_8446622_RNA45SN1_45S_50ntFlanks@12606.12626.21|ATGTGCTTGGCTGAGGAGCCA</td><td>>hg38_21_+_8433172_8446622_RNA45SN1_45S_50ntFlanks</td><td>12606</td><td>12626</td><td>21</td><td>ATGTGCTTGGCTGAGGAGCCA</td><td>376.4</td><td>-4.4</td><td>0.7</td><td>-6.5</td><td>1.1E-10</td><td>1.9E-10</td><td>BT20/MB468</td></tr>
<tr><td>>hg38_21_+_8433172_8446622_RNA45SN1_45S_50ntFlanks@10841.10859.19|CGTAACTTCGGGATAAGGA</td><td>>hg38_21_+_8433172_8446622_RNA45SN1_45S_50ntFlanks</td><td>10841</td><td>10859</td><td>19</td><td>CGTAACTTCGGGATAAGGA</td><td>796.6</td><td>-5.1</td><td>0.8</td><td>-6.5</td><td>9.5E-11</td><td>1.9E-10</td><td>MB231/MB468</td></tr>
<tr><td>>hg38_1_-_228634819_228635039_RNA5S12_5S_50ntFlanks@141.171.31|CCGCCTGGGAATACCGGGTGCTGTAGGCTTT</td><td>>hg38_1_-_228634819_228635039_RNA5S12_5S_50ntFlanks</td><td>141</td><td>171</td><td>31</td><td>CCGCCTGGGAATACCGGGTGCTGTAGGCTTT</td><td>5073.5</td><td>6.5</td><td>1.0</td><td>6.5</td><td>1.1E-10</td><td>2.0E-10</td><td>BT20/MB468</td></tr>
<tr><td>>hg38_21_+_8433172_8446622_RNA45SN1_45S_50ntFlanks@12477.12497.21|CACTAATAGGGAACGTGAGCT</td><td>>hg38_21_+_8433172_8446622_RNA45SN1_45S_50ntFlanks</td><td>12477</td><td>12497</td><td>21</td><td>CACTAATAGGGAACGTGAGCT</td><td>1405.7</td><td>2.3</td><td>0.4</td><td>6.5</td><td>1.1E-10</td><td>2.0E-10</td><td>BT20/MB468</td></tr>
<tr><td>>hg38_21_+_8433172_8446622_RNA45SN1_45S_50ntFlanks@10819.10837.19|AGGGAAGTCGGCAAGCCGG</td><td>>hg38_21_+_8433172_8446622_RNA45SN1_45S_50ntFlanks</td><td>10819</td><td>10837</td><td>19</td><td>AGGGAAGTCGGCAAGCCGG</td><td>285.1</td><td>-3.8</td><td>0.6</td><td>-6.5</td><td>1.1E-10</td><td>2.0E-10</td><td>BT20/MB468</td></tr>
<tr><td>>hg38_21_+_8433172_8446622_RNA45SN1_45S_50ntFlanks@10806.10830.25|AACAATGTAGGTAAGGGAAGTCGGC</td><td>>hg38_21_+_8433172_8446622_RNA45SN1_45S_50ntFlanks</td><td>10806</td><td>10830</td><td>25</td><td>AACAATGTAGGTAAGGGAAGTCGGC</td><td>88.1</td><td>-3.6</td><td>0.6</td><td>-6.5</td><td>1.1E-10</td><td>2.0E-10</td><td>BT20/MB468</td></tr>
<tr><td>>hg38_21_+_8433172_8446622_RNA45SN1_45S_50ntFlanks@12892.12933.42|CCCTCGCCCGTCACGCACCGCACGTTCGTGGGGAACCTGGCG</td><td>>hg38_21_+_8433172_8446622_RNA45SN1_45S_50ntFlanks</td><td>12892</td><td>12933</td><td>42</td><td>CCCTCGCCCGTCACGCACCGCACGTTCGTGGGGAACCTGGCG</td><td>120.9</td><td>-8.5</td><td>1.3</td><td>-6.5</td><td>9.7E-11</td><td>2.0E-10</td><td>MB231/MB468</td></tr>
<tr><td>>hg38_21_+_8433172_8446622_RNA45SN1_45S_50ntFlanks@4370.4389.20|TTAAAAAGCTCGTAGTTGGA</td><td>>hg38_21_+_8433172_8446622_RNA45SN1_45S_50ntFlanks</td><td>4370</td><td>4389</td><td>20</td><td>TTAAAAAGCTCGTAGTTGGA</td><td>196.9</td><td>-4.7</td><td>0.7</td><td>-6.5</td><td>9.7E-11</td><td>2.0E-10</td><td>MB231/MB468</td></tr>
<tr><td>>hg38_21_+_8433172_8446622_RNA45SN1_45S_50ntFlanks@11602.11619.18|TCCGACTGTTTAATTAAA</td><td>>hg38_21_+_8433172_8446622_RNA45SN1_45S_50ntFlanks</td><td>11602</td><td>11619</td><td>18</td><td>TCCGACTGTTTAATTAAA</td><td>724.3</td><td>-4.0</td><td>0.6</td><td>-6.4</td><td>1.1E-10</td><td>2.0E-10</td><td>BT20/MB468</td></tr>
<tr><td>>hg38_21_+_8433172_8446622_RNA45SN1_45S_50ntFlanks@4043.4063.21|ACCCATTCGAACGTCTGCCCT</td><td>>hg38_21_+_8433172_8446622_RNA45SN1_45S_50ntFlanks</td><td>4043</td><td>4063</td><td>21</td><td>ACCCATTCGAACGTCTGCCCT</td><td>89.1</td><td>-8.1</td><td>1.2</td><td>-6.5</td><td>9.7E-11</td><td>2.0E-10</td><td>MB231/MB468</td></tr>
<tr><td>>hg38_21_+_8433172_8446622_RNA45SN1_45S_50ntFlanks@10458.10490.33|ATGGCCTCCGTTGCCCTCGGCCGATCGAAAGGG</td><td>>hg38_21_+_8433172_8446622_RNA45SN1_45S_50ntFlanks</td><td>10458</td><td>10490</td><td>33</td><td>ATGGCCTCCGTTGCCCTCGGCCGATCGAAAGGG</td><td>109.8</td><td>-8.8</td><td>1.4</td><td>-6.4</td><td>1.1E-10</td><td>2.0E-10</td><td>BT20/MB468</td></tr>
<tr><td>>hg38_21_+_8433172_8446622_RNA45SN1_45S_50ntFlanks@12126.12151.26|CGACCCGCTCCGGGGACAGTGCCAGG</td><td>>hg38_21_+_8433172_8446622_RNA45SN1_45S_50ntFlanks</td><td>12126</td><td>12151</td><td>26</td><td>CGACCCGCTCCGGGGACAGTGCCAGG</td><td>98.4</td><td>-8.2</td><td>1.3</td><td>-6.5</td><td>9.8E-11</td><td>2.0E-10</td><td>MB231/MB468</td></tr>
<tr><td>>hg38_21_+_8433172_8446622_RNA45SN1_45S_50ntFlanks@10684.10705.22|CGGCGGCGTCCGGTGAGCTCTC</td><td>>hg38_21_+_8433172_8446622_RNA45SN1_45S_50ntFlanks</td><td>10684</td><td>10705</td><td>22</td><td>CGGCGGCGTCCGGTGAGCTCTC</td><td>164.5</td><td>-3.1</td><td>0.5</td><td>-6.5</td><td>9.9E-11</td><td>2.0E-10</td><td>MB231/MB468</td></tr>
<tr><td>>hg38_21_+_8433172_8446622_RNA45SN1_45S_50ntFlanks@12607.12624.18|TGTGCTTGGCTGAGGAGC</td><td>>hg38_21_+_8433172_8446622_RNA45SN1_45S_50ntFlanks</td><td>12607</td><td>12624</td><td>18</td><td>TGTGCTTGGCTGAGGAGC</td><td>838.6</td><td>4.0</td><td>0.6</td><td>6.5</td><td>9.9E-11</td><td>2.0E-10</td><td>MB231/MB468</td></tr>
<tr><td>>am_trna146_GlnCTG_6_-_27515531_27515602@1.29.29,trna1_GlnCTG_6_+_18836402_18836473@1.29.29,trna3_GlnCTG_17_+_8023070_8023141@1.29.29,trna42_GlnCTG_6_+_27263212_27263283@1.29.29,trna49_GlnCTG_6_+_27487308_27487379@1.29.29,trna7_GlnCTG_15_-_66161400_66161471@1.29.29,trna99_GlnCTG_6_-_28909378_28909449@1.29.29|GGTTCCATGGTGTAATGGTTAGCACTCTG</td><td>>am_trna146_GlnCTG_6_-_27515531_27515602</td><td>1</td><td>29</td><td>29</td><td>GGTTCCATGGTGTAATGGTTAGCACTCTG</td><td>10344.1</td><td>-7.4</td><td>1.1</td><td>-6.4</td><td>1.1E-10</td><td>2.0E-10</td><td>BT20/MB468</td></tr>
<tr><td>>hg38_21_+_8433172_8446622_RNA45SN1_45S_50ntFlanks@11315.11336.22|CCACCCCACGTCTCGTCGCGCG</td><td>>hg38_21_+_8433172_8446622_RNA45SN1_45S_50ntFlanks</td><td>11315</td><td>11336</td><td>22</td><td>CCACCCCACGTCTCGTCGCGCG</td><td>208.3</td><td>-7.4</td><td>1.1</td><td>-6.5</td><td>9.9E-11</td><td>2.0E-10</td><td>MB231/MB468</td></tr>
<tr><td>>hg38_21_+_8433172_8446622_RNA45SN1_45S_50ntFlanks@9736.9757.22|CCGAAACGATCTCAACCTATTC</td><td>>hg38_21_+_8433172_8446622_RNA45SN1_45S_50ntFlanks</td><td>9736</td><td>9757</td><td>22</td><td>CCGAAACGATCTCAACCTATTC</td><td>95.8</td><td>-8.2</td><td>1.3</td><td>-6.5</td><td>9.9E-11</td><td>2.0E-10</td><td>MB231/MB468</td></tr>
<tr><td>>hg38_21_+_8433172_8446622_RNA45SN1_45S_50ntFlanks@8191.8214.24|CAGCCCGTGGACGGTGTGAGGCCG</td><td>>hg38_21_+_8433172_8446622_RNA45SN1_45S_50ntFlanks</td><td>8191</td><td>8214</td><td>24</td><td>CAGCCCGTGGACGGTGTGAGGCCG</td><td>454.6</td><td>-3.6</td><td>0.6</td><td>-6.5</td><td>1.0E-10</td><td>2.0E-10</td><td>MB231/MB468</td></tr>
<tr><td>>MI0000285|hsa-mir-205&WithFlank&1|+|209432127|209432248@40.61.22(+1G)&[MIMAT0000266&hsa-miR-205-5p&offsets|0|0(+1G);m-351&1|+|209432166|209432187&offsets|0|0(+1G)]|TCCTTCATTCCACCGGAGTCTGG</td><td>>MI0000285|hsa-mir-205&WithFlank&1|+|209432127|209432248</td><td>40</td><td>61</td><td>22(+1G)</td><td>TCCTTCATTCCACCGGAGTCTGG</td><td>96.8</td><td>-8.2</td><td>1.3</td><td>-6.5</td><td>1.0E-10</td><td>2.0E-10</td><td>MB231/MB468</td></tr>
<tr><td>>hg38_21_+_8433172_8446622_RNA45SN1_45S_50ntFlanks@11718.11742.25|AACGGCGGGAGTAACTATGACTCTC</td><td>>hg38_21_+_8433172_8446622_RNA45SN1_45S_50ntFlanks</td><td>11718</td><td>11742</td><td>25</td><td>AACGGCGGGAGTAACTATGACTCTC</td><td>1100.0</td><td>-4.3</td><td>0.7</td><td>-6.5</td><td>1.0E-10</td><td>2.0E-10</td><td>MB231/MB468</td></tr>
<tr><td>>hg38_21_+_8433172_8446622_RNA45SN1_45S_50ntFlanks@4003.4022.20|ATAACCTCGGGCCGATCGCA</td><td>>hg38_21_+_8433172_8446622_RNA45SN1_45S_50ntFlanks</td><td>4003</td><td>4022</td><td>20</td><td>ATAACCTCGGGCCGATCGCA</td><td>369.3</td><td>-3.5</td><td>0.5</td><td>-6.5</td><td>1.0E-10</td><td>2.0E-10</td><td>MB231/MB468</td></tr>
<tr><td>>hg38_21_+_8433172_8446622_RNA45SN1_45S_50ntFlanks@7985.8006.22|GATCAGACGTGGCGACCCGCTG</td><td>>hg38_21_+_8433172_8446622_RNA45SN1_45S_50ntFlanks</td><td>7985</td><td>8006</td><td>22</td><td>GATCAGACGTGGCGACCCGCTG</td><td>1456.0</td><td>-7.0</td><td>1.1</td><td>-6.5</td><td>1.0E-10</td><td>2.0E-10</td><td>MB231/MB468</td></tr>
<tr><td>>hg38_21_+_8433172_8446622_RNA45SN1_45S_50ntFlanks@12490.12520.31|CGTGAGCTGGGTTTAGACCGTCGTGAGACAG</td><td>>hg38_21_+_8433172_8446622_RNA45SN1_45S_50ntFlanks</td><td>12490</td><td>12520</td><td>31</td><td>CGTGAGCTGGGTTTAGACCGTCGTGAGACAG</td><td>139.1</td><td>-5.2</td><td>0.8</td><td>-6.4</td><td>1.2E-10</td><td>2.1E-10</td><td>BT20/MB468</td></tr>
<tr><td>>hg38_21_+_8433172_8446622_RNA45SN1_45S_50ntFlanks@12436.12473.38|TCATTGTGAAGCAGAATTCACCAAGCGTTGGATTGTTC</td><td>>hg38_21_+_8433172_8446622_RNA45SN1_45S_50ntFlanks</td><td>12436</td><td>12473</td><td>38</td><td>TCATTGTGAAGCAGAATTCACCAAGCGTTGGATTGTTC</td><td>255.7</td><td>-4.2</td><td>0.6</td><td>-6.4</td><td>1.2E-10</td><td>2.1E-10</td><td>BT20/MB468</td></tr>
<tr><td>>trnaMT_PheGAA_MT_+_577_647@1.34.34|GTTTATGTAGCTTACCTCCTCAAAGCAATACACT</td><td>>trnaMT_PheGAA_MT_+_577_647</td><td>1</td><td>34</td><td>34</td><td>GTTTATGTAGCTTACCTCCTCAAAGCAATACACT</td><td>1243.7</td><td>9.2</td><td>1.4</td><td>6.4</td><td>1.2E-10</td><td>2.1E-10</td><td>BT20/MB468</td></tr>
<tr><td>>MI0000074|hsa-mir-19b-1&WithFlank&13|+|91351186|91351284@60.82.23;MI0000075|hsa-mir-19b-2&WithFlank&X|-|134169665|134169772@68.90.23&[MIMAT0000074&hsa-miR-19b-3p&offsets|0|0;m-122&13|+|91351245|91351267&offsets|0|0];[MIMAT0000074_1&hsa-miR-19b-3p&offsets|0|0;m-121&X|-|134169683|134169705&offsets|0|0]|TGTGCAAATCCATGCAAAACTGA</td><td>>MI0000074|hsa-mir-19b-1&WithFlank&13|+|91351186|91351284</td><td>60</td><td>82</td><td>23</td><td>TGTGCAAATCCATGCAAAACTGA</td><td>4426.1</td><td>7.7</td><td>1.2</td><td>6.4</td><td>1.2E-10</td><td>2.1E-10</td><td>BT20/MB468</td></tr>
<tr><td>>trnaMT_PheGAA_MT_+_577_647@39.62.24|ATGTTTAGACGGGCTCACATCACC</td><td>>trnaMT_PheGAA_MT_+_577_647</td><td>39</td><td>62</td><td>24</td><td>ATGTTTAGACGGGCTCACATCACC</td><td>548.9</td><td>8.0</td><td>1.2</td><td>6.4</td><td>1.2E-10</td><td>2.1E-10</td><td>BT20/MB468</td></tr>
<tr><td>>hg38_21_+_8433172_8446622_RNA45SN1_45S_50ntFlanks@5472.5514.43|ACGGCCCTGGCGGAGCGCTGAGAAGACGGTCGAACTTGACTAT</td><td>>hg38_21_+_8433172_8446622_RNA45SN1_45S_50ntFlanks</td><td>5472</td><td>5514</td><td>43</td><td>ACGGCCCTGGCGGAGCGCTGAGAAGACGGTCGAACTTGACTAT</td><td>3033.6</td><td>5.8</td><td>0.9</td><td>6.5</td><td>1.0E-10</td><td>2.1E-10</td><td>MB231/MB468</td></tr>
<tr><td>>hg38_21_+_8433172_8446622_RNA45SN1_45S_50ntFlanks@12324.12356.33|TTGGGTTTTAAGCAGGAGGTGTCAGAAAAGTTA</td><td>>hg38_21_+_8433172_8446622_RNA45SN1_45S_50ntFlanks</td><td>12324</td><td>12356</td><td>33</td><td>TTGGGTTTTAAGCAGGAGGTGTCAGAAAAGTTA</td><td>89.1</td><td>-8.1</td><td>1.2</td><td>-6.5</td><td>1.0E-10</td><td>2.1E-10</td><td>MB231/MB468</td></tr>
<tr><td>>hg38_21_+_8433172_8446622_RNA45SN1_45S_50ntFlanks@11785.11805.21|TGAATGGATGAACGAGATTCC</td><td>>hg38_21_+_8433172_8446622_RNA45SN1_45S_50ntFlanks</td><td>11785</td><td>11805</td><td>21</td><td>TGAATGGATGAACGAGATTCC</td><td>1041.1</td><td>-4.9</td><td>0.8</td><td>-6.4</td><td>1.2E-10</td><td>2.1E-10</td><td>BT20/MB468</td></tr>
<tr><td>>hg38_21_+_8433172_8446622_RNA45SN1_45S_50ntFlanks@10841.10861.21|CGTAACTTCGGGATAAGGATT</td><td>>hg38_21_+_8433172_8446622_RNA45SN1_45S_50ntFlanks</td><td>10841</td><td>10861</td><td>21</td><td>CGTAACTTCGGGATAAGGATT</td><td>1505.3</td><td>-3.7</td><td>0.6</td><td>-6.4</td><td>1.2E-10</td><td>2.1E-10</td><td>BT20/MB468</td></tr>
<tr><td>>hg38_21_+_8433172_8446622_RNA45SN1_45S_50ntFlanks@12939.12966.28|CCATTCGTAGACGACCTGCTTCTGGGTC</td><td>>hg38_21_+_8433172_8446622_RNA45SN1_45S_50ntFlanks</td><td>12939</td><td>12966</td><td>28</td><td>CCATTCGTAGACGACCTGCTTCTGGGTC</td><td>100.8</td><td>-5.7</td><td>0.9</td><td>-6.5</td><td>1.0E-10</td><td>2.1E-10</td><td>MB231/MB468</td></tr>
<tr><td>>hg38_21_+_8433172_8446622_RNA45SN1_45S_50ntFlanks@475.492.18|CTTCGTGATCGATGTGGT</td><td>>hg38_21_+_8433172_8446622_RNA45SN1_45S_50ntFlanks</td><td>475</td><td>492</td><td>18</td><td>CTTCGTGATCGATGTGGT</td><td>4511.9</td><td>5.5</td><td>0.9</td><td>6.4</td><td>1.2E-10</td><td>2.2E-10</td><td>BT20/MB468</td></tr>
<tr><td>>hg38_21_+_8433172_8446622_RNA45SN1_45S_50ntFlanks@8198.8223.26|TGGACGGTGTGAGGCCGGTAGCGGCC</td><td>>hg38_21_+_8433172_8446622_RNA45SN1_45S_50ntFlanks</td><td>8198</td><td>8223</td><td>26</td><td>TGGACGGTGTGAGGCCGGTAGCGGCC</td><td>296.9</td><td>-4.4</td><td>0.7</td><td>-6.4</td><td>1.2E-10</td><td>2.2E-10</td><td>BT20/MB468</td></tr>
<tr><td>>hg38_21_+_8433172_8446622_RNA45SN1_45S_50ntFlanks@4568.4589.22|AATGGAATAGGACCGCGGTTCT</td><td>>hg38_21_+_8433172_8446622_RNA45SN1_45S_50ntFlanks</td><td>4568</td><td>4589</td><td>22</td><td>AATGGAATAGGACCGCGGTTCT</td><td>821.6</td><td>4.9</td><td>0.8</td><td>6.5</td><td>1.1E-10</td><td>2.2E-10</td><td>MB231/MB468</td></tr>
<tr><td>>hg38_21_+_8433172_8446622_RNA45SN1_45S_50ntFlanks@11768.11785.18|CTAATTAGTGACGCGCAT</td><td>>hg38_21_+_8433172_8446622_RNA45SN1_45S_50ntFlanks</td><td>11768</td><td>11785</td><td>18</td><td>CTAATTAGTGACGCGCAT</td><td>121.7</td><td>-8.9</td><td>1.4</td><td>-6.4</td><td>1.2E-10</td><td>2.2E-10</td><td>BT20/MB468</td></tr>
<tr><td>>hg38_21_+_8433172_8446622_RNA45SN1_45S_50ntFlanks@11769.11791.23|TAATTAGTGACGCGCATGAATGG</td><td>>hg38_21_+_8433172_8446622_RNA45SN1_45S_50ntFlanks</td><td>11769</td><td>11791</td><td>23</td><td>TAATTAGTGACGCGCATGAATGG</td><td>920.7</td><td>-5.5</td><td>0.9</td><td>-6.4</td><td>1.3E-10</td><td>2.2E-10</td><td>BT20/MB468</td></tr>
<tr><td>>hg38_21_+_8433172_8446622_RNA45SN1_45S_50ntFlanks@11787.11806.20|AATGGATGAACGAGATTCCC</td><td>>hg38_21_+_8433172_8446622_RNA45SN1_45S_50ntFlanks</td><td>11787</td><td>11806</td><td>20</td><td>AATGGATGAACGAGATTCCC</td><td>191.3</td><td>-9.6</td><td>1.5</td><td>-6.4</td><td>1.3E-10</td><td>2.2E-10</td><td>BT20/MB468</td></tr>
<tr><td>>hg38_21_+_8433172_8446622_RNA45SN1_45S_50ntFlanks@11804.11821.18|CCCACTGTCCCTACCTAC</td><td>>hg38_21_+_8433172_8446622_RNA45SN1_45S_50ntFlanks</td><td>11804</td><td>11821</td><td>18</td><td>CCCACTGTCCCTACCTAC</td><td>109.2</td><td>-5.5</td><td>0.9</td><td>-6.4</td><td>1.3E-10</td><td>2.2E-10</td><td>BT20/MB468</td></tr>
<tr><td>>hg38_21_+_8433172_8446622_RNA45SN1_45S_50ntFlanks@8552.8579.28|CTCCCCGGAGGGGGCGGGCTCCGGCGGG</td><td>>hg38_21_+_8433172_8446622_RNA45SN1_45S_50ntFlanks</td><td>8552</td><td>8579</td><td>28</td><td>CTCCCCGGAGGGGGCGGGCTCCGGCGGG</td><td>76.7</td><td>-8.3</td><td>1.3</td><td>-6.4</td><td>1.3E-10</td><td>2.2E-10</td><td>BT20/MB468</td></tr>
<tr><td>>hg38_21_+_8433172_8446622_RNA45SN1_45S_50ntFlanks@9572.9598.27|CAAATCGGTCGTCCGACCTGGGTATAG</td><td>>hg38_21_+_8433172_8446622_RNA45SN1_45S_50ntFlanks</td><td>9572</td><td>9598</td><td>27</td><td>CAAATCGGTCGTCCGACCTGGGTATAG</td><td>144.0</td><td>-6.9</td><td>1.1</td><td>-6.5</td><td>1.1E-10</td><td>2.2E-10</td><td>MB231/MB468</td></tr>
</table>

| | | | | | | | | | | | | |
|---|---|---|---|---|---|---|---|---|---|---|---|---|
| >hg38_21_+_8433172_8446622_RNA45SN1_45S_50ntFlanks@6779.6802.24\|CCGGGGCTACGCCTGTCTGAGCGT | >hg38_21_+_8433172_8446622_RNA45SN1_45S_50ntFlanks | 6779 | 6802 | 24 | CCGGGGCTACGCCTGTCTGAGCGT | 711.0 | 4.3 | 0.7 | 6.5 | 1.1E-10 | 2.2E-10 | MB231/MB468 |
| >hg38_21_+_8433172_8446622_RNA45SN1_45S_50ntFlanks@11680.11701.22\|TGAATGTCAAAGTGAAGAAATT | >hg38_21_+_8433172_8446622_RNA45SN1_45S_50ntFlanks | 11680 | 11701 | 22 | TGAATGTCAAAGTGAAGAAATT | 126.2 | -9.0 | 1.4 | -6.4 | 1.3E-10 | 2.2E-10 | BT20/MB468 |
| >hg38_MT_+_1621_3279_RNR2_16S_50ntFlanks@1216.1248.33\|CAACCTCCGAGCAGTACATGCTAAGACTTCACC | >hg38_MT_+_1621_3279_RNR2_16S_50ntFlanks | 1216 | 1248 | 33 | CAACCTCCGAGCAGTACATGCTAAGACTTCACC | 571.7 | 8.5 | 1.3 | 6.4 | 1.3E-10 | 2.2E-10 | BT20/MB468 |
| >hg38_21_+_8433172_8446622_RNA45SN1_45S_50ntFlanks@10728.10755.28\|GAGAGGGTGTAAATCTCGCGCCGGGCCG | >hg38_21_+_8433172_8446622_RNA45SN1_45S_50ntFlanks | 10728 | 10755 | 28 | GAGAGGGTGTAAATCTCGCGCCGGGCCG | 108.4 | -8.3 | 1.3 | -6.5 | 1.1E-10 | 2.2E-10 | MB231/MB468 |
| >hg38_21_+_8433172_8446622_RNA45SN1_45S_50ntFlanks@10385.10406.22\|CATGTGAACAGCAGTTGAACAT | >hg38_21_+_8433172_8446622_RNA45SN1_45S_50ntFlanks | 10385 | 10406 | 22 | CATGTGAACAGCAGTTGAACAT | 90.9 | -8.5 | 1.3 | -6.4 | 1.3E-10 | 2.2E-10 | BT20/MB468 |
| >hg38_21_+_8433172_8446622_RNA45SN1_45S_50ntFlanks@11766.11784.19\|ATCTAATTAGTGACGCGCA | >hg38_21_+_8433172_8446622_RNA45SN1_45S_50ntFlanks | 11766 | 11784 | 19 | ATCTAATTAGTGACGCGCA | 195.6 | -9.6 | 1.5 | -6.4 | 1.3E-10 | 2.2E-10 | BT20/MB468 |
| >hg38_21_+_8433172_8446622_RNA45SN1_45S_50ntFlanks@11584.11611.28\|TGGTGCGGACCAGGGGAATCCGACTGTT | >hg38_21_+_8433172_8446622_RNA45SN1_45S_50ntFlanks | 11584 | 11611 | 28 | TGGTGCGGACCAGGGGAATCCGACTGTT | 478.8 | 4.1 | 0.6 | 6.4 | 1.3E-10 | 2.2E-10 | BT20/MB468 |
| >hg38_21_+_8433172_8446622_RNA45SN1_45S_50ntFlanks@10404.10430.27\|CATGGGTCAGTCGGTCCTGAGAGATGG | >hg38_21_+_8433172_8446622_RNA45SN1_45S_50ntFlanks | 10404 | 10430 | 27 | CATGGGTCAGTCGGTCCTGAGAGATGG | 301.5 | -5.8 | 0.9 | -6.4 | 1.3E-10 | 2.3E-10 | BT20/MB468 |
| >hg38_21_+_8433172_8446622_RNA45SN1_45S_50ntFlanks@3854.3882.29\|ACTGTGGTAATTCTAGAGCTAATACATGC | >hg38_21_+_8433172_8446622_RNA45SN1_45S_50ntFlanks | 3854 | 3882 | 29 | ACTGTGGTAATTCTAGAGCTAATACATGC | 184.1 | -3.6 | 0.6 | -6.4 | 1.3E-10 | 2.3E-10 | BT20/MB468 |
| >hg38_21_+_8433172_8446622_RNA45SN1_45S_50ntFlanks@11534.11551.18\|GTTCCGCGCGGCGCCTCG | >hg38_21_+_8433172_8446622_RNA45SN1_45S_50ntFlanks | 11534 | 11551 | 18 | GTTCCGCGCGGCGCCTCG | 826.2 | -5.7 | 0.9 | -6.4 | 1.1E-10 | 2.3E-10 | MB231/MB468 |
| >hg38_21_+_8433172_8446622_RNA45SN1_45S_50ntFlanks@12568.12587.20\|TCAGTACGAGAGGAACCGCA | >hg38_21_+_8433172_8446622_RNA45SN1_45S_50ntFlanks | 12568 | 12587 | 20 | TCAGTACGAGAGGAACCGCA | 114.9 | -4.9 | 0.8 | -6.4 | 1.1E-10 | 2.3E-10 | MB231/MB468 |
| >hg38_21_+_8433172_8446622_RNA45SN1_45S_50ntFlanks@12447.12475.29\|CAGAATTCACCAAGCGTTGGATTGTTCAC | >hg38_21_+_8433172_8446622_RNA45SN1_45S_50ntFlanks | 12447 | 12475 | 29 | CAGAATTCACCAAGCGTTGGATTGTTCAC | 156.4 | -9.3 | 1.4 | -6.4 | 1.3E-10 | 2.3E-10 | BT20/MB468 |
| >hg38_21_+_8433172_8446622_RNA45SN1_45S_50ntFlanks@10404.10435.32\|CATGGGTCAGTCGGTCCTGAGAGATGGGCGAG | >hg38_21_+_8433172_8446622_RNA45SN1_45S_50ntFlanks | 10404 | 10435 | 32 | CATGGGTCAGTCGGTCCTGAGAGATGGGCGAG | 92.0 | -8.1 | 1.3 | -6.4 | 1.1E-10 | 2.3E-10 | MB231/MB468 |
| >hg38_21_+_8433172_8446622_RNA45SN1_45S_50ntFlanks@10378.10407.30\|AGGGTTCCATGTGAACAGCAGTTGAACATG | >hg38_21_+_8433172_8446622_RNA45SN1_45S_50ntFlanks | 10378 | 10407 | 30 | AGGGTTCCATGTGAACAGCAGTTGAACATG | 109.5 | -4.1 | 0.6 | -6.4 | 1.1E-10 | 2.3E-10 | MB231/MB468 |
| >hg38_21_+_8433172_8446622_RNA45SN1_45S_50ntFlanks@4657.4688.32\|CTAGAGGTGAAATTCTTGGACCGGCGCAAGAC | >hg38_21_+_8433172_8446622_RNA45SN1_45S_50ntFlanks | 4657 | 4688 | 32 | CTAGAGGTGAAATTCTTGGACCGGCGCAAGAC | 161.7 | -8.9 | 1.4 | -6.4 | 1.2E-10 | 2.3E-10 | MB231/MB468 |
| >MI0000077\|hsa-mir-21&WithFlank&17\|+\|59841260\|59841343@14.31.18(+1C)&[MIMAT0000076&hsa-miR-21-5p&offsets\|0\|-4(+1C);m-1&17\|+\|59841273\|59841294&offsets\|0\|-4(+1C)]\|TAGCTTATCAGACTGATGC | >MI0000077\|hsa-mir-21&WithFlank&17\|+\|59841260\|59841343 | 14 | 31 | 18(+1C) | TAGCTTATCAGACTGATGC | 507.5 | 8.4 | 1.3 | 6.4 | 1.4E-10 | 2.4E-10 | BT20/MB468 |
| >hg38_21_+_8433172_8446622_RNA45SN1_45S_50ntFlanks@3984.4026.43\|GCGGCTTTGGTGACTCTAGATAACCTCGGGCCGATCGCACGCC | >hg38_21_+_8433172_8446622_RNA45SN1_45S_50ntFlanks | 3984 | 4026 | 43 | GCGGCTTTGGTGACTCTAGATAACCTCGGGCCGATCGCACGCC | 343.8 | -10.4 | 1.6 | -6.4 | 1.4E-10 | 2.4E-10 | BT20/MB468 |
| >hg38_21_+_8433172_8446622_RNA45SN1_45S_50ntFlanks@3782.3812.31\|CAGTGAAACTGCGAATGGCTCATTAAATCAG | >hg38_21_+_8433172_8446622_RNA45SN1_45S_50ntFlanks | 3782 | 3812 | 31 | CAGTGAAACTGCGAATGGCTCATTAAATCAG | 113.6 | -5.1 | 0.8 | -6.4 | 1.2E-10 | 2.4E-10 | MB231/MB468 |
| >am_trna162_MetCAT_6_-_26758550_26758622@31.51.21,trna164_MetCAT_6_-_26735574_26735646@31.51.21,trna20_MetCAT_16_+_71460396_71460468@31.51.21,trna21_MetCAT_6_+_26701712_26701784@31.51.21,trna27_MetCAT_6_+_26766444_26766516@31.51.21,trna75_MetCAT_6_+_28912352_28912424@31.51.21,trna97_MetCAT_6_-_28921042_28921114@31.51.21\|TCTCATAATCTGAAGGTCCTG | >am_trna162_MetCAT_6_-_26758550_26758622 | 31 | 51 | 21 | TCTCATAATCTGAAGGTCCTG | 364.2 | -7.1 | 1.1 | -6.4 | 1.2E-10 | 2.4E-10 | MB231/MB468 |
| >hg38_21_+_8433172_8446622_RNA45SN1_45S_50ntFlanks@11208.11233.26\|GCGGCGACTCTGGACGCGAGCCGGGC | >hg38_21_+_8433172_8446622_RNA45SN1_45S_50ntFlanks | 11208 | 11233 | 26 | GCGGCGACTCTGGACGCGAGCCGGGC | 1766.1 | -5.1 | 0.8 | -6.4 | 1.2E-10 | 2.4E-10 | MB231/MB468 |
| >hg38_21_+_8433172_8446622_RNA45SN1_45S_50ntFlanks@12785.12805.21\|CCCCGCGCGCGCGGGAGGGCG | >hg38_21_+_8433172_8446622_RNA45SN1_45S_50ntFlanks | 12785 | 12805 | 21 | CCCCGCGCGCGCGGGAGGGCG | 648.7 | -6.7 | 1.0 | -6.4 | 1.4E-10 | 2.4E-10 | BT20/MB468 |
| >hg38_21_+_8433172_8446622_RNA45SN1_45S_50ntFlanks@11566.11585.20\|TAGCAGCCGACTTAGAACTG | >hg38_21_+_8433172_8446622_RNA45SN1_45S_50ntFlanks | 11566 | 11585 | 20 | TAGCAGCCGACTTAGAACTG | 1115.8 | 4.0 | 0.6 | 6.4 | 1.2E-10 | 2.5E-10 | MB231/MB468 |
| >hg38_21_+_8433172_8446622_RNA45SN1_45S_50ntFlanks@9538.9563.26\|AACTCTGGTGGAGGTCCGTAGCGGTC | >hg38_21_+_8433172_8446622_RNA45SN1_45S_50ntFlanks | 9538 | 9563 | 26 | AACTCTGGTGGAGGTCCGTAGCGGTC | 99.4 | -8.6 | 1.3 | -6.4 | 1.4E-10 | 2.5E-10 | BT20/MB468 |
| >hg38_21_+_8433172_8446622_RNA45SN1_45S_50ntFlanks@5417.5443.27\|CTACCGATTGGATGGTTTAGTGAGGCC | >hg38_21_+_8433172_8446622_RNA45SN1_45S_50ntFlanks | 5417 | 5443 | 27 | CTACCGATTGGATGGTTTAGTGAGGCC | 106.5 | -8.3 | 1.3 | -6.4 | 1.2E-10 | 2.5E-10 | MB231/MB468 |
| >hg38_21_+_8433172_8446622_RNA45SN1_45S_50ntFlanks@9454.9503.50\|CCGCGCCGGGGAGGTGGAGCACGAGCGCACGTGTTAGGACCCGAAAGATG | >hg38_21_+_8433172_8446622_RNA45SN1_45S_50ntFlanks | 9454 | 9503 | 50 | CCGCGCCGGGGAGGTGGAGCACGAGCGCACGTGTTAGGACCCGAAAGATG | 157.4 | -8.9 | 1.4 | -6.4 | 1.2E-10 | 2.5E-10 | MB231/MB468 |
| >hg38_21_+_8433172_8446622_RNA45SN1_45S_50ntFlanks@10410.10440.31\|TCAGTCGGTCCTGAGAGATGGGCGAGCGCCG | >hg38_21_+_8433172_8446622_RNA45SN1_45S_50ntFlanks | 10410 | 10440 | 31 | TCAGTCGGTCCTGAGAGATGGGCGAGCGCCG | 139.5 | -4.0 | 0.6 | -6.4 | 1.4E-10 | 2.5E-10 | BT20/MB468 |
| >hg38_21_+_8433172_8446622_RNA45SN1_45S_50ntFlanks@12420.12451.32\|ATGTCGGCTCTTCCTATCATTGTGAAGCAGAA | >hg38_21_+_8433172_8446622_RNA45SN1_45S_50ntFlanks | 12420 | 12451 | 32 | ATGTCGGCTCTTCCTATCATTGTGAAGCAGAA | 160.8 | -2.2 | 0.3 | -6.4 | 1.4E-10 | 2.5E-10 | BT20/MB468 |
| >hg38_21_+_8433172_8446622_RNA45SN1_45S_50ntFlanks@8171.8188.18\|AAGTCCTTCTGATCGAGG | >hg38_21_+_8433172_8446622_RNA45SN1_45S_50ntFlanks | 8171 | 8188 | 18 | AAGTCCTTCTGATCGAGG | 251.8 | -10.0 | 1.6 | -6.4 | 1.4E-10 | 2.5E-10 | BT20/MB468 |
| >hg38_21_+_8433172_8446622_RNA45SN1_45S_50ntFlanks@10802.10822.21\|TTGGAACAATGTAGGTAAGGG | >hg38_21_+_8433172_8446622_RNA45SN1_45S_50ntFlanks | 10802 | 10822 | 21 | TTGGAACAATGTAGGTAAGGG | 105.8 | -4.6 | 0.7 | -6.4 | 1.2E-10 | 2.5E-10 | MB231/MB468 |
| >MI0000102\|hsa-mir-100&WithFlank&11\|-\|122152223\|122152314@19.39.21(+1A)&[MIMAT0000098&hsa-miR-100-5p&offsets\|0\|-1(+1A);m-27&11\|-\|122152276\|122152296&offsets\|0\|0(+1A)]\|AACCCGTAGATCCGAACTTGTA | >MI0000102\|hsa-mir-100&WithFlank&11\|-\|122152223\|122152314 | 19 | 39 | 21(+1A) | AACCCGTAGATCCGAACTTGTA | 316689.9 | 15.1 | 2.3 | 6.4 | 1.2E-10 | 2.5E-10 | MB231/MB468 |
| >hg38_21_+_8433172_8446622_RNA45SN1_45S_50ntFlanks@10410.10432.23\|TCAGTCGGTCCTGAGAGATGGGC | >hg38_21_+_8433172_8446622_RNA45SN1_45S_50ntFlanks | 10410 | 10432 | 23 | TCAGTCGGTCCTGAGAGATGGGC | 575.9 | -3.6 | 0.6 | -6.4 | 1.2E-10 | 2.5E-10 | MB231/MB468 |
| >hg38_21_+_8433172_8446622_RNA45SN1_45S_50ntFlanks@6725.6747.23\|GCAGGACACATTGATCATCGACA | >hg38_21_+_8433172_8446622_RNA45SN1_45S_50ntFlanks | 6725 | 6747 | 23 | GCAGGACACATTGATCATCGACA | 1153.5 | 6.0 | 0.9 | 6.4 | 1.2E-10 | 2.5E-10 | MB231/MB468 |
| >hg38_21_+_8433172_8446622_RNA45SN1_45S_50ntFlanks@12415.12438.24\|CTTCGATGTCGGCTCTTCCTATCA | >hg38_21_+_8433172_8446622_RNA45SN1_45S_50ntFlanks | 12415 | 12438 | 24 | CTTCGATGTCGGCTCTTCCTATCA | 852.7 | 4.3 | 0.7 | 6.4 | 1.5E-10 | 2.6E-10 | BT20/MB468 |
| >hg38_21_+_8433172_8446622_RNA45SN1_45S_50ntFlanks@8196.8225.30\|CGTGGACGGTGTGAGGCCGGTAGCGGCCCC | >hg38_21_+_8433172_8446622_RNA45SN1_45S_50ntFlanks | 8196 | 8225 | 30 | CGTGGACGGTGTGAGGCCGGTAGCGGCCCC | 310.3 | -3.1 | 0.5 | -6.4 | 1.5E-10 | 2.6E-10 | BT20/MB468 |
| >hg38_21_+_8433172_8446622_RNA45SN1_45S_50ntFlanks@3984.4005.22\|GCGGCTTTGGTGACTCTAGATA | >hg38_21_+_8433172_8446622_RNA45SN1_45S_50ntFlanks | 3984 | 4005 | 22 | GCGGCTTTGGTGACTCTAGATA | 113.9 | -5.6 | 0.9 | -6.4 | 1.5E-10 | 2.6E-10 | BT20/MB468 |
| >hg38_21_+_8433172_8446622_RNA45SN1_45S_50ntFlanks@11208.11230.23\|GCGGCGACTCTGGACGCGAGCCG | >hg38_21_+_8433172_8446622_RNA45SN1_45S_50ntFlanks | 11208 | 11230 | 23 | GCGGCGACTCTGGACGCGAGCCG | 519.5 | -4.6 | 0.7 | -6.4 | 1.3E-10 | 2.6E-10 | MB231/MB468 |
| >hg38_21_+_8433172_8446622_RNA45SN1_45S_50ntFlanks@12936.12962.27\|AAACCATTCGTAGACGACCTGCTTCTG | >hg38_21_+_8433172_8446622_RNA45SN1_45S_50ntFlanks | 12936 | 12962 | 27 | AAACCATTCGTAGACGACCTGCTTCTG | 141.2 | -4.4 | 0.7 | -6.4 | 1.5E-10 | 2.6E-10 | BT20/MB468 |
| >am_trna10_ValCAC_5_-_180649395_180649467@1.19.19,trna12_ValAAC_5_-_180645270_180645342@1.19.19,trna132_ValAAC_6_-_27721179_27721251@1.19.19,trna133_ValCAC_6_-_27696327_27696399@1.19.19,trna136_ValAAC_6_-_27648885_27648957@1.19.19,trna139_ValAAC_6_-_27618707_27618779@1.19.19,trna15_ValAAC_5_-_180615416_180615488@1.19.19,trna18_ValCAC_5_-_180529253_180529325@1.19.19,trna2_ValAAC_3_+_169490018_169490090@1.19.19,trna2_ValCAC_5_+_180524070_180524142@1.19.19,trna37_ValAAC_6_+_27203288_27203360@1.19.19,trna4_ValAAC_5_+_180591154_180591226@1.19.19,trna5_ValAAC_5_+_180596610_180596682@1.19.19,trna6_ValCAC_5_+_180600650_180600722@1.19.19,trna85_ValCAC_1_-_161369490_161369562@1.19.19,trna90_ValCAC_1_-_149684088_149684161@1.19.19,trna98_ValCAC_1_-_149298555_149298627@1.19.19,trna9_ValCAC_6_+_26538282_26538354@1.19.19\|GTTTCCGTAGTGTAGTGGT | >am_trna10_ValCAC_5_-_180649395_180649467 | 1 | 19 | 19 | GTTTCCGTAGTGTAGTGGT | 677.5 | 6.3 | 1.0 | 6.4 | 1.5E-10 | 2.6E-10 | BT20/MB468 |

| | | | | | | | | | | | | |
|---|---|---|---|---|---|---|---|---|---|---|---|---|
| >trna143_LysTTT_6_-_27559593_27559665@13.30.18,trna149_LysTTT_6_-_27302769_27302841@13.30.18,trna14_LysTTT_11_-_59327808_59327880@13.30.18,trna2_LysTTT_17_+_8022473_8022545@13.30.18,trna54_LysTTT_1_+_204475655_204475727@13.30.18,trna5_LysTTT_11_+_59323902_59323974@13.30.18,trna62_LysTTT_1_-_204476158_204476230@13.30.18,trna76_LysTTT_6_+_28918806_28918878@13.30.18\|CAGTCGGTAGAGCATCAG | >trna143_LysTTT_6_-_27559593_27559665 | 13 | 30 | 18 | CAGTCGGTAGAGCATCAG | 771.7 | -5.1 | 0.8 | -6.4 | 1.5E-10 | 2.6E-10 | BT20/MB468 |
| >MI0000298\|hsa-mir-221&WithFlank&X\|-\|45746151\|45746272@71.93.23&[MIMAT0000278&hsa-miR-221-3p&offsets\|0\|0;m-91&X\|-\|45746181\|45746202&offsets\|0\|+1]\|AGCTACATTGTCTGCTGGGTTTC | >MI0000298\|hsa-mir-221&WithFlank&X\|-\|45746151\|45746272 | 71 | 93 | 23 | AGCTACATTGTCTGCTGGGTTTC | 15100.4 | 4.6 | 0.7 | 6.4 | 1.5E-10 | 2.7E-10 | BT20/MB468 |
| >hg38_21_+_8433172_8446622_RNA45SN1_45S_50ntFlanks@5423.5443.21\|ATTGGATGGTTTAGTGAGGCC | >hg38_21_+_8433172_8446622_RNA45SN1_45S_50ntFlanks | 5423 | 5443 | 21 | ATTGGATGGTTTAGTGAGGCC | 93.2 | -5.6 | 0.9 | -6.4 | 1.3E-10 | 2.7E-10 | MB231/MB468 |
| >MI0000071\|hsa-mir-17&WithFlank&13\|+\|91350599\|91350694@20.42.23(+1A)&[MIMAT0000070&hsa-miR-17-5p&offsets\|0\|0(+1A);m-88&13\|+\|91350618\|91350640&offsets\|0\|0(+1A)]\|CAAAGTGCTTACAGTGCAGGTAGA | >MI0000071\|hsa-mir-17&WithFlank&13\|+\|91350599\|91350694 | 20 | 42 | 23(+1A) | CAAAGTGCTTACAGTGCAGGTAGA | 5558.4 | 4.9 | 0.8 | 6.4 | 1.5E-10 | 2.7E-10 | BT20/MB468 |
| >trna112_GlnCTG_1_-_147737382_147737453@31.50.20,trna131_GlnCTG_6_-_27759135_27759206@31.50.20,trna146_GlnCTG_6_-_27515531_27515602@31.50.20,trna15_GlnCTG_1_+_145963304_145963375@31.50.20,trna19_GlnCTG_1_+_147505038_147505109@31.50.20,trna1_GlnCTG_6_+_18836402_18836473@31.50.20,trna28_GlnCTG_1_+_149186125_149186196@31.50.20,trna3_GlnCTG_17_+_8023070_8023141@31.50.20,trna49_GlnCTG_6_+_27487308_27487379@31.50.20,trna7_GlnCTG_15_-_66161400_66161471@31.50.20,trna99_GlnCTG_6_-_28909378_28909449@31.50.20\|ACTCTGAATCCAGCGATCCG | >trna112_GlnCTG_1_-_147737382_147737453 | 31 | 50 | 20 | ACTCTGAATCCAGCGATCCG | 196.9 | -6.4 | 1.0 | -6.4 | 1.6E-10 | 2.7E-10 | BT20/MB468 |
| >MI0000070\|hsa-mir-16-1&WithFlank&13\|-\|50048967\|50049067@20.41.22(+1G);MI0000115\|hsa-mir-16-2&WithFlank&3\|+\|160404739\|160404831@16.37.22(+1G)&[MIMAT0000069&hsa-miR-16-5p&offsets\|0\|0(+1G);m-62&13\|-\|50049027\|50049048&offsets\|0\|0(+1G)];[MIMAT0000069_1&hsa-miR-16-5p&offsets\|0\|0(+1G);m-61&3\|+\|160404754\|160404775&offsets\|0\|0(+1G)]\|TAGCAGCACGTAAATATTGGCGG | >MI0000070\|hsa-mir-16-1&WithFlank&13\|-\|50048967\|50049067 | 20 | 41 | 22(+1G) | TAGCAGCACGTAAATATTGGCGG | 2337.6 | 9.7 | 1.5 | 6.4 | 1.6E-10 | 2.7E-10 | BT20/MB468 |
| >hg38_21_+_8433172_8446622_RNA45SN1_45S_50ntFlanks@10378.10406.29\|AGGGTTCCATGTGAACAGCAGTTGAACAT | >hg38_21_+_8433172_8446622_RNA45SN1_45S_50ntFlanks | 10378 | 10406 | 29 | AGGGTTCCATGTGAACAGCAGTTGAACAT | 112.1 | -5.7 | 0.9 | -6.4 | 1.6E-10 | 2.8E-10 | BT20/MB468 |
| >hg38_21_+_8433172_8446622_RNA45SN1_45S_50ntFlanks@12995.13014.20\|CTCGCTGCGATCTATTGAAA | >hg38_21_+_8433172_8446622_RNA45SN1_45S_50ntFlanks | 12995 | 13014 | 20 | CTCGCTGCGATCTATTGAAA | 828.2 | 3.8 | 0.6 | 6.4 | 1.4E-10 | 2.8E-10 | MB231/MB468 |
| >hg38_21_+_8433172_8446622_RNA45SN1_45S_50ntFlanks@12420.12447.28\|ATGTCGGCTCTTCCTATCATTGTGAAGC | >hg38_21_+_8433172_8446622_RNA45SN1_45S_50ntFlanks | 12420 | 12447 | 28 | ATGTCGGCTCTTCCTATCATTGTGAAGC | 1022.5 | 3.6 | 0.6 | 6.4 | 1.4E-10 | 2.8E-10 | MB231/MB468 |
| >hg38_21_+_8433172_8446622_RNA45SN1_45S_50ntFlanks@11768.11794.27\|CTAATTAGTGACGCGCATGAATGGATG | >hg38_21_+_8433172_8446622_RNA45SN1_45S_50ntFlanks | 11768 | 11794 | 27 | CTAATTAGTGACGCGCATGAATGGATG | 2220.4 | -3.7 | 0.6 | -6.4 | 1.4E-10 | 2.8E-10 | MB231/MB468 |
| >hg38_21_+_8433172_8446622_RNA45SN1_45S_50ntFlanks@6788.6807.20\|CGCCTGTCTGAGCGTCGCTT | >hg38_21_+_8433172_8446622_RNA45SN1_45S_50ntFlanks | 6788 | 6807 | 20 | CGCCTGTCTGAGCGTCGCTT | 1235.5 | 5.1 | 0.8 | 6.4 | 1.6E-10 | 2.8E-10 | BT20/MB468 |
| >hg38_21_+_8433172_8446622_RNA45SN1_45S_50ntFlanks@10842.10865.24\|GTAACTTCGGGATAAGGATTGGCT | >hg38_21_+_8433172_8446622_RNA45SN1_45S_50ntFlanks | 10842 | 10865 | 24 | GTAACTTCGGGATAAGGATTGGCT | 215.4 | -3.8 | 0.6 | -6.4 | 1.6E-10 | 2.8E-10 | BT20/MB468 |
| >hg38_21_+_8433172_8446622_RNA45SN1_45S_50ntFlanks@4570.4587.18\|TGGAATAGGACCGCGGTT | >hg38_21_+_8433172_8446622_RNA45SN1_45S_50ntFlanks | 4570 | 4587 | 18 | TGGAATAGGACCGCGGTT | 254.5 | -3.3 | 0.5 | -6.4 | 1.4E-10 | 2.8E-10 | MB231/MB468 |
| >hg38_21_+_8433172_8446622_RNA45SN1_45S_50ntFlanks@12488.12525.38\|AACGTGAGCTGGGTTTAGACCGTCGTGAGACAGGTTAG | >hg38_21_+_8433172_8446622_RNA45SN1_45S_50ntFlanks | 12488 | 12525 | 38 | AACGTGAGCTGGGTTTAGACCGTCGTGAGACAGGTTAG | 115.3 | -5.9 | 0.9 | -6.4 | 1.4E-10 | 2.8E-10 | MB231/MB468 |
| >hg38_1_-_228634819_228635039_RNA5S12_5S_50ntFlanks@148.170.23\|GGAATACCGGGTGCTGTAGGCTT | >hg38_1_-_228634819_228635039_RNA5S12_5S_50ntFlanks | 148 | 170 | 23 | GGAATACCGGGTGCTGTAGGCTT | 2402.2 | 3.4 | 0.5 | 6.4 | 1.6E-10 | 2.8E-10 | BT20/MB468 |
| >hg38_21_+_8433172_8446622_RNA45SN1_45S_50ntFlanks@12356.12375.20\|ACCACAGGGATAACTGGCTT | >hg38_21_+_8433172_8446622_RNA45SN1_45S_50ntFlanks | 12356 | 12375 | 20 | ACCACAGGGATAACTGGCTT | 98.2 | -8.2 | 1.3 | -6.4 | 1.4E-10 | 2.8E-10 | MB231/MB468 |
| >hg38_21_+_8433172_8446622_RNA45SN1_45S_50ntFlanks@10355.10388.34\|AACTTTGAAGGCCGAAGTGGAGAAGGGTTCCATG | >hg38_21_+_8433172_8446622_RNA45SN1_45S_50ntFlanks | 10355 | 10388 | 34 | AACTTTGAAGGCCGAAGTGGAGAAGGGTTCCATG | 366.2 | -5.5 | 0.9 | -6.4 | 1.4E-10 | 2.8E-10 | MB231/MB468 |
| >hg38_21_+_8433172_8446622_RNA45SN1_45S_50ntFlanks@6646.6692.47\|TCGTACGACTCTTAGCGGTGGATCACTCGGCTCGTGCGTCGATGAAG | >hg38_21_+_8433172_8446622_RNA45SN1_45S_50ntFlanks | 6646 | 6692 | 47 | TCGTACGACTCTTAGCGGTGGATCACTCGGCTCGTGCGTCGATGAAG | 621.2 | 4.7 | 0.7 | 6.4 | 1.4E-10 | 2.9E-10 | MB231/MB468 |
| >hg38_21_+_8433172_8446622_RNA45SN1_45S_50ntFlanks@11233.11251.19\|CCCTTCCCGTGGATCGCCC | >hg38_21_+_8433172_8446622_RNA45SN1_45S_50ntFlanks | 11233 | 11251 | 19 | CCCTTCCCGTGGATCGCCC | 104.5 | -8.7 | 1.4 | -6.4 | 1.6E-10 | 2.9E-10 | BT20/MB468 |
| >MI0000111\|hsa-mir-105-1&WithFlank&X\|-\|152392213\|152392305@19.41.23;MI0000112\|hsa-mir-105-2&WithFlank&X\|-\|152394406\|152394498@19.41.23&[MIMAT0000102&hsa-miR-105-5p&offsets\|0\|0;m-735&X\|-\|152392266\|152392287&offsets\|0\|+1];[MIMAT0000102_1&hsa-miR-105-5p&offsets\|0\|0;m-735&X\|-\|152394459\|152394480&offsets\|0\|+1]\|TCAAATGCTCAGACTCCTGTGGT | >MI0000111\|hsa-mir-105-1&WithFlank&X\|-\|152392213\|152392305 | 19 | 41 | 23 | TCAAATGCTCAGACTCCTGTGGT | 554.9 | 5.3 | 0.8 | 6.4 | 1.7E-10 | 2.9E-10 | BT20/MB468 |
| >hg38_21_+_8433172_8446622_RNA45SN1_45S_50ntFlanks@10498.10521.24\|TTCAGATCCCCGAATCCGGAGTGG | >hg38_21_+_8433172_8446622_RNA45SN1_45S_50ntFlanks | 10498 | 10521 | 24 | TTCAGATCCCCGAATCCGGAGTGG | 205.0 | -9.3 | 1.4 | -6.4 | 1.4E-10 | 2.9E-10 | MB231/MB468 |
| >hg38_21_+_8433172_8446622_RNA45SN1_45S_50ntFlanks@4445.4472.28\|CTTGCCTCTCGGCGCCCCCTCGATGCTC | >hg38_21_+_8433172_8446622_RNA45SN1_45S_50ntFlanks | 4445 | 4472 | 28 | CTTGCCTCTCGGCGCCCCCTCGATGCTC | 99.1 | -8.2 | 1.3 | -6.4 | 1.4E-10 | 2.9E-10 | MB231/MB468 |
| >hg38_21_+_8433172_8446622_RNA45SN1_45S_50ntFlanks@10348.10374.27\|AAACGAGAACTTTGAAGGCCGAAGTGG | >hg38_21_+_8433172_8446622_RNA45SN1_45S_50ntFlanks | 10348 | 10374 | 27 | AAACGAGAACTTTGAAGGCCGAAGTGG | 498.1 | -3.3 | 0.5 | -6.4 | 1.7E-10 | 2.9E-10 | BT20/MB468 |
| >hg38_21_+_8433172_8446622_RNA45SN1_45S_50ntFlanks@6741.6772.32\|ATCGACACTTCGAACGCACTTGCGGCCCCGGG | >hg38_21_+_8433172_8446622_RNA45SN1_45S_50ntFlanks | 6741 | 6772 | 32 | ATCGACACTTCGAACGCACTTGCGGCCCCGGG | 296.4 | -5.0 | 0.8 | -6.4 | 1.5E-10 | 2.9E-10 | MB231/MB468 |
| >am_trna162_MetCAT_6_-_26758550_26758622@31.52.22,trna164_MetCAT_6_-_26735574_26735646@31.52.22,trna20_MetCAT_16_+_71460396_71460468@31.52.22,trna21_MetCAT_6_+_26701712_26701784@31.52.22,trna27_MetCAT_6_+_26766444_26766516@31.52.22,trna75_MetCAT_6_+_28912352_28912424@31.52.22,trna97_MetCAT_6_-_28921042_28921114@31.52.22\|TCTCATAATCTGAAGGTCCTGA | >am_trna162_MetCAT_6_-_26758550_26758622 | 31 | 52 | 22 | TCTCATAATCTGAAGGTCCTGA | 97.2 | -8.2 | 1.3 | -6.4 | 1.5E-10 | 3.0E-10 | MB231/MB468 |
| >hg38_21_+_8433172_8446622_RNA45SN1_45S_50ntFlanks@11567.11590.24\|AGCAGCCGACTTAGAACTGGTGCG | >hg38_21_+_8433172_8446622_RNA45SN1_45S_50ntFlanks | 11567 | 11590 | 24 | AGCAGCCGACTTAGAACTGGTGCG | 376.2 | 3.6 | 0.6 | 6.4 | 1.5E-10 | 3.0E-10 | MB231/MB468 |
| >hg38_21_+_8433172_8446622_RNA45SN1_45S_50ntFlanks@5446.5467.22\|CGGATCGGCCCCGCCGGGGTCG | >hg38_21_+_8433172_8446622_RNA45SN1_45S_50ntFlanks | 5446 | 5467 | 22 | CGGATCGGCCCCGCCGGGGTCG | 111.5 | -8.4 | 1.3 | -6.4 | 1.5E-10 | 3.0E-10 | MB231/MB468 |
| >hg38_21_+_8433172_8446622_RNA45SN1_45S_50ntFlanks@12074.12095.22\|CGAGGGGCTCTCGCTTCTGGCG | >hg38_21_+_8433172_8446622_RNA45SN1_45S_50ntFlanks | 12074 | 12095 | 22 | CGAGGGGCTCTCGCTTCTGGCG | 583.6 | -4.8 | 0.8 | -6.4 | 1.7E-10 | 3.0E-10 | BT20/MB468 |
| >hg38_21_+_8433172_8446622_RNA45SN1_45S_50ntFlanks@6648.6668.21\|GTACGACTCTTAGCGGTGGAT | >hg38_21_+_8433172_8446622_RNA45SN1_45S_50ntFlanks | 6648 | 6668 | 21 | GTACGACTCTTAGCGGTGGAT | 852.2 | 3.9 | 0.6 | 6.4 | 1.5E-10 | 3.0E-10 | MB231/MB468 |
| >hg38_21_+_8433172_8446622_RNA45SN1_45S_50ntFlanks@10511.10533.23\|ATCCGGAGTGGCGGAGATGGGCG | >hg38_21_+_8433172_8446622_RNA45SN1_45S_50ntFlanks | 10511 | 10533 | 23 | ATCCGGAGTGGCGGAGATGGGCG | 110.8 | -8.4 | 1.3 | -6.4 | 1.5E-10 | 3.0E-10 | MB231/MB468 |
| >hg38_21_+_8433172_8446622_RNA45SN1_45S_50ntFlanks@12957.12980.24\|CTTCTGGGTCGGGGTTTCGTACGT | >hg38_21_+_8433172_8446622_RNA45SN1_45S_50ntFlanks | 12957 | 12980 | 24 | CTTCTGGGTCGGGGTTTCGTACGT | 89.9 | -8.1 | 1.3 | -6.4 | 1.5E-10 | 3.0E-10 | MB231/MB468 |
| >hg38_21_+_8433172_8446622_RNA45SN1_45S_50ntFlanks@12588.12614.27\|GGTTCAGACATTTGGTGTATGTGCTTG | >hg38_21_+_8433172_8446622_RNA45SN1_45S_50ntFlanks | 12588 | 12614 | 27 | GGTTCAGACATTTGGTGTATGTGCTTG | 112.8 | -8.4 | 1.3 | -6.4 | 1.5E-10 | 3.0E-10 | MB231/MB468 |
| >hg38_21_+_8433172_8446622_RNA45SN1_45S_50ntFlanks@13009.13027.19\|TTGAAAGTCAGCCCTCGAC | >hg38_21_+_8433172_8446622_RNA45SN1_45S_50ntFlanks | 13009 | 13027 | 19 | TTGAAAGTCAGCCCTCGAC | 333.4 | -6.2 | 1.0 | -6.4 | 1.5E-10 | 3.0E-10 | MB231/MB468 |
| >hg38_21_+_8433172_8446622_RNA45SN1_45S_50ntFlanks@6748.6771.24\|CTTCGAACGCACTTGCGGCCCCGG | >hg38_21_+_8433172_8446622_RNA45SN1_45S_50ntFlanks | 6748 | 6771 | 24 | CTTCGAACGCACTTGCGGCCCCGG | 130.2 | -5.1 | 0.8 | -6.4 | 1.8E-10 | 3.1E-10 | BT20/MB468 |
| >hg38_21_+_8433172_8446622_RNA45SN1_45S_50ntFlanks@13002.13029.28\|CGATCTATTGAAAGTCAGCCCTCGACAC | >hg38_21_+_8433172_8446622_RNA45SN1_45S_50ntFlanks | 13002 | 13029 | 28 | CGATCTATTGAAAGTCAGCCCTCGACAC | 1494.2 | -6.9 | 1.1 | -6.4 | 1.5E-10 | 3.1E-10 | MB231/MB468 |
| >hg38_21_+_8433172_8446622_RNA45SN1_45S_50ntFlanks@6651.6688.38\|CGACTCTTAGCGGTGGATCACTCGGCTCGTGCGTCGAT | >hg38_21_+_8433172_8446622_RNA45SN1_45S_50ntFlanks | 6651 | 6688 | 38 | CGACTCTTAGCGGTGGATCACTCGGCTCGTGCGTCGAT | 4111.6 | 4.0 | 0.6 | 6.4 | 1.6E-10 | 3.2E-10 | MB231/MB468 |
| >hg38_21_+_8433172_8446622_RNA45SN1_45S_50ntFlanks@9576.9597.22\|TCGGTCGTCCGACCTGGGTATA | >hg38_21_+_8433172_8446622_RNA45SN1_45S_50ntFlanks | 9576 | 9597 | 22 | TCGGTCGTCCGACCTGGGTATA | 162.2 | -4.0 | 0.6 | -6.4 | 1.6E-10 | 3.2E-10 | MB231/MB468 |
| >hg38_21_+_8433172_8446622_RNA45SN1_45S_50ntFlanks@5485.5510.26\|AGCGCTGAGAAGACGGTCGAACTTGA | >hg38_21_+_8433172_8446622_RNA45SN1_45S_50ntFlanks | 5485 | 5510 | 26 | AGCGCTGAGAAGACGGTCGAACTTGA | 92.4 | -8.1 | 1.3 | -6.4 | 1.6E-10 | 3.2E-10 | MB231/MB468 |
| >hg38_21_+_8433172_8446622_RNA45SN1_45S_50ntFlanks@10713.10736.24\|CTTGAAAATCCGGGGGAGAGGGTG | >hg38_21_+_8433172_8446622_RNA45SN1_45S_50ntFlanks | 10713 | 10736 | 24 | CTTGAAAATCCGGGGGAGAGGGTG | 139.9 | -3.6 | 0.6 | -6.4 | 1.8E-10 | 3.2E-10 | BT20/MB468 |

| | | | | | | | | | | | | |
|---|---|---|---|---|---|---|---|---|---|---|---|---|
| >hg38_21_+_8433172_8446622_RNA45SN1_45S_50ntFlanks@12324.12343.20\|TTGGGTTTTAAGCAGGAGGT | >hg38_21_+_8433172_8446622_RNA45SN1_45S_50ntFlanks | 12324 | 12343 | 20 | TTGGGTTTTAAGCAGGAGGT | 86.0 | -8.0 | 1.3 | -6.4 | 1.6E-10 | 3.2E-10 | MB231/MB468 |
| >hg38_21_+_8433172_8446622_RNA45SN1_45S_50ntFlanks@10681.10706.26\|TTCCGGCGGCGTCCGGTGAGCTCTCG | >hg38_21_+_8433172_8446622_RNA45SN1_45S_50ntFlanks | 10681 | 10706 | 26 | TTCCGGCGGCGTCCGGTGAGCTCTCG | 1895.0 | -4.7 | 0.7 | -6.4 | 1.8E-10 | 3.2E-10 | BT20/MB468 |
| >hg38_21_+_8433172_8446622_RNA45SN1_45S_50ntFlanks@10729.10752.24\|AGAGGGTGTAAATCTCGCGCCGGG | >hg38_21_+_8433172_8446622_RNA45SN1_45S_50ntFlanks | 10729 | 10752 | 24 | AGAGGGTGTAAATCTCGCGCCGGG | 98.6 | -8.2 | 1.3 | -6.4 | 1.6E-10 | 3.2E-10 | MB231/MB468 |
| >hg38_21_+_8433172_8446622_RNA45SN1_45S_50ntFlanks@4657.4689.33\|CTAGAGGTGAAATTCTTGGACCGGCGCAAGACG | >hg38_21_+_8433172_8446622_RNA45SN1_45S_50ntFlanks | 4657 | 4689 | 33 | CTAGAGGTGAAATTCTTGGACCGGCGCAAGACG | 2272.5 | -7.3 | 1.1 | -6.4 | 1.8E-10 | 3.2E-10 | BT20/MB468 |
| >trna10_LysCTT_16_+_3241501_3241573@1.30.30,trna13_LysCTT_14_-_58706613_58706685@1.30.30,trna2_LysCTT_15_+_79152904_79152976@1.30.30\|GCCCGGCTAGCTCAGTCGGTAGAGCATGGG | >trna10_LysCTT_16_+_3241501_3241573 | 1 | 30 | 30 | GCCCGGCTAGCTCAGTCGGTAGAGCATGGG | 166.1 | -5.9 | 0.9 | -6.4 | 1.8E-10 | 3.2E-10 | BT20/MB468 |
| >hg38_21_+_8433172_8446622_RNA45SN1_45S_50ntFlanks@10914.10946.33\|CGCCGCGGCTGGACGAGGCGCCGCCGCCCCCCC | >hg38_21_+_8433172_8446622_RNA45SN1_45S_50ntFlanks | 10914 | 10946 | 33 | CGCCGCGGCTGGACGAGGCGCCGCCGCCCCCCC | 95.2 | -8.1 | 1.3 | -6.4 | 1.6E-10 | 3.2E-10 | MB231/MB468 |
| >MI0000298\|hsa-mir-221&WithFlank&X\|-\|45746151\|45746272@71.94.24&[MIMAT0000278&hsa-miR-221-3p&offsets\|0\|+1;m-91&X\|-\|45746181\|45746202&offsets\|0\|+2]\|AGCTACATTGTCTGCTGGGTTTCA | >MI0000298\|hsa-mir-221&WithFlank&X\|-\|45746151\|45746272 | 71 | 94 | 24 | AGCTACATTGTCTGCTGGGTTTCA | 1779.4 | 4.8 | 0.7 | 6.4 | 1.9E-10 | 3.2E-10 | BT20/MB468 |
| >hg38_21_+_8433172_8446622_RNA45SN1_45S_50ntFlanks@7982.8008.27\|TCAGATCAGACGTGGCGACCCGCTGAA | >hg38_21_+_8433172_8446622_RNA45SN1_45S_50ntFlanks | 7982 | 8008 | 27 | TCAGATCAGACGTGGCGACCCGCTGAA | 493.5 | -3.8 | 0.6 | -6.4 | 1.9E-10 | 3.2E-10 | BT20/MB468 |
| >hg38_21_+_8433172_8446622_RNA45SN1_45S_50ntFlanks@8196.8214.19\|CGTGGACGGTGTGAGGCCG | >hg38_21_+_8433172_8446622_RNA45SN1_45S_50ntFlanks | 8196 | 8214 | 19 | CGTGGACGGTGTGAGGCCG | 993.5 | -4.0 | 0.6 | -6.4 | 1.9E-10 | 3.2E-10 | BT20/MB468 |
| >hg38_21_+_8433172_8446622_RNA45SN1_45S_50ntFlanks@10116.10134.19\|TGCGTCGGAGGGCGGCGGC | >hg38_21_+_8433172_8446622_RNA45SN1_45S_50ntFlanks | 10116 | 10134 | 19 | TGCGTCGGAGGGCGGCGGC | 95.5 | -8.2 | 1.3 | -6.4 | 1.6E-10 | 3.3E-10 | MB231/MB468 |
| >hg38_21_+_8433172_8446622_RNA45SN1_45S_50ntFlanks@4733.4765.33\|GAACGAAAGTCGGAGGTTCGAAGACGATCAGAT | >hg38_21_+_8433172_8446622_RNA45SN1_45S_50ntFlanks | 4733 | 4765 | 33 | GAACGAAAGTCGGAGGTTCGAAGACGATCAGAT | 299.9 | 7.3 | 1.1 | 6.4 | 1.9E-10 | 3.3E-10 | BT20/MB468 |
| >hg38_21_+_8433172_8446622_RNA45SN1_45S_50ntFlanks@12633.12659.27\|CGAAGCTACCATCTGTGGGATTATGAC | >hg38_21_+_8433172_8446622_RNA45SN1_45S_50ntFlanks | 12633 | 12659 | 27 | CGAAGCTACCATCTGTGGGATTATGAC | 955.3 | -7.1 | 1.1 | -6.4 | 1.9E-10 | 3.3E-10 | BT20/MB468 |
| >hg38_21_+_8433172_8446622_RNA45SN1_45S_50ntFlanks@10682.10711.30\|TCCGGCGGCGTCCGGTGAGCTCTCGCTGGC | >hg38_21_+_8433172_8446622_RNA45SN1_45S_50ntFlanks | 10682 | 10711 | 30 | TCCGGCGGCGTCCGGTGAGCTCTCGCTGGC | 147.2 | -4.6 | 0.7 | -6.4 | 1.9E-10 | 3.3E-10 | BT20/MB468 |
| >hg38_21_+_8433172_8446622_RNA45SN1_45S_50ntFlanks@10384.10403.20\|CCATGTGAACAGCAGTTGAA | >hg38_21_+_8433172_8446622_RNA45SN1_45S_50ntFlanks | 10384 | 10403 | 20 | CCATGTGAACAGCAGTTGAA | 199.3 | -7.3 | 1.1 | -6.4 | 1.6E-10 | 3.3E-10 | MB231/MB468 |
| >hg38_21_+_8433172_8446622_RNA45SN1_45S_50ntFlanks@8799.8823.25\|CCTCGGGAGGGCGCGCGGGTCGGGG | >hg38_21_+_8433172_8446622_RNA45SN1_45S_50ntFlanks | 8799 | 8823 | 25 | CCTCGGGAGGGCGCGCGGGTCGGGG | 85.1 | -8.0 | 1.3 | -6.4 | 1.7E-10 | 3.3E-10 | MB231/MB468 |
| >MI0000764\|hsa-mir-363&WithFlank&X\|-\|134169372\|134169458@56.77.22&[MIMAT0000707&hsa-miR-363-3p&offsets\|0\|0;m-178&X\|-\|134169381\|134169403&offsets\|0\|-1]\|AATTGCACGGTATCCATCTGTA | >MI0000764\|hsa-mir-363&WithFlank&X\|-\|134169372\|134169458 | 56 | 77 | 22 | AATTGCACGGTATCCATCTGTA | 4096.5 | 5.2 | 0.8 | 6.4 | 1.9E-10 | 3.3E-10 | BT20/MB468 |
| >MI0000270\|hsa-mir-181b-1&WithFlank&1\|-\|198858867\|198858988@42.65.24;MI0000683\|hsa-mir-181b-2&WithFlank&9\|+\|124693704\|124693804@22.45.24&[MIMAT0000257&hsa-miR-181b-5p&offsets\|0\|+1;m-73&1\|-\|198858925\|198858947&offsets\|0\|+1];[MIMAT0000257_1&hsa-miR-181b-5p&offsets\|0\|+1;m-72&9\|+\|124693725\|124693747&offsets\|0\|+1]\|AACATTCATTGCTGTCGGTGGGTT | >MI0000270\|hsa-mir-181b-1&WithFlank&1\|-\|198858867\|198858988 | 42 | 65 | 24 | AACATTCATTGCTGTCGGTGGGTT | 2658.9 | 5.8 | 0.9 | 6.4 | 1.9E-10 | 3.4E-10 | BT20/MB468 |
| >hg38_21_+_8433172_8446622_RNA45SN1_45S_50ntFlanks@12625.12657.33\|CAATGGGGCGAAGCTACCATCTGTGGGATTATG | >hg38_21_+_8433172_8446622_RNA45SN1_45S_50ntFlanks | 12625 | 12657 | 33 | CAATGGGGCGAAGCTACCATCTGTGGGATTATG | 159.0 | -3.1 | 0.5 | -6.4 | 1.9E-10 | 3.4E-10 | BT20/MB468 |
| >hg38_21_+_8433172_8446622_RNA45SN1_45S_50ntFlanks@5490.5512.23\|TGAGAAGACGGTCGAACTTGACT | >hg38_21_+_8433172_8446622_RNA45SN1_45S_50ntFlanks | 5490 | 5512 | 23 | TGAGAAGACGGTCGAACTTGACT | 507.5 | 8.0 | 1.3 | 6.4 | 2.0E-10 | 3.4E-10 | BT20/MB468 |
| >hg38_21_+_8433172_8446622_RNA45SN1_45S_50ntFlanks@13022.13039.18\|CTCGACACAAGGGTTTGT | >hg38_21_+_8433172_8446622_RNA45SN1_45S_50ntFlanks | 13022 | 13039 | 18 | CTCGACACAAGGGTTTGT | 1383.8 | 3.5 | 0.6 | 6.4 | 2.0E-10 | 3.4E-10 | BT20/MB468 |
| >hg38_21_+_8433172_8446622_RNA45SN1_45S_50ntFlanks@9033.9058.26\|CTCGTGGGGGGGCCGGGCCACCCCTC | >hg38_21_+_8433172_8446622_RNA45SN1_45S_50ntFlanks | 9033 | 9058 | 26 | CTCGTGGGGGGGCCGGGCCACCCCTC | 104.8 | -8.7 | 1.4 | -6.4 | 2.0E-10 | 3.5E-10 | BT20/MB468 |
| >hg38_21_+_8433172_8446622_RNA45SN1_45S_50ntFlanks@6745.6762.18\|ACACTTCGAACGCACTTG | >hg38_21_+_8433172_8446622_RNA45SN1_45S_50ntFlanks | 6745 | 6762 | 18 | ACACTTCGAACGCACTTG | 235.0 | -3.7 | 0.6 | -6.4 | 1.7E-10 | 3.5E-10 | MB231/MB468 |
| >hg38_21_+_8433172_8446622_RNA45SN1_45S_50ntFlanks@10678.10705.28\|CGGTTCCGGCGGCGTCCGGTGAGCTCTC | >hg38_21_+_8433172_8446622_RNA45SN1_45S_50ntFlanks | 10678 | 10705 | 28 | CGGTTCCGGCGGCGTCCGGTGAGCTCTC | 430.7 | -5.2 | 0.8 | -6.4 | 1.7E-10 | 3.5E-10 | MB231/MB468 |
| >MI0001733\|hsa-mir-452&WithFlank&X\|-\|151959622\|151959718@20.41.22&[MIMAT0001635&hsa-miR-452-5p&offsets\|0\|0;m-188&X\|-\|151959677\|151959699&offsets\|0\|-1]\|AACTGTTTGCAGAGGAAACTGA | >MI0001733\|hsa-mir-452&WithFlank&X\|-\|151959622\|151959718 | 20 | 41 | 22 | AACTGTTTGCAGAGGAAACTGA | 686.3 | 5.2 | 0.8 | 6.8 | 1.2E-11 | 3.5E-10 | BT20/MB231 |
| >hg38_21_+_8433172_8446622_RNA45SN1_45S_50ntFlanks@11778.11806.29\|ACGCGCATGAATGGATGAACGAGATTCCC | >hg38_21_+_8433172_8446622_RNA45SN1_45S_50ntFlanks | 11778 | 11806 | 29 | ACGCGCATGAATGGATGAACGAGATTCCC | 139.7 | -3.4 | 0.5 | -6.4 | 2.0E-10 | 3.5E-10 | BT20/MB468 |
| >hg38_21_+_8433172_8446622_RNA45SN1_45S_50ntFlanks@10351.10378.28\|CGAGAACTTTGAAGGCCGAAGTGGAGAA | >hg38_21_+_8433172_8446622_RNA45SN1_45S_50ntFlanks | 10351 | 10378 | 28 | CGAGAACTTTGAAGGCCGAAGTGGAGAA | 756.2 | -4.7 | 0.7 | -6.4 | 1.8E-10 | 3.5E-10 | MB231/MB468 |
| >hg38_21_+_8433172_8446622_RNA45SN1_45S_50ntFlanks@11212.11233.22\|CGACTCTGGACGCGAGCCGGGC | >hg38_21_+_8433172_8446622_RNA45SN1_45S_50ntFlanks | 11212 | 11233 | 22 | CGACTCTGGACGCGAGCCGGGC | 118.7 | -5.8 | 0.9 | -6.4 | 2.1E-10 | 3.6E-10 | BT20/MB468 |
| >trnaMT_GluTTC_MT_-_14674_14742@39.69.31,trnalookalike8_GluTTC_5_-_93905172_93905240@39.69.31\|ATTGGTCGTGGTTGTAGTCCGTGCGAGAATA | >trnaMT_GluTTC_MT_-_14674_14742 | 39 | 69 | 31 | ATTGGTCGTGGTTGTAGTCCGTGCGAGAATA | 297.2 | 7.3 | 1.1 | 6.4 | 2.1E-10 | 3.6E-10 | BT20/MB468 |
| >hg38_21_+_8433172_8446622_RNA45SN1_45S_50ntFlanks@10783.10801.19\|GTGAACAGCCTCTGGCATG | >hg38_21_+_8433172_8446622_RNA45SN1_45S_50ntFlanks | 10783 | 10801 | 19 | GTGAACAGCCTCTGGCATG | 209.7 | -3.5 | 0.5 | -6.4 | 1.8E-10 | 3.6E-10 | MB231/MB468 |
| >hg38_21_+_8433172_8446622_RNA45SN1_45S_50ntFlanks@3780.3801.22\|TACAGTGAAACTGCGAATGGCT | >hg38_21_+_8433172_8446622_RNA45SN1_45S_50ntFlanks | 3780 | 3801 | 22 | TACAGTGAAACTGCGAATGGCT | 175.9 | -3.3 | 0.5 | -6.4 | 1.8E-10 | 3.6E-10 | MB231/MB468 |
| >hg38_21_+_8433172_8446622_RNA45SN1_45S_50ntFlanks@8169.8188.20\|CCAAGTCCTTCTGATCGAGG | >hg38_21_+_8433172_8446622_RNA45SN1_45S_50ntFlanks | 8169 | 8188 | 20 | CCAAGTCCTTCTGATCGAGG | 207.1 | -9.7 | 1.5 | -6.4 | 2.1E-10 | 3.6E-10 | BT20/MB468 |
| >hg38_21_+_8433172_8446622_RNA45SN1_45S_50ntFlanks@10392.10431.40\|ACAGCAGTTGAACATGGGTCAGTCGGTCCTGAGAGATGGG | >hg38_21_+_8433172_8446622_RNA45SN1_45S_50ntFlanks | 10392 | 10431 | 40 | ACAGCAGTTGAACATGGGTCAGTCGGTCCTGAGAGATGGG | 170.0 | -9.4 | 1.5 | -6.4 | 2.1E-10 | 3.6E-10 | BT20/MB468 |
| >hg38_21_+_8433172_8446622_RNA45SN1_45S_50ntFlanks@11876.11904.29\|AAGACCCTGTTGAGCTTGACTCTAGTCTG | >hg38_21_+_8433172_8446622_RNA45SN1_45S_50ntFlanks | 11876 | 11904 | 29 | AAGACCCTGTTGAGCTTGACTCTAGTCTG | 123.0 | -8.5 | 1.3 | -6.4 | 1.8E-10 | 3.6E-10 | MB231/MB468 |
| >hg38_21_+_8433172_8446622_RNA45SN1_45S_50ntFlanks@6778.6802.25\|CCCGGGGCTACGCCTGTCTGAGCGT | >hg38_21_+_8433172_8446622_RNA45SN1_45S_50ntFlanks | 6778 | 6802 | 25 | CCCGGGGCTACGCCTGTCTGAGCGT | 1201.1 | 5.4 | 0.9 | 6.4 | 2.1E-10 | 3.7E-10 | BT20/MB468 |
| >hg38_21_+_8433172_8446622_RNA45SN1_45S_50ntFlanks@5472.5502.31\|ACGGCCCTGGCGGAGCGCTGAGAAGACGGTC | >hg38_21_+_8433172_8446622_RNA45SN1_45S_50ntFlanks | 5472 | 5502 | 31 | ACGGCCCTGGCGGAGCGCTGAGAAGACGGTC | 442.9 | 5.9 | 0.9 | 6.4 | 2.1E-10 | 3.7E-10 | BT20/MB468 |
| >hg38_21_+_8433172_8446622_RNA45SN1_45S_50ntFlanks@9573.9596.24\|AAATCGGTCGTCCGACCTGGGTAT | >hg38_21_+_8433172_8446622_RNA45SN1_45S_50ntFlanks | 9573 | 9596 | 24 | AAATCGGTCGTCCGACCTGGGTAT | 2376.5 | -4.7 | 0.7 | -6.4 | 1.8E-10 | 3.7E-10 | MB231/MB468 |
| >MI0000101\|hsa-mir-99a&WithFlank&21\|+\|16539083\|16539175@19.40.22&[MIMAT0000097&hsa-miR-99a-5p&offsets\|0\|0;m-64&21\|+\|16539101\|16539121&offsets\|0\|+1]\|AACCCGTAGATCCGATCTTGTG | >MI0000101\|hsa-mir-99a&WithFlank&21\|+\|16539083\|16539175 | 19 | 40 | 22 | AACCCGTAGATCCGATCTTGTG | 1460.1 | -6.9 | 1.0 | -6.8 | 1.3E-11 | 3.7E-10 | BT20/MB231 |
| >hg38_21_+_8433172_8446622_RNA45SN1_45S_50ntFlanks@3852.3881.30\|TAACTGTGGTAATTCTAGAGCTAATACATG | >hg38_21_+_8433172_8446622_RNA45SN1_45S_50ntFlanks | 3852 | 3881 | 30 | TAACTGTGGTAATTCTAGAGCTAATACATG | 142.8 | -3.3 | 0.5 | -6.4 | 2.1E-10 | 3.7E-10 | BT20/MB468 |
| >hg38_21_+_8433172_8446622_RNA45SN1_45S_50ntFlanks@12417.12436.20\|TCGATGTCGGCTCTTCCTAT | >hg38_21_+_8433172_8446622_RNA45SN1_45S_50ntFlanks | 12417 | 12436 | 20 | TCGATGTCGGCTCTTCCTAT | 4242.6 | -3.4 | 0.5 | -6.4 | 1.9E-10 | 3.7E-10 | MB231/MB468 |
| >hg38_21_+_8433172_8446622_RNA45SN1_45S_50ntFlanks@5526.5550.25\|AAAAGTCGTAACAAGGTTTCCGTAG | >hg38_21_+_8433172_8446622_RNA45SN1_45S_50ntFlanks | 5526 | 5550 | 25 | AAAAGTCGTAACAAGGTTTCCGTAG | 140.2 | -2.9 | 0.5 | -6.4 | 1.9E-10 | 3.8E-10 | MB231/MB468 |
| >hg38_21_+_8433172_8446622_RNA45SN1_45S_50ntFlanks@11567.11611.45\|AGCAGCCGACTTAGAACTGGTGCGGACCAGGGGAATCCGACTGTT | >hg38_21_+_8433172_8446622_RNA45SN1_45S_50ntFlanks | 11567 | 11611 | 45 | AGCAGCCGACTTAGAACTGGTGCGGACCAGGGGAATCCGACTGTT | 583.7 | 4.9 | 0.8 | 6.4 | 1.9E-10 | 3.8E-10 | MB231/MB468 |
| >hg38_21_+_8433172_8446622_RNA45SN1_45S_50ntFlanks@12941.12966.26\|ATTCGTAGACGACCTGCTTCTGGGTC | >hg38_21_+_8433172_8446622_RNA45SN1_45S_50ntFlanks | 12941 | 12966 | 26 | ATTCGTAGACGACCTGCTTCTGGGTC | 556.7 | -4.5 | 0.7 | -6.4 | 1.9E-10 | 3.8E-10 | MB231/MB468 |
| >hg38_21_+_8433172_8446622_RNA45SN1_45S_50ntFlanks@10404.10422.19\|CATGGGTCAGTCGGTCCTG | >hg38_21_+_8433172_8446622_RNA45SN1_45S_50ntFlanks | 10404 | 10422 | 19 | CATGGGTCAGTCGGTCCTG | 324.1 | -4.2 | 0.7 | -6.3 | 2.2E-10 | 3.8E-10 | BT20/MB468 |
| >hg38_21_+_8433172_8446622_RNA45SN1_45S_50ntFlanks@12479.12498.20\|CTAATAGGGAACGTGAGCTG | >hg38_21_+_8433172_8446622_RNA45SN1_45S_50ntFlanks | 12479 | 12498 | 20 | CTAATAGGGAACGTGAGCTG | 864.8 | -3.2 | 0.5 | -6.3 | 2.2E-10 | 3.8E-10 | BT20/MB468 |
| >hg38_21_+_8433172_8446622_RNA45SN1_45S_50ntFlanks@10679.10709.31\|GGTTCCGGCGGCGTCCGGTGAGCTCTCGCTG | >hg38_21_+_8433172_8446622_RNA45SN1_45S_50ntFlanks | 10679 | 10709 | 31 | GGTTCCGGCGGCGTCCGGTGAGCTCTCGCTG | 8394.2 | -5.9 | 0.9 | -6.3 | 2.2E-10 | 3.9E-10 | BT20/MB468 |
| >hg38_21_+_8433172_8446622_RNA45SN1_45S_50ntFlanks@10787.10815.29\|ACAGCCTCTGGCATGTTGGAACAATGTAG | >hg38_21_+_8433172_8446622_RNA45SN1_45S_50ntFlanks | 10787 | 10815 | 29 | ACAGCCTCTGGCATGTTGGAACAATGTAG | 168.2 | -3.1 | 0.5 | -6.3 | 2.3E-10 | 3.9E-10 | BT20/MB468 |

<table>
<tr><th></th><th></th><th></th><th></th><th></th><th></th><th></th><th></th><th></th><th></th><th></th><th></th><th></th></tr>
<tr><td>>hg38_21_+_8433172_8446622_RNA45SN1_45S_50ntFlanks@6721.6756.36|AATTGCAGGACACATTGATCATCGACACTTCGAACG</td><td>>hg38_21_+_8433172_8446622_RNA45SN1_45S_50ntFlanks</td><td>6721</td><td>6756</td><td>36</td><td>AATTGCAGGACACATTGATCATCGACACTTCGAACG</td><td>356.5</td><td>-3.3</td><td>0.5</td><td>-6.3</td><td>2.3E-10</td><td>3.9E-10</td><td>BT20/MB468</td></tr>
<tr><td>>hg38_21_+_8433172_8446622_RNA45SN1_45S_50ntFlanks@5472.5489.18|ACGGCCCTGGCGGAGCGC</td><td>>hg38_21_+_8433172_8446622_RNA45SN1_45S_50ntFlanks</td><td>5472</td><td>5489</td><td>18</td><td>ACGGCCCTGGCGGAGCGC</td><td>416.4</td><td>8.1</td><td>1.3</td><td>6.3</td><td>2.3E-10</td><td>3.9E-10</td><td>BT20/MB468</td></tr>
<tr><td>>hg38_21_+_8433172_8446622_RNA45SN1_45S_50ntFlanks@9180.9197.18|CCGTCGGGCCCGGGGGAG</td><td>>hg38_21_+_8433172_8446622_RNA45SN1_45S_50ntFlanks</td><td>9180</td><td>9197</td><td>18</td><td>CCGTCGGGCCCGGGGGAG</td><td>160.6</td><td>-7.0</td><td>1.1</td><td>-6.4</td><td>1.9E-10</td><td>3.9E-10</td><td>MB231/MB468</td></tr>
<tr><td>>hg38_21_+_8433172_8446622_RNA45SN1_45S_50ntFlanks@9569.9603.35|GTGCAAATCGGTCGTCCGACCTGGGTATAGGGGCG</td><td>>hg38_21_+_8433172_8446622_RNA45SN1_45S_50ntFlanks</td><td>9569</td><td>9603</td><td>35</td><td>GTGCAAATCGGTCGTCCGACCTGGGTATAGGGGCG</td><td>720.7</td><td>4.7</td><td>0.7</td><td>6.4</td><td>2.0E-10</td><td>3.9E-10</td><td>MB231/MB468</td></tr>
<tr><td>>hg38_21_+_8433172_8446622_RNA45SN1_45S_50ntFlanks@4657.4680.24|CTAGAGGTGAAATTCTTGGACCGG</td><td>>hg38_21_+_8433172_8446622_RNA45SN1_45S_50ntFlanks</td><td>4657</td><td>4680</td><td>24</td><td>CTAGAGGTGAAATTCTTGGACCGG</td><td>116.3</td><td>-8.9</td><td>1.4</td><td>-6.3</td><td>2.3E-10</td><td>3.9E-10</td><td>BT20/MB468</td></tr>
<tr><td>>hg38_21_+_8433172_8446622_RNA45SN1_45S_50ntFlanks@11873.11912.40|AAGAAGACCCTGTTGAGCTTGACTCTAGTCTGGCACGGTG</td><td>>hg38_21_+_8433172_8446622_RNA45SN1_45S_50ntFlanks</td><td>11873</td><td>11912</td><td>40</td><td>AAGAAGACCCTGTTGAGCTTGACTCTAGTCTGGCACGGTG</td><td>92.4</td><td>-8.1</td><td>1.3</td><td>-6.4</td><td>2.0E-10</td><td>4.0E-10</td><td>MB231/MB468</td></tr>
<tr><td>>hg38_21_+_8433172_8446622_RNA45SN1_45S_50ntFlanks@11575.11602.28|ACTTAGAACTGGTGCGGACCAGGGGAAT</td><td>>hg38_21_+_8433172_8446622_RNA45SN1_45S_50ntFlanks</td><td>11575</td><td>11602</td><td>28</td><td>ACTTAGAACTGGTGCGGACCAGGGGAAT</td><td>801.9</td><td>-4.2</td><td>0.7</td><td>-6.4</td><td>2.0E-10</td><td>4.0E-10</td><td>MB231/MB468</td></tr>
<tr><td>>hg38_21_+_8433172_8446622_RNA45SN1_45S_50ntFlanks@6663.6681.19|GTGGATCACTCGGCTCGTG</td><td>>hg38_21_+_8433172_8446622_RNA45SN1_45S_50ntFlanks</td><td>6663</td><td>6681</td><td>19</td><td>GTGGATCACTCGGCTCGTG</td><td>1349.7</td><td>-5.5</td><td>0.9</td><td>-6.3</td><td>2.3E-10</td><td>4.0E-10</td><td>BT20/MB468</td></tr>
<tr><td>>hg38_21_+_8433172_8446622_RNA45SN1_45S_50ntFlanks@9570.9597.28|TGCAAATCGGTCGTCCGACCTGGGTATA</td><td>>hg38_21_+_8433172_8446622_RNA45SN1_45S_50ntFlanks</td><td>9570</td><td>9597</td><td>28</td><td>TGCAAATCGGTCGTCCGACCTGGGTATA</td><td>2281.2</td><td>-5.5</td><td>0.9</td><td>-6.4</td><td>2.0E-10</td><td>4.0E-10</td><td>MB231/MB468</td></tr>
<tr><td>>hg38_21_+_8433172_8446622_RNA45SN1_45S_50ntFlanks@9198.9225.28|GTTCTCTCGGGGCCACGCGCGCGTCCCC</td><td>>hg38_21_+_8433172_8446622_RNA45SN1_45S_50ntFlanks</td><td>9198</td><td>9225</td><td>28</td><td>GTTCTCTCGGGGCCACGCGCGCGTCCCC</td><td>115.0</td><td>-8.4</td><td>1.3</td><td>-6.4</td><td>2.0E-10</td><td>4.0E-10</td><td>MB231/MB468</td></tr>
<tr><td>>hg38_21_+_8433172_8446622_RNA45SN1_45S_50ntFlanks@10414.10431.18|TCGGTCCTGAGAGATGGG</td><td>>hg38_21_+_8433172_8446622_RNA45SN1_45S_50ntFlanks</td><td>10414</td><td>10431</td><td>18</td><td>TCGGTCCTGAGAGATGGG</td><td>314.9</td><td>-4.3</td><td>0.7</td><td>-6.4</td><td>2.0E-10</td><td>4.1E-10</td><td>MB231/MB468</td></tr>
<tr><td>>hg38_21_+_8433172_8446622_RNA45SN1_45S_50ntFlanks@10717.10745.29|AAAATCCGGGGGAGAGGGTGTAAATCTCG</td><td>>hg38_21_+_8433172_8446622_RNA45SN1_45S_50ntFlanks</td><td>10717</td><td>10745</td><td>29</td><td>AAAATCCGGGGGAGAGGGTGTAAATCTCG</td><td>438.2</td><td>4.7</td><td>0.7</td><td>6.4</td><td>2.0E-10</td><td>4.1E-10</td><td>MB231/MB468</td></tr>
<tr><td>>hg38_21_+_8433172_8446622_RNA45SN1_45S_50ntFlanks@3705.3723.19|TACCTGGTTGATCCTGCCA</td><td>>hg38_21_+_8433172_8446622_RNA45SN1_45S_50ntFlanks</td><td>3705</td><td>3723</td><td>19</td><td>TACCTGGTTGATCCTGCCA</td><td>841.8</td><td>4.3</td><td>0.7</td><td>6.3</td><td>2.4E-10</td><td>4.1E-10</td><td>BT20/MB468</td></tr>
<tr><td>>hg38_21_+_8433172_8446622_RNA45SN1_45S_50ntFlanks@10839.10868.30|TCCGTAACTTCGGGATAAGGATTGGCTCTA</td><td>>hg38_21_+_8433172_8446622_RNA45SN1_45S_50ntFlanks</td><td>10839</td><td>10868</td><td>30</td><td>TCCGTAACTTCGGGATAAGGATTGGCTCTA</td><td>202.1</td><td>-4.7</td><td>0.7</td><td>-6.3</td><td>2.4E-10</td><td>4.1E-10</td><td>BT20/MB468</td></tr>
<tr><td>>hg38_21_+_8433172_8446622_RNA45SN1_45S_50ntFlanks@6671.6699.29|CTCGGCTCGTGCGTCGATGAAGAACGCAG</td><td>>hg38_21_+_8433172_8446622_RNA45SN1_45S_50ntFlanks</td><td>6671</td><td>6699</td><td>29</td><td>CTCGGCTCGTGCGTCGATGAAGAACGCAG</td><td>2065.1</td><td>-4.0</td><td>0.6</td><td>-6.4</td><td>2.1E-10</td><td>4.2E-10</td><td>MB231/MB468</td></tr>
<tr><td>>trna37_ValAAC_6_+_27203288_27203360@1.29.29|GTTTCCGTAGTGTAGTGGTTATCACGTTT</td><td>>trna37_ValAAC_6_+_27203288_27203360</td><td>1</td><td>29</td><td>29</td><td>GTTTCCGTAGTGTAGTGGTTATCACGTTT</td><td>986.0</td><td>-6.0</td><td>0.9</td><td>-6.3</td><td>2.4E-10</td><td>4.2E-10</td><td>BT20/MB468</td></tr>
<tr><td>>hg38_21_+_8433172_8446622_RNA45SN1_45S_50ntFlanks@10781.10818.38|AGGTGAACAGCCTCTGGCATGTTGGAACAATGTAGGTA</td><td>>hg38_21_+_8433172_8446622_RNA45SN1_45S_50ntFlanks</td><td>10781</td><td>10818</td><td>38</td><td>AGGTGAACAGCCTCTGGCATGTTGGAACAATGTAGGTA</td><td>1409.5</td><td>-4.4</td><td>0.7</td><td>-6.4</td><td>2.1E-10</td><td>4.2E-10</td><td>MB231/MB468</td></tr>
<tr><td>>MI0000071|hsa-mir-17&WithFlank&13|+|91350599|91350694@20.42.23(+1G)&[MIMAT0000070&hsa-miR-17-5p&offsets|0|0(+1G);m-88&13|+|91350618|91350640&offsets|0|0(+1G)]|CAAAGTGCTTACAGTGCAGGTAGG</td><td>>MI0000071|hsa-mir-17&WithFlank&13|+|91350599|91350694</td><td>20</td><td>42</td><td>23(+1G)</td><td>CAAAGTGCTTACAGTGCAGGTAGG</td><td>671.7</td><td>5.9</td><td>0.9</td><td>6.3</td><td>2.4E-10</td><td>4.2E-10</td><td>BT20/MB468</td></tr>
<tr><td>>hg38_21_+_8433172_8446622_RNA45SN1_45S_50ntFlanks@6652.6693.42|GACTCTTAGCGGTGGATCACTCGGCTCGTGCGTCGATGAAGA</td><td>>hg38_21_+_8433172_8446622_RNA45SN1_45S_50ntFlanks</td><td>6652</td><td>6693</td><td>42</td><td>GACTCTTAGCGGTGGATCACTCGGCTCGTGCGTCGATGAAGA</td><td>1096.0</td><td>5.8</td><td>0.9</td><td>6.3</td><td>2.4E-10</td><td>4.2E-10</td><td>BT20/MB468</td></tr>
<tr><td>>hg38_21_+_8433172_8446622_RNA45SN1_45S_50ntFlanks@10760.10785.26|CATATCCGCAGCAGGTCTCCAAGGTG</td><td>>hg38_21_+_8433172_8446622_RNA45SN1_45S_50ntFlanks</td><td>10760</td><td>10785</td><td>26</td><td>CATATCCGCAGCAGGTCTCCAAGGTG</td><td>90.1</td><td>-4.8</td><td>0.8</td><td>-6.3</td><td>2.5E-10</td><td>4.2E-10</td><td>BT20/MB468</td></tr>
<tr><td>>hg38_21_+_8433172_8446622_RNA45SN1_45S_50ntFlanks@8378.8400.23|TTCAAGAGGGCGTGAAACCGTTA</td><td>>hg38_21_+_8433172_8446622_RNA45SN1_45S_50ntFlanks</td><td>8378</td><td>8400</td><td>23</td><td>TTCAAGAGGGCGTGAAACCGTTA</td><td>109.2</td><td>-5.5</td><td>0.9</td><td>-6.3</td><td>2.5E-10</td><td>4.3E-10</td><td>BT20/MB468</td></tr>
<tr><td>>hg38_21_+_8433172_8446622_RNA45SN1_45S_50ntFlanks@12631.12653.23|GGCGAAGCTACCATCTGTGGGAT</td><td>>hg38_21_+_8433172_8446622_RNA45SN1_45S_50ntFlanks</td><td>12631</td><td>12653</td><td>23</td><td>GGCGAAGCTACCATCTGTGGGAT</td><td>111.7</td><td>-8.4</td><td>1.3</td><td>-6.3</td><td>2.2E-10</td><td>4.3E-10</td><td>MB231/MB468</td></tr>
<tr><td>>hg38_21_+_8433172_8446622_RNA45SN1_45S_50ntFlanks@11564.11592.29|CCTAGCAGCCGACTTAGAACTGGTGCGGA</td><td>>hg38_21_+_8433172_8446622_RNA45SN1_45S_50ntFlanks</td><td>11564</td><td>11592</td><td>29</td><td>CCTAGCAGCCGACTTAGAACTGGTGCGGA</td><td>342.0</td><td>-4.2</td><td>0.7</td><td>-6.3</td><td>2.2E-10</td><td>4.3E-10</td><td>MB231/MB468</td></tr>
<tr><td>>hg38_21_+_8433172_8446622_RNA45SN1_45S_50ntFlanks@12893.12933.41|CCTCGCCCGTCACGCACCGCACGTTCGTGGGGAACCTGGCG</td><td>>hg38_21_+_8433172_8446622_RNA45SN1_45S_50ntFlanks</td><td>12893</td><td>12933</td><td>41</td><td>CCTCGCCCGTCACGCACCGCACGTTCGTGGGGAACCTGGCG</td><td>279.6</td><td>-5.1</td><td>0.8</td><td>-6.3</td><td>2.2E-10</td><td>4.3E-10</td><td>MB231/MB468</td></tr>
<tr><td>>hg38_21_+_8433172_8446622_RNA45SN1_45S_50ntFlanks@9537.9563.27|AAACTCTGGTGGAGGTCCGTAGCGGTC</td><td>>hg38_21_+_8433172_8446622_RNA45SN1_45S_50ntFlanks</td><td>9537</td><td>9563</td><td>27</td><td>AAACTCTGGTGGAGGTCCGTAGCGGTC</td><td>112.0</td><td>-5.7</td><td>0.9</td><td>-6.3</td><td>2.5E-10</td><td>4.4E-10</td><td>BT20/MB468</td></tr>
<tr><td>>hg38_1_-_228634819_228635039_RNA5S12_5S_50ntFlanks@142.170.29|CGCCTGGGAATACCGGGTGCTGTAGGCTT</td><td>>hg38_1_-_228634819_228635039_RNA5S12_5S_50ntFlanks</td><td>142</td><td>170</td><td>29</td><td>CGCCTGGGAATACCGGGTGCTGTAGGCTT</td><td>1790.3</td><td>5.7</td><td>0.9</td><td>6.3</td><td>2.5E-10</td><td>4.4E-10</td><td>BT20/MB468</td></tr>
<tr><td>>am_trna111_HisGTG_1_-_147774845_147774916@-1G.29.30,trna118_HisGTG_1_-_145396881_145396952@-1G.29.30,trna16_HisGTG_1_+_146544773_146544844@-1G.29.30,trna1_HisGTG_15_+_45493349_45493420@-1G.29.30,trna21_HisGTG_1_+_147753471_147753542@-1G.29.30,trna33_HisGTG_6_+_27125906_27125977@-1G.29.30,trna7_HisGTG_9_-_14433938_14434009@-1G.29.30,trna8_HisGTG_15_-_45492611_45492682@-1G.29.30,trna9_HisGTG_15_-_45490804_45490875@-1G.29.30|GGCCGTGATCGTATAGTGGTTAGTACTCTG</td><td>>am_trna111_HisGTG_1_-_147774845_147774916</td><td>-1G</td><td>29</td><td>30</td><td>GGCCGTGATCGTATAGTGGTTAGTACTCTG</td><td>3996.1</td><td>-6.5</td><td>1.0</td><td>-6.3</td><td>2.6E-10</td><td>4.4E-10</td><td>BT20/MB468</td></tr>
<tr><td>>hg38_21_+_8433172_8446622_RNA45SN1_45S_50ntFlanks@11540.11584.45|CGCGGCGCCTCGCCTCGGCCGGCGCCTAGCAGCCGACTTAGAACT</td><td>>hg38_21_+_8433172_8446622_RNA45SN1_45S_50ntFlanks</td><td>11540</td><td>11584</td><td>45</td><td>CGCGGCGCCTCGCCTCGGCCGGCGCCTAGCAGCCGACTTAGAACT</td><td>381.7</td><td>7.5</td><td>1.2</td><td>6.3</td><td>2.6E-10</td><td>4.4E-10</td><td>BT20/MB468</td></tr>
<tr><td>>hg38_21_+_8433172_8446622_RNA45SN1_45S_50ntFlanks@11999.12043.45|CCCTGCGGGCCGCCGGTGAAATACCACTACTCTGATCGTTTTTTC</td><td>>hg38_21_+_8433172_8446622_RNA45SN1_45S_50ntFlanks</td><td>11999</td><td>12043</td><td>45</td><td>CCCTGCGGGCCGCCGGTGAAATACCACTACTCTGATCGTTTTTTC</td><td>153.8</td><td>-9.3</td><td>1.5</td><td>-6.3</td><td>2.6E-10</td><td>4.5E-10</td><td>BT20/MB468</td></tr>
<tr><td>>hg38_21_+_8433172_8446622_RNA45SN1_45S_50ntFlanks@10680.10708.29|GTTCCGGCGGCGTCCGGTGAGCTCTCGCT</td><td>>hg38_21_+_8433172_8446622_RNA45SN1_45S_50ntFlanks</td><td>10680</td><td>10708</td><td>29</td><td>GTTCCGGCGGCGTCCGGTGAGCTCTCGCT</td><td>15411.2</td><td>-5.3</td><td>0.8</td><td>-6.3</td><td>2.6E-10</td><td>4.5E-10</td><td>BT20/MB468</td></tr>
<tr><td>>hg38_21_+_8433172_8446622_RNA45SN1_45S_50ntFlanks@10342.10373.32|ATATTCAAACGAGAACTTTGAAGGCCGAAGTG</td><td>>hg38_21_+_8433172_8446622_RNA45SN1_45S_50ntFlanks</td><td>10342</td><td>10373</td><td>32</td><td>ATATTCAAACGAGAACTTTGAAGGCCGAAGTG</td><td>348.6</td><td>4.6</td><td>0.7</td><td>6.3</td><td>2.2E-10</td><td>4.5E-10</td><td>MB231/MB468</td></tr>
<tr><td>>hg38_21_+_8433172_8446622_RNA45SN1_45S_50ntFlanks@11573.11601.29|CGACTTAGAACTGGTGCGGACCAGGGGAA</td><td>>hg38_21_+_8433172_8446622_RNA45SN1_45S_50ntFlanks</td><td>11573</td><td>11601</td><td>29</td><td>CGACTTAGAACTGGTGCGGACCAGGGGAA</td><td>569.7</td><td>-3.7</td><td>0.6</td><td>-6.3</td><td>2.2E-10</td><td>4.5E-10</td><td>MB231/MB468</td></tr>
<tr><td>>hg38_21_+_8433172_8446622_RNA45SN1_45S_50ntFlanks@10678.10695.18|CGGTTCCGGCGGCGTCCG</td><td>>hg38_21_+_8433172_8446622_RNA45SN1_45S_50ntFlanks</td><td>10678</td><td>10695</td><td>18</td><td>CGGTTCCGGCGGCGTCCG</td><td>86.6</td><td>-8.0</td><td>1.3</td><td>-6.3</td><td>2.3E-10</td><td>4.5E-10</td><td>MB231/MB468</td></tr>
<tr><td>>hg38_21_+_8433172_8446622_RNA45SN1_45S_50ntFlanks@11569.11603.35|CAGCCGACTTAGAACTGGTGCGGACCAGGGGAATC</td><td>>hg38_21_+_8433172_8446622_RNA45SN1_45S_50ntFlanks</td><td>11569</td><td>11603</td><td>35</td><td>CAGCCGACTTAGAACTGGTGCGGACCAGGGGAATC</td><td>1886.5</td><td>-5.6</td><td>0.9</td><td>-6.3</td><td>2.6E-10</td><td>4.5E-10</td><td>BT20/MB468</td></tr>
<tr><td>>hg38_21_+_8433172_8446622_RNA45SN1_45S_50ntFlanks@10682.10712.31|TCCGGCGGCGTCCGGTGAGCTCTCGCTGGCC</td><td>>hg38_21_+_8433172_8446622_RNA45SN1_45S_50ntFlanks</td><td>10682</td><td>10712</td><td>31</td><td>TCCGGCGGCGTCCGGTGAGCTCTCGCTGGCC</td><td>216.6</td><td>-4.3</td><td>0.7</td><td>-6.3</td><td>2.6E-10</td><td>4.6E-10</td><td>BT20/MB468</td></tr>
<tr><td>>hg38_21_+_8433172_8446622_RNA45SN1_45S_50ntFlanks@10574.10592.19|AAGCCGGCGGGAGCCCCGG</td><td>>hg38_21_+_8433172_8446622_RNA45SN1_45S_50ntFlanks</td><td>10574</td><td>10592</td><td>19</td><td>AAGCCGGCGGGAGCCCCGG</td><td>100.3</td><td>-8.6</td><td>1.4</td><td>-6.3</td><td>2.6E-10</td><td>4.6E-10</td><td>BT20/MB468</td></tr>
<tr><td>>hg38_21_+_8433172_8446622_RNA45SN1_45S_50ntFlanks@10787.10822.36|ACAGCCTCTGGCATGTTGGAACAATGTAGGTAAGGG</td><td>>hg38_21_+_8433172_8446622_RNA45SN1_45S_50ntFlanks</td><td>10787</td><td>10822</td><td>36</td><td>ACAGCCTCTGGCATGTTGGAACAATGTAGGTAAGGG</td><td>110.3</td><td>-3.5</td><td>0.6</td><td>-6.3</td><td>2.7E-10</td><td>4.6E-10</td><td>BT20/MB468</td></tr>
<tr><td>>hg38_21_+_8433172_8446622_RNA45SN1_45S_50ntFlanks@485.505.21|GATGTGGTGACGTCGTGCTCT</td><td>>hg38_21_+_8433172_8446622_RNA45SN1_45S_50ntFlanks</td><td>485</td><td>505</td><td>21</td><td>GATGTGGTGACGTCGTGCTCT</td><td>47236.0</td><td>20.4</td><td>3.2</td><td>6.3</td><td>2.3E-10</td><td>4.6E-10</td><td>MB231/MB468</td></tr>
<tr><td>>hg38_21_+_8433172_8446622_RNA45SN1_45S_50ntFlanks@11918.11949.32|ACATGAGAGGTGTAGAATAAGTGGGAGGCCCC</td><td>>hg38_21_+_8433172_8446622_RNA45SN1_45S_50ntFlanks</td><td>11918</td><td>11949</td><td>32</td><td>ACATGAGAGGTGTAGAATAAGTGGGAGGCCCC</td><td>96.9</td><td>-8.6</td><td>1.4</td><td>-6.3</td><td>2.7E-10</td><td>4.6E-10</td><td>BT20/MB468</td></tr>
<tr><td>>hg38_21_+_8433172_8446622_RNA45SN1_45S_50ntFlanks@12686.12703.18|CCAGGCGGAACGATACGG</td><td>>hg38_21_+_8433172_8446622_RNA45SN1_45S_50ntFlanks</td><td>12686</td><td>12703</td><td>18</td><td>CCAGGCGGAACGATACGG</td><td>129.8</td><td>-5.6</td><td>0.9</td><td>-6.3</td><td>2.7E-10</td><td>4.6E-10</td><td>BT20/MB468</td></tr>
<tr><td>>hg38_21_+_8433172_8446622_RNA45SN1_45S_50ntFlanks@12997.13014.18|CGCTGCGATCTATTGAAA</td><td>>hg38_21_+_8433172_8446622_RNA45SN1_45S_50ntFlanks</td><td>12997</td><td>13014</td><td>18</td><td>CGCTGCGATCTATTGAAA</td><td>2006.3</td><td>3.3</td><td>0.5</td><td>6.3</td><td>2.7E-10</td><td>4.6E-10</td><td>BT20/MB468</td></tr>
<tr><td>>trna108_AlaAGC_6_-_28763741_28763812@58.75.18|TCCCCAGTACCTCCACCA</td><td>>trna108_AlaAGC_6_-_28763741_28763812</td><td>58</td><td>75</td><td>18</td><td>TCCCCAGTACCTCCACCA</td><td>930.4</td><td>6.0</td><td>0.9</td><td>6.3</td><td>2.3E-10</td><td>4.6E-10</td><td>MB231/MB468</td></tr>
<tr><td>>MI0000764|hsa-mir-363&WithFlank&X|-|134169372|134169458@56.77.22&[MIMAT0000707&hsa-miR-363-3p&offsets|0|0;m-178&X|-|134169381|134169403&offsets|0|-1]|AATTGCACGGTATCCATCTGTA</td><td>>MI0000764|hsa-mir-363&WithFlank&X|-|134169372|134169458</td><td>56</td><td>77</td><td>22</td><td>AATTGCACGGTATCCATCTGTA</td><td>117.2</td><td>-8.4</td><td>1.3</td><td>-6.3</td><td>2.3E-10</td><td>4.6E-10</td><td>MB231/MB468</td></tr>
<tr><td>>hg38_21_+_8433172_8446622_RNA45SN1_45S_50ntFlanks@12363.12383.21|GGATAACTGGCTTGTGGCGGC</td><td>>hg38_21_+_8433172_8446622_RNA45SN1_45S_50ntFlanks</td><td>12363</td><td>12383</td><td>21</td><td>GGATAACTGGCTTGTGGCGGC</td><td>99.9</td><td>-8.6</td><td>1.4</td><td>-6.3</td><td>2.7E-10</td><td>4.7E-10</td><td>BT20/MB468</td></tr>
<tr><td>>hg38_21_+_8433172_8446622_RNA45SN1_45S_50ntFlanks@10826.10865.40|TCGGCAAGCCGGATCCGTAACTTCGGGATAAGGATTGGCT</td><td>>hg38_21_+_8433172_8446622_RNA45SN1_45S_50ntFlanks</td><td>10826</td><td>10865</td><td>40</td><td>TCGGCAAGCCGGATCCGTAACTTCGGGATAAGGATTGGCT</td><td>144.0</td><td>-9.2</td><td>1.5</td><td>-6.3</td><td>2.7E-10</td><td>4.7E-10</td><td>BT20/MB468</td></tr>
<tr><td>>hg38_21_+_8433172_8446622_RNA45SN1_45S_50ntFlanks@10737.10757.21|TAAATCTCGCGCCGGGCCGTA</td><td>>hg38_21_+_8433172_8446622_RNA45SN1_45S_50ntFlanks</td><td>10737</td><td>10757</td><td>21</td><td>TAAATCTCGCGCCGGGCCGTA</td><td>110.2</td><td>-6.6</td><td>1.0</td><td>-6.3</td><td>2.4E-10</td><td>4.7E-10</td><td>MB231/MB468</td></tr>
<tr><td>>hg38_21_+_8433172_8446622_RNA45SN1_45S_50ntFlanks@3984.4005.22|GCGGCTTTGGTGACTCTAGATA</td><td>>hg38_21_+_8433172_8446622_RNA45SN1_45S_50ntFlanks</td><td>3984</td><td>4005</td><td>22</td><td>GCGGCTTTGGTGACTCTAGATA</td><td>92.7</td><td>-8.1</td><td>1.3</td><td>-6.3</td><td>2.4E-10</td><td>4.7E-10</td><td>MB231/MB468</td></tr>
<tr><td>>hg38_21_+_8433172_8446622_RNA45SN1_45S_50ntFlanks@11561.11602.42|GCGCCTAGCAGCCGACTTAGAACTGGTGCGGACCAGGGGAAT</td><td>>hg38_21_+_8433172_8446622_RNA45SN1_45S_50ntFlanks</td><td>11561</td><td>11602</td><td>42</td><td>GCGCCTAGCAGCCGACTTAGAACTGGTGCGGACCAGGGGAAT</td><td>266.9</td><td>-5.2</td><td>0.8</td><td>-6.3</td><td>2.8E-10</td><td>4.7E-10</td><td>BT20/MB468</td></tr>
</table>

| | | | | | | | | | | | | |
|---|---|---|---|---|---|---|---|---|---|---|---|---|
| >hg38_21_+_8433172_8446622_RNA45SN1_45S_50ntFlanks@11561.11582.22\|GCGCCTAGCAGCCGACTTAGAA | >hg38_21_+_8433172_8446622_RNA45SN1_45S_50ntFlanks | 11561 | 11582 | 22 | GCGCCTAGCAGCCGACTTAGAA | 198.9 | -5.8 | 0.9 | -6.3 | 2.4E-10 | 4.7E-10 | MB231/MB468 |
| >hg38_21_+_8433172_8446622_RNA45SN1_45S_50ntFlanks@12324.12342.19\|TTGGGTTTTAAGCAGGAGG | >hg38_21_+_8433172_8446622_RNA45SN1_45S_50ntFlanks | 12324 | 12342 | 19 | TTGGGTTTTAAGCAGGAGG | 94.2 | -8.1 | 1.3 | -6.3 | 2.4E-10 | 4.8E-10 | MB231/MB468 |
| >hg38_21_+_8433172_8446622_RNA45SN1_45S_50ntFlanks@12488.12519.32\|AACGTGAGCTGGGTTTAGACCGTCGTGAGACA | >hg38_21_+_8433172_8446622_RNA45SN1_45S_50ntFlanks | 12488 | 12519 | 32 | AACGTGAGCTGGGTTTAGACCGTCGTGAGACA | 184.4 | -3.8 | 0.6 | -6.3 | 2.8E-10 | 4.8E-10 | BT20/MB468 |
| >hg38_21_+_8433172_8446622_RNA45SN1_45S_50ntFlanks@10572.10596.25\|AGAAGCCGGCGGGAGCCCCGGGGAG | >hg38_21_+_8433172_8446622_RNA45SN1_45S_50ntFlanks | 10572 | 10596 | 25 | AGAAGCCGGCGGGAGCCCCGGGGAG | 119.5 | -6.7 | 1.1 | -6.3 | 2.4E-10 | 4.8E-10 | MB231/MB468 |
| >hg38_21_+_8433172_8446622_RNA45SN1_45S_50ntFlanks@12045.12070.26\|CTGACCCGGTGAGGCGGGGGGGCGAG | >hg38_21_+_8433172_8446622_RNA45SN1_45S_50ntFlanks | 12045 | 12070 | 26 | CTGACCCGGTGAGGCGGGGGGGCGAG | 85.9 | -8.0 | 1.3 | -6.3 | 2.4E-10 | 4.8E-10 | MB231/MB468 |
| >hg38_21_+_8433172_8446622_RNA45SN1_45S_50ntFlanks@8191.8211.21\|CAGCCCGTGGACGGTGTGAGG | >hg38_21_+_8433172_8446622_RNA45SN1_45S_50ntFlanks | 8191 | 8211 | 21 | CAGCCCGTGGACGGTGTGAGG | 131.5 | -5.2 | 0.8 | -6.3 | 2.8E-10 | 4.8E-10 | BT20/MB468 |
| >hg38_21_+_8433172_8446622_RNA45SN1_45S_50ntFlanks@12990.13039.50\|GCTCCCTCGCTGCGATCTATTGAAAGTCAGCCCTCGACACAAGGGTTTGT | >hg38_21_+_8433172_8446622_RNA45SN1_45S_50ntFlanks | 12990 | 13039 | 50 | GCTCCCTCGCTGCGATCTATTGAAAGTCAGCCCTCGACACAAGGGTTTGT | 858.1 | 6.4 | 1.0 | 6.3 | 2.4E-10 | 4.8E-10 | MB231/MB468 |
| >hg38_21_+_8433172_8446622_RNA45SN1_45S_50ntFlanks@4044.4063.20\|CCCATTCGAACGTCTGCCCT | >hg38_21_+_8433172_8446622_RNA45SN1_45S_50ntFlanks | 4044 | 4063 | 20 | CCCATTCGAACGTCTGCCCT | 83.4 | -8.0 | 1.3 | -6.3 | 2.4E-10 | 4.8E-10 | MB231/MB468 |
| >hg38_21_+_8433172_8446622_RNA45SN1_45S_50ntFlanks@6646.6682.37\|TCGTACGACTCTTAGCGGTGGATCACTCGGCTCGTGC | >hg38_21_+_8433172_8446622_RNA45SN1_45S_50ntFlanks | 6646 | 6682 | 37 | TCGTACGACTCTTAGCGGTGGATCACTCGGCTCGTGC | 411.0 | 6.9 | 1.1 | 6.3 | 2.8E-10 | 4.9E-10 | BT20/MB468 |
| >hg38_21_+_8433172_8446622_RNA45SN1_45S_50ntFlanks@4560.4579.20\|CTAGGAATAATGGAATAGGA | >hg38_21_+_8433172_8446622_RNA45SN1_45S_50ntFlanks | 4560 | 4579 | 20 | CTAGGAATAATGGAATAGGA | 214.3 | -4.0 | 0.6 | -6.3 | 2.5E-10 | 4.9E-10 | MB231/MB468 |
| >hg38_21_+_8433172_8446622_RNA45SN1_45S_50ntFlanks@12804.12821.18\|CGCGTGCCCCGCCGCGCG | >hg38_21_+_8433172_8446622_RNA45SN1_45S_50ntFlanks | 12804 | 12821 | 18 | CGCGTGCCCCGCCGCGCG | 81.8 | -7.9 | 1.3 | -6.3 | 2.5E-10 | 4.9E-10 | MB231/MB468 |
| >hg38_21_+_8433172_8446622_RNA45SN1_45S_50ntFlanks@8327.8356.30\|ATAGTCAACAAGTACCGTAAGGGAAAGTTG | >hg38_21_+_8433172_8446622_RNA45SN1_45S_50ntFlanks | 8327 | 8356 | 30 | ATAGTCAACAAGTACCGTAAGGGAAAGTTG | 123.4 | -8.9 | 1.4 | -6.3 | 2.9E-10 | 4.9E-10 | BT20/MB468 |
| >hg38_21_+_8433172_8446622_RNA45SN1_45S_50ntFlanks@6690.6719.30\|AAGAACGCAGCTAGCTGCGAGAATTAATGT | >hg38_21_+_8433172_8446622_RNA45SN1_45S_50ntFlanks | 6690 | 6719 | 30 | AAGAACGCAGCTAGCTGCGAGAATTAATGT | 84.6 | -8.4 | 1.3 | -6.3 | 2.9E-10 | 4.9E-10 | BT20/MB468 |
| >hg38_21_+_8433172_8446622_RNA45SN1_45S_50ntFlanks@12436.12454.19\|TCATTGTGAAGCAGAATTC | >hg38_21_+_8433172_8446622_RNA45SN1_45S_50ntFlanks | 12436 | 12454 | 19 | TCATTGTGAAGCAGAATTC | 996.6 | -4.4 | 0.7 | -6.3 | 2.5E-10 | 4.9E-10 | MB231/MB468 |
| >hg38_1_-_228634819_228635039_RNA5S12_5S_50ntFlanks@137.169.33\|GAGACCGCCTGGGAATACCGGGTGCTGTAGGCT | >hg38_1_-_228634819_228635039_RNA5S12_5S_50ntFlanks | 137 | 169 | 33 | GAGACCGCCTGGGAATACCGGGTGCTGTAGGCT | 533.2 | 8.3 | 1.3 | 6.3 | 2.9E-10 | 5.0E-10 | BT20/MB468 |
| >hg38_21_+_8433172_8446622_RNA45SN1_45S_50ntFlanks@12957.12987.31\|CTTCTGGGTCGGGGTTTCGTACGTAGCAGAG | >hg38_21_+_8433172_8446622_RNA45SN1_45S_50ntFlanks | 12957 | 12987 | 31 | CTTCTGGGTCGGGGTTTCGTACGTAGCAGAG | 253.9 | -5.7 | 0.9 | -6.3 | 2.9E-10 | 5.0E-10 | BT20/MB468 |
| >hg38_21_+_8433172_8446622_RNA45SN1_45S_50ntFlanks@10849.10868.20\|CGGGATAAGGATTGGCTCTA | >hg38_21_+_8433172_8446622_RNA45SN1_45S_50ntFlanks | 10849 | 10868 | 20 | CGGGATAAGGATTGGCTCTA | 649.4 | -7.5 | 1.2 | -6.3 | 2.9E-10 | 5.0E-10 | BT20/MB468 |
| >hg38_21_+_8433172_8446622_RNA45SN1_45S_50ntFlanks@12636.12657.22\|AGCTACCATCTGTGGGATTATG | >hg38_21_+_8433172_8446622_RNA45SN1_45S_50ntFlanks | 12636 | 12657 | 22 | AGCTACCATCTGTGGGATTATG | 229.3 | -5.0 | 0.8 | -6.3 | 2.5E-10 | 5.0E-10 | MB231/MB468 |
| >hg38_21_+_8433172_8446622_RNA45SN1_45S_50ntFlanks@10846.10863.18\|CTTCGGGATAAGGATTGG | >hg38_21_+_8433172_8446622_RNA45SN1_45S_50ntFlanks | 10846 | 10863 | 18 | CTTCGGGATAAGGATTGG | 300.6 | -6.5 | 1.0 | -6.3 | 2.9E-10 | 5.0E-10 | BT20/MB468 |
| >hg38_21_+_8433172_8446622_RNA45SN1_45S_50ntFlanks@11670.11702.33\|CCCAGTGCTCTGAATGTCAAAGTGAAGAAATTC | >hg38_21_+_8433172_8446622_RNA45SN1_45S_50ntFlanks | 11670 | 11702 | 33 | CCCAGTGCTCTGAATGTCAAAGTGAAGAAATTC | 1056.4 | -4.3 | 0.7 | -6.3 | 2.9E-10 | 5.0E-10 | BT20/MB468 |
| >hg38_21_+_8433172_8446622_RNA45SN1_45S_50ntFlanks@10386.10424.39\|ATGTGAACAGCAGTTGAACATGGGTCAGTCGGTCCTGAG | >hg38_21_+_8433172_8446622_RNA45SN1_45S_50ntFlanks | 10386 | 10424 | 39 | ATGTGAACAGCAGTTGAACATGGGTCAGTCGGTCCTGAG | 841.1 | 7.7 | 1.2 | 6.3 | 2.9E-10 | 5.0E-10 | BT20/MB468 |
| >hg38_21_+_8433172_8446622_RNA45SN1_45S_50ntFlanks@8171.8188.18\|AAGTCCTTCTGATCGAGG | >hg38_21_+_8433172_8446622_RNA45SN1_45S_50ntFlanks | 8171 | 8188 | 18 | AAGTCCTTCTGATCGAGG | 200.0 | -9.2 | 1.5 | -6.3 | 2.5E-10 | 5.0E-10 | MB231/MB468 |
| >hg38_21_+_8433172_8446622_RNA45SN1_45S_50ntFlanks@12083.12110.28\|CTCGCTTCTGGCGCCAAGCGCCCGGCCG | >hg38_21_+_8433172_8446622_RNA45SN1_45S_50ntFlanks | 12083 | 12110 | 28 | CTCGCTTCTGGCGCCAAGCGCCCGGCCG | 91.7 | -8.1 | 1.3 | -6.3 | 2.6E-10 | 5.0E-10 | MB231/MB468 |
| >hg38_21_+_8433172_8446622_RNA45SN1_45S_50ntFlanks@12033.12055.23\|ATCGTTTTTTCACTGACCCGGTG | >hg38_21_+_8433172_8446622_RNA45SN1_45S_50ntFlanks | 12033 | 12055 | 23 | ATCGTTTTTTCACTGACCCGGTG | 87.8 | -8.0 | 1.3 | -6.3 | 2.6E-10 | 5.1E-10 | MB231/MB468 |
| >hg38_21_+_8433172_8446622_RNA45SN1_45S_50ntFlanks@9306.9323.18\|CAAGGAGTCTAACACGTG | >hg38_21_+_8433172_8446622_RNA45SN1_45S_50ntFlanks | 9306 | 9323 | 18 | CAAGGAGTCTAACACGTG | 115.9 | -8.4 | 1.3 | -6.3 | 2.6E-10 | 5.1E-10 | MB231/MB468 |
| >hg38_21_+_8433172_8446622_RNA45SN1_45S_50ntFlanks@10677.10708.32\|GCGGTTCCGGCGGCGTCCGGTGAGCTCTCGCT | >hg38_21_+_8433172_8446622_RNA45SN1_45S_50ntFlanks | 10677 | 10708 | 32 | GCGGTTCCGGCGGCGTCCGGTGAGCTCTCGCT | 592.7 | 3.3 | 0.5 | 6.3 | 2.6E-10 | 5.2E-10 | MB231/MB468 |
| >hg38_21_+_8433172_8446622_RNA45SN1_45S_50ntFlanks@5472.5516.45\|ACGGCCCTGGCGGAGCGCTGAGAAGACGGTCGAACTTGACTATCT | >hg38_21_+_8433172_8446622_RNA45SN1_45S_50ntFlanks | 5472 | 5516 | 45 | ACGGCCCTGGCGGAGCGCTGAGAAGACGGTCGAACTTGACTATCT | 27230.5 | 13.6 | 2.2 | 6.3 | 2.6E-10 | 5.2E-10 | MB231/MB468 |
| >hg38_21_+_8433172_8446622_RNA45SN1_45S_50ntFlanks@10411.10440.30\|CAGTCGGTCCTGAGAGATGGGCGAGCGCCG | >hg38_21_+_8433172_8446622_RNA45SN1_45S_50ntFlanks | 10411 | 10440 | 30 | CAGTCGGTCCTGAGAGATGGGCGAGCGCCG | 151.5 | -4.5 | 0.7 | -6.3 | 2.6E-10 | 5.2E-10 | MB231/MB468 |
| >hg38_21_+_8433172_8446622_RNA45SN1_45S_50ntFlanks@12991.13010.20\|CTCCCTCGCTGCGATCTATT | >hg38_21_+_8433172_8446622_RNA45SN1_45S_50ntFlanks | 12991 | 13010 | 20 | CTCCCTCGCTGCGATCTATT | 355.6 | -3.6 | 0.6 | -6.3 | 2.6E-10 | 5.2E-10 | MB231/MB468 |
| >hg38_21_+_8433172_8446622_RNA45SN1_45S_50ntFlanks@6730.6770.41\|ACACATTGATCATCGACACTTCGAACGCACTTGCGGCCCCG | >hg38_21_+_8433172_8446622_RNA45SN1_45S_50ntFlanks | 6730 | 6770 | 41 | ACACATTGATCATCGACACTTCGAACGCACTTGCGGCCCCG | 205.3 | -5.8 | 0.9 | -6.3 | 3.1E-10 | 5.2E-10 | BT20/MB468 |
| >hg38_21_+_8433172_8446622_RNA45SN1_45S_50ntFlanks@11781.11807.27\|CGCATGAATGGATGAACGAGATTCCCA | >hg38_21_+_8433172_8446622_RNA45SN1_45S_50ntFlanks | 11781 | 11807 | 27 | CGCATGAATGGATGAACGAGATTCCCA | 83.4 | -8.0 | 1.3 | -6.3 | 2.7E-10 | 5.3E-10 | MB231/MB468 |
| >hg38_21_+_8433172_8446622_RNA45SN1_45S_50ntFlanks@12511.12538.28\|CGTGAGACAGGTTAGTTTTACCCTACTG | >hg38_21_+_8433172_8446622_RNA45SN1_45S_50ntFlanks | 12511 | 12538 | 28 | CGTGAGACAGGTTAGTTTTACCCTACTG | 85.5 | -8.0 | 1.3 | -6.3 | 2.7E-10 | 5.3E-10 | MB231/MB468 |
| >hg38_21_+_8433172_8446622_RNA45SN1_45S_50ntFlanks@10128.10149.22\|CGGCGGCGGCGGCGGCGGCGGG | >hg38_21_+_8433172_8446622_RNA45SN1_45S_50ntFlanks | 10128 | 10149 | 22 | CGGCGGCGGCGGCGGCGGCGGG | 73.7 | -8.2 | 1.3 | -6.3 | 3.1E-10 | 5.3E-10 | BT20/MB468 |
| >hg38_21_+_8433172_8446622_RNA45SN1_45S_50ntFlanks@8845.8866.22\|CGGCGGCGGCGGCGGCGGCGGG | >hg38_21_+_8433172_8446622_RNA45SN1_45S_50ntFlanks | 8845 | 8866 | 22 | CGGCGGCGGCGGCGGCGGCGGG | 73.7 | -8.2 | 1.3 | -6.3 | 3.1E-10 | 5.3E-10 | BT20/MB468 |
| >hg38_21_+_8433172_8446622_RNA45SN1_45S_50ntFlanks@11580.11600.21\|GAACTGGTGCGGACCAGGGGA | >hg38_21_+_8433172_8446622_RNA45SN1_45S_50ntFlanks | 11580 | 11600 | 21 | GAACTGGTGCGGACCAGGGGA | 333.5 | -4.0 | 0.6 | -6.3 | 3.1E-10 | 5.4E-10 | BT20/MB468 |
| >hg38_21_+_8433172_8446622_RNA45SN1_45S_50ntFlanks@4724.4758.35\|ATTAATCAAGAACGAAAGTCGGAGGTTCGAAGACG | >hg38_21_+_8433172_8446622_RNA45SN1_45S_50ntFlanks | 4724 | 4758 | 35 | ATTAATCAAGAACGAAAGTCGGAGGTTCGAAGACG | 4095.4 | -3.7 | 0.6 | -6.3 | 2.7E-10 | 5.4E-10 | MB231/MB468 |
| >MI0000076\|hsa-mir-20a&WithFlank&13\|+\|91351059\|91351141@14.36.23(+1A)&[MIMAT0000075&hsa-miR-20a-5p&offsets\|0\|0(+1A);m-96&13\|+\|91351072\|91351094&offsets\|0\|0(+1A)]\|\|TAAAGTGCTTATAGTGCAGGTAGA | >MI0000076\|hsa-mir-20a&WithFlank&13\|+\|91351059\|91351141 | 14 | 36 | 23(+1A) | TAAAGTGCTTATAGTGCAGGTAGA | 6972.0 | 5.6 | 0.9 | 6.3 | 3.2E-10 | 5.4E-10 | BT20/MB468 |
| >hg38_21_+_8433172_8446622_RNA45SN1_45S_50ntFlanks@11729.11761.33\|TAACTATGACTCTCTTAAGGTAGCCAAATGCCT | >hg38_21_+_8433172_8446622_RNA45SN1_45S_50ntFlanks | 11729 | 11761 | 33 | TAACTATGACTCTCTTAAGGTAGCCAAATGCCT | 342.7 | -4.1 | 0.7 | -6.3 | 3.2E-10 | 5.4E-10 | BT20/MB468 |
| >hg38_21_+_8433172_8446622_RNA45SN1_45S_50ntFlanks@10388.10407.20\|GTGAACAGCAGTTGAACATG | >hg38_21_+_8433172_8446622_RNA45SN1_45S_50ntFlanks | 10388 | 10407 | 20 | GTGAACAGCAGTTGAACATG | 212.7 | -6.3 | 1.0 | -6.3 | 3.2E-10 | 5.4E-10 | BT20/MB468 |
| >hg38_21_+_8433172_8446622_RNA45SN1_45S_50ntFlanks@5436.5466.31\|GTGAGGCCCTCGGATCGGCCCCGCCGGGGTC | >hg38_21_+_8433172_8446622_RNA45SN1_45S_50ntFlanks | 5436 | 5466 | 31 | GTGAGGCCCTCGGATCGGCCCCGCCGGGGTC | 92.1 | -8.1 | 1.3 | -6.3 | 2.8E-10 | 5.5E-10 | MB231/MB468 |
| >hg38_21_+_8433172_8446622_RNA45SN1_45S_50ntFlanks@10687.10711.25\|CGGCGTCCGGTGAGCTCTCGCTGGC | >hg38_21_+_8433172_8446622_RNA45SN1_45S_50ntFlanks | 10687 | 10711 | 25 | CGGCGTCCGGTGAGCTCTCGCTGGC | 2329.7 | -4.7 | 0.7 | -6.3 | 2.8E-10 | 5.5E-10 | MB231/MB468 |
| >hg38_21_+_8433172_8446622_RNA45SN1_45S_50ntFlanks@4762.4780.19\|AGATACCGTCGTAGTTCCG | >hg38_21_+_8433172_8446622_RNA45SN1_45S_50ntFlanks | 4762 | 4780 | 19 | AGATACCGTCGTAGTTCCG | 595.1 | -4.1 | 0.6 | -6.3 | 3.2E-10 | 5.5E-10 | BT20/MB468 |
| >hg38_21_+_8433172_8446622_RNA45SN1_45S_50ntFlanks@9198.9218.21\|GTTCTCTCGGGGCCACGCGCG | >hg38_21_+_8433172_8446622_RNA45SN1_45S_50ntFlanks | 9198 | 9218 | 21 | GTTCTCTCGGGGCCACGCGCG | 133.9 | -8.6 | 1.4 | -6.3 | 2.8E-10 | 5.5E-10 | MB231/MB468 |
| >hg38_21_+_8433172_8446622_RNA45SN1_45S_50ntFlanks@12619.12654.36\|AGGAGCCAATGGGGCGAAGCTACCATCTGTGGGATT | >hg38_21_+_8433172_8446622_RNA45SN1_45S_50ntFlanks | 12619 | 12654 | 36 | AGGAGCCAATGGGGCGAAGCTACCATCTGTGGGATT | 862.4 | -4.5 | 0.7 | -6.3 | 2.8E-10 | 5.5E-10 | MB231/MB468 |
| >hg38_21_+_8433172_8446622_RNA45SN1_45S_50ntFlanks@11115.11132.18\|CGGCAGGGGCCGGCGGCG | >hg38_21_+_8433172_8446622_RNA45SN1_45S_50ntFlanks | 11115 | 11132 | 18 | CGGCAGGGGCCGGCGGCG | 86.6 | -8.0 | 1.3 | -6.3 | 2.8E-10 | 5.5E-10 | MB231/MB468 |
| >hg38_21_+_8433172_8446622_RNA45SN1_45S_50ntFlanks@10766.10785.20\|CGCAGCAGGTCTCCAAGGTG | >hg38_21_+_8433172_8446622_RNA45SN1_45S_50ntFlanks | 10766 | 10785 | 20 | CGCAGCAGGTCTCCAAGGTG | 163.8 | -2.6 | 0.4 | -6.3 | 3.3E-10 | 5.6E-10 | BT20/MB468 |
| >hg38_21_+_8433172_8446622_RNA45SN1_45S_50ntFlanks@11760.11777.18\|CTCGTCATCTAATTAGTG | >hg38_21_+_8433172_8446622_RNA45SN1_45S_50ntFlanks | 11760 | 11777 | 18 | CTCGTCATCTAATTAGTG | 120.2 | -4.8 | 0.8 | -6.3 | 2.8E-10 | 5.6E-10 | MB231/MB468 |
| >hg38_21_+_8433172_8446622_RNA45SN1_45S_50ntFlanks@476.498.23\|TTCGTGATCGATGTGGTGACGTC | >hg38_21_+_8433172_8446622_RNA45SN1_45S_50ntFlanks | 476 | 498 | 23 | TTCGTGATCGATGTGGTGACGTC | 1710.1 | 5.9 | 0.9 | 6.3 | 3.3E-10 | 5.6E-10 | BT20/MB468 |
| >hg38_21_+_8433172_8446622_RNA45SN1_45S_50ntFlanks@8157.8178.22\|CTCGTGGGGGGCCCAAGTCCTT | >hg38_21_+_8433172_8446622_RNA45SN1_45S_50ntFlanks | 8157 | 8178 | 22 | CTCGTGGGGGGCCCAAGTCCTT | 198.5 | -6.3 | 1.0 | -6.3 | 3.3E-10 | 5.6E-10 | BT20/MB468 |
| >hg38_21_+_8433172_8446622_RNA45SN1_45S_50ntFlanks@11795.11820.26\|AACGAGATTCCCACTGTCCCTACCTA | >hg38_21_+_8433172_8446622_RNA45SN1_45S_50ntFlanks | 11795 | 11820 | 26 | AACGAGATTCCCACTGTCCCTACCTA | 83.5 | -8.0 | 1.3 | -6.3 | 2.9E-10 | 5.7E-10 | MB231/MB468 |

<table>
<tr><td>>hg38_21_+_8433172_8446622_RNA45SN1_45S_50ntFlanks@3940.3959.20|ACCCGGTCAGCCCCTCTCCG</td><td>>hg38_21_+_8433172_8446622_RNA45SN1_45S_50ntFlanks</td><td>3940</td><td>3959</td><td>20</td><td>ACCCGGTCAGCCCCTCTCCG</td><td>88.8</td><td>-8.0</td><td>1.3</td><td>-6.3</td><td>2.9E-10</td><td>5.7E-10</td><td>MB231/MB468</td></tr>
<tr><td>>hg38_21_+_8433172_8446622_RNA45SN1_45S_50ntFlanks@10384.10404.21|CCATGTGAACAGCAGTTGAAC</td><td>>hg38_21_+_8433172_8446622_RNA45SN1_45S_50ntFlanks</td><td>10384</td><td>10404</td><td>21</td><td>CCATGTGAACAGCAGTTGAAC</td><td>182.1</td><td>-2.6</td><td>0.4</td><td>-6.3</td><td>2.9E-10</td><td>5.7E-10</td><td>MB231/MB468</td></tr>
<tr><td>>hg38_21_+_8433172_8446622_RNA45SN1_45S_50ntFlanks@10513.10534.22|CCGGAGTGGCGGAGATGGGCGC</td><td>>hg38_21_+_8433172_8446622_RNA45SN1_45S_50ntFlanks</td><td>10513</td><td>10534</td><td>22</td><td>CCGGAGTGGCGGAGATGGGCGC</td><td>125.6</td><td>-5.2</td><td>0.8</td><td>-6.3</td><td>2.9E-10</td><td>5.7E-10</td><td>MB231/MB468</td></tr>
<tr><td>>hg38_21_+_8433172_8446622_RNA45SN1_45S_50ntFlanks@11561.11602.42|GCGCCTAGCAGCCGACTTAGAACTGGTGCGGACCAGGGGAAT</td><td>>hg38_21_+_8433172_8446622_RNA45SN1_45S_50ntFlanks</td><td>11561</td><td>11602</td><td>42</td><td>GCGCCTAGCAGCCGACTTAGAACTGGTGCGGACCAGGGGAAT</td><td>219.9</td><td>-4.5</td><td>0.7</td><td>-6.3</td><td>2.9E-10</td><td>5.8E-10</td><td>MB231/MB468</td></tr>
<tr><td>>hg38_21_+_8433172_8446622_RNA45SN1_45S_50ntFlanks@11759.11780.22|CCTCGTCATCTAATTAGTGACG</td><td>>hg38_21_+_8433172_8446622_RNA45SN1_45S_50ntFlanks</td><td>11759</td><td>11780</td><td>22</td><td>CCTCGTCATCTAATTAGTGACG</td><td>162.0</td><td>-8.9</td><td>1.4</td><td>-6.3</td><td>2.9E-10</td><td>5.8E-10</td><td>MB231/MB468</td></tr>
<tr><td>>hg38_21_+_8433172_8446622_RNA45SN1_45S_50ntFlanks@6661.6680.20|CGGTGGATCACTCGGCTCGT</td><td>>hg38_21_+_8433172_8446622_RNA45SN1_45S_50ntFlanks</td><td>6661</td><td>6680</td><td>20</td><td>CGGTGGATCACTCGGCTCGT</td><td>612.9</td><td>2.8</td><td>0.4</td><td>6.3</td><td>3.4E-10</td><td>5.8E-10</td><td>BT20/MB468</td></tr>
<tr><td>>hg38_21_+_8433172_8446622_RNA45SN1_45S_50ntFlanks@12416.12434.19|TTCGATGTCGGCTCTTCCT</td><td>>hg38_21_+_8433172_8446622_RNA45SN1_45S_50ntFlanks</td><td>12416</td><td>12434</td><td>19</td><td>TTCGATGTCGGCTCTTCCT</td><td>3187.7</td><td>3.9</td><td>0.6</td><td>6.3</td><td>3.0E-10</td><td>5.9E-10</td><td>MB231/MB468</td></tr>
<tr><td>>hg38_21_+_8433172_8446622_RNA45SN1_45S_50ntFlanks@4449.4472.24|CCTCTCGGCGCCCCCTCGATGCTC</td><td>>hg38_21_+_8433172_8446622_RNA45SN1_45S_50ntFlanks</td><td>4449</td><td>4472</td><td>24</td><td>CCTCTCGGCGCCCCCTCGATGCTC</td><td>221.4</td><td>-4.8</td><td>0.8</td><td>-6.3</td><td>3.0E-10</td><td>5.9E-10</td><td>MB231/MB468</td></tr>
<tr><td>>hg38_21_+_8433172_8446622_RNA45SN1_45S_50ntFlanks@10425.10457.33|AGATGGGCGAGCGCCGTTCCGAAGGGACGGGCG</td><td>>hg38_21_+_8433172_8446622_RNA45SN1_45S_50ntFlanks</td><td>10425</td><td>10457</td><td>33</td><td>AGATGGGCGAGCGCCGTTCCGAAGGGACGGGCG</td><td>90.6</td><td>-8.1</td><td>1.3</td><td>-6.3</td><td>3.0E-10</td><td>5.9E-10</td><td>MB231/MB468</td></tr>
<tr><td>>hg38_21_+_8433172_8446622_RNA45SN1_45S_50ntFlanks@10914.10932.19|CGCCGCGGCTGGACGAGGC</td><td>>hg38_21_+_8433172_8446622_RNA45SN1_45S_50ntFlanks</td><td>10914</td><td>10932</td><td>19</td><td>CGCCGCGGCTGGACGAGGC</td><td>759.0</td><td>-8.0</td><td>1.3</td><td>-6.3</td><td>3.5E-10</td><td>6.0E-10</td><td>BT20/MB468</td></tr>
<tr><td>>hg38_21_+_8433172_8446622_RNA45SN1_45S_50ntFlanks@11998.12032.35|GCCCTGCGGGCCGCCGGTGAAATACCACTACTCTG</td><td>>hg38_21_+_8433172_8446622_RNA45SN1_45S_50ntFlanks</td><td>11998</td><td>12032</td><td>35</td><td>GCCCTGCGGGCCGCCGGTGAAATACCACTACTCTG</td><td>83.3</td><td>-8.0</td><td>1.3</td><td>-6.3</td><td>3.1E-10</td><td>6.0E-10</td><td>MB231/MB468</td></tr>
<tr><td>>hg38_21_+_8433172_8446622_RNA45SN1_45S_50ntFlanks@11804.11821.18|CCCACTGTCCCTACCTAC</td><td>>hg38_21_+_8433172_8446622_RNA45SN1_45S_50ntFlanks</td><td>11804</td><td>11821</td><td>18</td><td>CCCACTGTCCCTACCTAC</td><td>89.6</td><td>-8.1</td><td>1.3</td><td>-6.3</td><td>3.1E-10</td><td>6.0E-10</td><td>MB231/MB468</td></tr>
<tr><td>>hg38_21_+_8433172_8446622_RNA45SN1_45S_50ntFlanks@4472.4492.21|CTTAGCTGAGTGTCCCGCGGG</td><td>>hg38_21_+_8433172_8446622_RNA45SN1_45S_50ntFlanks</td><td>4472</td><td>4492</td><td>21</td><td>CTTAGCTGAGTGTCCCGCGGG</td><td>147.2</td><td>-4.6</td><td>0.7</td><td>-6.3</td><td>3.1E-10</td><td>6.1E-10</td><td>MB231/MB468</td></tr>
<tr><td>>trna9_ArgTCT_1_+_94313129_94313213@1.29.29|GGCTCCGTGGCGCAATGGATAGCGCATTG</td><td>>trna9_ArgTCT_1_+_94313129_94313213</td><td>1</td><td>29</td><td>29</td><td>GGCTCCGTGGCGCAATGGATAGCGCATTG</td><td>110.3</td><td>-8.4</td><td>1.3</td><td>-6.3</td><td>3.1E-10</td><td>6.1E-10</td><td>MB231/MB468</td></tr>
<tr><td>>hg38_21_+_8433172_8446622_RNA45SN1_45S_50ntFlanks@11999.12033.35|CCCTGCGGGCCGCCGGTGAAATACCACTACTCTGA</td><td>>hg38_21_+_8433172_8446622_RNA45SN1_45S_50ntFlanks</td><td>11999</td><td>12033</td><td>35</td><td>CCCTGCGGGCCGCCGGTGAAATACCACTACTCTGA</td><td>140.2</td><td>-3.9</td><td>0.6</td><td>-6.3</td><td>3.6E-10</td><td>6.2E-10</td><td>BT20/MB468</td></tr>
<tr><td>>hg38_21_+_8433172_8446622_RNA45SN1_45S_50ntFlanks@12152.12180.29|TGGGGAGTTTGACTGGGGCGGTACACCTG</td><td>>hg38_21_+_8433172_8446622_RNA45SN1_45S_50ntFlanks</td><td>12152</td><td>12180</td><td>29</td><td>TGGGGAGTTTGACTGGGGCGGTACACCTG</td><td>86.6</td><td>-8.0</td><td>1.3</td><td>-6.3</td><td>3.2E-10</td><td>6.2E-10</td><td>MB231/MB468</td></tr>
<tr><td>>hg38_21_+_8433172_8446622_RNA45SN1_45S_50ntFlanks@12941.12964.24|ATTCGTAGACGACCTGCTTCTGGG</td><td>>hg38_21_+_8433172_8446622_RNA45SN1_45S_50ntFlanks</td><td>12941</td><td>12964</td><td>24</td><td>ATTCGTAGACGACCTGCTTCTGGG</td><td>122.7</td><td>-3.7</td><td>0.6</td><td>-6.3</td><td>3.6E-10</td><td>6.2E-10</td><td>BT20/MB468</td></tr>
<tr><td>>hg38_21_+_8433172_8446622_RNA45SN1_45S_50ntFlanks@11853.11872.20|CTTGGCGGAATCAGCGGGGA</td><td>>hg38_21_+_8433172_8446622_RNA45SN1_45S_50ntFlanks</td><td>11853</td><td>11872</td><td>20</td><td>CTTGGCGGAATCAGCGGGGA</td><td>102.4</td><td>-8.3</td><td>1.3</td><td>-6.3</td><td>3.2E-10</td><td>6.2E-10</td><td>MB231/MB468</td></tr>
<tr><td>>hg38_21_+_8433172_8446622_RNA45SN1_45S_50ntFlanks@8378.8400.23|TTCAAGAGGGCGTGAAACCGTTA</td><td>>hg38_21_+_8433172_8446622_RNA45SN1_45S_50ntFlanks</td><td>8378</td><td>8400</td><td>23</td><td>TTCAAGAGGGCGTGAAACCGTTA</td><td>88.4</td><td>-8.0</td><td>1.3</td><td>-6.3</td><td>3.2E-10</td><td>6.3E-10</td><td>MB231/MB468</td></tr>
<tr><td>>hg38_21_+_8433172_8446622_RNA45SN1_45S_50ntFlanks@12934.12962.29|CTAAACCATTCGTAGACGACCTGCTTCTG</td><td>>hg38_21_+_8433172_8446622_RNA45SN1_45S_50ntFlanks</td><td>12934</td><td>12962</td><td>29</td><td>CTAAACCATTCGTAGACGACCTGCTTCTG</td><td>241.8</td><td>-6.0</td><td>1.0</td><td>-6.3</td><td>3.7E-10</td><td>6.3E-10</td><td>BT20/MB468</td></tr>
<tr><td>>hg38_21_+_8433172_8446622_RNA45SN1_45S_50ntFlanks@10735.10757.23|TGTAAATCTCGCGCCGGGCCGTA</td><td>>hg38_21_+_8433172_8446622_RNA45SN1_45S_50ntFlanks</td><td>10735</td><td>10757</td><td>23</td><td>TGTAAATCTCGCGCCGGGCCGTA</td><td>103.4</td><td>-3.7</td><td>0.6</td><td>-6.3</td><td>3.2E-10</td><td>6.3E-10</td><td>MB231/MB468</td></tr>
<tr><td>>hg38_21_+_8433172_8446622_RNA45SN1_45S_50ntFlanks@12607.12626.20|TGTGCTTGGCTGAGGAGCCA</td><td>>hg38_21_+_8433172_8446622_RNA45SN1_45S_50ntFlanks</td><td>12607</td><td>12626</td><td>20</td><td>TGTGCTTGGCTGAGGAGCCA</td><td>219.8</td><td>-3.2</td><td>0.5</td><td>-6.3</td><td>3.7E-10</td><td>6.3E-10</td><td>BT20/MB468</td></tr>
<tr><td>>hg38_21_+_8433172_8446622_RNA45SN1_45S_50ntFlanks@8475.8492.18|CCCGGCGGATCTTTCCCG</td><td>>hg38_21_+_8433172_8446622_RNA45SN1_45S_50ntFlanks</td><td>8475</td><td>8492</td><td>18</td><td>CCCGGCGGATCTTTCCCG</td><td>99.6</td><td>-8.2</td><td>1.3</td><td>-6.3</td><td>3.2E-10</td><td>6.4E-10</td><td>MB231/MB468</td></tr>
<tr><td>>hg38_21_+_8433172_8446622_RNA45SN1_45S_50ntFlanks@7975.8017.43|CGCGACCTCAGATCAGACGTGGCGACCCGCTGAATTTAAGCAT</td><td>>hg38_21_+_8433172_8446622_RNA45SN1_45S_50ntFlanks</td><td>7975</td><td>8017</td><td>43</td><td>CGCGACCTCAGATCAGACGTGGCGACCCGCTGAATTTAAGCAT</td><td>4883.1</td><td>7.2</td><td>1.1</td><td>6.3</td><td>3.8E-10</td><td>6.4E-10</td><td>BT20/MB468</td></tr>
<tr><td>>hg38_21_+_8433172_8446622_RNA45SN1_45S_50ntFlanks@11786.11804.19|GAATGGATGAACGAGATTC</td><td>>hg38_21_+_8433172_8446622_RNA45SN1_45S_50ntFlanks</td><td>11786</td><td>11804</td><td>19</td><td>GAATGGATGAACGAGATTC</td><td>1245.5</td><td>-4.1</td><td>0.7</td><td>-6.3</td><td>3.3E-10</td><td>6.4E-10</td><td>MB231/MB468</td></tr>
<tr><td>>hg38_21_+_8433172_8446622_RNA45SN1_45S_50ntFlanks@6651.6685.35|CGACTCTTAGCGGTGGATCACTCGGCTCGTGCGTC</td><td>>hg38_21_+_8433172_8446622_RNA45SN1_45S_50ntFlanks</td><td>6651</td><td>6685</td><td>35</td><td>CGACTCTTAGCGGTGGATCACTCGGCTCGTGCGTC</td><td>2277.7</td><td>4.0</td><td>0.6</td><td>6.3</td><td>3.8E-10</td><td>6.5E-10</td><td>BT20/MB468</td></tr>
<tr><td>>hg38_21_+_8433172_8446622_RNA45SN1_45S_50ntFlanks@4160.4178.19|CCACATCCAAGGAAGGCAG</td><td>>hg38_21_+_8433172_8446622_RNA45SN1_45S_50ntFlanks</td><td>4160</td><td>4178</td><td>19</td><td>CCACATCCAAGGAAGGCAG</td><td>83.4</td><td>-8.0</td><td>1.3</td><td>-6.3</td><td>3.3E-10</td><td>6.5E-10</td><td>MB231/MB468</td></tr>
<tr><td>>hg38_21_+_8433172_8446622_RNA45SN1_45S_50ntFlanks@11211.11232.22|GCGACTCTGGACGCGAGCCGGG</td><td>>hg38_21_+_8433172_8446622_RNA45SN1_45S_50ntFlanks</td><td>11211</td><td>11232</td><td>22</td><td>GCGACTCTGGACGCGAGCCGGG</td><td>713.6</td><td>-4.9</td><td>0.8</td><td>-6.3</td><td>3.8E-10</td><td>6.5E-10</td><td>BT20/MB468</td></tr>
<tr><td>>trnaMT_GluTTC_MT_-_14674_14742@39.57.19,trnalookalike8_GluTTC_5_-_93905172_93905240@39.57.19|ATTGGTCGTGGTTGTAGTC</td><td>>trnaMT_GluTTC_MT_-_14674_14742</td><td>39</td><td>57</td><td>19</td><td>ATTGGTCGTGGTTGTAGTC</td><td>628.0</td><td>8.2</td><td>1.3</td><td>6.3</td><td>3.9E-10</td><td>6.6E-10</td><td>BT20/MB468</td></tr>
<tr><td>>hg38_21_+_8433172_8446622_RNA45SN1_45S_50ntFlanks@12072.12090.19|CCCGAGGGGCTCTCGCTTC</td><td>>hg38_21_+_8433172_8446622_RNA45SN1_45S_50ntFlanks</td><td>12072</td><td>12090</td><td>19</td><td>CCCGAGGGGCTCTCGCTTC</td><td>164.5</td><td>-2.9</td><td>0.5</td><td>-6.3</td><td>3.4E-10</td><td>6.6E-10</td><td>MB231/MB468</td></tr>
<tr><td>>hg38_21_+_8433172_8446622_RNA45SN1_45S_50ntFlanks@7990.8009.20|GACGTGGCGACCCGCTGAAT</td><td>>hg38_21_+_8433172_8446622_RNA45SN1_45S_50ntFlanks</td><td>7990</td><td>8009</td><td>20</td><td>GACGTGGCGACCCGCTGAAT</td><td>1816.2</td><td>5.2</td><td>0.8</td><td>6.3</td><td>3.4E-10</td><td>6.6E-10</td><td>MB231/MB468</td></tr>
<tr><td>>trnaMT_PheGAA_MT_+_577_647@36.53.18|AAAATGTTTAGACGGGCT</td><td>>trnaMT_PheGAA_MT_+_577_647</td><td>36</td><td>53</td><td>18</td><td>AAAATGTTTAGACGGGCT</td><td>503.2</td><td>4.9</td><td>0.8</td><td>6.3</td><td>3.9E-10</td><td>6.6E-10</td><td>BT20/MB468</td></tr>
<tr><td>>hg38_1_-_228634819_228635039_RNA5S12_5S_50ntFlanks@124.147.24|AGTACTTGGATGGGAGACCGCCTG</td><td>>hg38_1_-_228634819_228635039_RNA5S12_5S_50ntFlanks</td><td>124</td><td>147</td><td>24</td><td>AGTACTTGGATGGGAGACCGCCTG</td><td>201.9</td><td>-5.2</td><td>0.8</td><td>-6.3</td><td>3.4E-10</td><td>6.7E-10</td><td>MB231/MB468</td></tr>
<tr><td>>hg38_21_+_8433172_8446622_RNA45SN1_45S_50ntFlanks@5469.5499.31|CCCACGGCCCTGGCGGAGCGCTGAGAAGACG</td><td>>hg38_21_+_8433172_8446622_RNA45SN1_45S_50ntFlanks</td><td>5469</td><td>5499</td><td>31</td><td>CCCACGGCCCTGGCGGAGCGCTGAGAAGACG</td><td>140.2</td><td>-5.2</td><td>0.8</td><td>-6.3</td><td>3.5E-10</td><td>6.8E-10</td><td>MB231/MB468</td></tr>
<tr><td>>hg38_21_+_8433172_8446622_RNA45SN1_45S_50ntFlanks@10788.10807.20|CAGCCTCTGGCATGTTGGAA</td><td>>hg38_21_+_8433172_8446622_RNA45SN1_45S_50ntFlanks</td><td>10788</td><td>10807</td><td>20</td><td>CAGCCTCTGGCATGTTGGAA</td><td>143.1</td><td>-6.0</td><td>1.0</td><td>-6.3</td><td>4.0E-10</td><td>6.8E-10</td><td>BT20/MB468</td></tr>
<tr><td>>hg38_21_+_8433172_8446622_RNA45SN1_45S_50ntFlanks@10806.10829.24|AACAATGTAGGTAAGGGAAGTCGG</td><td>>hg38_21_+_8433172_8446622_RNA45SN1_45S_50ntFlanks</td><td>10806</td><td>10829</td><td>24</td><td>AACAATGTAGGTAAGGGAAGTCGG</td><td>83.7</td><td>-5.4</td><td>0.9</td><td>-6.3</td><td>3.5E-10</td><td>6.8E-10</td><td>MB231/MB468</td></tr>
<tr><td>>hg38_21_+_8433172_8446622_RNA45SN1_45S_50ntFlanks@11581.11603.23|AACTGGTGCGGACCAGGGGAATC</td><td>>hg38_21_+_8433172_8446622_RNA45SN1_45S_50ntFlanks</td><td>11581</td><td>11603</td><td>23</td><td>AACTGGTGCGGACCAGGGGAATC</td><td>3403.9</td><td>-4.1</td><td>0.6</td><td>-6.3</td><td>3.5E-10</td><td>6.8E-10</td><td>MB231/MB468</td></tr>
<tr><td>>trna10_AspGTC_12_-_125424193_125424264@32.70.39,trna12_AspGTC_12_-_125411891_125411962@32.70.39,trna144_AspGTC_6_-_27551236_27551307@32.70.39,trna38_AspGTC_17_-_8125556_8125627@32.70.39,trna45_AspGTC_6_+_27447453_27447524@32.70.39,trna48_AspGTC_6_+_27471523_27471594@32.70.39,trna4_AspGTC_12_+_96429799_96429870@32.70.39,trna69_AspGTC_1_-_161440205_161440276@32.70.39,trna72_AspGTC_1_-_161432824_161432895@32.70.39,trna75_AspGTC_1_-_161425414_161425485@32.70.39,trna78_AspGTC_1_-_161418033_161418104@32.70.39,trna81_AspGTC_1_-_161410615_161410686@32.70.39|CTGTCACGCGGGAGACCGGGGTTCGATTCCCCGACGGGG</td><td>>trna10_AspGTC_12_-_125424193_125424264</td><td>32</td><td>70</td><td>39</td><td>CTGTCACGCGGGAGACCGGGGTTCGATTCCCCGACGGGG</td><td>86.1</td><td>-8.0</td><td>1.3</td><td>-6.3</td><td>3.5E-10</td><td>6.8E-10</td><td>MB231/MB468</td></tr>
<tr><td>>hg38_21_+_8433172_8446622_RNA45SN1_45S_50ntFlanks@11672.11709.38|CAGTGCTCTGAATGTCAAAGTGAAGAAATTCAATGAAG</td><td>>hg38_21_+_8433172_8446622_RNA45SN1_45S_50ntFlanks</td><td>11672</td><td>11709</td><td>38</td><td>CAGTGCTCTGAATGTCAAAGTGAAGAAATTCAATGAAG</td><td>696.0</td><td>4.5</td><td>0.7</td><td>6.3</td><td>3.5E-10</td><td>6.8E-10</td><td>MB231/MB468</td></tr>
<tr><td>>hg38_21_+_8433172_8446622_RNA45SN1_45S_50ntFlanks@12628.12661.34|TGGGGCGAAGCTACCATCTGTGGGATTATGACTG</td><td>>hg38_21_+_8433172_8446622_RNA45SN1_45S_50ntFlanks</td><td>12628</td><td>12661</td><td>34</td><td>TGGGGCGAAGCTACCATCTGTGGGATTATGACTG</td><td>507.8</td><td>-6.5</td><td>1.0</td><td>-6.3</td><td>4.1E-10</td><td>6.9E-10</td><td>BT20/MB468</td></tr>
<tr><td>>MI0000285|hsa-mir-205&WithFlank&1|+|209432127|209432248@40.60.21(+1A)&[MIMAT0000266&hsa-miR-205-5p&offsets|0|-1(+1A);m-351&1|+|209432166|209432187&offsets|0|-1(+1A)]|TCCTTCATTCCACCGGAGTCTA</td><td>>MI0000285|hsa-mir-205&WithFlank&1|+|209432127|209432248</td><td>40</td><td>60</td><td>21(+1A)</td><td>TCCTTCATTCCACCGGAGTCTA</td><td>90.6</td><td>-8.1</td><td>1.3</td><td>-6.3</td><td>3.6E-10</td><td>6.9E-10</td><td>MB231/MB468</td></tr>
<tr><td>>hg38_21_+_8433172_8446622_RNA45SN1_45S_50ntFlanks@11564.11602.39|CCTAGCAGCCGACTTAGAACTGGTGCGGACCAGGGGAAT</td><td>>hg38_21_+_8433172_8446622_RNA45SN1_45S_50ntFlanks</td><td>11564</td><td>11602</td><td>39</td><td>CCTAGCAGCCGACTTAGAACTGGTGCGGACCAGGGGAAT</td><td>668.3</td><td>-3.8</td><td>0.6</td><td>-6.3</td><td>4.1E-10</td><td>7.0E-10</td><td>BT20/MB468</td></tr>
<tr><td>>hg38_21_+_8433172_8446622_RNA45SN1_45S_50ntFlanks@12154.12180.27|GGGAGTTTGACTGGGGCGGTACACCTG</td><td>>hg38_21_+_8433172_8446622_RNA45SN1_45S_50ntFlanks</td><td>12154</td><td>12180</td><td>27</td><td>GGGAGTTTGACTGGGGCGGTACACCTG</td><td>85.4</td><td>-8.0</td><td>1.3</td><td>-6.3</td><td>3.6E-10</td><td>7.1E-10</td><td>MB231/MB468</td></tr>
<tr><td>>hg38_21_+_8433172_8446622_RNA45SN1_45S_50ntFlanks@9027.9064.38|CGTCTCCTCGTGGGGGGGCCGGGCCACCCCTCCCACGG</td><td>>hg38_21_+_8433172_8446622_RNA45SN1_45S_50ntFlanks</td><td>9027</td><td>9064</td><td>38</td><td>CGTCTCCTCGTGGGGGGGCCGGGCCACCCCTCCCACGG</td><td>161.0</td><td>-5.7</td><td>0.9</td><td>-6.2</td><td>4.2E-10</td><td>7.1E-10</td><td>BT20/MB468</td></tr>
<tr><td>>hg38_21_+_8433172_8446622_RNA45SN1_45S_50ntFlanks@11255.11281.27|CTGCGGCGGGCGTCGCGGCCGCCCCCG</td><td>>hg38_21_+_8433172_8446622_RNA45SN1_45S_50ntFlanks</td><td>11255</td><td>11281</td><td>27</td><td>CTGCGGCGGGCGTCGCGGCCGCCCCCG</td><td>91.6</td><td>-8.1</td><td>1.3</td><td>-6.3</td><td>3.7E-10</td><td>7.1E-10</td><td>MB231/MB468</td></tr>
<tr><td>>hg38_21_+_8433172_8446622_RNA45SN1_45S_50ntFlanks@6744.6772.29|GACACTTCGAACGCACTTGCGGCCCCGGG</td><td>>hg38_21_+_8433172_8446622_RNA45SN1_45S_50ntFlanks</td><td>6744</td><td>6772</td><td>29</td><td>GACACTTCGAACGCACTTGCGGCCCCGGG</td><td>106.0</td><td>-8.3</td><td>1.3</td><td>-6.3</td><td>3.7E-10</td><td>7.1E-10</td><td>MB231/MB468</td></tr>
</table>

<table>
<tr><th></th><th></th><th></th><th></th><th></th><th></th><th></th><th></th><th></th><th></th><th></th><th></th><th></th></tr>
<tr><td>>hg38_21_+_8433172_8446622_RNA45SN1_45S_50ntFlanks@10921.10946.26|GCTGGACGAGGCGCCGCCGCCCCCCC</td><td>>hg38_21_+_8433172_8446622_RNA45SN1_45S_50ntFlanks</td><td>10921</td><td>10946</td><td>26</td><td>GCTGGACGAGGCGCCGCCGCCCCCCC</td><td>79.5</td><td>-7.9</td><td>1.3</td><td>-6.3</td><td>3.7E-10</td><td>7.1E-10</td><td>MB231/MB468</td></tr>
<tr><td>>hg38_21_+_8433172_8446622_RNA45SN1_45S_50ntFlanks@10841.10866.26|CGTAACTTCGGGATAAGGATTGGCTC</td><td>>hg38_21_+_8433172_8446622_RNA45SN1_45S_50ntFlanks</td><td>10841</td><td>10866</td><td>26</td><td>CGTAACTTCGGGATAAGGATTGGCTC</td><td>1035.5</td><td>-3.9</td><td>0.6</td><td>-6.2</td><td>4.2E-10</td><td>7.2E-10</td><td>BT20/MB468</td></tr>
<tr><td>>hg38_21_+_8433172_8446622_RNA45SN1_45S_50ntFlanks@6773.6793.21|TTCCTCCCGGGGCTACGCCTG</td><td>>hg38_21_+_8433172_8446622_RNA45SN1_45S_50ntFlanks</td><td>6773</td><td>6793</td><td>21</td><td>TTCCTCCCGGGGCTACGCCTG</td><td>354.7</td><td>-6.3</td><td>1.0</td><td>-6.2</td><td>4.3E-10</td><td>7.2E-10</td><td>BT20/MB468</td></tr>
<tr><td>>hg38_21_+_8433172_8446622_RNA45SN1_45S_50ntFlanks@12324.12347.24|TTGGGTTTTAAGCAGGAGGTGTCA</td><td>>hg38_21_+_8433172_8446622_RNA45SN1_45S_50ntFlanks</td><td>12324</td><td>12347</td><td>24</td><td>TTGGGTTTTAAGCAGGAGGTGTCA</td><td>79.7</td><td>-7.9</td><td>1.3</td><td>-6.3</td><td>3.7E-10</td><td>7.2E-10</td><td>MB231/MB468</td></tr>
<tr><td>>hg38_21_+_8433172_8446622_RNA45SN1_45S_50ntFlanks@11787.11805.19|AATGGATGAACGAGATTCC</td><td>>hg38_21_+_8433172_8446622_RNA45SN1_45S_50ntFlanks</td><td>11787</td><td>11805</td><td>19</td><td>AATGGATGAACGAGATTCC</td><td>139.8</td><td>-9.1</td><td>1.5</td><td>-6.2</td><td>4.3E-10</td><td>7.4E-10</td><td>BT20/MB468</td></tr>
<tr><td>>hg38_21_+_8433172_8446622_RNA45SN1_45S_50ntFlanks@12073.12110.38|CCGAGGGGCTCTCGCTTCTGGCGCCAAGCGCCCGGCCG</td><td>>hg38_21_+_8433172_8446622_RNA45SN1_45S_50ntFlanks</td><td>12073</td><td>12110</td><td>38</td><td>CCGAGGGGCTCTCGCTTCTGGCGCCAAGCGCCCGGCCG</td><td>222.9</td><td>-5.6</td><td>0.9</td><td>-6.2</td><td>4.5E-10</td><td>7.6E-10</td><td>BT20/MB468</td></tr>
<tr><td>>hg38_21_+_8433172_8446622_RNA45SN1_45S_50ntFlanks@9736.9758.23|CCGAAACGATCTCAACCTATTCT</td><td>>hg38_21_+_8433172_8446622_RNA45SN1_45S_50ntFlanks</td><td>9736</td><td>9758</td><td>23</td><td>CCGAAACGATCTCAACCTATTCT</td><td>109.9</td><td>-6.5</td><td>1.0</td><td>-6.3</td><td>3.9E-10</td><td>7.6E-10</td><td>MB231/MB468</td></tr>
<tr><td>>hg38_21_+_8433172_8446622_RNA45SN1_45S_50ntFlanks@12717.12742.26|CCTCGGTTGGCCTCGGATAGCCGGTC</td><td>>hg38_21_+_8433172_8446622_RNA45SN1_45S_50ntFlanks</td><td>12717</td><td>12742</td><td>26</td><td>CCTCGGTTGGCCTCGGATAGCCGGTC</td><td>76.9</td><td>-8.3</td><td>1.3</td><td>-6.2</td><td>4.5E-10</td><td>7.6E-10</td><td>BT20/MB468</td></tr>
<tr><td>>hg38_21_+_8433172_8446622_RNA45SN1_45S_50ntFlanks@8194.8212.19|CCCGTGGACGGTGTGAGGC</td><td>>hg38_21_+_8433172_8446622_RNA45SN1_45S_50ntFlanks</td><td>8194</td><td>8212</td><td>19</td><td>CCCGTGGACGGTGTGAGGC</td><td>486.2</td><td>-4.4</td><td>0.7</td><td>-6.2</td><td>4.5E-10</td><td>7.6E-10</td><td>BT20/MB468</td></tr>
<tr><td>>hg38_21_+_8433172_8446622_RNA45SN1_45S_50ntFlanks@4473.4494.22|TTAGCTGAGTGTCCCGCGGGGC</td><td>>hg38_21_+_8433172_8446622_RNA45SN1_45S_50ntFlanks</td><td>4473</td><td>4494</td><td>22</td><td>TTAGCTGAGTGTCCCGCGGGGC</td><td>77.3</td><td>-7.8</td><td>1.3</td><td>-6.3</td><td>3.9E-10</td><td>7.6E-10</td><td>MB231/MB468</td></tr>
<tr><td>>hg38_21_+_8433172_8446622_RNA45SN1_45S_50ntFlanks@11320.11338.19|CCACGTCTCGTCGCGCGCG</td><td>>hg38_21_+_8433172_8446622_RNA45SN1_45S_50ntFlanks</td><td>11320</td><td>11338</td><td>19</td><td>CCACGTCTCGTCGCGCGCG</td><td>84.5</td><td>-8.0</td><td>1.3</td><td>-6.3</td><td>3.9E-10</td><td>7.6E-10</td><td>MB231/MB468</td></tr>
<tr><td>>MI0000102|hsa-mir-100&WithFlank&11|-|122152223|122152314@21.40.20&[MIMAT0000098&hsa-miR-100-5p&offsets|+2|0;m-27&11|-|122152276|122152296&offsets|+2|+1]|CCCGTAGATCCGAACTTGTG</td><td>>MI0000102|hsa-mir-100&WithFlank&11|-|122152223|122152314</td><td>21</td><td>40</td><td>20</td><td>CCCGTAGATCCGAACTTGTG</td><td>249.2</td><td>-6.8</td><td>1.0</td><td>-6.7</td><td>2.7E-11</td><td>7.7E-10</td><td>BT20/MB231</td></tr>
<tr><td>>hg38_21_+_8433172_8446622_RNA45SN1_45S_50ntFlanks@12003.12029.27|GCGGGCCGCCGGTGAAATACCACTACT</td><td>>hg38_21_+_8433172_8446622_RNA45SN1_45S_50ntFlanks</td><td>12003</td><td>12029</td><td>27</td><td>GCGGGCCGCCGGTGAAATACCACTACT</td><td>1057.4</td><td>5.2</td><td>0.8</td><td>6.3</td><td>4.0E-10</td><td>7.7E-10</td><td>MB231/MB468</td></tr>
<tr><td>>trna20_GluTTC_2_-_131094701_131094772@1.33.33,trna5_GluTTC_13_-_41634874_41634945@1.33.33|TCCCATATGGTCTAGCGGTTAGGATTCCTGGTT</td><td>>trna20_GluTTC_2_-_131094701_131094772</td><td>1</td><td>33</td><td>33</td><td>TCCCATATGGTCTAGCGGTTAGGATTCCTGGTT</td><td>8536.8</td><td>14.1</td><td>2.3</td><td>6.3</td><td>4.0E-10</td><td>7.7E-10</td><td>MB231/MB468</td></tr>
<tr><td>>hg38_21_+_8433172_8446622_RNA45SN1_45S_50ntFlanks@11849.11872.24|CGGGCTTGGCGGAATCAGCGGGGA</td><td>>hg38_21_+_8433172_8446622_RNA45SN1_45S_50ntFlanks</td><td>11849</td><td>11872</td><td>24</td><td>CGGGCTTGGCGGAATCAGCGGGGA</td><td>256.6</td><td>-5.6</td><td>0.9</td><td>-6.2</td><td>4.6E-10</td><td>7.8E-10</td><td>BT20/MB468</td></tr>
<tr><td>>hg38_21_+_8433172_8446622_RNA45SN1_45S_50ntFlanks@12419.12449.31|GATGTCGGCTCTTCCTATCATTGTGAAGCAG</td><td>>hg38_21_+_8433172_8446622_RNA45SN1_45S_50ntFlanks</td><td>12419</td><td>12449</td><td>31</td><td>GATGTCGGCTCTTCCTATCATTGTGAAGCAG</td><td>326.6</td><td>3.2</td><td>0.5</td><td>6.2</td><td>4.6E-10</td><td>7.8E-10</td><td>BT20/MB468</td></tr>
<tr><td>>hg38_21_+_8433172_8446622_RNA45SN1_45S_50ntFlanks@12347.12366.20|AGAAAAGTTACCACAGGGAT</td><td>>hg38_21_+_8433172_8446622_RNA45SN1_45S_50ntFlanks</td><td>12347</td><td>12366</td><td>20</td><td>AGAAAAGTTACCACAGGGAT</td><td>79.9</td><td>-7.9</td><td>1.3</td><td>-6.3</td><td>4.0E-10</td><td>7.8E-10</td><td>MB231/MB468</td></tr>
<tr><td>>hg38_21_+_8433172_8446622_RNA45SN1_45S_50ntFlanks@12356.12376.21|ACCACAGGGATAACTGGCTTG</td><td>>hg38_21_+_8433172_8446622_RNA45SN1_45S_50ntFlanks</td><td>12356</td><td>12376</td><td>21</td><td>ACCACAGGGATAACTGGCTTG</td><td>314.4</td><td>-6.1</td><td>1.0</td><td>-6.2</td><td>4.6E-10</td><td>7.8E-10</td><td>BT20/MB468</td></tr>
<tr><td>>hg38_21_+_8433172_8446622_RNA45SN1_45S_50ntFlanks@10681.10712.32|TTCCGGCGGCGTCCGGTGAGCTCTCGCTGGCC</td><td>>hg38_21_+_8433172_8446622_RNA45SN1_45S_50ntFlanks</td><td>10681</td><td>10712</td><td>32</td><td>TTCCGGCGGCGTCCGGTGAGCTCTCGCTGGCC</td><td>1341.1</td><td>-5.4</td><td>0.9</td><td>-6.2</td><td>4.6E-10</td><td>7.8E-10</td><td>BT20/MB468</td></tr>
<tr><td>>hg38_21_+_8433172_8446622_RNA45SN1_45S_50ntFlanks@11573.11603.31|CGACTTAGAACTGGTGCGGACCAGGGGAATC</td><td>>hg38_21_+_8433172_8446622_RNA45SN1_45S_50ntFlanks</td><td>11573</td><td>11603</td><td>31</td><td>CGACTTAGAACTGGTGCGGACCAGGGGAATC</td><td>480.4</td><td>-3.2</td><td>0.5</td><td>-6.2</td><td>4.6E-10</td><td>7.8E-10</td><td>BT20/MB468</td></tr>
<tr><td>>hg38_21_+_8433172_8446622_RNA45SN1_45S_50ntFlanks@12994.13012.19|CCTCGCTGCGATCTATTGA</td><td>>hg38_21_+_8433172_8446622_RNA45SN1_45S_50ntFlanks</td><td>12994</td><td>13012</td><td>19</td><td>CCTCGCTGCGATCTATTGA</td><td>204.9</td><td>-2.8</td><td>0.4</td><td>-6.2</td><td>4.6E-10</td><td>7.8E-10</td><td>BT20/MB468</td></tr>
<tr><td>>hg38_21_+_8433172_8446622_RNA45SN1_45S_50ntFlanks@12957.12985.29|CTTCTGGGTCGGGGTTTCGTACGTAGCAG</td><td>>hg38_21_+_8433172_8446622_RNA45SN1_45S_50ntFlanks</td><td>12957</td><td>12985</td><td>29</td><td>CTTCTGGGTCGGGGTTTCGTACGTAGCAG</td><td>94.4</td><td>-5.0</td><td>0.8</td><td>-6.2</td><td>4.6E-10</td><td>7.8E-10</td><td>BT20/MB468</td></tr>
<tr><td>>hg38_21_+_8433172_8446622_RNA45SN1_45S_50ntFlanks@12995.13038.44|CTCGCTGCGATCTATTGAAAGTCAGCCCTCGACACAAGGGTTTG</td><td>>hg38_21_+_8433172_8446622_RNA45SN1_45S_50ntFlanks</td><td>12995</td><td>13038</td><td>44</td><td>CTCGCTGCGATCTATTGAAAGTCAGCCCTCGACACAAGGGTTTG</td><td>1543.0</td><td>9.8</td><td>1.6</td><td>6.2</td><td>4.6E-10</td><td>7.8E-10</td><td>BT20/MB468</td></tr>
<tr><td>>hg38_1_-_228634819_228635039_RNA5S12_5S_50ntFlanks@140.169.30|ACCGCCTGGGAATACCGGGTGCTGTAGGCT</td><td>>hg38_1_-_228634819_228635039_RNA5S12_5S_50ntFlanks</td><td>140</td><td>169</td><td>30</td><td>ACCGCCTGGGAATACCGGGTGCTGTAGGCT</td><td>8168.8</td><td>4.6</td><td>0.7</td><td>6.3</td><td>4.1E-10</td><td>7.9E-10</td><td>MB231/MB468</td></tr>
<tr><td>>hg38_21_+_8433172_8446622_RNA45SN1_45S_50ntFlanks@12457.12474.18|CAAGCGTTGGATTGTTCA</td><td>>hg38_21_+_8433172_8446622_RNA45SN1_45S_50ntFlanks</td><td>12457</td><td>12474</td><td>18</td><td>CAAGCGTTGGATTGTTCA</td><td>777.4</td><td>-4.9</td><td>0.8</td><td>-6.2</td><td>4.7E-10</td><td>7.9E-10</td><td>BT20/MB468</td></tr>
<tr><td>>hg38_21_+_8433172_8446622_RNA45SN1_45S_50ntFlanks@12626.12652.27|AATGGGGCGAAGCTACCATCTGTGGGA</td><td>>hg38_21_+_8433172_8446622_RNA45SN1_45S_50ntFlanks</td><td>12626</td><td>12652</td><td>27</td><td>AATGGGGCGAAGCTACCATCTGTGGGA</td><td>211.9</td><td>-3.3</td><td>0.5</td><td>-6.3</td><td>4.1E-10</td><td>7.9E-10</td><td>MB231/MB468</td></tr>
<tr><td>>hg38_21_+_8433172_8446622_RNA45SN1_45S_50ntFlanks@10510.10533.24|AATCCGGAGTGGCGGAGATGGGCG</td><td>>hg38_21_+_8433172_8446622_RNA45SN1_45S_50ntFlanks</td><td>10510</td><td>10533</td><td>24</td><td>AATCCGGAGTGGCGGAGATGGGCG</td><td>682.3</td><td>-6.9</td><td>1.1</td><td>-6.2</td><td>4.7E-10</td><td>8.0E-10</td><td>BT20/MB468</td></tr>
<tr><td>>hg38_21_+_8433172_8446622_RNA45SN1_45S_50ntFlanks@12475.12498.24|CCCACTAATAGGGAACGTGAGCTG</td><td>>hg38_21_+_8433172_8446622_RNA45SN1_45S_50ntFlanks</td><td>12475</td><td>12498</td><td>24</td><td>CCCACTAATAGGGAACGTGAGCTG</td><td>598.6</td><td>-4.8</td><td>0.8</td><td>-6.2</td><td>4.7E-10</td><td>8.0E-10</td><td>BT20/MB468</td></tr>
<tr><td>>hg38_21_+_8433172_8446622_RNA45SN1_45S_50ntFlanks@6725.6747.23|GCAGGACACATTGATCATCGACA</td><td>>hg38_21_+_8433172_8446622_RNA45SN1_45S_50ntFlanks</td><td>6725</td><td>6747</td><td>23</td><td>GCAGGACACATTGATCATCGACA</td><td>340.2</td><td>4.0</td><td>0.6</td><td>6.2</td><td>4.8E-10</td><td>8.1E-10</td><td>BT20/MB468</td></tr>
<tr><td>>hg38_MT_+_1621_3279_RNR2_16S_50ntFlanks@1370.1396.27|ATGTTGGATCAGGACATCCCGATGGTG</td><td>>hg38_MT_+_1621_3279_RNR2_16S_50ntFlanks</td><td>1370</td><td>1396</td><td>27</td><td>ATGTTGGATCAGGACATCCCGATGGTG</td><td>1514.6</td><td>8.7</td><td>1.4</td><td>6.2</td><td>4.2E-10</td><td>8.1E-10</td><td>MB231/MB468</td></tr>
<tr><td>>hg38_21_+_8433172_8446622_RNA45SN1_45S_50ntFlanks@12934.12965.32|CTAAACCATTCGTAGACGACCTGCTTCTGGGT</td><td>>hg38_21_+_8433172_8446622_RNA45SN1_45S_50ntFlanks</td><td>12934</td><td>12965</td><td>32</td><td>CTAAACCATTCGTAGACGACCTGCTTCTGGGT</td><td>361.5</td><td>-3.6</td><td>0.6</td><td>-6.2</td><td>4.2E-10</td><td>8.1E-10</td><td>MB231/MB468</td></tr>
<tr><td>>hg38_21_+_8433172_8446622_RNA45SN1_45S_50ntFlanks@6652.6697.46|GACTCTTAGCGGTGGATCACTCGGCTCGTGCGTCGATGAAGAACGC</td><td>>hg38_21_+_8433172_8446622_RNA45SN1_45S_50ntFlanks</td><td>6652</td><td>6697</td><td>46</td><td>GACTCTTAGCGGTGGATCACTCGGCTCGTGCGTCGATGAAGAACGC</td><td>2224.8</td><td>4.6</td><td>0.7</td><td>6.2</td><td>4.2E-10</td><td>8.2E-10</td><td>MB231/MB468</td></tr>
<tr><td>>hg38_21_+_8433172_8446622_RNA45SN1_45S_50ntFlanks@8325.8356.32|CGATAGTCAACAAGTACCGTAAGGGAAAGTTG</td><td>>hg38_21_+_8433172_8446622_RNA45SN1_45S_50ntFlanks</td><td>8325</td><td>8356</td><td>32</td><td>CGATAGTCAACAAGTACCGTAAGGGAAAGTTG</td><td>82.1</td><td>-7.9</td><td>1.3</td><td>-6.2</td><td>4.2E-10</td><td>8.2E-10</td><td>MB231/MB468</td></tr>
<tr><td>>MI0000447|hsa-mir-128-1&WithFlank&2|+|135665391|135665484@56.76.21(+2A);MI0000727|hsa-mir-128-2&WithFlank&3|+|35744470|35744565@58.78.21(+2A)&[MIMAT0000424&hsa-miR-128-3p&offsets|0|0(+2A);m-108&2|+|135665446|135665466&offsets|0|0(+2A)];[MIMAT0000424_1&hsa-miR-128-3p&offsets|0|0(+2A);m-117&3|+|35744527|35744547&offsets|0|0(+2A)]|TCACAGTGAACCGGTCTCTTTAA</td><td>>MI0000447|hsa-mir-128-1&WithFlank&2|+|135665391|135665484</td><td>56</td><td>76</td><td>21(+2A)</td><td>TCACAGTGAACCGGTCTCTTTAA</td><td>1225.8</td><td>4.7</td><td>0.7</td><td>6.2</td><td>4.2E-10</td><td>8.2E-10</td><td>MB231/MB468</td></tr>
<tr><td>>hg38_21_+_8433172_8446622_RNA45SN1_45S_50ntFlanks@10571.10589.19|GAGAAGCCGGCGGGAGCCC</td><td>>hg38_21_+_8433172_8446622_RNA45SN1_45S_50ntFlanks</td><td>10571</td><td>10589</td><td>19</td><td>GAGAAGCCGGCGGGAGCCC</td><td>87.2</td><td>-8.0</td><td>1.3</td><td>-6.2</td><td>4.2E-10</td><td>8.2E-10</td><td>MB231/MB468</td></tr>
<tr><td>>hg38_21_+_8433172_8446622_RNA45SN1_45S_50ntFlanks@10694.10711.18|CGGTGAGCTCTCGCTGGC</td><td>>hg38_21_+_8433172_8446622_RNA45SN1_45S_50ntFlanks</td><td>10694</td><td>10711</td><td>18</td><td>CGGTGAGCTCTCGCTGGC</td><td>412.3</td><td>-6.2</td><td>1.0</td><td>-6.2</td><td>4.9E-10</td><td>8.2E-10</td><td>BT20/MB468</td></tr>
<tr><td>>hg38_21_+_8433172_8446622_RNA45SN1_45S_50ntFlanks@12476.12496.21|CCACTAATAGGGAACGTGAGC</td><td>>hg38_21_+_8433172_8446622_RNA45SN1_45S_50ntFlanks</td><td>12476</td><td>12496</td><td>21</td><td>CCACTAATAGGGAACGTGAGC</td><td>1784.9</td><td>-3.8</td><td>0.6</td><td>-6.2</td><td>4.9E-10</td><td>8.2E-10</td><td>BT20/MB468</td></tr>
<tr><td>>hg38_21_+_8433172_8446622_RNA45SN1_45S_50ntFlanks@6735.6771.37|TTGATCATCGACACTTCGAACGCACTTGCGGCCCCGG</td><td>>hg38_21_+_8433172_8446622_RNA45SN1_45S_50ntFlanks</td><td>6735</td><td>6771</td><td>37</td><td>TTGATCATCGACACTTCGAACGCACTTGCGGCCCCGG</td><td>178.9</td><td>-3.6</td><td>0.6</td><td>-6.2</td><td>4.9E-10</td><td>8.2E-10</td><td>BT20/MB468</td></tr>
<tr><td>>hg38_21_+_8433172_8446622_RNA45SN1_45S_50ntFlanks@7987.8006.20|TCAGACGTGGCGACCCGCTG</td><td>>hg38_21_+_8433172_8446622_RNA45SN1_45S_50ntFlanks</td><td>7987</td><td>8006</td><td>20</td><td>TCAGACGTGGCGACCCGCTG</td><td>110.4</td><td>-5.5</td><td>0.9</td><td>-6.2</td><td>4.3E-10</td><td>8.2E-10</td><td>MB231/MB468</td></tr>
<tr><td>>hg38_21_+_8433172_8446622_RNA45SN1_45S_50ntFlanks@10099.10127.29|CGCGCGCGCGTGTGGTGTGCGTCGGAGGG</td><td>>hg38_21_+_8433172_8446622_RNA45SN1_45S_50ntFlanks</td><td>10099</td><td>10127</td><td>29</td><td>CGCGCGCGCGTGTGGTGTGCGTCGGAGGG</td><td>92.8</td><td>-8.1</td><td>1.3</td><td>-6.2</td><td>4.3E-10</td><td>8.4E-10</td><td>MB231/MB468</td></tr>
<tr><td>>hg38_21_+_8433172_8446622_RNA45SN1_45S_50ntFlanks@11880.11904.25|CCCTGTTGAGCTTGACTCTAGTCTG</td><td>>hg38_21_+_8433172_8446622_RNA45SN1_45S_50ntFlanks</td><td>11880</td><td>11904</td><td>25</td><td>CCCTGTTGAGCTTGACTCTAGTCTG</td><td>78.0</td><td>-7.9</td><td>1.3</td><td>-6.2</td><td>4.4E-10</td><td>8.5E-10</td><td>MB231/MB468</td></tr>
<tr><td>>hg38_21_+_8433172_8446622_RNA45SN1_45S_50ntFlanks@12893.12933.41|CCTCGCCCGTCACGCACCGCACGTTCGTGGGGAACCTGGCG</td><td>>hg38_21_+_8433172_8446622_RNA45SN1_45S_50ntFlanks</td><td>12893</td><td>12933</td><td>41</td><td>CCTCGCCCGTCACGCACCGCACGTTCGTGGGGAACCTGGCG</td><td>334.4</td><td>-6.2</td><td>1.0</td><td>-6.2</td><td>5.0E-10</td><td>8.5E-10</td><td>BT20/MB468</td></tr>
<tr><td>>trna15_ValAAC_5_-_180615416_180615488@1.29.29|GTTTCCGTAGTGTAGTGGTCATCACGTTC</td><td>>trna15_ValAAC_5_-_180615416_180615488</td><td>1</td><td>29</td><td>29</td><td>GTTTCCGTAGTGTAGTGGTCATCACGTTC</td><td>1724.2</td><td>-5.5</td><td>0.9</td><td>-6.2</td><td>4.4E-10</td><td>8.6E-10</td><td>MB231/MB468</td></tr>
<tr><td>>hg38_21_+_8433172_8446622_RNA45SN1_45S_50ntFlanks@3866.3898.33|CTAGAGCTAATACATGCCGACGGGCGCTGACCC</td><td>>hg38_21_+_8433172_8446622_RNA45SN1_45S_50ntFlanks</td><td>3866</td><td>3898</td><td>33</td><td>CTAGAGCTAATACATGCCGACGGGCGCTGACCC</td><td>137.8</td><td>-5.2</td><td>0.8</td><td>-6.2</td><td>4.4E-10</td><td>8.6E-10</td><td>MB231/MB468</td></tr>
<tr><td>>MI0000477|hsa-mir-146a&WithFlank&5|+|160485346|160485456@27.47.21&[MIMAT0000449&hsa-miR-146a-5p&offsets|0|-1;m-129&5|+|160485372|160485393&offsets|0|-1]|TGAGAACTGAATTCCATGGGT</td><td>>MI0000477|hsa-mir-146a&WithFlank&5|+|160485346|160485456</td><td>27</td><td>47</td><td>21</td><td>TGAGAACTGAATTCCATGGGT</td><td>6772.9</td><td>14.0</td><td>2.2</td><td>6.2</td><td>4.4E-10</td><td>8.6E-10</td><td>MB231/MB468</td></tr>
<tr><td>>hg38_21_+_8433172_8446622_RNA45SN1_45S_50ntFlanks@12128.12146.19|ACCCGCTCCGGGGACAGTG</td><td>>hg38_21_+_8433172_8446622_RNA45SN1_45S_50ntFlanks</td><td>12128</td><td>12146</td><td>19</td><td>ACCCGCTCCGGGGACAGTG</td><td>75.8</td><td>-7.8</td><td>1.3</td><td>-6.2</td><td>4.5E-10</td><td>8.6E-10</td><td>MB231/MB468</td></tr>
<tr><td>>hg38_21_+_8433172_8446622_RNA45SN1_45S_50ntFlanks@9568.9586.19|CGTGCAAATCGGTCGTCCG</td><td>>hg38_21_+_8433172_8446622_RNA45SN1_45S_50ntFlanks</td><td>9568</td><td>9586</td><td>19</td><td>CGTGCAAATCGGTCGTCCG</td><td>237.8</td><td>-5.8</td><td>0.9</td><td>-6.2</td><td>5.1E-10</td><td>8.7E-10</td><td>BT20/MB468</td></tr>
<tr><td>>hg38_21_+_8433172_8446622_RNA45SN1_45S_50ntFlanks@4560.4578.19|CTAGGAATAATGGAATAGG</td><td>>hg38_21_+_8433172_8446622_RNA45SN1_45S_50ntFlanks</td><td>4560</td><td>4578</td><td>19</td><td>CTAGGAATAATGGAATAGG</td><td>120.1</td><td>-4.8</td><td>0.8</td><td>-6.2</td><td>4.5E-10</td><td>8.7E-10</td><td>MB231/MB468</td></tr>
</table>

| | | | | | | | | | | | | |
|---|---|---|---|---|---|---|---|---|---|---|---|---|
| >hg38_21_+_8433172_8446622_RNA45SN1_45S_50ntFlanks@11161.11189.29\|CACGGTCCCCCGCGAGGGGGGCCCGGGCA | >hg38_21_+_8433172_8446622_RNA45SN1_45S_50ntFlanks | 11161 | 11189 | 29 | CACGGTCCCCCGCGAGGGGGGCCCGGGCA | 91.6 | -8.1 | 1.3 | -6.2 | 4.5E-10 | 8.7E-10 | MB231/MB468 |
| >hg38_1_-_228634819_228635039_RNA5S12_5S_50ntFlanks@141.169.29\|CCGCCTGGGAATACCGGGTGCTGTAGGCT | >hg38_1_-_228634819_228635039_RNA5S12_5S_50ntFlanks | 141 | 169 | 29 | CCGCCTGGGAATACCGGGTGCTGTAGGCT | 2335.2 | 5.0 | 0.8 | 6.2 | 5.2E-10 | 8.8E-10 | BT20/MB468 |
| >hg38_21_+_8433172_8446622_RNA45SN1_45S_50ntFlanks@3773.3792.20\|CGGCCGGTACAGTGAAACTG | >hg38_21_+_8433172_8446622_RNA45SN1_45S_50ntFlanks | 3773 | 3792 | 20 | CGGCCGGTACAGTGAAACTG | 204.7 | -3.1 | 0.5 | -6.2 | 4.5E-10 | 8.8E-10 | MB231/MB468 |
| >hg38_21_+_8433172_8446622_RNA45SN1_45S_50ntFlanks@4728.4747.20\|ATCAAGAACGAAAGTCGGAG | >hg38_21_+_8433172_8446622_RNA45SN1_45S_50ntFlanks | 4728 | 4747 | 20 | ATCAAGAACGAAAGTCGGAG | 70.4 | -8.1 | 1.3 | -6.2 | 5.2E-10 | 8.8E-10 | BT20/MB468 |
| >hg38_21_+_8433172_8446622_RNA45SN1_45S_50ntFlanks@4732.4761.30\|AGAACGAAAGTCGGAGGTTCGAAGACGATC | >hg38_21_+_8433172_8446622_RNA45SN1_45S_50ntFlanks | 4732 | 4761 | 30 | AGAACGAAAGTCGGAGGTTCGAAGACGATC | 380.1 | 4.2 | 0.7 | 6.2 | 5.2E-10 | 8.8E-10 | BT20/MB468 |
| >hg38_21_+_8433172_8446622_RNA45SN1_45S_50ntFlanks@11517.11539.23\|CCGCCCCCGGGGCCGCGGTTCCG | >hg38_21_+_8433172_8446622_RNA45SN1_45S_50ntFlanks | 11517 | 11539 | 23 | CCGCCCCCGGGGCCGCGGTTCCG | 79.1 | -7.9 | 1.3 | -6.2 | 4.6E-10 | 8.8E-10 | MB231/MB468 |
| >hg38_21_+_8433172_8446622_RNA45SN1_45S_50ntFlanks@10684.10709.26\|CGGCGGCGTCCGGTGAGCTCTCGCTG | >hg38_21_+_8433172_8446622_RNA45SN1_45S_50ntFlanks | 10684 | 10709 | 26 | CGGCGGCGTCCGGTGAGCTCTCGCTG | 2221.6 | -4.6 | 0.7 | -6.2 | 5.3E-10 | 8.9E-10 | BT20/MB468 |
| >hg38_21_+_8433172_8446622_RNA45SN1_45S_50ntFlanks@10819.10838.20\|AGGGAAGTCGGCAAGCCGGA | >hg38_21_+_8433172_8446622_RNA45SN1_45S_50ntFlanks | 10819 | 10838 | 20 | AGGGAAGTCGGCAAGCCGGA | 177.8 | -5.9 | 0.9 | -6.2 | 5.3E-10 | 8.9E-10 | BT20/MB468 |
| >hg38_21_+_8433172_8446622_RNA45SN1_45S_50ntFlanks@7986.8003.18\|ATCAGACGTGGCGACCCG | >hg38_21_+_8433172_8446622_RNA45SN1_45S_50ntFlanks | 7986 | 8003 | 18 | ATCAGACGTGGCGACCCG | 745.7 | -6.5 | 1.0 | -6.2 | 4.6E-10 | 9.0E-10 | MB231/MB468 |
| >hg38_21_+_8433172_8446622_RNA45SN1_45S_50ntFlanks@7986.8007.22\|ATCAGACGTGGCGACCCGCTGA | >hg38_21_+_8433172_8446622_RNA45SN1_45S_50ntFlanks | 7986 | 8007 | 22 | ATCAGACGTGGCGACCCGCTGA | 89.5 | -8.1 | 1.3 | -6.2 | 4.7E-10 | 9.0E-10 | MB231/MB468 |
| >hg38_21_+_8433172_8446622_RNA45SN1_45S_50ntFlanks@12970.12990.21\|GTTTCGTACGTAGCAGAGCAG | >hg38_21_+_8433172_8446622_RNA45SN1_45S_50ntFlanks | 12970 | 12990 | 21 | GTTTCGTACGTAGCAGAGCAG | 131.9 | -4.2 | 0.7 | -6.2 | 4.7E-10 | 9.0E-10 | MB231/MB468 |
| >hg38_21_+_8433172_8446622_RNA45SN1_45S_50ntFlanks@5423.5445.23\|ATTGGATGGTTTAGTGAGGCCCT | >hg38_21_+_8433172_8446622_RNA45SN1_45S_50ntFlanks | 5423 | 5445 | 23 | ATTGGATGGTTTAGTGAGGCCCT | 139.7 | -5.5 | 0.9 | -6.2 | 5.4E-10 | 9.1E-10 | BT20/MB468 |
| >hg38_21_+_8433172_8446622_RNA45SN1_45S_50ntFlanks@4657.4686.30\|CTAGAGGTGAAATTCTTGGACCGGCGCAAG | >hg38_21_+_8433172_8446622_RNA45SN1_45S_50ntFlanks | 4657 | 4686 | 30 | CTAGAGGTGAAATTCTTGGACCGGCGCAAG | 111.6 | -8.4 | 1.3 | -6.2 | 4.7E-10 | 9.1E-10 | MB231/MB468 |
| >hg38_21_+_8433172_8446622_RNA45SN1_45S_50ntFlanks@11703.11726.24\|AATGAAGCGCGGGTAAACGGCGGG | >hg38_21_+_8433172_8446622_RNA45SN1_45S_50ntFlanks | 11703 | 11726 | 24 | AATGAAGCGCGGGTAAACGGCGGG | 216.5 | -3.7 | 0.6 | -6.2 | 5.4E-10 | 9.1E-10 | BT20/MB468 |
| >hg38_MT_+_1621_3279_RNR2_16S_50ntFlanks@493.518.26\|GCTCTTTGGACACTAGGAAAAAACCT | >hg38_MT_+_1621_3279_RNR2_16S_50ntFlanks | 493 | 518 | 26 | GCTCTTTGGACACTAGGAAAAAACCT | 464.6 | 7.7 | 1.2 | 6.2 | 5.5E-10 | 9.2E-10 | BT20/MB468 |
| >MI0000447\|hsa-mir-128-1&WithFlank&2\|+\|135665391\|135665484@56.76.21(+1A);MI0000727\|hsa-mir-128-2&WithFlank&3\|+\|35744470\|35744565@58.78.21(+1A)&[MIMAT0000424&hsa-miR-128-3p&offsets\|0\|0(+1A);m-108&2\|+\|135665446\|135665466&offsets\|0\|0(+1A)];[MIMAT0000424_1&hsa-miR-128-3p&offsets\|0\|0(+1A);m-117&3\|+\|35744527\|35744547&offsets\|0\|0(+1A)]\|TCACAGTGAACCGGTCTCTTTA | >MI0000447\|hsa-mir-128-1&WithFlank&2\|+\|135665391\|135665484 | 56 | 76 | 21(+1A) | TCACAGTGAACCGGTCTCTTTA | 1714.4 | 4.5 | 0.7 | 6.2 | 4.8E-10 | 9.3E-10 | MB231/MB468 |
| >hg38_21_+_8433172_8446622_RNA45SN1_45S_50ntFlanks@10421.10457.37\|TGAGAGATGGGCGAGCGCCGTTCCGAAGGGACGGGCG | >hg38_21_+_8433172_8446622_RNA45SN1_45S_50ntFlanks | 10421 | 10457 | 37 | TGAGAGATGGGCGAGCGCCGTTCCGAAGGGACGGGCG | 109.9 | -6.5 | 1.1 | -6.2 | 4.9E-10 | 9.4E-10 | MB231/MB468 |
| >hg38_21_+_8433172_8446622_RNA45SN1_45S_50ntFlanks@9572.9592.21\|CAAATCGGTCGTCCGACCTGG | >hg38_21_+_8433172_8446622_RNA45SN1_45S_50ntFlanks | 9572 | 9592 | 21 | CAAATCGGTCGTCCGACCTGG | 156.2 | -6.2 | 1.0 | -6.2 | 4.9E-10 | 9.5E-10 | MB231/MB468 |
| >hg38_21_+_8433172_8446622_RNA45SN1_45S_50ntFlanks@11076.11103.28\|CGCCGCGTGGGGGCGGCGGCGGGGGGAG | >hg38_21_+_8433172_8446622_RNA45SN1_45S_50ntFlanks | 11076 | 11103 | 28 | CGCCGCGTGGGGGCGGCGGCGGGGGGAG | 102.2 | -8.3 | 1.3 | -6.2 | 4.9E-10 | 9.5E-10 | MB231/MB468 |
| >hg38_21_+_8433172_8446622_RNA45SN1_45S_50ntFlanks@11212.11230.19\|CGACTCTGGACGCGAGCCG | >hg38_21_+_8433172_8446622_RNA45SN1_45S_50ntFlanks | 11212 | 11230 | 19 | CGACTCTGGACGCGAGCCG | 129.8 | -6.7 | 1.1 | -6.2 | 4.9E-10 | 9.5E-10 | MB231/MB468 |
| >hg38_21_+_8433172_8446622_RNA45SN1_45S_50ntFlanks@3985.4026.42\|CGGCTTTGGTGACTCTAGATAACCTCGGGCCGATCGCACGCC | >hg38_21_+_8433172_8446622_RNA45SN1_45S_50ntFlanks | 3985 | 4026 | 42 | CGGCTTTGGTGACTCTAGATAACCTCGGGCCGATCGCACGCC | 511.7 | -8.4 | 1.4 | -6.2 | 5.0E-10 | 9.6E-10 | MB231/MB468 |
| >hg38_21_+_8433172_8446622_RNA45SN1_45S_50ntFlanks@10681.10708.28\|TTCCGGCGGCGTCCGGTGAGCTCTCGCT | >hg38_21_+_8433172_8446622_RNA45SN1_45S_50ntFlanks | 10681 | 10708 | 28 | TTCCGGCGGCGTCCGGTGAGCTCTCGCT | 417.3 | -4.0 | 0.6 | -6.2 | 5.7E-10 | 9.7E-10 | BT20/MB468 |
| >hg38_21_+_8433172_8446622_RNA45SN1_45S_50ntFlanks@7983.8003.21\|CAGATCAGACGTGGCGACCCG | >hg38_21_+_8433172_8446622_RNA45SN1_45S_50ntFlanks | 7983 | 8003 | 21 | CAGATCAGACGTGGCGACCCG | 271.5 | -3.9 | 0.6 | -6.2 | 5.8E-10 | 9.7E-10 | BT20/MB468 |
| >MI0000085\|hsa-mir-27a&WithFlank&19\|-\|13836434\|13836523@57.75.19(+1A)&[MIMAT0000084&hsa-miR-27a-3p&offsets\|0\|-2(+1A);m-43&19\|-\|13836447\|13836467&offsets\|0\|-2(+1A)]\|TTCACAGTGGCTAAGTTCCA | >MI0000085\|hsa-mir-27a&WithFlank&19\|-\|13836434\|13836523 | 57 | 75 | 19(+1A) | TTCACAGTGGCTAAGTTCCA | 4111.5 | 5.6 | 0.9 | 6.2 | 5.8E-10 | 9.8E-10 | BT20/MB468 |
| >hg38_21_+_8433172_8446622_RNA45SN1_45S_50ntFlanks@12654.12673.20\|TATGACTGAACGCCTCTAAG | >hg38_21_+_8433172_8446622_RNA45SN1_45S_50ntFlanks | 12654 | 12673 | 20 | TATGACTGAACGCCTCTAAG | 89.1 | -8.1 | 1.3 | -6.2 | 5.1E-10 | 9.9E-10 | MB231/MB468 |
| >hg38_21_+_8433172_8446622_RNA45SN1_45S_50ntFlanks@10727.10745.19\|GGAGAGGGTGTAAATCTCG | >hg38_21_+_8433172_8446622_RNA45SN1_45S_50ntFlanks | 10727 | 10745 | 19 | GGAGAGGGTGTAAATCTCG | 149.7 | -5.6 | 0.9 | -6.2 | 5.9E-10 | 9.9E-10 | BT20/MB468 |
| >hg38_21_+_8433172_8446622_RNA45SN1_45S_50ntFlanks@12081.12107.27\|CTCTCGCTTCTGGCGCCAAGCGCCCGG | >hg38_21_+_8433172_8446622_RNA45SN1_45S_50ntFlanks | 12081 | 12107 | 27 | CTCTCGCTTCTGGCGCCAAGCGCCCGG | 429.9 | -4.3 | 0.7 | -6.2 | 5.9E-10 | 9.9E-10 | BT20/MB468 |
| >hg38_21_+_8433172_8446622_RNA45SN1_45S_50ntFlanks@3738.3756.19\|TCTCAAAGATTAAGCCATG | >hg38_21_+_8433172_8446622_RNA45SN1_45S_50ntFlanks | 3738 | 3756 | 19 | TCTCAAAGATTAAGCCATG | 82.8 | -7.9 | 1.3 | -6.2 | 5.2E-10 | 9.9E-10 | MB231/MB468 |
| >hg38_21_+_8433172_8446622_RNA45SN1_45S_50ntFlanks@10834.10867.34\|CCGGATCCGTAACTTCGGGATAAGGATTGGCTCT | >hg38_21_+_8433172_8446622_RNA45SN1_45S_50ntFlanks | 10834 | 10867 | 34 | CCGGATCCGTAACTTCGGGATAAGGATTGGCTCT | 599.9 | 6.7 | 1.1 | 6.2 | 5.2E-10 | 9.9E-10 | MB231/MB468 |
| >MI0003686\|hsa-mir-542&WithFlank&X\|-\|134541335\|134541443@59.80.22&[MIMAT0003389&hsa-miR-542-3p&offsets\|0\|0;m-136&X\|-\|134541364\|134541385&offsets\|0\|0]\|TGTGACAGATTGATAACTGAAA | >MI0003686\|hsa-mir-542&WithFlank&X\|-\|134541335\|134541443 | 59 | 80 | 22 | TGTGACAGATTGATAACTGAAA | 3995.3 | 16.6 | 2.7 | 6.2 | 5.9E-10 | 1.0E-09 | BT20/MB468 |
| >hg38_21_+_8433172_8446622_RNA45SN1_45S_50ntFlanks@8169.8188.20\|CCAAGTCCTTCTGATCGAGG | >hg38_21_+_8433172_8446622_RNA45SN1_45S_50ntFlanks | 8169 | 8188 | 20 | CCAAGTCCTTCTGATCGAGG | 164.7 | -8.9 | 1.4 | -6.2 | 5.2E-10 | 1.0E-09 | MB231/MB468 |
| >trna116_GluCTC_1_-_145399233_145399304@37.73.37,trna134_GluTTC_1_-_16861774_16861845@37.73.37,trna71_GluCTC_1_-_161439189_161439260@37.73.37,trna74_GluCTC_1_-_161431809_161431880@37.73.37,trna77_GluCTC_1_-_161424398_161424469@37.73.37,trna77_GluCTC_6_+_28949976_28950047@37.73.37,trna80_GluCTC_1_-_161417018_161417089@37.73.37,trna84_GluTTC_1_-_161391883_161391954@37.73.37,trna87_GluCTC_6_-_126101393_126101464@37.73.37\|ACCGCCGCGGCCCGGGTTCGATTCCCGGTCAGGGAAC | >trna116_GluCTC_1_-_145399233_145399304 | 37 | 73 | 37 | ACCGCCGCGGCCCGGGTTCGATTCCCGGTCAGGGAAC | 2308.6 | 7.0 | 1.1 | 6.2 | 5.2E-10 | 1.0E-09 | MB231/MB468 |
| >hg38_21_+_8433172_8446622_RNA45SN1_45S_50ntFlanks@4367.4390.24\|CAGTTAAAAAGCTCGTAGTTGGAT | >hg38_21_+_8433172_8446622_RNA45SN1_45S_50ntFlanks | 4367 | 4390 | 24 | CAGTTAAAAAGCTCGTAGTTGGAT | 73.8 | -4.8 | 0.8 | -6.2 | 6.0E-10 | 1.0E-09 | BT20/MB468 |
| >hg38_21_+_8433172_8446622_RNA45SN1_45S_50ntFlanks@12510.12538.29\|TCGTGAGACAGGTTAGTTTTACCCTACTG | >hg38_21_+_8433172_8446622_RNA45SN1_45S_50ntFlanks | 12510 | 12538 | 29 | TCGTGAGACAGGTTAGTTTTACCCTACTG | 92.2 | -8.1 | 1.3 | -6.2 | 5.2E-10 | 1.0E-09 | MB231/MB468 |
| >hg38_21_+_8433172_8446622_RNA45SN1_45S_50ntFlanks@5420.5438.19\|CCGATTGGATGGTTTAGTG | >hg38_21_+_8433172_8446622_RNA45SN1_45S_50ntFlanks | 5420 | 5438 | 19 | CCGATTGGATGGTTTAGTG | 100.7 | -8.2 | 1.3 | -6.2 | 5.2E-10 | 1.0E-09 | MB231/MB468 |
| >hg38_21_+_8433172_8446622_RNA45SN1_45S_50ntFlanks@10388.10409.22\|GTGAACAGCAGTTGAACATGGG | >hg38_21_+_8433172_8446622_RNA45SN1_45S_50ntFlanks | 10388 | 10409 | 22 | GTGAACAGCAGTTGAACATGGG | 93.5 | -8.1 | 1.3 | -6.2 | 5.2E-10 | 1.0E-09 | MB231/MB468 |
| >hg38_21_+_8433172_8446622_RNA45SN1_45S_50ntFlanks@10690.10711.22\|CGTCCGGTGAGCTCTCGCTGGC | >hg38_21_+_8433172_8446622_RNA45SN1_45S_50ntFlanks | 10690 | 10711 | 22 | CGTCCGGTGAGCTCTCGCTGGC | 85.1 | -8.0 | 1.3 | -6.2 | 5.3E-10 | 1.0E-09 | MB231/MB468 |
| >hg38_21_+_8433172_8446622_RNA45SN1_45S_50ntFlanks@10402.10419.18\|AACATGGGTCAGTCGGTC | >hg38_21_+_8433172_8446622_RNA45SN1_45S_50ntFlanks | 10402 | 10419 | 18 | AACATGGGTCAGTCGGTC | 545.8 | -4.2 | 0.7 | -6.2 | 5.3E-10 | 1.0E-09 | MB231/MB468 |
| >hg38_21_+_8433172_8446622_RNA45SN1_45S_50ntFlanks@11581.11599.19\|AACTGGTGCGGACCAGGGG | >hg38_21_+_8433172_8446622_RNA45SN1_45S_50ntFlanks | 11581 | 11599 | 19 | AACTGGTGCGGACCAGGGG | 84.4 | -5.4 | 0.9 | -6.2 | 5.3E-10 | 1.0E-09 | MB231/MB468 |
| >hg38_21_+_8433172_8446622_RNA45SN1_45S_50ntFlanks@10689.10711.23\|GCGTCCGGTGAGCTCTCGCTGGC | >hg38_21_+_8433172_8446622_RNA45SN1_45S_50ntFlanks | 10689 | 10711 | 23 | GCGTCCGGTGAGCTCTCGCTGGC | 85.6 | -8.0 | 1.3 | -6.2 | 5.3E-10 | 1.0E-09 | MB231/MB468 |
| >hg38_21_+_8433172_8446622_RNA45SN1_45S_50ntFlanks@6666.6689.24\|GATCACTCGGCTCGTGCGTCGATG | >hg38_21_+_8433172_8446622_RNA45SN1_45S_50ntFlanks | 6666 | 6689 | 24 | GATCACTCGGCTCGTGCGTCGATG | 76.0 | -7.8 | 1.3 | -6.2 | 5.4E-10 | 1.0E-09 | MB231/MB468 |
| >hg38_21_+_8433172_8446622_RNA45SN1_45S_50ntFlanks@11452.11471.20\|CTCCTCGGCGCGCGGCGGCG | >hg38_21_+_8433172_8446622_RNA45SN1_45S_50ntFlanks | 11452 | 11471 | 20 | CTCCTCGGCGCGCGGCGGCG | 77.4 | -7.9 | 1.3 | -6.2 | 5.4E-10 | 1.0E-09 | MB231/MB468 |
| >hg38_21_+_8433172_8446622_RNA45SN1_45S_50ntFlanks@11848.11872.25\|ACGGGCTTGGCGGAATCAGCGGGGA | >hg38_21_+_8433172_8446622_RNA45SN1_45S_50ntFlanks | 11848 | 11872 | 25 | ACGGGCTTGGCGGAATCAGCGGGGA | 107.4 | -6.5 | 1.0 | -6.2 | 5.5E-10 | 1.0E-09 | MB231/MB468 |
| >hg38_21_+_8433172_8446622_RNA45SN1_45S_50ntFlanks@10841.10874.34\|CGTAACTTCGGGATAAGGATTGGCTCTAAGGGCT | >hg38_21_+_8433172_8446622_RNA45SN1_45S_50ntFlanks | 10841 | 10874 | 34 | CGTAACTTCGGGATAAGGATTGGCTCTAAGGGCT | 419.4 | -3.5 | 0.6 | -6.2 | 6.3E-10 | 1.1E-09 | BT20/MB468 |
| >hg38_21_+_8433172_8446622_RNA45SN1_45S_50ntFlanks@7985.8003.19\|GATCAGACGTGGCGACCCG | >hg38_21_+_8433172_8446622_RNA45SN1_45S_50ntFlanks | 7985 | 8003 | 19 | GATCAGACGTGGCGACCCG | 83.4 | -8.4 | 1.4 | -6.2 | 6.3E-10 | 1.1E-09 | BT20/MB468 |

| | | | | | | | | | | | | |
|---|---|---|---|---|---|---|---|---|---|---|---|---|
| >hg38_21_+_8433172_8446622_RNA45SN1_45S_50ntFlanks@7984.8004.21\|AGATCAGACGTGGCGACCCGC | >hg38_21_+_8433172_8446622_RNA45SN1_45S_50ntFlanks | 7984 | 8004 | 21 | AGATCAGACGTGGCGACCCGC | 219.2 | -5.2 | 0.8 | -6.2 | 6.4E-10 | 1.1E-09 | BT20/MB468 |
| >hg38_21_+_8433172_8446622_RNA45SN1_45S_50ntFlanks@10514.10533.20\|CGGAGTGGCGGAGATGGGCG | >hg38_21_+_8433172_8446622_RNA45SN1_45S_50ntFlanks | 10514 | 10533 | 20 | CGGAGTGGCGGAGATGGGCG | 169.1 | -5.4 | 0.9 | -6.2 | 5.6E-10 | 1.1E-09 | MB231/MB468 |
| >trnaMT_GluTTC_MT_-_14674_14742@40.61.22,trnalookalike8_GluTTC_5_-_93905172_93905240@40.61.22\|TTGGTCGTGGTTGTAGTCCGTG | >trnaMT_GluTTC_MT_-_14674_14742 | 40 | 61 | 22 | TTGGTCGTGGTTGTAGTCCGTG | 416.2 | 6.1 | 1.0 | 6.2 | 6.4E-10 | 1.1E-09 | BT20/MB468 |
| >hg38_21_+_8433172_8446622_RNA45SN1_45S_50ntFlanks@6645.6685.41\|CTCGTACGACTCTTAGCGGTGGATCACTCGGCTCGTGCGTC | >hg38_21_+_8433172_8446622_RNA45SN1_45S_50ntFlanks | 6645 | 6685 | 41 | CTCGTACGACTCTTAGCGGTGGATCACTCGGCTCGTGCGTC | 779.3 | 7.9 | 1.3 | 6.2 | 6.4E-10 | 1.1E-09 | BT20/MB468 |
| >am_trnaMT_GlyTCC_MT_+_9991_10058@28.48.21\|ACTTCCAATTAACTAGTTTTG | >am_trnaMT_GlyTCC_MT_+_9991_10058 | 28 | 48 | 21 | ACTTCCAATTAACTAGTTTTG | 99.2 | -8.2 | 1.3 | -6.2 | 5.6E-10 | 1.1E-09 | MB231/MB468 |
| >hg38_21_+_8433172_8446622_RNA45SN1_45S_50ntFlanks@10794.10817.24\|CTGGCATGTTGGAACAATGTAGGT | >hg38_21_+_8433172_8446622_RNA45SN1_45S_50ntFlanks | 10794 | 10817 | 24 | CTGGCATGTTGGAACAATGTAGGT | 571.7 | -3.9 | 0.6 | -6.2 | 5.6E-10 | 1.1E-09 | MB231/MB468 |
| >hg38_21_+_8433172_8446622_RNA45SN1_45S_50ntFlanks@11875.11893.19\|GAAGACCCTGTTGAGCTTG | >hg38_21_+_8433172_8446622_RNA45SN1_45S_50ntFlanks | 11875 | 11893 | 19 | GAAGACCCTGTTGAGCTTG | 624.5 | -4.2 | 0.7 | -6.2 | 5.6E-10 | 1.1E-09 | MB231/MB468 |
| >hg38_21_+_8433172_8446622_RNA45SN1_45S_50ntFlanks@10678.10708.31\|CGGTTCCGGCGGCGTCCGGTGAGCTCTCGCT | >hg38_21_+_8433172_8446622_RNA45SN1_45S_50ntFlanks | 10678 | 10708 | 31 | CGGTTCCGGCGGCGTCCGGTGAGCTCTCGCT | 9089.6 | -4.1 | 0.7 | -6.2 | 5.6E-10 | 1.1E-09 | MB231/MB468 |
| >hg38_21_+_8433172_8446622_RNA45SN1_45S_50ntFlanks@10905.10939.35\|CTGGGCGCGCGCCGCGGCTGGACGAGGCGCCGCCG | >hg38_21_+_8433172_8446622_RNA45SN1_45S_50ntFlanks | 10905 | 10939 | 35 | CTGGGCGCGCGCCGCGGCTGGACGAGGCGCCGCCG | 94.1 | -8.1 | 1.3 | -6.2 | 5.6E-10 | 1.1E-09 | MB231/MB468 |
| >hg38_21_+_8433172_8446622_RNA45SN1_45S_50ntFlanks@11660.11685.26\|GTGATTTCTGCCCAGTGCTCTGAATG | >hg38_21_+_8433172_8446622_RNA45SN1_45S_50ntFlanks | 11660 | 11685 | 26 | GTGATTTCTGCCCAGTGCTCTGAATG | 224.1 | -2.8 | 0.4 | -6.2 | 5.7E-10 | 1.1E-09 | MB231/MB468 |
| >hg38_21_+_8433172_8446622_RNA45SN1_45S_50ntFlanks@11731.11751.21\|ACTATGACTCTCTTAAGGTAG | >hg38_21_+_8433172_8446622_RNA45SN1_45S_50ntFlanks | 11731 | 11751 | 21 | ACTATGACTCTCTTAAGGTAG | 749.4 | -5.4 | 0.9 | -6.2 | 6.5E-10 | 1.1E-09 | BT20/MB468 |
| >hg38_21_+_8433172_8446622_RNA45SN1_45S_50ntFlanks@12161.12180.20\|TGACTGGGGCGGTACACCTG | >hg38_21_+_8433172_8446622_RNA45SN1_45S_50ntFlanks | 12161 | 12180 | 20 | TGACTGGGGCGGTACACCTG | 370.0 | -6.2 | 1.0 | -6.2 | 6.5E-10 | 1.1E-09 | BT20/MB468 |
| >hg38_21_+_8433172_8446622_RNA45SN1_45S_50ntFlanks@3991.4023.33\|TGGTGACTCTAGATAACCTCGGGCCGATCGCAC | >hg38_21_+_8433172_8446622_RNA45SN1_45S_50ntFlanks | 3991 | 4023 | 33 | TGGTGACTCTAGATAACCTCGGGCCGATCGCAC | 400.8 | 3.6 | 0.6 | 6.2 | 6.5E-10 | 1.1E-09 | BT20/MB468 |
| >hg38_21_+_8433172_8446622_RNA45SN1_45S_50ntFlanks@10922.10947.26\|CTGGACGAGGCGCCGCCGCCCCCCCC | >hg38_21_+_8433172_8446622_RNA45SN1_45S_50ntFlanks | 10922 | 10947 | 26 | CTGGACGAGGCGCCGCCGCCCCCCCC | 74.5 | -7.8 | 1.3 | -6.2 | 5.9E-10 | 1.1E-09 | MB231/MB468 |
| >hg38_21_+_8433172_8446622_RNA45SN1_45S_50ntFlanks@11317.11334.18\|ACCCCACGTCTCGTCGCG | >hg38_21_+_8433172_8446622_RNA45SN1_45S_50ntFlanks | 11317 | 11334 | 18 | ACCCCACGTCTCGTCGCG | 365.5 | -5.7 | 0.9 | -6.2 | 5.9E-10 | 1.1E-09 | MB231/MB468 |
| >hg38_21_+_8433172_8446622_RNA45SN1_45S_50ntFlanks@12157.12178.22\|AGTTTGACTGGGGCGGTACACC | >hg38_21_+_8433172_8446622_RNA45SN1_45S_50ntFlanks | 12157 | 12178 | 22 | AGTTTGACTGGGGCGGTACACC | 83.7 | -8.4 | 1.4 | -6.2 | 6.8E-10 | 1.1E-09 | BT20/MB468 |
| >hg38_21_+_8433172_8446622_RNA45SN1_45S_50ntFlanks@9546.9573.28\|TGGAGGTCCGTAGCGGTCCTGACGTGCA | >hg38_21_+_8433172_8446622_RNA45SN1_45S_50ntFlanks | 9546 | 9573 | 28 | TGGAGGTCCGTAGCGGTCCTGACGTGCA | 486.9 | 3.9 | 0.6 | 6.2 | 5.9E-10 | 1.1E-09 | MB231/MB468 |
| >hg38_21_+_8433172_8446622_RNA45SN1_45S_50ntFlanks@10866.10883.18\|CTAAGGGCTGGGTCGGTC | >hg38_21_+_8433172_8446622_RNA45SN1_45S_50ntFlanks | 10866 | 10883 | 18 | CTAAGGGCTGGGTCGGTC | 85.9 | -4.7 | 0.8 | -6.2 | 5.9E-10 | 1.1E-09 | MB231/MB468 |
| >hg38_21_+_8433172_8446622_RNA45SN1_45S_50ntFlanks@12941.12972.32\|ATTCGTAGACGACCTGCTTCTGGGTCGGGGTT | >hg38_21_+_8433172_8446622_RNA45SN1_45S_50ntFlanks | 12941 | 12972 | 32 | ATTCGTAGACGACCTGCTTCTGGGTCGGGGTT | 95.3 | -5.3 | 0.9 | -6.2 | 6.1E-10 | 1.2E-09 | MB231/MB468 |
| >hg38_21_+_8433172_8446622_RNA45SN1_45S_50ntFlanks@12418.12455.38\|CGATGTCGGCTCTTCCTATCATTGTGAAGCAGAATTCA | >hg38_21_+_8433172_8446622_RNA45SN1_45S_50ntFlanks | 12418 | 12455 | 38 | CGATGTCGGCTCTTCCTATCATTGTGAAGCAGAATTCA | 994.7 | -3.8 | 0.6 | -6.2 | 6.9E-10 | 1.2E-09 | BT20/MB468 |
| >hg38_21_+_8433172_8446622_RNA45SN1_45S_50ntFlanks@4257.4275.19\|TAATTGGAATGAGTCCACT | >hg38_21_+_8433172_8446622_RNA45SN1_45S_50ntFlanks | 4257 | 4275 | 19 | TAATTGGAATGAGTCCACT | 94.4 | -8.6 | 1.4 | -6.2 | 7.0E-10 | 1.2E-09 | BT20/MB468 |
| >MI0000683\|hsa-mir-181b-2&WithFlank&9\|+\|124693704\|124693804@22.46.25&[MIMAT0000257_1&hsa-miR-181b-5p&offsets\|0\|+2;m-72&9\|+\|124693725\|124693747&offsets\|0\|+2]\|AACATTCATTGCTGTCGGTGGGTTT | >MI0000683\|hsa-mir-181b-2&WithFlank&9\|+\|124693704\|124693804 | 22 | 46 | 25 | AACATTCATTGCTGTCGGTGGGTTT | 709.2 | 5.5 | 0.9 | 6.2 | 7.0E-10 | 1.2E-09 | BT20/MB468 |
| >hg38_21_+_8433172_8446622_RNA45SN1_45S_50ntFlanks@9571.9596.26\|GCAAATCGGTCGTCCGACCTGGGTAT | >hg38_21_+_8433172_8446622_RNA45SN1_45S_50ntFlanks | 9571 | 9596 | 26 | GCAAATCGGTCGTCCGACCTGGGTAT | 246.8 | -5.0 | 0.8 | -6.2 | 7.1E-10 | 1.2E-09 | BT20/MB468 |
| >hg38_21_+_8433172_8446622_RNA45SN1_45S_50ntFlanks@6652.6688.37\|GACTCTTAGCGGTGGATCACTCGGCTCGTGCGTCGAT | >hg38_21_+_8433172_8446622_RNA45SN1_45S_50ntFlanks | 6652 | 6688 | 37 | GACTCTTAGCGGTGGATCACTCGGCTCGTGCGTCGAT | 3244.8 | 4.2 | 0.7 | 6.2 | 6.2E-10 | 1.2E-09 | MB231/MB468 |
| >trna10_AspGTC_12_-_125424193_125424264@1.29.29,trna12_AspGTC_12_-_125411891_125411962@1.29.29,trna38_AspGTC_17_-_8125556_8125627@1.29.29,trna45_AspGTC_6_+_27447453_27447524@1.29.29,trna48_AspGTC_6_+_27471523_27471594@1.29.29,trna4_AspGTC_12_+_96429799_96429870@1.29.29,trna69_AspGTC_1_-_161440205_161440276@1.29.29,trna72_AspGTC_1_-_161432824_161432895@1.29.29,trna75_AspGTC_1_-_161425414_161425485@1.29.29,trna78_AspGTC_1_-_161418033_161418104@1.29.29,trna81_AspGTC_1_-_161410615_161410686@1.29.29\|TCCTCGTTAGTATAGTGGTGAGTATCCCC | >trna10_AspGTC_12_-_125424193_125424264 | 1 | 29 | 29 | TCCTCGTTAGTATAGTGGTGAGTATCCCC | 191.4 | -3.7 | 0.6 | -6.2 | 6.2E-10 | 1.2E-09 | MB231/MB468 |
| >hg38_21_+_8433172_8446622_RNA45SN1_45S_50ntFlanks@6651.6683.33\|CGACTCTTAGCGGTGGATCACTCGGCTCGTGCG | >hg38_21_+_8433172_8446622_RNA45SN1_45S_50ntFlanks | 6651 | 6683 | 33 | CGACTCTTAGCGGTGGATCACTCGGCTCGTGCG | 3762.6 | 3.7 | 0.6 | 6.2 | 7.1E-10 | 1.2E-09 | BT20/MB468 |
| >hg38_21_+_8433172_8446622_RNA45SN1_45S_50ntFlanks@10807.10840.34\|ACAATGTAGGTAAGGGAAGTCGGCAAGCCGGATC | >hg38_21_+_8433172_8446622_RNA45SN1_45S_50ntFlanks | 10807 | 10840 | 34 | ACAATGTAGGTAAGGGAAGTCGGCAAGCCGGATC | 134.1 | -2.6 | 0.4 | -6.2 | 6.3E-10 | 1.2E-09 | MB231/MB468 |
| >hg38_21_+_8433172_8446622_RNA45SN1_45S_50ntFlanks@10513.10534.22\|CCGGAGTGGCGGAGATGGGCGC | >hg38_21_+_8433172_8446622_RNA45SN1_45S_50ntFlanks | 10513 | 10534 | 22 | CCGGAGTGGCGGAGATGGGCGC | 150.5 | -5.6 | 0.9 | -6.2 | 7.2E-10 | 1.2E-09 | BT20/MB468 |
| >hg38_21_+_8433172_8446622_RNA45SN1_45S_50ntFlanks@13001.13027.27\|GCGATCTATTGAAAGTCAGCCCTCGAC | >hg38_21_+_8433172_8446622_RNA45SN1_45S_50ntFlanks | 13001 | 13027 | 27 | GCGATCTATTGAAAGTCAGCCCTCGAC | 266.2 | -6.1 | 1.0 | -6.2 | 7.2E-10 | 1.2E-09 | BT20/MB468 |
| >trna108_AlaAGC_6_-_28763741_28763812@58.75.18\|TCCCCAGTACCTCCACCA | >trna108_AlaAGC_6_-_28763741_28763812 | 58 | 75 | 18 | TCCCCAGTACCTCCACCA | 970.3 | 5.8 | 0.9 | 6.2 | 7.2E-10 | 1.2E-09 | BT20/MB468 |
| >hg38_21_+_8433172_8446622_RNA45SN1_45S_50ntFlanks@6744.6770.27\|GACACTTCGAACGCACTTGCGGCCCCG | >hg38_21_+_8433172_8446622_RNA45SN1_45S_50ntFlanks | 6744 | 6770 | 27 | GACACTTCGAACGCACTTGCGGCCCCG | 1124.7 | -5.1 | 0.8 | -6.2 | 6.3E-10 | 1.2E-09 | MB231/MB468 |
| >hg38_21_+_8433172_8446622_RNA45SN1_45S_50ntFlanks@476.496.21\|TTCGTGATCGATGTGGTGACG | >hg38_21_+_8433172_8446622_RNA45SN1_45S_50ntFlanks | 476 | 496 | 21 | TTCGTGATCGATGTGGTGACG | 887.6 | 5.4 | 0.9 | 6.2 | 7.3E-10 | 1.2E-09 | BT20/MB468 |
| >hg38_21_+_8433172_8446622_RNA45SN1_45S_50ntFlanks@4738.4767.30\|AAAGTCGGAGGTTCGAAGACGATCAGATAC | >hg38_21_+_8433172_8446622_RNA45SN1_45S_50ntFlanks | 4738 | 4767 | 30 | AAAGTCGGAGGTTCGAAGACGATCAGATAC | 469.2 | 7.7 | 1.2 | 6.2 | 7.3E-10 | 1.2E-09 | BT20/MB468 |
| >hg38_21_+_8433172_8446622_RNA45SN1_45S_50ntFlanks@12457.12485.29\|CAAGCGTTGGATTGTTCACCCACTAATAG | >hg38_21_+_8433172_8446622_RNA45SN1_45S_50ntFlanks | 12457 | 12485 | 29 | CAAGCGTTGGATTGTTCACCCACTAATAG | 286.6 | -5.7 | 0.9 | -6.2 | 7.3E-10 | 1.2E-09 | BT20/MB468 |
| >hg38_21_+_8433172_8446622_RNA45SN1_45S_50ntFlanks@9570.9594.25\|TGCAAATCGGTCGTCCGACCTGGGT | >hg38_21_+_8433172_8446622_RNA45SN1_45S_50ntFlanks | 9570 | 9594 | 25 | TGCAAATCGGTCGTCCGACCTGGGT | 330.8 | -3.7 | 0.6 | -6.2 | 7.4E-10 | 1.2E-09 | BT20/MB468 |
| >hg38_21_+_8433172_8446622_RNA45SN1_45S_50ntFlanks@5470.5499.30\|CCACGGCCCTGGCGGAGCGCTGAGAAGACG | >hg38_21_+_8433172_8446622_RNA45SN1_45S_50ntFlanks | 5470 | 5499 | 30 | CCACGGCCCTGGCGGAGCGCTGAGAAGACG | 262.5 | -3.7 | 0.6 | -6.2 | 6.4E-10 | 1.2E-09 | MB231/MB468 |
| >hg38_21_+_8433172_8446622_RNA45SN1_45S_50ntFlanks@6776.6802.27\|CTCCCGGGGCTACGCCTGTCTGAGCGT | >hg38_21_+_8433172_8446622_RNA45SN1_45S_50ntFlanks | 6776 | 6802 | 27 | CTCCCGGGGCTACGCCTGTCTGAGCGT | 903.3 | 3.9 | 0.6 | 6.2 | 6.5E-10 | 1.2E-09 | MB231/MB468 |
| >hg38_21_+_8433172_8446622_RNA45SN1_45S_50ntFlanks@6650.6696.47\|ACGACTCTTAGCGGTGGATCACTCGGCTCGTGCGTCGATGAAGAACG | >hg38_21_+_8433172_8446622_RNA45SN1_45S_50ntFlanks | 6650 | 6696 | 47 | ACGACTCTTAGCGGTGGATCACTCGGCTCGTGCGTCGATGAAGAACG | 460.7 | 5.3 | 0.9 | 6.2 | 7.4E-10 | 1.2E-09 | BT20/MB468 |
| >hg38_21_+_8433172_8446622_RNA45SN1_45S_50ntFlanks@6740.6772.33\|CATCGACACTTCGAACGCACTTGCGGCCCCGGG | >hg38_21_+_8433172_8446622_RNA45SN1_45S_50ntFlanks | 6740 | 6772 | 33 | CATCGACACTTCGAACGCACTTGCGGCCCCGGG | 340.3 | -4.9 | 0.8 | -6.2 | 7.4E-10 | 1.2E-09 | BT20/MB468 |
| >hg38_21_+_8433172_8446622_RNA45SN1_45S_50ntFlanks@10186.10208.23\|TCCTCCCCTCCTCCCGCCCACGC | >hg38_21_+_8433172_8446622_RNA45SN1_45S_50ntFlanks | 10186 | 10208 | 23 | TCCTCCCCTCCTCCCGCCCACGC | 79.3 | -7.9 | 1.3 | -6.2 | 6.5E-10 | 1.2E-09 | MB231/MB468 |
| >hg38_21_+_8433172_8446622_RNA45SN1_45S_50ntFlanks@476.514.39\|TTCGTGATCGATGTGGTGACGTCGTGCTCTCCCGGGCCG | >hg38_21_+_8433172_8446622_RNA45SN1_45S_50ntFlanks | 476 | 514 | 39 | TTCGTGATCGATGTGGTGACGTCGTGCTCTCCCGGGCCG | 466.1 | 8.2 | 1.3 | 6.2 | 7.4E-10 | 1.2E-09 | BT20/MB468 |
| >hg38_21_+_8433172_8446622_RNA45SN1_45S_50ntFlanks@11578.11613.36\|TAGAACTGGTGCGGACCAGGGGAATCCGACTGTTTA | >hg38_21_+_8433172_8446622_RNA45SN1_45S_50ntFlanks | 11578 | 11613 | 36 | TAGAACTGGTGCGGACCAGGGGAATCCGACTGTTTA | 501.9 | 5.3 | 0.9 | 6.2 | 7.4E-10 | 1.2E-09 | BT20/MB468 |
| >hg38_21_+_8433172_8446622_RNA45SN1_45S_50ntFlanks@3843.3860.18\|CTACTTGGATAACTGTGG | >hg38_21_+_8433172_8446622_RNA45SN1_45S_50ntFlanks | 3843 | 3860 | 18 | CTACTTGGATAACTGTGG | 222.6 | -2.9 | 0.5 | -6.2 | 6.5E-10 | 1.2E-09 | MB231/MB468 |
| >hg38_21_+_8433172_8446622_RNA45SN1_45S_50ntFlanks@8107.8143.37\|CGGCGGGGCGCGGGACATGTGGCGTACGGAAGACCCG | >hg38_21_+_8433172_8446622_RNA45SN1_45S_50ntFlanks | 8107 | 8143 | 37 | CGGCGGGGCGCGGGACATGTGGCGTACGGAAGACCCG | 172.8 | -6.1 | 1.0 | -6.2 | 7.5E-10 | 1.2E-09 | BT20/MB468 |
| >hg38_21_+_8433172_8446622_RNA45SN1_45S_50ntFlanks@4367.4393.27\|CAGTTAAAAAGCTCGTAGTTGGATCTT | >hg38_21_+_8433172_8446622_RNA45SN1_45S_50ntFlanks | 4367 | 4393 | 27 | CAGTTAAAAAGCTCGTAGTTGGATCTT | 119.8 | -4.5 | 0.7 | -6.2 | 6.6E-10 | 1.3E-09 | MB231/MB468 |

| | | | | | | | | | | | | |
|---|---|---|---|---|---|---|---|---|---|---|---|---|
| >hg38_21_+_8433172_8446622_RNA45SN1_45S_50ntFlanks@12632.12661.30\|GCGAAGCTACCATCTGTGGGATTATGACTG | >hg38_21_+_8433172_8446622_RNA45SN1_45S_50ntFlanks | 12632 | 12661 | 30 | GCGAAGCTACCATCTGTGGGATTATGACTG | 302.8 | -3.0 | 0.5 | -6.2 | 7.6E-10 | 1.3E-09 | BT20/MB468 |
| >hg38_21_+_8433172_8446622_RNA45SN1_45S_50ntFlanks@4657.4682.26\|CTAGAGGTGAAATTCTTGGACCGGCG | >hg38_21_+_8433172_8446622_RNA45SN1_45S_50ntFlanks | 4657 | 4682 | 26 | CTAGAGGTGAAATTCTTGGACCGGCG | 344.3 | -7.0 | 1.1 | -6.2 | 6.7E-10 | 1.3E-09 | MB231/MB468 |
| >hg38_21_+_8433172_8446622_RNA45SN1_45S_50ntFlanks@485.506.22\|GATGTGGTGACGTCGTGCTCTC | >hg38_21_+_8433172_8446622_RNA45SN1_45S_50ntFlanks | 485 | 506 | 22 | GATGTGGTGACGTCGTGCTCTC | 24052.1 | 19.5 | 3.2 | 6.2 | 6.7E-10 | 1.3E-09 | MB231/MB468 |
| >hg38_21_+_8433172_8446622_RNA45SN1_45S_50ntFlanks@4722.4751.30\|TCATTAATCAAGAACGAAAGTCGGAGGTTC | >hg38_21_+_8433172_8446622_RNA45SN1_45S_50ntFlanks | 4722 | 4751 | 30 | TCATTAATCAAGAACGAAAGTCGGAGGTTC | 389.0 | -3.8 | 0.6 | -6.2 | 6.7E-10 | 1.3E-09 | MB231/MB468 |
| >hg38_21_+_8433172_8446622_RNA45SN1_45S_50ntFlanks@12940.12972.33\|CATTCGTAGACGACCTGCTTCTGGGTCGGGGTT | >hg38_21_+_8433172_8446622_RNA45SN1_45S_50ntFlanks | 12940 | 12972 | 33 | CATTCGTAGACGACCTGCTTCTGGGTCGGGGTT | 264.1 | -5.3 | 0.9 | -6.2 | 7.7E-10 | 1.3E-09 | BT20/MB468 |
| >hg38_21_+_8433172_8446622_RNA45SN1_45S_50ntFlanks@12415.12440.26\|CTTCGATGTCGGCTCTTCCTATCATT | >hg38_21_+_8433172_8446622_RNA45SN1_45S_50ntFlanks | 12415 | 12440 | 26 | CTTCGATGTCGGCTCTTCCTATCATT | 678.3 | 3.8 | 0.6 | 6.2 | 6.8E-10 | 1.3E-09 | MB231/MB468 |
| >hg38_21_+_8433172_8446622_RNA45SN1_45S_50ntFlanks@3882.3899.18\|CCGACGGGCGCTGACCCC | >hg38_21_+_8433172_8446622_RNA45SN1_45S_50ntFlanks | 3882 | 3899 | 18 | CCGACGGGCGCTGACCCC | 1512.3 | -4.9 | 0.8 | -6.2 | 6.8E-10 | 1.3E-09 | MB231/MB468 |
| >hg38_21_+_8433172_8446622_RNA45SN1_45S_50ntFlanks@12935.12953.19\|TAAACCATTCGTAGACGAC | >hg38_21_+_8433172_8446622_RNA45SN1_45S_50ntFlanks | 12935 | 12953 | 19 | TAAACCATTCGTAGACGAC | 192.8 | -3.4 | 0.6 | -6.2 | 6.8E-10 | 1.3E-09 | MB231/MB468 |
| >hg38_21_+_8433172_8446622_RNA45SN1_45S_50ntFlanks@12957.12985.29\|CTTCTGGGTCGGGGTTTCGTACGTAGCAG | >hg38_21_+_8433172_8446622_RNA45SN1_45S_50ntFlanks | 12957 | 12985 | 29 | CTTCTGGGTCGGGGTTTCGTACGTAGCAG | 77.6 | -7.9 | 1.3 | -6.2 | 6.8E-10 | 1.3E-09 | MB231/MB468 |
| >MI0000102\|hsa-mir-100&WithFlank&11\|-\|122152223\|122152314@19.38.20(+1A)&[MIMAT0000098&hsa-miR-100-5p&offsets\|0\|-2(+1A);m-27&11\|-\|122152276\|122152296&offsets\|0\|-1(+1A)]]\|AACCCGTAGATCCGAACTTGA | >MI0000102\|hsa-mir-100&WithFlank&11\|-\|122152223\|122152314 | 19 | 38 | 20(+1A) | AACCCGTAGATCCGAACTTGA | 16127.1 | 14.5 | 2.4 | 6.2 | 6.8E-10 | 1.3E-09 | MB231/MB468 |
| >hg38_21_+_8433172_8446622_RNA45SN1_45S_50ntFlanks@11778.11804.27\|ACGCGCATGAATGGATGAACGAGATTC | >hg38_21_+_8433172_8446622_RNA45SN1_45S_50ntFlanks | 11778 | 11804 | 27 | ACGCGCATGAATGGATGAACGAGATTC | 411.0 | -4.9 | 0.8 | -6.1 | 7.9E-10 | 1.3E-09 | BT20/MB468 |
| >trna2_GlyGCC_21_-_18827107_18827177@1.32.32,trna35_GlyGCC_1_+_161413094_161413164@1.32.32,trna37_GlyGCC_1_+_161420467_161420537@1.32.32,trna39_GlyGCC_1_+_161427898_161427968@1.32.32,trna41_GlyGCC_1_+_161435258_161435328@1.32.32\|GCATGGGTGGTTCAGTGGTAGAATTCTCGCCT | >trna2_GlyGCC_21_-_18827107_18827177 | 1 | 32 | 32 | GCATGGGTGGTTCAGTGGTAGAATTCTCGCCT | 1256.6 | 5.5 | 0.9 | 6.2 | 7.0E-10 | 1.3E-09 | MB231/MB468 |
| >hg38_21_+_8433172_8446622_RNA45SN1_45S_50ntFlanks@7989.8009.21\|AGACGTGGCGACCCGCTGAAT | >hg38_21_+_8433172_8446622_RNA45SN1_45S_50ntFlanks | 7989 | 8009 | 21 | AGACGTGGCGACCCGCTGAAT | 10952.7 | 6.2 | 1.0 | 6.2 | 7.0E-10 | 1.3E-09 | MB231/MB468 |
| >hg38_21_+_8433172_8446622_RNA45SN1_45S_50ntFlanks@5442.5466.25\|CCCTCGGATCGGCCCCGCCGGGGTC | >hg38_21_+_8433172_8446622_RNA45SN1_45S_50ntFlanks | 5442 | 5466 | 25 | CCCTCGGATCGGCCCCGCCGGGGTC | 144.3 | -4.9 | 0.8 | -6.2 | 7.1E-10 | 1.3E-09 | MB231/MB468 |
| >hg38_21_+_8433172_8446622_RNA45SN1_45S_50ntFlanks@8155.8181.27\|CGCTCGTGGGGGGCCCAAGTCCTTCTG | >hg38_21_+_8433172_8446622_RNA45SN1_45S_50ntFlanks | 8155 | 8181 | 27 | CGCTCGTGGGGGGCCCAAGTCCTTCTG | 390.4 | -4.4 | 0.7 | -6.1 | 8.1E-10 | 1.3E-09 | BT20/MB468 |
| >hg38_21_+_8433172_8446622_RNA45SN1_45S_50ntFlanks@10421.10443.23\|TGAGAGATGGGCGAGCGCCGTTC | >hg38_21_+_8433172_8446622_RNA45SN1_45S_50ntFlanks | 10421 | 10443 | 23 | TGAGAGATGGGCGAGCGCCGTTC | 79.2 | -7.9 | 1.3 | -6.2 | 7.1E-10 | 1.3E-09 | MB231/MB468 |
| >hg38_21_+_8433172_8446622_RNA45SN1_45S_50ntFlanks@8194.8211.18\|CCCGTGGACGGTGTGAGG | >hg38_21_+_8433172_8446622_RNA45SN1_45S_50ntFlanks | 8194 | 8211 | 18 | CCCGTGGACGGTGTGAGG | 594.7 | -3.5 | 0.6 | -6.2 | 7.1E-10 | 1.4E-09 | MB231/MB468 |
| >hg38_21_+_8433172_8446622_RNA45SN1_45S_50ntFlanks@10427.10456.30\|ATGGGCGAGCGCCGTTCCGAAGGGACGGGC | >hg38_21_+_8433172_8446622_RNA45SN1_45S_50ntFlanks | 10427 | 10456 | 30 | ATGGGCGAGCGCCGTTCCGAAGGGACGGGC | 591.0 | -4.8 | 0.8 | -6.1 | 8.2E-10 | 1.4E-09 | BT20/MB468 |
| >hg38_21_+_8433172_8446622_RNA45SN1_45S_50ntFlanks@12489.12509.21\|ACGTGAGCTGGGTTTAGACCG | >hg38_21_+_8433172_8446622_RNA45SN1_45S_50ntFlanks | 12489 | 12509 | 21 | ACGTGAGCTGGGTTTAGACCG | 894.7 | -5.5 | 0.9 | -6.1 | 8.2E-10 | 1.4E-09 | BT20/MB468 |
| >hg38_21_+_8433172_8446622_RNA45SN1_45S_50ntFlanks@11207.11234.28\|GGCGGCGACTCTGGACGCGAGCCGGGCC | >hg38_21_+_8433172_8446622_RNA45SN1_45S_50ntFlanks | 11207 | 11234 | 28 | GGCGGCGACTCTGGACGCGAGCCGGGCC | 85.1 | -8.0 | 1.3 | -6.2 | 7.2E-10 | 1.4E-09 | MB231/MB468 |
| >hg38_21_+_8433172_8446622_RNA45SN1_45S_50ntFlanks@4723.4748.26\|CATTAATCAAGAACGAAAGTCGGAGG | >hg38_21_+_8433172_8446622_RNA45SN1_45S_50ntFlanks | 4723 | 4748 | 26 | CATTAATCAAGAACGAAAGTCGGAGG | 332.1 | -3.7 | 0.6 | -6.1 | 8.2E-10 | 1.4E-09 | BT20/MB468 |
| >hg38_21_+_8433172_8446622_RNA45SN1_45S_50ntFlanks@11038.11060.23\|TCTCCCCCGCTCCCCGTCCTCCC | >hg38_21_+_8433172_8446622_RNA45SN1_45S_50ntFlanks | 11038 | 11060 | 23 | TCTCCCCCGCTCCCCGTCCTCCC | 91.0 | -8.1 | 1.3 | -6.2 | 7.3E-10 | 1.4E-09 | MB231/MB468 |
| >hg38_21_+_8433172_8446622_RNA45SN1_45S_50ntFlanks@10866.10892.27\|CTAAGGGCTGGGTCGGTCGGGCTGGGG | >hg38_21_+_8433172_8446622_RNA45SN1_45S_50ntFlanks | 10866 | 10892 | 27 | CTAAGGGCTGGGTCGGTCGGGCTGGGG | 376.0 | -6.0 | 1.0 | -6.1 | 8.4E-10 | 1.4E-09 | BT20/MB468 |
| >hg38_21_+_8433172_8446622_RNA45SN1_45S_50ntFlanks@12499.12517.19\|GGTTTAGACCGTCGTGAGA | >hg38_21_+_8433172_8446622_RNA45SN1_45S_50ntFlanks | 12499 | 12517 | 19 | GGTTTAGACCGTCGTGAGA | 77.1 | -7.8 | 1.3 | -6.2 | 7.4E-10 | 1.4E-09 | MB231/MB468 |
| >hg38_21_+_8433172_8446622_RNA45SN1_45S_50ntFlanks@8378.8402.25\|TTCAAGAGGGCGTGAAACCGTTAAG | >hg38_21_+_8433172_8446622_RNA45SN1_45S_50ntFlanks | 8378 | 8402 | 25 | TTCAAGAGGGCGTGAAACCGTTAAG | 158.9 | -5.5 | 0.9 | -6.1 | 8.4E-10 | 1.4E-09 | BT20/MB468 |
| >hg38_21_+_8433172_8446622_RNA45SN1_45S_50ntFlanks@12461.12488.28\|CGTTGGATTGTTCACCCACTAATAGGGA | >hg38_21_+_8433172_8446622_RNA45SN1_45S_50ntFlanks | 12461 | 12488 | 28 | CGTTGGATTGTTCACCCACTAATAGGGA | 113.6 | -8.4 | 1.4 | -6.2 | 7.4E-10 | 1.4E-09 | MB231/MB468 |
| >hg38_21_+_8433172_8446622_RNA45SN1_45S_50ntFlanks@6785.6806.22\|CTACGCCTGTCTGAGCGTCGCT | >hg38_21_+_8433172_8446622_RNA45SN1_45S_50ntFlanks | 6785 | 6806 | 22 | CTACGCCTGTCTGAGCGTCGCT | 2878.7 | 4.2 | 0.7 | 6.2 | 7.4E-10 | 1.4E-09 | MB231/MB468 |
| >hg38_21_+_8433172_8446622_RNA45SN1_45S_50ntFlanks@4297.4321.25\|CATTGGAGGGCAAGTCTGGTGCCAG | >hg38_21_+_8433172_8446622_RNA45SN1_45S_50ntFlanks | 4297 | 4321 | 25 | CATTGGAGGGCAAGTCTGGTGCCAG | 191.9 | -4.8 | 0.8 | -6.2 | 7.5E-10 | 1.4E-09 | MB231/MB468 |
| >trna10_GlyTCC_17_+_8124866_8124937@1.30.30\|GCGTTGGTGGTATAGTGGTAAGCATAGCTG | >trna10_GlyTCC_17_+_8124866_8124937 | 1 | 30 | 30 | GCGTTGGTGGTATAGTGGTAAGCATAGCTG | 222.0 | -5.9 | 1.0 | -6.1 | 8.5E-10 | 1.4E-09 | BT20/MB468 |
| >hg38_21_+_8433172_8446622_RNA45SN1_45S_50ntFlanks@10830.10848.19\|CAAGCCGGATCCGTAACTT | >hg38_21_+_8433172_8446622_RNA45SN1_45S_50ntFlanks | 10830 | 10848 | 19 | CAAGCCGGATCCGTAACTT | 155.3 | -4.7 | 0.8 | -6.1 | 8.6E-10 | 1.4E-09 | BT20/MB468 |
| >hg38_21_+_8433172_8446622_RNA45SN1_45S_50ntFlanks@6652.6669.18\|GACTCTTAGCGGTGGATC | >hg38_21_+_8433172_8446622_RNA45SN1_45S_50ntFlanks | 6652 | 6669 | 18 | GACTCTTAGCGGTGGATC | 464.2 | 3.8 | 0.6 | 6.1 | 8.6E-10 | 1.4E-09 | BT20/MB468 |
| >hg38_21_+_8433172_8446622_RNA45SN1_45S_50ntFlanks@6699.6717.19\|GCTAGCTGCGAGAATTAAT | >hg38_21_+_8433172_8446622_RNA45SN1_45S_50ntFlanks | 6699 | 6717 | 19 | GCTAGCTGCGAGAATTAAT | 371.0 | -6.1 | 1.0 | -6.1 | 8.6E-10 | 1.4E-09 | BT20/MB468 |
| >trna119_LysCTT_1_-_145395522_145395594@1.30.30,trna11_LysCTT_5_-_180648979_180649051@1.30.30,trna13_LysCTT_6_+_26556774_26556846@1.30.30,trna32_LysCTT_16_-_3207406_3207478@1.30.30,trna7_LysCTT_16_+_3225692_3225764@1.30.30,trna9_LysCTT_5_+_180634755_180634827@1.30.30\|GCCCGGCTAGCTCAGTCGGTAGAGCATGAG | >trna119_LysCTT_1_-_145395522_145395594 | 1 | 30 | 30 | GCCCGGCTAGCTCAGTCGGTAGAGCATGAG | 11521.2 | -4.7 | 0.8 | -6.2 | 7.5E-10 | 1.4E-09 | MB231/MB468 |
| >hg38_21_+_8433172_8446622_RNA45SN1_45S_50ntFlanks@10350.10367.18\|ACGAGAACTTTGAAGGCC | >hg38_21_+_8433172_8446622_RNA45SN1_45S_50ntFlanks | 10350 | 10367 | 18 | ACGAGAACTTTGAAGGCC | 1212.4 | -3.1 | 0.5 | -6.1 | 8.6E-10 | 1.4E-09 | BT20/MB468 |
| >trna2_ArgCCT_16_+_3202901_3202973@13.30.18,trna3_ArgCCT_7_+_139025446_139025518@13.30.18\|TAATGGATAAGGCATTGG | >trna2_ArgCCT_16_+_3202901_3202973 | 13 | 30 | 18 | TAATGGATAAGGCATTGG | 98.6 | -8.2 | 1.3 | -6.2 | 7.6E-10 | 1.4E-09 | MB231/MB468 |
| >hg38_21_+_8433172_8446622_RNA45SN1_45S_50ntFlanks@11552.11591.40\|CCTCGGCCGGCGCCTAGCAGCCGACTTAGAACTGGTGCGG | >hg38_21_+_8433172_8446622_RNA45SN1_45S_50ntFlanks | 11552 | 11591 | 40 | CCTCGGCCGGCGCCTAGCAGCCGACTTAGAACTGGTGCGG | 154.9 | -3.3 | 0.5 | -6.1 | 8.7E-10 | 1.4E-09 | BT20/MB468 |
| >hg38_21_+_8433172_8446622_RNA45SN1_45S_50ntFlanks@12073.12093.21\|CCGAGGGGCTCTCGCTTCTGG | >hg38_21_+_8433172_8446622_RNA45SN1_45S_50ntFlanks | 12073 | 12093 | 21 | CCGAGGGGCTCTCGCTTCTGG | 96.4 | -5.1 | 0.8 | -6.2 | 7.7E-10 | 1.5E-09 | MB231/MB468 |
| >hg38_21_+_8433172_8446622_RNA45SN1_45S_50ntFlanks@9352.9371.20\|CCGTGGCGCAATGAAGGTGA | >hg38_21_+_8433172_8446622_RNA45SN1_45S_50ntFlanks | 9352 | 9371 | 20 | CCGTGGCGCAATGAAGGTGA | 98.1 | -8.6 | 1.4 | -6.1 | 8.8E-10 | 1.5E-09 | BT20/MB468 |
| >hg38_21_+_8433172_8446622_RNA45SN1_45S_50ntFlanks@10783.10801.19\|GTGAACAGCCTCTGGCATG | >hg38_21_+_8433172_8446622_RNA45SN1_45S_50ntFlanks | 10783 | 10801 | 19 | GTGAACAGCCTCTGGCATG | 246.8 | -5.1 | 0.8 | -6.1 | 8.8E-10 | 1.5E-09 | BT20/MB468 |
| >hg38_21_+_8433172_8446622_RNA45SN1_45S_50ntFlanks@11679.11702.24\|CTGAATGTCAAAGTGAAGAAATTC | >hg38_21_+_8433172_8446622_RNA45SN1_45S_50ntFlanks | 11679 | 11702 | 24 | CTGAATGTCAAAGTGAAGAAATTC | 104.3 | -3.9 | 0.6 | -6.1 | 8.8E-10 | 1.5E-09 | BT20/MB468 |
| >hg38_21_+_8433172_8446622_RNA45SN1_45S_50ntFlanks@10389.10411.23\|TGAACAGCAGTTGAACATGGGTC | >hg38_21_+_8433172_8446622_RNA45SN1_45S_50ntFlanks | 10389 | 10411 | 23 | TGAACAGCAGTTGAACATGGGTC | 127.8 | -3.7 | 0.6 | -6.1 | 8.9E-10 | 1.5E-09 | BT20/MB468 |
| >hg38_21_+_8433172_8446622_RNA45SN1_45S_50ntFlanks@12786.12805.20\|CCCGCGCGCGCGGGAGGGCG | >hg38_21_+_8433172_8446622_RNA45SN1_45S_50ntFlanks | 12786 | 12805 | 20 | CCCGCGCGCGCGGGAGGGCG | 71.3 | -7.7 | 1.3 | -6.1 | 7.9E-10 | 1.5E-09 | MB231/MB468 |
| >hg38_21_+_8433172_8446622_RNA45SN1_45S_50ntFlanks@4007.4024.18\|CCTCGGGCCGATCGCACG | >hg38_21_+_8433172_8446622_RNA45SN1_45S_50ntFlanks | 4007 | 4024 | 18 | CCTCGGGCCGATCGCACG | 415.0 | -2.8 | 0.5 | -6.1 | 9.0E-10 | 1.5E-09 | BT20/MB468 |
| >hg38_21_+_8433172_8446622_RNA45SN1_45S_50ntFlanks@8155.8181.27\|CGCTCGTGGGGGGCCCAAGTCCTTCTG | >hg38_21_+_8433172_8446622_RNA45SN1_45S_50ntFlanks | 8155 | 8181 | 27 | CGCTCGTGGGGGGCCCAAGTCCTTCTG | 345.7 | -3.0 | 0.5 | -6.1 | 7.9E-10 | 1.5E-09 | MB231/MB468 |
| >hg38_21_+_8433172_8446622_RNA45SN1_45S_50ntFlanks@10682.10710.29\|TCCGGCGGCGTCCGGTGAGCTCTCGCTGG | >hg38_21_+_8433172_8446622_RNA45SN1_45S_50ntFlanks | 10682 | 10710 | 29 | TCCGGCGGCGTCCGGTGAGCTCTCGCTGG | 170.7 | -6.1 | 1.0 | -6.1 | 9.1E-10 | 1.5E-09 | BT20/MB468 |
| >MI0000736\|hsa-mir-30c-1&WithFlank&1\|+\|40757278\|40757378@23.46.24(+1A);MI0000254\|hsa-mir-30c-2&WithFlank&6\|-\|71376954\|71377037@13.36.24(+1A)&[MIMAT0000244&hsa-miR-30c-5p&offsets\|0\|+1(+1A);m-57&1\|+\|40757300\|40757323&offsets\|0\|0(+1A)];[MIMAT0000244_1&hsa-miR-30c-5p&offsets\|0\|+1(+1A);m-56&6\|-\|71377002\|71377025&offsets\|0\|0(+1A)]]\|TGTAAACATCCTACACTCTCAGCTA | >MI0000736\|hsa-mir-30c-1&WithFlank&1\|+\|40757278\|40757378 | 23 | 46 | 24(+1A) | TGTAAACATCCTACACTCTCAGCTA | 526.4 | 5.2 | 0.8 | 6.1 | 9.3E-10 | 1.5E-09 | BT20/MB468 |
| >hg38_21_+_8433172_8446622_RNA45SN1_45S_50ntFlanks@10391.10411.21\|AACAGCAGTTGAACATGGGTC | >hg38_21_+_8433172_8446622_RNA45SN1_45S_50ntFlanks | 10391 | 10411 | 21 | AACAGCAGTTGAACATGGGTC | 166.8 | -3.3 | 0.5 | -6.1 | 9.3E-10 | 1.5E-09 | BT20/MB468 |

<table>
<tr><td>>hg38_21_+_8433172_8446622_RNA45SN1_45S_50ntFlanks@11873.11903.31|AAGAAGACCCTGTTGAGCTTGACTCTAGTCT</td><td>>hg38_21_+_8433172_8446622_RNA45SN1_45S_50ntFlanks</td><td>11873</td><td>11903</td><td>31</td><td>AAGAAGACCCTGTTGAGCTTGACTCTAGTCT</td><td>244.1</td><td>-4.6</td><td>0.8</td><td>-6.1</td><td>8.2E-10</td><td>1.6E-09</td><td>MB231/MB468</td></tr>
<tr><td>>hg38_21_+_8433172_8446622_RNA45SN1_45S_50ntFlanks@10408.10427.20|GGTCAGTCGGTCCTGAGAGA</td><td>>hg38_21_+_8433172_8446622_RNA45SN1_45S_50ntFlanks</td><td>10408</td><td>10427</td><td>20</td><td>GGTCAGTCGGTCCTGAGAGA</td><td>91.8</td><td>-8.1</td><td>1.3</td><td>-6.1</td><td>8.2E-10</td><td>1.6E-09</td><td>MB231/MB468</td></tr>
<tr><td>>hg38_21_+_8433172_8446622_RNA45SN1_45S_50ntFlanks@11212.11229.18|CGACTCTGGACGCGAGCC</td><td>>hg38_21_+_8433172_8446622_RNA45SN1_45S_50ntFlanks</td><td>11212</td><td>11229</td><td>18</td><td>CGACTCTGGACGCGAGCC</td><td>267.1</td><td>-3.8</td><td>0.6</td><td>-6.1</td><td>9.4E-10</td><td>1.6E-09</td><td>BT20/MB468</td></tr>
<tr><td>>hg38_21_+_8433172_8446622_RNA45SN1_45S_50ntFlanks@12160.12180.21|TTGACTGGGGCGGTACACCTG</td><td>>hg38_21_+_8433172_8446622_RNA45SN1_45S_50ntFlanks</td><td>12160</td><td>12180</td><td>21</td><td>TTGACTGGGGCGGTACACCTG</td><td>105.8</td><td>-5.0</td><td>0.8</td><td>-6.1</td><td>8.2E-10</td><td>1.6E-09</td><td>MB231/MB468</td></tr>
<tr><td>>hg38_21_+_8433172_8446622_RNA45SN1_45S_50ntFlanks@12712.12751.40|CGGAGCCTCGGTTGGCCTCGGATAGCCGGTCCCCCGCCTG</td><td>>hg38_21_+_8433172_8446622_RNA45SN1_45S_50ntFlanks</td><td>12712</td><td>12751</td><td>40</td><td>CGGAGCCTCGGTTGGCCTCGGATAGCCGGTCCCCCGCCTG</td><td>90.6</td><td>-8.1</td><td>1.3</td><td>-6.1</td><td>8.3E-10</td><td>1.6E-09</td><td>MB231/MB468</td></tr>
<tr><td>>hg38_21_+_8433172_8446622_RNA45SN1_45S_50ntFlanks@6719.6751.33|TGAATTGCAGGACACATTGATCATCGACACTTC</td><td>>hg38_21_+_8433172_8446622_RNA45SN1_45S_50ntFlanks</td><td>6719</td><td>6751</td><td>33</td><td>TGAATTGCAGGACACATTGATCATCGACACTTC</td><td>281.5</td><td>-4.5</td><td>0.7</td><td>-6.1</td><td>9.4E-10</td><td>1.6E-09</td><td>BT20/MB468</td></tr>
<tr><td>>hg38_21_+_8433172_8446622_RNA45SN1_45S_50ntFlanks@9343.9370.28|CGAAAGCCGCCGTGGCGCAATGAAGGTG</td><td>>hg38_21_+_8433172_8446622_RNA45SN1_45S_50ntFlanks</td><td>9343</td><td>9370</td><td>28</td><td>CGAAAGCCGCCGTGGCGCAATGAAGGTG</td><td>101.5</td><td>-8.2</td><td>1.3</td><td>-6.1</td><td>8.3E-10</td><td>1.6E-09</td><td>MB231/MB468</td></tr>
<tr><td>>trna111_HisGTG_1_-_147774845_147774916@-1T.26.27,trna118_HisGTG_1_-_145396881_145396952@-1T.26.27,trna16_HisGTG_1_+_146544773_146544844@-1T.26.27,trna1_HisGTG_15_+_45493349_45493420@-1T.26.27,trna21_HisGTG_1_+_147753471_147753542@-1T.26.27,trna33_HisGTG_6_+_27125906_27125977@-1T.26.27,trna7_HisGTG_9_-_14433938_14434009@-1T.26.27,trna8_HisGTG_15_-_45492611_45492682@-1T.26.27,trna9_HisGTG_15_-_45490804_45490875@-1T.26.27|TGCCGTGATCGTATAGTGGTTAGTACT</td><td>>trna111_HisGTG_1_-_147774845_147774916</td><td>-1T</td><td>26</td><td>27</td><td>TGCCGTGATCGTATAGTGGTTAGTACT</td><td>1029.4</td><td>4.3</td><td>0.7</td><td>6.1</td><td>9.5E-10</td><td>1.6E-09</td><td>BT20/MB468</td></tr>
<tr><td>>hg38_21_+_8433172_8446622_RNA45SN1_45S_50ntFlanks@10678.10696.19|CGGTTCCGGCGGCGTCCGG</td><td>>hg38_21_+_8433172_8446622_RNA45SN1_45S_50ntFlanks</td><td>10678</td><td>10696</td><td>19</td><td>CGGTTCCGGCGGCGTCCGG</td><td>69.1</td><td>-7.7</td><td>1.3</td><td>-6.1</td><td>8.3E-10</td><td>1.6E-09</td><td>MB231/MB468</td></tr>
<tr><td>>hg38_21_+_8433172_8446622_RNA45SN1_45S_50ntFlanks@6648.6680.33|GTACGACTCTTAGCGGTGGATCACTCGGCTCGT</td><td>>hg38_21_+_8433172_8446622_RNA45SN1_45S_50ntFlanks</td><td>6648</td><td>6680</td><td>33</td><td>GTACGACTCTTAGCGGTGGATCACTCGGCTCGT</td><td>1305.4</td><td>2.9</td><td>0.5</td><td>6.1</td><td>9.5E-10</td><td>1.6E-09</td><td>BT20/MB468</td></tr>
<tr><td>>hg38_21_+_8433172_8446622_RNA45SN1_45S_50ntFlanks@12934.12959.26|CTAAACCATTCGTAGACGACCTGCTT</td><td>>hg38_21_+_8433172_8446622_RNA45SN1_45S_50ntFlanks</td><td>12934</td><td>12959</td><td>26</td><td>CTAAACCATTCGTAGACGACCTGCTT</td><td>490.4</td><td>-3.2</td><td>0.5</td><td>-6.1</td><td>8.4E-10</td><td>1.6E-09</td><td>MB231/MB468</td></tr>
<tr><td>>hg38_21_+_8433172_8446622_RNA45SN1_45S_50ntFlanks@11537.11560.24|CCGCGCGGCGCCTCGCCTCGGCCG</td><td>>hg38_21_+_8433172_8446622_RNA45SN1_45S_50ntFlanks</td><td>11537</td><td>11560</td><td>24</td><td>CCGCGCGGCGCCTCGCCTCGGCCG</td><td>80.3</td><td>-7.9</td><td>1.3</td><td>-6.1</td><td>8.4E-10</td><td>1.6E-09</td><td>MB231/MB468</td></tr>
<tr><td>>hg38_21_+_8433172_8446622_RNA45SN1_45S_50ntFlanks@3870.3898.29|AGCTAATACATGCCGACGGGCGCTGACCC</td><td>>hg38_21_+_8433172_8446622_RNA45SN1_45S_50ntFlanks</td><td>3870</td><td>3898</td><td>29</td><td>AGCTAATACATGCCGACGGGCGCTGACCC</td><td>182.7</td><td>-5.0</td><td>0.8</td><td>-6.1</td><td>8.4E-10</td><td>1.6E-09</td><td>MB231/MB468</td></tr>
<tr><td>>MI0000102|hsa-mir-100&WithFlank&11|-|122152223|122152314@19.38.20(+1C)&[MIMAT0000098&hsa-miR-100-5p&offsets|0|-2(+1C);m-27&11|-|122152276|122152296&offsets|0|-1(+1C)]|AACCCGTAGATCCGAACTTGC</td><td>>MI0000102|hsa-mir-100&WithFlank&11|-|122152223|122152314</td><td>19</td><td>38</td><td>20(+1C)</td><td>AACCCGTAGATCCGAACTTGC</td><td>20275.9</td><td>-5.9</td><td>0.9</td><td>-6.6</td><td>5.7E-11</td><td>1.6E-09</td><td>BT20/MB231</td></tr>
<tr><td>>MI0000102|hsa-mir-100&WithFlank&11|-|122152223|122152314@20.40.21&[MIMAT0000098&hsa-miR-100-5p&offsets|+1|0;m-27&11|-|122152276|122152296&offsets|+1|+1]|ACCCGTAGATCCGAACTTGTG</td><td>>MI0000102|hsa-mir-100&WithFlank&11|-|122152223|122152314</td><td>20</td><td>40</td><td>21</td><td>ACCCGTAGATCCGAACTTGTG</td><td>1810.1</td><td>-6.8</td><td>1.0</td><td>-6.6</td><td>5.7E-11</td><td>1.6E-09</td><td>BT20/MB231</td></tr>
<tr><td>>hg38_21_+_8433172_8446622_RNA45SN1_45S_50ntFlanks@11583.11605.23|CTGGTGCGGACCAGGGGAATCCG</td><td>>hg38_21_+_8433172_8446622_RNA45SN1_45S_50ntFlanks</td><td>11583</td><td>11605</td><td>23</td><td>CTGGTGCGGACCAGGGGAATCCG</td><td>685.0</td><td>-5.3</td><td>0.9</td><td>-6.1</td><td>9.7E-10</td><td>1.6E-09</td><td>BT20/MB468</td></tr>
<tr><td>>hg38_21_+_8433172_8446622_RNA45SN1_45S_50ntFlanks@10905.10941.37|CTGGGCGCGCGCCGCGGCTGGACGAGGCGCCGCCGCC</td><td>>hg38_21_+_8433172_8446622_RNA45SN1_45S_50ntFlanks</td><td>10905</td><td>10941</td><td>37</td><td>CTGGGCGCGCGCCGCGGCTGGACGAGGCGCCGCCGCC</td><td>79.2</td><td>-7.9</td><td>1.3</td><td>-6.1</td><td>8.6E-10</td><td>1.6E-09</td><td>MB231/MB468</td></tr>
<tr><td>>hg38_21_+_8433172_8446622_RNA45SN1_45S_50ntFlanks@10786.10817.32|AACAGCCTCTGGCATGTTGGAACAATGTAGGT</td><td>>hg38_21_+_8433172_8446622_RNA45SN1_45S_50ntFlanks</td><td>10786</td><td>10817</td><td>32</td><td>AACAGCCTCTGGCATGTTGGAACAATGTAGGT</td><td>1260.1</td><td>-4.0</td><td>0.7</td><td>-6.1</td><td>9.8E-10</td><td>1.6E-09</td><td>BT20/MB468</td></tr>
<tr><td>>hg38_21_+_8433172_8446622_RNA45SN1_45S_50ntFlanks@3838.3859.22|CTCTCCTACTTGGATAACTGTG</td><td>>hg38_21_+_8433172_8446622_RNA45SN1_45S_50ntFlanks</td><td>3838</td><td>3859</td><td>22</td><td>CTCTCCTACTTGGATAACTGTG</td><td>93.6</td><td>-2.3</td><td>0.4</td><td>-6.1</td><td>9.9E-10</td><td>1.6E-09</td><td>BT20/MB468</td></tr>
<tr><td>>hg38_21_+_8433172_8446622_RNA45SN1_45S_50ntFlanks@4588.4617.30|CTATTTTGTTGGTTTTCGGAACTGAGGCCA</td><td>>hg38_21_+_8433172_8446622_RNA45SN1_45S_50ntFlanks</td><td>4588</td><td>4617</td><td>30</td><td>CTATTTTGTTGGTTTTCGGAACTGAGGCCA</td><td>72.2</td><td>-7.7</td><td>1.3</td><td>-6.1</td><td>8.7E-10</td><td>1.6E-09</td><td>MB231/MB468</td></tr>
<tr><td>>hg38_21_+_8433172_8446622_RNA45SN1_45S_50ntFlanks@10350.10374.25|ACGAGAACTTTGAAGGCCGAAGTGG</td><td>>hg38_21_+_8433172_8446622_RNA45SN1_45S_50ntFlanks</td><td>10350</td><td>10374</td><td>25</td><td>ACGAGAACTTTGAAGGCCGAAGTGG</td><td>1281.9</td><td>-3.6</td><td>0.6</td><td>-6.1</td><td>8.8E-10</td><td>1.7E-09</td><td>MB231/MB468</td></tr>
<tr><td>>hg38_21_+_8433172_8446622_RNA45SN1_45S_50ntFlanks@7988.8008.21|CAGACGTGGCGACCCGCTGAA</td><td>>hg38_21_+_8433172_8446622_RNA45SN1_45S_50ntFlanks</td><td>7988</td><td>8008</td><td>21</td><td>CAGACGTGGCGACCCGCTGAA</td><td>2473.3</td><td>-3.4</td><td>0.6</td><td>-6.1</td><td>1.0E-09</td><td>1.7E-09</td><td>BT20/MB468</td></tr>
<tr><td>>hg38_21_+_8433172_8446622_RNA45SN1_45S_50ntFlanks@4443.4470.28|CCCTTGCCTCTCGGCGCCCCCTCGATGC</td><td>>hg38_21_+_8433172_8446622_RNA45SN1_45S_50ntFlanks</td><td>4443</td><td>4470</td><td>28</td><td>CCCTTGCCTCTCGGCGCCCCCTCGATGC</td><td>83.2</td><td>-8.0</td><td>1.3</td><td>-6.1</td><td>8.9E-10</td><td>1.7E-09</td><td>MB231/MB468</td></tr>
<tr><td>>hg38_21_+_8433172_8446622_RNA45SN1_45S_50ntFlanks@13008.13039.32|ATTGAAAGTCAGCCCTCGACACAAGGGTTTGT</td><td>>hg38_21_+_8433172_8446622_RNA45SN1_45S_50ntFlanks</td><td>13008</td><td>13039</td><td>32</td><td>ATTGAAAGTCAGCCCTCGACACAAGGGTTTGT</td><td>3026.6</td><td>4.8</td><td>0.8</td><td>6.1</td><td>8.9E-10</td><td>1.7E-09</td><td>MB231/MB468</td></tr>
<tr><td>>hg38_21_+_8433172_8446622_RNA45SN1_45S_50ntFlanks@6719.6758.40|TGAATTGCAGGACACATTGATCATCGACACTTCGAACGCA</td><td>>hg38_21_+_8433172_8446622_RNA45SN1_45S_50ntFlanks</td><td>6719</td><td>6758</td><td>40</td><td>TGAATTGCAGGACACATTGATCATCGACACTTCGAACGCA</td><td>105.2</td><td>-3.9</td><td>0.6</td><td>-6.1</td><td>1.0E-09</td><td>1.7E-09</td><td>BT20/MB468</td></tr>
<tr><td>>hg38_21_+_8433172_8446622_RNA45SN1_45S_50ntFlanks@8552.8581.30|CTCCCCGGAGGGGGCGGGCTCCGGCGGGTG</td><td>>hg38_21_+_8433172_8446622_RNA45SN1_45S_50ntFlanks</td><td>8552</td><td>8581</td><td>30</td><td>CTCCCCGGAGGGGGCGGGCTCCGGCGGGTG</td><td>87.5</td><td>-8.0</td><td>1.3</td><td>-6.1</td><td>9.1E-10</td><td>1.7E-09</td><td>MB231/MB468</td></tr>
<tr><td>>hg38_21_+_8433172_8446622_RNA45SN1_45S_50ntFlanks@12156.12180.25|GAGTTTGACTGGGGCGGTACACCTG</td><td>>hg38_21_+_8433172_8446622_RNA45SN1_45S_50ntFlanks</td><td>12156</td><td>12180</td><td>25</td><td>GAGTTTGACTGGGGCGGTACACCTG</td><td>82.9</td><td>-8.0</td><td>1.3</td><td>-6.1</td><td>9.3E-10</td><td>1.7E-09</td><td>MB231/MB468</td></tr>
<tr><td>>hg38_21_+_8433172_8446622_RNA45SN1_45S_50ntFlanks@11104.11133.30|AAGGGTCGGGGCGGCAGGGGCCGGCGGCGG</td><td>>hg38_21_+_8433172_8446622_RNA45SN1_45S_50ntFlanks</td><td>11104</td><td>11133</td><td>30</td><td>AAGGGTCGGGGCGGCAGGGGCCGGCGGCGG</td><td>69.5</td><td>-7.7</td><td>1.3</td><td>-6.1</td><td>9.3E-10</td><td>1.8E-09</td><td>MB231/MB468</td></tr>
<tr><td>>trna111_HisGTG_1_-_147774845_147774916@-1T.23.24,trna118_HisGTG_1_-_145396881_145396952@-1T.23.24,trna16_HisGTG_1_+_146544773_146544844@-1T.23.24,trna1_HisGTG_15_+_45493349_45493420@-1T.23.24,trna21_HisGTG_1_+_147753471_147753542@-1T.23.24,trna33_HisGTG_6_+_27125906_27125977@-1T.23.24,trna7_HisGTG_9_-_14433938_14434009@-1T.23.24,trna8_HisGTG_15_-_45492611_45492682@-1T.23.24,trna9_HisGTG_15_-_45490804_45490875@-1T.23.24|TGCCGTGATCGTATAGTGGTTAGT</td><td>>trna111_HisGTG_1_-_147774845_147774916</td><td>-1T</td><td>23</td><td>24</td><td>TGCCGTGATCGTATAGTGGTTAGT</td><td>4899.1</td><td>-4.6</td><td>0.7</td><td>-6.5</td><td>6.4E-11</td><td>1.8E-09</td><td>BT20/MB231</td></tr>
<tr><td>>hg38_21_+_8433172_8446622_RNA45SN1_45S_50ntFlanks@12002.12043.42|TGCGGGCCGCCGGTGAAATACCACTACTCTGATCGTTTTTTC</td><td>>hg38_21_+_8433172_8446622_RNA45SN1_45S_50ntFlanks</td><td>12002</td><td>12043</td><td>42</td><td>TGCGGGCCGCCGGTGAAATACCACTACTCTGATCGTTTTTTC</td><td>155.2</td><td>-9.3</td><td>1.5</td><td>-6.1</td><td>1.1E-09</td><td>1.8E-09</td><td>BT20/MB468</td></tr>
<tr><td>>hg38_21_+_8433172_8446622_RNA45SN1_45S_50ntFlanks@4773.4793.21|TAGTTCCGACCATAAACGATG</td><td>>hg38_21_+_8433172_8446622_RNA45SN1_45S_50ntFlanks</td><td>4773</td><td>4793</td><td>21</td><td>TAGTTCCGACCATAAACGATG</td><td>1384.6</td><td>-4.1</td><td>0.7</td><td>-6.1</td><td>9.4E-10</td><td>1.8E-09</td><td>MB231/MB468</td></tr>
<tr><td>>hg38_21_+_8433172_8446622_RNA45SN1_45S_50ntFlanks@10461.10480.20|GCCTCCGTTGCCCTCGGCCG</td><td>>hg38_21_+_8433172_8446622_RNA45SN1_45S_50ntFlanks</td><td>10461</td><td>10480</td><td>20</td><td>GCCTCCGTTGCCCTCGGCCG</td><td>69.3</td><td>-7.7</td><td>1.3</td><td>-6.1</td><td>9.5E-10</td><td>1.8E-09</td><td>MB231/MB468</td></tr>
<tr><td>>trna116_GluCTC_1_-_145399233_145399304@32.72.41,trna71_GluCTC_1_-_161439189_161439260@32.72.41,trna74_GluCTC_1_-_161431809_161431880@32.72.41,trna77_GluCTC_1_-_161424398_161424469@32.72.41,trna77_GluCTC_6_+_28949976_28950047@32.72.41,trna80_GluCTC_1_-_161417018_161417089@32.72.41,trna87_GluCTC_6_-_126101393_126101464@32.72.41|CTCTCACCGCCGCGGCCCGGGTTCGATTCCCGGTCAGGGAA</td><td>>trna116_GluCTC_1_-_145399233_145399304</td><td>32</td><td>72</td><td>41</td><td>CTCTCACCGCCGCGGCCCGGGTTCGATTCCCGGTCAGGGAA</td><td>79.3</td><td>-7.9</td><td>1.3</td><td>-6.1</td><td>9.5E-10</td><td>1.8E-09</td><td>MB231/MB468</td></tr>
<tr><td>>hg38_21_+_8433172_8446622_RNA45SN1_45S_50ntFlanks@9739.9765.27|AAACGATCTCAACCTATTCTCAAACTT</td><td>>hg38_21_+_8433172_8446622_RNA45SN1_45S_50ntFlanks</td><td>9739</td><td>9765</td><td>27</td><td>AAACGATCTCAACCTATTCTCAAACTT</td><td>725.0</td><td>-4.1</td><td>0.7</td><td>-6.1</td><td>1.1E-09</td><td>1.8E-09</td><td>BT20/MB468</td></tr>
<tr><td>>hg38_21_+_8433172_8446622_RNA45SN1_45S_50ntFlanks@9534.9567.34|AGGAAACTCTGGTGGAGGTCCGTAGCGGTCCTGA</td><td>>hg38_21_+_8433172_8446622_RNA45SN1_45S_50ntFlanks</td><td>9534</td><td>9567</td><td>34</td><td>AGGAAACTCTGGTGGAGGTCCGTAGCGGTCCTGA</td><td>616.5</td><td>-4.8</td><td>0.8</td><td>-6.1</td><td>9.6E-10</td><td>1.8E-09</td><td>MB231/MB468</td></tr>
<tr><td>>hg38_21_+_8433172_8446622_RNA45SN1_45S_50ntFlanks@11319.11338.20|CCCACGTCTCGTCGCGCGCG</td><td>>hg38_21_+_8433172_8446622_RNA45SN1_45S_50ntFlanks</td><td>11319</td><td>11338</td><td>20</td><td>CCCACGTCTCGTCGCGCGCG</td><td>76.9</td><td>-7.8</td><td>1.3</td><td>-6.1</td><td>9.6E-10</td><td>1.8E-09</td><td>MB231/MB468</td></tr>
<tr><td>>MI0000782|hsa-mir-374a&WithFlank&X|-|74287280|74287363@18.38.21&[MIMAT0000727&hsa-miR-374a-5p&offsets|0|-1;m-184&X|-|74287325|74287346&offsets|0|-1]|TTATAATACAACCTGATAAGT</td><td>>MI0000782|hsa-mir-374a&WithFlank&X|-|74287280|74287363</td><td>18</td><td>38</td><td>21</td><td>TTATAATACAACCTGATAAGT</td><td>2230.3</td><td>6.5</td><td>1.1</td><td>6.1</td><td>1.1E-09</td><td>1.8E-09</td><td>BT20/MB468</td></tr>
</table>

<table>
<tr><th></th><th></th><th></th><th></th><th></th><th></th><th></th><th></th><th></th><th></th><th></th><th></th><th></th></tr>
<tr><td>>trna116_GluCTC_1_-_145399233_145399304@32.75.44,trna71_GluCTC_1_-_161439189_161439260@32.75.44,trna74_GluCTC_1_-_161431809_161431880@32.75.44,trna77_GluCTC_1_-_161424398_161424469@32.75.44,trna77_GluCTC_6_+_28949976_28950047@32.75.44,trna80_GluCTC_1_-_161417018_161417089@32.75.44,trna87_GluCTC_6_-_126101393_126101464@32.75.44|CTCTCACCGCCGCGGCCCGGGTTCGATTCCCGGTCAGGGAACCA</td><td>>trna116_GluCTC_1_-_145399233_145399304</td><td>32</td><td>75</td><td>44</td><td>CTCTCACCGCCGCGGCCCGGGTTCGATTCCCGGTCAGGGAACCA</td><td>103.7</td><td>-8.3</td><td>1.4</td><td>-6.1</td><td>9.6E-10</td><td>1.8E-09</td><td>MB231/MB468</td></tr>
<tr><td>>hg38_21_+_8433172_8446622_RNA45SN1_45S_50ntFlanks@10692.10709.18|TCCGGTGAGCTCTCGCTG</td><td>>hg38_21_+_8433172_8446622_RNA45SN1_45S_50ntFlanks</td><td>10692</td><td>10709</td><td>18</td><td>TCCGGTGAGCTCTCGCTG</td><td>145.5</td><td>-3.9</td><td>0.6</td><td>-6.1</td><td>1.1E-09</td><td>1.8E-09</td><td>BT20/MB468</td></tr>
<tr><td>>hg38_21_+_8433172_8446622_RNA45SN1_45S_50ntFlanks@9544.9563.20|GGTGGAGGTCCGTAGCGGTC</td><td>>hg38_21_+_8433172_8446622_RNA45SN1_45S_50ntFlanks</td><td>9544</td><td>9563</td><td>20</td><td>GGTGGAGGTCCGTAGCGGTC</td><td>142.9</td><td>-4.0</td><td>0.7</td><td>-6.1</td><td>1.1E-09</td><td>1.8E-09</td><td>BT20/MB468</td></tr>
<tr><td>>hg38_21_+_8433172_8446622_RNA45SN1_45S_50ntFlanks@9544.9566.23|GGTGGAGGTCCGTAGCGGTCCTG</td><td>>hg38_21_+_8433172_8446622_RNA45SN1_45S_50ntFlanks</td><td>9544</td><td>9566</td><td>23</td><td>GGTGGAGGTCCGTAGCGGTCCTG</td><td>118.4</td><td>-3.9</td><td>0.6</td><td>-6.1</td><td>1.1E-09</td><td>1.9E-09</td><td>BT20/MB468</td></tr>
<tr><td>>hg38_21_+_8433172_8446622_RNA45SN1_45S_50ntFlanks@12938.12956.19|ACCATTCGTAGACGACCTG</td><td>>hg38_21_+_8433172_8446622_RNA45SN1_45S_50ntFlanks</td><td>12938</td><td>12956</td><td>19</td><td>ACCATTCGTAGACGACCTG</td><td>1295.2</td><td>-3.9</td><td>0.6</td><td>-6.1</td><td>1.0E-09</td><td>1.9E-09</td><td>MB231/MB468</td></tr>
<tr><td>>hg38_21_+_8433172_8446622_RNA45SN1_45S_50ntFlanks@11564.11585.22|CCTAGCAGCCGACTTAGAACTG</td><td>>hg38_21_+_8433172_8446622_RNA45SN1_45S_50ntFlanks</td><td>11564</td><td>11585</td><td>22</td><td>CCTAGCAGCCGACTTAGAACTG</td><td>1363.7</td><td>-3.6</td><td>0.6</td><td>-6.1</td><td>1.0E-09</td><td>1.9E-09</td><td>MB231/MB468</td></tr>
<tr><td>>hg38_21_+_8433172_8446622_RNA45SN1_45S_50ntFlanks@12991.13011.21|CTCCCTCGCTGCGATCTATTG</td><td>>hg38_21_+_8433172_8446622_RNA45SN1_45S_50ntFlanks</td><td>12991</td><td>13011</td><td>21</td><td>CTCCCTCGCTGCGATCTATTG</td><td>1061.9</td><td>-4.9</td><td>0.8</td><td>-6.1</td><td>1.0E-09</td><td>1.9E-09</td><td>MB231/MB468</td></tr>
<tr><td>>am_trna2_ArgCCT_16_+_3202901_3202973@14.31.18,trna3_ArgCCT_7_+_139025446_139025518@14.31.18|AATGGATAAGGCATTGGC</td><td>>am_trna2_ArgCCT_16_+_3202901_3202973</td><td>14</td><td>31</td><td>18</td><td>AATGGATAAGGCATTGGC</td><td>234.0</td><td>-5.0</td><td>0.8</td><td>-6.1</td><td>1.0E-09</td><td>1.9E-09</td><td>MB231/MB468</td></tr>
<tr><td>>hg38_21_+_8433172_8446622_RNA45SN1_45S_50ntFlanks@11234.11252.19|CCTTCCCGTGGATCGCCCC</td><td>>hg38_21_+_8433172_8446622_RNA45SN1_45S_50ntFlanks</td><td>11234</td><td>11252</td><td>19</td><td>CCTTCCCGTGGATCGCCCC</td><td>76.6</td><td>-7.8</td><td>1.3</td><td>-6.1</td><td>1.0E-09</td><td>1.9E-09</td><td>MB231/MB468</td></tr>
<tr><td>>hg38_21_+_8433172_8446622_RNA45SN1_45S_50ntFlanks@12418.12453.36|CGATGTCGGCTCTTCCTATCATTGTGAAGCAGAATT</td><td>>hg38_21_+_8433172_8446622_RNA45SN1_45S_50ntFlanks</td><td>12418</td><td>12453</td><td>36</td><td>CGATGTCGGCTCTTCCTATCATTGTGAAGCAGAATT</td><td>431.6</td><td>4.1</td><td>0.7</td><td>6.1</td><td>1.0E-09</td><td>1.9E-09</td><td>MB231/MB468</td></tr>
<tr><td>>hg38_21_+_8433172_8446622_RNA45SN1_45S_50ntFlanks@12944.12966.23|CGTAGACGACCTGCTTCTGGGTC</td><td>>hg38_21_+_8433172_8446622_RNA45SN1_45S_50ntFlanks</td><td>12944</td><td>12966</td><td>23</td><td>CGTAGACGACCTGCTTCTGGGTC</td><td>131.9</td><td>-9.0</td><td>1.5</td><td>-6.1</td><td>1.2E-09</td><td>1.9E-09</td><td>BT20/MB468</td></tr>
<tr><td>>hg38_21_+_8433172_8446622_RNA45SN1_45S_50ntFlanks@4752.4769.18|GAAGACGATCAGATACCG</td><td>>hg38_21_+_8433172_8446622_RNA45SN1_45S_50ntFlanks</td><td>4752</td><td>4769</td><td>18</td><td>GAAGACGATCAGATACCG</td><td>73.5</td><td>-8.2</td><td>1.3</td><td>-6.1</td><td>1.2E-09</td><td>1.9E-09</td><td>BT20/MB468</td></tr>
<tr><td>>hg38_21_+_8433172_8446622_RNA45SN1_45S_50ntFlanks@4759.4777.19|ATCAGATACCGTCGTAGTT</td><td>>hg38_21_+_8433172_8446622_RNA45SN1_45S_50ntFlanks</td><td>4759</td><td>4777</td><td>19</td><td>ATCAGATACCGTCGTAGTT</td><td>143.8</td><td>-4.7</td><td>0.8</td><td>-6.1</td><td>1.0E-09</td><td>1.9E-09</td><td>MB231/MB468</td></tr>
<tr><td>>hg38_21_+_8433172_8446622_RNA45SN1_45S_50ntFlanks@10390.10424.35|GAACAGCAGTTGAACATGGGTCAGTCGGTCCTGAG</td><td>>hg38_21_+_8433172_8446622_RNA45SN1_45S_50ntFlanks</td><td>10390</td><td>10424</td><td>35</td><td>GAACAGCAGTTGAACATGGGTCAGTCGGTCCTGAG</td><td>450.9</td><td>7.2</td><td>1.2</td><td>6.1</td><td>1.2E-09</td><td>1.9E-09</td><td>BT20/MB468</td></tr>
<tr><td>>hg38_21_+_8433172_8446622_RNA45SN1_45S_50ntFlanks@10409.10429.21|GTCAGTCGGTCCTGAGAGATG</td><td>>hg38_21_+_8433172_8446622_RNA45SN1_45S_50ntFlanks</td><td>10409</td><td>10429</td><td>21</td><td>GTCAGTCGGTCCTGAGAGATG</td><td>896.8</td><td>-3.1</td><td>0.5</td><td>-6.1</td><td>1.2E-09</td><td>1.9E-09</td><td>BT20/MB468</td></tr>
<tr><td>>hg38_21_+_8433172_8446622_RNA45SN1_45S_50ntFlanks@9533.9554.22|GAGGAAACTCTGGTGGAGGTCC</td><td>>hg38_21_+_8433172_8446622_RNA45SN1_45S_50ntFlanks</td><td>9533</td><td>9554</td><td>22</td><td>GAGGAAACTCTGGTGGAGGTCC</td><td>222.2</td><td>-4.9</td><td>0.8</td><td>-6.1</td><td>1.2E-09</td><td>1.9E-09</td><td>BT20/MB468</td></tr>
<tr><td>>hg38_21_+_8433172_8446622_RNA45SN1_45S_50ntFlanks@12513.12538.26|TGAGACAGGTTAGTTTTACCCTACTG</td><td>>hg38_21_+_8433172_8446622_RNA45SN1_45S_50ntFlanks</td><td>12513</td><td>12538</td><td>26</td><td>TGAGACAGGTTAGTTTTACCCTACTG</td><td>100.8</td><td>-8.2</td><td>1.3</td><td>-6.1</td><td>1.0E-09</td><td>1.9E-09</td><td>MB231/MB468</td></tr>
<tr><td>>hg38_21_+_8433172_8446622_RNA45SN1_45S_50ntFlanks@10910.10932.23|CGCGCGCCGCGGCTGGACGAGGC</td><td>>hg38_21_+_8433172_8446622_RNA45SN1_45S_50ntFlanks</td><td>10910</td><td>10932</td><td>23</td><td>CGCGCGCCGCGGCTGGACGAGGC</td><td>103.5</td><td>-8.3</td><td>1.4</td><td>-6.1</td><td>1.0E-09</td><td>1.9E-09</td><td>MB231/MB468</td></tr>
<tr><td>>hg38_21_+_8433172_8446622_RNA45SN1_45S_50ntFlanks@8357.8387.31|AAAAGAACTTTGAAGAGAGAGTTCAAGAGGG</td><td>>hg38_21_+_8433172_8446622_RNA45SN1_45S_50ntFlanks</td><td>8357</td><td>8387</td><td>31</td><td>AAAAGAACTTTGAAGAGAGAGTTCAAGAGGG</td><td>128.8</td><td>-6.0</td><td>1.0</td><td>-6.1</td><td>1.0E-09</td><td>1.9E-09</td><td>MB231/MB468</td></tr>
<tr><td>>hg38_21_+_8433172_8446622_RNA45SN1_45S_50ntFlanks@12008.12039.32|CCGCCGGTGAAATACCACTACTCTGATCGTTT</td><td>>hg38_21_+_8433172_8446622_RNA45SN1_45S_50ntFlanks</td><td>12008</td><td>12039</td><td>32</td><td>CCGCCGGTGAAATACCACTACTCTGATCGTTT</td><td>99.5</td><td>-6.4</td><td>1.0</td><td>-6.1</td><td>1.0E-09</td><td>2.0E-09</td><td>MB231/MB468</td></tr>
<tr><td>>hg38_21_+_8433172_8446622_RNA45SN1_45S_50ntFlanks@10817.10840.24|TAAGGGAAGTCGGCAAGCCGGATC</td><td>>hg38_21_+_8433172_8446622_RNA45SN1_45S_50ntFlanks</td><td>10817</td><td>10840</td><td>24</td><td>TAAGGGAAGTCGGCAAGCCGGATC</td><td>162.3</td><td>-2.7</td><td>0.4</td><td>-6.1</td><td>1.0E-09</td><td>2.0E-09</td><td>MB231/MB468</td></tr>
<tr><td>>hg38_21_+_8433172_8446622_RNA45SN1_45S_50ntFlanks@485.514.30|GATGTGGTGACGTCGTGCTCTCCCGGGCCG</td><td>>hg38_21_+_8433172_8446622_RNA45SN1_45S_50ntFlanks</td><td>485</td><td>514</td><td>30</td><td>GATGTGGTGACGTCGTGCTCTCCCGGGCCG</td><td>2108.1</td><td>15.9</td><td>2.6</td><td>6.1</td><td>1.1E-09</td><td>2.0E-09</td><td>MB231/MB468</td></tr>
<tr><td>>hg38_21_+_8433172_8446622_RNA45SN1_45S_50ntFlanks@11876.11909.34|AAGACCCTGTTGAGCTTGACTCTAGTCTGGCACG</td><td>>hg38_21_+_8433172_8446622_RNA45SN1_45S_50ntFlanks</td><td>11876</td><td>11909</td><td>34</td><td>AAGACCCTGTTGAGCTTGACTCTAGTCTGGCACG</td><td>93.7</td><td>-3.2</td><td>0.5</td><td>-6.1</td><td>1.1E-09</td><td>2.0E-09</td><td>MB231/MB468</td></tr>
<tr><td>>hg38_21_+_8433172_8446622_RNA45SN1_45S_50ntFlanks@9562.9595.34|TCCTGACGTGCAAATCGGTCGTCCGACCTGGGTA</td><td>>hg38_21_+_8433172_8446622_RNA45SN1_45S_50ntFlanks</td><td>9562</td><td>9595</td><td>34</td><td>TCCTGACGTGCAAATCGGTCGTCCGACCTGGGTA</td><td>160.4</td><td>-3.1</td><td>0.5</td><td>-6.1</td><td>1.1E-09</td><td>2.0E-09</td><td>MB231/MB468</td></tr>
<tr><td>>hg38_21_+_8433172_8446622_RNA45SN1_45S_50ntFlanks@4163.4182.20|CATCCAAGGAAGGCAGCAGG</td><td>>hg38_21_+_8433172_8446622_RNA45SN1_45S_50ntFlanks</td><td>4163</td><td>4182</td><td>20</td><td>CATCCAAGGAAGGCAGCAGG</td><td>576.0</td><td>-5.4</td><td>0.9</td><td>-6.1</td><td>1.1E-09</td><td>2.0E-09</td><td>MB231/MB468</td></tr>
<tr><td>>hg38_21_+_8433172_8446622_RNA45SN1_45S_50ntFlanks@10361.10388.28|GAAGGCCGAAGTGGAGAAGGGTTCCATG</td><td>>hg38_21_+_8433172_8446622_RNA45SN1_45S_50ntFlanks</td><td>10361</td><td>10388</td><td>28</td><td>GAAGGCCGAAGTGGAGAAGGGTTCCATG</td><td>388.6</td><td>-5.6</td><td>0.9</td><td>-6.1</td><td>1.1E-09</td><td>2.0E-09</td><td>MB231/MB468</td></tr>
<tr><td>>hg38_21_+_8433172_8446622_RNA45SN1_45S_50ntFlanks@4558.4587.30|AGCTAGGAATAATGGAATAGGACCGCGGTT</td><td>>hg38_21_+_8433172_8446622_RNA45SN1_45S_50ntFlanks</td><td>4558</td><td>4587</td><td>30</td><td>AGCTAGGAATAATGGAATAGGACCGCGGTT</td><td>120.2</td><td>-5.2</td><td>0.9</td><td>-6.1</td><td>1.2E-09</td><td>2.0E-09</td><td>BT20/MB468</td></tr>
<tr><td>>hg38_21_+_8433172_8446622_RNA45SN1_45S_50ntFlanks@11881.11906.26|CCTGTTGAGCTTGACTCTAGTCTGGC</td><td>>hg38_21_+_8433172_8446622_RNA45SN1_45S_50ntFlanks</td><td>11881</td><td>11906</td><td>26</td><td>CCTGTTGAGCTTGACTCTAGTCTGGC</td><td>145.2</td><td>-6.1</td><td>1.0</td><td>-6.1</td><td>1.1E-09</td><td>2.0E-09</td><td>MB231/MB468</td></tr>
<tr><td>>MI0000073|hsa-mir-19a&WithFlank&13|+|91350885|91350978@55.76.22(+1U)&[MIMAT0000073&hsa-miR-19a-3p&offsets|0|-1(+1U);m-201&13|+|91350939|91350961&offsets|0|-1(+1U)]|TGTGCAAATCTATGCAAAACTGT</td><td>>MI0000073|hsa-mir-19a&WithFlank&13|+|91350885|91350978</td><td>55</td><td>76</td><td>22(+1U)</td><td>TGTGCAAATCTATGCAAAACTGT</td><td>493.7</td><td>5.0</td><td>0.8</td><td>6.1</td><td>1.2E-09</td><td>2.0E-09</td><td>BT20/MB468</td></tr>
<tr><td>>am_trnaMT_PheGAA_MT_+_577_647@41.62.22|GTTTAGACGGGCTCACATCACC</td><td>>am_trnaMT_PheGAA_MT_+_577_647</td><td>41</td><td>62</td><td>22</td><td>GTTTAGACGGGCTCACATCACC</td><td>904.4</td><td>8.2</td><td>1.3</td><td>6.1</td><td>1.3E-09</td><td>2.1E-09</td><td>BT20/MB468</td></tr>
<tr><td>>hg38_21_+_8433172_8446622_RNA45SN1_45S_50ntFlanks@10840.10868.29|CCGTAACTTCGGGATAAGGATTGGCTCTA</td><td>>hg38_21_+_8433172_8446622_RNA45SN1_45S_50ntFlanks</td><td>10840</td><td>10868</td><td>29</td><td>CCGTAACTTCGGGATAAGGATTGGCTCTA</td><td>473.4</td><td>-3.0</td><td>0.5</td><td>-6.1</td><td>1.1E-09</td><td>2.1E-09</td><td>MB231/MB468</td></tr>
<tr><td>>hg38_21_+_8433172_8446622_RNA45SN1_45S_50ntFlanks@9027.9059.33|CGTCTCCTCGTGGGGGGGCCGGGCCACCCCTCC</td><td>>hg38_21_+_8433172_8446622_RNA45SN1_45S_50ntFlanks</td><td>9027</td><td>9059</td><td>33</td><td>CGTCTCCTCGTGGGGGGGCCGGGCCACCCCTCC</td><td>81.1</td><td>-7.9</td><td>1.3</td><td>-6.1</td><td>1.1E-09</td><td>2.1E-09</td><td>MB231/MB468</td></tr>
<tr><td>>hg38_21_+_8433172_8446622_RNA45SN1_45S_50ntFlanks@10806.10840.35|AACAATGTAGGTAAGGGAAGTCGGCAAGCCGGATC</td><td>>hg38_21_+_8433172_8446622_RNA45SN1_45S_50ntFlanks</td><td>10806</td><td>10840</td><td>35</td><td>AACAATGTAGGTAAGGGAAGTCGGCAAGCCGGATC</td><td>230.3</td><td>-2.5</td><td>0.4</td><td>-6.1</td><td>1.1E-09</td><td>2.1E-09</td><td>MB231/MB468</td></tr>
<tr><td>>hg38_21_+_8433172_8446622_RNA45SN1_45S_50ntFlanks@11873.11892.20|AAGAAGACCCTGTTGAGCTT</td><td>>hg38_21_+_8433172_8446622_RNA45SN1_45S_50ntFlanks</td><td>11873</td><td>11892</td><td>20</td><td>AAGAAGACCCTGTTGAGCTT</td><td>858.4</td><td>-3.8</td><td>0.6</td><td>-6.1</td><td>1.3E-09</td><td>2.1E-09</td><td>BT20/MB468</td></tr>
<tr><td>>hg38_21_+_8433172_8446622_RNA45SN1_45S_50ntFlanks@10689.10706.18|GCGTCCGGTGAGCTCTCG</td><td>>hg38_21_+_8433172_8446622_RNA45SN1_45S_50ntFlanks</td><td>10689</td><td>10706</td><td>18</td><td>GCGTCCGGTGAGCTCTCG</td><td>1047.6</td><td>-7.4</td><td>1.2</td><td>-6.1</td><td>1.1E-09</td><td>2.1E-09</td><td>MB231/MB468</td></tr>
<tr><td>>hg38_21_+_8433172_8446622_RNA45SN1_45S_50ntFlanks@12347.12364.18|AGAAAAGTTACCACAGGG</td><td>>hg38_21_+_8433172_8446622_RNA45SN1_45S_50ntFlanks</td><td>12347</td><td>12364</td><td>18</td><td>AGAAAAGTTACCACAGGG</td><td>66.9</td><td>-7.6</td><td>1.3</td><td>-6.1</td><td>1.1E-09</td><td>2.1E-09</td><td>MB231/MB468</td></tr>
<tr><td>>hg38_21_+_8433172_8446622_RNA45SN1_45S_50ntFlanks@4005.4025.21|AACCTCGGGCCGATCGCACGC</td><td>>hg38_21_+_8433172_8446622_RNA45SN1_45S_50ntFlanks</td><td>4005</td><td>4025</td><td>21</td><td>AACCTCGGGCCGATCGCACGC</td><td>183.6</td><td>-3.0</td><td>0.5</td><td>-6.1</td><td>1.3E-09</td><td>2.1E-09</td><td>BT20/MB468</td></tr>
<tr><td>>hg38_21_+_8433172_8446622_RNA45SN1_45S_50ntFlanks@7984.8004.21|AGATCAGACGTGGCGACCCGC</td><td>>hg38_21_+_8433172_8446622_RNA45SN1_45S_50ntFlanks</td><td>7984</td><td>8004</td><td>21</td><td>AGATCAGACGTGGCGACCCGC</td><td>183.3</td><td>-5.8</td><td>1.0</td><td>-6.1</td><td>1.2E-09</td><td>2.2E-09</td><td>MB231/MB468</td></tr>
<tr><td>>hg38_21_+_8433172_8446622_RNA45SN1_45S_50ntFlanks@11755.11777.23|AATGCCTCGTCATCTAATTAGTG</td><td>>hg38_21_+_8433172_8446622_RNA45SN1_45S_50ntFlanks</td><td>11755</td><td>11777</td><td>23</td><td>AATGCCTCGTCATCTAATTAGTG</td><td>262.3</td><td>-5.7</td><td>0.9</td><td>-6.1</td><td>1.3E-09</td><td>2.2E-09</td><td>BT20/MB468</td></tr>
<tr><td>>hg38_21_+_8433172_8446622_RNA45SN1_45S_50ntFlanks@12617.12654.38|TGAGGAGCCAATGGGGCGAAGCTACCATCTGTGGGATT</td><td>>hg38_21_+_8433172_8446622_RNA45SN1_45S_50ntFlanks</td><td>12617</td><td>12654</td><td>38</td><td>TGAGGAGCCAATGGGGCGAAGCTACCATCTGTGGGATT</td><td>546.5</td><td>-3.4</td><td>0.6</td><td>-6.1</td><td>1.2E-09</td><td>2.2E-09</td><td>MB231/MB468</td></tr>
<tr><td>>hg38_21_+_8433172_8446622_RNA45SN1_45S_50ntFlanks@12415.12436.22|CTTCGATGTCGGCTCTTCCTAT</td><td>>hg38_21_+_8433172_8446622_RNA45SN1_45S_50ntFlanks</td><td>12415</td><td>12436</td><td>22</td><td>CTTCGATGTCGGCTCTTCCTAT</td><td>2284.7</td><td>-3.4</td><td>0.6</td><td>-6.1</td><td>1.2E-09</td><td>2.2E-09</td><td>MB231/MB468</td></tr>
<tr><td>>hg38_21_+_8433172_8446622_RNA45SN1_45S_50ntFlanks@12875.12892.18|CTGGAAAGGCGGCCGCCC</td><td>>hg38_21_+_8433172_8446622_RNA45SN1_45S_50ntFlanks</td><td>12875</td><td>12892</td><td>18</td><td>CTGGAAAGGCGGCCGCCC</td><td>80.9</td><td>-7.9</td><td>1.3</td><td>-6.1</td><td>1.2E-09</td><td>2.2E-09</td><td>MB231/MB468</td></tr>
<tr><td>>MI0003674|hsa-mir-653&WithFlank&7|-|93482754|93482861@22.43.22&[MIMAT0003328&hsa-miR-653-5p&offsets|+3|+4;m-368&7|-|93482819|93482840&offsets|0|0]|TTGAAACAATCTCTACTGAACC</td><td>>MI0003674|hsa-mir-653&WithFlank&7|-|93482754|93482861</td><td>22</td><td>43</td><td>22</td><td>TTGAAACAATCTCTACTGAACC</td><td>221.7</td><td>-6.3</td><td>1.0</td><td>-6.5</td><td>8.0E-11</td><td>2.2E-09</td><td>BT20/MB231</td></tr>
<tr><td>>hg38_21_+_8433172_8446622_RNA45SN1_45S_50ntFlanks@10786.10825.40|AACAGCCTCTGGCATGTTGGAACAATGTAGGTAAGGGAAG</td><td>>hg38_21_+_8433172_8446622_RNA45SN1_45S_50ntFlanks</td><td>10786</td><td>10825</td><td>40</td><td>AACAGCCTCTGGCATGTTGGAACAATGTAGGTAAGGGAAG</td><td>310.7</td><td>-4.8</td><td>0.8</td><td>-6.1</td><td>1.2E-09</td><td>2.2E-09</td><td>MB231/MB468</td></tr>
<tr><td>>hg38_21_+_8433172_8446622_RNA45SN1_45S_50ntFlanks@7973.8008.36|GACGCGACCTCAGATCAGACGTGGCGACCCGCTGAA</td><td>>hg38_21_+_8433172_8446622_RNA45SN1_45S_50ntFlanks</td><td>7973</td><td>8008</td><td>36</td><td>GACGCGACCTCAGATCAGACGTGGCGACCCGCTGAA</td><td>1670.8</td><td>4.6</td><td>0.8</td><td>6.1</td><td>1.2E-09</td><td>2.2E-09</td><td>MB231/MB468</td></tr>
<tr><td>>hg38_21_+_8433172_8446622_RNA45SN1_45S_50ntFlanks@10786.10817.32|AACAGCCTCTGGCATGTTGGAACAATGTAGGT</td><td>>hg38_21_+_8433172_8446622_RNA45SN1_45S_50ntFlanks</td><td>10786</td><td>10817</td><td>32</td><td>AACAGCCTCTGGCATGTTGGAACAATGTAGGT</td><td>1087.5</td><td>-3.0</td><td>0.5</td><td>-6.1</td><td>1.2E-09</td><td>2.2E-09</td><td>MB231/MB468</td></tr>
<tr><td>>MI0016003|hsa-mir-3613&WithFlank&13|-|49996409|49996507@22.43.22&[MIMAT0017990&hsa-miR-3613-5p&offsets|0|0;m-259&13|-|49996465|49996486&offsets|0|0]|TGTTGTACTTTTTTTTTTGTTC</td><td>>MI0016003|hsa-mir-3613&WithFlank&13|-|49996409|49996507</td><td>22</td><td>43</td><td>22</td><td>TGTTGTACTTTTTTTTTTGTTC</td><td>1007.8</td><td>5.6</td><td>0.9</td><td>6.1</td><td>1.4E-09</td><td>2.2E-09</td><td>BT20/MB468</td></tr>
<tr><td>>hg38_21_+_8433172_8446622_RNA45SN1_45S_50ntFlanks@9503.9526.24|GGTGAACTATGCCTGGGCAGGGCG</td><td>>hg38_21_+_8433172_8446622_RNA45SN1_45S_50ntFlanks</td><td>9503</td><td>9526</td><td>24</td><td>GGTGAACTATGCCTGGGCAGGGCG</td><td>83.0</td><td>-5.4</td><td>0.9</td><td>-6.1</td><td>1.2E-09</td><td>2.2E-09</td><td>MB231/MB468</td></tr>
</table>

| | | | | | | | | | | | | |
|---|---|---|---|---|---|---|---|---|---|---|---|---|
| >hg38_21_+_8433172_8446622_RNA45SN1_45S_50ntFlanks@11586.11611.26\|GTGCGGACCAGGGGAATCCGACTGTT | >hg38_21_+_8433172_8446622_RNA45SN1_45S_50ntFlanks | 11586 | 11611 | 26 | GTGCGGACCAGGGGAATCCGACTGTT | 2103.3 | -3.5 | 0.6 | -6.1 | 1.4E-09 | 2.2E-09 | BT20/MB468 |
| >hg38_21_+_8433172_8446622_RNA45SN1_45S_50ntFlanks@10922.10944.23\|CTGGACGAGGCGCCGCCGCCCCC | >hg38_21_+_8433172_8446622_RNA45SN1_45S_50ntFlanks | 10922 | 10944 | 23 | CTGGACGAGGCGCCGCCGCCCCC | 68.1 | -7.7 | 1.3 | -6.1 | 1.2E-09 | 2.2E-09 | MB231/MB468 |
| >hg38_21_+_8433172_8446622_RNA45SN1_45S_50ntFlanks@11999.12043.45\|CCCTGCGGGCCGCCGGTGAAATACCACTACTCTGATCGTTTTTTC | >hg38_21_+_8433172_8446622_RNA45SN1_45S_50ntFlanks | 11999 | 12043 | 45 | CCCTGCGGGCCGCCGGTGAAATACCACTACTCTGATCGTTTTTTC | 123.7 | -8.5 | 1.4 | -6.1 | 1.2E-09 | 2.3E-09 | MB231/MB468 |
| >hg38_21_+_8433172_8446622_RNA45SN1_45S_50ntFlanks@11768.11785.18\|CTAATTAGTGACGCGCAT | >hg38_21_+_8433172_8446622_RNA45SN1_45S_50ntFlanks | 11768 | 11785 | 18 | CTAATTAGTGACGCGCAT | 98.2 | -8.2 | 1.3 | -6.1 | 1.2E-09 | 2.3E-09 | MB231/MB468 |
| >MI0000061\|hsa-let-7a-2&WithFlank&11\|-\|122146516\|122146599@11.30.20;MI0000062\|hsa-let-7a-3&WithFlank&22\|+\|46112743\|46112828@10.29.20;MI0000060\|hsa-let-7a-1&WithFlank&9\|+\|94175951\|94176042@12.31.20&[MIMAT0000062&hsa-let-7a-5p&offsets\|0\|-2;m-6&11\|-\|122146568\|122146589&offsets\|0\|-2];[MIMAT0000062_1&hsa-let-7a-5p&offsets\|0\|-2;m-5&22\|+\|46112752\|46112773&offsets\|0\|-2];[MIMAT0000062_2&hsa-let-7a-5p&offsets\|0\|-2;m-4&9\|+\|94175962\|94175983&offsets\|0\|-2]\|TGAGGTAGTAGGTTGTATAG | >MI0000061\|hsa-let-7a-2&WithFlank&11\|-\|122146516\|122146599 | 11 | 30 | 20 | TGAGGTAGTAGGTTGTATAG | 418.5 | 8.0 | 1.3 | 6.1 | 1.4E-09 | 2.3E-09 | BT20/MB468 |
| >hg38_21_+_8433172_8446622_RNA45SN1_45S_50ntFlanks@12624.12652.29\|CCAATGGGGCGAAGCTACCATCTGTGGGA | >hg38_21_+_8433172_8446622_RNA45SN1_45S_50ntFlanks | 12624 | 12652 | 29 | CCAATGGGGCGAAGCTACCATCTGTGGGA | 95.5 | -5.6 | 0.9 | -6.1 | 1.2E-09 | 2.3E-09 | MB231/MB468 |
| >hg38_21_+_8433172_8446622_RNA45SN1_45S_50ntFlanks@12446.12480.35\|GCAGAATTCACCAAGCGTTGGATTGTTCACCCACT | >hg38_21_+_8433172_8446622_RNA45SN1_45S_50ntFlanks | 12446 | 12480 | 35 | GCAGAATTCACCAAGCGTTGGATTGTTCACCCACT | 388.3 | 6.0 | 1.0 | 6.1 | 1.2E-09 | 2.3E-09 | MB231/MB468 |
| >hg38_21_+_8433172_8446622_RNA45SN1_45S_50ntFlanks@12500.12518.19\|GTTTAGACCGTCGTGAGAC | >hg38_21_+_8433172_8446622_RNA45SN1_45S_50ntFlanks | 12500 | 12518 | 19 | GTTTAGACCGTCGTGAGAC | 93.7 | -8.1 | 1.3 | -6.1 | 1.2E-09 | 2.3E-09 | MB231/MB468 |
| >MI0000483\|hsa-mir-186&WithFlank&1\|-\|71067625\|71067722@21.44.24&[MIMAT0000456&hsa-miR-186-5p&offsets\|0\|+2;m-84&1\|-\|71067681\|71067702&offsets\|0\|+2]\|CAAAGAATTCTCCTTTTGGGCTTT | >MI0000483\|hsa-mir-186&WithFlank&1\|-\|71067625\|71067722 | 21 | 44 | 24 | CAAAGAATTCTCCTTTTGGGCTTT | 814.5 | 6.3 | 1.0 | 6.1 | 1.4E-09 | 2.3E-09 | BT20/MB468 |
| >MI0000071\|hsa-mir-17&WithFlank&13\|+\|91350599\|91350694@20.43.24&[MIMAT0000070&hsa-miR-17-5p&offsets\|0\|+1;m-88&13\|+\|91350618\|91350640&offsets\|0\|+1]\|CAAAGTGCTTACAGTGCAGGTAGT | >MI0000071\|hsa-mir-17&WithFlank&13\|+\|91350599\|91350694 | 20 | 43 | 24 | CAAAGTGCTTACAGTGCAGGTAGT | 3017.7 | 5.3 | 0.9 | 6.1 | 1.4E-09 | 2.3E-09 | BT20/MB468 |
| >hg38_21_+_8433172_8446622_RNA45SN1_45S_50ntFlanks@4564.4587.24\|GAATAATGGAATAGGACCGCGGTT | >hg38_21_+_8433172_8446622_RNA45SN1_45S_50ntFlanks | 4564 | 4587 | 24 | GAATAATGGAATAGGACCGCGGTT | 125.1 | -4.9 | 0.8 | -6.1 | 1.3E-09 | 2.3E-09 | MB231/MB468 |
| >hg38_21_+_8433172_8446622_RNA45SN1_45S_50ntFlanks@11572.11600.29\|CCGACTTAGAACTGGTGCGGACCAGGGGA | >hg38_21_+_8433172_8446622_RNA45SN1_45S_50ntFlanks | 11572 | 11600 | 29 | CCGACTTAGAACTGGTGCGGACCAGGGGA | 2059.9 | -3.5 | 0.6 | -6.1 | 1.3E-09 | 2.3E-09 | MB231/MB468 |
| >hg38_21_+_8433172_8446622_RNA45SN1_45S_50ntFlanks@6699.6717.19\|GCTAGCTGCGAGAATTAAT | >hg38_21_+_8433172_8446622_RNA45SN1_45S_50ntFlanks | 6699 | 6717 | 19 | GCTAGCTGCGAGAATTAAT | 322.5 | -3.5 | 0.6 | -6.1 | 1.3E-09 | 2.3E-09 | MB231/MB468 |
| >hg38_21_+_8433172_8446622_RNA45SN1_45S_50ntFlanks@6667.6689.23\|ATCACTCGGCTCGTGCGTCGATG | >hg38_21_+_8433172_8446622_RNA45SN1_45S_50ntFlanks | 6667 | 6689 | 23 | ATCACTCGGCTCGTGCGTCGATG | 74.2 | -7.8 | 1.3 | -6.1 | 1.3E-09 | 2.4E-09 | MB231/MB468 |
| >hg38_21_+_8433172_8446622_RNA45SN1_45S_50ntFlanks@10690.10709.20\|CGTCCGGTGAGCTCTCGCTG | >hg38_21_+_8433172_8446622_RNA45SN1_45S_50ntFlanks | 10690 | 10709 | 20 | CGTCCGGTGAGCTCTCGCTG | 1053.9 | -6.2 | 1.0 | -6.1 | 1.3E-09 | 2.4E-09 | MB231/MB468 |
| >hg38_21_+_8433172_8446622_RNA45SN1_45S_50ntFlanks@11569.11600.32\|CAGCCGACTTAGAACTGGTGCGGACCAGGGGA | >hg38_21_+_8433172_8446622_RNA45SN1_45S_50ntFlanks | 11569 | 11600 | 32 | CAGCCGACTTAGAACTGGTGCGGACCAGGGGA | 955.8 | -3.4 | 0.6 | -6.1 | 1.3E-09 | 2.4E-09 | MB231/MB468 |
| >hg38_21_+_8433172_8446622_RNA45SN1_45S_50ntFlanks@12457.12484.28\|CAAGCGTTGGATTGTTCACCCACTAATA | >hg38_21_+_8433172_8446622_RNA45SN1_45S_50ntFlanks | 12457 | 12484 | 28 | CAAGCGTTGGATTGTTCACCCACTAATA | 1076.5 | -6.9 | 1.1 | -6.0 | 1.5E-09 | 2.4E-09 | BT20/MB468 |
| >hg38_21_+_8433172_8446622_RNA45SN1_45S_50ntFlanks@10916.10935.20\|CCGCGGCTGGACGAGGCGCC | >hg38_21_+_8433172_8446622_RNA45SN1_45S_50ntFlanks | 10916 | 10935 | 20 | CCGCGGCTGGACGAGGCGCC | 87.2 | -8.0 | 1.3 | -6.1 | 1.3E-09 | 2.4E-09 | MB231/MB468 |
| >MI0000736\|hsa-mir-30c-1&WithFlank&1\|+\|40757278\|40757378@23.42.20;MI0000254\|hsa-mir-30c-2&WithFlank&6\|-\|71376954\|71377037@13.32.20&[MIMAT0000244&hsa-miR-30c-5p&offsets\|0\|-3;m-57&1\|+\|40757300\|40757323&offsets\|0\|-4];[MIMAT0000244_1&hsa-miR-30c-5p&offsets\|0\|-3;m-56&6\|-\|71377002\|71377025&offsets\|0\|-4]\|TGTAAACATCCTACACTCTC | >MI0000736\|hsa-mir-30c-1&WithFlank&1\|+\|40757278\|40757378 | 23 | 42 | 20 | TGTAAACATCCTACACTCTC | 1403.6 | 5.3 | 0.9 | 6.0 | 1.5E-09 | 2.4E-09 | BT20/MB468 |
| >MI0001519\|hsa-mir-20b&WithFlank&X\|-\|134169803\|134169883@12.34.23&[MIMAT0001413&hsa-miR-20b-5p&offsets\|0\|0;m-241&X\|-\|134169850\|134169872&offsets\|0\|0]\|CAAAGTGCTCATAGTGCAGGTAG | >MI0001519\|hsa-mir-20b&WithFlank&X\|-\|134169803\|134169883 | 12 | 34 | 23 | CAAAGTGCTCATAGTGCAGGTAG | 652.6 | 5.6 | 0.9 | 6.0 | 1.5E-09 | 2.4E-09 | BT20/MB468 |
| >hg38_21_+_8433172_8446622_RNA45SN1_45S_50ntFlanks@5487.5526.40\|CGCTGAGAAGACGGTCGAACTTGACTATCTAGAGGAAGTA | >hg38_21_+_8433172_8446622_RNA45SN1_45S_50ntFlanks | 5487 | 5526 | 40 | CGCTGAGAAGACGGTCGAACTTGACTATCTAGAGGAAGTA | 207.6 | -4.9 | 0.8 | -6.1 | 1.3E-09 | 2.4E-09 | MB231/MB468 |
| >hg38_21_+_8433172_8446622_RNA45SN1_45S_50ntFlanks@12934.12962.29\|CTAAACCATTCGTAGACGACCTGCTTCTG | >hg38_21_+_8433172_8446622_RNA45SN1_45S_50ntFlanks | 12934 | 12962 | 29 | CTAAACCATTCGTAGACGACCTGCTTCTG | 207.0 | -3.6 | 0.6 | -6.1 | 1.3E-09 | 2.4E-09 | MB231/MB468 |
| >hg38_21_+_8433172_8446622_RNA45SN1_45S_50ntFlanks@5420.5443.24\|CCGATTGGATGGTTTAGTGAGGCC | >hg38_21_+_8433172_8446622_RNA45SN1_45S_50ntFlanks | 5420 | 5443 | 24 | CCGATTGGATGGTTTAGTGAGGCC | 792.1 | -6.4 | 1.1 | -6.1 | 1.3E-09 | 2.4E-09 | MB231/MB468 |
| >hg38_21_+_8433172_8446622_RNA45SN1_45S_50ntFlanks@12626.12651.26\|AATGGGGCGAAGCTACCATCTGTGGG | >hg38_21_+_8433172_8446622_RNA45SN1_45S_50ntFlanks | 12626 | 12651 | 26 | AATGGGGCGAAGCTACCATCTGTGGG | 78.9 | -4.7 | 0.8 | -6.0 | 1.5E-09 | 2.4E-09 | BT20/MB468 |
| >hg38_21_+_8433172_8446622_RNA45SN1_45S_50ntFlanks@12540.12569.30\|TGATGTGTTGTTGCCATGGTAATCCTGCTC | >hg38_21_+_8433172_8446622_RNA45SN1_45S_50ntFlanks | 12540 | 12569 | 30 | TGATGTGTTGTTGCCATGGTAATCCTGCTC | 100.0 | -4.0 | 0.7 | -6.0 | 1.5E-09 | 2.5E-09 | BT20/MB468 |
| >hg38_21_+_8433172_8446622_RNA45SN1_45S_50ntFlanks@12709.12742.34\|CCGCGGAGCCTCGGTTGGCCTCGGATAGCCGGTC | >hg38_21_+_8433172_8446622_RNA45SN1_45S_50ntFlanks | 12709 | 12742 | 34 | CCGCGGAGCCTCGGTTGGCCTCGGATAGCCGGTC | 80.9 | -7.9 | 1.3 | -6.1 | 1.3E-09 | 2.5E-09 | MB231/MB468 |
| >hg38_21_+_8433172_8446622_RNA45SN1_45S_50ntFlanks@10687.10709.23\|CGGCGTCCGGTGAGCTCTCGCTG | >hg38_21_+_8433172_8446622_RNA45SN1_45S_50ntFlanks | 10687 | 10709 | 23 | CGGCGTCCGGTGAGCTCTCGCTG | 2914.9 | -4.8 | 0.8 | -6.0 | 1.5E-09 | 2.5E-09 | BT20/MB468 |
| >hg38_21_+_8433172_8446622_RNA45SN1_45S_50ntFlanks@11027.11061.35\|CTCTCTCTCTCTCTCCCCCGCTCCCCGTCCTCCCC | >hg38_21_+_8433172_8446622_RNA45SN1_45S_50ntFlanks | 11027 | 11061 | 35 | CTCTCTCTCTCTCTCCCCCGCTCCCCGTCCTCCCC | 75.3 | -7.8 | 1.3 | -6.1 | 1.4E-09 | 2.5E-09 | MB231/MB468 |
| >hg38_21_+_8433172_8446622_RNA45SN1_45S_50ntFlanks@10357.10388.32\|CTTTGAAGGCCGAAGTGGAGAAGGGTTCCATG | >hg38_21_+_8433172_8446622_RNA45SN1_45S_50ntFlanks | 10357 | 10388 | 32 | CTTTGAAGGCCGAAGTGGAGAAGGGTTCCATG | 346.3 | -4.1 | 0.7 | -6.1 | 1.4E-09 | 2.6E-09 | MB231/MB468 |
| >hg38_21_+_8433172_8446622_RNA45SN1_45S_50ntFlanks@11783.11807.25\|CATGAATGGATGAACGAGATTCCCA | >hg38_21_+_8433172_8446622_RNA45SN1_45S_50ntFlanks | 11783 | 11807 | 25 | CATGAATGGATGAACGAGATTCCCA | 233.5 | -3.7 | 0.6 | -6.1 | 1.4E-09 | 2.6E-09 | MB231/MB468 |
| >hg38_21_+_8433172_8446622_RNA45SN1_45S_50ntFlanks@12626.12661.36\|AATGGGGCGAAGCTACCATCTGTGGGATTATGACTG | >hg38_21_+_8433172_8446622_RNA45SN1_45S_50ntFlanks | 12626 | 12661 | 36 | AATGGGGCGAAGCTACCATCTGTGGGATTATGACTG | 1075.0 | -4.0 | 0.7 | -6.0 | 1.6E-09 | 2.6E-09 | BT20/MB468 |
| >hg38_21_+_8433172_8446622_RNA45SN1_45S_50ntFlanks@8472.8489.18\|CGGCCCGGCGGATCTTTC | >hg38_21_+_8433172_8446622_RNA45SN1_45S_50ntFlanks | 8472 | 8489 | 18 | CGGCCCGGCGGATCTTTC | 79.8 | -7.9 | 1.3 | -6.1 | 1.4E-09 | 2.6E-09 | MB231/MB468 |
| >hg38_21_+_8433172_8446622_RNA45SN1_45S_50ntFlanks@6646.6674.29\|TCGTACGACTCTTAGCGGTGGATCACTCG | >hg38_21_+_8433172_8446622_RNA45SN1_45S_50ntFlanks | 6646 | 6674 | 29 | TCGTACGACTCTTAGCGGTGGATCACTCG | 766.8 | 4.0 | 0.7 | 6.1 | 1.4E-09 | 2.6E-09 | MB231/MB468 |
| >hg38_21_+_8433172_8446622_RNA45SN1_45S_50ntFlanks@6652.6677.26\|GACTCTTAGCGGTGGATCACTCGGCT | >hg38_21_+_8433172_8446622_RNA45SN1_45S_50ntFlanks | 6652 | 6677 | 26 | GACTCTTAGCGGTGGATCACTCGGCT | 829.0 | 4.1 | 0.7 | 6.1 | 1.4E-09 | 2.6E-09 | MB231/MB468 |
| >hg38_21_+_8433172_8446622_RNA45SN1_45S_50ntFlanks@6719.6739.21\|TGAATTGCAGGACACATTGAT | >hg38_21_+_8433172_8446622_RNA45SN1_45S_50ntFlanks | 6719 | 6739 | 21 | TGAATTGCAGGACACATTGAT | 1395.2 | -6.1 | 1.0 | -6.0 | 1.6E-09 | 2.6E-09 | BT20/MB468 |
| >hg38_21_+_8433172_8446622_RNA45SN1_45S_50ntFlanks@11874.11899.26\|AGAAGACCCTGTTGAGCTTGACTCTA | >hg38_21_+_8433172_8446622_RNA45SN1_45S_50ntFlanks | 11874 | 11899 | 26 | AGAAGACCCTGTTGAGCTTGACTCTA | 358.9 | -3.9 | 0.6 | -6.0 | 1.6E-09 | 2.6E-09 | BT20/MB468 |
| >hg38_21_+_8433172_8446622_RNA45SN1_45S_50ntFlanks@10392.10411.20\|ACAGCAGTTGAACATGGGTC | >hg38_21_+_8433172_8446622_RNA45SN1_45S_50ntFlanks | 10392 | 10411 | 20 | ACAGCAGTTGAACATGGGTC | 319.1 | -4.5 | 0.8 | -6.0 | 1.6E-09 | 2.6E-09 | BT20/MB468 |
| >hg38_21_+_8433172_8446622_RNA45SN1_45S_50ntFlanks@6722.6739.18\|ATTGCAGGACACATTGAT | >hg38_21_+_8433172_8446622_RNA45SN1_45S_50ntFlanks | 6722 | 6739 | 18 | ATTGCAGGACACATTGAT | 361.7 | -4.0 | 0.7 | -6.0 | 1.6E-09 | 2.6E-09 | BT20/MB468 |
| >hg38_21_+_8433172_8446622_RNA45SN1_45S_50ntFlanks@8381.8404.24\|AAGAGGGCGTGAAACCGTTAAGAG | >hg38_21_+_8433172_8446622_RNA45SN1_45S_50ntFlanks | 8381 | 8404 | 24 | AAGAGGGCGTGAAACCGTTAAGAG | 170.7 | -5.6 | 0.9 | -6.0 | 1.6E-09 | 2.7E-09 | BT20/MB468 |
| >MI0003137\|hsa-mir-193b&WithFlank&16\|+\|14303961\|14304055@57.77.21(+1A)&[MIMAT0002819&hsa-miR-193b-3p&offsets\|0\|-1(+1A);m-166&16\|+\|14304017\|14304038&offsets\|0\|-1(+1A)]\|AACTGGCCCTCAAAGTCCCGCA | >MI0003137\|hsa-mir-193b&WithFlank&16\|+\|14303961\|14304055 | 57 | 77 | 21(+1A) | AACTGGCCCTCAAAGTCCCGCA | 922.4 | 6.6 | 1.0 | 6.5 | 9.8E-11 | 2.7E-09 | BT20/MB231 |
| >hg38_21_+_8433172_8446622_RNA45SN1_45S_50ntFlanks@10826.10865.40\|TCGGCAAGCCGGATCCGTAACTTCGGGATAAGGATTGGCT | >hg38_21_+_8433172_8446622_RNA45SN1_45S_50ntFlanks | 10826 | 10865 | 40 | TCGGCAAGCCGGATCCGTAACTTCGGGATAAGGATTGGCT | 115.5 | -8.4 | 1.4 | -6.0 | 1.5E-09 | 2.7E-09 | MB231/MB468 |
| >hg38_21_+_8433172_8446622_RNA45SN1_45S_50ntFlanks@8381.8399.19\|AAGAGGGCGTGAAACCGTT | >hg38_21_+_8433172_8446622_RNA45SN1_45S_50ntFlanks | 8381 | 8399 | 19 | AAGAGGGCGTGAAACCGTT | 250.3 | -3.6 | 0.6 | -6.0 | 1.5E-09 | 2.7E-09 | MB231/MB468 |
| >hg38_21_+_8433172_8446622_RNA45SN1_45S_50ntFlanks@10681.10708.28\|TTCCGGCGGCGTCCGGTGAGCTCTCGCT | >hg38_21_+_8433172_8446622_RNA45SN1_45S_50ntFlanks | 10681 | 10708 | 28 | TTCCGGCGGCGTCCGGTGAGCTCTCGCT | 338.1 | -4.3 | 0.7 | -6.0 | 1.5E-09 | 2.7E-09 | MB231/MB468 |
| >hg38_21_+_8433172_8446622_RNA45SN1_45S_50ntFlanks@10771.10790.20\|CAGGTCTCCAAGGTGAACAG | >hg38_21_+_8433172_8446622_RNA45SN1_45S_50ntFlanks | 10771 | 10790 | 20 | CAGGTCTCCAAGGTGAACAG | 185.9 | -4.4 | 0.7 | -6.0 | 1.5E-09 | 2.7E-09 | MB231/MB468 |
| >hg38_21_+_8433172_8446622_RNA45SN1_45S_50ntFlanks@10846.10863.18\|CTTCGGGATAAGGATTGG | >hg38_21_+_8433172_8446622_RNA45SN1_45S_50ntFlanks | 10846 | 10863 | 18 | CTTCGGGATAAGGATTGG | 255.2 | -3.8 | 0.6 | -6.0 | 1.5E-09 | 2.7E-09 | MB231/MB468 |

<table>
<tr><td>>hg38_21_+_8433172_8446622_RNA45SN1_45S_50ntFlanks@8552.8581.30|CTCCCCGGAGGGGGCGGGCTCCGGCGGGTG</td><td>>hg38_21_+_8433172_8446622_RNA45SN1_45S_50ntFlanks</td><td>8552</td><td>8581</td><td>30</td><td>CTCCCCGGAGGGGGCGGGCTCCGGCGGGTG</td><td>104.5</td><td>-5.4</td><td>0.9</td><td>-6.0</td><td>1.7E-09</td><td>2.7E-09</td><td>BT20/MB468</td></tr>
<tr><td>>hg38_21_+_8433172_8446622_RNA45SN1_45S_50ntFlanks@10386.10410.25|ATGTGAACAGCAGTTGAACATGGGT</td><td>>hg38_21_+_8433172_8446622_RNA45SN1_45S_50ntFlanks</td><td>10386</td><td>10410</td><td>25</td><td>ATGTGAACAGCAGTTGAACATGGGT</td><td>205.1</td><td>-3.1</td><td>0.5</td><td>-6.0</td><td>1.5E-09</td><td>2.7E-09</td><td>MB231/MB468</td></tr>
<tr><td>>hg38_21_+_8433172_8446622_RNA45SN1_45S_50ntFlanks@12685.12711.27|CCCAGGCGGAACGATACGGCAGCGCCG</td><td>>hg38_21_+_8433172_8446622_RNA45SN1_45S_50ntFlanks</td><td>12685</td><td>12711</td><td>27</td><td>CCCAGGCGGAACGATACGGCAGCGCCG</td><td>93.2</td><td>-8.1</td><td>1.3</td><td>-6.0</td><td>1.5E-09</td><td>2.7E-09</td><td>MB231/MB468</td></tr>
<tr><td>>hg38_21_+_8433172_8446622_RNA45SN1_45S_50ntFlanks@5419.5445.27|ACCGATTGGATGGTTTAGTGAGGCCCT</td><td>>hg38_21_+_8433172_8446622_RNA45SN1_45S_50ntFlanks</td><td>5419</td><td>5445</td><td>27</td><td>ACCGATTGGATGGTTTAGTGAGGCCCT</td><td>76.3</td><td>-7.8</td><td>1.3</td><td>-6.0</td><td>1.5E-09</td><td>2.8E-09</td><td>MB231/MB468</td></tr>
<tr><td>>trna16_GlnTTG_17_+_47269890_47269961@1.29.29,trna64_GlnTTG_6_+_28557156_28557227@1.29.29,trna84_GlnTTG_6_+_145503859_145503930@1.29.29|GGTCCCATGGTGTAATGGTTAGCACTCTG</td><td>>trna16_GlnTTG_17_+_47269890_47269961</td><td>1</td><td>29</td><td>29</td><td>GGTCCCATGGTGTAATGGTTAGCACTCTG</td><td>1998.3</td><td>-6.6</td><td>1.1</td><td>-6.0</td><td>1.7E-09</td><td>2.8E-09</td><td>BT20/MB468</td></tr>
<tr><td>>hg38_21_+_8433172_8446622_RNA45SN1_45S_50ntFlanks@6663.6689.27|GTGGATCACTCGGCTCGTGCGTCGATG</td><td>>hg38_21_+_8433172_8446622_RNA45SN1_45S_50ntFlanks</td><td>6663</td><td>6689</td><td>27</td><td>GTGGATCACTCGGCTCGTGCGTCGATG</td><td>1204.8</td><td>-3.7</td><td>0.6</td><td>-6.0</td><td>1.5E-09</td><td>2.8E-09</td><td>MB231/MB468</td></tr>
<tr><td>>hg38_21_+_8433172_8446622_RNA45SN1_45S_50ntFlanks@12607.12626.20|TGTGCTTGGCTGAGGAGCCA</td><td>>hg38_21_+_8433172_8446622_RNA45SN1_45S_50ntFlanks</td><td>12607</td><td>12626</td><td>20</td><td>TGTGCTTGGCTGAGGAGCCA</td><td>185.8</td><td>-2.8</td><td>0.5</td><td>-6.0</td><td>1.5E-09</td><td>2.8E-09</td><td>MB231/MB468</td></tr>
<tr><td>>hg38_21_+_8433172_8446622_RNA45SN1_45S_50ntFlanks@11579.11605.27|AGAACTGGTGCGGACCAGGGGAATCCG</td><td>>hg38_21_+_8433172_8446622_RNA45SN1_45S_50ntFlanks</td><td>11579</td><td>11605</td><td>27</td><td>AGAACTGGTGCGGACCAGGGGAATCCG</td><td>1151.5</td><td>-7.0</td><td>1.2</td><td>-6.0</td><td>1.7E-09</td><td>2.8E-09</td><td>BT20/MB468</td></tr>
<tr><td>>hg38_1_-_228634819_228635039_RNA5S12_5S_50ntFlanks@124.170.47|AGTACTTGGATGGGAGACCGCCTGGGAATACCGGGTGCTGTAGGCTT</td><td>>hg38_1_-_228634819_228635039_RNA5S12_5S_50ntFlanks</td><td>124</td><td>170</td><td>47</td><td>AGTACTTGGATGGGAGACCGCCTGGGAATACCGGGTGCTGTAGGCTT</td><td>746.0</td><td>7.7</td><td>1.3</td><td>6.0</td><td>1.5E-09</td><td>2.8E-09</td><td>MB231/MB468</td></tr>
<tr><td>>hg38_21_+_8433172_8446622_RNA45SN1_45S_50ntFlanks@3793.3814.22|CGAATGGCTCATTAAATCAGTT</td><td>>hg38_21_+_8433172_8446622_RNA45SN1_45S_50ntFlanks</td><td>3793</td><td>3814</td><td>22</td><td>CGAATGGCTCATTAAATCAGTT</td><td>76.1</td><td>-5.3</td><td>0.9</td><td>-6.0</td><td>1.5E-09</td><td>2.8E-09</td><td>MB231/MB468</td></tr>
<tr><td>>hg38_21_+_8433172_8446622_RNA45SN1_45S_50ntFlanks@6645.6663.19|CTCGTACGACTCTTAGCGG</td><td>>hg38_21_+_8433172_8446622_RNA45SN1_45S_50ntFlanks</td><td>6645</td><td>6663</td><td>19</td><td>CTCGTACGACTCTTAGCGG</td><td>2631.5</td><td>3.2</td><td>0.5</td><td>6.0</td><td>1.7E-09</td><td>2.9E-09</td><td>BT20/MB468</td></tr>
<tr><td>>hg38_21_+_8433172_8446622_RNA45SN1_45S_50ntFlanks@9554.9579.26|CGTAGCGGTCCTGACGTGCAAATCGG</td><td>>hg38_21_+_8433172_8446622_RNA45SN1_45S_50ntFlanks</td><td>9554</td><td>9579</td><td>26</td><td>CGTAGCGGTCCTGACGTGCAAATCGG</td><td>235.5</td><td>-4.8</td><td>0.8</td><td>-6.0</td><td>1.8E-09</td><td>2.9E-09</td><td>BT20/MB468</td></tr>
<tr><td>>MI0000100|hsa-mir-98&WithFlank&X|-|53556217|53556347@28.49.22&[MIMAT0000096&hsa-miR-98-5p&offsets|0|0;m-143&X|-|53556299|53556320&offsets|0|0]|TGAGGTAGTAAGTTGTATTGTT</td><td>>MI0000100|hsa-mir-98&WithFlank&X|-|53556217|53556347</td><td>28</td><td>49</td><td>22</td><td>TGAGGTAGTAAGTTGTATTGTT</td><td>10827.3</td><td>4.9</td><td>0.8</td><td>6.0</td><td>1.8E-09</td><td>2.9E-09</td><td>BT20/MB468</td></tr>
<tr><td>>hg38_21_+_8433172_8446622_RNA45SN1_45S_50ntFlanks@9621.9648.28|TCTAGTAGCTGGTTCCCTCCGAAGTTTC</td><td>>hg38_21_+_8433172_8446622_RNA45SN1_45S_50ntFlanks</td><td>9621</td><td>9648</td><td>28</td><td>TCTAGTAGCTGGTTCCCTCCGAAGTTTC</td><td>115.3</td><td>-4.0</td><td>0.7</td><td>-6.0</td><td>1.8E-09</td><td>2.9E-09</td><td>BT20/MB468</td></tr>
<tr><td>>hg38_21_+_8433172_8446622_RNA45SN1_45S_50ntFlanks@7988.8012.25|CAGACGTGGCGACCCGCTGAATTTA</td><td>>hg38_21_+_8433172_8446622_RNA45SN1_45S_50ntFlanks</td><td>7988</td><td>8012</td><td>25</td><td>CAGACGTGGCGACCCGCTGAATTTA</td><td>1616.8</td><td>-3.9</td><td>0.6</td><td>-6.0</td><td>1.8E-09</td><td>2.9E-09</td><td>BT20/MB468</td></tr>
<tr><td>>hg38_21_+_8433172_8446622_RNA45SN1_45S_50ntFlanks@6722.6739.18|ATTGCAGGACACATTGAT</td><td>>hg38_21_+_8433172_8446622_RNA45SN1_45S_50ntFlanks</td><td>6722</td><td>6739</td><td>18</td><td>ATTGCAGGACACATTGAT</td><td>298.7</td><td>-3.7</td><td>0.6</td><td>-6.0</td><td>1.6E-09</td><td>2.9E-09</td><td>MB231/MB468</td></tr>
<tr><td>>hg38_21_+_8433172_8446622_RNA45SN1_45S_50ntFlanks@11729.11746.18|TAACTATGACTCTCTTAA</td><td>>hg38_21_+_8433172_8446622_RNA45SN1_45S_50ntFlanks</td><td>11729</td><td>11746</td><td>18</td><td>TAACTATGACTCTCTTAA</td><td>182.4</td><td>-3.3</td><td>0.5</td><td>-6.0</td><td>1.8E-09</td><td>2.9E-09</td><td>BT20/MB468</td></tr>
<tr><td>>hg38_21_+_8433172_8446622_RNA45SN1_45S_50ntFlanks@8189.8212.24|CCCAGCCCGTGGACGGTGTGAGGC</td><td>>hg38_21_+_8433172_8446622_RNA45SN1_45S_50ntFlanks</td><td>8189</td><td>8212</td><td>24</td><td>CCCAGCCCGTGGACGGTGTGAGGC</td><td>264.5</td><td>-6.6</td><td>1.1</td><td>-6.0</td><td>1.8E-09</td><td>2.9E-09</td><td>BT20/MB468</td></tr>
<tr><td>>MI0000100|hsa-mir-98&WithFlank&X|-|53556217|53556347@28.48.21&[MIMAT0000096&hsa-miR-98-5p&offsets|0|-1;m-143&X|-|53556299|53556320&offsets|0|-1]|TGAGGTAGTAAGTTGTATTGT</td><td>>MI0000100|hsa-mir-98&WithFlank&X|-|53556217|53556347</td><td>28</td><td>48</td><td>21</td><td>TGAGGTAGTAAGTTGTATTGT</td><td>662.8</td><td>4.9</td><td>0.8</td><td>6.0</td><td>1.8E-09</td><td>2.9E-09</td><td>BT20/MB468</td></tr>
<tr><td>>hg38_21_+_8433172_8446622_RNA45SN1_45S_50ntFlanks@12625.12652.28|CAATGGGGCGAAGCTACCATCTGTGGGA</td><td>>hg38_21_+_8433172_8446622_RNA45SN1_45S_50ntFlanks</td><td>12625</td><td>12652</td><td>28</td><td>CAATGGGGCGAAGCTACCATCTGTGGGA</td><td>89.8</td><td>-4.8</td><td>0.8</td><td>-6.0</td><td>1.8E-09</td><td>2.9E-09</td><td>BT20/MB468</td></tr>
<tr><td>>hg38_21_+_8433172_8446622_RNA45SN1_45S_50ntFlanks@7974.8012.39|ACGCGACCTCAGATCAGACGTGGCGACCCGCTGAATTTA</td><td>>hg38_21_+_8433172_8446622_RNA45SN1_45S_50ntFlanks</td><td>7974</td><td>8012</td><td>39</td><td>ACGCGACCTCAGATCAGACGTGGCGACCCGCTGAATTTA</td><td>4250.0</td><td>4.6</td><td>0.8</td><td>6.0</td><td>1.6E-09</td><td>2.9E-09</td><td>MB231/MB468</td></tr>
<tr><td>>hg38_21_+_8433172_8446622_RNA45SN1_45S_50ntFlanks@8109.8143.35|GCGGGGCGCGGGACATGTGGCGTACGGAAGACCCG</td><td>>hg38_21_+_8433172_8446622_RNA45SN1_45S_50ntFlanks</td><td>8109</td><td>8143</td><td>35</td><td>GCGGGGCGCGGGACATGTGGCGTACGGAAGACCCG</td><td>71.8</td><td>-8.2</td><td>1.4</td><td>-6.0</td><td>1.8E-09</td><td>3.0E-09</td><td>BT20/MB468</td></tr>
<tr><td>>trna10_GlyTCC_17_+_8124866_8124937@1.29.29|GCGTTGGTGGTATAGTGGTAAGCATAGCT</td><td>>trna10_GlyTCC_17_+_8124866_8124937</td><td>1</td><td>29</td><td>29</td><td>GCGTTGGTGGTATAGTGGTAAGCATAGCT</td><td>89.4</td><td>-8.1</td><td>1.3</td><td>-6.0</td><td>1.6E-09</td><td>3.0E-09</td><td>MB231/MB468</td></tr>
<tr><td>>hg38_21_+_8433172_8446622_RNA45SN1_45S_50ntFlanks@12477.12496.20|CACTAATAGGGAACGTGAGC</td><td>>hg38_21_+_8433172_8446622_RNA45SN1_45S_50ntFlanks</td><td>12477</td><td>12496</td><td>20</td><td>CACTAATAGGGAACGTGAGC</td><td>2428.4</td><td>3.3</td><td>0.5</td><td>6.0</td><td>1.6E-09</td><td>3.0E-09</td><td>MB231/MB468</td></tr>
<tr><td>>hg38_21_+_8433172_8446622_RNA45SN1_45S_50ntFlanks@8106.8143.38|GCGGCGGGGCGCGGGACATGTGGCGTACGGAAGACCCG</td><td>>hg38_21_+_8433172_8446622_RNA45SN1_45S_50ntFlanks</td><td>8106</td><td>8143</td><td>38</td><td>GCGGCGGGGCGCGGGACATGTGGCGTACGGAAGACCCG</td><td>92.0</td><td>-8.1</td><td>1.3</td><td>-6.0</td><td>1.6E-09</td><td>3.0E-09</td><td>MB231/MB468</td></tr>
<tr><td>>hg38_21_+_8433172_8446622_RNA45SN1_45S_50ntFlanks@11561.11601.41|GCGCCTAGCAGCCGACTTAGAACTGGTGCGGACCAGGGGAA</td><td>>hg38_21_+_8433172_8446622_RNA45SN1_45S_50ntFlanks</td><td>11561</td><td>11601</td><td>41</td><td>GCGCCTAGCAGCCGACTTAGAACTGGTGCGGACCAGGGGAA</td><td>426.7</td><td>-5.3</td><td>0.9</td><td>-6.0</td><td>1.9E-09</td><td>3.0E-09</td><td>BT20/MB468</td></tr>
<tr><td>>MI0000749|hsa-mir-30e&WithFlank&1|+|40754349|40754452@65.83.19&[MIMAT0000693&hsa-miR-30e-3p&offsets|0|-3;m-25&1|+|40754413|40754434&offsets|0|-3]|CTTTCAGTCGGATGTTTAC</td><td>>MI0000749|hsa-mir-30e&WithFlank&1|+|40754349|40754452</td><td>65</td><td>83</td><td>19</td><td>CTTTCAGTCGGATGTTTAC</td><td>971.8</td><td>4.6</td><td>0.8</td><td>6.0</td><td>1.9E-09</td><td>3.1E-09</td><td>BT20/MB468</td></tr>
<tr><td>>hg38_21_+_8433172_8446622_RNA45SN1_45S_50ntFlanks@7983.8012.30|CAGATCAGACGTGGCGACCCGCTGAATTTA</td><td>>hg38_21_+_8433172_8446622_RNA45SN1_45S_50ntFlanks</td><td>7983</td><td>8012</td><td>30</td><td>CAGATCAGACGTGGCGACCCGCTGAATTTA</td><td>234.8</td><td>-4.0</td><td>0.7</td><td>-6.0</td><td>1.9E-09</td><td>3.1E-09</td><td>BT20/MB468</td></tr>
<tr><td>>hg38_21_+_8433172_8446622_RNA45SN1_45S_50ntFlanks@5445.5467.23|TCGGATCGGCCCCGCCGGGGTCG</td><td>>hg38_21_+_8433172_8446622_RNA45SN1_45S_50ntFlanks</td><td>5445</td><td>5467</td><td>23</td><td>TCGGATCGGCCCCGCCGGGGTCG</td><td>96.0</td><td>-4.6</td><td>0.8</td><td>-6.0</td><td>1.9E-09</td><td>3.1E-09</td><td>BT20/MB468</td></tr>
<tr><td>>am_trna117_GlyTCC_1_-_145397864_145397935@1.30.30,trna45_GlyTCC_1_+_161500903_161500974@1.30.30,trna70_GlyTCC_1_-_161439547_161439618@1.30.30,trna73_GlyTCC_1_-_161432166_161432237@1.30.30,trna76_GlyTCC_1_-_161424756_161424827@1.30.30,trna79_GlyTCC_1_-_161417375_161417446@1.30.30|GCGTTGGTGGTATAGTGGTGAGCATAGCTG</td><td>>am_trna117_GlyTCC_1_-_145397864_145397935</td><td>1</td><td>30</td><td>30</td><td>GCGTTGGTGGTATAGTGGTGAGCATAGCTG</td><td>1093.1</td><td>-6.1</td><td>1.0</td><td>-6.0</td><td>1.7E-09</td><td>3.1E-09</td><td>MB231/MB468</td></tr>
<tr><td>>hg38_21_+_8433172_8446622_RNA45SN1_45S_50ntFlanks@4064.4101.38|ATCAACTTTCGATGGTAGTCGCCGTGCCTACCATGGTG</td><td>>hg38_21_+_8433172_8446622_RNA45SN1_45S_50ntFlanks</td><td>4064</td><td>4101</td><td>38</td><td>ATCAACTTTCGATGGTAGTCGCCGTGCCTACCATGGTG</td><td>84.3</td><td>-8.0</td><td>1.3</td><td>-6.0</td><td>1.7E-09</td><td>3.1E-09</td><td>MB231/MB468</td></tr>
<tr><td>>hg38_21_+_8433172_8446622_RNA45SN1_45S_50ntFlanks@4003.4022.20|ATAACCTCGGGCCGATCGCA</td><td>>hg38_21_+_8433172_8446622_RNA45SN1_45S_50ntFlanks</td><td>4003</td><td>4022</td><td>20</td><td>ATAACCTCGGGCCGATCGCA</td><td>442.4</td><td>-3.4</td><td>0.6</td><td>-6.0</td><td>1.9E-09</td><td>3.1E-09</td><td>BT20/MB468</td></tr>
<tr><td>>hg38_21_+_8433172_8446622_RNA45SN1_45S_50ntFlanks@12073.12091.19|CCGAGGGGCTCTCGCTTCT</td><td>>hg38_21_+_8433172_8446622_RNA45SN1_45S_50ntFlanks</td><td>12073</td><td>12091</td><td>19</td><td>CCGAGGGGCTCTCGCTTCT</td><td>115.0</td><td>-5.2</td><td>0.9</td><td>-6.0</td><td>1.9E-09</td><td>3.1E-09</td><td>BT20/MB468</td></tr>
<tr><td>>hg38_21_+_8433172_8446622_RNA45SN1_45S_50ntFlanks@12358.12376.19|CACAGGGATAACTGGCTTG</td><td>>hg38_21_+_8433172_8446622_RNA45SN1_45S_50ntFlanks</td><td>12358</td><td>12376</td><td>19</td><td>CACAGGGATAACTGGCTTG</td><td>119.9</td><td>-6.6</td><td>1.1</td><td>-6.0</td><td>1.7E-09</td><td>3.1E-09</td><td>MB231/MB468</td></tr>
<tr><td>>hg38_21_+_8433172_8446622_RNA45SN1_45S_50ntFlanks@12476.12509.34|CCACTAATAGGGAACGTGAGCTGGGTTTAGACCG</td><td>>hg38_21_+_8433172_8446622_RNA45SN1_45S_50ntFlanks</td><td>12476</td><td>12509</td><td>34</td><td>CCACTAATAGGGAACGTGAGCTGGGTTTAGACCG</td><td>93.4</td><td>-8.1</td><td>1.3</td><td>-6.0</td><td>1.7E-09</td><td>3.1E-09</td><td>MB231/MB468</td></tr>
<tr><td>>trna23_ArgCCG_17_-_66016013_66016085@14.31.18|AATGGATAAGGCATCAGC</td><td>>trna23_ArgCCG_17_-_66016013_66016085</td><td>14</td><td>31</td><td>18</td><td>AATGGATAAGGCATCAGC</td><td>138.4</td><td>-6.8</td><td>1.1</td><td>-6.0</td><td>1.7E-09</td><td>3.1E-09</td><td>MB231/MB468</td></tr>
<tr><td>>hg38_21_+_8433172_8446622_RNA45SN1_45S_50ntFlanks@4372.4389.18|AAAAAGCTCGTAGTTGGA</td><td>>hg38_21_+_8433172_8446622_RNA45SN1_45S_50ntFlanks</td><td>4372</td><td>4389</td><td>18</td><td>AAAAAGCTCGTAGTTGGA</td><td>168.7</td><td>-3.4</td><td>0.6</td><td>-6.0</td><td>1.9E-09</td><td>3.2E-09</td><td>BT20/MB468</td></tr>
<tr><td>>trna59_GluCTC_1_+_249168447_249168518@32.70.39|CTCTCACCGCCGCGGCCCGGGTTCGATTCCCGGTCAGGA</td><td>>trna59_GluCTC_1_+_249168447_249168518</td><td>32</td><td>70</td><td>39</td><td>CTCTCACCGCCGCGGCCCGGGTTCGATTCCCGGTCAGGA</td><td>73.6</td><td>-7.8</td><td>1.3</td><td>-6.0</td><td>1.7E-09</td><td>3.2E-09</td><td>MB231/MB468</td></tr>
<tr><td>>hg38_21_+_8433172_8446622_RNA45SN1_45S_50ntFlanks@11876.11908.33|AAGACCCTGTTGAGCTTGACTCTAGTCTGGCAC</td><td>>hg38_21_+_8433172_8446622_RNA45SN1_45S_50ntFlanks</td><td>11876</td><td>11908</td><td>33</td><td>AAGACCCTGTTGAGCTTGACTCTAGTCTGGCAC</td><td>154.7</td><td>-5.2</td><td>0.9</td><td>-6.0</td><td>2.0E-09</td><td>3.2E-09</td><td>BT20/MB468</td></tr>
<tr><td>>hg38_21_+_8433172_8446622_RNA45SN1_45S_50ntFlanks@6751.6770.20|CGAACGCACTTGCGGCCCCG</td><td>>hg38_21_+_8433172_8446622_RNA45SN1_45S_50ntFlanks</td><td>6751</td><td>6770</td><td>20</td><td>CGAACGCACTTGCGGCCCCG</td><td>74.1</td><td>-5.2</td><td>0.9</td><td>-6.0</td><td>1.8E-09</td><td>3.2E-09</td><td>MB231/MB468</td></tr>
<tr><td>>hg38_21_+_8433172_8446622_RNA45SN1_45S_50ntFlanks@9621.9650.30|TCTAGTAGCTGGTTCCCTCCGAAGTTTCCC</td><td>>hg38_21_+_8433172_8446622_RNA45SN1_45S_50ntFlanks</td><td>9621</td><td>9650</td><td>30</td><td>TCTAGTAGCTGGTTCCCTCCGAAGTTTCCC</td><td>67.6</td><td>-8.1</td><td>1.3</td><td>-6.0</td><td>2.0E-09</td><td>3.2E-09</td><td>BT20/MB468</td></tr>
<tr><td>>hg38_21_+_8433172_8446622_RNA45SN1_45S_50ntFlanks@9027.9058.32|CGTCTCCTCGTGGGGGGGCCGGGCCACCCCTC</td><td>>hg38_21_+_8433172_8446622_RNA45SN1_45S_50ntFlanks</td><td>9027</td><td>9058</td><td>32</td><td>CGTCTCCTCGTGGGGGGGCCGGGCCACCCCTC</td><td>76.4</td><td>-7.8</td><td>1.3</td><td>-6.0</td><td>1.8E-09</td><td>3.2E-09</td><td>MB231/MB468</td></tr>
<tr><td>>hg38_21_+_8433172_8446622_RNA45SN1_45S_50ntFlanks@11619.11640.22|AACAAAGCATCGCGAAGGCCCG</td><td>>hg38_21_+_8433172_8446622_RNA45SN1_45S_50ntFlanks</td><td>11619</td><td>11640</td><td>22</td><td>AACAAAGCATCGCGAAGGCCCG</td><td>158.9</td><td>-4.6</td><td>0.8</td><td>-6.0</td><td>1.8E-09</td><td>3.3E-09</td><td>MB231/MB468</td></tr>
<tr><td>>hg38_21_+_8433172_8446622_RNA45SN1_45S_50ntFlanks@11210.11227.18|GGCGACTCTGGACGCGAG</td><td>>hg38_21_+_8433172_8446622_RNA45SN1_45S_50ntFlanks</td><td>11210</td><td>11227</td><td>18</td><td>GGCGACTCTGGACGCGAG</td><td>135.9</td><td>-5.9</td><td>1.0</td><td>-6.0</td><td>1.8E-09</td><td>3.3E-09</td><td>MB231/MB468</td></tr>
<tr><td>>hg38_21_+_8433172_8446622_RNA45SN1_45S_50ntFlanks@10798.10816.19|CATGTTGGAACAATGTAGG</td><td>>hg38_21_+_8433172_8446622_RNA45SN1_45S_50ntFlanks</td><td>10798</td><td>10816</td><td>19</td><td>CATGTTGGAACAATGTAGG</td><td>447.4</td><td>-3.6</td><td>0.6</td><td>-6.0</td><td>1.8E-09</td><td>3.3E-09</td><td>MB231/MB468</td></tr>
<tr><td>>hg38_21_+_8433172_8446622_RNA45SN1_45S_50ntFlanks@10832.10862.31|AGCCGGATCCGTAACTTCGGGATAAGGATTG</td><td>>hg38_21_+_8433172_8446622_RNA45SN1_45S_50ntFlanks</td><td>10832</td><td>10862</td><td>31</td><td>AGCCGGATCCGTAACTTCGGGATAAGGATTG</td><td>116.1</td><td>-5.0</td><td>0.8</td><td>-6.0</td><td>1.8E-09</td><td>3.3E-09</td><td>MB231/MB468</td></tr>
<tr><td>>hg38_21_+_8433172_8446622_RNA45SN1_45S_50ntFlanks@10819.10839.21|AGGGAAGTCGGCAAGCCGGAT</td><td>>hg38_21_+_8433172_8446622_RNA45SN1_45S_50ntFlanks</td><td>10819</td><td>10839</td><td>21</td><td>AGGGAAGTCGGCAAGCCGGAT</td><td>132.4</td><td>-3.0</td><td>0.5</td><td>-6.0</td><td>1.8E-09</td><td>3.3E-09</td><td>MB231/MB468</td></tr>
<tr><td>>hg38_21_+_8433172_8446622_RNA45SN1_45S_50ntFlanks@11874.11897.24|AGAAGACCCTGTTGAGCTTGACTC</td><td>>hg38_21_+_8433172_8446622_RNA45SN1_45S_50ntFlanks</td><td>11874</td><td>11897</td><td>24</td><td>AGAAGACCCTGTTGAGCTTGACTC</td><td>709.1</td><td>-3.9</td><td>0.7</td><td>-6.0</td><td>2.0E-09</td><td>3.3E-09</td><td>BT20/MB468</td></tr>
</table>

<table>
<tr><td>>hg38_21_+_8433172_8446622_RNA45SN1_45S_50ntFlanks@8115.8142.28|CGCGGGACATGTGGCGTACGGAAGACCC</td><td>>hg38_21_+_8433172_8446622_RNA45SN1_45S_50ntFlanks</td><td>8115</td><td>8142</td><td>28</td><td>CGCGGGACATGTGGCGTACGGAAGACCC</td><td>77.9</td><td>-7.9</td><td>1.3</td><td>-6.0</td><td>1.8E-09</td><td>3.4E-09</td><td>MB231/MB468</td></tr>
<tr><td>>hg38_21_+_8433172_8446622_RNA45SN1_45S_50ntFlanks@11316.11335.20|CACCCCACGTCTCGTCGCGC</td><td>>hg38_21_+_8433172_8446622_RNA45SN1_45S_50ntFlanks</td><td>11316</td><td>11335</td><td>20</td><td>CACCCCACGTCTCGTCGCGC</td><td>74.1</td><td>-7.8</td><td>1.3</td><td>-6.0</td><td>1.8E-09</td><td>3.4E-09</td><td>MB231/MB468</td></tr>
<tr><td>>hg38_21_+_8433172_8446622_RNA45SN1_45S_50ntFlanks@4161.4182.22|CACATCCAAGGAAGGCAGCAGG</td><td>>hg38_21_+_8433172_8446622_RNA45SN1_45S_50ntFlanks</td><td>4161</td><td>4182</td><td>22</td><td>CACATCCAAGGAAGGCAGCAGG</td><td>84.8</td><td>-5.2</td><td>0.9</td><td>-6.0</td><td>2.1E-09</td><td>3.4E-09</td><td>BT20/MB468</td></tr>
<tr><td>>hg38_21_+_8433172_8446622_RNA45SN1_45S_50ntFlanks@11703.11725.23|AATGAAGCGCGGGTAAACGGCGG</td><td>>hg38_21_+_8433172_8446622_RNA45SN1_45S_50ntFlanks</td><td>11703</td><td>11725</td><td>23</td><td>AATGAAGCGCGGGTAAACGGCGG</td><td>122.2</td><td>-5.3</td><td>0.9</td><td>-6.0</td><td>2.1E-09</td><td>3.4E-09</td><td>BT20/MB468</td></tr>
<tr><td>>hg38_MT_+_1621_3279_RNR2_16S_50ntFlanks@1428.1458.31|ATTAAAGTCCTACGTGATCTGAGTTCAGACC</td><td>>hg38_MT_+_1621_3279_RNR2_16S_50ntFlanks</td><td>1428</td><td>1458</td><td>31</td><td>ATTAAAGTCCTACGTGATCTGAGTTCAGACC</td><td>597.5</td><td>7.9</td><td>1.3</td><td>6.0</td><td>2.1E-09</td><td>3.4E-09</td><td>BT20/MB468</td></tr>
<tr><td>>hg38_21_+_8433172_8446622_RNA45SN1_45S_50ntFlanks@7974.8005.32|ACGCGACCTCAGATCAGACGTGGCGACCCGCT</td><td>>hg38_21_+_8433172_8446622_RNA45SN1_45S_50ntFlanks</td><td>7974</td><td>8005</td><td>32</td><td>ACGCGACCTCAGATCAGACGTGGCGACCCGCT</td><td>1772.7</td><td>4.2</td><td>0.7</td><td>6.0</td><td>1.9E-09</td><td>3.5E-09</td><td>MB231/MB468</td></tr>
<tr><td>>hg38_21_+_8433172_8446622_RNA45SN1_45S_50ntFlanks@12409.12448.40|TTGATCCTTCGATGTCGGCTCTTCCTATCATTGTGAAGCA</td><td>>hg38_21_+_8433172_8446622_RNA45SN1_45S_50ntFlanks</td><td>12409</td><td>12448</td><td>40</td><td>TTGATCCTTCGATGTCGGCTCTTCCTATCATTGTGAAGCA</td><td>456.6</td><td>10.9</td><td>1.8</td><td>6.0</td><td>1.9E-09</td><td>3.5E-09</td><td>MB231/MB468</td></tr>
<tr><td>>hg38_21_+_8433172_8446622_RNA45SN1_45S_50ntFlanks@3846.3869.24|CTTGGATAACTGTGGTAATTCTAG</td><td>>hg38_21_+_8433172_8446622_RNA45SN1_45S_50ntFlanks</td><td>3846</td><td>3869</td><td>24</td><td>CTTGGATAACTGTGGTAATTCTAG</td><td>306.0</td><td>-3.5</td><td>0.6</td><td>-6.0</td><td>2.2E-09</td><td>3.5E-09</td><td>BT20/MB468</td></tr>
<tr><td>>hg38_21_+_8433172_8446622_RNA45SN1_45S_50ntFlanks@11875.11906.32|GAAGACCCTGTTGAGCTTGACTCTAGTCTGGC</td><td>>hg38_21_+_8433172_8446622_RNA45SN1_45S_50ntFlanks</td><td>11875</td><td>11906</td><td>32</td><td>GAAGACCCTGTTGAGCTTGACTCTAGTCTGGC</td><td>116.3</td><td>-5.2</td><td>0.9</td><td>-6.0</td><td>2.2E-09</td><td>3.6E-09</td><td>BT20/MB468</td></tr>
<tr><td>>hg38_21_+_8433172_8446622_RNA45SN1_45S_50ntFlanks@5472.5516.45|ACGGCCCTGGCGGAGCGCTGAGAAGACGGTCGAACTTGACTATCT</td><td>>hg38_21_+_8433172_8446622_RNA45SN1_45S_50ntFlanks</td><td>5472</td><td>5516</td><td>45</td><td>ACGGCCCTGGCGGAGCGCTGAGAAGACGGTCGAACTTGACTATCT</td><td>1066.6</td><td>8.7</td><td>1.4</td><td>6.0</td><td>2.2E-09</td><td>3.6E-09</td><td>BT20/MB468</td></tr>
<tr><td>>hg38_21_+_8433172_8446622_RNA45SN1_45S_50ntFlanks@10394.10424.31|AGCAGTTGAACATGGGTCAGTCGGTCCTGAG</td><td>>hg38_21_+_8433172_8446622_RNA45SN1_45S_50ntFlanks</td><td>10394</td><td>10424</td><td>31</td><td>AGCAGTTGAACATGGGTCAGTCGGTCCTGAG</td><td>767.8</td><td>3.7</td><td>0.6</td><td>6.0</td><td>2.0E-09</td><td>3.6E-09</td><td>MB231/MB468</td></tr>
<tr><td>>hg38_21_+_8433172_8446622_RNA45SN1_45S_50ntFlanks@7988.8006.19|CAGACGTGGCGACCCGCTG</td><td>>hg38_21_+_8433172_8446622_RNA45SN1_45S_50ntFlanks</td><td>7988</td><td>8006</td><td>19</td><td>CAGACGTGGCGACCCGCTG</td><td>130.3</td><td>-5.0</td><td>0.8</td><td>-6.0</td><td>2.0E-09</td><td>3.7E-09</td><td>MB231/MB468</td></tr>
<tr><td>>MI0000749|hsa-mir-30e&WithFlank&1|+|40754349|40754452@23.46.24&[MIMAT0000692&hsa-miR-30e-5p&offsets|0|+2;m-24&1|+|40754371|40754394&offsets|0|0]|TGTAAACATCCTTGACTGGAAGCT</td><td>>MI0000749|hsa-mir-30e&WithFlank&1|+|40754349|40754452</td><td>23</td><td>46</td><td>24</td><td>TGTAAACATCCTTGACTGGAAGCT</td><td>16384.9</td><td>4.7</td><td>0.8</td><td>6.0</td><td>2.0E-09</td><td>3.7E-09</td><td>MB231/MB468</td></tr>
<tr><td>>hg38_21_+_8433172_8446622_RNA45SN1_45S_50ntFlanks@12396.12418.23|GCGACGTCGCTTTTTGATCCTTC</td><td>>hg38_21_+_8433172_8446622_RNA45SN1_45S_50ntFlanks</td><td>12396</td><td>12418</td><td>23</td><td>GCGACGTCGCTTTTTGATCCTTC</td><td>722.0</td><td>4.9</td><td>0.8</td><td>6.0</td><td>2.1E-09</td><td>3.7E-09</td><td>MB231/MB468</td></tr>
<tr><td>>hg38_21_+_8433172_8446622_RNA45SN1_45S_50ntFlanks@4657.4684.28|CTAGAGGTGAAATTCTTGGACCGGCGCA</td><td>>hg38_21_+_8433172_8446622_RNA45SN1_45S_50ntFlanks</td><td>4657</td><td>4684</td><td>28</td><td>CTAGAGGTGAAATTCTTGGACCGGCGCA</td><td>85.1</td><td>-8.0</td><td>1.3</td><td>-6.0</td><td>2.1E-09</td><td>3.7E-09</td><td>MB231/MB468</td></tr>
<tr><td>>hg38_21_+_8433172_8446622_RNA45SN1_45S_50ntFlanks@4444.4472.29|CCTTGCCTCTCGGCGCCCCCTCGATGCTC</td><td>>hg38_21_+_8433172_8446622_RNA45SN1_45S_50ntFlanks</td><td>4444</td><td>4472</td><td>29</td><td>CCTTGCCTCTCGGCGCCCCCTCGATGCTC</td><td>84.4</td><td>-8.4</td><td>1.4</td><td>-6.0</td><td>2.3E-09</td><td>3.7E-09</td><td>BT20/MB468</td></tr>
<tr><td>>hg38_21_+_8433172_8446622_RNA45SN1_45S_50ntFlanks@12655.12673.19|ATGACTGAACGCCTCTAAG</td><td>>hg38_21_+_8433172_8446622_RNA45SN1_45S_50ntFlanks</td><td>12655</td><td>12673</td><td>19</td><td>ATGACTGAACGCCTCTAAG</td><td>353.3</td><td>-4.3</td><td>0.7</td><td>-6.0</td><td>2.1E-09</td><td>3.8E-09</td><td>MB231/MB468</td></tr>
<tr><td>>hg38_21_+_8433172_8446622_RNA45SN1_45S_50ntFlanks@10534.10570.37|CCGCGAGGCGTCCAGTGCGGTAACGCGACCGATCCCG</td><td>>hg38_21_+_8433172_8446622_RNA45SN1_45S_50ntFlanks</td><td>10534</td><td>10570</td><td>37</td><td>CCGCGAGGCGTCCAGTGCGGTAACGCGACCGATCCCG</td><td>192.8</td><td>-7.2</td><td>1.2</td><td>-6.0</td><td>2.1E-09</td><td>3.8E-09</td><td>MB231/MB468</td></tr>
<tr><td>>hg38_21_+_8433172_8446622_RNA45SN1_45S_50ntFlanks@9504.9522.19|GTGAACTATGCCTGGGCAG</td><td>>hg38_21_+_8433172_8446622_RNA45SN1_45S_50ntFlanks</td><td>9504</td><td>9522</td><td>19</td><td>GTGAACTATGCCTGGGCAG</td><td>515.3</td><td>-4.3</td><td>0.7</td><td>-6.0</td><td>2.1E-09</td><td>3.8E-09</td><td>MB231/MB468</td></tr>
<tr><td>>hg38_21_+_8433172_8446622_RNA45SN1_45S_50ntFlanks@9198.9220.23|GTTCTCTCGGGGCCACGCGCGCG</td><td>>hg38_21_+_8433172_8446622_RNA45SN1_45S_50ntFlanks</td><td>9198</td><td>9220</td><td>23</td><td>GTTCTCTCGGGGCCACGCGCGCG</td><td>106.8</td><td>-8.3</td><td>1.4</td><td>-6.0</td><td>2.1E-09</td><td>3.8E-09</td><td>MB231/MB468</td></tr>
<tr><td>>MI0000081|hsa-mir-24-2&WithFlank&19|-|13836281|13836365@18.41.24&[MIMAT0004497&hsa-miR-24-2-5p&offsets|-1|+1;m-226&19|-|13836326|13836348&offsets|0|+1]|GTGCCTACTGAGCTGAAACACAGT</td><td>>MI0000081|hsa-mir-24-2&WithFlank&19|-|13836281|13836365</td><td>18</td><td>41</td><td>24</td><td>GTGCCTACTGAGCTGAAACACAGT</td><td>634.0</td><td>5.0</td><td>0.8</td><td>6.0</td><td>2.3E-09</td><td>3.8E-09</td><td>BT20/MB468</td></tr>
<tr><td>>hg38_21_+_8433172_8446622_RNA45SN1_45S_50ntFlanks@11592.11612.21|ACCAGGGGAATCCGACTGTTT</td><td>>hg38_21_+_8433172_8446622_RNA45SN1_45S_50ntFlanks</td><td>11592</td><td>11612</td><td>21</td><td>ACCAGGGGAATCCGACTGTTT</td><td>381.3</td><td>-4.5</td><td>0.8</td><td>-6.0</td><td>2.1E-09</td><td>3.9E-09</td><td>MB231/MB468</td></tr>
<tr><td>>hg38_21_+_8433172_8446622_RNA45SN1_45S_50ntFlanks@11694.11711.18|AAGAAATTCAATGAAGCG</td><td>>hg38_21_+_8433172_8446622_RNA45SN1_45S_50ntFlanks</td><td>11694</td><td>11711</td><td>18</td><td>AAGAAATTCAATGAAGCG</td><td>245.9</td><td>-5.4</td><td>0.9</td><td>-6.0</td><td>2.4E-09</td><td>3.9E-09</td><td>BT20/MB468</td></tr>
<tr><td>>hg38_21_+_8433172_8446622_RNA45SN1_45S_50ntFlanks@6740.6772.33|CATCGACACTTCGAACGCACTTGCGGCCCCGGG</td><td>>hg38_21_+_8433172_8446622_RNA45SN1_45S_50ntFlanks</td><td>6740</td><td>6772</td><td>33</td><td>CATCGACACTTCGAACGCACTTGCGGCCCCGGG</td><td>284.8</td><td>-4.7</td><td>0.8</td><td>-6.0</td><td>2.1E-09</td><td>3.9E-09</td><td>MB231/MB468</td></tr>
<tr><td>>hg38_21_+_8433172_8446622_RNA45SN1_45S_50ntFlanks@6645.6665.21|CTCGTACGACTCTTAGCGGTG</td><td>>hg38_21_+_8433172_8446622_RNA45SN1_45S_50ntFlanks</td><td>6645</td><td>6665</td><td>21</td><td>CTCGTACGACTCTTAGCGGTG</td><td>3934.1</td><td>4.5</td><td>0.7</td><td>6.0</td><td>2.1E-09</td><td>3.9E-09</td><td>MB231/MB468</td></tr>
<tr><td>>hg38_21_+_8433172_8446622_RNA45SN1_45S_50ntFlanks@11873.11910.38|AAGAAGACCCTGTTGAGCTTGACTCTAGTCTGGCACGG</td><td>>hg38_21_+_8433172_8446622_RNA45SN1_45S_50ntFlanks</td><td>11873</td><td>11910</td><td>38</td><td>AAGAAGACCCTGTTGAGCTTGACTCTAGTCTGGCACGG</td><td>96.7</td><td>-6.3</td><td>1.1</td><td>-6.0</td><td>2.2E-09</td><td>3.9E-09</td><td>MB231/MB468</td></tr>
<tr><td>>hg38_21_+_8433172_8446622_RNA45SN1_45S_50ntFlanks@11233.11251.19|CCCTTCCCGTGGATCGCCC</td><td>>hg38_21_+_8433172_8446622_RNA45SN1_45S_50ntFlanks</td><td>11233</td><td>11251</td><td>19</td><td>CCCTTCCCGTGGATCGCCC</td><td>84.8</td><td>-8.0</td><td>1.3</td><td>-6.0</td><td>2.2E-09</td><td>3.9E-09</td><td>MB231/MB468</td></tr>
<tr><td>>hg38_21_+_8433172_8446622_RNA45SN1_45S_50ntFlanks@8845.8867.23|CGGCGGCGGCGGCGGCGGCGGGA</td><td>>hg38_21_+_8433172_8446622_RNA45SN1_45S_50ntFlanks</td><td>8845</td><td>8867</td><td>23</td><td>CGGCGGCGGCGGCGGCGGCGGGA</td><td>73.2</td><td>-7.8</td><td>1.3</td><td>-6.0</td><td>2.2E-09</td><td>3.9E-09</td><td>MB231/MB468</td></tr>
<tr><td>>hg38_21_+_8433172_8446622_RNA45SN1_45S_50ntFlanks@7989.8016.28|AGACGTGGCGACCCGCTGAATTTAAGCA</td><td>>hg38_21_+_8433172_8446622_RNA45SN1_45S_50ntFlanks</td><td>7989</td><td>8016</td><td>28</td><td>AGACGTGGCGACCCGCTGAATTTAAGCA</td><td>764.7</td><td>4.2</td><td>0.7</td><td>6.0</td><td>2.4E-09</td><td>4.0E-09</td><td>BT20/MB468</td></tr>
<tr><td>>hg38_21_+_8433172_8446622_RNA45SN1_45S_50ntFlanks@9538.9563.26|AACTCTGGTGGAGGTCCGTAGCGGTC</td><td>>hg38_21_+_8433172_8446622_RNA45SN1_45S_50ntFlanks</td><td>9538</td><td>9563</td><td>26</td><td>AACTCTGGTGGAGGTCCGTAGCGGTC</td><td>80.7</td><td>-7.9</td><td>1.3</td><td>-6.0</td><td>2.2E-09</td><td>4.0E-09</td><td>MB231/MB468</td></tr>
<tr><td>>hg38_21_+_8433172_8446622_RNA45SN1_45S_50ntFlanks@8327.8356.30|ATAGTCAACAAGTACCGTAAGGGAAAGTTG</td><td>>hg38_21_+_8433172_8446622_RNA45SN1_45S_50ntFlanks</td><td>8327</td><td>8356</td><td>30</td><td>ATAGTCAACAAGTACCGTAAGGGAAAGTTG</td><td>99.7</td><td>-8.2</td><td>1.4</td><td>-6.0</td><td>2.2E-09</td><td>4.0E-09</td><td>MB231/MB468</td></tr>
<tr><td>>hg38_21_+_8433172_8446622_RNA45SN1_45S_50ntFlanks@12418.12446.29|CGATGTCGGCTCTTCCTATCATTGTGAAG</td><td>>hg38_21_+_8433172_8446622_RNA45SN1_45S_50ntFlanks</td><td>12418</td><td>12446</td><td>29</td><td>CGATGTCGGCTCTTCCTATCATTGTGAAG</td><td>1201.5</td><td>-3.6</td><td>0.6</td><td>-6.0</td><td>2.5E-09</td><td>4.0E-09</td><td>BT20/MB468</td></tr>
<tr><td>>hg38_21_+_8433172_8446622_RNA45SN1_45S_50ntFlanks@10756.10785.30|TACCCATATCCGCAGCAGGTCTCCAAGGTG</td><td>>hg38_21_+_8433172_8446622_RNA45SN1_45S_50ntFlanks</td><td>10756</td><td>10785</td><td>30</td><td>TACCCATATCCGCAGCAGGTCTCCAAGGTG</td><td>71.9</td><td>-7.7</td><td>1.3</td><td>-6.0</td><td>2.2E-09</td><td>4.0E-09</td><td>MB231/MB468</td></tr>
<tr><td>>hg38_21_+_8433172_8446622_RNA45SN1_45S_50ntFlanks@6698.6718.21|AGCTAGCTGCGAGAATTAATG</td><td>>hg38_21_+_8433172_8446622_RNA45SN1_45S_50ntFlanks</td><td>6698</td><td>6718</td><td>21</td><td>AGCTAGCTGCGAGAATTAATG</td><td>4423.1</td><td>-3.7</td><td>0.6</td><td>-6.0</td><td>2.2E-09</td><td>4.1E-09</td><td>MB231/MB468</td></tr>
<tr><td>>hg38_21_+_8433172_8446622_RNA45SN1_45S_50ntFlanks@10389.10408.20|TGAACAGCAGTTGAACATGG</td><td>>hg38_21_+_8433172_8446622_RNA45SN1_45S_50ntFlanks</td><td>10389</td><td>10408</td><td>20</td><td>TGAACAGCAGTTGAACATGG</td><td>233.2</td><td>-3.0</td><td>0.5</td><td>-6.0</td><td>2.2E-09</td><td>4.1E-09</td><td>MB231/MB468</td></tr>
<tr><td>>hg38_21_+_8433172_8446622_RNA45SN1_45S_50ntFlanks@12627.12651.25|ATGGGGCGAAGCTACCATCTGTGGG</td><td>>hg38_21_+_8433172_8446622_RNA45SN1_45S_50ntFlanks</td><td>12627</td><td>12651</td><td>25</td><td>ATGGGGCGAAGCTACCATCTGTGGG</td><td>63.3</td><td>-7.6</td><td>1.3</td><td>-6.0</td><td>2.2E-09</td><td>4.1E-09</td><td>MB231/MB468</td></tr>
<tr><td>>hg38_21_+_8433172_8446622_RNA45SN1_45S_50ntFlanks@10516.10533.18|GAGTGGCGGAGATGGGCG</td><td>>hg38_21_+_8433172_8446622_RNA45SN1_45S_50ntFlanks</td><td>10516</td><td>10533</td><td>18</td><td>GAGTGGCGGAGATGGGCG</td><td>309.1</td><td>-6.1</td><td>1.0</td><td>-6.0</td><td>2.5E-09</td><td>4.1E-09</td><td>BT20/MB468</td></tr>
<tr><td>>hg38_21_+_8433172_8446622_RNA45SN1_45S_50ntFlanks@12995.13015.21|CTCGCTGCGATCTATTGAAAG</td><td>>hg38_21_+_8433172_8446622_RNA45SN1_45S_50ntFlanks</td><td>12995</td><td>13015</td><td>21</td><td>CTCGCTGCGATCTATTGAAAG</td><td>1311.3</td><td>4.1</td><td>0.7</td><td>6.0</td><td>2.3E-09</td><td>4.1E-09</td><td>MB231/MB468</td></tr>
<tr><td>>hg38_21_+_8433172_8446622_RNA45SN1_45S_50ntFlanks@11894.11916.23|ACTCTAGTCTGGCACGGTGAAGA</td><td>>hg38_21_+_8433172_8446622_RNA45SN1_45S_50ntFlanks</td><td>11894</td><td>11916</td><td>23</td><td>ACTCTAGTCTGGCACGGTGAAGA</td><td>313.4</td><td>-5.1</td><td>0.9</td><td>-6.0</td><td>2.6E-09</td><td>4.2E-09</td><td>BT20/MB468</td></tr>
<tr><td>>hg38_21_+_8433172_8446622_RNA45SN1_45S_50ntFlanks@6723.6746.24|TTGCAGGACACATTGATCATCGAC</td><td>>hg38_21_+_8433172_8446622_RNA45SN1_45S_50ntFlanks</td><td>6723</td><td>6746</td><td>24</td><td>TTGCAGGACACATTGATCATCGAC</td><td>131.4</td><td>-3.7</td><td>0.6</td><td>-6.0</td><td>2.6E-09</td><td>4.2E-09</td><td>BT20/MB468</td></tr>
<tr><td>>am_trna111_HisGTG_1_-_147774845_147774916@-1G.23.24,trna118_HisGTG_1_-_145396881_145396952@-1G.23.24,trna16_HisGTG_1_+_146544773_146544844@-1G.23.24,trna1_HisGTG_15_+_45493349_45493420@-1G.23.24,trna21_HisGTG_1_+_147753471_147753542@-1G.23.24,trna33_HisGTG_6_+_27125906_27125977@-1G.23.24,trna7_HisGTG_9_-_14433938_14434009@-1G.23.24,trna8_HisGTG_15_-_45492611_45492682@-1G.23.24,trna9_HisGTG_15_-_45490804_45490875@-1G.23.24|GGCCGTGATCGTATAGTGGTTAGT</td><td>>am_trna111_HisGTG_1_-_147774845_147774916</td><td>-1G</td><td>23</td><td>24</td><td>GGCCGTGATCGTATAGTGGTTAGT</td><td>1021.3</td><td>-5.5</td><td>0.9</td><td>-6.4</td><td>1.6E-10</td><td>4.2E-09</td><td>BT20/MB231</td></tr>
<tr><td>>hg38_21_+_8433172_8446622_RNA45SN1_45S_50ntFlanks@12991.13014.24|CTCCCTCGCTGCGATCTATTGAAA</td><td>>hg38_21_+_8433172_8446622_RNA45SN1_45S_50ntFlanks</td><td>12991</td><td>13014</td><td>24</td><td>CTCCCTCGCTGCGATCTATTGAAA</td><td>1580.1</td><td>-4.5</td><td>0.8</td><td>-6.0</td><td>2.3E-09</td><td>4.2E-09</td><td>MB231/MB468</td></tr>
<tr><td>>MI0000270|hsa-mir-181b-1&WithFlank&1|-|198858867|198858988@42.64.23;MI0000683|hsa-mir-181b-2&WithFlank&9|+|124693704|124693804@22.44.23&[MIMAT0000257&hsa-miR-181b-5p&offsets|0|0;m-73&1|-|198858925|198858947&offsets|0|0];[MIMAT0000257_1&hsa-miR-181b-5p&offsets|0|0;m-72&9|+|124693725|124693747&offsets|0|0]|AACATTCATTGCTGTCGGTGGGT</td><td>>MI0000270|hsa-mir-181b-1&WithFlank&1|-|198858867|198858988</td><td>42</td><td>64</td><td>23</td><td>AACATTCATTGCTGTCGGTGGGT</td><td>2560.2</td><td>5.2</td><td>0.9</td><td>6.0</td><td>2.6E-09</td><td>4.2E-09</td><td>BT20/MB468</td></tr>
<tr><td>>hg38_21_+_8433172_8446622_RNA45SN1_45S_50ntFlanks@12874.12892.19|GCTGGAAAGGCGGCCGCCC</td><td>>hg38_21_+_8433172_8446622_RNA45SN1_45S_50ntFlanks</td><td>12874</td><td>12892</td><td>19</td><td>GCTGGAAAGGCGGCCGCCC</td><td>72.3</td><td>-7.8</td><td>1.3</td><td>-6.0</td><td>2.3E-09</td><td>4.2E-09</td><td>MB231/MB468</td></tr>
<tr><td>>MI0000749|hsa-mir-30e&WithFlank&1|+|40754349|40754452@24.46.23&[MIMAT0000692&hsa-miR-30e-5p&offsets|+1|+2;m-24&1|+|40754371|40754394&offsets|+1|0]|GTAAACATCCTTGACTGGAAGCT</td><td>>MI0000749|hsa-mir-30e&WithFlank&1|+|40754349|40754452</td><td>24</td><td>46</td><td>23</td><td>GTAAACATCCTTGACTGGAAGCT</td><td>1525.5</td><td>5.6</td><td>0.9</td><td>6.0</td><td>2.6E-09</td><td>4.2E-09</td><td>BT20/MB468</td></tr>
</table>

| | | | | | | | | | | | | |
|---|---|---|---|---|---|---|---|---|---|---|---|---|
| >hg38_21_+_8433172_8446622_RNA45SN1_45S_50ntFlanks@5419.5445.27\|ACCGATTGGATGGTTTAGTGAGGCCCT | >hg38_21_+_8433172_8446622_RNA45SN1_45S_50ntFlanks | 5419 | 5445 | 27 | ACCGATTGGATGGTTTAGTGAGGCCCT | 91.2 | -5.0 | 0.8 | -6.0 | 2.6E-09 | 4.3E-09 | BT20/MB468 |
| >hg38_21_+_8433172_8446622_RNA45SN1_45S_50ntFlanks@3872.3899.28\|CTAATACATGCCGACGGGCGCTGACCCC | >hg38_21_+_8433172_8446622_RNA45SN1_45S_50ntFlanks | 3872 | 3899 | 28 | CTAATACATGCCGACGGGCGCTGACCCC | 126.2 | -4.3 | 0.7 | -6.0 | 2.3E-09 | 4.3E-09 | MB231/MB468 |
| >hg38_21_+_8433172_8446622_RNA45SN1_45S_50ntFlanks@4243.4263.21\|CTTTCGAGGCCCTGTAATTGG | >hg38_21_+_8433172_8446622_RNA45SN1_45S_50ntFlanks | 4243 | 4263 | 21 | CTTTCGAGGCCCTGTAATTGG | 225.8 | -2.6 | 0.4 | -6.0 | 2.6E-09 | 4.3E-09 | BT20/MB468 |
| >hg38_21_+_8433172_8446622_RNA45SN1_45S_50ntFlanks@485.532.48\|GATGTGGTGACGTCGTGCTCTCCCGGGCCGGGTCCGAGCCGCGACGGG | >hg38_21_+_8433172_8446622_RNA45SN1_45S_50ntFlanks | 485 | 532 | 48 | GATGTGGTGACGTCGTGCTCTCCCGGGCCGGGTCCGAGCCGCGACGGG | 3520.5 | 16.4 | 2.8 | 6.0 | 2.6E-09 | 4.3E-09 | BT20/MB468 |
| >hg38_21_+_8433172_8446622_RNA45SN1_45S_50ntFlanks@11873.11909.37\|AAGAAGACCCTGTTGAGCTTGACTCTAGTCTGGCACG | >hg38_21_+_8433172_8446622_RNA45SN1_45S_50ntFlanks | 11873 | 11909 | 37 | AAGAAGACCCTGTTGAGCTTGACTCTAGTCTGGCACG | 120.6 | -5.9 | 1.0 | -6.0 | 2.4E-09 | 4.3E-09 | MB231/MB468 |
| >hg38_21_+_8433172_8446622_RNA45SN1_45S_50ntFlanks@9545.9572.28\|GTGGAGGTCCGTAGCGGTCCTGACGTGC | >hg38_21_+_8433172_8446622_RNA45SN1_45S_50ntFlanks | 9545 | 9572 | 28 | GTGGAGGTCCGTAGCGGTCCTGACGTGC | 905.0 | -3.3 | 0.6 | -5.9 | 2.7E-09 | 4.3E-09 | BT20/MB468 |
| >hg38_21_+_8433172_8446622_RNA45SN1_45S_50ntFlanks@6723.6740.18\|TTGCAGGACACATTGATC | >hg38_21_+_8433172_8446622_RNA45SN1_45S_50ntFlanks | 6723 | 6740 | 18 | TTGCAGGACACATTGATC | 612.2 | -3.9 | 0.6 | -5.9 | 2.7E-09 | 4.4E-09 | BT20/MB468 |
| >hg38_21_+_8433172_8446622_RNA45SN1_45S_50ntFlanks@11561.11600.40\|GCGCCTAGCAGCCGACTTAGAACTGGTGCGGACCAGGGGA | >hg38_21_+_8433172_8446622_RNA45SN1_45S_50ntFlanks | 11561 | 11600 | 40 | GCGCCTAGCAGCCGACTTAGAACTGGTGCGGACCAGGGGA | 162.5 | -4.3 | 0.7 | -6.0 | 2.4E-09 | 4.4E-09 | MB231/MB468 |
| >hg38_21_+_8433172_8446622_RNA45SN1_45S_50ntFlanks@10410.10440.31\|TCAGTCGGTCCTGAGAGATGGGCGAGCGCCG | >hg38_21_+_8433172_8446622_RNA45SN1_45S_50ntFlanks | 10410 | 10440 | 31 | TCAGTCGGTCCTGAGAGATGGGCGAGCGCCG | 117.3 | -3.4 | 0.6 | -6.0 | 2.4E-09 | 4.4E-09 | MB231/MB468 |
| >hg38_21_+_8433172_8446622_RNA45SN1_45S_50ntFlanks@6651.6696.46\|CGACTCTTAGCGGTGGATCACTCGGCTCGTGCGTCGATGAAGAACG | >hg38_21_+_8433172_8446622_RNA45SN1_45S_50ntFlanks | 6651 | 6696 | 46 | CGACTCTTAGCGGTGGATCACTCGGCTCGTGCGTCGATGAAGAACG | 5377.4 | 4.5 | 0.8 | 5.9 | 2.8E-09 | 4.5E-09 | BT20/MB468 |
| >hg38_21_+_8433172_8446622_RNA45SN1_45S_50ntFlanks@10781.10803.23\|AGGTGAACAGCCTCTGGCATGTT | >hg38_21_+_8433172_8446622_RNA45SN1_45S_50ntFlanks | 10781 | 10803 | 23 | AGGTGAACAGCCTCTGGCATGTT | 240.3 | -3.2 | 0.5 | -6.0 | 2.5E-09 | 4.5E-09 | MB231/MB468 |
| >hg38_21_+_8433172_8446622_RNA45SN1_45S_50ntFlanks@6651.6669.19\|CGACTCTTAGCGGTGGATC | >hg38_21_+_8433172_8446622_RNA45SN1_45S_50ntFlanks | 6651 | 6669 | 19 | CGACTCTTAGCGGTGGATC | 12131.1 | 3.1 | 0.5 | 5.9 | 2.8E-09 | 4.5E-09 | BT20/MB468 |
| >am_trna128_GlyGCC_6_-_27870686_27870756@1.34.34,trna18_GlyGCC_16_+_70822597_70822667@1.34.34,trna19_GlyGCC_16_+_70823410_70823480@1.34.34,trna19_GlyGCC_2_-_157257659_157257729@1.34.34,trna24_GlyGCC_16_-_70812942_70813012@1.34.34,trna25_GlyGCC_16_-_70812114_70812184@1.34.34,trna5_GlyGCC_17_+_8029064_8029134@1.34.34,trna68_GlyGCC_1_-_161493637_161493707@1.34.34\|GCATTGGTGGTTCAGTGGTAGAATTCTCGCCTGC | >am_trna128_GlyGCC_6_-_27870686_27870756 | 1 | 34 | 34 | GCATTGGTGGTTCAGTGGTAGAATTCTCGCCTGC | 458.2 | 4.8 | 0.8 | 5.9 | 2.8E-09 | 4.5E-09 | BT20/MB468 |
| >hg38_21_+_8433172_8446622_RNA45SN1_45S_50ntFlanks@10689.10708.20\|GCGTCCGGTGAGCTCTCGCT | >hg38_21_+_8433172_8446622_RNA45SN1_45S_50ntFlanks | 10689 | 10708 | 20 | GCGTCCGGTGAGCTCTCGCT | 637.5 | -3.8 | 0.6 | -6.0 | 2.5E-09 | 4.5E-09 | MB231/MB468 |
| >hg38_21_+_8433172_8446622_RNA45SN1_45S_50ntFlanks@11768.11806.39\|CTAATTAGTGACGCGCATGAATGGATGAACGAGATTCCC | >hg38_21_+_8433172_8446622_RNA45SN1_45S_50ntFlanks | 11768 | 11806 | 39 | CTAATTAGTGACGCGCATGAATGGATGAACGAGATTCCC | 107.3 | -5.1 | 0.9 | -5.9 | 2.8E-09 | 4.5E-09 | BT20/MB468 |
| >hg38_21_+_8433172_8446622_RNA45SN1_45S_50ntFlanks@10350.10379.30\|ACGAGAACTTTGAAGGCCGAAGTGGAGAAG | >hg38_21_+_8433172_8446622_RNA45SN1_45S_50ntFlanks | 10350 | 10379 | 30 | ACGAGAACTTTGAAGGCCGAAGTGGAGAAG | 299.5 | -2.6 | 0.4 | -5.9 | 2.8E-09 | 4.5E-09 | BT20/MB468 |
| >hg38_21_+_8433172_8446622_RNA45SN1_45S_50ntFlanks@10914.10932.19\|CGCCGCGGCTGGACGAGGC | >hg38_21_+_8433172_8446622_RNA45SN1_45S_50ntFlanks | 10914 | 10932 | 19 | CGCCGCGGCTGGACGAGGC | 607.6 | -7.6 | 1.3 | -6.0 | 2.5E-09 | 4.6E-09 | MB231/MB468 |
| >hg38_21_+_8433172_8446622_RNA45SN1_45S_50ntFlanks@9033.9058.26\|CTCGTGGGGGGGCCGGGCCACCCCTC | >hg38_21_+_8433172_8446622_RNA45SN1_45S_50ntFlanks | 9033 | 9058 | 26 | CTCGTGGGGGGGCCGGGCCACCCCTC | 85.3 | -8.0 | 1.3 | -6.0 | 2.5E-09 | 4.6E-09 | MB231/MB468 |
| >hg38_21_+_8433172_8446622_RNA45SN1_45S_50ntFlanks@9737.9765.29\|CGAAACGATCTCAACCTATTCTCAAACTT | >hg38_21_+_8433172_8446622_RNA45SN1_45S_50ntFlanks | 9737 | 9765 | 29 | CGAAACGATCTCAACCTATTCTCAAACTT | 1253.1 | 4.9 | 0.8 | 6.0 | 2.5E-09 | 4.6E-09 | MB231/MB468 |
| >hg38_21_+_8433172_8446622_RNA45SN1_45S_50ntFlanks@4817.4835.19\|ATTCCCATGACCCGCCGGG | >hg38_21_+_8433172_8446622_RNA45SN1_45S_50ntFlanks | 4817 | 4835 | 19 | ATTCCCATGACCCGCCGGG | 64.6 | -7.6 | 1.3 | -6.0 | 2.5E-09 | 4.6E-09 | MB231/MB468 |
| >hg38_21_+_8433172_8446622_RNA45SN1_45S_50ntFlanks@12971.12989.19\|TTTCGTACGTAGCAGAGCA | >hg38_21_+_8433172_8446622_RNA45SN1_45S_50ntFlanks | 12971 | 12989 | 19 | TTTCGTACGTAGCAGAGCA | 238.9 | -2.4 | 0.4 | -6.0 | 2.6E-09 | 4.6E-09 | MB231/MB468 |
| >hg38_21_+_8433172_8446622_RNA45SN1_45S_50ntFlanks@12656.12675.20\|TGACTGAACGCCTCTAAGTC | >hg38_21_+_8433172_8446622_RNA45SN1_45S_50ntFlanks | 12656 | 12675 | 20 | TGACTGAACGCCTCTAAGTC | 233.1 | -3.9 | 0.7 | -6.0 | 2.6E-09 | 4.7E-09 | MB231/MB468 |
| >hg38_21_+_8433172_8446622_RNA45SN1_45S_50ntFlanks@8169.8186.18\|CCAAGTCCTTCTGATCGA | >hg38_21_+_8433172_8446622_RNA45SN1_45S_50ntFlanks | 8169 | 8186 | 18 | CCAAGTCCTTCTGATCGA | 462.2 | -5.1 | 0.9 | -5.9 | 2.9E-09 | 4.7E-09 | BT20/MB468 |
| >hg38_21_+_8433172_8446622_RNA45SN1_45S_50ntFlanks@12935.12952.18\|TAAACCATTCGTAGACGA | >hg38_21_+_8433172_8446622_RNA45SN1_45S_50ntFlanks | 12935 | 12952 | 18 | TAAACCATTCGTAGACGA | 157.6 | -4.7 | 0.8 | -5.9 | 2.9E-09 | 4.7E-09 | BT20/MB468 |
| >hg38_21_+_8433172_8446622_RNA45SN1_45S_50ntFlanks@12074.12095.22\|CGAGGGGCTCTCGCTTCTGGCG | >hg38_21_+_8433172_8446622_RNA45SN1_45S_50ntFlanks | 12074 | 12095 | 22 | CGAGGGGCTCTCGCTTCTGGCG | 511.9 | -3.1 | 0.5 | -6.0 | 2.6E-09 | 4.7E-09 | MB231/MB468 |
| >trna10_AspGTC_12_-_125424193_125424264@37.74.38,trna12_AspGTC_12_-_125411891_125411962@37.74.38,trna144_AspGTC_6_-_27551236_27551307@37.74.38,trna38_AspGTC_17_-_8125556_8125627@37.74.38,trna45_AspGTC_6_+_27447453_27447524@37.74.38,trna48_AspGTC_6_+_27471523_27471594@37.74.38,trna4_AspGTC_12_+_96429799_96429870@37.74.38,trna69_AspGTC_1_-_161440205_161440276@37.74.38,trna72_AspGTC_1_-_161432824_161432895@37.74.38,trna75_AspGTC_1_-_161425414_161425485@37.74.38,trna78_AspGTC_1_-_161418033_161418104@37.74.38,trna81_AspGTC_1_-_161410615_161410686@37.74.38\|ACGCGGGAGACCGGGGTTCGATTCCCCGACGGGGAGCC | >trna10_AspGTC_12_-_125424193_125424264 | 37 | 74 | 38 | ACGCGGGAGACCGGGGTTCGATTCCCCGACGGGGAGCC | 4653.9 | 12.6 | 2.1 | 6.0 | 2.6E-09 | 4.7E-09 | MB231/MB468 |
| >trna119_LysCTT_1_-_145395522_145395594@1.32.32,trna11_LysCTT_5_-_180648979_180649051@1.32.32,trna13_LysCTT_6_+_26556774_26556846@1.32.32,trna32_LysCTT_16_-_3207406_3207478@1.32.32,trna7_LysCTT_16_+_3225692_3225764@1.32.32,trna9_LysCTT_5_+_180634755_180634827@1.32.32\|GCCCGGCTAGCTCAGTCGGTAGAGCATGAGAC | >trna119_LysCTT_1_-_145395522_145395594 | 1 | 32 | 32 | GCCCGGCTAGCTCAGTCGGTAGAGCATGAGAC | 7764.2 | 5.0 | 0.8 | 6.0 | 2.6E-09 | 4.7E-09 | MB231/MB468 |
| >hg38_21_+_8433172_8446622_RNA45SN1_45S_50ntFlanks@11315.11334.20\|CCACCCCACGTCTCGTCGCG | >hg38_21_+_8433172_8446622_RNA45SN1_45S_50ntFlanks | 11315 | 11334 | 20 | CCACCCCACGTCTCGTCGCG | 183.6 | -5.3 | 0.9 | -6.0 | 2.6E-09 | 4.7E-09 | MB231/MB468 |
| >hg38_1_-_228634819_228635039_RNA5S12_5S_50ntFlanks@150.170.21\|AATACCGGGTGCTGTAGGCTT | >hg38_1_-_228634819_228635039_RNA5S12_5S_50ntFlanks | 150 | 170 | 21 | AATACCGGGTGCTGTAGGCTT | 3142.3 | 5.0 | 0.8 | 6.0 | 2.6E-09 | 4.7E-09 | MB231/MB468 |
| >hg38_21_+_8433172_8446622_RNA45SN1_45S_50ntFlanks@10878.10901.24\|TCGGTCGGGCTGGGGCGCGAAGCG | >hg38_21_+_8433172_8446622_RNA45SN1_45S_50ntFlanks | 10878 | 10901 | 24 | TCGGTCGGGCTGGGGCGCGAAGCG | 178.0 | -3.7 | 0.6 | -5.9 | 3.0E-09 | 4.8E-09 | BT20/MB468 |
| >hg38_21_+_8433172_8446622_RNA45SN1_45S_50ntFlanks@4657.4680.24\|CTAGAGGTGAAATTCTTGGACCGG | >hg38_21_+_8433172_8446622_RNA45SN1_45S_50ntFlanks | 4657 | 4680 | 24 | CTAGAGGTGAAATTCTTGGACCGG | 100.3 | -8.2 | 1.4 | -6.0 | 2.6E-09 | 4.8E-09 | MB231/MB468 |
| >hg38_21_+_8433172_8446622_RNA45SN1_45S_50ntFlanks@12406.12436.31\|TTTTTGATCCTTCGATGTCGGCTCTTCCTAT | >hg38_21_+_8433172_8446622_RNA45SN1_45S_50ntFlanks | 12406 | 12436 | 31 | TTTTTGATCCTTCGATGTCGGCTCTTCCTAT | 191.8 | -3.0 | 0.5 | -6.0 | 2.7E-09 | 4.8E-09 | MB231/MB468 |
| >hg38_21_+_8433172_8446622_RNA45SN1_45S_50ntFlanks@10821.10840.20\|GGAAGTCGGCAAGCCGGATC | >hg38_21_+_8433172_8446622_RNA45SN1_45S_50ntFlanks | 10821 | 10840 | 20 | GGAAGTCGGCAAGCCGGATC | 1966.3 | -3.8 | 0.6 | -6.0 | 2.7E-09 | 4.8E-09 | MB231/MB468 |
| >hg38_21_+_8433172_8446622_RNA45SN1_45S_50ntFlanks@10905.10935.31\|CTGGGCGCGCGCCGCGGCTGGACGAGGCGCC | >hg38_21_+_8433172_8446622_RNA45SN1_45S_50ntFlanks | 10905 | 10935 | 31 | CTGGGCGCGCGCCGCGGCTGGACGAGGCGCC | 78.5 | -7.9 | 1.3 | -5.9 | 2.7E-09 | 4.8E-09 | MB231/MB468 |
| >hg38_21_+_8433172_8446622_RNA45SN1_45S_50ntFlanks@13005.13025.21\|TCTATTGAAAGTCAGCCCTCG | >hg38_21_+_8433172_8446622_RNA45SN1_45S_50ntFlanks | 13005 | 13025 | 21 | TCTATTGAAAGTCAGCCCTCG | 133.9 | -6.7 | 1.1 | -5.9 | 2.7E-09 | 4.8E-09 | MB231/MB468 |
| >hg38_21_+_8433172_8446622_RNA45SN1_45S_50ntFlanks@11235.11253.19\|CTTCCCGTGGATCGCCCCA | >hg38_21_+_8433172_8446622_RNA45SN1_45S_50ntFlanks | 11235 | 11253 | 19 | CTTCCCGTGGATCGCCCCA | 1072.1 | -6.3 | 1.1 | -5.9 | 3.0E-09 | 4.9E-09 | BT20/MB468 |
| >hg38_21_+_8433172_8446622_RNA45SN1_45S_50ntFlanks@10391.10426.36\|AACAGCAGTTGAACATGGGTCAGTCGGTCCTGAGAG | >hg38_21_+_8433172_8446622_RNA45SN1_45S_50ntFlanks | 10391 | 10426 | 36 | AACAGCAGTTGAACATGGGTCAGTCGGTCCTGAGAG | 184.1 | -6.1 | 1.0 | -5.9 | 3.1E-09 | 4.9E-09 | BT20/MB468 |
| >MI0000749\|hsa-mir-30e&WithFlank&1\|+\|40754349\|40754452@23.45.23(+1A)&[MIMAT0000692&hsa-miR-30e-5p&offsets\|0\|+1(+1A);m-24&1\|+\|40754371\|40754394&offsets\|0\|-1(+1A)]\|TGTAAACATCCTTGACTGGAAGCA | >MI0000749\|hsa-mir-30e&WithFlank&1\|+\|40754349\|40754452 | 23 | 45 | 23(+1A) | TGTAAACATCCTTGACTGGAAGCA | 3928.7 | 6.2 | 1.0 | 5.9 | 3.1E-09 | 4.9E-09 | BT20/MB468 |
| >hg38_21_+_8433172_8446622_RNA45SN1_45S_50ntFlanks@11741.11758.18\|TCTTAAGGTAGCCAAATG | >hg38_21_+_8433172_8446622_RNA45SN1_45S_50ntFlanks | 11741 | 11758 | 18 | TCTTAAGGTAGCCAAATG | 230.9 | -5.9 | 1.0 | -5.9 | 2.8E-09 | 5.0E-09 | MB231/MB468 |
| >MI0003129\|hsa-mir-146b&WithFlank&10\|+\|102436506\|102436590@15.38.24&[MIMAT0002809&hsa-miR-146b-5p&offsets\|0\|+1;m-70&10\|+\|102436520\|102436543&offsets\|0\|0]\|TGAGAACTGAATTCCATAGGCTGT | >MI0003129\|hsa-mir-146b&WithFlank&10\|+\|102436506\|102436590 | 15 | 38 | 24 | TGAGAACTGAATTCCATAGGCTGT | 703.9 | 4.9 | 0.8 | 5.9 | 3.1E-09 | 5.1E-09 | BT20/MB468 |
| >hg38_21_+_8433172_8446622_RNA45SN1_45S_50ntFlanks@6646.6663.18\|TCGTACGACTCTTAGCGG | >hg38_21_+_8433172_8446622_RNA45SN1_45S_50ntFlanks | 6646 | 6663 | 18 | TCGTACGACTCTTAGCGG | 17395.0 | 3.0 | 0.5 | 5.9 | 3.2E-09 | 5.1E-09 | BT20/MB468 |
| >hg38_21_+_8433172_8446622_RNA45SN1_45S_50ntFlanks@7983.8005.23\|CAGATCAGACGTGGCGACCCGCT | >hg38_21_+_8433172_8446622_RNA45SN1_45S_50ntFlanks | 7983 | 8005 | 23 | CAGATCAGACGTGGCGACCCGCT | 436.6 | -3.5 | 0.6 | -5.9 | 3.2E-09 | 5.1E-09 | BT20/MB468 |
| >hg38_21_+_8433172_8446622_RNA45SN1_45S_50ntFlanks@10829.10855.27\|GCAAGCCGGATCCGTAACTTCGGGATA | >hg38_21_+_8433172_8446622_RNA45SN1_45S_50ntFlanks | 10829 | 10855 | 27 | GCAAGCCGGATCCGTAACTTCGGGATA | 169.2 | -4.2 | 0.7 | -5.9 | 3.2E-09 | 5.1E-09 | BT20/MB468 |

<table>
<tr><td>>hg38_21_+_8433172_8446622_RNA45SN1_45S_50ntFlanks@11768.11806.39|CTAATTAGTGACGCGCATGAATGGATGAACGAGATTCCC</td><td>>hg38_21_+_8433172_8446622_RNA45SN1_45S_50ntFlanks</td><td>11768</td><td>11806</td><td>39</td><td>CTAATTAGTGACGCGCATGAATGGATGAACGAGATTCCC</td><td>88.6</td><td>-5.0</td><td>0.8</td><td>-5.9</td><td>2.8E-09</td><td>5.1E-09</td><td>MB231/MB468</td></tr>
<tr><td>>hg38_21_+_8433172_8446622_RNA45SN1_45S_50ntFlanks@4390.4408.19|TCTTGGGAGCGGGCGGGCG</td><td>>hg38_21_+_8433172_8446622_RNA45SN1_45S_50ntFlanks</td><td>4390</td><td>4408</td><td>19</td><td>TCTTGGGAGCGGGCGGGCG</td><td>76.2</td><td>-7.8</td><td>1.3</td><td>-5.9</td><td>2.9E-09</td><td>5.1E-09</td><td>MB231/MB468</td></tr>
<tr><td>>hg38_21_+_8433172_8446622_RNA45SN1_45S_50ntFlanks@11730.11751.22|AACTATGACTCTCTTAAGGTAG</td><td>>hg38_21_+_8433172_8446622_RNA45SN1_45S_50ntFlanks</td><td>11730</td><td>11751</td><td>22</td><td>AACTATGACTCTCTTAAGGTAG</td><td>430.6</td><td>-4.7</td><td>0.8</td><td>-5.9</td><td>3.2E-09</td><td>5.2E-09</td><td>BT20/MB468</td></tr>
<tr><td>>hg38_21_+_8433172_8446622_RNA45SN1_45S_50ntFlanks@8472.8490.19|CGGCCCGGCGGATCTTTCC</td><td>>hg38_21_+_8433172_8446622_RNA45SN1_45S_50ntFlanks</td><td>8472</td><td>8490</td><td>19</td><td>CGGCCCGGCGGATCTTTCC</td><td>117.0</td><td>-6.6</td><td>1.1</td><td>-5.9</td><td>2.9E-09</td><td>5.2E-09</td><td>MB231/MB468</td></tr>
<tr><td>>hg38_21_+_8433172_8446622_RNA45SN1_45S_50ntFlanks@4049.4066.18|TCGAACGTCTGCCCTATC</td><td>>hg38_21_+_8433172_8446622_RNA45SN1_45S_50ntFlanks</td><td>4049</td><td>4066</td><td>18</td><td>TCGAACGTCTGCCCTATC</td><td>455.5</td><td>3.1</td><td>0.5</td><td>5.9</td><td>2.9E-09</td><td>5.2E-09</td><td>MB231/MB468</td></tr>
<tr><td>>hg38_21_+_8433172_8446622_RNA45SN1_45S_50ntFlanks@9561.9596.36|GTCCTGACGTGCAAATCGGTCGTCCGACCTGGGTAT</td><td>>hg38_21_+_8433172_8446622_RNA45SN1_45S_50ntFlanks</td><td>9561</td><td>9596</td><td>36</td><td>GTCCTGACGTGCAAATCGGTCGTCCGACCTGGGTAT</td><td>114.2</td><td>-3.6</td><td>0.6</td><td>-5.9</td><td>3.2E-09</td><td>5.2E-09</td><td>BT20/MB468</td></tr>
<tr><td>>hg38_21_+_8433172_8446622_RNA45SN1_45S_50ntFlanks@3854.3871.18|ACTGTGGTAATTCTAGAG</td><td>>hg38_21_+_8433172_8446622_RNA45SN1_45S_50ntFlanks</td><td>3854</td><td>3871</td><td>18</td><td>ACTGTGGTAATTCTAGAG</td><td>403.3</td><td>-3.9</td><td>0.7</td><td>-5.9</td><td>3.2E-09</td><td>5.2E-09</td><td>BT20/MB468</td></tr>
<tr><td>>hg38_21_+_8433172_8446622_RNA45SN1_45S_50ntFlanks@12461.12487.27|CGTTGGATTGTTCACCCACTAATAGGG</td><td>>hg38_21_+_8433172_8446622_RNA45SN1_45S_50ntFlanks</td><td>12461</td><td>12487</td><td>27</td><td>CGTTGGATTGTTCACCCACTAATAGGG</td><td>215.8</td><td>-5.7</td><td>1.0</td><td>-5.9</td><td>3.2E-09</td><td>5.2E-09</td><td>BT20/MB468</td></tr>
<tr><td>>hg38_21_+_8433172_8446622_RNA45SN1_45S_50ntFlanks@3993.4022.30|GTGACTCTAGATAACCTCGGGCCGATCGCA</td><td>>hg38_21_+_8433172_8446622_RNA45SN1_45S_50ntFlanks</td><td>3993</td><td>4022</td><td>30</td><td>GTGACTCTAGATAACCTCGGGCCGATCGCA</td><td>352.3</td><td>3.5</td><td>0.6</td><td>5.9</td><td>2.9E-09</td><td>5.2E-09</td><td>MB231/MB468</td></tr>
<tr><td>>hg38_21_+_8433172_8446622_RNA45SN1_45S_50ntFlanks@9738.9760.23|GAAACGATCTCAACCTATTCTCA</td><td>>hg38_21_+_8433172_8446622_RNA45SN1_45S_50ntFlanks</td><td>9738</td><td>9760</td><td>23</td><td>GAAACGATCTCAACCTATTCTCA</td><td>92.3</td><td>-3.0</td><td>0.5</td><td>-5.9</td><td>3.3E-09</td><td>5.2E-09</td><td>BT20/MB468</td></tr>
<tr><td>>hg38_21_+_8433172_8446622_RNA45SN1_45S_50ntFlanks@12002.12030.29|TGCGGGCCGCCGGTGAAATACCACTACTC</td><td>>hg38_21_+_8433172_8446622_RNA45SN1_45S_50ntFlanks</td><td>12002</td><td>12030</td><td>29</td><td>TGCGGGCCGCCGGTGAAATACCACTACTC</td><td>267.2</td><td>-4.0</td><td>0.7</td><td>-5.9</td><td>3.3E-09</td><td>5.3E-09</td><td>BT20/MB468</td></tr>
<tr><td>>hg38_21_+_8433172_8446622_RNA45SN1_45S_50ntFlanks@12354.12376.23|TTACCACAGGGATAACTGGCTTG</td><td>>hg38_21_+_8433172_8446622_RNA45SN1_45S_50ntFlanks</td><td>12354</td><td>12376</td><td>23</td><td>TTACCACAGGGATAACTGGCTTG</td><td>104.9</td><td>-5.8</td><td>1.0</td><td>-5.9</td><td>3.0E-09</td><td>5.3E-09</td><td>MB231/MB468</td></tr>
<tr><td>>hg38_21_+_8433172_8446622_RNA45SN1_45S_50ntFlanks@10826.10858.33|TCGGCAAGCCGGATCCGTAACTTCGGGATAAGG</td><td>>hg38_21_+_8433172_8446622_RNA45SN1_45S_50ntFlanks</td><td>10826</td><td>10858</td><td>33</td><td>TCGGCAAGCCGGATCCGTAACTTCGGGATAAGG</td><td>331.0</td><td>-5.6</td><td>1.0</td><td>-5.9</td><td>3.0E-09</td><td>5.4E-09</td><td>MB231/MB468</td></tr>
<tr><td>>hg38_21_+_8433172_8446622_RNA45SN1_45S_50ntFlanks@4588.4611.24|CTATTTTGTTGGTTTTCGGAACTG</td><td>>hg38_21_+_8433172_8446622_RNA45SN1_45S_50ntFlanks</td><td>4588</td><td>4611</td><td>24</td><td>CTATTTTGTTGGTTTTCGGAACTG</td><td>93.8</td><td>-6.3</td><td>1.1</td><td>-5.9</td><td>3.0E-09</td><td>5.4E-09</td><td>MB231/MB468</td></tr>
<tr><td>>hg38_21_+_8433172_8446622_RNA45SN1_45S_50ntFlanks@3985.4003.19|CGGCTTTGGTGACTCTAGA</td><td>>hg38_21_+_8433172_8446622_RNA45SN1_45S_50ntFlanks</td><td>3985</td><td>4003</td><td>19</td><td>CGGCTTTGGTGACTCTAGA</td><td>68.8</td><td>-7.7</td><td>1.3</td><td>-5.9</td><td>3.0E-09</td><td>5.4E-09</td><td>MB231/MB468</td></tr>
<tr><td>>hg38_21_+_8433172_8446622_RNA45SN1_45S_50ntFlanks@9574.9595.22|AATCGGTCGTCCGACCTGGGTA</td><td>>hg38_21_+_8433172_8446622_RNA45SN1_45S_50ntFlanks</td><td>9574</td><td>9595</td><td>22</td><td>AATCGGTCGTCCGACCTGGGTA</td><td>2242.8</td><td>-4.0</td><td>0.7</td><td>-5.9</td><td>3.4E-09</td><td>5.4E-09</td><td>BT20/MB468</td></tr>
<tr><td>>hg38_21_+_8433172_8446622_RNA45SN1_45S_50ntFlanks@10349.10372.24|AACGAGAACTTTGAAGGCCGAAGT</td><td>>hg38_21_+_8433172_8446622_RNA45SN1_45S_50ntFlanks</td><td>10349</td><td>10372</td><td>24</td><td>AACGAGAACTTTGAAGGCCGAAGT</td><td>867.5</td><td>2.9</td><td>0.5</td><td>5.9</td><td>3.4E-09</td><td>5.4E-09</td><td>BT20/MB468</td></tr>
<tr><td>>MI0000102|hsa-mir-100&WithFlank&11|-|122152223|122152314@19.41.23&[MIMAT0000098&hsa-miR-100-5p&offsets|0|+1;m-27&11|-|122152276|122152296&offsets|0|+2]|AACCCGTAGATCCGAACTTGTGG</td><td>>MI0000102|hsa-mir-100&WithFlank&11|-|122152223|122152314</td><td>19</td><td>41</td><td>23</td><td>AACCCGTAGATCCGAACTTGTGG</td><td>6009.2</td><td>-5.8</td><td>0.9</td><td>-6.4</td><td>2.0E-10</td><td>5.4E-09</td><td>BT20/MB231</td></tr>
<tr><td>>hg38_21_+_8433172_8446622_RNA45SN1_45S_50ntFlanks@12936.12953.18|AAACCATTCGTAGACGAC</td><td>>hg38_21_+_8433172_8446622_RNA45SN1_45S_50ntFlanks</td><td>12936</td><td>12953</td><td>18</td><td>AAACCATTCGTAGACGAC</td><td>550.4</td><td>-3.4</td><td>0.6</td><td>-5.9</td><td>3.0E-09</td><td>5.4E-09</td><td>MB231/MB468</td></tr>
<tr><td>>hg38_21_+_8433172_8446622_RNA45SN1_45S_50ntFlanks@9556.9584.29|TAGCGGTCCTGACGTGCAAATCGGTCGTC</td><td>>hg38_21_+_8433172_8446622_RNA45SN1_45S_50ntFlanks</td><td>9556</td><td>9584</td><td>29</td><td>TAGCGGTCCTGACGTGCAAATCGGTCGTC</td><td>186.1</td><td>-6.1</td><td>1.0</td><td>-5.9</td><td>3.4E-09</td><td>5.5E-09</td><td>BT20/MB468</td></tr>
<tr><td>>MI0000746|hsa-mir-99b&WithFlank&19|+|51692606|51692687@13.32.20(+2G)&[MIMAT0000689&hsa-miR-99b-5p&offsets|0|-2(+2G);m-10&19|+|51692618|51692639&offsets|0|-2(+2G)]|CACCCGTAGAACCGACCTTGGG</td><td>>MI0000746|hsa-mir-99b&WithFlank&19|+|51692606|51692687</td><td>13</td><td>32</td><td>20(+2G)</td><td>CACCCGTAGAACCGACCTTGGG</td><td>1572.4</td><td>15.3</td><td>2.6</td><td>5.9</td><td>3.4E-09</td><td>5.5E-09</td><td>BT20/MB468</td></tr>
<tr><td>>MI0000782|hsa-mir-374a&WithFlank&X|-|74287280|74287363@18.39.22&[MIMAT0000727&hsa-miR-374a-5p&offsets|0|0;m-184&X|-|74287325|74287346&offsets|0|0]|TTATAATACAACCTGATAAGTG</td><td>>MI0000782|hsa-mir-374a&WithFlank&X|-|74287280|74287363</td><td>18</td><td>39</td><td>22</td><td>TTATAATACAACCTGATAAGTG</td><td>5146.7</td><td>6.7</td><td>1.1</td><td>5.9</td><td>3.4E-09</td><td>5.5E-09</td><td>BT20/MB468</td></tr>
<tr><td>>MI0003137|hsa-mir-193b&WithFlank&16|+|14303961|14304055@57.78.22&[MIMAT0002819&hsa-miR-193b-3p&offsets|0|0;m-166&16|+|14304017|14304038&offsets|0|0]|AACTGGCCCTCAAAGTCCCGCT</td><td>>MI0003137|hsa-mir-193b&WithFlank&16|+|14303961|14304055</td><td>57</td><td>78</td><td>22</td><td>AACTGGCCCTCAAAGTCCCGCT</td><td>5084.6</td><td>5.3</td><td>0.8</td><td>6.4</td><td>2.1E-10</td><td>5.5E-09</td><td>BT20/MB231</td></tr>
<tr><td>>hg38_21_+_8433172_8446622_RNA45SN1_45S_50ntFlanks@10348.10382.35|AAACGAGAACTTTGAAGGCCGAAGTGGAGAAGGGT</td><td>>hg38_21_+_8433172_8446622_RNA45SN1_45S_50ntFlanks</td><td>10348</td><td>10382</td><td>35</td><td>AAACGAGAACTTTGAAGGCCGAAGTGGAGAAGGGT</td><td>698.5</td><td>4.2</td><td>0.7</td><td>5.9</td><td>3.1E-09</td><td>5.6E-09</td><td>MB231/MB468</td></tr>
<tr><td>>MI0000732|hsa-mir-194-2&WithFlank&11|-|64891349|64891445@21.43.23&[MIMAT0000460_1&hsa-miR-194-5p&offsets|0|+1;m-126&11|-|64891404|64891425&offsets|0|+1]|TGTAACAGCAACTCCATGTGGAA</td><td>>MI0000732|hsa-mir-194-2&WithFlank&11|-|64891349|64891445</td><td>21</td><td>43</td><td>23</td><td>TGTAACAGCAACTCCATGTGGAA</td><td>609.3</td><td>5.1</td><td>0.9</td><td>5.9</td><td>3.5E-09</td><td>5.6E-09</td><td>BT20/MB468</td></tr>
<tr><td>>hg38_21_+_8433172_8446622_RNA45SN1_45S_50ntFlanks@12405.12430.26|CTTTTTGATCCTTCGATGTCGGCTCT</td><td>>hg38_21_+_8433172_8446622_RNA45SN1_45S_50ntFlanks</td><td>12405</td><td>12430</td><td>26</td><td>CTTTTTGATCCTTCGATGTCGGCTCT</td><td>648.4</td><td>-3.7</td><td>0.6</td><td>-5.9</td><td>3.2E-09</td><td>5.7E-09</td><td>MB231/MB468</td></tr>
<tr><td>>hg38_21_+_8433172_8446622_RNA45SN1_45S_50ntFlanks@7978.8011.34|GACCTCAGATCAGACGTGGCGACCCGCTGAATTT</td><td>>hg38_21_+_8433172_8446622_RNA45SN1_45S_50ntFlanks</td><td>7978</td><td>8011</td><td>34</td><td>GACCTCAGATCAGACGTGGCGACCCGCTGAATTT</td><td>349.4</td><td>6.0</td><td>1.0</td><td>5.9</td><td>3.5E-09</td><td>5.7E-09</td><td>BT20/MB468</td></tr>
<tr><td>>hg38_1_-_228634819_228635039_RNA5S12_5S_50ntFlanks@125.170.46|GTACTTGGATGGGAGACCGCCTGGGAATACCGGGTGCTGTAGGCTT</td><td>>hg38_1_-_228634819_228635039_RNA5S12_5S_50ntFlanks</td><td>125</td><td>170</td><td>46</td><td>GTACTTGGATGGGAGACCGCCTGGGAATACCGGGTGCTGTAGGCTT</td><td>712.7</td><td>12.3</td><td>2.1</td><td>5.9</td><td>3.2E-09</td><td>5.7E-09</td><td>MB231/MB468</td></tr>
<tr><td>>hg38_21_+_8433172_8446622_RNA45SN1_45S_50ntFlanks@485.507.23|GATGTGGTGACGTCGTGCTCTCC</td><td>>hg38_21_+_8433172_8446622_RNA45SN1_45S_50ntFlanks</td><td>485</td><td>507</td><td>23</td><td>GATGTGGTGACGTCGTGCTCTCC</td><td>15986.3</td><td>18.9</td><td>3.2</td><td>5.9</td><td>3.2E-09</td><td>5.7E-09</td><td>MB231/MB468</td></tr>
<tr><td>>hg38_21_+_8433172_8446622_RNA45SN1_45S_50ntFlanks@4588.4613.26|CTATTTTGTTGGTTTTCGGAACTGAG</td><td>>hg38_21_+_8433172_8446622_RNA45SN1_45S_50ntFlanks</td><td>4588</td><td>4613</td><td>26</td><td>CTATTTTGTTGGTTTTCGGAACTGAG</td><td>150.4</td><td>-6.1</td><td>1.0</td><td>-5.9</td><td>3.2E-09</td><td>5.8E-09</td><td>MB231/MB468</td></tr>
<tr><td>>hg38_21_+_8433172_8446622_RNA45SN1_45S_50ntFlanks@12499.12521.23|GGTTTAGACCGTCGTGAGACAGG</td><td>>hg38_21_+_8433172_8446622_RNA45SN1_45S_50ntFlanks</td><td>12499</td><td>12521</td><td>23</td><td>GGTTTAGACCGTCGTGAGACAGG</td><td>90.1</td><td>-6.2</td><td>1.1</td><td>-5.9</td><td>3.3E-09</td><td>5.8E-09</td><td>MB231/MB468</td></tr>
<tr><td>>hg38_21_+_8433172_8446622_RNA45SN1_45S_50ntFlanks@7979.8008.30|ACCTCAGATCAGACGTGGCGACCCGCTGAA</td><td>>hg38_21_+_8433172_8446622_RNA45SN1_45S_50ntFlanks</td><td>7979</td><td>8008</td><td>30</td><td>ACCTCAGATCAGACGTGGCGACCCGCTGAA</td><td>924.6</td><td>-3.8</td><td>0.7</td><td>-5.9</td><td>3.6E-09</td><td>5.8E-09</td><td>BT20/MB468</td></tr>
<tr><td>>hg38_21_+_8433172_8446622_RNA45SN1_45S_50ntFlanks@13006.13039.34|CTATTGAAAGTCAGCCCTCGACACAAGGGTTTGT</td><td>>hg38_21_+_8433172_8446622_RNA45SN1_45S_50ntFlanks</td><td>13006</td><td>13039</td><td>34</td><td>CTATTGAAAGTCAGCCCTCGACACAAGGGTTTGT</td><td>1608.2</td><td>4.6</td><td>0.8</td><td>5.9</td><td>3.3E-09</td><td>5.9E-09</td><td>MB231/MB468</td></tr>
<tr><td>>trna10_ValCAC_5_-_180649395_180649467@35.54.20,trna12_ValAAC_5_-_180645270_180645342@35.54.20,trna132_ValAAC_6_-_27721179_27721251@35.54.20,trna13_ValCAC_19_-_4724647_4724719@35.54.20,trna152_ValCAC_6_-_27248049_27248121@35.54.20,trna157_ValCAC_6_-_27173867_27173939@35.54.20,trna15_ValAAC_5_-_180615416_180615488@35.54.20,trna18_ValCAC_5_-_180529253_180529325@35.54.20,trna2_ValAAC_3_+_169490018_169490090@35.54.20,trna2_ValCAC_5_+_180524070_180524142@35.54.20,trna37_ValAAC_6_+_27203288_27203360@35.54.20,trna4_ValAAC_5_+_180591154_180591226@35.54.20,trna5_ValAAC_5_+_180596610_180596682@35.54.20,trna6_ValCAC_5_+_180600650_180600722@35.54.20,trna85_ValCAC_1_-_161369490_161369562@35.54.20,trna98_ValCAC_1_-_149298555_149298627@35.54.20,trna9_ValCAC_6_+_26538282_26538354@35.54.20|ACACGCGAAAGGTCCCCGGT</td><td>>trna10_ValCAC_5_-_180649395_180649467</td><td>35</td><td>54</td><td>20</td><td>ACACGCGAAAGGTCCCCGGT</td><td>73.1</td><td>-7.8</td><td>1.3</td><td>-5.9</td><td>3.3E-09</td><td>5.9E-09</td><td>MB231/MB468</td></tr>
<tr><td>>hg38_21_+_8433172_8446622_RNA45SN1_45S_50ntFlanks@9716.9737.22|AATGATTAGAGGTCTTGGGGCC</td><td>>hg38_21_+_8433172_8446622_RNA45SN1_45S_50ntFlanks</td><td>9716</td><td>9737</td><td>22</td><td>AATGATTAGAGGTCTTGGGGCC</td><td>229.1</td><td>-2.3</td><td>0.4</td><td>-5.9</td><td>3.3E-09</td><td>5.9E-09</td><td>MB231/MB468</td></tr>
<tr><td>>hg38_21_+_8433172_8446622_RNA45SN1_45S_50ntFlanks@10819.10839.21|AGGGAAGTCGGCAAGCCGGAT</td><td>>hg38_21_+_8433172_8446622_RNA45SN1_45S_50ntFlanks</td><td>10819</td><td>10839</td><td>21</td><td>AGGGAAGTCGGCAAGCCGGAT</td><td>147.7</td><td>-5.0</td><td>0.9</td><td>-5.9</td><td>3.7E-09</td><td>6.0E-09</td><td>BT20/MB468</td></tr>
<tr><td>>hg38_21_+_8433172_8446622_RNA45SN1_45S_50ntFlanks@11873.11898.26|AAGAAGACCCTGTTGAGCTTGACTCT</td><td>>hg38_21_+_8433172_8446622_RNA45SN1_45S_50ntFlanks</td><td>11873</td><td>11898</td><td>26</td><td>AAGAAGACCCTGTTGAGCTTGACTCT</td><td>949.7</td><td>-3.4</td><td>0.6</td><td>-5.9</td><td>3.4E-09</td><td>6.0E-09</td><td>MB231/MB468</td></tr>
<tr><td>>hg38_21_+_8433172_8446622_RNA45SN1_45S_50ntFlanks@10391.10430.40|AACAGCAGTTGAACATGGGTCAGTCGGTCCTGAGAGATGG</td><td>>hg38_21_+_8433172_8446622_RNA45SN1_45S_50ntFlanks</td><td>10391</td><td>10430</td><td>40</td><td>AACAGCAGTTGAACATGGGTCAGTCGGTCCTGAGAGATGG</td><td>279.5</td><td>-3.8</td><td>0.6</td><td>-5.9</td><td>3.4E-09</td><td>6.0E-09</td><td>MB231/MB468</td></tr>
<tr><td>>hg38_21_+_8433172_8446622_RNA45SN1_45S_50ntFlanks@11874.11908.35|AGAAGACCCTGTTGAGCTTGACTCTAGTCTGGCAC</td><td>>hg38_21_+_8433172_8446622_RNA45SN1_45S_50ntFlanks</td><td>11874</td><td>11908</td><td>35</td><td>AGAAGACCCTGTTGAGCTTGACTCTAGTCTGGCAC</td><td>344.6</td><td>-5.9</td><td>1.0</td><td>-5.9</td><td>3.8E-09</td><td>6.1E-09</td><td>BT20/MB468</td></tr>
<tr><td>>hg38_21_+_8433172_8446622_RNA45SN1_45S_50ntFlanks@13009.13028.20|TTGAAAGTCAGCCCTCGACA</td><td>>hg38_21_+_8433172_8446622_RNA45SN1_45S_50ntFlanks</td><td>13009</td><td>13028</td><td>20</td><td>TTGAAAGTCAGCCCTCGACA</td><td>152.2</td><td>-6.0</td><td>1.0</td><td>-5.9</td><td>3.4E-09</td><td>6.1E-09</td><td>MB231/MB468</td></tr>
<tr><td>>hg38_21_+_8433172_8446622_RNA45SN1_45S_50ntFlanks@11594.11612.19|CAGGGGAATCCGACTGTTT</td><td>>hg38_21_+_8433172_8446622_RNA45SN1_45S_50ntFlanks</td><td>11594</td><td>11612</td><td>19</td><td>CAGGGGAATCCGACTGTTT</td><td>760.8</td><td>-3.3</td><td>0.6</td><td>-5.9</td><td>3.8E-09</td><td>6.1E-09</td><td>BT20/MB468</td></tr>
<tr><td>>hg38_21_+_8433172_8446622_RNA45SN1_45S_50ntFlanks@10574.10592.19|AAGCCGGCGGGAGCCCCGG</td><td>>hg38_21_+_8433172_8446622_RNA45SN1_45S_50ntFlanks</td><td>10574</td><td>10592</td><td>19</td><td>AAGCCGGCGGGAGCCCCGG</td><td>81.7</td><td>-7.9</td><td>1.3</td><td>-5.9</td><td>3.4E-09</td><td>6.2E-09</td><td>MB231/MB468</td></tr>
</table>

<table>
<tr><td>>hg38_21_+_8433172_8446622_RNA45SN1_45S_50ntFlanks@9534.9573.40|AGGAAACTCTGGTGGAGGTCCGTAGCGGTCCTGACGTGCA</td><td>>hg38_21_+_8433172_8446622_RNA45SN1_45S_50ntFlanks</td><td>9534</td><td>9573</td><td>40</td><td>AGGAAACTCTGGTGGAGGTCCGTAGCGGTCCTGACGTGCA</td><td>479.7</td><td>-3.1</td><td>0.5</td><td>-5.9</td><td>3.9E-09</td><td>6.2E-09</td><td>BT20/MB468</td></tr>
<tr><td>>hg38_21_+_8433172_8446622_RNA45SN1_45S_50ntFlanks@11778.11798.21|ACGCGCATGAATGGATGAACG</td><td>>hg38_21_+_8433172_8446622_RNA45SN1_45S_50ntFlanks</td><td>11778</td><td>11798</td><td>21</td><td>ACGCGCATGAATGGATGAACG</td><td>192.9</td><td>-3.8</td><td>0.7</td><td>-5.9</td><td>3.9E-09</td><td>6.2E-09</td><td>BT20/MB468</td></tr>
<tr><td>>hg38_21_+_8433172_8446622_RNA45SN1_45S_50ntFlanks@10843.10861.19|TAACTTCGGGATAAGGATT</td><td>>hg38_21_+_8433172_8446622_RNA45SN1_45S_50ntFlanks</td><td>10843</td><td>10861</td><td>19</td><td>TAACTTCGGGATAAGGATT</td><td>200.9</td><td>-3.1</td><td>0.5</td><td>-5.9</td><td>3.9E-09</td><td>6.2E-09</td><td>BT20/MB468</td></tr>
<tr><td>>hg38_21_+_8433172_8446622_RNA45SN1_45S_50ntFlanks@12002.12043.42|TGCGGGCCGCCGGTGAAATACCACTACTCTGATCGTTTTTTC</td><td>>hg38_21_+_8433172_8446622_RNA45SN1_45S_50ntFlanks</td><td>12002</td><td>12043</td><td>42</td><td>TGCGGGCCGCCGGTGAAATACCACTACTCTGATCGTTTTTTC</td><td>124.0</td><td>-8.5</td><td>1.4</td><td>-5.9</td><td>3.5E-09</td><td>6.3E-09</td><td>MB231/MB468</td></tr>
<tr><td>>hg38_21_+_8433172_8446622_RNA45SN1_45S_50ntFlanks@4046.4065.20|CATTCGAACGTCTGCCCTAT</td><td>>hg38_21_+_8433172_8446622_RNA45SN1_45S_50ntFlanks</td><td>4046</td><td>4065</td><td>20</td><td>CATTCGAACGTCTGCCCTAT</td><td>140.6</td><td>-2.3</td><td>0.4</td><td>-5.9</td><td>3.9E-09</td><td>6.3E-09</td><td>BT20/MB468</td></tr>
<tr><td>>hg38_21_+_8433172_8446622_RNA45SN1_45S_50ntFlanks@12363.12383.21|GGATAACTGGCTTGTGGCGGC</td><td>>hg38_21_+_8433172_8446622_RNA45SN1_45S_50ntFlanks</td><td>12363</td><td>12383</td><td>21</td><td>GGATAACTGGCTTGTGGCGGC</td><td>81.3</td><td>-7.9</td><td>1.3</td><td>-5.9</td><td>3.5E-09</td><td>6.3E-09</td><td>MB231/MB468</td></tr>
<tr><td>>hg38_21_+_8433172_8446622_RNA45SN1_45S_50ntFlanks@10458.10484.27|ATGGCCTCCGTTGCCCTCGGCCGATCG</td><td>>hg38_21_+_8433172_8446622_RNA45SN1_45S_50ntFlanks</td><td>10458</td><td>10484</td><td>27</td><td>ATGGCCTCCGTTGCCCTCGGCCGATCG</td><td>228.0</td><td>-3.2</td><td>0.5</td><td>-5.9</td><td>3.5E-09</td><td>6.3E-09</td><td>MB231/MB468</td></tr>
<tr><td>>hg38_21_+_8433172_8446622_RNA45SN1_45S_50ntFlanks@10116.10134.19|TGCGTCGGAGGGCGGCGGC</td><td>>hg38_21_+_8433172_8446622_RNA45SN1_45S_50ntFlanks</td><td>10116</td><td>10134</td><td>19</td><td>TGCGTCGGAGGGCGGCGGC</td><td>118.2</td><td>-5.1</td><td>0.9</td><td>-5.9</td><td>4.0E-09</td><td>6.3E-09</td><td>BT20/MB468</td></tr>
<tr><td>>hg38_21_+_8433172_8446622_RNA45SN1_45S_50ntFlanks@475.523.49|CTTCGTGATCGATGTGGTGACGTCGTGCTCTCCCGGGCCGGGTCCGAGC</td><td>>hg38_21_+_8433172_8446622_RNA45SN1_45S_50ntFlanks</td><td>475</td><td>523</td><td>49</td><td>CTTCGTGATCGATGTGGTGACGTCGTGCTCTCCCGGGCCGGGTCCGAGC</td><td>497.5</td><td>8.7</td><td>1.5</td><td>5.9</td><td>4.0E-09</td><td>6.4E-09</td><td>BT20/MB468</td></tr>
<tr><td>>MI0003628|hsa-mir-615&WithFlank&12|+|54033944|54034051@67.86.20(+1C)&[MIMAT0003283&hsa-miR-615-3p&offsets|0|-2(+1C);m-495&12|+|54034010|54034030&offsets|0|-1(+1C)]|TCCGAGCCTGGGTCTCCCTCC</td><td>>MI0003628|hsa-mir-615&WithFlank&12|+|54033944|54034051</td><td>67</td><td>86</td><td>20(+1C)</td><td>TCCGAGCCTGGGTCTCCCTCC</td><td>284.0</td><td>5.1</td><td>0.8</td><td>6.3</td><td>2.5E-10</td><td>6.4E-09</td><td>BT20/MB231</td></tr>
<tr><td>>hg38_21_+_8433172_8446622_RNA45SN1_45S_50ntFlanks@9563.9595.33|CCTGACGTGCAAATCGGTCGTCCGACCTGGGTA</td><td>>hg38_21_+_8433172_8446622_RNA45SN1_45S_50ntFlanks</td><td>9563</td><td>9595</td><td>33</td><td>CCTGACGTGCAAATCGGTCGTCCGACCTGGGTA</td><td>102.0</td><td>-3.7</td><td>0.6</td><td>-5.9</td><td>4.0E-09</td><td>6.5E-09</td><td>BT20/MB468</td></tr>
<tr><td>>hg38_21_+_8433172_8446622_RNA45SN1_45S_50ntFlanks@9304.9338.35|ACCAAGGAGTCTAACACGTGCGCGAGTCGGGGGCT</td><td>>hg38_21_+_8433172_8446622_RNA45SN1_45S_50ntFlanks</td><td>9304</td><td>9338</td><td>35</td><td>ACCAAGGAGTCTAACACGTGCGCGAGTCGGGGGCT</td><td>80.6</td><td>-7.9</td><td>1.3</td><td>-5.9</td><td>3.6E-09</td><td>6.5E-09</td><td>MB231/MB468</td></tr>
<tr><td>>hg38_21_+_8433172_8446622_RNA45SN1_45S_50ntFlanks@6651.6688.38|CGACTCTTAGCGGTGGATCACTCGGCTCGTGCGTCGAT</td><td>>hg38_21_+_8433172_8446622_RNA45SN1_45S_50ntFlanks</td><td>6651</td><td>6688</td><td>38</td><td>CGACTCTTAGCGGTGGATCACTCGGCTCGTGCGTCGAT</td><td>6065.4</td><td>4.3</td><td>0.7</td><td>5.9</td><td>4.1E-09</td><td>6.5E-09</td><td>BT20/MB468</td></tr>
<tr><td>>hg38_21_+_8433172_8446622_RNA45SN1_45S_50ntFlanks@11918.11949.32|ACATGAGAGGTGTAGAATAAGTGGGAGGCCCC</td><td>>hg38_21_+_8433172_8446622_RNA45SN1_45S_50ntFlanks</td><td>11918</td><td>11949</td><td>32</td><td>ACATGAGAGGTGTAGAATAAGTGGGAGGCCCC</td><td>78.6</td><td>-7.9</td><td>1.3</td><td>-5.9</td><td>3.7E-09</td><td>6.5E-09</td><td>MB231/MB468</td></tr>
<tr><td>>hg38_21_+_8433172_8446622_RNA45SN1_45S_50ntFlanks@11576.11609.34|CTTAGAACTGGTGCGGACCAGGGGAATCCGACTG</td><td>>hg38_21_+_8433172_8446622_RNA45SN1_45S_50ntFlanks</td><td>11576</td><td>11609</td><td>34</td><td>CTTAGAACTGGTGCGGACCAGGGGAATCCGACTG</td><td>522.6</td><td>-3.4</td><td>0.6</td><td>-5.9</td><td>3.7E-09</td><td>6.6E-09</td><td>MB231/MB468</td></tr>
<tr><td>>hg38_21_+_8433172_8446622_RNA45SN1_45S_50ntFlanks@6745.6769.25|ACACTTCGAACGCACTTGCGGCCCC</td><td>>hg38_21_+_8433172_8446622_RNA45SN1_45S_50ntFlanks</td><td>6745</td><td>6769</td><td>25</td><td>ACACTTCGAACGCACTTGCGGCCCC</td><td>156.7</td><td>-3.3</td><td>0.6</td><td>-5.9</td><td>3.7E-09</td><td>6.6E-09</td><td>MB231/MB468</td></tr>
<tr><td>>hg38_21_+_8433172_8446622_RNA45SN1_45S_50ntFlanks@4005.4023.19|AACCTCGGGCCGATCGCAC</td><td>>hg38_21_+_8433172_8446622_RNA45SN1_45S_50ntFlanks</td><td>4005</td><td>4023</td><td>19</td><td>AACCTCGGGCCGATCGCAC</td><td>488.7</td><td>2.9</td><td>0.5</td><td>5.9</td><td>3.7E-09</td><td>6.6E-09</td><td>MB231/MB468</td></tr>
<tr><td>>hg38_21_+_8433172_8446622_RNA45SN1_45S_50ntFlanks@4722.4759.38|TCATTAATCAAGAACGAAAGTCGGAGGTTCGAAGACGA</td><td>>hg38_21_+_8433172_8446622_RNA45SN1_45S_50ntFlanks</td><td>4722</td><td>4759</td><td>38</td><td>TCATTAATCAAGAACGAAAGTCGGAGGTTCGAAGACGA</td><td>1033.7</td><td>-5.0</td><td>0.8</td><td>-5.9</td><td>4.2E-09</td><td>6.7E-09</td><td>BT20/MB468</td></tr>
<tr><td>>hg38_21_+_8433172_8446622_RNA45SN1_45S_50ntFlanks@3782.3805.24|CAGTGAAACTGCGAATGGCTCATT</td><td>>hg38_21_+_8433172_8446622_RNA45SN1_45S_50ntFlanks</td><td>3782</td><td>3805</td><td>24</td><td>CAGTGAAACTGCGAATGGCTCATT</td><td>329.7</td><td>-2.6</td><td>0.5</td><td>-5.9</td><td>4.2E-09</td><td>6.7E-09</td><td>BT20/MB468</td></tr>
<tr><td>>hg38_21_+_8433172_8446622_RNA45SN1_45S_50ntFlanks@8552.8577.26|CTCCCCGGAGGGGGCGGGCTCCGGCG</td><td>>hg38_21_+_8433172_8446622_RNA45SN1_45S_50ntFlanks</td><td>8552</td><td>8577</td><td>26</td><td>CTCCCCGGAGGGGGCGGGCTCCGGCG</td><td>194.8</td><td>-5.2</td><td>0.9</td><td>-5.9</td><td>4.2E-09</td><td>6.7E-09</td><td>BT20/MB468</td></tr>
<tr><td>>hg38_21_+_8433172_8446622_RNA45SN1_45S_50ntFlanks@11579.11605.27|AGAACTGGTGCGGACCAGGGGAATCCG</td><td>>hg38_21_+_8433172_8446622_RNA45SN1_45S_50ntFlanks</td><td>11579</td><td>11605</td><td>27</td><td>AGAACTGGTGCGGACCAGGGGAATCCG</td><td>964.8</td><td>-5.5</td><td>0.9</td><td>-5.9</td><td>3.8E-09</td><td>6.7E-09</td><td>MB231/MB468</td></tr>
<tr><td>>hg38_21_+_8433172_8446622_RNA45SN1_45S_50ntFlanks@11920.11951.32|ATGAGAGGTGTAGAATAAGTGGGAGGCCCCCG</td><td>>hg38_21_+_8433172_8446622_RNA45SN1_45S_50ntFlanks</td><td>11920</td><td>11951</td><td>32</td><td>ATGAGAGGTGTAGAATAAGTGGGAGGCCCCCG</td><td>98.1</td><td>-4.9</td><td>0.8</td><td>-5.9</td><td>4.2E-09</td><td>6.7E-09</td><td>BT20/MB468</td></tr>
<tr><td>>hg38_21_+_8433172_8446622_RNA45SN1_45S_50ntFlanks@10729.10752.24|AGAGGGTGTAAATCTCGCGCCGGG</td><td>>hg38_21_+_8433172_8446622_RNA45SN1_45S_50ntFlanks</td><td>10729</td><td>10752</td><td>24</td><td>AGAGGGTGTAAATCTCGCGCCGGG</td><td>121.1</td><td>-5.1</td><td>0.9</td><td>-5.9</td><td>4.2E-09</td><td>6.8E-09</td><td>BT20/MB468</td></tr>
<tr><td>>hg38_21_+_8433172_8446622_RNA45SN1_45S_50ntFlanks@4588.4608.21|CTATTTTGTTGGTTTTCGGAA</td><td>>hg38_21_+_8433172_8446622_RNA45SN1_45S_50ntFlanks</td><td>4588</td><td>4608</td><td>21</td><td>CTATTTTGTTGGTTTTCGGAA</td><td>152.0</td><td>-6.9</td><td>1.2</td><td>-5.9</td><td>3.8E-09</td><td>6.8E-09</td><td>MB231/MB468</td></tr>
<tr><td>>trna2_GlyGCC_21_-_18827107_18827177@1.32.32,trna35_GlyGCC_1_+_161413094_161413164@1.32.32,trna37_GlyGCC_1_+_161420467_161420537@1.32.32,trna39_GlyGCC_1_+_161427898_161427968@1.32.32,trna41_GlyGCC_1_+_161435258_161435328@1.32.32|GCATGGGTGGTTCAGTGGTAGAATTCTCGCCT</td><td>>trna2_GlyGCC_21_-_18827107_18827177</td><td>1</td><td>32</td><td>32</td><td>GCATGGGTGGTTCAGTGGTAGAATTCTCGCCT</td><td>7617.6</td><td>4.7</td><td>0.7</td><td>6.3</td><td>2.6E-10</td><td>6.8E-09</td><td>BT20/MB231</td></tr>
<tr><td>>hg38_21_+_8433172_8446622_RNA45SN1_45S_50ntFlanks@11670.11700.31|CCCAGTGCTCTGAATGTCAAAGTGAAGAAAT</td><td>>hg38_21_+_8433172_8446622_RNA45SN1_45S_50ntFlanks</td><td>11670</td><td>11700</td><td>31</td><td>CCCAGTGCTCTGAATGTCAAAGTGAAGAAAT</td><td>266.0</td><td>-4.0</td><td>0.7</td><td>-5.9</td><td>4.3E-09</td><td>6.9E-09</td><td>BT20/MB468</td></tr>
<tr><td>>hg38_21_+_8433172_8446622_RNA45SN1_45S_50ntFlanks@5471.5514.44|CACGGCCCTGGCGGAGCGCTGAGAAGACGGTCGAACTTGACTAT</td><td>>hg38_21_+_8433172_8446622_RNA45SN1_45S_50ntFlanks</td><td>5471</td><td>5514</td><td>44</td><td>CACGGCCCTGGCGGAGCGCTGAGAAGACGGTCGAACTTGACTAT</td><td>139.0</td><td>-4.6</td><td>0.8</td><td>-5.9</td><td>4.3E-09</td><td>6.9E-09</td><td>BT20/MB468</td></tr>
<tr><td>>hg38_21_+_8433172_8446622_RNA45SN1_45S_50ntFlanks@11875.11906.32|GAAGACCCTGTTGAGCTTGACTCTAGTCTGGC</td><td>>hg38_21_+_8433172_8446622_RNA45SN1_45S_50ntFlanks</td><td>11875</td><td>11906</td><td>32</td><td>GAAGACCCTGTTGAGCTTGACTCTAGTCTGGC</td><td>95.9</td><td>-4.8</td><td>0.8</td><td>-5.9</td><td>3.9E-09</td><td>7.0E-09</td><td>MB231/MB468</td></tr>
<tr><td>>hg38_21_+_8433172_8446622_RNA45SN1_45S_50ntFlanks@11873.11898.26|AAGAAGACCCTGTTGAGCTTGACTCT</td><td>>hg38_21_+_8433172_8446622_RNA45SN1_45S_50ntFlanks</td><td>11873</td><td>11898</td><td>26</td><td>AAGAAGACCCTGTTGAGCTTGACTCT</td><td>1136.5</td><td>-3.5</td><td>0.6</td><td>-5.9</td><td>4.5E-09</td><td>7.1E-09</td><td>BT20/MB468</td></tr>
<tr><td>>hg38_21_+_8433172_8446622_RNA45SN1_45S_50ntFlanks@6652.6696.45|GACTCTTAGCGGTGGATCACTCGGCTCGTGCGTCGATGAAGAACG</td><td>>hg38_21_+_8433172_8446622_RNA45SN1_45S_50ntFlanks</td><td>6652</td><td>6696</td><td>45</td><td>GACTCTTAGCGGTGGATCACTCGGCTCGTGCGTCGATGAAGAACG</td><td>986.5</td><td>4.6</td><td>0.8</td><td>5.9</td><td>4.5E-09</td><td>7.1E-09</td><td>BT20/MB468</td></tr>
<tr><td>>hg38_21_+_8433172_8446622_RNA45SN1_45S_50ntFlanks@11668.11702.35|TGCCCAGTGCTCTGAATGTCAAAGTGAAGAAATTC</td><td>>hg38_21_+_8433172_8446622_RNA45SN1_45S_50ntFlanks</td><td>11668</td><td>11702</td><td>35</td><td>TGCCCAGTGCTCTGAATGTCAAAGTGAAGAAATTC</td><td>2940.3</td><td>4.2</td><td>0.7</td><td>5.9</td><td>4.0E-09</td><td>7.2E-09</td><td>MB231/MB468</td></tr>
<tr><td>>hg38_21_+_8433172_8446622_RNA45SN1_45S_50ntFlanks@4243.4264.22|CTTTCGAGGCCCTGTAATTGGA</td><td>>hg38_21_+_8433172_8446622_RNA45SN1_45S_50ntFlanks</td><td>4243</td><td>4264</td><td>22</td><td>CTTTCGAGGCCCTGTAATTGGA</td><td>118.3</td><td>-2.5</td><td>0.4</td><td>-5.9</td><td>4.5E-09</td><td>7.2E-09</td><td>BT20/MB468</td></tr>
<tr><td>>hg38_21_+_8433172_8446622_RNA45SN1_45S_50ntFlanks@10684.10710.27|CGGCGGCGTCCGGTGAGCTCTCGCTGG</td><td>>hg38_21_+_8433172_8446622_RNA45SN1_45S_50ntFlanks</td><td>10684</td><td>10710</td><td>27</td><td>CGGCGGCGTCCGGTGAGCTCTCGCTGG</td><td>7257.9</td><td>-6.3</td><td>1.1</td><td>-5.9</td><td>4.5E-09</td><td>7.2E-09</td><td>BT20/MB468</td></tr>
<tr><td>>hg38_21_+_8433172_8446622_RNA45SN1_45S_50ntFlanks@8169.8187.19|CCAAGTCCTTCTGATCGAG</td><td>>hg38_21_+_8433172_8446622_RNA45SN1_45S_50ntFlanks</td><td>8169</td><td>8187</td><td>19</td><td>CCAAGTCCTTCTGATCGAG</td><td>178.0</td><td>-3.9</td><td>0.7</td><td>-5.9</td><td>4.0E-09</td><td>7.2E-09</td><td>MB231/MB468</td></tr>
<tr><td>>hg38_21_+_8433172_8446622_RNA45SN1_45S_50ntFlanks@5493.5516.24|GAAGACGGTCGAACTTGACTATCT</td><td>>hg38_21_+_8433172_8446622_RNA45SN1_45S_50ntFlanks</td><td>5493</td><td>5516</td><td>24</td><td>GAAGACGGTCGAACTTGACTATCT</td><td>607.4</td><td>8.2</td><td>1.4</td><td>5.9</td><td>4.5E-09</td><td>7.2E-09</td><td>BT20/MB468</td></tr>
<tr><td>>hg38_21_+_8433172_8446622_RNA45SN1_45S_50ntFlanks@12678.12703.26|AATCCCGCCCAGGCGGAACGATACGG</td><td>>hg38_21_+_8433172_8446622_RNA45SN1_45S_50ntFlanks</td><td>12678</td><td>12703</td><td>26</td><td>AATCCCGCCCAGGCGGAACGATACGG</td><td>256.2</td><td>-4.8</td><td>0.8</td><td>-5.9</td><td>4.1E-09</td><td>7.2E-09</td><td>MB231/MB468</td></tr>
<tr><td>>hg38_21_+_8433172_8446622_RNA45SN1_45S_50ntFlanks@9822.9850.29|CTAGTGGGCCACTTTTGGTAAGCAGAACT</td><td>>hg38_21_+_8433172_8446622_RNA45SN1_45S_50ntFlanks</td><td>9822</td><td>9850</td><td>29</td><td>CTAGTGGGCCACTTTTGGTAAGCAGAACT</td><td>272.3</td><td>-3.4</td><td>0.6</td><td>-5.9</td><td>4.6E-09</td><td>7.3E-09</td><td>BT20/MB468</td></tr>
<tr><td>>hg38_21_+_8433172_8446622_RNA45SN1_45S_50ntFlanks@6744.6772.29|GACACTTCGAACGCACTTGCGGCCCCGGG</td><td>>hg38_21_+_8433172_8446622_RNA45SN1_45S_50ntFlanks</td><td>6744</td><td>6772</td><td>29</td><td>GACACTTCGAACGCACTTGCGGCCCCGGG</td><td>132.0</td><td>-5.8</td><td>1.0</td><td>-5.9</td><td>4.6E-09</td><td>7.3E-09</td><td>BT20/MB468</td></tr>
<tr><td>>hg38_21_+_8433172_8446622_RNA45SN1_45S_50ntFlanks@11203.11225.23|CGGCGGCGGCGACTCTGGACGCG</td><td>>hg38_21_+_8433172_8446622_RNA45SN1_45S_50ntFlanks</td><td>11203</td><td>11225</td><td>23</td><td>CGGCGGCGGCGACTCTGGACGCG</td><td>128.0</td><td>-3.3</td><td>0.6</td><td>-5.9</td><td>4.1E-09</td><td>7.3E-09</td><td>MB231/MB468</td></tr>
<tr><td>>hg38_21_+_8433172_8446622_RNA45SN1_45S_50ntFlanks@8168.8186.19|CCCAAGTCCTTCTGATCGA</td><td>>hg38_21_+_8433172_8446622_RNA45SN1_45S_50ntFlanks</td><td>8168</td><td>8186</td><td>19</td><td>CCCAAGTCCTTCTGATCGA</td><td>1222.3</td><td>-7.6</td><td>1.3</td><td>-5.9</td><td>4.7E-09</td><td>7.4E-09</td><td>BT20/MB468</td></tr>
<tr><td>>hg38_21_+_8433172_8446622_RNA45SN1_45S_50ntFlanks@6731.6770.40|CACATTGATCATCGACACTTCGAACGCACTTGCGGCCCCG</td><td>>hg38_21_+_8433172_8446622_RNA45SN1_45S_50ntFlanks</td><td>6731</td><td>6770</td><td>40</td><td>CACATTGATCATCGACACTTCGAACGCACTTGCGGCCCCG</td><td>667.7</td><td>-6.6</td><td>1.1</td><td>-5.9</td><td>4.7E-09</td><td>7.4E-09</td><td>BT20/MB468</td></tr>
<tr><td>>MI0000102|hsa-mir-100&WithFlank&11|-|122152223|122152314@19.39.21(+1U)&[MIMAT0000098&hsa-miR-100-5p&offsets|0|-1(+1U);m-27&11|-|122152276|122152296&offsets|0|0(+1U)]|AACCCGTAGATCCGAACTTGTT</td><td>>MI0000102|hsa-mir-100&WithFlank&11|-|122152223|122152314</td><td>19</td><td>39</td><td>21(+1U)</td><td>AACCCGTAGATCCGAACTTGTT</td><td>29929.7</td><td>14.8</td><td>2.5</td><td>5.9</td><td>4.2E-09</td><td>7.5E-09</td><td>MB231/MB468</td></tr>
<tr><td>>hg38_21_+_8433172_8446622_RNA45SN1_45S_50ntFlanks@10782.10818.37|GGTGAACAGCCTCTGGCATGTTGGAACAATGTAGGTA</td><td>>hg38_21_+_8433172_8446622_RNA45SN1_45S_50ntFlanks</td><td>10782</td><td>10818</td><td>37</td><td>GGTGAACAGCCTCTGGCATGTTGGAACAATGTAGGTA</td><td>344.8</td><td>-6.3</td><td>1.1</td><td>-5.9</td><td>4.7E-09</td><td>7.5E-09</td><td>BT20/MB468</td></tr>
<tr><td>>hg38_21_+_8433172_8446622_RNA45SN1_45S_50ntFlanks@11894.11916.23|ACTCTAGTCTGGCACGGTGAAGA</td><td>>hg38_21_+_8433172_8446622_RNA45SN1_45S_50ntFlanks</td><td>11894</td><td>11916</td><td>23</td><td>ACTCTAGTCTGGCACGGTGAAGA</td><td>273.2</td><td>-4.1</td><td>0.7</td><td>-5.9</td><td>4.2E-09</td><td>7.5E-09</td><td>MB231/MB468</td></tr>
<tr><td>>trnaMT_GluTTC_MT_-_14674_14742@40.65.26,trnalookalike8_GluTTC_5_-_93905172_93905240@40.65.26|TTGGTCGTGGTTGTAGTCCGTGCGAG</td><td>>trnaMT_GluTTC_MT_-_14674_14742</td><td>40</td><td>65</td><td>26</td><td>TTGGTCGTGGTTGTAGTCCGTGCGAG</td><td>356.1</td><td>7.2</td><td>1.2</td><td>5.9</td><td>4.7E-09</td><td>7.5E-09</td><td>BT20/MB468</td></tr>
<tr><td>>hg38_21_+_8433172_8446622_RNA45SN1_45S_50ntFlanks@9565.9595.31|TGACGTGCAAATCGGTCGTCCGACCTGGGTA</td><td>>hg38_21_+_8433172_8446622_RNA45SN1_45S_50ntFlanks</td><td>9565</td><td>9595</td><td>31</td><td>TGACGTGCAAATCGGTCGTCCGACCTGGGTA</td><td>568.6</td><td>-4.0</td><td>0.7</td><td>-5.9</td><td>4.7E-09</td><td>7.5E-09</td><td>BT20/MB468</td></tr>
</table>

<table>
<tr><td>>hg38_1_-_228634819_228635039_RNA5S12_5S_50ntFlanks@142.171.30|CGCCTGGGAATACCGGGTGCTGTAGGCTTT</td><td>>hg38_1_-_228634819_228635039_RNA5S12_5S_50ntFlanks</td><td>142</td><td>171</td><td>30</td><td>CGCCTGGGAATACCGGGTGCTGTAGGCTTT</td><td>2428.8</td><td>5.7</td><td>1.0</td><td>5.9</td><td>4.7E-09</td><td>7.5E-09</td><td>BT20/MB468</td></tr>
<tr><td>>hg38_21_+_8433172_8446622_RNA45SN1_45S_50ntFlanks@3993.4013.21|GTGACTCTAGATAACCTCGGG</td><td>>hg38_21_+_8433172_8446622_RNA45SN1_45S_50ntFlanks</td><td>3993</td><td>4013</td><td>21</td><td>GTGACTCTAGATAACCTCGGG</td><td>351.8</td><td>-2.4</td><td>0.4</td><td>-5.9</td><td>4.8E-09</td><td>7.6E-09</td><td>BT20/MB468</td></tr>
<tr><td>>trnaMT_ValTAC_MT_+_1602_1670@41.72.32|GAGATTTCAACTTAACTTGACCGCTCTGACCA</td><td>>trnaMT_ValTAC_MT_+_1602_1670</td><td>41</td><td>72</td><td>32</td><td>GAGATTTCAACTTAACTTGACCGCTCTGACCA</td><td>554.6</td><td>5.2</td><td>0.9</td><td>5.9</td><td>4.9E-09</td><td>7.7E-09</td><td>BT20/MB468</td></tr>
<tr><td>>hg38_21_+_8433172_8446622_RNA45SN1_45S_50ntFlanks@7986.8009.24|ATCAGACGTGGCGACCCGCTGAAT</td><td>>hg38_21_+_8433172_8446622_RNA45SN1_45S_50ntFlanks</td><td>7986</td><td>8009</td><td>24</td><td>ATCAGACGTGGCGACCCGCTGAAT</td><td>67.1</td><td>-7.6</td><td>1.3</td><td>-5.9</td><td>4.4E-09</td><td>7.8E-09</td><td>MB231/MB468</td></tr>
<tr><td>>hg38_21_+_8433172_8446622_RNA45SN1_45S_50ntFlanks@12328.12346.19|GTTTTAAGCAGGAGGTGTC</td><td>>hg38_21_+_8433172_8446622_RNA45SN1_45S_50ntFlanks</td><td>12328</td><td>12346</td><td>19</td><td>GTTTTAAGCAGGAGGTGTC</td><td>126.8</td><td>-3.1</td><td>0.5</td><td>-5.9</td><td>4.4E-09</td><td>7.8E-09</td><td>MB231/MB468</td></tr>
<tr><td>>hg38_21_+_8433172_8446622_RNA45SN1_45S_50ntFlanks@6731.6771.41|CACATTGATCATCGACACTTCGAACGCACTTGCGGCCCCGG</td><td>>hg38_21_+_8433172_8446622_RNA45SN1_45S_50ntFlanks</td><td>6731</td><td>6771</td><td>41</td><td>CACATTGATCATCGACACTTCGAACGCACTTGCGGCCCCGG</td><td>397.0</td><td>-6.0</td><td>1.0</td><td>-5.9</td><td>4.9E-09</td><td>7.8E-09</td><td>BT20/MB468</td></tr>
<tr><td>>hg38_21_+_8433172_8446622_RNA45SN1_45S_50ntFlanks@11452.11478.27|CTCCTCGGCGCGCGGCGGCGGCGG</td><td>>hg38_21_+_8433172_8446622_RNA45SN1_45S_50ntFlanks</td><td>11452</td><td>11478</td><td>27</td><td>CTCCTCGGCGCGCGGCGGCGGCGG</td><td>182.0</td><td>-5.2</td><td>0.9</td><td>-5.8</td><td>5.0E-09</td><td>7.9E-09</td><td>BT20/MB468</td></tr>
<tr><td>>MI0000749|hsa-mir-30e&WithFlank&1|+|40754349|40754452@23.45.23(+1C)&[MIMAT0000692&hsa-miR-30e-5p&offsets|0|+1(+1C);m-24&1|+|40754371|40754394&offsets|0|-1(+1C)]|TGTAAACATCCTTGACTGGAAGCC</td><td>>MI0000749|hsa-mir-30e&WithFlank&1|+|40754349|40754452</td><td>23</td><td>45</td><td>23(+1C)</td><td>TGTAAACATCCTTGACTGGAAGCC</td><td>2169.6</td><td>4.6</td><td>0.8</td><td>5.9</td><td>4.5E-09</td><td>7.9E-09</td><td>MB231/MB468</td></tr>
<tr><td>>hg38_21_+_8433172_8446622_RNA45SN1_45S_50ntFlanks@7975.8005.31|CGCGACCTCAGATCAGACGTGGCGACCCGCT</td><td>>hg38_21_+_8433172_8446622_RNA45SN1_45S_50ntFlanks</td><td>7975</td><td>8005</td><td>31</td><td>CGCGACCTCAGATCAGACGTGGCGACCCGCT</td><td>16173.0</td><td>4.4</td><td>0.7</td><td>5.9</td><td>4.5E-09</td><td>8.0E-09</td><td>MB231/MB468</td></tr>
<tr><td>>MI0000072|hsa-mir-18a&WithFlank&13|+|91350745|91350827@12.34.23&[MIMAT0000072&hsa-miR-18a-5p&offsets|0|0;m-262&13|+|91350756|91350778&offsets|0|0]|TAAGGTGCATCTAGTGCAGATAG</td><td>>MI0000072|hsa-mir-18a&WithFlank&13|+|91350745|91350827</td><td>12</td><td>34</td><td>23</td><td>TAAGGTGCATCTAGTGCAGATAG</td><td>1039.7</td><td>5.8</td><td>1.0</td><td>5.8</td><td>5.1E-09</td><td>8.1E-09</td><td>BT20/MB468</td></tr>
<tr><td>>hg38_21_+_8433172_8446622_RNA45SN1_45S_50ntFlanks@6719.6738.20|TGAATTGCAGGACACATTGA</td><td>>hg38_21_+_8433172_8446622_RNA45SN1_45S_50ntFlanks</td><td>6719</td><td>6738</td><td>20</td><td>TGAATTGCAGGACACATTGA</td><td>163.1</td><td>-3.2</td><td>0.6</td><td>-5.9</td><td>4.5E-09</td><td>8.1E-09</td><td>MB231/MB468</td></tr>
<tr><td>>hg38_1_-_228634819_228635039_RNA5S12_5S_50ntFlanks@150.170.21|AATACCGGGTGCTGTAGGCTT</td><td>>hg38_1_-_228634819_228635039_RNA5S12_5S_50ntFlanks</td><td>150</td><td>170</td><td>21</td><td>AATACCGGGTGCTGTAGGCTT</td><td>2390.3</td><td>4.3</td><td>0.7</td><td>5.8</td><td>5.1E-09</td><td>8.1E-09</td><td>BT20/MB468</td></tr>
<tr><td>>hg38_21_+_8433172_8446622_RNA45SN1_45S_50ntFlanks@6784.6803.20|GCTACGCCTGTCTGAGCGTC</td><td>>hg38_21_+_8433172_8446622_RNA45SN1_45S_50ntFlanks</td><td>6784</td><td>6803</td><td>20</td><td>GCTACGCCTGTCTGAGCGTC</td><td>338.0</td><td>-6.1</td><td>1.1</td><td>-5.8</td><td>5.1E-09</td><td>8.1E-09</td><td>BT20/MB468</td></tr>
<tr><td>>hg38_21_+_8433172_8446622_RNA45SN1_45S_50ntFlanks@7986.8005.20|ATCAGACGTGGCGACCCGCT</td><td>>hg38_21_+_8433172_8446622_RNA45SN1_45S_50ntFlanks</td><td>7986</td><td>8005</td><td>20</td><td>ATCAGACGTGGCGACCCGCT</td><td>557.9</td><td>-5.8</td><td>1.0</td><td>-5.9</td><td>4.6E-09</td><td>8.1E-09</td><td>MB231/MB468</td></tr>
<tr><td>>hg38_21_+_8433172_8446622_RNA45SN1_45S_50ntFlanks@3985.4004.20|CGGCTTTGGTGACTCTAGAT</td><td>>hg38_21_+_8433172_8446622_RNA45SN1_45S_50ntFlanks</td><td>3985</td><td>4004</td><td>20</td><td>CGGCTTTGGTGACTCTAGAT</td><td>123.3</td><td>-5.9</td><td>1.0</td><td>-5.9</td><td>4.6E-09</td><td>8.1E-09</td><td>MB231/MB468</td></tr>
<tr><td>>hg38_21_+_8433172_8446622_RNA45SN1_45S_50ntFlanks@11670.11701.32|CCCAGTGCTCTGAATGTCAAAGTGAAGAAATT</td><td>>hg38_21_+_8433172_8446622_RNA45SN1_45S_50ntFlanks</td><td>11670</td><td>11701</td><td>32</td><td>CCCAGTGCTCTGAATGTCAAAGTGAAGAAATT</td><td>575.5</td><td>-3.8</td><td>0.6</td><td>-5.8</td><td>5.1E-09</td><td>8.2E-09</td><td>BT20/MB468</td></tr>
<tr><td>>hg38_21_+_8433172_8446622_RNA45SN1_45S_50ntFlanks@10866.10904.39|CTAAGGGCTGGGTCGGTCGGGCTGGGGCGCGAAGCGGGG</td><td>>hg38_21_+_8433172_8446622_RNA45SN1_45S_50ntFlanks</td><td>10866</td><td>10904</td><td>39</td><td>CTAAGGGCTGGGTCGGTCGGGCTGGGGCGCGAAGCGGGG</td><td>75.2</td><td>-5.1</td><td>0.9</td><td>-5.8</td><td>5.2E-09</td><td>8.2E-09</td><td>BT20/MB468</td></tr>
<tr><td>>hg38_21_+_8433172_8446622_RNA45SN1_45S_50ntFlanks@11583.11601.19|CTGGTGCGGACCAGGGGAA</td><td>>hg38_21_+_8433172_8446622_RNA45SN1_45S_50ntFlanks</td><td>11583</td><td>11601</td><td>19</td><td>CTGGTGCGGACCAGGGGAA</td><td>769.5</td><td>-4.1</td><td>0.7</td><td>-5.8</td><td>5.2E-09</td><td>8.2E-09</td><td>BT20/MB468</td></tr>
<tr><td>>hg38_21_+_8433172_8446622_RNA45SN1_45S_50ntFlanks@10869.10890.22|AGGGCTGGGTCGGTCGGGCTGG</td><td>>hg38_21_+_8433172_8446622_RNA45SN1_45S_50ntFlanks</td><td>10869</td><td>10890</td><td>22</td><td>AGGGCTGGGTCGGTCGGGCTGG</td><td>256.3</td><td>-3.9</td><td>0.7</td><td>-5.9</td><td>4.7E-09</td><td>8.3E-09</td><td>MB231/MB468</td></tr>
<tr><td>>am_trna128_GlyGCC_6_-_27870686_27870756@7.28.22,trna133_GlyCCC_1_-_16872434_16872504@7.28.22,trna17_GluTTC_2_-_204229131_204229201@7.28.22,trna18_GlyGCC_16_+_70822597_70822667@7.28.22,trna19_GlyGCC_16_+_70823410_70823480@7.28.22,trna19_GlyGCC_2_-_157257659_157257729@7.28.22,trna24_GlyGCC_16_-_70812942_70813012@7.28.22,trna25_GlyGCC_16_-_70812114_70812184@7.28.22,trna2_GlyGCC_21_-_18827107_18827177@7.28.22,trna35_GlyGCC_1_+_161413094_161413164@7.28.22,trna37_GlyGCC_1_+_161420467_161420537@7.28.22,trna39_GlyGCC_1_+_161427898_161427968@7.28.22,trna41_GlyGCC_1_+_161435258_161435328@7.28.22,trna4_GlyCCC_1_+_17188416_17188486@7.28.22,trna5_GlyGCC_17_+_8029064_8029134@7.28.22,trna68_GlyGCC_1_-_161493637_161493707@7.28.22,trna91_GlyCCC_1_-_149680210_149680280@7.28.22,trna99_ValCAC_1_-_149294666_149294736@7.28.22|GTGGTTCAGTGGTAGAATTCTC</td><td>>am_trna128_GlyGCC_6_-_27870686_27870756</td><td>7</td><td>28</td><td>22</td><td>GTGGTTCAGTGGTAGAATTCTC</td><td>150.4</td><td>-4.1</td><td>0.7</td><td>-5.9</td><td>4.7E-09</td><td>8.3E-09</td><td>MB231/MB468</td></tr>
<tr><td>>hg38_21_+_8433172_8446622_RNA45SN1_45S_50ntFlanks@4002.4026.25|GATAACCTCGGGCCGATCGCACGCC</td><td>>hg38_21_+_8433172_8446622_RNA45SN1_45S_50ntFlanks</td><td>4002</td><td>4026</td><td>25</td><td>GATAACCTCGGGCCGATCGCACGCC</td><td>112.9</td><td>-5.8</td><td>1.0</td><td>-5.9</td><td>4.7E-09</td><td>8.3E-09</td><td>MB231/MB468</td></tr>
<tr><td>>hg38_21_+_8433172_8446622_RNA45SN1_45S_50ntFlanks@7984.8005.22|AGATCAGACGTGGCGACCCGCT</td><td>>hg38_21_+_8433172_8446622_RNA45SN1_45S_50ntFlanks</td><td>7984</td><td>8005</td><td>22</td><td>AGATCAGACGTGGCGACCCGCT</td><td>216.9</td><td>-3.7</td><td>0.6</td><td>-5.8</td><td>5.3E-09</td><td>8.4E-09</td><td>BT20/MB468</td></tr>
<tr><td>>hg38_21_+_8433172_8446622_RNA45SN1_45S_50ntFlanks@12605.12623.19|TATGTGCTTGGCTGAGGAG</td><td>>hg38_21_+_8433172_8446622_RNA45SN1_45S_50ntFlanks</td><td>12605</td><td>12623</td><td>19</td><td>TATGTGCTTGGCTGAGGAG</td><td>262.9</td><td>-3.3</td><td>0.6</td><td>-5.9</td><td>4.8E-09</td><td>8.4E-09</td><td>MB231/MB468</td></tr>
<tr><td>>hg38_21_+_8433172_8446622_RNA45SN1_45S_50ntFlanks@11319.11336.18|CCCACGTCTCGTCGCGCG</td><td>>hg38_21_+_8433172_8446622_RNA45SN1_45S_50ntFlanks</td><td>11319</td><td>11336</td><td>18</td><td>CCCACGTCTCGTCGCGCG</td><td>83.1</td><td>-8.0</td><td>1.4</td><td>-5.9</td><td>4.8E-09</td><td>8.4E-09</td><td>MB231/MB468</td></tr>
<tr><td>>MI0000749|hsa-mir-30e&WithFlank&1|+|40754349|40754452@23.45.23(+1G)&[MIMAT0000692&hsa-miR-30e-5p&offsets|0|+1(+1G);m-24&1|+|40754371|40754394&offsets|0|-1(+1G)]|TGTAAACATCCTTGACTGGAAGCG</td><td>>MI0000749|hsa-mir-30e&WithFlank&1|+|40754349|40754452</td><td>23</td><td>45</td><td>23(+1G)</td><td>TGTAAACATCCTTGACTGGAAGCG</td><td>915.0</td><td>4.3</td><td>0.7</td><td>5.9</td><td>4.8E-09</td><td>8.5E-09</td><td>MB231/MB468</td></tr>
<tr><td>>hg38_21_+_8433172_8446622_RNA45SN1_45S_50ntFlanks@12936.12959.24|AAACCATTCGTAGACGACCTGCTT</td><td>>hg38_21_+_8433172_8446622_RNA45SN1_45S_50ntFlanks</td><td>12936</td><td>12959</td><td>24</td><td>AAACCATTCGTAGACGACCTGCTT</td><td>355.9</td><td>-2.5</td><td>0.4</td><td>-5.8</td><td>5.4E-09</td><td>8.5E-09</td><td>BT20/MB468</td></tr>
<tr><td>>hg38_21_+_8433172_8446622_RNA45SN1_45S_50ntFlanks@11581.11611.31|AACTGGTGCGGACCAGGGGAATCCGACTGTT</td><td>>hg38_21_+_8433172_8446622_RNA45SN1_45S_50ntFlanks</td><td>11581</td><td>11611</td><td>31</td><td>AACTGGTGCGGACCAGGGGAATCCGACTGTT</td><td>1797.6</td><td>-4.0</td><td>0.7</td><td>-5.8</td><td>5.4E-09</td><td>8.5E-09</td><td>BT20/MB468</td></tr>
<tr><td>>hg38_21_+_8433172_8446622_RNA45SN1_45S_50ntFlanks@9554.9582.29|CGTAGCGGTCCTGACGTGCAAATCGGTCG</td><td>>hg38_21_+_8433172_8446622_RNA45SN1_45S_50ntFlanks</td><td>9554</td><td>9582</td><td>29</td><td>CGTAGCGGTCCTGACGTGCAAATCGGTCG</td><td>505.1</td><td>-4.8</td><td>0.8</td><td>-5.8</td><td>5.4E-09</td><td>8.6E-09</td><td>BT20/MB468</td></tr>
<tr><td>>hg38_21_+_8433172_8446622_RNA45SN1_45S_50ntFlanks@12413.12436.24|TCCTTCGATGTCGGCTCTTCCTAT</td><td>>hg38_21_+_8433172_8446622_RNA45SN1_45S_50ntFlanks</td><td>12413</td><td>12436</td><td>24</td><td>TCCTTCGATGTCGGCTCTTCCTAT</td><td>150.2</td><td>-3.2</td><td>0.6</td><td>-5.9</td><td>4.9E-09</td><td>8.6E-09</td><td>MB231/MB468</td></tr>
<tr><td>>hg38_21_+_8433172_8446622_RNA45SN1_45S_50ntFlanks@12942.12973.32|TTCGTAGACGACCTGCTTCTGGGTCGGGGTTT</td><td>>hg38_21_+_8433172_8446622_RNA45SN1_45S_50ntFlanks</td><td>12942</td><td>12973</td><td>32</td><td>TTCGTAGACGACCTGCTTCTGGGTCGGGGTTT</td><td>170.1</td><td>-4.4</td><td>0.8</td><td>-5.9</td><td>4.9E-09</td><td>8.6E-09</td><td>MB231/MB468</td></tr>
<tr><td>>hg38_21_+_8433172_8446622_RNA45SN1_45S_50ntFlanks@6690.6718.29|AAGAACGCAGCTAGCTGCGAGAATTAATG</td><td>>hg38_21_+_8433172_8446622_RNA45SN1_45S_50ntFlanks</td><td>6690</td><td>6718</td><td>29</td><td>AAGAACGCAGCTAGCTGCGAGAATTAATG</td><td>94.8</td><td>-5.6</td><td>1.0</td><td>-5.9</td><td>4.9E-09</td><td>8.6E-09</td><td>MB231/MB468</td></tr>
<tr><td>>hg38_21_+_8433172_8446622_RNA45SN1_45S_50ntFlanks@4297.4314.18|CATTGGAGGGCAAGTCTG</td><td>>hg38_21_+_8433172_8446622_RNA45SN1_45S_50ntFlanks</td><td>4297</td><td>4314</td><td>18</td><td>CATTGGAGGGCAAGTCTG</td><td>424.9</td><td>-3.4</td><td>0.6</td><td>-5.8</td><td>5.5E-09</td><td>8.7E-09</td><td>BT20/MB468</td></tr>
<tr><td>>hg38_21_+_8433172_8446622_RNA45SN1_45S_50ntFlanks@9569.9596.28|GTGCAAATCGGTCGTCCGACCTGGGTAT</td><td>>hg38_21_+_8433172_8446622_RNA45SN1_45S_50ntFlanks</td><td>9569</td><td>9596</td><td>28</td><td>GTGCAAATCGGTCGTCCGACCTGGGTAT</td><td>393.1</td><td>-3.2</td><td>0.6</td><td>-5.8</td><td>4.9E-09</td><td>8.7E-09</td><td>MB231/MB468</td></tr>
<tr><td>>hg38_21_+_8433172_8446622_RNA45SN1_45S_50ntFlanks@12455.12487.33|ACCAAGCGTTGGATTGTTCACCCACTAATAGGG</td><td>>hg38_21_+_8433172_8446622_RNA45SN1_45S_50ntFlanks</td><td>12455</td><td>12487</td><td>33</td><td>ACCAAGCGTTGGATTGTTCACCCACTAATAGGG</td><td>308.1</td><td>-4.1</td><td>0.7</td><td>-5.8</td><td>5.5E-09</td><td>8.8E-09</td><td>BT20/MB468</td></tr>
<tr><td>>hg38_21_+_8433172_8446622_RNA45SN1_45S_50ntFlanks@9572.9594.23|CAAATCGGTCGTCCGACCTGGGT</td><td>>hg38_21_+_8433172_8446622_RNA45SN1_45S_50ntFlanks</td><td>9572</td><td>9594</td><td>23</td><td>CAAATCGGTCGTCCGACCTGGGT</td><td>628.4</td><td>-4.1</td><td>0.7</td><td>-5.8</td><td>5.6E-09</td><td>8.8E-09</td><td>BT20/MB468</td></tr>
<tr><td>>hg38_21_+_8433172_8446622_RNA45SN1_45S_50ntFlanks@12437.12474.38|CATTGTGAAGCAGAATTCACCAAGCGTTGGATTGTTCA</td><td>>hg38_21_+_8433172_8446622_RNA45SN1_45S_50ntFlanks</td><td>12437</td><td>12474</td><td>38</td><td>CATTGTGAAGCAGAATTCACCAAGCGTTGGATTGTTCA</td><td>241.9</td><td>-3.5</td><td>0.6</td><td>-5.8</td><td>5.6E-09</td><td>8.8E-09</td><td>BT20/MB468</td></tr>
<tr><td>>hg38_21_+_8433172_8446622_RNA45SN1_45S_50ntFlanks@10806.10828.23|AACAATGTAGGTAAGGGAAGTCG</td><td>>hg38_21_+_8433172_8446622_RNA45SN1_45S_50ntFlanks</td><td>10806</td><td>10828</td><td>23</td><td>AACAATGTAGGTAAGGGAAGTCG</td><td>183.7</td><td>-4.6</td><td>0.8</td><td>-5.8</td><td>5.0E-09</td><td>8.9E-09</td><td>MB231/MB468</td></tr>
<tr><td>>hg38_21_+_8433172_8446622_RNA45SN1_45S_50ntFlanks@4558.4587.30|AGCTAGGAATAATGGAATAGGACCGCGGTT</td><td>>hg38_21_+_8433172_8446622_RNA45SN1_45S_50ntFlanks</td><td>4558</td><td>4587</td><td>30</td><td>AGCTAGGAATAATGGAATAGGACCGCGGTT</td><td>101.3</td><td>-6.4</td><td>1.1</td><td>-5.8</td><td>5.0E-09</td><td>8.9E-09</td><td>MB231/MB468</td></tr>
<tr><td>>hg38_21_+_8433172_8446622_RNA45SN1_45S_50ntFlanks@11886.11904.19|TGAGCTTGACTCTAGTCTG</td><td>>hg38_21_+_8433172_8446622_RNA45SN1_45S_50ntFlanks</td><td>11886</td><td>11904</td><td>19</td><td>TGAGCTTGACTCTAGTCTG</td><td>802.4</td><td>-7.3</td><td>1.3</td><td>-5.8</td><td>5.6E-09</td><td>8.9E-09</td><td>BT20/MB468</td></tr>
<tr><td>>hg38_21_+_8433172_8446622_RNA45SN1_45S_50ntFlanks@7975.8007.33|CGCGACCTCAGATCAGACGTGGCGACCCGCTGA</td><td>>hg38_21_+_8433172_8446622_RNA45SN1_45S_50ntFlanks</td><td>7975</td><td>8007</td><td>33</td><td>CGCGACCTCAGATCAGACGTGGCGACCCGCTGA</td><td>9894.1</td><td>3.5</td><td>0.6</td><td>5.8</td><td>5.7E-09</td><td>8.9E-09</td><td>BT20/MB468</td></tr>
<tr><td>>hg38_21_+_8433172_8446622_RNA45SN1_45S_50ntFlanks@5527.5548.22|AAAGTCGTAACAAGGTTTCCGT</td><td>>hg38_21_+_8433172_8446622_RNA45SN1_45S_50ntFlanks</td><td>5527</td><td>5548</td><td>22</td><td>AAAGTCGTAACAAGGTTTCCGT</td><td>96.2</td><td>-4.6</td><td>0.8</td><td>-5.8</td><td>5.7E-09</td><td>9.0E-09</td><td>BT20/MB468</td></tr>
<tr><td>>hg38_21_+_8433172_8446622_RNA45SN1_45S_50ntFlanks@12991.13018.28|CTCCCTCGCTGCGATCTATTGAAAGTCA</td><td>>hg38_21_+_8433172_8446622_RNA45SN1_45S_50ntFlanks</td><td>12991</td><td>13018</td><td>28</td><td>CTCCCTCGCTGCGATCTATTGAAAGTCA</td><td>73.4</td><td>-4.9</td><td>0.8</td><td>-5.8</td><td>5.7E-09</td><td>9.1E-09</td><td>BT20/MB468</td></tr>
<tr><td>>hg38_21_+_8433172_8446622_RNA45SN1_45S_50ntFlanks@3835.3859.25|CTCCTCTCCTACTTGGATAACTGTG</td><td>>hg38_21_+_8433172_8446622_RNA45SN1_45S_50ntFlanks</td><td>3835</td><td>3859</td><td>25</td><td>CTCCTCTCCTACTTGGATAACTGTG</td><td>86.5</td><td>-5.0</td><td>0.9</td><td>-5.8</td><td>5.8E-09</td><td>9.1E-09</td><td>BT20/MB468</td></tr>
<tr><td>>hg38_21_+_8433172_8446622_RNA45SN1_45S_50ntFlanks@12639.12661.23|TACCATCTGTGGGATTATGACTG</td><td>>hg38_21_+_8433172_8446622_RNA45SN1_45S_50ntFlanks</td><td>12639</td><td>12661</td><td>23</td><td>TACCATCTGTGGGATTATGACTG</td><td>210.3</td><td>-2.3</td><td>0.4</td><td>-5.8</td><td>5.8E-09</td><td>9.1E-09</td><td>BT20/MB468</td></tr>
</table>

| | | | | | | | | | | | | |
|---|---|---|---|---|---|---|---|---|---|---|---|---|
| >hg38_21_+_8433172_8446622_RNA45SN1_45S_50ntFlanks@11671.11699.29\|CCAGTGCTCTGAATGTCAAAGTGAAGAAA | >hg38_21_+_8433172_8446622_RNA45SN1_45S_50ntFlanks | 11671 | 11699 | 29 | CCAGTGCTCTGAATGTCAAAGTGAAGAAA | 172.6 | -3.7 | 0.6 | -5.8 | 5.8E-09 | 9.2E-09 | BT20/MB468 |
| >hg38_21_+_8433172_8446622_RNA45SN1_45S_50ntFlanks@11577.11601.25\|TTAGAACTGGTGCGGACCAGGGGAA | >hg38_21_+_8433172_8446622_RNA45SN1_45S_50ntFlanks | 11577 | 11601 | 25 | TTAGAACTGGTGCGGACCAGGGGAA | 299.1 | -3.8 | 0.6 | -5.8 | 5.8E-09 | 9.2E-09 | BT20/MB468 |
| >hg38_21_+_8433172_8446622_RNA45SN1_45S_50ntFlanks@11200.11225.26\|CGGCGGCGGCGGCGACTCTGGACGCG | >hg38_21_+_8433172_8446622_RNA45SN1_45S_50ntFlanks | 11200 | 11225 | 26 | CGGCGGCGGCGGCGACTCTGGACGCG | 569.4 | -6.2 | 1.1 | -5.8 | 5.8E-09 | 9.2E-09 | BT20/MB468 |
| >trna119_LysCTT_1_-_145395522_145395594@3.33.31,trna11_LysCTT_5_-_180648979_180649051@3.33.31,trna13_LysCTT_6_+_26556774_26556846@3.33.31,trna7_LysCTT_16_+_3225692_3225764@3.33.31,trna9_LysCTT_5_+_180634755_180634827@3.33.31\|CCGGCTAGCTCAGTCGGTAGAGCATGAGACT | >trna119_LysCTT_1_-_145395522_145395594 | 3 | 33 | 31 | CCGGCTAGCTCAGTCGGTAGAGCATGAGACT | 2855.2 | 13.0 | 2.2 | 5.8 | 5.2E-09 | 9.2E-09 | MB231/MB468 |
| >hg38_21_+_8433172_8446622_RNA45SN1_45S_50ntFlanks@6733.6752.20\|CATTGATCATCGACACTTCG | >hg38_21_+_8433172_8446622_RNA45SN1_45S_50ntFlanks | 6733 | 6752 | 20 | CATTGATCATCGACACTTCG | 707.4 | -2.6 | 0.4 | -5.8 | 5.9E-09 | 9.3E-09 | BT20/MB468 |
| >hg38_21_+_8433172_8446622_RNA45SN1_45S_50ntFlanks@7984.8012.29\|AGATCAGACGTGGCGACCCGCTGAATTTA | >hg38_21_+_8433172_8446622_RNA45SN1_45S_50ntFlanks | 7984 | 8012 | 29 | AGATCAGACGTGGCGACCCGCTGAATTTA | 117.3 | -4.8 | 0.8 | -5.8 | 5.3E-09 | 9.3E-09 | MB231/MB468 |
| >hg38_21_+_8433172_8446622_RNA45SN1_45S_50ntFlanks@10796.10818.23\|GGCATGTTGGAACAATGTAGGTA | >hg38_21_+_8433172_8446622_RNA45SN1_45S_50ntFlanks | 10796 | 10818 | 23 | GGCATGTTGGAACAATGTAGGTA | 656.6 | -4.1 | 0.7 | -5.8 | 5.9E-09 | 9.3E-09 | BT20/MB468 |
| >hg38_21_+_8433172_8446622_RNA45SN1_45S_50ntFlanks@9534.9572.39\|AGGAAACTCTGGTGGAGGTCCGTAGCGGTCCTGACGTGC | >hg38_21_+_8433172_8446622_RNA45SN1_45S_50ntFlanks | 9534 | 9572 | 39 | AGGAAACTCTGGTGGAGGTCCGTAGCGGTCCTGACGTGC | 1038.7 | -3.6 | 0.6 | -5.8 | 5.9E-09 | 9.4E-09 | BT20/MB468 |
| >hg38_21_+_8433172_8446622_RNA45SN1_45S_50ntFlanks@11210.11230.21\|GGCGACTCTGGACGCGAGCCG | >hg38_21_+_8433172_8446622_RNA45SN1_45S_50ntFlanks | 11210 | 11230 | 21 | GGCGACTCTGGACGCGAGCCG | 78.7 | -7.9 | 1.3 | -5.8 | 5.3E-09 | 9.4E-09 | MB231/MB468 |
| >hg38_21_+_8433172_8446622_RNA45SN1_45S_50ntFlanks@8423.8442.20\|CGCAGTCCGCCCGGAGGATT | >hg38_21_+_8433172_8446622_RNA45SN1_45S_50ntFlanks | 8423 | 8442 | 20 | CGCAGTCCGCCCGGAGGATT | 167.0 | -3.4 | 0.6 | -5.8 | 5.3E-09 | 9.4E-09 | MB231/MB468 |
| >hg38_21_+_8433172_8446622_RNA45SN1_45S_50ntFlanks@11891.11910.20\|TTGACTCTAGTCTGGCACGG | >hg38_21_+_8433172_8446622_RNA45SN1_45S_50ntFlanks | 11891 | 11910 | 20 | TTGACTCTAGTCTGGCACGG | 178.0 | -3.2 | 0.5 | -5.8 | 6.0E-09 | 9.4E-09 | BT20/MB468 |
| >hg38_21_+_8433172_8446622_RNA45SN1_45S_50ntFlanks@11235.11252.18\|CTTCCCGTGGATCGCCCC | >hg38_21_+_8433172_8446622_RNA45SN1_45S_50ntFlanks | 11235 | 11252 | 18 | CTTCCCGTGGATCGCCCC | 986.6 | -5.7 | 1.0 | -5.8 | 5.4E-09 | 9.5E-09 | MB231/MB468 |
| >hg38_21_+_8433172_8446622_RNA45SN1_45S_50ntFlanks@12944.12966.23\|CGTAGACGACCTGCTTCTGGGTC | >hg38_21_+_8433172_8446622_RNA45SN1_45S_50ntFlanks | 12944 | 12966 | 23 | CGTAGACGACCTGCTTCTGGGTC | 106.5 | -8.3 | 1.4 | -5.8 | 5.4E-09 | 9.5E-09 | MB231/MB468 |
| >hg38_21_+_8433172_8446622_RNA45SN1_45S_50ntFlanks@8380.8406.27\|CAAGAGGGCGTGAAACCGTTAAGAGGT | >hg38_21_+_8433172_8446622_RNA45SN1_45S_50ntFlanks | 8380 | 8406 | 27 | CAAGAGGGCGTGAAACCGTTAAGAGGT | 345.4 | -3.2 | 0.6 | -5.8 | 6.1E-09 | 9.5E-09 | BT20/MB468 |
| >hg38_21_+_8433172_8446622_RNA45SN1_45S_50ntFlanks@8552.8579.28\|CTCCCCGGAGGGGGCGGGCTCCGGCGGG | >hg38_21_+_8433172_8446622_RNA45SN1_45S_50ntFlanks | 8552 | 8579 | 28 | CTCCCCGGAGGGGGCGGGCTCCGGCGGG | 65.0 | -7.6 | 1.3 | -5.8 | 5.4E-09 | 9.6E-09 | MB231/MB468 |
| >am_trna129_MetCAT_6_-_27870271_27870342@34.51.18,trna142_MetCAT_6_-_27560600_27560671@34.51.18,trna150_MetCAT_6_-_27300764_27300835@34.51.18,trna169_MetCAT_6_-_26330529_26330600@34.51.18,trna171_MetCAT_6_-_26313352_26313423@34.51.18,trna20_MetCAT_17_-_80452597_80452668@34.51.18,trna2_MetCAT_6_+_26286754_26286825@34.51.18,trna32_MetCAT_1_+_153643726_153643797@34.51.18,trna61_MetCAT_6_+_27745664_27745735@34.51.18\|ATAACCCAGAGGTCGATG | >am_trna129_MetCAT_6_-_27870271_27870342 | 34 | 51 | 18 | ATAACCCAGAGGTCGATG | 212.6 | -5.3 | 0.9 | -5.8 | 5.4E-09 | 9.6E-09 | MB231/MB468 |
| >hg38_21_+_8433172_8446622_RNA45SN1_45S_50ntFlanks@9566.9592.27\|GACGTGCAAATCGGTCGTCCGACCTGG | >hg38_21_+_8433172_8446622_RNA45SN1_45S_50ntFlanks | 9566 | 9592 | 27 | GACGTGCAAATCGGTCGTCCGACCTGG | 172.9 | -2.6 | 0.5 | -5.8 | 6.1E-09 | 9.6E-09 | BT20/MB468 |
| >hg38_21_+_8433172_8446622_RNA45SN1_45S_50ntFlanks@4995.5022.28\|ACAGATTGATAGCTCTTTCTCGATTCCG | >hg38_21_+_8433172_8446622_RNA45SN1_45S_50ntFlanks | 4995 | 5022 | 28 | ACAGATTGATAGCTCTTTCTCGATTCCG | 116.3 | -5.1 | 0.9 | -5.8 | 6.1E-09 | 9.6E-09 | BT20/MB468 |
| >hg38_21_+_8433172_8446622_RNA45SN1_45S_50ntFlanks@12992.13014.23\|TCCCTCGCTGCGATCTATTGAAA | >hg38_21_+_8433172_8446622_RNA45SN1_45S_50ntFlanks | 12992 | 13014 | 23 | TCCCTCGCTGCGATCTATTGAAA | 258.4 | -2.9 | 0.5 | -5.8 | 6.1E-09 | 9.6E-09 | BT20/MB468 |
| >hg38_21_+_8433172_8446622_RNA45SN1_45S_50ntFlanks@10350.10388.39\|ACGAGAACTTTGAAGGCCGAAGTGGAGAAGGGTTCCATG | >hg38_21_+_8433172_8446622_RNA45SN1_45S_50ntFlanks | 10350 | 10388 | 39 | ACGAGAACTTTGAAGGCCGAAGTGGAGAAGGGTTCCATG | 2480.7 | -5.6 | 1.0 | -5.8 | 6.2E-09 | 9.8E-09 | BT20/MB468 |
| >hg38_21_+_8433172_8446622_RNA45SN1_45S_50ntFlanks@9021.9039.19\|CGCCGGCGTCTCCTCGTGG | >hg38_21_+_8433172_8446622_RNA45SN1_45S_50ntFlanks | 9021 | 9039 | 19 | CGCCGGCGTCTCCTCGTGG | 77.7 | -4.5 | 0.8 | -5.8 | 5.5E-09 | 9.8E-09 | MB231/MB468 |
| >hg38_21_+_8433172_8446622_RNA45SN1_45S_50ntFlanks@11561.11586.26\|GCGCCTAGCAGCCGACTTAGAACTGG | >hg38_21_+_8433172_8446622_RNA45SN1_45S_50ntFlanks | 11561 | 11586 | 26 | GCGCCTAGCAGCCGACTTAGAACTGG | 470.3 | -2.8 | 0.5 | -5.8 | 5.6E-09 | 9.8E-09 | MB231/MB468 |
| >hg38_21_+_8433172_8446622_RNA45SN1_45S_50ntFlanks@8168.8185.18\|CCCAAGTCCTTCTGATCG | >hg38_21_+_8433172_8446622_RNA45SN1_45S_50ntFlanks | 8168 | 8185 | 18 | CCCAAGTCCTTCTGATCG | 485.9 | -4.0 | 0.7 | -5.8 | 6.3E-09 | 9.9E-09 | BT20/MB468 |
| >hg38_21_+_8433172_8446622_RNA45SN1_45S_50ntFlanks@3994.4013.20\|TGACTCTAGATAACCTCGGG | >hg38_21_+_8433172_8446622_RNA45SN1_45S_50ntFlanks | 3994 | 4013 | 20 | TGACTCTAGATAACCTCGGG | 131.1 | -2.9 | 0.5 | -5.8 | 6.3E-09 | 9.9E-09 | BT20/MB468 |
| >hg38_21_+_8433172_8446622_RNA45SN1_45S_50ntFlanks@12420.12453.34\|ATGTCGGCTCTTCCTATCATTGTGAAGCAGAATT | >hg38_21_+_8433172_8446622_RNA45SN1_45S_50ntFlanks | 12420 | 12453 | 34 | ATGTCGGCTCTTCCTATCATTGTGAAGCAGAATT | 672.3 | 3.6 | 0.6 | 5.8 | 5.6E-09 | 1.0E-08 | MB231/MB468 |
| >MI0000253\|hsa-mir-148a&WithFlank&7\|-\|25949913\|25949992@50.70.21(+1A)&[MIMAT0000243&hsa-miR-148a-3p&offsets\|0\|-1(+1A);m-7&7\|-\|25949922\|25949943&offsets\|0\|-1(+1A)]\|TCAGTGCACTACAGAACTTTGA | >MI0000253\|hsa-mir-148a&WithFlank&7\|-\|25949913\|25949992 | 50 | 70 | 21(+1A) | TCAGTGCACTACAGAACTTTGA | 1465.0 | 5.2 | 0.9 | 5.8 | 5.7E-09 | 1.0E-08 | MB231/MB468 |
| >MI0000289\|hsa-mir-181a-1&WithFlank&1\|-\|198859038\|198859159@30.51.22(+1C);MI0000269\|hsa-mir-181a-2&WithFlank&9\|+\|124692436\|124692557@45.66.22(+1C)&[MIMAT0000256&hsa-miR-181a-5p&offsets\|0\|-1(+1C);m-32&1\|-\|198859108\|198859130&offsets\|0\|-1(+1C)];[MIMAT0000256_1&hsa-miR-181a-5p&offsets\|0\|-1(+1C);m-32&9\|+\|124692480\|124692502&offsets\|0\|-1(+1C)]\|AACATTCAACGCTGTCGGTGAGC | >MI0000289\|hsa-mir-181a-1&WithFlank&1\|-\|198859038\|198859159 | 30 | 51 | 22(+1C) | AACATTCAACGCTGTCGGTGAGC | 454.4 | 4.7 | 0.8 | 5.8 | 6.4E-09 | 1.0E-08 | BT20/MB468 |
| >trnaMT_LeuTAA_MT_+_3230_3304@16.44.29\|CCCGGTAATCGCATAAAACTTAAAACTTT | >trnaMT_LeuTAA_MT_+_3230_3304 | 16 | 44 | 29 | CCCGGTAATCGCATAAAACTTAAAACTTT | 465.3 | -3.7 | 0.6 | -5.8 | 6.5E-09 | 1.0E-08 | BT20/MB468 |
| >hg38_21_+_8433172_8446622_RNA45SN1_45S_50ntFlanks@9574.9592.19\|AATCGGTCGTCCGACCTGG | >hg38_21_+_8433172_8446622_RNA45SN1_45S_50ntFlanks | 9574 | 9592 | 19 | AATCGGTCGTCCGACCTGG | 130.0 | -5.2 | 0.9 | -5.8 | 6.5E-09 | 1.0E-08 | BT20/MB468 |
| >hg38_21_+_8433172_8446622_RNA45SN1_45S_50ntFlanks@13008.13026.19\|ATTGAAAGTCAGCCCTCGA | >hg38_21_+_8433172_8446622_RNA45SN1_45S_50ntFlanks | 13008 | 13026 | 19 | ATTGAAAGTCAGCCCTCGA | 119.8 | -5.5 | 0.9 | -5.8 | 6.6E-09 | 1.0E-08 | BT20/MB468 |
| >hg38_21_+_8433172_8446622_RNA45SN1_45S_50ntFlanks@11641.11662.22\|CGGCGGGTGTTGACGCGATGTG | >hg38_21_+_8433172_8446622_RNA45SN1_45S_50ntFlanks | 11641 | 11662 | 22 | CGGCGGGTGTTGACGCGATGTG | 1663.7 | -7.2 | 1.2 | -5.8 | 6.6E-09 | 1.0E-08 | BT20/MB468 |
| >hg38_21_+_8433172_8446622_RNA45SN1_45S_50ntFlanks@12404.12436.33\|GCTTTTTGATCCTTCGATGTCGGCTCTTCCTAT | >hg38_21_+_8433172_8446622_RNA45SN1_45S_50ntFlanks | 12404 | 12436 | 33 | GCTTTTTGATCCTTCGATGTCGGCTCTTCCTAT | 109.5 | -3.3 | 0.6 | -5.8 | 6.6E-09 | 1.0E-08 | BT20/MB468 |
| >hg38_21_+_8433172_8446622_RNA45SN1_45S_50ntFlanks@10764.10785.22\|TCCGCAGCAGGTCTCCAAGGTG | >hg38_21_+_8433172_8446622_RNA45SN1_45S_50ntFlanks | 10764 | 10785 | 22 | TCCGCAGCAGGTCTCCAAGGTG | 71.2 | -4.9 | 0.8 | -5.8 | 5.9E-09 | 1.0E-08 | MB231/MB468 |
| >hg38_21_+_8433172_8446622_RNA45SN1_45S_50ntFlanks@11918.11948.31\|ACATGAGAGGTGTAGAATAAGTGGGAGGCCC | >hg38_21_+_8433172_8446622_RNA45SN1_45S_50ntFlanks | 11918 | 11948 | 31 | ACATGAGAGGTGTAGAATAAGTGGGAGGCCC | 165.1 | -6.3 | 1.1 | -5.8 | 6.7E-09 | 1.1E-08 | BT20/MB468 |
| >hg38_21_+_8433172_8446622_RNA45SN1_45S_50ntFlanks@10914.10931.18\|CGCCGCGGCTGGACGAGG | >hg38_21_+_8433172_8446622_RNA45SN1_45S_50ntFlanks | 10914 | 10931 | 18 | CGCCGCGGCTGGACGAGG | 730.1 | -5.9 | 1.0 | -5.8 | 6.1E-09 | 1.1E-08 | MB231/MB468 |
| >hg38_21_+_8433172_8446622_RNA45SN1_45S_50ntFlanks@11884.11904.21\|GTTGAGCTTGACTCTAGTCTG | >hg38_21_+_8433172_8446622_RNA45SN1_45S_50ntFlanks | 11884 | 11904 | 21 | GTTGAGCTTGACTCTAGTCTG | 143.7 | -6.0 | 1.0 | -5.8 | 6.1E-09 | 1.1E-08 | MB231/MB468 |
| >MI0000253\|hsa-mir-148a&WithFlank&7\|-\|25949913\|25949992@50.70.21(+1C)&[MIMAT0000243&hsa-miR-148a-3p&offsets\|0\|-1(+1C);m-7&7\|-\|25949922\|25949943&offsets\|0\|-1(+1C)]\|TCAGTGCACTACAGAACTTTGC | >MI0000253\|hsa-mir-148a&WithFlank&7\|-\|25949913\|25949992 | 50 | 70 | 21(+1C) | TCAGTGCACTACAGAACTTTGC | 7516.7 | 5.0 | 0.9 | 5.8 | 6.1E-09 | 1.1E-08 | MB231/MB468 |
| >hg38_21_+_8433172_8446622_RNA45SN1_45S_50ntFlanks@6731.6754.24\|CACATTGATCATCGACACTTCGAA | >hg38_21_+_8433172_8446622_RNA45SN1_45S_50ntFlanks | 6731 | 6754 | 24 | CACATTGATCATCGACACTTCGAA | 127.5 | -3.4 | 0.6 | -5.8 | 6.1E-09 | 1.1E-08 | MB231/MB468 |
| >hg38_21_+_8433172_8446622_RNA45SN1_45S_50ntFlanks@10800.10818.19\|TGTTGGAACAATGTAGGTA | >hg38_21_+_8433172_8446622_RNA45SN1_45S_50ntFlanks | 10800 | 10818 | 19 | TGTTGGAACAATGTAGGTA | 189.2 | -5.3 | 0.9 | -5.8 | 6.3E-09 | 1.1E-08 | MB231/MB468 |
| >hg38_21_+_8433172_8446622_RNA45SN1_45S_50ntFlanks@3835.3860.26\|CTCCTCTCCTACTTGGATAACTGTGG | >hg38_21_+_8433172_8446622_RNA45SN1_45S_50ntFlanks | 3835 | 3860 | 26 | CTCCTCTCCTACTTGGATAACTGTGG | 1160.7 | -4.2 | 0.7 | -5.8 | 7.0E-09 | 1.1E-08 | BT20/MB468 |
| >hg38_21_+_8433172_8446622_RNA45SN1_45S_50ntFlanks@6663.6680.18\|GTGGATCACTCGGCTCGT | >hg38_21_+_8433172_8446622_RNA45SN1_45S_50ntFlanks | 6663 | 6680 | 18 | GTGGATCACTCGGCTCGT | 74.1 | -5.3 | 0.9 | -5.8 | 6.3E-09 | 1.1E-08 | MB231/MB468 |
| >hg38_21_+_8433172_8446622_RNA45SN1_45S_50ntFlanks@6779.6806.28\|CCGGGGCTACGCCTGTCTGAGCGTCGCT | >hg38_21_+_8433172_8446622_RNA45SN1_45S_50ntFlanks | 6779 | 6806 | 28 | CCGGGGCTACGCCTGTCTGAGCGTCGCT | 784.6 | 3.6 | 0.6 | 5.8 | 7.1E-09 | 1.1E-08 | BT20/MB468 |
| >trna119_LysCTT_1_-_145395522_145395594@4.33.30,trna11_LysCTT_5_-_180648979_180649051@4.33.30,trna13_LysCTT_6_+_26556774_26556846@4.33.30,trna7_LysCTT_16_+_3225692_3225764@4.33.30,trna9_LysCTT_5_+_180634755_180634827@4.33.30\|CGGCTAGCTCAGTCGGTAGAGCATGAGACT | >trna119_LysCTT_1_-_145395522_145395594 | 4 | 33 | 30 | CGGCTAGCTCAGTCGGTAGAGCATGAGACT | 2699.1 | 13.0 | 2.2 | 5.8 | 6.3E-09 | 1.1E-08 | MB231/MB468 |
| >hg38_21_+_8433172_8446622_RNA45SN1_45S_50ntFlanks@12004.12035.32\|CGGGCCGCCGGTGAAATACCACTACTCTGATC | >hg38_21_+_8433172_8446622_RNA45SN1_45S_50ntFlanks | 12004 | 12035 | 32 | CGGGCCGCCGGTGAAATACCACTACTCTGATC | 594.3 | 5.1 | 0.8 | 6.2 | 4.3E-10 | 1.1E-08 | BT20/MB231 |

<table>
<tr><th></th><th></th><th></th><th></th><th></th><th></th><th></th><th></th><th></th><th></th><th></th><th></th><th></th></tr>
<tr><td>>hg38_21_+_8433172_8446622_RNA45SN1_45S_50ntFlanks@11583.11603.21|CTGGTGCGGACCAGGGGAATC</td><td>>hg38_21_+_8433172_8446622_RNA45SN1_45S_50ntFlanks</td><td>11583</td><td>11603</td><td>21</td><td>CTGGTGCGGACCAGGGGAATC</td><td>1473.6</td><td>-4.1</td><td>0.7</td><td>-5.8</td><td>7.3E-09</td><td>1.1E-08</td><td>BT20/MB468</td></tr>
<tr><td>>MI0000085|hsa-mir-27a&WithFlank&19|-|13836434|13836523@57.77.21&[MIMAT0000084&hsa-miR-27a-3p&offsets|0|0;m-43&19|-|13836447|13836467&offsets|0|0]|TTCACAGTGGCTAAGTTCCGC</td><td>>MI0000085|hsa-mir-27a&WithFlank&19|-|13836434|13836523</td><td>57</td><td>77</td><td>21</td><td>TTCACAGTGGCTAAGTTCCGC</td><td>6017.1</td><td>5.5</td><td>1.0</td><td>5.8</td><td>7.3E-09</td><td>1.2E-08</td><td>BT20/MB468</td></tr>
<tr><td>>hg38_21_+_8433172_8446622_RNA45SN1_45S_50ntFlanks@8179.8212.34|CTGATCGAGGCCCAGCCCGTGGACGGTGTGAGGC</td><td>>hg38_21_+_8433172_8446622_RNA45SN1_45S_50ntFlanks</td><td>8179</td><td>8212</td><td>34</td><td>CTGATCGAGGCCCAGCCCGTGGACGGTGTGAGGC</td><td>163.6</td><td>-6.3</td><td>1.1</td><td>-5.8</td><td>7.4E-09</td><td>1.2E-08</td><td>BT20/MB468</td></tr>
<tr><td>>hg38_21_+_8433172_8446622_RNA45SN1_45S_50ntFlanks@11617.11640.24|AAAACAAAGCATCGCGAAGGCCCG</td><td>>hg38_21_+_8433172_8446622_RNA45SN1_45S_50ntFlanks</td><td>11617</td><td>11640</td><td>24</td><td>AAAACAAAGCATCGCGAAGGCCCG</td><td>79.7</td><td>-5.4</td><td>0.9</td><td>-5.8</td><td>6.6E-09</td><td>1.2E-08</td><td>MB231/MB468</td></tr>
<tr><td>>hg38_21_+_8433172_8446622_RNA45SN1_45S_50ntFlanks@9576.9596.21|TCGGTCGTCCGACCTGGGTAT</td><td>>hg38_21_+_8433172_8446622_RNA45SN1_45S_50ntFlanks</td><td>9576</td><td>9596</td><td>21</td><td>TCGGTCGTCCGACCTGGGTAT</td><td>148.1</td><td>-4.8</td><td>0.8</td><td>-5.8</td><td>6.6E-09</td><td>1.2E-08</td><td>MB231/MB468</td></tr>
<tr><td>>hg38_21_+_8433172_8446622_RNA45SN1_45S_50ntFlanks@11729.11766.38|TAACTATGACTCTCTTAAGGTAGCCAAATGCCTCGTCA</td><td>>hg38_21_+_8433172_8446622_RNA45SN1_45S_50ntFlanks</td><td>11729</td><td>11766</td><td>38</td><td>TAACTATGACTCTCTTAAGGTAGCCAAATGCCTCGTCA</td><td>130.5</td><td>-3.6</td><td>0.6</td><td>-5.8</td><td>6.6E-09</td><td>1.2E-08</td><td>MB231/MB468</td></tr>
<tr><td>>hg38_21_+_8433172_8446622_RNA45SN1_45S_50ntFlanks@6690.6719.30|AAGAACGCAGCTAGCTGCGAGAATTAATGT</td><td>>hg38_21_+_8433172_8446622_RNA45SN1_45S_50ntFlanks</td><td>6690</td><td>6719</td><td>30</td><td>AAGAACGCAGCTAGCTGCGAGAATTAATGT</td><td>71.5</td><td>-7.7</td><td>1.3</td><td>-5.8</td><td>6.7E-09</td><td>1.2E-08</td><td>MB231/MB468</td></tr>
<tr><td>>am_trnaMT_ValTAC_MT_+_1602_1670@49.67.19|AACTTAACTTGACCGCTCT</td><td>>am_trnaMT_ValTAC_MT_+_1602_1670</td><td>49</td><td>67</td><td>19</td><td>AACTTAACTTGACCGCTCT</td><td>1193.0</td><td>5.5</td><td>1.0</td><td>5.8</td><td>6.7E-09</td><td>1.2E-08</td><td>MB231/MB468</td></tr>
<tr><td>>hg38_21_+_8433172_8446622_RNA45SN1_45S_50ntFlanks@11727.11752.26|AGTAACTATGACTCTCTTAAGGTAGC</td><td>>hg38_21_+_8433172_8446622_RNA45SN1_45S_50ntFlanks</td><td>11727</td><td>11752</td><td>26</td><td>AGTAACTATGACTCTCTTAAGGTAGC</td><td>493.1</td><td>3.4</td><td>0.6</td><td>5.8</td><td>6.7E-09</td><td>1.2E-08</td><td>MB231/MB468</td></tr>
<tr><td>>hg38_21_+_8433172_8446622_RNA45SN1_45S_50ntFlanks@12944.12972.29|CGTAGACGACCTGCTTCTGGGTCGGGGTT</td><td>>hg38_21_+_8433172_8446622_RNA45SN1_45S_50ntFlanks</td><td>12944</td><td>12972</td><td>29</td><td>CGTAGACGACCTGCTTCTGGGTCGGGGTT</td><td>182.9</td><td>-4.7</td><td>0.8</td><td>-5.8</td><td>7.6E-09</td><td>1.2E-08</td><td>BT20/MB468</td></tr>
<tr><td>>trna111_HisGTG_1_-_147774845_147774916@-1G.32.33,trna118_HisGTG_1_-_145396881_145396952@-1G.32.33,trna16_HisGTG_1_+_146544773_146544844@-1G.32.33,trna1_HisGTG_15_+_45493349_45493420@-1G.32.33,trna21_HisGTG_1_+_147753471_147753542@-1G.32.33,trna33_HisGTG_6_+_27125906_27125977@-1G.32.33,trna7_HisGTG_9_-_14433938_14434009@-1G.32.33,trna8_HisGTG_15_-_45492611_45492682@-1G.32.33,trna9_HisGTG_15_-_45490804_45490875@-1G.32.33|GGCCGTGATCGTATAGTGGTTAGTACTCTGCGT</td><td>>trna111_HisGTG_1_-_147774845_147774916</td><td>-1G</td><td>32</td><td>33</td><td>GGCCGTGATCGTATAGTGGTTAGTACTCTGCGT</td><td>7504.3</td><td>13.8</td><td>2.4</td><td>5.8</td><td>6.9E-09</td><td>1.2E-08</td><td>MB231/MB468</td></tr>
<tr><td>>hg38_21_+_8433172_8446622_RNA45SN1_45S_50ntFlanks@5482.5514.33|CGGAGCGCTGAGAAGACGGTCGAACTTGACTAT</td><td>>hg38_21_+_8433172_8446622_RNA45SN1_45S_50ntFlanks</td><td>5482</td><td>5514</td><td>33</td><td>CGGAGCGCTGAGAAGACGGTCGAACTTGACTAT</td><td>88.2</td><td>-4.6</td><td>0.8</td><td>-5.8</td><td>7.8E-09</td><td>1.2E-08</td><td>BT20/MB468</td></tr>
<tr><td>>hg38_21_+_8433172_8446622_RNA45SN1_45S_50ntFlanks@11199.11233.35|CCGGCGGCGGCGGCGACTCTGGACGCGAGCCGGGC</td><td>>hg38_21_+_8433172_8446622_RNA45SN1_45S_50ntFlanks</td><td>11199</td><td>11233</td><td>35</td><td>CCGGCGGCGGCGGCGACTCTGGACGCGAGCCGGGC</td><td>80.2</td><td>-3.6</td><td>0.6</td><td>-5.8</td><td>7.9E-09</td><td>1.2E-08</td><td>BT20/MB468</td></tr>
<tr><td>>hg38_21_+_8433172_8446622_RNA45SN1_45S_50ntFlanks@3782.3802.21|CAGTGAAACTGCGAATGGCTC</td><td>>hg38_21_+_8433172_8446622_RNA45SN1_45S_50ntFlanks</td><td>3782</td><td>3802</td><td>21</td><td>CAGTGAAACTGCGAATGGCTC</td><td>142.1</td><td>-2.6</td><td>0.4</td><td>-5.8</td><td>7.1E-09</td><td>1.2E-08</td><td>MB231/MB468</td></tr>
<tr><td>>MI0000102|hsa-mir-100&WithFlank&11|-|122152223|122152314@19.38.20(+1C)&[MIMAT0000098&hsa-miR-100-5p&offsets|0|-2(+1C);m-27&11|-|122152276|122152296&offsets|0|-1(+1C)]|AACCCGTAGATCCGAACTTGC</td><td>>MI0000102|hsa-mir-100&WithFlank&11|-|122152223|122152314</td><td>19</td><td>38</td><td>20(+1C)</td><td>AACCCGTAGATCCGAACTTGC</td><td>907.4</td><td>8.9</td><td>1.5</td><td>5.8</td><td>8.1E-09</td><td>1.3E-08</td><td>BT20/MB468</td></tr>
<tr><td>>hg38_21_+_8433172_8446622_RNA45SN1_45S_50ntFlanks@6645.6680.36|CTCGTACGACTCTTAGCGGTGGATCACTCGGCTCGT</td><td>>hg38_21_+_8433172_8446622_RNA45SN1_45S_50ntFlanks</td><td>6645</td><td>6680</td><td>36</td><td>CTCGTACGACTCTTAGCGGTGGATCACTCGGCTCGT</td><td>759.9</td><td>2.7</td><td>0.5</td><td>5.8</td><td>8.1E-09</td><td>1.3E-08</td><td>BT20/MB468</td></tr>
<tr><td>>hg38_21_+_8433172_8446622_RNA45SN1_45S_50ntFlanks@12941.12958.18|ATTCGTAGACGACCTGCT</td><td>>hg38_21_+_8433172_8446622_RNA45SN1_45S_50ntFlanks</td><td>12941</td><td>12958</td><td>18</td><td>ATTCGTAGACGACCTGCT</td><td>728.8</td><td>3.3</td><td>0.6</td><td>5.8</td><td>7.2E-09</td><td>1.3E-08</td><td>MB231/MB468</td></tr>
<tr><td>>hg38_21_+_8433172_8446622_RNA45SN1_45S_50ntFlanks@12933.12953.21|GCTAAACCATTCGTAGACGAC</td><td>>hg38_21_+_8433172_8446622_RNA45SN1_45S_50ntFlanks</td><td>12933</td><td>12953</td><td>21</td><td>GCTAAACCATTCGTAGACGAC</td><td>181.5</td><td>-4.3</td><td>0.7</td><td>-5.8</td><td>8.1E-09</td><td>1.3E-08</td><td>BT20/MB468</td></tr>
<tr><td>>hg38_21_+_8433172_8446622_RNA45SN1_45S_50ntFlanks@8113.8143.31|GGCGCGGGACATGTGGCGTACGGAAGACCCG</td><td>>hg38_21_+_8433172_8446622_RNA45SN1_45S_50ntFlanks</td><td>8113</td><td>8143</td><td>31</td><td>GGCGCGGGACATGTGGCGTACGGAAGACCCG</td><td>117.3</td><td>-6.5</td><td>1.1</td><td>-5.8</td><td>7.3E-09</td><td>1.3E-08</td><td>MB231/MB468</td></tr>
<tr><td>>hg38_21_+_8433172_8446622_RNA45SN1_45S_50ntFlanks@9564.9597.34|CTGACGTGCAAATCGGTCGTCCGACCTGGGTATA</td><td>>hg38_21_+_8433172_8446622_RNA45SN1_45S_50ntFlanks</td><td>9564</td><td>9597</td><td>34</td><td>CTGACGTGCAAATCGGTCGTCCGACCTGGGTATA</td><td>4300.0</td><td>-3.6</td><td>0.6</td><td>-5.8</td><td>7.3E-09</td><td>1.3E-08</td><td>MB231/MB468</td></tr>
<tr><td>>hg38_21_+_8433172_8446622_RNA45SN1_45S_50ntFlanks@11670.11700.31|CCCAGTGCTCTGAATGTCAAAGTGAAGAAAT</td><td>>hg38_21_+_8433172_8446622_RNA45SN1_45S_50ntFlanks</td><td>11670</td><td>11700</td><td>31</td><td>CCCAGTGCTCTGAATGTCAAAGTGAAGAAAT</td><td>235.1</td><td>-3.2</td><td>0.6</td><td>-5.8</td><td>7.4E-09</td><td>1.3E-08</td><td>MB231/MB468</td></tr>
<tr><td>>hg38_21_+_8433172_8446622_RNA45SN1_45S_50ntFlanks@9554.9597.44|CGTAGCGGTCCTGACGTGCAAATCGGTCGTCCGACCTGGGTATA</td><td>>hg38_21_+_8433172_8446622_RNA45SN1_45S_50ntFlanks</td><td>9554</td><td>9597</td><td>44</td><td>CGTAGCGGTCCTGACGTGCAAATCGGTCGTCCGACCTGGGTATA</td><td>234.2</td><td>-6.1</td><td>1.1</td><td>-5.8</td><td>8.3E-09</td><td>1.3E-08</td><td>BT20/MB468</td></tr>
<tr><td>>hg38_21_+_8433172_8446622_RNA45SN1_45S_50ntFlanks@6646.6680.35|TCGTACGACTCTTAGCGGTGGATCACTCGGCTCGT</td><td>>hg38_21_+_8433172_8446622_RNA45SN1_45S_50ntFlanks</td><td>6646</td><td>6680</td><td>35</td><td>TCGTACGACTCTTAGCGGTGGATCACTCGGCTCGT</td><td>6222.0</td><td>3.1</td><td>0.5</td><td>5.8</td><td>8.3E-09</td><td>1.3E-08</td><td>BT20/MB468</td></tr>
<tr><td>>MI0000253|hsa-mir-148a&WithFlank&7|-|25949913|25949992@50.71.22&[MIMAT0000243&hsa-miR-148a-3p&offsets|0|0;m-7&7|-|25949922|25949943&offsets|0|0]|TCAGTGCACTACAGAACTTTGT</td><td>>MI0000253|hsa-mir-148a&WithFlank&7|-|25949913|25949992</td><td>50</td><td>71</td><td>22</td><td>TCAGTGCACTACAGAACTTTGT</td><td>309619.0</td><td>4.2</td><td>0.7</td><td>6.2</td><td>5.1E-10</td><td>1.3E-08</td><td>BT20/MB231</td></tr>
<tr><td>>hg38_21_+_8433172_8446622_RNA45SN1_45S_50ntFlanks@11575.11602.28|ACTTAGAACTGGTGCGGACCAGGGGAAT</td><td>>hg38_21_+_8433172_8446622_RNA45SN1_45S_50ntFlanks</td><td>11575</td><td>11602</td><td>28</td><td>ACTTAGAACTGGTGCGGACCAGGGGAAT</td><td>972.6</td><td>-4.3</td><td>0.7</td><td>-5.8</td><td>8.3E-09</td><td>1.3E-08</td><td>BT20/MB468</td></tr>
<tr><td>>MI0000253|hsa-mir-148a&WithFlank&7|-|25949913|25949992@50.71.22&[MIMAT0000243&hsa-miR-148a-3p&offsets|0|0;m-7&7|-|25949922|25949943&offsets|0|0]|TCAGTGCACTACAGAACTTTGT</td><td>>MI0000253|hsa-mir-148a&WithFlank&7|-|25949913|25949992</td><td>50</td><td>71</td><td>22</td><td>TCAGTGCACTACAGAACTTTGT</td><td>70513.0</td><td>5.1</td><td>0.9</td><td>5.8</td><td>7.4E-09</td><td>1.3E-08</td><td>MB231/MB468</td></tr>
<tr><td>>hg38_21_+_8433172_8446622_RNA45SN1_45S_50ntFlanks@10843.10863.21|TAACTTCGGGATAAGGATTGG</td><td>>hg38_21_+_8433172_8446622_RNA45SN1_45S_50ntFlanks</td><td>10843</td><td>10863</td><td>21</td><td>TAACTTCGGGATAAGGATTGG</td><td>158.9</td><td>-4.6</td><td>0.8</td><td>-5.8</td><td>7.4E-09</td><td>1.3E-08</td><td>MB231/MB468</td></tr>
<tr><td>>hg38_21_+_8433172_8446622_RNA45SN1_45S_50ntFlanks@11573.11600.28|CGACTTAGAACTGGTGCGGACCAGGGGA</td><td>>hg38_21_+_8433172_8446622_RNA45SN1_45S_50ntFlanks</td><td>11573</td><td>11600</td><td>28</td><td>CGACTTAGAACTGGTGCGGACCAGGGGA</td><td>290.2</td><td>-2.5</td><td>0.4</td><td>-5.8</td><td>8.3E-09</td><td>1.3E-08</td><td>BT20/MB468</td></tr>
<tr><td>>hg38_21_+_8433172_8446622_RNA45SN1_45S_50ntFlanks@12656.12674.19|TGACTGAACGCCTCTAAGT</td><td>>hg38_21_+_8433172_8446622_RNA45SN1_45S_50ntFlanks</td><td>12656</td><td>12674</td><td>19</td><td>TGACTGAACGCCTCTAAGT</td><td>233.7</td><td>-2.5</td><td>0.4</td><td>-5.8</td><td>8.5E-09</td><td>1.3E-08</td><td>BT20/MB468</td></tr>
<tr><td>>hg38_21_+_8433172_8446622_RNA45SN1_45S_50ntFlanks@11641.11679.39|CGGCGGGTGTTGACGCGATGTGATTTCTGCCCAGTGCTC</td><td>>hg38_21_+_8433172_8446622_RNA45SN1_45S_50ntFlanks</td><td>11641</td><td>11679</td><td>39</td><td>CGGCGGGTGTTGACGCGATGTGATTTCTGCCCAGTGCTC</td><td>132.9</td><td>-5.2</td><td>0.9</td><td>-5.8</td><td>7.7E-09</td><td>1.3E-08</td><td>MB231/MB468</td></tr>
<tr><td>>hg38_21_+_8433172_8446622_RNA45SN1_45S_50ntFlanks@11873.11907.35|AAGAAGACCCTGTTGAGCTTGACTCTAGTCTGGCA</td><td>>hg38_21_+_8433172_8446622_RNA45SN1_45S_50ntFlanks</td><td>11873</td><td>11907</td><td>35</td><td>AAGAAGACCCTGTTGAGCTTGACTCTAGTCTGGCA</td><td>259.3</td><td>-6.1</td><td>1.1</td><td>-5.8</td><td>8.7E-09</td><td>1.4E-08</td><td>BT20/MB468</td></tr>
<tr><td>>hg38_21_+_8433172_8446622_RNA45SN1_45S_50ntFlanks@10866.10904.39|CTAAGGGCTGGGTCGGTCGGGCTGGGGCGCGAAGCGGGG</td><td>>hg38_21_+_8433172_8446622_RNA45SN1_45S_50ntFlanks</td><td>10866</td><td>10904</td><td>39</td><td>CTAAGGGCTGGGTCGGTCGGGCTGGGGCGCGAAGCGGGG</td><td>63.0</td><td>-7.6</td><td>1.3</td><td>-5.8</td><td>7.9E-09</td><td>1.4E-08</td><td>MB231/MB468</td></tr>
<tr><td>>hg38_21_+_8433172_8446622_RNA45SN1_45S_50ntFlanks@4257.4275.19|TAATTGGAATGAGTCCACT</td><td>>hg38_21_+_8433172_8446622_RNA45SN1_45S_50ntFlanks</td><td>4257</td><td>4275</td><td>19</td><td>TAATTGGAATGAGTCCACT</td><td>76.7</td><td>-7.8</td><td>1.4</td><td>-5.8</td><td>7.9E-09</td><td>1.4E-08</td><td>MB231/MB468</td></tr>
<tr><td>>hg38_21_+_8433172_8446622_RNA45SN1_45S_50ntFlanks@485.504.20|GATGTGGTGACGTCGTGCTC</td><td>>hg38_21_+_8433172_8446622_RNA45SN1_45S_50ntFlanks</td><td>485</td><td>504</td><td>20</td><td>GATGTGGTGACGTCGTGCTC</td><td>2294.9</td><td>16.1</td><td>2.8</td><td>5.8</td><td>7.9E-09</td><td>1.4E-08</td><td>MB231/MB468</td></tr>
<tr><td>>hg38_21_+_8433172_8446622_RNA45SN1_45S_50ntFlanks@6650.6669.20|ACGACTCTTAGCGGTGGATC</td><td>>hg38_21_+_8433172_8446622_RNA45SN1_45S_50ntFlanks</td><td>6650</td><td>6669</td><td>20</td><td>ACGACTCTTAGCGGTGGATC</td><td>1683.8</td><td>2.9</td><td>0.5</td><td>5.8</td><td>8.9E-09</td><td>1.4E-08</td><td>BT20/MB468</td></tr>
<tr><td>>MI0000283|hsa-mir-203a&WithFlank&14|+|104117399|104117520@71.92.22(+1U)&[MIMAT0000264&hsa-miR-203a-3p&offsets|0|0(+1U);m-79&14|+|104117469|104117490&offsets|0|0(+1U)]|GTGAAATGTTTAGGACCACTAGT</td><td>>MI0000283|hsa-mir-203a&WithFlank&14|+|104117399|104117520</td><td>71</td><td>92</td><td>22(+1U)</td><td>GTGAAATGTTTAGGACCACTAGT</td><td>547.0</td><td>8.1</td><td>1.4</td><td>5.8</td><td>8.9E-09</td><td>1.4E-08</td><td>BT20/MB468</td></tr>
<tr><td>>hg38_21_+_8433172_8446622_RNA45SN1_45S_50ntFlanks@11876.11897.22|AAGACCCTGTTGAGCTTGACTC</td><td>>hg38_21_+_8433172_8446622_RNA45SN1_45S_50ntFlanks</td><td>11876</td><td>11897</td><td>22</td><td>AAGACCCTGTTGAGCTTGACTC</td><td>408.0</td><td>-2.4</td><td>0.4</td><td>-5.8</td><td>8.9E-09</td><td>1.4E-08</td><td>BT20/MB468</td></tr>
<tr><td>>hg38_21_+_8433172_8446622_RNA45SN1_45S_50ntFlanks@4367.4388.22|CAGTTAAAAAGCTCGTAGTTGG</td><td>>hg38_21_+_8433172_8446622_RNA45SN1_45S_50ntFlanks</td><td>4367</td><td>4388</td><td>22</td><td>CAGTTAAAAAGCTCGTAGTTGG</td><td>201.3</td><td>-4.8</td><td>0.8</td><td>-5.8</td><td>8.0E-09</td><td>1.4E-08</td><td>MB231/MB468</td></tr>
<tr><td>>am_trnaMT_SerGCT_MT_+_12207_12265@39.58.20|TCTAACAACATGGCTTTCTC</td><td>>am_trnaMT_SerGCT_MT_+_12207_12265</td><td>39</td><td>58</td><td>20</td><td>TCTAACAACATGGCTTTCTC</td><td>65.2</td><td>-8.0</td><td>1.4</td><td>-5.7</td><td>9.0E-09</td><td>1.4E-08</td><td>BT20/MB468</td></tr>
<tr><td>>hg38_21_+_8433172_8446622_RNA45SN1_45S_50ntFlanks@11728.11751.24|GTAACTATGACTCTCTTAAGGTAG</td><td>>hg38_21_+_8433172_8446622_RNA45SN1_45S_50ntFlanks</td><td>11728</td><td>11751</td><td>24</td><td>GTAACTATGACTCTCTTAAGGTAG</td><td>1169.9</td><td>-5.9</td><td>1.0</td><td>-5.7</td><td>9.0E-09</td><td>1.4E-08</td><td>BT20/MB468</td></tr>
<tr><td>>hg38_21_+_8433172_8446622_RNA45SN1_45S_50ntFlanks@3854.3889.36|ACTGTGGTAATTCTAGAGCTAATACATGCCGACGGG</td><td>>hg38_21_+_8433172_8446622_RNA45SN1_45S_50ntFlanks</td><td>3854</td><td>3889</td><td>36</td><td>ACTGTGGTAATTCTAGAGCTAATACATGCCGACGGG</td><td>89.9</td><td>-4.6</td><td>0.8</td><td>-5.7</td><td>9.0E-09</td><td>1.4E-08</td><td>BT20/MB468</td></tr>
<tr><td>>hg38_21_+_8433172_8446622_RNA45SN1_45S_50ntFlanks@6653.6683.31|ACTCTTAGCGGTGGATCACTCGGCTCGTGCG</td><td>>hg38_21_+_8433172_8446622_RNA45SN1_45S_50ntFlanks</td><td>6653</td><td>6683</td><td>31</td><td>ACTCTTAGCGGTGGATCACTCGGCTCGTGCG</td><td>997.5</td><td>5.6</td><td>1.0</td><td>5.7</td><td>9.1E-09</td><td>1.4E-08</td><td>BT20/MB468</td></tr>
<tr><td>>hg38_21_+_8433172_8446622_RNA45SN1_45S_50ntFlanks@13008.13028.21|ATTGAAAGTCAGCCCTCGACA</td><td>>hg38_21_+_8433172_8446622_RNA45SN1_45S_50ntFlanks</td><td>13008</td><td>13028</td><td>21</td><td>ATTGAAAGTCAGCCCTCGACA</td><td>645.1</td><td>-5.0</td><td>0.9</td><td>-5.7</td><td>9.1E-09</td><td>1.4E-08</td><td>BT20/MB468</td></tr>
<tr><td>>hg38_21_+_8433172_8446622_RNA45SN1_45S_50ntFlanks@9534.9558.25|AGGAAACTCTGGTGGAGGTCCGTAG</td><td>>hg38_21_+_8433172_8446622_RNA45SN1_45S_50ntFlanks</td><td>9534</td><td>9558</td><td>25</td><td>AGGAAACTCTGGTGGAGGTCCGTAG</td><td>337.3</td><td>-5.4</td><td>0.9</td><td>-5.7</td><td>9.1E-09</td><td>1.4E-08</td><td>BT20/MB468</td></tr>
<tr><td>>hg38_21_+_8433172_8446622_RNA45SN1_45S_50ntFlanks@11767.11790.24|TCTAATTAGTGACGCGCATGAATG</td><td>>hg38_21_+_8433172_8446622_RNA45SN1_45S_50ntFlanks</td><td>11767</td><td>11790</td><td>24</td><td>TCTAATTAGTGACGCGCATGAATG</td><td>170.3</td><td>-3.8</td><td>0.7</td><td>-5.7</td><td>9.1E-09</td><td>1.4E-08</td><td>BT20/MB468</td></tr>
</table>

<table>
<tr><th></th><th></th><th></th><th></th><th></th><th></th><th></th><th></th><th></th><th></th><th></th><th></th><th></th></tr>
<tr><td>>trnaMT_GluTTC_MT_-_14674_14742@25.72.48,trnalookalike8_GluTTC_5_-_93905172_93905240@25.72.48|TGGTTTTTCATATCATTGGTCGTGGTTGTAGTCCGTGCGAGAATACCA</td><td>>trnaMT_GluTTC_MT_-_14674_14742</td><td>25</td><td>72</td><td>48</td><td>TGGTTTTTCATATCATTGGTCGTGGTTGTAGTCCGTGCGAGAATACCA</td><td>3389.3</td><td>13.5</td><td>2.3</td><td>5.8</td><td>8.1E-09</td><td>1.4E-08</td><td>MB231/MB468</td></tr>
<tr><td>>hg38_21_+_8433172_8446622_RNA45SN1_45S_50ntFlanks@11877.11906.30|AGACCCTGTTGAGCTTGACTCTAGTCTGGC</td><td>>hg38_21_+_8433172_8446622_RNA45SN1_45S_50ntFlanks</td><td>11877</td><td>11906</td><td>30</td><td>AGACCCTGTTGAGCTTGACTCTAGTCTGGC</td><td>81.8</td><td>-6.1</td><td>1.1</td><td>-5.8</td><td>8.3E-09</td><td>1.4E-08</td><td>MB231/MB468</td></tr>
<tr><td>>hg38_21_+_8433172_8446622_RNA45SN1_45S_50ntFlanks@11726.11751.26|GAGTAACTATGACTCTCTTAAGGTAG</td><td>>hg38_21_+_8433172_8446622_RNA45SN1_45S_50ntFlanks</td><td>11726</td><td>11751</td><td>26</td><td>GAGTAACTATGACTCTCTTAAGGTAG</td><td>129.2</td><td>-4.9</td><td>0.8</td><td>-5.7</td><td>9.3E-09</td><td>1.5E-08</td><td>BT20/MB468</td></tr>
<tr><td>>hg38_21_+_8433172_8446622_RNA45SN1_45S_50ntFlanks@12942.12962.21|TTCGTAGACGACCTGCTTCTG</td><td>>hg38_21_+_8433172_8446622_RNA45SN1_45S_50ntFlanks</td><td>12942</td><td>12962</td><td>21</td><td>TTCGTAGACGACCTGCTTCTG</td><td>161.0</td><td>-3.1</td><td>0.5</td><td>-5.7</td><td>9.3E-09</td><td>1.5E-08</td><td>BT20/MB468</td></tr>
<tr><td>>hg38_21_+_8433172_8446622_RNA45SN1_45S_50ntFlanks@12459.12488.30|AGCGTTGGATTGTTCACCCACTAATAGGGA</td><td>>hg38_21_+_8433172_8446622_RNA45SN1_45S_50ntFlanks</td><td>12459</td><td>12488</td><td>30</td><td>AGCGTTGGATTGTTCACCCACTAATAGGGA</td><td>127.0</td><td>-4.8</td><td>0.8</td><td>-5.8</td><td>8.4E-09</td><td>1.5E-08</td><td>MB231/MB468</td></tr>
<tr><td>>hg38_21_+_8433172_8446622_RNA45SN1_45S_50ntFlanks@10409.10426.18|GTCAGTCGGTCCTGAGAG</td><td>>hg38_21_+_8433172_8446622_RNA45SN1_45S_50ntFlanks</td><td>10409</td><td>10426</td><td>18</td><td>GTCAGTCGGTCCTGAGAG</td><td>119.5</td><td>-3.1</td><td>0.5</td><td>-5.7</td><td>9.4E-09</td><td>1.5E-08</td><td>BT20/MB468</td></tr>
<tr><td>>hg38_21_+_8433172_8446622_RNA45SN1_45S_50ntFlanks@8198.8223.26|TGGACGGTGTGAGGCCGGTAGCGGCC</td><td>>hg38_21_+_8433172_8446622_RNA45SN1_45S_50ntFlanks</td><td>8198</td><td>8223</td><td>26</td><td>TGGACGGTGTGAGGCCGGTAGCGGCC</td><td>265.2</td><td>-2.9</td><td>0.5</td><td>-5.8</td><td>8.4E-09</td><td>1.5E-08</td><td>MB231/MB468</td></tr>
<tr><td>>hg38_21_+_8433172_8446622_RNA45SN1_45S_50ntFlanks@12161.12181.21|TGACTGGGGCGGTACACCTGT</td><td>>hg38_21_+_8433172_8446622_RNA45SN1_45S_50ntFlanks</td><td>12161</td><td>12181</td><td>21</td><td>TGACTGGGGCGGTACACCTGT</td><td>183.3</td><td>-5.3</td><td>0.9</td><td>-5.7</td><td>9.5E-09</td><td>1.5E-08</td><td>BT20/MB468</td></tr>
<tr><td>>hg38_21_+_8433172_8446622_RNA45SN1_45S_50ntFlanks@4007.4026.20|CCTCGGGCCGATCGCACGCC</td><td>>hg38_21_+_8433172_8446622_RNA45SN1_45S_50ntFlanks</td><td>4007</td><td>4026</td><td>20</td><td>CCTCGGGCCGATCGCACGCC</td><td>106.7</td><td>-4.3</td><td>0.7</td><td>-5.7</td><td>9.6E-09</td><td>1.5E-08</td><td>BT20/MB468</td></tr>
<tr><td>>hg38_21_+_8433172_8446622_RNA45SN1_45S_50ntFlanks@10841.10875.35|CGTAACTTCGGGATAAGGATTGGCTCTAAGGGCTG</td><td>>hg38_21_+_8433172_8446622_RNA45SN1_45S_50ntFlanks</td><td>10841</td><td>10875</td><td>35</td><td>CGTAACTTCGGGATAAGGATTGGCTCTAAGGGCTG</td><td>245.1</td><td>-3.5</td><td>0.6</td><td>-5.8</td><td>8.6E-09</td><td>1.5E-08</td><td>MB231/MB468</td></tr>
<tr><td>>hg38_21_+_8433172_8446622_RNA45SN1_45S_50ntFlanks@10692.10709.18|TCCGGTGAGCTCTCGCTG</td><td>>hg38_21_+_8433172_8446622_RNA45SN1_45S_50ntFlanks</td><td>10692</td><td>10709</td><td>18</td><td>TCCGGTGAGCTCTCGCTG</td><td>118.8</td><td>-5.8</td><td>1.0</td><td>-5.8</td><td>8.6E-09</td><td>1.5E-08</td><td>MB231/MB468</td></tr>
<tr><td>>hg38_21_+_8433172_8446622_RNA45SN1_45S_50ntFlanks@11783.11806.24|CATGAATGGATGAACGAGATTCCC</td><td>>hg38_21_+_8433172_8446622_RNA45SN1_45S_50ntFlanks</td><td>11783</td><td>11806</td><td>24</td><td>CATGAATGGATGAACGAGATTCCC</td><td>2673.5</td><td>-3.7</td><td>0.6</td><td>-5.8</td><td>8.7E-09</td><td>1.5E-08</td><td>MB231/MB468</td></tr>
<tr><td>>hg38_21_+_8433172_8446622_RNA45SN1_45S_50ntFlanks@12934.12954.21|CTAAACCATTCGTAGACGACC</td><td>>hg38_21_+_8433172_8446622_RNA45SN1_45S_50ntFlanks</td><td>12934</td><td>12954</td><td>21</td><td>CTAAACCATTCGTAGACGACC</td><td>211.7</td><td>-3.7</td><td>0.6</td><td>-5.7</td><td>9.9E-09</td><td>1.5E-08</td><td>BT20/MB468</td></tr>
<tr><td>>hg38_21_+_8433172_8446622_RNA45SN1_45S_50ntFlanks@10347.10374.28|CAAACGAGAACTTTGAAGGCCGAAGTGG</td><td>>hg38_21_+_8433172_8446622_RNA45SN1_45S_50ntFlanks</td><td>10347</td><td>10374</td><td>28</td><td>CAAACGAGAACTTTGAAGGCCGAAGTGG</td><td>156.9</td><td>-2.6</td><td>0.5</td><td>-5.7</td><td>1.0E-08</td><td>1.6E-08</td><td>BT20/MB468</td></tr>
<tr><td>>hg38_21_+_8433172_8446622_RNA45SN1_45S_50ntFlanks@10588.10607.20|CCCGGGGAGAGTTCTCTTTT</td><td>>hg38_21_+_8433172_8446622_RNA45SN1_45S_50ntFlanks</td><td>10588</td><td>10607</td><td>20</td><td>CCCGGGGAGAGTTCTCTTTT</td><td>112.8</td><td>-4.9</td><td>0.8</td><td>-5.7</td><td>1.0E-08</td><td>1.6E-08</td><td>BT20/MB468</td></tr>
<tr><td>>hg38_21_+_8433172_8446622_RNA45SN1_45S_50ntFlanks@11603.11620.18|CCGACTGTTTAATTAAAA</td><td>>hg38_21_+_8433172_8446622_RNA45SN1_45S_50ntFlanks</td><td>11603</td><td>11620</td><td>18</td><td>CCGACTGTTTAATTAAAA</td><td>356.2</td><td>-3.1</td><td>0.5</td><td>-5.7</td><td>1.0E-08</td><td>1.6E-08</td><td>BT20/MB468</td></tr>
<tr><td>>hg38_21_+_8433172_8446622_RNA45SN1_45S_50ntFlanks@6652.6692.41|GACTCTTAGCGGTGGATCACTCGGCTCGTGCGTCGATGAAG</td><td>>hg38_21_+_8433172_8446622_RNA45SN1_45S_50ntFlanks</td><td>6652</td><td>6692</td><td>41</td><td>GACTCTTAGCGGTGGATCACTCGGCTCGTGCGTCGATGAAG</td><td>2607.6</td><td>4.2</td><td>0.7</td><td>5.7</td><td>9.0E-09</td><td>1.6E-08</td><td>MB231/MB468</td></tr>
<tr><td>>hg38_21_+_8433172_8446622_RNA45SN1_45S_50ntFlanks@10355.10388.34|AACTTTGAAGGCCGAAGTGGAGAAGGGTTCCATG</td><td>>hg38_21_+_8433172_8446622_RNA45SN1_45S_50ntFlanks</td><td>10355</td><td>10388</td><td>34</td><td>AACTTTGAAGGCCGAAGTGGAGAAGGGTTCCATG</td><td>453.5</td><td>-7.0</td><td>1.2</td><td>-5.7</td><td>1.0E-08</td><td>1.6E-08</td><td>BT20/MB468</td></tr>
<tr><td>>hg38_21_+_8433172_8446622_RNA45SN1_45S_50ntFlanks@11316.11343.28|CACCCCACGTCTCGTCGCGCGCGCGTCC</td><td>>hg38_21_+_8433172_8446622_RNA45SN1_45S_50ntFlanks</td><td>11316</td><td>11343</td><td>28</td><td>CACCCCACGTCTCGTCGCGCGCGCGTCC</td><td>406.5</td><td>-4.6</td><td>0.8</td><td>-5.7</td><td>1.0E-08</td><td>1.6E-08</td><td>BT20/MB468</td></tr>
<tr><td>>hg38_21_+_8433172_8446622_RNA45SN1_45S_50ntFlanks@10421.10453.33|TGAGAGATGGGCGAGCGCCGTTCCGAAGGGACG</td><td>>hg38_21_+_8433172_8446622_RNA45SN1_45S_50ntFlanks</td><td>10421</td><td>10453</td><td>33</td><td>TGAGAGATGGGCGAGCGCCGTTCCGAAGGGACG</td><td>200.5</td><td>-5.1</td><td>0.9</td><td>-5.7</td><td>1.0E-08</td><td>1.6E-08</td><td>BT20/MB468</td></tr>
<tr><td>>hg38_21_+_8433172_8446622_RNA45SN1_45S_50ntFlanks@11769.11804.36|TAATTAGTGACGCGCATGAATGGATGAACGAGATTC</td><td>>hg38_21_+_8433172_8446622_RNA45SN1_45S_50ntFlanks</td><td>11769</td><td>11804</td><td>36</td><td>TAATTAGTGACGCGCATGAATGGATGAACGAGATTC</td><td>367.7</td><td>-2.5</td><td>0.4</td><td>-5.7</td><td>9.1E-09</td><td>1.6E-08</td><td>MB231/MB468</td></tr>
<tr><td>>hg38_21_+_8433172_8446622_RNA45SN1_45S_50ntFlanks@11317.11344.28|ACCCCACGTCTCGTCGCGCGCGCGTCCG</td><td>>hg38_21_+_8433172_8446622_RNA45SN1_45S_50ntFlanks</td><td>11317</td><td>11344</td><td>28</td><td>ACCCCACGTCTCGTCGCGCGCGCGTCCG</td><td>3400.6</td><td>-7.5</td><td>1.3</td><td>-5.7</td><td>1.0E-08</td><td>1.6E-08</td><td>BT20/MB468</td></tr>
<tr><td>>hg38_21_+_8433172_8446622_RNA45SN1_45S_50ntFlanks@9305.9335.31|CCAAGGAGTCTAACACGTGCGCGAGTCGGGG</td><td>>hg38_21_+_8433172_8446622_RNA45SN1_45S_50ntFlanks</td><td>9305</td><td>9335</td><td>31</td><td>CCAAGGAGTCTAACACGTGCGCGAGTCGGGG</td><td>62.8</td><td>-7.5</td><td>1.3</td><td>-5.7</td><td>9.2E-09</td><td>1.6E-08</td><td>MB231/MB468</td></tr>
<tr><td>>hg38_1_-_228634819_228635039_RNA5S12_5S_50ntFlanks@124.147.24|AGTACTTGGATGGGAGACCGCCTG</td><td>>hg38_1_-_228634819_228635039_RNA5S12_5S_50ntFlanks</td><td>124</td><td>147</td><td>24</td><td>AGTACTTGGATGGGAGACCGCCTG</td><td>259.9</td><td>-3.4</td><td>0.6</td><td>-5.7</td><td>1.0E-08</td><td>1.6E-08</td><td>BT20/MB468</td></tr>
<tr><td>>hg38_21_+_8433172_8446622_RNA45SN1_45S_50ntFlanks@7992.8010.19|CGTGGCGACCCGCTGAATT</td><td>>hg38_21_+_8433172_8446622_RNA45SN1_45S_50ntFlanks</td><td>7992</td><td>8010</td><td>19</td><td>CGTGGCGACCCGCTGAATT</td><td>1526.5</td><td>5.3</td><td>0.9</td><td>5.7</td><td>9.2E-09</td><td>1.6E-08</td><td>MB231/MB468</td></tr>
<tr><td>>hg38_21_+_8433172_8446622_RNA45SN1_45S_50ntFlanks@4297.4320.24|CATTGGAGGGCAAGTCTGGTGCCA</td><td>>hg38_21_+_8433172_8446622_RNA45SN1_45S_50ntFlanks</td><td>4297</td><td>4320</td><td>24</td><td>CATTGGAGGGCAAGTCTGGTGCCA</td><td>133.7</td><td>-4.7</td><td>0.8</td><td>-5.7</td><td>9.4E-09</td><td>1.6E-08</td><td>MB231/MB468</td></tr>
<tr><td>>hg38_21_+_8433172_8446622_RNA45SN1_45S_50ntFlanks@9352.9371.20|CCGTGGCGCAATGAAGGTGA</td><td>>hg38_21_+_8433172_8446622_RNA45SN1_45S_50ntFlanks</td><td>9352</td><td>9371</td><td>20</td><td>CCGTGGCGCAATGAAGGTGA</td><td>82.6</td><td>-7.9</td><td>1.4</td><td>-5.7</td><td>9.4E-09</td><td>1.6E-08</td><td>MB231/MB468</td></tr>
<tr><td>>hg38_21_+_8433172_8446622_RNA45SN1_45S_50ntFlanks@11209.11254.46|CGGCGACTCTGGACGCGAGCCGGGCCCTTCCCGTGGATCGCCCCAG</td><td>>hg38_21_+_8433172_8446622_RNA45SN1_45S_50ntFlanks</td><td>11209</td><td>11254</td><td>46</td><td>CGGCGACTCTGGACGCGAGCCGGGCCCTTCCCGTGGATCGCCCCAG</td><td>1206.0</td><td>-4.3</td><td>0.7</td><td>-5.7</td><td>9.5E-09</td><td>1.7E-08</td><td>MB231/MB468</td></tr>
<tr><td>>hg38_21_+_8433172_8446622_RNA45SN1_45S_50ntFlanks@12539.12560.22|ATGATGTGTTGTTGCCATGGTA</td><td>>hg38_21_+_8433172_8446622_RNA45SN1_45S_50ntFlanks</td><td>12539</td><td>12560</td><td>22</td><td>ATGATGTGTTGTTGCCATGGTA</td><td>141.5</td><td>-4.3</td><td>0.8</td><td>-5.7</td><td>1.1E-08</td><td>1.7E-08</td><td>BT20/MB468</td></tr>
<tr><td>>hg38_21_+_8433172_8446622_RNA45SN1_45S_50ntFlanks@9545.9564.20|GTGGAGGTCCGTAGCGGTCC</td><td>>hg38_21_+_8433172_8446622_RNA45SN1_45S_50ntFlanks</td><td>9545</td><td>9564</td><td>20</td><td>GTGGAGGTCCGTAGCGGTCC</td><td>485.2</td><td>-3.9</td><td>0.7</td><td>-5.7</td><td>9.5E-09</td><td>1.7E-08</td><td>MB231/MB468</td></tr>
<tr><td>>hg38_21_+_8433172_8446622_RNA45SN1_45S_50ntFlanks@11561.11593.33|GCGCCTAGCAGCCGACTTAGAACTGGTGCGGAC</td><td>>hg38_21_+_8433172_8446622_RNA45SN1_45S_50ntFlanks</td><td>11561</td><td>11593</td><td>33</td><td>GCGCCTAGCAGCCGACTTAGAACTGGTGCGGAC</td><td>262.2</td><td>-3.7</td><td>0.7</td><td>-5.7</td><td>9.6E-09</td><td>1.7E-08</td><td>MB231/MB468</td></tr>
<tr><td>>MI0002468|hsa-mir-484&WithFlank&16|+|15643288|15643378@14.33.20(+2U)&[MIMAT0002174&hsa-miR-484&offsets|0|-2(+2U);m-127&16|+|15643301|15643321&offsets|0|-1(+2U)]|TCAGGCTCAGTCCCCTCCCGTT</td><td>>MI0002468|hsa-mir-484&WithFlank&16|+|15643288|15643378</td><td>14</td><td>33</td><td>20(+2U)</td><td>TCAGGCTCAGTCCCCTCCCGTT</td><td>595.4</td><td>8.1</td><td>1.4</td><td>5.7</td><td>1.1E-08</td><td>1.7E-08</td><td>BT20/MB468</td></tr>
<tr><td>>hg38_21_+_8433172_8446622_RNA45SN1_45S_50ntFlanks@12416.12447.32|TTCGATGTCGGCTCTTCCTATCATTGTGAAGC</td><td>>hg38_21_+_8433172_8446622_RNA45SN1_45S_50ntFlanks</td><td>12416</td><td>12447</td><td>32</td><td>TTCGATGTCGGCTCTTCCTATCATTGTGAAGC</td><td>309.0</td><td>5.9</td><td>1.0</td><td>5.7</td><td>1.1E-08</td><td>1.7E-08</td><td>BT20/MB468</td></tr>
<tr><td>>MI0000446|hsa-mir-125b-1&WithFlank&11|-|122099751|122099850@21.43.23&[MIMAT0000423&hsa-miR-125b-5p&offsets|0|+1;m-67&11|-|122099809|122099830&offsets|0|+1]|TCCCTGAGACCCTAACTTGTGAT</td><td>>MI0000446|hsa-mir-125b-1&WithFlank&11|-|122099751|122099850</td><td>21</td><td>43</td><td>23</td><td>TCCCTGAGACCCTAACTTGTGAT</td><td>2288.2</td><td>-6.5</td><td>1.1</td><td>-6.2</td><td>6.8E-10</td><td>1.7E-08</td><td>BT20/MB231</td></tr>
<tr><td>>hg38_21_+_8433172_8446622_RNA45SN1_45S_50ntFlanks@10788.10818.31|CAGCCTCTGGCATGTTGGAACAATGTAGGTA</td><td>>hg38_21_+_8433172_8446622_RNA45SN1_45S_50ntFlanks</td><td>10788</td><td>10818</td><td>31</td><td>CAGCCTCTGGCATGTTGGAACAATGTAGGTA</td><td>2188.0</td><td>-4.0</td><td>0.7</td><td>-5.7</td><td>1.0E-08</td><td>1.7E-08</td><td>MB231/MB468</td></tr>
<tr><td>>hg38_21_+_8433172_8446622_RNA45SN1_45S_50ntFlanks@6748.6771.24|CTTCGAACGCACTTGCGGCCCCGG</td><td>>hg38_21_+_8433172_8446622_RNA45SN1_45S_50ntFlanks</td><td>6748</td><td>6771</td><td>24</td><td>CTTCGAACGCACTTGCGGCCCCGG</td><td>110.9</td><td>-4.4</td><td>0.8</td><td>-5.7</td><td>1.0E-08</td><td>1.8E-08</td><td>MB231/MB468</td></tr>
<tr><td>>hg38_21_+_8433172_8446622_RNA45SN1_45S_50ntFlanks@10128.10149.22|CGGCGGCGGCGGCGGCGGCGGG</td><td>>hg38_21_+_8433172_8446622_RNA45SN1_45S_50ntFlanks</td><td>10128</td><td>10149</td><td>22</td><td>CGGCGGCGGCGGCGGCGGCGGG</td><td>62.2</td><td>-7.5</td><td>1.3</td><td>-5.7</td><td>1.0E-08</td><td>1.8E-08</td><td>MB231/MB468</td></tr>
<tr><td>>hg38_21_+_8433172_8446622_RNA45SN1_45S_50ntFlanks@8845.8866.22|CGGCGGCGGCGGCGGCGGCGGG</td><td>>hg38_21_+_8433172_8446622_RNA45SN1_45S_50ntFlanks</td><td>8845</td><td>8866</td><td>22</td><td>CGGCGGCGGCGGCGGCGGCGGG</td><td>62.2</td><td>-7.5</td><td>1.3</td><td>-5.7</td><td>1.0E-08</td><td>1.8E-08</td><td>MB231/MB468</td></tr>
<tr><td>>hg38_21_+_8433172_8446622_RNA45SN1_45S_50ntFlanks@10798.10816.19|CATGTTGGAACAATGTAGG</td><td>>hg38_21_+_8433172_8446622_RNA45SN1_45S_50ntFlanks</td><td>10798</td><td>10816</td><td>19</td><td>CATGTTGGAACAATGTAGG</td><td>541.7</td><td>-4.0</td><td>0.7</td><td>-5.7</td><td>1.1E-08</td><td>1.8E-08</td><td>BT20/MB468</td></tr>
<tr><td>>hg38_21_+_8433172_8446622_RNA45SN1_45S_50ntFlanks@10466.10497.32|CGTTGCCCTCGGCCGATCGAAAGGGAGTCGGG</td><td>>hg38_21_+_8433172_8446622_RNA45SN1_45S_50ntFlanks</td><td>10466</td><td>10497</td><td>32</td><td>CGTTGCCCTCGGCCGATCGAAAGGGAGTCGGG</td><td>465.4</td><td>-4.9</td><td>0.9</td><td>-5.7</td><td>1.0E-08</td><td>1.8E-08</td><td>MB231/MB468</td></tr>
<tr><td>>hg38_21_+_8433172_8446622_RNA45SN1_45S_50ntFlanks@11205.11232.28|GCGGCGGCGACTCTGGACGCGAGCCGGG</td><td>>hg38_21_+_8433172_8446622_RNA45SN1_45S_50ntFlanks</td><td>11205</td><td>11232</td><td>28</td><td>GCGGCGGCGACTCTGGACGCGAGCCGGG</td><td>318.7</td><td>-4.1</td><td>0.7</td><td>-5.7</td><td>1.0E-08</td><td>1.8E-08</td><td>MB231/MB468</td></tr>
<tr><td>>hg38_21_+_8433172_8446622_RNA45SN1_45S_50ntFlanks@11741.11759.19|TCTTAAGGTAGCCAAATGC</td><td>>hg38_21_+_8433172_8446622_RNA45SN1_45S_50ntFlanks</td><td>11741</td><td>11759</td><td>19</td><td>TCTTAAGGTAGCCAAATGC</td><td>145.8</td><td>-5.2</td><td>0.9</td><td>-5.7</td><td>1.0E-08</td><td>1.8E-08</td><td>MB231/MB468</td></tr>
<tr><td>>hg38_21_+_8433172_8446622_RNA45SN1_45S_50ntFlanks@7992.8011.20|CGTGGCGACCCGCTGAATTT</td><td>>hg38_21_+_8433172_8446622_RNA45SN1_45S_50ntFlanks</td><td>7992</td><td>8011</td><td>20</td><td>CGTGGCGACCCGCTGAATTT</td><td>3152.3</td><td>5.0</td><td>0.9</td><td>5.7</td><td>1.0E-08</td><td>1.8E-08</td><td>MB231/MB468</td></tr>
<tr><td>>am_MI0000815|hsa-mir-339&WithFlank&7|-|1022927|1023032@21.41.21&[MIMAT0000764&hsa-miR-339-5p&offsets|0|-2;m-163&7|-|1022990|1023012&offsets|0|-2]|TCCCTGTCCTCCAGGAGCTCA</td><td>>am_MI0000815|hsa-mir-339&WithFlank&7|-|1022927|1023032</td><td>21</td><td>41</td><td>21</td><td>TCCCTGTCCTCCAGGAGCTCA</td><td>928.3</td><td>9.0</td><td>1.6</td><td>5.7</td><td>1.2E-08</td><td>1.8E-08</td><td>BT20/MB468</td></tr>
<tr><td>>hg38_21_+_8433172_8446622_RNA45SN1_45S_50ntFlanks@9033.9059.27|CTCGTGGGGGGGCCGGGCCACCCCTCC</td><td>>hg38_21_+_8433172_8446622_RNA45SN1_45S_50ntFlanks</td><td>9033</td><td>9059</td><td>27</td><td>CTCGTGGGGGGGCCGGGCCACCCCTCC</td><td>170.8</td><td>-5.8</td><td>1.0</td><td>-5.7</td><td>1.2E-08</td><td>1.8E-08</td><td>BT20/MB468</td></tr>
<tr><td>>hg38_21_+_8433172_8446622_RNA45SN1_45S_50ntFlanks@9575.9594.20|ATCGGTCGTCCGACCTGGGT</td><td>>hg38_21_+_8433172_8446622_RNA45SN1_45S_50ntFlanks</td><td>9575</td><td>9594</td><td>20</td><td>ATCGGTCGTCCGACCTGGGT</td><td>125.8</td><td>-4.5</td><td>0.8</td><td>-5.7</td><td>1.1E-08</td><td>1.8E-08</td><td>MB231/MB468</td></tr>
<tr><td>>hg38_21_+_8433172_8446622_RNA45SN1_45S_50ntFlanks@12083.12107.25|CTCGCTTCTGGCGCCAAGCGCCCGG</td><td>>hg38_21_+_8433172_8446622_RNA45SN1_45S_50ntFlanks</td><td>12083</td><td>12107</td><td>25</td><td>CTCGCTTCTGGCGCCAAGCGCCCGG</td><td>265.2</td><td>-3.1</td><td>0.5</td><td>-5.7</td><td>1.1E-08</td><td>1.8E-08</td><td>MB231/MB468</td></tr>
<tr><td>>hg38_21_+_8433172_8446622_RNA45SN1_45S_50ntFlanks@5473.5516.44|CGGCCCTGGCGGAGCGCTGAGAAGACGGTCGAACTTGACTATCT</td><td>>hg38_21_+_8433172_8446622_RNA45SN1_45S_50ntFlanks</td><td>5473</td><td>5516</td><td>44</td><td>CGGCCCTGGCGGAGCGCTGAGAAGACGGTCGAACTTGACTATCT</td><td>597.7</td><td>8.3</td><td>1.5</td><td>5.7</td><td>1.2E-08</td><td>1.8E-08</td><td>BT20/MB468</td></tr>
<tr><td>>hg38_21_+_8433172_8446622_RNA45SN1_45S_50ntFlanks@11540.11580.41|CGCGGCGCCTCGCCTCGGCCGGCGCCTAGCAGCCGACTTAG</td><td>>hg38_21_+_8433172_8446622_RNA45SN1_45S_50ntFlanks</td><td>11540</td><td>11580</td><td>41</td><td>CGCGGCGCCTCGCCTCGGCCGGCGCCTAGCAGCCGACTTAG</td><td>626.9</td><td>3.5</td><td>0.6</td><td>5.7</td><td>1.1E-08</td><td>1.8E-08</td><td>MB231/MB468</td></tr>
<tr><td>>hg38_21_+_8433172_8446622_RNA45SN1_45S_50ntFlanks@12072.12095.24|CCCGAGGGGCTCTCGCTTCTGGCG</td><td>>hg38_21_+_8433172_8446622_RNA45SN1_45S_50ntFlanks</td><td>12072</td><td>12095</td><td>24</td><td>CCCGAGGGGCTCTCGCTTCTGGCG</td><td>838.1</td><td>-4.4</td><td>0.8</td><td>-5.7</td><td>1.1E-08</td><td>1.9E-08</td><td>MB231/MB468</td></tr>
</table>

<table>
<tr><td>>trnaMT_HisGTG_MT_+_12138_12206@32.61.30|TGAATCTGACAACAGAGGCTTACGACCCCT</td><td>>trnaMT_HisGTG_MT_+_12138_12206</td><td>32</td><td>61</td><td>30</td><td>TGAATCTGACAACAGAGGCTTACGACCCCT</td><td>332.2</td><td>5.1</td><td>0.9</td><td>5.7</td><td>1.2E-08</td><td>1.9E-08</td><td>BT20/MB468</td></tr>
<tr><td>>hg38_21_+_8433172_8446622_RNA45SN1_45S_50ntFlanks@11666.11687.22|TCTGCCCAGTGCTCTGAATGTC</td><td>>hg38_21_+_8433172_8446622_RNA45SN1_45S_50ntFlanks</td><td>11666</td><td>11687</td><td>22</td><td>TCTGCCCAGTGCTCTGAATGTC</td><td>416.3</td><td>3.1</td><td>0.6</td><td>5.7</td><td>1.2E-08</td><td>1.9E-08</td><td>BT20/MB468</td></tr>
<tr><td>>hg38_21_+_8433172_8446622_RNA45SN1_45S_50ntFlanks@6649.6698.50|TACGACTCTTAGCGGTGGATCACTCGGCTCGTGCGTCGATGAAGAACGCA</td><td>>hg38_21_+_8433172_8446622_RNA45SN1_45S_50ntFlanks</td><td>6649</td><td>6698</td><td>50</td><td>TACGACTCTTAGCGGTGGATCACTCGGCTCGTGCGTCGATGAAGAACGCA</td><td>847.9</td><td>4.8</td><td>0.8</td><td>5.7</td><td>1.1E-08</td><td>1.9E-08</td><td>MB231/MB468</td></tr>
<tr><td>>hg38_21_+_8433172_8446622_RNA45SN1_45S_50ntFlanks@12474.12497.24|ACCCACTAATAGGGAACGTGAGCT</td><td>>hg38_21_+_8433172_8446622_RNA45SN1_45S_50ntFlanks</td><td>12474</td><td>12497</td><td>24</td><td>ACCCACTAATAGGGAACGTGAGCT</td><td>1874.7</td><td>4.1</td><td>0.7</td><td>5.7</td><td>1.1E-08</td><td>1.9E-08</td><td>MB231/MB468</td></tr>
<tr><td>>hg38_21_+_8433172_8446622_RNA45SN1_45S_50ntFlanks@8198.8225.28|TGGACGGTGTGAGGCCGGTAGCGGCCCC</td><td>>hg38_21_+_8433172_8446622_RNA45SN1_45S_50ntFlanks</td><td>8198</td><td>8225</td><td>28</td><td>TGGACGGTGTGAGGCCGGTAGCGGCCCC</td><td>211.4</td><td>-2.6</td><td>0.5</td><td>-5.7</td><td>1.1E-08</td><td>1.9E-08</td><td>MB231/MB468</td></tr>
<tr><td>>hg38_21_+_8433172_8446622_RNA45SN1_45S_50ntFlanks@10841.10873.33|CGTAACTTCGGGATAAGGATTGGCTCTAAGGGC</td><td>>hg38_21_+_8433172_8446622_RNA45SN1_45S_50ntFlanks</td><td>10841</td><td>10873</td><td>33</td><td>CGTAACTTCGGGATAAGGATTGGCTCTAAGGGC</td><td>277.0</td><td>-3.0</td><td>0.5</td><td>-5.7</td><td>1.2E-08</td><td>1.9E-08</td><td>BT20/MB468</td></tr>
<tr><td>>hg38_21_+_8433172_8446622_RNA45SN1_45S_50ntFlanks@6650.6698.49|ACGACTCTTAGCGGTGGATCACTCGGCTCGTGCGTCGATGAAGAACGCA</td><td>>hg38_21_+_8433172_8446622_RNA45SN1_45S_50ntFlanks</td><td>6650</td><td>6698</td><td>49</td><td>ACGACTCTTAGCGGTGGATCACTCGGCTCGTGCGTCGATGAAGAACGCA</td><td>2245.6</td><td>4.6</td><td>0.8</td><td>5.7</td><td>1.1E-08</td><td>1.9E-08</td><td>MB231/MB468</td></tr>
<tr><td>>hg38_21_+_8433172_8446622_RNA45SN1_45S_50ntFlanks@4726.4752.27|TAATCAAGAACGAAAGTCGGAGGTTCG</td><td>>hg38_21_+_8433172_8446622_RNA45SN1_45S_50ntFlanks</td><td>4726</td><td>4752</td><td>27</td><td>TAATCAAGAACGAAAGTCGGAGGTTCG</td><td>200.2</td><td>-3.8</td><td>0.7</td><td>-5.7</td><td>1.1E-08</td><td>1.9E-08</td><td>MB231/MB468</td></tr>
<tr><td>>hg38_21_+_8433172_8446622_RNA45SN1_45S_50ntFlanks@12445.12480.36|AGCAGAATTCACCAAGCGTTGGATTGTTCACCCACT</td><td>>hg38_21_+_8433172_8446622_RNA45SN1_45S_50ntFlanks</td><td>12445</td><td>12480</td><td>36</td><td>AGCAGAATTCACCAAGCGTTGGATTGTTCACCCACT</td><td>389.0</td><td>3.6</td><td>0.6</td><td>5.7</td><td>1.1E-08</td><td>1.9E-08</td><td>MB231/MB468</td></tr>
<tr><td>>trnaMT_SerGCT_MT_+_12207_12265@1.38.38|GAGAAAGCTCACAAGAACTGCTAACTCATGCCCCCATG</td><td>>trnaMT_SerGCT_MT_+_12207_12265</td><td>1</td><td>38</td><td>38</td><td>GAGAAAGCTCACAAGAACTGCTAACTCATGCCCCCATG</td><td>808.6</td><td>4.8</td><td>0.9</td><td>5.7</td><td>1.2E-08</td><td>1.9E-08</td><td>BT20/MB468</td></tr>
<tr><td>>MI0000098|hsa-mir-96&WithFlank&7|-|129774686|129774775@15.36.22&[MIMAT0000095&hsa-miR-96-5p&offsets|0|-1;m-248&7|-|129774739|129774761&offsets|0|-1]|TTTGGCACTAGCACATTTTTGC</td><td>>MI0000098|hsa-mir-96&WithFlank&7|-|129774686|129774775</td><td>15</td><td>36</td><td>22</td><td>TTTGGCACTAGCACATTTTTGC</td><td>1015.7</td><td>9.0</td><td>1.6</td><td>5.7</td><td>1.3E-08</td><td>1.9E-08</td><td>BT20/MB468</td></tr>
<tr><td>>hg38_MT_+_1621_3279_RNR2_16S_50ntFlanks@1404.1421.18|TATTAAAGGTTCGTTTGT</td><td>>hg38_MT_+_1621_3279_RNR2_16S_50ntFlanks</td><td>1404</td><td>1421</td><td>18</td><td>TATTAAAGGTTCGTTTGT</td><td>1033.4</td><td>5.2</td><td>0.9</td><td>5.7</td><td>1.3E-08</td><td>1.9E-08</td><td>BT20/MB468</td></tr>
<tr><td>>MI0000085|hsa-mir-27a&WithFlank&19|-|13836434|13836523@57.76.20(+1A)&[MIMAT0000084&hsa-miR-27a-3p&offsets|0|-1(+1A);m-43&19|-|13836447|13836467&offsets|0|-1(+1A)]|TTCACAGTGGCTAAGTTCCGA</td><td>>MI0000085|hsa-mir-27a&WithFlank&19|-|13836434|13836523</td><td>57</td><td>76</td><td>20(+1A)</td><td>TTCACAGTGGCTAAGTTCCGA</td><td>4947.2</td><td>4.8</td><td>0.8</td><td>5.7</td><td>1.3E-08</td><td>2.0E-08</td><td>BT20/MB468</td></tr>
<tr><td>>hg38_21_+_8433172_8446622_RNA45SN1_45S_50ntFlanks@10846.10866.21|CTTCGGGATAAGGATTGGCTC</td><td>>hg38_21_+_8433172_8446622_RNA45SN1_45S_50ntFlanks</td><td>10846</td><td>10866</td><td>21</td><td>CTTCGGGATAAGGATTGGCTC</td><td>143.3</td><td>-3.7</td><td>0.7</td><td>-5.7</td><td>1.1E-08</td><td>2.0E-08</td><td>MB231/MB468</td></tr>
<tr><td>>hg38_21_+_8433172_8446622_RNA45SN1_45S_50ntFlanks@8180.8211.32|TGATCGAGGCCCAGCCCGTGGACGGTGTGAGG</td><td>>hg38_21_+_8433172_8446622_RNA45SN1_45S_50ntFlanks</td><td>8180</td><td>8211</td><td>32</td><td>TGATCGAGGCCCAGCCCGTGGACGGTGTGAGG</td><td>159.3</td><td>-5.4</td><td>0.9</td><td>-5.7</td><td>1.3E-08</td><td>2.0E-08</td><td>BT20/MB468</td></tr>
<tr><td>>hg38_21_+_8433172_8446622_RNA45SN1_45S_50ntFlanks@11876.11900.25|AAGACCCTGTTGAGCTTGACTCTAG</td><td>>hg38_21_+_8433172_8446622_RNA45SN1_45S_50ntFlanks</td><td>11876</td><td>11900</td><td>25</td><td>AAGACCCTGTTGAGCTTGACTCTAG</td><td>281.5</td><td>-3.5</td><td>0.6</td><td>-5.7</td><td>1.1E-08</td><td>2.0E-08</td><td>MB231/MB468</td></tr>
<tr><td>>hg38_21_+_8433172_8446622_RNA45SN1_45S_50ntFlanks@5467.5499.33|GGCCCACGGCCCTGGCGGAGCGCTGAGAAGACG</td><td>>hg38_21_+_8433172_8446622_RNA45SN1_45S_50ntFlanks</td><td>5467</td><td>5499</td><td>33</td><td>GGCCCACGGCCCTGGCGGAGCGCTGAGAAGACG</td><td>877.0</td><td>-4.3</td><td>0.8</td><td>-5.7</td><td>1.2E-08</td><td>2.0E-08</td><td>MB231/MB468</td></tr>
<tr><td>>hg38_21_+_8433172_8446622_RNA45SN1_45S_50ntFlanks@485.510.26|GATGTGGTGACGTCGTGCTCTCCCGG</td><td>>hg38_21_+_8433172_8446622_RNA45SN1_45S_50ntFlanks</td><td>485</td><td>510</td><td>26</td><td>GATGTGGTGACGTCGTGCTCTCCCGG</td><td>4494.4</td><td>17.0</td><td>3.0</td><td>5.7</td><td>1.2E-08</td><td>2.0E-08</td><td>MB231/MB468</td></tr>
<tr><td>>hg38_21_+_8433172_8446622_RNA45SN1_45S_50ntFlanks@10403.10430.28|ACATGGGTCAGTCGGTCCTGAGAGATGG</td><td>>hg38_21_+_8433172_8446622_RNA45SN1_45S_50ntFlanks</td><td>10403</td><td>10430</td><td>28</td><td>ACATGGGTCAGTCGGTCCTGAGAGATGG</td><td>136.3</td><td>-2.5</td><td>0.4</td><td>-5.7</td><td>1.2E-08</td><td>2.0E-08</td><td>MB231/MB468</td></tr>
<tr><td>>hg38_21_+_8433172_8446622_RNA45SN1_45S_50ntFlanks@11890.11907.18|CTTGACTCTAGTCTGGCA</td><td>>hg38_21_+_8433172_8446622_RNA45SN1_45S_50ntFlanks</td><td>11890</td><td>11907</td><td>18</td><td>CTTGACTCTAGTCTGGCA</td><td>560.1</td><td>-5.0</td><td>0.9</td><td>-5.7</td><td>1.2E-08</td><td>2.0E-08</td><td>MB231/MB468</td></tr>
<tr><td>>hg38_21_+_8433172_8446622_RNA45SN1_45S_50ntFlanks@12488.12520.33|AACGTGAGCTGGGTTTAGACCGTCGTGAGACAG</td><td>>hg38_21_+_8433172_8446622_RNA45SN1_45S_50ntFlanks</td><td>12488</td><td>12520</td><td>33</td><td>AACGTGAGCTGGGTTTAGACCGTCGTGAGACAG</td><td>321.8</td><td>-4.7</td><td>0.8</td><td>-5.7</td><td>1.2E-08</td><td>2.0E-08</td><td>MB231/MB468</td></tr>
<tr><td>>MI0000253|hsa-mir-148a&WithFlank&7|-|25949913|25949992@50.72.23&[MIMAT0000243&hsa-miR-148a-3p&offsets|0|+1;m-7&7|-|25949922|25949943&offsets|0|+1]|TCAGTGCACTACAGAACTTTGTC</td><td>>MI0000253|hsa-mir-148a&WithFlank&7|-|25949913|25949992</td><td>50</td><td>72</td><td>23</td><td>TCAGTGCACTACAGAACTTTGTC</td><td>4552.6</td><td>4.7</td><td>0.8</td><td>6.1</td><td>8.1E-10</td><td>2.0E-08</td><td>BT20/MB231</td></tr>
<tr><td>>MI0000253|hsa-mir-148a&WithFlank&7|-|25949913|25949992@50.70.21(+1C)&[MIMAT0000243&hsa-miR-148a-3p&offsets|0|-1(+1C);m-7&7|-|25949922|25949943&offsets|0|-1(+1C)]|TCAGTGCACTACAGAACTTTGC</td><td>>MI0000253|hsa-mir-148a&WithFlank&7|-|25949913|25949992</td><td>50</td><td>70</td><td>21(+1C)</td><td>TCAGTGCACTACAGAACTTTGC</td><td>39890.8</td><td>4.5</td><td>0.7</td><td>6.1</td><td>8.2E-10</td><td>2.0E-08</td><td>BT20/MB231</td></tr>
<tr><td>>hg38_21_+_8433172_8446622_RNA45SN1_45S_50ntFlanks@9573.9595.23|AAATCGGTCGTCCGACCTGGGTA</td><td>>hg38_21_+_8433172_8446622_RNA45SN1_45S_50ntFlanks</td><td>9573</td><td>9595</td><td>23</td><td>AAATCGGTCGTCCGACCTGGGTA</td><td>1332.5</td><td>-4.1</td><td>0.7</td><td>-5.7</td><td>1.3E-08</td><td>2.0E-08</td><td>BT20/MB468</td></tr>
<tr><td>>hg38_21_+_8433172_8446622_RNA45SN1_45S_50ntFlanks@10761.10785.25|ATATCCGCAGCAGGTCTCCAAGGTG</td><td>>hg38_21_+_8433172_8446622_RNA45SN1_45S_50ntFlanks</td><td>10761</td><td>10785</td><td>25</td><td>ATATCCGCAGCAGGTCTCCAAGGTG</td><td>104.3</td><td>-2.9</td><td>0.5</td><td>-5.7</td><td>1.2E-08</td><td>2.1E-08</td><td>MB231/MB468</td></tr>
<tr><td>>hg38_21_+_8433172_8446622_RNA45SN1_45S_50ntFlanks@11569.11591.23|CAGCCGACTTAGAACTGGTGCGG</td><td>>hg38_21_+_8433172_8446622_RNA45SN1_45S_50ntFlanks</td><td>11569</td><td>11591</td><td>23</td><td>CAGCCGACTTAGAACTGGTGCGG</td><td>222.1</td><td>-5.7</td><td>1.0</td><td>-5.7</td><td>1.3E-08</td><td>2.1E-08</td><td>BT20/MB468</td></tr>
<tr><td>>hg38_1_-_228634819_228635039_RNA5S12_5S_50ntFlanks@125.171.47|GTACTTGGATGGGAGACCGCCTGGGAATACCGGGTGCTGTAGGCTTT</td><td>>hg38_1_-_228634819_228635039_RNA5S12_5S_50ntFlanks</td><td>125</td><td>171</td><td>47</td><td>GTACTTGGATGGGAGACCGCCTGGGAATACCGGGTGCTGTAGGCTTT</td><td>733.5</td><td>12.3</td><td>2.2</td><td>5.7</td><td>1.2E-08</td><td>2.1E-08</td><td>MB231/MB468</td></tr>
<tr><td>>trna14_LysTTT_11_-_59327808_59327880@1.30.30,trna2_LysTTT_17_+_8022473_8022545@1.30.30,trna54_LysTTT_1_+_204475655_204475727@1.30.30,trna5_LysTTT_11_+_59323902_59323974@1.30.30,trna62_LysTTT_1_-_204476158_204476230@1.30.30,trna76_LysTTT_6_+_28918806_28918878@1.30.30|GCCCGGATAGCTCAGTCGGTAGAGCATCAG</td><td>>trna14_LysTTT_11_-_59327808_59327880</td><td>1</td><td>30</td><td>30</td><td>GCCCGGATAGCTCAGTCGGTAGAGCATCAG</td><td>774.2</td><td>-4.3</td><td>0.7</td><td>-5.7</td><td>1.2E-08</td><td>2.1E-08</td><td>MB231/MB468</td></tr>
<tr><td>>hg38_21_+_8433172_8446622_RNA45SN1_45S_50ntFlanks@4047.4081.35|ATTCGAACGTCTGCCCTATCAACTTTCGATGGTAG</td><td>>hg38_21_+_8433172_8446622_RNA45SN1_45S_50ntFlanks</td><td>4047</td><td>4081</td><td>35</td><td>ATTCGAACGTCTGCCCTATCAACTTTCGATGGTAG</td><td>80.8</td><td>-6.1</td><td>1.1</td><td>-5.7</td><td>1.2E-08</td><td>2.1E-08</td><td>MB231/MB468</td></tr>
<tr><td>>hg38_21_+_8433172_8446622_RNA45SN1_45S_50ntFlanks@10349.10384.36|AACGAGAACTTTGAAGGCCGAAGTGGAGAAGGGTTC</td><td>>hg38_21_+_8433172_8446622_RNA45SN1_45S_50ntFlanks</td><td>10349</td><td>10384</td><td>36</td><td>AACGAGAACTTTGAAGGCCGAAGTGGAGAAGGGTTC</td><td>535.0</td><td>-2.2</td><td>0.4</td><td>-5.7</td><td>1.3E-08</td><td>2.1E-08</td><td>BT20/MB468</td></tr>
<tr><td>>hg38_21_+_8433172_8446622_RNA45SN1_45S_50ntFlanks@9339.9370.32|CGCACGAAAGCCGCCGTGGCGCAATGAAGGTG</td><td>>hg38_21_+_8433172_8446622_RNA45SN1_45S_50ntFlanks</td><td>9339</td><td>9370</td><td>32</td><td>CGCACGAAAGCCGCCGTGGCGCAATGAAGGTG</td><td>70.7</td><td>-7.7</td><td>1.4</td><td>-5.7</td><td>1.2E-08</td><td>2.1E-08</td><td>MB231/MB468</td></tr>
<tr><td>>hg38_21_+_8433172_8446622_RNA45SN1_45S_50ntFlanks@10781.10808.28|AGGTGAACAGCCTCTGGCATGTTGGAAC</td><td>>hg38_21_+_8433172_8446622_RNA45SN1_45S_50ntFlanks</td><td>10781</td><td>10808</td><td>28</td><td>AGGTGAACAGCCTCTGGCATGTTGGAAC</td><td>97.6</td><td>-4.9</td><td>0.9</td><td>-5.7</td><td>1.2E-08</td><td>2.1E-08</td><td>MB231/MB468</td></tr>
<tr><td>>hg38_21_+_8433172_8446622_RNA45SN1_45S_50ntFlanks@8179.8208.30|CTGATCGAGGCCCAGCCCGTGGACGGTGTG</td><td>>hg38_21_+_8433172_8446622_RNA45SN1_45S_50ntFlanks</td><td>8179</td><td>8208</td><td>30</td><td>CTGATCGAGGCCCAGCCCGTGGACGGTGTG</td><td>120.4</td><td>-2.9</td><td>0.5</td><td>-5.7</td><td>1.2E-08</td><td>2.1E-08</td><td>MB231/MB468</td></tr>
<tr><td>>hg38_21_+_8433172_8446622_RNA45SN1_45S_50ntFlanks@11581.11601.21|AACTGGTGCGGACCAGGGGAA</td><td>>hg38_21_+_8433172_8446622_RNA45SN1_45S_50ntFlanks</td><td>11581</td><td>11601</td><td>21</td><td>AACTGGTGCGGACCAGGGGAA</td><td>975.2</td><td>-4.2</td><td>0.7</td><td>-5.7</td><td>1.2E-08</td><td>2.1E-08</td><td>MB231/MB468</td></tr>
<tr><td>>hg38_21_+_8433172_8446622_RNA45SN1_45S_50ntFlanks@10101.10125.25|CGCGCGCGTGTGGTGTGCGTCGGAG</td><td>>hg38_21_+_8433172_8446622_RNA45SN1_45S_50ntFlanks</td><td>10101</td><td>10125</td><td>25</td><td>CGCGCGCGTGTGGTGTGCGTCGGAG</td><td>327.1</td><td>-6.3</td><td>1.1</td><td>-5.7</td><td>1.4E-08</td><td>2.1E-08</td><td>BT20/MB468</td></tr>
<tr><td>>hg38_21_+_8433172_8446622_RNA45SN1_45S_50ntFlanks@3813.3837.25|TTATGGTTCCTTTGGTCGCTCGCTC</td><td>>hg38_21_+_8433172_8446622_RNA45SN1_45S_50ntFlanks</td><td>3813</td><td>3837</td><td>25</td><td>TTATGGTTCCTTTGGTCGCTCGCTC</td><td>90.6</td><td>-2.9</td><td>0.5</td><td>-5.7</td><td>1.4E-08</td><td>2.2E-08</td><td>BT20/MB468</td></tr>
<tr><td>>hg38_21_+_8433172_8446622_RNA45SN1_45S_50ntFlanks@6651.6690.40|CGACTCTTAGCGGTGGATCACTCGGCTCGTGCGTCGATGA</td><td>>hg38_21_+_8433172_8446622_RNA45SN1_45S_50ntFlanks</td><td>6651</td><td>6690</td><td>40</td><td>CGACTCTTAGCGGTGGATCACTCGGCTCGTGCGTCGATGA</td><td>2416.1</td><td>4.1</td><td>0.7</td><td>5.7</td><td>1.4E-08</td><td>2.2E-08</td><td>BT20/MB468</td></tr>
<tr><td>>hg38_21_+_8433172_8446622_RNA45SN1_45S_50ntFlanks@11885.11903.19|TTGAGCTTGACTCTAGTCT</td><td>>hg38_21_+_8433172_8446622_RNA45SN1_45S_50ntFlanks</td><td>11885</td><td>11903</td><td>19</td><td>TTGAGCTTGACTCTAGTCT</td><td>446.3</td><td>-6.8</td><td>1.2</td><td>-5.7</td><td>1.4E-08</td><td>2.2E-08</td><td>BT20/MB468</td></tr>
<tr><td>>MI0000471|hsa-mir-126&WithFlank&9|+|136670596|136670692@58.78.21&[MIMAT0000445&hsa-miR-126-3p&offsets|0|-1;m-40&9|+|136670653|136670674&offsets|0|-1]|TCGTACCGTGAGTAATAATGC</td><td>>MI0000471|hsa-mir-126&WithFlank&9|+|136670596|136670692</td><td>58</td><td>78</td><td>21</td><td>TCGTACCGTGAGTAATAATGC</td><td>5789.6</td><td>9.4</td><td>1.7</td><td>5.7</td><td>1.4E-08</td><td>2.2E-08</td><td>BT20/MB468</td></tr>
<tr><td>>hg38_21_+_8433172_8446622_RNA45SN1_45S_50ntFlanks@9821.9850.30|CCTAGTGGGCCACTTTTGGTAAGCAGAACT</td><td>>hg38_21_+_8433172_8446622_RNA45SN1_45S_50ntFlanks</td><td>9821</td><td>9850</td><td>30</td><td>CCTAGTGGGCCACTTTTGGTAAGCAGAACT</td><td>257.3</td><td>-3.3</td><td>0.6</td><td>-5.7</td><td>1.4E-08</td><td>2.2E-08</td><td>BT20/MB468</td></tr>
<tr><td>>hg38_21_+_8433172_8446622_RNA45SN1_45S_50ntFlanks@12447.12475.29|CAGAATTCACCAAGCGTTGGATTGTTCAC</td><td>>hg38_21_+_8433172_8446622_RNA45SN1_45S_50ntFlanks</td><td>12447</td><td>12475</td><td>29</td><td>CAGAATTCACCAAGCGTTGGATTGTTCAC</td><td>126.7</td><td>-6.1</td><td>1.1</td><td>-5.7</td><td>1.3E-08</td><td>2.2E-08</td><td>MB231/MB468</td></tr>
<tr><td>>hg38_21_+_8433172_8446622_RNA45SN1_45S_50ntFlanks@10463.10483.21|CTCCGTTGCCCTCGGCCGATC</td><td>>hg38_21_+_8433172_8446622_RNA45SN1_45S_50ntFlanks</td><td>10463</td><td>10483</td><td>21</td><td>CTCCGTTGCCCTCGGCCGATC</td><td>123.8</td><td>-3.3</td><td>0.6</td><td>-5.7</td><td>1.3E-08</td><td>2.2E-08</td><td>MB231/MB468</td></tr>
<tr><td>>MI0000736|hsa-mir-30c-1&WithFlank&1|+|40757278|40757378@23.46.24(+1U);MI0000254|hsa-mir-30c-2&WithFlank&6|-|71376954|71377037@13.36.24(+1U)&[MIMAT0000244&hsa-miR-30c-5p&offsets|0|+1(+1U);m-57&1|+|40757300|40757323&offsets|0|0(+1U)];[MIMAT0000244_1&hsa-miR-30c-5p&offsets|0|+1(+1U);m-56&6|-|71377002|71377025&offsets|0|0(+1U)]|TGTAAACATCCTACACTCTCAGCTT</td><td>>MI0000736|hsa-mir-30c-1&WithFlank&1|+|40757278|40757378</td><td>23</td><td>46</td><td>24(+1U)</td><td>TGTAAACATCCTACACTCTCAGCTT</td><td>727.0</td><td>4.9</td><td>0.9</td><td>5.7</td><td>1.5E-08</td><td>2.2E-08</td><td>BT20/MB468</td></tr>
<tr><td>>hg38_1_-_228634819_228635039_RNA5S12_5S_50ntFlanks@140.169.30|ACCGCCTGGGAATACCGGGTGCTGTAGGCT</td><td>>hg38_1_-_228634819_228635039_RNA5S12_5S_50ntFlanks</td><td>140</td><td>169</td><td>30</td><td>ACCGCCTGGGAATACCGGGTGCTGTAGGCT</td><td>7986.6</td><td>4.3</td><td>0.8</td><td>5.7</td><td>1.5E-08</td><td>2.3E-08</td><td>BT20/MB468</td></tr>
</table>

<table>
<tr><th></th><th></th><th></th><th></th><th></th><th></th><th></th><th></th><th></th><th></th><th></th><th></th><th></th><th></th></tr>
<tr><td>>hg38_21_+_8433172_8446622_RNA45SN1_45S_50ntFlanks@9381.9409.29|CGCTCGCCGGCCGAGGTGGGATCCCGAGG</td><td>>hg38_21_+_8433172_8446622_RNA45SN1_45S_50ntFlanks</td><td>9381</td><td>9409</td><td>29</td><td>CGCTCGCCGGCCGAGGTGGGATCCCGAGG</td><td>156.0</td><td>-4.7</td><td>0.8</td><td>-5.7</td><td>1.3E-08</td><td>2.3E-08</td><td>MB231/MB468</td></tr>
<tr><td>>hg38_21_+_8433172_8446622_RNA45SN1_45S_50ntFlanks@11234.11252.19|CCTTCCCGTGGATCGCCCC</td><td>>hg38_21_+_8433172_8446622_RNA45SN1_45S_50ntFlanks</td><td>11234</td><td>11252</td><td>19</td><td>CCTTCCCGTGGATCGCCCC</td><td>95.2</td><td>-4.9</td><td>0.9</td><td>-5.7</td><td>1.5E-08</td><td>2.3E-08</td><td>BT20/MB468</td></tr>
<tr><td>>hg38_21_+_8433172_8446622_RNA45SN1_45S_50ntFlanks@9738.9767.30|GAAACGATCTCAACCTATTCTCAAACTTTA</td><td>>hg38_21_+_8433172_8446622_RNA45SN1_45S_50ntFlanks</td><td>9738</td><td>9767</td><td>30</td><td>GAAACGATCTCAACCTATTCTCAAACTTTA</td><td>467.2</td><td>-3.1</td><td>0.5</td><td>-5.7</td><td>1.5E-08</td><td>2.3E-08</td><td>BT20/MB468</td></tr>
<tr><td>>hg38_1_-_228634819_228635039_RNA5S12_5S_50ntFlanks@132.169.38|GATGGGAGACCGCCTGGGAATACCGGGTGCTGTAGGCT</td><td>>hg38_1_-_228634819_228635039_RNA5S12_5S_50ntFlanks</td><td>132</td><td>169</td><td>38</td><td>GATGGGAGACCGCCTGGGAATACCGGGTGCTGTAGGCT</td><td>3923.0</td><td>4.4</td><td>0.8</td><td>5.7</td><td>1.5E-08</td><td>2.3E-08</td><td>BT20/MB468</td></tr>
<tr><td>>hg38_21_+_8433172_8446622_RNA45SN1_45S_50ntFlanks@11591.11611.21|GACCAGGGGAATCCGACTGTT</td><td>>hg38_21_+_8433172_8446622_RNA45SN1_45S_50ntFlanks</td><td>11591</td><td>11611</td><td>21</td><td>GACCAGGGGAATCCGACTGTT</td><td>140.8</td><td>-3.0</td><td>0.5</td><td>-5.7</td><td>1.5E-08</td><td>2.3E-08</td><td>BT20/MB468</td></tr>
<tr><td>>hg38_21_+_8433172_8446622_RNA45SN1_45S_50ntFlanks@12076.12095.20|AGGGGCTCTCGCTTCTGGCG</td><td>>hg38_21_+_8433172_8446622_RNA45SN1_45S_50ntFlanks</td><td>12076</td><td>12095</td><td>20</td><td>AGGGGCTCTCGCTTCTGGCG</td><td>2474.5</td><td>-3.2</td><td>0.6</td><td>-5.7</td><td>1.3E-08</td><td>2.3E-08</td><td>MB231/MB468</td></tr>
<tr><td>>hg38_1_-_228634819_228635039_RNA5S12_5S_50ntFlanks@132.170.39|GATGGGAGACCGCCTGGGAATACCGGGTGCTGTAGGCTT</td><td>>hg38_1_-_228634819_228635039_RNA5S12_5S_50ntFlanks</td><td>132</td><td>170</td><td>39</td><td>GATGGGAGACCGCCTGGGAATACCGGGTGCTGTAGGCTT</td><td>26534.5</td><td>4.4</td><td>0.8</td><td>5.7</td><td>1.3E-08</td><td>2.3E-08</td><td>MB231/MB468</td></tr>
<tr><td>>hg38_21_+_8433172_8446622_RNA45SN1_45S_50ntFlanks@6773.6803.31|TTCCTCCCGGGGCTACGCCTGTCTGAGCGTC</td><td>>hg38_21_+_8433172_8446622_RNA45SN1_45S_50ntFlanks</td><td>6773</td><td>6803</td><td>31</td><td>TTCCTCCCGGGGCTACGCCTGTCTGAGCGTC</td><td>1118.0</td><td>-4.3</td><td>0.8</td><td>-5.7</td><td>1.3E-08</td><td>2.3E-08</td><td>MB231/MB468</td></tr>
<tr><td>>hg38_21_+_8433172_8446622_RNA45SN1_45S_50ntFlanks@10391.10430.40|AACAGCAGTTGAACATGGGTCAGTCGGTCCTGAGAGATGG</td><td>>hg38_21_+_8433172_8446622_RNA45SN1_45S_50ntFlanks</td><td>10391</td><td>10430</td><td>40</td><td>AACAGCAGTTGAACATGGGTCAGTCGGTCCTGAGAGATGG</td><td>328.0</td><td>-6.1</td><td>1.1</td><td>-5.7</td><td>1.5E-08</td><td>2.3E-08</td><td>BT20/MB468</td></tr>
<tr><td>>hg38_21_+_8433172_8446622_RNA45SN1_45S_50ntFlanks@12416.12441.26|TTCGATGTCGGCTCTTCCTATCATTG</td><td>>hg38_21_+_8433172_8446622_RNA45SN1_45S_50ntFlanks</td><td>12416</td><td>12441</td><td>26</td><td>TTCGATGTCGGCTCTTCCTATCATTG</td><td>1909.3</td><td>4.1</td><td>0.7</td><td>5.7</td><td>1.4E-08</td><td>2.3E-08</td><td>MB231/MB468</td></tr>
<tr><td>>hg38_21_+_8433172_8446622_RNA45SN1_45S_50ntFlanks@6731.6754.24|CACATTGATCATCGACACTTCGAA</td><td>>hg38_21_+_8433172_8446622_RNA45SN1_45S_50ntFlanks</td><td>6731</td><td>6754</td><td>24</td><td>CACATTGATCATCGACACTTCGAA</td><td>157.9</td><td>-3.1</td><td>0.5</td><td>-5.7</td><td>1.5E-08</td><td>2.4E-08</td><td>BT20/MB468</td></tr>
<tr><td>>hg38_21_+_8433172_8446622_RNA45SN1_45S_50ntFlanks@11901.11920.20|TCTGGCACGGTGAAGAGACA</td><td>>hg38_21_+_8433172_8446622_RNA45SN1_45S_50ntFlanks</td><td>11901</td><td>11920</td><td>20</td><td>TCTGGCACGGTGAAGAGACA</td><td>92.9</td><td>-5.3</td><td>0.9</td><td>-5.7</td><td>1.6E-08</td><td>2.4E-08</td><td>BT20/MB468</td></tr>
<tr><td>>hg38_21_+_8433172_8446622_RNA45SN1_45S_50ntFlanks@9551.9571.21|GTCCGTAGCGGTCCTGACGTG</td><td>>hg38_21_+_8433172_8446622_RNA45SN1_45S_50ntFlanks</td><td>9551</td><td>9571</td><td>21</td><td>GTCCGTAGCGGTCCTGACGTG</td><td>581.4</td><td>2.9</td><td>0.5</td><td>5.7</td><td>1.4E-08</td><td>2.4E-08</td><td>MB231/MB468</td></tr>
<tr><td>>hg38_21_+_8433172_8446622_RNA45SN1_45S_50ntFlanks@12474.12498.25|ACCCACTAATAGGGAACGTGAGCTG</td><td>>hg38_21_+_8433172_8446622_RNA45SN1_45S_50ntFlanks</td><td>12474</td><td>12498</td><td>25</td><td>ACCCACTAATAGGGAACGTGAGCTG</td><td>575.3</td><td>-4.7</td><td>0.8</td><td>-5.7</td><td>1.6E-08</td><td>2.4E-08</td><td>BT20/MB468</td></tr>
<tr><td>>hg38_21_+_8433172_8446622_RNA45SN1_45S_50ntFlanks@8169.8187.19|CCAAGTCCTTCTGATCGAG</td><td>>hg38_21_+_8433172_8446622_RNA45SN1_45S_50ntFlanks</td><td>8169</td><td>8187</td><td>19</td><td>CCAAGTCCTTCTGATCGAG</td><td>212.9</td><td>-5.7</td><td>1.0</td><td>-5.7</td><td>1.6E-08</td><td>2.4E-08</td><td>BT20/MB468</td></tr>
<tr><td>>MI0000071|hsa-mir-17&WithFlank&13|+|91350599|91350694@20.41.22&[MIMAT0000070&hsa-miR-17-5p&offsets|0|-1;m-88&13|+|91350618|91350640&offsets|0|-1]|CAAAGTGCTTACAGTGCAGGTA</td><td>>MI0000071|hsa-mir-17&WithFlank&13|+|91350599|91350694</td><td>20</td><td>41</td><td>22</td><td>CAAAGTGCTTACAGTGCAGGTA</td><td>1465.4</td><td>5.5</td><td>1.0</td><td>5.6</td><td>1.6E-08</td><td>2.5E-08</td><td>BT20/MB468</td></tr>
<tr><td>>hg38_21_+_8433172_8446622_RNA45SN1_45S_50ntFlanks@10408.10437.30|GGTCAGTCGGTCCTGAGAGATGGGCGAGCG</td><td>>hg38_21_+_8433172_8446622_RNA45SN1_45S_50ntFlanks</td><td>10408</td><td>10437</td><td>30</td><td>GGTCAGTCGGTCCTGAGAGATGGGCGAGCG</td><td>98.0</td><td>-4.8</td><td>0.8</td><td>-5.6</td><td>1.6E-08</td><td>2.5E-08</td><td>BT20/MB468</td></tr>
<tr><td>>hg38_21_+_8433172_8446622_RNA45SN1_45S_50ntFlanks@12934.12969.36|CTAAACCATTCGTAGACGACCTGCTTCTGGGTCGGG</td><td>>hg38_21_+_8433172_8446622_RNA45SN1_45S_50ntFlanks</td><td>12934</td><td>12969</td><td>36</td><td>CTAAACCATTCGTAGACGACCTGCTTCTGGGTCGGG</td><td>168.2</td><td>-3.7</td><td>0.7</td><td>-5.7</td><td>1.5E-08</td><td>2.5E-08</td><td>MB231/MB468</td></tr>
<tr><td>>MI0025750|hsa-mir-7974&WithFlank&19|-|11495538|11495628@62.85.24&[MIMAT0031177&hsa-miR-7974&offsets|0|0;m-867&19|-|11495544|11495567&offsets|0|0]|AGGCTGTGATGCTCTCCTGAGCCC</td><td>>MI0025750|hsa-mir-7974&WithFlank&19|-|11495538|11495628</td><td>62</td><td>85</td><td>24</td><td>AGGCTGTGATGCTCTCCTGAGCCC</td><td>898.2</td><td>3.6</td><td>0.6</td><td>5.7</td><td>1.5E-08</td><td>2.5E-08</td><td>MB231/MB468</td></tr>
<tr><td>>hg38_21_+_8433172_8446622_RNA45SN1_45S_50ntFlanks@11641.11666.26|CGGCGGGTGTTGACGCGATGTGATTT</td><td>>hg38_21_+_8433172_8446622_RNA45SN1_45S_50ntFlanks</td><td>11641</td><td>11666</td><td>26</td><td>CGGCGGGTGTTGACGCGATGTGATTT</td><td>105.7</td><td>-5.2</td><td>0.9</td><td>-5.7</td><td>1.5E-08</td><td>2.5E-08</td><td>MB231/MB468</td></tr>
<tr><td>>hg38_21_+_8433172_8446622_RNA45SN1_45S_50ntFlanks@11774.11794.21|AGTGACGCGCATGAATGGATG</td><td>>hg38_21_+_8433172_8446622_RNA45SN1_45S_50ntFlanks</td><td>11774</td><td>11794</td><td>21</td><td>AGTGACGCGCATGAATGGATG</td><td>500.0</td><td>3.1</td><td>0.5</td><td>5.7</td><td>1.5E-08</td><td>2.5E-08</td><td>MB231/MB468</td></tr>
<tr><td>>hg38_21_+_8433172_8446622_RNA45SN1_45S_50ntFlanks@9621.9648.28|TCTAGTAGCTGGTTCCCTCCGAAGTTTC</td><td>>hg38_21_+_8433172_8446622_RNA45SN1_45S_50ntFlanks</td><td>9621</td><td>9648</td><td>28</td><td>TCTAGTAGCTGGTTCCCTCCGAAGTTTC</td><td>97.2</td><td>-4.8</td><td>0.8</td><td>-5.7</td><td>1.5E-08</td><td>2.5E-08</td><td>MB231/MB468</td></tr>
<tr><td>>hg38_21_+_8433172_8446622_RNA45SN1_45S_50ntFlanks@3846.3881.36|CTTGGATAACTGTGGTAATTCTAGAGCTAATACATG</td><td>>hg38_21_+_8433172_8446622_RNA45SN1_45S_50ntFlanks</td><td>3846</td><td>3881</td><td>36</td><td>CTTGGATAACTGTGGTAATTCTAGAGCTAATACATG</td><td>106.3</td><td>-3.4</td><td>0.6</td><td>-5.6</td><td>1.6E-08</td><td>2.5E-08</td><td>BT20/MB468</td></tr>
<tr><td>>hg38_21_+_8433172_8446622_RNA45SN1_45S_50ntFlanks@11670.11704.35|CCCAGTGCTCTGAATGTCAAAGTGAAGAAATTCAA</td><td>>hg38_21_+_8433172_8446622_RNA45SN1_45S_50ntFlanks</td><td>11670</td><td>11704</td><td>35</td><td>CCCAGTGCTCTGAATGTCAAAGTGAAGAAATTCAA</td><td>151.3</td><td>-2.6</td><td>0.5</td><td>-5.6</td><td>1.6E-08</td><td>2.5E-08</td><td>BT20/MB468</td></tr>
<tr><td>>hg38_21_+_8433172_8446622_RNA45SN1_45S_50ntFlanks@10873.10901.29|CTGGGTCGGTCGGGCTGGGGCGCGAAGCG</td><td>>hg38_21_+_8433172_8446622_RNA45SN1_45S_50ntFlanks</td><td>10873</td><td>10901</td><td>29</td><td>CTGGGTCGGTCGGGCTGGGGCGCGAAGCG</td><td>121.3</td><td>-4.7</td><td>0.8</td><td>-5.6</td><td>1.7E-08</td><td>2.5E-08</td><td>BT20/MB468</td></tr>
<tr><td>>hg38_21_+_8433172_8446622_RNA45SN1_45S_50ntFlanks@12405.12441.37|CTTTTTGATCCTTCGATGTCGGCTCTTCCTATCATTG</td><td>>hg38_21_+_8433172_8446622_RNA45SN1_45S_50ntFlanks</td><td>12405</td><td>12441</td><td>37</td><td>CTTTTTGATCCTTCGATGTCGGCTCTTCCTATCATTG</td><td>958.5</td><td>-3.9</td><td>0.7</td><td>-5.7</td><td>1.5E-08</td><td>2.6E-08</td><td>MB231/MB468</td></tr>
<tr><td>>hg38_21_+_8433172_8446622_RNA45SN1_45S_50ntFlanks@12081.12109.29|CTCTCGCTTCTGGCGCCAAGCGCCCGGCC</td><td>>hg38_21_+_8433172_8446622_RNA45SN1_45S_50ntFlanks</td><td>12081</td><td>12109</td><td>29</td><td>CTCTCGCTTCTGGCGCCAAGCGCCCGGCC</td><td>82.9</td><td>-4.9</td><td>0.9</td><td>-5.7</td><td>1.5E-08</td><td>2.6E-08</td><td>MB231/MB468</td></tr>
<tr><td>>hg38_21_+_8433172_8446622_RNA45SN1_45S_50ntFlanks@12685.12703.19|CCCAGGCGGAACGATACGG</td><td>>hg38_21_+_8433172_8446622_RNA45SN1_45S_50ntFlanks</td><td>12685</td><td>12703</td><td>19</td><td>CCCAGGCGGAACGATACGG</td><td>97.1</td><td>-4.7</td><td>0.8</td><td>-5.7</td><td>1.5E-08</td><td>2.6E-08</td><td>MB231/MB468</td></tr>
<tr><td>>hg38_21_+_8433172_8446622_RNA45SN1_45S_50ntFlanks@12073.12098.26|CCGAGGGGCTCTCGCTTCTGGCGCCA</td><td>>hg38_21_+_8433172_8446622_RNA45SN1_45S_50ntFlanks</td><td>12073</td><td>12098</td><td>26</td><td>CCGAGGGGCTCTCGCTTCTGGCGCCA</td><td>213.8</td><td>-3.8</td><td>0.7</td><td>-5.6</td><td>1.7E-08</td><td>2.6E-08</td><td>BT20/MB468</td></tr>
<tr><td>>hg38_21_+_8433172_8446622_RNA45SN1_45S_50ntFlanks@10794.10825.32|CTGGCATGTTGGAACAATGTAGGTAAGGGAAG</td><td>>hg38_21_+_8433172_8446622_RNA45SN1_45S_50ntFlanks</td><td>10794</td><td>10825</td><td>32</td><td>CTGGCATGTTGGAACAATGTAGGTAAGGGAAG</td><td>273.2</td><td>-3.9</td><td>0.7</td><td>-5.7</td><td>1.5E-08</td><td>2.6E-08</td><td>MB231/MB468</td></tr>
<tr><td>>hg38_1_-_228634819_228635039_RNA5S12_5S_50ntFlanks@148.171.24|GGAATACCGGGTGCTGTAGGCTTT</td><td>>hg38_1_-_228634819_228635039_RNA5S12_5S_50ntFlanks</td><td>148</td><td>171</td><td>24</td><td>GGAATACCGGGTGCTGTAGGCTTT</td><td>3372.0</td><td>4.2</td><td>0.7</td><td>5.7</td><td>1.5E-08</td><td>2.6E-08</td><td>MB231/MB468</td></tr>
<tr><td>>hg38_21_+_8433172_8446622_RNA45SN1_45S_50ntFlanks@3874.3899.26|AATACATGCCGACGGGCGCTGACCCC</td><td>>hg38_21_+_8433172_8446622_RNA45SN1_45S_50ntFlanks</td><td>3874</td><td>3899</td><td>26</td><td>AATACATGCCGACGGGCGCTGACCCC</td><td>445.2</td><td>-3.0</td><td>0.5</td><td>-5.7</td><td>1.5E-08</td><td>2.7E-08</td><td>MB231/MB468</td></tr>
<tr><td>>hg38_21_+_8433172_8446622_RNA45SN1_45S_50ntFlanks@12417.12453.37|TCGATGTCGGCTCTTCCTATCATTGTGAAGCAGAATT</td><td>>hg38_21_+_8433172_8446622_RNA45SN1_45S_50ntFlanks</td><td>12417</td><td>12453</td><td>37</td><td>TCGATGTCGGCTCTTCCTATCATTGTGAAGCAGAATT</td><td>459.4</td><td>5.8</td><td>1.0</td><td>5.7</td><td>1.6E-08</td><td>2.7E-08</td><td>MB231/MB468</td></tr>
<tr><td>>hg38_21_+_8433172_8446622_RNA45SN1_45S_50ntFlanks@6719.6739.21|TGAATTGCAGGACACATTGAT</td><td>>hg38_21_+_8433172_8446622_RNA45SN1_45S_50ntFlanks</td><td>6719</td><td>6739</td><td>21</td><td>TGAATTGCAGGACACATTGAT</td><td>1235.0</td><td>-3.4</td><td>0.6</td><td>-5.7</td><td>1.6E-08</td><td>2.7E-08</td><td>MB231/MB468</td></tr>
<tr><td>>hg38_21_+_8433172_8446622_RNA45SN1_45S_50ntFlanks@4738.4768.31|AAAGTCGGAGGTTCGAAGACGATCAGATACC</td><td>>hg38_21_+_8433172_8446622_RNA45SN1_45S_50ntFlanks</td><td>4738</td><td>4768</td><td>31</td><td>AAAGTCGGAGGTTCGAAGACGATCAGATACC</td><td>471.0</td><td>7.4</td><td>1.3</td><td>5.6</td><td>1.7E-08</td><td>2.7E-08</td><td>BT20/MB468</td></tr>
<tr><td>>hg38_21_+_8433172_8446622_RNA45SN1_45S_50ntFlanks@9201.9218.18|CTCTCGGGGCCACGCGCG</td><td>>hg38_21_+_8433172_8446622_RNA45SN1_45S_50ntFlanks</td><td>9201</td><td>9218</td><td>18</td><td>CTCTCGGGGCCACGCGCG</td><td>73.3</td><td>-7.8</td><td>1.4</td><td>-5.7</td><td>1.6E-08</td><td>2.7E-08</td><td>MB231/MB468</td></tr>
<tr><td>>hg38_21_+_8433172_8446622_RNA45SN1_45S_50ntFlanks@6772.6797.26|GTTCCTCCCGGGGCTACGCCTGTCTG</td><td>>hg38_21_+_8433172_8446622_RNA45SN1_45S_50ntFlanks</td><td>6772</td><td>6797</td><td>26</td><td>GTTCCTCCCGGGGCTACGCCTGTCTG</td><td>203.0</td><td>-5.3</td><td>0.9</td><td>-5.6</td><td>1.8E-08</td><td>2.7E-08</td><td>BT20/MB468</td></tr>
<tr><td>>hg38_21_+_8433172_8446622_RNA45SN1_45S_50ntFlanks@10351.10390.40|CGAGAACTTTGAAGGCCGAAGTGGAGAAGGGTTCCATGTG</td><td>>hg38_21_+_8433172_8446622_RNA45SN1_45S_50ntFlanks</td><td>10351</td><td>10390</td><td>40</td><td>CGAGAACTTTGAAGGCCGAAGTGGAGAAGGGTTCCATGTG</td><td>4047.7</td><td>-3.9</td><td>0.7</td><td>-5.7</td><td>1.6E-08</td><td>2.7E-08</td><td>MB231/MB468</td></tr>
<tr><td>>hg38_21_+_8433172_8446622_RNA45SN1_45S_50ntFlanks@10905.10933.29|CTGGGCGCGCGCCGCGGCTGGACGAGGCG</td><td>>hg38_21_+_8433172_8446622_RNA45SN1_45S_50ntFlanks</td><td>10905</td><td>10933</td><td>29</td><td>CTGGGCGCGCGCCGCGGCTGGACGAGGCG</td><td>116.5</td><td>-6.5</td><td>1.2</td><td>-5.7</td><td>1.6E-08</td><td>2.7E-08</td><td>MB231/MB468</td></tr>
<tr><td>>hg38_21_+_8433172_8446622_RNA45SN1_45S_50ntFlanks@12157.12183.27|AGTTTGACTGGGGCGGTACACCTGTCA</td><td>>hg38_21_+_8433172_8446622_RNA45SN1_45S_50ntFlanks</td><td>12157</td><td>12183</td><td>27</td><td>AGTTTGACTGGGGCGGTACACCTGTCA</td><td>91.7</td><td>-3.3</td><td>0.6</td><td>-5.6</td><td>1.6E-08</td><td>2.8E-08</td><td>MB231/MB468</td></tr>
<tr><td>>hg38_21_+_8433172_8446622_RNA45SN1_45S_50ntFlanks@4297.4319.23|CATTGGAGGGCAAGTCTGGTGCC</td><td>>hg38_21_+_8433172_8446622_RNA45SN1_45S_50ntFlanks</td><td>4297</td><td>4319</td><td>23</td><td>CATTGGAGGGCAAGTCTGGTGCC</td><td>149.5</td><td>-3.4</td><td>0.6</td><td>-5.6</td><td>1.8E-08</td><td>2.8E-08</td><td>BT20/MB468</td></tr>
<tr><td>>hg38_21_+_8433172_8446622_RNA45SN1_45S_50ntFlanks@12415.12440.26|CTTCGATGTCGGCTCTTCCTATCATT</td><td>>hg38_21_+_8433172_8446622_RNA45SN1_45S_50ntFlanks</td><td>12415</td><td>12440</td><td>26</td><td>CTTCGATGTCGGCTCTTCCTATCATT</td><td>450.5</td><td>2.9</td><td>0.5</td><td>5.6</td><td>1.8E-08</td><td>2.8E-08</td><td>BT20/MB468</td></tr>
<tr><td>>hg38_21_+_8433172_8446622_RNA45SN1_45S_50ntFlanks@4102.4121.20|ACCACGGGTGACGGGGAATC</td><td>>hg38_21_+_8433172_8446622_RNA45SN1_45S_50ntFlanks</td><td>4102</td><td>4121</td><td>20</td><td>ACCACGGGTGACGGGGAATC</td><td>482.7</td><td>-3.5</td><td>0.6</td><td>-5.6</td><td>1.7E-08</td><td>2.9E-08</td><td>MB231/MB468</td></tr>
<tr><td>>hg38_21_+_8433172_8446622_RNA45SN1_45S_50ntFlanks@6737.6770.34|GATCATCGACACTTCGAACGCACTTGCGGCCCCG</td><td>>hg38_21_+_8433172_8446622_RNA45SN1_45S_50ntFlanks</td><td>6737</td><td>6770</td><td>34</td><td>GATCATCGACACTTCGAACGCACTTGCGGCCCCG</td><td>379.9</td><td>-5.3</td><td>0.9</td><td>-5.6</td><td>1.7E-08</td><td>2.9E-08</td><td>MB231/MB468</td></tr>
<tr><td>>hg38_21_+_8433172_8446622_RNA45SN1_45S_50ntFlanks@4380.4408.29|CGTAGTTGGATCTTGGGAGCGGGCGGGCG</td><td>>hg38_21_+_8433172_8446622_RNA45SN1_45S_50ntFlanks</td><td>4380</td><td>4408</td><td>29</td><td>CGTAGTTGGATCTTGGGAGCGGGCGGGCG</td><td>77.5</td><td>-7.9</td><td>1.4</td><td>-5.6</td><td>1.7E-08</td><td>2.9E-08</td><td>MB231/MB468</td></tr>
<tr><td>>hg38_21_+_8433172_8446622_RNA45SN1_45S_50ntFlanks@10766.10797.32|CGCAGCAGGTCTCCAAGGTGAACAGCCTCTGG</td><td>>hg38_21_+_8433172_8446622_RNA45SN1_45S_50ntFlanks</td><td>10766</td><td>10797</td><td>32</td><td>CGCAGCAGGTCTCCAAGGTGAACAGCCTCTGG</td><td>102.3</td><td>-3.1</td><td>0.6</td><td>-5.6</td><td>1.7E-08</td><td>2.9E-08</td><td>MB231/MB468</td></tr>
<tr><td>>hg38_21_+_8433172_8446622_RNA45SN1_45S_50ntFlanks@11318.11344.27|CCCCACGTCTCGTCGCGCGCGCGTCCG</td><td>>hg38_21_+_8433172_8446622_RNA45SN1_45S_50ntFlanks</td><td>11318</td><td>11344</td><td>27</td><td>CCCCACGTCTCGTCGCGCGCGCGTCCG</td><td>2754.7</td><td>-7.3</td><td>1.3</td><td>-5.6</td><td>1.9E-08</td><td>2.9E-08</td><td>BT20/MB468</td></tr>
</table>

<table>
<tr><td>>hg38_21_+_8433172_8446622_RNA45SN1_45S_50ntFlanks@9569.9592.24|GTGCAAATCGGTCGTCCGACCTGG</td><td>>hg38_21_+_8433172_8446622_RNA45SN1_45S_50ntFlanks</td><td>9569</td><td>9592</td><td>24</td><td>GTGCAAATCGGTCGTCCGACCTGG</td><td>145.8</td><td>-4.7</td><td>0.8</td><td>-5.6</td><td>1.9E-08</td><td>2.9E-08</td><td>BT20/MB468</td></tr>
<tr><td>>hg38_21_+_8433172_8446622_RNA45SN1_45S_50ntFlanks@10351.10373.23|CGAGAACTTTGAAGGCCGAAGTG</td><td>>hg38_21_+_8433172_8446622_RNA45SN1_45S_50ntFlanks</td><td>10351</td><td>10373</td><td>23</td><td>CGAGAACTTTGAAGGCCGAAGTG</td><td>630.4</td><td>-3.1</td><td>0.6</td><td>-5.6</td><td>1.9E-08</td><td>2.9E-08</td><td>BT20/MB468</td></tr>
<tr><td>>trnaMT_HisGTG_MT_+_12138_12206@32.63.32|TGAATCTGACAACAGAGGCTTACGACCCCTTA</td><td>>trnaMT_HisGTG_MT_+_12138_12206</td><td>32</td><td>63</td><td>32</td><td>TGAATCTGACAACAGAGGCTTACGACCCCTTA</td><td>915.3</td><td>5.9</td><td>1.0</td><td>5.6</td><td>1.9E-08</td><td>2.9E-08</td><td>BT20/MB468</td></tr>
<tr><td>>hg38_21_+_8433172_8446622_RNA45SN1_45S_50ntFlanks@11918.11948.31|ACATGAGAGGTGTAGAATAAGTGGGAGGCCC</td><td>>hg38_21_+_8433172_8446622_RNA45SN1_45S_50ntFlanks</td><td>11918</td><td>11948</td><td>31</td><td>ACATGAGAGGTGTAGAATAAGTGGGAGGCCC</td><td>134.4</td><td>-6.7</td><td>1.2</td><td>-5.6</td><td>1.7E-08</td><td>2.9E-08</td><td>MB231/MB468</td></tr>
<tr><td>>am_trnaMT_HisGTG_MT_+_12138_12206@28.45.18|ATTGTGAATCTGACAACA</td><td>>am_trnaMT_HisGTG_MT_+_12138_12206</td><td>28</td><td>45</td><td>18</td><td>ATTGTGAATCTGACAACA</td><td>531.6</td><td>6.9</td><td>1.2</td><td>5.6</td><td>1.9E-08</td><td>3.0E-08</td><td>BT20/MB468</td></tr>
<tr><td>>hg38_21_+_8433172_8446622_RNA45SN1_45S_50ntFlanks@3873.3899.27|TAATACATGCCGACGGGCGCTGACCCC</td><td>>hg38_21_+_8433172_8446622_RNA45SN1_45S_50ntFlanks</td><td>3873</td><td>3899</td><td>27</td><td>TAATACATGCCGACGGGCGCTGACCCC</td><td>317.0</td><td>-5.7</td><td>1.0</td><td>-5.6</td><td>1.9E-08</td><td>3.0E-08</td><td>BT20/MB468</td></tr>
<tr><td>>hg38_21_+_8433172_8446622_RNA45SN1_45S_50ntFlanks@10481.10498.18|ATCGAAAGGGAGTCGGGT</td><td>>hg38_21_+_8433172_8446622_RNA45SN1_45S_50ntFlanks</td><td>10481</td><td>10498</td><td>18</td><td>ATCGAAAGGGAGTCGGGT</td><td>99.3</td><td>-4.5</td><td>0.8</td><td>-5.6</td><td>1.8E-08</td><td>3.0E-08</td><td>MB231/MB468</td></tr>
<tr><td>>hg38_21_+_8433172_8446622_RNA45SN1_45S_50ntFlanks@12526.12546.21|TTTTACCCTACTGATGATGTG</td><td>>hg38_21_+_8433172_8446622_RNA45SN1_45S_50ntFlanks</td><td>12526</td><td>12546</td><td>21</td><td>TTTTACCCTACTGATGATGTG</td><td>59.2</td><td>-7.5</td><td>1.3</td><td>-5.6</td><td>1.8E-08</td><td>3.0E-08</td><td>MB231/MB468</td></tr>
<tr><td>>trnaMT_SerGCT_MT_+_12207_12265@8.38.31|CTCACAAGAACTGCTAACTCATGCCCCCATG</td><td>>trnaMT_SerGCT_MT_+_12207_12265</td><td>8</td><td>38</td><td>31</td><td>CTCACAAGAACTGCTAACTCATGCCCCCATG</td><td>207.1</td><td>-6.0</td><td>1.1</td><td>-5.6</td><td>1.8E-08</td><td>3.0E-08</td><td>MB231/MB468</td></tr>
<tr><td>>hg38_21_+_8433172_8446622_RNA45SN1_45S_50ntFlanks@8175.8203.29|CCTTCTGATCGAGGCCCAGCCCGTGGACG</td><td>>hg38_21_+_8433172_8446622_RNA45SN1_45S_50ntFlanks</td><td>8175</td><td>8203</td><td>29</td><td>CCTTCTGATCGAGGCCCAGCCCGTGGACG</td><td>365.0</td><td>4.0</td><td>0.7</td><td>5.6</td><td>1.8E-08</td><td>3.0E-08</td><td>MB231/MB468</td></tr>
<tr><td>>hg38_21_+_8433172_8446622_RNA45SN1_45S_50ntFlanks@11323.11341.19|CGTCTCGTCGCGCGCGCGT</td><td>>hg38_21_+_8433172_8446622_RNA45SN1_45S_50ntFlanks</td><td>11323</td><td>11341</td><td>19</td><td>CGTCTCGTCGCGCGCGCGT</td><td>80.2</td><td>-6.0</td><td>1.1</td><td>-5.6</td><td>1.8E-08</td><td>3.0E-08</td><td>MB231/MB468</td></tr>
<tr><td>>hg38_21_+_8433172_8446622_RNA45SN1_45S_50ntFlanks@9556.9573.18|TAGCGGTCCTGACGTGCA</td><td>>hg38_21_+_8433172_8446622_RNA45SN1_45S_50ntFlanks</td><td>9556</td><td>9573</td><td>18</td><td>TAGCGGTCCTGACGTGCA</td><td>410.7</td><td>-3.4</td><td>0.6</td><td>-5.6</td><td>2.0E-08</td><td>3.0E-08</td><td>BT20/MB468</td></tr>
<tr><td>>hg38_21_+_8433172_8446622_RNA45SN1_45S_50ntFlanks@12957.12989.33|CTTCTGGGTCGGGGTTTCGTACGTAGCAGAGCA</td><td>>hg38_21_+_8433172_8446622_RNA45SN1_45S_50ntFlanks</td><td>12957</td><td>12989</td><td>33</td><td>CTTCTGGGTCGGGGTTTCGTACGTAGCAGAGCA</td><td>110.9</td><td>-4.7</td><td>0.8</td><td>-5.6</td><td>2.0E-08</td><td>3.1E-08</td><td>BT20/MB468</td></tr>
<tr><td>>hg38_21_+_8433172_8446622_RNA45SN1_45S_50ntFlanks@12459.12476.18|AGCGTTGGATTGTTCACC</td><td>>hg38_21_+_8433172_8446622_RNA45SN1_45S_50ntFlanks</td><td>12459</td><td>12476</td><td>18</td><td>AGCGTTGGATTGTTCACC</td><td>2185.0</td><td>-2.9</td><td>0.5</td><td>-5.6</td><td>1.8E-08</td><td>3.1E-08</td><td>MB231/MB468</td></tr>
<tr><td>>hg38_21_+_8433172_8446622_RNA45SN1_45S_50ntFlanks@10768.10785.18|CAGCAGGTCTCCAAGGTG</td><td>>hg38_21_+_8433172_8446622_RNA45SN1_45S_50ntFlanks</td><td>10768</td><td>10785</td><td>18</td><td>CAGCAGGTCTCCAAGGTG</td><td>610.0</td><td>-3.7</td><td>0.7</td><td>-5.6</td><td>1.8E-08</td><td>3.1E-08</td><td>MB231/MB468</td></tr>
<tr><td>>hg38_21_+_8433172_8446622_RNA45SN1_45S_50ntFlanks@6722.6740.19|ATTGCAGGACACATTGATC</td><td>>hg38_21_+_8433172_8446622_RNA45SN1_45S_50ntFlanks</td><td>6722</td><td>6740</td><td>19</td><td>ATTGCAGGACACATTGATC</td><td>1017.7</td><td>-3.2</td><td>0.6</td><td>-5.6</td><td>1.8E-08</td><td>3.1E-08</td><td>MB231/MB468</td></tr>
<tr><td>>hg38_21_+_8433172_8446622_RNA45SN1_45S_50ntFlanks@6719.6742.24|TGAATTGCAGGACACATTGATCAT</td><td>>hg38_21_+_8433172_8446622_RNA45SN1_45S_50ntFlanks</td><td>6719</td><td>6742</td><td>24</td><td>TGAATTGCAGGACACATTGATCAT</td><td>553.6</td><td>-4.2</td><td>0.7</td><td>-5.6</td><td>1.8E-08</td><td>3.1E-08</td><td>MB231/MB468</td></tr>
<tr><td>>hg38_21_+_8433172_8446622_RNA45SN1_45S_50ntFlanks@10682.10709.28|TCCGGCGGCGTCCGGTGAGCTCTCGCTG</td><td>>hg38_21_+_8433172_8446622_RNA45SN1_45S_50ntFlanks</td><td>10682</td><td>10709</td><td>28</td><td>TCCGGCGGCGTCCGGTGAGCTCTCGCTG</td><td>109.9</td><td>-4.0</td><td>0.7</td><td>-5.6</td><td>2.1E-08</td><td>3.1E-08</td><td>BT20/MB468</td></tr>
<tr><td>>hg38_21_+_8433172_8446622_RNA45SN1_45S_50ntFlanks@12626.12652.27|AATGGGGCGAAGCTACCATCTGTGGGA</td><td>>hg38_21_+_8433172_8446622_RNA45SN1_45S_50ntFlanks</td><td>12626</td><td>12652</td><td>27</td><td>AATGGGGCGAAGCTACCATCTGTGGGA</td><td>235.3</td><td>-5.2</td><td>0.9</td><td>-5.6</td><td>2.1E-08</td><td>3.1E-08</td><td>BT20/MB468</td></tr>
<tr><td>>hg38_21_+_8433172_8446622_RNA45SN1_45S_50ntFlanks@10823.10844.22|AAGTCGGCAAGCCGGATCCGTA</td><td>>hg38_21_+_8433172_8446622_RNA45SN1_45S_50ntFlanks</td><td>10823</td><td>10844</td><td>22</td><td>AAGTCGGCAAGCCGGATCCGTA</td><td>179.9</td><td>-3.8</td><td>0.7</td><td>-5.6</td><td>2.1E-08</td><td>3.1E-08</td><td>BT20/MB468</td></tr>
<tr><td>>hg38_21_+_8433172_8446622_RNA45SN1_45S_50ntFlanks@12416.12455.40|TTCGATGTCGGCTCTTCCTATCATTGTGAAGCAGAATTCA</td><td>>hg38_21_+_8433172_8446622_RNA45SN1_45S_50ntFlanks</td><td>12416</td><td>12455</td><td>40</td><td>TTCGATGTCGGCTCTTCCTATCATTGTGAAGCAGAATTCA</td><td>624.0</td><td>3.6</td><td>0.6</td><td>5.6</td><td>1.8E-08</td><td>3.1E-08</td><td>MB231/MB468</td></tr>
<tr><td>>hg38_21_+_8433172_8446622_RNA45SN1_45S_50ntFlanks@6653.6688.36|ACTCTTAGCGGTGGATCACTCGGCTCGTGCGTCGAT</td><td>>hg38_21_+_8433172_8446622_RNA45SN1_45S_50ntFlanks</td><td>6653</td><td>6688</td><td>36</td><td>ACTCTTAGCGGTGGATCACTCGGCTCGTGCGTCGAT</td><td>1000.2</td><td>3.9</td><td>0.7</td><td>5.6</td><td>1.8E-08</td><td>3.1E-08</td><td>MB231/MB468</td></tr>
<tr><td>>hg38_21_+_8433172_8446622_RNA45SN1_45S_50ntFlanks@11573.11613.41|CGACTTAGAACTGGTGCGGACCAGGGGAATCCGACTGTTTA</td><td>>hg38_21_+_8433172_8446622_RNA45SN1_45S_50ntFlanks</td><td>11573</td><td>11613</td><td>41</td><td>CGACTTAGAACTGGTGCGGACCAGGGGAATCCGACTGTTTA</td><td>498.3</td><td>6.2</td><td>1.1</td><td>5.6</td><td>1.9E-08</td><td>3.2E-08</td><td>MB231/MB468</td></tr>
<tr><td>>hg38_21_+_8433172_8446622_RNA45SN1_45S_50ntFlanks@12588.12616.29|GGTTCAGACATTTGGTGTATGTGCTTGGC</td><td>>hg38_21_+_8433172_8446622_RNA45SN1_45S_50ntFlanks</td><td>12588</td><td>12616</td><td>29</td><td>GGTTCAGACATTTGGTGTATGTGCTTGGC</td><td>87.8</td><td>-4.5</td><td>0.8</td><td>-5.6</td><td>1.9E-08</td><td>3.2E-08</td><td>MB231/MB468</td></tr>
<tr><td>>hg38_21_+_8433172_8446622_RNA45SN1_45S_50ntFlanks@10411.10442.32|CAGTCGGTCCTGAGAGATGGGCGAGCGCCGTT</td><td>>hg38_21_+_8433172_8446622_RNA45SN1_45S_50ntFlanks</td><td>10411</td><td>10442</td><td>32</td><td>CAGTCGGTCCTGAGAGATGGGCGAGCGCCGTT</td><td>459.5</td><td>-3.5</td><td>0.6</td><td>-5.6</td><td>2.1E-08</td><td>3.2E-08</td><td>BT20/MB468</td></tr>
<tr><td>>hg38_21_+_8433172_8446622_RNA45SN1_45S_50ntFlanks@12616.12654.39|CTGAGGAGCCAATGGGGCGAAGCTACCATCTGTGGGATT</td><td>>hg38_21_+_8433172_8446622_RNA45SN1_45S_50ntFlanks</td><td>12616</td><td>12654</td><td>39</td><td>CTGAGGAGCCAATGGGGCGAAGCTACCATCTGTGGGATT</td><td>437.2</td><td>-3.1</td><td>0.6</td><td>-5.6</td><td>1.9E-08</td><td>3.2E-08</td><td>MB231/MB468</td></tr>
<tr><td>>MI0000253|hsa-mir-148a&WithFlank&7|-|25949913|25949992@50.73.24&[MIMAT0000243&hsa-miR-148a-3p&offsets|0|+2;m-7&7|-|25949922|25949943&offsets|0|+2]|TCAGTGCACTACAGAACTTTGTCT</td><td>>MI0000253|hsa-mir-148a&WithFlank&7|-|25949913|25949992</td><td>50</td><td>73</td><td>24</td><td>TCAGTGCACTACAGAACTTTGTCT</td><td>840.3</td><td>4.9</td><td>0.8</td><td>6.1</td><td>1.3E-09</td><td>3.3E-08</td><td>BT20/MB231</td></tr>
<tr><td>>hg38_21_+_8433172_8446622_RNA45SN1_45S_50ntFlanks@11781.11798.18|CGCATGAATGGATGAACG</td><td>>hg38_21_+_8433172_8446622_RNA45SN1_45S_50ntFlanks</td><td>11781</td><td>11798</td><td>18</td><td>CGCATGAATGGATGAACG</td><td>652.3</td><td>-3.7</td><td>0.7</td><td>-5.6</td><td>1.9E-08</td><td>3.3E-08</td><td>MB231/MB468</td></tr>
<tr><td>>hg38_21_+_8433172_8446622_RNA45SN1_45S_50ntFlanks@11729.11764.36|TAACTATGACTCTCTTAAGGTAGCCAAATGCCTCGT</td><td>>hg38_21_+_8433172_8446622_RNA45SN1_45S_50ntFlanks</td><td>11729</td><td>11764</td><td>36</td><td>TAACTATGACTCTCTTAAGGTAGCCAAATGCCTCGT</td><td>385.1</td><td>-3.6</td><td>0.6</td><td>-5.6</td><td>2.2E-08</td><td>3.3E-08</td><td>BT20/MB468</td></tr>
<tr><td>>hg38_21_+_8433172_8446622_RNA45SN1_45S_50ntFlanks@10735.10757.23|TGTAAATCTCGCGCCGGGCCGTA</td><td>>hg38_21_+_8433172_8446622_RNA45SN1_45S_50ntFlanks</td><td>10735</td><td>10757</td><td>23</td><td>TGTAAATCTCGCGCCGGGCCGTA</td><td>126.9</td><td>-3.3</td><td>0.6</td><td>-5.6</td><td>2.2E-08</td><td>3.3E-08</td><td>BT20/MB468</td></tr>
<tr><td>>hg38_21_+_8433172_8446622_RNA45SN1_45S_50ntFlanks@4002.4025.24|GATAACCTCGGGCCGATCGCACGC</td><td>>hg38_21_+_8433172_8446622_RNA45SN1_45S_50ntFlanks</td><td>4002</td><td>4025</td><td>24</td><td>GATAACCTCGGGCCGATCGCACGC</td><td>175.0</td><td>-3.5</td><td>0.6</td><td>-5.6</td><td>2.0E-08</td><td>3.3E-08</td><td>MB231/MB468</td></tr>
<tr><td>>hg38_21_+_8433172_8446622_RNA45SN1_45S_50ntFlanks@11894.11920.27|ACTCTAGTCTGGCACGGTGAAGAGACA</td><td>>hg38_21_+_8433172_8446622_RNA45SN1_45S_50ntFlanks</td><td>11894</td><td>11920</td><td>27</td><td>ACTCTAGTCTGGCACGGTGAAGAGACA</td><td>124.9</td><td>-3.6</td><td>0.6</td><td>-5.6</td><td>2.0E-08</td><td>3.3E-08</td><td>MB231/MB468</td></tr>
<tr><td>>hg38_21_+_8433172_8446622_RNA45SN1_45S_50ntFlanks@3785.3810.26|TGAAACTGCGAATGGCTCATTAAATC</td><td>>hg38_21_+_8433172_8446622_RNA45SN1_45S_50ntFlanks</td><td>3785</td><td>3810</td><td>26</td><td>TGAAACTGCGAATGGCTCATTAAATC</td><td>167.3</td><td>-3.3</td><td>0.6</td><td>-5.6</td><td>2.2E-08</td><td>3.3E-08</td><td>BT20/MB468</td></tr>
<tr><td>>hg38_21_+_8433172_8446622_RNA45SN1_45S_50ntFlanks@9564.9598.35|CTGACGTGCAAATCGGTCGTCCGACCTGGGTATAG</td><td>>hg38_21_+_8433172_8446622_RNA45SN1_45S_50ntFlanks</td><td>9564</td><td>9598</td><td>35</td><td>CTGACGTGCAAATCGGTCGTCCGACCTGGGTATAG</td><td>97.2</td><td>-4.6</td><td>0.8</td><td>-5.6</td><td>2.2E-08</td><td>3.3E-08</td><td>BT20/MB468</td></tr>
<tr><td>>hg38_21_+_8433172_8446622_RNA45SN1_45S_50ntFlanks@9554.9577.24|CGTAGCGGTCCTGACGTGCAAATC</td><td>>hg38_21_+_8433172_8446622_RNA45SN1_45S_50ntFlanks</td><td>9554</td><td>9577</td><td>24</td><td>CGTAGCGGTCCTGACGTGCAAATC</td><td>2617.4</td><td>3.4</td><td>0.6</td><td>5.6</td><td>2.0E-08</td><td>3.4E-08</td><td>MB231/MB468</td></tr>
<tr><td>>hg38_21_+_8433172_8446622_RNA45SN1_45S_50ntFlanks@9305.9336.32|CCAAGGAGTCTAACACGTGCGCGAGTCGGGGG</td><td>>hg38_21_+_8433172_8446622_RNA45SN1_45S_50ntFlanks</td><td>9305</td><td>9336</td><td>32</td><td>CCAAGGAGTCTAACACGTGCGCGAGTCGGGGG</td><td>99.4</td><td>-4.3</td><td>0.8</td><td>-5.6</td><td>2.2E-08</td><td>3.4E-08</td><td>BT20/MB468</td></tr>
<tr><td>>hg38_21_+_8433172_8446622_RNA45SN1_45S_50ntFlanks@9571.9595.25|GCAAATCGGTCGTCCGACCTGGGTA</td><td>>hg38_21_+_8433172_8446622_RNA45SN1_45S_50ntFlanks</td><td>9571</td><td>9595</td><td>25</td><td>GCAAATCGGTCGTCCGACCTGGGTA</td><td>144.2</td><td>-2.6</td><td>0.5</td><td>-5.6</td><td>2.2E-08</td><td>3.4E-08</td><td>BT20/MB468</td></tr>
<tr><td>>MI0003684|hsa-mir-660&WithFlank&X|+|50013235|50013343@22.44.23&[MIMAT0003338&hsa-miR-660-5p&offsets|0|+1;m-157&X|+|50013256|50013278&offsets|0|0]|TACCCATTGCATATCGGAGTTGT</td><td>>MI0003684|hsa-mir-660&WithFlank&X|+|50013235|50013343</td><td>22</td><td>44</td><td>23</td><td>TACCCATTGCATATCGGAGTTGT</td><td>886.5</td><td>5.2</td><td>0.9</td><td>5.6</td><td>2.3E-08</td><td>3.4E-08</td><td>BT20/MB468</td></tr>
<tr><td>>hg38_21_+_8433172_8446622_RNA45SN1_45S_50ntFlanks@12407.12425.19|TTTTGATCCTTCGATGTCG</td><td>>hg38_21_+_8433172_8446622_RNA45SN1_45S_50ntFlanks</td><td>12407</td><td>12425</td><td>19</td><td>TTTTGATCCTTCGATGTCG</td><td>119.5</td><td>-5.4</td><td>1.0</td><td>-5.6</td><td>2.3E-08</td><td>3.4E-08</td><td>BT20/MB468</td></tr>
<tr><td>>hg38_21_+_8433172_8446622_RNA45SN1_45S_50ntFlanks@9563.9596.34|CCTGACGTGCAAATCGGTCGTCCGACCTGGGTAT</td><td>>hg38_21_+_8433172_8446622_RNA45SN1_45S_50ntFlanks</td><td>9563</td><td>9596</td><td>34</td><td>CCTGACGTGCAAATCGGTCGTCCGACCTGGGTAT</td><td>876.4</td><td>-3.5</td><td>0.6</td><td>-5.6</td><td>2.0E-08</td><td>3.4E-08</td><td>MB231/MB468</td></tr>
<tr><td>>hg38_21_+_8433172_8446622_RNA45SN1_45S_50ntFlanks@7984.8003.20|AGATCAGACGTGGCGACCCG</td><td>>hg38_21_+_8433172_8446622_RNA45SN1_45S_50ntFlanks</td><td>7984</td><td>8003</td><td>20</td><td>AGATCAGACGTGGCGACCCG</td><td>137.3</td><td>-5.2</td><td>0.9</td><td>-5.6</td><td>2.0E-08</td><td>3.5E-08</td><td>MB231/MB468</td></tr>
<tr><td>>hg38_21_+_8433172_8446622_RNA45SN1_45S_50ntFlanks@10392.10435.44|ACAGCAGTTGAACATGGGTCAGTCGGTCCTGAGAGATGGGCGAG</td><td>>hg38_21_+_8433172_8446622_RNA45SN1_45S_50ntFlanks</td><td>10392</td><td>10435</td><td>44</td><td>ACAGCAGTTGAACATGGGTCAGTCGGTCCTGAGAGATGGGCGAG</td><td>339.3</td><td>-6.3</td><td>1.1</td><td>-5.6</td><td>2.1E-08</td><td>3.5E-08</td><td>MB231/MB468</td></tr>
<tr><td>>hg38_21_+_8433172_8446622_RNA45SN1_45S_50ntFlanks@12685.12711.27|CCCAGGCGGAACGATACGGCAGCGCCG</td><td>>hg38_21_+_8433172_8446622_RNA45SN1_45S_50ntFlanks</td><td>12685</td><td>12711</td><td>27</td><td>CCCAGGCGGAACGATACGGCAGCGCCG</td><td>113.8</td><td>-5.6</td><td>1.0</td><td>-5.6</td><td>2.3E-08</td><td>3.5E-08</td><td>BT20/MB468</td></tr>
<tr><td>>hg38_21_+_8433172_8446622_RNA45SN1_45S_50ntFlanks@12935.12955.21|TAAACCATTCGTAGACGACCT</td><td>>hg38_21_+_8433172_8446622_RNA45SN1_45S_50ntFlanks</td><td>12935</td><td>12955</td><td>21</td><td>TAAACCATTCGTAGACGACCT</td><td>90.5</td><td>-3.2</td><td>0.6</td><td>-5.6</td><td>2.1E-08</td><td>3.5E-08</td><td>MB231/MB468</td></tr>
<tr><td>>hg38_21_+_8433172_8446622_RNA45SN1_45S_50ntFlanks@10788.10815.28|CAGCCTCTGGCATGTTGGAACAATGTAG</td><td>>hg38_21_+_8433172_8446622_RNA45SN1_45S_50ntFlanks</td><td>10788</td><td>10815</td><td>28</td><td>CAGCCTCTGGCATGTTGGAACAATGTAG</td><td>251.2</td><td>-2.5</td><td>0.5</td><td>-5.6</td><td>2.1E-08</td><td>3.5E-08</td><td>MB231/MB468</td></tr>
<tr><td>>hg38_21_+_8433172_8446622_RNA45SN1_45S_50ntFlanks@4732.4768.37|AGAACGAAAGTCGGAGGTTCGAAGACGATCAGATACC</td><td>>hg38_21_+_8433172_8446622_RNA45SN1_45S_50ntFlanks</td><td>4732</td><td>4768</td><td>37</td><td>AGAACGAAAGTCGGAGGTTCGAAGACGATCAGATACC</td><td>843.4</td><td>11.5</td><td>2.1</td><td>5.6</td><td>2.1E-08</td><td>3.5E-08</td><td>MB231/MB468</td></tr>
<tr><td>>hg38_21_+_8433172_8446622_RNA45SN1_45S_50ntFlanks@12489.12518.30|ACGTGAGCTGGGTTTAGACCGTCGTGAGAC</td><td>>hg38_21_+_8433172_8446622_RNA45SN1_45S_50ntFlanks</td><td>12489</td><td>12518</td><td>30</td><td>ACGTGAGCTGGGTTTAGACCGTCGTGAGAC</td><td>97.4</td><td>-3.1</td><td>0.6</td><td>-5.6</td><td>2.3E-08</td><td>3.5E-08</td><td>BT20/MB468</td></tr>
<tr><td>>hg38_21_+_8433172_8446622_RNA45SN1_45S_50ntFlanks@4753.4782.30|AAGACGATCAGATACCGTCGTAGTTCCGAC</td><td>>hg38_21_+_8433172_8446622_RNA45SN1_45S_50ntFlanks</td><td>4753</td><td>4782</td><td>30</td><td>AAGACGATCAGATACCGTCGTAGTTCCGAC</td><td>248.3</td><td>-5.3</td><td>1.0</td><td>-5.6</td><td>2.3E-08</td><td>3.5E-08</td><td>BT20/MB468</td></tr>
<tr><td>>MI0000102|hsa-mir-100&WithFlank&11|-|122152223|122152314@19.40.22(+3A)&[MIMAT0000098&hsa-miR-100-5p&offsets|0|0(+3A);m-27&11|-|122152276|122152296&offsets|0|+1(+3A)]|AACCCGTAGATCCGAACTTGTGAAA</td><td>>MI0000102|hsa-mir-100&WithFlank&11|-|122152223|122152314</td><td>19</td><td>40</td><td>22(+3A)</td><td>AACCCGTAGATCCGAACTTGTGAAA</td><td>234.9</td><td>-5.7</td><td>0.9</td><td>-6.1</td><td>1.4E-09</td><td>3.5E-08</td><td>BT20/MB231</td></tr>
<tr><td>>hg38_21_+_8433172_8446622_RNA45SN1_45S_50ntFlanks@12928.12956.29|CTGGCGCTAAACCATTCGTAGACGACCTG</td><td>>hg38_21_+_8433172_8446622_RNA45SN1_45S_50ntFlanks</td><td>12928</td><td>12956</td><td>29</td><td>CTGGCGCTAAACCATTCGTAGACGACCTG</td><td>88.6</td><td>-4.4</td><td>0.8</td><td>-5.6</td><td>2.1E-08</td><td>3.6E-08</td><td>MB231/MB468</td></tr>
<tr><td>>MI0000085|hsa-mir-27a&WithFlank&19|-|13836434|13836523@16.37.22&[MIMAT0004501&hsa-miR-27a-5p&offsets|0|0;m-270&19|-|13836487|13836508&offsets|0|0]|AGGGCTTAGCTGCTTGTGAGCA</td><td>>MI0000085|hsa-mir-27a&WithFlank&19|-|13836434|13836523</td><td>16</td><td>37</td><td>22</td><td>AGGGCTTAGCTGCTTGTGAGCA</td><td>690.3</td><td>5.6</td><td>1.0</td><td>5.6</td><td>2.4E-08</td><td>3.6E-08</td><td>BT20/MB468</td></tr>
</table>

<table>
<tr><th></th><th></th><th></th><th></th><th></th><th></th><th></th><th></th><th></th><th></th><th></th><th></th><th></th></tr>
<tr><td>>hg38_21_+_8433172_8446622_RNA45SN1_45S_50ntFlanks@12458.12487.30|AAGCGTTGGATTGTTCACCCACTAATAGGG</td><td>>hg38_21_+_8433172_8446622_RNA45SN1_45S_50ntFlanks</td><td>12458</td><td>12487</td><td>30</td><td>AAGCGTTGGATTGTTCACCCACTAATAGGG</td><td>572.3</td><td>-2.6</td><td>0.5</td><td>-5.6</td><td>2.1E-08</td><td>3.6E-08</td><td>MB231/MB468</td></tr>
<tr><td>>hg38_21_+_8433172_8446622_RNA45SN1_45S_50ntFlanks@4554.4573.20|CCGCAGCTAGGAATAATGGA</td><td>>hg38_21_+_8433172_8446622_RNA45SN1_45S_50ntFlanks</td><td>4554</td><td>4573</td><td>20</td><td>CCGCAGCTAGGAATAATGGA</td><td>82.9</td><td>-4.0</td><td>0.7</td><td>-5.6</td><td>2.4E-08</td><td>3.7E-08</td><td>BT20/MB468</td></tr>
<tr><td>>hg38_21_+_8433172_8446622_RNA45SN1_45S_50ntFlanks@11573.11591.19|CGACTTAGAACTGGTGCGG</td><td>>hg38_21_+_8433172_8446622_RNA45SN1_45S_50ntFlanks</td><td>11573</td><td>11591</td><td>19</td><td>CGACTTAGAACTGGTGCGG</td><td>339.3</td><td>-3.2</td><td>0.6</td><td>-5.6</td><td>2.2E-08</td><td>3.7E-08</td><td>MB231/MB468</td></tr>
<tr><td>>hg38_21_+_8433172_8446622_RNA45SN1_45S_50ntFlanks@11592.11613.22|ACCAGGGGAATCCGACTGTTTA</td><td>>hg38_21_+_8433172_8446622_RNA45SN1_45S_50ntFlanks</td><td>11592</td><td>11613</td><td>22</td><td>ACCAGGGGAATCCGACTGTTTA</td><td>215.5</td><td>-3.6</td><td>0.6</td><td>-5.6</td><td>2.4E-08</td><td>3.7E-08</td><td>BT20/MB468</td></tr>
<tr><td>>hg38_21_+_8433172_8446622_RNA45SN1_45S_50ntFlanks@4729.4758.30|TCAAGAACGAAAGTCGGAGGTTCGAAGACG</td><td>>hg38_21_+_8433172_8446622_RNA45SN1_45S_50ntFlanks</td><td>4729</td><td>4758</td><td>30</td><td>TCAAGAACGAAAGTCGGAGGTTCGAAGACG</td><td>730.1</td><td>-2.7</td><td>0.5</td><td>-5.6</td><td>2.4E-08</td><td>3.7E-08</td><td>BT20/MB468</td></tr>
<tr><td>>hg38_21_+_8433172_8446622_RNA45SN1_45S_50ntFlanks@4566.4587.22|ATAATGGAATAGGACCGCGGTT</td><td>>hg38_21_+_8433172_8446622_RNA45SN1_45S_50ntFlanks</td><td>4566</td><td>4587</td><td>22</td><td>ATAATGGAATAGGACCGCGGTT</td><td>254.2</td><td>-4.2</td><td>0.8</td><td>-5.6</td><td>2.4E-08</td><td>3.7E-08</td><td>BT20/MB468</td></tr>
<tr><td>>hg38_21_+_8433172_8446622_RNA45SN1_45S_50ntFlanks@3775.3793.19|GCCGGTACAGTGAAACTGC</td><td>>hg38_21_+_8433172_8446622_RNA45SN1_45S_50ntFlanks</td><td>3775</td><td>3793</td><td>19</td><td>GCCGGTACAGTGAAACTGC</td><td>86.9</td><td>-2.4</td><td>0.4</td><td>-5.6</td><td>2.5E-08</td><td>3.8E-08</td><td>BT20/MB468</td></tr>
<tr><td>>hg38_21_+_8433172_8446622_RNA45SN1_45S_50ntFlanks@9622.9648.27|CTAGTAGCTGGTTCCCTCCGAAGTTTC</td><td>>hg38_21_+_8433172_8446622_RNA45SN1_45S_50ntFlanks</td><td>9622</td><td>9648</td><td>27</td><td>CTAGTAGCTGGTTCCCTCCGAAGTTTC</td><td>76.4</td><td>-5.3</td><td>0.9</td><td>-5.6</td><td>2.2E-08</td><td>3.8E-08</td><td>MB231/MB468</td></tr>
<tr><td>>hg38_21_+_8433172_8446622_RNA45SN1_45S_50ntFlanks@10905.10936.32|CTGGGCGCGCGCCGCGGCTGGACGAGGCGCCG</td><td>>hg38_21_+_8433172_8446622_RNA45SN1_45S_50ntFlanks</td><td>10905</td><td>10936</td><td>32</td><td>CTGGGCGCGCGCCGCGGCTGGACGAGGCGCCG</td><td>188.2</td><td>-5.5</td><td>1.0</td><td>-5.6</td><td>2.5E-08</td><td>3.8E-08</td><td>BT20/MB468</td></tr>
<tr><td>>hg38_21_+_8433172_8446622_RNA45SN1_45S_50ntFlanks@4069.4089.21|CTTTCGATGGTAGTCGCCGTG</td><td>>hg38_21_+_8433172_8446622_RNA45SN1_45S_50ntFlanks</td><td>4069</td><td>4089</td><td>21</td><td>CTTTCGATGGTAGTCGCCGTG</td><td>147.1</td><td>-4.6</td><td>0.8</td><td>-5.6</td><td>2.3E-08</td><td>3.8E-08</td><td>MB231/MB468</td></tr>
<tr><td>>hg38_21_+_8433172_8446622_RNA45SN1_45S_50ntFlanks@12073.12091.19|CCGAGGGGCTCTCGCTTCT</td><td>>hg38_21_+_8433172_8446622_RNA45SN1_45S_50ntFlanks</td><td>12073</td><td>12091</td><td>19</td><td>CCGAGGGGCTCTCGCTTCT</td><td>99.5</td><td>-3.6</td><td>0.6</td><td>-5.6</td><td>2.3E-08</td><td>3.9E-08</td><td>MB231/MB468</td></tr>
<tr><td>>hg38_21_+_8433172_8446622_RNA45SN1_45S_50ntFlanks@11778.11803.26|ACGCGCATGAATGGATGAACGAGATT</td><td>>hg38_21_+_8433172_8446622_RNA45SN1_45S_50ntFlanks</td><td>11778</td><td>11803</td><td>26</td><td>ACGCGCATGAATGGATGAACGAGATT</td><td>690.4</td><td>-3.8</td><td>0.7</td><td>-5.6</td><td>2.6E-08</td><td>3.9E-08</td><td>BT20/MB468</td></tr>
<tr><td>>MI0000253|hsa-mir-148a&WithFlank&7|-|25949913|25949992@50.71.22(+1U)&[MIMAT0000243&hsa-miR-148a-3p&offsets|0|0(+1U);m-7&7|-|25949922|25949943&offsets|0|0(+1U)]|TCAGTGCACTACAGAACTTTGTT</td><td>>MI0000253|hsa-mir-148a&WithFlank&7|-|25949913|25949992</td><td>50</td><td>71</td><td>22(+1U)</td><td>TCAGTGCACTACAGAACTTTGTT</td><td>2026.1</td><td>4.9</td><td>0.8</td><td>6.0</td><td>1.6E-09</td><td>4.0E-08</td><td>BT20/MB231</td></tr>
<tr><td>>hg38_21_+_8433172_8446622_RNA45SN1_45S_50ntFlanks@10878.10902.25|TCGGTCGGGCTGGGGCGCGAAGCGG</td><td>>hg38_21_+_8433172_8446622_RNA45SN1_45S_50ntFlanks</td><td>10878</td><td>10902</td><td>25</td><td>TCGGTCGGGCTGGGGCGCGAAGCGG</td><td>201.1</td><td>-4.1</td><td>0.7</td><td>-5.6</td><td>2.6E-08</td><td>4.0E-08</td><td>BT20/MB468</td></tr>
<tr><td>>hg38_21_+_8433172_8446622_RNA45SN1_45S_50ntFlanks@10820.10840.21|GGGAAGTCGGCAAGCCGGATC</td><td>>hg38_21_+_8433172_8446622_RNA45SN1_45S_50ntFlanks</td><td>10820</td><td>10840</td><td>21</td><td>GGGAAGTCGGCAAGCCGGATC</td><td>880.1</td><td>-3.2</td><td>0.6</td><td>-5.6</td><td>2.4E-08</td><td>4.0E-08</td><td>MB231/MB468</td></tr>
<tr><td>>hg38_21_+_8433172_8446622_RNA45SN1_45S_50ntFlanks@4444.4472.29|CCTTGCCTCTCGGCGCCCCCTCGATGCTC</td><td>>hg38_21_+_8433172_8446622_RNA45SN1_45S_50ntFlanks</td><td>4444</td><td>4472</td><td>29</td><td>CCTTGCCTCTCGGCGCCCCCTCGATGCTC</td><td>68.4</td><td>-7.7</td><td>1.4</td><td>-5.6</td><td>2.4E-08</td><td>4.1E-08</td><td>MB231/MB468</td></tr>
<tr><td>>hg38_21_+_8433172_8446622_RNA45SN1_45S_50ntFlanks@6652.6688.37|GACTCTTAGCGGTGGATCACTCGGCTCGTGCGTCGAT</td><td>>hg38_21_+_8433172_8446622_RNA45SN1_45S_50ntFlanks</td><td>6652</td><td>6688</td><td>37</td><td>GACTCTTAGCGGTGGATCACTCGGCTCGTGCGTCGAT</td><td>2265.9</td><td>3.3</td><td>0.6</td><td>5.6</td><td>2.7E-08</td><td>4.1E-08</td><td>BT20/MB468</td></tr>
<tr><td>>hg38_21_+_8433172_8446622_RNA45SN1_45S_50ntFlanks@12656.12675.20|TGACTGAACGCCTCTAAGTC</td><td>>hg38_21_+_8433172_8446622_RNA45SN1_45S_50ntFlanks</td><td>12656</td><td>12675</td><td>20</td><td>TGACTGAACGCCTCTAAGTC</td><td>285.8</td><td>-3.9</td><td>0.7</td><td>-5.6</td><td>2.7E-08</td><td>4.1E-08</td><td>BT20/MB468</td></tr>
<tr><td>>hg38_21_+_8433172_8446622_RNA45SN1_45S_50ntFlanks@12332.12349.18|TAAGCAGGAGGTGTCAGA</td><td>>hg38_21_+_8433172_8446622_RNA45SN1_45S_50ntFlanks</td><td>12332</td><td>12349</td><td>18</td><td>TAAGCAGGAGGTGTCAGA</td><td>87.2</td><td>-5.0</td><td>0.9</td><td>-5.6</td><td>2.7E-08</td><td>4.1E-08</td><td>BT20/MB468</td></tr>
<tr><td>>hg38_21_+_8433172_8446622_RNA45SN1_45S_50ntFlanks@11682.11702.21|AATGTCAAAGTGAAGAAATTC</td><td>>hg38_21_+_8433172_8446622_RNA45SN1_45S_50ntFlanks</td><td>11682</td><td>11702</td><td>21</td><td>AATGTCAAAGTGAAGAAATTC</td><td>411.5</td><td>-4.8</td><td>0.9</td><td>-5.6</td><td>2.7E-08</td><td>4.1E-08</td><td>BT20/MB468</td></tr>
<tr><td>>MI0000266|hsa-mir-10a&WithFlank&17|-|48579832|48579953@28.49.22&[MIMAT0000253&hsa-miR-10a-5p&offsets|0|-1;m-18&17|-|48579905|48579926&offsets|0|0]|TACCCTGTAGATCCGAATTTGT</td><td>>MI0000266|hsa-mir-10a&WithFlank&17|-|48579832|48579953</td><td>28</td><td>49</td><td>22</td><td>TACCCTGTAGATCCGAATTTGT</td><td>27491.4</td><td>12.3</td><td>2.2</td><td>5.6</td><td>2.4E-08</td><td>4.1E-08</td><td>MB231/MB468</td></tr>
<tr><td>>hg38_21_+_8433172_8446622_RNA45SN1_45S_50ntFlanks@12539.12558.20|ATGATGTGTTGTTGCCATGG</td><td>>hg38_21_+_8433172_8446622_RNA45SN1_45S_50ntFlanks</td><td>12539</td><td>12558</td><td>20</td><td>ATGATGTGTTGTTGCCATGG</td><td>80.2</td><td>-6.0</td><td>1.1</td><td>-5.6</td><td>2.5E-08</td><td>4.2E-08</td><td>MB231/MB468</td></tr>
<tr><td>>hg38_21_+_8433172_8446622_RNA45SN1_45S_50ntFlanks@7983.8004.22|CAGATCAGACGTGGCGACCCGC</td><td>>hg38_21_+_8433172_8446622_RNA45SN1_45S_50ntFlanks</td><td>7983</td><td>8004</td><td>22</td><td>CAGATCAGACGTGGCGACCCGC</td><td>485.1</td><td>-4.9</td><td>0.9</td><td>-5.6</td><td>2.8E-08</td><td>4.2E-08</td><td>BT20/MB468</td></tr>
<tr><td>>hg38_21_+_8433172_8446622_RNA45SN1_45S_50ntFlanks@11161.11178.18|CACGGTCCCCCGCGAGGG</td><td>>hg38_21_+_8433172_8446622_RNA45SN1_45S_50ntFlanks</td><td>11161</td><td>11178</td><td>18</td><td>CACGGTCCCCCGCGAGGG</td><td>1374.3</td><td>-6.8</td><td>1.2</td><td>-5.6</td><td>2.8E-08</td><td>4.2E-08</td><td>BT20/MB468</td></tr>
<tr><td>>hg38_21_+_8433172_8446622_RNA45SN1_45S_50ntFlanks@8356.8378.23|GAAAAGAACTTTGAAGAGAGAGT</td><td>>hg38_21_+_8433172_8446622_RNA45SN1_45S_50ntFlanks</td><td>8356</td><td>8378</td><td>23</td><td>GAAAAGAACTTTGAAGAGAGAGT</td><td>4154.8</td><td>-3.1</td><td>0.6</td><td>-5.6</td><td>2.5E-08</td><td>4.2E-08</td><td>MB231/MB468</td></tr>
<tr><td>>hg38_21_+_8433172_8446622_RNA45SN1_45S_50ntFlanks@10798.10823.26|CATGTTGGAACAATGTAGGTAAGGGA</td><td>>hg38_21_+_8433172_8446622_RNA45SN1_45S_50ntFlanks</td><td>10798</td><td>10823</td><td>26</td><td>CATGTTGGAACAATGTAGGTAAGGGA</td><td>107.0</td><td>-3.1</td><td>0.5</td><td>-5.6</td><td>2.5E-08</td><td>4.2E-08</td><td>MB231/MB468</td></tr>
<tr><td>>hg38_21_+_8433172_8446622_RNA45SN1_45S_50ntFlanks@7975.8012.38|CGCGACCTCAGATCAGACGTGGCGACCCGCTGAATTTA</td><td>>hg38_21_+_8433172_8446622_RNA45SN1_45S_50ntFlanks</td><td>7975</td><td>8012</td><td>38</td><td>CGCGACCTCAGATCAGACGTGGCGACCCGCTGAATTTA</td><td>27471.9</td><td>4.4</td><td>0.8</td><td>5.6</td><td>2.5E-08</td><td>4.3E-08</td><td>MB231/MB468</td></tr>
<tr><td>>hg38_21_+_8433172_8446622_RNA45SN1_45S_50ntFlanks@10786.10815.30|AACAGCCTCTGGCATGTTGGAACAATGTAG</td><td>>hg38_21_+_8433172_8446622_RNA45SN1_45S_50ntFlanks</td><td>10786</td><td>10815</td><td>30</td><td>AACAGCCTCTGGCATGTTGGAACAATGTAG</td><td>188.4</td><td>-3.4</td><td>0.6</td><td>-5.6</td><td>2.5E-08</td><td>4.3E-08</td><td>MB231/MB468</td></tr>
<tr><td>>hg38_21_+_8433172_8446622_RNA45SN1_45S_50ntFlanks@11580.11611.32|GAACTGGTGCGGACCAGGGGAATCCGACTGTT</td><td>>hg38_21_+_8433172_8446622_RNA45SN1_45S_50ntFlanks</td><td>11580</td><td>11611</td><td>32</td><td>GAACTGGTGCGGACCAGGGGAATCCGACTGTT</td><td>1887.0</td><td>-3.3</td><td>0.6</td><td>-5.6</td><td>2.8E-08</td><td>4.3E-08</td><td>BT20/MB468</td></tr>
<tr><td>>hg38_21_+_8433172_8446622_RNA45SN1_45S_50ntFlanks@485.511.27|GATGTGGTGACGTCGTGCTCTCCCGGG</td><td>>hg38_21_+_8433172_8446622_RNA45SN1_45S_50ntFlanks</td><td>485</td><td>511</td><td>27</td><td>GATGTGGTGACGTCGTGCTCTCCCGGG</td><td>2650.3</td><td>16.3</td><td>2.9</td><td>5.6</td><td>2.6E-08</td><td>4.3E-08</td><td>MB231/MB468</td></tr>
<tr><td>>hg38_21_+_8433172_8446622_RNA45SN1_45S_50ntFlanks@11766.11786.21|ATCTAATTAGTGACGCGCATG</td><td>>hg38_21_+_8433172_8446622_RNA45SN1_45S_50ntFlanks</td><td>11766</td><td>11786</td><td>21</td><td>ATCTAATTAGTGACGCGCATG</td><td>93.4</td><td>-4.6</td><td>0.8</td><td>-5.6</td><td>2.6E-08</td><td>4.4E-08</td><td>MB231/MB468</td></tr>
<tr><td>>hg38_21_+_8433172_8446622_RNA45SN1_45S_50ntFlanks@3841.3868.28|TCCTACTTGGATAACTGTGGTAATTCTA</td><td>>hg38_21_+_8433172_8446622_RNA45SN1_45S_50ntFlanks</td><td>3841</td><td>3868</td><td>28</td><td>TCCTACTTGGATAACTGTGGTAATTCTA</td><td>336.3</td><td>2.1</td><td>0.4</td><td>5.5</td><td>2.9E-08</td><td>4.4E-08</td><td>BT20/MB468</td></tr>
<tr><td>>hg38_21_+_8433172_8446622_RNA45SN1_45S_50ntFlanks@4723.4758.36|CATTAATCAAGAACGAAAGTCGGAGGTTCGAAGACG</td><td>>hg38_21_+_8433172_8446622_RNA45SN1_45S_50ntFlanks</td><td>4723</td><td>4758</td><td>36</td><td>CATTAATCAAGAACGAAAGTCGGAGGTTCGAAGACG</td><td>2554.9</td><td>-3.2</td><td>0.6</td><td>-5.6</td><td>2.6E-08</td><td>4.4E-08</td><td>MB231/MB468</td></tr>
<tr><td>>trnaMT_GlnTTG_MT_-_4329_4400@1.30.30,trnalookalike2_GlnTTG_1_-_564879_564950@1.30.30|TAGGATGGGGTGTGATAGGTGGCACGGAGA</td><td>>trnaMT_GlnTTG_MT_-_4329_4400</td><td>1</td><td>30</td><td>30</td><td>TAGGATGGGGTGTGATAGGTGGCACGGAGA</td><td>1008.4</td><td>5.3</td><td>0.9</td><td>5.5</td><td>2.9E-08</td><td>4.4E-08</td><td>BT20/MB468</td></tr>
<tr><td>>hg38_21_+_8433172_8446622_RNA45SN1_45S_50ntFlanks@6784.6803.20|GCTACGCCTGTCTGAGCGTC</td><td>>hg38_21_+_8433172_8446622_RNA45SN1_45S_50ntFlanks</td><td>6784</td><td>6803</td><td>20</td><td>GCTACGCCTGTCTGAGCGTC</td><td>269.8</td><td>-6.1</td><td>1.1</td><td>-5.6</td><td>2.6E-08</td><td>4.4E-08</td><td>MB231/MB468</td></tr>
<tr><td>>hg38_21_+_8433172_8446622_RNA45SN1_45S_50ntFlanks@12624.12657.34|CCAATGGGGCGAAGCTACCATCTGTGGGATTATG</td><td>>hg38_21_+_8433172_8446622_RNA45SN1_45S_50ntFlanks</td><td>12624</td><td>12657</td><td>34</td><td>CCAATGGGGCGAAGCTACCATCTGTGGGATTATG</td><td>520.6</td><td>-5.4</td><td>1.0</td><td>-5.5</td><td>3.0E-08</td><td>4.5E-08</td><td>BT20/MB468</td></tr>
<tr><td>>hg38_21_+_8433172_8446622_RNA45SN1_45S_50ntFlanks@6666.6686.21|GATCACTCGGCTCGTGCGTCG</td><td>>hg38_21_+_8433172_8446622_RNA45SN1_45S_50ntFlanks</td><td>6666</td><td>6686</td><td>21</td><td>GATCACTCGGCTCGTGCGTCG</td><td>89.1</td><td>-4.7</td><td>0.9</td><td>-5.6</td><td>2.7E-08</td><td>4.5E-08</td><td>MB231/MB468</td></tr>
<tr><td>>hg38_21_+_8433172_8446622_RNA45SN1_45S_50ntFlanks@7979.7996.18|ACCTCAGATCAGACGTGG</td><td>>hg38_21_+_8433172_8446622_RNA45SN1_45S_50ntFlanks</td><td>7979</td><td>7996</td><td>18</td><td>ACCTCAGATCAGACGTGG</td><td>87.8</td><td>-4.4</td><td>0.8</td><td>-5.6</td><td>2.7E-08</td><td>4.5E-08</td><td>MB231/MB468</td></tr>
<tr><td>>hg38_21_+_8433172_8446622_RNA45SN1_45S_50ntFlanks@6721.6743.23|AATTGCAGGACACATTGATCATC</td><td>>hg38_21_+_8433172_8446622_RNA45SN1_45S_50ntFlanks</td><td>6721</td><td>6743</td><td>23</td><td>AATTGCAGGACACATTGATCATC</td><td>5604.7</td><td>-3.7</td><td>0.7</td><td>-5.6</td><td>2.7E-08</td><td>4.5E-08</td><td>MB231/MB468</td></tr>
<tr><td>>hg38_21_+_8433172_8446622_RNA45SN1_45S_50ntFlanks@10845.10872.28|ACTTCGGGATAAGGATTGGCTCTAAGGG</td><td>>hg38_21_+_8433172_8446622_RNA45SN1_45S_50ntFlanks</td><td>10845</td><td>10872</td><td>28</td><td>ACTTCGGGATAAGGATTGGCTCTAAGGG</td><td>108.0</td><td>-3.9</td><td>0.7</td><td>-5.5</td><td>3.0E-08</td><td>4.6E-08</td><td>BT20/MB468</td></tr>
<tr><td>>hg38_1_-_228634819_228635039_RNA5S12_5S_50ntFlanks@127.171.45|ACTTGGATGGGAGACCGCCTGGGAATACCGGGTGCTGTAGGCTTT</td><td>>hg38_1_-_228634819_228635039_RNA5S12_5S_50ntFlanks</td><td>127</td><td>171</td><td>45</td><td>ACTTGGATGGGAGACCGCCTGGGAATACCGGGTGCTGTAGGCTTT</td><td>2178.9</td><td>12.7</td><td>2.3</td><td>5.6</td><td>2.8E-08</td><td>4.6E-08</td><td>MB231/MB468</td></tr>
<tr><td>>hg38_21_+_8433172_8446622_RNA45SN1_45S_50ntFlanks@9738.9762.25|GAAACGATCTCAACCTATTCTCAAA</td><td>>hg38_21_+_8433172_8446622_RNA45SN1_45S_50ntFlanks</td><td>9738</td><td>9762</td><td>25</td><td>GAAACGATCTCAACCTATTCTCAAA</td><td>119.8</td><td>-4.8</td><td>0.9</td><td>-5.5</td><td>3.1E-08</td><td>4.6E-08</td><td>BT20/MB468</td></tr>
<tr><td>>hg38_21_+_8433172_8446622_RNA45SN1_45S_50ntFlanks@12002.12031.30|TGCGGGCCGCCGGTGAAATACCACTACTCT</td><td>>hg38_21_+_8433172_8446622_RNA45SN1_45S_50ntFlanks</td><td>12002</td><td>12031</td><td>30</td><td>TGCGGGCCGCCGGTGAAATACCACTACTCT</td><td>531.9</td><td>-2.8</td><td>0.5</td><td>-5.6</td><td>2.8E-08</td><td>4.7E-08</td><td>MB231/MB468</td></tr>
<tr><td>>trna10_GluCTC_3_-_103879524_103879595@16.42.27,trna116_GluCTC_1_-_145399233_145399304@16.42.27,trna59_GluCTC_1_+_249168447_249168518@16.42.27,trna71_GluCTC_1_-_161439189_161439260@16.42.27,trna74_GluCTC_1_-_161431809_161431880@16.42.27,trna77_GluCTC_1_-_161424398_161424469@16.42.27,trna77_GluCTC_6_+_28949976_28950047@16.42.27,trna80_GluCTC_1_-_161417018_161417089@16.42.27,trna87_GluCTC_6_-_126101393_126101464@16.42.27|TGGTTAGGATTCGGCGCTCTCACCGCC</td><td>>trna10_GluCTC_3_-_103879524_103879595</td><td>16</td><td>42</td><td>27</td><td>TGGTTAGGATTCGGCGCTCTCACCGCC</td><td>828.2</td><td>4.8</td><td>0.9</td><td>5.6</td><td>2.8E-08</td><td>4.7E-08</td><td>MB231/MB468</td></tr>
<tr><td>>MI0000462|hsa-mir-152&WithFlank&17|-|48037155|48037253@60.81.22&[MIMAT0000438&hsa-miR-152-3p&offsets|0|+1;m-97&17|-|48037174|48037194&offsets|0|+1]|TCAGTGCATGACAGAACTTGGG</td><td>>MI0000462|hsa-mir-152&WithFlank&17|-|48037155|48037253</td><td>60</td><td>81</td><td>22</td><td>TCAGTGCATGACAGAACTTGGG</td><td>1703.3</td><td>12.0</td><td>2.2</td><td>5.6</td><td>2.8E-08</td><td>4.7E-08</td><td>MB231/MB468</td></tr>
<tr><td>>hg38_21_+_8433172_8446622_RNA45SN1_45S_50ntFlanks@485.516.32|GATGTGGTGACGTCGTGCTCTCCCGGGCCGGG</td><td>>hg38_21_+_8433172_8446622_RNA45SN1_45S_50ntFlanks</td><td>485</td><td>516</td><td>32</td><td>GATGTGGTGACGTCGTGCTCTCCCGGGCCGGG</td><td>3474.5</td><td>16.7</td><td>3.0</td><td>5.6</td><td>2.8E-08</td><td>4.7E-08</td><td>MB231/MB468</td></tr>
</table>

| | | | | | | | | | | | | |
|---|---|---|---|---|---|---|---|---|---|---|---|---|
| >hg38_21_+_8433172_8446622_RNA45SN1_45S_50ntFlanks@10843.10867.25\|TAACTTCGGGATAAGGATTGGCTCT | >hg38_21_+_8433172_8446622_RNA45SN1_45S_50ntFlanks | 10843 | 10867 | 25 | TAACTTCGGGATAAGGATTGGCTCT | 229.7 | -3.3 | 0.6 | -5.5 | 3.1E-08 | 4.7E-08 | BT20/MB468 |
| >hg38_21_+_8433172_8446622_RNA45SN1_45S_50ntFlanks@3802.3829.28\|CATTAAATCAGTTATGGTTCCTTTGGTC | >hg38_21_+_8433172_8446622_RNA45SN1_45S_50ntFlanks | 3802 | 3829 | 28 | CATTAAATCAGTTATGGTTCCTTTGGTC | 157.2 | -3.0 | 0.5 | -5.6 | 2.8E-08 | 4.7E-08 | MB231/MB468 |
| >hg38_21_+_8433172_8446622_RNA45SN1_45S_50ntFlanks@7980.7998.19\|CCTCAGATCAGACGTGGCG | >hg38_21_+_8433172_8446622_RNA45SN1_45S_50ntFlanks | 7980 | 7998 | 19 | CCTCAGATCAGACGTGGCG | 104.9 | -5.0 | 0.9 | -5.6 | 2.8E-08 | 4.7E-08 | MB231/MB468 |
| >hg38_21_+_8433172_8446622_RNA45SN1_45S_50ntFlanks@10351.10380.30\|CGAGAACTTTGAAGGCCGAAGTGGAGAAGG | >hg38_21_+_8433172_8446622_RNA45SN1_45S_50ntFlanks | 10351 | 10380 | 30 | CGAGAACTTTGAAGGCCGAAGTGGAGAAGG | 343.0 | -2.8 | 0.5 | -5.5 | 3.1E-08 | 4.7E-08 | BT20/MB468 |
| >hg38_21_+_8433172_8446622_RNA45SN1_45S_50ntFlanks@9305.9335.31\|CCAAGGAGTCTAACACGTGCGCGAGTCGGGG | >hg38_21_+_8433172_8446622_RNA45SN1_45S_50ntFlanks | 9305 | 9335 | 31 | CCAAGGAGTCTAACACGTGCGCGAGTCGGGG | 75.7 | -4.9 | 0.9 | -5.5 | 3.1E-08 | 4.8E-08 | BT20/MB468 |
| >hg38_21_+_8433172_8446622_RNA45SN1_45S_50ntFlanks@10689.10709.21\|GCGTCCGGTGAGCTCTCGCTG | >hg38_21_+_8433172_8446622_RNA45SN1_45S_50ntFlanks | 10689 | 10709 | 21 | GCGTCCGGTGAGCTCTCGCTG | 484.7 | -6.0 | 1.1 | -5.5 | 3.2E-08 | 4.8E-08 | BT20/MB468 |
| >hg38_21_+_8433172_8446622_RNA45SN1_45S_50ntFlanks@7975.7998.24\|CGCGACCTCAGATCAGACGTGGCG | >hg38_21_+_8433172_8446622_RNA45SN1_45S_50ntFlanks | 7975 | 7998 | 24 | CGCGACCTCAGATCAGACGTGGCG | 875.5 | 4.9 | 0.9 | 5.5 | 3.2E-08 | 4.8E-08 | BT20/MB468 |
| >trna10_LysCTT_16_+_3241501_3241573@3.33.31,trna13_LysCTT_14_-_58706613_58706685@3.33.31,trna2_LysCTT_15_+_79152904_79152976@3.33.31\|CCGGCTAGCTCAGTCGGTAGAGCATGGGACT | >trna10_LysCTT_16_+_3241501_3241573 | 3 | 33 | 31 | CCGGCTAGCTCAGTCGGTAGAGCATGGGACT | 2035.2 | 12.4 | 2.2 | 5.6 | 2.9E-08 | 4.8E-08 | MB231/MB468 |
| >hg38_21_+_8433172_8446622_RNA45SN1_45S_50ntFlanks@12457.12484.28\|CAAGCGTTGGATTGTTCACCCACTAATA | >hg38_21_+_8433172_8446622_RNA45SN1_45S_50ntFlanks | 12457 | 12484 | 28 | CAAGCGTTGGATTGTTCACCCACTAATA | 882.1 | -4.7 | 0.8 | -5.5 | 2.9E-08 | 4.8E-08 | MB231/MB468 |
| >trna12_ArgCCT_16_+_3243918_3243990@34.51.18,trna18_ArgCCT_17_+_73030001_73030073@34.51.18,trna21_ArgCCT_17_-_73030526_73030598@34.51.18,trna2_ArgCCT_16_+_3202901_3202973@34.51.18,trna3_ArgCCT_7_+_139025446_139025518@34.51.18\|CCTAAGCCAGGGATTGTG | >trna12_ArgCCT_16_+_3243918_3243990 | 34 | 51 | 18 | CCTAAGCCAGGGATTGTG | 88.4 | -6.1 | 1.1 | -5.5 | 2.9E-08 | 4.8E-08 | MB231/MB468 |
| >hg38_21_+_8433172_8446622_RNA45SN1_45S_50ntFlanks@7989.8011.23\|AGACGTGGCGACCCGCTGAATTT | >hg38_21_+_8433172_8446622_RNA45SN1_45S_50ntFlanks | 7989 | 8011 | 23 | AGACGTGGCGACCCGCTGAATTT | 2908.9 | 3.9 | 0.7 | 5.5 | 3.2E-08 | 4.9E-08 | BT20/MB468 |
| >hg38_21_+_8433172_8446622_RNA45SN1_45S_50ntFlanks@10392.10433.42\|ACAGCAGTTGAACATGGGTCAGTCGGTCCTGAGAGATGGGCG | >hg38_21_+_8433172_8446622_RNA45SN1_45S_50ntFlanks | 10392 | 10433 | 42 | ACAGCAGTTGAACATGGGTCAGTCGGTCCTGAGAGATGGGCG | 322.9 | -5.8 | 1.0 | -5.5 | 3.3E-08 | 5.0E-08 | BT20/MB468 |
| >MI0000462\|hsa-mir-152&WithFlank&17\|-\|48037155\|48037253@60.80.21&[MIMAT0000438&hsa-miR-152-3p&offsets\|0\|0;m-97&17\|-\|48037174\|48037194&offsets\|0\|0]\|TCAGTGCATGACAGAACTTGG | >MI0000462\|hsa-mir-152&WithFlank&17\|-\|48037155\|48037253 | 60 | 80 | 21 | TCAGTGCATGACAGAACTTGG | 2509.6 | 12.4 | 2.2 | 5.5 | 3.0E-08 | 5.1E-08 | MB231/MB468 |
| >hg38_21_+_8433172_8446622_RNA45SN1_45S_50ntFlanks@7984.8003.20\|AGATCAGACGTGGCGACCCG | >hg38_21_+_8433172_8446622_RNA45SN1_45S_50ntFlanks | 7984 | 8003 | 20 | AGATCAGACGTGGCGACCCG | 164.4 | -4.4 | 0.8 | -5.5 | 3.4E-08 | 5.2E-08 | BT20/MB468 |
| >hg38_21_+_8433172_8446622_RNA45SN1_45S_50ntFlanks@11770.11794.25\|AATTAGTGACGCGCATGAATGGATG | >hg38_21_+_8433172_8446622_RNA45SN1_45S_50ntFlanks | 11770 | 11794 | 25 | AATTAGTGACGCGCATGAATGGATG | 841.1 | 3.1 | 0.6 | 5.5 | 3.1E-08 | 5.2E-08 | MB231/MB468 |
| >hg38_21_+_8433172_8446622_RNA45SN1_45S_50ntFlanks@11671.11698.28\|CCAGTGCTCTGAATGTCAAAGTGAAGAA | >hg38_21_+_8433172_8446622_RNA45SN1_45S_50ntFlanks | 11671 | 11698 | 28 | CCAGTGCTCTGAATGTCAAAGTGAAGAA | 126.9 | -3.1 | 0.6 | -5.5 | 3.5E-08 | 5.2E-08 | BT20/MB468 |
| >hg38_21_+_8433172_8446622_RNA45SN1_45S_50ntFlanks@4045.4062.18\|CCATTCGAACGTCTGCCC | >hg38_21_+_8433172_8446622_RNA45SN1_45S_50ntFlanks | 4045 | 4062 | 18 | CCATTCGAACGTCTGCCC | 262.5 | -2.6 | 0.5 | -5.5 | 3.1E-08 | 5.2E-08 | MB231/MB468 |
| >hg38_21_+_8433172_8446622_RNA45SN1_45S_50ntFlanks@11315.11343.29\|CCACCCCACGTCTCGTCGCGCGCGCGTCC | >hg38_21_+_8433172_8446622_RNA45SN1_45S_50ntFlanks | 11315 | 11343 | 29 | CCACCCCACGTCTCGTCGCGCGCGCGTCC | 85.2 | -6.1 | 1.1 | -5.5 | 3.2E-08 | 5.3E-08 | MB231/MB468 |
| >MI0000289\|hsa-mir-181a-1&WithFlank&1\|-\|198859038\|198859159@30.53.24;MI0000269\|hsa-mir-181a-2&WithFlank&9\|+\|124692436\|124692557@45.68.24&[MIMAT0000256&hsa-miR-181a-5p&offsets\|0\|+1;m-32&1\|-\|198859108\|198859130&offsets\|0\|+1];[MIMAT0000256_1&hsa-miR-181a-5p&offsets\|0\|+1;m-32&9\|+\|124692480\|124692502&offsets\|0\|+1]\|AACATTCAACGCTGTCGGTGAGTT | >MI0000289\|hsa-mir-181a-1&WithFlank&1\|-\|198859038\|198859159 | 30 | 53 | 24 | AACATTCAACGCTGTCGGTGAGTT | 1019.9 | 5.2 | 0.9 | 5.5 | 3.5E-08 | 5.3E-08 | BT20/MB468 |
| >hg38_21_+_8433172_8446622_RNA45SN1_45S_50ntFlanks@10684.10711.28\|CGGCGGCGTCCGGTGAGCTCTCGCTGGC | >hg38_21_+_8433172_8446622_RNA45SN1_45S_50ntFlanks | 10684 | 10711 | 28 | CGGCGGCGTCCGGTGAGCTCTCGCTGGC | 2409.2 | -3.0 | 0.5 | -5.5 | 3.2E-08 | 5.4E-08 | MB231/MB468 |
| >hg38_21_+_8433172_8446622_RNA45SN1_45S_50ntFlanks@9005.9026.22\|CTCCCCCGCGGGGGCGCGCCGG | >hg38_21_+_8433172_8446622_RNA45SN1_45S_50ntFlanks | 9005 | 9026 | 22 | CTCCCCCGCGGGGGCGCGCCGG | 95.7 | -4.8 | 0.9 | -5.5 | 3.2E-08 | 5.4E-08 | MB231/MB468 |
| >MI0000266\|hsa-mir-10a&WithFlank&17\|-\|48579832\|48579953@28.50.23(+1A)&[MIMAT0000253&hsa-miR-10a-5p&offsets\|0\|0(+1A);m-18&17\|-\|48579905\|48579926&offsets\|0\|+1(+1A)]\|TACCCTGTAGATCCGAATTTGTGA | >MI0000266\|hsa-mir-10a&WithFlank&17\|-\|48579832\|48579953 | 28 | 50 | 23(+1A) | TACCCTGTAGATCCGAATTTGTGA | 2001.6 | 12.1 | 2.2 | 5.5 | 3.2E-08 | 5.4E-08 | MB231/MB468 |
| >MI0000076\|hsa-mir-20a&WithFlank&13\|+\|91351059\|91351141@14.36.23&[MIMAT0000075&hsa-miR-20a-5p&offsets\|0\|0;m-96&13\|+\|91351072\|91351094&offsets\|0\|0]\|TAAAGTGCTTATAGTGCAGGTAG | >MI0000076\|hsa-mir-20a&WithFlank&13\|+\|91351059\|91351141 | 14 | 36 | 23 | TAAAGTGCTTATAGTGCAGGTAG | 31250.6 | 5.1 | 0.9 | 5.5 | 3.7E-08 | 5.5E-08 | BT20/MB468 |
| >hg38_21_+_8433172_8446622_RNA45SN1_45S_50ntFlanks@9565.9596.32\|TGACGTGCAAATCGGTCGTCCGACCTGGGTAT | >hg38_21_+_8433172_8446622_RNA45SN1_45S_50ntFlanks | 9565 | 9596 | 32 | TGACGTGCAAATCGGTCGTCCGACCTGGGTAT | 4359.9 | -3.5 | 0.6 | -5.5 | 3.3E-08 | 5.6E-08 | MB231/MB468 |
| >am_trnaMT_ValTAC_MT_+_1602_1670@30.50.21\|CTTACACTTAGGAGATTTCAA | >am_trnaMT_ValTAC_MT_+_1602_1670 | 30 | 50 | 21 | CTTACACTTAGGAGATTTCAA | 63.2 | -8.0 | 1.4 | -5.5 | 3.7E-08 | 5.6E-08 | BT20/MB468 |
| >hg38_21_+_8433172_8446622_RNA45SN1_45S_50ntFlanks@7975.8018.44\|CGCGACCTCAGATCAGACGTGGCGACCCGCTGAATTTAAGCATA | >hg38_21_+_8433172_8446622_RNA45SN1_45S_50ntFlanks | 7975 | 8018 | 44 | CGCGACCTCAGATCAGACGTGGCGACCCGCTGAATTTAAGCATA | 2269.7 | 7.5 | 1.4 | 5.5 | 3.7E-08 | 5.6E-08 | BT20/MB468 |
| >hg38_21_+_8433172_8446622_RNA45SN1_45S_50ntFlanks@13000.13039.40\|TGCGATCTATTGAAAGTCAGCCCTCGACACAAGGGTTTGT | >hg38_21_+_8433172_8446622_RNA45SN1_45S_50ntFlanks | 13000 | 13039 | 40 | TGCGATCTATTGAAAGTCAGCCCTCGACACAAGGGTTTGT | 528.6 | 5.6 | 1.0 | 5.5 | 3.7E-08 | 5.6E-08 | BT20/MB468 |
| >hg38_21_+_8433172_8446622_RNA45SN1_45S_50ntFlanks@11752.11777.26\|CCAAATGCCTCGTCATCTAATTAGTG | >hg38_21_+_8433172_8446622_RNA45SN1_45S_50ntFlanks | 11752 | 11777 | 26 | CCAAATGCCTCGTCATCTAATTAGTG | 78.9 | -4.2 | 0.8 | -5.5 | 3.4E-08 | 5.6E-08 | MB231/MB468 |
| >hg38_21_+_8433172_8446622_RNA45SN1_45S_50ntFlanks@3831.3860.30\|CTCGCTCCTCTCCTACTTGGATAACTGTGG | >hg38_21_+_8433172_8446622_RNA45SN1_45S_50ntFlanks | 3831 | 3860 | 30 | CTCGCTCCTCTCCTACTTGGATAACTGTGG | 191.9 | -3.3 | 0.6 | -5.5 | 3.7E-08 | 5.7E-08 | BT20/MB468 |
| >hg38_21_+_8433172_8446622_RNA45SN1_45S_50ntFlanks@10508.10533.26\|CGAATCCGGAGTGGCGGAGATGGGCG | >hg38_21_+_8433172_8446622_RNA45SN1_45S_50ntFlanks | 10508 | 10533 | 26 | CGAATCCGGAGTGGCGGAGATGGGCG | 143.9 | -5.1 | 0.9 | -5.5 | 3.8E-08 | 5.7E-08 | BT20/MB468 |
| >MI0000102\|hsa-mir-100&WithFlank&11\|-\|122152223\|122152314@19.40.22(+1A)&[MIMAT0000098&hsa-miR-100-5p&offsets\|0\|0(+1A);m-27&11\|-\|122152276\|122152296&offsets\|0\|+1(+1A)]\|AACCCGTAGATCCGAACTTGTGA | >MI0000102\|hsa-mir-100&WithFlank&11\|-\|122152223\|122152314 | 19 | 40 | 22(+1A) | AACCCGTAGATCCGAACTTGTGA | 33755.5 | -5.4 | 0.9 | -6.0 | 2.4E-09 | 5.7E-08 | BT20/MB231 |
| >hg38_21_+_8433172_8446622_RNA45SN1_45S_50ntFlanks@12456.12480.25\|CCAAGCGTTGGATTGTTCACCCACT | >hg38_21_+_8433172_8446622_RNA45SN1_45S_50ntFlanks | 12456 | 12480 | 25 | CCAAGCGTTGGATTGTTCACCCACT | 1902.9 | -3.8 | 0.7 | -5.5 | 3.4E-08 | 5.8E-08 | MB231/MB468 |
| >hg38_21_+_8433172_8446622_RNA45SN1_45S_50ntFlanks@12160.12179.20\|TTGACTGGGGCGGTACACCT | >hg38_21_+_8433172_8446622_RNA45SN1_45S_50ntFlanks | 12160 | 12179 | 20 | TTGACTGGGGCGGTACACCT | 124.0 | -4.6 | 0.8 | -5.5 | 3.5E-08 | 5.8E-08 | MB231/MB468 |
| >hg38_21_+_8433172_8446622_RNA45SN1_45S_50ntFlanks@12633.12651.19\|CGAAGCTACCATCTGTGGG | >hg38_21_+_8433172_8446622_RNA45SN1_45S_50ntFlanks | 12633 | 12651 | 19 | CGAAGCTACCATCTGTGGG | 597.4 | -5.0 | 0.9 | -5.5 | 3.5E-08 | 5.8E-08 | MB231/MB468 |
| >hg38_21_+_8433172_8446622_RNA45SN1_45S_50ntFlanks@12926.12956.31\|ACCTGGCGCTAAACCATTCGTAGACGACCTG | >hg38_21_+_8433172_8446622_RNA45SN1_45S_50ntFlanks | 12926 | 12956 | 31 | ACCTGGCGCTAAACCATTCGTAGACGACCTG | 109.5 | -5.6 | 1.0 | -5.5 | 3.9E-08 | 5.8E-08 | BT20/MB468 |
| >hg38_21_+_8433172_8446622_RNA45SN1_45S_50ntFlanks@4724.4748.25\|ATTAATCAAGAACGAAAGTCGGAGG | >hg38_21_+_8433172_8446622_RNA45SN1_45S_50ntFlanks | 4724 | 4748 | 25 | ATTAATCAAGAACGAAAGTCGGAGG | 356.4 | -3.3 | 0.6 | -5.5 | 3.9E-08 | 5.8E-08 | BT20/MB468 |
| >hg38_21_+_8433172_8446622_RNA45SN1_45S_50ntFlanks@5487.5526.40\|CGCTGAGAAGACGGTCGAACTTGACTATCTAGAGGAAGTA | >hg38_21_+_8433172_8446622_RNA45SN1_45S_50ntFlanks | 5487 | 5526 | 40 | CGCTGAGAAGACGGTCGAACTTGACTATCTAGAGGAAGTA | 258.8 | -4.8 | 0.9 | -5.5 | 3.9E-08 | 5.8E-08 | BT20/MB468 |
| >trna111_HisGTG_1_-_147774845_147774916@-1A.23.24,trna118_HisGTG_1_-_145396881_145396952@-1A.23.24,trna16_HisGTG_1_+_146544773_146544844@-1A.23.24,trna1_HisGTG_15_+_45493349_45493420@-1A.23.24,trna21_HisGTG_1_+_147753471_147753542@-1A.23.24,trna33_HisGTG_6_+_27125906_27125977@-1A.23.24,trna7_HisGTG_9_-_14433938_14434009@-1A.23.24,trna8_HisGTG_15_-_45492611_45492682@-1A.23.24,trna9_HisGTG_15_-_45490804_45490875@-1A.23.24\|AGCCGTGATCGTATAGTGGTTAGT | >trna111_HisGTG_1_-_147774845_147774916 | -1A | 23 | 24 | AGCCGTGATCGTATAGTGGTTAGT | 577.7 | -4.6 | 0.8 | -6.0 | 2.5E-09 | 5.9E-08 | BT20/MB231 |
| >hg38_21_+_8433172_8446622_RNA45SN1_45S_50ntFlanks@10798.10822.25\|CATGTTGGAACAATGTAGGTAAGGG | >hg38_21_+_8433172_8446622_RNA45SN1_45S_50ntFlanks | 10798 | 10822 | 25 | CATGTTGGAACAATGTAGGTAAGGG | 154.4 | -3.1 | 0.6 | -5.5 | 3.5E-08 | 5.9E-08 | MB231/MB468 |
| >hg38_21_+_8433172_8446622_RNA45SN1_45S_50ntFlanks@7977.7995.19\|CGACCTCAGATCAGACGTG | >hg38_21_+_8433172_8446622_RNA45SN1_45S_50ntFlanks | 7977 | 7995 | 19 | CGACCTCAGATCAGACGTG | 179.2 | -3.9 | 0.7 | -5.5 | 3.9E-08 | 5.9E-08 | BT20/MB468 |
| >hg38_21_+_8433172_8446622_RNA45SN1_45S_50ntFlanks@4370.4398.29\|TTAAAAAGCTCGTAGTTGGATCTTGGGAG | >hg38_21_+_8433172_8446622_RNA45SN1_45S_50ntFlanks | 4370 | 4398 | 29 | TTAAAAAGCTCGTAGTTGGATCTTGGGAG | 156.1 | -5.0 | 0.9 | -5.5 | 3.9E-08 | 5.9E-08 | BT20/MB468 |
| >hg38_21_+_8433172_8446622_RNA45SN1_45S_50ntFlanks@8198.8215.18\|TGGACGGTGTGAGGCCGG | >hg38_21_+_8433172_8446622_RNA45SN1_45S_50ntFlanks | 8198 | 8215 | 18 | TGGACGGTGTGAGGCCGG | 1834.8 | -4.9 | 0.9 | -5.5 | 3.6E-08 | 6.0E-08 | MB231/MB468 |
| >MI0000440\|hsa-mir-27b&WithFlank&9\|+\|95085439\|95085547@70.87.18&[MIMAT0000419&hsa-miR-27b-3p&offsets\|+3\|0;m-39&9\|+\|95085505\|95085525&offsets\|+3\|0]\|ACAGTGGCTAAGTTCTGC | >MI0000440\|hsa-mir-27b&WithFlank&9\|+\|95085439\|95085547 | 70 | 87 | 18 | ACAGTGGCTAAGTTCTGC | 1858.7 | 15.8 | 2.9 | 5.5 | 3.6E-08 | 6.0E-08 | MB231/MB468 |

<table>
<tr><td>>hg38_21_+_8433172_8446622_RNA45SN1_45S_50ntFlanks@12081.12106.26|CTCTCGCTTCTGGCGCCAAGCGCCCG</td><td>>hg38_21_+_8433172_8446622_RNA45SN1_45S_50ntFlanks</td><td>12081</td><td>12106</td><td>26</td><td>CTCTCGCTTCTGGCGCCAAGCGCCCG</td><td>437.8</td><td>-4.9</td><td>0.9</td><td>-5.5</td><td>4.0E-08</td><td>6.0E-08</td><td>BT20/MB468</td></tr>
<tr><td>>hg38_21_+_8433172_8446622_RNA45SN1_45S_50ntFlanks@12003.12033.31|GCGGGCCGCCGGTGAAATACCACTACTCTGA</td><td>>hg38_21_+_8433172_8446622_RNA45SN1_45S_50ntFlanks</td><td>12003</td><td>12033</td><td>31</td><td>GCGGGCCGCCGGTGAAATACCACTACTCTGA</td><td>606.0</td><td>4.3</td><td>0.8</td><td>5.5</td><td>4.0E-08</td><td>6.0E-08</td><td>BT20/MB468</td></tr>
<tr><td>>hg38_21_+_8433172_8446622_RNA45SN1_45S_50ntFlanks@12637.12657.21|GCTACCATCTGTGGGATTATG</td><td>>hg38_21_+_8433172_8446622_RNA45SN1_45S_50ntFlanks</td><td>12637</td><td>12657</td><td>21</td><td>GCTACCATCTGTGGGATTATG</td><td>130.1</td><td>-3.1</td><td>0.6</td><td>-5.5</td><td>4.0E-08</td><td>6.0E-08</td><td>BT20/MB468</td></tr>
<tr><td>>MI0000253|hsa-mir-148a&WithFlank&7|-|25949913|25949992@12.33.22&[MIMAT0004549&hsa-miR-148a-5p&offsets|0|0;m-189&7|-|25949960|25949981&offsets|0|0]|AAAGTTCTGAGACACTCCGACT</td><td>>MI0000253|hsa-mir-148a&WithFlank&7|-|25949913|25949992</td><td>12</td><td>33</td><td>22</td><td>AAAGTTCTGAGACACTCCGACT</td><td>726.9</td><td>4.8</td><td>0.8</td><td>6.0</td><td>2.6E-09</td><td>6.1E-08</td><td>BT20/MB231</td></tr>
<tr><td>>hg38_21_+_8433172_8446622_RNA45SN1_45S_50ntFlanks@9505.9523.19|TGAACTATGCCTGGGCAGG</td><td>>hg38_21_+_8433172_8446622_RNA45SN1_45S_50ntFlanks</td><td>9505</td><td>9523</td><td>19</td><td>TGAACTATGCCTGGGCAGG</td><td>107.7</td><td>-4.2</td><td>0.8</td><td>-5.5</td><td>3.6E-08</td><td>6.1E-08</td><td>MB231/MB468</td></tr>
<tr><td>>hg38_21_+_8433172_8446622_RNA45SN1_45S_50ntFlanks@12417.12439.23|TCGATGTCGGCTCTTCCTATCAT</td><td>>hg38_21_+_8433172_8446622_RNA45SN1_45S_50ntFlanks</td><td>12417</td><td>12439</td><td>23</td><td>TCGATGTCGGCTCTTCCTATCAT</td><td>731.7</td><td>3.2</td><td>0.6</td><td>5.5</td><td>3.6E-08</td><td>6.1E-08</td><td>MB231/MB468</td></tr>
<tr><td>>MI0000076|hsa-mir-20a&WithFlank&13|+|91351059|91351141@14.37.24&[MIMAT0000075&hsa-miR-20a-5p&offsets|0|+1;m-96&13|+|91351072|91351094&offsets|0|+1]|TAAAGTGCTTATAGTGCAGGTAGT</td><td>>MI0000076|hsa-mir-20a&WithFlank&13|+|91351059|91351141</td><td>14</td><td>37</td><td>24</td><td>TAAAGTGCTTATAGTGCAGGTAGT</td><td>2157.6</td><td>5.3</td><td>1.0</td><td>5.5</td><td>4.0E-08</td><td>6.1E-08</td><td>BT20/MB468</td></tr>
<tr><td>>hg38_21_+_8433172_8446622_RNA45SN1_45S_50ntFlanks@9533.9561.29|GAGGAAACTCTGGTGGAGGTCCGTAGCGG</td><td>>hg38_21_+_8433172_8446622_RNA45SN1_45S_50ntFlanks</td><td>9533</td><td>9561</td><td>29</td><td>GAGGAAACTCTGGTGGAGGTCCGTAGCGG</td><td>172.1</td><td>-3.4</td><td>0.6</td><td>-5.5</td><td>3.7E-08</td><td>6.1E-08</td><td>MB231/MB468</td></tr>
<tr><td>>hg38_21_+_8433172_8446622_RNA45SN1_45S_50ntFlanks@4325.4346.22|CCGCGGTAATTCCAGCTCCAAT</td><td>>hg38_21_+_8433172_8446622_RNA45SN1_45S_50ntFlanks</td><td>4325</td><td>4346</td><td>22</td><td>CCGCGGTAATTCCAGCTCCAAT</td><td>100.4</td><td>-4.9</td><td>0.9</td><td>-5.5</td><td>3.7E-08</td><td>6.1E-08</td><td>MB231/MB468</td></tr>
<tr><td>>hg38_21_+_8433172_8446622_RNA45SN1_45S_50ntFlanks@11882.11904.23|CTGTTGAGCTTGACTCTAGTCTG</td><td>>hg38_21_+_8433172_8446622_RNA45SN1_45S_50ntFlanks</td><td>11882</td><td>11904</td><td>23</td><td>CTGTTGAGCTTGACTCTAGTCTG</td><td>876.7</td><td>-5.5</td><td>1.0</td><td>-5.5</td><td>3.7E-08</td><td>6.2E-08</td><td>MB231/MB468</td></tr>
<tr><td>>am_trnaMT_ValTAC_MT_+_1602_1670@50.67.18|ACTTAACTTGACCGCTCT</td><td>>am_trnaMT_ValTAC_MT_+_1602_1670</td><td>50</td><td>67</td><td>18</td><td>ACTTAACTTGACCGCTCT</td><td>5829.2</td><td>5.5</td><td>1.0</td><td>5.5</td><td>4.1E-08</td><td>6.2E-08</td><td>BT20/MB468</td></tr>
<tr><td>>hg38_21_+_8433172_8446622_RNA45SN1_45S_50ntFlanks@11561.11600.40|GCGCCTAGCAGCCGACTTAGAACTGGTGCGGACCAGGGGA</td><td>>hg38_21_+_8433172_8446622_RNA45SN1_45S_50ntFlanks</td><td>11561</td><td>11600</td><td>40</td><td>GCGCCTAGCAGCCGACTTAGAACTGGTGCGGACCAGGGGA</td><td>201.2</td><td>-4.1</td><td>0.7</td><td>-5.5</td><td>4.1E-08</td><td>6.2E-08</td><td>BT20/MB468</td></tr>
<tr><td>>hg38_21_+_8433172_8446622_RNA45SN1_45S_50ntFlanks@13006.13039.34|CTATTGAAAGTCAGCCCTCGACACAAGGGTTTGT</td><td>>hg38_21_+_8433172_8446622_RNA45SN1_45S_50ntFlanks</td><td>13006</td><td>13039</td><td>34</td><td>CTATTGAAAGTCAGCCCTCGACACAAGGGTTTGT</td><td>1221.9</td><td>3.8</td><td>0.7</td><td>5.5</td><td>4.1E-08</td><td>6.2E-08</td><td>BT20/MB468</td></tr>
<tr><td>>MI0000299|hsa-mir-222&WithFlank&X|-|45747009|45747130@75.99.25(+4U)&[MIMAT0000279&hsa-miR-222-3p&offsets|0|+4(+4U);m-140&X|-|45747033|45747056&offsets|0|+1(+4U)]|AGCTACATCTGGCTACTGGGTCTCTTTTT</td><td>>MI0000299|hsa-mir-222&WithFlank&X|-|45747009|45747130</td><td>75</td><td>99</td><td>25(+4U)</td><td>AGCTACATCTGGCTACTGGGTCTCTTTTT</td><td>1821.1</td><td>11.9</td><td>2.2</td><td>5.5</td><td>3.7E-08</td><td>6.2E-08</td><td>MB231/MB468</td></tr>
<tr><td>>hg38_21_+_8433172_8446622_RNA45SN1_45S_50ntFlanks@10806.10830.25|AACAATGTAGGTAAGGGAAGTCGGC</td><td>>hg38_21_+_8433172_8446622_RNA45SN1_45S_50ntFlanks</td><td>10806</td><td>10830</td><td>25</td><td>AACAATGTAGGTAAGGGAAGTCGGC</td><td>72.9</td><td>-5.9</td><td>1.1</td><td>-5.5</td><td>3.7E-08</td><td>6.2E-08</td><td>MB231/MB468</td></tr>
<tr><td>>hg38_21_+_8433172_8446622_RNA45SN1_45S_50ntFlanks@10351.10385.35|CGAGAACTTTGAAGGCCGAAGTGGAGAAGGGTTCC</td><td>>hg38_21_+_8433172_8446622_RNA45SN1_45S_50ntFlanks</td><td>10351</td><td>10385</td><td>35</td><td>CGAGAACTTTGAAGGCCGAAGTGGAGAAGGGTTCC</td><td>161.3</td><td>-3.1</td><td>0.6</td><td>-5.5</td><td>4.2E-08</td><td>6.3E-08</td><td>BT20/MB468</td></tr>
<tr><td>>MI0000289|hsa-mir-181a-1&WithFlank&1|-|198859038|198859159@30.52.23;MI0000269|hsa-mir-181a-2&WithFlank&9|+|124692436|124692557@45.67.23&[MIMAT0000256&hsa-miR-181a-5p&offsets|0|0;m-32&1|-|198859108|198859130&offsets|0|0];[MIMAT0000256_1&hsa-miR-181a-5p&offsets|0|0;m-32&9|+|124692480|124692502&offsets|0|0]|AACATTCAACGCTGTCGGTGAGT</td><td>>MI0000289|hsa-mir-181a-1&WithFlank&1|-|198859038|198859159</td><td>30</td><td>52</td><td>23</td><td>AACATTCAACGCTGTCGGTGAGT</td><td>3624.5</td><td>4.6</td><td>0.8</td><td>5.5</td><td>4.2E-08</td><td>6.3E-08</td><td>BT20/MB468</td></tr>
<tr><td>>hg38_21_+_8433172_8446622_RNA45SN1_45S_50ntFlanks@11729.11765.37|TAACTATGACTCTCTTAAGGTAGCCAAATGCCTCGTC</td><td>>hg38_21_+_8433172_8446622_RNA45SN1_45S_50ntFlanks</td><td>11729</td><td>11765</td><td>37</td><td>TAACTATGACTCTCTTAAGGTAGCCAAATGCCTCGTC</td><td>2551.2</td><td>-4.0</td><td>0.7</td><td>-5.5</td><td>3.9E-08</td><td>6.5E-08</td><td>MB231/MB468</td></tr>
<tr><td>>hg38_21_+_8433172_8446622_RNA45SN1_45S_50ntFlanks@11580.11601.22|GAACTGGTGCGGACCAGGGGAA</td><td>>hg38_21_+_8433172_8446622_RNA45SN1_45S_50ntFlanks</td><td>11580</td><td>11601</td><td>22</td><td>GAACTGGTGCGGACCAGGGGAA</td><td>670.8</td><td>-3.6</td><td>0.7</td><td>-5.5</td><td>3.9E-08</td><td>6.5E-08</td><td>MB231/MB468</td></tr>
<tr><td>>hg38_21_+_8433172_8446622_RNA45SN1_45S_50ntFlanks@12083.12108.26|CTCGCTTCTGGCGCCAAGCGCCCGGC</td><td>>hg38_21_+_8433172_8446622_RNA45SN1_45S_50ntFlanks</td><td>12083</td><td>12108</td><td>26</td><td>CTCGCTTCTGGCGCCAAGCGCCCGGC</td><td>195.6</td><td>-3.8</td><td>0.7</td><td>-5.5</td><td>3.9E-08</td><td>6.6E-08</td><td>MB231/MB468</td></tr>
<tr><td>>MI0000471|hsa-mir-126&WithFlank&9|+|136670596|136670692@58.77.20&[MIMAT0000445&hsa-miR-126-3p&offsets|0|-2;m-40&9|+|136670653|136670674&offsets|0|-2]|TCGTACCGTGAGTAATAATG</td><td>>MI0000471|hsa-mir-126&WithFlank&9|+|136670596|136670692</td><td>58</td><td>77</td><td>20</td><td>TCGTACCGTGAGTAATAATG</td><td>1337.5</td><td>8.7</td><td>1.6</td><td>5.5</td><td>4.4E-08</td><td>6.6E-08</td><td>BT20/MB468</td></tr>
<tr><td>>hg38_1_-_228634819_228635039_RNA5S12_5S_50ntFlanks@140.171.32|ACCGCCTGGGAATACCGGGTGCTGTAGGCTTT</td><td>>hg38_1_-_228634819_228635039_RNA5S12_5S_50ntFlanks</td><td>140</td><td>171</td><td>32</td><td>ACCGCCTGGGAATACCGGGTGCTGTAGGCTTT</td><td>6512.7</td><td>4.9</td><td>0.9</td><td>5.5</td><td>4.4E-08</td><td>6.7E-08</td><td>BT20/MB468</td></tr>
<tr><td>>MI0001733|hsa-mir-452&WithFlank&X|-|151959622|151959718@20.41.22(+1A)&[MIMAT0001635&hsa-miR-452-5p&offsets|0|0(+1A);m-188&X|-|151959677|151959699&offsets|0|-1(+1A)]|AACTGTTTGCAGAGGAAACTGAA</td><td>>MI0001733|hsa-mir-452&WithFlank&X|-|151959622|151959718</td><td>20</td><td>41</td><td>22(+1A)</td><td>AACTGTTTGCAGAGGAAACTGAA</td><td>244.5</td><td>4.5</td><td>0.7</td><td>5.9</td><td>2.8E-09</td><td>6.7E-08</td><td>BT20/MB231</td></tr>
<tr><td>>hg38_21_+_8433172_8446622_RNA45SN1_45S_50ntFlanks@12475.12496.22|CCCACTAATAGGGAACGTGAGC</td><td>>hg38_21_+_8433172_8446622_RNA45SN1_45S_50ntFlanks</td><td>12475</td><td>12496</td><td>22</td><td>CCCACTAATAGGGAACGTGAGC</td><td>1451.6</td><td>3.2</td><td>0.6</td><td>5.5</td><td>4.0E-08</td><td>6.7E-08</td><td>MB231/MB468</td></tr>
<tr><td>>am_trnaMT_ArgTCG_MT_+_10405_10469@31.48.18|TCGACTCATTAAATTATG</td><td>>am_trnaMT_ArgTCG_MT_+_10405_10469</td><td>31</td><td>48</td><td>18</td><td>TCGACTCATTAAATTATG</td><td>145.6</td><td>-4.4</td><td>0.8</td><td>-5.5</td><td>4.5E-08</td><td>6.7E-08</td><td>BT20/MB468</td></tr>
<tr><td>>hg38_21_+_8433172_8446622_RNA45SN1_45S_50ntFlanks@10816.10840.25|GTAAGGGAAGTCGGCAAGCCGGATC</td><td>>hg38_21_+_8433172_8446622_RNA45SN1_45S_50ntFlanks</td><td>10816</td><td>10840</td><td>25</td><td>GTAAGGGAAGTCGGCAAGCCGGATC</td><td>139.2</td><td>-2.8</td><td>0.5</td><td>-5.5</td><td>4.0E-08</td><td>6.8E-08</td><td>MB231/MB468</td></tr>
<tr><td>>hg38_21_+_8433172_8446622_RNA45SN1_45S_50ntFlanks@10116.10135.20|TGCGTCGGAGGGCGGCGGCG</td><td>>hg38_21_+_8433172_8446622_RNA45SN1_45S_50ntFlanks</td><td>10116</td><td>10135</td><td>20</td><td>TGCGTCGGAGGGCGGCGGCG</td><td>159.9</td><td>-4.8</td><td>0.9</td><td>-5.5</td><td>4.5E-08</td><td>6.8E-08</td><td>BT20/MB468</td></tr>
<tr><td>>hg38_21_+_8433172_8446622_RNA45SN1_45S_50ntFlanks@7979.7999.21|ACCTCAGATCAGACGTGGCGA</td><td>>hg38_21_+_8433172_8446622_RNA45SN1_45S_50ntFlanks</td><td>7979</td><td>7999</td><td>21</td><td>ACCTCAGATCAGACGTGGCGA</td><td>101.2</td><td>-6.3</td><td>1.1</td><td>-5.5</td><td>4.1E-08</td><td>6.8E-08</td><td>MB231/MB468</td></tr>
<tr><td>>hg38_21_+_8433172_8446622_RNA45SN1_45S_50ntFlanks@4730.4752.23|CAAGAACGAAAGTCGGAGGTTCG</td><td>>hg38_21_+_8433172_8446622_RNA45SN1_45S_50ntFlanks</td><td>4730</td><td>4752</td><td>23</td><td>CAAGAACGAAAGTCGGAGGTTCG</td><td>237.8</td><td>-3.1</td><td>0.6</td><td>-5.5</td><td>4.5E-08</td><td>6.8E-08</td><td>BT20/MB468</td></tr>
<tr><td>>trna10_LysCTT_16_+_3241501_3241573@1.29.29,trna13_LysCTT_14_-_58706613_58706685@1.29.29,trna2_LysCTT_15_+_79152904_79152976@1.29.29|GCCCGGCTAGCTCAGTCGGTAGAGCATGG</td><td>>trna10_LysCTT_16_+_3241501_3241573</td><td>1</td><td>29</td><td>29</td><td>GCCCGGCTAGCTCAGTCGGTAGAGCATGG</td><td>515.1</td><td>-3.7</td><td>0.7</td><td>-5.5</td><td>4.1E-08</td><td>6.9E-08</td><td>MB231/MB468</td></tr>
<tr><td>>hg38_21_+_8433172_8446622_RNA45SN1_45S_50ntFlanks@10761.10780.20|ATATCCGCAGCAGGTCTCCA</td><td>>hg38_21_+_8433172_8446622_RNA45SN1_45S_50ntFlanks</td><td>10761</td><td>10780</td><td>20</td><td>ATATCCGCAGCAGGTCTCCA</td><td>96.6</td><td>-6.2</td><td>1.1</td><td>-5.5</td><td>4.1E-08</td><td>6.9E-08</td><td>MB231/MB468</td></tr>
<tr><td>>hg38_21_+_8433172_8446622_RNA45SN1_45S_50ntFlanks@11592.11613.22|ACCAGGGGAATCCGACTGTTTA</td><td>>hg38_21_+_8433172_8446622_RNA45SN1_45S_50ntFlanks</td><td>11592</td><td>11613</td><td>22</td><td>ACCAGGGGAATCCGACTGTTTA</td><td>177.5</td><td>-3.3</td><td>0.6</td><td>-5.5</td><td>4.1E-08</td><td>6.9E-08</td><td>MB231/MB468</td></tr>
<tr><td>>MI0000298|hsa-mir-221&WithFlank&X|-|45746151|45746272@31.56.26(+1U)&[MIMAT0004568&hsa-miR-221-5p&offsets|0|+4(+1U);m-405&X|-|45746220|45746242&offsets|0|+3(+1U)]|ACCTGGCATACAATGTAGATTTCTGTT</td><td>>MI0000298|hsa-mir-221&WithFlank&X|-|45746151|45746272</td><td>31</td><td>56</td><td>26(+1U)</td><td>ACCTGGCATACAATGTAGATTTCTGTT</td><td>679.4</td><td>11.0</td><td>2.0</td><td>5.5</td><td>4.1E-08</td><td>6.9E-08</td><td>MB231/MB468</td></tr>
<tr><td>>hg38_21_+_8433172_8446622_RNA45SN1_45S_50ntFlanks@3840.3873.34|CTCCTACTTGGATAACTGTGGTAATTCTAGAGCT</td><td>>hg38_21_+_8433172_8446622_RNA45SN1_45S_50ntFlanks</td><td>3840</td><td>3873</td><td>34</td><td>CTCCTACTTGGATAACTGTGGTAATTCTAGAGCT</td><td>677.0</td><td>4.5</td><td>0.8</td><td>5.5</td><td>4.6E-08</td><td>6.9E-08</td><td>BT20/MB468</td></tr>
<tr><td>>hg38_21_+_8433172_8446622_RNA45SN1_45S_50ntFlanks@9621.9650.30|TCTAGTAGCTGGTTCCCTCCGAAGTTTCCC</td><td>>hg38_21_+_8433172_8446622_RNA45SN1_45S_50ntFlanks</td><td>9621</td><td>9650</td><td>30</td><td>TCTAGTAGCTGGTTCCCTCCGAAGTTTCCC</td><td>57.0</td><td>-7.4</td><td>1.4</td><td>-5.5</td><td>4.2E-08</td><td>7.0E-08</td><td>MB231/MB468</td></tr>
<tr><td>>MI0000294|hsa-mir-218-1&WithFlank&4|+|20528269|20528390@31.51.21;MI0000295|hsa-mir-218-2&WithFlank&5|-|168768140|168768261@31.51.21&[MIMAT0000275&hsa-miR-218-5p&offsets|0|0;m-185&4|+|20528299|20528319&offsets|0|0];[MIMAT0000275_1&hsa-miR-218-5p&offsets|0|0;m-187&5|-|168768211|168768231&offsets|0|0]|TTGTGCTTGATCTAACCATGT</td><td>>MI0000294|hsa-mir-218-1&WithFlank&4|+|20528269|20528390</td><td>31</td><td>51</td><td>21</td><td>TTGTGCTTGATCTAACCATGT</td><td>1808.2</td><td>11.9</td><td>2.2</td><td>5.5</td><td>4.2E-08</td><td>7.0E-08</td><td>MB231/MB468</td></tr>
<tr><td>>hg38_21_+_8433172_8446622_RNA45SN1_45S_50ntFlanks@8356.8375.20|GAAAAGAACTTTGAAGAGAG</td><td>>hg38_21_+_8433172_8446622_RNA45SN1_45S_50ntFlanks</td><td>8356</td><td>8375</td><td>20</td><td>GAAAAGAACTTTGAAGAGAG</td><td>894.3</td><td>-4.1</td><td>0.8</td><td>-5.5</td><td>4.7E-08</td><td>7.0E-08</td><td>BT20/MB468</td></tr>
<tr><td>>MI0000071|hsa-mir-17&WithFlank&13|+|91350599|91350694@20.42.23&[MIMAT0000070&hsa-miR-17-5p&offsets|0|0;m-88&13|+|91350618|91350640&offsets|0|0]|CAAAGTGCTTACAGTGCAGGTAG</td><td>>MI0000071|hsa-mir-17&WithFlank&13|+|91350599|91350694</td><td>20</td><td>42</td><td>23</td><td>CAAAGTGCTTACAGTGCAGGTAG</td><td>14244.5</td><td>4.6</td><td>0.8</td><td>5.5</td><td>4.7E-08</td><td>7.0E-08</td><td>BT20/MB468</td></tr>
<tr><td>>hg38_21_+_8433172_8446622_RNA45SN1_45S_50ntFlanks@11576.11609.34|CTTAGAACTGGTGCGGACCAGGGGAATCCGACTG</td><td>>hg38_21_+_8433172_8446622_RNA45SN1_45S_50ntFlanks</td><td>11576</td><td>11609</td><td>34</td><td>CTTAGAACTGGTGCGGACCAGGGGAATCCGACTG</td><td>637.4</td><td>-3.5</td><td>0.6</td><td>-5.5</td><td>4.7E-08</td><td>7.1E-08</td><td>BT20/MB468</td></tr>
<tr><td>>hg38_21_+_8433172_8446622_RNA45SN1_45S_50ntFlanks@6646.6677.32|TCGTACGACTCTTAGCGGTGGATCACTCGGCT</td><td>>hg38_21_+_8433172_8446622_RNA45SN1_45S_50ntFlanks</td><td>6646</td><td>6677</td><td>32</td><td>TCGTACGACTCTTAGCGGTGGATCACTCGGCT</td><td>1425.1</td><td>4.1</td><td>0.7</td><td>5.5</td><td>4.3E-08</td><td>7.2E-08</td><td>MB231/MB468</td></tr>
<tr><td>>hg38_21_+_8433172_8446622_RNA45SN1_45S_50ntFlanks@10916.10936.21|CCGCGGCTGGACGAGGCGCCG</td><td>>hg38_21_+_8433172_8446622_RNA45SN1_45S_50ntFlanks</td><td>10916</td><td>10936</td><td>21</td><td>CCGCGGCTGGACGAGGCGCCG</td><td>231.1</td><td>-5.7</td><td>1.0</td><td>-5.5</td><td>4.8E-08</td><td>7.2E-08</td><td>BT20/MB468</td></tr>
<tr><td>>hg38_21_+_8433172_8446622_RNA45SN1_45S_50ntFlanks@4804.4822.19|ATGCGGCGGCGTTATTCCC</td><td>>hg38_21_+_8433172_8446622_RNA45SN1_45S_50ntFlanks</td><td>4804</td><td>4822</td><td>19</td><td>ATGCGGCGGCGTTATTCCC</td><td>290.4</td><td>3.6</td><td>0.7</td><td>5.5</td><td>4.3E-08</td><td>7.2E-08</td><td>MB231/MB468</td></tr>
<tr><td>>hg38_21_+_8433172_8446622_RNA45SN1_45S_50ntFlanks@9534.9567.34|AGGAAACTCTGGTGGAGGTCCGTAGCGGTCCTGA</td><td>>hg38_21_+_8433172_8446622_RNA45SN1_45S_50ntFlanks</td><td>9534</td><td>9567</td><td>34</td><td>AGGAAACTCTGGTGGAGGTCCGTAGCGGTCCTGA</td><td>757.3</td><td>-5.5</td><td>1.0</td><td>-5.5</td><td>4.8E-08</td><td>7.2E-08</td><td>BT20/MB468</td></tr>
<tr><td>>hg38_21_+_8433172_8446622_RNA45SN1_45S_50ntFlanks@5467.5491.25|GGCCCACGGCCCTGGCGGAGCGCTG</td><td>>hg38_21_+_8433172_8446622_RNA45SN1_45S_50ntFlanks</td><td>5467</td><td>5491</td><td>25</td><td>GGCCCACGGCCCTGGCGGAGCGCTG</td><td>120.9</td><td>-4.7</td><td>0.9</td><td>-5.5</td><td>4.3E-08</td><td>7.2E-08</td><td>MB231/MB468</td></tr>
</table>

| | | | | | | | | | | | | |
|---|---|---|---|---|---|---|---|---|---|---|---|---|
| >hg38_21_+_8433172_8446622_RNA45SN1_45S_50ntFlanks@11671.11701.31\|CCAGTGCTCTGAATGTCAAAGTGAAGAAATT | >hg38_21_+_8433172_8446622_RNA45SN1_45S_50ntFlanks | 11671 | 11701 | 31 | CCAGTGCTCTGAATGTCAAAGTGAAGAAATT | 370.8 | -2.9 | 0.5 | -5.5 | 4.9E-08 | 7.3E-08 | BT20/MB468 |
| >hg38_21_+_8433172_8446622_RNA45SN1_45S_50ntFlanks@11672.11707.36\|CAGTGCTCTGAATGTCAAAGTGAAGAAATTCAATGA | >hg38_21_+_8433172_8446622_RNA45SN1_45S_50ntFlanks | 11672 | 11707 | 36 | CAGTGCTCTGAATGTCAAAGTGAAGAAATTCAATGA | 613.9 | 4.3 | 0.8 | 5.5 | 4.4E-08 | 7.3E-08 | MB231/MB468 |
| >hg38_21_+_8433172_8446622_RNA45SN1_45S_50ntFlanks@12630.12667.38\|GGGCGAAGCTACCATCTGTGGGATTATGACTGAACGCC | >hg38_21_+_8433172_8446622_RNA45SN1_45S_50ntFlanks | 12630 | 12667 | 38 | GGGCGAAGCTACCATCTGTGGGATTATGACTGAACGCC | 284.9 | 9.9 | 1.8 | 5.5 | 4.4E-08 | 7.3E-08 | MB231/MB468 |
| >hg38_21_+_8433172_8446622_RNA45SN1_45S_50ntFlanks@10350.10378.29\|ACGAGAACTTTGAAGGCCGAAGTGGAGAA | >hg38_21_+_8433172_8446622_RNA45SN1_45S_50ntFlanks | 10350 | 10378 | 29 | ACGAGAACTTTGAAGGCCGAAGTGGAGAA | 342.6 | -2.3 | 0.4 | -5.5 | 4.4E-08 | 7.3E-08 | MB231/MB468 |
| >trna20_GluTTC_2_-_131094701_131094772@1.31.31,trna5_GluTTC_13_-_41634874_41634945@1.31.31\|TCCCATATGGTCTAGCGGTTAGGATTCCTGG | >trna20_GluTTC_2_-_131094701_131094772 | 1 | 31 | 31 | TCCCATATGGTCTAGCGGTTAGGATTCCTGG | 255.0 | -2.8 | 0.5 | -5.5 | 4.9E-08 | 7.3E-08 | BT20/MB468 |
| >hg38_21_+_8433172_8446622_RNA45SN1_45S_50ntFlanks@10794.10814.21\|CTGGCATGTTGGAACAATGTA | >hg38_21_+_8433172_8446622_RNA45SN1_45S_50ntFlanks | 10794 | 10814 | 21 | CTGGCATGTTGGAACAATGTA | 202.3 | -3.9 | 0.7 | -5.5 | 4.4E-08 | 7.4E-08 | MB231/MB468 |
| >hg38_21_+_8433172_8446622_RNA45SN1_45S_50ntFlanks@10808.10831.24\|CAATGTAGGTAAGGGAAGTCGGCA | >hg38_21_+_8433172_8446622_RNA45SN1_45S_50ntFlanks | 10808 | 10831 | 24 | CAATGTAGGTAAGGGAAGTCGGCA | 98.9 | -2.9 | 0.5 | -5.5 | 4.4E-08 | 7.4E-08 | MB231/MB468 |
| >trna14_LysTTT_11_-_59327808_59327880@1.24.24,trna2_LysTTT_17_+_8022473_8022545@1.24.24,trna54_LysTTT_1_+_204475655_204475727@1.24.24,trna5_LysTTT_11_+_59323902_59323974@1.24.24,trna62_LysTTT_1_-_204476158_204476230@1.24.24,trna76_LysTTT_6_+_28918806_28918878@1.24.24\|GCCCGGATAGCTCAGTCGGTAGAG | >trna14_LysTTT_11_-_59327808_59327880 | 1 | 24 | 24 | GCCCGGATAGCTCAGTCGGTAGAG | 475.9 | -4.2 | 0.8 | -5.5 | 4.9E-08 | 7.4E-08 | BT20/MB468 |
| >MI0000102\|hsa-mir-100&WithFlank&11\|-\|122152223\|122152314@19.39.21&[MIMAT0000098&hsa-miR-100-5p&offsets\|0\|-1;m-27&11\|-\|122152276\|122152296&offsets\|0\|0]\|AACCCGTAGATCCGAACTTGT | >MI0000102\|hsa-mir-100&WithFlank&11\|-\|122152223\|122152314 | 19 | 39 | 21 | AACCCGTAGATCCGAACTTGT | 6736.2 | 8.5 | 1.6 | 5.5 | 5.0E-08 | 7.5E-08 | BT20/MB468 |
| >hg38_21_+_8433172_8446622_RNA45SN1_45S_50ntFlanks@10392.10422.31\|ACAGCAGTTGAACATGGGTCAGTCGGTCCTG | >hg38_21_+_8433172_8446622_RNA45SN1_45S_50ntFlanks | 10392 | 10422 | 31 | ACAGCAGTTGAACATGGGTCAGTCGGTCCTG | 603.1 | -4.4 | 0.8 | -5.5 | 5.0E-08 | 7.5E-08 | BT20/MB468 |
| >MI0003591\|hsa-mir-584&WithFlank&5\|-\|149062307\|149062415@22.42.21(+1U)&[MIMAT0003249&hsa-miR-584-5p&offsets\|0\|-1(+1U);m-167&5\|-\|149062374\|149062394&offsets\|0\|0(+1U)]\|TTATGGTTTGCCTGGGACTGAT | >MI0003591\|hsa-mir-584&WithFlank&5\|-\|149062307\|149062415 | 22 | 42 | 21(+1U) | TTATGGTTTGCCTGGGACTGAT | 1675.4 | 12.0 | 2.2 | 5.5 | 4.5E-08 | 7.6E-08 | MB231/MB468 |
| >hg38_21_+_8433172_8446622_RNA45SN1_45S_50ntFlanks@11210.11254.45\|GGCGACTCTGGACGCGAGCCGGGCCCTTCCCGTGGATCGCCCCAG | >hg38_21_+_8433172_8446622_RNA45SN1_45S_50ntFlanks | 11210 | 11254 | 45 | GGCGACTCTGGACGCGAGCCGGGCCCTTCCCGTGGATCGCCCCAG | 154.0 | -5.8 | 1.1 | -5.4 | 5.1E-08 | 7.6E-08 | BT20/MB468 |
| >hg38_21_+_8433172_8446622_RNA45SN1_45S_50ntFlanks@6753.6770.18\|AACGCACTTGCGGCCCCG | >hg38_21_+_8433172_8446622_RNA45SN1_45S_50ntFlanks | 6753 | 6770 | 18 | AACGCACTTGCGGCCCCG | 76.2 | -4.3 | 0.8 | -5.5 | 4.6E-08 | 7.7E-08 | MB231/MB468 |
| >MI0000102\|hsa-mir-100&WithFlank&11\|-\|122152223\|122152314@19.39.21(+1C)&[MIMAT0000098&hsa-miR-100-5p&offsets\|0\|-1(+1C);m-27&11\|-\|122152276\|122152296&offsets\|0\|0(+1C)]\|AACCCGTAGATCCGAACTTGTC | >MI0000102\|hsa-mir-100&WithFlank&11\|-\|122152223\|122152314 | 19 | 39 | 21(+1C) | AACCCGTAGATCCGAACTTGTC | 10152.7 | 13.5 | 2.5 | 5.5 | 4.7E-08 | 7.8E-08 | MB231/MB468 |
| >hg38_21_+_8433172_8446622_RNA45SN1_45S_50ntFlanks@7980.7998.19\|CCTCAGATCAGACGTGGCG | >hg38_21_+_8433172_8446622_RNA45SN1_45S_50ntFlanks | 7980 | 7998 | 19 | CCTCAGATCAGACGTGGCG | 124.5 | -5.2 | 1.0 | -5.4 | 5.2E-08 | 7.8E-08 | BT20/MB468 |
| >hg38_21_+_8433172_8446622_RNA45SN1_45S_50ntFlanks@12073.12106.34\|CCGAGGGGCTCTCGCTTCTGGCGCCAAGCGCCCG | >hg38_21_+_8433172_8446622_RNA45SN1_45S_50ntFlanks | 12073 | 12106 | 34 | CCGAGGGGCTCTCGCTTCTGGCGCCAAGCGCCCG | 237.7 | -3.6 | 0.7 | -5.5 | 4.7E-08 | 7.8E-08 | MB231/MB468 |
| >hg38_21_+_8433172_8446622_RNA45SN1_45S_50ntFlanks@11576.11602.27\|CTTAGAACTGGTGCGGACCAGGGGAAT | >hg38_21_+_8433172_8446622_RNA45SN1_45S_50ntFlanks | 11576 | 11602 | 27 | CTTAGAACTGGTGCGGACCAGGGGAAT | 744.8 | -4.4 | 0.8 | -5.5 | 4.7E-08 | 7.8E-08 | MB231/MB468 |
| >hg38_21_+_8433172_8446622_RNA45SN1_45S_50ntFlanks@9534.9570.37\|AGGAAACTCTGGTGGAGGTCCGTAGCGGTCCTGACGT | >hg38_21_+_8433172_8446622_RNA45SN1_45S_50ntFlanks | 9534 | 9570 | 37 | AGGAAACTCTGGTGGAGGTCCGTAGCGGTCCTGACGT | 689.9 | -2.9 | 0.5 | -5.4 | 5.3E-08 | 7.9E-08 | BT20/MB468 |
| >hg38_21_+_8433172_8446622_RNA45SN1_45S_50ntFlanks@10401.10424.24\|GAACATGGGTCAGTCGGTCCTGAG | >hg38_21_+_8433172_8446622_RNA45SN1_45S_50ntFlanks | 10401 | 10424 | 24 | GAACATGGGTCAGTCGGTCCTGAG | 1583.0 | 4.3 | 0.8 | 5.4 | 5.3E-08 | 7.9E-08 | BT20/MB468 |
| >hg38_21_+_8433172_8446622_RNA45SN1_45S_50ntFlanks@3984.4024.41\|GCGGCTTTGGTGACTCTAGATAACCTCGGGCCGATCGCACG | >hg38_21_+_8433172_8446622_RNA45SN1_45S_50ntFlanks | 3984 | 4024 | 41 | GCGGCTTTGGTGACTCTAGATAACCTCGGGCCGATCGCACG | 310.0 | -7.3 | 1.3 | -5.4 | 5.3E-08 | 7.9E-08 | BT20/MB468 |
| >hg38_21_+_8433172_8446622_RNA45SN1_45S_50ntFlanks@10686.10706.21\|GCGGCGTCCGGTGAGCTCTCG | >hg38_21_+_8433172_8446622_RNA45SN1_45S_50ntFlanks | 10686 | 10706 | 21 | GCGGCGTCCGGTGAGCTCTCG | 602.0 | -3.6 | 0.7 | -5.5 | 4.8E-08 | 8.0E-08 | MB231/MB468 |
| >hg38_21_+_8433172_8446622_RNA45SN1_45S_50ntFlanks@12433.12454.22\|CTATCATTGTGAAGCAGAATTC | >hg38_21_+_8433172_8446622_RNA45SN1_45S_50ntFlanks | 12433 | 12454 | 22 | CTATCATTGTGAAGCAGAATTC | 169.2 | -5.4 | 1.0 | -5.4 | 5.4E-08 | 8.0E-08 | BT20/MB468 |
| >hg38_21_+_8433172_8446622_RNA45SN1_45S_50ntFlanks@12416.12437.22\|TTCGATGTCGGCTCTTCCTATC | >hg38_21_+_8433172_8446622_RNA45SN1_45S_50ntFlanks | 12416 | 12437 | 22 | TTCGATGTCGGCTCTTCCTATC | 1115.2 | 2.6 | 0.5 | 5.4 | 5.4E-08 | 8.1E-08 | BT20/MB468 |
| >hg38_21_+_8433172_8446622_RNA45SN1_45S_50ntFlanks@10797.10823.27\|GCATGTTGGAACAATGTAGGTAAGGGA | >hg38_21_+_8433172_8446622_RNA45SN1_45S_50ntFlanks | 10797 | 10823 | 27 | GCATGTTGGAACAATGTAGGTAAGGGA | 101.7 | -2.8 | 0.5 | -5.5 | 4.9E-08 | 8.1E-08 | MB231/MB468 |
| >MI0000253\|hsa-mir-148a&WithFlank&7\|-\|25949913\|25949992@50.72.23(+1A)&[MIMAT0000243&hsa-miR-148a-3p&offsets\|0\|+1(+1A);m-7&7\|-\|25949922\|25949943&offsets\|0\|+1(+1A)]\|TCAGTGCACTACAGAACTTTGTCA | >MI0000253\|hsa-mir-148a&WithFlank&7\|-\|25949913\|25949992 | 50 | 72 | 23(+1A) | TCAGTGCACTACAGAACTTTGTCA | 311.4 | 6.1 | 1.0 | 5.9 | 3.5E-09 | 8.1E-08 | BT20/MB231 |
| >hg38_21_+_8433172_8446622_RNA45SN1_45S_50ntFlanks@12008.12033.26\|CCGCCGGTGAAATACCACTACTCTGA | >hg38_21_+_8433172_8446622_RNA45SN1_45S_50ntFlanks | 12008 | 12033 | 26 | CCGCCGGTGAAATACCACTACTCTGA | 301.8 | -5.0 | 0.9 | -5.4 | 5.4E-08 | 8.1E-08 | BT20/MB468 |
| >MI0000286\|hsa-mir-210&WithFlank&11\|-\|568083\|568204@72.93.22(+1G)&[MIMAT0000267&hsa-miR-210-3p&offsets\|0\|0(+1G);m-102&11\|-\|568112\|568133&offsets\|0\|0(+1G)]\|CTGTGCGTGTGACAGCGGCTGAG | >MI0000286\|hsa-mir-210&WithFlank&11\|-\|568083\|568204 | 72 | 93 | 22(+1G) | CTGTGCGTGTGACAGCGGCTGAG | 2076.7 | 15.7 | 2.9 | 5.4 | 5.5E-08 | 8.2E-08 | BT20/MB468 |
| >hg38_21_+_8433172_8446622_RNA45SN1_45S_50ntFlanks@8158.8188.31\|TCGTGGGGGGCCCAAGTCCTTCTGATCGAGG | >hg38_21_+_8433172_8446622_RNA45SN1_45S_50ntFlanks | 8158 | 8188 | 31 | TCGTGGGGGGCCCAAGTCCTTCTGATCGAGG | 180.6 | -4.5 | 0.8 | -5.4 | 5.6E-08 | 8.3E-08 | BT20/MB468 |
| >hg38_21_+_8433172_8446622_RNA45SN1_45S_50ntFlanks@12941.12960.20\|ATTCGTAGACGACCTGCTTC | >hg38_21_+_8433172_8446622_RNA45SN1_45S_50ntFlanks | 12941 | 12960 | 20 | ATTCGTAGACGACCTGCTTC | 460.6 | 2.7 | 0.5 | 5.4 | 5.6E-08 | 8.4E-08 | BT20/MB468 |
| >hg38_21_+_8433172_8446622_RNA45SN1_45S_50ntFlanks@4547.4565.19\|CTGGATACCGCAGCTAGGA | >hg38_21_+_8433172_8446622_RNA45SN1_45S_50ntFlanks | 4547 | 4565 | 19 | CTGGATACCGCAGCTAGGA | 185.9 | -5.4 | 1.0 | -5.4 | 5.6E-08 | 8.4E-08 | BT20/MB468 |
| >hg38_21_+_8433172_8446622_RNA45SN1_45S_50ntFlanks@10781.10802.22\|AGGTGAACAGCCTCTGGCATGT | >hg38_21_+_8433172_8446622_RNA45SN1_45S_50ntFlanks | 10781 | 10802 | 22 | AGGTGAACAGCCTCTGGCATGT | 209.3 | -2.3 | 0.4 | -5.4 | 5.1E-08 | 8.4E-08 | MB231/MB468 |
| >trna143_LysTTT_6_-_27559593_27559665@13.30.18,trna149_LysTTT_6_-_27302769_27302841@13.30.18,trna14_LysTTT_11_-_59327808_59327880@13.30.18,trna2_LysTTT_17_+_8022473_8022545@13.30.18,trna54_LysTTT_1_+_204475655_204475727@13.30.18,trna5_LysTTT_11_+_59323902_59323974@13.30.18,trna62_LysTTT_1_-_204476158_204476230@13.30.18,trna76_LysTTT_6_+_28918806_28918878@13.30.18\|CAGTCGGTAGAGCATCAG | >trna143_LysTTT_6_-_27559593_27559665 | 13 | 30 | 18 | CAGTCGGTAGAGCATCAG | 641.2 | -4.1 | 0.7 | -5.4 | 5.1E-08 | 8.5E-08 | MB231/MB468 |
| >hg38_21_+_8433172_8446622_RNA45SN1_45S_50ntFlanks@12954.12973.20\|CTGCTTCTGGGTCGGGGTTT | >hg38_21_+_8433172_8446622_RNA45SN1_45S_50ntFlanks | 12954 | 12973 | 20 | CTGCTTCTGGGTCGGGGTTT | 132.8 | -4.7 | 0.9 | -5.4 | 5.1E-08 | 8.5E-08 | MB231/MB468 |
| >am_trna128_GlyGCC_6_-_27870686_27870756@1.30.30,trna133_GlyCCC_1_-_16872434_16872504@1.30.30,trna18_GlyGCC_16_+_70822597_70822667@1.30.30,trna19_GlyGCC_16_+_70823410_70823480@1.30.30,trna19_GlyGCC_2_-_157257659_157257729@1.30.30,trna24_GlyGCC_16_-_70812942_70813012@1.30.30,trna25_GlyGCC_16_-_70812114_70812184@1.30.30,trna4_GlyCCC_1_+_17188416_17188486@1.30.30,trna5_GlyGCC_17_+_8029064_8029134@1.30.30,trna68_GlyGCC_1_-_161493637_161493707@1.30.30\|GCATTGGTGGTTCAGTGGTAGAATTCTCGC | >am_trna128_GlyGCC_6_-_27870686_27870756 | 1 | 30 | 30 | GCATTGGTGGTTCAGTGGTAGAATTCTCGC | 8887.2 | -2.9 | 0.5 | -5.4 | 5.8E-08 | 8.6E-08 | BT20/MB468 |
| >hg38_21_+_8433172_8446622_RNA45SN1_45S_50ntFlanks@12686.12703.18\|CCAGGCGGAACGATACGG | >hg38_21_+_8433172_8446622_RNA45SN1_45S_50ntFlanks | 12686 | 12703 | 18 | CCAGGCGGAACGATACGG | 112.0 | -4.2 | 0.8 | -5.4 | 5.2E-08 | 8.7E-08 | MB231/MB468 |
| >hg38_21_+_8433172_8446622_RNA45SN1_45S_50ntFlanks@10403.10429.27\|ACATGGGTCAGTCGGTCCTGAGAGATG | >hg38_21_+_8433172_8446622_RNA45SN1_45S_50ntFlanks | 10403 | 10429 | 27 | ACATGGGTCAGTCGGTCCTGAGAGATG | 401.1 | -2.2 | 0.4 | -5.4 | 5.9E-08 | 8.8E-08 | BT20/MB468 |
| >hg38_1_-_228634819_228635039_RNA5S12_5S_50ntFlanks@148.169.22\|GGAATACCGGGTGCTGTAGGCT | >hg38_1_-_228634819_228635039_RNA5S12_5S_50ntFlanks | 148 | 169 | 22 | GGAATACCGGGTGCTGTAGGCT | 1145.7 | 3.4 | 0.6 | 5.4 | 5.9E-08 | 8.8E-08 | BT20/MB468 |
| >hg38_21_+_8433172_8446622_RNA45SN1_45S_50ntFlanks@10822.10840.19\|GAAGTCGGCAAGCCGGATC | >hg38_21_+_8433172_8446622_RNA45SN1_45S_50ntFlanks | 10822 | 10840 | 19 | GAAGTCGGCAAGCCGGATC | 240.3 | -3.2 | 0.6 | -5.4 | 5.3E-08 | 8.8E-08 | MB231/MB468 |
| >hg38_21_+_8433172_8446622_RNA45SN1_45S_50ntFlanks@9343.9370.28\|CGAAAGCCGCCGTGGCGCAATGAAGGTG | >hg38_21_+_8433172_8446622_RNA45SN1_45S_50ntFlanks | 9343 | 9370 | 28 | CGAAAGCCGCCGTGGCGCAATGAAGGTG | 121.6 | -5.3 | 1.0 | -5.4 | 6.0E-08 | 8.9E-08 | BT20/MB468 |
| >hg38_21_+_8433172_8446622_RNA45SN1_45S_50ntFlanks@11577.11594.18\|TTAGAACTGGTGCGGACC | >hg38_21_+_8433172_8446622_RNA45SN1_45S_50ntFlanks | 11577 | 11594 | 18 | TTAGAACTGGTGCGGACC | 217.1 | -2.1 | 0.4 | -5.4 | 6.0E-08 | 8.9E-08 | BT20/MB468 |
| >hg38_21_+_8433172_8446622_RNA45SN1_45S_50ntFlanks@11894.11911.18\|ACTCTAGTCTGGCACGGT | >hg38_21_+_8433172_8446622_RNA45SN1_45S_50ntFlanks | 11894 | 11911 | 18 | ACTCTAGTCTGGCACGGT | 97.8 | -4.5 | 0.8 | -5.4 | 6.0E-08 | 8.9E-08 | BT20/MB468 |

| | | | | | | | | | | | | |
|---|---|---|---|---|---|---|---|---|---|---|---|---|
| >hg38_21_+_8433172_8446622_RNA45SN1_45S_50ntFlanks@5152.5185.34\|AGGGACAAGTGGCGTTCAGCCACCCGAGATTGAG | >hg38_21_+_8433172_8446622_RNA45SN1_45S_50ntFlanks | 5152 | 5185 | 34 | AGGGACAAGTGGCGTTCAGCCACCCGAGATTGAG | 533.7 | -5.1 | 0.9 | -5.4 | 5.4E-08 | 9.0E-08 | MB231/MB468 |
| >hg38_21_+_8433172_8446622_RNA45SN1_45S_50ntFlanks@11641.11667.27\|CGGCGGGTGTTGACGCGATGTGATTTC | >hg38_21_+_8433172_8446622_RNA45SN1_45S_50ntFlanks | 11641 | 11667 | 27 | CGGCGGGTGTTGACGCGATGTGATTTC | 394.5 | -3.4 | 0.6 | -5.4 | 5.4E-08 | 9.0E-08 | MB231/MB468 |
| >hg38_21_+_8433172_8446622_RNA45SN1_45S_50ntFlanks@9575.9597.23\|ATCGGTCGTCCGACCTGGGTATA | >hg38_21_+_8433172_8446622_RNA45SN1_45S_50ntFlanks | 9575 | 9597 | 23 | ATCGGTCGTCCGACCTGGGTATA | 1864.3 | -6.5 | 1.2 | -5.4 | 5.4E-08 | 9.0E-08 | MB231/MB468 |
| >hg38_21_+_8433172_8446622_RNA45SN1_45S_50ntFlanks@10788.10818.31\|CAGCCTCTGGCATGTTGGAACAATGTAGGTA | >hg38_21_+_8433172_8446622_RNA45SN1_45S_50ntFlanks | 10788 | 10818 | 31 | CAGCCTCTGGCATGTTGGAACAATGTAGGTA | 2656.1 | -4.6 | 0.9 | -5.4 | 6.1E-08 | 9.1E-08 | BT20/MB468 |
| >hg38_21_+_8433172_8446622_RNA45SN1_45S_50ntFlanks@4160.4184.25\|CCACATCCAAGGAAGGCAGCAGGCG | >hg38_21_+_8433172_8446622_RNA45SN1_45S_50ntFlanks | 4160 | 4184 | 25 | CCACATCCAAGGAAGGCAGCAGGCG | 101.8 | -4.2 | 0.8 | -5.4 | 5.5E-08 | 9.2E-08 | MB231/MB468 |
| >hg38_21_+_8433172_8446622_RNA45SN1_45S_50ntFlanks@12072.12092.21\|CCCGAGGGGCTCTCGCTTCTG | >hg38_21_+_8433172_8446622_RNA45SN1_45S_50ntFlanks | 12072 | 12092 | 21 | CCCGAGGGGCTCTCGCTTCTG | 167.9 | -3.4 | 0.6 | -5.4 | 6.2E-08 | 9.2E-08 | BT20/MB468 |
| >hg38_21_+_8433172_8446622_RNA45SN1_45S_50ntFlanks@11199.11220.22\|CCGGCGGCGGCGGCGACTCTGG | >hg38_21_+_8433172_8446622_RNA45SN1_45S_50ntFlanks | 11199 | 11220 | 22 | CCGGCGGCGGCGGCGACTCTGG | 243.0 | -4.1 | 0.8 | -5.4 | 6.2E-08 | 9.2E-08 | BT20/MB468 |
| >hg38_21_+_8433172_8446622_RNA45SN1_45S_50ntFlanks@12475.12498.24\|CCCACTAATAGGGAACGTGAGCTG | >hg38_21_+_8433172_8446622_RNA45SN1_45S_50ntFlanks | 12475 | 12498 | 24 | CCCACTAATAGGGAACGTGAGCTG | 531.8 | -3.1 | 0.6 | -5.4 | 5.7E-08 | 9.4E-08 | MB231/MB468 |
| >hg38_21_+_8433172_8446622_RNA45SN1_45S_50ntFlanks@12626.12651.26\|AATGGGGCGAAGCTACCATCTGTGGG | >hg38_21_+_8433172_8446622_RNA45SN1_45S_50ntFlanks | 12626 | 12651 | 26 | AATGGGGCGAAGCTACCATCTGTGGG | 68.2 | -4.3 | 0.8 | -5.4 | 5.7E-08 | 9.4E-08 | MB231/MB468 |
| >hg38_21_+_8433172_8446622_RNA45SN1_45S_50ntFlanks@11781.11803.23\|CGCATGAATGGATGAACGAGATT | >hg38_21_+_8433172_8446622_RNA45SN1_45S_50ntFlanks | 11781 | 11803 | 23 | CGCATGAATGGATGAACGAGATT | 1398.5 | -4.0 | 0.7 | -5.4 | 5.8E-08 | 9.5E-08 | MB231/MB468 |
| >hg38_21_+_8433172_8446622_RNA45SN1_45S_50ntFlanks@7986.8014.29\|ATCAGACGTGGCGACCCGCTGAATTTAAG | >hg38_21_+_8433172_8446622_RNA45SN1_45S_50ntFlanks | 7986 | 8014 | 29 | ATCAGACGTGGCGACCCGCTGAATTTAAG | 111.8 | -4.4 | 0.8 | -5.4 | 5.8E-08 | 9.5E-08 | MB231/MB468 |
| >hg38_21_+_8433172_8446622_RNA45SN1_45S_50ntFlanks@10798.10825.28\|CATGTTGGAACAATGTAGGTAAGGGAAG | >hg38_21_+_8433172_8446622_RNA45SN1_45S_50ntFlanks | 10798 | 10825 | 28 | CATGTTGGAACAATGTAGGTAAGGGAAG | 202.9 | -3.2 | 0.6 | -5.4 | 6.4E-08 | 9.5E-08 | BT20/MB468 |
| >hg38_21_+_8433172_8446622_RNA45SN1_45S_50ntFlanks@11876.11899.24\|AAGACCCTGTTGAGCTTGACTCTA | >hg38_21_+_8433172_8446622_RNA45SN1_45S_50ntFlanks | 11876 | 11899 | 24 | AAGACCCTGTTGAGCTTGACTCTA | 226.3 | -2.6 | 0.5 | -5.4 | 5.8E-08 | 9.6E-08 | MB231/MB468 |
| >hg38_21_+_8433172_8446622_RNA45SN1_45S_50ntFlanks@12416.12444.29\|TTCGATGTCGGCTCTTCCTATCATTGTGA | >hg38_21_+_8433172_8446622_RNA45SN1_45S_50ntFlanks | 12416 | 12444 | 29 | TTCGATGTCGGCTCTTCCTATCATTGTGA | 595.8 | 3.0 | 0.6 | 5.4 | 6.5E-08 | 9.7E-08 | BT20/MB468 |
| >hg38_21_+_8433172_8446622_RNA45SN1_45S_50ntFlanks@4004.4024.21\|TAACCTCGGGCCGATCGCACG | >hg38_21_+_8433172_8446622_RNA45SN1_45S_50ntFlanks | 4004 | 4024 | 21 | TAACCTCGGGCCGATCGCACG | 89.4 | -3.1 | 0.6 | -5.4 | 6.5E-08 | 9.7E-08 | BT20/MB468 |
| >hg38_21_+_8433172_8446622_RNA45SN1_45S_50ntFlanks@11730.11751.22\|AACTATGACTCTCTTAAGGTAG | >hg38_21_+_8433172_8446622_RNA45SN1_45S_50ntFlanks | 11730 | 11751 | 22 | AACTATGACTCTCTTAAGGTAG | 367.1 | -3.4 | 0.6 | -5.4 | 5.9E-08 | 9.7E-08 | MB231/MB468 |
| >hg38_21_+_8433172_8446622_RNA45SN1_45S_50ntFlanks@12942.12964.23\|TTCGTAGACGACCTGCTTCTGGG | >hg38_21_+_8433172_8446622_RNA45SN1_45S_50ntFlanks | 12942 | 12964 | 23 | TTCGTAGACGACCTGCTTCTGGG | 190.3 | -5.7 | 1.1 | -5.4 | 6.5E-08 | 9.7E-08 | BT20/MB468 |
| >hg38_21_+_8433172_8446622_RNA45SN1_45S_50ntFlanks@13007.13028.22\|TATTGAAAGTCAGCCCTCGACA | >hg38_21_+_8433172_8446622_RNA45SN1_45S_50ntFlanks | 13007 | 13028 | 22 | TATTGAAAGTCAGCCCTCGACA | 583.6 | -5.0 | 0.9 | -5.4 | 6.5E-08 | 9.7E-08 | BT20/MB468 |
| >hg38_21_+_8433172_8446622_RNA45SN1_45S_50ntFlanks@10912.10931.20\|CGCGCCGCGGCTGGACGAGG | >hg38_21_+_8433172_8446622_RNA45SN1_45S_50ntFlanks | 10912 | 10931 | 20 | CGCGCCGCGGCTGGACGAGG | 183.7 | -4.7 | 0.9 | -5.4 | 5.9E-08 | 9.8E-08 | MB231/MB468 |
| >hg38_21_+_8433172_8446622_RNA45SN1_45S_50ntFlanks@10385.10406.22\|CATGTGAACAGCAGTTGAACAT | >hg38_21_+_8433172_8446622_RNA45SN1_45S_50ntFlanks | 10385 | 10406 | 22 | CATGTGAACAGCAGTTGAACAT | 76.0 | -5.0 | 0.9 | -5.4 | 5.9E-08 | 9.8E-08 | MB231/MB468 |
| >hg38_21_+_8433172_8446622_RNA45SN1_45S_50ntFlanks@9562.9592.31\|TCCTGACGTGCAAATCGGTCGTCCGACCTGG | >hg38_21_+_8433172_8446622_RNA45SN1_45S_50ntFlanks | 9562 | 9592 | 31 | TCCTGACGTGCAAATCGGTCGTCCGACCTGG | 108.7 | -3.9 | 0.7 | -5.4 | 5.9E-08 | 9.8E-08 | MB231/MB468 |
| >hg38_21_+_8433172_8446622_RNA45SN1_45S_50ntFlanks@10409.10430.22\|GTCAGTCGGTCCTGAGAGATGG | >hg38_21_+_8433172_8446622_RNA45SN1_45S_50ntFlanks | 10409 | 10430 | 22 | GTCAGTCGGTCCTGAGAGATGG | 450.0 | -4.9 | 0.9 | -5.4 | 6.6E-08 | 9.8E-08 | BT20/MB468 |
| >trna10_AspGTC_12_-_125424193_125424264@1.29.29,trna12_AspGTC_12_-_125411891_125411962@1.29.29,trna38_AspGTC_17_-_8125556_8125627@1.29.29,trna45_AspGTC_6_+_27447453_27447524@1.29.29,trna48_AspGTC_6_+_27471523_27471594@1.29.29,trna4_AspGTC_12_+_96429799_96429870@1.29.29,trna69_AspGTC_1_-_161440205_161440276@1.29.29,trna72_AspGTC_1_-_161432824_161432895@1.29.29,trna75_AspGTC_1_-_161425414_161425485@1.29.29,trna78_AspGTC_1_-_161418033_161418104@1.29.29,trna81_AspGTC_1_-_161410615_161410686@1.29.29\|TCCTCGTTAGTATAGTGGTGAGTATCCCC | >trna10_AspGTC_12_-_125424193_125424264 | 1 | 29 | 29 | TCCTCGTTAGTATAGTGGTGAGTATCCCC | 225.1 | -5.4 | 1.0 | -5.4 | 6.6E-08 | 9.9E-08 | BT20/MB468 |
| >hg38_21_+_8433172_8446622_RNA45SN1_45S_50ntFlanks@10821.10839.19\|GGAAGTCGGCAAGCCGGAT | >hg38_21_+_8433172_8446622_RNA45SN1_45S_50ntFlanks | 10821 | 10839 | 19 | GGAAGTCGGCAAGCCGGAT | 168.2 | -3.0 | 0.6 | -5.4 | 6.6E-08 | 9.9E-08 | BT20/MB468 |
| >hg38_21_+_8433172_8446622_RNA45SN1_45S_50ntFlanks@11592.11612.21\|ACCAGGGGAATCCGACTGTTT | >hg38_21_+_8433172_8446622_RNA45SN1_45S_50ntFlanks | 11592 | 11612 | 21 | ACCAGGGGAATCCGACTGTTT | 475.7 | -4.4 | 0.8 | -5.4 | 6.6E-08 | 9.9E-08 | BT20/MB468 |
| >am_trna2_GlyGCC_21_-_18827107_18827177@1.27.27,trna35_GlyGCC_1_+_161413094_161413164@1.27.27,trna37_GlyGCC_1_+_161420467_161420537@1.27.27,trna39_GlyGCC_1_+_161427898_161427968@1.27.27,trna41_GlyGCC_1_+_161435258_161435328@1.27.27\|GCATGGGTGGTTCAGTGGTAGAATTCT | >am_trna2_GlyGCC_21_-_18827107_18827177 | 1 | 27 | 27 | GCATGGGTGGTTCAGTGGTAGAATTCT | 334.3 | -3.5 | 0.6 | -5.4 | 6.0E-08 | 9.9E-08 | MB231/MB468 |
| >hg38_21_+_8433172_8446622_RNA45SN1_45S_50ntFlanks@4995.5024.30\|ACAGATTGATAGCTCTTTCTCGATTCCGTG | >hg38_21_+_8433172_8446622_RNA45SN1_45S_50ntFlanks | 4995 | 5024 | 30 | ACAGATTGATAGCTCTTTCTCGATTCCGTG | 260.2 | -3.2 | 0.6 | -5.4 | 6.0E-08 | 9.9E-08 | MB231/MB468 |
| >hg38_21_+_8433172_8446622_RNA45SN1_45S_50ntFlanks@10101.10127.27\|CGCGCGCGTGTGGTGTGCGTCGGAGGG | >hg38_21_+_8433172_8446622_RNA45SN1_45S_50ntFlanks | 10101 | 10127 | 27 | CGCGCGCGTGTGGTGTGCGTCGGAGGG | 474.6 | -6.1 | 1.1 | -5.4 | 6.8E-08 | 1.0E-07 | BT20/MB468 |
| >hg38_MT_+_1621_3279_RNR2_16S_50ntFlanks@322.340.19\|CACACCCGTCTATGTAGCA | >hg38_MT_+_1621_3279_RNR2_16S_50ntFlanks | 322 | 340 | 19 | CACACCCGTCTATGTAGCA | 886.5 | 11.1 | 2.0 | 5.4 | 6.1E-08 | 1.0E-07 | MB231/MB468 |
| >MI0005756\|hsa-mir-934&WithFlank&X\|+\|136550872\|136550966@21.41.21(+1A)&[MIMAT0004977&hsa-miR-934&offsets\|0\|-1(+1A);m-943&X\|+\|136550892\|136550913&offsets\|0\|-1(+1A)]\|TGTCTACTACTGGAGACACTGA | >MI0005756\|hsa-mir-934&WithFlank&X\|+\|136550872\|136550966 | 21 | 41 | 21(+1A) | TGTCTACTACTGGAGACACTGA | 2931.2 | 15.0 | 2.6 | 5.9 | 4.4E-09 | 1.0E-07 | BT20/MB231 |
| >trna14_LysTTT_11_-_59327808_59327880@1.29.29,trna2_LysTTT_17_+_8022473_8022545@1.29.29,trna54_LysTTT_1_+_204475655_204475727@1.29.29,trna5_LysTTT_11_+_59323902_59323974@1.29.29,trna62_LysTTT_1_-_204476158_204476230@1.29.29,trna76_LysTTT_6_+_28918806_28918878@1.29.29\|GCCCGGATAGCTCAGTCGGTAGAGCATCA | >trna14_LysTTT_11_-_59327808_59327880 | 1 | 29 | 29 | GCCCGGATAGCTCAGTCGGTAGAGCATCA | 237.8 | -6.7 | 1.2 | -5.4 | 6.9E-08 | 1.0E-07 | BT20/MB468 |
| >hg38_21_+_8433172_8446622_RNA45SN1_45S_50ntFlanks@10780.10801.22\|AAGGTGAACAGCCTCTGGCATG | >hg38_21_+_8433172_8446622_RNA45SN1_45S_50ntFlanks | 10780 | 10801 | 22 | AAGGTGAACAGCCTCTGGCATG | 232.3 | -2.3 | 0.4 | -5.4 | 6.9E-08 | 1.0E-07 | BT20/MB468 |
| >hg38_21_+_8433172_8446622_RNA45SN1_45S_50ntFlanks@10788.10822.35\|CAGCCTCTGGCATGTTGGAACAATGTAGGTAAGGG | >hg38_21_+_8433172_8446622_RNA45SN1_45S_50ntFlanks | 10788 | 10822 | 35 | CAGCCTCTGGCATGTTGGAACAATGTAGGTAAGGG | 133.1 | -3.6 | 0.7 | -5.4 | 6.9E-08 | 1.0E-07 | BT20/MB468 |
| >hg38_21_+_8433172_8446622_RNA45SN1_45S_50ntFlanks@4731.4768.38\|AAGAACGAAAGTCGGAGGTTCGAAGACGATCAGATACC | >hg38_21_+_8433172_8446622_RNA45SN1_45S_50ntFlanks | 4731 | 4768 | 38 | AAGAACGAAAGTCGGAGGTTCGAAGACGATCAGATACC | 1279.9 | 11.6 | 2.1 | 5.4 | 6.3E-08 | 1.0E-07 | MB231/MB468 |
| >hg38_21_+_8433172_8446622_RNA45SN1_45S_50ntFlanks@5071.5098.28\|TTAATTCCGATAACGAACGAGACTCTGG | >hg38_21_+_8433172_8446622_RNA45SN1_45S_50ntFlanks | 5071 | 5098 | 28 | TTAATTCCGATAACGAACGAGACTCTGG | 184.6 | -3.4 | 0.6 | -5.4 | 7.0E-08 | 1.0E-07 | BT20/MB468 |
| >hg38_21_+_8433172_8446622_RNA45SN1_45S_50ntFlanks@10346.10374.29\|TCAAACGAGAACTTTGAAGGCCGAAGTGG | >hg38_21_+_8433172_8446622_RNA45SN1_45S_50ntFlanks | 10346 | 10374 | 29 | TCAAACGAGAACTTTGAAGGCCGAAGTGG | 563.1 | -3.8 | 0.7 | -5.4 | 7.1E-08 | 1.0E-07 | BT20/MB468 |
| >hg38_21_+_8433172_8446622_RNA45SN1_45S_50ntFlanks@6739.6771.33\|TCATCGACACTTCGAACGCACTTGCGGCCCCGG | >hg38_21_+_8433172_8446622_RNA45SN1_45S_50ntFlanks | 6739 | 6771 | 33 | TCATCGACACTTCGAACGCACTTGCGGCCCCGG | 305.7 | -4.2 | 0.8 | -5.4 | 6.5E-08 | 1.1E-07 | MB231/MB468 |
| >hg38_21_+_8433172_8446622_RNA45SN1_45S_50ntFlanks@10355.10383.29\|AACTTTGAAGGCCGAAGTGGAGAAGGGTT | >hg38_21_+_8433172_8446622_RNA45SN1_45S_50ntFlanks | 10355 | 10383 | 29 | AACTTTGAAGGCCGAAGTGGAGAAGGGTT | 160.8 | -2.3 | 0.4 | -5.4 | 6.5E-08 | 1.1E-07 | MB231/MB468 |
| >hg38_21_+_8433172_8446622_RNA45SN1_45S_50ntFlanks@10361.10390.30\|GAAGGCCGAAGTGGAGAAGGGTTCCATGTG | >hg38_21_+_8433172_8446622_RNA45SN1_45S_50ntFlanks | 10361 | 10390 | 30 | GAAGGCCGAAGTGGAGAAGGGTTCCATGTG | 223.2 | -3.5 | 0.6 | -5.4 | 6.5E-08 | 1.1E-07 | MB231/MB468 |
| >hg38_21_+_8433172_8446622_RNA45SN1_45S_50ntFlanks@11208.11254.47\|GCGGCGACTCTGGACGCGAGCCGGGCCCTTCCCGTGGATCGCCCCAG | >hg38_21_+_8433172_8446622_RNA45SN1_45S_50ntFlanks | 11208 | 11254 | 47 | GCGGCGACTCTGGACGCGAGCCGGGCCCTTCCCGTGGATCGCCCCAG | 1098.6 | -5.9 | 1.1 | -5.4 | 7.3E-08 | 1.1E-07 | BT20/MB468 |

<table>
<tr><td></td><td></td><td></td><td></td><td></td><td></td><td></td><td></td><td></td><td></td><td></td><td></td><td></td></tr>
<tr><td>>hg38_21_+_8433172_8446622_RNA45SN1_45S_50ntFlanks@10829.10852.24|GCAAGCCGGATCCGTAACTTCGGG</td><td>>hg38_21_+_8433172_8446622_RNA45SN1_45S_50ntFlanks</td><td>10829</td><td>10852</td><td>24</td><td>GCAAGCCGGATCCGTAACTTCGGG</td><td>273.5</td><td>-3.4</td><td>0.6</td><td>-5.4</td><td>6.7E-08</td><td>1.1E-07</td><td>MB231/MB468</td></tr>
<tr><td>>hg38_21_+_8433172_8446622_RNA45SN1_45S_50ntFlanks@11573.11602.30|CGACTTAGAACTGGTGCGGACCAGGGGAAT</td><td>>hg38_21_+_8433172_8446622_RNA45SN1_45S_50ntFlanks</td><td>11573</td><td>11602</td><td>30</td><td>CGACTTAGAACTGGTGCGGACCAGGGGAAT</td><td>401.9</td><td>-2.8</td><td>0.5</td><td>-5.4</td><td>7.4E-08</td><td>1.1E-07</td><td>BT20/MB468</td></tr>
<tr><td>>hg38_21_+_8433172_8446622_RNA45SN1_45S_50ntFlanks@10827.10844.18|CGGCAAGCCGGATCCGTA</td><td>>hg38_21_+_8433172_8446622_RNA45SN1_45S_50ntFlanks</td><td>10827</td><td>10844</td><td>18</td><td>CGGCAAGCCGGATCCGTA</td><td>114.3</td><td>-4.5</td><td>0.8</td><td>-5.4</td><td>6.7E-08</td><td>1.1E-07</td><td>MB231/MB468</td></tr>
<tr><td>>hg38_21_+_8433172_8446622_RNA45SN1_45S_50ntFlanks@6721.6749.29|AATTGCAGGACACATTGATCATCGACACT</td><td>>hg38_21_+_8433172_8446622_RNA45SN1_45S_50ntFlanks</td><td>6721</td><td>6749</td><td>29</td><td>AATTGCAGGACACATTGATCATCGACACT</td><td>185.3</td><td>-1.9</td><td>0.3</td><td>-5.4</td><td>7.5E-08</td><td>1.1E-07</td><td>BT20/MB468</td></tr>
<tr><td>>hg38_21_+_8433172_8446622_RNA45SN1_45S_50ntFlanks@13008.13025.18|ATTGAAAGTCAGCCCTCG</td><td>>hg38_21_+_8433172_8446622_RNA45SN1_45S_50ntFlanks</td><td>13008</td><td>13025</td><td>18</td><td>ATTGAAAGTCAGCCCTCG</td><td>1083.8</td><td>-3.9</td><td>0.7</td><td>-5.4</td><td>6.8E-08</td><td>1.1E-07</td><td>MB231/MB468</td></tr>
<tr><td>>hg38_21_+_8433172_8446622_RNA45SN1_45S_50ntFlanks@10843.10864.22|TAACTTCGGGATAAGGATTGGC</td><td>>hg38_21_+_8433172_8446622_RNA45SN1_45S_50ntFlanks</td><td>10843</td><td>10864</td><td>22</td><td>TAACTTCGGGATAAGGATTGGC</td><td>121.6</td><td>-5.1</td><td>1.0</td><td>-5.4</td><td>7.6E-08</td><td>1.1E-07</td><td>BT20/MB468</td></tr>
<tr><td>>hg38_21_+_8433172_8446622_RNA45SN1_45S_50ntFlanks@11569.11586.18|CAGCCGACTTAGAACTGG</td><td>>hg38_21_+_8433172_8446622_RNA45SN1_45S_50ntFlanks</td><td>11569</td><td>11586</td><td>18</td><td>CAGCCGACTTAGAACTGG</td><td>1281.5</td><td>-2.8</td><td>0.5</td><td>-5.4</td><td>7.6E-08</td><td>1.1E-07</td><td>BT20/MB468</td></tr>
<tr><td>>hg38_21_+_8433172_8446622_RNA45SN1_45S_50ntFlanks@6651.6674.24|CGACTCTTAGCGGTGGATCACTCG</td><td>>hg38_21_+_8433172_8446622_RNA45SN1_45S_50ntFlanks</td><td>6651</td><td>6674</td><td>24</td><td>CGACTCTTAGCGGTGGATCACTCG</td><td>957.0</td><td>3.7</td><td>0.7</td><td>5.4</td><td>6.8E-08</td><td>1.1E-07</td><td>MB231/MB468</td></tr>
<tr><td>>hg38_21_+_8433172_8446622_RNA45SN1_45S_50ntFlanks@3996.4024.29|ACTCTAGATAACCTCGGGCCGATCGCACG</td><td>>hg38_21_+_8433172_8446622_RNA45SN1_45S_50ntFlanks</td><td>3996</td><td>4024</td><td>29</td><td>ACTCTAGATAACCTCGGGCCGATCGCACG</td><td>130.6</td><td>-3.4</td><td>0.6</td><td>-5.4</td><td>7.6E-08</td><td>1.1E-07</td><td>BT20/MB468</td></tr>
<tr><td>>hg38_21_+_8433172_8446622_RNA45SN1_45S_50ntFlanks@4002.4022.21|GATAACCTCGGGCCGATCGCA</td><td>>hg38_21_+_8433172_8446622_RNA45SN1_45S_50ntFlanks</td><td>4002</td><td>4022</td><td>21</td><td>GATAACCTCGGGCCGATCGCA</td><td>235.8</td><td>-2.6</td><td>0.5</td><td>-5.4</td><td>7.7E-08</td><td>1.1E-07</td><td>BT20/MB468</td></tr>
<tr><td>>hg38_21_+_8433172_8446622_RNA45SN1_45S_50ntFlanks@3775.3799.25|GCCGGTACAGTGAAACTGCGAATGG</td><td>>hg38_21_+_8433172_8446622_RNA45SN1_45S_50ntFlanks</td><td>3775</td><td>3799</td><td>25</td><td>GCCGGTACAGTGAAACTGCGAATGG</td><td>84.2</td><td>-4.5</td><td>0.8</td><td>-5.4</td><td>6.9E-08</td><td>1.1E-07</td><td>MB231/MB468</td></tr>
<tr><td>>hg38_21_+_8433172_8446622_RNA45SN1_45S_50ntFlanks@13008.13027.20|ATTGAAAGTCAGCCCTCGAC</td><td>>hg38_21_+_8433172_8446622_RNA45SN1_45S_50ntFlanks</td><td>13008</td><td>13027</td><td>20</td><td>ATTGAAAGTCAGCCCTCGAC</td><td>571.3</td><td>-4.2</td><td>0.8</td><td>-5.4</td><td>7.0E-08</td><td>1.1E-07</td><td>MB231/MB468</td></tr>
<tr><td>>hg38_21_+_8433172_8446622_RNA45SN1_45S_50ntFlanks@12658.12675.18|ACTGAACGCCTCTAAGTC</td><td>>hg38_21_+_8433172_8446622_RNA45SN1_45S_50ntFlanks</td><td>12658</td><td>12675</td><td>18</td><td>ACTGAACGCCTCTAAGTC</td><td>601.7</td><td>-4.2</td><td>0.8</td><td>-5.4</td><td>7.0E-08</td><td>1.1E-07</td><td>MB231/MB468</td></tr>
<tr><td>>hg38_21_+_8433172_8446622_RNA45SN1_45S_50ntFlanks@11568.11592.25|GCAGCCGACTTAGAACTGGTGCGGA</td><td>>hg38_21_+_8433172_8446622_RNA45SN1_45S_50ntFlanks</td><td>11568</td><td>11592</td><td>25</td><td>GCAGCCGACTTAGAACTGGTGCGGA</td><td>226.5</td><td>-2.4</td><td>0.4</td><td>-5.4</td><td>7.0E-08</td><td>1.2E-07</td><td>MB231/MB468</td></tr>
<tr><td>>hg38_21_+_8433172_8446622_RNA45SN1_45S_50ntFlanks@11874.11900.27|AGAAGACCCTGTTGAGCTTGACTCTAG</td><td>>hg38_21_+_8433172_8446622_RNA45SN1_45S_50ntFlanks</td><td>11874</td><td>11900</td><td>27</td><td>AGAAGACCCTGTTGAGCTTGACTCTAG</td><td>695.5</td><td>-5.6</td><td>1.0</td><td>-5.4</td><td>7.8E-08</td><td>1.2E-07</td><td>BT20/MB468</td></tr>
<tr><td>>hg38_21_+_8433172_8446622_RNA45SN1_45S_50ntFlanks@9574.9592.19|AATCGGTCGTCCGACCTGG</td><td>>hg38_21_+_8433172_8446622_RNA45SN1_45S_50ntFlanks</td><td>9574</td><td>9592</td><td>19</td><td>AATCGGTCGTCCGACCTGG</td><td>109.0</td><td>-5.6</td><td>1.0</td><td>-5.4</td><td>7.0E-08</td><td>1.2E-07</td><td>MB231/MB468</td></tr>
<tr><td>>MI0000749|hsa-mir-30e&WithFlank&1|+|40754349|40754452@23.45.23&[MIMAT0000692&hsa-miR-30e-5p&offsets|0|+1;m-24&1|+|40754371|40754394&offsets|0|-1]|TGTAAACATCCTTGACTGGAAGC</td><td>>MI0000749|hsa-mir-30e&WithFlank&1|+|40754349|40754452</td><td>23</td><td>45</td><td>23</td><td>TGTAAACATCCTTGACTGGAAGC</td><td>566.2</td><td>4.1</td><td>0.8</td><td>5.4</td><td>7.1E-08</td><td>1.2E-07</td><td>MB231/MB468</td></tr>
<tr><td>>MI0000101|hsa-mir-99a&WithFlank&21|+|16539083|16539175@19.40.22(+1U)&[MIMAT0000097&hsa-miR-99a-5p&offsets|0|0(+1U);m-64&21|+|16539101|16539121&offsets|0|+1(+1U)]|AACCCGTAGATCCGATCTTGTGT</td><td>>MI0000101|hsa-mir-99a&WithFlank&21|+|16539083|16539175</td><td>19</td><td>40</td><td>22(+1U)</td><td>AACCCGTAGATCCGATCTTGTGT</td><td>102.9</td><td>-4.7</td><td>0.9</td><td>-5.4</td><td>7.9E-08</td><td>1.2E-07</td><td>BT20/MB468</td></tr>
<tr><td>>hg38_21_+_8433172_8446622_RNA45SN1_45S_50ntFlanks@11779.11803.25|CGCGCATGAATGGATGAACGAGATT</td><td>>hg38_21_+_8433172_8446622_RNA45SN1_45S_50ntFlanks</td><td>11779</td><td>11803</td><td>25</td><td>CGCGCATGAATGGATGAACGAGATT</td><td>314.4</td><td>2.5</td><td>0.5</td><td>5.4</td><td>7.1E-08</td><td>1.2E-07</td><td>MB231/MB468</td></tr>
<tr><td>>hg38_21_+_8433172_8446622_RNA45SN1_45S_50ntFlanks@11671.11702.32|CCAGTGCTCTGAATGTCAAAGTGAAGAAATTC</td><td>>hg38_21_+_8433172_8446622_RNA45SN1_45S_50ntFlanks</td><td>11671</td><td>11702</td><td>32</td><td>CCAGTGCTCTGAATGTCAAAGTGAAGAAATTC</td><td>913.4</td><td>-3.3</td><td>0.6</td><td>-5.4</td><td>7.9E-08</td><td>1.2E-07</td><td>BT20/MB468</td></tr>
<tr><td>>hg38_21_+_8433172_8446622_RNA45SN1_45S_50ntFlanks@12712.12742.31|CGGAGCCTCGGTTGGCCTCGGATAGCCGGTC</td><td>>hg38_21_+_8433172_8446622_RNA45SN1_45S_50ntFlanks</td><td>12712</td><td>12742</td><td>31</td><td>CGGAGCCTCGGTTGGCCTCGGATAGCCGGTC</td><td>78.1</td><td>-4.8</td><td>0.9</td><td>-5.4</td><td>7.2E-08</td><td>1.2E-07</td><td>MB231/MB468</td></tr>
<tr><td>>hg38_21_+_8433172_8446622_RNA45SN1_45S_50ntFlanks@10680.10709.30|GTTCCGGCGGCGTCCGGTGAGCTCTCGCTG</td><td>>hg38_21_+_8433172_8446622_RNA45SN1_45S_50ntFlanks</td><td>10680</td><td>10709</td><td>30</td><td>GTTCCGGCGGCGTCCGGTGAGCTCTCGCTG</td><td>4951.7</td><td>-4.8</td><td>0.9</td><td>-5.4</td><td>8.1E-08</td><td>1.2E-07</td><td>BT20/MB468</td></tr>
<tr><td>>hg38_21_+_8433172_8446622_RNA45SN1_45S_50ntFlanks@12416.12443.28|TTCGATGTCGGCTCTTCCTATCATTGTG</td><td>>hg38_21_+_8433172_8446622_RNA45SN1_45S_50ntFlanks</td><td>12416</td><td>12443</td><td>28</td><td>TTCGATGTCGGCTCTTCCTATCATTGTG</td><td>2596.4</td><td>3.8</td><td>0.7</td><td>5.4</td><td>7.3E-08</td><td>1.2E-07</td><td>MB231/MB468</td></tr>
<tr><td>>hg38_21_+_8433172_8446622_RNA45SN1_45S_50ntFlanks@6730.6771.42|ACACATTGATCATCGACACTTCGAACGCACTTGCGGCCCCGG</td><td>>hg38_21_+_8433172_8446622_RNA45SN1_45S_50ntFlanks</td><td>6730</td><td>6771</td><td>42</td><td>ACACATTGATCATCGACACTTCGAACGCACTTGCGGCCCCGG</td><td>144.9</td><td>-4.9</td><td>0.9</td><td>-5.4</td><td>8.2E-08</td><td>1.2E-07</td><td>BT20/MB468</td></tr>
<tr><td>>MI0000462|hsa-mir-152&WithFlank&17|-|48037155|48037253@60.80.21&[MIMAT0000438&hsa-miR-152-3p&offsets|0|0;m-97&17|-|48037174|48037194&offsets|0|0]|TCAGTGCATGACAGAACTTGG</td><td>>MI0000462|hsa-mir-152&WithFlank&17|-|48037155|48037253</td><td>60</td><td>80</td><td>21</td><td>TCAGTGCATGACAGAACTTGG</td><td>3974.0</td><td>12.7</td><td>2.4</td><td>5.4</td><td>8.3E-08</td><td>1.2E-07</td><td>BT20/MB468</td></tr>
<tr><td>>hg38_21_+_8433172_8446622_RNA45SN1_45S_50ntFlanks@7974.7994.21|ACGCGACCTCAGATCAGACGT</td><td>>hg38_21_+_8433172_8446622_RNA45SN1_45S_50ntFlanks</td><td>7974</td><td>7994</td><td>21</td><td>ACGCGACCTCAGATCAGACGT</td><td>2478.9</td><td>3.5</td><td>0.7</td><td>5.4</td><td>8.3E-08</td><td>1.2E-07</td><td>BT20/MB468</td></tr>
<tr><td>>MI0000478|hsa-mir-149&WithFlank&2|+|240455995|240456095@21.42.22(+1A)&[MIMAT0000450&hsa-miR-149-5p&offsets|0|-1(+1A);m-195&2|+|240456015|240456037&offsets|0|-1(+1A)]|TCTGGCTCCGTGTCTTCACTCCA</td><td>>MI0000478|hsa-mir-149&WithFlank&2|+|240455995|240456095</td><td>21</td><td>42</td><td>22(+1A)</td><td>TCTGGCTCCGTGTCTTCACTCCA</td><td>526.4</td><td>4.8</td><td>0.8</td><td>5.8</td><td>5.4E-09</td><td>1.2E-07</td><td>BT20/MB231</td></tr>
<tr><td>>MI0000810|hsa-mir-135b&WithFlank&1|-|205448296|205448404@22.44.23&[MIMAT0000758&hsa-miR-135b-5p&offsets|0|0;m-321&1|-|205448361|205448383&offsets|0|0]|TATGGCTTTTCATTCCTATGTGA</td><td>>MI0000810|hsa-mir-135b&WithFlank&1|-|205448296|205448404</td><td>22</td><td>44</td><td>23</td><td>TATGGCTTTTCATTCCTATGTGA</td><td>3343.4</td><td>4.9</td><td>0.9</td><td>5.4</td><td>8.4E-08</td><td>1.2E-07</td><td>BT20/MB468</td></tr>
<tr><td>>hg38_21_+_8433172_8446622_RNA45SN1_45S_50ntFlanks@11718.11751.34|AACGGCGGGAGTAACTATGACTCTCTTAAGGTAG</td><td>>hg38_21_+_8433172_8446622_RNA45SN1_45S_50ntFlanks</td><td>11718</td><td>11751</td><td>34</td><td>AACGGCGGGAGTAACTATGACTCTCTTAAGGTAG</td><td>581.3</td><td>-4.0</td><td>0.7</td><td>-5.4</td><td>7.6E-08</td><td>1.2E-07</td><td>MB231/MB468</td></tr>
<tr><td>>hg38_21_+_8433172_8446622_RNA45SN1_45S_50ntFlanks@5482.5499.18|CGGAGCGCTGAGAAGACG</td><td>>hg38_21_+_8433172_8446622_RNA45SN1_45S_50ntFlanks</td><td>5482</td><td>5499</td><td>18</td><td>CGGAGCGCTGAGAAGACG</td><td>129.2</td><td>-2.1</td><td>0.4</td><td>-5.4</td><td>8.4E-08</td><td>1.2E-07</td><td>BT20/MB468</td></tr>
<tr><td>>hg38_21_+_8433172_8446622_RNA45SN1_45S_50ntFlanks@10801.10818.18|GTTGGAACAATGTAGGTA</td><td>>hg38_21_+_8433172_8446622_RNA45SN1_45S_50ntFlanks</td><td>10801</td><td>10818</td><td>18</td><td>GTTGGAACAATGTAGGTA</td><td>296.2</td><td>-4.5</td><td>0.8</td><td>-5.4</td><td>7.6E-08</td><td>1.2E-07</td><td>MB231/MB468</td></tr>
<tr><td>>hg38_21_+_8433172_8446622_RNA45SN1_45S_50ntFlanks@4729.4751.23|TCAAGAACGAAAGTCGGAGGTTC</td><td>>hg38_21_+_8433172_8446622_RNA45SN1_45S_50ntFlanks</td><td>4729</td><td>4751</td><td>23</td><td>TCAAGAACGAAAGTCGGAGGTTC</td><td>240.9</td><td>-2.7</td><td>0.5</td><td>-5.4</td><td>8.5E-08</td><td>1.3E-07</td><td>BT20/MB468</td></tr>
<tr><td>>hg38_21_+_8433172_8446622_RNA45SN1_45S_50ntFlanks@10345.10373.29|TTCAAACGAGAACTTTGAAGGCCGAAGTG</td><td>>hg38_21_+_8433172_8446622_RNA45SN1_45S_50ntFlanks</td><td>10345</td><td>10373</td><td>29</td><td>TTCAAACGAGAACTTTGAAGGCCGAAGTG</td><td>582.5</td><td>5.9</td><td>1.1</td><td>5.4</td><td>7.6E-08</td><td>1.3E-07</td><td>MB231/MB468</td></tr>
<tr><td>>hg38_21_+_8433172_8446622_RNA45SN1_45S_50ntFlanks@6646.6665.20|TCGTACGACTCTTAGCGGTG</td><td>>hg38_21_+_8433172_8446622_RNA45SN1_45S_50ntFlanks</td><td>6646</td><td>6665</td><td>20</td><td>TCGTACGACTCTTAGCGGTG</td><td>10998.1</td><td>4.0</td><td>0.7</td><td>5.4</td><td>7.7E-08</td><td>1.3E-07</td><td>MB231/MB468</td></tr>
<tr><td>>hg38_21_+_8433172_8446622_RNA45SN1_45S_50ntFlanks@6725.6744.20|GCAGGACACATTGATCATCG</td><td>>hg38_21_+_8433172_8446622_RNA45SN1_45S_50ntFlanks</td><td>6725</td><td>6744</td><td>20</td><td>GCAGGACACATTGATCATCG</td><td>408.6</td><td>-2.4</td><td>0.4</td><td>-5.4</td><td>8.6E-08</td><td>1.3E-07</td><td>BT20/MB468</td></tr>
<tr><td>>hg38_21_+_8433172_8446622_RNA45SN1_45S_50ntFlanks@11322.11344.23|ACGTCTCGTCGCGCGCGCGTCCG</td><td>>hg38_21_+_8433172_8446622_RNA45SN1_45S_50ntFlanks</td><td>11322</td><td>11344</td><td>23</td><td>ACGTCTCGTCGCGCGCGCGTCCG</td><td>3877.5</td><td>-4.5</td><td>0.8</td><td>-5.4</td><td>7.8E-08</td><td>1.3E-07</td><td>MB231/MB468</td></tr>
<tr><td>>hg38_21_+_8433172_8446622_RNA45SN1_45S_50ntFlanks@11575.11609.35|ACTTAGAACTGGTGCGGACCAGGGGAATCCGACTG</td><td>>hg38_21_+_8433172_8446622_RNA45SN1_45S_50ntFlanks</td><td>11575</td><td>11609</td><td>35</td><td>ACTTAGAACTGGTGCGGACCAGGGGAATCCGACTG</td><td>445.6</td><td>-3.0</td><td>0.6</td><td>-5.3</td><td>8.8E-08</td><td>1.3E-07</td><td>BT20/MB468</td></tr>
<tr><td>>hg38_21_+_8433172_8446622_RNA45SN1_45S_50ntFlanks@10393.10419.27|CAGCAGTTGAACATGGGTCAGTCGGTC</td><td>>hg38_21_+_8433172_8446622_RNA45SN1_45S_50ntFlanks</td><td>10393</td><td>10419</td><td>27</td><td>CAGCAGTTGAACATGGGTCAGTCGGTC</td><td>231.4</td><td>-2.5</td><td>0.5</td><td>-5.4</td><td>8.0E-08</td><td>1.3E-07</td><td>MB231/MB468</td></tr>
<tr><td>>hg38_21_+_8433172_8446622_RNA45SN1_45S_50ntFlanks@11785.11803.19|TGAATGGATGAACGAGATT</td><td>>hg38_21_+_8433172_8446622_RNA45SN1_45S_50ntFlanks</td><td>11785</td><td>11803</td><td>19</td><td>TGAATGGATGAACGAGATT</td><td>2965.8</td><td>-3.5</td><td>0.7</td><td>-5.4</td><td>8.1E-08</td><td>1.3E-07</td><td>MB231/MB468</td></tr>
<tr><td>>MI0000071|hsa-mir-17&WithFlank&13|+|91350599|91350694@20.40.21&[MIMAT0000070&hsa-miR-17-5p&offsets|0|-2;m-88&13|+|91350618|91350640&offsets|0|-2]|CAAAGTGCTTACAGTGCAGGT</td><td>>MI0000071|hsa-mir-17&WithFlank&13|+|91350599|91350694</td><td>20</td><td>40</td><td>21</td><td>CAAAGTGCTTACAGTGCAGGT</td><td>1140.7</td><td>4.9</td><td>0.9</td><td>5.3</td><td>9.0E-08</td><td>1.3E-07</td><td>BT20/MB468</td></tr>
<tr><td>>hg38_21_+_8433172_8446622_RNA45SN1_45S_50ntFlanks@11718.11737.20|AACGGCGGGAGTAACTATGA</td><td>>hg38_21_+_8433172_8446622_RNA45SN1_45S_50ntFlanks</td><td>11718</td><td>11737</td><td>20</td><td>AACGGCGGGAGTAACTATGA</td><td>161.8</td><td>-3.7</td><td>0.7</td><td>-5.4</td><td>8.2E-08</td><td>1.3E-07</td><td>MB231/MB468</td></tr>
<tr><td>>hg38_21_+_8433172_8446622_RNA45SN1_45S_50ntFlanks@11208.11254.47|GCGGCGACTCTGGACGCGAGCCGGGCCCTTCCCGTGGATCGCCCCAG</td><td>>hg38_21_+_8433172_8446622_RNA45SN1_45S_50ntFlanks</td><td>11208</td><td>11254</td><td>47</td><td>GCGGCGACTCTGGACGCGAGCCGGGCCCTTCCCGTGGATCGCCCCAG</td><td>896.4</td><td>-5.7</td><td>1.1</td><td>-5.4</td><td>8.3E-08</td><td>1.4E-07</td><td>MB231/MB468</td></tr>
<tr><td>>hg38_21_+_8433172_8446622_RNA45SN1_45S_50ntFlanks@3990.4024.35|TTGGTGACTCTAGATAACCTCGGGCCGATCGCACG</td><td>>hg38_21_+_8433172_8446622_RNA45SN1_45S_50ntFlanks</td><td>3990</td><td>4024</td><td>35</td><td>TTGGTGACTCTAGATAACCTCGGGCCGATCGCACG</td><td>380.8</td><td>2.9</td><td>0.5</td><td>5.4</td><td>8.3E-08</td><td>1.4E-07</td><td>MB231/MB468</td></tr>
<tr><td>>hg38_21_+_8433172_8446622_RNA45SN1_45S_50ntFlanks@11578.11603.26|TAGAACTGGTGCGGACCAGGGGAATC</td><td>>hg38_21_+_8433172_8446622_RNA45SN1_45S_50ntFlanks</td><td>11578</td><td>11603</td><td>26</td><td>TAGAACTGGTGCGGACCAGGGGAATC</td><td>679.8</td><td>-3.5</td><td>0.7</td><td>-5.3</td><td>9.2E-08</td><td>1.4E-07</td><td>BT20/MB468</td></tr>
<tr><td>>hg38_21_+_8433172_8446622_RNA45SN1_45S_50ntFlanks@5425.5450.26|TGGATGGTTTAGTGAGGCCCTCGGAT</td><td>>hg38_21_+_8433172_8446622_RNA45SN1_45S_50ntFlanks</td><td>5425</td><td>5450</td><td>26</td><td>TGGATGGTTTAGTGAGGCCCTCGGAT</td><td>385.5</td><td>10.2</td><td>1.9</td><td>5.4</td><td>8.3E-08</td><td>1.4E-07</td><td>MB231/MB468</td></tr>
<tr><td>>hg38_21_+_8433172_8446622_RNA45SN1_45S_50ntFlanks@11680.11701.22|TGAATGTCAAAGTGAAGAAATT</td><td>>hg38_21_+_8433172_8446622_RNA45SN1_45S_50ntFlanks</td><td>11680</td><td>11701</td><td>22</td><td>TGAATGTCAAAGTGAAGAAATT</td><td>103.5</td><td>-5.2</td><td>1.0</td><td>-5.4</td><td>8.3E-08</td><td>1.4E-07</td><td>MB231/MB468</td></tr>
<tr><td>>hg38_21_+_8433172_8446622_RNA45SN1_45S_50ntFlanks@11883.11904.22|TGTTGAGCTTGACTCTAGTCTG</td><td>>hg38_21_+_8433172_8446622_RNA45SN1_45S_50ntFlanks</td><td>11883</td><td>11904</td><td>22</td><td>TGTTGAGCTTGACTCTAGTCTG</td><td>123.1</td><td>-3.4</td><td>0.6</td><td>-5.3</td><td>9.3E-08</td><td>1.4E-07</td><td>BT20/MB468</td></tr>
<tr><td>>hg38_21_+_8433172_8446622_RNA45SN1_45S_50ntFlanks@6650.6699.50|ACGACTCTTAGCGGTGGATCACTCGGCTCGTGCGTCGATGAAGAACGCAG</td><td>>hg38_21_+_8433172_8446622_RNA45SN1_45S_50ntFlanks</td><td>6650</td><td>6699</td><td>50</td><td>ACGACTCTTAGCGGTGGATCACTCGGCTCGTGCGTCGATGAAGAACGCAG</td><td>2057.0</td><td>5.2</td><td>1.0</td><td>5.4</td><td>8.4E-08</td><td>1.4E-07</td><td>MB231/MB468</td></tr>
</table>

| | | | | | | | | | | | | |
|---|---|---|---|---|---|---|---|---|---|---|---|---|
| >hg38_21_+_8433172_8446622_RNA45SN1_45S_50ntFlanks@9568.9603.36\|CGTGCAAATCGGTCGTCCGACCTGGGTATAGGGGCG | >hg38_21_+_8433172_8446622_RNA45SN1_45S_50ntFlanks | 9568 | 9603 | 36 | CGTGCAAATCGGTCGTCCGACCTGGGTATAGGGGCG | 319.3 | -3.7 | 0.7 | -5.3 | 9.3E-08 | 1.4E-07 | BT20/MB468 |
| >hg38_21_+_8433172_8446622_RNA45SN1_45S_50ntFlanks@10482.10512.31\|TCGAAAGGGAGTCGGGTTCAGATCCCCGAAT | >hg38_21_+_8433172_8446622_RNA45SN1_45S_50ntFlanks | 10482 | 10512 | 31 | TCGAAAGGGAGTCGGGTTCAGATCCCCGAAT | 140.4 | -3.7 | 0.7 | -5.4 | 8.4E-08 | 1.4E-07 | MB231/MB468 |
| >hg38_21_+_8433172_8446622_RNA45SN1_45S_50ntFlanks@9539.9571.33\|ACTCTGGTGGAGGTCCGTAGCGGTCCTGACGTG | >hg38_21_+_8433172_8446622_RNA45SN1_45S_50ntFlanks | 9539 | 9571 | 33 | ACTCTGGTGGAGGTCCGTAGCGGTCCTGACGTG | 557.0 | -3.0 | 0.6 | -5.3 | 9.4E-08 | 1.4E-07 | BT20/MB468 |
| >hg38_21_+_8433172_8446622_RNA45SN1_45S_50ntFlanks@12641.12659.19\|CCATCTGTGGGATTATGAC | >hg38_21_+_8433172_8446622_RNA45SN1_45S_50ntFlanks | 12641 | 12659 | 19 | CCATCTGTGGGATTATGAC | 605.5 | -3.7 | 0.7 | -5.4 | 8.5E-08 | 1.4E-07 | MB231/MB468 |
| >hg38_21_+_8433172_8446622_RNA45SN1_45S_50ntFlanks@12635.12661.27\|AAGCTACCATCTGTGGGATTATGACTG | >hg38_21_+_8433172_8446622_RNA45SN1_45S_50ntFlanks | 12635 | 12661 | 27 | AAGCTACCATCTGTGGGATTATGACTG | 2688.2 | -3.2 | 0.6 | -5.4 | 8.6E-08 | 1.4E-07 | MB231/MB468 |
| >hg38_21_+_8433172_8446622_RNA45SN1_45S_50ntFlanks@11208.11225.18\|GCGGCGACTCTGGACGCG | >hg38_21_+_8433172_8446622_RNA45SN1_45S_50ntFlanks | 11208 | 11225 | 18 | GCGGCGACTCTGGACGCG | 3262.3 | -4.9 | 0.9 | -5.3 | 9.5E-08 | 1.4E-07 | BT20/MB468 |
| >hg38_21_+_8433172_8446622_RNA45SN1_45S_50ntFlanks@11843.11872.30\|AGGGAACGGGCTTGGCGGAATCAGCGGGGA | >hg38_21_+_8433172_8446622_RNA45SN1_45S_50ntFlanks | 11843 | 11872 | 30 | AGGGAACGGGCTTGGCGGAATCAGCGGGGA | 137.7 | -4.7 | 0.9 | -5.3 | 9.5E-08 | 1.4E-07 | BT20/MB468 |
| >hg38_21_+_8433172_8446622_RNA45SN1_45S_50ntFlanks@11641.11685.45\|CGGCGGGTGTTGACGCGATGTGATTTCTGCCCAGTGCTCTGAATG | >hg38_21_+_8433172_8446622_RNA45SN1_45S_50ntFlanks | 11641 | 11685 | 45 | CGGCGGGTGTTGACGCGATGTGATTTCTGCCCAGTGCTCTGAATG | 292.3 | -5.1 | 1.0 | -5.3 | 9.6E-08 | 1.4E-07 | BT20/MB468 |
| >am_trnaMT_SerGCT_MT_+_12207_12265@13.31.19\|AAGAACTGCTAACTCATGC | >am_trnaMT_SerGCT_MT_+_12207_12265 | 13 | 31 | 19 | AAGAACTGCTAACTCATGC | 458.2 | 4.3 | 0.8 | 5.3 | 9.6E-08 | 1.4E-07 | BT20/MB468 |
| >hg38_21_+_8433172_8446622_RNA45SN1_45S_50ntFlanks@8176.8206.31\|CTTCTGATCGAGGCCCAGCCCGTGGACGGTG | >hg38_21_+_8433172_8446622_RNA45SN1_45S_50ntFlanks | 8176 | 8206 | 31 | CTTCTGATCGAGGCCCAGCCCGTGGACGGTG | 441.8 | -2.9 | 0.5 | -5.3 | 9.6E-08 | 1.4E-07 | BT20/MB468 |
| >hg38_21_+_8433172_8446622_RNA45SN1_45S_50ntFlanks@483.529.47\|TCGATGTGGTGACGTCGTGCTCTCCCGGGCCGGGTCCGAGCCGCGAC | >hg38_21_+_8433172_8446622_RNA45SN1_45S_50ntFlanks | 483 | 529 | 47 | TCGATGTGGTGACGTCGTGCTCTCCCGGGCCGGGTCCGAGCCGCGAC | 732.7 | 11.3 | 2.1 | 5.4 | 8.8E-08 | 1.4E-07 | MB231/MB468 |
| >hg38_21_+_8433172_8446622_RNA45SN1_45S_50ntFlanks@10922.10946.25\|CTGGACGAGGCGCCGCCGCCCCCCC | >hg38_21_+_8433172_8446622_RNA45SN1_45S_50ntFlanks | 10922 | 10946 | 25 | CTGGACGAGGCGCCGCCGCCCCCCC | 451.8 | -5.8 | 1.1 | -5.3 | 9.7E-08 | 1.4E-07 | BT20/MB468 |
| >hg38_21_+_8433172_8446622_RNA45SN1_45S_50ntFlanks@12457.12474.18\|CAAGCGTTGGATTGTTCA | >hg38_21_+_8433172_8446622_RNA45SN1_45S_50ntFlanks | 12457 | 12474 | 18 | CAAGCGTTGGATTGTTCA | 653.3 | -3.6 | 0.7 | -5.3 | 8.8E-08 | 1.4E-07 | MB231/MB468 |
| >hg38_21_+_8433172_8446622_RNA45SN1_45S_50ntFlanks@4724.4751.28\|ATTAATCAAGAACGAAAGTCGGAGGTTC | >hg38_21_+_8433172_8446622_RNA45SN1_45S_50ntFlanks | 4724 | 4751 | 28 | ATTAATCAAGAACGAAAGTCGGAGGTTC | 443.9 | -3.2 | 0.6 | -5.3 | 8.9E-08 | 1.5E-07 | MB231/MB468 |
| >hg38_21_+_8433172_8446622_RNA45SN1_45S_50ntFlanks@4560.4577.18\|CTAGGAATAATGGAATAG | >hg38_21_+_8433172_8446622_RNA45SN1_45S_50ntFlanks | 4560 | 4577 | 18 | CTAGGAATAATGGAATAG | 145.2 | -4.2 | 0.8 | -5.3 | 8.9E-08 | 1.5E-07 | MB231/MB468 |
| >hg38_21_+_8433172_8446622_RNA45SN1_45S_50ntFlanks@10349.10369.21\|AACGAGAACTTTGAAGGCCGA | >hg38_21_+_8433172_8446622_RNA45SN1_45S_50ntFlanks | 10349 | 10369 | 21 | AACGAGAACTTTGAAGGCCGA | 1430.9 | 3.3 | 0.6 | 5.3 | 8.9E-08 | 1.5E-07 | MB231/MB468 |
| >hg38_21_+_8433172_8446622_RNA45SN1_45S_50ntFlanks@9535.9553.19\|GGAAACTCTGGTGGAGGTC | >hg38_21_+_8433172_8446622_RNA45SN1_45S_50ntFlanks | 9535 | 9553 | 19 | GGAAACTCTGGTGGAGGTC | 151.3 | -3.1 | 0.6 | -5.3 | 9.0E-08 | 1.5E-07 | MB231/MB468 |
| >hg38_21_+_8433172_8446622_RNA45SN1_45S_50ntFlanks@12938.12966.29\|ACCATTCGTAGACGACCTGCTTCTGGGTC | >hg38_21_+_8433172_8446622_RNA45SN1_45S_50ntFlanks | 12938 | 12966 | 29 | ACCATTCGTAGACGACCTGCTTCTGGGTC | 103.1 | -3.5 | 0.7 | -5.3 | 9.0E-08 | 1.5E-07 | MB231/MB468 |
| >hg38_21_+_8433172_8446622_RNA45SN1_45S_50ntFlanks@11561.11588.28\|GCGCCTAGCAGCCGACTTAGAACTGGTG | >hg38_21_+_8433172_8446622_RNA45SN1_45S_50ntFlanks | 11561 | 11588 | 28 | GCGCCTAGCAGCCGACTTAGAACTGGTG | 553.9 | -4.4 | 0.8 | -5.3 | 9.9E-08 | 1.5E-07 | BT20/MB468 |
| >hg38_21_+_8433172_8446622_RNA45SN1_45S_50ntFlanks@8175.8216.42\|CCTTCTGATCGAGGCCCAGCCCGTGGACGGTGTGAGGCCGGT | >hg38_21_+_8433172_8446622_RNA45SN1_45S_50ntFlanks | 8175 | 8216 | 42 | CCTTCTGATCGAGGCCCAGCCCGTGGACGGTGTGAGGCCGGT | 926.2 | 11.4 | 2.1 | 5.3 | 9.0E-08 | 1.5E-07 | MB231/MB468 |
| >hg38_21_+_8433172_8446622_RNA45SN1_45S_50ntFlanks@9534.9574.41\|AGGAAACTCTGGTGGAGGTCCGTAGCGGTCCTGACGTGCAA | >hg38_21_+_8433172_8446622_RNA45SN1_45S_50ntFlanks | 9534 | 9574 | 41 | AGGAAACTCTGGTGGAGGTCCGTAGCGGTCCTGACGTGCAA | 185.5 | -3.8 | 0.7 | -5.3 | 1.0E-07 | 1.5E-07 | BT20/MB468 |
| >hg38_21_+_8433172_8446622_RNA45SN1_45S_50ntFlanks@12073.12094.22\|CCGAGGGGCTCTCGCTTCTGGC | >hg38_21_+_8433172_8446622_RNA45SN1_45S_50ntFlanks | 12073 | 12094 | 22 | CCGAGGGGCTCTCGCTTCTGGC | 295.8 | -3.1 | 0.6 | -5.3 | 1.0E-07 | 1.5E-07 | BT20/MB468 |
| >hg38_21_+_8433172_8446622_RNA45SN1_45S_50ntFlanks@12988.13013.26\|CAGCTCCCTCGCTGCGATCTATTGAA | >hg38_21_+_8433172_8446622_RNA45SN1_45S_50ntFlanks | 12988 | 13013 | 26 | CAGCTCCCTCGCTGCGATCTATTGAA | 717.5 | 2.8 | 0.5 | 5.3 | 9.2E-08 | 1.5E-07 | MB231/MB468 |
| >hg38_21_+_8433172_8446622_RNA45SN1_45S_50ntFlanks@11873.11897.25\|AAGAAGACCCTGTTGAGCTTGACTC | >hg38_21_+_8433172_8446622_RNA45SN1_45S_50ntFlanks | 11873 | 11897 | 25 | AAGAAGACCCTGTTGAGCTTGACTC | 626.3 | -3.9 | 0.7 | -5.3 | 1.0E-07 | 1.5E-07 | BT20/MB468 |
| >hg38_21_+_8433172_8446622_RNA45SN1_45S_50ntFlanks@7974.7991.18\|ACGCGACCTCAGATCAGA | >hg38_21_+_8433172_8446622_RNA45SN1_45S_50ntFlanks | 7974 | 7991 | 18 | ACGCGACCTCAGATCAGA | 2158.1 | 4.5 | 0.8 | 5.3 | 1.0E-07 | 1.5E-07 | BT20/MB468 |
| >hg38_21_+_8433172_8446622_RNA45SN1_45S_50ntFlanks@12367.12384.18\|AACTGGCTTGTGGCGGCC | >hg38_21_+_8433172_8446622_RNA45SN1_45S_50ntFlanks | 12367 | 12384 | 18 | AACTGGCTTGTGGCGGCC | 142.7 | -2.8 | 0.5 | -5.3 | 9.5E-08 | 1.6E-07 | MB231/MB468 |
| >hg38_21_+_8433172_8446622_RNA45SN1_45S_50ntFlanks@9198.9225.28\|GTTCTCTCGGGGCCACGCGCGCGTCCCC | >hg38_21_+_8433172_8446622_RNA45SN1_45S_50ntFlanks | 9198 | 9225 | 28 | GTTCTCTCGGGGCCACGCGCGCGTCCCC | 143.6 | -5.3 | 1.0 | -5.3 | 1.1E-07 | 1.6E-07 | BT20/MB468 |
| >hg38_1_-_228634819_228635039_RNA5S12_5S_50ntFlanks@130.171.42\|TGGATGGGAGACCGCCTGGGAATACCGGGTGCTGTAGGCTTT | >hg38_1_-_228634819_228635039_RNA5S12_5S_50ntFlanks | 130 | 171 | 42 | TGGATGGGAGACCGCCTGGGAATACCGGGTGCTGTAGGCTTT | 2784.2 | 12.2 | 2.3 | 5.3 | 9.6E-08 | 1.6E-07 | MB231/MB468 |
| >hg38_21_+_8433172_8446622_RNA45SN1_45S_50ntFlanks@7983.8012.30\|CAGATCAGACGTGGCGACCCGCTGAATTTA | >hg38_21_+_8433172_8446622_RNA45SN1_45S_50ntFlanks | 7983 | 8012 | 30 | CAGATCAGACGTGGCGACCCGCTGAATTTA | 209.4 | -2.9 | 0.5 | -5.3 | 9.6E-08 | 1.6E-07 | MB231/MB468 |
| >trna128_GlyGCC_6_-_27870686_27870756@19.38.20,trna19_GlyGCC_16_+_70823410_70823480@19.38.20,trna19_GlyGCC_2_-_157257659_157257729@19.38.20,trna24_GlyGCC_16_-_70812942_70813012@19.38.20,trna25_GlyGCC_16_-_70812114_70812184@19.38.20,trna2_GlyGCC_21_-_18827107_18827177@19.38.20,trna35_GlyGCC_1_+_161413094_161413164@19.38.20,trna37_GlyGCC_1_+_161420467_161420537@19.38.20,trna39_GlyGCC_1_+_161427898_161427968@19.38.20,trna41_GlyGCC_1_+_161435258_161435328@19.38.20,trna5_GlyGCC_17_+_8029064_8029134@19.38.20,trna68_GlyGCC_1_-_161493637_161493707@19.38.20\|TAGAATTCTCGCCTGCCACG | >trna128_GlyGCC_6_-_27870686_27870756 | 19 | 38 | 20 | TAGAATTCTCGCCTGCCACG | 825.9 | 6.7 | 1.3 | 5.3 | 1.1E-07 | 1.6E-07 | BT20/MB468 |
| >hg38_21_+_8433172_8446622_RNA45SN1_45S_50ntFlanks@3837.3860.24\|CCTCTCCTACTTGGATAACTGTGG | >hg38_21_+_8433172_8446622_RNA45SN1_45S_50ntFlanks | 3837 | 3860 | 24 | CCTCTCCTACTTGGATAACTGTGG | 175.6 | -3.7 | 0.7 | -5.3 | 1.1E-07 | 1.6E-07 | BT20/MB468 |
| >hg38_21_+_8433172_8446622_RNA45SN1_45S_50ntFlanks@11570.11603.34\|AGCCGACTTAGAACTGGTGCGGACCAGGGGAATC | >hg38_21_+_8433172_8446622_RNA45SN1_45S_50ntFlanks | 11570 | 11603 | 34 | AGCCGACTTAGAACTGGTGCGGACCAGGGGAATC | 379.6 | -2.7 | 0.5 | -5.3 | 9.9E-08 | 1.6E-07 | MB231/MB468 |
| >hg38_21_+_8433172_8446622_RNA45SN1_45S_50ntFlanks@6723.6745.23\|TTGCAGGACACATTGATCATCGA | >hg38_21_+_8433172_8446622_RNA45SN1_45S_50ntFlanks | 6723 | 6745 | 23 | TTGCAGGACACATTGATCATCGA | 97.2 | -2.8 | 0.5 | -5.3 | 9.9E-08 | 1.6E-07 | MB231/MB468 |
| >hg38_21_+_8433172_8446622_RNA45SN1_45S_50ntFlanks@11670.11687.18\|CCCAGTGCTCTGAATGTC | >hg38_21_+_8433172_8446622_RNA45SN1_45S_50ntFlanks | 11670 | 11687 | 18 | CCCAGTGCTCTGAATGTC | 725.0 | -2.7 | 0.5 | -5.3 | 1.1E-07 | 1.6E-07 | BT20/MB468 |
| >hg38_21_+_8433172_8446622_RNA45SN1_45S_50ntFlanks@3854.3881.28\|ACTGTGGTAATTCTAGAGCTAATACATG | >hg38_21_+_8433172_8446622_RNA45SN1_45S_50ntFlanks | 3854 | 3881 | 28 | ACTGTGGTAATTCTAGAGCTAATACATG | 1491.7 | -3.8 | 0.7 | -5.3 | 1.0E-07 | 1.6E-07 | MB231/MB468 |
| >MI0003628\|hsa-mir-615&WithFlank&12\|+\|54033944\|54034051@67.88.22&[MIMAT0003283&hsa-miR-615-3p&offsets\|0\|0;m-495&12\|+\|54034010\|54034030&offsets\|0\|+1]\|TCCGAGCCTGGGTCTCCCTCTT | >MI0003628\|hsa-mir-615&WithFlank&12\|+\|54033944\|54034051 | 67 | 88 | 22 | TCCGAGCCTGGGTCTCCCTCTT | 1243.5 | 4.8 | 0.8 | 5.8 | 7.2E-09 | 1.6E-07 | BT20/MB231 |
| >hg38_21_+_8433172_8446622_RNA45SN1_45S_50ntFlanks@11874.11910.37\|AGAAGACCCTGTTGAGCTTGACTCTAGTCTGGCACGG | >hg38_21_+_8433172_8446622_RNA45SN1_45S_50ntFlanks | 11874 | 11910 | 37 | AGAAGACCCTGTTGAGCTTGACTCTAGTCTGGCACGG | 179.6 | -5.6 | 1.1 | -5.3 | 1.1E-07 | 1.6E-07 | BT20/MB468 |
| >hg38_21_+_8433172_8446622_RNA45SN1_45S_50ntFlanks@12500.12518.19\|GTTTAGACCGTCGTGAGAC | >hg38_21_+_8433172_8446622_RNA45SN1_45S_50ntFlanks | 12500 | 12518 | 19 | GTTTAGACCGTCGTGAGAC | 118.0 | -4.2 | 0.8 | -5.3 | 1.1E-07 | 1.6E-07 | BT20/MB468 |
| >hg38_21_+_8433172_8446622_RNA45SN1_45S_50ntFlanks@7975.7998.24\|CGCGACCTCAGATCAGACGTGGCG | >hg38_21_+_8433172_8446622_RNA45SN1_45S_50ntFlanks | 7975 | 7998 | 24 | CGCGACCTCAGATCAGACGTGGCG | 504.8 | 4.5 | 0.8 | 5.8 | 7.3E-09 | 1.7E-07 | BT20/MB231 |
| >hg38_21_+_8433172_8446622_RNA45SN1_45S_50ntFlanks@12249.12272.24\|CAAAAGCTCGCTTGATCTTGATTT | >hg38_21_+_8433172_8446622_RNA45SN1_45S_50ntFlanks | 12249 | 12272 | 24 | CAAAAGCTCGCTTGATCTTGATTT | 101.4 | -3.1 | 0.6 | -5.3 | 1.1E-07 | 1.7E-07 | BT20/MB468 |
| >hg38_21_+_8433172_8446622_RNA45SN1_45S_50ntFlanks@12892.12932.41\|CCCTCGCCCGTCACGCACCGCACGTTCGTGGGGAACCTGGC | >hg38_21_+_8433172_8446622_RNA45SN1_45S_50ntFlanks | 12892 | 12932 | 41 | CCCTCGCCCGTCACGCACCGCACGTTCGTGGGGAACCTGGC | 414.3 | -4.8 | 0.9 | -5.3 | 1.2E-07 | 1.7E-07 | BT20/MB468 |
| >hg38_21_+_8433172_8446622_RNA45SN1_45S_50ntFlanks@12605.12622.18\|TATGTGCTTGGCTGAGGA | >hg38_21_+_8433172_8446622_RNA45SN1_45S_50ntFlanks | 12605 | 12622 | 18 | TATGTGCTTGGCTGAGGA | 204.7 | -3.8 | 0.7 | -5.3 | 1.2E-07 | 1.7E-07 | BT20/MB468 |
| >hg38_21_+_8433172_8446622_RNA45SN1_45S_50ntFlanks@3705.3722.18\|TACCTGGTTGATCCTGCC | >hg38_21_+_8433172_8446622_RNA45SN1_45S_50ntFlanks | 3705 | 3722 | 18 | TACCTGGTTGATCCTGCC | 4767.8 | 3.8 | 0.7 | 5.3 | 1.1E-07 | 1.7E-07 | MB231/MB468 |
| >hg38_21_+_8433172_8446622_RNA45SN1_45S_50ntFlanks@10873.10902.30\|CTGGGTCGGTCGGGCTGGGGCGCGAAGCGG | >hg38_21_+_8433172_8446622_RNA45SN1_45S_50ntFlanks | 10873 | 10902 | 30 | CTGGGTCGGTCGGGCTGGGGCGCGAAGCGG | 319.3 | -5.5 | 1.0 | -5.3 | 1.2E-07 | 1.7E-07 | BT20/MB468 |
| >hg38_21_+_8433172_8446622_RNA45SN1_45S_50ntFlanks@4588.4613.26\|CTATTTTGTTGGTTTTCGGAACTGAG | >hg38_21_+_8433172_8446622_RNA45SN1_45S_50ntFlanks | 4588 | 4613 | 26 | CTATTTTGTTGGTTTTCGGAACTGAG | 185.1 | -4.7 | 0.9 | -5.3 | 1.2E-07 | 1.7E-07 | BT20/MB468 |
| >hg38_21_+_8433172_8446622_RNA45SN1_45S_50ntFlanks@7981.8008.28\|CTCAGATCAGACGTGGCGACCCGCTGAA | >hg38_21_+_8433172_8446622_RNA45SN1_45S_50ntFlanks | 7981 | 8008 | 28 | CTCAGATCAGACGTGGCGACCCGCTGAA | 463.7 | -3.6 | 0.7 | -5.3 | 1.1E-07 | 1.7E-07 | MB231/MB468 |
| >hg38_21_+_8433172_8446622_RNA45SN1_45S_50ntFlanks@6723.6743.21\|TTGCAGGACACATTGATCATC | >hg38_21_+_8433172_8446622_RNA45SN1_45S_50ntFlanks | 6723 | 6743 | 21 | TTGCAGGACACATTGATCATC | 599.4 | -3.3 | 0.6 | -5.3 | 1.2E-07 | 1.7E-07 | BT20/MB468 |
| >hg38_21_+_8433172_8446622_RNA45SN1_45S_50ntFlanks@7980.8007.28\|CCTCAGATCAGACGTGGCGACCCGCTGA | >hg38_21_+_8433172_8446622_RNA45SN1_45S_50ntFlanks | 7980 | 8007 | 28 | CCTCAGATCAGACGTGGCGACCCGCTGA | 135.2 | -3.5 | 0.7 | -5.3 | 1.2E-07 | 1.7E-07 | BT20/MB468 |

| | | | | | | | | | | | | |
|---|---|---|---|---|---|---|---|---|---|---|---|---|
| >hg38_21_+_8433172_8446622_RNA45SN1_45S_50ntFlanks@11593.11611.19\|CCAGGGGAATCCGACTGTT | >hg38_21_+_8433172_8446622_RNA45SN1_45S_50ntFlanks | 11593 | 11611 | 19 | CCAGGGGAATCCGACTGTT | 2251.4 | -3.7 | 0.7 | -5.3 | 1.1E-07 | 1.7E-07 | MB231/MB468 |
| >hg38_21_+_8433172_8446622_RNA45SN1_45S_50ntFlanks@9178.9197.20\|CGCCGTCGGGCCCGGGGGAG | >hg38_21_+_8433172_8446622_RNA45SN1_45S_50ntFlanks | 9178 | 9197 | 20 | CGCCGTCGGGCCCGGGGGAG | 82.7 | -6.0 | 1.1 | -5.3 | 1.1E-07 | 1.8E-07 | MB231/MB468 |
| >hg38_MT_+_1621_3279_RNR2_16S_50ntFlanks@1370.1396.27\|ATGTTGGATCAGGACATCCCGATGGTG | >hg38_MT_+_1621_3279_RNR2_16S_50ntFlanks | 1370 | 1396 | 27 | ATGTTGGATCAGGACATCCCGATGGTG | 628.1 | 7.3 | 1.4 | 5.3 | 1.2E-07 | 1.8E-07 | BT20/MB468 |
| >hg38_21_+_8433172_8446622_RNA45SN1_45S_50ntFlanks@6709.6734.26\|AGAATTAATGTGAATTGCAGGACACA | >hg38_21_+_8433172_8446622_RNA45SN1_45S_50ntFlanks | 6709 | 6734 | 26 | AGAATTAATGTGAATTGCAGGACACA | 77.7 | -3.5 | 0.7 | -5.3 | 1.1E-07 | 1.8E-07 | MB231/MB468 |
| >hg38_21_+_8433172_8446622_RNA45SN1_45S_50ntFlanks@6724.6744.21\|TGCAGGACACATTGATCATCG | >hg38_21_+_8433172_8446622_RNA45SN1_45S_50ntFlanks | 6724 | 6744 | 21 | TGCAGGACACATTGATCATCG | 173.1 | -2.6 | 0.5 | -5.3 | 1.2E-07 | 1.8E-07 | BT20/MB468 |
| >hg38_21_+_8433172_8446622_RNA45SN1_45S_50ntFlanks@6719.6741.23\|TGAATTGCAGGACACATTGATCA | >hg38_21_+_8433172_8446622_RNA45SN1_45S_50ntFlanks | 6719 | 6741 | 23 | TGAATTGCAGGACACATTGATCA | 424.0 | -5.1 | 1.0 | -5.3 | 1.2E-07 | 1.8E-07 | BT20/MB468 |
| >trnaMT_LeuTAA_MT_+_3230_3304@16.44.29\|CCCGGTAATCGCATAAAACTTAAAACTTT | >trnaMT_LeuTAA_MT_+_3230_3304 | 16 | 44 | 29 | CCCGGTAATCGCATAAAACTTAAAACTTT | 405.6 | -3.4 | 0.6 | -5.3 | 1.1E-07 | 1.8E-07 | MB231/MB468 |
| >hg38_21_+_8433172_8446622_RNA45SN1_45S_50ntFlanks@12634.12661.28\|GAAGCTACCATCTGTGGGATTATGACTG | >hg38_21_+_8433172_8446622_RNA45SN1_45S_50ntFlanks | 12634 | 12661 | 28 | GAAGCTACCATCTGTGGGATTATGACTG | 184.0 | -2.1 | 0.4 | -5.3 | 1.2E-07 | 1.8E-07 | BT20/MB468 |
| >hg38_21_+_8433172_8446622_RNA45SN1_45S_50ntFlanks@10509.10533.25\|GAATCCGGAGTGGCGGAGATGGGCG | >hg38_21_+_8433172_8446622_RNA45SN1_45S_50ntFlanks | 10509 | 10533 | 25 | GAATCCGGAGTGGCGGAGATGGGCG | 89.6 | -5.0 | 0.9 | -5.3 | 1.1E-07 | 1.8E-07 | MB231/MB468 |
| >hg38_MT_+_598_1651_RNR1_12S_50ntFlanks@428.462.35\|AAAGTGGCTTTAACATATCTGAACACACAATAGCT | >hg38_MT_+_598_1651_RNR1_12S_50ntFlanks | 428 | 462 | 35 | AAAGTGGCTTTAACATATCTGAACACACAATAGCT | 1665.6 | 12.0 | 2.3 | 5.3 | 1.1E-07 | 1.8E-07 | MB231/MB468 |
| >hg38_21_+_8433172_8446622_RNA45SN1_45S_50ntFlanks@12420.12443.24\|ATGTCGGCTCTTCCTATCATTGTG | >hg38_21_+_8433172_8446622_RNA45SN1_45S_50ntFlanks | 12420 | 12443 | 24 | ATGTCGGCTCTTCCTATCATTGTG | 2831.4 | -3.3 | 0.6 | -5.3 | 1.2E-07 | 1.8E-07 | BT20/MB468 |
| >hg38_21_+_8433172_8446622_RNA45SN1_45S_50ntFlanks@12431.12455.25\|TCCTATCATTGTGAAGCAGAATTCA | >hg38_21_+_8433172_8446622_RNA45SN1_45S_50ntFlanks | 12431 | 12455 | 25 | TCCTATCATTGTGAAGCAGAATTCA | 83.8 | -3.4 | 0.6 | -5.3 | 1.2E-07 | 1.8E-07 | BT20/MB468 |
| >hg38_21_+_8433172_8446622_RNA45SN1_45S_50ntFlanks@4783.4800.18\|CATAAACGATGCCGACCG | >hg38_21_+_8433172_8446622_RNA45SN1_45S_50ntFlanks | 4783 | 4800 | 18 | CATAAACGATGCCGACCG | 70.5 | -4.6 | 0.9 | -5.3 | 1.1E-07 | 1.8E-07 | MB231/MB468 |
| >hg38_21_+_8433172_8446622_RNA45SN1_45S_50ntFlanks@12405.12443.39\|CTTTTTGATCCTTCGATGTCGGCTCTTCCTATCATTGTG | >hg38_21_+_8433172_8446622_RNA45SN1_45S_50ntFlanks | 12405 | 12443 | 39 | CTTTTTGATCCTTCGATGTCGGCTCTTCCTATCATTGTG | 1805.7 | -3.4 | 0.6 | -5.3 | 1.1E-07 | 1.8E-07 | MB231/MB468 |
| >hg38_21_+_8433172_8446622_RNA45SN1_45S_50ntFlanks@11671.11700.30\|CCAGTGCTCTGAATGTCAAAGTGAAGAAAT | >hg38_21_+_8433172_8446622_RNA45SN1_45S_50ntFlanks | 11671 | 11700 | 30 | CCAGTGCTCTGAATGTCAAAGTGAAGAAAT | 222.0 | -3.5 | 0.7 | -5.3 | 1.3E-07 | 1.8E-07 | BT20/MB468 |
| >MI0003628\|hsa-mir-615&WithFlank&12\|+\|54033944\|54034051@67.87.21&[MIMAT0003283&hsa-miR-615-3p&offsets\|0\|-1;m-495&12\|+\|54034010\|54034030&offsets\|0\|0]\|TCCGAGCCTGGGTCTCCCTCT | >MI0003628\|hsa-mir-615&WithFlank&12\|+\|54033944\|54034051 | 67 | 87 | 21 | TCCGAGCCTGGGTCTCCCTCT | 1911.0 | 5.1 | 0.9 | 5.8 | 8.2E-09 | 1.8E-07 | BT20/MB231 |
| >hg38_21_+_8433172_8446622_RNA45SN1_45S_50ntFlanks@12478.12496.19\|ACTAATAGGGAACGTGAGC | >hg38_21_+_8433172_8446622_RNA45SN1_45S_50ntFlanks | 12478 | 12496 | 19 | ACTAATAGGGAACGTGAGC | 1246.8 | 2.9 | 0.5 | 5.3 | 1.1E-07 | 1.8E-07 | MB231/MB468 |
| >hg38_21_+_8433172_8446622_RNA45SN1_45S_50ntFlanks@10410.10443.34\|TCAGTCGGTCCTGAGAGATGGGCGAGCGCCGTTC | >hg38_21_+_8433172_8446622_RNA45SN1_45S_50ntFlanks | 10410 | 10443 | 34 | TCAGTCGGTCCTGAGAGATGGGCGAGCGCCGTTC | 211.1 | -2.6 | 0.5 | -5.3 | 1.1E-07 | 1.9E-07 | MB231/MB468 |
| >hg38_21_+_8433172_8446622_RNA45SN1_45S_50ntFlanks@12574.12604.31\|CGAGAGGAACCGCAGGTTCAGACATTTGGTG | >hg38_21_+_8433172_8446622_RNA45SN1_45S_50ntFlanks | 12574 | 12604 | 31 | CGAGAGGAACCGCAGGTTCAGACATTTGGTG | 92.7 | -5.4 | 1.0 | -5.3 | 1.2E-07 | 1.9E-07 | MB231/MB468 |
| >hg38_21_+_8433172_8446622_RNA45SN1_45S_50ntFlanks@10392.10426.35\|ACAGCAGTTGAACATGGGTCAGTCGGTCCTGAGAG | >hg38_21_+_8433172_8446622_RNA45SN1_45S_50ntFlanks | 10392 | 10426 | 35 | ACAGCAGTTGAACATGGGTCAGTCGGTCCTGAGAG | 408.6 | -5.2 | 1.0 | -5.3 | 1.3E-07 | 1.9E-07 | BT20/MB468 |
| >MI0000446\|hsa-mir-125b-1&WithFlank&11\|-\|122099751\|122099850@22.42.21;MI0000470\|hsa-mir-125b-2&WithFlank&21\|+\|16590231\|16590331@24.44.21&[MIMAT0000423&hsa-miR-125b-5p&offsets\|+1\|0;m-67&11\|-\|122099809\|122099830&offsets\|+1\|0];[MIMAT0000423_1&hsa-miR-125b-5p&offsets\|+1\|0;m-68&21\|+\|16590253\|16590274&offsets\|+1\|0]\|CCCTGAGACCCTAACTTGTGA | >MI0000446\|hsa-mir-125b-1&WithFlank&11\|-\|122099751\|122099850 | 22 | 42 | 21 | CCCTGAGACCCTAACTTGTGA | 197.9 | -6.3 | 1.1 | -5.8 | 8.5E-09 | 1.9E-07 | BT20/MB231 |
| >hg38_21_+_8433172_8446622_RNA45SN1_45S_50ntFlanks@10401.10420.20\|GAACATGGGTCAGTCGGTCC | >hg38_21_+_8433172_8446622_RNA45SN1_45S_50ntFlanks | 10401 | 10420 | 20 | GAACATGGGTCAGTCGGTCC | 133.5 | -2.8 | 0.5 | -5.3 | 1.3E-07 | 1.9E-07 | BT20/MB468 |
| >hg38_21_+_8433172_8446622_RNA45SN1_45S_50ntFlanks@10353.10383.31\|AGAACTTTGAAGGCCGAAGTGGAGAAGGGTT | >hg38_21_+_8433172_8446622_RNA45SN1_45S_50ntFlanks | 10353 | 10383 | 31 | AGAACTTTGAAGGCCGAAGTGGAGAAGGGTT | 223.5 | -1.9 | 0.4 | -5.3 | 1.4E-07 | 2.0E-07 | BT20/MB468 |
| >hg38_21_+_8433172_8446622_RNA45SN1_45S_50ntFlanks@12434.12455.22\|TATCATTGTGAAGCAGAATTCA | >hg38_21_+_8433172_8446622_RNA45SN1_45S_50ntFlanks | 12434 | 12455 | 22 | TATCATTGTGAAGCAGAATTCA | 260.8 | -5.3 | 1.0 | -5.3 | 1.4E-07 | 2.0E-07 | BT20/MB468 |
| >hg38_21_+_8433172_8446622_RNA45SN1_45S_50ntFlanks@10458.10488.31\|ATGGCCTCCGTTGCCCTCGGCCGATCGAAAG | >hg38_21_+_8433172_8446622_RNA45SN1_45S_50ntFlanks | 10458 | 10488 | 31 | ATGGCCTCCGTTGCCCTCGGCCGATCGAAAG | 695.4 | -5.2 | 1.0 | -5.3 | 1.4E-07 | 2.0E-07 | BT20/MB468 |
| >hg38_21_+_8433172_8446622_RNA45SN1_45S_50ntFlanks@11579.11600.22\|AGAACTGGTGCGGACCAGGGGA | >hg38_21_+_8433172_8446622_RNA45SN1_45S_50ntFlanks | 11579 | 11600 | 22 | AGAACTGGTGCGGACCAGGGGA | 160.5 | -3.4 | 0.6 | -5.3 | 1.2E-07 | 2.0E-07 | MB231/MB468 |
| >hg38_21_+_8433172_8446622_RNA45SN1_45S_50ntFlanks@8189.8208.20\|CCCAGCCCGTGGACGGTGTG | >hg38_21_+_8433172_8446622_RNA45SN1_45S_50ntFlanks | 8189 | 8208 | 20 | CCCAGCCCGTGGACGGTGTG | 148.8 | -5.8 | 1.1 | -5.3 | 1.4E-07 | 2.0E-07 | BT20/MB468 |
| >hg38_21_+_8433172_8446622_RNA45SN1_45S_50ntFlanks@8170.8215.46\|CAAGTCCTTCTGATCGAGGCCCAGCCCGTGGACGGTGTGAGGCCGG | >hg38_21_+_8433172_8446622_RNA45SN1_45S_50ntFlanks | 8170 | 8215 | 46 | CAAGTCCTTCTGATCGAGGCCCAGCCCGTGGACGGTGTGAGGCCGG | 140.9 | -3.6 | 0.7 | -5.3 | 1.2E-07 | 2.0E-07 | MB231/MB468 |
| >trna111_HisGTG_1_-_147774845_147774916@-1A.29.30,trna118_HisGTG_1_-_145396881_145396952@-1A.29.30,trna16_HisGTG_1_+_146544773_146544844@-1A.29.30,trna1_HisGTG_15_+_45493349_45493420@-1A.29.30,trna21_HisGTG_1_+_147753471_147753542@-1A.29.30,trna33_HisGTG_6_+_27125906_27125977@-1A.29.30,trna7_HisGTG_9_-_14433938_14434009@-1A.29.30,trna8_HisGTG_15_-_45492611_45492682@-1A.29.30,trna9_HisGTG_15_-_45490804_45490875@-1A.29.30\|AGCCGTGATCGTATAGTGGTTAGTACTCTG | >trna111_HisGTG_1_-_147774845_147774916 | -1A | 29 | 30 | AGCCGTGATCGTATAGTGGTTAGTACTCTG | 290.1 | -3.7 | 0.7 | -5.3 | 1.3E-07 | 2.1E-07 | MB231/MB468 |
| >am_trnaMT_SerGCT_MT_+_12207_12265@39.58.20\|TCTAACAACATGGCTTTCTC | >am_trnaMT_SerGCT_MT_+_12207_12265 | 39 | 58 | 20 | TCTAACAACATGGCTTTCTC | 55.8 | -7.4 | 1.4 | -5.3 | 1.3E-07 | 2.1E-07 | MB231/MB468 |
| >hg38_21_+_8433172_8446622_RNA45SN1_45S_50ntFlanks@10468.10497.30\|TTGCCCTCGGCCGATCGAAAGGGAGTCGGG | >hg38_21_+_8433172_8446622_RNA45SN1_45S_50ntFlanks | 10468 | 10497 | 30 | TTGCCCTCGGCCGATCGAAAGGGAGTCGGG | 247.3 | -4.2 | 0.8 | -5.3 | 1.3E-07 | 2.1E-07 | MB231/MB468 |
| >hg38_21_+_8433172_8446622_RNA45SN1_45S_50ntFlanks@8179.8203.25\|CTGATCGAGGCCCAGCCCGTGGACG | >hg38_21_+_8433172_8446622_RNA45SN1_45S_50ntFlanks | 8179 | 8203 | 25 | CTGATCGAGGCCCAGCCCGTGGACG | 160.6 | -3.4 | 0.6 | -5.3 | 1.4E-07 | 2.1E-07 | BT20/MB468 |
| >hg38_21_+_8433172_8446622_RNA45SN1_45S_50ntFlanks@10380.10424.45\|GGTTCCATGTGAACAGCAGTTGAACATGGGTCAGTCGGTCCTGAG | >hg38_21_+_8433172_8446622_RNA45SN1_45S_50ntFlanks | 10380 | 10424 | 45 | GGTTCCATGTGAACAGCAGTTGAACATGGGTCAGTCGGTCCTGAG | 440.8 | 10.2 | 1.9 | 5.3 | 1.3E-07 | 2.1E-07 | MB231/MB468 |
| >hg38_MT_+_1621_3279_RNR2_16S_50ntFlanks@51.70.20\|GCTAAACCTAGCCCCAAACC | >hg38_MT_+_1621_3279_RNR2_16S_50ntFlanks | 51 | 70 | 20 | GCTAAACCTAGCCCCAAACC | 869.4 | 11.3 | 2.1 | 5.3 | 1.3E-07 | 2.1E-07 | MB231/MB468 |
| >hg38_1_-_228634819_228635039_RNA5S12_5S_50ntFlanks@129.170.42\|TTGGATGGGAGACCGCCTGGGAATACCGGGTGCTGTAGGCTT | >hg38_1_-_228634819_228635039_RNA5S12_5S_50ntFlanks | 129 | 170 | 42 | TTGGATGGGAGACCGCCTGGGAATACCGGGTGCTGTAGGCTT | 2946.8 | 12.2 | 2.3 | 5.3 | 1.3E-07 | 2.1E-07 | MB231/MB468 |
| >hg38_21_+_8433172_8446622_RNA45SN1_45S_50ntFlanks@7980.8012.33\|CCTCAGATCAGACGTGGCGACCCGCTGAATTTA | >hg38_21_+_8433172_8446622_RNA45SN1_45S_50ntFlanks | 7980 | 8012 | 33 | CCTCAGATCAGACGTGGCGACCCGCTGAATTTA | 183.2 | -3.3 | 0.6 | -5.3 | 1.3E-07 | 2.1E-07 | MB231/MB468 |
| >trnaMT_PheGAA_MT_+_577_647@35.56.22\|GAAAATGTTTAGACGGGCTCAC | >trnaMT_PheGAA_MT_+_577_647 | 35 | 56 | 22 | GAAAATGTTTAGACGGGCTCAC | 1143.1 | 11.4 | 2.2 | 5.3 | 1.3E-07 | 2.1E-07 | MB231/MB468 |
| >hg38_21_+_8433172_8446622_RNA45SN1_45S_50ntFlanks@12640.12669.30\|ACCATCTGTGGGATTATGACTGAACGCCTC | >hg38_21_+_8433172_8446622_RNA45SN1_45S_50ntFlanks | 12640 | 12669 | 30 | ACCATCTGTGGGATTATGACTGAACGCCTC | 251.0 | 6.4 | 1.2 | 5.3 | 1.5E-07 | 2.1E-07 | BT20/MB468 |
| >hg38_21_+_8433172_8446622_RNA45SN1_45S_50ntFlanks@11564.11588.25\|CCTAGCAGCCGACTTAGAACTGGTG | >hg38_21_+_8433172_8446622_RNA45SN1_45S_50ntFlanks | 11564 | 11588 | 25 | CCTAGCAGCCGACTTAGAACTGGTG | 864.4 | -3.3 | 0.6 | -5.3 | 1.3E-07 | 2.2E-07 | MB231/MB468 |
| >hg38_21_+_8433172_8446622_RNA45SN1_45S_50ntFlanks@12405.12437.33\|CTTTTTGATCCTTCGATGTCGGCTCTTCCTATC | >hg38_21_+_8433172_8446622_RNA45SN1_45S_50ntFlanks | 12405 | 12437 | 33 | CTTTTTGATCCTTCGATGTCGGCTCTTCCTATC | 1555.3 | -3.6 | 0.7 | -5.3 | 1.3E-07 | 2.2E-07 | MB231/MB468 |
| >MI0000749\|hsa-mir-30e&WithFlank&1\|+\|40754349\|40754452@23.45.23(+1G)&[MIMAT0000692&hsa-miR-30e-5p&offsets\|0\|+1(+1G);m-24&1\|+\|40754371\|40754394&offsets\|0\|-1(+1G)]\|TGTAAACATCCTTGACTGGAAGCG | >MI0000749\|hsa-mir-30e&WithFlank&1\|+\|40754349\|40754452 | 23 | 45 | 23(+1G) | TGTAAACATCCTTGACTGGAAGCG | 2852.7 | 5.7 | 1.1 | 5.3 | 1.5E-07 | 2.2E-07 | BT20/MB468 |
| >hg38_21_+_8433172_8446622_RNA45SN1_45S_50ntFlanks@7975.8008.34\|CGCGACCTCAGATCAGACGTGGCGACCCGCTGAA | >hg38_21_+_8433172_8446622_RNA45SN1_45S_50ntFlanks | 7975 | 8008 | 34 | CGCGACCTCAGATCAGACGTGGCGACCCGCTGAA | 53994.3 | 4.3 | 0.8 | 5.3 | 1.3E-07 | 2.2E-07 | MB231/MB468 |
| >hg38_21_+_8433172_8446622_RNA45SN1_45S_50ntFlanks@12420.12441.22\|ATGTCGGCTCTTCCTATCATTG | >hg38_21_+_8433172_8446622_RNA45SN1_45S_50ntFlanks | 12420 | 12441 | 22 | ATGTCGGCTCTTCCTATCATTG | 1649.2 | -3.3 | 0.6 | -5.3 | 1.5E-07 | 2.2E-07 | BT20/MB468 |
| >hg38_21_+_8433172_8446622_RNA45SN1_45S_50ntFlanks@12071.12093.23\|CCCCGAGGGGCTCTCGCTTCTGG | >hg38_21_+_8433172_8446622_RNA45SN1_45S_50ntFlanks | 12071 | 12093 | 23 | CCCCGAGGGGCTCTCGCTTCTGG | 113.9 | -4.6 | 0.9 | -5.3 | 1.5E-07 | 2.2E-07 | BT20/MB468 |
| >hg38_21_+_8433172_8446622_RNA45SN1_45S_50ntFlanks@10737.10757.21\|TAAATCTCGCGCCGGGCCGTA | >hg38_21_+_8433172_8446622_RNA45SN1_45S_50ntFlanks | 10737 | 10757 | 21 | TAAATCTCGCGCCGGGCCGTA | 140.0 | -2.9 | 0.6 | -5.2 | 1.5E-07 | 2.2E-07 | BT20/MB468 |

| | | | | | | | | | | | | |
|---|---|---|---|---|---|---|---|---|---|---|---|---|
| >hg38_21_+_8433172_8446622_RNA45SN1_45S_50ntFlanks@8168.8186.19\|CCCAAGTCCTTCTGATCGA | >hg38_21_+_8433172_8446622_RNA45SN1_45S_50ntFlanks | 8168 | 8186 | 19 | CCCAAGTCCTTCTGATCGA | 1010.7 | -4.4 | 0.8 | -5.3 | 1.4E-07 | 2.2E-07 | MB231/MB468 |
| >hg38_21_+_8433172_8446622_RNA45SN1_45S_50ntFlanks@12444.12477.34\|AAGCAGAATTCACCAAGCGTTGGATTGTTCACCC | >hg38_21_+_8433172_8446622_RNA45SN1_45S_50ntFlanks | 12444 | 12477 | 34 | AAGCAGAATTCACCAAGCGTTGGATTGTTCACCC | 668.6 | 3.5 | 0.7 | 5.3 | 1.4E-07 | 2.2E-07 | MB231/MB468 |
| >hg38_21_+_8433172_8446622_RNA45SN1_45S_50ntFlanks@10116.10136.21\|TGCGTCGGAGGGCGGCGGCGG | >hg38_21_+_8433172_8446622_RNA45SN1_45S_50ntFlanks | 10116 | 10136 | 21 | TGCGTCGGAGGGCGGCGGCGG | 181.5 | -4.8 | 0.9 | -5.2 | 1.5E-07 | 2.2E-07 | BT20/MB468 |
| >hg38_21_+_8433172_8446622_RNA45SN1_45S_50ntFlanks@12164.12182.19\|CTGGGGCGGTACACCTGTC | >hg38_21_+_8433172_8446622_RNA45SN1_45S_50ntFlanks | 12164 | 12182 | 19 | CTGGGGCGGTACACCTGTC | 118.6 | -4.2 | 0.8 | -5.3 | 1.4E-07 | 2.3E-07 | MB231/MB468 |
| >hg38_21_+_8433172_8446622_RNA45SN1_45S_50ntFlanks@10771.10806.36\|CAGGTCTCCAAGGTGAACAGCCTCTGGCATGTTGGA | >hg38_21_+_8433172_8446622_RNA45SN1_45S_50ntFlanks | 10771 | 10806 | 36 | CAGGTCTCCAAGGTGAACAGCCTCTGGCATGTTGGA | 451.4 | -4.6 | 0.9 | -5.3 | 1.4E-07 | 2.3E-07 | MB231/MB468 |
| >hg38_21_+_8433172_8446622_RNA45SN1_45S_50ntFlanks@8194.8225.32\|CCCGTGGACGGTGTGAGGCCGGTAGCGGCCCC | >hg38_21_+_8433172_8446622_RNA45SN1_45S_50ntFlanks | 8194 | 8225 | 32 | CCCGTGGACGGTGTGAGGCCGGTAGCGGCCCC | 114.3 | -2.8 | 0.5 | -5.3 | 1.4E-07 | 2.3E-07 | MB231/MB468 |
| >hg38_21_+_8433172_8446622_RNA45SN1_45S_50ntFlanks@5485.5526.42\|AGCGCTGAGAAGACGGTCGAACTTGACTATCTAGAGGAAGTA | >hg38_21_+_8433172_8446622_RNA45SN1_45S_50ntFlanks | 5485 | 5526 | 42 | AGCGCTGAGAAGACGGTCGAACTTGACTATCTAGAGGAAGTA | 136.1 | -5.2 | 1.0 | -5.3 | 1.4E-07 | 2.3E-07 | MB231/MB468 |
| >trna14_LysTTT_11_-_59327808_59327880@1.24.24,trna2_LysTTT_17_+_8022473_8022545@1.24.24,trna54_LysTTT_1_+_204475655_204475727@1.24.24,trna5_LysTTT_11_+_59323902_59323974@1.24.24,trna62_LysTTT_1_-_204476158_204476230@1.24.24,trna76_LysTTT_6_+_28918806_28918878@1.24.24\|GCCCGGATAGCTCAGTCGGTAGAG | >trna14_LysTTT_11_-_59327808_59327880 | 1 | 24 | 24 | GCCCGGATAGCTCAGTCGGTAGAG | 388.9 | -4.5 | 0.8 | -5.3 | 1.4E-07 | 2.3E-07 | MB231/MB468 |
| >hg38_21_+_8433172_8446622_RNA45SN1_45S_50ntFlanks@12632.12659.28\|GCGAAGCTACCATCTGTGGGATTATGAC | >hg38_21_+_8433172_8446622_RNA45SN1_45S_50ntFlanks | 12632 | 12659 | 28 | GCGAAGCTACCATCTGTGGGATTATGAC | 219.2 | -2.7 | 0.5 | -5.3 | 1.4E-07 | 2.3E-07 | MB231/MB468 |
| >hg38_21_+_8433172_8446622_RNA45SN1_45S_50ntFlanks@12404.12432.29\|GCTTTTTGATCCTTCGATGTCGGCTCTTC | >hg38_21_+_8433172_8446622_RNA45SN1_45S_50ntFlanks | 12404 | 12432 | 29 | GCTTTTTGATCCTTCGATGTCGGCTCTTC | 168.0 | -2.5 | 0.5 | -5.2 | 1.6E-07 | 2.3E-07 | BT20/MB468 |
| >hg38_21_+_8433172_8446622_RNA45SN1_45S_50ntFlanks@12163.12183.21\|ACTGGGGCGGTACACCTGTCA | >hg38_21_+_8433172_8446622_RNA45SN1_45S_50ntFlanks | 12163 | 12183 | 21 | ACTGGGGCGGTACACCTGTCA | 472.9 | -6.0 | 1.1 | -5.2 | 1.6E-07 | 2.3E-07 | BT20/MB468 |
| >hg38_21_+_8433172_8446622_RNA45SN1_45S_50ntFlanks@6722.6746.25\|ATTGCAGGACACATTGATCATCGAC | >hg38_21_+_8433172_8446622_RNA45SN1_45S_50ntFlanks | 6722 | 6746 | 25 | ATTGCAGGACACATTGATCATCGAC | 222.3 | -2.3 | 0.4 | -5.3 | 1.4E-07 | 2.3E-07 | MB231/MB468 |
| >hg38_1_-_228634819_228635039_RNA5S12_5S_50ntFlanks@132.170.39\|GATGGGAGACCGCCTGGGAATACCGGGTGCTGTAGGCTT | >hg38_1_-_228634819_228635039_RNA5S12_5S_50ntFlanks | 132 | 170 | 39 | GATGGGAGACCGCCTGGGAATACCGGGTGCTGTAGGCTT | 19070.6 | 3.7 | 0.7 | 5.2 | 1.6E-07 | 2.3E-07 | BT20/MB468 |
| >hg38_21_+_8433172_8446622_RNA45SN1_45S_50ntFlanks@11683.11702.20\|ATGTCAAAGTGAAGAAATTC | >hg38_21_+_8433172_8446622_RNA45SN1_45S_50ntFlanks | 11683 | 11702 | 20 | ATGTCAAAGTGAAGAAATTC | 222.5 | -3.4 | 0.6 | -5.3 | 1.4E-07 | 2.3E-07 | MB231/MB468 |
| >hg38_21_+_8433172_8446622_RNA45SN1_45S_50ntFlanks@12083.12106.24\|CTCGCTTCTGGCGCCAAGCGCCCG | >hg38_21_+_8433172_8446622_RNA45SN1_45S_50ntFlanks | 12083 | 12106 | 24 | CTCGCTTCTGGCGCCAAGCGCCCG | 320.5 | -4.0 | 0.8 | -5.3 | 1.5E-07 | 2.4E-07 | MB231/MB468 |
| >hg38_21_+_8433172_8446622_RNA45SN1_45S_50ntFlanks@8379.8400.22\|TCAAGAGGGCGTGAAACCGTTA | >hg38_21_+_8433172_8446622_RNA45SN1_45S_50ntFlanks | 8379 | 8400 | 22 | TCAAGAGGGCGTGAAACCGTTA | 164.7 | -4.5 | 0.9 | -5.3 | 1.5E-07 | 2.4E-07 | MB231/MB468 |
| >hg38_21_+_8433172_8446622_RNA45SN1_45S_50ntFlanks@12007.12043.37\|GCCGCCGGTGAAATACCACTACTCTGATCGTTTTTTC | >hg38_21_+_8433172_8446622_RNA45SN1_45S_50ntFlanks | 12007 | 12043 | 37 | GCCGCCGGTGAAATACCACTACTCTGATCGTTTTTTC | 227.2 | -4.4 | 0.8 | -5.2 | 1.6E-07 | 2.4E-07 | BT20/MB468 |
| >hg38_21_+_8433172_8446622_RNA45SN1_45S_50ntFlanks@9352.9377.26\|CCGTGGCGCAATGAAGGTGAAGGCCG | >hg38_21_+_8433172_8446622_RNA45SN1_45S_50ntFlanks | 9352 | 9377 | 26 | CCGTGGCGCAATGAAGGTGAAGGCCG | 78.2 | -5.9 | 1.1 | -5.3 | 1.5E-07 | 2.4E-07 | MB231/MB468 |
| >hg38_21_+_8433172_8446622_RNA45SN1_45S_50ntFlanks@11324.11344.21\|GTCTCGTCGCGCGCGCGTCCG | >hg38_21_+_8433172_8446622_RNA45SN1_45S_50ntFlanks | 11324 | 11344 | 21 | GTCTCGTCGCGCGCGCGTCCG | 110.4 | -3.8 | 0.7 | -5.3 | 1.5E-07 | 2.4E-07 | MB231/MB468 |
| >hg38_21_+_8433172_8446622_RNA45SN1_45S_50ntFlanks@4753.4782.30\|AAGACGATCAGATACCGTCGTAGTTCCGAC | >hg38_21_+_8433172_8446622_RNA45SN1_45S_50ntFlanks | 4753 | 4782 | 30 | AAGACGATCAGATACCGTCGTAGTTCCGAC | 225.8 | -2.8 | 0.5 | -5.3 | 1.5E-07 | 2.4E-07 | MB231/MB468 |
| >hg38_21_+_8433172_8446622_RNA45SN1_45S_50ntFlanks@12397.12419.23\|CGACGTCGCTTTTTGATCCTTCG | >hg38_21_+_8433172_8446622_RNA45SN1_45S_50ntFlanks | 12397 | 12419 | 23 | CGACGTCGCTTTTTGATCCTTCG | 99.6 | -2.8 | 0.5 | -5.2 | 1.7E-07 | 2.4E-07 | BT20/MB468 |
| >hg38_21_+_8433172_8446622_RNA45SN1_45S_50ntFlanks@10830.10848.19\|CAAGCCGGATCCGTAACTT | >hg38_21_+_8433172_8446622_RNA45SN1_45S_50ntFlanks | 10830 | 10848 | 19 | CAAGCCGGATCCGTAACTT | 132.2 | -3.4 | 0.6 | -5.2 | 1.5E-07 | 2.5E-07 | MB231/MB468 |
| >hg38_21_+_8433172_8446622_RNA45SN1_45S_50ntFlanks@6733.6758.26\|CATTGATCATCGACACTTCGAACGCA | >hg38_21_+_8433172_8446622_RNA45SN1_45S_50ntFlanks | 6733 | 6758 | 26 | CATTGATCATCGACACTTCGAACGCA | 356.4 | -2.4 | 0.5 | -5.2 | 1.7E-07 | 2.5E-07 | BT20/MB468 |
| >hg38_21_+_8433172_8446622_RNA45SN1_45S_50ntFlanks@11570.11603.34\|AGCCGACTTAGAACTGGTGCGGACCAGGGGAATC | >hg38_21_+_8433172_8446622_RNA45SN1_45S_50ntFlanks | 11570 | 11603 | 34 | AGCCGACTTAGAACTGGTGCGGACCAGGGGAATC | 424.3 | -3.9 | 0.8 | -5.2 | 1.7E-07 | 2.5E-07 | BT20/MB468 |
| >hg38_21_+_8433172_8446622_RNA45SN1_45S_50ntFlanks@12008.12031.24\|CCGCCGGTGAAATACCACTACTCT | >hg38_21_+_8433172_8446622_RNA45SN1_45S_50ntFlanks | 12008 | 12031 | 24 | CCGCCGGTGAAATACCACTACTCT | 1457.1 | -4.5 | 0.9 | -5.2 | 1.7E-07 | 2.5E-07 | BT20/MB468 |
| >MI0005756\|hsa-mir-934&WithFlank&X\|+\|136550872\|136550966@21.42.22(+1G)&[MIMAT0004977&hsa-miR-934&offsets\|0\|0(+1G);m-943&X\|+\|136550892\|136550913&offsets\|0\|0(+1G)]\|TGTCTACTACTGGAGACACTGGG | >MI0005756\|hsa-mir-934&WithFlank&X\|+\|136550872\|136550966 | 21 | 42 | 22(+1G) | TGTCTACTACTGGAGACACTGGG | 724.1 | 13.0 | 2.3 | 5.7 | 1.1E-08 | 2.5E-07 | BT20/MB231 |
| >hg38_21_+_8433172_8446622_RNA45SN1_45S_50ntFlanks@12073.12093.21\|CCGAGGGGCTCTCGCTTCTGG | >hg38_21_+_8433172_8446622_RNA45SN1_45S_50ntFlanks | 12073 | 12093 | 21 | CCGAGGGGCTCTCGCTTCTGG | 116.9 | -4.6 | 0.9 | -5.2 | 1.7E-07 | 2.5E-07 | BT20/MB468 |
| >hg38_21_+_8433172_8446622_RNA45SN1_45S_50ntFlanks@9545.9570.26\|GTGGAGGTCCGTAGCGGTCCTGACGT | >hg38_21_+_8433172_8446622_RNA45SN1_45S_50ntFlanks | 9545 | 9570 | 26 | GTGGAGGTCCGTAGCGGTCCTGACGT | 645.0 | -2.6 | 0.5 | -5.2 | 1.7E-07 | 2.5E-07 | BT20/MB468 |
| >hg38_21_+_8433172_8446622_RNA45SN1_45S_50ntFlanks@9567.9586.20\|ACGTGCAAATCGGTCGTCCG | >hg38_21_+_8433172_8446622_RNA45SN1_45S_50ntFlanks | 9567 | 9586 | 20 | ACGTGCAAATCGGTCGTCCG | 357.6 | -3.1 | 0.6 | -5.2 | 1.6E-07 | 2.6E-07 | MB231/MB468 |
| >hg38_21_+_8433172_8446622_RNA45SN1_45S_50ntFlanks@6656.6681.26\|CTTAGCGGTGGATCACTCGGCTCGTG | >hg38_21_+_8433172_8446622_RNA45SN1_45S_50ntFlanks | 6656 | 6681 | 26 | CTTAGCGGTGGATCACTCGGCTCGTG | 1606.0 | -3.5 | 0.7 | -5.2 | 1.6E-07 | 2.6E-07 | MB231/MB468 |
| >hg38_21_+_8433172_8446622_RNA45SN1_45S_50ntFlanks@6721.6742.22\|AATTGCAGGACACATTGATCAT | >hg38_21_+_8433172_8446622_RNA45SN1_45S_50ntFlanks | 6721 | 6742 | 22 | AATTGCAGGACACATTGATCAT | 593.8 | -4.2 | 0.8 | -5.2 | 1.6E-07 | 2.6E-07 | MB231/MB468 |
| >hg38_21_+_8433172_8446622_RNA45SN1_45S_50ntFlanks@12627.12661.35\|ATGGGGCGAAGCTACCATCTGTGGGATTATGACTG | >hg38_21_+_8433172_8446622_RNA45SN1_45S_50ntFlanks | 12627 | 12661 | 35 | ATGGGGCGAAGCTACCATCTGTGGGATTATGACTG | 888.4 | -6.4 | 1.2 | -5.2 | 1.8E-07 | 2.6E-07 | BT20/MB468 |
| >hg38_21_+_8433172_8446622_RNA45SN1_45S_50ntFlanks@8170.8214.45\|CAAGTCCTTCTGATCGAGGCCCAGCCCGTGGACGGTGTGAGGCCG | >hg38_21_+_8433172_8446622_RNA45SN1_45S_50ntFlanks | 8170 | 8214 | 45 | CAAGTCCTTCTGATCGAGGCCCAGCCCGTGGACGGTGTGAGGCCG | 308.7 | -3.5 | 0.7 | -5.2 | 1.8E-07 | 2.6E-07 | BT20/MB468 |
| >hg38_21_+_8433172_8446622_RNA45SN1_45S_50ntFlanks@6661.6706.46\|CGGTGGATCACTCGGCTCGTGCGTCGATGAAGAACGCAGCTAGCTG | >hg38_21_+_8433172_8446622_RNA45SN1_45S_50ntFlanks | 6661 | 6706 | 46 | CGGTGGATCACTCGGCTCGTGCGTCGATGAAGAACGCAGCTAGCTG | 660.1 | 3.8 | 0.7 | 5.2 | 1.6E-07 | 2.6E-07 | MB231/MB468 |
| >hg38_21_+_8433172_8446622_RNA45SN1_45S_50ntFlanks@6658.6681.24\|TAGCGGTGGATCACTCGGCTCGTG | >hg38_21_+_8433172_8446622_RNA45SN1_45S_50ntFlanks | 6658 | 6681 | 24 | TAGCGGTGGATCACTCGGCTCGTG | 152.6 | -2.1 | 0.4 | -5.2 | 1.8E-07 | 2.6E-07 | BT20/MB468 |
| >hg38_21_+_8433172_8446622_RNA45SN1_45S_50ntFlanks@7975.7993.19\|CGCGACCTCAGATCAGACG | >hg38_21_+_8433172_8446622_RNA45SN1_45S_50ntFlanks | 7975 | 7993 | 19 | CGCGACCTCAGATCAGACG | 20268.4 | 4.1 | 0.8 | 5.2 | 1.8E-07 | 2.6E-07 | BT20/MB468 |
| >hg38_21_+_8433172_8446622_RNA45SN1_45S_50ntFlanks@12406.12432.27\|TTTTTGATCCTTCGATGTCGGCTCTTC | >hg38_21_+_8433172_8446622_RNA45SN1_45S_50ntFlanks | 12406 | 12432 | 27 | TTTTTGATCCTTCGATGTCGGCTCTTC | 147.6 | -3.7 | 0.7 | -5.2 | 1.8E-07 | 2.7E-07 | BT20/MB468 |
| >hg38_21_+_8433172_8446622_RNA45SN1_45S_50ntFlanks@12488.12519.32\|AACGTGAGCTGGGTTTAGACCGTCGTGAGACA | >hg38_21_+_8433172_8446622_RNA45SN1_45S_50ntFlanks | 12488 | 12519 | 32 | AACGTGAGCTGGGTTTAGACCGTCGTGAGACA | 168.5 | -2.7 | 0.5 | -5.2 | 1.7E-07 | 2.7E-07 | MB231/MB468 |
| >hg38_21_+_8433172_8446622_RNA45SN1_45S_50ntFlanks@3769.3792.24\|CGCACGGCCGGTACAGTGAAACTG | >hg38_21_+_8433172_8446622_RNA45SN1_45S_50ntFlanks | 3769 | 3792 | 24 | CGCACGGCCGGTACAGTGAAACTG | 171.0 | -3.1 | 0.6 | -5.2 | 1.7E-07 | 2.7E-07 | MB231/MB468 |
| >hg38_21_+_8433172_8446622_RNA45SN1_45S_50ntFlanks@4047.4066.20\|ATTCGAACGTCTGCCCTATC | >hg38_21_+_8433172_8446622_RNA45SN1_45S_50ntFlanks | 4047 | 4066 | 20 | ATTCGAACGTCTGCCCTATC | 973.4 | -2.5 | 0.5 | -5.2 | 1.8E-07 | 2.7E-07 | BT20/MB468 |
| >hg38_21_+_8433172_8446622_RNA45SN1_45S_50ntFlanks@4722.4758.37\|TCATTAATCAAGAACGAAAGTCGGAGGTTCGAAGACG | >hg38_21_+_8433172_8446622_RNA45SN1_45S_50ntFlanks | 4722 | 4758 | 37 | TCATTAATCAAGAACGAAAGTCGGAGGTTCGAAGACG | 2556.7 | -3.5 | 0.7 | -5.2 | 1.7E-07 | 2.7E-07 | MB231/MB468 |
| >hg38_21_+_8433172_8446622_RNA45SN1_45S_50ntFlanks@10770.10797.28\|GCAGGTCTCCAAGGTGAACAGCCTCTGG | >hg38_21_+_8433172_8446622_RNA45SN1_45S_50ntFlanks | 10770 | 10797 | 28 | GCAGGTCTCCAAGGTGAACAGCCTCTGG | 95.9 | -2.8 | 0.5 | -5.2 | 1.7E-07 | 2.7E-07 | MB231/MB468 |
| >hg38_21_+_8433172_8446622_RNA45SN1_45S_50ntFlanks@11573.11592.20\|CGACTTAGAACTGGTGCGGA | >hg38_21_+_8433172_8446622_RNA45SN1_45S_50ntFlanks | 11573 | 11592 | 20 | CGACTTAGAACTGGTGCGGA | 455.9 | -2.9 | 0.6 | -5.2 | 1.7E-07 | 2.7E-07 | MB231/MB468 |
| >hg38_21_+_8433172_8446622_RNA45SN1_45S_50ntFlanks@10411.10442.32\|CAGTCGGTCCTGAGAGATGGGCGAGCGCCGTT | >hg38_21_+_8433172_8446622_RNA45SN1_45S_50ntFlanks | 10411 | 10442 | 32 | CAGTCGGTCCTGAGAGATGGGCGAGCGCCGTT | 399.9 | -3.1 | 0.6 | -5.2 | 1.7E-07 | 2.7E-07 | MB231/MB468 |
| >hg38_21_+_8433172_8446622_RNA45SN1_45S_50ntFlanks@11575.11607.33\|ACTTAGAACTGGTGCGGACCAGGGGAATCCGAC | >hg38_21_+_8433172_8446622_RNA45SN1_45S_50ntFlanks | 11575 | 11607 | 33 | ACTTAGAACTGGTGCGGACCAGGGGAATCCGAC | 323.2 | -2.6 | 0.5 | -5.2 | 1.9E-07 | 2.7E-07 | BT20/MB468 |

| | | | | | | | | | | | | |
|---|---|---|---|---|---|---|---|---|---|---|---|---|
| >hg38_21_+_8433172_8446622_RNA45SN1_45S_50ntFlanks@11618.11640.23\|AAACAAAGCATCGCGAAGGCCCG | >hg38_21_+_8433172_8446622_RNA45SN1_45S_50ntFlanks | 11618 | 11640 | 23 | AAACAAAGCATCGCGAAGGCCCG | 127.9 | -4.4 | 0.8 | -5.2 | 1.7E-07 | 2.7E-07 | MB231/MB468 |
| >hg38_21_+_8433172_8446622_RNA45SN1_45S_50ntFlanks@5527.5548.22\|AAAGTCGTAACAAGGTTTCCGT | >hg38_21_+_8433172_8446622_RNA45SN1_45S_50ntFlanks | 5527 | 5548 | 22 | AAAGTCGTAACAAGGTTTCCGT | 82.1 | -5.2 | 1.0 | -5.2 | 1.7E-07 | 2.7E-07 | MB231/MB468 |
| >hg38_21_+_8433172_8446622_RNA45SN1_45S_50ntFlanks@10409.10435.27\|GTCAGTCGGTCCTGAGAGATGGGCGAG | >hg38_21_+_8433172_8446622_RNA45SN1_45S_50ntFlanks | 10409 | 10435 | 27 | GTCAGTCGGTCCTGAGAGATGGGCGAG | 192.9 | -4.0 | 0.8 | -5.2 | 1.9E-07 | 2.7E-07 | BT20/MB468 |
| >hg38_21_+_8433172_8446622_RNA45SN1_45S_50ntFlanks@12407.12431.25\|TTTTGATCCTTCGATGTCGGCTCTT | >hg38_21_+_8433172_8446622_RNA45SN1_45S_50ntFlanks | 12407 | 12431 | 25 | TTTTGATCCTTCGATGTCGGCTCTT | 804.3 | -3.2 | 0.6 | -5.2 | 1.9E-07 | 2.7E-07 | BT20/MB468 |
| >hg38_21_+_8433172_8446622_RNA45SN1_45S_50ntFlanks@12419.12441.23\|GATGTCGGCTCTTCCTATCATTG | >hg38_21_+_8433172_8446622_RNA45SN1_45S_50ntFlanks | 12419 | 12441 | 23 | GATGTCGGCTCTTCCTATCATTG | 1719.8 | 3.7 | 0.7 | 5.2 | 1.7E-07 | 2.7E-07 | MB231/MB468 |
| >hg38_21_+_8433172_8446622_RNA45SN1_45S_50ntFlanks@9554.9581.28\|CGTAGCGGTCCTGACGTGCAAATCGGTC | >hg38_21_+_8433172_8446622_RNA45SN1_45S_50ntFlanks | 9554 | 9581 | 28 | CGTAGCGGTCCTGACGTGCAAATCGGTC | 1254.0 | 3.0 | 0.6 | 5.2 | 1.7E-07 | 2.7E-07 | MB231/MB468 |
| >MI0000286\|hsa-mir-210&WithFlank&11\|-\|568083\|568204@72.92.21(+1G)&[MIMAT0000267&hsa-miR-210-3p&offsets\|0\|-1(+1G);m-102&11\|-\|568112\|568133&offsets\|0\|-1(+1G)]\|CTGTGCGTGTGACAGCGGCTGG | >MI0000286\|hsa-mir-210&WithFlank&11\|-\|568083\|568204 | 72 | 92 | 21(+1G) | CTGTGCGTGTGACAGCGGCTGG | 2508.5 | 15.9 | 3.1 | 5.2 | 1.9E-07 | 2.7E-07 | BT20/MB468 |
| >hg38_21_+_8433172_8446622_RNA45SN1_45S_50ntFlanks@12934.12960.27\|CTAAACCATTCGTAGACGACCTGCTTC | >hg38_21_+_8433172_8446622_RNA45SN1_45S_50ntFlanks | 12934 | 12960 | 27 | CTAAACCATTCGTAGACGACCTGCTTC | 380.8 | -3.1 | 0.6 | -5.2 | 1.9E-07 | 2.7E-07 | BT20/MB468 |
| >MI0016887\|hsa-mir-4521&WithFlank&17\|+\|8186939\|8187010@10.35.26&[MIMAT0019058&hsa-miR-4521&offsets\|0\|+4;m-765&17\|+\|8186948\|8186969&offsets\|0\|+4]\|GCTAAGGAAGTCCTGTGCTCAGTTTT | >MI0016887\|hsa-mir-4521&WithFlank&17\|+\|8186939\|8187010 | 10 | 35 | 26 | GCTAAGGAAGTCCTGTGCTCAGTTTT | 3103.7 | 5.9 | 1.1 | 5.2 | 1.9E-07 | 2.7E-07 | BT20/MB468 |
| >hg38_21_+_8433172_8446622_RNA45SN1_45S_50ntFlanks@12420.12437.18\|ATGTCGGCTCTTCCTATC | >hg38_21_+_8433172_8446622_RNA45SN1_45S_50ntFlanks | 12420 | 12437 | 18 | ATGTCGGCTCTTCCTATC | 4915.4 | -2.6 | 0.5 | -5.2 | 1.9E-07 | 2.8E-07 | BT20/MB468 |
| >hg38_21_+_8433172_8446622_RNA45SN1_45S_50ntFlanks@4241.4258.18\|CTCTTTCGAGGCCCTGTA | >hg38_21_+_8433172_8446622_RNA45SN1_45S_50ntFlanks | 4241 | 4258 | 18 | CTCTTTCGAGGCCCTGTA | 468.4 | -6.3 | 1.2 | -5.2 | 1.9E-07 | 2.8E-07 | BT20/MB468 |
| >MI0000749\|hsa-mir-30e&WithFlank&1\|+\|40754349\|40754452@23.45.23(+1C)&[MIMAT0000692&hsa-miR-30e-5p&offsets\|0\|+1(+1C);m-24&1\|+\|40754371\|40754394&offsets\|0\|-1(+1C)]\|TGTAAACATCCTTGACTGGAAGCC | >MI0000749\|hsa-mir-30e&WithFlank&1\|+\|40754349\|40754452 | 23 | 45 | 23(+1C) | TGTAAACATCCTTGACTGGAAGCC | 3505.4 | 5.0 | 1.0 | 5.2 | 1.9E-07 | 2.8E-07 | BT20/MB468 |
| >hg38_21_+_8433172_8446622_RNA45SN1_45S_50ntFlanks@6734.6770.37\|ATTGATCATCGACACTTCGAACGCACTTGCGGCCCCG | >hg38_21_+_8433172_8446622_RNA45SN1_45S_50ntFlanks | 6734 | 6770 | 37 | ATTGATCATCGACACTTCGAACGCACTTGCGGCCCCG | 482.5 | -2.2 | 0.4 | -5.2 | 1.9E-07 | 2.8E-07 | BT20/MB468 |
| >hg38_21_+_8433172_8446622_RNA45SN1_45S_50ntFlanks@6650.6683.34\|ACGACTCTTAGCGGTGGATCACTCGGCTCGTGCG | >hg38_21_+_8433172_8446622_RNA45SN1_45S_50ntFlanks | 6650 | 6683 | 34 | ACGACTCTTAGCGGTGGATCACTCGGCTCGTGCG | 1003.0 | 4.1 | 0.8 | 5.2 | 2.0E-07 | 2.8E-07 | BT20/MB468 |
| >hg38_21_+_8433172_8446622_RNA45SN1_45S_50ntFlanks@9563.9597.35\|CCTGACGTGCAAATCGGTCGTCCGACCTGGGTATA | >hg38_21_+_8433172_8446622_RNA45SN1_45S_50ntFlanks | 9563 | 9597 | 35 | CCTGACGTGCAAATCGGTCGTCCGACCTGGGTATA | 377.3 | -5.6 | 1.1 | -5.2 | 2.0E-07 | 2.8E-07 | BT20/MB468 |
| >hg38_21_+_8433172_8446622_RNA45SN1_45S_50ntFlanks@9544.9566.23\|GGTGGAGGTCCGTAGCGGTCCTG | >hg38_21_+_8433172_8446622_RNA45SN1_45S_50ntFlanks | 9544 | 9566 | 23 | GGTGGAGGTCCGTAGCGGTCCTG | 100.1 | -3.1 | 0.6 | -5.2 | 1.8E-07 | 2.9E-07 | MB231/MB468 |
| >hg38_21_+_8433172_8446622_RNA45SN1_45S_50ntFlanks@10844.10865.22\|AACTTCGGGATAAGGATTGGCT | >hg38_21_+_8433172_8446622_RNA45SN1_45S_50ntFlanks | 10844 | 10865 | 22 | AACTTCGGGATAAGGATTGGCT | 350.8 | -5.0 | 1.0 | -5.2 | 2.0E-07 | 2.9E-07 | BT20/MB468 |
| >MI0000088\|hsa-mir-30a&WithFlank&6\|-\|71403545\|71403627@12.35.24(+3U)&[MIMAT0000087&hsa-miR-30a-5p&offsets\|0\|+2(+3U);m-15&6\|-\|71403595\|71403616&offsets\|0\|+2(+3U)]\|TGTAAACATCCTCGACTGGAAGCTTTT | >MI0000088\|hsa-mir-30a&WithFlank&6\|-\|71403545\|71403627 | 12 | 35 | 24(+3U) | TGTAAACATCCTCGACTGGAAGCTTTT | 1279.7 | 11.4 | 2.2 | 5.2 | 1.8E-07 | 2.9E-07 | MB231/MB468 |
| >hg38_21_+_8433172_8446622_RNA45SN1_45S_50ntFlanks@10685.10706.22\|GGCGGCGTCCGGTGAGCTCTCG | >hg38_21_+_8433172_8446622_RNA45SN1_45S_50ntFlanks | 10685 | 10706 | 22 | GGCGGCGTCCGGTGAGCTCTCG | 152.4 | -2.9 | 0.5 | -5.2 | 1.8E-07 | 2.9E-07 | MB231/MB468 |
| >hg38_21_+_8433172_8446622_RNA45SN1_45S_50ntFlanks@10781.10817.37\|AGGTGAACAGCCTCTGGCATGTTGGAACAATGTAGGT | >hg38_21_+_8433172_8446622_RNA45SN1_45S_50ntFlanks | 10781 | 10817 | 37 | AGGTGAACAGCCTCTGGCATGTTGGAACAATGTAGGT | 332.2 | -3.6 | 0.7 | -5.2 | 2.0E-07 | 2.9E-07 | BT20/MB468 |
| >hg38_21_+_8433172_8446622_RNA45SN1_45S_50ntFlanks@4002.4042.41\|GATAACCTCGGGCCGATCGCACGCCCCCCGTGGCGGCGACG | >hg38_21_+_8433172_8446622_RNA45SN1_45S_50ntFlanks | 4002 | 4042 | 41 | GATAACCTCGGGCCGATCGCACGCCCCCCGTGGCGGCGACG | 128.9 | -3.1 | 0.6 | -5.2 | 1.8E-07 | 2.9E-07 | MB231/MB468 |
| >hg38_21_+_8433172_8446622_RNA45SN1_45S_50ntFlanks@11573.11592.20\|CGACTTAGAACTGGTGCGGA | >hg38_21_+_8433172_8446622_RNA45SN1_45S_50ntFlanks | 11573 | 11592 | 20 | CGACTTAGAACTGGTGCGGA | 540.7 | -3.3 | 0.6 | -5.2 | 2.0E-07 | 2.9E-07 | BT20/MB468 |
| >hg38_21_+_8433172_8446622_RNA45SN1_45S_50ntFlanks@6663.6689.27\|GTGGATCACTCGGCTCGTGCGTCGATG | >hg38_21_+_8433172_8446622_RNA45SN1_45S_50ntFlanks | 6663 | 6689 | 27 | GTGGATCACTCGGCTCGTGCGTCGATG | 1408.9 | -3.9 | 0.8 | -5.2 | 2.0E-07 | 2.9E-07 | BT20/MB468 |
| >hg38_21_+_8433172_8446622_RNA45SN1_45S_50ntFlanks@9545.9569.25\|GTGGAGGTCCGTAGCGGTCCTGACG | >hg38_21_+_8433172_8446622_RNA45SN1_45S_50ntFlanks | 9545 | 9569 | 25 | GTGGAGGTCCGTAGCGGTCCTGACG | 409.9 | -3.3 | 0.6 | -5.2 | 2.0E-07 | 2.9E-07 | BT20/MB468 |
| >hg38_21_+_8433172_8446622_RNA45SN1_45S_50ntFlanks@9339.9370.32\|CGCACGAAAGCCGCCGTGGCGCAATGAAGGTG | >hg38_21_+_8433172_8446622_RNA45SN1_45S_50ntFlanks | 9339 | 9370 | 32 | CGCACGAAAGCCGCCGTGGCGCAATGAAGGTG | 84.6 | -5.1 | 1.0 | -5.2 | 2.0E-07 | 2.9E-07 | BT20/MB468 |
| >MI0000253\|hsa-mir-148a&WithFlank&7\|-\|25949913\|25949992@50.70.21(+1G)&[MIMAT0000243&hsa-miR-148a-3p&offsets\|0\|-1(+1G);m-7&7\|-\|25949922\|25949943&offsets\|0\|-1(+1G)]\|TCAGTGCACTACAGAACTTTGG | >MI0000253\|hsa-mir-148a&WithFlank&7\|-\|25949913\|25949992 | 50 | 70 | 21(+1G) | TCAGTGCACTACAGAACTTTGG | 2761.5 | 4.8 | 0.9 | 5.2 | 1.8E-07 | 2.9E-07 | MB231/MB468 |
| >hg38_1_-_228634819_228635039_RNA5S12_5S_50ntFlanks@130.170.41\|TGGATGGGAGACCGCCTGGGAATACCGGGTGCTGTAGGCTT | >hg38_1_-_228634819_228635039_RNA5S12_5S_50ntFlanks | 130 | 170 | 41 | TGGATGGGAGACCGCCTGGGAATACCGGGTGCTGTAGGCTT | 2896.4 | 12.2 | 2.3 | 5.2 | 1.8E-07 | 3.0E-07 | MB231/MB468 |
| >hg38_21_+_8433172_8446622_RNA45SN1_45S_50ntFlanks@7989.8010.22\|AGACGTGGCGACCCGCTGAATT | >hg38_21_+_8433172_8446622_RNA45SN1_45S_50ntFlanks | 7989 | 8010 | 22 | AGACGTGGCGACCCGCTGAATT | 1535.6 | 4.0 | 0.8 | 5.2 | 2.1E-07 | 3.0E-07 | BT20/MB468 |
| >hg38_21_+_8433172_8446622_RNA45SN1_45S_50ntFlanks@7974.8016.43\|ACGCGACCTCAGATCAGACGTGGCGACCCGCTGAATTTAAGCA | >hg38_21_+_8433172_8446622_RNA45SN1_45S_50ntFlanks | 7974 | 8016 | 43 | ACGCGACCTCAGATCAGACGTGGCGACCCGCTGAATTTAAGCA | 638.1 | 6.4 | 1.2 | 5.2 | 2.1E-07 | 3.0E-07 | BT20/MB468 |
| >hg38_21_+_8433172_8446622_RNA45SN1_45S_50ntFlanks@3853.3872.20\|AACTGTGGTAATTCTAGAGC | >hg38_21_+_8433172_8446622_RNA45SN1_45S_50ntFlanks | 3853 | 3872 | 20 | AACTGTGGTAATTCTAGAGC | 265.9 | -2.3 | 0.4 | -5.2 | 2.1E-07 | 3.0E-07 | BT20/MB468 |
| >hg38_21_+_8433172_8446622_RNA45SN1_45S_50ntFlanks@10355.10390.36\|AACTTTGAAGGCCGAAGTGGAGAAGGGTTCCATGTG | >hg38_21_+_8433172_8446622_RNA45SN1_45S_50ntFlanks | 10355 | 10390 | 36 | AACTTTGAAGGCCGAAGTGGAGAAGGGTTCCATGTG | 182.9 | -3.3 | 0.6 | -5.2 | 2.1E-07 | 3.1E-07 | BT20/MB468 |
| >hg38_21_+_8433172_8446622_RNA45SN1_45S_50ntFlanks@10798.10817.20\|CATGTTGGAACAATGTAGGT | >hg38_21_+_8433172_8446622_RNA45SN1_45S_50ntFlanks | 10798 | 10817 | 20 | CATGTTGGAACAATGTAGGT | 700.3 | -3.8 | 0.7 | -5.2 | 2.1E-07 | 3.1E-07 | BT20/MB468 |
| >hg38_21_+_8433172_8446622_RNA45SN1_45S_50ntFlanks@10869.10904.36\|AGGGCTGGGTCGGTCGGGCTGGGGCGCGAAGCGGGG | >hg38_21_+_8433172_8446622_RNA45SN1_45S_50ntFlanks | 10869 | 10904 | 36 | AGGGCTGGGTCGGTCGGGCTGGGGCGCGAAGCGGGG | 88.4 | -4.3 | 0.8 | -5.2 | 2.1E-07 | 3.1E-07 | BT20/MB468 |
| >hg38_21_+_8433172_8446622_RNA45SN1_45S_50ntFlanks@11578.11603.26\|TAGAACTGGTGCGGACCAGGGGAATC | >hg38_21_+_8433172_8446622_RNA45SN1_45S_50ntFlanks | 11578 | 11603 | 26 | TAGAACTGGTGCGGACCAGGGGAATC | 584.2 | -2.9 | 0.6 | -5.2 | 1.9E-07 | 3.1E-07 | MB231/MB468 |
| >hg38_21_+_8433172_8446622_RNA45SN1_45S_50ntFlanks@12458.12478.21\|AAGCGTTGGATTGTTCACCCA | >hg38_21_+_8433172_8446622_RNA45SN1_45S_50ntFlanks | 12458 | 12478 | 21 | AAGCGTTGGATTGTTCACCCA | 1032.6 | -2.5 | 0.5 | -5.2 | 2.2E-07 | 3.1E-07 | BT20/MB468 |
| >hg38_21_+_8433172_8446622_RNA45SN1_45S_50ntFlanks@9623.9648.26\|TAGTAGCTGGTTCCCTCCGAAGTTTC | >hg38_21_+_8433172_8446622_RNA45SN1_45S_50ntFlanks | 9623 | 9648 | 26 | TAGTAGCTGGTTCCCTCCGAAGTTTC | 158.5 | -3.2 | 0.6 | -5.2 | 1.9E-07 | 3.1E-07 | MB231/MB468 |
| >hg38_21_+_8433172_8446622_RNA45SN1_45S_50ntFlanks@4068.4089.22\|ACTTTCGATGGTAGTCGCCGTG | >hg38_21_+_8433172_8446622_RNA45SN1_45S_50ntFlanks | 4068 | 4089 | 22 | ACTTTCGATGGTAGTCGCCGTG | 87.5 | -3.5 | 0.7 | -5.2 | 2.2E-07 | 3.2E-07 | BT20/MB468 |
| >hg38_MT_+_1621_3279_RNR2_16S_50ntFlanks@51.77.27\|GCTAAACCTAGCCCCAAACCCACTCCA | >hg38_MT_+_1621_3279_RNR2_16S_50ntFlanks | 51 | 77 | 27 | GCTAAACCTAGCCCCAAACCCACTCCA | 719.1 | 10.9 | 2.1 | 5.2 | 2.0E-07 | 3.2E-07 | MB231/MB468 |
| >hg38_21_+_8433172_8446622_RNA45SN1_45S_50ntFlanks@11874.11892.19\|AGAAGACCCTGTTGAGCTT | >hg38_21_+_8433172_8446622_RNA45SN1_45S_50ntFlanks | 11874 | 11892 | 19 | AGAAGACCCTGTTGAGCTT | 682.9 | -3.8 | 0.7 | -5.2 | 2.2E-07 | 3.2E-07 | BT20/MB468 |
| >hg38_21_+_8433172_8446622_RNA45SN1_45S_50ntFlanks@12478.12498.21\|ACTAATAGGGAACGTGAGCTG | >hg38_21_+_8433172_8446622_RNA45SN1_45S_50ntFlanks | 12478 | 12498 | 21 | ACTAATAGGGAACGTGAGCTG | 553.1 | -2.4 | 0.5 | -5.2 | 2.2E-07 | 3.2E-07 | BT20/MB468 |
| >hg38_21_+_8433172_8446622_RNA45SN1_45S_50ntFlanks@11727.11753.27\|AGTAACTATGACTCTCTTAAGGTAGCC | >hg38_21_+_8433172_8446622_RNA45SN1_45S_50ntFlanks | 11727 | 11753 | 27 | AGTAACTATGACTCTCTTAAGGTAGCC | 308.7 | -4.7 | 0.9 | -5.2 | 2.2E-07 | 3.2E-07 | BT20/MB468 |
| >hg38_1_-_228634819_228635039_RNA5S12_5S_50ntFlanks@150.171.22\|AATACCGGGTGCTGTAGGCTTT | >hg38_1_-_228634819_228635039_RNA5S12_5S_50ntFlanks | 150 | 171 | 22 | AATACCGGGTGCTGTAGGCTTT | 2161.8 | 3.9 | 0.8 | 5.2 | 2.2E-07 | 3.2E-07 | BT20/MB468 |
| >trnaMT_GluTTC_MT_-_14674_14742@39.65.27,trnalookalike8_GluTTC_5_-_93905172_93905240@39.65.27\|ATTGGTCGTGGTTGTAGTCCGTGCGAG | >trnaMT_GluTTC_MT_-_14674_14742 | 39 | 65 | 27 | ATTGGTCGTGGTTGTAGTCCGTGCGAG | 427.2 | 6.9 | 1.3 | 5.2 | 2.2E-07 | 3.2E-07 | BT20/MB468 |
| >hg38_21_+_8433172_8446622_RNA45SN1_45S_50ntFlanks@9739.9758.20\|AAACGATCTCAACCTATTCT | >hg38_21_+_8433172_8446622_RNA45SN1_45S_50ntFlanks | 9739 | 9758 | 20 | AAACGATCTCAACCTATTCT | 1820.5 | -2.3 | 0.4 | -5.2 | 2.0E-07 | 3.2E-07 | MB231/MB468 |
| >hg38_21_+_8433172_8446622_RNA45SN1_45S_50ntFlanks@4588.4615.28\|CTATTTTGTTGGTTTTCGGAACTGAGGC | >hg38_21_+_8433172_8446622_RNA45SN1_45S_50ntFlanks | 4588 | 4615 | 28 | CTATTTTGTTGGTTTTCGGAACTGAGGC | 139.3 | -3.3 | 0.6 | -5.2 | 2.0E-07 | 3.3E-07 | MB231/MB468 |
| >hg38_21_+_8433172_8446622_RNA45SN1_45S_50ntFlanks@6661.6688.28\|CGGTGGATCACTCGGCTCGTGCGTCGAT | >hg38_21_+_8433172_8446622_RNA45SN1_45S_50ntFlanks | 6661 | 6688 | 28 | CGGTGGATCACTCGGCTCGTGCGTCGAT | 2793.0 | 4.0 | 0.8 | 5.2 | 2.3E-07 | 3.3E-07 | BT20/MB468 |
| >hg38_21_+_8433172_8446622_RNA45SN1_45S_50ntFlanks@11579.11601.23\|AGAACTGGTGCGGACCAGGGGAA | >hg38_21_+_8433172_8446622_RNA45SN1_45S_50ntFlanks | 11579 | 11601 | 23 | AGAACTGGTGCGGACCAGGGGAA | 696.7 | -4.0 | 0.8 | -5.2 | 2.3E-07 | 3.3E-07 | BT20/MB468 |
| >hg38_21_+_8433172_8446622_RNA45SN1_45S_50ntFlanks@4555.4572.18\|CGCAGCTAGGAATAATGG | >hg38_21_+_8433172_8446622_RNA45SN1_45S_50ntFlanks | 4555 | 4572 | 18 | CGCAGCTAGGAATAATGG | 361.4 | -3.1 | 0.6 | -5.2 | 2.1E-07 | 3.3E-07 | MB231/MB468 |

<table>
<tr><td>>hg38_21_+_8433172_8446622_RNA45SN1_45S_50ntFlanks@12932.12953.22|CGCTAAACCATTCGTAGACGAC</td><td>>hg38_21_+_8433172_8446622_RNA45SN1_45S_50ntFlanks</td><td>12932</td><td>12953</td><td>22</td><td>CGCTAAACCATTCGTAGACGAC</td><td>102.8</td><td>-3.4</td><td>0.7</td><td>-5.2</td><td>2.3E-07</td><td>3.3E-07</td><td>BT20/MB468</td></tr>
<tr><td>>hg38_21_+_8433172_8446622_RNA45SN1_45S_50ntFlanks@9305.9336.32|CCAAGGAGTCTAACACGTGCGCGAGTCGGGGG</td><td>>hg38_21_+_8433172_8446622_RNA45SN1_45S_50ntFlanks</td><td>9305</td><td>9336</td><td>32</td><td>CCAAGGAGTCTAACACGTGCGCGAGTCGGGGG</td><td>83.9</td><td>-6.0</td><td>1.2</td><td>-5.2</td><td>2.1E-07</td><td>3.3E-07</td><td>MB231/MB468</td></tr>
<tr><td>>MI0000446|hsa-mir-125b-1&WithFlank&11|-|122099751|122099850@21.42.22;MI0000470|hsa-mir-125b-2&WithFlank&21|+|16590231|16590331@23.44.22&[MIMAT0000423&hsa-miR-125b-5p&offsets|0|0;m-67&11|-|122099809|122099830&offsets|0|0];[MIMAT0000423_1&hsa-miR-125b-5p&offsets|0|0;m-68&21|+|16590253|16590274&offsets|0|0]|TCCCTGAGACCCTAACTTGTGA</td><td>>MI0000446|hsa-mir-125b-1&WithFlank&11|-|122099751|122099850</td><td>21</td><td>42</td><td>22</td><td>TCCCTGAGACCCTAACTTGTGA</td><td>12565.9</td><td>-5.4</td><td>1.0</td><td>-5.7</td><td>1.5E-08</td><td>3.4E-07</td><td>BT20/MB231</td></tr>
<tr><td>>hg38_21_+_8433172_8446622_RNA45SN1_45S_50ntFlanks@11578.11601.24|TAGAACTGGTGCGGACCAGGGGAA</td><td>>hg38_21_+_8433172_8446622_RNA45SN1_45S_50ntFlanks</td><td>11578</td><td>11601</td><td>24</td><td>TAGAACTGGTGCGGACCAGGGGAA</td><td>180.8</td><td>-2.8</td><td>0.5</td><td>-5.2</td><td>2.1E-07</td><td>3.4E-07</td><td>MB231/MB468</td></tr>
<tr><td>>hg38_21_+_8433172_8446622_RNA45SN1_45S_50ntFlanks@10411.10441.31|CAGTCGGTCCTGAGAGATGGGCGAGCGCCGT</td><td>>hg38_21_+_8433172_8446622_RNA45SN1_45S_50ntFlanks</td><td>10411</td><td>10441</td><td>31</td><td>CAGTCGGTCCTGAGAGATGGGCGAGCGCCGT</td><td>103.6</td><td>-2.7</td><td>0.5</td><td>-5.2</td><td>2.1E-07</td><td>3.4E-07</td><td>MB231/MB468</td></tr>
<tr><td>>hg38_21_+_8433172_8446622_RNA45SN1_45S_50ntFlanks@11766.11784.19|ATCTAATTAGTGACGCGCA</td><td>>hg38_21_+_8433172_8446622_RNA45SN1_45S_50ntFlanks</td><td>11766</td><td>11784</td><td>19</td><td>ATCTAATTAGTGACGCGCA</td><td>158.2</td><td>-5.4</td><td>1.0</td><td>-5.2</td><td>2.1E-07</td><td>3.4E-07</td><td>MB231/MB468</td></tr>
<tr><td>>hg38_21_+_8433172_8446622_RNA45SN1_45S_50ntFlanks@10393.10422.30|CAGCAGTTGAACATGGGTCAGTCGGTCCTG</td><td>>hg38_21_+_8433172_8446622_RNA45SN1_45S_50ntFlanks</td><td>10393</td><td>10422</td><td>30</td><td>CAGCAGTTGAACATGGGTCAGTCGGTCCTG</td><td>400.2</td><td>-3.7</td><td>0.7</td><td>-5.2</td><td>2.4E-07</td><td>3.4E-07</td><td>BT20/MB468</td></tr>
<tr><td>>hg38_21_+_8433172_8446622_RNA45SN1_45S_50ntFlanks@6733.6754.22|CATTGATCATCGACACTTCGAA</td><td>>hg38_21_+_8433172_8446622_RNA45SN1_45S_50ntFlanks</td><td>6733</td><td>6754</td><td>22</td><td>CATTGATCATCGACACTTCGAA</td><td>134.6</td><td>-2.6</td><td>0.5</td><td>-5.2</td><td>2.2E-07</td><td>3.4E-07</td><td>MB231/MB468</td></tr>
<tr><td>>hg38_21_+_8433172_8446622_RNA45SN1_45S_50ntFlanks@10458.10490.33|ATGGCCTCCGTTGCCCTCGGCCGATCGAAAGGG</td><td>>hg38_21_+_8433172_8446622_RNA45SN1_45S_50ntFlanks</td><td>10458</td><td>10490</td><td>33</td><td>ATGGCCTCCGTTGCCCTCGGCCGATCGAAAGGG</td><td>91.0</td><td>-4.8</td><td>0.9</td><td>-5.2</td><td>2.2E-07</td><td>3.5E-07</td><td>MB231/MB468</td></tr>
<tr><td>>hg38_21_+_8433172_8446622_RNA45SN1_45S_50ntFlanks@11672.11699.28|CAGTGCTCTGAATGTCAAAGTGAAGAAA</td><td>>hg38_21_+_8433172_8446622_RNA45SN1_45S_50ntFlanks</td><td>11672</td><td>11699</td><td>28</td><td>CAGTGCTCTGAATGTCAAAGTGAAGAAA</td><td>144.3</td><td>-2.4</td><td>0.5</td><td>-5.2</td><td>2.4E-07</td><td>3.5E-07</td><td>BT20/MB468</td></tr>
<tr><td>>hg38_21_+_8433172_8446622_RNA45SN1_45S_50ntFlanks@10772.10805.34|AGGTCTCCAAGGTGAACAGCCTCTGGCATGTTGG</td><td>>hg38_21_+_8433172_8446622_RNA45SN1_45S_50ntFlanks</td><td>10772</td><td>10805</td><td>34</td><td>AGGTCTCCAAGGTGAACAGCCTCTGGCATGTTGG</td><td>93.3</td><td>-2.6</td><td>0.5</td><td>-5.2</td><td>2.2E-07</td><td>3.5E-07</td><td>MB231/MB468</td></tr>
<tr><td>>hg38_21_+_8433172_8446622_RNA45SN1_45S_50ntFlanks@3705.3746.42|TACCTGGTTGATCCTGCCAGTAGCATATGCTTGTCTCAAAGA</td><td>>hg38_21_+_8433172_8446622_RNA45SN1_45S_50ntFlanks</td><td>3705</td><td>3746</td><td>42</td><td>TACCTGGTTGATCCTGCCAGTAGCATATGCTTGTCTCAAAGA</td><td>13647.7</td><td>14.6</td><td>2.8</td><td>5.2</td><td>2.4E-07</td><td>3.5E-07</td><td>BT20/MB468</td></tr>
<tr><td>>MI0003628|hsa-mir-615&WithFlank&12|+|54033944|54034051@67.86.20&[MIMAT0003283&hsa-miR-615-3p&offsets|0|-2;m-495&12|+|54034010|54034030&offsets|0|-1]|TCCGAGCCTGGGTCTCCCTC</td><td>>MI0003628|hsa-mir-615&WithFlank&12|+|54033944|54034051</td><td>67</td><td>86</td><td>20</td><td>TCCGAGCCTGGGTCTCCCTC</td><td>510.8</td><td>5.0</td><td>0.9</td><td>5.7</td><td>1.6E-08</td><td>3.5E-07</td><td>BT20/MB231</td></tr>
<tr><td>>hg38_21_+_8433172_8446622_RNA45SN1_45S_50ntFlanks@9571.9597.27|GCAAATCGGTCGTCCGACCTGGGTATA</td><td>>hg38_21_+_8433172_8446622_RNA45SN1_45S_50ntFlanks</td><td>9571</td><td>9597</td><td>27</td><td>GCAAATCGGTCGTCCGACCTGGGTATA</td><td>252.3</td><td>-4.7</td><td>0.9</td><td>-5.2</td><td>2.4E-07</td><td>3.5E-07</td><td>BT20/MB468</td></tr>
<tr><td>>trnaMT_AsnGTT_MT_-_5657_5729@1.40.40,trnalookalike5_AsnGTT_1_-_566207_566279@1.40.40|TAGATTGAAGCCAGTTGATTAGGGTGCTTAGCTGTTAACT</td><td>>trnaMT_AsnGTT_MT_-_5657_5729</td><td>1</td><td>40</td><td>40</td><td>TAGATTGAAGCCAGTTGATTAGGGTGCTTAGCTGTTAACT</td><td>508.6</td><td>10.2</td><td>2.0</td><td>5.2</td><td>2.2E-07</td><td>3.6E-07</td><td>MB231/MB468</td></tr>
<tr><td>>MI0000272|hsa-mir-182&WithFlank&7|-|129770377|129770498@29.51.23(+1A)&[MIMAT0000259&hsa-miR-182-5p&offsets|0|-1(+1A);m-22&7|-|129770449|129770470&offsets|0|+1(+1A)]|TTTGGCAATGGTAGAACTCACACA</td><td>>MI0000272|hsa-mir-182&WithFlank&7|-|129770377|129770498</td><td>29</td><td>51</td><td>23(+1A)</td><td>TTTGGCAATGGTAGAACTCACACA</td><td>7860.0</td><td>4.4</td><td>0.8</td><td>5.6</td><td>1.6E-08</td><td>3.6E-07</td><td>BT20/MB231</td></tr>
<tr><td>>hg38_21_+_8433172_8446622_RNA45SN1_45S_50ntFlanks@4760.4777.18|TCAGATACCGTCGTAGTT</td><td>>hg38_21_+_8433172_8446622_RNA45SN1_45S_50ntFlanks</td><td>4760</td><td>4777</td><td>18</td><td>TCAGATACCGTCGTAGTT</td><td>121.0</td><td>-2.8</td><td>0.5</td><td>-5.2</td><td>2.3E-07</td><td>3.6E-07</td><td>MB231/MB468</td></tr>
<tr><td>>trna2_GlyGCC_21_-_18827107_18827177@1.33.33,trna35_GlyGCC_1_+_161413094_161413164@1.33.33,trna37_GlyGCC_1_+_161420467_161420537@1.33.33,trna39_GlyGCC_1_+_161427898_161427968@1.33.33,trna41_GlyGCC_1_+_161435258_161435328@1.33.33|GCATGGGTGGTTCAGTGGTAGAATTCTCGCCTG</td><td>>trna2_GlyGCC_21_-_18827107_18827177</td><td>1</td><td>33</td><td>33</td><td>GCATGGGTGGTTCAGTGGTAGAATTCTCGCCTG</td><td>1239.4</td><td>4.8</td><td>0.8</td><td>5.6</td><td>1.7E-08</td><td>3.6E-07</td><td>BT20/MB231</td></tr>
<tr><td>>hg38_21_+_8433172_8446622_RNA45SN1_45S_50ntFlanks@12957.12989.33|CTTCTGGGTCGGGGTTTCGTACGTAGCAGAGCA</td><td>>hg38_21_+_8433172_8446622_RNA45SN1_45S_50ntFlanks</td><td>12957</td><td>12989</td><td>33</td><td>CTTCTGGGTCGGGGTTTCGTACGTAGCAGAGCA</td><td>93.8</td><td>-4.2</td><td>0.8</td><td>-5.2</td><td>2.3E-07</td><td>3.6E-07</td><td>MB231/MB468</td></tr>
<tr><td>>hg38_21_+_8433172_8446622_RNA45SN1_45S_50ntFlanks@9567.9603.37|ACGTGCAAATCGGTCGTCCGACCTGGGTATAGGGGCG</td><td>>hg38_21_+_8433172_8446622_RNA45SN1_45S_50ntFlanks</td><td>9567</td><td>9603</td><td>37</td><td>ACGTGCAAATCGGTCGTCCGACCTGGGTATAGGGGCG</td><td>241.2</td><td>-2.5</td><td>0.5</td><td>-5.2</td><td>2.5E-07</td><td>3.6E-07</td><td>BT20/MB468</td></tr>
<tr><td>>hg38_21_+_8433172_8446622_RNA45SN1_45S_50ntFlanks@12356.12375.20|ACCACAGGGATAACTGGCTT</td><td>>hg38_21_+_8433172_8446622_RNA45SN1_45S_50ntFlanks</td><td>12356</td><td>12375</td><td>20</td><td>ACCACAGGGATAACTGGCTT</td><td>122.2</td><td>-4.7</td><td>0.9</td><td>-5.2</td><td>2.5E-07</td><td>3.6E-07</td><td>BT20/MB468</td></tr>
<tr><td>>hg38_21_+_8433172_8446622_RNA45SN1_45S_50ntFlanks@11906.11929.24|CACGGTGAAGAGACATGAGAGGTG</td><td>>hg38_21_+_8433172_8446622_RNA45SN1_45S_50ntFlanks</td><td>11906</td><td>11929</td><td>24</td><td>CACGGTGAAGAGACATGAGAGGTG</td><td>86.2</td><td>-4.1</td><td>0.8</td><td>-5.2</td><td>2.3E-07</td><td>3.6E-07</td><td>MB231/MB468</td></tr>
<tr><td>>hg38_21_+_8433172_8446622_RNA45SN1_45S_50ntFlanks@3885.3915.31|ACGGGCGCTGACCCCCTTCGCGGGGGGGATG</td><td>>hg38_21_+_8433172_8446622_RNA45SN1_45S_50ntFlanks</td><td>3885</td><td>3915</td><td>31</td><td>ACGGGCGCTGACCCCCTTCGCGGGGGGGATG</td><td>150.6</td><td>-4.0</td><td>0.8</td><td>-5.2</td><td>2.5E-07</td><td>3.6E-07</td><td>BT20/MB468</td></tr>
<tr><td>>hg38_21_+_8433172_8446622_RNA45SN1_45S_50ntFlanks@10869.10889.21|AGGGCTGGGTCGGTCGGGCTG</td><td>>hg38_21_+_8433172_8446622_RNA45SN1_45S_50ntFlanks</td><td>10869</td><td>10889</td><td>21</td><td>AGGGCTGGGTCGGTCGGGCTG</td><td>326.5</td><td>-4.4</td><td>0.8</td><td>-5.2</td><td>2.5E-07</td><td>3.7E-07</td><td>BT20/MB468</td></tr>
<tr><td>>hg38_21_+_8433172_8446622_RNA45SN1_45S_50ntFlanks@10683.10705.23|CCGGCGGCGTCCGGTGAGCTCTC</td><td>>hg38_21_+_8433172_8446622_RNA45SN1_45S_50ntFlanks</td><td>10683</td><td>10705</td><td>23</td><td>CCGGCGGCGTCCGGTGAGCTCTC</td><td>343.4</td><td>-2.9</td><td>0.6</td><td>-5.2</td><td>2.3E-07</td><td>3.7E-07</td><td>MB231/MB468</td></tr>
<tr><td>>hg38_21_+_8433172_8446622_RNA45SN1_45S_50ntFlanks@12406.12435.30|TTTTTGATCCTTCGATGTCGGCTCTTCCTA</td><td>>hg38_21_+_8433172_8446622_RNA45SN1_45S_50ntFlanks</td><td>12406</td><td>12435</td><td>30</td><td>TTTTTGATCCTTCGATGTCGGCTCTTCCTA</td><td>93.1</td><td>-3.4</td><td>0.6</td><td>-5.2</td><td>2.3E-07</td><td>3.7E-07</td><td>MB231/MB468</td></tr>
<tr><td>>hg38_21_+_8433172_8446622_RNA45SN1_45S_50ntFlanks@11199.11233.35|CCGGCGGCGGCGGCGACTCTGGACGCGAGCCGGGC</td><td>>hg38_21_+_8433172_8446622_RNA45SN1_45S_50ntFlanks</td><td>11199</td><td>11233</td><td>35</td><td>CCGGCGGCGGCGGCGACTCTGGACGCGAGCCGGGC</td><td>67.4</td><td>-5.7</td><td>1.1</td><td>-5.2</td><td>2.3E-07</td><td>3.7E-07</td><td>MB231/MB468</td></tr>
<tr><td>>hg38_21_+_8433172_8446622_RNA45SN1_45S_50ntFlanks@6729.6770.42|GACACATTGATCATCGACACTTCGAACGCACTTGCGGCCCCG</td><td>>hg38_21_+_8433172_8446622_RNA45SN1_45S_50ntFlanks</td><td>6729</td><td>6770</td><td>42</td><td>GACACATTGATCATCGACACTTCGAACGCACTTGCGGCCCCG</td><td>173.6</td><td>-3.8</td><td>0.7</td><td>-5.2</td><td>2.6E-07</td><td>3.7E-07</td><td>BT20/MB468</td></tr>
<tr><td>>hg38_21_+_8433172_8446622_RNA45SN1_45S_50ntFlanks@12994.13039.46|CCTCGCTGCGATCTATTGAAAGTCAGCCCTCGACACAAGGGTTTGT</td><td>>hg38_21_+_8433172_8446622_RNA45SN1_45S_50ntFlanks</td><td>12994</td><td>13039</td><td>46</td><td>CCTCGCTGCGATCTATTGAAAGTCAGCCCTCGACACAAGGGTTTGT</td><td>3604.0</td><td>5.5</td><td>1.1</td><td>5.2</td><td>2.6E-07</td><td>3.7E-07</td><td>BT20/MB468</td></tr>
<tr><td>>hg38_21_+_8433172_8446622_RNA45SN1_45S_50ntFlanks@4163.4186.24|CATCCAAGGAAGGCAGCAGGCGCG</td><td>>hg38_21_+_8433172_8446622_RNA45SN1_45S_50ntFlanks</td><td>4163</td><td>4186</td><td>24</td><td>CATCCAAGGAAGGCAGCAGGCGCG</td><td>179.9</td><td>-3.8</td><td>0.7</td><td>-5.2</td><td>2.6E-07</td><td>3.7E-07</td><td>BT20/MB468</td></tr>
<tr><td>>hg38_21_+_8433172_8446622_RNA45SN1_45S_50ntFlanks@12641.12661.21|CCATCTGTGGGATTATGACTG</td><td>>hg38_21_+_8433172_8446622_RNA45SN1_45S_50ntFlanks</td><td>12641</td><td>12661</td><td>21</td><td>CCATCTGTGGGATTATGACTG</td><td>406.1</td><td>-2.9</td><td>0.6</td><td>-5.2</td><td>2.6E-07</td><td>3.7E-07</td><td>BT20/MB468</td></tr>
<tr><td>>hg38_21_+_8433172_8446622_RNA45SN1_45S_50ntFlanks@10357.10388.32|CTTTGAAGGCCGAAGTGGAGAAGGGTTCCATG</td><td>>hg38_21_+_8433172_8446622_RNA45SN1_45S_50ntFlanks</td><td>10357</td><td>10388</td><td>32</td><td>CTTTGAAGGCCGAAGTGGAGAAGGGTTCCATG</td><td>419.0</td><td>-5.6</td><td>1.1</td><td>-5.2</td><td>2.6E-07</td><td>3.7E-07</td><td>BT20/MB468</td></tr>
<tr><td>>hg38_21_+_8433172_8446622_RNA45SN1_45S_50ntFlanks@12007.12032.26|GCCGCCGGTGAAATACCACTACTCTG</td><td>>hg38_21_+_8433172_8446622_RNA45SN1_45S_50ntFlanks</td><td>12007</td><td>12032</td><td>26</td><td>GCCGCCGGTGAAATACCACTACTCTG</td><td>1768.6</td><td>-3.9</td><td>0.8</td><td>-5.1</td><td>2.6E-07</td><td>3.8E-07</td><td>BT20/MB468</td></tr>
<tr><td>>hg38_21_+_8433172_8446622_RNA45SN1_45S_50ntFlanks@11783.11810.28|CATGAATGGATGAACGAGATTCCCACTG</td><td>>hg38_21_+_8433172_8446622_RNA45SN1_45S_50ntFlanks</td><td>11783</td><td>11810</td><td>28</td><td>CATGAATGGATGAACGAGATTCCCACTG</td><td>100.4</td><td>-3.8</td><td>0.7</td><td>-5.2</td><td>2.4E-07</td><td>3.8E-07</td><td>MB231/MB468</td></tr>
<tr><td>>hg38_21_+_8433172_8446622_RNA45SN1_45S_50ntFlanks@7989.8017.29|AGACGTGGCGACCCGCTGAATTTAAGCAT</td><td>>hg38_21_+_8433172_8446622_RNA45SN1_45S_50ntFlanks</td><td>7989</td><td>8017</td><td>29</td><td>AGACGTGGCGACCCGCTGAATTTAAGCAT</td><td>441.0</td><td>3.3</td><td>0.6</td><td>5.1</td><td>2.6E-07</td><td>3.8E-07</td><td>BT20/MB468</td></tr>
<tr><td>>MI0000272|hsa-mir-182&WithFlank&7|-|129770377|129770498@29.50.22(+1U)&[MIMAT0000259&hsa-miR-182-5p&offsets|0|-2(+1U);m-22&7|-|129770449|129770470&offsets|0|0(+1U)]|TTTGGCAATGGTAGAACTCACAT</td><td>>MI0000272|hsa-mir-182&WithFlank&7|-|129770377|129770498</td><td>29</td><td>50</td><td>22(+1U)</td><td>TTTGGCAATGGTAGAACTCACAT</td><td>836.9</td><td>4.5</td><td>0.8</td><td>5.6</td><td>1.8E-08</td><td>3.8E-07</td><td>BT20/MB231</td></tr>
<tr><td>>hg38_21_+_8433172_8446622_RNA45SN1_45S_50ntFlanks@10787.10822.36|ACAGCCTCTGGCATGTTGGAACAATGTAGGTAAGGG</td><td>>hg38_21_+_8433172_8446622_RNA45SN1_45S_50ntFlanks</td><td>10787</td><td>10822</td><td>36</td><td>ACAGCCTCTGGCATGTTGGAACAATGTAGGTAAGGG</td><td>97.8</td><td>-2.6</td><td>0.5</td><td>-5.2</td><td>2.4E-07</td><td>3.8E-07</td><td>MB231/MB468</td></tr>
<tr><td>>MI0000102|hsa-mir-100&WithFlank&11|-|122152223|122152314@19.38.20(+1G)&[MIMAT0000098&hsa-miR-100-5p&offsets|0|-2(+1G);m-27&11|-|122152276|122152296&offsets|0|-1(+1G)]|AACCCGTAGATCCGAACTTGG</td><td>>MI0000102|hsa-mir-100&WithFlank&11|-|122152223|122152314</td><td>19</td><td>38</td><td>20(+1G)</td><td>AACCCGTAGATCCGAACTTGG</td><td>9012.0</td><td>-5.8</td><td>1.0</td><td>-5.6</td><td>1.8E-08</td><td>3.9E-07</td><td>BT20/MB231</td></tr>
<tr><td>>hg38_21_+_8433172_8446622_RNA45SN1_45S_50ntFlanks@10798.10823.26|CATGTTGGAACAATGTAGGTAAGGGA</td><td>>hg38_21_+_8433172_8446622_RNA45SN1_45S_50ntFlanks</td><td>10798</td><td>10823</td><td>26</td><td>CATGTTGGAACAATGTAGGTAAGGGA</td><td>129.9</td><td>-2.9</td><td>0.6</td><td>-5.1</td><td>2.7E-07</td><td>3.9E-07</td><td>BT20/MB468</td></tr>
<tr><td>>hg38_21_+_8433172_8446622_RNA45SN1_45S_50ntFlanks@12625.12652.28|CAATGGGGCGAAGCTACCATCTGTGGGA</td><td>>hg38_21_+_8433172_8446622_RNA45SN1_45S_50ntFlanks</td><td>12625</td><td>12652</td><td>28</td><td>CAATGGGGCGAAGCTACCATCTGTGGGA</td><td>79.1</td><td>-3.2</td><td>0.6</td><td>-5.2</td><td>2.5E-07</td><td>3.9E-07</td><td>MB231/MB468</td></tr>
<tr><td>>hg38_21_+_8433172_8446622_RNA45SN1_45S_50ntFlanks@10186.10207.22|TCCTCCCCTCCTCCCGCCCACG</td><td>>hg38_21_+_8433172_8446622_RNA45SN1_45S_50ntFlanks</td><td>10186</td><td>10207</td><td>22</td><td>TCCTCCCCTCCTCCCGCCCACG</td><td>168.7</td><td>-4.8</td><td>0.9</td><td>-5.1</td><td>2.8E-07</td><td>4.0E-07</td><td>BT20/MB468</td></tr>
<tr><td>>hg38_21_+_8433172_8446622_RNA45SN1_45S_50ntFlanks@8379.8400.22|TCAAGAGGGCGTGAAACCGTTA</td><td>>hg38_21_+_8433172_8446622_RNA45SN1_45S_50ntFlanks</td><td>8379</td><td>8400</td><td>22</td><td>TCAAGAGGGCGTGAAACCGTTA</td><td>199.3</td><td>-5.7</td><td>1.1</td><td>-5.1</td><td>2.8E-07</td><td>4.0E-07</td><td>BT20/MB468</td></tr>
</table>

<table>
<tr><td>>trna10_AspGTC_12_-_125424193_125424264@31.72.42,trna12_AspGTC_12_-_125411891_125411962@31.72.42,trna38_AspGTC_17_-_8125556_8125627@31.72.42,trna45_AspGTC_6_+_27447453_27447524@31.72.42,trna48_AspGTC_6_+_27471523_27471594@31.72.42,trna4_AspGTC_12_+_96429799_96429870@31.72.42,trna69_AspGTC_1_-_161440205_161440276@31.72.42,trna72_AspGTC_1_-_161432824_161432895@31.72.42,trna75_AspGTC_1_-_161425414_161425485@31.72.42,trna78_AspGTC_1_-_161418033_161418104@31.72.42,trna81_AspGTC_1_-_161410615_161410686@31.72.42|CCTGTCACGCGGGAGACCGGGGTTCGATTCCCCGACGGGGAG</td><td>>trna10_AspGTC_12_-_125424193_125424264</td><td>31</td><td>72</td><td>42</td><td>CCTGTCACGCGGGAGACCGGGGTTCGATTCCCCGACGGGGAG</td><td>244.0</td><td>-5.7</td><td>1.1</td><td>-5.1</td><td>2.8E-07</td><td>4.0E-07</td><td>BT20/MB468</td></tr>
<tr><td>>hg38_21_+_8433172_8446622_RNA45SN1_45S_50ntFlanks@4367.4388.22|CAGTTAAAAAGCTCGTAGTTGG</td><td>>hg38_21_+_8433172_8446622_RNA45SN1_45S_50ntFlanks</td><td>4367</td><td>4388</td><td>22</td><td>CAGTTAAAAAGCTCGTAGTTGG</td><td>237.7</td><td>-4.6</td><td>0.9</td><td>-5.1</td><td>2.8E-07</td><td>4.0E-07</td><td>BT20/MB468</td></tr>
<tr><td>>hg38_21_+_8433172_8446622_RNA45SN1_45S_50ntFlanks@4736.4758.23|CGAAAGTCGGAGGTTCGAAGACG</td><td>>hg38_21_+_8433172_8446622_RNA45SN1_45S_50ntFlanks</td><td>4736</td><td>4758</td><td>23</td><td>CGAAAGTCGGAGGTTCGAAGACG</td><td>192.6</td><td>-2.0</td><td>0.4</td><td>-5.1</td><td>2.8E-07</td><td>4.1E-07</td><td>BT20/MB468</td></tr>
<tr><td>>hg38_21_+_8433172_8446622_RNA45SN1_45S_50ntFlanks@12076.12095.20|AGGGGCTCTCGCTTCTGGCG</td><td>>hg38_21_+_8433172_8446622_RNA45SN1_45S_50ntFlanks</td><td>12076</td><td>12095</td><td>20</td><td>AGGGGCTCTCGCTTCTGGCG</td><td>2896.9</td><td>-4.1</td><td>0.8</td><td>-5.1</td><td>2.9E-07</td><td>4.1E-07</td><td>BT20/MB468</td></tr>
<tr><td>>hg38_21_+_8433172_8446622_RNA45SN1_45S_50ntFlanks@10403.10430.28|ACATGGGTCAGTCGGTCCTGAGAGATGG</td><td>>hg38_21_+_8433172_8446622_RNA45SN1_45S_50ntFlanks</td><td>10403</td><td>10430</td><td>28</td><td>ACATGGGTCAGTCGGTCCTGAGAGATGG</td><td>148.1</td><td>-4.6</td><td>0.9</td><td>-5.1</td><td>2.9E-07</td><td>4.2E-07</td><td>BT20/MB468</td></tr>
<tr><td>>hg38_21_+_8433172_8446622_RNA45SN1_45S_50ntFlanks@4068.4089.22|ACTTTCGATGGTAGTCGCCGTG</td><td>>hg38_21_+_8433172_8446622_RNA45SN1_45S_50ntFlanks</td><td>4068</td><td>4089</td><td>22</td><td>ACTTTCGATGGTAGTCGCCGTG</td><td>72.6</td><td>-5.8</td><td>1.1</td><td>-5.1</td><td>2.6E-07</td><td>4.2E-07</td><td>MB231/MB468</td></tr>
<tr><td>>MI0000085|hsa-mir-27a&WithFlank&19|-|13836434|13836523@57.76.20(+1G)&[MIMAT0000084&hsa-miR-27a-3p&offsets|0|-1(+1G);m-43&19|-|13836447|13836467&offsets|0|-1(+1G)]|TTCACAGTGGCTAAGTTCCGG</td><td>>MI0000085|hsa-mir-27a&WithFlank&19|-|13836434|13836523</td><td>57</td><td>76</td><td>20(+1G)</td><td>TTCACAGTGGCTAAGTTCCGG</td><td>843.0</td><td>4.9</td><td>1.0</td><td>5.1</td><td>2.9E-07</td><td>4.2E-07</td><td>BT20/MB468</td></tr>
<tr><td>>hg38_21_+_8433172_8446622_RNA45SN1_45S_50ntFlanks@8176.8197.22|CTTCTGATCGAGGCCCAGCCCG</td><td>>hg38_21_+_8433172_8446622_RNA45SN1_45S_50ntFlanks</td><td>8176</td><td>8197</td><td>22</td><td>CTTCTGATCGAGGCCCAGCCCG</td><td>475.9</td><td>-2.1</td><td>0.4</td><td>-5.1</td><td>2.6E-07</td><td>4.2E-07</td><td>MB231/MB468</td></tr>
<tr><td>>hg38_21_+_8433172_8446622_RNA45SN1_45S_50ntFlanks@12072.12089.18|CCCGAGGGGCTCTCGCTT</td><td>>hg38_21_+_8433172_8446622_RNA45SN1_45S_50ntFlanks</td><td>12072</td><td>12089</td><td>18</td><td>CCCGAGGGGCTCTCGCTT</td><td>98.5</td><td>-3.7</td><td>0.7</td><td>-5.1</td><td>2.6E-07</td><td>4.2E-07</td><td>MB231/MB468</td></tr>
<tr><td>>hg38_21_+_8433172_8446622_RNA45SN1_45S_50ntFlanks@11727.11745.19|AGTAACTATGACTCTCTTA</td><td>>hg38_21_+_8433172_8446622_RNA45SN1_45S_50ntFlanks</td><td>11727</td><td>11745</td><td>19</td><td>AGTAACTATGACTCTCTTA</td><td>594.1</td><td>3.4</td><td>0.7</td><td>5.1</td><td>2.7E-07</td><td>4.3E-07</td><td>MB231/MB468</td></tr>
<tr><td>>MI0000114|hsa-mir-107&WithFlank&10|-|89592741|89592833@56.78.23&[MIMAT0000104&hsa-miR-107&offsets|0|0;m-78&10|-|89592758|89592778&offsets|0|+2]|AGCAGCATTGTACAGGGCTATCA</td><td>>MI0000114|hsa-mir-107&WithFlank&10|-|89592741|89592833</td><td>56</td><td>78</td><td>23</td><td>AGCAGCATTGTACAGGGCTATCA</td><td>775.0</td><td>4.6</td><td>0.9</td><td>5.1</td><td>3.0E-07</td><td>4.3E-07</td><td>BT20/MB468</td></tr>
<tr><td>>hg38_21_+_8433172_8446622_RNA45SN1_45S_50ntFlanks@11573.11594.22|CGACTTAGAACTGGTGCGGACC</td><td>>hg38_21_+_8433172_8446622_RNA45SN1_45S_50ntFlanks</td><td>11573</td><td>11594</td><td>22</td><td>CGACTTAGAACTGGTGCGGACC</td><td>357.6</td><td>2.3</td><td>0.4</td><td>5.1</td><td>2.7E-07</td><td>4.3E-07</td><td>MB231/MB468</td></tr>
<tr><td>>hg38_21_+_8433172_8446622_RNA45SN1_45S_50ntFlanks@2959.2979.21|CGACGTGTGGCGTGGGTCGAC</td><td>>hg38_21_+_8433172_8446622_RNA45SN1_45S_50ntFlanks</td><td>2959</td><td>2979</td><td>21</td><td>CGACGTGTGGCGTGGGTCGAC</td><td>121.3</td><td>-3.9</td><td>0.8</td><td>-5.1</td><td>2.7E-07</td><td>4.3E-07</td><td>MB231/MB468</td></tr>
<tr><td>>hg38_21_+_8433172_8446622_RNA45SN1_45S_50ntFlanks@12407.12436.30|TTTTGATCCTTCGATGTCGGCTCTTCCTAT</td><td>>hg38_21_+_8433172_8446622_RNA45SN1_45S_50ntFlanks</td><td>12407</td><td>12436</td><td>30</td><td>TTTTGATCCTTCGATGTCGGCTCTTCCTAT</td><td>817.0</td><td>-2.8</td><td>0.5</td><td>-5.1</td><td>2.7E-07</td><td>4.4E-07</td><td>MB231/MB468</td></tr>
<tr><td>>hg38_21_+_8433172_8446622_RNA45SN1_45S_50ntFlanks@11570.11591.22|AGCCGACTTAGAACTGGTGCGG</td><td>>hg38_21_+_8433172_8446622_RNA45SN1_45S_50ntFlanks</td><td>11570</td><td>11591</td><td>22</td><td>AGCCGACTTAGAACTGGTGCGG</td><td>552.0</td><td>-3.9</td><td>0.8</td><td>-5.1</td><td>3.1E-07</td><td>4.4E-07</td><td>BT20/MB468</td></tr>
<tr><td>>hg38_21_+_8433172_8446622_RNA45SN1_45S_50ntFlanks@6722.6758.37|ATTGCAGGACACATTGATCATCGACACTTCGAACGCA</td><td>>hg38_21_+_8433172_8446622_RNA45SN1_45S_50ntFlanks</td><td>6722</td><td>6758</td><td>37</td><td>ATTGCAGGACACATTGATCATCGACACTTCGAACGCA</td><td>111.1</td><td>-2.4</td><td>0.5</td><td>-5.1</td><td>3.1E-07</td><td>4.4E-07</td><td>BT20/MB468</td></tr>
<tr><td>>MI0000085|hsa-mir-27a&WithFlank&19|-|13836434|13836523@57.76.20&[MIMAT0000084&hsa-miR-27a-3p&offsets|0|-1;m-43&19|-|13836447|13836467&offsets|0|-1]|TTCACAGTGGCTAAGTTCCG</td><td>>MI0000085|hsa-mir-27a&WithFlank&19|-|13836434|13836523</td><td>57</td><td>76</td><td>20</td><td>TTCACAGTGGCTAAGTTCCG</td><td>14221.7</td><td>4.4</td><td>0.9</td><td>5.1</td><td>3.1E-07</td><td>4.4E-07</td><td>BT20/MB468</td></tr>
<tr><td>>hg38_21_+_8433172_8446622_RNA45SN1_45S_50ntFlanks@4128.4158.31|CGATTCCGGAGAGGGAGCCTGAGAAACGGCT</td><td>>hg38_21_+_8433172_8446622_RNA45SN1_45S_50ntFlanks</td><td>4128</td><td>4158</td><td>31</td><td>CGATTCCGGAGAGGGAGCCTGAGAAACGGCT</td><td>79.0</td><td>-2.9</td><td>0.6</td><td>-5.1</td><td>2.8E-07</td><td>4.5E-07</td><td>MB231/MB468</td></tr>
<tr><td>>hg38_21_+_8433172_8446622_RNA45SN1_45S_50ntFlanks@3875.3899.25|ATACATGCCGACGGGCGCTGACCCC</td><td>>hg38_21_+_8433172_8446622_RNA45SN1_45S_50ntFlanks</td><td>3875</td><td>3899</td><td>25</td><td>ATACATGCCGACGGGCGCTGACCCC</td><td>516.4</td><td>-2.9</td><td>0.6</td><td>-5.1</td><td>2.8E-07</td><td>4.5E-07</td><td>MB231/MB468</td></tr>
<tr><td>>hg38_21_+_8433172_8446622_RNA45SN1_45S_50ntFlanks@10770.10805.36|GCAGGTCTCCAAGGTGAACAGCCTCTGGCATGTTGG</td><td>>hg38_21_+_8433172_8446622_RNA45SN1_45S_50ntFlanks</td><td>10770</td><td>10805</td><td>36</td><td>GCAGGTCTCCAAGGTGAACAGCCTCTGGCATGTTGG</td><td>120.2</td><td>-2.5</td><td>0.5</td><td>-5.1</td><td>2.8E-07</td><td>4.5E-07</td><td>MB231/MB468</td></tr>
<tr><td>>hg38_21_+_8433172_8446622_RNA45SN1_45S_50ntFlanks@10396.10424.29|CAGTTGAACATGGGTCAGTCGGTCCTGAG</td><td>>hg38_21_+_8433172_8446622_RNA45SN1_45S_50ntFlanks</td><td>10396</td><td>10424</td><td>29</td><td>CAGTTGAACATGGGTCAGTCGGTCCTGAG</td><td>2000.4</td><td>4.0</td><td>0.8</td><td>5.1</td><td>3.2E-07</td><td>4.6E-07</td><td>BT20/MB468</td></tr>
<tr><td>>hg38_21_+_8433172_8446622_RNA45SN1_45S_50ntFlanks@8195.8213.19|CCGTGGACGGTGTGAGGCC</td><td>>hg38_21_+_8433172_8446622_RNA45SN1_45S_50ntFlanks</td><td>8195</td><td>8213</td><td>19</td><td>CCGTGGACGGTGTGAGGCC</td><td>387.1</td><td>3.3</td><td>0.6</td><td>5.1</td><td>3.2E-07</td><td>4.6E-07</td><td>BT20/MB468</td></tr>
<tr><td>>hg38_21_+_8433172_8446622_RNA45SN1_45S_50ntFlanks@10402.10432.31|AACATGGGTCAGTCGGTCCTGAGAGATGGGC</td><td>>hg38_21_+_8433172_8446622_RNA45SN1_45S_50ntFlanks</td><td>10402</td><td>10432</td><td>31</td><td>AACATGGGTCAGTCGGTCCTGAGAGATGGGC</td><td>139.4</td><td>-3.3</td><td>0.7</td><td>-5.1</td><td>3.2E-07</td><td>4.6E-07</td><td>BT20/MB468</td></tr>
<tr><td>>hg38_21_+_8433172_8446622_RNA45SN1_45S_50ntFlanks@8356.8373.18|GAAAAGAACTTTGAAGAG</td><td>>hg38_21_+_8433172_8446622_RNA45SN1_45S_50ntFlanks</td><td>8356</td><td>8373</td><td>18</td><td>GAAAAGAACTTTGAAGAG</td><td>175.7</td><td>-3.2</td><td>0.6</td><td>-5.1</td><td>2.9E-07</td><td>4.7E-07</td><td>MB231/MB468</td></tr>
<tr><td>>hg38_21_+_8433172_8446622_RNA45SN1_45S_50ntFlanks@13010.13027.18|TGAAAGTCAGCCCTCGAC</td><td>>hg38_21_+_8433172_8446622_RNA45SN1_45S_50ntFlanks</td><td>13010</td><td>13027</td><td>18</td><td>TGAAAGTCAGCCCTCGAC</td><td>134.7</td><td>-4.2</td><td>0.8</td><td>-5.1</td><td>3.0E-07</td><td>4.7E-07</td><td>MB231/MB468</td></tr>
<tr><td>>hg38_21_+_8433172_8446622_RNA45SN1_45S_50ntFlanks@11317.11340.24|ACCCCACGTCTCGTCGCGCGCGCG</td><td>>hg38_21_+_8433172_8446622_RNA45SN1_45S_50ntFlanks</td><td>11317</td><td>11340</td><td>24</td><td>ACCCCACGTCTCGTCGCGCGCGCG</td><td>98.4</td><td>-5.4</td><td>1.1</td><td>-5.1</td><td>3.0E-07</td><td>4.8E-07</td><td>MB231/MB468</td></tr>
<tr><td>>hg38_1_-_228634819_228635039_RNA5S12_5S_50ntFlanks@148.171.24|GGAATACCGGGTGCTGTAGGCTTT</td><td>>hg38_1_-_228634819_228635039_RNA5S12_5S_50ntFlanks</td><td>148</td><td>171</td><td>24</td><td>GGAATACCGGGTGCTGTAGGCTTT</td><td>1994.6</td><td>3.1</td><td>0.6</td><td>5.1</td><td>3.4E-07</td><td>4.8E-07</td><td>BT20/MB468</td></tr>
<tr><td>>hg38_21_+_8433172_8446622_RNA45SN1_45S_50ntFlanks@3984.4026.43|GCGGCTTTGGTGACTCTAGATAACCTCGGGCCGATCGCACGCC</td><td>>hg38_21_+_8433172_8446622_RNA45SN1_45S_50ntFlanks</td><td>3984</td><td>4026</td><td>43</td><td>GCGGCTTTGGTGACTCTAGATAACCTCGGGCCGATCGCACGCC</td><td>273.1</td><td>-9.7</td><td>1.9</td><td>-5.1</td><td>3.0E-07</td><td>4.8E-07</td><td>MB231/MB468</td></tr>
<tr><td>>hg38_21_+_8433172_8446622_RNA45SN1_45S_50ntFlanks@12605.12622.18|TATGTGCTTGGCTGAGGA</td><td>>hg38_21_+_8433172_8446622_RNA45SN1_45S_50ntFlanks</td><td>12605</td><td>12622</td><td>18</td><td>TATGTGCTTGGCTGAGGA</td><td>176.8</td><td>-3.0</td><td>0.6</td><td>-5.1</td><td>3.1E-07</td><td>4.8E-07</td><td>MB231/MB468</td></tr>
<tr><td>>hg38_21_+_8433172_8446622_RNA45SN1_45S_50ntFlanks@9545.9562.18|GTGGAGGTCCGTAGCGGT</td><td>>hg38_21_+_8433172_8446622_RNA45SN1_45S_50ntFlanks</td><td>9545</td><td>9562</td><td>18</td><td>GTGGAGGTCCGTAGCGGT</td><td>150.7</td><td>-2.7</td><td>0.5</td><td>-5.1</td><td>3.1E-07</td><td>4.9E-07</td><td>MB231/MB468</td></tr>
<tr><td>>hg38_21_+_8433172_8446622_RNA45SN1_45S_50ntFlanks@10848.10865.18|TCGGGATAAGGATTGGCT</td><td>>hg38_21_+_8433172_8446622_RNA45SN1_45S_50ntFlanks</td><td>10848</td><td>10865</td><td>18</td><td>TCGGGATAAGGATTGGCT</td><td>124.5</td><td>-2.8</td><td>0.6</td><td>-5.1</td><td>3.5E-07</td><td>5.0E-07</td><td>BT20/MB468</td></tr>
<tr><td>>hg38_21_+_8433172_8446622_RNA45SN1_45S_50ntFlanks@10830.10852.23|CAAGCCGGATCCGTAACTTCGGG</td><td>>hg38_21_+_8433172_8446622_RNA45SN1_45S_50ntFlanks</td><td>10830</td><td>10852</td><td>23</td><td>CAAGCCGGATCCGTAACTTCGGG</td><td>266.3</td><td>-3.9</td><td>0.8</td><td>-5.1</td><td>3.5E-07</td><td>5.0E-07</td><td>BT20/MB468</td></tr>
<tr><td>>hg38_21_+_8433172_8446622_RNA45SN1_45S_50ntFlanks@11570.11591.22|AGCCGACTTAGAACTGGTGCGG</td><td>>hg38_21_+_8433172_8446622_RNA45SN1_45S_50ntFlanks</td><td>11570</td><td>11591</td><td>22</td><td>AGCCGACTTAGAACTGGTGCGG</td><td>470.5</td><td>-3.1</td><td>0.6</td><td>-5.1</td><td>3.2E-07</td><td>5.0E-07</td><td>MB231/MB468</td></tr>
<tr><td>>hg38_21_+_8433172_8446622_RNA45SN1_45S_50ntFlanks@10408.10437.30|GGTCAGTCGGTCCTGAGAGATGGGCGAGCG</td><td>>hg38_21_+_8433172_8446622_RNA45SN1_45S_50ntFlanks</td><td>10408</td><td>10437</td><td>30</td><td>GGTCAGTCGGTCCTGAGAGATGGGCGAGCG</td><td>84.7</td><td>-3.1</td><td>0.6</td><td>-5.1</td><td>3.2E-07</td><td>5.0E-07</td><td>MB231/MB468</td></tr>
<tr><td>>MI0000489|hsa-mir-195&WithFlank&17|-|7017609|7017707@21.42.22&[MIMAT0000461&hsa-miR-195-5p&offsets|0|+1;m-191&17|-|7017666|7017687&offsets|0|0]|TAGCAGCACAGAAATATTGGCA</td><td>>MI0000489|hsa-mir-195&WithFlank&17|-|7017609|7017707</td><td>21</td><td>42</td><td>22</td><td>TAGCAGCACAGAAATATTGGCA</td><td>428.4</td><td>4.9</td><td>0.9</td><td>5.6</td><td>2.4E-08</td><td>5.1E-07</td><td>BT20/MB231</td></tr>
<tr><td>>hg38_21_+_8433172_8446622_RNA45SN1_45S_50ntFlanks@12083.12107.25|CTCGCTTCTGGCGCCAAGCGCCCGG</td><td>>hg38_21_+_8433172_8446622_RNA45SN1_45S_50ntFlanks</td><td>12083</td><td>12107</td><td>25</td><td>CTCGCTTCTGGCGCCAAGCGCCCGG</td><td>304.5</td><td>-5.5</td><td>1.1</td><td>-5.1</td><td>3.6E-07</td><td>5.1E-07</td><td>BT20/MB468</td></tr>
<tr><td>>hg38_21_+_8433172_8446622_RNA45SN1_45S_50ntFlanks@9553.9571.19|CCGTAGCGGTCCTGACGTG</td><td>>hg38_21_+_8433172_8446622_RNA45SN1_45S_50ntFlanks</td><td>9553</td><td>9571</td><td>19</td><td>CCGTAGCGGTCCTGACGTG</td><td>188.8</td><td>-2.1</td><td>0.4</td><td>-5.1</td><td>3.6E-07</td><td>5.1E-07</td><td>BT20/MB468</td></tr>
<tr><td>>hg38_21_+_8433172_8446622_RNA45SN1_45S_50ntFlanks@12357.12376.20|CCACAGGGATAACTGGCTTG</td><td>>hg38_21_+_8433172_8446622_RNA45SN1_45S_50ntFlanks</td><td>12357</td><td>12376</td><td>20</td><td>CCACAGGGATAACTGGCTTG</td><td>1021.1</td><td>-6.4</td><td>1.3</td><td>-5.1</td><td>3.6E-07</td><td>5.2E-07</td><td>BT20/MB468</td></tr>
<tr><td>>hg38_21_+_8433172_8446622_RNA45SN1_45S_50ntFlanks@8189.8208.20|CCCAGCCCGTGGACGGTGTG</td><td>>hg38_21_+_8433172_8446622_RNA45SN1_45S_50ntFlanks</td><td>8189</td><td>8208</td><td>20</td><td>CCCAGCCCGTGGACGGTGTG</td><td>120.3</td><td>-5.1</td><td>1.0</td><td>-5.1</td><td>3.3E-07</td><td>5.2E-07</td><td>MB231/MB468</td></tr>
<tr><td>>hg38_21_+_8433172_8446622_RNA45SN1_45S_50ntFlanks@3777.3801.25|CGGTACAGTGAAACTGCGAATGGCT</td><td>>hg38_21_+_8433172_8446622_RNA45SN1_45S_50ntFlanks</td><td>3777</td><td>3801</td><td>25</td><td>CGGTACAGTGAAACTGCGAATGGCT</td><td>414.1</td><td>-2.7</td><td>0.5</td><td>-5.1</td><td>3.3E-07</td><td>5.2E-07</td><td>MB231/MB468</td></tr>
<tr><td>>am_trna128_GlyGCC_6_-_27870686_27870756@7.28.22,trna133_GlyCCC_1_-_16872434_16872504@7.28.22,trna17_GluTTC_2_-_204229131_204229201@7.28.22,trna18_GlyGCC_16_+_70822597_70822667@7.28.22,trna19_GlyGCC_16_+_70823410_70823480@7.28.22,trna19_GlyGCC_2_-_157257659_157257729@7.28.22,trna24_GlyGCC_16_-_70812942_70813012@7.28.22,trna25_GlyGCC_16_-_70812114_70812184@7.28.22,trna2_GlyGCC_21_-_18827107_18827177@7.28.22,trna35_GlyGCC_1_+_161413094_161413164@7.28.22,trna37_GlyGCC_1_+_161420467_161420537@7.28.22,trna39_GlyGCC_1_+_161427898_161427968@7.28.22,trna41_GlyGCC_1_+_161435258_161435328@7.28.22,trna4_GlyCCC_1_+_17188416_17188486@7.28.22,trna5_GlyGCC_17_+_8029064_8029134@7.28.22,trna68_GlyGCC_1_-_161493637_161493707@7.28.22,trna91_GlyCCC_1_-_149680210_149680280@7.28.22,trna99_ValCAC_1_-_149294666_149294736@7.28.22|GTGGTTCAGTGGTAGAATTCTC</td><td>>am_trna128_GlyGCC_6_-_27870686_27870756</td><td>7</td><td>28</td><td>22</td><td>GTGGTTCAGTGGTAGAATTCTC</td><td>192.8</td><td>-3.3</td><td>0.6</td><td>-5.1</td><td>3.7E-07</td><td>5.2E-07</td><td>BT20/MB468</td></tr>
</table>

| | | | | | | | | | | | | |
|---|---|---|---|---|---|---|---|---|---|---|---|---|
| >trna134_GluTTC_1_-_16861774_16861845@1.31.31,trna5_GluTTC_1_+_17199078_17199149@1.31.31,trna84_GluTTC_1_-_161391883_161391954@1.31.31,trna94_GluTTC_1_-_149664355_149664427@1.31.31\|TCCCTGGTGGTCTAGTGGCTAGGATTCGGCG | >trna134_GluTTC_1_-_16861774_16861845 | 1 | 31 | 31 | TCCCTGGTGGTCTAGTGGCTAGGATTCGGCG | 1417.3 | -3.6 | 0.7 | -5.1 | 3.7E-07 | 5.3E-07 | BT20/MB468 |
| >hg38_21_+_8433172_8446622_RNA45SN1_45S_50ntFlanks@11879.11904.26\|ACCCTGTTGAGCTTGACTCTAGTCTG | >hg38_21_+_8433172_8446622_RNA45SN1_45S_50ntFlanks | 11879 | 11904 | 26 | ACCCTGTTGAGCTTGACTCTAGTCTG | 157.8 | -3.3 | 0.6 | -5.1 | 3.7E-07 | 5.3E-07 | BT20/MB468 |
| >hg38_21_+_8433172_8446622_RNA45SN1_45S_50ntFlanks@10473.10490.18\|CTCGGCCGATCGAAAGGG | >hg38_21_+_8433172_8446622_RNA45SN1_45S_50ntFlanks | 10473 | 10490 | 18 | CTCGGCCGATCGAAAGGG | 127.8 | -2.9 | 0.6 | -5.1 | 3.7E-07 | 5.3E-07 | BT20/MB468 |
| >trnaMT_SerGCT_MT_+_12207_12265@14.45.32\|AGAACTGCTAACTCATGCCCCCATGTCTAACA | >trnaMT_SerGCT_MT_+_12207_12265 | 14 | 45 | 32 | AGAACTGCTAACTCATGCCCCCATGTCTAACA | 417.1 | 7.0 | 1.4 | 5.1 | 3.7E-07 | 5.3E-07 | BT20/MB468 |
| >hg38_21_+_8433172_8446622_RNA45SN1_45S_50ntFlanks@11573.11605.33\|CGACTTAGAACTGGTGCGGACCAGGGGAATCCG | >hg38_21_+_8433172_8446622_RNA45SN1_45S_50ntFlanks | 11573 | 11605 | 33 | CGACTTAGAACTGGTGCGGACCAGGGGAATCCG | 195.1 | -2.2 | 0.4 | -5.1 | 3.4E-07 | 5.3E-07 | MB231/MB468 |
| >hg38_21_+_8433172_8446622_RNA45SN1_45S_50ntFlanks@5492.5510.19\|AGAAGACGGTCGAACTTGA | >hg38_21_+_8433172_8446622_RNA45SN1_45S_50ntFlanks | 5492 | 5510 | 19 | AGAAGACGGTCGAACTTGA | 222.4 | -3.2 | 0.6 | -5.1 | 3.4E-07 | 5.3E-07 | MB231/MB468 |
| >am_trna11_ProAGG_16_+_3241989_3242060@1.27.27,trna12_ProAGG_6_+_26555498_26555569@1.27.27,trna12_ProTGG_11_-_75946869_75946940@1.27.27,trna14_ProTGG_5_-_180615854_180615925@1.27.27,trna22_ProAGG_14_-_21081560_21081631@1.27.27,trna23_ProAGG_14_-_21077495_21077566@1.27.27,trna28_ProTGG_16_-_3234133_3234204@1.27.27,trna29_ProAGG_16_-_3232635_3232706@1.27.27,trna2_ProAGG_7_+_128423504_128423575@1.27.27,trna30_ProCGG_6_+_27059521_27059592@1.27.27,trna37_ProCGG_17_-_8126151_8126222@1.27.27,trna3_ProTGG_16_+_3208923_3208994@1.27.27,trna52_ProCGG_1_+_167683962_167684033@1.27.27,trna65_ProAGG_1_-_167684725_167684796@1.27.27,trna6_ProCGG_16_+_3222049_3222120@1.27.27,trna6_ProTGG_14_+_21152175_21152246@1.27.27,trna8_ProTGG_16_+_3238094_3238165@1.27.27,trna9_ProAGG_11_+_75946557_75946628@1.27.27,trna9_ProAGG_16_+_3239634_3239705@1.27.27\|GGCTCGTTGGTCTAGGGGTATGATTCT | >am_trna11_ProAGG_16_+_3241989_3242060 | 1 | 27 | 27 | GGCTCGTTGGTCTAGGGGTATGATTCT | 94.5 | -4.1 | 0.8 | -5.1 | 3.4E-07 | 5.3E-07 | MB231/MB468 |
| >hg38_21_+_8433172_8446622_RNA45SN1_45S_50ntFlanks@3705.3746.42\|TACCTGGTTGATCCTGCCAGTAGCATATGCTTGTCTCAAAGA | >hg38_21_+_8433172_8446622_RNA45SN1_45S_50ntFlanks | 3705 | 3746 | 42 | TACCTGGTTGATCCTGCCAGTAGCATATGCTTGTCTCAAAGA | 719.9 | 10.5 | 2.1 | 5.1 | 3.4E-07 | 5.4E-07 | MB231/MB468 |
| >hg38_21_+_8433172_8446622_RNA45SN1_45S_50ntFlanks@10713.10736.24\|CTTGAAAATCCGGGGGAGAGGGTG | >hg38_21_+_8433172_8446622_RNA45SN1_45S_50ntFlanks | 10713 | 10736 | 24 | CTTGAAAATCCGGGGGAGAGGGTG | 122.3 | -2.6 | 0.5 | -5.1 | 3.4E-07 | 5.4E-07 | MB231/MB468 |
| >hg38_21_+_8433172_8446622_RNA45SN1_45S_50ntFlanks@4006.4026.21\|ACCTCGGGCCGATCGCACGCC | >hg38_21_+_8433172_8446622_RNA45SN1_45S_50ntFlanks | 4006 | 4026 | 21 | ACCTCGGGCCGATCGCACGCC | 130.9 | -4.2 | 0.8 | -5.1 | 3.4E-07 | 5.4E-07 | MB231/MB468 |
| >hg38_21_+_8433172_8446622_RNA45SN1_45S_50ntFlanks@6785.6806.22\|CTACGCCTGTCTGAGCGTCGCT | >hg38_21_+_8433172_8446622_RNA45SN1_45S_50ntFlanks | 6785 | 6806 | 22 | CTACGCCTGTCTGAGCGTCGCT | 3312.9 | 4.1 | 0.8 | 5.1 | 3.8E-07 | 5.5E-07 | BT20/MB468 |
| >MI0000272\|hsa-mir-182&WithFlank&7\|-\|129770377\|129770498@29.51.23&[MIMAT0000259&hsa-miR-182-5p&offsets\|0\|-1;m-22&7\|-\|129770449\|129770470&offsets\|0\|+1]\|TTTGGCAATGGTAGAACTCACAC | >MI0000272\|hsa-mir-182&WithFlank&7\|-\|129770377\|129770498 | 29 | 51 | 23 | TTTGGCAATGGTAGAACTCACAC | 3303.5 | 3.9 | 0.7 | 5.6 | 2.6E-08 | 5.5E-07 | BT20/MB231 |
| >MI0003137\|hsa-mir-193b&WithFlank&16\|+\|14303961\|14304055@57.77.21(+1C)&[MIMAT0002819&hsa-miR-193b-3p&offsets\|0\|-1(+1C);m-166&16\|+\|14304017\|14304038&offsets\|0\|-1(+1C)]\|AACTGGCCCTCAAAGTCCCGCC | >MI0003137\|hsa-mir-193b&WithFlank&16\|+\|14303961\|14304055 | 57 | 77 | 21(+1C) | AACTGGCCCTCAAAGTCCCGCC | 824.3 | 5.0 | 0.9 | 5.6 | 2.6E-08 | 5.5E-07 | BT20/MB231 |
| >hg38_21_+_8433172_8446622_RNA45SN1_45S_50ntFlanks@9533.9566.34\|GAGGAAACTCTGGTGGAGGTCCGTAGCGGTCCTG | >hg38_21_+_8433172_8446622_RNA45SN1_45S_50ntFlanks | 9533 | 9566 | 34 | GAGGAAACTCTGGTGGAGGTCCGTAGCGGTCCTG | 116.5 | -3.8 | 0.8 | -5.1 | 3.9E-07 | 5.5E-07 | BT20/MB468 |
| >am_trnaMT_ValTAC_MT_+_1602_1670@30.50.21\|CTTACACTTAGGAGATTTCAA | >am_trnaMT_ValTAC_MT_+_1602_1670 | 30 | 50 | 21 | CTTACACTTAGGAGATTTCAA | 53.8 | -7.3 | 1.4 | -5.1 | 3.5E-07 | 5.6E-07 | MB231/MB468 |
| >hg38_21_+_8433172_8446622_RNA45SN1_45S_50ntFlanks@3900.3919.20\|CTTCGCGGGGGGGATGCGTG | >hg38_21_+_8433172_8446622_RNA45SN1_45S_50ntFlanks | 3900 | 3919 | 20 | CTTCGCGGGGGGGATGCGTG | 165.5 | -2.6 | 0.5 | -5.1 | 4.0E-07 | 5.7E-07 | BT20/MB468 |
| >trna111_HisGTG_1_-_147774845_147774916@-1T.34.35,trna118_HisGTG_1_-_145396881_145396952@-1T.34.35,trna16_HisGTG_1_+_146544773_146544844@-1T.34.35,trna1_HisGTG_15_+_45493349_45493420@-1T.34.35,trna21_HisGTG_1_+_147753471_147753542@-1T.34.35,trna33_HisGTG_6_+_27125906_27125977@-1T.34.35,trna7_HisGTG_9_-_14433938_14434009@-1T.34.35,trna8_HisGTG_15_-_45492611_45492682@-1T.34.35,trna9_HisGTG_15_-_45490804_45490875@-1T.34.35\|TGCCGTGATCGTATAGTGGTTAGTACTCTGCGTTG | >trna111_HisGTG_1_-_147774845_147774916 | -1T | 34 | 35 | TGCCGTGATCGTATAGTGGTTAGTACTCTGCGTTG | 1192.3 | 5.0 | 1.0 | 5.1 | 4.0E-07 | 5.7E-07 | BT20/MB468 |
| >hg38_21_+_8433172_8446622_RNA45SN1_45S_50ntFlanks@4659.4678.20\|AGAGGTGAAATTCTTGGACC | >hg38_21_+_8433172_8446622_RNA45SN1_45S_50ntFlanks | 4659 | 4678 | 20 | AGAGGTGAAATTCTTGGACC | 319.2 | 10.0 | 2.0 | 5.1 | 3.6E-07 | 5.7E-07 | MB231/MB468 |
| >hg38_21_+_8433172_8446622_RNA45SN1_45S_50ntFlanks@12476.12497.22\|CCACTAATAGGGAACGTGAGCT | >hg38_21_+_8433172_8446622_RNA45SN1_45S_50ntFlanks | 12476 | 12497 | 22 | CCACTAATAGGGAACGTGAGCT | 1574.1 | -3.0 | 0.6 | -5.1 | 4.0E-07 | 5.7E-07 | BT20/MB468 |
| >hg38_21_+_8433172_8446622_RNA45SN1_45S_50ntFlanks@6722.6742.21\|ATTGCAGGACACATTGATCAT | >hg38_21_+_8433172_8446622_RNA45SN1_45S_50ntFlanks | 6722 | 6742 | 21 | ATTGCAGGACACATTGATCAT | 120.0 | -4.0 | 0.8 | -5.1 | 4.0E-07 | 5.8E-07 | BT20/MB468 |
| >hg38_21_+_8433172_8446622_RNA45SN1_45S_50ntFlanks@9567.9588.22\|ACGTGCAAATCGGTCGTCCGAC | >hg38_21_+_8433172_8446622_RNA45SN1_45S_50ntFlanks | 9567 | 9588 | 22 | ACGTGCAAATCGGTCGTCCGAC | 117.0 | -3.3 | 0.7 | -5.1 | 3.7E-07 | 5.8E-07 | MB231/MB468 |
| >hg38_21_+_8433172_8446622_RNA45SN1_45S_50ntFlanks@11581.11600.20\|AACTGGTGCGGACCAGGGGA | >hg38_21_+_8433172_8446622_RNA45SN1_45S_50ntFlanks | 11581 | 11600 | 20 | AACTGGTGCGGACCAGGGGA | 524.5 | -3.3 | 0.6 | -5.1 | 4.1E-07 | 5.8E-07 | BT20/MB468 |
| >trna10_LysCTT_16_+_3241501_3241573@14.31.18,trna13_LysCTT_14_-_58706613_58706685@14.31.18,trna2_LysCTT_15_+_79152904_79152976@14.31.18,trna4_LysCTT_18_-_43669271_43669343@14.31.18\|AGTCGGTAGAGCATGGGA | >trna10_LysCTT_16_+_3241501_3241573 | 14 | 31 | 18 | AGTCGGTAGAGCATGGGA | 887.4 | -4.2 | 0.8 | -5.1 | 4.1E-07 | 5.8E-07 | BT20/MB468 |
| >hg38_21_+_8433172_8446622_RNA45SN1_45S_50ntFlanks@476.523.48\|TTCGTGATCGATGTGGTGACGTCGTGCTCTCCCGGGCCGGGTCCGAGC | >hg38_21_+_8433172_8446622_RNA45SN1_45S_50ntFlanks | 476 | 523 | 48 | TTCGTGATCGATGTGGTGACGTCGTGCTCTCCCGGGCCGGGTCCGAGC | 3522.0 | 12.2 | 2.4 | 5.1 | 3.7E-07 | 5.8E-07 | MB231/MB468 |
| >hg38_21_+_8433172_8446622_RNA45SN1_45S_50ntFlanks@4372.4389.18\|AAAAAGCTCGTAGTTGGA | >hg38_21_+_8433172_8446622_RNA45SN1_45S_50ntFlanks | 4372 | 4389 | 18 | AAAAAGCTCGTAGTTGGA | 145.2 | -2.9 | 0.6 | -5.1 | 3.7E-07 | 5.8E-07 | MB231/MB468 |
| >hg38_21_+_8433172_8446622_RNA45SN1_45S_50ntFlanks@9567.9585.19\|ACGTGCAAATCGGTCGTCC | >hg38_21_+_8433172_8446622_RNA45SN1_45S_50ntFlanks | 9567 | 9585 | 19 | ACGTGCAAATCGGTCGTCC | 167.6 | -5.2 | 1.0 | -5.1 | 4.1E-07 | 5.9E-07 | BT20/MB468 |
| >hg38_21_+_8433172_8446622_RNA45SN1_45S_50ntFlanks@7974.8008.35\|ACGCGACCTCAGATCAGACGTGGCGACCCGCTGAA | >hg38_21_+_8433172_8446622_RNA45SN1_45S_50ntFlanks | 7974 | 8008 | 35 | ACGCGACCTCAGATCAGACGTGGCGACCCGCTGAA | 8617.4 | 3.9 | 0.8 | 5.1 | 3.7E-07 | 5.9E-07 | MB231/MB468 |
| >hg38_21_+_8433172_8446622_RNA45SN1_45S_50ntFlanks@12405.12428.24\|CTTTTTGATCCTTCGATGTCGGCT | >hg38_21_+_8433172_8446622_RNA45SN1_45S_50ntFlanks | 12405 | 12428 | 24 | CTTTTTGATCCTTCGATGTCGGCT | 108.2 | -4.0 | 0.8 | -5.1 | 3.8E-07 | 5.9E-07 | MB231/MB468 |
| >hg38_1_-_228634819_228635039_RNA5S12_5S_50ntFlanks@84.102.19\|CTCGTCTGATCTCGGAAGC | >hg38_1_-_228634819_228635039_RNA5S12_5S_50ntFlanks | 84 | 102 | 19 | CTCGTCTGATCTCGGAAGC | 398.0 | -4.8 | 0.9 | -5.1 | 4.3E-07 | 6.1E-07 | BT20/MB468 |
| >hg38_21_+_8433172_8446622_RNA45SN1_45S_50ntFlanks@11576.11602.27\|CTTAGAACTGGTGCGGACCAGGGGAAT | >hg38_21_+_8433172_8446622_RNA45SN1_45S_50ntFlanks | 11576 | 11602 | 27 | CTTAGAACTGGTGCGGACCAGGGGAAT | 908.6 | -4.6 | 0.9 | -5.1 | 4.3E-07 | 6.1E-07 | BT20/MB468 |
| >hg38_21_+_8433172_8446622_RNA45SN1_45S_50ntFlanks@10458.10484.27\|ATGGCCTCCGTTGCCCTCGGCCGATCG | >hg38_21_+_8433172_8446622_RNA45SN1_45S_50ntFlanks | 10458 | 10484 | 27 | ATGGCCTCCGTTGCCCTCGGCCGATCG | 261.0 | -5.2 | 1.0 | -5.1 | 4.3E-07 | 6.1E-07 | BT20/MB468 |
| >hg38_21_+_8433172_8446622_RNA45SN1_45S_50ntFlanks@11660.11685.26\|GTGATTTCTGCCCAGTGCTCTGAATG | >hg38_21_+_8433172_8446622_RNA45SN1_45S_50ntFlanks | 11660 | 11685 | 26 | GTGATTTCTGCCCAGTGCTCTGAATG | 278.4 | -2.2 | 0.4 | -5.1 | 4.3E-07 | 6.1E-07 | BT20/MB468 |
| >hg38_21_+_8433172_8446622_RNA45SN1_45S_50ntFlanks@9554.9576.23\|CGTAGCGGTCCTGACGTGCAAAT | >hg38_21_+_8433172_8446622_RNA45SN1_45S_50ntFlanks | 9554 | 9576 | 23 | CGTAGCGGTCCTGACGTGCAAAT | 652.8 | 1.8 | 0.4 | 5.1 | 4.3E-07 | 6.1E-07 | BT20/MB468 |
| >hg38_21_+_8433172_8446622_RNA45SN1_45S_50ntFlanks@11580.11603.24\|GAACTGGTGCGGACCAGGGGAATC | >hg38_21_+_8433172_8446622_RNA45SN1_45S_50ntFlanks | 11580 | 11603 | 24 | GAACTGGTGCGGACCAGGGGAATC | 2577.4 | -3.4 | 0.7 | -5.1 | 3.9E-07 | 6.2E-07 | MB231/MB468 |
| >hg38_21_+_8433172_8446622_RNA45SN1_45S_50ntFlanks@4047.4064.18\|ATTCGAACGTCTGCCCTA | >hg38_21_+_8433172_8446622_RNA45SN1_45S_50ntFlanks | 4047 | 4064 | 18 | ATTCGAACGTCTGCCCTA | 169.8 | -2.9 | 0.6 | -5.1 | 4.0E-07 | 6.2E-07 | MB231/MB468 |
| >hg38_21_+_8433172_8446622_RNA45SN1_45S_50ntFlanks@8380.8407.28\|CAAGAGGGCGTGAAACCGTTAAGAGGTA | >hg38_21_+_8433172_8446622_RNA45SN1_45S_50ntFlanks | 8380 | 8407 | 28 | CAAGAGGGCGTGAAACCGTTAAGAGGTA | 130.4 | -2.6 | 0.5 | -5.1 | 4.4E-07 | 6.3E-07 | BT20/MB468 |
| >hg38_21_+_8433172_8446622_RNA45SN1_45S_50ntFlanks@10351.10369.19\|CGAGAACTTTGAAGGCCGA | >hg38_21_+_8433172_8446622_RNA45SN1_45S_50ntFlanks | 10351 | 10369 | 19 | CGAGAACTTTGAAGGCCGA | 800.5 | -3.2 | 0.6 | -5.1 | 4.4E-07 | 6.3E-07 | BT20/MB468 |
| >hg38_21_+_8433172_8446622_RNA45SN1_45S_50ntFlanks@12164.12182.19\|CTGGGGCGGTACACCTGTC | >hg38_21_+_8433172_8446622_RNA45SN1_45S_50ntFlanks | 12164 | 12182 | 19 | CTGGGGCGGTACACCTGTC | 144.4 | -3.6 | 0.7 | -5.0 | 4.5E-07 | 6.3E-07 | BT20/MB468 |
| >hg38_21_+_8433172_8446622_RNA45SN1_45S_50ntFlanks@11670.11698.29\|CCCAGTGCTCTGAATGTCAAAGTGAAGAA | >hg38_21_+_8433172_8446622_RNA45SN1_45S_50ntFlanks | 11670 | 11698 | 29 | CCCAGTGCTCTGAATGTCAAAGTGAAGAA | 235.5 | -3.3 | 0.7 | -5.0 | 4.5E-07 | 6.4E-07 | BT20/MB468 |
| >hg38_21_+_8433172_8446622_RNA45SN1_45S_50ntFlanks@12488.12522.35\|AACGTGAGCTGGGTTTAGACCGTCGTGAGACAGGT | >hg38_21_+_8433172_8446622_RNA45SN1_45S_50ntFlanks | 12488 | 12522 | 35 | AACGTGAGCTGGGTTTAGACCGTCGTGAGACAGGT | 178.5 | -3.4 | 0.7 | -5.1 | 4.2E-07 | 6.6E-07 | MB231/MB468 |
| >hg38_21_+_8433172_8446622_RNA45SN1_45S_50ntFlanks@10362.10382.21\|AAGGCCGAAGTGGAGAAGGGT | >hg38_21_+_8433172_8446622_RNA45SN1_45S_50ntFlanks | 10362 | 10382 | 21 | AAGGCCGAAGTGGAGAAGGGT | 181.6 | -2.5 | 0.5 | -5.0 | 4.7E-07 | 6.6E-07 | BT20/MB468 |

<table>
<tr><td></td><td></td><td></td><td></td><td></td><td></td><td></td><td></td><td></td><td></td><td></td><td></td><td></td></tr>
<tr><td>>hg38_21_+_8433172_8446622_RNA45SN1_45S_50ntFlanks@8168.8195.28|CCCAAGTCCTTCTGATCGAGGCCCAGCC</td><td>>hg38_21_+_8433172_8446622_RNA45SN1_45S_50ntFlanks</td><td>8168</td><td>8195</td><td>28</td><td>CCCAAGTCCTTCTGATCGAGGCCCAGCC</td><td>145.4</td><td>-3.2</td><td>0.6</td><td>-5.0</td><td>4.7E-07</td><td>6.6E-07</td><td>BT20/MB468</td></tr>
<tr><td>>hg38_21_+_8433172_8446622_RNA45SN1_45S_50ntFlanks@11562.11588.27|CGCCTAGCAGCCGACTTAGAACTGGTG</td><td>>hg38_21_+_8433172_8446622_RNA45SN1_45S_50ntFlanks</td><td>11562</td><td>11588</td><td>27</td><td>CGCCTAGCAGCCGACTTAGAACTGGTG</td><td>348.1</td><td>-2.8</td><td>0.6</td><td>-5.1</td><td>4.2E-07</td><td>6.7E-07</td><td>MB231/MB468</td></tr>
<tr><td>>hg38_21_+_8433172_8446622_RNA45SN1_45S_50ntFlanks@10869.10894.26|AGGGCTGGGTCGGTCGGGCTGGGGCG</td><td>>hg38_21_+_8433172_8446622_RNA45SN1_45S_50ntFlanks</td><td>10869</td><td>10894</td><td>26</td><td>AGGGCTGGGTCGGTCGGGCTGGGGCG</td><td>150.5</td><td>-2.9</td><td>0.6</td><td>-5.1</td><td>4.2E-07</td><td>6.7E-07</td><td>MB231/MB468</td></tr>
<tr><td>>hg38_21_+_8433172_8446622_RNA45SN1_45S_50ntFlanks@10873.10892.20|CTGGGTCGGTCGGGCTGGGG</td><td>>hg38_21_+_8433172_8446622_RNA45SN1_45S_50ntFlanks</td><td>10873</td><td>10892</td><td>20</td><td>CTGGGTCGGTCGGGCTGGGG</td><td>295.1</td><td>-4.3</td><td>0.9</td><td>-5.0</td><td>4.7E-07</td><td>6.7E-07</td><td>BT20/MB468</td></tr>
<tr><td>>hg38_21_+_8433172_8446622_RNA45SN1_45S_50ntFlanks@13001.13028.28|GCGATCTATTGAAAGTCAGCCCTCGACA</td><td>>hg38_21_+_8433172_8446622_RNA45SN1_45S_50ntFlanks</td><td>13001</td><td>13028</td><td>28</td><td>GCGATCTATTGAAAGTCAGCCCTCGACA</td><td>103.9</td><td>-4.8</td><td>0.9</td><td>-5.0</td><td>4.7E-07</td><td>6.7E-07</td><td>BT20/MB468</td></tr>
<tr><td>>hg38_21_+_8433172_8446622_RNA45SN1_45S_50ntFlanks@11568.11587.20|GCAGCCGACTTAGAACTGGT</td><td>>hg38_21_+_8433172_8446622_RNA45SN1_45S_50ntFlanks</td><td>11568</td><td>11587</td><td>20</td><td>GCAGCCGACTTAGAACTGGT</td><td>532.8</td><td>2.3</td><td>0.5</td><td>5.1</td><td>4.3E-07</td><td>6.7E-07</td><td>MB231/MB468</td></tr>
<tr><td>>trna10_AspGTC_12_-_125424193_125424264@1.31.31,trna12_AspGTC_12_-_125411891_125411962@1.31.31,trna38_AspGTC_17_-_8125556_8125627@1.31.31,trna45_AspGTC_6_+_27447453_27447524@1.31.31,trna48_AspGTC_6_+_27471523_27471594@1.31.31,trna4_AspGTC_12_+_96429799_96429870@1.31.31,trna69_AspGTC_1_-_161440205_161440276@1.31.31,trna72_AspGTC_1_-_161432824_161432895@1.31.31,trna75_AspGTC_1_-_161425414_161425485@1.31.31,trna78_AspGTC_1_-_161418033_161418104@1.31.31,trna81_AspGTC_1_-_161410615_161410686@1.31.31|TCCTCGTTAGTATAGTGGTGAGTATCCCCGC</td><td>>trna10_AspGTC_12_-_125424193_125424264</td><td>1</td><td>31</td><td>31</td><td>TCCTCGTTAGTATAGTGGTGAGTATCCCCGC</td><td>153.2</td><td>-4.9</td><td>1.0</td><td>-5.0</td><td>4.8E-07</td><td>6.8E-07</td><td>BT20/MB468</td></tr>
<tr><td>>hg38_21_+_8433172_8446622_RNA45SN1_45S_50ntFlanks@12419.12460.42|GATGTCGGCTCTTCCTATCATTGTGAAGCAGAATTCACCAAG</td><td>>hg38_21_+_8433172_8446622_RNA45SN1_45S_50ntFlanks</td><td>12419</td><td>12460</td><td>42</td><td>GATGTCGGCTCTTCCTATCATTGTGAAGCAGAATTCACCAAG</td><td>371.8</td><td>9.7</td><td>1.9</td><td>5.1</td><td>4.3E-07</td><td>6.8E-07</td><td>MB231/MB468</td></tr>
<tr><td>>hg38_21_+_8433172_8446622_RNA45SN1_45S_50ntFlanks@12960.12982.23|CTGGGTCGGGGTTTCGTACGTAG</td><td>>hg38_21_+_8433172_8446622_RNA45SN1_45S_50ntFlanks</td><td>12960</td><td>12982</td><td>23</td><td>CTGGGTCGGGGTTTCGTACGTAG</td><td>116.3</td><td>-4.0</td><td>0.8</td><td>-5.1</td><td>4.3E-07</td><td>6.8E-07</td><td>MB231/MB468</td></tr>
<tr><td>>hg38_21_+_8433172_8446622_RNA45SN1_45S_50ntFlanks@5444.5467.24|CTCGGATCGGCCCCGCCGGGGTCG</td><td>>hg38_21_+_8433172_8446622_RNA45SN1_45S_50ntFlanks</td><td>5444</td><td>5467</td><td>24</td><td>CTCGGATCGGCCCCGCCGGGGTCG</td><td>349.8</td><td>-3.7</td><td>0.7</td><td>-5.1</td><td>4.3E-07</td><td>6.8E-07</td><td>MB231/MB468</td></tr>
<tr><td>>hg38_21_+_8433172_8446622_RNA45SN1_45S_50ntFlanks@5470.5499.30|CCACGGCCCTGGCGGAGCGCTGAGAAGACG</td><td>>hg38_21_+_8433172_8446622_RNA45SN1_45S_50ntFlanks</td><td>5470</td><td>5499</td><td>30</td><td>CCACGGCCCTGGCGGAGCGCTGAGAAGACG</td><td>309.9</td><td>-4.3</td><td>0.8</td><td>-5.0</td><td>4.8E-07</td><td>6.8E-07</td><td>BT20/MB468</td></tr>
<tr><td>>hg38_21_+_8433172_8446622_RNA45SN1_45S_50ntFlanks@4569.4587.19|ATGGAATAGGACCGCGGTT</td><td>>hg38_21_+_8433172_8446622_RNA45SN1_45S_50ntFlanks</td><td>4569</td><td>4587</td><td>19</td><td>ATGGAATAGGACCGCGGTT</td><td>258.6</td><td>-3.6</td><td>0.7</td><td>-5.0</td><td>4.8E-07</td><td>6.8E-07</td><td>BT20/MB468</td></tr>
<tr><td>>hg38_21_+_8433172_8446622_RNA45SN1_45S_50ntFlanks@10393.10435.43|CAGCAGTTGAACATGGGTCAGTCGGTCCTGAGAGATGGGCGAG</td><td>>hg38_21_+_8433172_8446622_RNA45SN1_45S_50ntFlanks</td><td>10393</td><td>10435</td><td>43</td><td>CAGCAGTTGAACATGGGTCAGTCGGTCCTGAGAGATGGGCGAG</td><td>231.8</td><td>-6.3</td><td>1.3</td><td>-5.0</td><td>4.9E-07</td><td>6.9E-07</td><td>BT20/MB468</td></tr>
<tr><td>>MI0000253|hsa-mir-148a&WithFlank&7|-|25949913|25949992@50.70.21&[MIMAT0000243&hsa-miR-148a-3p&offsets|0|-1;m-7&7|-|25949922|25949943&offsets|0|-1]|TCAGTGCACTACAGAACTTTG</td><td>>MI0000253|hsa-mir-148a&WithFlank&7|-|25949913|25949992</td><td>50</td><td>70</td><td>21</td><td>TCAGTGCACTACAGAACTTTG</td><td>979.1</td><td>5.2</td><td>0.9</td><td>5.5</td><td>3.3E-08</td><td>6.9E-07</td><td>BT20/MB231</td></tr>
<tr><td>>hg38_21_+_8433172_8446622_RNA45SN1_45S_50ntFlanks@11702.11728.27|CAATGAAGCGCGGGTAAACGGCGGGAG</td><td>>hg38_21_+_8433172_8446622_RNA45SN1_45S_50ntFlanks</td><td>11702</td><td>11728</td><td>27</td><td>CAATGAAGCGCGGGTAAACGGCGGGAG</td><td>94.1</td><td>-3.7</td><td>0.7</td><td>-5.1</td><td>4.4E-07</td><td>6.9E-07</td><td>MB231/MB468</td></tr>
<tr><td>>hg38_21_+_8433172_8446622_RNA45SN1_45S_50ntFlanks@10402.10429.28|AACATGGGTCAGTCGGTCCTGAGAGATG</td><td>>hg38_21_+_8433172_8446622_RNA45SN1_45S_50ntFlanks</td><td>10402</td><td>10429</td><td>28</td><td>AACATGGGTCAGTCGGTCCTGAGAGATG</td><td>2903.1</td><td>-2.9</td><td>0.6</td><td>-5.1</td><td>4.4E-07</td><td>6.9E-07</td><td>MB231/MB468</td></tr>
<tr><td>>hg38_21_+_8433172_8446622_RNA45SN1_45S_50ntFlanks@6731.6756.26|CACATTGATCATCGACACTTCGAACG</td><td>>hg38_21_+_8433172_8446622_RNA45SN1_45S_50ntFlanks</td><td>6731</td><td>6756</td><td>26</td><td>CACATTGATCATCGACACTTCGAACG</td><td>658.2</td><td>-2.9</td><td>0.6</td><td>-5.0</td><td>4.9E-07</td><td>7.0E-07</td><td>BT20/MB468</td></tr>
<tr><td>>am_trnaMT_SerGCT_MT_+_12207_12265@14.31.18|AGAACTGCTAACTCATGC</td><td>>am_trnaMT_SerGCT_MT_+_12207_12265</td><td>14</td><td>31</td><td>18</td><td>AGAACTGCTAACTCATGC</td><td>1039.4</td><td>7.4</td><td>1.5</td><td>5.0</td><td>4.9E-07</td><td>7.0E-07</td><td>BT20/MB468</td></tr>
<tr><td>>hg38_21_+_8433172_8446622_RNA45SN1_45S_50ntFlanks@11572.11589.18|CCGACTTAGAACTGGTGC</td><td>>hg38_21_+_8433172_8446622_RNA45SN1_45S_50ntFlanks</td><td>11572</td><td>11589</td><td>18</td><td>CCGACTTAGAACTGGTGC</td><td>315.9</td><td>-2.8</td><td>0.5</td><td>-5.0</td><td>4.9E-07</td><td>7.0E-07</td><td>BT20/MB468</td></tr>
<tr><td>>hg38_21_+_8433172_8446622_RNA45SN1_45S_50ntFlanks@9305.9323.19|CCAAGGAGTCTAACACGTG</td><td>>hg38_21_+_8433172_8446622_RNA45SN1_45S_50ntFlanks</td><td>9305</td><td>9323</td><td>19</td><td>CCAAGGAGTCTAACACGTG</td><td>81.6</td><td>-5.9</td><td>1.2</td><td>-5.0</td><td>4.5E-07</td><td>7.0E-07</td><td>MB231/MB468</td></tr>
<tr><td>>hg38_21_+_8433172_8446622_RNA45SN1_45S_50ntFlanks@11200.11233.34|CGGCGGCGGCGGCGACTCTGGACGCGAGCCGGGC</td><td>>hg38_21_+_8433172_8446622_RNA45SN1_45S_50ntFlanks</td><td>11200</td><td>11233</td><td>34</td><td>CGGCGGCGGCGGCGACTCTGGACGCGAGCCGGGC</td><td>86.3</td><td>-4.4</td><td>0.9</td><td>-5.0</td><td>5.0E-07</td><td>7.1E-07</td><td>BT20/MB468</td></tr>
<tr><td>>hg38_1_-_228634819_228635039_RNA5S12_5S_50ntFlanks@122.147.26|TTAGTACTTGGATGGGAGACCGCCTG</td><td>>hg38_1_-_228634819_228635039_RNA5S12_5S_50ntFlanks</td><td>122</td><td>147</td><td>26</td><td>TTAGTACTTGGATGGGAGACCGCCTG</td><td>135.2</td><td>-4.1</td><td>0.8</td><td>-5.0</td><td>5.0E-07</td><td>7.1E-07</td><td>BT20/MB468</td></tr>
<tr><td>>hg38_21_+_8433172_8446622_RNA45SN1_45S_50ntFlanks@10392.10430.39|ACAGCAGTTGAACATGGGTCAGTCGGTCCTGAGAGATGG</td><td>>hg38_21_+_8433172_8446622_RNA45SN1_45S_50ntFlanks</td><td>10392</td><td>10430</td><td>39</td><td>ACAGCAGTTGAACATGGGTCAGTCGGTCCTGAGAGATGG</td><td>552.1</td><td>-3.9</td><td>0.8</td><td>-5.0</td><td>4.5E-07</td><td>7.1E-07</td><td>MB231/MB468</td></tr>
<tr><td>>trnaMT_SerGCT_MT_+_12207_12265@16.62.47|AACTGCTAACTCATGCCCCCATGTCTAACAACATGGCTTTCTCACCA</td><td>>trnaMT_SerGCT_MT_+_12207_12265</td><td>16</td><td>62</td><td>47</td><td>AACTGCTAACTCATGCCCCCATGTCTAACAACATGGCTTTCTCACCA</td><td>594.4</td><td>10.4</td><td>2.1</td><td>5.0</td><td>4.5E-07</td><td>7.1E-07</td><td>MB231/MB468</td></tr>
<tr><td>>MI0000478|hsa-mir-149&WithFlank&2|+|240455995|240456095@21.42.22(+1U)&[MIMAT0000450&hsa-miR-149-5p&offsets|0|-1(+1U);m-195&2|+|240456015|240456037&offsets|0|-1(+1U)]|TCTGGCTCCGTGTCTTCACTCCT</td><td>>MI0000478|hsa-mir-149&WithFlank&2|+|240455995|240456095</td><td>21</td><td>42</td><td>22(+1U)</td><td>TCTGGCTCCGTGTCTTCACTCCT</td><td>1114.5</td><td>4.3</td><td>0.8</td><td>5.5</td><td>3.5E-08</td><td>7.1E-07</td><td>BT20/MB231</td></tr>
<tr><td>>hg38_21_+_8433172_8446622_RNA45SN1_45S_50ntFlanks@11769.11786.18|TAATTAGTGACGCGCATG</td><td>>hg38_21_+_8433172_8446622_RNA45SN1_45S_50ntFlanks</td><td>11769</td><td>11786</td><td>18</td><td>TAATTAGTGACGCGCATG</td><td>319.0</td><td>-2.7</td><td>0.5</td><td>-5.0</td><td>4.5E-07</td><td>7.1E-07</td><td>MB231/MB468</td></tr>
<tr><td>>hg38_21_+_8433172_8446622_RNA45SN1_45S_50ntFlanks@12274.12293.20|CAGTACGAATACAGACCGTG</td><td>>hg38_21_+_8433172_8446622_RNA45SN1_45S_50ntFlanks</td><td>12274</td><td>12293</td><td>20</td><td>CAGTACGAATACAGACCGTG</td><td>109.7</td><td>-3.4</td><td>0.7</td><td>-5.0</td><td>5.0E-07</td><td>7.1E-07</td><td>BT20/MB468</td></tr>
<tr><td>>hg38_21_+_8433172_8446622_RNA45SN1_45S_50ntFlanks@10350.10373.24|ACGAGAACTTTGAAGGCCGAAGTG</td><td>>hg38_21_+_8433172_8446622_RNA45SN1_45S_50ntFlanks</td><td>10350</td><td>10373</td><td>24</td><td>ACGAGAACTTTGAAGGCCGAAGTG</td><td>432.5</td><td>-2.0</td><td>0.4</td><td>-5.0</td><td>5.1E-07</td><td>7.2E-07</td><td>BT20/MB468</td></tr>
<tr><td>>hg38_21_+_8433172_8446622_RNA45SN1_45S_50ntFlanks@12893.12932.40|CCTCGCCCGTCACGCACCGCACGTTCGTGGGGAACCTGGC</td><td>>hg38_21_+_8433172_8446622_RNA45SN1_45S_50ntFlanks</td><td>12893</td><td>12932</td><td>40</td><td>CCTCGCCCGTCACGCACCGCACGTTCGTGGGGAACCTGGC</td><td>408.9</td><td>-3.7</td><td>0.7</td><td>-5.0</td><td>5.1E-07</td><td>7.2E-07</td><td>BT20/MB468</td></tr>
<tr><td>>hg38_21_+_8433172_8446622_RNA45SN1_45S_50ntFlanks@9575.9595.21|ATCGGTCGTCCGACCTGGGTA</td><td>>hg38_21_+_8433172_8446622_RNA45SN1_45S_50ntFlanks</td><td>9575</td><td>9595</td><td>21</td><td>ATCGGTCGTCCGACCTGGGTA</td><td>619.7</td><td>-5.3</td><td>1.0</td><td>-5.0</td><td>4.6E-07</td><td>7.2E-07</td><td>MB231/MB468</td></tr>
<tr><td>>am_MI0039740|hsa-mir-12136&WithFlank&1|-|632609|632691@7.32.26&[MIMAT0049032&hsa-miR-12136&offsets|0|+8]|GAAAAAGTCATGGAGGCCATGGGGTT</td><td>>am_MI0039740|hsa-mir-12136&WithFlank&1|-|632609|632691</td><td>7</td><td>32</td><td>26</td><td>GAAAAAGTCATGGAGGCCATGGGGTT</td><td>716.8</td><td>6.2</td><td>1.2</td><td>5.0</td><td>5.1E-07</td><td>7.2E-07</td><td>BT20/MB468</td></tr>
<tr><td>>trnaMT_SerTGA_MT_-_7446_7514@1.26.26,trnalookalike6_SerTGA_1_-_567997_568065@1.26.26|GAAAAAGTCATGGAGGCCATGGGGTT</td><td>>trnaMT_SerTGA_MT_-_7446_7514</td><td>1</td><td>26</td><td>26</td><td>GAAAAAGTCATGGAGGCCATGGGGTT</td><td>716.8</td><td>6.2</td><td>1.2</td><td>5.0</td><td>5.1E-07</td><td>7.2E-07</td><td>BT20/MB468</td></tr>
<tr><td>>MI0000102|hsa-mir-100&WithFlank&11|-|122152223|122152314@19.40.22&[MIMAT0000098&hsa-miR-100-5p&offsets|0|0;m-27&11|-|122152276|122152296&offsets|0|+1]|AACCCGTAGATCCGAACTTGTG</td><td>>MI0000102|hsa-mir-100&WithFlank&11|-|122152223|122152314</td><td>19</td><td>40</td><td>22</td><td>AACCCGTAGATCCGAACTTGTG</td><td>35476.4</td><td>8.7</td><td>1.7</td><td>5.0</td><td>5.1E-07</td><td>7.2E-07</td><td>BT20/MB468</td></tr>
<tr><td>>hg38_21_+_8433172_8446622_RNA45SN1_45S_50ntFlanks@11211.11232.22|GCGACTCTGGACGCGAGCCGGG</td><td>>hg38_21_+_8433172_8446622_RNA45SN1_45S_50ntFlanks</td><td>11211</td><td>11232</td><td>22</td><td>GCGACTCTGGACGCGAGCCGGG</td><td>577.6</td><td>-5.9</td><td>1.2</td><td>-5.0</td><td>4.6E-07</td><td>7.2E-07</td><td>MB231/MB468</td></tr>
<tr><td>>MI0000102|hsa-mir-100&WithFlank&11|-|122152223|122152314@19.39.21(+1C)&[MIMAT0000098&hsa-miR-100-5p&offsets|0|-1(+1C);m-27&11|-|122152276|122152296&offsets|0|0(+1C)]|AACCCGTAGATCCGAACTTGTC</td><td>>MI0000102|hsa-mir-100&WithFlank&11|-|122152223|122152314</td><td>19</td><td>39</td><td>21(+1C)</td><td>AACCCGTAGATCCGAACTTGTC</td><td>2916.3</td><td>-5.3</td><td>1.0</td><td>-5.5</td><td>3.6E-08</td><td>7.2E-07</td><td>BT20/MB231</td></tr>
<tr><td>>hg38_21_+_8433172_8446622_RNA45SN1_45S_50ntFlanks@6731.6771.41|CACATTGATCATCGACACTTCGAACGCACTTGCGGCCCCGG</td><td>>hg38_21_+_8433172_8446622_RNA45SN1_45S_50ntFlanks</td><td>6731</td><td>6771</td><td>41</td><td>CACATTGATCATCGACACTTCGAACGCACTTGCGGCCCCGG</td><td>351.7</td><td>-3.3</td><td>0.6</td><td>-5.0</td><td>4.6E-07</td><td>7.3E-07</td><td>MB231/MB468</td></tr>
<tr><td>>hg38_21_+_8433172_8446622_RNA45SN1_45S_50ntFlanks@7980.8005.26|CCTCAGATCAGACGTGGCGACCCGCT</td><td>>hg38_21_+_8433172_8446622_RNA45SN1_45S_50ntFlanks</td><td>7980</td><td>8005</td><td>26</td><td>CCTCAGATCAGACGTGGCGACCCGCT</td><td>150.2</td><td>-2.7</td><td>0.5</td><td>-5.0</td><td>4.6E-07</td><td>7.3E-07</td><td>MB231/MB468</td></tr>
<tr><td>>hg38_21_+_8433172_8446622_RNA45SN1_45S_50ntFlanks@6670.6696.27|ACTCGGCTCGTGCGTCGATGAAGAACG</td><td>>hg38_21_+_8433172_8446622_RNA45SN1_45S_50ntFlanks</td><td>6670</td><td>6696</td><td>27</td><td>ACTCGGCTCGTGCGTCGATGAAGAACG</td><td>381.0</td><td>2.2</td><td>0.4</td><td>5.0</td><td>4.7E-07</td><td>7.3E-07</td><td>MB231/MB468</td></tr>
</table>

| | | | | | | | | | | | | |
|---|---|---|---|---|---|---|---|---|---|---|---|---|
| >am_trna11_ProAGG_16_+_3241989_3242060@1.28.28,trna12_ProAGG_6_+_26555498_26555569@1.28.28,trna12_ProTGG_11_-_75946869_75946940@1.28.28,trna14_ProTGG_5_-_180615854_180615925@1.28.28,trna22_ProAGG_14_-_21081560_21081631@1.28.28,trna23_ProAGG_14_-_21077495_21077566@1.28.28,trna28_ProTGG_16_-_3234133_3234204@1.28.28,trna29_ProAGG_16_-_3232635_3232706@1.28.28,trna2_ProAGG_7_+_128423504_128423575@1.28.28,trna30_ProCGG_6_+_27059521_27059592@1.28.28,trna37_ProCGG_17_-_8126151_8126222@1.28.28,trna3_ProTGG_16_+_3208923_3208994@1.28.28,trna52_ProCGG_1_+_167683962_167684033@1.28.28,trna65_ProAGG_1_-_167684725_167684796@1.28.28,trna6_ProCGG_16_+_3222049_3222120@1.28.28,trna6_ProTGG_14_+_21152175_21152246@1.28.28,trna8_ProTGG_16_+_3238094_3238165@1.28.28,trna9_ProAGG_11_+_75946557_75946628@1.28.28,trna9_ProAGG_16_+_3239634_3239705@1.28.28\|GGCTCGTTGGTCTAGGGGTATGATTCTC | >am_trna11_ProAGG_16_+_3241989_3242060 | 1 | 28 | 28 | GGCTCGTTGGTCTAGGGGTATGATTCTC | 615.7 | -6.0 | 1.2 | -5.0 | 5.2E-07 | 7.4E-07 | BT20/MB468 |
| >hg38_21_+_8433172_8446622_RNA45SN1_45S_50ntFlanks@10458.10486.29\|ATGGCCTCCGTTGCCCTCGGCCGATCGAA | >hg38_21_+_8433172_8446622_RNA45SN1_45S_50ntFlanks | 10458 | 10486 | 29 | ATGGCCTCCGTTGCCCTCGGCCGATCGAA | 154.3 | -3.7 | 0.7 | -5.0 | 5.2E-07 | 7.4E-07 | BT20/MB468 |
| >MI0000749\|hsa-mir-30e&WithFlank&1\|+\|40754349\|40754452@23.45.23&[MIMAT0000692&hsa-miR-30e-5p&offsets\|0\|+1;m-24&1\|+\|40754371\|40754394&offsets\|0\|-1]\|TGTAAACATCCTTGACTGGAAGC | >MI0000749\|hsa-mir-30e&WithFlank&1\|+\|40754349\|40754452 | 23 | 45 | 23 | TGTAAACATCCTTGACTGGAAGC | 1675.1 | 5.5 | 1.1 | 5.0 | 5.2E-07 | 7.4E-07 | BT20/MB468 |
| >hg38_21_+_8433172_8446622_RNA45SN1_45S_50ntFlanks@4006.4042.37\|ACCTCGGGCCGATCGCACGCCCCCCGTGGCGGCGACG | >hg38_21_+_8433172_8446622_RNA45SN1_45S_50ntFlanks | 4006 | 4042 | 37 | ACCTCGGGCCGATCGCACGCCCCCCGTGGCGGCGACG | 191.6 | -2.4 | 0.5 | -5.0 | 4.7E-07 | 7.4E-07 | MB231/MB468 |
| >hg38_21_+_8433172_8446622_RNA45SN1_45S_50ntFlanks@8195.8215.21\|CCGTGGACGGTGTGAGGCCGG | >hg38_21_+_8433172_8446622_RNA45SN1_45S_50ntFlanks | 8195 | 8215 | 21 | CCGTGGACGGTGTGAGGCCGG | 127.7 | -2.7 | 0.5 | -5.0 | 5.3E-07 | 7.5E-07 | BT20/MB468 |
| >hg38_21_+_8433172_8446622_RNA45SN1_45S_50ntFlanks@4730.4759.30\|CAAGAACGAAAGTCGGAGGTTCGAAGACGA | >hg38_21_+_8433172_8446622_RNA45SN1_45S_50ntFlanks | 4730 | 4759 | 30 | CAAGAACGAAAGTCGGAGGTTCGAAGACGA | 305.7 | -3.0 | 0.6 | -5.0 | 5.3E-07 | 7.5E-07 | BT20/MB468 |
| >hg38_21_+_8433172_8446622_RNA45SN1_45S_50ntFlanks@4731.4759.29\|AAGAACGAAAGTCGGAGGTTCGAAGACGA | >hg38_21_+_8433172_8446622_RNA45SN1_45S_50ntFlanks | 4731 | 4759 | 29 | AAGAACGAAAGTCGGAGGTTCGAAGACGA | 842.7 | -2.6 | 0.5 | -5.0 | 5.3E-07 | 7.6E-07 | BT20/MB468 |
| >hg38_21_+_8433172_8446622_RNA45SN1_45S_50ntFlanks@11703.11726.24\|AATGAAGCGCGGGTAAACGGCGGG | >hg38_21_+_8433172_8446622_RNA45SN1_45S_50ntFlanks | 11703 | 11726 | 24 | AATGAAGCGCGGGTAAACGGCGGG | 200.3 | -2.1 | 0.4 | -5.0 | 4.8E-07 | 7.6E-07 | MB231/MB468 |
| >hg38_21_+_8433172_8446622_RNA45SN1_45S_50ntFlanks@11580.11601.22\|GAACTGGTGCGGACCAGGGGAA | >hg38_21_+_8433172_8446622_RNA45SN1_45S_50ntFlanks | 11580 | 11601 | 22 | GAACTGGTGCGGACCAGGGGAA | 811.0 | -3.8 | 0.8 | -5.0 | 5.4E-07 | 7.6E-07 | BT20/MB468 |
| >hg38_21_+_8433172_8446622_RNA45SN1_45S_50ntFlanks@475.519.45\|CTTCGTGATCGATGTGGTGACGTCGTGCTCTCCCGGGCCGGGTCC | >hg38_21_+_8433172_8446622_RNA45SN1_45S_50ntFlanks | 475 | 519 | 45 | CTTCGTGATCGATGTGGTGACGTCGTGCTCTCCCGGGCCGGGTCC | 1061.6 | 11.1 | 2.2 | 5.0 | 4.8E-07 | 7.6E-07 | MB231/MB468 |
| >hg38_21_+_8433172_8446622_RNA45SN1_45S_50ntFlanks@10786.10826.41\|AACAGCCTCTGGCATGTTGGAACAATGTAGGTAAGGGAAGT | >hg38_21_+_8433172_8446622_RNA45SN1_45S_50ntFlanks | 10786 | 10826 | 41 | AACAGCCTCTGGCATGTTGGAACAATGTAGGTAAGGGAAGT | 366.7 | -2.7 | 0.5 | -5.0 | 4.9E-07 | 7.7E-07 | MB231/MB468 |
| >hg38_21_+_8433172_8446622_RNA45SN1_45S_50ntFlanks@3783.3801.19\|AGTGAAACTGCGAATGGCT | >hg38_21_+_8433172_8446622_RNA45SN1_45S_50ntFlanks | 3783 | 3801 | 19 | AGTGAAACTGCGAATGGCT | 298.6 | -3.0 | 0.6 | -5.0 | 5.5E-07 | 7.7E-07 | BT20/MB468 |
| >hg38_21_+_8433172_8446622_RNA45SN1_45S_50ntFlanks@5492.5514.23\|AGAAGACGGTCGAACTTGACTAT | >hg38_21_+_8433172_8446622_RNA45SN1_45S_50ntFlanks | 5492 | 5514 | 23 | AGAAGACGGTCGAACTTGACTAT | 1026.9 | -3.9 | 0.8 | -5.0 | 5.0E-07 | 7.8E-07 | MB231/MB468 |
| >MI0000749\|hsa-mir-30e&WithFlank&1\|+\|40754349\|40754452@23.46.24&[MIMAT0000692&hsa-miR-30e-5p&offsets\|0\|+2;m-24&1\|+\|40754371\|40754394&offsets\|0\|0]\|TGTAAACATCCTTGACTGGAAGCT | >MI0000749\|hsa-mir-30e&WithFlank&1\|+\|40754349\|40754452 | 23 | 46 | 24 | TGTAAACATCCTTGACTGGAAGCT | 21943.2 | 4.8 | 1.0 | 5.0 | 5.6E-07 | 7.9E-07 | BT20/MB468 |
| >MI0000289\|hsa-mir-181a-1&WithFlank&1\|-\|198859038\|198859159@30.54.25;MI0000269\|hsa-mir-181a-2&WithFlank&9\|+\|124692436\|124692557@45.69.25&[MIMAT0000256&hsa-miR-181a-5p&offsets\|0\|+2;m-32&1\|-\|198859108\|198859130&offsets\|0\|+2];[MIMAT0000256_1&hsa-miR-181a-5p&offsets\|0\|+2;m-32&9\|+\|124692480\|124692502&offsets\|0\|+2]\|AACATTCAACGCTGTCGGTGAGTTT | >MI0000289\|hsa-mir-181a-1&WithFlank&1\|-\|198859038\|198859159 | 30 | 54 | 25 | AACATTCAACGCTGTCGGTGAGTTT | 888.9 | 5.0 | 1.0 | 5.0 | 5.6E-07 | 7.9E-07 | BT20/MB468 |
| >hg38_21_+_8433172_8446622_RNA45SN1_45S_50ntFlanks@10361.10383.23\|GAAGGCCGAAGTGGAGAAGGGTT | >hg38_21_+_8433172_8446622_RNA45SN1_45S_50ntFlanks | 10361 | 10383 | 23 | GAAGGCCGAAGTGGAGAAGGGTT | 296.1 | -2.1 | 0.4 | -5.0 | 5.1E-07 | 8.0E-07 | MB231/MB468 |
| >hg38_21_+_8433172_8446622_RNA45SN1_45S_50ntFlanks@7975.7992.18\|CGCGACCTCAGATCAGAC | >hg38_21_+_8433172_8446622_RNA45SN1_45S_50ntFlanks | 7975 | 7992 | 18 | CGCGACCTCAGATCAGAC | 12232.0 | 4.4 | 0.9 | 5.0 | 5.6E-07 | 8.0E-07 | BT20/MB468 |
| >hg38_21_+_8433172_8446622_RNA45SN1_45S_50ntFlanks@10350.10390.41\|ACGAGAACTTTGAAGGCCGAAGTGGAGAAGGGTTCCATGTG | >hg38_21_+_8433172_8446622_RNA45SN1_45S_50ntFlanks | 10350 | 10390 | 41 | ACGAGAACTTTGAAGGCCGAAGTGGAGAAGGGTTCCATGTG | 1485.9 | -4.1 | 0.8 | -5.0 | 5.7E-07 | 8.0E-07 | BT20/MB468 |
| >hg38_21_+_8433172_8446622_RNA45SN1_45S_50ntFlanks@8197.8215.19\|GTGGACGGTGTGAGGCCGG | >hg38_21_+_8433172_8446622_RNA45SN1_45S_50ntFlanks | 8197 | 8215 | 19 | GTGGACGGTGTGAGGCCGG | 162.0 | -3.1 | 0.6 | -5.0 | 5.7E-07 | 8.0E-07 | BT20/MB468 |
| >hg38_21_+_8433172_8446622_RNA45SN1_45S_50ntFlanks@13007.13025.19\|TATTGAAAGTCAGCCCTCG | >hg38_21_+_8433172_8446622_RNA45SN1_45S_50ntFlanks | 13007 | 13025 | 19 | TATTGAAAGTCAGCCCTCG | 617.2 | -3.6 | 0.7 | -5.0 | 5.1E-07 | 8.0E-07 | MB231/MB468 |
| >hg38_21_+_8433172_8446622_RNA45SN1_45S_50ntFlanks@10829.10853.25\|GCAAGCCGGATCCGTAACTTCGGGA | >hg38_21_+_8433172_8446622_RNA45SN1_45S_50ntFlanks | 10829 | 10853 | 25 | GCAAGCCGGATCCGTAACTTCGGGA | 173.5 | -3.1 | 0.6 | -5.0 | 5.7E-07 | 8.1E-07 | BT20/MB468 |
| >hg38_21_+_8433172_8446622_RNA45SN1_45S_50ntFlanks@11314.11344.31\|CCCACCCCACGTCTCGTCGCGCGCGCGTCCG | >hg38_21_+_8433172_8446622_RNA45SN1_45S_50ntFlanks | 11314 | 11344 | 31 | CCCACCCCACGTCTCGTCGCGCGCGCGTCCG | 263.0 | -5.1 | 1.0 | -5.0 | 5.8E-07 | 8.1E-07 | BT20/MB468 |
| >hg38_21_+_8433172_8446622_RNA45SN1_45S_50ntFlanks@5515.5535.21\|CTAGAGGAAGTAAAAGTCGTA | >hg38_21_+_8433172_8446622_RNA45SN1_45S_50ntFlanks | 5515 | 5535 | 21 | CTAGAGGAAGTAAAAGTCGTA | 300.6 | -3.7 | 0.7 | -5.0 | 5.8E-07 | 8.2E-07 | BT20/MB468 |
| >MI0000253\|hsa-mir-148a&WithFlank&7\|-\|25949913\|25949992@50.70.21(+1G)&[MIMAT0000243&hsa-miR-148a-3p&offsets\|0\|-1(+1G);m-7&7\|-\|25949922\|25949943&offsets\|0\|-1(+1G)]\|TCAGTGCACTACAGAACTTTGG | >MI0000253\|hsa-mir-148a&WithFlank&7\|-\|25949913\|25949992 | 50 | 70 | 21(+1G) | TCAGTGCACTACAGAACTTTGG | 22421.7 | 5.0 | 0.9 | 5.5 | 4.1E-08 | 8.3E-07 | BT20/MB231 |
| >hg38_21_+_8433172_8446622_RNA45SN1_45S_50ntFlanks@11587.11605.19\|TGCGGACCAGGGGAATCCG | >hg38_21_+_8433172_8446622_RNA45SN1_45S_50ntFlanks | 11587 | 11605 | 19 | TGCGGACCAGGGGAATCCG | 120.4 | -4.2 | 0.8 | -5.0 | 5.3E-07 | 8.3E-07 | MB231/MB468 |
| >hg38_21_+_8433172_8446622_RNA45SN1_45S_50ntFlanks@12934.12961.28\|CTAAACCATTCGTAGACGACCTGCTTCT | >hg38_21_+_8433172_8446622_RNA45SN1_45S_50ntFlanks | 12934 | 12961 | 28 | CTAAACCATTCGTAGACGACCTGCTTCT | 379.4 | -2.8 | 0.6 | -5.0 | 5.3E-07 | 8.4E-07 | MB231/MB468 |
| >hg38_21_+_8433172_8446622_RNA45SN1_45S_50ntFlanks@4759.4779.21\|ATCAGATACCGTCGTAGTTCC | >hg38_21_+_8433172_8446622_RNA45SN1_45S_50ntFlanks | 4759 | 4779 | 21 | ATCAGATACCGTCGTAGTTCC | 505.4 | -5.1 | 1.0 | -5.0 | 5.4E-07 | 8.4E-07 | MB231/MB468 |
| >hg38_21_+_8433172_8446622_RNA45SN1_45S_50ntFlanks@12419.12436.18\|GATGTCGGCTCTTCCTAT | >hg38_21_+_8433172_8446622_RNA45SN1_45S_50ntFlanks | 12419 | 12436 | 18 | GATGTCGGCTCTTCCTAT | 1859.2 | -2.9 | 0.6 | -5.0 | 6.0E-07 | 8.4E-07 | BT20/MB468 |
| >hg38_21_+_8433172_8446622_RNA45SN1_45S_50ntFlanks@7991.8012.22\|ACGTGGCGACCCGCTGAATTTA | >hg38_21_+_8433172_8446622_RNA45SN1_45S_50ntFlanks | 7991 | 8012 | 22 | ACGTGGCGACCCGCTGAATTTA | 2989.7 | -3.8 | 0.8 | -5.0 | 6.0E-07 | 8.5E-07 | BT20/MB468 |
| >am_trna10_LysCTT_16_+_3241501_3241573@11.28.18,trna119_LysCTT_1_-_145395522_145395594@11.28.18,trna11_LysCTT_5_-_180648979_180649051@11.28.18,trna128_LysCTT_1_-_55423542_55423614@11.28.18,trna13_LysCTT_14_-_58706613_58706685@11.28.18,trna13_LysCTT_6_+_26556774_26556846@11.28.18,trna2_LysCTT_15_+_79152904_79152976@11.28.18,trna32_LysCTT_16_-_3207406_3207478@11.28.18,trna4_LysCTT_18_-_43669271_43669343@11.28.18,trna6_LysCTT_19_-_52425396_52425468@11.28.18,trna7_LysCTT_16_+_3225692_3225764@11.28.18,trna9_LysCTT_5_+_180634755_180634827@11.28.18\|CTCAGTCGGTAGAGCATG | >am_trna10_LysCTT_16_+_3241501_3241573 | 11 | 28 | 18 | CTCAGTCGGTAGAGCATG | 488.5 | -3.3 | 0.7 | -5.0 | 6.0E-07 | 8.5E-07 | BT20/MB468 |
| >hg38_21_+_8433172_8446622_RNA45SN1_45S_50ntFlanks@9534.9558.25\|AGGAAACTCTGGTGGAGGTCCGTAG | >hg38_21_+_8433172_8446622_RNA45SN1_45S_50ntFlanks | 9534 | 9558 | 25 | AGGAAACTCTGGTGGAGGTCCGTAG | 289.8 | -3.3 | 0.7 | -5.0 | 5.5E-07 | 8.6E-07 | MB231/MB468 |
| >hg38_21_+_8433172_8446622_RNA45SN1_45S_50ntFlanks@12540.12569.30\|TGATGTGTTGTTGCCATGGTAATCCTGCTC | >hg38_21_+_8433172_8446622_RNA45SN1_45S_50ntFlanks | 12540 | 12569 | 30 | TGATGTGTTGTTGCCATGGTAATCCTGCTC | 83.6 | -4.3 | 0.9 | -5.0 | 5.5E-07 | 8.7E-07 | MB231/MB468 |
| >hg38_21_+_8433172_8446622_RNA45SN1_45S_50ntFlanks@12407.12432.26\|TTTTGATCCTTCGATGTCGGCTCTTC | >hg38_21_+_8433172_8446622_RNA45SN1_45S_50ntFlanks | 12407 | 12432 | 26 | TTTTGATCCTTCGATGTCGGCTCTTC | 325.1 | -3.0 | 0.6 | -5.0 | 6.1E-07 | 8.7E-07 | BT20/MB468 |
| >hg38_21_+_8433172_8446622_RNA45SN1_45S_50ntFlanks@3985.4024.40\|CGGCTTTGGTGACTCTAGATAACCTCGGGCCGATCGCACG | >hg38_21_+_8433172_8446622_RNA45SN1_45S_50ntFlanks | 3985 | 4024 | 40 | CGGCTTTGGTGACTCTAGATAACCTCGGGCCGATCGCACG | 349.8 | -6.5 | 1.3 | -5.0 | 6.2E-07 | 8.7E-07 | BT20/MB468 |
| >trnaMT_ValTAC_MT_+_1602_1670@46.72.27\|TTCAACTTAACTTGACCGCTCTGACCA | >trnaMT_ValTAC_MT_+_1602_1670 | 46 | 72 | 27 | TTCAACTTAACTTGACCGCTCTGACCA | 593.2 | 10.3 | 2.0 | 5.0 | 5.6E-07 | 8.7E-07 | MB231/MB468 |
| >hg38_21_+_8433172_8446622_RNA45SN1_45S_50ntFlanks@10798.10822.25\|CATGTTGGAACAATGTAGGTAAGGG | >hg38_21_+_8433172_8446622_RNA45SN1_45S_50ntFlanks | 10798 | 10822 | 25 | CATGTTGGAACAATGTAGGTAAGGG | 194.5 | -2.5 | 0.5 | -5.0 | 6.2E-07 | 8.7E-07 | BT20/MB468 |
| >hg38_21_+_8433172_8446622_RNA45SN1_45S_50ntFlanks@4730.4751.22\|CAAGAACGAAAGTCGGAGGTTC | >hg38_21_+_8433172_8446622_RNA45SN1_45S_50ntFlanks | 4730 | 4751 | 22 | CAAGAACGAAAGTCGGAGGTTC | 396.6 | -2.0 | 0.4 | -5.0 | 6.3E-07 | 8.8E-07 | BT20/MB468 |
| >hg38_21_+_8433172_8446622_RNA45SN1_45S_50ntFlanks@11671.11703.33\|CCAGTGCTCTGAATGTCAAAGTGAAGAAATTCA | >hg38_21_+_8433172_8446622_RNA45SN1_45S_50ntFlanks | 11671 | 11703 | 33 | CCAGTGCTCTGAATGTCAAAGTGAAGAAATTCA | 501.2 | -3.4 | 0.7 | -5.0 | 6.3E-07 | 8.8E-07 | BT20/MB468 |
| >hg38_21_+_8433172_8446622_RNA45SN1_45S_50ntFlanks@12158.12180.23\|GTTTGACTGGGGCGGTACACCTG | >hg38_21_+_8433172_8446622_RNA45SN1_45S_50ntFlanks | 12158 | 12180 | 23 | GTTTGACTGGGGCGGTACACCTG | 93.0 | -5.3 | 1.1 | -5.0 | 5.6E-07 | 8.8E-07 | MB231/MB468 |

<table>
<tr><td>>MI0000085|hsa-mir-27a&WithFlank&19|-|13836434|13836523@57.76.20(+1U)&[MIMAT0000084&hsa-miR-27a-3p&offsets|0|-1(+1U);m-43&19|-|13836447|13836467&offsets|0|-1(+1U)]|TTCACAGTGGCTAAGTTCCGT</td><td>>MI0000085|hsa-mir-27a&WithFlank&19|-|13836434|13836523</td><td>57</td><td>76</td><td>20(+1U)</td><td>TTCACAGTGGCTAAGTTCCGT</td><td>8654.1</td><td>4.5</td><td>0.9</td><td>5.0</td><td>6.4E-07</td><td>8.9E-07</td><td>BT20/MB468</td></tr>
<tr><td>>hg38_21_+_8433172_8446622_RNA45SN1_45S_50ntFlanks@7977.8012.36|CGACCTCAGATCAGACGTGGCGACCCGCTGAATTTA</td><td>>hg38_21_+_8433172_8446622_RNA45SN1_45S_50ntFlanks</td><td>7977</td><td>8012</td><td>36</td><td>CGACCTCAGATCAGACGTGGCGACCCGCTGAATTTA</td><td>734.5</td><td>4.0</td><td>0.8</td><td>5.0</td><td>5.7E-07</td><td>9.0E-07</td><td>MB231/MB468</td></tr>
<tr><td>>hg38_21_+_8433172_8446622_RNA45SN1_45S_50ntFlanks@13007.13027.21|TATTGAAAGTCAGCCCTCGAC</td><td>>hg38_21_+_8433172_8446622_RNA45SN1_45S_50ntFlanks</td><td>13007</td><td>13027</td><td>21</td><td>TATTGAAAGTCAGCCCTCGAC</td><td>512.8</td><td>-4.7</td><td>1.0</td><td>-5.0</td><td>6.4E-07</td><td>9.0E-07</td><td>BT20/MB468</td></tr>
<tr><td>>hg38_21_+_8433172_8446622_RNA45SN1_45S_50ntFlanks@9574.9597.24|AATCGGTCGTCCGACCTGGGTATA</td><td>>hg38_21_+_8433172_8446622_RNA45SN1_45S_50ntFlanks</td><td>9574</td><td>9597</td><td>24</td><td>AATCGGTCGTCCGACCTGGGTATA</td><td>4794.8</td><td>-3.7</td><td>0.7</td><td>-5.0</td><td>5.8E-07</td><td>9.0E-07</td><td>MB231/MB468</td></tr>
<tr><td>>hg38_21_+_8433172_8446622_RNA45SN1_45S_50ntFlanks@10466.10488.23|CGTTGCCCTCGGCCGATCGAAAG</td><td>>hg38_21_+_8433172_8446622_RNA45SN1_45S_50ntFlanks</td><td>10466</td><td>10488</td><td>23</td><td>CGTTGCCCTCGGCCGATCGAAAG</td><td>209.8</td><td>-4.7</td><td>0.9</td><td>-5.0</td><td>6.5E-07</td><td>9.1E-07</td><td>BT20/MB468</td></tr>
<tr><td>>hg38_21_+_8433172_8446622_RNA45SN1_45S_50ntFlanks@11729.11750.22|TAACTATGACTCTCTTAAGGTA</td><td>>hg38_21_+_8433172_8446622_RNA45SN1_45S_50ntFlanks</td><td>11729</td><td>11750</td><td>22</td><td>TAACTATGACTCTCTTAAGGTA</td><td>1165.0</td><td>-2.8</td><td>0.6</td><td>-5.0</td><td>5.8E-07</td><td>9.1E-07</td><td>MB231/MB468</td></tr>
<tr><td>>hg38_21_+_8433172_8446622_RNA45SN1_45S_50ntFlanks@10679.10706.28|GGTTCCGGCGGCGTCCGGTGAGCTCTCG</td><td>>hg38_21_+_8433172_8446622_RNA45SN1_45S_50ntFlanks</td><td>10679</td><td>10706</td><td>28</td><td>GGTTCCGGCGGCGTCCGGTGAGCTCTCG</td><td>4855.3</td><td>-5.1</td><td>1.0</td><td>-5.0</td><td>6.5E-07</td><td>9.2E-07</td><td>BT20/MB468</td></tr>
<tr><td>>hg38_21_+_8433172_8446622_RNA45SN1_45S_50ntFlanks@4772.4793.22|GTAGTTCCGACCATAAACGATG</td><td>>hg38_21_+_8433172_8446622_RNA45SN1_45S_50ntFlanks</td><td>4772</td><td>4793</td><td>22</td><td>GTAGTTCCGACCATAAACGATG</td><td>192.7</td><td>-3.4</td><td>0.7</td><td>-5.0</td><td>6.6E-07</td><td>9.3E-07</td><td>BT20/MB468</td></tr>
<tr><td>>hg38_21_+_8433172_8446622_RNA45SN1_45S_50ntFlanks@11315.11344.30|CCACCCCACGTCTCGTCGCGCGCGCGTCCG</td><td>>hg38_21_+_8433172_8446622_RNA45SN1_45S_50ntFlanks</td><td>11315</td><td>11344</td><td>30</td><td>CCACCCCACGTCTCGTCGCGCGCGCGTCCG</td><td>1146.9</td><td>-6.3</td><td>1.3</td><td>-5.0</td><td>6.6E-07</td><td>9.3E-07</td><td>BT20/MB468</td></tr>
<tr><td>>hg38_21_+_8433172_8446622_RNA45SN1_45S_50ntFlanks@12475.12497.23|CCCACTAATAGGGAACGTGAGCT</td><td>>hg38_21_+_8433172_8446622_RNA45SN1_45S_50ntFlanks</td><td>12475</td><td>12497</td><td>23</td><td>CCCACTAATAGGGAACGTGAGCT</td><td>1011.2</td><td>3.2</td><td>0.6</td><td>5.0</td><td>6.0E-07</td><td>9.4E-07</td><td>MB231/MB468</td></tr>
<tr><td>>trna10_GlyTCC_17_+_8124866_8124937@1.29.29|GCGTTGGTGGTATAGTGGTAAGCATAGCT</td><td>>trna10_GlyTCC_17_+_8124866_8124937</td><td>1</td><td>29</td><td>29</td><td>GCGTTGGTGGTATAGTGGTAAGCATAGCT</td><td>112.1</td><td>-3.7</td><td>0.8</td><td>-5.0</td><td>6.8E-07</td><td>9.5E-07</td><td>BT20/MB468</td></tr>
<tr><td>>hg38_21_+_8433172_8446622_RNA45SN1_45S_50ntFlanks@11781.11805.25|CGCATGAATGGATGAACGAGATTCC</td><td>>hg38_21_+_8433172_8446622_RNA45SN1_45S_50ntFlanks</td><td>11781</td><td>11805</td><td>25</td><td>CGCATGAATGGATGAACGAGATTCC</td><td>421.7</td><td>-3.5</td><td>0.7</td><td>-5.0</td><td>6.1E-07</td><td>9.5E-07</td><td>MB231/MB468</td></tr>
<tr><td>>MI0003628|hsa-mir-615&WithFlank&12|+|54033944|54034051@67.85.19&[MIMAT0003283&hsa-miR-615-3p&offsets|0|-3;m-495&12|+|54034010|54034030&offsets|0|-2]|TCCGAGCCTGGGTCTCCCT</td><td>>MI0003628|hsa-mir-615&WithFlank&12|+|54033944|54034051</td><td>67</td><td>85</td><td>19</td><td>TCCGAGCCTGGGTCTCCCT</td><td>278.2</td><td>4.9</td><td>0.9</td><td>5.5</td><td>4.8E-08</td><td>9.6E-07</td><td>BT20/MB231</td></tr>
<tr><td>>hg38_21_+_8433172_8446622_RNA45SN1_45S_50ntFlanks@6721.6760.40|AATTGCAGGACACATTGATCATCGACACTTCGAACGCACT</td><td>>hg38_21_+_8433172_8446622_RNA45SN1_45S_50ntFlanks</td><td>6721</td><td>6760</td><td>40</td><td>AATTGCAGGACACATTGATCATCGACACTTCGAACGCACT</td><td>211.2</td><td>-1.8</td><td>0.4</td><td>-5.0</td><td>6.8E-07</td><td>9.6E-07</td><td>BT20/MB468</td></tr>
<tr><td>>hg38_21_+_8433172_8446622_RNA45SN1_45S_50ntFlanks@9551.9568.18|GTCCGTAGCGGTCCTGAC</td><td>>hg38_21_+_8433172_8446622_RNA45SN1_45S_50ntFlanks</td><td>9551</td><td>9568</td><td>18</td><td>GTCCGTAGCGGTCCTGAC</td><td>277.8</td><td>8.0</td><td>1.6</td><td>5.0</td><td>6.1E-07</td><td>9.6E-07</td><td>MB231/MB468</td></tr>
<tr><td>>hg38_1_-_228634819_228635039_RNA5S12_5S_50ntFlanks@132.169.38|GATGGGAGACCGCCTGGGAATACCGGGTGCTGTAGGCT</td><td>>hg38_1_-_228634819_228635039_RNA5S12_5S_50ntFlanks</td><td>132</td><td>169</td><td>38</td><td>GATGGGAGACCGCCTGGGAATACCGGGTGCTGTAGGCT</td><td>2435.2</td><td>3.9</td><td>0.8</td><td>5.0</td><td>6.2E-07</td><td>9.6E-07</td><td>MB231/MB468</td></tr>
<tr><td>>MI0000102|hsa-mir-100&WithFlank&11|-|122152223|122152314@19.39.21(+1A)&[MIMAT0000098&hsa-miR-100-5p&offsets|0|-1(+1A);m-27&11|-|122152276|122152296&offsets|0|0(+1A)]|AACCCGTAGATCCGAACTTGTA</td><td>>MI0000102|hsa-mir-100&WithFlank&11|-|122152223|122152314</td><td>19</td><td>39</td><td>21(+1A)</td><td>AACCCGTAGATCCGAACTTGTA</td><td>91838.1</td><td>-5.0</td><td>0.9</td><td>-5.5</td><td>4.9E-08</td><td>9.8E-07</td><td>BT20/MB231</td></tr>
<tr><td>>hg38_21_+_8433172_8446622_RNA45SN1_45S_50ntFlanks@8170.8195.26|CAAGTCCTTCTGATCGAGGCCCAGCC</td><td>>hg38_21_+_8433172_8446622_RNA45SN1_45S_50ntFlanks</td><td>8170</td><td>8195</td><td>26</td><td>CAAGTCCTTCTGATCGAGGCCCAGCC</td><td>739.4</td><td>-3.2</td><td>0.6</td><td>-5.0</td><td>6.3E-07</td><td>9.8E-07</td><td>MB231/MB468</td></tr>
<tr><td>>hg38_21_+_8433172_8446622_RNA45SN1_45S_50ntFlanks@8176.8194.19|CTTCTGATCGAGGCCCAGC</td><td>>hg38_21_+_8433172_8446622_RNA45SN1_45S_50ntFlanks</td><td>8176</td><td>8194</td><td>19</td><td>CTTCTGATCGAGGCCCAGC</td><td>455.7</td><td>3.2</td><td>0.6</td><td>5.0</td><td>6.3E-07</td><td>9.8E-07</td><td>MB231/MB468</td></tr>
<tr><td>>hg38_21_+_8433172_8446622_RNA45SN1_45S_50ntFlanks@6733.6774.42|CATTGATCATCGACACTTCGAACGCACTTGCGGCCCCGGGTT</td><td>>hg38_21_+_8433172_8446622_RNA45SN1_45S_50ntFlanks</td><td>6733</td><td>6774</td><td>42</td><td>CATTGATCATCGACACTTCGAACGCACTTGCGGCCCCGGGTT</td><td>797.8</td><td>10.8</td><td>2.2</td><td>5.0</td><td>6.4E-07</td><td>9.9E-07</td><td>MB231/MB468</td></tr>
<tr><td>>hg38_21_+_8433172_8446622_RNA45SN1_45S_50ntFlanks@6661.6706.46|CGGTGGATCACTCGGCTCGTGCGTCGATGAAGAACGCAGCTAGCTG</td><td>>hg38_21_+_8433172_8446622_RNA45SN1_45S_50ntFlanks</td><td>6661</td><td>6706</td><td>46</td><td>CGGTGGATCACTCGGCTCGTGCGTCGATGAAGAACGCAGCTAGCTG</td><td>1119.3</td><td>4.3</td><td>0.9</td><td>5.0</td><td>7.1E-07</td><td>9.9E-07</td><td>BT20/MB468</td></tr>
<tr><td>>hg38_21_+_8433172_8446622_RNA45SN1_45S_50ntFlanks@12274.12293.20|CAGTACGAATACAGACCGTG</td><td>>hg38_21_+_8433172_8446622_RNA45SN1_45S_50ntFlanks</td><td>12274</td><td>12293</td><td>20</td><td>CAGTACGAATACAGACCGTG</td><td>93.0</td><td>-3.6</td><td>0.7</td><td>-5.0</td><td>6.4E-07</td><td>1.0E-06</td><td>MB231/MB468</td></tr>
<tr><td>>hg38_21_+_8433172_8446622_RNA45SN1_45S_50ntFlanks@6664.6689.26|TGGATCACTCGGCTCGTGCGTCGATG</td><td>>hg38_21_+_8433172_8446622_RNA45SN1_45S_50ntFlanks</td><td>6664</td><td>6689</td><td>26</td><td>TGGATCACTCGGCTCGTGCGTCGATG</td><td>193.9</td><td>-2.8</td><td>0.6</td><td>-5.0</td><td>6.5E-07</td><td>1.0E-06</td><td>MB231/MB468</td></tr>
<tr><td>>hg38_1_-_228634819_228635039_RNA5S12_5S_50ntFlanks@148.170.23|GGAATACCGGGTGCTGTAGGCTT</td><td>>hg38_1_-_228634819_228635039_RNA5S12_5S_50ntFlanks</td><td>148</td><td>170</td><td>23</td><td>GGAATACCGGGTGCTGTAGGCTT</td><td>2566.1</td><td>3.8</td><td>0.8</td><td>5.0</td><td>6.5E-07</td><td>1.0E-06</td><td>MB231/MB468</td></tr>
<tr><td>>hg38_21_+_8433172_8446622_RNA45SN1_45S_50ntFlanks@6731.6752.22|CACATTGATCATCGACACTTCG</td><td>>hg38_21_+_8433172_8446622_RNA45SN1_45S_50ntFlanks</td><td>6731</td><td>6752</td><td>22</td><td>CACATTGATCATCGACACTTCG</td><td>425.0</td><td>-3.4</td><td>0.7</td><td>-5.0</td><td>7.2E-07</td><td>1.0E-06</td><td>BT20/MB468</td></tr>
<tr><td>>hg38_21_+_8433172_8446622_RNA45SN1_45S_50ntFlanks@10849.10868.20|CGGGATAAGGATTGGCTCTA</td><td>>hg38_21_+_8433172_8446622_RNA45SN1_45S_50ntFlanks</td><td>10849</td><td>10868</td><td>20</td><td>CGGGATAAGGATTGGCTCTA</td><td>531.9</td><td>-5.0</td><td>1.0</td><td>-5.0</td><td>6.5E-07</td><td>1.0E-06</td><td>MB231/MB468</td></tr>
<tr><td>>hg38_21_+_8433172_8446622_RNA45SN1_45S_50ntFlanks@12445.12472.28|AGCAGAATTCACCAAGCGTTGGATTGTT</td><td>>hg38_21_+_8433172_8446622_RNA45SN1_45S_50ntFlanks</td><td>12445</td><td>12472</td><td>28</td><td>AGCAGAATTCACCAAGCGTTGGATTGTT</td><td>114.1</td><td>-3.9</td><td>0.8</td><td>-5.0</td><td>7.3E-07</td><td>1.0E-06</td><td>BT20/MB468</td></tr>
<tr><td>>hg38_21_+_8433172_8446622_RNA45SN1_45S_50ntFlanks@12627.12661.35|ATGGGGCGAAGCTACCATCTGTGGGATTATGACTG</td><td>>hg38_21_+_8433172_8446622_RNA45SN1_45S_50ntFlanks</td><td>12627</td><td>12661</td><td>35</td><td>ATGGGGCGAAGCTACCATCTGTGGGATTATGACTG</td><td>779.8</td><td>-3.3</td><td>0.7</td><td>-5.0</td><td>6.6E-07</td><td>1.0E-06</td><td>MB231/MB468</td></tr>
<tr><td>>hg38_21_+_8433172_8446622_RNA45SN1_45S_50ntFlanks@12163.12183.21|ACTGGGGCGGTACACCTGTCA</td><td>>hg38_21_+_8433172_8446622_RNA45SN1_45S_50ntFlanks</td><td>12163</td><td>12183</td><td>21</td><td>ACTGGGGCGGTACACCTGTCA</td><td>403.1</td><td>-3.7</td><td>0.7</td><td>-5.0</td><td>6.6E-07</td><td>1.0E-06</td><td>MB231/MB468</td></tr>
<tr><td>>hg38_21_+_8433172_8446622_RNA45SN1_45S_50ntFlanks@12992.13015.24|TCCCTCGCTGCGATCTATTGAAAG</td><td>>hg38_21_+_8433172_8446622_RNA45SN1_45S_50ntFlanks</td><td>12992</td><td>13015</td><td>24</td><td>TCCCTCGCTGCGATCTATTGAAAG</td><td>350.3</td><td>-4.4</td><td>0.9</td><td>-5.0</td><td>7.4E-07</td><td>1.0E-06</td><td>BT20/MB468</td></tr>
<tr><td>>hg38_21_+_8433172_8446622_RNA45SN1_45S_50ntFlanks@9622.9648.27|CTAGTAGCTGGTTCCCTCCGAAGTTTC</td><td>>hg38_21_+_8433172_8446622_RNA45SN1_45S_50ntFlanks</td><td>9622</td><td>9648</td><td>27</td><td>CTAGTAGCTGGTTCCCTCCGAAGTTTC</td><td>91.9</td><td>-4.6</td><td>0.9</td><td>-5.0</td><td>7.4E-07</td><td>1.0E-06</td><td>BT20/MB468</td></tr>
<tr><td>>hg38_21_+_8433172_8446622_RNA45SN1_45S_50ntFlanks@12942.12966.25|TTCGTAGACGACCTGCTTCTGGGTC</td><td>>hg38_21_+_8433172_8446622_RNA45SN1_45S_50ntFlanks</td><td>12942</td><td>12966</td><td>25</td><td>TTCGTAGACGACCTGCTTCTGGGTC</td><td>516.6</td><td>-5.3</td><td>1.1</td><td>-5.0</td><td>6.6E-07</td><td>1.0E-06</td><td>MB231/MB468</td></tr>
<tr><td>>hg38_21_+_8433172_8446622_RNA45SN1_45S_50ntFlanks@10846.10868.23|CTTCGGGATAAGGATTGGCTCTA</td><td>>hg38_21_+_8433172_8446622_RNA45SN1_45S_50ntFlanks</td><td>10846</td><td>10868</td><td>23</td><td>CTTCGGGATAAGGATTGGCTCTA</td><td>717.6</td><td>-5.2</td><td>1.0</td><td>-5.0</td><td>6.7E-07</td><td>1.0E-06</td><td>MB231/MB468</td></tr>
<tr><td>>hg38_21_+_8433172_8446622_RNA45SN1_45S_50ntFlanks@12459.12484.26|AGCGTTGGATTGTTCACCCACTAATA</td><td>>hg38_21_+_8433172_8446622_RNA45SN1_45S_50ntFlanks</td><td>12459</td><td>12484</td><td>26</td><td>AGCGTTGGATTGTTCACCCACTAATA</td><td>147.3</td><td>-4.5</td><td>0.9</td><td>-5.0</td><td>6.7E-07</td><td>1.0E-06</td><td>MB231/MB468</td></tr>
<tr><td>>hg38_21_+_8433172_8446622_RNA45SN1_45S_50ntFlanks@3872.3899.28|CTAATACATGCCGACGGGCGCTGACCCC</td><td>>hg38_21_+_8433172_8446622_RNA45SN1_45S_50ntFlanks</td><td>3872</td><td>3899</td><td>28</td><td>CTAATACATGCCGACGGGCGCTGACCCC</td><td>147.6</td><td>-4.6</td><td>0.9</td><td>-5.0</td><td>7.4E-07</td><td>1.0E-06</td><td>BT20/MB468</td></tr>
<tr><td>>hg38_21_+_8433172_8446622_RNA45SN1_45S_50ntFlanks@12083.12108.26|CTCGCTTCTGGCGCCAAGCGCCCGGC</td><td>>hg38_21_+_8433172_8446622_RNA45SN1_45S_50ntFlanks</td><td>12083</td><td>12108</td><td>26</td><td>CTCGCTTCTGGCGCCAAGCGCCCGGC</td><td>239.1</td><td>-4.2</td><td>0.9</td><td>-4.9</td><td>7.5E-07</td><td>1.0E-06</td><td>BT20/MB468</td></tr>
<tr><td>>hg38_21_+_8433172_8446622_RNA45SN1_45S_50ntFlanks@6703.6720.18|GCTGCGAGAATTAATGTG</td><td>>hg38_21_+_8433172_8446622_RNA45SN1_45S_50ntFlanks</td><td>6703</td><td>6720</td><td>18</td><td>GCTGCGAGAATTAATGTG</td><td>187.9</td><td>-3.4</td><td>0.7</td><td>-4.9</td><td>7.5E-07</td><td>1.1E-06</td><td>BT20/MB468</td></tr>
<tr><td>>hg38_21_+_8433172_8446622_RNA45SN1_45S_50ntFlanks@4128.4158.31|CGATTCCGGAGAGGGAGCCTGAGAAACGGCT</td><td>>hg38_21_+_8433172_8446622_RNA45SN1_45S_50ntFlanks</td><td>4128</td><td>4158</td><td>31</td><td>CGATTCCGGAGAGGGAGCCTGAGAAACGGCT</td><td>90.9</td><td>-4.3</td><td>0.9</td><td>-4.9</td><td>7.5E-07</td><td>1.1E-06</td><td>BT20/MB468</td></tr>
<tr><td>>hg38_21_+_8433172_8446622_RNA45SN1_45S_50ntFlanks@11671.11688.18|CCAGTGCTCTGAATGTCA</td><td>>hg38_21_+_8433172_8446622_RNA45SN1_45S_50ntFlanks</td><td>11671</td><td>11688</td><td>18</td><td>CCAGTGCTCTGAATGTCA</td><td>681.4</td><td>-4.7</td><td>0.9</td><td>-4.9</td><td>7.5E-07</td><td>1.1E-06</td><td>BT20/MB468</td></tr>
<tr><td>>trna116_GluCTC_1_-_145399233_145399304@32.69.38,trna59_GluCTC_1_+_249168447_249168518@32.69.38,trna71_GluCTC_1_-_161439189_161439260@32.69.38,trna74_GluCTC_1_-_161431809_161431880@32.69.38,trna77_GluCTC_1_-_161424398_161424469@32.69.38,trna77_GluCTC_6_+_28949976_28950047@32.69.38,trna80_GluCTC_1_-_161417018_161417089@32.69.38,trna87_GluCTC_6_-_126101393_126101464@32.69.38|CTCTCACCGCCGCGGCCCGGGTTCGATTCCCGGTCAGG</td><td>>trna116_GluCTC_1_-_145399233_145399304</td><td>32</td><td>69</td><td>38</td><td>CTCTCACCGCCGCGGCCCGGGTTCGATTCCCGGTCAGG</td><td>153.1</td><td>-3.7</td><td>0.7</td><td>-4.9</td><td>7.6E-07</td><td>1.1E-06</td><td>BT20/MB468</td></tr>
<tr><td>>hg38_21_+_8433172_8446622_RNA45SN1_45S_50ntFlanks@12937.12955.19|AACCATTCGTAGACGACCT</td><td>>hg38_21_+_8433172_8446622_RNA45SN1_45S_50ntFlanks</td><td>12937</td><td>12955</td><td>19</td><td>AACCATTCGTAGACGACCT</td><td>244.3</td><td>-4.2</td><td>0.8</td><td>-4.9</td><td>7.6E-07</td><td>1.1E-06</td><td>BT20/MB468</td></tr>
<tr><td>>hg38_21_+_8433172_8446622_RNA45SN1_45S_50ntFlanks@3705.3725.21|TACCTGGTTGATCCTGCCAGT</td><td>>hg38_21_+_8433172_8446622_RNA45SN1_45S_50ntFlanks</td><td>3705</td><td>3725</td><td>21</td><td>TACCTGGTTGATCCTGCCAGT</td><td>8850.4</td><td>4.4</td><td>0.8</td><td>5.4</td><td>5.4E-08</td><td>1.1E-06</td><td>BT20/MB231</td></tr>
<tr><td>>hg38_21_+_8433172_8446622_RNA45SN1_45S_50ntFlanks@10411.10435.25|CAGTCGGTCCTGAGAGATGGGCGAG</td><td>>hg38_21_+_8433172_8446622_RNA45SN1_45S_50ntFlanks</td><td>10411</td><td>10435</td><td>25</td><td>CAGTCGGTCCTGAGAGATGGGCGAG</td><td>498.4</td><td>-6.3</td><td>1.3</td><td>-4.9</td><td>7.7E-07</td><td>1.1E-06</td><td>BT20/MB468</td></tr>
<tr><td>>hg38_21_+_8433172_8446622_RNA45SN1_45S_50ntFlanks@10412.10429.18|AGTCGGTCCTGAGAGATG</td><td>>hg38_21_+_8433172_8446622_RNA45SN1_45S_50ntFlanks</td><td>10412</td><td>10429</td><td>18</td><td>AGTCGGTCCTGAGAGATG</td><td>188.8</td><td>-4.7</td><td>1.0</td><td>-4.9</td><td>7.7E-07</td><td>1.1E-06</td><td>BT20/MB468</td></tr>
<tr><td>>hg38_21_+_8433172_8446622_RNA45SN1_45S_50ntFlanks@12893.12932.40|CCTCGCCCGTCACGCACCGCACGTTCGTGGGGAACCTGGC</td><td>>hg38_21_+_8433172_8446622_RNA45SN1_45S_50ntFlanks</td><td>12893</td><td>12932</td><td>40</td><td>CCTCGCCCGTCACGCACCGCACGTTCGTGGGGAACCTGGC</td><td>342.9</td><td>-3.2</td><td>0.6</td><td>-5.0</td><td>7.1E-07</td><td>1.1E-06</td><td>MB231/MB468</td></tr>
<tr><td>>hg38_21_+_8433172_8446622_RNA45SN1_45S_50ntFlanks@8196.8226.31|CGTGGACGGTGTGAGGCCGGTAGCGGCCCCC</td><td>>hg38_21_+_8433172_8446622_RNA45SN1_45S_50ntFlanks</td><td>8196</td><td>8226</td><td>31</td><td>CGTGGACGGTGTGAGGCCGGTAGCGGCCCCC</td><td>175.0</td><td>-3.1</td><td>0.6</td><td>-4.9</td><td>7.9E-07</td><td>1.1E-06</td><td>BT20/MB468</td></tr>
</table>

<table>
<tr><td>>hg38_21_+_8433172_8446622_RNA45SN1_45S_50ntFlanks@3782.3800.19|CAGTGAAACTGCGAATGGC</td><td>>hg38_21_+_8433172_8446622_RNA45SN1_45S_50ntFlanks</td><td>3782</td><td>3800</td><td>19</td><td>CAGTGAAACTGCGAATGGC</td><td>327.3</td><td>-3.6</td><td>0.7</td><td>-4.9</td><td>7.9E-07</td><td>1.1E-06</td><td>BT20/MB468</td></tr>
<tr><td>>hg38_21_+_8433172_8446622_RNA45SN1_45S_50ntFlanks@11564.11590.27|CCTAGCAGCCGACTTAGAACTGGTGCG</td><td>>hg38_21_+_8433172_8446622_RNA45SN1_45S_50ntFlanks</td><td>11564</td><td>11590</td><td>27</td><td>CCTAGCAGCCGACTTAGAACTGGTGCG</td><td>267.5</td><td>-3.2</td><td>0.7</td><td>-4.9</td><td>7.9E-07</td><td>1.1E-06</td><td>BT20/MB468</td></tr>
<tr><td>>hg38_21_+_8433172_8446622_RNA45SN1_45S_50ntFlanks@6651.6698.48|CGACTCTTAGCGGTGGATCACTCGGCTCGTGCGTCGATGAAGAACGCA</td><td>>hg38_21_+_8433172_8446622_RNA45SN1_45S_50ntFlanks</td><td>6651</td><td>6698</td><td>48</td><td>CGACTCTTAGCGGTGGATCACTCGGCTCGTGCGTCGATGAAGAACGCA</td><td>12852.6</td><td>4.0</td><td>0.8</td><td>5.0</td><td>7.2E-07</td><td>1.1E-06</td><td>MB231/MB468</td></tr>
<tr><td>>hg38_21_+_8433172_8446622_RNA45SN1_45S_50ntFlanks@11784.11806.23|ATGAATGGATGAACGAGATTCCC</td><td>>hg38_21_+_8433172_8446622_RNA45SN1_45S_50ntFlanks</td><td>11784</td><td>11806</td><td>23</td><td>ATGAATGGATGAACGAGATTCCC</td><td>890.9</td><td>-3.1</td><td>0.6</td><td>-5.0</td><td>7.2E-07</td><td>1.1E-06</td><td>MB231/MB468</td></tr>
<tr><td>>hg38_21_+_8433172_8446622_RNA45SN1_45S_50ntFlanks@10679.10708.30|GGTTCCGGCGGCGTCCGGTGAGCTCTCGCT</td><td>>hg38_21_+_8433172_8446622_RNA45SN1_45S_50ntFlanks</td><td>10679</td><td>10708</td><td>30</td><td>GGTTCCGGCGGCGTCCGGTGAGCTCTCGCT</td><td>5429.1</td><td>-4.5</td><td>0.9</td><td>-4.9</td><td>8.1E-07</td><td>1.1E-06</td><td>BT20/MB468</td></tr>
<tr><td>>hg38_21_+_8433172_8446622_RNA45SN1_45S_50ntFlanks@6726.6743.18|CAGGACACATTGATCATC</td><td>>hg38_21_+_8433172_8446622_RNA45SN1_45S_50ntFlanks</td><td>6726</td><td>6743</td><td>18</td><td>CAGGACACATTGATCATC</td><td>266.5</td><td>-4.5</td><td>0.9</td><td>-4.9</td><td>8.2E-07</td><td>1.1E-06</td><td>BT20/MB468</td></tr>
<tr><td>>hg38_21_+_8433172_8446622_RNA45SN1_45S_50ntFlanks@4724.4759.36|ATTAATCAAGAACGAAAGTCGGAGGTTCGAAGACGA</td><td>>hg38_21_+_8433172_8446622_RNA45SN1_45S_50ntFlanks</td><td>4724</td><td>4759</td><td>36</td><td>ATTAATCAAGAACGAAAGTCGGAGGTTCGAAGACGA</td><td>1108.0</td><td>-3.7</td><td>0.7</td><td>-5.0</td><td>7.4E-07</td><td>1.1E-06</td><td>MB231/MB468</td></tr>
<tr><td>>hg38_21_+_8433172_8446622_RNA45SN1_45S_50ntFlanks@11873.11904.32|AAGAAGACCCTGTTGAGCTTGACTCTAGTCTG</td><td>>hg38_21_+_8433172_8446622_RNA45SN1_45S_50ntFlanks</td><td>11873</td><td>11904</td><td>32</td><td>AAGAAGACCCTGTTGAGCTTGACTCTAGTCTG</td><td>654.2</td><td>-6.5</td><td>1.3</td><td>-4.9</td><td>8.2E-07</td><td>1.2E-06</td><td>BT20/MB468</td></tr>
<tr><td>>hg38_21_+_8433172_8446622_RNA45SN1_45S_50ntFlanks@12081.12109.29|CTCTCGCTTCTGGCGCCAAGCGCCCGGCC</td><td>>hg38_21_+_8433172_8446622_RNA45SN1_45S_50ntFlanks</td><td>12081</td><td>12109</td><td>29</td><td>CTCTCGCTTCTGGCGCCAAGCGCCCGGCC</td><td>102.0</td><td>-4.5</td><td>0.9</td><td>-4.9</td><td>8.3E-07</td><td>1.2E-06</td><td>BT20/MB468</td></tr>
<tr><td>>hg38_21_+_8433172_8446622_RNA45SN1_45S_50ntFlanks@9623.9648.26|TAGTAGCTGGTTCCCTCCGAAGTTTC</td><td>>hg38_21_+_8433172_8446622_RNA45SN1_45S_50ntFlanks</td><td>9623</td><td>9648</td><td>26</td><td>TAGTAGCTGGTTCCCTCCGAAGTTTC</td><td>184.3</td><td>-3.6</td><td>0.7</td><td>-4.9</td><td>8.3E-07</td><td>1.2E-06</td><td>BT20/MB468</td></tr>
<tr><td>>hg38_21_+_8433172_8446622_RNA45SN1_45S_50ntFlanks@6661.6679.19|CGGTGGATCACTCGGCTCG</td><td>>hg38_21_+_8433172_8446622_RNA45SN1_45S_50ntFlanks</td><td>6661</td><td>6679</td><td>19</td><td>CGGTGGATCACTCGGCTCG</td><td>447.9</td><td>2.2</td><td>0.4</td><td>4.9</td><td>8.3E-07</td><td>1.2E-06</td><td>BT20/MB468</td></tr>
<tr><td>>hg38_21_+_8433172_8446622_RNA45SN1_45S_50ntFlanks@8176.8193.18|CTTCTGATCGAGGCCCAG</td><td>>hg38_21_+_8433172_8446622_RNA45SN1_45S_50ntFlanks</td><td>8176</td><td>8193</td><td>18</td><td>CTTCTGATCGAGGCCCAG</td><td>144.7</td><td>-2.6</td><td>0.5</td><td>-4.9</td><td>8.3E-07</td><td>1.2E-06</td><td>BT20/MB468</td></tr>
<tr><td>>hg38_21_+_8433172_8446622_RNA45SN1_45S_50ntFlanks@10905.10933.29|CTGGGCGCGCGCCGCGGCTGGACGAGGCG</td><td>>hg38_21_+_8433172_8446622_RNA45SN1_45S_50ntFlanks</td><td>10905</td><td>10933</td><td>29</td><td>CTGGGCGCGCGCCGCGGCTGGACGAGGCG</td><td>145.6</td><td>-4.0</td><td>0.8</td><td>-4.9</td><td>8.3E-07</td><td>1.2E-06</td><td>BT20/MB468</td></tr>
<tr><td>>MI0000810|hsa-mir-135b&WithFlank&1|-|205448296|205448404@22.44.23&[MIMAT0000758&hsa-miR-135b-5p&offsets|0|0;m-321&1|-|205448361|205448383&offsets|0|0]|TATGGCTTTTCATTCCTATGTGA</td><td>>MI0000810|hsa-mir-135b&WithFlank&1|-|205448296|205448404</td><td>22</td><td>44</td><td>23</td><td>TATGGCTTTTCATTCCTATGTGA</td><td>1643.4</td><td>4.1</td><td>0.8</td><td>4.9</td><td>7.5E-07</td><td>1.2E-06</td><td>MB231/MB468</td></tr>
<tr><td>>hg38_21_+_8433172_8446622_RNA45SN1_45S_50ntFlanks@11583.11603.21|CTGGTGCGGACCAGGGGAATC</td><td>>hg38_21_+_8433172_8446622_RNA45SN1_45S_50ntFlanks</td><td>11583</td><td>11603</td><td>21</td><td>CTGGTGCGGACCAGGGGAATC</td><td>1232.6</td><td>-3.5</td><td>0.7</td><td>-4.9</td><td>7.5E-07</td><td>1.2E-06</td><td>MB231/MB468</td></tr>
<tr><td>>hg38_21_+_8433172_8446622_RNA45SN1_45S_50ntFlanks@10781.10809.29|AGGTGAACAGCCTCTGGCATGTTGGAACA</td><td>>hg38_21_+_8433172_8446622_RNA45SN1_45S_50ntFlanks</td><td>10781</td><td>10809</td><td>29</td><td>AGGTGAACAGCCTCTGGCATGTTGGAACA</td><td>123.1</td><td>-2.4</td><td>0.5</td><td>-4.9</td><td>8.4E-07</td><td>1.2E-06</td><td>BT20/MB468</td></tr>
<tr><td>>hg38_21_+_8433172_8446622_RNA45SN1_45S_50ntFlanks@6738.6756.19|ATCATCGACACTTCGAACG</td><td>>hg38_21_+_8433172_8446622_RNA45SN1_45S_50ntFlanks</td><td>6738</td><td>6756</td><td>19</td><td>ATCATCGACACTTCGAACG</td><td>516.7</td><td>-5.4</td><td>1.1</td><td>-4.9</td><td>7.6E-07</td><td>1.2E-06</td><td>MB231/MB468</td></tr>
<tr><td>>hg38_21_+_8433172_8446622_RNA45SN1_45S_50ntFlanks@7981.8012.32|CTCAGATCAGACGTGGCGACCCGCTGAATTTA</td><td>>hg38_21_+_8433172_8446622_RNA45SN1_45S_50ntFlanks</td><td>7981</td><td>8012</td><td>32</td><td>CTCAGATCAGACGTGGCGACCCGCTGAATTTA</td><td>131.6</td><td>-3.3</td><td>0.7</td><td>-4.9</td><td>8.5E-07</td><td>1.2E-06</td><td>BT20/MB468</td></tr>
<tr><td>>MI0000076|hsa-mir-20a&WithFlank&13|+|91351059|91351141@14.34.21&[MIMAT0000075&hsa-miR-20a-5p&offsets|0|-2;m-96&13|+|91351072|91351094&offsets|0|-2]|TAAAGTGCTTATAGTGCAGGT</td><td>>MI0000076|hsa-mir-20a&WithFlank&13|+|91351059|91351141</td><td>14</td><td>34</td><td>21</td><td>TAAAGTGCTTATAGTGCAGGT</td><td>1245.3</td><td>4.9</td><td>1.0</td><td>4.9</td><td>8.5E-07</td><td>1.2E-06</td><td>BT20/MB468</td></tr>
<tr><td>>am_trna116_GluCTC_1_-_145399233_145399304@1.28.28,trna59_GluCTC_1_+_249168447_249168518@1.28.28,trna71_GluCTC_1_-_161439189_161439260@1.28.28,trna74_GluCTC_1_-_161431809_161431880@1.28.28,trna77_GluCTC_1_-_161424398_161424469@1.28.28,trna77_GluCTC_6_+_28949976_28950047@1.28.28,trna80_GluCTC_1_-_161417018_161417089@1.28.28,trna87_GluCTC_6_-_126101393_126101464@1.28.28|TCCCTGGTGGTCTAGTGGTTAGGATTCG</td><td>>am_trna116_GluCTC_1_-_145399233_145399304</td><td>1</td><td>28</td><td>28</td><td>TCCCTGGTGGTCTAGTGGTTAGGATTCG</td><td>20048.0</td><td>-3.6</td><td>0.7</td><td>-4.9</td><td>8.6E-07</td><td>1.2E-06</td><td>BT20/MB468</td></tr>
<tr><td>>hg38_21_+_8433172_8446622_RNA45SN1_45S_50ntFlanks@11784.11805.22|ATGAATGGATGAACGAGATTCC</td><td>>hg38_21_+_8433172_8446622_RNA45SN1_45S_50ntFlanks</td><td>11784</td><td>11805</td><td>22</td><td>ATGAATGGATGAACGAGATTCC</td><td>722.1</td><td>-4.7</td><td>0.9</td><td>-4.9</td><td>8.6E-07</td><td>1.2E-06</td><td>BT20/MB468</td></tr>
<tr><td>>hg38_21_+_8433172_8446622_RNA45SN1_45S_50ntFlanks@10798.10817.20|CATGTTGGAACAATGTAGGT</td><td>>hg38_21_+_8433172_8446622_RNA45SN1_45S_50ntFlanks</td><td>10798</td><td>10817</td><td>20</td><td>CATGTTGGAACAATGTAGGT</td><td>596.1</td><td>-2.9</td><td>0.6</td><td>-4.9</td><td>7.7E-07</td><td>1.2E-06</td><td>MB231/MB468</td></tr>
<tr><td>>hg38_21_+_8433172_8446622_RNA45SN1_45S_50ntFlanks@4739.4758.20|AAGTCGGAGGTTCGAAGACG</td><td>>hg38_21_+_8433172_8446622_RNA45SN1_45S_50ntFlanks</td><td>4739</td><td>4758</td><td>20</td><td>AAGTCGGAGGTTCGAAGACG</td><td>135.2</td><td>-2.2</td><td>0.4</td><td>-4.9</td><td>8.8E-07</td><td>1.2E-06</td><td>BT20/MB468</td></tr>
<tr><td>>hg38_21_+_8433172_8446622_RNA45SN1_45S_50ntFlanks@9005.9026.22|CTCCCCCGCGGGGGCGCGCCGG</td><td>>hg38_21_+_8433172_8446622_RNA45SN1_45S_50ntFlanks</td><td>9005</td><td>9026</td><td>22</td><td>CTCCCCCGCGGGGGCGCGCCGG</td><td>113.3</td><td>-4.7</td><td>1.0</td><td>-4.9</td><td>8.9E-07</td><td>1.2E-06</td><td>BT20/MB468</td></tr>
<tr><td>>hg38_21_+_8433172_8446622_RNA45SN1_45S_50ntFlanks@9027.9065.39|CGTCTCCTCGTGGGGGGGCCGGGCCACCCCTCCCACGGC</td><td>>hg38_21_+_8433172_8446622_RNA45SN1_45S_50ntFlanks</td><td>9027</td><td>9065</td><td>39</td><td>CGTCTCCTCGTGGGGGGGCCGGGCCACCCCTCCCACGGC</td><td>95.8</td><td>-5.3</td><td>1.1</td><td>-4.9</td><td>8.1E-07</td><td>1.3E-06</td><td>MB231/MB468</td></tr>
<tr><td>>hg38_21_+_8433172_8446622_RNA45SN1_45S_50ntFlanks@11561.11593.33|GCGCCTAGCAGCCGACTTAGAACTGGTGCGGAC</td><td>>hg38_21_+_8433172_8446622_RNA45SN1_45S_50ntFlanks</td><td>11561</td><td>11593</td><td>33</td><td>GCGCCTAGCAGCCGACTTAGAACTGGTGCGGAC</td><td>309.0</td><td>-4.8</td><td>1.0</td><td>-4.9</td><td>9.0E-07</td><td>1.3E-06</td><td>BT20/MB468</td></tr>
<tr><td>>hg38_21_+_8433172_8446622_RNA45SN1_45S_50ntFlanks@12488.12522.35|AACGTGAGCTGGGTTTAGACCGTCGTGAGACAGGT</td><td>>hg38_21_+_8433172_8446622_RNA45SN1_45S_50ntFlanks</td><td>12488</td><td>12522</td><td>35</td><td>AACGTGAGCTGGGTTTAGACCGTCGTGAGACAGGT</td><td>201.2</td><td>-4.0</td><td>0.8</td><td>-4.9</td><td>9.0E-07</td><td>1.3E-06</td><td>BT20/MB468</td></tr>
<tr><td>>hg38_21_+_8433172_8446622_RNA45SN1_45S_50ntFlanks@10392.10429.38|ACAGCAGTTGAACATGGGTCAGTCGGTCCTGAGAGATG</td><td>>hg38_21_+_8433172_8446622_RNA45SN1_45S_50ntFlanks</td><td>10392</td><td>10429</td><td>38</td><td>ACAGCAGTTGAACATGGGTCAGTCGGTCCTGAGAGATG</td><td>1755.6</td><td>-4.3</td><td>0.9</td><td>-4.9</td><td>9.0E-07</td><td>1.3E-06</td><td>BT20/MB468</td></tr>
<tr><td>>hg38_21_+_8433172_8446622_RNA45SN1_45S_50ntFlanks@11874.11898.25|AGAAGACCCTGTTGAGCTTGACTCT</td><td>>hg38_21_+_8433172_8446622_RNA45SN1_45S_50ntFlanks</td><td>11874</td><td>11898</td><td>25</td><td>AGAAGACCCTGTTGAGCTTGACTCT</td><td>1772.9</td><td>-3.5</td><td>0.7</td><td>-4.9</td><td>9.1E-07</td><td>1.3E-06</td><td>BT20/MB468</td></tr>
<tr><td>>MI0000090|hsa-mir-32&WithFlank&9|-|109046223|109046304@12.33.22&[MIMAT0000090&hsa-miR-32-5p&offsets|0|0;m-208&9|-|109046273|109046293&offsets|0|+1]|TATTGCACATTACTAAGTTGCA</td><td>>MI0000090|hsa-mir-32&WithFlank&9|-|109046223|109046304</td><td>12</td><td>33</td><td>22</td><td>TATTGCACATTACTAAGTTGCA</td><td>1562.8</td><td>5.0</td><td>1.0</td><td>4.9</td><td>9.1E-07</td><td>1.3E-06</td><td>BT20/MB468</td></tr>
<tr><td>>hg38_21_+_8433172_8446622_RNA45SN1_45S_50ntFlanks@8378.8402.25|TTCAAGAGGGCGTGAAACCGTTAAG</td><td>>hg38_21_+_8433172_8446622_RNA45SN1_45S_50ntFlanks</td><td>8378</td><td>8402</td><td>25</td><td>TTCAAGAGGGCGTGAAACCGTTAAG</td><td>145.0</td><td>-2.5</td><td>0.5</td><td>-4.9</td><td>8.2E-07</td><td>1.3E-06</td><td>MB231/MB468</td></tr>
<tr><td>>hg38_21_+_8433172_8446622_RNA45SN1_45S_50ntFlanks@9574.9609.36|AATCGGTCGTCCGACCTGGGTATAGGGGCGAAAGAC</td><td>>hg38_21_+_8433172_8446622_RNA45SN1_45S_50ntFlanks</td><td>9574</td><td>9609</td><td>36</td><td>AATCGGTCGTCCGACCTGGGTATAGGGGCGAAAGAC</td><td>434.8</td><td>8.6</td><td>1.7</td><td>4.9</td><td>8.2E-07</td><td>1.3E-06</td><td>MB231/MB468</td></tr>
<tr><td>>hg38_21_+_8433172_8446622_RNA45SN1_45S_50ntFlanks@10829.10858.30|GCAAGCCGGATCCGTAACTTCGGGATAAGG</td><td>>hg38_21_+_8433172_8446622_RNA45SN1_45S_50ntFlanks</td><td>10829</td><td>10858</td><td>30</td><td>GCAAGCCGGATCCGTAACTTCGGGATAAGG</td><td>173.9</td><td>-3.3</td><td>0.7</td><td>-4.9</td><td>8.3E-07</td><td>1.3E-06</td><td>MB231/MB468</td></tr>
<tr><td>>hg38_21_+_8433172_8446622_RNA45SN1_45S_50ntFlanks@10421.10453.33|TGAGAGATGGGCGAGCGCCGTTCCGAAGGGACG</td><td>>hg38_21_+_8433172_8446622_RNA45SN1_45S_50ntFlanks</td><td>10421</td><td>10453</td><td>33</td><td>TGAGAGATGGGCGAGCGCCGTTCCGAAGGGACG</td><td>194.2</td><td>-2.1</td><td>0.4</td><td>-4.9</td><td>8.4E-07</td><td>1.3E-06</td><td>MB231/MB468</td></tr>
<tr><td>>hg38_21_+_8433172_8446622_RNA45SN1_45S_50ntFlanks@9545.9575.31|GTGGAGGTCCGTAGCGGTCCTGACGTGCAAA</td><td>>hg38_21_+_8433172_8446622_RNA45SN1_45S_50ntFlanks</td><td>9545</td><td>9575</td><td>31</td><td>GTGGAGGTCCGTAGCGGTCCTGACGTGCAAA</td><td>100.1</td><td>-4.3</td><td>0.9</td><td>-4.9</td><td>9.4E-07</td><td>1.3E-06</td><td>BT20/MB468</td></tr>
<tr><td>>hg38_21_+_8433172_8446622_RNA45SN1_45S_50ntFlanks@10788.10807.20|CAGCCTCTGGCATGTTGGAA</td><td>>hg38_21_+_8433172_8446622_RNA45SN1_45S_50ntFlanks</td><td>10788</td><td>10807</td><td>20</td><td>CAGCCTCTGGCATGTTGGAA</td><td>122.0</td><td>-4.0</td><td>0.8</td><td>-4.9</td><td>8.4E-07</td><td>1.3E-06</td><td>MB231/MB468</td></tr>
<tr><td>>MI0000086|hsa-mir-28&WithFlank&3|+|188688775|188688872@60.81.22(+1U)&[MIMAT0004502&hsa-miR-28-3p&offsets|0|0(+1U);m-28&3|+|188688834|188688855&offsets|0|0(+1U)]|CACTAGATTGTGAGCTCCTGGAT</td><td>>MI0000086|hsa-mir-28&WithFlank&3|+|188688775|188688872</td><td>60</td><td>81</td><td>22(+1U)</td><td>CACTAGATTGTGAGCTCCTGGAT</td><td>1915.3</td><td>11.6</td><td>2.3</td><td>4.9</td><td>8.5E-07</td><td>1.3E-06</td><td>MB231/MB468</td></tr>
<tr><td>>hg38_21_+_8433172_8446622_RNA45SN1_45S_50ntFlanks@7980.7999.20|CCTCAGATCAGACGTGGCGA</td><td>>hg38_21_+_8433172_8446622_RNA45SN1_45S_50ntFlanks</td><td>7980</td><td>7999</td><td>20</td><td>CCTCAGATCAGACGTGGCGA</td><td>75.0</td><td>-4.2</td><td>0.9</td><td>-4.9</td><td>8.5E-07</td><td>1.3E-06</td><td>MB231/MB468</td></tr>
<tr><td>>hg38_21_+_8433172_8446622_RNA45SN1_45S_50ntFlanks@6649.6683.35|TACGACTCTTAGCGGTGGATCACTCGGCTCGTGCG</td><td>>hg38_21_+_8433172_8446622_RNA45SN1_45S_50ntFlanks</td><td>6649</td><td>6683</td><td>35</td><td>TACGACTCTTAGCGGTGGATCACTCGGCTCGTGCG</td><td>568.5</td><td>3.6</td><td>0.7</td><td>4.9</td><td>9.5E-07</td><td>1.3E-06</td><td>BT20/MB468</td></tr>
<tr><td>>hg38_21_+_8433172_8446622_RNA45SN1_45S_50ntFlanks@6702.6720.19|AGCTGCGAGAATTAATGTG</td><td>>hg38_21_+_8433172_8446622_RNA45SN1_45S_50ntFlanks</td><td>6702</td><td>6720</td><td>19</td><td>AGCTGCGAGAATTAATGTG</td><td>506.8</td><td>-2.2</td><td>0.5</td><td>-4.9</td><td>9.5E-07</td><td>1.3E-06</td><td>BT20/MB468</td></tr>
<tr><td>>hg38_21_+_8433172_8446622_RNA45SN1_45S_50ntFlanks@11564.11602.39|CCTAGCAGCCGACTTAGAACTGGTGCGGACCAGGGGAAT</td><td>>hg38_21_+_8433172_8446622_RNA45SN1_45S_50ntFlanks</td><td>11564</td><td>11602</td><td>39</td><td>CCTAGCAGCCGACTTAGAACTGGTGCGGACCAGGGGAAT</td><td>564.8</td><td>-3.2</td><td>0.6</td><td>-4.9</td><td>8.5E-07</td><td>1.3E-06</td><td>MB231/MB468</td></tr>
<tr><td>>hg38_21_+_8433172_8446622_RNA45SN1_45S_50ntFlanks@3782.3800.19|CAGTGAAACTGCGAATGGC</td><td>>hg38_21_+_8433172_8446622_RNA45SN1_45S_50ntFlanks</td><td>3782</td><td>3800</td><td>19</td><td>CAGTGAAACTGCGAATGGC</td><td>295.8</td><td>-2.4</td><td>0.5</td><td>-4.9</td><td>8.5E-07</td><td>1.3E-06</td><td>MB231/MB468</td></tr>
<tr><td>>hg38_21_+_8433172_8446622_RNA45SN1_45S_50ntFlanks@6652.6683.32|GACTCTTAGCGGTGGATCACTCGGCTCGTGCG</td><td>>hg38_21_+_8433172_8446622_RNA45SN1_45S_50ntFlanks</td><td>6652</td><td>6683</td><td>32</td><td>GACTCTTAGCGGTGGATCACTCGGCTCGTGCG</td><td>2526.3</td><td>4.7</td><td>1.0</td><td>4.9</td><td>9.5E-07</td><td>1.3E-06</td><td>BT20/MB468</td></tr>
<tr><td>>hg38_21_+_8433172_8446622_RNA45SN1_45S_50ntFlanks@12423.12455.33|TCGGCTCTTCCTATCATTGTGAAGCAGAATTCA</td><td>>hg38_21_+_8433172_8446622_RNA45SN1_45S_50ntFlanks</td><td>12423</td><td>12455</td><td>33</td><td>TCGGCTCTTCCTATCATTGTGAAGCAGAATTCA</td><td>106.1</td><td>-3.8</td><td>0.8</td><td>-4.9</td><td>9.6E-07</td><td>1.3E-06</td><td>BT20/MB468</td></tr>
<tr><td>>hg38_21_+_8433172_8446622_RNA45SN1_45S_50ntFlanks@9571.9597.27|GCAAATCGGTCGTCCGACCTGGGTATA</td><td>>hg38_21_+_8433172_8446622_RNA45SN1_45S_50ntFlanks</td><td>9571</td><td>9597</td><td>27</td><td>GCAAATCGGTCGTCCGACCTGGGTATA</td><td>226.0</td><td>-2.6</td><td>0.5</td><td>-4.9</td><td>8.6E-07</td><td>1.3E-06</td><td>MB231/MB468</td></tr>
<tr><td>>hg38_21_+_8433172_8446622_RNA45SN1_45S_50ntFlanks@4560.4589.30|CTAGGAATAATGGAATAGGACCGCGGTTCT</td><td>>hg38_21_+_8433172_8446622_RNA45SN1_45S_50ntFlanks</td><td>4560</td><td>4589</td><td>30</td><td>CTAGGAATAATGGAATAGGACCGCGGTTCT</td><td>131.1</td><td>-3.0</td><td>0.6</td><td>-4.9</td><td>9.6E-07</td><td>1.3E-06</td><td>BT20/MB468</td></tr>
</table>

| | | | | | | | | | | | | |
|---|---|---|---|---|---|---|---|---|---|---|---|---|
| >hg38_21_+_8433172_8446622_RNA45SN1_45S_50ntFlanks@6719.6756.38\|TGAATTGCAGGACACATTGATCATCGACACTTCGAACG | >hg38_21_+_8433172_8446622_RNA45SN1_45S_50ntFlanks | 6719 | 6756 | 38 | TGAATTGCAGGACACATTGATCATCGACACTTCGAACG | 100.8 | -3.2 | 0.7 | -4.9 | 9.6E-07 | 1.3E-06 | BT20/MB468 |
| >hg38_21_+_8433172_8446622_RNA45SN1_45S_50ntFlanks@11703.11727.25\|AATGAAGCGCGGGTAAACGGCGGGA | >hg38_21_+_8433172_8446622_RNA45SN1_45S_50ntFlanks | 11703 | 11727 | 25 | AATGAAGCGCGGGTAAACGGCGGGA | 148.8 | -2.3 | 0.5 | -4.9 | 8.7E-07 | 1.4E-06 | MB231/MB468 |
| >hg38_21_+_8433172_8446622_RNA45SN1_45S_50ntFlanks@12083.12106.24\|CTCGCTTCTGGCGCCAAGCGCCCG | >hg38_21_+_8433172_8446622_RNA45SN1_45S_50ntFlanks | 12083 | 12106 | 24 | CTCGCTTCTGGCGCCAAGCGCCCG | 384.8 | -5.8 | 1.2 | -4.9 | 9.8E-07 | 1.4E-06 | BT20/MB468 |
| >hg38_21_+_8433172_8446622_RNA45SN1_45S_50ntFlanks@11581.11599.19\|AACTGGTGCGGACCAGGGG | >hg38_21_+_8433172_8446622_RNA45SN1_45S_50ntFlanks | 11581 | 11599 | 19 | AACTGGTGCGGACCAGGGG | 106.7 | -3.7 | 0.7 | -4.9 | 9.9E-07 | 1.4E-06 | BT20/MB468 |
| >hg38_21_+_8433172_8446622_RNA45SN1_45S_50ntFlanks@11920.11950.31\|ATGAGAGGTGTAGAATAAGTGGGAGGCCCCC | >hg38_21_+_8433172_8446622_RNA45SN1_45S_50ntFlanks | 11920 | 11950 | 31 | ATGAGAGGTGTAGAATAAGTGGGAGGCCCCC | 103.4 | -3.0 | 0.6 | -4.9 | 1.0E-06 | 1.4E-06 | BT20/MB468 |
| >hg38_21_+_8433172_8446622_RNA45SN1_45S_50ntFlanks@12076.12096.21\|AGGGGCTCTCGCTTCTGGCGC | >hg38_21_+_8433172_8446622_RNA45SN1_45S_50ntFlanks | 12076 | 12096 | 21 | AGGGGCTCTCGCTTCTGGCGC | 186.0 | -2.4 | 0.5 | -4.9 | 9.1E-07 | 1.4E-06 | MB231/MB468 |
| >hg38_21_+_8433172_8446622_RNA45SN1_45S_50ntFlanks@10866.10889.24\|CTAAGGGCTGGGTCGGTCGGGCTG | >hg38_21_+_8433172_8446622_RNA45SN1_45S_50ntFlanks | 10866 | 10889 | 24 | CTAAGGGCTGGGTCGGTCGGGCTG | 222.9 | -4.9 | 1.0 | -4.9 | 1.0E-06 | 1.4E-06 | BT20/MB468 |
| >MI0000286\|hsa-mir-210&WithFlank&11\|-\|568083\|568204@72.92.21(+1U)&[MIMAT0000267&hsa-miR-210-3p&offsets\|0\|-1(+1U);m-102&11\|-\|568112\|568133&offsets\|0\|-1(+1U)]\|CTGTGCGTGTGACAGCGGCTGT | >MI0000286\|hsa-mir-210&WithFlank&11\|-\|568083\|568204 | 72 | 92 | 21(+1U) | CTGTGCGTGTGACAGCGGCTGT | 6052.9 | 13.5 | 2.8 | 4.9 | 1.0E-06 | 1.4E-06 | BT20/MB468 |
| >hg38_21_+_8433172_8446622_RNA45SN1_45S_50ntFlanks@4732.4765.34\|AGAACGAAAGTCGGAGGTTCGAAGACGATCAGAT | >hg38_21_+_8433172_8446622_RNA45SN1_45S_50ntFlanks | 4732 | 4765 | 34 | AGAACGAAAGTCGGAGGTTCGAAGACGATCAGAT | 346.7 | 6.4 | 1.3 | 4.9 | 1.0E-06 | 1.4E-06 | BT20/MB468 |
| >hg38_21_+_8433172_8446622_RNA45SN1_45S_50ntFlanks@11784.11803.20\|ATGAATGGATGAACGAGATT | >hg38_21_+_8433172_8446622_RNA45SN1_45S_50ntFlanks | 11784 | 11803 | 20 | ATGAATGGATGAACGAGATT | 2223.5 | -3.1 | 0.6 | -4.9 | 9.2E-07 | 1.4E-06 | MB231/MB468 |
| >hg38_21_+_8433172_8446622_RNA45SN1_45S_50ntFlanks@6732.6770.39\|ACATTGATCATCGACACTTCGAACGCACTTGCGGCCCCG | >hg38_21_+_8433172_8446622_RNA45SN1_45S_50ntFlanks | 6732 | 6770 | 39 | ACATTGATCATCGACACTTCGAACGCACTTGCGGCCCCG | 598.8 | -2.7 | 0.6 | -4.9 | 1.0E-06 | 1.4E-06 | BT20/MB468 |
| >hg38_21_+_8433172_8446622_RNA45SN1_45S_50ntFlanks@6676.6696.21\|CTCGTGCGTCGATGAAGAACG | >hg38_21_+_8433172_8446622_RNA45SN1_45S_50ntFlanks | 6676 | 6696 | 21 | CTCGTGCGTCGATGAAGAACG | 184.8 | -2.4 | 0.5 | -4.9 | 1.0E-06 | 1.4E-06 | BT20/MB468 |
| >hg38_21_+_8433172_8446622_RNA45SN1_45S_50ntFlanks@11778.11804.27\|ACGCGCATGAATGGATGAACGAGATTC | >hg38_21_+_8433172_8446622_RNA45SN1_45S_50ntFlanks | 11778 | 11804 | 27 | ACGCGCATGAATGGATGAACGAGATTC | 370.4 | -2.6 | 0.5 | -4.9 | 9.3E-07 | 1.4E-06 | MB231/MB468 |
| >MI0000266\|hsa-mir-10a&WithFlank&17\|-\|48579832\|48579953@29.50.22(+1A)&[MIMAT0000253&hsa-miR-10a-5p&offsets\|+1\|0(+1A);m-18&17\|-\|48579905\|48579926&offsets\|+1\|+1(+1A)]\|ACCCTGTAGATCCGAATTTGTGA | >MI0000266\|hsa-mir-10a&WithFlank&17\|-\|48579832\|48579953 | 29 | 50 | 22(+1A) | ACCCTGTAGATCCGAATTTGTGA | 2855.8 | 11.6 | 2.4 | 4.9 | 1.0E-06 | 1.4E-06 | BT20/MB468 |
| >hg38_21_+_8433172_8446622_RNA45SN1_45S_50ntFlanks@10409.10433.25\|GTCAGTCGGTCCTGAGAGATGGGCG | >hg38_21_+_8433172_8446622_RNA45SN1_45S_50ntFlanks | 10409 | 10433 | 25 | GTCAGTCGGTCCTGAGAGATGGGCG | 194.4 | -2.4 | 0.5 | -4.9 | 9.4E-07 | 1.5E-06 | MB231/MB468 |
| >MI0000102\|hsa-mir-100&WithFlank&11\|-\|122152223\|122152314@19.38.20(+2G)&[MIMAT0000098&hsa-miR-100-5p&offsets\|0\|-2(+2G);m-27&11\|-\|122152276\|122152296&offsets\|0\|-1(+2G)]\|AACCCGTAGATCCGAACTTGGG | >MI0000102\|hsa-mir-100&WithFlank&11\|-\|122152223\|122152314 | 19 | 38 | 20(+2G) | AACCCGTAGATCCGAACTTGGG | 2206.9 | -5.6 | 1.0 | -5.4 | 7.5E-08 | 1.5E-06 | BT20/MB231 |
| >MI0000746\|hsa-mir-99b&WithFlank&19\|+\|51692606\|51692687@13.32.20(+1U)&[MIMAT0000689&hsa-miR-99b-5p&offsets\|0\|-2(+1U);m-10&19\|+\|51692618\|51692639&offsets\|0\|-2(+1U)]\|CACCCGTAGAACCGACCTTGT | >MI0000746\|hsa-mir-99b&WithFlank&19\|+\|51692606\|51692687 | 13 | 32 | 20(+1U) | CACCCGTAGAACCGACCTTGT | 13217.1 | 3.6 | 0.7 | 5.4 | 7.5E-08 | 1.5E-06 | BT20/MB231 |
| >hg38_21_+_8433172_8446622_RNA45SN1_45S_50ntFlanks@9738.9766.29\|GAAACGATCTCAACCTATTCTCAAACTTT | >hg38_21_+_8433172_8446622_RNA45SN1_45S_50ntFlanks | 9738 | 9766 | 29 | GAAACGATCTCAACCTATTCTCAAACTTT | 89.6 | -3.0 | 0.6 | -4.9 | 1.1E-06 | 1.5E-06 | BT20/MB468 |
| >hg38_21_+_8433172_8446622_RNA45SN1_45S_50ntFlanks@10686.10710.25\|GCGGCGTCCGGTGAGCTCTCGCTGG | >hg38_21_+_8433172_8446622_RNA45SN1_45S_50ntFlanks | 10686 | 10710 | 25 | GCGGCGTCCGGTGAGCTCTCGCTGG | 1966.9 | -3.0 | 0.6 | -4.9 | 9.5E-07 | 1.5E-06 | MB231/MB468 |
| >hg38_21_+_8433172_8446622_RNA45SN1_45S_50ntFlanks@5494.5514.21\|AAGACGGTCGAACTTGACTAT | >hg38_21_+_8433172_8446622_RNA45SN1_45S_50ntFlanks | 5494 | 5514 | 21 | AAGACGGTCGAACTTGACTAT | 186.2 | -3.1 | 0.6 | -4.9 | 9.6E-07 | 1.5E-06 | MB231/MB468 |
| >MI0000077\|hsa-mir-21&WithFlank&17\|+\|59841260\|59841343@14.34.21(+3C)&[MIMAT0000076&hsa-miR-21-5p&offsets\|0\|-1(+3C);m-1&17\|+\|59841273\|59841294&offsets\|0\|-1(+3C)]\|TAGCTTATCAGACTGATGTTGCCC | >MI0000077\|hsa-mir-21&WithFlank&17\|+\|59841260\|59841343 | 14 | 34 | 21(+3C) | TAGCTTATCAGACTGATGTTGCCC | 1028.4 | 11.0 | 2.2 | 4.9 | 9.6E-07 | 1.5E-06 | MB231/MB468 |
| >hg38_21_+_8433172_8446622_RNA45SN1_45S_50ntFlanks@12974.12992.19\|CGTACGTAGCAGAGCAGCT | >hg38_21_+_8433172_8446622_RNA45SN1_45S_50ntFlanks | 12974 | 12992 | 19 | CGTACGTAGCAGAGCAGCT | 437.6 | -2.0 | 0.4 | -4.9 | 9.7E-07 | 1.5E-06 | MB231/MB468 |
| >am_trna128_GlyGCC_6_-_27870686_27870756@3.35.33,trna18_GlyGCC_16_+_70822597_70822667@3.35.33,trna19_GlyGCC_16_+_70823410_70823480@3.35.33,trna19_GlyGCC_2_-_157257659_157257729@3.35.33,trna24_GlyGCC_16_-_70812942_70813012@3.35.33,trna25_GlyGCC_16_-_70812114_70812184@3.35.33,trna5_GlyGCC_17_+_8029064_8029134@3.35.33,trna68_GlyGCC_1_-_161493637_161493707@3.35.33\|ATTGGTGGTTCAGTGGTAGAATTCTCGCCTGCC | >am_trna128_GlyGCC_6_-_27870686_27870756 | 3 | 35 | 33 | ATTGGTGGTTCAGTGGTAGAATTCTCGCCTGCC | 656.0 | -4.1 | 0.8 | -5.4 | 7.7E-08 | 1.5E-06 | BT20/MB231 |
| >hg38_21_+_8433172_8446622_RNA45SN1_45S_50ntFlanks@4000.4024.25\|TAGATAACCTCGGGCCGATCGCACG | >hg38_21_+_8433172_8446622_RNA45SN1_45S_50ntFlanks | 4000 | 4024 | 25 | TAGATAACCTCGGGCCGATCGCACG | 266.2 | -2.0 | 0.4 | -4.9 | 9.8E-07 | 1.5E-06 | MB231/MB468 |
| >hg38_21_+_8433172_8446622_RNA45SN1_45S_50ntFlanks@8190.8212.23\|CCAGCCCGTGGACGGTGTGAGGC | >hg38_21_+_8433172_8446622_RNA45SN1_45S_50ntFlanks | 8190 | 8212 | 23 | CCAGCCCGTGGACGGTGTGAGGC | 458.3 | -2.7 | 0.5 | -4.9 | 9.9E-07 | 1.5E-06 | MB231/MB468 |
| >hg38_21_+_8433172_8446622_RNA45SN1_45S_50ntFlanks@4122.4145.24\|AGGGTTCGATTCCGGAGAGGGAGC | >hg38_21_+_8433172_8446622_RNA45SN1_45S_50ntFlanks | 4122 | 4145 | 24 | AGGGTTCGATTCCGGAGAGGGAGC | 94.7 | -4.4 | 0.9 | -4.9 | 1.1E-06 | 1.5E-06 | BT20/MB468 |
| >MI0000102\|hsa-mir-100&WithFlank&11\|-\|122152223\|122152314@19.40.22(+1U)&[MIMAT0000098&hsa-miR-100-5p&offsets\|0\|0(+1U);m-27&11\|-\|122152276\|122152296&offsets\|0\|+1(+1U)]\|AACCCGTAGATCCGAACTTGTGT | >MI0000102\|hsa-mir-100&WithFlank&11\|-\|122152223\|122152314 | 19 | 40 | 22(+1U) | AACCCGTAGATCCGAACTTGTGT | 1572.0 | 8.2 | 1.7 | 4.9 | 1.1E-06 | 1.6E-06 | BT20/MB468 |
| >hg38_21_+_8433172_8446622_RNA45SN1_45S_50ntFlanks@11581.11600.20\|AACTGGTGCGGACCAGGGGA | >hg38_21_+_8433172_8446622_RNA45SN1_45S_50ntFlanks | 11581 | 11600 | 20 | AACTGGTGCGGACCAGGGGA | 443.9 | -2.8 | 0.6 | -4.9 | 1.0E-06 | 1.6E-06 | MB231/MB468 |
| >hg38_21_+_8433172_8446622_RNA45SN1_45S_50ntFlanks@10186.10208.23\|TCCTCCCCTCCTCCCGCCCACGC | >hg38_21_+_8433172_8446622_RNA45SN1_45S_50ntFlanks | 10186 | 10208 | 23 | TCCTCCCCTCCTCCCGCCCACGC | 99.4 | -4.4 | 0.9 | -4.9 | 1.1E-06 | 1.6E-06 | BT20/MB468 |
| >hg38_21_+_8433172_8446622_RNA45SN1_45S_50ntFlanks@12996.13015.20\|TCGCTGCGATCTATTGAAAG | >hg38_21_+_8433172_8446622_RNA45SN1_45S_50ntFlanks | 12996 | 13015 | 20 | TCGCTGCGATCTATTGAAAG | 1058.5 | 4.3 | 0.9 | 4.9 | 1.1E-06 | 1.6E-06 | BT20/MB468 |
| >hg38_21_+_8433172_8446622_RNA45SN1_45S_50ntFlanks@6647.6665.19\|CGTACGACTCTTAGCGGTG | >hg38_21_+_8433172_8446622_RNA45SN1_45S_50ntFlanks | 6647 | 6665 | 19 | CGTACGACTCTTAGCGGTG | 2128.5 | 3.8 | 0.8 | 4.9 | 1.0E-06 | 1.6E-06 | MB231/MB468 |
| >trna27_GlyCCC_2_-_70476123_70476193@1.32.32,trna34_GlyCCC_16_-_686736_686806@1.32.32\|GCGCCGCTGGTGTAGTGGTATCATGCAAGATT | >trna27_GlyCCC_2_-_70476123_70476193 | 1 | 32 | 32 | GCGCCGCTGGTGTAGTGGTATCATGCAAGATT | 941.2 | 11.7 | 2.4 | 4.9 | 1.0E-06 | 1.6E-06 | MB231/MB468 |
| >hg38_21_+_8433172_8446622_RNA45SN1_45S_50ntFlanks@4723.4751.29\|CATTAATCAAGAACGAAAGTCGGAGGTTC | >hg38_21_+_8433172_8446622_RNA45SN1_45S_50ntFlanks | 4723 | 4751 | 29 | CATTAATCAAGAACGAAAGTCGGAGGTTC | 409.6 | -2.8 | 0.6 | -4.9 | 1.0E-06 | 1.6E-06 | MB231/MB468 |
| >hg38_21_+_8433172_8446622_RNA45SN1_45S_50ntFlanks@3793.3810.18\|CGAATGGCTCATTAAATC | >hg38_21_+_8433172_8446622_RNA45SN1_45S_50ntFlanks | 3793 | 3810 | 18 | CGAATGGCTCATTAAATC | 143.4 | -2.2 | 0.5 | -4.9 | 1.0E-06 | 1.6E-06 | MB231/MB468 |
| >hg38_21_+_8433172_8446622_RNA45SN1_45S_50ntFlanks@9738.9762.25\|GAAACGATCTCAACCTATTCTCAAA | >hg38_21_+_8433172_8446622_RNA45SN1_45S_50ntFlanks | 9738 | 9762 | 25 | GAAACGATCTCAACCTATTCTCAAA | 110.0 | -2.5 | 0.5 | -4.9 | 1.0E-06 | 1.6E-06 | MB231/MB468 |
| >hg38_21_+_8433172_8446622_RNA45SN1_45S_50ntFlanks@9736.9767.32\|CCGAAACGATCTCAACCTATTCTCAAACTTTA | >hg38_21_+_8433172_8446622_RNA45SN1_45S_50ntFlanks | 9736 | 9767 | 32 | CCGAAACGATCTCAACCTATTCTCAAACTTTA | 82.3 | -4.1 | 0.8 | -4.9 | 1.1E-06 | 1.6E-06 | MB231/MB468 |
| >hg38_21_+_8433172_8446622_RNA45SN1_45S_50ntFlanks@11581.11612.32\|AACTGGTGCGGACCAGGGGAATCCGACTGTTT | >hg38_21_+_8433172_8446622_RNA45SN1_45S_50ntFlanks | 11581 | 11612 | 32 | AACTGGTGCGGACCAGGGGAATCCGACTGTTT | 2034.0 | -3.0 | 0.6 | -4.9 | 1.2E-06 | 1.6E-06 | BT20/MB468 |
| >MI0000487\|hsa-mir-193a&WithFlank&17\|+\|31559990\|31560089@27.48.22&[MIMAT0004614&hsa-miR-193a-5p&offsets\|0\|0;m-123&17\|+\|31560016\|31560037&offsets\|0\|0]\|TGGGTCTTTGCGGGCGAGATGA | >MI0000487\|hsa-mir-193a&WithFlank&17\|+\|31559990\|31560089 | 27 | 48 | 22 | TGGGTCTTTGCGGGCGAGATGA | 346.9 | 4.2 | 0.8 | 5.4 | 8.5E-08 | 1.6E-06 | BT20/MB231 |
| >hg38_21_+_8433172_8446622_RNA45SN1_45S_50ntFlanks@4726.4759.34\|TAATCAAGAACGAAAGTCGGAGGTTCGAAGACGA | >hg38_21_+_8433172_8446622_RNA45SN1_45S_50ntFlanks | 4726 | 4759 | 34 | TAATCAAGAACGAAAGTCGGAGGTTCGAAGACGA | 562.3 | -3.5 | 0.7 | -4.9 | 1.1E-06 | 1.7E-06 | MB231/MB468 |
| >hg38_21_+_8433172_8446622_RNA45SN1_45S_50ntFlanks@11873.11910.38\|AAGAAGACCCTGTTGAGCTTGACTCTAGTCTGGCACGG | >hg38_21_+_8433172_8446622_RNA45SN1_45S_50ntFlanks | 11873 | 11910 | 38 | AAGAAGACCCTGTTGAGCTTGACTCTAGTCTGGCACGG | 115.7 | -5.0 | 1.0 | -4.9 | 1.2E-06 | 1.7E-06 | BT20/MB468 |

<table>
<tr><th></th><th></th><th></th><th></th><th></th><th></th><th></th><th></th><th></th><th></th><th></th><th></th><th></th></tr>
<tr><td>>MI0000446|hsa-mir-125b-1&WithFlank&11|-|122099751|122099850@21.42.22(+1A);MI0000470|hsa-mir-125b-2&WithFlank&21|+|16590231|16590331@23.44.22(+1A)&[MIMAT0000423&hsa-miR-125b-5p&offsets|0|0(+1A);m-67&11|-|122099809|122099830&offsets|0|0(+1A)];[MIMAT0000423_1&hsa-miR-125b-5p&offsets|0|0(+1A);m-68&21|+|16590253|16590274&offsets|0|0(+1A)]|TCCCTGAGACCCTAACTTGTGAA</td><td>>MI0000446|hsa-mir-125b-1&WithFlank&11|-|122099751|122099850</td><td>21</td><td>42</td><td>22(+1A)</td><td>TCCCTGAGACCCTAACTTGTGAA</td><td>4588.8</td><td>-5.0</td><td>0.9</td><td>-5.4</td><td>8.7E-08</td><td>1.7E-06</td><td>BT20/MB231</td></tr>
<tr><td>>hg38_21_+_8433172_8446622_RNA45SN1_45S_50ntFlanks@12932.12960.29|CGCTAAACCATTCGTAGACGACCTGCTTC</td><td>>hg38_21_+_8433172_8446622_RNA45SN1_45S_50ntFlanks</td><td>12932</td><td>12960</td><td>29</td><td>CGCTAAACCATTCGTAGACGACCTGCTTC</td><td>353.8</td><td>2.7</td><td>0.6</td><td>4.9</td><td>1.2E-06</td><td>1.7E-06</td><td>BT20/MB468</td></tr>
<tr><td>>hg38_21_+_8433172_8446622_RNA45SN1_45S_50ntFlanks@9568.9585.18|CGTGCAAATCGGTCGTCC</td><td>>hg38_21_+_8433172_8446622_RNA45SN1_45S_50ntFlanks</td><td>9568</td><td>9585</td><td>18</td><td>CGTGCAAATCGGTCGTCC</td><td>98.8</td><td>-3.7</td><td>0.8</td><td>-4.9</td><td>1.2E-06</td><td>1.7E-06</td><td>BT20/MB468</td></tr>
<tr><td>>hg38_21_+_8433172_8446622_RNA45SN1_45S_50ntFlanks@6738.6758.21|ATCATCGACACTTCGAACGCA</td><td>>hg38_21_+_8433172_8446622_RNA45SN1_45S_50ntFlanks</td><td>6738</td><td>6758</td><td>21</td><td>ATCATCGACACTTCGAACGCA</td><td>155.2</td><td>-4.4</td><td>0.9</td><td>-4.9</td><td>1.1E-06</td><td>1.7E-06</td><td>MB231/MB468</td></tr>
<tr><td>>hg38_21_+_8433172_8446622_RNA45SN1_45S_50ntFlanks@12941.12959.19|ATTCGTAGACGACCTGCTT</td><td>>hg38_21_+_8433172_8446622_RNA45SN1_45S_50ntFlanks</td><td>12941</td><td>12959</td><td>19</td><td>ATTCGTAGACGACCTGCTT</td><td>668.5</td><td>3.0</td><td>0.6</td><td>4.9</td><td>1.1E-06</td><td>1.7E-06</td><td>MB231/MB468</td></tr>
<tr><td>>hg38_21_+_8433172_8446622_RNA45SN1_45S_50ntFlanks@3999.4024.26|CTAGATAACCTCGGGCCGATCGCACG</td><td>>hg38_21_+_8433172_8446622_RNA45SN1_45S_50ntFlanks</td><td>3999</td><td>4024</td><td>26</td><td>CTAGATAACCTCGGGCCGATCGCACG</td><td>105.1</td><td>-3.2</td><td>0.7</td><td>-4.8</td><td>1.3E-06</td><td>1.7E-06</td><td>BT20/MB468</td></tr>
<tr><td>>hg38_21_+_8433172_8446622_RNA45SN1_45S_50ntFlanks@8388.8405.18|CGTGAAACCGTTAAGAGG</td><td>>hg38_21_+_8433172_8446622_RNA45SN1_45S_50ntFlanks</td><td>8388</td><td>8405</td><td>18</td><td>CGTGAAACCGTTAAGAGG</td><td>138.2</td><td>-3.7</td><td>0.8</td><td>-4.9</td><td>1.1E-06</td><td>1.7E-06</td><td>MB231/MB468</td></tr>
<tr><td>>MI0000272|hsa-mir-182&WithFlank&7|-|129770377|129770498@29.51.23(+1C)&[MIMAT0000259&hsa-miR-182-5p&offsets|0|-1(+1C);m-22&7|-|129770449|129770470&offsets|0|+1(+1C)]|TTTGGCAATGGTAGAACTCACACC</td><td>>MI0000272|hsa-mir-182&WithFlank&7|-|129770377|129770498</td><td>29</td><td>51</td><td>23(+1C)</td><td>TTTGGCAATGGTAGAACTCACACC</td><td>10068.6</td><td>3.7</td><td>0.7</td><td>5.3</td><td>9.1E-08</td><td>1.7E-06</td><td>BT20/MB231</td></tr>
<tr><td>>am_trna10_ValCAC_5_-_180649395_180649467@1.29.29,trna12_ValAAC_5_-_180645270_180645342@1.29.29,trna132_ValAAC_6_-_27721179_27721251@1.29.29,trna136_ValAAC_6_-_27648885_27648957@1.29.29,trna139_ValAAC_6_-_27618707_27618779@1.29.29,trna18_ValCAC_5_-_180529253_180529325@1.29.29,trna2_ValAAC_3_+_169490018_169490090@1.29.29,trna2_ValCAC_5_+_180524070_180524142@1.29.29,trna4_ValAAC_5_+_180591154_180591226@1.29.29,trna5_ValAAC_5_+_180596610_180596682@1.29.29,trna6_ValCAC_5_+_180600650_180600722@1.29.29,trna85_ValCAC_1_-_161369490_161369562@1.29.29,trna90_ValCAC_1_-_149684088_149684161@1.29.29,trna98_ValCAC_1_-_149298555_149298627@1.29.29,trna9_ValCAC_6_+_26538282_26538354@1.29.29|GTTTCCGTAGTGTAGTGGTTATCACGTTC</td><td>>am_trna10_ValCAC_5_-_180649395_180649467</td><td>1</td><td>29</td><td>29</td><td>GTTTCCGTAGTGTAGTGGTTATCACGTTC</td><td>14764.5</td><td>-4.3</td><td>0.9</td><td>-4.9</td><td>1.1E-06</td><td>1.7E-06</td><td>MB231/MB468</td></tr>
<tr><td>>hg38_21_+_8433172_8446622_RNA45SN1_45S_50ntFlanks@10384.10424.41|CCATGTGAACAGCAGTTGAACATGGGTCAGTCGGTCCTGAG</td><td>>hg38_21_+_8433172_8446622_RNA45SN1_45S_50ntFlanks</td><td>10384</td><td>10424</td><td>41</td><td>CCATGTGAACAGCAGTTGAACATGGGTCAGTCGGTCCTGAG</td><td>775.9</td><td>3.8</td><td>0.8</td><td>4.9</td><td>1.1E-06</td><td>1.8E-06</td><td>MB231/MB468</td></tr>
<tr><td>>hg38_21_+_8433172_8446622_RNA45SN1_45S_50ntFlanks@10404.10424.21|CATGGGTCAGTCGGTCCTGAG</td><td>>hg38_21_+_8433172_8446622_RNA45SN1_45S_50ntFlanks</td><td>10404</td><td>10424</td><td>21</td><td>CATGGGTCAGTCGGTCCTGAG</td><td>1058.5</td><td>2.9</td><td>0.6</td><td>4.8</td><td>1.3E-06</td><td>1.8E-06</td><td>BT20/MB468</td></tr>
<tr><td>>hg38_21_+_8433172_8446622_RNA45SN1_45S_50ntFlanks@7981.8007.27|CTCAGATCAGACGTGGCGACCCGCTGA</td><td>>hg38_21_+_8433172_8446622_RNA45SN1_45S_50ntFlanks</td><td>7981</td><td>8007</td><td>27</td><td>CTCAGATCAGACGTGGCGACCCGCTGA</td><td>87.6</td><td>-5.2</td><td>1.1</td><td>-4.9</td><td>1.2E-06</td><td>1.8E-06</td><td>MB231/MB468</td></tr>
<tr><td>>hg38_21_+_8433172_8446622_RNA45SN1_45S_50ntFlanks@7975.8008.34|CGCGACCTCAGATCAGACGTGGCGACCCGCTGAA</td><td>>hg38_21_+_8433172_8446622_RNA45SN1_45S_50ntFlanks</td><td>7975</td><td>8008</td><td>34</td><td>CGCGACCTCAGATCAGACGTGGCGACCCGCTGAA</td><td>25546.8</td><td>2.8</td><td>0.6</td><td>4.8</td><td>1.3E-06</td><td>1.8E-06</td><td>BT20/MB468</td></tr>
<tr><td>>hg38_21_+_8433172_8446622_RNA45SN1_45S_50ntFlanks@10444.10462.19|CGAAGGGACGGGCGATGGC</td><td>>hg38_21_+_8433172_8446622_RNA45SN1_45S_50ntFlanks</td><td>10444</td><td>10462</td><td>19</td><td>CGAAGGGACGGGCGATGGC</td><td>141.2</td><td>-2.6</td><td>0.5</td><td>-4.9</td><td>1.2E-06</td><td>1.8E-06</td><td>MB231/MB468</td></tr>
<tr><td>>hg38_1_-_228634819_228635039_RNA5S12_5S_50ntFlanks@128.171.44|CTTGGATGGGAGACCGCCTGGGAATACCGGGTGCTGTAGGCTTT</td><td>>hg38_1_-_228634819_228635039_RNA5S12_5S_50ntFlanks</td><td>128</td><td>171</td><td>44</td><td>CTTGGATGGGAGACCGCCTGGGAATACCGGGTGCTGTAGGCTTT</td><td>1736.9</td><td>11.7</td><td>2.4</td><td>4.9</td><td>1.2E-06</td><td>1.8E-06</td><td>MB231/MB468</td></tr>
<tr><td>>hg38_21_+_8433172_8446622_RNA45SN1_45S_50ntFlanks@7991.8016.26|ACGTGGCGACCCGCTGAATTTAAGCA</td><td>>hg38_21_+_8433172_8446622_RNA45SN1_45S_50ntFlanks</td><td>7991</td><td>8016</td><td>26</td><td>ACGTGGCGACCCGCTGAATTTAAGCA</td><td>987.3</td><td>2.9</td><td>0.6</td><td>4.9</td><td>1.2E-06</td><td>1.8E-06</td><td>MB231/MB468</td></tr>
<tr><td>>hg38_21_+_8433172_8446622_RNA45SN1_45S_50ntFlanks@12417.12440.24|TCGATGTCGGCTCTTCCTATCATT</td><td>>hg38_21_+_8433172_8446622_RNA45SN1_45S_50ntFlanks</td><td>12417</td><td>12440</td><td>24</td><td>TCGATGTCGGCTCTTCCTATCATT</td><td>986.6</td><td>3.3</td><td>0.7</td><td>4.9</td><td>1.2E-06</td><td>1.8E-06</td><td>MB231/MB468</td></tr>
<tr><td>>hg38_21_+_8433172_8446622_RNA45SN1_45S_50ntFlanks@11561.11590.30|GCGCCTAGCAGCCGACTTAGAACTGGTGCG</td><td>>hg38_21_+_8433172_8446622_RNA45SN1_45S_50ntFlanks</td><td>11561</td><td>11590</td><td>30</td><td>GCGCCTAGCAGCCGACTTAGAACTGGTGCG</td><td>317.2</td><td>-2.8</td><td>0.6</td><td>-4.9</td><td>1.2E-06</td><td>1.8E-06</td><td>MB231/MB468</td></tr>
<tr><td>>hg38_21_+_8433172_8446622_RNA45SN1_45S_50ntFlanks@8198.8224.27|TGGACGGTGTGAGGCCGGTAGCGGCCC</td><td>>hg38_21_+_8433172_8446622_RNA45SN1_45S_50ntFlanks</td><td>8198</td><td>8224</td><td>27</td><td>TGGACGGTGTGAGGCCGGTAGCGGCCC</td><td>173.6</td><td>-2.2</td><td>0.5</td><td>-4.9</td><td>1.2E-06</td><td>1.8E-06</td><td>MB231/MB468</td></tr>
<tr><td>>MI0000102|hsa-mir-100&WithFlank&11|-|122152223|122152314@20.39.20(+1A)&[MIMAT0000098&hsa-miR-100-5p&offsets|+1|-1(+1A);m-27&11|-|122152276|122152296&offsets|+1|0(+1A)]|ACCCGTAGATCCGAACTTGTA</td><td>>MI0000102|hsa-mir-100&WithFlank&11|-|122152223|122152314</td><td>20</td><td>39</td><td>20(+1A)</td><td>ACCCGTAGATCCGAACTTGTA</td><td>194.3</td><td>-5.5</td><td>1.0</td><td>-5.3</td><td>9.8E-08</td><td>1.9E-06</td><td>BT20/MB231</td></tr>
<tr><td>>hg38_21_+_8433172_8446622_RNA45SN1_45S_50ntFlanks@11577.11600.24|TTAGAACTGGTGCGGACCAGGGGA</td><td>>hg38_21_+_8433172_8446622_RNA45SN1_45S_50ntFlanks</td><td>11577</td><td>11600</td><td>24</td><td>TTAGAACTGGTGCGGACCAGGGGA</td><td>103.6</td><td>-3.8</td><td>0.8</td><td>-4.9</td><td>1.2E-06</td><td>1.9E-06</td><td>MB231/MB468</td></tr>
<tr><td>>hg38_21_+_8433172_8446622_RNA45SN1_45S_50ntFlanks@12071.12097.27|CCCCGAGGGGCTCTCGCTTCTGGCGCC</td><td>>hg38_21_+_8433172_8446622_RNA45SN1_45S_50ntFlanks</td><td>12071</td><td>12097</td><td>27</td><td>CCCCGAGGGGCTCTCGCTTCTGGCGCC</td><td>236.6</td><td>-3.8</td><td>0.8</td><td>-4.8</td><td>1.3E-06</td><td>1.9E-06</td><td>BT20/MB468</td></tr>
<tr><td>>hg38_21_+_8433172_8446622_RNA45SN1_45S_50ntFlanks@9740.9757.18|AACGATCTCAACCTATTC</td><td>>hg38_21_+_8433172_8446622_RNA45SN1_45S_50ntFlanks</td><td>9740</td><td>9757</td><td>18</td><td>AACGATCTCAACCTATTC</td><td>139.5</td><td>-3.5</td><td>0.7</td><td>-4.8</td><td>1.3E-06</td><td>1.9E-06</td><td>BT20/MB468</td></tr>
<tr><td>>hg38_21_+_8433172_8446622_RNA45SN1_45S_50ntFlanks@10794.10821.28|CTGGCATGTTGGAACAATGTAGGTAAGG</td><td>>hg38_21_+_8433172_8446622_RNA45SN1_45S_50ntFlanks</td><td>10794</td><td>10821</td><td>28</td><td>CTGGCATGTTGGAACAATGTAGGTAAGG</td><td>159.4</td><td>-2.7</td><td>0.6</td><td>-4.8</td><td>1.3E-06</td><td>1.9E-06</td><td>BT20/MB468</td></tr>
<tr><td>>hg38_21_+_8433172_8446622_RNA45SN1_45S_50ntFlanks@10348.10365.18|AAACGAGAACTTTGAAGG</td><td>>hg38_21_+_8433172_8446622_RNA45SN1_45S_50ntFlanks</td><td>10348</td><td>10365</td><td>18</td><td>AAACGAGAACTTTGAAGG</td><td>288.6</td><td>-2.1</td><td>0.4</td><td>-4.8</td><td>1.4E-06</td><td>1.9E-06</td><td>BT20/MB468</td></tr>
<tr><td>>hg38_21_+_8433172_8446622_RNA45SN1_45S_50ntFlanks@10678.10706.29|CGGTTCCGGCGGCGTCCGGTGAGCTCTCG</td><td>>hg38_21_+_8433172_8446622_RNA45SN1_45S_50ntFlanks</td><td>10678</td><td>10706</td><td>29</td><td>CGGTTCCGGCGGCGTCCGGTGAGCTCTCG</td><td>19303.6</td><td>-5.3</td><td>1.1</td><td>-4.8</td><td>1.4E-06</td><td>1.9E-06</td><td>BT20/MB468</td></tr>
<tr><td>>hg38_21_+_8433172_8446622_RNA45SN1_45S_50ntFlanks@9602.9622.21|CGAAAGACTAATCGAACCATC</td><td>>hg38_21_+_8433172_8446622_RNA45SN1_45S_50ntFlanks</td><td>9602</td><td>9622</td><td>21</td><td>CGAAAGACTAATCGAACCATC</td><td>141.9</td><td>-3.7</td><td>0.8</td><td>-4.9</td><td>1.2E-06</td><td>1.9E-06</td><td>MB231/MB468</td></tr>
<tr><td>>hg38_21_+_8433172_8446622_RNA45SN1_45S_50ntFlanks@12625.12661.37|CAATGGGGCGAAGCTACCATCTGTGGGATTATGACTG</td><td>>hg38_21_+_8433172_8446622_RNA45SN1_45S_50ntFlanks</td><td>12625</td><td>12661</td><td>37</td><td>CAATGGGGCGAAGCTACCATCTGTGGGATTATGACTG</td><td>244.0</td><td>-2.7</td><td>0.6</td><td>-4.8</td><td>1.4E-06</td><td>1.9E-06</td><td>BT20/MB468</td></tr>
<tr><td>>hg38_21_+_8433172_8446622_RNA45SN1_45S_50ntFlanks@10795.10817.23|TGGCATGTTGGAACAATGTAGGT</td><td>>hg38_21_+_8433172_8446622_RNA45SN1_45S_50ntFlanks</td><td>10795</td><td>10817</td><td>23</td><td>TGGCATGTTGGAACAATGTAGGT</td><td>197.7</td><td>-2.4</td><td>0.5</td><td>-4.8</td><td>1.2E-06</td><td>1.9E-06</td><td>MB231/MB468</td></tr>
<tr><td>>hg38_21_+_8433172_8446622_RNA45SN1_45S_50ntFlanks@4729.4752.24|TCAAGAACGAAAGTCGGAGGTTCG</td><td>>hg38_21_+_8433172_8446622_RNA45SN1_45S_50ntFlanks</td><td>4729</td><td>4752</td><td>24</td><td>TCAAGAACGAAAGTCGGAGGTTCG</td><td>143.7</td><td>-3.8</td><td>0.8</td><td>-4.8</td><td>1.4E-06</td><td>1.9E-06</td><td>BT20/MB468</td></tr>
<tr><td>>hg38_21_+_8433172_8446622_RNA45SN1_45S_50ntFlanks@10348.10372.25|AAACGAGAACTTTGAAGGCCGAAGT</td><td>>hg38_21_+_8433172_8446622_RNA45SN1_45S_50ntFlanks</td><td>10348</td><td>10372</td><td>25</td><td>AAACGAGAACTTTGAAGGCCGAAGT</td><td>572.3</td><td>2.4</td><td>0.5</td><td>4.8</td><td>1.4E-06</td><td>1.9E-06</td><td>BT20/MB468</td></tr>
<tr><td>>hg38_21_+_8433172_8446622_RNA45SN1_45S_50ntFlanks@6656.6680.25|CTTAGCGGTGGATCACTCGGCTCGT</td><td>>hg38_21_+_8433172_8446622_RNA45SN1_45S_50ntFlanks</td><td>6656</td><td>6680</td><td>25</td><td>CTTAGCGGTGGATCACTCGGCTCGT</td><td>108.0</td><td>-3.1</td><td>0.6</td><td>-4.8</td><td>1.4E-06</td><td>1.9E-06</td><td>BT20/MB468</td></tr>
<tr><td>>trna152_ValCAC_6_-_27248049_27248121@1.28.28|GCTTCTGTAGTGTAGTGGTTATCACGTT</td><td>>trna152_ValCAC_6_-_27248049_27248121</td><td>1</td><td>28</td><td>28</td><td>GCTTCTGTAGTGTAGTGGTTATCACGTT</td><td>210.9</td><td>-3.6</td><td>0.7</td><td>-4.8</td><td>1.4E-06</td><td>2.0E-06</td><td>BT20/MB468</td></tr>
<tr><td>>hg38_21_+_8433172_8446622_RNA45SN1_45S_50ntFlanks@10402.10422.21|AACATGGGTCAGTCGGTCCTG</td><td>>hg38_21_+_8433172_8446622_RNA45SN1_45S_50ntFlanks</td><td>10402</td><td>10422</td><td>21</td><td>AACATGGGTCAGTCGGTCCTG</td><td>313.5</td><td>-3.1</td><td>0.6</td><td>-4.8</td><td>1.4E-06</td><td>2.0E-06</td><td>BT20/MB468</td></tr>
<tr><td>>hg38_21_+_8433172_8446622_RNA45SN1_45S_50ntFlanks@6731.6770.40|CACATTGATCATCGACACTTCGAACGCACTTGCGGCCCCG</td><td>>hg38_21_+_8433172_8446622_RNA45SN1_45S_50ntFlanks</td><td>6731</td><td>6770</td><td>40</td><td>CACATTGATCATCGACACTTCGAACGCACTTGCGGCCCCG</td><td>578.3</td><td>-3.7</td><td>0.8</td><td>-4.8</td><td>1.3E-06</td><td>2.0E-06</td><td>MB231/MB468</td></tr>
<tr><td>>hg38_21_+_8433172_8446622_RNA45SN1_45S_50ntFlanks@10794.10817.24|CTGGCATGTTGGAACAATGTAGGT</td><td>>hg38_21_+_8433172_8446622_RNA45SN1_45S_50ntFlanks</td><td>10794</td><td>10817</td><td>24</td><td>CTGGCATGTTGGAACAATGTAGGT</td><td>699.8</td><td>-4.0</td><td>0.8</td><td>-4.8</td><td>1.4E-06</td><td>2.0E-06</td><td>BT20/MB468</td></tr>
<tr><td>>hg38_21_+_8433172_8446622_RNA45SN1_45S_50ntFlanks@6733.6756.24|CATTGATCATCGACACTTCGAACG</td><td>>hg38_21_+_8433172_8446622_RNA45SN1_45S_50ntFlanks</td><td>6733</td><td>6756</td><td>24</td><td>CATTGATCATCGACACTTCGAACG</td><td>891.1</td><td>-2.3</td><td>0.5</td><td>-4.8</td><td>1.4E-06</td><td>2.0E-06</td><td>BT20/MB468</td></tr>
<tr><td>>hg38_21_+_8433172_8446622_RNA45SN1_45S_50ntFlanks@8380.8403.24|CAAGAGGGCGTGAAACCGTTAAGA</td><td>>hg38_21_+_8433172_8446622_RNA45SN1_45S_50ntFlanks</td><td>8380</td><td>8403</td><td>24</td><td>CAAGAGGGCGTGAAACCGTTAAGA</td><td>523.6</td><td>-2.8</td><td>0.6</td><td>-4.8</td><td>1.3E-06</td><td>2.0E-06</td><td>MB231/MB468</td></tr>
<tr><td>>hg38_21_+_8433172_8446622_RNA45SN1_45S_50ntFlanks@10395.10424.30|GCAGTTGAACATGGGTCAGTCGGTCCTGAG</td><td>>hg38_21_+_8433172_8446622_RNA45SN1_45S_50ntFlanks</td><td>10395</td><td>10424</td><td>30</td><td>GCAGTTGAACATGGGTCAGTCGGTCCTGAG</td><td>954.4</td><td>4.6</td><td>0.9</td><td>4.8</td><td>1.4E-06</td><td>2.0E-06</td><td>BT20/MB468</td></tr>
<tr><td>>MI0000767|hsa-mir-365a&WithFlank&16|+|14309279|14309377@62.81.20;MI0000769|hsa-mir-365b&WithFlank&17|+|31575405|31575527@74.93.20&[MIMAT0000710&hsa-miR-365a-3p&offsets|0|-2;m-176&16|+|14309340|14309361&offsets|0|-2];[MIMAT0022834&hsa-miR-365b-3p&offsets|0|-2;m-175&17|+|31575478|31575499&offsets|0|-2]|TAATGCCCCTAAAAATCCTT</td><td>>MI0000767|hsa-mir-365a&WithFlank&16|+|14309279|14309377</td><td>62</td><td>81</td><td>20</td><td>TAATGCCCCTAAAAATCCTT</td><td>298.2</td><td>4.4</td><td>0.8</td><td>5.3</td><td>1.1E-07</td><td>2.0E-06</td><td>BT20/MB231</td></tr>
</table>

| | | | | | | | | | | | | |
|---|---|---|---|---|---|---|---|---|---|---|---|---|
| >MI0003776\|hsa-mir-320b-1&WithFlank&1\|+\|116671740\|116671823@48.66.19(+1A);MI0003839\|hsa-mir-320b-2&WithFlank&1\|-\|224257034\|224257116@47.65.19(+1A);MI0003778\|hsa-mir-320c-1&WithFlank&18\|+\|21683512\|21683595@48.66.19(+1A);MI0008191\|hsa-mir-320c-2&WithFlank&18\|+\|24321669\|24321752@48.66.19(+1A);MI0000542\|hsa-mir-320a&WithFlank&8\|-\|22244960\|22245043@48.66.19(+1A)&[MIMAT0005792&hsa-miR-320b&offsets\|0\|-3(+1A);m-183&1\|+\|116671787\|116671806&offsets\|0\|-1(+1A)];[MIMAT0005792_1&hsa-miR-320b&offsets\|0\|-3(+1A);m-183&1\|-\|224257051\|224257070&offsets\|0\|-1(+1A)];[MIMAT0005793&hsa-miR-320c&offsets\|0\|-1(+1A);m-222&18\|+\|21683559\|21683578&offsets\|0\|-1(+1A)];[MIMAT0005793_1&hsa-miR-320c&offsets\|0\|-1(+1A);m-219&18\|+\|24321716\|24321735&offsets\|0\|-1(+1A)];[MIMAT0000510&hsa-miR-320a-3p&offsets\|0\|-3(+1A);m-60&8\|-\|22244975\|22244996&offsets\|0\|-3(+1A)]\|AAAAGCTGGGTTGAGAGGGA | >MI0003776\|hsa-mir-320b-1&WithFlank&1\|+\|116671740\|116671823 | 48 | 66 | 19(+1A) | AAAAGCTGGGTTGAGAGGGA | 311.9 | 3.8 | 0.7 | 5.3 | 1.1E-07 | 2.0E-06 | BT20/MB231 |
| >trna14_LysTTT_11_-_59327808_59327880@1.30.30,trna2_LysTTT_17_+_8022473_8022545@1.30.30,trna54_LysTTT_1_+_204475655_204475727@1.30.30,trna5_LysTTT_11_+_59323902_59323974@1.30.30,trna62_LysTTT_1_-_204476158_204476230@1.30.30,trna76_LysTTT_6_+_28918806_28918878@1.30.30\|GCCCGGATAGCTCAGTCGGTAGAGCATCAG | >trna14_LysTTT_11_-_59327808_59327880 | 1 | 30 | 30 | GCCCGGATAGCTCAGTCGGTAGAGCATCAG | 923.7 | -6.6 | 1.4 | -4.8 | 1.4E-06 | 2.0E-06 | BT20/MB468 |
| >hg38_21_+_8433172_8446622_RNA45SN1_45S_50ntFlanks@8357.8376.20\|AAAAGAACTTTGAAGAGAGA | >hg38_21_+_8433172_8446622_RNA45SN1_45S_50ntFlanks | 8357 | 8376 | 20 | AAAAGAACTTTGAAGAGAGA | 958.1 | -3.2 | 0.7 | -4.8 | 1.5E-06 | 2.0E-06 | BT20/MB468 |
| >hg38_21_+_8433172_8446622_RNA45SN1_45S_50ntFlanks@3876.3898.23\|TACATGCCGACGGGCGCTGACCC | >hg38_21_+_8433172_8446622_RNA45SN1_45S_50ntFlanks | 3876 | 3898 | 23 | TACATGCCGACGGGCGCTGACCC | 667.4 | -2.7 | 0.6 | -4.8 | 1.3E-06 | 2.0E-06 | MB231/MB468 |
| >hg38_21_+_8433172_8446622_RNA45SN1_45S_50ntFlanks@6661.6696.36\|CGGTGGATCACTCGGCTCGTGCGTCGATGAAGAACG | >hg38_21_+_8433172_8446622_RNA45SN1_45S_50ntFlanks | 6661 | 6696 | 36 | CGGTGGATCACTCGGCTCGTGCGTCGATGAAGAACG | 917.9 | 3.3 | 0.7 | 4.8 | 1.5E-06 | 2.0E-06 | BT20/MB468 |
| >hg38_21_+_8433172_8446622_RNA45SN1_45S_50ntFlanks@7979.7999.21\|ACCTCAGATCAGACGTGGCGA | >hg38_21_+_8433172_8446622_RNA45SN1_45S_50ntFlanks | 7979 | 7999 | 21 | ACCTCAGATCAGACGTGGCGA | 121.5 | -4.8 | 1.0 | -4.8 | 1.5E-06 | 2.0E-06 | BT20/MB468 |
| >hg38_21_+_8433172_8446622_RNA45SN1_45S_50ntFlanks@11206.11225.20\|CGGCGGCGACTCTGGACGCG | >hg38_21_+_8433172_8446622_RNA45SN1_45S_50ntFlanks | 11206 | 11225 | 20 | CGGCGGCGACTCTGGACGCG | 133.1 | -4.0 | 0.8 | -4.8 | 1.3E-06 | 2.1E-06 | MB231/MB468 |
| >hg38_21_+_8433172_8446622_RNA45SN1_45S_50ntFlanks@11565.11585.21\|CTAGCAGCCGACTTAGAACTG | >hg38_21_+_8433172_8446622_RNA45SN1_45S_50ntFlanks | 11565 | 11585 | 21 | CTAGCAGCCGACTTAGAACTG | 229.1 | -2.0 | 0.4 | -4.8 | 1.5E-06 | 2.1E-06 | BT20/MB468 |
| >hg38_21_+_8433172_8446622_RNA45SN1_45S_50ntFlanks@7981.8003.23\|CTCAGATCAGACGTGGCGACCCG | >hg38_21_+_8433172_8446622_RNA45SN1_45S_50ntFlanks | 7981 | 8003 | 23 | CTCAGATCAGACGTGGCGACCCG | 75.5 | -4.2 | 0.9 | -4.8 | 1.4E-06 | 2.1E-06 | MB231/MB468 |
| >hg38_21_+_8433172_8446622_RNA45SN1_45S_50ntFlanks@10348.10371.24\|AAACGAGAACTTTGAAGGCCGAAG | >hg38_21_+_8433172_8446622_RNA45SN1_45S_50ntFlanks | 10348 | 10371 | 24 | AAACGAGAACTTTGAAGGCCGAAG | 1365.4 | 3.2 | 0.7 | 4.8 | 1.4E-06 | 2.1E-06 | MB231/MB468 |
| >hg38_21_+_8433172_8446622_RNA45SN1_45S_50ntFlanks@5486.5516.31\|GCGCTGAGAAGACGGTCGAACTTGACTATCT | >hg38_21_+_8433172_8446622_RNA45SN1_45S_50ntFlanks | 5486 | 5516 | 31 | GCGCTGAGAAGACGGTCGAACTTGACTATCT | 596.0 | 4.7 | 1.0 | 4.8 | 1.5E-06 | 2.1E-06 | BT20/MB468 |
| >hg38_21_+_8433172_8446622_RNA45SN1_45S_50ntFlanks@3843.3869.27\|CTACTTGGATAACTGTGGTAATTCTAG | >hg38_21_+_8433172_8446622_RNA45SN1_45S_50ntFlanks | 3843 | 3869 | 27 | CTACTTGGATAACTGTGGTAATTCTAG | 128.8 | -2.2 | 0.5 | -4.8 | 1.4E-06 | 2.1E-06 | MB231/MB468 |
| >hg38_21_+_8433172_8446622_RNA45SN1_45S_50ntFlanks@12394.12418.25\|TAGCGACGTCGCTTTTTGATCCTTC | >hg38_21_+_8433172_8446622_RNA45SN1_45S_50ntFlanks | 12394 | 12418 | 25 | TAGCGACGTCGCTTTTTGATCCTTC | 474.7 | 3.6 | 0.7 | 4.8 | 1.4E-06 | 2.1E-06 | MB231/MB468 |
| >hg38_21_+_8433172_8446622_RNA45SN1_45S_50ntFlanks@11579.11600.22\|AGAACTGGTGCGGACCAGGGGA | >hg38_21_+_8433172_8446622_RNA45SN1_45S_50ntFlanks | 11579 | 11600 | 22 | AGAACTGGTGCGGACCAGGGGA | 190.2 | -4.5 | 0.9 | -4.8 | 1.6E-06 | 2.2E-06 | BT20/MB468 |
| >am_trna130_GlnTTG_6_-_27763640_27763711@13.30.18,trna146_GlnCTG_6_-_27515531_27515602@13.30.18,trna16_GlnTTG_17_+_47269890_47269961@13.30.18,trna173_GlnTTG_6_-_26311975_26312046@13.30.18,trna174_GlnTTG_6_-_26311424_26311495@13.30.18,trna1_GlnCTG_6_+_18836402_18836473@13.30.18,trna3_GlnCTG_17_+_8023070_8023141@13.30.18,trna42_GlnCTG_6_+_27263212_27263283@13.30.18,trna49_GlnCTG_6_+_27487308_27487379@13.30.18,trna64_GlnTTG_6_+_28557156_28557227@13.30.18,trna7_GlnCTG_15_-_66161400_66161471@13.30.18,trna84_GlnTTG_6_+_145503859_145503930@13.30.18,trna99_GlnCTG_6_-_28909378_28909449@13.30.18\|TAATGGTTAGCACTCTGG | >am_trna130_GlnTTG_6_-_27763640_27763711 | 13 | 30 | 18 | TAATGGTTAGCACTCTGG | 159.9 | -3.6 | 0.8 | -4.8 | 1.4E-06 | 2.2E-06 | MB231/MB468 |
| >hg38_21_+_8433172_8446622_RNA45SN1_45S_50ntFlanks@11718.11744.27\|AACGGCGGGAGTAACTATGACTCTCTT | >hg38_21_+_8433172_8446622_RNA45SN1_45S_50ntFlanks | 11718 | 11744 | 27 | AACGGCGGGAGTAACTATGACTCTCTT | 472.7 | -3.7 | 0.8 | -4.8 | 1.4E-06 | 2.2E-06 | MB231/MB468 |
| >hg38_21_+_8433172_8446622_RNA45SN1_45S_50ntFlanks@10409.10431.23\|GTCAGTCGGTCCTGAGAGATGGG | >hg38_21_+_8433172_8446622_RNA45SN1_45S_50ntFlanks | 10409 | 10431 | 23 | GTCAGTCGGTCCTGAGAGATGGG | 178.0 | -4.6 | 1.0 | -4.8 | 1.6E-06 | 2.2E-06 | BT20/MB468 |
| >hg38_21_+_8433172_8446622_RNA45SN1_45S_50ntFlanks@12940.12965.26\|CATTCGTAGACGACCTGCTTCTGGGT | >hg38_21_+_8433172_8446622_RNA45SN1_45S_50ntFlanks | 12940 | 12965 | 26 | CATTCGTAGACGACCTGCTTCTGGGT | 150.9 | -2.7 | 0.6 | -4.8 | 1.6E-06 | 2.2E-06 | BT20/MB468 |
| >hg38_21_+_8433172_8446622_RNA45SN1_45S_50ntFlanks@12936.12962.27\|AAACCATTCGTAGACGACCTGCTTCTG | >hg38_21_+_8433172_8446622_RNA45SN1_45S_50ntFlanks | 12936 | 12962 | 27 | AAACCATTCGTAGACGACCTGCTTCTG | 121.4 | -3.0 | 0.6 | -4.8 | 1.5E-06 | 2.2E-06 | MB231/MB468 |
| >hg38_21_+_8433172_8446622_RNA45SN1_45S_50ntFlanks@10873.10890.18\|CTGGGTCGGTCGGGCTGG | >hg38_21_+_8433172_8446622_RNA45SN1_45S_50ntFlanks | 10873 | 10890 | 18 | CTGGGTCGGTCGGGCTGG | 167.0 | -2.9 | 0.6 | -4.8 | 1.5E-06 | 2.2E-06 | MB231/MB468 |
| >hg38_21_+_8433172_8446622_RNA45SN1_45S_50ntFlanks@5489.5516.28\|CTGAGAAGACGGTCGAACTTGACTATCT | >hg38_21_+_8433172_8446622_RNA45SN1_45S_50ntFlanks | 5489 | 5516 | 28 | CTGAGAAGACGGTCGAACTTGACTATCT | 4165.0 | 4.9 | 1.0 | 4.8 | 1.6E-06 | 2.2E-06 | BT20/MB468 |
| >hg38_21_+_8433172_8446622_RNA45SN1_45S_50ntFlanks@10411.10440.30\|CAGTCGGTCCTGAGAGATGGGCGAGCGCCG | >hg38_21_+_8433172_8446622_RNA45SN1_45S_50ntFlanks | 10411 | 10440 | 30 | CAGTCGGTCCTGAGAGATGGGCGAGCGCCG | 180.8 | -4.7 | 1.0 | -4.8 | 1.6E-06 | 2.2E-06 | BT20/MB468 |
| >hg38_21_+_8433172_8446622_RNA45SN1_45S_50ntFlanks@4122.4145.24\|AGGGTTCGATTCCGGAGAGGGAGC | >hg38_21_+_8433172_8446622_RNA45SN1_45S_50ntFlanks | 4122 | 4145 | 24 | AGGGTTCGATTCCGGAGAGGGAGC | 83.1 | -3.7 | 0.8 | -4.8 | 1.5E-06 | 2.3E-06 | MB231/MB468 |
| >MI0000342\|hsa-mir-200b&WithFlank&1\|+\|1167098\|1167204@64.85.22(+1U)&[MIMAT0000318&hsa-miR-200b-3p&offsets\|+1\|+1(+1U);m-93&1\|+\|1167160\|1167182&offsets\|+1\|0(+1U)]\|AATACTGCCTGGTAATGATGACT | >MI0000342\|hsa-mir-200b&WithFlank&1\|+\|1167098\|1167204 | 64 | 85 | 22(+1U) | AATACTGCCTGGTAATGATGACT | 211.7 | -2.4 | 0.5 | -4.8 | 1.5E-06 | 2.3E-06 | MB231/MB468 |
| >hg38_21_+_8433172_8446622_RNA45SN1_45S_50ntFlanks@11787.11806.20\|AATGGATGAACGAGATTCCC | >hg38_21_+_8433172_8446622_RNA45SN1_45S_50ntFlanks | 11787 | 11806 | 20 | AATGGATGAACGAGATTCCC | 158.1 | -4.7 | 1.0 | -4.8 | 1.5E-06 | 2.3E-06 | MB231/MB468 |
| >hg38_21_+_8433172_8446622_RNA45SN1_45S_50ntFlanks@11569.11599.31\|CAGCCGACTTAGAACTGGTGCGGACCAGGGG | >hg38_21_+_8433172_8446622_RNA45SN1_45S_50ntFlanks | 11569 | 11599 | 31 | CAGCCGACTTAGAACTGGTGCGGACCAGGGG | 212.2 | -2.1 | 0.4 | -4.8 | 1.7E-06 | 2.3E-06 | BT20/MB468 |
| >hg38_21_+_8433172_8446622_RNA45SN1_45S_50ntFlanks@9740.9758.19\|AACGATCTCAACCTATTCT | >hg38_21_+_8433172_8446622_RNA45SN1_45S_50ntFlanks | 9740 | 9758 | 19 | AACGATCTCAACCTATTCT | 296.0 | -2.8 | 0.6 | -4.8 | 1.7E-06 | 2.3E-06 | BT20/MB468 |
| >hg38_21_+_8433172_8446622_RNA45SN1_45S_50ntFlanks@4007.4026.20\|CCTCGGGCCGATCGCACGCC | >hg38_21_+_8433172_8446622_RNA45SN1_45S_50ntFlanks | 4007 | 4026 | 20 | CCTCGGGCCGATCGCACGCC | 87.1 | -4.3 | 0.9 | -4.8 | 1.5E-06 | 2.3E-06 | MB231/MB468 |
| >hg38_21_+_8433172_8446622_RNA45SN1_45S_50ntFlanks@12624.12657.34\|CCAATGGGGCGAAGCTACCATCTGTGGGATTATG | >hg38_21_+_8433172_8446622_RNA45SN1_45S_50ntFlanks | 12624 | 12657 | 34 | CCAATGGGGCGAAGCTACCATCTGTGGGATTATG | 477.4 | -3.1 | 0.6 | -4.8 | 1.5E-06 | 2.3E-06 | MB231/MB468 |
| >hg38_MT_+_598_1651_RNR1_12S_50ntFlanks@428.455.28\|AAAGTGGCTTTAACATATCTGAACACAC | >hg38_MT_+_598_1651_RNR1_12S_50ntFlanks | 428 | 455 | 28 | AAAGTGGCTTTAACATATCTGAACACAC | 1704.6 | 11.2 | 2.3 | 4.8 | 1.5E-06 | 2.3E-06 | MB231/MB468 |
| >am_trna128_GlyGCC_6_-_27870686_27870756@1.34.34,trna18_GlyGCC_16_+_70822597_70822667@1.34.34,trna19_GlyGCC_16_+_70823410_70823480@1.34.34,trna19_GlyGCC_2_-_157257659_157257729@1.34.34,trna24_GlyGCC_16_-_70812942_70813012@1.34.34,trna25_GlyGCC_16_-_70812114_70812184@1.34.34,trna5_GlyGCC_17_+_8029064_8029134@1.34.34,trna68_GlyGCC_1_-_161493637_161493707@1.34.34\|GCATTGGTGGTTCAGTGGTAGAATTCTCGCCTGC | >am_trna128_GlyGCC_6_-_27870686_27870756 | 1 | 34 | 34 | GCATTGGTGGTTCAGTGGTAGAATTCTCGCCTGC | 3843.6 | -4.3 | 0.8 | -5.3 | 1.2E-07 | 2.3E-06 | BT20/MB231 |
| >hg38_21_+_8433172_8446622_RNA45SN1_45S_50ntFlanks@10798.10825.28\|CATGTTGGAACAATGTAGGTAAGGGAAG | >hg38_21_+_8433172_8446622_RNA45SN1_45S_50ntFlanks | 10798 | 10825 | 28 | CATGTTGGAACAATGTAGGTAAGGGAAG | 161.8 | -3.8 | 0.8 | -4.8 | 1.5E-06 | 2.3E-06 | MB231/MB468 |
| >hg38_21_+_8433172_8446622_RNA45SN1_45S_50ntFlanks@11669.11702.34\|GCCCAGTGCTCTGAATGTCAAAGTGAAGAAATTC | >hg38_21_+_8433172_8446622_RNA45SN1_45S_50ntFlanks | 11669 | 11702 | 34 | GCCCAGTGCTCTGAATGTCAAAGTGAAGAAATTC | 449.0 | 4.0 | 0.8 | 4.8 | 1.7E-06 | 2.3E-06 | BT20/MB468 |
| >hg38_21_+_8433172_8446622_RNA45SN1_45S_50ntFlanks@11592.11611.20\|ACCAGGGGAATCCGACTGTT | >hg38_21_+_8433172_8446622_RNA45SN1_45S_50ntFlanks | 11592 | 11611 | 20 | ACCAGGGGAATCCGACTGTT | 514.7 | -3.9 | 0.8 | -4.8 | 1.7E-06 | 2.3E-06 | BT20/MB468 |
| >MI0000102\|hsa-mir-100&WithFlank&11\|-\|122152223\|122152314@19.38.20(+1A)&[MIMAT0000098&hsa-miR-100-5p&offsets\|0\|-2(+1A);m-27&11\|-\|122152276\|122152296&offsets\|0\|-1(+1A)]\|AACCCGTAGATCCGAACTTGA | >MI0000102\|hsa-mir-100&WithFlank&11\|-\|122152223\|122152314 | 19 | 38 | 20(+1A) | AACCCGTAGATCCGAACTTGA | 4549.8 | -5.0 | 0.9 | -5.3 | 1.3E-07 | 2.4E-06 | BT20/MB231 |

<table>
<thead>
<tr><th></th><th></th><th></th><th></th><th></th><th></th><th></th><th></th><th></th><th></th><th></th><th></th><th></th></tr>
</thead>
<tbody>
<tr><td>>trna111_HisGTG_1_-_147774845_147774916@-1G.34.35,trna118_HisGTG_1_-_145396881_145396952@-1G.34.35,trna16_HisGTG_1_+_146544773_146544844@-1G.34.35,trna1_HisGTG_15_+_45493349_45493420@-1G.34.35,trna21_HisGTG_1_+_147753471_147753542@-1G.34.35,trna33_HisGTG_6_+_27125906_27125977@-1G.34.35,trna7_HisGTG_9_-_14433938_14434009@-1G.34.35,trna8_HisGTG_15_-_45492611_45492682@-1G.34.35,trna9_HisGTG_15_-_45490804_45490875@-1G.34.35|GGCCGTGATCGTATAGTGGTTAGTACTCTGCGTTG</td><td>>trna111_HisGTG_1_-_147774845_147774916</td><td>-1G</td><td>34</td><td>35</td><td>GGCCGTGATCGTATAGTGGTTAGTACTCTGCGTTG</td><td>4181.2</td><td>5.6</td><td>1.2</td><td>4.8</td><td>1.7E-06</td><td>2.4E-06</td><td>BT20/MB468</td></tr>
<tr><td>>MI0000101|hsa-mir-99a&WithFlank&21|+|16539083|16539175@19.38.20(+1C)&[MIMAT0000097&hsa-miR-99a-5p&offsets|0|-2(+1C);m-64&21|+|16539101|16539121&offsets|0|-1(+1C)]|AACCCGTAGATCCGATCTTGC</td><td>>MI0000101|hsa-mir-99a&WithFlank&21|+|16539083|16539175</td><td>19</td><td>38</td><td>20(+1C)</td><td>AACCCGTAGATCCGATCTTGC</td><td>126.2</td><td>-4.5</td><td>0.9</td><td>-4.8</td><td>1.8E-06</td><td>2.4E-06</td><td>BT20/MB468</td></tr>
<tr><td>>hg38_21_+_8433172_8446622_RNA45SN1_45S_50ntFlanks@10668.10706.39|GAAAGCGTCGCGGTTCCGGCGGCGTCCGGTGAGCTCTCG</td><td>>hg38_21_+_8433172_8446622_RNA45SN1_45S_50ntFlanks</td><td>10668</td><td>10706</td><td>39</td><td>GAAAGCGTCGCGGTTCCGGCGGCGTCCGGTGAGCTCTCG</td><td>207.2</td><td>-4.9</td><td>1.0</td><td>-4.8</td><td>1.6E-06</td><td>2.4E-06</td><td>MB231/MB468</td></tr>
<tr><td>>hg38_21_+_8433172_8446622_RNA45SN1_45S_50ntFlanks@12202.12220.19|CTAAGGCGAGCTCAGGGAG</td><td>>hg38_21_+_8433172_8446622_RNA45SN1_45S_50ntFlanks</td><td>12202</td><td>12220</td><td>19</td><td>CTAAGGCGAGCTCAGGGAG</td><td>110.6</td><td>-3.4</td><td>0.7</td><td>-4.8</td><td>1.8E-06</td><td>2.4E-06</td><td>BT20/MB468</td></tr>
<tr><td>>MI0000085|hsa-mir-27a&WithFlank&19|-|13836434|13836523@57.76.20(+3U)&[MIMAT0000084&hsa-miR-27a-3p&offsets|0|-1(+3U);m-43&19|-|13836447|13836467&offsets|0|-1(+3U)]|TTCACAGTGGCTAAGTTCCGTTT</td><td>>MI0000085|hsa-mir-27a&WithFlank&19|-|13836434|13836523</td><td>57</td><td>76</td><td>20(+3U)</td><td>TTCACAGTGGCTAAGTTCCGTTT</td><td>1005.6</td><td>4.7</td><td>1.0</td><td>4.8</td><td>1.8E-06</td><td>2.4E-06</td><td>BT20/MB468</td></tr>
<tr><td>>hg38_21_+_8433172_8446622_RNA45SN1_45S_50ntFlanks@10391.10422.32|AACAGCAGTTGAACATGGGTCAGTCGGTCCTG</td><td>>hg38_21_+_8433172_8446622_RNA45SN1_45S_50ntFlanks</td><td>10391</td><td>10422</td><td>32</td><td>AACAGCAGTTGAACATGGGTCAGTCGGTCCTG</td><td>251.6</td><td>-3.0</td><td>0.6</td><td>-4.8</td><td>1.8E-06</td><td>2.4E-06</td><td>BT20/MB468</td></tr>
<tr><td>>hg38_21_+_8433172_8446622_RNA45SN1_45S_50ntFlanks@6776.6806.31|CTCCCGGGGCTACGCCTGTCTGAGCGTCGCT</td><td>>hg38_21_+_8433172_8446622_RNA45SN1_45S_50ntFlanks</td><td>6776</td><td>6806</td><td>31</td><td>CTCCCGGGGCTACGCCTGTCTGAGCGTCGCT</td><td>2618.1</td><td>3.3</td><td>0.7</td><td>4.8</td><td>1.8E-06</td><td>2.4E-06</td><td>BT20/MB468</td></tr>
<tr><td>>hg38_21_+_8433172_8446622_RNA45SN1_45S_50ntFlanks@12637.12657.21|GCTACCATCTGTGGGATTATG</td><td>>hg38_21_+_8433172_8446622_RNA45SN1_45S_50ntFlanks</td><td>12637</td><td>12657</td><td>21</td><td>GCTACCATCTGTGGGATTATG</td><td>109.4</td><td>-2.7</td><td>0.6</td><td>-4.8</td><td>1.6E-06</td><td>2.4E-06</td><td>MB231/MB468</td></tr>
<tr><td>>hg38_1_-_228634819_228635039_RNA5S12_5S_50ntFlanks@140.170.31|ACCGCCTGGGAATACCGGGTGCTGTAGGCTT</td><td>>hg38_1_-_228634819_228635039_RNA5S12_5S_50ntFlanks</td><td>140</td><td>170</td><td>31</td><td>ACCGCCTGGGAATACCGGGTGCTGTAGGCTT</td><td>16178.9</td><td>3.7</td><td>0.8</td><td>4.8</td><td>1.8E-06</td><td>2.5E-06</td><td>BT20/MB468</td></tr>
<tr><td>>hg38_21_+_8433172_8446622_RNA45SN1_45S_50ntFlanks@11209.11254.46|CGGCGACTCTGGACGCGAGCCGGGCCCTTCCCGTGGATCGCCCCAG</td><td>>hg38_21_+_8433172_8446622_RNA45SN1_45S_50ntFlanks</td><td>11209</td><td>11254</td><td>46</td><td>CGGCGACTCTGGACGCGAGCCGGGCCCTTCCCGTGGATCGCCCCAG</td><td>1485.7</td><td>-4.2</td><td>0.9</td><td>-4.8</td><td>1.8E-06</td><td>2.5E-06</td><td>BT20/MB468</td></tr>
<tr><td>>hg38_21_+_8433172_8446622_RNA45SN1_45S_50ntFlanks@3852.3872.21|TAACTGTGGTAATTCTAGAGC</td><td>>hg38_21_+_8433172_8446622_RNA45SN1_45S_50ntFlanks</td><td>3852</td><td>3872</td><td>21</td><td>TAACTGTGGTAATTCTAGAGC</td><td>146.6</td><td>-2.1</td><td>0.4</td><td>-4.8</td><td>1.8E-06</td><td>2.5E-06</td><td>BT20/MB468</td></tr>
<tr><td>>hg38_21_+_8433172_8446622_RNA45SN1_45S_50ntFlanks@9305.9323.19|CCAAGGAGTCTAACACGTG</td><td>>hg38_21_+_8433172_8446622_RNA45SN1_45S_50ntFlanks</td><td>9305</td><td>9323</td><td>19</td><td>CCAAGGAGTCTAACACGTG</td><td>97.6</td><td>-3.7</td><td>0.8</td><td>-4.8</td><td>1.8E-06</td><td>2.5E-06</td><td>BT20/MB468</td></tr>
<tr><td>>hg38_21_+_8433172_8446622_RNA45SN1_45S_50ntFlanks@11670.11701.32|CCCAGTGCTCTGAATGTCAAAGTGAAGAAATT</td><td>>hg38_21_+_8433172_8446622_RNA45SN1_45S_50ntFlanks</td><td>11670</td><td>11701</td><td>32</td><td>CCCAGTGCTCTGAATGTCAAAGTGAAGAAATT</td><td>522.3</td><td>-2.7</td><td>0.6</td><td>-4.8</td><td>1.7E-06</td><td>2.5E-06</td><td>MB231/MB468</td></tr>
<tr><td>>hg38_21_+_8433172_8446622_RNA45SN1_45S_50ntFlanks@12488.12508.21|AACGTGAGCTGGGTTTAGACC</td><td>>hg38_21_+_8433172_8446622_RNA45SN1_45S_50ntFlanks</td><td>12488</td><td>12508</td><td>21</td><td>AACGTGAGCTGGGTTTAGACC</td><td>141.3</td><td>-4.0</td><td>0.8</td><td>-4.8</td><td>1.9E-06</td><td>2.6E-06</td><td>BT20/MB468</td></tr>
<tr><td>>hg38_21_+_8433172_8446622_RNA45SN1_45S_50ntFlanks@11718.11738.21|AACGGCGGGAGTAACTATGAC</td><td>>hg38_21_+_8433172_8446622_RNA45SN1_45S_50ntFlanks</td><td>11718</td><td>11738</td><td>21</td><td>AACGGCGGGAGTAACTATGAC</td><td>618.6</td><td>-3.9</td><td>0.8</td><td>-4.8</td><td>1.7E-06</td><td>2.6E-06</td><td>MB231/MB468</td></tr>
<tr><td>>hg38_21_+_8433172_8446622_RNA45SN1_45S_50ntFlanks@12642.12659.18|CATCTGTGGGATTATGAC</td><td>>hg38_21_+_8433172_8446622_RNA45SN1_45S_50ntFlanks</td><td>12642</td><td>12659</td><td>18</td><td>CATCTGTGGGATTATGAC</td><td>179.9</td><td>-2.8</td><td>0.6</td><td>-4.8</td><td>1.7E-06</td><td>2.6E-06</td><td>MB231/MB468</td></tr>
<tr><td>>hg38_21_+_8433172_8446622_RNA45SN1_45S_50ntFlanks@11321.11343.23|CACGTCTCGTCGCGCGCGCGTCC</td><td>>hg38_21_+_8433172_8446622_RNA45SN1_45S_50ntFlanks</td><td>11321</td><td>11343</td><td>23</td><td>CACGTCTCGTCGCGCGCGCGTCC</td><td>338.0</td><td>-4.0</td><td>0.8</td><td>-4.8</td><td>1.7E-06</td><td>2.6E-06</td><td>MB231/MB468</td></tr>
<tr><td>>hg38_21_+_8433172_8446622_RNA45SN1_45S_50ntFlanks@12994.13014.21|CCTCGCTGCGATCTATTGAAA</td><td>>hg38_21_+_8433172_8446622_RNA45SN1_45S_50ntFlanks</td><td>12994</td><td>13014</td><td>21</td><td>CCTCGCTGCGATCTATTGAAA</td><td>189.9</td><td>-3.3</td><td>0.7</td><td>-4.8</td><td>1.9E-06</td><td>2.6E-06</td><td>BT20/MB468</td></tr>
<tr><td>>hg38_21_+_8433172_8446622_RNA45SN1_45S_50ntFlanks@10350.10384.35|ACGAGAACTTTGAAGGCCGAAGTGGAGAAGGGTTC</td><td>>hg38_21_+_8433172_8446622_RNA45SN1_45S_50ntFlanks</td><td>10350</td><td>10384</td><td>35</td><td>ACGAGAACTTTGAAGGCCGAAGTGGAGAAGGGTTC</td><td>1066.3</td><td>-2.9</td><td>0.6</td><td>-4.8</td><td>1.7E-06</td><td>2.6E-06</td><td>MB231/MB468</td></tr>
<tr><td>>hg38_21_+_8433172_8446622_RNA45SN1_45S_50ntFlanks@12459.12478.20|AGCGTTGGATTGTTCACCCA</td><td>>hg38_21_+_8433172_8446622_RNA45SN1_45S_50ntFlanks</td><td>12459</td><td>12478</td><td>20</td><td>AGCGTTGGATTGTTCACCCA</td><td>719.2</td><td>-4.7</td><td>1.0</td><td>-4.8</td><td>1.9E-06</td><td>2.6E-06</td><td>BT20/MB468</td></tr>
<tr><td>>hg38_21_+_8433172_8446622_RNA45SN1_45S_50ntFlanks@12934.12958.25|CTAAACCATTCGTAGACGACCTGCT</td><td>>hg38_21_+_8433172_8446622_RNA45SN1_45S_50ntFlanks</td><td>12934</td><td>12958</td><td>25</td><td>CTAAACCATTCGTAGACGACCTGCT</td><td>302.0</td><td>-3.1</td><td>0.6</td><td>-4.8</td><td>1.9E-06</td><td>2.6E-06</td><td>BT20/MB468</td></tr>
<tr><td>>hg38_21_+_8433172_8446622_RNA45SN1_45S_50ntFlanks@9569.9592.24|GTGCAAATCGGTCGTCCGACCTGG</td><td>>hg38_21_+_8433172_8446622_RNA45SN1_45S_50ntFlanks</td><td>9569</td><td>9592</td><td>24</td><td>GTGCAAATCGGTCGTCCGACCTGG</td><td>125.2</td><td>-2.9</td><td>0.6</td><td>-4.8</td><td>1.7E-06</td><td>2.6E-06</td><td>MB231/MB468</td></tr>
<tr><td>>hg38_21_+_8433172_8446622_RNA45SN1_45S_50ntFlanks@9554.9571.18|CGTAGCGGTCCTGACGTG</td><td>>hg38_21_+_8433172_8446622_RNA45SN1_45S_50ntFlanks</td><td>9554</td><td>9571</td><td>18</td><td>CGTAGCGGTCCTGACGTG</td><td>101835.9</td><td>-3.2</td><td>0.7</td><td>-4.8</td><td>1.8E-06</td><td>2.7E-06</td><td>MB231/MB468</td></tr>
<tr><td>>hg38_21_+_8433172_8446622_RNA45SN1_45S_50ntFlanks@7990.8009.20|GACGTGGCGACCCGCTGAAT</td><td>>hg38_21_+_8433172_8446622_RNA45SN1_45S_50ntFlanks</td><td>7990</td><td>8009</td><td>20</td><td>GACGTGGCGACCCGCTGAAT</td><td>737.5</td><td>3.6</td><td>0.8</td><td>4.8</td><td>2.0E-06</td><td>2.7E-06</td><td>BT20/MB468</td></tr>
<tr><td>>hg38_21_+_8433172_8446622_RNA45SN1_45S_50ntFlanks@6732.6771.40|ACATTGATCATCGACACTTCGAACGCACTTGCGGCCCCGG</td><td>>hg38_21_+_8433172_8446622_RNA45SN1_45S_50ntFlanks</td><td>6732</td><td>6771</td><td>40</td><td>ACATTGATCATCGACACTTCGAACGCACTTGCGGCCCCGG</td><td>489.2</td><td>-3.0</td><td>0.6</td><td>-4.8</td><td>2.0E-06</td><td>2.7E-06</td><td>BT20/MB468</td></tr>
<tr><td>>MI0000273|hsa-mir-183&WithFlank&7|-|129774899|129775020@34.54.21(+1A)&[MIMAT0000261&hsa-miR-183-5p&offsets|+1|0(+1A);m-41&7|-|129774966|129774987&offsets|0|-1(+1A)]|ATGGCACTGGTAGAATTCACTA</td><td>>MI0000273|hsa-mir-183&WithFlank&7|-|129774899|129775020</td><td>34</td><td>54</td><td>21(+1A)</td><td>ATGGCACTGGTAGAATTCACTA</td><td>798.1</td><td>4.0</td><td>0.8</td><td>5.3</td><td>1.5E-07</td><td>2.7E-06</td><td>BT20/MB231</td></tr>
<tr><td>>hg38_21_+_8433172_8446622_RNA45SN1_45S_50ntFlanks@4726.4758.33|TAATCAAGAACGAAAGTCGGAGGTTCGAAGACG</td><td>>hg38_21_+_8433172_8446622_RNA45SN1_45S_50ntFlanks</td><td>4726</td><td>4758</td><td>33</td><td>TAATCAAGAACGAAAGTCGGAGGTTCGAAGACG</td><td>1476.6</td><td>-3.0</td><td>0.6</td><td>-4.8</td><td>1.8E-06</td><td>2.7E-06</td><td>MB231/MB468</td></tr>
<tr><td>>hg38_21_+_8433172_8446622_RNA45SN1_45S_50ntFlanks@6651.6699.49|CGACTCTTAGCGGTGGATCACTCGGCTCGTGCGTCGATGAAGAACGCAG</td><td>>hg38_21_+_8433172_8446622_RNA45SN1_45S_50ntFlanks</td><td>6651</td><td>6699</td><td>49</td><td>CGACTCTTAGCGGTGGATCACTCGGCTCGTGCGTCGATGAAGAACGCAG</td><td>25917.3</td><td>4.5</td><td>0.9</td><td>4.8</td><td>1.8E-06</td><td>2.7E-06</td><td>MB231/MB468</td></tr>
<tr><td>>hg38_21_+_8433172_8446622_RNA45SN1_45S_50ntFlanks@7980.8012.33|CCTCAGATCAGACGTGGCGACCCGCTGAATTTA</td><td>>hg38_21_+_8433172_8446622_RNA45SN1_45S_50ntFlanks</td><td>7980</td><td>8012</td><td>33</td><td>CCTCAGATCAGACGTGGCGACCCGCTGAATTTA</td><td>216.3</td><td>-3.6</td><td>0.8</td><td>-4.8</td><td>2.0E-06</td><td>2.7E-06</td><td>BT20/MB468</td></tr>
<tr><td>>hg38_21_+_8433172_8446622_RNA45SN1_45S_50ntFlanks@6661.6678.18|CGGTGGATCACTCGGCTC</td><td>>hg38_21_+_8433172_8446622_RNA45SN1_45S_50ntFlanks</td><td>6661</td><td>6678</td><td>18</td><td>CGGTGGATCACTCGGCTC</td><td>777.4</td><td>4.2</td><td>0.8</td><td>5.3</td><td>1.5E-07</td><td>2.7E-06</td><td>BT20/MB231</td></tr>
<tr><td>>hg38_21_+_8433172_8446622_RNA45SN1_45S_50ntFlanks@8175.8202.28|CCTTCTGATCGAGGCCCAGCCCGTGGAC</td><td>>hg38_21_+_8433172_8446622_RNA45SN1_45S_50ntFlanks</td><td>8175</td><td>8202</td><td>28</td><td>CCTTCTGATCGAGGCCCAGCCCGTGGAC</td><td>517.6</td><td>8.8</td><td>1.8</td><td>4.8</td><td>1.8E-06</td><td>2.7E-06</td><td>MB231/MB468</td></tr>
<tr><td>>hg38_21_+_8433172_8446622_RNA45SN1_45S_50ntFlanks@6699.6719.21|GCTAGCTGCGAGAATTAATGT</td><td>>hg38_21_+_8433172_8446622_RNA45SN1_45S_50ntFlanks</td><td>6699</td><td>6719</td><td>21</td><td>GCTAGCTGCGAGAATTAATGT</td><td>1073.4</td><td>-2.3</td><td>0.5</td><td>-4.8</td><td>1.8E-06</td><td>2.7E-06</td><td>MB231/MB468</td></tr>
<tr><td>>trnaMT_SerGCT_MT_+_12207_12265@8.38.31|CTCACAAGAACTGCTAACTCATGCCCCCATG</td><td>>trnaMT_SerGCT_MT_+_12207_12265</td><td>8</td><td>38</td><td>31</td><td>CTCACAAGAACTGCTAACTCATGCCCCCATG</td><td>240.7</td><td>-5.7</td><td>1.2</td><td>-4.8</td><td>2.0E-06</td><td>2.7E-06</td><td>BT20/MB468</td></tr>
<tr><td>>hg38_21_+_8433172_8446622_RNA45SN1_45S_50ntFlanks@11785.11806.22|TGAATGGATGAACGAGATTCCC</td><td>>hg38_21_+_8433172_8446622_RNA45SN1_45S_50ntFlanks</td><td>11785</td><td>11806</td><td>22</td><td>TGAATGGATGAACGAGATTCCC</td><td>899.4</td><td>-3.7</td><td>0.8</td><td>-4.8</td><td>1.8E-06</td><td>2.8E-06</td><td>MB231/MB468</td></tr>
<tr><td>>hg38_21_+_8433172_8446622_RNA45SN1_45S_50ntFlanks@3813.3834.22|TTATGGTTCCTTTGGTCGCTCG</td><td>>hg38_21_+_8433172_8446622_RNA45SN1_45S_50ntFlanks</td><td>3813</td><td>3834</td><td>22</td><td>TTATGGTTCCTTTGGTCGCTCG</td><td>111.0</td><td>-2.6</td><td>0.5</td><td>-4.8</td><td>1.8E-06</td><td>2.8E-06</td><td>MB231/MB468</td></tr>
<tr><td>>hg38_21_+_8433172_8446622_RNA45SN1_45S_50ntFlanks@7981.7998.18|CTCAGATCAGACGTGGCG</td><td>>hg38_21_+_8433172_8446622_RNA45SN1_45S_50ntFlanks</td><td>7981</td><td>7998</td><td>18</td><td>CTCAGATCAGACGTGGCG</td><td>67.9</td><td>-5.7</td><td>1.2</td><td>-4.8</td><td>1.8E-06</td><td>2.8E-06</td><td>MB231/MB468</td></tr>
<tr><td>>hg38_21_+_8433172_8446622_RNA45SN1_45S_50ntFlanks@11710.11728.19|CGCGGGTAAACGGCGGGAG</td><td>>hg38_21_+_8433172_8446622_RNA45SN1_45S_50ntFlanks</td><td>11710</td><td>11728</td><td>19</td><td>CGCGGGTAAACGGCGGGAG</td><td>326.4</td><td>-4.2</td><td>0.9</td><td>-4.7</td><td>2.0E-06</td><td>2.8E-06</td><td>BT20/MB468</td></tr>
<tr><td>>hg38_21_+_8433172_8446622_RNA45SN1_45S_50ntFlanks@9574.9602.29|AATCGGTCGTCCGACCTGGGTATAGGGGC</td><td>>hg38_21_+_8433172_8446622_RNA45SN1_45S_50ntFlanks</td><td>9574</td><td>9602</td><td>29</td><td>AATCGGTCGTCCGACCTGGGTATAGGGGC</td><td>392.0</td><td>2.1</td><td>0.4</td><td>4.7</td><td>2.0E-06</td><td>2.8E-06</td><td>BT20/MB468</td></tr>
<tr><td>>hg38_21_+_8433172_8446622_RNA45SN1_45S_50ntFlanks@4588.4611.24|CTATTTTGTTGGTTTTCGGAACTG</td><td>>hg38_21_+_8433172_8446622_RNA45SN1_45S_50ntFlanks</td><td>4588</td><td>4611</td><td>24</td><td>CTATTTTGTTGGTTTTCGGAACTG</td><td>113.9</td><td>-5.0</td><td>1.0</td><td>-4.7</td><td>2.1E-06</td><td>2.8E-06</td><td>BT20/MB468</td></tr>
<tr><td>>hg38_21_+_8433172_8446622_RNA45SN1_45S_50ntFlanks@12007.12043.37|GCCGCCGGTGAAATACCACTACTCTGATCGTTTTTTC</td><td>>hg38_21_+_8433172_8446622_RNA45SN1_45S_50ntFlanks</td><td>12007</td><td>12043</td><td>37</td><td>GCCGCCGGTGAAATACCACTACTCTGATCGTTTTTTC</td><td>193.4</td><td>-3.2</td><td>0.7</td><td>-4.8</td><td>1.9E-06</td><td>2.8E-06</td><td>MB231/MB468</td></tr>
<tr><td>>hg38_21_+_8433172_8446622_RNA45SN1_45S_50ntFlanks@4762.4780.19|AGATACCGTCGTAGTTCCG</td><td>>hg38_21_+_8433172_8446622_RNA45SN1_45S_50ntFlanks</td><td>4762</td><td>4780</td><td>19</td><td>AGATACCGTCGTAGTTCCG</td><td>511.5</td><td>-3.1</td><td>0.7</td><td>-4.8</td><td>1.9E-06</td><td>2.8E-06</td><td>MB231/MB468</td></tr>
<tr><td>>hg38_21_+_8433172_8446622_RNA45SN1_45S_50ntFlanks@11200.11232.33|CGGCGGCGGCGGCGACTCTGGACGCGAGCCGGG</td><td>>hg38_21_+_8433172_8446622_RNA45SN1_45S_50ntFlanks</td><td>11200</td><td>11232</td><td>33</td><td>CGGCGGCGGCGGCGACTCTGGACGCGAGCCGGG</td><td>1253.1</td><td>-4.8</td><td>1.0</td><td>-4.7</td><td>2.1E-06</td><td>2.8E-06</td><td>BT20/MB468</td></tr>
<tr><td>>hg38_21_+_8433172_8446622_RNA45SN1_45S_50ntFlanks@10680.10706.27|GTTCCGGCGGCGTCCGGTGAGCTCTCG</td><td>>hg38_21_+_8433172_8446622_RNA45SN1_45S_50ntFlanks</td><td>10680</td><td>10706</td><td>27</td><td>GTTCCGGCGGCGTCCGGTGAGCTCTCG</td><td>8557.9</td><td>-4.8</td><td>1.0</td><td>-4.7</td><td>2.1E-06</td><td>2.8E-06</td><td>BT20/MB468</td></tr>
<tr><td>>hg38_21_+_8433172_8446622_RNA45SN1_45S_50ntFlanks@10771.10791.21|CAGGTCTCCAAGGTGAACAGC</td><td>>hg38_21_+_8433172_8446622_RNA45SN1_45S_50ntFlanks</td><td>10771</td><td>10791</td><td>21</td><td>CAGGTCTCCAAGGTGAACAGC</td><td>414.8</td><td>-3.2</td><td>0.7</td><td>-4.7</td><td>2.1E-06</td><td>2.9E-06</td><td>BT20/MB468</td></tr>
<tr><td>>hg38_21_+_8433172_8446622_RNA45SN1_45S_50ntFlanks@8342.8364.23|CGTAAGGGAAAGTTGAAAAGAAC</td><td>>hg38_21_+_8433172_8446622_RNA45SN1_45S_50ntFlanks</td><td>8342</td><td>8364</td><td>23</td><td>CGTAAGGGAAAGTTGAAAAGAAC</td><td>437.7</td><td>8.5</td><td>1.8</td><td>4.8</td><td>1.9E-06</td><td>2.9E-06</td><td>MB231/MB468</td></tr>
</tbody>
</table>

| | | | | | | | | | | | | |
|---|---|---|---|---|---|---|---|---|---|---|---|---|
| >hg38_21_+_8433172_8446622_RNA45SN1_45S_50ntFlanks@11569.11592.24\|CAGCCGACTTAGAACTGGTGCGGA | >hg38_21_+_8433172_8446622_RNA45SN1_45S_50ntFlanks | 11569 | 11592 | 24 | CAGCCGACTTAGAACTGGTGCGGA | 552.8 | -3.9 | 0.8 | -4.7 | 2.1E-06 | 2.9E-06 | BT20/MB468 |
| >hg38_21_+_8433172_8446622_RNA45SN1_45S_50ntFlanks@4569.4587.19\|ATGGAATAGGACCGCGGTT | >hg38_21_+_8433172_8446622_RNA45SN1_45S_50ntFlanks | 4569 | 4587 | 19 | ATGGAATAGGACCGCGGTT | 235.0 | -2.5 | 0.5 | -4.8 | 1.9E-06 | 2.9E-06 | MB231/MB468 |
| >hg38_21_+_8433172_8446622_RNA45SN1_45S_50ntFlanks@10818.10840.23\|AAGGGAAGTCGGCAAGCCGGATC | >hg38_21_+_8433172_8446622_RNA45SN1_45S_50ntFlanks | 10818 | 10840 | 23 | AAGGGAAGTCGGCAAGCCGGATC | 105.5 | -2.9 | 0.6 | -4.7 | 2.1E-06 | 2.9E-06 | BT20/MB468 |
| >hg38_21_+_8433172_8446622_RNA45SN1_45S_50ntFlanks@3854.3880.27\|ACTGTGGTAATTCTAGAGCTAATACAT | >hg38_21_+_8433172_8446622_RNA45SN1_45S_50ntFlanks | 3854 | 3880 | 27 | ACTGTGGTAATTCTAGAGCTAATACAT | 342.2 | -2.7 | 0.6 | -4.7 | 2.1E-06 | 2.9E-06 | BT20/MB468 |
| >hg38_21_+_8433172_8446622_RNA45SN1_45S_50ntFlanks@4738.4761.24\|AAAGTCGGAGGTTCGAAGACGATC | >hg38_21_+_8433172_8446622_RNA45SN1_45S_50ntFlanks | 4738 | 4761 | 24 | AAAGTCGGAGGTTCGAAGACGATC | 382.8 | 2.6 | 0.6 | 4.7 | 2.1E-06 | 2.9E-06 | BT20/MB468 |
| >hg38_21_+_8433172_8446622_RNA45SN1_45S_50ntFlanks@12002.12031.30\|TGCGGGCCGCCGGTGAAATACCACTACTCT | >hg38_21_+_8433172_8446622_RNA45SN1_45S_50ntFlanks | 12002 | 12031 | 30 | TGCGGGCCGCCGGTGAAATACCACTACTCT | 605.2 | -4.2 | 0.9 | -4.7 | 2.2E-06 | 2.9E-06 | BT20/MB468 |
| >hg38_21_+_8433172_8446622_RNA45SN1_45S_50ntFlanks@12954.12973.20\|CTGCTTCTGGGTCGGGGTTT | >hg38_21_+_8433172_8446622_RNA45SN1_45S_50ntFlanks | 12954 | 12973 | 20 | CTGCTTCTGGGTCGGGGTTT | 163.2 | -5.4 | 1.1 | -4.7 | 2.2E-06 | 3.0E-06 | BT20/MB468 |
| >hg38_21_+_8433172_8446622_RNA45SN1_45S_50ntFlanks@10350.10371.22\|ACGAGAACTTTGAAGGCCGAAG | >hg38_21_+_8433172_8446622_RNA45SN1_45S_50ntFlanks | 10350 | 10371 | 22 | ACGAGAACTTTGAAGGCCGAAG | 1139.5 | -2.6 | 0.5 | -4.7 | 2.2E-06 | 3.0E-06 | BT20/MB468 |
| >hg38_21_+_8433172_8446622_RNA45SN1_45S_50ntFlanks@10768.10797.30\|CAGCAGGTCTCCAAGGTGAACAGCCTCTGG | >hg38_21_+_8433172_8446622_RNA45SN1_45S_50ntFlanks | 10768 | 10797 | 30 | CAGCAGGTCTCCAAGGTGAACAGCCTCTGG | 171.4 | -5.2 | 1.1 | -4.7 | 2.2E-06 | 3.0E-06 | BT20/MB468 |
| >MI0000102\|hsa-mir-100&WithFlank&11\|-\|122152223\|122152314@19.38.20&[MIMAT0000098&hsa-miR-100-5p&offsets\|0\|-2;m-27&11\|-\|122152276\|122152296&offsets\|0\|-1]\|AACCCGTAGATCCGAACTTG | >MI0000102\|hsa-mir-100&WithFlank&11\|-\|122152223\|122152314 | 19 | 38 | 20 | AACCCGTAGATCCGAACTTG | 20512.7 | -5.6 | 1.1 | -5.2 | 1.6E-07 | 3.0E-06 | BT20/MB231 |
| >MI0000272\|hsa-mir-182&WithFlank&7\|-\|129770377\|129770498@29.52.24&[MIMAT0000259&hsa-miR-182-5p&offsets\|0\|0;m-22&7\|-\|129770449\|129770470&offsets\|0\|+2]\|TTTGGCAATGGTAGAACTCACACT | >MI0000272\|hsa-mir-182&WithFlank&7\|-\|129770377\|129770498 | 29 | 52 | 24 | TTTGGCAATGGTAGAACTCACACT | 70970.2 | 3.4 | 0.7 | 5.2 | 1.6E-07 | 3.0E-06 | BT20/MB231 |
| >hg38_21_+_8433172_8446622_RNA45SN1_45S_50ntFlanks@12072.12097.26\|CCCGAGGGGCTCTCGCTTCTGGCGCC | >hg38_21_+_8433172_8446622_RNA45SN1_45S_50ntFlanks | 12072 | 12097 | 26 | CCCGAGGGGCTCTCGCTTCTGGCGCC | 349.1 | -3.9 | 0.8 | -4.7 | 2.2E-06 | 3.0E-06 | BT20/MB468 |
| >hg38_21_+_8433172_8446622_RNA45SN1_45S_50ntFlanks@11768.11798.31\|CTAATTAGTGACGCGCATGAATGGATGAACG | >hg38_21_+_8433172_8446622_RNA45SN1_45S_50ntFlanks | 11768 | 11798 | 31 | CTAATTAGTGACGCGCATGAATGGATGAACG | 83.0 | -3.6 | 0.8 | -4.8 | 2.0E-06 | 3.0E-06 | MB231/MB468 |
| >hg38_21_+_8433172_8446622_RNA45SN1_45S_50ntFlanks@11576.11600.25\|CTTAGAACTGGTGCGGACCAGGGGA | >hg38_21_+_8433172_8446622_RNA45SN1_45S_50ntFlanks | 11576 | 11600 | 25 | CTTAGAACTGGTGCGGACCAGGGGA | 604.9 | -5.2 | 1.1 | -4.7 | 2.2E-06 | 3.0E-06 | BT20/MB468 |
| >hg38_21_+_8433172_8446622_RNA45SN1_45S_50ntFlanks@12458.12488.31\|AAGCGTTGGATTGTTCACCCACTAATAGGGA | >hg38_21_+_8433172_8446622_RNA45SN1_45S_50ntFlanks | 12458 | 12488 | 31 | AAGCGTTGGATTGTTCACCCACTAATAGGGA | 139.4 | -3.2 | 0.7 | -4.7 | 2.2E-06 | 3.0E-06 | BT20/MB468 |
| >hg38_21_+_8433172_8446622_RNA45SN1_45S_50ntFlanks@5444.5467.24\|CTCGGATCGGCCCCGCCGGGGTCG | >hg38_21_+_8433172_8446622_RNA45SN1_45S_50ntFlanks | 5444 | 5467 | 24 | CTCGGATCGGCCCCGCCGGGGTCG | 389.8 | -5.4 | 1.1 | -4.7 | 2.3E-06 | 3.1E-06 | BT20/MB468 |
| >hg38_21_+_8433172_8446622_RNA45SN1_45S_50ntFlanks@4202.4219.18\|CGACCCGGGGAGGTAGTG | >hg38_21_+_8433172_8446622_RNA45SN1_45S_50ntFlanks | 4202 | 4219 | 18 | CGACCCGGGGAGGTAGTG | 76.4 | -4.1 | 0.9 | -4.7 | 2.3E-06 | 3.1E-06 | BT20/MB468 |
| >hg38_21_+_8433172_8446622_RNA45SN1_45S_50ntFlanks@10846.10867.22\|CTTCGGGATAAGGATTGGCTCT | >hg38_21_+_8433172_8446622_RNA45SN1_45S_50ntFlanks | 10846 | 10867 | 22 | CTTCGGGATAAGGATTGGCTCT | 280.8 | -3.0 | 0.6 | -4.7 | 2.3E-06 | 3.1E-06 | BT20/MB468 |
| >hg38_21_+_8433172_8446622_RNA45SN1_45S_50ntFlanks@10797.10823.27\|GCATGTTGGAACAATGTAGGTAAGGGA | >hg38_21_+_8433172_8446622_RNA45SN1_45S_50ntFlanks | 10797 | 10823 | 27 | GCATGTTGGAACAATGTAGGTAAGGGA | 115.6 | -4.3 | 0.9 | -4.7 | 2.3E-06 | 3.1E-06 | BT20/MB468 |
| >hg38_21_+_8433172_8446622_RNA45SN1_45S_50ntFlanks@12419.12443.25\|GATGTCGGCTCTTCCTATCATTGTG | >hg38_21_+_8433172_8446622_RNA45SN1_45S_50ntFlanks | 12419 | 12443 | 25 | GATGTCGGCTCTTCCTATCATTGTG | 1911.6 | 3.3 | 0.7 | 4.8 | 2.0E-06 | 3.1E-06 | MB231/MB468 |
| >hg38_21_+_8433172_8446622_RNA45SN1_45S_50ntFlanks@10534.10570.37\|CCGCGAGGCGTCCAGTGCGGTAACGCGACCGATCCCG | >hg38_21_+_8433172_8446622_RNA45SN1_45S_50ntFlanks | 10534 | 10570 | 37 | CCGCGAGGCGTCCAGTGCGGTAACGCGACCGATCCCG | 240.7 | -5.5 | 1.2 | -4.7 | 2.3E-06 | 3.1E-06 | BT20/MB468 |
| >hg38_21_+_8433172_8446622_RNA45SN1_45S_50ntFlanks@8169.8186.18\|CCAAGTCCTTCTGATCGA | >hg38_21_+_8433172_8446622_RNA45SN1_45S_50ntFlanks | 8169 | 8186 | 18 | CCAAGTCCTTCTGATCGA | 383.5 | -3.7 | 0.8 | -4.7 | 2.0E-06 | 3.1E-06 | MB231/MB468 |
| >hg38_21_+_8433172_8446622_RNA45SN1_45S_50ntFlanks@10393.10411.19\|CAGCAGTTGAACATGGGTC | >hg38_21_+_8433172_8446622_RNA45SN1_45S_50ntFlanks | 10393 | 10411 | 19 | CAGCAGTTGAACATGGGTC | 117.9 | -2.9 | 0.6 | -4.7 | 2.3E-06 | 3.1E-06 | BT20/MB468 |
| >hg38_21_+_8433172_8446622_RNA45SN1_45S_50ntFlanks@11729.11747.19\|TAACTATGACTCTCTTAAG | >hg38_21_+_8433172_8446622_RNA45SN1_45S_50ntFlanks | 11729 | 11747 | 19 | TAACTATGACTCTCTTAAG | 3064.3 | -2.9 | 0.6 | -4.7 | 2.1E-06 | 3.1E-06 | MB231/MB468 |
| >hg38_1_-_228634819_228635039_RNA5S12_5S_50ntFlanks@128.170.43\|CTTGGATGGGAGACCGCCTGGGAATACCGGGTGCTGTAGGCTT | >hg38_1_-_228634819_228635039_RNA5S12_5S_50ntFlanks | 128 | 170 | 43 | CTTGGATGGGAGACCGCCTGGGAATACCGGGTGCTGTAGGCTT | 1484.6 | 11.3 | 2.4 | 4.7 | 2.1E-06 | 3.1E-06 | MB231/MB468 |
| >hg38_21_+_8433172_8446622_RNA45SN1_45S_50ntFlanks@11572.11592.21\|CCGACTTAGAACTGGTGCGGA | >hg38_21_+_8433172_8446622_RNA45SN1_45S_50ntFlanks | 11572 | 11592 | 21 | CCGACTTAGAACTGGTGCGGA | 983.0 | -4.0 | 0.8 | -4.7 | 2.3E-06 | 3.1E-06 | BT20/MB468 |
| >hg38_21_+_8433172_8446622_RNA45SN1_45S_50ntFlanks@11873.11892.20\|AAGAAGACCCTGTTGAGCTT | >hg38_21_+_8433172_8446622_RNA45SN1_45S_50ntFlanks | 11873 | 11892 | 20 | AAGAAGACCCTGTTGAGCTT | 700.5 | -4.5 | 0.9 | -4.7 | 2.1E-06 | 3.2E-06 | MB231/MB468 |
| >hg38_21_+_8433172_8446622_RNA45SN1_45S_50ntFlanks@11209.11234.26\|CGGCGACTCTGGACGCGAGCCGGGCC | >hg38_21_+_8433172_8446622_RNA45SN1_45S_50ntFlanks | 11209 | 11234 | 26 | CGGCGACTCTGGACGCGAGCCGGGCC | 1787.3 | -4.9 | 1.0 | -4.7 | 2.3E-06 | 3.2E-06 | BT20/MB468 |
| >hg38_21_+_8433172_8446622_RNA45SN1_45S_50ntFlanks@4370.4391.22\|TTAAAAAGCTCGTAGTTGGATC | >hg38_21_+_8433172_8446622_RNA45SN1_45S_50ntFlanks | 4370 | 4391 | 22 | TTAAAAAGCTCGTAGTTGGATC | 204.8 | -2.4 | 0.5 | -4.7 | 2.1E-06 | 3.2E-06 | MB231/MB468 |
| >MI0000478\|hsa-mir-149&WithFlank&2\|+\|240455995\|240456095@21.41.21(+1A)&[MIMAT0000450&hsa-miR-149-5p&offsets\|0\|-2(+1A);m-195&2\|+\|240456015\|240456037&offsets\|0\|-2(+1A)]\|TCTGGCTCCGTGTCTTCACTCA | >MI0000478\|hsa-mir-149&WithFlank&2\|+\|240455995\|240456095 | 21 | 41 | 21(+1A) | TCTGGCTCCGTGTCTTCACTCA | 351.8 | 4.4 | 0.8 | 5.2 | 1.8E-07 | 3.2E-06 | BT20/MB231 |
| >hg38_21_+_8433172_8446622_RNA45SN1_45S_50ntFlanks@12445.12476.32\|AGCAGAATTCACCAAGCGTTGGATTGTTCACC | >hg38_21_+_8433172_8446622_RNA45SN1_45S_50ntFlanks | 12445 | 12476 | 32 | AGCAGAATTCACCAAGCGTTGGATTGTTCACC | 569.8 | 3.1 | 0.7 | 4.7 | 2.1E-06 | 3.2E-06 | MB231/MB468 |
| >hg38_21_+_8433172_8446622_RNA45SN1_45S_50ntFlanks@11883.11904.22\|TGTTGAGCTTGACTCTAGTCTG | >hg38_21_+_8433172_8446622_RNA45SN1_45S_50ntFlanks | 11883 | 11904 | 22 | TGTTGAGCTTGACTCTAGTCTG | 99.5 | -4.1 | 0.9 | -4.7 | 2.1E-06 | 3.2E-06 | MB231/MB468 |
| >hg38_21_+_8433172_8446622_RNA45SN1_45S_50ntFlanks@12073.12092.20\|CCGAGGGGCTCTCGCTTCTG | >hg38_21_+_8433172_8446622_RNA45SN1_45S_50ntFlanks | 12073 | 12092 | 20 | CCGAGGGGCTCTCGCTTCTG | 276.6 | -2.6 | 0.5 | -4.7 | 2.1E-06 | 3.2E-06 | MB231/MB468 |
| >hg38_1_-_228634819_228635039_RNA5S12_5S_50ntFlanks@126.170.45\|TACTTGGATGGGAGACCGCCTGGGAATACCGGGTGCTGTAGGCTT | >hg38_1_-_228634819_228635039_RNA5S12_5S_50ntFlanks | 126 | 170 | 45 | TACTTGGATGGGAGACCGCCTGGGAATACCGGGTGCTGTAGGCTT | 846.7 | 7.2 | 1.5 | 4.7 | 2.4E-06 | 3.3E-06 | BT20/MB468 |
| >hg38_21_+_8433172_8446622_RNA45SN1_45S_50ntFlanks@3899.3927.29\|CCTTCGCGGGGGGGATGCGTGCATTTATC | >hg38_21_+_8433172_8446622_RNA45SN1_45S_50ntFlanks | 3899 | 3927 | 29 | CCTTCGCGGGGGGGATGCGTGCATTTATC | 191.0 | -4.1 | 0.9 | -4.7 | 2.1E-06 | 3.3E-06 | MB231/MB468 |
| >hg38_21_+_8433172_8446622_RNA45SN1_45S_50ntFlanks@9564.9598.35\|CTGACGTGCAAATCGGTCGTCCGACCTGGGTATAG | >hg38_21_+_8433172_8446622_RNA45SN1_45S_50ntFlanks | 9564 | 9598 | 35 | CTGACGTGCAAATCGGTCGTCCGACCTGGGTATAG | 90.0 | -2.3 | 0.5 | -4.7 | 2.2E-06 | 3.3E-06 | MB231/MB468 |
| >MI0000266\|hsa-mir-10a&WithFlank&17\|-\|48579832\|48579953@29.50.22(+1A)&[MIMAT0000253&hsa-miR-10a-5p&offsets\|+1\|0(+1A);m-18&17\|-\|48579905\|48579926&offsets\|+1\|+1(+1A)]\|ACCCTGTAGATCCGAATTTGTGA | >MI0000266\|hsa-mir-10a&WithFlank&17\|-\|48579832\|48579953 | 29 | 50 | 22(+1A) | ACCCTGTAGATCCGAATTTGTGA | 1617.5 | 11.2 | 2.4 | 4.7 | 2.2E-06 | 3.3E-06 | MB231/MB468 |
| >hg38_21_+_8433172_8446622_RNA45SN1_45S_50ntFlanks@5445.5467.23\|TCGGATCGGCCCCGCCGGGGTCG | >hg38_21_+_8433172_8446622_RNA45SN1_45S_50ntFlanks | 5445 | 5467 | 23 | TCGGATCGGCCCCGCCGGGGTCG | 89.9 | -2.8 | 0.6 | -4.7 | 2.2E-06 | 3.3E-06 | MB231/MB468 |
| >hg38_21_+_8433172_8446622_RNA45SN1_45S_50ntFlanks@10791.10818.28\|CCTCTGGCATGTTGGAACAATGTAGGTA | >hg38_21_+_8433172_8446622_RNA45SN1_45S_50ntFlanks | 10791 | 10818 | 28 | CCTCTGGCATGTTGGAACAATGTAGGTA | 247.3 | -3.1 | 0.7 | -4.7 | 2.2E-06 | 3.3E-06 | MB231/MB468 |
| >hg38_21_+_8433172_8446622_RNA45SN1_45S_50ntFlanks@11872.11893.22\|AAAGAAGACCCTGTTGAGCTTG | >hg38_21_+_8433172_8446622_RNA45SN1_45S_50ntFlanks | 11872 | 11893 | 22 | AAAGAAGACCCTGTTGAGCTTG | 210.1 | -2.8 | 0.6 | -4.7 | 2.2E-06 | 3.4E-06 | MB231/MB468 |
| >hg38_21_+_8433172_8446622_RNA45SN1_45S_50ntFlanks@11881.11903.23\|CCTGTTGAGCTTGACTCTAGTCT | >hg38_21_+_8433172_8446622_RNA45SN1_45S_50ntFlanks | 11881 | 11903 | 23 | CCTGTTGAGCTTGACTCTAGTCT | 357.2 | -2.9 | 0.6 | -4.7 | 2.5E-06 | 3.4E-06 | BT20/MB468 |
| >hg38_21_+_8433172_8446622_RNA45SN1_45S_50ntFlanks@12418.12456.39\|CGATGTCGGCTCTTCCTATCATTGTGAAGCAGAATTCAC | >hg38_21_+_8433172_8446622_RNA45SN1_45S_50ntFlanks | 12418 | 12456 | 39 | CGATGTCGGCTCTTCCTATCATTGTGAAGCAGAATTCAC | 246.7 | -2.7 | 0.6 | -4.7 | 2.5E-06 | 3.4E-06 | BT20/MB468 |
| >hg38_21_+_8433172_8446622_RNA45SN1_45S_50ntFlanks@6779.6802.24\|CCGGGGCTACGCCTGTCTGAGCGT | >hg38_21_+_8433172_8446622_RNA45SN1_45S_50ntFlanks | 6779 | 6802 | 24 | CCGGGGCTACGCCTGTCTGAGCGT | 456.6 | 3.4 | 0.7 | 4.7 | 2.5E-06 | 3.4E-06 | BT20/MB468 |
| >hg38_21_+_8433172_8446622_RNA45SN1_45S_50ntFlanks@12624.12661.38\|CCAATGGGGCGAAGCTACCATCTGTGGGATTATGACTG | >hg38_21_+_8433172_8446622_RNA45SN1_45S_50ntFlanks | 12624 | 12661 | 38 | CCAATGGGGCGAAGCTACCATCTGTGGGATTATGACTG | 714.0 | -2.5 | 0.5 | -4.7 | 2.3E-06 | 3.5E-06 | MB231/MB468 |
| >hg38_21_+_8433172_8446622_RNA45SN1_45S_50ntFlanks@7981.8003.23\|CTCAGATCAGACGTGGCGACCCG | >hg38_21_+_8433172_8446622_RNA45SN1_45S_50ntFlanks | 7981 | 8003 | 23 | CTCAGATCAGACGTGGCGACCCG | 89.8 | -3.4 | 0.7 | -4.7 | 2.6E-06 | 3.5E-06 | BT20/MB468 |

<table>
<tr><th></th><th></th><th></th><th></th><th></th><th></th><th></th><th></th><th></th><th></th><th></th><th></th><th></th></tr>
<tr><td>>hg38_21_+_8433172_8446622_RNA45SN1_45S_50ntFlanks@6733.6770.38|CATTGATCATCGACACTTCGAACGCACTTGCGGCCCCG</td><td>>hg38_21_+_8433172_8446622_RNA45SN1_45S_50ntFlanks</td><td>6733</td><td>6770</td><td>38</td><td>CATTGATCATCGACACTTCGAACGCACTTGCGGCCCCG</td><td>851.7</td><td>-2.6</td><td>0.6</td><td>-4.7</td><td>2.3E-06</td><td>3.5E-06</td><td>MB231/MB468</td></tr>
<tr><td>>hg38_21_+_8433172_8446622_RNA45SN1_45S_50ntFlanks@11778.11805.28|ACGCGCATGAATGGATGAACGAGATTCC</td><td>>hg38_21_+_8433172_8446622_RNA45SN1_45S_50ntFlanks</td><td>11778</td><td>11805</td><td>28</td><td>ACGCGCATGAATGGATGAACGAGATTCC</td><td>118.8</td><td>-2.0</td><td>0.4</td><td>-4.7</td><td>2.6E-06</td><td>3.5E-06</td><td>BT20/MB468</td></tr>
<tr><td>>hg38_21_+_8433172_8446622_RNA45SN1_45S_50ntFlanks@10458.10487.30|ATGGCCTCCGTTGCCCTCGGCCGATCGAAA</td><td>>hg38_21_+_8433172_8446622_RNA45SN1_45S_50ntFlanks</td><td>10458</td><td>10487</td><td>30</td><td>ATGGCCTCCGTTGCCCTCGGCCGATCGAAA</td><td>366.7</td><td>-4.3</td><td>0.9</td><td>-4.7</td><td>2.6E-06</td><td>3.5E-06</td><td>BT20/MB468</td></tr>
<tr><td>>hg38_21_+_8433172_8446622_RNA45SN1_45S_50ntFlanks@12944.12962.19|CGTAGACGACCTGCTTCTG</td><td>>hg38_21_+_8433172_8446622_RNA45SN1_45S_50ntFlanks</td><td>12944</td><td>12962</td><td>19</td><td>CGTAGACGACCTGCTTCTG</td><td>146.4</td><td>-2.4</td><td>0.5</td><td>-4.7</td><td>2.6E-06</td><td>3.5E-06</td><td>BT20/MB468</td></tr>
<tr><td>>MI0000446|hsa-mir-125b-1&WithFlank&11|-|122099751|122099850@21.41.21;MI0000470|hsa-mir-125b-2&WithFlank&21|+|16590231|16590331@23.43.21&[MIMAT0000423&hsa-miR-125b-5p&offsets|0|-1;m-67&11|-|122099809|122099830&offsets|0|-1];[MIMAT0000423_1&hsa-miR-125b-5p&offsets|0|-1;m-68&21|+|16590253|16590274&offsets|0|-1]|TCCCTGAGACCCTAACTTGTG</td><td>>MI0000446|hsa-mir-125b-1&WithFlank&11|-|122099751|122099850</td><td>21</td><td>41</td><td>21</td><td>TCCCTGAGACCCTAACTTGTG</td><td>7939.3</td><td>-5.1</td><td>1.0</td><td>-5.2</td><td>2.0E-07</td><td>3.5E-06</td><td>BT20/MB231</td></tr>
<tr><td>>hg38_21_+_8433172_8446622_RNA45SN1_45S_50ntFlanks@9572.9605.34|CAAATCGGTCGTCCGACCTGGGTATAGGGGCGAA</td><td>>hg38_21_+_8433172_8446622_RNA45SN1_45S_50ntFlanks</td><td>9572</td><td>9605</td><td>34</td><td>CAAATCGGTCGTCCGACCTGGGTATAGGGGCGAA</td><td>334.4</td><td>-2.9</td><td>0.6</td><td>-4.7</td><td>2.6E-06</td><td>3.6E-06</td><td>BT20/MB468</td></tr>
<tr><td>>hg38_21_+_8433172_8446622_RNA45SN1_45S_50ntFlanks@8113.8140.28|GGCGCGGGACATGTGGCGTACGGAAGAC</td><td>>hg38_21_+_8433172_8446622_RNA45SN1_45S_50ntFlanks</td><td>8113</td><td>8140</td><td>28</td><td>GGCGCGGGACATGTGGCGTACGGAAGAC</td><td>122.8</td><td>-4.8</td><td>1.0</td><td>-4.7</td><td>2.7E-06</td><td>3.6E-06</td><td>BT20/MB468</td></tr>
<tr><td>>hg38_21_+_8433172_8446622_RNA45SN1_45S_50ntFlanks@7989.8006.18|AGACGTGGCGACCCGCTG</td><td>>hg38_21_+_8433172_8446622_RNA45SN1_45S_50ntFlanks</td><td>7989</td><td>8006</td><td>18</td><td>AGACGTGGCGACCCGCTG</td><td>1599.7</td><td>-3.8</td><td>0.8</td><td>-4.7</td><td>2.4E-06</td><td>3.7E-06</td><td>MB231/MB468</td></tr>
<tr><td>>MI0000746|hsa-mir-99b&WithFlank&19|+|51692606|51692687@13.32.20(+1G)&[MIMAT0000689&hsa-miR-99b-5p&offsets|0|-2(+1G);m-10&19|+|51692618|51692639&offsets|0|-2(+1G)]|CACCCGTAGAACCGACCTTGG</td><td>>MI0000746|hsa-mir-99b&WithFlank&19|+|51692606|51692687</td><td>13</td><td>32</td><td>20(+1G)</td><td>CACCCGTAGAACCGACCTTGG</td><td>1030.2</td><td>5.3</td><td>1.0</td><td>5.2</td><td>2.1E-07</td><td>3.7E-06</td><td>BT20/MB231</td></tr>
<tr><td>>hg38_21_+_8433172_8446622_RNA45SN1_45S_50ntFlanks@7985.8004.20|GATCAGACGTGGCGACCCGC</td><td>>hg38_21_+_8433172_8446622_RNA45SN1_45S_50ntFlanks</td><td>7985</td><td>8004</td><td>20</td><td>GATCAGACGTGGCGACCCGC</td><td>117.8</td><td>-4.4</td><td>0.9</td><td>-4.7</td><td>2.4E-06</td><td>3.7E-06</td><td>MB231/MB468</td></tr>
<tr><td>>hg38_21_+_8433172_8446622_RNA45SN1_45S_50ntFlanks@6652.6698.47|GACTCTTAGCGGTGGATCACTCGGCTCGTGCGTCGATGAAGAACGCA</td><td>>hg38_21_+_8433172_8446622_RNA45SN1_45S_50ntFlanks</td><td>6652</td><td>6698</td><td>47</td><td>GACTCTTAGCGGTGGATCACTCGGCTCGTGCGTCGATGAAGAACGCA</td><td>6504.3</td><td>3.8</td><td>0.8</td><td>4.7</td><td>2.5E-06</td><td>3.7E-06</td><td>MB231/MB468</td></tr>
<tr><td>>hg38_21_+_8433172_8446622_RNA45SN1_45S_50ntFlanks@8196.8219.24|CGTGGACGGTGTGAGGCCGGTAGC</td><td>>hg38_21_+_8433172_8446622_RNA45SN1_45S_50ntFlanks</td><td>8196</td><td>8219</td><td>24</td><td>CGTGGACGGTGTGAGGCCGGTAGC</td><td>401.3</td><td>3.4</td><td>0.7</td><td>4.7</td><td>2.5E-06</td><td>3.8E-06</td><td>MB231/MB468</td></tr>
<tr><td>>hg38_21_+_8433172_8446622_RNA45SN1_45S_50ntFlanks@10678.10709.32|CGGTTCCGGCGGCGTCCGGTGAGCTCTCGCTG</td><td>>hg38_21_+_8433172_8446622_RNA45SN1_45S_50ntFlanks</td><td>10678</td><td>10709</td><td>32</td><td>CGGTTCCGGCGGCGTCCGGTGAGCTCTCGCTG</td><td>21220.1</td><td>-4.8</td><td>1.0</td><td>-4.7</td><td>2.8E-06</td><td>3.8E-06</td><td>BT20/MB468</td></tr>
<tr><td>>hg38_21_+_8433172_8446622_RNA45SN1_45S_50ntFlanks@10404.10424.21|CATGGGTCAGTCGGTCCTGAG</td><td>>hg38_21_+_8433172_8446622_RNA45SN1_45S_50ntFlanks</td><td>10404</td><td>10424</td><td>21</td><td>CATGGGTCAGTCGGTCCTGAG</td><td>857.7</td><td>2.8</td><td>0.6</td><td>4.7</td><td>2.5E-06</td><td>3.8E-06</td><td>MB231/MB468</td></tr>
<tr><td>>hg38_21_+_8433172_8446622_RNA45SN1_45S_50ntFlanks@4283.4314.32|CTTTAACGAGGATCCATTGGAGGGCAAGTCTG</td><td>>hg38_21_+_8433172_8446622_RNA45SN1_45S_50ntFlanks</td><td>4283</td><td>4314</td><td>32</td><td>CTTTAACGAGGATCCATTGGAGGGCAAGTCTG</td><td>158.3</td><td>-2.4</td><td>0.5</td><td>-4.7</td><td>2.5E-06</td><td>3.8E-06</td><td>MB231/MB468</td></tr>
<tr><td>>hg38_21_+_8433172_8446622_RNA45SN1_45S_50ntFlanks@10588.10607.20|CCCGGGGAGAGTTCTCTTTT</td><td>>hg38_21_+_8433172_8446622_RNA45SN1_45S_50ntFlanks</td><td>10588</td><td>10607</td><td>20</td><td>CCCGGGGAGAGTTCTCTTTT</td><td>96.4</td><td>-4.7</td><td>1.0</td><td>-4.7</td><td>2.5E-06</td><td>3.8E-06</td><td>MB231/MB468</td></tr>
<tr><td>>hg38_21_+_8433172_8446622_RNA45SN1_45S_50ntFlanks@6647.6680.34|CGTACGACTCTTAGCGGTGGATCACTCGGCTCGT</td><td>>hg38_21_+_8433172_8446622_RNA45SN1_45S_50ntFlanks</td><td>6647</td><td>6680</td><td>34</td><td>CGTACGACTCTTAGCGGTGGATCACTCGGCTCGT</td><td>700.3</td><td>2.8</td><td>0.6</td><td>4.7</td><td>2.5E-06</td><td>3.8E-06</td><td>MB231/MB468</td></tr>
<tr><td>>hg38_21_+_8433172_8446622_RNA45SN1_45S_50ntFlanks@9737.9754.18|CGAAACGATCTCAACCTA</td><td>>hg38_21_+_8433172_8446622_RNA45SN1_45S_50ntFlanks</td><td>9737</td><td>9754</td><td>18</td><td>CGAAACGATCTCAACCTA</td><td>89.1</td><td>-2.7</td><td>0.6</td><td>-4.7</td><td>2.5E-06</td><td>3.8E-06</td><td>MB231/MB468</td></tr>
<tr><td>>hg38_21_+_8433172_8446622_RNA45SN1_45S_50ntFlanks@12933.12952.20|GCTAAACCATTCGTAGACGA</td><td>>hg38_21_+_8433172_8446622_RNA45SN1_45S_50ntFlanks</td><td>12933</td><td>12952</td><td>20</td><td>GCTAAACCATTCGTAGACGA</td><td>111.0</td><td>-4.5</td><td>1.0</td><td>-4.7</td><td>2.8E-06</td><td>3.8E-06</td><td>BT20/MB468</td></tr>
<tr><td>>hg38_21_+_8433172_8446622_RNA45SN1_45S_50ntFlanks@12926.12956.31|ACCTGGCGCTAAACCATTCGTAGACGACCTG</td><td>>hg38_21_+_8433172_8446622_RNA45SN1_45S_50ntFlanks</td><td>12926</td><td>12956</td><td>31</td><td>ACCTGGCGCTAAACCATTCGTAGACGACCTG</td><td>97.2</td><td>-3.7</td><td>0.8</td><td>-4.7</td><td>2.5E-06</td><td>3.8E-06</td><td>MB231/MB468</td></tr>
<tr><td>>hg38_21_+_8433172_8446622_RNA45SN1_45S_50ntFlanks@12072.12093.22|CCCGAGGGGCTCTCGCTTCTGG</td><td>>hg38_21_+_8433172_8446622_RNA45SN1_45S_50ntFlanks</td><td>12072</td><td>12093</td><td>22</td><td>CCCGAGGGGCTCTCGCTTCTGG</td><td>91.5</td><td>-2.6</td><td>0.6</td><td>-4.7</td><td>2.5E-06</td><td>3.9E-06</td><td>MB231/MB468</td></tr>
<tr><td>>hg38_21_+_8433172_8446622_RNA45SN1_45S_50ntFlanks@4735.4766.32|ACGAAAGTCGGAGGTTCGAAGACGATCAGATA</td><td>>hg38_21_+_8433172_8446622_RNA45SN1_45S_50ntFlanks</td><td>4735</td><td>4766</td><td>32</td><td>ACGAAAGTCGGAGGTTCGAAGACGATCAGATA</td><td>412.8</td><td>8.1</td><td>1.7</td><td>4.7</td><td>2.6E-06</td><td>3.9E-06</td><td>MB231/MB468</td></tr>
<tr><td>>hg38_21_+_8433172_8446622_RNA45SN1_45S_50ntFlanks@12996.13013.18|TCGCTGCGATCTATTGAA</td><td>>hg38_21_+_8433172_8446622_RNA45SN1_45S_50ntFlanks</td><td>12996</td><td>13013</td><td>18</td><td>TCGCTGCGATCTATTGAA</td><td>1699.1</td><td>2.7</td><td>0.6</td><td>4.7</td><td>2.8E-06</td><td>3.9E-06</td><td>BT20/MB468</td></tr>
<tr><td>>hg38_21_+_8433172_8446622_RNA45SN1_45S_50ntFlanks@11583.11601.19|CTGGTGCGGACCAGGGGAA</td><td>>hg38_21_+_8433172_8446622_RNA45SN1_45S_50ntFlanks</td><td>11583</td><td>11601</td><td>19</td><td>CTGGTGCGGACCAGGGGAA</td><td>658.2</td><td>-3.0</td><td>0.6</td><td>-4.7</td><td>2.6E-06</td><td>3.9E-06</td><td>MB231/MB468</td></tr>
<tr><td>>hg38_21_+_8433172_8446622_RNA45SN1_45S_50ntFlanks@11874.11892.19|AGAAGACCCTGTTGAGCTT</td><td>>hg38_21_+_8433172_8446622_RNA45SN1_45S_50ntFlanks</td><td>11874</td><td>11892</td><td>19</td><td>AGAAGACCCTGTTGAGCTT</td><td>579.2</td><td>-3.4</td><td>0.7</td><td>-4.7</td><td>2.6E-06</td><td>3.9E-06</td><td>MB231/MB468</td></tr>
<tr><td>>hg38_21_+_8433172_8446622_RNA45SN1_45S_50ntFlanks@6722.6744.23|ATTGCAGGACACATTGATCATCG</td><td>>hg38_21_+_8433172_8446622_RNA45SN1_45S_50ntFlanks</td><td>6722</td><td>6744</td><td>23</td><td>ATTGCAGGACACATTGATCATCG</td><td>2196.8</td><td>-3.3</td><td>0.7</td><td>-4.7</td><td>2.6E-06</td><td>3.9E-06</td><td>MB231/MB468</td></tr>
<tr><td>>hg38_21_+_8433172_8446622_RNA45SN1_45S_50ntFlanks@6730.6770.41|ACACATTGATCATCGACACTTCGAACGCACTTGCGGCCCCG</td><td>>hg38_21_+_8433172_8446622_RNA45SN1_45S_50ntFlanks</td><td>6730</td><td>6770</td><td>41</td><td>ACACATTGATCATCGACACTTCGAACGCACTTGCGGCCCCG</td><td>192.1</td><td>-2.3</td><td>0.5</td><td>-4.7</td><td>2.6E-06</td><td>3.9E-06</td><td>MB231/MB468</td></tr>
<tr><td>>hg38_21_+_8433172_8446622_RNA45SN1_45S_50ntFlanks@11576.11607.32|CTTAGAACTGGTGCGGACCAGGGGAATCCGAC</td><td>>hg38_21_+_8433172_8446622_RNA45SN1_45S_50ntFlanks</td><td>11576</td><td>11607</td><td>32</td><td>CTTAGAACTGGTGCGGACCAGGGGAATCCGAC</td><td>300.7</td><td>-2.9</td><td>0.6</td><td>-4.7</td><td>2.9E-06</td><td>3.9E-06</td><td>BT20/MB468</td></tr>
<tr><td>>hg38_21_+_8433172_8446622_RNA45SN1_45S_50ntFlanks@12959.12982.24|TCTGGGTCGGGGTTTCGTACGTAG</td><td>>hg38_21_+_8433172_8446622_RNA45SN1_45S_50ntFlanks</td><td>12959</td><td>12982</td><td>24</td><td>TCTGGGTCGGGGTTTCGTACGTAG</td><td>121.7</td><td>-2.4</td><td>0.5</td><td>-4.7</td><td>2.6E-06</td><td>4.0E-06</td><td>MB231/MB468</td></tr>
<tr><td>>hg38_21_+_8433172_8446622_RNA45SN1_45S_50ntFlanks@11575.11604.30|ACTTAGAACTGGTGCGGACCAGGGGAATCC</td><td>>hg38_21_+_8433172_8446622_RNA45SN1_45S_50ntFlanks</td><td>11575</td><td>11604</td><td>30</td><td>ACTTAGAACTGGTGCGGACCAGGGGAATCC</td><td>121.4</td><td>-2.5</td><td>0.5</td><td>-4.7</td><td>3.0E-06</td><td>4.0E-06</td><td>BT20/MB468</td></tr>
<tr><td>>hg38_21_+_8433172_8446622_RNA45SN1_45S_50ntFlanks@11581.11601.21|AACTGGTGCGGACCAGGGGAA</td><td>>hg38_21_+_8433172_8446622_RNA45SN1_45S_50ntFlanks</td><td>11581</td><td>11601</td><td>21</td><td>AACTGGTGCGGACCAGGGGAA</td><td>1204.7</td><td>-3.8</td><td>0.8</td><td>-4.7</td><td>3.0E-06</td><td>4.1E-06</td><td>BT20/MB468</td></tr>
<tr><td>>hg38_21_+_8433172_8446622_RNA45SN1_45S_50ntFlanks@4005.4023.19|AACCTCGGGCCGATCGCAC</td><td>>hg38_21_+_8433172_8446622_RNA45SN1_45S_50ntFlanks</td><td>4005</td><td>4023</td><td>19</td><td>AACCTCGGGCCGATCGCAC</td><td>397.6</td><td>2.2</td><td>0.5</td><td>4.7</td><td>3.0E-06</td><td>4.1E-06</td><td>BT20/MB468</td></tr>
<tr><td>>MI0000102|hsa-mir-100&WithFlank&11|-|122152223|122152314@19.37.19(+1A)&[MIMAT0000098&hsa-miR-100-5p&offsets|0|-3(+1A);m-27&11|-|122152276|122152296&offsets|0|-2(+1A)]|AACCCGTAGATCCGAACTTA</td><td>>MI0000102|hsa-mir-100&WithFlank&11|-|122152223|122152314</td><td>19</td><td>37</td><td>19(+1A)</td><td>AACCCGTAGATCCGAACTTA</td><td>1908.3</td><td>-5.1</td><td>1.0</td><td>-5.2</td><td>2.3E-07</td><td>4.1E-06</td><td>BT20/MB231</td></tr>
<tr><td>>hg38_21_+_8433172_8446622_RNA45SN1_45S_50ntFlanks@3854.3874.21|ACTGTGGTAATTCTAGAGCTA</td><td>>hg38_21_+_8433172_8446622_RNA45SN1_45S_50ntFlanks</td><td>3854</td><td>3874</td><td>21</td><td>ACTGTGGTAATTCTAGAGCTA</td><td>389.1</td><td>-2.5</td><td>0.5</td><td>-4.7</td><td>2.7E-06</td><td>4.1E-06</td><td>MB231/MB468</td></tr>
<tr><td>>hg38_21_+_8433172_8446622_RNA45SN1_45S_50ntFlanks@8176.8202.27|CTTCTGATCGAGGCCCAGCCCGTGGAC</td><td>>hg38_21_+_8433172_8446622_RNA45SN1_45S_50ntFlanks</td><td>8176</td><td>8202</td><td>27</td><td>CTTCTGATCGAGGCCCAGCCCGTGGAC</td><td>2238.4</td><td>3.8</td><td>0.8</td><td>4.7</td><td>2.7E-06</td><td>4.1E-06</td><td>MB231/MB468</td></tr>
<tr><td>>hg38_21_+_8433172_8446622_RNA45SN1_45S_50ntFlanks@11781.11807.27|CGCATGAATGGATGAACGAGATTCCCA</td><td>>hg38_21_+_8433172_8446622_RNA45SN1_45S_50ntFlanks</td><td>11781</td><td>11807</td><td>27</td><td>CGCATGAATGGATGAACGAGATTCCCA</td><td>104.0</td><td>-4.8</td><td>1.0</td><td>-4.7</td><td>3.1E-06</td><td>4.2E-06</td><td>BT20/MB468</td></tr>
<tr><td>>hg38_21_+_8433172_8446622_RNA45SN1_45S_50ntFlanks@12912.12933.22|CACGTTCGTGGGGAACCTGGCG</td><td>>hg38_21_+_8433172_8446622_RNA45SN1_45S_50ntFlanks</td><td>12912</td><td>12933</td><td>22</td><td>CACGTTCGTGGGGAACCTGGCG</td><td>460.3</td><td>-5.1</td><td>1.1</td><td>-4.7</td><td>3.1E-06</td><td>4.2E-06</td><td>BT20/MB468</td></tr>
<tr><td>>hg38_21_+_8433172_8446622_RNA45SN1_45S_50ntFlanks@11212.11229.18|CGACTCTGGACGCGAGCC</td><td>>hg38_21_+_8433172_8446622_RNA45SN1_45S_50ntFlanks</td><td>11212</td><td>11229</td><td>18</td><td>CGACTCTGGACGCGAGCC</td><td>218.8</td><td>-3.9</td><td>0.8</td><td>-4.7</td><td>2.8E-06</td><td>4.2E-06</td><td>MB231/MB468</td></tr>
<tr><td>>hg38_21_+_8433172_8446622_RNA45SN1_45S_50ntFlanks@3848.3866.19|TGGATAACTGTGGTAATTC</td><td>>hg38_21_+_8433172_8446622_RNA45SN1_45S_50ntFlanks</td><td>3848</td><td>3866</td><td>19</td><td>TGGATAACTGTGGTAATTC</td><td>382.1</td><td>8.0</td><td>1.7</td><td>4.7</td><td>2.8E-06</td><td>4.2E-06</td><td>MB231/MB468</td></tr>
<tr><td>>hg38_21_+_8433172_8446622_RNA45SN1_45S_50ntFlanks@10349.10382.34|AACGAGAACTTTGAAGGCCGAAGTGGAGAAGGGT</td><td>>hg38_21_+_8433172_8446622_RNA45SN1_45S_50ntFlanks</td><td>10349</td><td>10382</td><td>34</td><td>AACGAGAACTTTGAAGGCCGAAGTGGAGAAGGGT</td><td>1164.1</td><td>3.4</td><td>0.7</td><td>4.7</td><td>2.8E-06</td><td>4.2E-06</td><td>MB231/MB468</td></tr>
<tr><td>>hg38_21_+_8433172_8446622_RNA45SN1_45S_50ntFlanks@10350.10372.23|ACGAGAACTTTGAAGGCCGAAGT</td><td>>hg38_21_+_8433172_8446622_RNA45SN1_45S_50ntFlanks</td><td>10350</td><td>10372</td><td>23</td><td>ACGAGAACTTTGAAGGCCGAAGT</td><td>1251.6</td><td>2.7</td><td>0.6</td><td>4.7</td><td>2.8E-06</td><td>4.2E-06</td><td>MB231/MB468</td></tr>
<tr><td>>hg38_21_+_8433172_8446622_RNA45SN1_45S_50ntFlanks@11199.11229.31|CCGGCGGCGGCGGCGACTCTGGACGCGAGCC</td><td>>hg38_21_+_8433172_8446622_RNA45SN1_45S_50ntFlanks</td><td>11199</td><td>11229</td><td>31</td><td>CCGGCGGCGGCGGCGACTCTGGACGCGAGCC</td><td>64.6</td><td>-3.9</td><td>0.8</td><td>-4.7</td><td>2.8E-06</td><td>4.3E-06</td><td>MB231/MB468</td></tr>
<tr><td>>hg38_21_+_8433172_8446622_RNA45SN1_45S_50ntFlanks@9565.9592.28|TGACGTGCAAATCGGTCGTCCGACCTGG</td><td>>hg38_21_+_8433172_8446622_RNA45SN1_45S_50ntFlanks</td><td>9565</td><td>9592</td><td>28</td><td>TGACGTGCAAATCGGTCGTCCGACCTGG</td><td>445.0</td><td>-2.3</td><td>0.5</td><td>-4.7</td><td>2.9E-06</td><td>4.3E-06</td><td>MB231/MB468</td></tr>
<tr><td>>hg38_21_+_8433172_8446622_RNA45SN1_45S_50ntFlanks@12971.12992.22|TTTCGTACGTAGCAGAGCAGCT</td><td>>hg38_21_+_8433172_8446622_RNA45SN1_45S_50ntFlanks</td><td>12971</td><td>12992</td><td>22</td><td>TTTCGTACGTAGCAGAGCAGCT</td><td>122.3</td><td>-2.2</td><td>0.5</td><td>-4.7</td><td>3.2E-06</td><td>4.4E-06</td><td>BT20/MB468</td></tr>
<tr><td>>hg38_21_+_8433172_8446622_RNA45SN1_45S_50ntFlanks@7988.8016.29|CAGACGTGGCGACCCGCTGAATTTAAGCA</td><td>>hg38_21_+_8433172_8446622_RNA45SN1_45S_50ntFlanks</td><td>7988</td><td>8016</td><td>29</td><td>CAGACGTGGCGACCCGCTGAATTTAAGCA</td><td>734.7</td><td>2.6</td><td>0.6</td><td>4.7</td><td>2.9E-06</td><td>4.4E-06</td><td>MB231/MB468</td></tr>
<tr><td>>hg38_21_+_8433172_8446622_RNA45SN1_45S_50ntFlanks@11207.11227.21|GGCGGCGACTCTGGACGCGAG</td><td>>hg38_21_+_8433172_8446622_RNA45SN1_45S_50ntFlanks</td><td>11207</td><td>11227</td><td>21</td><td>GGCGGCGACTCTGGACGCGAG</td><td>85.7</td><td>-3.7</td><td>0.8</td><td>-4.7</td><td>2.9E-06</td><td>4.4E-06</td><td>MB231/MB468</td></tr>
<tr><td>>hg38_21_+_8433172_8446622_RNA45SN1_45S_50ntFlanks@9353.9370.18|CGTGGCGCAATGAAGGTG</td><td>>hg38_21_+_8433172_8446622_RNA45SN1_45S_50ntFlanks</td><td>9353</td><td>9370</td><td>18</td><td>CGTGGCGCAATGAAGGTG</td><td>79.2</td><td>-4.9</td><td>1.0</td><td>-4.7</td><td>3.2E-06</td><td>4.4E-06</td><td>BT20/MB468</td></tr>
<tr><td>>hg38_21_+_8433172_8446622_RNA45SN1_45S_50ntFlanks@7979.8003.25|ACCTCAGATCAGACGTGGCGACCCG</td><td>>hg38_21_+_8433172_8446622_RNA45SN1_45S_50ntFlanks</td><td>7979</td><td>8003</td><td>25</td><td>ACCTCAGATCAGACGTGGCGACCCG</td><td>324.9</td><td>-2.9</td><td>0.6</td><td>-4.7</td><td>2.9E-06</td><td>4.4E-06</td><td>MB231/MB468</td></tr>
<tr><td>>hg38_21_+_8433172_8446622_RNA45SN1_45S_50ntFlanks@10404.10422.19|CATGGGTCAGTCGGTCCTG</td><td>>hg38_21_+_8433172_8446622_RNA45SN1_45S_50ntFlanks</td><td>10404</td><td>10422</td><td>19</td><td>CATGGGTCAGTCGGTCCTG</td><td>282.1</td><td>-2.8</td><td>0.6</td><td>-4.7</td><td>3.0E-06</td><td>4.5E-06</td><td>MB231/MB468</td></tr>
</table>

| | | | | | | | | | | | | |
|---|---|---|---|---|---|---|---|---|---|---|---|---|
| >hg38_21_+_8433172_8446622_RNA45SN1_45S_50ntFlanks@11718.11739.22\|AACGGCGGGAGTAACTATGACT | >hg38_21_+_8433172_8446622_RNA45SN1_45S_50ntFlanks | 11718 | 11739 | 22 | AACGGCGGGAGTAACTATGACT | 255.1 | -4.3 | 0.9 | -4.7 | 3.3E-06 | 4.5E-06 | BT20/MB468 |
| >trnaMT_MetCAT_MT_+_4402_4469@46.71.26\|TTGGTTATACCCTTCCCGTACTACCA | >trnaMT_MetCAT_MT_+_4402_4469 | 46 | 71 | 26 | TTGGTTATACCCTTCCCGTACTACCA | 1401.3 | 11.2 | 2.4 | 4.7 | 3.0E-06 | 4.5E-06 | MB231/MB468 |
| >hg38_21_+_8433172_8446622_RNA45SN1_45S_50ntFlanks@10849.10866.18\|CGGGATAAGGATTGGCTC | >hg38_21_+_8433172_8446622_RNA45SN1_45S_50ntFlanks | 10849 | 10866 | 18 | CGGGATAAGGATTGGCTC | 162.2 | -3.8 | 0.8 | -4.6 | 3.3E-06 | 4.5E-06 | BT20/MB468 |
| >hg38_21_+_8433172_8446622_RNA45SN1_45S_50ntFlanks@12626.12659.34\|AATGGGGCGAAGCTACCATCTGTGGGATTATGAC | >hg38_21_+_8433172_8446622_RNA45SN1_45S_50ntFlanks | 12626 | 12659 | 34 | AATGGGGCGAAGCTACCATCTGTGGGATTATGAC | 432.2 | -2.8 | 0.6 | -4.7 | 3.1E-06 | 4.6E-06 | MB231/MB468 |
| >hg38_21_+_8433172_8446622_RNA45SN1_45S_50ntFlanks@9534.9565.32\|AGGAAACTCTGGTGGAGGTCCGTAGCGGTCCT | >hg38_21_+_8433172_8446622_RNA45SN1_45S_50ntFlanks | 9534 | 9565 | 32 | AGGAAACTCTGGTGGAGGTCCGTAGCGGTCCT | 948.8 | -2.2 | 0.5 | -4.7 | 3.1E-06 | 4.6E-06 | MB231/MB468 |
| >hg38_21_+_8433172_8446622_RNA45SN1_45S_50ntFlanks@10411.10453.43\|CAGTCGGTCCTGAGAGATGGGCGAGCGCCGTTCCGAAGGGACG | >hg38_21_+_8433172_8446622_RNA45SN1_45S_50ntFlanks | 10411 | 10453 | 43 | CAGTCGGTCCTGAGAGATGGGCGAGCGCCGTTCCGAAGGGACG | 300.2 | -1.9 | 0.4 | -4.7 | 3.1E-06 | 4.7E-06 | MB231/MB468 |
| >MI0000342\|hsa-mir-200b&WithFlank&1\|+\|1167098\|1167204@63.83.21&[MIMAT0000318&hsa-miR-200b-3p&offsets\|0\|-1;m-93&1\|+\|1167160\|1167182&offsets\|0\|-2]\|TAATACTGCCTGGTAATGATG | >MI0000342\|hsa-mir-200b&WithFlank&1\|+\|1167098\|1167204 | 63 | 83 | 21 | TAATACTGCCTGGTAATGATG | 289.6 | -3.1 | 0.7 | -4.7 | 3.1E-06 | 4.7E-06 | MB231/MB468 |
| >hg38_21_+_8433172_8446622_RNA45SN1_45S_50ntFlanks@12487.12509.23\|GAACGTGAGCTGGGTTTAGACCG | >hg38_21_+_8433172_8446622_RNA45SN1_45S_50ntFlanks | 12487 | 12509 | 23 | GAACGTGAGCTGGGTTTAGACCG | 230.3 | -3.1 | 0.7 | -4.6 | 3.5E-06 | 4.7E-06 | BT20/MB468 |
| >hg38_21_+_8433172_8446622_RNA45SN1_45S_50ntFlanks@12442.12473.32\|TGAAGCAGAATTCACCAAGCGTTGGATTGTTC | >hg38_21_+_8433172_8446622_RNA45SN1_45S_50ntFlanks | 12442 | 12473 | 32 | TGAAGCAGAATTCACCAAGCGTTGGATTGTTC | 196.6 | -2.8 | 0.6 | -4.6 | 3.5E-06 | 4.7E-06 | BT20/MB468 |
| >MI0000101\|hsa-mir-99a&WithFlank&21\|+\|16539083\|16539175@19.40.22(+1A)&[MIMAT0000097&hsa-miR-99a-5p&offsets\|0\|0(+1A);m-64&21\|+\|16539101\|16539121&offsets\|0\|+1(+1A)]\|AACCCGTAGATCCGATCTTGTGA | >MI0000101\|hsa-mir-99a&WithFlank&21\|+\|16539083\|16539175 | 19 | 40 | 22(+1A) | AACCCGTAGATCCGATCTTGTGA | 318.1 | -5.2 | 1.0 | -5.1 | 2.7E-07 | 4.7E-06 | BT20/MB231 |
| >hg38_21_+_8433172_8446622_RNA45SN1_45S_50ntFlanks@4367.4400.34\|CAGTTAAAAAGCTCGTAGTTGGATCTTGGGAGCG | >hg38_21_+_8433172_8446622_RNA45SN1_45S_50ntFlanks | 4367 | 4400 | 34 | CAGTTAAAAAGCTCGTAGTTGGATCTTGGGAGCG | 76.8 | -3.4 | 0.7 | -4.7 | 3.2E-06 | 4.8E-06 | MB231/MB468 |
| >hg38_21_+_8433172_8446622_RNA45SN1_45S_50ntFlanks@3813.3837.25\|TTATGGTTCCTTTGGTCGCTCGCTC | >hg38_21_+_8433172_8446622_RNA45SN1_45S_50ntFlanks | 3813 | 3837 | 25 | TTATGGTTCCTTTGGTCGCTCGCTC | 77.2 | -2.5 | 0.5 | -4.7 | 3.2E-06 | 4.8E-06 | MB231/MB468 |
| >hg38_21_+_8433172_8446622_RNA45SN1_45S_50ntFlanks@11199.11227.29\|CCGGCGGCGGCGGCGACTCTGGACGCGAG | >hg38_21_+_8433172_8446622_RNA45SN1_45S_50ntFlanks | 11199 | 11227 | 29 | CCGGCGGCGGCGGCGACTCTGGACGCGAG | 318.6 | -5.1 | 1.1 | -4.6 | 3.6E-06 | 4.8E-06 | BT20/MB468 |
| >trna152_ValCAC_6_-_27248049_27248121@1.30.30\|GCTTCTGTAGTGTAGTGGTTATCACGTTCG | >trna152_ValCAC_6_-_27248049_27248121 | 1 | 30 | 30 | GCTTCTGTAGTGTAGTGGTTATCACGTTCG | 155.5 | -5.0 | 1.1 | -4.6 | 3.6E-06 | 4.8E-06 | BT20/MB468 |
| >hg38_21_+_8433172_8446622_RNA45SN1_45S_50ntFlanks@10784.10818.35\|TGAACAGCCTCTGGCATGTTGGAACAATGTAGGTA | >hg38_21_+_8433172_8446622_RNA45SN1_45S_50ntFlanks | 10784 | 10818 | 35 | TGAACAGCCTCTGGCATGTTGGAACAATGTAGGTA | 398.4 | -2.1 | 0.5 | -4.7 | 3.2E-06 | 4.8E-06 | MB231/MB468 |
| >hg38_21_+_8433172_8446622_RNA45SN1_45S_50ntFlanks@12404.12431.28\|GCTTTTTGATCCTTCGATGTCGGCTCTT | >hg38_21_+_8433172_8446622_RNA45SN1_45S_50ntFlanks | 12404 | 12431 | 28 | GCTTTTTGATCCTTCGATGTCGGCTCTT | 193.0 | -2.7 | 0.6 | -4.6 | 3.6E-06 | 4.8E-06 | BT20/MB468 |
| >trna111_HisGTG_1_-_147774845_147774916@-1T.32.33,trna118_HisGTG_1_-_145396881_145396952@-1T.32.33,trna16_HisGTG_1_+_146544773_146544844@-1T.32.33,trna1_HisGTG_15_+_45493349_45493420@-1T.32.33,trna21_HisGTG_1_+_147753471_147753542@-1T.32.33,trna33_HisGTG_6_+_27125906_27125977@-1T.32.33,trna7_HisGTG_9_-_14433938_14434009@-1T.32.33,trna8_HisGTG_15_-_45492611_45492682@-1T.32.33,trna9_HisGTG_15_-_45490804_45490875@-1T.32.33\|TGCCGTGATCGTATAGTGGTTAGTACTCTGCGT | >trna111_HisGTG_1_-_147774845_147774916 | -1T | 32 | 33 | TGCCGTGATCGTATAGTGGTTAGTACTCTGCGT | 1086.1 | 11.1 | 2.4 | 4.7 | 3.2E-06 | 4.9E-06 | MB231/MB468 |
| >hg38_21_+_8433172_8446622_RNA45SN1_45S_50ntFlanks@5515.5535.21\|CTAGAGGAAGTAAAAGTCGTA | >hg38_21_+_8433172_8446622_RNA45SN1_45S_50ntFlanks | 5515 | 5535 | 21 | CTAGAGGAAGTAAAAGTCGTA | 263.3 | -2.5 | 0.5 | -4.7 | 3.3E-06 | 4.9E-06 | MB231/MB468 |
| >hg38_21_+_8433172_8446622_RNA45SN1_45S_50ntFlanks@10830.10853.24\|CAAGCCGGATCCGTAACTTCGGGA | >hg38_21_+_8433172_8446622_RNA45SN1_45S_50ntFlanks | 10830 | 10853 | 24 | CAAGCCGGATCCGTAACTTCGGGA | 156.5 | -3.3 | 0.7 | -4.7 | 3.3E-06 | 4.9E-06 | MB231/MB468 |
| >hg38_21_+_8433172_8446622_RNA45SN1_45S_50ntFlanks@10387.10407.21\|TGTGAACAGCAGTTGAACATG | >hg38_21_+_8433172_8446622_RNA45SN1_45S_50ntFlanks | 10387 | 10407 | 21 | TGTGAACAGCAGTTGAACATG | 189.9 | -3.3 | 0.7 | -4.6 | 3.6E-06 | 4.9E-06 | BT20/MB468 |
| >hg38_21_+_8433172_8446622_RNA45SN1_45S_50ntFlanks@7979.8011.33\|ACCTCAGATCAGACGTGGCGACCCGCTGAATTT | >hg38_21_+_8433172_8446622_RNA45SN1_45S_50ntFlanks | 7979 | 8011 | 33 | ACCTCAGATCAGACGTGGCGACCCGCTGAATTT | 409.0 | 2.6 | 0.6 | 4.7 | 3.3E-06 | 4.9E-06 | MB231/MB468 |
| >hg38_21_+_8433172_8446622_RNA45SN1_45S_50ntFlanks@12409.12434.26\|TTGATCCTTCGATGTCGGCTCTTCCT | >hg38_21_+_8433172_8446622_RNA45SN1_45S_50ntFlanks | 12409 | 12434 | 26 | TTGATCCTTCGATGTCGGCTCTTCCT | 479.6 | 3.1 | 0.7 | 4.6 | 3.3E-06 | 5.0E-06 | MB231/MB468 |
| >hg38_21_+_8433172_8446622_RNA45SN1_45S_50ntFlanks@5485.5518.34\|AGCGCTGAGAAGACGGTCGAACTTGACTATCTAG | >hg38_21_+_8433172_8446622_RNA45SN1_45S_50ntFlanks | 5485 | 5518 | 34 | AGCGCTGAGAAGACGGTCGAACTTGACTATCTAG | 416.1 | -4.1 | 0.9 | -4.6 | 3.7E-06 | 5.0E-06 | BT20/MB468 |
| >hg38_21_+_8433172_8446622_RNA45SN1_45S_50ntFlanks@11874.11904.31\|AGAAGACCCTGTTGAGCTTGACTCTAGTCTG | >hg38_21_+_8433172_8446622_RNA45SN1_45S_50ntFlanks | 11874 | 11904 | 31 | AGAAGACCCTGTTGAGCTTGACTCTAGTCTG | 529.5 | -6.0 | 1.3 | -4.6 | 3.7E-06 | 5.0E-06 | BT20/MB468 |
| >am_trna10_ValCAC_5_-_180649395_180649467@1.30.30,trna12_ValAAC_5_-_180645270_180645342@1.30.30,trna132_ValAAC_6_-_27721179_27721251@1.30.30,trna136_ValAAC_6_-_27648885_27648957@1.30.30,trna139_ValAAC_6_-_27618707_27618779@1.30.30,trna18_ValCAC_5_-_180529253_180529325@1.30.30,trna2_ValAAC_3_+_169490018_169490090@1.30.30,trna2_ValCAC_5_+_180524070_180524142@1.30.30,trna4_ValAAC_5_+_180591154_180591226@1.30.30,trna5_ValAAC_5_+_180596610_180596682@1.30.30,trna6_ValCAC_5_+_180600650_180600722@1.30.30,trna85_ValCAC_1_-_161369490_161369562@1.30.30,trna90_ValCAC_1_-_149684088_149684161@1.30.30,trna98_ValCAC_1_-_149298555_149298627@1.30.30,trna9_ValCAC_6_+_26538282_26538354@1.30.30\|GTTTCCGTAGTGTAGTGGTTATCACGTTCG | >am_trna10_ValCAC_5_-_180649395_180649467 | 1 | 30 | 30 | GTTTCCGTAGTGTAGTGGTTATCACGTTCG | 182.7 | -5.3 | 1.1 | -4.6 | 3.7E-06 | 5.0E-06 | BT20/MB468 |
| >hg38_21_+_8433172_8446622_RNA45SN1_45S_50ntFlanks@9623.9650.28\|TAGTAGCTGGTTCCCTCCGAAGTTTCCC | >hg38_21_+_8433172_8446622_RNA45SN1_45S_50ntFlanks | 9623 | 9650 | 28 | TAGTAGCTGGTTCCCTCCGAAGTTTCCC | 123.3 | -4.1 | 0.9 | -4.6 | 3.7E-06 | 5.0E-06 | BT20/MB468 |
| >hg38_21_+_8433172_8446622_RNA45SN1_45S_50ntFlanks@9533.9563.31\|GAGGAAACTCTGGTGGAGGTCCGTAGCGGTC | >hg38_21_+_8433172_8446622_RNA45SN1_45S_50ntFlanks | 9533 | 9563 | 31 | GAGGAAACTCTGGTGGAGGTCCGTAGCGGTC | 945.1 | -2.8 | 0.6 | -4.6 | 3.4E-06 | 5.1E-06 | MB231/MB468 |
| >hg38_21_+_8433172_8446622_RNA45SN1_45S_50ntFlanks@4725.4759.35\|TTAATCAAGAACGAAAGTCGGAGGTTCGAAGACGA | >hg38_21_+_8433172_8446622_RNA45SN1_45S_50ntFlanks | 4725 | 4759 | 35 | TTAATCAAGAACGAAAGTCGGAGGTTCGAAGACGA | 150.7 | -3.1 | 0.7 | -4.6 | 3.8E-06 | 5.1E-06 | BT20/MB468 |
| >hg38_21_+_8433172_8446622_RNA45SN1_45S_50ntFlanks@8912.8936.25\|CGCCGAATCCCGGGGCCGAGGGAGC | >hg38_21_+_8433172_8446622_RNA45SN1_45S_50ntFlanks | 8912 | 8936 | 25 | CGCCGAATCCCGGGGCCGAGGGAGC | 911.3 | 9.6 | 2.1 | 4.6 | 3.4E-06 | 5.1E-06 | MB231/MB468 |
| >hg38_21_+_8433172_8446622_RNA45SN1_45S_50ntFlanks@12490.12509.20\|CGTGAGCTGGGTTTAGACCG | >hg38_21_+_8433172_8446622_RNA45SN1_45S_50ntFlanks | 12490 | 12509 | 20 | CGTGAGCTGGGTTTAGACCG | 127.6 | -4.3 | 0.9 | -4.6 | 3.8E-06 | 5.2E-06 | BT20/MB468 |
| >hg38_21_+_8433172_8446622_RNA45SN1_45S_50ntFlanks@10393.10426.34\|CAGCAGTTGAACATGGGTCAGTCGGTCCTGAGAG | >hg38_21_+_8433172_8446622_RNA45SN1_45S_50ntFlanks | 10393 | 10426 | 34 | CAGCAGTTGAACATGGGTCAGTCGGTCCTGAGAG | 223.6 | -4.1 | 0.9 | -4.6 | 3.9E-06 | 5.2E-06 | BT20/MB468 |
| >hg38_21_+_8433172_8446622_RNA45SN1_45S_50ntFlanks@10905.10939.35\|CTGGGCGCGCGCCGCGGCTGGACGAGGCGCCGCCG | >hg38_21_+_8433172_8446622_RNA45SN1_45S_50ntFlanks | 10905 | 10939 | 35 | CTGGGCGCGCGCCGCGGCTGGACGAGGCGCCGCCG | 118.9 | -4.6 | 1.0 | -4.6 | 3.9E-06 | 5.2E-06 | BT20/MB468 |
| >hg38_21_+_8433172_8446622_RNA45SN1_45S_50ntFlanks@11874.11898.25\|AGAAGACCCTGTTGAGCTTGACTCT | >hg38_21_+_8433172_8446622_RNA45SN1_45S_50ntFlanks | 11874 | 11898 | 25 | AGAAGACCCTGTTGAGCTTGACTCT | 1520.4 | -3.1 | 0.7 | -4.6 | 3.5E-06 | 5.3E-06 | MB231/MB468 |
| >hg38_21_+_8433172_8446622_RNA45SN1_45S_50ntFlanks@10351.10370.20\|CGAGAACTTTGAAGGCCGAA | >hg38_21_+_8433172_8446622_RNA45SN1_45S_50ntFlanks | 10351 | 10370 | 20 | CGAGAACTTTGAAGGCCGAA | 388.2 | -2.8 | 0.6 | -4.6 | 3.9E-06 | 5.3E-06 | BT20/MB468 |
| >MI0000286\|hsa-mir-210&WithFlank&11\|-\|568083\|568204@72.93.22(+1A)&[MIMAT0000267&hsa-miR-210-3p&offsets\|0\|0(+1A);m-102&11\|-\|568112\|568133&offsets\|0\|0(+1A)]\|CTGTGCGTGTGACAGCGGCTGAA | >MI0000286\|hsa-mir-210&WithFlank&11\|-\|568083\|568204 | 72 | 93 | 22(+1A) | CTGTGCGTGTGACAGCGGCTGAA | 590.8 | 8.5 | 1.8 | 4.6 | 3.6E-06 | 5.3E-06 | MB231/MB468 |
| >hg38_21_+_8433172_8446622_RNA45SN1_45S_50ntFlanks@12933.12951.19\|GCTAAACCATTCGTAGACG | >hg38_21_+_8433172_8446622_RNA45SN1_45S_50ntFlanks | 12933 | 12951 | 19 | GCTAAACCATTCGTAGACG | 137.2 | -2.7 | 0.6 | -4.6 | 4.0E-06 | 5.4E-06 | BT20/MB468 |
| >hg38_21_+_8433172_8446622_RNA45SN1_45S_50ntFlanks@11873.11909.37\|AAGAAGACCCTGTTGAGCTTGACTCTAGTCTGGCACG | >hg38_21_+_8433172_8446622_RNA45SN1_45S_50ntFlanks | 11873 | 11909 | 37 | AAGAAGACCCTGTTGAGCTTGACTCTAGTCTGGCACG | 144.2 | -4.9 | 1.1 | -4.6 | 4.0E-06 | 5.4E-06 | BT20/MB468 |
| >hg38_21_+_8433172_8446622_RNA45SN1_45S_50ntFlanks@11663.11685.23\|ATTTCTGCCCAGTGCTCTGAATG | >hg38_21_+_8433172_8446622_RNA45SN1_45S_50ntFlanks | 11663 | 11685 | 23 | ATTTCTGCCCAGTGCTCTGAATG | 517.5 | -2.3 | 0.5 | -4.6 | 4.0E-06 | 5.4E-06 | BT20/MB468 |
| >hg38_21_+_8433172_8446622_RNA45SN1_45S_50ntFlanks@9566.9596.31\|GACGTGCAAATCGGTCGTCCGACCTGGGTAT | >hg38_21_+_8433172_8446622_RNA45SN1_45S_50ntFlanks | 9566 | 9596 | 31 | GACGTGCAAATCGGTCGTCCGACCTGGGTAT | 1776.8 | -2.7 | 0.6 | -4.6 | 3.6E-06 | 5.4E-06 | MB231/MB468 |
| >MI0000085\|hsa-mir-27a&WithFlank&19\|-\|13836434\|13836523@57.76.20(+2U)&[MIMAT0000084&hsa-miR-27a-3p&offsets\|0\|-1(+2U);m-43&19\|-\|13836447\|13836467&offsets\|0\|-1(+2U)]\|TTCACAGTGGCTAAGTTCCGTT | >MI0000085\|hsa-mir-27a&WithFlank&19\|-\|13836434\|13836523 | 57 | 76 | 20(+2U) | TTCACAGTGGCTAAGTTCCGTT | 4055.4 | 4.4 | 1.0 | 4.6 | 4.0E-06 | 5.4E-06 | BT20/MB468 |

<table>
<tr><td>>hg38_21_+_8433172_8446622_RNA45SN1_45S_50ntFlanks@11675.11702.28|TGCTCTGAATGTCAAAGTGAAGAAATTC</td><td>>hg38_21_+_8433172_8446622_RNA45SN1_45S_50ntFlanks</td><td>11675</td><td>11702</td><td>28</td><td>TGCTCTGAATGTCAAAGTGAAGAAATTC</td><td>482.9</td><td>2.1</td><td>0.5</td><td>4.6</td><td>4.0E-06</td><td>5.4E-06</td><td>BT20/MB468</td></tr>
<tr><td>>hg38_21_+_8433172_8446622_RNA45SN1_45S_50ntFlanks@4322.4339.18|CAGCCGCGGTAATTCCAG</td><td>>hg38_21_+_8433172_8446622_RNA45SN1_45S_50ntFlanks</td><td>4322</td><td>4339</td><td>18</td><td>CAGCCGCGGTAATTCCAG</td><td>107.5</td><td>-3.9</td><td>0.8</td><td>-4.6</td><td>3.7E-06</td><td>5.5E-06</td><td>MB231/MB468</td></tr>
<tr><td>>hg38_21_+_8433172_8446622_RNA45SN1_45S_50ntFlanks@11561.11594.34|GCGCCTAGCAGCCGACTTAGAACTGGTGCGGACC</td><td>>hg38_21_+_8433172_8446622_RNA45SN1_45S_50ntFlanks</td><td>11561</td><td>11594</td><td>34</td><td>GCGCCTAGCAGCCGACTTAGAACTGGTGCGGACC</td><td>235.5</td><td>-2.8</td><td>0.6</td><td>-4.6</td><td>4.1E-06</td><td>5.5E-06</td><td>BT20/MB468</td></tr>
<tr><td>>MI0000737|hsa-mir-200a&WithFlank&1|+|1167857|1167958@60.82.23&[MIMAT0000682&hsa-miR-200a-3p&offsets|0|+1;m-142&1|+|1167916|1167938&offsets|0|0]|TAACACTGTCTGGTAACGATGTT</td><td>>MI0000737|hsa-mir-200a&WithFlank&1|+|1167857|1167958</td><td>60</td><td>82</td><td>23</td><td>TAACACTGTCTGGTAACGATGTT</td><td>1439.8</td><td>-3.6</td><td>0.8</td><td>-4.6</td><td>3.7E-06</td><td>5.6E-06</td><td>MB231/MB468</td></tr>
<tr><td>>hg38_21_+_8433172_8446622_RNA45SN1_45S_50ntFlanks@13008.13028.21|ATTGAAAGTCAGCCCTCGACA</td><td>>hg38_21_+_8433172_8446622_RNA45SN1_45S_50ntFlanks</td><td>13008</td><td>13028</td><td>21</td><td>ATTGAAAGTCAGCCCTCGACA</td><td>534.6</td><td>-3.7</td><td>0.8</td><td>-4.6</td><td>3.7E-06</td><td>5.6E-06</td><td>MB231/MB468</td></tr>
<tr><td>>MI0000767|hsa-mir-365a&WithFlank&16|+|14309279|14309377@62.82.21;MI0000769|hsa-mir-365b&WithFlank&17|+|31575405|31575527@74.94.21&[MIMAT0000710&hsa-miR-365a-3p&offsets|0|-1;m-176&16|+|14309340|14309361&offsets|0|-1];[MIMAT0022834&hsa-miR-365b-3p&offsets|0|-1;m-175&17|+|31575478|31575499&offsets|0|-1]|TAATGCCCCTAAAAATCCTTA</td><td>>MI0000767|hsa-mir-365a&WithFlank&16|+|14309279|14309377</td><td>62</td><td>82</td><td>21</td><td>TAATGCCCCTAAAAATCCTTA</td><td>607.2</td><td>4.5</td><td>0.9</td><td>5.1</td><td>3.2E-07</td><td>5.6E-06</td><td>BT20/MB231</td></tr>
<tr><td>>hg38_21_+_8433172_8446622_RNA45SN1_45S_50ntFlanks@10684.10705.22|CGGCGGCGTCCGGTGAGCTCTC</td><td>>hg38_21_+_8433172_8446622_RNA45SN1_45S_50ntFlanks</td><td>10684</td><td>10705</td><td>22</td><td>CGGCGGCGTCCGGTGAGCTCTC</td><td>191.0</td><td>-3.3</td><td>0.7</td><td>-4.6</td><td>4.2E-06</td><td>5.6E-06</td><td>BT20/MB468</td></tr>
<tr><td>>hg38_21_+_8433172_8446622_RNA45SN1_45S_50ntFlanks@7974.7992.19|ACGCGACCTCAGATCAGAC</td><td>>hg38_21_+_8433172_8446622_RNA45SN1_45S_50ntFlanks</td><td>7974</td><td>7992</td><td>19</td><td>ACGCGACCTCAGATCAGAC</td><td>1649.8</td><td>4.1</td><td>0.9</td><td>4.6</td><td>4.2E-06</td><td>5.7E-06</td><td>BT20/MB468</td></tr>
<tr><td>>hg38_21_+_8433172_8446622_RNA45SN1_45S_50ntFlanks@11671.11693.23|CCAGTGCTCTGAATGTCAAAGTG</td><td>>hg38_21_+_8433172_8446622_RNA45SN1_45S_50ntFlanks</td><td>11671</td><td>11693</td><td>23</td><td>CCAGTGCTCTGAATGTCAAAGTG</td><td>1009.7</td><td>-4.1</td><td>0.9</td><td>-4.6</td><td>4.3E-06</td><td>5.7E-06</td><td>BT20/MB468</td></tr>
<tr><td>>hg38_21_+_8433172_8446622_RNA45SN1_45S_50ntFlanks@7982.8008.27|TCAGATCAGACGTGGCGACCCGCTGAA</td><td>>hg38_21_+_8433172_8446622_RNA45SN1_45S_50ntFlanks</td><td>7982</td><td>8008</td><td>27</td><td>TCAGATCAGACGTGGCGACCCGCTGAA</td><td>440.7</td><td>-2.8</td><td>0.6</td><td>-4.6</td><td>3.8E-06</td><td>5.8E-06</td><td>MB231/MB468</td></tr>
<tr><td>>hg38_21_+_8433172_8446622_RNA45SN1_45S_50ntFlanks@3845.3868.24|ACTTGGATAACTGTGGTAATTCTA</td><td>>hg38_21_+_8433172_8446622_RNA45SN1_45S_50ntFlanks</td><td>3845</td><td>3868</td><td>24</td><td>ACTTGGATAACTGTGGTAATTCTA</td><td>509.8</td><td>2.5</td><td>0.5</td><td>4.6</td><td>3.9E-06</td><td>5.8E-06</td><td>MB231/MB468</td></tr>
<tr><td>>hg38_21_+_8433172_8446622_RNA45SN1_45S_50ntFlanks@9539.9563.25|ACTCTGGTGGAGGTCCGTAGCGGTC</td><td>>hg38_21_+_8433172_8446622_RNA45SN1_45S_50ntFlanks</td><td>9539</td><td>9563</td><td>25</td><td>ACTCTGGTGGAGGTCCGTAGCGGTC</td><td>126.9</td><td>-3.8</td><td>0.8</td><td>-4.6</td><td>4.0E-06</td><td>6.0E-06</td><td>MB231/MB468</td></tr>
<tr><td>>hg38_21_+_8433172_8446622_RNA45SN1_45S_50ntFlanks@5485.5514.30|AGCGCTGAGAAGACGGTCGAACTTGACTAT</td><td>>hg38_21_+_8433172_8446622_RNA45SN1_45S_50ntFlanks</td><td>5485</td><td>5514</td><td>30</td><td>AGCGCTGAGAAGACGGTCGAACTTGACTAT</td><td>1930.3</td><td>-3.9</td><td>0.8</td><td>-4.6</td><td>4.5E-06</td><td>6.0E-06</td><td>BT20/MB468</td></tr>
<tr><td>>hg38_21_+_8433172_8446622_RNA45SN1_45S_50ntFlanks@4759.4776.18|ATCAGATACCGTCGTAGT</td><td>>hg38_21_+_8433172_8446622_RNA45SN1_45S_50ntFlanks</td><td>4759</td><td>4776</td><td>18</td><td>ATCAGATACCGTCGTAGT</td><td>149.9</td><td>-4.2</td><td>0.9</td><td>-4.6</td><td>4.1E-06</td><td>6.1E-06</td><td>MB231/MB468</td></tr>
<tr><td>>MI0000102|hsa-mir-100&WithFlank&11|-|122152223|122152314@22.40.19&[MIMAT0000098&hsa-miR-100-5p&offsets|+3|0;m-27&11|-|122152276|122152296&offsets|+3|+1]|CCGTAGATCCGAACTTGTG</td><td>>MI0000102|hsa-mir-100&WithFlank&11|-|122152223|122152314</td><td>22</td><td>40</td><td>19</td><td>CCGTAGATCCGAACTTGTG</td><td>918.6</td><td>-6.2</td><td>1.2</td><td>-5.1</td><td>3.6E-07</td><td>6.2E-06</td><td>BT20/MB231</td></tr>
<tr><td>>hg38_21_+_8433172_8446622_RNA45SN1_45S_50ntFlanks@12393.12414.22|ATAGCGACGTCGCTTTTTGATC</td><td>>hg38_21_+_8433172_8446622_RNA45SN1_45S_50ntFlanks</td><td>12393</td><td>12414</td><td>22</td><td>ATAGCGACGTCGCTTTTTGATC</td><td>606.0</td><td>2.8</td><td>0.6</td><td>4.6</td><td>4.2E-06</td><td>6.2E-06</td><td>MB231/MB468</td></tr>
<tr><td>>hg38_21_+_8433172_8446622_RNA45SN1_45S_50ntFlanks@7974.7993.20|ACGCGACCTCAGATCAGACG</td><td>>hg38_21_+_8433172_8446622_RNA45SN1_45S_50ntFlanks</td><td>7974</td><td>7993</td><td>20</td><td>ACGCGACCTCAGATCAGACG</td><td>3690.8</td><td>3.8</td><td>0.8</td><td>4.6</td><td>4.6E-06</td><td>6.2E-06</td><td>BT20/MB468</td></tr>
<tr><td>>hg38_21_+_8433172_8446622_RNA45SN1_45S_50ntFlanks@12397.12418.22|CGACGTCGCTTTTTGATCCTTC</td><td>>hg38_21_+_8433172_8446622_RNA45SN1_45S_50ntFlanks</td><td>12397</td><td>12418</td><td>22</td><td>CGACGTCGCTTTTTGATCCTTC</td><td>1297.6</td><td>3.4</td><td>0.7</td><td>4.6</td><td>4.2E-06</td><td>6.3E-06</td><td>MB231/MB468</td></tr>
<tr><td>>hg38_21_+_8433172_8446622_RNA45SN1_45S_50ntFlanks@6697.6718.22|CAGCTAGCTGCGAGAATTAATG</td><td>>hg38_21_+_8433172_8446622_RNA45SN1_45S_50ntFlanks</td><td>6697</td><td>6718</td><td>22</td><td>CAGCTAGCTGCGAGAATTAATG</td><td>371.8</td><td>-2.5</td><td>0.6</td><td>-4.6</td><td>4.7E-06</td><td>6.3E-06</td><td>BT20/MB468</td></tr>
<tr><td>>hg38_21_+_8433172_8446622_RNA45SN1_45S_50ntFlanks@6645.6666.22|CTCGTACGACTCTTAGCGGTGG</td><td>>hg38_21_+_8433172_8446622_RNA45SN1_45S_50ntFlanks</td><td>6645</td><td>6666</td><td>22</td><td>CTCGTACGACTCTTAGCGGTGG</td><td>3358.0</td><td>3.3</td><td>0.7</td><td>4.6</td><td>4.7E-06</td><td>6.3E-06</td><td>BT20/MB468</td></tr>
<tr><td>>hg38_21_+_8433172_8446622_RNA45SN1_45S_50ntFlanks@10680.10705.26|GTTCCGGCGGCGTCCGGTGAGCTCTC</td><td>>hg38_21_+_8433172_8446622_RNA45SN1_45S_50ntFlanks</td><td>10680</td><td>10705</td><td>26</td><td>GTTCCGGCGGCGTCCGGTGAGCTCTC</td><td>294.7</td><td>-2.8</td><td>0.6</td><td>-4.6</td><td>4.3E-06</td><td>6.4E-06</td><td>MB231/MB468</td></tr>
<tr><td>>hg38_21_+_8433172_8446622_RNA45SN1_45S_50ntFlanks@10378.10406.29|AGGGTTCCATGTGAACAGCAGTTGAACAT</td><td>>hg38_21_+_8433172_8446622_RNA45SN1_45S_50ntFlanks</td><td>10378</td><td>10406</td><td>29</td><td>AGGGTTCCATGTGAACAGCAGTTGAACAT</td><td>99.0</td><td>-2.8</td><td>0.6</td><td>-4.6</td><td>4.3E-06</td><td>6.4E-06</td><td>MB231/MB468</td></tr>
<tr><td>>MI0000272|hsa-mir-182&WithFlank&7|-|129770377|129770498@29.50.22&[MIMAT0000259&hsa-miR-182-5p&offsets|0|-2;m-22&7|-|129770449|129770470&offsets|0|0]|TTTGGCAATGGTAGAACTCACA</td><td>>MI0000272|hsa-mir-182&WithFlank&7|-|129770377|129770498</td><td>29</td><td>50</td><td>22</td><td>TTTGGCAATGGTAGAACTCACA</td><td>5393.7</td><td>3.9</td><td>0.8</td><td>5.1</td><td>3.8E-07</td><td>6.5E-06</td><td>BT20/MB231</td></tr>
<tr><td>>hg38_21_+_8433172_8446622_RNA45SN1_45S_50ntFlanks@10830.10855.26|CAAGCCGGATCCGTAACTTCGGGATA</td><td>>hg38_21_+_8433172_8446622_RNA45SN1_45S_50ntFlanks</td><td>10830</td><td>10855</td><td>26</td><td>CAAGCCGGATCCGTAACTTCGGGATA</td><td>116.3</td><td>-4.0</td><td>0.9</td><td>-4.6</td><td>4.4E-06</td><td>6.6E-06</td><td>MB231/MB468</td></tr>
<tr><td>>MI0000471|hsa-mir-126&WithFlank&9|+|136670596|136670692@58.78.21&[MIMAT0000445&hsa-miR-126-3p&offsets|0|-1;m-40&9|+|136670653|136670674&offsets|0|-1]|TCGTACCGTGAGTAATAATGC</td><td>>MI0000471|hsa-mir-126&WithFlank&9|+|136670596|136670692</td><td>58</td><td>78</td><td>21</td><td>TCGTACCGTGAGTAATAATGC</td><td>8698.5</td><td>10.3</td><td>2.3</td><td>4.6</td><td>4.4E-06</td><td>6.6E-06</td><td>MB231/MB468</td></tr>
<tr><td>>hg38_21_+_8433172_8446622_RNA45SN1_45S_50ntFlanks@10795.10817.23|TGGCATGTTGGAACAATGTAGGT</td><td>>hg38_21_+_8433172_8446622_RNA45SN1_45S_50ntFlanks</td><td>10795</td><td>10817</td><td>23</td><td>TGGCATGTTGGAACAATGTAGGT</td><td>236.0</td><td>-2.7</td><td>0.6</td><td>-4.6</td><td>5.0E-06</td><td>6.7E-06</td><td>BT20/MB468</td></tr>
<tr><td>>hg38_21_+_8433172_8446622_RNA45SN1_45S_50ntFlanks@12332.12349.18|TAAGCAGGAGGTGTCAGA</td><td>>hg38_21_+_8433172_8446622_RNA45SN1_45S_50ntFlanks</td><td>12332</td><td>12349</td><td>18</td><td>TAAGCAGGAGGTGTCAGA</td><td>79.3</td><td>-2.8</td><td>0.6</td><td>-4.6</td><td>4.5E-06</td><td>6.7E-06</td><td>MB231/MB468</td></tr>
<tr><td>>hg38_21_+_8433172_8446622_RNA45SN1_45S_50ntFlanks@8191.8211.21|CAGCCCGTGGACGGTGTGAGG</td><td>>hg38_21_+_8433172_8446622_RNA45SN1_45S_50ntFlanks</td><td>8191</td><td>8211</td><td>21</td><td>CAGCCCGTGGACGGTGTGAGG</td><td>111.7</td><td>-3.8</td><td>0.8</td><td>-4.6</td><td>4.5E-06</td><td>6.7E-06</td><td>MB231/MB468</td></tr>
<tr><td>>hg38_21_+_8433172_8446622_RNA45SN1_45S_50ntFlanks@11569.11593.25|CAGCCGACTTAGAACTGGTGCGGAC</td><td>>hg38_21_+_8433172_8446622_RNA45SN1_45S_50ntFlanks</td><td>11569</td><td>11593</td><td>25</td><td>CAGCCGACTTAGAACTGGTGCGGAC</td><td>169.6</td><td>-3.7</td><td>0.8</td><td>-4.6</td><td>4.6E-06</td><td>6.8E-06</td><td>MB231/MB468</td></tr>
<tr><td>>hg38_21_+_8433172_8446622_RNA45SN1_45S_50ntFlanks@9465.9503.39|AGGTGGAGCACGAGCGCACGTGTTAGGACCCGAAAGATG</td><td>>hg38_21_+_8433172_8446622_RNA45SN1_45S_50ntFlanks</td><td>9465</td><td>9503</td><td>39</td><td>AGGTGGAGCACGAGCGCACGTGTTAGGACCCGAAAGATG</td><td>69.4</td><td>-4.8</td><td>1.1</td><td>-4.6</td><td>4.6E-06</td><td>6.8E-06</td><td>MB231/MB468</td></tr>
<tr><td>>hg38_21_+_8433172_8446622_RNA45SN1_45S_50ntFlanks@10411.10445.35|CAGTCGGTCCTGAGAGATGGGCGAGCGCCGTTCCG</td><td>>hg38_21_+_8433172_8446622_RNA45SN1_45S_50ntFlanks</td><td>10411</td><td>10445</td><td>35</td><td>CAGTCGGTCCTGAGAGATGGGCGAGCGCCGTTCCG</td><td>93.1</td><td>-4.0</td><td>0.9</td><td>-4.6</td><td>5.1E-06</td><td>6.9E-06</td><td>BT20/MB468</td></tr>
<tr><td>>hg38_21_+_8433172_8446622_RNA45SN1_45S_50ntFlanks@11581.11611.31|AACTGGTGCGGACCAGGGGAATCCGACTGTT</td><td>>hg38_21_+_8433172_8446622_RNA45SN1_45S_50ntFlanks</td><td>11581</td><td>11611</td><td>31</td><td>AACTGGTGCGGACCAGGGGAATCCGACTGTT</td><td>1577.0</td><td>-2.6</td><td>0.6</td><td>-4.6</td><td>4.6E-06</td><td>6.9E-06</td><td>MB231/MB468</td></tr>
<tr><td>>hg38_21_+_8433172_8446622_RNA45SN1_45S_50ntFlanks@10392.10411.20|ACAGCAGTTGAACATGGGTC</td><td>>hg38_21_+_8433172_8446622_RNA45SN1_45S_50ntFlanks</td><td>10392</td><td>10411</td><td>20</td><td>ACAGCAGTTGAACATGGGTC</td><td>279.9</td><td>-2.8</td><td>0.6</td><td>-4.6</td><td>4.6E-06</td><td>6.9E-06</td><td>MB231/MB468</td></tr>
<tr><td>>hg38_21_+_8433172_8446622_RNA45SN1_45S_50ntFlanks@3769.3794.26|CGCACGGCCGGTACAGTGAAACTGCG</td><td>>hg38_21_+_8433172_8446622_RNA45SN1_45S_50ntFlanks</td><td>3769</td><td>3794</td><td>26</td><td>CGCACGGCCGGTACAGTGAAACTGCG</td><td>196.4</td><td>-3.2</td><td>0.7</td><td>-4.6</td><td>5.2E-06</td><td>6.9E-06</td><td>BT20/MB468</td></tr>
<tr><td>>hg38_21_+_8433172_8446622_RNA45SN1_45S_50ntFlanks@10797.10825.29|GCATGTTGGAACAATGTAGGTAAGGGAAG</td><td>>hg38_21_+_8433172_8446622_RNA45SN1_45S_50ntFlanks</td><td>10797</td><td>10825</td><td>29</td><td>GCATGTTGGAACAATGTAGGTAAGGGAAG</td><td>146.7</td><td>-3.2</td><td>0.7</td><td>-4.6</td><td>4.7E-06</td><td>6.9E-06</td><td>MB231/MB468</td></tr>
<tr><td>>hg38_21_+_8433172_8446622_RNA45SN1_45S_50ntFlanks@7981.8005.25|CTCAGATCAGACGTGGCGACCCGCT</td><td>>hg38_21_+_8433172_8446622_RNA45SN1_45S_50ntFlanks</td><td>7981</td><td>8005</td><td>25</td><td>CTCAGATCAGACGTGGCGACCCGCT</td><td>165.2</td><td>-2.1</td><td>0.5</td><td>-4.6</td><td>5.2E-06</td><td>7.0E-06</td><td>BT20/MB468</td></tr>
<tr><td>>hg38_21_+_8433172_8446622_RNA45SN1_45S_50ntFlanks@11581.11606.26|AACTGGTGCGGACCAGGGGAATCCGA</td><td>>hg38_21_+_8433172_8446622_RNA45SN1_45S_50ntFlanks</td><td>11581</td><td>11606</td><td>26</td><td>AACTGGTGCGGACCAGGGGAATCCGA</td><td>234.6</td><td>-2.0</td><td>0.4</td><td>-4.6</td><td>4.7E-06</td><td>7.0E-06</td><td>MB231/MB468</td></tr>
<tr><td>>trna12_ArgCCT_16_+_3243918_3243990@35.52.18,trna18_ArgCCT_17_+_73030001_73030073@35.52.18,trna21_ArgCCT_17_-_73030526_73030598@35.52.18,trna2_ArgCCT_16_+_3202901_3202973@35.52.18,trna3_ArgCCT_7_+_139025446_139025518@35.52.18|CTAAGCCAGGGATTGTGG</td><td>>trna12_ArgCCT_16_+_3243918_3243990</td><td>35</td><td>52</td><td>18</td><td>CTAAGCCAGGGATTGTGG</td><td>92.2</td><td>-3.8</td><td>0.8</td><td>-4.6</td><td>4.7E-06</td><td>7.0E-06</td><td>MB231/MB468</td></tr>
<tr><td>>MI0000446|hsa-mir-125b-1&WithFlank&11|-|122099751|122099850@21.39.19(+1C);MI0000470|hsa-mir-125b-2&WithFlank&21|+|16590231|16590331@23.41.19(+1C)&[MIMAT0000423&hsa-miR-125b-5p&offsets|0|-3(+1C);m-67&11|-|122099809|122099830&offsets|0|-3(+1C)];[MIMAT0000423_1&hsa-miR-125b-5p&offsets|0|-3(+1C);m-68&21|+|16590253|16590274&offsets|0|-3(+1C)]|TCCCTGAGACCCTAACTTGC</td><td>>MI0000446|hsa-mir-125b-1&WithFlank&11|-|122099751|122099850</td><td>21</td><td>39</td><td>19(+1C)</td><td>TCCCTGAGACCCTAACTTGC</td><td>158.4</td><td>-5.1</td><td>1.0</td><td>-5.1</td><td>4.2E-07</td><td>7.0E-06</td><td>BT20/MB231</td></tr>
<tr><td>>trna119_LysCTT_1_-_145395522_145395594@3.33.31,trna11_LysCTT_5_-_180648979_180649051@3.33.31,trna13_LysCTT_6_+_26556774_26556846@3.33.31,trna7_LysCTT_16_+_3225692_3225764@3.33.31,trna9_LysCTT_5_+_180634755_180634827@3.33.31|CCGGCTAGCTCAGTCGGTAGAGCATGAGACT</td><td>>trna119_LysCTT_1_-_145395522_145395594</td><td>3</td><td>33</td><td>31</td><td>CCGGCTAGCTCAGTCGGTAGAGCATGAGACT</td><td>699.2</td><td>-4.6</td><td>0.9</td><td>-5.1</td><td>4.2E-07</td><td>7.0E-06</td><td>BT20/MB231</td></tr>
<tr><td>>trna20_GluTTC_2_-_131094701_131094772@1.31.31,trna5_GluTTC_13_-_41634874_41634945@1.31.31|TCCCATATGGTCTAGCGGTTAGGATTCCTGG</td><td>>trna20_GluTTC_2_-_131094701_131094772</td><td>1</td><td>31</td><td>31</td><td>TCCCATATGGTCTAGCGGTTAGGATTCCTGG</td><td>303.8</td><td>-4.4</td><td>0.9</td><td>-5.1</td><td>4.2E-07</td><td>7.0E-06</td><td>BT20/MB231</td></tr>
<tr><td>>hg38_21_+_8433172_8446622_RNA45SN1_45S_50ntFlanks@12907.12933.27|CACCGCACGTTCGTGGGGAACCTGGCG</td><td>>hg38_21_+_8433172_8446622_RNA45SN1_45S_50ntFlanks</td><td>12907</td><td>12933</td><td>27</td><td>CACCGCACGTTCGTGGGGAACCTGGCG</td><td>556.7</td><td>-5.1</td><td>1.1</td><td>-4.6</td><td>5.3E-06</td><td>7.0E-06</td><td>BT20/MB468</td></tr>
<tr><td>>hg38_21_+_8433172_8446622_RNA45SN1_45S_50ntFlanks@12415.12442.28|CTTCGATGTCGGCTCTTCCTATCATTGT</td><td>>hg38_21_+_8433172_8446622_RNA45SN1_45S_50ntFlanks</td><td>12415</td><td>12442</td><td>28</td><td>CTTCGATGTCGGCTCTTCCTATCATTGT</td><td>282.6</td><td>2.7</td><td>0.6</td><td>4.6</td><td>5.3E-06</td><td>7.1E-06</td><td>BT20/MB468</td></tr>
<tr><td>>hg38_21_+_8433172_8446622_RNA45SN1_45S_50ntFlanks@11325.11344.20|TCTCGTCGCGCGCGCGTCCG</td><td>>hg38_21_+_8433172_8446622_RNA45SN1_45S_50ntFlanks</td><td>11325</td><td>11344</td><td>20</td><td>TCTCGTCGCGCGCGCGTCCG</td><td>279.0</td><td>-3.2</td><td>0.7</td><td>-4.6</td><td>4.8E-06</td><td>7.1E-06</td><td>MB231/MB468</td></tr>
</table>

<table>
<tr><td>>hg38_21_+_8433172_8446622_RNA45SN1_45S_50ntFlanks@8170.8196.27|CAAGTCCTTCTGATCGAGGCCCAGCCC</td><td>>hg38_21_+_8433172_8446622_RNA45SN1_45S_50ntFlanks</td><td>8170</td><td>8196</td><td>27</td><td>CAAGTCCTTCTGATCGAGGCCCAGCCC</td><td>82.9</td><td>-2.3</td><td>0.5</td><td>-4.6</td><td>4.8E-06</td><td>7.1E-06</td><td>MB231/MB468</td></tr>
<tr><td>>MI0000102|hsa-mir-100&WithFlank&11|-|122152223|122152314@19.39.21(+1U)&[MIMAT0000098&hsa-miR-100-5p&offsets|0|-1(+1U);m-27&11|-|122152276|122152296&offsets|0|0(+1U)]|AACCCGTAGATCCGAACTTGTT</td><td>>MI0000102|hsa-mir-100&WithFlank&11|-|122152223|122152314</td><td>19</td><td>39</td><td>21(+1U)</td><td>AACCCGTAGATCCGAACTTGTT</td><td>8629.3</td><td>-5.0</td><td>1.0</td><td>-5.1</td><td>4.3E-07</td><td>7.2E-06</td><td>BT20/MB231</td></tr>
<tr><td>>hg38_21_+_8433172_8446622_RNA45SN1_45S_50ntFlanks@9545.9571.27|GTGGAGGTCCGTAGCGGTCCTGACGTG</td><td>>hg38_21_+_8433172_8446622_RNA45SN1_45S_50ntFlanks</td><td>9545</td><td>9571</td><td>27</td><td>GTGGAGGTCCGTAGCGGTCCTGACGTG</td><td>6436.5</td><td>-2.9</td><td>0.6</td><td>-4.6</td><td>4.8E-06</td><td>7.2E-06</td><td>MB231/MB468</td></tr>
<tr><td>>hg38_21_+_8433172_8446622_RNA45SN1_45S_50ntFlanks@10869.10891.23|AGGGCTGGGTCGGTCGGGCTGGG</td><td>>hg38_21_+_8433172_8446622_RNA45SN1_45S_50ntFlanks</td><td>10869</td><td>10891</td><td>23</td><td>AGGGCTGGGTCGGTCGGGCTGGG</td><td>333.6</td><td>-3.5</td><td>0.8</td><td>-4.6</td><td>4.8E-06</td><td>7.2E-06</td><td>MB231/MB468</td></tr>
<tr><td>>hg38_21_+_8433172_8446622_RNA45SN1_45S_50ntFlanks@3811.3834.24|AGTTATGGTTCCTTTGGTCGCTCG</td><td>>hg38_21_+_8433172_8446622_RNA45SN1_45S_50ntFlanks</td><td>3811</td><td>3834</td><td>24</td><td>AGTTATGGTTCCTTTGGTCGCTCG</td><td>100.4</td><td>-2.8</td><td>0.6</td><td>-4.5</td><td>5.4E-06</td><td>7.2E-06</td><td>BT20/MB468</td></tr>
<tr><td>>hg38_21_+_8433172_8446622_RNA45SN1_45S_50ntFlanks@10768.10793.26|CAGCAGGTCTCCAAGGTGAACAGCCT</td><td>>hg38_21_+_8433172_8446622_RNA45SN1_45S_50ntFlanks</td><td>10768</td><td>10793</td><td>26</td><td>CAGCAGGTCTCCAAGGTGAACAGCCT</td><td>124.4</td><td>-4.6</td><td>1.0</td><td>-4.5</td><td>5.4E-06</td><td>7.2E-06</td><td>BT20/MB468</td></tr>
<tr><td>>hg38_21_+_8433172_8446622_RNA45SN1_45S_50ntFlanks@10387.10410.24|TGTGAACAGCAGTTGAACATGGGT</td><td>>hg38_21_+_8433172_8446622_RNA45SN1_45S_50ntFlanks</td><td>10387</td><td>10410</td><td>24</td><td>TGTGAACAGCAGTTGAACATGGGT</td><td>266.5</td><td>-2.9</td><td>0.6</td><td>-4.6</td><td>4.9E-06</td><td>7.2E-06</td><td>MB231/MB468</td></tr>
<tr><td>>hg38_21_+_8433172_8446622_RNA45SN1_45S_50ntFlanks@12431.12455.25|TCCTATCATTGTGAAGCAGAATTCA</td><td>>hg38_21_+_8433172_8446622_RNA45SN1_45S_50ntFlanks</td><td>12431</td><td>12455</td><td>25</td><td>TCCTATCATTGTGAAGCAGAATTCA</td><td>74.4</td><td>-2.6</td><td>0.6</td><td>-4.6</td><td>4.9E-06</td><td>7.2E-06</td><td>MB231/MB468</td></tr>
<tr><td>>hg38_21_+_8433172_8446622_RNA45SN1_45S_50ntFlanks@5515.5534.20|CTAGAGGAAGTAAAAGTCGT</td><td>>hg38_21_+_8433172_8446622_RNA45SN1_45S_50ntFlanks</td><td>5515</td><td>5534</td><td>20</td><td>CTAGAGGAAGTAAAAGTCGT</td><td>87.7</td><td>-2.8</td><td>0.6</td><td>-4.6</td><td>4.9E-06</td><td>7.3E-06</td><td>MB231/MB468</td></tr>
<tr><td>>hg38_21_+_8433172_8446622_RNA45SN1_45S_50ntFlanks@10471.10488.18|CCCTCGGCCGATCGAAAG</td><td>>hg38_21_+_8433172_8446622_RNA45SN1_45S_50ntFlanks</td><td>10471</td><td>10488</td><td>18</td><td>CCCTCGGCCGATCGAAAG</td><td>449.6</td><td>-2.5</td><td>0.5</td><td>-4.6</td><td>4.9E-06</td><td>7.3E-06</td><td>MB231/MB468</td></tr>
<tr><td>>hg38_21_+_8433172_8446622_RNA45SN1_45S_50ntFlanks@10362.10379.18|AAGGCCGAAGTGGAGAAG</td><td>>hg38_21_+_8433172_8446622_RNA45SN1_45S_50ntFlanks</td><td>10362</td><td>10379</td><td>18</td><td>AAGGCCGAAGTGGAGAAG</td><td>152.6</td><td>-3.5</td><td>0.8</td><td>-4.5</td><td>5.4E-06</td><td>7.3E-06</td><td>BT20/MB468</td></tr>
<tr><td>>MI0000102|hsa-mir-100&WithFlank&11|-|122152223|122152314@19.40.22(+2A)&[MIMAT0000098&hsa-miR-100-5p&offsets|0|0(+2A);m-27&11|-|122152276|122152296&offsets|0|+1(+2A)]|AACCCGTAGATCCGAACTTGTGAA</td><td>>MI0000102|hsa-mir-100&WithFlank&11|-|122152223|122152314</td><td>19</td><td>40</td><td>22(+2A)</td><td>AACCCGTAGATCCGAACTTGTGAA</td><td>1423.2</td><td>-5.6</td><td>1.1</td><td>-5.0</td><td>4.4E-07</td><td>7.4E-06</td><td>BT20/MB231</td></tr>
<tr><td>>hg38_21_+_8433172_8446622_RNA45SN1_45S_50ntFlanks@6644.6683.40|CCTCGTACGACTCTTAGCGGTGGATCACTCGGCTCGTGCG</td><td>>hg38_21_+_8433172_8446622_RNA45SN1_45S_50ntFlanks</td><td>6644</td><td>6683</td><td>40</td><td>CCTCGTACGACTCTTAGCGGTGGATCACTCGGCTCGTGCG</td><td>1432.8</td><td>10.9</td><td>2.4</td><td>4.6</td><td>5.0E-06</td><td>7.5E-06</td><td>MB231/MB468</td></tr>
<tr><td>>am_trna2_GlyTCC_19_+_4724082_4724153@1.29.29|GCGTTGGTGGTATAGTGGTTAGCATAGCT</td><td>>am_trna2_GlyTCC_19_+_4724082_4724153</td><td>1</td><td>29</td><td>29</td><td>GCGTTGGTGGTATAGTGGTTAGCATAGCT</td><td>99.4</td><td>-3.9</td><td>0.9</td><td>-4.6</td><td>5.1E-06</td><td>7.5E-06</td><td>MB231/MB468</td></tr>
<tr><td>>hg38_21_+_8433172_8446622_RNA45SN1_45S_50ntFlanks@10843.10864.22|TAACTTCGGGATAAGGATTGGC</td><td>>hg38_21_+_8433172_8446622_RNA45SN1_45S_50ntFlanks</td><td>10843</td><td>10864</td><td>22</td><td>TAACTTCGGGATAAGGATTGGC</td><td>109.1</td><td>-2.8</td><td>0.6</td><td>-4.6</td><td>5.1E-06</td><td>7.6E-06</td><td>MB231/MB468</td></tr>
<tr><td>>MI0003513|hsa-mir-455&WithFlank&9|+|114209428|114209535@60.80.21(+1C)&[MIMAT0004784&hsa-miR-455-3p&offsets|0|0(+1C);m-135&9|+|114209487|114209508&offsets|0|-1(+1C)]|GCAGTCCATGGGCATATACACC</td><td>>MI0003513|hsa-mir-455&WithFlank&9|+|114209428|114209535</td><td>60</td><td>80</td><td>21(+1C)</td><td>GCAGTCCATGGGCATATACACC</td><td>275.1</td><td>-3.7</td><td>0.7</td><td>-5.0</td><td>4.6E-07</td><td>7.6E-06</td><td>BT20/MB231</td></tr>
<tr><td>>hg38_21_+_8433172_8446622_RNA45SN1_45S_50ntFlanks@4002.4022.21|GATAACCTCGGGCCGATCGCA</td><td>>hg38_21_+_8433172_8446622_RNA45SN1_45S_50ntFlanks</td><td>4002</td><td>4022</td><td>21</td><td>GATAACCTCGGGCCGATCGCA</td><td>207.9</td><td>-2.0</td><td>0.4</td><td>-4.6</td><td>5.2E-06</td><td>7.7E-06</td><td>MB231/MB468</td></tr>
<tr><td>>trna12_ArgCCT_16_+_3243918_3243990@35.52.18,trna18_ArgCCT_17_+_73030001_73030073@35.52.18,trna21_ArgCCT_17_-_73030526_73030598@35.52.18,trna2_ArgCCT_16_+_3202901_3202973@35.52.18,trna3_ArgCCT_7_+_139025446_139025518@35.52.18|CTAAGCCAGGGATTGTGG</td><td>>trna12_ArgCCT_16_+_3243918_3243990</td><td>35</td><td>52</td><td>18</td><td>CTAAGCCAGGGATTGTGG</td><td>111.0</td><td>-3.9</td><td>0.9</td><td>-4.5</td><td>5.8E-06</td><td>7.7E-06</td><td>BT20/MB468</td></tr>
<tr><td>>hg38_21_+_8433172_8446622_RNA45SN1_45S_50ntFlanks@6646.6678.33|TCGTACGACTCTTAGCGGTGGATCACTCGGCTC</td><td>>hg38_21_+_8433172_8446622_RNA45SN1_45S_50ntFlanks</td><td>6646</td><td>6678</td><td>33</td><td>TCGTACGACTCTTAGCGGTGGATCACTCGGCTC</td><td>880.7</td><td>3.7</td><td>0.8</td><td>4.6</td><td>5.3E-06</td><td>7.8E-06</td><td>MB231/MB468</td></tr>
<tr><td>>hg38_21_+_8433172_8446622_RNA45SN1_45S_50ntFlanks@8114.8140.27|GCGCGGGACATGTGGCGTACGGAAGAC</td><td>>hg38_21_+_8433172_8446622_RNA45SN1_45S_50ntFlanks</td><td>8114</td><td>8140</td><td>27</td><td>GCGCGGGACATGTGGCGTACGGAAGAC</td><td>393.0</td><td>-4.2</td><td>0.9</td><td>-4.5</td><td>5.9E-06</td><td>7.8E-06</td><td>BT20/MB468</td></tr>
<tr><td>>am_trnaMT_SerGCT_MT_+_12207_12265@12.32.21|CAAGAACTGCTAACTCATGCC</td><td>>am_trnaMT_SerGCT_MT_+_12207_12265</td><td>12</td><td>32</td><td>21</td><td>CAAGAACTGCTAACTCATGCC</td><td>596.3</td><td>6.6</td><td>1.4</td><td>4.5</td><td>5.9E-06</td><td>7.9E-06</td><td>BT20/MB468</td></tr>
<tr><td>>hg38_21_+_8433172_8446622_RNA45SN1_45S_50ntFlanks@8358.8379.22|AAAGAACTTTGAAGAGAGAGTT</td><td>>hg38_21_+_8433172_8446622_RNA45SN1_45S_50ntFlanks</td><td>8358</td><td>8379</td><td>22</td><td>AAAGAACTTTGAAGAGAGAGTT</td><td>1146.3</td><td>-2.8</td><td>0.6</td><td>-4.6</td><td>5.3E-06</td><td>7.9E-06</td><td>MB231/MB468</td></tr>
<tr><td>>hg38_21_+_8433172_8446622_RNA45SN1_45S_50ntFlanks@4995.5013.19|ACAGATTGATAGCTCTTTC</td><td>>hg38_21_+_8433172_8446622_RNA45SN1_45S_50ntFlanks</td><td>4995</td><td>5013</td><td>19</td><td>ACAGATTGATAGCTCTTTC</td><td>128.2</td><td>-2.8</td><td>0.6</td><td>-4.6</td><td>5.4E-06</td><td>7.9E-06</td><td>MB231/MB468</td></tr>
<tr><td>>MI0000273|hsa-mir-183&WithFlank&7|-|129774899|129775020@33.54.22(+1A)&[MIMAT0000261&hsa-miR-183-5p&offsets|0|0(+1A);m-41&7|-|129774966|129774987&offsets|-1|-1(+1A)]|TATGGCACTGGTAGAATTCACTA</td><td>>MI0000273|hsa-mir-183&WithFlank&7|-|129774899|129775020</td><td>33</td><td>54</td><td>22(+1A)</td><td>TATGGCACTGGTAGAATTCACTA</td><td>997.0</td><td>3.9</td><td>0.8</td><td>5.0</td><td>4.8E-07</td><td>8.0E-06</td><td>BT20/MB231</td></tr>
<tr><td>>hg38_21_+_8433172_8446622_RNA45SN1_45S_50ntFlanks@12936.12961.26|AAACCATTCGTAGACGACCTGCTTCT</td><td>>hg38_21_+_8433172_8446622_RNA45SN1_45S_50ntFlanks</td><td>12936</td><td>12961</td><td>26</td><td>AAACCATTCGTAGACGACCTGCTTCT</td><td>291.9</td><td>-2.5</td><td>0.6</td><td>-4.5</td><td>6.0E-06</td><td>8.0E-06</td><td>BT20/MB468</td></tr>
<tr><td>>hg38_21_+_8433172_8446622_RNA45SN1_45S_50ntFlanks@10357.10383.27|CTTTGAAGGCCGAAGTGGAGAAGGGTT</td><td>>hg38_21_+_8433172_8446622_RNA45SN1_45S_50ntFlanks</td><td>10357</td><td>10383</td><td>27</td><td>CTTTGAAGGCCGAAGTGGAGAAGGGTT</td><td>279.3</td><td>-2.2</td><td>0.5</td><td>-4.5</td><td>5.4E-06</td><td>8.0E-06</td><td>MB231/MB468</td></tr>
<tr><td>>hg38_21_+_8433172_8446622_RNA45SN1_45S_50ntFlanks@10349.10382.34|AACGAGAACTTTGAAGGCCGAAGTGGAGAAGGGT</td><td>>hg38_21_+_8433172_8446622_RNA45SN1_45S_50ntFlanks</td><td>10349</td><td>10382</td><td>34</td><td>AACGAGAACTTTGAAGGCCGAAGTGGAGAAGGGT</td><td>554.2</td><td>1.8</td><td>0.4</td><td>4.5</td><td>6.0E-06</td><td>8.0E-06</td><td>BT20/MB468</td></tr>
<tr><td>>hg38_21_+_8433172_8446622_RNA45SN1_45S_50ntFlanks@9736.9753.18|CCGAAACGATCTCAACCT</td><td>>hg38_21_+_8433172_8446622_RNA45SN1_45S_50ntFlanks</td><td>9736</td><td>9753</td><td>18</td><td>CCGAAACGATCTCAACCT</td><td>161.2</td><td>-4.9</td><td>1.1</td><td>-4.5</td><td>6.1E-06</td><td>8.1E-06</td><td>BT20/MB468</td></tr>
<tr><td>>MI0000272|hsa-mir-182&WithFlank&7|-|129770377|129770498@29.49.21&[MIMAT0000259&hsa-miR-182-5p&offsets|0|-3;m-22&7|-|129770449|129770470&offsets|0|-1]|TTTGGCAATGGTAGAACTCAC</td><td>>MI0000272|hsa-mir-182&WithFlank&7|-|129770377|129770498</td><td>29</td><td>49</td><td>21</td><td>TTTGGCAATGGTAGAACTCAC</td><td>668.4</td><td>4.0</td><td>0.8</td><td>5.0</td><td>5.0E-07</td><td>8.2E-06</td><td>BT20/MB231</td></tr>
<tr><td>>MI0003591|hsa-mir-584&WithFlank&5|-|149062307|149062415@22.43.22&[MIMAT0003249&hsa-miR-584-5p&offsets|0|0;m-167&5|-|149062374|149062394&offsets|0|+1]|TTATGGTTTGCCTGGGACTGAG</td><td>>MI0003591|hsa-mir-584&WithFlank&5|-|149062307|149062415</td><td>22</td><td>43</td><td>22</td><td>TTATGGTTTGCCTGGGACTGAG</td><td>525.6</td><td>10.0</td><td>2.2</td><td>4.5</td><td>5.5E-06</td><td>8.2E-06</td><td>MB231/MB468</td></tr>
<tr><td>>hg38_21_+_8433172_8446622_RNA45SN1_45S_50ntFlanks@10514.10533.20|CGGAGTGGCGGAGATGGGCG</td><td>>hg38_21_+_8433172_8446622_RNA45SN1_45S_50ntFlanks</td><td>10514</td><td>10533</td><td>20</td><td>CGGAGTGGCGGAGATGGGCG</td><td>213.6</td><td>-4.1</td><td>0.9</td><td>-4.5</td><td>6.1E-06</td><td>8.2E-06</td><td>BT20/MB468</td></tr>
<tr><td>>hg38_21_+_8433172_8446622_RNA45SN1_45S_50ntFlanks@10679.10712.34|GGTTCCGGCGGCGTCCGGTGAGCTCTCGCTGGCC</td><td>>hg38_21_+_8433172_8446622_RNA45SN1_45S_50ntFlanks</td><td>10679</td><td>10712</td><td>34</td><td>GGTTCCGGCGGCGTCCGGTGAGCTCTCGCTGGCC</td><td>282.9</td><td>-5.0</td><td>1.1</td><td>-4.5</td><td>6.1E-06</td><td>8.2E-06</td><td>BT20/MB468</td></tr>
<tr><td>>hg38_21_+_8433172_8446622_RNA45SN1_45S_50ntFlanks@5472.5499.28|ACGGCCCTGGCGGAGCGCTGAGAAGACG</td><td>>hg38_21_+_8433172_8446622_RNA45SN1_45S_50ntFlanks</td><td>5472</td><td>5499</td><td>28</td><td>ACGGCCCTGGCGGAGCGCTGAGAAGACG</td><td>2046.9</td><td>2.5</td><td>0.5</td><td>4.5</td><td>5.5E-06</td><td>8.2E-06</td><td>MB231/MB468</td></tr>
<tr><td>>hg38_21_+_8433172_8446622_RNA45SN1_45S_50ntFlanks@12417.12435.19|TCGATGTCGGCTCTTCCTA</td><td>>hg38_21_+_8433172_8446622_RNA45SN1_45S_50ntFlanks</td><td>12417</td><td>12435</td><td>19</td><td>TCGATGTCGGCTCTTCCTA</td><td>3104.6</td><td>-4.0</td><td>0.9</td><td>-4.5</td><td>6.1E-06</td><td>8.2E-06</td><td>BT20/MB468</td></tr>
<tr><td>>hg38_21_+_8433172_8446622_RNA45SN1_45S_50ntFlanks@8194.8212.19|CCCGTGGACGGTGTGAGGC</td><td>>hg38_21_+_8433172_8446622_RNA45SN1_45S_50ntFlanks</td><td>8194</td><td>8212</td><td>19</td><td>CCCGTGGACGGTGTGAGGC</td><td>450.2</td><td>-2.3</td><td>0.5</td><td>-4.5</td><td>5.6E-06</td><td>8.3E-06</td><td>MB231/MB468</td></tr>
<tr><td>>hg38_21_+_8433172_8446622_RNA45SN1_45S_50ntFlanks@12405.12440.36|CTTTTTGATCCTTCGATGTCGGCTCTTCCTATCATT</td><td>>hg38_21_+_8433172_8446622_RNA45SN1_45S_50ntFlanks</td><td>12405</td><td>12440</td><td>36</td><td>CTTTTTGATCCTTCGATGTCGGCTCTTCCTATCATT</td><td>164.6</td><td>-2.2</td><td>0.5</td><td>-4.5</td><td>6.2E-06</td><td>8.3E-06</td><td>BT20/MB468</td></tr>
<tr><td>>hg38_21_+_8433172_8446622_RNA45SN1_45S_50ntFlanks@12008.12034.27|CCGCCGGTGAAATACCACTACTCTGAT</td><td>>hg38_21_+_8433172_8446622_RNA45SN1_45S_50ntFlanks</td><td>12008</td><td>12034</td><td>27</td><td>CCGCCGGTGAAATACCACTACTCTGAT</td><td>183.8</td><td>-3.8</td><td>0.8</td><td>-4.5</td><td>5.6E-06</td><td>8.3E-06</td><td>MB231/MB468</td></tr>
<tr><td>>trna14_LysTTT_11_-_59327808_59327880@1.36.36,trna2_LysTTT_17_+_8022473_8022545@1.36.36,trna54_LysTTT_1_+_204475655_204475727@1.36.36,trna5_LysTTT_11_+_59323902_59323974@1.36.36,trna62_LysTTT_1_-_204476158_204476230@1.36.36,trna76_LysTTT_6_+_28918806_28918878@1.36.36|GCCCGGATAGCTCAGTCGGTAGAGCATCAGACTTTT</td><td>>trna14_LysTTT_11_-_59327808_59327880</td><td>1</td><td>36</td><td>36</td><td>GCCCGGATAGCTCAGTCGGTAGAGCATCAGACTTTT</td><td>431.6</td><td>7.9</td><td>1.7</td><td>4.5</td><td>5.6E-06</td><td>8.4E-06</td><td>MB231/MB468</td></tr>
<tr><td>>hg38_21_+_8433172_8446622_RNA45SN1_45S_50ntFlanks@6723.6740.18|TTGCAGGACACATTGATC</td><td>>hg38_21_+_8433172_8446622_RNA45SN1_45S_50ntFlanks</td><td>6723</td><td>6740</td><td>18</td><td>TTGCAGGACACATTGATC</td><td>533.3</td><td>-2.6</td><td>0.6</td><td>-4.5</td><td>5.6E-06</td><td>8.4E-06</td><td>MB231/MB468</td></tr>
<tr><td>>hg38_21_+_8433172_8446622_RNA45SN1_45S_50ntFlanks@11778.11798.21|ACGCGCATGAATGGATGAACG</td><td>>hg38_21_+_8433172_8446622_RNA45SN1_45S_50ntFlanks</td><td>11778</td><td>11798</td><td>21</td><td>ACGCGCATGAATGGATGAACG</td><td>165.0</td><td>-2.9</td><td>0.6</td><td>-4.5</td><td>5.7E-06</td><td>8.4E-06</td><td>MB231/MB468</td></tr>
<tr><td>>MI0000098|hsa-mir-96&WithFlank&7|-|129774686|129774775@15.36.22(+1A)&[MIMAT0000095&hsa-miR-96-5p&offsets|0|-1(+1A);m-248&7|-|129774739|129774761&offsets|0|-1(+1A)]|TTTGGCACTAGCACATTTTTGCA</td><td>>MI0000098|hsa-mir-96&WithFlank&7|-|129774686|129774775</td><td>15</td><td>36</td><td>22(+1A)</td><td>TTTGGCACTAGCACATTTTTGCA</td><td>2320.8</td><td>4.3</td><td>0.9</td><td>5.0</td><td>5.1E-07</td><td>8.4E-06</td><td>BT20/MB231</td></tr>
<tr><td>>MI0001733|hsa-mir-452&WithFlank&X|-|151959622|151959718@20.42.23&[MIMAT0001635&hsa-miR-452-5p&offsets|0|+1;m-188&X|-|151959677|151959699&offsets|0|0]|AACTGTTTGCAGAGGAAACTGAG</td><td>>MI0001733|hsa-mir-452&WithFlank&X|-|151959622|151959718</td><td>20</td><td>42</td><td>23</td><td>AACTGTTTGCAGAGGAAACTGAG</td><td>1601.7</td><td>3.5</td><td>0.7</td><td>5.0</td><td>5.2E-07</td><td>8.4E-06</td><td>BT20/MB231</td></tr>
<tr><td>>MI0000239|hsa-mir-197&WithFlank&1|+|109598887|109598973@54.74.21(+1A)&[MIMAT0000227&hsa-miR-197-3p&offsets|0|-1(+1A);m-99&1|+|109598940|109598961&offsets|0|-1(+1A)]|TTCACCACCTTCTCCACCCAGA</td><td>>MI0000239|hsa-mir-197&WithFlank&1|+|109598887|109598973</td><td>54</td><td>74</td><td>21(+1A)</td><td>TTCACCACCTTCTCCACCCAGA</td><td>924.2</td><td>4.4</td><td>0.9</td><td>5.0</td><td>5.2E-07</td><td>8.4E-06</td><td>BT20/MB231</td></tr>
<tr><td>>hg38_21_+_8433172_8446622_RNA45SN1_45S_50ntFlanks@4724.4748.25|ATTAATCAAGAACGAAAGTCGGAGG</td><td>>hg38_21_+_8433172_8446622_RNA45SN1_45S_50ntFlanks</td><td>4724</td><td>4748</td><td>25</td><td>ATTAATCAAGAACGAAAGTCGGAGG</td><td>315.6</td><td>-2.8</td><td>0.6</td><td>-4.5</td><td>5.7E-06</td><td>8.5E-06</td><td>MB231/MB468</td></tr>
</table>

<table>
<tr><td>>hg38_21_+_8433172_8446622_RNA45SN1_45S_50ntFlanks@10727.10745.19|GGAGAGGGTGTAAATCTCG</td><td>>hg38_21_+_8433172_8446622_RNA45SN1_45S_50ntFlanks</td><td>10727</td><td>10745</td><td>19</td><td>GGAGAGGGTGTAAATCTCG</td><td>132.3</td><td>-2.8</td><td>0.6</td><td>-4.5</td><td>5.8E-06</td><td>8.5E-06</td><td>MB231/MB468</td></tr>
<tr><td>>hg38_21_+_8433172_8446622_RNA45SN1_45S_50ntFlanks@10349.10370.22|AACGAGAACTTTGAAGGCCGAA</td><td>>hg38_21_+_8433172_8446622_RNA45SN1_45S_50ntFlanks</td><td>10349</td><td>10370</td><td>22</td><td>AACGAGAACTTTGAAGGCCGAA</td><td>638.5</td><td>2.8</td><td>0.6</td><td>4.5</td><td>5.8E-06</td><td>8.5E-06</td><td>MB231/MB468</td></tr>
<tr><td>>hg38_21_+_8433172_8446622_RNA45SN1_45S_50ntFlanks@7979.8003.25|ACCTCAGATCAGACGTGGCGACCCG</td><td>>hg38_21_+_8433172_8446622_RNA45SN1_45S_50ntFlanks</td><td>7979</td><td>8003</td><td>25</td><td>ACCTCAGATCAGACGTGGCGACCCG</td><td>365.8</td><td>-3.9</td><td>0.9</td><td>-4.5</td><td>6.4E-06</td><td>8.6E-06</td><td>BT20/MB468</td></tr>
<tr><td>>hg38_21_+_8433172_8446622_RNA45SN1_45S_50ntFlanks@6652.6699.48|GACTCTTAGCGGTGGATCACTCGGCTCGTGCGTCGATGAAGAACGCAG</td><td>>hg38_21_+_8433172_8446622_RNA45SN1_45S_50ntFlanks</td><td>6652</td><td>6699</td><td>48</td><td>GACTCTTAGCGGTGGATCACTCGGCTCGTGCGTCGATGAAGAACGCAG</td><td>4511.5</td><td>4.2</td><td>0.9</td><td>4.5</td><td>5.8E-06</td><td>8.6E-06</td><td>MB231/MB468</td></tr>
<tr><td>>MI0000266|hsa-mir-10a&WithFlank&17|-|48579832|48579953@29.49.21&[MIMAT0000253&hsa-miR-10a-5p&offsets|+1|-1;m-18&17|-|48579905|48579926&offsets|+1|0]|ACCCTGTAGATCCGAATTTGT</td><td>>MI0000266|hsa-mir-10a&WithFlank&17|-|48579832|48579953</td><td>29</td><td>49</td><td>21</td><td>ACCCTGTAGATCCGAATTTGT</td><td>5198.3</td><td>11.7</td><td>2.6</td><td>4.5</td><td>6.4E-06</td><td>8.6E-06</td><td>BT20/MB468</td></tr>
<tr><td>>hg38_21_+_8433172_8446622_RNA45SN1_45S_50ntFlanks@5446.5466.21|CGGATCGGCCCCGCCGGGGTC</td><td>>hg38_21_+_8433172_8446622_RNA45SN1_45S_50ntFlanks</td><td>5446</td><td>5466</td><td>21</td><td>CGGATCGGCCCCGCCGGGGTC</td><td>3621.9</td><td>-6.6</td><td>1.5</td><td>-4.5</td><td>6.4E-06</td><td>8.6E-06</td><td>BT20/MB468</td></tr>
<tr><td>>hg38_21_+_8433172_8446622_RNA45SN1_45S_50ntFlanks@8176.8201.26|CTTCTGATCGAGGCCCAGCCCGTGGA</td><td>>hg38_21_+_8433172_8446622_RNA45SN1_45S_50ntFlanks</td><td>8176</td><td>8201</td><td>26</td><td>CTTCTGATCGAGGCCCAGCCCGTGGA</td><td>285.5</td><td>-3.2</td><td>0.7</td><td>-4.5</td><td>6.5E-06</td><td>8.6E-06</td><td>BT20/MB468</td></tr>
<tr><td>>hg38_21_+_8433172_8446622_RNA45SN1_45S_50ntFlanks@9504.9531.28|GTGAACTATGCCTGGGCAGGGCGAAGCC</td><td>>hg38_21_+_8433172_8446622_RNA45SN1_45S_50ntFlanks</td><td>9504</td><td>9531</td><td>28</td><td>GTGAACTATGCCTGGGCAGGGCGAAGCC</td><td>97.6</td><td>-3.2</td><td>0.7</td><td>-4.5</td><td>5.8E-06</td><td>8.6E-06</td><td>MB231/MB468</td></tr>
<tr><td>>hg38_21_+_8433172_8446622_RNA45SN1_45S_50ntFlanks@12002.12030.29|TGCGGGCCGCCGGTGAAATACCACTACTC</td><td>>hg38_21_+_8433172_8446622_RNA45SN1_45S_50ntFlanks</td><td>12002</td><td>12030</td><td>29</td><td>TGCGGGCCGCCGGTGAAATACCACTACTC</td><td>237.4</td><td>-2.8</td><td>0.6</td><td>-4.5</td><td>5.9E-06</td><td>8.7E-06</td><td>MB231/MB468</td></tr>
<tr><td>>hg38_21_+_8433172_8446622_RNA45SN1_45S_50ntFlanks@11643.11660.18|GCGGGTGTTGACGCGATG</td><td>>hg38_21_+_8433172_8446622_RNA45SN1_45S_50ntFlanks</td><td>11643</td><td>11660</td><td>18</td><td>GCGGGTGTTGACGCGATG</td><td>235.3</td><td>-3.1</td><td>0.7</td><td>-4.5</td><td>6.5E-06</td><td>8.7E-06</td><td>BT20/MB468</td></tr>
<tr><td>>hg38_21_+_8433172_8446622_RNA45SN1_45S_50ntFlanks@10778.10818.41|CCAAGGTGAACAGCCTCTGGCATGTTGGAACAATGTAGGTA</td><td>>hg38_21_+_8433172_8446622_RNA45SN1_45S_50ntFlanks</td><td>10778</td><td>10818</td><td>41</td><td>CCAAGGTGAACAGCCTCTGGCATGTTGGAACAATGTAGGTA</td><td>173.3</td><td>-3.6</td><td>0.8</td><td>-4.5</td><td>6.5E-06</td><td>8.7E-06</td><td>BT20/MB468</td></tr>
<tr><td>>trna10_MetCAT_8_-_124169470_124169542@34.51.18,trna22_MetCAT_16_-_87417628_87417700@34.51.18|CATAATCTGAAGGTCGTG</td><td>>trna10_MetCAT_8_-_124169470_124169542</td><td>34</td><td>51</td><td>18</td><td>CATAATCTGAAGGTCGTG</td><td>158.8</td><td>-2.5</td><td>0.6</td><td>-4.5</td><td>6.5E-06</td><td>8.7E-06</td><td>BT20/MB468</td></tr>
<tr><td>>hg38_21_+_8433172_8446622_RNA45SN1_45S_50ntFlanks@11891.11909.19|TTGACTCTAGTCTGGCACG</td><td>>hg38_21_+_8433172_8446622_RNA45SN1_45S_50ntFlanks</td><td>11891</td><td>11909</td><td>19</td><td>TTGACTCTAGTCTGGCACG</td><td>155.6</td><td>-2.6</td><td>0.6</td><td>-4.5</td><td>6.5E-06</td><td>8.7E-06</td><td>BT20/MB468</td></tr>
<tr><td>>trna10_AspGTC_12_-_125424193_125424264@37.74.38,trna12_AspGTC_12_-_125411891_125411962@37.74.38,trna144_AspGTC_6_-_27551236_27551307@37.74.38,trna38_AspGTC_17_-_8125556_8125627@37.74.38,trna45_AspGTC_6_+_27447453_27447524@37.74.38,trna48_AspGTC_6_+_27471523_27471594@37.74.38,trna4_AspGTC_12_+_96429799_96429870@37.74.38,trna69_AspGTC_1_-_161440205_161440276@37.74.38,trna72_AspGTC_1_-_161432824_161432895@37.74.38,trna75_AspGTC_1_-_161425414_161425485@37.74.38,trna78_AspGTC_1_-_161418033_161418104@37.74.38,trna81_AspGTC_1_-_161410615_161410686@37.74.38|ACGCGGGAGACCGGGGTTCGATTCCCCGACGGGGAGCC</td><td>>trna10_AspGTC_12_-_125424193_125424264</td><td>37</td><td>74</td><td>38</td><td>ACGCGGGAGACCGGGGTTCGATTCCCCGACGGGGAGCC</td><td>1013.8</td><td>-5.4</td><td>1.1</td><td>-5.0</td><td>5.4E-07</td><td>8.8E-06</td><td>BT20/MB231</td></tr>
<tr><td>>hg38_21_+_8433172_8446622_RNA45SN1_45S_50ntFlanks@11209.11232.24|CGGCGACTCTGGACGCGAGCCGGG</td><td>>hg38_21_+_8433172_8446622_RNA45SN1_45S_50ntFlanks</td><td>11209</td><td>11232</td><td>24</td><td>CGGCGACTCTGGACGCGAGCCGGG</td><td>10676.5</td><td>-4.9</td><td>1.1</td><td>-4.5</td><td>6.6E-06</td><td>8.8E-06</td><td>BT20/MB468</td></tr>
<tr><td>>hg38_21_+_8433172_8446622_RNA45SN1_45S_50ntFlanks@10664.10706.43|CTTGGAAAGCGTCGCGGTTCCGGCGGCGTCCGGTGAGCTCTCG</td><td>>hg38_21_+_8433172_8446622_RNA45SN1_45S_50ntFlanks</td><td>10664</td><td>10706</td><td>43</td><td>CTTGGAAAGCGTCGCGGTTCCGGCGGCGTCCGGTGAGCTCTCG</td><td>231.5</td><td>-5.3</td><td>1.2</td><td>-4.5</td><td>6.6E-06</td><td>8.8E-06</td><td>BT20/MB468</td></tr>
<tr><td>>hg38_21_+_8433172_8446622_RNA45SN1_45S_50ntFlanks@12456.12480.25|CCAAGCGTTGGATTGTTCACCCACT</td><td>>hg38_21_+_8433172_8446622_RNA45SN1_45S_50ntFlanks</td><td>12456</td><td>12480</td><td>25</td><td>CCAAGCGTTGGATTGTTCACCCACT</td><td>2307.0</td><td>-3.9</td><td>0.9</td><td>-4.5</td><td>6.6E-06</td><td>8.8E-06</td><td>BT20/MB468</td></tr>
<tr><td>>hg38_21_+_8433172_8446622_RNA45SN1_45S_50ntFlanks@12935.12966.32|TAAACCATTCGTAGACGACCTGCTTCTGGGTC</td><td>>hg38_21_+_8433172_8446622_RNA45SN1_45S_50ntFlanks</td><td>12935</td><td>12966</td><td>32</td><td>TAAACCATTCGTAGACGACCTGCTTCTGGGTC</td><td>80.0</td><td>-3.7</td><td>0.8</td><td>-4.5</td><td>6.0E-06</td><td>8.8E-06</td><td>MB231/MB468</td></tr>
<tr><td>>hg38_21_+_8433172_8446622_RNA45SN1_45S_50ntFlanks@4322.4346.25|CAGCCGCGGTAATTCCAGCTCCAAT</td><td>>hg38_21_+_8433172_8446622_RNA45SN1_45S_50ntFlanks</td><td>4322</td><td>4346</td><td>25</td><td>CAGCCGCGGTAATTCCAGCTCCAAT</td><td>161.6</td><td>-4.2</td><td>0.9</td><td>-4.5</td><td>6.0E-06</td><td>8.8E-06</td><td>MB231/MB468</td></tr>
<tr><td>>hg38_21_+_8433172_8446622_RNA45SN1_45S_50ntFlanks@10410.10442.33|TCAGTCGGTCCTGAGAGATGGGCGAGCGCCGTT</td><td>>hg38_21_+_8433172_8446622_RNA45SN1_45S_50ntFlanks</td><td>10410</td><td>10442</td><td>33</td><td>TCAGTCGGTCCTGAGAGATGGGCGAGCGCCGTT</td><td>232.9</td><td>-2.5</td><td>0.6</td><td>-4.5</td><td>6.6E-06</td><td>8.8E-06</td><td>BT20/MB468</td></tr>
<tr><td>>hg38_21_+_8433172_8446622_RNA45SN1_45S_50ntFlanks@9739.9767.29|AAACGATCTCAACCTATTCTCAAACTTTA</td><td>>hg38_21_+_8433172_8446622_RNA45SN1_45S_50ntFlanks</td><td>9739</td><td>9767</td><td>29</td><td>AAACGATCTCAACCTATTCTCAAACTTTA</td><td>240.2</td><td>-2.5</td><td>0.6</td><td>-4.5</td><td>6.7E-06</td><td>8.9E-06</td><td>BT20/MB468</td></tr>
<tr><td>>hg38_21_+_8433172_8446622_RNA45SN1_45S_50ntFlanks@4006.4024.19|ACCTCGGGCCGATCGCACG</td><td>>hg38_21_+_8433172_8446622_RNA45SN1_45S_50ntFlanks</td><td>4006</td><td>4024</td><td>19</td><td>ACCTCGGGCCGATCGCACG</td><td>544.7</td><td>-2.7</td><td>0.6</td><td>-4.5</td><td>6.1E-06</td><td>9.0E-06</td><td>MB231/MB468</td></tr>
<tr><td>>hg38_21_+_8433172_8446622_RNA45SN1_45S_50ntFlanks@11767.11803.37|TCTAATTAGTGACGCGCATGAATGGATGAACGAGATT</td><td>>hg38_21_+_8433172_8446622_RNA45SN1_45S_50ntFlanks</td><td>11767</td><td>11803</td><td>37</td><td>TCTAATTAGTGACGCGCATGAATGGATGAACGAGATT</td><td>427.3</td><td>-1.9</td><td>0.4</td><td>-4.5</td><td>6.2E-06</td><td>9.1E-06</td><td>MB231/MB468</td></tr>
<tr><td>>hg38_21_+_8433172_8446622_RNA45SN1_45S_50ntFlanks@12478.12496.19|ACTAATAGGGAACGTGAGC</td><td>>hg38_21_+_8433172_8446622_RNA45SN1_45S_50ntFlanks</td><td>12478</td><td>12496</td><td>19</td><td>ACTAATAGGGAACGTGAGC</td><td>770.6</td><td>1.7</td><td>0.4</td><td>4.5</td><td>6.9E-06</td><td>9.3E-06</td><td>BT20/MB468</td></tr>
<tr><td>>am_trnaMT_SerGCT_MT_+_12207_12265@13.32.20|AAGAACTGCTAACTCATGCC</td><td>>am_trnaMT_SerGCT_MT_+_12207_12265</td><td>13</td><td>32</td><td>20</td><td>AAGAACTGCTAACTCATGCC</td><td>420.6</td><td>6.3</td><td>1.4</td><td>4.5</td><td>7.0E-06</td><td>9.4E-06</td><td>BT20/MB468</td></tr>
<tr><td>>hg38_21_+_8433172_8446622_RNA45SN1_45S_50ntFlanks@4130.4162.33|ATTCCGGAGAGGGAGCCTGAGAAACGGCTACCA</td><td>>hg38_21_+_8433172_8446622_RNA45SN1_45S_50ntFlanks</td><td>4130</td><td>4162</td><td>33</td><td>ATTCCGGAGAGGGAGCCTGAGAAACGGCTACCA</td><td>367.9</td><td>-3.2</td><td>0.7</td><td>-4.5</td><td>6.4E-06</td><td>9.4E-06</td><td>MB231/MB468</td></tr>
<tr><td>>am_trna128_GlyGCC_6_-_27870686_27870756@1.28.28,trna133_GlyCCC_1_-_16872434_16872504@1.28.28,trna18_GlyGCC_16_+_70822597_70822667@1.28.28,trna19_GlyGCC_16_+_70823410_70823480@1.28.28,trna19_GlyGCC_2_-_157257659_157257729@1.28.28,trna24_GlyGCC_16_-_70812942_70813012@1.28.28,trna25_GlyGCC_16_-_70812114_70812184@1.28.28,trna4_GlyCCC_1_+_17188416_17188486@1.28.28,trna5_GlyGCC_17_+_8029064_8029134@1.28.28,trna68_GlyGCC_1_-_161493637_161493707@1.28.28|GCATTGGTGGTTCAGTGGTAGAATTCTC</td><td>>am_trna128_GlyGCC_6_-_27870686_27870756</td><td>1</td><td>28</td><td>28</td><td>GCATTGGTGGTTCAGTGGTAGAATTCTC</td><td>30518.5</td><td>-3.3</td><td>0.7</td><td>-4.5</td><td>6.5E-06</td><td>9.6E-06</td><td>MB231/MB468</td></tr>
<tr><td>>hg38_21_+_8433172_8446622_RNA45SN1_45S_50ntFlanks@10668.10706.39|GAAAGCGTCGCGGTTCCGGCGGCGTCCGGTGAGCTCTCG</td><td>>hg38_21_+_8433172_8446622_RNA45SN1_45S_50ntFlanks</td><td>10668</td><td>10706</td><td>39</td><td>GAAAGCGTCGCGGTTCCGGCGGCGTCCGGTGAGCTCTCG</td><td>264.4</td><td>-4.0</td><td>0.9</td><td>-4.5</td><td>7.2E-06</td><td>9.6E-06</td><td>BT20/MB468</td></tr>
<tr><td>>hg38_21_+_8433172_8446622_RNA45SN1_45S_50ntFlanks@10795.10816.22|TGGCATGTTGGAACAATGTAGG</td><td>>hg38_21_+_8433172_8446622_RNA45SN1_45S_50ntFlanks</td><td>10795</td><td>10816</td><td>22</td><td>TGGCATGTTGGAACAATGTAGG</td><td>120.8</td><td>-2.6</td><td>0.6</td><td>-4.5</td><td>6.6E-06</td><td>9.7E-06</td><td>MB231/MB468</td></tr>
<tr><td>>hg38_21_+_8433172_8446622_RNA45SN1_45S_50ntFlanks@11317.11343.27|ACCCCACGTCTCGTCGCGCGCGCGTCC</td><td>>hg38_21_+_8433172_8446622_RNA45SN1_45S_50ntFlanks</td><td>11317</td><td>11343</td><td>27</td><td>ACCCCACGTCTCGTCGCGCGCGCGTCC</td><td>274.8</td><td>-3.8</td><td>0.9</td><td>-4.5</td><td>6.6E-06</td><td>9.7E-06</td><td>MB231/MB468</td></tr>
<tr><td>>hg38_21_+_8433172_8446622_RNA45SN1_45S_50ntFlanks@9533.9566.34|GAGGAAACTCTGGTGGAGGTCCGTAGCGGTCCTG</td><td>>hg38_21_+_8433172_8446622_RNA45SN1_45S_50ntFlanks</td><td>9533</td><td>9566</td><td>34</td><td>GAGGAAACTCTGGTGGAGGTCCGTAGCGGTCCTG</td><td>97.8</td><td>-3.0</td><td>0.7</td><td>-4.5</td><td>6.6E-06</td><td>9.8E-06</td><td>MB231/MB468</td></tr>
<tr><td>>hg38_21_+_8433172_8446622_RNA45SN1_45S_50ntFlanks@12162.12181.20|GACTGGGGCGGTACACCTGT</td><td>>hg38_21_+_8433172_8446622_RNA45SN1_45S_50ntFlanks</td><td>12162</td><td>12181</td><td>20</td><td>GACTGGGGCGGTACACCTGT</td><td>146.5</td><td>-3.7</td><td>0.8</td><td>-4.5</td><td>6.6E-06</td><td>9.8E-06</td><td>MB231/MB468</td></tr>
<tr><td>>hg38_21_+_8433172_8446622_RNA45SN1_45S_50ntFlanks@6649.6670.22|TACGACTCTTAGCGGTGGATCA</td><td>>hg38_21_+_8433172_8446622_RNA45SN1_45S_50ntFlanks</td><td>6649</td><td>6670</td><td>22</td><td>TACGACTCTTAGCGGTGGATCA</td><td>91.2</td><td>-2.7</td><td>0.6</td><td>-4.5</td><td>6.7E-06</td><td>9.8E-06</td><td>MB231/MB468</td></tr>
<tr><td>>hg38_21_+_8433172_8446622_RNA45SN1_45S_50ntFlanks@12454.12476.23|CACCAAGCGTTGGATTGTTCACC</td><td>>hg38_21_+_8433172_8446622_RNA45SN1_45S_50ntFlanks</td><td>12454</td><td>12476</td><td>23</td><td>CACCAAGCGTTGGATTGTTCACC</td><td>749.6</td><td>2.4</td><td>0.5</td><td>4.5</td><td>6.7E-06</td><td>9.9E-06</td><td>MB231/MB468</td></tr>
<tr><td>>hg38_21_+_8433172_8446622_RNA45SN1_45S_50ntFlanks@12944.12965.22|CGTAGACGACCTGCTTCTGGGT</td><td>>hg38_21_+_8433172_8446622_RNA45SN1_45S_50ntFlanks</td><td>12944</td><td>12965</td><td>22</td><td>CGTAGACGACCTGCTTCTGGGT</td><td>137.4</td><td>-3.2</td><td>0.7</td><td>-4.5</td><td>7.5E-06</td><td>1.0E-05</td><td>BT20/MB468</td></tr>
<tr><td>>MI0000273|hsa-mir-183&WithFlank&7|-|129774899|129775020@34.53.20(+1A)&[MIMAT0000261&hsa-miR-183-5p&offsets|+1|-1(+1A);m-41&7|-|129774966|129774987&offsets|0|-2(+1A)]|ATGGCACTGGTAGAATTCACA</td><td>>MI0000273|hsa-mir-183&WithFlank&7|-|129774899|129775020</td><td>34</td><td>53</td><td>20(+1A)</td><td>ATGGCACTGGTAGAATTCACA</td><td>408.6</td><td>3.7</td><td>0.8</td><td>5.0</td><td>6.3E-07</td><td>1.0E-05</td><td>BT20/MB231</td></tr>
<tr><td>>MI0000301|hsa-mir-224&WithFlank&X|-|151958572|151958664@14.34.21(+1A)&[MIMAT0000281&hsa-miR-224-5p&offsets|+1|-3(+1A);m-238&X|-|151958630|151958651&offsets|0|-1(+1A)]|CAAGTCACTAGTGGTTCCGTTA</td><td>>MI0000301|hsa-mir-224&WithFlank&X|-|151958572|151958664</td><td>14</td><td>34</td><td>21(+1A)</td><td>CAAGTCACTAGTGGTTCCGTTA</td><td>475.6</td><td>3.6</td><td>0.7</td><td>5.0</td><td>6.3E-07</td><td>1.0E-05</td><td>BT20/MB231</td></tr>
<tr><td>>hg38_21_+_8433172_8446622_RNA45SN1_45S_50ntFlanks@12417.12436.20|TCGATGTCGGCTCTTCCTAT</td><td>>hg38_21_+_8433172_8446622_RNA45SN1_45S_50ntFlanks</td><td>12417</td><td>12436</td><td>20</td><td>TCGATGTCGGCTCTTCCTAT</td><td>4927.4</td><td>-4.1</td><td>0.9</td><td>-4.5</td><td>7.6E-06</td><td>1.0E-05</td><td>BT20/MB468</td></tr>
<tr><td>>hg38_21_+_8433172_8446622_RNA45SN1_45S_50ntFlanks@3881.3898.18|GCCGACGGGCGCTGACCC</td><td>>hg38_21_+_8433172_8446622_RNA45SN1_45S_50ntFlanks</td><td>3881</td><td>3898</td><td>18</td><td>GCCGACGGGCGCTGACCC</td><td>211.9</td><td>-4.1</td><td>0.9</td><td>-4.5</td><td>7.7E-06</td><td>1.0E-05</td><td>BT20/MB468</td></tr>
<tr><td>>hg38_21_+_8433172_8446622_RNA45SN1_45S_50ntFlanks@8384.8402.19|AGGGCGTGAAACCGTTAAG</td><td>>hg38_21_+_8433172_8446622_RNA45SN1_45S_50ntFlanks</td><td>8384</td><td>8402</td><td>19</td><td>AGGGCGTGAAACCGTTAAG</td><td>126.5</td><td>-3.2</td><td>0.7</td><td>-4.5</td><td>7.0E-06</td><td>1.0E-05</td><td>MB231/MB468</td></tr>
<tr><td>>hg38_21_+_8433172_8446622_RNA45SN1_45S_50ntFlanks@12071.12093.23|CCCCGAGGGGCTCTCGCTTCTGG</td><td>>hg38_21_+_8433172_8446622_RNA45SN1_45S_50ntFlanks</td><td>12071</td><td>12093</td><td>23</td><td>CCCCGAGGGGCTCTCGCTTCTGG</td><td>108.0</td><td>-2.1</td><td>0.5</td><td>-4.5</td><td>7.0E-06</td><td>1.0E-05</td><td>MB231/MB468</td></tr>
</table>

| | | | | | | | | | | | | |
|---|---|---|---|---|---|---|---|---|---|---|---|---|
| >hg38_21_+_8433172_8446622_RNA45SN1_45S_50ntFlanks@8192.8215.24\|AGCCCGTGGACGGTGTGAGGCCGG | >hg38_21_+_8433172_8446622_RNA45SN1_45S_50ntFlanks | 8192 | 8215 | 24 | AGCCCGTGGACGGTGTGAGGCCGG | 141.8 | -2.7 | 0.6 | -4.5 | 7.1E-06 | 1.0E-05 | MB231/MB468 |
| >hg38_21_+_8433172_8446622_RNA45SN1_45S_50ntFlanks@12419.12444.26\|GATGTCGGCTCTTCCTATCATTGTGA | >hg38_21_+_8433172_8446622_RNA45SN1_45S_50ntFlanks | 12419 | 12444 | 26 | GATGTCGGCTCTTCCTATCATTGTGA | 426.1 | 2.6 | 0.6 | 4.5 | 7.8E-06 | 1.0E-05 | BT20/MB468 |
| >hg38_21_+_8433172_8446622_RNA45SN1_45S_50ntFlanks@12035.12053.19\|CGTTTTTTCACTGACCCGG | >hg38_21_+_8433172_8446622_RNA45SN1_45S_50ntFlanks | 12035 | 12053 | 19 | CGTTTTTTCACTGACCCGG | 112.7 | -3.6 | 0.8 | -4.5 | 7.1E-06 | 1.0E-05 | MB231/MB468 |
| >hg38_21_+_8433172_8446622_RNA45SN1_45S_50ntFlanks@10798.10815.18\|CATGTTGGAACAATGTAG | >hg38_21_+_8433172_8446622_RNA45SN1_45S_50ntFlanks | 10798 | 10815 | 18 | CATGTTGGAACAATGTAG | 106.3 | -3.4 | 0.8 | -4.5 | 7.1E-06 | 1.0E-05 | MB231/MB468 |
| >hg38_21_+_8433172_8446622_RNA45SN1_45S_50ntFlanks@4297.4317.21\|CATTGGAGGGCAAGTCTGGTG | >hg38_21_+_8433172_8446622_RNA45SN1_45S_50ntFlanks | 4297 | 4317 | 21 | CATTGGAGGGCAAGTCTGGTG | 420.1 | -3.9 | 0.9 | -4.5 | 7.9E-06 | 1.0E-05 | BT20/MB468 |
| >am_trnaMT_ValTAC_MT_+_1602_1670@50.68.19\|ACTTAACTTGACCGCTCTG | >am_trnaMT_ValTAC_MT_+_1602_1670 | 50 | 68 | 19 | ACTTAACTTGACCGCTCTG | 2683.7 | -4.1 | 0.9 | -4.5 | 7.1E-06 | 1.1E-05 | MB231/MB468 |
| >hg38_21_+_8433172_8446622_RNA45SN1_45S_50ntFlanks@9566.9595.30\|GACGTGCAAATCGGTCGTCCGACCTGGGTA | >hg38_21_+_8433172_8446622_RNA45SN1_45S_50ntFlanks | 9566 | 9595 | 30 | GACGTGCAAATCGGTCGTCCGACCTGGGTA | 223.2 | -2.8 | 0.6 | -4.5 | 7.9E-06 | 1.1E-05 | BT20/MB468 |
| >trna37_ValAAC_6_+_27203288_27203360@1.29.29\|GTTTCCGTAGTGTAGTGGTTATCACGTTT | >trna37_ValAAC_6_+_27203288_27203360 | 1 | 29 | 29 | GTTTCCGTAGTGTAGTGGTTATCACGTTT | 878.5 | -3.9 | 0.9 | -4.5 | 7.2E-06 | 1.1E-05 | MB231/MB468 |
| >hg38_21_+_8433172_8446622_RNA45SN1_45S_50ntFlanks@7985.8003.19\|GATCAGACGTGGCGACCCG | >hg38_21_+_8433172_8446622_RNA45SN1_45S_50ntFlanks | 7985 | 8003 | 19 | GATCAGACGTGGCGACCCG | 72.7 | -4.3 | 1.0 | -4.5 | 7.2E-06 | 1.1E-05 | MB231/MB468 |
| >hg38_21_+_8433172_8446622_RNA45SN1_45S_50ntFlanks@11027.11050.24\|CTCTCTCTCTCTCTCCCCCGCTCC | >hg38_21_+_8433172_8446622_RNA45SN1_45S_50ntFlanks | 11027 | 11050 | 24 | CTCTCTCTCTCTCTCCCCCGCTCC | 541.5 | -5.4 | 1.2 | -4.5 | 8.0E-06 | 1.1E-05 | BT20/MB468 |
| >hg38_21_+_8433172_8446622_RNA45SN1_45S_50ntFlanks@11569.11599.31\|CAGCCGACTTAGAACTGGTGCGGACCAGGGG | >hg38_21_+_8433172_8446622_RNA45SN1_45S_50ntFlanks | 11569 | 11599 | 31 | CAGCCGACTTAGAACTGGTGCGGACCAGGGG | 175.0 | -2.4 | 0.5 | -4.5 | 7.2E-06 | 1.1E-05 | MB231/MB468 |
| >trna11_GluTTC_15_-_26327381_26327452@1.31.31,trna3_GluTTC_13_-_45492062_45492133@1.31.31\|TCCCACATGGTCTAGCGGTTAGGATTCCTGG | >trna11_GluTTC_15_-_26327381_26327452 | 1 | 31 | 31 | TCCCACATGGTCTAGCGGTTAGGATTCCTGG | 1147.4 | -2.6 | 0.6 | -4.5 | 8.0E-06 | 1.1E-05 | BT20/MB468 |
| >hg38_21_+_8433172_8446622_RNA45SN1_45S_50ntFlanks@12417.12446.30\|TCGATGTCGGCTCTTCCTATCATTGTGAAG | >hg38_21_+_8433172_8446622_RNA45SN1_45S_50ntFlanks | 12417 | 12446 | 30 | TCGATGTCGGCTCTTCCTATCATTGTGAAG | 612.4 | -2.5 | 0.6 | -4.5 | 8.1E-06 | 1.1E-05 | BT20/MB468 |
| >hg38_21_+_8433172_8446622_RNA45SN1_45S_50ntFlanks@11561.11590.30\|GCGCCTAGCAGCCGACTTAGAACTGGTGCG | >hg38_21_+_8433172_8446622_RNA45SN1_45S_50ntFlanks | 11561 | 11590 | 30 | GCGCCTAGCAGCCGACTTAGAACTGGTGCG | 364.8 | -4.6 | 1.0 | -4.5 | 8.1E-06 | 1.1E-05 | BT20/MB468 |
| >hg38_21_+_8433172_8446622_RNA45SN1_45S_50ntFlanks@9604.9621.18\|AAAGACTAATCGAACCAT | >hg38_21_+_8433172_8446622_RNA45SN1_45S_50ntFlanks | 9604 | 9621 | 18 | AAAGACTAATCGAACCAT | 275.0 | -2.5 | 0.5 | -4.5 | 7.4E-06 | 1.1E-05 | MB231/MB468 |
| >hg38_21_+_8433172_8446622_RNA45SN1_45S_50ntFlanks@10389.10429.41\|TGAACAGCAGTTGAACATGGGTCAGTCGGTCCTGAGAGATG | >hg38_21_+_8433172_8446622_RNA45SN1_45S_50ntFlanks | 10389 | 10429 | 41 | TGAACAGCAGTTGAACATGGGTCAGTCGGTCCTGAGAGATG | 308.9 | -3.8 | 0.8 | -4.5 | 8.1E-06 | 1.1E-05 | BT20/MB468 |
| >hg38_21_+_8433172_8446622_RNA45SN1_45S_50ntFlanks@8381.8403.23\|AAGAGGGCGTGAAACCGTTAAGA | >hg38_21_+_8433172_8446622_RNA45SN1_45S_50ntFlanks | 8381 | 8403 | 23 | AAGAGGGCGTGAAACCGTTAAGA | 231.9 | -2.2 | 0.5 | -4.5 | 7.4E-06 | 1.1E-05 | MB231/MB468 |
| >MI0000266\|hsa-mir-10a&WithFlank&17\|-\|48579832\|48579953@29.49.21&[MIMAT0000253&hsa-miR-10a-5p&offsets\|+1\|-1;m-18&17\|-\|48579905\|48579926&offsets\|+1\|0]\|ACCCTGTAGATCCGAATTTGT | >MI0000266\|hsa-mir-10a&WithFlank&17\|-\|48579832\|48579953 | 29 | 49 | 21 | ACCCTGTAGATCCGAATTTGT | 3812.6 | 11.6 | 2.6 | 4.5 | 7.4E-06 | 1.1E-05 | MB231/MB468 |
| >MI0000342\|hsa-mir-200b&WithFlank&1\|+\|1167098\|1167204@63.85.23(+1A)&[MIMAT0000318&hsa-miR-200b-3p&offsets\|0\|+1(+1A);m-93&1\|+\|1167160\|1167182&offsets\|0\|0(+1A)]\|TAATACTGCCTGGTAATGATGACA | >MI0000342\|hsa-mir-200b&WithFlank&1\|+\|1167098\|1167204 | 63 | 85 | 23(+1A) | TAATACTGCCTGGTAATGATGACA | 665.2 | -2.8 | 0.6 | -4.5 | 7.5E-06 | 1.1E-05 | MB231/MB468 |
| >hg38_21_+_8433172_8446622_RNA45SN1_45S_50ntFlanks@10764.10785.22\|TCCGCAGCAGGTCTCCAAGGTG | >hg38_21_+_8433172_8446622_RNA45SN1_45S_50ntFlanks | 10764 | 10785 | 22 | TCCGCAGCAGGTCTCCAAGGTG | 86.6 | -4.6 | 1.0 | -4.5 | 8.3E-06 | 1.1E-05 | BT20/MB468 |
| >hg38_21_+_8433172_8446622_RNA45SN1_45S_50ntFlanks@10797.10822.26\|GCATGTTGGAACAATGTAGGTAAGGG | >hg38_21_+_8433172_8446622_RNA45SN1_45S_50ntFlanks | 10797 | 10822 | 26 | GCATGTTGGAACAATGTAGGTAAGGG | 187.8 | -3.2 | 0.7 | -4.5 | 8.3E-06 | 1.1E-05 | BT20/MB468 |
| >hg38_21_+_8433172_8446622_RNA45SN1_45S_50ntFlanks@12946.12964.19\|TAGACGACCTGCTTCTGGG | >hg38_21_+_8433172_8446622_RNA45SN1_45S_50ntFlanks | 12946 | 12964 | 19 | TAGACGACCTGCTTCTGGG | 170.1 | -2.5 | 0.6 | -4.5 | 7.5E-06 | 1.1E-05 | MB231/MB468 |
| >hg38_21_+_8433172_8446622_RNA45SN1_45S_50ntFlanks@9552.9571.20\|TCCGTAGCGGTCCTGACGTG | >hg38_21_+_8433172_8446622_RNA45SN1_45S_50ntFlanks | 9552 | 9571 | 20 | TCCGTAGCGGTCCTGACGTG | 562.4 | -1.9 | 0.4 | -4.5 | 7.5E-06 | 1.1E-05 | MB231/MB468 |
| >MI0000101\|hsa-mir-99a&WithFlank&21\|+\|16539083\|16539175@19.40.22&[MIMAT0000097&hsa-miR-99a-5p&offsets\|0\|0;m-64&21\|+\|16539101\|16539121&offsets\|0\|+1]\|AACCCGTAGATCCGATCTTGTG | >MI0000101\|hsa-mir-99a&WithFlank&21\|+\|16539083\|16539175 | 19 | 40 | 22 | AACCCGTAGATCCGATCTTGTG | 5926.6 | 3.4 | 0.8 | 4.5 | 7.5E-06 | 1.1E-05 | MB231/MB468 |
| >hg38_21_+_8433172_8446622_RNA45SN1_45S_50ntFlanks@9806.9830.25\|CGTGGAATGCGAGTGCCTAGTGGGC | >hg38_21_+_8433172_8446622_RNA45SN1_45S_50ntFlanks | 9806 | 9830 | 25 | CGTGGAATGCGAGTGCCTAGTGGGC | 85.6 | -3.0 | 0.7 | -4.5 | 8.3E-06 | 1.1E-05 | BT20/MB468 |
| >hg38_21_+_8433172_8446622_RNA45SN1_45S_50ntFlanks@10830.10851.22\|CAAGCCGGATCCGTAACTTCGG | >hg38_21_+_8433172_8446622_RNA45SN1_45S_50ntFlanks | 10830 | 10851 | 22 | CAAGCCGGATCCGTAACTTCGG | 231.4 | -2.2 | 0.5 | -4.5 | 8.4E-06 | 1.1E-05 | BT20/MB468 |
| >hg38_21_+_8433172_8446622_RNA45SN1_45S_50ntFlanks@6651.6677.27\|CGACTCTTAGCGGTGGATCACTCGGCT | >hg38_21_+_8433172_8446622_RNA45SN1_45S_50ntFlanks | 6651 | 6677 | 27 | CGACTCTTAGCGGTGGATCACTCGGCT | 3204.6 | 3.1 | 0.7 | 4.5 | 7.6E-06 | 1.1E-05 | MB231/MB468 |
| >hg38_21_+_8433172_8446622_RNA45SN1_45S_50ntFlanks@12416.12435.20\|TTCGATGTCGGCTCTTCCTA | >hg38_21_+_8433172_8446622_RNA45SN1_45S_50ntFlanks | 12416 | 12435 | 20 | TTCGATGTCGGCTCTTCCTA | 550.2 | -2.1 | 0.5 | -4.5 | 8.5E-06 | 1.1E-05 | BT20/MB468 |
| >hg38_21_+_8433172_8446622_RNA45SN1_45S_50ntFlanks@12941.12965.25\|ATTCGTAGACGACCTGCTTCTGGGT | >hg38_21_+_8433172_8446622_RNA45SN1_45S_50ntFlanks | 12941 | 12965 | 25 | ATTCGTAGACGACCTGCTTCTGGGT | 258.3 | -2.5 | 0.6 | -4.4 | 8.6E-06 | 1.1E-05 | BT20/MB468 |
| >hg38_21_+_8433172_8446622_RNA45SN1_45S_50ntFlanks@11594.11611.18\|CAGGGGAATCCGACTGTT | >hg38_21_+_8433172_8446622_RNA45SN1_45S_50ntFlanks | 11594 | 11611 | 18 | CAGGGGAATCCGACTGTT | 1589.2 | -2.4 | 0.5 | -4.5 | 7.8E-06 | 1.1E-05 | MB231/MB468 |
| >hg38_21_+_8433172_8446622_RNA45SN1_45S_50ntFlanks@12394.12417.24\|TAGCGACGTCGCTTTTTGATCCTT | >hg38_21_+_8433172_8446622_RNA45SN1_45S_50ntFlanks | 12394 | 12417 | 24 | TAGCGACGTCGCTTTTTGATCCTT | 118.4 | -2.8 | 0.6 | -4.4 | 8.7E-06 | 1.2E-05 | BT20/MB468 |
| >hg38_21_+_8433172_8446622_RNA45SN1_45S_50ntFlanks@7974.8007.34\|ACGCGACCTCAGATCAGACGTGGCGACCCGCTGA | >hg38_21_+_8433172_8446622_RNA45SN1_45S_50ntFlanks | 7974 | 8007 | 34 | ACGCGACCTCAGATCAGACGTGGCGACCCGCTGA | 1153.4 | 2.9 | 0.6 | 4.4 | 8.7E-06 | 1.2E-05 | BT20/MB468 |
| >hg38_21_+_8433172_8446622_RNA45SN1_45S_50ntFlanks@8170.8212.43\|CAAGTCCTTCTGATCGAGGCCCAGCCCGTGGACGGTGTGAGGC | >hg38_21_+_8433172_8446622_RNA45SN1_45S_50ntFlanks | 8170 | 8212 | 43 | CAAGTCCTTCTGATCGAGGCCCAGCCCGTGGACGGTGTGAGGC | 388.2 | -2.1 | 0.5 | -4.5 | 7.9E-06 | 1.2E-05 | MB231/MB468 |
| >MI0000478\|hsa-mir-149&WithFlank&2\|+\|240455995\|240456095@21.43.23&[MIMAT0000450&hsa-miR-149-5p&offsets\|0\|0;m-195&2\|+\|240456015\|240456037&offsets\|0\|0]\|TCTGGCTCCGTGTCTTCACTCCC | >MI0000478\|hsa-mir-149&WithFlank&2\|+\|240455995\|240456095 | 21 | 43 | 23 | TCTGGCTCCGTGTCTTCACTCCC | 1609.1 | 3.8 | 0.8 | 5.0 | 7.3E-07 | 1.2E-05 | BT20/MB231 |
| >hg38_21_+_8433172_8446622_RNA45SN1_45S_50ntFlanks@6648.6680.33\|GTACGACTCTTAGCGGTGGATCACTCGGCTCGT | >hg38_21_+_8433172_8446622_RNA45SN1_45S_50ntFlanks | 6648 | 6680 | 33 | GTACGACTCTTAGCGGTGGATCACTCGGCTCGT | 959.0 | 2.7 | 0.6 | 4.5 | 8.0E-06 | 1.2E-05 | MB231/MB468 |
| >MI0000239\|hsa-mir-197&WithFlank&1\|+\|109598887\|109598973@54.74.21(+1U)&[MIMAT0000227&hsa-miR-197-3p&offsets\|0\|-1(+1U);m-99&1\|+\|109598940\|109598961&offsets\|0\|-1(+1U)]\|TTCACCACCTTCTCCACCCAGT | >MI0000239\|hsa-mir-197&WithFlank&1\|+\|109598887\|109598973 | 54 | 74 | 21(+1U) | TTCACCACCTTCTCCACCCAGT | 3023.9 | 3.9 | 0.8 | 4.9 | 7.5E-07 | 1.2E-05 | BT20/MB231 |
| >MI0000273\|hsa-mir-183&WithFlank&7\|-\|129774899\|129775020@34.55.22(+1A)&[MIMAT0000261&hsa-miR-183-5p&offsets\|+1\|+1(+1A);m-41&7\|-\|129774966\|129774987&offsets\|0\|0(+1A)]\|ATGGCACTGGTAGAATTCACTGA | >MI0000273\|hsa-mir-183&WithFlank&7\|-\|129774899\|129775020 | 34 | 55 | 22(+1A) | ATGGCACTGGTAGAATTCACTGA | 186.4 | 3.8 | 0.8 | 4.9 | 7.4E-07 | 1.2E-05 | BT20/MB231 |
| >hg38_21_+_8433172_8446622_RNA45SN1_45S_50ntFlanks@10351.10375.25\|CGAGAACTTTGAAGGCCGAAGTGGA | >hg38_21_+_8433172_8446622_RNA45SN1_45S_50ntFlanks | 10351 | 10375 | 25 | CGAGAACTTTGAAGGCCGAAGTGGA | 150.3 | -2.8 | 0.6 | -4.5 | 8.0E-06 | 1.2E-05 | MB231/MB468 |
| >hg38_21_+_8433172_8446622_RNA45SN1_45S_50ntFlanks@10350.10388.39\|ACGAGAACTTTGAAGGCCGAAGTGGAGAAGGGTTCCATG | >hg38_21_+_8433172_8446622_RNA45SN1_45S_50ntFlanks | 10350 | 10388 | 39 | ACGAGAACTTTGAAGGCCGAAGTGGAGAAGGGTTCCATG | 2046.0 | -3.8 | 0.9 | -4.5 | 8.0E-06 | 1.2E-05 | MB231/MB468 |
| >hg38_21_+_8433172_8446622_RNA45SN1_45S_50ntFlanks@11318.11343.26\|CCCCACGTCTCGTCGCGCGCGCGTCC | >hg38_21_+_8433172_8446622_RNA45SN1_45S_50ntFlanks | 11318 | 11343 | 26 | CCCCACGTCTCGTCGCGCGCGCGTCC | 311.8 | -3.8 | 0.9 | -4.4 | 8.9E-06 | 1.2E-05 | BT20/MB468 |
| >hg38_21_+_8433172_8446622_RNA45SN1_45S_50ntFlanks@12444.12475.32\|AAGCAGAATTCACCAAGCGTTGGATTGTTCAC | >hg38_21_+_8433172_8446622_RNA45SN1_45S_50ntFlanks | 12444 | 12475 | 32 | AAGCAGAATTCACCAAGCGTTGGATTGTTCAC | 137.7 | -4.3 | 1.0 | -4.4 | 8.9E-06 | 1.2E-05 | BT20/MB468 |
| >hg38_21_+_8433172_8446622_RNA45SN1_45S_50ntFlanks@4726.4751.26\|TAATCAAGAACGAAAGTCGGAGGTTC | >hg38_21_+_8433172_8446622_RNA45SN1_45S_50ntFlanks | 4726 | 4751 | 26 | TAATCAAGAACGAAAGTCGGAGGTTC | 150.2 | -2.6 | 0.6 | -4.5 | 8.1E-06 | 1.2E-05 | MB231/MB468 |
| >hg38_21_+_8433172_8446622_RNA45SN1_45S_50ntFlanks@10840.10864.25\|CCGTAACTTCGGGATAAGGATTGGC | >hg38_21_+_8433172_8446622_RNA45SN1_45S_50ntFlanks | 10840 | 10864 | 25 | CCGTAACTTCGGGATAAGGATTGGC | 91.7 | -4.6 | 1.0 | -4.4 | 8.9E-06 | 1.2E-05 | BT20/MB468 |
| >hg38_21_+_8433172_8446622_RNA45SN1_45S_50ntFlanks@11782.11804.23\|GCATGAATGGATGAACGAGATTC | >hg38_21_+_8433172_8446622_RNA45SN1_45S_50ntFlanks | 11782 | 11804 | 23 | GCATGAATGGATGAACGAGATTC | 159.6 | -2.0 | 0.4 | -4.5 | 8.1E-06 | 1.2E-05 | MB231/MB468 |
| >hg38_21_+_8433172_8446622_RNA45SN1_45S_50ntFlanks@12396.12417.22\|GCGACGTCGCTTTTTGATCCTT | >hg38_21_+_8433172_8446622_RNA45SN1_45S_50ntFlanks | 12396 | 12417 | 22 | GCGACGTCGCTTTTTGATCCTT | 455.6 | -3.8 | 0.9 | -4.4 | 9.1E-06 | 1.2E-05 | BT20/MB468 |
| >hg38_21_+_8433172_8446622_RNA45SN1_45S_50ntFlanks@6671.6696.26\|CTCGGCTCGTGCGTCGATGAAGAACG | >hg38_21_+_8433172_8446622_RNA45SN1_45S_50ntFlanks | 6671 | 6696 | 26 | CTCGGCTCGTGCGTCGATGAAGAACG | 625.1 | -2.4 | 0.5 | -4.4 | 9.1E-06 | 1.2E-05 | BT20/MB468 |
| >hg38_21_+_8433172_8446622_RNA45SN1_45S_50ntFlanks@4367.4400.34\|CAGTTAAAAAGCTCGTAGTTGGATCTTGGGAGCG | >hg38_21_+_8433172_8446622_RNA45SN1_45S_50ntFlanks | 4367 | 4400 | 34 | CAGTTAAAAAGCTCGTAGTTGGATCTTGGGAGCG | 88.3 | -4.6 | 1.0 | -4.4 | 9.1E-06 | 1.2E-05 | BT20/MB468 |
| >hg38_21_+_8433172_8446622_RNA45SN1_45S_50ntFlanks@10771.10788.18\|CAGGTCTCCAAGGTGAAC | >hg38_21_+_8433172_8446622_RNA45SN1_45S_50ntFlanks | 10771 | 10788 | 18 | CAGGTCTCCAAGGTGAAC | 125.4 | -2.5 | 0.6 | -4.4 | 9.1E-06 | 1.2E-05 | BT20/MB468 |
| >hg38_21_+_8433172_8446622_RNA45SN1_45S_50ntFlanks@9623.9650.28\|TAGTAGCTGGTTCCCTCCGAAGTTTCCC | >hg38_21_+_8433172_8446622_RNA45SN1_45S_50ntFlanks | 9623 | 9650 | 28 | TAGTAGCTGGTTCCCTCCGAAGTTTCCC | 109.9 | -2.9 | 0.7 | -4.5 | 8.2E-06 | 1.2E-05 | MB231/MB468 |

<table>
<tr><td>>hg38_21_+_8433172_8446622_RNA45SN1_45S_50ntFlanks@11562.11591.30|CGCCTAGCAGCCGACTTAGAACTGGTGCGG</td><td>>hg38_21_+_8433172_8446622_RNA45SN1_45S_50ntFlanks</td><td>11562</td><td>11591</td><td>30</td><td>CGCCTAGCAGCCGACTTAGAACTGGTGCGG</td><td>138.9</td><td>-3.6</td><td>0.8</td><td>-4.5</td><td>8.3E-06</td><td>1.2E-05</td><td>MB231/MB468</td></tr>
<tr><td>>trnaMT_GluTTC_MT_-_14674_14742@39.61.23,trnalookalike8_GluTTC_5_-_93905172_93905240@39.61.23|ATTGGTCGTGGTTGTAGTCCGTG</td><td>>trnaMT_GluTTC_MT_-_14674_14742</td><td>39</td><td>61</td><td>23</td><td>ATTGGTCGTGGTTGTAGTCCGTG</td><td>558.9</td><td>6.5</td><td>1.5</td><td>4.4</td><td>9.2E-06</td><td>1.2E-05</td><td>BT20/MB468</td></tr>
<tr><td>>MI0000737|hsa-mir-200a&WithFlank&1|+|1167857|1167958@60.81.22(+1C)&[MIMAT0000682&hsa-miR-200a-3p&offsets|0|0(+1C);m-142&1|+|1167916|1167938&offsets|0|-1(+1C)]|TAACACTGTCTGGTAACGATGTC</td><td>>MI0000737|hsa-mir-200a&WithFlank&1|+|1167857|1167958</td><td>60</td><td>81</td><td>22(+1C)</td><td>TAACACTGTCTGGTAACGATGTC</td><td>151.3</td><td>-3.5</td><td>0.8</td><td>-4.5</td><td>8.4E-06</td><td>1.2E-05</td><td>MB231/MB468</td></tr>
<tr><td>>hg38_21_+_8433172_8446622_RNA45SN1_45S_50ntFlanks@11580.11600.21|GAACTGGTGCGGACCAGGGGA</td><td>>hg38_21_+_8433172_8446622_RNA45SN1_45S_50ntFlanks</td><td>11580</td><td>11600</td><td>21</td><td>GAACTGGTGCGGACCAGGGGA</td><td>297.3</td><td>-2.4</td><td>0.5</td><td>-4.5</td><td>8.4E-06</td><td>1.2E-05</td><td>MB231/MB468</td></tr>
<tr><td>>hg38_21_+_8433172_8446622_RNA45SN1_45S_50ntFlanks@4547.4565.19|CTGGATACCGCAGCTAGGA</td><td>>hg38_21_+_8433172_8446622_RNA45SN1_45S_50ntFlanks</td><td>4547</td><td>4565</td><td>19</td><td>CTGGATACCGCAGCTAGGA</td><td>171.5</td><td>-2.9</td><td>0.7</td><td>-4.5</td><td>8.4E-06</td><td>1.2E-05</td><td>MB231/MB468</td></tr>
<tr><td>>hg38_21_+_8433172_8446622_RNA45SN1_45S_50ntFlanks@10792.10818.27|CTCTGGCATGTTGGAACAATGTAGGTA</td><td>>hg38_21_+_8433172_8446622_RNA45SN1_45S_50ntFlanks</td><td>10792</td><td>10818</td><td>27</td><td>CTCTGGCATGTTGGAACAATGTAGGTA</td><td>489.9</td><td>-3.6</td><td>0.8</td><td>-4.4</td><td>9.3E-06</td><td>1.2E-05</td><td>BT20/MB468</td></tr>
<tr><td>>hg38_21_+_8433172_8446622_RNA45SN1_45S_50ntFlanks@10841.10859.19|CGTAACTTCGGGATAAGGA</td><td>>hg38_21_+_8433172_8446622_RNA45SN1_45S_50ntFlanks</td><td>10841</td><td>10859</td><td>19</td><td>CGTAACTTCGGGATAAGGA</td><td>985.0</td><td>-5.3</td><td>1.2</td><td>-4.4</td><td>9.3E-06</td><td>1.2E-05</td><td>BT20/MB468</td></tr>
<tr><td>>hg38_21_+_8433172_8446622_RNA45SN1_45S_50ntFlanks@11682.11706.25|AATGTCAAAGTGAAGAAATTCAATG</td><td>>hg38_21_+_8433172_8446622_RNA45SN1_45S_50ntFlanks</td><td>11682</td><td>11706</td><td>25</td><td>AATGTCAAAGTGAAGAAATTCAATG</td><td>629.5</td><td>-4.9</td><td>1.1</td><td>-4.4</td><td>9.3E-06</td><td>1.2E-05</td><td>BT20/MB468</td></tr>
<tr><td>>MI0003513|hsa-mir-455&WithFlank&9|+|114209428|114209535@60.81.22&[MIMAT0004784&hsa-miR-455-3p&offsets|0|+1;m-135&9|+|114209487|114209508&offsets|0|0]|GCAGTCCATGGGCATATACACT</td><td>>MI0003513|hsa-mir-455&WithFlank&9|+|114209428|114209535</td><td>60</td><td>81</td><td>22</td><td>GCAGTCCATGGGCATATACACT</td><td>308.8</td><td>-3.7</td><td>0.8</td><td>-4.9</td><td>8.0E-07</td><td>1.2E-05</td><td>BT20/MB231</td></tr>
<tr><td>>hg38_21_+_8433172_8446622_RNA45SN1_45S_50ntFlanks@9534.9572.39|AGGAAACTCTGGTGGAGGTCCGTAGCGGTCCTGACGTGC</td><td>>hg38_21_+_8433172_8446622_RNA45SN1_45S_50ntFlanks</td><td>9534</td><td>9572</td><td>39</td><td>AGGAAACTCTGGTGGAGGTCCGTAGCGGTCCTGACGTGC</td><td>921.4</td><td>-2.4</td><td>0.5</td><td>-4.5</td><td>8.5E-06</td><td>1.2E-05</td><td>MB231/MB468</td></tr>
<tr><td>>hg38_21_+_8433172_8446622_RNA45SN1_45S_50ntFlanks@11671.11699.29|CCAGTGCTCTGAATGTCAAAGTGAAGAAA</td><td>>hg38_21_+_8433172_8446622_RNA45SN1_45S_50ntFlanks</td><td>11671</td><td>11699</td><td>29</td><td>CCAGTGCTCTGAATGTCAAAGTGAAGAAA</td><td>156.0</td><td>-2.4</td><td>0.5</td><td>-4.5</td><td>8.5E-06</td><td>1.2E-05</td><td>MB231/MB468</td></tr>
<tr><td>>hg38_21_+_8433172_8446622_RNA45SN1_45S_50ntFlanks@7989.8009.21|AGACGTGGCGACCCGCTGAAT</td><td>>hg38_21_+_8433172_8446622_RNA45SN1_45S_50ntFlanks</td><td>7989</td><td>8009</td><td>21</td><td>AGACGTGGCGACCCGCTGAAT</td><td>1544.5</td><td>2.9</td><td>0.7</td><td>4.4</td><td>9.4E-06</td><td>1.2E-05</td><td>BT20/MB468</td></tr>
<tr><td>>hg38_21_+_8433172_8446622_RNA45SN1_45S_50ntFlanks@10788.10816.29|CAGCCTCTGGCATGTTGGAACAATGTAGG</td><td>>hg38_21_+_8433172_8446622_RNA45SN1_45S_50ntFlanks</td><td>10788</td><td>10816</td><td>29</td><td>CAGCCTCTGGCATGTTGGAACAATGTAGG</td><td>439.4</td><td>-2.2</td><td>0.5</td><td>-4.5</td><td>8.6E-06</td><td>1.3E-05</td><td>MB231/MB468</td></tr>
<tr><td>>hg38_21_+_8433172_8446622_RNA45SN1_45S_50ntFlanks@12419.12446.28|GATGTCGGCTCTTCCTATCATTGTGAAG</td><td>>hg38_21_+_8433172_8446622_RNA45SN1_45S_50ntFlanks</td><td>12419</td><td>12446</td><td>28</td><td>GATGTCGGCTCTTCCTATCATTGTGAAG</td><td>1453.2</td><td>3.2</td><td>0.7</td><td>4.5</td><td>8.6E-06</td><td>1.3E-05</td><td>MB231/MB468</td></tr>
<tr><td>>MI0000289|hsa-mir-181a-1&WithFlank&1|-|198859038|198859159@30.54.25(+1U);MI0000269|hsa-mir-181a-2&WithFlank&9|+|124692436|124692557@45.69.25(+1U)&[MIMAT0000256&hsa-miR-181a-5p&offsets|0|+2(+1U);m-32&1|-|198859108|198859130&offsets|0|+2(+1U)];[MIMAT0000256_1&hsa-miR-181a-5p&offsets|0|+2(+1U);m-32&9|+|124692480|124692502&offsets|0|+2(+1U)]|AACATTCAACGCTGTCGGTGAGTTTT</td><td>>MI0000289|hsa-mir-181a-1&WithFlank&1|-|198859038|198859159</td><td>30</td><td>54</td><td>25(+1U)</td><td>AACATTCAACGCTGTCGGTGAGTTTT</td><td>637.5</td><td>-3.9</td><td>0.8</td><td>-4.9</td><td>8.1E-07</td><td>1.3E-05</td><td>BT20/MB231</td></tr>
<tr><td>>hg38_21_+_8433172_8446622_RNA45SN1_45S_50ntFlanks@6699.6720.22|GCTAGCTGCGAGAATTAATGTG</td><td>>hg38_21_+_8433172_8446622_RNA45SN1_45S_50ntFlanks</td><td>6699</td><td>6720</td><td>22</td><td>GCTAGCTGCGAGAATTAATGTG</td><td>22208.8</td><td>-3.1</td><td>0.7</td><td>-4.4</td><td>8.6E-06</td><td>1.3E-05</td><td>MB231/MB468</td></tr>
<tr><td>>trna10_LysCTT_16_+_3241501_3241573@1.35.35,trna13_LysCTT_14_-_58706613_58706685@1.35.35,trna2_LysCTT_15_+_79152904_79152976@1.35.35|GCCCGGCTAGCTCAGTCGGTAGAGCATGGGACTCT</td><td>>trna10_LysCTT_16_+_3241501_3241573</td><td>1</td><td>35</td><td>35</td><td>GCCCGGCTAGCTCAGTCGGTAGAGCATGGGACTCT</td><td>168438.6</td><td>4.7</td><td>0.9</td><td>4.9</td><td>8.3E-07</td><td>1.3E-05</td><td>BT20/MB231</td></tr>
<tr><td>>hg38_21_+_8433172_8446622_RNA45SN1_45S_50ntFlanks@6652.6678.27|GACTCTTAGCGGTGGATCACTCGGCTC</td><td>>hg38_21_+_8433172_8446622_RNA45SN1_45S_50ntFlanks</td><td>6652</td><td>6678</td><td>27</td><td>GACTCTTAGCGGTGGATCACTCGGCTC</td><td>1099.0</td><td>3.7</td><td>0.8</td><td>4.4</td><td>8.7E-06</td><td>1.3E-05</td><td>MB231/MB468</td></tr>
<tr><td>>hg38_21_+_8433172_8446622_RNA45SN1_45S_50ntFlanks@12892.12932.41|CCCTCGCCCGTCACGCACCGCACGTTCGTGGGGAACCTGGC</td><td>>hg38_21_+_8433172_8446622_RNA45SN1_45S_50ntFlanks</td><td>12892</td><td>12932</td><td>41</td><td>CCCTCGCCCGTCACGCACCGCACGTTCGTGGGGAACCTGGC</td><td>356.6</td><td>-3.1</td><td>0.7</td><td>-4.4</td><td>8.7E-06</td><td>1.3E-05</td><td>MB231/MB468</td></tr>
<tr><td>>hg38_21_+_8433172_8446622_RNA45SN1_45S_50ntFlanks@4224.4242.19|AAAATAACAATACAGGACT</td><td>>hg38_21_+_8433172_8446622_RNA45SN1_45S_50ntFlanks</td><td>4224</td><td>4242</td><td>19</td><td>AAAATAACAATACAGGACT</td><td>553.4</td><td>-3.3</td><td>0.7</td><td>-4.4</td><td>9.7E-06</td><td>1.3E-05</td><td>BT20/MB468</td></tr>
<tr><td>>am_trna2_GlyTCC_19_+_4724082_4724153@1.29.29|GCGTTGGTGGTATAGTGGTTAGCATAGCT</td><td>>am_trna2_GlyTCC_19_+_4724082_4724153</td><td>1</td><td>29</td><td>29</td><td>GCGTTGGTGGTATAGTGGTTAGCATAGCT</td><td>124.0</td><td>-2.8</td><td>0.6</td><td>-4.4</td><td>9.7E-06</td><td>1.3E-05</td><td>BT20/MB468</td></tr>
<tr><td>>hg38_21_+_8433172_8446622_RNA45SN1_45S_50ntFlanks@8182.8211.30|ATCGAGGCCCAGCCCGTGGACGGTGTGAGG</td><td>>hg38_21_+_8433172_8446622_RNA45SN1_45S_50ntFlanks</td><td>8182</td><td>8211</td><td>30</td><td>ATCGAGGCCCAGCCCGTGGACGGTGTGAGG</td><td>167.9</td><td>-3.9</td><td>0.9</td><td>-4.4</td><td>8.8E-06</td><td>1.3E-05</td><td>MB231/MB468</td></tr>
<tr><td>>hg38_21_+_8433172_8446622_RNA45SN1_45S_50ntFlanks@10882.10902.21|TCGGGCTGGGGCGCGAAGCGG</td><td>>hg38_21_+_8433172_8446622_RNA45SN1_45S_50ntFlanks</td><td>10882</td><td>10902</td><td>21</td><td>TCGGGCTGGGGCGCGAAGCGG</td><td>177.6</td><td>-3.2</td><td>0.7</td><td>-4.4</td><td>9.8E-06</td><td>1.3E-05</td><td>BT20/MB468</td></tr>
<tr><td>>hg38_21_+_8433172_8446622_RNA45SN1_45S_50ntFlanks@12652.12675.24|ATTATGACTGAACGCCTCTAAGTC</td><td>>hg38_21_+_8433172_8446622_RNA45SN1_45S_50ntFlanks</td><td>12652</td><td>12675</td><td>24</td><td>ATTATGACTGAACGCCTCTAAGTC</td><td>96.8</td><td>-3.5</td><td>0.8</td><td>-4.4</td><td>8.9E-06</td><td>1.3E-05</td><td>MB231/MB468</td></tr>
<tr><td>>hg38_21_+_8433172_8446622_RNA45SN1_45S_50ntFlanks@4102.4120.19|ACCACGGGTGACGGGGAAT</td><td>>hg38_21_+_8433172_8446622_RNA45SN1_45S_50ntFlanks</td><td>4102</td><td>4120</td><td>19</td><td>ACCACGGGTGACGGGGAAT</td><td>144.6</td><td>-2.4</td><td>0.5</td><td>-4.4</td><td>9.9E-06</td><td>1.3E-05</td><td>BT20/MB468</td></tr>
<tr><td>>trna111_HisGTG_1_-_147774845_147774916@-1G.30.31,trna118_HisGTG_1_-_145396881_145396952@-1G.30.31,trna16_HisGTG_1_+_146544773_146544844@-1G.30.31,trna1_HisGTG_15_+_45493349_45493420@-1G.30.31,trna21_HisGTG_1_+_147753471_147753542@-1G.30.31,trna33_HisGTG_6_+_27125906_27125977@-1G.30.31,trna7_HisGTG_9_-_14433938_14434009@-1G.30.31,trna8_HisGTG_15_-_45492611_45492682@-1G.30.31,trna9_HisGTG_15_-_45490804_45490875@-1G.30.31|GGCCGTGATCGTATAGTGGTTAGTACTCTGC</td><td>>trna111_HisGTG_1_-_147774845_147774916</td><td>-1G</td><td>30</td><td>31</td><td>GGCCGTGATCGTATAGTGGTTAGTACTCTGC</td><td>6039.6</td><td>4.0</td><td>0.9</td><td>4.4</td><td>9.0E-06</td><td>1.3E-05</td><td>MB231/MB468</td></tr>
<tr><td>>hg38_21_+_8433172_8446622_RNA45SN1_45S_50ntFlanks@6733.6771.39|CATTGATCATCGACACTTCGAACGCACTTGCGGCCCCGG</td><td>>hg38_21_+_8433172_8446622_RNA45SN1_45S_50ntFlanks</td><td>6733</td><td>6771</td><td>39</td><td>CATTGATCATCGACACTTCGAACGCACTTGCGGCCCCGG</td><td>729.7</td><td>-2.3</td><td>0.5</td><td>-4.4</td><td>9.0E-06</td><td>1.3E-05</td><td>MB231/MB468</td></tr>
<tr><td>>hg38_21_+_8433172_8446622_RNA45SN1_45S_50ntFlanks@5084.5110.27|CGAACGAGACTCTGGCATGCTAACTAG</td><td>>hg38_21_+_8433172_8446622_RNA45SN1_45S_50ntFlanks</td><td>5084</td><td>5110</td><td>27</td><td>CGAACGAGACTCTGGCATGCTAACTAG</td><td>148.1</td><td>-3.0</td><td>0.7</td><td>-4.4</td><td>1.0E-05</td><td>1.3E-05</td><td>BT20/MB468</td></tr>
<tr><td>>hg38_21_+_8433172_8446622_RNA45SN1_45S_50ntFlanks@12420.12438.19|ATGTCGGCTCTTCCTATCA</td><td>>hg38_21_+_8433172_8446622_RNA45SN1_45S_50ntFlanks</td><td>12420</td><td>12438</td><td>19</td><td>ATGTCGGCTCTTCCTATCA</td><td>1273.5</td><td>2.8</td><td>0.6</td><td>4.4</td><td>9.1E-06</td><td>1.3E-05</td><td>MB231/MB468</td></tr>
<tr><td>>hg38_21_+_8433172_8446622_RNA45SN1_45S_50ntFlanks@12655.12674.20|ATGACTGAACGCCTCTAAGT</td><td>>hg38_21_+_8433172_8446622_RNA45SN1_45S_50ntFlanks</td><td>12655</td><td>12674</td><td>20</td><td>ATGACTGAACGCCTCTAAGT</td><td>3678.7</td><td>-4.0</td><td>0.9</td><td>-4.4</td><td>9.1E-06</td><td>1.3E-05</td><td>MB231/MB468</td></tr>
<tr><td>>hg38_21_+_8433172_8446622_RNA45SN1_45S_50ntFlanks@10869.10892.24|AGGGCTGGGTCGGTCGGGCTGGGG</td><td>>hg38_21_+_8433172_8446622_RNA45SN1_45S_50ntFlanks</td><td>10869</td><td>10892</td><td>24</td><td>AGGGCTGGGTCGGTCGGGCTGGGG</td><td>345.3</td><td>-5.3</td><td>1.2</td><td>-4.4</td><td>1.0E-05</td><td>1.3E-05</td><td>BT20/MB468</td></tr>
<tr><td>>hg38_21_+_8433172_8446622_RNA45SN1_45S_50ntFlanks@11728.11751.24|GTAACTATGACTCTCTTAAGGTAG</td><td>>hg38_21_+_8433172_8446622_RNA45SN1_45S_50ntFlanks</td><td>11728</td><td>11751</td><td>24</td><td>GTAACTATGACTCTCTTAAGGTAG</td><td>1006.9</td><td>-3.6</td><td>0.8</td><td>-4.4</td><td>9.1E-06</td><td>1.3E-05</td><td>MB231/MB468</td></tr>
<tr><td>>MI0000825|hsa-mir-345&WithFlank&14|+|100307853|100307962@24.45.22(+1U)&[MIMAT0000772&hsa-miR-345-5p&offsets|0|0(+1U);m-193&14|+|100307876|100307896&offsets|0|+1(+1U)]|GCTGACTCCTAGTCCAGGGCTCT</td><td>>MI0000825|hsa-mir-345&WithFlank&14|+|100307853|100307962</td><td>24</td><td>45</td><td>22(+1U)</td><td>GCTGACTCCTAGTCCAGGGCTCT</td><td>732.3</td><td>8.5</td><td>1.9</td><td>4.4</td><td>9.2E-06</td><td>1.3E-05</td><td>MB231/MB468</td></tr>
<tr><td>>hg38_21_+_8433172_8446622_RNA45SN1_45S_50ntFlanks@10393.10429.37|CAGCAGTTGAACATGGGTCAGTCGGTCCTGAGAGATG</td><td>>hg38_21_+_8433172_8446622_RNA45SN1_45S_50ntFlanks</td><td>10393</td><td>10429</td><td>37</td><td>CAGCAGTTGAACATGGGTCAGTCGGTCCTGAGAGATG</td><td>1021.6</td><td>-3.3</td><td>0.7</td><td>-4.4</td><td>1.0E-05</td><td>1.3E-05</td><td>BT20/MB468</td></tr>
<tr><td>>hg38_21_+_8433172_8446622_RNA45SN1_45S_50ntFlanks@10349.10373.25|AACGAGAACTTTGAAGGCCGAAGTG</td><td>>hg38_21_+_8433172_8446622_RNA45SN1_45S_50ntFlanks</td><td>10349</td><td>10373</td><td>25</td><td>AACGAGAACTTTGAAGGCCGAAGTG</td><td>1051.3</td><td>3.3</td><td>0.8</td><td>4.4</td><td>9.3E-06</td><td>1.4E-05</td><td>MB231/MB468</td></tr>
<tr><td>>hg38_21_+_8433172_8446622_RNA45SN1_45S_50ntFlanks@10840.10864.25|CCGTAACTTCGGGATAAGGATTGGC</td><td>>hg38_21_+_8433172_8446622_RNA45SN1_45S_50ntFlanks</td><td>10840</td><td>10864</td><td>25</td><td>CCGTAACTTCGGGATAAGGATTGGC</td><td>80.8</td><td>-2.6</td><td>0.6</td><td>-4.4</td><td>9.4E-06</td><td>1.4E-05</td><td>MB231/MB468</td></tr>
<tr><td>>hg38_21_+_8433172_8446622_RNA45SN1_45S_50ntFlanks@4770.4793.24|TCGTAGTTCCGACCATAAACGATG</td><td>>hg38_21_+_8433172_8446622_RNA45SN1_45S_50ntFlanks</td><td>4770</td><td>4793</td><td>24</td><td>TCGTAGTTCCGACCATAAACGATG</td><td>207.1</td><td>-2.9</td><td>0.7</td><td>-4.4</td><td>1.0E-05</td><td>1.4E-05</td><td>BT20/MB468</td></tr>
<tr><td>>hg38_21_+_8433172_8446622_RNA45SN1_45S_50ntFlanks@12357.12375.19|CCACAGGGATAACTGGCTT</td><td>>hg38_21_+_8433172_8446622_RNA45SN1_45S_50ntFlanks</td><td>12357</td><td>12375</td><td>19</td><td>CCACAGGGATAACTGGCTT</td><td>510.5</td><td>-5.3</td><td>1.2</td><td>-4.4</td><td>1.0E-05</td><td>1.4E-05</td><td>BT20/MB468</td></tr>
<tr><td>>hg38_21_+_8433172_8446622_RNA45SN1_45S_50ntFlanks@12071.12094.24|CCCCGAGGGGCTCTCGCTTCTGGC</td><td>>hg38_21_+_8433172_8446622_RNA45SN1_45S_50ntFlanks</td><td>12071</td><td>12094</td><td>24</td><td>CCCCGAGGGGCTCTCGCTTCTGGC</td><td>220.6</td><td>-3.2</td><td>0.7</td><td>-4.4</td><td>1.0E-05</td><td>1.4E-05</td><td>BT20/MB468</td></tr>
<tr><td>>hg38_21_+_8433172_8446622_RNA45SN1_45S_50ntFlanks@12957.12986.30|CTTCTGGGTCGGGGTTTCGTACGTAGCAGA</td><td>>hg38_21_+_8433172_8446622_RNA45SN1_45S_50ntFlanks</td><td>12957</td><td>12986</td><td>30</td><td>CTTCTGGGTCGGGGTTTCGTACGTAGCAGA</td><td>156.6</td><td>-3.2</td><td>0.7</td><td>-4.4</td><td>1.1E-05</td><td>1.4E-05</td><td>BT20/MB468</td></tr>
<tr><td>>hg38_21_+_8433172_8446622_RNA45SN1_45S_50ntFlanks@11572.11593.22|CCGACTTAGAACTGGTGCGGAC</td><td>>hg38_21_+_8433172_8446622_RNA45SN1_45S_50ntFlanks</td><td>11572</td><td>11593</td><td>22</td><td>CCGACTTAGAACTGGTGCGGAC</td><td>556.2</td><td>-3.7</td><td>0.8</td><td>-4.4</td><td>1.1E-05</td><td>1.4E-05</td><td>BT20/MB468</td></tr>
<tr><td>>hg38_21_+_8433172_8446622_RNA45SN1_45S_50ntFlanks@7979.8007.29|ACCTCAGATCAGACGTGGCGACCCGCTGA</td><td>>hg38_21_+_8433172_8446622_RNA45SN1_45S_50ntFlanks</td><td>7979</td><td>8007</td><td>29</td><td>ACCTCAGATCAGACGTGGCGACCCGCTGA</td><td>162.9</td><td>-2.8</td><td>0.6</td><td>-4.4</td><td>9.6E-06</td><td>1.4E-05</td><td>MB231/MB468</td></tr>
</table>

<table>
<tr><th></th><th></th><th></th><th></th><th></th><th></th><th></th><th></th><th></th><th></th><th></th><th></th><th></th></tr>
<tr><td>>hg38_1_-_228634819_228635039_RNA5S12_5S_50ntFlanks@126.171.46|TACTTGGATGGGAGACCGCCTGGGAATACCGGGTGCTGTAGGCTTT</td><td>>hg38_1_-_228634819_228635039_RNA5S12_5S_50ntFlanks</td><td>126</td><td>171</td><td>46</td><td>TACTTGGATGGGAGACCGCCTGGGAATACCGGGTGCTGTAGGCTTT</td><td>2025.5</td><td>9.8</td><td>2.2</td><td>4.4</td><td>9.6E-06</td><td>1.4E-05</td><td>MB231/MB468</td></tr>
<tr><td>>hg38_21_+_8433172_8446622_RNA45SN1_45S_50ntFlanks@9306.9338.33|CAAGGAGTCTAACACGTGCGCGAGTCGGGGGCT</td><td>>hg38_21_+_8433172_8446622_RNA45SN1_45S_50ntFlanks</td><td>9306</td><td>9338</td><td>33</td><td>CAAGGAGTCTAACACGTGCGCGAGTCGGGGGCT</td><td>121.1</td><td>-3.7</td><td>0.8</td><td>-4.4</td><td>9.7E-06</td><td>1.4E-05</td><td>MB231/MB468</td></tr>
<tr><td>>hg38_21_+_8433172_8446622_RNA45SN1_45S_50ntFlanks@12406.12431.26|TTTTTGATCCTTCGATGTCGGCTCTT</td><td>>hg38_21_+_8433172_8446622_RNA45SN1_45S_50ntFlanks</td><td>12406</td><td>12431</td><td>26</td><td>TTTTTGATCCTTCGATGTCGGCTCTT</td><td>246.9</td><td>-2.4</td><td>0.5</td><td>-4.4</td><td>9.7E-06</td><td>1.4E-05</td><td>MB231/MB468</td></tr>
<tr><td>>hg38_21_+_8433172_8446622_RNA45SN1_45S_50ntFlanks@7980.7999.20|CCTCAGATCAGACGTGGCGA</td><td>>hg38_21_+_8433172_8446622_RNA45SN1_45S_50ntFlanks</td><td>7980</td><td>7999</td><td>20</td><td>CCTCAGATCAGACGTGGCGA</td><td>87.1</td><td>-4.2</td><td>1.0</td><td>-4.4</td><td>1.1E-05</td><td>1.4E-05</td><td>BT20/MB468</td></tr>
<tr><td>>hg38_21_+_8433172_8446622_RNA45SN1_45S_50ntFlanks@12073.12097.25|CCGAGGGGCTCTCGCTTCTGGCGCC</td><td>>hg38_21_+_8433172_8446622_RNA45SN1_45S_50ntFlanks</td><td>12073</td><td>12097</td><td>25</td><td>CCGAGGGGCTCTCGCTTCTGGCGCC</td><td>1340.0</td><td>-4.3</td><td>1.0</td><td>-4.4</td><td>1.1E-05</td><td>1.4E-05</td><td>BT20/MB468</td></tr>
<tr><td>>hg38_1_-_228634819_228635039_RNA5S12_5S_50ntFlanks@128.170.43|CTTGGATGGGAGACCGCCTGGGAATACCGGGTGCTGTAGGCTT</td><td>>hg38_1_-_228634819_228635039_RNA5S12_5S_50ntFlanks</td><td>128</td><td>170</td><td>43</td><td>CTTGGATGGGAGACCGCCTGGGAATACCGGGTGCTGTAGGCTT</td><td>1149.8</td><td>10.7</td><td>2.4</td><td>4.4</td><td>1.1E-05</td><td>1.5E-05</td><td>BT20/MB468</td></tr>
<tr><td>>hg38_21_+_8433172_8446622_RNA45SN1_45S_50ntFlanks@8381.8404.24|AAGAGGGCGTGAAACCGTTAAGAG</td><td>>hg38_21_+_8433172_8446622_RNA45SN1_45S_50ntFlanks</td><td>8381</td><td>8404</td><td>24</td><td>AAGAGGGCGTGAAACCGTTAAGAG</td><td>154.0</td><td>-3.0</td><td>0.7</td><td>-4.4</td><td>1.0E-05</td><td>1.5E-05</td><td>MB231/MB468</td></tr>
<tr><td>>am_trnaMT_ValTAC_MT_+_1602_1670@50.67.18|ACTTAACTTGACCGCTCT</td><td>>am_trnaMT_ValTAC_MT_+_1602_1670</td><td>50</td><td>67</td><td>18</td><td>ACTTAACTTGACCGCTCT</td><td>1911.6</td><td>4.0</td><td>0.9</td><td>4.4</td><td>1.0E-05</td><td>1.5E-05</td><td>MB231/MB468</td></tr>
<tr><td>>hg38_21_+_8433172_8446622_RNA45SN1_45S_50ntFlanks@12397.12417.21|CGACGTCGCTTTTTGATCCTT</td><td>>hg38_21_+_8433172_8446622_RNA45SN1_45S_50ntFlanks</td><td>12397</td><td>12417</td><td>21</td><td>CGACGTCGCTTTTTGATCCTT</td><td>583.5</td><td>-3.1</td><td>0.7</td><td>-4.4</td><td>1.1E-05</td><td>1.5E-05</td><td>BT20/MB468</td></tr>
<tr><td>>hg38_21_+_8433172_8446622_RNA45SN1_45S_50ntFlanks@11873.11903.31|AAGAAGACCCTGTTGAGCTTGACTCTAGTCT</td><td>>hg38_21_+_8433172_8446622_RNA45SN1_45S_50ntFlanks</td><td>11873</td><td>11903</td><td>31</td><td>AAGAAGACCCTGTTGAGCTTGACTCTAGTCT</td><td>308.1</td><td>-3.5</td><td>0.8</td><td>-4.4</td><td>1.1E-05</td><td>1.5E-05</td><td>BT20/MB468</td></tr>
<tr><td>>hg38_21_+_8433172_8446622_RNA45SN1_45S_50ntFlanks@9740.9758.19|AACGATCTCAACCTATTCT</td><td>>hg38_21_+_8433172_8446622_RNA45SN1_45S_50ntFlanks</td><td>9740</td><td>9758</td><td>19</td><td>AACGATCTCAACCTATTCT</td><td>271.7</td><td>-1.8</td><td>0.4</td><td>-4.4</td><td>1.0E-05</td><td>1.5E-05</td><td>MB231/MB468</td></tr>
<tr><td>>hg38_21_+_8433172_8446622_RNA45SN1_45S_50ntFlanks@8194.8213.20|CCCGTGGACGGTGTGAGGCC</td><td>>hg38_21_+_8433172_8446622_RNA45SN1_45S_50ntFlanks</td><td>8194</td><td>8213</td><td>20</td><td>CCCGTGGACGGTGTGAGGCC</td><td>774.5</td><td>-4.0</td><td>0.9</td><td>-4.4</td><td>1.1E-05</td><td>1.5E-05</td><td>BT20/MB468</td></tr>
<tr><td>>hg38_21_+_8433172_8446622_RNA45SN1_45S_50ntFlanks@10797.10818.22|GCATGTTGGAACAATGTAGGTA</td><td>>hg38_21_+_8433172_8446622_RNA45SN1_45S_50ntFlanks</td><td>10797</td><td>10818</td><td>22</td><td>GCATGTTGGAACAATGTAGGTA</td><td>2904.6</td><td>-3.3</td><td>0.7</td><td>-4.4</td><td>1.0E-05</td><td>1.5E-05</td><td>MB231/MB468</td></tr>
<tr><td>>hg38_21_+_8433172_8446622_RNA45SN1_45S_50ntFlanks@12489.12518.30|ACGTGAGCTGGGTTTAGACCGTCGTGAGAC</td><td>>hg38_21_+_8433172_8446622_RNA45SN1_45S_50ntFlanks</td><td>12489</td><td>12518</td><td>30</td><td>ACGTGAGCTGGGTTTAGACCGTCGTGAGAC</td><td>89.1</td><td>-2.2</td><td>0.5</td><td>-4.4</td><td>1.0E-05</td><td>1.5E-05</td><td>MB231/MB468</td></tr>
<tr><td>>hg38_21_+_8433172_8446622_RNA45SN1_45S_50ntFlanks@10819.10837.19|AGGGAAGTCGGCAAGCCGG</td><td>>hg38_21_+_8433172_8446622_RNA45SN1_45S_50ntFlanks</td><td>10819</td><td>10837</td><td>19</td><td>AGGGAAGTCGGCAAGCCGG</td><td>253.5</td><td>-2.3</td><td>0.5</td><td>-4.4</td><td>1.0E-05</td><td>1.5E-05</td><td>MB231/MB468</td></tr>
<tr><td>>MI0000268|hsa-mir-34a&WithFlank&1|-|9151662|9151783@28.50.23&[MIMAT0000255&hsa-miR-34a-5p&offsets|0|+1;m-119&1|-|9151735|9151756&offsets|0|+1]|TGGCAGTGTCTTAGCTGGTTGTT</td><td>>MI0000268|hsa-mir-34a&WithFlank&1|-|9151662|9151783</td><td>28</td><td>50</td><td>23</td><td>TGGCAGTGTCTTAGCTGGTTGTT</td><td>430.4</td><td>4.4</td><td>0.9</td><td>4.9</td><td>9.8E-07</td><td>1.5E-05</td><td>BT20/MB231</td></tr>
<tr><td>>hg38_21_+_8433172_8446622_RNA45SN1_45S_50ntFlanks@12912.12933.22|CACGTTCGTGGGGAACCTGGCG</td><td>>hg38_21_+_8433172_8446622_RNA45SN1_45S_50ntFlanks</td><td>12912</td><td>12933</td><td>22</td><td>CACGTTCGTGGGGAACCTGGCG</td><td>411.8</td><td>-4.0</td><td>0.9</td><td>-4.4</td><td>1.0E-05</td><td>1.5E-05</td><td>MB231/MB468</td></tr>
<tr><td>>hg38_21_+_8433172_8446622_RNA45SN1_45S_50ntFlanks@5487.5509.23|CGCTGAGAAGACGGTCGAACTTG</td><td>>hg38_21_+_8433172_8446622_RNA45SN1_45S_50ntFlanks</td><td>5487</td><td>5509</td><td>23</td><td>CGCTGAGAAGACGGTCGAACTTG</td><td>88.4</td><td>-2.6</td><td>0.6</td><td>-4.4</td><td>1.1E-05</td><td>1.5E-05</td><td>MB231/MB468</td></tr>
<tr><td>>hg38_21_+_8433172_8446622_RNA45SN1_45S_50ntFlanks@4297.4317.21|CATTGGAGGGCAAGTCTGGTG</td><td>>hg38_21_+_8433172_8446622_RNA45SN1_45S_50ntFlanks</td><td>4297</td><td>4317</td><td>21</td><td>CATTGGAGGGCAAGTCTGGTG</td><td>358.1</td><td>-3.9</td><td>0.9</td><td>-4.4</td><td>1.1E-05</td><td>1.5E-05</td><td>MB231/MB468</td></tr>
<tr><td>>trna111_HisGTG_1_-_147774845_147774916@1.31.31,trna118_HisGTG_1_-_145396881_145396952@1.31.31,trna16_HisGTG_1_+_146544773_146544844@1.31.31,trna1_HisGTG_15_+_45493349_45493420@1.31.31,trna21_HisGTG_1_+_147753471_147753542@1.31.31,trna33_HisGTG_6_+_27125906_27125977@1.31.31,trna7_HisGTG_9_-_14433938_14434009@1.31.31,trna8_HisGTG_15_-_45492611_45492682@1.31.31,trna9_HisGTG_15_-_45490804_45490875@1.31.31|GCCGTGATCGTATAGTGGTTAGTACTCTGCG</td><td>>trna111_HisGTG_1_-_147774845_147774916</td><td>1</td><td>31</td><td>31</td><td>GCCGTGATCGTATAGTGGTTAGTACTCTGCG</td><td>8167.5</td><td>-2.4</td><td>0.6</td><td>-4.4</td><td>1.2E-05</td><td>1.5E-05</td><td>BT20/MB468</td></tr>
<tr><td>>hg38_21_+_8433172_8446622_RNA45SN1_45S_50ntFlanks@3838.3860.23|CTCTCCTACTTGGATAACTGTGG</td><td>>hg38_21_+_8433172_8446622_RNA45SN1_45S_50ntFlanks</td><td>3838</td><td>3860</td><td>23</td><td>CTCTCCTACTTGGATAACTGTGG</td><td>546.4</td><td>-2.4</td><td>0.5</td><td>-4.4</td><td>1.1E-05</td><td>1.5E-05</td><td>MB231/MB468</td></tr>
<tr><td>>hg38_21_+_8433172_8446622_RNA45SN1_45S_50ntFlanks@10829.10855.27|GCAAGCCGGATCCGTAACTTCGGGATA</td><td>>hg38_21_+_8433172_8446622_RNA45SN1_45S_50ntFlanks</td><td>10829</td><td>10855</td><td>27</td><td>GCAAGCCGGATCCGTAACTTCGGGATA</td><td>150.7</td><td>-2.5</td><td>0.6</td><td>-4.4</td><td>1.1E-05</td><td>1.6E-05</td><td>MB231/MB468</td></tr>
<tr><td>>hg38_21_+_8433172_8446622_RNA45SN1_45S_50ntFlanks@5515.5534.20|CTAGAGGAAGTAAAAGTCGT</td><td>>hg38_21_+_8433172_8446622_RNA45SN1_45S_50ntFlanks</td><td>5515</td><td>5534</td><td>20</td><td>CTAGAGGAAGTAAAAGTCGT</td><td>102.4</td><td>-3.7</td><td>0.9</td><td>-4.4</td><td>1.2E-05</td><td>1.6E-05</td><td>BT20/MB468</td></tr>
<tr><td>>hg38_21_+_8433172_8446622_RNA45SN1_45S_50ntFlanks@4005.4042.38|AACCTCGGGCCGATCGCACGCCCCCCGTGGCGGCGACG</td><td>>hg38_21_+_8433172_8446622_RNA45SN1_45S_50ntFlanks</td><td>4005</td><td>4042</td><td>38</td><td>AACCTCGGGCCGATCGCACGCCCCCCGTGGCGGCGACG</td><td>160.5</td><td>-2.5</td><td>0.6</td><td>-4.4</td><td>1.1E-05</td><td>1.6E-05</td><td>MB231/MB468</td></tr>
<tr><td>>hg38_21_+_8433172_8446622_RNA45SN1_45S_50ntFlanks@8196.8223.28|CGTGGACGGTGTGAGGCCGGTAGCGGCC</td><td>>hg38_21_+_8433172_8446622_RNA45SN1_45S_50ntFlanks</td><td>8196</td><td>8223</td><td>28</td><td>CGTGGACGGTGTGAGGCCGGTAGCGGCC</td><td>399.7</td><td>-2.3</td><td>0.5</td><td>-4.4</td><td>1.2E-05</td><td>1.6E-05</td><td>BT20/MB468</td></tr>
<tr><td>>hg38_21_+_8433172_8446622_RNA45SN1_45S_50ntFlanks@11581.11609.29|AACTGGTGCGGACCAGGGGAATCCGACTG</td><td>>hg38_21_+_8433172_8446622_RNA45SN1_45S_50ntFlanks</td><td>11581</td><td>11609</td><td>29</td><td>AACTGGTGCGGACCAGGGGAATCCGACTG</td><td>1683.7</td><td>-2.6</td><td>0.6</td><td>-4.4</td><td>1.2E-05</td><td>1.6E-05</td><td>BT20/MB468</td></tr>
<tr><td>>hg38_21_+_8433172_8446622_RNA45SN1_45S_50ntFlanks@12157.12178.22|AGTTTGACTGGGGCGGTACACC</td><td>>hg38_21_+_8433172_8446622_RNA45SN1_45S_50ntFlanks</td><td>12157</td><td>12178</td><td>22</td><td>AGTTTGACTGGGGCGGTACACC</td><td>74.7</td><td>-3.4</td><td>0.8</td><td>-4.4</td><td>1.1E-05</td><td>1.6E-05</td><td>MB231/MB468</td></tr>
<tr><td>>hg38_21_+_8433172_8446622_RNA45SN1_45S_50ntFlanks@11573.11603.31|CGACTTAGAACTGGTGCGGACCAGGGGAATC</td><td>>hg38_21_+_8433172_8446622_RNA45SN1_45S_50ntFlanks</td><td>11573</td><td>11603</td><td>31</td><td>CGACTTAGAACTGGTGCGGACCAGGGGAATC</td><td>421.5</td><td>-2.4</td><td>0.5</td><td>-4.4</td><td>1.1E-05</td><td>1.6E-05</td><td>MB231/MB468</td></tr>
<tr><td>>hg38_21_+_8433172_8446622_RNA45SN1_45S_50ntFlanks@5515.5533.19|CTAGAGGAAGTAAAAGTCG</td><td>>hg38_21_+_8433172_8446622_RNA45SN1_45S_50ntFlanks</td><td>5515</td><td>5533</td><td>19</td><td>CTAGAGGAAGTAAAAGTCG</td><td>94.7</td><td>-2.6</td><td>0.6</td><td>-4.4</td><td>1.1E-05</td><td>1.6E-05</td><td>MB231/MB468</td></tr>
<tr><td>>hg38_21_+_8433172_8446622_RNA45SN1_45S_50ntFlanks@12418.12449.32|CGATGTCGGCTCTTCCTATCATTGTGAAGCAG</td><td>>hg38_21_+_8433172_8446622_RNA45SN1_45S_50ntFlanks</td><td>12418</td><td>12449</td><td>32</td><td>CGATGTCGGCTCTTCCTATCATTGTGAAGCAG</td><td>810.9</td><td>2.6</td><td>0.6</td><td>4.4</td><td>1.1E-05</td><td>1.6E-05</td><td>MB231/MB468</td></tr>
<tr><td>>hg38_21_+_8433172_8446622_RNA45SN1_45S_50ntFlanks@6776.6803.28|CTCCCGGGGCTACGCCTGTCTGAGCGTC</td><td>>hg38_21_+_8433172_8446622_RNA45SN1_45S_50ntFlanks</td><td>6776</td><td>6803</td><td>28</td><td>CTCCCGGGGCTACGCCTGTCTGAGCGTC</td><td>672.9</td><td>-2.9</td><td>0.7</td><td>-4.4</td><td>1.2E-05</td><td>1.6E-05</td><td>BT20/MB468</td></tr>
<tr><td>>hg38_21_+_8433172_8446622_RNA45SN1_45S_50ntFlanks@3793.3814.22|CGAATGGCTCATTAAATCAGTT</td><td>>hg38_21_+_8433172_8446622_RNA45SN1_45S_50ntFlanks</td><td>3793</td><td>3814</td><td>22</td><td>CGAATGGCTCATTAAATCAGTT</td><td>93.1</td><td>-3.9</td><td>0.9</td><td>-4.4</td><td>1.2E-05</td><td>1.6E-05</td><td>BT20/MB468</td></tr>
<tr><td>>MI0000289|hsa-mir-181a-1&WithFlank&1|-|198859038|198859159@70.91.22&[MIMAT0000270&hsa-miR-181a-3p&offsets|0|0;m-125&1|-|198859069|198859090&offsets|0|0]|ACCATCGACCGTTGATTGTACC</td><td>>MI0000289|hsa-mir-181a-1&WithFlank&1|-|198859038|198859159</td><td>70</td><td>91</td><td>22</td><td>ACCATCGACCGTTGATTGTACC</td><td>1494.1</td><td>-9.5</td><td>2.0</td><td>-4.9</td><td>1.1E-06</td><td>1.6E-05</td><td>BT20/MB231</td></tr>
<tr><td>>hg38_21_+_8433172_8446622_RNA45SN1_45S_50ntFlanks@11873.11902.30|AAGAAGACCCTGTTGAGCTTGACTCTAGTC</td><td>>hg38_21_+_8433172_8446622_RNA45SN1_45S_50ntFlanks</td><td>11873</td><td>11902</td><td>30</td><td>AAGAAGACCCTGTTGAGCTTGACTCTAGTC</td><td>215.0</td><td>-2.4</td><td>0.5</td><td>-4.4</td><td>1.1E-05</td><td>1.6E-05</td><td>MB231/MB468</td></tr>
<tr><td>>hg38_21_+_8433172_8446622_RNA45SN1_45S_50ntFlanks@12479.12509.31|CTAATAGGGAACGTGAGCTGGGTTTAGACCG</td><td>>hg38_21_+_8433172_8446622_RNA45SN1_45S_50ntFlanks</td><td>12479</td><td>12509</td><td>31</td><td>CTAATAGGGAACGTGAGCTGGGTTTAGACCG</td><td>128.1</td><td>-4.0</td><td>0.9</td><td>-4.4</td><td>1.1E-05</td><td>1.6E-05</td><td>MB231/MB468</td></tr>
<tr><td>>hg38_21_+_8433172_8446622_RNA45SN1_45S_50ntFlanks@11208.11238.31|GCGGCGACTCTGGACGCGAGCCGGGCCCTTC</td><td>>hg38_21_+_8433172_8446622_RNA45SN1_45S_50ntFlanks</td><td>11208</td><td>11238</td><td>31</td><td>GCGGCGACTCTGGACGCGAGCCGGGCCCTTC</td><td>83.5</td><td>-4.4</td><td>1.0</td><td>-4.4</td><td>1.1E-05</td><td>1.6E-05</td><td>MB231/MB468</td></tr>
<tr><td>>hg38_21_+_8433172_8446622_RNA45SN1_45S_50ntFlanks@11452.11471.20|CTCCTCGGCGCGCGGCGGCG</td><td>>hg38_21_+_8433172_8446622_RNA45SN1_45S_50ntFlanks</td><td>11452</td><td>11471</td><td>20</td><td>CTCCTCGGCGCGCGGCGGCG</td><td>97.0</td><td>-4.0</td><td>0.9</td><td>-4.4</td><td>1.2E-05</td><td>1.6E-05</td><td>BT20/MB468</td></tr>
<tr><td>>hg38_21_+_8433172_8446622_RNA45SN1_45S_50ntFlanks@3846.3869.24|CTTGGATAACTGTGGTAATTCTAG</td><td>>hg38_21_+_8433172_8446622_RNA45SN1_45S_50ntFlanks</td><td>3846</td><td>3869</td><td>24</td><td>CTTGGATAACTGTGGTAATTCTAG</td><td>274.9</td><td>-2.6</td><td>0.6</td><td>-4.4</td><td>1.1E-05</td><td>1.6E-05</td><td>MB231/MB468</td></tr>
<tr><td>>hg38_21_+_8433172_8446622_RNA45SN1_45S_50ntFlanks@6649.6681.33|TACGACTCTTAGCGGTGGATCACTCGGCTCGTG</td><td>>hg38_21_+_8433172_8446622_RNA45SN1_45S_50ntFlanks</td><td>6649</td><td>6681</td><td>33</td><td>TACGACTCTTAGCGGTGGATCACTCGGCTCGTG</td><td>652.5</td><td>-1.8</td><td>0.4</td><td>-4.4</td><td>1.2E-05</td><td>1.6E-05</td><td>BT20/MB468</td></tr>
<tr><td>>hg38_21_+_8433172_8446622_RNA45SN1_45S_50ntFlanks@10869.10886.18|AGGGCTGGGTCGGTCGGG</td><td>>hg38_21_+_8433172_8446622_RNA45SN1_45S_50ntFlanks</td><td>10869</td><td>10886</td><td>18</td><td>AGGGCTGGGTCGGTCGGG</td><td>223.5</td><td>-5.0</td><td>1.1</td><td>-4.4</td><td>1.2E-05</td><td>1.6E-05</td><td>BT20/MB468</td></tr>
<tr><td>>hg38_21_+_8433172_8446622_RNA45SN1_45S_50ntFlanks@11661.11680.20|TGATTTCTGCCCAGTGCTCT</td><td>>hg38_21_+_8433172_8446622_RNA45SN1_45S_50ntFlanks</td><td>11661</td><td>11680</td><td>20</td><td>TGATTTCTGCCCAGTGCTCT</td><td>589.1</td><td>-3.5</td><td>0.8</td><td>-4.4</td><td>1.3E-05</td><td>1.6E-05</td><td>BT20/MB468</td></tr>
<tr><td>>hg38_21_+_8433172_8446622_RNA45SN1_45S_50ntFlanks@12073.12092.20|CCGAGGGGCTCTCGCTTCTG</td><td>>hg38_21_+_8433172_8446622_RNA45SN1_45S_50ntFlanks</td><td>12073</td><td>12092</td><td>20</td><td>CCGAGGGGCTCTCGCTTCTG</td><td>305.1</td><td>-4.8</td><td>1.1</td><td>-4.4</td><td>1.3E-05</td><td>1.7E-05</td><td>BT20/MB468</td></tr>
<tr><td>>hg38_21_+_8433172_8446622_RNA45SN1_45S_50ntFlanks@7981.8005.25|CTCAGATCAGACGTGGCGACCCGCT</td><td>>hg38_21_+_8433172_8446622_RNA45SN1_45S_50ntFlanks</td><td>7981</td><td>8005</td><td>25</td><td>CTCAGATCAGACGTGGCGACCCGCT</td><td>135.4</td><td>-2.5</td><td>0.6</td><td>-4.4</td><td>1.1E-05</td><td>1.7E-05</td><td>MB231/MB468</td></tr>
<tr><td>>hg38_21_+_8433172_8446622_RNA45SN1_45S_50ntFlanks@11562.11585.24|CGCCTAGCAGCCGACTTAGAACTG</td><td>>hg38_21_+_8433172_8446622_RNA45SN1_45S_50ntFlanks</td><td>11562</td><td>11585</td><td>24</td><td>CGCCTAGCAGCCGACTTAGAACTG</td><td>92.9</td><td>-2.6</td><td>0.6</td><td>-4.4</td><td>1.1E-05</td><td>1.7E-05</td><td>MB231/MB468</td></tr>
</table>

| | | | | | | | | | | | | |
|---|---|---|---|---|---|---|---|---|---|---|---|---|
| >am_trna116_GluCTC_1_-_145399233_145399304@1.28.28,trna59_GluCTC_1_+_249168447_249168518@1.28.28,trna71_GluCTC_1_-_161439189_161439260@1.28.28,trna74_GluCTC_1_-_161431809_161431880@1.28.28,trna77_GluCTC_1_-_161424398_161424469@1.28.28,trna77_GluCTC_6_+_28949976_28950047@1.28.28,trna80_GluCTC_1_-_161417018_161417089@1.28.28,trna87_GluCTC_6_-_126101393_126101464@1.28.28\|TCCCTGGTGGTCTAGTGGTTAGGATTCG | >am_trna116_GluCTC_1_-_145399233_145399304 | 1 | 28 | 28 | TCCCTGGTGGTCTAGTGGTTAGGATTCG | 16611.1 | -3.2 | 0.7 | -4.4 | 1.2E-05 | 1.7E-05 | MB231/MB468 |
| >MI0000736\|hsa-mir-30c-1&WithFlank&1\|+\|40757278\|40757378@23.46.24(+2U);MI0000254\|hsa-mir-30c-2&WithFlank&6\|-\|71376954\|71377037@13.36.24(+2U)&[MIMAT0000244&hsa-miR-30c-5p&offsets\|0\|+1(+2U);m-57&1\|+\|40757300\|40757323&offsets\|0\|0(+2U)];[MIMAT0000244_1&hsa-miR-30c-5p&offsets\|0\|+1(+2U);m-56&6\|-\|71377002\|71377025&offsets\|0\|0(+2U)]\|TGTAAACATCCTACACTCTCAGCTTT | >MI0000736\|hsa-mir-30c-1&WithFlank&1\|+\|40757278\|40757378 | 23 | 46 | 24(+2U) | TGTAAACATCCTACACTCTCAGCTTT | 375.5 | -3.6 | 0.7 | -4.9 | 1.1E-06 | 1.7E-05 | BT20/MB231 |
| >hg38_21_+_8433172_8446622_RNA45SN1_45S_50ntFlanks@12994.13013.20\|CCTCGCTGCGATCTATTGAA | >hg38_21_+_8433172_8446622_RNA45SN1_45S_50ntFlanks | 12994 | 13013 | 20 | CCTCGCTGCGATCTATTGAA | 647.6 | -2.3 | 0.5 | -4.4 | 1.2E-05 | 1.7E-05 | MB231/MB468 |
| >hg38_21_+_8433172_8446622_RNA45SN1_45S_50ntFlanks@12992.13012.21\|TCCCTCGCTGCGATCTATTGA | >hg38_21_+_8433172_8446622_RNA45SN1_45S_50ntFlanks | 12992 | 13012 | 21 | TCCCTCGCTGCGATCTATTGA | 279.3 | -3.8 | 0.9 | -4.4 | 1.3E-05 | 1.7E-05 | BT20/MB468 |
| >hg38_21_+_8433172_8446622_RNA45SN1_45S_50ntFlanks@12997.13015.19\|CGCTGCGATCTATTGAAAG | >hg38_21_+_8433172_8446622_RNA45SN1_45S_50ntFlanks | 12997 | 13015 | 19 | CGCTGCGATCTATTGAAAG | 1032.0 | 3.4 | 0.8 | 4.4 | 1.3E-05 | 1.7E-05 | BT20/MB468 |
| >MI0000466\|hsa-mir-9-1&WithFlank&1\|-\|156420335\|156420435@22.44.23;MI0000468\|hsa-mir-9-3&WithFlank&15\|+\|89368011\|89368112@22.44.23;MI0000467\|hsa-mir-9-2&WithFlank&5\|-\|88666847\|88666945@22.44.23&[MIMAT0000441&hsa-miR-9-5p&offsets\|0\|0;m-104&1\|-\|156420392\|156420414&offsets\|0\|0];[MIMAT0000441_1&hsa-miR-9-5p&offsets\|0\|0;m-105&15\|+\|89368032\|89368054&offsets\|0\|0];[MIMAT0000441_2&hsa-miR-9-5p&offsets\|0\|0;m-104&5\|-\|88666902\|88666924&offsets\|0\|0]\|TCTTTGGTTATCTAGCTGTATGA | >MI0000466\|hsa-mir-9-1&WithFlank&1\|-\|156420335\|156420435 | 22 | 44 | 23 | TCTTTGGTTATCTAGCTGTATGA | 3913.8 | 3.4 | 0.7 | 4.9 | 1.1E-06 | 1.7E-05 | BT20/MB231 |
| >hg38_21_+_8433172_8446622_RNA45SN1_45S_50ntFlanks@6663.6686.24\|GTGGATCACTCGGCTCGTGCGTCG | >hg38_21_+_8433172_8446622_RNA45SN1_45S_50ntFlanks | 6663 | 6686 | 24 | GTGGATCACTCGGCTCGTGCGTCG | 337.2 | -2.6 | 0.6 | -4.4 | 1.2E-05 | 1.7E-05 | MB231/MB468 |
| >hg38_21_+_8433172_8446622_RNA45SN1_45S_50ntFlanks@11872.11893.22\|AAAGAAGACCCTGTTGAGCTTG | >hg38_21_+_8433172_8446622_RNA45SN1_45S_50ntFlanks | 11872 | 11893 | 22 | AAAGAAGACCCTGTTGAGCTTG | 253.3 | -2.5 | 0.6 | -4.4 | 1.3E-05 | 1.7E-05 | BT20/MB468 |
| >hg38_21_+_8433172_8446622_RNA45SN1_45S_50ntFlanks@9552.9570.19\|TCCGTAGCGGTCCTGACGT | >hg38_21_+_8433172_8446622_RNA45SN1_45S_50ntFlanks | 9552 | 9570 | 19 | TCCGTAGCGGTCCTGACGT | 289.0 | 2.0 | 0.5 | 4.4 | 1.2E-05 | 1.7E-05 | MB231/MB468 |
| >trnaMT_ProTGG_MT_-_15956_16023@41.67.27\|ATGGTGGAGTTAAAGACTTTTTCTCTG | >trnaMT_ProTGG_MT_-_15956_16023 | 41 | 67 | 27 | ATGGTGGAGTTAAAGACTTTTTCTCTG | 84.2 | -4.2 | 1.0 | -4.4 | 1.2E-05 | 1.7E-05 | MB231/MB468 |
| >MI0000767\|hsa-mir-365a&WithFlank&16\|+\|14309279\|14309377@62.83.22;MI0000769\|hsa-mir-365b&WithFlank&17\|+\|31575405\|31575527@74.95.22&[MIMAT0000710&hsa-miR-365a-3p&offsets\|0\|0;m-176&16\|+\|14309340\|14309361&offsets\|0\|0];[MIMAT0022834&hsa-miR-365b-3p&offsets\|0\|0;m-175&17\|+\|31575478\|31575499&offsets\|0\|0]\|TAATGCCCCTAAAAATCCTTAT | >MI0000767\|hsa-mir-365a&WithFlank&16\|+\|14309279\|14309377 | 62 | 83 | 22 | TAATGCCCCTAAAAATCCTTAT | 2688.1 | 4.0 | 0.8 | 4.9 | 1.2E-06 | 1.7E-05 | BT20/MB231 |
| >hg38_21_+_8433172_8446622_RNA45SN1_45S_50ntFlanks@4591.4616.26\|TTTTGTTGGTTTTCGGAACTGAGGCC | >hg38_21_+_8433172_8446622_RNA45SN1_45S_50ntFlanks | 4591 | 4616 | 26 | TTTTGTTGGTTTTCGGAACTGAGGCC | 164.1 | -2.3 | 0.5 | -4.4 | 1.3E-05 | 1.7E-05 | BT20/MB468 |
| >hg38_21_+_8433172_8446622_RNA45SN1_45S_50ntFlanks@3931.3959.29\|TCAAAACCAACCCGGTCAGCCCCTCTCCG | >hg38_21_+_8433172_8446622_RNA45SN1_45S_50ntFlanks | 3931 | 3959 | 29 | TCAAAACCAACCCGGTCAGCCCCTCTCCG | 121.3 | -4.1 | 0.9 | -4.4 | 1.3E-05 | 1.7E-05 | BT20/MB468 |
| >hg38_21_+_8433172_8446622_RNA45SN1_45S_50ntFlanks@12934.12960.27\|CTAAACCATTCGTAGACGACCTGCTTC | >hg38_21_+_8433172_8446622_RNA45SN1_45S_50ntFlanks | 12934 | 12960 | 27 | CTAAACCATTCGTAGACGACCTGCTTC | 327.8 | -2.3 | 0.5 | -4.4 | 1.2E-05 | 1.8E-05 | MB231/MB468 |
| >hg38_21_+_8433172_8446622_RNA45SN1_45S_50ntFlanks@11027.11051.25\|CTCTCTCTCTCTCTCCCCCGCTCCC | >hg38_21_+_8433172_8446622_RNA45SN1_45S_50ntFlanks | 11027 | 11051 | 25 | CTCTCTCTCTCTCTCCCCCGCTCCC | 557.1 | -5.3 | 1.2 | -4.4 | 1.3E-05 | 1.8E-05 | BT20/MB468 |
| >hg38_21_+_8433172_8446622_RNA45SN1_45S_50ntFlanks@9552.9569.18\|TCCGTAGCGGTCCTGACG | >hg38_21_+_8433172_8446622_RNA45SN1_45S_50ntFlanks | 9552 | 9569 | 18 | TCCGTAGCGGTCCTGACG | 664.0 | 1.9 | 0.4 | 4.4 | 1.2E-05 | 1.8E-05 | MB231/MB468 |
| >hg38_21_+_8433172_8446622_RNA45SN1_45S_50ntFlanks@7991.8012.22\|ACGTGGCGACCCGCTGAATTTA | >hg38_21_+_8433172_8446622_RNA45SN1_45S_50ntFlanks | 7991 | 8012 | 22 | ACGTGGCGACCCGCTGAATTTA | 2700.1 | -2.7 | 0.6 | -4.4 | 1.2E-05 | 1.8E-05 | MB231/MB468 |
| >MI0000101\|hsa-mir-99a&WithFlank&21\|+\|16539083\|16539175@19.39.21&[MIMAT0000097&hsa-miR-99a-5p&offsets\|0\|-1;m-64&21\|+\|16539101\|16539121&offsets\|0\|0]\|AACCCGTAGATCCGATCTTGT | >MI0000101\|hsa-mir-99a&WithFlank&21\|+\|16539083\|16539175 | 19 | 39 | 21 | AACCCGTAGATCCGATCTTGT | 970.8 | -5.6 | 1.2 | -4.9 | 1.2E-06 | 1.8E-05 | BT20/MB231 |
| >hg38_21_+_8433172_8446622_RNA45SN1_45S_50ntFlanks@9545.9574.30\|GTGGAGGTCCGTAGCGGTCCTGACGTGCAA | >hg38_21_+_8433172_8446622_RNA45SN1_45S_50ntFlanks | 9545 | 9574 | 30 | GTGGAGGTCCGTAGCGGTCCTGACGTGCAA | 570.5 | -2.6 | 0.6 | -4.4 | 1.3E-05 | 1.9E-05 | MB231/MB468 |
| >hg38_21_+_8433172_8446622_RNA45SN1_45S_50ntFlanks@12474.12499.26\|ACCCACTAATAGGGAACGTGAGCTGG | >hg38_21_+_8433172_8446622_RNA45SN1_45S_50ntFlanks | 12474 | 12499 | 26 | ACCCACTAATAGGGAACGTGAGCTGG | 143.0 | -3.0 | 0.7 | -4.3 | 1.4E-05 | 1.9E-05 | BT20/MB468 |
| >MI0000064\|hsa-let-7c&WithFlank&21\|+\|16539822\|16539917@17.38.22&[MIMAT0000064&hsa-let-7c-5p&offsets\|0\|0;m-37&21\|+\|16539838\|16539859&offsets\|0\|0]\|TGAGGTAGTAGGTTGTATGGTT | >MI0000064\|hsa-let-7c&WithFlank&21\|+\|16539822\|16539917 | 17 | 38 | 22 | TGAGGTAGTAGGTTGTATGGTT | 1555.0 | 3.3 | 0.8 | 4.4 | 1.3E-05 | 1.9E-05 | MB231/MB468 |
| >hg38_21_+_8433172_8446622_RNA45SN1_45S_50ntFlanks@10411.10445.35\|CAGTCGGTCCTGAGAGATGGGCGAGCGCCGTTCCG | >hg38_21_+_8433172_8446622_RNA45SN1_45S_50ntFlanks | 10411 | 10445 | 35 | CAGTCGGTCCTGAGAGATGGGCGAGCGCCGTTCCG | 80.6 | -4.4 | 1.0 | -4.4 | 1.3E-05 | 1.9E-05 | MB231/MB468 |
| >hg38_21_+_8433172_8446622_RNA45SN1_45S_50ntFlanks@4730.4758.29\|CAAGAACGAAAGTCGGAGGTTCGAAGACG | >hg38_21_+_8433172_8446622_RNA45SN1_45S_50ntFlanks | 4730 | 4758 | 29 | CAAGAACGAAAGTCGGAGGTTCGAAGACG | 1100.2 | -2.1 | 0.5 | -4.3 | 1.4E-05 | 1.9E-05 | BT20/MB468 |
| >hg38_21_+_8433172_8446622_RNA45SN1_45S_50ntFlanks@4760.4779.20\|TCAGATACCGTCGTAGTTCC | >hg38_21_+_8433172_8446622_RNA45SN1_45S_50ntFlanks | 4760 | 4779 | 20 | TCAGATACCGTCGTAGTTCC | 306.7 | -5.2 | 1.2 | -4.3 | 1.4E-05 | 1.9E-05 | BT20/MB468 |
| >hg38_21_+_8433172_8446622_RNA45SN1_45S_50ntFlanks@12418.12437.20\|CGATGTCGGCTCTTCCTATC | >hg38_21_+_8433172_8446622_RNA45SN1_45S_50ntFlanks | 12418 | 12437 | 20 | CGATGTCGGCTCTTCCTATC | 3845.8 | -2.5 | 0.6 | -4.3 | 1.5E-05 | 1.9E-05 | BT20/MB468 |
| >hg38_21_+_8433172_8446622_RNA45SN1_45S_50ntFlanks@12997.13039.43\|CGCTGCGATCTATTGAAAGTCAGCCCTCGACACAAGGGTTTGT | >hg38_21_+_8433172_8446622_RNA45SN1_45S_50ntFlanks | 12997 | 13039 | 43 | CGCTGCGATCTATTGAAAGTCAGCCCTCGACACAAGGGTTTGT | 301221.1 | -3.3 | 0.7 | -4.8 | 1.3E-06 | 1.9E-05 | BT20/MB231 |
| >hg38_21_+_8433172_8446622_RNA45SN1_45S_50ntFlanks@10458.10495.38\|ATGGCCTCCGTTGCCCTCGGCCGATCGAAAGGGAGTCG | >hg38_21_+_8433172_8446622_RNA45SN1_45S_50ntFlanks | 10458 | 10495 | 38 | ATGGCCTCCGTTGCCCTCGGCCGATCGAAAGGGAGTCG | 211.3 | -3.2 | 0.7 | -4.4 | 1.3E-05 | 1.9E-05 | MB231/MB468 |
| >hg38_21_+_8433172_8446622_RNA45SN1_45S_50ntFlanks@10823.10844.22\|AAGTCGGCAAGCCGGATCCGTA | >hg38_21_+_8433172_8446622_RNA45SN1_45S_50ntFlanks | 10823 | 10844 | 22 | AAGTCGGCAAGCCGGATCCGTA | 154.7 | -2.8 | 0.6 | -4.4 | 1.4E-05 | 2.0E-05 | MB231/MB468 |
| >hg38_21_+_8433172_8446622_RNA45SN1_45S_50ntFlanks@9822.9850.29\|CTAGTGGGCCACTTTTGGTAAGCAGAACT | >hg38_21_+_8433172_8446622_RNA45SN1_45S_50ntFlanks | 9822 | 9850 | 29 | CTAGTGGGCCACTTTTGGTAAGCAGAACT | 246.9 | -2.5 | 0.6 | -4.3 | 1.4E-05 | 2.0E-05 | MB231/MB468 |
| >hg38_21_+_8433172_8446622_RNA45SN1_45S_50ntFlanks@12526.12546.21\|TTTTACCCTACTGATGATGTG | >hg38_21_+_8433172_8446622_RNA45SN1_45S_50ntFlanks | 12526 | 12546 | 21 | TTTTACCCTACTGATGATGTG | 72.0 | -4.1 | 0.9 | -4.3 | 1.5E-05 | 2.0E-05 | BT20/MB468 |
| >MI0000273\|hsa-mir-183&WithFlank&7\|-\|129774899\|129775020@34.55.22&[MIMAT0000261&hsa-miR-183-5p&offsets\|+1\|+1;m-41&7\|-\|129774966\|129774987&offsets\|0\|0]\|ATGGCACTGGTAGAATTCACTG | >MI0000273\|hsa-mir-183&WithFlank&7\|-\|129774899\|129775020 | 34 | 55 | 22 | ATGGCACTGGTAGAATTCACTG | 3962.5 | 3.1 | 0.6 | 4.8 | 1.3E-06 | 2.0E-05 | BT20/MB231 |
| >hg38_1_-_228634819_228635039_RNA5S12_5S_50ntFlanks@134.173.40\|TGGGAGACCGCCTGGGAATACCGGGTGCTGTAGGCTTTTT | >hg38_1_-_228634819_228635039_RNA5S12_5S_50ntFlanks | 134 | 173 | 40 | TGGGAGACCGCCTGGGAATACCGGGTGCTGTAGGCTTTTT | 221.9 | -4.3 | 0.9 | -4.8 | 1.3E-06 | 2.0E-05 | BT20/MB231 |
| >hg38_21_+_8433172_8446622_RNA45SN1_45S_50ntFlanks@4297.4318.22\|CATTGGAGGGCAAGTCTGGTGC | >hg38_21_+_8433172_8446622_RNA45SN1_45S_50ntFlanks | 4297 | 4318 | 22 | CATTGGAGGGCAAGTCTGGTGC | 100.0 | -2.9 | 0.7 | -4.3 | 1.5E-05 | 2.0E-05 | BT20/MB468 |
| >hg38_21_+_8433172_8446622_RNA45SN1_45S_50ntFlanks@4555.4573.19\|CGCAGCTAGGAATAATGGA | >hg38_21_+_8433172_8446622_RNA45SN1_45S_50ntFlanks | 4555 | 4573 | 19 | CGCAGCTAGGAATAATGGA | 196.8 | -4.2 | 1.0 | -4.3 | 1.5E-05 | 2.0E-05 | BT20/MB468 |
| >hg38_21_+_8433172_8446622_RNA45SN1_45S_50ntFlanks@10391.10411.21\|AACAGCAGTTGAACATGGGTC | >hg38_21_+_8433172_8446622_RNA45SN1_45S_50ntFlanks | 10391 | 10411 | 21 | AACAGCAGTTGAACATGGGTC | 152.9 | -1.9 | 0.4 | -4.3 | 1.4E-05 | 2.0E-05 | MB231/MB468 |
| >hg38_21_+_8433172_8446622_RNA45SN1_45S_50ntFlanks@4752.4782.31\|GAAGACGATCAGATACCGTCGTAGTTCCGAC | >hg38_21_+_8433172_8446622_RNA45SN1_45S_50ntFlanks | 4752 | 4782 | 31 | GAAGACGATCAGATACCGTCGTAGTTCCGAC | 106.0 | -2.0 | 0.5 | -4.3 | 1.4E-05 | 2.0E-05 | MB231/MB468 |
| >hg38_21_+_8433172_8446622_RNA45SN1_45S_50ntFlanks@6650.6693.44\|ACGACTCTTAGCGGTGGATCACTCGGCTCGTGCGTCGATGAAGA | >hg38_21_+_8433172_8446622_RNA45SN1_45S_50ntFlanks | 6650 | 6693 | 44 | ACGACTCTTAGCGGTGGATCACTCGGCTCGTGCGTCGATGAAGA | 458.3 | 7.8 | 1.8 | 4.3 | 1.4E-05 | 2.0E-05 | MB231/MB468 |
| >hg38_21_+_8433172_8446622_RNA45SN1_45S_50ntFlanks@12605.12634.30\|TATGTGCTTGGCTGAGGAGCCAATGGGGCG | >hg38_21_+_8433172_8446622_RNA45SN1_45S_50ntFlanks | 12605 | 12634 | 30 | TATGTGCTTGGCTGAGGAGCCAATGGGGCG | 88.9 | -4.3 | 1.0 | -4.3 | 1.5E-05 | 2.0E-05 | BT20/MB468 |
| >hg38_21_+_8433172_8446622_RNA45SN1_45S_50ntFlanks@11567.11585.19\|AGCAGCCGACTTAGAACTG | >hg38_21_+_8433172_8446622_RNA45SN1_45S_50ntFlanks | 11567 | 11585 | 19 | AGCAGCCGACTTAGAACTG | 4168.8 | 2.6 | 0.6 | 4.3 | 1.4E-05 | 2.0E-05 | MB231/MB468 |
| >MI0000749\|hsa-mir-30e&WithFlank&1\|+\|40754349\|40754452@23.46.24(+1U)&[MIMAT0000692&hsa-miR-30e-5p&offsets\|0\|+2(+1U);m-24&1\|+\|40754371\|40754394&offsets\|0\|0(+1U)]\|TGTAAACATCCTTGACTGGAAGCTT | >MI0000749\|hsa-mir-30e&WithFlank&1\|+\|40754349\|40754452 | 23 | 46 | 24(+1U) | TGTAAACATCCTTGACTGGAAGCTT | 661.7 | 4.1 | 0.9 | 4.3 | 1.5E-05 | 2.0E-05 | BT20/MB468 |
| >hg38_21_+_8433172_8446622_RNA45SN1_45S_50ntFlanks@11570.11602.33\|AGCCGACTTAGAACTGGTGCGGACCAGGGGAAT | >hg38_21_+_8433172_8446622_RNA45SN1_45S_50ntFlanks | 11570 | 11602 | 33 | AGCCGACTTAGAACTGGTGCGGACCAGGGGAAT | 346.7 | -2.2 | 0.5 | -4.3 | 1.5E-05 | 2.0E-05 | BT20/MB468 |

| | | | | | | | | | | | | |
|---|---|---|---|---|---|---|---|---|---|---|---|---|
| >hg38_21_+_8433172_8446622_RNA45SN1_45S_50ntFlanks@6649.6669.21\|TACGACTCTTAGCGGTGGATC | >hg38_21_+_8433172_8446622_RNA45SN1_45S_50ntFlanks | 6649 | 6669 | 21 | TACGACTCTTAGCGGTGGATC | 1037.7 | 2.2 | 0.5 | 4.3 | 1.5E-05 | 2.0E-05 | BT20/MB468 |
| >trnaMT_ProTGG_MT_-_15956_16023@41.67.27\|ATGGTGGAGTTAAAGACTTTTTCTCTG | >trnaMT_ProTGG_MT_-_15956_16023 | 41 | 67 | 27 | ATGGTGGAGTTAAAGACTTTTTCTCTG | 100.5 | -3.4 | 0.8 | -4.3 | 1.5E-05 | 2.0E-05 | BT20/MB468 |
| >hg38_21_+_8433172_8446622_RNA45SN1_45S_50ntFlanks@9308.9338.31\|AGGAGTCTAACACGTGCGCGAGTCGGGGGCT | >hg38_21_+_8433172_8446622_RNA45SN1_45S_50ntFlanks | 9308 | 9338 | 31 | AGGAGTCTAACACGTGCGCGAGTCGGGGGCT | 92.2 | -4.2 | 1.0 | -4.3 | 1.6E-05 | 2.0E-05 | BT20/MB468 |
| >hg38_21_+_8433172_8446622_RNA45SN1_45S_50ntFlanks@8360.8379.20\|AGAACTTTGAAGAGAGAGTT | >hg38_21_+_8433172_8446622_RNA45SN1_45S_50ntFlanks | 8360 | 8379 | 20 | AGAACTTTGAAGAGAGAGTT | 115.8 | -3.1 | 0.7 | -4.3 | 1.6E-05 | 2.0E-05 | BT20/MB468 |
| >MI0000286\|hsa-mir-210&WithFlank&11\|-\|568083\|568204@72.93.22(+1A)&[MIMAT0000267&hsa-miR-210-3p&offsets\|0\|0(+1A);m-102&11\|-\|568112\|568133&offsets\|0\|0(+1A)]\|CTGTGCGTGTGACAGCGGCTGAA | >MI0000286\|hsa-mir-210&WithFlank&11\|-\|568083\|568204 | 72 | 93 | 22(+1A) | CTGTGCGTGTGACAGCGGCTGAA | 4686.1 | 4.7 | 1.0 | 4.8 | 1.4E-06 | 2.0E-05 | BT20/MB231 |
| >hg38_21_+_8433172_8446622_RNA45SN1_45S_50ntFlanks@4369.4389.21\|GTTAAAAAGCTCGTAGTTGGA | >hg38_21_+_8433172_8446622_RNA45SN1_45S_50ntFlanks | 4369 | 4389 | 21 | GTTAAAAAGCTCGTAGTTGGA | 78.0 | -3.4 | 0.8 | -4.3 | 1.4E-05 | 2.1E-05 | MB231/MB468 |
| >hg38_21_+_8433172_8446622_RNA45SN1_45S_50ntFlanks@9736.9767.32\|CCGAAACGATCTCAACCTATTCTCAAACTTTA | >hg38_21_+_8433172_8446622_RNA45SN1_45S_50ntFlanks | 9736 | 9767 | 32 | CCGAAACGATCTCAACCTATTCTCAAACTTTA | 97.3 | -4.1 | 0.9 | -4.3 | 1.6E-05 | 2.1E-05 | BT20/MB468 |
| >hg38_21_+_8433172_8446622_RNA45SN1_45S_50ntFlanks@11317.11343.27\|ACCCCACGTCTCGTCGCGCGCGCGTCC | >hg38_21_+_8433172_8446622_RNA45SN1_45S_50ntFlanks | 11317 | 11343 | 27 | ACCCCACGTCTCGTCGCGCGCGCGTCC | 326.4 | -4.0 | 0.9 | -4.3 | 1.6E-05 | 2.1E-05 | BT20/MB468 |
| >hg38_21_+_8433172_8446622_RNA45SN1_45S_50ntFlanks@11208.11235.28\|GCGGCGACTCTGGACGCGAGCCGGGCCC | >hg38_21_+_8433172_8446622_RNA45SN1_45S_50ntFlanks | 11208 | 11235 | 28 | GCGGCGACTCTGGACGCGAGCCGGGCCC | 77.6 | -3.4 | 0.8 | -4.3 | 1.5E-05 | 2.1E-05 | MB231/MB468 |
| >hg38_21_+_8433172_8446622_RNA45SN1_45S_50ntFlanks@11317.11342.26\|ACCCCACGTCTCGTCGCGCGCGCGTC | >hg38_21_+_8433172_8446622_RNA45SN1_45S_50ntFlanks | 11317 | 11342 | 26 | ACCCCACGTCTCGTCGCGCGCGCGTC | 834.8 | -4.7 | 1.1 | -4.3 | 1.6E-05 | 2.1E-05 | BT20/MB468 |
| >hg38_21_+_8433172_8446622_RNA45SN1_45S_50ntFlanks@11104.11133.30\|AAGGGTCGGGGCGGCAGGGGCCGGCGGCGG | >hg38_21_+_8433172_8446622_RNA45SN1_45S_50ntFlanks | 11104 | 11133 | 30 | AAGGGTCGGGGCGGCAGGGGCCGGCGGCGG | 86.6 | -3.8 | 0.9 | -4.3 | 1.6E-05 | 2.1E-05 | BT20/MB468 |
| >hg38_21_+_8433172_8446622_RNA45SN1_45S_50ntFlanks@11586.11609.24\|GTGCGGACCAGGGGAATCCGACTG | >hg38_21_+_8433172_8446622_RNA45SN1_45S_50ntFlanks | 11586 | 11609 | 24 | GTGCGGACCAGGGGAATCCGACTG | 521.2 | -2.9 | 0.7 | -4.3 | 1.6E-05 | 2.1E-05 | BT20/MB468 |
| >hg38_21_+_8433172_8446622_RNA45SN1_45S_50ntFlanks@9533.9553.21\|GAGGAAACTCTGGTGGAGGTC | >hg38_21_+_8433172_8446622_RNA45SN1_45S_50ntFlanks | 9533 | 9553 | 21 | GAGGAAACTCTGGTGGAGGTC | 5591.5 | -2.8 | 0.7 | -4.3 | 1.5E-05 | 2.2E-05 | MB231/MB468 |
| >hg38_21_+_8433172_8446622_RNA45SN1_45S_50ntFlanks@12072.12093.22\|CCCGAGGGGCTCTCGCTTCTGG | >hg38_21_+_8433172_8446622_RNA45SN1_45S_50ntFlanks | 12072 | 12093 | 22 | CCCGAGGGGCTCTCGCTTCTGG | 101.2 | -4.0 | 0.9 | -4.3 | 1.7E-05 | 2.2E-05 | BT20/MB468 |
| >hg38_21_+_8433172_8446622_RNA45SN1_45S_50ntFlanks@7981.7998.18\|CTCAGATCAGACGTGGCG | >hg38_21_+_8433172_8446622_RNA45SN1_45S_50ntFlanks | 7981 | 7998 | 18 | CTCAGATCAGACGTGGCG | 79.1 | -4.4 | 1.0 | -4.3 | 1.7E-05 | 2.2E-05 | BT20/MB468 |
| >hg38_21_+_8433172_8446622_RNA45SN1_45S_50ntFlanks@5495.5516.22\|AGACGGTCGAACTTGACTATCT | >hg38_21_+_8433172_8446622_RNA45SN1_45S_50ntFlanks | 5495 | 5516 | 22 | AGACGGTCGAACTTGACTATCT | 703.5 | 10.4 | 2.4 | 4.3 | 1.7E-05 | 2.2E-05 | BT20/MB468 |
| >trna128_GlyGCC_6_-_27870686_27870756@19.38.20,trna19_GlyGCC_16_+_70823410_70823480@19.38.20,trna19_GlyGCC_2_-_157257659_157257729@19.38.20,trna24_GlyGCC_16_-_70812942_70813012@19.38.20,trna25_GlyGCC_16_-_70812114_70812184@19.38.20,trna2_GlyGCC_21_-_18827107_18827177@19.38.20,trna35_GlyGCC_1_+_161413094_161413164@19.38.20,trna37_GlyGCC_1_+_161420467_161420537@19.38.20,trna39_GlyGCC_1_+_161427898_161427968@19.38.20,trna41_GlyGCC_1_+_161435258_161435328@19.38.20,trna5_GlyGCC_17_+_8029064_8029134@19.38.20,trna68_GlyGCC_1_-_161493637_161493707@19.38.20\|TAGAATTCTCGCCTGCCACG | >trna128_GlyGCC_6_-_27870686_27870756 | 19 | 38 | 20 | TAGAATTCTCGCCTGCCACG | 404.7 | 3.9 | 0.8 | 4.8 | 1.5E-06 | 2.2E-05 | BT20/MB231 |
| >hg38_21_+_8433172_8446622_RNA45SN1_45S_50ntFlanks@9567.9584.18\|ACGTGCAAATCGGTCGTC | >hg38_21_+_8433172_8446622_RNA45SN1_45S_50ntFlanks | 9567 | 9584 | 18 | ACGTGCAAATCGGTCGTC | 312.0 | -2.8 | 0.6 | -4.3 | 1.5E-05 | 2.2E-05 | MB231/MB468 |
| >hg38_21_+_8433172_8446622_RNA45SN1_45S_50ntFlanks@10677.10708.32\|GCGGTTCCGGCGGCGTCCGGTGAGCTCTCGCT | >hg38_21_+_8433172_8446622_RNA45SN1_45S_50ntFlanks | 10677 | 10708 | 32 | GCGGTTCCGGCGGCGTCCGGTGAGCTCTCGCT | 1236.1 | 4.2 | 1.0 | 4.3 | 1.7E-05 | 2.2E-05 | BT20/MB468 |
| >hg38_21_+_8433172_8446622_RNA45SN1_45S_50ntFlanks@10402.10432.31\|AACATGGGTCAGTCGGTCCTGAGAGATGGGC | >hg38_21_+_8433172_8446622_RNA45SN1_45S_50ntFlanks | 10402 | 10432 | 31 | AACATGGGTCAGTCGGTCCTGAGAGATGGGC | 130.4 | -1.9 | 0.4 | -4.3 | 1.5E-05 | 2.2E-05 | MB231/MB468 |
| >hg38_21_+_8433172_8446622_RNA45SN1_45S_50ntFlanks@11583.11600.18\|CTGGTGCGGACCAGGGGA | >hg38_21_+_8433172_8446622_RNA45SN1_45S_50ntFlanks | 11583 | 11600 | 18 | CTGGTGCGGACCAGGGGA | 382.5 | -2.8 | 0.7 | -4.3 | 1.7E-05 | 2.2E-05 | BT20/MB468 |
| >hg38_21_+_8433172_8446622_RNA45SN1_45S_50ntFlanks@3771.3799.29\|CACGGCCGGTACAGTGAAACTGCGAATGG | >hg38_21_+_8433172_8446622_RNA45SN1_45S_50ntFlanks | 3771 | 3799 | 29 | CACGGCCGGTACAGTGAAACTGCGAATGG | 81.7 | -2.2 | 0.5 | -4.3 | 1.6E-05 | 2.2E-05 | MB231/MB468 |
| >hg38_21_+_8433172_8446622_RNA45SN1_45S_50ntFlanks@3939.3958.20\|AACCCGGTCAGCCCCTCTCC | >hg38_21_+_8433172_8446622_RNA45SN1_45S_50ntFlanks | 3939 | 3958 | 20 | AACCCGGTCAGCCCCTCTCC | 114.8 | -2.9 | 0.7 | -4.3 | 1.6E-05 | 2.2E-05 | MB231/MB468 |
| >hg38_21_+_8433172_8446622_RNA45SN1_45S_50ntFlanks@10842.10865.24\|GTAACTTCGGGATAAGGATTGGCT | >hg38_21_+_8433172_8446622_RNA45SN1_45S_50ntFlanks | 10842 | 10865 | 24 | GTAACTTCGGGATAAGGATTGGCT | 188.2 | -2.5 | 0.6 | -4.3 | 1.6E-05 | 2.3E-05 | MB231/MB468 |
| >hg38_21_+_8433172_8446622_RNA45SN1_45S_50ntFlanks@7979.8008.30\|ACCTCAGATCAGACGTGGCGACCCGCTGAA | >hg38_21_+_8433172_8446622_RNA45SN1_45S_50ntFlanks | 7979 | 8008 | 30 | ACCTCAGATCAGACGTGGCGACCCGCTGAA | 816.8 | -2.7 | 0.6 | -4.3 | 1.6E-05 | 2.3E-05 | MB231/MB468 |
| >hg38_21_+_8433172_8446622_RNA45SN1_45S_50ntFlanks@9546.9571.26\|TGGAGGTCCGTAGCGGTCCTGACGTG | >hg38_21_+_8433172_8446622_RNA45SN1_45S_50ntFlanks | 9546 | 9571 | 26 | TGGAGGTCCGTAGCGGTCCTGACGTG | 1022.0 | 2.6 | 0.6 | 4.3 | 1.6E-05 | 2.3E-05 | MB231/MB468 |
| >hg38_21_+_8433172_8446622_RNA45SN1_45S_50ntFlanks@12934.12957.24\|CTAAACCATTCGTAGACGACCTGC | >hg38_21_+_8433172_8446622_RNA45SN1_45S_50ntFlanks | 12934 | 12957 | 24 | CTAAACCATTCGTAGACGACCTGC | 223.7 | -1.9 | 0.4 | -4.3 | 1.6E-05 | 2.3E-05 | MB231/MB468 |
| >hg38_21_+_8433172_8446622_RNA45SN1_45S_50ntFlanks@12637.12661.25\|GCTACCATCTGTGGGATTATGACTG | >hg38_21_+_8433172_8446622_RNA45SN1_45S_50ntFlanks | 12637 | 12661 | 25 | GCTACCATCTGTGGGATTATGACTG | 401.5 | -2.0 | 0.5 | -4.3 | 1.6E-05 | 2.3E-05 | MB231/MB468 |
| >hg38_21_+_8433172_8446622_RNA45SN1_45S_50ntFlanks@10346.10374.29\|TCAAACGAGAACTTTGAAGGCCGAAGTGG | >hg38_21_+_8433172_8446622_RNA45SN1_45S_50ntFlanks | 10346 | 10374 | 29 | TCAAACGAGAACTTTGAAGGCCGAAGTGG | 507.9 | -2.5 | 0.6 | -4.3 | 1.6E-05 | 2.3E-05 | MB231/MB468 |
| >MI0000737\|hsa-mir-200a&WithFlank&1\|+\|1167857\|1167958@60.81.22&[MIMAT0000682&hsa-miR-200a-3p&offsets\|0\|0;m-142&1\|+\|1167916\|1167938&offsets\|0\|-1]\|TAACACTGTCTGGTAACGATGT | >MI0000737\|hsa-mir-200a&WithFlank&1\|+\|1167857\|1167958 | 60 | 81 | 22 | TAACACTGTCTGGTAACGATGT | 268.3 | -2.9 | 0.7 | -4.3 | 1.6E-05 | 2.3E-05 | MB231/MB468 |
| >hg38_21_+_8433172_8446622_RNA45SN1_45S_50ntFlanks@12418.12454.37\|CGATGTCGGCTCTTCCTATCATTGTGAAGCAGAATTC | >hg38_21_+_8433172_8446622_RNA45SN1_45S_50ntFlanks | 12418 | 12454 | 37 | CGATGTCGGCTCTTCCTATCATTGTGAAGCAGAATTC | 413.0 | 2.5 | 0.6 | 4.3 | 1.6E-05 | 2.3E-05 | MB231/MB468 |
| >trna119_LysCTT_1_-_145395522_145395594@4.36.33,trna11_LysCTT_5_-_180648979_180649051@4.36.33,trna13_LysCTT_6_+_26556774_26556846@4.36.33,trna7_LysCTT_16_+_3225692_3225764@4.36.33,trna9_LysCTT_5_+_180634755_180634827@4.36.33\|CGGCTAGCTCAGTCGGTAGAGCATGAGACTCTT | >trna119_LysCTT_1_-_145395522_145395594 | 4 | 36 | 33 | CGGCTAGCTCAGTCGGTAGAGCATGAGACTCTT | 683.8 | -4.1 | 0.8 | -4.8 | 1.6E-06 | 2.4E-05 | BT20/MB231 |
| >hg38_21_+_8433172_8446622_RNA45SN1_45S_50ntFlanks@7980.8005.26\|CCTCAGATCAGACGTGGCGACCCGCT | >hg38_21_+_8433172_8446622_RNA45SN1_45S_50ntFlanks | 7980 | 8005 | 26 | CCTCAGATCAGACGTGGCGACCCGCT | 181.6 | -2.3 | 0.5 | -4.3 | 1.8E-05 | 2.4E-05 | BT20/MB468 |
| >am_trna162_MetCAT_6_-_26758550_26758622@36.54.19,trna164_MetCAT_6_-_26735574_26735646@36.54.19,trna20_MetCAT_16_+_71460396_71460468@36.54.19,trna21_MetCAT_6_+_26701712_26701784@36.54.19,trna27_MetCAT_6_+_26766444_26766516@36.54.19,trna75_MetCAT_6_+_28912352_28912424@36.54.19,trna97_MetCAT_6_-_28921042_28921114@36.54.19\|TAATCTGAAGGTCCTGAGT | >am_trna162_MetCAT_6_-_26758550_26758622 | 36 | 54 | 19 | TAATCTGAAGGTCCTGAGT | 864.0 | 3.9 | 0.9 | 4.3 | 1.6E-05 | 2.4E-05 | MB231/MB468 |
| >hg38_21_+_8433172_8446622_RNA45SN1_45S_50ntFlanks@11579.11601.23\|AGAACTGGTGCGGACCAGGGGAA | >hg38_21_+_8433172_8446622_RNA45SN1_45S_50ntFlanks | 11579 | 11601 | 23 | AGAACTGGTGCGGACCAGGGGAA | 581.8 | -3.5 | 0.8 | -4.3 | 1.6E-05 | 2.4E-05 | MB231/MB468 |
| >hg38_21_+_8433172_8446622_RNA45SN1_45S_50ntFlanks@3839.3860.22\|TCTCCTACTTGGATAACTGTGG | >hg38_21_+_8433172_8446622_RNA45SN1_45S_50ntFlanks | 3839 | 3860 | 22 | TCTCCTACTTGGATAACTGTGG | 237.0 | 1.8 | 0.4 | 4.3 | 1.7E-05 | 2.4E-05 | MB231/MB468 |
| >hg38_21_+_8433172_8446622_RNA45SN1_45S_50ntFlanks@6652.6686.35\|GACTCTTAGCGGTGGATCACTCGGCTCGTGCGTCG | >hg38_21_+_8433172_8446622_RNA45SN1_45S_50ntFlanks | 6652 | 6686 | 35 | GACTCTTAGCGGTGGATCACTCGGCTCGTGCGTCG | 2625.3 | 2.8 | 0.6 | 4.3 | 1.7E-05 | 2.4E-05 | MB231/MB468 |
| >hg38_21_+_8433172_8446622_RNA45SN1_45S_50ntFlanks@12994.13039.46\|CCTCGCTGCGATCTATTGAAAGTCAGCCCTCGACACAAGGGTTTGT | >hg38_21_+_8433172_8446622_RNA45SN1_45S_50ntFlanks | 12994 | 13039 | 46 | CCTCGCTGCGATCTATTGAAAGTCAGCCCTCGACACAAGGGTTTGT | 36812.5 | -4.7 | 1.0 | -4.8 | 1.7E-06 | 2.4E-05 | BT20/MB231 |
| >MI0000272\|hsa-mir-182&WithFlank&7\|-\|129770377\|129770498@29.52.24(+1A)&[MIMAT0000259&hsa-miR-182-5p&offsets\|0\|0(+1A);m-22&7\|-\|129770449\|129770470&offsets\|0\|+2(+1A)]\|TTTGGCAATGGTAGAACTCACACTA | >MI0000272\|hsa-mir-182&WithFlank&7\|-\|129770377\|129770498 | 29 | 52 | 24(+1A) | TTTGGCAATGGTAGAACTCACACTA | 2670.2 | 3.5 | 0.7 | 4.8 | 1.7E-06 | 2.4E-05 | BT20/MB231 |
| >hg38_21_+_8433172_8446622_RNA45SN1_45S_50ntFlanks@9304.9338.35\|ACCAAGGAGTCTAACACGTGCGCGAGTCGGGGGCT | >hg38_21_+_8433172_8446622_RNA45SN1_45S_50ntFlanks | 9304 | 9338 | 35 | ACCAAGGAGTCTAACACGTGCGCGAGTCGGGGGCT | 98.7 | -4.3 | 1.0 | -4.3 | 1.9E-05 | 2.5E-05 | BT20/MB468 |
| >hg38_21_+_8433172_8446622_RNA45SN1_45S_50ntFlanks@10846.10866.21\|CTTCGGGATAAGGATTGGCTC | >hg38_21_+_8433172_8446622_RNA45SN1_45S_50ntFlanks | 10846 | 10866 | 21 | CTTCGGGATAAGGATTGGCTC | 171.8 | -4.6 | 1.1 | -4.3 | 1.9E-05 | 2.5E-05 | BT20/MB468 |

<table>
<tr><td>>MI0000446|hsa-mir-125b-1&WithFlank&11|-|122099751|122099850@21.41.21(+1G);MI0000470|hsa-mir-125b-2&WithFlank&21|+|16590231|16590331@23.43.21(+1G)&[MIMAT0000423&hsa-miR-125b-5p&offsets|0|-1(+1G);m-67&11|-|122099809|122099830&offsets|0|-1(+1G)];[MIMAT0000423_1&hsa-miR-125b-5p&offsets|0|-1(+1G);m-68&21|+|16590253|16590274&offsets|0|-1(+1G)]|TCCCTGAGACCCTAACTTGTGG</td><td>>MI0000446|hsa-mir-125b-1&WithFlank&11|-|122099751|122099850</td><td>21</td><td>41</td><td>21(+1G)</td><td>TCCCTGAGACCCTAACTTGTGG</td><td>623.5</td><td>-5.3</td><td>1.1</td><td>-4.8</td><td>1.7E-06</td><td>2.5E-05</td><td>BT20/MB231</td></tr>
<tr><td>>hg38_21_+_8433172_8446622_RNA45SN1_45S_50ntFlanks@11727.11751.25|AGTAACTATGACTCTCTTAAGGTAG</td><td>>hg38_21_+_8433172_8446622_RNA45SN1_45S_50ntFlanks</td><td>11727</td><td>11751</td><td>25</td><td>AGTAACTATGACTCTCTTAAGGTAG</td><td>809.2</td><td>-3.3</td><td>0.8</td><td>-4.3</td><td>1.9E-05</td><td>2.5E-05</td><td>BT20/MB468</td></tr>
<tr><td>>MI0000255|hsa-mir-30d&WithFlank&8|-|134804870|134804951@12.35.24(+4U)&[MIMAT0000245&hsa-miR-30d-5p&offsets|0|+2(+4U);m-21&8|-|134804919|134804940&offsets|0|+2(+4U)]|TGTAAACATCCCCGACTGGAAGCTTTTT</td><td>>MI0000255|hsa-mir-30d&WithFlank&8|-|134804870|134804951</td><td>12</td><td>35</td><td>24(+4U)</td><td>TGTAAACATCCCCGACTGGAAGCTTTTT</td><td>952.9</td><td>10.4</td><td>2.4</td><td>4.3</td><td>1.7E-05</td><td>2.5E-05</td><td>MB231/MB468</td></tr>
<tr><td>>hg38_21_+_8433172_8446622_RNA45SN1_45S_50ntFlanks@12957.12975.19|CTTCTGGGTCGGGGTTTCG</td><td>>hg38_21_+_8433172_8446622_RNA45SN1_45S_50ntFlanks</td><td>12957</td><td>12975</td><td>19</td><td>CTTCTGGGTCGGGGTTTCG</td><td>109.5</td><td>-4.0</td><td>0.9</td><td>-4.3</td><td>1.9E-05</td><td>2.5E-05</td><td>BT20/MB468</td></tr>
<tr><td>>hg38_21_+_8433172_8446622_RNA45SN1_45S_50ntFlanks@10781.10817.37|AGGTGAACAGCCTCTGGCATGTTGGAACAATGTAGGT</td><td>>hg38_21_+_8433172_8446622_RNA45SN1_45S_50ntFlanks</td><td>10781</td><td>10817</td><td>37</td><td>AGGTGAACAGCCTCTGGCATGTTGGAACAATGTAGGT</td><td>293.0</td><td>-2.4</td><td>0.6</td><td>-4.3</td><td>1.7E-05</td><td>2.5E-05</td><td>MB231/MB468</td></tr>
<tr><td>>hg38_21_+_8433172_8446622_RNA45SN1_45S_50ntFlanks@9737.9754.18|CGAAACGATCTCAACCTA</td><td>>hg38_21_+_8433172_8446622_RNA45SN1_45S_50ntFlanks</td><td>9737</td><td>9754</td><td>18</td><td>CGAAACGATCTCAACCTA</td><td>97.6</td><td>-4.0</td><td>0.9</td><td>-4.3</td><td>1.9E-05</td><td>2.5E-05</td><td>BT20/MB468</td></tr>
<tr><td>>hg38_21_+_8433172_8446622_RNA45SN1_45S_50ntFlanks@11573.11591.19|CGACTTAGAACTGGTGCGG</td><td>>hg38_21_+_8433172_8446622_RNA45SN1_45S_50ntFlanks</td><td>11573</td><td>11591</td><td>19</td><td>CGACTTAGAACTGGTGCGG</td><td>396.2</td><td>-4.2</td><td>1.0</td><td>-4.3</td><td>1.9E-05</td><td>2.5E-05</td><td>BT20/MB468</td></tr>
<tr><td>>hg38_21_+_8433172_8446622_RNA45SN1_45S_50ntFlanks@10389.10406.18|TGAACAGCAGTTGAACAT</td><td>>hg38_21_+_8433172_8446622_RNA45SN1_45S_50ntFlanks</td><td>10389</td><td>10406</td><td>18</td><td>TGAACAGCAGTTGAACAT</td><td>104.6</td><td>-2.7</td><td>0.6</td><td>-4.3</td><td>1.8E-05</td><td>2.5E-05</td><td>MB231/MB468</td></tr>
<tr><td>>hg38_21_+_8433172_8446622_RNA45SN1_45S_50ntFlanks@8196.8217.22|CGTGGACGGTGTGAGGCCGGTA</td><td>>hg38_21_+_8433172_8446622_RNA45SN1_45S_50ntFlanks</td><td>8196</td><td>8217</td><td>22</td><td>CGTGGACGGTGTGAGGCCGGTA</td><td>323.9</td><td>2.9</td><td>0.7</td><td>4.3</td><td>1.8E-05</td><td>2.5E-05</td><td>MB231/MB468</td></tr>
<tr><td>>hg38_21_+_8433172_8446622_RNA45SN1_45S_50ntFlanks@10792.10816.25|CTCTGGCATGTTGGAACAATGTAGG</td><td>>hg38_21_+_8433172_8446622_RNA45SN1_45S_50ntFlanks</td><td>10792</td><td>10816</td><td>25</td><td>CTCTGGCATGTTGGAACAATGTAGG</td><td>235.1</td><td>-2.2</td><td>0.5</td><td>-4.3</td><td>2.0E-05</td><td>2.6E-05</td><td>BT20/MB468</td></tr>
<tr><td>>hg38_21_+_8433172_8446622_RNA45SN1_45S_50ntFlanks@10797.10822.26|GCATGTTGGAACAATGTAGGTAAGGG</td><td>>hg38_21_+_8433172_8446622_RNA45SN1_45S_50ntFlanks</td><td>10797</td><td>10822</td><td>26</td><td>GCATGTTGGAACAATGTAGGTAAGGG</td><td>158.1</td><td>-2.6</td><td>0.6</td><td>-4.3</td><td>1.8E-05</td><td>2.6E-05</td><td>MB231/MB468</td></tr>
<tr><td>>hg38_21_+_8433172_8446622_RNA45SN1_45S_50ntFlanks@6729.6771.43|GACACATTGATCATCGACACTTCGAACGCACTTGCGGCCCCGG</td><td>>hg38_21_+_8433172_8446622_RNA45SN1_45S_50ntFlanks</td><td>6729</td><td>6771</td><td>43</td><td>GACACATTGATCATCGACACTTCGAACGCACTTGCGGCCCCGG</td><td>115.7</td><td>-3.0</td><td>0.7</td><td>-4.3</td><td>2.0E-05</td><td>2.6E-05</td><td>BT20/MB468</td></tr>
<tr><td>>hg38_21_+_8433172_8446622_RNA45SN1_45S_50ntFlanks@10808.10837.30|CAATGTAGGTAAGGGAAGTCGGCAAGCCGG</td><td>>hg38_21_+_8433172_8446622_RNA45SN1_45S_50ntFlanks</td><td>10808</td><td>10837</td><td>30</td><td>CAATGTAGGTAAGGGAAGTCGGCAAGCCGG</td><td>110.3</td><td>-1.9</td><td>0.4</td><td>-4.3</td><td>1.8E-05</td><td>2.6E-05</td><td>MB231/MB468</td></tr>
<tr><td>>hg38_21_+_8433172_8446622_RNA45SN1_45S_50ntFlanks@12618.12654.37|GAGGAGCCAATGGGGCGAAGCTACCATCTGTGGGATT</td><td>>hg38_21_+_8433172_8446622_RNA45SN1_45S_50ntFlanks</td><td>12618</td><td>12654</td><td>37</td><td>GAGGAGCCAATGGGGCGAAGCTACCATCTGTGGGATT</td><td>104.3</td><td>-2.5</td><td>0.6</td><td>-4.3</td><td>2.0E-05</td><td>2.7E-05</td><td>BT20/MB468</td></tr>
<tr><td>>hg38_21_+_8433172_8446622_RNA45SN1_45S_50ntFlanks@12479.12498.20|CTAATAGGGAACGTGAGCTG</td><td>>hg38_21_+_8433172_8446622_RNA45SN1_45S_50ntFlanks</td><td>12479</td><td>12498</td><td>20</td><td>CTAATAGGGAACGTGAGCTG</td><td>759.3</td><td>-2.5</td><td>0.6</td><td>-4.3</td><td>1.9E-05</td><td>2.7E-05</td><td>MB231/MB468</td></tr>
<tr><td>>hg38_21_+_8433172_8446622_RNA45SN1_45S_50ntFlanks@10391.10426.36|AACAGCAGTTGAACATGGGTCAGTCGGTCCTGAGAG</td><td>>hg38_21_+_8433172_8446622_RNA45SN1_45S_50ntFlanks</td><td>10391</td><td>10426</td><td>36</td><td>AACAGCAGTTGAACATGGGTCAGTCGGTCCTGAGAG</td><td>163.0</td><td>-2.8</td><td>0.6</td><td>-4.3</td><td>1.9E-05</td><td>2.7E-05</td><td>MB231/MB468</td></tr>
<tr><td>>hg38_21_+_8433172_8446622_RNA45SN1_45S_50ntFlanks@8356.8375.20|GAAAAGAACTTTGAAGAGAG</td><td>>hg38_21_+_8433172_8446622_RNA45SN1_45S_50ntFlanks</td><td>8356</td><td>8375</td><td>20</td><td>GAAAAGAACTTTGAAGAGAG</td><td>764.8</td><td>-3.2</td><td>0.7</td><td>-4.3</td><td>1.9E-05</td><td>2.7E-05</td><td>MB231/MB468</td></tr>
<tr><td>>hg38_21_+_8433172_8446622_RNA45SN1_45S_50ntFlanks@10342.10361.20|ATATTCAAACGAGAACTTTG</td><td>>hg38_21_+_8433172_8446622_RNA45SN1_45S_50ntFlanks</td><td>10342</td><td>10361</td><td>20</td><td>ATATTCAAACGAGAACTTTG</td><td>684.4</td><td>-2.1</td><td>0.5</td><td>-4.3</td><td>1.9E-05</td><td>2.7E-05</td><td>MB231/MB468</td></tr>
<tr><td>>hg38_21_+_8433172_8446622_RNA45SN1_45S_50ntFlanks@9546.9563.18|TGGAGGTCCGTAGCGGTC</td><td>>hg38_21_+_8433172_8446622_RNA45SN1_45S_50ntFlanks</td><td>9546</td><td>9563</td><td>18</td><td>TGGAGGTCCGTAGCGGTC</td><td>117.6</td><td>-2.6</td><td>0.6</td><td>-4.3</td><td>2.1E-05</td><td>2.7E-05</td><td>BT20/MB468</td></tr>
<tr><td>>hg38_21_+_8433172_8446622_RNA45SN1_45S_50ntFlanks@8170.8215.46|CAAGTCCTTCTGATCGAGGCCCAGCCCGTGGACGGTGTGAGGCCGG</td><td>>hg38_21_+_8433172_8446622_RNA45SN1_45S_50ntFlanks</td><td>8170</td><td>8215</td><td>46</td><td>CAAGTCCTTCTGATCGAGGCCCAGCCCGTGGACGGTGTGAGGCCGG</td><td>166.6</td><td>-5.1</td><td>1.2</td><td>-4.3</td><td>2.1E-05</td><td>2.7E-05</td><td>BT20/MB468</td></tr>
<tr><td>>MI0000477|hsa-mir-146a&WithFlank&5|+|160485346|160485456@27.49.23&[MIMAT0000449&hsa-miR-146a-5p&offsets|0|+1;m-129&5|+|160485372|160485393&offsets|0|+1]|TGAGAACTGAATTCCATGGGTTG</td><td>>MI0000477|hsa-mir-146a&WithFlank&5|+|160485346|160485456</td><td>27</td><td>49</td><td>23</td><td>TGAGAACTGAATTCCATGGGTTG</td><td>1269.2</td><td>-9.7</td><td>2.0</td><td>-4.8</td><td>1.9E-06</td><td>2.7E-05</td><td>BT20/MB231</td></tr>
<tr><td>>hg38_21_+_8433172_8446622_RNA45SN1_45S_50ntFlanks@12932.12953.22|CGCTAAACCATTCGTAGACGAC</td><td>>hg38_21_+_8433172_8446622_RNA45SN1_45S_50ntFlanks</td><td>12932</td><td>12953</td><td>22</td><td>CGCTAAACCATTCGTAGACGAC</td><td>91.5</td><td>-2.4</td><td>0.6</td><td>-4.3</td><td>1.9E-05</td><td>2.7E-05</td><td>MB231/MB468</td></tr>
<tr><td>>hg38_21_+_8433172_8446622_RNA45SN1_45S_50ntFlanks@10393.10435.43|CAGCAGTTGAACATGGGTCAGTCGGTCCTGAGAGATGGGCGAG</td><td>>hg38_21_+_8433172_8446622_RNA45SN1_45S_50ntFlanks</td><td>10393</td><td>10435</td><td>43</td><td>CAGCAGTTGAACATGGGTCAGTCGGTCCTGAGAGATGGGCGAG</td><td>194.1</td><td>-3.6</td><td>0.8</td><td>-4.3</td><td>1.9E-05</td><td>2.7E-05</td><td>MB231/MB468</td></tr>
<tr><td>>MI0003834|hsa-mir-769&WithFlank&19|+|46018926|46019055@36.57.22(+2U)&[MIMAT0003886&hsa-miR-769-5p&offsets|0|0(+2U);m-216&19|+|46018961|46018982&offsets|0|0(+2U)]|TGAGACCTCTGGGTTCTGAGCTTT</td><td>>MI0003834|hsa-mir-769&WithFlank&19|+|46018926|46019055</td><td>36</td><td>57</td><td>22(+2U)</td><td>TGAGACCTCTGGGTTCTGAGCTTT</td><td>564.6</td><td>10.1</td><td>2.4</td><td>4.3</td><td>2.1E-05</td><td>2.7E-05</td><td>BT20/MB468</td></tr>
<tr><td>>hg38_21_+_8433172_8446622_RNA45SN1_45S_50ntFlanks@6719.6752.34|TGAATTGCAGGACACATTGATCATCGACACTTCG</td><td>>hg38_21_+_8433172_8446622_RNA45SN1_45S_50ntFlanks</td><td>6719</td><td>6752</td><td>34</td><td>TGAATTGCAGGACACATTGATCATCGACACTTCG</td><td>219.4</td><td>-1.9</td><td>0.4</td><td>-4.3</td><td>1.9E-05</td><td>2.7E-05</td><td>MB231/MB468</td></tr>
<tr><td>>hg38_21_+_8433172_8446622_RNA45SN1_45S_50ntFlanks@5492.5526.35|AGAAGACGGTCGAACTTGACTATCTAGAGGAAGTA</td><td>>hg38_21_+_8433172_8446622_RNA45SN1_45S_50ntFlanks</td><td>5492</td><td>5526</td><td>35</td><td>AGAAGACGGTCGAACTTGACTATCTAGAGGAAGTA</td><td>303.2</td><td>-2.6</td><td>0.6</td><td>-4.3</td><td>1.9E-05</td><td>2.7E-05</td><td>MB231/MB468</td></tr>
<tr><td>>MI0000446|hsa-mir-125b-1&WithFlank&11|-|122099751|122099850@21.41.21(+1U);MI0000470|hsa-mir-125b-2&WithFlank&21|+|16590231|16590331@23.43.21(+1U)&[MIMAT0000423&hsa-miR-125b-5p&offsets|0|-1(+1U);m-67&11|-|122099809|122099830&offsets|0|-1(+1U)];[MIMAT0000423_1&hsa-miR-125b-5p&offsets|0|-1(+1U);m-68&21|+|16590253|16590274&offsets|0|-1(+1U)]|TCCCTGAGACCCTAACTTGTGT</td><td>>MI0000446|hsa-mir-125b-1&WithFlank&11|-|122099751|122099850</td><td>21</td><td>41</td><td>21(+1U)</td><td>TCCCTGAGACCCTAACTTGTGT</td><td>849.4</td><td>-5.1</td><td>1.1</td><td>-4.8</td><td>1.9E-06</td><td>2.7E-05</td><td>BT20/MB231</td></tr>
<tr><td>>MI0000101|hsa-mir-99a&WithFlank&21|+|16539083|16539175@19.40.22(+1A)&[MIMAT0000097&hsa-miR-99a-5p&offsets|0|0(+1A);m-64&21|+|16539101|16539121&offsets|0|+1(+1A)]|AACCCGTAGATCCGATCTTGTGA</td><td>>MI0000101|hsa-mir-99a&WithFlank&21|+|16539083|16539175</td><td>19</td><td>40</td><td>22(+1A)</td><td>AACCCGTAGATCCGATCTTGTGA</td><td>594.9</td><td>-4.6</td><td>1.1</td><td>-4.3</td><td>2.1E-05</td><td>2.8E-05</td><td>BT20/MB468</td></tr>
<tr><td>>hg38_21_+_8433172_8446622_RNA45SN1_45S_50ntFlanks@10783.10817.35|GTGAACAGCCTCTGGCATGTTGGAACAATGTAGGT</td><td>>hg38_21_+_8433172_8446622_RNA45SN1_45S_50ntFlanks</td><td>10783</td><td>10817</td><td>35</td><td>GTGAACAGCCTCTGGCATGTTGGAACAATGTAGGT</td><td>246.9</td><td>-2.2</td><td>0.5</td><td>-4.3</td><td>2.0E-05</td><td>2.8E-05</td><td>MB231/MB468</td></tr>
<tr><td>>hg38_21_+_8433172_8446622_RNA45SN1_45S_50ntFlanks@10787.10816.30|ACAGCCTCTGGCATGTTGGAACAATGTAGG</td><td>>hg38_21_+_8433172_8446622_RNA45SN1_45S_50ntFlanks</td><td>10787</td><td>10816</td><td>30</td><td>ACAGCCTCTGGCATGTTGGAACAATGTAGG</td><td>291.8</td><td>-2.4</td><td>0.6</td><td>-4.3</td><td>2.0E-05</td><td>2.8E-05</td><td>MB231/MB468</td></tr>
<tr><td>>hg38_21_+_8433172_8446622_RNA45SN1_45S_50ntFlanks@8191.8212.22|CAGCCCGTGGACGGTGTGAGGC</td><td>>hg38_21_+_8433172_8446622_RNA45SN1_45S_50ntFlanks</td><td>8191</td><td>8212</td><td>22</td><td>CAGCCCGTGGACGGTGTGAGGC</td><td>503.2</td><td>-3.3</td><td>0.8</td><td>-4.3</td><td>2.0E-05</td><td>2.8E-05</td><td>MB231/MB468</td></tr>
<tr><td>>hg38_21_+_8433172_8446622_RNA45SN1_45S_50ntFlanks@12652.12675.24|ATTATGACTGAACGCCTCTAAGTC</td><td>>hg38_21_+_8433172_8446622_RNA45SN1_45S_50ntFlanks</td><td>12652</td><td>12675</td><td>24</td><td>ATTATGACTGAACGCCTCTAAGTC</td><td>114.6</td><td>-3.0</td><td>0.7</td><td>-4.2</td><td>2.2E-05</td><td>2.9E-05</td><td>BT20/MB468</td></tr>
<tr><td>>hg38_21_+_8433172_8446622_RNA45SN1_45S_50ntFlanks@4588.4619.32|CTATTTTGTTGGTTTTCGGAACTGAGGCCATG</td><td>>hg38_21_+_8433172_8446622_RNA45SN1_45S_50ntFlanks</td><td>4588</td><td>4619</td><td>32</td><td>CTATTTTGTTGGTTTTCGGAACTGAGGCCATG</td><td>187.4</td><td>-2.4</td><td>0.6</td><td>-4.2</td><td>2.2E-05</td><td>2.9E-05</td><td>BT20/MB468</td></tr>
<tr><td>>hg38_21_+_8433172_8446622_RNA45SN1_45S_50ntFlanks@11235.11268.34|CTTCCCGTGGATCGCCCCAGCTGCGGCGGGCGTC</td><td>>hg38_21_+_8433172_8446622_RNA45SN1_45S_50ntFlanks</td><td>11235</td><td>11268</td><td>34</td><td>CTTCCCGTGGATCGCCCCAGCTGCGGCGGGCGTC</td><td>119.4</td><td>-3.3</td><td>0.8</td><td>-4.3</td><td>2.1E-05</td><td>3.0E-05</td><td>MB231/MB468</td></tr>
<tr><td>>trnaMT_PheGAA_MT_+_577_647@42.74.33|TTTAGACGGGCTCACATCACCCCATAAACACCA</td><td>>trnaMT_PheGAA_MT_+_577_647</td><td>42</td><td>74</td><td>33</td><td>TTTAGACGGGCTCACATCACCCCATAAACACCA</td><td>579.8</td><td>9.6</td><td>2.3</td><td>4.3</td><td>2.1E-05</td><td>3.0E-05</td><td>MB231/MB468</td></tr>
<tr><td>>hg38_21_+_8433172_8446622_RNA45SN1_45S_50ntFlanks@3985.4003.19|CGGCTTTGGTGACTCTAGA</td><td>>hg38_21_+_8433172_8446622_RNA45SN1_45S_50ntFlanks</td><td>3985</td><td>4003</td><td>19</td><td>CGGCTTTGGTGACTCTAGA</td><td>92.7</td><td>-2.6</td><td>0.6</td><td>-4.2</td><td>2.3E-05</td><td>3.0E-05</td><td>BT20/MB468</td></tr>
<tr><td>>hg38_21_+_8433172_8446622_RNA45SN1_45S_50ntFlanks@5496.5516.21|GACGGTCGAACTTGACTATCT</td><td>>hg38_21_+_8433172_8446622_RNA45SN1_45S_50ntFlanks</td><td>5496</td><td>5516</td><td>21</td><td>GACGGTCGAACTTGACTATCT</td><td>401.0</td><td>10.0</td><td>2.4</td><td>4.2</td><td>2.3E-05</td><td>3.0E-05</td><td>BT20/MB468</td></tr>
<tr><td>>hg38_21_+_8433172_8446622_RNA45SN1_45S_50ntFlanks@3861.3881.21|TAATTCTAGAGCTAATACATG</td><td>>hg38_21_+_8433172_8446622_RNA45SN1_45S_50ntFlanks</td><td>3861</td><td>3881</td><td>21</td><td>TAATTCTAGAGCTAATACATG</td><td>169.4</td><td>-3.4</td><td>0.8</td><td>-4.3</td><td>2.1E-05</td><td>3.0E-05</td><td>MB231/MB468</td></tr>
<tr><td>>MI0000101|hsa-mir-99a&WithFlank&21|+|16539083|16539175@19.40.22(+1U)&[MIMAT0000097&hsa-miR-99a-5p&offsets|0|0(+1U);m-64&21|+|16539101|16539121&offsets|0|+1(+1U)]|AACCCGTAGATCCGATCTTGTGT</td><td>>MI0000101|hsa-mir-99a&WithFlank&21|+|16539083|16539175</td><td>19</td><td>40</td><td>22(+1U)</td><td>AACCCGTAGATCCGATCTTGTGT</td><td>640.8</td><td>3.2</td><td>0.7</td><td>4.3</td><td>2.1E-05</td><td>3.0E-05</td><td>MB231/MB468</td></tr>
<tr><td>>hg38_21_+_8433172_8446622_RNA45SN1_45S_50ntFlanks@12405.12439.35|CTTTTTGATCCTTCGATGTCGGCTCTTCCTATCAT</td><td>>hg38_21_+_8433172_8446622_RNA45SN1_45S_50ntFlanks</td><td>12405</td><td>12439</td><td>35</td><td>CTTTTTGATCCTTCGATGTCGGCTCTTCCTATCAT</td><td>110.4</td><td>-2.0</td><td>0.5</td><td>-4.2</td><td>2.3E-05</td><td>3.0E-05</td><td>BT20/MB468</td></tr>
<tr><td>>hg38_21_+_8433172_8446622_RNA45SN1_45S_50ntFlanks@10819.10838.20|AGGGAAGTCGGCAAGCCGGA</td><td>>hg38_21_+_8433172_8446622_RNA45SN1_45S_50ntFlanks</td><td>10819</td><td>10838</td><td>20</td><td>AGGGAAGTCGGCAAGCCGGA</td><td>154.2</td><td>-3.6</td><td>0.9</td><td>-4.3</td><td>2.1E-05</td><td>3.0E-05</td><td>MB231/MB468</td></tr>
<tr><td>>hg38_21_+_8433172_8446622_RNA45SN1_45S_50ntFlanks@11876.11908.33|AAGACCCTGTTGAGCTTGACTCTAGTCTGGCAC</td><td>>hg38_21_+_8433172_8446622_RNA45SN1_45S_50ntFlanks</td><td>11876</td><td>11908</td><td>33</td><td>AAGACCCTGTTGAGCTTGACTCTAGTCTGGCAC</td><td>138.3</td><td>-3.3</td><td>0.8</td><td>-4.3</td><td>2.1E-05</td><td>3.0E-05</td><td>MB231/MB468</td></tr>
<tr><td>>trna2_GlyGCC_21_-_18827107_18827177@1.31.31,trna35_GlyGCC_1_+_161413094_161413164@1.31.31,trna37_GlyGCC_1_+_161420467_161420537@1.31.31,trna39_GlyGCC_1_+_161427898_161427968@1.31.31,trna41_GlyGCC_1_+_161435258_161435328@1.31.31|GCATGGGTGGTTCAGTGGTAGAATTCTCGCC</td><td>>trna2_GlyGCC_21_-_18827107_18827177</td><td>1</td><td>31</td><td>31</td><td>GCATGGGTGGTTCAGTGGTAGAATTCTCGCC</td><td>1739.2</td><td>3.6</td><td>0.9</td><td>4.2</td><td>2.2E-05</td><td>3.1E-05</td><td>MB231/MB468</td></tr>
</table>

<table>
<tr><th></th><th></th><th></th><th></th><th></th><th></th><th></th><th></th><th></th><th></th><th></th><th></th><th></th></tr>
<tr><td>>hg38_21_+_8433172_8446622_RNA45SN1_45S_50ntFlanks@8171.8195.25|AAGTCCTTCTGATCGAGGCCCAGCC</td><td>>hg38_21_+_8433172_8446622_RNA45SN1_45S_50ntFlanks</td><td>8171</td><td>8195</td><td>25</td><td>AAGTCCTTCTGATCGAGGCCCAGCC</td><td>132.5</td><td>-2.3</td><td>0.5</td><td>-4.2</td><td>2.4E-05</td><td>3.1E-05</td><td>BT20/MB468</td></tr>
<tr><td>>hg38_21_+_8433172_8446622_RNA45SN1_45S_50ntFlanks@9554.9584.31|CGTAGCGGTCCTGACGTGCAAATCGGTCGTC</td><td>>hg38_21_+_8433172_8446622_RNA45SN1_45S_50ntFlanks</td><td>9554</td><td>9584</td><td>31</td><td>CGTAGCGGTCCTGACGTGCAAATCGGTCGTC</td><td>1503.9</td><td>-2.4</td><td>0.6</td><td>-4.2</td><td>2.2E-05</td><td>3.1E-05</td><td>MB231/MB468</td></tr>
<tr><td>>hg38_21_+_8433172_8446622_RNA45SN1_45S_50ntFlanks@6738.6769.32|ATCATCGACACTTCGAACGCACTTGCGGCCCC</td><td>>hg38_21_+_8433172_8446622_RNA45SN1_45S_50ntFlanks</td><td>6738</td><td>6769</td><td>32</td><td>ATCATCGACACTTCGAACGCACTTGCGGCCCC</td><td>100.5</td><td>-3.6</td><td>0.9</td><td>-4.2</td><td>2.2E-05</td><td>3.1E-05</td><td>MB231/MB468</td></tr>
<tr><td>>hg38_21_+_8433172_8446622_RNA45SN1_45S_50ntFlanks@10805.10840.36|GAACAATGTAGGTAAGGGAAGTCGGCAAGCCGGATC</td><td>>hg38_21_+_8433172_8446622_RNA45SN1_45S_50ntFlanks</td><td>10805</td><td>10840</td><td>36</td><td>GAACAATGTAGGTAAGGGAAGTCGGCAAGCCGGATC</td><td>109.8</td><td>-1.9</td><td>0.5</td><td>-4.2</td><td>2.2E-05</td><td>3.2E-05</td><td>MB231/MB468</td></tr>
<tr><td>>hg38_21_+_8433172_8446622_RNA45SN1_45S_50ntFlanks@4368.4389.22|AGTTAAAAAGCTCGTAGTTGGA</td><td>>hg38_21_+_8433172_8446622_RNA45SN1_45S_50ntFlanks</td><td>4368</td><td>4389</td><td>22</td><td>AGTTAAAAAGCTCGTAGTTGGA</td><td>88.2</td><td>-2.7</td><td>0.6</td><td>-4.2</td><td>2.2E-05</td><td>3.2E-05</td><td>MB231/MB468</td></tr>
<tr><td>>hg38_21_+_8433172_8446622_RNA45SN1_45S_50ntFlanks@7982.8004.23|TCAGATCAGACGTGGCGACCCGC</td><td>>hg38_21_+_8433172_8446622_RNA45SN1_45S_50ntFlanks</td><td>7982</td><td>8004</td><td>23</td><td>TCAGATCAGACGTGGCGACCCGC</td><td>128.5</td><td>-3.2</td><td>0.8</td><td>-4.2</td><td>2.5E-05</td><td>3.2E-05</td><td>BT20/MB468</td></tr>
<tr><td>>hg38_21_+_8433172_8446622_RNA45SN1_45S_50ntFlanks@6726.6744.19|CAGGACACATTGATCATCG</td><td>>hg38_21_+_8433172_8446622_RNA45SN1_45S_50ntFlanks</td><td>6726</td><td>6744</td><td>19</td><td>CAGGACACATTGATCATCG</td><td>260.5</td><td>-2.9</td><td>0.7</td><td>-4.2</td><td>2.3E-05</td><td>3.2E-05</td><td>MB231/MB468</td></tr>
<tr><td>>hg38_21_+_8433172_8446622_RNA45SN1_45S_50ntFlanks@12640.12669.30|ACCATCTGTGGGATTATGACTGAACGCCTC</td><td>>hg38_21_+_8433172_8446622_RNA45SN1_45S_50ntFlanks</td><td>12640</td><td>12669</td><td>30</td><td>ACCATCTGTGGGATTATGACTGAACGCCTC</td><td>603.6</td><td>7.9</td><td>1.9</td><td>4.2</td><td>2.3E-05</td><td>3.2E-05</td><td>MB231/MB468</td></tr>
<tr><td>>hg38_21_+_8433172_8446622_RNA45SN1_45S_50ntFlanks@10783.10817.35|GTGAACAGCCTCTGGCATGTTGGAACAATGTAGGT</td><td>>hg38_21_+_8433172_8446622_RNA45SN1_45S_50ntFlanks</td><td>10783</td><td>10817</td><td>35</td><td>GTGAACAGCCTCTGGCATGTTGGAACAATGTAGGT</td><td>281.4</td><td>-3.1</td><td>0.7</td><td>-4.2</td><td>2.5E-05</td><td>3.2E-05</td><td>BT20/MB468</td></tr>
<tr><td>>hg38_21_+_8433172_8446622_RNA45SN1_45S_50ntFlanks@10348.10373.26|AAACGAGAACTTTGAAGGCCGAAGTG</td><td>>hg38_21_+_8433172_8446622_RNA45SN1_45S_50ntFlanks</td><td>10348</td><td>10373</td><td>26</td><td>AAACGAGAACTTTGAAGGCCGAAGTG</td><td>703.5</td><td>3.5</td><td>0.8</td><td>4.2</td><td>2.3E-05</td><td>3.3E-05</td><td>MB231/MB468</td></tr>
<tr><td>>hg38_21_+_8433172_8446622_RNA45SN1_45S_50ntFlanks@10466.10487.22|CGTTGCCCTCGGCCGATCGAAA</td><td>>hg38_21_+_8433172_8446622_RNA45SN1_45S_50ntFlanks</td><td>10466</td><td>10487</td><td>22</td><td>CGTTGCCCTCGGCCGATCGAAA</td><td>136.9</td><td>-3.2</td><td>0.8</td><td>-4.2</td><td>2.5E-05</td><td>3.3E-05</td><td>BT20/MB468</td></tr>
<tr><td>>hg38_21_+_8433172_8446622_RNA45SN1_45S_50ntFlanks@4069.4101.33|CTTTCGATGGTAGTCGCCGTGCCTACCATGGTG</td><td>>hg38_21_+_8433172_8446622_RNA45SN1_45S_50ntFlanks</td><td>4069</td><td>4101</td><td>33</td><td>CTTTCGATGGTAGTCGCCGTGCCTACCATGGTG</td><td>201.6</td><td>-2.5</td><td>0.6</td><td>-4.2</td><td>2.6E-05</td><td>3.3E-05</td><td>BT20/MB468</td></tr>
<tr><td>>hg38_21_+_8433172_8446622_RNA45SN1_45S_50ntFlanks@10802.10826.25|TTGGAACAATGTAGGTAAGGGAAGT</td><td>>hg38_21_+_8433172_8446622_RNA45SN1_45S_50ntFlanks</td><td>10802</td><td>10826</td><td>25</td><td>TTGGAACAATGTAGGTAAGGGAAGT</td><td>89.1</td><td>-2.2</td><td>0.5</td><td>-4.2</td><td>2.4E-05</td><td>3.4E-05</td><td>MB231/MB468</td></tr>
<tr><td>>hg38_21_+_8433172_8446622_RNA45SN1_45S_50ntFlanks@10806.10827.22|AACAATGTAGGTAAGGGAAGTC</td><td>>hg38_21_+_8433172_8446622_RNA45SN1_45S_50ntFlanks</td><td>10806</td><td>10827</td><td>22</td><td>AACAATGTAGGTAAGGGAAGTC</td><td>107.9</td><td>-2.1</td><td>0.5</td><td>-4.2</td><td>2.4E-05</td><td>3.4E-05</td><td>MB231/MB468</td></tr>
<tr><td>>am_trna2_GlyGCC_21_-_18827107_18827177@1.25.25,trna35_GlyGCC_1_+_161413094_161413164@1.25.25,trna37_GlyGCC_1_+_161420467_161420537@1.25.25,trna39_GlyGCC_1_+_161427898_161427968@1.25.25,trna41_GlyGCC_1_+_161435258_161435328@1.25.25|GCATGGGTGGTTCAGTGGTAGAATT</td><td>>am_trna2_GlyGCC_21_-_18827107_18827177</td><td>1</td><td>25</td><td>25</td><td>GCATGGGTGGTTCAGTGGTAGAATT</td><td>642.8</td><td>3.2</td><td>0.8</td><td>4.2</td><td>2.6E-05</td><td>3.4E-05</td><td>BT20/MB468</td></tr>
<tr><td>>hg38_21_+_8433172_8446622_RNA45SN1_45S_50ntFlanks@4723.4748.26|CATTAATCAAGAACGAAAGTCGGAGG</td><td>>hg38_21_+_8433172_8446622_RNA45SN1_45S_50ntFlanks</td><td>4723</td><td>4748</td><td>26</td><td>CATTAATCAAGAACGAAAGTCGGAGG</td><td>305.7</td><td>-2.5</td><td>0.6</td><td>-4.2</td><td>2.4E-05</td><td>3.4E-05</td><td>MB231/MB468</td></tr>
<tr><td>>MI0000478|hsa-mir-149&WithFlank&2|+|240455995|240456095@21.41.21(+1U)&[MIMAT0000450&hsa-miR-149-5p&offsets|0|-2(+1U);m-195&2|+|240456015|240456037&offsets|0|-2(+1U)]|TCTGGCTCCGTGTCTTCACTCT</td><td>>MI0000478|hsa-mir-149&WithFlank&2|+|240455995|240456095</td><td>21</td><td>41</td><td>21(+1U)</td><td>TCTGGCTCCGTGTCTTCACTCT</td><td>698.0</td><td>3.6</td><td>0.8</td><td>4.7</td><td>2.4E-06</td><td>3.4E-05</td><td>BT20/MB231</td></tr>
<tr><td>>hg38_21_+_8433172_8446622_RNA45SN1_45S_50ntFlanks@10808.10837.30|CAATGTAGGTAAGGGAAGTCGGCAAGCCGG</td><td>>hg38_21_+_8433172_8446622_RNA45SN1_45S_50ntFlanks</td><td>10808</td><td>10837</td><td>30</td><td>CAATGTAGGTAAGGGAAGTCGGCAAGCCGG</td><td>116.3</td><td>-4.0</td><td>1.0</td><td>-4.2</td><td>2.6E-05</td><td>3.4E-05</td><td>BT20/MB468</td></tr>
<tr><td>>hg38_21_+_8433172_8446622_RNA45SN1_45S_50ntFlanks@10351.10375.25|CGAGAACTTTGAAGGCCGAAGTGGA</td><td>>hg38_21_+_8433172_8446622_RNA45SN1_45S_50ntFlanks</td><td>10351</td><td>10375</td><td>25</td><td>CGAGAACTTTGAAGGCCGAAGTGGA</td><td>172.2</td><td>-4.8</td><td>1.2</td><td>-4.2</td><td>2.7E-05</td><td>3.5E-05</td><td>BT20/MB468</td></tr>
<tr><td>>hg38_21_+_8433172_8446622_RNA45SN1_45S_50ntFlanks@11572.11606.35|CCGACTTAGAACTGGTGCGGACCAGGGGAATCCGA</td><td>>hg38_21_+_8433172_8446622_RNA45SN1_45S_50ntFlanks</td><td>11572</td><td>11606</td><td>35</td><td>CCGACTTAGAACTGGTGCGGACCAGGGGAATCCGA</td><td>154.8</td><td>-2.6</td><td>0.6</td><td>-4.2</td><td>2.7E-05</td><td>3.5E-05</td><td>BT20/MB468</td></tr>
<tr><td>>hg38_21_+_8433172_8446622_RNA45SN1_45S_50ntFlanks@6646.6666.21|TCGTACGACTCTTAGCGGTGG</td><td>>hg38_21_+_8433172_8446622_RNA45SN1_45S_50ntFlanks</td><td>6646</td><td>6666</td><td>21</td><td>TCGTACGACTCTTAGCGGTGG</td><td>17373.9</td><td>3.0</td><td>0.7</td><td>4.2</td><td>2.7E-05</td><td>3.5E-05</td><td>BT20/MB468</td></tr>
<tr><td>>hg38_21_+_8433172_8446622_RNA45SN1_45S_50ntFlanks@6731.6756.26|CACATTGATCATCGACACTTCGAACG</td><td>>hg38_21_+_8433172_8446622_RNA45SN1_45S_50ntFlanks</td><td>6731</td><td>6756</td><td>26</td><td>CACATTGATCATCGACACTTCGAACG</td><td>593.5</td><td>-2.1</td><td>0.5</td><td>-4.2</td><td>2.5E-05</td><td>3.5E-05</td><td>MB231/MB468</td></tr>
<tr><td>>hg38_21_+_8433172_8446622_RNA45SN1_45S_50ntFlanks@11668.11709.42|TGCCCAGTGCTCTGAATGTCAAAGTGAAGAAATTCAATGAAG</td><td>>hg38_21_+_8433172_8446622_RNA45SN1_45S_50ntFlanks</td><td>11668</td><td>11709</td><td>42</td><td>TGCCCAGTGCTCTGAATGTCAAAGTGAAGAAATTCAATGAAG</td><td>156.0</td><td>-3.9</td><td>0.8</td><td>-4.7</td><td>2.5E-06</td><td>3.5E-05</td><td>BT20/MB231</td></tr>
<tr><td>>hg38_MT_+_598_1651_RNR1_12S_50ntFlanks@972.1004.33|GGTAAGTGTACTGGAAAGTGCACTTGGACGAAC</td><td>>hg38_MT_+_598_1651_RNR1_12S_50ntFlanks</td><td>972</td><td>1004</td><td>33</td><td>GGTAAGTGTACTGGAAAGTGCACTTGGACGAAC</td><td>334.8</td><td>-3.2</td><td>0.7</td><td>-4.7</td><td>2.5E-06</td><td>3.5E-05</td><td>BT20/MB231</td></tr>
<tr><td>>hg38_21_+_8433172_8446622_RNA45SN1_45S_50ntFlanks@3776.3801.26|CCGGTACAGTGAAACTGCGAATGGCT</td><td>>hg38_21_+_8433172_8446622_RNA45SN1_45S_50ntFlanks</td><td>3776</td><td>3801</td><td>26</td><td>CCGGTACAGTGAAACTGCGAATGGCT</td><td>301.9</td><td>-2.8</td><td>0.7</td><td>-4.2</td><td>2.5E-05</td><td>3.6E-05</td><td>MB231/MB468</td></tr>
<tr><td>>hg38_21_+_8433172_8446622_RNA45SN1_45S_50ntFlanks@4567.4589.23|TAATGGAATAGGACCGCGGTTCT</td><td>>hg38_21_+_8433172_8446622_RNA45SN1_45S_50ntFlanks</td><td>4567</td><td>4589</td><td>23</td><td>TAATGGAATAGGACCGCGGTTCT</td><td>178.4</td><td>-2.2</td><td>0.5</td><td>-4.2</td><td>2.5E-05</td><td>3.6E-05</td><td>MB231/MB468</td></tr>
<tr><td>>hg38_21_+_8433172_8446622_RNA45SN1_45S_50ntFlanks@12941.12964.24|ATTCGTAGACGACCTGCTTCTGGG</td><td>>hg38_21_+_8433172_8446622_RNA45SN1_45S_50ntFlanks</td><td>12941</td><td>12964</td><td>24</td><td>ATTCGTAGACGACCTGCTTCTGGG</td><td>116.6</td><td>-1.8</td><td>0.4</td><td>-4.2</td><td>2.6E-05</td><td>3.6E-05</td><td>MB231/MB468</td></tr>
<tr><td>>hg38_21_+_8433172_8446622_RNA45SN1_45S_50ntFlanks@11320.11344.25|CCACGTCTCGTCGCGCGCGCGTCCG</td><td>>hg38_21_+_8433172_8446622_RNA45SN1_45S_50ntFlanks</td><td>11320</td><td>11344</td><td>25</td><td>CCACGTCTCGTCGCGCGCGCGTCCG</td><td>1938.7</td><td>-5.0</td><td>1.2</td><td>-4.2</td><td>2.8E-05</td><td>3.6E-05</td><td>BT20/MB468</td></tr>
<tr><td>>hg38_21_+_8433172_8446622_RNA45SN1_45S_50ntFlanks@7977.7995.19|CGACCTCAGATCAGACGTG</td><td>>hg38_21_+_8433172_8446622_RNA45SN1_45S_50ntFlanks</td><td>7977</td><td>7995</td><td>19</td><td>CGACCTCAGATCAGACGTG</td><td>156.8</td><td>-2.7</td><td>0.6</td><td>-4.2</td><td>2.6E-05</td><td>3.7E-05</td><td>MB231/MB468</td></tr>
<tr><td>>hg38_21_+_8433172_8446622_RNA45SN1_45S_50ntFlanks@11574.11603.30|GACTTAGAACTGGTGCGGACCAGGGGAATC</td><td>>hg38_21_+_8433172_8446622_RNA45SN1_45S_50ntFlanks</td><td>11574</td><td>11603</td><td>30</td><td>GACTTAGAACTGGTGCGGACCAGGGGAATC</td><td>123.4</td><td>-3.6</td><td>0.8</td><td>-4.2</td><td>2.8E-05</td><td>3.7E-05</td><td>BT20/MB468</td></tr>
<tr><td>>hg38_21_+_8433172_8446622_RNA45SN1_45S_50ntFlanks@10342.10361.20|ATATTCAAACGAGAACTTTG</td><td>>hg38_21_+_8433172_8446622_RNA45SN1_45S_50ntFlanks</td><td>10342</td><td>10361</td><td>20</td><td>ATATTCAAACGAGAACTTTG</td><td>791.0</td><td>-2.6</td><td>0.6</td><td>-4.2</td><td>2.8E-05</td><td>3.7E-05</td><td>BT20/MB468</td></tr>
<tr><td>>am_trna116_GluCTC_1_-_145399233_145399304@1.29.29,trna59_GluCTC_1_+_249168447_249168518@1.29.29,trna71_GluCTC_1_-_161439189_161439260@1.29.29,trna74_GluCTC_1_-_161431809_161431880@1.29.29,trna77_GluCTC_1_-_161424398_161424469@1.29.29,trna77_GluCTC_6_+_28949976_28950047@1.29.29,trna80_GluCTC_1_-_161417018_161417089@1.29.29,trna87_GluCTC_6_-_126101393_126101464@1.29.29|TCCCTGGTGGTCTAGTGGTTAGGATTCGG</td><td>>am_trna116_GluCTC_1_-_145399233_145399304</td><td>1</td><td>29</td><td>29</td><td>TCCCTGGTGGTCTAGTGGTTAGGATTCGG</td><td>28614.1</td><td>-3.7</td><td>0.9</td><td>-4.2</td><td>2.9E-05</td><td>3.7E-05</td><td>BT20/MB468</td></tr>
<tr><td>>hg38_21_+_8433172_8446622_RNA45SN1_45S_50ntFlanks@11586.11612.27|GTGCGGACCAGGGGAATCCGACTGTTT</td><td>>hg38_21_+_8433172_8446622_RNA45SN1_45S_50ntFlanks</td><td>11586</td><td>11612</td><td>27</td><td>GTGCGGACCAGGGGAATCCGACTGTTT</td><td>1056.0</td><td>-3.1</td><td>0.7</td><td>-4.2</td><td>2.9E-05</td><td>3.7E-05</td><td>BT20/MB468</td></tr>
<tr><td>>hg38_21_+_8433172_8446622_RNA45SN1_45S_50ntFlanks@10681.10712.32|TTCCGGCGGCGTCCGGTGAGCTCTCGCTGGCC</td><td>>hg38_21_+_8433172_8446622_RNA45SN1_45S_50ntFlanks</td><td>10681</td><td>10712</td><td>32</td><td>TTCCGGCGGCGTCCGGTGAGCTCTCGCTGGCC</td><td>1186.9</td><td>-3.0</td><td>0.7</td><td>-4.2</td><td>2.6E-05</td><td>3.7E-05</td><td>MB231/MB468</td></tr>
<tr><td>>hg38_21_+_8433172_8446622_RNA45SN1_45S_50ntFlanks@10348.10376.29|AAACGAGAACTTTGAAGGCCGAAGTGGAG</td><td>>hg38_21_+_8433172_8446622_RNA45SN1_45S_50ntFlanks</td><td>10348</td><td>10376</td><td>29</td><td>AAACGAGAACTTTGAAGGCCGAAGTGGAG</td><td>140.3</td><td>-2.0</td><td>0.5</td><td>-4.2</td><td>2.9E-05</td><td>3.7E-05</td><td>BT20/MB468</td></tr>
<tr><td>>hg38_21_+_8433172_8446622_RNA45SN1_45S_50ntFlanks@3777.3794.18|CGGTACAGTGAAACTGCG</td><td>>hg38_21_+_8433172_8446622_RNA45SN1_45S_50ntFlanks</td><td>3777</td><td>3794</td><td>18</td><td>CGGTACAGTGAAACTGCG</td><td>137.6</td><td>-4.3</td><td>1.0</td><td>-4.2</td><td>2.9E-05</td><td>3.7E-05</td><td>BT20/MB468</td></tr>
<tr><td>>hg38_21_+_8433172_8446622_RNA45SN1_45S_50ntFlanks@10772.10796.25|AGGTCTCCAAGGTGAACAGCCTCTG</td><td>>hg38_21_+_8433172_8446622_RNA45SN1_45S_50ntFlanks</td><td>10772</td><td>10796</td><td>25</td><td>AGGTCTCCAAGGTGAACAGCCTCTG</td><td>422.1</td><td>-2.6</td><td>0.6</td><td>-4.2</td><td>2.9E-05</td><td>3.8E-05</td><td>BT20/MB468</td></tr>
<tr><td>>hg38_21_+_8433172_8446622_RNA45SN1_45S_50ntFlanks@6671.6697.27|CTCGGCTCGTGCGTCGATGAAGAACGC</td><td>>hg38_21_+_8433172_8446622_RNA45SN1_45S_50ntFlanks</td><td>6671</td><td>6697</td><td>27</td><td>CTCGGCTCGTGCGTCGATGAAGAACGC</td><td>414.1</td><td>-1.8</td><td>0.4</td><td>-4.2</td><td>2.7E-05</td><td>3.8E-05</td><td>MB231/MB468</td></tr>
<tr><td>>am_trna128_GlyGCC_6_-_27870686_27870756@3.34.32,trna18_GlyGCC_16_+_70822597_70822667@3.34.32,trna19_GlyGCC_16_+_70823410_70823480@3.34.32,trna19_GlyGCC_2_-_157257659_157257729@3.34.32,trna24_GlyGCC_16_-_70812942_70813012@3.34.32,trna25_GlyGCC_16_-_70812114_70812184@3.34.32,trna5_GlyGCC_17_+_8029064_8029134@3.34.32,trna68_GlyGCC_1_-_161493637_161493707@3.34.32|ATTGGTGGTTCAGTGGTAGAATTCTCGCCTGC</td><td>>am_trna128_GlyGCC_6_-_27870686_27870756</td><td>3</td><td>34</td><td>32</td><td>ATTGGTGGTTCAGTGGTAGAATTCTCGCCTGC</td><td>286.0</td><td>-4.0</td><td>0.8</td><td>-4.7</td><td>2.7E-06</td><td>3.8E-05</td><td>BT20/MB231</td></tr>
<tr><td>>hg38_21_+_8433172_8446622_RNA45SN1_45S_50ntFlanks@11571.11588.18|GCCGACTTAGAACTGGTG</td><td>>hg38_21_+_8433172_8446622_RNA45SN1_45S_50ntFlanks</td><td>11571</td><td>11588</td><td>18</td><td>GCCGACTTAGAACTGGTG</td><td>410.9</td><td>2.6</td><td>0.6</td><td>4.2</td><td>2.7E-05</td><td>3.8E-05</td><td>MB231/MB468</td></tr>
<tr><td>>hg38_21_+_8433172_8446622_RNA45SN1_45S_50ntFlanks@12456.12491.36|CCAAGCGTTGGATTGTTCACCCACTAATAGGGAACG</td><td>>hg38_21_+_8433172_8446622_RNA45SN1_45S_50ntFlanks</td><td>12456</td><td>12491</td><td>36</td><td>CCAAGCGTTGGATTGTTCACCCACTAATAGGGAACG</td><td>97.8</td><td>-3.8</td><td>0.9</td><td>-4.2</td><td>2.9E-05</td><td>3.8E-05</td><td>BT20/MB468</td></tr>
</table>

<table>
<tr><td>>hg38_21_+_8433172_8446622_RNA45SN1_45S_50ntFlanks@11577.11602.26|TTAGAACTGGTGCGGACCAGGGGAAT</td><td>>hg38_21_+_8433172_8446622_RNA45SN1_45S_50ntFlanks</td><td>11577</td><td>11602</td><td>26</td><td>TTAGAACTGGTGCGGACCAGGGGAAT</td><td>172.0</td><td>-2.5</td><td>0.6</td><td>-4.2</td><td>2.7E-05</td><td>3.9E-05</td><td>MB231/MB468</td></tr>
<tr><td>>trna119_LysCTT_1_-_145395522_145395594@1.29.29,trna11_LysCTT_5_-_180648979_180649051@1.29.29,trna13_LysCTT_6_+_26556774_26556846@1.29.29,trna32_LysCTT_16_-_3207406_3207478@1.29.29,trna7_LysCTT_16_+_3225692_3225764@1.29.29,trna9_LysCTT_5_+_180634755_180634827@1.29.29|GCCCGGCTAGCTCAGTCGGTAGAGCATGA</td><td>>trna119_LysCTT_1_-_145395522_145395594</td><td>1</td><td>29</td><td>29</td><td>GCCCGGCTAGCTCAGTCGGTAGAGCATGA</td><td>1597.9</td><td>-3.1</td><td>0.7</td><td>-4.2</td><td>2.7E-05</td><td>3.9E-05</td><td>MB231/MB468</td></tr>
<tr><td>>hg38_21_+_8433172_8446622_RNA45SN1_45S_50ntFlanks@12418.12445.28|CGATGTCGGCTCTTCCTATCATTGTGAA</td><td>>hg38_21_+_8433172_8446622_RNA45SN1_45S_50ntFlanks</td><td>12418</td><td>12445</td><td>28</td><td>CGATGTCGGCTCTTCCTATCATTGTGAA</td><td>419.3</td><td>-2.2</td><td>0.5</td><td>-4.2</td><td>3.0E-05</td><td>3.9E-05</td><td>BT20/MB468</td></tr>
<tr><td>>hg38_21_+_8433172_8446622_RNA45SN1_45S_50ntFlanks@11694.11717.24|AAGAAATTCAATGAAGCGCGGGTA</td><td>>hg38_21_+_8433172_8446622_RNA45SN1_45S_50ntFlanks</td><td>11694</td><td>11717</td><td>24</td><td>AAGAAATTCAATGAAGCGCGGGTA</td><td>229.4</td><td>-2.9</td><td>0.7</td><td>-4.2</td><td>2.8E-05</td><td>3.9E-05</td><td>MB231/MB468</td></tr>
<tr><td>>hg38_21_+_8433172_8446622_RNA45SN1_45S_50ntFlanks@10411.10441.31|CAGTCGGTCCTGAGAGATGGGCGAGCGCCGT</td><td>>hg38_21_+_8433172_8446622_RNA45SN1_45S_50ntFlanks</td><td>10411</td><td>10441</td><td>31</td><td>CAGTCGGTCCTGAGAGATGGGCGAGCGCCGT</td><td>118.8</td><td>-3.3</td><td>0.8</td><td>-4.2</td><td>3.0E-05</td><td>3.9E-05</td><td>BT20/MB468</td></tr>
<tr><td>>hg38_21_+_8433172_8446622_RNA45SN1_45S_50ntFlanks@11718.11736.19|AACGGCGGGAGTAACTATG</td><td>>hg38_21_+_8433172_8446622_RNA45SN1_45S_50ntFlanks</td><td>11718</td><td>11736</td><td>19</td><td>AACGGCGGGAGTAACTATG</td><td>1188.2</td><td>-3.9</td><td>0.9</td><td>-4.2</td><td>3.0E-05</td><td>3.9E-05</td><td>BT20/MB468</td></tr>
<tr><td>>hg38_21_+_8433172_8446622_RNA45SN1_45S_50ntFlanks@10853.10873.21|ATAAGGATTGGCTCTAAGGGC</td><td>>hg38_21_+_8433172_8446622_RNA45SN1_45S_50ntFlanks</td><td>10853</td><td>10873</td><td>21</td><td>ATAAGGATTGGCTCTAAGGGC</td><td>162.5</td><td>-2.5</td><td>0.6</td><td>-4.2</td><td>3.0E-05</td><td>3.9E-05</td><td>BT20/MB468</td></tr>
<tr><td>>hg38_21_+_8433172_8446622_RNA45SN1_45S_50ntFlanks@6647.6689.43|CGTACGACTCTTAGCGGTGGATCACTCGGCTCGTGCGTCGATG</td><td>>hg38_21_+_8433172_8446622_RNA45SN1_45S_50ntFlanks</td><td>6647</td><td>6689</td><td>43</td><td>CGTACGACTCTTAGCGGTGGATCACTCGGCTCGTGCGTCGATG</td><td>408.9</td><td>2.2</td><td>0.5</td><td>4.2</td><td>2.8E-05</td><td>4.0E-05</td><td>MB231/MB468</td></tr>
<tr><td>>MI0000471|hsa-mir-126&WithFlank&9|+|136670596|136670692@58.77.20(+1A)&[MIMAT0000445&hsa-miR-126-3p&offsets|0|-2(+1A);m-40&9|+|136670653|136670674&offsets|0|-2(+1A)]|TCGTACCGTGAGTAATAATGA</td><td>>MI0000471|hsa-mir-126&WithFlank&9|+|136670596|136670692</td><td>58</td><td>77</td><td>20(+1A)</td><td>TCGTACCGTGAGTAATAATGA</td><td>597.7</td><td>9.8</td><td>2.3</td><td>4.2</td><td>3.1E-05</td><td>4.0E-05</td><td>BT20/MB468</td></tr>
<tr><td>>trnaMT_AsnGTT_MT_-_5657_5729@1.40.40,trnalookalike5_AsnGTT_1_-_566207_566279@1.40.40|TAGATTGAAGCCAGTTGATTAGGGTGCTTAGCTGTTAACT</td><td>>trnaMT_AsnGTT_MT_-_5657_5729</td><td>1</td><td>40</td><td>40</td><td>TAGATTGAAGCCAGTTGATTAGGGTGCTTAGCTGTTAACT</td><td>441.1</td><td>9.8</td><td>2.3</td><td>4.2</td><td>3.1E-05</td><td>4.1E-05</td><td>BT20/MB468</td></tr>
<tr><td>>hg38_21_+_8433172_8446622_RNA45SN1_45S_50ntFlanks@3835.3861.27|CTCCTCTCCTACTTGGATAACTGTGGT</td><td>>hg38_21_+_8433172_8446622_RNA45SN1_45S_50ntFlanks</td><td>3835</td><td>3861</td><td>27</td><td>CTCCTCTCCTACTTGGATAACTGTGGT</td><td>190.8</td><td>-2.7</td><td>0.6</td><td>-4.2</td><td>3.1E-05</td><td>4.1E-05</td><td>BT20/MB468</td></tr>
<tr><td>>hg38_21_+_8433172_8446622_RNA45SN1_45S_50ntFlanks@11564.11590.27|CCTAGCAGCCGACTTAGAACTGGTGCG</td><td>>hg38_21_+_8433172_8446622_RNA45SN1_45S_50ntFlanks</td><td>11564</td><td>11590</td><td>27</td><td>CCTAGCAGCCGACTTAGAACTGGTGCG</td><td>233.1</td><td>-2.3</td><td>0.6</td><td>-4.2</td><td>2.9E-05</td><td>4.1E-05</td><td>MB231/MB468</td></tr>
<tr><td>>hg38_21_+_8433172_8446622_RNA45SN1_45S_50ntFlanks@11570.11600.31|AGCCGACTTAGAACTGGTGCGGACCAGGGGA</td><td>>hg38_21_+_8433172_8446622_RNA45SN1_45S_50ntFlanks</td><td>11570</td><td>11600</td><td>31</td><td>AGCCGACTTAGAACTGGTGCGGACCAGGGGA</td><td>229.4</td><td>-1.9</td><td>0.5</td><td>-4.2</td><td>3.2E-05</td><td>4.1E-05</td><td>BT20/MB468</td></tr>
<tr><td>>hg38_21_+_8433172_8446622_RNA45SN1_45S_50ntFlanks@11572.11590.19|CCGACTTAGAACTGGTGCG</td><td>>hg38_21_+_8433172_8446622_RNA45SN1_45S_50ntFlanks</td><td>11572</td><td>11590</td><td>19</td><td>CCGACTTAGAACTGGTGCG</td><td>2078.5</td><td>-2.2</td><td>0.5</td><td>-4.2</td><td>2.9E-05</td><td>4.1E-05</td><td>MB231/MB468</td></tr>
<tr><td>>hg38_21_+_8433172_8446622_RNA45SN1_45S_50ntFlanks@10809.10826.18|AATGTAGGTAAGGGAAGT</td><td>>hg38_21_+_8433172_8446622_RNA45SN1_45S_50ntFlanks</td><td>10809</td><td>10826</td><td>18</td><td>AATGTAGGTAAGGGAAGT</td><td>945.6</td><td>3.9</td><td>0.9</td><td>4.2</td><td>2.9E-05</td><td>4.2E-05</td><td>MB231/MB468</td></tr>
<tr><td>>hg38_21_+_8433172_8446622_RNA45SN1_45S_50ntFlanks@11578.11601.24|TAGAACTGGTGCGGACCAGGGGAA</td><td>>hg38_21_+_8433172_8446622_RNA45SN1_45S_50ntFlanks</td><td>11578</td><td>11601</td><td>24</td><td>TAGAACTGGTGCGGACCAGGGGAA</td><td>221.9</td><td>-2.6</td><td>0.6</td><td>-4.2</td><td>3.2E-05</td><td>4.2E-05</td><td>BT20/MB468</td></tr>
<tr><td>>MI0000061|hsa-let-7a-2&WithFlank&11|-|122146516|122146599@11.30.20(+1A);MI0000062|hsa-let-7a-3&WithFlank&22|+|46112743|46112828@10.29.20(+1A);MI0000060|hsa-let-7a-1&WithFlank&9|+|94175951|94176042@12.31.20(+1A)&[MIMAT0000062&hsa-let-7a-5p&offsets|0|-2(+1A);m-6&11|-|122146568|122146589&offsets|0|-2(+1A)];[MIMAT0000062_1&hsa-let-7a-5p&offsets|0|-2(+1A);m-5&22|+|46112752|46112773&offsets|0|-2(+1A)];[MIMAT0000062_2&hsa-let-7a-5p&offsets|0|-2(+1A);m-4&9|+|94175962|94175983&offsets|0|-2(+1A)]|TGAGGTAGTAGGTTGTATAGA</td><td>>MI0000061|hsa-let-7a-2&WithFlank&11|-|122146516|122146599</td><td>11</td><td>30</td><td>20(+1A)</td><td>TGAGGTAGTAGGTTGTATAGA</td><td>739.3</td><td>10.1</td><td>2.4</td><td>4.2</td><td>3.2E-05</td><td>4.2E-05</td><td>BT20/MB468</td></tr>
<tr><td>>hg38_21_+_8433172_8446622_RNA45SN1_45S_50ntFlanks@6737.6756.20|GATCATCGACACTTCGAACG</td><td>>hg38_21_+_8433172_8446622_RNA45SN1_45S_50ntFlanks</td><td>6737</td><td>6756</td><td>20</td><td>GATCATCGACACTTCGAACG</td><td>107.1</td><td>-3.6</td><td>0.9</td><td>-4.2</td><td>3.0E-05</td><td>4.2E-05</td><td>MB231/MB468</td></tr>
<tr><td>>hg38_1_-_228634819_228635039_RNA5S12_5S_50ntFlanks@84.102.19|CTCGTCTGATCTCGGAAGC</td><td>>hg38_1_-_228634819_228635039_RNA5S12_5S_50ntFlanks</td><td>84</td><td>102</td><td>19</td><td>CTCGTCTGATCTCGGAAGC</td><td>344.5</td><td>-3.7</td><td>0.9</td><td>-4.2</td><td>3.0E-05</td><td>4.3E-05</td><td>MB231/MB468</td></tr>
<tr><td>>hg38_21_+_8433172_8446622_RNA45SN1_45S_50ntFlanks@8176.8212.37|CTTCTGATCGAGGCCCAGCCCGTGGACGGTGTGAGGC</td><td>>hg38_21_+_8433172_8446622_RNA45SN1_45S_50ntFlanks</td><td>8176</td><td>8212</td><td>37</td><td>CTTCTGATCGAGGCCCAGCCCGTGGACGGTGTGAGGC</td><td>561.3</td><td>2.7</td><td>0.6</td><td>4.2</td><td>3.0E-05</td><td>4.3E-05</td><td>MB231/MB468</td></tr>
<tr><td>>hg38_21_+_8433172_8446622_RNA45SN1_45S_50ntFlanks@4555.4573.19|CGCAGCTAGGAATAATGGA</td><td>>hg38_21_+_8433172_8446622_RNA45SN1_45S_50ntFlanks</td><td>4555</td><td>4573</td><td>19</td><td>CGCAGCTAGGAATAATGGA</td><td>179.2</td><td>-2.6</td><td>0.6</td><td>-4.2</td><td>3.0E-05</td><td>4.3E-05</td><td>MB231/MB468</td></tr>
<tr><td>>hg38_21_+_8433172_8446622_RNA45SN1_45S_50ntFlanks@10357.10376.20|CTTTGAAGGCCGAAGTGGAG</td><td>>hg38_21_+_8433172_8446622_RNA45SN1_45S_50ntFlanks</td><td>10357</td><td>10376</td><td>20</td><td>CTTTGAAGGCCGAAGTGGAG</td><td>94.9</td><td>-2.4</td><td>0.6</td><td>-4.2</td><td>3.1E-05</td><td>4.3E-05</td><td>MB231/MB468</td></tr>
<tr><td>>hg38_21_+_8433172_8446622_RNA45SN1_45S_50ntFlanks@6669.6686.18|CACTCGGCTCGTGCGTCG</td><td>>hg38_21_+_8433172_8446622_RNA45SN1_45S_50ntFlanks</td><td>6669</td><td>6686</td><td>18</td><td>CACTCGGCTCGTGCGTCG</td><td>130.7</td><td>-3.0</td><td>0.7</td><td>-4.1</td><td>3.3E-05</td><td>4.3E-05</td><td>BT20/MB468</td></tr>
<tr><td>>MI0001445|hsa-mir-423&WithFlank&17|+|30117073|30117178@58.80.23(+1G)&[MIMAT0001340&hsa-miR-423-3p&offsets|-1|-1(+1G);m-100&17|+|30117131|30117153&offsets|-1|-1(+1G)]|AAGCTCGGTCTGAGGCCCCTCAGG</td><td>>MI0001445|hsa-mir-423&WithFlank&17|+|30117073|30117178</td><td>58</td><td>80</td><td>23(+1G)</td><td>AAGCTCGGTCTGAGGCCCCTCAGG</td><td>402.7</td><td>9.4</td><td>2.3</td><td>4.1</td><td>3.4E-05</td><td>4.3E-05</td><td>BT20/MB468</td></tr>
<tr><td>>hg38_21_+_8433172_8446622_RNA45SN1_45S_50ntFlanks@10682.10709.28|TCCGGCGGCGTCCGGTGAGCTCTCGCTG</td><td>>hg38_21_+_8433172_8446622_RNA45SN1_45S_50ntFlanks</td><td>10682</td><td>10709</td><td>28</td><td>TCCGGCGGCGTCCGGTGAGCTCTCGCTG</td><td>92.9</td><td>-3.1</td><td>0.7</td><td>-4.2</td><td>3.1E-05</td><td>4.3E-05</td><td>MB231/MB468</td></tr>
<tr><td>>hg38_21_+_8433172_8446622_RNA45SN1_45S_50ntFlanks@11602.11620.19|TCCGACTGTTTAATTAAAA</td><td>>hg38_21_+_8433172_8446622_RNA45SN1_45S_50ntFlanks</td><td>11602</td><td>11620</td><td>19</td><td>TCCGACTGTTTAATTAAAA</td><td>559.7</td><td>-2.1</td><td>0.5</td><td>-4.2</td><td>3.1E-05</td><td>4.4E-05</td><td>MB231/MB468</td></tr>
<tr><td>>hg38_21_+_8433172_8446622_RNA45SN1_45S_50ntFlanks@12407.12448.42|TTTTGATCCTTCGATGTCGGCTCTTCCTATCATTGTGAAGCA</td><td>>hg38_21_+_8433172_8446622_RNA45SN1_45S_50ntFlanks</td><td>12407</td><td>12448</td><td>42</td><td>TTTTGATCCTTCGATGTCGGCTCTTCCTATCATTGTGAAGCA</td><td>534.3</td><td>9.5</td><td>2.3</td><td>4.2</td><td>3.1E-05</td><td>4.4E-05</td><td>MB231/MB468</td></tr>
<tr><td>>hg38_21_+_8433172_8446622_RNA45SN1_45S_50ntFlanks@8153.8181.29|GCCGCTCGTGGGGGGCCCAAGTCCTTCTG</td><td>>hg38_21_+_8433172_8446622_RNA45SN1_45S_50ntFlanks</td><td>8153</td><td>8181</td><td>29</td><td>GCCGCTCGTGGGGGGCCCAAGTCCTTCTG</td><td>150.7</td><td>-2.7</td><td>0.7</td><td>-4.1</td><td>3.4E-05</td><td>4.4E-05</td><td>BT20/MB468</td></tr>
<tr><td>>hg38_21_+_8433172_8446622_RNA45SN1_45S_50ntFlanks@12479.12497.19|CTAATAGGGAACGTGAGCT</td><td>>hg38_21_+_8433172_8446622_RNA45SN1_45S_50ntFlanks</td><td>12479</td><td>12497</td><td>19</td><td>CTAATAGGGAACGTGAGCT</td><td>1579.0</td><td>1.5</td><td>0.4</td><td>4.1</td><td>3.4E-05</td><td>4.4E-05</td><td>BT20/MB468</td></tr>
<tr><td>>trna111_HisGTG_1_-_147774845_147774916@-1T.22.23,trna118_HisGTG_1_-_145396881_145396952@-1T.22.23,trna16_HisGTG_1_+_146544773_146544844@-1T.22.23,trna1_HisGTG_15_+_45493349_45493420@-1T.22.23,trna21_HisGTG_1_+_147753471_147753542@-1T.22.23,trna33_HisGTG_6_+_27125906_27125977@-1T.22.23,trna7_HisGTG_9_-_14433938_14434009@-1T.22.23,trna8_HisGTG_15_-_45492611_45492682@-1T.22.23,trna9_HisGTG_15_-_45490804_45490875@-1T.22.23|TGCCGTGATCGTATAGTGGTTAG</td><td>>trna111_HisGTG_1_-_147774845_147774916</td><td>-1T</td><td>22</td><td>23</td><td>TGCCGTGATCGTATAGTGGTTAG</td><td>251.3</td><td>-3.5</td><td>0.7</td><td>-4.7</td><td>3.2E-06</td><td>4.4E-05</td><td>BT20/MB231</td></tr>
<tr><td>>hg38_21_+_8433172_8446622_RNA45SN1_45S_50ntFlanks@12992.13039.48|TCCCTCGCTGCGATCTATTGAAAGTCAGCCCTCGACACAAGGGTTTGT</td><td>>hg38_21_+_8433172_8446622_RNA45SN1_45S_50ntFlanks</td><td>12992</td><td>13039</td><td>48</td><td>TCCCTCGCTGCGATCTATTGAAAGTCAGCCCTCGACACAAGGGTTTGT</td><td>2336.6</td><td>4.0</td><td>1.0</td><td>4.2</td><td>3.1E-05</td><td>4.4E-05</td><td>MB231/MB468</td></tr>
<tr><td>>hg38_21_+_8433172_8446622_RNA45SN1_45S_50ntFlanks@12420.12445.26|ATGTCGGCTCTTCCTATCATTGTGAA</td><td>>hg38_21_+_8433172_8446622_RNA45SN1_45S_50ntFlanks</td><td>12420</td><td>12445</td><td>26</td><td>ATGTCGGCTCTTCCTATCATTGTGAA</td><td>744.8</td><td>-2.3</td><td>0.6</td><td>-4.1</td><td>3.4E-05</td><td>4.4E-05</td><td>BT20/MB468</td></tr>
<tr><td>>hg38_21_+_8433172_8446622_RNA45SN1_45S_50ntFlanks@8384.8402.19|AGGGCGTGAAACCGTTAAG</td><td>>hg38_21_+_8433172_8446622_RNA45SN1_45S_50ntFlanks</td><td>8384</td><td>8402</td><td>19</td><td>AGGGCGTGAAACCGTTAAG</td><td>146.8</td><td>-4.6</td><td>1.1</td><td>-4.1</td><td>3.4E-05</td><td>4.5E-05</td><td>BT20/MB468</td></tr>
<tr><td>>hg38_21_+_8433172_8446622_RNA45SN1_45S_50ntFlanks@12479.12497.19|CTAATAGGGAACGTGAGCT</td><td>>hg38_21_+_8433172_8446622_RNA45SN1_45S_50ntFlanks</td><td>12479</td><td>12497</td><td>19</td><td>CTAATAGGGAACGTGAGCT</td><td>2309.9</td><td>2.6</td><td>0.6</td><td>4.2</td><td>3.2E-05</td><td>4.5E-05</td><td>MB231/MB468</td></tr>
<tr><td>>trna116_GluCTC_1_-_145399233_145399304@36.74.39,trna134_GluTTC_1_-_16861774_16861845@36.74.39,trna71_GluCTC_1_-_161439189_161439260@36.74.39,trna74_GluCTC_1_-_161431809_161431880@36.74.39,trna77_GluCTC_1_-_161424398_161424469@36.74.39,trna77_GluCTC_6_+_28949976_28950047@36.74.39,trna80_GluCTC_1_-_161417018_161417089@36.74.39,trna84_GluTTC_1_-_161391883_161391954@36.74.39,trna87_GluCTC_6_-_126101393_126101464@36.74.39|CACCGCCGCGGCCCGGGTTCGATTCCCGGTCAGGGAACC</td><td>>trna116_GluCTC_1_-_145399233_145399304</td><td>36</td><td>74</td><td>39</td><td>CACCGCCGCGGCCCGGGTTCGATTCCCGGTCAGGGAACC</td><td>1160.7</td><td>8.7</td><td>2.1</td><td>4.2</td><td>3.2E-05</td><td>4.5E-05</td><td>MB231/MB468</td></tr>
<tr><td>>hg38_21_+_8433172_8446622_RNA45SN1_45S_50ntFlanks@6661.6688.28|CGGTGGATCACTCGGCTCGTGCGTCGAT</td><td>>hg38_21_+_8433172_8446622_RNA45SN1_45S_50ntFlanks</td><td>6661</td><td>6688</td><td>28</td><td>CGGTGGATCACTCGGCTCGTGCGTCGAT</td><td>997.3</td><td>2.7</td><td>0.6</td><td>4.2</td><td>3.2E-05</td><td>4.5E-05</td><td>MB231/MB468</td></tr>
<tr><td>>hg38_21_+_8433172_8446622_RNA45SN1_45S_50ntFlanks@9534.9569.36|AGGAAACTCTGGTGGAGGTCCGTAGCGGTCCTGACG</td><td>>hg38_21_+_8433172_8446622_RNA45SN1_45S_50ntFlanks</td><td>9534</td><td>9569</td><td>36</td><td>AGGAAACTCTGGTGGAGGTCCGTAGCGGTCCTGACG</td><td>1682.3</td><td>-2.9</td><td>0.7</td><td>-4.2</td><td>3.2E-05</td><td>4.5E-05</td><td>MB231/MB468</td></tr>
<tr><td>>MI0000299|hsa-mir-222&WithFlank&X|-|45747009|45747130@75.99.25(+4U)&[MIMAT0000279&hsa-miR-222-3p&offsets|0|+4(+4U);m-140&X|-|45747033|45747056&offsets|0|+1(+4U)]|AGCTACATCTGGCTACTGGGTCTCTTTTT</td><td>>MI0000299|hsa-mir-222&WithFlank&X|-|45747009|45747130</td><td>75</td><td>99</td><td>25(+4U)</td><td>AGCTACATCTGGCTACTGGGTCTCTTTTT</td><td>546.7</td><td>-3.8</td><td>0.8</td><td>-4.7</td><td>3.3E-06</td><td>4.5E-05</td><td>BT20/MB231</td></tr>
</table>

<table>
<tr><th></th><th></th><th></th><th></th><th></th><th></th><th></th><th></th><th></th><th></th><th></th><th></th><th></th></tr>
<tr><td>>MI0000466|hsa-mir-9-1&WithFlank&1|-|156420335|156420435@22.42.21(+1A);MI0000468|hsa-mir-9-3&WithFlank&15|+|89368011|89368112@22.42.21(+1A);MI0000467|hsa-mir-9-2&WithFlank&5|-|88666847|88666945@22.42.21(+1A)&[MIMAT0000441&hsa-miR-9-5p&offsets|0|-2(+1A);m-104&1|-|156420392|156420414&offsets|0|-2(+1A)];[MIMAT0000441_1&hsa-miR-9-5p&offsets|0|-2(+1A);m-105&15|+|89368032|89368054&offsets|0|-2(+1A)];[MIMAT0000441_2&hsa-miR-9-5p&offsets|0|-2(+1A);m-104&5|-|88666902|88666924&offsets|0|-2(+1A)]|TCTTTGGTTATCTAGCTGTATA</td><td>>MI0000466|hsa-mir-9-1&WithFlank&1|-|156420335|156420435</td><td>22</td><td>42</td><td>21(+1A)</td><td>TCTTTGGTTATCTAGCTGTATA</td><td>504.2</td><td>4.0</td><td>0.8</td><td>4.7</td><td>3.3E-06</td><td>4.5E-05</td><td>BT20/MB231</td></tr>
<tr><td>>hg38_21_+_8433172_8446622_RNA45SN1_45S_50ntFlanks@6663.6680.18|GTGGATCACTCGGCTCGT</td><td>>hg38_21_+_8433172_8446622_RNA45SN1_45S_50ntFlanks</td><td>6663</td><td>6680</td><td>18</td><td>GTGGATCACTCGGCTCGT</td><td>94.9</td><td>-2.8</td><td>0.7</td><td>-4.1</td><td>3.5E-05</td><td>4.5E-05</td><td>BT20/MB468</td></tr>
<tr><td>>am_MI0003639|hsa-mir-625&WithFlank&14|+|65471096|65471192@58.80.23&[MIMAT0004808&hsa-miR-625-3p&offsets|0|+1;m-137&14|+|65471153|65471174&offsets|0|+1]|GACTATAGAACTTTCCCCCTCAT</td><td>>am_MI0003639|hsa-mir-625&WithFlank&14|+|65471096|65471192</td><td>58</td><td>80</td><td>23</td><td>GACTATAGAACTTTCCCCCTCAT</td><td>428.8</td><td>-3.7</td><td>0.8</td><td>-4.6</td><td>3.3E-06</td><td>4.5E-05</td><td>BT20/MB231</td></tr>
<tr><td>>MI0000102|hsa-mir-100&WithFlank&11|-|122152223|122152314@19.37.19&[MIMAT0000098&hsa-miR-100-5p&offsets|0|-3;m-27&11|-|122152276|122152296&offsets|0|-2]|AACCCGTAGATCCGAACTT</td><td>>MI0000102|hsa-mir-100&WithFlank&11|-|122152223|122152314</td><td>19</td><td>37</td><td>19</td><td>AACCCGTAGATCCGAACTT</td><td>355.3</td><td>-4.7</td><td>1.0</td><td>-4.6</td><td>3.3E-06</td><td>4.5E-05</td><td>BT20/MB231</td></tr>
<tr><td>>MI0000273|hsa-mir-183&WithFlank&7|-|129774899|129775020@34.54.21&[MIMAT0000261&hsa-miR-183-5p&offsets|+1|0;m-41&7|-|129774966|129774987&offsets|0|-1]|ATGGCACTGGTAGAATTCACT</td><td>>MI0000273|hsa-mir-183&WithFlank&7|-|129774899|129775020</td><td>34</td><td>54</td><td>21</td><td>ATGGCACTGGTAGAATTCACT</td><td>4992.9</td><td>3.0</td><td>0.6</td><td>4.6</td><td>3.3E-06</td><td>4.5E-05</td><td>BT20/MB231</td></tr>
<tr><td>>hg38_21_+_8433172_8446622_RNA45SN1_45S_50ntFlanks@11670.11698.29|CCCAGTGCTCTGAATGTCAAAGTGAAGAA</td><td>>hg38_21_+_8433172_8446622_RNA45SN1_45S_50ntFlanks</td><td>11670</td><td>11698</td><td>29</td><td>CCCAGTGCTCTGAATGTCAAAGTGAAGAA</td><td>209.5</td><td>-2.4</td><td>0.6</td><td>-4.2</td><td>3.2E-05</td><td>4.6E-05</td><td>MB231/MB468</td></tr>
<tr><td>>hg38_21_+_8433172_8446622_RNA45SN1_45S_50ntFlanks@12640.12668.29|ACCATCTGTGGGATTATGACTGAACGCCT</td><td>>hg38_21_+_8433172_8446622_RNA45SN1_45S_50ntFlanks</td><td>12640</td><td>12668</td><td>29</td><td>ACCATCTGTGGGATTATGACTGAACGCCT</td><td>403.5</td><td>7.7</td><td>1.8</td><td>4.2</td><td>3.2E-05</td><td>4.6E-05</td><td>MB231/MB468</td></tr>
<tr><td>>hg38_21_+_8433172_8446622_RNA45SN1_45S_50ntFlanks@11572.11597.26|CCGACTTAGAACTGGTGCGGACCAGG</td><td>>hg38_21_+_8433172_8446622_RNA45SN1_45S_50ntFlanks</td><td>11572</td><td>11597</td><td>26</td><td>CCGACTTAGAACTGGTGCGGACCAGG</td><td>313.4</td><td>-2.6</td><td>0.6</td><td>-4.1</td><td>3.6E-05</td><td>4.6E-05</td><td>BT20/MB468</td></tr>
<tr><td>>hg38_21_+_8433172_8446622_RNA45SN1_45S_50ntFlanks@12459.12488.30|AGCGTTGGATTGTTCACCCACTAATAGGGA</td><td>>hg38_21_+_8433172_8446622_RNA45SN1_45S_50ntFlanks</td><td>12459</td><td>12488</td><td>30</td><td>AGCGTTGGATTGTTCACCCACTAATAGGGA</td><td>156.3</td><td>-4.3</td><td>1.0</td><td>-4.1</td><td>3.6E-05</td><td>4.6E-05</td><td>BT20/MB468</td></tr>
<tr><td>>hg38_21_+_8433172_8446622_RNA45SN1_45S_50ntFlanks@6699.6721.23|GCTAGCTGCGAGAATTAATGTGA</td><td>>hg38_21_+_8433172_8446622_RNA45SN1_45S_50ntFlanks</td><td>6699</td><td>6721</td><td>23</td><td>GCTAGCTGCGAGAATTAATGTGA</td><td>221.4</td><td>-1.8</td><td>0.4</td><td>-4.1</td><td>3.6E-05</td><td>4.6E-05</td><td>BT20/MB468</td></tr>
<tr><td>>MI0000286|hsa-mir-210&WithFlank&11|-|568083|568204@72.93.22&[MIMAT0000267&hsa-miR-210-3p&offsets|0|0;m-102&11|-|568112|568133&offsets|0|0]|CTGTGCGTGTGACAGCGGCTGA</td><td>>MI0000286|hsa-mir-210&WithFlank&11|-|568083|568204</td><td>72</td><td>93</td><td>22</td><td>CTGTGCGTGTGACAGCGGCTGA</td><td>6461.3</td><td>4.7</td><td>1.0</td><td>4.6</td><td>3.4E-06</td><td>4.6E-05</td><td>BT20/MB231</td></tr>
<tr><td>>hg38_21_+_8433172_8446622_RNA45SN1_45S_50ntFlanks@10393.10433.41|CAGCAGTTGAACATGGGTCAGTCGGTCCTGAGAGATGGGCG</td><td>>hg38_21_+_8433172_8446622_RNA45SN1_45S_50ntFlanks</td><td>10393</td><td>10433</td><td>41</td><td>CAGCAGTTGAACATGGGTCAGTCGGTCCTGAGAGATGGGCG</td><td>176.6</td><td>-4.0</td><td>1.0</td><td>-4.1</td><td>3.6E-05</td><td>4.6E-05</td><td>BT20/MB468</td></tr>
<tr><td>>MI0000737|hsa-mir-200a&WithFlank&1|+|1167857|1167958@22.43.22&[MIMAT0001620&hsa-miR-200a-5p&offsets|0|0;m-154&1|+|1167878|1167899&offsets|0|0]|CATCTTACCGGACAGTGCTGGA</td><td>>MI0000737|hsa-mir-200a&WithFlank&1|+|1167857|1167958</td><td>22</td><td>43</td><td>22</td><td>CATCTTACCGGACAGTGCTGGA</td><td>198.1</td><td>-2.2</td><td>0.5</td><td>-4.1</td><td>3.6E-05</td><td>4.7E-05</td><td>BT20/MB468</td></tr>
<tr><td>>hg38_21_+_8433172_8446622_RNA45SN1_45S_50ntFlanks@3782.3805.24|CAGTGAAACTGCGAATGGCTCATT</td><td>>hg38_21_+_8433172_8446622_RNA45SN1_45S_50ntFlanks</td><td>3782</td><td>3805</td><td>24</td><td>CAGTGAAACTGCGAATGGCTCATT</td><td>303.1</td><td>-1.7</td><td>0.4</td><td>-4.2</td><td>3.3E-05</td><td>4.7E-05</td><td>MB231/MB468</td></tr>
<tr><td>>hg38_21_+_8433172_8446622_RNA45SN1_45S_50ntFlanks@10403.10420.18|ACATGGGTCAGTCGGTCC</td><td>>hg38_21_+_8433172_8446622_RNA45SN1_45S_50ntFlanks</td><td>10403</td><td>10420</td><td>18</td><td>ACATGGGTCAGTCGGTCC</td><td>121.3</td><td>-2.5</td><td>0.6</td><td>-4.2</td><td>3.3E-05</td><td>4.7E-05</td><td>MB231/MB468</td></tr>
<tr><td>>hg38_21_+_8433172_8446622_RNA45SN1_45S_50ntFlanks@11873.11901.29|AAGAAGACCCTGTTGAGCTTGACTCTAGT</td><td>>hg38_21_+_8433172_8446622_RNA45SN1_45S_50ntFlanks</td><td>11873</td><td>11901</td><td>29</td><td>AAGAAGACCCTGTTGAGCTTGACTCTAGT</td><td>142.1</td><td>-1.4</td><td>0.3</td><td>-4.1</td><td>3.7E-05</td><td>4.7E-05</td><td>BT20/MB468</td></tr>
<tr><td>>hg38_21_+_8433172_8446622_RNA45SN1_45S_50ntFlanks@12444.12480.37|AAGCAGAATTCACCAAGCGTTGGATTGTTCACCCACT</td><td>>hg38_21_+_8433172_8446622_RNA45SN1_45S_50ntFlanks</td><td>12444</td><td>12480</td><td>37</td><td>AAGCAGAATTCACCAAGCGTTGGATTGTTCACCCACT</td><td>599.6</td><td>2.8</td><td>0.7</td><td>4.1</td><td>3.3E-05</td><td>4.7E-05</td><td>MB231/MB468</td></tr>
<tr><td>>hg38_21_+_8433172_8446622_RNA45SN1_45S_50ntFlanks@12076.12097.22|AGGGGCTCTCGCTTCTGGCGCC</td><td>>hg38_21_+_8433172_8446622_RNA45SN1_45S_50ntFlanks</td><td>12076</td><td>12097</td><td>22</td><td>AGGGGCTCTCGCTTCTGGCGCC</td><td>192.7</td><td>-2.1</td><td>0.5</td><td>-4.1</td><td>3.4E-05</td><td>4.8E-05</td><td>MB231/MB468</td></tr>
<tr><td>>hg38_21_+_8433172_8446622_RNA45SN1_45S_50ntFlanks@4367.4393.27|CAGTTAAAAAGCTCGTAGTTGGATCTT</td><td>>hg38_21_+_8433172_8446622_RNA45SN1_45S_50ntFlanks</td><td>4367</td><td>4393</td><td>27</td><td>CAGTTAAAAAGCTCGTAGTTGGATCTT</td><td>142.4</td><td>-4.6</td><td>1.1</td><td>-4.1</td><td>3.7E-05</td><td>4.8E-05</td><td>BT20/MB468</td></tr>
<tr><td>>hg38_21_+_8433172_8446622_RNA45SN1_45S_50ntFlanks@6656.6680.25|CTTAGCGGTGGATCACTCGGCTCGT</td><td>>hg38_21_+_8433172_8446622_RNA45SN1_45S_50ntFlanks</td><td>6656</td><td>6680</td><td>25</td><td>CTTAGCGGTGGATCACTCGGCTCGT</td><td>98.6</td><td>-2.5</td><td>0.6</td><td>-4.1</td><td>3.4E-05</td><td>4.9E-05</td><td>MB231/MB468</td></tr>
<tr><td>>hg38_21_+_8433172_8446622_RNA45SN1_45S_50ntFlanks@4727.4758.32|AATCAAGAACGAAAGTCGGAGGTTCGAAGACG</td><td>>hg38_21_+_8433172_8446622_RNA45SN1_45S_50ntFlanks</td><td>4727</td><td>4758</td><td>32</td><td>AATCAAGAACGAAAGTCGGAGGTTCGAAGACG</td><td>1087.0</td><td>2.8</td><td>0.7</td><td>4.1</td><td>3.5E-05</td><td>4.9E-05</td><td>MB231/MB468</td></tr>
<tr><td>>hg38_21_+_8433172_8446622_RNA45SN1_45S_50ntFlanks@10669.10706.38|AAAGCGTCGCGGTTCCGGCGGCGTCCGGTGAGCTCTCG</td><td>>hg38_21_+_8433172_8446622_RNA45SN1_45S_50ntFlanks</td><td>10669</td><td>10706</td><td>38</td><td>AAAGCGTCGCGGTTCCGGCGGCGTCCGGTGAGCTCTCG</td><td>228.2</td><td>-4.4</td><td>1.1</td><td>-4.1</td><td>3.8E-05</td><td>4.9E-05</td><td>BT20/MB468</td></tr>
<tr><td>>hg38_21_+_8433172_8446622_RNA45SN1_45S_50ntFlanks@6661.6698.38|CGGTGGATCACTCGGCTCGTGCGTCGATGAAGAACGCA</td><td>>hg38_21_+_8433172_8446622_RNA45SN1_45S_50ntFlanks</td><td>6661</td><td>6698</td><td>38</td><td>CGGTGGATCACTCGGCTCGTGCGTCGATGAAGAACGCA</td><td>592.4</td><td>2.0</td><td>0.5</td><td>4.1</td><td>3.8E-05</td><td>4.9E-05</td><td>BT20/MB468</td></tr>
<tr><td>>MI0000439|hsa-mir-23b&WithFlank&9|+|95085202|95085310@67.87.21&[MIMAT0000418&hsa-miR-23b-3p&offsets|+3|+1;m-42&9|+|95085265|95085285&offsets|+3|+3]|ACATTGCCAGGGATTACCACG</td><td>>MI0000439|hsa-mir-23b&WithFlank&9|+|95085202|95085310</td><td>67</td><td>87</td><td>21</td><td>ACATTGCCAGGGATTACCACG</td><td>1113.2</td><td>15.0</td><td>3.6</td><td>4.1</td><td>3.5E-05</td><td>4.9E-05</td><td>MB231/MB468</td></tr>
<tr><td>>hg38_21_+_8433172_8446622_RNA45SN1_45S_50ntFlanks@12259.12284.26|CTTGATCTTGATTTTCAGTACGAATA</td><td>>hg38_21_+_8433172_8446622_RNA45SN1_45S_50ntFlanks</td><td>12259</td><td>12284</td><td>26</td><td>CTTGATCTTGATTTTCAGTACGAATA</td><td>295.9</td><td>-2.7</td><td>0.6</td><td>-4.1</td><td>3.8E-05</td><td>4.9E-05</td><td>BT20/MB468</td></tr>
<tr><td>>hg38_21_+_8433172_8446622_RNA45SN1_45S_50ntFlanks@3775.3798.24|GCCGGTACAGTGAAACTGCGAATG</td><td>>hg38_21_+_8433172_8446622_RNA45SN1_45S_50ntFlanks</td><td>3775</td><td>3798</td><td>24</td><td>GCCGGTACAGTGAAACTGCGAATG</td><td>163.2</td><td>-2.0</td><td>0.5</td><td>-4.1</td><td>3.8E-05</td><td>5.0E-05</td><td>BT20/MB468</td></tr>
<tr><td>>hg38_21_+_8433172_8446622_RNA45SN1_45S_50ntFlanks@12605.12634.30|TATGTGCTTGGCTGAGGAGCCAATGGGGCG</td><td>>hg38_21_+_8433172_8446622_RNA45SN1_45S_50ntFlanks</td><td>12605</td><td>12634</td><td>30</td><td>TATGTGCTTGGCTGAGGAGCCAATGGGGCG</td><td>77.1</td><td>-4.0</td><td>1.0</td><td>-4.1</td><td>3.5E-05</td><td>5.0E-05</td><td>MB231/MB468</td></tr>
<tr><td>>hg38_21_+_8433172_8446622_RNA45SN1_45S_50ntFlanks@10866.10883.18|CTAAGGGCTGGGTCGGTC</td><td>>hg38_21_+_8433172_8446622_RNA45SN1_45S_50ntFlanks</td><td>10866</td><td>10883</td><td>18</td><td>CTAAGGGCTGGGTCGGTC</td><td>109.1</td><td>-2.7</td><td>0.7</td><td>-4.1</td><td>3.9E-05</td><td>5.0E-05</td><td>BT20/MB468</td></tr>
<tr><td>>hg38_21_+_8433172_8446622_RNA45SN1_45S_50ntFlanks@11562.11585.24|CGCCTAGCAGCCGACTTAGAACTG</td><td>>hg38_21_+_8433172_8446622_RNA45SN1_45S_50ntFlanks</td><td>11562</td><td>11585</td><td>24</td><td>CGCCTAGCAGCCGACTTAGAACTG</td><td>117.1</td><td>-2.3</td><td>0.6</td><td>-4.1</td><td>3.9E-05</td><td>5.0E-05</td><td>BT20/MB468</td></tr>
<tr><td>>trnaMT_MetCAT_MT_+_4402_4469@40.71.32|AAAATGTTGGTTATACCCTTCCCGTACTACCA</td><td>>trnaMT_MetCAT_MT_+_4402_4469</td><td>40</td><td>71</td><td>32</td><td>AAAATGTTGGTTATACCCTTCCCGTACTACCA</td><td>468.2</td><td>2.9</td><td>0.7</td><td>4.1</td><td>3.9E-05</td><td>5.0E-05</td><td>BT20/MB468</td></tr>
<tr><td>>hg38_21_+_8433172_8446622_RNA45SN1_45S_50ntFlanks@9306.9338.33|CAAGGAGTCTAACACGTGCGCGAGTCGGGGGCT</td><td>>hg38_21_+_8433172_8446622_RNA45SN1_45S_50ntFlanks</td><td>9306</td><td>9338</td><td>33</td><td>CAAGGAGTCTAACACGTGCGCGAGTCGGGGGCT</td><td>138.5</td><td>-4.3</td><td>1.0</td><td>-4.1</td><td>3.9E-05</td><td>5.0E-05</td><td>BT20/MB468</td></tr>
<tr><td>>hg38_21_+_8433172_8446622_RNA45SN1_45S_50ntFlanks@3813.3834.22|TTATGGTTCCTTTGGTCGCTCG</td><td>>hg38_21_+_8433172_8446622_RNA45SN1_45S_50ntFlanks</td><td>3813</td><td>3834</td><td>22</td><td>TTATGGTTCCTTTGGTCGCTCG</td><td>134.2</td><td>-2.3</td><td>0.6</td><td>-4.1</td><td>3.9E-05</td><td>5.0E-05</td><td>BT20/MB468</td></tr>
<tr><td>>hg38_21_+_8433172_8446622_RNA45SN1_45S_50ntFlanks@9738.9763.26|GAAACGATCTCAACCTATTCTCAAAC</td><td>>hg38_21_+_8433172_8446622_RNA45SN1_45S_50ntFlanks</td><td>9738</td><td>9763</td><td>26</td><td>GAAACGATCTCAACCTATTCTCAAAC</td><td>153.9</td><td>-2.2</td><td>0.5</td><td>-4.1</td><td>3.9E-05</td><td>5.1E-05</td><td>BT20/MB468</td></tr>
<tr><td>>hg38_21_+_8433172_8446622_RNA45SN1_45S_50ntFlanks@12488.12524.37|AACGTGAGCTGGGTTTAGACCGTCGTGAGACAGGTTA</td><td>>hg38_21_+_8433172_8446622_RNA45SN1_45S_50ntFlanks</td><td>12488</td><td>12524</td><td>37</td><td>AACGTGAGCTGGGTTTAGACCGTCGTGAGACAGGTTA</td><td>124.4</td><td>-2.9</td><td>0.7</td><td>-4.1</td><td>3.6E-05</td><td>5.1E-05</td><td>MB231/MB468</td></tr>
<tr><td>>trna119_LysCTT_1_-_145395522_145395594@2.33.32,trna11_LysCTT_5_-_180648979_180649051@2.33.32,trna13_LysCTT_6_+_26556774_26556846@2.33.32,trna7_LysCTT_16_+_3225692_3225764@2.33.32,trna9_LysCTT_5_+_180634755_180634827@2.33.32|CCCGGCTAGCTCAGTCGGTAGAGCATGAGACT</td><td>>trna119_LysCTT_1_-_145395522_145395594</td><td>2</td><td>33</td><td>32</td><td>CCCGGCTAGCTCAGTCGGTAGAGCATGAGACT</td><td>357.0</td><td>-5.0</td><td>1.1</td><td>-4.6</td><td>3.8E-06</td><td>5.1E-05</td><td>BT20/MB231</td></tr>
<tr><td>>hg38_21_+_8433172_8446622_RNA45SN1_45S_50ntFlanks@11315.11332.18|CCACCCCACGTCTCGTCG</td><td>>hg38_21_+_8433172_8446622_RNA45SN1_45S_50ntFlanks</td><td>11315</td><td>11332</td><td>18</td><td>CCACCCCACGTCTCGTCG</td><td>363.6</td><td>-5.3</td><td>1.3</td><td>-4.1</td><td>4.0E-05</td><td>5.1E-05</td><td>BT20/MB468</td></tr>
<tr><td>>hg38_21_+_8433172_8446622_RNA45SN1_45S_50ntFlanks@11734.11751.18|ATGACTCTCTTAAGGTAG</td><td>>hg38_21_+_8433172_8446622_RNA45SN1_45S_50ntFlanks</td><td>11734</td><td>11751</td><td>18</td><td>ATGACTCTCTTAAGGTAG</td><td>122.9</td><td>-3.2</td><td>0.8</td><td>-4.1</td><td>4.0E-05</td><td>5.2E-05</td><td>BT20/MB468</td></tr>
<tr><td>>am_trnaMT_HisGTG_MT_+_12138_12206@26.45.20|AGATTGTGAATCTGACAACA</td><td>>am_trnaMT_HisGTG_MT_+_12138_12206</td><td>26</td><td>45</td><td>20</td><td>AGATTGTGAATCTGACAACA</td><td>2859.1</td><td>11.4</td><td>2.8</td><td>4.1</td><td>4.0E-05</td><td>5.2E-05</td><td>BT20/MB468</td></tr>
<tr><td>>hg38_21_+_8433172_8446622_RNA45SN1_45S_50ntFlanks@12937.12961.25|AACCATTCGTAGACGACCTGCTTCT</td><td>>hg38_21_+_8433172_8446622_RNA45SN1_45S_50ntFlanks</td><td>12937</td><td>12961</td><td>25</td><td>AACCATTCGTAGACGACCTGCTTCT</td><td>231.5</td><td>-2.5</td><td>0.6</td><td>-4.1</td><td>4.1E-05</td><td>5.2E-05</td><td>BT20/MB468</td></tr>
<tr><td>>hg38_21_+_8433172_8446622_RNA45SN1_45S_50ntFlanks@12081.12108.28|CTCTCGCTTCTGGCGCCAAGCGCCCGGC</td><td>>hg38_21_+_8433172_8446622_RNA45SN1_45S_50ntFlanks</td><td>12081</td><td>12108</td><td>28</td><td>CTCTCGCTTCTGGCGCCAAGCGCCCGGC</td><td>466.4</td><td>-5.1</td><td>1.3</td><td>-4.1</td><td>4.1E-05</td><td>5.3E-05</td><td>BT20/MB468</td></tr>
<tr><td>>hg38_21_+_8433172_8446622_RNA45SN1_45S_50ntFlanks@7982.8007.26|TCAGATCAGACGTGGCGACCCGCTGA</td><td>>hg38_21_+_8433172_8446622_RNA45SN1_45S_50ntFlanks</td><td>7982</td><td>8007</td><td>26</td><td>TCAGATCAGACGTGGCGACCCGCTGA</td><td>102.5</td><td>-2.8</td><td>0.7</td><td>-4.1</td><td>3.7E-05</td><td>5.3E-05</td><td>MB231/MB468</td></tr>
<tr><td>>hg38_21_+_8433172_8446622_RNA45SN1_45S_50ntFlanks@10353.10388.36|AGAACTTTGAAGGCCGAAGTGGAGAAGGGTTCCATG</td><td>>hg38_21_+_8433172_8446622_RNA45SN1_45S_50ntFlanks</td><td>10353</td><td>10388</td><td>36</td><td>AGAACTTTGAAGGCCGAAGTGGAGAAGGGTTCCATG</td><td>310.1</td><td>-3.4</td><td>0.8</td><td>-4.1</td><td>4.1E-05</td><td>5.3E-05</td><td>BT20/MB468</td></tr>
<tr><td>>hg38_21_+_8433172_8446622_RNA45SN1_45S_50ntFlanks@11874.11902.29|AGAAGACCCTGTTGAGCTTGACTCTAGTC</td><td>>hg38_21_+_8433172_8446622_RNA45SN1_45S_50ntFlanks</td><td>11874</td><td>11902</td><td>29</td><td>AGAAGACCCTGTTGAGCTTGACTCTAGTC</td><td>240.1</td><td>-3.3</td><td>0.8</td><td>-4.1</td><td>3.8E-05</td><td>5.3E-05</td><td>MB231/MB468</td></tr>
<tr><td>>hg38_21_+_8433172_8446622_RNA45SN1_45S_50ntFlanks@12073.12097.25|CCGAGGGGCTCTCGCTTCTGGCGCC</td><td>>hg38_21_+_8433172_8446622_RNA45SN1_45S_50ntFlanks</td><td>12073</td><td>12097</td><td>25</td><td>CCGAGGGGCTCTCGCTTCTGGCGCC</td><td>1183.5</td><td>-2.7</td><td>0.7</td><td>-4.1</td><td>3.8E-05</td><td>5.4E-05</td><td>MB231/MB468</td></tr>
<tr><td>>MI0000286|hsa-mir-210&WithFlank&11|-|568083|568204@72.92.21(+1G)&[MIMAT0000267&hsa-miR-210-3p&offsets|0|-1(+1G);m-102&11|-|568112|568133&offsets|0|-1(+1G)]|CTGTGCGTGTGACAGCGGCTGG</td><td>>MI0000286|hsa-mir-210&WithFlank&11|-|568083|568204</td><td>72</td><td>92</td><td>21(+1G)</td><td>CTGTGCGTGTGACAGCGGCTGG</td><td>821.7</td><td>5.8</td><td>1.3</td><td>4.6</td><td>4.0E-06</td><td>5.4E-05</td><td>BT20/MB231</td></tr>
<tr><td>>hg38_21_+_8433172_8446622_RNA45SN1_45S_50ntFlanks@7975.7994.20|CGCGACCTCAGATCAGACGT</td><td>>hg38_21_+_8433172_8446622_RNA45SN1_45S_50ntFlanks</td><td>7975</td><td>7994</td><td>20</td><td>CGCGACCTCAGATCAGACGT</td><td>63525.2</td><td>3.1</td><td>0.8</td><td>4.1</td><td>4.2E-05</td><td>5.4E-05</td><td>BT20/MB468</td></tr>
</table>

| | | | | | | | | | | | | |
|---|---|---|---|---|---|---|---|---|---|---|---|---|
| >hg38_21_+_8433172_8446622_RNA45SN1_45S_50ntFlanks@10866.10886.21\|CTAAGGGCTGGGTCGGTCGGG | >hg38_21_+_8433172_8446622_RNA45SN1_45S_50ntFlanks | 10866 | 10886 | 21 | CTAAGGGCTGGGTCGGTCGGG | 177.4 | -4.3 | 1.0 | -4.1 | 4.3E-05 | 5.5E-05 | BT20/MB468 |
| >hg38_21_+_8433172_8446622_RNA45SN1_45S_50ntFlanks@12626.12657.32\|AATGGGGCGAAGCTACCATCTGTGGGATTATG | >hg38_21_+_8433172_8446622_RNA45SN1_45S_50ntFlanks | 12626 | 12657 | 32 | AATGGGGCGAAGCTACCATCTGTGGGATTATG | 336.3 | -2.0 | 0.5 | -4.1 | 3.9E-05 | 5.5E-05 | MB231/MB468 |
| >hg38_21_+_8433172_8446622_RNA45SN1_45S_50ntFlanks@12419.12455.37\|GATGTCGGCTCTTCCTATCATTGTGAAGCAGAATTCA | >hg38_21_+_8433172_8446622_RNA45SN1_45S_50ntFlanks | 12419 | 12455 | 37 | GATGTCGGCTCTTCCTATCATTGTGAAGCAGAATTCA | 734.8 | 2.8 | 0.7 | 4.1 | 3.9E-05 | 5.6E-05 | MB231/MB468 |
| >hg38_21_+_8433172_8446622_RNA45SN1_45S_50ntFlanks@12418.12438.21\|CGATGTCGGCTCTTCCTATCA | >hg38_21_+_8433172_8446622_RNA45SN1_45S_50ntFlanks | 12418 | 12438 | 21 | CGATGTCGGCTCTTCCTATCA | 869.8 | 2.5 | 0.6 | 4.1 | 3.9E-05 | 5.6E-05 | MB231/MB468 |
| >hg38_21_+_8433172_8446622_RNA45SN1_45S_50ntFlanks@6722.6741.20\|ATTGCAGGACACATTGATCA | >hg38_21_+_8433172_8446622_RNA45SN1_45S_50ntFlanks | 6722 | 6741 | 20 | ATTGCAGGACACATTGATCA | 120.0 | -4.6 | 1.1 | -4.1 | 4.3E-05 | 5.6E-05 | BT20/MB468 |
| >hg38_21_+_8433172_8446622_RNA45SN1_45S_50ntFlanks@6651.6686.36\|CGACTCTTAGCGGTGGATCACTCGGCTCGTGCGTCG | >hg38_21_+_8433172_8446622_RNA45SN1_45S_50ntFlanks | 6651 | 6686 | 36 | CGACTCTTAGCGGTGGATCACTCGGCTCGTGCGTCG | 1455.3 | 2.9 | 0.7 | 4.1 | 4.0E-05 | 5.6E-05 | MB231/MB468 |
| >hg38_21_+_8433172_8446622_RNA45SN1_45S_50ntFlanks@9736.9753.18\|CCGAAACGATCTCAACCT | >hg38_21_+_8433172_8446622_RNA45SN1_45S_50ntFlanks | 9736 | 9753 | 18 | CCGAAACGATCTCAACCT | 149.6 | -3.2 | 0.8 | -4.1 | 4.0E-05 | 5.6E-05 | MB231/MB468 |
| >hg38_21_+_8433172_8446622_RNA45SN1_45S_50ntFlanks@12913.12933.21\|ACGTTCGTGGGGAACCTGGCG | >hg38_21_+_8433172_8446622_RNA45SN1_45S_50ntFlanks | 12913 | 12933 | 21 | ACGTTCGTGGGGAACCTGGCG | 153.5 | -4.2 | 1.0 | -4.1 | 4.4E-05 | 5.6E-05 | BT20/MB468 |
| >hg38_21_+_8433172_8446622_RNA45SN1_45S_50ntFlanks@7974.7991.18\|ACGCGACCTCAGATCAGA | >hg38_21_+_8433172_8446622_RNA45SN1_45S_50ntFlanks | 7974 | 7991 | 18 | ACGCGACCTCAGATCAGA | 989.4 | 3.5 | 0.9 | 4.1 | 4.0E-05 | 5.6E-05 | MB231/MB468 |
| >hg38_21_+_8433172_8446622_RNA45SN1_45S_50ntFlanks@11783.11805.23\|CATGAATGGATGAACGAGATTCC | >hg38_21_+_8433172_8446622_RNA45SN1_45S_50ntFlanks | 11783 | 11805 | 23 | CATGAATGGATGAACGAGATTCC | 2178.8 | -2.4 | 0.6 | -4.1 | 4.0E-05 | 5.7E-05 | MB231/MB468 |
| >MI0000063\|hsa-let-7b&WithFlank&22\|+\|46113680\|46113774@12.32.21(+1A)&[MIMAT0000063&hsa-let-7b-5p&offsets\|0\|-1(+1A);m-14&22\|+\|46113691\|46113712&offsets\|0\|-1(+1A)]\|TGAGGTAGTAGGTTGTGTGGTA | >MI0000063\|hsa-let-7b&WithFlank&22\|+\|46113680\|46113774 | 12 | 32 | 21(+1A) | TGAGGTAGTAGGTTGTGTGGTA | 3413.6 | 3.3 | 0.7 | 4.6 | 4.3E-06 | 5.7E-05 | BT20/MB231 |
| >hg38_21_+_8433172_8446622_RNA45SN1_45S_50ntFlanks@10916.10933.18\|CCGCGGCTGGACGAGGCG | >hg38_21_+_8433172_8446622_RNA45SN1_45S_50ntFlanks | 10916 | 10933 | 18 | CCGCGGCTGGACGAGGCG | 275.4 | -3.4 | 0.8 | -4.1 | 4.5E-05 | 5.7E-05 | BT20/MB468 |
| >hg38_21_+_8433172_8446622_RNA45SN1_45S_50ntFlanks@9604.9622.19\|AAAGACTAATCGAACCATC | >hg38_21_+_8433172_8446622_RNA45SN1_45S_50ntFlanks | 9604 | 9622 | 19 | AAAGACTAATCGAACCATC | 862.6 | -2.5 | 0.6 | -4.1 | 4.1E-05 | 5.7E-05 | MB231/MB468 |
| >hg38_21_+_8433172_8446622_RNA45SN1_45S_50ntFlanks@13007.13028.22\|TATTGAAAGTCAGCCCTCGACA | >hg38_21_+_8433172_8446622_RNA45SN1_45S_50ntFlanks | 13007 | 13028 | 22 | TATTGAAAGTCAGCCCTCGACA | 509.2 | -2.8 | 0.7 | -4.1 | 4.1E-05 | 5.8E-05 | MB231/MB468 |
| >trna152_ValCAC_6_-_27248049_27248121@1.29.29\|GCTTCTGTAGTGTAGTGGTTATCACGTTC | >trna152_ValCAC_6_-_27248049_27248121 | 1 | 29 | 29 | GCTTCTGTAGTGTAGTGGTTATCACGTTC | 8614.1 | -3.8 | 0.9 | -4.1 | 4.1E-05 | 5.8E-05 | MB231/MB468 |
| >trna134_GluTTC_1_-_16861774_16861845@1.33.33,trna5_GluTTC_1_+_17199078_17199149@1.33.33,trna84_GluTTC_1_-_161391883_161391954@1.33.33,trna94_GluTTC_1_-_149664355_149664427@1.33.33\|TCCCTGGTGGTCTAGTGGCTAGGATTCGGCGCT | >trna134_GluTTC_1_-_16861774_16861845 | 1 | 33 | 33 | TCCCTGGTGGTCTAGTGGCTAGGATTCGGCGCT | 1657.1 | 3.9 | 0.8 | 4.6 | 4.4E-06 | 5.8E-05 | BT20/MB231 |
| >hg38_21_+_8433172_8446622_RNA45SN1_45S_50ntFlanks@10869.10894.26\|AGGGCTGGGTCGGTCGGGCTGGGGCG | >hg38_21_+_8433172_8446622_RNA45SN1_45S_50ntFlanks | 10869 | 10894 | 26 | AGGGCTGGGTCGGTCGGGCTGGGGCG | 189.6 | -2.1 | 0.5 | -4.1 | 4.5E-05 | 5.8E-05 | BT20/MB468 |
| >hg38_21_+_8433172_8446622_RNA45SN1_45S_50ntFlanks@11877.11906.30\|AGACCCTGTTGAGCTTGACTCTAGTCTGGC | >hg38_21_+_8433172_8446622_RNA45SN1_45S_50ntFlanks | 11877 | 11906 | 30 | AGACCCTGTTGAGCTTGACTCTAGTCTGGC | 99.5 | -4.4 | 1.1 | -4.1 | 4.6E-05 | 5.9E-05 | BT20/MB468 |
| >hg38_21_+_8433172_8446622_RNA45SN1_45S_50ntFlanks@4224.4242.19\|AAAATAACAATACAGGACT | >hg38_21_+_8433172_8446622_RNA45SN1_45S_50ntFlanks | 4224 | 4242 | 19 | AAAATAACAATACAGGACT | 490.3 | -2.3 | 0.6 | -4.1 | 4.2E-05 | 5.9E-05 | MB231/MB468 |
| >hg38_21_+_8433172_8446622_RNA45SN1_45S_50ntFlanks@12420.12454.35\|ATGTCGGCTCTTCCTATCATTGTGAAGCAGAATTC | >hg38_21_+_8433172_8446622_RNA45SN1_45S_50ntFlanks | 12420 | 12454 | 35 | ATGTCGGCTCTTCCTATCATTGTGAAGCAGAATTC | 577.7 | 2.3 | 0.6 | 4.1 | 4.2E-05 | 5.9E-05 | MB231/MB468 |
| >hg38_21_+_8433172_8446622_RNA45SN1_45S_50ntFlanks@11718.11743.26\|AACGGCGGGAGTAACTATGACTCTCT | >hg38_21_+_8433172_8446622_RNA45SN1_45S_50ntFlanks | 11718 | 11743 | 26 | AACGGCGGGAGTAACTATGACTCTCT | 283.1 | -2.9 | 0.7 | -4.1 | 4.2E-05 | 5.9E-05 | MB231/MB468 |
| >hg38_21_+_8433172_8446622_RNA45SN1_45S_50ntFlanks@5485.5506.22\|AGCGCTGAGAAGACGGTCGAAC | >hg38_21_+_8433172_8446622_RNA45SN1_45S_50ntFlanks | 5485 | 5506 | 22 | AGCGCTGAGAAGACGGTCGAAC | 69.9 | -2.4 | 0.6 | -4.1 | 4.2E-05 | 6.0E-05 | MB231/MB468 |
| >hg38_21_+_8433172_8446622_RNA45SN1_45S_50ntFlanks@11728.11749.22\|GTAACTATGACTCTCTTAAGGT | >hg38_21_+_8433172_8446622_RNA45SN1_45S_50ntFlanks | 11728 | 11749 | 22 | GTAACTATGACTCTCTTAAGGT | 132.2 | -3.0 | 0.7 | -4.1 | 4.7E-05 | 6.0E-05 | BT20/MB468 |
| >hg38_21_+_8433172_8446622_RNA45SN1_45S_50ntFlanks@10683.10712.30\|CCGGCGGCGTCCGGTGAGCTCTCGCTGGCC | >hg38_21_+_8433172_8446622_RNA45SN1_45S_50ntFlanks | 10683 | 10712 | 30 | CCGGCGGCGTCCGGTGAGCTCTCGCTGGCC | 1240.8 | -2.3 | 0.6 | -4.1 | 4.3E-05 | 6.0E-05 | MB231/MB468 |
| >hg38_21_+_8433172_8446622_RNA45SN1_45S_50ntFlanks@12488.12508.21\|AACGTGAGCTGGGTTTAGACC | >hg38_21_+_8433172_8446622_RNA45SN1_45S_50ntFlanks | 12488 | 12508 | 21 | AACGTGAGCTGGGTTTAGACC | 124.1 | -3.6 | 0.9 | -4.1 | 4.3E-05 | 6.0E-05 | MB231/MB468 |
| >MI0000767\|hsa-mir-365a&WithFlank&16\|+\|14309279\|14309377@62.81.20(+2U);MI0000769\|hsa-mir-365b&WithFlank&17\|+\|31575405\|31575527@74.93.20(+2U)&[MIMAT0000710&hsa-miR-365a-3p&offsets\|0\|-2(+2U);m-176&16\|+\|14309340\|14309361&offsets\|0\|-2(+2U)];[MIMAT0022834&hsa-miR-365b-3p&offsets\|0\|-2(+2U);m-175&17\|+\|31575478\|31575499&offsets\|0\|-2(+2U)]\|TAATGCCCCTAAAAATCCTTTT | >MI0000767\|hsa-mir-365a&WithFlank&16\|+\|14309279\|14309377 | 62 | 81 | 20(+2U) | TAATGCCCCTAAAAATCCTTTT | 438.9 | 3.7 | 0.8 | 4.6 | 4.6E-06 | 6.0E-05 | BT20/MB231 |
| >hg38_21_+_8433172_8446622_RNA45SN1_45S_50ntFlanks@8359.8379.21\|AAGAACTTTGAAGAGAGAGTT | >hg38_21_+_8433172_8446622_RNA45SN1_45S_50ntFlanks | 8359 | 8379 | 21 | AAGAACTTTGAAGAGAGAGTT | 1207.6 | -3.5 | 0.9 | -4.1 | 4.7E-05 | 6.0E-05 | BT20/MB468 |
| >hg38_21_+_8433172_8446622_RNA45SN1_45S_50ntFlanks@12444.12475.32\|AAGCAGAATTCACCAAGCGTTGGATTGTTCAC | >hg38_21_+_8433172_8446622_RNA45SN1_45S_50ntFlanks | 12444 | 12475 | 32 | AAGCAGAATTCACCAAGCGTTGGATTGTTCAC | 115.1 | -3.0 | 0.7 | -4.1 | 4.3E-05 | 6.0E-05 | MB231/MB468 |
| >hg38_21_+_8433172_8446622_RNA45SN1_45S_50ntFlanks@12539.12568.30\|ATGATGTGTTGTTGCCATGGTAATCCTGCT | >hg38_21_+_8433172_8446622_RNA45SN1_45S_50ntFlanks | 12539 | 12568 | 30 | ATGATGTGTTGTTGCCATGGTAATCCTGCT | 113.7 | -3.4 | 0.8 | -4.1 | 4.7E-05 | 6.0E-05 | BT20/MB468 |
| >hg38_21_+_8433172_8446622_RNA45SN1_45S_50ntFlanks@12437.12474.38\|CATTGTGAAGCAGAATTCACCAAGCGTTGGATTGTTCA | >hg38_21_+_8433172_8446622_RNA45SN1_45S_50ntFlanks | 12437 | 12474 | 38 | CATTGTGAAGCAGAATTCACCAAGCGTTGGATTGTTCA | 226.8 | -2.0 | 0.5 | -4.1 | 4.3E-05 | 6.0E-05 | MB231/MB468 |
| >trnaMT_PheGAA_MT_+_577_647@35.57.23\|GAAAATGTTTAGACGGGCTCACA | >trnaMT_PheGAA_MT_+_577_647 | 35 | 57 | 23 | GAAAATGTTTAGACGGGCTCACA | 3144.3 | 3.1 | 0.7 | 4.6 | 4.6E-06 | 6.1E-05 | BT20/MB231 |
| >MI0000255\|hsa-mir-30d&WithFlank&8\|-\|134804870\|134804951@12.35.24(+4U)&[MIMAT0000245&hsa-miR-30d-5p&offsets\|0\|+2(+4U);m-21&8\|-\|134804919\|134804940&offsets\|0\|+2(+4U)]\|TGTAAACATCCCCGACTGGAAGCTTTTT | >MI0000255\|hsa-mir-30d&WithFlank&8\|-\|134804870\|134804951 | 12 | 35 | 24(+4U) | TGTAAACATCCCCGACTGGAAGCTTTTT | 267.9 | -3.5 | 0.8 | -4.6 | 4.7E-06 | 6.1E-05 | BT20/MB231 |
| >hg38_21_+_8433172_8446622_RNA45SN1_45S_50ntFlanks@6776.6807.32\|CTCCCGGGGCTACGCCTGTCTGAGCGTCGCTT | >hg38_21_+_8433172_8446622_RNA45SN1_45S_50ntFlanks | 6776 | 6807 | 32 | CTCCCGGGGCTACGCCTGTCTGAGCGTCGCTT | 398.6 | 2.3 | 0.6 | 4.1 | 4.4E-05 | 6.1E-05 | MB231/MB468 |
| >hg38_21_+_8433172_8446622_RNA45SN1_45S_50ntFlanks@10681.10709.29\|TTCCGGCGGCGTCCGGTGAGCTCTCGCTG | >hg38_21_+_8433172_8446622_RNA45SN1_45S_50ntFlanks | 10681 | 10709 | 29 | TTCCGGCGGCGTCCGGTGAGCTCTCGCTG | 486.3 | -5.0 | 1.2 | -4.1 | 4.8E-05 | 6.1E-05 | BT20/MB468 |
| >hg38_21_+_8433172_8446622_RNA45SN1_45S_50ntFlanks@12934.12958.25\|CTAAACCATTCGTAGACGACCTGCT | >hg38_21_+_8433172_8446622_RNA45SN1_45S_50ntFlanks | 12934 | 12958 | 25 | CTAAACCATTCGTAGACGACCTGCT | 274.2 | -1.9 | 0.5 | -4.1 | 4.4E-05 | 6.1E-05 | MB231/MB468 |
| >hg38_21_+_8433172_8446622_RNA45SN1_45S_50ntFlanks@6651.6678.28\|CGACTCTTAGCGGTGGATCACTCGGCTC | >hg38_21_+_8433172_8446622_RNA45SN1_45S_50ntFlanks | 6651 | 6678 | 28 | CGACTCTTAGCGGTGGATCACTCGGCTC | 4352.1 | 3.4 | 0.8 | 4.1 | 4.4E-05 | 6.2E-05 | MB231/MB468 |
| >hg38_21_+_8433172_8446622_RNA45SN1_45S_50ntFlanks@6664.6689.26\|TGGATCACTCGGCTCGTGCGTCGATG | >hg38_21_+_8433172_8446622_RNA45SN1_45S_50ntFlanks | 6664 | 6689 | 26 | TGGATCACTCGGCTCGTGCGTCGATG | 234.7 | -2.5 | 0.6 | -4.1 | 4.8E-05 | 6.2E-05 | BT20/MB468 |
| >hg38_21_+_8433172_8446622_RNA45SN1_45S_50ntFlanks@12477.12496.20\|CACTAATAGGGAACGTGAGC | >hg38_21_+_8433172_8446622_RNA45SN1_45S_50ntFlanks | 12477 | 12496 | 20 | CACTAATAGGGAACGTGAGC | 1078.5 | 1.5 | 0.4 | 4.1 | 4.8E-05 | 6.2E-05 | BT20/MB468 |
| >hg38_21_+_8433172_8446622_RNA45SN1_45S_50ntFlanks@9471.9503.33\|AGCACGAGCGCACGTGTTAGGACCCGAAAGATG | >hg38_21_+_8433172_8446622_RNA45SN1_45S_50ntFlanks | 9471 | 9503 | 33 | AGCACGAGCGCACGTGTTAGGACCCGAAAGATG | 79.4 | -2.5 | 0.6 | -4.1 | 4.8E-05 | 6.2E-05 | BT20/MB468 |
| >hg38_21_+_8433172_8446622_RNA45SN1_45S_50ntFlanks@4995.5022.28\|ACAGATTGATAGCTCTTTCTCGATTCCG | >hg38_21_+_8433172_8446622_RNA45SN1_45S_50ntFlanks | 4995 | 5022 | 28 | ACAGATTGATAGCTCTTTCTCGATTCCG | 102.8 | -3.5 | 0.9 | -4.1 | 4.5E-05 | 6.3E-05 | MB231/MB468 |
| >hg38_21_+_8433172_8446622_RNA45SN1_45S_50ntFlanks@11564.11587.24\|CCTAGCAGCCGACTTAGAACTGGT | >hg38_21_+_8433172_8446622_RNA45SN1_45S_50ntFlanks | 11564 | 11587 | 24 | CCTAGCAGCCGACTTAGAACTGGT | 219.4 | -1.9 | 0.5 | -4.1 | 4.9E-05 | 6.3E-05 | BT20/MB468 |
| >hg38_21_+_8433172_8446622_RNA45SN1_45S_50ntFlanks@9537.9563.27\|AAACTCTGGTGGAGGTCCGTAGCGGTC | >hg38_21_+_8433172_8446622_RNA45SN1_45S_50ntFlanks | 9537 | 9563 | 27 | AAACTCTGGTGGAGGTCCGTAGCGGTC | 102.5 | -2.4 | 0.6 | -4.1 | 4.5E-05 | 6.3E-05 | MB231/MB468 |
| >hg38_21_+_8433172_8446622_RNA45SN1_45S_50ntFlanks@4045.4066.22\|CCATTCGAACGTCTGCCCTATC | >hg38_21_+_8433172_8446622_RNA45SN1_45S_50ntFlanks | 4045 | 4066 | 22 | CCATTCGAACGTCTGCCCTATC | 288.5 | -2.0 | 0.5 | -4.1 | 4.9E-05 | 6.3E-05 | BT20/MB468 |
| >hg38_21_+_8433172_8446622_RNA45SN1_45S_50ntFlanks@4725.4758.34\|TTAATCAAGAACGAAAGTCGGAGGTTCGAAGACG | >hg38_21_+_8433172_8446622_RNA45SN1_45S_50ntFlanks | 4725 | 4758 | 34 | TTAATCAAGAACGAAAGTCGGAGGTTCGAAGACG | 508.4 | -2.6 | 0.6 | -4.1 | 5.0E-05 | 6.3E-05 | BT20/MB468 |
| >hg38_21_+_8433172_8446622_RNA45SN1_45S_50ntFlanks@10845.10872.28\|ACTTCGGGATAAGGATTGGCTCTAAGGG | >hg38_21_+_8433172_8446622_RNA45SN1_45S_50ntFlanks | 10845 | 10872 | 28 | ACTTCGGGATAAGGATTGGCTCTAAGGG | 95.7 | -2.4 | 0.6 | -4.1 | 4.5E-05 | 6.4E-05 | MB231/MB468 |
| >hg38_21_+_8433172_8446622_RNA45SN1_45S_50ntFlanks@11210.11232.23\|GGCGACTCTGGACGCGAGCCGGG | >hg38_21_+_8433172_8446622_RNA45SN1_45S_50ntFlanks | 11210 | 11232 | 23 | GGCGACTCTGGACGCGAGCCGGG | 811.3 | -5.5 | 1.4 | -4.1 | 5.0E-05 | 6.4E-05 | BT20/MB468 |

<table>
<tr><th></th><th></th><th></th><th></th><th></th><th></th><th></th><th></th><th></th><th></th><th></th><th></th><th></th></tr>
<tr><td>>hg38_21_+_8433172_8446622_RNA45SN1_45S_50ntFlanks@12940.12965.26|CATTCGTAGACGACCTGCTTCTGGGT</td><td>>hg38_21_+_8433172_8446622_RNA45SN1_45S_50ntFlanks</td><td>12940</td><td>12965</td><td>26</td><td>CATTCGTAGACGACCTGCTTCTGGGT</td><td>139.4</td><td>-1.6</td><td>0.4</td><td>-4.1</td><td>4.6E-05</td><td>6.4E-05</td><td>MB231/MB468</td></tr>
<tr><td>>MI0000273|hsa-mir-183&WithFlank&7|-|129774899|129775020@33.53.21(+1A)&[MIMAT0000261&hsa-miR-183-5p&offsets|0|-1(+1A);m-41&7|-|129774966|129774987&offsets|-1|-2(+1A)]|TATGGCACTGGTAGAATTCACA</td><td>>MI0000273|hsa-mir-183&WithFlank&7|-|129774899|129775020</td><td>33</td><td>53</td><td>21(+1A)</td><td>TATGGCACTGGTAGAATTCACA</td><td>1539.6</td><td>3.5</td><td>0.8</td><td>4.6</td><td>5.0E-06</td><td>6.4E-05</td><td>BT20/MB231</td></tr>
<tr><td>>trna2_GlyGCC_21_-_18827107_18827177@1.29.29,trna35_GlyGCC_1_+_161413094_161413164@1.29.29,trna37_GlyGCC_1_+_161420467_161420537@1.29.29,trna39_GlyGCC_1_+_161427898_161427968@1.29.29,trna41_GlyGCC_1_+_161435258_161435328@1.29.29|GCATGGGTGGTTCAGTGGTAGAATTCTCG</td><td>>trna2_GlyGCC_21_-_18827107_18827177</td><td>1</td><td>29</td><td>29</td><td>GCATGGGTGGTTCAGTGGTAGAATTCTCG</td><td>361.6</td><td>5.0</td><td>1.1</td><td>4.6</td><td>5.0E-06</td><td>6.4E-05</td><td>BT20/MB231</td></tr>
<tr><td>>hg38_21_+_8433172_8446622_RNA45SN1_45S_50ntFlanks@7981.8007.27|CTCAGATCAGACGTGGCGACCCGCTGA</td><td>>hg38_21_+_8433172_8446622_RNA45SN1_45S_50ntFlanks</td><td>7981</td><td>8007</td><td>27</td><td>CTCAGATCAGACGTGGCGACCCGCTGA</td><td>108.4</td><td>-2.9</td><td>0.7</td><td>-4.1</td><td>5.0E-05</td><td>6.4E-05</td><td>BT20/MB468</td></tr>
<tr><td>>hg38_21_+_8433172_8446622_RNA45SN1_45S_50ntFlanks@10348.10367.20|AAACGAGAACTTTGAAGGCC</td><td>>hg38_21_+_8433172_8446622_RNA45SN1_45S_50ntFlanks</td><td>10348</td><td>10367</td><td>20</td><td>AAACGAGAACTTTGAAGGCC</td><td>997.3</td><td>2.8</td><td>0.7</td><td>4.1</td><td>4.6E-05</td><td>6.5E-05</td><td>MB231/MB468</td></tr>
<tr><td>>hg38_21_+_8433172_8446622_RNA45SN1_45S_50ntFlanks@9738.9764.27|GAAACGATCTCAACCTATTCTCAAACT</td><td>>hg38_21_+_8433172_8446622_RNA45SN1_45S_50ntFlanks</td><td>9738</td><td>9764</td><td>27</td><td>GAAACGATCTCAACCTATTCTCAAACT</td><td>167.9</td><td>-2.2</td><td>0.5</td><td>-4.1</td><td>5.0E-05</td><td>6.5E-05</td><td>BT20/MB468</td></tr>
<tr><td>>hg38_21_+_8433172_8446622_RNA45SN1_45S_50ntFlanks@10392.10431.40|ACAGCAGTTGAACATGGGTCAGTCGGTCCTGAGAGATGGG</td><td>>hg38_21_+_8433172_8446622_RNA45SN1_45S_50ntFlanks</td><td>10392</td><td>10431</td><td>40</td><td>ACAGCAGTTGAACATGGGTCAGTCGGTCCTGAGAGATGGG</td><td>144.2</td><td>-3.4</td><td>0.8</td><td>-4.1</td><td>4.6E-05</td><td>6.5E-05</td><td>MB231/MB468</td></tr>
<tr><td>>hg38_21_+_8433172_8446622_RNA45SN1_45S_50ntFlanks@4220.4243.24|ACGAAAAATAACAATACAGGACTC</td><td>>hg38_21_+_8433172_8446622_RNA45SN1_45S_50ntFlanks</td><td>4220</td><td>4243</td><td>24</td><td>ACGAAAAATAACAATACAGGACTC</td><td>111.2</td><td>-2.2</td><td>0.6</td><td>-4.1</td><td>5.1E-05</td><td>6.5E-05</td><td>BT20/MB468</td></tr>
<tr><td>>hg38_21_+_8433172_8446622_RNA45SN1_45S_50ntFlanks@12633.12656.24|CGAAGCTACCATCTGTGGGATTAT</td><td>>hg38_21_+_8433172_8446622_RNA45SN1_45S_50ntFlanks</td><td>12633</td><td>12656</td><td>24</td><td>CGAAGCTACCATCTGTGGGATTAT</td><td>124.1</td><td>-2.4</td><td>0.6</td><td>-4.1</td><td>4.6E-05</td><td>6.5E-05</td><td>MB231/MB468</td></tr>
<tr><td>>hg38_21_+_8433172_8446622_RNA45SN1_45S_50ntFlanks@3855.3881.27|CTGTGGTAATTCTAGAGCTAATACATG</td><td>>hg38_21_+_8433172_8446622_RNA45SN1_45S_50ntFlanks</td><td>3855</td><td>3881</td><td>27</td><td>CTGTGGTAATTCTAGAGCTAATACATG</td><td>243.3</td><td>-2.9</td><td>0.7</td><td>-4.1</td><td>5.1E-05</td><td>6.5E-05</td><td>BT20/MB468</td></tr>
<tr><td>>hg38_21_+_8433172_8446622_RNA45SN1_45S_50ntFlanks@4726.4748.23|TAATCAAGAACGAAAGTCGGAGG</td><td>>hg38_21_+_8433172_8446622_RNA45SN1_45S_50ntFlanks</td><td>4726</td><td>4748</td><td>23</td><td>TAATCAAGAACGAAAGTCGGAGG</td><td>135.9</td><td>-2.3</td><td>0.6</td><td>-4.1</td><td>4.7E-05</td><td>6.6E-05</td><td>MB231/MB468</td></tr>
<tr><td>>hg38_21_+_8433172_8446622_RNA45SN1_45S_50ntFlanks@6779.6803.25|CCGGGGCTACGCCTGTCTGAGCGTC</td><td>>hg38_21_+_8433172_8446622_RNA45SN1_45S_50ntFlanks</td><td>6779</td><td>6803</td><td>25</td><td>CCGGGGCTACGCCTGTCTGAGCGTC</td><td>278.3</td><td>-1.6</td><td>0.4</td><td>-4.0</td><td>5.1E-05</td><td>6.6E-05</td><td>BT20/MB468</td></tr>
<tr><td>>hg38_21_+_8433172_8446622_RNA45SN1_45S_50ntFlanks@10798.10826.29|CATGTTGGAACAATGTAGGTAAGGGAAGT</td><td>>hg38_21_+_8433172_8446622_RNA45SN1_45S_50ntFlanks</td><td>10798</td><td>10826</td><td>29</td><td>CATGTTGGAACAATGTAGGTAAGGGAAGT</td><td>172.7</td><td>-2.0</td><td>0.5</td><td>-4.1</td><td>4.7E-05</td><td>6.6E-05</td><td>MB231/MB468</td></tr>
<tr><td>>am_trnaMT_GluTTC_MT_-_14674_14742@27.55.29,trnalookalike8_GluTTC_5_-_93905172_93905240@27.55.29|GTTTTTCATATCATTGGTCGTGGTTGTAG</td><td>>am_trnaMT_GluTTC_MT_-_14674_14742</td><td>27</td><td>55</td><td>29</td><td>GTTTTTCATATCATTGGTCGTGGTTGTAG</td><td>94.2</td><td>-4.3</td><td>1.1</td><td>-4.0</td><td>5.2E-05</td><td>6.6E-05</td><td>BT20/MB468</td></tr>
<tr><td>>MI0000273|hsa-mir-183&WithFlank&7|-|129774899|129775020@34.53.20(+1C)&[MIMAT0000261&hsa-miR-183-5p&offsets|+1|-1(+1C);m-41&7|-|129774966|129774987&offsets|0|-2(+1C)]|ATGGCACTGGTAGAATTCACC</td><td>>MI0000273|hsa-mir-183&WithFlank&7|-|129774899|129775020</td><td>34</td><td>53</td><td>20(+1C)</td><td>ATGGCACTGGTAGAATTCACC</td><td>703.5</td><td>3.3</td><td>0.7</td><td>4.6</td><td>5.1E-06</td><td>6.6E-05</td><td>BT20/MB231</td></tr>
<tr><td>>am_trna128_GlyGCC_6_-_27870686_27870756@1.30.30,trna133_GlyCCC_1_-_16872434_16872504@1.30.30,trna18_GlyGCC_16_+_70822597_70822667@1.30.30,trna19_GlyGCC_16_+_70823410_70823480@1.30.30,trna19_GlyGCC_2_-_157257659_157257729@1.30.30,trna24_GlyGCC_16_-_70812942_70813012@1.30.30,trna25_GlyGCC_16_-_70812114_70812184@1.30.30,trna4_GlyCCC_1_+_17188416_17188486@1.30.30,trna5_GlyGCC_17_+_8029064_8029134@1.30.30,trna68_GlyGCC_1_-_161493637_161493707@1.30.30|GCATTGGTGGTTCAGTGGTAGAATTCTCGC</td><td>>am_trna128_GlyGCC_6_-_27870686_27870756</td><td>1</td><td>30</td><td>30</td><td>GCATTGGTGGTTCAGTGGTAGAATTCTCGC</td><td>6002.5</td><td>-3.7</td><td>0.8</td><td>-4.6</td><td>5.1E-06</td><td>6.6E-05</td><td>BT20/MB231</td></tr>
<tr><td>>hg38_21_+_8433172_8446622_RNA45SN1_45S_50ntFlanks@7980.8007.28|CCTCAGATCAGACGTGGCGACCCGCTGA</td><td>>hg38_21_+_8433172_8446622_RNA45SN1_45S_50ntFlanks</td><td>7980</td><td>8007</td><td>28</td><td>CCTCAGATCAGACGTGGCGACCCGCTGA</td><td>116.8</td><td>-3.3</td><td>0.8</td><td>-4.1</td><td>4.7E-05</td><td>6.6E-05</td><td>MB231/MB468</td></tr>
<tr><td>>hg38_21_+_8433172_8446622_RNA45SN1_45S_50ntFlanks@6730.6756.27|ACACATTGATCATCGACACTTCGAACG</td><td>>hg38_21_+_8433172_8446622_RNA45SN1_45S_50ntFlanks</td><td>6730</td><td>6756</td><td>27</td><td>ACACATTGATCATCGACACTTCGAACG</td><td>176.3</td><td>-1.9</td><td>0.5</td><td>-4.0</td><td>5.2E-05</td><td>6.7E-05</td><td>BT20/MB468</td></tr>
<tr><td>>hg38_21_+_8433172_8446622_RNA45SN1_45S_50ntFlanks@12644.12661.18|TCTGTGGGATTATGACTG</td><td>>hg38_21_+_8433172_8446622_RNA45SN1_45S_50ntFlanks</td><td>12644</td><td>12661</td><td>18</td><td>TCTGTGGGATTATGACTG</td><td>343.1</td><td>-2.7</td><td>0.7</td><td>-4.0</td><td>5.2E-05</td><td>6.7E-05</td><td>BT20/MB468</td></tr>
<tr><td>>hg38_21_+_8433172_8446622_RNA45SN1_45S_50ntFlanks@12076.12097.22|AGGGGCTCTCGCTTCTGGCGCC</td><td>>hg38_21_+_8433172_8446622_RNA45SN1_45S_50ntFlanks</td><td>12076</td><td>12097</td><td>22</td><td>AGGGGCTCTCGCTTCTGGCGCC</td><td>216.9</td><td>-3.1</td><td>0.8</td><td>-4.0</td><td>5.2E-05</td><td>6.7E-05</td><td>BT20/MB468</td></tr>
<tr><td>>hg38_21_+_8433172_8446622_RNA45SN1_45S_50ntFlanks@12457.12485.29|CAAGCGTTGGATTGTTCACCCACTAATAG</td><td>>hg38_21_+_8433172_8446622_RNA45SN1_45S_50ntFlanks</td><td>12457</td><td>12485</td><td>29</td><td>CAAGCGTTGGATTGTTCACCCACTAATAG</td><td>252.1</td><td>-2.8</td><td>0.7</td><td>-4.1</td><td>4.8E-05</td><td>6.7E-05</td><td>MB231/MB468</td></tr>
<tr><td>>hg38_21_+_8433172_8446622_RNA45SN1_45S_50ntFlanks@12632.12657.26|GCGAAGCTACCATCTGTGGGATTATG</td><td>>hg38_21_+_8433172_8446622_RNA45SN1_45S_50ntFlanks</td><td>12632</td><td>12657</td><td>26</td><td>GCGAAGCTACCATCTGTGGGATTATG</td><td>103.9</td><td>-2.2</td><td>0.5</td><td>-4.1</td><td>4.9E-05</td><td>6.8E-05</td><td>MB231/MB468</td></tr>
<tr><td>>MI0000446|hsa-mir-125b-1&WithFlank&11|-|122099751|122099850@21.42.22(+2A);MI0000470|hsa-mir-125b-2&WithFlank&21|+|16590231|16590331@23.44.22(+2A)&[MIMAT0000423&hsa-miR-125b-5p&offsets|0|0(+2A);m-67&11|-|122099809|122099830&offsets|0|0(+2A)];[MIMAT0000423_1&hsa-miR-125b-5p&offsets|0|0(+2A);m-68&21|+|16590253|16590274&offsets|0|0(+2A)]|TCCCTGAGACCCTAACTTGTGAAA</td><td>>MI0000446|hsa-mir-125b-1&WithFlank&11|-|122099751|122099850</td><td>21</td><td>42</td><td>22(+2A)</td><td>TCCCTGAGACCCTAACTTGTGAAA</td><td>867.3</td><td>-4.9</td><td>1.1</td><td>-4.6</td><td>5.3E-06</td><td>6.8E-05</td><td>BT20/MB231</td></tr>
<tr><td>>hg38_21_+_8433172_8446622_RNA45SN1_45S_50ntFlanks@12933.12952.20|GCTAAACCATTCGTAGACGA</td><td>>hg38_21_+_8433172_8446622_RNA45SN1_45S_50ntFlanks</td><td>12933</td><td>12952</td><td>20</td><td>GCTAAACCATTCGTAGACGA</td><td>99.6</td><td>-2.5</td><td>0.6</td><td>-4.1</td><td>4.9E-05</td><td>6.8E-05</td><td>MB231/MB468</td></tr>
<tr><td>>trna111_HisGTG_1_-_147774845_147774916@-1T.31.32,trna118_HisGTG_1_-_145396881_145396952@-1T.31.32,trna16_HisGTG_1_+_146544773_146544844@-1T.31.32,trna1_HisGTG_15_+_45493349_45493420@-1T.31.32,trna21_HisGTG_1_+_147753471_147753542@-1T.31.32,trna33_HisGTG_6_+_27125906_27125977@-1T.31.32,trna7_HisGTG_9_-_14433938_14434009@-1T.31.32,trna8_HisGTG_15_-_45492611_45492682@-1T.31.32,trna9_HisGTG_15_-_45490804_45490875@-1T.31.32|TGCCGTGATCGTATAGTGGTTAGTACTCTGCG</td><td>>trna111_HisGTG_1_-_147774845_147774916</td><td>-1T</td><td>31</td><td>32</td><td>TGCCGTGATCGTATAGTGGTTAGTACTCTGCG</td><td>5005.7</td><td>-3.3</td><td>0.8</td><td>-4.0</td><td>5.4E-05</td><td>6.9E-05</td><td>BT20/MB468</td></tr>
<tr><td>>hg38_21_+_8433172_8446622_RNA45SN1_45S_50ntFlanks@12165.12182.18|TGGGGCGGTACACCTGTC</td><td>>hg38_21_+_8433172_8446622_RNA45SN1_45S_50ntFlanks</td><td>12165</td><td>12182</td><td>18</td><td>TGGGGCGGTACACCTGTC</td><td>116.5</td><td>-3.1</td><td>0.8</td><td>-4.0</td><td>5.4E-05</td><td>6.9E-05</td><td>BT20/MB468</td></tr>
<tr><td>>hg38_21_+_8433172_8446622_RNA45SN1_45S_50ntFlanks@4007.4025.19|CCTCGGGCCGATCGCACGC</td><td>>hg38_21_+_8433172_8446622_RNA45SN1_45S_50ntFlanks</td><td>4007</td><td>4025</td><td>19</td><td>CCTCGGGCCGATCGCACGC</td><td>135.7</td><td>-2.6</td><td>0.6</td><td>-4.0</td><td>5.4E-05</td><td>6.9E-05</td><td>BT20/MB468</td></tr>
<tr><td>>hg38_21_+_8433172_8446622_RNA45SN1_45S_50ntFlanks@11881.11906.26|CCTGTTGAGCTTGACTCTAGTCTGGC</td><td>>hg38_21_+_8433172_8446622_RNA45SN1_45S_50ntFlanks</td><td>11881</td><td>11906</td><td>26</td><td>CCTGTTGAGCTTGACTCTAGTCTGGC</td><td>181.1</td><td>-3.8</td><td>0.9</td><td>-4.0</td><td>5.4E-05</td><td>6.9E-05</td><td>BT20/MB468</td></tr>
<tr><td>>hg38_21_+_8433172_8446622_RNA45SN1_45S_50ntFlanks@12416.12458.43|TTCGATGTCGGCTCTTCCTATCATTGTGAAGCAGAATTCACCA</td><td>>hg38_21_+_8433172_8446622_RNA45SN1_45S_50ntFlanks</td><td>12416</td><td>12458</td><td>43</td><td>TTCGATGTCGGCTCTTCCTATCATTGTGAAGCAGAATTCACCA</td><td>256.4</td><td>-3.4</td><td>0.8</td><td>-4.5</td><td>5.4E-06</td><td>6.9E-05</td><td>BT20/MB231</td></tr>
<tr><td>>hg38_21_+_8433172_8446622_RNA45SN1_45S_50ntFlanks@3785.3802.18|TGAAACTGCGAATGGCTC</td><td>>hg38_21_+_8433172_8446622_RNA45SN1_45S_50ntFlanks</td><td>3785</td><td>3802</td><td>18</td><td>TGAAACTGCGAATGGCTC</td><td>138.9</td><td>-1.4</td><td>0.3</td><td>-4.0</td><td>5.5E-05</td><td>7.0E-05</td><td>BT20/MB468</td></tr>
<tr><td>>hg38_21_+_8433172_8446622_RNA45SN1_45S_50ntFlanks@10347.10368.22|CAAACGAGAACTTTGAAGGCCG</td><td>>hg38_21_+_8433172_8446622_RNA45SN1_45S_50ntFlanks</td><td>10347</td><td>10368</td><td>22</td><td>CAAACGAGAACTTTGAAGGCCG</td><td>486.3</td><td>-2.3</td><td>0.6</td><td>-4.0</td><td>5.5E-05</td><td>7.0E-05</td><td>BT20/MB468</td></tr>
<tr><td>>MI0000471|hsa-mir-126&WithFlank&9|+|136670596|136670692@58.77.20&[MIMAT0000445&hsa-miR-126-3p&offsets|0|-2;m-40&9|+|136670653|136670674&offsets|0|-2]|TCGTACCGTGAGTAATAATG</td><td>>MI0000471|hsa-mir-126&WithFlank&9|+|136670596|136670692</td><td>58</td><td>77</td><td>20</td><td>TCGTACCGTGAGTAATAATG</td><td>1042.5</td><td>8.7</td><td>2.2</td><td>4.1</td><td>5.0E-05</td><td>7.0E-05</td><td>MB231/MB468</td></tr>
<tr><td>>hg38_21_+_8433172_8446622_RNA45SN1_45S_50ntFlanks@9825.9850.26|GTGGGCCACTTTTGGTAAGCAGAACT</td><td>>hg38_21_+_8433172_8446622_RNA45SN1_45S_50ntFlanks</td><td>9825</td><td>9850</td><td>26</td><td>GTGGGCCACTTTTGGTAAGCAGAACT</td><td>76.5</td><td>-3.1</td><td>0.8</td><td>-4.0</td><td>5.5E-05</td><td>7.0E-05</td><td>BT20/MB468</td></tr>
<tr><td>>hg38_21_+_8433172_8446622_RNA45SN1_45S_50ntFlanks@6738.6760.23|ATCATCGACACTTCGAACGCACT</td><td>>hg38_21_+_8433172_8446622_RNA45SN1_45S_50ntFlanks</td><td>6738</td><td>6760</td><td>23</td><td>ATCATCGACACTTCGAACGCACT</td><td>98.4</td><td>-3.7</td><td>0.9</td><td>-4.1</td><td>5.1E-05</td><td>7.2E-05</td><td>MB231/MB468</td></tr>
<tr><td>>hg38_21_+_8433172_8446622_RNA45SN1_45S_50ntFlanks@4129.4162.34|GATTCCGGAGAGGGAGCCTGAGAAACGGCTACCA</td><td>>hg38_21_+_8433172_8446622_RNA45SN1_45S_50ntFlanks</td><td>4129</td><td>4162</td><td>34</td><td>GATTCCGGAGAGGGAGCCTGAGAAACGGCTACCA</td><td>137.2</td><td>-2.7</td><td>0.7</td><td>-4.0</td><td>5.1E-05</td><td>7.2E-05</td><td>MB231/MB468</td></tr>
<tr><td>>am_trna2_GlyGCC_21_-_18827107_18827177@1.27.27,trna35_GlyGCC_1_+_161413094_161413164@1.27.27,trna37_GlyGCC_1_+_161420467_161420537@1.27.27,trna39_GlyGCC_1_+_161427898_161427968@1.27.27,trna41_GlyGCC_1_+_161435258_161435328@1.27.27|GCATGGGTGGTTCAGTGGTAGAATTCT</td><td>>am_trna2_GlyGCC_21_-_18827107_18827177</td><td>1</td><td>27</td><td>27</td><td>GCATGGGTGGTTCAGTGGTAGAATTCT</td><td>1990.2</td><td>2.5</td><td>0.6</td><td>4.0</td><td>5.6E-05</td><td>7.2E-05</td><td>BT20/MB468</td></tr>
<tr><td>>hg38_21_+_8433172_8446622_RNA45SN1_45S_50ntFlanks@11571.11591.21|GCCGACTTAGAACTGGTGCGG</td><td>>hg38_21_+_8433172_8446622_RNA45SN1_45S_50ntFlanks</td><td>11571</td><td>11591</td><td>21</td><td>GCCGACTTAGAACTGGTGCGG</td><td>276.9</td><td>-2.7</td><td>0.7</td><td>-4.0</td><td>5.6E-05</td><td>7.2E-05</td><td>BT20/MB468</td></tr>
<tr><td>>hg38_21_+_8433172_8446622_RNA45SN1_45S_50ntFlanks@12423.12441.19|TCGGCTCTTCCTATCATTG</td><td>>hg38_21_+_8433172_8446622_RNA45SN1_45S_50ntFlanks</td><td>12423</td><td>12441</td><td>19</td><td>TCGGCTCTTCCTATCATTG</td><td>135.2</td><td>-2.8</td><td>0.7</td><td>-4.0</td><td>5.6E-05</td><td>7.2E-05</td><td>BT20/MB468</td></tr>
<tr><td>>hg38_21_+_8433172_8446622_RNA45SN1_45S_50ntFlanks@8157.8181.25|CTCGTGGGGGGCCCAAGTCCTTCTG</td><td>>hg38_21_+_8433172_8446622_RNA45SN1_45S_50ntFlanks</td><td>8157</td><td>8181</td><td>25</td><td>CTCGTGGGGGGCCCAAGTCCTTCTG</td><td>258.0</td><td>-3.2</td><td>0.8</td><td>-4.0</td><td>5.7E-05</td><td>7.2E-05</td><td>BT20/MB468</td></tr>
</table>

<table>
<tr><td>>hg38_21_+_8433172_8446622_RNA45SN1_45S_50ntFlanks@7977.8006.30|CGACCTCAGATCAGACGTGGCGACCCGCTG</td><td>>hg38_21_+_8433172_8446622_RNA45SN1_45S_50ntFlanks</td><td>7977</td><td>8006</td><td>30</td><td>CGACCTCAGATCAGACGTGGCGACCCGCTG</td><td>811.3</td><td>-3.9</td><td>1.0</td><td>-4.0</td><td>5.7E-05</td><td>7.2E-05</td><td>BT20/MB468</td></tr>
<tr><td>>hg38_21_+_8433172_8446622_RNA45SN1_45S_50ntFlanks@7982.8007.26|TCAGATCAGACGTGGCGACCCGCTGA</td><td>>hg38_21_+_8433172_8446622_RNA45SN1_45S_50ntFlanks</td><td>7982</td><td>8007</td><td>26</td><td>TCAGATCAGACGTGGCGACCCGCTGA</td><td>125.1</td><td>-2.4</td><td>0.6</td><td>-4.0</td><td>5.7E-05</td><td>7.3E-05</td><td>BT20/MB468</td></tr>
<tr><td>>hg38_21_+_8433172_8446622_RNA45SN1_45S_50ntFlanks@10841.10872.32|CGTAACTTCGGGATAAGGATTGGCTCTAAGGG</td><td>>hg38_21_+_8433172_8446622_RNA45SN1_45S_50ntFlanks</td><td>10841</td><td>10872</td><td>32</td><td>CGTAACTTCGGGATAAGGATTGGCTCTAAGGG</td><td>386.0</td><td>-3.4</td><td>0.9</td><td>-4.0</td><td>5.7E-05</td><td>7.3E-05</td><td>BT20/MB468</td></tr>
<tr><td>>hg38_21_+_8433172_8446622_RNA45SN1_45S_50ntFlanks@12635.12660.26|AAGCTACCATCTGTGGGATTATGACT</td><td>>hg38_21_+_8433172_8446622_RNA45SN1_45S_50ntFlanks</td><td>12635</td><td>12660</td><td>26</td><td>AAGCTACCATCTGTGGGATTATGACT</td><td>144.2</td><td>-2.2</td><td>0.5</td><td>-4.0</td><td>5.7E-05</td><td>7.3E-05</td><td>BT20/MB468</td></tr>
<tr><td>>hg38_21_+_8433172_8446622_RNA45SN1_45S_50ntFlanks@6759.6806.48|CTTGCGGCCCCGGGTTCCTCCCGGGGCTACGCCTGTCTGAGCGTCGCT</td><td>>hg38_21_+_8433172_8446622_RNA45SN1_45S_50ntFlanks</td><td>6759</td><td>6806</td><td>48</td><td>CTTGCGGCCCCGGGTTCCTCCCGGGGCTACGCCTGTCTGAGCGTCGCT</td><td>1480.4</td><td>10.5</td><td>2.6</td><td>4.0</td><td>5.2E-05</td><td>7.3E-05</td><td>MB231/MB468</td></tr>
<tr><td>>hg38_21_+_8433172_8446622_RNA45SN1_45S_50ntFlanks@11319.11344.26|CCCACGTCTCGTCGCGCGCGCGTCCG</td><td>>hg38_21_+_8433172_8446622_RNA45SN1_45S_50ntFlanks</td><td>11319</td><td>11344</td><td>26</td><td>CCCACGTCTCGTCGCGCGCGCGTCCG</td><td>1062.6</td><td>-5.5</td><td>1.4</td><td>-4.0</td><td>5.8E-05</td><td>7.4E-05</td><td>BT20/MB468</td></tr>
<tr><td>>hg38_21_+_8433172_8446622_RNA45SN1_45S_50ntFlanks@11561.11589.29|GCGCCTAGCAGCCGACTTAGAACTGGTGC</td><td>>hg38_21_+_8433172_8446622_RNA45SN1_45S_50ntFlanks</td><td>11561</td><td>11589</td><td>29</td><td>GCGCCTAGCAGCCGACTTAGAACTGGTGC</td><td>184.8</td><td>-2.8</td><td>0.7</td><td>-4.0</td><td>5.8E-05</td><td>7.4E-05</td><td>BT20/MB468</td></tr>
<tr><td>>hg38_21_+_8433172_8446622_RNA45SN1_45S_50ntFlanks@10866.10890.25|CTAAGGGCTGGGTCGGTCGGGCTGG</td><td>>hg38_21_+_8433172_8446622_RNA45SN1_45S_50ntFlanks</td><td>10866</td><td>10890</td><td>25</td><td>CTAAGGGCTGGGTCGGTCGGGCTGG</td><td>331.5</td><td>-3.6</td><td>0.9</td><td>-4.0</td><td>5.8E-05</td><td>7.4E-05</td><td>BT20/MB468</td></tr>
<tr><td>>hg38_21_+_8433172_8446622_RNA45SN1_45S_50ntFlanks@10351.10385.35|CGAGAACTTTGAAGGCCGAAGTGGAGAAGGGTTCC</td><td>>hg38_21_+_8433172_8446622_RNA45SN1_45S_50ntFlanks</td><td>10351</td><td>10385</td><td>35</td><td>CGAGAACTTTGAAGGCCGAAGTGGAGAAGGGTTCC</td><td>142.8</td><td>-2.1</td><td>0.5</td><td>-4.0</td><td>5.4E-05</td><td>7.5E-05</td><td>MB231/MB468</td></tr>
<tr><td>>hg38_21_+_8433172_8446622_RNA45SN1_45S_50ntFlanks@12405.12446.42|CTTTTTGATCCTTCGATGTCGGCTCTTCCTATCATTGTGAAG</td><td>>hg38_21_+_8433172_8446622_RNA45SN1_45S_50ntFlanks</td><td>12405</td><td>12446</td><td>42</td><td>CTTTTTGATCCTTCGATGTCGGCTCTTCCTATCATTGTGAAG</td><td>394.8</td><td>-1.8</td><td>0.5</td><td>-4.0</td><td>5.4E-05</td><td>7.5E-05</td><td>MB231/MB468</td></tr>
<tr><td>>hg38_21_+_8433172_8446622_RNA45SN1_45S_50ntFlanks@11703.11725.23|AATGAAGCGCGGGTAAACGGCGG</td><td>>hg38_21_+_8433172_8446622_RNA45SN1_45S_50ntFlanks</td><td>11703</td><td>11725</td><td>23</td><td>AATGAAGCGCGGGTAAACGGCGG</td><td>109.3</td><td>-2.6</td><td>0.6</td><td>-4.0</td><td>5.4E-05</td><td>7.6E-05</td><td>MB231/MB468</td></tr>
<tr><td>>hg38_21_+_8433172_8446622_RNA45SN1_45S_50ntFlanks@11568.11603.36|GCAGCCGACTTAGAACTGGTGCGGACCAGGGGAATC</td><td>>hg38_21_+_8433172_8446622_RNA45SN1_45S_50ntFlanks</td><td>11568</td><td>11603</td><td>36</td><td>GCAGCCGACTTAGAACTGGTGCGGACCAGGGGAATC</td><td>265.2</td><td>-2.1</td><td>0.5</td><td>-4.0</td><td>5.4E-05</td><td>7.6E-05</td><td>MB231/MB468</td></tr>
<tr><td>>hg38_21_+_8433172_8446622_RNA45SN1_45S_50ntFlanks@10349.10388.40|AACGAGAACTTTGAAGGCCGAAGTGGAGAAGGGTTCCATG</td><td>>hg38_21_+_8433172_8446622_RNA45SN1_45S_50ntFlanks</td><td>10349</td><td>10388</td><td>40</td><td>AACGAGAACTTTGAAGGCCGAAGTGGAGAAGGGTTCCATG</td><td>767.7</td><td>-3.4</td><td>0.9</td><td>-4.0</td><td>6.0E-05</td><td>7.6E-05</td><td>BT20/MB468</td></tr>
<tr><td>>hg38_21_+_8433172_8446622_RNA45SN1_45S_50ntFlanks@3782.3810.29|CAGTGAAACTGCGAATGGCTCATTAAATC</td><td>>hg38_21_+_8433172_8446622_RNA45SN1_45S_50ntFlanks</td><td>3782</td><td>3810</td><td>29</td><td>CAGTGAAACTGCGAATGGCTCATTAAATC</td><td>132.9</td><td>-2.9</td><td>0.7</td><td>-4.0</td><td>6.0E-05</td><td>7.6E-05</td><td>BT20/MB468</td></tr>
<tr><td>>hg38_21_+_8433172_8446622_RNA45SN1_45S_50ntFlanks@6758.6806.49|ACTTGCGGCCCCGGGTTCCTCCCGGGGCTACGCCTGTCTGAGCGTCGCT</td><td>>hg38_21_+_8433172_8446622_RNA45SN1_45S_50ntFlanks</td><td>6758</td><td>6806</td><td>49</td><td>ACTTGCGGCCCCGGGTTCCTCCCGGGGCTACGCCTGTCTGAGCGTCGCT</td><td>1205.7</td><td>10.6</td><td>2.6</td><td>4.0</td><td>5.5E-05</td><td>7.6E-05</td><td>MB231/MB468</td></tr>
<tr><td>>hg38_21_+_8433172_8446622_RNA45SN1_45S_50ntFlanks@12388.12433.46|CGTTCATAGCGACGTCGCTTTTTGATCCTTCGATGTCGGCTCTTCC</td><td>>hg38_21_+_8433172_8446622_RNA45SN1_45S_50ntFlanks</td><td>12388</td><td>12433</td><td>46</td><td>CGTTCATAGCGACGTCGCTTTTTGATCCTTCGATGTCGGCTCTTCC</td><td>251.1</td><td>-3.8</td><td>0.8</td><td>-4.5</td><td>6.0E-06</td><td>7.6E-05</td><td>BT20/MB231</td></tr>
<tr><td>>hg38_21_+_8433172_8446622_RNA45SN1_45S_50ntFlanks@12995.13039.45|CTCGCTGCGATCTATTGAAAGTCAGCCCTCGACACAAGGGTTTGT</td><td>>hg38_21_+_8433172_8446622_RNA45SN1_45S_50ntFlanks</td><td>12995</td><td>13039</td><td>45</td><td>CTCGCTGCGATCTATTGAAAGTCAGCCCTCGACACAAGGGTTTGT</td><td>601406.2</td><td>-3.7</td><td>0.8</td><td>-4.5</td><td>6.1E-06</td><td>7.7E-05</td><td>BT20/MB231</td></tr>
<tr><td>>hg38_21_+_8433172_8446622_RNA45SN1_45S_50ntFlanks@11539.11584.46|GCGCGGCGCCTCGCCTCGGCCGGCGCCTAGCAGCCGACTTAGAACT</td><td>>hg38_21_+_8433172_8446622_RNA45SN1_45S_50ntFlanks</td><td>11539</td><td>11584</td><td>46</td><td>GCGCGGCGCCTCGCCTCGGCCGGCGCCTAGCAGCCGACTTAGAACT</td><td>457.9</td><td>-3.5</td><td>0.8</td><td>-4.5</td><td>6.1E-06</td><td>7.7E-05</td><td>BT20/MB231</td></tr>
<tr><td>>hg38_21_+_8433172_8446622_RNA45SN1_45S_50ntFlanks@8359.8379.21|AAGAACTTTGAAGAGAGAGTT</td><td>>hg38_21_+_8433172_8446622_RNA45SN1_45S_50ntFlanks</td><td>8359</td><td>8379</td><td>21</td><td>AAGAACTTTGAAGAGAGAGTT</td><td>1016.2</td><td>-2.8</td><td>0.7</td><td>-4.0</td><td>5.5E-05</td><td>7.7E-05</td><td>MB231/MB468</td></tr>
<tr><td>>hg38_21_+_8433172_8446622_RNA45SN1_45S_50ntFlanks@11728.11753.26|GTAACTATGACTCTCTTAAGGTAGCC</td><td>>hg38_21_+_8433172_8446622_RNA45SN1_45S_50ntFlanks</td><td>11728</td><td>11753</td><td>26</td><td>GTAACTATGACTCTCTTAAGGTAGCC</td><td>169.3</td><td>-3.2</td><td>0.8</td><td>-4.0</td><td>5.6E-05</td><td>7.7E-05</td><td>MB231/MB468</td></tr>
<tr><td>>hg38_21_+_8433172_8446622_RNA45SN1_45S_50ntFlanks@6650.6675.26|ACGACTCTTAGCGGTGGATCACTCGG</td><td>>hg38_21_+_8433172_8446622_RNA45SN1_45S_50ntFlanks</td><td>6650</td><td>6675</td><td>26</td><td>ACGACTCTTAGCGGTGGATCACTCGG</td><td>390.6</td><td>2.6</td><td>0.6</td><td>4.0</td><td>6.1E-05</td><td>7.8E-05</td><td>BT20/MB468</td></tr>
<tr><td>>hg38_21_+_8433172_8446622_RNA45SN1_45S_50ntFlanks@11672.11692.21|CAGTGCTCTGAATGTCAAAGT</td><td>>hg38_21_+_8433172_8446622_RNA45SN1_45S_50ntFlanks</td><td>11672</td><td>11692</td><td>21</td><td>CAGTGCTCTGAATGTCAAAGT</td><td>786.2</td><td>2.8</td><td>0.7</td><td>4.0</td><td>5.6E-05</td><td>7.9E-05</td><td>MB231/MB468</td></tr>
<tr><td>>trna13_ValCAC_19_-_4724647_4724719@1.34.34|GTTTCCGTAGTGTAGCGGTTATCACATTCGCCTC</td><td>>trna13_ValCAC_19_-_4724647_4724719</td><td>1</td><td>34</td><td>34</td><td>GTTTCCGTAGTGTAGCGGTTATCACATTCGCCTC</td><td>424.1</td><td>-4.2</td><td>0.9</td><td>-4.5</td><td>6.3E-06</td><td>7.9E-05</td><td>BT20/MB231</td></tr>
<tr><td>>hg38_21_+_8433172_8446622_RNA45SN1_45S_50ntFlanks@11873.11905.33|AAGAAGACCCTGTTGAGCTTGACTCTAGTCTGG</td><td>>hg38_21_+_8433172_8446622_RNA45SN1_45S_50ntFlanks</td><td>11873</td><td>11905</td><td>33</td><td>AAGAAGACCCTGTTGAGCTTGACTCTAGTCTGG</td><td>593.8</td><td>-5.5</td><td>1.4</td><td>-4.0</td><td>6.3E-05</td><td>7.9E-05</td><td>BT20/MB468</td></tr>
<tr><td>>hg38_21_+_8433172_8446622_RNA45SN1_45S_50ntFlanks@12992.13015.24|TCCCTCGCTGCGATCTATTGAAAG</td><td>>hg38_21_+_8433172_8446622_RNA45SN1_45S_50ntFlanks</td><td>12992</td><td>13015</td><td>24</td><td>TCCCTCGCTGCGATCTATTGAAAG</td><td>336.4</td><td>-1.8</td><td>0.5</td><td>-4.0</td><td>5.7E-05</td><td>8.0E-05</td><td>MB231/MB468</td></tr>
<tr><td>>hg38_21_+_8433172_8446622_RNA45SN1_45S_50ntFlanks@5473.5499.27|CGGCCCTGGCGGAGCGCTGAGAAGACG</td><td>>hg38_21_+_8433172_8446622_RNA45SN1_45S_50ntFlanks</td><td>5473</td><td>5499</td><td>27</td><td>CGGCCCTGGCGGAGCGCTGAGAAGACG</td><td>1395.1</td><td>2.6</td><td>0.7</td><td>4.0</td><td>5.7E-05</td><td>8.0E-05</td><td>MB231/MB468</td></tr>
<tr><td>>hg38_21_+_8433172_8446622_RNA45SN1_45S_50ntFlanks@5186.5205.20|CAATAACAGGTCTGTGATGC</td><td>>hg38_21_+_8433172_8446622_RNA45SN1_45S_50ntFlanks</td><td>5186</td><td>5205</td><td>20</td><td>CAATAACAGGTCTGTGATGC</td><td>125.8</td><td>-1.8</td><td>0.5</td><td>-4.0</td><td>6.3E-05</td><td>8.0E-05</td><td>BT20/MB468</td></tr>
<tr><td>>hg38_21_+_8433172_8446622_RNA45SN1_45S_50ntFlanks@3835.3862.28|CTCCTCTCCTACTTGGATAACTGTGGTA</td><td>>hg38_21_+_8433172_8446622_RNA45SN1_45S_50ntFlanks</td><td>3835</td><td>3862</td><td>28</td><td>CTCCTCTCCTACTTGGATAACTGTGGTA</td><td>122.0</td><td>-2.2</td><td>0.6</td><td>-4.0</td><td>6.4E-05</td><td>8.1E-05</td><td>BT20/MB468</td></tr>
<tr><td>>MI0000283|hsa-mir-203a&WithFlank&14|+|104117399|104117520@71.91.21(+1A)&[MIMAT0000264&hsa-miR-203a-3p&offsets|0|-1(+1A);m-79&14|+|104117469|104117490&offsets|0|-1(+1A)]|GTGAAATGTTTAGGACCACTAA</td><td>>MI0000283|hsa-mir-203a&WithFlank&14|+|104117399|104117520</td><td>71</td><td>91</td><td>21(+1A)</td><td>GTGAAATGTTTAGGACCACTAA</td><td>1211.1</td><td>10.3</td><td>2.6</td><td>4.0</td><td>6.4E-05</td><td>8.1E-05</td><td>BT20/MB468</td></tr>
<tr><td>>hg38_21_+_8433172_8446622_RNA45SN1_45S_50ntFlanks@9740.9757.18|AACGATCTCAACCTATTC</td><td>>hg38_21_+_8433172_8446622_RNA45SN1_45S_50ntFlanks</td><td>9740</td><td>9757</td><td>18</td><td>AACGATCTCAACCTATTC</td><td>119.1</td><td>-3.4</td><td>0.8</td><td>-4.0</td><td>5.8E-05</td><td>8.1E-05</td><td>MB231/MB468</td></tr>
<tr><td>>hg38_21_+_8433172_8446622_RNA45SN1_45S_50ntFlanks@11579.11609.31|AGAACTGGTGCGGACCAGGGGAATCCGACTG</td><td>>hg38_21_+_8433172_8446622_RNA45SN1_45S_50ntFlanks</td><td>11579</td><td>11609</td><td>31</td><td>AGAACTGGTGCGGACCAGGGGAATCCGACTG</td><td>573.0</td><td>-2.6</td><td>0.6</td><td>-4.0</td><td>6.4E-05</td><td>8.1E-05</td><td>BT20/MB468</td></tr>
<tr><td>>trnaMT_PheGAA_MT_+_577_647@54.74.21|CACATCACCCCATAAACACCA</td><td>>trnaMT_PheGAA_MT_+_577_647</td><td>54</td><td>74</td><td>21</td><td>CACATCACCCCATAAACACCA</td><td>605.4</td><td>2.8</td><td>0.7</td><td>4.0</td><td>6.4E-05</td><td>8.2E-05</td><td>BT20/MB468</td></tr>
<tr><td>>hg38_21_+_8433172_8446622_RNA45SN1_45S_50ntFlanks@4995.5013.19|ACAGATTGATAGCTCTTTC</td><td>>hg38_21_+_8433172_8446622_RNA45SN1_45S_50ntFlanks</td><td>4995</td><td>5013</td><td>19</td><td>ACAGATTGATAGCTCTTTC</td><td>152.2</td><td>-2.4</td><td>0.6</td><td>-4.0</td><td>6.5E-05</td><td>8.2E-05</td><td>BT20/MB468</td></tr>
<tr><td>>hg38_21_+_8433172_8446622_RNA45SN1_45S_50ntFlanks@6721.6747.27|AATTGCAGGACACATTGATCATCGACA</td><td>>hg38_21_+_8433172_8446622_RNA45SN1_45S_50ntFlanks</td><td>6721</td><td>6747</td><td>27</td><td>AATTGCAGGACACATTGATCATCGACA</td><td>2183.6</td><td>-2.3</td><td>0.6</td><td>-4.0</td><td>5.9E-05</td><td>8.2E-05</td><td>MB231/MB468</td></tr>
<tr><td>>hg38_21_+_8433172_8446622_RNA45SN1_45S_50ntFlanks@11562.11592.31|CGCCTAGCAGCCGACTTAGAACTGGTGCGGA</td><td>>hg38_21_+_8433172_8446622_RNA45SN1_45S_50ntFlanks</td><td>11562</td><td>11592</td><td>31</td><td>CGCCTAGCAGCCGACTTAGAACTGGTGCGGA</td><td>969.2</td><td>-3.2</td><td>0.8</td><td>-4.0</td><td>6.5E-05</td><td>8.2E-05</td><td>BT20/MB468</td></tr>
<tr><td>>hg38_21_+_8433172_8446622_RNA45SN1_45S_50ntFlanks@10808.10831.24|CAATGTAGGTAAGGGAAGTCGGCA</td><td>>hg38_21_+_8433172_8446622_RNA45SN1_45S_50ntFlanks</td><td>10808</td><td>10831</td><td>24</td><td>CAATGTAGGTAAGGGAAGTCGGCA</td><td>118.1</td><td>-3.2</td><td>0.8</td><td>-4.0</td><td>6.5E-05</td><td>8.3E-05</td><td>BT20/MB468</td></tr>
<tr><td>>hg38_21_+_8433172_8446622_RNA45SN1_45S_50ntFlanks@11668.11701.34|TGCCCAGTGCTCTGAATGTCAAAGTGAAGAAATT</td><td>>hg38_21_+_8433172_8446622_RNA45SN1_45S_50ntFlanks</td><td>11668</td><td>11701</td><td>34</td><td>TGCCCAGTGCTCTGAATGTCAAAGTGAAGAAATT</td><td>440.7</td><td>2.3</td><td>0.6</td><td>4.0</td><td>6.0E-05</td><td>8.3E-05</td><td>MB231/MB468</td></tr>
<tr><td>>MI0000286|hsa-mir-210&WithFlank&11|-|568083|568204@72.92.21(+1U)&[MIMAT0000267&hsa-miR-210-3p&offsets|0|-1(+1U);m-102&11|-|568112|568133&offsets|0|-1(+1U)]|CTGTGCGTGTGACAGCGGCTGT</td><td>>MI0000286|hsa-mir-210&WithFlank&11|-|568083|568204</td><td>72</td><td>92</td><td>21(+1U)</td><td>CTGTGCGTGTGACAGCGGCTGT</td><td>2036.9</td><td>4.9</td><td>1.1</td><td>4.5</td><td>6.7E-06</td><td>8.4E-05</td><td>BT20/MB231</td></tr>
<tr><td>>hg38_21_+_8433172_8446622_RNA45SN1_45S_50ntFlanks@10839.10868.30|TCCGTAACTTCGGGATAAGGATTGGCTCTA</td><td>>hg38_21_+_8433172_8446622_RNA45SN1_45S_50ntFlanks</td><td>10839</td><td>10868</td><td>30</td><td>TCCGTAACTTCGGGATAAGGATTGGCTCTA</td><td>185.1</td><td>-2.3</td><td>0.6</td><td>-4.0</td><td>6.0E-05</td><td>8.4E-05</td><td>MB231/MB468</td></tr>
<tr><td>>hg38_21_+_8433172_8446622_RNA45SN1_45S_50ntFlanks@5485.5509.25|AGCGCTGAGAAGACGGTCGAACTTG</td><td>>hg38_21_+_8433172_8446622_RNA45SN1_45S_50ntFlanks</td><td>5485</td><td>5509</td><td>25</td><td>AGCGCTGAGAAGACGGTCGAACTTG</td><td>172.4</td><td>-2.4</td><td>0.6</td><td>-4.0</td><td>6.1E-05</td><td>8.5E-05</td><td>MB231/MB468</td></tr>
<tr><td>>hg38_21_+_8433172_8446622_RNA45SN1_45S_50ntFlanks@10351.10391.41|CGAGAACTTTGAAGGCCGAAGTGGAGAAGGGTTCCATGTGA</td><td>>hg38_21_+_8433172_8446622_RNA45SN1_45S_50ntFlanks</td><td>10351</td><td>10391</td><td>41</td><td>CGAGAACTTTGAAGGCCGAAGTGGAGAAGGGTTCCATGTGA</td><td>369.1</td><td>-3.8</td><td>0.9</td><td>-4.0</td><td>6.7E-05</td><td>8.5E-05</td><td>BT20/MB468</td></tr>
<tr><td>>hg38_21_+_8433172_8446622_RNA45SN1_45S_50ntFlanks@11569.11593.25|CAGCCGACTTAGAACTGGTGCGGAC</td><td>>hg38_21_+_8433172_8446622_RNA45SN1_45S_50ntFlanks</td><td>11569</td><td>11593</td><td>25</td><td>CAGCCGACTTAGAACTGGTGCGGAC</td><td>207.0</td><td>-3.8</td><td>1.0</td><td>-4.0</td><td>6.7E-05</td><td>8.5E-05</td><td>BT20/MB468</td></tr>
<tr><td>>MI0000101|hsa-mir-99a&WithFlank&21|+|16539083|16539175@19.40.22&[MIMAT0000097&hsa-miR-99a-5p&offsets|0|0;m-64&21|+|16539101|16539121&offsets|0|+1]|AACCCGTAGATCCGATCTTGTG</td><td>>MI0000101|hsa-mir-99a&WithFlank&21|+|16539083|16539175</td><td>19</td><td>40</td><td>22</td><td>AACCCGTAGATCCGATCTTGTG</td><td>839.1</td><td>-4.5</td><td>1.1</td><td>-4.0</td><td>6.8E-05</td><td>8.6E-05</td><td>BT20/MB468</td></tr>
<tr><td>>hg38_21_+_8433172_8446622_RNA45SN1_45S_50ntFlanks@11210.11231.22|GGCGACTCTGGACGCGAGCCGG</td><td>>hg38_21_+_8433172_8446622_RNA45SN1_45S_50ntFlanks</td><td>11210</td><td>11231</td><td>22</td><td>GGCGACTCTGGACGCGAGCCGG</td><td>143.0</td><td>-3.3</td><td>0.8</td><td>-4.0</td><td>6.2E-05</td><td>8.7E-05</td><td>MB231/MB468</td></tr>
<tr><td>>hg38_21_+_8433172_8446622_RNA45SN1_45S_50ntFlanks@7979.8007.29|ACCTCAGATCAGACGTGGCGACCCGCTGA</td><td>>hg38_21_+_8433172_8446622_RNA45SN1_45S_50ntFlanks</td><td>7979</td><td>8007</td><td>29</td><td>ACCTCAGATCAGACGTGGCGACCCGCTGA</td><td>190.4</td><td>-3.1</td><td>0.8</td><td>-4.0</td><td>6.8E-05</td><td>8.7E-05</td><td>BT20/MB468</td></tr>
<tr><td>>MI0000255|hsa-mir-30d&WithFlank&8|-|134804870|134804951@12.32.21(+1A)&[MIMAT0000245&hsa-miR-30d-5p&offsets|0|-1(+1A);m-21&8|-|134804919|134804940&offsets|0|-1(+1A)]|TGTAAACATCCCCGACTGGAAA</td><td>>MI0000255|hsa-mir-30d&WithFlank&8|-|134804870|134804951</td><td>12</td><td>32</td><td>21(+1A)</td><td>TGTAAACATCCCCGACTGGAAA</td><td>1685.0</td><td>3.5</td><td>0.8</td><td>4.5</td><td>7.0E-06</td><td>8.7E-05</td><td>BT20/MB231</td></tr>
</table>

<table>
<tr><td>>hg38_21_+_8433172_8446622_RNA45SN1_45S_50ntFlanks@11569.11590.22|CAGCCGACTTAGAACTGGTGCG</td><td>>hg38_21_+_8433172_8446622_RNA45SN1_45S_50ntFlanks</td><td>11569</td><td>11590</td><td>22</td><td>CAGCCGACTTAGAACTGGTGCG</td><td>179.2</td><td>-2.5</td><td>0.6</td><td>-4.0</td><td>6.9E-05</td><td>8.7E-05</td><td>BT20/MB468</td></tr>
<tr><td>>hg38_21_+_8433172_8446622_RNA45SN1_45S_50ntFlanks@10392.10426.35|ACAGCAGTTGAACATGGGTCAGTCGGTCCTGAGAG</td><td>>hg38_21_+_8433172_8446622_RNA45SN1_45S_50ntFlanks</td><td>10392</td><td>10426</td><td>35</td><td>ACAGCAGTTGAACATGGGTCAGTCGGTCCTGAGAG</td><td>351.8</td><td>-2.9</td><td>0.7</td><td>-4.0</td><td>6.2E-05</td><td>8.7E-05</td><td>MB231/MB468</td></tr>
<tr><td>>hg38_21_+_8433172_8446622_RNA45SN1_45S_50ntFlanks@8174.8203.30|TCCTTCTGATCGAGGCCCAGCCCGTGGACG</td><td>>hg38_21_+_8433172_8446622_RNA45SN1_45S_50ntFlanks</td><td>8174</td><td>8203</td><td>30</td><td>TCCTTCTGATCGAGGCCCAGCCCGTGGACG</td><td>506.2</td><td>3.2</td><td>0.8</td><td>4.0</td><td>6.3E-05</td><td>8.7E-05</td><td>MB231/MB468</td></tr>
<tr><td>>hg38_21_+_8433172_8446622_RNA45SN1_45S_50ntFlanks@12941.12972.32|ATTCGTAGACGACCTGCTTCTGGGTCGGGGTT</td><td>>hg38_21_+_8433172_8446622_RNA45SN1_45S_50ntFlanks</td><td>12941</td><td>12972</td><td>32</td><td>ATTCGTAGACGACCTGCTTCTGGGTCGGGGTT</td><td>122.3</td><td>-3.5</td><td>0.9</td><td>-4.0</td><td>6.9E-05</td><td>8.8E-05</td><td>BT20/MB468</td></tr>
<tr><td>>hg38_21_+_8433172_8446622_RNA45SN1_45S_50ntFlanks@10873.10891.19|CTGGGTCGGTCGGGCTGGG</td><td>>hg38_21_+_8433172_8446622_RNA45SN1_45S_50ntFlanks</td><td>10873</td><td>10891</td><td>19</td><td>CTGGGTCGGTCGGGCTGGG</td><td>264.5</td><td>-4.6</td><td>1.2</td><td>-4.0</td><td>7.0E-05</td><td>8.9E-05</td><td>BT20/MB468</td></tr>
<tr><td>>hg38_21_+_8433172_8446622_RNA45SN1_45S_50ntFlanks@10350.10381.32|ACGAGAACTTTGAAGGCCGAAGTGGAGAAGGG</td><td>>hg38_21_+_8433172_8446622_RNA45SN1_45S_50ntFlanks</td><td>10350</td><td>10381</td><td>32</td><td>ACGAGAACTTTGAAGGCCGAAGTGGAGAAGGG</td><td>180.6</td><td>-1.8</td><td>0.5</td><td>-4.0</td><td>6.4E-05</td><td>8.9E-05</td><td>MB231/MB468</td></tr>
<tr><td>>hg38_21_+_8433172_8446622_RNA45SN1_45S_50ntFlanks@13008.13026.19|ATTGAAAGTCAGCCCTCGA</td><td>>hg38_21_+_8433172_8446622_RNA45SN1_45S_50ntFlanks</td><td>13008</td><td>13026</td><td>19</td><td>ATTGAAAGTCAGCCCTCGA</td><td>101.2</td><td>-3.6</td><td>0.9</td><td>-4.0</td><td>6.4E-05</td><td>8.9E-05</td><td>MB231/MB468</td></tr>
<tr><td>>hg38_21_+_8433172_8446622_RNA45SN1_45S_50ntFlanks@4005.4024.20|AACCTCGGGCCGATCGCACG</td><td>>hg38_21_+_8433172_8446622_RNA45SN1_45S_50ntFlanks</td><td>4005</td><td>4024</td><td>20</td><td>AACCTCGGGCCGATCGCACG</td><td>524.8</td><td>-1.8</td><td>0.5</td><td>-4.0</td><td>6.4E-05</td><td>8.9E-05</td><td>MB231/MB468</td></tr>
<tr><td>>hg38_21_+_8433172_8446622_RNA45SN1_45S_50ntFlanks@8192.8211.20|AGCCCGTGGACGGTGTGAGG</td><td>>hg38_21_+_8433172_8446622_RNA45SN1_45S_50ntFlanks</td><td>8192</td><td>8211</td><td>20</td><td>AGCCCGTGGACGGTGTGAGG</td><td>166.6</td><td>-3.3</td><td>0.8</td><td>-4.0</td><td>7.1E-05</td><td>9.0E-05</td><td>BT20/MB468</td></tr>
<tr><td>>hg38_21_+_8433172_8446622_RNA45SN1_45S_50ntFlanks@6719.6747.29|TGAATTGCAGGACACATTGATCATCGACA</td><td>>hg38_21_+_8433172_8446622_RNA45SN1_45S_50ntFlanks</td><td>6719</td><td>6747</td><td>29</td><td>TGAATTGCAGGACACATTGATCATCGACA</td><td>948.1</td><td>-2.2</td><td>0.6</td><td>-4.0</td><td>6.6E-05</td><td>9.1E-05</td><td>MB231/MB468</td></tr>
<tr><td>>hg38_21_+_8433172_8446622_RNA45SN1_45S_50ntFlanks@12957.12975.19|CTTCTGGGTCGGGGTTTCG</td><td>>hg38_21_+_8433172_8446622_RNA45SN1_45S_50ntFlanks</td><td>12957</td><td>12975</td><td>19</td><td>CTTCTGGGTCGGGGTTTCG</td><td>104.4</td><td>-2.1</td><td>0.5</td><td>-4.0</td><td>6.6E-05</td><td>9.1E-05</td><td>MB231/MB468</td></tr>
<tr><td>>hg38_21_+_8433172_8446622_RNA45SN1_45S_50ntFlanks@11906.11929.24|CACGGTGAAGAGACATGAGAGGTG</td><td>>hg38_21_+_8433172_8446622_RNA45SN1_45S_50ntFlanks</td><td>11906</td><td>11929</td><td>24</td><td>CACGGTGAAGAGACATGAGAGGTG</td><td>100.6</td><td>-4.4</td><td>1.1</td><td>-4.0</td><td>7.2E-05</td><td>9.2E-05</td><td>BT20/MB468</td></tr>
<tr><td>>hg38_21_+_8433172_8446622_RNA45SN1_45S_50ntFlanks@7975.7995.21|CGCGACCTCAGATCAGACGTG</td><td>>hg38_21_+_8433172_8446622_RNA45SN1_45S_50ntFlanks</td><td>7975</td><td>7995</td><td>21</td><td>CGCGACCTCAGATCAGACGTG</td><td>10593.6</td><td>2.3</td><td>0.6</td><td>4.0</td><td>7.3E-05</td><td>9.2E-05</td><td>BT20/MB468</td></tr>
<tr><td>>hg38_21_+_8433172_8446622_RNA45SN1_45S_50ntFlanks@11565.11582.18|CTAGCAGCCGACTTAGAA</td><td>>hg38_21_+_8433172_8446622_RNA45SN1_45S_50ntFlanks</td><td>11565</td><td>11582</td><td>18</td><td>CTAGCAGCCGACTTAGAA</td><td>123.7</td><td>-1.8</td><td>0.5</td><td>-4.0</td><td>6.7E-05</td><td>9.3E-05</td><td>MB231/MB468</td></tr>
<tr><td>>hg38_21_+_8433172_8446622_RNA45SN1_45S_50ntFlanks@12072.12106.35|CCCGAGGGGCTCTCGCTTCTGGCGCCAAGCGCCCG</td><td>>hg38_21_+_8433172_8446622_RNA45SN1_45S_50ntFlanks</td><td>12072</td><td>12106</td><td>35</td><td>CCCGAGGGGCTCTCGCTTCTGGCGCCAAGCGCCCG</td><td>152.4</td><td>-2.5</td><td>0.6</td><td>-4.0</td><td>6.7E-05</td><td>9.3E-05</td><td>MB231/MB468</td></tr>
<tr><td>>hg38_21_+_8433172_8446622_RNA45SN1_45S_50ntFlanks@6661.6697.37|CGGTGGATCACTCGGCTCGTGCGTCGATGAAGAACGC</td><td>>hg38_21_+_8433172_8446622_RNA45SN1_45S_50ntFlanks</td><td>6661</td><td>6697</td><td>37</td><td>CGGTGGATCACTCGGCTCGTGCGTCGATGAAGAACGC</td><td>1023.7</td><td>2.1</td><td>0.5</td><td>4.0</td><td>7.4E-05</td><td>9.4E-05</td><td>BT20/MB468</td></tr>
<tr><td>>hg38_21_+_8433172_8446622_RNA45SN1_45S_50ntFlanks@11552.11592.41|CCTCGGCCGGCGCCTAGCAGCCGACTTAGAACTGGTGCGGA</td><td>>hg38_21_+_8433172_8446622_RNA45SN1_45S_50ntFlanks</td><td>11552</td><td>11592</td><td>41</td><td>CCTCGGCCGGCGCCTAGCAGCCGACTTAGAACTGGTGCGGA</td><td>102.2</td><td>-2.2</td><td>0.5</td><td>-4.0</td><td>7.4E-05</td><td>9.4E-05</td><td>BT20/MB468</td></tr>
<tr><td>>hg38_21_+_8433172_8446622_RNA45SN1_45S_50ntFlanks@12073.12098.26|CCGAGGGGCTCTCGCTTCTGGCGCCA</td><td>>hg38_21_+_8433172_8446622_RNA45SN1_45S_50ntFlanks</td><td>12073</td><td>12098</td><td>26</td><td>CCGAGGGGCTCTCGCTTCTGGCGCCA</td><td>197.0</td><td>-2.0</td><td>0.5</td><td>-4.0</td><td>6.8E-05</td><td>9.4E-05</td><td>MB231/MB468</td></tr>
<tr><td>>trnaMT_MetCAT_MT_+_4402_4469@46.71.26|TTGGTTATACCCTTCCCGTACTACCA</td><td>>trnaMT_MetCAT_MT_+_4402_4469</td><td>46</td><td>71</td><td>26</td><td>TTGGTTATACCCTTCCCGTACTACCA</td><td>491.9</td><td>9.3</td><td>2.4</td><td>4.0</td><td>7.5E-05</td><td>9.4E-05</td><td>BT20/MB468</td></tr>
<tr><td>>hg38_21_+_8433172_8446622_RNA45SN1_45S_50ntFlanks@12072.12106.35|CCCGAGGGGCTCTCGCTTCTGGCGCCAAGCGCCCG</td><td>>hg38_21_+_8433172_8446622_RNA45SN1_45S_50ntFlanks</td><td>12072</td><td>12106</td><td>35</td><td>CCCGAGGGGCTCTCGCTTCTGGCGCCAAGCGCCCG</td><td>177.1</td><td>-3.3</td><td>0.8</td><td>-4.0</td><td>7.5E-05</td><td>9.5E-05</td><td>BT20/MB468</td></tr>
<tr><td>>hg38_21_+_8433172_8446622_RNA45SN1_45S_50ntFlanks@6655.6681.27|TCTTAGCGGTGGATCACTCGGCTCGTG</td><td>>hg38_21_+_8433172_8446622_RNA45SN1_45S_50ntFlanks</td><td>6655</td><td>6681</td><td>27</td><td>TCTTAGCGGTGGATCACTCGGCTCGTG</td><td>1611.3</td><td>-2.6</td><td>0.6</td><td>-4.0</td><td>7.5E-05</td><td>9.5E-05</td><td>BT20/MB468</td></tr>
<tr><td>>hg38_21_+_8433172_8446622_RNA45SN1_45S_50ntFlanks@3789.3810.22|ACTGCGAATGGCTCATTAAATC</td><td>>hg38_21_+_8433172_8446622_RNA45SN1_45S_50ntFlanks</td><td>3789</td><td>3810</td><td>22</td><td>ACTGCGAATGGCTCATTAAATC</td><td>256.8</td><td>-3.3</td><td>0.8</td><td>-4.0</td><td>7.6E-05</td><td>9.6E-05</td><td>BT20/MB468</td></tr>
<tr><td>>hg38_21_+_8433172_8446622_RNA45SN1_45S_50ntFlanks@11876.11909.34|AAGACCCTGTTGAGCTTGACTCTAGTCTGGCACG</td><td>>hg38_21_+_8433172_8446622_RNA45SN1_45S_50ntFlanks</td><td>11876</td><td>11909</td><td>34</td><td>AAGACCCTGTTGAGCTTGACTCTAGTCTGGCACG</td><td>111.7</td><td>-3.2</td><td>0.8</td><td>-4.0</td><td>7.6E-05</td><td>9.6E-05</td><td>BT20/MB468</td></tr>
<tr><td>>hg38_21_+_8433172_8446622_RNA45SN1_45S_50ntFlanks@12417.12438.22|TCGATGTCGGCTCTTCCTATCA</td><td>>hg38_21_+_8433172_8446622_RNA45SN1_45S_50ntFlanks</td><td>12417</td><td>12438</td><td>22</td><td>TCGATGTCGGCTCTTCCTATCA</td><td>615.0</td><td>2.6</td><td>0.7</td><td>4.0</td><td>7.7E-05</td><td>9.7E-05</td><td>BT20/MB468</td></tr>
<tr><td>>hg38_21_+_8433172_8446622_RNA45SN1_45S_50ntFlanks@12407.12433.27|TTTTGATCCTTCGATGTCGGCTCTTCC</td><td>>hg38_21_+_8433172_8446622_RNA45SN1_45S_50ntFlanks</td><td>12407</td><td>12433</td><td>27</td><td>TTTTGATCCTTCGATGTCGGCTCTTCC</td><td>1009.8</td><td>2.7</td><td>0.7</td><td>4.0</td><td>7.0E-05</td><td>9.7E-05</td><td>MB231/MB468</td></tr>
<tr><td>>hg38_21_+_8433172_8446622_RNA45SN1_45S_50ntFlanks@10677.10712.36|GCGGTTCCGGCGGCGTCCGGTGAGCTCTCGCTGGCC</td><td>>hg38_21_+_8433172_8446622_RNA45SN1_45S_50ntFlanks</td><td>10677</td><td>10712</td><td>36</td><td>GCGGTTCCGGCGGCGTCCGGTGAGCTCTCGCTGGCC</td><td>697.1</td><td>2.3</td><td>0.6</td><td>4.0</td><td>7.0E-05</td><td>9.7E-05</td><td>MB231/MB468</td></tr>
<tr><td>>MI0000272|hsa-mir-182&WithFlank&7|-|129770377|129770498@29.49.21(+1G)&[MIMAT0000259&hsa-miR-182-5p&offsets|0|-3(+1G);m-22&7|-|129770449|129770470&offsets|0|-1(+1G)]|TTTGGCAATGGTAGAACTCACG</td><td>>MI0000272|hsa-mir-182&WithFlank&7|-|129770377|129770498</td><td>29</td><td>49</td><td>21(+1G)</td><td>TTTGGCAATGGTAGAACTCACG</td><td>533.8</td><td>4.6</td><td>1.0</td><td>4.5</td><td>7.9E-06</td><td>9.8E-05</td><td>BT20/MB231</td></tr>
<tr><td>>hg38_21_+_8433172_8446622_RNA45SN1_45S_50ntFlanks@10853.10873.21|ATAAGGATTGGCTCTAAGGGC</td><td>>hg38_21_+_8433172_8446622_RNA45SN1_45S_50ntFlanks</td><td>10853</td><td>10873</td><td>21</td><td>ATAAGGATTGGCTCTAAGGGC</td><td>138.9</td><td>-2.2</td><td>0.6</td><td>-4.0</td><td>7.1E-05</td><td>9.9E-05</td><td>MB231/MB468</td></tr>
<tr><td>>MI0000273|hsa-mir-183&WithFlank&7|-|129774899|129775020@33.55.23&[MIMAT0000261&hsa-miR-183-5p&offsets|0|+1;m-41&7|-|129774966|129774987&offsets|-1|0]|TATGGCACTGGTAGAATTCACTG</td><td>>MI0000273|hsa-mir-183&WithFlank&7|-|129774899|129775020</td><td>33</td><td>55</td><td>23</td><td>TATGGCACTGGTAGAATTCACTG</td><td>4977.6</td><td>2.8</td><td>0.6</td><td>4.5</td><td>8.0E-06</td><td>9.9E-05</td><td>BT20/MB231</td></tr>
<tr><td>>hg38_21_+_8433172_8446622_RNA45SN1_45S_50ntFlanks@7975.8014.40|CGCGACCTCAGATCAGACGTGGCGACCCGCTGAATTTAAG</td><td>>hg38_21_+_8433172_8446622_RNA45SN1_45S_50ntFlanks</td><td>7975</td><td>8014</td><td>40</td><td>CGCGACCTCAGATCAGACGTGGCGACCCGCTGAATTTAAG</td><td>3249.7</td><td>3.5</td><td>0.9</td><td>3.9</td><td>7.8E-05</td><td>9.9E-05</td><td>BT20/MB468</td></tr>
<tr><td>>hg38_1_-_228634819_228635039_RNA5S12_5S_50ntFlanks@86.103.18|CGTCTGATCTCGGAAGCT</td><td>>hg38_1_-_228634819_228635039_RNA5S12_5S_50ntFlanks</td><td>86</td><td>103</td><td>18</td><td>CGTCTGATCTCGGAAGCT</td><td>196.5</td><td>-2.1</td><td>0.5</td><td>-3.9</td><td>8.0E-05</td><td>1.0E-04</td><td>BT20/MB468</td></tr>
<tr><td>>hg38_21_+_8433172_8446622_RNA45SN1_45S_50ntFlanks@6718.6744.27|GTGAATTGCAGGACACATTGATCATCG</td><td>>hg38_21_+_8433172_8446622_RNA45SN1_45S_50ntFlanks</td><td>6718</td><td>6744</td><td>27</td><td>GTGAATTGCAGGACACATTGATCATCG</td><td>115.0</td><td>-1.9</td><td>0.5</td><td>-3.9</td><td>8.0E-05</td><td>1.0E-04</td><td>BT20/MB468</td></tr>
<tr><td>>MI0000478|hsa-mir-149&WithFlank&2|+|240455995|240456095@21.42.22&[MIMAT0000450&hsa-miR-149-5p&offsets|0|-1;m-195&2|+|240456015|240456037&offsets|0|-1]|TCTGGCTCCGTGTCTTCACTCC</td><td>>MI0000478|hsa-mir-149&WithFlank&2|+|240455995|240456095</td><td>21</td><td>42</td><td>22</td><td>TCTGGCTCCGTGTCTTCACTCC</td><td>1199.6</td><td>3.3</td><td>0.7</td><td>4.5</td><td>8.2E-06</td><td>1.0E-04</td><td>BT20/MB231</td></tr>
<tr><td>>hg38_21_+_8433172_8446622_RNA45SN1_45S_50ntFlanks@8356.8377.22|GAAAAGAACTTTGAAGAGAGAG</td><td>>hg38_21_+_8433172_8446622_RNA45SN1_45S_50ntFlanks</td><td>8356</td><td>8377</td><td>22</td><td>GAAAAGAACTTTGAAGAGAGAG</td><td>964.4</td><td>-2.6</td><td>0.7</td><td>-4.0</td><td>7.3E-05</td><td>1.0E-04</td><td>MB231/MB468</td></tr>
<tr><td>>hg38_21_+_8433172_8446622_RNA45SN1_45S_50ntFlanks@7975.8007.33|CGCGACCTCAGATCAGACGTGGCGACCCGCTGA</td><td>>hg38_21_+_8433172_8446622_RNA45SN1_45S_50ntFlanks</td><td>7975</td><td>8007</td><td>33</td><td>CGCGACCTCAGATCAGACGTGGCGACCCGCTGA</td><td>7106.6</td><td>3.2</td><td>0.8</td><td>4.0</td><td>7.3E-05</td><td>1.0E-04</td><td>MB231/MB468</td></tr>
<tr><td>>hg38_21_+_8433172_8446622_RNA45SN1_45S_50ntFlanks@12433.12452.20|CTATCATTGTGAAGCAGAAT</td><td>>hg38_21_+_8433172_8446622_RNA45SN1_45S_50ntFlanks</td><td>12433</td><td>12452</td><td>20</td><td>CTATCATTGTGAAGCAGAAT</td><td>109.8</td><td>-2.8</td><td>0.7</td><td>-3.9</td><td>8.0E-05</td><td>1.0E-04</td><td>BT20/MB468</td></tr>
<tr><td>>hg38_21_+_8433172_8446622_RNA45SN1_45S_50ntFlanks@7992.8019.28|CGTGGCGACCCGCTGAATTTAAGCATAT</td><td>>hg38_21_+_8433172_8446622_RNA45SN1_45S_50ntFlanks</td><td>7992</td><td>8019</td><td>28</td><td>CGTGGCGACCCGCTGAATTTAAGCATAT</td><td>402.1</td><td>7.7</td><td>2.0</td><td>4.0</td><td>7.3E-05</td><td>1.0E-04</td><td>MB231/MB468</td></tr>
<tr><td>>hg38_21_+_8433172_8446622_RNA45SN1_45S_50ntFlanks@12631.12661.31|GGCGAAGCTACCATCTGTGGGATTATGACTG</td><td>>hg38_21_+_8433172_8446622_RNA45SN1_45S_50ntFlanks</td><td>12631</td><td>12661</td><td>31</td><td>GGCGAAGCTACCATCTGTGGGATTATGACTG</td><td>109.7</td><td>-2.5</td><td>0.6</td><td>-3.9</td><td>8.0E-05</td><td>1.0E-04</td><td>BT20/MB468</td></tr>
<tr><td>>MI0000088|hsa-mir-30a&WithFlank&6|-|71403545|71403627@12.35.24(+2U)&[MIMAT0000087&hsa-miR-30a-5p&offsets|0|+2(+2U);m-15&6|-|71403595|71403616&offsets|0|+2(+2U)]|TGTAAACATCCTCGACTGGAAGCTTT</td><td>>MI0000088|hsa-mir-30a&WithFlank&6|-|71403545|71403627</td><td>12</td><td>35</td><td>24(+2U)</td><td>TGTAAACATCCTCGACTGGAAGCTTT</td><td>3001.2</td><td>-3.2</td><td>0.7</td><td>-4.5</td><td>8.3E-06</td><td>1.0E-04</td><td>BT20/MB231</td></tr>
<tr><td>>am_trna134_GluTTC_1_-_16861774_16861845@1.28.28,trna5_GluTTC_1_+_17199078_17199149@1.28.28,trna84_GluTTC_1_-_161391883_161391954@1.28.28,trna94_GluTTC_1_-_149664355_149664427@1.28.28|TCCCTGGTGGTCTAGTGGCTAGGATTCG</td><td>>am_trna134_GluTTC_1_-_16861774_16861845</td><td>1</td><td>28</td><td>28</td><td>TCCCTGGTGGTCTAGTGGCTAGGATTCG</td><td>446.7</td><td>-2.4</td><td>0.6</td><td>-3.9</td><td>8.1E-05</td><td>1.0E-04</td><td>BT20/MB468</td></tr>
<tr><td>>hg38_21_+_8433172_8446622_RNA45SN1_45S_50ntFlanks@8171.8212.42|AAGTCCTTCTGATCGAGGCCCAGCCCGTGGACGGTGTGAGGC</td><td>>hg38_21_+_8433172_8446622_RNA45SN1_45S_50ntFlanks</td><td>8171</td><td>8212</td><td>42</td><td>AAGTCCTTCTGATCGAGGCCCAGCCCGTGGACGGTGTGAGGC</td><td>175.4</td><td>-2.6</td><td>0.7</td><td>-3.9</td><td>8.1E-05</td><td>1.0E-04</td><td>BT20/MB468</td></tr>
<tr><td>>hg38_21_+_8433172_8446622_RNA45SN1_45S_50ntFlanks@5482.5514.33|CGGAGCGCTGAGAAGACGGTCGAACTTGACTAT</td><td>>hg38_21_+_8433172_8446622_RNA45SN1_45S_50ntFlanks</td><td>5482</td><td>5514</td><td>33</td><td>CGGAGCGCTGAGAAGACGGTCGAACTTGACTAT</td><td>85.5</td><td>-2.3</td><td>0.6</td><td>-4.0</td><td>7.4E-05</td><td>1.0E-04</td><td>MB231/MB468</td></tr>
<tr><td>>hg38_21_+_8433172_8446622_RNA45SN1_45S_50ntFlanks@4367.4396.30|CAGTTAAAAAGCTCGTAGTTGGATCTTGGG</td><td>>hg38_21_+_8433172_8446622_RNA45SN1_45S_50ntFlanks</td><td>4367</td><td>4396</td><td>30</td><td>CAGTTAAAAAGCTCGTAGTTGGATCTTGGG</td><td>123.6</td><td>-4.4</td><td>1.1</td><td>-3.9</td><td>8.2E-05</td><td>1.0E-04</td><td>BT20/MB468</td></tr>
<tr><td>>hg38_21_+_8433172_8446622_RNA45SN1_45S_50ntFlanks@12992.13013.22|TCCCTCGCTGCGATCTATTGAA</td><td>>hg38_21_+_8433172_8446622_RNA45SN1_45S_50ntFlanks</td><td>12992</td><td>13013</td><td>22</td><td>TCCCTCGCTGCGATCTATTGAA</td><td>823.9</td><td>-2.2</td><td>0.6</td><td>-4.0</td><td>7.5E-05</td><td>1.0E-04</td><td>MB231/MB468</td></tr>
<tr><td>>hg38_21_+_8433172_8446622_RNA45SN1_45S_50ntFlanks@10349.10378.30|AACGAGAACTTTGAAGGCCGAAGTGGAGAA</td><td>>hg38_21_+_8433172_8446622_RNA45SN1_45S_50ntFlanks</td><td>10349</td><td>10378</td><td>30</td><td>AACGAGAACTTTGAAGGCCGAAGTGGAGAA</td><td>192.6</td><td>-1.4</td><td>0.3</td><td>-3.9</td><td>8.2E-05</td><td>1.0E-04</td><td>BT20/MB468</td></tr>
<tr><td>>hg38_21_+_8433172_8446622_RNA45SN1_45S_50ntFlanks@12416.12451.36|TTCGATGTCGGCTCTTCCTATCATTGTGAAGCAGAA</td><td>>hg38_21_+_8433172_8446622_RNA45SN1_45S_50ntFlanks</td><td>12416</td><td>12451</td><td>36</td><td>TTCGATGTCGGCTCTTCCTATCATTGTGAAGCAGAA</td><td>295.8</td><td>7.1</td><td>1.8</td><td>4.0</td><td>7.5E-05</td><td>1.0E-04</td><td>MB231/MB468</td></tr>
</table>

<table>
<tr><th></th><th></th><th></th><th></th><th></th><th></th><th></th><th></th><th></th><th></th><th></th><th></th><th></th></tr>
<tr><td>>hg38_21_+_8433172_8446622_RNA45SN1_45S_50ntFlanks@11208.11229.22|GCGGCGACTCTGGACGCGAGCC</td><td>>hg38_21_+_8433172_8446622_RNA45SN1_45S_50ntFlanks</td><td>11208</td><td>11229</td><td>22</td><td>GCGGCGACTCTGGACGCGAGCC</td><td>374.8</td><td>-2.2</td><td>0.5</td><td>-4.0</td><td>7.5E-05</td><td>1.0E-04</td><td>MB231/MB468</td></tr>
<tr><td>>trna5_AspGTC_12_+_98897281_98897352@35.75.41|TCACGCGGGAGACCGGGGTTCAATTCCCCGACGGGGAGCCA</td><td>>trna5_AspGTC_12_+_98897281_98897352</td><td>35</td><td>75</td><td>41</td><td>TCACGCGGGAGACCGGGGTTCAATTCCCCGACGGGGAGCCA</td><td>893.0</td><td>10.3</td><td>2.6</td><td>4.0</td><td>7.6E-05</td><td>1.0E-04</td><td>MB231/MB468</td></tr>
<tr><td>>hg38_21_+_8433172_8446622_RNA45SN1_45S_50ntFlanks@12259.12284.26|CTTGATCTTGATTTTCAGTACGAATA</td><td>>hg38_21_+_8433172_8446622_RNA45SN1_45S_50ntFlanks</td><td>12259</td><td>12284</td><td>26</td><td>CTTGATCTTGATTTTCAGTACGAATA</td><td>260.8</td><td>-2.3</td><td>0.6</td><td>-4.0</td><td>7.6E-05</td><td>1.0E-04</td><td>MB231/MB468</td></tr>
<tr><td>>hg38_21_+_8433172_8446622_RNA45SN1_45S_50ntFlanks@11874.11894.21|AGAAGACCCTGTTGAGCTTGA</td><td>>hg38_21_+_8433172_8446622_RNA45SN1_45S_50ntFlanks</td><td>11874</td><td>11894</td><td>21</td><td>AGAAGACCCTGTTGAGCTTGA</td><td>143.5</td><td>-3.6</td><td>0.9</td><td>-3.9</td><td>8.3E-05</td><td>1.0E-04</td><td>BT20/MB468</td></tr>
<tr><td>>hg38_21_+_8433172_8446622_RNA45SN1_45S_50ntFlanks@12992.13011.20|TCCCTCGCTGCGATCTATTG</td><td>>hg38_21_+_8433172_8446622_RNA45SN1_45S_50ntFlanks</td><td>12992</td><td>13011</td><td>20</td><td>TCCCTCGCTGCGATCTATTG</td><td>205.2</td><td>-2.1</td><td>0.5</td><td>-4.0</td><td>7.6E-05</td><td>1.1E-04</td><td>MB231/MB468</td></tr>
<tr><td>>hg38_21_+_8433172_8446622_RNA45SN1_45S_50ntFlanks@10357.10377.21|CTTTGAAGGCCGAAGTGGAGA</td><td>>hg38_21_+_8433172_8446622_RNA45SN1_45S_50ntFlanks</td><td>10357</td><td>10377</td><td>21</td><td>CTTTGAAGGCCGAAGTGGAGA</td><td>203.1</td><td>-2.1</td><td>0.5</td><td>-4.0</td><td>7.6E-05</td><td>1.1E-04</td><td>MB231/MB468</td></tr>
<tr><td>>hg38_21_+_8433172_8446622_RNA45SN1_45S_50ntFlanks@10845.10867.23|ACTTCGGGATAAGGATTGGCTCT</td><td>>hg38_21_+_8433172_8446622_RNA45SN1_45S_50ntFlanks</td><td>10845</td><td>10867</td><td>23</td><td>ACTTCGGGATAAGGATTGGCTCT</td><td>345.5</td><td>-2.1</td><td>0.5</td><td>-4.0</td><td>7.6E-05</td><td>1.1E-04</td><td>MB231/MB468</td></tr>
<tr><td>>hg38_21_+_8433172_8446622_RNA45SN1_45S_50ntFlanks@11208.11232.25|GCGGCGACTCTGGACGCGAGCCGGG</td><td>>hg38_21_+_8433172_8446622_RNA45SN1_45S_50ntFlanks</td><td>11208</td><td>11232</td><td>25</td><td>GCGGCGACTCTGGACGCGAGCCGGG</td><td>6457.1</td><td>-5.1</td><td>1.3</td><td>-3.9</td><td>8.4E-05</td><td>1.1E-04</td><td>BT20/MB468</td></tr>
<tr><td>>hg38_21_+_8433172_8446622_RNA45SN1_45S_50ntFlanks@10768.10793.26|CAGCAGGTCTCCAAGGTGAACAGCCT</td><td>>hg38_21_+_8433172_8446622_RNA45SN1_45S_50ntFlanks</td><td>10768</td><td>10793</td><td>26</td><td>CAGCAGGTCTCCAAGGTGAACAGCCT</td><td>106.0</td><td>-3.3</td><td>0.8</td><td>-4.0</td><td>7.7E-05</td><td>1.1E-04</td><td>MB231/MB468</td></tr>
<tr><td>>hg38_21_+_8433172_8446622_RNA45SN1_45S_50ntFlanks@11562.11588.27|CGCCTAGCAGCCGACTTAGAACTGGTG</td><td>>hg38_21_+_8433172_8446622_RNA45SN1_45S_50ntFlanks</td><td>11562</td><td>11588</td><td>27</td><td>CGCCTAGCAGCCGACTTAGAACTGGTG</td><td>422.8</td><td>-2.7</td><td>0.7</td><td>-3.9</td><td>8.5E-05</td><td>1.1E-04</td><td>BT20/MB468</td></tr>
<tr><td>>am_trnaMT_SerGCT_MT_+_12207_12265@16.38.23|AACTGCTAACTCATGCCCCCATG</td><td>>am_trnaMT_SerGCT_MT_+_12207_12265</td><td>16</td><td>38</td><td>23</td><td>AACTGCTAACTCATGCCCCCATG</td><td>182.1</td><td>-3.2</td><td>0.8</td><td>-3.9</td><td>8.6E-05</td><td>1.1E-04</td><td>BT20/MB468</td></tr>
<tr><td>>hg38_21_+_8433172_8446622_RNA45SN1_45S_50ntFlanks@5492.5510.19|AGAAGACGGTCGAACTTGA</td><td>>hg38_21_+_8433172_8446622_RNA45SN1_45S_50ntFlanks</td><td>5492</td><td>5510</td><td>19</td><td>AGAAGACGGTCGAACTTGA</td><td>270.7</td><td>-2.4</td><td>0.6</td><td>-3.9</td><td>8.6E-05</td><td>1.1E-04</td><td>BT20/MB468</td></tr>
<tr><td>>hg38_21_+_8433172_8446622_RNA45SN1_45S_50ntFlanks@6653.6688.36|ACTCTTAGCGGTGGATCACTCGGCTCGTGCGTCGAT</td><td>>hg38_21_+_8433172_8446622_RNA45SN1_45S_50ntFlanks</td><td>6653</td><td>6688</td><td>36</td><td>ACTCTTAGCGGTGGATCACTCGGCTCGTGCGTCGAT</td><td>812.6</td><td>3.3</td><td>0.8</td><td>3.9</td><td>8.6E-05</td><td>1.1E-04</td><td>BT20/MB468</td></tr>
<tr><td>>hg38_21_+_8433172_8446622_RNA45SN1_45S_50ntFlanks@11568.11588.21|GCAGCCGACTTAGAACTGGTG</td><td>>hg38_21_+_8433172_8446622_RNA45SN1_45S_50ntFlanks</td><td>11568</td><td>11588</td><td>21</td><td>GCAGCCGACTTAGAACTGGTG</td><td>678.5</td><td>-1.7</td><td>0.4</td><td>-3.9</td><td>8.7E-05</td><td>1.1E-04</td><td>BT20/MB468</td></tr>
<tr><td>>hg38_21_+_8433172_8446622_RNA45SN1_45S_50ntFlanks@8356.8373.18|GAAAAGAACTTTGAAGAG</td><td>>hg38_21_+_8433172_8446622_RNA45SN1_45S_50ntFlanks</td><td>8356</td><td>8373</td><td>18</td><td>GAAAAGAACTTTGAAGAG</td><td>218.1</td><td>-2.9</td><td>0.7</td><td>-3.9</td><td>8.8E-05</td><td>1.1E-04</td><td>BT20/MB468</td></tr>
<tr><td>>hg38_21_+_8433172_8446622_RNA45SN1_45S_50ntFlanks@10404.10429.26|CATGGGTCAGTCGGTCCTGAGAGATG</td><td>>hg38_21_+_8433172_8446622_RNA45SN1_45S_50ntFlanks</td><td>10404</td><td>10429</td><td>26</td><td>CATGGGTCAGTCGGTCCTGAGAGATG</td><td>948.5</td><td>-1.9</td><td>0.5</td><td>-3.9</td><td>8.2E-05</td><td>1.1E-04</td><td>MB231/MB468</td></tr>
<tr><td>>hg38_21_+_8433172_8446622_RNA45SN1_45S_50ntFlanks@6730.6771.42|ACACATTGATCATCGACACTTCGAACGCACTTGCGGCCCCGG</td><td>>hg38_21_+_8433172_8446622_RNA45SN1_45S_50ntFlanks</td><td>6730</td><td>6771</td><td>42</td><td>ACACATTGATCATCGACACTTCGAACGCACTTGCGGCCCCGG</td><td>132.4</td><td>-2.4</td><td>0.6</td><td>-3.9</td><td>8.2E-05</td><td>1.1E-04</td><td>MB231/MB468</td></tr>
<tr><td>>hg38_21_+_8433172_8446622_RNA45SN1_45S_50ntFlanks@12455.12475.21|ACCAAGCGTTGGATTGTTCAC</td><td>>hg38_21_+_8433172_8446622_RNA45SN1_45S_50ntFlanks</td><td>12455</td><td>12475</td><td>21</td><td>ACCAAGCGTTGGATTGTTCAC</td><td>166.2</td><td>-3.8</td><td>1.0</td><td>-3.9</td><td>9.1E-05</td><td>1.1E-04</td><td>BT20/MB468</td></tr>
<tr><td>>hg38_21_+_8433172_8446622_RNA45SN1_45S_50ntFlanks@10849.10867.19|CGGGATAAGGATTGGCTCT</td><td>>hg38_21_+_8433172_8446622_RNA45SN1_45S_50ntFlanks</td><td>10849</td><td>10867</td><td>19</td><td>CGGGATAAGGATTGGCTCT</td><td>243.1</td><td>-2.9</td><td>0.7</td><td>-3.9</td><td>9.1E-05</td><td>1.1E-04</td><td>BT20/MB468</td></tr>
<tr><td>>hg38_21_+_8433172_8446622_RNA45SN1_45S_50ntFlanks@11785.11802.18|TGAATGGATGAACGAGAT</td><td>>hg38_21_+_8433172_8446622_RNA45SN1_45S_50ntFlanks</td><td>11785</td><td>11802</td><td>18</td><td>TGAATGGATGAACGAGAT</td><td>144.5</td><td>-1.8</td><td>0.5</td><td>-3.9</td><td>9.1E-05</td><td>1.2E-04</td><td>BT20/MB468</td></tr>
<tr><td>>hg38_21_+_8433172_8446622_RNA45SN1_45S_50ntFlanks@5484.5509.26|GAGCGCTGAGAAGACGGTCGAACTTG</td><td>>hg38_21_+_8433172_8446622_RNA45SN1_45S_50ntFlanks</td><td>5484</td><td>5509</td><td>26</td><td>GAGCGCTGAGAAGACGGTCGAACTTG</td><td>102.9</td><td>-1.9</td><td>0.5</td><td>-3.9</td><td>9.2E-05</td><td>1.2E-04</td><td>BT20/MB468</td></tr>
<tr><td>>hg38_21_+_8433172_8446622_RNA45SN1_45S_50ntFlanks@11731.11751.21|ACTATGACTCTCTTAAGGTAG</td><td>>hg38_21_+_8433172_8446622_RNA45SN1_45S_50ntFlanks</td><td>11731</td><td>11751</td><td>21</td><td>ACTATGACTCTCTTAAGGTAG</td><td>670.3</td><td>-2.9</td><td>0.7</td><td>-3.9</td><td>8.4E-05</td><td>1.2E-04</td><td>MB231/MB468</td></tr>
<tr><td>>MI0000239|hsa-mir-197&WithFlank&1|+|109598887|109598973@54.76.23&[MIMAT0000227&hsa-miR-197-3p&offsets|0|+1;m-99&1|+|109598940|109598961&offsets|0|+1]|TTCACCACCTTCTCCACCCAGCA</td><td>>MI0000239|hsa-mir-197&WithFlank&1|+|109598887|109598973</td><td>54</td><td>76</td><td>23</td><td>TTCACCACCTTCTCCACCCAGCA</td><td>242.5</td><td>3.7</td><td>0.8</td><td>4.4</td><td>9.5E-06</td><td>1.2E-04</td><td>BT20/MB231</td></tr>
<tr><td>>hg38_21_+_8433172_8446622_RNA45SN1_45S_50ntFlanks@4726.4748.23|TAATCAAGAACGAAAGTCGGAGG</td><td>>hg38_21_+_8433172_8446622_RNA45SN1_45S_50ntFlanks</td><td>4726</td><td>4748</td><td>23</td><td>TAATCAAGAACGAAAGTCGGAGG</td><td>162.4</td><td>-2.1</td><td>0.5</td><td>-3.9</td><td>9.3E-05</td><td>1.2E-04</td><td>BT20/MB468</td></tr>
<tr><td>>hg38_21_+_8433172_8446622_RNA45SN1_45S_50ntFlanks@4738.4759.22|AAAGTCGGAGGTTCGAAGACGA</td><td>>hg38_21_+_8433172_8446622_RNA45SN1_45S_50ntFlanks</td><td>4738</td><td>4759</td><td>22</td><td>AAAGTCGGAGGTTCGAAGACGA</td><td>260.2</td><td>-2.4</td><td>0.6</td><td>-3.9</td><td>9.3E-05</td><td>1.2E-04</td><td>BT20/MB468</td></tr>
<tr><td>>hg38_21_+_8433172_8446622_RNA45SN1_45S_50ntFlanks@10393.10430.38|CAGCAGTTGAACATGGGTCAGTCGGTCCTGAGAGATGG</td><td>>hg38_21_+_8433172_8446622_RNA45SN1_45S_50ntFlanks</td><td>10393</td><td>10430</td><td>38</td><td>CAGCAGTTGAACATGGGTCAGTCGGTCCTGAGAGATGG</td><td>349.5</td><td>-2.6</td><td>0.7</td><td>-3.9</td><td>8.5E-05</td><td>1.2E-04</td><td>MB231/MB468</td></tr>
<tr><td>>MI0000079|hsa-mir-23a&WithFlank&19|-|13836581|13836665@54.72.19(+1G)&[MIMAT0000078&hsa-miR-23a-3p&offsets|+3|+1(+1G);m-31&19|-|13836595|13836615&offsets|+3|+1(+1G)]|ACATTGCCAGGGATTTCCAG</td><td>>MI0000079|hsa-mir-23a&WithFlank&19|-|13836581|13836665</td><td>54</td><td>72</td><td>19(+1G)</td><td>ACATTGCCAGGGATTTCCAG</td><td>655.3</td><td>14.3</td><td>3.6</td><td>3.9</td><td>8.6E-05</td><td>1.2E-04</td><td>MB231/MB468</td></tr>
<tr><td>>hg38_21_+_8433172_8446622_RNA45SN1_45S_50ntFlanks@6659.6681.23|AGCGGTGGATCACTCGGCTCGTG</td><td>>hg38_21_+_8433172_8446622_RNA45SN1_45S_50ntFlanks</td><td>6659</td><td>6681</td><td>23</td><td>AGCGGTGGATCACTCGGCTCGTG</td><td>663.5</td><td>-2.7</td><td>0.7</td><td>-3.9</td><td>9.5E-05</td><td>1.2E-04</td><td>BT20/MB468</td></tr>
<tr><td>>hg38_21_+_8433172_8446622_RNA45SN1_45S_50ntFlanks@5073.5098.26|AATTCCGATAACGAACGAGACTCTGG</td><td>>hg38_21_+_8433172_8446622_RNA45SN1_45S_50ntFlanks</td><td>5073</td><td>5098</td><td>26</td><td>AATTCCGATAACGAACGAGACTCTGG</td><td>298.6</td><td>-2.3</td><td>0.6</td><td>-3.9</td><td>8.6E-05</td><td>1.2E-04</td><td>MB231/MB468</td></tr>
<tr><td>>hg38_21_+_8433172_8446622_RNA45SN1_45S_50ntFlanks@4322.4346.25|CAGCCGCGGTAATTCCAGCTCCAAT</td><td>>hg38_21_+_8433172_8446622_RNA45SN1_45S_50ntFlanks</td><td>4322</td><td>4346</td><td>25</td><td>CAGCCGCGGTAATTCCAGCTCCAAT</td><td>202.8</td><td>-4.0</td><td>1.0</td><td>-3.9</td><td>9.5E-05</td><td>1.2E-04</td><td>BT20/MB468</td></tr>
<tr><td>>hg38_21_+_8433172_8446622_RNA45SN1_45S_50ntFlanks@4729.4759.31|TCAAGAACGAAAGTCGGAGGTTCGAAGACGA</td><td>>hg38_21_+_8433172_8446622_RNA45SN1_45S_50ntFlanks</td><td>4729</td><td>4759</td><td>31</td><td>TCAAGAACGAAAGTCGGAGGTTCGAAGACGA</td><td>257.6</td><td>-2.3</td><td>0.6</td><td>-3.9</td><td>9.6E-05</td><td>1.2E-04</td><td>BT20/MB468</td></tr>
<tr><td>>hg38_21_+_8433172_8446622_RNA45SN1_45S_50ntFlanks@12072.12096.25|CCCGAGGGGCTCTCGCTTCTGGCGC</td><td>>hg38_21_+_8433172_8446622_RNA45SN1_45S_50ntFlanks</td><td>12072</td><td>12096</td><td>25</td><td>CCCGAGGGGCTCTCGCTTCTGGCGC</td><td>259.3</td><td>-2.6</td><td>0.7</td><td>-3.9</td><td>8.7E-05</td><td>1.2E-04</td><td>MB231/MB468</td></tr>
<tr><td>>hg38_21_+_8433172_8446622_RNA45SN1_45S_50ntFlanks@12717.12742.26|CCTCGGTTGGCCTCGGATAGCCGGTC</td><td>>hg38_21_+_8433172_8446622_RNA45SN1_45S_50ntFlanks</td><td>12717</td><td>12742</td><td>26</td><td>CCTCGGTTGGCCTCGGATAGCCGGTC</td><td>67.5</td><td>-3.6</td><td>0.9</td><td>-3.9</td><td>8.7E-05</td><td>1.2E-04</td><td>MB231/MB468</td></tr>
<tr><td>>hg38_21_+_8433172_8446622_RNA45SN1_45S_50ntFlanks@6731.6758.28|CACATTGATCATCGACACTTCGAACGCA</td><td>>hg38_21_+_8433172_8446622_RNA45SN1_45S_50ntFlanks</td><td>6731</td><td>6758</td><td>28</td><td>CACATTGATCATCGACACTTCGAACGCA</td><td>194.7</td><td>-1.8</td><td>0.5</td><td>-3.9</td><td>8.7E-05</td><td>1.2E-04</td><td>MB231/MB468</td></tr>
<tr><td>>hg38_21_+_8433172_8446622_RNA45SN1_45S_50ntFlanks@6646.6675.30|TCGTACGACTCTTAGCGGTGGATCACTCGG</td><td>>hg38_21_+_8433172_8446622_RNA45SN1_45S_50ntFlanks</td><td>6646</td><td>6675</td><td>30</td><td>TCGTACGACTCTTAGCGGTGGATCACTCGG</td><td>1722.6</td><td>2.3</td><td>0.6</td><td>3.9</td><td>9.7E-05</td><td>1.2E-04</td><td>BT20/MB468</td></tr>
<tr><td>>hg38_21_+_8433172_8446622_RNA45SN1_45S_50ntFlanks@10402.10426.25|AACATGGGTCAGTCGGTCCTGAGAG</td><td>>hg38_21_+_8433172_8446622_RNA45SN1_45S_50ntFlanks</td><td>10402</td><td>10426</td><td>25</td><td>AACATGGGTCAGTCGGTCCTGAGAG</td><td>404.8</td><td>-2.5</td><td>0.6</td><td>-3.9</td><td>8.8E-05</td><td>1.2E-04</td><td>MB231/MB468</td></tr>
<tr><td>>MI0001733|hsa-mir-452&WithFlank&X|-|151959622|151959718@23.43.21&[MIMAT0001635&hsa-miR-452-5p&offsets|+3|+2;m-188&X|-|151959677|151959699&offsets|+3|+1]|TGTTTGCAGAGGAAACTGAGA</td><td>>MI0001733|hsa-mir-452&WithFlank&X|-|151959622|151959718</td><td>23</td><td>43</td><td>21</td><td>TGTTTGCAGAGGAAACTGAGA</td><td>993.7</td><td>3.1</td><td>0.7</td><td>4.4</td><td>1.0E-05</td><td>1.2E-04</td><td>BT20/MB231</td></tr>
<tr><td>>MI0003513|hsa-mir-455&WithFlank&9|+|114209428|114209535@22.44.23&[MIMAT0003150&hsa-miR-455-5p&offsets|0|+1;m-232&9|+|114209449|114209470&offsets|0|+1]|TATGTGCCTTTGGACTACATCGT</td><td>>MI0003513|hsa-mir-455&WithFlank&9|+|114209428|114209535</td><td>22</td><td>44</td><td>23</td><td>TATGTGCCTTTGGACTACATCGT</td><td>1421.4</td><td>-3.1</td><td>0.7</td><td>-4.4</td><td>1.0E-05</td><td>1.2E-04</td><td>BT20/MB231</td></tr>
<tr><td>>hg38_1_-_228634819_228635039_RNA5S12_5S_50ntFlanks@148.169.22|GGAATACCGGGTGCTGTAGGCT</td><td>>hg38_1_-_228634819_228635039_RNA5S12_5S_50ntFlanks</td><td>148</td><td>169</td><td>22</td><td>GGAATACCGGGTGCTGTAGGCT</td><td>699.1</td><td>2.9</td><td>0.7</td><td>3.9</td><td>9.0E-05</td><td>1.2E-04</td><td>MB231/MB468</td></tr>
<tr><td>>hg38_21_+_8433172_8446622_RNA45SN1_45S_50ntFlanks@11206.11232.27|CGGCGGCGACTCTGGACGCGAGCCGGG</td><td>>hg38_21_+_8433172_8446622_RNA45SN1_45S_50ntFlanks</td><td>11206</td><td>11232</td><td>27</td><td>CGGCGGCGACTCTGGACGCGAGCCGGG</td><td>295.6</td><td>-2.8</td><td>0.7</td><td>-3.9</td><td>9.0E-05</td><td>1.2E-04</td><td>MB231/MB468</td></tr>
<tr><td>>hg38_21_+_8433172_8446622_RNA45SN1_45S_50ntFlanks@12071.12096.26|CCCCGAGGGGCTCTCGCTTCTGGCGC</td><td>>hg38_21_+_8433172_8446622_RNA45SN1_45S_50ntFlanks</td><td>12071</td><td>12096</td><td>26</td><td>CCCCGAGGGGCTCTCGCTTCTGGCGC</td><td>388.4</td><td>-2.3</td><td>0.6</td><td>-3.9</td><td>9.1E-05</td><td>1.3E-04</td><td>MB231/MB468</td></tr>
<tr><td>>hg38_21_+_8433172_8446622_RNA45SN1_45S_50ntFlanks@9574.9595.22|AATCGGTCGTCCGACCTGGGTA</td><td>>hg38_21_+_8433172_8446622_RNA45SN1_45S_50ntFlanks</td><td>9574</td><td>9595</td><td>22</td><td>AATCGGTCGTCCGACCTGGGTA</td><td>2024.3</td><td>-2.5</td><td>0.6</td><td>-3.9</td><td>9.2E-05</td><td>1.3E-04</td><td>MB231/MB468</td></tr>
<tr><td>>hg38_21_+_8433172_8446622_RNA45SN1_45S_50ntFlanks@3855.3872.18|CTGTGGTAATTCTAGAGC</td><td>>hg38_21_+_8433172_8446622_RNA45SN1_45S_50ntFlanks</td><td>3855</td><td>3872</td><td>18</td><td>CTGTGGTAATTCTAGAGC</td><td>138.1</td><td>-1.8</td><td>0.5</td><td>-3.9</td><td>1.0E-04</td><td>1.3E-04</td><td>BT20/MB468</td></tr>
<tr><td>>hg38_21_+_8433172_8446622_RNA45SN1_45S_50ntFlanks@10760.10785.26|CATATCCGCAGCAGGTCTCCAAGGTG</td><td>>hg38_21_+_8433172_8446622_RNA45SN1_45S_50ntFlanks</td><td>10760</td><td>10785</td><td>26</td><td>CATATCCGCAGCAGGTCTCCAAGGTG</td><td>83.1</td><td>-2.3</td><td>0.6</td><td>-3.9</td><td>9.2E-05</td><td>1.3E-04</td><td>MB231/MB468</td></tr>
<tr><td>>hg38_21_+_8433172_8446622_RNA45SN1_45S_50ntFlanks@12973.12994.22|TCGTACGTAGCAGAGCAGCTCC</td><td>>hg38_21_+_8433172_8446622_RNA45SN1_45S_50ntFlanks</td><td>12973</td><td>12994</td><td>22</td><td>TCGTACGTAGCAGAGCAGCTCC</td><td>100.1</td><td>-2.3</td><td>0.6</td><td>-3.9</td><td>1.0E-04</td><td>1.3E-04</td><td>BT20/MB468</td></tr>
<tr><td>>hg38_21_+_8433172_8446622_RNA45SN1_45S_50ntFlanks@12072.12094.23|CCCGAGGGGCTCTCGCTTCTGGC</td><td>>hg38_21_+_8433172_8446622_RNA45SN1_45S_50ntFlanks</td><td>12072</td><td>12094</td><td>23</td><td>CCCGAGGGGCTCTCGCTTCTGGC</td><td>148.4</td><td>-2.0</td><td>0.5</td><td>-3.9</td><td>1.0E-04</td><td>1.3E-04</td><td>BT20/MB468</td></tr>
<tr><td>>MI0000470|hsa-mir-125b-2&WithFlank&21|+|16590231|16590331@23.45.23&[MIMAT0000423_1&hsa-miR-125b-5p&offsets|0|+1;m-68&21|+|16590253|16590274&offsets|0|+1]|TCCCTGAGACCCTAACTTGTGAG</td><td>>MI0000470|hsa-mir-125b-2&WithFlank&21|+|16590231|16590331</td><td>23</td><td>45</td><td>23</td><td>TCCCTGAGACCCTAACTTGTGAG</td><td>303.2</td><td>-4.9</td><td>1.1</td><td>-4.4</td><td>1.1E-05</td><td>1.3E-04</td><td>BT20/MB231</td></tr>
</table>

<table>
<tr><td>>hg38_21_+_8433172_8446622_RNA45SN1_45S_50ntFlanks@7975.8002.28|CGCGACCTCAGATCAGACGTGGCGACCC</td><td>>hg38_21_+_8433172_8446622_RNA45SN1_45S_50ntFlanks</td><td>7975</td><td>8002</td><td>28</td><td>CGCGACCTCAGATCAGACGTGGCGACCC</td><td>1684.2</td><td>3.2</td><td>0.7</td><td>4.4</td><td>1.1E-05</td><td>1.3E-04</td><td>BT20/MB231</td></tr>
<tr><td>>hg38_21_+_8433172_8446622_RNA45SN1_45S_50ntFlanks@4722.4748.27|TCATTAATCAAGAACGAAAGTCGGAGG</td><td>>hg38_21_+_8433172_8446622_RNA45SN1_45S_50ntFlanks</td><td>4722</td><td>4748</td><td>27</td><td>TCATTAATCAAGAACGAAAGTCGGAGG</td><td>343.7</td><td>-2.9</td><td>0.7</td><td>-3.9</td><td>9.5E-05</td><td>1.3E-04</td><td>MB231/MB468</td></tr>
<tr><td>>MI0000301|hsa-mir-224&WithFlank&X|-|151958572|151958664@14.34.21(+1C)&[MIMAT0000281&hsa-miR-224-5p&offsets|+1|-3(+1C);m-238&X|-|151958630|151958651&offsets|0|-1(+1C)]|CAAGTCACTAGTGGTTCCGTTC</td><td>>MI0000301|hsa-mir-224&WithFlank&X|-|151958572|151958664</td><td>14</td><td>34</td><td>21(+1C)</td><td>CAAGTCACTAGTGGTTCCGTTC</td><td>1450.2</td><td>3.1</td><td>0.7</td><td>4.4</td><td>1.1E-05</td><td>1.3E-04</td><td>BT20/MB231</td></tr>
<tr><td>>trna116_GluCTC_1_-_145399233_145399304@37.74.38,trna134_GluTTC_1_-_16861774_16861845@37.74.38,trna71_GluCTC_1_-_161439189_161439260@37.74.38,trna74_GluCTC_1_-_161431809_161431880@37.74.38,trna77_GluCTC_1_-_161424398_161424469@37.74.38,trna77_GluCTC_6_+_28949976_28950047@37.74.38,trna80_GluCTC_1_-_161417018_161417089@37.74.38,trna84_GluTTC_1_-_161391883_161391954@37.74.38,trna87_GluCTC_6_-_126101393_126101464@37.74.38|ACCGCCGCGGCCCGGGTTCGATTCCCGGTCAGGGAACC</td><td>>trna116_GluCTC_1_-_145399233_145399304</td><td>37</td><td>74</td><td>38</td><td>ACCGCCGCGGCCCGGGTTCGATTCCCGGTCAGGGAACC</td><td>2263.1</td><td>-4.6</td><td>1.0</td><td>-4.4</td><td>1.1E-05</td><td>1.3E-04</td><td>BT20/MB231</td></tr>
<tr><td>>hg38_21_+_8433172_8446622_RNA45SN1_45S_50ntFlanks@12258.12284.27|GCTTGATCTTGATTTTCAGTACGAATA</td><td>>hg38_21_+_8433172_8446622_RNA45SN1_45S_50ntFlanks</td><td>12258</td><td>12284</td><td>27</td><td>GCTTGATCTTGATTTTCAGTACGAATA</td><td>153.2</td><td>-3.1</td><td>0.8</td><td>-3.9</td><td>1.1E-04</td><td>1.3E-04</td><td>BT20/MB468</td></tr>
<tr><td>>hg38_21_+_8433172_8446622_RNA45SN1_45S_50ntFlanks@7975.8001.27|CGCGACCTCAGATCAGACGTGGCGACC</td><td>>hg38_21_+_8433172_8446622_RNA45SN1_45S_50ntFlanks</td><td>7975</td><td>8001</td><td>27</td><td>CGCGACCTCAGATCAGACGTGGCGACC</td><td>792.5</td><td>3.2</td><td>0.7</td><td>4.4</td><td>1.1E-05</td><td>1.3E-04</td><td>BT20/MB231</td></tr>
<tr><td>>hg38_21_+_8433172_8446622_RNA45SN1_45S_50ntFlanks@12420.12442.23|ATGTCGGCTCTTCCTATCATTGT</td><td>>hg38_21_+_8433172_8446622_RNA45SN1_45S_50ntFlanks</td><td>12420</td><td>12442</td><td>23</td><td>ATGTCGGCTCTTCCTATCATTGT</td><td>559.1</td><td>2.5</td><td>0.6</td><td>3.9</td><td>9.7E-05</td><td>1.3E-04</td><td>MB231/MB468</td></tr>
<tr><td>>hg38_21_+_8433172_8446622_RNA45SN1_45S_50ntFlanks@11881.11900.20|CCTGTTGAGCTTGACTCTAG</td><td>>hg38_21_+_8433172_8446622_RNA45SN1_45S_50ntFlanks</td><td>11881</td><td>11900</td><td>20</td><td>CCTGTTGAGCTTGACTCTAG</td><td>166.8</td><td>-3.8</td><td>1.0</td><td>-3.9</td><td>1.1E-04</td><td>1.3E-04</td><td>BT20/MB468</td></tr>
<tr><td>>trna152_ValCAC_6_-_27248049_27248121@1.31.31|GCTTCTGTAGTGTAGTGGTTATCACGTTCGC</td><td>>trna152_ValCAC_6_-_27248049_27248121</td><td>1</td><td>31</td><td>31</td><td>GCTTCTGTAGTGTAGTGGTTATCACGTTCGC</td><td>92.2</td><td>-2.7</td><td>0.7</td><td>-3.9</td><td>1.1E-04</td><td>1.3E-04</td><td>BT20/MB468</td></tr>
<tr><td>>hg38_21_+_8433172_8446622_RNA45SN1_45S_50ntFlanks@5492.5514.23|AGAAGACGGTCGAACTTGACTAT</td><td>>hg38_21_+_8433172_8446622_RNA45SN1_45S_50ntFlanks</td><td>5492</td><td>5514</td><td>23</td><td>AGAAGACGGTCGAACTTGACTAT</td><td>1228.1</td><td>-3.2</td><td>0.8</td><td>-3.9</td><td>1.1E-04</td><td>1.4E-04</td><td>BT20/MB468</td></tr>
<tr><td>>hg38_21_+_8433172_8446622_RNA45SN1_45S_50ntFlanks@8191.8215.25|CAGCCCGTGGACGGTGTGAGGCCGG</td><td>>hg38_21_+_8433172_8446622_RNA45SN1_45S_50ntFlanks</td><td>8191</td><td>8215</td><td>25</td><td>CAGCCCGTGGACGGTGTGAGGCCGG</td><td>917.3</td><td>-4.5</td><td>1.2</td><td>-3.9</td><td>1.1E-04</td><td>1.4E-04</td><td>BT20/MB468</td></tr>
<tr><td>>MI0000272|hsa-mir-182&WithFlank&7|-|129770377|129770498@29.51.23(+1G)&[MIMAT0000259&hsa-miR-182-5p&offsets|0|-1(+1G);m-22&7|-|129770449|129770470&offsets|0|+1(+1G)]|TTTGGCAATGGTAGAACTCACACG</td><td>>MI0000272|hsa-mir-182&WithFlank&7|-|129770377|129770498</td><td>29</td><td>51</td><td>23(+1G)</td><td>TTTGGCAATGGTAGAACTCACACG</td><td>7028.1</td><td>4.2</td><td>1.0</td><td>4.4</td><td>1.2E-05</td><td>1.4E-04</td><td>BT20/MB231</td></tr>
<tr><td>>hg38_21_+_8433172_8446622_RNA45SN1_45S_50ntFlanks@4726.4751.26|TAATCAAGAACGAAAGTCGGAGGTTC</td><td>>hg38_21_+_8433172_8446622_RNA45SN1_45S_50ntFlanks</td><td>4726</td><td>4751</td><td>26</td><td>TAATCAAGAACGAAAGTCGGAGGTTC</td><td>161.6</td><td>-4.5</td><td>1.2</td><td>-3.9</td><td>1.1E-04</td><td>1.4E-04</td><td>BT20/MB468</td></tr>
<tr><td>>hg38_21_+_8433172_8446622_RNA45SN1_45S_50ntFlanks@3985.4009.25|CGGCTTTGGTGACTCTAGATAACCT</td><td>>hg38_21_+_8433172_8446622_RNA45SN1_45S_50ntFlanks</td><td>3985</td><td>4009</td><td>25</td><td>CGGCTTTGGTGACTCTAGATAACCT</td><td>69.4</td><td>-3.2</td><td>0.8</td><td>-3.9</td><td>1.0E-04</td><td>1.4E-04</td><td>MB231/MB468</td></tr>
<tr><td>>hg38_21_+_8433172_8446622_RNA45SN1_45S_50ntFlanks@6722.6747.26|ATTGCAGGACACATTGATCATCGACA</td><td>>hg38_21_+_8433172_8446622_RNA45SN1_45S_50ntFlanks</td><td>6722</td><td>6747</td><td>26</td><td>ATTGCAGGACACATTGATCATCGACA</td><td>271.6</td><td>-1.5</td><td>0.4</td><td>-3.9</td><td>1.1E-04</td><td>1.4E-04</td><td>BT20/MB468</td></tr>
<tr><td>>hg38_21_+_8433172_8446622_RNA45SN1_45S_50ntFlanks@10478.10512.35|CCGATCGAAAGGGAGTCGGGTTCAGATCCCCGAAT</td><td>>hg38_21_+_8433172_8446622_RNA45SN1_45S_50ntFlanks</td><td>10478</td><td>10512</td><td>35</td><td>CCGATCGAAAGGGAGTCGGGTTCAGATCCCCGAAT</td><td>85.8</td><td>-3.4</td><td>0.9</td><td>-3.9</td><td>1.0E-04</td><td>1.4E-04</td><td>MB231/MB468</td></tr>
<tr><td>>hg38_21_+_8433172_8446622_RNA45SN1_45S_50ntFlanks@3775.3794.20|GCCGGTACAGTGAAACTGCG</td><td>>hg38_21_+_8433172_8446622_RNA45SN1_45S_50ntFlanks</td><td>3775</td><td>3794</td><td>20</td><td>GCCGGTACAGTGAAACTGCG</td><td>178.4</td><td>-1.8</td><td>0.5</td><td>-3.9</td><td>1.0E-04</td><td>1.4E-04</td><td>MB231/MB468</td></tr>
<tr><td>>hg38_21_+_8433172_8446622_RNA45SN1_45S_50ntFlanks@4566.4587.22|ATAATGGAATAGGACCGCGGTT</td><td>>hg38_21_+_8433172_8446622_RNA45SN1_45S_50ntFlanks</td><td>4566</td><td>4587</td><td>22</td><td>ATAATGGAATAGGACCGCGGTT</td><td>235.4</td><td>-2.4</td><td>0.6</td><td>-3.9</td><td>1.0E-04</td><td>1.4E-04</td><td>MB231/MB468</td></tr>
<tr><td>>hg38_21_+_8433172_8446622_RNA45SN1_45S_50ntFlanks@6734.6756.23|ATTGATCATCGACACTTCGAACG</td><td>>hg38_21_+_8433172_8446622_RNA45SN1_45S_50ntFlanks</td><td>6734</td><td>6756</td><td>23</td><td>ATTGATCATCGACACTTCGAACG</td><td>435.6</td><td>1.5</td><td>0.4</td><td>3.9</td><td>1.0E-04</td><td>1.4E-04</td><td>MB231/MB468</td></tr>
<tr><td>>hg38_21_+_8433172_8446622_RNA45SN1_45S_50ntFlanks@10922.10939.18|CTGGACGAGGCGCCGCCG</td><td>>hg38_21_+_8433172_8446622_RNA45SN1_45S_50ntFlanks</td><td>10922</td><td>10939</td><td>18</td><td>CTGGACGAGGCGCCGCCG</td><td>955.7</td><td>-4.6</td><td>1.2</td><td>-3.9</td><td>1.1E-04</td><td>1.4E-04</td><td>BT20/MB468</td></tr>
<tr><td>>hg38_21_+_8433172_8446622_RNA45SN1_45S_50ntFlanks@11027.11061.35|CTCTCTCTCTCTCTCCCCCGCTCCCCGTCCTCCCC</td><td>>hg38_21_+_8433172_8446622_RNA45SN1_45S_50ntFlanks</td><td>11027</td><td>11061</td><td>35</td><td>CTCTCTCTCTCTCTCCCCCGCTCCCCGTCCTCCCC</td><td>94.1</td><td>-3.8</td><td>1.0</td><td>-3.9</td><td>1.1E-04</td><td>1.4E-04</td><td>BT20/MB468</td></tr>
<tr><td>>MI0000098|hsa-mir-96&WithFlank&7|-|129774686|129774775@15.37.23&[MIMAT0000095&hsa-miR-96-5p&offsets|0|0;m-248&7|-|129774739|129774761&offsets|0|0]|TTTGGCACTAGCACATTTTTGCT</td><td>>MI0000098|hsa-mir-96&WithFlank&7|-|129774686|129774775</td><td>15</td><td>37</td><td>23</td><td>TTTGGCACTAGCACATTTTTGCT</td><td>13216.5</td><td>3.2</td><td>0.7</td><td>4.4</td><td>1.2E-05</td><td>1.4E-04</td><td>BT20/MB231</td></tr>
<tr><td>>trna119_LysCTT_1_-_145395522_145395594@3.34.32,trna11_LysCTT_5_-_180648979_180649051@3.34.32,trna13_LysCTT_6_+_26556774_26556846@3.34.32,trna7_LysCTT_16_+_3225692_3225764@3.34.32,trna9_LysCTT_5_+_180634755_180634827@3.34.32|CCGGCTAGCTCAGTCGGTAGAGCATGAGACTC</td><td>>trna119_LysCTT_1_-_145395522_145395594</td><td>3</td><td>34</td><td>32</td><td>CCGGCTAGCTCAGTCGGTAGAGCATGAGACTC</td><td>203.9</td><td>-3.5</td><td>0.8</td><td>-4.4</td><td>1.2E-05</td><td>1.4E-04</td><td>BT20/MB231</td></tr>
<tr><td>>hg38_21_+_8433172_8446622_RNA45SN1_45S_50ntFlanks@4328.4346.19|CGGTAATTCCAGCTCCAAT</td><td>>hg38_21_+_8433172_8446622_RNA45SN1_45S_50ntFlanks</td><td>4328</td><td>4346</td><td>19</td><td>CGGTAATTCCAGCTCCAAT</td><td>149.5</td><td>-2.7</td><td>0.7</td><td>-3.9</td><td>1.1E-04</td><td>1.4E-04</td><td>BT20/MB468</td></tr>
<tr><td>>hg38_21_+_8433172_8446622_RNA45SN1_45S_50ntFlanks@11683.11701.19|ATGTCAAAGTGAAGAAATT</td><td>>hg38_21_+_8433172_8446622_RNA45SN1_45S_50ntFlanks</td><td>11683</td><td>11701</td><td>19</td><td>ATGTCAAAGTGAAGAAATT</td><td>103.0</td><td>-2.8</td><td>0.7</td><td>-3.9</td><td>1.1E-04</td><td>1.5E-04</td><td>MB231/MB468</td></tr>
<tr><td>>hg38_21_+_8433172_8446622_RNA45SN1_45S_50ntFlanks@12202.12221.20|CTAAGGCGAGCTCAGGGAGG</td><td>>hg38_21_+_8433172_8446622_RNA45SN1_45S_50ntFlanks</td><td>12202</td><td>12221</td><td>20</td><td>CTAAGGCGAGCTCAGGGAGG</td><td>90.7</td><td>-2.7</td><td>0.7</td><td>-3.9</td><td>1.2E-04</td><td>1.5E-04</td><td>BT20/MB468</td></tr>
<tr><td>>hg38_21_+_8433172_8446622_RNA45SN1_45S_50ntFlanks@8104.8142.39|CCGCGGCGGGGCGCGGGACATGTGGCGTACGGAAGACCC</td><td>>hg38_21_+_8433172_8446622_RNA45SN1_45S_50ntFlanks</td><td>8104</td><td>8142</td><td>39</td><td>CCGCGGCGGGGCGCGGGACATGTGGCGTACGGAAGACCC</td><td>89.2</td><td>-2.9</td><td>0.8</td><td>-3.9</td><td>1.2E-04</td><td>1.5E-04</td><td>BT20/MB468</td></tr>
<tr><td>>hg38_21_+_8433172_8446622_RNA45SN1_45S_50ntFlanks@10914.10931.18|CGCCGCGGCTGGACGAGG</td><td>>hg38_21_+_8433172_8446622_RNA45SN1_45S_50ntFlanks</td><td>10914</td><td>10931</td><td>18</td><td>CGCCGCGGCTGGACGAGG</td><td>906.4</td><td>-5.5</td><td>1.4</td><td>-3.9</td><td>1.2E-04</td><td>1.5E-04</td><td>BT20/MB468</td></tr>
<tr><td>>hg38_21_+_8433172_8446622_RNA45SN1_45S_50ntFlanks@11322.11342.21|ACGTCTCGTCGCGCGCGCGTC</td><td>>hg38_21_+_8433172_8446622_RNA45SN1_45S_50ntFlanks</td><td>11322</td><td>11342</td><td>21</td><td>ACGTCTCGTCGCGCGCGCGTC</td><td>814.1</td><td>-3.5</td><td>0.9</td><td>-3.9</td><td>1.1E-04</td><td>1.5E-04</td><td>MB231/MB468</td></tr>
<tr><td>>hg38_21_+_8433172_8446622_RNA45SN1_45S_50ntFlanks@5527.5549.23|AAAGTCGTAACAAGGTTTCCGTA</td><td>>hg38_21_+_8433172_8446622_RNA45SN1_45S_50ntFlanks</td><td>5527</td><td>5549</td><td>23</td><td>AAAGTCGTAACAAGGTTTCCGTA</td><td>287.0</td><td>-4.8</td><td>1.2</td><td>-3.9</td><td>1.2E-04</td><td>1.5E-04</td><td>BT20/MB468</td></tr>
<tr><td>>hg38_21_+_8433172_8446622_RNA45SN1_45S_50ntFlanks@4047.4065.19|ATTCGAACGTCTGCCCTAT</td><td>>hg38_21_+_8433172_8446622_RNA45SN1_45S_50ntFlanks</td><td>4047</td><td>4065</td><td>19</td><td>ATTCGAACGTCTGCCCTAT</td><td>253.5</td><td>-1.8</td><td>0.5</td><td>-3.9</td><td>1.1E-04</td><td>1.5E-04</td><td>MB231/MB468</td></tr>
<tr><td>>hg38_21_+_8433172_8446622_RNA45SN1_45S_50ntFlanks@4760.4779.20|TCAGATACCGTCGTAGTTCC</td><td>>hg38_21_+_8433172_8446622_RNA45SN1_45S_50ntFlanks</td><td>4760</td><td>4779</td><td>20</td><td>TCAGATACCGTCGTAGTTCC</td><td>287.6</td><td>-2.3</td><td>0.6</td><td>-3.9</td><td>1.1E-04</td><td>1.5E-04</td><td>MB231/MB468</td></tr>
<tr><td>>MI0000098|hsa-mir-96&WithFlank&7|-|129774686|129774775@15.36.22(+1C)&[MIMAT0000095&hsa-miR-96-5p&offsets|0|-1(+1C);m-248&7|-|129774739|129774761&offsets|0|-1(+1C)]|TTTGGCACTAGCACATTTTTGCC</td><td>>MI0000098|hsa-mir-96&WithFlank&7|-|129774686|129774775</td><td>15</td><td>36</td><td>22(+1C)</td><td>TTTGGCACTAGCACATTTTTGCC</td><td>2053.8</td><td>3.4</td><td>0.8</td><td>4.4</td><td>1.3E-05</td><td>1.5E-04</td><td>BT20/MB231</td></tr>
<tr><td>>hg38_21_+_8433172_8446622_RNA45SN1_45S_50ntFlanks@12459.12486.28|AGCGTTGGATTGTTCACCCACTAATAGG</td><td>>hg38_21_+_8433172_8446622_RNA45SN1_45S_50ntFlanks</td><td>12459</td><td>12486</td><td>28</td><td>AGCGTTGGATTGTTCACCCACTAATAGG</td><td>134.3</td><td>-1.9</td><td>0.5</td><td>-3.9</td><td>1.1E-04</td><td>1.5E-04</td><td>MB231/MB468</td></tr>
<tr><td>>hg38_21_+_8433172_8446622_RNA45SN1_45S_50ntFlanks@6726.6743.18|CAGGACACATTGATCATC</td><td>>hg38_21_+_8433172_8446622_RNA45SN1_45S_50ntFlanks</td><td>6726</td><td>6743</td><td>18</td><td>CAGGACACATTGATCATC</td><td>225.4</td><td>-3.0</td><td>0.8</td><td>-3.9</td><td>1.1E-04</td><td>1.5E-04</td><td>MB231/MB468</td></tr>
<tr><td>>hg38_21_+_8433172_8446622_RNA45SN1_45S_50ntFlanks@11880.11904.25|CCCTGTTGAGCTTGACTCTAGTCTG</td><td>>hg38_21_+_8433172_8446622_RNA45SN1_45S_50ntFlanks</td><td>11880</td><td>11904</td><td>25</td><td>CCCTGTTGAGCTTGACTCTAGTCTG</td><td>107.2</td><td>-2.7</td><td>0.7</td><td>-3.8</td><td>1.2E-04</td><td>1.5E-04</td><td>BT20/MB468</td></tr>
<tr><td>>hg38_21_+_8433172_8446622_RNA45SN1_45S_50ntFlanks@8180.8211.32|TGATCGAGGCCCAGCCCGTGGACGGTGTGAGG</td><td>>hg38_21_+_8433172_8446622_RNA45SN1_45S_50ntFlanks</td><td>8180</td><td>8211</td><td>32</td><td>TGATCGAGGCCCAGCCCGTGGACGGTGTGAGG</td><td>142.6</td><td>-2.6</td><td>0.7</td><td>-3.9</td><td>1.1E-04</td><td>1.5E-04</td><td>MB231/MB468</td></tr>
<tr><td>>hg38_21_+_8433172_8446622_RNA45SN1_45S_50ntFlanks@7990.8012.23|GACGTGGCGACCCGCTGAATTTA</td><td>>hg38_21_+_8433172_8446622_RNA45SN1_45S_50ntFlanks</td><td>7990</td><td>8012</td><td>23</td><td>GACGTGGCGACCCGCTGAATTTA</td><td>1021.2</td><td>2.6</td><td>0.7</td><td>3.9</td><td>1.1E-04</td><td>1.5E-04</td><td>MB231/MB468</td></tr>
<tr><td>>hg38_21_+_8433172_8446622_RNA45SN1_45S_50ntFlanks@3854.3874.21|ACTGTGGTAATTCTAGAGCTA</td><td>>hg38_21_+_8433172_8446622_RNA45SN1_45S_50ntFlanks</td><td>3854</td><td>3874</td><td>21</td><td>ACTGTGGTAATTCTAGAGCTA</td><td>464.3</td><td>-2.8</td><td>0.7</td><td>-3.8</td><td>1.2E-04</td><td>1.5E-04</td><td>BT20/MB468</td></tr>
<tr><td>>hg38_21_+_8433172_8446622_RNA45SN1_45S_50ntFlanks@12405.12433.29|CTTTTTGATCCTTCGATGTCGGCTCTTCC</td><td>>hg38_21_+_8433172_8446622_RNA45SN1_45S_50ntFlanks</td><td>12405</td><td>12433</td><td>29</td><td>CTTTTTGATCCTTCGATGTCGGCTCTTCC</td><td>1767.9</td><td>-2.7</td><td>0.7</td><td>-3.9</td><td>1.1E-04</td><td>1.5E-04</td><td>MB231/MB468</td></tr>
<tr><td>>hg38_21_+_8433172_8446622_RNA45SN1_45S_50ntFlanks@10392.10422.31|ACAGCAGTTGAACATGGGTCAGTCGGTCCTG</td><td>>hg38_21_+_8433172_8446622_RNA45SN1_45S_50ntFlanks</td><td>10392</td><td>10422</td><td>31</td><td>ACAGCAGTTGAACATGGGTCAGTCGGTCCTG</td><td>513.9</td><td>-2.9</td><td>0.8</td><td>-3.9</td><td>1.1E-04</td><td>1.5E-04</td><td>MB231/MB468</td></tr>
<tr><td>>hg38_21_+_8433172_8446622_RNA45SN1_45S_50ntFlanks@4683.4707.25|CAAGACGGACCAGAGCGAAAGCATT</td><td>>hg38_21_+_8433172_8446622_RNA45SN1_45S_50ntFlanks</td><td>4683</td><td>4707</td><td>25</td><td>CAAGACGGACCAGAGCGAAAGCATT</td><td>226.5</td><td>-2.9</td><td>0.8</td><td>-3.8</td><td>1.2E-04</td><td>1.5E-04</td><td>BT20/MB468</td></tr>
<tr><td>>hg38_21_+_8433172_8446622_RNA45SN1_45S_50ntFlanks@8176.8202.27|CTTCTGATCGAGGCCCAGCCCGTGGAC</td><td>>hg38_21_+_8433172_8446622_RNA45SN1_45S_50ntFlanks</td><td>8176</td><td>8202</td><td>27</td><td>CTTCTGATCGAGGCCCAGCCCGTGGAC</td><td>1146.8</td><td>2.4</td><td>0.6</td><td>3.8</td><td>1.2E-04</td><td>1.5E-04</td><td>BT20/MB468</td></tr>
<tr><td>>hg38_21_+_8433172_8446622_RNA45SN1_45S_50ntFlanks@9564.9603.40|CTGACGTGCAAATCGGTCGTCCGACCTGGGTATAGGGGCG</td><td>>hg38_21_+_8433172_8446622_RNA45SN1_45S_50ntFlanks</td><td>9564</td><td>9603</td><td>40</td><td>CTGACGTGCAAATCGGTCGTCCGACCTGGGTATAGGGGCG</td><td>476.6</td><td>2.6</td><td>0.7</td><td>3.9</td><td>1.1E-04</td><td>1.5E-04</td><td>MB231/MB468</td></tr>
<tr><td>>hg38_21_+_8433172_8446622_RNA45SN1_45S_50ntFlanks@9534.9568.35|AGGAAACTCTGGTGGAGGTCCGTAGCGGTCCTGAC</td><td>>hg38_21_+_8433172_8446622_RNA45SN1_45S_50ntFlanks</td><td>9534</td><td>9568</td><td>35</td><td>AGGAAACTCTGGTGGAGGTCCGTAGCGGTCCTGAC</td><td>247.0</td><td>-3.8</td><td>1.0</td><td>-3.8</td><td>1.2E-04</td><td>1.5E-04</td><td>BT20/MB468</td></tr>
</table>

<table>
<tr><td>>hg38_21_+_8433172_8446622_RNA45SN1_45S_50ntFlanks@3785.3810.26|TGAAACTGCGAATGGCTCATTAAATC</td><td>>hg38_21_+_8433172_8446622_RNA45SN1_45S_50ntFlanks</td><td>3785</td><td>3810</td><td>26</td><td>TGAAACTGCGAATGGCTCATTAAATC</td><td>163.4</td><td>-1.6</td><td>0.4</td><td>-3.9</td><td>1.1E-04</td><td>1.5E-04</td><td>MB231/MB468</td></tr>
<tr><td>>hg38_21_+_8433172_8446622_RNA45SN1_45S_50ntFlanks@12936.12961.26|AAACCATTCGTAGACGACCTGCTTCT</td><td>>hg38_21_+_8433172_8446622_RNA45SN1_45S_50ntFlanks</td><td>12936</td><td>12961</td><td>26</td><td>AAACCATTCGTAGACGACCTGCTTCT</td><td>261.2</td><td>-1.7</td><td>0.4</td><td>-3.9</td><td>1.1E-04</td><td>1.6E-04</td><td>MB231/MB468</td></tr>
<tr><td>>hg38_21_+_8433172_8446622_RNA45SN1_45S_50ntFlanks@10869.10891.23|AGGGCTGGGTCGGTCGGGCTGGG</td><td>>hg38_21_+_8433172_8446622_RNA45SN1_45S_50ntFlanks</td><td>10869</td><td>10891</td><td>23</td><td>AGGGCTGGGTCGGTCGGGCTGGG</td><td>407.6</td><td>-3.6</td><td>0.9</td><td>-3.8</td><td>1.3E-04</td><td>1.6E-04</td><td>BT20/MB468</td></tr>
<tr><td>>hg38_21_+_8433172_8446622_RNA45SN1_45S_50ntFlanks@11767.11804.38|TCTAATTAGTGACGCGCATGAATGGATGAACGAGATTC</td><td>>hg38_21_+_8433172_8446622_RNA45SN1_45S_50ntFlanks</td><td>11767</td><td>11804</td><td>38</td><td>TCTAATTAGTGACGCGCATGAATGGATGAACGAGATTC</td><td>98.6</td><td>-1.8</td><td>0.5</td><td>-3.9</td><td>1.1E-04</td><td>1.6E-04</td><td>MB231/MB468</td></tr>
<tr><td>>MI0000746|hsa-mir-99b&WithFlank&19|+|51692606|51692687@13.32.20(+2G)&[MIMAT0000689&hsa-miR-99b-5p&offsets|0|-2(+2G);m-10&19|+|51692618|51692639&offsets|0|-2(+2G)]|CACCCGTAGAACCGACCTTGGG</td><td>>MI0000746|hsa-mir-99b&WithFlank&19|+|51692606|51692687</td><td>13</td><td>32</td><td>20(+2G)</td><td>CACCCGTAGAACCGACCTTGGG</td><td>540.4</td><td>4.5</td><td>1.0</td><td>4.4</td><td>1.4E-05</td><td>1.6E-04</td><td>BT20/MB231</td></tr>
<tr><td>>hg38_21_+_8433172_8446622_RNA45SN1_45S_50ntFlanks@3940.3959.20|ACCCGGTCAGCCCCTCTCCG</td><td>>hg38_21_+_8433172_8446622_RNA45SN1_45S_50ntFlanks</td><td>3940</td><td>3959</td><td>20</td><td>ACCCGGTCAGCCCCTCTCCG</td><td>113.1</td><td>-3.9</td><td>1.0</td><td>-3.8</td><td>1.3E-04</td><td>1.6E-04</td><td>BT20/MB468</td></tr>
<tr><td>>MI0000299|hsa-mir-222&WithFlank&X|-|45747009|45747130@75.97.23(+3U)&[MIMAT0000279&hsa-miR-222-3p&offsets|0|+2(+3U);m-140&X|-|45747033|45747056&offsets|0|-1(+3U)]|AGCTACATCTGGCTACTGGGTCTTTT</td><td>>MI0000299|hsa-mir-222&WithFlank&X|-|45747009|45747130</td><td>75</td><td>97</td><td>23(+3U)</td><td>AGCTACATCTGGCTACTGGGTCTTTT</td><td>319.9</td><td>-3.4</td><td>0.8</td><td>-4.3</td><td>1.4E-05</td><td>1.6E-04</td><td>BT20/MB231</td></tr>
<tr><td>>hg38_21_+_8433172_8446622_RNA45SN1_45S_50ntFlanks@6646.6665.20|TCGTACGACTCTTAGCGGTG</td><td>>hg38_21_+_8433172_8446622_RNA45SN1_45S_50ntFlanks</td><td>6646</td><td>6665</td><td>20</td><td>TCGTACGACTCTTAGCGGTG</td><td>38309.2</td><td>3.3</td><td>0.8</td><td>4.3</td><td>1.4E-05</td><td>1.6E-04</td><td>BT20/MB231</td></tr>
<tr><td>>hg38_21_+_8433172_8446622_RNA45SN1_45S_50ntFlanks@11671.11700.30|CCAGTGCTCTGAATGTCAAAGTGAAGAAAT</td><td>>hg38_21_+_8433172_8446622_RNA45SN1_45S_50ntFlanks</td><td>11671</td><td>11700</td><td>30</td><td>CCAGTGCTCTGAATGTCAAAGTGAAGAAAT</td><td>214.4</td><td>-1.7</td><td>0.4</td><td>-3.8</td><td>1.2E-04</td><td>1.6E-04</td><td>MB231/MB468</td></tr>
<tr><td>>hg38_21_+_8433172_8446622_RNA45SN1_45S_50ntFlanks@10792.10818.27|CTCTGGCATGTTGGAACAATGTAGGTA</td><td>>hg38_21_+_8433172_8446622_RNA45SN1_45S_50ntFlanks</td><td>10792</td><td>10818</td><td>27</td><td>CTCTGGCATGTTGGAACAATGTAGGTA</td><td>420.1</td><td>-2.5</td><td>0.7</td><td>-3.8</td><td>1.2E-04</td><td>1.6E-04</td><td>MB231/MB468</td></tr>
<tr><td>>hg38_21_+_8433172_8446622_RNA45SN1_45S_50ntFlanks@6730.6752.23|ACACATTGATCATCGACACTTCG</td><td>>hg38_21_+_8433172_8446622_RNA45SN1_45S_50ntFlanks</td><td>6730</td><td>6752</td><td>23</td><td>ACACATTGATCATCGACACTTCG</td><td>130.6</td><td>-2.9</td><td>0.8</td><td>-3.8</td><td>1.3E-04</td><td>1.6E-04</td><td>BT20/MB468</td></tr>
<tr><td>>MI0000239|hsa-mir-197&WithFlank&1|+|109598887|109598973@54.75.22&[MIMAT0000227&hsa-miR-197-3p&offsets|0|0;m-99&1|+|109598940|109598961&offsets|0|0]|TTCACCACCTTCTCCACCCAGC</td><td>>MI0000239|hsa-mir-197&WithFlank&1|+|109598887|109598973</td><td>54</td><td>75</td><td>22</td><td>TTCACCACCTTCTCCACCCAGC</td><td>7198.1</td><td>3.0</td><td>0.7</td><td>4.3</td><td>1.4E-05</td><td>1.7E-04</td><td>BT20/MB231</td></tr>
<tr><td>>am_trnaMT_ArgTCG_MT_+_10405_10469@29.48.20|TTTCGACTCATTAAATTATG</td><td>>am_trnaMT_ArgTCG_MT_+_10405_10469</td><td>29</td><td>48</td><td>20</td><td>TTTCGACTCATTAAATTATG</td><td>146.5</td><td>-3.0</td><td>0.8</td><td>-3.8</td><td>1.3E-04</td><td>1.7E-04</td><td>BT20/MB468</td></tr>
<tr><td>>hg38_21_+_8433172_8446622_RNA45SN1_45S_50ntFlanks@7977.8019.43|CGACCTCAGATCAGACGTGGCGACCCGCTGAATTTAAGCATAT</td><td>>hg38_21_+_8433172_8446622_RNA45SN1_45S_50ntFlanks</td><td>7977</td><td>8019</td><td>43</td><td>CGACCTCAGATCAGACGTGGCGACCCGCTGAATTTAAGCATAT</td><td>290.2</td><td>-4.4</td><td>1.0</td><td>-4.3</td><td>1.5E-05</td><td>1.7E-04</td><td>BT20/MB231</td></tr>
<tr><td>>hg38_21_+_8433172_8446622_RNA45SN1_45S_50ntFlanks@8176.8215.40|CTTCTGATCGAGGCCCAGCCCGTGGACGGTGTGAGGCCGG</td><td>>hg38_21_+_8433172_8446622_RNA45SN1_45S_50ntFlanks</td><td>8176</td><td>8215</td><td>40</td><td>CTTCTGATCGAGGCCCAGCCCGTGGACGGTGTGAGGCCGG</td><td>232.8</td><td>-3.7</td><td>1.0</td><td>-3.8</td><td>1.3E-04</td><td>1.7E-04</td><td>BT20/MB468</td></tr>
<tr><td>>hg38_21_+_8433172_8446622_RNA45SN1_45S_50ntFlanks@12995.13037.43|CTCGCTGCGATCTATTGAAAGTCAGCCCTCGACACAAGGGTTT</td><td>>hg38_21_+_8433172_8446622_RNA45SN1_45S_50ntFlanks</td><td>12995</td><td>13037</td><td>43</td><td>CTCGCTGCGATCTATTGAAAGTCAGCCCTCGACACAAGGGTTT</td><td>950.4</td><td>-4.1</td><td>0.9</td><td>-4.3</td><td>1.5E-05</td><td>1.7E-04</td><td>BT20/MB231</td></tr>
<tr><td>>hg38_21_+_8433172_8446622_RNA45SN1_45S_50ntFlanks@12936.12954.19|AAACCATTCGTAGACGACC</td><td>>hg38_21_+_8433172_8446622_RNA45SN1_45S_50ntFlanks</td><td>12936</td><td>12954</td><td>19</td><td>AAACCATTCGTAGACGACC</td><td>182.6</td><td>-2.2</td><td>0.6</td><td>-3.8</td><td>1.3E-04</td><td>1.7E-04</td><td>BT20/MB468</td></tr>
<tr><td>>hg38_21_+_8433172_8446622_RNA45SN1_45S_50ntFlanks@12935.12961.27|TAAACCATTCGTAGACGACCTGCTTCT</td><td>>hg38_21_+_8433172_8446622_RNA45SN1_45S_50ntFlanks</td><td>12935</td><td>12961</td><td>27</td><td>TAAACCATTCGTAGACGACCTGCTTCT</td><td>122.9</td><td>-2.2</td><td>0.6</td><td>-3.8</td><td>1.4E-04</td><td>1.7E-04</td><td>BT20/MB468</td></tr>
<tr><td>>hg38_21_+_8433172_8446622_RNA45SN1_45S_50ntFlanks@9548.9573.26|GAGGTCCGTAGCGGTCCTGACGTGCA</td><td>>hg38_21_+_8433172_8446622_RNA45SN1_45S_50ntFlanks</td><td>9548</td><td>9573</td><td>26</td><td>GAGGTCCGTAGCGGTCCTGACGTGCA</td><td>478.0</td><td>2.3</td><td>0.6</td><td>3.8</td><td>1.2E-04</td><td>1.7E-04</td><td>MB231/MB468</td></tr>
<tr><td>>hg38_21_+_8433172_8446622_RNA45SN1_45S_50ntFlanks@10771.10796.26|CAGGTCTCCAAGGTGAACAGCCTCTG</td><td>>hg38_21_+_8433172_8446622_RNA45SN1_45S_50ntFlanks</td><td>10771</td><td>10796</td><td>26</td><td>CAGGTCTCCAAGGTGAACAGCCTCTG</td><td>582.5</td><td>-2.7</td><td>0.7</td><td>-3.8</td><td>1.3E-04</td><td>1.7E-04</td><td>MB231/MB468</td></tr>
<tr><td>>hg38_21_+_8433172_8446622_RNA45SN1_45S_50ntFlanks@3835.3871.37|CTCCTCTCCTACTTGGATAACTGTGGTAATTCTAGAG</td><td>>hg38_21_+_8433172_8446622_RNA45SN1_45S_50ntFlanks</td><td>3835</td><td>3871</td><td>37</td><td>CTCCTCTCCTACTTGGATAACTGTGGTAATTCTAGAG</td><td>152.8</td><td>-3.0</td><td>0.8</td><td>-3.8</td><td>1.4E-04</td><td>1.7E-04</td><td>BT20/MB468</td></tr>
<tr><td>>hg38_21_+_8433172_8446622_RNA45SN1_45S_50ntFlanks@8194.8216.23|CCCGTGGACGGTGTGAGGCCGGT</td><td>>hg38_21_+_8433172_8446622_RNA45SN1_45S_50ntFlanks</td><td>8194</td><td>8216</td><td>23</td><td>CCCGTGGACGGTGTGAGGCCGGT</td><td>132.2</td><td>-2.0</td><td>0.5</td><td>-3.8</td><td>1.3E-04</td><td>1.7E-04</td><td>MB231/MB468</td></tr>
<tr><td>>hg38_21_+_8433172_8446622_RNA45SN1_45S_50ntFlanks@8105.8142.38|CGCGGCGGGGCGCGGGACATGTGGCGTACGGAAGACCC</td><td>>hg38_21_+_8433172_8446622_RNA45SN1_45S_50ntFlanks</td><td>8105</td><td>8142</td><td>38</td><td>CGCGGCGGGGCGCGGGACATGTGGCGTACGGAAGACCC</td><td>80.7</td><td>-2.9</td><td>0.8</td><td>-3.8</td><td>1.4E-04</td><td>1.8E-04</td><td>BT20/MB468</td></tr>
<tr><td>>hg38_21_+_8433172_8446622_RNA45SN1_45S_50ntFlanks@11540.11586.47|CGCGGCGCCTCGCCTCGGCCGGCGCCTAGCAGCCGACTTAGAACTGG</td><td>>hg38_21_+_8433172_8446622_RNA45SN1_45S_50ntFlanks</td><td>11540</td><td>11586</td><td>47</td><td>CGCGGCGCCTCGCCTCGGCCGGCGCCTAGCAGCCGACTTAGAACTGG</td><td>329.3</td><td>-3.6</td><td>0.8</td><td>-4.3</td><td>1.6E-05</td><td>1.8E-04</td><td>BT20/MB231</td></tr>
<tr><td>>hg38_21_+_8433172_8446622_RNA45SN1_45S_50ntFlanks@4243.4263.21|CTTTCGAGGCCCTGTAATTGG</td><td>>hg38_21_+_8433172_8446622_RNA45SN1_45S_50ntFlanks</td><td>4243</td><td>4263</td><td>21</td><td>CTTTCGAGGCCCTGTAATTGG</td><td>199.6</td><td>-2.0</td><td>0.5</td><td>-3.8</td><td>1.3E-04</td><td>1.8E-04</td><td>MB231/MB468</td></tr>
<tr><td>>MI0000272|hsa-mir-182&WithFlank&7|-|129770377|129770498@29.52.24(+1U)&[MIMAT0000259&hsa-miR-182-5p&offsets|0|0(+1U);m-22&7|-|129770449|129770470&offsets|0|+2(+1U)]|TTTGGCAATGGTAGAACTCACACTT</td><td>>MI0000272|hsa-mir-182&WithFlank&7|-|129770377|129770498</td><td>29</td><td>52</td><td>24(+1U)</td><td>TTTGGCAATGGTAGAACTCACACTT</td><td>484.6</td><td>3.2</td><td>0.7</td><td>4.3</td><td>1.6E-05</td><td>1.8E-04</td><td>BT20/MB231</td></tr>
<tr><td>>hg38_21_+_8433172_8446622_RNA45SN1_45S_50ntFlanks@12993.13015.23|CCCTCGCTGCGATCTATTGAAAG</td><td>>hg38_21_+_8433172_8446622_RNA45SN1_45S_50ntFlanks</td><td>12993</td><td>13015</td><td>23</td><td>CCCTCGCTGCGATCTATTGAAAG</td><td>132.3</td><td>-1.9</td><td>0.5</td><td>-3.8</td><td>1.4E-04</td><td>1.8E-04</td><td>BT20/MB468</td></tr>
<tr><td>>MI0000298|hsa-mir-221&WithFlank&X|-|45746151|45746272@31.56.26(+1U)&[MIMAT0004568&hsa-miR-221-5p&offsets|0|+4(+1U);m-405&X|-|45746220|45746242&offsets|0|+3(+1U)]|ACCTGGCATACAATGTAGATTTCTGTT</td><td>>MI0000298|hsa-mir-221&WithFlank&X|-|45746151|45746272</td><td>31</td><td>56</td><td>26(+1U)</td><td>ACCTGGCATACAATGTAGATTTCTGTT</td><td>201.4</td><td>-3.6</td><td>0.8</td><td>-4.3</td><td>1.6E-05</td><td>1.8E-04</td><td>BT20/MB231</td></tr>
<tr><td>>hg38_21_+_8433172_8446622_RNA45SN1_45S_50ntFlanks@6719.6749.31|TGAATTGCAGGACACATTGATCATCGACACT</td><td>>hg38_21_+_8433172_8446622_RNA45SN1_45S_50ntFlanks</td><td>6719</td><td>6749</td><td>31</td><td>TGAATTGCAGGACACATTGATCATCGACACT</td><td>287.0</td><td>-2.1</td><td>0.5</td><td>-3.8</td><td>1.3E-04</td><td>1.8E-04</td><td>MB231/MB468</td></tr>
<tr><td>>hg38_21_+_8433172_8446622_RNA45SN1_45S_50ntFlanks@8379.8402.24|TCAAGAGGGCGTGAAACCGTTAAG</td><td>>hg38_21_+_8433172_8446622_RNA45SN1_45S_50ntFlanks</td><td>8379</td><td>8402</td><td>24</td><td>TCAAGAGGGCGTGAAACCGTTAAG</td><td>180.3</td><td>-2.7</td><td>0.7</td><td>-3.8</td><td>1.3E-04</td><td>1.8E-04</td><td>MB231/MB468</td></tr>
<tr><td>>hg38_21_+_8433172_8446622_RNA45SN1_45S_50ntFlanks@4732.4765.34|AGAACGAAAGTCGGAGGTTCGAAGACGATCAGAT</td><td>>hg38_21_+_8433172_8446622_RNA45SN1_45S_50ntFlanks</td><td>4732</td><td>4765</td><td>34</td><td>AGAACGAAAGTCGGAGGTTCGAAGACGATCAGAT</td><td>632.0</td><td>7.5</td><td>2.0</td><td>3.8</td><td>1.3E-04</td><td>1.8E-04</td><td>MB231/MB468</td></tr>
<tr><td>>hg38_21_+_8433172_8446622_RNA45SN1_45S_50ntFlanks@11729.11748.20|TAACTATGACTCTCTTAAGG</td><td>>hg38_21_+_8433172_8446622_RNA45SN1_45S_50ntFlanks</td><td>11729</td><td>11748</td><td>20</td><td>TAACTATGACTCTCTTAAGG</td><td>2780.5</td><td>-2.0</td><td>0.5</td><td>-3.8</td><td>1.3E-04</td><td>1.8E-04</td><td>MB231/MB468</td></tr>
<tr><td>>hg38_21_+_8433172_8446622_RNA45SN1_45S_50ntFlanks@11208.11233.26|GCGGCGACTCTGGACGCGAGCCGGGC</td><td>>hg38_21_+_8433172_8446622_RNA45SN1_45S_50ntFlanks</td><td>11208</td><td>11233</td><td>26</td><td>GCGGCGACTCTGGACGCGAGCCGGGC</td><td>2220.6</td><td>-3.6</td><td>0.9</td><td>-3.8</td><td>1.5E-04</td><td>1.8E-04</td><td>BT20/MB468</td></tr>
<tr><td>>hg38_21_+_8433172_8446622_RNA45SN1_45S_50ntFlanks@10394.10424.31|AGCAGTTGAACATGGGTCAGTCGGTCCTGAG</td><td>>hg38_21_+_8433172_8446622_RNA45SN1_45S_50ntFlanks</td><td>10394</td><td>10424</td><td>31</td><td>AGCAGTTGAACATGGGTCAGTCGGTCCTGAG</td><td>855.5</td><td>3.6</td><td>1.0</td><td>3.8</td><td>1.5E-04</td><td>1.8E-04</td><td>BT20/MB468</td></tr>
<tr><td>>trna111_HisGTG_1_-_147774845_147774916@-1T.29.30,trna118_HisGTG_1_-_145396881_145396952@-1T.29.30,trna16_HisGTG_1_+_146544773_146544844@-1T.29.30,trna1_HisGTG_15_+_45493349_45493420@-1T.29.30,trna21_HisGTG_1_+_147753471_147753542@-1T.29.30,trna33_HisGTG_6_+_27125906_27125977@-1T.29.30,trna7_HisGTG_9_-_14433938_14434009@-1T.29.30,trna8_HisGTG_15_-_45492611_45492682@-1T.29.30,trna9_HisGTG_15_-_45490804_45490875@-1T.29.30|TGCCGTGATCGTATAGTGGTTAGTACTCTG</td><td>>trna111_HisGTG_1_-_147774845_147774916</td><td>-1T</td><td>29</td><td>30</td><td>TGCCGTGATCGTATAGTGGTTAGTACTCTG</td><td>2353.2</td><td>-3.2</td><td>0.8</td><td>-3.8</td><td>1.3E-04</td><td>1.8E-04</td><td>MB231/MB468</td></tr>
<tr><td>>MI0000299|hsa-mir-222&WithFlank&X|-|45747009|45747130@75.99.25(+3U)&[MIMAT0000279&hsa-miR-222-3p&offsets|0|+4(+3U);m-140&X|-|45747033|45747056&offsets|0|+1(+3U)]|AGCTACATCTGGCTACTGGGTCTCTTTT</td><td>>MI0000299|hsa-mir-222&WithFlank&X|-|45747009|45747130</td><td>75</td><td>99</td><td>25(+3U)</td><td>AGCTACATCTGGCTACTGGGTCTCTTTT</td><td>2887.8</td><td>-3.5</td><td>0.8</td><td>-4.3</td><td>1.6E-05</td><td>1.9E-04</td><td>BT20/MB231</td></tr>
<tr><td>>hg38_21_+_8433172_8446622_RNA45SN1_45S_50ntFlanks@9623.9649.27|TAGTAGCTGGTTCCCTCCGAAGTTTCC</td><td>>hg38_21_+_8433172_8446622_RNA45SN1_45S_50ntFlanks</td><td>9623</td><td>9649</td><td>27</td><td>TAGTAGCTGGTTCCCTCCGAAGTTTCC</td><td>84.8</td><td>-3.0</td><td>0.8</td><td>-3.8</td><td>1.5E-04</td><td>1.9E-04</td><td>BT20/MB468</td></tr>
<tr><td>>hg38_21_+_8433172_8446622_RNA45SN1_45S_50ntFlanks@3771.3794.24|CACGGCCGGTACAGTGAAACTGCG</td><td>>hg38_21_+_8433172_8446622_RNA45SN1_45S_50ntFlanks</td><td>3771</td><td>3794</td><td>24</td><td>CACGGCCGGTACAGTGAAACTGCG</td><td>156.5</td><td>-2.6</td><td>0.7</td><td>-3.8</td><td>1.5E-04</td><td>1.9E-04</td><td>BT20/MB468</td></tr>
<tr><td>>MI0003776|hsa-mir-320b-1&WithFlank&1|+|116671740|116671823@48.66.19;MI0003839|hsa-mir-320b-2&WithFlank&1|-|224257034|224257116@47.65.19;MI0003778|hsa-mir-320c-1&WithFlank&18|+|21683512|21683595@48.66.19;MI0008191|hsa-mir-320c-2&WithFlank&18|+|24321669|24321752@48.66.19;MI0000542|hsa-mir-320a&WithFlank&8|-|22244960|22245043@48.66.19&[MIMAT0005792&hsa-miR-320b&offsets|0|-3;m-183&1|+|116671787|116671806&offsets|0|-1];[MIMAT0005792_1&hsa-miR-320b&offsets|0|-3;m-183&1|-|224257051|224257070&offsets|0|-1];[MIMAT0005793&hsa-miR-320c&offsets|0|-1;m-222&18|+|21683559|21683578&offsets|0|-1];[MIMAT0005793_1&hsa-miR-320c&offsets|0|-1;m-219&18|+|24321716|24321735&offsets|0|-1];[MIMAT0000510&hsa-miR-320a-3p&offsets|0|-3;m-60&8|-|22244975|22244996&offsets|0|-3]|AAAAGCTGGGTTGAGAGGG</td><td>>MI0003776|hsa-mir-320b-1&WithFlank&1|+|116671740|116671823</td><td>48</td><td>66</td><td>19</td><td>AAAAGCTGGGTTGAGAGGG</td><td>315.6</td><td>3.1</td><td>0.7</td><td>4.3</td><td>1.7E-05</td><td>1.9E-04</td><td>BT20/MB231</td></tr>
<tr><td>>hg38_21_+_8433172_8446622_RNA45SN1_45S_50ntFlanks@9554.9582.29|CGTAGCGGTCCTGACGTGCAAATCGGTCG</td><td>>hg38_21_+_8433172_8446622_RNA45SN1_45S_50ntFlanks</td><td>9554</td><td>9582</td><td>29</td><td>CGTAGCGGTCCTGACGTGCAAATCGGTCG</td><td>470.2</td><td>-2.1</td><td>0.6</td><td>-3.8</td><td>1.4E-04</td><td>1.9E-04</td><td>MB231/MB468</td></tr>
</table>

| | | | | | | | | | | | | |
|---|---|---|---|---|---|---|---|---|---|---|---|---|
| >hg38_21_+_8433172_8446622_RNA45SN1_45S_50ntFlanks@11317.11338.22\|ACCCCACGTCTCGTCGCGCGCG | >hg38_21_+_8433172_8446622_RNA45SN1_45S_50ntFlanks | 11317 | 11338 | 22 | ACCCCACGTCTCGTCGCGCGCG | 624.8 | -4.2 | 1.1 | -3.8 | 1.5E-04 | 1.9E-04 | BT20/MB468 |
| >hg38_1_-_228634819_228635039_RNA5S12_5S_50ntFlanks@72.92.21\|AACGCGCCCGATCTCGTCTGA | >hg38_1_-_228634819_228635039_RNA5S12_5S_50ntFlanks | 72 | 92 | 21 | AACGCGCCCGATCTCGTCTGA | 80.8 | -2.5 | 0.7 | -3.8 | 1.5E-04 | 1.9E-04 | BT20/MB468 |
| >hg38_21_+_8433172_8446622_RNA45SN1_45S_50ntFlanks@4657.4684.28\|CTAGAGGTGAAATTCTTGGACCGGCGCA | >hg38_21_+_8433172_8446622_RNA45SN1_45S_50ntFlanks | 4657 | 4684 | 28 | CTAGAGGTGAAATTCTTGGACCGGCGCA | 107.5 | -3.5 | 0.9 | -3.8 | 1.5E-04 | 1.9E-04 | BT20/MB468 |
| >hg38_21_+_8433172_8446622_RNA45SN1_45S_50ntFlanks@11575.11606.32\|ACTTAGAACTGGTGCGGACCAGGGGAATCCGA | >hg38_21_+_8433172_8446622_RNA45SN1_45S_50ntFlanks | 11575 | 11606 | 32 | ACTTAGAACTGGTGCGGACCAGGGGAATCCGA | 147.5 | -3.0 | 0.8 | -3.8 | 1.4E-04 | 1.9E-04 | MB231/MB468 |
| >hg38_21_+_8433172_8446622_RNA45SN1_45S_50ntFlanks@4161.4182.22\|CACATCCAAGGAAGGCAGCAGG | >hg38_21_+_8433172_8446622_RNA45SN1_45S_50ntFlanks | 4161 | 4182 | 22 | CACATCCAAGGAAGGCAGCAGG | 75.4 | -3.4 | 0.9 | -3.8 | 1.4E-04 | 1.9E-04 | MB231/MB468 |
| >hg38_21_+_8433172_8446622_RNA45SN1_45S_50ntFlanks@11874.11905.32\|AGAAGACCCTGTTGAGCTTGACTCTAGTCTGG | >hg38_21_+_8433172_8446622_RNA45SN1_45S_50ntFlanks | 11874 | 11905 | 32 | AGAAGACCCTGTTGAGCTTGACTCTAGTCTGG | 525.7 | -5.2 | 1.4 | -3.8 | 1.5E-04 | 1.9E-04 | BT20/MB468 |
| >hg38_21_+_8433172_8446622_RNA45SN1_45S_50ntFlanks@10797.10816.20\|GCATGTTGGAACAATGTAGG | >hg38_21_+_8433172_8446622_RNA45SN1_45S_50ntFlanks | 10797 | 10816 | 20 | GCATGTTGGAACAATGTAGG | 263.2 | -3.1 | 0.8 | -3.8 | 1.6E-04 | 1.9E-04 | BT20/MB468 |
| >hg38_21_+_8433172_8446622_RNA45SN1_45S_50ntFlanks@11710.11740.31\|CGCGGGTAAACGGCGGGAGTAACTATGACTC | >hg38_21_+_8433172_8446622_RNA45SN1_45S_50ntFlanks | 11710 | 11740 | 31 | CGCGGGTAAACGGCGGGAGTAACTATGACTC | 276.5 | -4.0 | 1.1 | -3.8 | 1.6E-04 | 2.0E-04 | BT20/MB468 |
| >MI0000298\|hsa-mir-221&WithFlank&X\|-\|45746151\|45746272@31.56.26&[MIMAT0004568&hsa-miR-221-5p&offsets\|0\|+4;m-405&X\|-\|45746220\|45746242&offsets\|0\|+3]\|ACCTGGCATACAATGTAGATTTCTGT | >MI0000298\|hsa-mir-221&WithFlank&X\|-\|45746151\|45746272 | 31 | 56 | 26 | ACCTGGCATACAATGTAGATTTCTGT | 2851.0 | -2.9 | 0.7 | -4.3 | 1.8E-05 | 2.0E-04 | BT20/MB231 |
| >hg38_1_-_228634819_228635039_RNA5S12_5S_50ntFlanks@84.111.28\|CTCGTCTGATCTCGGAAGCTAAGCAGGG | >hg38_1_-_228634819_228635039_RNA5S12_5S_50ntFlanks | 84 | 111 | 28 | CTCGTCTGATCTCGGAAGCTAAGCAGGG | 93.2 | -3.1 | 0.8 | -3.8 | 1.6E-04 | 2.0E-04 | BT20/MB468 |
| >hg38_21_+_8433172_8446622_RNA45SN1_45S_50ntFlanks@8176.8205.30\|CTTCTGATCGAGGCCCAGCCCGTGGACGGT | >hg38_21_+_8433172_8446622_RNA45SN1_45S_50ntFlanks | 8176 | 8205 | 30 | CTTCTGATCGAGGCCCAGCCCGTGGACGGT | 253.8 | 1.7 | 0.4 | 3.8 | 1.4E-04 | 2.0E-04 | MB231/MB468 |
| >hg38_21_+_8433172_8446622_RNA45SN1_45S_50ntFlanks@12944.12965.22\|CGTAGACGACCTGCTTCTGGGT | >hg38_21_+_8433172_8446622_RNA45SN1_45S_50ntFlanks | 12944 | 12965 | 22 | CGTAGACGACCTGCTTCTGGGT | 126.5 | -2.0 | 0.5 | -3.8 | 1.4E-04 | 2.0E-04 | MB231/MB468 |
| >hg38_21_+_8433172_8446622_RNA45SN1_45S_50ntFlanks@5486.5514.29\|GCGCTGAGAAGACGGTCGAACTTGACTAT | >hg38_21_+_8433172_8446622_RNA45SN1_45S_50ntFlanks | 5486 | 5514 | 29 | GCGCTGAGAAGACGGTCGAACTTGACTAT | 344.3 | -2.4 | 0.6 | -3.8 | 1.4E-04 | 2.0E-04 | MB231/MB468 |
| >am_trna134_GluTTC_1_-_16861774_16861845@1.29.29,trna5_GluTTC_1_+_17199078_17199149@1.29.29,trna84_GluTTC_1_-_161391883_161391954@1.29.29,trna94_GluTTC_1_-_149664355_149664427@1.29.29\|TCCCTGGTGGTCTAGTGGCTAGGATTCGG | >am_trna134_GluTTC_1_-_16861774_16861845 | 1 | 29 | 29 | TCCCTGGTGGTCTAGTGGCTAGGATTCGG | 1023.2 | -3.9 | 1.0 | -3.8 | 1.6E-04 | 2.0E-04 | BT20/MB468 |
| >hg38_21_+_8433172_8446622_RNA45SN1_45S_50ntFlanks@3777.3799.23\|CGGTACAGTGAAACTGCGAATGG | >hg38_21_+_8433172_8446622_RNA45SN1_45S_50ntFlanks | 3777 | 3799 | 23 | CGGTACAGTGAAACTGCGAATGG | 75.2 | -2.0 | 0.5 | -3.8 | 1.4E-04 | 2.0E-04 | MB231/MB468 |
| >hg38_1_-_228634819_228635039_RNA5S12_5S_50ntFlanks@122.170.49\|TTAGTACTTGGATGGGAGACCGCCTGGGAATACCGGGTGCTGTAGGCTT | >hg38_1_-_228634819_228635039_RNA5S12_5S_50ntFlanks | 122 | 170 | 49 | TTAGTACTTGGATGGGAGACCGCCTGGGAATACCGGGTGCTGTAGGCTT | 1007.9 | 8.9 | 2.3 | 3.8 | 1.4E-04 | 2.0E-04 | MB231/MB468 |
| >hg38_21_+_8433172_8446622_RNA45SN1_45S_50ntFlanks@10346.10367.22\|TCAAACGAGAACTTTGAAGGCC | >hg38_21_+_8433172_8446622_RNA45SN1_45S_50ntFlanks | 10346 | 10367 | 22 | TCAAACGAGAACTTTGAAGGCC | 478.0 | 2.5 | 0.7 | 3.8 | 1.5E-04 | 2.0E-04 | MB231/MB468 |
| >trna152_ValCAC_6_-_27248049_27248121@1.23.23\|GCTTCTGTAGTGTAGTGGTTATC | >trna152_ValCAC_6_-_27248049_27248121 | 1 | 23 | 23 | GCTTCTGTAGTGTAGTGGTTATC | 828.5 | 2.7 | 0.7 | 3.8 | 1.5E-04 | 2.0E-04 | MB231/MB468 |
| >am_trnaMT_PheGAA_MT_+_577_647@45.62.18\|AGACGGGCTCACATCACC | >am_trnaMT_PheGAA_MT_+_577_647 | 45 | 62 | 18 | AGACGGGCTCACATCACC | 697.1 | 3.5 | 0.9 | 3.8 | 1.6E-04 | 2.0E-04 | BT20/MB468 |
| >hg38_21_+_8433172_8446622_RNA45SN1_45S_50ntFlanks@11755.11777.23\|AATGCCTCGTCATCTAATTAGTG | >hg38_21_+_8433172_8446622_RNA45SN1_45S_50ntFlanks | 11755 | 11777 | 23 | AATGCCTCGTCATCTAATTAGTG | 257.9 | -1.8 | 0.5 | -3.8 | 1.5E-04 | 2.0E-04 | MB231/MB468 |
| >hg38_21_+_8433172_8446622_RNA45SN1_45S_50ntFlanks@3854.3884.31\|ACTGTGGTAATTCTAGAGCTAATACATGCCG | >hg38_21_+_8433172_8446622_RNA45SN1_45S_50ntFlanks | 3854 | 3884 | 31 | ACTGTGGTAATTCTAGAGCTAATACATGCCG | 155.7 | -2.6 | 0.7 | -3.8 | 1.6E-04 | 2.0E-04 | BT20/MB468 |
| >hg38_21_+_8433172_8446622_RNA45SN1_45S_50ntFlanks@3855.3881.27\|CTGTGGTAATTCTAGAGCTAATACATG | >hg38_21_+_8433172_8446622_RNA45SN1_45S_50ntFlanks | 3855 | 3881 | 27 | CTGTGGTAATTCTAGAGCTAATACATG | 208.8 | -2.3 | 0.6 | -3.8 | 1.5E-04 | 2.0E-04 | MB231/MB468 |
| >hg38_21_+_8433172_8446622_RNA45SN1_45S_50ntFlanks@9554.9575.22\|CGTAGCGGTCCTGACGTGCAAA | >hg38_21_+_8433172_8446622_RNA45SN1_45S_50ntFlanks | 9554 | 9575 | 22 | CGTAGCGGTCCTGACGTGCAAA | 615.9 | -2.1 | 0.6 | -3.8 | 1.6E-04 | 2.0E-04 | BT20/MB468 |
| >hg38_21_+_8433172_8446622_RNA45SN1_45S_50ntFlanks@11642.11660.19\|GGCGGGTGTTGACGCGATG | >hg38_21_+_8433172_8446622_RNA45SN1_45S_50ntFlanks | 11642 | 11660 | 19 | GGCGGGTGTTGACGCGATG | 90.8 | -2.6 | 0.7 | -3.8 | 1.6E-04 | 2.0E-04 | BT20/MB468 |
| >hg38_21_+_8433172_8446622_RNA45SN1_45S_50ntFlanks@8176.8195.20\|CTTCTGATCGAGGCCCAGCC | >hg38_21_+_8433172_8446622_RNA45SN1_45S_50ntFlanks | 8176 | 8195 | 20 | CTTCTGATCGAGGCCCAGCC | 3635.9 | 2.7 | 0.7 | 3.8 | 1.5E-04 | 2.1E-04 | MB231/MB468 |
| >hg38_21_+_8433172_8446622_RNA45SN1_45S_50ntFlanks@12959.12982.24\|TCTGGGTCGGGGTTTCGTACGTAG | >hg38_21_+_8433172_8446622_RNA45SN1_45S_50ntFlanks | 12959 | 12982 | 24 | TCTGGGTCGGGGTTTCGTACGTAG | 142.6 | -2.6 | 0.7 | -3.8 | 1.7E-04 | 2.1E-04 | BT20/MB468 |
| >hg38_21_+_8433172_8446622_RNA45SN1_45S_50ntFlanks@10441.10460.20\|TTCCGAAGGGACGGGCGATG | >hg38_21_+_8433172_8446622_RNA45SN1_45S_50ntFlanks | 10441 | 10460 | 20 | TTCCGAAGGGACGGGCGATG | 245.6 | -2.4 | 0.7 | -3.8 | 1.7E-04 | 2.1E-04 | BT20/MB468 |
| >hg38_21_+_8433172_8446622_RNA45SN1_45S_50ntFlanks@12995.13013.19\|CTCGCTGCGATCTATTGAA | >hg38_21_+_8433172_8446622_RNA45SN1_45S_50ntFlanks | 12995 | 13013 | 19 | CTCGCTGCGATCTATTGAA | 1209.9 | 2.2 | 0.6 | 3.8 | 1.5E-04 | 2.1E-04 | MB231/MB468 |
| >MI0000098\|hsa-mir-96&WithFlank&7\|-\|129774686\|129774775@15.36.22(+1G)&[MIMAT0000095&hsa-miR-96-5p&offsets\|0\|-1(+1G);m-248&7\|-\|129774739\|129774761&offsets\|0\|-1(+1G)]\|TTTGGCACTAGCACATTTTTGCG | >MI0000098\|hsa-mir-96&WithFlank&7\|-\|129774686\|129774775 | 15 | 36 | 22(+1G) | TTTGGCACTAGCACATTTTTGCG | 1736.4 | 4.2 | 1.0 | 4.3 | 1.9E-05 | 2.1E-04 | BT20/MB231 |
| >hg38_21_+_8433172_8446622_RNA45SN1_45S_50ntFlanks@5485.5513.29\|AGCGCTGAGAAGACGGTCGAACTTGACTA | >hg38_21_+_8433172_8446622_RNA45SN1_45S_50ntFlanks | 5485 | 5513 | 29 | AGCGCTGAGAAGACGGTCGAACTTGACTA | 84.9 | -3.0 | 0.8 | -3.8 | 1.6E-04 | 2.1E-04 | MB231/MB468 |
| >hg38_21_+_8433172_8446622_RNA45SN1_45S_50ntFlanks@4760.4793.34\|TCAGATACCGTCGTAGTTCCGACCATAAACGATG | >hg38_21_+_8433172_8446622_RNA45SN1_45S_50ntFlanks | 4760 | 4793 | 34 | TCAGATACCGTCGTAGTTCCGACCATAAACGATG | 129.2 | -1.9 | 0.5 | -3.8 | 1.6E-04 | 2.1E-04 | MB231/MB468 |
| >hg38_21_+_8433172_8446622_RNA45SN1_45S_50ntFlanks@10869.10890.22\|AGGGCTGGGTCGGTCGGGCTGG | >hg38_21_+_8433172_8446622_RNA45SN1_45S_50ntFlanks | 10869 | 10890 | 22 | AGGGCTGGGTCGGTCGGGCTGG | 317.9 | -3.6 | 1.0 | -3.8 | 1.7E-04 | 2.1E-04 | BT20/MB468 |
| >hg38_21_+_8433172_8446622_RNA45SN1_45S_50ntFlanks@9574.9598.25\|AATCGGTCGTCCGACCTGGGTATAG | >hg38_21_+_8433172_8446622_RNA45SN1_45S_50ntFlanks | 9574 | 9598 | 25 | AATCGGTCGTCCGACCTGGGTATAG | 131.6 | -2.1 | 0.5 | -3.8 | 1.6E-04 | 2.1E-04 | MB231/MB468 |
| >hg38_21_+_8433172_8446622_RNA45SN1_45S_50ntFlanks@9806.9830.25\|CGTGGAATGCGAGTGCCTAGTGGGC | >hg38_21_+_8433172_8446622_RNA45SN1_45S_50ntFlanks | 9806 | 9830 | 25 | CGTGGAATGCGAGTGCCTAGTGGGC | 77.9 | -2.5 | 0.7 | -3.8 | 1.6E-04 | 2.1E-04 | MB231/MB468 |
| >hg38_21_+_8433172_8446622_RNA45SN1_45S_50ntFlanks@11566.11585.20\|TAGCAGCCGACTTAGAACTG | >hg38_21_+_8433172_8446622_RNA45SN1_45S_50ntFlanks | 11566 | 11585 | 20 | TAGCAGCCGACTTAGAACTG | 320.3 | 1.5 | 0.4 | 3.8 | 1.7E-04 | 2.2E-04 | BT20/MB468 |
| >hg38_21_+_8433172_8446622_RNA45SN1_45S_50ntFlanks@10788.10822.35\|CAGCCTCTGGCATGTTGGAACAATGTAGGTAAGGG | >hg38_21_+_8433172_8446622_RNA45SN1_45S_50ntFlanks | 10788 | 10822 | 35 | CAGCCTCTGGCATGTTGGAACAATGTAGGTAAGGG | 121.5 | -2.0 | 0.5 | -3.8 | 1.6E-04 | 2.2E-04 | MB231/MB468 |
| >hg38_21_+_8433172_8446622_RNA45SN1_45S_50ntFlanks@7974.8007.34\|ACGCGACCTCAGATCAGACGTGGCGACCCGCTGA | >hg38_21_+_8433172_8446622_RNA45SN1_45S_50ntFlanks | 7974 | 8007 | 34 | ACGCGACCTCAGATCAGACGTGGCGACCCGCTGA | 1015.2 | 2.9 | 0.8 | 3.8 | 1.6E-04 | 2.2E-04 | MB231/MB468 |
| >hg38_21_+_8433172_8446622_RNA45SN1_45S_50ntFlanks@12413.12432.20\|TCCTTCGATGTCGGCTCTTC | >hg38_21_+_8433172_8446622_RNA45SN1_45S_50ntFlanks | 12413 | 12432 | 20 | TCCTTCGATGTCGGCTCTTC | 334.2 | -2.1 | 0.6 | -3.8 | 1.6E-04 | 2.2E-04 | MB231/MB468 |
| >hg38_21_+_8433172_8446622_RNA45SN1_45S_50ntFlanks@10389.10411.23\|TGAACAGCAGTTGAACATGGGTC | >hg38_21_+_8433172_8446622_RNA45SN1_45S_50ntFlanks | 10389 | 10411 | 23 | TGAACAGCAGTTGAACATGGGTC | 123.6 | -1.7 | 0.5 | -3.8 | 1.6E-04 | 2.2E-04 | MB231/MB468 |
| >hg38_21_+_8433172_8446622_RNA45SN1_45S_50ntFlanks@6651.6692.42\|CGACTCTTAGCGGTGGATCACTCGGCTCGTGCGTCGATGAAG | >hg38_21_+_8433172_8446622_RNA45SN1_45S_50ntFlanks | 6651 | 6692 | 42 | CGACTCTTAGCGGTGGATCACTCGGCTCGTGCGTCGATGAAG | 4106.5 | 2.4 | 0.6 | 3.7 | 1.8E-04 | 2.2E-04 | BT20/MB468 |
| >hg38_21_+_8433172_8446622_RNA45SN1_45S_50ntFlanks@12420.12460.41\|ATGTCGGCTCTTCCTATCATTGTGAAGCAGAATTCACCAAG | >hg38_21_+_8433172_8446622_RNA45SN1_45S_50ntFlanks | 12420 | 12460 | 41 | ATGTCGGCTCTTCCTATCATTGTGAAGCAGAATTCACCAAG | 246.4 | -3.2 | 0.8 | -4.3 | 2.0E-05 | 2.2E-04 | BT20/MB231 |
| >hg38_21_+_8433172_8446622_RNA45SN1_45S_50ntFlanks@10793.10818.26\|TCTGGCATGTTGGAACAATGTAGGTA | >hg38_21_+_8433172_8446622_RNA45SN1_45S_50ntFlanks | 10793 | 10818 | 26 | TCTGGCATGTTGGAACAATGTAGGTA | 486.6 | -3.3 | 0.9 | -3.7 | 1.8E-04 | 2.2E-04 | BT20/MB468 |
| >hg38_21_+_8433172_8446622_RNA45SN1_45S_50ntFlanks@12932.12972.41\|CGCTAAACCATTCGTAGACGACCTGCTTCTGGGTCGGGGTT | >hg38_21_+_8433172_8446622_RNA45SN1_45S_50ntFlanks | 12932 | 12972 | 41 | CGCTAAACCATTCGTAGACGACCTGCTTCTGGGTCGGGGTT | 199.1 | -2.7 | 0.7 | -3.8 | 1.6E-04 | 2.2E-04 | MB231/MB468 |
| >MI0000286\|hsa-mir-210&WithFlank&11\|-\|568083\|568204@72.93.22(+1G)&[MIMAT0000267&hsa-miR-210-3p&offsets\|0\|0(+1G);m-102&11\|-\|568112\|568133&offsets\|0\|0(+1G)]\|CTGTGCGTGTGACAGCGGCTGAG | >MI0000286\|hsa-mir-210&WithFlank&11\|-\|568083\|568204 | 72 | 93 | 22(+1G) | CTGTGCGTGTGACAGCGGCTGAG | 684.5 | 5.1 | 1.2 | 4.3 | 2.1E-05 | 2.3E-04 | BT20/MB231 |
| >hg38_21_+_8433172_8446622_RNA45SN1_45S_50ntFlanks@4047.4073.27\|ATTCGAACGTCTGCCCTATCAACTTTC | >hg38_21_+_8433172_8446622_RNA45SN1_45S_50ntFlanks | 4047 | 4073 | 27 | ATTCGAACGTCTGCCCTATCAACTTTC | 102.6 | -2.1 | 0.6 | -3.7 | 1.8E-04 | 2.3E-04 | BT20/MB468 |
| >hg38_21_+_8433172_8446622_RNA45SN1_45S_50ntFlanks@8189.8212.24\|CCCAGCCCGTGGACGGTGTGAGGC | >hg38_21_+_8433172_8446622_RNA45SN1_45S_50ntFlanks | 8189 | 8212 | 24 | CCCAGCCCGTGGACGGTGTGAGGC | 230.5 | -2.9 | 0.8 | -3.8 | 1.7E-04 | 2.3E-04 | MB231/MB468 |

<table>
<tr><td>>hg38_21_+_8433172_8446622_RNA45SN1_45S_50ntFlanks@3851.3881.31|ATAACTGTGGTAATTCTAGAGCTAATACATG</td><td>>hg38_21_+_8433172_8446622_RNA45SN1_45S_50ntFlanks</td><td>3851</td><td>3881</td><td>31</td><td>ATAACTGTGGTAATTCTAGAGCTAATACATG</td><td>118.8</td><td>-2.2</td><td>0.6</td><td>-3.7</td><td>1.8E-04</td><td>2.3E-04</td><td>BT20/MB468</td></tr>
<tr><td>>hg38_21_+_8433172_8446622_RNA45SN1_45S_50ntFlanks@4047.4064.18|ATTCGAACGTCTGCCCTA</td><td>>hg38_21_+_8433172_8446622_RNA45SN1_45S_50ntFlanks</td><td>4047</td><td>4064</td><td>18</td><td>ATTCGAACGTCTGCCCTA</td><td>197.9</td><td>-3.7</td><td>1.0</td><td>-3.7</td><td>1.9E-04</td><td>2.3E-04</td><td>BT20/MB468</td></tr>
<tr><td>>hg38_21_+_8433172_8446622_RNA45SN1_45S_50ntFlanks@10797.10825.29|GCATGTTGGAACAATGTAGGTAAGGGAAG</td><td>>hg38_21_+_8433172_8446622_RNA45SN1_45S_50ntFlanks</td><td>10797</td><td>10825</td><td>29</td><td>GCATGTTGGAACAATGTAGGTAAGGGAAG</td><td>175.2</td><td>-4.2</td><td>1.1</td><td>-3.7</td><td>1.9E-04</td><td>2.3E-04</td><td>BT20/MB468</td></tr>
<tr><td>>hg38_21_+_8433172_8446622_RNA45SN1_45S_50ntFlanks@10425.10453.29|AGATGGGCGAGCGCCGTTCCGAAGGGACG</td><td>>hg38_21_+_8433172_8446622_RNA45SN1_45S_50ntFlanks</td><td>10425</td><td>10453</td><td>29</td><td>AGATGGGCGAGCGCCGTTCCGAAGGGACG</td><td>174.9</td><td>-1.5</td><td>0.4</td><td>-3.8</td><td>1.7E-04</td><td>2.3E-04</td><td>MB231/MB468</td></tr>
<tr><td>>MI0000446|hsa-mir-125b-1&WithFlank&11|-|122099751|122099850@21.40.20;MI0000470|hsa-mir-125b-2&WithFlank&21|+|16590231|16590331@23.42.20&[MIMAT0000423&hsa-miR-125b-5p&offsets|0|-2;m-67&11|-|122099809|122099830&offsets|0|-2];[MIMAT0000423_1&hsa-miR-125b-5p&offsets|0|-2;m-68&21|+|16590253|16590274&offsets|0|-2]|TCCCTGAGACCCTAACTTGT</td><td>>MI0000446|hsa-mir-125b-1&WithFlank&11|-|122099751|122099850</td><td>21</td><td>40</td><td>20</td><td>TCCCTGAGACCCTAACTTGT</td><td>1421.9</td><td>-5.0</td><td>1.2</td><td>-4.3</td><td>2.1E-05</td><td>2.3E-04</td><td>BT20/MB231</td></tr>
<tr><td>>hg38_21_+_8433172_8446622_RNA45SN1_45S_50ntFlanks@12413.12432.20|TCCTTCGATGTCGGCTCTTC</td><td>>hg38_21_+_8433172_8446622_RNA45SN1_45S_50ntFlanks</td><td>12413</td><td>12432</td><td>20</td><td>TCCTTCGATGTCGGCTCTTC</td><td>369.6</td><td>-3.2</td><td>0.9</td><td>-3.7</td><td>1.9E-04</td><td>2.3E-04</td><td>BT20/MB468</td></tr>
<tr><td>>MI0000301|hsa-mir-224&WithFlank&X|-|151958572|151958664@14.35.22&[MIMAT0000281&hsa-miR-224-5p&offsets|+1|-2;m-238&X|-|151958630|151958651&offsets|0|0]|CAAGTCACTAGTGGTTCCGTTT</td><td>>MI0000301|hsa-mir-224&WithFlank&X|-|151958572|151958664</td><td>14</td><td>35</td><td>22</td><td>CAAGTCACTAGTGGTTCCGTTT</td><td>12788.0</td><td>2.7</td><td>0.6</td><td>4.3</td><td>2.1E-05</td><td>2.3E-04</td><td>BT20/MB231</td></tr>
<tr><td>>hg38_21_+_8433172_8446622_RNA45SN1_45S_50ntFlanks@10775.10796.22|TCTCCAAGGTGAACAGCCTCTG</td><td>>hg38_21_+_8433172_8446622_RNA45SN1_45S_50ntFlanks</td><td>10775</td><td>10796</td><td>22</td><td>TCTCCAAGGTGAACAGCCTCTG</td><td>109.9</td><td>-3.0</td><td>0.8</td><td>-3.7</td><td>1.9E-04</td><td>2.3E-04</td><td>BT20/MB468</td></tr>
<tr><td>>MI0000446|hsa-mir-125b-1&WithFlank&11|-|122099751|122099850@21.40.20(+1A);MI0000470|hsa-mir-125b-2&WithFlank&21|+|16590231|16590331@23.42.20(+1A)&[MIMAT0000423&hsa-miR-125b-5p&offsets|0|-2(+1A);m-67&11|-|122099809|122099830&offsets|0|-2(+1A)];[MIMAT0000423_1&hsa-miR-125b-5p&offsets|0|-2(+1A);m-68&21|+|16590253|16590274&offsets|0|-2(+1A)]|TCCCTGAGACCCTAACTTGTA</td><td>>MI0000446|hsa-mir-125b-1&WithFlank&11|-|122099751|122099850</td><td>21</td><td>40</td><td>20(+1A)</td><td>TCCCTGAGACCCTAACTTGTA</td><td>841.7</td><td>-4.6</td><td>1.1</td><td>-4.2</td><td>2.2E-05</td><td>2.4E-04</td><td>BT20/MB231</td></tr>
<tr><td>>hg38_21_+_8433172_8446622_RNA45SN1_45S_50ntFlanks@8358.8379.22|AAAGAACTTTGAAGAGAGAGTT</td><td>>hg38_21_+_8433172_8446622_RNA45SN1_45S_50ntFlanks</td><td>8358</td><td>8379</td><td>22</td><td>AAAGAACTTTGAAGAGAGAGTT</td><td>1372.9</td><td>-3.1</td><td>0.8</td><td>-3.7</td><td>1.9E-04</td><td>2.4E-04</td><td>BT20/MB468</td></tr>
<tr><td>>hg38_21_+_8433172_8446622_RNA45SN1_45S_50ntFlanks@12458.12484.27|AAGCGTTGGATTGTTCACCCACTAATA</td><td>>hg38_21_+_8433172_8446622_RNA45SN1_45S_50ntFlanks</td><td>12458</td><td>12484</td><td>27</td><td>AAGCGTTGGATTGTTCACCCACTAATA</td><td>298.8</td><td>-2.6</td><td>0.7</td><td>-3.7</td><td>1.9E-04</td><td>2.4E-04</td><td>BT20/MB468</td></tr>
<tr><td>>hg38_21_+_8433172_8446622_RNA45SN1_45S_50ntFlanks@12914.12933.20|CGTTCGTGGGGAACCTGGCG</td><td>>hg38_21_+_8433172_8446622_RNA45SN1_45S_50ntFlanks</td><td>12914</td><td>12933</td><td>20</td><td>CGTTCGTGGGGAACCTGGCG</td><td>211.1</td><td>-3.6</td><td>1.0</td><td>-3.7</td><td>1.9E-04</td><td>2.4E-04</td><td>BT20/MB468</td></tr>
<tr><td>>hg38_21_+_8433172_8446622_RNA45SN1_45S_50ntFlanks@10402.10428.27|AACATGGGTCAGTCGGTCCTGAGAGAT</td><td>>hg38_21_+_8433172_8446622_RNA45SN1_45S_50ntFlanks</td><td>10402</td><td>10428</td><td>27</td><td>AACATGGGTCAGTCGGTCCTGAGAGAT</td><td>176.1</td><td>-1.7</td><td>0.5</td><td>-3.7</td><td>1.9E-04</td><td>2.4E-04</td><td>BT20/MB468</td></tr>
<tr><td>>hg38_21_+_8433172_8446622_RNA45SN1_45S_50ntFlanks@11572.11593.22|CCGACTTAGAACTGGTGCGGAC</td><td>>hg38_21_+_8433172_8446622_RNA45SN1_45S_50ntFlanks</td><td>11572</td><td>11593</td><td>22</td><td>CCGACTTAGAACTGGTGCGGAC</td><td>495.9</td><td>-2.2</td><td>0.6</td><td>-3.8</td><td>1.7E-04</td><td>2.4E-04</td><td>MB231/MB468</td></tr>
<tr><td>>hg38_21_+_8433172_8446622_RNA45SN1_45S_50ntFlanks@4367.4396.30|CAGTTAAAAAGCTCGTAGTTGGATCTTGGG</td><td>>hg38_21_+_8433172_8446622_RNA45SN1_45S_50ntFlanks</td><td>4367</td><td>4396</td><td>30</td><td>CAGTTAAAAAGCTCGTAGTTGGATCTTGGG</td><td>110.9</td><td>-3.1</td><td>0.8</td><td>-3.8</td><td>1.7E-04</td><td>2.4E-04</td><td>MB231/MB468</td></tr>
<tr><td>>MI0001733|hsa-mir-452&WithFlank&X|-|151959622|151959718@20.43.24&[MIMAT0001635&hsa-miR-452-5p&offsets|0|+2;m-188&X|-|151959677|151959699&offsets|0|+1]|AACTGTTTGCAGAGGAAACTGAGA</td><td>>MI0001733|hsa-mir-452&WithFlank&X|-|151959622|151959718</td><td>20</td><td>43</td><td>24</td><td>AACTGTTTGCAGAGGAAACTGAGA</td><td>1493.4</td><td>2.9</td><td>0.7</td><td>4.2</td><td>2.2E-05</td><td>2.4E-04</td><td>BT20/MB231</td></tr>
<tr><td>>hg38_21_+_8433172_8446622_RNA45SN1_45S_50ntFlanks@10794.10813.20|CTGGCATGTTGGAACAATGT</td><td>>hg38_21_+_8433172_8446622_RNA45SN1_45S_50ntFlanks</td><td>10794</td><td>10813</td><td>20</td><td>CTGGCATGTTGGAACAATGT</td><td>110.9</td><td>-2.7</td><td>0.7</td><td>-3.8</td><td>1.8E-04</td><td>2.4E-04</td><td>MB231/MB468</td></tr>
<tr><td>>hg38_21_+_8433172_8446622_RNA45SN1_45S_50ntFlanks@8194.8216.23|CCCGTGGACGGTGTGAGGCCGGT</td><td>>hg38_21_+_8433172_8446622_RNA45SN1_45S_50ntFlanks</td><td>8194</td><td>8216</td><td>23</td><td>CCCGTGGACGGTGTGAGGCCGGT</td><td>152.1</td><td>-2.4</td><td>0.6</td><td>-3.7</td><td>1.9E-04</td><td>2.4E-04</td><td>BT20/MB468</td></tr>
<tr><td>>hg38_21_+_8433172_8446622_RNA45SN1_45S_50ntFlanks@11552.11569.18|CCTCGGCCGGCGCCTAGC</td><td>>hg38_21_+_8433172_8446622_RNA45SN1_45S_50ntFlanks</td><td>11552</td><td>11569</td><td>18</td><td>CCTCGGCCGGCGCCTAGC</td><td>278.6</td><td>-2.1</td><td>0.6</td><td>-3.7</td><td>2.0E-04</td><td>2.4E-04</td><td>BT20/MB468</td></tr>
<tr><td>>trna119_LysCTT_1_-_145395522_145395594@1.33.33,trna11_LysCTT_5_-_180648979_180649051@1.33.33,trna13_LysCTT_6_+_26556774_26556846@1.33.33,trna7_LysCTT_16_+_3225692_3225764@1.33.33,trna9_LysCTT_5_+_180634755_180634827@1.33.33|GCCCGGCTAGCTCAGTCGGTAGAGCATGAGACT</td><td>>trna119_LysCTT_1_-_145395522_145395594</td><td>1</td><td>33</td><td>33</td><td>GCCCGGCTAGCTCAGTCGGTAGAGCATGAGACT</td><td>15283.8</td><td>-3.6</td><td>0.8</td><td>-4.2</td><td>2.2E-05</td><td>2.4E-04</td><td>BT20/MB231</td></tr>
<tr><td>>hg38_21_+_8433172_8446622_RNA45SN1_45S_50ntFlanks@11767.11794.28|TCTAATTAGTGACGCGCATGAATGGATG</td><td>>hg38_21_+_8433172_8446622_RNA45SN1_45S_50ntFlanks</td><td>11767</td><td>11794</td><td>28</td><td>TCTAATTAGTGACGCGCATGAATGGATG</td><td>1120.7</td><td>-2.1</td><td>0.6</td><td>-3.7</td><td>1.8E-04</td><td>2.4E-04</td><td>MB231/MB468</td></tr>
<tr><td>>hg38_1_-_228634819_228635039_RNA5S12_5S_50ntFlanks@84.111.28|CTCGTCTGATCTCGGAAGCTAAGCAGGG</td><td>>hg38_1_-_228634819_228635039_RNA5S12_5S_50ntFlanks</td><td>84</td><td>111</td><td>28</td><td>CTCGTCTGATCTCGGAAGCTAAGCAGGG</td><td>76.1</td><td>-3.3</td><td>0.9</td><td>-3.7</td><td>1.8E-04</td><td>2.4E-04</td><td>MB231/MB468</td></tr>
<tr><td>>hg38_21_+_8433172_8446622_RNA45SN1_45S_50ntFlanks@12405.12444.40|CTTTTTGATCCTTCGATGTCGGCTCTTCCTATCATTGTGA</td><td>>hg38_21_+_8433172_8446622_RNA45SN1_45S_50ntFlanks</td><td>12405</td><td>12444</td><td>40</td><td>CTTTTTGATCCTTCGATGTCGGCTCTTCCTATCATTGTGA</td><td>260.4</td><td>-3.6</td><td>1.0</td><td>-3.7</td><td>2.0E-04</td><td>2.4E-04</td><td>BT20/MB468</td></tr>
<tr><td>>hg38_21_+_8433172_8446622_RNA45SN1_45S_50ntFlanks@9198.9222.25|GTTCTCTCGGGGCCACGCGCGCGTC</td><td>>hg38_21_+_8433172_8446622_RNA45SN1_45S_50ntFlanks</td><td>9198</td><td>9222</td><td>25</td><td>GTTCTCTCGGGGCCACGCGCGCGTC</td><td>385.3</td><td>-3.9</td><td>1.1</td><td>-3.7</td><td>2.0E-04</td><td>2.4E-04</td><td>BT20/MB468</td></tr>
<tr><td>>hg38_21_+_8433172_8446622_RNA45SN1_45S_50ntFlanks@12997.13040.44|CGCTGCGATCTATTGAAAGTCAGCCCTCGACACAAGGGTTTGTC</td><td>>hg38_21_+_8433172_8446622_RNA45SN1_45S_50ntFlanks</td><td>12997</td><td>13040</td><td>44</td><td>CGCTGCGATCTATTGAAAGTCAGCCCTCGACACAAGGGTTTGTC</td><td>532.2</td><td>-4.5</td><td>1.1</td><td>-4.2</td><td>2.3E-05</td><td>2.5E-04</td><td>BT20/MB231</td></tr>
<tr><td>>hg38_21_+_8433172_8446622_RNA45SN1_45S_50ntFlanks@3811.3834.24|AGTTATGGTTCCTTTGGTCGCTCG</td><td>>hg38_21_+_8433172_8446622_RNA45SN1_45S_50ntFlanks</td><td>3811</td><td>3834</td><td>24</td><td>AGTTATGGTTCCTTTGGTCGCTCG</td><td>92.4</td><td>-2.0</td><td>0.5</td><td>-3.7</td><td>1.8E-04</td><td>2.5E-04</td><td>MB231/MB468</td></tr>
<tr><td>>hg38_21_+_8433172_8446622_RNA45SN1_45S_50ntFlanks@11767.11790.24|TCTAATTAGTGACGCGCATGAATG</td><td>>hg38_21_+_8433172_8446622_RNA45SN1_45S_50ntFlanks</td><td>11767</td><td>11790</td><td>24</td><td>TCTAATTAGTGACGCGCATGAATG</td><td>171.8</td><td>-1.6</td><td>0.4</td><td>-3.7</td><td>1.8E-04</td><td>2.5E-04</td><td>MB231/MB468</td></tr>
<tr><td>>MI0000650|hsa-mir-200c&WithFlank&12|+|6963693|6963772@50.71.22&[MIMAT0000617&hsa-miR-200c-3p&offsets|0|-1;m-54&12|+|6963742|6963764&offsets|0|-1]|TAATACTGCCGGGTAATGATGG</td><td>>MI0000650|hsa-mir-200c&WithFlank&12|+|6963693|6963772</td><td>50</td><td>71</td><td>22</td><td>TAATACTGCCGGGTAATGATGG</td><td>209.9</td><td>-2.2</td><td>0.6</td><td>-3.7</td><td>1.8E-04</td><td>2.5E-04</td><td>MB231/MB468</td></tr>
<tr><td>>hg38_21_+_8433172_8446622_RNA45SN1_45S_50ntFlanks@12990.13013.24|GCTCCCTCGCTGCGATCTATTGAA</td><td>>hg38_21_+_8433172_8446622_RNA45SN1_45S_50ntFlanks</td><td>12990</td><td>13013</td><td>24</td><td>GCTCCCTCGCTGCGATCTATTGAA</td><td>1165.4</td><td>-3.0</td><td>0.8</td><td>-3.7</td><td>2.0E-04</td><td>2.5E-04</td><td>BT20/MB468</td></tr>
<tr><td>>hg38_21_+_8433172_8446622_RNA45SN1_45S_50ntFlanks@10687.10708.22|CGGCGTCCGGTGAGCTCTCGCT</td><td>>hg38_21_+_8433172_8446622_RNA45SN1_45S_50ntFlanks</td><td>10687</td><td>10708</td><td>22</td><td>CGGCGTCCGGTGAGCTCTCGCT</td><td>1272.5</td><td>-2.4</td><td>0.7</td><td>-3.7</td><td>1.8E-04</td><td>2.5E-04</td><td>MB231/MB468</td></tr>
<tr><td>>hg38_21_+_8433172_8446622_RNA45SN1_45S_50ntFlanks@6647.6666.20|CGTACGACTCTTAGCGGTGG</td><td>>hg38_21_+_8433172_8446622_RNA45SN1_45S_50ntFlanks</td><td>6647</td><td>6666</td><td>20</td><td>CGTACGACTCTTAGCGGTGG</td><td>2341.6</td><td>2.7</td><td>0.7</td><td>3.7</td><td>2.0E-04</td><td>2.5E-04</td><td>BT20/MB468</td></tr>
<tr><td>>MI0002470|hsa-mir-486-1&WithFlank&8|-|41660435|41660514@10.31.22;MI0023622|hsa-mir-486-2&WithFlank&8|+|41660438|41660513@7.28.22&[MIMAT0002177&hsa-miR-486-5p&offsets|0|0;m-107&8|-|41660484|41660505&offsets|0|0];[MIMAT0002177_1&hsa-miR-486-5p&offsets|0|0;TJU_CMC_MD2.ID03138.5p-miR&offsets|0|-2;m-106&8|+|41660444|41660465&offsets|0|0]|TCCTGTACTGAGCTGCCCCGAG</td><td>>MI0002470|hsa-mir-486-1&WithFlank&8|-|41660435|41660514</td><td>10</td><td>31</td><td>22</td><td>TCCTGTACTGAGCTGCCCCGAG</td><td>679.6</td><td>9.3</td><td>2.5</td><td>3.7</td><td>1.9E-04</td><td>2.5E-04</td><td>MB231/MB468</td></tr>
<tr><td>>MI0000063|hsa-let-7b&WithFlank&22|+|46113680|46113774@12.32.21(+1C)&[MIMAT0000063&hsa-let-7b-5p&offsets|0|-1(+1C);m-14&22|+|46113691|46113712&offsets|0|-1(+1C)]|TGAGGTAGTAGGTTGTGTGGTC</td><td>>MI0000063|hsa-let-7b&WithFlank&22|+|46113680|46113774</td><td>12</td><td>32</td><td>21(+1C)</td><td>TGAGGTAGTAGGTTGTGTGGTC</td><td>6752.6</td><td>2.6</td><td>0.6</td><td>4.2</td><td>2.4E-05</td><td>2.5E-04</td><td>BT20/MB231</td></tr>
<tr><td>>hg38_21_+_8433172_8446622_RNA45SN1_45S_50ntFlanks@11672.11709.38|CAGTGCTCTGAATGTCAAAGTGAAGAAATTCAATGAAG</td><td>>hg38_21_+_8433172_8446622_RNA45SN1_45S_50ntFlanks</td><td>11672</td><td>11709</td><td>38</td><td>CAGTGCTCTGAATGTCAAAGTGAAGAAATTCAATGAAG</td><td>311.2</td><td>-3.6</td><td>0.8</td><td>-4.2</td><td>2.4E-05</td><td>2.5E-04</td><td>BT20/MB231</td></tr>
<tr><td>>hg38_21_+_8433172_8446622_RNA45SN1_45S_50ntFlanks@11671.11704.34|CCAGTGCTCTGAATGTCAAAGTGAAGAAATTCAA</td><td>>hg38_21_+_8433172_8446622_RNA45SN1_45S_50ntFlanks</td><td>11671</td><td>11704</td><td>34</td><td>CCAGTGCTCTGAATGTCAAAGTGAAGAAATTCAA</td><td>314.7</td><td>-3.6</td><td>1.0</td><td>-3.7</td><td>2.1E-04</td><td>2.5E-04</td><td>BT20/MB468</td></tr>
<tr><td>>MI0003778|hsa-mir-320c-1&WithFlank&18|+|21683512|21683595@48.67.20;MI0008191|hsa-mir-320c-2&WithFlank&18|+|24321669|24321752@48.67.20&[MIMAT0005793&hsa-miR-320c&offsets|0|0;m-222&18|+|21683559|21683578&offsets|0|0];[MIMAT0005793_1&hsa-miR-320c&offsets|0|0;m-219&18|+|24321716|24321735&offsets|0|0]|AAAAGCTGGGTTGAGAGGGT</td><td>>MI0003778|hsa-mir-320c-1&WithFlank&18|+|21683512|21683595</td><td>48</td><td>67</td><td>20</td><td>AAAAGCTGGGTTGAGAGGGT</td><td>625.7</td><td>3.0</td><td>0.7</td><td>4.2</td><td>2.4E-05</td><td>2.6E-04</td><td>BT20/MB231</td></tr>
<tr><td>>hg38_21_+_8433172_8446622_RNA45SN1_45S_50ntFlanks@12933.12965.33|GCTAAACCATTCGTAGACGACCTGCTTCTGGGT</td><td>>hg38_21_+_8433172_8446622_RNA45SN1_45S_50ntFlanks</td><td>12933</td><td>12965</td><td>33</td><td>GCTAAACCATTCGTAGACGACCTGCTTCTGGGT</td><td>112.1</td><td>-2.6</td><td>0.7</td><td>-3.7</td><td>2.1E-04</td><td>2.6E-04</td><td>BT20/MB468</td></tr>
<tr><td>>hg38_21_+_8433172_8446622_RNA45SN1_45S_50ntFlanks@10509.10533.25|GAATCCGGAGTGGCGGAGATGGGCG</td><td>>hg38_21_+_8433172_8446622_RNA45SN1_45S_50ntFlanks</td><td>10509</td><td>10533</td><td>25</td><td>GAATCCGGAGTGGCGGAGATGGGCG</td><td>116.5</td><td>-3.2</td><td>0.9</td><td>-3.7</td><td>2.1E-04</td><td>2.6E-04</td><td>BT20/MB468</td></tr>
<tr><td>>hg38_21_+_8433172_8446622_RNA45SN1_45S_50ntFlanks@11316.11336.21|CACCCCACGTCTCGTCGCGCG</td><td>>hg38_21_+_8433172_8446622_RNA45SN1_45S_50ntFlanks</td><td>11316</td><td>11336</td><td>21</td><td>CACCCCACGTCTCGTCGCGCG</td><td>1134.3</td><td>-4.4</td><td>1.2</td><td>-3.7</td><td>2.1E-04</td><td>2.6E-04</td><td>BT20/MB468</td></tr>
<tr><td>>MI0000102|hsa-mir-100&WithFlank&11|-|122152223|122152314@19.39.21(+1A)&[MIMAT0000098&hsa-miR-100-5p&offsets|0|-1(+1A);m-27&11|-|122152276|122152296&offsets|0|0(+1A)]|AACCCGTAGATCCGAACTTGTA</td><td>>MI0000102|hsa-mir-100&WithFlank&11|-|122152223|122152314</td><td>19</td><td>39</td><td>21(+1A)</td><td>AACCCGTAGATCCGAACTTGTA</td><td>7830.8</td><td>9.5</td><td>2.6</td><td>3.7</td><td>2.1E-04</td><td>2.6E-04</td><td>BT20/MB468</td></tr>
<tr><td>>hg38_21_+_8433172_8446622_RNA45SN1_45S_50ntFlanks@10664.10706.43|CTTGGAAAGCGTCGCGGTTCCGGCGGCGTCCGGTGAGCTCTCG</td><td>>hg38_21_+_8433172_8446622_RNA45SN1_45S_50ntFlanks</td><td>10664</td><td>10706</td><td>43</td><td>CTTGGAAAGCGTCGCGGTTCCGGCGGCGTCCGGTGAGCTCTCG</td><td>196.3</td><td>-3.5</td><td>0.9</td><td>-3.7</td><td>1.9E-04</td><td>2.6E-04</td><td>MB231/MB468</td></tr>
</table>

<table>
<tr><th></th><th></th><th></th><th></th><th></th><th></th><th></th><th></th><th></th><th></th><th></th><th></th><th></th></tr>
<tr><td>>hg38_21_+_8433172_8446622_RNA45SN1_45S_50ntFlanks@12937.12961.25|AACCATTCGTAGACGACCTGCTTCT</td><td>>hg38_21_+_8433172_8446622_RNA45SN1_45S_50ntFlanks</td><td>12937</td><td>12961</td><td>25</td><td>AACCATTCGTAGACGACCTGCTTCT</td><td>208.3</td><td>-1.6</td><td>0.4</td><td>-3.7</td><td>1.9E-04</td><td>2.6E-04</td><td>MB231/MB468</td></tr>
<tr><td>>hg38_21_+_8433172_8446622_RNA45SN1_45S_50ntFlanks@6661.6697.37|CGGTGGATCACTCGGCTCGTGCGTCGATGAAGAACGC</td><td>>hg38_21_+_8433172_8446622_RNA45SN1_45S_50ntFlanks</td><td>6661</td><td>6697</td><td>37</td><td>CGGTGGATCACTCGGCTCGTGCGTCGATGAAGAACGC</td><td>733.8</td><td>1.8</td><td>0.5</td><td>3.7</td><td>1.9E-04</td><td>2.6E-04</td><td>MB231/MB468</td></tr>
<tr><td>>hg38_21_+_8433172_8446622_RNA45SN1_45S_50ntFlanks@4220.4242.23|ACGAAAAATAACAATACAGGACT</td><td>>hg38_21_+_8433172_8446622_RNA45SN1_45S_50ntFlanks</td><td>4220</td><td>4242</td><td>23</td><td>ACGAAAAATAACAATACAGGACT</td><td>478.9</td><td>-2.8</td><td>0.7</td><td>-3.7</td><td>2.2E-04</td><td>2.7E-04</td><td>BT20/MB468</td></tr>
<tr><td>>hg38_21_+_8433172_8446622_RNA45SN1_45S_50ntFlanks@6651.6670.20|CGACTCTTAGCGGTGGATCA</td><td>>hg38_21_+_8433172_8446622_RNA45SN1_45S_50ntFlanks</td><td>6651</td><td>6670</td><td>20</td><td>CGACTCTTAGCGGTGGATCA</td><td>901.7</td><td>-2.0</td><td>0.5</td><td>-3.7</td><td>2.0E-04</td><td>2.7E-04</td><td>MB231/MB468</td></tr>
<tr><td>>hg38_21_+_8433172_8446622_RNA45SN1_45S_50ntFlanks@11319.11338.20|CCCACGTCTCGTCGCGCGCG</td><td>>hg38_21_+_8433172_8446622_RNA45SN1_45S_50ntFlanks</td><td>11319</td><td>11338</td><td>20</td><td>CCCACGTCTCGTCGCGCGCG</td><td>98.5</td><td>-3.7</td><td>1.0</td><td>-3.7</td><td>2.2E-04</td><td>2.7E-04</td><td>BT20/MB468</td></tr>
<tr><td>>hg38_21_+_8433172_8446622_RNA45SN1_45S_50ntFlanks@8359.8378.20|AAGAACTTTGAAGAGAGAGT</td><td>>hg38_21_+_8433172_8446622_RNA45SN1_45S_50ntFlanks</td><td>8359</td><td>8378</td><td>20</td><td>AAGAACTTTGAAGAGAGAGT</td><td>602.3</td><td>-2.2</td><td>0.6</td><td>-3.7</td><td>2.2E-04</td><td>2.7E-04</td><td>BT20/MB468</td></tr>
<tr><td>>hg38_21_+_8433172_8446622_RNA45SN1_45S_50ntFlanks@12628.12652.25|TGGGGCGAAGCTACCATCTGTGGGA</td><td>>hg38_21_+_8433172_8446622_RNA45SN1_45S_50ntFlanks</td><td>12628</td><td>12652</td><td>25</td><td>TGGGGCGAAGCTACCATCTGTGGGA</td><td>82.7</td><td>-2.8</td><td>0.8</td><td>-3.7</td><td>2.0E-04</td><td>2.7E-04</td><td>MB231/MB468</td></tr>
<tr><td>>MI0000446|hsa-mir-125b-1&WithFlank&11|-|122099751|122099850@21.39.19;MI0000470|hsa-mir-125b-2&WithFlank&21|+|16590231|16590331@23.41.19&[MIMAT0000423&hsa-miR-125b-5p&offsets|0|-3;m-67&11|-|122099809|122099830&offsets|0|-3];[MIMAT0000423_1&hsa-miR-125b-5p&offsets|0|-3;m-68&21|+|16590253|16590274&offsets|0|-3]|TCCCTGAGACCCTAACTTG</td><td>>MI0000446|hsa-mir-125b-1&WithFlank&11|-|122099751|122099850</td><td>21</td><td>39</td><td>19</td><td>TCCCTGAGACCCTAACTTG</td><td>1033.8</td><td>-5.1</td><td>1.2</td><td>-4.2</td><td>2.5E-05</td><td>2.7E-04</td><td>BT20/MB231</td></tr>
<tr><td>>hg38_21_+_8433172_8446622_RNA45SN1_45S_50ntFlanks@12420.12449.30|ATGTCGGCTCTTCCTATCATTGTGAAGCAG</td><td>>hg38_21_+_8433172_8446622_RNA45SN1_45S_50ntFlanks</td><td>12420</td><td>12449</td><td>30</td><td>ATGTCGGCTCTTCCTATCATTGTGAAGCAG</td><td>605.1</td><td>-1.6</td><td>0.4</td><td>-3.7</td><td>2.2E-04</td><td>2.7E-04</td><td>BT20/MB468</td></tr>
<tr><td>>hg38_21_+_8433172_8446622_RNA45SN1_45S_50ntFlanks@10771.10818.48|CAGGTCTCCAAGGTGAACAGCCTCTGGCATGTTGGAACAATGTAGGTA</td><td>>hg38_21_+_8433172_8446622_RNA45SN1_45S_50ntFlanks</td><td>10771</td><td>10818</td><td>48</td><td>CAGGTCTCCAAGGTGAACAGCCTCTGGCATGTTGGAACAATGTAGGTA</td><td>183.4</td><td>-3.5</td><td>0.9</td><td>-3.7</td><td>2.0E-04</td><td>2.7E-04</td><td>MB231/MB468</td></tr>
<tr><td>>hg38_21_+_8433172_8446622_RNA45SN1_45S_50ntFlanks@8176.8204.29|CTTCTGATCGAGGCCCAGCCCGTGGACGG</td><td>>hg38_21_+_8433172_8446622_RNA45SN1_45S_50ntFlanks</td><td>8176</td><td>8204</td><td>29</td><td>CTTCTGATCGAGGCCCAGCCCGTGGACGG</td><td>751.4</td><td>-1.9</td><td>0.5</td><td>-3.7</td><td>2.0E-04</td><td>2.7E-04</td><td>MB231/MB468</td></tr>
<tr><td>>hg38_21_+_8433172_8446622_RNA45SN1_45S_50ntFlanks@7981.8012.32|CTCAGATCAGACGTGGCGACCCGCTGAATTTA</td><td>>hg38_21_+_8433172_8446622_RNA45SN1_45S_50ntFlanks</td><td>7981</td><td>8012</td><td>32</td><td>CTCAGATCAGACGTGGCGACCCGCTGAATTTA</td><td>119.9</td><td>-2.2</td><td>0.6</td><td>-3.7</td><td>2.0E-04</td><td>2.7E-04</td><td>MB231/MB468</td></tr>
<tr><td>>MI0003776|hsa-mir-320b-1&WithFlank&1|+|116671740|116671823@48.68.21;MI0003839|hsa-mir-320b-2&WithFlank&1|-|224257034|224257116@47.67.21&[MIMAT0005792&hsa-miR-320b&offsets|0|-1;m-183&1|+|116671787|116671806&offsets|0|+1];[MIMAT0005792_1&hsa-miR-320b&offsets|0|-1;m-183&1|-|224257051|224257070&offsets|0|+1]|AAAAGCTGGGTTGAGAGGGCA</td><td>>MI0003776|hsa-mir-320b-1&WithFlank&1|+|116671740|116671823</td><td>48</td><td>68</td><td>21</td><td>AAAAGCTGGGTTGAGAGGGCA</td><td>1374.8</td><td>3.4</td><td>0.8</td><td>4.2</td><td>2.6E-05</td><td>2.8E-04</td><td>BT20/MB231</td></tr>
<tr><td>>am_trna116_GluCTC_1_-_145399233_145399304@1.29.29,trna59_GluCTC_1_+_249168447_249168518@1.29.29,trna71_GluCTC_1_-_161439189_161439260@1.29.29,trna74_GluCTC_1_-_161431809_161431880@1.29.29,trna77_GluCTC_1_-_161424398_161424469@1.29.29,trna77_GluCTC_6_+_28949976_28950047@1.29.29,trna80_GluCTC_1_-_161417018_161417089@1.29.29,trna87_GluCTC_6_-_126101393_126101464@1.29.29|TCCCTGGTGGTCTAGTGGTTAGGATTCGG</td><td>>am_trna116_GluCTC_1_-_145399233_145399304</td><td>1</td><td>29</td><td>29</td><td>TCCCTGGTGGTCTAGTGGTTAGGATTCGG</td><td>24444.0</td><td>-3.0</td><td>0.8</td><td>-3.7</td><td>2.0E-04</td><td>2.8E-04</td><td>MB231/MB468</td></tr>
<tr><td>>hg38_21_+_8433172_8446622_RNA45SN1_45S_50ntFlanks@7992.8012.21|CGTGGCGACCCGCTGAATTTA</td><td>>hg38_21_+_8433172_8446622_RNA45SN1_45S_50ntFlanks</td><td>7992</td><td>8012</td><td>21</td><td>CGTGGCGACCCGCTGAATTTA</td><td>403.6</td><td>-2.4</td><td>0.6</td><td>-3.7</td><td>2.2E-04</td><td>2.8E-04</td><td>BT20/MB468</td></tr>
<tr><td>>hg38_21_+_8433172_8446622_RNA45SN1_45S_50ntFlanks@7991.8017.27|ACGTGGCGACCCGCTGAATTTAAGCAT</td><td>>hg38_21_+_8433172_8446622_RNA45SN1_45S_50ntFlanks</td><td>7991</td><td>8017</td><td>27</td><td>ACGTGGCGACCCGCTGAATTTAAGCAT</td><td>518.7</td><td>1.8</td><td>0.5</td><td>3.7</td><td>2.0E-04</td><td>2.8E-04</td><td>MB231/MB468</td></tr>
<tr><td>>hg38_21_+_8433172_8446622_RNA45SN1_45S_50ntFlanks@9567.9585.19|ACGTGCAAATCGGTCGTCC</td><td>>hg38_21_+_8433172_8446622_RNA45SN1_45S_50ntFlanks</td><td>9567</td><td>9585</td><td>19</td><td>ACGTGCAAATCGGTCGTCC</td><td>154.6</td><td>-2.2</td><td>0.6</td><td>-3.7</td><td>2.0E-04</td><td>2.8E-04</td><td>MB231/MB468</td></tr>
<tr><td>>hg38_21_+_8433172_8446622_RNA45SN1_45S_50ntFlanks@9569.9597.29|GTGCAAATCGGTCGTCCGACCTGGGTATA</td><td>>hg38_21_+_8433172_8446622_RNA45SN1_45S_50ntFlanks</td><td>9569</td><td>9597</td><td>29</td><td>GTGCAAATCGGTCGTCCGACCTGGGTATA</td><td>786.6</td><td>-2.3</td><td>0.6</td><td>-3.7</td><td>2.1E-04</td><td>2.8E-04</td><td>MB231/MB468</td></tr>
<tr><td>>hg38_21_+_8433172_8446622_RNA45SN1_45S_50ntFlanks@10686.10711.26|GCGGCGTCCGGTGAGCTCTCGCTGGC</td><td>>hg38_21_+_8433172_8446622_RNA45SN1_45S_50ntFlanks</td><td>10686</td><td>10711</td><td>26</td><td>GCGGCGTCCGGTGAGCTCTCGCTGGC</td><td>427.7</td><td>-3.4</td><td>0.9</td><td>-3.7</td><td>2.3E-04</td><td>2.8E-04</td><td>BT20/MB468</td></tr>
<tr><td>>trna134_GluTTC_1_-_16861774_16861845@1.31.31,trna5_GluTTC_1_+_17199078_17199149@1.31.31,trna84_GluTTC_1_-_161391883_161391954@1.31.31,trna94_GluTTC_1_-_149664355_149664427@1.31.31|TCCCTGGTGGTCTAGTGGCTAGGATTCGGCG</td><td>>trna134_GluTTC_1_-_16861774_16861845</td><td>1</td><td>31</td><td>31</td><td>TCCCTGGTGGTCTAGTGGCTAGGATTCGGCG</td><td>1171.8</td><td>-4.1</td><td>1.1</td><td>-3.7</td><td>2.1E-04</td><td>2.8E-04</td><td>MB231/MB468</td></tr>
<tr><td>>hg38_21_+_8433172_8446622_RNA45SN1_45S_50ntFlanks@12607.12625.19|TGTGCTTGGCTGAGGAGCC</td><td>>hg38_21_+_8433172_8446622_RNA45SN1_45S_50ntFlanks</td><td>12607</td><td>12625</td><td>19</td><td>TGTGCTTGGCTGAGGAGCC</td><td>345.5</td><td>-1.6</td><td>0.4</td><td>-3.7</td><td>2.3E-04</td><td>2.8E-04</td><td>BT20/MB468</td></tr>
<tr><td>>am_trnaMT_ValTAC_MT_+_1602_1670@48.67.20|CAACTTAACTTGACCGCTCT</td><td>>am_trnaMT_ValTAC_MT_+_1602_1670</td><td>48</td><td>67</td><td>20</td><td>CAACTTAACTTGACCGCTCT</td><td>7842.0</td><td>8.6</td><td>2.3</td><td>3.7</td><td>2.3E-04</td><td>2.8E-04</td><td>BT20/MB468</td></tr>
<tr><td>>hg38_21_+_8433172_8446622_RNA45SN1_45S_50ntFlanks@3777.3805.29|CGGTACAGTGAAACTGCGAATGGCTCATT</td><td>>hg38_21_+_8433172_8446622_RNA45SN1_45S_50ntFlanks</td><td>3777</td><td>3805</td><td>29</td><td>CGGTACAGTGAAACTGCGAATGGCTCATT</td><td>137.8</td><td>-2.1</td><td>0.6</td><td>-3.7</td><td>2.3E-04</td><td>2.8E-04</td><td>BT20/MB468</td></tr>
<tr><td>>hg38_21_+_8433172_8446622_RNA45SN1_45S_50ntFlanks@10841.10872.32|CGTAACTTCGGGATAAGGATTGGCTCTAAGGG</td><td>>hg38_21_+_8433172_8446622_RNA45SN1_45S_50ntFlanks</td><td>10841</td><td>10872</td><td>32</td><td>CGTAACTTCGGGATAAGGATTGGCTCTAAGGG</td><td>346.1</td><td>-2.2</td><td>0.6</td><td>-3.7</td><td>2.1E-04</td><td>2.8E-04</td><td>MB231/MB468</td></tr>
<tr><td>>hg38_21_+_8433172_8446622_RNA45SN1_45S_50ntFlanks@6731.6753.23|CACATTGATCATCGACACTTCGA</td><td>>hg38_21_+_8433172_8446622_RNA45SN1_45S_50ntFlanks</td><td>6731</td><td>6753</td><td>23</td><td>CACATTGATCATCGACACTTCGA</td><td>283.0</td><td>-2.2</td><td>0.6</td><td>-3.7</td><td>2.1E-04</td><td>2.8E-04</td><td>MB231/MB468</td></tr>
<tr><td>>hg38_21_+_8433172_8446622_RNA45SN1_45S_50ntFlanks@10320.10341.22|CAGATCTTGGTGGTAGTAGCAA</td><td>>hg38_21_+_8433172_8446622_RNA45SN1_45S_50ntFlanks</td><td>10320</td><td>10341</td><td>22</td><td>CAGATCTTGGTGGTAGTAGCAA</td><td>81.0</td><td>-2.6</td><td>0.7</td><td>-3.7</td><td>2.1E-04</td><td>2.8E-04</td><td>MB231/MB468</td></tr>
<tr><td>>hg38_21_+_8433172_8446622_RNA45SN1_45S_50ntFlanks@10916.10935.20|CCGCGGCTGGACGAGGCGCC</td><td>>hg38_21_+_8433172_8446622_RNA45SN1_45S_50ntFlanks</td><td>10916</td><td>10935</td><td>20</td><td>CCGCGGCTGGACGAGGCGCC</td><td>116.8</td><td>-2.9</td><td>0.8</td><td>-3.7</td><td>2.3E-04</td><td>2.8E-04</td><td>BT20/MB468</td></tr>
<tr><td>>hg38_21_+_8433172_8446622_RNA45SN1_45S_50ntFlanks@8196.8213.18|CGTGGACGGTGTGAGGCC</td><td>>hg38_21_+_8433172_8446622_RNA45SN1_45S_50ntFlanks</td><td>8196</td><td>8213</td><td>18</td><td>CGTGGACGGTGTGAGGCC</td><td>2474.8</td><td>2.4</td><td>0.7</td><td>3.7</td><td>2.1E-04</td><td>2.9E-04</td><td>MB231/MB468</td></tr>
<tr><td>>hg38_21_+_8433172_8446622_RNA45SN1_45S_50ntFlanks@7981.8004.24|CTCAGATCAGACGTGGCGACCCGC</td><td>>hg38_21_+_8433172_8446622_RNA45SN1_45S_50ntFlanks</td><td>7981</td><td>8004</td><td>24</td><td>CTCAGATCAGACGTGGCGACCCGC</td><td>103.9</td><td>-2.2</td><td>0.6</td><td>-3.7</td><td>2.3E-04</td><td>2.9E-04</td><td>BT20/MB468</td></tr>
<tr><td>>hg38_21_+_8433172_8446622_RNA45SN1_45S_50ntFlanks@12433.12454.22|CTATCATTGTGAAGCAGAATTC</td><td>>hg38_21_+_8433172_8446622_RNA45SN1_45S_50ntFlanks</td><td>12433</td><td>12454</td><td>22</td><td>CTATCATTGTGAAGCAGAATTC</td><td>160.2</td><td>-2.1</td><td>0.6</td><td>-3.7</td><td>2.1E-04</td><td>2.9E-04</td><td>MB231/MB468</td></tr>
<tr><td>>hg38_21_+_8433172_8446622_RNA45SN1_45S_50ntFlanks@10684.10702.19|CGGCGGCGTCCGGTGAGCT</td><td>>hg38_21_+_8433172_8446622_RNA45SN1_45S_50ntFlanks</td><td>10684</td><td>10702</td><td>19</td><td>CGGCGGCGTCCGGTGAGCT</td><td>188.7</td><td>-3.4</td><td>0.9</td><td>-3.7</td><td>2.3E-04</td><td>2.9E-04</td><td>BT20/MB468</td></tr>
<tr><td>>hg38_21_+_8433172_8446622_RNA45SN1_45S_50ntFlanks@7977.7994.18|CGACCTCAGATCAGACGT</td><td>>hg38_21_+_8433172_8446622_RNA45SN1_45S_50ntFlanks</td><td>7977</td><td>7994</td><td>18</td><td>CGACCTCAGATCAGACGT</td><td>203.1</td><td>-2.2</td><td>0.6</td><td>-3.7</td><td>2.3E-04</td><td>2.9E-04</td><td>BT20/MB468</td></tr>
<tr><td>>hg38_21_+_8433172_8446622_RNA45SN1_45S_50ntFlanks@3853.3881.29|AACTGTGGTAATTCTAGAGCTAATACATG</td><td>>hg38_21_+_8433172_8446622_RNA45SN1_45S_50ntFlanks</td><td>3853</td><td>3881</td><td>29</td><td>AACTGTGGTAATTCTAGAGCTAATACATG</td><td>514.2</td><td>-2.1</td><td>0.6</td><td>-3.7</td><td>2.1E-04</td><td>2.9E-04</td><td>MB231/MB468</td></tr>
<tr><td>>hg38_21_+_8433172_8446622_RNA45SN1_45S_50ntFlanks@11641.11680.40|CGGCGGGTGTTGACGCGATGTGATTTCTGCCCAGTGCTCT</td><td>>hg38_21_+_8433172_8446622_RNA45SN1_45S_50ntFlanks</td><td>11641</td><td>11680</td><td>40</td><td>CGGCGGGTGTTGACGCGATGTGATTTCTGCCCAGTGCTCT</td><td>207.2</td><td>-3.7</td><td>1.0</td><td>-3.7</td><td>2.4E-04</td><td>2.9E-04</td><td>BT20/MB468</td></tr>
<tr><td>>hg38_21_+_8433172_8446622_RNA45SN1_45S_50ntFlanks@7985.8007.23|GATCAGACGTGGCGACCCGCTGA</td><td>>hg38_21_+_8433172_8446622_RNA45SN1_45S_50ntFlanks</td><td>7985</td><td>8007</td><td>23</td><td>GATCAGACGTGGCGACCCGCTGA</td><td>90.5</td><td>-3.3</td><td>0.9</td><td>-3.7</td><td>2.2E-04</td><td>2.9E-04</td><td>MB231/MB468</td></tr>
<tr><td>>hg38_21_+_8433172_8446622_RNA45SN1_45S_50ntFlanks@11569.11588.20|CAGCCGACTTAGAACTGGTG</td><td>>hg38_21_+_8433172_8446622_RNA45SN1_45S_50ntFlanks</td><td>11569</td><td>11588</td><td>20</td><td>CAGCCGACTTAGAACTGGTG</td><td>129.3</td><td>-2.7</td><td>0.7</td><td>-3.7</td><td>2.2E-04</td><td>2.9E-04</td><td>MB231/MB468</td></tr>
<tr><td>>hg38_21_+_8433172_8446622_RNA45SN1_45S_50ntFlanks@4733.4758.26|GAACGAAAGTCGGAGGTTCGAAGACG</td><td>>hg38_21_+_8433172_8446622_RNA45SN1_45S_50ntFlanks</td><td>4733</td><td>4758</td><td>26</td><td>GAACGAAAGTCGGAGGTTCGAAGACG</td><td>824.7</td><td>1.9</td><td>0.5</td><td>3.7</td><td>2.2E-04</td><td>2.9E-04</td><td>MB231/MB468</td></tr>
<tr><td>>trnaMT_PheGAA_MT_+_577_647@54.74.21|CACATCACCCCATAAACACCA</td><td>>trnaMT_PheGAA_MT_+_577_647</td><td>54</td><td>74</td><td>21</td><td>CACATCACCCCATAAACACCA</td><td>443.1</td><td>2.5</td><td>0.7</td><td>3.7</td><td>2.2E-04</td><td>2.9E-04</td><td>MB231/MB468</td></tr>
<tr><td>>MI0000737|hsa-mir-200a&WithFlank&1|+|1167857|1167958@60.82.23(+1U)&[MIMAT0000682&hsa-miR-200a-3p&offsets|0|+1(+1U);m-142&1|+|1167916|1167938&offsets|0|0(+1U)]|TAACACTGTCTGGTAACGATGTTT</td><td>>MI0000737|hsa-mir-200a&WithFlank&1|+|1167857|1167958</td><td>60</td><td>82</td><td>23(+1U)</td><td>TAACACTGTCTGGTAACGATGTTT</td><td>118.1</td><td>-2.9</td><td>0.8</td><td>-3.7</td><td>2.2E-04</td><td>2.9E-04</td><td>MB231/MB468</td></tr>
<tr><td>>hg38_21_+_8433172_8446622_RNA45SN1_45S_50ntFlanks@11316.11342.27|CACCCCACGTCTCGTCGCGCGCGCGTC</td><td>>hg38_21_+_8433172_8446622_RNA45SN1_45S_50ntFlanks</td><td>11316</td><td>11342</td><td>27</td><td>CACCCCACGTCTCGTCGCGCGCGCGTC</td><td>1765.1</td><td>-5.4</td><td>1.5</td><td>-3.7</td><td>2.4E-04</td><td>3.0E-04</td><td>BT20/MB468</td></tr>
<tr><td>>hg38_1_-_228634819_228635039_RNA5S12_5S_50ntFlanks@84.109.26|CTCGTCTGATCTCGGAAGCTAAGCAG</td><td>>hg38_1_-_228634819_228635039_RNA5S12_5S_50ntFlanks</td><td>84</td><td>109</td><td>26</td><td>CTCGTCTGATCTCGGAAGCTAAGCAG</td><td>143.2</td><td>-2.3</td><td>0.6</td><td>-3.7</td><td>2.4E-04</td><td>3.0E-04</td><td>BT20/MB468</td></tr>
<tr><td>>hg38_21_+_8433172_8446622_RNA45SN1_45S_50ntFlanks@10352.10371.20|GAGAACTTTGAAGGCCGAAG</td><td>>hg38_21_+_8433172_8446622_RNA45SN1_45S_50ntFlanks</td><td>10352</td><td>10371</td><td>20</td><td>GAGAACTTTGAAGGCCGAAG</td><td>237.2</td><td>-3.3</td><td>0.8</td><td>-4.2</td><td>2.8E-05</td><td>3.0E-04</td><td>BT20/MB231</td></tr>
<tr><td>>hg38_21_+_8433172_8446622_RNA45SN1_45S_50ntFlanks@7975.8000.26|CGCGACCTCAGATCAGACGTGGCGAC</td><td>>hg38_21_+_8433172_8446622_RNA45SN1_45S_50ntFlanks</td><td>7975</td><td>8000</td><td>26</td><td>CGCGACCTCAGATCAGACGTGGCGAC</td><td>365.9</td><td>3.0</td><td>0.7</td><td>4.2</td><td>2.8E-05</td><td>3.0E-04</td><td>BT20/MB231</td></tr>
<tr><td>>trnaMT_PheGAA_MT_+_577_647@36.57.22|AAAATGTTTAGACGGGCTCACA</td><td>>trnaMT_PheGAA_MT_+_577_647</td><td>36</td><td>57</td><td>22</td><td>AAAATGTTTAGACGGGCTCACA</td><td>1896.4</td><td>3.1</td><td>0.7</td><td>4.2</td><td>2.8E-05</td><td>3.0E-04</td><td>BT20/MB231</td></tr>
</table>

| | | | | | | | | | | | | |
|---|---|---|---|---|---|---|---|---|---|---|---|---|
| >MI0000542\|hsa-mir-320a&WithFlank&8\|-\|22244960\|22245043@49.71.23&[MIMAT0000510&hsa-miR-320a-3p&offsets\|+1\|+2;m-60&8\|-\|22244975\|22244996&offsets\|+1\|+2]\|AAAGCTGGGTTGAGAGGGCGAAA | >MI0000542\|hsa-mir-320a&WithFlank&8\|-\|22244960\|22245043 | 49 | 71 | 23 | AAAGCTGGGTTGAGAGGGCGAAA | 297.3 | 3.1 | 0.7 | 4.2 | 2.8E-05 | 3.0E-04 | BT20/MB231 |
| >MI0003513\|hsa-mir-455&WithFlank&9\|+\|114209428\|114209535@22.45.24&[MIMAT0003150&hsa-miR-455-5p&offsets\|0\|+2;m-232&9\|+\|114209449\|114209470&offsets\|0\|+2]\|TATGTGCCTTTGGACTACATCGTG | >MI0003513\|hsa-mir-455&WithFlank&9\|+\|114209428\|114209535 | 22 | 45 | 24 | TATGTGCCTTTGGACTACATCGTG | 346.1 | -2.9 | 0.7 | -4.2 | 2.9E-05 | 3.0E-04 | BT20/MB231 |
| >am_trnaMT_SerGCT_MT_+_12207_12265@14.31.18\|AGAACTGCTAACTCATGC | >am_trnaMT_SerGCT_MT_+_12207_12265 | 14 | 31 | 18 | AGAACTGCTAACTCATGC | 1178.6 | 7.7 | 2.1 | 3.7 | 2.2E-04 | 3.0E-04 | MB231/MB468 |
| >hg38_21_+_8433172_8446622_RNA45SN1_45S_50ntFlanks@10421.10456.36\|TGAGAGATGGGCGAGCGCCGTTCCGAAGGGACGGGC | >hg38_21_+_8433172_8446622_RNA45SN1_45S_50ntFlanks | 10421 | 10456 | 36 | TGAGAGATGGGCGAGCGCCGTTCCGAAGGGACGGGC | 408.5 | -1.9 | 0.5 | -3.7 | 2.2E-04 | 3.0E-04 | MB231/MB468 |
| >hg38_21_+_8433172_8446622_RNA45SN1_45S_50ntFlanks@12459.12477.19\|AGCGTTGGATTGTTCACCC | >hg38_21_+_8433172_8446622_RNA45SN1_45S_50ntFlanks | 12459 | 12477 | 19 | AGCGTTGGATTGTTCACCC | 642.1 | -2.0 | 0.5 | -3.7 | 2.2E-04 | 3.0E-04 | MB231/MB468 |
| >hg38_21_+_8433172_8446622_RNA45SN1_45S_50ntFlanks@10686.10711.26\|GCGGCGTCCGGTGAGCTCTCGCTGGC | >hg38_21_+_8433172_8446622_RNA45SN1_45S_50ntFlanks | 10686 | 10711 | 26 | GCGGCGTCCGGTGAGCTCTCGCTGGC | 396.4 | -2.0 | 0.5 | -3.7 | 2.2E-04 | 3.0E-04 | MB231/MB468 |
| >hg38_21_+_8433172_8446622_RNA45SN1_45S_50ntFlanks@5485.5518.34\|AGCGCTGAGAAGACGGTCGAACTTGACTATCTAG | >hg38_21_+_8433172_8446622_RNA45SN1_45S_50ntFlanks | 5485 | 5518 | 34 | AGCGCTGAGAAGACGGTCGAACTTGACTATCTAG | 324.5 | -6.3 | 1.7 | -3.7 | 2.2E-04 | 3.0E-04 | MB231/MB468 |
| >hg38_21_+_8433172_8446622_RNA45SN1_45S_50ntFlanks@10787.10815.29\|ACAGCCTCTGGCATGTTGGAACAATGTAG | >hg38_21_+_8433172_8446622_RNA45SN1_45S_50ntFlanks | 10787 | 10815 | 29 | ACAGCCTCTGGCATGTTGGAACAATGTAG | 150.1 | -2.3 | 0.6 | -3.7 | 2.2E-04 | 3.0E-04 | MB231/MB468 |
| >hg38_21_+_8433172_8446622_RNA45SN1_45S_50ntFlanks@12202.12220.19\|CTAAGGCGAGCTCAGGGAG | >hg38_21_+_8433172_8446622_RNA45SN1_45S_50ntFlanks | 12202 | 12220 | 19 | CTAAGGCGAGCTCAGGGAG | 102.4 | -2.6 | 0.7 | -3.7 | 2.2E-04 | 3.0E-04 | MB231/MB468 |
| >MI0000298\|hsa-mir-221&WithFlank&X\|-\|45746151\|45746272@71.93.23(+1U)&[MIMAT0000278&hsa-miR-221-3p&offsets\|0\|0(+1U);m-91&X\|-\|45746181\|45746202&offsets\|0\|+1(+1U)]\|AGCTACATTGTCTGCTGGGTTTCT | >MI0000298\|hsa-mir-221&WithFlank&X\|-\|45746151\|45746272 | 71 | 93 | 23(+1U) | AGCTACATTGTCTGCTGGGTTTCT | 19201.9 | -2.6 | 0.6 | -4.2 | 2.9E-05 | 3.0E-04 | BT20/MB231 |
| >hg38_21_+_8433172_8446622_RNA45SN1_45S_50ntFlanks@5443.5466.24\|CCTCGGATCGGCCCCGCCGGGGTC | >hg38_21_+_8433172_8446622_RNA45SN1_45S_50ntFlanks | 5443 | 5466 | 24 | CCTCGGATCGGCCCCGCCGGGGTC | 357.1 | -1.8 | 0.5 | -3.7 | 2.2E-04 | 3.0E-04 | MB231/MB468 |
| >hg38_21_+_8433172_8446622_RNA45SN1_45S_50ntFlanks@10683.10705.23\|CCGGCGGCGTCCGGTGAGCTCTC | >hg38_21_+_8433172_8446622_RNA45SN1_45S_50ntFlanks | 10683 | 10705 | 23 | CCGGCGGCGTCCGGTGAGCTCTC | 396.1 | -3.2 | 0.9 | -3.7 | 2.5E-04 | 3.0E-04 | BT20/MB468 |
| >hg38_21_+_8433172_8446622_RNA45SN1_45S_50ntFlanks@12416.12456.41\|TTCGATGTCGGCTCTTCCTATCATTGTGAAGCAGAATTCAC | >hg38_21_+_8433172_8446622_RNA45SN1_45S_50ntFlanks | 12416 | 12456 | 41 | TTCGATGTCGGCTCTTCCTATCATTGTGAAGCAGAATTCAC | 242.1 | -3.1 | 0.7 | -4.2 | 2.9E-05 | 3.0E-04 | BT20/MB231 |
| >MI0000477\|hsa-mir-146a&WithFlank&5\|+\|160485346\|160485456@27.47.21(+1G)&[MIMAT0000449&hsa-miR-146a-5p&offsets\|0\|-1(+1G);m-129&5\|+\|160485372\|160485393&offsets\|0\|-1(+1G)]\|TGAGAACTGAATTCCATGGGTG | >MI0000477\|hsa-mir-146a&WithFlank&5\|+\|160485346\|160485456 | 27 | 47 | 21(+1G) | TGAGAACTGAATTCCATGGGTG | 810.2 | -9.5 | 2.3 | -4.2 | 2.9E-05 | 3.0E-04 | BT20/MB231 |
| >hg38_21_+_8433172_8446622_RNA45SN1_45S_50ntFlanks@3854.3872.19\|ACTGTGGTAATTCTAGAGC | >hg38_21_+_8433172_8446622_RNA45SN1_45S_50ntFlanks | 3854 | 3872 | 19 | ACTGTGGTAATTCTAGAGC | 627.1 | -1.8 | 0.5 | -3.7 | 2.3E-04 | 3.0E-04 | MB231/MB468 |
| >hg38_21_+_8433172_8446622_RNA45SN1_45S_50ntFlanks@9559.9584.26\|CGGTCCTGACGTGCAAATCGGTCGTC | >hg38_21_+_8433172_8446622_RNA45SN1_45S_50ntFlanks | 9559 | 9584 | 26 | CGGTCCTGACGTGCAAATCGGTCGTC | 277.9 | 1.4 | 0.4 | 3.7 | 2.3E-04 | 3.0E-04 | MB231/MB468 |
| >hg38_21_+_8433172_8446622_RNA45SN1_45S_50ntFlanks@6721.6751.31\|AATTGCAGGACACATTGATCATCGACACTTC | >hg38_21_+_8433172_8446622_RNA45SN1_45S_50ntFlanks | 6721 | 6751 | 31 | AATTGCAGGACACATTGATCATCGACACTTC | 108.1 | -1.7 | 0.5 | -3.7 | 2.5E-04 | 3.1E-04 | BT20/MB468 |
| >hg38_21_+_8433172_8446622_RNA45SN1_45S_50ntFlanks@12909.12933.25\|CCGCACGTTCGTGGGGAACCTGGCG | >hg38_21_+_8433172_8446622_RNA45SN1_45S_50ntFlanks | 12909 | 12933 | 25 | CCGCACGTTCGTGGGGAACCTGGCG | 132.2 | -2.9 | 0.8 | -3.7 | 2.5E-04 | 3.1E-04 | BT20/MB468 |
| >hg38_21_+_8433172_8446622_RNA45SN1_45S_50ntFlanks@11571.11600.30\|GCCGACTTAGAACTGGTGCGGACCAGGGGA | >hg38_21_+_8433172_8446622_RNA45SN1_45S_50ntFlanks | 11571 | 11600 | 30 | GCCGACTTAGAACTGGTGCGGACCAGGGGA | 341.6 | 1.8 | 0.5 | 3.7 | 2.3E-04 | 3.1E-04 | MB231/MB468 |
| >hg38_21_+_8433172_8446622_RNA45SN1_45S_50ntFlanks@11580.11606.27\|GAACTGGTGCGGACCAGGGGAATCCGA | >hg38_21_+_8433172_8446622_RNA45SN1_45S_50ntFlanks | 11580 | 11606 | 27 | GAACTGGTGCGGACCAGGGGAATCCGA | 133.8 | -2.2 | 0.6 | -3.7 | 2.3E-04 | 3.1E-04 | MB231/MB468 |
| >hg38_21_+_8433172_8446622_RNA45SN1_45S_50ntFlanks@4325.4346.22\|CCGCGGTAATTCCAGCTCCAAT | >hg38_21_+_8433172_8446622_RNA45SN1_45S_50ntFlanks | 4325 | 4346 | 22 | CCGCGGTAATTCCAGCTCCAAT | 130.1 | -3.4 | 0.9 | -3.7 | 2.5E-04 | 3.1E-04 | BT20/MB468 |
| >hg38_21_+_8433172_8446622_RNA45SN1_45S_50ntFlanks@12999.13039.41\|CTGCGATCTATTGAAAGTCAGCCCTCGACACAAGGGTTTGT | >hg38_21_+_8433172_8446622_RNA45SN1_45S_50ntFlanks | 12999 | 13039 | 41 | CTGCGATCTATTGAAAGTCAGCCCTCGACACAAGGGTTTGT | 9123.5 | -2.5 | 0.6 | -4.2 | 3.0E-05 | 3.1E-04 | BT20/MB231 |
| >hg38_21_+_8433172_8446622_RNA45SN1_45S_50ntFlanks@5485.5515.31\|AGCGCTGAGAAGACGGTCGAACTTGACTATC | >hg38_21_+_8433172_8446622_RNA45SN1_45S_50ntFlanks | 5485 | 5515 | 31 | AGCGCTGAGAAGACGGTCGAACTTGACTATC | 118.2 | -1.9 | 0.5 | -3.7 | 2.3E-04 | 3.1E-04 | MB231/MB468 |
| >hg38_21_+_8433172_8446622_RNA45SN1_45S_50ntFlanks@11581.11606.26\|AACTGGTGCGGACCAGGGGAATCCGA | >hg38_21_+_8433172_8446622_RNA45SN1_45S_50ntFlanks | 11581 | 11606 | 26 | AACTGGTGCGGACCAGGGGAATCCGA | 283.0 | -1.9 | 0.5 | -3.7 | 2.5E-04 | 3.1E-04 | BT20/MB468 |
| >hg38_21_+_8433172_8446622_RNA45SN1_45S_50ntFlanks@6653.6686.34\|ACTCTTAGCGGTGGATCACTCGGCTCGTGCGTCG | >hg38_21_+_8433172_8446622_RNA45SN1_45S_50ntFlanks | 6653 | 6686 | 34 | ACTCTTAGCGGTGGATCACTCGGCTCGTGCGTCG | 547.3 | 2.5 | 0.7 | 3.7 | 2.3E-04 | 3.1E-04 | MB231/MB468 |
| >hg38_21_+_8433172_8446622_RNA45SN1_45S_50ntFlanks@8197.8215.19\|GTGGACGGTGTGAGGCCGG | >hg38_21_+_8433172_8446622_RNA45SN1_45S_50ntFlanks | 8197 | 8215 | 19 | GTGGACGGTGTGAGGCCGG | 142.5 | -2.1 | 0.6 | -3.7 | 2.3E-04 | 3.1E-04 | MB231/MB468 |
| >hg38_21_+_8433172_8446622_RNA45SN1_45S_50ntFlanks@11670.11703.34\|CCCAGTGCTCTGAATGTCAAAGTGAAGAAATTCA | >hg38_21_+_8433172_8446622_RNA45SN1_45S_50ntFlanks | 11670 | 11703 | 34 | CCCAGTGCTCTGAATGTCAAAGTGAAGAAATTCA | 298.1 | -3.0 | 0.8 | -3.7 | 2.6E-04 | 3.1E-04 | BT20/MB468 |
| >hg38_21_+_8433172_8446622_RNA45SN1_45S_50ntFlanks@12997.13037.41\|CGCTGCGATCTATTGAAAGTCAGCCCTCGACACAAGGGTTT | >hg38_21_+_8433172_8446622_RNA45SN1_45S_50ntFlanks | 12997 | 13037 | 41 | CGCTGCGATCTATTGAAAGTCAGCCCTCGACACAAGGGTTT | 542.6 | -3.4 | 0.8 | -4.2 | 3.1E-05 | 3.1E-04 | BT20/MB231 |
| >hg38_21_+_8433172_8446622_RNA45SN1_45S_50ntFlanks@3939.3959.21\|AACCCGGTCAGCCCCTCTCCG | >hg38_21_+_8433172_8446622_RNA45SN1_45S_50ntFlanks | 3939 | 3959 | 21 | AACCCGGTCAGCCCCTCTCCG | 537.4 | -4.2 | 1.1 | -3.7 | 2.6E-04 | 3.2E-04 | BT20/MB468 |
| >hg38_21_+_8433172_8446622_RNA45SN1_45S_50ntFlanks@3842.3861.20\|CCTACTTGGATAACTGTGGT | >hg38_21_+_8433172_8446622_RNA45SN1_45S_50ntFlanks | 3842 | 3861 | 20 | CCTACTTGGATAACTGTGGT | 269.2 | 1.5 | 0.4 | 3.7 | 2.3E-04 | 3.2E-04 | MB231/MB468 |
| >hg38_21_+_8433172_8446622_RNA45SN1_45S_50ntFlanks@10689.10708.20\|GCGTCCGGTGAGCTCTCGCT | >hg38_21_+_8433172_8446622_RNA45SN1_45S_50ntFlanks | 10689 | 10708 | 20 | GCGTCCGGTGAGCTCTCGCT | 765.6 | -3.8 | 1.0 | -3.7 | 2.6E-04 | 3.2E-04 | BT20/MB468 |
| >hg38_21_+_8433172_8446622_RNA45SN1_45S_50ntFlanks@12993.13013.21\|CCCTCGCTGCGATCTATTGAA | >hg38_21_+_8433172_8446622_RNA45SN1_45S_50ntFlanks | 12993 | 13013 | 21 | CCCTCGCTGCGATCTATTGAA | 347.5 | -3.1 | 0.8 | -3.7 | 2.6E-04 | 3.2E-04 | BT20/MB468 |
| >am_trna111_HisGTG_1_-_147774845_147774916@3.29.27,trna118_HisGTG_1_-_145396881_145396952@3.29.27,trna16_HisGTG_1_+_146544773_146544844@3.29.27,trna1_HisGTG_15_+_45493349_45493420@3.29.27,trna21_HisGTG_1_+_147753471_147753542@3.29.27,trna33_HisGTG_6_+_27125906_27125977@3.29.27,trna7_HisGTG_9_-_14433938_14434009@3.29.27,trna8_HisGTG_15_-_45492611_45492682@3.29.27,trna9_HisGTG_15_-_45490804_45490875@3.29.27\|CGTGATCGTATAGTGGTTAGTACTCTG | >am_trna111_HisGTG_1_-_147774845_147774916 | 3 | 29 | 27 | CGTGATCGTATAGTGGTTAGTACTCTG | 84.9 | -3.0 | 0.8 | -3.7 | 2.4E-04 | 3.2E-04 | MB231/MB468 |
| >hg38_21_+_8433172_8446622_RNA45SN1_45S_50ntFlanks@6700.6721.22\|CTAGCTGCGAGAATTAATGTGA | >hg38_21_+_8433172_8446622_RNA45SN1_45S_50ntFlanks | 6700 | 6721 | 22 | CTAGCTGCGAGAATTAATGTGA | 127.6 | -1.7 | 0.5 | -3.7 | 2.6E-04 | 3.2E-04 | BT20/MB468 |
| >hg38_21_+_8433172_8446622_RNA45SN1_45S_50ntFlanks@10787.10817.31\|ACAGCCTCTGGCATGTTGGAACAATGTAGGT | >hg38_21_+_8433172_8446622_RNA45SN1_45S_50ntFlanks | 10787 | 10817 | 31 | ACAGCCTCTGGCATGTTGGAACAATGTAGGT | 296.0 | -2.5 | 0.7 | -3.7 | 2.6E-04 | 3.2E-04 | BT20/MB468 |
| >hg38_21_+_8433172_8446622_RNA45SN1_45S_50ntFlanks@12077.12095.19\|GGGGCTCTCGCTTCTGGCG | >hg38_21_+_8433172_8446622_RNA45SN1_45S_50ntFlanks | 12077 | 12095 | 19 | GGGGCTCTCGCTTCTGGCG | 220.2 | -2.8 | 0.8 | -3.7 | 2.6E-04 | 3.2E-04 | BT20/MB468 |
| >hg38_21_+_8433172_8446622_RNA45SN1_45S_50ntFlanks@7974.7992.19\|ACGCGACCTCAGATCAGAC | >hg38_21_+_8433172_8446622_RNA45SN1_45S_50ntFlanks | 7974 | 7992 | 19 | ACGCGACCTCAGATCAGAC | 732.0 | 3.1 | 0.8 | 3.7 | 2.4E-04 | 3.2E-04 | MB231/MB468 |
| >hg38_21_+_8433172_8446622_RNA45SN1_45S_50ntFlanks@8176.8220.45\|CTTCTGATCGAGGCCCAGCCCGTGGACGGTGTGAGGCCGGTAGCG | >hg38_21_+_8433172_8446622_RNA45SN1_45S_50ntFlanks | 8176 | 8220 | 45 | CTTCTGATCGAGGCCCAGCCCGTGGACGGTGTGAGGCCGGTAGCG | 229.4 | 1.7 | 0.5 | 3.7 | 2.4E-04 | 3.3E-04 | MB231/MB468 |
| >hg38_21_+_8433172_8446622_RNA45SN1_45S_50ntFlanks@4367.4390.24\|CAGTTAAAAAGCTCGTAGTTGGAT | >hg38_21_+_8433172_8446622_RNA45SN1_45S_50ntFlanks | 4367 | 4390 | 24 | CAGTTAAAAAGCTCGTAGTTGGAT | 65.4 | -3.2 | 0.9 | -3.7 | 2.4E-04 | 3.3E-04 | MB231/MB468 |
| >hg38_21_+_8433172_8446622_RNA45SN1_45S_50ntFlanks@9533.9551.19\|GAGGAAACTCTGGTGGAGG | >hg38_21_+_8433172_8446622_RNA45SN1_45S_50ntFlanks | 9533 | 9551 | 19 | GAGGAAACTCTGGTGGAGG | 228.6 | -2.8 | 0.8 | -3.6 | 2.7E-04 | 3.3E-04 | BT20/MB468 |
| >hg38_21_+_8433172_8446622_RNA45SN1_45S_50ntFlanks@3777.3798.22\|CGGTACAGTGAAACTGCGAATG | >hg38_21_+_8433172_8446622_RNA45SN1_45S_50ntFlanks | 3777 | 3798 | 22 | CGGTACAGTGAAACTGCGAATG | 90.3 | -1.8 | 0.5 | -3.6 | 2.7E-04 | 3.3E-04 | BT20/MB468 |
| >hg38_21_+_8433172_8446622_RNA45SN1_45S_50ntFlanks@8380.8406.27\|CAAGAGGGCGTGAAACCGTTAAGAGGT | >hg38_21_+_8433172_8446622_RNA45SN1_45S_50ntFlanks | 8380 | 8406 | 27 | CAAGAGGGCGTGAAACCGTTAAGAGGT | 324.5 | -1.9 | 0.5 | -3.7 | 2.4E-04 | 3.3E-04 | MB231/MB468 |
| >am_trna2_GlyGCC_21_-_18827107_18827177@1.25.25,trna35_GlyGCC_1_+_161413094_161413164@1.25.25,trna37_GlyGCC_1_+_161420467_161420537@1.25.25,trna39_GlyGCC_1_+_161427898_161427968@1.25.25,trna41_GlyGCC_1_+_161435258_161435328@1.25.25\|GCATGGGTGGTTCAGTGGTAGAATT | >am_trna2_GlyGCC_21_-_18827107_18827177 | 1 | 25 | 25 | GCATGGGTGGTTCAGTGGTAGAATT | 296.8 | 8.0 | 1.9 | 4.2 | 3.3E-05 | 3.3E-04 | BT20/MB231 |
| >hg38_21_+_8433172_8446622_RNA45SN1_45S_50ntFlanks@6646.6685.40\|TCGTACGACTCTTAGCGGTGGATCACTCGGCTCGTGCGTC | >hg38_21_+_8433172_8446622_RNA45SN1_45S_50ntFlanks | 6646 | 6685 | 40 | TCGTACGACTCTTAGCGGTGGATCACTCGGCTCGTGCGTC | 5774.0 | 3.3 | 0.8 | 4.2 | 3.3E-05 | 3.3E-04 | BT20/MB231 |
| >hg38_21_+_8433172_8446622_RNA45SN1_45S_50ntFlanks@10686.10708.23\|GCGGCGTCCGGTGAGCTCTCGCT | >hg38_21_+_8433172_8446622_RNA45SN1_45S_50ntFlanks | 10686 | 10708 | 23 | GCGGCGTCCGGTGAGCTCTCGCT | 1245.9 | 2.1 | 0.6 | 3.7 | 2.5E-04 | 3.3E-04 | MB231/MB468 |

<table>
<tr><td>>MI0001641|hsa-mir-429&WithFlank&1|+|1168999|1169093@57.77.21(+1C)&[MIMAT0001536&hsa-miR-429&offsets|0|-1(+1C);m-207&1|+|1169055|1169076&offsets|0|-1(+1C)]|TAATACTGTCTGGTAAAACCGC</td><td>>MI0001641|hsa-mir-429&WithFlank&1|+|1168999|1169093</td><td>57</td><td>77</td><td>21(+1C)</td><td>TAATACTGTCTGGTAAAACCGC</td><td>90.2</td><td>-3.2</td><td>0.9</td><td>-3.7</td><td>2.5E-04</td><td>3.4E-04</td><td>MB231/MB468</td></tr>
<tr><td>>hg38_21_+_8433172_8446622_RNA45SN1_45S_50ntFlanks@10683.10708.26|CCGGCGGCGTCCGGTGAGCTCTCGCT</td><td>>hg38_21_+_8433172_8446622_RNA45SN1_45S_50ntFlanks</td><td>10683</td><td>10708</td><td>26</td><td>CCGGCGGCGTCCGGTGAGCTCTCGCT</td><td>230.6</td><td>-3.1</td><td>0.8</td><td>-3.6</td><td>2.8E-04</td><td>3.4E-04</td><td>BT20/MB468</td></tr>
<tr><td>>trna12_ArgCCT_16_+_3243918_3243990@35.54.20,trna18_ArgCCT_17_+_73030001_73030073@35.54.20,trna21_ArgCCT_17_-_73030526_73030598@35.54.20,trna2_ArgCCT_16_+_3202901_3202973@35.54.20,trna3_ArgCCT_7_+_139025446_139025518@35.54.20|CTAAGCCAGGGATTGTGGGT</td><td>>trna12_ArgCCT_16_+_3243918_3243990</td><td>35</td><td>54</td><td>20</td><td>CTAAGCCAGGGATTGTGGGT</td><td>1361.1</td><td>3.1</td><td>0.8</td><td>3.7</td><td>2.5E-04</td><td>3.4E-04</td><td>MB231/MB468</td></tr>
<tr><td>>hg38_21_+_8433172_8446622_RNA45SN1_45S_50ntFlanks@11570.11592.23|AGCCGACTTAGAACTGGTGCGGA</td><td>>hg38_21_+_8433172_8446622_RNA45SN1_45S_50ntFlanks</td><td>11570</td><td>11592</td><td>23</td><td>AGCCGACTTAGAACTGGTGCGGA</td><td>330.9</td><td>-2.3</td><td>0.6</td><td>-3.7</td><td>2.5E-04</td><td>3.4E-04</td><td>MB231/MB468</td></tr>
<tr><td>>hg38_21_+_8433172_8446622_RNA45SN1_45S_50ntFlanks@12420.12440.21|ATGTCGGCTCTTCCTATCATT</td><td>>hg38_21_+_8433172_8446622_RNA45SN1_45S_50ntFlanks</td><td>12420</td><td>12440</td><td>21</td><td>ATGTCGGCTCTTCCTATCATT</td><td>1345.0</td><td>2.5</td><td>0.7</td><td>3.7</td><td>2.5E-04</td><td>3.4E-04</td><td>MB231/MB468</td></tr>
<tr><td>>hg38_21_+_8433172_8446622_RNA45SN1_45S_50ntFlanks@12418.12442.25|CGATGTCGGCTCTTCCTATCATTGT</td><td>>hg38_21_+_8433172_8446622_RNA45SN1_45S_50ntFlanks</td><td>12418</td><td>12442</td><td>25</td><td>CGATGTCGGCTCTTCCTATCATTGT</td><td>529.0</td><td>2.5</td><td>0.7</td><td>3.7</td><td>2.6E-04</td><td>3.4E-04</td><td>MB231/MB468</td></tr>
<tr><td>>hg38_21_+_8433172_8446622_RNA45SN1_45S_50ntFlanks@9564.9594.31|CTGACGTGCAAATCGGTCGTCCGACCTGGGT</td><td>>hg38_21_+_8433172_8446622_RNA45SN1_45S_50ntFlanks</td><td>9564</td><td>9594</td><td>31</td><td>CTGACGTGCAAATCGGTCGTCCGACCTGGGT</td><td>359.8</td><td>-1.9</td><td>0.5</td><td>-3.7</td><td>2.6E-04</td><td>3.4E-04</td><td>MB231/MB468</td></tr>
<tr><td>>hg38_21_+_8433172_8446622_RNA45SN1_45S_50ntFlanks@7982.8004.23|TCAGATCAGACGTGGCGACCCGC</td><td>>hg38_21_+_8433172_8446622_RNA45SN1_45S_50ntFlanks</td><td>7982</td><td>8004</td><td>23</td><td>TCAGATCAGACGTGGCGACCCGC</td><td>116.1</td><td>-2.5</td><td>0.7</td><td>-3.7</td><td>2.6E-04</td><td>3.4E-04</td><td>MB231/MB468</td></tr>
<tr><td>>hg38_1_-_228634819_228635039_RNA5S12_5S_50ntFlanks@134.172.39|TGGGAGACCGCCTGGGAATACCGGGTGCTGTAGGCTTTT</td><td>>hg38_1_-_228634819_228635039_RNA5S12_5S_50ntFlanks</td><td>134</td><td>172</td><td>39</td><td>TGGGAGACCGCCTGGGAATACCGGGTGCTGTAGGCTTTT</td><td>1128.3</td><td>-3.8</td><td>0.9</td><td>-4.1</td><td>3.4E-05</td><td>3.4E-04</td><td>BT20/MB231</td></tr>
<tr><td>>hg38_21_+_8433172_8446622_RNA45SN1_45S_50ntFlanks@3854.3889.36|ACTGTGGTAATTCTAGAGCTAATACATGCCGACGGG</td><td>>hg38_21_+_8433172_8446622_RNA45SN1_45S_50ntFlanks</td><td>3854</td><td>3889</td><td>36</td><td>ACTGTGGTAATTCTAGAGCTAATACATGCCGACGGG</td><td>90.4</td><td>-2.0</td><td>0.5</td><td>-3.7</td><td>2.6E-04</td><td>3.5E-04</td><td>MB231/MB468</td></tr>
<tr><td>>hg38_21_+_8433172_8446622_RNA45SN1_45S_50ntFlanks@4752.4769.18|GAAGACGATCAGATACCG</td><td>>hg38_21_+_8433172_8446622_RNA45SN1_45S_50ntFlanks</td><td>4752</td><td>4769</td><td>18</td><td>GAAGACGATCAGATACCG</td><td>68.3</td><td>-2.7</td><td>0.7</td><td>-3.7</td><td>2.6E-04</td><td>3.5E-04</td><td>MB231/MB468</td></tr>
<tr><td>>trna119_LysCTT_1_-_145395522_145395594@4.33.30,trna11_LysCTT_5_-_180648979_180649051@4.33.30,trna13_LysCTT_6_+_26556774_26556846@4.33.30,trna7_LysCTT_16_+_3225692_3225764@4.33.30,trna9_LysCTT_5_+_180634755_180634827@4.33.30|CGGCTAGCTCAGTCGGTAGAGCATGAGACT</td><td>>trna119_LysCTT_1_-_145395522_145395594</td><td>4</td><td>33</td><td>30</td><td>CGGCTAGCTCAGTCGGTAGAGCATGAGACT</td><td>687.1</td><td>-3.9</td><td>0.9</td><td>-4.1</td><td>3.4E-05</td><td>3.5E-04</td><td>BT20/MB231</td></tr>
<tr><td>>hg38_21_+_8433172_8446622_RNA45SN1_45S_50ntFlanks@12932.12972.41|CGCTAAACCATTCGTAGACGACCTGCTTCTGGGTCGGGGTT</td><td>>hg38_21_+_8433172_8446622_RNA45SN1_45S_50ntFlanks</td><td>12932</td><td>12972</td><td>41</td><td>CGCTAAACCATTCGTAGACGACCTGCTTCTGGGTCGGGGTT</td><td>245.7</td><td>-2.9</td><td>0.8</td><td>-3.6</td><td>2.8E-04</td><td>3.5E-04</td><td>BT20/MB468</td></tr>
<tr><td>>hg38_21_+_8433172_8446622_RNA45SN1_45S_50ntFlanks@12408.12434.27|TTTGATCCTTCGATGTCGGCTCTTCCT</td><td>>hg38_21_+_8433172_8446622_RNA45SN1_45S_50ntFlanks</td><td>12408</td><td>12434</td><td>27</td><td>TTTGATCCTTCGATGTCGGCTCTTCCT</td><td>409.4</td><td>2.1</td><td>0.6</td><td>3.7</td><td>2.6E-04</td><td>3.5E-04</td><td>MB231/MB468</td></tr>
<tr><td>>hg38_21_+_8433172_8446622_RNA45SN1_45S_50ntFlanks@10349.10379.31|AACGAGAACTTTGAAGGCCGAAGTGGAGAAG</td><td>>hg38_21_+_8433172_8446622_RNA45SN1_45S_50ntFlanks</td><td>10349</td><td>10379</td><td>31</td><td>AACGAGAACTTTGAAGGCCGAAGTGGAGAAG</td><td>457.6</td><td>2.4</td><td>0.7</td><td>3.7</td><td>2.6E-04</td><td>3.5E-04</td><td>MB231/MB468</td></tr>
<tr><td>>hg38_21_+_8433172_8446622_RNA45SN1_45S_50ntFlanks@8104.8140.37|CCGCGGCGGGGCGCGGGACATGTGGCGTACGGAAGAC</td><td>>hg38_21_+_8433172_8446622_RNA45SN1_45S_50ntFlanks</td><td>8104</td><td>8140</td><td>37</td><td>CCGCGGCGGGGCGCGGGACATGTGGCGTACGGAAGAC</td><td>321.5</td><td>-2.7</td><td>0.7</td><td>-3.7</td><td>2.6E-04</td><td>3.5E-04</td><td>MB231/MB468</td></tr>
<tr><td>>hg38_21_+_8433172_8446622_RNA45SN1_45S_50ntFlanks@7975.7992.18|CGCGACCTCAGATCAGAC</td><td>>hg38_21_+_8433172_8446622_RNA45SN1_45S_50ntFlanks</td><td>7975</td><td>7992</td><td>18</td><td>CGCGACCTCAGATCAGAC</td><td>4629.6</td><td>3.2</td><td>0.9</td><td>3.6</td><td>2.6E-04</td><td>3.5E-04</td><td>MB231/MB468</td></tr>
<tr><td>>hg38_21_+_8433172_8446622_RNA45SN1_45S_50ntFlanks@12993.13039.47|CCCTCGCTGCGATCTATTGAAAGTCAGCCCTCGACACAAGGGTTTGT</td><td>>hg38_21_+_8433172_8446622_RNA45SN1_45S_50ntFlanks</td><td>12993</td><td>13039</td><td>47</td><td>CCCTCGCTGCGATCTATTGAAAGTCAGCCCTCGACACAAGGGTTTGT</td><td>1514.4</td><td>3.8</td><td>1.0</td><td>3.6</td><td>2.9E-04</td><td>3.5E-04</td><td>BT20/MB468</td></tr>
<tr><td>>hg38_21_+_8433172_8446622_RNA45SN1_45S_50ntFlanks@5472.5514.43|ACGGCCCTGGCGGAGCGCTGAGAAGACGGTCGAACTTGACTAT</td><td>>hg38_21_+_8433172_8446622_RNA45SN1_45S_50ntFlanks</td><td>5472</td><td>5514</td><td>43</td><td>ACGGCCCTGGCGGAGCGCTGAGAAGACGGTCGAACTTGACTAT</td><td>788.5</td><td>-3.7</td><td>0.9</td><td>-4.1</td><td>3.5E-05</td><td>3.5E-04</td><td>BT20/MB231</td></tr>
<tr><td>>hg38_21_+_8433172_8446622_RNA45SN1_45S_50ntFlanks@8114.8138.25|GCGCGGGACATGTGGCGTACGGAAG</td><td>>hg38_21_+_8433172_8446622_RNA45SN1_45S_50ntFlanks</td><td>8114</td><td>8138</td><td>25</td><td>GCGCGGGACATGTGGCGTACGGAAG</td><td>426.9</td><td>-3.8</td><td>1.0</td><td>-3.6</td><td>2.9E-04</td><td>3.6E-04</td><td>BT20/MB468</td></tr>
<tr><td>>hg38_21_+_8433172_8446622_RNA45SN1_45S_50ntFlanks@12405.12438.34|CTTTTTGATCCTTCGATGTCGGCTCTTCCTATCA</td><td>>hg38_21_+_8433172_8446622_RNA45SN1_45S_50ntFlanks</td><td>12405</td><td>12438</td><td>34</td><td>CTTTTTGATCCTTCGATGTCGGCTCTTCCTATCA</td><td>116.1</td><td>-1.6</td><td>0.4</td><td>-3.6</td><td>2.9E-04</td><td>3.6E-04</td><td>BT20/MB468</td></tr>
<tr><td>>MI0000272|hsa-mir-182&WithFlank&7|-|129770377|129770498@29.53.25&[MIMAT0000259&hsa-miR-182-5p&offsets|0|+1;m-22&7|-|129770449|129770470&offsets|0|+3]|TTTGGCAATGGTAGAACTCACACTG</td><td>>MI0000272|hsa-mir-182&WithFlank&7|-|129770377|129770498</td><td>29</td><td>53</td><td>25</td><td>TTTGGCAATGGTAGAACTCACACTG</td><td>13106.7</td><td>2.6</td><td>0.6</td><td>4.1</td><td>3.6E-05</td><td>3.6E-04</td><td>BT20/MB231</td></tr>
<tr><td>>hg38_21_+_8433172_8446622_RNA45SN1_45S_50ntFlanks@7988.8012.25|CAGACGTGGCGACCCGCTGAATTTA</td><td>>hg38_21_+_8433172_8446622_RNA45SN1_45S_50ntFlanks</td><td>7988</td><td>8012</td><td>25</td><td>CAGACGTGGCGACCCGCTGAATTTA</td><td>1507.3</td><td>-2.3</td><td>0.6</td><td>-3.6</td><td>2.7E-04</td><td>3.7E-04</td><td>MB231/MB468</td></tr>
<tr><td>>hg38_21_+_8433172_8446622_RNA45SN1_45S_50ntFlanks@12488.12512.25|AACGTGAGCTGGGTTTAGACCGTCG</td><td>>hg38_21_+_8433172_8446622_RNA45SN1_45S_50ntFlanks</td><td>12488</td><td>12512</td><td>25</td><td>AACGTGAGCTGGGTTTAGACCGTCG</td><td>232.7</td><td>-3.2</td><td>0.9</td><td>-3.6</td><td>3.0E-04</td><td>3.7E-04</td><td>BT20/MB468</td></tr>
<tr><td>>trna116_GluCTC_1_-_145399233_145399304@35.74.40,trna134_GluTTC_1_-_16861774_16861845@35.74.40,trna71_GluCTC_1_-_161439189_161439260@35.74.40,trna74_GluCTC_1_-_161431809_161431880@35.74.40,trna77_GluCTC_1_-_161424398_161424469@35.74.40,trna77_GluCTC_6_+_28949976_28950047@35.74.40,trna80_GluCTC_1_-_161417018_161417089@35.74.40,trna84_GluTTC_1_-_161391883_161391954@35.74.40,trna87_GluCTC_6_-_126101393_126101464@35.74.40|TCACCGCCGCGGCCCGGGTTCGATTCCCGGTCAGGGAACC</td><td>>trna116_GluCTC_1_-_145399233_145399304</td><td>35</td><td>74</td><td>40</td><td>TCACCGCCGCGGCCCGGGTTCGATTCCCGGTCAGGGAACC</td><td>779.7</td><td>9.4</td><td>2.6</td><td>3.6</td><td>2.7E-04</td><td>3.7E-04</td><td>MB231/MB468</td></tr>
<tr><td>>hg38_21_+_8433172_8446622_RNA45SN1_45S_50ntFlanks@10912.10932.21|CGCGCCGCGGCTGGACGAGGC</td><td>>hg38_21_+_8433172_8446622_RNA45SN1_45S_50ntFlanks</td><td>10912</td><td>10932</td><td>21</td><td>CGCGCCGCGGCTGGACGAGGC</td><td>305.7</td><td>-4.7</td><td>1.3</td><td>-3.6</td><td>3.0E-04</td><td>3.7E-04</td><td>BT20/MB468</td></tr>
<tr><td>>hg38_21_+_8433172_8446622_RNA45SN1_45S_50ntFlanks@11564.11592.29|CCTAGCAGCCGACTTAGAACTGGTGCGGA</td><td>>hg38_21_+_8433172_8446622_RNA45SN1_45S_50ntFlanks</td><td>11564</td><td>11592</td><td>29</td><td>CCTAGCAGCCGACTTAGAACTGGTGCGGA</td><td>431.3</td><td>-3.3</td><td>0.9</td><td>-3.6</td><td>3.0E-04</td><td>3.7E-04</td><td>BT20/MB468</td></tr>
<tr><td>>trna10_AspGTC_12_-_125424193_125424264@37.73.37,trna12_AspGTC_12_-_125411891_125411962@37.73.37,trna144_AspGTC_6_-_27551236_27551307@37.73.37,trna38_AspGTC_17_-_8125556_8125627@37.73.37,trna45_AspGTC_6_+_27447453_27447524@37.73.37,trna48_AspGTC_6_+_27471523_27471594@37.73.37,trna4_AspGTC_12_+_96429799_96429870@37.73.37,trna69_AspGTC_1_-_161440205_161440276@37.73.37,trna72_AspGTC_1_-_161432824_161432895@37.73.37,trna75_AspGTC_1_-_161425414_161425485@37.73.37,trna78_AspGTC_1_-_161418033_161418104@37.73.37,trna81_AspGTC_1_-_161410615_161410686@37.73.37|ACGCGGGAGACCGGGGTTCGATTCCCCGACGGGGAGC</td><td>>trna10_AspGTC_12_-_125424193_125424264</td><td>37</td><td>73</td><td>37</td><td>ACGCGGGAGACCGGGGTTCGATTCCCCGACGGGGAGC</td><td>1105.7</td><td>-4.3</td><td>1.0</td><td>-4.1</td><td>3.7E-05</td><td>3.7E-04</td><td>BT20/MB231</td></tr>
<tr><td>>hg38_21_+_8433172_8446622_RNA45SN1_45S_50ntFlanks@7977.8018.42|CGACCTCAGATCAGACGTGGCGACCCGCTGAATTTAAGCATA</td><td>>hg38_21_+_8433172_8446622_RNA45SN1_45S_50ntFlanks</td><td>7977</td><td>8018</td><td>42</td><td>CGACCTCAGATCAGACGTGGCGACCCGCTGAATTTAAGCATA</td><td>173.1</td><td>-4.2</td><td>1.0</td><td>-4.1</td><td>3.7E-05</td><td>3.7E-04</td><td>BT20/MB231</td></tr>
<tr><td>>hg38_21_+_8433172_8446622_RNA45SN1_45S_50ntFlanks@11320.11338.19|CCACGTCTCGTCGCGCGCG</td><td>>hg38_21_+_8433172_8446622_RNA45SN1_45S_50ntFlanks</td><td>11320</td><td>11338</td><td>19</td><td>CCACGTCTCGTCGCGCGCG</td><td>108.3</td><td>-3.7</td><td>1.0</td><td>-3.6</td><td>3.1E-04</td><td>3.7E-04</td><td>BT20/MB468</td></tr>
<tr><td>>hg38_21_+_8433172_8446622_RNA45SN1_45S_50ntFlanks@12457.12477.21|CAAGCGTTGGATTGTTCACCC</td><td>>hg38_21_+_8433172_8446622_RNA45SN1_45S_50ntFlanks</td><td>12457</td><td>12477</td><td>21</td><td>CAAGCGTTGGATTGTTCACCC</td><td>2714.2</td><td>-2.6</td><td>0.7</td><td>-3.6</td><td>2.8E-04</td><td>3.8E-04</td><td>MB231/MB468</td></tr>
<tr><td>>hg38_21_+_8433172_8446622_RNA45SN1_45S_50ntFlanks@8196.8222.27|CGTGGACGGTGTGAGGCCGGTAGCGGC</td><td>>hg38_21_+_8433172_8446622_RNA45SN1_45S_50ntFlanks</td><td>8196</td><td>8222</td><td>27</td><td>CGTGGACGGTGTGAGGCCGGTAGCGGC</td><td>203.5</td><td>-2.0</td><td>0.5</td><td>-3.6</td><td>3.1E-04</td><td>3.8E-04</td><td>BT20/MB468</td></tr>
<tr><td>>hg38_21_+_8433172_8446622_RNA45SN1_45S_50ntFlanks@12418.12444.27|CGATGTCGGCTCTTCCTATCATTGTGA</td><td>>hg38_21_+_8433172_8446622_RNA45SN1_45S_50ntFlanks</td><td>12418</td><td>12444</td><td>27</td><td>CGATGTCGGCTCTTCCTATCATTGTGA</td><td>765.9</td><td>-2.1</td><td>0.6</td><td>-3.6</td><td>3.1E-04</td><td>3.8E-04</td><td>BT20/MB468</td></tr>
<tr><td>>hg38_21_+_8433172_8446622_RNA45SN1_45S_50ntFlanks@6645.6669.25|CTCGTACGACTCTTAGCGGTGGATC</td><td>>hg38_21_+_8433172_8446622_RNA45SN1_45S_50ntFlanks</td><td>6645</td><td>6669</td><td>25</td><td>CTCGTACGACTCTTAGCGGTGGATC</td><td>2689.9</td><td>3.0</td><td>0.8</td><td>3.6</td><td>2.8E-04</td><td>3.8E-04</td><td>MB231/MB468</td></tr>
<tr><td>>hg38_21_+_8433172_8446622_RNA45SN1_45S_50ntFlanks@9739.9759.21|AAACGATCTCAACCTATTCTC</td><td>>hg38_21_+_8433172_8446622_RNA45SN1_45S_50ntFlanks</td><td>9739</td><td>9759</td><td>21</td><td>AAACGATCTCAACCTATTCTC</td><td>407.9</td><td>-1.8</td><td>0.5</td><td>-3.6</td><td>2.9E-04</td><td>3.8E-04</td><td>MB231/MB468</td></tr>
<tr><td>>hg38_21_+_8433172_8446622_RNA45SN1_45S_50ntFlanks@11576.11607.32|CTTAGAACTGGTGCGGACCAGGGGAATCCGAC</td><td>>hg38_21_+_8433172_8446622_RNA45SN1_45S_50ntFlanks</td><td>11576</td><td>11607</td><td>32</td><td>CTTAGAACTGGTGCGGACCAGGGGAATCCGAC</td><td>262.7</td><td>-2.0</td><td>0.6</td><td>-3.6</td><td>2.9E-04</td><td>3.8E-04</td><td>MB231/MB468</td></tr>
<tr><td>>hg38_21_+_8433172_8446622_RNA45SN1_45S_50ntFlanks@6649.6698.50|TACGACTCTTAGCGGTGGATCACTCGGCTCGTGCGTCGATGAAGAACGCA</td><td>>hg38_21_+_8433172_8446622_RNA45SN1_45S_50ntFlanks</td><td>6649</td><td>6698</td><td>50</td><td>TACGACTCTTAGCGGTGGATCACTCGGCTCGTGCGTCGATGAAGAACGCA</td><td>269.0</td><td>-3.4</td><td>0.8</td><td>-4.1</td><td>3.9E-05</td><td>3.8E-04</td><td>BT20/MB231</td></tr>
<tr><td>>hg38_21_+_8433172_8446622_RNA45SN1_45S_50ntFlanks@6701.6720.20|TAGCTGCGAGAATTAATGTG</td><td>>hg38_21_+_8433172_8446622_RNA45SN1_45S_50ntFlanks</td><td>6701</td><td>6720</td><td>20</td><td>TAGCTGCGAGAATTAATGTG</td><td>102.2</td><td>-1.4</td><td>0.4</td><td>-3.6</td><td>3.1E-04</td><td>3.8E-04</td><td>BT20/MB468</td></tr>
</table>

| | | | | | | | | | | | | |
|---|---|---|---|---|---|---|---|---|---|---|---|---|
| >hg38_21_+_8433172_8446622_RNA45SN1_45S_50ntFlanks@12995.13015.21\|CTCGCTGCGATCTATTGAAAG | >hg38_21_+_8433172_8446622_RNA45SN1_45S_50ntFlanks | 12995 | 13015 | 21 | CTCGCTGCGATCTATTGAAAG | 1248.0 | 3.7 | 1.0 | 3.6 | 3.2E-04 | 3.9E-04 | BT20/MB468 |
| >hg38_21_+_8433172_8446622_RNA45SN1_45S_50ntFlanks@11580.11609.30\|GAACTGGTGCGGACCAGGGGAATCCGACTG | >hg38_21_+_8433172_8446622_RNA45SN1_45S_50ntFlanks | 11580 | 11609 | 30 | GAACTGGTGCGGACCAGGGGAATCCGACTG | 850.2 | -2.4 | 0.7 | -3.6 | 3.2E-04 | 3.9E-04 | BT20/MB468 |
| >hg38_21_+_8433172_8446622_RNA45SN1_45S_50ntFlanks@7991.8010.20\|ACGTGGCGACCCGCTGAATT | >hg38_21_+_8433172_8446622_RNA45SN1_45S_50ntFlanks | 7991 | 8010 | 20 | ACGTGGCGACCCGCTGAATT | 2457.7 | 3.3 | 0.9 | 3.6 | 2.9E-04 | 3.9E-04 | MB231/MB468 |
| >hg38_21_+_8433172_8446622_RNA45SN1_45S_50ntFlanks@9555.9584.30\|GTAGCGGTCCTGACGTGCAAATCGGTCGTC | >hg38_21_+_8433172_8446622_RNA45SN1_45S_50ntFlanks | 9555 | 9584 | 30 | GTAGCGGTCCTGACGTGCAAATCGGTCGTC | 198.1 | -2.2 | 0.6 | -3.6 | 3.2E-04 | 3.9E-04 | BT20/MB468 |
| >MI0000063\|hsa-let-7b&WithFlank&22\|+\|46113680\|46113774@12.33.22(+1A)&[MIMAT0000063&hsa-let-7b-5p&offsets\|0\|0(+1A);m-14&22\|+\|46113691\|46113712&offsets\|0\|0(+1A)]\|TGAGGTAGTAGGTTGTGTGGTTA | >MI0000063\|hsa-let-7b&WithFlank&22\|+\|46113680\|46113774 | 12 | 33 | 22(+1A) | TGAGGTAGTAGGTTGTGTGGTTA | 3437.8 | 2.7 | 0.7 | 4.1 | 4.0E-05 | 3.9E-04 | BT20/MB231 |
| >hg38_21_+_8433172_8446622_RNA45SN1_45S_50ntFlanks@10683.10708.26\|CCGGCGGCGTCCGGTGAGCTCTCGCT | >hg38_21_+_8433172_8446622_RNA45SN1_45S_50ntFlanks | 10683 | 10708 | 26 | CCGGCGGCGTCCGGTGAGCTCTCGCT | 208.3 | -1.9 | 0.5 | -3.6 | 2.9E-04 | 3.9E-04 | MB231/MB468 |
| >hg38_21_+_8433172_8446622_RNA45SN1_45S_50ntFlanks@3844.3868.25\|TACTTGGATAACTGTGGTAATTCTA | >hg38_21_+_8433172_8446622_RNA45SN1_45S_50ntFlanks | 3844 | 3868 | 25 | TACTTGGATAACTGTGGTAATTCTA | 403.5 | 2.2 | 0.6 | 3.6 | 3.0E-04 | 4.0E-04 | MB231/MB468 |
| >hg38_21_+_8433172_8446622_RNA45SN1_45S_50ntFlanks@10409.10432.24\|GTCAGTCGGTCCTGAGAGATGGGC | >hg38_21_+_8433172_8446622_RNA45SN1_45S_50ntFlanks | 10409 | 10432 | 24 | GTCAGTCGGTCCTGAGAGATGGGC | 159.1 | -1.5 | 0.4 | -3.6 | 3.3E-04 | 4.0E-04 | BT20/MB468 |
| >hg38_21_+_8433172_8446622_RNA45SN1_45S_50ntFlanks@3837.3860.24\|CCTCTCCTACTTGGATAACTGTGG | >hg38_21_+_8433172_8446622_RNA45SN1_45S_50ntFlanks | 3837 | 3860 | 24 | CCTCTCCTACTTGGATAACTGTGG | 172.0 | -1.8 | 0.5 | -3.6 | 3.0E-04 | 4.0E-04 | MB231/MB468 |
| >MI0000063\|hsa-let-7b&WithFlank&22\|+\|46113680\|46113774@12.31.20(+1A)&[MIMAT0000063&hsa-let-7b-5p&offsets\|0\|-2(+1A);m-14&22\|+\|46113691\|46113712&offsets\|0\|-2(+1A)]\|TGAGGTAGTAGGTTGTGTGGA | >MI0000063\|hsa-let-7b&WithFlank&22\|+\|46113680\|46113774 | 12 | 31 | 20(+1A) | TGAGGTAGTAGGTTGTGTGGA | 338.3 | 3.2 | 0.8 | 4.1 | 4.1E-05 | 4.0E-04 | BT20/MB231 |
| >hg38_21_+_8433172_8446622_RNA45SN1_45S_50ntFlanks@10353.10390.38\|AGAACTTTGAAGGCCGAAGTGGAGAAGGGTTCCATGTG | >hg38_21_+_8433172_8446622_RNA45SN1_45S_50ntFlanks | 10353 | 10390 | 38 | AGAACTTTGAAGGCCGAAGTGGAGAAGGGTTCCATGTG | 695.3 | 2.4 | 0.7 | 3.6 | 3.0E-04 | 4.0E-04 | MB231/MB468 |
| >MI0001641\|hsa-mir-429&WithFlank&1\|+\|1168999\|1169093@57.78.22&[MIMAT0001536&hsa-miR-429&offsets\|0\|0;m-207&1\|+\|1169055\|1169076&offsets\|0\|0]\|TAATACTGTCTGGTAAAACCGT | >MI0001641\|hsa-mir-429&WithFlank&1\|+\|1168999\|1169093 | 57 | 78 | 22 | TAATACTGTCTGGTAAAACCGT | 789.4 | -3.0 | 0.8 | -3.6 | 3.0E-04 | 4.0E-04 | MB231/MB468 |
| >MI0000272\|hsa-mir-182&WithFlank&7\|-\|129770377\|129770498@29.54.26&[MIMAT0000259&hsa-miR-182-5p&offsets\|0\|+2;m-22&7\|-\|129770449\|129770470&offsets\|0\|+4]\|TTTGGCAATGGTAGAACTCACACTGG | >MI0000272\|hsa-mir-182&WithFlank&7\|-\|129770377\|129770498 | 29 | 54 | 26 | TTTGGCAATGGTAGAACTCACACTGG | 6283.7 | 2.7 | 0.7 | 4.1 | 4.1E-05 | 4.1E-04 | BT20/MB231 |
| >hg38_21_+_8433172_8446622_RNA45SN1_45S_50ntFlanks@12418.12441.24\|CGATGTCGGCTCTTCCTATCATTG | >hg38_21_+_8433172_8446622_RNA45SN1_45S_50ntFlanks | 12418 | 12441 | 24 | CGATGTCGGCTCTTCCTATCATTG | 812.9 | -2.4 | 0.7 | -3.6 | 3.0E-04 | 4.1E-04 | MB231/MB468 |
| >hg38_21_+_8433172_8446622_RNA45SN1_45S_50ntFlanks@11319.11336.18\|CCCACGTCTCGTCGCGCG | >hg38_21_+_8433172_8446622_RNA45SN1_45S_50ntFlanks | 11319 | 11336 | 18 | CCCACGTCTCGTCGCGCG | 107.3 | -4.0 | 1.1 | -3.6 | 3.3E-04 | 4.1E-04 | BT20/MB468 |
| >MI0000542\|hsa-mir-320a&WithFlank&8\|-\|22244960\|22245043@48.69.22&[MIMAT0000510&hsa-miR-320a-3p&offsets\|0\|0;m-60&8\|-\|22244975\|22244996&offsets\|0\|0]\|AAAAGCTGGGTTGAGAGGGCGA | >MI0000542\|hsa-mir-320a&WithFlank&8\|-\|22244960\|22245043 | 48 | 69 | 22 | AAAAGCTGGGTTGAGAGGGCGA | 16520.3 | 2.9 | 0.7 | 4.1 | 4.2E-05 | 4.1E-04 | BT20/MB231 |
| >hg38_21_+_8433172_8446622_RNA45SN1_45S_50ntFlanks@11316.11335.20\|CACCCCACGTCTCGTCGCGC | >hg38_21_+_8433172_8446622_RNA45SN1_45S_50ntFlanks | 11316 | 11335 | 20 | CACCCCACGTCTCGTCGCGC | 95.3 | -3.7 | 1.0 | -3.6 | 3.4E-04 | 4.1E-04 | BT20/MB468 |
| >hg38_21_+_8433172_8446622_RNA45SN1_45S_50ntFlanks@12481.12500.20\|AATAGGGAACGTGAGCTGGG | >hg38_21_+_8433172_8446622_RNA45SN1_45S_50ntFlanks | 12481 | 12500 | 20 | AATAGGGAACGTGAGCTGGG | 231.4 | 1.5 | 0.4 | 3.6 | 3.1E-04 | 4.2E-04 | MB231/MB468 |
| >hg38_21_+_8433172_8446622_RNA45SN1_45S_50ntFlanks@12478.12498.21\|ACTAATAGGGAACGTGAGCTG | >hg38_21_+_8433172_8446622_RNA45SN1_45S_50ntFlanks | 12478 | 12498 | 21 | ACTAATAGGGAACGTGAGCTG | 496.4 | -1.7 | 0.5 | -3.6 | 3.1E-04 | 4.2E-04 | MB231/MB468 |
| >hg38_21_+_8433172_8446622_RNA45SN1_45S_50ntFlanks@4730.4752.23\|CAAGAACGAAAGTCGGAGGTTCG | >hg38_21_+_8433172_8446622_RNA45SN1_45S_50ntFlanks | 4730 | 4752 | 23 | CAAGAACGAAAGTCGGAGGTTCG | 222.3 | -1.7 | 0.5 | -3.6 | 3.1E-04 | 4.2E-04 | MB231/MB468 |
| >hg38_21_+_8433172_8446622_RNA45SN1_45S_50ntFlanks@11565.11582.18\|CTAGCAGCCGACTTAGAA | >hg38_21_+_8433172_8446622_RNA45SN1_45S_50ntFlanks | 11565 | 11582 | 18 | CTAGCAGCCGACTTAGAA | 137.6 | -2.7 | 0.7 | -3.6 | 3.4E-04 | 4.2E-04 | BT20/MB468 |
| >hg38_21_+_8433172_8446622_RNA45SN1_45S_50ntFlanks@10841.10873.33\|CGTAACTTCGGGATAAGGATTGGCTCTAAGGGC | >hg38_21_+_8433172_8446622_RNA45SN1_45S_50ntFlanks | 10841 | 10873 | 33 | CGTAACTTCGGGATAAGGATTGGCTCTAAGGGC | 259.6 | -1.6 | 0.5 | -3.6 | 3.1E-04 | 4.2E-04 | MB231/MB468 |
| >hg38_21_+_8433172_8446622_RNA45SN1_45S_50ntFlanks@4760.4786.27\|TCAGATACCGTCGTAGTTCCGACCATA | >hg38_21_+_8433172_8446622_RNA45SN1_45S_50ntFlanks | 4760 | 4786 | 27 | TCAGATACCGTCGTAGTTCCGACCATA | 353.0 | -1.8 | 0.5 | -3.6 | 3.1E-04 | 4.2E-04 | MB231/MB468 |
| >hg38_21_+_8433172_8446622_RNA45SN1_45S_50ntFlanks@13002.13030.29\|CGATCTATTGAAAGTCAGCCCTCGACACA | >hg38_21_+_8433172_8446622_RNA45SN1_45S_50ntFlanks | 13002 | 13030 | 29 | CGATCTATTGAAAGTCAGCCCTCGACACA | 115.4 | -3.3 | 0.9 | -3.6 | 3.1E-04 | 4.2E-04 | MB231/MB468 |
| >hg38_21_+_8433172_8446622_RNA45SN1_45S_50ntFlanks@4328.4346.19\|CGGTAATTCCAGCTCCAAT | >hg38_21_+_8433172_8446622_RNA45SN1_45S_50ntFlanks | 4328 | 4346 | 19 | CGGTAATTCCAGCTCCAAT | 126.3 | -2.1 | 0.6 | -3.6 | 3.1E-04 | 4.2E-04 | MB231/MB468 |
| >hg38_21_+_8433172_8446622_RNA45SN1_45S_50ntFlanks@10830.10852.23\|CAAGCCGGATCCGTAACTTCGGG | >hg38_21_+_8433172_8446622_RNA45SN1_45S_50ntFlanks | 10830 | 10852 | 23 | CAAGCCGGATCCGTAACTTCGGG | 228.4 | -2.9 | 0.8 | -3.6 | 3.2E-04 | 4.2E-04 | MB231/MB468 |
| >hg38_21_+_8433172_8446622_RNA45SN1_45S_50ntFlanks@10791.10818.28\|CCTCTGGCATGTTGGAACAATGTAGGTA | >hg38_21_+_8433172_8446622_RNA45SN1_45S_50ntFlanks | 10791 | 10818 | 28 | CCTCTGGCATGTTGGAACAATGTAGGTA | 306.9 | -2.9 | 0.8 | -3.6 | 3.5E-04 | 4.2E-04 | BT20/MB468 |
| >hg38_21_+_8433172_8446622_RNA45SN1_45S_50ntFlanks@3853.3871.19\|AACTGTGGTAATTCTAGAG | >hg38_21_+_8433172_8446622_RNA45SN1_45S_50ntFlanks | 3853 | 3871 | 19 | AACTGTGGTAATTCTAGAG | 221.7 | -2.4 | 0.7 | -3.6 | 3.2E-04 | 4.2E-04 | MB231/MB468 |
| >hg38_21_+_8433172_8446622_RNA45SN1_45S_50ntFlanks@10385.10405.21\|CATGTGAACAGCAGTTGAACA | >hg38_21_+_8433172_8446622_RNA45SN1_45S_50ntFlanks | 10385 | 10405 | 21 | CATGTGAACAGCAGTTGAACA | 123.5 | -1.9 | 0.5 | -3.6 | 3.2E-04 | 4.3E-04 | MB231/MB468 |
| >hg38_21_+_8433172_8446622_RNA45SN1_45S_50ntFlanks@12996.13040.45\|TCGCTGCGATCTATTGAAAGTCAGCCCTCGACACAAGGGTTTGTC | >hg38_21_+_8433172_8446622_RNA45SN1_45S_50ntFlanks | 12996 | 13040 | 45 | TCGCTGCGATCTATTGAAAGTCAGCCCTCGACACAAGGGTTTGTC | 781.7 | -3.9 | 1.0 | -4.1 | 4.4E-05 | 4.3E-04 | BT20/MB231 |
| >hg38_21_+_8433172_8446622_RNA45SN1_45S_50ntFlanks@476.523.48\|TTCGTGATCGATGTGGTGACGTCGTGCTCTCCCGGGCCGGGTCCGAGC | >hg38_21_+_8433172_8446622_RNA45SN1_45S_50ntFlanks | 476 | 523 | 48 | TTCGTGATCGATGTGGTGACGTCGTGCTCTCCCGGGCCGGGTCCGAGC | 422.4 | 9.0 | 2.5 | 3.6 | 3.5E-04 | 4.3E-04 | BT20/MB468 |
| >Antisense_hg38_21_-_8433172_8446622_RNA45SN1_45S_50ntFlanks@7674.7693.20\|CTCTCTCTCTCTCTCCCCCG | >Antisense_hg38_21_-_8433172_8446622_RNA45SN1_45S_50ntFlanks | 7674 | 7693 | 20 | CTCTCTCTCTCTCTCCCCCG | 207.3 | -5.7 | 1.6 | -3.6 | 3.2E-04 | 4.3E-04 | MB231/MB468 |
| >hg38_21_+_8433172_8446622_RNA45SN1_45S_50ntFlanks@11027.11046.20\|CTCTCTCTCTCTCTCCCCCG | >hg38_21_+_8433172_8446622_RNA45SN1_45S_50ntFlanks | 11027 | 11046 | 20 | CTCTCTCTCTCTCTCCCCCG | 207.3 | -5.7 | 1.6 | -3.6 | 3.2E-04 | 4.3E-04 | MB231/MB468 |
| >hg38_21_+_8433172_8446622_RNA45SN1_45S_50ntFlanks@12991.13017.27\|CTCCCTCGCTGCGATCTATTGAAAGTC | >hg38_21_+_8433172_8446622_RNA45SN1_45S_50ntFlanks | 12991 | 13017 | 27 | CTCCCTCGCTGCGATCTATTGAAAGTC | 291.5 | -2.6 | 0.7 | -3.6 | 3.5E-04 | 4.3E-04 | BT20/MB468 |
| >hg38_21_+_8433172_8446622_RNA45SN1_45S_50ntFlanks@5125.5145.21\|AGCGGTCGGCGTCCCCCAACT | >hg38_21_+_8433172_8446622_RNA45SN1_45S_50ntFlanks | 5125 | 5145 | 21 | AGCGGTCGGCGTCCCCCAACT | 213.5 | 3.6 | 0.9 | 4.1 | 4.5E-05 | 4.3E-04 | BT20/MB231 |
| >hg38_21_+_8433172_8446622_RNA45SN1_45S_50ntFlanks@6645.6680.36\|CTCGTACGACTCTTAGCGGTGGATCACTCGGCTCGT | >hg38_21_+_8433172_8446622_RNA45SN1_45S_50ntFlanks | 6645 | 6680 | 36 | CTCGTACGACTCTTAGCGGTGGATCACTCGGCTCGT | 551.5 | 2.5 | 0.7 | 3.6 | 3.3E-04 | 4.4E-04 | MB231/MB468 |
| >hg38_21_+_8433172_8446622_RNA45SN1_45S_50ntFlanks@8342.8365.24\|CGTAAGGGAAAGTTGAAAAGAACT | >hg38_21_+_8433172_8446622_RNA45SN1_45S_50ntFlanks | 8342 | 8365 | 24 | CGTAAGGGAAAGTTGAAAAGAACT | 678.7 | 5.7 | 1.6 | 3.6 | 3.6E-04 | 4.4E-04 | BT20/MB468 |
| >hg38_21_+_8433172_8446622_RNA45SN1_45S_50ntFlanks@6733.6754.22\|CATTGATCATCGACACTTCGAA | >hg38_21_+_8433172_8446622_RNA45SN1_45S_50ntFlanks | 6733 | 6754 | 22 | CATTGATCATCGACACTTCGAA | 173.9 | -2.0 | 0.6 | -3.6 | 3.6E-04 | 4.4E-04 | BT20/MB468 |
| >MI0000272\|hsa-mir-182&WithFlank&7\|-\|129770377\|129770498@29.53.25(+1A)&[MIMAT0000259&hsa-miR-182-5p&offsets\|0\|+1(+1A);m-22&7\|-\|129770449\|129770470&offsets\|0\|+3(+1A)]\|TTTGGCAATGGTAGAACTCACACTGA | >MI0000272\|hsa-mir-182&WithFlank&7\|-\|129770377\|129770498 | 29 | 53 | 25(+1A) | TTTGGCAATGGTAGAACTCACACTGA | 1678.6 | 3.3 | 0.8 | 4.1 | 4.6E-05 | 4.4E-04 | BT20/MB231 |
| >hg38_21_+_8433172_8446622_RNA45SN1_45S_50ntFlanks@12938.12955.18\|ACCATTCGTAGACGACCT | >hg38_21_+_8433172_8446622_RNA45SN1_45S_50ntFlanks | 12938 | 12955 | 18 | ACCATTCGTAGACGACCT | 98.0 | -1.6 | 0.5 | -3.6 | 3.3E-04 | 4.4E-04 | MB231/MB468 |
| >hg38_21_+_8433172_8446622_RNA45SN1_45S_50ntFlanks@12073.12106.34\|CCGAGGGGCTCTCGCTTCTGGCGCCAAGCGCCCG | >hg38_21_+_8433172_8446622_RNA45SN1_45S_50ntFlanks | 12073 | 12106 | 34 | CCGAGGGGCTCTCGCTTCTGGCGCCAAGCGCCCG | 286.9 | -4.4 | 1.2 | -3.6 | 3.6E-04 | 4.5E-04 | BT20/MB468 |
| >hg38_21_+_8433172_8446622_RNA45SN1_45S_50ntFlanks@10686.10710.25\|GCGGCGTCCGGTGAGCTCTCGCTGG | >hg38_21_+_8433172_8446622_RNA45SN1_45S_50ntFlanks | 10686 | 10710 | 25 | GCGGCGTCCGGTGAGCTCTCGCTGG | 2331.7 | -3.2 | 0.9 | -3.6 | 3.6E-04 | 4.5E-04 | BT20/MB468 |
| >hg38_21_+_8433172_8446622_RNA45SN1_45S_50ntFlanks@6776.6802.27\|CTCCCGGGGCTACGCCTGTCTGAGCGT | >hg38_21_+_8433172_8446622_RNA45SN1_45S_50ntFlanks | 6776 | 6802 | 27 | CTCCCGGGGCTACGCCTGTCTGAGCGT | 414.5 | 2.4 | 0.7 | 3.6 | 3.7E-04 | 4.5E-04 | BT20/MB468 |
| >hg38_21_+_8433172_8446622_RNA45SN1_45S_50ntFlanks@4729.4752.24\|TCAAGAACGAAAGTCGGAGGTTCG | >hg38_21_+_8433172_8446622_RNA45SN1_45S_50ntFlanks | 4729 | 4752 | 24 | TCAAGAACGAAAGTCGGAGGTTCG | 130.8 | -2.1 | 0.6 | -3.6 | 3.4E-04 | 4.5E-04 | MB231/MB468 |
| >hg38_1_-_228634819_228635039_RNA5S12_5S_50ntFlanks@72.92.21\|AACGCGCCCGATCTCGTCTGA | >hg38_1_-_228634819_228635039_RNA5S12_5S_50ntFlanks | 72 | 92 | 21 | AACGCGCCCGATCTCGTCTGA | 71.7 | -2.2 | 0.6 | -3.6 | 3.4E-04 | 4.5E-04 | MB231/MB468 |
| >hg38_21_+_8433172_8446622_RNA45SN1_45S_50ntFlanks@12994.13015.22\|CCTCGCTGCGATCTATTGAAAG | >hg38_21_+_8433172_8446622_RNA45SN1_45S_50ntFlanks | 12994 | 13015 | 22 | CCTCGCTGCGATCTATTGAAAG | 279.7 | -2.4 | 0.7 | -3.6 | 3.7E-04 | 4.5E-04 | BT20/MB468 |
| >hg38_21_+_8433172_8446622_RNA45SN1_45S_50ntFlanks@6689.6720.32\|GAAGAACGCAGCTAGCTGCGAGAATTAATGTG | >hg38_21_+_8433172_8446622_RNA45SN1_45S_50ntFlanks | 6689 | 6720 | 32 | GAAGAACGCAGCTAGCTGCGAGAATTAATGTG | 127.8 | -1.8 | 0.5 | -3.6 | 3.4E-04 | 4.6E-04 | MB231/MB468 |

| | | | | | | | | | | | | |
|---|---|---|---|---|---|---|---|---|---|---|---|---|
| >MI0003776\|hsa-mir-320b-1&WithFlank&1\|+\|116671740\|116671823@48.67.20(+1U);MI0003839\|hsa-mir-320b-2&WithFlank&1\|-\|224257034\|224257116@47.66.20(+1U);MI0000542\|hsa-mir-320a&WithFlank&8\|-\|22244960\|22245043@48.67.20(+1U)&[MIMAT0005792&hsa-miR-320b&offsets\|0\|-2(+1U);m-183&1\|+\|116671787\|116671806&offsets\|0\|0(+1U)];[MIMAT0005792_1&hsa-miR-320b&offsets\|0\|-2(+1U);m-183&1\|-\|224257051\|224257070&offsets\|0\|0(+1U)];[MIMAT0000510&hsa-miR-320a-3p&offsets\|0\|-2(+1U);m-60&8\|-\|22244975\|22244996&offsets\|0\|-2(+1U)]\|AAAAGCTGGGTTGAGAGGGCT | >MI0003776\|hsa-mir-320b-1&WithFlank&1\|+\|116671740\|116671823 | 48 | 67 | 20(+1U) | AAAAGCTGGGTTGAGAGGGCT | 467.6 | 3.0 | 0.7 | 4.1 | 4.7E-05 | 4.6E-04 | BT20/MB231 |
| >hg38_21_+_8433172_8446622_RNA45SN1_45S_50ntFlanks@5472.5515.44\|ACGGCCCTGGCGGAGCGCTGAGAAGACGGTCGAACTTGACTATC | >hg38_21_+_8433172_8446622_RNA45SN1_45S_50ntFlanks | 5472 | 5515 | 44 | ACGGCCCTGGCGGAGCGCTGAGAAGACGGTCGAACTTGACTATC | 239.0 | -3.1 | 0.8 | -4.1 | 4.7E-05 | 4.6E-04 | BT20/MB231 |
| >hg38_21_+_8433172_8446622_RNA45SN1_45S_50ntFlanks@11552.11574.23\|CCTCGGCCGGCGCCTAGCAGCCG | >hg38_21_+_8433172_8446622_RNA45SN1_45S_50ntFlanks | 11552 | 11574 | 23 | CCTCGGCCGGCGCCTAGCAGCCG | 98.0 | -1.7 | 0.5 | -3.6 | 3.4E-04 | 4.6E-04 | MB231/MB468 |
| >hg38_21_+_8433172_8446622_RNA45SN1_45S_50ntFlanks@11663.11685.23\|ATTTCTGCCCAGTGCTCTGAATG | >hg38_21_+_8433172_8446622_RNA45SN1_45S_50ntFlanks | 11663 | 11685 | 23 | ATTTCTGCCCAGTGCTCTGAATG | 452.3 | -1.9 | 0.5 | -3.6 | 3.5E-04 | 4.6E-04 | MB231/MB468 |
| >hg38_21_+_8433172_8446622_RNA45SN1_45S_50ntFlanks@10778.10818.41\|CCAAGGTGAACAGCCTCTGGCATGTTGGAACAATGTAGGTA | >hg38_21_+_8433172_8446622_RNA45SN1_45S_50ntFlanks | 10778 | 10818 | 41 | CCAAGGTGAACAGCCTCTGGCATGTTGGAACAATGTAGGTA | 150.9 | -2.3 | 0.6 | -3.6 | 3.5E-04 | 4.6E-04 | MB231/MB468 |
| >hg38_21_+_8433172_8446622_RNA45SN1_45S_50ntFlanks@11881.11899.19\|CCTGTTGAGCTTGACTCTA | >hg38_21_+_8433172_8446622_RNA45SN1_45S_50ntFlanks | 11881 | 11899 | 19 | CCTGTTGAGCTTGACTCTA | 99.6 | -3.2 | 0.9 | -3.6 | 3.5E-04 | 4.7E-04 | MB231/MB468 |
| >hg38_21_+_8433172_8446622_RNA45SN1_45S_50ntFlanks@4565.4589.25\|AATAATGGAATAGGACCGCGGTTCT | >hg38_21_+_8433172_8446622_RNA45SN1_45S_50ntFlanks | 4565 | 4589 | 25 | AATAATGGAATAGGACCGCGGTTCT | 278.1 | 2.4 | 0.7 | 3.6 | 3.5E-04 | 4.7E-04 | MB231/MB468 |
| >hg38_21_+_8433172_8446622_RNA45SN1_45S_50ntFlanks@9353.9370.18\|CGTGGCGCAATGAAGGTG | >hg38_21_+_8433172_8446622_RNA45SN1_45S_50ntFlanks | 9353 | 9370 | 18 | CGTGGCGCAATGAAGGTG | 71.1 | -2.9 | 0.8 | -3.6 | 3.5E-04 | 4.7E-04 | MB231/MB468 |
| >hg38_21_+_8433172_8446622_RNA45SN1_45S_50ntFlanks@11769.11790.22\|TAATTAGTGACGCGCATGAATG | >hg38_21_+_8433172_8446622_RNA45SN1_45S_50ntFlanks | 11769 | 11790 | 22 | TAATTAGTGACGCGCATGAATG | 508.0 | -1.9 | 0.5 | -3.6 | 3.5E-04 | 4.7E-04 | MB231/MB468 |
| >hg38_21_+_8433172_8446622_RNA45SN1_45S_50ntFlanks@4370.4391.22\|TTAAAAAGCTCGTAGTTGGATC | >hg38_21_+_8433172_8446622_RNA45SN1_45S_50ntFlanks | 4370 | 4391 | 22 | TTAAAAAGCTCGTAGTTGGATC | 247.5 | -2.4 | 0.7 | -3.6 | 3.8E-04 | 4.7E-04 | BT20/MB468 |
| >hg38_21_+_8433172_8446622_RNA45SN1_45S_50ntFlanks@10099.10117.19\|CGCGCGCGCGTGTGGTGTG | >hg38_21_+_8433172_8446622_RNA45SN1_45S_50ntFlanks | 10099 | 10117 | 19 | CGCGCGCGCGTGTGGTGTG | 403.2 | -3.6 | 1.0 | -3.6 | 3.8E-04 | 4.7E-04 | BT20/MB468 |
| >hg38_21_+_8433172_8446622_RNA45SN1_45S_50ntFlanks@3878.3898.21\|CATGCCGACGGGCGCTGACCC | >hg38_21_+_8433172_8446622_RNA45SN1_45S_50ntFlanks | 3878 | 3898 | 21 | CATGCCGACGGGCGCTGACCC | 191.5 | -2.0 | 0.6 | -3.6 | 3.8E-04 | 4.7E-04 | BT20/MB468 |
| >MI0000286\|hsa-mir-210&WithFlank&11\|-\|568083\|568204@72.93.22(+2A)&[MIMAT0000267&hsa-miR-210-3p&offsets\|0\|0(+2A);m-102&11\|-\|568112\|568133&offsets\|0\|0(+2A)]\|CTGTGCGTGTGACAGCGGCTGAAA | >MI0000286\|hsa-mir-210&WithFlank&11\|-\|568083\|568204 | 72 | 93 | 22(+2A) | CTGTGCGTGTGACAGCGGCTGAAA | 4790.1 | 3.0 | 0.7 | 4.1 | 4.9E-05 | 4.7E-04 | BT20/MB231 |
| >hg38_21_+_8433172_8446622_RNA45SN1_45S_50ntFlanks@9739.9762.24\|AAACGATCTCAACCTATTCTCAAA | >hg38_21_+_8433172_8446622_RNA45SN1_45S_50ntFlanks | 9739 | 9762 | 24 | AAACGATCTCAACCTATTCTCAAA | 81.5 | -2.4 | 0.7 | -3.5 | 3.9E-04 | 4.7E-04 | BT20/MB468 |
| >hg38_21_+_8433172_8446622_RNA45SN1_45S_50ntFlanks@8182.8223.42\|ATCGAGGCCCAGCCCGTGGACGGTGTGAGGCCGGTAGCGGCC | >hg38_21_+_8433172_8446622_RNA45SN1_45S_50ntFlanks | 8182 | 8223 | 42 | ATCGAGGCCCAGCCCGTGGACGGTGTGAGGCCGGTAGCGGCC | 97.1 | -3.2 | 0.9 | -3.6 | 3.6E-04 | 4.8E-04 | MB231/MB468 |
| >hg38_21_+_8433172_8446622_RNA45SN1_45S_50ntFlanks@9564.9595.32\|CTGACGTGCAAATCGGTCGTCCGACCTGGGTA | >hg38_21_+_8433172_8446622_RNA45SN1_45S_50ntFlanks | 9564 | 9595 | 32 | CTGACGTGCAAATCGGTCGTCCGACCTGGGTA | 1173.7 | -2.8 | 0.8 | -3.6 | 3.6E-04 | 4.8E-04 | MB231/MB468 |
| >hg38_21_+_8433172_8446622_RNA45SN1_45S_50ntFlanks@10922.10945.24\|CTGGACGAGGCGCCGCCGCCCCCC | >hg38_21_+_8433172_8446622_RNA45SN1_45S_50ntFlanks | 10922 | 10945 | 24 | CTGGACGAGGCGCCGCCGCCCCCC | 262.8 | -5.6 | 1.6 | -3.6 | 3.6E-04 | 4.8E-04 | MB231/MB468 |
| >hg38_21_+_8433172_8446622_RNA45SN1_45S_50ntFlanks@11323.11344.22\|CGTCTCGTCGCGCGCGCGTCCG | >hg38_21_+_8433172_8446622_RNA45SN1_45S_50ntFlanks | 11323 | 11344 | 22 | CGTCTCGTCGCGCGCGCGTCCG | 7022.8 | -5.1 | 1.4 | -3.5 | 4.0E-04 | 4.8E-04 | BT20/MB468 |
| >hg38_21_+_8433172_8446622_RNA45SN1_45S_50ntFlanks@4048.4066.19\|TTCGAACGTCTGCCCTATC | >hg38_21_+_8433172_8446622_RNA45SN1_45S_50ntFlanks | 4048 | 4066 | 19 | TTCGAACGTCTGCCCTATC | 458.5 | 2.0 | 0.6 | 3.6 | 3.6E-04 | 4.8E-04 | MB231/MB468 |
| >MI0000298\|hsa-mir-221&WithFlank&X\|-\|45746151\|45746272@71.92.22(+2U)&[MIMAT0000278&hsa-miR-221-3p&offsets\|0\|-1(+2U);m-91&X\|-\|45746181\|45746202&offsets\|0\|0(+2U)]\|AGCTACATTGTCTGCTGGGTTTTT | >MI0000298\|hsa-mir-221&WithFlank&X\|-\|45746151\|45746272 | 71 | 92 | 22(+2U) | AGCTACATTGTCTGCTGGGTTTTT | 565.8 | -2.9 | 0.7 | -4.1 | 5.1E-05 | 4.8E-04 | BT20/MB231 |
| >hg38_21_+_8433172_8446622_RNA45SN1_45S_50ntFlanks@6775.6803.29\|CCTCCCGGGGCTACGCCTGTCTGAGCGTC | >hg38_21_+_8433172_8446622_RNA45SN1_45S_50ntFlanks | 6775 | 6803 | 29 | CCTCCCGGGGCTACGCCTGTCTGAGCGTC | 274.5 | -3.0 | 0.8 | -3.5 | 4.0E-04 | 4.9E-04 | BT20/MB468 |
| >MI0000273\|hsa-mir-183&WithFlank&7\|-\|129774899\|129775020@33.53.21(+1C)&[MIMAT0000261&hsa-miR-183-5p&offsets\|0\|-1(+1C);m-41&7\|-\|129774966\|129774987&offsets\|-1\|-2(+1C)]\|TATGGCACTGGTAGAATTCACC | >MI0000273\|hsa-mir-183&WithFlank&7\|-\|129774899\|129775020 | 33 | 53 | 21(+1C) | TATGGCACTGGTAGAATTCACC | 2648.8 | 2.7 | 0.7 | 4.0 | 5.2E-05 | 4.9E-04 | BT20/MB231 |
| >hg38_21_+_8433172_8446622_RNA45SN1_45S_50ntFlanks@9533.9552.20\|GAGGAAACTCTGGTGGAGGT | >hg38_21_+_8433172_8446622_RNA45SN1_45S_50ntFlanks | 9533 | 9552 | 20 | GAGGAAACTCTGGTGGAGGT | 227.8 | -2.1 | 0.6 | -3.5 | 4.1E-04 | 4.9E-04 | BT20/MB468 |
| >trnaMT_SerGCT_MT_+_12207_12265@10.38.29\|CACAAGAACTGCTAACTCATGCCCCCATG | >trnaMT_SerGCT_MT_+_12207_12265 | 10 | 38 | 29 | CACAAGAACTGCTAACTCATGCCCCCATG | 170.1 | -4.4 | 1.3 | -3.5 | 4.1E-04 | 5.0E-04 | BT20/MB468 |
| >hg38_21_+_8433172_8446622_RNA45SN1_45S_50ntFlanks@10391.10422.32\|AACAGCAGTTGAACATGGGTCAGTCGGTCCTG | >hg38_21_+_8433172_8446622_RNA45SN1_45S_50ntFlanks | 10391 | 10422 | 32 | AACAGCAGTTGAACATGGGTCAGTCGGTCCTG | 221.4 | -2.0 | 0.6 | -3.6 | 3.7E-04 | 5.0E-04 | MB231/MB468 |
| >hg38_21_+_8433172_8446622_RNA45SN1_45S_50ntFlanks@11576.11611.36\|CTTAGAACTGGTGCGGACCAGGGGAATCCGACTGTT | >hg38_21_+_8433172_8446622_RNA45SN1_45S_50ntFlanks | 11576 | 11611 | 36 | CTTAGAACTGGTGCGGACCAGGGGAATCCGACTGTT | 387.5 | -2.7 | 0.8 | -3.5 | 4.1E-04 | 5.0E-04 | BT20/MB468 |
| >hg38_21_+_8433172_8446622_RNA45SN1_45S_50ntFlanks@4728.4747.20\|ATCAAGAACGAAAGTCGGAG | >hg38_21_+_8433172_8446622_RNA45SN1_45S_50ntFlanks | 4728 | 4747 | 20 | ATCAAGAACGAAAGTCGGAG | 63.4 | -3.3 | 0.9 | -3.6 | 3.8E-04 | 5.0E-04 | MB231/MB468 |
| >hg38_21_+_8433172_8446622_RNA45SN1_45S_50ntFlanks@12997.13038.42\|CGCTGCGATCTATTGAAAGTCAGCCCTCGACACAAGGGTTTG | >hg38_21_+_8433172_8446622_RNA45SN1_45S_50ntFlanks | 12997 | 13038 | 42 | CGCTGCGATCTATTGAAAGTCAGCCCTCGACACAAGGGTTTG | 4144.5 | -3.2 | 0.8 | -4.0 | 5.3E-05 | 5.1E-04 | BT20/MB231 |
| >hg38_21_+_8433172_8446622_RNA45SN1_45S_50ntFlanks@6646.6680.35\|TCGTACGACTCTTAGCGGTGGATCACTCGGCTCGT | >hg38_21_+_8433172_8446622_RNA45SN1_45S_50ntFlanks | 6646 | 6680 | 35 | TCGTACGACTCTTAGCGGTGGATCACTCGGCTCGT | 3064.0 | 2.2 | 0.6 | 3.6 | 3.8E-04 | 5.1E-04 | MB231/MB468 |
| >hg38_21_+_8433172_8446622_RNA45SN1_45S_50ntFlanks@11567.11603.37\|AGCAGCCGACTTAGAACTGGTGCGGACCAGGGGAATC | >hg38_21_+_8433172_8446622_RNA45SN1_45S_50ntFlanks | 11567 | 11603 | 37 | AGCAGCCGACTTAGAACTGGTGCGGACCAGGGGAATC | 172.2 | -1.6 | 0.5 | -3.5 | 4.2E-04 | 5.1E-04 | BT20/MB468 |
| >hg38_21_+_8433172_8446622_RNA45SN1_45S_50ntFlanks@12477.12499.23\|CACTAATAGGGAACGTGAGCTGG | >hg38_21_+_8433172_8446622_RNA45SN1_45S_50ntFlanks | 12477 | 12499 | 23 | CACTAATAGGGAACGTGAGCTGG | 257.8 | -1.6 | 0.4 | -3.5 | 4.2E-04 | 5.1E-04 | BT20/MB468 |
| >hg38_21_+_8433172_8446622_RNA45SN1_45S_50ntFlanks@12162.12180.19\|GACTGGGGCGGTACACCTG | >hg38_21_+_8433172_8446622_RNA45SN1_45S_50ntFlanks | 12162 | 12180 | 19 | GACTGGGGCGGTACACCTG | 312.2 | -5.8 | 1.6 | -3.5 | 3.9E-04 | 5.1E-04 | MB231/MB468 |
| >hg38_21_+_8433172_8446622_RNA45SN1_45S_50ntFlanks@9623.9652.30\|TAGTAGCTGGTTCCCTCCGAAGTTTCCCTC | >hg38_21_+_8433172_8446622_RNA45SN1_45S_50ntFlanks | 9623 | 9652 | 30 | TAGTAGCTGGTTCCCTCCGAAGTTTCCCTC | 77.3 | -2.2 | 0.6 | -3.5 | 4.2E-04 | 5.2E-04 | BT20/MB468 |
| >trna20_GluTTC_2_-_131094701_131094772@1.31.31,trna5_GluTTC_13_-_41634874_41634945@1.31.31\|TCCCATATGGTCTAGCGGTTAGGATTCCTGG | >trna20_GluTTC_2_-_131094701_131094772 | 1 | 31 | 31 | TCCCATATGGTCTAGCGGTTAGGATTCCTGG | 1356.0 | 3.0 | 0.8 | 3.5 | 3.9E-04 | 5.2E-04 | MB231/MB468 |
| >hg38_21_+_8433172_8446622_RNA45SN1_45S_50ntFlanks@10355.10390.36\|AACTTTGAAGGCCGAAGTGGAGAAGGGTTCCATGTG | >hg38_21_+_8433172_8446622_RNA45SN1_45S_50ntFlanks | 10355 | 10390 | 36 | AACTTTGAAGGCCGAAGTGGAGAAGGGTTCCATGTG | 164.1 | -2.0 | 0.6 | -3.5 | 3.9E-04 | 5.2E-04 | MB231/MB468 |
| >hg38_21_+_8433172_8446622_RNA45SN1_45S_50ntFlanks@8319.8356.38\|CGAGACCGATAGTCAACAAGTACCGTAAGGGAAAGTTG | >hg38_21_+_8433172_8446622_RNA45SN1_45S_50ntFlanks | 8319 | 8356 | 38 | CGAGACCGATAGTCAACAAGTACCGTAAGGGAAAGTTG | 131.3 | -2.1 | 0.6 | -3.5 | 3.9E-04 | 5.2E-04 | MB231/MB468 |
| >hg38_21_+_8433172_8446622_RNA45SN1_45S_50ntFlanks@11200.11223.24\|CGGCGGCGGCGGCGACTCTGGACG | >hg38_21_+_8433172_8446622_RNA45SN1_45S_50ntFlanks | 11200 | 11223 | 24 | CGGCGGCGGCGGCGACTCTGGACG | 130.0 | -3.6 | 1.0 | -3.5 | 4.2E-04 | 5.2E-04 | BT20/MB468 |
| >hg38_21_+_8433172_8446622_RNA45SN1_45S_50ntFlanks@5494.5514.21\|AAGACGGTCGAACTTGACTAT | >hg38_21_+_8433172_8446622_RNA45SN1_45S_50ntFlanks | 5494 | 5514 | 21 | AAGACGGTCGAACTTGACTAT | 217.9 | -3.6 | 1.0 | -3.5 | 4.3E-04 | 5.2E-04 | BT20/MB468 |
| >MI0000478\|hsa-mir-149&WithFlank&2\|+\|240455995\|240456095@21.41.21&[MIMAT0000450&hsa-miR-149-5p&offsets\|0\|-2;m-195&2\|+\|240456015\|240456037&offsets\|0\|-2]\|TCTGGCTCCGTGTCTTCACTC | >MI0000478\|hsa-mir-149&WithFlank&2\|+\|240455995\|240456095 | 21 | 41 | 21 | TCTGGCTCCGTGTCTTCACTC | 613.4 | 2.9 | 0.7 | 4.0 | 5.5E-05 | 5.2E-04 | BT20/MB231 |
| >trna116_GluCTC_1_-_145399233_145399304@35.74.40,trna134_GluTTC_1_-_16861774_16861845@35.74.40,trna71_GluCTC_1_-_161439189_161439260@35.74.40,trna74_GluCTC_1_-_161431809_161431880@35.74.40,trna77_GluCTC_1_-_161424398_161424469@35.74.40,trna77_GluCTC_6_+_28949976_28950047@35.74.40,trna80_GluCTC_1_-_161417018_161417089@35.74.40,trna84_GluTTC_1_-_161391883_161391954@35.74.40,trna87_GluCTC_6_-_126101393_126101464@35.74.40\|TCACCGCCGCGGCCCGGGTTCGATTCCCGGTCAGGGAACC | >trna116_GluCTC_1_-_145399233_145399304 | 35 | 74 | 40 | TCACCGCCGCGGCCCGGGTTCGATTCCCGGTCAGGGAACC | 216.6 | -4.3 | 1.1 | -4.0 | 5.5E-05 | 5.2E-04 | BT20/MB231 |
| >hg38_21_+_8433172_8446622_RNA45SN1_45S_50ntFlanks@12420.12450.31\|ATGTCGGCTCTTCCTATCATTGTGAAGCAGA | >hg38_21_+_8433172_8446622_RNA45SN1_45S_50ntFlanks | 12420 | 12450 | 31 | ATGTCGGCTCTTCCTATCATTGTGAAGCAGA | 559.0 | 1.8 | 0.5 | 3.5 | 4.0E-04 | 5.2E-04 | MB231/MB468 |
| >hg38_21_+_8433172_8446622_RNA45SN1_45S_50ntFlanks@3984.4024.41\|GCGGCTTTGGTGACTCTAGATAACCTCGGGCCGATCGCACG | >hg38_21_+_8433172_8446622_RNA45SN1_45S_50ntFlanks | 3984 | 4024 | 41 | GCGGCTTTGGTGACTCTAGATAACCTCGGGCCGATCGCACG | 247.8 | -6.7 | 1.9 | -3.5 | 4.0E-04 | 5.3E-04 | MB231/MB468 |

| | | | | | | | | | | | | |
|---|---|---|---|---|---|---|---|---|---|---|---|---|
| >hg38_21_+_8433172_8446622_RNA45SN1_45S_50ntFlanks@11668.11685.18\|TGCCCAGTGCTCTGAATG | >hg38_21_+_8433172_8446622_RNA45SN1_45S_50ntFlanks | 11668 | 11685 | 18 | TGCCCAGTGCTCTGAATG | 1367.2 | 2.2 | 0.6 | 3.5 | 4.0E-04 | 5.3E-04 | MB231/MB468 |
| >hg38_21_+_8433172_8446622_RNA45SN1_45S_50ntFlanks@8381.8406.26\|AAGAGGGCGTGAAACCGTTAAGAGGT | >hg38_21_+_8433172_8446622_RNA45SN1_45S_50ntFlanks | 8381 | 8406 | 26 | AAGAGGGCGTGAAACCGTTAAGAGGT | 600.7 | 2.9 | 0.8 | 3.5 | 4.0E-04 | 5.3E-04 | MB231/MB468 |
| >trna111_HisGTG_1_-_147774845_147774916@-1T.28.29,trna118_HisGTG_1_-_145396881_145396952@-1T.28.29,trna16_HisGTG_1_+_146544773_146544844@-1T.28.29,trna1_HisGTG_15_+_45493349_45493420@-1T.28.29,trna21_HisGTG_1_+_147753471_147753542@-1T.28.29,trna33_HisGTG_6_+_27125906_27125977@-1T.28.29,trna7_HisGTG_9_-_14433938_14434009@-1T.28.29,trna8_HisGTG_15_-_45492611_45492682@-1T.28.29,trna9_HisGTG_15_-_45490804_45490875@-1T.28.29\|TGCCGTGATCGTATAGTGGTTAGTACTCT | >trna111_HisGTG_1_-_147774845_147774916 | -1T | 28 | 29 | TGCCGTGATCGTATAGTGGTTAGTACTCT | 2584.5 | 3.0 | 0.8 | 3.5 | 4.0E-04 | 5.3E-04 | MB231/MB468 |
| >hg38_21_+_8433172_8446622_RNA45SN1_45S_50ntFlanks@12444.12467.24\|AAGCAGAATTCACCAAGCGTTGGA | >hg38_21_+_8433172_8446622_RNA45SN1_45S_50ntFlanks | 12444 | 12467 | 24 | AAGCAGAATTCACCAAGCGTTGGA | 143.3 | -2.4 | 0.7 | -3.5 | 4.0E-04 | 5.3E-04 | MB231/MB468 |
| >trna14_LysTTT_11_-_59327808_59327880@1.35.35,trna2_LysTTT_17_+_8022473_8022545@1.35.35,trna54_LysTTT_1_+_204475655_204475727@1.35.35,trna5_LysTTT_11_+_59323902_59323974@1.35.35,trna62_LysTTT_1_-_204476158_204476230@1.35.35,trna76_LysTTT_6_+_28918806_28918878@1.35.35\|GCCCGGATAGCTCAGTCGGTAGAGCATCAGACTTT | >trna14_LysTTT_11_-_59327808_59327880 | 1 | 35 | 35 | GCCCGGATAGCTCAGTCGGTAGAGCATCAGACTTT | 1076.9 | 3.9 | 1.0 | 4.0 | 5.6E-05 | 5.3E-04 | BT20/MB231 |
| >hg38_21_+_8433172_8446622_RNA45SN1_45S_50ntFlanks@6661.6680.20\|CGGTGGATCACTCGGCTCGT | >hg38_21_+_8433172_8446622_RNA45SN1_45S_50ntFlanks | 6661 | 6680 | 20 | CGGTGGATCACTCGGCTCGT | 302.3 | 2.6 | 0.7 | 4.0 | 5.7E-05 | 5.3E-04 | BT20/MB231 |
| >hg38_21_+_8433172_8446622_RNA45SN1_45S_50ntFlanks@12076.12096.21\|AGGGGCTCTCGCTTCTGGCGC | >hg38_21_+_8433172_8446622_RNA45SN1_45S_50ntFlanks | 12076 | 12096 | 21 | AGGGGCTCTCGCTTCTGGCGC | 220.9 | -2.8 | 0.8 | -3.5 | 4.4E-04 | 5.3E-04 | BT20/MB468 |
| >hg38_21_+_8433172_8446622_RNA45SN1_45S_50ntFlanks@3882.3915.34\|CCGACGGGCGCTGACCCCCTTCGCGGGGGGGATG | >hg38_21_+_8433172_8446622_RNA45SN1_45S_50ntFlanks | 3882 | 3915 | 34 | CCGACGGGCGCTGACCCCCTTCGCGGGGGGGATG | 91.5 | -2.6 | 0.8 | -3.5 | 4.5E-04 | 5.4E-04 | BT20/MB468 |
| >hg38_21_+_8433172_8446622_RNA45SN1_45S_50ntFlanks@11663.11681.19\|ATTTCTGCCCAGTGCTCTG | >hg38_21_+_8433172_8446622_RNA45SN1_45S_50ntFlanks | 11663 | 11681 | 19 | ATTTCTGCCCAGTGCTCTG | 140.9 | -1.6 | 0.5 | -3.5 | 4.1E-04 | 5.4E-04 | MB231/MB468 |
| >hg38_21_+_8433172_8446622_RNA45SN1_45S_50ntFlanks@11572.11609.38\|CCGACTTAGAACTGGTGCGGACCAGGGGAATCCGACTG | >hg38_21_+_8433172_8446622_RNA45SN1_45S_50ntFlanks | 11572 | 11609 | 38 | CCGACTTAGAACTGGTGCGGACCAGGGGAATCCGACTG | 224.7 | -2.1 | 0.6 | -3.5 | 4.5E-04 | 5.4E-04 | BT20/MB468 |
| >hg38_21_+_8433172_8446622_RNA45SN1_45S_50ntFlanks@10856.10873.18\|AGGATTGGCTCTAAGGGC | >hg38_21_+_8433172_8446622_RNA45SN1_45S_50ntFlanks | 10856 | 10873 | 18 | AGGATTGGCTCTAAGGGC | 121.1 | -1.5 | 0.4 | -3.5 | 4.5E-04 | 5.5E-04 | BT20/MB468 |
| >hg38_21_+_8433172_8446622_RNA45SN1_45S_50ntFlanks@10807.10825.19\|ACAATGTAGGTAAGGGAAG | >hg38_21_+_8433172_8446622_RNA45SN1_45S_50ntFlanks | 10807 | 10825 | 19 | ACAATGTAGGTAAGGGAAG | 166.6 | -5.4 | 1.5 | -3.5 | 4.1E-04 | 5.5E-04 | MB231/MB468 |
| >hg38_21_+_8433172_8446622_RNA45SN1_45S_50ntFlanks@10468.10487.20\|TTGCCCTCGGCCGATCGAAA | >hg38_21_+_8433172_8446622_RNA45SN1_45S_50ntFlanks | 10468 | 10487 | 20 | TTGCCCTCGGCCGATCGAAA | 401.0 | 2.1 | 0.6 | 3.5 | 4.1E-04 | 5.5E-04 | MB231/MB468 |
| >hg38_21_+_8433172_8446622_RNA45SN1_45S_50ntFlanks@11568.11601.34\|GCAGCCGACTTAGAACTGGTGCGGACCAGGGGAA | >hg38_21_+_8433172_8446622_RNA45SN1_45S_50ntFlanks | 11568 | 11601 | 34 | GCAGCCGACTTAGAACTGGTGCGGACCAGGGGAA | 274.9 | -1.6 | 0.4 | -3.5 | 4.2E-04 | 5.5E-04 | MB231/MB468 |
| >hg38_21_+_8433172_8446622_RNA45SN1_45S_50ntFlanks@10797.10826.30\|GCATGTTGGAACAATGTAGGTAAGGGAAGT | >hg38_21_+_8433172_8446622_RNA45SN1_45S_50ntFlanks | 10797 | 10826 | 30 | GCATGTTGGAACAATGTAGGTAAGGGAAGT | 184.0 | -2.0 | 0.6 | -3.5 | 4.5E-04 | 5.5E-04 | BT20/MB468 |
| >hg38_MT_+_1621_3279_RNR2_16S_50ntFlanks@1347.1389.43\|CAACAATAGGGTTTACGACCTCGATGTTGGATCAGGACATCCC | >hg38_MT_+_1621_3279_RNR2_16S_50ntFlanks | 1347 | 1389 | 43 | CAACAATAGGGTTTACGACCTCGATGTTGGATCAGGACATCCC | 425.3 | -3.5 | 0.9 | -4.0 | 5.9E-05 | 5.5E-04 | BT20/MB231 |
| >hg38_21_+_8433172_8446622_RNA45SN1_45S_50ntFlanks@3705.3748.44\|TACCTGGTTGATCCTGCCAGTAGCATATGCTTGTCTCAAAGATT | >hg38_21_+_8433172_8446622_RNA45SN1_45S_50ntFlanks | 3705 | 3748 | 44 | TACCTGGTTGATCCTGCCAGTAGCATATGCTTGTCTCAAAGATT | 6175.9 | 4.3 | 1.1 | 4.0 | 5.9E-05 | 5.5E-04 | BT20/MB231 |
| >MI0000273\|hsa-mir-183&WithFlank&7\|-\|129774899\|129775020@34.53.20(+1G)&[MIMAT0000261&hsa-miR-183-5p&offsets\|+1\|-1(+1G);m-41&7\|-\|129774966\|129774987&offsets\|0\|-2(+1G)]\|ATGGCACTGGTAGAATTCACG | >MI0000273\|hsa-mir-183&WithFlank&7\|-\|129774899\|129775020 | 34 | 53 | 20(+1G) | ATGGCACTGGTAGAATTCACG | 380.4 | 3.6 | 0.9 | 4.0 | 5.9E-05 | 5.5E-04 | BT20/MB231 |
| >hg38_21_+_8433172_8446622_RNA45SN1_45S_50ntFlanks@7988.8017.30\|CAGACGTGGCGACCCGCTGAATTTAAGCAT | >hg38_21_+_8433172_8446622_RNA45SN1_45S_50ntFlanks | 7988 | 8017 | 30 | CAGACGTGGCGACCCGCTGAATTTAAGCAT | 476.2 | 1.9 | 0.5 | 3.5 | 4.2E-04 | 5.6E-04 | MB231/MB468 |
| >hg38_21_+_8433172_8446622_RNA45SN1_45S_50ntFlanks@12907.12933.27\|CACCGCACGTTCGTGGGGAACCTGGCG | >hg38_21_+_8433172_8446622_RNA45SN1_45S_50ntFlanks | 12907 | 12933 | 27 | CACCGCACGTTCGTGGGGAACCTGGCG | 510.0 | -3.3 | 0.9 | -3.5 | 4.2E-04 | 5.6E-04 | MB231/MB468 |
| >hg38_21_+_8433172_8446622_RNA45SN1_45S_50ntFlanks@8168.8185.18\|CCCAAGTCCTTCTGATCG | >hg38_21_+_8433172_8446622_RNA45SN1_45S_50ntFlanks | 8168 | 8185 | 18 | CCCAAGTCCTTCTGATCG | 424.0 | -2.5 | 0.7 | -3.5 | 4.3E-04 | 5.6E-04 | MB231/MB468 |
| >hg38_21_+_8433172_8446622_RNA45SN1_45S_50ntFlanks@5436.5466.31\|GTGAGGCCCTCGGATCGGCCCCGCCGGGGTC | >hg38_21_+_8433172_8446622_RNA45SN1_45S_50ntFlanks | 5436 | 5466 | 31 | GTGAGGCCCTCGGATCGGCCCCGCCGGGGTC | 119.6 | -2.8 | 0.8 | -3.5 | 4.7E-04 | 5.7E-04 | BT20/MB468 |
| >hg38_21_+_8433172_8446622_RNA45SN1_45S_50ntFlanks@11672.11703.32\|CAGTGCTCTGAATGTCAAAGTGAAGAAATTCA | >hg38_21_+_8433172_8446622_RNA45SN1_45S_50ntFlanks | 11672 | 11703 | 32 | CAGTGCTCTGAATGTCAAAGTGAAGAAATTCA | 812.0 | 2.0 | 0.6 | 3.5 | 4.3E-04 | 5.7E-04 | MB231/MB468 |
| >hg38_21_+_8433172_8446622_RNA45SN1_45S_50ntFlanks@5494.5526.33\|AAGACGGTCGAACTTGACTATCTAGAGGAAGTA | >hg38_21_+_8433172_8446622_RNA45SN1_45S_50ntFlanks | 5494 | 5526 | 33 | AAGACGGTCGAACTTGACTATCTAGAGGAAGTA | 144.5 | -2.2 | 0.6 | -3.5 | 4.7E-04 | 5.7E-04 | BT20/MB468 |
| >hg38_21_+_8433172_8446622_RNA45SN1_45S_50ntFlanks@11318.11336.19\|CCCCACGTCTCGTCGCGCG | >hg38_21_+_8433172_8446622_RNA45SN1_45S_50ntFlanks | 11318 | 11336 | 19 | CCCCACGTCTCGTCGCGCG | 400.7 | -4.6 | 1.3 | -3.5 | 4.7E-04 | 5.7E-04 | BT20/MB468 |
| >MI0001733\|hsa-mir-452&WithFlank&X\|-\|151959622\|151959718@23.44.22&[MIMAT0001635&hsa-miR-452-5p&offsets\|+3\|+3;m-188&X\|-\|151959677\|151959699&offsets\|+3\|+2]\|TGTTTGCAGAGGAAACTGAGAC | >MI0001733\|hsa-mir-452&WithFlank&X\|-\|151959622\|151959718 | 23 | 44 | 22 | TGTTTGCAGAGGAAACTGAGAC | 540.4 | 2.8 | 0.7 | 4.0 | 6.1E-05 | 5.7E-04 | BT20/MB231 |
| >hg38_21_+_8433172_8446622_RNA45SN1_45S_50ntFlanks@10458.10486.29\|ATGGCCTCCGTTGCCCTCGGCCGATCGAA | >hg38_21_+_8433172_8446622_RNA45SN1_45S_50ntFlanks | 10458 | 10486 | 29 | ATGGCCTCCGTTGCCCTCGGCCGATCGAA | 138.2 | -2.2 | 0.6 | -3.5 | 4.3E-04 | 5.7E-04 | MB231/MB468 |
| >hg38_21_+_8433172_8446622_RNA45SN1_45S_50ntFlanks@12487.12509.23\|GAACGTGAGCTGGGTTTAGACCG | >hg38_21_+_8433172_8446622_RNA45SN1_45S_50ntFlanks | 12487 | 12509 | 23 | GAACGTGAGCTGGGTTTAGACCG | 222.8 | -1.6 | 0.5 | -3.5 | 4.3E-04 | 5.7E-04 | MB231/MB468 |
| >hg38_21_+_8433172_8446622_RNA45SN1_45S_50ntFlanks@12416.12434.19\|TTCGATGTCGGCTCTTCCT | >hg38_21_+_8433172_8446622_RNA45SN1_45S_50ntFlanks | 12416 | 12434 | 19 | TTCGATGTCGGCTCTTCCT | 1169.7 | 1.8 | 0.5 | 3.5 | 4.8E-04 | 5.8E-04 | BT20/MB468 |
| >hg38_21_+_8433172_8446622_RNA45SN1_45S_50ntFlanks@7975.8012.38\|CGCGACCTCAGATCAGACGTGGCGACCCGCTGAATTTA | >hg38_21_+_8433172_8446622_RNA45SN1_45S_50ntFlanks | 7975 | 8012 | 38 | CGCGACCTCAGATCAGACGTGGCGACCCGCTGAATTTA | 12081.7 | 2.8 | 0.8 | 3.5 | 4.8E-04 | 5.8E-04 | BT20/MB468 |
| >hg38_21_+_8433172_8446622_RNA45SN1_45S_50ntFlanks@11577.11609.33\|TTAGAACTGGTGCGGACCAGGGGAATCCGACTG | >hg38_21_+_8433172_8446622_RNA45SN1_45S_50ntFlanks | 11577 | 11609 | 33 | TTAGAACTGGTGCGGACCAGGGGAATCCGACTG | 207.7 | -1.9 | 0.6 | -3.5 | 4.8E-04 | 5.8E-04 | BT20/MB468 |
| >hg38_21_+_8433172_8446622_RNA45SN1_45S_50ntFlanks@9535.9553.19\|GGAAACTCTGGTGGAGGTC | >hg38_21_+_8433172_8446622_RNA45SN1_45S_50ntFlanks | 9535 | 9553 | 19 | GGAAACTCTGGTGGAGGTC | 178.7 | -4.3 | 1.2 | -3.5 | 4.8E-04 | 5.8E-04 | BT20/MB468 |
| >trna119_LysCTT_1_-_145395522_145395594@4.34.31,trna11_LysCTT_5_-_180648979_180649051@4.34.31,trna13_LysCTT_6_+_26556774_26556846@4.34.31,trna7_LysCTT_16_+_3225692_3225764@4.34.31,trna9_LysCTT_5_+_180634755_180634827@4.34.31\|CGGCTAGCTCAGTCGGTAGAGCATGAGACTC | >trna119_LysCTT_1_-_145395522_145395594 | 4 | 34 | 31 | CGGCTAGCTCAGTCGGTAGAGCATGAGACTC | 228.1 | -3.3 | 0.8 | -4.0 | 6.3E-05 | 5.9E-04 | BT20/MB231 |
| >hg38_21_+_8433172_8446622_RNA45SN1_45S_50ntFlanks@12488.12524.37\|AACGTGAGCTGGGTTTAGACCGTCGTGAGACAGGTTA | >hg38_21_+_8433172_8446622_RNA45SN1_45S_50ntFlanks | 12488 | 12524 | 37 | AACGTGAGCTGGGTTTAGACCGTCGTGAGACAGGTTA | 145.5 | -2.9 | 0.8 | -3.5 | 4.8E-04 | 5.9E-04 | BT20/MB468 |
| >hg38_21_+_8433172_8446622_RNA45SN1_45S_50ntFlanks@4554.4572.19\|CCGCAGCTAGGAATAATGG | >hg38_21_+_8433172_8446622_RNA45SN1_45S_50ntFlanks | 4554 | 4572 | 19 | CCGCAGCTAGGAATAATGG | 106.9 | -2.2 | 0.6 | -3.5 | 4.5E-04 | 5.9E-04 | MB231/MB468 |
| >hg38_21_+_8433172_8446622_RNA45SN1_45S_50ntFlanks@8113.8138.26\|GGCGCGGGACATGTGGCGTACGGAAG | >hg38_21_+_8433172_8446622_RNA45SN1_45S_50ntFlanks | 8113 | 8138 | 26 | GGCGCGGGACATGTGGCGTACGGAAG | 168.6 | -4.1 | 1.2 | -3.5 | 4.9E-04 | 5.9E-04 | BT20/MB468 |
| >hg38_21_+_8433172_8446622_RNA45SN1_45S_50ntFlanks@11641.11685.45\|CGGCGGGTGTTGACGCGATGTGATTTCTGCCCAGTGCTCTGAATG | >hg38_21_+_8433172_8446622_RNA45SN1_45S_50ntFlanks | 11641 | 11685 | 45 | CGGCGGGTGTTGACGCGATGTGATTTCTGCCCAGTGCTCTGAATG | 270.6 | -2.2 | 0.6 | -3.5 | 4.5E-04 | 5.9E-04 | MB231/MB468 |
| >trna152_ValCAC_6_-_27248049_27248121@1.24.24\|GCTTCTGTAGTGTAGTGGTTATCA | >trna152_ValCAC_6_-_27248049_27248121 | 1 | 24 | 24 | GCTTCTGTAGTGTAGTGGTTATCA | 118.9 | -4.1 | 1.2 | -3.5 | 4.9E-04 | 6.0E-04 | BT20/MB468 |
| >hg38_21_+_8433172_8446622_RNA45SN1_45S_50ntFlanks@11317.11334.18\|ACCCCACGTCTCGTCGCG | >hg38_21_+_8433172_8446622_RNA45SN1_45S_50ntFlanks | 11317 | 11334 | 18 | ACCCCACGTCTCGTCGCG | 465.1 | -4.0 | 1.2 | -3.5 | 4.9E-04 | 6.0E-04 | BT20/MB468 |
| >hg38_21_+_8433172_8446622_RNA45SN1_45S_50ntFlanks@10351.10380.30\|CGAGAACTTTGAAGGCCGAAGTGGAGAAGG | >hg38_21_+_8433172_8446622_RNA45SN1_45S_50ntFlanks | 10351 | 10380 | 30 | CGAGAACTTTGAAGGCCGAAGTGGAGAAGG | 314.7 | -1.8 | 0.5 | -3.5 | 4.5E-04 | 6.0E-04 | MB231/MB468 |
| >hg38_21_+_8433172_8446622_RNA45SN1_45S_50ntFlanks@12415.12444.30\|CTTCGATGTCGGCTCTTCCTATCATTGTGA | >hg38_21_+_8433172_8446622_RNA45SN1_45S_50ntFlanks | 12415 | 12444 | 30 | CTTCGATGTCGGCTCTTCCTATCATTGTGA | 1098.4 | 2.2 | 0.6 | 3.5 | 4.6E-04 | 6.1E-04 | MB231/MB468 |
| >hg38_21_+_8433172_8446622_RNA45SN1_45S_50ntFlanks@12419.12460.42\|GATGTCGGCTCTTCCTATCATTGTGAAGCAGAATTCACCAAG | >hg38_21_+_8433172_8446622_RNA45SN1_45S_50ntFlanks | 12419 | 12460 | 42 | GATGTCGGCTCTTCCTATCATTGTGAAGCAGAATTCACCAAG | 156.0 | -3.0 | 0.7 | -4.0 | 6.6E-05 | 6.1E-04 | BT20/MB231 |
| >hg38_21_+_8433172_8446622_RNA45SN1_45S_50ntFlanks@11572.11597.26\|CCGACTTAGAACTGGTGCGGACCAGG | >hg38_21_+_8433172_8446622_RNA45SN1_45S_50ntFlanks | 11572 | 11597 | 26 | CCGACTTAGAACTGGTGCGGACCAGG | 262.8 | -2.5 | 0.7 | -3.5 | 4.6E-04 | 6.1E-04 | MB231/MB468 |
| >hg38_21_+_8433172_8446622_RNA45SN1_45S_50ntFlanks@8115.8142.28\|CGCGGGACATGTGGCGTACGGAAGACCC | >hg38_21_+_8433172_8446622_RNA45SN1_45S_50ntFlanks | 8115 | 8142 | 28 | CGCGGGACATGTGGCGTACGGAAGACCC | 103.1 | -2.7 | 0.8 | -3.5 | 5.1E-04 | 6.1E-04 | BT20/MB468 |
| >hg38_21_+_8433172_8446622_RNA45SN1_45S_50ntFlanks@12001.12032.32\|CTGCGGGCCGCCGGTGAAATACCACTACTCTG | >hg38_21_+_8433172_8446622_RNA45SN1_45S_50ntFlanks | 12001 | 12032 | 32 | CTGCGGGCCGCCGGTGAAATACCACTACTCTG | 1404.8 | -3.5 | 1.0 | -3.5 | 4.7E-04 | 6.1E-04 | MB231/MB468 |
| >hg38_21_+_8433172_8446622_RNA45SN1_45S_50ntFlanks@10409.10429.21\|GTCAGTCGGTCCTGAGAGATG | >hg38_21_+_8433172_8446622_RNA45SN1_45S_50ntFlanks | 10409 | 10429 | 21 | GTCAGTCGGTCCTGAGAGATG | 856.9 | -1.6 | 0.5 | -3.5 | 4.7E-04 | 6.2E-04 | MB231/MB468 |

<table>
<tr><th></th><th></th><th></th><th></th><th></th><th></th><th></th><th></th><th></th><th></th><th></th><th></th><th></th><th></th></tr>
<tr><td>>hg38_21_+_8433172_8446622_RNA45SN1_45S_50ntFlanks@6723.6746.24|TTGCAGGACACATTGATCATCGAC</td><td>>hg38_21_+_8433172_8446622_RNA45SN1_45S_50ntFlanks</td><td>6723</td><td>6746</td><td>24</td><td>TTGCAGGACACATTGATCATCGAC</td><td>126.0</td><td>-1.7</td><td>0.5</td><td>-3.5</td><td>4.7E-04</td><td>6.2E-04</td><td>MB231/MB468</td></tr>
<tr><td>>hg38_21_+_8433172_8446622_RNA45SN1_45S_50ntFlanks@12420.12456.37|ATGTCGGCTCTTCCTATCATTGTGAAGCAGAATTCAC</td><td>>hg38_21_+_8433172_8446622_RNA45SN1_45S_50ntFlanks</td><td>12420</td><td>12456</td><td>37</td><td>ATGTCGGCTCTTCCTATCATTGTGAAGCAGAATTCAC</td><td>487.0</td><td>-2.3</td><td>0.7</td><td>-3.5</td><td>5.1E-04</td><td>6.2E-04</td><td>BT20/MB468</td></tr>
<tr><td>>am_trnaMT_PheGAA_MT_+_577_647@45.62.18|AGACGGGCTCACATCACC</td><td>>am_trnaMT_PheGAA_MT_+_577_647</td><td>45</td><td>62</td><td>18</td><td>AGACGGGCTCACATCACC</td><td>381.7</td><td>2.7</td><td>0.8</td><td>3.5</td><td>4.7E-04</td><td>6.2E-04</td><td>MB231/MB468</td></tr>
<tr><td>>MI0000063|hsa-let-7b&WithFlank&22|+|46113680|46113774@12.33.22&[MIMAT0000063&hsa-let-7b-5p&offsets|0|0;m-14&22|+|46113691|46113712&offsets|0|0]|TGAGGTAGTAGGTTGTGTGGTT</td><td>>MI0000063|hsa-let-7b&WithFlank&22|+|46113680|46113774</td><td>12</td><td>33</td><td>22</td><td>TGAGGTAGTAGGTTGTGTGGTT</td><td>48329.1</td><td>2.4</td><td>0.6</td><td>4.0</td><td>6.8E-05</td><td>6.2E-04</td><td>BT20/MB231</td></tr>
<tr><td>>MI0000477|hsa-mir-146a&WithFlank&5|+|160485346|160485456@27.46.20(+1C)&[MIMAT0000449&hsa-miR-146a-5p&offsets|0|-2(+1C);m-129&5|+|160485372|160485393&offsets|0|-2(+1C)]|TGAGAACTGAATTCCATGGGC</td><td>>MI0000477|hsa-mir-146a&WithFlank&5|+|160485346|160485456</td><td>27</td><td>46</td><td>20(+1C)</td><td>TGAGAACTGAATTCCATGGGC</td><td>242.7</td><td>-7.8</td><td>2.0</td><td>-4.0</td><td>6.8E-05</td><td>6.2E-04</td><td>BT20/MB231</td></tr>
<tr><td>>hg38_21_+_8433172_8446622_RNA45SN1_45S_50ntFlanks@13007.13027.21|TATTGAAAGTCAGCCCTCGAC</td><td>>hg38_21_+_8433172_8446622_RNA45SN1_45S_50ntFlanks</td><td>13007</td><td>13027</td><td>21</td><td>TATTGAAAGTCAGCCCTCGAC</td><td>457.5</td><td>-2.4</td><td>0.7</td><td>-3.5</td><td>4.7E-04</td><td>6.3E-04</td><td>MB231/MB468</td></tr>
<tr><td>>MI0000102|hsa-mir-100&WithFlank&11|-|122152223|122152314@19.40.22(+1A)&[MIMAT0000098&hsa-miR-100-5p&offsets|0|0(+1A);m-27&11|-|122152276|122152296&offsets|0|+1(+1A)]|AACCCGTAGATCCGAACTTGTGA</td><td>>MI0000102|hsa-mir-100&WithFlank&11|-|122152223|122152314</td><td>19</td><td>40</td><td>22(+1A)</td><td>AACCCGTAGATCCGAACTTGTGA</td><td>1897.8</td><td>8.2</td><td>2.4</td><td>3.5</td><td>5.2E-04</td><td>6.3E-04</td><td>BT20/MB468</td></tr>
<tr><td>>MI0000477|hsa-mir-146a&WithFlank&5|+|160485346|160485456@27.47.21&[MIMAT0000449&hsa-miR-146a-5p&offsets|0|-1;m-129&5|+|160485372|160485393&offsets|0|-1]|TGAGAACTGAATTCCATGGGT</td><td>>MI0000477|hsa-mir-146a&WithFlank&5|+|160485346|160485456</td><td>27</td><td>47</td><td>21</td><td>TGAGAACTGAATTCCATGGGT</td><td>2155.4</td><td>-7.5</td><td>1.9</td><td>-4.0</td><td>6.8E-05</td><td>6.3E-04</td><td>BT20/MB231</td></tr>
<tr><td>>hg38_21_+_8433172_8446622_RNA45SN1_45S_50ntFlanks@11683.11702.20|ATGTCAAAGTGAAGAAATTC</td><td>>hg38_21_+_8433172_8446622_RNA45SN1_45S_50ntFlanks</td><td>11683</td><td>11702</td><td>20</td><td>ATGTCAAAGTGAAGAAATTC</td><td>269.6</td><td>-3.7</td><td>1.1</td><td>-3.5</td><td>5.2E-04</td><td>6.3E-04</td><td>BT20/MB468</td></tr>
<tr><td>>hg38_21_+_8433172_8446622_RNA45SN1_45S_50ntFlanks@8342.8365.24|CGTAAGGGAAAGTTGAAAAGAACT</td><td>>hg38_21_+_8433172_8446622_RNA45SN1_45S_50ntFlanks</td><td>8342</td><td>8365</td><td>24</td><td>CGTAAGGGAAAGTTGAAAAGAACT</td><td>2847.9</td><td>7.9</td><td>2.3</td><td>3.5</td><td>4.8E-04</td><td>6.3E-04</td><td>MB231/MB468</td></tr>
<tr><td>>hg38_21_+_8433172_8446622_RNA45SN1_45S_50ntFlanks@4369.4389.21|GTTAAAAAGCTCGTAGTTGGA</td><td>>hg38_21_+_8433172_8446622_RNA45SN1_45S_50ntFlanks</td><td>4369</td><td>4389</td><td>21</td><td>GTTAAAAAGCTCGTAGTTGGA</td><td>90.2</td><td>-3.9</td><td>1.1</td><td>-3.5</td><td>5.2E-04</td><td>6.3E-04</td><td>BT20/MB468</td></tr>
<tr><td>>hg38_21_+_8433172_8446622_RNA45SN1_45S_50ntFlanks@11574.11603.30|GACTTAGAACTGGTGCGGACCAGGGGAATC</td><td>>hg38_21_+_8433172_8446622_RNA45SN1_45S_50ntFlanks</td><td>11574</td><td>11603</td><td>30</td><td>GACTTAGAACTGGTGCGGACCAGGGGAATC</td><td>115.2</td><td>-1.8</td><td>0.5</td><td>-3.5</td><td>4.8E-04</td><td>6.3E-04</td><td>MB231/MB468</td></tr>
<tr><td>>hg38_21_+_8433172_8446622_RNA45SN1_45S_50ntFlanks@10787.10817.31|ACAGCCTCTGGCATGTTGGAACAATGTAGGT</td><td>>hg38_21_+_8433172_8446622_RNA45SN1_45S_50ntFlanks</td><td>10787</td><td>10817</td><td>31</td><td>ACAGCCTCTGGCATGTTGGAACAATGTAGGT</td><td>258.3</td><td>-2.0</td><td>0.6</td><td>-3.5</td><td>4.8E-04</td><td>6.3E-04</td><td>MB231/MB468</td></tr>
<tr><td>>hg38_21_+_8433172_8446622_RNA45SN1_45S_50ntFlanks@6784.6801.18|GCTACGCCTGTCTGAGCG</td><td>>hg38_21_+_8433172_8446622_RNA45SN1_45S_50ntFlanks</td><td>6784</td><td>6801</td><td>18</td><td>GCTACGCCTGTCTGAGCG</td><td>138.8</td><td>-2.9</td><td>0.8</td><td>-3.5</td><td>5.2E-04</td><td>6.4E-04</td><td>BT20/MB468</td></tr>
<tr><td>>hg38_21_+_8433172_8446622_RNA45SN1_45S_50ntFlanks@9533.9561.29|GAGGAAACTCTGGTGGAGGTCCGTAGCGG</td><td>>hg38_21_+_8433172_8446622_RNA45SN1_45S_50ntFlanks</td><td>9533</td><td>9561</td><td>29</td><td>GAGGAAACTCTGGTGGAGGTCCGTAGCGG</td><td>203.9</td><td>-4.0</td><td>1.2</td><td>-3.5</td><td>5.2E-04</td><td>6.4E-04</td><td>BT20/MB468</td></tr>
<tr><td>>hg38_21_+_8433172_8446622_RNA45SN1_45S_50ntFlanks@9535.9571.37|GGAAACTCTGGTGGAGGTCCGTAGCGGTCCTGACGTG</td><td>>hg38_21_+_8433172_8446622_RNA45SN1_45S_50ntFlanks</td><td>9535</td><td>9571</td><td>37</td><td>GGAAACTCTGGTGGAGGTCCGTAGCGGTCCTGACGTG</td><td>555.1</td><td>2.2</td><td>0.6</td><td>3.5</td><td>4.9E-04</td><td>6.4E-04</td><td>MB231/MB468</td></tr>
<tr><td>>hg38_21_+_8433172_8446622_RNA45SN1_45S_50ntFlanks@12990.13015.26|GCTCCCTCGCTGCGATCTATTGAAAG</td><td>>hg38_21_+_8433172_8446622_RNA45SN1_45S_50ntFlanks</td><td>12990</td><td>13015</td><td>26</td><td>GCTCCCTCGCTGCGATCTATTGAAAG</td><td>195.6</td><td>-3.1</td><td>0.9</td><td>-3.5</td><td>5.3E-04</td><td>6.4E-04</td><td>BT20/MB468</td></tr>
<tr><td>>MI0001733|hsa-mir-452&WithFlank&X|-|151959622|151959718@59.80.22&[MIMAT0001636&hsa-miR-452-3p&offsets|-5|-5;m-457&X|-|151959639|151959660&offsets|0|0]|TCAGTCTCATCTGCAAAGAAGT</td><td>>MI0001733|hsa-mir-452&WithFlank&X|-|151959622|151959718</td><td>59</td><td>80</td><td>22</td><td>TCAGTCTCATCTGCAAAGAAGT</td><td>435.0</td><td>3.0</td><td>0.8</td><td>4.0</td><td>7.1E-05</td><td>6.5E-04</td><td>BT20/MB231</td></tr>
<tr><td>>hg38_21_+_8433172_8446622_RNA45SN1_45S_50ntFlanks@12996.13039.44|TCGCTGCGATCTATTGAAAGTCAGCCCTCGACACAAGGGTTTGT</td><td>>hg38_21_+_8433172_8446622_RNA45SN1_45S_50ntFlanks</td><td>12996</td><td>13039</td><td>44</td><td>TCGCTGCGATCTATTGAAAGTCAGCCCTCGACACAAGGGTTTGT</td><td>244337.2</td><td>-2.9</td><td>0.7</td><td>-4.0</td><td>7.2E-05</td><td>6.5E-04</td><td>BT20/MB231</td></tr>
<tr><td>>hg38_21_+_8433172_8446622_RNA45SN1_45S_50ntFlanks@6648.6696.49|GTACGACTCTTAGCGGTGGATCACTCGGCTCGTGCGTCGATGAAGAACG</td><td>>hg38_21_+_8433172_8446622_RNA45SN1_45S_50ntFlanks</td><td>6648</td><td>6696</td><td>49</td><td>GTACGACTCTTAGCGGTGGATCACTCGGCTCGTGCGTCGATGAAGAACG</td><td>1210.0</td><td>-3.4</td><td>0.8</td><td>-4.0</td><td>7.2E-05</td><td>6.5E-04</td><td>BT20/MB231</td></tr>
<tr><td>>hg38_21_+_8433172_8446622_RNA45SN1_45S_50ntFlanks@10792.10822.31|CTCTGGCATGTTGGAACAATGTAGGTAAGGG</td><td>>hg38_21_+_8433172_8446622_RNA45SN1_45S_50ntFlanks</td><td>10792</td><td>10822</td><td>31</td><td>CTCTGGCATGTTGGAACAATGTAGGTAAGGG</td><td>173.2</td><td>-2.0</td><td>0.6</td><td>-3.5</td><td>5.4E-04</td><td>6.5E-04</td><td>BT20/MB468</td></tr>
<tr><td>>hg38_21_+_8433172_8446622_RNA45SN1_45S_50ntFlanks@9537.9571.35|AAACTCTGGTGGAGGTCCGTAGCGGTCCTGACGTG</td><td>>hg38_21_+_8433172_8446622_RNA45SN1_45S_50ntFlanks</td><td>9537</td><td>9571</td><td>35</td><td>AAACTCTGGTGGAGGTCCGTAGCGGTCCTGACGTG</td><td>176.6</td><td>-1.5</td><td>0.4</td><td>-3.5</td><td>5.4E-04</td><td>6.5E-04</td><td>BT20/MB468</td></tr>
<tr><td>>hg38_21_+_8433172_8446622_RNA45SN1_45S_50ntFlanks@11571.11592.22|GCCGACTTAGAACTGGTGCGGA</td><td>>hg38_21_+_8433172_8446622_RNA45SN1_45S_50ntFlanks</td><td>11571</td><td>11592</td><td>22</td><td>GCCGACTTAGAACTGGTGCGGA</td><td>349.8</td><td>-2.3</td><td>0.7</td><td>-3.5</td><td>5.4E-04</td><td>6.6E-04</td><td>BT20/MB468</td></tr>
<tr><td>>hg38_21_+_8433172_8446622_RNA45SN1_45S_50ntFlanks@11210.11233.24|GGCGACTCTGGACGCGAGCCGGGC</td><td>>hg38_21_+_8433172_8446622_RNA45SN1_45S_50ntFlanks</td><td>11210</td><td>11233</td><td>24</td><td>GGCGACTCTGGACGCGAGCCGGGC</td><td>266.9</td><td>-3.5</td><td>1.0</td><td>-3.5</td><td>5.4E-04</td><td>6.6E-04</td><td>BT20/MB468</td></tr>
<tr><td>>hg38_21_+_8433172_8446622_RNA45SN1_45S_50ntFlanks@11568.11586.19|GCAGCCGACTTAGAACTGG</td><td>>hg38_21_+_8433172_8446622_RNA45SN1_45S_50ntFlanks</td><td>11568</td><td>11586</td><td>19</td><td>GCAGCCGACTTAGAACTGG</td><td>2102.0</td><td>2.0</td><td>0.6</td><td>3.5</td><td>5.0E-04</td><td>6.6E-04</td><td>MB231/MB468</td></tr>
<tr><td>>am_trnaMT_ValTAC_MT_+_1602_1670@48.68.21|CAACTTAACTTGACCGCTCTG</td><td>>am_trnaMT_ValTAC_MT_+_1602_1670</td><td>48</td><td>68</td><td>21</td><td>CAACTTAACTTGACCGCTCTG</td><td>619.5</td><td>2.9</td><td>0.8</td><td>3.5</td><td>5.5E-04</td><td>6.6E-04</td><td>BT20/MB468</td></tr>
<tr><td>>hg38_21_+_8433172_8446622_RNA45SN1_45S_50ntFlanks@11316.11338.23|CACCCCACGTCTCGTCGCGCGCG</td><td>>hg38_21_+_8433172_8446622_RNA45SN1_45S_50ntFlanks</td><td>11316</td><td>11338</td><td>23</td><td>CACCCCACGTCTCGTCGCGCGCG</td><td>1263.0</td><td>-5.0</td><td>1.5</td><td>-3.5</td><td>5.5E-04</td><td>6.6E-04</td><td>BT20/MB468</td></tr>
<tr><td>>hg38_21_+_8433172_8446622_RNA45SN1_45S_50ntFlanks@6663.6688.26|GTGGATCACTCGGCTCGTGCGTCGAT</td><td>>hg38_21_+_8433172_8446622_RNA45SN1_45S_50ntFlanks</td><td>6663</td><td>6688</td><td>26</td><td>GTGGATCACTCGGCTCGTGCGTCGAT</td><td>176.7</td><td>-1.8</td><td>0.5</td><td>-3.5</td><td>5.0E-04</td><td>6.6E-04</td><td>MB231/MB468</td></tr>
<tr><td>>hg38_21_+_8433172_8446622_RNA45SN1_45S_50ntFlanks@11210.11227.18|GGCGACTCTGGACGCGAG</td><td>>hg38_21_+_8433172_8446622_RNA45SN1_45S_50ntFlanks</td><td>11210</td><td>11227</td><td>18</td><td>GGCGACTCTGGACGCGAG</td><td>173.7</td><td>-4.1</td><td>1.2</td><td>-3.5</td><td>5.5E-04</td><td>6.7E-04</td><td>BT20/MB468</td></tr>
<tr><td>>hg38_21_+_8433172_8446622_RNA45SN1_45S_50ntFlanks@11565.11586.22|CTAGCAGCCGACTTAGAACTGG</td><td>>hg38_21_+_8433172_8446622_RNA45SN1_45S_50ntFlanks</td><td>11565</td><td>11586</td><td>22</td><td>CTAGCAGCCGACTTAGAACTGG</td><td>115.7</td><td>-1.4</td><td>0.4</td><td>-3.5</td><td>5.5E-04</td><td>6.7E-04</td><td>BT20/MB468</td></tr>
<tr><td>>MI0000286|hsa-mir-210&WithFlank&11|-|568083|568204@72.94.23&[MIMAT0000267&hsa-miR-210-3p&offsets|0|+1;m-102&11|-|568112|568133&offsets|0|+1]|CTGTGCGTGTGACAGCGGCTGAT</td><td>>MI0000286|hsa-mir-210&WithFlank&11|-|568083|568204</td><td>72</td><td>94</td><td>23</td><td>CTGTGCGTGTGACAGCGGCTGAT</td><td>2981.3</td><td>3.9</td><td>1.0</td><td>4.0</td><td>7.4E-05</td><td>6.7E-04</td><td>BT20/MB231</td></tr>
<tr><td>>MI0000101|hsa-mir-99a&WithFlank&21|+|16539083|16539175@19.39.21&[MIMAT0000097&hsa-miR-99a-5p&offsets|0|-1;m-64&21|+|16539101|16539121&offsets|0|0]|AACCCGTAGATCCGATCTTGT</td><td>>MI0000101|hsa-mir-99a&WithFlank&21|+|16539083|16539175</td><td>19</td><td>39</td><td>21</td><td>AACCCGTAGATCCGATCTTGT</td><td>1064.2</td><td>-4.2</td><td>1.2</td><td>-3.5</td><td>5.6E-04</td><td>6.7E-04</td><td>BT20/MB468</td></tr>
<tr><td>>hg38_21_+_8433172_8446622_RNA45SN1_45S_50ntFlanks@9198.9218.21|GTTCTCTCGGGGCCACGCGCG</td><td>>hg38_21_+_8433172_8446622_RNA45SN1_45S_50ntFlanks</td><td>9198</td><td>9218</td><td>21</td><td>GTTCTCTCGGGGCCACGCGCG</td><td>172.1</td><td>-4.2</td><td>1.2</td><td>-3.5</td><td>5.6E-04</td><td>6.7E-04</td><td>BT20/MB468</td></tr>
<tr><td>>hg38_21_+_8433172_8446622_RNA45SN1_45S_50ntFlanks@6719.6751.33|TGAATTGCAGGACACATTGATCATCGACACTTC</td><td>>hg38_21_+_8433172_8446622_RNA45SN1_45S_50ntFlanks</td><td>6719</td><td>6751</td><td>33</td><td>TGAATTGCAGGACACATTGATCATCGACACTTC</td><td>286.5</td><td>-1.6</td><td>0.5</td><td>-3.5</td><td>5.1E-04</td><td>6.8E-04</td><td>MB231/MB468</td></tr>
<tr><td>>hg38_21_+_8433172_8446622_RNA45SN1_45S_50ntFlanks@12991.13018.28|CTCCCTCGCTGCGATCTATTGAAAGTCA</td><td>>hg38_21_+_8433172_8446622_RNA45SN1_45S_50ntFlanks</td><td>12991</td><td>13018</td><td>28</td><td>CTCCCTCGCTGCGATCTATTGAAAGTCA</td><td>68.5</td><td>-2.3</td><td>0.7</td><td>-3.5</td><td>5.1E-04</td><td>6.8E-04</td><td>MB231/MB468</td></tr>
<tr><td>>hg38_21_+_8433172_8446622_RNA45SN1_45S_50ntFlanks@4661.4689.29|AGGTGAAATTCTTGGACCGGCGCAAGACG</td><td>>hg38_21_+_8433172_8446622_RNA45SN1_45S_50ntFlanks</td><td>4661</td><td>4689</td><td>29</td><td>AGGTGAAATTCTTGGACCGGCGCAAGACG</td><td>108.0</td><td>-3.0</td><td>0.9</td><td>-3.4</td><td>5.6E-04</td><td>6.8E-04</td><td>BT20/MB468</td></tr>
<tr><td>>hg38_21_+_8433172_8446622_RNA45SN1_45S_50ntFlanks@5471.5499.29|CACGGCCCTGGCGGAGCGCTGAGAAGACG</td><td>>hg38_21_+_8433172_8446622_RNA45SN1_45S_50ntFlanks</td><td>5471</td><td>5499</td><td>29</td><td>CACGGCCCTGGCGGAGCGCTGAGAAGACG</td><td>757.7</td><td>-2.0</td><td>0.6</td><td>-3.5</td><td>5.2E-04</td><td>6.8E-04</td><td>MB231/MB468</td></tr>
<tr><td>>hg38_21_+_8433172_8446622_RNA45SN1_45S_50ntFlanks@9554.9579.26|CGTAGCGGTCCTGACGTGCAAATCGG</td><td>>hg38_21_+_8433172_8446622_RNA45SN1_45S_50ntFlanks</td><td>9554</td><td>9579</td><td>26</td><td>CGTAGCGGTCCTGACGTGCAAATCGG</td><td>223.8</td><td>-1.9</td><td>0.6</td><td>-3.5</td><td>5.2E-04</td><td>6.9E-04</td><td>MB231/MB468</td></tr>
<tr><td>>hg38_21_+_8433172_8446622_RNA45SN1_45S_50ntFlanks@3785.3802.18|TGAAACTGCGAATGGCTC</td><td>>hg38_21_+_8433172_8446622_RNA45SN1_45S_50ntFlanks</td><td>3785</td><td>3802</td><td>18</td><td>TGAAACTGCGAATGGCTC</td><td>116.9</td><td>-1.5</td><td>0.4</td><td>-3.5</td><td>5.3E-04</td><td>6.9E-04</td><td>MB231/MB468</td></tr>
<tr><td>>hg38_21_+_8433172_8446622_RNA45SN1_45S_50ntFlanks@9536.9571.36|GAAACTCTGGTGGAGGTCCGTAGCGGTCCTGACGTG</td><td>>hg38_21_+_8433172_8446622_RNA45SN1_45S_50ntFlanks</td><td>9536</td><td>9571</td><td>36</td><td>GAAACTCTGGTGGAGGTCCGTAGCGGTCCTGACGTG</td><td>170.9</td><td>-1.5</td><td>0.4</td><td>-3.4</td><td>5.7E-04</td><td>6.9E-04</td><td>BT20/MB468</td></tr>
<tr><td>>MI0003776|hsa-mir-320b-1&WithFlank&1|+|116671740|116671823@48.67.20;MI0003839|hsa-mir-320b-2&WithFlank&1|-|224257034|224257116@47.66.20;MI0000542|hsa-mir-320a&WithFlank&8|-|22244960|22245043@48.67.20&[MIMAT0005792&hsa-miR-320b&offsets|0|-2;m-183&1|+|116671787|116671806&offsets|0|0];[MIMAT0005792_1&hsa-miR-320b&offsets|0|-2;m-183&1|-|224257051|224257070&offsets|0|0];[MIMAT0000510&hsa-miR-320a-3p&offsets|0|-2;m-60&8|-|22244975|22244996&offsets|0|-2]|AAAAGCTGGGTTGAGAGGGC</td><td>>MI0003776|hsa-mir-320b-1&WithFlank&1|+|116671740|116671823</td><td>48</td><td>67</td><td>20</td><td>AAAAGCTGGGTTGAGAGGGC</td><td>1411.3</td><td>2.8</td><td>0.7</td><td>4.0</td><td>7.7E-05</td><td>7.0E-04</td><td>BT20/MB231</td></tr>
<tr><td>>hg38_21_+_8433172_8446622_RNA45SN1_45S_50ntFlanks@11670.11688.19|CCCAGTGCTCTGAATGTCA</td><td>>hg38_21_+_8433172_8446622_RNA45SN1_45S_50ntFlanks</td><td>11670</td><td>11688</td><td>19</td><td>CCCAGTGCTCTGAATGTCA</td><td>604.3</td><td>-2.9</td><td>0.8</td><td>-3.4</td><td>5.8E-04</td><td>7.0E-04</td><td>BT20/MB468</td></tr>
<tr><td>>hg38_21_+_8433172_8446622_RNA45SN1_45S_50ntFlanks@6696.6720.25|GCAGCTAGCTGCGAGAATTAATGTG</td><td>>hg38_21_+_8433172_8446622_RNA45SN1_45S_50ntFlanks</td><td>6696</td><td>6720</td><td>25</td><td>GCAGCTAGCTGCGAGAATTAATGTG</td><td>305.3</td><td>5.3</td><td>1.5</td><td>3.5</td><td>5.4E-04</td><td>7.1E-04</td><td>MB231/MB468</td></tr>
</table>

<table>
<tr><td>>trna116_GluCTC_1_-_145399233_145399304@37.75.39,trna134_GluTTC_1_-_16861774_16861845@37.75.39,trna71_GluCTC_1_-_161439189_161439260@37.75.39,trna74_GluCTC_1_-_161431809_161431880@37.75.39,trna77_GluCTC_1_-_161424398_161424469@37.75.39,trna77_GluCTC_6_+_28949976_28950047@37.75.39,trna80_GluCTC_1_-_161417018_161417089@37.75.39,trna84_GluTTC_1_-_161391883_161391954@37.75.39,trna87_GluCTC_6_-_126101393_126101464@37.75.39|ACCGCCGCGGCCCGGGTTCGATTCCCGGTCAGGGAACCA</td><td>>trna116_GluCTC_1_-_145399233_145399304</td><td>37</td><td>75</td><td>39</td><td>ACCGCCGCGGCCCGGGTTCGATTCCCGGTCAGGGAACCA</td><td>2229.8</td><td>-4.1</td><td>1.0</td><td>-3.9</td><td>8.0E-05</td><td>7.2E-04</td><td>BT20/MB231</td></tr>
<tr><td>>hg38_21_+_8433172_8446622_RNA45SN1_45S_50ntFlanks@4590.4614.25|ATTTTGTTGGTTTTCGGAACTGAGG</td><td>>hg38_21_+_8433172_8446622_RNA45SN1_45S_50ntFlanks</td><td>4590</td><td>4614</td><td>25</td><td>ATTTTGTTGGTTTTCGGAACTGAGG</td><td>123.8</td><td>-3.0</td><td>0.9</td><td>-3.5</td><td>5.5E-04</td><td>7.2E-04</td><td>MB231/MB468</td></tr>
<tr><td>>hg38_21_+_8433172_8446622_RNA45SN1_45S_50ntFlanks@11318.11338.21|CCCCACGTCTCGTCGCGCGCG</td><td>>hg38_21_+_8433172_8446622_RNA45SN1_45S_50ntFlanks</td><td>11318</td><td>11338</td><td>21</td><td>CCCCACGTCTCGTCGCGCGCG</td><td>386.8</td><td>-4.4</td><td>1.3</td><td>-3.4</td><td>6.0E-04</td><td>7.2E-04</td><td>BT20/MB468</td></tr>
<tr><td>>hg38_21_+_8433172_8446622_RNA45SN1_45S_50ntFlanks@6734.6752.19|ATTGATCATCGACACTTCG</td><td>>hg38_21_+_8433172_8446622_RNA45SN1_45S_50ntFlanks</td><td>6734</td><td>6752</td><td>19</td><td>ATTGATCATCGACACTTCG</td><td>359.0</td><td>1.8</td><td>0.5</td><td>3.5</td><td>5.6E-04</td><td>7.3E-04</td><td>MB231/MB468</td></tr>
<tr><td>>MI0000101|hsa-mir-99a&WithFlank&21|+|16539083|16539175@19.38.20(+1C)&[MIMAT0000097&hsa-miR-99a-5p&offsets|0|-2(+1C);m-64&21|+|16539101|16539121&offsets|0|-1(+1C)]|AACCCGTAGATCCGATCTTGC</td><td>>MI0000101|hsa-mir-99a&WithFlank&21|+|16539083|16539175</td><td>19</td><td>38</td><td>20(+1C)</td><td>AACCCGTAGATCCGATCTTGC</td><td>495.4</td><td>2.4</td><td>0.7</td><td>3.5</td><td>5.6E-04</td><td>7.3E-04</td><td>MB231/MB468</td></tr>
<tr><td>>hg38_1_-_228634819_228635039_RNA5S12_5S_50ntFlanks@86.103.18|CGTCTGATCTCGGAAGCT</td><td>>hg38_1_-_228634819_228635039_RNA5S12_5S_50ntFlanks</td><td>86</td><td>103</td><td>18</td><td>CGTCTGATCTCGGAAGCT</td><td>166.3</td><td>-2.3</td><td>0.7</td><td>-3.5</td><td>5.6E-04</td><td>7.3E-04</td><td>MB231/MB468</td></tr>
<tr><td>>hg38_21_+_8433172_8446622_RNA45SN1_45S_50ntFlanks@12006.12032.27|GGCCGCCGGTGAAATACCACTACTCTG</td><td>>hg38_21_+_8433172_8446622_RNA45SN1_45S_50ntFlanks</td><td>12006</td><td>12032</td><td>27</td><td>GGCCGCCGGTGAAATACCACTACTCTG</td><td>244.6</td><td>-1.8</td><td>0.5</td><td>-3.4</td><td>6.1E-04</td><td>7.3E-04</td><td>BT20/MB468</td></tr>
<tr><td>>hg38_21_+_8433172_8446622_RNA45SN1_45S_50ntFlanks@8177.8203.27|TTCTGATCGAGGCCCAGCCCGTGGACG</td><td>>hg38_21_+_8433172_8446622_RNA45SN1_45S_50ntFlanks</td><td>8177</td><td>8203</td><td>27</td><td>TTCTGATCGAGGCCCAGCCCGTGGACG</td><td>832.7</td><td>2.4</td><td>0.7</td><td>3.4</td><td>5.6E-04</td><td>7.4E-04</td><td>MB231/MB468</td></tr>
<tr><td>>MI0000253|hsa-mir-148a&WithFlank&7|-|25949913|25949992@50.69.20(+2U)&[MIMAT0000243&hsa-miR-148a-3p&offsets|0|-2(+2U);m-7&7|-|25949922|25949943&offsets|0|-2(+2U)]|TCAGTGCACTACAGAACTTTTT</td><td>>MI0000253|hsa-mir-148a&WithFlank&7|-|25949913|25949992</td><td>50</td><td>69</td><td>20(+2U)</td><td>TCAGTGCACTACAGAACTTTTT</td><td>234.8</td><td>3.7</td><td>0.9</td><td>3.9</td><td>8.3E-05</td><td>7.4E-04</td><td>BT20/MB231</td></tr>
<tr><td>>hg38_21_+_8433172_8446622_RNA45SN1_45S_50ntFlanks@3841.3868.28|TCCTACTTGGATAACTGTGGTAATTCTA</td><td>>hg38_21_+_8433172_8446622_RNA45SN1_45S_50ntFlanks</td><td>3841</td><td>3868</td><td>28</td><td>TCCTACTTGGATAACTGTGGTAATTCTA</td><td>235.8</td><td>1.8</td><td>0.5</td><td>3.4</td><td>5.7E-04</td><td>7.4E-04</td><td>MB231/MB468</td></tr>
<tr><td>>hg38_21_+_8433172_8446622_RNA45SN1_45S_50ntFlanks@10796.10817.22|GGCATGTTGGAACAATGTAGGT</td><td>>hg38_21_+_8433172_8446622_RNA45SN1_45S_50ntFlanks</td><td>10796</td><td>10817</td><td>22</td><td>GGCATGTTGGAACAATGTAGGT</td><td>139.9</td><td>-2.2</td><td>0.6</td><td>-3.4</td><td>6.1E-04</td><td>7.4E-04</td><td>BT20/MB468</td></tr>
<tr><td>>hg38_21_+_8433172_8446622_RNA45SN1_45S_50ntFlanks@9305.9338.34|CCAAGGAGTCTAACACGTGCGCGAGTCGGGGGCT</td><td>>hg38_21_+_8433172_8446622_RNA45SN1_45S_50ntFlanks</td><td>9305</td><td>9338</td><td>34</td><td>CCAAGGAGTCTAACACGTGCGCGAGTCGGGGGCT</td><td>271.6</td><td>-3.2</td><td>0.9</td><td>-3.4</td><td>5.7E-04</td><td>7.4E-04</td><td>MB231/MB468</td></tr>
<tr><td>>hg38_21_+_8433172_8446622_RNA45SN1_45S_50ntFlanks@11703.11740.38|AATGAAGCGCGGGTAAACGGCGGGAGTAACTATGACTC</td><td>>hg38_21_+_8433172_8446622_RNA45SN1_45S_50ntFlanks</td><td>11703</td><td>11740</td><td>38</td><td>AATGAAGCGCGGGTAAACGGCGGGAGTAACTATGACTC</td><td>140.7</td><td>-3.0</td><td>0.9</td><td>-3.4</td><td>6.2E-04</td><td>7.4E-04</td><td>BT20/MB468</td></tr>
<tr><td>>hg38_21_+_8433172_8446622_RNA45SN1_45S_50ntFlanks@12419.12462.44|GATGTCGGCTCTTCCTATCATTGTGAAGCAGAATTCACCAAGCG</td><td>>hg38_21_+_8433172_8446622_RNA45SN1_45S_50ntFlanks</td><td>12419</td><td>12462</td><td>44</td><td>GATGTCGGCTCTTCCTATCATTGTGAAGCAGAATTCACCAAGCG</td><td>161.8</td><td>-3.2</td><td>0.8</td><td>-3.9</td><td>8.4E-05</td><td>7.5E-04</td><td>BT20/MB231</td></tr>
<tr><td>>MI0000542|hsa-mir-320a&WithFlank&8|-|22244960|22245043@49.68.20(+2U)&[MIMAT0000510&hsa-miR-320a-3p&offsets|+1|-1(+2U);m-60&8|-|22244975|22244996&offsets|+1|-1(+2U)]|AAAGCTGGGTTGAGAGGGCGTT</td><td>>MI0000542|hsa-mir-320a&WithFlank&8|-|22244960|22245043</td><td>49</td><td>68</td><td>20(+2U)</td><td>AAAGCTGGGTTGAGAGGGCGTT</td><td>238.7</td><td>2.9</td><td>0.7</td><td>3.9</td><td>8.4E-05</td><td>7.5E-04</td><td>BT20/MB231</td></tr>
<tr><td>>hg38_21_+_8433172_8446622_RNA45SN1_45S_50ntFlanks@9554.9580.27|CGTAGCGGTCCTGACGTGCAAATCGGT</td><td>>hg38_21_+_8433172_8446622_RNA45SN1_45S_50ntFlanks</td><td>9554</td><td>9580</td><td>27</td><td>CGTAGCGGTCCTGACGTGCAAATCGGT</td><td>251.6</td><td>-1.6</td><td>0.5</td><td>-3.4</td><td>6.2E-04</td><td>7.5E-04</td><td>BT20/MB468</td></tr>
<tr><td>>hg38_21_+_8433172_8446622_RNA45SN1_45S_50ntFlanks@4760.4783.24|TCAGATACCGTCGTAGTTCCGACC</td><td>>hg38_21_+_8433172_8446622_RNA45SN1_45S_50ntFlanks</td><td>4760</td><td>4783</td><td>24</td><td>TCAGATACCGTCGTAGTTCCGACC</td><td>534.7</td><td>-2.0</td><td>0.6</td><td>-3.4</td><td>5.7E-04</td><td>7.5E-04</td><td>MB231/MB468</td></tr>
<tr><td>>hg38_21_+_8433172_8446622_RNA45SN1_45S_50ntFlanks@11672.11701.30|CAGTGCTCTGAATGTCAAAGTGAAGAAATT</td><td>>hg38_21_+_8433172_8446622_RNA45SN1_45S_50ntFlanks</td><td>11672</td><td>11701</td><td>30</td><td>CAGTGCTCTGAATGTCAAAGTGAAGAAATT</td><td>659.2</td><td>1.8</td><td>0.5</td><td>3.4</td><td>5.8E-04</td><td>7.6E-04</td><td>MB231/MB468</td></tr>
<tr><td>>hg38_21_+_8433172_8446622_RNA45SN1_45S_50ntFlanks@11235.11268.34|CTTCCCGTGGATCGCCCCAGCTGCGGCGGGCGTC</td><td>>hg38_21_+_8433172_8446622_RNA45SN1_45S_50ntFlanks</td><td>11235</td><td>11268</td><td>34</td><td>CTTCCCGTGGATCGCCCCAGCTGCGGCGGGCGTC</td><td>137.4</td><td>-3.6</td><td>1.1</td><td>-3.4</td><td>6.3E-04</td><td>7.6E-04</td><td>BT20/MB468</td></tr>
<tr><td>>hg38_21_+_8433172_8446622_RNA45SN1_45S_50ntFlanks@11737.11768.32|ACTCTCTTAAGGTAGCCAAATGCCTCGTCATC</td><td>>hg38_21_+_8433172_8446622_RNA45SN1_45S_50ntFlanks</td><td>11737</td><td>11768</td><td>32</td><td>ACTCTCTTAAGGTAGCCAAATGCCTCGTCATC</td><td>94.4</td><td>-2.0</td><td>0.6</td><td>-3.4</td><td>5.8E-04</td><td>7.6E-04</td><td>MB231/MB468</td></tr>
<tr><td>>MI0003686|hsa-mir-542&WithFlank&X|-|134541335|134541443@59.80.22&[MIMAT0003389&hsa-miR-542-3p&offsets|0|0;m-136&X|-|134541364|134541385&offsets|0|0]|TGTGACAGATTGATAACTGAAA</td><td>>MI0003686|hsa-mir-542&WithFlank&X|-|134541335|134541443</td><td>59</td><td>80</td><td>22</td><td>TGTGACAGATTGATAACTGAAA</td><td>1352.3</td><td>4.1</td><td>1.1</td><td>3.9</td><td>8.6E-05</td><td>7.7E-04</td><td>BT20/MB231</td></tr>
<tr><td>>trna10_AspGTC_12_-_125424193_125424264@36.74.39,trna12_AspGTC_12_-_125411891_125411962@36.74.39,trna144_AspGTC_6_-_27551236_27551307@36.74.39,trna38_AspGTC_17_-_8125556_8125627@36.74.39,trna45_AspGTC_6_+_27447453_27447524@36.74.39,trna48_AspGTC_6_+_27471523_27471594@36.74.39,trna4_AspGTC_12_+_96429799_96429870@36.74.39,trna69_AspGTC_1_-_161440205_161440276@36.74.39,trna72_AspGTC_1_-_161432824_161432895@36.74.39,trna75_AspGTC_1_-_161425414_161425485@36.74.39,trna78_AspGTC_1_-_161418033_161418104@36.74.39,trna81_AspGTC_1_-_161410615_161410686@36.74.39|CACGCGGGAGACCGGGGTTCGATTCCCCGACGGGGAGCC</td><td>>trna10_AspGTC_12_-_125424193_125424264</td><td>36</td><td>74</td><td>39</td><td>CACGCGGGAGACCGGGGTTCGATTCCCCGACGGGGAGCC</td><td>444.5</td><td>-3.7</td><td>1.0</td><td>-3.9</td><td>8.6E-05</td><td>7.7E-04</td><td>BT20/MB231</td></tr>
<tr><td>>hg38_21_+_8433172_8446622_RNA45SN1_45S_50ntFlanks@12007.12029.23|GCCGCCGGTGAAATACCACTACT</td><td>>hg38_21_+_8433172_8446622_RNA45SN1_45S_50ntFlanks</td><td>12007</td><td>12029</td><td>23</td><td>GCCGCCGGTGAAATACCACTACT</td><td>760.9</td><td>3.6</td><td>1.0</td><td>3.4</td><td>6.4E-04</td><td>7.7E-04</td><td>BT20/MB468</td></tr>
<tr><td>>trna111_HisGTG_1_-_147774845_147774916@1.30.30,trna118_HisGTG_1_-_145396881_145396952@1.30.30,trna16_HisGTG_1_+_146544773_146544844@1.30.30,trna1_HisGTG_15_+_45493349_45493420@1.30.30,trna21_HisGTG_1_+_147753471_147753542@1.30.30,trna33_HisGTG_6_+_27125906_27125977@1.30.30,trna7_HisGTG_9_-_14433938_14434009@1.30.30,trna8_HisGTG_15_-_45492611_45492682@1.30.30,trna9_HisGTG_15_-_45490804_45490875@1.30.30|GCCGTGATCGTATAGTGGTTAGTACTCTGC</td><td>>trna111_HisGTG_1_-_147774845_147774916</td><td>1</td><td>30</td><td>30</td><td>GCCGTGATCGTATAGTGGTTAGTACTCTGC</td><td>3453.5</td><td>3.5</td><td>1.0</td><td>3.4</td><td>5.9E-04</td><td>7.7E-04</td><td>MB231/MB468</td></tr>
<tr><td>>hg38_21_+_8433172_8446622_RNA45SN1_45S_50ntFlanks@3835.3859.25|CTCCTCTCCTACTTGGATAACTGTG</td><td>>hg38_21_+_8433172_8446622_RNA45SN1_45S_50ntFlanks</td><td>3835</td><td>3859</td><td>25</td><td>CTCCTCTCCTACTTGGATAACTGTG</td><td>85.2</td><td>-2.1</td><td>0.6</td><td>-3.4</td><td>5.9E-04</td><td>7.7E-04</td><td>MB231/MB468</td></tr>
<tr><td>>hg38_21_+_8433172_8446622_RNA45SN1_45S_50ntFlanks@8114.8141.28|GCGCGGGACATGTGGCGTACGGAAGACC</td><td>>hg38_21_+_8433172_8446622_RNA45SN1_45S_50ntFlanks</td><td>8114</td><td>8141</td><td>28</td><td>GCGCGGGACATGTGGCGTACGGAAGACC</td><td>185.4</td><td>-3.1</td><td>0.9</td><td>-3.4</td><td>5.9E-04</td><td>7.7E-04</td><td>MB231/MB468</td></tr>
<tr><td>>hg38_21_+_8433172_8446622_RNA45SN1_45S_50ntFlanks@11561.11587.27|GCGCCTAGCAGCCGACTTAGAACTGGT</td><td>>hg38_21_+_8433172_8446622_RNA45SN1_45S_50ntFlanks</td><td>11561</td><td>11587</td><td>27</td><td>GCGCCTAGCAGCCGACTTAGAACTGGT</td><td>133.7</td><td>-2.1</td><td>0.6</td><td>-3.4</td><td>6.4E-04</td><td>7.7E-04</td><td>BT20/MB468</td></tr>
<tr><td>>hg38_21_+_8433172_8446622_RNA45SN1_45S_50ntFlanks@5468.5499.32|GCCCACGGCCCTGGCGGAGCGCTGAGAAGACG</td><td>>hg38_21_+_8433172_8446622_RNA45SN1_45S_50ntFlanks</td><td>5468</td><td>5499</td><td>32</td><td>GCCCACGGCCCTGGCGGAGCGCTGAGAAGACG</td><td>96.0</td><td>-2.2</td><td>0.6</td><td>-3.4</td><td>5.9E-04</td><td>7.7E-04</td><td>MB231/MB468</td></tr>
<tr><td>>hg38_21_+_8433172_8446622_RNA45SN1_45S_50ntFlanks@11579.11606.28|AGAACTGGTGCGGACCAGGGGAATCCGA</td><td>>hg38_21_+_8433172_8446622_RNA45SN1_45S_50ntFlanks</td><td>11579</td><td>11606</td><td>28</td><td>AGAACTGGTGCGGACCAGGGGAATCCGA</td><td>144.4</td><td>-1.9</td><td>0.5</td><td>-3.4</td><td>5.9E-04</td><td>7.8E-04</td><td>MB231/MB468</td></tr>
<tr><td>>Antisense_hg38_21_-_8433172_8446622_RNA45SN1_45S_50ntFlanks@7674.7693.20|CTCTCTCTCTCTCTCCCCCG</td><td>>Antisense_hg38_21_-_8433172_8446622_RNA45SN1_45S_50ntFlanks</td><td>7674</td><td>7693</td><td>20</td><td>CTCTCTCTCTCTCTCCCCCG</td><td>260.9</td><td>-4.2</td><td>1.2</td><td>-3.4</td><td>6.5E-04</td><td>7.8E-04</td><td>BT20/MB468</td></tr>
<tr><td>>hg38_21_+_8433172_8446622_RNA45SN1_45S_50ntFlanks@11027.11046.20|CTCTCTCTCTCTCTCCCCCG</td><td>>hg38_21_+_8433172_8446622_RNA45SN1_45S_50ntFlanks</td><td>11027</td><td>11046</td><td>20</td><td>CTCTCTCTCTCTCTCCCCCG</td><td>260.9</td><td>-4.2</td><td>1.2</td><td>-3.4</td><td>6.5E-04</td><td>7.8E-04</td><td>BT20/MB468</td></tr>
<tr><td>>hg38_21_+_8433172_8446622_RNA45SN1_45S_50ntFlanks@6648.6689.42|GTACGACTCTTAGCGGTGGATCACTCGGCTCGTGCGTCGATG</td><td>>hg38_21_+_8433172_8446622_RNA45SN1_45S_50ntFlanks</td><td>6648</td><td>6689</td><td>42</td><td>GTACGACTCTTAGCGGTGGATCACTCGGCTCGTGCGTCGATG</td><td>1941.2</td><td>1.9</td><td>0.6</td><td>3.4</td><td>6.0E-04</td><td>7.8E-04</td><td>MB231/MB468</td></tr>
<tr><td>>hg38_21_+_8433172_8446622_RNA45SN1_45S_50ntFlanks@8356.8376.21|GAAAAGAACTTTGAAGAGAGA</td><td>>hg38_21_+_8433172_8446622_RNA45SN1_45S_50ntFlanks</td><td>8356</td><td>8376</td><td>21</td><td>GAAAAGAACTTTGAAGAGAGA</td><td>222.9</td><td>-2.3</td><td>0.7</td><td>-3.4</td><td>6.5E-04</td><td>7.8E-04</td><td>BT20/MB468</td></tr>
<tr><td>>hg38_21_+_8433172_8446622_RNA45SN1_45S_50ntFlanks@9198.9220.23|GTTCTCTCGGGGCCACGCGCGCG</td><td>>hg38_21_+_8433172_8446622_RNA45SN1_45S_50ntFlanks</td><td>9198</td><td>9220</td><td>23</td><td>GTTCTCTCGGGGCCACGCGCGCG</td><td>137.7</td><td>-4.1</td><td>1.2</td><td>-3.4</td><td>6.5E-04</td><td>7.9E-04</td><td>BT20/MB468</td></tr>
<tr><td>>hg38_21_+_8433172_8446622_RNA45SN1_45S_50ntFlanks@6652.6680.29|GACTCTTAGCGGTGGATCACTCGGCTCGT</td><td>>hg38_21_+_8433172_8446622_RNA45SN1_45S_50ntFlanks</td><td>6652</td><td>6680</td><td>29</td><td>GACTCTTAGCGGTGGATCACTCGGCTCGT</td><td>531.8</td><td>2.0</td><td>0.6</td><td>3.4</td><td>6.0E-04</td><td>7.9E-04</td><td>MB231/MB468</td></tr>
<tr><td>>hg38_21_+_8433172_8446622_RNA45SN1_45S_50ntFlanks@11879.11898.20|ACCCTGTTGAGCTTGACTCT</td><td>>hg38_21_+_8433172_8446622_RNA45SN1_45S_50ntFlanks</td><td>11879</td><td>11898</td><td>20</td><td>ACCCTGTTGAGCTTGACTCT</td><td>1047.9</td><td>3.8</td><td>1.1</td><td>3.4</td><td>6.5E-04</td><td>7.9E-04</td><td>BT20/MB468</td></tr>
<tr><td>>hg38_21_+_8433172_8446622_RNA45SN1_45S_50ntFlanks@11217.11234.18|CTGGACGCGAGCCGGGCC</td><td>>hg38_21_+_8433172_8446622_RNA45SN1_45S_50ntFlanks</td><td>11217</td><td>11234</td><td>18</td><td>CTGGACGCGAGCCGGGCC</td><td>990.9</td><td>-3.4</td><td>1.0</td><td>-3.4</td><td>6.6E-04</td><td>7.9E-04</td><td>BT20/MB468</td></tr>
<tr><td>>hg38_21_+_8433172_8446622_RNA45SN1_45S_50ntFlanks@12474.12498.25|ACCCACTAATAGGGAACGTGAGCTG</td><td>>hg38_21_+_8433172_8446622_RNA45SN1_45S_50ntFlanks</td><td>12474</td><td>12498</td><td>25</td><td>ACCCACTAATAGGGAACGTGAGCTG</td><td>535.6</td><td>-2.4</td><td>0.7</td><td>-3.4</td><td>6.0E-04</td><td>7.9E-04</td><td>MB231/MB468</td></tr>
</table>

| | | | | | | | | | | | | |
|---|---|---|---|---|---|---|---|---|---|---|---|---|
| >hg38_21_+_8433172_8446622_RNA45SN1_45S_50ntFlanks@4723.4759.37\|CATTAATCAAGAACGAAAGTCGGAGGTTCGAAGACGA | >hg38_21_+_8433172_8446622_RNA45SN1_45S_50ntFlanks | 4723 | 4759 | 37 | CATTAATCAAGAACGAAAGTCGGAGGTTCGAAGACGA | 653.3 | -2.4 | 0.7 | -3.4 | 6.0E-04 | 7.9E-04 | MB231/MB468 |
| >MI0000298\|hsa-mir-221&WithFlank&X\|-\|45746151\|45746272@31.55.25&[MIMAT0004568&hsa-miR-221-5p&offsets\|0\|+3;m-405&X\|-\|45746220\|45746242&offsets\|0\|+2]\|ACCTGGCATACAATGTAGATTTCTG | >MI0000298\|hsa-mir-221&WithFlank&X\|-\|45746151\|45746272 | 31 | 55 | 25 | ACCTGGCATACAATGTAGATTTCTG | 469.8 | -2.8 | 0.7 | -3.9 | 9.0E-05 | 7.9E-04 | BT20/MB231 |
| >hg38_21_+_8433172_8446622_RNA45SN1_45S_50ntFlanks@12416.12457.42\|TTCGATGTCGGCTCTTCCTATCATTGTGAAGCAGAATTCACC | >hg38_21_+_8433172_8446622_RNA45SN1_45S_50ntFlanks | 12416 | 12457 | 42 | TTCGATGTCGGCTCTTCCTATCATTGTGAAGCAGAATTCACC | 904.1 | -3.1 | 0.8 | -3.9 | 9.1E-05 | 8.0E-04 | BT20/MB231 |
| >hg38_21_+_8433172_8446622_RNA45SN1_45S_50ntFlanks@6667.6699.33\|ATCACTCGGCTCGTGCGTCGATGAAGAACGCAG | >hg38_21_+_8433172_8446622_RNA45SN1_45S_50ntFlanks | 6667 | 6699 | 33 | ATCACTCGGCTCGTGCGTCGATGAAGAACGCAG | 111.4 | -3.2 | 0.9 | -3.4 | 6.1E-04 | 8.0E-04 | MB231/MB468 |
| >hg38_21_+_8433172_8446622_RNA45SN1_45S_50ntFlanks@11670.11688.19\|CCCAGTGCTCTGAATGTCA | >hg38_21_+_8433172_8446622_RNA45SN1_45S_50ntFlanks | 11670 | 11688 | 19 | CCCAGTGCTCTGAATGTCA | 523.2 | -2.1 | 0.6 | -3.4 | 6.2E-04 | 8.1E-04 | MB231/MB468 |
| >hg38_21_+_8433172_8446622_RNA45SN1_45S_50ntFlanks@9533.9562.30\|GAGGAAACTCTGGTGGAGGTCCGTAGCGGT | >hg38_21_+_8433172_8446622_RNA45SN1_45S_50ntFlanks | 9533 | 9562 | 30 | GAGGAAACTCTGGTGGAGGTCCGTAGCGGT | 161.7 | -1.9 | 0.5 | -3.4 | 6.7E-04 | 8.1E-04 | BT20/MB468 |
| >hg38_21_+_8433172_8446622_RNA45SN1_45S_50ntFlanks@6722.6741.20\|ATTGCAGGACACATTGATCA | >hg38_21_+_8433172_8446622_RNA45SN1_45S_50ntFlanks | 6722 | 6741 | 20 | ATTGCAGGACACATTGATCA | 107.9 | -2.2 | 0.6 | -3.4 | 6.2E-04 | 8.1E-04 | MB231/MB468 |
| >hg38_21_+_8433172_8446622_RNA45SN1_45S_50ntFlanks@12003.12030.28\|GCGGGCCGCCGGTGAAATACCACTACTC | >hg38_21_+_8433172_8446622_RNA45SN1_45S_50ntFlanks | 12003 | 12030 | 28 | GCGGGCCGCCGGTGAAATACCACTACTC | 176.6 | -2.9 | 0.8 | -3.4 | 6.7E-04 | 8.1E-04 | BT20/MB468 |
| >hg38_21_+_8433172_8446622_RNA45SN1_45S_50ntFlanks@12914.12933.20\|CGTTCGTGGGGAACCTGGCG | >hg38_21_+_8433172_8446622_RNA45SN1_45S_50ntFlanks | 12914 | 12933 | 20 | CGTTCGTGGGGAACCTGGCG | 195.6 | -2.7 | 0.8 | -3.4 | 6.2E-04 | 8.1E-04 | MB231/MB468 |
| >MI0000286\|hsa-mir-210&WithFlank&11\|-\|568083\|568204@72.93.22(+1C)&[MIMAT0000267&hsa-miR-210-3p&offsets\|0\|0(+1C);m-102&11\|-\|568112\|568133&offsets\|0\|0(+1C)]\|CTGTGCGTGTGACAGCGGCTGAC | >MI0000286\|hsa-mir-210&WithFlank&11\|-\|568083\|568204 | 72 | 93 | 22(+1C) | CTGTGCGTGTGACAGCGGCTGAC | 571.0 | 3.7 | 1.0 | 3.9 | 9.3E-05 | 8.2E-04 | BT20/MB231 |
| >hg38_21_+_8433172_8446622_RNA45SN1_45S_50ntFlanks@12458.12476.19\|AAGCGTTGGATTGTTCACC | >hg38_21_+_8433172_8446622_RNA45SN1_45S_50ntFlanks | 12458 | 12476 | 19 | AAGCGTTGGATTGTTCACC | 2584.6 | -1.6 | 0.5 | -3.4 | 6.8E-04 | 8.2E-04 | BT20/MB468 |
| >hg38_21_+_8433172_8446622_RNA45SN1_45S_50ntFlanks@8319.8356.38\|CGAGACCGATAGTCAACAAGTACCGTAAGGGAAAGTTG | >hg38_21_+_8433172_8446622_RNA45SN1_45S_50ntFlanks | 8319 | 8356 | 38 | CGAGACCGATAGTCAACAAGTACCGTAAGGGAAAGTTG | 141.2 | -3.0 | 0.9 | -3.4 | 6.8E-04 | 8.2E-04 | BT20/MB468 |
| >hg38_21_+_8433172_8446622_RNA45SN1_45S_50ntFlanks@12003.12029.27\|GCGGGCCGCCGGTGAAATACCACTACT | >hg38_21_+_8433172_8446622_RNA45SN1_45S_50ntFlanks | 12003 | 12029 | 27 | GCGGGCCGCCGGTGAAATACCACTACT | 307.9 | -3.2 | 0.8 | -3.9 | 9.4E-05 | 8.2E-04 | BT20/MB231 |
| >hg38_21_+_8433172_8446622_RNA45SN1_45S_50ntFlanks@3705.3746.42\|TACCTGGTTGATCCTGCCAGTAGCATATGCTTGTCTCAAAGA | >hg38_21_+_8433172_8446622_RNA45SN1_45S_50ntFlanks | 3705 | 3746 | 42 | TACCTGGTTGATCCTGCCAGTAGCATATGCTTGTCTCAAAGA | 4403.6 | 4.2 | 1.1 | 3.9 | 9.5E-05 | 8.3E-04 | BT20/MB231 |
| >hg38_21_+_8433172_8446622_RNA45SN1_45S_50ntFlanks@9539.9571.33\|ACTCTGGTGGAGGTCCGTAGCGGTCCTGACGTG | >hg38_21_+_8433172_8446622_RNA45SN1_45S_50ntFlanks | 9539 | 9571 | 33 | ACTCTGGTGGAGGTCCGTAGCGGTCCTGACGTG | 538.2 | -1.5 | 0.4 | -3.4 | 6.3E-04 | 8.3E-04 | MB231/MB468 |
| >hg38_21_+_8433172_8446622_RNA45SN1_45S_50ntFlanks@3843.3871.29\|CTACTTGGATAACTGTGGTAATTCTAGAG | >hg38_21_+_8433172_8446622_RNA45SN1_45S_50ntFlanks | 3843 | 3871 | 29 | CTACTTGGATAACTGTGGTAATTCTAGAG | 358.2 | -2.3 | 0.7 | -3.4 | 6.9E-04 | 8.3E-04 | BT20/MB468 |
| >hg38_21_+_8433172_8446622_RNA45SN1_45S_50ntFlanks@10388.10407.20\|GTGAACAGCAGTTGAACATG | >hg38_21_+_8433172_8446622_RNA45SN1_45S_50ntFlanks | 10388 | 10407 | 20 | GTGAACAGCAGTTGAACATG | 199.7 | -2.1 | 0.6 | -3.4 | 6.3E-04 | 8.3E-04 | MB231/MB468 |
| >hg38_21_+_8433172_8446622_RNA45SN1_45S_50ntFlanks@12404.12431.28\|GCTTTTTGATCCTTCGATGTCGGCTCTT | >hg38_21_+_8433172_8446622_RNA45SN1_45S_50ntFlanks | 12404 | 12431 | 28 | GCTTTTTGATCCTTCGATGTCGGCTCTT | 175.9 | -1.7 | 0.5 | -3.4 | 6.4E-04 | 8.3E-04 | MB231/MB468 |
| >MI0003667\|hsa-mir-652&WithFlank&X\|+\|110055323\|110055432@67.87.21&[MIMAT0003322&hsa-miR-652-3p&offsets\|0\|0;m-148&X\|+\|110055389\|110055410&offsets\|0\|-1]\|AATGGCGCCACTAGGGTTGTG | >MI0003667\|hsa-mir-652&WithFlank&X\|+\|110055323\|110055432 | 67 | 87 | 21 | AATGGCGCCACTAGGGTTGTG | 266.7 | 3.0 | 0.8 | 3.9 | 9.6E-05 | 8.3E-04 | BT20/MB231 |
| >hg38_MT_+_598_1651_RNR1_12S_50ntFlanks@966.1004.39\|TAACATGGTAAGTGTACTGGAAAGTGCACTTGGACGAAC | >hg38_MT_+_598_1651_RNR1_12S_50ntFlanks | 966 | 1004 | 39 | TAACATGGTAAGTGTACTGGAAAGTGCACTTGGACGAAC | 552.7 | -3.1 | 0.8 | -3.9 | 9.6E-05 | 8.3E-04 | BT20/MB231 |
| >hg38_21_+_8433172_8446622_RNA45SN1_45S_50ntFlanks@12071.12092.22\|CCCCGAGGGGCTCTCGCTTCTG | >hg38_21_+_8433172_8446622_RNA45SN1_45S_50ntFlanks | 12071 | 12092 | 22 | CCCCGAGGGGCTCTCGCTTCTG | 134.1 | -2.5 | 0.7 | -3.4 | 6.9E-04 | 8.4E-04 | BT20/MB468 |
| >hg38_21_+_8433172_8446622_RNA45SN1_45S_50ntFlanks@6701.6720.20\|TAGCTGCGAGAATTAATGTG | >hg38_21_+_8433172_8446622_RNA45SN1_45S_50ntFlanks | 6701 | 6720 | 20 | TAGCTGCGAGAATTAATGTG | 255.4 | 2.0 | 0.6 | 3.4 | 6.4E-04 | 8.4E-04 | MB231/MB468 |
| >hg38_21_+_8433172_8446622_RNA45SN1_45S_50ntFlanks@9539.9569.31\|ACTCTGGTGGAGGTCCGTAGCGGTCCTGACG | >hg38_21_+_8433172_8446622_RNA45SN1_45S_50ntFlanks | 9539 | 9569 | 31 | ACTCTGGTGGAGGTCCGTAGCGGTCCTGACG | 129.4 | -2.9 | 0.8 | -3.4 | 7.0E-04 | 8.4E-04 | BT20/MB468 |
| >hg38_21_+_8433172_8446622_RNA45SN1_45S_50ntFlanks@9573.9595.23\|AAATCGGTCGTCCGACCTGGGTA | >hg38_21_+_8433172_8446622_RNA45SN1_45S_50ntFlanks | 9573 | 9595 | 23 | AAATCGGTCGTCCGACCTGGGTA | 1227.0 | -2.4 | 0.7 | -3.4 | 6.4E-04 | 8.4E-04 | MB231/MB468 |
| >hg38_21_+_8433172_8446622_RNA45SN1_45S_50ntFlanks@3985.4009.25\|CGGCTTTGGTGACTCTAGATAACCT | >hg38_21_+_8433172_8446622_RNA45SN1_45S_50ntFlanks | 3985 | 4009 | 25 | CGGCTTTGGTGACTCTAGATAACCT | 82.3 | -2.9 | 0.9 | -3.4 | 7.0E-04 | 8.5E-04 | BT20/MB468 |
| >hg38_21_+_8433172_8446622_RNA45SN1_45S_50ntFlanks@11208.11234.27\|GCGGCGACTCTGGACGCGAGCCGGGCC | >hg38_21_+_8433172_8446622_RNA45SN1_45S_50ntFlanks | 11208 | 11234 | 27 | GCGGCGACTCTGGACGCGAGCCGGGCC | 2358.8 | -5.3 | 1.6 | -3.4 | 7.1E-04 | 8.5E-04 | BT20/MB468 |
| >hg38_21_+_8433172_8446622_RNA45SN1_45S_50ntFlanks@11561.11594.34\|GCGCCTAGCAGCCGACTTAGAACTGGTGCGGACC | >hg38_21_+_8433172_8446622_RNA45SN1_45S_50ntFlanks | 11561 | 11594 | 34 | GCGCCTAGCAGCCGACTTAGAACTGGTGCGGACC | 211.6 | -1.9 | 0.6 | -3.4 | 6.6E-04 | 8.6E-04 | MB231/MB468 |
| >MI0000810\|hsa-mir-135b&WithFlank&1\|-\|205448296\|205448404@22.43.22&[MIMAT0000758&hsa-miR-135b-5p&offsets\|0\|-1;m-321&1\|-\|205448361\|205448383&offsets\|0\|-1]\|TATGGCTTTTCATTCCTATGTG | >MI0000810\|hsa-mir-135b&WithFlank&1\|-\|205448296\|205448404 | 22 | 43 | 22 | TATGGCTTTTCATTCCTATGTG | 317.3 | 2.9 | 0.7 | 3.9 | 1.0E-04 | 8.7E-04 | BT20/MB231 |
| >hg38_21_+_8433172_8446622_RNA45SN1_45S_50ntFlanks@11661.11680.20\|TGATTTCTGCCCAGTGCTCT | >hg38_21_+_8433172_8446622_RNA45SN1_45S_50ntFlanks | 11661 | 11680 | 20 | TGATTTCTGCCCAGTGCTCT | 504.1 | -2.4 | 0.7 | -3.4 | 6.7E-04 | 8.8E-04 | MB231/MB468 |
| >hg38_21_+_8433172_8446622_RNA45SN1_45S_50ntFlanks@7975.7993.19\|CGCGACCTCAGATCAGACG | >hg38_21_+_8433172_8446622_RNA45SN1_45S_50ntFlanks | 7975 | 7993 | 19 | CGCGACCTCAGATCAGACG | 8406.5 | 3.0 | 0.9 | 3.4 | 6.7E-04 | 8.8E-04 | MB231/MB468 |
| >hg38_21_+_8433172_8446622_RNA45SN1_45S_50ntFlanks@11564.11586.23\|CCTAGCAGCCGACTTAGAACTGG | >hg38_21_+_8433172_8446622_RNA45SN1_45S_50ntFlanks | 11564 | 11586 | 23 | CCTAGCAGCCGACTTAGAACTGG | 567.9 | -1.6 | 0.5 | -3.4 | 6.7E-04 | 8.8E-04 | MB231/MB468 |
| >hg38_21_+_8433172_8446622_RNA45SN1_45S_50ntFlanks@12003.12031.29\|GCGGGCCGCCGGTGAAATACCACTACTCT | >hg38_21_+_8433172_8446622_RNA45SN1_45S_50ntFlanks | 12003 | 12031 | 29 | GCGGGCCGCCGGTGAAATACCACTACTCT | 360.3 | -2.0 | 0.6 | -3.4 | 7.4E-04 | 8.9E-04 | BT20/MB468 |
| >hg38_21_+_8433172_8446622_RNA45SN1_45S_50ntFlanks@11894.11911.18\|ACTCTAGTCTGGCACGGT | >hg38_21_+_8433172_8446622_RNA45SN1_45S_50ntFlanks | 11894 | 11911 | 18 | ACTCTAGTCTGGCACGGT | 90.8 | -2.8 | 0.8 | -3.4 | 6.9E-04 | 9.0E-04 | MB231/MB468 |
| >hg38_21_+_8433172_8446622_RNA45SN1_45S_50ntFlanks@7982.8005.24\|TCAGATCAGACGTGGCGACCCGCT | >hg38_21_+_8433172_8446622_RNA45SN1_45S_50ntFlanks | 7982 | 8005 | 24 | TCAGATCAGACGTGGCGACCCGCT | 133.9 | -1.9 | 0.6 | -3.4 | 6.9E-04 | 9.1E-04 | MB231/MB468 |
| >MI0000301\|hsa-mir-224&WithFlank&X\|-\|151958572\|151958664@14.35.22(+1U)&[MIMAT0000281&hsa-miR-224-5p&offsets\|+1\|-2(+1U);m-238&X\|-\|151958630\|151958651&offsets\|0\|0(+1U)]\|CAAGTCACTAGTGGTTCCGTTTT | >MI0000301\|hsa-mir-224&WithFlank&X\|-\|151958572\|151958664 | 14 | 35 | 22(+1U) | CAAGTCACTAGTGGTTCCGTTTT | 286.2 | 2.7 | 0.7 | 3.9 | 1.1E-04 | 9.1E-04 | BT20/MB231 |
| >MI0000542\|hsa-mir-320a&WithFlank&8\|-\|22244960\|22245043@48.68.21&[MIMAT0000510&hsa-miR-320a-3p&offsets\|0\|-1;m-60&8\|-\|22244975\|22244996&offsets\|0\|-1]\|AAAAGCTGGGTTGAGAGGGCG | >MI0000542\|hsa-mir-320a&WithFlank&8\|-\|22244960\|22245043 | 48 | 68 | 21 | AAAAGCTGGGTTGAGAGGGCG | 4587.2 | 2.7 | 0.7 | 3.9 | 1.1E-04 | 9.1E-04 | BT20/MB231 |
| >am_trnaMT_ValTAC_MT_+_1602_1670@48.67.20\|CAACTTAACTTGACCGCTCT | >am_trnaMT_ValTAC_MT_+_1602_1670 | 48 | 67 | 20 | CAACTTAACTTGACCGCTCT | 3956.2 | 7.8 | 2.3 | 3.4 | 7.1E-04 | 9.2E-04 | MB231/MB468 |
| >hg38_21_+_8433172_8446622_RNA45SN1_45S_50ntFlanks@4764.4781.18\|ATACCGTCGTAGTTCCGA | >hg38_21_+_8433172_8446622_RNA45SN1_45S_50ntFlanks | 4764 | 4781 | 18 | ATACCGTCGTAGTTCCGA | 101.5 | -2.8 | 0.8 | -3.4 | 7.1E-04 | 9.3E-04 | MB231/MB468 |
| >hg38_21_+_8433172_8446622_RNA45SN1_45S_50ntFlanks@12996.13037.42\|TCGCTGCGATCTATTGAAAGTCAGCCCTCGACACAAGGGTTT | >hg38_21_+_8433172_8446622_RNA45SN1_45S_50ntFlanks | 12996 | 13037 | 42 | TCGCTGCGATCTATTGAAAGTCAGCCCTCGACACAAGGGTTT | 353.3 | -2.9 | 0.7 | -3.9 | 1.1E-04 | 9.3E-04 | BT20/MB231 |
| >MI0000542\|hsa-mir-320a&WithFlank&8\|-\|22244960\|22245043@48.71.24&[MIMAT0000510&hsa-miR-320a-3p&offsets\|0\|+2;m-60&8\|-\|22244975\|22244996&offsets\|0\|+2]\|AAAAGCTGGGTTGAGAGGGCGAAA | >MI0000542\|hsa-mir-320a&WithFlank&8\|-\|22244960\|22245043 | 48 | 71 | 24 | AAAAGCTGGGTTGAGAGGGCGAAA | 2859.8 | 2.8 | 0.7 | 3.9 | 1.1E-04 | 9.3E-04 | BT20/MB231 |
| >hg38_21_+_8433172_8446622_RNA45SN1_45S_50ntFlanks@7991.8011.21\|ACGTGGCGACCCGCTGAATTT | >hg38_21_+_8433172_8446622_RNA45SN1_45S_50ntFlanks | 7991 | 8011 | 21 | ACGTGGCGACCCGCTGAATTT | 5214.3 | 2.8 | 0.8 | 3.4 | 7.1E-04 | 9.3E-04 | MB231/MB468 |
| >hg38_21_+_8433172_8446622_RNA45SN1_45S_50ntFlanks@12456.12490.35\|CCAAGCGTTGGATTGTTCACCCACTAATAGGGAAC | >hg38_21_+_8433172_8446622_RNA45SN1_45S_50ntFlanks | 12456 | 12490 | 35 | CCAAGCGTTGGATTGTTCACCCACTAATAGGGAAC | 526.3 | -2.0 | 0.6 | -3.4 | 7.3E-04 | 9.5E-04 | MB231/MB468 |
| >hg38_21_+_8433172_8446622_RNA45SN1_45S_50ntFlanks@8105.8142.38\|CGCGGCGGGGCGCGGGACATGTGGCGTACGGAAGACCC | >hg38_21_+_8433172_8446622_RNA45SN1_45S_50ntFlanks | 8105 | 8142 | 38 | CGCGGCGGGGCGCGGGACATGTGGCGTACGGAAGACCC | 69.8 | -2.9 | 0.9 | -3.4 | 7.3E-04 | 9.6E-04 | MB231/MB468 |
| >hg38_21_+_8433172_8446622_RNA45SN1_45S_50ntFlanks@6656.6689.34\|CTTAGCGGTGGATCACTCGGCTCGTGCGTCGATG | >hg38_21_+_8433172_8446622_RNA45SN1_45S_50ntFlanks | 6656 | 6689 | 34 | CTTAGCGGTGGATCACTCGGCTCGTGCGTCGATG | 273.8 | -1.7 | 0.5 | -3.4 | 7.4E-04 | 9.6E-04 | MB231/MB468 |
| >hg38_21_+_8433172_8446622_RNA45SN1_45S_50ntFlanks@6648.6670.23\|GTACGACTCTTAGCGGTGGATCA | >hg38_21_+_8433172_8446622_RNA45SN1_45S_50ntFlanks | 6648 | 6670 | 23 | GTACGACTCTTAGCGGTGGATCA | 104.0 | -1.8 | 0.5 | -3.4 | 7.4E-04 | 9.6E-04 | MB231/MB468 |
| >hg38_21_+_8433172_8446622_RNA45SN1_45S_50ntFlanks@12418.12439.22\|CGATGTCGGCTCTTCCTATCAT | >hg38_21_+_8433172_8446622_RNA45SN1_45S_50ntFlanks | 12418 | 12439 | 22 | CGATGTCGGCTCTTCCTATCAT | 680.4 | 1.9 | 0.5 | 3.4 | 7.4E-04 | 9.6E-04 | MB231/MB468 |
| >hg38_21_+_8433172_8446622_RNA45SN1_45S_50ntFlanks@10841.10868.28\|CGTAACTTCGGGATAAGGATTGGCTCTA | >hg38_21_+_8433172_8446622_RNA45SN1_45S_50ntFlanks | 10841 | 10868 | 28 | CGTAACTTCGGGATAAGGATTGGCTCTA | 5752.2 | -3.2 | 1.0 | -3.4 | 7.4E-04 | 9.6E-04 | MB231/MB468 |
| >hg38_21_+_8433172_8446622_RNA45SN1_45S_50ntFlanks@12940.12959.20\|CATTCGTAGACGACCTGCTT | >hg38_21_+_8433172_8446622_RNA45SN1_45S_50ntFlanks | 12940 | 12959 | 20 | CATTCGTAGACGACCTGCTT | 137.5 | -1.6 | 0.5 | -3.4 | 8.1E-04 | 9.7E-04 | BT20/MB468 |

<table>
<tr><td></td><td></td><td></td><td></td><td></td><td></td><td></td><td></td><td></td><td></td><td></td><td></td><td></td><td></td></tr>
<tr><td>>hg38_21_+_8433172_8446622_RNA45SN1_45S_50ntFlanks@12935.12959.25|TAAACCATTCGTAGACGACCTGCTT</td><td>>hg38_21_+_8433172_8446622_RNA45SN1_45S_50ntFlanks</td><td>12935</td><td>12959</td><td>25</td><td>TAAACCATTCGTAGACGACCTGCTT</td><td>140.3</td><td>-1.9</td><td>0.6</td><td>-3.3</td><td>8.1E-04</td><td>9.7E-04</td><td>BT20/MB468</td></tr>
<tr><td>>hg38_21_+_8433172_8446622_RNA45SN1_45S_50ntFlanks@12539.12567.29|ATGATGTGTTGTTGCCATGGTAATCCTGC</td><td>>hg38_21_+_8433172_8446622_RNA45SN1_45S_50ntFlanks</td><td>12539</td><td>12567</td><td>29</td><td>ATGATGTGTTGTTGCCATGGTAATCCTGC</td><td>425.3</td><td>-5.8</td><td>1.7</td><td>-3.4</td><td>7.5E-04</td><td>9.7E-04</td><td>MB231/MB468</td></tr>
<tr><td>>hg38_21_+_8433172_8446622_RNA45SN1_45S_50ntFlanks@12007.12032.26|GCCGCCGGTGAAATACCACTACTCTG</td><td>>hg38_21_+_8433172_8446622_RNA45SN1_45S_50ntFlanks</td><td>12007</td><td>12032</td><td>26</td><td>GCCGCCGGTGAAATACCACTACTCTG</td><td>1566.4</td><td>-2.7</td><td>0.8</td><td>-3.4</td><td>7.5E-04</td><td>9.8E-04</td><td>MB231/MB468</td></tr>
<tr><td>>hg38_21_+_8433172_8446622_RNA45SN1_45S_50ntFlanks@5527.5553.27|AAAGTCGTAACAAGGTTTCCGTAGGTG</td><td>>hg38_21_+_8433172_8446622_RNA45SN1_45S_50ntFlanks</td><td>5527</td><td>5553</td><td>27</td><td>AAAGTCGTAACAAGGTTTCCGTAGGTG</td><td>76.7</td><td>-2.6</td><td>0.8</td><td>-3.4</td><td>7.5E-04</td><td>9.8E-04</td><td>MB231/MB468</td></tr>
<tr><td>>hg38_21_+_8433172_8446622_RNA45SN1_45S_50ntFlanks@5485.5516.32|AGCGCTGAGAAGACGGTCGAACTTGACTATCT</td><td>>hg38_21_+_8433172_8446622_RNA45SN1_45S_50ntFlanks</td><td>5485</td><td>5516</td><td>32</td><td>AGCGCTGAGAAGACGGTCGAACTTGACTATCT</td><td>4414.9</td><td>3.9</td><td>1.2</td><td>3.3</td><td>8.2E-04</td><td>9.8E-04</td><td>BT20/MB468</td></tr>
<tr><td>>hg38_21_+_8433172_8446622_RNA45SN1_45S_50ntFlanks@5487.5516.30|CGCTGAGAAGACGGTCGAACTTGACTATCT</td><td>>hg38_21_+_8433172_8446622_RNA45SN1_45S_50ntFlanks</td><td>5487</td><td>5516</td><td>30</td><td>CGCTGAGAAGACGGTCGAACTTGACTATCT</td><td>1704.6</td><td>2.8</td><td>0.8</td><td>3.4</td><td>7.6E-04</td><td>9.9E-04</td><td>MB231/MB468</td></tr>
<tr><td>>hg38_21_+_8433172_8446622_RNA45SN1_45S_50ntFlanks@11317.11336.20|ACCCCACGTCTCGTCGCGCG</td><td>>hg38_21_+_8433172_8446622_RNA45SN1_45S_50ntFlanks</td><td>11317</td><td>11336</td><td>20</td><td>ACCCCACGTCTCGTCGCGCG</td><td>558.6</td><td>-3.9</td><td>1.2</td><td>-3.3</td><td>8.2E-04</td><td>9.9E-04</td><td>BT20/MB468</td></tr>
<tr><td>>hg38_21_+_8433172_8446622_RNA45SN1_45S_50ntFlanks@6772.6807.36|GTTCCTCCCGGGGCTACGCCTGTCTGAGCGTCGCTT</td><td>>hg38_21_+_8433172_8446622_RNA45SN1_45S_50ntFlanks</td><td>6772</td><td>6807</td><td>36</td><td>GTTCCTCCCGGGGCTACGCCTGTCTGAGCGTCGCTT</td><td>418.9</td><td>-1.9</td><td>0.6</td><td>-3.3</td><td>8.2E-04</td><td>9.9E-04</td><td>BT20/MB468</td></tr>
<tr><td>>hg38_21_+_8433172_8446622_RNA45SN1_45S_50ntFlanks@6731.6753.23|CACATTGATCATCGACACTTCGA</td><td>>hg38_21_+_8433172_8446622_RNA45SN1_45S_50ntFlanks</td><td>6731</td><td>6753</td><td>23</td><td>CACATTGATCATCGACACTTCGA</td><td>333.0</td><td>-2.3</td><td>0.7</td><td>-3.3</td><td>8.3E-04</td><td>9.9E-04</td><td>BT20/MB468</td></tr>
<tr><td>>hg38_21_+_8433172_8446622_RNA45SN1_45S_50ntFlanks@3783.3801.19|AGTGAAACTGCGAATGGCT</td><td>>hg38_21_+_8433172_8446622_RNA45SN1_45S_50ntFlanks</td><td>3783</td><td>3801</td><td>19</td><td>AGTGAAACTGCGAATGGCT</td><td>282.3</td><td>-1.6</td><td>0.5</td><td>-3.4</td><td>7.6E-04</td><td>1.0E-03</td><td>MB231/MB468</td></tr>
<tr><td>>hg38_21_+_8433172_8446622_RNA45SN1_45S_50ntFlanks@6725.6743.19|GCAGGACACATTGATCATC</td><td>>hg38_21_+_8433172_8446622_RNA45SN1_45S_50ntFlanks</td><td>6725</td><td>6743</td><td>19</td><td>GCAGGACACATTGATCATC</td><td>1357.1</td><td>2.8</td><td>0.8</td><td>3.4</td><td>7.7E-04</td><td>1.0E-03</td><td>MB231/MB468</td></tr>
<tr><td>>hg38_21_+_8433172_8446622_RNA45SN1_45S_50ntFlanks@12416.12455.40|TTCGATGTCGGCTCTTCCTATCATTGTGAAGCAGAATTCA</td><td>>hg38_21_+_8433172_8446622_RNA45SN1_45S_50ntFlanks</td><td>12416</td><td>12455</td><td>40</td><td>TTCGATGTCGGCTCTTCCTATCATTGTGAAGCAGAATTCA</td><td>273.7</td><td>-3.1</td><td>0.8</td><td>-3.8</td><td>1.2E-04</td><td>1.0E-03</td><td>BT20/MB231</td></tr>
<tr><td>>hg38_21_+_8433172_8446622_RNA45SN1_45S_50ntFlanks@10101.10126.26|CGCGCGCGTGTGGTGTGCGTCGGAGG</td><td>>hg38_21_+_8433172_8446622_RNA45SN1_45S_50ntFlanks</td><td>10101</td><td>10126</td><td>26</td><td>CGCGCGCGTGTGGTGTGCGTCGGAGG</td><td>331.7</td><td>-4.3</td><td>1.3</td><td>-3.3</td><td>8.4E-04</td><td>1.0E-03</td><td>BT20/MB468</td></tr>
<tr><td>>hg38_21_+_8433172_8446622_RNA45SN1_45S_50ntFlanks@10873.10892.20|CTGGGTCGGTCGGGCTGGGG</td><td>>hg38_21_+_8433172_8446622_RNA45SN1_45S_50ntFlanks</td><td>10873</td><td>10892</td><td>20</td><td>CTGGGTCGGTCGGGCTGGGG</td><td>238.7</td><td>-5.4</td><td>1.6</td><td>-3.4</td><td>7.8E-04</td><td>1.0E-03</td><td>MB231/MB468</td></tr>
<tr><td>>hg38_21_+_8433172_8446622_RNA45SN1_45S_50ntFlanks@11672.11691.20|CAGTGCTCTGAATGTCAAAG</td><td>>hg38_21_+_8433172_8446622_RNA45SN1_45S_50ntFlanks</td><td>11672</td><td>11691</td><td>20</td><td>CAGTGCTCTGAATGTCAAAG</td><td>613.7</td><td>1.8</td><td>0.5</td><td>3.4</td><td>7.8E-04</td><td>1.0E-03</td><td>MB231/MB468</td></tr>
<tr><td>>hg38_21_+_8433172_8446622_RNA45SN1_45S_50ntFlanks@6648.6666.19|GTACGACTCTTAGCGGTGG</td><td>>hg38_21_+_8433172_8446622_RNA45SN1_45S_50ntFlanks</td><td>6648</td><td>6666</td><td>19</td><td>GTACGACTCTTAGCGGTGG</td><td>490.6</td><td>2.4</td><td>0.7</td><td>3.3</td><td>8.6E-04</td><td>1.0E-03</td><td>BT20/MB468</td></tr>
<tr><td>>MI0000101|hsa-mir-99a&WithFlank&21|+|16539083|16539175@19.39.21&[MIMAT0000097&hsa-miR-99a-5p&offsets|0|-1;m-64&21|+|16539101|16539121&offsets|0|0]|AACCCGTAGATCCGATCTTGT</td><td>>MI0000101|hsa-mir-99a&WithFlank&21|+|16539083|16539175</td><td>19</td><td>39</td><td>21</td><td>AACCCGTAGATCCGATCTTGT</td><td>4382.9</td><td>2.5</td><td>0.7</td><td>3.4</td><td>8.0E-04</td><td>1.0E-03</td><td>MB231/MB468</td></tr>
<tr><td>>hg38_21_+_8433172_8446622_RNA45SN1_45S_50ntFlanks@4370.4389.20|TTAAAAAGCTCGTAGTTGGA</td><td>>hg38_21_+_8433172_8446622_RNA45SN1_45S_50ntFlanks</td><td>4370</td><td>4389</td><td>20</td><td>TTAAAAAGCTCGTAGTTGGA</td><td>237.6</td><td>-4.2</td><td>1.3</td><td>-3.3</td><td>8.7E-04</td><td>1.0E-03</td><td>BT20/MB468</td></tr>
<tr><td>>hg38_21_+_8433172_8446622_RNA45SN1_45S_50ntFlanks@7979.8005.27|ACCTCAGATCAGACGTGGCGACCCGCT</td><td>>hg38_21_+_8433172_8446622_RNA45SN1_45S_50ntFlanks</td><td>7979</td><td>8005</td><td>27</td><td>ACCTCAGATCAGACGTGGCGACCCGCT</td><td>185.3</td><td>-1.8</td><td>0.5</td><td>-3.3</td><td>8.7E-04</td><td>1.0E-03</td><td>BT20/MB468</td></tr>
<tr><td>>hg38_21_+_8433172_8446622_RNA45SN1_45S_50ntFlanks@12418.12440.23|CGATGTCGGCTCTTCCTATCATT</td><td>>hg38_21_+_8433172_8446622_RNA45SN1_45S_50ntFlanks</td><td>12418</td><td>12440</td><td>23</td><td>CGATGTCGGCTCTTCCTATCATT</td><td>761.0</td><td>2.2</td><td>0.7</td><td>3.4</td><td>8.1E-04</td><td>1.0E-03</td><td>MB231/MB468</td></tr>
<tr><td>>hg38_21_+_8433172_8446622_RNA45SN1_45S_50ntFlanks@10320.10341.22|CAGATCTTGGTGGTAGTAGCAA</td><td>>hg38_21_+_8433172_8446622_RNA45SN1_45S_50ntFlanks</td><td>10320</td><td>10341</td><td>22</td><td>CAGATCTTGGTGGTAGTAGCAA</td><td>93.2</td><td>-2.5</td><td>0.8</td><td>-3.3</td><td>8.7E-04</td><td>1.0E-03</td><td>BT20/MB468</td></tr>
<tr><td>>hg38_21_+_8433172_8446622_RNA45SN1_45S_50ntFlanks@10793.10816.24|TCTGGCATGTTGGAACAATGTAGG</td><td>>hg38_21_+_8433172_8446622_RNA45SN1_45S_50ntFlanks</td><td>10793</td><td>10816</td><td>24</td><td>TCTGGCATGTTGGAACAATGTAGG</td><td>190.4</td><td>-1.9</td><td>0.6</td><td>-3.3</td><td>8.8E-04</td><td>1.1E-03</td><td>BT20/MB468</td></tr>
<tr><td>>hg38_21_+_8433172_8446622_RNA45SN1_45S_50ntFlanks@3993.4024.32|GTGACTCTAGATAACCTCGGGCCGATCGCACG</td><td>>hg38_21_+_8433172_8446622_RNA45SN1_45S_50ntFlanks</td><td>3993</td><td>4024</td><td>32</td><td>GTGACTCTAGATAACCTCGGGCCGATCGCACG</td><td>682.8</td><td>2.3</td><td>0.7</td><td>3.3</td><td>8.1E-04</td><td>1.1E-03</td><td>MB231/MB468</td></tr>
<tr><td>>hg38_21_+_8433172_8446622_RNA45SN1_45S_50ntFlanks@12417.12457.41|TCGATGTCGGCTCTTCCTATCATTGTGAAGCAGAATTCACC</td><td>>hg38_21_+_8433172_8446622_RNA45SN1_45S_50ntFlanks</td><td>12417</td><td>12457</td><td>41</td><td>TCGATGTCGGCTCTTCCTATCATTGTGAAGCAGAATTCACC</td><td>295.1</td><td>-2.9</td><td>0.8</td><td>-3.8</td><td>1.2E-04</td><td>1.1E-03</td><td>BT20/MB231</td></tr>
<tr><td>>hg38_21_+_8433172_8446622_RNA45SN1_45S_50ntFlanks@4220.4243.24|ACGAAAAATAACAATACAGGACTC</td><td>>hg38_21_+_8433172_8446622_RNA45SN1_45S_50ntFlanks</td><td>4220</td><td>4243</td><td>24</td><td>ACGAAAAATAACAATACAGGACTC</td><td>92.5</td><td>-1.9</td><td>0.6</td><td>-3.3</td><td>8.1E-04</td><td>1.1E-03</td><td>MB231/MB468</td></tr>
<tr><td>>hg38_21_+_8433172_8446622_RNA45SN1_45S_50ntFlanks@12420.12455.36|ATGTCGGCTCTTCCTATCATTGTGAAGCAGAATTCA</td><td>>hg38_21_+_8433172_8446622_RNA45SN1_45S_50ntFlanks</td><td>12420</td><td>12455</td><td>36</td><td>ATGTCGGCTCTTCCTATCATTGTGAAGCAGAATTCA</td><td>1812.2</td><td>-2.0</td><td>0.6</td><td>-3.3</td><td>8.1E-04</td><td>1.1E-03</td><td>MB231/MB468</td></tr>
<tr><td>>MI0000446|hsa-mir-125b-1&WithFlank&11|-|122099751|122099850@21.42.22;MI0000470|hsa-mir-125b-2&WithFlank&21|+|16590231|16590331@23.44.22&[MIMAT0000423&hsa-miR-125b-5p&offsets|0|0;m-67&11|-|122099809|122099830&offsets|0|0];[MIMAT0000423_1&hsa-miR-125b-5p&offsets|0|0;m-68&21|+|16590253|16590274&offsets|0|0]|TCCCTGAGACCCTAACTTGTGA</td><td>>MI0000446|hsa-mir-125b-1&WithFlank&11|-|122099751|122099850</td><td>21</td><td>42</td><td>22</td><td>TCCCTGAGACCCTAACTTGTGA</td><td>779.1</td><td>3.5</td><td>1.1</td><td>3.3</td><td>8.8E-04</td><td>1.1E-03</td><td>BT20/MB468</td></tr>
<tr><td>>hg38_21_+_8433172_8446622_RNA45SN1_45S_50ntFlanks@7982.8012.31|TCAGATCAGACGTGGCGACCCGCTGAATTTA</td><td>>hg38_21_+_8433172_8446622_RNA45SN1_45S_50ntFlanks</td><td>7982</td><td>8012</td><td>31</td><td>TCAGATCAGACGTGGCGACCCGCTGAATTTA</td><td>103.7</td><td>-1.9</td><td>0.6</td><td>-3.3</td><td>8.9E-04</td><td>1.1E-03</td><td>BT20/MB468</td></tr>
<tr><td>>am_trna134_GluTTC_1_-_16861774_16861845@1.30.30,trna5_GluTTC_1_+_17199078_17199149@1.30.30,trna84_GluTTC_1_-_161391883_161391954@1.30.30,trna94_GluTTC_1_-_149664355_149664427@1.30.30|TCCCTGGTGGTCTAGTGGCTAGGATTCGGC</td><td>>am_trna134_GluTTC_1_-_16861774_16861845</td><td>1</td><td>30</td><td>30</td><td>TCCCTGGTGGTCTAGTGGCTAGGATTCGGC</td><td>342.8</td><td>-2.3</td><td>0.7</td><td>-3.3</td><td>8.2E-04</td><td>1.1E-03</td><td>MB231/MB468</td></tr>
<tr><td>>hg38_21_+_8433172_8446622_RNA45SN1_45S_50ntFlanks@8360.8379.20|AGAACTTTGAAGAGAGAGTT</td><td>>hg38_21_+_8433172_8446622_RNA45SN1_45S_50ntFlanks</td><td>8360</td><td>8379</td><td>20</td><td>AGAACTTTGAAGAGAGAGTT</td><td>94.6</td><td>-2.9</td><td>0.9</td><td>-3.3</td><td>8.2E-04</td><td>1.1E-03</td><td>MB231/MB468</td></tr>
<tr><td>>hg38_21_+_8433172_8446622_RNA45SN1_45S_50ntFlanks@11552.11588.37|CCTCGGCCGGCGCCTAGCAGCCGACTTAGAACTGGTG</td><td>>hg38_21_+_8433172_8446622_RNA45SN1_45S_50ntFlanks</td><td>11552</td><td>11588</td><td>37</td><td>CCTCGGCCGGCGCCTAGCAGCCGACTTAGAACTGGTG</td><td>311.0</td><td>1.6</td><td>0.5</td><td>3.3</td><td>8.3E-04</td><td>1.1E-03</td><td>MB231/MB468</td></tr>
<tr><td>>hg38_21_+_8433172_8446622_RNA45SN1_45S_50ntFlanks@11771.11794.24|ATTAGTGACGCGCATGAATGGATG</td><td>>hg38_21_+_8433172_8446622_RNA45SN1_45S_50ntFlanks</td><td>11771</td><td>11794</td><td>24</td><td>ATTAGTGACGCGCATGAATGGATG</td><td>300.5</td><td>-1.2</td><td>0.4</td><td>-3.3</td><td>9.0E-04</td><td>1.1E-03</td><td>BT20/MB468</td></tr>
<tr><td>>hg38_21_+_8433172_8446622_RNA45SN1_45S_50ntFlanks@11586.11613.28|GTGCGGACCAGGGGAATCCGACTGTTTA</td><td>>hg38_21_+_8433172_8446622_RNA45SN1_45S_50ntFlanks</td><td>11586</td><td>11613</td><td>28</td><td>GTGCGGACCAGGGGAATCCGACTGTTTA</td><td>170.9</td><td>-1.9</td><td>0.6</td><td>-3.3</td><td>9.0E-04</td><td>1.1E-03</td><td>BT20/MB468</td></tr>
<tr><td>>hg38_21_+_8433172_8446622_RNA45SN1_45S_50ntFlanks@12417.12439.23|TCGATGTCGGCTCTTCCTATCAT</td><td>>hg38_21_+_8433172_8446622_RNA45SN1_45S_50ntFlanks</td><td>12417</td><td>12439</td><td>23</td><td>TCGATGTCGGCTCTTCCTATCAT</td><td>503.9</td><td>2.2</td><td>0.7</td><td>3.3</td><td>9.0E-04</td><td>1.1E-03</td><td>BT20/MB468</td></tr>
<tr><td>>MI0000542|hsa-mir-320a&WithFlank&8|-|22244960|22245043@48.72.25&[MIMAT0000510&hsa-miR-320a-3p&offsets|0|+3;m-60&8|-|22244975|22244996&offsets|0|+3]|AAAAGCTGGGTTGAGAGGGCGAAAA</td><td>>MI0000542|hsa-mir-320a&WithFlank&8|-|22244960|22245043</td><td>48</td><td>72</td><td>25</td><td>AAAAGCTGGGTTGAGAGGGCGAAAA</td><td>248.4</td><td>3.1</td><td>0.8</td><td>3.8</td><td>1.3E-04</td><td>1.1E-03</td><td>BT20/MB231</td></tr>
<tr><td>>hg38_21_+_8433172_8446622_RNA45SN1_45S_50ntFlanks@10802.10826.25|TTGGAACAATGTAGGTAAGGGAAGT</td><td>>hg38_21_+_8433172_8446622_RNA45SN1_45S_50ntFlanks</td><td>10802</td><td>10826</td><td>25</td><td>TTGGAACAATGTAGGTAAGGGAAGT</td><td>104.5</td><td>-2.8</td><td>0.9</td><td>-3.3</td><td>9.1E-04</td><td>1.1E-03</td><td>BT20/MB468</td></tr>
<tr><td>>hg38_21_+_8433172_8446622_RNA45SN1_45S_50ntFlanks@10366.10383.18|CCGAAGTGGAGAAGGGTT</td><td>>hg38_21_+_8433172_8446622_RNA45SN1_45S_50ntFlanks</td><td>10366</td><td>10383</td><td>18</td><td>CCGAAGTGGAGAAGGGTT</td><td>92.8</td><td>-2.5</td><td>0.8</td><td>-3.3</td><td>8.4E-04</td><td>1.1E-03</td><td>MB231/MB468</td></tr>
<tr><td>>hg38_21_+_8433172_8446622_RNA45SN1_45S_50ntFlanks@3852.3874.23|TAACTGTGGTAATTCTAGAGCTA</td><td>>hg38_21_+_8433172_8446622_RNA45SN1_45S_50ntFlanks</td><td>3852</td><td>3874</td><td>23</td><td>TAACTGTGGTAATTCTAGAGCTA</td><td>154.3</td><td>-1.9</td><td>0.6</td><td>-3.3</td><td>8.5E-04</td><td>1.1E-03</td><td>MB231/MB468</td></tr>
<tr><td>>hg38_21_+_8433172_8446622_RNA45SN1_45S_50ntFlanks@11567.11601.35|AGCAGCCGACTTAGAACTGGTGCGGACCAGGGGAA</td><td>>hg38_21_+_8433172_8446622_RNA45SN1_45S_50ntFlanks</td><td>11567</td><td>11601</td><td>35</td><td>AGCAGCCGACTTAGAACTGGTGCGGACCAGGGGAA</td><td>199.5</td><td>-1.7</td><td>0.5</td><td>-3.3</td><td>9.2E-04</td><td>1.1E-03</td><td>BT20/MB468</td></tr>
<tr><td>>hg38_21_+_8433172_8446622_RNA45SN1_45S_50ntFlanks@3705.3735.31|TACCTGGTTGATCCTGCCAGTAGCATATGCT</td><td>>hg38_21_+_8433172_8446622_RNA45SN1_45S_50ntFlanks</td><td>3705</td><td>3735</td><td>31</td><td>TACCTGGTTGATCCTGCCAGTAGCATATGCT</td><td>650.5</td><td>3.1</td><td>0.8</td><td>3.8</td><td>1.3E-04</td><td>1.1E-03</td><td>BT20/MB231</td></tr>
<tr><td>>MI0000102|hsa-mir-100&WithFlank&11|-|122152223|122152314@19.39.21(+1U)&[MIMAT0000098&hsa-miR-100-5p&offsets|0|-1(+1U);m-27&11|-|122152276|122152296&offsets|0|0(+1U)]|AACCCGTAGATCCGAACTTGTT</td><td>>MI0000102|hsa-mir-100&WithFlank&11|-|122152223|122152314</td><td>19</td><td>39</td><td>21(+1U)</td><td>AACCCGTAGATCCGAACTTGTT</td><td>768.2</td><td>9.3</td><td>2.8</td><td>3.3</td><td>9.3E-04</td><td>1.1E-03</td><td>BT20/MB468</td></tr>
<tr><td>>hg38_21_+_8433172_8446622_RNA45SN1_45S_50ntFlanks@6647.6681.35|CGTACGACTCTTAGCGGTGGATCACTCGGCTCGTG</td><td>>hg38_21_+_8433172_8446622_RNA45SN1_45S_50ntFlanks</td><td>6647</td><td>6681</td><td>35</td><td>CGTACGACTCTTAGCGGTGGATCACTCGGCTCGTG</td><td>6036.2</td><td>2.2</td><td>0.7</td><td>3.3</td><td>8.5E-04</td><td>1.1E-03</td><td>MB231/MB468</td></tr>
<tr><td>>hg38_21_+_8433172_8446622_RNA45SN1_45S_50ntFlanks@12941.12965.25|ATTCGTAGACGACCTGCTTCTGGGT</td><td>>hg38_21_+_8433172_8446622_RNA45SN1_45S_50ntFlanks</td><td>12941</td><td>12965</td><td>25</td><td>ATTCGTAGACGACCTGCTTCTGGGT</td><td>249.2</td><td>-1.2</td><td>0.4</td><td>-3.3</td><td>8.6E-04</td><td>1.1E-03</td><td>MB231/MB468</td></tr>
<tr><td>>hg38_21_+_8433172_8446622_RNA45SN1_45S_50ntFlanks@10789.10818.30|AGCCTCTGGCATGTTGGAACAATGTAGGTA</td><td>>hg38_21_+_8433172_8446622_RNA45SN1_45S_50ntFlanks</td><td>10789</td><td>10818</td><td>30</td><td>AGCCTCTGGCATGTTGGAACAATGTAGGTA</td><td>450.8</td><td>-2.0</td><td>0.6</td><td>-3.3</td><td>8.6E-04</td><td>1.1E-03</td><td>MB231/MB468</td></tr>
<tr><td>>hg38_21_+_8433172_8446622_RNA45SN1_45S_50ntFlanks@11315.11342.28|CCACCCCACGTCTCGTCGCGCGCGTC</td><td>>hg38_21_+_8433172_8446622_RNA45SN1_45S_50ntFlanks</td><td>11315</td><td>11342</td><td>28</td><td>CCACCCCACGTCTCGTCGCGCGCGTC</td><td>283.3</td><td>-3.1</td><td>1.0</td><td>-3.3</td><td>9.3E-04</td><td>1.1E-03</td><td>BT20/MB468</td></tr>
<tr><td>>hg38_21_+_8433172_8446622_RNA45SN1_45S_50ntFlanks@9821.9850.30|CCTAGTGGGCCACTTTTGGTAAGCAGAACT</td><td>>hg38_21_+_8433172_8446622_RNA45SN1_45S_50ntFlanks</td><td>9821</td><td>9850</td><td>30</td><td>CCTAGTGGGCCACTTTTGGTAAGCAGAACT</td><td>245.0</td><td>-1.9</td><td>0.6</td><td>-3.3</td><td>8.7E-04</td><td>1.1E-03</td><td>MB231/MB468</td></tr>
</table>

| | | | | | | | | | | | | |
|---|---|---|---|---|---|---|---|---|---|---|---|---|
| >hg38_21_+_8433172_8446622_RNA45SN1_45S_50ntFlanks@12957.12986.30\|CTTCTGGGTCGGGGTTTCGTACGTAGCAGA | >hg38_21_+_8433172_8446622_RNA45SN1_45S_50ntFlanks | 12957 | 12986 | 30 | CTTCTGGGTCGGGGTTTCGTACGTAGCAGA | 149.7 | -1.9 | 0.6 | -3.3 | 8.7E-04 | 1.1E-03 | MB231/MB468 |
| >hg38_21_+_8433172_8446622_RNA45SN1_45S_50ntFlanks@11569.11588.20\|CAGCCGACTTAGAACTGGTG | >hg38_21_+_8433172_8446622_RNA45SN1_45S_50ntFlanks | 11569 | 11588 | 20 | CAGCCGACTTAGAACTGGTG | 156.4 | -3.3 | 1.0 | -3.3 | 9.4E-04 | 1.1E-03 | BT20/MB468 |
| >MI0000273\|hsa-mir-183&WithFlank&7\|-\|129774899\|129775020@33.54.22&[MIMAT0000261&hsa-miR-183-5p&offsets\|0\|0;m-41&7\|-\|129774966\|129774987&offsets\|-1\|-1]\|TATGGCACTGGTAGAATTCACT | >MI0000273\|hsa-mir-183&WithFlank&7\|-\|129774899\|129775020 | 33 | 54 | 22 | TATGGCACTGGTAGAATTCACT | 19108.4 | 2.4 | 0.6 | 3.8 | 1.3E-04 | 1.1E-03 | BT20/MB231 |
| >hg38_21_+_8433172_8446622_RNA45SN1_45S_50ntFlanks@6647.6679.33\|CGTACGACTCTTAGCGGTGGATCACTCGGCTCG | >hg38_21_+_8433172_8446622_RNA45SN1_45S_50ntFlanks | 6647 | 6679 | 33 | CGTACGACTCTTAGCGGTGGATCACTCGGCTCG | 372.5 | 1.8 | 0.5 | 3.3 | 8.7E-04 | 1.1E-03 | MB231/MB468 |
| >hg38_21_+_8433172_8446622_RNA45SN1_45S_50ntFlanks@9738.9760.23\|GAAACGATCTCAACCTATTCTCA | >hg38_21_+_8433172_8446622_RNA45SN1_45S_50ntFlanks | 9738 | 9760 | 23 | GAAACGATCTCAACCTATTCTCA | 86.0 | -1.7 | 0.5 | -3.3 | 8.7E-04 | 1.1E-03 | MB231/MB468 |
| >hg38_21_+_8433172_8446622_RNA45SN1_45S_50ntFlanks@11027.11053.27\|CTCTCTCTCTCTCTCCCCCGCTCCCCG | >hg38_21_+_8433172_8446622_RNA45SN1_45S_50ntFlanks | 11027 | 11053 | 27 | CTCTCTCTCTCTCTCCCCCGCTCCCCG | 406.5 | -4.3 | 1.3 | -3.3 | 9.5E-04 | 1.1E-03 | BT20/MB468 |
| >hg38_21_+_8433172_8446622_RNA45SN1_45S_50ntFlanks@6734.6753.20\|ATTGATCATCGACACTTCGA | >hg38_21_+_8433172_8446622_RNA45SN1_45S_50ntFlanks | 6734 | 6753 | 20 | ATTGATCATCGACACTTCGA | 254.0 | 1.7 | 0.5 | 3.3 | 8.7E-04 | 1.1E-03 | MB231/MB468 |
| >am_trnaMT_ArgTCG_MT_+_10405_10469@31.48.18\|TCGACTCATTAAATTATG | >am_trnaMT_ArgTCG_MT_+_10405_10469 | 31 | 48 | 18 | TCGACTCATTAAATTATG | 133.1 | -2.6 | 0.8 | -3.3 | 8.8E-04 | 1.1E-03 | MB231/MB468 |
| >hg38_21_+_8433172_8446622_RNA45SN1_45S_50ntFlanks@12941.12958.18\|ATTCGTAGACGACCTGCT | >hg38_21_+_8433172_8446622_RNA45SN1_45S_50ntFlanks | 12941 | 12958 | 18 | ATTCGTAGACGACCTGCT | 417.8 | 2.0 | 0.6 | 3.3 | 9.6E-04 | 1.1E-03 | BT20/MB468 |
| >hg38_21_+_8433172_8446622_RNA45SN1_45S_50ntFlanks@6698.6720.23\|AGCTAGCTGCGAGAATTAATGTG | >hg38_21_+_8433172_8446622_RNA45SN1_45S_50ntFlanks | 6698 | 6720 | 23 | AGCTAGCTGCGAGAATTAATGTG | 11961.0 | -2.2 | 0.7 | -3.3 | 8.9E-04 | 1.2E-03 | MB231/MB468 |
| >hg38_21_+_8433172_8446622_RNA45SN1_45S_50ntFlanks@10770.10796.27\|GCAGGTCTCCAAGGTGAACAGCCTCTG | >hg38_21_+_8433172_8446622_RNA45SN1_45S_50ntFlanks | 10770 | 10796 | 27 | GCAGGTCTCCAAGGTGAACAGCCTCTG | 594.9 | -2.1 | 0.6 | -3.3 | 9.7E-04 | 1.2E-03 | BT20/MB468 |
| >MI0000477\|hsa-mir-146a&WithFlank&5\|+\|160485346\|160485456@27.48.22(+1A)&[MIMAT0000449&hsa-miR-146a-5p&offsets\|0\|0(+1A);m-129&5\|+\|160485372\|160485393&offsets\|0\|0(+1A)]\|TGAGAACTGAATTCCATGGGTTA | >MI0000477\|hsa-mir-146a&WithFlank&5\|+\|160485346\|160485456 | 27 | 48 | 22(+1A) | TGAGAACTGAATTCCATGGGTTA | 361.1 | -7.9 | 2.1 | -3.8 | 1.4E-04 | 1.2E-03 | BT20/MB231 |
| >hg38_21_+_8433172_8446622_RNA45SN1_45S_50ntFlanks@3853.3871.19\|AACTGTGGTAATTCTAGAG | >hg38_21_+_8433172_8446622_RNA45SN1_45S_50ntFlanks | 3853 | 3871 | 19 | AACTGTGGTAATTCTAGAG | 245.3 | -3.1 | 0.9 | -3.3 | 9.7E-04 | 1.2E-03 | BT20/MB468 |
| >MI0000088\|hsa-mir-30a&WithFlank&6\|-\|71403545\|71403627@12.35.24(+3U)&[MIMAT0000087&hsa-miR-30a-5p&offsets\|0\|+2(+3U);m-15&6\|-\|71403595\|71403616&offsets\|0\|+2(+3U)]\|TGTAAACATCCTCGACTGGAAGCTTTT | >MI0000088\|hsa-mir-30a&WithFlank&6\|-\|71403545\|71403627 | 12 | 35 | 24(+3U) | TGTAAACATCCTCGACTGGAAGCTTTT | 387.3 | -3.4 | 0.9 | -3.8 | 1.4E-04 | 1.2E-03 | BT20/MB231 |
| >hg38_21_+_8433172_8446622_RNA45SN1_45S_50ntFlanks@11027.11049.23\|CTCTCTCTCTCTCTCCCCCGCTC | >hg38_21_+_8433172_8446622_RNA45SN1_45S_50ntFlanks | 11027 | 11049 | 23 | CTCTCTCTCTCTCTCCCCCGCTC | 368.2 | -4.2 | 1.3 | -3.3 | 9.8E-04 | 1.2E-03 | BT20/MB468 |
| >hg38_21_+_8433172_8446622_RNA45SN1_45S_50ntFlanks@9566.9592.27\|GACGTGCAAATCGGTCGTCCGACCTGG | >hg38_21_+_8433172_8446622_RNA45SN1_45S_50ntFlanks | 9566 | 9592 | 27 | GACGTGCAAATCGGTCGTCCGACCTGG | 167.5 | -1.4 | 0.4 | -3.3 | 9.1E-04 | 1.2E-03 | MB231/MB468 |
| >hg38_21_+_8433172_8446622_RNA45SN1_45S_50ntFlanks@4005.4025.21\|AACCTCGGGCCGATCGCACGC | >hg38_21_+_8433172_8446622_RNA45SN1_45S_50ntFlanks | 4005 | 4025 | 21 | AACCTCGGGCCGATCGCACGC | 177.0 | -1.7 | 0.5 | -3.3 | 9.1E-04 | 1.2E-03 | MB231/MB468 |
| >hg38_21_+_8433172_8446622_RNA45SN1_45S_50ntFlanks@12420.12459.40\|ATGTCGGCTCTTCCTATCATTGTGAAGCAGAATTCACCAA | >hg38_21_+_8433172_8446622_RNA45SN1_45S_50ntFlanks | 12420 | 12459 | 40 | ATGTCGGCTCTTCCTATCATTGTGAAGCAGAATTCACCAA | 259.5 | -2.8 | 0.7 | -3.8 | 1.4E-04 | 1.2E-03 | BT20/MB231 |
| >hg38_21_+_8433172_8446622_RNA45SN1_45S_50ntFlanks@10420.10453.34\|CTGAGAGATGGGCGAGCGCCGTTCCGAAGGGACG | >hg38_21_+_8433172_8446622_RNA45SN1_45S_50ntFlanks | 10420 | 10453 | 34 | CTGAGAGATGGGCGAGCGCCGTTCCGAAGGGACG | 493.8 | -2.1 | 0.6 | -3.3 | 9.2E-04 | 1.2E-03 | MB231/MB468 |
| >hg38_21_+_8433172_8446622_RNA45SN1_45S_50ntFlanks@9552.9572.21\|TCCGTAGCGGTCCTGACGTGC | >hg38_21_+_8433172_8446622_RNA45SN1_45S_50ntFlanks | 9552 | 9572 | 21 | TCCGTAGCGGTCCTGACGTGC | 213.0 | -1.7 | 0.5 | -3.3 | 1.0E-03 | 1.2E-03 | BT20/MB468 |
| >hg38_21_+_8433172_8446622_RNA45SN1_45S_50ntFlanks@12640.12661.22\|ACCATCTGTGGGATTATGACTG | >hg38_21_+_8433172_8446622_RNA45SN1_45S_50ntFlanks | 12640 | 12661 | 22 | ACCATCTGTGGGATTATGACTG | 376.7 | -2.1 | 0.6 | -3.3 | 1.0E-03 | 1.2E-03 | BT20/MB468 |
| >MI0000466\|hsa-mir-9-1&WithFlank&1\|-\|156420335\|156420435@22.43.22;MI0000468\|hsa-mir-9-3&WithFlank&15\|+\|89368011\|89368112@22.43.22;MI0000467\|hsa-mir-9-2&WithFlank&5\|-\|88666847\|88666945@22.43.22&[MIMAT0000441&hsa-miR-9-5p&offsets\|0\|-1;m-104&1\|-\|156420392\|156420414&offsets\|0\|-1];[MIMAT0000441_1&hsa-miR-9-5p&offsets\|0\|-1;m-105&15\|+\|89368032\|89368054&offsets\|0\|-1];[MIMAT0000441_2&hsa-miR-9-5p&offsets\|0\|-1;m-104&5\|-\|88666902\|88666924&offsets\|0\|-1]\|TCTTTGGTTATCTAGCTGTATG | >MI0000466\|hsa-mir-9-1&WithFlank&1\|-\|156420335\|156420435 | 22 | 43 | 22 | TCTTTGGTTATCTAGCTGTATG | 3232.6 | 2.7 | 0.7 | 3.8 | 1.4E-04 | 1.2E-03 | BT20/MB231 |
| >trnaMT_GlnTTG_MT_-_4329_4400@1.33.33,trnalookalike2_GlnTTG_1_-_564879_564950@1.33.33\|TAGGATGGGGTGTGATAGGTGGCACGGAGAATT | >trnaMT_GlnTTG_MT_-_4329_4400 | 1 | 33 | 33 | TAGGATGGGGTGTGATAGGTGGCACGGAGAATT | 570.9 | 3.1 | 0.8 | 3.8 | 1.4E-04 | 1.2E-03 | BT20/MB231 |
| >hg38_21_+_8433172_8446622_RNA45SN1_45S_50ntFlanks@12937.12957.21\|AACCATTCGTAGACGACCTGC | >hg38_21_+_8433172_8446622_RNA45SN1_45S_50ntFlanks | 12937 | 12957 | 21 | AACCATTCGTAGACGACCTGC | 428.0 | 1.9 | 0.6 | 3.3 | 9.3E-04 | 1.2E-03 | MB231/MB468 |
| >hg38_21_+_8433172_8446622_RNA45SN1_45S_50ntFlanks@9533.9550.18\|GAGGAAACTCTGGTGGAG | >hg38_21_+_8433172_8446622_RNA45SN1_45S_50ntFlanks | 9533 | 9550 | 18 | GAGGAAACTCTGGTGGAG | 177.7 | -1.9 | 0.6 | -3.3 | 1.0E-03 | 1.2E-03 | BT20/MB468 |
| >hg38_21_+_8433172_8446622_RNA45SN1_45S_50ntFlanks@11539.11583.45\|GCGCGGCGCCTCGCCTCGGCCGGCGCCTAGCAGCCGACTTAGAAC | >hg38_21_+_8433172_8446622_RNA45SN1_45S_50ntFlanks | 11539 | 11583 | 45 | GCGCGGCGCCTCGCCTCGGCCGGCGCCTAGCAGCCGACTTAGAAC | 233.6 | -3.2 | 0.9 | -3.8 | 1.5E-04 | 1.2E-03 | BT20/MB231 |
| >hg38_21_+_8433172_8446622_RNA45SN1_45S_50ntFlanks@12417.12437.21\|TCGATGTCGGCTCTTCCTATC | >hg38_21_+_8433172_8446622_RNA45SN1_45S_50ntFlanks | 12417 | 12437 | 21 | TCGATGTCGGCTCTTCCTATC | 1724.5 | -1.6 | 0.5 | -3.3 | 1.0E-03 | 1.2E-03 | BT20/MB468 |
| >MI0000255\|hsa-mir-30d&WithFlank&8\|-\|134804870\|134804951@12.33.22(+1A)&[MIMAT0000245&hsa-miR-30d-5p&offsets\|0\|0(+1A);m-21&8\|-\|134804919\|134804940&offsets\|0\|0(+1A)]\|TGTAAACATCCCCGACTGGAAGA | >MI0000255\|hsa-mir-30d&WithFlank&8\|-\|134804870\|134804951 | 12 | 33 | 22(+1A) | TGTAAACATCCCCGACTGGAAGA | 4432.9 | 2.3 | 0.6 | 3.8 | 1.5E-04 | 1.2E-03 | BT20/MB231 |
| >MI0001641\|hsa-mir-429&WithFlank&1\|+\|1168999\|1169093@57.77.21(+1C)&[MIMAT0001536&hsa-miR-429&offsets\|0\|-1(+1C);m-207&1\|+\|1169055\|1169076&offsets\|0\|-1(+1C)]\|TAATACTGTCTGGTAAAACCGC | >MI0001641\|hsa-mir-429&WithFlank&1\|+\|1168999\|1169093 | 57 | 77 | 21(+1C) | TAATACTGTCTGGTAAAACCGC | 109.5 | -3.2 | 1.0 | -3.3 | 1.0E-03 | 1.2E-03 | BT20/MB468 |
| >MI0000542\|hsa-mir-320a&WithFlank&8\|-\|22244960\|22245043@48.68.21(+2U)&[MIMAT0000510&hsa-miR-320a-3p&offsets\|0\|-1(+2U);m-60&8\|-\|22244975\|22244996&offsets\|0\|-1(+2U)]\|AAAAGCTGGGTTGAGAGGGCGTT | >MI0000542\|hsa-mir-320a&WithFlank&8\|-\|22244960\|22245043 | 48 | 68 | 21(+2U) | AAAAGCTGGGTTGAGAGGGCGTT | 3245.4 | 2.7 | 0.7 | 3.8 | 1.5E-04 | 1.2E-03 | BT20/MB231 |
| >hg38_21_+_8433172_8446622_RNA45SN1_45S_50ntFlanks@10240.10259.20\|CTACGCCGCGACGAGTAGGA | >hg38_21_+_8433172_8446622_RNA45SN1_45S_50ntFlanks | 10240 | 10259 | 20 | CTACGCCGCGACGAGTAGGA | 221.3 | 2.7 | 0.7 | 3.8 | 1.5E-04 | 1.2E-03 | BT20/MB231 |
| >hg38_21_+_8433172_8446622_RNA45SN1_45S_50ntFlanks@11728.11745.18\|GTAACTATGACTCTCTTA | >hg38_21_+_8433172_8446622_RNA45SN1_45S_50ntFlanks | 11728 | 11745 | 18 | GTAACTATGACTCTCTTA | 368.0 | 1.9 | 0.6 | 3.3 | 9.7E-04 | 1.2E-03 | MB231/MB468 |
| >hg38_21_+_8433172_8446622_RNA45SN1_45S_50ntFlanks@4733.4751.19\|GAACGAAAGTCGGAGGTTC | >hg38_21_+_8433172_8446622_RNA45SN1_45S_50ntFlanks | 4733 | 4751 | 19 | GAACGAAAGTCGGAGGTTC | 491.6 | 1.9 | 0.6 | 3.3 | 9.7E-04 | 1.3E-03 | MB231/MB468 |
| >hg38_21_+_8433172_8446622_RNA45SN1_45S_50ntFlanks@9563.9597.35\|CCTGACGTGCAAATCGGTCGTCCGACCTGGGTATA | >hg38_21_+_8433172_8446622_RNA45SN1_45S_50ntFlanks | 9563 | 9597 | 35 | CCTGACGTGCAAATCGGTCGTCCGACCTGGGTATA | 362.7 | -2.0 | 0.6 | -3.3 | 9.7E-04 | 1.3E-03 | MB231/MB468 |
| >hg38_21_+_8433172_8446622_RNA45SN1_45S_50ntFlanks@12006.12033.28\|GGCCGCCGGTGAAATACCACTACTCTGA | >hg38_21_+_8433172_8446622_RNA45SN1_45S_50ntFlanks | 12006 | 12033 | 28 | GGCCGCCGGTGAAATACCACTACTCTGA | 241.4 | 2.7 | 0.7 | 3.8 | 1.5E-04 | 1.3E-03 | BT20/MB231 |
| >am_trnaMT_ValTAC_MT_+_1602_1670@51.69.19\|CTTAACTTGACCGCTCTGA | >am_trnaMT_ValTAC_MT_+_1602_1670 | 51 | 69 | 19 | CTTAACTTGACCGCTCTGA | 124.5 | -3.2 | 1.0 | -3.3 | 1.1E-03 | 1.3E-03 | BT20/MB468 |
| >hg38_21_+_8433172_8446622_RNA45SN1_45S_50ntFlanks@10350.10382.33\|ACGAGAACTTTGAAGGCCGAAGTGGAGAAGGGT | >hg38_21_+_8433172_8446622_RNA45SN1_45S_50ntFlanks | 10350 | 10382 | 33 | ACGAGAACTTTGAAGGCCGAAGTGGAGAAGGGT | 500.4 | -1.5 | 0.5 | -3.3 | 1.1E-03 | 1.3E-03 | BT20/MB468 |
| >hg38_21_+_8433172_8446622_RNA45SN1_45S_50ntFlanks@10427.10453.27\|ATGGGCGAGCGCCGTTCCGAAGGGACG | >hg38_21_+_8433172_8446622_RNA45SN1_45S_50ntFlanks | 10427 | 10453 | 27 | ATGGGCGAGCGCCGTTCCGAAGGGACG | 376.2 | -2.0 | 0.6 | -3.3 | 9.9E-04 | 1.3E-03 | MB231/MB468 |
| >trnaMT_LeuTAA_MT_+_3230_3304@18.44.27\|CGGTAATCGCATAAAACTTAAAACTTT | >trnaMT_LeuTAA_MT_+_3230_3304 | 18 | 44 | 27 | CGGTAATCGCATAAAACTTAAAACTTT | 618.9 | 1.8 | 0.5 | 3.3 | 1.1E-03 | 1.3E-03 | BT20/MB468 |
| >hg38_21_+_8433172_8446622_RNA45SN1_45S_50ntFlanks@4602.4619.18\|TTCGGAACTGAGGCCATG | >hg38_21_+_8433172_8446622_RNA45SN1_45S_50ntFlanks | 4602 | 4619 | 18 | TTCGGAACTGAGGCCATG | 122.0 | -1.9 | 0.6 | -3.3 | 1.1E-03 | 1.3E-03 | BT20/MB468 |
| >hg38_21_+_8433172_8446622_RNA45SN1_45S_50ntFlanks@12420.12444.25\|ATGTCGGCTCTTCCTATCATTGTGA | >hg38_21_+_8433172_8446622_RNA45SN1_45S_50ntFlanks | 12420 | 12444 | 25 | ATGTCGGCTCTTCCTATCATTGTGA | 1013.9 | -1.9 | 0.6 | -3.3 | 1.1E-03 | 1.3E-03 | BT20/MB468 |
| >hg38_21_+_8433172_8446622_RNA45SN1_45S_50ntFlanks@12456.12493.38\|CCAAGCGTTGGATTGTTCACCCACTAATAGGGAACGTG | >hg38_21_+_8433172_8446622_RNA45SN1_45S_50ntFlanks | 12456 | 12493 | 38 | CCAAGCGTTGGATTGTTCACCCACTAATAGGGAACGTG | 261.4 | -2.0 | 0.6 | -3.3 | 1.1E-03 | 1.3E-03 | BT20/MB468 |
| >hg38_21_+_8433172_8446622_RNA45SN1_45S_50ntFlanks@10389.10424.36\|TGAACAGCAGTTGAACATGGGTCAGTCGGTCCTGAG | >hg38_21_+_8433172_8446622_RNA45SN1_45S_50ntFlanks | 10389 | 10424 | 36 | TGAACAGCAGTTGAACATGGGTCAGTCGGTCCTGAG | 1595.9 | 2.4 | 0.7 | 3.3 | 1.0E-03 | 1.3E-03 | MB231/MB468 |
| >hg38_21_+_8433172_8446622_RNA45SN1_45S_50ntFlanks@4223.4242.20\|AAAAATAACAATACAGGACT | >hg38_21_+_8433172_8446622_RNA45SN1_45S_50ntFlanks | 4223 | 4242 | 20 | AAAAATAACAATACAGGACT | 172.0 | -2.0 | 0.6 | -3.3 | 1.0E-03 | 1.3E-03 | MB231/MB468 |

| | | | | | | | | | | | | |
|---|---|---|---|---|---|---|---|---|---|---|---|---|
| >trna10_LysCTT_16_+_3241501_3241573@1.26.26,trna119_LysCTT_1_-_145395522_145395594@1.26.26,trna11_LysCTT_5_-_180648979_180649051@1.26.26,trna13_LysCTT_14_-_58706613_58706685@1.26.26,trna13_LysCTT_6_+_26556774_26556846@1.26.26,trna2_LysCTT_15_+_79152904_79152976@1.26.26,trna32_LysCTT_16_-_3207406_3207478@1.26.26,trna7_LysCTT_16_+_3225692_3225764@1.26.26,trna9_LysCTT_5_+_180634755_180634827@1.26.26\|GCCCGGCTAGCTCAGTCGGTAGAGCA | >trna10_LysCTT_16_+_3241501_3241573 | 1 | 26 | 26 | GCCCGGCTAGCTCAGTCGGTAGAGCA | 85.7 | -1.8 | 0.6 | -3.3 | 1.0E-03 | 1.3E-03 | MB231/MB468 |
| >trna2_GlyGCC_21_-_18827107_18827177@1.28.28,trna35_GlyGCC_1_+_161413094_161413164@1.28.28,trna37_GlyGCC_1_+_161420467_161420537@1.28.28,trna39_GlyGCC_1_+_161427898_161427968@1.28.28,trna41_GlyGCC_1_+_161435258_161435328@1.28.28\|GCATGGGTGGTTCAGTGGTAGAATTCTC | >trna2_GlyGCC_21_-_18827107_18827177 | 1 | 28 | 28 | GCATGGGTGGTTCAGTGGTAGAATTCTC | 1543.7 | 7.2 | 1.9 | 3.8 | 1.7E-04 | 1.4E-03 | BT20/MB231 |
| >trna119_LysCTT_1_-_145395522_145395594@1.35.35,trna11_LysCTT_5_-_180648979_180649051@1.35.35,trna13_LysCTT_6_+_26556774_26556846@1.35.35,trna7_LysCTT_16_+_3225692_3225764@1.35.35,trna9_LysCTT_5_+_180634755_180634827@1.35.35\|GCCCGGCTAGCTCAGTCGGTAGAGCATGAGACTCT | >trna119_LysCTT_1_-_145395522_145395594 | 1 | 35 | 35 | GCCCGGCTAGCTCAGTCGGTAGAGCATGAGACTCT | 16535.0 | 2.8 | 0.8 | 3.8 | 1.7E-04 | 1.4E-03 | BT20/MB231 |
| >hg38_MT_+_598_1651_RNR1_12S_50ntFlanks@967.1004.38\|AACATGGTAAGTGTACTGGAAAGTGCACTTGGACGAAC | >hg38_MT_+_598_1651_RNR1_12S_50ntFlanks | 967 | 1004 | 38 | AACATGGTAAGTGTACTGGAAAGTGCACTTGGACGAAC | 279.2 | -2.9 | 0.8 | -3.8 | 1.7E-04 | 1.4E-03 | BT20/MB231 |
| >trna20_GluTTC_2_-_131094701_131094772@1.36.36,trna5_GluTTC_13_-_41634874_41634945@1.36.36\|TCCCATATGGTCTAGCGGTTAGGATTCCTGGTTTTC | >trna20_GluTTC_2_-_131094701_131094772 | 1 | 36 | 36 | TCCCATATGGTCTAGCGGTTAGGATTCCTGGTTTTC | 321.8 | -3.5 | 0.9 | -3.8 | 1.7E-04 | 1.4E-03 | BT20/MB231 |
| >hg38_21_+_8433172_8446622_RNA45SN1_45S_50ntFlanks@9572.9601.30\|CAAATCGGTCGTCCGACCTGGGTATAGGGG | >hg38_21_+_8433172_8446622_RNA45SN1_45S_50ntFlanks | 9572 | 9601 | 30 | CAAATCGGTCGTCCGACCTGGGTATAGGGG | 300.6 | -1.7 | 0.5 | -3.3 | 1.1E-03 | 1.4E-03 | MB231/MB468 |
| >hg38_21_+_8433172_8446622_RNA45SN1_45S_50ntFlanks@11885.11903.19\|TTGAGCTTGACTCTAGTCT | >hg38_21_+_8433172_8446622_RNA45SN1_45S_50ntFlanks | 11885 | 11903 | 19 | TTGAGCTTGACTCTAGTCT | 368.5 | -5.6 | 1.7 | -3.3 | 1.1E-03 | 1.4E-03 | MB231/MB468 |
| >hg38_21_+_8433172_8446622_RNA45SN1_45S_50ntFlanks@10351.10379.29\|CGAGAACTTTGAAGGCCGAAGTGGAGAAG | >hg38_21_+_8433172_8446622_RNA45SN1_45S_50ntFlanks | 10351 | 10379 | 29 | CGAGAACTTTGAAGGCCGAAGTGGAGAAG | 546.2 | -1.9 | 0.6 | -3.3 | 1.1E-03 | 1.4E-03 | MB231/MB468 |
| >hg38_21_+_8433172_8446622_RNA45SN1_45S_50ntFlanks@12423.12445.23\|TCGGCTCTTCCTATCATTGTGAA | >hg38_21_+_8433172_8446622_RNA45SN1_45S_50ntFlanks | 12423 | 12445 | 23 | TCGGCTCTTCCTATCATTGTGAA | 109.1 | -2.3 | 0.7 | -3.2 | 1.2E-03 | 1.4E-03 | BT20/MB468 |
| >hg38_21_+_8433172_8446622_RNA45SN1_45S_50ntFlanks@11771.11791.21\|ATTAGTGACGCGCATGAATGG | >hg38_21_+_8433172_8446622_RNA45SN1_45S_50ntFlanks | 11771 | 11791 | 21 | ATTAGTGACGCGCATGAATGG | 157.4 | -2.4 | 0.7 | -3.2 | 1.2E-03 | 1.4E-03 | BT20/MB468 |
| >MI0000299\|hsa-mir-222&WithFlank&X\|-\|45747009\|45747130@75.99.25(+2U)&[MIMAT0000279&hsa-miR-222-3p&offsets\|0\|+4(+2U);m-140&X\|-\|45747033\|45747056&offsets\|0\|+1(+2U)]\|AGCTACATCTGGCTACTGGGTCTCTTT | >MI0000299\|hsa-mir-222&WithFlank&X\|-\|45747009\|45747130 | 75 | 99 | 25(+2U) | AGCTACATCTGGCTACTGGGTCTCTTT | 3437.2 | -2.9 | 0.8 | -3.8 | 1.7E-04 | 1.4E-03 | BT20/MB231 |
| >hg38_21_+_8433172_8446622_RNA45SN1_45S_50ntFlanks@6645.6662.18\|CTCGTACGACTCTTAGCG | >hg38_21_+_8433172_8446622_RNA45SN1_45S_50ntFlanks | 6645 | 6662 | 18 | CTCGTACGACTCTTAGCG | 2078.4 | 2.4 | 0.7 | 3.3 | 1.1E-03 | 1.4E-03 | MB231/MB468 |
| >hg38_21_+_8433172_8446622_RNA45SN1_45S_50ntFlanks@9568.9585.18\|CGTGCAAATCGGTCGTCC | >hg38_21_+_8433172_8446622_RNA45SN1_45S_50ntFlanks | 9568 | 9585 | 18 | CGTGCAAATCGGTCGTCC | 92.4 | -1.9 | 0.6 | -3.3 | 1.1E-03 | 1.4E-03 | MB231/MB468 |
| >trna32_LysCTT_16_-_3207406_3207478@1.33.33\|GCCCGGCTAGCTCAGTCGGTAGAGCATGAGACC | >trna32_LysCTT_16_-_3207406_3207478 | 1 | 33 | 33 | GCCCGGCTAGCTCAGTCGGTAGAGCATGAGACC | 2064.2 | -3.4 | 0.9 | -3.8 | 1.8E-04 | 1.4E-03 | BT20/MB231 |
| >hg38_21_+_8433172_8446622_RNA45SN1_45S_50ntFlanks@12935.12959.25\|TAAACCATTCGTAGACGACCTGCTT | >hg38_21_+_8433172_8446622_RNA45SN1_45S_50ntFlanks | 12935 | 12959 | 25 | TAAACCATTCGTAGACGACCTGCTT | 119.8 | -1.5 | 0.5 | -3.3 | 1.1E-03 | 1.4E-03 | MB231/MB468 |
| >hg38_21_+_8433172_8446622_RNA45SN1_45S_50ntFlanks@11572.11599.28\|CCGACTTAGAACTGGTGCGGACCAGGGG | >hg38_21_+_8433172_8446622_RNA45SN1_45S_50ntFlanks | 11572 | 11599 | 28 | CCGACTTAGAACTGGTGCGGACCAGGGG | 303.1 | -2.2 | 0.7 | -3.3 | 1.1E-03 | 1.4E-03 | MB231/MB468 |
| >hg38_21_+_8433172_8446622_RNA45SN1_45S_50ntFlanks@7992.8011.20\|CGTGGCGACCCGCTGAATTT | >hg38_21_+_8433172_8446622_RNA45SN1_45S_50ntFlanks | 7992 | 8011 | 20 | CGTGGCGACCCGCTGAATTT | 636.8 | 2.2 | 0.7 | 3.2 | 1.2E-03 | 1.4E-03 | BT20/MB468 |
| >hg38_21_+_8433172_8446622_RNA45SN1_45S_50ntFlanks@9544.9571.28\|GGTGGAGGTCCGTAGCGGTCCTGACGTG | >hg38_21_+_8433172_8446622_RNA45SN1_45S_50ntFlanks | 9544 | 9571 | 28 | GGTGGAGGTCCGTAGCGGTCCTGACGTG | 1224.7 | -2.7 | 0.8 | -3.2 | 1.2E-03 | 1.4E-03 | BT20/MB468 |
| >hg38_21_+_8433172_8446622_RNA45SN1_45S_50ntFlanks@4735.4757.23\|ACGAAAGTCGGAGGTTCGAAGAC | >hg38_21_+_8433172_8446622_RNA45SN1_45S_50ntFlanks | 4735 | 4757 | 23 | ACGAAAGTCGGAGGTTCGAAGAC | 289.3 | 6.3 | 1.9 | 3.3 | 1.1E-03 | 1.5E-03 | MB231/MB468 |
| >hg38_21_+_8433172_8446622_RNA45SN1_45S_50ntFlanks@10781.10800.20\|AGGTGAACAGCCTCTGGCAT | >hg38_21_+_8433172_8446622_RNA45SN1_45S_50ntFlanks | 10781 | 10800 | 20 | AGGTGAACAGCCTCTGGCAT | 126.7 | -2.1 | 0.7 | -3.2 | 1.2E-03 | 1.5E-03 | BT20/MB468 |
| >hg38_21_+_8433172_8446622_RNA45SN1_45S_50ntFlanks@3705.3739.35\|TACCTGGTTGATCCTGCCAGTAGCATATGCTTGTC | >hg38_21_+_8433172_8446622_RNA45SN1_45S_50ntFlanks | 3705 | 3739 | 35 | TACCTGGTTGATCCTGCCAGTAGCATATGCTTGTC | 1285.1 | 2.7 | 0.7 | 3.7 | 1.8E-04 | 1.5E-03 | BT20/MB231 |
| >hg38_21_+_8433172_8446622_RNA45SN1_45S_50ntFlanks@12394.12419.26\|TAGCGACGTCGCTTTTTGATCCTTCG | >hg38_21_+_8433172_8446622_RNA45SN1_45S_50ntFlanks | 12394 | 12419 | 26 | TAGCGACGTCGCTTTTTGATCCTTCG | 421.7 | -3.0 | 0.9 | -3.2 | 1.2E-03 | 1.5E-03 | BT20/MB468 |
| >hg38_21_+_8433172_8446622_RNA45SN1_45S_50ntFlanks@10684.10712.29\|CGGCGGCGTCCGGTGAGCTCTCGCTGGCC | >hg38_21_+_8433172_8446622_RNA45SN1_45S_50ntFlanks | 10684 | 10712 | 29 | CGGCGGCGTCCGGTGAGCTCTCGCTGGCC | 311.1 | -1.5 | 0.5 | -3.2 | 1.2E-03 | 1.5E-03 | MB231/MB468 |
| >hg38_21_+_8433172_8446622_RNA45SN1_45S_50ntFlanks@4220.4242.23\|ACGAAAAATAACAATACAGGACT | >hg38_21_+_8433172_8446622_RNA45SN1_45S_50ntFlanks | 4220 | 4242 | 23 | ACGAAAAATAACAATACAGGACT | 421.6 | -2.0 | 0.6 | -3.2 | 1.2E-03 | 1.5E-03 | MB231/MB468 |
| >hg38_21_+_8433172_8446622_RNA45SN1_45S_50ntFlanks@11200.11227.28\|CGGCGGCGGCGGCGACTCTGGACGCGAG | >hg38_21_+_8433172_8446622_RNA45SN1_45S_50ntFlanks | 11200 | 11227 | 28 | CGGCGGCGGCGGCGACTCTGGACGCGAG | 131.5 | -3.4 | 1.1 | -3.2 | 1.3E-03 | 1.5E-03 | BT20/MB468 |
| >hg38_21_+_8433172_8446622_RNA45SN1_45S_50ntFlanks@11778.11803.26\|ACGCGCATGAATGGATGAACGAGATT | >hg38_21_+_8433172_8446622_RNA45SN1_45S_50ntFlanks | 11778 | 11803 | 26 | ACGCGCATGAATGGATGAACGAGATT | 652.5 | -1.8 | 0.6 | -3.2 | 1.2E-03 | 1.5E-03 | MB231/MB468 |
| >hg38_21_+_8433172_8446622_RNA45SN1_45S_50ntFlanks@10796.10818.23\|GGCATGTTGGAACAATGTAGGTA | >hg38_21_+_8433172_8446622_RNA45SN1_45S_50ntFlanks | 10796 | 10818 | 23 | GGCATGTTGGAACAATGTAGGTA | 595.0 | -2.2 | 0.7 | -3.2 | 1.2E-03 | 1.5E-03 | MB231/MB468 |
| >hg38_21_+_8433172_8446622_RNA45SN1_45S_50ntFlanks@11575.11609.35\|ACTTAGAACTGGTGCGGACCAGGGGAATCCGACTG | >hg38_21_+_8433172_8446622_RNA45SN1_45S_50ntFlanks | 11575 | 11609 | 35 | ACTTAGAACTGGTGCGGACCAGGGGAATCCGACTG | 409.4 | -1.7 | 0.5 | -3.2 | 1.2E-03 | 1.5E-03 | MB231/MB468 |
| >hg38_21_+_8433172_8446622_RNA45SN1_45S_50ntFlanks@10771.10790.20\|CAGGTCTCCAAGGTGAACAG | >hg38_21_+_8433172_8446622_RNA45SN1_45S_50ntFlanks | 10771 | 10790 | 20 | CAGGTCTCCAAGGTGAACAG | 230.6 | -4.0 | 1.2 | -3.2 | 1.3E-03 | 1.5E-03 | BT20/MB468 |
| >hg38_21_+_8433172_8446622_RNA45SN1_45S_50ntFlanks@6719.6750.32\|TGAATTGCAGGACACATTGATCATCGACACTT | >hg38_21_+_8433172_8446622_RNA45SN1_45S_50ntFlanks | 6719 | 6750 | 32 | TGAATTGCAGGACACATTGATCATCGACACTT | 117.7 | -1.4 | 0.4 | -3.2 | 1.2E-03 | 1.5E-03 | MB231/MB468 |
| >hg38_21_+_8433172_8446622_RNA45SN1_45S_50ntFlanks@11873.11902.30\|AAGAAGACCCTGTTGAGCTTGACTCTAGTC | >hg38_21_+_8433172_8446622_RNA45SN1_45S_50ntFlanks | 11873 | 11902 | 30 | AAGAAGACCCTGTTGAGCTTGACTCTAGTC | 260.1 | -2.4 | 0.8 | -3.2 | 1.3E-03 | 1.6E-03 | BT20/MB468 |
| >MI0000299\|hsa-mir-222&WithFlank&X\|-\|45747009\|45747130@75.99.25&[MIMAT0000279&hsa-miR-222-3p&offsets\|0\|+4;m-140&X\|-\|45747033\|45747056&offsets\|0\|+1]\|AGCTACATCTGGCTACTGGGTCTCT | >MI0000299\|hsa-mir-222&WithFlank&X\|-\|45747009\|45747130 | 75 | 99 | 25 | AGCTACATCTGGCTACTGGGTCTCT | 53759.7 | -2.3 | 0.6 | -3.7 | 1.9E-04 | 1.6E-03 | BT20/MB231 |
| >hg38_21_+_8433172_8446622_RNA45SN1_45S_50ntFlanks@5444.5465.22\|CTCGGATCGGCCCCGCCGGGGT | >hg38_21_+_8433172_8446622_RNA45SN1_45S_50ntFlanks | 5444 | 5465 | 22 | CTCGGATCGGCCCCGCCGGGGT | 130.7 | -2.2 | 0.7 | -3.2 | 1.2E-03 | 1.6E-03 | MB231/MB468 |
| >hg38_21_+_8433172_8446622_RNA45SN1_45S_50ntFlanks@4008.4025.18\|CTCGGGCCGATCGCACGC | >hg38_21_+_8433172_8446622_RNA45SN1_45S_50ntFlanks | 4008 | 4025 | 18 | CTCGGGCCGATCGCACGC | 176.3 | -1.7 | 0.5 | -3.2 | 1.3E-03 | 1.6E-03 | BT20/MB468 |
| >hg38_21_+_8433172_8446622_RNA45SN1_45S_50ntFlanks@9554.9578.25\|CGTAGCGGTCCTGACGTGCAAATCG | >hg38_21_+_8433172_8446622_RNA45SN1_45S_50ntFlanks | 9554 | 9578 | 25 | CGTAGCGGTCCTGACGTGCAAATCG | 156.1 | -1.6 | 0.5 | -3.2 | 1.3E-03 | 1.6E-03 | BT20/MB468 |
| >hg38_21_+_8433172_8446622_RNA45SN1_45S_50ntFlanks@12004.12033.30\|CGGGCCGCCGGTGAAATACCACTACTCTGA | >hg38_21_+_8433172_8446622_RNA45SN1_45S_50ntFlanks | 12004 | 12033 | 30 | CGGGCCGCCGGTGAAATACCACTACTCTGA | 1300.7 | 3.5 | 0.9 | 3.7 | 1.9E-04 | 1.6E-03 | BT20/MB231 |
| >am_trnaMT_ArgTCG_MT_+_10405_10469@29.48.20\|TTTCGACTCATTAAATTATG | >am_trnaMT_ArgTCG_MT_+_10405_10469 | 29 | 48 | 20 | TTTCGACTCATTAAATTATG | 135.9 | -2.1 | 0.6 | -3.2 | 1.2E-03 | 1.6E-03 | MB231/MB468 |
| >hg38_21_+_8433172_8446622_RNA45SN1_45S_50ntFlanks@10794.10821.28\|CTGGCATGTTGGAACAATGTAGGTAAGG | >hg38_21_+_8433172_8446622_RNA45SN1_45S_50ntFlanks | 10794 | 10821 | 28 | CTGGCATGTTGGAACAATGTAGGTAAGG | 147.8 | -1.6 | 0.5 | -3.2 | 1.2E-03 | 1.6E-03 | MB231/MB468 |
| >hg38_21_+_8433172_8446622_RNA45SN1_45S_50ntFlanks@7980.8003.24\|CCTCAGATCAGACGTGGCGACCCG | >hg38_21_+_8433172_8446622_RNA45SN1_45S_50ntFlanks | 7980 | 8003 | 24 | CCTCAGATCAGACGTGGCGACCCG | 206.9 | -2.4 | 0.8 | -3.2 | 1.2E-03 | 1.6E-03 | MB231/MB468 |
| >hg38_21_+_8433172_8446622_RNA45SN1_45S_50ntFlanks@7980.8008.29\|CCTCAGATCAGACGTGGCGACCCGCTGAA | >hg38_21_+_8433172_8446622_RNA45SN1_45S_50ntFlanks | 7980 | 8008 | 29 | CCTCAGATCAGACGTGGCGACCCGCTGAA | 642.1 | -2.5 | 0.8 | -3.2 | 1.2E-03 | 1.6E-03 | MB231/MB468 |
| >hg38_21_+_8433172_8446622_RNA45SN1_45S_50ntFlanks@9544.9563.20\|GGTGGAGGTCCGTAGCGGTC | >hg38_21_+_8433172_8446622_RNA45SN1_45S_50ntFlanks | 9544 | 9563 | 20 | GGTGGAGGTCCGTAGCGGTC | 136.5 | -1.8 | 0.6 | -3.2 | 1.2E-03 | 1.6E-03 | MB231/MB468 |
| >hg38_21_+_8433172_8446622_RNA45SN1_45S_50ntFlanks@4661.4689.29\|AGGTGAAATTCTTGGACCGGCGCAAGACG | >hg38_21_+_8433172_8446622_RNA45SN1_45S_50ntFlanks | 4661 | 4689 | 29 | AGGTGAAATTCTTGGACCGGCGCAAGACG | 96.6 | -2.7 | 0.8 | -3.2 | 1.2E-03 | 1.6E-03 | MB231/MB468 |
| >hg38_21_+_8433172_8446622_RNA45SN1_45S_50ntFlanks@9533.9554.22\|GAGGAAACTCTGGTGGAGGTCC | >hg38_21_+_8433172_8446622_RNA45SN1_45S_50ntFlanks | 9533 | 9554 | 22 | GAGGAAACTCTGGTGGAGGTCC | 208.4 | -2.0 | 0.6 | -3.2 | 1.2E-03 | 1.6E-03 | MB231/MB468 |
| >hg38_21_+_8433172_8446622_RNA45SN1_45S_50ntFlanks@3881.3899.19\|GCCGACGGGCGCTGACCCC | >hg38_21_+_8433172_8446622_RNA45SN1_45S_50ntFlanks | 3881 | 3899 | 19 | GCCGACGGGCGCTGACCCC | 119.0 | -2.5 | 0.8 | -3.2 | 1.3E-03 | 1.6E-03 | BT20/MB468 |
| >hg38_21_+_8433172_8446622_RNA45SN1_45S_50ntFlanks@12438.12473.36\|ATTGTGAAGCAGAATTCACCAAGCGTTGGATTGTTC | >hg38_21_+_8433172_8446622_RNA45SN1_45S_50ntFlanks | 12438 | 12473 | 36 | ATTGTGAAGCAGAATTCACCAAGCGTTGGATTGTTC | 1091.8 | 2.0 | 0.6 | 3.2 | 1.2E-03 | 1.6E-03 | MB231/MB468 |

| | | | | | | | | | | | | |
|---|---|---|---|---|---|---|---|---|---|---|---|---|
| >hg38_21_+_8433172_8446622_RNA45SN1_45S_50ntFlanks@10425.10456.32\|AGATGGGCGAGCGCCGTTCCGAAGGGACGGGC | >hg38_21_+_8433172_8446622_RNA45SN1_45S_50ntFlanks | 10425 | 10456 | 32 | AGATGGGCGAGCGCCGTTCCGAAGGGACGGGC | 237.4 | -1.2 | 0.4 | -3.2 | 1.2E-03 | 1.6E-03 | MB231/MB468 |
| >hg38_21_+_8433172_8446622_RNA45SN1_45S_50ntFlanks@5467.5499.33\|GGCCCACGGCCCTGGCGGAGCGCTGAGAAGACG | >hg38_21_+_8433172_8446622_RNA45SN1_45S_50ntFlanks | 5467 | 5499 | 33 | GGCCCACGGCCCTGGCGGAGCGCTGAGAAGACG | 1070.9 | -3.6 | 1.1 | -3.2 | 1.4E-03 | 1.6E-03 | BT20/MB468 |
| >hg38_21_+_8433172_8446622_RNA45SN1_45S_50ntFlanks@9555.9573.19\|GTAGCGGTCCTGACGTGCA | >hg38_21_+_8433172_8446622_RNA45SN1_45S_50ntFlanks | 9555 | 9573 | 19 | GTAGCGGTCCTGACGTGCA | 456.2 | 1.8 | 0.6 | 3.2 | 1.3E-03 | 1.6E-03 | MB231/MB468 |
| >hg38_21_+_8433172_8446622_RNA45SN1_45S_50ntFlanks@4067.4089.23\|AACTTTCGATGGTAGTCGCCGTG | >hg38_21_+_8433172_8446622_RNA45SN1_45S_50ntFlanks | 4067 | 4089 | 23 | AACTTTCGATGGTAGTCGCCGTG | 95.8 | -1.6 | 0.5 | -3.2 | 1.4E-03 | 1.6E-03 | BT20/MB468 |
| >hg38_21_+_8433172_8446622_RNA45SN1_45S_50ntFlanks@12416.12436.21\|TTCGATGTCGGCTCTTCCTAT | >hg38_21_+_8433172_8446622_RNA45SN1_45S_50ntFlanks | 12416 | 12436 | 21 | TTCGATGTCGGCTCTTCCTAT | 781.4 | -1.8 | 0.5 | -3.2 | 1.4E-03 | 1.6E-03 | BT20/MB468 |
| >hg38_21_+_8433172_8446622_RNA45SN1_45S_50ntFlanks@6721.6758.38\|AATTGCAGGACACATTGATCATCGACACTTCGAACGCA | >hg38_21_+_8433172_8446622_RNA45SN1_45S_50ntFlanks | 6721 | 6758 | 38 | AATTGCAGGACACATTGATCATCGACACTTCGAACGCA | 1084.3 | -2.1 | 0.7 | -3.2 | 1.3E-03 | 1.6E-03 | MB231/MB468 |
| >hg38_21_+_8433172_8446622_RNA45SN1_45S_50ntFlanks@11564.11584.21\|CCTAGCAGCCGACTTAGAACT | >hg38_21_+_8433172_8446622_RNA45SN1_45S_50ntFlanks | 11564 | 11584 | 21 | CCTAGCAGCCGACTTAGAACT | 371.9 | 1.8 | 0.6 | 3.2 | 1.3E-03 | 1.6E-03 | MB231/MB468 |
| >hg38_21_+_8433172_8446622_RNA45SN1_45S_50ntFlanks@11318.11342.25\|CCCCACGTCTCGTCGCGCGCGCGTC | >hg38_21_+_8433172_8446622_RNA45SN1_45S_50ntFlanks | 11318 | 11342 | 25 | CCCCACGTCTCGTCGCGCGCGCGTC | 653.5 | -4.4 | 1.4 | -3.2 | 1.4E-03 | 1.6E-03 | BT20/MB468 |
| >hg38_21_+_8433172_8446622_RNA45SN1_45S_50ntFlanks@11316.11334.19\|CACCCCACGTCTCGTCGCG | >hg38_21_+_8433172_8446622_RNA45SN1_45S_50ntFlanks | 11316 | 11334 | 19 | CACCCCACGTCTCGTCGCG | 952.5 | -4.7 | 1.5 | -3.2 | 1.4E-03 | 1.6E-03 | BT20/MB468 |
| >hg38_21_+_8433172_8446622_RNA45SN1_45S_50ntFlanks@6646.6693.48\|TCGTACGACTCTTAGCGGTGGATCACTCGGCTCGTGCGTCGATGAAGA | >hg38_21_+_8433172_8446622_RNA45SN1_45S_50ntFlanks | 6646 | 6693 | 48 | TCGTACGACTCTTAGCGGTGGATCACTCGGCTCGTGCGTCGATGAAGA | 552.2 | 7.3 | 2.3 | 3.2 | 1.4E-03 | 1.6E-03 | BT20/MB468 |
| >hg38_21_+_8433172_8446622_RNA45SN1_45S_50ntFlanks@11027.11060.34\|CTCTCTCTCTCTCTCCCCCGCTCCCCGTCCTCCC | >hg38_21_+_8433172_8446622_RNA45SN1_45S_50ntFlanks | 11027 | 11060 | 34 | CTCTCTCTCTCTCTCCCCCGCTCCCCGTCCTCCC | 328.9 | -4.0 | 1.3 | -3.2 | 1.4E-03 | 1.7E-03 | BT20/MB468 |
| >hg38_21_+_8433172_8446622_RNA45SN1_45S_50ntFlanks@7992.8009.18\|CGTGGCGACCCGCTGAAT | >hg38_21_+_8433172_8446622_RNA45SN1_45S_50ntFlanks | 7992 | 8009 | 18 | CGTGGCGACCCGCTGAAT | 1335.2 | 3.1 | 1.0 | 3.2 | 1.3E-03 | 1.7E-03 | MB231/MB468 |
| >hg38_21_+_8433172_8446622_RNA45SN1_45S_50ntFlanks@10349.10371.23\|AACGAGAACTTTGAAGGCCGAAG | >hg38_21_+_8433172_8446622_RNA45SN1_45S_50ntFlanks | 10349 | 10371 | 23 | AACGAGAACTTTGAAGGCCGAAG | 2019.5 | 2.2 | 0.7 | 3.2 | 1.3E-03 | 1.7E-03 | MB231/MB468 |
| >hg38_21_+_8433172_8446622_RNA45SN1_45S_50ntFlanks@12420.12456.37\|ATGTCGGCTCTTCCTATCATTGTGAAGCAGAATTCAC | >hg38_21_+_8433172_8446622_RNA45SN1_45S_50ntFlanks | 12420 | 12456 | 37 | ATGTCGGCTCTTCCTATCATTGTGAAGCAGAATTCAC | 293.9 | -2.9 | 0.8 | -3.7 | 2.1E-04 | 1.7E-03 | BT20/MB231 |
| >hg38_21_+_8433172_8446622_RNA45SN1_45S_50ntFlanks@6721.6753.33\|AATTGCAGGACACATTGATCATCGACACTTCGA | >hg38_21_+_8433172_8446622_RNA45SN1_45S_50ntFlanks | 6721 | 6753 | 33 | AATTGCAGGACACATTGATCATCGACACTTCGA | 121.6 | -1.7 | 0.5 | -3.2 | 1.3E-03 | 1.7E-03 | MB231/MB468 |
| >hg38_21_+_8433172_8446622_RNA45SN1_45S_50ntFlanks@12481.12499.19\|AATAGGGAACGTGAGCTGG | >hg38_21_+_8433172_8446622_RNA45SN1_45S_50ntFlanks | 12481 | 12499 | 19 | AATAGGGAACGTGAGCTGG | 259.8 | 1.4 | 0.4 | 3.2 | 1.3E-03 | 1.7E-03 | MB231/MB468 |
| >hg38_21_+_8433172_8446622_RNA45SN1_45S_50ntFlanks@10873.10891.19\|CTGGGTCGGTCGGGCTGGG | >hg38_21_+_8433172_8446622_RNA45SN1_45S_50ntFlanks | 10873 | 10891 | 19 | CTGGGTCGGTCGGGCTGGG | 214.7 | -5.2 | 1.6 | -3.2 | 1.3E-03 | 1.7E-03 | MB231/MB468 |
| >hg38_21_+_8433172_8446622_RNA45SN1_45S_50ntFlanks@7973.8008.36\|GACGCGACCTCAGATCAGACGTGGCGACCCGCTGAA | >hg38_21_+_8433172_8446622_RNA45SN1_45S_50ntFlanks | 7973 | 8008 | 36 | GACGCGACCTCAGATCAGACGTGGCGACCCGCTGAA | 504.7 | 2.4 | 0.7 | 3.2 | 1.4E-03 | 1.7E-03 | BT20/MB468 |
| >hg38_21_+_8433172_8446622_RNA45SN1_45S_50ntFlanks@12463.12487.25\|TTGGATTGTTCACCCACTAATAGGG | >hg38_21_+_8433172_8446622_RNA45SN1_45S_50ntFlanks | 12463 | 12487 | 25 | TTGGATTGTTCACCCACTAATAGGG | 189.2 | -2.1 | 0.7 | -3.2 | 1.4E-03 | 1.7E-03 | BT20/MB468 |
| >hg38_21_+_8433172_8446622_RNA45SN1_45S_50ntFlanks@3850.3871.22\|GATAACTGTGGTAATTCTAGAG | >hg38_21_+_8433172_8446622_RNA45SN1_45S_50ntFlanks | 3850 | 3871 | 22 | GATAACTGTGGTAATTCTAGAG | 142.6 | -3.2 | 1.0 | -3.2 | 1.4E-03 | 1.7E-03 | BT20/MB468 |
| >hg38_21_+_8433172_8446622_RNA45SN1_45S_50ntFlanks@9533.9571.39\|GAGGAAACTCTGGTGGAGGTCCGTAGCGGTCCTGACGTG | >hg38_21_+_8433172_8446622_RNA45SN1_45S_50ntFlanks | 9533 | 9571 | 39 | GAGGAAACTCTGGTGGAGGTCCGTAGCGGTCCTGACGTG | 723.2 | -2.1 | 0.7 | -3.2 | 1.4E-03 | 1.7E-03 | BT20/MB468 |
| >hg38_21_+_8433172_8446622_RNA45SN1_45S_50ntFlanks@3835.3860.26\|CTCCTCTCCTACTTGGATAACTGTGG | >hg38_21_+_8433172_8446622_RNA45SN1_45S_50ntFlanks | 3835 | 3860 | 26 | CTCCTCTCCTACTTGGATAACTGTGG | 1105.7 | -2.4 | 0.7 | -3.2 | 1.3E-03 | 1.7E-03 | MB231/MB468 |
| >hg38_21_+_8433172_8446622_RNA45SN1_45S_50ntFlanks@6646.6689.44\|TCGTACGACTCTTAGCGGTGGATCACTCGGCTCGTGCGTCGATG | >hg38_21_+_8433172_8446622_RNA45SN1_45S_50ntFlanks | 6646 | 6689 | 44 | TCGTACGACTCTTAGCGGTGGATCACTCGGCTCGTGCGTCGATG | 3651.4 | 1.9 | 0.6 | 3.2 | 1.3E-03 | 1.7E-03 | MB231/MB468 |
| >MI0000342\|hsa-mir-200b&WithFlank&1\|+\|1167098\|1167204@63.86.24&[MIMAT0000318&hsa-miR-200b-3p&offsets\|0\|+2;m-93&1\|+\|1167160\|1167182&offsets\|0\|+1]\|TAATACTGCCTGGTAATGATGACG | >MI0000342\|hsa-mir-200b&WithFlank&1\|+\|1167098\|1167204 | 63 | 86 | 24 | TAATACTGCCTGGTAATGATGACG | 167.8 | -2.1 | 0.7 | -3.2 | 1.5E-03 | 1.7E-03 | BT20/MB468 |
| >hg38_21_+_8433172_8446622_RNA45SN1_45S_50ntFlanks@12418.12443.26\|CGATGTCGGCTCTTCCTATCATTGTG | >hg38_21_+_8433172_8446622_RNA45SN1_45S_50ntFlanks | 12418 | 12443 | 26 | CGATGTCGGCTCTTCCTATCATTGTG | 3574.7 | -2.1 | 0.7 | -3.2 | 1.3E-03 | 1.7E-03 | MB231/MB468 |
| >hg38_21_+_8433172_8446622_RNA45SN1_45S_50ntFlanks@12415.12433.19\|CTTCGATGTCGGCTCTTCC | >hg38_21_+_8433172_8446622_RNA45SN1_45S_50ntFlanks | 12415 | 12433 | 19 | CTTCGATGTCGGCTCTTCC | 1468.2 | 2.0 | 0.6 | 3.2 | 1.4E-03 | 1.7E-03 | MB231/MB468 |
| >hg38_21_+_8433172_8446622_RNA45SN1_45S_50ntFlanks@4499.4517.19\|AGCGTTTACTTTGAAAAAA | >hg38_21_+_8433172_8446622_RNA45SN1_45S_50ntFlanks | 4499 | 4517 | 19 | AGCGTTTACTTTGAAAAAA | 126.1 | -2.0 | 0.6 | -3.2 | 1.4E-03 | 1.7E-03 | MB231/MB468 |
| >hg38_21_+_8433172_8446622_RNA45SN1_45S_50ntFlanks@12407.12446.40\|TTTTGATCCTTCGATGTCGGCTCTTCCTATCATTGTGAAG | >hg38_21_+_8433172_8446622_RNA45SN1_45S_50ntFlanks | 12407 | 12446 | 40 | TTTTGATCCTTCGATGTCGGCTCTTCCTATCATTGTGAAG | 338.1 | 1.9 | 0.6 | 3.2 | 1.4E-03 | 1.8E-03 | MB231/MB468 |
| >am_trna128_GlyGCC_6_-_27870686_27870756@1.29.29,trna133_GlyCCC_1_-_16872434_16872504@1.29.29,trna18_GlyGCC_16_+_70822597_70822667@1.29.29,trna19_GlyGCC_16_+_70823410_70823480@1.29.29,trna19_GlyGCC_2_-_157257659_157257729@1.29.29,trna24_GlyGCC_16_-_70812942_70813012@1.29.29,trna25_GlyGCC_16_-_70812114_70812184@1.29.29,trna4_GlyCCC_1_+_17188416_17188486@1.29.29,trna5_GlyGCC_17_+_8029064_8029134@1.29.29,trna68_GlyGCC_1_-_161493637_161493707@1.29.29\|GCATTGGTGGTTCAGTGGTAGAATTCTCG | >am_trna128_GlyGCC_6_-_27870686_27870756 | 1 | 29 | 29 | GCATTGGTGGTTCAGTGGTAGAATTCTCG | 20616.2 | -2.8 | 0.9 | -3.2 | 1.4E-03 | 1.8E-03 | MB231/MB468 |
| >hg38_21_+_8433172_8446622_RNA45SN1_45S_50ntFlanks@10829.10851.23\|GCAAGCCGGATCCGTAACTTCGG | >hg38_21_+_8433172_8446622_RNA45SN1_45S_50ntFlanks | 10829 | 10851 | 23 | GCAAGCCGGATCCGTAACTTCGG | 189.7 | -1.3 | 0.4 | -3.2 | 1.5E-03 | 1.8E-03 | BT20/MB468 |
| >hg38_21_+_8433172_8446622_RNA45SN1_45S_50ntFlanks@7974.8008.35\|ACGCGACCTCAGATCAGACGTGGCGACCCGCTGAA | >hg38_21_+_8433172_8446622_RNA45SN1_45S_50ntFlanks | 7974 | 8008 | 35 | ACGCGACCTCAGATCAGACGTGGCGACCCGCTGAA | 3190.0 | 1.8 | 0.6 | 3.2 | 1.5E-03 | 1.8E-03 | BT20/MB468 |
| >hg38_21_+_8433172_8446622_RNA45SN1_45S_50ntFlanks@7974.7993.20\|ACGCGACCTCAGATCAGACG | >hg38_21_+_8433172_8446622_RNA45SN1_45S_50ntFlanks | 7974 | 7993 | 20 | ACGCGACCTCAGATCAGACG | 1764.6 | 2.8 | 0.9 | 3.2 | 1.4E-03 | 1.8E-03 | MB231/MB468 |
| >hg38_21_+_8433172_8446622_RNA45SN1_45S_50ntFlanks@4590.4616.27\|ATTTTGTTGGTTTTCGGAACTGAGGCC | >hg38_21_+_8433172_8446622_RNA45SN1_45S_50ntFlanks | 4590 | 4616 | 27 | ATTTTGTTGGTTTTCGGAACTGAGGCC | 278.3 | -2.1 | 0.7 | -3.2 | 1.5E-03 | 1.8E-03 | BT20/MB468 |
| >hg38_21_+_8433172_8446622_RNA45SN1_45S_50ntFlanks@11672.11706.35\|CAGTGCTCTGAATGTCAAAGTGAAGAAATTCAATG | >hg38_21_+_8433172_8446622_RNA45SN1_45S_50ntFlanks | 11672 | 11706 | 35 | CAGTGCTCTGAATGTCAAAGTGAAGAAATTCAATG | 895.5 | 2.5 | 0.8 | 3.2 | 1.4E-03 | 1.8E-03 | MB231/MB468 |
| >hg38_21_+_8433172_8446622_RNA45SN1_45S_50ntFlanks@11785.11805.21\|TGAATGGATGAACGAGATTCC | >hg38_21_+_8433172_8446622_RNA45SN1_45S_50ntFlanks | 11785 | 11805 | 21 | TGAATGGATGAACGAGATTCC | 931.3 | -2.4 | 0.8 | -3.2 | 1.4E-03 | 1.8E-03 | MB231/MB468 |
| >hg38_MT_+_598_1651_RNR1_12S_50ntFlanks@719.744.26\|TAAAGACGTTAGGTCAAGGTGTAGCC | >hg38_MT_+_598_1651_RNR1_12S_50ntFlanks | 719 | 744 | 26 | TAAAGACGTTAGGTCAAGGTGTAGCC | 797.4 | -3.3 | 0.9 | -3.7 | 2.2E-04 | 1.8E-03 | BT20/MB231 |
| >hg38_21_+_8433172_8446622_RNA45SN1_45S_50ntFlanks@10410.10442.33\|TCAGTCGGTCCTGAGAGATGGGCGAGCGCCGTT | >hg38_21_+_8433172_8446622_RNA45SN1_45S_50ntFlanks | 10410 | 10442 | 33 | TCAGTCGGTCCTGAGAGATGGGCGAGCGCCGTT | 215.4 | -1.8 | 0.6 | -3.2 | 1.4E-03 | 1.8E-03 | MB231/MB468 |
| >hg38_21_+_8433172_8446622_RNA45SN1_45S_50ntFlanks@12241.12272.32\|CAGAAGGGCAAAAGCTCGCTTGATCTTGATTT | >hg38_21_+_8433172_8446622_RNA45SN1_45S_50ntFlanks | 12241 | 12272 | 32 | CAGAAGGGCAAAAGCTCGCTTGATCTTGATTT | 311.5 | 2.2 | 0.7 | 3.2 | 1.4E-03 | 1.8E-03 | MB231/MB468 |
| >hg38_21_+_8433172_8446622_RNA45SN1_45S_50ntFlanks@12416.12454.39\|TTCGATGTCGGCTCTTCCTATCATTGTGAAGCAGAATTC | >hg38_21_+_8433172_8446622_RNA45SN1_45S_50ntFlanks | 12416 | 12454 | 39 | TTCGATGTCGGCTCTTCCTATCATTGTGAAGCAGAATTC | 442.5 | -2.9 | 0.8 | -3.7 | 2.3E-04 | 1.8E-03 | BT20/MB231 |
| >MI0000542\|hsa-mir-320a&WithFlank&8\|-\|22244960\|22245043@48.70.23(+1G)&[MIMAT0000510&hsa-miR-320a-3p&offsets\|0\|+1(+1G);m-60&8\|-\|22244975\|22244996&offsets\|0\|+1(+1G)]\|AAAAGCTGGGTTGAGAGGGCGAAG | >MI0000542\|hsa-mir-320a&WithFlank&8\|-\|22244960\|22245043 | 48 | 70 | 23(+1G) | AAAAGCTGGGTTGAGAGGGCGAAG | 848.6 | 2.9 | 0.8 | 3.7 | 2.3E-04 | 1.8E-03 | BT20/MB231 |
| >MI0000746\|hsa-mir-99b&WithFlank&19\|+\|51692606\|51692687@13.31.19(+1A)&[MIMAT0000689&hsa-miR-99b-5p&offsets\|0\|-3(+1A);m-10&19\|+\|51692618\|51692639&offsets\|0\|-3(+1A)]\|CACCCGTAGAACCGACCTTA | >MI0000746\|hsa-mir-99b&WithFlank&19\|+\|51692606\|51692687 | 13 | 31 | 19(+1A) | CACCCGTAGAACCGACCTTA | 284.3 | 2.9 | 0.8 | 3.7 | 2.3E-04 | 1.8E-03 | BT20/MB231 |
| >MI0033423\|hsa-mir-10399&WithFlank&7\|-\|139044094\|139044163@43.63.21(+1U)&[MIMAT0041630&hsa-miR-10399-3p&offsets\|-1\|-1(+1U);TJU_CMC_MD2.ID03061.3p-miR&offsets\|+3\|+2(+1U);m-2657&7\|-\|139044103\|139044124&offsets\|+3\|+2(+1U)]\|TCTCTCGGACAAGCTGTAGGTT | >MI0033423\|hsa-mir-10399&WithFlank&7\|-\|139044094\|139044163 | 43 | 63 | 21(+1U) | TCTCTCGGACAAGCTGTAGGTT | 252.9 | -2.7 | 0.7 | -3.7 | 2.3E-04 | 1.8E-03 | BT20/MB231 |
| >hg38_21_+_8433172_8446622_RNA45SN1_45S_50ntFlanks@11891.11909.19\|TTGACTCTAGTCTGGCACG | >hg38_21_+_8433172_8446622_RNA45SN1_45S_50ntFlanks | 11891 | 11909 | 19 | TTGACTCTAGTCTGGCACG | 129.1 | -2.9 | 0.9 | -3.2 | 1.4E-03 | 1.8E-03 | MB231/MB468 |
| >hg38_21_+_8433172_8446622_RNA45SN1_45S_50ntFlanks@6649.6696.48\|TACGACTCTTAGCGGTGGATCACTCGGCTCGTGCGTCGATGAAGAACG | >hg38_21_+_8433172_8446622_RNA45SN1_45S_50ntFlanks | 6649 | 6696 | 48 | TACGACTCTTAGCGGTGGATCACTCGGCTCGTGCGTCGATGAAGAACG | 369.7 | -2.6 | 0.7 | -3.7 | 2.3E-04 | 1.8E-03 | BT20/MB231 |
| >MI0000063\|hsa-let-7b&WithFlank&22\|+\|46113680\|46113774@12.31.20&[MIMAT0000063&hsa-let-7b-5p&offsets\|0\|-2;m-14&22\|+\|46113691\|46113712&offsets\|0\|-2]\|TGAGGTAGTAGGTTGTGTGG | >MI0000063\|hsa-let-7b&WithFlank&22\|+\|46113680\|46113774 | 12 | 31 | 20 | TGAGGTAGTAGGTTGTGTGG | 634.8 | 2.6 | 0.7 | 3.7 | 2.3E-04 | 1.8E-03 | BT20/MB231 |

<table>
<tr><td>>hg38_21_+_8433172_8446622_RNA45SN1_45S_50ntFlanks@11575.11611.37|ACTTAGAACTGGTGCGGACCAGGGGAATCCGACTGTT</td><td>>hg38_21_+_8433172_8446622_RNA45SN1_45S_50ntFlanks</td><td>11575</td><td>11611</td><td>37</td><td>ACTTAGAACTGGTGCGGACCAGGGGAATCCGACTGTT</td><td>309.1</td><td>-2.3</td><td>0.7</td><td>-3.2</td><td>1.5E-03</td><td>1.8E-03</td><td>BT20/MB468</td></tr>
<tr><td>>hg38_21_+_8433172_8446622_RNA45SN1_45S_50ntFlanks@3854.3877.24|ACTGTGGTAATTCTAGAGCTAATA</td><td>>hg38_21_+_8433172_8446622_RNA45SN1_45S_50ntFlanks</td><td>3854</td><td>3877</td><td>24</td><td>ACTGTGGTAATTCTAGAGCTAATA</td><td>68.0</td><td>-2.6</td><td>0.8</td><td>-3.2</td><td>1.4E-03</td><td>1.8E-03</td><td>MB231/MB468</td></tr>
<tr><td>>hg38_21_+_8433172_8446622_RNA45SN1_45S_50ntFlanks@11587.11611.25|TGCGGACCAGGGGAATCCGACTGTT</td><td>>hg38_21_+_8433172_8446622_RNA45SN1_45S_50ntFlanks</td><td>11587</td><td>11611</td><td>25</td><td>TGCGGACCAGGGGAATCCGACTGTT</td><td>557.6</td><td>-1.6</td><td>0.5</td><td>-3.2</td><td>1.5E-03</td><td>1.8E-03</td><td>BT20/MB468</td></tr>
<tr><td>>hg38_21_+_8433172_8446622_RNA45SN1_45S_50ntFlanks@6643.6683.41|ACCTCGTACGACTCTTAGCGGTGGATCACTCGGCTCGTGCG</td><td>>hg38_21_+_8433172_8446622_RNA45SN1_45S_50ntFlanks</td><td>6643</td><td>6683</td><td>41</td><td>ACCTCGTACGACTCTTAGCGGTGGATCACTCGGCTCGTGCG</td><td>282.6</td><td>-2.8</td><td>0.8</td><td>-3.7</td><td>2.3E-04</td><td>1.8E-03</td><td>BT20/MB231</td></tr>
<tr><td>>MI0000804|hsa-mir-328&WithFlank&16|-|67202315|67202401@54.75.22&[MIMAT0000752&hsa-miR-328-3p&offsets|0|0;m-160&16|-|67202327|67202348&offsets|0|0]|CTGGCCCTCTCTGCCCTTCCGT</td><td>>MI0000804|hsa-mir-328&WithFlank&16|-|67202315|67202401</td><td>54</td><td>75</td><td>22</td><td>CTGGCCCTCTCTGCCCTTCCGT</td><td>899.3</td><td>2.7</td><td>0.7</td><td>3.7</td><td>2.3E-04</td><td>1.8E-03</td><td>BT20/MB231</td></tr>
<tr><td>>hg38_21_+_8433172_8446622_RNA45SN1_45S_50ntFlanks@10843.10867.25|TAACTTCGGGATAAGGATTGGCTCT</td><td>>hg38_21_+_8433172_8446622_RNA45SN1_45S_50ntFlanks</td><td>10843</td><td>10867</td><td>25</td><td>TAACTTCGGGATAAGGATTGGCTCT</td><td>214.2</td><td>-1.7</td><td>0.5</td><td>-3.2</td><td>1.4E-03</td><td>1.8E-03</td><td>MB231/MB468</td></tr>
<tr><td>>hg38_21_+_8433172_8446622_RNA45SN1_45S_50ntFlanks@12418.12436.19|CGATGTCGGCTCTTCCTAT</td><td>>hg38_21_+_8433172_8446622_RNA45SN1_45S_50ntFlanks</td><td>12418</td><td>12436</td><td>19</td><td>CGATGTCGGCTCTTCCTAT</td><td>12954.3</td><td>-3.8</td><td>1.2</td><td>-3.2</td><td>1.5E-03</td><td>1.8E-03</td><td>BT20/MB468</td></tr>
<tr><td>>hg38_21_+_8433172_8446622_RNA45SN1_45S_50ntFlanks@11594.11612.19|CAGGGGAATCCGACTGTTT</td><td>>hg38_21_+_8433172_8446622_RNA45SN1_45S_50ntFlanks</td><td>11594</td><td>11612</td><td>19</td><td>CAGGGGAATCCGACTGTTT</td><td>693.4</td><td>-1.9</td><td>0.6</td><td>-3.2</td><td>1.4E-03</td><td>1.8E-03</td><td>MB231/MB468</td></tr>
<tr><td>>hg38_21_+_8433172_8446622_RNA45SN1_45S_50ntFlanks@12907.12932.26|CACCGCACGTTCGTGGGGAACCTGGC</td><td>>hg38_21_+_8433172_8446622_RNA45SN1_45S_50ntFlanks</td><td>12907</td><td>12932</td><td>26</td><td>CACCGCACGTTCGTGGGGAACCTGGC</td><td>123.6</td><td>-2.5</td><td>0.8</td><td>-3.2</td><td>1.5E-03</td><td>1.8E-03</td><td>BT20/MB468</td></tr>
<tr><td>>hg38_21_+_8433172_8446622_RNA45SN1_45S_50ntFlanks@12479.12496.18|CTAATAGGGAACGTGAGC</td><td>>hg38_21_+_8433172_8446622_RNA45SN1_45S_50ntFlanks</td><td>12479</td><td>12496</td><td>18</td><td>CTAATAGGGAACGTGAGC</td><td>1468.4</td><td>2.0</td><td>0.6</td><td>3.2</td><td>1.4E-03</td><td>1.8E-03</td><td>MB231/MB468</td></tr>
<tr><td>>hg38_1_-_228634819_228635039_RNA5S12_5S_50ntFlanks@122.147.26|TTAGTACTTGGATGGGAGACCGCCTG</td><td>>hg38_1_-_228634819_228635039_RNA5S12_5S_50ntFlanks</td><td>122</td><td>147</td><td>26</td><td>TTAGTACTTGGATGGGAGACCGCCTG</td><td>118.4</td><td>-2.7</td><td>0.9</td><td>-3.2</td><td>1.4E-03</td><td>1.8E-03</td><td>MB231/MB468</td></tr>
<tr><td>>hg38_21_+_8433172_8446622_RNA45SN1_45S_50ntFlanks@9178.9197.20|CGCCGTCGGGCCCGGGGGAG</td><td>>hg38_21_+_8433172_8446622_RNA45SN1_45S_50ntFlanks</td><td>9178</td><td>9197</td><td>20</td><td>CGCCGTCGGGCCCGGGGGAG</td><td>104.2</td><td>-3.5</td><td>1.1</td><td>-3.2</td><td>1.6E-03</td><td>1.9E-03</td><td>BT20/MB468</td></tr>
<tr><td>>am_trna119_LysCTT_1_-_145395522_145395594@16.33.18,trna11_LysCTT_5_-_180648979_180649051@16.33.18,trna128_LysCTT_1_-_55423542_55423614@16.33.18,trna13_LysCTT_6_+_26556774_26556846@16.33.18,trna6_LysCTT_19_-_52425396_52425468@16.33.18,trna7_LysCTT_16_+_3225692_3225764@16.33.18,trna9_LysCTT_5_+_180634755_180634827@16.33.18|TCGGTAGAGCATGAGACT</td><td>>am_trna119_LysCTT_1_-_145395522_145395594</td><td>16</td><td>33</td><td>18</td><td>TCGGTAGAGCATGAGACT</td><td>416.0</td><td>-3.8</td><td>1.0</td><td>-3.7</td><td>2.4E-04</td><td>1.9E-03</td><td>BT20/MB231</td></tr>
<tr><td>>MI0000301|hsa-mir-224&WithFlank&X|-|151958572|151958664@14.34.21(+1G)&[MIMAT0000281&hsa-miR-224-5p&offsets|+1|-3(+1G);m-238&X|-|151958630|151958651&offsets|0|-1(+1G)]|CAAGTCACTAGTGGTTCCGTTG</td><td>>MI0000301|hsa-mir-224&WithFlank&X|-|151958572|151958664</td><td>14</td><td>34</td><td>21(+1G)</td><td>CAAGTCACTAGTGGTTCCGTTG</td><td>418.4</td><td>3.3</td><td>0.9</td><td>3.7</td><td>2.4E-04</td><td>1.9E-03</td><td>BT20/MB231</td></tr>
<tr><td>>MI0000301|hsa-mir-224&WithFlank&X|-|151958572|151958664@14.36.23&[MIMAT0000281&hsa-miR-224-5p&offsets|+1|-1;m-238&X|-|151958630|151958651&offsets|0|+1]|CAAGTCACTAGTGGTTCCGTTTA</td><td>>MI0000301|hsa-mir-224&WithFlank&X|-|151958572|151958664</td><td>14</td><td>36</td><td>23</td><td>CAAGTCACTAGTGGTTCCGTTTA</td><td>2179.3</td><td>2.5</td><td>0.7</td><td>3.7</td><td>2.4E-04</td><td>1.9E-03</td><td>BT20/MB231</td></tr>
<tr><td>>hg38_21_+_8433172_8446622_RNA45SN1_45S_50ntFlanks@11784.11805.22|ATGAATGGATGAACGAGATTCC</td><td>>hg38_21_+_8433172_8446622_RNA45SN1_45S_50ntFlanks</td><td>11784</td><td>11805</td><td>22</td><td>ATGAATGGATGAACGAGATTCC</td><td>675.0</td><td>-2.0</td><td>0.6</td><td>-3.2</td><td>1.5E-03</td><td>1.9E-03</td><td>MB231/MB468</td></tr>
<tr><td>>hg38_21_+_8433172_8446622_RNA45SN1_45S_50ntFlanks@10473.10490.18|CTCGGCCGATCGAAAGGG</td><td>>hg38_21_+_8433172_8446622_RNA45SN1_45S_50ntFlanks</td><td>10473</td><td>10490</td><td>18</td><td>CTCGGCCGATCGAAAGGG</td><td>113.1</td><td>-2.1</td><td>0.7</td><td>-3.2</td><td>1.5E-03</td><td>1.9E-03</td><td>MB231/MB468</td></tr>
<tr><td>>hg38_21_+_8433172_8446622_RNA45SN1_45S_50ntFlanks@4735.4752.18|ACGAAAGTCGGAGGTTCG</td><td>>hg38_21_+_8433172_8446622_RNA45SN1_45S_50ntFlanks</td><td>4735</td><td>4752</td><td>18</td><td>ACGAAAGTCGGAGGTTCG</td><td>175.8</td><td>-1.8</td><td>0.6</td><td>-3.2</td><td>1.6E-03</td><td>1.9E-03</td><td>BT20/MB468</td></tr>
<tr><td>>hg38_21_+_8433172_8446622_RNA45SN1_45S_50ntFlanks@12419.12458.40|GATGTCGGCTCTTCCTATCATTGTGAAGCAGAATTCACCA</td><td>>hg38_21_+_8433172_8446622_RNA45SN1_45S_50ntFlanks</td><td>12419</td><td>12458</td><td>40</td><td>GATGTCGGCTCTTCCTATCATTGTGAAGCAGAATTCACCA</td><td>390.7</td><td>-2.7</td><td>0.7</td><td>-3.7</td><td>2.4E-04</td><td>1.9E-03</td><td>BT20/MB231</td></tr>
<tr><td>>hg38_21_+_8433172_8446622_RNA45SN1_45S_50ntFlanks@11561.11589.29|GCGCCTAGCAGCCGACTTAGAACTGGTGC</td><td>>hg38_21_+_8433172_8446622_RNA45SN1_45S_50ntFlanks</td><td>11561</td><td>11589</td><td>29</td><td>GCGCCTAGCAGCCGACTTAGAACTGGTGC</td><td>172.0</td><td>-1.6</td><td>0.5</td><td>-3.2</td><td>1.5E-03</td><td>1.9E-03</td><td>MB231/MB468</td></tr>
<tr><td>>hg38_21_+_8433172_8446622_RNA45SN1_45S_50ntFlanks@6698.6717.20|AGCTAGCTGCGAGAATTAAT</td><td>>hg38_21_+_8433172_8446622_RNA45SN1_45S_50ntFlanks</td><td>6698</td><td>6717</td><td>20</td><td>AGCTAGCTGCGAGAATTAAT</td><td>166.4</td><td>-1.6</td><td>0.5</td><td>-3.2</td><td>1.5E-03</td><td>1.9E-03</td><td>MB231/MB468</td></tr>
<tr><td>>MI0000298|hsa-mir-221&WithFlank&X|-|45746151|45746272@71.93.23&[MIMAT0000278&hsa-miR-221-3p&offsets|0|0;m-91&X|-|45746181|45746202&offsets|0|+1]|AGCTACATTGTCTGCTGGGTTTC</td><td>>MI0000298|hsa-mir-221&WithFlank&X|-|45746151|45746272</td><td>71</td><td>93</td><td>23</td><td>AGCTACATTGTCTGCTGGGTTTC</td><td>41039.8</td><td>-2.4</td><td>0.6</td><td>-3.7</td><td>2.5E-04</td><td>1.9E-03</td><td>BT20/MB231</td></tr>
<tr><td>>trnaMT_SerGCT_MT_+_12207_12265@8.31.24|CTCACAAGAACTGCTAACTCATGC</td><td>>trnaMT_SerGCT_MT_+_12207_12265</td><td>8</td><td>31</td><td>24</td><td>CTCACAAGAACTGCTAACTCATGC</td><td>91.7</td><td>-2.9</td><td>0.9</td><td>-3.2</td><td>1.6E-03</td><td>1.9E-03</td><td>BT20/MB468</td></tr>
<tr><td>>hg38_21_+_8433172_8446622_RNA45SN1_45S_50ntFlanks@12475.12499.25|CCCACTAATAGGGAACGTGAGCTGG</td><td>>hg38_21_+_8433172_8446622_RNA45SN1_45S_50ntFlanks</td><td>12475</td><td>12499</td><td>25</td><td>CCCACTAATAGGGAACGTGAGCTGG</td><td>121.3</td><td>-2.2</td><td>0.7</td><td>-3.2</td><td>1.6E-03</td><td>1.9E-03</td><td>BT20/MB468</td></tr>
<tr><td>>hg38_21_+_8433172_8446622_RNA45SN1_45S_50ntFlanks@7977.8017.41|CGACCTCAGATCAGACGTGGCGACCCGCTGAATTTAAGCAT</td><td>>hg38_21_+_8433172_8446622_RNA45SN1_45S_50ntFlanks</td><td>7977</td><td>8017</td><td>41</td><td>CGACCTCAGATCAGACGTGGCGACCCGCTGAATTTAAGCAT</td><td>375.7</td><td>-3.5</td><td>1.0</td><td>-3.7</td><td>2.5E-04</td><td>1.9E-03</td><td>BT20/MB231</td></tr>
<tr><td>>MI0000102|hsa-mir-100&WithFlank&11|-|122152223|122152314@19.36.18&[MIMAT0000098&hsa-miR-100-5p&offsets|0|-4;m-27&11|-|122152276|122152296&offsets|0|-3]|AACCCGTAGATCCGAACT</td><td>>MI0000102|hsa-mir-100&WithFlank&11|-|122152223|122152314</td><td>19</td><td>36</td><td>18</td><td>AACCCGTAGATCCGAACT</td><td>377.2</td><td>-3.8</td><td>1.0</td><td>-3.7</td><td>2.5E-04</td><td>1.9E-03</td><td>BT20/MB231</td></tr>
<tr><td>>hg38_21_+_8433172_8446622_RNA45SN1_45S_50ntFlanks@3840.3862.23|CTCCTACTTGGATAACTGTGGTA</td><td>>hg38_21_+_8433172_8446622_RNA45SN1_45S_50ntFlanks</td><td>3840</td><td>3862</td><td>23</td><td>CTCCTACTTGGATAACTGTGGTA</td><td>334.1</td><td>3.0</td><td>0.8</td><td>3.7</td><td>2.5E-04</td><td>1.9E-03</td><td>BT20/MB231</td></tr>
<tr><td>>trna10_LysCTT_16_+_3241501_3241573@3.35.33,trna13_LysCTT_14_-_58706613_58706685@3.35.33,trna2_LysCTT_15_+_79152904_79152976@3.35.33|CCGGCTAGCTCAGTCGGTAGAGCATGGGACTCT</td><td>>trna10_LysCTT_16_+_3241501_3241573</td><td>3</td><td>35</td><td>33</td><td>CCGGCTAGCTCAGTCGGTAGAGCATGGGACTCT</td><td>403.8</td><td>-2.6</td><td>0.7</td><td>-3.7</td><td>2.5E-04</td><td>1.9E-03</td><td>BT20/MB231</td></tr>
<tr><td>>hg38_21_+_8433172_8446622_RNA45SN1_45S_50ntFlanks@4592.4616.25|TTTGTTGGTTTTCGGAACTGAGGCC</td><td>>hg38_21_+_8433172_8446622_RNA45SN1_45S_50ntFlanks</td><td>4592</td><td>4616</td><td>25</td><td>TTTGTTGGTTTTCGGAACTGAGGCC</td><td>123.0</td><td>-1.6</td><td>0.5</td><td>-3.2</td><td>1.5E-03</td><td>1.9E-03</td><td>MB231/MB468</td></tr>
<tr><td>>hg38_21_+_8433172_8446622_RNA45SN1_45S_50ntFlanks@3843.3871.29|CTACTTGGATAACTGTGGTAATTCTAGAG</td><td>>hg38_21_+_8433172_8446622_RNA45SN1_45S_50ntFlanks</td><td>3843</td><td>3871</td><td>29</td><td>CTACTTGGATAACTGTGGTAATTCTAGAG</td><td>317.2</td><td>-2.1</td><td>0.7</td><td>-3.2</td><td>1.5E-03</td><td>1.9E-03</td><td>MB231/MB468</td></tr>
<tr><td>>trna119_LysCTT_1_-_145395522_145395594@1.31.31,trna11_LysCTT_5_-_180648979_180649051@1.31.31,trna13_LysCTT_6_+_26556774_26556846@1.31.31,trna32_LysCTT_16_-_3207406_3207478@1.31.31,trna7_LysCTT_16_+_3225692_3225764@1.31.31,trna9_LysCTT_5_+_180634755_180634827@1.31.31|GCCCGGCTAGCTCAGTCGGTAGAGCATGAGA</td><td>>trna119_LysCTT_1_-_145395522_145395594</td><td>1</td><td>31</td><td>31</td><td>GCCCGGCTAGCTCAGTCGGTAGAGCATGAGA</td><td>389.4</td><td>-2.9</td><td>0.8</td><td>-3.7</td><td>2.6E-04</td><td>2.0E-03</td><td>BT20/MB231</td></tr>
<tr><td>>MI0000737|hsa-mir-200a&WithFlank&1|+|1167857|1167958@60.81.22(+1C)&[MIMAT0000682&hsa-miR-200a-3p&offsets|0|0(+1C);m-142&1|+|1167916|1167938&offsets|0|-1(+1C)]|TAACACTGTCTGGTAACGATGTC</td><td>>MI0000737|hsa-mir-200a&WithFlank&1|+|1167857|1167958</td><td>60</td><td>81</td><td>22(+1C)</td><td>TAACACTGTCTGGTAACGATGTC</td><td>195.5</td><td>-2.6</td><td>0.8</td><td>-3.1</td><td>1.6E-03</td><td>2.0E-03</td><td>BT20/MB468</td></tr>
<tr><td>>hg38_21_+_8433172_8446622_RNA45SN1_45S_50ntFlanks@12154.12182.29|GGGAGTTTGACTGGGGCGGTACACCTGTC</td><td>>hg38_21_+_8433172_8446622_RNA45SN1_45S_50ntFlanks</td><td>12154</td><td>12182</td><td>29</td><td>GGGAGTTTGACTGGGGCGGTACACCTGTC</td><td>104.5</td><td>-1.9</td><td>0.6</td><td>-3.2</td><td>1.5E-03</td><td>2.0E-03</td><td>MB231/MB468</td></tr>
<tr><td>>hg38_21_+_8433172_8446622_RNA45SN1_45S_50ntFlanks@12971.12988.18|TTTCGTACGTAGCAGAGC</td><td>>hg38_21_+_8433172_8446622_RNA45SN1_45S_50ntFlanks</td><td>12971</td><td>12988</td><td>18</td><td>TTTCGTACGTAGCAGAGC</td><td>113.5</td><td>-1.3</td><td>0.4</td><td>-3.2</td><td>1.5E-03</td><td>2.0E-03</td><td>MB231/MB468</td></tr>
<tr><td>>hg38_21_+_8433172_8446622_RNA45SN1_45S_50ntFlanks@12569.12587.19|CAGTACGAGAGGAACCGCA</td><td>>hg38_21_+_8433172_8446622_RNA45SN1_45S_50ntFlanks</td><td>12569</td><td>12587</td><td>19</td><td>CAGTACGAGAGGAACCGCA</td><td>133.6</td><td>-1.9</td><td>0.6</td><td>-3.2</td><td>1.5E-03</td><td>2.0E-03</td><td>MB231/MB468</td></tr>
<tr><td>>hg38_21_+_8433172_8446622_RNA45SN1_45S_50ntFlanks@10402.10422.21|AACATGGGTCAGTCGGTCCTG</td><td>>hg38_21_+_8433172_8446622_RNA45SN1_45S_50ntFlanks</td><td>10402</td><td>10422</td><td>21</td><td>AACATGGGTCAGTCGGTCCTG</td><td>278.0</td><td>-2.0</td><td>0.6</td><td>-3.2</td><td>1.5E-03</td><td>2.0E-03</td><td>MB231/MB468</td></tr>
<tr><td>>hg38_21_+_8433172_8446622_RNA45SN1_45S_50ntFlanks@12077.12095.19|GGGGCTCTCGCTTCTGGCG</td><td>>hg38_21_+_8433172_8446622_RNA45SN1_45S_50ntFlanks</td><td>12077</td><td>12095</td><td>19</td><td>GGGGCTCTCGCTTCTGGCG</td><td>203.8</td><td>-1.6</td><td>0.5</td><td>-3.2</td><td>1.5E-03</td><td>2.0E-03</td><td>MB231/MB468</td></tr>
<tr><td>>MI0000542|hsa-mir-320a&WithFlank&8|-|22244960|22245043@49.69.21&[MIMAT0000510&hsa-miR-320a-3p&offsets|+1|0;m-60&8|-|22244975|22244996&offsets|+1|0]|AAAGCTGGGTTGAGAGGGCGA</td><td>>MI0000542|hsa-mir-320a&WithFlank&8|-|22244960|22245043</td><td>49</td><td>69</td><td>21</td><td>AAAGCTGGGTTGAGAGGGCGA</td><td>880.8</td><td>2.8</td><td>0.8</td><td>3.7</td><td>2.6E-04</td><td>2.0E-03</td><td>BT20/MB231</td></tr>
<tr><td>>hg38_21_+_8433172_8446622_RNA45SN1_45S_50ntFlanks@12006.12032.27|GGCCGCCGGTGAAATACCACTACTCTG</td><td>>hg38_21_+_8433172_8446622_RNA45SN1_45S_50ntFlanks</td><td>12006</td><td>12032</td><td>27</td><td>GGCCGCCGGTGAAATACCACTACTCTG</td><td>198.4</td><td>-2.0</td><td>0.6</td><td>-3.2</td><td>1.6E-03</td><td>2.0E-03</td><td>MB231/MB468</td></tr>
<tr><td>>hg38_21_+_8433172_8446622_RNA45SN1_45S_50ntFlanks@7988.8018.31|CAGACGTGGCGACCCGCTGAATTTAAGCATA</td><td>>hg38_21_+_8433172_8446622_RNA45SN1_45S_50ntFlanks</td><td>7988</td><td>8018</td><td>31</td><td>CAGACGTGGCGACCCGCTGAATTTAAGCATA</td><td>220.8</td><td>1.5</td><td>0.5</td><td>3.2</td><td>1.6E-03</td><td>2.0E-03</td><td>MB231/MB468</td></tr>
<tr><td>>hg38_21_+_8433172_8446622_RNA45SN1_45S_50ntFlanks@11540.11584.45|CGCGGCGCCTCGCCTCGGCCGGCGCCTAGCAGCCGACTTAGAACT</td><td>>hg38_21_+_8433172_8446622_RNA45SN1_45S_50ntFlanks</td><td>11540</td><td>11584</td><td>45</td><td>CGCGGCGCCTCGCCTCGGCCGGCGCCTAGCAGCCGACTTAGAACT</td><td>1842.2</td><td>-3.0</td><td>0.8</td><td>-3.6</td><td>2.6E-04</td><td>2.0E-03</td><td>BT20/MB231</td></tr>
<tr><td>>hg38_21_+_8433172_8446622_RNA45SN1_45S_50ntFlanks@7983.8009.27|CAGATCAGACGTGGCGACCCGCTGAAT</td><td>>hg38_21_+_8433172_8446622_RNA45SN1_45S_50ntFlanks</td><td>7983</td><td>8009</td><td>27</td><td>CAGATCAGACGTGGCGACCCGCTGAAT</td><td>253.8</td><td>-1.6</td><td>0.5</td><td>-3.2</td><td>1.6E-03</td><td>2.0E-03</td><td>MB231/MB468</td></tr>
<tr><td>>hg38_21_+_8433172_8446622_RNA45SN1_45S_50ntFlanks@6657.6681.25|TTAGCGGTGGATCACTCGGCTCGTG</td><td>>hg38_21_+_8433172_8446622_RNA45SN1_45S_50ntFlanks</td><td>6657</td><td>6681</td><td>25</td><td>TTAGCGGTGGATCACTCGGCTCGTG</td><td>476.3</td><td>-1.7</td><td>0.6</td><td>-3.2</td><td>1.6E-03</td><td>2.0E-03</td><td>MB231/MB468</td></tr>
<tr><td>>MI0000477|hsa-mir-146a&WithFlank&5|+|160485346|160485456@27.47.21(+1C)&[MIMAT0000449&hsa-miR-146a-5p&offsets|0|-1(+1C);m-129&5|+|160485372|160485393&offsets|0|-1(+1C)]|TGAGAACTGAATTCCATGGGTC</td><td>>MI0000477|hsa-mir-146a&WithFlank&5|+|160485346|160485456</td><td>27</td><td>47</td><td>21(+1C)</td><td>TGAGAACTGAATTCCATGGGTC</td><td>1714.0</td><td>-8.6</td><td>2.4</td><td>-3.6</td><td>2.7E-04</td><td>2.0E-03</td><td>BT20/MB231</td></tr>
<tr><td>>trnaMT_MetCAT_MT_+_4402_4469@36.71.36|CCCGAAAATGTTGGTTATACCCTTCCCGTACTACCA</td><td>>trnaMT_MetCAT_MT_+_4402_4469</td><td>36</td><td>71</td><td>36</td><td>CCCGAAAATGTTGGTTATACCCTTCCCGTACTACCA</td><td>275.4</td><td>-2.7</td><td>0.7</td><td>-3.6</td><td>2.7E-04</td><td>2.0E-03</td><td>BT20/MB231</td></tr>
</table>

<table>
<tr><th></th><th></th><th></th><th></th><th></th><th></th><th></th><th></th><th></th><th></th><th></th><th></th><th></th></tr>
<tr><td>>hg38_21_+_8433172_8446622_RNA45SN1_45S_50ntFlanks@6641.6683.43|CGACCTCGTACGACTCTTAGCGGTGGATCACTCGGCTCGTGCG</td><td>>hg38_21_+_8433172_8446622_RNA45SN1_45S_50ntFlanks</td><td>6641</td><td>6683</td><td>43</td><td>CGACCTCGTACGACTCTTAGCGGTGGATCACTCGGCTCGTGCG</td><td>167.0</td><td>-3.8</td><td>1.0</td><td>-3.6</td><td>2.7E-04</td><td>2.0E-03</td><td>BT20/MB231</td></tr>
<tr><td>>hg38_21_+_8433172_8446622_RNA45SN1_45S_50ntFlanks@9021.9039.19|CGCCGGCGTCTCCTCGTGG</td><td>>hg38_21_+_8433172_8446622_RNA45SN1_45S_50ntFlanks</td><td>9021</td><td>9039</td><td>19</td><td>CGCCGGCGTCTCCTCGTGG</td><td>97.9</td><td>-3.2</td><td>1.0</td><td>-3.1</td><td>1.7E-03</td><td>2.0E-03</td><td>BT20/MB468</td></tr>
<tr><td>>hg38_21_+_8433172_8446622_RNA45SN1_45S_50ntFlanks@11581.11613.33|AACTGGTGCGGACCAGGGGAATCCGACTGTTTA</td><td>>hg38_21_+_8433172_8446622_RNA45SN1_45S_50ntFlanks</td><td>11581</td><td>11613</td><td>33</td><td>AACTGGTGCGGACCAGGGGAATCCGACTGTTTA</td><td>282.1</td><td>-1.4</td><td>0.4</td><td>-3.1</td><td>1.7E-03</td><td>2.0E-03</td><td>BT20/MB468</td></tr>
<tr><td>>hg38_21_+_8433172_8446622_RNA45SN1_45S_50ntFlanks@11203.11232.30|CGGCGGCGGCGACTCTGGACGCGAGCCGGG</td><td>>hg38_21_+_8433172_8446622_RNA45SN1_45S_50ntFlanks</td><td>11203</td><td>11232</td><td>30</td><td>CGGCGGCGGCGACTCTGGACGCGAGCCGGG</td><td>162.0</td><td>-2.0</td><td>0.6</td><td>-3.2</td><td>1.6E-03</td><td>2.1E-03</td><td>MB231/MB468</td></tr>
<tr><td>>hg38_21_+_8433172_8446622_RNA45SN1_45S_50ntFlanks@11568.11585.18|GCAGCCGACTTAGAACTG</td><td>>hg38_21_+_8433172_8446622_RNA45SN1_45S_50ntFlanks</td><td>11568</td><td>11585</td><td>18</td><td>GCAGCCGACTTAGAACTG</td><td>2023.0</td><td>-1.3</td><td>0.4</td><td>-3.1</td><td>1.7E-03</td><td>2.1E-03</td><td>BT20/MB468</td></tr>
<tr><td>>hg38_21_+_8433172_8446622_RNA45SN1_45S_50ntFlanks@12360.12384.25|CAGGGATAACTGGCTTGTGGCGGCC</td><td>>hg38_21_+_8433172_8446622_RNA45SN1_45S_50ntFlanks</td><td>12360</td><td>12384</td><td>25</td><td>CAGGGATAACTGGCTTGTGGCGGCC</td><td>142.8</td><td>-1.5</td><td>0.5</td><td>-3.2</td><td>1.6E-03</td><td>2.1E-03</td><td>MB231/MB468</td></tr>
<tr><td>>hg38_21_+_8433172_8446622_RNA45SN1_45S_50ntFlanks@3705.3728.24|TACCTGGTTGATCCTGCCAGTAGC</td><td>>hg38_21_+_8433172_8446622_RNA45SN1_45S_50ntFlanks</td><td>3705</td><td>3728</td><td>24</td><td>TACCTGGTTGATCCTGCCAGTAGC</td><td>2201.7</td><td>3.0</td><td>0.8</td><td>3.6</td><td>2.8E-04</td><td>2.1E-03</td><td>BT20/MB231</td></tr>
<tr><td>>MI0001446|hsa-mir-424&WithFlank&X|-|134546608|134546717@17.37.21&[MIMAT0001341&hsa-miR-424-5p&offsets|0|-1;m-152&X|-|134546681|134546701&offsets|0|0]|CAGCAGCAATTCATGTTTTGA</td><td>>MI0001446|hsa-mir-424&WithFlank&X|-|134546608|134546717</td><td>17</td><td>37</td><td>21</td><td>CAGCAGCAATTCATGTTTTGA</td><td>1159.4</td><td>3.7</td><td>1.0</td><td>3.6</td><td>2.7E-04</td><td>2.1E-03</td><td>BT20/MB231</td></tr>
<tr><td>>hg38_21_+_8433172_8446622_RNA45SN1_45S_50ntFlanks@12258.12284.27|GCTTGATCTTGATTTTCAGTACGAATA</td><td>>hg38_21_+_8433172_8446622_RNA45SN1_45S_50ntFlanks</td><td>12258</td><td>12284</td><td>27</td><td>GCTTGATCTTGATTTTCAGTACGAATA</td><td>146.8</td><td>-1.9</td><td>0.6</td><td>-3.1</td><td>1.6E-03</td><td>2.1E-03</td><td>MB231/MB468</td></tr>
<tr><td>>hg38_21_+_8433172_8446622_RNA45SN1_45S_50ntFlanks@10392.10418.27|ACAGCAGTTGAACATGGGTCAGTCGGT</td><td>>hg38_21_+_8433172_8446622_RNA45SN1_45S_50ntFlanks</td><td>10392</td><td>10418</td><td>27</td><td>ACAGCAGTTGAACATGGGTCAGTCGGT</td><td>146.1</td><td>-1.3</td><td>0.4</td><td>-3.1</td><td>1.7E-03</td><td>2.1E-03</td><td>MB231/MB468</td></tr>
<tr><td>>hg38_21_+_8433172_8446622_RNA45SN1_45S_50ntFlanks@11876.11900.25|AAGACCCTGTTGAGCTTGACTCTAG</td><td>>hg38_21_+_8433172_8446622_RNA45SN1_45S_50ntFlanks</td><td>11876</td><td>11900</td><td>25</td><td>AAGACCCTGTTGAGCTTGACTCTAG</td><td>366.5</td><td>-2.4</td><td>0.8</td><td>-3.1</td><td>1.8E-03</td><td>2.1E-03</td><td>BT20/MB468</td></tr>
<tr><td>>hg38_21_+_8433172_8446622_RNA45SN1_45S_50ntFlanks@11672.11702.31|CAGTGCTCTGAATGTCAAAGTGAAGAAATTC</td><td>>hg38_21_+_8433172_8446622_RNA45SN1_45S_50ntFlanks</td><td>11672</td><td>11702</td><td>31</td><td>CAGTGCTCTGAATGTCAAAGTGAAGAAATTC</td><td>748.4</td><td>-1.6</td><td>0.5</td><td>-3.1</td><td>1.8E-03</td><td>2.1E-03</td><td>BT20/MB468</td></tr>
<tr><td>>hg38_21_+_8433172_8446622_RNA45SN1_45S_50ntFlanks@11568.11592.25|GCAGCCGACTTAGAACTGGTGCGGA</td><td>>hg38_21_+_8433172_8446622_RNA45SN1_45S_50ntFlanks</td><td>11568</td><td>11592</td><td>25</td><td>GCAGCCGACTTAGAACTGGTGCGGA</td><td>299.6</td><td>-1.6</td><td>0.5</td><td>-3.1</td><td>1.8E-03</td><td>2.1E-03</td><td>BT20/MB468</td></tr>
<tr><td>>hg38_21_+_8433172_8446622_RNA45SN1_45S_50ntFlanks@10427.10456.30|ATGGGCGAGCGCCGTTCCGAAGGGACGGGC</td><td>>hg38_21_+_8433172_8446622_RNA45SN1_45S_50ntFlanks</td><td>10427</td><td>10456</td><td>30</td><td>ATGGGCGAGCGCCGTTCCGAAGGGACGGGC</td><td>607.4</td><td>-1.4</td><td>0.5</td><td>-3.1</td><td>1.7E-03</td><td>2.1E-03</td><td>MB231/MB468</td></tr>
<tr><td>>hg38_21_+_8433172_8446622_RNA45SN1_45S_50ntFlanks@8176.8196.21|CTTCTGATCGAGGCCCAGCCC</td><td>>hg38_21_+_8433172_8446622_RNA45SN1_45S_50ntFlanks</td><td>8176</td><td>8196</td><td>21</td><td>CTTCTGATCGAGGCCCAGCCC</td><td>447.9</td><td>1.8</td><td>0.6</td><td>3.1</td><td>1.7E-03</td><td>2.1E-03</td><td>MB231/MB468</td></tr>
<tr><td>>hg38_21_+_8433172_8446622_RNA45SN1_45S_50ntFlanks@4006.4023.18|ACCTCGGGCCGATCGCAC</td><td>>hg38_21_+_8433172_8446622_RNA45SN1_45S_50ntFlanks</td><td>4006</td><td>4023</td><td>18</td><td>ACCTCGGGCCGATCGCAC</td><td>450.2</td><td>1.9</td><td>0.6</td><td>3.1</td><td>1.7E-03</td><td>2.1E-03</td><td>MB231/MB468</td></tr>
<tr><td>>trna116_GluCTC_1_-_145399233_145399304@37.73.37,trna134_GluTTC_1_-_16861774_16861845@37.73.37,trna71_GluCTC_1_-_161439189_161439260@37.73.37,trna74_GluCTC_1_-_161431809_161431880@37.73.37,trna77_GluCTC_1_-_161424398_161424469@37.73.37,trna77_GluCTC_6_+_28949976_28950047@37.73.37,trna80_GluCTC_1_-_161417018_161417089@37.73.37,trna84_GluTTC_1_-_161391883_161391954@37.73.37,trna87_GluCTC_6_-_126101393_126101464@37.73.37|ACCGCCGCGGCCCGGGTTCGATTCCCGGTCAGGGAAC</td><td>>trna116_GluCTC_1_-_145399233_145399304</td><td>37</td><td>73</td><td>37</td><td>ACCGCCGCGGCCCGGGTTCGATTCCCGGTCAGGGAAC</td><td>548.9</td><td>-3.6</td><td>1.0</td><td>-3.6</td><td>2.9E-04</td><td>2.2E-03</td><td>BT20/MB231</td></tr>
<tr><td>>hg38_21_+_8433172_8446622_RNA45SN1_45S_50ntFlanks@6646.6675.30|TCGTACGACTCTTAGCGGTGGATCACTCGG</td><td>>hg38_21_+_8433172_8446622_RNA45SN1_45S_50ntFlanks</td><td>6646</td><td>6675</td><td>30</td><td>TCGTACGACTCTTAGCGGTGGATCACTCGG</td><td>1357.2</td><td>2.1</td><td>0.7</td><td>3.1</td><td>1.7E-03</td><td>2.2E-03</td><td>MB231/MB468</td></tr>
<tr><td>>hg38_21_+_8433172_8446622_RNA45SN1_45S_50ntFlanks@8176.8220.45|CTTCTGATCGAGGCCCAGCCCGTGGACGGTGTGAGGCCGGTAGCG</td><td>>hg38_21_+_8433172_8446622_RNA45SN1_45S_50ntFlanks</td><td>8176</td><td>8220</td><td>45</td><td>CTTCTGATCGAGGCCCAGCCCGTGGACGGTGTGAGGCCGGTAGCG</td><td>97.8</td><td>-1.6</td><td>0.5</td><td>-3.1</td><td>1.8E-03</td><td>2.2E-03</td><td>BT20/MB468</td></tr>
<tr><td>>MI0000239|hsa-mir-197&WithFlank&1|+|109598887|109598973@54.74.21&[MIMAT0000227&hsa-miR-197-3p&offsets|0|-1;m-99&1|+|109598940|109598961&offsets|0|-1]|TTCACCACCTTCTCCACCCAG</td><td>>MI0000239|hsa-mir-197&WithFlank&1|+|109598887|109598973</td><td>54</td><td>74</td><td>21</td><td>TTCACCACCTTCTCCACCCAG</td><td>310.7</td><td>2.7</td><td>0.8</td><td>3.6</td><td>2.9E-04</td><td>2.2E-03</td><td>BT20/MB231</td></tr>
<tr><td>>hg38_21_+_8433172_8446622_RNA45SN1_45S_50ntFlanks@5496.5516.21|GACGGTCGAACTTGACTATCT</td><td>>hg38_21_+_8433172_8446622_RNA45SN1_45S_50ntFlanks</td><td>5496</td><td>5516</td><td>21</td><td>GACGGTCGAACTTGACTATCT</td><td>354.6</td><td>4.0</td><td>1.1</td><td>3.6</td><td>2.9E-04</td><td>2.2E-03</td><td>BT20/MB231</td></tr>
<tr><td>>hg38_21_+_8433172_8446622_RNA45SN1_45S_50ntFlanks@12458.12475.18|AAGCGTTGGATTGTTCAC</td><td>>hg38_21_+_8433172_8446622_RNA45SN1_45S_50ntFlanks</td><td>12458</td><td>12475</td><td>18</td><td>AAGCGTTGGATTGTTCAC</td><td>444.4</td><td>-1.8</td><td>0.6</td><td>-3.1</td><td>1.7E-03</td><td>2.2E-03</td><td>MB231/MB468</td></tr>
<tr><td>>MI0000070|hsa-mir-16-1&WithFlank&13|-|50048967|50049067@20.40.21(+1A);MI0000115|hsa-mir-16-2&WithFlank&3|+|160404739|160404831@16.36.21(+1A)&[MIMAT0000069&hsa-miR-16-5p&offsets|0|-1(+1A);m-62&13|-|50049027|50049048&offsets|0|-1(+1A)];[MIMAT0000069_1&hsa-miR-16-5p&offsets|0|-1(+1A);m-61&3|+|160404754|160404775&offsets|0|-1(+1A)]|TAGCAGCACGTAAATATTGGCA</td><td>>MI0000070|hsa-mir-16-1&WithFlank&13|-|50048967|50049067</td><td>20</td><td>40</td><td>21(+1A)</td><td>TAGCAGCACGTAAATATTGGCA</td><td>5032.1</td><td>3.7</td><td>1.0</td><td>3.6</td><td>3.0E-04</td><td>2.2E-03</td><td>BT20/MB231</td></tr>
<tr><td>>hg38_21_+_8433172_8446622_RNA45SN1_45S_50ntFlanks@12444.12476.33|AAGCAGAATTCACCAAGCGTTGGATTGTTCACC</td><td>>hg38_21_+_8433172_8446622_RNA45SN1_45S_50ntFlanks</td><td>12444</td><td>12476</td><td>33</td><td>AAGCAGAATTCACCAAGCGTTGGATTGTTCACC</td><td>1103.9</td><td>2.1</td><td>0.7</td><td>3.1</td><td>1.8E-03</td><td>2.2E-03</td><td>MB231/MB468</td></tr>
<tr><td>>hg38_21_+_8433172_8446622_RNA45SN1_45S_50ntFlanks@11718.11751.34|AACGGCGGGAGTAACTATGACTCTCTTAAGGTAG</td><td>>hg38_21_+_8433172_8446622_RNA45SN1_45S_50ntFlanks</td><td>11718</td><td>11751</td><td>34</td><td>AACGGCGGGAGTAACTATGACTCTCTTAAGGTAG</td><td>703.6</td><td>-4.5</td><td>1.5</td><td>-3.1</td><td>1.9E-03</td><td>2.2E-03</td><td>BT20/MB468</td></tr>
<tr><td>>hg38_21_+_8433172_8446622_RNA45SN1_45S_50ntFlanks@12405.12444.40|CTTTTTGATCCTTCGATGTCGGCTCTTCCTATCATTGTGA</td><td>>hg38_21_+_8433172_8446622_RNA45SN1_45S_50ntFlanks</td><td>12405</td><td>12444</td><td>40</td><td>CTTTTTGATCCTTCGATGTCGGCTCTTCCTATCATTGTGA</td><td>256.7</td><td>-1.5</td><td>0.5</td><td>-3.1</td><td>1.8E-03</td><td>2.2E-03</td><td>MB231/MB468</td></tr>
<tr><td>>hg38_21_+_8433172_8446622_RNA45SN1_45S_50ntFlanks@11671.11698.28|CCAGTGCTCTGAATGTCAAAGTGAAGAA</td><td>>hg38_21_+_8433172_8446622_RNA45SN1_45S_50ntFlanks</td><td>11671</td><td>11698</td><td>28</td><td>CCAGTGCTCTGAATGTCAAAGTGAAGAA</td><td>124.2</td><td>-1.4</td><td>0.5</td><td>-3.1</td><td>1.8E-03</td><td>2.2E-03</td><td>MB231/MB468</td></tr>
<tr><td>>MI0000273|hsa-mir-183&WithFlank&7|-|129774899|129775020@33.53.21(+1G)&[MIMAT0000261&hsa-miR-183-5p&offsets|0|-1(+1G);m-41&7|-|129774966|129774987&offsets|-1|-2(+1G)]|TATGGCACTGGTAGAATTCACG</td><td>>MI0000273|hsa-mir-183&WithFlank&7|-|129774899|129775020</td><td>33</td><td>53</td><td>21(+1G)</td><td>TATGGCACTGGTAGAATTCACG</td><td>1408.5</td><td>3.2</td><td>0.9</td><td>3.6</td><td>3.0E-04</td><td>2.2E-03</td><td>BT20/MB231</td></tr>
<tr><td>>MI0000294|hsa-mir-218-1&WithFlank&4|+|20528269|20528390@31.52.22;MI0000295|hsa-mir-218-2&WithFlank&5|-|168768140|168768261@31.52.22&[MIMAT0000275&hsa-miR-218-5p&offsets|0|+1;m-185&4|+|20528299|20528319&offsets|0|+1];[MIMAT0000275_1&hsa-miR-218-5p&offsets|0|+1;m-187&5|-|168768211|168768231&offsets|0|+1]|TTGTGCTTGATCTAACCATGTG</td><td>>MI0000294|hsa-mir-218-1&WithFlank&4|+|20528269|20528390</td><td>31</td><td>52</td><td>22</td><td>TTGTGCTTGATCTAACCATGTG</td><td>255.3</td><td>-6.8</td><td>1.9</td><td>-3.6</td><td>3.0E-04</td><td>2.2E-03</td><td>BT20/MB231</td></tr>
<tr><td>>hg38_21_+_8433172_8446622_RNA45SN1_45S_50ntFlanks@3705.3736.32|TACCTGGTTGATCCTGCCAGTAGCATATGCTT</td><td>>hg38_21_+_8433172_8446622_RNA45SN1_45S_50ntFlanks</td><td>3705</td><td>3736</td><td>32</td><td>TACCTGGTTGATCCTGCCAGTAGCATATGCTT</td><td>1719.3</td><td>2.9</td><td>0.8</td><td>3.6</td><td>3.1E-04</td><td>2.2E-03</td><td>BT20/MB231</td></tr>
<tr><td>>trna20_GluTTC_2_-_131094701_131094772@1.32.32,trna5_GluTTC_13_-_41634874_41634945@1.32.32|TCCCATATGGTCTAGCGGTTAGGATTCCTGGT</td><td>>trna20_GluTTC_2_-_131094701_131094772</td><td>1</td><td>32</td><td>32</td><td>TCCCATATGGTCTAGCGGTTAGGATTCCTGGT</td><td>239.7</td><td>-2.8</td><td>0.8</td><td>-3.6</td><td>3.1E-04</td><td>2.2E-03</td><td>BT20/MB231</td></tr>
<tr><td>>hg38_21_+_8433172_8446622_RNA45SN1_45S_50ntFlanks@4367.4397.31|CAGTTAAAAAGCTCGTAGTTGGATCTTGGGA</td><td>>hg38_21_+_8433172_8446622_RNA45SN1_45S_50ntFlanks</td><td>4367</td><td>4397</td><td>31</td><td>CAGTTAAAAAGCTCGTAGTTGGATCTTGGGA</td><td>76.7</td><td>-2.5</td><td>0.8</td><td>-3.1</td><td>1.8E-03</td><td>2.2E-03</td><td>MB231/MB468</td></tr>
<tr><td>>hg38_21_+_8433172_8446622_RNA45SN1_45S_50ntFlanks@11207.11232.26|GGCGGCGACTCTGGACGCGAGCCGGG</td><td>>hg38_21_+_8433172_8446622_RNA45SN1_45S_50ntFlanks</td><td>11207</td><td>11232</td><td>26</td><td>GGCGGCGACTCTGGACGCGAGCCGGG</td><td>271.0</td><td>-5.2</td><td>1.7</td><td>-3.1</td><td>1.8E-03</td><td>2.3E-03</td><td>MB231/MB468</td></tr>
<tr><td>>hg38_21_+_8433172_8446622_RNA45SN1_45S_50ntFlanks@11567.11587.21|AGCAGCCGACTTAGAACTGGT</td><td>>hg38_21_+_8433172_8446622_RNA45SN1_45S_50ntFlanks</td><td>11567</td><td>11587</td><td>21</td><td>AGCAGCCGACTTAGAACTGGT</td><td>299.6</td><td>1.3</td><td>0.4</td><td>3.1</td><td>1.9E-03</td><td>2.3E-03</td><td>BT20/MB468</td></tr>
<tr><td>>hg38_21_+_8433172_8446622_RNA45SN1_45S_50ntFlanks@11580.11607.28|GAACTGGTGCGGACCAGGGGAATCCGAC</td><td>>hg38_21_+_8433172_8446622_RNA45SN1_45S_50ntFlanks</td><td>11580</td><td>11607</td><td>28</td><td>GAACTGGTGCGGACCAGGGGAATCCGAC</td><td>341.9</td><td>-1.7</td><td>0.5</td><td>-3.1</td><td>1.9E-03</td><td>2.3E-03</td><td>BT20/MB468</td></tr>
<tr><td>>hg38_21_+_8433172_8446622_RNA45SN1_45S_50ntFlanks@12396.12416.21|GCGACGTCGCTTTTTGATCCT</td><td>>hg38_21_+_8433172_8446622_RNA45SN1_45S_50ntFlanks</td><td>12396</td><td>12416</td><td>21</td><td>GCGACGTCGCTTTTTGATCCT</td><td>211.4</td><td>-1.9</td><td>0.6</td><td>-3.1</td><td>1.9E-03</td><td>2.3E-03</td><td>BT20/MB468</td></tr>
<tr><td>>hg38_21_+_8433172_8446622_RNA45SN1_45S_50ntFlanks@9306.9323.18|CAAGGAGTCTAACACGTG</td><td>>hg38_21_+_8433172_8446622_RNA45SN1_45S_50ntFlanks</td><td>9306</td><td>9323</td><td>18</td><td>CAAGGAGTCTAACACGTG</td><td>144.3</td><td>-3.6</td><td>1.2</td><td>-3.1</td><td>1.9E-03</td><td>2.3E-03</td><td>BT20/MB468</td></tr>
<tr><td>>MI0000342|hsa-mir-200b&WithFlank&1|+|1167098|1167204@64.85.22(+1U)&[MIMAT0000318&hsa-miR-200b-3p&offsets|+1|+1(+1U);m-93&1|+|1167160|1167182&offsets|+1|0(+1U)]|AATACTGCCTGGTAATGATGACT</td><td>>MI0000342|hsa-mir-200b&WithFlank&1|+|1167098|1167204</td><td>64</td><td>85</td><td>22(+1U)</td><td>AATACTGCCTGGTAATGATGACT</td><td>256.5</td><td>-2.4</td><td>0.8</td><td>-3.1</td><td>1.9E-03</td><td>2.3E-03</td><td>BT20/MB468</td></tr>
<tr><td>>hg38_21_+_8433172_8446622_RNA45SN1_45S_50ntFlanks@12627.12667.41|ATGGGGCGAAGCTACCATCTGTGGGATTATGACTGAACGCC</td><td>>hg38_21_+_8433172_8446622_RNA45SN1_45S_50ntFlanks</td><td>12627</td><td>12667</td><td>41</td><td>ATGGGGCGAAGCTACCATCTGTGGGATTATGACTGAACGCC</td><td>377.3</td><td>-3.1</td><td>0.9</td><td>-3.6</td><td>3.2E-04</td><td>2.3E-03</td><td>BT20/MB231</td></tr>
<tr><td>>hg38_21_+_8433172_8446622_RNA45SN1_45S_50ntFlanks@6647.6670.24|CGTACGACTCTTAGCGGTGGATCA</td><td>>hg38_21_+_8433172_8446622_RNA45SN1_45S_50ntFlanks</td><td>6647</td><td>6670</td><td>24</td><td>CGTACGACTCTTAGCGGTGGATCA</td><td>343.0</td><td>-1.8</td><td>0.6</td><td>-3.1</td><td>1.8E-03</td><td>2.3E-03</td><td>MB231/MB468</td></tr>
<tr><td>>hg38_21_+_8433172_8446622_RNA45SN1_45S_50ntFlanks@10686.10706.21|GCGGCGTCCGGTGAGCTCTCG</td><td>>hg38_21_+_8433172_8446622_RNA45SN1_45S_50ntFlanks</td><td>10686</td><td>10706</td><td>21</td><td>GCGGCGTCCGGTGAGCTCTCG</td><td>771.7</td><td>-2.6</td><td>0.8</td><td>-3.1</td><td>2.0E-03</td><td>2.3E-03</td><td>BT20/MB468</td></tr>
<tr><td>>hg38_21_+_8433172_8446622_RNA45SN1_45S_50ntFlanks@11573.11600.28|CGACTTAGAACTGGTGCGGACCAGGGGA</td><td>>hg38_21_+_8433172_8446622_RNA45SN1_45S_50ntFlanks</td><td>11573</td><td>11600</td><td>28</td><td>CGACTTAGAACTGGTGCGGACCAGGGGA</td><td>267.7</td><td>-1.5</td><td>0.5</td><td>-3.1</td><td>1.8E-03</td><td>2.3E-03</td><td>MB231/MB468</td></tr>
<tr><td>>hg38_21_+_8433172_8446622_RNA45SN1_45S_50ntFlanks@12641.12661.21|CCATCTGTGGGATTATGACTG</td><td>>hg38_21_+_8433172_8446622_RNA45SN1_45S_50ntFlanks</td><td>12641</td><td>12661</td><td>21</td><td>CCATCTGTGGGATTATGACTG</td><td>379.4</td><td>-1.9</td><td>0.6</td><td>-3.1</td><td>1.8E-03</td><td>2.3E-03</td><td>MB231/MB468</td></tr>
</table>

<table>
<tr><td>>hg38_21_+_8433172_8446622_RNA45SN1_45S_50ntFlanks@10841.10874.34|CGTAACTTCGGGATAAGGATTGGCTCTAAGGGCT</td><td>>hg38_21_+_8433172_8446622_RNA45SN1_45S_50ntFlanks</td><td>10841</td><td>10874</td><td>34</td><td>CGTAACTTCGGGATAAGGATTGGCTCTAAGGGCT</td><td>401.6</td><td>-1.6</td><td>0.5</td><td>-3.1</td><td>1.8E-03</td><td>2.3E-03</td><td>MB231/MB468</td></tr>
<tr><td>>hg38_21_+_8433172_8446622_RNA45SN1_45S_50ntFlanks@9555.9573.19|GTAGCGGTCCTGACGTGCA</td><td>>hg38_21_+_8433172_8446622_RNA45SN1_45S_50ntFlanks</td><td>9555</td><td>9573</td><td>19</td><td>GTAGCGGTCCTGACGTGCA</td><td>196.1</td><td>-1.3</td><td>0.4</td><td>-3.1</td><td>2.0E-03</td><td>2.4E-03</td><td>BT20/MB468</td></tr>
<tr><td>>hg38_21_+_8433172_8446622_RNA45SN1_45S_50ntFlanks@10841.10860.20|CGTAACTTCGGGATAAGGAT</td><td>>hg38_21_+_8433172_8446622_RNA45SN1_45S_50ntFlanks</td><td>10841</td><td>10860</td><td>20</td><td>CGTAACTTCGGGATAAGGAT</td><td>271.0</td><td>-2.4</td><td>0.8</td><td>-3.1</td><td>2.0E-03</td><td>2.4E-03</td><td>BT20/MB468</td></tr>
<tr><td>>hg38_21_+_8433172_8446622_RNA45SN1_45S_50ntFlanks@4163.4186.24|CATCCAAGGAAGGCAGCAGGCGCG</td><td>>hg38_21_+_8433172_8446622_RNA45SN1_45S_50ntFlanks</td><td>4163</td><td>4186</td><td>24</td><td>CATCCAAGGAAGGCAGCAGGCGCG</td><td>175.4</td><td>-2.0</td><td>0.6</td><td>-3.1</td><td>1.9E-03</td><td>2.4E-03</td><td>MB231/MB468</td></tr>
<tr><td>>hg38_21_+_8433172_8446622_RNA45SN1_45S_50ntFlanks@10905.10941.37|CTGGGCGCGCGCCGCGGCTGGACGAGGCGCCGCCGCC</td><td>>hg38_21_+_8433172_8446622_RNA45SN1_45S_50ntFlanks</td><td>10905</td><td>10941</td><td>37</td><td>CTGGGCGCGCGCCGCGGCTGGACGAGGCGCCGCCGCC</td><td>103.0</td><td>-3.3</td><td>1.1</td><td>-3.1</td><td>2.0E-03</td><td>2.4E-03</td><td>BT20/MB468</td></tr>
<tr><td>>hg38_21_+_8433172_8446622_RNA45SN1_45S_50ntFlanks@12437.12473.37|CATTGTGAAGCAGAATTCACCAAGCGTTGGATTGTTC</td><td>>hg38_21_+_8433172_8446622_RNA45SN1_45S_50ntFlanks</td><td>12437</td><td>12473</td><td>37</td><td>CATTGTGAAGCAGAATTCACCAAGCGTTGGATTGTTC</td><td>1349.7</td><td>-2.1</td><td>0.7</td><td>-3.1</td><td>1.9E-03</td><td>2.4E-03</td><td>MB231/MB468</td></tr>
<tr><td>>hg38_21_+_8433172_8446622_RNA45SN1_45S_50ntFlanks@11217.11234.18|CTGGACGCGAGCCGGGCC</td><td>>hg38_21_+_8433172_8446622_RNA45SN1_45S_50ntFlanks</td><td>11217</td><td>11234</td><td>18</td><td>CTGGACGCGAGCCGGGCC</td><td>855.9</td><td>-2.3</td><td>0.7</td><td>-3.1</td><td>1.9E-03</td><td>2.4E-03</td><td>MB231/MB468</td></tr>
<tr><td>>hg38_21_+_8433172_8446622_RNA45SN1_45S_50ntFlanks@9553.9572.20|CCGTAGCGGTCCTGACGTGC</td><td>>hg38_21_+_8433172_8446622_RNA45SN1_45S_50ntFlanks</td><td>9553</td><td>9572</td><td>20</td><td>CCGTAGCGGTCCTGACGTGC</td><td>348.3</td><td>1.4</td><td>0.5</td><td>3.1</td><td>1.9E-03</td><td>2.4E-03</td><td>MB231/MB468</td></tr>
<tr><td>>hg38_21_+_8433172_8446622_RNA45SN1_45S_50ntFlanks@8198.8216.19|TGGACGGTGTGAGGCCGGT</td><td>>hg38_21_+_8433172_8446622_RNA45SN1_45S_50ntFlanks</td><td>8198</td><td>8216</td><td>19</td><td>TGGACGGTGTGAGGCCGGT</td><td>116.2</td><td>-2.4</td><td>0.8</td><td>-3.1</td><td>2.0E-03</td><td>2.4E-03</td><td>BT20/MB468</td></tr>
<tr><td>>hg38_21_+_8433172_8446622_RNA45SN1_45S_50ntFlanks@4565.4587.23|AATAATGGAATAGGACCGCGGTT</td><td>>hg38_21_+_8433172_8446622_RNA45SN1_45S_50ntFlanks</td><td>4565</td><td>4587</td><td>23</td><td>AATAATGGAATAGGACCGCGGTT</td><td>519.7</td><td>-2.0</td><td>0.6</td><td>-3.1</td><td>2.0E-03</td><td>2.4E-03</td><td>BT20/MB468</td></tr>
<tr><td>>hg38_21_+_8433172_8446622_RNA45SN1_45S_50ntFlanks@11208.11228.21|GCGGCGACTCTGGACGCGAGC</td><td>>hg38_21_+_8433172_8446622_RNA45SN1_45S_50ntFlanks</td><td>11208</td><td>11228</td><td>21</td><td>GCGGCGACTCTGGACGCGAGC</td><td>281.5</td><td>-1.8</td><td>0.6</td><td>-3.1</td><td>1.9E-03</td><td>2.4E-03</td><td>MB231/MB468</td></tr>
<tr><td>>hg38_21_+_8433172_8446622_RNA45SN1_45S_50ntFlanks@10401.10430.30|GAACATGGGTCAGTCGGTCCTGAGAGATGG</td><td>>hg38_21_+_8433172_8446622_RNA45SN1_45S_50ntFlanks</td><td>10401</td><td>10430</td><td>30</td><td>GAACATGGGTCAGTCGGTCCTGAGAGATGG</td><td>174.9</td><td>-2.9</td><td>0.9</td><td>-3.1</td><td>2.0E-03</td><td>2.4E-03</td><td>BT20/MB468</td></tr>
<tr><td>>hg38_21_+_8433172_8446622_RNA45SN1_45S_50ntFlanks@9556.9584.29|TAGCGGTCCTGACGTGCAAATCGGTCGTC</td><td>>hg38_21_+_8433172_8446622_RNA45SN1_45S_50ntFlanks</td><td>9556</td><td>9584</td><td>29</td><td>TAGCGGTCCTGACGTGCAAATCGGTCGTC</td><td>174.2</td><td>-2.1</td><td>0.7</td><td>-3.1</td><td>1.9E-03</td><td>2.4E-03</td><td>MB231/MB468</td></tr>
<tr><td>>MI0000342|hsa-mir-200b&WithFlank&1|+|1167098|1167204@63.84.22&[MIMAT0000318&hsa-miR-200b-3p&offsets|0|0;m-93&1|+|1167160|1167182&offsets|0|-1]|TAATACTGCCTGGTAATGATGA</td><td>>MI0000342|hsa-mir-200b&WithFlank&1|+|1167098|1167204</td><td>63</td><td>84</td><td>22</td><td>TAATACTGCCTGGTAATGATGA</td><td>745.5</td><td>-2.2</td><td>0.7</td><td>-3.1</td><td>1.9E-03</td><td>2.4E-03</td><td>MB231/MB468</td></tr>
<tr><td>>hg38_21_+_8433172_8446622_RNA45SN1_45S_50ntFlanks@11573.11602.30|CGACTTAGAACTGGTGCGGACCAGGGGAAT</td><td>>hg38_21_+_8433172_8446622_RNA45SN1_45S_50ntFlanks</td><td>11573</td><td>11602</td><td>30</td><td>CGACTTAGAACTGGTGCGGACCAGGGGAAT</td><td>375.8</td><td>-1.6</td><td>0.5</td><td>-3.1</td><td>1.9E-03</td><td>2.4E-03</td><td>MB231/MB468</td></tr>
<tr><td>>hg38_21_+_8433172_8446622_RNA45SN1_45S_50ntFlanks@11205.11225.21|GCGGCGGCGACTCTGGACGCG</td><td>>hg38_21_+_8433172_8446622_RNA45SN1_45S_50ntFlanks</td><td>11205</td><td>11225</td><td>21</td><td>GCGGCGGCGACTCTGGACGCG</td><td>204.9</td><td>-5.1</td><td>1.7</td><td>-3.1</td><td>1.9E-03</td><td>2.4E-03</td><td>MB231/MB468</td></tr>
<tr><td>>hg38_21_+_8433172_8446622_RNA45SN1_45S_50ntFlanks@11726.11751.26|GAGTAACTATGACTCTCTTAAGGTAG</td><td>>hg38_21_+_8433172_8446622_RNA45SN1_45S_50ntFlanks</td><td>11726</td><td>11751</td><td>26</td><td>GAGTAACTATGACTCTCTTAAGGTAG</td><td>116.2</td><td>-2.3</td><td>0.7</td><td>-3.1</td><td>1.9E-03</td><td>2.4E-03</td><td>MB231/MB468</td></tr>
<tr><td>>hg38_21_+_8433172_8446622_RNA45SN1_45S_50ntFlanks@3787.3810.24|AAACTGCGAATGGCTCATTAAATC</td><td>>hg38_21_+_8433172_8446622_RNA45SN1_45S_50ntFlanks</td><td>3787</td><td>3810</td><td>24</td><td>AAACTGCGAATGGCTCATTAAATC</td><td>459.4</td><td>-2.2</td><td>0.7</td><td>-3.1</td><td>2.1E-03</td><td>2.4E-03</td><td>BT20/MB468</td></tr>
<tr><td>>MI0000298|hsa-mir-221&WithFlank&X|-|45746151|45746272@31.55.25(+1C)&[MIMAT0004568&hsa-miR-221-5p&offsets|0|+3(+1C);m-405&X|-|45746220|45746242&offsets|0|+2(+1C)]|ACCTGGCATACAATGTAGATTTCTGC</td><td>>MI0000298|hsa-mir-221&WithFlank&X|-|45746151|45746272</td><td>31</td><td>55</td><td>25(+1C)</td><td>ACCTGGCATACAATGTAGATTTCTGC</td><td>347.3</td><td>-2.6</td><td>0.7</td><td>-3.6</td><td>3.3E-04</td><td>2.4E-03</td><td>BT20/MB231</td></tr>
<tr><td>>hg38_21_+_8433172_8446622_RNA45SN1_45S_50ntFlanks@12419.12456.38|GATGTCGGCTCTTCCTATCATTGTGAAGCAGAATTCAC</td><td>>hg38_21_+_8433172_8446622_RNA45SN1_45S_50ntFlanks</td><td>12419</td><td>12456</td><td>38</td><td>GATGTCGGCTCTTCCTATCATTGTGAAGCAGAATTCAC</td><td>380.0</td><td>-2.6</td><td>0.7</td><td>-3.6</td><td>3.4E-04</td><td>2.4E-03</td><td>BT20/MB231</td></tr>
<tr><td>>hg38_21_+_8433172_8446622_RNA45SN1_45S_50ntFlanks@11879.11904.26|ACCCTGTTGAGCTTGACTCTAGTCTG</td><td>>hg38_21_+_8433172_8446622_RNA45SN1_45S_50ntFlanks</td><td>11879</td><td>11904</td><td>26</td><td>ACCCTGTTGAGCTTGACTCTAGTCTG</td><td>135.4</td><td>-2.5</td><td>0.8</td><td>-3.1</td><td>1.9E-03</td><td>2.4E-03</td><td>MB231/MB468</td></tr>
<tr><td>>hg38_21_+_8433172_8446622_RNA45SN1_45S_50ntFlanks@10795.10816.22|TGGCATGTTGGAACAATGTAGG</td><td>>hg38_21_+_8433172_8446622_RNA45SN1_45S_50ntFlanks</td><td>10795</td><td>10816</td><td>22</td><td>TGGCATGTTGGAACAATGTAGG</td><td>142.7</td><td>-3.1</td><td>1.0</td><td>-3.1</td><td>2.1E-03</td><td>2.5E-03</td><td>BT20/MB468</td></tr>
<tr><td>>hg38_21_+_8433172_8446622_RNA45SN1_45S_50ntFlanks@6650.6679.30|ACGACTCTTAGCGGTGGATCACTCGGCTCG</td><td>>hg38_21_+_8433172_8446622_RNA45SN1_45S_50ntFlanks</td><td>6650</td><td>6679</td><td>30</td><td>ACGACTCTTAGCGGTGGATCACTCGGCTCG</td><td>1260.3</td><td>1.9</td><td>0.6</td><td>3.1</td><td>1.9E-03</td><td>2.5E-03</td><td>MB231/MB468</td></tr>
<tr><td>>hg38_21_+_8433172_8446622_RNA45SN1_45S_50ntFlanks@9554.9572.19|CGTAGCGGTCCTGACGTGC</td><td>>hg38_21_+_8433172_8446622_RNA45SN1_45S_50ntFlanks</td><td>9554</td><td>9572</td><td>19</td><td>CGTAGCGGTCCTGACGTGC</td><td>10944.0</td><td>-2.0</td><td>0.6</td><td>-3.1</td><td>1.9E-03</td><td>2.5E-03</td><td>MB231/MB468</td></tr>
<tr><td>>hg38_21_+_8433172_8446622_RNA45SN1_45S_50ntFlanks@10792.10822.31|CTCTGGCATGTTGGAACAATGTAGGTAAGGG</td><td>>hg38_21_+_8433172_8446622_RNA45SN1_45S_50ntFlanks</td><td>10792</td><td>10822</td><td>31</td><td>CTCTGGCATGTTGGAACAATGTAGGTAAGGG</td><td>151.1</td><td>-1.4</td><td>0.5</td><td>-3.1</td><td>2.0E-03</td><td>2.5E-03</td><td>MB231/MB468</td></tr>
<tr><td>>hg38_21_+_8433172_8446622_RNA45SN1_45S_50ntFlanks@10343.10361.19|TATTCAAACGAGAACTTTG</td><td>>hg38_21_+_8433172_8446622_RNA45SN1_45S_50ntFlanks</td><td>10343</td><td>10361</td><td>19</td><td>TATTCAAACGAGAACTTTG</td><td>252.8</td><td>-2.0</td><td>0.7</td><td>-3.1</td><td>2.1E-03</td><td>2.5E-03</td><td>BT20/MB468</td></tr>
<tr><td>>hg38_21_+_8433172_8446622_RNA45SN1_45S_50ntFlanks@11577.11609.33|TTAGAACTGGTGCGGACCAGGGGAATCCGACTG</td><td>>hg38_21_+_8433172_8446622_RNA45SN1_45S_50ntFlanks</td><td>11577</td><td>11609</td><td>33</td><td>TTAGAACTGGTGCGGACCAGGGGAATCCGACTG</td><td>171.8</td><td>-1.8</td><td>0.6</td><td>-3.1</td><td>2.0E-03</td><td>2.5E-03</td><td>MB231/MB468</td></tr>
<tr><td>>MI0000299|hsa-mir-222&WithFlank&X|-|45747009|45747130@75.99.25(+1U)&[MIMAT0000279&hsa-miR-222-3p&offsets|0|+4(+1U);m-140&X|-|45747033|45747056&offsets|0|+1(+1U)]|AGCTACATCTGGCTACTGGGTCTCTT</td><td>>MI0000299|hsa-mir-222&WithFlank&X|-|45747009|45747130</td><td>75</td><td>99</td><td>25(+1U)</td><td>AGCTACATCTGGCTACTGGGTCTCTT</td><td>6606.8</td><td>-2.6</td><td>0.7</td><td>-3.6</td><td>3.5E-04</td><td>2.5E-03</td><td>BT20/MB231</td></tr>
<tr><td>>am_trna119_LysCTT_1_-_145395522_145395594@14.31.18,trna11_LysCTT_5_-_180648979_180649051@14.31.18,trna128_LysCTT_1_-_55423542_55423614@14.31.18,trna13_LysCTT_6_+_26556774_26556846@14.31.18,trna32_LysCTT_16_-_3207406_3207478@14.31.18,trna6_LysCTT_19_-_52425396_52425468@14.31.18,trna7_LysCTT_16_+_3225692_3225764@14.31.18,trna9_LysCTT_5_+_180634755_180634827@14.31.18|AGTCGGTAGAGCATGAGA</td><td>>am_trna119_LysCTT_1_-_145395522_145395594</td><td>14</td><td>31</td><td>18</td><td>AGTCGGTAGAGCATGAGA</td><td>396.4</td><td>-2.1</td><td>0.7</td><td>-3.1</td><td>2.1E-03</td><td>2.5E-03</td><td>BT20/MB468</td></tr>
<tr><td>>MI0000298|hsa-mir-221&WithFlank&X|-|45746151|45746272@71.93.23(+1C)&[MIMAT0000278&hsa-miR-221-3p&offsets|0|0(+1C);m-91&X|-|45746181|45746202&offsets|0|+1(+1C)]|AGCTACATTGTCTGCTGGGTTTCC</td><td>>MI0000298|hsa-mir-221&WithFlank&X|-|45746151|45746272</td><td>71</td><td>93</td><td>23(+1C)</td><td>AGCTACATTGTCTGCTGGGTTTCC</td><td>2525.6</td><td>-2.3</td><td>0.7</td><td>-3.6</td><td>3.5E-04</td><td>2.5E-03</td><td>BT20/MB231</td></tr>
<tr><td>>MI0000301|hsa-mir-224&WithFlank&X|-|151958572|151958664@14.34.21&[MIMAT0000281&hsa-miR-224-5p&offsets|+1|-3;m-238&X|-|151958630|151958651&offsets|0|-1]|CAAGTCACTAGTGGTTCCGTT</td><td>>MI0000301|hsa-mir-224&WithFlank&X|-|151958572|151958664</td><td>14</td><td>34</td><td>21</td><td>CAAGTCACTAGTGGTTCCGTT</td><td>1763.5</td><td>2.4</td><td>0.7</td><td>3.6</td><td>3.5E-04</td><td>2.5E-03</td><td>BT20/MB231</td></tr>
<tr><td>>MI0000542|hsa-mir-320a&WithFlank&8|-|22244960|22245043@48.70.23&[MIMAT0000510&hsa-miR-320a-3p&offsets|0|+1;m-60&8|-|22244975|22244996&offsets|0|+1]|AAAAGCTGGGTTGAGAGGGCGAA</td><td>>MI0000542|hsa-mir-320a&WithFlank&8|-|22244960|22245043</td><td>48</td><td>70</td><td>23</td><td>AAAAGCTGGGTTGAGAGGGCGAA</td><td>9020.3</td><td>2.5</td><td>0.7</td><td>3.6</td><td>3.5E-04</td><td>2.5E-03</td><td>BT20/MB231</td></tr>
<tr><td>>MI0003643|hsa-mir-629&WithFlank&15|-|70079366|70079474@28.48.21(+1C)&[MIMAT0004810&hsa-miR-629-5p&offsets|0|0(+1C);m-150&15|-|70079427|70079447&offsets|0|0(+1C)]|TGGGTTTACGTTGGGAGAACTC</td><td>>MI0003643|hsa-mir-629&WithFlank&15|-|70079366|70079474</td><td>28</td><td>48</td><td>21(+1C)</td><td>TGGGTTTACGTTGGGAGAACTC</td><td>314.3</td><td>2.4</td><td>0.7</td><td>3.6</td><td>3.5E-04</td><td>2.5E-03</td><td>BT20/MB231</td></tr>
<tr><td>>hg38_21_+_8433172_8446622_RNA45SN1_45S_50ntFlanks@7988.8014.27|CAGACGTGGCGACCCGCTGAATTTAAG</td><td>>hg38_21_+_8433172_8446622_RNA45SN1_45S_50ntFlanks</td><td>7988</td><td>8014</td><td>27</td><td>CAGACGTGGCGACCCGCTGAATTTAAG</td><td>319.8</td><td>-1.8</td><td>0.6</td><td>-3.1</td><td>2.2E-03</td><td>2.5E-03</td><td>BT20/MB468</td></tr>
<tr><td>>hg38_21_+_8433172_8446622_RNA45SN1_45S_50ntFlanks@11602.11619.18|TCCGACTGTTTAATTAAA</td><td>>hg38_21_+_8433172_8446622_RNA45SN1_45S_50ntFlanks</td><td>11602</td><td>11619</td><td>18</td><td>TCCGACTGTTTAATTAAA</td><td>704.0</td><td>-1.6</td><td>0.5</td><td>-3.1</td><td>2.0E-03</td><td>2.6E-03</td><td>MB231/MB468</td></tr>
<tr><td>>hg38_21_+_8433172_8446622_RNA45SN1_45S_50ntFlanks@12419.12437.19|GATGTCGGCTCTTCCTATC</td><td>>hg38_21_+_8433172_8446622_RNA45SN1_45S_50ntFlanks</td><td>12419</td><td>12437</td><td>19</td><td>GATGTCGGCTCTTCCTATC</td><td>1964.8</td><td>1.6</td><td>0.5</td><td>3.1</td><td>2.2E-03</td><td>2.6E-03</td><td>BT20/MB468</td></tr>
<tr><td>>hg38_21_+_8433172_8446622_RNA45SN1_45S_50ntFlanks@10788.10817.30|CAGCCTCTGGCATGTTGGAACAATGTAGGT</td><td>>hg38_21_+_8433172_8446622_RNA45SN1_45S_50ntFlanks</td><td>10788</td><td>10817</td><td>30</td><td>CAGCCTCTGGCATGTTGGAACAATGTAGGT</td><td>253.3</td><td>-1.9</td><td>0.6</td><td>-3.1</td><td>2.0E-03</td><td>2.6E-03</td><td>MB231/MB468</td></tr>
<tr><td>>hg38_21_+_8433172_8446622_RNA45SN1_45S_50ntFlanks@12973.13000.28|TCGTACGTAGCAGAGCAGCTCCCTCGCT</td><td>>hg38_21_+_8433172_8446622_RNA45SN1_45S_50ntFlanks</td><td>12973</td><td>13000</td><td>28</td><td>TCGTACGTAGCAGAGCAGCTCCCTCGCT</td><td>105.4</td><td>-1.3</td><td>0.4</td><td>-3.1</td><td>2.0E-03</td><td>2.6E-03</td><td>MB231/MB468</td></tr>
<tr><td>>am_MI0003776|hsa-mir-320b-1&WithFlank&1|+|116671740|116671823@48.65.18;MI0003839|hsa-mir-320b-2&WithFlank&1|-|224257034|224257116@47.64.18;MI0008190|hsa-mir-320d-1&WithFlank&13|-|40727810|40727893@48.65.18;MI0003778|hsa-mir-320c-1&WithFlank&18|+|21683512|21683595@48.65.18;MI0008191|hsa-mir-320c-2&WithFlank&18|+|24321669|24321752@48.65.18;MI0000542|hsa-mir-320a&WithFlank&8|-|22244960|22245043@48.65.18;MI0008192|hsa-mir-320d-2&WithFlank&X|-|140926154|140926237@48.65.18&[MIMAT0005792&hsa-miR-320b&offsets|0|-4;m-183&1|+|116671787|116671806&offsets|0|-2];[MIMAT0005792_1&hsa-miR-320b&offsets|0|-4;m-183&1|-|224257051|224257070&offsets|0|-2];[MIMAT0006764&hsa-miR-320d&offsets|0|-1;m-246&13|-|40727829|40727846&offsets|0|0];[MIMAT0005793&hsa-miR-320c&offsets|0|-2;m-222&18|+|21683559|21683578&offsets|0|-2];[MIMAT0005793_1&hsa-miR-320c&offsets|0|-2;m-219&18|+|24321716|24321735&offsets|0|-2];[MIMAT0000510&hsa-miR-320a-3p&offsets|0|-4;m-60&8|-|22244975|22244996&offsets|0|-4];[MIMAT0006764_1&hsa-miR-320d&offsets|0|-1;m-253&X|-|140926173|140926190&offsets|0|0]|AAAAGCTGGGTTGAGAGG</td><td>>am_MI0003776|hsa-mir-320b-1&WithFlank&1|+|116671740|116671823</td><td>48</td><td>65</td><td>18</td><td>AAAAGCTGGGTTGAGAGG</td><td>379.9</td><td>2.5</td><td>0.7</td><td>3.6</td><td>3.7E-04</td><td>2.6E-03</td><td>BT20/MB231</td></tr>
<tr><td>>hg38_21_+_8433172_8446622_RNA45SN1_45S_50ntFlanks@11729.11746.18|TAACTATGACTCTCTTAA</td><td>>hg38_21_+_8433172_8446622_RNA45SN1_45S_50ntFlanks</td><td>11729</td><td>11746</td><td>18</td><td>TAACTATGACTCTCTTAA</td><td>164.2</td><td>-2.0</td><td>0.6</td><td>-3.1</td><td>2.1E-03</td><td>2.7E-03</td><td>MB231/MB468</td></tr>
<tr><td>>hg38_21_+_8433172_8446622_RNA45SN1_45S_50ntFlanks@12415.12457.43|CTTCGATGTCGGCTCTTCCTATCATTGTGAAGCAGAATTCACC</td><td>>hg38_21_+_8433172_8446622_RNA45SN1_45S_50ntFlanks</td><td>12415</td><td>12457</td><td>43</td><td>CTTCGATGTCGGCTCTTCCTATCATTGTGAAGCAGAATTCACC</td><td>214.5</td><td>-2.7</td><td>0.8</td><td>-3.6</td><td>3.7E-04</td><td>2.7E-03</td><td>BT20/MB231</td></tr>
</table>

| | | | | | | | | | | | | |
|---|---|---|---|---|---|---|---|---|---|---|---|---|
| >hg38_21_+_8433172_8446622_RNA45SN1_45S_50ntFlanks@3881.3898.18\|GCCGACGGGCGCTGACCC | >hg38_21_+_8433172_8446622_RNA45SN1_45S_50ntFlanks | 3881 | 3898 | 18 | GCCGACGGGCGCTGACCC | 193.9 | -2.2 | 0.7 | -3.1 | 2.1E-03 | 2.7E-03 | MB231/MB468 |
| >hg38_21_+_8433172_8446622_RNA45SN1_45S_50ntFlanks@6647.6669.23\|CGTACGACTCTTAGCGGTGGATC | >hg38_21_+_8433172_8446622_RNA45SN1_45S_50ntFlanks | 6647 | 6669 | 23 | CGTACGACTCTTAGCGGTGGATC | 2063.3 | 2.5 | 0.8 | 3.1 | 2.1E-03 | 2.7E-03 | MB231/MB468 |
| >hg38_21_+_8433172_8446622_RNA45SN1_45S_50ntFlanks@11579.11607.29\|AGAACTGGTGCGGACCAGGGGAATCCGAC | >hg38_21_+_8433172_8446622_RNA45SN1_45S_50ntFlanks | 11579 | 11607 | 29 | AGAACTGGTGCGGACCAGGGGAATCCGAC | 402.8 | -1.6 | 0.5 | -3.1 | 2.3E-03 | 2.7E-03 | BT20/MB468 |
| >hg38_21_+_8433172_8446622_RNA45SN1_45S_50ntFlanks@8176.8199.24\|CTTCTGATCGAGGCCCAGCCCGTG | >hg38_21_+_8433172_8446622_RNA45SN1_45S_50ntFlanks | 8176 | 8199 | 24 | CTTCTGATCGAGGCCCAGCCCGTG | 159.7 | -1.4 | 0.5 | -3.1 | 2.3E-03 | 2.7E-03 | BT20/MB468 |
| >hg38_21_+_8433172_8446622_RNA45SN1_45S_50ntFlanks@11104.11123.20\|AAGGGTCGGGGCGGCAGGGG | >hg38_21_+_8433172_8446622_RNA45SN1_45S_50ntFlanks | 11104 | 11123 | 20 | AAGGGTCGGGGCGGCAGGGG | 170.4 | -3.5 | 1.2 | -3.1 | 2.3E-03 | 2.7E-03 | BT20/MB468 |
| >hg38_21_+_8433172_8446622_RNA45SN1_45S_50ntFlanks@10353.10390.38\|AGAACTTTGAAGGCCGAAGTGGAGAAGGGTTCCATGTG | >hg38_21_+_8433172_8446622_RNA45SN1_45S_50ntFlanks | 10353 | 10390 | 38 | AGAACTTTGAAGGCCGAAGTGGAGAAGGGTTCCATGTG | 284.5 | -3.1 | 0.9 | -3.6 | 3.8E-04 | 2.7E-03 | BT20/MB231 |
| >hg38_21_+_8433172_8446622_RNA45SN1_45S_50ntFlanks@12072.12092.21\|CCCGAGGGGCTCTCGCTTCTG | >hg38_21_+_8433172_8446622_RNA45SN1_45S_50ntFlanks | 12072 | 12092 | 21 | CCCGAGGGGCTCTCGCTTCTG | 162.3 | -1.6 | 0.5 | -3.1 | 2.2E-03 | 2.7E-03 | MB231/MB468 |
| >hg38_21_+_8433172_8446622_RNA45SN1_45S_50ntFlanks@3840.3865.26\|CTCCTACTTGGATAACTGTGGTAATT | >hg38_21_+_8433172_8446622_RNA45SN1_45S_50ntFlanks | 3840 | 3865 | 26 | CTCCTACTTGGATAACTGTGGTAATT | 252.4 | 2.6 | 0.7 | 3.6 | 3.8E-04 | 2.7E-03 | BT20/MB231 |
| >MI0000737\|hsa-mir-200a&WithFlank&1\|+\|1167857\|1167958@60.82.23&[MIMAT0000682&hsa-miR-200a-3p&offsets\|0\|+1;m-142&1\|+\|1167916\|1167938&offsets\|0\|0]\|TAACACTGTCTGGTAACGATGTT | >MI0000737\|hsa-mir-200a&WithFlank&1\|+\|1167857\|1167958 | 60 | 82 | 23 | TAACACTGTCTGGTAACGATGTT | 1837.3 | -3.0 | 1.0 | -3.0 | 2.3E-03 | 2.7E-03 | BT20/MB468 |
| >hg38_21_+_8433172_8446622_RNA45SN1_45S_50ntFlanks@11569.11597.29\|CAGCCGACTTAGAACTGGTGCGGACCAGG | >hg38_21_+_8433172_8446622_RNA45SN1_45S_50ntFlanks | 11569 | 11597 | 29 | CAGCCGACTTAGAACTGGTGCGGACCAGG | 195.4 | -2.0 | 0.7 | -3.0 | 2.3E-03 | 2.7E-03 | BT20/MB468 |
| >hg38_21_+_8433172_8446622_RNA45SN1_45S_50ntFlanks@10907.10930.24\|GGGCGCGCGCCGCGGCTGGACGAG | >hg38_21_+_8433172_8446622_RNA45SN1_45S_50ntFlanks | 10907 | 10930 | 24 | GGGCGCGCGCCGCGGCTGGACGAG | 373.3 | 10.4 | 3.4 | 3.0 | 2.3E-03 | 2.8E-03 | BT20/MB468 |
| >hg38_21_+_8433172_8446622_RNA45SN1_45S_50ntFlanks@6660.6681.22\|GCGGTGGATCACTCGGCTCGTG | >hg38_21_+_8433172_8446622_RNA45SN1_45S_50ntFlanks | 6660 | 6681 | 22 | GCGGTGGATCACTCGGCTCGTG | 524.2 | -2.0 | 0.6 | -3.0 | 2.3E-03 | 2.8E-03 | BT20/MB468 |
| >MI0000273\|hsa-mir-183&WithFlank&7\|-\|129774899\|129775020@33.53.21&[MIMAT0000261&hsa-miR-183-5p&offsets\|0\|-1;m-41&7\|-\|129774966\|129774987&offsets\|-1\|-2]\|TATGGCACTGGTAGAATTCAC | >MI0000273\|hsa-mir-183&WithFlank&7\|-\|129774899\|129775020 | 33 | 53 | 21 | TATGGCACTGGTAGAATTCAC | 255.9 | 2.6 | 0.7 | 3.5 | 3.9E-04 | 2.8E-03 | BT20/MB231 |
| >trna10_AspGTC_12_-_125424193_125424264@37.75.39,trna12_AspGTC_12_-_125411891_125411962@37.75.39,trna144_AspGTC_6_-_27551236_27551307@37.75.39,trna38_AspGTC_17_-_8125556_8125627@37.75.39,trna45_AspGTC_6_+_27447453_27447524@37.75.39,trna48_AspGTC_6_+_27471523_27471594@37.75.39,trna4_AspGTC_12_+_96429799_96429870@37.75.39,trna69_AspGTC_1_-_161440205_161440276@37.75.39,trna72_AspGTC_1_-_161432824_161432895@37.75.39,trna75_AspGTC_1_-_161425414_161425485@37.75.39,trna78_AspGTC_1_-_161418033_161418104@37.75.39,trna81_AspGTC_1_-_161410615_161410686@37.75.39\|ACGCGGGAGACCGGGGTTCGATTCCCCGACGGGGAGCCA | >trna10_AspGTC_12_-_125424193_125424264 | 37 | 75 | 39 | ACGCGGGAGACCGGGGTTCGATTCCCCGACGGGGAGCCA | 709.7 | -3.7 | 1.1 | -3.5 | 3.9E-04 | 2.8E-03 | BT20/MB231 |
| >hg38_21_+_8433172_8446622_RNA45SN1_45S_50ntFlanks@11200.11232.33\|CGGCGGCGGCGGCGACTCTGGACGCGAGCCGGG | >hg38_21_+_8433172_8446622_RNA45SN1_45S_50ntFlanks | 11200 | 11232 | 33 | CGGCGGCGGCGGCGACTCTGGACGCGAGCCGGG | 1124.4 | -3.0 | 1.0 | -3.1 | 2.2E-03 | 2.8E-03 | MB231/MB468 |
| >MI0000542\|hsa-mir-320a&WithFlank&8\|-\|22244960\|22245043@48.68.21(+3U)&[MIMAT0000510&hsa-miR-320a-3p&offsets\|0\|-1(+3U);m-60&8\|-\|22244975\|22244996&offsets\|0\|-1(+3U)]\|AAAAGCTGGGTTGAGAGGGCGTTT | >MI0000542\|hsa-mir-320a&WithFlank&8\|-\|22244960\|22245043 | 48 | 68 | 21(+3U) | AAAAGCTGGGTTGAGAGGGCGTTT | 877.1 | 2.5 | 0.7 | 3.5 | 3.9E-04 | 2.8E-03 | BT20/MB231 |
| >hg38_21_+_8433172_8446622_RNA45SN1_45S_50ntFlanks@11577.11611.35\|TTAGAACTGGTGCGGACCAGGGGAATCCGACTGTT | >hg38_21_+_8433172_8446622_RNA45SN1_45S_50ntFlanks | 11577 | 11611 | 35 | TTAGAACTGGTGCGGACCAGGGGAATCCGACTGTT | 290.8 | -1.7 | 0.6 | -3.0 | 2.4E-03 | 2.8E-03 | BT20/MB468 |
| >hg38_21_+_8433172_8446622_RNA45SN1_45S_50ntFlanks@7978.8019.42\|GACCTCAGATCAGACGTGGCGACCCGCTGAATTTAAGCATAT | >hg38_21_+_8433172_8446622_RNA45SN1_45S_50ntFlanks | 7978 | 8019 | 42 | GACCTCAGATCAGACGTGGCGACCCGCTGAATTTAAGCATAT | 157.1 | -4.3 | 1.2 | -3.5 | 4.0E-04 | 2.8E-03 | BT20/MB231 |
| >hg38_21_+_8433172_8446622_RNA45SN1_45S_50ntFlanks@12676.12703.28\|AGAATCCCGCCCAGGCGGAACGATACGG | >hg38_21_+_8433172_8446622_RNA45SN1_45S_50ntFlanks | 12676 | 12703 | 28 | AGAATCCCGCCCAGGCGGAACGATACGG | 358.0 | -5.3 | 1.7 | -3.1 | 2.2E-03 | 2.8E-03 | MB231/MB468 |
| >hg38_21_+_8433172_8446622_RNA45SN1_45S_50ntFlanks@11209.11231.23\|CGGCGACTCTGGACGCGAGCCGG | >hg38_21_+_8433172_8446622_RNA45SN1_45S_50ntFlanks | 11209 | 11231 | 23 | CGGCGACTCTGGACGCGAGCCGG | 894.1 | -2.8 | 0.9 | -3.0 | 2.4E-03 | 2.8E-03 | BT20/MB468 |
| >hg38_21_+_8433172_8446622_RNA45SN1_45S_50ntFlanks@10351.10372.22\|CGAGAACTTTGAAGGCCGAAGT | >hg38_21_+_8433172_8446622_RNA45SN1_45S_50ntFlanks | 10351 | 10372 | 22 | CGAGAACTTTGAAGGCCGAAGT | 369.8 | -1.4 | 0.5 | -3.0 | 2.4E-03 | 2.8E-03 | BT20/MB468 |
| >MI0000063\|hsa-let-7b&WithFlank&22\|+\|46113680\|46113774@12.32.21&[MIMAT0000063&hsa-let-7b-5p&offsets\|0\|-1;m-14&22\|+\|46113691\|46113712&offsets\|0\|-1]\|TGAGGTAGTAGGTTGTGTGGT | >MI0000063\|hsa-let-7b&WithFlank&22\|+\|46113680\|46113774 | 12 | 32 | 21 | TGAGGTAGTAGGTTGTGTGGT | 5833.1 | 2.2 | 0.6 | 3.5 | 4.0E-04 | 2.8E-03 | BT20/MB231 |
| >hg38_21_+_8433172_8446622_RNA45SN1_45S_50ntFlanks@12420.12450.31\|ATGTCGGCTCTTCCTATCATTGTGAAGCAGA | >hg38_21_+_8433172_8446622_RNA45SN1_45S_50ntFlanks | 12420 | 12450 | 31 | ATGTCGGCTCTTCCTATCATTGTGAAGCAGA | 233.4 | -1.3 | 0.4 | -3.0 | 2.4E-03 | 2.9E-03 | BT20/MB468 |
| >MI0000542\|hsa-mir-320a&WithFlank&8\|-\|22244960\|22245043@49.70.22&[MIMAT0000510&hsa-miR-320a-3p&offsets\|+1\|+1;m-60&8\|-\|22244975\|22244996&offsets\|+1\|+1]\|AAAGCTGGGTTGAGAGGGCGAA | >MI0000542\|hsa-mir-320a&WithFlank&8\|-\|22244960\|22245043 | 49 | 70 | 22 | AAAGCTGGGTTGAGAGGGCGAA | 589.0 | 2.6 | 0.7 | 3.5 | 4.1E-04 | 2.9E-03 | BT20/MB231 |
| >hg38_21_+_8433172_8446622_RNA45SN1_45S_50ntFlanks@6645.6681.37\|CTCGTACGACTCTTAGCGGTGGATCACTCGGCTCGTG | >hg38_21_+_8433172_8446622_RNA45SN1_45S_50ntFlanks | 6645 | 6681 | 37 | CTCGTACGACTCTTAGCGGTGGATCACTCGGCTCGTG | 1247.2 | -1.4 | 0.5 | -3.0 | 2.4E-03 | 2.9E-03 | BT20/MB468 |
| >hg38_21_+_8433172_8446622_RNA45SN1_45S_50ntFlanks@9571.9595.25\|GCAAATCGGTCGTCCGACCTGGGTA | >hg38_21_+_8433172_8446622_RNA45SN1_45S_50ntFlanks | 9571 | 9595 | 25 | GCAAATCGGTCGTCCGACCTGGGTA | 136.7 | -1.5 | 0.5 | -3.1 | 2.3E-03 | 2.9E-03 | MB231/MB468 |
| >MI0000298\|hsa-mir-221&WithFlank&X\|-\|45746151\|45746272@31.54.24&[MIMAT0004568&hsa-miR-221-5p&offsets\|0\|+2;m-405&X\|-\|45746220\|45746242&offsets\|0\|+1]\|ACCTGGCATACAATGTAGATTTCT | >MI0000298\|hsa-mir-221&WithFlank&X\|-\|45746151\|45746272 | 31 | 54 | 24 | ACCTGGCATACAATGTAGATTTCT | 3025.5 | -2.4 | 0.7 | -3.5 | 4.1E-04 | 2.9E-03 | BT20/MB231 |
| >MI0000750\|hsa-mir-26a-2&WithFlank&12\|-\|57824603\|57824698@20.41.22(+3U);MI0000083\|hsa-mir-26a-1&WithFlank&3\|+\|37969398\|37969486@16.37.22(+3U)&[MIMAT0000082&hsa-miR-26a-5p&offsets\|0\|0(+3U);m-33&12\|-\|57824658\|57824679&offsets\|0\|0(+3U)];[MIMAT0000082_1&hsa-miR-26a-5p&offsets\|0\|0(+3U);m-34&3\|+\|37969413\|37969434&offsets\|0\|0(+3U)]\|TTCAAGTAATCCAGGATAGGCTTTT | >MI0000750\|hsa-mir-26a-2&WithFlank&12\|-\|57824603\|57824698 | 20 | 41 | 22(+3U) | TTCAAGTAATCCAGGATAGGCTTTT | 249.7 | -2.5 | 0.7 | -3.5 | 4.1E-04 | 2.9E-03 | BT20/MB231 |
| >trna111_HisGTG_1_-_147774845_147774916@-1T.24.25,trna118_HisGTG_1_-_145396881_145396952@-1T.24.25,trna16_HisGTG_1_+_146544773_146544844@-1T.24.25,trna1_HisGTG_15_+_45493349_45493420@-1T.24.25,trna21_HisGTG_1_+_147753471_147753542@-1T.24.25,trna33_HisGTG_6_+_27125906_27125977@-1T.24.25,trna7_HisGTG_9_-_14433938_14434009@-1T.24.25,trna8_HisGTG_15_-_45492611_45492682@-1T.24.25,trna9_HisGTG_15_-_45490804_45490875@-1T.24.25\|TGCCGTGATCGTATAGTGGTTAGTA | >trna111_HisGTG_1_-_147774845_147774916 | -1T | 24 | 25 | TGCCGTGATCGTATAGTGGTTAGTA | 227.9 | -3.7 | 1.1 | -3.5 | 4.1E-04 | 2.9E-03 | BT20/MB231 |
| >hg38_21_+_8433172_8446622_RNA45SN1_45S_50ntFlanks@11580.11613.34\|GAACTGGTGCGGACCAGGGGAATCCGACTGTTTA | >hg38_21_+_8433172_8446622_RNA45SN1_45S_50ntFlanks | 11580 | 11613 | 34 | GAACTGGTGCGGACCAGGGGAATCCGACTGTTTA | 1534.0 | 1.7 | 0.6 | 3.0 | 2.4E-03 | 2.9E-03 | BT20/MB468 |
| >hg38_21_+_8433172_8446622_RNA45SN1_45S_50ntFlanks@3843.3868.26\|CTACTTGGATAACTGTGGTAATTCTA | >hg38_21_+_8433172_8446622_RNA45SN1_45S_50ntFlanks | 3843 | 3868 | 26 | CTACTTGGATAACTGTGGTAATTCTA | 582.1 | -1.5 | 0.5 | -3.0 | 2.4E-03 | 2.9E-03 | BT20/MB468 |
| >hg38_21_+_8433172_8446622_RNA45SN1_45S_50ntFlanks@9534.9560.27\|AGGAAACTCTGGTGGAGGTCCGTAGCG | >hg38_21_+_8433172_8446622_RNA45SN1_45S_50ntFlanks | 9534 | 9560 | 27 | AGGAAACTCTGGTGGAGGTCCGTAGCG | 388.9 | -1.7 | 0.6 | -3.1 | 2.3E-03 | 2.9E-03 | MB231/MB468 |
| >hg38_21_+_8433172_8446622_RNA45SN1_45S_50ntFlanks@11583.11602.20\|CTGGTGCGGACCAGGGGAAT | >hg38_21_+_8433172_8446622_RNA45SN1_45S_50ntFlanks | 11583 | 11602 | 20 | CTGGTGCGGACCAGGGGAAT | 658.3 | -2.4 | 0.8 | -3.0 | 2.5E-03 | 2.9E-03 | BT20/MB468 |
| >hg38_21_+_8433172_8446622_RNA45SN1_45S_50ntFlanks@4046.4067.22\|CATTCGAACGTCTGCCCTATCA | >hg38_21_+_8433172_8446622_RNA45SN1_45S_50ntFlanks | 4046 | 4067 | 22 | CATTCGAACGTCTGCCCTATCA | 143.9 | -1.4 | 0.4 | -3.0 | 2.5E-03 | 2.9E-03 | BT20/MB468 |
| >hg38_21_+_8433172_8446622_RNA45SN1_45S_50ntFlanks@4364.4391.28\|CTGCAGTTAAAAAGCTCGTAGTTGGATC | >hg38_21_+_8433172_8446622_RNA45SN1_45S_50ntFlanks | 4364 | 4391 | 28 | CTGCAGTTAAAAAGCTCGTAGTTGGATC | 124.6 | -1.9 | 0.6 | -3.0 | 2.5E-03 | 2.9E-03 | BT20/MB468 |
| >hg38_21_+_8433172_8446622_RNA45SN1_45S_50ntFlanks@10797.10817.21\|GCATGTTGGAACAATGTAGGT | >hg38_21_+_8433172_8446622_RNA45SN1_45S_50ntFlanks | 10797 | 10817 | 21 | GCATGTTGGAACAATGTAGGT | 666.9 | -2.6 | 0.9 | -3.0 | 2.5E-03 | 2.9E-03 | BT20/MB468 |
| >hg38_21_+_8433172_8446622_RNA45SN1_45S_50ntFlanks@11567.11600.34\|AGCAGCCGACTTAGAACTGGTGCGGACCAGGGGA | >hg38_21_+_8433172_8446622_RNA45SN1_45S_50ntFlanks | 11567 | 11600 | 34 | AGCAGCCGACTTAGAACTGGTGCGGACCAGGGGA | 199.2 | 1.2 | 0.4 | 3.0 | 2.3E-03 | 2.9E-03 | MB231/MB468 |
| >hg38_21_+_8433172_8446622_RNA45SN1_45S_50ntFlanks@11235.11255.21\|CTTCCCGTGGATCGCCCCAGC | >hg38_21_+_8433172_8446622_RNA45SN1_45S_50ntFlanks | 11235 | 11255 | 21 | CTTCCCGTGGATCGCCCCAGC | 245.9 | -1.8 | 0.6 | -3.0 | 2.3E-03 | 2.9E-03 | MB231/MB468 |

<table>
<tr><th></th><th></th><th></th><th></th><th></th><th></th><th></th><th></th><th></th><th></th><th></th><th></th><th></th></tr>
<tr><td>>MI0000063|hsa-let-7b&WithFlank&22|+|46113680|46113774@12.33.22(+1G)&[MIMAT0000063&hsa-let-7b-5p&offsets|0|0(+1G);m-14&22|+|46113691|46113712&offsets|0|0(+1G)]|TGAGGTAGTAGGTTGTGTGGTTG</td><td>>MI0000063|hsa-let-7b&WithFlank&22|+|46113680|46113774</td><td>12</td><td>33</td><td>22(+1G)</td><td>TGAGGTAGTAGGTTGTGTGGTTG</td><td>582.2</td><td>2.5</td><td>0.7</td><td>3.5</td><td>4.2E-04</td><td>2.9E-03</td><td>BT20/MB231</td></tr>
<tr><td>>hg38_21_+_8433172_8446622_RNA45SN1_45S_50ntFlanks@9574.9594.21|AATCGGTCGTCCGACCTGGGT</td><td>>hg38_21_+_8433172_8446622_RNA45SN1_45S_50ntFlanks</td><td>9574</td><td>9594</td><td>21</td><td>AATCGGTCGTCCGACCTGGGT</td><td>469.2</td><td>-1.9</td><td>0.6</td><td>-3.0</td><td>2.5E-03</td><td>2.9E-03</td><td>BT20/MB468</td></tr>
<tr><td>>am_trnaMT_GluTTC_MT_-_14674_14742@27.48.22,trnalookalike8_GluTTC_5_-_93905172_93905240@27.48.22|GTTTTTCATATCATTGGTCGTG</td><td>>am_trnaMT_GluTTC_MT_-_14674_14742</td><td>27</td><td>48</td><td>22</td><td>GTTTTTCATATCATTGGTCGTG</td><td>119.3</td><td>-1.9</td><td>0.6</td><td>-3.0</td><td>2.3E-03</td><td>2.9E-03</td><td>MB231/MB468</td></tr>
<tr><td>>hg38_21_+_8433172_8446622_RNA45SN1_45S_50ntFlanks@6723.6744.22|TTGCAGGACACATTGATCATCG</td><td>>hg38_21_+_8433172_8446622_RNA45SN1_45S_50ntFlanks</td><td>6723</td><td>6744</td><td>22</td><td>TTGCAGGACACATTGATCATCG</td><td>1146.1</td><td>-2.3</td><td>0.8</td><td>-3.0</td><td>2.3E-03</td><td>2.9E-03</td><td>MB231/MB468</td></tr>
<tr><td>>MI0000542|hsa-mir-320a&WithFlank&8|-|22244960|22245043@48.69.22(+1G)&[MIMAT0000510&hsa-miR-320a-3p&offsets|0|0(+1G);m-60&8|-|22244975|22244996&offsets|0|0(+1G)]|AAAAGCTGGGTTGAGAGGGCGAG</td><td>>MI0000542|hsa-mir-320a&WithFlank&8|-|22244960|22245043</td><td>48</td><td>69</td><td>22(+1G)</td><td>AAAAGCTGGGTTGAGAGGGCGAG</td><td>1230.0</td><td>3.1</td><td>0.9</td><td>3.5</td><td>4.3E-04</td><td>3.0E-03</td><td>BT20/MB231</td></tr>
<tr><td>>trnaMT_SerGCT_MT_+_12207_12265@8.30.23|CTCACAAGAACTGCTAACTCATG</td><td>>trnaMT_SerGCT_MT_+_12207_12265</td><td>8</td><td>30</td><td>23</td><td>CTCACAAGAACTGCTAACTCATG</td><td>94.5</td><td>-2.4</td><td>0.8</td><td>-3.0</td><td>2.3E-03</td><td>3.0E-03</td><td>MB231/MB468</td></tr>
<tr><td>>hg38_21_+_8433172_8446622_RNA45SN1_45S_50ntFlanks@4731.4758.28|AAGAACGAAAGTCGGAGGTTCGAAGACG</td><td>>hg38_21_+_8433172_8446622_RNA45SN1_45S_50ntFlanks</td><td>4731</td><td>4758</td><td>28</td><td>AAGAACGAAAGTCGGAGGTTCGAAGACG</td><td>2786.6</td><td>-1.4</td><td>0.5</td><td>-3.0</td><td>2.5E-03</td><td>3.0E-03</td><td>BT20/MB468</td></tr>
<tr><td>>hg38_21_+_8433172_8446622_RNA45SN1_45S_50ntFlanks@10793.10818.26|TCTGGCATGTTGGAACAATGTAGGTA</td><td>>hg38_21_+_8433172_8446622_RNA45SN1_45S_50ntFlanks</td><td>10793</td><td>10818</td><td>26</td><td>TCTGGCATGTTGGAACAATGTAGGTA</td><td>423.4</td><td>-2.2</td><td>0.7</td><td>-3.0</td><td>2.3E-03</td><td>3.0E-03</td><td>MB231/MB468</td></tr>
<tr><td>>hg38_21_+_8433172_8446622_RNA45SN1_45S_50ntFlanks@11572.11611.40|CCGACTTAGAACTGGTGCGGACCAGGGGAATCCGACTGTT</td><td>>hg38_21_+_8433172_8446622_RNA45SN1_45S_50ntFlanks</td><td>11572</td><td>11611</td><td>40</td><td>CCGACTTAGAACTGGTGCGGACCAGGGGAATCCGACTGTT</td><td>416.9</td><td>-2.6</td><td>0.9</td><td>-3.0</td><td>2.5E-03</td><td>3.0E-03</td><td>BT20/MB468</td></tr>
<tr><td>>hg38_21_+_8433172_8446622_RNA45SN1_45S_50ntFlanks@4735.4758.24|ACGAAAGTCGGAGGTTCGAAGACG</td><td>>hg38_21_+_8433172_8446622_RNA45SN1_45S_50ntFlanks</td><td>4735</td><td>4758</td><td>24</td><td>ACGAAAGTCGGAGGTTCGAAGACG</td><td>1401.8</td><td>2.3</td><td>0.8</td><td>3.0</td><td>2.4E-03</td><td>3.0E-03</td><td>MB231/MB468</td></tr>
<tr><td>>hg38_21_+_8433172_8446622_RNA45SN1_45S_50ntFlanks@12418.12451.34|CGATGTCGGCTCTTCCTATCATTGTGAAGCAGAA</td><td>>hg38_21_+_8433172_8446622_RNA45SN1_45S_50ntFlanks</td><td>12418</td><td>12451</td><td>34</td><td>CGATGTCGGCTCTTCCTATCATTGTGAAGCAGAA</td><td>198.0</td><td>1.4</td><td>0.5</td><td>3.0</td><td>2.4E-03</td><td>3.0E-03</td><td>MB231/MB468</td></tr>
<tr><td>>MI0000767|hsa-mir-365a&WithFlank&16|+|14309279|14309377@62.84.23;MI0000769|hsa-mir-365b&WithFlank&17|+|31575405|31575527@74.96.23&[MIMAT0000710&hsa-miR-365a-3p&offsets|0|+1;m-176&16|+|14309340|14309361&offsets|0|+1];[MIMAT0022834&hsa-miR-365b-3p&offsets|0|+1;m-175&17|+|31575478|31575499&offsets|0|+1]|TAATGCCCCTAAAAATCCTTATT</td><td>>MI0000767|hsa-mir-365a&WithFlank&16|+|14309279|14309377</td><td>62</td><td>84</td><td>23</td><td>TAATGCCCCTAAAAATCCTTATT</td><td>1004.5</td><td>3.0</td><td>0.9</td><td>3.5</td><td>4.3E-04</td><td>3.0E-03</td><td>BT20/MB231</td></tr>
<tr><td>>hg38_21_+_8433172_8446622_RNA45SN1_45S_50ntFlanks@12418.12457.40|CGATGTCGGCTCTTCCTATCATTGTGAAGCAGAATTCACC</td><td>>hg38_21_+_8433172_8446622_RNA45SN1_45S_50ntFlanks</td><td>12418</td><td>12457</td><td>40</td><td>CGATGTCGGCTCTTCCTATCATTGTGAAGCAGAATTCACC</td><td>293.4</td><td>-2.6</td><td>0.7</td><td>-3.5</td><td>4.3E-04</td><td>3.0E-03</td><td>BT20/MB231</td></tr>
<tr><td>>hg38_21_+_8433172_8446622_RNA45SN1_45S_50ntFlanks@3840.3860.21|CTCCTACTTGGATAACTGTGG</td><td>>hg38_21_+_8433172_8446622_RNA45SN1_45S_50ntFlanks</td><td>3840</td><td>3860</td><td>21</td><td>CTCCTACTTGGATAACTGTGG</td><td>625.9</td><td>1.3</td><td>0.4</td><td>3.0</td><td>2.6E-03</td><td>3.0E-03</td><td>BT20/MB468</td></tr>
<tr><td>>hg38_21_+_8433172_8446622_RNA45SN1_45S_50ntFlanks@9738.9767.30|GAAACGATCTCAACCTATTCTCAAACTTTA</td><td>>hg38_21_+_8433172_8446622_RNA45SN1_45S_50ntFlanks</td><td>9738</td><td>9767</td><td>30</td><td>GAAACGATCTCAACCTATTCTCAAACTTTA</td><td>456.2</td><td>-1.5</td><td>0.5</td><td>-3.0</td><td>2.4E-03</td><td>3.0E-03</td><td>MB231/MB468</td></tr>
<tr><td>>hg38_21_+_8433172_8446622_RNA45SN1_45S_50ntFlanks@6651.6696.46|CGACTCTTAGCGGTGGATCACTCGGCTCGTGCGTCGATGAAGAACG</td><td>>hg38_21_+_8433172_8446622_RNA45SN1_45S_50ntFlanks</td><td>6651</td><td>6696</td><td>46</td><td>CGACTCTTAGCGGTGGATCACTCGGCTCGTGCGTCGATGAAGAACG</td><td>13094.3</td><td>-2.3</td><td>0.7</td><td>-3.5</td><td>4.4E-04</td><td>3.0E-03</td><td>BT20/MB231</td></tr>
<tr><td>>MI0000255|hsa-mir-30d&WithFlank&8|-|134804870|134804951@12.32.21&[MIMAT0000245&hsa-miR-30d-5p&offsets|0|-1;m-21&8|-|134804919|134804940&offsets|0|-1]|TGTAAACATCCCCGACTGGAA</td><td>>MI0000255|hsa-mir-30d&WithFlank&8|-|134804870|134804951</td><td>12</td><td>32</td><td>21</td><td>TGTAAACATCCCCGACTGGAA</td><td>1621.1</td><td>2.4</td><td>0.7</td><td>3.5</td><td>4.4E-04</td><td>3.0E-03</td><td>BT20/MB231</td></tr>
<tr><td>>hg38_21_+_8433172_8446622_RNA45SN1_45S_50ntFlanks@4753.4783.31|AAGACGATCAGATACCGTCGTAGTTCCGACC</td><td>>hg38_21_+_8433172_8446622_RNA45SN1_45S_50ntFlanks</td><td>4753</td><td>4783</td><td>31</td><td>AAGACGATCAGATACCGTCGTAGTTCCGACC</td><td>217.5</td><td>-1.5</td><td>0.5</td><td>-3.0</td><td>2.6E-03</td><td>3.0E-03</td><td>BT20/MB468</td></tr>
<tr><td>>hg38_21_+_8433172_8446622_RNA45SN1_45S_50ntFlanks@6733.6758.26|CATTGATCATCGACACTTCGAACGCA</td><td>>hg38_21_+_8433172_8446622_RNA45SN1_45S_50ntFlanks</td><td>6733</td><td>6758</td><td>26</td><td>CATTGATCATCGACACTTCGAACGCA</td><td>340.3</td><td>-1.3</td><td>0.4</td><td>-3.0</td><td>2.4E-03</td><td>3.0E-03</td><td>MB231/MB468</td></tr>
<tr><td>>hg38_21_+_8433172_8446622_RNA45SN1_45S_50ntFlanks@4297.4319.23|CATTGGAGGGCAAGTCTGGTGCC</td><td>>hg38_21_+_8433172_8446622_RNA45SN1_45S_50ntFlanks</td><td>4297</td><td>4319</td><td>23</td><td>CATTGGAGGGCAAGTCTGGTGCC</td><td>131.6</td><td>-2.6</td><td>0.8</td><td>-3.0</td><td>2.4E-03</td><td>3.0E-03</td><td>MB231/MB468</td></tr>
<tr><td>>hg38_21_+_8433172_8446622_RNA45SN1_45S_50ntFlanks@7974.8002.29|ACGCGACCTCAGATCAGACGTGGCGACCC</td><td>>hg38_21_+_8433172_8446622_RNA45SN1_45S_50ntFlanks</td><td>7974</td><td>8002</td><td>29</td><td>ACGCGACCTCAGATCAGACGTGGCGACCC</td><td>217.2</td><td>2.6</td><td>0.7</td><td>3.5</td><td>4.4E-04</td><td>3.0E-03</td><td>BT20/MB231</td></tr>
<tr><td>>MI0016887|hsa-mir-4521&WithFlank&17|+|8186939|8187010@14.34.21&[MIMAT0019058&hsa-miR-4521&offsets|+4|+3;m-765&17|+|8186948|8186969&offsets|+4|+3]|AGGAAGTCCTGTGCTCAGTTT</td><td>>MI0016887|hsa-mir-4521&WithFlank&17|+|8186939|8187010</td><td>14</td><td>34</td><td>21</td><td>AGGAAGTCCTGTGCTCAGTTT</td><td>274.9</td><td>2.7</td><td>0.8</td><td>3.5</td><td>4.4E-04</td><td>3.0E-03</td><td>BT20/MB231</td></tr>
<tr><td>>trnaMT_CysGCA_MT_-_5761_5826@47.69.23|CTTCAAACCTGCCGGGGCTTCCA</td><td>>trnaMT_CysGCA_MT_-_5761_5826</td><td>47</td><td>69</td><td>23</td><td>CTTCAAACCTGCCGGGGCTTCCA</td><td>151.4</td><td>-1.9</td><td>0.6</td><td>-3.0</td><td>2.4E-03</td><td>3.1E-03</td><td>MB231/MB468</td></tr>
<tr><td>>hg38_21_+_8433172_8446622_RNA45SN1_45S_50ntFlanks@6646.6672.27|TCGTACGACTCTTAGCGGTGGATCACT</td><td>>hg38_21_+_8433172_8446622_RNA45SN1_45S_50ntFlanks</td><td>6646</td><td>6672</td><td>27</td><td>TCGTACGACTCTTAGCGGTGGATCACT</td><td>6481.1</td><td>2.7</td><td>0.8</td><td>3.5</td><td>4.5E-04</td><td>3.1E-03</td><td>BT20/MB231</td></tr>
<tr><td>>hg38_21_+_8433172_8446622_RNA45SN1_45S_50ntFlanks@11876.11893.18|AAGACCCTGTTGAGCTTG</td><td>>hg38_21_+_8433172_8446622_RNA45SN1_45S_50ntFlanks</td><td>11876</td><td>11893</td><td>18</td><td>AAGACCCTGTTGAGCTTG</td><td>2420.2</td><td>-2.9</td><td>1.0</td><td>-3.0</td><td>2.6E-03</td><td>3.1E-03</td><td>BT20/MB468</td></tr>
<tr><td>>hg38_21_+_8433172_8446622_RNA45SN1_45S_50ntFlanks@4729.4748.20|TCAAGAACGAAAGTCGGAGG</td><td>>hg38_21_+_8433172_8446622_RNA45SN1_45S_50ntFlanks</td><td>4729</td><td>4748</td><td>20</td><td>TCAAGAACGAAAGTCGGAGG</td><td>129.9</td><td>-1.5</td><td>0.5</td><td>-3.0</td><td>2.6E-03</td><td>3.1E-03</td><td>BT20/MB468</td></tr>
<tr><td>>hg38_21_+_8433172_8446622_RNA45SN1_45S_50ntFlanks@10905.10935.31|CTGGGCGCGCGCCGCGGCTGGACGAGGCGCC</td><td>>hg38_21_+_8433172_8446622_RNA45SN1_45S_50ntFlanks</td><td>10905</td><td>10935</td><td>31</td><td>CTGGGCGCGCGCCGCGGCTGGACGAGGCGCC</td><td>103.0</td><td>-3.4</td><td>1.1</td><td>-3.0</td><td>2.6E-03</td><td>3.1E-03</td><td>BT20/MB468</td></tr>
<tr><td>>hg38_21_+_8433172_8446622_RNA45SN1_45S_50ntFlanks@7974.8012.39|ACGCGACCTCAGATCAGACGTGGCGACCCGCTGAATTTA</td><td>>hg38_21_+_8433172_8446622_RNA45SN1_45S_50ntFlanks</td><td>7974</td><td>8012</td><td>39</td><td>ACGCGACCTCAGATCAGACGTGGCGACCCGCTGAATTTA</td><td>1333.8</td><td>2.4</td><td>0.8</td><td>3.0</td><td>2.6E-03</td><td>3.1E-03</td><td>BT20/MB468</td></tr>
<tr><td>>hg38_21_+_8433172_8446622_RNA45SN1_45S_50ntFlanks@12419.12455.37|GATGTCGGCTCTTCCTATCATTGTGAAGCAGAATTCA</td><td>>hg38_21_+_8433172_8446622_RNA45SN1_45S_50ntFlanks</td><td>12419</td><td>12455</td><td>37</td><td>GATGTCGGCTCTTCCTATCATTGTGAAGCAGAATTCA</td><td>287.6</td><td>-2.6</td><td>0.7</td><td>-3.5</td><td>4.6E-04</td><td>3.1E-03</td><td>BT20/MB231</td></tr>
<tr><td>>hg38_21_+_8433172_8446622_RNA45SN1_45S_50ntFlanks@13000.13027.28|TGCGATCTATTGAAAGTCAGCCCTCGAC</td><td>>hg38_21_+_8433172_8446622_RNA45SN1_45S_50ntFlanks</td><td>13000</td><td>13027</td><td>28</td><td>TGCGATCTATTGAAAGTCAGCCCTCGAC</td><td>286.3</td><td>-3.9</td><td>1.1</td><td>-3.5</td><td>4.6E-04</td><td>3.1E-03</td><td>BT20/MB231</td></tr>
<tr><td>>MI0000746|hsa-mir-99b&WithFlank&19|+|51692606|51692687@13.31.19&[MIMAT0000689&hsa-miR-99b-5p&offsets|0|-3;m-10&19|+|51692618|51692639&offsets|0|-3]|CACCCGTAGAACCGACCTT</td><td>>MI0000746|hsa-mir-99b&WithFlank&19|+|51692606|51692687</td><td>13</td><td>31</td><td>19</td><td>CACCCGTAGAACCGACCTT</td><td>203.6</td><td>2.7</td><td>0.8</td><td>3.5</td><td>4.6E-04</td><td>3.1E-03</td><td>BT20/MB231</td></tr>
<tr><td>>hg38_21_+_8433172_8446622_RNA45SN1_45S_50ntFlanks@11874.11897.24|AGAAGACCCTGTTGAGCTTGACTC</td><td>>hg38_21_+_8433172_8446622_RNA45SN1_45S_50ntFlanks</td><td>11874</td><td>11897</td><td>24</td><td>AGAAGACCCTGTTGAGCTTGACTC</td><td>645.1</td><td>-2.6</td><td>0.9</td><td>-3.0</td><td>2.5E-03</td><td>3.1E-03</td><td>MB231/MB468</td></tr>
<tr><td>>hg38_21_+_8433172_8446622_RNA45SN1_45S_50ntFlanks@6657.6681.25|TTAGCGGTGGATCACTCGGCTCGTG</td><td>>hg38_21_+_8433172_8446622_RNA45SN1_45S_50ntFlanks</td><td>6657</td><td>6681</td><td>25</td><td>TTAGCGGTGGATCACTCGGCTCGTG</td><td>504.6</td><td>-2.8</td><td>0.9</td><td>-3.0</td><td>2.7E-03</td><td>3.1E-03</td><td>BT20/MB468</td></tr>
<tr><td>>trnaMT_ValTAC_MT_+_1602_1670@49.70.22|AACTTAACTTGACCGCTCTGAC</td><td>>trnaMT_ValTAC_MT_+_1602_1670</td><td>49</td><td>70</td><td>22</td><td>AACTTAACTTGACCGCTCTGAC</td><td>180.0</td><td>2.9</td><td>0.8</td><td>3.5</td><td>4.6E-04</td><td>3.1E-03</td><td>BT20/MB231</td></tr>
<tr><td>>hg38_21_+_8433172_8446622_RNA45SN1_45S_50ntFlanks@4399.4423.25|CGGGCGGGCGGTCCGCCGCGAGGCG</td><td>>hg38_21_+_8433172_8446622_RNA45SN1_45S_50ntFlanks</td><td>4399</td><td>4423</td><td>25</td><td>CGGGCGGGCGGTCCGCCGCGAGGCG</td><td>105.8</td><td>-2.5</td><td>0.8</td><td>-3.0</td><td>2.5E-03</td><td>3.1E-03</td><td>MB231/MB468</td></tr>
<tr><td>>hg38_21_+_8433172_8446622_RNA45SN1_45S_50ntFlanks@9544.9573.30|GGTGGAGGTCCGTAGCGGTCCTGACGTGCA</td><td>>hg38_21_+_8433172_8446622_RNA45SN1_45S_50ntFlanks</td><td>9544</td><td>9573</td><td>30</td><td>GGTGGAGGTCCGTAGCGGTCCTGACGTGCA</td><td>359.3</td><td>1.8</td><td>0.6</td><td>3.0</td><td>2.5E-03</td><td>3.1E-03</td><td>MB231/MB468</td></tr>
<tr><td>>am_trnaMT_GluTTC_MT_-_14674_14742@27.55.29,trnalookalike8_GluTTC_5_-_93905172_93905240@27.55.29|GTTTTTCATATCATTGGTCGTGGTTGTAG</td><td>>am_trnaMT_GluTTC_MT_-_14674_14742</td><td>27</td><td>55</td><td>29</td><td>GTTTTTCATATCATTGGTCGTGGTTGTAG</td><td>87.9</td><td>-2.7</td><td>0.9</td><td>-3.0</td><td>2.5E-03</td><td>3.2E-03</td><td>MB231/MB468</td></tr>
<tr><td>>hg38_21_+_8433172_8446622_RNA45SN1_45S_50ntFlanks@3841.3879.39|TCCTACTTGGATAACTGTGGTAATTCTAGAGCTAATACA</td><td>>hg38_21_+_8433172_8446622_RNA45SN1_45S_50ntFlanks</td><td>3841</td><td>3879</td><td>39</td><td>TCCTACTTGGATAACTGTGGTAATTCTAGAGCTAATACA</td><td>209.8</td><td>2.7</td><td>0.8</td><td>3.5</td><td>4.7E-04</td><td>3.2E-03</td><td>BT20/MB231</td></tr>
<tr><td>>hg38_21_+_8433172_8446622_RNA45SN1_45S_50ntFlanks@12457.12483.27|CAAGCGTTGGATTGTTCACCCACTAAT</td><td>>hg38_21_+_8433172_8446622_RNA45SN1_45S_50ntFlanks</td><td>12457</td><td>12483</td><td>27</td><td>CAAGCGTTGGATTGTTCACCCACTAAT</td><td>150.4</td><td>-3.2</td><td>1.1</td><td>-3.0</td><td>2.7E-03</td><td>3.2E-03</td><td>BT20/MB468</td></tr>
<tr><td>>hg38_21_+_8433172_8446622_RNA45SN1_45S_50ntFlanks@6735.6770.36|TTGATCATCGACACTTCGAACGCACTTGCGGCCCCG</td><td>>hg38_21_+_8433172_8446622_RNA45SN1_45S_50ntFlanks</td><td>6735</td><td>6770</td><td>36</td><td>TTGATCATCGACACTTCGAACGCACTTGCGGCCCCG</td><td>221.0</td><td>-2.0</td><td>0.7</td><td>-3.0</td><td>2.6E-03</td><td>3.2E-03</td><td>MB231/MB468</td></tr>
<tr><td>>hg38_21_+_8433172_8446622_RNA45SN1_45S_50ntFlanks@6645.6666.22|CTCGTACGACTCTTAGCGGTGG</td><td>>hg38_21_+_8433172_8446622_RNA45SN1_45S_50ntFlanks</td><td>6645</td><td>6666</td><td>22</td><td>CTCGTACGACTCTTAGCGGTGG</td><td>1931.5</td><td>2.6</td><td>0.9</td><td>3.0</td><td>2.6E-03</td><td>3.2E-03</td><td>MB231/MB468</td></tr>
<tr><td>>hg38_21_+_8433172_8446622_RNA45SN1_45S_50ntFlanks@11669.11709.41|GCCCAGTGCTCTGAATGTCAAAGTGAAGAAATTCAATGAAG</td><td>>hg38_21_+_8433172_8446622_RNA45SN1_45S_50ntFlanks</td><td>11669</td><td>11709</td><td>41</td><td>GCCCAGTGCTCTGAATGTCAAAGTGAAGAAATTCAATGAAG</td><td>174.3</td><td>-2.8</td><td>0.8</td><td>-3.5</td><td>4.8E-04</td><td>3.3E-03</td><td>BT20/MB231</td></tr>
<tr><td>>hg38_21_+_8433172_8446622_RNA45SN1_45S_50ntFlanks@9266.9285.20|CGACGTCGGCTACCCACCCG</td><td>>hg38_21_+_8433172_8446622_RNA45SN1_45S_50ntFlanks</td><td>9266</td><td>9285</td><td>20</td><td>CGACGTCGGCTACCCACCCG</td><td>143.4</td><td>-2.8</td><td>0.9</td><td>-3.0</td><td>2.8E-03</td><td>3.3E-03</td><td>BT20/MB468</td></tr>
<tr><td>>trna10_LysCTT_16_+_3241501_3241573@4.35.32,trna13_LysCTT_14_-_58706613_58706685@4.35.32,trna2_LysCTT_15_+_79152904_79152976@4.35.32|CGGCTAGCTCAGTCGGTAGAGCATGGGACTCT</td><td>>trna10_LysCTT_16_+_3241501_3241573</td><td>4</td><td>35</td><td>32</td><td>CGGCTAGCTCAGTCGGTAGAGCATGGGACTCT</td><td>421.7</td><td>-2.8</td><td>0.8</td><td>-3.5</td><td>4.9E-04</td><td>3.3E-03</td><td>BT20/MB231</td></tr>
<tr><td>>hg38_21_+_8433172_8446622_RNA45SN1_45S_50ntFlanks@6654.6681.28|CTCTTAGCGGTGGATCACTCGGCTCGTG</td><td>>hg38_21_+_8433172_8446622_RNA45SN1_45S_50ntFlanks</td><td>6654</td><td>6681</td><td>28</td><td>CTCTTAGCGGTGGATCACTCGGCTCGTG</td><td>1016.5</td><td>-2.4</td><td>0.8</td><td>-3.0</td><td>2.8E-03</td><td>3.3E-03</td><td>BT20/MB468</td></tr>
<tr><td>>hg38_21_+_8433172_8446622_RNA45SN1_45S_50ntFlanks@12419.12459.41|GATGTCGGCTCTTCCTATCATTGTGAAGCAGAATTCACCAA</td><td>>hg38_21_+_8433172_8446622_RNA45SN1_45S_50ntFlanks</td><td>12419</td><td>12459</td><td>41</td><td>GATGTCGGCTCTTCCTATCATTGTGAAGCAGAATTCACCAA</td><td>163.9</td><td>-2.6</td><td>0.7</td><td>-3.5</td><td>4.9E-04</td><td>3.3E-03</td><td>BT20/MB231</td></tr>
</table>

| | | | | | | | | | | | | |
|---|---|---|---|---|---|---|---|---|---|---|---|---|
| >hg38_21_+_8433172_8446622_RNA45SN1_45S_50ntFlanks@7979.8012.34\|ACCTCAGATCAGACGTGGCGACCCGCTGAATTTA | >hg38_21_+_8433172_8446622_RNA45SN1_45S_50ntFlanks | 7979 | 8012 | 34 | ACCTCAGATCAGACGTGGCGACCCGCTGAATTTA | 300.5 | -2.6 | 0.9 | -3.0 | 2.8E-03 | 3.3E-03 | BT20/MB468 |
| >hg38_21_+_8433172_8446622_RNA45SN1_45S_50ntFlanks@8176.8208.33\|CTTCTGATCGAGGCCCAGCCCGTGGACGGTGTG | >hg38_21_+_8433172_8446622_RNA45SN1_45S_50ntFlanks | 8176 | 8208 | 33 | CTTCTGATCGAGGCCCAGCCCGTGGACGGTGTG | 305.5 | -1.8 | 0.6 | -3.0 | 2.8E-03 | 3.3E-03 | BT20/MB468 |
| >hg38_21_+_8433172_8446622_RNA45SN1_45S_50ntFlanks@11315.11338.24\|CCACCCCACGTCTCGTCGCGCGCG | >hg38_21_+_8433172_8446622_RNA45SN1_45S_50ntFlanks | 11315 | 11338 | 24 | CCACCCCACGTCTCGTCGCGCGCG | 297.4 | -3.9 | 1.3 | -3.0 | 2.9E-03 | 3.3E-03 | BT20/MB468 |
| >hg38_21_+_8433172_8446622_RNA45SN1_45S_50ntFlanks@10349.10367.19\|AACGAGAACTTTGAAGGCC | >hg38_21_+_8433172_8446622_RNA45SN1_45S_50ntFlanks | 10349 | 10367 | 19 | AACGAGAACTTTGAAGGCC | 1432.7 | 2.0 | 0.7 | 3.0 | 2.7E-03 | 3.4E-03 | MB231/MB468 |
| >hg38_21_+_8433172_8446622_RNA45SN1_45S_50ntFlanks@6651.6697.47\|CGACTCTTAGCGGTGGATCACTCGGCTCGTGCGTCGATGAAGAACGC | >hg38_21_+_8433172_8446622_RNA45SN1_45S_50ntFlanks | 6651 | 6697 | 47 | CGACTCTTAGCGGTGGATCACTCGGCTCGTGCGTCGATGAAGAACGC | 2275.6 | 2.7 | 0.9 | 3.0 | 2.9E-03 | 3.4E-03 | BT20/MB468 |
| >hg38_21_+_8433172_8446622_RNA45SN1_45S_50ntFlanks@9737.9765.29\|CGAAACGATCTCAACCTATTCTCAAACTT | >hg38_21_+_8433172_8446622_RNA45SN1_45S_50ntFlanks | 9737 | 9765 | 29 | CGAAACGATCTCAACCTATTCTCAAACTT | 333.6 | -3.3 | 1.0 | -3.5 | 5.1E-04 | 3.4E-03 | BT20/MB231 |
| >hg38_21_+_8433172_8446622_RNA45SN1_45S_50ntFlanks@6721.6770.50\|AATTGCAGGACACATTGATCATCGACACTTCGAACGCACTTGCGGCCCCG | >hg38_21_+_8433172_8446622_RNA45SN1_45S_50ntFlanks | 6721 | 6770 | 50 | AATTGCAGGACACATTGATCATCGACACTTCGAACGCACTTGCGGCCCCG | 191.4 | -2.3 | 0.8 | -3.0 | 2.7E-03 | 3.4E-03 | MB231/MB468 |
| >hg38_21_+_8433172_8446622_RNA45SN1_45S_50ntFlanks@6700.6720.21\|CTAGCTGCGAGAATTAATGTG | >hg38_21_+_8433172_8446622_RNA45SN1_45S_50ntFlanks | 6700 | 6720 | 21 | CTAGCTGCGAGAATTAATGTG | 11936.4 | -2.3 | 0.8 | -3.0 | 2.7E-03 | 3.4E-03 | MB231/MB468 |
| >hg38_21_+_8433172_8446622_RNA45SN1_45S_50ntFlanks@6723.6747.25\|TTGCAGGACACATTGATCATCGACA | >hg38_21_+_8433172_8446622_RNA45SN1_45S_50ntFlanks | 6723 | 6747 | 25 | TTGCAGGACACATTGATCATCGACA | 253.7 | 1.3 | 0.4 | 3.0 | 2.7E-03 | 3.4E-03 | MB231/MB468 |
| >hg38_21_+_8433172_8446622_RNA45SN1_45S_50ntFlanks@9536.9571.36\|GAAACTCTGGTGGAGGTCCGTAGCGGTCCTGACGTG | >hg38_21_+_8433172_8446622_RNA45SN1_45S_50ntFlanks | 9536 | 9571 | 36 | GAAACTCTGGTGGAGGTCCGTAGCGGTCCTGACGTG | 307.4 | 1.1 | 0.4 | 3.0 | 2.7E-03 | 3.4E-03 | MB231/MB468 |
| >hg38_21_+_8433172_8446622_RNA45SN1_45S_50ntFlanks@9565.9597.33\|TGACGTGCAAATCGGTCGTCCGACCTGGGTATA | >hg38_21_+_8433172_8446622_RNA45SN1_45S_50ntFlanks | 9565 | 9597 | 33 | TGACGTGCAAATCGGTCGTCCGACCTGGGTATA | 1754.6 | -2.0 | 0.7 | -3.0 | 2.7E-03 | 3.4E-03 | MB231/MB468 |
| >hg38_21_+_8433172_8446622_RNA45SN1_45S_50ntFlanks@6721.6757.37\|AATTGCAGGACACATTGATCATCGACACTTCGAACGC | >hg38_21_+_8433172_8446622_RNA45SN1_45S_50ntFlanks | 6721 | 6757 | 37 | AATTGCAGGACACATTGATCATCGACACTTCGAACGC | 159.6 | -1.8 | 0.6 | -3.0 | 2.7E-03 | 3.4E-03 | MB231/MB468 |
| >MI0000469\|hsa-mir-125a&WithFlank&19\|+\|51693248\|51693345@21.42.22(+1A)&[MIMAT0000443&hsa-miR-125a-5p&offsets\|0\|-2(+1A);m-63&19\|+\|51693268\|51693289&offsets\|0\|0(+1A)]\|TCCCTGAGACCCTTTAACCTGTA | >MI0000469\|hsa-mir-125a&WithFlank&19\|+\|51693248\|51693345 | 21 | 42 | 22(+1A) | TCCCTGAGACCCTTTAACCTGTA | 4450.9 | 2.3 | 0.7 | 3.5 | 5.1E-04 | 3.4E-03 | BT20/MB231 |
| >hg38_21_+_8433172_8446622_RNA45SN1_45S_50ntFlanks@11694.11711.18\|AAGAAATTCAATGAAGCG | >hg38_21_+_8433172_8446622_RNA45SN1_45S_50ntFlanks | 11694 | 11711 | 18 | AAGAAATTCAATGAAGCG | 222.9 | -2.3 | 0.8 | -3.0 | 2.7E-03 | 3.4E-03 | MB231/MB468 |
| >hg38_21_+_8433172_8446622_RNA45SN1_45S_50ntFlanks@4733.4752.20\|GAACGAAAGTCGGAGGTTCG | >hg38_21_+_8433172_8446622_RNA45SN1_45S_50ntFlanks | 4733 | 4752 | 20 | GAACGAAAGTCGGAGGTTCG | 246.0 | 1.4 | 0.5 | 3.0 | 2.7E-03 | 3.4E-03 | MB231/MB468 |
| >hg38_21_+_8433172_8446622_RNA45SN1_45S_50ntFlanks@4297.4314.18\|CATTGGAGGGCAAGTCTG | >hg38_21_+_8433172_8446622_RNA45SN1_45S_50ntFlanks | 4297 | 4314 | 18 | CATTGGAGGGCAAGTCTG | 396.6 | -2.2 | 0.7 | -3.0 | 2.7E-03 | 3.5E-03 | MB231/MB468 |
| >hg38_21_+_8433172_8446622_RNA45SN1_45S_50ntFlanks@10841.10861.21\|CGTAACTTCGGGATAAGGATT | >hg38_21_+_8433172_8446622_RNA45SN1_45S_50ntFlanks | 10841 | 10861 | 21 | CGTAACTTCGGGATAAGGATT | 1392.7 | -2.0 | 0.7 | -3.0 | 2.8E-03 | 3.5E-03 | MB231/MB468 |
| >hg38_21_+_8433172_8446622_RNA45SN1_45S_50ntFlanks@6647.6665.19\|CGTACGACTCTTAGCGGTG | >hg38_21_+_8433172_8446622_RNA45SN1_45S_50ntFlanks | 6647 | 6665 | 19 | CGTACGACTCTTAGCGGTG | 5485.1 | 2.8 | 0.8 | 3.5 | 5.2E-04 | 3.5E-03 | BT20/MB231 |
| >hg38_21_+_8433172_8446622_RNA45SN1_45S_50ntFlanks@12396.12414.19\|GCGACGTCGCTTTTTGATC | >hg38_21_+_8433172_8446622_RNA45SN1_45S_50ntFlanks | 12396 | 12414 | 19 | GCGACGTCGCTTTTTGATC | 2528.7 | 1.5 | 0.5 | 3.0 | 2.8E-03 | 3.5E-03 | MB231/MB468 |
| >hg38_21_+_8433172_8446622_RNA45SN1_45S_50ntFlanks@3776.3805.30\|CCGGTACAGTGAAACTGCGAATGGCTCATT | >hg38_21_+_8433172_8446622_RNA45SN1_45S_50ntFlanks | 3776 | 3805 | 30 | CCGGTACAGTGAAACTGCGAATGGCTCATT | 209.5 | -1.4 | 0.5 | -3.0 | 3.0E-03 | 3.5E-03 | BT20/MB468 |
| >hg38_21_+_8433172_8446622_RNA45SN1_45S_50ntFlanks@12418.12451.34\|CGATGTCGGCTCTTCCTATCATTGTGAAGCAGAA | >hg38_21_+_8433172_8446622_RNA45SN1_45S_50ntFlanks | 12418 | 12451 | 34 | CGATGTCGGCTCTTCCTATCATTGTGAAGCAGAA | 100.9 | -1.4 | 0.5 | -3.0 | 3.0E-03 | 3.5E-03 | BT20/MB468 |
| >hg38_21_+_8433172_8446622_RNA45SN1_45S_50ntFlanks@12416.12441.26\|TTCGATGTCGGCTCTTCCTATCATTG | >hg38_21_+_8433172_8446622_RNA45SN1_45S_50ntFlanks | 12416 | 12441 | 26 | TTCGATGTCGGCTCTTCCTATCATTG | 486.4 | 1.4 | 0.5 | 3.0 | 3.0E-03 | 3.5E-03 | BT20/MB468 |
| >hg38_21_+_8433172_8446622_RNA45SN1_45S_50ntFlanks@4223.4242.20\|AAAAATAACAATACAGGACT | >hg38_21_+_8433172_8446622_RNA45SN1_45S_50ntFlanks | 4223 | 4242 | 20 | AAAAATAACAATACAGGACT | 199.5 | -2.1 | 0.7 | -3.0 | 3.0E-03 | 3.5E-03 | BT20/MB468 |
| >hg38_21_+_8433172_8446622_RNA45SN1_45S_50ntFlanks@4588.4619.32\|CTATTTTGTTGGTTTTCGGAACTGAGGCCATG | >hg38_21_+_8433172_8446622_RNA45SN1_45S_50ntFlanks | 4588 | 4619 | 32 | CTATTTTGTTGGTTTTCGGAACTGAGGCCATG | 172.6 | -1.6 | 0.5 | -3.0 | 2.8E-03 | 3.5E-03 | MB231/MB468 |
| >hg38_21_+_8433172_8446622_RNA45SN1_45S_50ntFlanks@11570.11592.23\|AGCCGACTTAGAACTGGTGCGGA | >hg38_21_+_8433172_8446622_RNA45SN1_45S_50ntFlanks | 11570 | 11592 | 23 | AGCCGACTTAGAACTGGTGCGGA | 404.4 | -2.2 | 0.7 | -3.0 | 3.0E-03 | 3.5E-03 | BT20/MB468 |
| >hg38_21_+_8433172_8446622_RNA45SN1_45S_50ntFlanks@10682.10711.30\|TCCGGCGGCGTCCGGTGAGCTCTCGCTGGC | >hg38_21_+_8433172_8446622_RNA45SN1_45S_50ntFlanks | 10682 | 10711 | 30 | TCCGGCGGCGTCCGGTGAGCTCTCGCTGGC | 139.5 | -2.0 | 0.7 | -3.0 | 2.8E-03 | 3.5E-03 | MB231/MB468 |
| >hg38_21_+_8433172_8446622_RNA45SN1_45S_50ntFlanks@10392.10429.38\|ACAGCAGTTGAACATGGGTCAGTCGGTCCTGAGAGATG | >hg38_21_+_8433172_8446622_RNA45SN1_45S_50ntFlanks | 10392 | 10429 | 38 | ACAGCAGTTGAACATGGGTCAGTCGGTCCTGAGAGATG | 1551.0 | -2.4 | 0.8 | -3.0 | 2.8E-03 | 3.5E-03 | MB231/MB468 |
| >hg38_21_+_8433172_8446622_RNA45SN1_45S_50ntFlanks@10346.10383.38\|TCAAACGAGAACTTTGAAGGCCGAAGTGGAGAAGGGTT | >hg38_21_+_8433172_8446622_RNA45SN1_45S_50ntFlanks | 10346 | 10383 | 38 | TCAAACGAGAACTTTGAAGGCCGAAGTGGAGAAGGGTT | 185.0 | 1.2 | 0.4 | 3.0 | 2.8E-03 | 3.6E-03 | MB231/MB468 |
| >hg38_21_+_8433172_8446622_RNA45SN1_45S_50ntFlanks@4732.4759.28\|AGAACGAAAGTCGGAGGTTCGAAGACGA | >hg38_21_+_8433172_8446622_RNA45SN1_45S_50ntFlanks | 4732 | 4759 | 28 | AGAACGAAAGTCGGAGGTTCGAAGACGA | 228.6 | -1.6 | 0.5 | -3.0 | 3.1E-03 | 3.6E-03 | BT20/MB468 |
| >trnaMT_ValTAC_MT_+_1602_1670@51.71.21\|CTTAACTTGACCGCTCTGACC | >trnaMT_ValTAC_MT_+_1602_1670 | 51 | 71 | 21 | CTTAACTTGACCGCTCTGACC | 218.7 | 1.5 | 0.5 | 3.0 | 2.9E-03 | 3.6E-03 | MB231/MB468 |
| >hg38_21_+_8433172_8446622_RNA45SN1_45S_50ntFlanks@10829.10853.25\|GCAAGCCGGATCCGTAACTTCGGGA | >hg38_21_+_8433172_8446622_RNA45SN1_45S_50ntFlanks | 10829 | 10853 | 25 | GCAAGCCGGATCCGTAACTTCGGGA | 161.3 | -1.8 | 0.6 | -3.0 | 2.9E-03 | 3.6E-03 | MB231/MB468 |
| >hg38_21_+_8433172_8446622_RNA45SN1_45S_50ntFlanks@10348.10369.22\|AAACGAGAACTTTGAAGGCCGA | >hg38_21_+_8433172_8446622_RNA45SN1_45S_50ntFlanks | 10348 | 10369 | 22 | AAACGAGAACTTTGAAGGCCGA | 337.0 | 1.8 | 0.6 | 3.0 | 3.1E-03 | 3.6E-03 | BT20/MB468 |
| >hg38_21_+_8433172_8446622_RNA45SN1_45S_50ntFlanks@9602.9622.21\|CGAAAGACTAATCGAACCATC | >hg38_21_+_8433172_8446622_RNA45SN1_45S_50ntFlanks | 9602 | 9622 | 21 | CGAAAGACTAATCGAACCATC | 174.3 | -2.9 | 1.0 | -3.0 | 3.1E-03 | 3.6E-03 | BT20/MB468 |
| >hg38_21_+_8433172_8446622_RNA45SN1_45S_50ntFlanks@11729.11752.24\|TAACTATGACTCTCTTAAGGTAGC | >hg38_21_+_8433172_8446622_RNA45SN1_45S_50ntFlanks | 11729 | 11752 | 24 | TAACTATGACTCTCTTAAGGTAGC | 1879.1 | -2.3 | 0.8 | -3.0 | 2.9E-03 | 3.7E-03 | MB231/MB468 |
| >hg38_21_+_8433172_8446622_RNA45SN1_45S_50ntFlanks@11579.11604.26\|AGAACTGGTGCGGACCAGGGGAATCC | >hg38_21_+_8433172_8446622_RNA45SN1_45S_50ntFlanks | 11579 | 11604 | 26 | AGAACTGGTGCGGACCAGGGGAATCC | 131.5 | -1.7 | 0.6 | -3.0 | 3.1E-03 | 3.7E-03 | BT20/MB468 |
| >hg38_21_+_8433172_8446622_RNA45SN1_45S_50ntFlanks@10830.10851.22\|CAAGCCGGATCCGTAACTTCGG | >hg38_21_+_8433172_8446622_RNA45SN1_45S_50ntFlanks | 10830 | 10851 | 22 | CAAGCCGGATCCGTAACTTCGG | 202.6 | -1.9 | 0.6 | -3.0 | 2.9E-03 | 3.7E-03 | MB231/MB468 |
| >hg38_21_+_8433172_8446622_RNA45SN1_45S_50ntFlanks@7975.7995.21\|CGCGACCTCAGATCAGACGTG | >hg38_21_+_8433172_8446622_RNA45SN1_45S_50ntFlanks | 7975 | 7995 | 21 | CGCGACCTCAGATCAGACGTG | 5248.8 | 2.7 | 0.8 | 3.5 | 5.5E-04 | 3.7E-03 | BT20/MB231 |
| >hg38_21_+_8433172_8446622_RNA45SN1_45S_50ntFlanks@6651.6686.36\|CGACTCTTAGCGGTGGATCACTCGGCTCGTGCGTCG | >hg38_21_+_8433172_8446622_RNA45SN1_45S_50ntFlanks | 6651 | 6686 | 36 | CGACTCTTAGCGGTGGATCACTCGGCTCGTGCGTCG | 1042.7 | 2.0 | 0.7 | 3.0 | 3.1E-03 | 3.7E-03 | BT20/MB468 |
| >hg38_21_+_8433172_8446622_RNA45SN1_45S_50ntFlanks@12415.12455.41\|CTTCGATGTCGGCTCTTCCTATCATTGTGAAGCAGAATTCA | >hg38_21_+_8433172_8446622_RNA45SN1_45S_50ntFlanks | 12415 | 12455 | 41 | CTTCGATGTCGGCTCTTCCTATCATTGTGAAGCAGAATTCA | 190.1 | -1.8 | 0.6 | -3.0 | 3.2E-03 | 3.7E-03 | BT20/MB468 |
| >hg38_21_+_8433172_8446622_RNA45SN1_45S_50ntFlanks@12933.12953.21\|GCTAAACCATTCGTAGACGAC | >hg38_21_+_8433172_8446622_RNA45SN1_45S_50ntFlanks | 12933 | 12953 | 21 | GCTAAACCATTCGTAGACGAC | 171.5 | -1.9 | 0.6 | -3.0 | 2.9E-03 | 3.7E-03 | MB231/MB468 |
| >hg38_21_+_8433172_8446622_RNA45SN1_45S_50ntFlanks@9554.9597.44\|CGTAGCGGTCCTGACGTGCAAATCGGTCGTCCGACCTGGGTATA | >hg38_21_+_8433172_8446622_RNA45SN1_45S_50ntFlanks | 9554 | 9597 | 44 | CGTAGCGGTCCTGACGTGCAAATCGGTCGTCCGACCTGGGTATA | 225.0 | -1.9 | 0.6 | -3.0 | 3.0E-03 | 3.7E-03 | MB231/MB468 |
| >hg38_21_+_8433172_8446622_RNA45SN1_45S_50ntFlanks@6664.6686.23\|TGGATCACTCGGCTCGTGCGTCG | >hg38_21_+_8433172_8446622_RNA45SN1_45S_50ntFlanks | 6664 | 6686 | 23 | TGGATCACTCGGCTCGTGCGTCG | 182.0 | -1.9 | 0.7 | -3.0 | 3.0E-03 | 3.7E-03 | MB231/MB468 |
| >hg38_21_+_8433172_8446622_RNA45SN1_45S_50ntFlanks@11571.11602.32\|GCCGACTTAGAACTGGTGCGGACCAGGGGAAT | >hg38_21_+_8433172_8446622_RNA45SN1_45S_50ntFlanks | 11571 | 11602 | 32 | GCCGACTTAGAACTGGTGCGGACCAGGGGAAT | 306.6 | 1.5 | 0.5 | 3.0 | 3.0E-03 | 3.7E-03 | MB231/MB468 |
| >hg38_21_+_8433172_8446622_RNA45SN1_45S_50ntFlanks@12457.12476.20\|CAAGCGTTGGATTGTTCACC | >hg38_21_+_8433172_8446622_RNA45SN1_45S_50ntFlanks | 12457 | 12476 | 20 | CAAGCGTTGGATTGTTCACC | 5193.2 | -2.1 | 0.7 | -3.0 | 3.0E-03 | 3.7E-03 | MB231/MB468 |
| >MI0000063\|hsa-let-7b&WithFlank&22\|+\|46113680\|46113774@12.31.20(+1C)&[MIMAT0000063&hsa-let-7b-5p&offsets\|0\|-2(+1C);m-14&22\|+\|46113691\|46113712&offsets\|0\|-2(+1C)]\|TGAGGTAGTAGGTTGTGTGGC | >MI0000063\|hsa-let-7b&WithFlank&22\|+\|46113680\|46113774 | 12 | 31 | 20(+1C) | TGAGGTAGTAGGTTGTGTGGC | 743.6 | 2.4 | 0.7 | 3.4 | 5.7E-04 | 3.8E-03 | BT20/MB231 |
| >hg38_21_+_8433172_8446622_RNA45SN1_45S_50ntFlanks@3843.3861.19\|CTACTTGGATAACTGTGGT | >hg38_21_+_8433172_8446622_RNA45SN1_45S_50ntFlanks | 3843 | 3861 | 19 | CTACTTGGATAACTGTGGT | 822.6 | -1.6 | 0.6 | -2.9 | 3.3E-03 | 3.8E-03 | BT20/MB468 |
| >hg38_21_+_8433172_8446622_RNA45SN1_45S_50ntFlanks@11579.11611.33\|AGAACTGGTGCGGACCAGGGGAATCCGACTGTT | >hg38_21_+_8433172_8446622_RNA45SN1_45S_50ntFlanks | 11579 | 11611 | 33 | AGAACTGGTGCGGACCAGGGGAATCCGACTGTT | 944.8 | -2.1 | 0.7 | -2.9 | 3.3E-03 | 3.8E-03 | BT20/MB468 |
| >hg38_21_+_8433172_8446622_RNA45SN1_45S_50ntFlanks@12415.12441.27\|CTTCGATGTCGGCTCTTCCTATCATTG | >hg38_21_+_8433172_8446622_RNA45SN1_45S_50ntFlanks | 12415 | 12441 | 27 | CTTCGATGTCGGCTCTTCCTATCATTG | 856.2 | -1.5 | 0.5 | -2.9 | 3.3E-03 | 3.8E-03 | BT20/MB468 |

| | | | | | | | | | | | | |
|---|---|---|---|---|---|---|---|---|---|---|---|---|
| >hg38_21_+_8433172_8446622_RNA45SN1_45S_50ntFlanks@11210.11254.45\|GGCGACTCTGGACGCGAGCCGGGCCCTTCCCGTGGATCGCCCCAG | >hg38_21_+_8433172_8446622_RNA45SN1_45S_50ntFlanks | 11210 | 11254 | 45 | GGCGACTCTGGACGCGAGCCGGGCCCTTCCCGTGGATCGCCCCAG | 126.9 | -5.0 | 1.7 | -3.0 | 3.1E-03 | 3.9E-03 | MB231/MB468 |
| >hg38_21_+_8433172_8446622_RNA45SN1_45S_50ntFlanks@10420.10456.37\|CTGAGAGATGGGCGAGCGCCGTTCCGAAGGGACGGGC | >hg38_21_+_8433172_8446622_RNA45SN1_45S_50ntFlanks | 10420 | 10456 | 37 | CTGAGAGATGGGCGAGCGCCGTTCCGAAGGGACGGGC | 813.7 | -2.0 | 0.7 | -3.0 | 3.1E-03 | 3.9E-03 | MB231/MB468 |
| >hg38_21_+_8433172_8446622_RNA45SN1_45S_50ntFlanks@5492.5513.22\|AGAAGACGGTCGAACTTGACTA | >hg38_21_+_8433172_8446622_RNA45SN1_45S_50ntFlanks | 5492 | 5513 | 22 | AGAAGACGGTCGAACTTGACTA | 143.3 | -2.1 | 0.7 | -3.0 | 3.1E-03 | 3.9E-03 | MB231/MB468 |
| >hg38_21_+_8433172_8446622_RNA45SN1_45S_50ntFlanks@10684.10702.19\|CGGCGGCGTCCGGTGAGCT | >hg38_21_+_8433172_8446622_RNA45SN1_45S_50ntFlanks | 10684 | 10702 | 19 | CGGCGGCGTCCGGTGAGCT | 168.0 | -2.0 | 0.7 | -3.0 | 3.1E-03 | 3.9E-03 | MB231/MB468 |
| >MI0000477\|hsa-mir-146a&WithFlank&5\|+\|160485346\|160485456@27.47.21(+1A)&[MIMAT0000449&hsa-miR-146a-5p&offsets\|0\|-1(+1A);m-129&5\|+\|160485372\|160485393&offsets\|0\|-1(+1A)]\|TGAGAACTGAATTCCATGGGTA | >MI0000477\|hsa-mir-146a&WithFlank&5\|+\|160485346\|160485456 | 27 | 47 | 21(+1A) | TGAGAACTGAATTCCATGGGTA | 577.1 | -7.7 | 2.3 | -3.4 | 5.9E-04 | 3.9E-03 | BT20/MB231 |
| >hg38_21_+_8433172_8446622_RNA45SN1_45S_50ntFlanks@8179.8203.25\|CTGATCGAGGCCCAGCCCGTGGACG | >hg38_21_+_8433172_8446622_RNA45SN1_45S_50ntFlanks | 8179 | 8203 | 25 | CTGATCGAGGCCCAGCCCGTGGACG | 153.8 | -1.5 | 0.5 | -3.0 | 3.1E-03 | 3.9E-03 | MB231/MB468 |
| >MI0000077\|hsa-mir-21&WithFlank&17\|+\|59841260\|59841343@14.32.19(+1C)&[MIMAT0000076&hsa-miR-21-5p&offsets\|0\|-3(+1C);m-1&17\|+\|59841273\|59841294&offsets\|0\|-3(+1C)]\|TAGCTTATCAGACTGATGTC | >MI0000077\|hsa-mir-21&WithFlank&17\|+\|59841260\|59841343 | 14 | 32 | 19(+1C) | TAGCTTATCAGACTGATGTC | 348.6 | 2.4 | 0.7 | 3.4 | 5.9E-04 | 3.9E-03 | BT20/MB231 |
| >hg38_21_+_8433172_8446622_RNA45SN1_45S_50ntFlanks@7977.8006.30\|CGACCTCAGATCAGACGTGGCGACCCGCTG | >hg38_21_+_8433172_8446622_RNA45SN1_45S_50ntFlanks | 7977 | 8006 | 30 | CGACCTCAGATCAGACGTGGCGACCCGCTG | 742.3 | -2.3 | 0.8 | -3.0 | 3.1E-03 | 3.9E-03 | MB231/MB468 |
| >MI0000286\|hsa-mir-210&WithFlank&11\|-\|568083\|568204@72.93.22(+3A)&[MIMAT0000267&hsa-miR-210-3p&offsets\|0\|0(+3A);m-102&11\|-\|568112\|568133&offsets\|0\|0(+3A)]\|CTGTGCGTGTGACAGCGGCTGAAAA | >MI0000286\|hsa-mir-210&WithFlank&11\|-\|568083\|568204 | 72 | 93 | 22(+3A) | CTGTGCGTGTGACAGCGGCTGAAAA | 344.3 | 2.7 | 0.8 | 3.4 | 5.9E-04 | 3.9E-03 | BT20/MB231 |
| >hg38_MT_+_598_1651_RNR1_12S_50ntFlanks@968.1004.37\|ACATGGTAAGTGTACTGGAAAGTGCACTTGGACGAAC | >hg38_MT_+_598_1651_RNR1_12S_50ntFlanks | 968 | 1004 | 37 | ACATGGTAAGTGTACTGGAAAGTGCACTTGGACGAAC | 455.8 | -2.6 | 0.8 | -3.4 | 6.0E-04 | 3.9E-03 | BT20/MB231 |
| >hg38_21_+_8433172_8446622_RNA45SN1_45S_50ntFlanks@3842.3860.19\|CCTACTTGGATAACTGTGG | >hg38_21_+_8433172_8446622_RNA45SN1_45S_50ntFlanks | 3842 | 3860 | 19 | CCTACTTGGATAACTGTGG | 151.9 | -1.6 | 0.5 | -2.9 | 3.4E-03 | 3.9E-03 | BT20/MB468 |
| >hg38_21_+_8433172_8446622_RNA45SN1_45S_50ntFlanks@6656.6689.34\|CTTAGCGGTGGATCACTCGGCTCGTGCGTCGATG | >hg38_21_+_8433172_8446622_RNA45SN1_45S_50ntFlanks | 6656 | 6689 | 34 | CTTAGCGGTGGATCACTCGGCTCGTGCGTCGATG | 319.6 | -1.9 | 0.6 | -2.9 | 3.4E-03 | 4.0E-03 | BT20/MB468 |
| >am_trnaMT_SerGCT_MT_+_12207_12265@12.31.20\|CAAGAACTGCTAACTCATGC | >am_trnaMT_SerGCT_MT_+_12207_12265 | 12 | 31 | 20 | CAAGAACTGCTAACTCATGC | 458.8 | 3.5 | 1.2 | 2.9 | 3.4E-03 | 4.0E-03 | BT20/MB468 |
| >hg38_21_+_8433172_8446622_RNA45SN1_45S_50ntFlanks@8357.8391.35\|AAAAGAACTTTGAAGAGAGAGTTCAAGAGGGCGTG | >hg38_21_+_8433172_8446622_RNA45SN1_45S_50ntFlanks | 8357 | 8391 | 35 | AAAAGAACTTTGAAGAGAGAGTTCAAGAGGGCGTG | 275.2 | -1.7 | 0.6 | -2.9 | 3.4E-03 | 4.0E-03 | BT20/MB468 |
| >hg38_21_+_8433172_8446622_RNA45SN1_45S_50ntFlanks@11570.11602.33\|AGCCGACTTAGAACTGGTGCGGACCAGGGGAAT | >hg38_21_+_8433172_8446622_RNA45SN1_45S_50ntFlanks | 11570 | 11602 | 33 | AGCCGACTTAGAACTGGTGCGGACCAGGGGAAT | 319.0 | -1.4 | 0.5 | -3.0 | 3.2E-03 | 4.0E-03 | MB231/MB468 |
| >am_trna111_HisGTG_1_-_147774845_147774916@2.29.28,trna118_HisGTG_1_-_145396881_145396952@2.29.28,trna16_HisGTG_1_+_146544773_146544844@2.29.28,trna1_HisGTG_15_+_45493349_45493420@2.29.28,trna21_HisGTG_1_+_147753471_147753542@2.29.28,trna33_HisGTG_6_+_27125906_27125977@2.29.28,trna7_HisGTG_9_-_14433938_14434009@2.29.28,trna8_HisGTG_15_-_45492611_45492682@2.29.28,trna9_HisGTG_15_-_45490804_45490875@2.29.28\|CCGTGATCGTATAGTGGTTAGTACTCTG | >am_trna111_HisGTG_1_-_147774845_147774916 | 2 | 29 | 28 | CCGTGATCGTATAGTGGTTAGTACTCTG | 115.0 | -1.4 | 0.5 | -2.9 | 3.4E-03 | 4.0E-03 | BT20/MB468 |
| >hg38_21_+_8433172_8446622_RNA45SN1_45S_50ntFlanks@12444.12478.35\|AAGCAGAATTCACCAAGCGTTGGATTGTTCACCCA | >hg38_21_+_8433172_8446622_RNA45SN1_45S_50ntFlanks | 12444 | 12478 | 35 | AAGCAGAATTCACCAAGCGTTGGATTGTTCACCCA | 180.2 | -1.7 | 0.6 | -2.9 | 3.4E-03 | 4.0E-03 | BT20/MB468 |
| >hg38_21_+_8433172_8446622_RNA45SN1_45S_50ntFlanks@11210.11230.21\|GGCGACTCTGGACGCGAGCCG | >hg38_21_+_8433172_8446622_RNA45SN1_45S_50ntFlanks | 11210 | 11230 | 21 | GGCGACTCTGGACGCGAGCCG | 103.6 | -3.4 | 1.2 | -2.9 | 3.4E-03 | 4.0E-03 | BT20/MB468 |
| >hg38_21_+_8433172_8446622_RNA45SN1_45S_50ntFlanks@10792.10815.24\|CTCTGGCATGTTGGAACAATGTAG | >hg38_21_+_8433172_8446622_RNA45SN1_45S_50ntFlanks | 10792 | 10815 | 24 | CTCTGGCATGTTGGAACAATGTAG | 220.0 | -1.7 | 0.6 | -2.9 | 3.4E-03 | 4.0E-03 | BT20/MB468 |
| >hg38_21_+_8433172_8446622_RNA45SN1_45S_50ntFlanks@12480.12498.19\|TAATAGGGAACGTGAGCTG | >hg38_21_+_8433172_8446622_RNA45SN1_45S_50ntFlanks | 12480 | 12498 | 19 | TAATAGGGAACGTGAGCTG | 203.1 | -1.1 | 0.4 | -2.9 | 3.4E-03 | 4.0E-03 | BT20/MB468 |
| >hg38_21_+_8433172_8446622_RNA45SN1_45S_50ntFlanks@12073.12094.22\|CCGAGGGGCTCTCGCTTCTGGC | >hg38_21_+_8433172_8446622_RNA45SN1_45S_50ntFlanks | 12073 | 12094 | 22 | CCGAGGGGCTCTCGCTTCTGGC | 285.7 | -1.6 | 0.5 | -2.9 | 3.2E-03 | 4.0E-03 | MB231/MB468 |
| >hg38_21_+_8433172_8446622_RNA45SN1_45S_50ntFlanks@10794.10812.19\|CTGGCATGTTGGAACAATG | >hg38_21_+_8433172_8446622_RNA45SN1_45S_50ntFlanks | 10794 | 10812 | 19 | CTGGCATGTTGGAACAATG | 111.7 | -2.0 | 0.7 | -2.9 | 3.2E-03 | 4.0E-03 | MB231/MB468 |
| >hg38_21_+_8433172_8446622_RNA45SN1_45S_50ntFlanks@9534.9574.41\|AGGAAACTCTGGTGGAGGTCCGTAGCGGTCCTGACGTGCAA | >hg38_21_+_8433172_8446622_RNA45SN1_45S_50ntFlanks | 9534 | 9574 | 41 | AGGAAACTCTGGTGGAGGTCCGTAGCGGTCCTGACGTGCAA | 188.9 | -1.3 | 0.4 | -2.9 | 3.2E-03 | 4.0E-03 | MB231/MB468 |
| >hg38_21_+_8433172_8446622_RNA45SN1_45S_50ntFlanks@12165.12182.18\|TGGGGCGGTACACCTGTC | >hg38_21_+_8433172_8446622_RNA45SN1_45S_50ntFlanks | 12165 | 12182 | 18 | TGGGGCGGTACACCTGTC | 108.1 | -1.8 | 0.6 | -2.9 | 3.2E-03 | 4.0E-03 | MB231/MB468 |
| >hg38_21_+_8433172_8446622_RNA45SN1_45S_50ntFlanks@8192.8214.23\|AGCCCGTGGACGGTGTGAGGCCG | >hg38_21_+_8433172_8446622_RNA45SN1_45S_50ntFlanks | 8192 | 8214 | 23 | AGCCCGTGGACGGTGTGAGGCCG | 133.2 | -1.9 | 0.6 | -2.9 | 3.4E-03 | 4.0E-03 | BT20/MB468 |
| >hg38_21_+_8433172_8446622_RNA45SN1_45S_50ntFlanks@5071.5098.28\|TTAATTCCGATAACGAACGAGACTCTGG | >hg38_21_+_8433172_8446622_RNA45SN1_45S_50ntFlanks | 5071 | 5098 | 28 | TTAATTCCGATAACGAACGAGACTCTGG | 181.5 | -1.7 | 0.6 | -2.9 | 3.2E-03 | 4.0E-03 | MB231/MB468 |
| >hg38_21_+_8433172_8446622_RNA45SN1_45S_50ntFlanks@10395.10429.35\|GCAGTTGAACATGGGTCAGTCGGTCCTGAGAGATG | >hg38_21_+_8433172_8446622_RNA45SN1_45S_50ntFlanks | 10395 | 10429 | 35 | GCAGTTGAACATGGGTCAGTCGGTCCTGAGAGATG | 444.0 | 2.2 | 0.7 | 2.9 | 3.2E-03 | 4.0E-03 | MB231/MB468 |
| >MI0000086\|hsa-mir-28&WithFlank&3\|+\|188688775\|188688872@62.82.21&[MIMAT0004502&hsa-miR-28-3p&offsets\|+2\|+1;m-28&3\|+\|188688834\|188688855&offsets\|+2\|+1]\|CTAGATTGTGAGCTCCTGGAG | >MI0000086\|hsa-mir-28&WithFlank&3\|+\|188688775\|188688872 | 62 | 82 | 21 | CTAGATTGTGAGCTCCTGGAG | 288.2 | 2.5 | 0.7 | 3.4 | 6.1E-04 | 4.0E-03 | BT20/MB231 |
| >hg38_21_+_8433172_8446622_RNA45SN1_45S_50ntFlanks@10349.10366.18\|AACGAGAACTTTGAAGGC | >hg38_21_+_8433172_8446622_RNA45SN1_45S_50ntFlanks | 10349 | 10366 | 18 | AACGAGAACTTTGAAGGC | 1307.5 | -1.3 | 0.5 | -2.9 | 3.5E-03 | 4.0E-03 | BT20/MB468 |
| >hg38_21_+_8433172_8446622_RNA45SN1_45S_50ntFlanks@6652.6692.41\|GACTCTTAGCGGTGGATCACTCGGCTCGTGCGTCGATGAAG | >hg38_21_+_8433172_8446622_RNA45SN1_45S_50ntFlanks | 6652 | 6692 | 41 | GACTCTTAGCGGTGGATCACTCGGCTCGTGCGTCGATGAAG | 1125.1 | 2.5 | 0.9 | 2.9 | 3.5E-03 | 4.0E-03 | BT20/MB468 |
| >hg38_21_+_8433172_8446622_RNA45SN1_45S_50ntFlanks@12942.12959.18\|TTCGTAGACGACCTGCTT | >hg38_21_+_8433172_8446622_RNA45SN1_45S_50ntFlanks | 12942 | 12959 | 18 | TTCGTAGACGACCTGCTT | 670.5 | -1.8 | 0.6 | -2.9 | 3.5E-03 | 4.0E-03 | BT20/MB468 |
| >hg38_21_+_8433172_8446622_RNA45SN1_45S_50ntFlanks@12458.12486.29\|AAGCGTTGGATTGTTCACCCACTAATAGG | >hg38_21_+_8433172_8446622_RNA45SN1_45S_50ntFlanks | 12458 | 12486 | 29 | AAGCGTTGGATTGTTCACCCACTAATAGG | 357.0 | 1.3 | 0.5 | 2.9 | 3.2E-03 | 4.1E-03 | MB231/MB468 |
| >hg38_21_+_8433172_8446622_RNA45SN1_45S_50ntFlanks@4592.4616.25\|TTTGTTGGTTTTCGGAACTGAGGCC | >hg38_21_+_8433172_8446622_RNA45SN1_45S_50ntFlanks | 4592 | 4616 | 25 | TTTGTTGGTTTTCGGAACTGAGGCC | 143.2 | -1.7 | 0.6 | -2.9 | 3.5E-03 | 4.1E-03 | BT20/MB468 |
| >hg38_21_+_8433172_8446622_RNA45SN1_45S_50ntFlanks@11787.11805.19\|AATGGATGAACGAGATTCC | >hg38_21_+_8433172_8446622_RNA45SN1_45S_50ntFlanks | 11787 | 11805 | 19 | AATGGATGAACGAGATTCC | 128.8 | -2.3 | 0.8 | -2.9 | 3.3E-03 | 4.1E-03 | MB231/MB468 |
| >trnaMT_SerGCT_MT_+_12207_12265@10.38.29\|CACAAGAACTGCTAACTCATGCCCCCATG | >trnaMT_SerGCT_MT_+_12207_12265 | 10 | 38 | 29 | CACAAGAACTGCTAACTCATGCCCCCATG | 164.0 | -2.4 | 0.8 | -2.9 | 3.3E-03 | 4.1E-03 | MB231/MB468 |
| >MI0000063\|hsa-let-7b&WithFlank&22\|+\|46113680\|46113774@12.32.21(+1G)&[MIMAT0000063&hsa-let-7b-5p&offsets\|0\|-1(+1G);m-14&22\|+\|46113691\|46113712&offsets\|0\|-1(+1G)]\|TGAGGTAGTAGGTTGTGTGGTG | >MI0000063\|hsa-let-7b&WithFlank&22\|+\|46113680\|46113774 | 12 | 32 | 21(+1G) | TGAGGTAGTAGGTTGTGTGGTG | 2757.7 | 3.0 | 0.9 | 3.4 | 6.2E-04 | 4.1E-03 | BT20/MB231 |
| >trna10_LysCTT_16_+_3241501_3241573@1.28.28,trna119_LysCTT_1_-_145395522_145395594@1.28.28,trna11_LysCTT_5_-_180648979_180649051@1.28.28,trna13_LysCTT_14_-_58706613_58706685@1.28.28,trna13_LysCTT_6_+_26556774_26556846@1.28.28,trna2_LysCTT_15_+_79152904_79152976@1.28.28,trna32_LysCTT_16_-_3207406_3207478@1.28.28,trna7_LysCTT_16_+_3225692_3225764@1.28.28,trna9_LysCTT_5_+_180634755_180634827@1.28.28\|GCCCGGCTAGCTCAGTCGGTAGAGCATG | >trna10_LysCTT_16_+_3241501_3241573 | 1 | 28 | 28 | GCCCGGCTAGCTCAGTCGGTAGAGCATG | 3112.7 | -2.3 | 0.8 | -2.9 | 3.3E-03 | 4.1E-03 | MB231/MB468 |
| >hg38_21_+_8433172_8446622_RNA45SN1_45S_50ntFlanks@9572.9605.34\|CAAATCGGTCGTCCGACCTGGGTATAGGGGCGAA | >hg38_21_+_8433172_8446622_RNA45SN1_45S_50ntFlanks | 9572 | 9605 | 34 | CAAATCGGTCGTCCGACCTGGGTATAGGGGCGAA | 310.4 | -1.6 | 0.5 | -2.9 | 3.3E-03 | 4.1E-03 | MB231/MB468 |
| >hg38_21_+_8433172_8446622_RNA45SN1_45S_50ntFlanks@12071.12106.36\|CCCCGAGGGGCTCTCGCTTCTGGCGCCAAGCGCCCG | >hg38_21_+_8433172_8446622_RNA45SN1_45S_50ntFlanks | 12071 | 12106 | 36 | CCCCGAGGGGCTCTCGCTTCTGGCGCCAAGCGCCCG | 224.8 | -2.6 | 0.9 | -2.9 | 3.5E-03 | 4.1E-03 | BT20/MB468 |
| >hg38_21_+_8433172_8446622_RNA45SN1_45S_50ntFlanks@12640.12659.20\|ACCATCTGTGGGATTATGAC | >hg38_21_+_8433172_8446622_RNA45SN1_45S_50ntFlanks | 12640 | 12659 | 20 | ACCATCTGTGGGATTATGAC | 326.8 | -1.1 | 0.4 | -2.9 | 3.3E-03 | 4.1E-03 | MB231/MB468 |
| >hg38_21_+_8433172_8446622_RNA45SN1_45S_50ntFlanks@9570.9603.34\|TGCAAATCGGTCGTCCGACCTGGGTATAGGGGCG | >hg38_21_+_8433172_8446622_RNA45SN1_45S_50ntFlanks | 9570 | 9603 | 34 | TGCAAATCGGTCGTCCGACCTGGGTATAGGGGCG | 348.5 | -1.3 | 0.4 | -2.9 | 3.6E-03 | 4.2E-03 | BT20/MB468 |
| >hg38_21_+_8433172_8446622_RNA45SN1_45S_50ntFlanks@10797.10820.24\|GCATGTTGGAACAATGTAGGTAAG | >hg38_21_+_8433172_8446622_RNA45SN1_45S_50ntFlanks | 10797 | 10820 | 24 | GCATGTTGGAACAATGTAGGTAAG | 296.9 | 1.9 | 0.7 | 2.9 | 3.4E-03 | 4.2E-03 | MB231/MB468 |

| | | | | | | | | | | | | |
|---|---|---|---|---|---|---|---|---|---|---|---|---|
| >hg38_21_+_8433172_8446622_RNA45SN1_45S_50ntFlanks@11586.11611.26\|GTGCGGACCAGGGGAATCCGACTGTT | >hg38_21_+_8433172_8446622_RNA45SN1_45S_50ntFlanks | 11586 | 11611 | 26 | GTGCGGACCAGGGGAATCCGACTGTT | 1959.7 | -1.8 | 0.6 | -2.9 | 3.4E-03 | 4.2E-03 | MB231/MB468 |
| >hg38_21_+_8433172_8446622_RNA45SN1_45S_50ntFlanks@12606.12624.19\|ATGTGCTTGGCTGAGGAGC | >hg38_21_+_8433172_8446622_RNA45SN1_45S_50ntFlanks | 12606 | 12624 | 19 | ATGTGCTTGGCTGAGGAGC | 393.8 | 1.5 | 0.5 | 2.9 | 3.4E-03 | 4.2E-03 | MB231/MB468 |
| >hg38_21_+_8433172_8446622_RNA45SN1_45S_50ntFlanks@12415.12445.31\|CTTCGATGTCGGCTCTTCCTATCATTGTGAA | >hg38_21_+_8433172_8446622_RNA45SN1_45S_50ntFlanks | 12415 | 12445 | 31 | CTTCGATGTCGGCTCTTCCTATCATTGTGAA | 397.9 | 1.5 | 0.5 | 2.9 | 3.4E-03 | 4.2E-03 | MB231/MB468 |
| >hg38_21_+_8433172_8446622_RNA45SN1_45S_50ntFlanks@3899.3927.29\|CCTTCGCGGGGGGGATGCGTGCATTTATC | >hg38_21_+_8433172_8446622_RNA45SN1_45S_50ntFlanks | 3899 | 3927 | 29 | CCTTCGCGGGGGGGATGCGTGCATTTATC | 249.9 | -2.8 | 1.0 | -2.9 | 3.6E-03 | 4.2E-03 | BT20/MB468 |
| >hg38_21_+_8433172_8446622_RNA45SN1_45S_50ntFlanks@4008.4042.35\|CTCGGGCCGATCGCACGCCCCCCGTGGCGGCGACG | >hg38_21_+_8433172_8446622_RNA45SN1_45S_50ntFlanks | 4008 | 4042 | 35 | CTCGGGCCGATCGCACGCCCCCCGTGGCGGCGACG | 180.5 | -1.5 | 0.5 | -2.9 | 3.4E-03 | 4.2E-03 | MB231/MB468 |
| >hg38_21_+_8433172_8446622_RNA45SN1_45S_50ntFlanks@12607.12624.18\|TGTGCTTGGCTGAGGAGC | >hg38_21_+_8433172_8446622_RNA45SN1_45S_50ntFlanks | 12607 | 12624 | 18 | TGTGCTTGGCTGAGGAGC | 233.5 | 1.5 | 0.5 | 2.9 | 3.6E-03 | 4.2E-03 | BT20/MB468 |
| >hg38_21_+_8433172_8446622_RNA45SN1_45S_50ntFlanks@9533.9551.19\|GAGGAAACTCTGGTGGAGG | >hg38_21_+_8433172_8446622_RNA45SN1_45S_50ntFlanks | 9533 | 9551 | 19 | GAGGAAACTCTGGTGGAGG | 225.3 | -1.2 | 0.4 | -2.9 | 3.4E-03 | 4.2E-03 | MB231/MB468 |
| >hg38_21_+_8433172_8446622_RNA45SN1_45S_50ntFlanks@12406.12434.29\|TTTTTGATCCTTCGATGTCGGCTCTTCCT | >hg38_21_+_8433172_8446622_RNA45SN1_45S_50ntFlanks | 12406 | 12434 | 29 | TTTTTGATCCTTCGATGTCGGCTCTTCCT | 313.6 | -1.4 | 0.5 | -2.9 | 3.7E-03 | 4.3E-03 | BT20/MB468 |
| >hg38_21_+_8433172_8446622_RNA45SN1_45S_50ntFlanks@12995.13038.44\|CTCGCTGCGATCTATTGAAAGTCAGCCCTCGACACAAGGGTTTG | >hg38_21_+_8433172_8446622_RNA45SN1_45S_50ntFlanks | 12995 | 13038 | 44 | CTCGCTGCGATCTATTGAAAGTCAGCCCTCGACACAAGGGTTTG | 5430.8 | -3.2 | 1.0 | -3.4 | 6.6E-04 | 4.3E-03 | BT20/MB231 |
| >hg38_21_+_8433172_8446622_RNA45SN1_45S_50ntFlanks@5527.5553.27\|AAAGTCGTAACAAGGTTTCCGTAGGTG | >hg38_21_+_8433172_8446622_RNA45SN1_45S_50ntFlanks | 5527 | 5553 | 27 | AAAGTCGTAACAAGGTTTCCGTAGGTG | 88.8 | -3.0 | 1.0 | -2.9 | 3.7E-03 | 4.3E-03 | BT20/MB468 |
| >hg38_21_+_8433172_8446622_RNA45SN1_45S_50ntFlanks@9548.9574.27\|GAGGTCCGTAGCGGTCCTGACGTGCAA | >hg38_21_+_8433172_8446622_RNA45SN1_45S_50ntFlanks | 9548 | 9574 | 27 | GAGGTCCGTAGCGGTCCTGACGTGCAA | 202.9 | 1.6 | 0.6 | 2.9 | 3.5E-03 | 4.3E-03 | MB231/MB468 |
| >hg38_21_+_8433172_8446622_RNA45SN1_45S_50ntFlanks@5485.5526.42\|AGCGCTGAGAAGACGGTCGAACTTGACTATCTAGAGGAAGTA | >hg38_21_+_8433172_8446622_RNA45SN1_45S_50ntFlanks | 5485 | 5526 | 42 | AGCGCTGAGAAGACGGTCGAACTTGACTATCTAGAGGAAGTA | 180.2 | -3.0 | 1.0 | -2.9 | 3.7E-03 | 4.3E-03 | BT20/MB468 |
| >hg38_21_+_8433172_8446622_RNA45SN1_45S_50ntFlanks@10798.10826.29\|CATGTTGGAACAATGTAGGTAAGGGAAGT | >hg38_21_+_8433172_8446622_RNA45SN1_45S_50ntFlanks | 10798 | 10826 | 29 | CATGTTGGAACAATGTAGGTAAGGGAAGT | 222.8 | -1.6 | 0.6 | -2.9 | 3.7E-03 | 4.3E-03 | BT20/MB468 |
| >hg38_21_+_8433172_8446622_RNA45SN1_45S_50ntFlanks@12456.12496.41\|CCAAGCGTTGGATTGTTCACCCACTAATAGGGAACGTGAGC | >hg38_21_+_8433172_8446622_RNA45SN1_45S_50ntFlanks | 12456 | 12496 | 41 | CCAAGCGTTGGATTGTTCACCCACTAATAGGGAACGTGAGC | 309.5 | 1.6 | 0.6 | 2.9 | 3.5E-03 | 4.3E-03 | MB231/MB468 |
| >hg38_21_+_8433172_8446622_RNA45SN1_45S_50ntFlanks@8359.8377.19\|AAGAACTTTGAAGAGAGAG | >hg38_21_+_8433172_8446622_RNA45SN1_45S_50ntFlanks | 8359 | 8377 | 19 | AAGAACTTTGAAGAGAGAG | 213.1 | -1.8 | 0.6 | -2.9 | 3.7E-03 | 4.3E-03 | BT20/MB468 |
| >hg38_21_+_8433172_8446622_RNA45SN1_45S_50ntFlanks@11205.11225.21\|GCGGCGGCGACTCTGGACGCG | >hg38_21_+_8433172_8446622_RNA45SN1_45S_50ntFlanks | 11205 | 11225 | 21 | GCGGCGGCGACTCTGGACGCG | 269.2 | -2.9 | 1.0 | -2.9 | 3.7E-03 | 4.3E-03 | BT20/MB468 |
| >hg38_21_+_8433172_8446622_RNA45SN1_45S_50ntFlanks@6651.6675.25\|CGACTCTTAGCGGTGGATCACTCGG | >hg38_21_+_8433172_8446622_RNA45SN1_45S_50ntFlanks | 6651 | 6675 | 25 | CGACTCTTAGCGGTGGATCACTCGG | 1855.5 | 1.9 | 0.6 | 2.9 | 3.8E-03 | 4.4E-03 | BT20/MB468 |
| >hg38_21_+_8433172_8446622_RNA45SN1_45S_50ntFlanks@3858.3881.24\|TGGTAATTCTAGAGCTAATACATG | >hg38_21_+_8433172_8446622_RNA45SN1_45S_50ntFlanks | 3858 | 3881 | 24 | TGGTAATTCTAGAGCTAATACATG | 141.3 | -3.0 | 1.0 | -2.9 | 3.8E-03 | 4.4E-03 | BT20/MB468 |
| >hg38_21_+_8433172_8446622_RNA45SN1_45S_50ntFlanks@11882.11903.22\|CTGTTGAGCTTGACTCTAGTCT | >hg38_21_+_8433172_8446622_RNA45SN1_45S_50ntFlanks | 11882 | 11903 | 22 | CTGTTGAGCTTGACTCTAGTCT | 105.2 | -2.5 | 0.8 | -2.9 | 3.5E-03 | 4.4E-03 | MB231/MB468 |
| >hg38_21_+_8433172_8446622_RNA45SN1_45S_50ntFlanks@11772.11794.23\|TTAGTGACGCGCATGAATGGATG | >hg38_21_+_8433172_8446622_RNA45SN1_45S_50ntFlanks | 11772 | 11794 | 23 | TTAGTGACGCGCATGAATGGATG | 234.4 | -1.5 | 0.5 | -2.9 | 3.8E-03 | 4.4E-03 | BT20/MB468 |
| >hg38_21_+_8433172_8446622_RNA45SN1_45S_50ntFlanks@11208.11231.24\|GCGGCGACTCTGGACGCGAGCCGG | >hg38_21_+_8433172_8446622_RNA45SN1_45S_50ntFlanks | 11208 | 11231 | 24 | GCGGCGACTCTGGACGCGAGCCGG | 982.2 | -2.7 | 0.9 | -2.9 | 3.8E-03 | 4.4E-03 | BT20/MB468 |
| >hg38_21_+_8433172_8446622_RNA45SN1_45S_50ntFlanks@4685.4707.23\|AGACGGACCAGAGCGAAAGCATT | >hg38_21_+_8433172_8446622_RNA45SN1_45S_50ntFlanks | 4685 | 4707 | 23 | AGACGGACCAGAGCGAAAGCATT | 148.2 | -2.1 | 0.7 | -2.9 | 3.8E-03 | 4.4E-03 | BT20/MB468 |
| >hg38_21_+_8433172_8446622_RNA45SN1_45S_50ntFlanks@11729.11764.36\|TAACTATGACTCTCTTAAGGTAGCCAAATGCCTCGT | >hg38_21_+_8433172_8446622_RNA45SN1_45S_50ntFlanks | 11729 | 11764 | 36 | TAACTATGACTCTCTTAAGGTAGCCAAATGCCTCGT | 382.5 | -1.7 | 0.6 | -2.9 | 3.6E-03 | 4.4E-03 | MB231/MB468 |
| >hg38_21_+_8433172_8446622_RNA45SN1_45S_50ntFlanks@10409.10426.18\|GTCAGTCGGTCCTGAGAG | >hg38_21_+_8433172_8446622_RNA45SN1_45S_50ntFlanks | 10409 | 10426 | 18 | GTCAGTCGGTCCTGAGAG | 116.0 | -1.4 | 0.5 | -2.9 | 3.6E-03 | 4.4E-03 | MB231/MB468 |
| >hg38_21_+_8433172_8446622_RNA45SN1_45S_50ntFlanks@6651.6698.48\|CGACTCTTAGCGGTGGATCACTCGGCTCGTGCGTCGATGAAGAACGCA | >hg38_21_+_8433172_8446622_RNA45SN1_45S_50ntFlanks | 6651 | 6698 | 48 | CGACTCTTAGCGGTGGATCACTCGGCTCGTGCGTCGATGAAGAACGCA | 4248.4 | -2.5 | 0.7 | -3.4 | 6.8E-04 | 4.4E-03 | BT20/MB231 |
| >hg38_21_+_8433172_8446622_RNA45SN1_45S_50ntFlanks@10771.10792.22\|CAGGTCTCCAAGGTGAACAGCC | >hg38_21_+_8433172_8446622_RNA45SN1_45S_50ntFlanks | 10771 | 10792 | 22 | CAGGTCTCCAAGGTGAACAGCC | 167.7 | -1.7 | 0.6 | -2.9 | 3.6E-03 | 4.5E-03 | MB231/MB468 |
| >hg38_21_+_8433172_8446622_RNA45SN1_45S_50ntFlanks@10786.10828.43\|AACAGCCTCTGGCATGTTGGAACAATGTAGGTAAGGGAAGTCG | >hg38_21_+_8433172_8446622_RNA45SN1_45S_50ntFlanks | 10786 | 10828 | 43 | AACAGCCTCTGGCATGTTGGAACAATGTAGGTAAGGGAAGTCG | 103.6 | -1.5 | 0.5 | -2.9 | 3.6E-03 | 4.5E-03 | MB231/MB468 |
| >hg38_21_+_8433172_8446622_RNA45SN1_45S_50ntFlanks@3788.3810.23\|AACTGCGAATGGCTCATTAAATC | >hg38_21_+_8433172_8446622_RNA45SN1_45S_50ntFlanks | 3788 | 3810 | 23 | AACTGCGAATGGCTCATTAAATC | 138.1 | -2.3 | 0.8 | -2.9 | 3.9E-03 | 4.5E-03 | BT20/MB468 |
| >hg38_21_+_8433172_8446622_RNA45SN1_45S_50ntFlanks@12937.12954.18\|AACCATTCGTAGACGACC | >hg38_21_+_8433172_8446622_RNA45SN1_45S_50ntFlanks | 12937 | 12954 | 18 | AACCATTCGTAGACGACC | 139.8 | -2.3 | 0.8 | -2.9 | 3.9E-03 | 4.5E-03 | BT20/MB468 |
| >hg38_MT_+_598_1651_RNR1_12S_50ntFlanks@970.1004.35\|ATGGTAAGTGTACTGGAAAGTGCACTTGGACGAAC | >hg38_MT_+_598_1651_RNR1_12S_50ntFlanks | 970 | 1004 | 35 | ATGGTAAGTGTACTGGAAAGTGCACTTGGACGAAC | 329.8 | -2.3 | 0.7 | -3.4 | 6.9E-04 | 4.5E-03 | BT20/MB231 |
| >hg38_21_+_8433172_8446622_RNA45SN1_45S_50ntFlanks@13000.13027.28\|TGCGATCTATTGAAAGTCAGCCCTCGAC | >hg38_21_+_8433172_8446622_RNA45SN1_45S_50ntFlanks | 13000 | 13027 | 28 | TGCGATCTATTGAAAGTCAGCCCTCGAC | 296.4 | -2.7 | 0.9 | -2.9 | 3.9E-03 | 4.5E-03 | BT20/MB468 |
| >hg38_21_+_8433172_8446622_RNA45SN1_45S_50ntFlanks@11207.11225.19\|GGCGGCGACTCTGGACGCG | >hg38_21_+_8433172_8446622_RNA45SN1_45S_50ntFlanks | 11207 | 11225 | 19 | GGCGGCGACTCTGGACGCG | 381.0 | -2.6 | 0.9 | -2.9 | 3.9E-03 | 4.5E-03 | BT20/MB468 |
| >hg38_21_+_8433172_8446622_RNA45SN1_45S_50ntFlanks@3850.3872.23\|GATAACTGTGGTAATTCTAGAGC | >hg38_21_+_8433172_8446622_RNA45SN1_45S_50ntFlanks | 3850 | 3872 | 23 | GATAACTGTGGTAATTCTAGAGC | 424.0 | 1.7 | 0.6 | 2.9 | 3.7E-03 | 4.5E-03 | MB231/MB468 |
| >hg38_21_+_8433172_8446622_RNA45SN1_45S_50ntFlanks@12413.12431.19\|TCCTTCGATGTCGGCTCTT | >hg38_21_+_8433172_8446622_RNA45SN1_45S_50ntFlanks | 12413 | 12431 | 19 | TCCTTCGATGTCGGCTCTT | 168.7 | -1.7 | 0.6 | -2.9 | 3.7E-03 | 4.6E-03 | MB231/MB468 |
| >hg38_21_+_8433172_8446622_RNA45SN1_45S_50ntFlanks@11874.11902.29\|AGAAGACCCTGTTGAGCTTGACTCTAGTC | >hg38_21_+_8433172_8446622_RNA45SN1_45S_50ntFlanks | 11874 | 11902 | 29 | AGAAGACCCTGTTGAGCTTGACTCTAGTC | 309.5 | -2.3 | 0.8 | -2.9 | 3.9E-03 | 4.6E-03 | BT20/MB468 |
| >hg38_21_+_8433172_8446622_RNA45SN1_45S_50ntFlanks@8192.8214.23\|AGCCCGTGGACGGTGTGAGGCCG | >hg38_21_+_8433172_8446622_RNA45SN1_45S_50ntFlanks | 8192 | 8214 | 23 | AGCCCGTGGACGGTGTGAGGCCG | 109.8 | -1.7 | 0.6 | -2.9 | 3.7E-03 | 4.6E-03 | MB231/MB468 |
| >hg38_21_+_8433172_8446622_RNA45SN1_45S_50ntFlanks@4243.4264.22\|CTTTCGAGGCCCTGTAATTGGA | >hg38_21_+_8433172_8446622_RNA45SN1_45S_50ntFlanks | 4243 | 4264 | 22 | CTTTCGAGGCCCTGTAATTGGA | 115.2 | -1.3 | 0.5 | -2.9 | 3.7E-03 | 4.6E-03 | MB231/MB468 |
| >hg38_21_+_8433172_8446622_RNA45SN1_45S_50ntFlanks@6646.6669.24\|TCGTACGACTCTTAGCGGTGGATC | >hg38_21_+_8433172_8446622_RNA45SN1_45S_50ntFlanks | 6646 | 6669 | 24 | TCGTACGACTCTTAGCGGTGGATC | 11044.1 | 2.3 | 0.8 | 2.9 | 3.7E-03 | 4.6E-03 | MB231/MB468 |
| >MI0000255\|hsa-mir-30d&WithFlank&8\|-\|134804870\|134804951@12.33.22(+1U)&[MIMAT0000245&hsa-miR-30d-5p&offsets\|0\|0(+1U);m-21&8\|-\|134804919\|134804940&offsets\|0\|0(+1U)]\|TGTAAACATCCCCGACTGGAAGT | >MI0000255\|hsa-mir-30d&WithFlank&8\|-\|134804870\|134804951 | 12 | 33 | 22(+1U) | TGTAAACATCCCCGACTGGAAGT | 17335.8 | 2.0 | 0.6 | 3.4 | 7.2E-04 | 4.7E-03 | BT20/MB231 |
| >hg38_21_+_8433172_8446622_RNA45SN1_45S_50ntFlanks@8171.8197.27\|AAGTCCTTCTGATCGAGGCCCAGCCCG | >hg38_21_+_8433172_8446622_RNA45SN1_45S_50ntFlanks | 8171 | 8197 | 27 | AAGTCCTTCTGATCGAGGCCCAGCCCG | 192.5 | -1.6 | 0.6 | -2.9 | 3.8E-03 | 4.7E-03 | MB231/MB468 |
| >hg38_21_+_8433172_8446622_RNA45SN1_45S_50ntFlanks@11576.11611.36\|CTTAGAACTGGTGCGGACCAGGGGAATCCGACTGTT | >hg38_21_+_8433172_8446622_RNA45SN1_45S_50ntFlanks | 11576 | 11611 | 36 | CTTAGAACTGGTGCGGACCAGGGGAATCCGACTGTT | 335.1 | -1.9 | 0.7 | -2.9 | 3.8E-03 | 4.7E-03 | MB231/MB468 |
| >hg38_21_+_8433172_8446622_RNA45SN1_45S_50ntFlanks@8178.8203.26\|TCTGATCGAGGCCCAGCCCGTGGACG | >hg38_21_+_8433172_8446622_RNA45SN1_45S_50ntFlanks | 8178 | 8203 | 26 | TCTGATCGAGGCCCAGCCCGTGGACG | 465.3 | 1.8 | 0.6 | 2.9 | 3.8E-03 | 4.7E-03 | MB231/MB468 |
| >hg38_21_+_8433172_8446622_RNA45SN1_45S_50ntFlanks@9556.9581.26\|TAGCGGTCCTGACGTGCAAATCGGTC | >hg38_21_+_8433172_8446622_RNA45SN1_45S_50ntFlanks | 9556 | 9581 | 26 | TAGCGGTCCTGACGTGCAAATCGGTC | 137.1 | -2.4 | 0.8 | -2.9 | 4.0E-03 | 4.7E-03 | BT20/MB468 |
| >hg38_21_+_8433172_8446622_RNA45SN1_45S_50ntFlanks@6731.6752.22\|CACATTGATCATCGACACTTCG | >hg38_21_+_8433172_8446622_RNA45SN1_45S_50ntFlanks | 6731 | 6752 | 22 | CACATTGATCATCGACACTTCG | 418.6 | -1.5 | 0.5 | -2.9 | 3.8E-03 | 4.7E-03 | MB231/MB468 |
| >MI0000482\|hsa-mir-185&WithFlank&22\|+\|20033133\|20033226@21.42.22(+1A)&[MIMAT0000455&hsa-miR-185-5p&offsets\|0\|0(+1A);m-145&22\|+\|20033153\|20033174&offsets\|0\|0(+1A)]\|TGGAGAGAAAGGCAGTTCCTGAA | >MI0000482\|hsa-mir-185&WithFlank&22\|+\|20033133\|20033226 | 21 | 42 | 22(+1A) | TGGAGAGAAAGGCAGTTCCTGAA | 335.6 | 2.2 | 0.6 | 3.4 | 7.3E-04 | 4.7E-03 | BT20/MB231 |
| >hg38_21_+_8433172_8446622_RNA45SN1_45S_50ntFlanks@8179.8210.32\|CTGATCGAGGCCCAGCCCGTGGACGGTGTGAG | >hg38_21_+_8433172_8446622_RNA45SN1_45S_50ntFlanks | 8179 | 8210 | 32 | CTGATCGAGGCCCAGCCCGTGGACGGTGTGAG | 131.6 | -2.0 | 0.7 | -2.9 | 3.8E-03 | 4.7E-03 | MB231/MB468 |
| >hg38_21_+_8433172_8446622_RNA45SN1_45S_50ntFlanks@12656.12674.19\|TGACTGAACGCCTCTAAGT | >hg38_21_+_8433172_8446622_RNA45SN1_45S_50ntFlanks | 12656 | 12674 | 19 | TGACTGAACGCCTCTAAGT | 225.5 | -1.4 | 0.5 | -2.9 | 3.8E-03 | 4.7E-03 | MB231/MB468 |
| >hg38_21_+_8433172_8446622_RNA45SN1_45S_50ntFlanks@11319.11342.24\|CCCACGTCTCGTCGCGCGCGCGTC | >hg38_21_+_8433172_8446622_RNA45SN1_45S_50ntFlanks | 11319 | 11342 | 24 | CCCACGTCTCGTCGCGCGCGCGTC | 322.6 | -3.0 | 1.0 | -2.9 | 4.1E-03 | 4.8E-03 | BT20/MB468 |

| | | | | | | | | | | | | |
|---|---|---|---|---|---|---|---|---|---|---|---|---|
| >am_trna111_HisGTG_1_-_147774845_147774916@-1G.29.30,trna118_HisGTG_1_-_145396881_145396952@-1G.29.30,trna16_HisGTG_1_+_146544773_146544844@-1G.29.30,trna1_HisGTG_15_+_45493349_45493420@-1G.29.30,trna21_HisGTG_1_+_147753471_147753542@-1G.29.30,trna33_HisGTG_6_+_27125906_27125977@-1G.29.30,trna7_HisGTG_9_-_14433938_14434009@-1G.29.30,trna8_HisGTG_15_-_45492611_45492682@-1G.29.30,trna9_HisGTG_15_-_45490804_45490875@-1G.29.30\|GGCCGTGATCGTATAGTGGTTAGTACTCTG | >am_trna111_HisGTG_1_-_147774845_147774916 | -1G | 29 | 30 | GGCCGTGATCGTATAGTGGTTAGTACTCTG | 3791.9 | -2.3 | 0.8 | -2.9 | 3.8E-03 | 4.8E-03 | MB231/MB468 |
| >hg38_21_+_8433172_8446622_RNA45SN1_45S_50ntFlanks@12420.12447.28\|ATGTCGGCTCTTCCTATCATTGTGAAGC | >hg38_21_+_8433172_8446622_RNA45SN1_45S_50ntFlanks | 12420 | 12447 | 28 | ATGTCGGCTCTTCCTATCATTGTGAAGC | 442.5 | -2.8 | 0.8 | -3.4 | 7.4E-04 | 4.8E-03 | BT20/MB231 |
| >hg38_21_+_8433172_8446622_RNA45SN1_45S_50ntFlanks@11661.11684.24\|TGATTTCTGCCCAGTGCTCTGAAT | >hg38_21_+_8433172_8446622_RNA45SN1_45S_50ntFlanks | 11661 | 11684 | 24 | TGATTTCTGCCCAGTGCTCTGAAT | 137.2 | -1.8 | 0.6 | -2.9 | 3.8E-03 | 4.8E-03 | MB231/MB468 |
| >hg38_21_+_8433172_8446622_RNA45SN1_45S_50ntFlanks@11568.11603.36\|GCAGCCGACTTAGAACTGGTGCGGACCAGGGGAATC | >hg38_21_+_8433172_8446622_RNA45SN1_45S_50ntFlanks | 11568 | 11603 | 36 | GCAGCCGACTTAGAACTGGTGCGGACCAGGGGAATC | 328.9 | -1.7 | 0.6 | -2.9 | 4.1E-03 | 4.8E-03 | BT20/MB468 |
| >trna152_ValCAC_6_-_27248049_27248121@1.32.32\|GCTTCTGTAGTGTAGTGGTTATCACGTTCGCC | >trna152_ValCAC_6_-_27248049_27248121 | 1 | 32 | 32 | GCTTCTGTAGTGTAGTGGTTATCACGTTCGCC | 2503.1 | -2.8 | 0.8 | -3.4 | 7.4E-04 | 4.8E-03 | BT20/MB231 |
| >hg38_21_+_8433172_8446622_RNA45SN1_45S_50ntFlanks@10678.10708.31\|CGGTTCCGGCGGCGTCCGGTGAGCTCTCGCT | >hg38_21_+_8433172_8446622_RNA45SN1_45S_50ntFlanks | 10678 | 10708 | 31 | CGGTTCCGGCGGCGTCCGGTGAGCTCTCGCT | 11409.9 | -2.8 | 1.0 | -2.9 | 4.1E-03 | 4.8E-03 | BT20/MB468 |
| >hg38_21_+_8433172_8446622_RNA45SN1_45S_50ntFlanks@10669.10706.38\|AAAGCGTCGCGGTTCCGGCGGCGTCCGGTGAGCTCTCG | >hg38_21_+_8433172_8446622_RNA45SN1_45S_50ntFlanks | 10669 | 10706 | 38 | AAAGCGTCGCGGTTCCGGCGGCGTCCGGTGAGCTCTCG | 179.0 | -5.4 | 1.9 | -2.9 | 3.9E-03 | 4.8E-03 | MB231/MB468 |
| >hg38_21_+_8433172_8446622_RNA45SN1_45S_50ntFlanks@10468.10495.28\|TTGCCCTCGGCCGATCGAAAGGGAGTCG | >hg38_21_+_8433172_8446622_RNA45SN1_45S_50ntFlanks | 10468 | 10495 | 28 | TTGCCCTCGGCCGATCGAAAGGGAGTCG | 176.8 | -2.8 | 1.0 | -2.9 | 4.2E-03 | 4.8E-03 | BT20/MB468 |
| >am_trna162_MetCAT_6_-_26758550_26758622@31.54.24,trna164_MetCAT_6_-_26735574_26735646@31.54.24,trna20_MetCAT_16_+_71460396_71460468@31.54.24,trna21_MetCAT_6_+_26701712_26701784@31.54.24,trna27_MetCAT_6_+_26766444_26766516@31.54.24,trna75_MetCAT_6_+_28912352_28912424@31.54.24,trna97_MetCAT_6_-_28921042_28921114@31.54.24\|TCTCATAATCTGAAGGTCCTGAGT | >am_trna162_MetCAT_6_-_26758550_26758622 | 31 | 54 | 24 | TCTCATAATCTGAAGGTCCTGAGT | 594.8 | -2.4 | 0.8 | -2.9 | 4.2E-03 | 4.9E-03 | BT20/MB468 |
| >hg38_21_+_8433172_8446622_RNA45SN1_45S_50ntFlanks@10346.10373.28\|TCAAACGAGAACTTTGAAGGCCGAAGTG | >hg38_21_+_8433172_8446622_RNA45SN1_45S_50ntFlanks | 10346 | 10373 | 28 | TCAAACGAGAACTTTGAAGGCCGAAGTG | 556.9 | 2.0 | 0.7 | 2.9 | 3.9E-03 | 4.9E-03 | MB231/MB468 |
| >hg38_21_+_8433172_8446622_RNA45SN1_45S_50ntFlanks@6669.6699.31\|CACTCGGCTCGTGCGTCGATGAAGAACGCAG | >hg38_21_+_8433172_8446622_RNA45SN1_45S_50ntFlanks | 6669 | 6699 | 31 | CACTCGGCTCGTGCGTCGATGAAGAACGCAG | 133.6 | -1.2 | 0.4 | -2.9 | 4.2E-03 | 4.9E-03 | BT20/MB468 |
| >hg38_21_+_8433172_8446622_RNA45SN1_45S_50ntFlanks@10781.10814.34\|AGGTGAACAGCCTCTGGCATGTTGGAACAATGTA | >hg38_21_+_8433172_8446622_RNA45SN1_45S_50ntFlanks | 10781 | 10814 | 34 | AGGTGAACAGCCTCTGGCATGTTGGAACAATGTA | 141.9 | -1.7 | 0.6 | -2.9 | 4.3E-03 | 4.9E-03 | BT20/MB468 |
| >hg38_21_+_8433172_8446622_RNA45SN1_45S_50ntFlanks@4759.4783.25\|ATCAGATACCGTCGTAGTTCCGACC | >hg38_21_+_8433172_8446622_RNA45SN1_45S_50ntFlanks | 4759 | 4783 | 25 | ATCAGATACCGTCGTAGTTCCGACC | 841.5 | -3.8 | 1.3 | -2.9 | 4.0E-03 | 4.9E-03 | MB231/MB468 |
| >hg38_21_+_8433172_8446622_RNA45SN1_45S_50ntFlanks@6661.6698.38\|CGGTGGATCACTCGGCTCGTGCGTCGATGAAGAACGCA | >hg38_21_+_8433172_8446622_RNA45SN1_45S_50ntFlanks | 6661 | 6698 | 38 | CGGTGGATCACTCGGCTCGTGCGTCGATGAAGAACGCA | 381.7 | 1.5 | 0.5 | 2.9 | 4.0E-03 | 5.0E-03 | MB231/MB468 |
| >hg38_21_+_8433172_8446622_RNA45SN1_45S_50ntFlanks@11581.11607.27\|AACTGGTGCGGACCAGGGGAATCCGAC | >hg38_21_+_8433172_8446622_RNA45SN1_45S_50ntFlanks | 11581 | 11607 | 27 | AACTGGTGCGGACCAGGGGAATCCGAC | 367.5 | -1.6 | 0.5 | -2.9 | 4.3E-03 | 5.0E-03 | BT20/MB468 |
| >hg38_1_-_228634819_228635039_RNA5S12_5S_50ntFlanks@84.109.26\|CTCGTCTGATCTCGGAAGCTAAGCAG | >hg38_1_-_228634819_228635039_RNA5S12_5S_50ntFlanks | 84 | 109 | 26 | CTCGTCTGATCTCGGAAGCTAAGCAG | 117.1 | -2.5 | 0.9 | -2.9 | 4.0E-03 | 5.0E-03 | MB231/MB468 |
| >hg38_21_+_8433172_8446622_RNA45SN1_45S_50ntFlanks@5492.5509.18\|AGAAGACGGTCGAACTTG | >hg38_21_+_8433172_8446622_RNA45SN1_45S_50ntFlanks | 5492 | 5509 | 18 | AGAAGACGGTCGAACTTG | 320.4 | -2.5 | 0.9 | -2.9 | 4.1E-03 | 5.0E-03 | MB231/MB468 |
| >hg38_21_+_8433172_8446622_RNA45SN1_45S_50ntFlanks@11668.11707.40\|TGCCCAGTGCTCTGAATGTCAAAGTGAAGAAATTCAATGA | >hg38_21_+_8433172_8446622_RNA45SN1_45S_50ntFlanks | 11668 | 11707 | 40 | TGCCCAGTGCTCTGAATGTCAAAGTGAAGAAATTCAATGA | 395.4 | -2.7 | 0.8 | -3.4 | 7.8E-04 | 5.0E-03 | BT20/MB231 |
| >hg38_21_+_8433172_8446622_RNA45SN1_45S_50ntFlanks@3853.3872.20\|AACTGTGGTAATTCTAGAGC | >hg38_21_+_8433172_8446622_RNA45SN1_45S_50ntFlanks | 3853 | 3872 | 20 | AACTGTGGTAATTCTAGAGC | 255.2 | -1.2 | 0.4 | -2.9 | 4.1E-03 | 5.0E-03 | MB231/MB468 |
| >MI0000299\|hsa-mir-222&WithFlank&X\|-\|45747009\|45747130@75.98.24(+1C)&[MIMAT0000279&hsa-miR-222-3p&offsets\|0\|+3(+1C);m-140&X\|-\|45747033\|45747056&offsets\|0\|0(+1C)]\|AGCTACATCTGGCTACTGGGTCTCC | >MI0000299\|hsa-mir-222&WithFlank&X\|-\|45747009\|45747130 | 75 | 98 | 24(+1C) | AGCTACATCTGGCTACTGGGTCTCC | 10417.8 | -2.3 | 0.7 | -3.4 | 7.9E-04 | 5.0E-03 | BT20/MB231 |
| >hg38_21_+_8433172_8446622_RNA45SN1_45S_50ntFlanks@3876.3899.24\|TACATGCCGACGGGCGCTGACCCC | >hg38_21_+_8433172_8446622_RNA45SN1_45S_50ntFlanks | 3876 | 3899 | 24 | TACATGCCGACGGGCGCTGACCCC | 201.2 | -1.2 | 0.4 | -2.9 | 4.1E-03 | 5.1E-03 | MB231/MB468 |
| >hg38_21_+_8433172_8446622_RNA45SN1_45S_50ntFlanks@9573.9603.31\|AAATCGGTCGTCCGACCTGGGTATAGGGGCG | >hg38_21_+_8433172_8446622_RNA45SN1_45S_50ntFlanks | 9573 | 9603 | 31 | AAATCGGTCGTCCGACCTGGGTATAGGGGCG | 807.9 | -1.6 | 0.6 | -2.8 | 4.4E-03 | 5.1E-03 | BT20/MB468 |
| >hg38_21_+_8433172_8446622_RNA45SN1_45S_50ntFlanks@11321.11344.24\|CACGTCTCGTCGCGCGCGCGTCCG | >hg38_21_+_8433172_8446622_RNA45SN1_45S_50ntFlanks | 11321 | 11344 | 24 | CACGTCTCGTCGCGCGCGCGTCCG | 3629.4 | -3.8 | 1.3 | -2.8 | 4.4E-03 | 5.1E-03 | BT20/MB468 |
| >am_MI0000809\|hsa-mir-151a&WithFlank&8\|-\|140732558\|140732659@53.71.19(+1A)&[MIMAT0000757&hsa-miR-151a-3p&offsets\|0\|-2(+1A);m-30&8\|-\|140732587\|140732607&offsets\|0\|-2(+1A)]\|CTAGACTGAAGCTCCTTGAA | >am_MI0000809\|hsa-mir-151a&WithFlank&8\|-\|140732558\|140732659 | 53 | 71 | 19(+1A) | CTAGACTGAAGCTCCTTGAA | 871.5 | 2.6 | 0.8 | 3.4 | 8.0E-04 | 5.1E-03 | BT20/MB231 |
| >am_MI0003772\|hsa-mir-151b&WithFlank&14\|-\|100109413\|100109520@66.85.20(+1C);MI0000809\|hsa-mir-151a&WithFlank&8\|-\|140732558\|140732659@17.36.20(+1C)&[MIMAT0010214&hsa-miR-151b&offsets\|0\|+2(+1C);m-179&14\|-\|100109436\|100109455&offsets\|0\|0(+1C)];[MIMAT0004697&hsa-miR-151a-5p&offsets\|0\|-1(+1C);m-75&8\|-\|140732623\|140732643&offsets\|0\|-1(+1C)]\|TCGAGGAGCTCACAGTCTAGC | >am_MI0003772\|hsa-mir-151b&WithFlank&14\|-\|100109413\|100109520 | 66 | 85 | 20(+1C) | TCGAGGAGCTCACAGTCTAGC | 489.2 | 2.9 | 0.9 | 3.4 | 8.0E-04 | 5.1E-03 | BT20/MB231 |
| >hg38_21_+_8433172_8446622_RNA45SN1_45S_50ntFlanks@7977.8016.40\|CGACCTCAGATCAGACGTGGCGACCCGCTGAATTTAAGCA | >hg38_21_+_8433172_8446622_RNA45SN1_45S_50ntFlanks | 7977 | 8016 | 40 | CGACCTCAGATCAGACGTGGCGACCCGCTGAATTTAAGCA | 965.3 | -3.4 | 1.0 | -3.4 | 8.0E-04 | 5.1E-03 | BT20/MB231 |
| >hg38_21_+_8433172_8446622_RNA45SN1_45S_50ntFlanks@3839.3860.22\|TCTCCTACTTGGATAACTGTGG | >hg38_21_+_8433172_8446622_RNA45SN1_45S_50ntFlanks | 3839 | 3860 | 22 | TCTCCTACTTGGATAACTGTGG | 312.9 | 2.0 | 0.7 | 2.8 | 4.4E-03 | 5.1E-03 | BT20/MB468 |
| >hg38_21_+_8433172_8446622_RNA45SN1_45S_50ntFlanks@10678.10707.30\|CGGTTCCGGCGGCGTCCGGTGAGCTCTCGC | >hg38_21_+_8433172_8446622_RNA45SN1_45S_50ntFlanks | 10678 | 10707 | 30 | CGGTTCCGGCGGCGTCCGGTGAGCTCTCGC | 298.6 | -1.7 | 0.6 | -2.9 | 4.1E-03 | 5.1E-03 | MB231/MB468 |
| >hg38_21_+_8433172_8446622_RNA45SN1_45S_50ntFlanks@8113.8140.28\|GGCGCGGGACATGTGGCGTACGGAAGAC | >hg38_21_+_8433172_8446622_RNA45SN1_45S_50ntFlanks | 8113 | 8140 | 28 | GGCGCGGGACATGTGGCGTACGGAAGAC | 103.4 | -4.7 | 1.6 | -2.9 | 4.2E-03 | 5.2E-03 | MB231/MB468 |
| >hg38_21_+_8433172_8446622_RNA45SN1_45S_50ntFlanks@3781.3801.21\|ACAGTGAAACTGCGAATGGCT | >hg38_21_+_8433172_8446622_RNA45SN1_45S_50ntFlanks | 3781 | 3801 | 21 | ACAGTGAAACTGCGAATGGCT | 135.8 | -2.3 | 0.8 | -2.8 | 4.5E-03 | 5.2E-03 | BT20/MB468 |
| >hg38_21_+_8433172_8446622_RNA45SN1_45S_50ntFlanks@6663.6699.37\|GTGGATCACTCGGCTCGTGCGTCGATGAAGAACGCAG | >hg38_21_+_8433172_8446622_RNA45SN1_45S_50ntFlanks | 6663 | 6699 | 37 | GTGGATCACTCGGCTCGTGCGTCGATGAAGAACGCAG | 154.4 | -1.4 | 0.5 | -2.9 | 4.2E-03 | 5.2E-03 | MB231/MB468 |
| >trna2_GlyGCC_21_-_18827107_18827177@1.30.30,trna35_GlyGCC_1_+_161413094_161413164@1.30.30,trna37_GlyGCC_1_+_161420467_161420537@1.30.30,trna39_GlyGCC_1_+_161427898_161427968@1.30.30,trna41_GlyGCC_1_+_161435258_161435328@1.30.30\|GCATGGGTGGTTCAGTGGTAGAATTCTCGC | >trna2_GlyGCC_21_-_18827107_18827177 | 1 | 30 | 30 | GCATGGGTGGTTCAGTGGTAGAATTCTCGC | 3485.5 | -2.9 | 1.0 | -2.9 | 4.2E-03 | 5.2E-03 | MB231/MB468 |
| >MI0000737\|hsa-mir-200a&WithFlank&1\|+\|1167857\|1167958@22.43.22&[MIMAT0001620&hsa-miR-200a-5p&offsets\|0\|0;m-154&1\|+\|1167878\|1167899&offsets\|0\|0]\|CATCTTACCGGACAGTGCTGGA | >MI0000737\|hsa-mir-200a&WithFlank&1\|+\|1167857\|1167958 | 22 | 43 | 22 | CATCTTACCGGACAGTGCTGGA | 172.5 | -1.8 | 0.6 | -2.9 | 4.2E-03 | 5.2E-03 | MB231/MB468 |
| >hg38_21_+_8433172_8446622_RNA45SN1_45S_50ntFlanks@10797.10816.20\|GCATGTTGGAACAATGTAGG | >hg38_21_+_8433172_8446622_RNA45SN1_45S_50ntFlanks | 10797 | 10816 | 20 | GCATGTTGGAACAATGTAGG | 233.7 | -1.9 | 0.7 | -2.9 | 4.2E-03 | 5.2E-03 | MB231/MB468 |
| >hg38_21_+_8433172_8446622_RNA45SN1_45S_50ntFlanks@12446.12476.31\|GCAGAATTCACCAAGCGTTGGATTGTTCACC | >hg38_21_+_8433172_8446622_RNA45SN1_45S_50ntFlanks | 12446 | 12476 | 31 | GCAGAATTCACCAAGCGTTGGATTGTTCACC | 421.0 | -3.1 | 0.9 | -3.3 | 8.2E-04 | 5.2E-03 | BT20/MB231 |
| >hg38_21_+_8433172_8446622_RNA45SN1_45S_50ntFlanks@4005.4026.22\|AACCTCGGGCCGATCGCACGCC | >hg38_21_+_8433172_8446622_RNA45SN1_45S_50ntFlanks | 4005 | 4026 | 22 | AACCTCGGGCCGATCGCACGCC | 128.2 | -2.5 | 0.9 | -2.9 | 4.2E-03 | 5.2E-03 | MB231/MB468 |
| >hg38_21_+_8433172_8446622_RNA45SN1_45S_50ntFlanks@5471.5499.29\|CACGGCCCTGGCGGAGCGCTGAGAAGACG | >hg38_21_+_8433172_8446622_RNA45SN1_45S_50ntFlanks | 5471 | 5499 | 29 | CACGGCCCTGGCGGAGCGCTGAGAAGACG | 833.4 | -3.1 | 1.1 | -2.8 | 4.5E-03 | 5.2E-03 | BT20/MB468 |
| >hg38_21_+_8433172_8446622_RNA45SN1_45S_50ntFlanks@11322.11344.23\|ACGTCTCGTCGCGCGCGCGTCCG | >hg38_21_+_8433172_8446622_RNA45SN1_45S_50ntFlanks | 11322 | 11344 | 23 | ACGTCTCGTCGCGCGCGCGTCCG | 4799.6 | -3.8 | 1.4 | -2.8 | 4.6E-03 | 5.3E-03 | BT20/MB468 |
| >hg38_21_+_8433172_8446622_RNA45SN1_45S_50ntFlanks@12001.12031.31\|CTGCGGGCCGCCGGTGAAATACCACTACTCT | >hg38_21_+_8433172_8446622_RNA45SN1_45S_50ntFlanks | 12001 | 12031 | 31 | CTGCGGGCCGCCGGTGAAATACCACTACTCT | 244.7 | -1.8 | 0.6 | -2.9 | 4.3E-03 | 5.3E-03 | MB231/MB468 |
| >trnaMT_ValTAC_MT_+_1602_1670@51.71.21\|CTTAACTTGACCGCTCTGACC | >trnaMT_ValTAC_MT_+_1602_1670 | 51 | 71 | 21 | CTTAACTTGACCGCTCTGACC | 223.0 | 1.2 | 0.4 | 2.8 | 4.6E-03 | 5.3E-03 | BT20/MB468 |
| >hg38_21_+_8433172_8446622_RNA45SN1_45S_50ntFlanks@5489.5514.26\|CTGAGAAGACGGTCGAACTTGACTAT | >hg38_21_+_8433172_8446622_RNA45SN1_45S_50ntFlanks | 5489 | 5514 | 26 | CTGAGAAGACGGTCGAACTTGACTAT | 462.0 | -2.4 | 0.8 | -2.8 | 4.6E-03 | 5.3E-03 | BT20/MB468 |

| | | | | | | | | | | | | |
|---|---|---|---|---|---|---|---|---|---|---|---|---|
| >MI0000103\|hsa-mir-101-1&WithFlank&1\|-\|65058428\|65058514@53.73.21(+1A);MI0000739\|hsa-mir-101-2&WithFlank&9\|+\|4850291\|4850381@55.75.21(+1A)&[MIMAT0000099&hsa-miR-101-3p&offsets\|0\|0(+1A);m-17&1\|-\|65058443\|65058462&offsets\|0\|+1(+1A)];[MIMAT0000099_1&hsa-miR-101-3p&offsets\|0\|0(+1A);m-16&9\|+\|4850345\|4850364&offsets\|0\|+1(+1A)]\|TACAGTACTGTGATAACTGAAA | >MI0000103\|hsa-mir-101-1&WithFlank&1\|-\|65058428\|65058514 | 53 | 73 | 21(+1A) | TACAGTACTGTGATAACTGAAA | 3720.5 | 3.5 | 1.0 | 3.3 | 8.4E-04 | 5.3E-03 | BT20/MB231 |
| >trnaMT_GluTTC_MT_-_14674_14742@35.71.37,trnalookalike8_GluTTC_5_-_93905172_93905240@35.71.37\|TATCATTGGTCGTGGTTGTAGTCCGTGCGAGAATACC | >trnaMT_GluTTC_MT_-_14674_14742 | 35 | 71 | 37 | TATCATTGGTCGTGGTTGTAGTCCGTGCGAGAATACC | 560.7 | -2.2 | 0.7 | -3.3 | 8.4E-04 | 5.3E-03 | BT20/MB231 |
| >hg38_21_+_8433172_8446622_RNA45SN1_45S_50ntFlanks@11873.11908.36\|AAGAAGACCCTGTTGAGCTTGACTCTAGTCTGGCAC | >hg38_21_+_8433172_8446622_RNA45SN1_45S_50ntFlanks | 11873 | 11908 | 36 | AAGAAGACCCTGTTGAGCTTGACTCTAGTCTGGCAC | 198.7 | -4.7 | 1.6 | -2.9 | 4.3E-03 | 5.3E-03 | MB231/MB468 |
| >hg38_21_+_8433172_8446622_RNA45SN1_45S_50ntFlanks@11661.11684.24\|TGATTTCTGCCCAGTGCTCTGAAT | >hg38_21_+_8433172_8446622_RNA45SN1_45S_50ntFlanks | 11661 | 11684 | 24 | TGATTTCTGCCCAGTGCTCTGAAT | 164.4 | -1.8 | 0.6 | -2.8 | 4.6E-03 | 5.3E-03 | BT20/MB468 |
| >hg38_21_+_8433172_8446622_RNA45SN1_45S_50ntFlanks@10687.10705.19\|CGGCGTCCGGTGAGCTCTC | >hg38_21_+_8433172_8446622_RNA45SN1_45S_50ntFlanks | 10687 | 10705 | 19 | CGGCGTCCGGTGAGCTCTC | 126.4 | -4.5 | 1.6 | -2.9 | 4.4E-03 | 5.4E-03 | MB231/MB468 |
| >hg38_21_+_8433172_8446622_RNA45SN1_45S_50ntFlanks@10349.10376.28\|AACGAGAACTTTGAAGGCCGAAGTGGAG | >hg38_21_+_8433172_8446622_RNA45SN1_45S_50ntFlanks | 10349 | 10376 | 28 | AACGAGAACTTTGAAGGCCGAAGTGGAG | 549.1 | -1.5 | 0.5 | -2.9 | 4.4E-03 | 5.4E-03 | MB231/MB468 |
| >hg38_21_+_8433172_8446622_RNA45SN1_45S_50ntFlanks@10873.10890.18\|CTGGGTCGGTCGGGCTGG | >hg38_21_+_8433172_8446622_RNA45SN1_45S_50ntFlanks | 10873 | 10890 | 18 | CTGGGTCGGTCGGGCTGG | 203.6 | -2.9 | 1.0 | -2.8 | 4.7E-03 | 5.4E-03 | BT20/MB468 |
| >hg38_21_+_8433172_8446622_RNA45SN1_45S_50ntFlanks@10922.10945.24\|CTGGACGAGGCGCCGCCGCCCCCC | >hg38_21_+_8433172_8446622_RNA45SN1_45S_50ntFlanks | 10922 | 10945 | 24 | CTGGACGAGGCGCCGCCGCCCCCC | 329.0 | -3.7 | 1.3 | -2.8 | 4.7E-03 | 5.4E-03 | BT20/MB468 |
| >hg38_21_+_8433172_8446622_RNA45SN1_45S_50ntFlanks@10350.10370.21\|ACGAGAACTTTGAAGGCCGAA | >hg38_21_+_8433172_8446622_RNA45SN1_45S_50ntFlanks | 10350 | 10370 | 21 | ACGAGAACTTTGAAGGCCGAA | 285.6 | -1.5 | 0.5 | -2.8 | 4.7E-03 | 5.4E-03 | BT20/MB468 |
| >hg38_21_+_8433172_8446622_RNA45SN1_45S_50ntFlanks@3993.4013.21\|GTGACTCTAGATAACCTCGGG | >hg38_21_+_8433172_8446622_RNA45SN1_45S_50ntFlanks | 3993 | 4013 | 21 | GTGACTCTAGATAACCTCGGG | 341.7 | -1.3 | 0.4 | -2.8 | 4.4E-03 | 5.5E-03 | MB231/MB468 |
| >MI0000255\|hsa-mir-30d&WithFlank&8\|-\|134804870\|134804951@12.35.24(+3U)&[MIMAT0000245&hsa-miR-30d-5p&offsets\|0\|+2(+3U);m-21&8\|-\|134804919\|134804940&offsets\|0\|+2(+3U)]\|TGTAAACATCCCCGACTGGAAGCTTTT | >MI0000255\|hsa-mir-30d&WithFlank&8\|-\|134804870\|134804951 | 12 | 35 | 24(+3U) | TGTAAACATCCCCGACTGGAAGCTTTT | 504.8 | -2.4 | 0.7 | -3.3 | 8.7E-04 | 5.5E-03 | BT20/MB231 |
| >hg38_21_+_8433172_8446622_RNA45SN1_45S_50ntFlanks@12417.12445.29\|TCGATGTCGGCTCTTCCTATCATTGTGAA | >hg38_21_+_8433172_8446622_RNA45SN1_45S_50ntFlanks | 12417 | 12445 | 29 | TCGATGTCGGCTCTTCCTATCATTGTGAA | 290.5 | -1.2 | 0.4 | -2.8 | 4.8E-03 | 5.5E-03 | BT20/MB468 |
| >hg38_21_+_8433172_8446622_RNA45SN1_45S_50ntFlanks@10794.10820.27\|CTGGCATGTTGGAACAATGTAGGTAAG | >hg38_21_+_8433172_8446622_RNA45SN1_45S_50ntFlanks | 10794 | 10820 | 27 | CTGGCATGTTGGAACAATGTAGGTAAG | 143.7 | -2.7 | 1.0 | -2.8 | 4.8E-03 | 5.5E-03 | BT20/MB468 |
| >hg38_21_+_8433172_8446622_RNA45SN1_45S_50ntFlanks@5527.5549.23\|AAAGTCGTAACAAGGTTTCCGTA | >hg38_21_+_8433172_8446622_RNA45SN1_45S_50ntFlanks | 5527 | 5549 | 23 | AAAGTCGTAACAAGGTTTCCGTA | 242.8 | -4.2 | 1.5 | -2.8 | 4.5E-03 | 5.5E-03 | MB231/MB468 |
| >hg38_21_+_8433172_8446622_RNA45SN1_45S_50ntFlanks@9564.9603.40\|CTGACGTGCAAATCGGTCGTCCGACCTGGGTATAGGGGCG | >hg38_21_+_8433172_8446622_RNA45SN1_45S_50ntFlanks | 9564 | 9603 | 40 | CTGACGTGCAAATCGGTCGTCCGACCTGGGTATAGGGGCG | 205.2 | -3.2 | 1.0 | -3.3 | 8.8E-04 | 5.6E-03 | BT20/MB231 |
| >hg38_21_+_8433172_8446622_RNA45SN1_45S_50ntFlanks@9313.9338.26\|TCTAACACGTGCGCGAGTCGGGGGCT | >hg38_21_+_8433172_8446622_RNA45SN1_45S_50ntFlanks | 9313 | 9338 | 26 | TCTAACACGTGCGCGAGTCGGGGGCT | 101.7 | -2.4 | 0.9 | -2.8 | 4.8E-03 | 5.6E-03 | BT20/MB468 |
| >hg38_21_+_8433172_8446622_RNA45SN1_45S_50ntFlanks@12477.12498.22\|CACTAATAGGGAACGTGAGCTG | >hg38_21_+_8433172_8446622_RNA45SN1_45S_50ntFlanks | 12477 | 12498 | 22 | CACTAATAGGGAACGTGAGCTG | 1030.6 | -1.7 | 0.6 | -2.8 | 4.5E-03 | 5.6E-03 | MB231/MB468 |
| >MI0000542\|hsa-mir-320a&WithFlank&8\|-\|22244960\|22245043@50.70.21&[MIMAT0000510&hsa-miR-320a-3p&offsets\|+2\|+1;m-60&8\|-\|22244975\|22244996&offsets\|+2\|+1]\|AAGCTGGGTTGAGAGGGCGAA | >MI0000542\|hsa-mir-320a&WithFlank&8\|-\|22244960\|22245043 | 50 | 70 | 21 | AAGCTGGGTTGAGAGGGCGAA | 199.0 | 2.6 | 0.8 | 3.3 | 8.9E-04 | 5.6E-03 | BT20/MB231 |
| >hg38_21_+_8433172_8446622_RNA45SN1_45S_50ntFlanks@11672.11704.33\|CAGTGCTCTGAATGTCAAAGTGAAGAAATTCAA | >hg38_21_+_8433172_8446622_RNA45SN1_45S_50ntFlanks | 11672 | 11704 | 33 | CAGTGCTCTGAATGTCAAAGTGAAGAAATTCAA | 289.6 | -1.7 | 0.6 | -2.8 | 4.8E-03 | 5.6E-03 | BT20/MB468 |
| >am_trna116_GluCTC_1_-_145399233_145399304@7.28.22,trna59_GluCTC_1_+_249168447_249168518@7.28.22,trna71_GluCTC_1_-_161439189_161439260@7.28.22,trna74_GluCTC_1_-_161431809_161431880@7.28.22,trna77_GluCTC_1_-_161424398_161424469@7.28.22,trna77_GluCTC_6_+_28949976_28950047@7.28.22,trna80_GluCTC_1_-_161417018_161417089@7.28.22,trna87_GluCTC_6_-_126101393_126101464@7.28.22\|GTGGTCTAGTGGTTAGGATTCG | >am_trna116_GluCTC_1_-_145399233_145399304 | 7 | 28 | 22 | GTGGTCTAGTGGTTAGGATTCG | 158.6 | -2.6 | 0.9 | -2.8 | 4.9E-03 | 5.6E-03 | BT20/MB468 |
| >hg38_21_+_8433172_8446622_RNA45SN1_45S_50ntFlanks@12418.12435.18\|CGATGTCGGCTCTTCCTA | >hg38_21_+_8433172_8446622_RNA45SN1_45S_50ntFlanks | 12418 | 12435 | 18 | CGATGTCGGCTCTTCCTA | 8571.3 | -3.4 | 1.2 | -2.8 | 4.9E-03 | 5.6E-03 | BT20/MB468 |
| >hg38_21_+_8433172_8446622_RNA45SN1_45S_50ntFlanks@9561.9597.37\|GTCCTGACGTGCAAATCGGTCGTCCGACCTGGGTATA | >hg38_21_+_8433172_8446622_RNA45SN1_45S_50ntFlanks | 9561 | 9597 | 37 | GTCCTGACGTGCAAATCGGTCGTCCGACCTGGGTATA | 122.0 | -3.2 | 1.1 | -2.8 | 4.9E-03 | 5.6E-03 | BT20/MB468 |
| >hg38_21_+_8433172_8446622_RNA45SN1_45S_50ntFlanks@11719.11742.24\|ACGGCGGGAGTAACTATGACTCTC | >hg38_21_+_8433172_8446622_RNA45SN1_45S_50ntFlanks | 11719 | 11742 | 24 | ACGGCGGGAGTAACTATGACTCTC | 116.1 | -2.2 | 0.8 | -2.8 | 4.9E-03 | 5.6E-03 | BT20/MB468 |
| >hg38_21_+_8433172_8446622_RNA45SN1_45S_50ntFlanks@11577.11600.24\|TTAGAACTGGTGCGGACCAGGGGA | >hg38_21_+_8433172_8446622_RNA45SN1_45S_50ntFlanks | 11577 | 11600 | 24 | TTAGAACTGGTGCGGACCAGGGGA | 132.0 | -2.8 | 1.0 | -2.8 | 4.9E-03 | 5.6E-03 | BT20/MB468 |
| >hg38_21_+_8433172_8446622_RNA45SN1_45S_50ntFlanks@11891.11910.20\|TTGACTCTAGTCTGGCACGG | >hg38_21_+_8433172_8446622_RNA45SN1_45S_50ntFlanks | 11891 | 11910 | 20 | TTGACTCTAGTCTGGCACGG | 169.8 | -1.9 | 0.7 | -2.8 | 4.6E-03 | 5.6E-03 | MB231/MB468 |
| >hg38_21_+_8433172_8446622_RNA45SN1_45S_50ntFlanks@4368.4389.22\|AGTTAAAAAGCTCGTAGTTGGA | >hg38_21_+_8433172_8446622_RNA45SN1_45S_50ntFlanks | 4368 | 4389 | 22 | AGTTAAAAAGCTCGTAGTTGGA | 99.4 | -3.4 | 1.2 | -2.8 | 4.9E-03 | 5.7E-03 | BT20/MB468 |
| >hg38_21_+_8433172_8446622_RNA45SN1_45S_50ntFlanks@11573.11590.18\|CGACTTAGAACTGGTGCG | >hg38_21_+_8433172_8446622_RNA45SN1_45S_50ntFlanks | 11573 | 11590 | 18 | CGACTTAGAACTGGTGCG | 359.1 | -1.4 | 0.5 | -2.8 | 4.9E-03 | 5.7E-03 | BT20/MB468 |
| >hg38_21_+_8433172_8446622_RNA45SN1_45S_50ntFlanks@12420.12457.38\|ATGTCGGCTCTTCCTATCATTGTGAAGCAGAATTCACC | >hg38_21_+_8433172_8446622_RNA45SN1_45S_50ntFlanks | 12420 | 12457 | 38 | ATGTCGGCTCTTCCTATCATTGTGAAGCAGAATTCACC | 861.5 | -2.6 | 0.8 | -3.3 | 9.1E-04 | 5.7E-03 | BT20/MB231 |
| >hg38_21_+_8433172_8446622_RNA45SN1_45S_50ntFlanks@5125.5158.34\|AGCGGTCGGCGTCCCCCAACTTCTTAGAGGGACA | >hg38_21_+_8433172_8446622_RNA45SN1_45S_50ntFlanks | 5125 | 5158 | 34 | AGCGGTCGGCGTCCCCCAACTTCTTAGAGGGACA | 304.3 | 2.8 | 0.8 | 3.3 | 9.1E-04 | 5.7E-03 | BT20/MB231 |
| >hg38_21_+_8433172_8446622_RNA45SN1_45S_50ntFlanks@6661.6686.26\|CGGTGGATCACTCGGCTCGTGCGTCG | >hg38_21_+_8433172_8446622_RNA45SN1_45S_50ntFlanks | 6661 | 6686 | 26 | CGGTGGATCACTCGGCTCGTGCGTCG | 1112.4 | 1.8 | 0.6 | 2.8 | 4.6E-03 | 5.7E-03 | MB231/MB468 |
| >hg38_21_+_8433172_8446622_RNA45SN1_45S_50ntFlanks@8179.8210.32\|CTGATCGAGGCCCAGCCCGTGGACGGTGTGAG | >hg38_21_+_8433172_8446622_RNA45SN1_45S_50ntFlanks | 8179 | 8210 | 32 | CTGATCGAGGCCCAGCCCGTGGACGGTGTGAG | 152.9 | -2.6 | 0.9 | -2.8 | 5.0E-03 | 5.7E-03 | BT20/MB468 |
| >trna20_GluTTC_2_-_131094701_131094772@1.29.29,trna5_GluTTC_13_-_41634874_41634945@1.29.29\|TCCCATATGGTCTAGCGGTTAGGATTCCT | >trna20_GluTTC_2_-_131094701_131094772 | 1 | 29 | 29 | TCCCATATGGTCTAGCGGTTAGGATTCCT | 329.2 | -2.7 | 1.0 | -2.8 | 5.0E-03 | 5.7E-03 | BT20/MB468 |
| >hg38_21_+_8433172_8446622_RNA45SN1_45S_50ntFlanks@10840.10867.28\|CCGTAACTTCGGGATAAGGATTGGCTCT | >hg38_21_+_8433172_8446622_RNA45SN1_45S_50ntFlanks | 10840 | 10867 | 28 | CCGTAACTTCGGGATAAGGATTGGCTCT | 214.0 | -1.6 | 0.6 | -2.8 | 5.0E-03 | 5.7E-03 | BT20/MB468 |
| >hg38_21_+_8433172_8446622_RNA45SN1_45S_50ntFlanks@6661.6686.26\|CGGTGGATCACTCGGCTCGTGCGTCG | >hg38_21_+_8433172_8446622_RNA45SN1_45S_50ntFlanks | 6661 | 6686 | 26 | CGGTGGATCACTCGGCTCGTGCGTCG | 1542.2 | 2.1 | 0.7 | 2.8 | 5.0E-03 | 5.7E-03 | BT20/MB468 |
| >hg38_21_+_8433172_8446622_RNA45SN1_45S_50ntFlanks@6650.6698.49\|ACGACTCTTAGCGGTGGATCACTCGGCTCGTGCGTCGATGAAGAACGCA | >hg38_21_+_8433172_8446622_RNA45SN1_45S_50ntFlanks | 6650 | 6698 | 49 | ACGACTCTTAGCGGTGGATCACTCGGCTCGTGCGTCGATGAAGAACGCA | 770.3 | -2.8 | 0.8 | -3.3 | 9.2E-04 | 5.7E-03 | BT20/MB231 |
| >am_trnaMT_ValTAC_MT_+_1602_1670@28.47.20\|AACTTACACTTAGGAGATTT | >am_trnaMT_ValTAC_MT_+_1602_1670 | 28 | 47 | 20 | AACTTACACTTAGGAGATTT | 351.1 | 3.2 | 1.0 | 3.3 | 9.2E-04 | 5.7E-03 | BT20/MB231 |
| >hg38_21_+_8433172_8446622_RNA45SN1_45S_50ntFlanks@6647.6689.43\|CGTACGACTCTTAGCGGTGGATCACTCGGCTCGTGCGTCGATG | >hg38_21_+_8433172_8446622_RNA45SN1_45S_50ntFlanks | 6647 | 6689 | 43 | CGTACGACTCTTAGCGGTGGATCACTCGGCTCGTGCGTCGATG | 422.3 | 1.9 | 0.7 | 2.8 | 5.0E-03 | 5.8E-03 | BT20/MB468 |
| >hg38_21_+_8433172_8446622_RNA45SN1_45S_50ntFlanks@10679.10707.29\|GGTTCCGGCGGCGTCCGGTGAGCTCTCGC | >hg38_21_+_8433172_8446622_RNA45SN1_45S_50ntFlanks | 10679 | 10707 | 29 | GGTTCCGGCGGCGTCCGGTGAGCTCTCGC | 163.5 | -1.3 | 0.5 | -2.8 | 4.7E-03 | 5.8E-03 | MB231/MB468 |
| >hg38_21_+_8433172_8446622_RNA45SN1_45S_50ntFlanks@4742.4759.18\|TCGGAGGTTCGAAGACGA | >hg38_21_+_8433172_8446622_RNA45SN1_45S_50ntFlanks | 4742 | 4759 | 18 | TCGGAGGTTCGAAGACGA | 643.3 | 2.3 | 0.8 | 2.8 | 4.7E-03 | 5.8E-03 | MB231/MB468 |
| >hg38_21_+_8433172_8446622_RNA45SN1_45S_50ntFlanks@12474.12496.23\|ACCCACTAATAGGGAACGTGAGC | >hg38_21_+_8433172_8446622_RNA45SN1_45S_50ntFlanks | 12474 | 12496 | 23 | ACCCACTAATAGGGAACGTGAGC | 684.8 | 1.1 | 0.4 | 2.8 | 5.1E-03 | 5.8E-03 | BT20/MB468 |
| >MI0001641\|hsa-mir-429&WithFlank&1\|+\|1168999\|1169093@57.78.22&[MIMAT0001536&hsa-miR-429&offsets\|0\|0;m-207&1\|+\|1169055\|1169076&offsets\|0\|0]\|TAATACTGTCTGGTAAAACCGT | >MI0001641\|hsa-mir-429&WithFlank&1\|+\|1168999\|1169093 | 57 | 78 | 22 | TAATACTGTCTGGTAAAACCGT | 1002.8 | -2.5 | 0.9 | -2.8 | 5.1E-03 | 5.9E-03 | BT20/MB468 |
| >hg38_21_+_8433172_8446622_RNA45SN1_45S_50ntFlanks@6651.6699.49\|CGACTCTTAGCGGTGGATCACTCGGCTCGTGCGTCGATGAAGAACGCAG | >hg38_21_+_8433172_8446622_RNA45SN1_45S_50ntFlanks | 6651 | 6699 | 49 | CGACTCTTAGCGGTGGATCACTCGGCTCGTGCGTCGATGAAGAACGCAG | 6761.1 | -2.8 | 0.8 | -3.3 | 9.4E-04 | 5.9E-03 | BT20/MB231 |
| >hg38_21_+_8433172_8446622_RNA45SN1_45S_50ntFlanks@6648.6669.22\|GTACGACTCTTAGCGGTGGATC | >hg38_21_+_8433172_8446622_RNA45SN1_45S_50ntFlanks | 6648 | 6669 | 22 | GTACGACTCTTAGCGGTGGATC | 760.5 | 2.1 | 0.7 | 2.8 | 4.8E-03 | 5.9E-03 | MB231/MB468 |
| >hg38_21_+_8433172_8446622_RNA45SN1_45S_50ntFlanks@7974.7994.21\|ACGCGACCTCAGATCAGACGT | >hg38_21_+_8433172_8446622_RNA45SN1_45S_50ntFlanks | 7974 | 7994 | 21 | ACGCGACCTCAGATCAGACGT | 1178.3 | 2.8 | 0.8 | 3.3 | 9.5E-04 | 5.9E-03 | BT20/MB231 |

<table>
<tr><td></td><td></td><td></td><td></td><td></td><td></td><td></td><td></td><td></td><td></td><td></td><td></td><td></td></tr>
<tr><td>>hg38_21_+_8433172_8446622_RNA45SN1_45S_50ntFlanks@6650.6689.40|ACGACTCTTAGCGGTGGATCACTCGGCTCGTGCGTCGATG</td><td>>hg38_21_+_8433172_8446622_RNA45SN1_45S_50ntFlanks</td><td>6650</td><td>6689</td><td>40</td><td>ACGACTCTTAGCGGTGGATCACTCGGCTCGTGCGTCGATG</td><td>1282.5</td><td>1.4</td><td>0.5</td><td>2.8</td><td>4.8E-03</td><td>5.9E-03</td><td>MB231/MB468</td></tr>
<tr><td>>MI0000272|hsa-mir-182&WithFlank&7|-|129770377|129770498@29.54.26(+1A)&[MIMAT0000259&hsa-miR-182-5p&offsets|0|+2(+1A);m-22&7|-|129770449|129770470&offsets|0|+4(+1A)]|TTTGGCAATGGTAGAACTCACACTGGA</td><td>>MI0000272|hsa-mir-182&WithFlank&7|-|129770377|129770498</td><td>29</td><td>54</td><td>26(+1A)</td><td>TTTGGCAATGGTAGAACTCACACTGGA</td><td>232.9</td><td>2.5</td><td>0.8</td><td>3.3</td><td>9.5E-04</td><td>5.9E-03</td><td>BT20/MB231</td></tr>
<tr><td>>hg38_21_+_8433172_8446622_RNA45SN1_45S_50ntFlanks@12416.12446.31|TTCGATGTCGGCTCTTCCTATCATTGTGAAG</td><td>>hg38_21_+_8433172_8446622_RNA45SN1_45S_50ntFlanks</td><td>12416</td><td>12446</td><td>31</td><td>TTCGATGTCGGCTCTTCCTATCATTGTGAAG</td><td>335.8</td><td>1.6</td><td>0.6</td><td>2.8</td><td>5.2E-03</td><td>5.9E-03</td><td>BT20/MB468</td></tr>
<tr><td>>trna14_LysTTT_11_-_59327808_59327880@1.29.29,trna2_LysTTT_17_+_8022473_8022545@1.29.29,trna54_LysTTT_1_+_204475655_204475727@1.29.29,trna5_LysTTT_11_+_59323902_59323974@1.29.29,trna62_LysTTT_1_-_204476158_204476230@1.29.29,trna76_LysTTT_6_+_28918806_28918878@1.29.29|GCCCGGATAGCTCAGTCGGTAGAGCATCA</td><td>>trna14_LysTTT_11_-_59327808_59327880</td><td>1</td><td>29</td><td>29</td><td>GCCCGGATAGCTCAGTCGGTAGAGCATCA</td><td>192.5</td><td>-4.8</td><td>1.7</td><td>-2.8</td><td>4.8E-03</td><td>6.0E-03</td><td>MB231/MB468</td></tr>
<tr><td>>hg38_21_+_8433172_8446622_RNA45SN1_45S_50ntFlanks@11580.11602.23|GAACTGGTGCGGACCAGGGGAAT</td><td>>hg38_21_+_8433172_8446622_RNA45SN1_45S_50ntFlanks</td><td>11580</td><td>11602</td><td>23</td><td>GAACTGGTGCGGACCAGGGGAAT</td><td>678.0</td><td>-2.5</td><td>0.9</td><td>-2.8</td><td>5.2E-03</td><td>6.0E-03</td><td>BT20/MB468</td></tr>
<tr><td>>hg38_21_+_8433172_8446622_RNA45SN1_45S_50ntFlanks@10349.10373.25|AACGAGAACTTTGAAGGCCGAAGTG</td><td>>hg38_21_+_8433172_8446622_RNA45SN1_45S_50ntFlanks</td><td>10349</td><td>10373</td><td>25</td><td>AACGAGAACTTTGAAGGCCGAAGTG</td><td>390.2</td><td>1.1</td><td>0.4</td><td>2.8</td><td>5.2E-03</td><td>6.0E-03</td><td>BT20/MB468</td></tr>
<tr><td>>hg38_21_+_8433172_8446622_RNA45SN1_45S_50ntFlanks@11874.11894.21|AGAAGACCCTGTTGAGCTTGA</td><td>>hg38_21_+_8433172_8446622_RNA45SN1_45S_50ntFlanks</td><td>11874</td><td>11894</td><td>21</td><td>AGAAGACCCTGTTGAGCTTGA</td><td>116.8</td><td>-4.4</td><td>1.5</td><td>-2.8</td><td>4.9E-03</td><td>6.0E-03</td><td>MB231/MB468</td></tr>
<tr><td>>hg38_21_+_8433172_8446622_RNA45SN1_45S_50ntFlanks@8115.8138.24|CGCGGGACATGTGGCGTACGGAAG</td><td>>hg38_21_+_8433172_8446622_RNA45SN1_45S_50ntFlanks</td><td>8115</td><td>8138</td><td>24</td><td>CGCGGGACATGTGGCGTACGGAAG</td><td>167.7</td><td>-4.8</td><td>1.7</td><td>-2.8</td><td>4.9E-03</td><td>6.0E-03</td><td>MB231/MB468</td></tr>
<tr><td>>hg38_21_+_8433172_8446622_RNA45SN1_45S_50ntFlanks@10789.10818.30|AGCCTCTGGCATGTTGGAACAATGTAGGTA</td><td>>hg38_21_+_8433172_8446622_RNA45SN1_45S_50ntFlanks</td><td>10789</td><td>10818</td><td>30</td><td>AGCCTCTGGCATGTTGGAACAATGTAGGTA</td><td>556.0</td><td>-2.0</td><td>0.7</td><td>-2.8</td><td>5.2E-03</td><td>6.0E-03</td><td>BT20/MB468</td></tr>
<tr><td>>hg38_21_+_8433172_8446622_RNA45SN1_45S_50ntFlanks@12606.12625.20|ATGTGCTTGGCTGAGGAGCC</td><td>>hg38_21_+_8433172_8446622_RNA45SN1_45S_50ntFlanks</td><td>12606</td><td>12625</td><td>20</td><td>ATGTGCTTGGCTGAGGAGCC</td><td>622.4</td><td>-1.5</td><td>0.5</td><td>-2.8</td><td>4.9E-03</td><td>6.1E-03</td><td>MB231/MB468</td></tr>
<tr><td>>hg38_21_+_8433172_8446622_RNA45SN1_45S_50ntFlanks@12475.12499.25|CCCACTAATAGGGAACGTGAGCTGG</td><td>>hg38_21_+_8433172_8446622_RNA45SN1_45S_50ntFlanks</td><td>12475</td><td>12499</td><td>25</td><td>CCCACTAATAGGGAACGTGAGCTGG</td><td>108.1</td><td>-1.6</td><td>0.6</td><td>-2.8</td><td>4.9E-03</td><td>6.1E-03</td><td>MB231/MB468</td></tr>
<tr><td>>hg38_21_+_8433172_8446622_RNA45SN1_45S_50ntFlanks@7973.7995.23|GACGCGACCTCAGATCAGACGTG</td><td>>hg38_21_+_8433172_8446622_RNA45SN1_45S_50ntFlanks</td><td>7973</td><td>7995</td><td>23</td><td>GACGCGACCTCAGATCAGACGTG</td><td>399.5</td><td>1.9</td><td>0.7</td><td>2.8</td><td>5.3E-03</td><td>6.1E-03</td><td>BT20/MB468</td></tr>
<tr><td>>hg38_21_+_8433172_8446622_RNA45SN1_45S_50ntFlanks@7979.8011.33|ACCTCAGATCAGACGTGGCGACCCGCTGAATTT</td><td>>hg38_21_+_8433172_8446622_RNA45SN1_45S_50ntFlanks</td><td>7979</td><td>8011</td><td>33</td><td>ACCTCAGATCAGACGTGGCGACCCGCTGAATTT</td><td>296.6</td><td>1.7</td><td>0.6</td><td>2.8</td><td>5.3E-03</td><td>6.1E-03</td><td>BT20/MB468</td></tr>
<tr><td>>hg38_21_+_8433172_8446622_RNA45SN1_45S_50ntFlanks@11572.11606.35|CCGACTTAGAACTGGTGCGGACCAGGGGAATCCGA</td><td>>hg38_21_+_8433172_8446622_RNA45SN1_45S_50ntFlanks</td><td>11572</td><td>11606</td><td>35</td><td>CCGACTTAGAACTGGTGCGGACCAGGGGAATCCGA</td><td>141.2</td><td>-1.5</td><td>0.5</td><td>-2.8</td><td>5.0E-03</td><td>6.1E-03</td><td>MB231/MB468</td></tr>
<tr><td>>hg38_21_+_8433172_8446622_RNA45SN1_45S_50ntFlanks@10408.10432.25|GGTCAGTCGGTCCTGAGAGATGGGC</td><td>>hg38_21_+_8433172_8446622_RNA45SN1_45S_50ntFlanks</td><td>10408</td><td>10432</td><td>25</td><td>GGTCAGTCGGTCCTGAGAGATGGGC</td><td>299.6</td><td>-4.4</td><td>1.6</td><td>-2.8</td><td>5.0E-03</td><td>6.2E-03</td><td>MB231/MB468</td></tr>
<tr><td>>hg38_21_+_8433172_8446622_RNA45SN1_45S_50ntFlanks@10677.10709.33|GCGGTTCCGGCGGCGTCCGGTGAGCTCTCGCTG</td><td>>hg38_21_+_8433172_8446622_RNA45SN1_45S_50ntFlanks</td><td>10677</td><td>10709</td><td>33</td><td>GCGGTTCCGGCGGCGTCCGGTGAGCTCTCGCTG</td><td>1365.9</td><td>2.9</td><td>1.1</td><td>2.8</td><td>5.4E-03</td><td>6.2E-03</td><td>BT20/MB468</td></tr>
<tr><td>>hg38_21_+_8433172_8446622_RNA45SN1_45S_50ntFlanks@12420.12448.29|ATGTCGGCTCTTCCTATCATTGTGAAGCA</td><td>>hg38_21_+_8433172_8446622_RNA45SN1_45S_50ntFlanks</td><td>12420</td><td>12448</td><td>29</td><td>ATGTCGGCTCTTCCTATCATTGTGAAGCA</td><td>441.3</td><td>1.4</td><td>0.5</td><td>2.8</td><td>5.4E-03</td><td>6.2E-03</td><td>BT20/MB468</td></tr>
<tr><td>>hg38_21_+_8433172_8446622_RNA45SN1_45S_50ntFlanks@7989.8014.26|AGACGTGGCGACCCGCTGAATTTAAG</td><td>>hg38_21_+_8433172_8446622_RNA45SN1_45S_50ntFlanks</td><td>7989</td><td>8014</td><td>26</td><td>AGACGTGGCGACCCGCTGAATTTAAG</td><td>514.0</td><td>2.0</td><td>0.7</td><td>2.8</td><td>5.1E-03</td><td>6.3E-03</td><td>MB231/MB468</td></tr>
<tr><td>>hg38_21_+_8433172_8446622_RNA45SN1_45S_50ntFlanks@8109.8143.35|GCGGGGCGCGGGACATGTGGCGTACGGAAGACCCG</td><td>>hg38_21_+_8433172_8446622_RNA45SN1_45S_50ntFlanks</td><td>8109</td><td>8143</td><td>35</td><td>GCGGGGCGCGGGACATGTGGCGTACGGAAGACCCG</td><td>72.1</td><td>-2.1</td><td>0.7</td><td>-2.8</td><td>5.1E-03</td><td>6.3E-03</td><td>MB231/MB468</td></tr>
<tr><td>>am_trna111_HisGTG_1_-_147774845_147774916@1.29.29,trna118_HisGTG_1_-_145396881_145396952@1.29.29,trna16_HisGTG_1_+_146544773_146544844@1.29.29,trna1_HisGTG_15_+_45493349_45493420@1.29.29,trna21_HisGTG_1_+_147753471_147753542@1.29.29,trna33_HisGTG_6_+_27125906_27125977@1.29.29,trna7_HisGTG_9_-_14433938_14434009@1.29.29,trna8_HisGTG_15_-_45492611_45492682@1.29.29,trna9_HisGTG_15_-_45490804_45490875@1.29.29|GCCGTGATCGTATAGTGGTTAGTACTCTG</td><td>>am_trna111_HisGTG_1_-_147774845_147774916</td><td>1</td><td>29</td><td>29</td><td>GCCGTGATCGTATAGTGGTTAGTACTCTG</td><td>4073.7</td><td>-2.6</td><td>0.9</td><td>-2.8</td><td>5.1E-03</td><td>6.3E-03</td><td>MB231/MB468</td></tr>
<tr><td>>hg38_21_+_8433172_8446622_RNA45SN1_45S_50ntFlanks@10389.10429.41|TGAACAGCAGTTGAACATGGGTCAGTCGGTCCTGAGAGATG</td><td>>hg38_21_+_8433172_8446622_RNA45SN1_45S_50ntFlanks</td><td>10389</td><td>10429</td><td>41</td><td>TGAACAGCAGTTGAACATGGGTCAGTCGGTCCTGAGAGATG</td><td>275.1</td><td>-2.1</td><td>0.7</td><td>-2.8</td><td>5.1E-03</td><td>6.3E-03</td><td>MB231/MB468</td></tr>
<tr><td>>am_trna119_LysCTT_1_-_145395522_145395594@13.30.18,trna11_LysCTT_5_-_180648979_180649051@13.30.18,trna128_LysCTT_1_-_55423542_55423614@13.30.18,trna13_LysCTT_6_+_26556774_26556846@13.30.18,trna32_LysCTT_16_-_3207406_3207478@13.30.18,trna6_LysCTT_19_-_52425396_52425468@13.30.18,trna7_LysCTT_16_+_3225692_3225764@13.30.18,trna9_LysCTT_5_+_180634755_180634827@13.30.18|CAGTCGGTAGAGCATGAG</td><td>>am_trna119_LysCTT_1_-_145395522_145395594</td><td>13</td><td>30</td><td>18</td><td>CAGTCGGTAGAGCATGAG</td><td>177.5</td><td>-1.5</td><td>0.5</td><td>-2.8</td><td>5.5E-03</td><td>6.3E-03</td><td>BT20/MB468</td></tr>
<tr><td>>hg38_21_+_8433172_8446622_RNA45SN1_45S_50ntFlanks@11682.11702.21|AATGTCAAAGTGAAGAAATTC</td><td>>hg38_21_+_8433172_8446622_RNA45SN1_45S_50ntFlanks</td><td>11682</td><td>11702</td><td>21</td><td>AATGTCAAAGTGAAGAAATTC</td><td>375.5</td><td>-2.1</td><td>0.7</td><td>-2.8</td><td>5.2E-03</td><td>6.4E-03</td><td>MB231/MB468</td></tr>
<tr><td>>hg38_21_+_8433172_8446622_RNA45SN1_45S_50ntFlanks@8179.8212.34|CTGATCGAGGCCCAGCCCGTGGACGGTGTGAGGC</td><td>>hg38_21_+_8433172_8446622_RNA45SN1_45S_50ntFlanks</td><td>8179</td><td>8212</td><td>34</td><td>CTGATCGAGGCCCAGCCCGTGGACGGTGTGAGGC</td><td>135.6</td><td>-4.3</td><td>1.5</td><td>-2.8</td><td>5.2E-03</td><td>6.4E-03</td><td>MB231/MB468</td></tr>
<tr><td>>hg38_21_+_8433172_8446622_RNA45SN1_45S_50ntFlanks@12420.12439.20|ATGTCGGCTCTTCCTATCAT</td><td>>hg38_21_+_8433172_8446622_RNA45SN1_45S_50ntFlanks</td><td>12420</td><td>12439</td><td>20</td><td>ATGTCGGCTCTTCCTATCAT</td><td>1648.8</td><td>1.7</td><td>0.6</td><td>2.8</td><td>5.2E-03</td><td>6.4E-03</td><td>MB231/MB468</td></tr>
<tr><td>>hg38_21_+_8433172_8446622_RNA45SN1_45S_50ntFlanks@10679.10705.27|GGTTCCGGCGGCGTCCGGTGAGCTCTC</td><td>>hg38_21_+_8433172_8446622_RNA45SN1_45S_50ntFlanks</td><td>10679</td><td>10705</td><td>27</td><td>GGTTCCGGCGGCGTCCGGTGAGCTCTC</td><td>292.6</td><td>-2.8</td><td>1.0</td><td>-2.8</td><td>5.6E-03</td><td>6.4E-03</td><td>BT20/MB468</td></tr>
<tr><td>>hg38_21_+_8433172_8446622_RNA45SN1_45S_50ntFlanks@10458.10487.30|ATGGCCTCCGTTGCCCTCGGCCGATCGAAA</td><td>>hg38_21_+_8433172_8446622_RNA45SN1_45S_50ntFlanks</td><td>10458</td><td>10487</td><td>30</td><td>ATGGCCTCCGTTGCCCTCGGCCGATCGAAA</td><td>342.1</td><td>-1.9</td><td>0.7</td><td>-2.8</td><td>5.2E-03</td><td>6.4E-03</td><td>MB231/MB468</td></tr>
<tr><td>>hg38_21_+_8433172_8446622_RNA45SN1_45S_50ntFlanks@12365.12383.19|ATAACTGGCTTGTGGCGGC</td><td>>hg38_21_+_8433172_8446622_RNA45SN1_45S_50ntFlanks</td><td>12365</td><td>12383</td><td>19</td><td>ATAACTGGCTTGTGGCGGC</td><td>183.2</td><td>-4.6</td><td>1.6</td><td>-2.8</td><td>5.2E-03</td><td>6.4E-03</td><td>MB231/MB468</td></tr>
<tr><td>>MI0000542|hsa-mir-320a&WithFlank&8|-|22244960|22245043@48.68.21(+1G)&[MIMAT0000510&hsa-miR-320a-3p&offsets|0|-1(+1G);m-60&8|-|22244975|22244996&offsets|0|-1(+1G)]|AAAAGCTGGGTTGAGAGGGCGG</td><td>>MI0000542|hsa-mir-320a&WithFlank&8|-|22244960|22245043</td><td>48</td><td>68</td><td>21(+1G)</td><td>AAAAGCTGGGTTGAGAGGGCGG</td><td>1434.6</td><td>3.1</td><td>1.0</td><td>3.3</td><td>1.0E-03</td><td>6.4E-03</td><td>BT20/MB231</td></tr>
<tr><td>>hg38_21_+_8433172_8446622_RNA45SN1_45S_50ntFlanks@9538.9571.34|AACTCTGGTGGAGGTCCGTAGCGGTCCTGACGTG</td><td>>hg38_21_+_8433172_8446622_RNA45SN1_45S_50ntFlanks</td><td>9538</td><td>9571</td><td>34</td><td>AACTCTGGTGGAGGTCCGTAGCGGTCCTGACGTG</td><td>289.3</td><td>-2.4</td><td>0.9</td><td>-2.8</td><td>5.6E-03</td><td>6.5E-03</td><td>BT20/MB468</td></tr>
<tr><td>>hg38_21_+_8433172_8446622_RNA45SN1_45S_50ntFlanks@9545.9572.28|GTGGAGGTCCGTAGCGGTCCTGACGTGC</td><td>>hg38_21_+_8433172_8446622_RNA45SN1_45S_50ntFlanks</td><td>9545</td><td>9572</td><td>28</td><td>GTGGAGGTCCGTAGCGGTCCTGACGTGC</td><td>874.4</td><td>-1.7</td><td>0.6</td><td>-2.8</td><td>5.3E-03</td><td>6.5E-03</td><td>MB231/MB468</td></tr>
<tr><td>>hg38_21_+_8433172_8446622_RNA45SN1_45S_50ntFlanks@6734.6753.20|ATTGATCATCGACACTTCGA</td><td>>hg38_21_+_8433172_8446622_RNA45SN1_45S_50ntFlanks</td><td>6734</td><td>6753</td><td>20</td><td>ATTGATCATCGACACTTCGA</td><td>272.4</td><td>1.5</td><td>0.5</td><td>2.8</td><td>5.7E-03</td><td>6.5E-03</td><td>BT20/MB468</td></tr>
<tr><td>>hg38_21_+_8433172_8446622_RNA45SN1_45S_50ntFlanks@11207.11232.26|GGCGGCGACTCTGGACGCGAGCCGGG</td><td>>hg38_21_+_8433172_8446622_RNA45SN1_45S_50ntFlanks</td><td>11207</td><td>11232</td><td>26</td><td>GGCGGCGACTCTGGACGCGAGCCGGG</td><td>368.8</td><td>-2.3</td><td>0.8</td><td>-2.8</td><td>5.7E-03</td><td>6.6E-03</td><td>BT20/MB468</td></tr>
<tr><td>>hg38_21_+_8433172_8446622_RNA45SN1_45S_50ntFlanks@4589.4616.28|TATTTTGTTGGTTTTCGGAACTGAGGCC</td><td>>hg38_21_+_8433172_8446622_RNA45SN1_45S_50ntFlanks</td><td>4589</td><td>4616</td><td>28</td><td>TATTTTGTTGGTTTTCGGAACTGAGGCC</td><td>89.8</td><td>-1.9</td><td>0.7</td><td>-2.8</td><td>5.7E-03</td><td>6.6E-03</td><td>BT20/MB468</td></tr>
<tr><td>>hg38_21_+_8433172_8446622_RNA45SN1_45S_50ntFlanks@4724.4757.34|ATTAATCAAGAACGAAAGTCGGAGGTTCGAAGAC</td><td>>hg38_21_+_8433172_8446622_RNA45SN1_45S_50ntFlanks</td><td>4724</td><td>4757</td><td>34</td><td>ATTAATCAAGAACGAAAGTCGGAGGTTCGAAGAC</td><td>89.7</td><td>-2.6</td><td>0.9</td><td>-2.8</td><td>5.7E-03</td><td>6.6E-03</td><td>BT20/MB468</td></tr>
<tr><td>>hg38_21_+_8433172_8446622_RNA45SN1_45S_50ntFlanks@7988.8011.24|CAGACGTGGCGACCCGCTGAATTT</td><td>>hg38_21_+_8433172_8446622_RNA45SN1_45S_50ntFlanks</td><td>7988</td><td>8011</td><td>24</td><td>CAGACGTGGCGACCCGCTGAATTT</td><td>968.4</td><td>1.8</td><td>0.6</td><td>2.8</td><td>5.4E-03</td><td>6.6E-03</td><td>MB231/MB468</td></tr>
<tr><td>>hg38_21_+_8433172_8446622_RNA45SN1_45S_50ntFlanks@12420.12449.30|ATGTCGGCTCTTCCTATCATTGTGAAGCAG</td><td>>hg38_21_+_8433172_8446622_RNA45SN1_45S_50ntFlanks</td><td>12420</td><td>12449</td><td>30</td><td>ATGTCGGCTCTTCCTATCATTGTGAAGCAG</td><td>1404.5</td><td>1.7</td><td>0.6</td><td>2.8</td><td>5.4E-03</td><td>6.6E-03</td><td>MB231/MB468</td></tr>
<tr><td>>MI0000542|hsa-mir-320a&WithFlank&8|-|22244960|22245043@49.68.20&[MIMAT0000510&hsa-miR-320a-3p&offsets|+1|-1;m-60&8|-|22244975|22244996&offsets|+1|-1]|AAAGCTGGGTTGAGAGGGCG</td><td>>MI0000542|hsa-mir-320a&WithFlank&8|-|22244960|22245043</td><td>49</td><td>68</td><td>20</td><td>AAAGCTGGGTTGAGAGGGCG</td><td>288.2</td><td>2.4</td><td>0.7</td><td>3.3</td><td>1.1E-03</td><td>6.6E-03</td><td>BT20/MB231</td></tr>
<tr><td>>hg38_21_+_8433172_8446622_RNA45SN1_45S_50ntFlanks@11671.11701.31|CCAGTGCTCTGAATGTCAAAGTGAAGAAATT</td><td>>hg38_21_+_8433172_8446622_RNA45SN1_45S_50ntFlanks</td><td>11671</td><td>11701</td><td>31</td><td>CCAGTGCTCTGAATGTCAAAGTGAAGAAATT</td><td>371.1</td><td>-1.2</td><td>0.4</td><td>-2.8</td><td>5.4E-03</td><td>6.7E-03</td><td>MB231/MB468</td></tr>
<tr><td>>hg38_21_+_8433172_8446622_RNA45SN1_45S_50ntFlanks@11319.11342.24|CCCACGTCTCGTCGCGCGCGCGTC</td><td>>hg38_21_+_8433172_8446622_RNA45SN1_45S_50ntFlanks</td><td>11319</td><td>11342</td><td>24</td><td>CCCACGTCTCGTCGCGCGCGCGTC</td><td>256.9</td><td>-4.8</td><td>1.7</td><td>-2.8</td><td>5.4E-03</td><td>6.7E-03</td><td>MB231/MB468</td></tr>
</table>

<table>
<tr><th></th><th></th><th></th><th></th><th></th><th></th><th></th><th></th><th></th><th></th><th></th><th></th><th></th></tr>
<tr><td>>am_MI0000809|hsa-mir-151a&WithFlank&8|-|140732558|140732659@17.38.22&[MIMAT0004697&hsa-miR-151a-5p&offsets|0|+1;m-75&8|-|140732623|140732643&offsets|0|+1]|TCGAGGAGCTCACAGTCTAGTA</td><td>>am_MI0000809|hsa-mir-151a&WithFlank&8|-|140732558|140732659</td><td>17</td><td>38</td><td>22</td><td>TCGAGGAGCTCACAGTCTAGTA</td><td>1789.6</td><td>2.7</td><td>0.8</td><td>3.3</td><td>1.1E-03</td><td>6.7E-03</td><td>BT20/MB231</td></tr>
<tr><td>>MI0000749|hsa-mir-30e&WithFlank&1|+|40754349|40754452@23.46.24(+2U)&[MIMAT0000692&hsa-miR-30e-5p&offsets|0|+2(+2U);m-24&1|+|40754371|40754394&offsets|0|0(+2U)]|TGTAAACATCCTTGACTGGAAGCTTT</td><td>>MI0000749|hsa-mir-30e&WithFlank&1|+|40754349|40754452</td><td>23</td><td>46</td><td>24(+2U)</td><td>TGTAAACATCCTTGACTGGAAGCTTT</td><td>256.2</td><td>-2.4</td><td>0.7</td><td>-3.3</td><td>1.1E-03</td><td>6.7E-03</td><td>BT20/MB231</td></tr>
<tr><td>>trna152_ValCAC_6_-_27248049_27248121@1.33.33|GCTTCTGTAGTGTAGTGGTTATCACGTTCGCCT</td><td>>trna152_ValCAC_6_-_27248049_27248121</td><td>1</td><td>33</td><td>33</td><td>GCTTCTGTAGTGTAGTGGTTATCACGTTCGCCT</td><td>1743.2</td><td>-3.1</td><td>0.9</td><td>-3.3</td><td>1.1E-03</td><td>6.7E-03</td><td>BT20/MB231</td></tr>
<tr><td>>hg38_21_+_8433172_8446622_RNA45SN1_45S_50ntFlanks@4683.4707.25|CAAGACGGACCAGAGCGAAAGCATT</td><td>>hg38_21_+_8433172_8446622_RNA45SN1_45S_50ntFlanks</td><td>4683</td><td>4707</td><td>25</td><td>CAAGACGGACCAGAGCGAAAGCATT</td><td>209.1</td><td>-2.2</td><td>0.8</td><td>-2.8</td><td>5.5E-03</td><td>6.7E-03</td><td>MB231/MB468</td></tr>
<tr><td>>hg38_21_+_8433172_8446622_RNA45SN1_45S_50ntFlanks@8176.8213.38|CTTCTGATCGAGGCCCAGCCCGTGGACGGTGTGAGGCC</td><td>>hg38_21_+_8433172_8446622_RNA45SN1_45S_50ntFlanks</td><td>8176</td><td>8213</td><td>38</td><td>CTTCTGATCGAGGCCCAGCCCGTGGACGGTGTGAGGCC</td><td>506.5</td><td>2.2</td><td>0.8</td><td>2.8</td><td>5.5E-03</td><td>6.7E-03</td><td>MB231/MB468</td></tr>
<tr><td>>MI0000746|hsa-mir-99b&WithFlank&19|+|51692606|51692687@51.72.22(+1A)&[MIMAT0004678&hsa-miR-99b-3p&offsets|0|0(+1A);m-151&19|+|51692656|51692678&offsets|0|-1(+1A)]|CAAGCTCGTGTCTGTGGGTCCGA</td><td>>MI0000746|hsa-mir-99b&WithFlank&19|+|51692606|51692687</td><td>51</td><td>72</td><td>22(+1A)</td><td>CAAGCTCGTGTCTGTGGGTCCGA</td><td>1036.3</td><td>2.5</td><td>0.8</td><td>3.3</td><td>1.1E-03</td><td>6.8E-03</td><td>BT20/MB231</td></tr>
<tr><td>>trna119_LysCTT_1_-_145395522_145395594@4.35.32,trna11_LysCTT_5_-_180648979_180649051@4.35.32,trna13_LysCTT_6_+_26556774_26556846@4.35.32,trna7_LysCTT_16_+_3225692_3225764@4.35.32,trna9_LysCTT_5_+_180634755_180634827@4.35.32|CGGCTAGCTCAGTCGGTAGAGCATGAGACTCT</td><td>>trna119_LysCTT_1_-_145395522_145395594</td><td>4</td><td>35</td><td>32</td><td>CGGCTAGCTCAGTCGGTAGAGCATGAGACTCT</td><td>1225.3</td><td>-2.9</td><td>0.9</td><td>-3.3</td><td>1.1E-03</td><td>6.8E-03</td><td>BT20/MB231</td></tr>
<tr><td>>hg38_21_+_8433172_8446622_RNA45SN1_45S_50ntFlanks@11037.11060.24|CTCTCCCCCGCTCCCCGTCCTCCC</td><td>>hg38_21_+_8433172_8446622_RNA45SN1_45S_50ntFlanks</td><td>11037</td><td>11060</td><td>24</td><td>CTCTCCCCCGCTCCCCGTCCTCCC</td><td>120.6</td><td>-4.4</td><td>1.6</td><td>-2.8</td><td>5.5E-03</td><td>6.8E-03</td><td>MB231/MB468</td></tr>
<tr><td>>MI0003643|hsa-mir-629&WithFlank&15|-|70079366|70079474@28.49.22&[MIMAT0004810&hsa-miR-629-5p&offsets|0|+1;m-150&15|-|70079427|70079447&offsets|0|+1]|TGGGTTTACGTTGGGAGAACTT</td><td>>MI0003643|hsa-mir-629&WithFlank&15|-|70079366|70079474</td><td>28</td><td>49</td><td>22</td><td>TGGGTTTACGTTGGGAGAACTT</td><td>2569.6</td><td>2.0</td><td>0.6</td><td>3.3</td><td>1.1E-03</td><td>6.8E-03</td><td>BT20/MB231</td></tr>
<tr><td>>hg38_21_+_8433172_8446622_RNA45SN1_45S_50ntFlanks@11682.11706.25|AATGTCAAAGTGAAGAAATTCAATG</td><td>>hg38_21_+_8433172_8446622_RNA45SN1_45S_50ntFlanks</td><td>11682</td><td>11706</td><td>25</td><td>AATGTCAAAGTGAAGAAATTCAATG</td><td>538.5</td><td>-2.7</td><td>1.0</td><td>-2.8</td><td>5.5E-03</td><td>6.8E-03</td><td>MB231/MB468</td></tr>
<tr><td>>hg38_21_+_8433172_8446622_RNA45SN1_45S_50ntFlanks@11567.11588.22|AGCAGCCGACTTAGAACTGGTG</td><td>>hg38_21_+_8433172_8446622_RNA45SN1_45S_50ntFlanks</td><td>11567</td><td>11588</td><td>22</td><td>AGCAGCCGACTTAGAACTGGTG</td><td>559.8</td><td>-1.5</td><td>0.5</td><td>-2.8</td><td>5.9E-03</td><td>6.8E-03</td><td>BT20/MB468</td></tr>
<tr><td>>hg38_21_+_8433172_8446622_RNA45SN1_45S_50ntFlanks@6645.6663.19|CTCGTACGACTCTTAGCGG</td><td>>hg38_21_+_8433172_8446622_RNA45SN1_45S_50ntFlanks</td><td>6645</td><td>6663</td><td>19</td><td>CTCGTACGACTCTTAGCGG</td><td>1231.0</td><td>2.1</td><td>0.8</td><td>2.8</td><td>5.5E-03</td><td>6.8E-03</td><td>MB231/MB468</td></tr>
<tr><td>>hg38_21_+_8433172_8446622_RNA45SN1_45S_50ntFlanks@8114.8141.28|GCGCGGGACATGTGGCGTACGGAAGACC</td><td>>hg38_21_+_8433172_8446622_RNA45SN1_45S_50ntFlanks</td><td>8114</td><td>8141</td><td>28</td><td>GCGCGGGACATGTGGCGTACGGAAGACC</td><td>228.8</td><td>-2.5</td><td>0.9</td><td>-2.7</td><td>6.0E-03</td><td>6.8E-03</td><td>BT20/MB468</td></tr>
<tr><td>>hg38_21_+_8433172_8446622_RNA45SN1_45S_50ntFlanks@10821.10839.19|GGAAGTCGGCAAGCCGGAT</td><td>>hg38_21_+_8433172_8446622_RNA45SN1_45S_50ntFlanks</td><td>10821</td><td>10839</td><td>19</td><td>GGAAGTCGGCAAGCCGGAT</td><td>162.9</td><td>-1.4</td><td>0.5</td><td>-2.8</td><td>5.6E-03</td><td>6.8E-03</td><td>MB231/MB468</td></tr>
<tr><td>>hg38_21_+_8433172_8446622_RNA45SN1_45S_50ntFlanks@12407.12425.19|TTTTGATCCTTCGATGTCG</td><td>>hg38_21_+_8433172_8446622_RNA45SN1_45S_50ntFlanks</td><td>12407</td><td>12425</td><td>19</td><td>TTTTGATCCTTCGATGTCG</td><td>112.8</td><td>-1.9</td><td>0.7</td><td>-2.8</td><td>5.6E-03</td><td>6.8E-03</td><td>MB231/MB468</td></tr>
<tr><td>>MI0000736|hsa-mir-30c-1&WithFlank&1|+|40757278|40757378@23.48.26;MI0000254|hsa-mir-30c-2&WithFlank&6|-|71376954|71377037@13.38.26&[MIMAT0000244&hsa-miR-30c-5p&offsets|0|+3;m-57&1|+|40757300|40757323&offsets|0|+2];[MIMAT0000244_1&hsa-miR-30c-5p&offsets|0|+3;m-56&6|-|71377002|71377025&offsets|0|+2]|TGTAAACATCCTACACTCTCAGCTGT</td><td>>MI0000736|hsa-mir-30c-1&WithFlank&1|+|40757278|40757378</td><td>23</td><td>48</td><td>26</td><td>TGTAAACATCCTACACTCTCAGCTGT</td><td>304.0</td><td>-2.8</td><td>0.9</td><td>-3.3</td><td>1.1E-03</td><td>6.9E-03</td><td>BT20/MB231</td></tr>
<tr><td>>hg38_21_+_8433172_8446622_RNA45SN1_45S_50ntFlanks@6644.6683.40|CCTCGTACGACTCTTAGCGGTGGATCACTCGGCTCGTGCG</td><td>>hg38_21_+_8433172_8446622_RNA45SN1_45S_50ntFlanks</td><td>6644</td><td>6683</td><td>40</td><td>CCTCGTACGACTCTTAGCGGTGGATCACTCGGCTCGTGCG</td><td>463.7</td><td>-2.5</td><td>0.8</td><td>-3.3</td><td>1.1E-03</td><td>6.9E-03</td><td>BT20/MB231</td></tr>
<tr><td>>hg38_21_+_8433172_8446622_RNA45SN1_45S_50ntFlanks@3852.3874.23|TAACTGTGGTAATTCTAGAGCTA</td><td>>hg38_21_+_8433172_8446622_RNA45SN1_45S_50ntFlanks</td><td>3852</td><td>3874</td><td>23</td><td>TAACTGTGGTAATTCTAGAGCTA</td><td>182.7</td><td>-1.9</td><td>0.7</td><td>-2.7</td><td>6.0E-03</td><td>6.9E-03</td><td>BT20/MB468</td></tr>
<tr><td>>hg38_21_+_8433172_8446622_RNA45SN1_45S_50ntFlanks@12481.12500.20|AATAGGGAACGTGAGCTGGG</td><td>>hg38_21_+_8433172_8446622_RNA45SN1_45S_50ntFlanks</td><td>12481</td><td>12500</td><td>20</td><td>AATAGGGAACGTGAGCTGGG</td><td>523.1</td><td>2.7</td><td>1.0</td><td>2.7</td><td>6.1E-03</td><td>7.0E-03</td><td>BT20/MB468</td></tr>
<tr><td>>am_trnaMT_ValTAC_MT_+_1602_1670@49.68.20|AACTTAACTTGACCGCTCTG</td><td>>am_trnaMT_ValTAC_MT_+_1602_1670</td><td>49</td><td>68</td><td>20</td><td>AACTTAACTTGACCGCTCTG</td><td>205.6</td><td>2.7</td><td>0.8</td><td>3.3</td><td>1.1E-03</td><td>7.0E-03</td><td>BT20/MB231</td></tr>
<tr><td>>MI0000746|hsa-mir-99b&WithFlank&19|+|51692606|51692687@51.71.21(+1A)&[MIMAT0004678&hsa-miR-99b-3p&offsets|0|-1(+1A);m-151&19|+|51692656|51692678&offsets|0|-2(+1A)]|CAAGCTCGTGTCTGTGGGTCCA</td><td>>MI0000746|hsa-mir-99b&WithFlank&19|+|51692606|51692687</td><td>51</td><td>71</td><td>21(+1A)</td><td>CAAGCTCGTGTCTGTGGGTCCA</td><td>729.9</td><td>2.5</td><td>0.8</td><td>3.3</td><td>1.1E-03</td><td>7.0E-03</td><td>BT20/MB231</td></tr>
<tr><td>>am_trnaMT_ValTAC_MT_+_1602_1670@49.67.19|AACTTAACTTGACCGCTCT</td><td>>am_trnaMT_ValTAC_MT_+_1602_1670</td><td>49</td><td>67</td><td>19</td><td>AACTTAACTTGACCGCTCT</td><td>2958.5</td><td>2.6</td><td>0.8</td><td>3.2</td><td>1.2E-03</td><td>7.1E-03</td><td>BT20/MB231</td></tr>
<tr><td>>trnaMT_ValTAC_MT_+_1602_1670@50.70.21|ACTTAACTTGACCGCTCTGAC</td><td>>trnaMT_ValTAC_MT_+_1602_1670</td><td>50</td><td>70</td><td>21</td><td>ACTTAACTTGACCGCTCTGAC</td><td>272.0</td><td>2.6</td><td>0.8</td><td>3.2</td><td>1.2E-03</td><td>7.1E-03</td><td>BT20/MB231</td></tr>
<tr><td>>hg38_21_+_8433172_8446622_RNA45SN1_45S_50ntFlanks@4590.4616.27|ATTTTGTTGGTTTTCGGAACTGAGGCC</td><td>>hg38_21_+_8433172_8446622_RNA45SN1_45S_50ntFlanks</td><td>4590</td><td>4616</td><td>27</td><td>ATTTTGTTGGTTTTCGGAACTGAGGCC</td><td>246.8</td><td>-1.9</td><td>0.7</td><td>-2.8</td><td>5.8E-03</td><td>7.1E-03</td><td>MB231/MB468</td></tr>
<tr><td>>am_MI0039740|hsa-mir-12136&WithFlank&1|-|632609|632691@7.32.26&[MIMAT0049032&hsa-miR-12136&offsets|0|+8]|GAAAAAGTCATGGAGGCCATGGGGTT</td><td>>am_MI0039740|hsa-mir-12136&WithFlank&1|-|632609|632691</td><td>7</td><td>32</td><td>26</td><td>GAAAAAGTCATGGAGGCCATGGGGTT</td><td>391.2</td><td>2.1</td><td>0.6</td><td>3.2</td><td>1.2E-03</td><td>7.1E-03</td><td>BT20/MB231</td></tr>
<tr><td>>hg38_21_+_8433172_8446622_RNA45SN1_45S_50ntFlanks@11540.11583.44|CGCGGCGCCTCGCCTCGGCCGGCGCCTAGCAGCCGACTTAGAAC</td><td>>hg38_21_+_8433172_8446622_RNA45SN1_45S_50ntFlanks</td><td>11540</td><td>11583</td><td>44</td><td>CGCGGCGCCTCGCCTCGGCCGGCGCCTAGCAGCCGACTTAGAAC</td><td>780.6</td><td>-3.0</td><td>0.9</td><td>-3.2</td><td>1.2E-03</td><td>7.1E-03</td><td>BT20/MB231</td></tr>
<tr><td>>hg38_21_+_8433172_8446622_RNA45SN1_45S_50ntFlanks@12420.12458.39|ATGTCGGCTCTTCCTATCATTGTGAAGCAGAATTCACCA</td><td>>hg38_21_+_8433172_8446622_RNA45SN1_45S_50ntFlanks</td><td>12420</td><td>12458</td><td>39</td><td>ATGTCGGCTCTTCCTATCATTGTGAAGCAGAATTCACCA</td><td>381.9</td><td>-2.5</td><td>0.8</td><td>-3.2</td><td>1.2E-03</td><td>7.1E-03</td><td>BT20/MB231</td></tr>
<tr><td>>MI0000098|hsa-mir-96&WithFlank&7|-|129774686|129774775@15.36.22&[MIMAT0000095&hsa-miR-96-5p&offsets|0|-1;m-248&7|-|129774739|129774761&offsets|0|-1]|TTTGGCACTAGCACATTTTTGC</td><td>>MI0000098|hsa-mir-96&WithFlank&7|-|129774686|129774775</td><td>15</td><td>36</td><td>22</td><td>TTTGGCACTAGCACATTTTTGC</td><td>392.2</td><td>2.9</td><td>0.9</td><td>3.2</td><td>1.2E-03</td><td>7.1E-03</td><td>BT20/MB231</td></tr>
<tr><td>>trnaMT_SerTGA_MT_-_7446_7514@1.26.26,trnalookalike6_SerTGA_1_-_567997_568065@1.26.26|GAAAAAGTCATGGAGGCCATGGGGTT</td><td>>trnaMT_SerTGA_MT_-_7446_7514</td><td>1</td><td>26</td><td>26</td><td>GAAAAAGTCATGGAGGCCATGGGGTT</td><td>391.2</td><td>2.1</td><td>0.6</td><td>3.2</td><td>1.2E-03</td><td>7.1E-03</td><td>BT20/MB231</td></tr>
<tr><td>>hg38_21_+_8433172_8446622_RNA45SN1_45S_50ntFlanks@12476.12496.21|CCACTAATAGGGAACGTGAGC</td><td>>hg38_21_+_8433172_8446622_RNA45SN1_45S_50ntFlanks</td><td>12476</td><td>12496</td><td>21</td><td>CCACTAATAGGGAACGTGAGC</td><td>1797.6</td><td>-1.6</td><td>0.6</td><td>-2.8</td><td>5.8E-03</td><td>7.1E-03</td><td>MB231/MB468</td></tr>
<tr><td>>hg38_21_+_8433172_8446622_RNA45SN1_45S_50ntFlanks@6778.6803.26|CCCGGGGCTACGCCTGTCTGAGCGTC</td><td>>hg38_21_+_8433172_8446622_RNA45SN1_45S_50ntFlanks</td><td>6778</td><td>6803</td><td>26</td><td>CCCGGGGCTACGCCTGTCTGAGCGTC</td><td>554.6</td><td>2.2</td><td>0.8</td><td>2.8</td><td>5.8E-03</td><td>7.1E-03</td><td>MB231/MB468</td></tr>
<tr><td>>MI0000478|hsa-mir-149&WithFlank&2|+|240455995|240456095@21.40.20&[MIMAT0000450&hsa-miR-149-5p&offsets|0|-3;m-195&2|+|240456015|240456037&offsets|0|-3]|TCTGGCTCCGTGTCTTCACT</td><td>>MI0000478|hsa-mir-149&WithFlank&2|+|240455995|240456095</td><td>21</td><td>40</td><td>20</td><td>TCTGGCTCCGTGTCTTCACT</td><td>501.2</td><td>2.9</td><td>0.9</td><td>3.2</td><td>1.2E-03</td><td>7.1E-03</td><td>BT20/MB231</td></tr>
<tr><td>>trna2_GlyGCC_21_-_18827107_18827177@1.30.30,trna35_GlyGCC_1_+_161413094_161413164@1.30.30,trna37_GlyGCC_1_+_161420467_161420537@1.30.30,trna39_GlyGCC_1_+_161427898_161427968@1.30.30,trna41_GlyGCC_1_+_161435258_161435328@1.30.30|GCATGGGTGGTTCAGTGGTAGAATTCTCGC</td><td>>trna2_GlyGCC_21_-_18827107_18827177</td><td>1</td><td>30</td><td>30</td><td>GCATGGGTGGTTCAGTGGTAGAATTCTCGC</td><td>811.1</td><td>3.2</td><td>1.0</td><td>3.2</td><td>1.2E-03</td><td>7.1E-03</td><td>BT20/MB231</td></tr>
<tr><td>>hg38_21_+_8433172_8446622_RNA45SN1_45S_50ntFlanks@11718.11736.19|AACGGCGGGAGTAACTATG</td><td>>hg38_21_+_8433172_8446622_RNA45SN1_45S_50ntFlanks</td><td>11718</td><td>11736</td><td>19</td><td>AACGGCGGGAGTAACTATG</td><td>1039.7</td><td>-2.3</td><td>0.8</td><td>-2.8</td><td>5.8E-03</td><td>7.1E-03</td><td>MB231/MB468</td></tr>
<tr><td>>hg38_21_+_8433172_8446622_RNA45SN1_45S_50ntFlanks@13000.13027.28|TGCGATCTATTGAAAGTCAGCCCTCGAC</td><td>>hg38_21_+_8433172_8446622_RNA45SN1_45S_50ntFlanks</td><td>13000</td><td>13027</td><td>28</td><td>TGCGATCTATTGAAAGTCAGCCCTCGAC</td><td>851.3</td><td>2.0</td><td>0.7</td><td>2.8</td><td>5.8E-03</td><td>7.1E-03</td><td>MB231/MB468</td></tr>
<tr><td>>hg38_21_+_8433172_8446622_RNA45SN1_45S_50ntFlanks@10468.10488.21|TTGCCCTCGGCCGATCGAAAG</td><td>>hg38_21_+_8433172_8446622_RNA45SN1_45S_50ntFlanks</td><td>10468</td><td>10488</td><td>21</td><td>TTGCCCTCGGCCGATCGAAAG</td><td>214.6</td><td>-1.9</td><td>0.7</td><td>-2.7</td><td>6.2E-03</td><td>7.2E-03</td><td>BT20/MB468</td></tr>
<tr><td>>am_trnaMT_HisGTG_MT_+_12138_12206@26.45.20|AGATTGTGAATCTGACAACA</td><td>>am_trnaMT_HisGTG_MT_+_12138_12206</td><td>26</td><td>45</td><td>20</td><td>AGATTGTGAATCTGACAACA</td><td>1041.1</td><td>3.2</td><td>1.0</td><td>3.2</td><td>1.2E-03</td><td>7.2E-03</td><td>BT20/MB231</td></tr>
<tr><td>>hg38_21_+_8433172_8446622_RNA45SN1_45S_50ntFlanks@6646.6670.25|TCGTACGACTCTTAGCGGTGGATCA</td><td>>hg38_21_+_8433172_8446622_RNA45SN1_45S_50ntFlanks</td><td>6646</td><td>6670</td><td>25</td><td>TCGTACGACTCTTAGCGGTGGATCA</td><td>2510.7</td><td>-1.8</td><td>0.6</td><td>-2.8</td><td>5.9E-03</td><td>7.3E-03</td><td>MB231/MB468</td></tr>
<tr><td>>hg38_21_+_8433172_8446622_RNA45SN1_45S_50ntFlanks@4565.4589.25|AATAATGGAATAGGACCGCGGTTCT</td><td>>hg38_21_+_8433172_8446622_RNA45SN1_45S_50ntFlanks</td><td>4565</td><td>4589</td><td>25</td><td>AATAATGGAATAGGACCGCGGTTCT</td><td>324.9</td><td>2.4</td><td>0.9</td><td>2.7</td><td>6.4E-03</td><td>7.3E-03</td><td>BT20/MB468</td></tr>
<tr><td>>hg38_21_+_8433172_8446622_RNA45SN1_45S_50ntFlanks@3769.3791.23|CGCACGGCCGGTACAGTGAAACT</td><td>>hg38_21_+_8433172_8446622_RNA45SN1_45S_50ntFlanks</td><td>3769</td><td>3791</td><td>23</td><td>CGCACGGCCGGTACAGTGAAACT</td><td>205.2</td><td>1.9</td><td>0.7</td><td>2.8</td><td>6.0E-03</td><td>7.3E-03</td><td>MB231/MB468</td></tr>
<tr><td>>MI0000298|hsa-mir-221&WithFlank&X|-|45746151|45746272@71.92.22(+1U)&[MIMAT0000278&hsa-miR-221-3p&offsets|0|-1(+1U);m-91&X|-|45746181|45746202&offsets|0|0(+1U)]|AGCTACATTGTCTGCTGGGTTTT</td><td>>MI0000298|hsa-mir-221&WithFlank&X|-|45746151|45746272</td><td>71</td><td>92</td><td>22(+1U)</td><td>AGCTACATTGTCTGCTGGGTTTT</td><td>5639.2</td><td>-2.1</td><td>0.6</td><td>-3.2</td><td>1.2E-03</td><td>7.3E-03</td><td>BT20/MB231</td></tr>
<tr><td>>hg38_21_+_8433172_8446622_RNA45SN1_45S_50ntFlanks@5472.5516.45|ACGGCCCTGGCGGAGCGCTGAGAAGACGGTCGAACTTGACTATCT</td><td>>hg38_21_+_8433172_8446622_RNA45SN1_45S_50ntFlanks</td><td>5472</td><td>5516</td><td>45</td><td>ACGGCCCTGGCGGAGCGCTGAGAAGACGGTCGAACTTGACTATCT</td><td>6708.3</td><td>-3.2</td><td>1.0</td><td>-3.2</td><td>1.2E-03</td><td>7.3E-03</td><td>BT20/MB231</td></tr>
<tr><td>>trna119_LysCTT_1_-_145395522_145395594@1.32.32,trna11_LysCTT_5_-_180648979_180649051@1.32.32,trna13_LysCTT_6_+_26556774_26556846@1.32.32,trna32_LysCTT_16_-_3207406_3207478@1.32.32,trna7_LysCTT_16_+_3225692_3225764@1.32.32,trna9_LysCTT_5_+_180634755_180634827@1.32.32|GCCCGGCTAGCTCAGTCGGTAGAGCATGAGAC</td><td>>trna119_LysCTT_1_-_145395522_145395594</td><td>1</td><td>32</td><td>32</td><td>GCCCGGCTAGCTCAGTCGGTAGAGCATGAGAC</td><td>2102.0</td><td>-2.6</td><td>0.8</td><td>-3.2</td><td>1.2E-03</td><td>7.3E-03</td><td>BT20/MB231</td></tr>
</table>

| | | | | | | | | | | | | |
|---|---|---|---|---|---|---|---|---|---|---|---|---|
| >MI0000542\|hsa-mir-320a&WithFlank&8\|-\|22244960\|22245043@48.68.21(+1U)&[MIMAT0000510&hsa-miR-320a-3p&offsets\|0\|-1(+1U);m-60&8\|-\|22244975\|22244996&offsets\|0\|-1(+1U)]\|AAAAGCTGGGTTGAGAGGGCGT | >MI0000542\|hsa-mir-320a&WithFlank&8\|-\|22244960\|22245043 | 48 | 68 | 21(+1U) | AAAAGCTGGGTTGAGAGGGCGT | 5904.1 | 2.2 | 0.7 | 3.2 | 1.2E-03 | 7.3E-03 | BT20/MB231 |
| >MI0000266\|hsa-mir-10a&WithFlank&17\|-\|48579832\|48579953@28.48.21(+1A)&[MIMAT0000253&hsa-miR-10a-5p&offsets\|0\|-2(+1A);m-18&17\|-\|48579905\|48579926&offsets\|0\|-1(+1A)]\|TACCCTGTAGATCCGAATTTGA | >MI0000266\|hsa-mir-10a&WithFlank&17\|-\|48579832\|48579953 | 28 | 48 | 21(+1A) | TACCCTGTAGATCCGAATTTGA | 1117.6 | 2.5 | 0.8 | 3.2 | 1.2E-03 | 7.3E-03 | BT20/MB231 |
| >hg38_21_+_8433172_8446622_RNA45SN1_45S_50ntFlanks@10443.10460.18\|CCGAAGGGACGGGCGATG | >hg38_21_+_8433172_8446622_RNA45SN1_45S_50ntFlanks | 10443 | 10460 | 18 | CCGAAGGGACGGGCGATG | 125.2 | -2.0 | 0.7 | -2.7 | 6.4E-03 | 7.4E-03 | BT20/MB468 |
| >hg38_21_+_8433172_8446622_RNA45SN1_45S_50ntFlanks@4734.4751.18\|AACGAAAGTCGGAGGTTC | >hg38_21_+_8433172_8446622_RNA45SN1_45S_50ntFlanks | 4734 | 4751 | 18 | AACGAAAGTCGGAGGTTC | 531.6 | -1.2 | 0.5 | -2.7 | 6.4E-03 | 7.4E-03 | BT20/MB468 |
| >hg38_21_+_8433172_8446622_RNA45SN1_45S_50ntFlanks@6652.6698.47\|GACTCTTAGCGGTGGATCACTCGGCTCGTGCGTCGATGAAGAACGCA | >hg38_21_+_8433172_8446622_RNA45SN1_45S_50ntFlanks | 6652 | 6698 | 47 | GACTCTTAGCGGTGGATCACTCGGCTCGTGCGTCGATGAAGAACGCA | 2265.9 | -2.6 | 0.8 | -3.2 | 1.3E-03 | 7.4E-03 | BT20/MB231 |
| >trna14_LysTTT_11_-_59327808_59327880@1.36.36,trna2_LysTTT_17_+_8022473_8022545@1.36.36,trna54_LysTTT_1_+_204475655_204475727@1.36.36,trna5_LysTTT_11_+_59323902_59323974@1.36.36,trna62_LysTTT_1_-_204476158_204476230@1.36.36,trna76_LysTTT_6_+_28918806_28918878@1.36.36\|GCCCGGATAGCTCAGTCGGTAGAGCATCAGACTTTT | >trna14_LysTTT_11_-_59327808_59327880 | 1 | 36 | 36 | GCCCGGATAGCTCAGTCGGTAGAGCATCAGACTTTT | 1362.4 | 3.0 | 0.9 | 3.2 | 1.3E-03 | 7.4E-03 | BT20/MB231 |
| >hg38_21_+_8433172_8446622_RNA45SN1_45S_50ntFlanks@12419.12452.34\|GATGTCGGCTCTTCCTATCATTGTGAAGCAGAAT | >hg38_21_+_8433172_8446622_RNA45SN1_45S_50ntFlanks | 12419 | 12452 | 34 | GATGTCGGCTCTTCCTATCATTGTGAAGCAGAAT | 1030.8 | -2.6 | 0.8 | -3.2 | 1.3E-03 | 7.4E-03 | BT20/MB231 |
| >hg38_21_+_8433172_8446622_RNA45SN1_45S_50ntFlanks@6646.6663.18\|TCGTACGACTCTTAGCGG | >hg38_21_+_8433172_8446622_RNA45SN1_45S_50ntFlanks | 6646 | 6663 | 18 | TCGTACGACTCTTAGCGG | 10129.5 | 2.5 | 0.8 | 3.2 | 1.3E-03 | 7.4E-03 | BT20/MB231 |
| >MI0006273\|hsa-mir-1180&WithFlank&17\|-\|19344500\|19344580@47.66.20&[MIMAT0005825&hsa-miR-1180-3p&offsets\|0\|-2;m-214&17\|-\|19344513\|19344534&offsets\|0\|-2]\|TTTCCGGCTCGCGTGGGTGT | >MI0006273\|hsa-mir-1180&WithFlank&17\|-\|19344500\|19344580 | 47 | 66 | 20 | TTTCCGGCTCGCGTGGGTGT | 421.2 | 2.2 | 0.7 | 3.2 | 1.3E-03 | 7.4E-03 | BT20/MB231 |
| >trnaMT_GlnTTG_MT_-_4329_4400@1.30.30,trnalookalike2_GlnTTG_1_-_564879_564950@1.30.30\|TAGGATGGGGTGTGATAGGTGGCACGGAGA | >trnaMT_GlnTTG_MT_-_4329_4400 | 1 | 30 | 30 | TAGGATGGGGTGTGATAGGTGGCACGGAGA | 564.7 | 2.3 | 0.7 | 3.2 | 1.3E-03 | 7.4E-03 | BT20/MB231 |
| >hg38_21_+_8433172_8446622_RNA45SN1_45S_50ntFlanks@10346.10369.24\|TCAAACGAGAACTTTGAAGGCCGA | >hg38_21_+_8433172_8446622_RNA45SN1_45S_50ntFlanks | 10346 | 10369 | 24 | TCAAACGAGAACTTTGAAGGCCGA | 362.2 | -1.4 | 0.5 | -2.7 | 6.5E-03 | 7.5E-03 | BT20/MB468 |
| >hg38_21_+_8433172_8446622_RNA45SN1_45S_50ntFlanks@11200.11223.24\|CGGCGGCGGCGGCGACTCTGGACG | >hg38_21_+_8433172_8446622_RNA45SN1_45S_50ntFlanks | 11200 | 11223 | 24 | CGGCGGCGGCGGCGACTCTGGACG | 108.3 | -4.2 | 1.5 | -2.7 | 6.1E-03 | 7.5E-03 | MB231/MB468 |
| >hg38_21_+_8433172_8446622_RNA45SN1_45S_50ntFlanks@12625.12659.35\|CAATGGGGCGAAGCTACCATCTGTGGGATTATGAC | >hg38_21_+_8433172_8446622_RNA45SN1_45S_50ntFlanks | 12625 | 12659 | 35 | CAATGGGGCGAAGCTACCATCTGTGGGATTATGAC | 134.4 | -1.2 | 0.4 | -2.7 | 6.1E-03 | 7.5E-03 | MB231/MB468 |
| >hg38_21_+_8433172_8446622_RNA45SN1_45S_50ntFlanks@6669.6699.31\|CACTCGGCTCGTGCGTCGATGAAGAACGCAG | >hg38_21_+_8433172_8446622_RNA45SN1_45S_50ntFlanks | 6669 | 6699 | 31 | CACTCGGCTCGTGCGTCGATGAAGAACGCAG | 243.8 | 1.3 | 0.5 | 2.7 | 6.1E-03 | 7.5E-03 | MB231/MB468 |
| >hg38_21_+_8433172_8446622_RNA45SN1_45S_50ntFlanks@6651.6675.25\|CGACTCTTAGCGGTGGATCACTCGG | >hg38_21_+_8433172_8446622_RNA45SN1_45S_50ntFlanks | 6651 | 6675 | 25 | CGACTCTTAGCGGTGGATCACTCGG | 1604.5 | 1.9 | 0.7 | 2.7 | 6.1E-03 | 7.5E-03 | MB231/MB468 |
| >hg38_21_+_8433172_8446622_RNA45SN1_45S_50ntFlanks@11570.11588.19\|AGCCGACTTAGAACTGGTG | >hg38_21_+_8433172_8446622_RNA45SN1_45S_50ntFlanks | 11570 | 11588 | 19 | AGCCGACTTAGAACTGGTG | 192.2 | -1.9 | 0.7 | -2.7 | 6.6E-03 | 7.5E-03 | BT20/MB468 |
| >hg38_21_+_8433172_8446622_RNA45SN1_45S_50ntFlanks@7975.7994.20\|CGCGACCTCAGATCAGACGT | >hg38_21_+_8433172_8446622_RNA45SN1_45S_50ntFlanks | 7975 | 7994 | 20 | CGCGACCTCAGATCAGACGT | 33388.5 | 2.3 | 0.8 | 2.7 | 6.2E-03 | 7.6E-03 | MB231/MB468 |
| >hg38_21_+_8433172_8446622_RNA45SN1_45S_50ntFlanks@5237.5255.19\|ACTGACTGGCTCAGCGTGT | >hg38_21_+_8433172_8446622_RNA45SN1_45S_50ntFlanks | 5237 | 5255 | 19 | ACTGACTGGCTCAGCGTGT | 274.1 | 2.5 | 0.8 | 3.2 | 1.3E-03 | 7.6E-03 | BT20/MB231 |
| >hg38_21_+_8433172_8446622_RNA45SN1_45S_50ntFlanks@12944.12962.19\|CGTAGACGACCTGCTTCTG | >hg38_21_+_8433172_8446622_RNA45SN1_45S_50ntFlanks | 12944 | 12962 | 19 | CGTAGACGACCTGCTTCTG | 136.1 | -1.3 | 0.5 | -2.7 | 6.2E-03 | 7.6E-03 | MB231/MB468 |
| >MI0000469\|hsa-mir-125a&WithFlank&19\|+\|51693248\|51693345@21.41.21(+1A)&[MIMAT0000443&hsa-miR-125a-5p&offsets\|0\|-3(+1A);m-63&19\|+\|51693268\|51693289&offsets\|0\|-1(+1A)]\|TCCCTGAGACCCTTTAACCTGA | >MI0000469\|hsa-mir-125a&WithFlank&19\|+\|51693248\|51693345 | 21 | 41 | 21(+1A) | TCCCTGAGACCCTTTAACCTGA | 229.9 | 2.5 | 0.8 | 3.2 | 1.3E-03 | 7.6E-03 | BT20/MB231 |
| >hg38_21_+_8433172_8446622_RNA45SN1_45S_50ntFlanks@11207.11234.28\|GGCGGCGACTCTGGACGCGAGCCGGGCC | >hg38_21_+_8433172_8446622_RNA45SN1_45S_50ntFlanks | 11207 | 11234 | 28 | GGCGGCGACTCTGGACGCGAGCCGGGCC | 112.7 | -3.1 | 1.1 | -2.7 | 6.7E-03 | 7.6E-03 | BT20/MB468 |
| >hg38_21_+_8433172_8446622_RNA45SN1_45S_50ntFlanks@11683.11701.19\|ATGTCAAAGTGAAGAAATT | >hg38_21_+_8433172_8446622_RNA45SN1_45S_50ntFlanks | 11683 | 11701 | 19 | ATGTCAAAGTGAAGAAATT | 138.5 | -2.0 | 0.7 | -2.7 | 6.7E-03 | 7.6E-03 | BT20/MB468 |
| >hg38_21_+_8433172_8446622_RNA45SN1_45S_50ntFlanks@12999.13027.29\|CTGCGATCTATTGAAAGTCAGCCCTCGAC | >hg38_21_+_8433172_8446622_RNA45SN1_45S_50ntFlanks | 12999 | 13027 | 29 | CTGCGATCTATTGAAAGTCAGCCCTCGAC | 459.8 | -2.2 | 0.8 | -2.7 | 6.8E-03 | 7.7E-03 | BT20/MB468 |
| >hg38_21_+_8433172_8446622_RNA45SN1_45S_50ntFlanks@12459.12480.22\|AGCGTTGGATTGTTCACCCACT | >hg38_21_+_8433172_8446622_RNA45SN1_45S_50ntFlanks | 12459 | 12480 | 22 | AGCGTTGGATTGTTCACCCACT | 244.7 | -2.2 | 0.8 | -2.7 | 6.3E-03 | 7.7E-03 | MB231/MB468 |
| >hg38_21_+_8433172_8446622_RNA45SN1_45S_50ntFlanks@11316.11340.25\|CACCCCACGTCTCGTCGCGCGCGCG | >hg38_21_+_8433172_8446622_RNA45SN1_45S_50ntFlanks | 11316 | 11340 | 25 | CACCCCACGTCTCGTCGCGCGCGCG | 262.6 | -2.9 | 1.1 | -2.7 | 6.8E-03 | 7.8E-03 | BT20/MB468 |
| >hg38_21_+_8433172_8446622_RNA45SN1_45S_50ntFlanks@3838.3859.22\|CTCTCCTACTTGGATAACTGTG | >hg38_21_+_8433172_8446622_RNA45SN1_45S_50ntFlanks | 3838 | 3859 | 22 | CTCTCCTACTTGGATAACTGTG | 83.1 | -1.5 | 0.5 | -2.7 | 6.4E-03 | 7.8E-03 | MB231/MB468 |
| >hg38_21_+_8433172_8446622_RNA45SN1_45S_50ntFlanks@9538.9571.34\|AACTCTGGTGGAGGTCCGTAGCGGTCCTGACGTG | >hg38_21_+_8433172_8446622_RNA45SN1_45S_50ntFlanks | 9538 | 9571 | 34 | AACTCTGGTGGAGGTCCGTAGCGGTCCTGACGTG | 276.3 | -1.4 | 0.5 | -2.7 | 6.4E-03 | 7.8E-03 | MB231/MB468 |
| >hg38_21_+_8433172_8446622_RNA45SN1_45S_50ntFlanks@11568.11591.24\|GCAGCCGACTTAGAACTGGTGCGG | >hg38_21_+_8433172_8446622_RNA45SN1_45S_50ntFlanks | 11568 | 11591 | 24 | GCAGCCGACTTAGAACTGGTGCGG | 105.4 | -1.5 | 0.6 | -2.7 | 6.4E-03 | 7.8E-03 | MB231/MB468 |
| >hg38_21_+_8433172_8446622_RNA45SN1_45S_50ntFlanks@10458.10488.31\|ATGGCCTCCGTTGCCCTCGGCCGATCGAAAG | >hg38_21_+_8433172_8446622_RNA45SN1_45S_50ntFlanks | 10458 | 10488 | 31 | ATGGCCTCCGTTGCCCTCGGCCGATCGAAAG | 612.6 | -2.7 | 1.0 | -2.7 | 6.4E-03 | 7.8E-03 | MB231/MB468 |
| >hg38_21_+_8433172_8446622_RNA45SN1_45S_50ntFlanks@12413.12434.22\|TCCTTCGATGTCGGCTCTTCCT | >hg38_21_+_8433172_8446622_RNA45SN1_45S_50ntFlanks | 12413 | 12434 | 22 | TCCTTCGATGTCGGCTCTTCCT | 1408.4 | 3.2 | 1.2 | 2.7 | 6.8E-03 | 7.8E-03 | BT20/MB468 |
| >trna116_GluCTC_1_-_145399233_145399304@35.75.41,trna134_GluTTC_1_-_16861774_16861845@35.75.41,trna71_GluCTC_1_-_161439189_161439260@35.75.41,trna74_GluCTC_1_-_161431809_161431880@35.75.41,trna77_GluCTC_1_-_161424398_161424469@35.75.41,trna77_GluCTC_6_+_28949976_28950047@35.75.41,trna80_GluCTC_1_-_161417018_161417089@35.75.41,trna84_GluTTC_1_-_161391883_161391954@35.75.41,trna87_GluCTC_6_-_126101393_126101464@35.75.41\|TCACCGCCGCGGCCCGGGTTCGATTCCCGGTCAGGGAACCA | >trna116_GluCTC_1_-_145399233_145399304 | 35 | 75 | 41 | TCACCGCCGCGGCCCGGGTTCGATTCCCGGTCAGGGAACCA | 299.4 | -3.6 | 1.1 | -3.2 | 1.4E-03 | 7.9E-03 | BT20/MB231 |
| >hg38_21_+_8433172_8446622_RNA45SN1_45S_50ntFlanks@10684.10708.25\|CGGCGGCGTCCGGTGAGCTCTCGCT | >hg38_21_+_8433172_8446622_RNA45SN1_45S_50ntFlanks | 10684 | 10708 | 25 | CGGCGGCGTCCGGTGAGCTCTCGCT | 721.0 | -1.6 | 0.6 | -2.7 | 6.9E-03 | 7.9E-03 | BT20/MB468 |
| >hg38_21_+_8433172_8446622_RNA45SN1_45S_50ntFlanks@9569.9586.18\|GTGCAAATCGGTCGTCCG | >hg38_21_+_8433172_8446622_RNA45SN1_45S_50ntFlanks | 9569 | 9586 | 18 | GTGCAAATCGGTCGTCCG | 129.8 | -1.6 | 0.6 | -2.7 | 6.9E-03 | 7.9E-03 | BT20/MB468 |
| >am_trnaMT_ValTAC_MT_+_1602_1670@50.68.19\|ACTTAACTTGACCGCTCTG | >am_trnaMT_ValTAC_MT_+_1602_1670 | 50 | 68 | 19 | ACTTAACTTGACCGCTCTG | 3388.1 | -2.4 | 0.9 | -2.7 | 6.9E-03 | 7.9E-03 | BT20/MB468 |
| >hg38_21_+_8433172_8446622_RNA45SN1_45S_50ntFlanks@6733.6756.24\|CATTGATCATCGACACTTCGAACG | >hg38_21_+_8433172_8446622_RNA45SN1_45S_50ntFlanks | 6733 | 6756 | 24 | CATTGATCATCGACACTTCGAACG | 843.8 | -1.4 | 0.5 | -2.7 | 6.5E-03 | 7.9E-03 | MB231/MB468 |
| >hg38_21_+_8433172_8446622_RNA45SN1_45S_50ntFlanks@10392.10433.42\|ACAGCAGTTGAACATGGGTCAGTCGGTCCTGAGAGATGGGCG | >hg38_21_+_8433172_8446622_RNA45SN1_45S_50ntFlanks | 10392 | 10433 | 42 | ACAGCAGTTGAACATGGGTCAGTCGGTCCTGAGAGATGGGCG | 291.4 | -2.3 | 0.8 | -2.7 | 6.5E-03 | 7.9E-03 | MB231/MB468 |
| >hg38_21_+_8433172_8446622_RNA45SN1_45S_50ntFlanks@12417.12454.38\|TCGATGTCGGCTCTTCCTATCATTGTGAAGCAGAATTC | >hg38_21_+_8433172_8446622_RNA45SN1_45S_50ntFlanks | 12417 | 12454 | 38 | TCGATGTCGGCTCTTCCTATCATTGTGAAGCAGAATTC | 172.9 | -2.6 | 0.8 | -3.2 | 1.4E-03 | 8.0E-03 | BT20/MB231 |
| >hg38_21_+_8433172_8446622_RNA45SN1_45S_50ntFlanks@6646.6666.21\|TCGTACGACTCTTAGCGGTGG | >hg38_21_+_8433172_8446622_RNA45SN1_45S_50ntFlanks | 6646 | 6666 | 21 | TCGTACGACTCTTAGCGGTGG | 9286.0 | 2.5 | 0.8 | 3.2 | 1.4E-03 | 8.0E-03 | BT20/MB231 |
| >hg38_21_+_8433172_8446622_RNA45SN1_45S_50ntFlanks@476.524.49\|TTCGTGATCGATGTGGTGACGTCGTGCTCTCCCGGGCCGGGTCCGAGCC | >hg38_21_+_8433172_8446622_RNA45SN1_45S_50ntFlanks | 476 | 524 | 49 | TTCGTGATCGATGTGGTGACGTCGTGCTCTCCCGGGCCGGGTCCGAGCC | 700.3 | -3.0 | 1.0 | -3.2 | 1.4E-03 | 8.0E-03 | BT20/MB231 |
| >hg38_21_+_8433172_8446622_RNA45SN1_45S_50ntFlanks@11568.11587.20\|GCAGCCGACTTAGAACTGGT | >hg38_21_+_8433172_8446622_RNA45SN1_45S_50ntFlanks | 11568 | 11587 | 20 | GCAGCCGACTTAGAACTGGT | 356.3 | 1.1 | 0.4 | 2.7 | 7.0E-03 | 8.1E-03 | BT20/MB468 |
| >hg38_21_+_8433172_8446622_RNA45SN1_45S_50ntFlanks@11540.11580.41\|CGCGGCGCCTCGCCTCGGCCGGCGCCTAGCAGCCGACTTAG | >hg38_21_+_8433172_8446622_RNA45SN1_45S_50ntFlanks | 11540 | 11580 | 41 | CGCGGCGCCTCGCCTCGGCCGGCGCCTAGCAGCCGACTTAG | 288.6 | -2.9 | 0.9 | -3.2 | 1.4E-03 | 8.1E-03 | BT20/MB231 |
| >hg38_21_+_8433172_8446622_RNA45SN1_45S_50ntFlanks@8358.8378.21\|AAAGAACTTTGAAGAGAGAGT | >hg38_21_+_8433172_8446622_RNA45SN1_45S_50ntFlanks | 8358 | 8378 | 21 | AAAGAACTTTGAAGAGAGAGT | 594.2 | -1.8 | 0.7 | -2.7 | 7.1E-03 | 8.1E-03 | BT20/MB468 |
| >hg38_21_+_8433172_8446622_RNA45SN1_45S_50ntFlanks@12478.12499.22\|ACTAATAGGGAACGTGAGCTGG | >hg38_21_+_8433172_8446622_RNA45SN1_45S_50ntFlanks | 12478 | 12499 | 22 | ACTAATAGGGAACGTGAGCTGG | 255.5 | -1.5 | 0.6 | -2.7 | 7.1E-03 | 8.1E-03 | BT20/MB468 |

<table>
<tr><td>>MI0000266|hsa-mir-10a&WithFlank&17|-|48579832|48579953@28.49.22(+1A)&[MIMAT0000253&hsa-miR-10a-5p&offsets|0|-1(+1A);m-18&17|-|48579905|48579926&offsets|0|0(+1A)]|TACCCTGTAGATCCGAATTTGTA</td><td>>MI0000266|hsa-mir-10a&WithFlank&17|-|48579832|48579953</td><td>28</td><td>49</td><td>22(+1A)</td><td>TACCCTGTAGATCCGAATTTGTA</td><td>3337.3</td><td>2.2</td><td>0.7</td><td>3.2</td><td>1.4E-03</td><td>8.1E-03</td><td>BT20/MB231</td></tr>
<tr><td>>hg38_21_+_8433172_8446622_RNA45SN1_45S_50ntFlanks@6645.6665.21|CTCGTACGACTCTTAGCGGTG</td><td>>hg38_21_+_8433172_8446622_RNA45SN1_45S_50ntFlanks</td><td>6645</td><td>6665</td><td>21</td><td>CTCGTACGACTCTTAGCGGTG</td><td>8705.8</td><td>2.5</td><td>0.8</td><td>3.2</td><td>1.4E-03</td><td>8.1E-03</td><td>BT20/MB231</td></tr>
<tr><td>>hg38_MT_+_1621_3279_RNR2_16S_50ntFlanks@460.491.32|TTGTAAATTTAACTGTTAGTCCAAAGAGGAAC</td><td>>hg38_MT_+_1621_3279_RNR2_16S_50ntFlanks</td><td>460</td><td>491</td><td>32</td><td>TTGTAAATTTAACTGTTAGTCCAAAGAGGAAC</td><td>235.2</td><td>-2.3</td><td>0.7</td><td>-3.2</td><td>1.4E-03</td><td>8.1E-03</td><td>BT20/MB231</td></tr>
<tr><td>>MI0000070|hsa-mir-16-1&WithFlank&13|-|50048967|50049067@20.40.21(+1U);MI0000115|hsa-mir-16-2&WithFlank&3|+|160404739|160404831@16.36.21(+1U)&[MIMAT0000069&hsa-miR-16-5p&offsets|0|-1(+1U);m-62&13|-|50049027|50049048&offsets|0|-1(+1U)];[MIMAT0000069_1&hsa-miR-16-5p&offsets|0|-1(+1U);m-61&3|+|160404754|160404775&offsets|0|-1(+1U)]|TAGCAGCACGTAAATATTGGCT</td><td>>MI0000070|hsa-mir-16-1&WithFlank&13|-|50048967|50049067</td><td>20</td><td>40</td><td>21(+1U)</td><td>TAGCAGCACGTAAATATTGGCT</td><td>1154.9</td><td>3.7</td><td>1.2</td><td>3.2</td><td>1.4E-03</td><td>8.1E-03</td><td>BT20/MB231</td></tr>
<tr><td>>trnaMT_GluTTC_MT_-_14674_14742@37.71.35,trnalookalike8_GluTTC_5_-_93905172_93905240@37.71.35|TCATTGGTCGTGGTTGTAGTCCGTGCGAGAATACC</td><td>>trnaMT_GluTTC_MT_-_14674_14742</td><td>37</td><td>71</td><td>35</td><td>TCATTGGTCGTGGTTGTAGTCCGTGCGAGAATACC</td><td>366.3</td><td>-2.4</td><td>0.8</td><td>-3.2</td><td>1.4E-03</td><td>8.1E-03</td><td>BT20/MB231</td></tr>
<tr><td>>hg38_21_+_8433172_8446622_RNA45SN1_45S_50ntFlanks@9572.9603.32|CAAATCGGTCGTCCGACCTGGGTATAGGGGCG</td><td>>hg38_21_+_8433172_8446622_RNA45SN1_45S_50ntFlanks</td><td>9572</td><td>9603</td><td>32</td><td>CAAATCGGTCGTCCGACCTGGGTATAGGGGCG</td><td>1650.5</td><td>-1.6</td><td>0.6</td><td>-2.7</td><td>6.7E-03</td><td>8.1E-03</td><td>MB231/MB468</td></tr>
<tr><td>>hg38_21_+_8433172_8446622_RNA45SN1_45S_50ntFlanks@6651.6681.31|CGACTCTTAGCGGTGGATCACTCGGCTCGTG</td><td>>hg38_21_+_8433172_8446622_RNA45SN1_45S_50ntFlanks</td><td>6651</td><td>6681</td><td>31</td><td>CGACTCTTAGCGGTGGATCACTCGGCTCGTG</td><td>31408.6</td><td>-1.4</td><td>0.5</td><td>-2.7</td><td>7.2E-03</td><td>8.2E-03</td><td>BT20/MB468</td></tr>
<tr><td>>hg38_21_+_8433172_8446622_RNA45SN1_45S_50ntFlanks@10349.10390.42|AACGAGAACTTTGAAGGCCGAAGTGGAGAAGGGTTCCATGTG</td><td>>hg38_21_+_8433172_8446622_RNA45SN1_45S_50ntFlanks</td><td>10349</td><td>10390</td><td>42</td><td>AACGAGAACTTTGAAGGCCGAAGTGGAGAAGGGTTCCATGTG</td><td>528.0</td><td>-1.7</td><td>0.6</td><td>-2.7</td><td>7.2E-03</td><td>8.3E-03</td><td>BT20/MB468</td></tr>
<tr><td>>hg38_MT_+_1621_3279_RNR2_16S_50ntFlanks@1216.1262.47|CAACCTCCGAGCAGTACATGCTAAGACTTCACCAGTCAAAGCGAACT</td><td>>hg38_MT_+_1621_3279_RNR2_16S_50ntFlanks</td><td>1216</td><td>1262</td><td>47</td><td>CAACCTCCGAGCAGTACATGCTAAGACTTCACCAGTCAAAGCGAACT</td><td>1276.0</td><td>-2.4</td><td>0.7</td><td>-3.2</td><td>1.5E-03</td><td>8.3E-03</td><td>BT20/MB231</td></tr>
<tr><td>>hg38_21_+_8433172_8446622_RNA45SN1_45S_50ntFlanks@10677.10709.33|GCGGTTCCGGCGGCGTCCGGTGAGCTCTCGCTG</td><td>>hg38_21_+_8433172_8446622_RNA45SN1_45S_50ntFlanks</td><td>10677</td><td>10709</td><td>33</td><td>GCGGTTCCGGCGGCGTCCGGTGAGCTCTCGCTG</td><td>496.0</td><td>1.3</td><td>0.5</td><td>2.7</td><td>6.8E-03</td><td>8.3E-03</td><td>MB231/MB468</td></tr>
<tr><td>>hg38_21_+_8433172_8446622_RNA45SN1_45S_50ntFlanks@12626.12661.36|AATGGGGCGAAGCTACCATCTGTGGGATTATGACTG</td><td>>hg38_21_+_8433172_8446622_RNA45SN1_45S_50ntFlanks</td><td>12626</td><td>12661</td><td>36</td><td>AATGGGGCGAAGCTACCATCTGTGGGATTATGACTG</td><td>1045.7</td><td>-1.7</td><td>0.6</td><td>-2.7</td><td>6.8E-03</td><td>8.3E-03</td><td>MB231/MB468</td></tr>
<tr><td>>MI0000802|hsa-mir-340&WithFlank&5|-|180015297|180015403@22.42.21(+1C)&[MIMAT0004692&hsa-miR-340-5p&offsets|0|-1(+1C);m-220&5|-|180015361|180015382&offsets|0|-1(+1C)]|TTATAAAGCAATGAGACTGATC</td><td>>MI0000802|hsa-mir-340&WithFlank&5|-|180015297|180015403</td><td>22</td><td>42</td><td>21(+1C)</td><td>TTATAAAGCAATGAGACTGATC</td><td>1019.0</td><td>3.0</td><td>0.9</td><td>3.2</td><td>1.5E-03</td><td>8.3E-03</td><td>BT20/MB231</td></tr>
<tr><td>>hg38_21_+_8433172_8446622_RNA45SN1_45S_50ntFlanks@3843.3872.30|CTACTTGGATAACTGTGGTAATTCTAGAGC</td><td>>hg38_21_+_8433172_8446622_RNA45SN1_45S_50ntFlanks</td><td>3843</td><td>3872</td><td>30</td><td>CTACTTGGATAACTGTGGTAATTCTAGAGC</td><td>213.9</td><td>1.1</td><td>0.4</td><td>2.7</td><td>6.8E-03</td><td>8.3E-03</td><td>MB231/MB468</td></tr>
<tr><td>>trna152_ValCAC_6_-_27248049_27248121@1.31.31|GCTTCTGTAGTGTAGTGGTTATCACGTTCGC</td><td>>trna152_ValCAC_6_-_27248049_27248121</td><td>1</td><td>31</td><td>31</td><td>GCTTCTGTAGTGTAGTGGTTATCACGTTCGC</td><td>410.7</td><td>2.7</td><td>1.0</td><td>2.7</td><td>6.8E-03</td><td>8.3E-03</td><td>MB231/MB468</td></tr>
<tr><td>>hg38_21_+_8433172_8446622_RNA45SN1_45S_50ntFlanks@11718.11739.22|AACGGCGGGAGTAACTATGACT</td><td>>hg38_21_+_8433172_8446622_RNA45SN1_45S_50ntFlanks</td><td>11718</td><td>11739</td><td>22</td><td>AACGGCGGGAGTAACTATGACT</td><td>233.4</td><td>-2.0</td><td>0.7</td><td>-2.7</td><td>6.8E-03</td><td>8.4E-03</td><td>MB231/MB468</td></tr>
<tr><td>>hg38_21_+_8433172_8446622_RNA45SN1_45S_50ntFlanks@4380.4408.29|CGTAGTTGGATCTTGGGAGCGGGCGGGCG</td><td>>hg38_21_+_8433172_8446622_RNA45SN1_45S_50ntFlanks</td><td>4380</td><td>4408</td><td>29</td><td>CGTAGTTGGATCTTGGGAGCGGGCGGGCG</td><td>96.7</td><td>-3.2</td><td>1.2</td><td>-2.7</td><td>7.4E-03</td><td>8.4E-03</td><td>BT20/MB468</td></tr>
<tr><td>>hg38_21_+_8433172_8446622_RNA45SN1_45S_50ntFlanks@6697.6720.24|CAGCTAGCTGCGAGAATTAATGTG</td><td>>hg38_21_+_8433172_8446622_RNA45SN1_45S_50ntFlanks</td><td>6697</td><td>6720</td><td>24</td><td>CAGCTAGCTGCGAGAATTAATGTG</td><td>1704.6</td><td>1.7</td><td>0.6</td><td>2.7</td><td>6.9E-03</td><td>8.4E-03</td><td>MB231/MB468</td></tr>
<tr><td>>hg38_21_+_8433172_8446622_RNA45SN1_45S_50ntFlanks@5494.5526.33|AAGACGGTCGAACTTGACTATCTAGAGGAAGTA</td><td>>hg38_21_+_8433172_8446622_RNA45SN1_45S_50ntFlanks</td><td>5494</td><td>5526</td><td>33</td><td>AAGACGGTCGAACTTGACTATCTAGAGGAAGTA</td><td>132.7</td><td>-1.6</td><td>0.6</td><td>-2.7</td><td>7.0E-03</td><td>8.5E-03</td><td>MB231/MB468</td></tr>
<tr><td>>MI0000063|hsa-let-7b&WithFlank&22|+|46113680|46113774@12.31.20(+1G)&[MIMAT0000063&hsa-let-7b-5p&offsets|0|-2(+1G);m-14&22|+|46113691|46113712&offsets|0|-2(+1G)]|TGAGGTAGTAGGTTGTGTGGG</td><td>>MI0000063|hsa-let-7b&WithFlank&22|+|46113680|46113774</td><td>12</td><td>31</td><td>20(+1G)</td><td>TGAGGTAGTAGGTTGTGTGGG</td><td>403.9</td><td>2.8</td><td>0.9</td><td>3.2</td><td>1.5E-03</td><td>8.5E-03</td><td>BT20/MB231</td></tr>
<tr><td>>hg38_21_+_8433172_8446622_RNA45SN1_45S_50ntFlanks@7975.8005.31|CGCGACCTCAGATCAGACGTGGCGACCCGCT</td><td>>hg38_21_+_8433172_8446622_RNA45SN1_45S_50ntFlanks</td><td>7975</td><td>8005</td><td>31</td><td>CGCGACCTCAGATCAGACGTGGCGACCCGCT</td><td>38720.3</td><td>2.5</td><td>0.8</td><td>3.2</td><td>1.5E-03</td><td>8.6E-03</td><td>BT20/MB231</td></tr>
<tr><td>>MI0000255|hsa-mir-30d&WithFlank&8|-|134804870|134804951@12.33.22&[MIMAT0000245&hsa-miR-30d-5p&offsets|0|0;m-21&8|-|134804919|134804940&offsets|0|0]|TGTAAACATCCCCGACTGGAAG</td><td>>MI0000255|hsa-mir-30d&WithFlank&8|-|134804870|134804951</td><td>12</td><td>33</td><td>22</td><td>TGTAAACATCCCCGACTGGAAG</td><td>10406.8</td><td>2.2</td><td>0.7</td><td>3.2</td><td>1.5E-03</td><td>8.6E-03</td><td>BT20/MB231</td></tr>
<tr><td>>MI0000466|hsa-mir-9-1&WithFlank&1|-|156420335|156420435@22.43.22(+1U);MI0000468|hsa-mir-9-3&WithFlank&15|+|89368011|89368112@22.43.22(+1U);MI0000467|hsa-mir-9-2&WithFlank&5|-|88666847|88666945@22.43.22(+1U)&[MIMAT0000441&hsa-miR-9-5p&offsets|0|-1(+1U);m-104&1|-|156420392|156420414&offsets|0|-1(+1U)];[MIMAT0000441_1&hsa-miR-9-5p&offsets|0|-1(+1U);m-105&15|+|89368032|89368054&offsets|0|-1(+1U)];[MIMAT0000441_2&hsa-miR-9-5p&offsets|0|-1(+1U);m-104&5|-|88666902|88666924&offsets|0|-1(+1U)]|TCTTTGGTTATCTAGCTGTATGT</td><td>>MI0000466|hsa-mir-9-1&WithFlank&1|-|156420335|156420435</td><td>22</td><td>43</td><td>22(+1U)</td><td>TCTTTGGTTATCTAGCTGTATGT</td><td>329.5</td><td>2.7</td><td>0.8</td><td>3.2</td><td>1.5E-03</td><td>8.6E-03</td><td>BT20/MB231</td></tr>
<tr><td>>hg38_21_+_8433172_8446622_RNA45SN1_45S_50ntFlanks@9535.9571.37|GGAAACTCTGGTGGAGGTCCGTAGCGGTCCTGACGTG</td><td>>hg38_21_+_8433172_8446622_RNA45SN1_45S_50ntFlanks</td><td>9535</td><td>9571</td><td>37</td><td>GGAAACTCTGGTGGAGGTCCGTAGCGGTCCTGACGTG</td><td>182.8</td><td>-1.4</td><td>0.5</td><td>-2.7</td><td>7.6E-03</td><td>8.6E-03</td><td>BT20/MB468</td></tr>
<tr><td>>hg38_21_+_8433172_8446622_RNA45SN1_45S_50ntFlanks@11579.11602.24|AGAACTGGTGCGGACCAGGGGAAT</td><td>>hg38_21_+_8433172_8446622_RNA45SN1_45S_50ntFlanks</td><td>11579</td><td>11602</td><td>24</td><td>AGAACTGGTGCGGACCAGGGGAAT</td><td>544.9</td><td>-2.4</td><td>0.9</td><td>-2.7</td><td>7.6E-03</td><td>8.7E-03</td><td>BT20/MB468</td></tr>
<tr><td>>hg38_21_+_8433172_8446622_RNA45SN1_45S_50ntFlanks@5445.5466.22|TCGGATCGGCCCCGCCGGGGTC</td><td>>hg38_21_+_8433172_8446622_RNA45SN1_45S_50ntFlanks</td><td>5445</td><td>5466</td><td>22</td><td>TCGGATCGGCCCCGCCGGGGTC</td><td>2237.4</td><td>-3.6</td><td>1.3</td><td>-2.7</td><td>7.6E-03</td><td>8.7E-03</td><td>BT20/MB468</td></tr>
<tr><td>>hg38_21_+_8433172_8446622_RNA45SN1_45S_50ntFlanks@12417.12450.34|TCGATGTCGGCTCTTCCTATCATTGTGAAGCAGA</td><td>>hg38_21_+_8433172_8446622_RNA45SN1_45S_50ntFlanks</td><td>12417</td><td>12450</td><td>34</td><td>TCGATGTCGGCTCTTCCTATCATTGTGAAGCAGA</td><td>151.8</td><td>-2.7</td><td>0.9</td><td>-3.2</td><td>1.6E-03</td><td>8.7E-03</td><td>BT20/MB231</td></tr>
<tr><td>>hg38_21_+_8433172_8446622_RNA45SN1_45S_50ntFlanks@4734.4758.25|AACGAAAGTCGGAGGTTCGAAGACG</td><td>>hg38_21_+_8433172_8446622_RNA45SN1_45S_50ntFlanks</td><td>4734</td><td>4758</td><td>25</td><td>AACGAAAGTCGGAGGTTCGAAGACG</td><td>1675.6</td><td>1.6</td><td>0.6</td><td>2.7</td><td>7.2E-03</td><td>8.8E-03</td><td>MB231/MB468</td></tr>
<tr><td>>MI0000650|hsa-mir-200c&WithFlank&12|+|6963693|6963772@50.72.23(+1A)&[MIMAT0000617&hsa-miR-200c-3p&offsets|0|0(+1A);m-54&12|+|6963742|6963764&offsets|0|0(+1A)]|TAATACTGCCGGGTAATGATGGAA</td><td>>MI0000650|hsa-mir-200c&WithFlank&12|+|6963693|6963772</td><td>50</td><td>72</td><td>23(+1A)</td><td>TAATACTGCCGGGTAATGATGGAA</td><td>109.5</td><td>-1.6</td><td>0.6</td><td>-2.7</td><td>7.2E-03</td><td>8.8E-03</td><td>MB231/MB468</td></tr>
<tr><td>>hg38_21_+_8433172_8446622_RNA45SN1_45S_50ntFlanks@6700.6720.21|CTAGCTGCGAGAATTAATGTG</td><td>>hg38_21_+_8433172_8446622_RNA45SN1_45S_50ntFlanks</td><td>6700</td><td>6720</td><td>21</td><td>CTAGCTGCGAGAATTAATGTG</td><td>863.9</td><td>-3.2</td><td>1.0</td><td>-3.2</td><td>1.6E-03</td><td>8.8E-03</td><td>BT20/MB231</td></tr>
<tr><td>>hg38_21_+_8433172_8446622_RNA45SN1_45S_50ntFlanks@9606.9623.18|AGACTAATCGAACCATCT</td><td>>hg38_21_+_8433172_8446622_RNA45SN1_45S_50ntFlanks</td><td>9606</td><td>9623</td><td>18</td><td>AGACTAATCGAACCATCT</td><td>117.3</td><td>-1.3</td><td>0.5</td><td>-2.7</td><td>7.2E-03</td><td>8.8E-03</td><td>MB231/MB468</td></tr>
<tr><td>>hg38_21_+_8433172_8446622_RNA45SN1_45S_50ntFlanks@11209.11227.19|CGGCGACTCTGGACGCGAG</td><td>>hg38_21_+_8433172_8446622_RNA45SN1_45S_50ntFlanks</td><td>11209</td><td>11227</td><td>19</td><td>CGGCGACTCTGGACGCGAG</td><td>921.7</td><td>-2.7</td><td>1.0</td><td>-2.7</td><td>7.8E-03</td><td>8.9E-03</td><td>BT20/MB468</td></tr>
<tr><td>>trna111_HisGTG_1_-_147774845_147774916@-1A.31.32,trna118_HisGTG_1_-_145396881_145396952@-1A.31.32,trna16_HisGTG_1_+_146544773_146544844@-1A.31.32,trna1_HisGTG_15_+_45493349_45493420@-1A.31.32,trna21_HisGTG_1_+_147753471_147753542@-1A.31.32,trna33_HisGTG_6_+_27125906_27125977@-1A.31.32,trna7_HisGTG_9_-_14433938_14434009@-1A.31.32,trna8_HisGTG_15_-_45492611_45492682@-1A.31.32,trna9_HisGTG_15_-_45490804_45490875@-1A.31.32|AGCCGTGATCGTATAGTGGTTAGTACTCTGCG</td><td>>trna111_HisGTG_1_-_147774845_147774916</td><td>-1A</td><td>31</td><td>32</td><td>AGCCGTGATCGTATAGTGGTTAGTACTCTGCG</td><td>833.9</td><td>-3.1</td><td>1.1</td><td>-2.7</td><td>7.8E-03</td><td>8.9E-03</td><td>BT20/MB468</td></tr>
<tr><td>>am_MI0000809|hsa-mir-151a&WithFlank&8|-|140732558|140732659@17.37.21&[MIMAT0004697&hsa-miR-151a-5p&offsets|0|0;m-75&8|-|140732623|140732643&offsets|0|0]|TCGAGGAGCTCACAGTCTAGT</td><td>>am_MI0000809|hsa-mir-151a&WithFlank&8|-|140732558|140732659</td><td>17</td><td>37</td><td>21</td><td>TCGAGGAGCTCACAGTCTAGT</td><td>4022.0</td><td>2.6</td><td>0.8</td><td>3.2</td><td>1.6E-03</td><td>8.9E-03</td><td>BT20/MB231</td></tr>
<tr><td>>MI0000294|hsa-mir-218-1&WithFlank&4|+|20528269|20528390@31.51.21;MI0000295|hsa-mir-218-2&WithFlank&5|-|168768140|168768261@31.51.21&[MIMAT0000275&hsa-miR-218-5p&offsets|0|0;m-185&4|+|20528299|20528319&offsets|0|0];[MIMAT0000275_1&hsa-miR-218-5p&offsets|0|0;m-187&5|-|168768211|168768231&offsets|0|0]|TTGTGCTTGATCTAACCATGT</td><td>>MI0000294|hsa-mir-218-1&WithFlank&4|+|20528269|20528390</td><td>31</td><td>51</td><td>21</td><td>TTGTGCTTGATCTAACCATGT</td><td>510.0</td><td>-6.9</td><td>2.2</td><td>-3.2</td><td>1.6E-03</td><td>9.0E-03</td><td>BT20/MB231</td></tr>
</table>

<table>
<tr><th></th><th></th><th></th><th></th><th></th><th></th><th></th><th></th><th></th><th></th><th></th><th></th><th></th></tr>
<tr><td>>am_trna10_ValCAC_5_-_180649395_180649467@1.31.31,trna12_ValAAC_5_-_180645270_180645342@1.31.31,trna132_ValAAC_6_-_27721179_27721251@1.31.31,trna136_ValAAC_6_-_27648885_27648957@1.31.31,trna139_ValAAC_6_-_27618707_27618779@1.31.31,trna18_ValCAC_5_-_180529253_180529325@1.31.31,trna2_ValAAC_3_+_169490018_169490090@1.31.31,trna2_ValCAC_5_+_180524070_180524142@1.31.31,trna4_ValAAC_5_+_180591154_180591226@1.31.31,trna5_ValAAC_5_+_180596610_180596682@1.31.31,trna6_ValCAC_5_+_180600650_180600722@1.31.31,trna85_ValCAC_1_-_161369490_161369562@1.31.31,trna90_ValCAC_1_-_149684088_149684161@1.31.31,trna98_ValCAC_1_-_149298555_149298627@1.31.31,trna9_ValCAC_6_+_26538282_26538354@1.31.31|GTTTCCGTAGTGTAGTGGTTATCACGTTCGC</td><td>>am_trna10_ValCAC_5_-_180649395_180649467</td><td>1</td><td>31</td><td>31</td><td>GTTTCCGTAGTGTAGTGGTTATCACGTTCGC</td><td>313.2</td><td>-3.1</td><td>1.0</td><td>-3.2</td><td>1.6E-03</td><td>9.0E-03</td><td>BT20/MB231</td></tr>
<tr><td>>hg38_21_+_8433172_8446622_RNA45SN1_45S_50ntFlanks@6646.6689.44|TCGTACGACTCTTAGCGGTGGATCACTCGGCTCGTGCGTCGATG</td><td>>hg38_21_+_8433172_8446622_RNA45SN1_45S_50ntFlanks</td><td>6646</td><td>6689</td><td>44</td><td>TCGTACGACTCTTAGCGGTGGATCACTCGGCTCGTGCGTCGATG</td><td>5278.9</td><td>2.2</td><td>0.8</td><td>2.7</td><td>7.9E-03</td><td>9.0E-03</td><td>BT20/MB468</td></tr>
<tr><td>>trna10_LysCTT_16_+_3241501_3241573@14.31.18,trna13_LysCTT_14_-_58706613_58706685@14.31.18,trna2_LysCTT_15_+_79152904_79152976@14.31.18,trna4_LysCTT_18_-_43669271_43669343@14.31.18|AGTCGGTAGAGCATGGGA</td><td>>trna10_LysCTT_16_+_3241501_3241573</td><td>14</td><td>31</td><td>18</td><td>AGTCGGTAGAGCATGGGA</td><td>801.3</td><td>-2.3</td><td>0.8</td><td>-2.7</td><td>7.4E-03</td><td>9.0E-03</td><td>MB231/MB468</td></tr>
<tr><td>>hg38_21_+_8433172_8446622_RNA45SN1_45S_50ntFlanks@11319.11343.25|CCCACGTCTCGTCGCGCGCGTCC</td><td>>hg38_21_+_8433172_8446622_RNA45SN1_45S_50ntFlanks</td><td>11319</td><td>11343</td><td>25</td><td>CCCACGTCTCGTCGCGCGCGTCC</td><td>119.3</td><td>-2.1</td><td>0.8</td><td>-2.7</td><td>8.0E-03</td><td>9.1E-03</td><td>BT20/MB468</td></tr>
<tr><td>>hg38_21_+_8433172_8446622_RNA45SN1_45S_50ntFlanks@11209.11233.25|CGGCGACTCTGGACGCGAGCCGGGC</td><td>>hg38_21_+_8433172_8446622_RNA45SN1_45S_50ntFlanks</td><td>11209</td><td>11233</td><td>25</td><td>CGGCGACTCTGGACGCGAGCCGGGC</td><td>1066.7</td><td>-2.7</td><td>1.0</td><td>-2.7</td><td>8.0E-03</td><td>9.1E-03</td><td>BT20/MB468</td></tr>
<tr><td>>hg38_21_+_8433172_8446622_RNA45SN1_45S_50ntFlanks@12003.12034.32|GCGGGCCGCCGGTGAAATACCACTACTCTGAT</td><td>>hg38_21_+_8433172_8446622_RNA45SN1_45S_50ntFlanks</td><td>12003</td><td>12034</td><td>32</td><td>GCGGGCCGCCGGTGAAATACCACTACTCTGAT</td><td>625.9</td><td>-2.6</td><td>0.8</td><td>-3.1</td><td>1.7E-03</td><td>9.1E-03</td><td>BT20/MB231</td></tr>
<tr><td>>hg38_21_+_8433172_8446622_RNA45SN1_45S_50ntFlanks@12419.12449.31|GATGTCGGCTCTTCCTATCATTGTGAAGCAG</td><td>>hg38_21_+_8433172_8446622_RNA45SN1_45S_50ntFlanks</td><td>12419</td><td>12449</td><td>31</td><td>GATGTCGGCTCTTCCTATCATTGTGAAGCAG</td><td>1277.6</td><td>-2.8</td><td>0.9</td><td>-3.1</td><td>1.7E-03</td><td>9.1E-03</td><td>BT20/MB231</td></tr>
<tr><td>>MI0000289|hsa-mir-181a-1&WithFlank&1|-|198859038|198859159@30.49.20;MI0000269|hsa-mir-181a-2&WithFlank&9|+|124692436|124692557@45.64.20&[MIMAT0000256&hsa-miR-181a-5p&offsets|0|-3;m-32&1|-|198859108|198859130&offsets|0|-3];[MIMAT0000256_1&hsa-miR-181a-5p&offsets|0|-3;m-32&9|+|124692480|124692502&offsets|0|-3]|AACATTCAACGCTGTCGGTG</td><td>>MI0000289|hsa-mir-181a-1&WithFlank&1|-|198859038|198859159</td><td>30</td><td>49</td><td>20</td><td>AACATTCAACGCTGTCGGTG</td><td>554.2</td><td>-2.3</td><td>0.7</td><td>-3.1</td><td>1.6E-03</td><td>9.1E-03</td><td>BT20/MB231</td></tr>
<tr><td>>MI0000683|hsa-mir-181b-2&WithFlank&9|+|124693704|124693804@22.46.25(+1U)&[MIMAT0000257_1&hsa-miR-181b-5p&offsets|0|+2(+1U);m-72&9|+|124693725|124693747&offsets|0|+2(+1U)]|AACATTCATTGCTGTCGGTGGGTTTT</td><td>>MI0000683|hsa-mir-181b-2&WithFlank&9|+|124693704|124693804</td><td>22</td><td>46</td><td>25(+1U)</td><td>AACATTCATTGCTGTCGGTGGGTTTT</td><td>492.7</td><td>-2.4</td><td>0.8</td><td>-3.1</td><td>1.7E-03</td><td>9.1E-03</td><td>BT20/MB231</td></tr>
<tr><td>>trna10_AspGTC_12_-_125424193_125424264@35.75.41,trna12_AspGTC_12_-_125411891_125411962@35.75.41,trna144_AspGTC_6_-_27551236_27551307@35.75.41,trna38_AspGTC_17_-_8125556_8125627@35.75.41,trna45_AspGTC_6_+_27447453_27447524@35.75.41,trna48_AspGTC_6_+_27471523_27471594@35.75.41,trna4_AspGTC_12_+_96429799_96429870@35.75.41,trna69_AspGTC_1_-_161440205_161440276@35.75.41,trna72_AspGTC_1_-_161432824_161432895@35.75.41,trna75_AspGTC_1_-_161425414_161425485@35.75.41,trna78_AspGTC_1_-_161418033_161418104@35.75.41,trna81_AspGTC_1_-_161410615_161410686@35.75.41|TCACGCGGGAGACCGGGGTTCGATTCCCCGACGGGGAGCCA</td><td>>trna10_AspGTC_12_-_125424193_125424264</td><td>35</td><td>75</td><td>41</td><td>TCACGCGGGAGACCGGGGTTCGATTCCCCGACGGGGAGCCA</td><td>1426.1</td><td>-3.8</td><td>1.2</td><td>-3.1</td><td>1.7E-03</td><td>9.1E-03</td><td>BT20/MB231</td></tr>
<tr><td>>trnaMT_PheGAA_MT_+_577_647@38.57.20|AATGTTTAGACGGGCTCACA</td><td>>trnaMT_PheGAA_MT_+_577_647</td><td>38</td><td>57</td><td>20</td><td>AATGTTTAGACGGGCTCACA</td><td>891.4</td><td>2.5</td><td>0.8</td><td>3.1</td><td>1.7E-03</td><td>9.1E-03</td><td>BT20/MB231</td></tr>
<tr><td>>trna11_GluTTC_15_-_26327381_26327452@1.31.31,trna3_GluTTC_13_-_45492062_45492133@1.31.31|TCCCACATGGTCTAGCGGTTAGGATTCCTGG</td><td>>trna11_GluTTC_15_-_26327381_26327452</td><td>1</td><td>31</td><td>31</td><td>TCCCACATGGTCTAGCGGTTAGGATTCCTGG</td><td>505.0</td><td>-2.6</td><td>0.8</td><td>-3.1</td><td>1.7E-03</td><td>9.1E-03</td><td>BT20/MB231</td></tr>
<tr><td>>hg38_21_+_8433172_8446622_RNA45SN1_45S_50ntFlanks@6653.6680.28|ACTCTTAGCGGTGGATCACTCGGCTCGT</td><td>>hg38_21_+_8433172_8446622_RNA45SN1_45S_50ntFlanks</td><td>6653</td><td>6680</td><td>28</td><td>ACTCTTAGCGGTGGATCACTCGGCTCGT</td><td>287.1</td><td>1.3</td><td>0.5</td><td>2.7</td><td>7.5E-03</td><td>9.1E-03</td><td>MB231/MB468</td></tr>
<tr><td>>MI0000342|hsa-mir-200b&WithFlank&1|+|1167098|1167204@63.85.23(+1U)&[MIMAT0000318&hsa-miR-200b-3p&offsets|0|+1(+1U);m-93&1|+|1167160|1167182&offsets|0|0(+1U)]|TAATACTGCCTGGTAATGATGACT</td><td>>MI0000342|hsa-mir-200b&WithFlank&1|+|1167098|1167204</td><td>63</td><td>85</td><td>23(+1U)</td><td>TAATACTGCCTGGTAATGATGACT</td><td>1440.8</td><td>-2.0</td><td>0.8</td><td>-2.6</td><td>8.1E-03</td><td>9.2E-03</td><td>BT20/MB468</td></tr>
<tr><td>>trnaMT_GluTTC_MT_-_14674_14742@31.72.42,trnalookalike8_GluTTC_5_-_93905172_93905240@31.72.42|TTCATATCATTGGTCGTGGTTGTAGTCCGTGCGAGAATACCA</td><td>>trnaMT_GluTTC_MT_-_14674_14742</td><td>31</td><td>72</td><td>42</td><td>TTCATATCATTGGTCGTGGTTGTAGTCCGTGCGAGAATACCA</td><td>788.1</td><td>-2.4</td><td>0.8</td><td>-3.1</td><td>1.7E-03</td><td>9.2E-03</td><td>BT20/MB231</td></tr>
<tr><td>>hg38_21_+_8433172_8446622_RNA45SN1_45S_50ntFlanks@12394.12414.21|TAGCGACGTCGCTTTTTGATC</td><td>>hg38_21_+_8433172_8446622_RNA45SN1_45S_50ntFlanks</td><td>12394</td><td>12414</td><td>21</td><td>TAGCGACGTCGCTTTTTGATC</td><td>876.4</td><td>-1.5</td><td>0.6</td><td>-2.6</td><td>8.1E-03</td><td>9.3E-03</td><td>BT20/MB468</td></tr>
<tr><td>>hg38_21_+_8433172_8446622_RNA45SN1_45S_50ntFlanks@8192.8215.24|AGCCCGTGGACGGTGTGAGGCCGG</td><td>>hg38_21_+_8433172_8446622_RNA45SN1_45S_50ntFlanks</td><td>8192</td><td>8215</td><td>24</td><td>AGCCCGTGGACGGTGTGAGGCCGG</td><td>172.0</td><td>-2.7</td><td>1.0</td><td>-2.6</td><td>8.1E-03</td><td>9.3E-03</td><td>BT20/MB468</td></tr>
<tr><td>>hg38_21_+_8433172_8446622_RNA45SN1_45S_50ntFlanks@12473.12496.24|CACCCACTAATAGGGAACGTGAGC</td><td>>hg38_21_+_8433172_8446622_RNA45SN1_45S_50ntFlanks</td><td>12473</td><td>12496</td><td>24</td><td>CACCCACTAATAGGGAACGTGAGC</td><td>432.0</td><td>-2.5</td><td>0.8</td><td>-3.1</td><td>1.7E-03</td><td>9.3E-03</td><td>BT20/MB231</td></tr>
<tr><td>>hg38_21_+_8433172_8446622_RNA45SN1_45S_50ntFlanks@6733.6753.21|CATTGATCATCGACACTTCGA</td><td>>hg38_21_+_8433172_8446622_RNA45SN1_45S_50ntFlanks</td><td>6733</td><td>6753</td><td>21</td><td>CATTGATCATCGACACTTCGA</td><td>389.3</td><td>-1.5</td><td>0.6</td><td>-2.7</td><td>7.7E-03</td><td>9.3E-03</td><td>MB231/MB468</td></tr>
<tr><td>>hg38_21_+_8433172_8446622_RNA45SN1_45S_50ntFlanks@12893.12930.38|CCTCGCCCGTCACGCACCGCACGTTCGTGGGGAACCTG</td><td>>hg38_21_+_8433172_8446622_RNA45SN1_45S_50ntFlanks</td><td>12893</td><td>12930</td><td>38</td><td>CCTCGCCCGTCACGCACCGCACGTTCGTGGGGAACCTG</td><td>330.9</td><td>-2.3</td><td>0.9</td><td>-2.6</td><td>8.2E-03</td><td>9.4E-03</td><td>BT20/MB468</td></tr>
<tr><td>>hg38_21_+_8433172_8446622_RNA45SN1_45S_50ntFlanks@11671.11688.18|CCAGTGCTCTGAATGTCA</td><td>>hg38_21_+_8433172_8446622_RNA45SN1_45S_50ntFlanks</td><td>11671</td><td>11688</td><td>18</td><td>CCAGTGCTCTGAATGTCA</td><td>605.8</td><td>-2.3</td><td>0.9</td><td>-2.7</td><td>7.7E-03</td><td>9.4E-03</td><td>MB231/MB468</td></tr>
<tr><td>>MI0001733|hsa-mir-452&WithFlank&X|-|151959622|151959718@20.42.23(+1U)&[MIMAT0001635&hsa-miR-452-5p&offsets|0|+1(+1U);m-188&X|-|151959677|151959699&offsets|0|0(+1U)]|AACTGTTTGCAGAGGAAACTGAGT</td><td>>MI0001733|hsa-mir-452&WithFlank&X|-|151959622|151959718</td><td>20</td><td>42</td><td>23(+1U)</td><td>AACTGTTTGCAGAGGAAACTGAGT</td><td>449.8</td><td>2.2</td><td>0.7</td><td>3.1</td><td>1.7E-03</td><td>9.4E-03</td><td>BT20/MB231</td></tr>
<tr><td>>hg38_21_+_8433172_8446622_RNA45SN1_45S_50ntFlanks@4735.4767.33|ACGAAAGTCGGAGGTTCGAAGACGATCAGATAC</td><td>>hg38_21_+_8433172_8446622_RNA45SN1_45S_50ntFlanks</td><td>4735</td><td>4767</td><td>33</td><td>ACGAAAGTCGGAGGTTCGAAGACGATCAGATAC</td><td>489.6</td><td>-3.3</td><td>1.0</td><td>-3.1</td><td>1.7E-03</td><td>9.4E-03</td><td>BT20/MB231</td></tr>
<tr><td>>MI0003667|hsa-mir-652&WithFlank&X|+|110055323|110055432@67.87.21(+1U)&[MIMAT0003322&hsa-miR-652-3p&offsets|0|0(+1U);m-148&X|+|110055389|110055410&offsets|0|-1(+1U)]|AATGGCGCCACTAGGGTTGTGT</td><td>>MI0003667|hsa-mir-652&WithFlank&X|+|110055323|110055432</td><td>67</td><td>87</td><td>21(+1U)</td><td>AATGGCGCCACTAGGGTTGTGT</td><td>944.5</td><td>2.8</td><td>0.9</td><td>3.1</td><td>1.7E-03</td><td>9.4E-03</td><td>BT20/MB231</td></tr>
<tr><td>>hg38_21_+_8433172_8446622_RNA45SN1_45S_50ntFlanks@11672.11700.29|CAGTGCTCTGAATGTCAAAGTGAAGAAAT</td><td>>hg38_21_+_8433172_8446622_RNA45SN1_45S_50ntFlanks</td><td>11672</td><td>11700</td><td>29</td><td>CAGTGCTCTGAATGTCAAAGTGAAGAAAT</td><td>340.3</td><td>1.3</td><td>0.5</td><td>2.7</td><td>7.7E-03</td><td>9.4E-03</td><td>MB231/MB468</td></tr>
<tr><td>>hg38_21_+_8433172_8446622_RNA45SN1_45S_50ntFlanks@4588.4621.34|CTATTTTGTTGGTTTTCGGAACTGAGGCCATGAT</td><td>>hg38_21_+_8433172_8446622_RNA45SN1_45S_50ntFlanks</td><td>4588</td><td>4621</td><td>34</td><td>CTATTTTGTTGGTTTTCGGAACTGAGGCCATGAT</td><td>129.7</td><td>-1.6</td><td>0.6</td><td>-2.6</td><td>8.3E-03</td><td>9.5E-03</td><td>BT20/MB468</td></tr>
<tr><td>>hg38_21_+_8433172_8446622_RNA45SN1_45S_50ntFlanks@12418.12454.37|CGATGTCGGCTCTTCCTATCATTGTGAAGCAGAATTC</td><td>>hg38_21_+_8433172_8446622_RNA45SN1_45S_50ntFlanks</td><td>12418</td><td>12454</td><td>37</td><td>CGATGTCGGCTCTTCCTATCATTGTGAAGCAGAATTC</td><td>155.3</td><td>-2.5</td><td>0.8</td><td>-3.1</td><td>1.7E-03</td><td>9.5E-03</td><td>BT20/MB231</td></tr>
<tr><td>>hg38_21_+_8433172_8446622_RNA45SN1_45S_50ntFlanks@7993.8016.24|GTGGCGACCCGCTGAATTTAAGCA</td><td>>hg38_21_+_8433172_8446622_RNA45SN1_45S_50ntFlanks</td><td>7993</td><td>8016</td><td>24</td><td>GTGGCGACCCGCTGAATTTAAGCA</td><td>263.1</td><td>-3.1</td><td>1.0</td><td>-3.1</td><td>1.8E-03</td><td>9.5E-03</td><td>BT20/MB231</td></tr>
<tr><td>>trnaMT_PheGAA_MT_+_577_647@37.57.21|AAATGTTTAGACGGGCTCACA</td><td>>trnaMT_PheGAA_MT_+_577_647</td><td>37</td><td>57</td><td>21</td><td>AAATGTTTAGACGGGCTCACA</td><td>280.5</td><td>2.2</td><td>0.7</td><td>3.1</td><td>1.8E-03</td><td>9.5E-03</td><td>BT20/MB231</td></tr>
<tr><td>>hg38_21_+_8433172_8446622_RNA45SN1_45S_50ntFlanks@8197.8214.18|GTGGACGGTGTGAGGCCG</td><td>>hg38_21_+_8433172_8446622_RNA45SN1_45S_50ntFlanks</td><td>8197</td><td>8214</td><td>18</td><td>GTGGACGGTGTGAGGCCG</td><td>133.4</td><td>-1.4</td><td>0.5</td><td>-2.6</td><td>8.4E-03</td><td>9.5E-03</td><td>BT20/MB468</td></tr>
<tr><td>>hg38_21_+_8433172_8446622_RNA45SN1_45S_50ntFlanks@7983.8009.27|CAGATCAGACGTGGCGACCCGCTGAAT</td><td>>hg38_21_+_8433172_8446622_RNA45SN1_45S_50ntFlanks</td><td>7983</td><td>8009</td><td>27</td><td>CAGATCAGACGTGGCGACCCGCTGAAT</td><td>311.5</td><td>-1.3</td><td>0.5</td><td>-2.6</td><td>8.4E-03</td><td>9.6E-03</td><td>BT20/MB468</td></tr>
<tr><td>>hg38_21_+_8433172_8446622_RNA45SN1_45S_50ntFlanks@10769.10796.28|AGCAGGTCTCCAAGGTGAACAGCCTCTG</td><td>>hg38_21_+_8433172_8446622_RNA45SN1_45S_50ntFlanks</td><td>10769</td><td>10796</td><td>28</td><td>AGCAGGTCTCCAAGGTGAACAGCCTCTG</td><td>1017.6</td><td>2.1</td><td>0.8</td><td>2.7</td><td>7.9E-03</td><td>9.6E-03</td><td>MB231/MB468</td></tr>
<tr><td>>hg38_21_+_8433172_8446622_RNA45SN1_45S_50ntFlanks@4734.4752.19|AACGAAAGTCGGAGGTTCG</td><td>>hg38_21_+_8433172_8446622_RNA45SN1_45S_50ntFlanks</td><td>4734</td><td>4752</td><td>19</td><td>AACGAAAGTCGGAGGTTCG</td><td>515.4</td><td>-1.6</td><td>0.6</td><td>-2.6</td><td>8.5E-03</td><td>9.6E-03</td><td>BT20/MB468</td></tr>
<tr><td>>hg38_21_+_8433172_8446622_RNA45SN1_45S_50ntFlanks@11569.11594.26|CAGCCGACTTAGAACTGGTGCGGACC</td><td>>hg38_21_+_8433172_8446622_RNA45SN1_45S_50ntFlanks</td><td>11569</td><td>11594</td><td>26</td><td>CAGCCGACTTAGAACTGGTGCGGACC</td><td>357.7</td><td>-2.0</td><td>0.7</td><td>-2.6</td><td>8.6E-03</td><td>9.7E-03</td><td>BT20/MB468</td></tr>
<tr><td>>trnaMT_LeuTAA_MT_+_3230_3304@18.44.27|CGGTAATCGCATAAAACTTAAAACTTT</td><td>>trnaMT_LeuTAA_MT_+_3230_3304</td><td>18</td><td>44</td><td>27</td><td>CGGTAATCGCATAAAACTTAAAACTTT</td><td>535.9</td><td>1.8</td><td>0.7</td><td>2.7</td><td>8.0E-03</td><td>9.8E-03</td><td>MB231/MB468</td></tr>
<tr><td>>hg38_21_+_8433172_8446622_RNA45SN1_45S_50ntFlanks@11727.11752.26|AGTAACTATGACTCTCTTAAGGTAGC</td><td>>hg38_21_+_8433172_8446622_RNA45SN1_45S_50ntFlanks</td><td>11727</td><td>11752</td><td>26</td><td>AGTAACTATGACTCTCTTAAGGTAGC</td><td>207.7</td><td>-2.8</td><td>0.9</td><td>-3.1</td><td>1.8E-03</td><td>9.8E-03</td><td>BT20/MB231</td></tr>
<tr><td>>hg38_21_+_8433172_8446622_RNA45SN1_45S_50ntFlanks@4731.4750.20|AAGAACGAAAGTCGGAGGTT</td><td>>hg38_21_+_8433172_8446622_RNA45SN1_45S_50ntFlanks</td><td>4731</td><td>4750</td><td>20</td><td>AAGAACGAAAGTCGGAGGTT</td><td>298.2</td><td>1.9</td><td>0.7</td><td>2.7</td><td>8.0E-03</td><td>9.8E-03</td><td>MB231/MB468</td></tr>
</table>

| | | | | | | | | | | | | |
|---|---|---|---|---|---|---|---|---|---|---|---|---|
| >hg38_21_+_8433172_8446622_RNA45SN1_45S_50ntFlanks@7979.8004.26\|ACCTCAGATCAGACGTGGCGACCCGC | >hg38_21_+_8433172_8446622_RNA45SN1_45S_50ntFlanks | 7979 | 8004 | 26 | ACCTCAGATCAGACGTGGCGACCCGC | 87.7 | -2.1 | 0.8 | -2.6 | 8.6E-03 | 9.8E-03 | BT20/MB468 |
| >hg38_21_+_8433172_8446622_RNA45SN1_45S_50ntFlanks@12907.12931.25\|CACCGCACGTTCGTGGGGAACCTGG | >hg38_21_+_8433172_8446622_RNA45SN1_45S_50ntFlanks | 12907 | 12931 | 25 | CACCGCACGTTCGTGGGGAACCTGG | 92.9 | -1.9 | 0.7 | -2.6 | 8.6E-03 | 9.8E-03 | BT20/MB468 |
| >hg38_21_+_8433172_8446622_RNA45SN1_45S_50ntFlanks@11672.11693.22\|CAGTGCTCTGAATGTCAAAGTG | >hg38_21_+_8433172_8446622_RNA45SN1_45S_50ntFlanks | 11672 | 11693 | 22 | CAGTGCTCTGAATGTCAAAGTG | 453.2 | -2.2 | 0.8 | -2.6 | 8.6E-03 | 9.8E-03 | BT20/MB468 |
| >hg38_21_+_8433172_8446622_RNA45SN1_45S_50ntFlanks@12407.12431.25\|TTTTGATCCTTCGATGTCGGCTCTT | >hg38_21_+_8433172_8446622_RNA45SN1_45S_50ntFlanks | 12407 | 12431 | 25 | TTTTGATCCTTCGATGTCGGCTCTT | 763.1 | -1.6 | 0.6 | -2.6 | 8.1E-03 | 9.8E-03 | MB231/MB468 |
| >hg38_21_+_8433172_8446622_RNA45SN1_45S_50ntFlanks@12481.12499.19\|AATAGGGAACGTGAGCTGG | >hg38_21_+_8433172_8446622_RNA45SN1_45S_50ntFlanks | 12481 | 12499 | 19 | AATAGGGAACGTGAGCTGG | 578.9 | 2.6 | 1.0 | 2.6 | 8.7E-03 | 9.8E-03 | BT20/MB468 |
| >hg38_21_+_8433172_8446622_RNA45SN1_45S_50ntFlanks@10774.10796.23\|GTCTCCAAGGTGAACAGCCTCTG | >hg38_21_+_8433172_8446622_RNA45SN1_45S_50ntFlanks | 10774 | 10796 | 23 | GTCTCCAAGGTGAACAGCCTCTG | 138.4 | -2.2 | 0.8 | -2.6 | 8.7E-03 | 9.9E-03 | BT20/MB468 |
| >hg38_21_+_8433172_8446622_RNA45SN1_45S_50ntFlanks@8153.8181.29\|GCCGCTCGTGGGGGGCCCAAGTCCTTCTG | >hg38_21_+_8433172_8446622_RNA45SN1_45S_50ntFlanks | 8153 | 8181 | 29 | GCCGCTCGTGGGGGGCCCAAGTCCTTCTG | 134.6 | -1.9 | 0.7 | -2.6 | 8.1E-03 | 9.9E-03 | MB231/MB468 |
| >hg38_21_+_8433172_8446622_RNA45SN1_45S_50ntFlanks@11576.11594.19\|CTTAGAACTGGTGCGGACC | >hg38_21_+_8433172_8446622_RNA45SN1_45S_50ntFlanks | 11576 | 11594 | 19 | CTTAGAACTGGTGCGGACC | 824.3 | -1.3 | 0.5 | -2.6 | 8.1E-03 | 9.9E-03 | MB231/MB468 |
| >am_trnaMT_GluTTC_MT_-_14674_14742@27.48.22,trnalookalike8_GluTTC_5_-_93905172_93905240@27.48.22\|GTTTTTCATATCATTGGTCGTG | >am_trnaMT_GluTTC_MT_-_14674_14742 | 27 | 48 | 22 | GTTTTTCATATCATTGGTCGTG | 126.3 | -3.2 | 1.2 | -2.6 | 8.7E-03 | 9.9E-03 | BT20/MB468 |
| >hg38_21_+_8433172_8446622_RNA45SN1_45S_50ntFlanks@12419.12450.32\|GATGTCGGCTCTTCCTATCATTGTGAAGCAGA | >hg38_21_+_8433172_8446622_RNA45SN1_45S_50ntFlanks | 12419 | 12450 | 32 | GATGTCGGCTCTTCCTATCATTGTGAAGCAGA | 389.7 | -2.4 | 0.8 | -3.1 | 1.8E-03 | 9.9E-03 | BT20/MB231 |
| >hg38_21_+_8433172_8446622_RNA45SN1_45S_50ntFlanks@11670.11702.33\|CCCAGTGCTCTGAATGTCAAAGTGAAGAAATTC | >hg38_21_+_8433172_8446622_RNA45SN1_45S_50ntFlanks | 11670 | 11702 | 33 | CCCAGTGCTCTGAATGTCAAAGTGAAGAAATTC | 1065.2 | -1.8 | 0.7 | -2.6 | 8.2E-03 | 1.0E-02 | MB231/MB468 |
| >hg38_21_+_8433172_8446622_RNA45SN1_45S_50ntFlanks@9574.9603.30\|AATCGGTCGTCCGACCTGGGTATAGGGGCG | >hg38_21_+_8433172_8446622_RNA45SN1_45S_50ntFlanks | 9574 | 9603 | 30 | AATCGGTCGTCCGACCTGGGTATAGGGGCG | 3567.9 | 1.7 | 0.7 | 2.6 | 8.3E-03 | 1.0E-02 | MB231/MB468 |
| >hg38_21_+_8433172_8446622_RNA45SN1_45S_50ntFlanks@8114.8140.27\|GCGCGGGACATGTGGCGTACGGAAGAC | >hg38_21_+_8433172_8446622_RNA45SN1_45S_50ntFlanks | 8114 | 8140 | 27 | GCGCGGGACATGTGGCGTACGGAAGAC | 320.6 | -4.8 | 1.8 | -2.6 | 8.3E-03 | 1.0E-02 | MB231/MB468 |
| >hg38_21_+_8433172_8446622_RNA45SN1_45S_50ntFlanks@10794.10814.21\|CTGGCATGTTGGAACAATGTA | >hg38_21_+_8433172_8446622_RNA45SN1_45S_50ntFlanks | 10794 | 10814 | 21 | CTGGCATGTTGGAACAATGTA | 279.9 | -2.1 | 0.8 | -2.6 | 8.9E-03 | 1.0E-02 | BT20/MB468 |
| >hg38_21_+_8433172_8446622_RNA45SN1_45S_50ntFlanks@12071.12106.36\|CCCCGAGGGGCTCTCGCTTCTGGCGCCAAGCGCCCG | >hg38_21_+_8433172_8446622_RNA45SN1_45S_50ntFlanks | 12071 | 12106 | 36 | CCCCGAGGGGCTCTCGCTTCTGGCGCCAAGCGCCCG | 200.4 | -1.7 | 0.6 | -2.6 | 8.4E-03 | 1.0E-02 | MB231/MB468 |
| >hg38_21_+_8433172_8446622_RNA45SN1_45S_50ntFlanks@9572.9602.31\|CAAATCGGTCGTCCGACCTGGGTATAGGGGC | >hg38_21_+_8433172_8446622_RNA45SN1_45S_50ntFlanks | 9572 | 9602 | 31 | CAAATCGGTCGTCCGACCTGGGTATAGGGGC | 150.2 | -1.1 | 0.4 | -2.6 | 9.0E-03 | 1.0E-02 | BT20/MB468 |
| >am_trna116_GluCTC_1_-_145399233_145399304@1.27.27,trna59_GluCTC_1_+_249168447_249168518@1.27.27,trna71_GluCTC_1_-_161439189_161439260@1.27.27,trna74_GluCTC_1_-_161431809_161431880@1.27.27,trna77_GluCTC_1_-_161424398_161424469@1.27.27,trna77_GluCTC_6_+_28949976_28950047@1.27.27,trna80_GluCTC_1_-_161417018_161417089@1.27.27,trna87_GluCTC_6_-_126101393_126101464@1.27.27\|TCCCTGGTGGTCTAGTGGTTAGGATTC | >am_trna116_GluCTC_1_-_145399233_145399304 | 1 | 27 | 27 | TCCCTGGTGGTCTAGTGGTTAGGATTC | 3318.8 | -2.0 | 0.8 | -2.6 | 8.4E-03 | 1.0E-02 | MB231/MB468 |
| >trna152_ValCAC_6_-_27248049_27248121@1.34.34\|GCTTCTGTAGTGTAGTGGTTATCACGTTCGCCTC | >trna152_ValCAC_6_-_27248049_27248121 | 1 | 34 | 34 | GCTTCTGTAGTGTAGTGGTTATCACGTTCGCCTC | 3452.6 | -3.0 | 1.0 | -3.1 | 1.9E-03 | 1.0E-02 | BT20/MB231 |
| >hg38_21_+_8433172_8446622_RNA45SN1_45S_50ntFlanks@12418.12438.21\|CGATGTCGGCTCTTCCTATCA | >hg38_21_+_8433172_8446622_RNA45SN1_45S_50ntFlanks | 12418 | 12438 | 21 | CGATGTCGGCTCTTCCTATCA | 961.1 | 2.4 | 0.9 | 2.6 | 9.1E-03 | 1.0E-02 | BT20/MB468 |
| >hg38_21_+_8433172_8446622_RNA45SN1_45S_50ntFlanks@6645.6679.35\|CTCGTACGACTCTTAGCGGTGGATCACTCGGCTCG | >hg38_21_+_8433172_8446622_RNA45SN1_45S_50ntFlanks | 6645 | 6679 | 35 | CTCGTACGACTCTTAGCGGTGGATCACTCGGCTCG | 686.7 | 1.8 | 0.7 | 2.6 | 8.5E-03 | 1.0E-02 | MB231/MB468 |
| >MI0000298\|hsa-mir-221&WithFlank&X\|-\|45746151\|45746272@71.94.24&[MIMAT0000278&hsa-miR-221-3p&offsets\|0\|+1;m-91&X\|-\|45746181\|45746202&offsets\|0\|+2]\|AGCTACATTGTCTGCTGGGTTTCA | >MI0000298\|hsa-mir-221&WithFlank&X\|-\|45746151\|45746272 | 71 | 94 | 24 | AGCTACATTGTCTGCTGGGTTTCA | 3762.8 | -2.1 | 0.7 | -3.1 | 1.9E-03 | 1.0E-02 | BT20/MB231 |
| >hg38_21_+_8433172_8446622_RNA45SN1_45S_50ntFlanks@12419.12457.39\|GATGTCGGCTCTTCCTATCATTGTGAAGCAGAATTCACC | >hg38_21_+_8433172_8446622_RNA45SN1_45S_50ntFlanks | 12419 | 12457 | 39 | GATGTCGGCTCTTCCTATCATTGTGAAGCAGAATTCACC | 970.4 | -2.5 | 0.8 | -3.1 | 1.9E-03 | 1.0E-02 | BT20/MB231 |
| >MI0000746\|hsa-mir-99b&WithFlank&19\|+\|51692606\|51692687@51.71.21(+1U)&[MIMAT0004678&hsa-miR-99b-3p&offsets\|0\|-1(+1U);m-151&19\|+\|51692656\|51692678&offsets\|0\|-2(+1U)]\|CAAGCTCGTGTCTGTGGGTCCT | >MI0000746\|hsa-mir-99b&WithFlank&19\|+\|51692606\|51692687 | 51 | 71 | 21(+1U) | CAAGCTCGTGTCTGTGGGTCCT | 243.5 | 2.3 | 0.7 | 3.1 | 1.9E-03 | 1.0E-02 | BT20/MB231 |
| >hg38_21_+_8433172_8446622_RNA45SN1_45S_50ntFlanks@12419.12454.36\|GATGTCGGCTCTTCCTATCATTGTGAAGCAGAATTC | >hg38_21_+_8433172_8446622_RNA45SN1_45S_50ntFlanks | 12419 | 12454 | 36 | GATGTCGGCTCTTCCTATCATTGTGAAGCAGAATTC | 617.8 | -2.5 | 0.8 | -3.1 | 1.9E-03 | 1.0E-02 | BT20/MB231 |
| >hg38_21_+_8433172_8446622_RNA45SN1_45S_50ntFlanks@11771.11794.24\|ATTAGTGACGCGCATGAATGGATG | >hg38_21_+_8433172_8446622_RNA45SN1_45S_50ntFlanks | 11771 | 11794 | 24 | ATTAGTGACGCGCATGAATGGATG | 537.7 | 1.3 | 0.5 | 2.6 | 8.6E-03 | 1.0E-02 | MB231/MB468 |
| >hg38_21_+_8433172_8446622_RNA45SN1_45S_50ntFlanks@6733.6753.21\|CATTGATCATCGACACTTCGA | >hg38_21_+_8433172_8446622_RNA45SN1_45S_50ntFlanks | 6733 | 6753 | 21 | CATTGATCATCGACACTTCGA | 466.1 | -1.4 | 0.6 | -2.6 | 9.2E-03 | 1.0E-02 | BT20/MB468 |
| >hg38_21_+_8433172_8446622_RNA45SN1_45S_50ntFlanks@6648.6679.32\|GTACGACTCTTAGCGGTGGATCACTCGGCTCG | >hg38_21_+_8433172_8446622_RNA45SN1_45S_50ntFlanks | 6648 | 6679 | 32 | GTACGACTCTTAGCGGTGGATCACTCGGCTCG | 1087.1 | 2.0 | 0.8 | 2.6 | 8.7E-03 | 1.1E-02 | MB231/MB468 |
| >hg38_21_+_8433172_8446622_RNA45SN1_45S_50ntFlanks@3845.3868.24\|ACTTGGATAACTGTGGTAATTCTA | >hg38_21_+_8433172_8446622_RNA45SN1_45S_50ntFlanks | 3845 | 3868 | 24 | ACTTGGATAACTGTGGTAATTCTA | 308.2 | 1.2 | 0.4 | 2.6 | 9.3E-03 | 1.1E-02 | BT20/MB468 |
| >hg38_21_+_8433172_8446622_RNA45SN1_45S_50ntFlanks@12453.12477.25\|TCACCAAGCGTTGGATTGTTCACCC | >hg38_21_+_8433172_8446622_RNA45SN1_45S_50ntFlanks | 12453 | 12477 | 25 | TCACCAAGCGTTGGATTGTTCACCC | 246.9 | -2.5 | 0.8 | -3.1 | 2.0E-03 | 1.1E-02 | BT20/MB231 |
| >hg38_21_+_8433172_8446622_RNA45SN1_45S_50ntFlanks@6650.6696.47\|ACGACTCTTAGCGGTGGATCACTCGGCTCGTGCGTCGATGAAGAACG | >hg38_21_+_8433172_8446622_RNA45SN1_45S_50ntFlanks | 6650 | 6696 | 47 | ACGACTCTTAGCGGTGGATCACTCGGCTCGTGCGTCGATGAAGAACG | 1069.6 | -2.3 | 0.7 | -3.1 | 2.0E-03 | 1.1E-02 | BT20/MB231 |
| >hg38_21_+_8433172_8446622_RNA45SN1_45S_50ntFlanks@6661.6700.40\|CGGTGGATCACTCGGCTCGTGCGTCGATGAAGAACGCAGC | >hg38_21_+_8433172_8446622_RNA45SN1_45S_50ntFlanks | 6661 | 6700 | 40 | CGGTGGATCACTCGGCTCGTGCGTCGATGAAGAACGCAGC | 665.0 | 2.1 | 0.7 | 3.1 | 2.0E-03 | 1.1E-02 | BT20/MB231 |
| >hg38_21_+_8433172_8446622_RNA45SN1_45S_50ntFlanks@11881.11899.19\|CCTGTTGAGCTTGACTCTA | >hg38_21_+_8433172_8446622_RNA45SN1_45S_50ntFlanks | 11881 | 11899 | 19 | CCTGTTGAGCTTGACTCTA | 128.6 | -2.0 | 0.8 | -2.6 | 9.3E-03 | 1.1E-02 | BT20/MB468 |
| >am_trna111_HisGTG_1_-_147774845_147774916@-1G.28.29,trna118_HisGTG_1_-_145396881_145396952@-1G.28.29,trna16_HisGTG_1_+_146544773_146544844@-1G.28.29,trna1_HisGTG_15_+_45493349_45493420@-1G.28.29,trna21_HisGTG_1_+_147753471_147753542@-1G.28.29,trna33_HisGTG_6_+_27125906_27125977@-1G.28.29,trna7_HisGTG_9_-_14433938_14434009@-1G.28.29,trna8_HisGTG_15_-_45492611_45492682@-1G.28.29,trna9_HisGTG_15_-_45490804_45490875@-1G.28.29\|GGCCGTGATCGTATAGTGGTTAGTACTCT | >am_trna111_HisGTG_1_-_147774845_147774916 | -1G | 28 | 29 | GGCCGTGATCGTATAGTGGTTAGTACTCT | 2110.2 | 2.0 | 0.8 | 2.6 | 8.7E-03 | 1.1E-02 | MB231/MB468 |
| >hg38_21_+_8433172_8446622_RNA45SN1_45S_50ntFlanks@3728.3748.21\|CATATGCTTGTCTCAAAGATT | >hg38_21_+_8433172_8446622_RNA45SN1_45S_50ntFlanks | 3728 | 3748 | 21 | CATATGCTTGTCTCAAAGATT | 106.0 | -1.5 | 0.6 | -2.6 | 8.8E-03 | 1.1E-02 | MB231/MB468 |
| >hg38_21_+_8433172_8446622_RNA45SN1_45S_50ntFlanks@7982.8005.24\|TCAGATCAGACGTGGCGACCCGCT | >hg38_21_+_8433172_8446622_RNA45SN1_45S_50ntFlanks | 7982 | 8005 | 24 | TCAGATCAGACGTGGCGACCCGCT | 167.6 | -1.5 | 0.6 | -2.6 | 9.4E-03 | 1.1E-02 | BT20/MB468 |
| >hg38_21_+_8433172_8446622_RNA45SN1_45S_50ntFlanks@8357.8376.20\|AAAAGAACTTTGAAGAGAGA | >hg38_21_+_8433172_8446622_RNA45SN1_45S_50ntFlanks | 8357 | 8376 | 20 | AAAAGAACTTTGAAGAGAGA | 889.7 | -1.7 | 0.7 | -2.6 | 8.8E-03 | 1.1E-02 | MB231/MB468 |
| >hg38_21_+_8433172_8446622_RNA45SN1_45S_50ntFlanks@3840.3861.22\|CTCCTACTTGGATAACTGTGGT | >hg38_21_+_8433172_8446622_RNA45SN1_45S_50ntFlanks | 3840 | 3861 | 22 | CTCCTACTTGGATAACTGTGGT | 319.8 | 2.4 | 0.8 | 3.1 | 2.0E-03 | 1.1E-02 | BT20/MB231 |
| >hg38_21_+_8433172_8446622_RNA45SN1_45S_50ntFlanks@12457.12483.27\|CAAGCGTTGGATTGTTCACCCACTAAT | >hg38_21_+_8433172_8446622_RNA45SN1_45S_50ntFlanks | 12457 | 12483 | 27 | CAAGCGTTGGATTGTTCACCCACTAAT | 137.5 | -1.7 | 0.6 | -2.6 | 8.9E-03 | 1.1E-02 | MB231/MB468 |

<table>
<tr><th></th><th></th><th></th><th></th><th></th><th></th><th></th><th></th><th></th><th></th><th></th><th></th><th></th></tr>
<tr><td>>am_trna10_ValCAC_5_-_180649395_180649467@1.31.31,trna12_ValAAC_5_-_180645270_180645342@1.31.31,trna132_ValAAC_6_-_27721179_27721251@1.31.31,trna136_ValAAC_6_-_27648885_27648957@1.31.31,trna139_ValAAC_6_-_27618707_27618779@1.31.31,trna18_ValCAC_5_-_180529253_180529325@1.31.31,trna2_ValAAC_3_+_169490018_169490090@1.31.31,trna2_ValCAC_5_+_180524070_180524142@1.31.31,trna4_ValAAC_5_+_180591154_180591226@1.31.31,trna5_ValAAC_5_+_180596610_180596682@1.31.31,trna6_ValCAC_5_+_180600650_180600722@1.31.31,trna85_ValCAC_1_-_161369490_161369562@1.31.31,trna90_ValCAC_1_-_149684088_149684161@1.31.31,trna98_ValCAC_1_-_149298555_149298627@1.31.31,trna9_ValCAC_6_+_26538282_26538354@1.31.31|GTTTCCGTAGTGTAGTGGTTATCACGTTCGC</td><td>>am_trna10_ValCAC_5_-_180649395_180649467</td><td>1</td><td>31</td><td>31</td><td>GTTTCCGTAGTGTAGTGGTTATCACGTTCGC</td><td>445.7</td><td>-2.1</td><td>0.8</td><td>-2.6</td><td>9.5E-03</td><td>1.1E-02</td><td>BT20/MB468</td></tr>
<tr><td>>hg38_21_+_8433172_8446622_RNA45SN1_45S_50ntFlanks@7978.8006.29|GACCTCAGATCAGACGTGGCGACCCGCTG</td><td>>hg38_21_+_8433172_8446622_RNA45SN1_45S_50ntFlanks</td><td>7978</td><td>8006</td><td>29</td><td>GACCTCAGATCAGACGTGGCGACCCGCTG</td><td>175.6</td><td>-2.2</td><td>0.9</td><td>-2.6</td><td>9.5E-03</td><td>1.1E-02</td><td>BT20/MB468</td></tr>
<tr><td>>hg38_21_+_8433172_8446622_RNA45SN1_45S_50ntFlanks@8176.8217.42|CTTCTGATCGAGGCCCAGCCCGTGGACGGTGTGAGGCCGGTA</td><td>>hg38_21_+_8433172_8446622_RNA45SN1_45S_50ntFlanks</td><td>8176</td><td>8217</td><td>42</td><td>CTTCTGATCGAGGCCCAGCCCGTGGACGGTGTGAGGCCGGTA</td><td>201.6</td><td>-2.6</td><td>0.8</td><td>-3.1</td><td>2.0E-03</td><td>1.1E-02</td><td>BT20/MB231</td></tr>
<tr><td>>trnaMT_ValTAC_MT_+_1602_1670@49.72.24|AACTTAACTTGACCGCTCTGACCA</td><td>>trnaMT_ValTAC_MT_+_1602_1670</td><td>49</td><td>72</td><td>24</td><td>AACTTAACTTGACCGCTCTGACCA</td><td>1550.2</td><td>2.4</td><td>0.8</td><td>3.1</td><td>2.0E-03</td><td>1.1E-02</td><td>BT20/MB231</td></tr>
<tr><td>>hg38_21_+_8433172_8446622_RNA45SN1_45S_50ntFlanks@7977.8003.27|CGACCTCAGATCAGACGTGGCGACCCG</td><td>>hg38_21_+_8433172_8446622_RNA45SN1_45S_50ntFlanks</td><td>7977</td><td>8003</td><td>27</td><td>CGACCTCAGATCAGACGTGGCGACCCG</td><td>108.3</td><td>-2.1</td><td>0.8</td><td>-2.6</td><td>9.6E-03</td><td>1.1E-02</td><td>BT20/MB468</td></tr>
<tr><td>>hg38_21_+_8433172_8446622_RNA45SN1_45S_50ntFlanks@10350.10369.20|ACGAGAACTTTGAAGGCCGA</td><td>>hg38_21_+_8433172_8446622_RNA45SN1_45S_50ntFlanks</td><td>10350</td><td>10369</td><td>20</td><td>ACGAGAACTTTGAAGGCCGA</td><td>561.4</td><td>-1.6</td><td>0.6</td><td>-2.6</td><td>9.6E-03</td><td>1.1E-02</td><td>BT20/MB468</td></tr>
<tr><td>>hg38_21_+_8433172_8446622_RNA45SN1_45S_50ntFlanks@9544.9570.27|GGTGGAGGTCCGTAGCGGTCCTGACGT</td><td>>hg38_21_+_8433172_8446622_RNA45SN1_45S_50ntFlanks</td><td>9544</td><td>9570</td><td>27</td><td>GGTGGAGGTCCGTAGCGGTCCTGACGT</td><td>243.9</td><td>1.1</td><td>0.4</td><td>2.6</td><td>9.0E-03</td><td>1.1E-02</td><td>MB231/MB468</td></tr>
<tr><td>>hg38_21_+_8433172_8446622_RNA45SN1_45S_50ntFlanks@7991.8018.28|ACGTGGCGACCCGCTGAATTTAAGCATA</td><td>>hg38_21_+_8433172_8446622_RNA45SN1_45S_50ntFlanks</td><td>7991</td><td>8018</td><td>28</td><td>ACGTGGCGACCCGCTGAATTTAAGCATA</td><td>145.2</td><td>-1.5</td><td>0.6</td><td>-2.6</td><td>9.6E-03</td><td>1.1E-02</td><td>BT20/MB468</td></tr>
<tr><td>>hg38_21_+_8433172_8446622_RNA45SN1_45S_50ntFlanks@475.492.18|CTTCGTGATCGATGTGGT</td><td>>hg38_21_+_8433172_8446622_RNA45SN1_45S_50ntFlanks</td><td>475</td><td>492</td><td>18</td><td>CTTCGTGATCGATGTGGT</td><td>2437.1</td><td>2.6</td><td>0.8</td><td>3.1</td><td>2.1E-03</td><td>1.1E-02</td><td>BT20/MB231</td></tr>
<tr><td>>MI0000289|hsa-mir-181a-1&WithFlank&1|-|198859038|198859159@30.51.22;MI0000269|hsa-mir-181a-2&WithFlank&9|+|124692436|124692557@45.66.22&[MIMAT0000256&hsa-miR-181a-5p&offsets|0|-1;m-32&1|-|198859108|198859130&offsets|0|-1];[MIMAT0000256_1&hsa-miR-181a-5p&offsets|0|-1;m-32&9|+|124692480|124692502&offsets|0|-1]|AACATTCAACGCTGTCGGTGAG</td><td>>MI0000289|hsa-mir-181a-1&WithFlank&1|-|198859038|198859159</td><td>30</td><td>51</td><td>22</td><td>AACATTCAACGCTGTCGGTGAG</td><td>604.6</td><td>-2.2</td><td>0.7</td><td>-3.1</td><td>2.1E-03</td><td>1.1E-02</td><td>BT20/MB231</td></tr>
<tr><td>>hg38_21_+_8433172_8446622_RNA45SN1_45S_50ntFlanks@10350.10380.31|ACGAGAACTTTGAAGGCCGAAGTGGAGAAGG</td><td>>hg38_21_+_8433172_8446622_RNA45SN1_45S_50ntFlanks</td><td>10350</td><td>10380</td><td>31</td><td>ACGAGAACTTTGAAGGCCGAAGTGGAGAAGG</td><td>162.7</td><td>-1.3</td><td>0.5</td><td>-2.6</td><td>9.7E-03</td><td>1.1E-02</td><td>BT20/MB468</td></tr>
<tr><td>>trnaMT_GluTTC_MT_-_14674_14742@25.72.48,trnalookalike8_GluTTC_5_-_93905172_93905240@25.72.48|TGGTTTTTCATATCATTGGTCGTGGTTGTAGTCCGTGCGAGAATACCA</td><td>>trnaMT_GluTTC_MT_-_14674_14742</td><td>25</td><td>72</td><td>48</td><td>TGGTTTTTCATATCATTGGTCGTGGTTGTAGTCCGTGCGAGAATACCA</td><td>818.0</td><td>-3.8</td><td>1.2</td><td>-3.1</td><td>2.1E-03</td><td>1.1E-02</td><td>BT20/MB231</td></tr>
<tr><td>>hg38_21_+_8433172_8446622_RNA45SN1_45S_50ntFlanks@12912.12935.24|CACGTTCGTGGGGAACCTGGCGCT</td><td>>hg38_21_+_8433172_8446622_RNA45SN1_45S_50ntFlanks</td><td>12912</td><td>12935</td><td>24</td><td>CACGTTCGTGGGGAACCTGGCGCT</td><td>142.5</td><td>-2.4</td><td>0.9</td><td>-2.6</td><td>9.8E-03</td><td>1.1E-02</td><td>BT20/MB468</td></tr>
<tr><td>>hg38_21_+_8433172_8446622_RNA45SN1_45S_50ntFlanks@10351.10391.41|CGAGAACTTTGAAGGCCGAAGTGGAGAAGGGTTCCATGTGA</td><td>>hg38_21_+_8433172_8446622_RNA45SN1_45S_50ntFlanks</td><td>10351</td><td>10391</td><td>41</td><td>CGAGAACTTTGAAGGCCGAAGTGGAGAAGGGTTCCATGTGA</td><td>330.8</td><td>-2.0</td><td>0.8</td><td>-2.6</td><td>9.2E-03</td><td>1.1E-02</td><td>MB231/MB468</td></tr>
<tr><td>>hg38_21_+_8433172_8446622_RNA45SN1_45S_50ntFlanks@8799.8823.25|CCTCGGGAGGGCGCGCGGGTCGGGG</td><td>>hg38_21_+_8433172_8446622_RNA45SN1_45S_50ntFlanks</td><td>8799</td><td>8823</td><td>25</td><td>CCTCGGGAGGGCGCGCGGGTCGGGG</td><td>111.6</td><td>-2.9</td><td>1.1</td><td>-2.6</td><td>9.9E-03</td><td>1.1E-02</td><td>BT20/MB468</td></tr>
<tr><td>>hg38_21_+_8433172_8446622_RNA45SN1_45S_50ntFlanks@5482.5499.18|CGGAGCGCTGAGAAGACG</td><td>>hg38_21_+_8433172_8446622_RNA45SN1_45S_50ntFlanks</td><td>5482</td><td>5499</td><td>18</td><td>CGGAGCGCTGAGAAGACG</td><td>120.2</td><td>-1.2</td><td>0.5</td><td>-2.6</td><td>9.3E-03</td><td>1.1E-02</td><td>MB231/MB468</td></tr>
<tr><td>>hg38_21_+_8433172_8446622_RNA45SN1_45S_50ntFlanks@8171.8214.44|AAGTCCTTCTGATCGAGGCCCAGCCCGTGGACGGTGTGAGGCCG</td><td>>hg38_21_+_8433172_8446622_RNA45SN1_45S_50ntFlanks</td><td>8171</td><td>8214</td><td>44</td><td>AAGTCCTTCTGATCGAGGCCCAGCCCGTGGACGGTGTGAGGCCG</td><td>165.7</td><td>-1.4</td><td>0.5</td><td>-2.6</td><td>1.0E-02</td><td>1.1E-02</td><td>BT20/MB468</td></tr>
<tr><td>>hg38_21_+_8433172_8446622_RNA45SN1_45S_50ntFlanks@11672.11700.29|CAGTGCTCTGAATGTCAAAGTGAAGAAAT</td><td>>hg38_21_+_8433172_8446622_RNA45SN1_45S_50ntFlanks</td><td>11672</td><td>11700</td><td>29</td><td>CAGTGCTCTGAATGTCAAAGTGAAGAAAT</td><td>196.2</td><td>-1.0</td><td>0.4</td><td>-2.6</td><td>1.0E-02</td><td>1.1E-02</td><td>BT20/MB468</td></tr>
<tr><td>>hg38_21_+_8433172_8446622_RNA45SN1_45S_50ntFlanks@3854.3883.30|ACTGTGGTAATTCTAGAGCTAATACATGCC</td><td>>hg38_21_+_8433172_8446622_RNA45SN1_45S_50ntFlanks</td><td>3854</td><td>3883</td><td>30</td><td>ACTGTGGTAATTCTAGAGCTAATACATGCC</td><td>284.9</td><td>1.1</td><td>0.4</td><td>2.6</td><td>9.4E-03</td><td>1.1E-02</td><td>MB231/MB468</td></tr>
<tr><td>>MI0000469|hsa-mir-125a&WithFlank&19|+|51693248|51693345@21.42.22(+1U)&[MIMAT0000443&hsa-miR-125a-5p&offsets|0|-2(+1U);m-63&19|+|51693268|51693289&offsets|0|0(+1U)]|TCCCTGAGACCCTTTAACCTGTT</td><td>>MI0000469|hsa-mir-125a&WithFlank&19|+|51693248|51693345</td><td>21</td><td>42</td><td>22(+1U)</td><td>TCCCTGAGACCCTTTAACCTGTT</td><td>452.6</td><td>2.2</td><td>0.7</td><td>3.1</td><td>2.2E-03</td><td>1.1E-02</td><td>BT20/MB231</td></tr>
<tr><td>>trnaMT_SerGCT_MT_+_12207_12265@8.30.23|CTCACAAGAACTGCTAACTCATG</td><td>>trnaMT_SerGCT_MT_+_12207_12265</td><td>8</td><td>30</td><td>23</td><td>CTCACAAGAACTGCTAACTCATG</td><td>115.9</td><td>-1.9</td><td>0.7</td><td>-2.6</td><td>1.0E-02</td><td>1.1E-02</td><td>BT20/MB468</td></tr>
<tr><td>>MI0000063|hsa-let-7b&WithFlank&22|+|46113680|46113774@12.33.22(+1C)&[MIMAT0000063&hsa-let-7b-5p&offsets|0|0(+1C);m-14&22|+|46113691|46113712&offsets|0|0(+1C)]|TGAGGTAGTAGGTTGTGTGGTTC</td><td>>MI0000063|hsa-let-7b&WithFlank&22|+|46113680|46113774</td><td>12</td><td>33</td><td>22(+1C)</td><td>TGAGGTAGTAGGTTGTGTGGTTC</td><td>979.2</td><td>2.1</td><td>0.7</td><td>3.1</td><td>2.2E-03</td><td>1.1E-02</td><td>BT20/MB231</td></tr>
<tr><td>>hg38_21_+_8433172_8446622_RNA45SN1_45S_50ntFlanks@12419.12447.29|GATGTCGGCTCTTCCTATCATTGTGAAGC</td><td>>hg38_21_+_8433172_8446622_RNA45SN1_45S_50ntFlanks</td><td>12419</td><td>12447</td><td>29</td><td>GATGTCGGCTCTTCCTATCATTGTGAAGC</td><td>914.6</td><td>-2.5</td><td>0.8</td><td>-3.1</td><td>2.2E-03</td><td>1.1E-02</td><td>BT20/MB231</td></tr>
<tr><td>>hg38_21_+_8433172_8446622_RNA45SN1_45S_50ntFlanks@11785.11802.18|TGAATGGATGAACGAGAT</td><td>>hg38_21_+_8433172_8446622_RNA45SN1_45S_50ntFlanks</td><td>11785</td><td>11802</td><td>18</td><td>TGAATGGATGAACGAGAT</td><td>128.8</td><td>-1.4</td><td>0.6</td><td>-2.6</td><td>9.5E-03</td><td>1.2E-02</td><td>MB231/MB468</td></tr>
<tr><td>>MI0000298|hsa-mir-221&WithFlank&X|-|45746151|45746272@71.92.22&[MIMAT0000278&hsa-miR-221-3p&offsets|0|-1;m-91&X|-|45746181|45746202&offsets|0|0]|AGCTACATTGTCTGCTGGGTTT</td><td>>MI0000298|hsa-mir-221&WithFlank&X|-|45746151|45746272</td><td>71</td><td>92</td><td>22</td><td>AGCTACATTGTCTGCTGGGTTT</td><td>27306.5</td><td>-2.0</td><td>0.6</td><td>-3.1</td><td>2.2E-03</td><td>1.2E-02</td><td>BT20/MB231</td></tr>
<tr><td>>hg38_21_+_8433172_8446622_RNA45SN1_45S_50ntFlanks@6698.6719.22|AGCTAGCTGCGAGAATTAATGT</td><td>>hg38_21_+_8433172_8446622_RNA45SN1_45S_50ntFlanks</td><td>6698</td><td>6719</td><td>22</td><td>AGCTAGCTGCGAGAATTAATGT</td><td>584.8</td><td>-1.5</td><td>0.6</td><td>-2.6</td><td>9.6E-03</td><td>1.2E-02</td><td>MB231/MB468</td></tr>
<tr><td>>hg38_21_+_8433172_8446622_RNA45SN1_45S_50ntFlanks@12417.12449.33|TCGATGTCGGCTCTTCCTATCATTGTGAAGCAG</td><td>>hg38_21_+_8433172_8446622_RNA45SN1_45S_50ntFlanks</td><td>12417</td><td>12449</td><td>33</td><td>TCGATGTCGGCTCTTCCTATCATTGTGAAGCAG</td><td>209.9</td><td>-2.8</td><td>0.9</td><td>-3.1</td><td>2.2E-03</td><td>1.2E-02</td><td>BT20/MB231</td></tr>
<tr><td>>MI0025750|hsa-mir-7974&WithFlank&19|-|11495538|11495628@62.85.24&[MIMAT0031177&hsa-miR-7974&offsets|0|0;m-867&19|-|11495544|11495567&offsets|0|0]|AGGCTGTGATGCTCTCCTGAGCCC</td><td>>MI0025750|hsa-mir-7974&WithFlank&19|-|11495538|11495628</td><td>62</td><td>85</td><td>24</td><td>AGGCTGTGATGCTCTCCTGAGCCC</td><td>519.0</td><td>2.4</td><td>0.9</td><td>2.6</td><td>1.0E-02</td><td>1.2E-02</td><td>BT20/MB468</td></tr>
<tr><td>>hg38_21_+_8433172_8446622_RNA45SN1_45S_50ntFlanks@3854.3877.24|ACTGTGGTAATTCTAGAGCTAATA</td><td>>hg38_21_+_8433172_8446622_RNA45SN1_45S_50ntFlanks</td><td>3854</td><td>3877</td><td>24</td><td>ACTGTGGTAATTCTAGAGCTAATA</td><td>91.0</td><td>-1.5</td><td>0.6</td><td>-2.6</td><td>1.0E-02</td><td>1.2E-02</td><td>BT20/MB468</td></tr>
<tr><td>>hg38_21_+_8433172_8446622_RNA45SN1_45S_50ntFlanks@6650.6699.50|ACGACTCTTAGCGGTGGATCACTCGGCTCGTGCGTCGATGAAGAACGCAG</td><td>>hg38_21_+_8433172_8446622_RNA45SN1_45S_50ntFlanks</td><td>6650</td><td>6699</td><td>50</td><td>ACGACTCTTAGCGGTGGATCACTCGGCTCGTGCGTCGATGAAGAACGCAG</td><td>544.6</td><td>-2.8</td><td>0.9</td><td>-3.1</td><td>2.3E-03</td><td>1.2E-02</td><td>BT20/MB231</td></tr>
<tr><td>>am_trna10_ValCAC_5_-_180649395_180649467@5.23.19,trna12_ValAAC_5_-_180645270_180645342@5.23.19,trna132_ValAAC_6_-_27721179_27721251@5.23.19,trna136_ValAAC_6_-_27648885_27648957@5.23.19,trna139_ValAAC_6_-_27618707_27618779@5.23.19,trna18_ValCAC_5_-_180529253_180529325@5.23.19,trna2_ValAAC_3_+_169490018_169490090@5.23.19,trna2_ValCAC_5_+_180524070_180524142@5.23.19,trna37_ValAAC_6_+_27203288_27203360@5.23.19,trna4_ValAAC_5_+_180591154_180591226@5.23.19,trna5_ValAAC_5_+_180596610_180596682@5.23.19,trna6_ValCAC_5_+_180600650_180600722@5.23.19,trna85_ValCAC_1_-_161369490_161369562@5.23.19,trna90_ValCAC_1_-_149684088_149684161@5.23.19,trna98_ValCAC_1_-_149298555_149298627@5.23.19,trna9_ValCAC_6_+_26538282_26538354@5.23.19|CCGTAGTGTAGTGGTTATC</td><td>>am_trna10_ValCAC_5_-_180649395_180649467</td><td>5</td><td>23</td><td>19</td><td>CCGTAGTGTAGTGGTTATC</td><td>452.2</td><td>2.6</td><td>1.0</td><td>2.6</td><td>1.0E-02</td><td>1.2E-02</td><td>BT20/MB468</td></tr>
<tr><td>>hg38_21_+_8433172_8446622_RNA45SN1_45S_50ntFlanks@12419.12451.33|GATGTCGGCTCTTCCTATCATTGTGAAGCAGAA</td><td>>hg38_21_+_8433172_8446622_RNA45SN1_45S_50ntFlanks</td><td>12419</td><td>12451</td><td>33</td><td>GATGTCGGCTCTTCCTATCATTGTGAAGCAGAA</td><td>182.2</td><td>-2.5</td><td>0.8</td><td>-3.1</td><td>2.3E-03</td><td>1.2E-02</td><td>BT20/MB231</td></tr>
<tr><td>>hg38_21_+_8433172_8446622_RNA45SN1_45S_50ntFlanks@12415.12434.20|CTTCGATGTCGGCTCTTCCT</td><td>>hg38_21_+_8433172_8446622_RNA45SN1_45S_50ntFlanks</td><td>12415</td><td>12434</td><td>20</td><td>CTTCGATGTCGGCTCTTCCT</td><td>4695.8</td><td>3.4</td><td>1.1</td><td>3.0</td><td>2.3E-03</td><td>1.2E-02</td><td>BT20/MB231</td></tr>
<tr><td>>hg38_21_+_8433172_8446622_RNA45SN1_45S_50ntFlanks@12458.12488.31|AAGCGTTGGATTGTTCACCCACTAATAGGGA</td><td>>hg38_21_+_8433172_8446622_RNA45SN1_45S_50ntFlanks</td><td>12458</td><td>12488</td><td>31</td><td>AAGCGTTGGATTGTTCACCCACTAATAGGGA</td><td>139.3</td><td>-1.3</td><td>0.5</td><td>-2.6</td><td>9.9E-03</td><td>1.2E-02</td><td>MB231/MB468</td></tr>
<tr><td>>hg38_21_+_8433172_8446622_RNA45SN1_45S_50ntFlanks@6649.6689.41|TACGACTCTTAGCGGTGGATCACTCGGCTCGTGCGTCGATG</td><td>>hg38_21_+_8433172_8446622_RNA45SN1_45S_50ntFlanks</td><td>6649</td><td>6689</td><td>41</td><td>TACGACTCTTAGCGGTGGATCACTCGGCTCGTGCGTCGATG</td><td>853.8</td><td>1.3</td><td>0.5</td><td>2.6</td><td>9.9E-03</td><td>1.2E-02</td><td>MB231/MB468</td></tr>
<tr><td>>hg38_21_+_8433172_8446622_RNA45SN1_45S_50ntFlanks@13000.13028.29|TGCGATCTATTGAAAGTCAGCCCTCGACA</td><td>>hg38_21_+_8433172_8446622_RNA45SN1_45S_50ntFlanks</td><td>13000</td><td>13028</td><td>29</td><td>TGCGATCTATTGAAAGTCAGCCCTCGACA</td><td>131.9</td><td>-3.1</td><td>1.2</td><td>-2.6</td><td>1.1E-02</td><td>1.2E-02</td><td>BT20/MB468</td></tr>
<tr><td>>hg38_21_+_8433172_8446622_RNA45SN1_45S_50ntFlanks@6646.6667.22|TCGTACGACTCTTAGCGGTGGA</td><td>>hg38_21_+_8433172_8446622_RNA45SN1_45S_50ntFlanks</td><td>6646</td><td>6667</td><td>22</td><td>TCGTACGACTCTTAGCGGTGGA</td><td>9020.6</td><td>2.3</td><td>0.8</td><td>3.0</td><td>2.3E-03</td><td>1.2E-02</td><td>BT20/MB231</td></tr>
</table>

| | | | | | | | | | | | | |
|---|---|---|---|---|---|---|---|---|---|---|---|---|
| >hg38_21_+_8433172_8446622_RNA45SN1_45S_50ntFlanks@8381.8405.25\|AAGAGGGCGTGAAACCGTTAAGAGG | >hg38_21_+_8433172_8446622_RNA45SN1_45S_50ntFlanks | 8381 | 8405 | 25 | AAGAGGGCGTGAAACCGTTAAGAGG | 283.0 | -1.7 | 0.7 | -2.6 | 1.0E-02 | 1.2E-02 | MB231/MB468 |
| >hg38_21_+_8433172_8446622_RNA45SN1_45S_50ntFlanks@7986.8004.19\|ATCAGACGTGGCGACCCGC | >hg38_21_+_8433172_8446622_RNA45SN1_45S_50ntFlanks | 7986 | 8004 | 19 | ATCAGACGTGGCGACCCGC | 215.0 | -4.1 | 1.6 | -2.6 | 1.0E-02 | 1.2E-02 | MB231/MB468 |
| >hg38_21_+_8433172_8446622_RNA45SN1_45S_50ntFlanks@7979.8005.27\|ACCTCAGATCAGACGTGGCGACCCGCT | >hg38_21_+_8433172_8446622_RNA45SN1_45S_50ntFlanks | 7979 | 8005 | 27 | ACCTCAGATCAGACGTGGCGACCCGCT | 171.6 | -1.2 | 0.5 | -2.6 | 1.0E-02 | 1.2E-02 | MB231/MB468 |
| >hg38_21_+_8433172_8446622_RNA45SN1_45S_50ntFlanks@11727.11753.27\|AGTAACTATGACTCTCTTAAGGTAGCC | >hg38_21_+_8433172_8446622_RNA45SN1_45S_50ntFlanks | 11727 | 11753 | 27 | AGTAACTATGACTCTCTTAAGGTAGCC | 290.9 | -1.8 | 0.7 | -2.6 | 1.0E-02 | 1.2E-02 | MB231/MB468 |
| >hg38_1_-_228634819_228635039_RNA5S12_5S_50ntFlanks@51.94.44\|GTCTACGGCCATACCACCCTGAACGCGCCCGATCTCGTCTGATC | >hg38_1_-_228634819_228635039_RNA5S12_5S_50ntFlanks | 51 | 94 | 44 | GTCTACGGCCATACCACCCTGAACGCGCCCGATCTCGTCTGATC | 455.0 | 3.3 | 1.1 | 3.0 | 2.4E-03 | 1.2E-02 | BT20/MB231 |
| >MI0003137\|hsa-mir-193b&WithFlank&16\|+\|14303961\|14304055@57.77.21(+1G)&[MIMAT0002819&hsa-miR-193b-3p&offsets\|0\|-1(+1G);m-166&16\|+\|14304017\|14304038&offsets\|0\|-1(+1G)]\|AACTGGCCCTCAAAGTCCCGCG | >MI0003137\|hsa-mir-193b&WithFlank&16\|+\|14303961\|14304055 | 57 | 77 | 21(+1G) | AACTGGCCCTCAAAGTCCCGCG | 685.9 | 6.1 | 2.0 | 3.0 | 2.4E-03 | 1.2E-02 | BT20/MB231 |
| >trna20_GluTTC_2_-_131094701_131094772@1.33.33,trna5_GluTTC_13_-_41634874_41634945@1.33.33\|TCCCATATGGTCTAGCGGTTAGGATTCCTGGTT | >trna20_GluTTC_2_-_131094701_131094772 | 1 | 33 | 33 | TCCCATATGGTCTAGCGGTTAGGATTCCTGGTT | 2641.0 | -2.2 | 0.7 | -3.0 | 2.4E-03 | 1.2E-02 | BT20/MB231 |
| >trnaMT_MetCAT_MT_+_4402_4469@34.71.38\|ACCCCGAAAATGTTGGTTATACCCTTCCCGTACTACCA | >trnaMT_MetCAT_MT_+_4402_4469 | 34 | 71 | 38 | ACCCCGAAAATGTTGGTTATACCCTTCCCGTACTACCA | 279.8 | -2.8 | 0.9 | -3.0 | 2.4E-03 | 1.2E-02 | BT20/MB231 |
| >hg38_21_+_8433172_8446622_RNA45SN1_45S_50ntFlanks@11581.11602.22\|AACTGGTGCGGACCAGGGGAAT | >hg38_21_+_8433172_8446622_RNA45SN1_45S_50ntFlanks | 11581 | 11602 | 22 | AACTGGTGCGGACCAGGGGAAT | 867.9 | -2.0 | 0.8 | -2.6 | 1.0E-02 | 1.2E-02 | MB231/MB468 |
| >hg38_21_+_8433172_8446622_RNA45SN1_45S_50ntFlanks@5487.5516.30\|CGCTGAGAAGACGGTCGAACTTGACTATCT | >hg38_21_+_8433172_8446622_RNA45SN1_45S_50ntFlanks | 5487 | 5516 | 30 | CGCTGAGAAGACGGTCGAACTTGACTATCT | 1776.0 | 2.5 | 1.0 | 2.5 | 1.1E-02 | 1.2E-02 | BT20/MB468 |
| >MI0000077\|hsa-mir-21&WithFlank&17\|+\|59841260\|59841343@14.39.26&[MIMAT0000076&hsa-miR-21-5p&offsets\|0\|+4;m-1&17\|+\|59841273\|59841294&offsets\|0\|+4]\|TAGCTTATCAGACTGATGTTGACTGT | >MI0000077\|hsa-mir-21&WithFlank&17\|+\|59841260\|59841343 | 14 | 39 | 26 | TAGCTTATCAGACTGATGTTGACTGT | 678.9 | 2.7 | 0.9 | 3.0 | 2.4E-03 | 1.2E-02 | BT20/MB231 |
| >MI0000298\|hsa-mir-221&WithFlank&X\|-\|45746151\|45746272@71.93.23(+2U)&[MIMAT0000278&hsa-miR-221-3p&offsets\|0\|0(+2U);m-91&X\|-\|45746181\|45746202&offsets\|0\|+1(+2U)]\|AGCTACATTGTCTGCTGGGTTTCTT | >MI0000298\|hsa-mir-221&WithFlank&X\|-\|45746151\|45746272 | 71 | 93 | 23(+2U) | AGCTACATTGTCTGCTGGGTTTCTT | 286.0 | -2.2 | 0.7 | -3.0 | 2.4E-03 | 1.2E-02 | BT20/MB231 |
| >hg38_21_+_8433172_8446622_RNA45SN1_45S_50ntFlanks@12419.12453.35\|GATGTCGGCTCTTCCTATCATTGTGAAGCAGAATT | >hg38_21_+_8433172_8446622_RNA45SN1_45S_50ntFlanks | 12419 | 12453 | 35 | GATGTCGGCTCTTCCTATCATTGTGAAGCAGAATT | 693.7 | -2.4 | 0.8 | -3.0 | 2.4E-03 | 1.2E-02 | BT20/MB231 |
| >hg38_21_+_8433172_8446622_RNA45SN1_45S_50ntFlanks@6651.6670.20\|CGACTCTTAGCGGTGGATCA | >hg38_21_+_8433172_8446622_RNA45SN1_45S_50ntFlanks | 6651 | 6670 | 20 | CGACTCTTAGCGGTGGATCA | 330.0 | 2.3 | 0.8 | 3.0 | 2.4E-03 | 1.2E-02 | BT20/MB231 |
| >trna10_AspGTC_12_-_125424193_125424264@36.75.40,trna12_AspGTC_12_-_125411891_125411962@36.75.40,trna144_AspGTC_6_-_27551236_27551307@36.75.40,trna38_AspGTC_17_-_8125556_8125627@36.75.40,trna45_AspGTC_6_+_27447453_27447524@36.75.40,trna48_AspGTC_6_+_27471523_27471594@36.75.40,trna4_AspGTC_12_+_96429799_96429870@36.75.40,trna69_AspGTC_1_-_161440205_161440276@36.75.40,trna72_AspGTC_1_-_161432824_161432895@36.75.40,trna75_AspGTC_1_-_161425414_161425485@36.75.40,trna78_AspGTC_1_-_161418033_161418104@36.75.40,trna81_AspGTC_1_-_161410615_161410686@36.75.40\|CACGCGGGAGACCGGGGTTCGATTCCCCGACGGGGAGCCA | >trna10_AspGTC_12_-_125424193_125424264 | 36 | 75 | 40 | CACGCGGGAGACCGGGGTTCGATTCCCCGACGGGGAGCCA | 631.5 | -2.6 | 0.9 | -3.0 | 2.4E-03 | 1.2E-02 | BT20/MB231 |
| >hg38_21_+_8433172_8446622_RNA45SN1_45S_50ntFlanks@10835.10858.24\|CGGATCCGTAACTTCGGGATAAGG | >hg38_21_+_8433172_8446622_RNA45SN1_45S_50ntFlanks | 10835 | 10858 | 24 | CGGATCCGTAACTTCGGGATAAGG | 274.2 | -1.2 | 0.5 | -2.5 | 1.1E-02 | 1.2E-02 | BT20/MB468 |
| >hg38_21_+_8433172_8446622_RNA45SN1_45S_50ntFlanks@10848.10865.18\|TCGGGATAAGGATTGGCT | >hg38_21_+_8433172_8446622_RNA45SN1_45S_50ntFlanks | 10848 | 10865 | 18 | TCGGGATAAGGATTGGCT | 119.6 | -1.3 | 0.5 | -2.6 | 1.0E-02 | 1.2E-02 | MB231/MB468 |
| >trna2_GlyGCC_21_-_18827107_18827177@1.28.28,trna35_GlyGCC_1_+_161413094_161413164@1.28.28,trna37_GlyGCC_1_+_161420467_161420537@1.28.28,trna39_GlyGCC_1_+_161427898_161427968@1.28.28,trna41_GlyGCC_1_+_161435258_161435328@1.28.28\|GCATGGGTGGTTCAGTGGTAGAATTCTC | >trna2_GlyGCC_21_-_18827107_18827177 | 1 | 28 | 28 | GCATGGGTGGTTCAGTGGTAGAATTCTC | 12243.2 | -2.0 | 0.8 | -2.5 | 1.1E-02 | 1.2E-02 | BT20/MB468 |
| >hg38_21_+_8433172_8446622_RNA45SN1_45S_50ntFlanks@11783.11800.18\|CATGAATGGATGAACGAG | >hg38_21_+_8433172_8446622_RNA45SN1_45S_50ntFlanks | 11783 | 11800 | 18 | CATGAATGGATGAACGAG | 405.3 | -1.8 | 0.7 | -2.6 | 1.0E-02 | 1.2E-02 | MB231/MB468 |
| >hg38_21_+_8433172_8446622_RNA45SN1_45S_50ntFlanks@11671.11691.21\|CCAGTGCTCTGAATGTCAAAG | >hg38_21_+_8433172_8446622_RNA45SN1_45S_50ntFlanks | 11671 | 11691 | 21 | CCAGTGCTCTGAATGTCAAAG | 422.4 | -1.6 | 0.6 | -2.5 | 1.1E-02 | 1.2E-02 | BT20/MB468 |
| >hg38_21_+_8433172_8446622_RNA45SN1_45S_50ntFlanks@12993.13012.20\|CCCTCGCTGCGATCTATTGA | >hg38_21_+_8433172_8446622_RNA45SN1_45S_50ntFlanks | 12993 | 13012 | 20 | CCCTCGCTGCGATCTATTGA | 124.3 | -1.8 | 0.7 | -2.5 | 1.1E-02 | 1.2E-02 | BT20/MB468 |
| >hg38_21_+_8433172_8446622_RNA45SN1_45S_50ntFlanks@4735.4759.25\|ACGAAAGTCGGAGGTTCGAAGACGA | >hg38_21_+_8433172_8446622_RNA45SN1_45S_50ntFlanks | 4735 | 4759 | 25 | ACGAAAGTCGGAGGTTCGAAGACGA | 609.4 | 2.1 | 0.8 | 2.6 | 1.0E-02 | 1.2E-02 | MB231/MB468 |
| >hg38_21_+_8433172_8446622_RNA45SN1_45S_50ntFlanks@11552.11591.40\|CCTCGGCCGGCGCCTAGCAGCCGACTTAGAACTGGTGCGG | >hg38_21_+_8433172_8446622_RNA45SN1_45S_50ntFlanks | 11552 | 11591 | 40 | CCTCGGCCGGCGCCTAGCAGCCGACTTAGAACTGGTGCGG | 157.4 | -1.1 | 0.4 | -2.6 | 1.0E-02 | 1.3E-02 | MB231/MB468 |
| >hg38_21_+_8433172_8446622_RNA45SN1_45S_50ntFlanks@11561.11584.24\|GCGCCTAGCAGCCGACTTAGAACT | >hg38_21_+_8433172_8446622_RNA45SN1_45S_50ntFlanks | 11561 | 11584 | 24 | GCGCCTAGCAGCCGACTTAGAACT | 217.1 | 1.2 | 0.5 | 2.6 | 1.0E-02 | 1.3E-02 | MB231/MB468 |
| >hg38_21_+_8433172_8446622_RNA45SN1_45S_50ntFlanks@12934.12954.21\|CTAAACCATTCGTAGACGACC | >hg38_21_+_8433172_8446622_RNA45SN1_45S_50ntFlanks | 12934 | 12954 | 21 | CTAAACCATTCGTAGACGACC | 202.1 | -1.7 | 0.7 | -2.6 | 1.0E-02 | 1.3E-02 | MB231/MB468 |
| >hg38_21_+_8433172_8446622_RNA45SN1_45S_50ntFlanks@11668.11706.39\|TGCCCAGTGCTCTGAATGTCAAAGTGAAGAAATTCAATG | >hg38_21_+_8433172_8446622_RNA45SN1_45S_50ntFlanks | 11668 | 11706 | 39 | TGCCCAGTGCTCTGAATGTCAAAGTGAAGAAATTCAATG | 643.6 | -2.5 | 0.8 | -3.0 | 2.5E-03 | 1.3E-02 | BT20/MB231 |
| >MI0000299\|hsa-mir-222&WithFlank&X\|-\|45747009\|45747130@75.98.24&[MIMAT0000279&hsa-miR-222-3p&offsets\|0\|+3;m-140&X\|-\|45747033\|45747056&offsets\|0\|0]\|AGCTACATCTGGCTACTGGGTCTC | >MI0000299\|hsa-mir-222&WithFlank&X\|-\|45747009\|45747130 | 75 | 98 | 24 | AGCTACATCTGGCTACTGGGTCTC | 25569.1 | -2.2 | 0.7 | -3.0 | 2.5E-03 | 1.3E-02 | BT20/MB231 |
| >trnaMT_PheGAA_MT_+_577_647@36.55.20\|AAAATGTTTAGACGGGCTCA | >trnaMT_PheGAA_MT_+_577_647 | 36 | 55 | 20 | AAAATGTTTAGACGGGCTCA | 729.7 | 2.3 | 0.8 | 3.0 | 2.5E-03 | 1.3E-02 | BT20/MB231 |
| >hg38_21_+_8433172_8446622_RNA45SN1_45S_50ntFlanks@12396.12417.22\|GCGACGTCGCTTTTTGATCCTT | >hg38_21_+_8433172_8446622_RNA45SN1_45S_50ntFlanks | 12396 | 12417 | 22 | GCGACGTCGCTTTTTGATCCTT | 464.6 | -1.4 | 0.6 | -2.6 | 1.1E-02 | 1.3E-02 | MB231/MB468 |
| >MI0000086\|hsa-mir-28&WithFlank&3\|+\|188688775\|188688872@20.41.22(+1A)&[MIMAT0000085&hsa-miR-28-5p&offsets\|0\|0(+1A);m-133&3\|+\|188688794\|188688815&offsets\|0\|0(+1A)]\|AAGGAGCTCACAGTCTATTGAGA | >MI0000086\|hsa-mir-28&WithFlank&3\|+\|188688775\|188688872 | 20 | 41 | 22(+1A) | AAGGAGCTCACAGTCTATTGAGA | 452.7 | 2.3 | 0.8 | 3.0 | 2.5E-03 | 1.3E-02 | BT20/MB231 |
| >hg38_21_+_8433172_8446622_RNA45SN1_45S_50ntFlanks@11589.11609.21\|CGGACCAGGGGAATCCGACTG | >hg38_21_+_8433172_8446622_RNA45SN1_45S_50ntFlanks | 11589 | 11609 | 21 | CGGACCAGGGGAATCCGACTG | 459.3 | 1.5 | 0.6 | 2.6 | 1.1E-02 | 1.3E-02 | MB231/MB468 |
| >hg38_21_+_8433172_8446622_RNA45SN1_45S_50ntFlanks@3705.3741.37\|TACCTGGTTGATCCTGCCAGTAGCATATGCTTGTCTC | >hg38_21_+_8433172_8446622_RNA45SN1_45S_50ntFlanks | 3705 | 3741 | 37 | TACCTGGTTGATCCTGCCAGTAGCATATGCTTGTCTC | 2496.8 | 2.4 | 0.8 | 3.0 | 2.5E-03 | 1.3E-02 | BT20/MB231 |
| >MI0000542\|hsa-mir-320a&WithFlank&8\|-\|22244960\|22245043@48.68.21(+1C)&[MIMAT0000510&hsa-miR-320a-3p&offsets\|0\|-1(+1C);m-60&8\|-\|22244975\|22244996&offsets\|0\|-1(+1C)]\|AAAAGCTGGGTTGAGAGGGCGC | >MI0000542\|hsa-mir-320a&WithFlank&8\|-\|22244960\|22245043 | 48 | 68 | 21(+1C) | AAAAGCTGGGTTGAGAGGGCGC | 757.4 | 2.3 | 0.8 | 3.0 | 2.6E-03 | 1.3E-02 | BT20/MB231 |
| >trna11_GluTTC_15_-_26327381_26327452@1.27.27,trna3_GluTTC_13_-_45492062_45492133@1.27.27\|TCCCACATGGTCTAGCGGTTAGGATTC | >trna11_GluTTC_15_-_26327381_26327452 | 1 | 27 | 27 | TCCCACATGGTCTAGCGGTTAGGATTC | 273.8 | -1.9 | 0.7 | -2.5 | 1.1E-02 | 1.3E-02 | BT20/MB468 |
| >hg38_21_+_8433172_8446622_RNA45SN1_45S_50ntFlanks@10393.10424.32\|CAGCAGTTGAACATGGGTCAGTCGGTCCTGAG | >hg38_21_+_8433172_8446622_RNA45SN1_45S_50ntFlanks | 10393 | 10424 | 32 | CAGCAGTTGAACATGGGTCAGTCGGTCCTGAG | 1368.9 | 1.6 | 0.6 | 2.6 | 1.1E-02 | 1.3E-02 | MB231/MB468 |
| >hg38_21_+_8433172_8446622_RNA45SN1_45S_50ntFlanks@4007.4025.19\|CCTCGGGCCGATCGCACGC | >hg38_21_+_8433172_8446622_RNA45SN1_45S_50ntFlanks | 4007 | 4025 | 19 | CCTCGGGCCGATCGCACGC | 129.8 | -1.3 | 0.5 | -2.6 | 1.1E-02 | 1.3E-02 | MB231/MB468 |
| >am_trna128_GlyGCC_6_-_27870686_27870756@3.31.29,trna133_GlyCCC_1_-_16872434_16872504@3.31.29,trna18_GlyGCC_16_+_70822597_70822667@3.31.29,trna19_GlyGCC_16_+_70823410_70823480@3.31.29,trna19_GlyGCC_2_-_157257659_157257729@3.31.29,trna24_GlyGCC_16_-_70812942_70813012@3.31.29,trna25_GlyGCC_16_-_70812114_70812184@3.31.29,trna4_GlyCCC_1_+_17188416_17188486@3.31.29,trna5_GlyGCC_17_+_8029064_8029134@3.31.29,trna68_GlyGCC_1_-_161493637_161493707@3.31.29\|ATTGGTGGTTCAGTGGTAGAATTCTCGCC | >am_trna128_GlyGCC_6_-_27870686_27870756 | 3 | 31 | 29 | ATTGGTGGTTCAGTGGTAGAATTCTCGCC | 235.9 | -2.4 | 0.8 | -3.0 | 2.6E-03 | 1.3E-02 | BT20/MB231 |
| >MI0000342\|hsa-mir-200b&WithFlank&1\|+\|1167098\|1167204@63.83.21&[MIMAT0000318&hsa-miR-200b-3p&offsets\|0\|-1;m-93&1\|+\|1167160\|1167182&offsets\|0\|-2]\|TAATACTGCCTGGTAATGATG | >MI0000342\|hsa-mir-200b&WithFlank&1\|+\|1167098\|1167204 | 63 | 83 | 21 | TAATACTGCCTGGTAATGATG | 384.4 | -2.1 | 0.8 | -2.5 | 1.1E-02 | 1.3E-02 | BT20/MB468 |
| >hg38_21_+_8433172_8446622_RNA45SN1_45S_50ntFlanks@7991.8013.23\|ACGTGGCGACCCGCTGAATTTAA | >hg38_21_+_8433172_8446622_RNA45SN1_45S_50ntFlanks | 7991 | 8013 | 23 | ACGTGGCGACCCGCTGAATTTAA | 362.0 | 1.3 | 0.5 | 2.5 | 1.1E-02 | 1.3E-02 | MB231/MB468 |

<table>
<tr><td>>hg38_21_+_8433172_8446622_RNA45SN1_45S_50ntFlanks@10686.10712.27|GCGGCGTCCGGTGAGCTCTCGCTGGCC</td><td>>hg38_21_+_8433172_8446622_RNA45SN1_45S_50ntFlanks</td><td>10686</td><td>10712</td><td>27</td><td>GCGGCGTCCGGTGAGCTCTCGCTGGCC</td><td>231.6</td><td>-2.2</td><td>0.9</td><td>-2.5</td><td>1.2E-02</td><td>1.3E-02</td><td>BT20/MB468</td></tr>
<tr><td>>hg38_21_+_8433172_8446622_RNA45SN1_45S_50ntFlanks@6661.6679.19|CGGTGGATCACTCGGCTCG</td><td>>hg38_21_+_8433172_8446622_RNA45SN1_45S_50ntFlanks</td><td>6661</td><td>6679</td><td>19</td><td>CGGTGGATCACTCGGCTCG</td><td>214.7</td><td>2.1</td><td>0.7</td><td>3.0</td><td>2.6E-03</td><td>1.3E-02</td><td>BT20/MB231</td></tr>
<tr><td>>hg38_21_+_8433172_8446622_RNA45SN1_45S_50ntFlanks@12415.12432.18|CTTCGATGTCGGCTCTTC</td><td>>hg38_21_+_8433172_8446622_RNA45SN1_45S_50ntFlanks</td><td>12415</td><td>12432</td><td>18</td><td>CTTCGATGTCGGCTCTTC</td><td>2925.3</td><td>-1.5</td><td>0.6</td><td>-2.5</td><td>1.1E-02</td><td>1.3E-02</td><td>MB231/MB468</td></tr>
<tr><td>>hg38_21_+_8433172_8446622_RNA45SN1_45S_50ntFlanks@12417.12444.28|TCGATGTCGGCTCTTCCTATCATTGTGA</td><td>>hg38_21_+_8433172_8446622_RNA45SN1_45S_50ntFlanks</td><td>12417</td><td>12444</td><td>28</td><td>TCGATGTCGGCTCTTCCTATCATTGTGA</td><td>643.9</td><td>-1.2</td><td>0.5</td><td>-2.5</td><td>1.2E-02</td><td>1.3E-02</td><td>BT20/MB468</td></tr>
<tr><td>>hg38_21_+_8433172_8446622_RNA45SN1_45S_50ntFlanks@10353.10390.38|AGAACTTTGAAGGCCGAAGTGGAGAAGGGTTCCATGTG</td><td>>hg38_21_+_8433172_8446622_RNA45SN1_45S_50ntFlanks</td><td>10353</td><td>10390</td><td>38</td><td>AGAACTTTGAAGGCCGAAGTGGAGAAGGGTTCCATGTG</td><td>222.5</td><td>-1.3</td><td>0.5</td><td>-2.5</td><td>1.2E-02</td><td>1.3E-02</td><td>BT20/MB468</td></tr>
<tr><td>>hg38_21_+_8433172_8446622_RNA45SN1_45S_50ntFlanks@6722.6742.21|ATTGCAGGACACATTGATCAT</td><td>>hg38_21_+_8433172_8446622_RNA45SN1_45S_50ntFlanks</td><td>6722</td><td>6742</td><td>21</td><td>ATTGCAGGACACATTGATCAT</td><td>109.7</td><td>-2.0</td><td>0.8</td><td>-2.5</td><td>1.1E-02</td><td>1.3E-02</td><td>MB231/MB468</td></tr>
<tr><td>>hg38_21_+_8433172_8446622_RNA45SN1_45S_50ntFlanks@8189.8213.25|CCCAGCCCGTGGACGGTGTGAGGCC</td><td>>hg38_21_+_8433172_8446622_RNA45SN1_45S_50ntFlanks</td><td>8189</td><td>8213</td><td>25</td><td>CCCAGCCCGTGGACGGTGTGAGGCC</td><td>179.1</td><td>-2.1</td><td>0.8</td><td>-2.5</td><td>1.2E-02</td><td>1.3E-02</td><td>BT20/MB468</td></tr>
<tr><td>>hg38_21_+_8433172_8446622_RNA45SN1_45S_50ntFlanks@11212.11233.22|CGACTCTGGACGCGAGCCGGGC</td><td>>hg38_21_+_8433172_8446622_RNA45SN1_45S_50ntFlanks</td><td>11212</td><td>11233</td><td>22</td><td>CGACTCTGGACGCGAGCCGGGC</td><td>101.5</td><td>-4.1</td><td>1.6</td><td>-2.5</td><td>1.1E-02</td><td>1.3E-02</td><td>MB231/MB468</td></tr>
<tr><td>>hg38_21_+_8433172_8446622_RNA45SN1_45S_50ntFlanks@6708.6731.24|GAGAATTAATGTGAATTGCAGGAC</td><td>>hg38_21_+_8433172_8446622_RNA45SN1_45S_50ntFlanks</td><td>6708</td><td>6731</td><td>24</td><td>GAGAATTAATGTGAATTGCAGGAC</td><td>289.1</td><td>-2.5</td><td>0.8</td><td>-3.0</td><td>2.7E-03</td><td>1.3E-02</td><td>BT20/MB231</td></tr>
<tr><td>>hg38_21_+_8433172_8446622_RNA45SN1_45S_50ntFlanks@12993.13039.47|CCCTCGCTGCGATCTATTGAAAGTCAGCCCTCGACACAAGGGTTTGT</td><td>>hg38_21_+_8433172_8446622_RNA45SN1_45S_50ntFlanks</td><td>12993</td><td>13039</td><td>47</td><td>CCCTCGCTGCGATCTATTGAAAGTCAGCCCTCGACACAAGGGTTTGT</td><td>3688.5</td><td>-2.7</td><td>0.9</td><td>-3.0</td><td>2.7E-03</td><td>1.3E-02</td><td>BT20/MB231</td></tr>
<tr><td>>hg38_21_+_8433172_8446622_RNA45SN1_45S_50ntFlanks@11577.11611.35|TTAGAACTGGTGCGGACCAGGGGAATCCGACTGTT</td><td>>hg38_21_+_8433172_8446622_RNA45SN1_45S_50ntFlanks</td><td>11577</td><td>11611</td><td>35</td><td>TTAGAACTGGTGCGGACCAGGGGAATCCGACTGTT</td><td>251.6</td><td>-1.3</td><td>0.5</td><td>-2.5</td><td>1.1E-02</td><td>1.3E-02</td><td>MB231/MB468</td></tr>
<tr><td>>hg38_21_+_8433172_8446622_RNA45SN1_45S_50ntFlanks@3840.3879.40|CTCCTACTTGGATAACTGTGGTAATTCTAGAGCTAATACA</td><td>>hg38_21_+_8433172_8446622_RNA45SN1_45S_50ntFlanks</td><td>3840</td><td>3879</td><td>40</td><td>CTCCTACTTGGATAACTGTGGTAATTCTAGAGCTAATACA</td><td>290.9</td><td>2.3</td><td>0.8</td><td>3.0</td><td>2.7E-03</td><td>1.3E-02</td><td>BT20/MB231</td></tr>
<tr><td>>MI0000063|hsa-let-7b&WithFlank&22|+|46113680|46113774@12.34.23&[MIMAT0000063&hsa-let-7b-5p&offsets|0|+1;m-14&22|+|46113691|46113712&offsets|0|+1]|TGAGGTAGTAGGTTGTGTGGTTT</td><td>>MI0000063|hsa-let-7b&WithFlank&22|+|46113680|46113774</td><td>12</td><td>34</td><td>23</td><td>TGAGGTAGTAGGTTGTGTGGTTT</td><td>7609.9</td><td>1.8</td><td>0.6</td><td>3.0</td><td>2.7E-03</td><td>1.3E-02</td><td>BT20/MB231</td></tr>
<tr><td>>hg38_21_+_8433172_8446622_RNA45SN1_45S_50ntFlanks@11565.11585.21|CTAGCAGCCGACTTAGAACTG</td><td>>hg38_21_+_8433172_8446622_RNA45SN1_45S_50ntFlanks</td><td>11565</td><td>11585</td><td>21</td><td>CTAGCAGCCGACTTAGAACTG</td><td>214.1</td><td>-1.1</td><td>0.4</td><td>-2.5</td><td>1.1E-02</td><td>1.4E-02</td><td>MB231/MB468</td></tr>
<tr><td>>hg38_21_+_8433172_8446622_RNA45SN1_45S_50ntFlanks@8157.8181.25|CTCGTGGGGGGCCCAAGTCCTTCTG</td><td>>hg38_21_+_8433172_8446622_RNA45SN1_45S_50ntFlanks</td><td>8157</td><td>8181</td><td>25</td><td>CTCGTGGGGGGCCCAAGTCCTTCTG</td><td>244.7</td><td>-1.6</td><td>0.6</td><td>-2.5</td><td>1.1E-02</td><td>1.4E-02</td><td>MB231/MB468</td></tr>
<tr><td>>hg38_21_+_8433172_8446622_RNA45SN1_45S_50ntFlanks@10362.10379.18|AAGGCCGAAGTGGAGAAG</td><td>>hg38_21_+_8433172_8446622_RNA45SN1_45S_50ntFlanks</td><td>10362</td><td>10379</td><td>18</td><td>AAGGCCGAAGTGGAGAAG</td><td>141.6</td><td>-1.8</td><td>0.7</td><td>-2.5</td><td>1.1E-02</td><td>1.4E-02</td><td>MB231/MB468</td></tr>
<tr><td>>am_trna162_MetCAT_6_-_26758550_26758622@31.54.24,trna164_MetCAT_6_-_26735574_26735646@31.54.24,trna20_MetCAT_16_+_71460396_71460468@31.54.24,trna21_MetCAT_6_+_26701712_26701784@31.54.24,trna27_MetCAT_6_+_26766444_26766516@31.54.24,trna75_MetCAT_6_+_28912352_28912424@31.54.24,trna97_MetCAT_6_-_28921042_28921114@31.54.24|TCTCATAATCTGAAGGTCCTGAGT</td><td>>am_trna162_MetCAT_6_-_26758550_26758622</td><td>31</td><td>54</td><td>24</td><td>TCTCATAATCTGAAGGTCCTGAGT</td><td>519.0</td><td>-1.8</td><td>0.7</td><td>-2.5</td><td>1.1E-02</td><td>1.4E-02</td><td>MB231/MB468</td></tr>
<tr><td>>hg38_21_+_8433172_8446622_RNA45SN1_45S_50ntFlanks@11679.11702.24|CTGAATGTCAAAGTGAAGAAATTC</td><td>>hg38_21_+_8433172_8446622_RNA45SN1_45S_50ntFlanks</td><td>11679</td><td>11702</td><td>24</td><td>CTGAATGTCAAAGTGAAGAAATTC</td><td>105.7</td><td>-1.3</td><td>0.5</td><td>-2.5</td><td>1.1E-02</td><td>1.4E-02</td><td>MB231/MB468</td></tr>
<tr><td>>hg38_21_+_8433172_8446622_RNA45SN1_45S_50ntFlanks@10466.10488.23|CGTTGCCCTCGGCCGATCGAAAG</td><td>>hg38_21_+_8433172_8446622_RNA45SN1_45S_50ntFlanks</td><td>10466</td><td>10488</td><td>23</td><td>CGTTGCCCTCGGCCGATCGAAAG</td><td>189.7</td><td>-2.2</td><td>0.9</td><td>-2.5</td><td>1.1E-02</td><td>1.4E-02</td><td>MB231/MB468</td></tr>
<tr><td>>hg38_21_+_8433172_8446622_RNA45SN1_45S_50ntFlanks@11670.11693.24|CCCAGTGCTCTGAATGTCAAAGTG</td><td>>hg38_21_+_8433172_8446622_RNA45SN1_45S_50ntFlanks</td><td>11670</td><td>11693</td><td>24</td><td>CCCAGTGCTCTGAATGTCAAAGTG</td><td>534.6</td><td>-1.9</td><td>0.7</td><td>-2.5</td><td>1.2E-02</td><td>1.4E-02</td><td>BT20/MB468</td></tr>
<tr><td>>hg38_21_+_8433172_8446622_RNA45SN1_45S_50ntFlanks@6646.6670.25|TCGTACGACTCTTAGCGGTGGATCA</td><td>>hg38_21_+_8433172_8446622_RNA45SN1_45S_50ntFlanks</td><td>6646</td><td>6670</td><td>25</td><td>TCGTACGACTCTTAGCGGTGGATCA</td><td>1045.8</td><td>2.4</td><td>0.8</td><td>3.0</td><td>2.8E-03</td><td>1.4E-02</td><td>BT20/MB231</td></tr>
<tr><td>>hg38_21_+_8433172_8446622_RNA45SN1_45S_50ntFlanks@7974.8005.32|ACGCGACCTCAGATCAGACGTGGCGACCCGCT</td><td>>hg38_21_+_8433172_8446622_RNA45SN1_45S_50ntFlanks</td><td>7974</td><td>8005</td><td>32</td><td>ACGCGACCTCAGATCAGACGTGGCGACCCGCT</td><td>4035.0</td><td>2.4</td><td>0.8</td><td>3.0</td><td>2.8E-03</td><td>1.4E-02</td><td>BT20/MB231</td></tr>
<tr><td>>hg38_21_+_8433172_8446622_RNA45SN1_45S_50ntFlanks@5492.5526.35|AGAAGACGGTCGAACTTGACTATCTAGAGGAAGTA</td><td>>hg38_21_+_8433172_8446622_RNA45SN1_45S_50ntFlanks</td><td>5492</td><td>5526</td><td>35</td><td>AGAAGACGGTCGAACTTGACTATCTAGAGGAAGTA</td><td>378.5</td><td>-1.8</td><td>0.7</td><td>-2.5</td><td>1.2E-02</td><td>1.4E-02</td><td>BT20/MB468</td></tr>
<tr><td>>MI0000077|hsa-mir-21&WithFlank&17|+|59841260|59841343@14.34.21(+3C)&[MIMAT0000076&hsa-miR-21-5p&offsets|0|-1(+3C);m-1&17|+|59841273|59841294&offsets|0|-1(+3C)]|TAGCTTATCAGACTGATGTTGCCC</td><td>>MI0000077|hsa-mir-21&WithFlank&17|+|59841260|59841343</td><td>14</td><td>34</td><td>21(+3C)</td><td>TAGCTTATCAGACTGATGTTGCCC</td><td>325.8</td><td>-2.4</td><td>0.8</td><td>-3.0</td><td>2.8E-03</td><td>1.4E-02</td><td>BT20/MB231</td></tr>
<tr><td>>trnaMT_ValTAC_MT_+_1602_1670@50.72.23|ACTTAACTTGACCGCTCTGACCA</td><td>>trnaMT_ValTAC_MT_+_1602_1670</td><td>50</td><td>72</td><td>23</td><td>ACTTAACTTGACCGCTCTGACCA</td><td>2687.1</td><td>2.2</td><td>0.7</td><td>3.0</td><td>2.8E-03</td><td>1.4E-02</td><td>BT20/MB231</td></tr>
<tr><td>>hg38_21_+_8433172_8446622_RNA45SN1_45S_50ntFlanks@11875.11893.19|GAAGACCCTGTTGAGCTTG</td><td>>hg38_21_+_8433172_8446622_RNA45SN1_45S_50ntFlanks</td><td>11875</td><td>11893</td><td>19</td><td>GAAGACCCTGTTGAGCTTG</td><td>834.6</td><td>-2.2</td><td>0.9</td><td>-2.5</td><td>1.2E-02</td><td>1.4E-02</td><td>BT20/MB468</td></tr>
<tr><td>>hg38_21_+_8433172_8446622_RNA45SN1_45S_50ntFlanks@6658.6681.24|TAGCGGTGGATCACTCGGCTCGTG</td><td>>hg38_21_+_8433172_8446622_RNA45SN1_45S_50ntFlanks</td><td>6658</td><td>6681</td><td>24</td><td>TAGCGGTGGATCACTCGGCTCGTG</td><td>148.3</td><td>-1.2</td><td>0.5</td><td>-2.5</td><td>1.2E-02</td><td>1.4E-02</td><td>MB231/MB468</td></tr>
<tr><td>>hg38_21_+_8433172_8446622_RNA45SN1_45S_50ntFlanks@12071.12094.24|CCCCGAGGGGCTCTCGCTTCTGGC</td><td>>hg38_21_+_8433172_8446622_RNA45SN1_45S_50ntFlanks</td><td>12071</td><td>12094</td><td>24</td><td>CCCCGAGGGGCTCTCGCTTCTGGC</td><td>226.0</td><td>-1.2</td><td>0.5</td><td>-2.5</td><td>1.2E-02</td><td>1.4E-02</td><td>MB231/MB468</td></tr>
<tr><td>>trna111_HisGTG_1_-_147774845_147774916@-1G.33.34,trna118_HisGTG_1_-_145396881_145396952@-1G.33.34,trna16_HisGTG_1_+_146544773_146544844@-1G.33.34,trna1_HisGTG_15_+_45493349_45493420@-1G.33.34,trna21_HisGTG_1_+_147753471_147753542@-1G.33.34,trna33_HisGTG_6_+_27125906_27125977@-1G.33.34,trna7_HisGTG_9_-_14433938_14434009@-1G.33.34,trna8_HisGTG_15_-_45492611_45492682@-1G.33.34,trna9_HisGTG_15_-_45490804_45490875@-1G.33.34|GGCCGTGATCGTATAGTGGTTAGTACTCTGCGTT</td><td>>trna111_HisGTG_1_-_147774845_147774916</td><td>-1G</td><td>33</td><td>34</td><td>GGCCGTGATCGTATAGTGGTTAGTACTCTGCGTT</td><td>27853.9</td><td>-3.1</td><td>1.0</td><td>-3.0</td><td>2.8E-03</td><td>1.4E-02</td><td>BT20/MB231</td></tr>
<tr><td>>hg38_21_+_8433172_8446622_RNA45SN1_45S_50ntFlanks@10351.10382.32|CGAGAACTTTGAAGGCCGAAGTGGAGAAGGGT</td><td>>hg38_21_+_8433172_8446622_RNA45SN1_45S_50ntFlanks</td><td>10351</td><td>10382</td><td>32</td><td>CGAGAACTTTGAAGGCCGAAGTGGAGAAGGGT</td><td>854.9</td><td>-1.3</td><td>0.5</td><td>-2.5</td><td>1.2E-02</td><td>1.4E-02</td><td>MB231/MB468</td></tr>
<tr><td>>hg38_21_+_8433172_8446622_RNA45SN1_45S_50ntFlanks@12008.12029.22|CCGCCGGTGAAATACCACTACT</td><td>>hg38_21_+_8433172_8446622_RNA45SN1_45S_50ntFlanks</td><td>12008</td><td>12029</td><td>22</td><td>CCGCCGGTGAAATACCACTACT</td><td>144.1</td><td>-4.2</td><td>1.7</td><td>-2.5</td><td>1.2E-02</td><td>1.4E-02</td><td>MB231/MB468</td></tr>
<tr><td>>hg38_21_+_8433172_8446622_RNA45SN1_45S_50ntFlanks@10775.10796.22|TCTCCAAGGTGAACAGCCTCTG</td><td>>hg38_21_+_8433172_8446622_RNA45SN1_45S_50ntFlanks</td><td>10775</td><td>10796</td><td>22</td><td>TCTCCAAGGTGAACAGCCTCTG</td><td>101.4</td><td>-1.7</td><td>0.7</td><td>-2.5</td><td>1.2E-02</td><td>1.4E-02</td><td>MB231/MB468</td></tr>
<tr><td>>trna11_GluTTC_15_-_26327381_26327452@1.27.27,trna3_GluTTC_13_-_45492062_45492133@1.27.27|TCCCACATGGTCTAGCGGTTAGGATTC</td><td>>trna11_GluTTC_15_-_26327381_26327452</td><td>1</td><td>27</td><td>27</td><td>TCCCACATGGTCTAGCGGTTAGGATTC</td><td>218.7</td><td>-2.0</td><td>0.8</td><td>-2.5</td><td>1.2E-02</td><td>1.4E-02</td><td>MB231/MB468</td></tr>
<tr><td>>hg38_21_+_8433172_8446622_RNA45SN1_45S_50ntFlanks@6664.6686.23|TGGATCACTCGGCTCGTGCGTCG</td><td>>hg38_21_+_8433172_8446622_RNA45SN1_45S_50ntFlanks</td><td>6664</td><td>6686</td><td>23</td><td>TGGATCACTCGGCTCGTGCGTCG</td><td>211.6</td><td>-2.2</td><td>0.9</td><td>-2.5</td><td>1.3E-02</td><td>1.4E-02</td><td>BT20/MB468</td></tr>
<tr><td>>hg38_21_+_8433172_8446622_RNA45SN1_45S_50ntFlanks@6728.6752.25|GGACACATTGATCATCGACACTTCG</td><td>>hg38_21_+_8433172_8446622_RNA45SN1_45S_50ntFlanks</td><td>6728</td><td>6752</td><td>25</td><td>GGACACATTGATCATCGACACTTCG</td><td>148.9</td><td>-3.1</td><td>1.0</td><td>-3.0</td><td>2.9E-03</td><td>1.4E-02</td><td>BT20/MB231</td></tr>
<tr><td>>hg38_21_+_8433172_8446622_RNA45SN1_45S_50ntFlanks@10846.10867.22|CTTCGGGATAAGGATTGGCTCT</td><td>>hg38_21_+_8433172_8446622_RNA45SN1_45S_50ntFlanks</td><td>10846</td><td>10867</td><td>22</td><td>CTTCGGGATAAGGATTGGCTCT</td><td>265.2</td><td>-1.5</td><td>0.6</td><td>-2.5</td><td>1.2E-02</td><td>1.4E-02</td><td>MB231/MB468</td></tr>
<tr><td>>am_trnaMT_ValTAC_MT_+_1602_1670@28.46.19|AACTTACACTTAGGAGATT</td><td>>am_trnaMT_ValTAC_MT_+_1602_1670</td><td>28</td><td>46</td><td>19</td><td>AACTTACACTTAGGAGATT</td><td>218.2</td><td>2.8</td><td>0.9</td><td>3.0</td><td>2.9E-03</td><td>1.4E-02</td><td>BT20/MB231</td></tr>
<tr><td>>hg38_21_+_8433172_8446622_RNA45SN1_45S_50ntFlanks@3843.3869.27|CTACTTGGATAACTGTGGTAATTCTAG</td><td>>hg38_21_+_8433172_8446622_RNA45SN1_45S_50ntFlanks</td><td>3843</td><td>3869</td><td>27</td><td>CTACTTGGATAACTGTGGTAATTCTAG</td><td>149.7</td><td>-2.5</td><td>1.0</td><td>-2.5</td><td>1.3E-02</td><td>1.4E-02</td><td>BT20/MB468</td></tr>
</table>

| | | | | | | | | | | | | |
|---|---|---|---|---|---|---|---|---|---|---|---|---|
| >am_trna11_ProAGG_16_+_3241989_3242060@1.27.27,trna12_ProAGG_6_+_26555498_26555569@1.27.27,trna12_ProTGG_11_-_75946869_75946940@1.27.27,trna14_ProTGG_5_-_180615854_180615925@1.27.27,trna22_ProAGG_14_-_21081560_21081631@1.27.27,trna23_ProAGG_14_-_21077495_21077566@1.27.27,trna28_ProTGG_16_-_3234133_3234204@1.27.27,trna29_ProAGG_16_-_3232635_3232706@1.27.27,trna2_ProAGG_7_+_128423504_128423575@1.27.27,trna30_ProCGG_6_+_27059521_27059592@1.27.27,trna37_ProCGG_17_-_8126151_8126222@1.27.27,trna3_ProTGG_16_+_3208923_3208994@1.27.27,trna52_ProCGG_1_+_167683962_167684033@1.27.27,trna65_ProAGG_1_-_167684725_167684796@1.27.27,trna6_ProCGG_16_+_3222049_3222120@1.27.27,trna6_ProTGG_14_+_21152175_21152246@1.27.27,trna8_ProTGG_16_+_3238094_3238165@1.27.27,trna9_ProAGG_11_+_75946557_75946628@1.27.27,trna9_ProAGG_16_+_3239634_3239705@1.27.27\|GGCTCGTTGGTCTAGGGGTATGATTCT | >am_trna11_ProAGG_16_+_3241989_3242060 | 1 | 27 | 27 | GGCTCGTTGGTCTAGGGGTATGATTCT | 127.1 | -2.5 | 1.0 | -2.5 | 1.3E-02 | 1.4E-02 | BT20/MB468 |
| >hg38_21_+_8433172_8446622_RNA45SN1_45S_50ntFlanks@3705.3742.38\|TACCTGGTTGATCCTGCCAGTAGCATATGCTTGTCTCA | >hg38_21_+_8433172_8446622_RNA45SN1_45S_50ntFlanks | 3705 | 3742 | 38 | TACCTGGTTGATCCTGCCAGTAGCATATGCTTGTCTCA | 2193.7 | 2.4 | 0.8 | 3.0 | 2.9E-03 | 1.5E-02 | BT20/MB231 |
| >hg38_21_+_8433172_8446622_RNA45SN1_45S_50ntFlanks@12075.12095.21\|GAGGGGCTCTCGCTTCTGGCG | >hg38_21_+_8433172_8446622_RNA45SN1_45S_50ntFlanks | 12075 | 12095 | 21 | GAGGGGCTCTCGCTTCTGGCG | 180.2 | -2.0 | 0.8 | -2.5 | 1.3E-02 | 1.5E-02 | BT20/MB468 |
| >hg38_21_+_8433172_8446622_RNA45SN1_45S_50ntFlanks@12455.12477.23\|ACCAAGCGTTGGATTGTTCACCC | >hg38_21_+_8433172_8446622_RNA45SN1_45S_50ntFlanks | 12455 | 12477 | 23 | ACCAAGCGTTGGATTGTTCACCC | 545.5 | 1.8 | 0.7 | 2.5 | 1.2E-02 | 1.5E-02 | MB231/MB468 |
| >hg38_21_+_8433172_8446622_RNA45SN1_45S_50ntFlanks@3996.4024.29\|ACTCTAGATAACCTCGGGCCGATCGCACG | >hg38_21_+_8433172_8446622_RNA45SN1_45S_50ntFlanks | 3996 | 4024 | 29 | ACTCTAGATAACCTCGGGCCGATCGCACG | 121.6 | -1.8 | 0.7 | -2.5 | 1.2E-02 | 1.5E-02 | MB231/MB468 |
| >hg38_21_+_8433172_8446622_RNA45SN1_45S_50ntFlanks@12418.12439.22\|CGATGTCGGCTCTTCCTATCAT | >hg38_21_+_8433172_8446622_RNA45SN1_45S_50ntFlanks | 12418 | 12439 | 22 | CGATGTCGGCTCTTCCTATCAT | 989.5 | 2.2 | 0.9 | 2.5 | 1.3E-02 | 1.5E-02 | BT20/MB468 |
| >hg38_21_+_8433172_8446622_RNA45SN1_45S_50ntFlanks@7988.8010.23\|CAGACGTGGCGACCCGCTGAATT | >hg38_21_+_8433172_8446622_RNA45SN1_45S_50ntFlanks | 7988 | 8010 | 23 | CAGACGTGGCGACCCGCTGAATT | 418.4 | 1.5 | 0.6 | 2.5 | 1.2E-02 | 1.5E-02 | MB231/MB468 |
| >hg38_21_+_8433172_8446622_RNA45SN1_45S_50ntFlanks@12416.12446.31\|TTCGATGTCGGCTCTTCCTATCATTGTGAAG | >hg38_21_+_8433172_8446622_RNA45SN1_45S_50ntFlanks | 12416 | 12446 | 31 | TTCGATGTCGGCTCTTCCTATCATTGTGAAG | 1287.6 | -3.0 | 1.0 | -3.0 | 3.0E-03 | 1.5E-02 | BT20/MB231 |
| >MI0000447\|hsa-mir-128-1&WithFlank&2\|+\|135665391\|135665484@56.75.20(+1G);MI0000727\|hsa-mir-128-2&WithFlank&3\|+\|35744470\|35744565@58.77.20(+1G)&[MIMAT0000424&hsa-miR-128-3p&offsets\|0\|-1(+1G);m-108&2\|+\|135665446\|135665466&offsets\|0\|-1(+1G)];[MIMAT0000424_1&hsa-miR-128-3p&offsets\|0\|-1(+1G);m-117&3\|+\|35744527\|35744547&offsets\|0\|-1(+1G)]\|TCACAGTGAACCGGTCTCTTG | >MI0000447\|hsa-mir-128-1&WithFlank&2\|+\|135665391\|135665484 | 56 | 75 | 20(+1G) | TCACAGTGAACCGGTCTCTTG | 376.2 | 2.3 | 0.8 | 3.0 | 3.0E-03 | 1.5E-02 | BT20/MB231 |
| >hg38_21_+_8433172_8446622_RNA45SN1_45S_50ntFlanks@6702.6720.19\|AGCTGCGAGAATTAATGTG | >hg38_21_+_8433172_8446622_RNA45SN1_45S_50ntFlanks | 6702 | 6720 | 19 | AGCTGCGAGAATTAATGTG | 300.7 | -2.6 | 0.9 | -3.0 | 3.0E-03 | 1.5E-02 | BT20/MB231 |
| >hg38_21_+_8433172_8446622_RNA45SN1_45S_50ntFlanks@6646.6671.26\|TCGTACGACTCTTAGCGGTGGATCAC | >hg38_21_+_8433172_8446622_RNA45SN1_45S_50ntFlanks | 6646 | 6671 | 26 | TCGTACGACTCTTAGCGGTGGATCAC | 347.5 | 2.1 | 0.7 | 3.0 | 3.0E-03 | 1.5E-02 | BT20/MB231 |
| >MI0000483\|hsa-mir-186&WithFlank&1\|-\|71067625\|71067722@21.41.21(+1A)&[MIMAT0000456&hsa-miR-186-5p&offsets\|0\|-1(+1A);m-84&1\|-\|71067681\|71067702&offsets\|0\|-1(+1A)]\|CAAAGAATTCTCCTTTTGGGCA | >MI0000483\|hsa-mir-186&WithFlank&1\|-\|71067625\|71067722 | 21 | 41 | 21(+1A) | CAAAGAATTCTCCTTTTGGGCA | 540.9 | 2.4 | 0.8 | 3.0 | 3.0E-03 | 1.5E-02 | BT20/MB231 |
| >MI0000065\|hsa-let-7d&WithFlank&9\|+\|94178828\|94178926@14.34.21(+1A)&[MIMAT0000065&hsa-let-7d-5p&offsets\|0\|-1(+1A);m-90&9\|+\|94178841\|94178862&offsets\|0\|-1(+1A)]\|AGAGGTAGTAGGTTGCATAGTA | >MI0000065\|hsa-let-7d&WithFlank&9\|+\|94178828\|94178926 | 14 | 34 | 21(+1A) | AGAGGTAGTAGGTTGCATAGTA | 649.2 | 2.5 | 0.9 | 3.0 | 3.0E-03 | 1.5E-02 | BT20/MB231 |
| >MI0000298\|hsa-mir-221&WithFlank&X\|-\|45746151\|45746272@31.52.22&[MIMAT0004568&hsa-miR-221-5p&offsets\|0\|0;m-405&X\|-\|45746220\|45746242&offsets\|0\|-1]\|ACCTGGCATACAATGTAGATTT | >MI0000298\|hsa-mir-221&WithFlank&X\|-\|45746151\|45746272 | 31 | 52 | 22 | ACCTGGCATACAATGTAGATTT | 254.8 | -2.2 | 0.8 | -3.0 | 3.0E-03 | 1.5E-02 | BT20/MB231 |
| >hg38_21_+_8433172_8446622_RNA45SN1_45S_50ntFlanks@11208.11252.45\|GCGGCGACTCTGGACGCGAGCCGGGCCCTTCCCGTGGATCGCCCC | >hg38_21_+_8433172_8446622_RNA45SN1_45S_50ntFlanks | 11208 | 11252 | 45 | GCGGCGACTCTGGACGCGAGCCGGGCCCTTCCCGTGGATCGCCCC | 98.9 | -2.3 | 0.9 | -2.5 | 1.2E-02 | 1.5E-02 | MB231/MB468 |
| >hg38_21_+_8433172_8446622_RNA45SN1_45S_50ntFlanks@12539.12568.30\|ATGATGTGTTGTTGCCATGGTAATCCTGCT | >hg38_21_+_8433172_8446622_RNA45SN1_45S_50ntFlanks | 12539 | 12568 | 30 | ATGATGTGTTGTTGCCATGGTAATCCTGCT | 93.4 | -4.0 | 1.6 | -2.5 | 1.2E-02 | 1.5E-02 | MB231/MB468 |
| >hg38_21_+_8433172_8446622_RNA45SN1_45S_50ntFlanks@11734.11751.18\|ATGACTCTCTTAAGGTAG | >hg38_21_+_8433172_8446622_RNA45SN1_45S_50ntFlanks | 11734 | 11751 | 18 | ATGACTCTCTTAAGGTAG | 118.8 | -1.4 | 0.6 | -2.5 | 1.2E-02 | 1.5E-02 | MB231/MB468 |
| >hg38_21_+_8433172_8446622_RNA45SN1_45S_50ntFlanks@12420.12449.30\|ATGTCGGCTCTTCCTATCATTGTGAAGCAG | >hg38_21_+_8433172_8446622_RNA45SN1_45S_50ntFlanks | 12420 | 12449 | 30 | ATGTCGGCTCTTCCTATCATTGTGAAGCAG | 513.5 | -2.6 | 0.9 | -3.0 | 3.1E-03 | 1.5E-02 | BT20/MB231 |
| >hg38_21_+_8433172_8446622_RNA45SN1_45S_50ntFlanks@11577.11602.26\|TTAGAACTGGTGCGGACCAGGGGAAT | >hg38_21_+_8433172_8446622_RNA45SN1_45S_50ntFlanks | 11577 | 11602 | 26 | TTAGAACTGGTGCGGACCAGGGGAAT | 217.8 | -2.0 | 0.8 | -2.5 | 1.3E-02 | 1.5E-02 | BT20/MB468 |
| >hg38_21_+_8433172_8446622_RNA45SN1_45S_50ntFlanks@11564.11587.24\|CCTAGCAGCCGACTTAGAACTGGT | >hg38_21_+_8433172_8446622_RNA45SN1_45S_50ntFlanks | 11564 | 11587 | 24 | CCTAGCAGCCGACTTAGAACTGGT | 209.3 | -1.0 | 0.4 | -2.5 | 1.3E-02 | 1.5E-02 | MB231/MB468 |
| >hg38_21_+_8433172_8446622_RNA45SN1_45S_50ntFlanks@6646.6664.19\|TCGTACGACTCTTAGCGGT | >hg38_21_+_8433172_8446622_RNA45SN1_45S_50ntFlanks | 6646 | 6664 | 19 | TCGTACGACTCTTAGCGGT | 24093.0 | 2.0 | 0.7 | 3.0 | 3.1E-03 | 1.5E-02 | BT20/MB231 |
| >hg38_21_+_8433172_8446622_RNA45SN1_45S_50ntFlanks@482.505.24\|ATCGATGTGGTGACGTCGTGCTCT | >hg38_21_+_8433172_8446622_RNA45SN1_45S_50ntFlanks | 482 | 505 | 24 | ATCGATGTGGTGACGTCGTGCTCT | 8705.8 | 6.1 | 2.4 | 2.5 | 1.3E-02 | 1.5E-02 | MB231/MB468 |
| >hg38_21_+_8433172_8446622_RNA45SN1_45S_50ntFlanks@9739.9760.22\|AAACGATCTCAACCTATTCTCA | >hg38_21_+_8433172_8446622_RNA45SN1_45S_50ntFlanks | 9739 | 9760 | 22 | AAACGATCTCAACCTATTCTCA | 99.2 | -1.8 | 0.7 | -2.5 | 1.4E-02 | 1.5E-02 | BT20/MB468 |
| >hg38_21_+_8433172_8446622_RNA45SN1_45S_50ntFlanks@10352.10369.18\|GAGAACTTTGAAGGCCGA | >hg38_21_+_8433172_8446622_RNA45SN1_45S_50ntFlanks | 10352 | 10369 | 18 | GAGAACTTTGAAGGCCGA | 371.4 | -2.4 | 0.8 | -3.0 | 3.2E-03 | 1.5E-02 | BT20/MB231 |
| >hg38_21_+_8433172_8446622_RNA45SN1_45S_50ntFlanks@12415.12446.32\|CTTCGATGTCGGCTCTTCCTATCATTGTGAAG | >hg38_21_+_8433172_8446622_RNA45SN1_45S_50ntFlanks | 12415 | 12446 | 32 | CTTCGATGTCGGCTCTTCCTATCATTGTGAAG | 1026.1 | 1.8 | 0.7 | 2.5 | 1.3E-02 | 1.5E-02 | MB231/MB468 |
| >hg38_21_+_8433172_8446622_RNA45SN1_45S_50ntFlanks@6648.6681.34\|GTACGACTCTTAGCGGTGGATCACTCGGCTCGTG | >hg38_21_+_8433172_8446622_RNA45SN1_45S_50ntFlanks | 6648 | 6681 | 34 | GTACGACTCTTAGCGGTGGATCACTCGGCTCGTG | 5764.8 | 1.6 | 0.7 | 2.5 | 1.3E-02 | 1.5E-02 | MB231/MB468 |
| >hg38_21_+_8433172_8446622_RNA45SN1_45S_50ntFlanks@11208.11227.20\|GCGGCGACTCTGGACGCGAG | >hg38_21_+_8433172_8446622_RNA45SN1_45S_50ntFlanks | 11208 | 11227 | 20 | GCGGCGACTCTGGACGCGAG | 1256.0 | -2.9 | 1.2 | -2.5 | 1.4E-02 | 1.5E-02 | BT20/MB468 |
| >hg38_21_+_8433172_8446622_RNA45SN1_45S_50ntFlanks@4727.4759.33\|AATCAAGAACGAAAGTCGGAGGTTCGAAGACGA | >hg38_21_+_8433172_8446622_RNA45SN1_45S_50ntFlanks | 4727 | 4759 | 33 | AATCAAGAACGAAAGTCGGAGGTTCGAAGACGA | 292.2 | 1.8 | 0.7 | 2.5 | 1.3E-02 | 1.5E-02 | MB231/MB468 |
| >hg38_21_+_8433172_8446622_RNA45SN1_45S_50ntFlanks@6697.6720.24\|CAGCTAGCTGCGAGAATTAATGTG | >hg38_21_+_8433172_8446622_RNA45SN1_45S_50ntFlanks | 6697 | 6720 | 24 | CAGCTAGCTGCGAGAATTAATGTG | 805.8 | -1.1 | 0.5 | -2.5 | 1.4E-02 | 1.5E-02 | BT20/MB468 |
| >hg38_21_+_8433172_8446622_RNA45SN1_45S_50ntFlanks@6646.6668.23\|TCGTACGACTCTTAGCGGTGGAT | >hg38_21_+_8433172_8446622_RNA45SN1_45S_50ntFlanks | 6646 | 6668 | 23 | TCGTACGACTCTTAGCGGTGGAT | 9131.5 | 2.0 | 0.7 | 2.9 | 3.2E-03 | 1.5E-02 | BT20/MB231 |
| >hg38_21_+_8433172_8446622_RNA45SN1_45S_50ntFlanks@12417.12440.24\|TCGATGTCGGCTCTTCCTATCATT | >hg38_21_+_8433172_8446622_RNA45SN1_45S_50ntFlanks | 12417 | 12440 | 24 | TCGATGTCGGCTCTTCCTATCATT | 369.0 | 1.1 | 0.5 | 2.5 | 1.4E-02 | 1.5E-02 | BT20/MB468 |
| >hg38_21_+_8433172_8446622_RNA45SN1_45S_50ntFlanks@3831.3860.30\|CTCGCTCCTCTCCTACTTGGATAACTGTGG | >hg38_21_+_8433172_8446622_RNA45SN1_45S_50ntFlanks | 3831 | 3860 | 30 | CTCGCTCCTCTCCTACTTGGATAACTGTGG | 189.8 | -1.6 | 0.6 | -2.5 | 1.3E-02 | 1.6E-02 | MB231/MB468 |
| >hg38_21_+_8433172_8446622_RNA45SN1_45S_50ntFlanks@11209.11230.22\|CGGCGACTCTGGACGCGAGCCG | >hg38_21_+_8433172_8446622_RNA45SN1_45S_50ntFlanks | 11209 | 11230 | 22 | CGGCGACTCTGGACGCGAGCCG | 734.7 | -2.3 | 0.9 | -2.5 | 1.4E-02 | 1.6E-02 | BT20/MB468 |
| >MI0000441\|hsa-mir-30b&WithFlank&8\|-\|134800514\|134800613@23.43.21(+1A)&[MIMAT0000420&hsa-miR-30b-5p&offsets\|0\|-1(+1A);m-76&8\|-\|134800570\|134800591&offsets\|0\|-1(+1A)]\|TGTAAACATCCTACACTCAGCA | >MI0000441\|hsa-mir-30b&WithFlank&8\|-\|134800514\|134800613 | 23 | 43 | 21(+1A) | TGTAAACATCCTACACTCAGCA | 4353.7 | 3.1 | 1.1 | 2.9 | 3.2E-03 | 1.6E-02 | BT20/MB231 |
| >hg38_21_+_8433172_8446622_RNA45SN1_45S_50ntFlanks@9554.9581.28\|CGTAGCGGTCCTGACGTGCAAATCGGTC | >hg38_21_+_8433172_8446622_RNA45SN1_45S_50ntFlanks | 9554 | 9581 | 28 | CGTAGCGGTCCTGACGTGCAAATCGGTC | 522.4 | -2.7 | 0.9 | -2.9 | 3.2E-03 | 1.6E-02 | BT20/MB231 |
| >hg38_21_+_8433172_8446622_RNA45SN1_45S_50ntFlanks@11583.11600.18\|CTGGTGCGGACCAGGGGA | >hg38_21_+_8433172_8446622_RNA45SN1_45S_50ntFlanks | 11583 | 11600 | 18 | CTGGTGCGGACCAGGGGA | 363.4 | -1.4 | 0.6 | -2.5 | 1.3E-02 | 1.6E-02 | MB231/MB468 |
| >MI0000342\|hsa-mir-200b&WithFlank&1\|+\|1167098\|1167204@63.85.23(+1C)&[MIMAT0000318&hsa-miR-200b-3p&offsets\|0\|+1(+1C);m-93&1\|+\|1167160\|1167182&offsets\|0\|0(+1C)]\|TAATACTGCCTGGTAATGATGACC | >MI0000342\|hsa-mir-200b&WithFlank&1\|+\|1167098\|1167204 | 63 | 85 | 23(+1C) | TAATACTGCCTGGTAATGATGACC | 253.0 | -1.9 | 0.8 | -2.5 | 1.4E-02 | 1.6E-02 | BT20/MB468 |
| >hg38_21_+_8433172_8446622_RNA45SN1_45S_50ntFlanks@12419.12446.28\|GATGTCGGCTCTTCCTATCATTGTGAAG | >hg38_21_+_8433172_8446622_RNA45SN1_45S_50ntFlanks | 12419 | 12446 | 28 | GATGTCGGCTCTTCCTATCATTGTGAAG | 674.4 | -2.7 | 0.9 | -2.9 | 3.3E-03 | 1.6E-02 | BT20/MB231 |
| >hg38_21_+_8433172_8446622_RNA45SN1_45S_50ntFlanks@12423.12457.35\|TCGGCTCTTCCTATCATTGTGAAGCAGAATTCACC | >hg38_21_+_8433172_8446622_RNA45SN1_45S_50ntFlanks | 12423 | 12457 | 35 | TCGGCTCTTCCTATCATTGTGAAGCAGAATTCACC | 279.4 | -2.5 | 0.8 | -2.9 | 3.3E-03 | 1.6E-02 | BT20/MB231 |
| >hg38_21_+_8433172_8446622_RNA45SN1_45S_50ntFlanks@8317.8356.40\|CACGAGACCGATAGTCAACAAGTACCGTAAGGGAAAGTTG | >hg38_21_+_8433172_8446622_RNA45SN1_45S_50ntFlanks | 8317 | 8356 | 40 | CACGAGACCGATAGTCAACAAGTACCGTAAGGGAAAGTTG | 123.4 | -1.8 | 0.7 | -2.5 | 1.4E-02 | 1.6E-02 | BT20/MB468 |

<table>
<tr><td>>MI0000299|hsa-mir-222&WithFlank&X|-|45747009|45747130@75.100.26&[MIMAT0000279&hsa-miR-222-3p&offsets|0|+5;m-140&X|-|45747033|45747056&offsets|0|+2]|AGCTACATCTGGCTACTGGGTCTCTG</td><td>>MI0000299|hsa-mir-222&WithFlank&X|-|45747009|45747130</td><td>75</td><td>100</td><td>26</td><td>AGCTACATCTGGCTACTGGGTCTCTG</td><td>246.2</td><td>-2.3</td><td>0.8</td><td>-2.9</td><td>3.3E-03</td><td>1.6E-02</td><td>BT20/MB231</td></tr>
<tr><td>>hg38_21_+_8433172_8446622_RNA45SN1_45S_50ntFlanks@12415.12443.29|CTTCGATGTCGGCTCTTCCTATCATTGTG</td><td>>hg38_21_+_8433172_8446622_RNA45SN1_45S_50ntFlanks</td><td>12415</td><td>12443</td><td>29</td><td>CTTCGATGTCGGCTCTTCCTATCATTGTG</td><td>1502.0</td><td>-1.4</td><td>0.6</td><td>-2.5</td><td>1.4E-02</td><td>1.6E-02</td><td>BT20/MB468</td></tr>
<tr><td>>MI0000064|hsa-let-7c&WithFlank&21|+|16539822|16539917@17.38.22&[MIMAT0000064&hsa-let-7c-5p&offsets|0|0;m-37&21|+|16539838|16539859&offsets|0|0]|TGAGGTAGTAGGTTGTATGGTT</td><td>>MI0000064|hsa-let-7c&WithFlank&21|+|16539822|16539917</td><td>17</td><td>38</td><td>22</td><td>TGAGGTAGTAGGTTGTATGGTT</td><td>802.5</td><td>1.7</td><td>0.7</td><td>2.5</td><td>1.4E-02</td><td>1.6E-02</td><td>BT20/MB468</td></tr>
<tr><td>>MI0000542|hsa-mir-320a&WithFlank&8|-|22244960|22245043@48.69.22(+1C)&[MIMAT0000510&hsa-miR-320a-3p&offsets|0|0(+1C);m-60&8|-|22244975|22244996&offsets|0|0(+1C)]|AAAAGCTGGGTTGAGAGGGCGAC</td><td>>MI0000542|hsa-mir-320a&WithFlank&8|-|22244960|22245043</td><td>48</td><td>69</td><td>22(+1C)</td><td>AAAAGCTGGGTTGAGAGGGCGAC</td><td>1394.2</td><td>2.4</td><td>0.8</td><td>2.9</td><td>3.3E-03</td><td>1.6E-02</td><td>BT20/MB231</td></tr>
<tr><td>>hg38_21_+_8433172_8446622_RNA45SN1_45S_50ntFlanks@12482.12499.18|ATAGGGAACGTGAGCTGG</td><td>>hg38_21_+_8433172_8446622_RNA45SN1_45S_50ntFlanks</td><td>12482</td><td>12499</td><td>18</td><td>ATAGGGAACGTGAGCTGG</td><td>776.6</td><td>3.2</td><td>1.3</td><td>2.5</td><td>1.4E-02</td><td>1.6E-02</td><td>BT20/MB468</td></tr>
<tr><td>>MI0000299|hsa-mir-222&WithFlank&X|-|45747009|45747130@75.99.25(+1A)&[MIMAT0000279&hsa-miR-222-3p&offsets|0|+4(+1A);m-140&X|-|45747033|45747056&offsets|0|+1(+1A)]|AGCTACATCTGGCTACTGGGTCTCTA</td><td>>MI0000299|hsa-mir-222&WithFlank&X|-|45747009|45747130</td><td>75</td><td>99</td><td>25(+1A)</td><td>AGCTACATCTGGCTACTGGGTCTCTA</td><td>502.1</td><td>-2.1</td><td>0.7</td><td>-2.9</td><td>3.3E-03</td><td>1.6E-02</td><td>BT20/MB231</td></tr>
<tr><td>>hg38_21_+_8433172_8446622_RNA45SN1_45S_50ntFlanks@12453.12476.24|TCACCAAGCGTTGGATTGTTCACC</td><td>>hg38_21_+_8433172_8446622_RNA45SN1_45S_50ntFlanks</td><td>12453</td><td>12476</td><td>24</td><td>TCACCAAGCGTTGGATTGTTCACC</td><td>413.4</td><td>-2.3</td><td>0.8</td><td>-2.9</td><td>3.3E-03</td><td>1.6E-02</td><td>BT20/MB231</td></tr>
<tr><td>>hg38_21_+_8433172_8446622_RNA45SN1_45S_50ntFlanks@12457.12486.30|CAAGCGTTGGATTGTTCACCCACTAATAGG</td><td>>hg38_21_+_8433172_8446622_RNA45SN1_45S_50ntFlanks</td><td>12457</td><td>12486</td><td>30</td><td>CAAGCGTTGGATTGTTCACCCACTAATAGG</td><td>483.4</td><td>-1.6</td><td>0.6</td><td>-2.5</td><td>1.3E-02</td><td>1.6E-02</td><td>MB231/MB468</td></tr>
<tr><td>>hg38_21_+_8433172_8446622_RNA45SN1_45S_50ntFlanks@12420.12454.35|ATGTCGGCTCTTCCTATCATTGTGAAGCAGAATTC</td><td>>hg38_21_+_8433172_8446622_RNA45SN1_45S_50ntFlanks</td><td>12420</td><td>12454</td><td>35</td><td>ATGTCGGCTCTTCCTATCATTGTGAAGCAGAATTC</td><td>219.4</td><td>-2.3</td><td>0.8</td><td>-2.9</td><td>3.3E-03</td><td>1.6E-02</td><td>BT20/MB231</td></tr>
<tr><td>>hg38_21_+_8433172_8446622_RNA45SN1_45S_50ntFlanks@12478.12500.23|ACTAATAGGGAACGTGAGCTGGG</td><td>>hg38_21_+_8433172_8446622_RNA45SN1_45S_50ntFlanks</td><td>12478</td><td>12500</td><td>23</td><td>ACTAATAGGGAACGTGAGCTGGG</td><td>157.5</td><td>-1.2</td><td>0.5</td><td>-2.5</td><td>1.4E-02</td><td>1.6E-02</td><td>BT20/MB468</td></tr>
<tr><td>>MI0000077|hsa-mir-21&WithFlank&17|+|59841260|59841343@14.33.20&[MIMAT0000076&hsa-miR-21-5p&offsets|0|-2;m-1&17|+|59841273|59841294&offsets|0|-2]|TAGCTTATCAGACTGATGTT</td><td>>MI0000077|hsa-mir-21&WithFlank&17|+|59841260|59841343</td><td>14</td><td>33</td><td>20</td><td>TAGCTTATCAGACTGATGTT</td><td>2690.5</td><td>2.0</td><td>0.7</td><td>2.9</td><td>3.3E-03</td><td>1.6E-02</td><td>BT20/MB231</td></tr>
<tr><td>>hg38_21_+_8433172_8446622_RNA45SN1_45S_50ntFlanks@8114.8138.25|GCGCGGGACATGTGGCGTACGGAAG</td><td>>hg38_21_+_8433172_8446622_RNA45SN1_45S_50ntFlanks</td><td>8114</td><td>8138</td><td>25</td><td>GCGCGGGACATGTGGCGTACGGAAG</td><td>346.9</td><td>-4.7</td><td>1.9</td><td>-2.5</td><td>1.3E-02</td><td>1.6E-02</td><td>MB231/MB468</td></tr>
<tr><td>>hg38_21_+_8433172_8446622_RNA45SN1_45S_50ntFlanks@9554.9580.27|CGTAGCGGTCCTGACGTGCAAATCGGT</td><td>>hg38_21_+_8433172_8446622_RNA45SN1_45S_50ntFlanks</td><td>9554</td><td>9580</td><td>27</td><td>CGTAGCGGTCCTGACGTGCAAATCGGT</td><td>442.3</td><td>1.1</td><td>0.5</td><td>2.5</td><td>1.3E-02</td><td>1.6E-02</td><td>MB231/MB468</td></tr>
<tr><td>>hg38_21_+_8433172_8446622_RNA45SN1_45S_50ntFlanks@10349.10368.20|AACGAGAACTTTGAAGGCCG</td><td>>hg38_21_+_8433172_8446622_RNA45SN1_45S_50ntFlanks</td><td>10349</td><td>10368</td><td>20</td><td>AACGAGAACTTTGAAGGCCG</td><td>3341.5</td><td>-1.8</td><td>0.7</td><td>-2.5</td><td>1.3E-02</td><td>1.6E-02</td><td>MB231/MB468</td></tr>
<tr><td>>hg38_21_+_8433172_8446622_RNA45SN1_45S_50ntFlanks@11567.11586.20|AGCAGCCGACTTAGAACTGG</td><td>>hg38_21_+_8433172_8446622_RNA45SN1_45S_50ntFlanks</td><td>11567</td><td>11586</td><td>20</td><td>AGCAGCCGACTTAGAACTGG</td><td>1004.3</td><td>1.1</td><td>0.5</td><td>2.4</td><td>1.4E-02</td><td>1.6E-02</td><td>BT20/MB468</td></tr>
<tr><td>>am_trnaMT_ValTAC_MT_+_1602_1670@38.57.20|TAGGAGATTTCAACTTAACT</td><td>>am_trnaMT_ValTAC_MT_+_1602_1670</td><td>38</td><td>57</td><td>20</td><td>TAGGAGATTTCAACTTAACT</td><td>394.8</td><td>2.2</td><td>0.7</td><td>2.9</td><td>3.4E-03</td><td>1.6E-02</td><td>BT20/MB231</td></tr>
<tr><td>>hg38_21_+_8433172_8446622_RNA45SN1_45S_50ntFlanks@12418.12452.35|CGATGTCGGCTCTTCCTATCATTGTGAAGCAGAAT</td><td>>hg38_21_+_8433172_8446622_RNA45SN1_45S_50ntFlanks</td><td>12418</td><td>12452</td><td>35</td><td>CGATGTCGGCTCTTCCTATCATTGTGAAGCAGAAT</td><td>255.7</td><td>-2.2</td><td>0.7</td><td>-2.9</td><td>3.4E-03</td><td>1.6E-02</td><td>BT20/MB231</td></tr>
<tr><td>>MI0000063|hsa-let-7b&WithFlank&22|+|46113680|46113774@12.33.22(+2A)&[MIMAT0000063&hsa-let-7b-5p&offsets|0|0(+2A);m-14&22|+|46113691|46113712&offsets|0|0(+2A)]|TGAGGTAGTAGGTTGTGTGGTTAA</td><td>>MI0000063|hsa-let-7b&WithFlank&22|+|46113680|46113774</td><td>12</td><td>33</td><td>22(+2A)</td><td>TGAGGTAGTAGGTTGTGTGGTTAA</td><td>304.6</td><td>2.2</td><td>0.8</td><td>2.9</td><td>3.4E-03</td><td>1.6E-02</td><td>BT20/MB231</td></tr>
<tr><td>>hg38_21_+_8433172_8446622_RNA45SN1_45S_50ntFlanks@11208.11239.32|GCGGCGACTCTGGACGCGAGCCGGGCCCTTCC</td><td>>hg38_21_+_8433172_8446622_RNA45SN1_45S_50ntFlanks</td><td>11208</td><td>11239</td><td>32</td><td>GCGGCGACTCTGGACGCGAGCCGGGCCCTTCC</td><td>133.5</td><td>-4.1</td><td>1.7</td><td>-2.5</td><td>1.4E-02</td><td>1.6E-02</td><td>MB231/MB468</td></tr>
<tr><td>>hg38_21_+_8433172_8446622_RNA45SN1_45S_50ntFlanks@12988.13013.26|CAGCTCCCTCGCTGCGATCTATTGAA</td><td>>hg38_21_+_8433172_8446622_RNA45SN1_45S_50ntFlanks</td><td>12988</td><td>13013</td><td>26</td><td>CAGCTCCCTCGCTGCGATCTATTGAA</td><td>290.5</td><td>-2.4</td><td>0.8</td><td>-2.9</td><td>3.4E-03</td><td>1.6E-02</td><td>BT20/MB231</td></tr>
<tr><td>>hg38_21_+_8433172_8446622_RNA45SN1_45S_50ntFlanks@12416.12447.32|TTCGATGTCGGCTCTTCCTATCATTGTGAAGC</td><td>>hg38_21_+_8433172_8446622_RNA45SN1_45S_50ntFlanks</td><td>12416</td><td>12447</td><td>32</td><td>TTCGATGTCGGCTCTTCCTATCATTGTGAAGC</td><td>1290.2</td><td>-2.7</td><td>0.9</td><td>-2.9</td><td>3.4E-03</td><td>1.6E-02</td><td>BT20/MB231</td></tr>
<tr><td>>hg38_21_+_8433172_8446622_RNA45SN1_45S_50ntFlanks@6646.6669.24|TCGTACGACTCTTAGCGGTGGATC</td><td>>hg38_21_+_8433172_8446622_RNA45SN1_45S_50ntFlanks</td><td>6646</td><td>6669</td><td>24</td><td>TCGTACGACTCTTAGCGGTGGATC</td><td>20516.0</td><td>2.4</td><td>0.8</td><td>2.9</td><td>3.5E-03</td><td>1.6E-02</td><td>BT20/MB231</td></tr>
<tr><td>>hg38_21_+_8433172_8446622_RNA45SN1_45S_50ntFlanks@10687.10708.22|CGGCGTCCGGTGAGCTCTCGCT</td><td>>hg38_21_+_8433172_8446622_RNA45SN1_45S_50ntFlanks</td><td>10687</td><td>10708</td><td>22</td><td>CGGCGTCCGGTGAGCTCTCGCT</td><td>1535.6</td><td>-2.2</td><td>0.9</td><td>-2.4</td><td>1.5E-02</td><td>1.6E-02</td><td>BT20/MB468</td></tr>
<tr><td>>hg38_21_+_8433172_8446622_RNA45SN1_45S_50ntFlanks@11582.11602.21|ACTGGTGCGGACCAGGGGAAT</td><td>>hg38_21_+_8433172_8446622_RNA45SN1_45S_50ntFlanks</td><td>11582</td><td>11602</td><td>21</td><td>ACTGGTGCGGACCAGGGGAAT</td><td>910.9</td><td>2.0</td><td>0.8</td><td>2.4</td><td>1.5E-02</td><td>1.6E-02</td><td>BT20/MB468</td></tr>
<tr><td>>MI0000466|hsa-mir-9-1&WithFlank&1|-|156420335|156420435@22.42.21;MI0000468|hsa-mir-9-3&WithFlank&15|+|89368011|89368112@22.42.21;MI0000467|hsa-mir-9-2&WithFlank&5|-|88666847|88666945@22.42.21&[MIMAT0000441&hsa-miR-9-5p&offsets|0|-2;m-104&1|-|156420392|156420414&offsets|0|-2];[MIMAT0000441_1&hsa-miR-9-5p&offsets|0|-2;m-105&15|+|89368032|89368054&offsets|0|-2];[MIMAT0000441_2&hsa-miR-9-5p&offsets|0|-2;m-104&5|-|88666902|88666924&offsets|0|-2]|TCTTTGGTTATCTAGCTGTAT</td><td>>MI0000466|hsa-mir-9-1&WithFlank&1|-|156420335|156420435</td><td>22</td><td>42</td><td>21</td><td>TCTTTGGTTATCTAGCTGTAT</td><td>284.1</td><td>2.2</td><td>0.8</td><td>2.9</td><td>3.5E-03</td><td>1.6E-02</td><td>BT20/MB231</td></tr>
<tr><td>>MI0000255|hsa-mir-30d&WithFlank&8|-|134804870|134804951@12.33.22(+1G)&[MIMAT0000245&hsa-miR-30d-5p&offsets|0|0(+1G);m-21&8|-|134804919|134804940&offsets|0|0(+1G)]|TGTAAACATCCCCGACTGGAAGG</td><td>>MI0000255|hsa-mir-30d&WithFlank&8|-|134804870|134804951</td><td>12</td><td>33</td><td>22(+1G)</td><td>TGTAAACATCCCCGACTGGAAGG</td><td>674.6</td><td>2.6</td><td>0.9</td><td>2.9</td><td>3.5E-03</td><td>1.6E-02</td><td>BT20/MB231</td></tr>
<tr><td>>am_trnaMT_ValTAC_MT_+_1602_1670@48.68.21|CAACTTAACTTGACCGCTCTG</td><td>>am_trnaMT_ValTAC_MT_+_1602_1670</td><td>48</td><td>68</td><td>21</td><td>CAACTTAACTTGACCGCTCTG</td><td>303.3</td><td>1.9</td><td>0.8</td><td>2.5</td><td>1.4E-02</td><td>1.6E-02</td><td>MB231/MB468</td></tr>
<tr><td>>hg38_21_+_8433172_8446622_RNA45SN1_45S_50ntFlanks@11766.11790.25|ATCTAATTAGTGACGCGCATGAATG</td><td>>hg38_21_+_8433172_8446622_RNA45SN1_45S_50ntFlanks</td><td>11766</td><td>11790</td><td>25</td><td>ATCTAATTAGTGACGCGCATGAATG</td><td>187.0</td><td>-3.8</td><td>1.5</td><td>-2.5</td><td>1.4E-02</td><td>1.6E-02</td><td>MB231/MB468</td></tr>
<tr><td>>hg38_21_+_8433172_8446622_RNA45SN1_45S_50ntFlanks@10796.10816.21|GGCATGTTGGAACAATGTAGG</td><td>>hg38_21_+_8433172_8446622_RNA45SN1_45S_50ntFlanks</td><td>10796</td><td>10816</td><td>21</td><td>GGCATGTTGGAACAATGTAGG</td><td>100.3</td><td>-2.4</td><td>1.0</td><td>-2.4</td><td>1.5E-02</td><td>1.6E-02</td><td>BT20/MB468</td></tr>
<tr><td>>MI0000289|hsa-mir-181a-1&WithFlank&1|-|198859038|198859159@30.54.25;MI0000269|hsa-mir-181a-2&WithFlank&9|+|124692436|124692557@45.69.25&[MIMAT0000256&hsa-miR-181a-5p&offsets|0|+2;m-32&1|-|198859108|198859130&offsets|0|+2];[MIMAT0000256_1&hsa-miR-181a-5p&offsets|0|+2;m-32&9|+|124692480|124692502&offsets|0|+2]|AACATTCAACGCTGTCGGTGAGTTT</td><td>>MI0000289|hsa-mir-181a-1&WithFlank&1|-|198859038|198859159</td><td>30</td><td>54</td><td>25</td><td>AACATTCAACGCTGTCGGTGAGTTT</td><td>1850.9</td><td>-2.3</td><td>0.8</td><td>-2.9</td><td>3.5E-03</td><td>1.7E-02</td><td>BT20/MB231</td></tr>
<tr><td>>hg38_21_+_8433172_8446622_RNA45SN1_45S_50ntFlanks@11571.11612.42|GCCGACTTAGAACTGGTGCGGACCAGGGGAATCCGACTGTTT</td><td>>hg38_21_+_8433172_8446622_RNA45SN1_45S_50ntFlanks</td><td>11571</td><td>11612</td><td>42</td><td>GCCGACTTAGAACTGGTGCGGACCAGGGGAATCCGACTGTTT</td><td>159.2</td><td>-2.2</td><td>0.8</td><td>-2.9</td><td>3.5E-03</td><td>1.7E-02</td><td>BT20/MB231</td></tr>
<tr><td>>hg38_21_+_8433172_8446622_RNA45SN1_45S_50ntFlanks@12626.12667.42|AATGGGGCGAAGCTACCATCTGTGGGATTATGACTGAACGCC</td><td>>hg38_21_+_8433172_8446622_RNA45SN1_45S_50ntFlanks</td><td>12626</td><td>12667</td><td>42</td><td>AATGGGGCGAAGCTACCATCTGTGGGATTATGACTGAACGCC</td><td>982.1</td><td>-2.5</td><td>0.9</td><td>-2.9</td><td>3.5E-03</td><td>1.7E-02</td><td>BT20/MB231</td></tr>
<tr><td>>hg38_21_+_8433172_8446622_RNA45SN1_45S_50ntFlanks@11572.11612.41|CCGACTTAGAACTGGTGCGGACCAGGGGAATCCGACTGTTT</td><td>>hg38_21_+_8433172_8446622_RNA45SN1_45S_50ntFlanks</td><td>11572</td><td>11612</td><td>41</td><td>CCGACTTAGAACTGGTGCGGACCAGGGGAATCCGACTGTTT</td><td>591.3</td><td>-2.3</td><td>1.0</td><td>-2.4</td><td>1.5E-02</td><td>1.7E-02</td><td>BT20/MB468</td></tr>
<tr><td>>hg38_21_+_8433172_8446622_RNA45SN1_45S_50ntFlanks@11540.11582.43|CGCGGCGCCTCGCCTCGGCCGGCGCCTAGCAGCCGACTTAGAA</td><td>>hg38_21_+_8433172_8446622_RNA45SN1_45S_50ntFlanks</td><td>11540</td><td>11582</td><td>43</td><td>CGCGGCGCCTCGCCTCGGCCGGCGCCTAGCAGCCGACTTAGAA</td><td>518.6</td><td>-2.8</td><td>0.9</td><td>-2.9</td><td>3.6E-03</td><td>1.7E-02</td><td>BT20/MB231</td></tr>
<tr><td>>am_trnaMT_ValTAC_MT_+_1602_1670@50.68.19|ACTTAACTTGACCGCTCTG</td><td>>am_trnaMT_ValTAC_MT_+_1602_1670</td><td>50</td><td>68</td><td>19</td><td>ACTTAACTTGACCGCTCTG</td><td>274.9</td><td>2.4</td><td>0.8</td><td>2.9</td><td>3.6E-03</td><td>1.7E-02</td><td>BT20/MB231</td></tr>
<tr><td>>MI0000811|hsa-mir-148b&WithFlank&12|+|54337210|54337320@69.90.22(+3U)&[MIMAT0000759&hsa-miR-148b-3p&offsets|0|0(+3U);m-77&12|+|54337278|54337299&offsets|0|0(+3U)]|TCAGTGCATCACAGAACTTTGTTTT</td><td>>MI0000811|hsa-mir-148b&WithFlank&12|+|54337210|54337320</td><td>69</td><td>90</td><td>22(+3U)</td><td>TCAGTGCATCACAGAACTTTGTTTT</td><td>383.9</td><td>-2.0</td><td>0.7</td><td>-2.9</td><td>3.6E-03</td><td>1.7E-02</td><td>BT20/MB231</td></tr>
<tr><td>>hg38_21_+_8433172_8446622_RNA45SN1_45S_50ntFlanks@12458.12477.20|AAGCGTTGGATTGTTCACCC</td><td>>hg38_21_+_8433172_8446622_RNA45SN1_45S_50ntFlanks</td><td>12458</td><td>12477</td><td>20</td><td>AAGCGTTGGATTGTTCACCC</td><td>985.1</td><td>-1.3</td><td>0.5</td><td>-2.4</td><td>1.5E-02</td><td>1.7E-02</td><td>BT20/MB468</td></tr>
<tr><td>>hg38_21_+_8433172_8446622_RNA45SN1_45S_50ntFlanks@12416.12452.37|TTCGATGTCGGCTCTTCCTATCATTGTGAAGCAGAAT</td><td>>hg38_21_+_8433172_8446622_RNA45SN1_45S_50ntFlanks</td><td>12416</td><td>12452</td><td>37</td><td>TTCGATGTCGGCTCTTCCTATCATTGTGAAGCAGAAT</td><td>668.2</td><td>-2.4</td><td>0.8</td><td>-2.9</td><td>3.6E-03</td><td>1.7E-02</td><td>BT20/MB231</td></tr>
<tr><td>>hg38_21_+_8433172_8446622_RNA45SN1_45S_50ntFlanks@5444.5466.23|CTCGGATCGGCCCCGCCGGGGTC</td><td>>hg38_21_+_8433172_8446622_RNA45SN1_45S_50ntFlanks</td><td>5444</td><td>5466</td><td>23</td><td>CTCGGATCGGCCCCGCCGGGGTC</td><td>8725.8</td><td>-3.0</td><td>1.2</td><td>-2.4</td><td>1.5E-02</td><td>1.7E-02</td><td>BT20/MB468</td></tr>
<tr><td>>hg38_21_+_8433172_8446622_RNA45SN1_45S_50ntFlanks@12420.12452.33|ATGTCGGCTCTTCCTATCATTGTGAAGCAGAAT</td><td>>hg38_21_+_8433172_8446622_RNA45SN1_45S_50ntFlanks</td><td>12420</td><td>12452</td><td>33</td><td>ATGTCGGCTCTTCCTATCATTGTGAAGCAGAAT</td><td>332.5</td><td>-2.3</td><td>0.8</td><td>-2.9</td><td>3.7E-03</td><td>1.7E-02</td><td>BT20/MB231</td></tr>
<tr><td>>hg38_21_+_8433172_8446622_RNA45SN1_45S_50ntFlanks@11675.11709.35|TGCTCTGAATGTCAAAGTGAAGAAATTCAATGAAG</td><td>>hg38_21_+_8433172_8446622_RNA45SN1_45S_50ntFlanks</td><td>11675</td><td>11709</td><td>35</td><td>TGCTCTGAATGTCAAAGTGAAGAAATTCAATGAAG</td><td>351.8</td><td>-2.5</td><td>0.9</td><td>-2.9</td><td>3.7E-03</td><td>1.7E-02</td><td>BT20/MB231</td></tr>
<tr><td>>hg38_21_+_8433172_8446622_RNA45SN1_45S_50ntFlanks@11670.11704.35|CCCAGTGCTCTGAATGTCAAAGTGAAGAAATTCAA</td><td>>hg38_21_+_8433172_8446622_RNA45SN1_45S_50ntFlanks</td><td>11670</td><td>11704</td><td>35</td><td>CCCAGTGCTCTGAATGTCAAAGTGAAGAAATTCAA</td><td>149.3</td><td>-1.2</td><td>0.5</td><td>-2.5</td><td>1.4E-02</td><td>1.7E-02</td><td>MB231/MB468</td></tr>
</table>

| | | | | | | | | | | | | |
|---|---|---|---|---|---|---|---|---|---|---|---|---|
| >hg38_21_+_8433172_8446622_RNA45SN1_45S_50ntFlanks@6652.6699.48\|GACTCTTAGCGGTGGATCACTCGGCTCGTGCGTCGATGAAGAACGCAG | >hg38_21_+_8433172_8446622_RNA45SN1_45S_50ntFlanks | 6652 | 6699 | 48 | GACTCTTAGCGGTGGATCACTCGGCTCGTGCGTCGATGAAGAACGCAG | 1232.0 | -2.6 | 0.9 | -2.9 | 3.7E-03 | 1.7E-02 | BT20/MB231 |
| >hg38_21_+_8433172_8446622_RNA45SN1_45S_50ntFlanks@5484.5509.26\|GAGCGCTGAGAAGACGGTCGAACTTG | >hg38_21_+_8433172_8446622_RNA45SN1_45S_50ntFlanks | 5484 | 5509 | 26 | GAGCGCTGAGAAGACGGTCGAACTTG | 94.9 | -1.4 | 0.6 | -2.5 | 1.4E-02 | 1.7E-02 | MB231/MB468 |
| >hg38_21_+_8433172_8446622_RNA45SN1_45S_50ntFlanks@11772.11794.23\|TTAGTGACGCGCATGAATGGATG | >hg38_21_+_8433172_8446622_RNA45SN1_45S_50ntFlanks | 11772 | 11794 | 23 | TTAGTGACGCGCATGAATGGATG | 422.0 | 1.2 | 0.5 | 2.5 | 1.4E-02 | 1.7E-02 | MB231/MB468 |
| >hg38_21_+_8433172_8446622_RNA45SN1_45S_50ntFlanks@6722.6743.22\|ATTGCAGGACACATTGATCATC | >hg38_21_+_8433172_8446622_RNA45SN1_45S_50ntFlanks | 6722 | 6743 | 22 | ATTGCAGGACACATTGATCATC | 1039.7 | -1.6 | 0.7 | -2.4 | 1.4E-02 | 1.7E-02 | MB231/MB468 |
| >hg38_21_+_8433172_8446622_RNA45SN1_45S_50ntFlanks@6649.6679.31\|TACGACTCTTAGCGGTGGATCACTCGGCTCG | >hg38_21_+_8433172_8446622_RNA45SN1_45S_50ntFlanks | 6649 | 6679 | 31 | TACGACTCTTAGCGGTGGATCACTCGGCTCG | 855.1 | 1.6 | 0.7 | 2.4 | 1.4E-02 | 1.7E-02 | MB231/MB468 |
| >hg38_21_+_8433172_8446622_RNA45SN1_45S_50ntFlanks@12479.12500.22\|CTAATAGGGAACGTGAGCTGGG | >hg38_21_+_8433172_8446622_RNA45SN1_45S_50ntFlanks | 12479 | 12500 | 22 | CTAATAGGGAACGTGAGCTGGG | 308.7 | -1.2 | 0.5 | -2.4 | 1.4E-02 | 1.7E-02 | MB231/MB468 |
| >hg38_21_+_8433172_8446622_RNA45SN1_45S_50ntFlanks@9555.9573.19\|GTAGCGGTCCTGACGTGCA | >hg38_21_+_8433172_8446622_RNA45SN1_45S_50ntFlanks | 9555 | 9573 | 19 | GTAGCGGTCCTGACGTGCA | 174.6 | -2.6 | 0.9 | -2.9 | 3.7E-03 | 1.7E-02 | BT20/MB231 |
| >hg38_21_+_8433172_8446622_RNA45SN1_45S_50ntFlanks@12479.12500.22\|CTAATAGGGAACGTGAGCTGGG | >hg38_21_+_8433172_8446622_RNA45SN1_45S_50ntFlanks | 12479 | 12500 | 22 | CTAATAGGGAACGTGAGCTGGG | 347.7 | -1.5 | 0.6 | -2.4 | 1.5E-02 | 1.7E-02 | BT20/MB468 |
| >trna152_ValCAC_6_-_27248049_27248121@1.36.36\|GCTTCTGTAGTGTAGTGGTTATCACGTTCGCCTCAC | >trna152_ValCAC_6_-_27248049_27248121 | 1 | 36 | 36 | GCTTCTGTAGTGTAGTGGTTATCACGTTCGCCTCAC | 435.5 | -2.5 | 0.9 | -2.9 | 3.7E-03 | 1.7E-02 | BT20/MB231 |
| >hg38_21_+_8433172_8446622_RNA45SN1_45S_50ntFlanks@12992.13012.21\|TCCCTCGCTGCGATCTATTGA | >hg38_21_+_8433172_8446622_RNA45SN1_45S_50ntFlanks | 12992 | 13012 | 21 | TCCCTCGCTGCGATCTATTGA | 298.0 | -1.0 | 0.4 | -2.4 | 1.4E-02 | 1.7E-02 | MB231/MB468 |
| >hg38_21_+_8433172_8446622_RNA45SN1_45S_50ntFlanks@9565.9597.33\|TGACGTGCAAATCGGTCGTCCGACCTGGGTATA | >hg38_21_+_8433172_8446622_RNA45SN1_45S_50ntFlanks | 9565 | 9597 | 33 | TGACGTGCAAATCGGTCGTCCGACCTGGGTATA | 141.2 | -2.8 | 1.0 | -2.9 | 3.8E-03 | 1.7E-02 | BT20/MB231 |
| >hg38_21_+_8433172_8446622_RNA45SN1_45S_50ntFlanks@5469.5499.31\|CCCACGGCCCTGGCGGAGCGCTGAGAAGACG | >hg38_21_+_8433172_8446622_RNA45SN1_45S_50ntFlanks | 5469 | 5499 | 31 | CCCACGGCCCTGGCGGAGCGCTGAGAAGACG | 189.8 | -2.2 | 0.9 | -2.4 | 1.5E-02 | 1.7E-02 | BT20/MB468 |
| >hg38_21_+_8433172_8446622_RNA45SN1_45S_50ntFlanks@10357.10390.34\|CTTTGAAGGCCGAAGTGGAGAAGGGTTCCATGTG | >hg38_21_+_8433172_8446622_RNA45SN1_45S_50ntFlanks | 10357 | 10390 | 34 | CTTTGAAGGCCGAAGTGGAGAAGGGTTCCATGTG | 231.8 | -1.2 | 0.5 | -2.4 | 1.5E-02 | 1.8E-02 | MB231/MB468 |
| >hg38_21_+_8433172_8446622_RNA45SN1_45S_50ntFlanks@12416.12453.38\|TTCGATGTCGGCTCTTCCTATCATTGTGAAGCAGAATT | >hg38_21_+_8433172_8446622_RNA45SN1_45S_50ntFlanks | 12416 | 12453 | 38 | TTCGATGTCGGCTCTTCCTATCATTGTGAAGCAGAATT | 578.7 | -2.4 | 0.8 | -2.9 | 3.8E-03 | 1.8E-02 | BT20/MB231 |
| >trna116_GluCTC_1_-_145399233_145399304@39.74.36,trna134_GluTTC_1_-_16861774_16861845@39.74.36,trna71_GluCTC_1_-_161439189_161439260@39.74.36,trna74_GluCTC_1_-_161431809_161431880@39.74.36,trna77_GluCTC_1_-_161424398_161424469@39.74.36,trna77_GluCTC_6_+_28949976_28950047@39.74.36,trna80_GluCTC_1_-_161417018_161417089@39.74.36,trna84_GluTTC_1_-_161391883_161391954@39.74.36,trna87_GluCTC_6_-_126101393_126101464@39.74.36\|CGCCGCGGCCCGGGTTCGATTCCCGGTCAGGGAACC | >trna116_GluCTC_1_-_145399233_145399304 | 39 | 74 | 36 | CGCCGCGGCCCGGGTTCGATTCCCGGTCAGGGAACC | 242.8 | -2.6 | 0.9 | -2.9 | 3.8E-03 | 1.8E-02 | BT20/MB231 |
| >hg38_21_+_8433172_8446622_RNA45SN1_45S_50ntFlanks@12397.12416.20\|CGACGTCGCTTTTTGATCCT | >hg38_21_+_8433172_8446622_RNA45SN1_45S_50ntFlanks | 12397 | 12416 | 20 | CGACGTCGCTTTTTGATCCT | 165.7 | -1.3 | 0.5 | -2.4 | 1.6E-02 | 1.8E-02 | BT20/MB468 |
| >hg38_21_+_8433172_8446622_RNA45SN1_45S_50ntFlanks@11583.11602.20\|CTGGTGCGGACCAGGGGAAT | >hg38_21_+_8433172_8446622_RNA45SN1_45S_50ntFlanks | 11583 | 11602 | 20 | CTGGTGCGGACCAGGGGAAT | 563.7 | -1.8 | 0.8 | -2.4 | 1.5E-02 | 1.8E-02 | MB231/MB468 |
| >MI0000298\|hsa-mir-221&WithFlank&X\|-\|45746151\|45746272@31.53.23(+1C)&[MIMAT0004568&hsa-miR-221-5p&offsets\|0\|+1(+1C);m-405&X\|-\|45746220\|45746242&offsets\|0\|0(+1C)]\|ACCTGGCATACAATGTAGATTTCC | >MI0000298\|hsa-mir-221&WithFlank&X\|-\|45746151\|45746272 | 31 | 53 | 23(+1C) | ACCTGGCATACAATGTAGATTTCC | 345.8 | -2.1 | 0.7 | -2.9 | 3.9E-03 | 1.8E-02 | BT20/MB231 |
| >hg38_21_+_8433172_8446622_RNA45SN1_45S_50ntFlanks@8176.8214.39\|CTTCTGATCGAGGCCCAGCCCGTGGACGGTGTGAGGCCG | >hg38_21_+_8433172_8446622_RNA45SN1_45S_50ntFlanks | 8176 | 8214 | 39 | CTTCTGATCGAGGCCCAGCCCGTGGACGGTGTGAGGCCG | 410.0 | -1.8 | 0.8 | -2.4 | 1.6E-02 | 1.8E-02 | BT20/MB468 |
| >hg38_21_+_8433172_8446622_RNA45SN1_45S_50ntFlanks@9471.9503.33\|AGCACGAGCGCACGTGTTAGGACCCGAAAGATG | >hg38_21_+_8433172_8446622_RNA45SN1_45S_50ntFlanks | 9471 | 9503 | 33 | AGCACGAGCGCACGTGTTAGGACCCGAAAGATG | 72.9 | -2.1 | 0.8 | -2.4 | 1.5E-02 | 1.8E-02 | MB231/MB468 |
| >MI0000746\|hsa-mir-99b&WithFlank&19\|+\|51692606\|51692687@13.32.20(+1A)&[MIMAT0000689&hsa-miR-99b-5p&offsets\|0\|-2(+1A);m-10&19\|+\|51692618\|51692639&offsets\|0\|-2(+1A)]\|CACCCGTAGAACCGACCTTGA | >MI0000746\|hsa-mir-99b&WithFlank&19\|+\|51692606\|51692687 | 13 | 32 | 20(+1A) | CACCCGTAGAACCGACCTTGA | 1549.7 | 2.3 | 0.8 | 2.9 | 3.9E-03 | 1.8E-02 | BT20/MB231 |
| >MI0000746\|hsa-mir-99b&WithFlank&19\|+\|51692606\|51692687@51.70.20(+1U)&[MIMAT0004678&hsa-miR-99b-3p&offsets\|0\|-2(+1U);m-151&19\|+\|51692656\|51692678&offsets\|0\|-3(+1U)]\|CAAGCTCGTGTCTGTGGGTCT | >MI0000746\|hsa-mir-99b&WithFlank&19\|+\|51692606\|51692687 | 51 | 70 | 20(+1U) | CAAGCTCGTGTCTGTGGGTCT | 557.2 | 2.3 | 0.8 | 2.9 | 3.9E-03 | 1.8E-02 | BT20/MB231 |
| >trnaMT_MetCAT_MT_+_4402_4469@52.70.19\|ATACCCTTCCCGTACTACC | >trnaMT_MetCAT_MT_+_4402_4469 | 52 | 70 | 19 | ATACCCTTCCCGTACTACC | 509.6 | -2.9 | 1.0 | -2.9 | 4.0E-03 | 1.8E-02 | BT20/MB231 |
| >MI0000802\|hsa-mir-340&WithFlank&5\|-\|180015297\|180015403@22.43.22&[MIMAT0004692&hsa-miR-340-5p&offsets\|0\|0;m-220&5\|-\|180015361\|180015382&offsets\|0\|0]\|TTATAAAGCAATGAGACTGATT | >MI0000802\|hsa-mir-340&WithFlank&5\|-\|180015297\|180015403 | 22 | 43 | 22 | TTATAAAGCAATGAGACTGATT | 10578.7 | 2.5 | 0.9 | 2.9 | 4.0E-03 | 1.8E-02 | BT20/MB231 |
| >trnaMT_PheGAA_MT_+_577_647@39.57.19\|ATGTTTAGACGGGCTCACA | >trnaMT_PheGAA_MT_+_577_647 | 39 | 57 | 19 | ATGTTTAGACGGGCTCACA | 1700.5 | 1.9 | 0.7 | 2.9 | 4.0E-03 | 1.8E-02 | BT20/MB231 |
| >hg38_21_+_8433172_8446622_RNA45SN1_45S_50ntFlanks@4069.4101.33\|CTTTCGATGGTAGTCGCCGTGCCTACCATGGTG | >hg38_21_+_8433172_8446622_RNA45SN1_45S_50ntFlanks | 4069 | 4101 | 33 | CTTTCGATGGTAGTCGCCGTGCCTACCATGGTG | 190.3 | -1.3 | 0.5 | -2.4 | 1.5E-02 | 1.8E-02 | MB231/MB468 |
| >hg38_21_+_8433172_8446622_RNA45SN1_45S_50ntFlanks@9545.9570.26\|GTGGAGGTCCGTAGCGGTCCTGACGT | >hg38_21_+_8433172_8446622_RNA45SN1_45S_50ntFlanks | 9545 | 9570 | 26 | GTGGAGGTCCGTAGCGGTCCTGACGT | 631.6 | -1.3 | 0.5 | -2.4 | 1.5E-02 | 1.8E-02 | MB231/MB468 |
| >hg38_21_+_8433172_8446622_RNA45SN1_45S_50ntFlanks@12420.12446.27\|ATGTCGGCTCTTCCTATCATTGTGAAG | >hg38_21_+_8433172_8446622_RNA45SN1_45S_50ntFlanks | 12420 | 12446 | 27 | ATGTCGGCTCTTCCTATCATTGTGAAG | 447.6 | -2.7 | 0.9 | -2.9 | 4.0E-03 | 1.8E-02 | BT20/MB231 |
| >hg38_21_+_8433172_8446622_RNA45SN1_45S_50ntFlanks@11565.11588.24\|CTAGCAGCCGACTTAGAACTGGTG | >hg38_21_+_8433172_8446622_RNA45SN1_45S_50ntFlanks | 11565 | 11588 | 24 | CTAGCAGCCGACTTAGAACTGGTG | 169.0 | -1.4 | 0.6 | -2.4 | 1.6E-02 | 1.8E-02 | BT20/MB468 |
| >hg38_21_+_8433172_8446622_RNA45SN1_45S_50ntFlanks@11672.11702.31\|CAGTGCTCTGAATGTCAAAGTGAAGAAATTC | >hg38_21_+_8433172_8446622_RNA45SN1_45S_50ntFlanks | 11672 | 11702 | 31 | CAGTGCTCTGAATGTCAAAGTGAAGAAATTC | 1468.8 | 1.4 | 0.6 | 2.4 | 1.5E-02 | 1.8E-02 | MB231/MB468 |
| >hg38_21_+_8433172_8446622_RNA45SN1_45S_50ntFlanks@10800.10818.19\|TGTTGGAACAATGTAGGTA | >hg38_21_+_8433172_8446622_RNA45SN1_45S_50ntFlanks | 10800 | 10818 | 19 | TGTTGGAACAATGTAGGTA | 236.0 | -4.6 | 1.9 | -2.4 | 1.7E-02 | 1.9E-02 | BT20/MB468 |
| >hg38_21_+_8433172_8446622_RNA45SN1_45S_50ntFlanks@6663.6699.37\|GTGGATCACTCGGCTCGTGCGTCGATGAAGAACGCAG | >hg38_21_+_8433172_8446622_RNA45SN1_45S_50ntFlanks | 6663 | 6699 | 37 | GTGGATCACTCGGCTCGTGCGTCGATGAAGAACGCAG | 167.6 | -2.1 | 0.9 | -2.4 | 1.7E-02 | 1.9E-02 | BT20/MB468 |
| >hg38_21_+_8433172_8446622_RNA45SN1_45S_50ntFlanks@8352.8379.28\|AGTTGAAAAGAACTTTGAAGAGAGAGTT | >hg38_21_+_8433172_8446622_RNA45SN1_45S_50ntFlanks | 8352 | 8379 | 28 | AGTTGAAAAGAACTTTGAAGAGAGAGTT | 199.0 | -1.6 | 0.7 | -2.4 | 1.7E-02 | 1.9E-02 | BT20/MB468 |
| >hg38_21_+_8433172_8446622_RNA45SN1_45S_50ntFlanks@6648.6665.18\|GTACGACTCTTAGCGGTG | >hg38_21_+_8433172_8446622_RNA45SN1_45S_50ntFlanks | 6648 | 6665 | 18 | GTACGACTCTTAGCGGTG | 730.6 | 2.5 | 0.9 | 2.9 | 4.1E-03 | 1.9E-02 | BT20/MB231 |
| >hg38_21_+_8433172_8446622_RNA45SN1_45S_50ntFlanks@12397.12414.18\|CGACGTCGCTTTTTGATC | >hg38_21_+_8433172_8446622_RNA45SN1_45S_50ntFlanks | 12397 | 12414 | 18 | CGACGTCGCTTTTTGATC | 1154.9 | -1.5 | 0.6 | -2.4 | 1.7E-02 | 1.9E-02 | BT20/MB468 |
| >hg38_21_+_8433172_8446622_RNA45SN1_45S_50ntFlanks@7987.8008.22\|TCAGACGTGGCGACCCGCTGAA | >hg38_21_+_8433172_8446622_RNA45SN1_45S_50ntFlanks | 7987 | 8008 | 22 | TCAGACGTGGCGACCCGCTGAA | 89.6 | -2.8 | 1.1 | -2.4 | 1.7E-02 | 1.9E-02 | BT20/MB468 |
| >hg38_21_+_8433172_8446622_RNA45SN1_45S_50ntFlanks@10349.10390.42\|AACGAGAACTTTGAAGGCCGAAGTGGAGAAGGGTTCCATGTG | >hg38_21_+_8433172_8446622_RNA45SN1_45S_50ntFlanks | 10349 | 10390 | 42 | AACGAGAACTTTGAAGGCCGAAGTGGAGAAGGGTTCCATGTG | 359.5 | -2.5 | 0.9 | -2.9 | 4.1E-03 | 1.9E-02 | BT20/MB231 |
| >trnaMT_TrpTCA_MT_+_5512_5579@53.71.19,trnalookalike3_TrpTCA_1_+_566062_566129@53.71.19\|CAATACTTAATTTCTGCCA | >trnaMT_TrpTCA_MT_+_5512_5579 | 53 | 71 | 19 | CAATACTTAATTTCTGCCA | 384.7 | 1.6 | 0.7 | 2.4 | 1.6E-02 | 1.9E-02 | MB231/MB468 |
| >MI0000299\|hsa-mir-222&WithFlank&X\|-\|45747009\|45747130@75.97.23(+2U)&[MIMAT0000279&hsa-miR-222-3p&offsets\|0\|+2(+2U);m-140&X\|-\|45747033\|45747056&offsets\|0\|-1(+2U)]\|AGCTACATCTGGCTACTGGGTCTTT | >MI0000299\|hsa-mir-222&WithFlank&X\|-\|45747009\|45747130 | 75 | 97 | 23(+2U) | AGCTACATCTGGCTACTGGGTCTTT | 1438.9 | -2.2 | 0.8 | -2.9 | 4.1E-03 | 1.9E-02 | BT20/MB231 |
| >hg38_21_+_8433172_8446622_RNA45SN1_45S_50ntFlanks@10921.10939.19\|GCTGGACGAGGCGCCGCCG | >hg38_21_+_8433172_8446622_RNA45SN1_45S_50ntFlanks | 10921 | 10939 | 19 | GCTGGACGAGGCGCCGCCG | 143.7 | -4.2 | 1.7 | -2.4 | 1.6E-02 | 1.9E-02 | MB231/MB468 |
| >hg38_21_+_8433172_8446622_RNA45SN1_45S_50ntFlanks@4735.4768.34\|ACGAAAGTCGGAGGTTCGAAGACGATCAGATACC | >hg38_21_+_8433172_8446622_RNA45SN1_45S_50ntFlanks | 4735 | 4768 | 34 | ACGAAAGTCGGAGGTTCGAAGACGATCAGATACC | 400.3 | -2.9 | 1.0 | -2.9 | 4.2E-03 | 1.9E-02 | BT20/MB231 |
| >hg38_21_+_8433172_8446622_RNA45SN1_45S_50ntFlanks@12417.12447.31\|TCGATGTCGGCTCTTCCTATCATTGTGAAGC | >hg38_21_+_8433172_8446622_RNA45SN1_45S_50ntFlanks | 12417 | 12447 | 31 | TCGATGTCGGCTCTTCCTATCATTGTGAAGC | 328.8 | -2.4 | 0.8 | -2.9 | 4.2E-03 | 1.9E-02 | BT20/MB231 |
| >hg38_21_+_8433172_8446622_RNA45SN1_45S_50ntFlanks@12625.12667.43\|CAATGGGGCGAAGCTACCATCTGTGGGATTATGACTGAACGCC | >hg38_21_+_8433172_8446622_RNA45SN1_45S_50ntFlanks | 12625 | 12667 | 43 | CAATGGGGCGAAGCTACCATCTGTGGGATTATGACTGAACGCC | 450.8 | -2.5 | 0.9 | -2.9 | 4.2E-03 | 1.9E-02 | BT20/MB231 |

| | | | | | | | | | | | | |
|---|---|---|---|---|---|---|---|---|---|---|---|---|
| >trna111_HisGTG_1_-_147774845_147774916@1.35.35,trna118_HisGTG_1_-_145396881_145396952@1.35.35,trna16_HisGTG_1_+_146544773_146544844@1.35.35,trna1_HisGTG_15_+_45493349_45493420@1.35.35,trna21_HisGTG_1_+_147753471_147753542@1.35.35,trna33_HisGTG_6_+_27125906_27125977@1.35.35,trna7_HisGTG_9_-_14433938_14434009@1.35.35,trna8_HisGTG_15_-_45492611_45492682@1.35.35,trna9_HisGTG_15_-_45490804_45490875@1.35.35\|GCCGTGATCGTATAGTGGTTAGTACTCTGCGTTGT | >trna111_HisGTG_1_-_147774845_147774916 | 1 | 35 | 35 | GCCGTGATCGTATAGTGGTTAGTACTCTGCGTTGT | 1062.4 | 3.0 | 1.0 | 2.9 | 4.2E-03 | 1.9E-02 | BT20/MB231 |
| >hg38_21_+_8433172_8446622_RNA45SN1_45S_50ntFlanks@3850.3873.24\|GATAACTGTGGTAATTCTAGAGCT | >hg38_21_+_8433172_8446622_RNA45SN1_45S_50ntFlanks | 3850 | 3873 | 24 | GATAACTGTGGTAATTCTAGAGCT | 564.9 | 1.7 | 0.7 | 2.4 | 1.6E-02 | 1.9E-02 | MB231/MB468 |
| >hg38_21_+_8433172_8446622_RNA45SN1_45S_50ntFlanks@11670.11693.24\|CCCAGTGCTCTGAATGTCAAAGTG | >hg38_21_+_8433172_8446622_RNA45SN1_45S_50ntFlanks | 11670 | 11693 | 24 | CCCAGTGCTCTGAATGTCAAAGTG | 475.1 | -1.3 | 0.5 | -2.4 | 1.6E-02 | 1.9E-02 | MB231/MB468 |
| >hg38_21_+_8433172_8446622_RNA45SN1_45S_50ntFlanks@11541.11584.44\|GCGGCGCCTCGCCTCGGCCGGCGCCTAGCAGCCGACTTAGAACT | >hg38_21_+_8433172_8446622_RNA45SN1_45S_50ntFlanks | 11541 | 11584 | 44 | GCGGCGCCTCGCCTCGGCCGGCGCCTAGCAGCCGACTTAGAACT | 239.6 | -2.4 | 0.8 | -2.9 | 4.2E-03 | 1.9E-02 | BT20/MB231 |
| >trna111_HisGTG_1_-_147774845_147774916@-1T.26.27,trna118_HisGTG_1_-_145396881_145396952@-1T.26.27,trna16_HisGTG_1_+_146544773_146544844@-1T.26.27,trna1_HisGTG_15_+_45493349_45493420@-1T.26.27,trna21_HisGTG_1_+_147753471_147753542@-1T.26.27,trna33_HisGTG_6_+_27125906_27125977@-1T.26.27,trna7_HisGTG_9_-_14433938_14434009@-1T.26.27,trna8_HisGTG_15_-_45492611_45492682@-1T.26.27,trna9_HisGTG_15_-_45490804_45490875@-1T.26.27\|TGCCGTGATCGTATAGTGGTTAGTACT | >trna111_HisGTG_1_-_147774845_147774916 | -1T | 26 | 27 | TGCCGTGATCGTATAGTGGTTAGTACT | 5028.0 | -2.7 | 1.0 | -2.9 | 4.2E-03 | 1.9E-02 | BT20/MB231 |
| >MI0006273\|hsa-mir-1180&WithFlank&17\|-\|19344500\|19344580@47.67.21(+1C)&[MIMAT0005825&hsa-miR-1180-3p&offsets\|0\|-1(+1C);m-214&17\|-\|19344513\|19344534&offsets\|0\|-1(+1C)]\|TTTCCGGCTCGCGTGGGTGTGC | >MI0006273\|hsa-mir-1180&WithFlank&17\|-\|19344500\|19344580 | 47 | 67 | 21(+1C) | TTTCCGGCTCGCGTGGGTGTGC | 275.1 | 1.9 | 0.7 | 2.9 | 4.3E-03 | 1.9E-02 | BT20/MB231 |
| >hg38_21_+_8433172_8446622_RNA45SN1_45S_50ntFlanks@10394.10429.36\|AGCAGTTGAACATGGGTCAGTCGGTCCTGAGAGATG | >hg38_21_+_8433172_8446622_RNA45SN1_45S_50ntFlanks | 10394 | 10429 | 36 | AGCAGTTGAACATGGGTCAGTCGGTCCTGAGAGATG | 439.0 | 1.5 | 0.6 | 2.4 | 1.6E-02 | 1.9E-02 | MB231/MB468 |
| >hg38_21_+_8433172_8446622_RNA45SN1_45S_50ntFlanks@6648.6695.48\|GTACGACTCTTAGCGGTGGATCACTCGGCTCGTGCGTCGATGAAGAAC | >hg38_21_+_8433172_8446622_RNA45SN1_45S_50ntFlanks | 6648 | 6695 | 48 | GTACGACTCTTAGCGGTGGATCACTCGGCTCGTGCGTCGATGAAGAAC | 765.2 | -2.3 | 0.8 | -2.9 | 4.3E-03 | 1.9E-02 | BT20/MB231 |
| >hg38_21_+_8433172_8446622_RNA45SN1_45S_50ntFlanks@11315.11336.22\|CCACCCCACGTCTCGTCGCGCG | >hg38_21_+_8433172_8446622_RNA45SN1_45S_50ntFlanks | 11315 | 11336 | 22 | CCACCCCACGTCTCGTCGCGCG | 282.4 | -2.7 | 1.1 | -2.4 | 1.7E-02 | 1.9E-02 | BT20/MB468 |
| >trna17_ValTAC_11_-_59318102_59318174@1.30.30,trna4_ValTAC_X_-_18693029_18693101@1.30.30\|GGTTCCATAGTGTAGTGGTTATCACGTCTG | >trna17_ValTAC_11_-_59318102_59318174 | 1 | 30 | 30 | GGTTCCATAGTGTAGTGGTTATCACGTCTG | 284.2 | -4.6 | 1.9 | -2.4 | 1.6E-02 | 1.9E-02 | MB231/MB468 |
| >hg38_21_+_8433172_8446622_RNA45SN1_45S_50ntFlanks@12408.12434.27\|TTTGATCCTTCGATGTCGGCTCTTCCT | >hg38_21_+_8433172_8446622_RNA45SN1_45S_50ntFlanks | 12408 | 12434 | 27 | TTTGATCCTTCGATGTCGGCTCTTCCT | 459.1 | 2.0 | 0.8 | 2.4 | 1.7E-02 | 1.9E-02 | BT20/MB468 |
| >hg38_21_+_8433172_8446622_RNA45SN1_45S_50ntFlanks@6652.6679.28\|GACTCTTAGCGGTGGATCACTCGGCTCG | >hg38_21_+_8433172_8446622_RNA45SN1_45S_50ntFlanks | 6652 | 6679 | 28 | GACTCTTAGCGGTGGATCACTCGGCTCG | 1264.2 | 1.7 | 0.7 | 2.4 | 1.6E-02 | 1.9E-02 | MB231/MB468 |
| >hg38_21_+_8433172_8446622_RNA45SN1_45S_50ntFlanks@3835.3862.28\|CTCCTCTCCTACTTGGATAACTGTGGTA | >hg38_21_+_8433172_8446622_RNA45SN1_45S_50ntFlanks | 3835 | 3862 | 28 | CTCCTCTCCTACTTGGATAACTGTGGTA | 112.4 | -1.5 | 0.6 | -2.4 | 1.6E-02 | 1.9E-02 | MB231/MB468 |
| >hg38_21_+_8433172_8446622_RNA45SN1_45S_50ntFlanks@9623.9649.27\|TAGTAGCTGGTTCCCTCCGAAGTTTCC | >hg38_21_+_8433172_8446622_RNA45SN1_45S_50ntFlanks | 9623 | 9649 | 27 | TAGTAGCTGGTTCCCTCCGAAGTTTCC | 78.4 | -1.9 | 0.8 | -2.4 | 1.6E-02 | 1.9E-02 | MB231/MB468 |
| >hg38_21_+_8433172_8446622_RNA45SN1_45S_50ntFlanks@4729.4751.23\|TCAAGAACGAAAGTCGGAGGTTC | >hg38_21_+_8433172_8446622_RNA45SN1_45S_50ntFlanks | 4729 | 4751 | 23 | TCAAGAACGAAAGTCGGAGGTTC | 240.6 | -1.2 | 0.5 | -2.4 | 1.6E-02 | 2.0E-02 | MB231/MB468 |
| >hg38_21_+_8433172_8446622_RNA45SN1_45S_50ntFlanks@8176.8215.40\|CTTCTGATCGAGGCCCAGCCCGTGGACGGTGTGAGGCCGG | >hg38_21_+_8433172_8446622_RNA45SN1_45S_50ntFlanks | 8176 | 8215 | 40 | CTTCTGATCGAGGCCCAGCCCGTGGACGGTGTGAGGCCGG | 231.7 | -1.4 | 0.6 | -2.4 | 1.6E-02 | 2.0E-02 | MB231/MB468 |
| >hg38_21_+_8433172_8446622_RNA45SN1_45S_50ntFlanks@4498.4520.23\|AAGCGTTTACTTTGAAAAAATTA | >hg38_21_+_8433172_8446622_RNA45SN1_45S_50ntFlanks | 4498 | 4520 | 23 | AAGCGTTTACTTTGAAAAAATTA | 308.3 | 1.4 | 0.6 | 2.4 | 1.8E-02 | 2.0E-02 | BT20/MB468 |
| >hg38_21_+_8433172_8446622_RNA45SN1_45S_50ntFlanks@11578.11613.36\|TAGAACTGGTGCGGACCAGGGGAATCCGACTGTTTA | >hg38_21_+_8433172_8446622_RNA45SN1_45S_50ntFlanks | 11578 | 11613 | 36 | TAGAACTGGTGCGGACCAGGGGAATCCGACTGTTTA | 304.0 | 2.8 | 1.0 | 2.8 | 4.4E-03 | 2.0E-02 | BT20/MB231 |
| >hg38_21_+_8433172_8446622_RNA45SN1_45S_50ntFlanks@6651.6697.47\|CGACTCTTAGCGGTGGATCACTCGGCTCGTGCGTCGATGAAGAACGC | >hg38_21_+_8433172_8446622_RNA45SN1_45S_50ntFlanks | 6651 | 6697 | 47 | CGACTCTTAGCGGTGGATCACTCGGCTCGTGCGTCGATGAAGAACGC | 3652.5 | -2.0 | 0.7 | -2.8 | 4.4E-03 | 2.0E-02 | BT20/MB231 |
| >hg38_MT_+_1621_3279_RNR2_16S_50ntFlanks@493.517.25\|GCTCTTTGGACACTAGGAAAAAACC | >hg38_MT_+_1621_3279_RNR2_16S_50ntFlanks | 493 | 517 | 25 | GCTCTTTGGACACTAGGAAAAAACC | 794.5 | -2.2 | 0.8 | -2.8 | 4.4E-03 | 2.0E-02 | BT20/MB231 |
| >trna111_HisGTG_1_-_147774845_147774916@-1G.35.36,trna118_HisGTG_1_-_145396881_145396952@-1G.35.36,trna16_HisGTG_1_+_146544773_146544844@-1G.35.36,trna1_HisGTG_15_+_45493349_45493420@-1G.35.36,trna21_HisGTG_1_+_147753471_147753542@-1G.35.36,trna33_HisGTG_6_+_27125906_27125977@-1G.35.36,trna7_HisGTG_9_-_14433938_14434009@-1G.35.36,trna8_HisGTG_15_-_45492611_45492682@-1G.35.36,trna9_HisGTG_15_-_45490804_45490875@-1G.35.36\|GGCCGTGATCGTATAGTGGTTAGTACTCTGCGTTGT | >trna111_HisGTG_1_-_147774845_147774916 | -1G | 35 | 36 | GGCCGTGATCGTATAGTGGTTAGTACTCTGCGTTGT | 1832.0 | -2.5 | 0.9 | -2.8 | 4.4E-03 | 2.0E-02 | BT20/MB231 |
| >hg38_21_+_8433172_8446622_RNA45SN1_45S_50ntFlanks@10818.10840.23\|AAGGGAAGTCGGCAAGCCGGATC | >hg38_21_+_8433172_8446622_RNA45SN1_45S_50ntFlanks | 10818 | 10840 | 23 | AAGGGAAGTCGGCAAGCCGGATC | 109.4 | -1.1 | 0.5 | -2.4 | 1.7E-02 | 2.0E-02 | MB231/MB468 |
| >hg38_21_+_8433172_8446622_RNA45SN1_45S_50ntFlanks@12935.12961.27\|TAAACCATTCGTAGACGACCTGCTTCT | >hg38_21_+_8433172_8446622_RNA45SN1_45S_50ntFlanks | 12935 | 12961 | 27 | TAAACCATTCGTAGACGACCTGCTTCT | 119.1 | -1.0 | 0.4 | -2.4 | 1.7E-02 | 2.0E-02 | MB231/MB468 |
| >hg38_21_+_8433172_8446622_RNA45SN1_45S_50ntFlanks@11887.11904.18\|GAGCTTGACTCTAGTCTG | >hg38_21_+_8433172_8446622_RNA45SN1_45S_50ntFlanks | 11887 | 11904 | 18 | GAGCTTGACTCTAGTCTG | 87.7 | -3.9 | 1.6 | -2.4 | 1.7E-02 | 2.0E-02 | MB231/MB468 |
| >hg38_21_+_8433172_8446622_RNA45SN1_45S_50ntFlanks@12458.12496.39\|AAGCGTTGGATTGTTCACCCACTAATAGGGAACGTGAGC | >hg38_21_+_8433172_8446622_RNA45SN1_45S_50ntFlanks | 12458 | 12496 | 39 | AAGCGTTGGATTGTTCACCCACTAATAGGGAACGTGAGC | 327.0 | -2.1 | 0.7 | -2.8 | 4.5E-03 | 2.0E-02 | BT20/MB231 |
| >hg38_21_+_8433172_8446622_RNA45SN1_45S_50ntFlanks@10682.10710.29\|TCCGGCGGCGTCCGGTGAGCTCTCGCTGG | >hg38_21_+_8433172_8446622_RNA45SN1_45S_50ntFlanks | 10682 | 10710 | 29 | TCCGGCGGCGTCCGGTGAGCTCTCGCTGG | 143.5 | -4.2 | 1.8 | -2.4 | 1.7E-02 | 2.0E-02 | MB231/MB468 |
| >trnaMT_MetCAT_MT_+_4402_4469@35.71.37\|CCCCGAAAATGTTGGTTATACCCTTCCCGTACTACCA | >trnaMT_MetCAT_MT_+_4402_4469 | 35 | 71 | 37 | CCCCGAAAATGTTGGTTATACCCTTCCCGTACTACCA | 213.4 | -2.2 | 0.8 | -2.8 | 4.5E-03 | 2.0E-02 | BT20/MB231 |
| >hg38_21_+_8433172_8446622_RNA45SN1_45S_50ntFlanks@12937.12959.23\|AACCATTCGTAGACGACCTGCTT | >hg38_21_+_8433172_8446622_RNA45SN1_45S_50ntFlanks | 12937 | 12959 | 23 | AACCATTCGTAGACGACCTGCTT | 343.6 | -1.6 | 0.7 | -2.4 | 1.8E-02 | 2.0E-02 | BT20/MB468 |
| >hg38_21_+_8433172_8446622_RNA45SN1_45S_50ntFlanks@6773.6806.34\|TTCCTCCCGGGGCTACGCCTGTCTGAGCGTCGCT | >hg38_21_+_8433172_8446622_RNA45SN1_45S_50ntFlanks | 6773 | 6806 | 34 | TTCCTCCCGGGGCTACGCCTGTCTGAGCGTCGCT | 1714.2 | 1.6 | 0.7 | 2.4 | 1.7E-02 | 2.0E-02 | MB231/MB468 |
| >hg38_21_+_8433172_8446622_RNA45SN1_45S_50ntFlanks@8196.8214.19\|CGTGGACGGTGTGAGGCCG | >hg38_21_+_8433172_8446622_RNA45SN1_45S_50ntFlanks | 8196 | 8214 | 19 | CGTGGACGGTGTGAGGCCG | 989.4 | -1.8 | 0.8 | -2.4 | 1.7E-02 | 2.0E-02 | MB231/MB468 |
| >hg38_21_+_8433172_8446622_RNA45SN1_45S_50ntFlanks@10830.10853.24\|CAAGCCGGATCCGTAACTTCGGGA | >hg38_21_+_8433172_8446622_RNA45SN1_45S_50ntFlanks | 10830 | 10853 | 24 | CAAGCCGGATCCGTAACTTCGGGA | 197.8 | -2.8 | 1.2 | -2.4 | 1.8E-02 | 2.0E-02 | BT20/MB468 |
| >hg38_21_+_8433172_8446622_RNA45SN1_45S_50ntFlanks@12413.12443.31\|TCCTTCGATGTCGGCTCTTCCTATCATTGTG | >hg38_21_+_8433172_8446622_RNA45SN1_45S_50ntFlanks | 12413 | 12443 | 31 | TCCTTCGATGTCGGCTCTTCCTATCATTGTG | 180.7 | -1.4 | 0.6 | -2.4 | 1.8E-02 | 2.0E-02 | BT20/MB468 |
| >hg38_21_+_8433172_8446622_RNA45SN1_45S_50ntFlanks@7975.8003.29\|CGCGACCTCAGATCAGACGTGGCGACCCG | >hg38_21_+_8433172_8446622_RNA45SN1_45S_50ntFlanks | 7975 | 8003 | 29 | CGCGACCTCAGATCAGACGTGGCGACCCG | 1908.2 | 1.4 | 0.6 | 2.4 | 1.8E-02 | 2.0E-02 | BT20/MB468 |
| >hg38_21_+_8433172_8446622_RNA45SN1_45S_50ntFlanks@12479.12499.21\|CTAATAGGGAACGTGAGCTGG | >hg38_21_+_8433172_8446622_RNA45SN1_45S_50ntFlanks | 12479 | 12499 | 21 | CTAATAGGGAACGTGAGCTGG | 327.7 | -1.1 | 0.5 | -2.4 | 1.7E-02 | 2.0E-02 | MB231/MB468 |
| >hg38_21_+_8433172_8446622_RNA45SN1_45S_50ntFlanks@3705.3744.40\|TACCTGGTTGATCCTGCCAGTAGCATATGCTTGTCTCAAA | >hg38_21_+_8433172_8446622_RNA45SN1_45S_50ntFlanks | 3705 | 3744 | 40 | TACCTGGTTGATCCTGCCAGTAGCATATGCTTGTCTCAAA | 932.8 | 2.5 | 0.9 | 2.8 | 4.6E-03 | 2.0E-02 | BT20/MB231 |
| >hg38_21_+_8433172_8446622_RNA45SN1_45S_50ntFlanks@11315.11343.29\|CCACCCCACGTCTCGTCGCGCGCGCGTCC | >hg38_21_+_8433172_8446622_RNA45SN1_45S_50ntFlanks | 11315 | 11343 | 29 | CCACCCCACGTCTCGTCGCGCGCGCGTCC | 110.6 | -2.8 | 1.2 | -2.4 | 1.8E-02 | 2.0E-02 | BT20/MB468 |
| >hg38_21_+_8433172_8446622_RNA45SN1_45S_50ntFlanks@11786.11803.18\|GAATGGATGAACGAGATT | >hg38_21_+_8433172_8446622_RNA45SN1_45S_50ntFlanks | 11786 | 11803 | 18 | GAATGGATGAACGAGATT | 1158.6 | -1.6 | 0.7 | -2.4 | 1.7E-02 | 2.0E-02 | MB231/MB468 |
| >hg38_21_+_8433172_8446622_RNA45SN1_45S_50ntFlanks@10389.10424.36\|TGAACAGCAGTTGAACATGGGTCAGTCGGTCCTGAG | >hg38_21_+_8433172_8446622_RNA45SN1_45S_50ntFlanks | 10389 | 10424 | 36 | TGAACAGCAGTTGAACATGGGTCAGTCGGTCCTGAG | 1529.4 | 1.9 | 0.8 | 2.4 | 1.8E-02 | 2.0E-02 | BT20/MB468 |
| >hg38_21_+_8433172_8446622_RNA45SN1_45S_50ntFlanks@3854.3873.20\|ACTGTGGTAATTCTAGAGCT | >hg38_21_+_8433172_8446622_RNA45SN1_45S_50ntFlanks | 3854 | 3873 | 20 | ACTGTGGTAATTCTAGAGCT | 1002.0 | -1.7 | 0.7 | -2.4 | 1.7E-02 | 2.0E-02 | MB231/MB468 |
| >hg38_21_+_8433172_8446622_RNA45SN1_45S_50ntFlanks@4367.4397.31\|CAGTTAAAAAGCTCGTAGTTGGATCTTGGGA | >hg38_21_+_8433172_8446622_RNA45SN1_45S_50ntFlanks | 4367 | 4397 | 31 | CAGTTAAAAAGCTCGTAGTTGGATCTTGGGA | 91.3 | -2.3 | 1.0 | -2.4 | 1.8E-02 | 2.0E-02 | BT20/MB468 |
| >hg38_21_+_8433172_8446622_RNA45SN1_45S_50ntFlanks@10850.10868.19\|GGGATAAGGATTGGCTCTA | >hg38_21_+_8433172_8446622_RNA45SN1_45S_50ntFlanks | 10850 | 10868 | 19 | GGGATAAGGATTGGCTCTA | 118.0 | -1.6 | 0.7 | -2.4 | 1.7E-02 | 2.1E-02 | MB231/MB468 |

<table>
<tr><td>>hg38_21_+_8433172_8446622_RNA45SN1_45S_50ntFlanks@485.503.19|GATGTGGTGACGTCGTGCT</td><td>>hg38_21_+_8433172_8446622_RNA45SN1_45S_50ntFlanks</td><td>485</td><td>503</td><td>19</td><td>GATGTGGTGACGTCGTGCT</td><td>205.5</td><td>2.7</td><td>0.9</td><td>2.8</td><td>4.7E-03</td><td>2.1E-02</td><td>BT20/MB231</td></tr>
<tr><td>>hg38_21_+_8433172_8446622_RNA45SN1_45S_50ntFlanks@4069.4094.26|CTTTCGATGGTAGTCGCCGTGCCTAC</td><td>>hg38_21_+_8433172_8446622_RNA45SN1_45S_50ntFlanks</td><td>4069</td><td>4094</td><td>26</td><td>CTTTCGATGGTAGTCGCCGTGCCTAC</td><td>110.3</td><td>-1.3</td><td>0.5</td><td>-2.4</td><td>1.7E-02</td><td>2.1E-02</td><td>MB231/MB468</td></tr>
<tr><td>>hg38_21_+_8433172_8446622_RNA45SN1_45S_50ntFlanks@8192.8212.21|AGCCCGTGGACGGTGTGAGGC</td><td>>hg38_21_+_8433172_8446622_RNA45SN1_45S_50ntFlanks</td><td>8192</td><td>8212</td><td>21</td><td>AGCCCGTGGACGGTGTGAGGC</td><td>144.4</td><td>-1.8</td><td>0.8</td><td>-2.4</td><td>1.8E-02</td><td>2.1E-02</td><td>BT20/MB468</td></tr>
<tr><td>>hg38_21_+_8433172_8446622_RNA45SN1_45S_50ntFlanks@3835.3871.37|CTCCTCTCCTACTTGGATAACTGTGGTAATTCTAGAG</td><td>>hg38_21_+_8433172_8446622_RNA45SN1_45S_50ntFlanks</td><td>3835</td><td>3871</td><td>37</td><td>CTCCTCTCCTACTTGGATAACTGTGGTAATTCTAGAG</td><td>148.3</td><td>-1.8</td><td>0.8</td><td>-2.4</td><td>1.7E-02</td><td>2.1E-02</td><td>MB231/MB468</td></tr>
<tr><td>>hg38_21_+_8433172_8446622_RNA45SN1_45S_50ntFlanks@3844.3868.25|TACTTGGATAACTGTGGTAATTCTA</td><td>>hg38_21_+_8433172_8446622_RNA45SN1_45S_50ntFlanks</td><td>3844</td><td>3868</td><td>25</td><td>TACTTGGATAACTGTGGTAATTCTA</td><td>287.7</td><td>1.1</td><td>0.5</td><td>2.4</td><td>1.9E-02</td><td>2.1E-02</td><td>BT20/MB468</td></tr>
<tr><td>>MI0000103|hsa-mir-101-1&WithFlank&1|-|65058428|65058514@52.72.21;MI0000739|hsa-mir-101-2&WithFlank&9|+|4850291|4850381@54.74.21&[MIMAT0000099&hsa-miR-101-3p&offsets|-1|-1;m-17&1|-|65058443|65058462&offsets|-1|0];[MIMAT0000099_1&hsa-miR-101-3p&offsets|-1|-1;m-16&9|+|4850345|4850364&offsets|-1|0]|GTACAGTACTGTGATAACTGA</td><td>>MI0000103|hsa-mir-101-1&WithFlank&1|-|65058428|65058514</td><td>52</td><td>72</td><td>21</td><td>GTACAGTACTGTGATAACTGA</td><td>4753.2</td><td>2.7</td><td>1.0</td><td>2.8</td><td>4.7E-03</td><td>2.1E-02</td><td>BT20/MB231</td></tr>
<tr><td>>hg38_21_+_8433172_8446622_RNA45SN1_45S_50ntFlanks@12399.12419.21|ACGTCGCTTTTTGATCCTTCG</td><td>>hg38_21_+_8433172_8446622_RNA45SN1_45S_50ntFlanks</td><td>12399</td><td>12419</td><td>21</td><td>ACGTCGCTTTTTGATCCTTCG</td><td>180.0</td><td>-1.2</td><td>0.5</td><td>-2.4</td><td>1.8E-02</td><td>2.1E-02</td><td>MB231/MB468</td></tr>
<tr><td>>hg38_21_+_8433172_8446622_RNA45SN1_45S_50ntFlanks@6647.6696.50|CGTACGACTCTTAGCGGTGGATCACTCGGCTCGTGCGTCGATGAAGAACG</td><td>>hg38_21_+_8433172_8446622_RNA45SN1_45S_50ntFlanks</td><td>6647</td><td>6696</td><td>50</td><td>CGTACGACTCTTAGCGGTGGATCACTCGGCTCGTGCGTCGATGAAGAACG</td><td>446.3</td><td>-2.6</td><td>0.9</td><td>-2.8</td><td>4.7E-03</td><td>2.1E-02</td><td>BT20/MB231</td></tr>
<tr><td>>hg38_21_+_8433172_8446622_RNA45SN1_45S_50ntFlanks@11552.11569.18|CCTCGGCCGGCGCCTAGC</td><td>>hg38_21_+_8433172_8446622_RNA45SN1_45S_50ntFlanks</td><td>11552</td><td>11569</td><td>18</td><td>CCTCGGCCGGCGCCTAGC</td><td>146.7</td><td>-2.5</td><td>0.9</td><td>-2.8</td><td>4.8E-03</td><td>2.1E-02</td><td>BT20/MB231</td></tr>
<tr><td>>hg38_21_+_8433172_8446622_RNA45SN1_45S_50ntFlanks@11580.11602.23|GAACTGGTGCGGACCAGGGGAAT</td><td>>hg38_21_+_8433172_8446622_RNA45SN1_45S_50ntFlanks</td><td>11580</td><td>11602</td><td>23</td><td>GAACTGGTGCGGACCAGGGGAAT</td><td>599.0</td><td>-1.7</td><td>0.7</td><td>-2.4</td><td>1.8E-02</td><td>2.1E-02</td><td>MB231/MB468</td></tr>
<tr><td>>hg38_21_+_8433172_8446622_RNA45SN1_45S_50ntFlanks@6662.6681.20|GGTGGATCACTCGGCTCGTG</td><td>>hg38_21_+_8433172_8446622_RNA45SN1_45S_50ntFlanks</td><td>6662</td><td>6681</td><td>20</td><td>GGTGGATCACTCGGCTCGTG</td><td>93.7</td><td>-1.3</td><td>0.5</td><td>-2.3</td><td>1.9E-02</td><td>2.1E-02</td><td>BT20/MB468</td></tr>
<tr><td>>hg38_21_+_8433172_8446622_RNA45SN1_45S_50ntFlanks@12407.12429.23|TTTTGATCCTTCGATGTCGGCTC</td><td>>hg38_21_+_8433172_8446622_RNA45SN1_45S_50ntFlanks</td><td>12407</td><td>12429</td><td>23</td><td>TTTTGATCCTTCGATGTCGGCTC</td><td>348.6</td><td>1.1</td><td>0.5</td><td>2.4</td><td>1.8E-02</td><td>2.1E-02</td><td>MB231/MB468</td></tr>
<tr><td>>hg38_21_+_8433172_8446622_RNA45SN1_45S_50ntFlanks@12438.12474.37|ATTGTGAAGCAGAATTCACCAAGCGTTGGATTGTTCA</td><td>>hg38_21_+_8433172_8446622_RNA45SN1_45S_50ntFlanks</td><td>12438</td><td>12474</td><td>37</td><td>ATTGTGAAGCAGAATTCACCAAGCGTTGGATTGTTCA</td><td>199.3</td><td>1.1</td><td>0.5</td><td>2.4</td><td>1.8E-02</td><td>2.1E-02</td><td>MB231/MB468</td></tr>
<tr><td>>hg38_21_+_8433172_8446622_RNA45SN1_45S_50ntFlanks@11571.11590.20|GCCGACTTAGAACTGGTGCG</td><td>>hg38_21_+_8433172_8446622_RNA45SN1_45S_50ntFlanks</td><td>11571</td><td>11590</td><td>20</td><td>GCCGACTTAGAACTGGTGCG</td><td>276.1</td><td>-2.4</td><td>0.8</td><td>-2.8</td><td>4.8E-03</td><td>2.1E-02</td><td>BT20/MB231</td></tr>
<tr><td>>hg38_21_+_8433172_8446622_RNA45SN1_45S_50ntFlanks@10420.10455.36|CTGAGAGATGGGCGAGCGCCGTTCCGAAGGGACGGG</td><td>>hg38_21_+_8433172_8446622_RNA45SN1_45S_50ntFlanks</td><td>10420</td><td>10455</td><td>36</td><td>CTGAGAGATGGGCGAGCGCCGTTCCGAAGGGACGGG</td><td>108.4</td><td>-1.5</td><td>0.6</td><td>-2.4</td><td>1.8E-02</td><td>2.1E-02</td><td>MB231/MB468</td></tr>
<tr><td>>hg38_21_+_8433172_8446622_RNA45SN1_45S_50ntFlanks@8352.8379.28|AGTTGAAAAGAACTTTGAAGAGAGAGTT</td><td>>hg38_21_+_8433172_8446622_RNA45SN1_45S_50ntFlanks</td><td>8352</td><td>8379</td><td>28</td><td>AGTTGAAAAGAACTTTGAAGAGAGAGTT</td><td>172.2</td><td>-1.2</td><td>0.5</td><td>-2.4</td><td>1.8E-02</td><td>2.1E-02</td><td>MB231/MB468</td></tr>
<tr><td>>am_trna128_GlyGCC_6_-_27870686_27870756@1.30.30,trna133_GlyCCC_1_-_16872434_16872504@1.30.30,trna18_GlyGCC_16_+_70822597_70822667@1.30.30,trna19_GlyGCC_16_+_70823410_70823480@1.30.30,trna19_GlyGCC_2_-_157257659_157257729@1.30.30,trna24_GlyGCC_16_-_70812942_70813012@1.30.30,trna25_GlyGCC_16_-_70812114_70812184@1.30.30,trna4_GlyCCC_1_+_17188416_17188486@1.30.30,trna5_GlyGCC_17_+_8029064_8029134@1.30.30,trna68_GlyGCC_1_-_161493637_161493707@1.30.30|GCATTGGTGGTTCAGTGGTAGAATTCTCGC</td><td>>am_trna128_GlyGCC_6_-_27870686_27870756</td><td>1</td><td>30</td><td>30</td><td>GCATTGGTGGTTCAGTGGTAGAATTCTCGC</td><td>31054.8</td><td>2.2</td><td>0.9</td><td>2.4</td><td>1.8E-02</td><td>2.1E-02</td><td>MB231/MB468</td></tr>
<tr><td>>hg38_21_+_8433172_8446622_RNA45SN1_45S_50ntFlanks@10791.10816.26|CCTCTGGCATGTTGGAACAATGTAGG</td><td>>hg38_21_+_8433172_8446622_RNA45SN1_45S_50ntFlanks</td><td>10791</td><td>10816</td><td>26</td><td>CCTCTGGCATGTTGGAACAATGTAGG</td><td>142.3</td><td>-1.3</td><td>0.5</td><td>-2.3</td><td>1.9E-02</td><td>2.1E-02</td><td>BT20/MB468</td></tr>
<tr><td>>hg38_MT_+_1621_3279_RNR2_16S_50ntFlanks@1217.1262.46|AACCTCCGAGCAGTACATGCTAAGACTTCACCAGTCAAAGCGAACT</td><td>>hg38_MT_+_1621_3279_RNR2_16S_50ntFlanks</td><td>1217</td><td>1262</td><td>46</td><td>AACCTCCGAGCAGTACATGCTAAGACTTCACCAGTCAAAGCGAACT</td><td>221.7</td><td>-2.1</td><td>0.7</td><td>-2.8</td><td>4.9E-03</td><td>2.1E-02</td><td>BT20/MB231</td></tr>
<tr><td>>hg38_21_+_8433172_8446622_RNA45SN1_45S_50ntFlanks@9548.9571.24|GAGGTCCGTAGCGGTCCTGACGTG</td><td>>hg38_21_+_8433172_8446622_RNA45SN1_45S_50ntFlanks</td><td>9548</td><td>9571</td><td>24</td><td>GAGGTCCGTAGCGGTCCTGACGTG</td><td>446.2</td><td>-1.3</td><td>0.6</td><td>-2.3</td><td>1.9E-02</td><td>2.1E-02</td><td>BT20/MB468</td></tr>
<tr><td>>hg38_21_+_8433172_8446622_RNA45SN1_45S_50ntFlanks@11703.11736.34|AATGAAGCGCGGGTAAACGGCGGGAGTAACTATG</td><td>>hg38_21_+_8433172_8446622_RNA45SN1_45S_50ntFlanks</td><td>11703</td><td>11736</td><td>34</td><td>AATGAAGCGCGGGTAAACGGCGGGAGTAACTATG</td><td>410.7</td><td>1.5</td><td>0.6</td><td>2.4</td><td>1.8E-02</td><td>2.1E-02</td><td>MB231/MB468</td></tr>
<tr><td>>hg38_21_+_8433172_8446622_RNA45SN1_45S_50ntFlanks@3851.3872.22|ATAACTGTGGTAATTCTAGAGC</td><td>>hg38_21_+_8433172_8446622_RNA45SN1_45S_50ntFlanks</td><td>3851</td><td>3872</td><td>22</td><td>ATAACTGTGGTAATTCTAGAGC</td><td>191.1</td><td>1.0</td><td>0.4</td><td>2.4</td><td>1.8E-02</td><td>2.1E-02</td><td>MB231/MB468</td></tr>
<tr><td>>am_trna128_GlyGCC_6_-_27870686_27870756@1.36.36,trna18_GlyGCC_16_+_70822597_70822667@1.36.36,trna19_GlyGCC_16_+_70823410_70823480@1.36.36,trna19_GlyGCC_2_-_157257659_157257729@1.36.36,trna24_GlyGCC_16_-_70812942_70813012@1.36.36,trna25_GlyGCC_16_-_70812114_70812184@1.36.36,trna5_GlyGCC_17_+_8029064_8029134@1.36.36,trna68_GlyGCC_1_-_161493637_161493707@1.36.36|GCATTGGTGGTTCAGTGGTAGAATTCTCGCCTGCCA</td><td>>am_trna128_GlyGCC_6_-_27870686_27870756</td><td>1</td><td>36</td><td>36</td><td>GCATTGGTGGTTCAGTGGTAGAATTCTCGCCTGCCA</td><td>957.8</td><td>-2.0</td><td>0.7</td><td>-2.8</td><td>4.9E-03</td><td>2.2E-02</td><td>BT20/MB231</td></tr>
<tr><td>>MI0000542|hsa-mir-320a&WithFlank&8|-|22244960|22245043@48.70.23(+1C)&[MIMAT0000510&hsa-miR-320a-3p&offsets|0|+1(+1C);m-60&8|-|22244975|22244996&offsets|0|+1(+1C)]|AAAAGCTGGGTTGAGAGGGCGAAC</td><td>>MI0000542|hsa-mir-320a&WithFlank&8|-|22244960|22245043</td><td>48</td><td>70</td><td>23(+1C)</td><td>AAAAGCTGGGTTGAGAGGGCGAAC</td><td>965.0</td><td>2.1</td><td>0.7</td><td>2.8</td><td>4.9E-03</td><td>2.2E-02</td><td>BT20/MB231</td></tr>
<tr><td>>MI0000077|hsa-mir-21&WithFlank&17|+|59841260|59841343@14.31.18(+1C)&[MIMAT0000076&hsa-miR-21-5p&offsets|0|-4(+1C);m-1&17|+|59841273|59841294&offsets|0|-4(+1C)]|TAGCTTATCAGACTGATGC</td><td>>MI0000077|hsa-mir-21&WithFlank&17|+|59841260|59841343</td><td>14</td><td>31</td><td>18(+1C)</td><td>TAGCTTATCAGACTGATGC</td><td>273.3</td><td>1.9</td><td>0.7</td><td>2.8</td><td>4.9E-03</td><td>2.2E-02</td><td>BT20/MB231</td></tr>
<tr><td>>hg38_21_+_8433172_8446622_RNA45SN1_45S_50ntFlanks@4725.4767.43|TTAATCAAGAACGAAAGTCGGAGGTTCGAAGACGATCAGATAC</td><td>>hg38_21_+_8433172_8446622_RNA45SN1_45S_50ntFlanks</td><td>4725</td><td>4767</td><td>43</td><td>TTAATCAAGAACGAAAGTCGGAGGTTCGAAGACGATCAGATAC</td><td>137.8</td><td>-2.7</td><td>0.9</td><td>-2.8</td><td>4.9E-03</td><td>2.2E-02</td><td>BT20/MB231</td></tr>
<tr><td>>hg38_21_+_8433172_8446622_RNA45SN1_45S_50ntFlanks@11199.11229.31|CCGGCGGCGGCGGCGACTCTGGACGCGAGCC</td><td>>hg38_21_+_8433172_8446622_RNA45SN1_45S_50ntFlanks</td><td>11199</td><td>11229</td><td>31</td><td>CCGGCGGCGGCGGCGACTCTGGACGCGAGCC</td><td>87.9</td><td>-1.8</td><td>0.8</td><td>-2.3</td><td>1.9E-02</td><td>2.2E-02</td><td>BT20/MB468</td></tr>
<tr><td>>MI0000446|hsa-mir-125b-1&WithFlank&11|-|122099751|122099850@21.42.22(+1C);MI0000470|hsa-mir-125b-2&WithFlank&21|+|16590231|16590331@23.44.22(+1C)&[MIMAT0000423&hsa-miR-125b-5p&offsets|0|0(+1C);m-67&11|-|122099809|122099830&offsets|0|0(+1C)];[MIMAT0000423_1&hsa-miR-125b-5p&offsets|0|0(+1C);m-68&21|+|16590253|16590274&offsets|0|0(+1C)]|TCCCTGAGACCCTAACTTGTGAC</td><td>>MI0000446|hsa-mir-125b-1&WithFlank&11|-|122099751|122099850</td><td>21</td><td>42</td><td>22(+1C)</td><td>TCCCTGAGACCCTAACTTGTGAC</td><td>324.2</td><td>-6.1</td><td>2.2</td><td>-2.8</td><td>5.0E-03</td><td>2.2E-02</td><td>BT20/MB231</td></tr>
<tr><td>>hg38_21_+_8433172_8446622_RNA45SN1_45S_50ntFlanks@12996.13038.43|TCGCTGCGATCTATTGAAAGTCAGCCCTCGACACAAGGGTTTG</td><td>>hg38_21_+_8433172_8446622_RNA45SN1_45S_50ntFlanks</td><td>12996</td><td>13038</td><td>43</td><td>TCGCTGCGATCTATTGAAAGTCAGCCCTCGACACAAGGGTTTG</td><td>2787.6</td><td>-2.4</td><td>0.9</td><td>-2.8</td><td>5.0E-03</td><td>2.2E-02</td><td>BT20/MB231</td></tr>
<tr><td>>hg38_21_+_8433172_8446622_RNA45SN1_45S_50ntFlanks@12445.12472.28|AGCAGAATTCACCAAGCGTTGGATTGTT</td><td>>hg38_21_+_8433172_8446622_RNA45SN1_45S_50ntFlanks</td><td>12445</td><td>12472</td><td>28</td><td>AGCAGAATTCACCAAGCGTTGGATTGTT</td><td>106.6</td><td>-1.8</td><td>0.8</td><td>-2.4</td><td>1.8E-02</td><td>2.2E-02</td><td>MB231/MB468</td></tr>
<tr><td>>hg38_21_+_8433172_8446622_RNA45SN1_45S_50ntFlanks@7991.8008.18|ACGTGGCGACCCGCTGAA</td><td>>hg38_21_+_8433172_8446622_RNA45SN1_45S_50ntFlanks</td><td>7991</td><td>8008</td><td>18</td><td>ACGTGGCGACCCGCTGAA</td><td>5957.3</td><td>-2.1</td><td>0.9</td><td>-2.4</td><td>1.8E-02</td><td>2.2E-02</td><td>MB231/MB468</td></tr>
<tr><td>>am_trnaMT_ValTAC_MT_+_1602_1670@44.68.25|ATTTCAACTTAACTTGACCGCTCTG</td><td>>am_trnaMT_ValTAC_MT_+_1602_1670</td><td>44</td><td>68</td><td>25</td><td>ATTTCAACTTAACTTGACCGCTCTG</td><td>125.8</td><td>-2.0</td><td>0.9</td><td>-2.4</td><td>1.8E-02</td><td>2.2E-02</td><td>MB231/MB468</td></tr>
<tr><td>>hg38_21_+_8433172_8446622_RNA45SN1_45S_50ntFlanks@4687.4707.21|ACGGACCAGAGCGAAAGCATT</td><td>>hg38_21_+_8433172_8446622_RNA45SN1_45S_50ntFlanks</td><td>4687</td><td>4707</td><td>21</td><td>ACGGACCAGAGCGAAAGCATT</td><td>133.0</td><td>-2.0</td><td>0.9</td><td>-2.3</td><td>2.0E-02</td><td>2.2E-02</td><td>BT20/MB468</td></tr>
<tr><td>>hg38_21_+_8433172_8446622_RNA45SN1_45S_50ntFlanks@11671.11704.34|CCAGTGCTCTGAATGTCAAAGTGAAGAAATTCAA</td><td>>hg38_21_+_8433172_8446622_RNA45SN1_45S_50ntFlanks</td><td>11671</td><td>11704</td><td>34</td><td>CCAGTGCTCTGAATGTCAAAGTGAAGAAATTCAA</td><td>298.8</td><td>-1.6</td><td>0.7</td><td>-2.4</td><td>1.8E-02</td><td>2.2E-02</td><td>MB231/MB468</td></tr>
<tr><td>>MI0000301|hsa-mir-224&WithFlank&X|-|151958572|151958664@14.36.23(+1A)&[MIMAT0000281&hsa-miR-224-5p&offsets|+1|-1(+1A);m-238&X|-|151958630|151958651&offsets|0|+1(+1A)]|CAAGTCACTAGTGGTTCCGTTTAA</td><td>>MI0000301|hsa-mir-224&WithFlank&X|-|151958572|151958664</td><td>14</td><td>36</td><td>23(+1A)</td><td>CAAGTCACTAGTGGTTCCGTTTAA</td><td>467.5</td><td>1.9</td><td>0.7</td><td>2.8</td><td>5.0E-03</td><td>2.2E-02</td><td>BT20/MB231</td></tr>
<tr><td>>hg38_21_+_8433172_8446622_RNA45SN1_45S_50ntFlanks@10841.10866.26|CGTAACTTCGGGATAAGGATTGGCTC</td><td>>hg38_21_+_8433172_8446622_RNA45SN1_45S_50ntFlanks</td><td>10841</td><td>10866</td><td>26</td><td>CGTAACTTCGGGATAAGGATTGGCTC</td><td>967.3</td><td>-1.9</td><td>0.8</td><td>-2.4</td><td>1.9E-02</td><td>2.2E-02</td><td>MB231/MB468</td></tr>
<tr><td>>hg38_21_+_8433172_8446622_RNA45SN1_45S_50ntFlanks@10348.10373.26|AAACGAGAACTTTGAAGGCCGAAGTG</td><td>>hg38_21_+_8433172_8446622_RNA45SN1_45S_50ntFlanks</td><td>10348</td><td>10373</td><td>26</td><td>AAACGAGAACTTTGAAGGCCGAAGTG</td><td>244.9</td><td>1.2</td><td>0.5</td><td>2.3</td><td>2.0E-02</td><td>2.2E-02</td><td>BT20/MB468</td></tr>
<tr><td>>hg38_21_+_8433172_8446622_RNA45SN1_45S_50ntFlanks@11603.11620.18|CCGACTGTTTAATTAAAA</td><td>>hg38_21_+_8433172_8446622_RNA45SN1_45S_50ntFlanks</td><td>11603</td><td>11620</td><td>18</td><td>CCGACTGTTTAATTAAAA</td><td>366.6</td><td>-1.0</td><td>0.4</td><td>-2.4</td><td>1.9E-02</td><td>2.2E-02</td><td>MB231/MB468</td></tr>
<tr><td>>hg38_21_+_8433172_8446622_RNA45SN1_45S_50ntFlanks@7991.8014.24|ACGTGGCGACCCGCTGAATTTAAG</td><td>>hg38_21_+_8433172_8446622_RNA45SN1_45S_50ntFlanks</td><td>7991</td><td>8014</td><td>24</td><td>ACGTGGCGACCCGCTGAATTTAAG</td><td>582.2</td><td>-1.5</td><td>0.6</td><td>-2.3</td><td>2.0E-02</td><td>2.2E-02</td><td>BT20/MB468</td></tr>
<tr><td>>hg38_21_+_8433172_8446622_RNA45SN1_45S_50ntFlanks@10866.10894.29|CTAAGGGCTGGGTCGGTCGGGCTGGGGCG</td><td>>hg38_21_+_8433172_8446622_RNA45SN1_45S_50ntFlanks</td><td>10866</td><td>10894</td><td>29</td><td>CTAAGGGCTGGGTCGGTCGGGCTGGGGCG</td><td>209.5</td><td>-4.2</td><td>1.8</td><td>-2.4</td><td>1.9E-02</td><td>2.2E-02</td><td>MB231/MB468</td></tr>
</table>

<table>
<tr><td>>hg38_21_+_8433172_8446622_RNA45SN1_45S_50ntFlanks@11586.11612.27|GTGCGGACCAGGGGAATCCGACTGTTT</td><td>>hg38_21_+_8433172_8446622_RNA45SN1_45S_50ntFlanks</td><td>11586</td><td>11612</td><td>27</td><td>GTGCGGACCAGGGGAATCCGACTGTTT</td><td>973.9</td><td>-1.6</td><td>0.7</td><td>-2.4</td><td>1.9E-02</td><td>2.2E-02</td><td>MB231/MB468</td></tr>
<tr><td>>MI0000469|hsa-mir-125a&WithFlank&19|+|51693248|51693345@21.41.21(+1C)&[MIMAT0000443&hsa-miR-125a-5p&offsets|0|-3(+1C);m-63&19|+|51693268|51693289&offsets|0|-1(+1C)]|TCCCTGAGACCCTTTAACCTGC</td><td>>MI0000469|hsa-mir-125a&WithFlank&19|+|51693248|51693345</td><td>21</td><td>41</td><td>21(+1C)</td><td>TCCCTGAGACCCTTTAACCTGC</td><td>757.1</td><td>2.0</td><td>0.7</td><td>2.8</td><td>5.1E-03</td><td>2.2E-02</td><td>BT20/MB231</td></tr>
<tr><td>>hg38_21_+_8433172_8446622_RNA45SN1_45S_50ntFlanks@6651.6669.19|CGACTCTTAGCGGTGGATC</td><td>>hg38_21_+_8433172_8446622_RNA45SN1_45S_50ntFlanks</td><td>6651</td><td>6669</td><td>19</td><td>CGACTCTTAGCGGTGGATC</td><td>4717.8</td><td>1.7</td><td>0.7</td><td>2.3</td><td>1.9E-02</td><td>2.2E-02</td><td>MB231/MB468</td></tr>
<tr><td>>hg38_21_+_8433172_8446622_RNA45SN1_45S_50ntFlanks@10346.10368.23|TCAAACGAGAACTTTGAAGGCCG</td><td>>hg38_21_+_8433172_8446622_RNA45SN1_45S_50ntFlanks</td><td>10346</td><td>10368</td><td>23</td><td>TCAAACGAGAACTTTGAAGGCCG</td><td>2008.5</td><td>-1.5</td><td>0.6</td><td>-2.3</td><td>1.9E-02</td><td>2.2E-02</td><td>MB231/MB468</td></tr>
<tr><td>>hg38_21_+_8433172_8446622_RNA45SN1_45S_50ntFlanks@12936.12957.22|AAACCATTCGTAGACGACCTGC</td><td>>hg38_21_+_8433172_8446622_RNA45SN1_45S_50ntFlanks</td><td>12936</td><td>12957</td><td>22</td><td>AAACCATTCGTAGACGACCTGC</td><td>456.8</td><td>1.4</td><td>0.6</td><td>2.3</td><td>1.9E-02</td><td>2.3E-02</td><td>MB231/MB468</td></tr>
<tr><td>>hg38_21_+_8433172_8446622_RNA45SN1_45S_50ntFlanks@12417.12452.36|TCGATGTCGGCTCTTCCTATCATTGTGAAGCAGAAT</td><td>>hg38_21_+_8433172_8446622_RNA45SN1_45S_50ntFlanks</td><td>12417</td><td>12452</td><td>36</td><td>TCGATGTCGGCTCTTCCTATCATTGTGAAGCAGAAT</td><td>270.2</td><td>-2.1</td><td>0.8</td><td>-2.8</td><td>5.2E-03</td><td>2.3E-02</td><td>BT20/MB231</td></tr>
<tr><td>>trnaMT_ValTAC_MT_+_1602_1670@55.72.18|ACTTGACCGCTCTGACCA</td><td>>trnaMT_ValTAC_MT_+_1602_1670</td><td>55</td><td>72</td><td>18</td><td>ACTTGACCGCTCTGACCA</td><td>3124.1</td><td>2.0</td><td>0.7</td><td>2.8</td><td>5.2E-03</td><td>2.3E-02</td><td>BT20/MB231</td></tr>
<tr><td>>hg38_21_+_8433172_8446622_RNA45SN1_45S_50ntFlanks@11580.11613.34|GAACTGGTGCGGACCAGGGGAATCCGACTGTTTA</td><td>>hg38_21_+_8433172_8446622_RNA45SN1_45S_50ntFlanks</td><td>11580</td><td>11613</td><td>34</td><td>GAACTGGTGCGGACCAGGGGAATCCGACTGTTTA</td><td>1368.1</td><td>1.9</td><td>0.8</td><td>2.3</td><td>1.9E-02</td><td>2.3E-02</td><td>MB231/MB468</td></tr>
<tr><td>>trnaMT_GluTTC_MT_-_14674_14742@41.71.31,trnalookalike8_GluTTC_5_-_93905172_93905240@41.71.31|TGGTCGTGGTTGTAGTCCGTGCGAGAATACC</td><td>>trnaMT_GluTTC_MT_-_14674_14742</td><td>41</td><td>71</td><td>31</td><td>TGGTCGTGGTTGTAGTCCGTGCGAGAATACC</td><td>399.2</td><td>-1.8</td><td>0.7</td><td>-2.8</td><td>5.3E-03</td><td>2.3E-02</td><td>BT20/MB231</td></tr>
<tr><td>>hg38_21_+_8433172_8446622_RNA45SN1_45S_50ntFlanks@12419.12448.30|GATGTCGGCTCTTCCTATCATTGTGAAGCA</td><td>>hg38_21_+_8433172_8446622_RNA45SN1_45S_50ntFlanks</td><td>12419</td><td>12448</td><td>30</td><td>GATGTCGGCTCTTCCTATCATTGTGAAGCA</td><td>2469.8</td><td>-2.3</td><td>0.8</td><td>-2.8</td><td>5.3E-03</td><td>2.3E-02</td><td>BT20/MB231</td></tr>
<tr><td>>MI0000342|hsa-mir-200b&WithFlank&1|+|1167098|1167204@63.85.23(+1A)&[MIMAT0000318&hsa-miR-200b-3p&offsets|0|+1(+1A);m-93&1|+|1167160|1167182&offsets|0|0(+1A)]|TAATACTGCCTGGTAATGATGACA</td><td>>MI0000342|hsa-mir-200b&WithFlank&1|+|1167098|1167204</td><td>63</td><td>85</td><td>23(+1A)</td><td>TAATACTGCCTGGTAATGATGACA</td><td>882.1</td><td>-1.8</td><td>0.8</td><td>-2.3</td><td>2.0E-02</td><td>2.3E-02</td><td>BT20/MB468</td></tr>
<tr><td>>hg38_21_+_8433172_8446622_RNA45SN1_45S_50ntFlanks@9573.9601.29|AAATCGGTCGTCCGACCTGGGTATAGGGG</td><td>>hg38_21_+_8433172_8446622_RNA45SN1_45S_50ntFlanks</td><td>9573</td><td>9601</td><td>29</td><td>AAATCGGTCGTCCGACCTGGGTATAGGGG</td><td>151.7</td><td>-1.0</td><td>0.5</td><td>-2.3</td><td>2.0E-02</td><td>2.3E-02</td><td>BT20/MB468</td></tr>
<tr><td>>hg38_21_+_8433172_8446622_RNA45SN1_45S_50ntFlanks@12479.12499.21|CTAATAGGGAACGTGAGCTGG</td><td>>hg38_21_+_8433172_8446622_RNA45SN1_45S_50ntFlanks</td><td>12479</td><td>12499</td><td>21</td><td>CTAATAGGGAACGTGAGCTGG</td><td>368.6</td><td>-1.5</td><td>0.7</td><td>-2.3</td><td>2.1E-02</td><td>2.3E-02</td><td>BT20/MB468</td></tr>
<tr><td>>hg38_21_+_8433172_8446622_RNA45SN1_45S_50ntFlanks@12420.12448.29|ATGTCGGCTCTTCCTATCATTGTGAAGCA</td><td>>hg38_21_+_8433172_8446622_RNA45SN1_45S_50ntFlanks</td><td>12420</td><td>12448</td><td>29</td><td>ATGTCGGCTCTTCCTATCATTGTGAAGCA</td><td>1170.6</td><td>-2.5</td><td>0.9</td><td>-2.8</td><td>5.3E-03</td><td>2.3E-02</td><td>BT20/MB231</td></tr>
<tr><td>>MI0000439|hsa-mir-23b&WithFlank&9|+|95085202|95085310@64.86.23(+1U)&[MIMAT0000418&hsa-miR-23b-3p&offsets|0|0(+1U);m-42&9|+|95085265|95085285&offsets|0|+2(+1U)]|ATCACATTGCCAGGGATTACCACT</td><td>>MI0000439|hsa-mir-23b&WithFlank&9|+|95085202|95085310</td><td>64</td><td>86</td><td>23(+1U)</td><td>ATCACATTGCCAGGGATTACCACT</td><td>2174.3</td><td>5.3</td><td>2.3</td><td>2.3</td><td>1.9E-02</td><td>2.3E-02</td><td>MB231/MB468</td></tr>
<tr><td>>hg38_21_+_8433172_8446622_RNA45SN1_45S_50ntFlanks@8191.8213.23|CAGCCCGTGGACGGTGTGAGGCC</td><td>>hg38_21_+_8433172_8446622_RNA45SN1_45S_50ntFlanks</td><td>8191</td><td>8213</td><td>23</td><td>CAGCCCGTGGACGGTGTGAGGCC</td><td>536.0</td><td>-1.5</td><td>0.6</td><td>-2.3</td><td>1.9E-02</td><td>2.3E-02</td><td>MB231/MB468</td></tr>
<tr><td>>hg38_21_+_8433172_8446622_RNA45SN1_45S_50ntFlanks@476.523.48|TTCGTGATCGATGTGGTGACGTCGTGCTCTCCCGGGCCGGGTCCGAGC</td><td>>hg38_21_+_8433172_8446622_RNA45SN1_45S_50ntFlanks</td><td>476</td><td>523</td><td>48</td><td>TTCGTGATCGATGTGGTGACGTCGTGCTCTCCCGGGCCGGGTCCGAGC</td><td>1127.1</td><td>-2.6</td><td>0.9</td><td>-2.8</td><td>5.3E-03</td><td>2.3E-02</td><td>BT20/MB231</td></tr>
<tr><td>>MI0000542|hsa-mir-320a&WithFlank&8|-|22244960|22245043@49.70.22(+1U)&[MIMAT0000510&hsa-miR-320a-3p&offsets|+1|+1(+1U);m-60&8|-|22244975|22244996&offsets|+1|+1(+1U)]|AAAGCTGGGTTGAGAGGGCGAAT</td><td>>MI0000542|hsa-mir-320a&WithFlank&8|-|22244960|22245043</td><td>49</td><td>70</td><td>22(+1U)</td><td>AAAGCTGGGTTGAGAGGGCGAAT</td><td>662.1</td><td>1.9</td><td>0.7</td><td>2.8</td><td>5.3E-03</td><td>2.3E-02</td><td>BT20/MB231</td></tr>
<tr><td>>hg38_21_+_8433172_8446622_RNA45SN1_45S_50ntFlanks@12459.12480.22|AGCGTTGGATTGTTCACCCACT</td><td>>hg38_21_+_8433172_8446622_RNA45SN1_45S_50ntFlanks</td><td>12459</td><td>12480</td><td>22</td><td>AGCGTTGGATTGTTCACCCACT</td><td>308.0</td><td>-2.2</td><td>0.9</td><td>-2.3</td><td>2.1E-02</td><td>2.3E-02</td><td>BT20/MB468</td></tr>
<tr><td>>hg38_21_+_8433172_8446622_RNA45SN1_45S_50ntFlanks@11671.11693.23|CCAGTGCTCTGAATGTCAAAGTG</td><td>>hg38_21_+_8433172_8446622_RNA45SN1_45S_50ntFlanks</td><td>11671</td><td>11693</td><td>23</td><td>CCAGTGCTCTGAATGTCAAAGTG</td><td>914.4</td><td>-2.0</td><td>0.9</td><td>-2.3</td><td>1.9E-02</td><td>2.3E-02</td><td>MB231/MB468</td></tr>
<tr><td>>MI0006273|hsa-mir-1180&WithFlank&17|-|19344500|19344580@47.68.22&[MIMAT0005825&hsa-miR-1180-3p&offsets|0|0;m-214&17|-|19344513|19344534&offsets|0|0]|TTTCCGGCTCGCGTGGGTGTGT</td><td>>MI0006273|hsa-mir-1180&WithFlank&17|-|19344500|19344580</td><td>47</td><td>68</td><td>22</td><td>TTTCCGGCTCGCGTGGGTGTGT</td><td>1840.8</td><td>1.8</td><td>0.7</td><td>2.8</td><td>5.4E-03</td><td>2.3E-02</td><td>BT20/MB231</td></tr>
<tr><td>>trna10_LysCTT_16_+_3241501_3241573@1.36.36,trna13_LysCTT_14_-_58706613_58706685@1.36.36,trna2_LysCTT_15_+_79152904_79152976@1.36.36|GCCCGGCTAGCTCAGTCGGTAGAGCATGGGACTCTT</td><td>>trna10_LysCTT_16_+_3241501_3241573</td><td>1</td><td>36</td><td>36</td><td>GCCCGGCTAGCTCAGTCGGTAGAGCATGGGACTCTT</td><td>7393.3</td><td>2.6</td><td>0.9</td><td>2.8</td><td>5.4E-03</td><td>2.3E-02</td><td>BT20/MB231</td></tr>
<tr><td>>hg38_21_+_8433172_8446622_RNA45SN1_45S_50ntFlanks@6651.6680.30|CGACTCTTAGCGGTGGATCACTCGGCTCGT</td><td>>hg38_21_+_8433172_8446622_RNA45SN1_45S_50ntFlanks</td><td>6651</td><td>6680</td><td>30</td><td>CGACTCTTAGCGGTGGATCACTCGGCTCGT</td><td>3916.9</td><td>1.4</td><td>0.6</td><td>2.3</td><td>2.0E-02</td><td>2.3E-02</td><td>MB231/MB468</td></tr>
<tr><td>>hg38_21_+_8433172_8446622_RNA45SN1_45S_50ntFlanks@12482.12500.19|ATAGGGAACGTGAGCTGGG</td><td>>hg38_21_+_8433172_8446622_RNA45SN1_45S_50ntFlanks</td><td>12482</td><td>12500</td><td>19</td><td>ATAGGGAACGTGAGCTGGG</td><td>811.3</td><td>2.9</td><td>1.3</td><td>2.3</td><td>2.1E-02</td><td>2.3E-02</td><td>BT20/MB468</td></tr>
<tr><td>>trna111_HisGTG_1_-_147774845_147774916@-1T.30.31,trna118_HisGTG_1_-_145396881_145396952@-1T.30.31,trna16_HisGTG_1_+_146544773_146544844@-1T.30.31,trna1_HisGTG_15_+_45493349_45493420@-1T.30.31,trna21_HisGTG_1_+_147753471_147753542@-1T.30.31,trna33_HisGTG_6_+_27125906_27125977@-1T.30.31,trna7_HisGTG_9_-_14433938_14434009@-1T.30.31,trna8_HisGTG_15_-_45492611_45492682@-1T.30.31,trna9_HisGTG_15_-_45490804_45490875@-1T.30.31|TGCCGTGATCGTATAGTGGTTAGTACTCTGC</td><td>>trna111_HisGTG_1_-_147774845_147774916</td><td>-1T</td><td>30</td><td>31</td><td>TGCCGTGATCGTATAGTGGTTAGTACTCTGC</td><td>1048.1</td><td>2.2</td><td>0.9</td><td>2.3</td><td>2.0E-02</td><td>2.3E-02</td><td>MB231/MB468</td></tr>
<tr><td>>hg38_21_+_8433172_8446622_RNA45SN1_45S_50ntFlanks@12474.12496.23|ACCCACTAATAGGGAACGTGAGC</td><td>>hg38_21_+_8433172_8446622_RNA45SN1_45S_50ntFlanks</td><td>12474</td><td>12496</td><td>23</td><td>ACCCACTAATAGGGAACGTGAGC</td><td>1408.2</td><td>-2.3</td><td>0.8</td><td>-2.8</td><td>5.4E-03</td><td>2.3E-02</td><td>BT20/MB231</td></tr>
<tr><td>>hg38_1_-_228634819_228635039_RNA5S12_5S_50ntFlanks@134.171.38|TGGGAGACCGCCTGGGAATACCGGGTGCTGTAGGCTTT</td><td>>hg38_1_-_228634819_228635039_RNA5S12_5S_50ntFlanks</td><td>134</td><td>171</td><td>38</td><td>TGGGAGACCGCCTGGGAATACCGGGTGCTGTAGGCTTT</td><td>46719.7</td><td>-2.2</td><td>0.8</td><td>-2.8</td><td>5.5E-03</td><td>2.3E-02</td><td>BT20/MB231</td></tr>
<tr><td>>hg38_21_+_8433172_8446622_RNA45SN1_45S_50ntFlanks@12909.12933.25|CCGCACGTTCGTGGGGAACCTGGCG</td><td>>hg38_21_+_8433172_8446622_RNA45SN1_45S_50ntFlanks</td><td>12909</td><td>12933</td><td>25</td><td>CCGCACGTTCGTGGGGAACCTGGCG</td><td>126.2</td><td>-1.8</td><td>0.8</td><td>-2.3</td><td>2.0E-02</td><td>2.3E-02</td><td>MB231/MB468</td></tr>
<tr><td>>MI0000067|hsa-let-7f-1&WithFlank&9|+|94176341|94176439@13.31.19;MI0000068|hsa-let-7f-2&WithFlank&X|-|53557186|53557280@14.32.19&[MIMAT0000067&hsa-let-7f-5p&offsets|0|-3;m-20&9|+|94176353|94176374&offsets|0|-3];[MIMAT0000067_1&hsa-let-7f-5p&offsets|0|-3;m-19&X|-|53557246|53557267&offsets|0|-3]|TGAGGTAGTAGATTGTATA</td><td>>MI0000067|hsa-let-7f-1&WithFlank&9|+|94176341|94176439</td><td>13</td><td>31</td><td>19</td><td>TGAGGTAGTAGATTGTATA</td><td>1256.9</td><td>2.2</td><td>0.8</td><td>2.8</td><td>5.5E-03</td><td>2.4E-02</td><td>BT20/MB231</td></tr>
<tr><td>>MI0000746|hsa-mir-99b&WithFlank&19|+|51692606|51692687@13.32.20&[MIMAT0000689&hsa-miR-99b-5p&offsets|0|-2;m-10&19|+|51692618|51692639&offsets|0|-2]|CACCCGTAGAACCGACCTTG</td><td>>MI0000746|hsa-mir-99b&WithFlank&19|+|51692606|51692687</td><td>13</td><td>32</td><td>20</td><td>CACCCGTAGAACCGACCTTG</td><td>1929.1</td><td>2.1</td><td>0.8</td><td>2.8</td><td>5.5E-03</td><td>2.4E-02</td><td>BT20/MB231</td></tr>
<tr><td>>hg38_21_+_8433172_8446622_RNA45SN1_45S_50ntFlanks@6648.6692.45|GTACGACTCTTAGCGGTGGATCACTCGGCTCGTGCGTCGATGAAG</td><td>>hg38_21_+_8433172_8446622_RNA45SN1_45S_50ntFlanks</td><td>6648</td><td>6692</td><td>45</td><td>GTACGACTCTTAGCGGTGGATCACTCGGCTCGTGCGTCGATGAAG</td><td>418.2</td><td>-2.1</td><td>0.8</td><td>-2.8</td><td>5.6E-03</td><td>2.4E-02</td><td>BT20/MB231</td></tr>
<tr><td>>hg38_21_+_8433172_8446622_RNA45SN1_45S_50ntFlanks@12644.12661.18|TCTGTGGGATTATGACTG</td><td>>hg38_21_+_8433172_8446622_RNA45SN1_45S_50ntFlanks</td><td>12644</td><td>12661</td><td>18</td><td>TCTGTGGGATTATGACTG</td><td>331.4</td><td>-1.6</td><td>0.7</td><td>-2.3</td><td>2.0E-02</td><td>2.4E-02</td><td>MB231/MB468</td></tr>
<tr><td>>am_trnaMT_ValTAC_MT_+_1602_1670@50.67.18|ACTTAACTTGACCGCTCT</td><td>>am_trnaMT_ValTAC_MT_+_1602_1670</td><td>50</td><td>67</td><td>18</td><td>ACTTAACTTGACCGCTCT</td><td>3511.3</td><td>2.2</td><td>0.8</td><td>2.8</td><td>5.6E-03</td><td>2.4E-02</td><td>BT20/MB231</td></tr>
<tr><td>>hg38_MT_+_1621_3279_RNR2_16S_50ntFlanks@1370.1414.45|ATGTTGGATCAGGACATCCCGATGGTGCAGCCGCTATTAAAGGTT</td><td>>hg38_MT_+_1621_3279_RNR2_16S_50ntFlanks</td><td>1370</td><td>1414</td><td>45</td><td>ATGTTGGATCAGGACATCCCGATGGTGCAGCCGCTATTAAAGGTT</td><td>275.6</td><td>-2.1</td><td>0.8</td><td>-2.8</td><td>5.6E-03</td><td>2.4E-02</td><td>BT20/MB231</td></tr>
<tr><td>>hg38_21_+_8433172_8446622_RNA45SN1_45S_50ntFlanks@7985.8008.24|GATCAGACGTGGCGACCCGCTGAA</td><td>>hg38_21_+_8433172_8446622_RNA45SN1_45S_50ntFlanks</td><td>7985</td><td>8008</td><td>24</td><td>GATCAGACGTGGCGACCCGCTGAA</td><td>285.6</td><td>-3.9</td><td>1.7</td><td>-2.3</td><td>2.0E-02</td><td>2.4E-02</td><td>MB231/MB468</td></tr>
<tr><td>>trna11_GluTTC_15_-_26327381_26327452@1.29.29,trna3_GluTTC_13_-_45492062_45492133@1.29.29|TCCCACATGGTCTAGCGGTTAGGATTCCT</td><td>>trna11_GluTTC_15_-_26327381_26327452</td><td>1</td><td>29</td><td>29</td><td>TCCCACATGGTCTAGCGGTTAGGATTCCT</td><td>2764.3</td><td>-1.8</td><td>0.8</td><td>-2.3</td><td>2.1E-02</td><td>2.4E-02</td><td>BT20/MB468</td></tr>
<tr><td>>hg38_21_+_8433172_8446622_RNA45SN1_45S_50ntFlanks@7978.8017.40|GACCTCAGATCAGACGTGGCGACCCGCTGAATTTAAGCAT</td><td>>hg38_21_+_8433172_8446622_RNA45SN1_45S_50ntFlanks</td><td>7978</td><td>8017</td><td>40</td><td>GACCTCAGATCAGACGTGGCGACCCGCTGAATTTAAGCAT</td><td>247.3</td><td>-2.8</td><td>1.0</td><td>-2.8</td><td>5.7E-03</td><td>2.4E-02</td><td>BT20/MB231</td></tr>
<tr><td>>hg38_21_+_8433172_8446622_RNA45SN1_45S_50ntFlanks@9545.9575.31|GTGGAGGTCCGTAGCGGTCCTGACGTGCAAA</td><td>>hg38_21_+_8433172_8446622_RNA45SN1_45S_50ntFlanks</td><td>9545</td><td>9575</td><td>31</td><td>GTGGAGGTCCGTAGCGGTCCTGACGTGCAAA</td><td>105.6</td><td>-1.2</td><td>0.5</td><td>-2.3</td><td>2.0E-02</td><td>2.4E-02</td><td>MB231/MB468</td></tr>
<tr><td>>hg38_21_+_8433172_8446622_RNA45SN1_45S_50ntFlanks@6661.6701.41|CGGTGGATCACTCGGCTCGTGCGTCGATGAAGAACGCAGCT</td><td>>hg38_21_+_8433172_8446622_RNA45SN1_45S_50ntFlanks</td><td>6661</td><td>6701</td><td>41</td><td>CGGTGGATCACTCGGCTCGTGCGTCGATGAAGAACGCAGCT</td><td>1053.9</td><td>2.0</td><td>0.7</td><td>2.8</td><td>5.7E-03</td><td>2.4E-02</td><td>BT20/MB231</td></tr>
<tr><td>>MI0000447|hsa-mir-128-1&WithFlank&2|+|135665391|135665484@56.75.20(+1A);MI0000727|hsa-mir-128-2&WithFlank&3|+|35744470|35744565@58.77.20(+1A)&[MIMAT0000424&hsa-miR-128-3p&offsets|0|-1(+1A);m-108&2|+|135665446|135665466&offsets|0|-1(+1A)];[MIMAT0000424_1&hsa-miR-128-3p&offsets|0|-1(+1A);m-117&3|+|35744527|35744547&offsets|0|-1(+1A)]|TCACAGTGAACCGGTCTCTTA</td><td>>MI0000447|hsa-mir-128-1&WithFlank&2|+|135665391|135665484</td><td>56</td><td>75</td><td>20(+1A)</td><td>TCACAGTGAACCGGTCTCTTA</td><td>608.7</td><td>2.1</td><td>0.8</td><td>2.8</td><td>5.7E-03</td><td>2.4E-02</td><td>BT20/MB231</td></tr>
</table>

<table>
<tr><td>>hg38_MT_+_1621_3279_RNR2_16S_50ntFlanks@493.518.26|GCTCTTTGGACACTAGGAAAAAACCT</td><td>>hg38_MT_+_1621_3279_RNR2_16S_50ntFlanks</td><td>493</td><td>518</td><td>26</td><td>GCTCTTTGGACACTAGGAAAAAACCT</td><td>1214.5</td><td>-2.1</td><td>0.8</td><td>-2.8</td><td>5.7E-03</td><td>2.4E-02</td><td>BT20/MB231</td></tr>
<tr><td>>hg38_21_+_8433172_8446622_RNA45SN1_45S_50ntFlanks@8176.8206.31|CTTCTGATCGAGGCCCAGCCCGTGGACGGTG</td><td>>hg38_21_+_8433172_8446622_RNA45SN1_45S_50ntFlanks</td><td>8176</td><td>8206</td><td>31</td><td>CTTCTGATCGAGGCCCAGCCCGTGGACGGTG</td><td>450.7</td><td>-1.0</td><td>0.4</td><td>-2.3</td><td>2.1E-02</td><td>2.4E-02</td><td>MB231/MB468</td></tr>
<tr><td>>am_trnaMT_ValTAC_MT_+_1602_1670@49.68.20|AACTTAACTTGACCGCTCTG</td><td>>am_trnaMT_ValTAC_MT_+_1602_1670</td><td>49</td><td>68</td><td>20</td><td>AACTTAACTTGACCGCTCTG</td><td>415.8</td><td>-1.9</td><td>0.8</td><td>-2.3</td><td>2.1E-02</td><td>2.4E-02</td><td>MB231/MB468</td></tr>
<tr><td>>hg38_21_+_8433172_8446622_RNA45SN1_45S_50ntFlanks@6650.6669.20|ACGACTCTTAGCGGTGGATC</td><td>>hg38_21_+_8433172_8446622_RNA45SN1_45S_50ntFlanks</td><td>6650</td><td>6669</td><td>20</td><td>ACGACTCTTAGCGGTGGATC</td><td>833.2</td><td>2.3</td><td>0.8</td><td>2.8</td><td>5.8E-03</td><td>2.4E-02</td><td>BT20/MB231</td></tr>
<tr><td>>MI0000070|hsa-mir-16-1&WithFlank&13|-|50048967|50049067@20.41.22;MI0000115|hsa-mir-16-2&WithFlank&3|+|160404739|160404831@16.37.22&[MIMAT0000069&hsa-miR-16-5p&offsets|0|0;m-62&13|-|50049027|50049048&offsets|0|0];[MIMAT0000069_1&hsa-miR-16-5p&offsets|0|0;m-61&3|+|160404754|160404775&offsets|0|0]|TAGCAGCACGTAAATATTGGCG</td><td>>MI0000070|hsa-mir-16-1&WithFlank&13|-|50048967|50049067</td><td>20</td><td>41</td><td>22</td><td>TAGCAGCACGTAAATATTGGCG</td><td>13095.6</td><td>2.6</td><td>0.9</td><td>2.8</td><td>5.8E-03</td><td>2.4E-02</td><td>BT20/MB231</td></tr>
<tr><td>>hg38_21_+_8433172_8446622_RNA45SN1_45S_50ntFlanks@4007.4042.36|CCTCGGGCCGATCGCACGCCCCCCGTGGCGGCGACG</td><td>>hg38_21_+_8433172_8446622_RNA45SN1_45S_50ntFlanks</td><td>4007</td><td>4042</td><td>36</td><td>CCTCGGGCCGATCGCACGCCCCCCGTGGCGGCGACG</td><td>153.6</td><td>-1.2</td><td>0.5</td><td>-2.3</td><td>2.1E-02</td><td>2.5E-02</td><td>MB231/MB468</td></tr>
<tr><td>>hg38_21_+_8433172_8446622_RNA45SN1_45S_50ntFlanks@11315.11334.20|CCACCCCACGTCTCGTCGCG</td><td>>hg38_21_+_8433172_8446622_RNA45SN1_45S_50ntFlanks</td><td>11315</td><td>11334</td><td>20</td><td>CCACCCCACGTCTCGTCGCG</td><td>258.2</td><td>-2.2</td><td>1.0</td><td>-2.3</td><td>2.2E-02</td><td>2.5E-02</td><td>BT20/MB468</td></tr>
<tr><td>>hg38_21_+_8433172_8446622_RNA45SN1_45S_50ntFlanks@7975.7996.22|CGCGACCTCAGATCAGACGTGG</td><td>>hg38_21_+_8433172_8446622_RNA45SN1_45S_50ntFlanks</td><td>7975</td><td>7996</td><td>22</td><td>CGCGACCTCAGATCAGACGTGG</td><td>3396.8</td><td>1.6</td><td>0.7</td><td>2.3</td><td>2.2E-02</td><td>2.5E-02</td><td>BT20/MB468</td></tr>
<tr><td>>hg38_21_+_8433172_8446622_RNA45SN1_45S_50ntFlanks@6647.6672.26|CGTACGACTCTTAGCGGTGGATCACT</td><td>>hg38_21_+_8433172_8446622_RNA45SN1_45S_50ntFlanks</td><td>6647</td><td>6672</td><td>26</td><td>CGTACGACTCTTAGCGGTGGATCACT</td><td>1070.3</td><td>2.3</td><td>0.8</td><td>2.8</td><td>5.9E-03</td><td>2.5E-02</td><td>BT20/MB231</td></tr>
<tr><td>>hg38_21_+_8433172_8446622_RNA45SN1_45S_50ntFlanks@5473.5501.29|CGGCCCTGGCGGAGCGCTGAGAAGACGGT</td><td>>hg38_21_+_8433172_8446622_RNA45SN1_45S_50ntFlanks</td><td>5473</td><td>5501</td><td>29</td><td>CGGCCCTGGCGGAGCGCTGAGAAGACGGT</td><td>667.5</td><td>3.2</td><td>1.2</td><td>2.8</td><td>5.9E-03</td><td>2.5E-02</td><td>BT20/MB231</td></tr>
<tr><td>>hg38_21_+_8433172_8446622_RNA45SN1_45S_50ntFlanks@10781.10809.29|AGGTGAACAGCCTCTGGCATGTTGGAACA</td><td>>hg38_21_+_8433172_8446622_RNA45SN1_45S_50ntFlanks</td><td>10781</td><td>10809</td><td>29</td><td>AGGTGAACAGCCTCTGGCATGTTGGAACA</td><td>121.5</td><td>-1.0</td><td>0.4</td><td>-2.3</td><td>2.1E-02</td><td>2.5E-02</td><td>MB231/MB468</td></tr>
<tr><td>>hg38_21_+_8433172_8446622_RNA45SN1_45S_50ntFlanks@12419.12437.19|GATGTCGGCTCTTCCTATC</td><td>>hg38_21_+_8433172_8446622_RNA45SN1_45S_50ntFlanks</td><td>12419</td><td>12437</td><td>19</td><td>GATGTCGGCTCTTCCTATC</td><td>4154.6</td><td>-2.1</td><td>0.8</td><td>-2.8</td><td>5.9E-03</td><td>2.5E-02</td><td>BT20/MB231</td></tr>
<tr><td>>hg38_21_+_8433172_8446622_RNA45SN1_45S_50ntFlanks@12441.12473.33|GTGAAGCAGAATTCACCAAGCGTTGGATTGTTC</td><td>>hg38_21_+_8433172_8446622_RNA45SN1_45S_50ntFlanks</td><td>12441</td><td>12473</td><td>33</td><td>GTGAAGCAGAATTCACCAAGCGTTGGATTGTTC</td><td>146.9</td><td>-2.4</td><td>0.9</td><td>-2.8</td><td>6.0E-03</td><td>2.5E-02</td><td>BT20/MB231</td></tr>
<tr><td>>MI0000299|hsa-mir-222&WithFlank&X|-|45747009|45747130@75.98.24(+1A)&[MIMAT0000279&hsa-miR-222-3p&offsets|0|+3(+1A);m-140&X|-|45747033|45747056&offsets|0|0(+1A)]|AGCTACATCTGGCTACTGGGTCTCA</td><td>>MI0000299|hsa-mir-222&WithFlank&X|-|45747009|45747130</td><td>75</td><td>98</td><td>24(+1A)</td><td>AGCTACATCTGGCTACTGGGTCTCA</td><td>6305.8</td><td>-1.8</td><td>0.6</td><td>-2.8</td><td>5.9E-03</td><td>2.5E-02</td><td>BT20/MB231</td></tr>
<tr><td>>MI0000434|hsa-let-7i&WithFlank&12|+|62603680|62603775@12.33.22(+2U)&[MIMAT0000415&hsa-let-7i-5p&offsets|0|0(+2U);m-49&12|+|62603691|62603712&offsets|0|0(+2U)]|TGAGGTAGTAGTTTGTGCTGTTTT</td><td>>MI0000434|hsa-let-7i&WithFlank&12|+|62603680|62603775</td><td>12</td><td>33</td><td>22(+2U)</td><td>TGAGGTAGTAGTTTGTGCTGTTTT</td><td>408.7</td><td>-1.8</td><td>0.7</td><td>-2.8</td><td>5.9E-03</td><td>2.5E-02</td><td>BT20/MB231</td></tr>
<tr><td>>hg38_21_+_8433172_8446622_RNA45SN1_45S_50ntFlanks@12942.12960.19|TTCGTAGACGACCTGCTTC</td><td>>hg38_21_+_8433172_8446622_RNA45SN1_45S_50ntFlanks</td><td>12942</td><td>12960</td><td>19</td><td>TTCGTAGACGACCTGCTTC</td><td>703.0</td><td>1.4</td><td>0.6</td><td>2.3</td><td>2.1E-02</td><td>2.5E-02</td><td>MB231/MB468</td></tr>
<tr><td>>hg38_21_+_8433172_8446622_RNA45SN1_45S_50ntFlanks@3777.3794.18|CGGTACAGTGAAACTGCG</td><td>>hg38_21_+_8433172_8446622_RNA45SN1_45S_50ntFlanks</td><td>3777</td><td>3794</td><td>18</td><td>CGGTACAGTGAAACTGCG</td><td>132.6</td><td>-1.8</td><td>0.8</td><td>-2.3</td><td>2.1E-02</td><td>2.5E-02</td><td>MB231/MB468</td></tr>
<tr><td>>hg38_21_+_8433172_8446622_RNA45SN1_45S_50ntFlanks@8105.8140.36|CGCGGCGGGGCGCGGGACATGTGGCGTACGGAAGAC</td><td>>hg38_21_+_8433172_8446622_RNA45SN1_45S_50ntFlanks</td><td>8105</td><td>8140</td><td>36</td><td>CGCGGCGGGGCGCGGGACATGTGGCGTACGGAAGAC</td><td>263.1</td><td>-4.3</td><td>1.9</td><td>-2.3</td><td>2.1E-02</td><td>2.5E-02</td><td>MB231/MB468</td></tr>
<tr><td>>hg38_21_+_8433172_8446622_RNA45SN1_45S_50ntFlanks@7982.8012.31|TCAGATCAGACGTGGCGACCCGCTGAATTTA</td><td>>hg38_21_+_8433172_8446622_RNA45SN1_45S_50ntFlanks</td><td>7982</td><td>8012</td><td>31</td><td>TCAGATCAGACGTGGCGACCCGCTGAATTTA</td><td>92.3</td><td>-1.4</td><td>0.6</td><td>-2.3</td><td>2.1E-02</td><td>2.5E-02</td><td>MB231/MB468</td></tr>
<tr><td>>hg38_21_+_8433172_8446622_RNA45SN1_45S_50ntFlanks@6646.6677.32|TCGTACGACTCTTAGCGGTGGATCACTCGGCT</td><td>>hg38_21_+_8433172_8446622_RNA45SN1_45S_50ntFlanks</td><td>6646</td><td>6677</td><td>32</td><td>TCGTACGACTCTTAGCGGTGGATCACTCGGCT</td><td>2794.6</td><td>2.2</td><td>0.8</td><td>2.7</td><td>6.0E-03</td><td>2.5E-02</td><td>BT20/MB231</td></tr>
<tr><td>>hg38_21_+_8433172_8446622_RNA45SN1_45S_50ntFlanks@12995.13040.46|CTCGCTGCGATCTATTGAAAGTCAGCCCTCGACACAAGGGTTTGTC</td><td>>hg38_21_+_8433172_8446622_RNA45SN1_45S_50ntFlanks</td><td>12995</td><td>13040</td><td>46</td><td>CTCGCTGCGATCTATTGAAAGTCAGCCCTCGACACAAGGGTTTGTC</td><td>2939.4</td><td>-5.5</td><td>2.0</td><td>-2.7</td><td>6.0E-03</td><td>2.5E-02</td><td>BT20/MB231</td></tr>
<tr><td>>hg38_21_+_8433172_8446622_RNA45SN1_45S_50ntFlanks@11672.11703.32|CAGTGCTCTGAATGTCAAAGTGAAGAAATTCA</td><td>>hg38_21_+_8433172_8446622_RNA45SN1_45S_50ntFlanks</td><td>11672</td><td>11703</td><td>32</td><td>CAGTGCTCTGAATGTCAAAGTGAAGAAATTCA</td><td>297.5</td><td>-2.2</td><td>0.8</td><td>-2.7</td><td>6.0E-03</td><td>2.5E-02</td><td>BT20/MB231</td></tr>
<tr><td>>hg38_21_+_8433172_8446622_RNA45SN1_45S_50ntFlanks@10350.10372.23|ACGAGAACTTTGAAGGCCGAAGT</td><td>>hg38_21_+_8433172_8446622_RNA45SN1_45S_50ntFlanks</td><td>10350</td><td>10372</td><td>23</td><td>ACGAGAACTTTGAAGGCCGAAGT</td><td>559.9</td><td>-2.2</td><td>0.8</td><td>-2.7</td><td>6.1E-03</td><td>2.5E-02</td><td>BT20/MB231</td></tr>
<tr><td>>hg38_21_+_8433172_8446622_RNA45SN1_45S_50ntFlanks@11672.11698.27|CAGTGCTCTGAATGTCAAAGTGAAGAA</td><td>>hg38_21_+_8433172_8446622_RNA45SN1_45S_50ntFlanks</td><td>11672</td><td>11698</td><td>27</td><td>CAGTGCTCTGAATGTCAAAGTGAAGAA</td><td>159.0</td><td>1.1</td><td>0.5</td><td>2.3</td><td>2.1E-02</td><td>2.5E-02</td><td>MB231/MB468</td></tr>
<tr><td>>hg38_21_+_8433172_8446622_RNA45SN1_45S_50ntFlanks@6650.6672.23|ACGACTCTTAGCGGTGGATCACT</td><td>>hg38_21_+_8433172_8446622_RNA45SN1_45S_50ntFlanks</td><td>6650</td><td>6672</td><td>23</td><td>ACGACTCTTAGCGGTGGATCACT</td><td>390.8</td><td>2.2</td><td>0.8</td><td>2.7</td><td>6.1E-03</td><td>2.5E-02</td><td>BT20/MB231</td></tr>
<tr><td>>MI0000066|hsa-let-7e&WithFlank&19|+|51692780|51692870@14.34.21(+1A)&[MIMAT0000066&hsa-let-7e-5p&offsets|0|-1(+1A);m-50&19|+|51692793|51692814&offsets|0|-1(+1A)]|TGAGGTAGGAGGTTGTATAGTA</td><td>>MI0000066|hsa-let-7e&WithFlank&19|+|51692780|51692870</td><td>14</td><td>34</td><td>21(+1A)</td><td>TGAGGTAGGAGGTTGTATAGTA</td><td>813.7</td><td>2.2</td><td>0.8</td><td>2.7</td><td>6.1E-03</td><td>2.5E-02</td><td>BT20/MB231</td></tr>
<tr><td>>MI0000103|hsa-mir-101-1&WithFlank&1|-|65058428|65058514@52.73.22(+1A);MI0000739|hsa-mir-101-2&WithFlank&9|+|4850291|4850381@54.75.22(+1A)&[MIMAT0000099&hsa-miR-101-3p&offsets|-1|0(+1A);m-17&1|-|65058443|65058462&offsets|-1|+1(+1A)];[MIMAT0000099_1&hsa-miR-101-3p&offsets|-1|0(+1A);m-16&9|+|4850345|4850364&offsets|-1|+1(+1A)]|GTACAGTACTGTGATAACTGAAA</td><td>>MI0000103|hsa-mir-101-1&WithFlank&1|-|65058428|65058514</td><td>52</td><td>73</td><td>22(+1A)</td><td>GTACAGTACTGTGATAACTGAAA</td><td>2494.0</td><td>2.8</td><td>1.0</td><td>2.7</td><td>6.1E-03</td><td>2.5E-02</td><td>BT20/MB231</td></tr>
<tr><td>>MI0000746|hsa-mir-99b&WithFlank&19|+|51692606|51692687@51.69.19&[MIMAT0004678&hsa-miR-99b-3p&offsets|0|-3;m-151&19|+|51692656|51692678&offsets|0|-4]|CAAGCTCGTGTCTGTGGGT</td><td>>MI0000746|hsa-mir-99b&WithFlank&19|+|51692606|51692687</td><td>51</td><td>69</td><td>19</td><td>CAAGCTCGTGTCTGTGGGT</td><td>426.2</td><td>2.3</td><td>0.8</td><td>2.7</td><td>6.1E-03</td><td>2.5E-02</td><td>BT20/MB231</td></tr>
<tr><td>>hg38_21_+_8433172_8446622_RNA45SN1_45S_50ntFlanks@12417.12453.37|TCGATGTCGGCTCTTCCTATCATTGTGAAGCAGAATT</td><td>>hg38_21_+_8433172_8446622_RNA45SN1_45S_50ntFlanks</td><td>12417</td><td>12453</td><td>37</td><td>TCGATGTCGGCTCTTCCTATCATTGTGAAGCAGAATT</td><td>231.6</td><td>-2.2</td><td>0.8</td><td>-2.7</td><td>6.2E-03</td><td>2.6E-02</td><td>BT20/MB231</td></tr>
<tr><td>>MI0003643|hsa-mir-629&WithFlank&15|-|70079366|70079474@28.48.21&[MIMAT0004810&hsa-miR-629-5p&offsets|0|0;m-150&15|-|70079427|70079447&offsets|0|0]|TGGGTTTACGTTGGGAGAACT</td><td>>MI0003643|hsa-mir-629&WithFlank&15|-|70079366|70079474</td><td>28</td><td>48</td><td>21</td><td>TGGGTTTACGTTGGGAGAACT</td><td>623.1</td><td>1.8</td><td>0.7</td><td>2.7</td><td>6.2E-03</td><td>2.6E-02</td><td>BT20/MB231</td></tr>
<tr><td>>hg38_21_+_8433172_8446622_RNA45SN1_45S_50ntFlanks@12415.12435.21|CTTCGATGTCGGCTCTTCCTA</td><td>>hg38_21_+_8433172_8446622_RNA45SN1_45S_50ntFlanks</td><td>12415</td><td>12435</td><td>21</td><td>CTTCGATGTCGGCTCTTCCTA</td><td>1716.0</td><td>-2.0</td><td>0.9</td><td>-2.3</td><td>2.3E-02</td><td>2.6E-02</td><td>BT20/MB468</td></tr>
<tr><td>>hg38_21_+_8433172_8446622_RNA45SN1_45S_50ntFlanks@10793.10816.24|TCTGGCATGTTGGAACAATGTAGG</td><td>>hg38_21_+_8433172_8446622_RNA45SN1_45S_50ntFlanks</td><td>10793</td><td>10816</td><td>24</td><td>TCTGGCATGTTGGAACAATGTAGG</td><td>172.3</td><td>-1.2</td><td>0.5</td><td>-2.3</td><td>2.2E-02</td><td>2.6E-02</td><td>MB231/MB468</td></tr>
<tr><td>>hg38_21_+_8433172_8446622_RNA45SN1_45S_50ntFlanks@11320.11343.24|CCACGTCTCGTCGCGCGCGCGTCC</td><td>>hg38_21_+_8433172_8446622_RNA45SN1_45S_50ntFlanks</td><td>11320</td><td>11343</td><td>24</td><td>CCACGTCTCGTCGCGCGCGCGTCC</td><td>146.3</td><td>-3.9</td><td>1.7</td><td>-2.3</td><td>2.2E-02</td><td>2.6E-02</td><td>MB231/MB468</td></tr>
<tr><td>>hg38_21_+_8433172_8446622_RNA45SN1_45S_50ntFlanks@4565.4587.23|AATAATGGAATAGGACCGCGGTT</td><td>>hg38_21_+_8433172_8446622_RNA45SN1_45S_50ntFlanks</td><td>4565</td><td>4587</td><td>23</td><td>AATAATGGAATAGGACCGCGGTT</td><td>467.3</td><td>-1.5</td><td>0.7</td><td>-2.3</td><td>2.2E-02</td><td>2.6E-02</td><td>MB231/MB468</td></tr>
<tr><td>>am_trnaMT_HisGTG_MT_+_12138_12206@20.51.32|AACATCAGATTGTGAATCTGACAACAGAGGCT</td><td>>am_trnaMT_HisGTG_MT_+_12138_12206</td><td>20</td><td>51</td><td>32</td><td>AACATCAGATTGTGAATCTGACAACAGAGGCT</td><td>328.2</td><td>2.3</td><td>0.9</td><td>2.7</td><td>6.3E-03</td><td>2.6E-02</td><td>BT20/MB231</td></tr>
<tr><td>>hg38_21_+_8433172_8446622_RNA45SN1_45S_50ntFlanks@11668.11688.21|TGCCCAGTGCTCTGAATGTCA</td><td>>hg38_21_+_8433172_8446622_RNA45SN1_45S_50ntFlanks</td><td>11668</td><td>11688</td><td>21</td><td>TGCCCAGTGCTCTGAATGTCA</td><td>681.0</td><td>-2.4</td><td>0.9</td><td>-2.7</td><td>6.3E-03</td><td>2.6E-02</td><td>BT20/MB231</td></tr>
<tr><td>>hg38_21_+_8433172_8446622_RNA45SN1_45S_50ntFlanks@7991.8018.28|ACGTGGCGACCCGCTGAATTTAAGCATA</td><td>>hg38_21_+_8433172_8446622_RNA45SN1_45S_50ntFlanks</td><td>7991</td><td>8018</td><td>28</td><td>ACGTGGCGACCCGCTGAATTTAAGCATA</td><td>315.0</td><td>1.6</td><td>0.7</td><td>2.3</td><td>2.2E-02</td><td>2.6E-02</td><td>MB231/MB468</td></tr>
<tr><td>>hg38_21_+_8433172_8446622_RNA45SN1_45S_50ntFlanks@12412.12435.24|ATCCTTCGATGTCGGCTCTTCCTA</td><td>>hg38_21_+_8433172_8446622_RNA45SN1_45S_50ntFlanks</td><td>12412</td><td>12435</td><td>24</td><td>ATCCTTCGATGTCGGCTCTTCCTA</td><td>162.4</td><td>-3.6</td><td>1.6</td><td>-2.3</td><td>2.2E-02</td><td>2.6E-02</td><td>MB231/MB468</td></tr>
<tr><td>>hg38_21_+_8433172_8446622_RNA45SN1_45S_50ntFlanks@7975.8006.32|CGCGACCTCAGATCAGACGTGGCGACCCGCTG</td><td>>hg38_21_+_8433172_8446622_RNA45SN1_45S_50ntFlanks</td><td>7975</td><td>8006</td><td>32</td><td>CGCGACCTCAGATCAGACGTGGCGACCCGCTG</td><td>10920.5</td><td>2.2</td><td>0.8</td><td>2.7</td><td>6.3E-03</td><td>2.6E-02</td><td>BT20/MB231</td></tr>
<tr><td>>hg38_21_+_8433172_8446622_RNA45SN1_45S_50ntFlanks@12416.12449.34|TTCGATGTCGGCTCTTCCTATCATTGTGAAGCAG</td><td>>hg38_21_+_8433172_8446622_RNA45SN1_45S_50ntFlanks</td><td>12416</td><td>12449</td><td>34</td><td>TTCGATGTCGGCTCTTCCTATCATTGTGAAGCAG</td><td>1502.7</td><td>-2.5</td><td>0.9</td><td>-2.7</td><td>6.4E-03</td><td>2.6E-02</td><td>BT20/MB231</td></tr>
<tr><td>>hg38_21_+_8433172_8446622_RNA45SN1_45S_50ntFlanks@9739.9764.26|AAACGATCTCAACCTATTCTCAAACT</td><td>>hg38_21_+_8433172_8446622_RNA45SN1_45S_50ntFlanks</td><td>9739</td><td>9764</td><td>26</td><td>AAACGATCTCAACCTATTCTCAAACT</td><td>195.9</td><td>1.4</td><td>0.6</td><td>2.3</td><td>2.2E-02</td><td>2.6E-02</td><td>MB231/MB468</td></tr>
<tr><td>>trnaMT_SerGCT_MT_+_12207_12265@8.31.24|CTCACAAGAACTGCTAACTCATGC</td><td>>trnaMT_SerGCT_MT_+_12207_12265</td><td>8</td><td>31</td><td>24</td><td>CTCACAAGAACTGCTAACTCATGC</td><td>89.4</td><td>-1.8</td><td>0.8</td><td>-2.3</td><td>2.2E-02</td><td>2.6E-02</td><td>MB231/MB468</td></tr>
<tr><td>>hg38_21_+_8433172_8446622_RNA45SN1_45S_50ntFlanks@6663.6686.24|GTGGATCACTCGGCTCGTGCGTCG</td><td>>hg38_21_+_8433172_8446622_RNA45SN1_45S_50ntFlanks</td><td>6663</td><td>6686</td><td>24</td><td>GTGGATCACTCGGCTCGTGCGTCG</td><td>411.8</td><td>-2.2</td><td>1.0</td><td>-2.3</td><td>2.4E-02</td><td>2.6E-02</td><td>BT20/MB468</td></tr>
<tr><td>>hg38_21_+_8433172_8446622_RNA45SN1_45S_50ntFlanks@12001.12030.30|CTGCGGGCCGCCGGTGAAATACCACTACTC</td><td>>hg38_21_+_8433172_8446622_RNA45SN1_45S_50ntFlanks</td><td>12001</td><td>12030</td><td>30</td><td>CTGCGGGCCGCCGGTGAAATACCACTACTC</td><td>103.0</td><td>-2.2</td><td>0.9</td><td>-2.3</td><td>2.2E-02</td><td>2.6E-02</td><td>MB231/MB468</td></tr>
<tr><td>>hg38_21_+_8433172_8446622_RNA45SN1_45S_50ntFlanks@10738.10757.20|AAATCTCGCGCCGGGCCGTA</td><td>>hg38_21_+_8433172_8446622_RNA45SN1_45S_50ntFlanks</td><td>10738</td><td>10757</td><td>20</td><td>AAATCTCGCGCCGGGCCGTA</td><td>96.8</td><td>-1.4</td><td>0.6</td><td>-2.3</td><td>2.2E-02</td><td>2.6E-02</td><td>MB231/MB468</td></tr>
<tr><td>>hg38_21_+_8433172_8446622_RNA45SN1_45S_50ntFlanks@10849.10866.18|CGGGATAAGGATTGGCTC</td><td>>hg38_21_+_8433172_8446622_RNA45SN1_45S_50ntFlanks</td><td>10849</td><td>10866</td><td>18</td><td>CGGGATAAGGATTGGCTC</td><td>156.6</td><td>-1.7</td><td>0.7</td><td>-2.3</td><td>2.3E-02</td><td>2.7E-02</td><td>MB231/MB468</td></tr>
<tr><td>>hg38_21_+_8433172_8446622_RNA45SN1_45S_50ntFlanks@3769.3786.18|CGCACGGCCGGTACAGTG</td><td>>hg38_21_+_8433172_8446622_RNA45SN1_45S_50ntFlanks</td><td>3769</td><td>3786</td><td>18</td><td>CGCACGGCCGGTACAGTG</td><td>160.0</td><td>-1.1</td><td>0.5</td><td>-2.3</td><td>2.3E-02</td><td>2.7E-02</td><td>MB231/MB468</td></tr>
</table>

| | | | | | | | | | | | | |
|---|---|---|---|---|---|---|---|---|---|---|---|---|
| >am_trna10_ValCAC_5_-_180649395_180649467@1.31.31,trna12_ValAAC_5_-_180645270_180645342@1.31.31,trna132_ValAAC_6_-_27721179_27721251@1.31.31,trna136_ValAAC_6_-_27648885_27648957@1.31.31,trna139_ValAAC_6_-_27618707_27618779@1.31.31,trna18_ValCAC_5_-_180529253_180529325@1.31.31,trna2_ValAAC_3_+_169490018_169490090@1.31.31,trna2_ValCAC_5_+_180524070_180524142@1.31.31,trna4_ValAAC_5_+_180591154_180591226@1.31.31,trna5_ValAAC_5_+_180596610_180596682@1.31.31,trna6_ValCAC_5_+_180600650_180600722@1.31.31,trna85_ValCAC_1_-_161369490_161369562@1.31.31,trna90_ValCAC_1_-_149684088_149684161@1.31.31,trna98_ValCAC_1_-_149298555_149298627@1.31.31,trna9_ValCAC_6_+_26538282_26538354@1.31.31\|GTTTCCGTAGTGTAGTGGTTATCACGTTCGC | >am_trna10_ValCAC_5_-_180649395_180649467 | 1 | 31 | 31 | GTTTCCGTAGTGTAGTGGTTATCACGTTCGC | 1600.6 | 2.4 | 1.0 | 2.3 | 2.3E-02 | 2.7E-02 | MB231/MB468 |
| >MI0000750\|hsa-mir-26a-2&WithFlank&12\|-\|57824603\|57824698@20.41.22(+2U);MI0000083\|hsa-mir-26a-1&WithFlank&3\|+\|37969398\|37969486@16.37.22(+2U)&[MIMAT0000082&hsa-miR-26a-5p&offsets\|0\|0(+2U);m-33&12\|-\|57824658\|57824679&offsets\|0\|0(+2U)];[MIMAT0000082_1&hsa-miR-26a-5p&offsets\|0\|0(+2U);m-34&3\|+\|37969413\|37969434&offsets\|0\|0(+2U)]\|TTCAAGTAATCCAGGATAGGCTTT | >MI0000750\|hsa-mir-26a-2&WithFlank&12\|-\|57824603\|57824698 | 20 | 41 | 22(+2U) | TTCAAGTAATCCAGGATAGGCTTT | 6270.9 | -1.8 | 0.7 | -2.7 | 6.5E-03 | 2.7E-02 | BT20/MB231 |
| >am_trnaMT_SerGCT_MT_+_12207_12265@16.38.23\|AACTGCTAACTCATGCCCCCATG | >am_trnaMT_SerGCT_MT_+_12207_12265 | 16 | 38 | 23 | AACTGCTAACTCATGCCCCCATG | 174.1 | -1.8 | 0.8 | -2.3 | 2.3E-02 | 2.7E-02 | MB231/MB468 |
| >trna111_HisGTG_1_-_147774845_147774916@-1G.31.32,trna118_HisGTG_1_-_145396881_145396952@-1G.31.32,trna16_HisGTG_1_+_146544773_146544844@-1G.31.32,trna1_HisGTG_15_+_45493349_45493420@-1G.31.32,trna21_HisGTG_1_+_147753471_147753542@-1G.31.32,trna33_HisGTG_6_+_27125906_27125977@-1G.31.32,trna7_HisGTG_9_-_14433938_14434009@-1G.31.32,trna8_HisGTG_15_-_45492611_45492682@-1G.31.32,trna9_HisGTG_15_-_45490804_45490875@-1G.31.32\|GGCCGTGATCGTATAGTGGTTAGTACTCTGCG | >trna111_HisGTG_1_-_147774845_147774916 | -1G | 31 | 32 | GGCCGTGATCGTATAGTGGTTAGTACTCTGCG | 9320.3 | -2.3 | 1.0 | -2.3 | 2.4E-02 | 2.7E-02 | BT20/MB468 |
| >hg38_21_+_8433172_8446622_RNA45SN1_45S_50ntFlanks@7978.8016.39\|GACCTCAGATCAGACGTGGCGACCCGCTGAATTTAAGCA | >hg38_21_+_8433172_8446622_RNA45SN1_45S_50ntFlanks | 7978 | 8016 | 39 | GACCTCAGATCAGACGTGGCGACCCGCTGAATTTAAGCA | 353.5 | -2.9 | 1.1 | -2.7 | 6.6E-03 | 2.7E-02 | BT20/MB231 |
| >hg38_21_+_8433172_8446622_RNA45SN1_45S_50ntFlanks@8170.8214.45\|CAAGTCCTTCTGATCGAGGCCCAGCCCGTGGACGGTGTGAGGCCG | >hg38_21_+_8433172_8446622_RNA45SN1_45S_50ntFlanks | 8170 | 8214 | 45 | CAAGTCCTTCTGATCGAGGCCCAGCCCGTGGACGGTGTGAGGCCG | 315.0 | -1.4 | 0.6 | -2.3 | 2.3E-02 | 2.7E-02 | MB231/MB468 |
| >MI0000088\|hsa-mir-30a&WithFlank&6\|-\|71403545\|71403627@12.32.21(+1A)&[MIMAT0000087&hsa-miR-30a-5p&offsets\|0\|-1(+1A);m-15&6\|-\|71403595\|71403616&offsets\|0\|-1(+1A)]\|TGTAAACATCCTCGACTGGAAA | >MI0000088\|hsa-mir-30a&WithFlank&6\|-\|71403545\|71403627 | 12 | 32 | 21(+1A) | TGTAAACATCCTCGACTGGAAA | 2113.3 | 2.7 | 1.0 | 2.7 | 6.6E-03 | 2.7E-02 | BT20/MB231 |
| >MI0000542\|hsa-mir-320a&WithFlank&8\|-\|22244960\|22245043@48.69.22(+1U)&[MIMAT0000510&hsa-miR-320a-3p&offsets\|0\|0(+1U);m-60&8\|-\|22244975\|22244996&offsets\|0\|0(+1U)]\|AAAAGCTGGGTTGAGAGGGCGAT | >MI0000542\|hsa-mir-320a&WithFlank&8\|-\|22244960\|22245043 | 48 | 69 | 22(+1U) | AAAAGCTGGGTTGAGAGGGCGAT | 11285.0 | 2.0 | 0.7 | 2.7 | 6.7E-03 | 2.7E-02 | BT20/MB231 |
| >hg38_21_+_8433172_8446622_RNA45SN1_45S_50ntFlanks@10387.10407.21\|TGTGAACAGCAGTTGAACATG | >hg38_21_+_8433172_8446622_RNA45SN1_45S_50ntFlanks | 10387 | 10407 | 21 | TGTGAACAGCAGTTGAACATG | 185.7 | -1.3 | 0.6 | -2.3 | 2.3E-02 | 2.7E-02 | MB231/MB468 |
| >hg38_21_+_8433172_8446622_RNA45SN1_45S_50ntFlanks@12991.13016.26\|CTCCCTCGCTGCGATCTATTGAAAGT | >hg38_21_+_8433172_8446622_RNA45SN1_45S_50ntFlanks | 12991 | 13016 | 26 | CTCCCTCGCTGCGATCTATTGAAAGT | 422.8 | -1.1 | 0.5 | -2.3 | 2.3E-02 | 2.7E-02 | MB231/MB468 |
| >hg38_21_+_8433172_8446622_RNA45SN1_45S_50ntFlanks@3985.4010.26\|CGGCTTTGGTGACTCTAGATAACCTC | >hg38_21_+_8433172_8446622_RNA45SN1_45S_50ntFlanks | 3985 | 4010 | 26 | CGGCTTTGGTGACTCTAGATAACCTC | 99.0 | -1.8 | 0.8 | -2.2 | 2.5E-02 | 2.7E-02 | BT20/MB468 |
| >hg38_21_+_8433172_8446622_RNA45SN1_45S_50ntFlanks@3852.3881.30\|TAACTGTGGTAATTCTAGAGCTAATACATG | >hg38_21_+_8433172_8446622_RNA45SN1_45S_50ntFlanks | 3852 | 3881 | 30 | TAACTGTGGTAATTCTAGAGCTAATACATG | 148.1 | -1.2 | 0.5 | -2.3 | 2.3E-02 | 2.7E-02 | MB231/MB468 |
| >hg38_21_+_8433172_8446622_RNA45SN1_45S_50ntFlanks@10807.10828.22\|ACAATGTAGGTAAGGGAAGTCG | >hg38_21_+_8433172_8446622_RNA45SN1_45S_50ntFlanks | 10807 | 10828 | 22 | ACAATGTAGGTAAGGGAAGTCG | 144.1 | -1.4 | 0.6 | -2.3 | 2.3E-02 | 2.7E-02 | MB231/MB468 |
| >hg38_21_+_8433172_8446622_RNA45SN1_45S_50ntFlanks@10677.10706.30\|GCGGTTCCGGCGGCGTCCGGTGAGCTCTCG | >hg38_21_+_8433172_8446622_RNA45SN1_45S_50ntFlanks | 10677 | 10706 | 30 | GCGGTTCCGGCGGCGTCCGGTGAGCTCTCG | 874.8 | 2.5 | 1.1 | 2.2 | 2.5E-02 | 2.7E-02 | BT20/MB468 |
| >hg38_21_+_8433172_8446622_RNA45SN1_45S_50ntFlanks@482.506.25\|ATCGATGTGGTGACGTCGTGCTCTC | >hg38_21_+_8433172_8446622_RNA45SN1_45S_50ntFlanks | 482 | 506 | 25 | ATCGATGTGGTGACGTCGTGCTCTC | 3852.8 | 5.3 | 2.4 | 2.3 | 2.3E-02 | 2.8E-02 | MB231/MB468 |
| >hg38_21_+_8433172_8446622_RNA45SN1_45S_50ntFlanks@3789.3810.22\|ACTGCGAATGGCTCATTAAATC | >hg38_21_+_8433172_8446622_RNA45SN1_45S_50ntFlanks | 3789 | 3810 | 22 | ACTGCGAATGGCTCATTAAATC | 249.6 | -1.6 | 0.7 | -2.3 | 2.3E-02 | 2.8E-02 | MB231/MB468 |
| >hg38_21_+_8433172_8446622_RNA45SN1_45S_50ntFlanks@12475.12496.22\|CCCACTAATAGGGAACGTGAGC | >hg38_21_+_8433172_8446622_RNA45SN1_45S_50ntFlanks | 12475 | 12496 | 22 | CCCACTAATAGGGAACGTGAGC | 664.9 | -2.2 | 0.8 | -2.7 | 6.8E-03 | 2.8E-02 | BT20/MB231 |
| >MI0000298\|hsa-mir-221&WithFlank&X\|-\|45746151\|45746272@71.91.21(+2C)&[MIMAT0000278&hsa-miR-221-3p&offsets\|0\|-2(+2C);m-91&X\|-\|45746181\|45746202&offsets\|0\|-1(+2C)]\|AGCTACATTGTCTGCTGGGTTCC | >MI0000298\|hsa-mir-221&WithFlank&X\|-\|45746151\|45746272 | 71 | 91 | 21(+2C) | AGCTACATTGTCTGCTGGGTTCC | 226.7 | -2.1 | 0.8 | -2.7 | 6.8E-03 | 2.8E-02 | BT20/MB231 |
| >hg38_21_+_8433172_8446622_RNA45SN1_45S_50ntFlanks@5485.5516.32\|AGCGCTGAGAAGACGGTCGAACTTGACTATCT | >hg38_21_+_8433172_8446622_RNA45SN1_45S_50ntFlanks | 5485 | 5516 | 32 | AGCGCTGAGAAGACGGTCGAACTTGACTATCT | 3600.0 | 4.7 | 1.7 | 2.7 | 6.8E-03 | 2.8E-02 | BT20/MB231 |
| >hg38_21_+_8433172_8446622_RNA45SN1_45S_50ntFlanks@6700.6717.18\|CTAGCTGCGAGAATTAAT | >hg38_21_+_8433172_8446622_RNA45SN1_45S_50ntFlanks | 6700 | 6717 | 18 | CTAGCTGCGAGAATTAAT | 189.5 | -1.2 | 0.5 | -2.3 | 2.4E-02 | 2.8E-02 | MB231/MB468 |
| >hg38_21_+_8433172_8446622_RNA45SN1_45S_50ntFlanks@3838.3871.34\|CTCTCCTACTTGGATAACTGTGGTAATTCTAGAG | >hg38_21_+_8433172_8446622_RNA45SN1_45S_50ntFlanks | 3838 | 3871 | 34 | CTCTCCTACTTGGATAACTGTGGTAATTCTAGAG | 195.3 | -1.5 | 0.7 | -2.2 | 2.5E-02 | 2.8E-02 | BT20/MB468 |
| >MI0000101\|hsa-mir-99a&WithFlank&21\|+\|16539083\|16539175@19.40.22(+1A)&[MIMAT0000097&hsa-miR-99a-5p&offsets\|0\|0(+1A);m-64&21\|+\|16539101\|16539121&offsets\|0\|+1(+1A)]\|AACCCGTAGATCCGATCTTGTGA | >MI0000101\|hsa-mir-99a&WithFlank&21\|+\|16539083\|16539175 | 19 | 40 | 22(+1A) | AACCCGTAGATCCGATCTTGTGA | 1626.4 | 1.7 | 0.7 | 2.3 | 2.4E-02 | 2.8E-02 | MB231/MB468 |
| >hg38_21_+_8433172_8446622_RNA45SN1_45S_50ntFlanks@12455.12478.24\|ACCAAGCGTTGGATTGTTCACCCA | >hg38_21_+_8433172_8446622_RNA45SN1_45S_50ntFlanks | 12455 | 12478 | 24 | ACCAAGCGTTGGATTGTTCACCCA | 223.4 | -1.7 | 0.8 | -2.2 | 2.5E-02 | 2.8E-02 | BT20/MB468 |
| >hg38_21_+_8433172_8446622_RNA45SN1_45S_50ntFlanks@10350.10390.41\|ACGAGAACTTTGAAGGCCGAAGTGGAGAAGGGTTCCATGTG | >hg38_21_+_8433172_8446622_RNA45SN1_45S_50ntFlanks | 10350 | 10390 | 41 | ACGAGAACTTTGAAGGCCGAAGTGGAGAAGGGTTCCATGTG | 1442.9 | -1.6 | 0.7 | -2.3 | 2.4E-02 | 2.8E-02 | MB231/MB468 |
| >hg38_21_+_8433172_8446622_RNA45SN1_45S_50ntFlanks@10766.10785.20\|CGCAGCAGGTCTCCAAGGTG | >hg38_21_+_8433172_8446622_RNA45SN1_45S_50ntFlanks | 10766 | 10785 | 20 | CGCAGCAGGTCTCCAAGGTG | 162.1 | -1.1 | 0.5 | -2.3 | 2.4E-02 | 2.8E-02 | MB231/MB468 |
| >MI0000081\|hsa-mir-24-2&WithFlank&19\|-\|13836281\|13836365@56.77.22(+1A);MI0000080\|hsa-mir-24-1&WithFlank&9\|+\|95086015\|95086094@50.71.22(+1A)&[MIMAT0000080&hsa-miR-24-3p&offsets\|0\|0(+1A);m-36&19\|-\|13836288\|13836310&offsets\|0\|-1(+1A)];[MIMAT0000080_1&hsa-miR-24-3p&offsets\|0\|0(+1A);m-35&9\|+\|95086064\|95086086&offsets\|0\|-1(+1A)]\|TGGCTCAGTTCAGCAGGAACAGA | >MI0000081\|hsa-mir-24-2&WithFlank&19\|-\|13836281\|13836365 | 56 | 77 | 22(+1A) | TGGCTCAGTTCAGCAGGAACAGA | 1446.7 | 2.3 | 0.9 | 2.7 | 7.0E-03 | 2.8E-02 | BT20/MB231 |
| >hg38_MT_+_598_1651_RNR1_12S_50ntFlanks@428.468.41\|AAAGTGGCTTTAACATATCTGAACACACAATAGCTAAGACC | >hg38_MT_+_598_1651_RNR1_12S_50ntFlanks | 428 | 468 | 41 | AAAGTGGCTTTAACATATCTGAACACACAATAGCTAAGACC | 1544.0 | -2.5 | 0.9 | -2.7 | 7.0E-03 | 2.8E-02 | BT20/MB231 |
| >MI0000081\|hsa-mir-24-2&WithFlank&19\|-\|13836281\|13836365@56.73.18;MI0000080\|hsa-mir-24-1&WithFlank&9\|+\|95086015\|95086094@50.67.18&[MIMAT0000080&hsa-miR-24-3p&offsets\|0\|-4;m-36&19\|-\|13836288\|13836310&offsets\|0\|-5];[MIMAT0000080_1&hsa-miR-24-3p&offsets\|0\|-4;m-35&9\|+\|95086064\|95086086&offsets\|0\|-5]\|TGGCTCAGTTCAGCAGGA | >MI0000081\|hsa-mir-24-2&WithFlank&19\|-\|13836281\|13836365 | 56 | 73 | 18 | TGGCTCAGTTCAGCAGGA | 201.5 | 2.0 | 0.7 | 2.7 | 7.0E-03 | 2.8E-02 | BT20/MB231 |
| >MI0000542\|hsa-mir-320a&WithFlank&8\|-\|22244960\|22245043@49.68.20(+1U)&[MIMAT0000510&hsa-miR-320a-3p&offsets\|+1\|-1(+1U);m-60&8\|-\|22244975\|22244996&offsets\|+1\|-1(+1U)]\|AAAGCTGGGTTGAGAGGGCGT | >MI0000542\|hsa-mir-320a&WithFlank&8\|-\|22244960\|22245043 | 49 | 68 | 20(+1U) | AAAGCTGGGTTGAGAGGGCGT | 308.4 | 2.0 | 0.7 | 2.7 | 7.0E-03 | 2.8E-02 | BT20/MB231 |
| >hg38_21_+_8433172_8446622_RNA45SN1_45S_50ntFlanks@9308.9338.31\|AGGAGTCTAACACGTGCGCGAGTCGGGGGCT | >hg38_21_+_8433172_8446622_RNA45SN1_45S_50ntFlanks | 9308 | 9338 | 31 | AGGAGTCTAACACGTGCGCGAGTCGGGGGCT | 80.1 | -3.7 | 1.7 | -2.3 | 2.4E-02 | 2.8E-02 | MB231/MB468 |
| >hg38_21_+_8433172_8446622_RNA45SN1_45S_50ntFlanks@6784.6801.18\|GCTACGCCTGTCTGAGCG | >hg38_21_+_8433172_8446622_RNA45SN1_45S_50ntFlanks | 6784 | 6801 | 18 | GCTACGCCTGTCTGAGCG | 109.5 | -3.4 | 1.5 | -2.3 | 2.4E-02 | 2.8E-02 | MB231/MB468 |
| >hg38_21_+_8433172_8446622_RNA45SN1_45S_50ntFlanks@475.524.50\|CTTCGTGATCGATGTGGTGACGTCGTGCTCTCCCGGGCCGGGTCCGAGCC | >hg38_21_+_8433172_8446622_RNA45SN1_45S_50ntFlanks | 475 | 524 | 50 | CTTCGTGATCGATGTGGTGACGTCGTGCTCTCCCGGGCCGGGTCCGAGCC | 731.2 | -2.7 | 1.0 | -2.7 | 7.1E-03 | 2.8E-02 | BT20/MB231 |
| >hg38_21_+_8433172_8446622_RNA45SN1_45S_50ntFlanks@12417.12446.30\|TCGATGTCGGCTCTTCCTATCATTGTGAAG | >hg38_21_+_8433172_8446622_RNA45SN1_45S_50ntFlanks | 12417 | 12446 | 30 | TCGATGTCGGCTCTTCCTATCATTGTGAAG | 317.5 | -2.6 | 1.0 | -2.7 | 7.1E-03 | 2.9E-02 | BT20/MB231 |
| >hg38_21_+_8433172_8446622_RNA45SN1_45S_50ntFlanks@12415.12436.22\|CTTCGATGTCGGCTCTTCCTAT | >hg38_21_+_8433172_8446622_RNA45SN1_45S_50ntFlanks | 12415 | 12436 | 22 | CTTCGATGTCGGCTCTTCCTAT | 3029.1 | -2.0 | 0.9 | -2.2 | 2.6E-02 | 2.9E-02 | BT20/MB468 |

<table>
<tr><td>>hg38_21_+_8433172_8446622_RNA45SN1_45S_50ntFlanks@6646.6682.37|TCGTACGACTCTTAGCGGTGGATCACTCGGCTCGTGC</td><td>>hg38_21_+_8433172_8446622_RNA45SN1_45S_50ntFlanks</td><td>6646</td><td>6682</td><td>37</td><td>TCGTACGACTCTTAGCGGTGGATCACTCGGCTCGTGC</td><td>216.0</td><td>1.9</td><td>0.7</td><td>2.7</td><td>7.1E-03</td><td>2.9E-02</td><td>BT20/MB231</td></tr>
<tr><td>>hg38_21_+_8433172_8446622_RNA45SN1_45S_50ntFlanks@11582.11611.30|ACTGGTGCGGACCAGGGGAATCCGACTGTT</td><td>>hg38_21_+_8433172_8446622_RNA45SN1_45S_50ntFlanks</td><td>11582</td><td>11611</td><td>30</td><td>ACTGGTGCGGACCAGGGGAATCCGACTGTT</td><td>1036.7</td><td>-1.3</td><td>0.6</td><td>-2.2</td><td>2.6E-02</td><td>2.9E-02</td><td>BT20/MB468</td></tr>
<tr><td>>MI0000299|hsa-mir-222&WithFlank&X|-|45747009|45747130@75.99.25(+1C)&[MIMAT0000279&hsa-miR-222-3p&offsets|0|+4(+1C);m-140&X|-|45747033|45747056&offsets|0|+1(+1C)]|AGCTACATCTGGCTACTGGGTCTCTC</td><td>>MI0000299|hsa-mir-222&WithFlank&X|-|45747009|45747130</td><td>75</td><td>99</td><td>25(+1C)</td><td>AGCTACATCTGGCTACTGGGTCTCTC</td><td>781.3</td><td>-2.1</td><td>0.8</td><td>-2.7</td><td>7.1E-03</td><td>2.9E-02</td><td>BT20/MB231</td></tr>
<tr><td>>hg38_21_+_8433172_8446622_RNA45SN1_45S_50ntFlanks@12993.13013.21|CCCTCGCTGCGATCTATTGAA</td><td>>hg38_21_+_8433172_8446622_RNA45SN1_45S_50ntFlanks</td><td>12993</td><td>13013</td><td>21</td><td>CCCTCGCTGCGATCTATTGAA</td><td>344.6</td><td>-1.2</td><td>0.5</td><td>-2.2</td><td>2.4E-02</td><td>2.9E-02</td><td>MB231/MB468</td></tr>
<tr><td>>am_MI0003639|hsa-mir-625&WithFlank&14|+|65471096|65471192@58.78.21(+1U)&[MIMAT0004808&hsa-miR-625-3p&offsets|0|-1(+1U);m-137&14|+|65471153|65471174&offsets|0|-1(+1U)]|GACTATAGAACTTTCCCCCTCT</td><td>>am_MI0003639|hsa-mir-625&WithFlank&14|+|65471096|65471192</td><td>58</td><td>78</td><td>21(+1U)</td><td>GACTATAGAACTTTCCCCCTCT</td><td>212.1</td><td>-2.1</td><td>0.8</td><td>-2.7</td><td>7.2E-03</td><td>2.9E-02</td><td>BT20/MB231</td></tr>
<tr><td>>hg38_21_+_8433172_8446622_RNA45SN1_45S_50ntFlanks@6670.6698.29|ACTCGGCTCGTGCGTCGATGAAGAACGCA</td><td>>hg38_21_+_8433172_8446622_RNA45SN1_45S_50ntFlanks</td><td>6670</td><td>6698</td><td>29</td><td>ACTCGGCTCGTGCGTCGATGAAGAACGCA</td><td>247.7</td><td>-1.0</td><td>0.4</td><td>-2.2</td><td>2.5E-02</td><td>2.9E-02</td><td>MB231/MB468</td></tr>
<tr><td>>hg38_21_+_8433172_8446622_RNA45SN1_45S_50ntFlanks@11666.11706.41|TCTGCCCAGTGCTCTGAATGTCAAAGTGAAGAAATTCAATG</td><td>>hg38_21_+_8433172_8446622_RNA45SN1_45S_50ntFlanks</td><td>11666</td><td>11706</td><td>41</td><td>TCTGCCCAGTGCTCTGAATGTCAAAGTGAAGAAATTCAATG</td><td>204.1</td><td>-2.2</td><td>0.8</td><td>-2.7</td><td>7.2E-03</td><td>2.9E-02</td><td>BT20/MB231</td></tr>
<tr><td>>hg38_21_+_8433172_8446622_RNA45SN1_45S_50ntFlanks@12912.12935.24|CACGTTCGTGGGGAACCTGGCGCT</td><td>>hg38_21_+_8433172_8446622_RNA45SN1_45S_50ntFlanks</td><td>12912</td><td>12935</td><td>24</td><td>CACGTTCGTGGGGAACCTGGCGCT</td><td>130.4</td><td>-2.0</td><td>0.9</td><td>-2.2</td><td>2.5E-02</td><td>2.9E-02</td><td>MB231/MB468</td></tr>
<tr><td>>hg38_21_+_8433172_8446622_RNA45SN1_45S_50ntFlanks@6645.6685.41|CTCGTACGACTCTTAGCGGTGGATCACTCGGCTCGTGCGTC</td><td>>hg38_21_+_8433172_8446622_RNA45SN1_45S_50ntFlanks</td><td>6645</td><td>6685</td><td>41</td><td>CTCGTACGACTCTTAGCGGTGGATCACTCGGCTCGTGCGTC</td><td>333.2</td><td>2.4</td><td>0.9</td><td>2.7</td><td>7.2E-03</td><td>2.9E-02</td><td>BT20/MB231</td></tr>
<tr><td>>hg38_21_+_8433172_8446622_RNA45SN1_45S_50ntFlanks@6718.6743.26|GTGAATTGCAGGACACATTGATCATC</td><td>>hg38_21_+_8433172_8446622_RNA45SN1_45S_50ntFlanks</td><td>6718</td><td>6743</td><td>26</td><td>GTGAATTGCAGGACACATTGATCATC</td><td>378.3</td><td>1.4</td><td>0.6</td><td>2.2</td><td>2.5E-02</td><td>2.9E-02</td><td>MB231/MB468</td></tr>
<tr><td>>MI0000273|hsa-mir-183&WithFlank&7|-|129774899|129775020@34.56.23&[MIMAT0000261&hsa-miR-183-5p&offsets|+1|+2;m-41&7|-|129774966|129774987&offsets|0|+1]|ATGGCACTGGTAGAATTCACTGT</td><td>>MI0000273|hsa-mir-183&WithFlank&7|-|129774899|129775020</td><td>34</td><td>56</td><td>23</td><td>ATGGCACTGGTAGAATTCACTGT</td><td>1299.2</td><td>1.9</td><td>0.7</td><td>2.7</td><td>7.3E-03</td><td>2.9E-02</td><td>BT20/MB231</td></tr>
<tr><td>>MI0000266|hsa-mir-10a&WithFlank&17|-|48579832|48579953@29.49.21(+1A)&[MIMAT0000253&hsa-miR-10a-5p&offsets|+1|-1(+1A);m-18&17|-|48579905|48579926&offsets|+1|0(+1A)]|ACCCTGTAGATCCGAATTTGTA</td><td>>MI0000266|hsa-mir-10a&WithFlank&17|-|48579832|48579953</td><td>29</td><td>49</td><td>21(+1A)</td><td>ACCCTGTAGATCCGAATTTGTA</td><td>2253.6</td><td>2.0</td><td>0.7</td><td>2.7</td><td>7.3E-03</td><td>2.9E-02</td><td>BT20/MB231</td></tr>
<tr><td>>hg38_21_+_8433172_8446622_RNA45SN1_45S_50ntFlanks@11571.11591.21|GCCGACTTAGAACTGGTGCGG</td><td>>hg38_21_+_8433172_8446622_RNA45SN1_45S_50ntFlanks</td><td>11571</td><td>11591</td><td>21</td><td>GCCGACTTAGAACTGGTGCGG</td><td>262.8</td><td>-1.3</td><td>0.6</td><td>-2.2</td><td>2.5E-02</td><td>2.9E-02</td><td>MB231/MB468</td></tr>
<tr><td>>MI0000264|hsa-mir-7-2&WithFlank&15|+|88611819|88611940@38.60.23(+1A);MI0000265|hsa-mir-7-3&WithFlank&19|+|4770664|4770785@37.59.23(+1A);MI0000263|hsa-mir-7-1&WithFlank&9|-|83969742|83969863@30.52.23(+1A)&[MIMAT0000252&hsa-miR-7-5p&offsets|0|-1(+1A);m-400&15|+|88611856|88611879&offsets|0|-1(+1A)];[MIMAT0000252_1&hsa-miR-7-5p&offsets|0|-1(+1A);m-402&19|+|4770700|4770723&offsets|0|-1(+1A)];[MIMAT0000252_2&hsa-miR-7-5p&offsets|0|-1(+1A);m-401&9|-|83969811|83969834&offsets|0|-1(+1A)]|TGGAAGACTAGTGATTTTGTTGTA</td><td>>MI0000264|hsa-mir-7-2&WithFlank&15|+|88611819|88611940</td><td>38</td><td>60</td><td>23(+1A)</td><td>TGGAAGACTAGTGATTTTGTTGTA</td><td>10719.5</td><td>2.2</td><td>0.8</td><td>2.7</td><td>7.3E-03</td><td>2.9E-02</td><td>BT20/MB231</td></tr>
<tr><td>>hg38_21_+_8433172_8446622_RNA45SN1_45S_50ntFlanks@11570.11600.31|AGCCGACTTAGAACTGGTGCGGACCAGGGGA</td><td>>hg38_21_+_8433172_8446622_RNA45SN1_45S_50ntFlanks</td><td>11570</td><td>11600</td><td>31</td><td>AGCCGACTTAGAACTGGTGCGGACCAGGGGA</td><td>219.7</td><td>-1.0</td><td>0.5</td><td>-2.2</td><td>2.5E-02</td><td>2.9E-02</td><td>MB231/MB468</td></tr>
<tr><td>>hg38_21_+_8433172_8446622_RNA45SN1_45S_50ntFlanks@4759.4793.35|ATCAGATACCGTCGTAGTTCCGACCATAAACGATG</td><td>>hg38_21_+_8433172_8446622_RNA45SN1_45S_50ntFlanks</td><td>4759</td><td>4793</td><td>35</td><td>ATCAGATACCGTCGTAGTTCCGACCATAAACGATG</td><td>157.3</td><td>-3.5</td><td>1.5</td><td>-2.2</td><td>2.5E-02</td><td>2.9E-02</td><td>MB231/MB468</td></tr>
<tr><td>>hg38_21_+_8433172_8446622_RNA45SN1_45S_50ntFlanks@12418.12447.30|CGATGTCGGCTCTTCCTATCATTGTGAAGC</td><td>>hg38_21_+_8433172_8446622_RNA45SN1_45S_50ntFlanks</td><td>12418</td><td>12447</td><td>30</td><td>CGATGTCGGCTCTTCCTATCATTGTGAAGC</td><td>246.9</td><td>-2.2</td><td>0.8</td><td>-2.7</td><td>7.3E-03</td><td>2.9E-02</td><td>BT20/MB231</td></tr>
<tr><td>>hg38_21_+_8433172_8446622_RNA45SN1_45S_50ntFlanks@11668.11686.19|TGCCCAGTGCTCTGAATGT</td><td>>hg38_21_+_8433172_8446622_RNA45SN1_45S_50ntFlanks</td><td>11668</td><td>11686</td><td>19</td><td>TGCCCAGTGCTCTGAATGT</td><td>282.4</td><td>-2.3</td><td>0.9</td><td>-2.7</td><td>7.4E-03</td><td>2.9E-02</td><td>BT20/MB231</td></tr>
<tr><td>>hg38_21_+_8433172_8446622_RNA45SN1_45S_50ntFlanks@11672.11707.36|CAGTGCTCTGAATGTCAAAGTGAAGAAATTCAATGA</td><td>>hg38_21_+_8433172_8446622_RNA45SN1_45S_50ntFlanks</td><td>11672</td><td>11707</td><td>36</td><td>CAGTGCTCTGAATGTCAAAGTGAAGAAATTCAATGA</td><td>307.4</td><td>-2.3</td><td>0.8</td><td>-2.7</td><td>7.4E-03</td><td>2.9E-02</td><td>BT20/MB231</td></tr>
<tr><td>>hg38_21_+_8433172_8446622_RNA45SN1_45S_50ntFlanks@12420.12450.31|ATGTCGGCTCTTCCTATCATTGTGAAGCAGA</td><td>>hg38_21_+_8433172_8446622_RNA45SN1_45S_50ntFlanks</td><td>12420</td><td>12450</td><td>31</td><td>ATGTCGGCTCTTCCTATCATTGTGAAGCAGA</td><td>205.0</td><td>-2.3</td><td>0.9</td><td>-2.7</td><td>7.4E-03</td><td>2.9E-02</td><td>BT20/MB231</td></tr>
<tr><td>>hg38_21_+_8433172_8446622_RNA45SN1_45S_50ntFlanks@6699.6720.22|GCTAGCTGCGAGAATTAATGTG</td><td>>hg38_21_+_8433172_8446622_RNA45SN1_45S_50ntFlanks</td><td>6699</td><td>6720</td><td>22</td><td>GCTAGCTGCGAGAATTAATGTG</td><td>953.8</td><td>-2.6</td><td>1.0</td><td>-2.7</td><td>7.4E-03</td><td>2.9E-02</td><td>BT20/MB231</td></tr>
<tr><td>>hg38_21_+_8433172_8446622_RNA45SN1_45S_50ntFlanks@9551.9568.18|GTCCGTAGCGGTCCTGAC</td><td>>hg38_21_+_8433172_8446622_RNA45SN1_45S_50ntFlanks</td><td>9551</td><td>9568</td><td>18</td><td>GTCCGTAGCGGTCCTGAC</td><td>145.0</td><td>-2.5</td><td>1.0</td><td>-2.7</td><td>7.4E-03</td><td>2.9E-02</td><td>BT20/MB231</td></tr>
<tr><td>>hg38_21_+_8433172_8446622_RNA45SN1_45S_50ntFlanks@9569.9610.42|GTGCAAATCGGTCGTCCGACCTGGGTATAGGGGCGAAAGACT</td><td>>hg38_21_+_8433172_8446622_RNA45SN1_45S_50ntFlanks</td><td>9569</td><td>9610</td><td>42</td><td>GTGCAAATCGGTCGTCCGACCTGGGTATAGGGGCGAAAGACT</td><td>157.3</td><td>-2.5</td><td>1.0</td><td>-2.7</td><td>7.4E-03</td><td>2.9E-02</td><td>BT20/MB231</td></tr>
<tr><td>>hg38_21_+_8433172_8446622_RNA45SN1_45S_50ntFlanks@10348.10366.19|AAACGAGAACTTTGAAGGC</td><td>>hg38_21_+_8433172_8446622_RNA45SN1_45S_50ntFlanks</td><td>10348</td><td>10366</td><td>19</td><td>AAACGAGAACTTTGAAGGC</td><td>1364.6</td><td>1.4</td><td>0.6</td><td>2.2</td><td>2.5E-02</td><td>2.9E-02</td><td>MB231/MB468</td></tr>
<tr><td>>MI0000289|hsa-mir-181a-1&WithFlank&1|-|198859038|198859159@30.50.21;MI0000269|hsa-mir-181a-2&WithFlank&9|+|124692436|124692557@45.65.21&[MIMAT0000256&hsa-miR-181a-5p&offsets|0|-2;m-32&1|-|198859108|198859130&offsets|0|-2];[MIMAT0000256_1&hsa-miR-181a-5p&offsets|0|-2;m-32&9|+|124692480|124692502&offsets|0|-2]|AACATTCAACGCTGTCGGTGA</td><td>>MI0000289|hsa-mir-181a-1&WithFlank&1|-|198859038|198859159</td><td>30</td><td>50</td><td>21</td><td>AACATTCAACGCTGTCGGTGA</td><td>561.3</td><td>-1.9</td><td>0.7</td><td>-2.7</td><td>7.5E-03</td><td>2.9E-02</td><td>BT20/MB231</td></tr>
<tr><td>>MI0000739|hsa-mir-101-2&WithFlank&9|+|4850291|4850381@55.77.23&[MIMAT0000099_1&hsa-miR-101-3p&offsets|0|+2;m-16&9|+|4850345|4850364&offsets|0|+3]|TACAGTACTGTGATAACTGAAGA</td><td>>MI0000739|hsa-mir-101-2&WithFlank&9|+|4850291|4850381</td><td>55</td><td>77</td><td>23</td><td>TACAGTACTGTGATAACTGAAGA</td><td>869.2</td><td>2.7</td><td>1.0</td><td>2.7</td><td>7.5E-03</td><td>2.9E-02</td><td>BT20/MB231</td></tr>
<tr><td>>am_trna116_GluCTC_1_-_145399233_145399304@1.27.27,trna59_GluCTC_1_+_249168447_249168518@1.27.27,trna71_GluCTC_1_-_161439189_161439260@1.27.27,trna74_GluCTC_1_-_161431809_161431880@1.27.27,trna77_GluCTC_1_-_161424398_161424469@1.27.27,trna77_GluCTC_6_+_28949976_28950047@1.27.27,trna80_GluCTC_1_-_161417018_161417089@1.27.27,trna87_GluCTC_6_-_126101393_126101464@1.27.27|TCCCTGGTGGTCTAGTGGTTAGGATTC</td><td>>am_trna116_GluCTC_1_-_145399233_145399304</td><td>1</td><td>27</td><td>27</td><td>TCCCTGGTGGTCTAGTGGTTAGGATTC</td><td>4268.0</td><td>-1.6</td><td>0.7</td><td>-2.2</td><td>2.6E-02</td><td>2.9E-02</td><td>BT20/MB468</td></tr>
<tr><td>>hg38_21_+_8433172_8446622_RNA45SN1_45S_50ntFlanks@6776.6803.28|CTCCCGGGGCTACGCCTGTCTGAGCGTC</td><td>>hg38_21_+_8433172_8446622_RNA45SN1_45S_50ntFlanks</td><td>6776</td><td>6803</td><td>28</td><td>CTCCCGGGGCTACGCCTGTCTGAGCGTC</td><td>652.3</td><td>-1.3</td><td>0.6</td><td>-2.2</td><td>2.5E-02</td><td>3.0E-02</td><td>MB231/MB468</td></tr>
<tr><td>>hg38_21_+_8433172_8446622_RNA45SN1_45S_50ntFlanks@6652.6669.18|GACTCTTAGCGGTGGATC</td><td>>hg38_21_+_8433172_8446622_RNA45SN1_45S_50ntFlanks</td><td>6652</td><td>6669</td><td>18</td><td>GACTCTTAGCGGTGGATC</td><td>239.6</td><td>2.1</td><td>0.8</td><td>2.7</td><td>7.6E-03</td><td>3.0E-02</td><td>BT20/MB231</td></tr>
<tr><td>>hg38_21_+_8433172_8446622_RNA45SN1_45S_50ntFlanks@3843.3862.20|CTACTTGGATAACTGTGGTA</td><td>>hg38_21_+_8433172_8446622_RNA45SN1_45S_50ntFlanks</td><td>3843</td><td>3862</td><td>20</td><td>CTACTTGGATAACTGTGGTA</td><td>166.6</td><td>-1.1</td><td>0.5</td><td>-2.2</td><td>2.5E-02</td><td>3.0E-02</td><td>MB231/MB468</td></tr>
<tr><td>>hg38_21_+_8433172_8446622_RNA45SN1_45S_50ntFlanks@12451.12476.26|ATTCACCAAGCGTTGGATTGTTCACC</td><td>>hg38_21_+_8433172_8446622_RNA45SN1_45S_50ntFlanks</td><td>12451</td><td>12476</td><td>26</td><td>ATTCACCAAGCGTTGGATTGTTCACC</td><td>394.1</td><td>-2.2</td><td>0.8</td><td>-2.7</td><td>7.6E-03</td><td>3.0E-02</td><td>BT20/MB231</td></tr>
<tr><td>>MI0000077|hsa-mir-21&WithFlank&17|+|59841260|59841343@14.31.18&[MIMAT0000076&hsa-miR-21-5p&offsets|0|-4;m-1&17|+|59841273|59841294&offsets|0|-4]|TAGCTTATCAGACTGATG</td><td>>MI0000077|hsa-mir-21&WithFlank&17|+|59841260|59841343</td><td>14</td><td>31</td><td>18</td><td>TAGCTTATCAGACTGATG</td><td>2605.7</td><td>1.8</td><td>0.7</td><td>2.7</td><td>7.6E-03</td><td>3.0E-02</td><td>BT20/MB231</td></tr>
<tr><td>>hg38_21_+_8433172_8446622_RNA45SN1_45S_50ntFlanks@4735.4757.23|ACGAAAGTCGGAGGTTCGAAGAC</td><td>>hg38_21_+_8433172_8446622_RNA45SN1_45S_50ntFlanks</td><td>4735</td><td>4757</td><td>23</td><td>ACGAAAGTCGGAGGTTCGAAGAC</td><td>148.7</td><td>-2.4</td><td>0.9</td><td>-2.7</td><td>7.6E-03</td><td>3.0E-02</td><td>BT20/MB231</td></tr>
<tr><td>>hg38_21_+_8433172_8446622_RNA45SN1_45S_50ntFlanks@3705.3723.19|TACCTGGTTGATCCTGCCA</td><td>>hg38_21_+_8433172_8446622_RNA45SN1_45S_50ntFlanks</td><td>3705</td><td>3723</td><td>19</td><td>TACCTGGTTGATCCTGCCA</td><td>396.4</td><td>2.2</td><td>0.8</td><td>2.7</td><td>7.7E-03</td><td>3.0E-02</td><td>BT20/MB231</td></tr>
<tr><td>>hg38_21_+_8433172_8446622_RNA45SN1_45S_50ntFlanks@8354.8379.26|TTGAAAAGAACTTTGAAGAGAGAGTT</td><td>>hg38_21_+_8433172_8446622_RNA45SN1_45S_50ntFlanks</td><td>8354</td><td>8379</td><td>26</td><td>TTGAAAAGAACTTTGAAGAGAGAGTT</td><td>150.0</td><td>-1.2</td><td>0.5</td><td>-2.2</td><td>2.6E-02</td><td>3.0E-02</td><td>MB231/MB468</td></tr>
<tr><td>>hg38_21_+_8433172_8446622_RNA45SN1_45S_50ntFlanks@10738.10755.18|AAATCTCGCGCCGGGCCG</td><td>>hg38_21_+_8433172_8446622_RNA45SN1_45S_50ntFlanks</td><td>10738</td><td>10755</td><td>18</td><td>AAATCTCGCGCCGGGCCG</td><td>307.7</td><td>-2.4</td><td>1.1</td><td>-2.2</td><td>2.7E-02</td><td>3.0E-02</td><td>BT20/MB468</td></tr>
<tr><td>>hg38_21_+_8433172_8446622_RNA45SN1_45S_50ntFlanks@12420.12446.27|ATGTCGGCTCTTCCTATCATTGTGAAG</td><td>>hg38_21_+_8433172_8446622_RNA45SN1_45S_50ntFlanks</td><td>12420</td><td>12446</td><td>27</td><td>ATGTCGGCTCTTCCTATCATTGTGAAG</td><td>4136.9</td><td>-1.6</td><td>0.7</td><td>-2.2</td><td>2.6E-02</td><td>3.0E-02</td><td>MB231/MB468</td></tr>
<tr><td>>hg38_21_+_8433172_8446622_RNA45SN1_45S_50ntFlanks@7992.8014.23|CGTGGCGACCCGCTGAATTTAAG</td><td>>hg38_21_+_8433172_8446622_RNA45SN1_45S_50ntFlanks</td><td>7992</td><td>8014</td><td>23</td><td>CGTGGCGACCCGCTGAATTTAAG</td><td>295.2</td><td>1.4</td><td>0.6</td><td>2.2</td><td>2.6E-02</td><td>3.1E-02</td><td>MB231/MB468</td></tr>
<tr><td>>hg38_21_+_8433172_8446622_RNA45SN1_45S_50ntFlanks@11876.11897.22|AAGACCCTGTTGAGCTTGACTC</td><td>>hg38_21_+_8433172_8446622_RNA45SN1_45S_50ntFlanks</td><td>11876</td><td>11897</td><td>22</td><td>AAGACCCTGTTGAGCTTGACTC</td><td>381.1</td><td>-1.5</td><td>0.7</td><td>-2.2</td><td>2.6E-02</td><td>3.1E-02</td><td>MB231/MB468</td></tr>
<tr><td>>hg38_21_+_8433172_8446622_RNA45SN1_45S_50ntFlanks@3853.3874.22|AACTGTGGTAATTCTAGAGCTA</td><td>>hg38_21_+_8433172_8446622_RNA45SN1_45S_50ntFlanks</td><td>3853</td><td>3874</td><td>22</td><td>AACTGTGGTAATTCTAGAGCTA</td><td>139.9</td><td>-1.3</td><td>0.6</td><td>-2.2</td><td>2.6E-02</td><td>3.1E-02</td><td>MB231/MB468</td></tr>
<tr><td>>hg38_21_+_8433172_8446622_RNA45SN1_45S_50ntFlanks@10794.10820.27|CTGGCATGTTGGAACAATGTAGGTAAG</td><td>>hg38_21_+_8433172_8446622_RNA45SN1_45S_50ntFlanks</td><td>10794</td><td>10820</td><td>27</td><td>CTGGCATGTTGGAACAATGTAGGTAAG</td><td>137.3</td><td>-1.3</td><td>0.6</td><td>-2.2</td><td>2.6E-02</td><td>3.1E-02</td><td>MB231/MB468</td></tr>
</table>

<table>
<tr><th></th><th></th><th></th><th></th><th></th><th></th><th></th><th></th><th></th><th></th><th></th><th></th><th></th></tr>
<tr><td>>hg38_21_+_8433172_8446622_RNA45SN1_45S_50ntFlanks@12435.12454.20|ATCATTGTGAAGCAGAATTC</td><td>>hg38_21_+_8433172_8446622_RNA45SN1_45S_50ntFlanks</td><td>12435</td><td>12454</td><td>20</td><td>ATCATTGTGAAGCAGAATTC</td><td>357.3</td><td>-4.1</td><td>1.8</td><td>-2.2</td><td>2.6E-02</td><td>3.1E-02</td><td>MB231/MB468</td></tr>
<tr><td>>hg38_21_+_8433172_8446622_RNA45SN1_45S_50ntFlanks@4732.4751.20|AGAACGAAAGTCGGAGGTTC</td><td>>hg38_21_+_8433172_8446622_RNA45SN1_45S_50ntFlanks</td><td>4732</td><td>4751</td><td>20</td><td>AGAACGAAAGTCGGAGGTTC</td><td>507.2</td><td>1.5</td><td>0.7</td><td>2.2</td><td>2.6E-02</td><td>3.1E-02</td><td>MB231/MB468</td></tr>
<tr><td>>hg38_1_-_228634819_228635039_RNA5S12_5S_50ntFlanks@51.70.20|GTCTACGGCCATACCACCCT</td><td>>hg38_1_-_228634819_228635039_RNA5S12_5S_50ntFlanks</td><td>51</td><td>70</td><td>20</td><td>GTCTACGGCCATACCACCCT</td><td>286.1</td><td>1.5</td><td>0.7</td><td>2.2</td><td>2.8E-02</td><td>3.1E-02</td><td>BT20/MB468</td></tr>
<tr><td>>hg38_21_+_8433172_8446622_RNA45SN1_45S_50ntFlanks@11568.11591.24|GCAGCCGACTTAGAACTGGTG CGG</td><td>>hg38_21_+_8433172_8446622_RNA45SN1_45S_50ntFlanks</td><td>11568</td><td>11591</td><td>24</td><td>GCAGCCGACTTAGAACTGGTGCGG</td><td>128.0</td><td>-1.4</td><td>0.7</td><td>-2.2</td><td>2.8E-02</td><td>3.1E-02</td><td>BT20/MB468</td></tr>
<tr><td>>am_trna2_GlyTCC_19_+_4724082_4724153@1.30.30|GCGTTGGTGGTATAGTGGTTAGCATAGCTG</td><td>>am_trna2_GlyTCC_19_+_4724082_4724153</td><td>1</td><td>30</td><td>30</td><td>GCGTTGGTGGTATAGTGGTTAGCATAGCTG</td><td>195.2</td><td>-4.1</td><td>1.8</td><td>-2.2</td><td>2.6E-02</td><td>3.1E-02</td><td>MB231/MB468</td></tr>
<tr><td>>trna111_HisGTG_1_-_147774845_147774916@-1G.31.32,trna118_HisGTG_1_-_145396881_145396952@-1G.31.32,trna16_HisGTG_1_+_146544773_146544844@-1G.31.32,trna1_HisGTG_15_+_45493349_45493420@-1G.31.32,trna21_HisGTG_1_+_147753471_147753542@-1G.31.32,trna33_HisGTG_6_+_27125906_27125977@-1G.31.32,trna7_HisGTG_9_-_14433938_14434009@-1G.31.32,trna8_HisGTG_15_-_45492611_45492682@-1G.31.32,trna9_HisGTG_15_-_45490804_45490875@-1G.31.32|GGCCGTGATCGTATAGTGGTTAGTACTCTGCG</td><td>>trna111_HisGTG_1_-_147774845_147774916</td><td>-1G</td><td>31</td><td>32</td><td>GGCCGTGATCGTATAGTGGTTAGTACTCTGCG</td><td>5287.0</td><td>-2.9</td><td>1.1</td><td>-2.7</td><td>7.9E-03</td><td>3.1E-02</td><td>BT20/MB231</td></tr>
<tr><td>>MI0000084|hsa-mir-26b&WithFlank&2|+|218402640|218402728@18.38.21(+1A)&[MIMAT0000083&hsa-miR-26b-5p&offsets|0|0(+1A);m-53&2|+|218402657|218402678&offsets|0|-1(+1A)]|TTCAAGTAATTCAGGATAGGTA</td><td>>MI0000084|hsa-mir-26b&WithFlank&2|+|218402640|218402728</td><td>18</td><td>38</td><td>21(+1A)</td><td>TTCAAGTAATTCAGGATAGGTA</td><td>1198.3</td><td>2.6</td><td>1.0</td><td>2.7</td><td>7.9E-03</td><td>3.1E-02</td><td>BT20/MB231</td></tr>
<tr><td>>hg38_21_+_8433172_8446622_RNA45SN1_45S_50ntFlanks@10391.10424.34|AACAGCAGTTGAACATGGGTC AGTCGGTCCTGAG</td><td>>hg38_21_+_8433172_8446622_RNA45SN1_45S_50ntFlanks</td><td>10391</td><td>10424</td><td>34</td><td>AACAGCAGTTGAACATGGGTCAGTCGGTCCTGAG</td><td>2530.8</td><td>2.1</td><td>1.0</td><td>2.2</td><td>2.8E-02</td><td>3.1E-02</td><td>BT20/MB468</td></tr>
<tr><td>>hg38_21_+_8433172_8446622_RNA45SN1_45S_50ntFlanks@11769.11803.35|TAATTAGTGACGCGCATGAAT GGATGAACGAGATT</td><td>>hg38_21_+_8433172_8446622_RNA45SN1_45S_50ntFlanks</td><td>11769</td><td>11803</td><td>35</td><td>TAATTAGTGACGCGCATGAATGGATGAACGAGATT</td><td>615.1</td><td>-1.3</td><td>0.6</td><td>-2.2</td><td>2.7E-02</td><td>3.1E-02</td><td>MB231/MB468</td></tr>
<tr><td>>hg38_21_+_8433172_8446622_RNA45SN1_45S_50ntFlanks@9740.9765.26|AACGATCTCAACCTATTCTCAAAC TT</td><td>>hg38_21_+_8433172_8446622_RNA45SN1_45S_50ntFlanks</td><td>9740</td><td>9765</td><td>26</td><td>AACGATCTCAACCTATTCTCAAACTT</td><td>220.4</td><td>1.5</td><td>0.7</td><td>2.2</td><td>2.7E-02</td><td>3.1E-02</td><td>MB231/MB468</td></tr>
<tr><td>>trnaMT_GluTTC_MT_-_14674_14742@34.71.38,trnalookalike8_GluTTC_5_-_93905172_93905240@34.71.38|ATATCATTGGTCGTGGTTGTAGTCCGTGCGAGAATACC</td><td>>trnaMT_GluTTC_MT_-_14674_14742</td><td>34</td><td>71</td><td>38</td><td>ATATCATTGGTCGTGGTTGTAGTCCGTGCGAGAATACC</td><td>695.1</td><td>-1.9</td><td>0.7</td><td>-2.7</td><td>8.0E-03</td><td>3.1E-02</td><td>BT20/MB231</td></tr>
<tr><td>>hg38_21_+_8433172_8446622_RNA45SN1_45S_50ntFlanks@11581.11609.29|AACTGGTGCGGACCAGGGGA ATCCGACTG</td><td>>hg38_21_+_8433172_8446622_RNA45SN1_45S_50ntFlanks</td><td>11581</td><td>11609</td><td>29</td><td>AACTGGTGCGGACCAGGGGAATCCGACTG</td><td>1603.6</td><td>-1.3</td><td>0.6</td><td>-2.2</td><td>2.7E-02</td><td>3.1E-02</td><td>MB231/MB468</td></tr>
<tr><td>>MI0016887|hsa-mir-4521&WithFlank&17|+|8186939|8187010@13.34.22&[MIMAT0019058&hsa-miR-4521&offsets|+3|+3;m-765&17|+|8186948|8186969&offsets|+3|+3]|AAGGAAGTCCTGTGCTCAGTTT</td><td>>MI0016887|hsa-mir-4521&WithFlank&17|+|8186939|8187010</td><td>13</td><td>34</td><td>22</td><td>AAGGAAGTCCTGTGCTCAGTTT</td><td>429.1</td><td>2.2</td><td>0.8</td><td>2.6</td><td>8.1E-03</td><td>3.1E-02</td><td>BT20/MB231</td></tr>
<tr><td>>hg38_21_+_8433172_8446622_RNA45SN1_45S_50ntFlanks@10686.10708.23|GCGGCGTCCGGTGAGCTCTCG CT</td><td>>hg38_21_+_8433172_8446622_RNA45SN1_45S_50ntFlanks</td><td>10686</td><td>10708</td><td>23</td><td>GCGGCGTCCGGTGAGCTCTCGCT</td><td>1450.9</td><td>2.0</td><td>0.9</td><td>2.2</td><td>2.8E-02</td><td>3.1E-02</td><td>BT20/MB468</td></tr>
<tr><td>>MI0000483|hsa-mir-186&WithFlank&1|-|71067625|71067722@21.43.23&[MIMAT0000456&hsa-miR-186-5p&offsets|0|+1;m-84&1|-|71067681|71067702&offsets|0|+1]|CAAAGAATTCTCCTTTTGGGCTT</td><td>>MI0000483|hsa-mir-186&WithFlank&1|-|71067625|71067722</td><td>21</td><td>43</td><td>23</td><td>CAAAGAATTCTCCTTTTGGGCTT</td><td>1754.3</td><td>2.1</td><td>0.8</td><td>2.6</td><td>8.1E-03</td><td>3.1E-02</td><td>BT20/MB231</td></tr>
<tr><td>>hg38_21_+_8433172_8446622_RNA45SN1_45S_50ntFlanks@6648.6681.34|GTACGACTCTTAGCGGTGGATCA CTCGGCTCGTG</td><td>>hg38_21_+_8433172_8446622_RNA45SN1_45S_50ntFlanks</td><td>6648</td><td>6681</td><td>34</td><td>GTACGACTCTTAGCGGTGGATCACTCGGCTCGTG</td><td>2646.8</td><td>-1.4</td><td>0.7</td><td>-2.2</td><td>2.8E-02</td><td>3.1E-02</td><td>BT20/MB468</td></tr>
<tr><td>>hg38_21_+_8433172_8446622_RNA45SN1_45S_50ntFlanks@12458.12477.20|AAGCGTTGGATTGTTCACCC</td><td>>hg38_21_+_8433172_8446622_RNA45SN1_45S_50ntFlanks</td><td>12458</td><td>12477</td><td>20</td><td>AAGCGTTGGATTGTTCACCC</td><td>1741.5</td><td>1.3</td><td>0.6</td><td>2.2</td><td>2.7E-02</td><td>3.2E-02</td><td>MB231/MB468</td></tr>
<tr><td>>MI0000342|hsa-mir-200b&WithFlank&1|+|1167098|1167204@63.85.23&[MIMAT0000318&hsa-miR-200b-3p&offsets|0|+1;m-93&1|+|1167160|1167182&offsets|0|0]|TAATACTGCCTGGTAATGATGAC</td><td>>MI0000342|hsa-mir-200b&WithFlank&1|+|1167098|1167204</td><td>63</td><td>85</td><td>23</td><td>TAATACTGCCTGGTAATGATGAC</td><td>3020.3</td><td>-1.6</td><td>0.7</td><td>-2.2</td><td>2.8E-02</td><td>3.2E-02</td><td>BT20/MB468</td></tr>
<tr><td>>hg38_21_+_8433172_8446622_RNA45SN1_45S_50ntFlanks@12942.12961.20|TTCGTAGACGACCTGCTTCT</td><td>>hg38_21_+_8433172_8446622_RNA45SN1_45S_50ntFlanks</td><td>12942</td><td>12961</td><td>20</td><td>TTCGTAGACGACCTGCTTCT</td><td>602.1</td><td>1.3</td><td>0.6</td><td>2.2</td><td>2.7E-02</td><td>3.2E-02</td><td>MB231/MB468</td></tr>
<tr><td>>trnaMT_CysGCA_MT_-_5761_5826@47.69.23|CTTCAAACCTGCCGGGGCTTCCA</td><td>>trnaMT_CysGCA_MT_-_5761_5826</td><td>47</td><td>69</td><td>23</td><td>CTTCAAACCTGCCGGGGCTTCCA</td><td>172.9</td><td>-2.4</td><td>1.1</td><td>-2.2</td><td>2.8E-02</td><td>3.2E-02</td><td>BT20/MB468</td></tr>
<tr><td>>hg38_21_+_8433172_8446622_RNA45SN1_45S_50ntFlanks@6735.6771.37|TTGATCATCGACACTTCGAACGC ACTTGCGGCCCCGG</td><td>>hg38_21_+_8433172_8446622_RNA45SN1_45S_50ntFlanks</td><td>6735</td><td>6771</td><td>37</td><td>TTGATCATCGACACTTCGAACGCACTTGCGGCCCCGG</td><td>173.9</td><td>-1.6</td><td>0.7</td><td>-2.2</td><td>2.7E-02</td><td>3.2E-02</td><td>MB231/MB468</td></tr>
<tr><td>>trnaMT_PheGAA_MT_+_577_647@35.55.21|GAAAATGTTTAGACGGGCTCA</td><td>>trnaMT_PheGAA_MT_+_577_647</td><td>35</td><td>55</td><td>21</td><td>GAAAATGTTTAGACGGGCTCA</td><td>1097.1</td><td>2.0</td><td>0.8</td><td>2.6</td><td>8.2E-03</td><td>3.2E-02</td><td>BT20/MB231</td></tr>
<tr><td>>hg38_21_+_8433172_8446622_RNA45SN1_45S_50ntFlanks@12990.13039.50|GCTCCCTCGCTGCGATCTATTG AAAGTCAGCCCTCGACACAAGGGTTTGT</td><td>>hg38_21_+_8433172_8446622_RNA45SN1_45S_50ntFlanks</td><td>12990</td><td>13039</td><td>50</td><td>GCTCCCTCGCTGCGATCTATTGAAAGTCAGCCCTCGACACAAGGGTTTGT</td><td>236.7</td><td>-3.1</td><td>1.2</td><td>-2.6</td><td>8.2E-03</td><td>3.2E-02</td><td>BT20/MB231</td></tr>
<tr><td>>hg38_MT_+_1621_3279_RNR2_16S_50ntFlanks@51.90.40|GCTAAACCTAGCCCCAAACCCACTCCACCTTACTACC AGA</td><td>>hg38_MT_+_1621_3279_RNR2_16S_50ntFlanks</td><td>51</td><td>90</td><td>40</td><td>GCTAAACCTAGCCCCAAACCCACTCCACCTTACTACCAGA</td><td>590.9</td><td>-2.0</td><td>0.8</td><td>-2.6</td><td>8.2E-03</td><td>3.2E-02</td><td>BT20/MB231</td></tr>
<tr><td>>MI0000469|hsa-mir-125a&WithFlank&19|+|51693248|51693345@21.42.22&[MIMAT0000443&hsa-miR-125a-5p&offsets|0|-2;m-63&19|+|51693268|51693289&offsets|0|0]|TCCCTGAGACCCTTTAACCTGT</td><td>>MI0000469|hsa-mir-125a&WithFlank&19|+|51693248|51693345</td><td>21</td><td>42</td><td>22</td><td>TCCCTGAGACCCTTTAACCTGT</td><td>6090.6</td><td>1.8</td><td>0.7</td><td>2.6</td><td>8.2E-03</td><td>3.2E-02</td><td>BT20/MB231</td></tr>
<tr><td>>MI0000736|hsa-mir-30c-1&WithFlank&1|+|40757278|40757378@23.46.24(+1U);MI0000254|hsa-mir-30c-2&WithFlank&6|-|71376954|71377037@13.36.24(+1U)&[MIMAT0000244&hsa-miR-30c-5p&offsets|0|+1(+1U);m-57&1|+|40757300|40757323&offsets|0|0(+1U)];[MIMAT0000244_1&hsa-miR-30c-5p&offsets|0|+1(+1U);m-56&6|-|71377002|71377025&offsets|0|0(+1U)]|TGTAAACATCCTACACTCTCAGCTT</td><td>>MI0000736|hsa-mir-30c-1&WithFlank&1|+|40757278|40757378</td><td>23</td><td>46</td><td>24(+1U)</td><td>TGTAAACATCCTACACTCTCAGCTT</td><td>1394.2</td><td>-2.0</td><td>0.8</td><td>-2.6</td><td>8.3E-03</td><td>3.2E-02</td><td>BT20/MB231</td></tr>
<tr><td>>hg38_21_+_8433172_8446622_RNA45SN1_45S_50ntFlanks@474.497.24|TCTTCGTGATCGATGTGGTGACGT</td><td>>hg38_21_+_8433172_8446622_RNA45SN1_45S_50ntFlanks</td><td>474</td><td>497</td><td>24</td><td>TCTTCGTGATCGATGTGGTGACGT</td><td>995.7</td><td>-2.3</td><td>0.9</td><td>-2.6</td><td>8.3E-03</td><td>3.2E-02</td><td>BT20/MB231</td></tr>
<tr><td>>hg38_21_+_8433172_8446622_RNA45SN1_45S_50ntFlanks@4067.4089.23|AACTTTCGATGGTAGTCGCCGTG</td><td>>hg38_21_+_8433172_8446622_RNA45SN1_45S_50ntFlanks</td><td>4067</td><td>4089</td><td>23</td><td>AACTTTCGATGGTAGTCGCCGTG</td><td>87.3</td><td>-1.4</td><td>0.6</td><td>-2.2</td><td>2.7E-02</td><td>3.2E-02</td><td>MB231/MB468</td></tr>
<tr><td>>trnaMT_ValTAC_MT_+_1602_1670@53.72.20|TAACTTGACCGCTCTGACCA</td><td>>trnaMT_ValTAC_MT_+_1602_1670</td><td>53</td><td>72</td><td>20</td><td>TAACTTGACCGCTCTGACCA</td><td>2462.1</td><td>2.0</td><td>0.7</td><td>2.6</td><td>8.3E-03</td><td>3.2E-02</td><td>BT20/MB231</td></tr>
<tr><td>>hg38_21_+_8433172_8446622_RNA45SN1_45S_50ntFlanks@8104.8142.39|CCGCGGCGGGGCGCGGGACATG TGGCGTACGGAAGACCC</td><td>>hg38_21_+_8433172_8446622_RNA45SN1_45S_50ntFlanks</td><td>8104</td><td>8142</td><td>39</td><td>CCGCGGCGGGGCGCGGGACATGTGGCGTACGGAAGACCC</td><td>83.1</td><td>-1.9</td><td>0.9</td><td>-2.2</td><td>2.8E-02</td><td>3.2E-02</td><td>MB231/MB468</td></tr>
<tr><td>>hg38_21_+_8433172_8446622_RNA45SN1_45S_50ntFlanks@3850.3871.22|GATAACTGTGGTAATTCTAGAG</td><td>>hg38_21_+_8433172_8446622_RNA45SN1_45S_50ntFlanks</td><td>3850</td><td>3871</td><td>22</td><td>GATAACTGTGGTAATTCTAGAG</td><td>145.1</td><td>-1.5</td><td>0.7</td><td>-2.2</td><td>2.8E-02</td><td>3.2E-02</td><td>MB231/MB468</td></tr>
<tr><td>>hg38_21_+_8433172_8446622_RNA45SN1_45S_50ntFlanks@3837.3868.32|CCTCTCCTACTTGGATAACTGTGG TAATTCTA</td><td>>hg38_21_+_8433172_8446622_RNA45SN1_45S_50ntFlanks</td><td>3837</td><td>3868</td><td>32</td><td>CCTCTCCTACTTGGATAACTGTGGTAATTCTA</td><td>115.2</td><td>-1.4</td><td>0.7</td><td>-2.2</td><td>2.9E-02</td><td>3.2E-02</td><td>BT20/MB468</td></tr>
<tr><td>>hg38_21_+_8433172_8446622_RNA45SN1_45S_50ntFlanks@6661.6680.20|CGGTGGATCACTCGGCTCGT</td><td>>hg38_21_+_8433172_8446622_RNA45SN1_45S_50ntFlanks</td><td>6661</td><td>6680</td><td>20</td><td>CGGTGGATCACTCGGCTCGT</td><td>220.1</td><td>1.1</td><td>0.5</td><td>2.2</td><td>2.8E-02</td><td>3.2E-02</td><td>MB231/MB468</td></tr>
<tr><td>>hg38_21_+_8433172_8446622_RNA45SN1_45S_50ntFlanks@6652.6697.46|GACTCTTAGCGGTGGATCACTCG GCTCGTGCGTCGATGAAGAACGC</td><td>>hg38_21_+_8433172_8446622_RNA45SN1_45S_50ntFlanks</td><td>6652</td><td>6697</td><td>46</td><td>GACTCTTAGCGGTGGATCACTCGGCTCGTGCGTCGATGAAGAACGC</td><td>525.3</td><td>1.9</td><td>0.9</td><td>2.2</td><td>2.9E-02</td><td>3.2E-02</td><td>BT20/MB468</td></tr>
<tr><td>>hg38_21_+_8433172_8446622_RNA45SN1_45S_50ntFlanks@12936.12958.23|AAACCATTCGTAGACGACCTG CT</td><td>>hg38_21_+_8433172_8446622_RNA45SN1_45S_50ntFlanks</td><td>12936</td><td>12958</td><td>23</td><td>AAACCATTCGTAGACGACCTGCT</td><td>240.8</td><td>-1.1</td><td>0.5</td><td>-2.2</td><td>2.9E-02</td><td>3.2E-02</td><td>BT20/MB468</td></tr>
<tr><td>>hg38_21_+_8433172_8446622_RNA45SN1_45S_50ntFlanks@12418.12449.32|CGATGTCGGCTCTTCCTATCAT TGTGAAGCAG</td><td>>hg38_21_+_8433172_8446622_RNA45SN1_45S_50ntFlanks</td><td>12418</td><td>12449</td><td>32</td><td>CGATGTCGGCTCTTCCTATCATTGTGAAGCAG</td><td>343.8</td><td>-2.4</td><td>0.9</td><td>-2.6</td><td>8.4E-03</td><td>3.2E-02</td><td>BT20/MB231</td></tr>
<tr><td>>hg38_21_+_8433172_8446622_RNA45SN1_45S_50ntFlanks@5489.5514.26|CTGAGAAGACGGTCGAACTTGA CTAT</td><td>>hg38_21_+_8433172_8446622_RNA45SN1_45S_50ntFlanks</td><td>5489</td><td>5514</td><td>26</td><td>CTGAGAAGACGGTCGAACTTGACTAT</td><td>414.1</td><td>-1.5</td><td>0.7</td><td>-2.2</td><td>2.8E-02</td><td>3.2E-02</td><td>MB231/MB468</td></tr>
<tr><td>>hg38_21_+_8433172_8446622_RNA45SN1_45S_50ntFlanks@11668.11702.35|TGCCCAGTGCTCTGAATGTCA AAGTGAAGAAATTC</td><td>>hg38_21_+_8433172_8446622_RNA45SN1_45S_50ntFlanks</td><td>11668</td><td>11702</td><td>35</td><td>TGCCCAGTGCTCTGAATGTCAAAGTGAAGAAATTC</td><td>737.0</td><td>1.6</td><td>0.7</td><td>2.2</td><td>2.9E-02</td><td>3.3E-02</td><td>BT20/MB468</td></tr>
<tr><td>>hg38_21_+_8433172_8446622_RNA45SN1_45S_50ntFlanks@11667.11706.40|CTGCCCAGTGCTCTGAATGTCA AAGTGAAGAAATTCAATG</td><td>>hg38_21_+_8433172_8446622_RNA45SN1_45S_50ntFlanks</td><td>11667</td><td>11706</td><td>40</td><td>CTGCCCAGTGCTCTGAATGTCAAAGTGAAGAAATTCAATG</td><td>174.4</td><td>-2.0</td><td>0.8</td><td>-2.6</td><td>8.5E-03</td><td>3.3E-02</td><td>BT20/MB231</td></tr>
<tr><td>>hg38_21_+_8433172_8446622_RNA45SN1_45S_50ntFlanks@10771.10788.18|CAGGTCTCCAAGGTGAAC</td><td>>hg38_21_+_8433172_8446622_RNA45SN1_45S_50ntFlanks</td><td>10771</td><td>10788</td><td>18</td><td>CAGGTCTCCAAGGTGAAC</td><td>119.9</td><td>-1.2</td><td>0.5</td><td>-2.2</td><td>2.8E-02</td><td>3.3E-02</td><td>MB231/MB468</td></tr>
<tr><td>>hg38_21_+_8433172_8446622_RNA45SN1_45S_50ntFlanks@10781.10814.34|AGGTGAACAGCCTCTGGCATG TTGGAACAATGTA</td><td>>hg38_21_+_8433172_8446622_RNA45SN1_45S_50ntFlanks</td><td>10781</td><td>10814</td><td>34</td><td>AGGTGAACAGCCTCTGGCATGTTGGAACAATGTA</td><td>126.1</td><td>-1.3</td><td>0.6</td><td>-2.2</td><td>2.8E-02</td><td>3.3E-02</td><td>MB231/MB468</td></tr>
<tr><td>>hg38_21_+_8433172_8446622_RNA45SN1_45S_50ntFlanks@4560.4589.30|CTAGGAATAATGGAATAGGACC GCGGTTCT</td><td>>hg38_21_+_8433172_8446622_RNA45SN1_45S_50ntFlanks</td><td>4560</td><td>4589</td><td>30</td><td>CTAGGAATAATGGAATAGGACCGCGGTTCT</td><td>128.2</td><td>-1.6</td><td>0.7</td><td>-2.2</td><td>2.8E-02</td><td>3.3E-02</td><td>MB231/MB468</td></tr>
<tr><td>>MI0000342|hsa-mir-200b&WithFlank&1|+|1167098|1167204@63.86.24&[MIMAT0000318&hsa-miR-200b-3p&offsets|0|+2;m-93&1|+|1167160|1167182&offsets|0|+1]|TAATACTGCCTGGTAATGATGACG</td><td>>MI0000342|hsa-mir-200b&WithFlank&1|+|1167098|1167204</td><td>63</td><td>86</td><td>24</td><td>TAATACTGCCTGGTAATGATGACG</td><td>141.7</td><td>-1.9</td><td>0.9</td><td>-2.2</td><td>2.8E-02</td><td>3.3E-02</td><td>MB231/MB468</td></tr>
<tr><td>>hg38_21_+_8433172_8446622_RNA45SN1_45S_50ntFlanks@6648.6694.47|GTACGACTCTTAGCGGTGGATCA CTCGGCTCGTGCGTCGATGAAGAA</td><td>>hg38_21_+_8433172_8446622_RNA45SN1_45S_50ntFlanks</td><td>6648</td><td>6694</td><td>47</td><td>GTACGACTCTTAGCGGTGGATCACTCGGCTCGTGCGTCGATGAAGAA</td><td>196.2</td><td>-2.2</td><td>0.8</td><td>-2.6</td><td>8.6E-03</td><td>3.3E-02</td><td>BT20/MB231</td></tr>
</table>

<table>
<tr><th></th><th></th><th></th><th></th><th></th><th></th><th></th><th></th><th></th><th></th><th></th><th></th><th></th></tr>
<tr><td>>hg38_21_+_8433172_8446622_RNA45SN1_45S_50ntFlanks@7992.8019.28|CGTGGCGACCCGCTGAATTTAAGCATAT</td><td>>hg38_21_+_8433172_8446622_RNA45SN1_45S_50ntFlanks</td><td>7992</td><td>8019</td><td>28</td><td>CGTGGCGACCCGCTGAATTTAAGCATAT</td><td>210.1</td><td>-2.9</td><td>1.1</td><td>-2.6</td><td>8.6E-03</td><td>3.3E-02</td><td>BT20/MB231</td></tr>
<tr><td>>MI0000065|hsa-let-7d&WithFlank&9|+|94178828|94178926@14.35.22(+1A)&[MIMAT0000065&hsa-let-7d-5p&offsets|0|0(+1A);m-90&9|+|94178841|94178862&offsets|0|0(+1A)]|AGAGGTAGTAGGTTGCATAGTTA</td><td>>MI0000065|hsa-let-7d&WithFlank&9|+|94178828|94178926</td><td>14</td><td>35</td><td>22(+1A)</td><td>AGAGGTAGTAGGTTGCATAGTTA</td><td>329.6</td><td>2.0</td><td>0.8</td><td>2.6</td><td>8.6E-03</td><td>3.3E-02</td><td>BT20/MB231</td></tr>
<tr><td>>MI0000542|hsa-mir-320a&WithFlank&8|-|22244960|22245043@48.69.22(+3U)&[MIMAT0000510&hsa-miR-320a-3p&offsets|0|0(+3U);m-60&8|-|22244975|22244996&offsets|0|0(+3U)]|AAAAGCTGGGTTGAGAGGGCGATTT</td><td>>MI0000542|hsa-mir-320a&WithFlank&8|-|22244960|22245043</td><td>48</td><td>69</td><td>22(+3U)</td><td>AAAAGCTGGGTTGAGAGGGCGATTT</td><td>335.4</td><td>2.0</td><td>0.7</td><td>2.6</td><td>8.7E-03</td><td>3.3E-02</td><td>BT20/MB231</td></tr>
<tr><td>>MI0000081|hsa-mir-24-2&WithFlank&19|-|13836281|13836365@56.74.19(+1A);MI0000080|hsa-mir-24-1&WithFlank&9|+|95086015|95086094@50.68.19(+1A)&[MIMAT0000080&hsa-miR-24-3p&offsets|0|-3(+1A);m-36&19|-|13836288|13836310&offsets|0|-4(+1A)];[MIMAT0000080_1&hsa-miR-24-3p&offsets|0|-3(+1A);m-35&9|+|95086064|95086086&offsets|0|-4(+1A)]|TGGCTCAGTTCAGCAGGAAA</td><td>>MI0000081|hsa-mir-24-2&WithFlank&19|-|13836281|13836365</td><td>56</td><td>74</td><td>19(+1A)</td><td>TGGCTCAGTTCAGCAGGAAA</td><td>430.3</td><td>2.1</td><td>0.8</td><td>2.6</td><td>8.7E-03</td><td>3.3E-02</td><td>BT20/MB231</td></tr>
<tr><td>>hg38_21_+_8433172_8446622_RNA45SN1_45S_50ntFlanks@4554.4573.20|CCGCAGCTAGGAATAATGGA</td><td>>hg38_21_+_8433172_8446622_RNA45SN1_45S_50ntFlanks</td><td>4554</td><td>4573</td><td>20</td><td>CCGCAGCTAGGAATAATGGA</td><td>89.4</td><td>-1.4</td><td>0.6</td><td>-2.2</td><td>2.9E-02</td><td>3.3E-02</td><td>MB231/MB468</td></tr>
<tr><td>>hg38_21_+_8433172_8446622_RNA45SN1_45S_50ntFlanks@8190.8213.24|CCAGCCCGTGGACGGTGTGAGGCC</td><td>>hg38_21_+_8433172_8446622_RNA45SN1_45S_50ntFlanks</td><td>8190</td><td>8213</td><td>24</td><td>CCAGCCCGTGGACGGTGTGAGGCC</td><td>372.8</td><td>1.5</td><td>0.7</td><td>2.2</td><td>2.9E-02</td><td>3.3E-02</td><td>MB231/MB468</td></tr>
<tr><td>>hg38_21_+_8433172_8446622_RNA45SN1_45S_50ntFlanks@12999.13028.30|CTGCGATCTATTGAAAGTCAGCCCTCGACA</td><td>>hg38_21_+_8433172_8446622_RNA45SN1_45S_50ntFlanks</td><td>12999</td><td>13028</td><td>30</td><td>CTGCGATCTATTGAAAGTCAGCCCTCGACA</td><td>283.7</td><td>-2.2</td><td>1.0</td><td>-2.2</td><td>3.0E-02</td><td>3.4E-02</td><td>BT20/MB468</td></tr>
<tr><td>>trnaMT_PheGAA_MT_+_577_647@35.60.26|GAAAATGTTTAGACGGGCTCACATCA</td><td>>trnaMT_PheGAA_MT_+_577_647</td><td>35</td><td>60</td><td>26</td><td>GAAAATGTTTAGACGGGCTCACATCA</td><td>629.3</td><td>2.4</td><td>0.9</td><td>2.6</td><td>8.8E-03</td><td>3.4E-02</td><td>BT20/MB231</td></tr>
<tr><td>>hg38_21_+_8433172_8446622_RNA45SN1_45S_50ntFlanks@12416.12451.36|TTCGATGTCGGCTCTTCCTATCATTGTGAAGCAGAA</td><td>>hg38_21_+_8433172_8446622_RNA45SN1_45S_50ntFlanks</td><td>12416</td><td>12451</td><td>36</td><td>TTCGATGTCGGCTCTTCCTATCATTGTGAAGCAGAA</td><td>148.1</td><td>-2.4</td><td>0.9</td><td>-2.6</td><td>8.9E-03</td><td>3.4E-02</td><td>BT20/MB231</td></tr>
<tr><td>>MI0000469|hsa-mir-125a&WithFlank&19|+|51693248|51693345@21.41.21&[MIMAT0000443&hsa-miR-125a-5p&offsets|0|-3;m-63&19|+|51693268|51693289&offsets|0|-1]|TCCCTGAGACCCTTTAACCTG</td><td>>MI0000469|hsa-mir-125a&WithFlank&19|+|51693248|51693345</td><td>21</td><td>41</td><td>21</td><td>TCCCTGAGACCCTTTAACCTG</td><td>588.5</td><td>2.3</td><td>0.9</td><td>2.6</td><td>8.9E-03</td><td>3.4E-02</td><td>BT20/MB231</td></tr>
<tr><td>>hg38_21_+_8433172_8446622_RNA45SN1_45S_50ntFlanks@3842.3860.19|CCTACTTGGATAACTGTGG</td><td>>hg38_21_+_8433172_8446622_RNA45SN1_45S_50ntFlanks</td><td>3842</td><td>3860</td><td>19</td><td>CCTACTTGGATAACTGTGG</td><td>134.5</td><td>-1.0</td><td>0.5</td><td>-2.2</td><td>2.9E-02</td><td>3.4E-02</td><td>MB231/MB468</td></tr>
<tr><td>>MI0000542|hsa-mir-320a&WithFlank&8|-|22244960|22245043@48.71.24(+1U)&[MIMAT0000510&hsa-miR-320a-3p&offsets|0|+2(+1U);m-60&8|-|22244975|22244996&offsets|0|+2(+1U)]|AAAAGCTGGGTTGAGAGGGCGAAAT</td><td>>MI0000542|hsa-mir-320a&WithFlank&8|-|22244960|22245043</td><td>48</td><td>71</td><td>24(+1U)</td><td>AAAAGCTGGGTTGAGAGGGCGAAAT</td><td>830.4</td><td>2.0</td><td>0.8</td><td>2.6</td><td>9.0E-03</td><td>3.4E-02</td><td>BT20/MB231</td></tr>
<tr><td>>hg38_21_+_8433172_8446622_RNA45SN1_45S_50ntFlanks@10349.10377.29|AACGAGAACTTTGAAGGCCGAAGTGGAGA</td><td>>hg38_21_+_8433172_8446622_RNA45SN1_45S_50ntFlanks</td><td>10349</td><td>10377</td><td>29</td><td>AACGAGAACTTTGAAGGCCGAAGTGGAGA</td><td>942.8</td><td>-1.3</td><td>0.6</td><td>-2.2</td><td>2.9E-02</td><td>3.4E-02</td><td>MB231/MB468</td></tr>
<tr><td>>trna111_HisGTG_1_-_147774845_147774916@-1G.30.31,trna118_HisGTG_1_-_145396881_145396952@-1G.30.31,trna16_HisGTG_1_+_146544773_146544844@-1G.30.31,trna1_HisGTG_15_+_45493349_45493420@-1G.30.31,trna21_HisGTG_1_+_147753471_147753542@-1G.30.31,trna33_HisGTG_6_+_27125906_27125977@-1G.30.31,trna7_HisGTG_9_-_14433938_14434009@-1G.30.31,trna8_HisGTG_15_-_45492611_45492682@-1G.30.31,trna9_HisGTG_15_-_45490804_45490875@-1G.30.31|GGCCGTGATCGTATAGTGGTTAGTACTCTGC</td><td>>trna111_HisGTG_1_-_147774845_147774916</td><td>-1G</td><td>30</td><td>31</td><td>GGCCGTGATCGTATAGTGGTTAGTACTCTGC</td><td>1634.3</td><td>-2.9</td><td>1.1</td><td>-2.6</td><td>9.0E-03</td><td>3.4E-02</td><td>BT20/MB231</td></tr>
<tr><td>>hg38_21_+_8433172_8446622_RNA45SN1_45S_50ntFlanks@12416.12448.33|TTCGATGTCGGCTCTTCCTATCATTGTGAAGCA</td><td>>hg38_21_+_8433172_8446622_RNA45SN1_45S_50ntFlanks</td><td>12416</td><td>12448</td><td>33</td><td>TTCGATGTCGGCTCTTCCTATCATTGTGAAGCA</td><td>3552.1</td><td>-2.3</td><td>0.9</td><td>-2.6</td><td>9.0E-03</td><td>3.4E-02</td><td>BT20/MB231</td></tr>
<tr><td>>MI0000261|hsa-mir-139&WithFlank&11|-|72615057|72615136@13.35.23&[MIMAT0000250&hsa-miR-139-5p&offsets|0|0;m-186&11|-|72615102|72615124&offsets|0|0]|TCTACAGTGCACGTGTCTCCAGT</td><td>>MI0000261|hsa-mir-139&WithFlank&11|-|72615057|72615136</td><td>13</td><td>35</td><td>23</td><td>TCTACAGTGCACGTGTCTCCAGT</td><td>1645.4</td><td>-2.0</td><td>0.8</td><td>-2.6</td><td>9.0E-03</td><td>3.4E-02</td><td>BT20/MB231</td></tr>
<tr><td>>hg38_21_+_8433172_8446622_RNA45SN1_45S_50ntFlanks@11581.11612.32|AACTGGTGCGGACCAGGGGAATCCGACTGTTT</td><td>>hg38_21_+_8433172_8446622_RNA45SN1_45S_50ntFlanks</td><td>11581</td><td>11612</td><td>32</td><td>AACTGGTGCGGACCAGGGGAATCCGACTGTTT</td><td>1956.8</td><td>-1.4</td><td>0.6</td><td>-2.2</td><td>2.9E-02</td><td>3.4E-02</td><td>MB231/MB468</td></tr>
<tr><td>>hg38_21_+_8433172_8446622_RNA45SN1_45S_50ntFlanks@10351.10371.21|CGAGAACTTTGAAGGCCGAAG</td><td>>hg38_21_+_8433172_8446622_RNA45SN1_45S_50ntFlanks</td><td>10351</td><td>10371</td><td>21</td><td>CGAGAACTTTGAAGGCCGAAG</td><td>905.4</td><td>-1.3</td><td>0.6</td><td>-2.2</td><td>2.9E-02</td><td>3.4E-02</td><td>MB231/MB468</td></tr>
<tr><td>>hg38_21_+_8433172_8446622_RNA45SN1_45S_50ntFlanks@3850.3869.20|GATAACTGTGGTAATTCTAG</td><td>>hg38_21_+_8433172_8446622_RNA45SN1_45S_50ntFlanks</td><td>3850</td><td>3869</td><td>20</td><td>GATAACTGTGGTAATTCTAG</td><td>100.1</td><td>-1.1</td><td>0.5</td><td>-2.2</td><td>2.9E-02</td><td>3.4E-02</td><td>MB231/MB468</td></tr>
<tr><td>>hg38_21_+_8433172_8446622_RNA45SN1_45S_50ntFlanks@12455.12476.22|ACCAAGCGTTGGATTGTTCACC</td><td>>hg38_21_+_8433172_8446622_RNA45SN1_45S_50ntFlanks</td><td>12455</td><td>12476</td><td>22</td><td>ACCAAGCGTTGGATTGTTCACC</td><td>428.5</td><td>-1.9</td><td>0.7</td><td>-2.6</td><td>9.0E-03</td><td>3.4E-02</td><td>BT20/MB231</td></tr>
<tr><td>>hg38_21_+_8433172_8446622_RNA45SN1_45S_50ntFlanks@12420.12453.34|ATGTCGGCTCTTCCTATCATTGTGAAGCAGAATT</td><td>>hg38_21_+_8433172_8446622_RNA45SN1_45S_50ntFlanks</td><td>12420</td><td>12453</td><td>34</td><td>ATGTCGGCTCTTCCTATCATTGTGAAGCAGAATT</td><td>315.3</td><td>-2.0</td><td>0.8</td><td>-2.6</td><td>9.1E-03</td><td>3.4E-02</td><td>BT20/MB231</td></tr>
<tr><td>>hg38_21_+_8433172_8446622_RNA45SN1_45S_50ntFlanks@4727.4768.42|AATCAAGAACGAAAGTCGGAGGTTCGAAGACGATCAGATACC</td><td>>hg38_21_+_8433172_8446622_RNA45SN1_45S_50ntFlanks</td><td>4727</td><td>4768</td><td>42</td><td>AATCAAGAACGAAAGTCGGAGGTTCGAAGACGATCAGATACC</td><td>362.4</td><td>-2.3</td><td>0.9</td><td>-2.6</td><td>9.1E-03</td><td>3.4E-02</td><td>BT20/MB231</td></tr>
<tr><td>>hg38_21_+_8433172_8446622_RNA45SN1_45S_50ntFlanks@6651.6669.19|CGACTCTTAGCGGTGGATC</td><td>>hg38_21_+_8433172_8446622_RNA45SN1_45S_50ntFlanks</td><td>6651</td><td>6669</td><td>19</td><td>CGACTCTTAGCGGTGGATC</td><td>6253.4</td><td>2.1</td><td>0.8</td><td>2.6</td><td>9.1E-03</td><td>3.4E-02</td><td>BT20/MB231</td></tr>
<tr><td>>hg38_21_+_8433172_8446622_RNA45SN1_45S_50ntFlanks@6652.6697.46|GACTCTTAGCGGTGGATCACTCGGCTCGTGCGTCGATGAAGAACGC</td><td>>hg38_21_+_8433172_8446622_RNA45SN1_45S_50ntFlanks</td><td>6652</td><td>6697</td><td>46</td><td>GACTCTTAGCGGTGGATCACTCGGCTCGTGCGTCGATGAAGAACGC</td><td>912.7</td><td>-2.2</td><td>0.8</td><td>-2.6</td><td>9.1E-03</td><td>3.4E-02</td><td>BT20/MB231</td></tr>
<tr><td>>hg38_21_+_8433172_8446622_RNA45SN1_45S_50ntFlanks@9554.9578.25|CGTAGCGGTCCTGACGTGCAAATCG</td><td>>hg38_21_+_8433172_8446622_RNA45SN1_45S_50ntFlanks</td><td>9554</td><td>9578</td><td>25</td><td>CGTAGCGGTCCTGACGTGCAAATCG</td><td>274.9</td><td>1.1</td><td>0.5</td><td>2.2</td><td>2.9E-02</td><td>3.4E-02</td><td>MB231/MB468</td></tr>
<tr><td>>hg38_21_+_8433172_8446622_RNA45SN1_45S_50ntFlanks@10835.10858.24|CGGATCCGTAACTTCGGGATAAGG</td><td>>hg38_21_+_8433172_8446622_RNA45SN1_45S_50ntFlanks</td><td>10835</td><td>10858</td><td>24</td><td>CGGATCCGTAACTTCGGGATAAGG</td><td>440.0</td><td>1.1</td><td>0.5</td><td>2.2</td><td>3.0E-02</td><td>3.4E-02</td><td>MB231/MB468</td></tr>
<tr><td>>MI0000077|hsa-mir-21&WithFlank&17|+|59841260|59841343@15.36.22(+1A)&[MIMAT0000076&hsa-miR-21-5p&offsets|+1|+1(+1A);m-1&17|+|59841273|59841294&offsets|+1|+1(+1A)]|AGCTTATCAGACTGATGTTGACA</td><td>>MI0000077|hsa-mir-21&WithFlank&17|+|59841260|59841343</td><td>15</td><td>36</td><td>22(+1A)</td><td>AGCTTATCAGACTGATGTTGACA</td><td>771.4</td><td>1.9</td><td>0.7</td><td>2.6</td><td>9.2E-03</td><td>3.4E-02</td><td>BT20/MB231</td></tr>
<tr><td>>MI0000093|hsa-mir-92a-1&WithFlank&13|+|91351308|91351397@54.76.23(+1U)&[MIMAT0000092&hsa-miR-92a-3p&offsets|0|+1(+1U);m-9&13|+|91351361|91351382&offsets|0|+1(+1U)]|TATTGCACTTGTCCCGGCCTGTTT</td><td>>MI0000093|hsa-mir-92a-1&WithFlank&13|+|91351308|91351397</td><td>54</td><td>76</td><td>23(+1U)</td><td>TATTGCACTTGTCCCGGCCTGTTT</td><td>588.2</td><td>-1.8</td><td>0.7</td><td>-2.6</td><td>9.2E-03</td><td>3.4E-02</td><td>BT20/MB231</td></tr>
<tr><td>>MI0005531|hsa-mir-450b&WithFlank&X|-|134540179|134540268@17.37.21&[MIMAT0004909&hsa-miR-450b-5p&offsets|0|-1;m-266&X|-|134540232|134540252&offsets|0|0]|TTTTGCAATATGTTCCTGAAT</td><td>>MI0005531|hsa-mir-450b&WithFlank&X|-|134540179|134540268</td><td>17</td><td>37</td><td>21</td><td>TTTTGCAATATGTTCCTGAAT</td><td>1558.1</td><td>2.5</td><td>1.0</td><td>2.6</td><td>9.2E-03</td><td>3.4E-02</td><td>BT20/MB231</td></tr>
<tr><td>>hg38_21_+_8433172_8446622_RNA45SN1_45S_50ntFlanks@11587.11612.26|TGCGGACCAGGGGAATCCGACTGTTT</td><td>>hg38_21_+_8433172_8446622_RNA45SN1_45S_50ntFlanks</td><td>11587</td><td>11612</td><td>26</td><td>TGCGGACCAGGGGAATCCGACTGTTT</td><td>219.2</td><td>-1.2</td><td>0.5</td><td>-2.2</td><td>3.1E-02</td><td>3.5E-02</td><td>BT20/MB468</td></tr>
<tr><td>>hg38_21_+_8433172_8446622_RNA45SN1_45S_50ntFlanks@4006.4023.18|ACCTCGGGCCGATCGCAC</td><td>>hg38_21_+_8433172_8446622_RNA45SN1_45S_50ntFlanks</td><td>4006</td><td>4023</td><td>18</td><td>ACCTCGGGCCGATCGCAC</td><td>504.5</td><td>1.7</td><td>0.8</td><td>2.2</td><td>3.1E-02</td><td>3.5E-02</td><td>BT20/MB468</td></tr>
<tr><td>>hg38_21_+_8433172_8446622_RNA45SN1_45S_50ntFlanks@12893.12930.38|CCTCGCCCGTCACGCACCGCACGTTCGTGGGGAACCTG</td><td>>hg38_21_+_8433172_8446622_RNA45SN1_45S_50ntFlanks</td><td>12893</td><td>12930</td><td>38</td><td>CCTCGCCCGTCACGCACCGCACGTTCGTGGGGAACCTG</td><td>244.5</td><td>-4.2</td><td>1.9</td><td>-2.2</td><td>3.0E-02</td><td>3.5E-02</td><td>MB231/MB468</td></tr>
<tr><td>>trna12_ArgCCT_16_+_3243918_3243990@36.54.19,trna18_ArgCCT_17_+_73030001_73030073@36.54.19,trna21_ArgCCT_17_-_73030526_73030598@36.54.19,trna2_ArgCCT_16_+_3202901_3202973@36.54.19,trna3_ArgCCT_7_+_139025446_139025518@36.54.19|TAAGCCAGGGATTGTGGGT</td><td>>trna12_ArgCCT_16_+_3243918_3243990</td><td>36</td><td>54</td><td>19</td><td>TAAGCCAGGGATTGTGGGT</td><td>564.9</td><td>1.7</td><td>0.8</td><td>2.2</td><td>3.0E-02</td><td>3.5E-02</td><td>MB231/MB468</td></tr>
<tr><td>>hg38_21_+_8433172_8446622_RNA45SN1_45S_50ntFlanks@8192.8211.20|AGCCCGTGGACGGTGTGAGG</td><td>>hg38_21_+_8433172_8446622_RNA45SN1_45S_50ntFlanks</td><td>8192</td><td>8211</td><td>20</td><td>AGCCCGTGGACGGTGTGAGG</td><td>164.5</td><td>-1.3</td><td>0.6</td><td>-2.2</td><td>3.0E-02</td><td>3.5E-02</td><td>MB231/MB468</td></tr>
<tr><td>>hg38_21_+_8433172_8446622_RNA45SN1_45S_50ntFlanks@6661.6688.28|CGGTGGATCACTCGGCTCGTGCGTCGAT</td><td>>hg38_21_+_8433172_8446622_RNA45SN1_45S_50ntFlanks</td><td>6661</td><td>6688</td><td>28</td><td>CGGTGGATCACTCGGCTCGTGCGTCGAT</td><td>1601.5</td><td>2.3</td><td>0.9</td><td>2.6</td><td>9.4E-03</td><td>3.5E-02</td><td>BT20/MB231</td></tr>
<tr><td>>hg38_21_+_8433172_8446622_RNA45SN1_45S_50ntFlanks@11568.11611.44|GCAGCCGACTTAGAACTGGTGCGGACCAGGGGAATCCGACTGTT</td><td>>hg38_21_+_8433172_8446622_RNA45SN1_45S_50ntFlanks</td><td>11568</td><td>11611</td><td>44</td><td>GCAGCCGACTTAGAACTGGTGCGGACCAGGGGAATCCGACTGTT</td><td>1678.8</td><td>2.1</td><td>0.8</td><td>2.6</td><td>9.4E-03</td><td>3.5E-02</td><td>BT20/MB231</td></tr>
<tr><td>>hg38_21_+_8433172_8446622_RNA45SN1_45S_50ntFlanks@6653.6696.44|ACTCTTAGCGGTGGATCACTCGGCTCGTGCGTCGATGAAGAACG</td><td>>hg38_21_+_8433172_8446622_RNA45SN1_45S_50ntFlanks</td><td>6653</td><td>6696</td><td>44</td><td>ACTCTTAGCGGTGGATCACTCGGCTCGTGCGTCGATGAAGAACG</td><td>291.5</td><td>-1.9</td><td>0.7</td><td>-2.6</td><td>9.4E-03</td><td>3.5E-02</td><td>BT20/MB231</td></tr>
<tr><td>>hg38_21_+_8433172_8446622_RNA45SN1_45S_50ntFlanks@4567.4589.23|TAATGGAATAGGACCGCGGTTCT</td><td>>hg38_21_+_8433172_8446622_RNA45SN1_45S_50ntFlanks</td><td>4567</td><td>4589</td><td>23</td><td>TAATGGAATAGGACCGCGGTTCT</td><td>215.8</td><td>-1.9</td><td>0.9</td><td>-2.1</td><td>3.2E-02</td><td>3.5E-02</td><td>BT20/MB468</td></tr>
<tr><td>>hg38_21_+_8433172_8446622_RNA45SN1_45S_50ntFlanks@11566.11586.21|TAGCAGCCGACTTAGAACTGG</td><td>>hg38_21_+_8433172_8446622_RNA45SN1_45S_50ntFlanks</td><td>11566</td><td>11586</td><td>21</td><td>TAGCAGCCGACTTAGAACTGG</td><td>547.5</td><td>-2.2</td><td>0.9</td><td>-2.6</td><td>9.4E-03</td><td>3.5E-02</td><td>BT20/MB231</td></tr>
<tr><td>>hg38_21_+_8433172_8446622_RNA45SN1_45S_50ntFlanks@6646.6678.33|TCGTACGACTCTTAGCGGTGGATCACTCGGCTC</td><td>>hg38_21_+_8433172_8446622_RNA45SN1_45S_50ntFlanks</td><td>6646</td><td>6678</td><td>33</td><td>TCGTACGACTCTTAGCGGTGGATCACTCGGCTC</td><td>1822.5</td><td>2.3</td><td>0.9</td><td>2.6</td><td>9.5E-03</td><td>3.5E-02</td><td>BT20/MB231</td></tr>
</table>

| | | | | | | | | | | | | |
|---|---|---|---|---|---|---|---|---|---|---|---|---|
| >am_trnaMT_LysTTT_MT_+_8295_8364@30.52.23,trnalookalike7_LysTTT_1_+_568844_568913@30.52.23\|TTAAGTTAAAGATTAAGAGAACC | >am_trnaMT_LysTTT_MT_+_8295_8364 | 30 | 52 | 23 | TTAAGTTAAAGATTAAGAGAACC | 522.7 | 2.1 | 0.8 | 2.6 | 9.5E-03 | 3.5E-02 | BT20/MB231 |
| >MI0000542\|hsa-mir-320a&WithFlank&8\|-\|22244960\|22245043@49.69.21(+1U)&[MIMAT0000510&hsa-miR-320a-3p&offsets\|+1\|0(+1U);m-60&8\|-\|22244975\|22244996&offsets\|+1\|0(+1U)]\|AAAGCTGGGTTGAGAGGGCGAT | >MI0000542\|hsa-mir-320a&WithFlank&8\|-\|22244960\|22245043 | 49 | 69 | 21(+1U) | AAAGCTGGGTTGAGAGGGCGAT | 688.3 | 2.1 | 0.8 | 2.6 | 9.5E-03 | 3.5E-02 | BT20/MB231 |
| >hg38_21_+_8433172_8446622_RNA45SN1_45S_50ntFlanks@6646.6676.31\|TCGTACGACTCTTAGCGGTGGATCACTCGGC | >hg38_21_+_8433172_8446622_RNA45SN1_45S_50ntFlanks | 6646 | 6676 | 31 | TCGTACGACTCTTAGCGGTGGATCACTCGGC | 1677.0 | 2.1 | 0.8 | 2.6 | 9.6E-03 | 3.6E-02 | BT20/MB231 |
| >hg38_21_+_8433172_8446622_RNA45SN1_45S_50ntFlanks@12417.12445.29\|TCGATGTCGGCTCTTCCTATCATTGTGAA | >hg38_21_+_8433172_8446622_RNA45SN1_45S_50ntFlanks | 12417 | 12445 | 29 | TCGATGTCGGCTCTTCCTATCATTGTGAA | 480.9 | 1.1 | 0.5 | 2.2 | 3.1E-02 | 3.6E-02 | MB231/MB468 |
| >hg38_21_+_8433172_8446622_RNA45SN1_45S_50ntFlanks@7977.7994.18\|CGACCTCAGATCAGACGT | >hg38_21_+_8433172_8446622_RNA45SN1_45S_50ntFlanks | 7977 | 7994 | 18 | CGACCTCAGATCAGACGT | 196.8 | -1.1 | 0.5 | -2.2 | 3.1E-02 | 3.6E-02 | MB231/MB468 |
| >hg38_21_+_8433172_8446622_RNA45SN1_45S_50ntFlanks@3991.4024.34\|TGGTGACTCTAGATAACCTCGGGCCGATCGCACG | >hg38_21_+_8433172_8446622_RNA45SN1_45S_50ntFlanks | 3991 | 4024 | 34 | TGGTGACTCTAGATAACCTCGGGCCGATCGCACG | 223.4 | 1.1 | 0.5 | 2.2 | 3.1E-02 | 3.6E-02 | MB231/MB468 |
| >MI0000734\|hsa-mir-106b&WithFlank&7\|-\|100093987\|100094080@18.39.22&[MIMAT0000680&hsa-miR-106b-5p&offsets\|0\|+1;m-103&7\|-\|100094043\|100094063&offsets\|0\|+1]\|TAAAGTGCTGACAGTGCAGATA | >MI0000734\|hsa-mir-106b&WithFlank&7\|-\|100093987\|100094080 | 18 | 39 | 22 | TAAAGTGCTGACAGTGCAGATA | 6121.7 | 2.4 | 0.9 | 2.6 | 9.7E-03 | 3.6E-02 | BT20/MB231 |
| >hg38_21_+_8433172_8446622_RNA45SN1_45S_50ntFlanks@12990.13012.23\|GCTCCCTCGCTGCGATCTATTGA | >hg38_21_+_8433172_8446622_RNA45SN1_45S_50ntFlanks | 12990 | 13012 | 23 | GCTCCCTCGCTGCGATCTATTGA | 284.7 | -1.7 | 0.8 | -2.1 | 3.2E-02 | 3.6E-02 | BT20/MB468 |
| >hg38_21_+_8433172_8446622_RNA45SN1_45S_50ntFlanks@11670.11690.21\|CCCAGTGCTCTGAATGTCAAA | >hg38_21_+_8433172_8446622_RNA45SN1_45S_50ntFlanks | 11670 | 11690 | 21 | CCCAGTGCTCTGAATGTCAAA | 104.5 | -1.1 | 0.5 | -2.2 | 3.1E-02 | 3.6E-02 | MB231/MB468 |
| >hg38_21_+_8433172_8446622_RNA45SN1_45S_50ntFlanks@11582.11602.21\|ACTGGTGCGGACCAGGGGAAT | >hg38_21_+_8433172_8446622_RNA45SN1_45S_50ntFlanks | 11582 | 11602 | 21 | ACTGGTGCGGACCAGGGGAAT | 652.6 | 1.7 | 0.8 | 2.2 | 3.1E-02 | 3.6E-02 | MB231/MB468 |
| >hg38_21_+_8433172_8446622_RNA45SN1_45S_50ntFlanks@12626.12669.44\|AATGGGGCGAAGCTACCATCTGTGGGATTATGACTGAACGCCTC | >hg38_21_+_8433172_8446622_RNA45SN1_45S_50ntFlanks | 12626 | 12669 | 44 | AATGGGGCGAAGCTACCATCTGTGGGATTATGACTGAACGCCTC | 174.2 | -2.4 | 0.9 | -2.6 | 9.7E-03 | 3.6E-02 | BT20/MB231 |
| >hg38_21_+_8433172_8446622_RNA45SN1_45S_50ntFlanks@12907.12935.29\|CACCGCACGTTCGTGGGGAACCTGGCGCT | >hg38_21_+_8433172_8446622_RNA45SN1_45S_50ntFlanks | 12907 | 12935 | 29 | CACCGCACGTTCGTGGGGAACCTGGCGCT | 168.3 | -1.9 | 0.9 | -2.1 | 3.3E-02 | 3.6E-02 | BT20/MB468 |
| >hg38_21_+_8433172_8446622_RNA45SN1_45S_50ntFlanks@10781.10800.20\|AGGTGAACAGCCTCTGGCAT | >hg38_21_+_8433172_8446622_RNA45SN1_45S_50ntFlanks | 10781 | 10800 | 20 | AGGTGAACAGCCTCTGGCAT | 118.5 | -1.1 | 0.5 | -2.2 | 3.1E-02 | 3.6E-02 | MB231/MB468 |
| >hg38_21_+_8433172_8446622_RNA45SN1_45S_50ntFlanks@3837.3871.35\|CCTCTCCTACTTGGATAACTGTGGTAATTCTAGAG | >hg38_21_+_8433172_8446622_RNA45SN1_45S_50ntFlanks | 3837 | 3871 | 35 | CCTCTCCTACTTGGATAACTGTGGTAATTCTAGAG | 102.2 | -1.4 | 0.7 | -2.2 | 3.1E-02 | 3.6E-02 | MB231/MB468 |
| >MI0000066\|hsa-let-7e&WithFlank&19\|+\|51692780\|51692870@14.35.22(+1A)&[MIMAT0000066&hsa-let-7e-5p&offsets\|0\|0(+1A);m-50&19\|+\|51692793\|51692814&offsets\|0\|0(+1A)]\|TGAGGTAGGAGGTTGTATAGTTA | >MI0000066\|hsa-let-7e&WithFlank&19\|+\|51692780\|51692870 | 14 | 35 | 22(+1A) | TGAGGTAGGAGGTTGTATAGTTA | 584.6 | 1.8 | 0.7 | 2.6 | 9.9E-03 | 3.6E-02 | BT20/MB231 |
| >hg38_21_+_8433172_8446622_RNA45SN1_45S_50ntFlanks@6661.6685.25\|CGGTGGATCACTCGGCTCGTGCGTC | >hg38_21_+_8433172_8446622_RNA45SN1_45S_50ntFlanks | 6661 | 6685 | 25 | CGGTGGATCACTCGGCTCGTGCGTC | 460.5 | 2.0 | 0.8 | 2.6 | 9.9E-03 | 3.6E-02 | BT20/MB231 |
| >trna14_LysTTT_11_-_59327808_59327880@1.31.31,trna2_LysTTT_17_+_8022473_8022545@1.31.31,trna54_LysTTT_1_+_204475655_204475727@1.31.31,trna5_LysTTT_11_+_59323902_59323974@1.31.31,trna62_LysTTT_1_-_204476158_204476230@1.31.31,trna76_LysTTT_6_+_28918806_28918878@1.31.31\|GCCCGGATAGCTCAGTCGGTAGAGCATCAGA | >trna14_LysTTT_11_-_59327808_59327880 | 1 | 31 | 31 | GCCCGGATAGCTCAGTCGGTAGAGCATCAGA | 274.0 | -2.4 | 1.1 | -2.1 | 3.3E-02 | 3.7E-02 | BT20/MB468 |
| >hg38_21_+_8433172_8446622_RNA45SN1_45S_50ntFlanks@11670.11691.22\|CCCAGTGCTCTGAATGTCAAAG | >hg38_21_+_8433172_8446622_RNA45SN1_45S_50ntFlanks | 11670 | 11691 | 22 | CCCAGTGCTCTGAATGTCAAAG | 449.8 | -1.3 | 0.6 | -2.1 | 3.3E-02 | 3.7E-02 | BT20/MB468 |
| >hg38_21_+_8433172_8446622_RNA45SN1_45S_50ntFlanks@9266.9285.20\|CGACGTCGGCTACCCACCCG | >hg38_21_+_8433172_8446622_RNA45SN1_45S_50ntFlanks | 9266 | 9285 | 20 | CGACGTCGGCTACCCACCCG | 136.6 | -1.9 | 0.9 | -2.2 | 3.2E-02 | 3.7E-02 | MB231/MB468 |
| >hg38_21_+_8433172_8446622_RNA45SN1_45S_50ntFlanks@10829.10851.23\|GCAAGCCGGATCCGTAACTTCGG | >hg38_21_+_8433172_8446622_RNA45SN1_45S_50ntFlanks | 10829 | 10851 | 23 | GCAAGCCGGATCCGTAACTTCGG | 170.2 | -1.0 | 0.5 | -2.1 | 3.2E-02 | 3.7E-02 | MB231/MB468 |
| >MI0000447\|hsa-mir-128-1&WithFlank&2\|+\|135665391\|135665484@56.76.21(+1A);MI0000727\|hsa-mir-128-2&WithFlank&3\|+\|35744470\|35744565@58.78.21(+1A)&[MIMAT0000424&hsa-miR-128-3p&offsets\|0\|0(+1A);m-108&2\|+\|135665446\|135665466&offsets\|0\|0(+1A)];[MIMAT0000424_1&hsa-miR-128-3p&offsets\|0\|0(+1A);m-117&3\|+\|35744527\|35744547&offsets\|0\|0(+1A)]\|TCACAGTGAACCGGTCTCTTTA | >MI0000447\|hsa-mir-128-1&WithFlank&2\|+\|135665391\|135665484 | 56 | 76 | 21(+1A) | TCACAGTGAACCGGTCTCTTTA | 2311.2 | 1.9 | 0.7 | 2.6 | 1.0E-02 | 3.7E-02 | BT20/MB231 |
| >hg38_21_+_8433172_8446622_RNA45SN1_45S_50ntFlanks@3846.3868.23\|CTTGGATAACTGTGGTAATTCTA | >hg38_21_+_8433172_8446622_RNA45SN1_45S_50ntFlanks | 3846 | 3868 | 23 | CTTGGATAACTGTGGTAATTCTA | 401.9 | -1.0 | 0.5 | -2.1 | 3.3E-02 | 3.7E-02 | BT20/MB468 |
| >hg38_21_+_8433172_8446622_RNA45SN1_45S_50ntFlanks@12439.12473.35\|TTGTGAAGCAGAATTCACCAAGCGTTGGATTGTTC | >hg38_21_+_8433172_8446622_RNA45SN1_45S_50ntFlanks | 12439 | 12473 | 35 | TTGTGAAGCAGAATTCACCAAGCGTTGGATTGTTC | 529.6 | -2.2 | 0.8 | -2.6 | 1.0E-02 | 3.7E-02 | BT20/MB231 |
| >hg38_21_+_8433172_8446622_RNA45SN1_45S_50ntFlanks@12408.12431.24\|TTTGATCCTTCGATGTCGGCTCTT | >hg38_21_+_8433172_8446622_RNA45SN1_45S_50ntFlanks | 12408 | 12431 | 24 | TTTGATCCTTCGATGTCGGCTCTT | 309.8 | -1.3 | 0.6 | -2.1 | 3.2E-02 | 3.7E-02 | MB231/MB468 |
| >hg38_21_+_8433172_8446622_RNA45SN1_45S_50ntFlanks@12455.12487.33\|ACCAAGCGTTGGATTGTTCACCCACTAATAGGG | >hg38_21_+_8433172_8446622_RNA45SN1_45S_50ntFlanks | 12455 | 12487 | 33 | ACCAAGCGTTGGATTGTTCACCCACTAATAGGG | 312.8 | -1.3 | 0.6 | -2.1 | 3.2E-02 | 3.7E-02 | MB231/MB468 |
| >hg38_21_+_8433172_8446622_RNA45SN1_45S_50ntFlanks@12997.13014.18\|CGCTGCGATCTATTGAAA | >hg38_21_+_8433172_8446622_RNA45SN1_45S_50ntFlanks | 12997 | 13014 | 18 | CGCTGCGATCTATTGAAA | 4824.1 | -2.0 | 0.8 | -2.6 | 1.0E-02 | 3.7E-02 | BT20/MB231 |
| >hg38_21_+_8433172_8446622_RNA45SN1_45S_50ntFlanks@8357.8391.35\|AAAAGAACTTTGAAGAGAGAGTTCAAGAGGGCGTG | >hg38_21_+_8433172_8446622_RNA45SN1_45S_50ntFlanks | 8357 | 8391 | 35 | AAAAGAACTTTGAAGAGAGAGTTCAAGAGGGCGTG | 249.8 | -1.4 | 0.6 | -2.1 | 3.2E-02 | 3.7E-02 | MB231/MB468 |
| >hg38_21_+_8433172_8446622_RNA45SN1_45S_50ntFlanks@10346.10373.28\|TCAAACGAGAACTTTGAAGGCCGAAGTG | >hg38_21_+_8433172_8446622_RNA45SN1_45S_50ntFlanks | 10346 | 10373 | 28 | TCAAACGAGAACTTTGAAGGCCGAAGTG | 226.1 | -2.2 | 0.9 | -2.6 | 1.0E-02 | 3.7E-02 | BT20/MB231 |
| >hg38_21_+_8433172_8446622_RNA45SN1_45S_50ntFlanks@5492.5509.18\|AGAAGACGGTCGAACTTG | >hg38_21_+_8433172_8446622_RNA45SN1_45S_50ntFlanks | 5492 | 5509 | 18 | AGAAGACGGTCGAACTTG | 402.7 | -1.7 | 0.8 | -2.1 | 3.4E-02 | 3.7E-02 | BT20/MB468 |
| >hg38_21_+_8433172_8446622_RNA45SN1_45S_50ntFlanks@12412.12436.25\|ATCCTTCGATGTCGGCTCTTCCTAT | >hg38_21_+_8433172_8446622_RNA45SN1_45S_50ntFlanks | 12412 | 12436 | 25 | ATCCTTCGATGTCGGCTCTTCCTAT | 290.1 | -3.7 | 1.7 | -2.1 | 3.2E-02 | 3.7E-02 | MB231/MB468 |
| >hg38_21_+_8433172_8446622_RNA45SN1_45S_50ntFlanks@11664.11685.22\|TTTCTGCCCAGTGCTCTGAATG | >hg38_21_+_8433172_8446622_RNA45SN1_45S_50ntFlanks | 11664 | 11685 | 22 | TTTCTGCCCAGTGCTCTGAATG | 177.4 | -1.1 | 0.5 | -2.1 | 3.2E-02 | 3.7E-02 | MB231/MB468 |
| >am_MI0000809\|hsa-mir-151a&WithFlank&8\|-\|140732558\|140732659@53.72.20(+1A)&[MIMAT0000757&hsa-miR-151a-3p&offsets\|0\|-1(+1A);m-30&8\|-\|140732587\|140732607&offsets\|0\|-1(+1A)]\|CTAGACTGAAGCTCCTTGAGA | >am_MI0000809\|hsa-mir-151a&WithFlank&8\|-\|140732558\|140732659 | 53 | 72 | 20(+1A) | CTAGACTGAAGCTCCTTGAGA | 18851.0 | 1.6 | 0.6 | 2.6 | 1.0E-02 | 3.8E-02 | BT20/MB231 |
| >am_trnaMT_ValTAC_MT_+_1602_1670@28.48.21\|AACTTACACTTAGGAGATTTC | >am_trnaMT_ValTAC_MT_+_1602_1670 | 28 | 48 | 21 | AACTTACACTTAGGAGATTTC | 257.3 | 2.4 | 0.9 | 2.6 | 1.0E-02 | 3.8E-02 | BT20/MB231 |
| >MI0000746\|hsa-mir-99b&WithFlank&19\|+\|51692606\|51692687@51.72.22&[MIMAT0004678&hsa-miR-99b-3p&offsets\|0\|0;m-151&19\|+\|51692656\|51692678&offsets\|0\|-1]\|CAAGCTCGTGTCTGTGGGTCCG | >MI0000746\|hsa-mir-99b&WithFlank&19\|+\|51692606\|51692687 | 51 | 72 | 22 | CAAGCTCGTGTCTGTGGGTCCG | 2305.9 | 1.6 | 0.6 | 2.6 | 1.0E-02 | 3.8E-02 | BT20/MB231 |
| >hg38_21_+_8433172_8446622_RNA45SN1_45S_50ntFlanks@8182.8206.25\|ATCGAGGCCCAGCCCGTGGACGGTG | >hg38_21_+_8433172_8446622_RNA45SN1_45S_50ntFlanks | 8182 | 8206 | 25 | ATCGAGGCCCAGCCCGTGGACGGTG | 173.6 | -3.4 | 1.6 | -2.1 | 3.3E-02 | 3.8E-02 | MB231/MB468 |
| >hg38_21_+_8433172_8446622_RNA45SN1_45S_50ntFlanks@6649.6670.22\|TACGACTCTTAGCGGTGGATCA | >hg38_21_+_8433172_8446622_RNA45SN1_45S_50ntFlanks | 6649 | 6670 | 22 | TACGACTCTTAGCGGTGGATCA | 135.2 | -1.2 | 0.6 | -2.1 | 3.4E-02 | 3.8E-02 | BT20/MB468 |
| >hg38_21_+_8433172_8446622_RNA45SN1_45S_50ntFlanks@9623.9652.30\|TAGTAGCTGGTTCCCTCCGAAGTTTCCCTC | >hg38_21_+_8433172_8446622_RNA45SN1_45S_50ntFlanks | 9623 | 9652 | 30 | TAGTAGCTGGTTCCCTCCGAAGTTTCCCTC | 72.7 | -1.4 | 0.7 | -2.1 | 3.3E-02 | 3.8E-02 | MB231/MB468 |
| >hg38_21_+_8433172_8446622_RNA45SN1_45S_50ntFlanks@11575.11613.39\|ACTTAGAACTGGTGCGGACCAGGGGAATCCGACTGTTTA | >hg38_21_+_8433172_8446622_RNA45SN1_45S_50ntFlanks | 11575 | 11613 | 39 | ACTTAGAACTGGTGCGGACCAGGGGAATCCGACTGTTTA | 403.5 | 1.5 | 0.7 | 2.1 | 3.3E-02 | 3.8E-02 | MB231/MB468 |
| >am_trnaMT_MetCAT_MT_+_4402_4469@1.22.22\|AGTAAGGTCAGCTAAATAAGCT | >am_trnaMT_MetCAT_MT_+_4402_4469 | 1 | 22 | 22 | AGTAAGGTCAGCTAAATAAGCT | 1397.0 | 2.0 | 0.8 | 2.6 | 1.0E-02 | 3.8E-02 | BT20/MB231 |
| >hg38_21_+_8433172_8446622_RNA45SN1_45S_50ntFlanks@6725.6744.20\|GCAGGACACATTGATCATCG | >hg38_21_+_8433172_8446622_RNA45SN1_45S_50ntFlanks | 6725 | 6744 | 20 | GCAGGACACATTGATCATCG | 171.3 | -2.4 | 0.9 | -2.6 | 1.0E-02 | 3.8E-02 | BT20/MB231 |
| >hg38_21_+_8433172_8446622_RNA45SN1_45S_50ntFlanks@6655.6681.27\|TCTTAGCGGTGGATCACTCGGCTCGTG | >hg38_21_+_8433172_8446622_RNA45SN1_45S_50ntFlanks | 6655 | 6681 | 27 | TCTTAGCGGTGGATCACTCGGCTCGTG | 1583.3 | -1.2 | 0.6 | -2.1 | 3.3E-02 | 3.8E-02 | MB231/MB468 |
| >hg38_21_+_8433172_8446622_RNA45SN1_45S_50ntFlanks@3842.3861.20\|CCTACTTGGATAACTGTGGT | >hg38_21_+_8433172_8446622_RNA45SN1_45S_50ntFlanks | 3842 | 3861 | 20 | CCTACTTGGATAACTGTGGT | 267.6 | 1.1 | 0.5 | 2.1 | 3.4E-02 | 3.8E-02 | BT20/MB468 |
| >hg38_21_+_8433172_8446622_RNA45SN1_45S_50ntFlanks@6645.6662.18\|CTCGTACGACTCTTAGCG | >hg38_21_+_8433172_8446622_RNA45SN1_45S_50ntFlanks | 6645 | 6662 | 18 | CTCGTACGACTCTTAGCG | 3044.3 | 2.0 | 0.8 | 2.6 | 1.0E-02 | 3.8E-02 | BT20/MB231 |
| >MI0000825\|hsa-mir-345&WithFlank&14\|+\|100307853\|100307962@24.45.22&[MIMAT0000772&hsa-miR-345-5p&offsets\|0\|0;m-193&14\|+\|100307876\|100307896&offsets\|0\|+1]\|GCTGACTCCTAGTCCAGGGCTC | >MI0000825\|hsa-mir-345&WithFlank&14\|+\|100307853\|100307962 | 24 | 45 | 22 | GCTGACTCCTAGTCCAGGGCTC | 349.2 | -1.8 | 0.7 | -2.6 | 1.0E-02 | 3.8E-02 | BT20/MB231 |
| >hg38_21_+_8433172_8446622_RNA45SN1_45S_50ntFlanks@7980.8004.25\|CCTCAGATCAGACGTGGCGACCCGC | >hg38_21_+_8433172_8446622_RNA45SN1_45S_50ntFlanks | 7980 | 8004 | 25 | CCTCAGATCAGACGTGGCGACCCGC | 94.2 | -2.5 | 1.2 | -2.1 | 3.5E-02 | 3.8E-02 | BT20/MB468 |

<table>
<tr><td>>hg38_21_+_8433172_8446622_RNA45SN1_45S_50ntFlanks@6721.6750.30|AATTGCAGGACACATTGATCATCGACACTT</td><td>>hg38_21_+_8433172_8446622_RNA45SN1_45S_50ntFlanks</td><td>6721</td><td>6750</td><td>30</td><td>AATTGCAGGACACATTGATCATCGACACTT</td><td>332.6</td><td>-1.2</td><td>0.6</td><td>-2.1</td><td>3.3E-02</td><td>3.8E-02</td><td>MB231/MB468</td></tr>
<tr><td>>MI0000087|hsa-mir-29a&WithFlank&7|-|130876741|130876816@47.70.24&[MIMAT0000086&hsa-miR-29a-3p&offsets|-1|+1;m-26&7|-|130876749|130876769&offsets|-1|+2]|CTAGCACCATCTGAAATCGGTTAT</td><td>>MI0000087|hsa-mir-29a&WithFlank&7|-|130876741|130876816</td><td>47</td><td>70</td><td>24</td><td>CTAGCACCATCTGAAATCGGTTAT</td><td>224.4</td><td>-2.1</td><td>0.8</td><td>-2.6</td><td>1.1E-02</td><td>3.8E-02</td><td>BT20/MB231</td></tr>
<tr><td>>hg38_21_+_8433172_8446622_RNA45SN1_45S_50ntFlanks@4728.4752.25|ATCAAGAACGAAAGTCGGAGGTTCG</td><td>>hg38_21_+_8433172_8446622_RNA45SN1_45S_50ntFlanks</td><td>4728</td><td>4752</td><td>25</td><td>ATCAAGAACGAAAGTCGGAGGTTCG</td><td>323.8</td><td>-3.7</td><td>1.7</td><td>-2.1</td><td>3.3E-02</td><td>3.8E-02</td><td>MB231/MB468</td></tr>
<tr><td>>hg38_21_+_8433172_8446622_RNA45SN1_45S_50ntFlanks@9565.9603.39|TGACGTGCAAATCGGTCGTCCGACCTGGGTATAGGGGCG</td><td>>hg38_21_+_8433172_8446622_RNA45SN1_45S_50ntFlanks</td><td>9565</td><td>9603</td><td>39</td><td>TGACGTGCAAATCGGTCGTCCGACCTGGGTATAGGGGCG</td><td>144.0</td><td>-2.6</td><td>1.0</td><td>-2.6</td><td>1.1E-02</td><td>3.8E-02</td><td>BT20/MB231</td></tr>
<tr><td>>MI0000802|hsa-mir-340&WithFlank&5|-|180015297|180015403@22.42.21&[MIMAT0004692&hsa-miR-340-5p&offsets|0|-1;m-220&5|-|180015361|180015382&offsets|0|-1]|TTATAAAGCAATGAGACTGAT</td><td>>MI0000802|hsa-mir-340&WithFlank&5|-|180015297|180015403</td><td>22</td><td>42</td><td>21</td><td>TTATAAAGCAATGAGACTGAT</td><td>1006.1</td><td>2.4</td><td>0.9</td><td>2.6</td><td>1.1E-02</td><td>3.8E-02</td><td>BT20/MB231</td></tr>
<tr><td>>hg38_21_+_8433172_8446622_RNA45SN1_45S_50ntFlanks@8198.8216.19|TGGACGGTGTGAGGCCGGT</td><td>>hg38_21_+_8433172_8446622_RNA45SN1_45S_50ntFlanks</td><td>8198</td><td>8216</td><td>19</td><td>TGGACGGTGTGAGGCCGGT</td><td>106.6</td><td>-1.4</td><td>0.6</td><td>-2.1</td><td>3.3E-02</td><td>3.8E-02</td><td>MB231/MB468</td></tr>
<tr><td>>MI0000264|hsa-mir-7-2&WithFlank&15|+|88611819|88611940@38.59.22(+1A);MI0000265|hsa-mir-7-3&WithFlank&19|+|4770664|4770785@37.58.22(+1A);MI0000263|hsa-mir-7-1&WithFlank&9|-|83969742|83969863@30.51.22(+1A)&[MIMAT0000252&hsa-miR-7-5p&offsets|0|-2(+1A);m-400&15|+|88611856|88611879&offsets|0|-2(+1A)];[MIMAT0000252_1&hsa-miR-7-5p&offsets|0|-2(+1A);m-402&19|+|4770700|4770723&offsets|0|-2(+1A)];[MIMAT0000252_2&hsa-miR-7-5p&offsets|0|-2(+1A);m-401&9|-|83969811|83969834&offsets|0|-2(+1A)]|TGGAAGACTAGTGATTTTGTTGA</td><td>>MI0000264|hsa-mir-7-2&WithFlank&15|+|88611819|88611940</td><td>38</td><td>59</td><td>22(+1A)</td><td>TGGAAGACTAGTGATTTTGTTGA</td><td>1787.1</td><td>2.4</td><td>0.9</td><td>2.6</td><td>1.1E-02</td><td>3.8E-02</td><td>BT20/MB231</td></tr>
<tr><td>>MI0000298|hsa-mir-221&WithFlank&X|-|45746151|45746272@71.93.23(+1G)&[MIMAT0000278&hsa-miR-221-3p&offsets|0|0(+1G);m-91&X|-|45746181|45746202&offsets|0|+1(+1G)]|AGCTACATTGTCTGCTGGGTTTCG</td><td>>MI0000298|hsa-mir-221&WithFlank&X|-|45746151|45746272</td><td>71</td><td>93</td><td>23(+1G)</td><td>AGCTACATTGTCTGCTGGGTTTCG</td><td>1428.3</td><td>-2.0</td><td>0.8</td><td>-2.6</td><td>1.1E-02</td><td>3.9E-02</td><td>BT20/MB231</td></tr>
<tr><td>>hg38_21_+_8433172_8446622_RNA45SN1_45S_50ntFlanks@10685.10711.27|GGCGGCGTCCGGTGAGCTCTCGCTGGC</td><td>>hg38_21_+_8433172_8446622_RNA45SN1_45S_50ntFlanks</td><td>10685</td><td>10711</td><td>27</td><td>GGCGGCGTCCGGTGAGCTCTCGCTGGC</td><td>146.0</td><td>-1.8</td><td>0.8</td><td>-2.1</td><td>3.5E-02</td><td>3.9E-02</td><td>BT20/MB468</td></tr>
<tr><td>>hg38_21_+_8433172_8446622_RNA45SN1_45S_50ntFlanks@5237.5254.18|ACTGACTGGCTCAGCGTG</td><td>>hg38_21_+_8433172_8446622_RNA45SN1_45S_50ntFlanks</td><td>5237</td><td>5254</td><td>18</td><td>ACTGACTGGCTCAGCGTG</td><td>249.1</td><td>1.6</td><td>0.7</td><td>2.1</td><td>3.3E-02</td><td>3.9E-02</td><td>MB231/MB468</td></tr>
<tr><td>>hg38_21_+_8433172_8446622_RNA45SN1_45S_50ntFlanks@11581.11602.22|AACTGGTGCGGACCAGGGGAAT</td><td>>hg38_21_+_8433172_8446622_RNA45SN1_45S_50ntFlanks</td><td>11581</td><td>11602</td><td>22</td><td>AACTGGTGCGGACCAGGGGAAT</td><td>1084.0</td><td>-1.8</td><td>0.8</td><td>-2.1</td><td>3.5E-02</td><td>3.9E-02</td><td>BT20/MB468</td></tr>
<tr><td>>hg38_21_+_8433172_8446622_RNA45SN1_45S_50ntFlanks@8195.8215.21|CCGTGGACGGTGTGAGGCCGG</td><td>>hg38_21_+_8433172_8446622_RNA45SN1_45S_50ntFlanks</td><td>8195</td><td>8215</td><td>21</td><td>CCGTGGACGGTGTGAGGCCGG</td><td>127.4</td><td>-1.2</td><td>0.6</td><td>-2.1</td><td>3.4E-02</td><td>3.9E-02</td><td>MB231/MB468</td></tr>
<tr><td>>hg38_21_+_8433172_8446622_RNA45SN1_45S_50ntFlanks@12451.12477.27|ATTCACCAAGCGTTGGATTGTTCACCC</td><td>>hg38_21_+_8433172_8446622_RNA45SN1_45S_50ntFlanks</td><td>12451</td><td>12477</td><td>27</td><td>ATTCACCAAGCGTTGGATTGTTCACCC</td><td>262.1</td><td>-2.0</td><td>0.8</td><td>-2.5</td><td>1.1E-02</td><td>3.9E-02</td><td>BT20/MB231</td></tr>
<tr><td>>MI0000266|hsa-mir-10a&WithFlank&17|-|48579832|48579953@28.49.22(+1U)&[MIMAT0000253&hsa-miR-10a-5p&offsets|0|-1(+1U);m-18&17|-|48579905|48579926&offsets|0|0(+1U)]|TACCCTGTAGATCCGAATTTGTT</td><td>>MI0000266|hsa-mir-10a&WithFlank&17|-|48579832|48579953</td><td>28</td><td>49</td><td>22(+1U)</td><td>TACCCTGTAGATCCGAATTTGTT</td><td>411.0</td><td>1.9</td><td>0.7</td><td>2.5</td><td>1.1E-02</td><td>3.9E-02</td><td>BT20/MB231</td></tr>
<tr><td>>hg38_21_+_8433172_8446622_RNA45SN1_45S_50ntFlanks@10401.10439.39|GAACATGGGTCAGTCGGTCCTGAGAGATGGGCGAGCGCC</td><td>>hg38_21_+_8433172_8446622_RNA45SN1_45S_50ntFlanks</td><td>10401</td><td>10439</td><td>39</td><td>GAACATGGGTCAGTCGGTCCTGAGAGATGGGCGAGCGCC</td><td>287.4</td><td>-2.4</td><td>0.9</td><td>-2.5</td><td>1.1E-02</td><td>3.9E-02</td><td>BT20/MB231</td></tr>
<tr><td>>hg38_21_+_8433172_8446622_RNA45SN1_45S_50ntFlanks@12004.12034.31|CGGGCCGCCGGTGAAATACCACTACTCTGAT</td><td>>hg38_21_+_8433172_8446622_RNA45SN1_45S_50ntFlanks</td><td>12004</td><td>12034</td><td>31</td><td>CGGGCCGCCGGTGAAATACCACTACTCTGAT</td><td>321.0</td><td>2.2</td><td>0.9</td><td>2.5</td><td>1.1E-02</td><td>3.9E-02</td><td>BT20/MB231</td></tr>
<tr><td>>MI0003560|hsa-mir-92b&WithFlank&1|+|155195171|155195278@67.88.22(+1A)&[MIMAT0003218&hsa-miR-92b-3p&offsets|0|0(+1A);m-115&1|+|155195237|155195258&offsets|0|0(+1A)]|TATTGCACTCGTCCCGGCCTCCA</td><td>>MI0003560|hsa-mir-92b&WithFlank&1|+|155195171|155195278</td><td>67</td><td>88</td><td>22(+1A)</td><td>TATTGCACTCGTCCCGGCCTCCA</td><td>466.2</td><td>1.8</td><td>0.7</td><td>2.5</td><td>1.1E-02</td><td>3.9E-02</td><td>BT20/MB231</td></tr>
<tr><td>>hg38_21_+_8433172_8446622_RNA45SN1_45S_50ntFlanks@10344.10373.30|ATTCAAACGAGAACTTTGAAGGCCGAAGTG</td><td>>hg38_21_+_8433172_8446622_RNA45SN1_45S_50ntFlanks</td><td>10344</td><td>10373</td><td>30</td><td>ATTCAAACGAGAACTTTGAAGGCCGAAGTG</td><td>256.1</td><td>-2.1</td><td>0.8</td><td>-2.5</td><td>1.1E-02</td><td>3.9E-02</td><td>BT20/MB231</td></tr>
<tr><td>>hg38_21_+_8433172_8446622_RNA45SN1_45S_50ntFlanks@10738.10756.19|AAATCTCGCGCCGGGCCGT</td><td>>hg38_21_+_8433172_8446622_RNA45SN1_45S_50ntFlanks</td><td>10738</td><td>10756</td><td>19</td><td>AAATCTCGCGCCGGGCCGT</td><td>327.6</td><td>-2.2</td><td>0.9</td><td>-2.5</td><td>1.1E-02</td><td>3.9E-02</td><td>BT20/MB231</td></tr>
<tr><td>>hg38_21_+_8433172_8446622_RNA45SN1_45S_50ntFlanks@10341.10361.21|AATATTCAAACGAGAACTTTG</td><td>>hg38_21_+_8433172_8446622_RNA45SN1_45S_50ntFlanks</td><td>10341</td><td>10361</td><td>21</td><td>AATATTCAAACGAGAACTTTG</td><td>104.1</td><td>-1.1</td><td>0.5</td><td>-2.1</td><td>3.4E-02</td><td>3.9E-02</td><td>MB231/MB468</td></tr>
<tr><td>>hg38_21_+_8433172_8446622_RNA45SN1_45S_50ntFlanks@4591.4616.26|TTTTGTTGGTTTTCGGAACTGAGGCC</td><td>>hg38_21_+_8433172_8446622_RNA45SN1_45S_50ntFlanks</td><td>4591</td><td>4616</td><td>26</td><td>TTTTGTTGGTTTTCGGAACTGAGGCC</td><td>157.9</td><td>-1.5</td><td>0.7</td><td>-2.1</td><td>3.4E-02</td><td>3.9E-02</td><td>MB231/MB468</td></tr>
<tr><td>>hg38_21_+_8433172_8446622_RNA45SN1_45S_50ntFlanks@10393.10422.30|CAGCAGTTGAACATGGGTCAGTCGGTCCTG</td><td>>hg38_21_+_8433172_8446622_RNA45SN1_45S_50ntFlanks</td><td>10393</td><td>10422</td><td>30</td><td>CAGCAGTTGAACATGGGTCAGTCGGTCCTG</td><td>383.3</td><td>-1.5</td><td>0.7</td><td>-2.1</td><td>3.4E-02</td><td>3.9E-02</td><td>MB231/MB468</td></tr>
<tr><td>>hg38_21_+_8433172_8446622_RNA45SN1_45S_50ntFlanks@11567.11592.26|AGCAGCCGACTTAGAACTGGTGCGGA</td><td>>hg38_21_+_8433172_8446622_RNA45SN1_45S_50ntFlanks</td><td>11567</td><td>11592</td><td>26</td><td>AGCAGCCGACTTAGAACTGGTGCGGA</td><td>181.4</td><td>-1.5</td><td>0.7</td><td>-2.1</td><td>3.6E-02</td><td>4.0E-02</td><td>BT20/MB468</td></tr>
<tr><td>>hg38_21_+_8433172_8446622_RNA45SN1_45S_50ntFlanks@9565.9594.30|TGACGTGCAAATCGGTCGTCCGACCTGGGT</td><td>>hg38_21_+_8433172_8446622_RNA45SN1_45S_50ntFlanks</td><td>9565</td><td>9594</td><td>30</td><td>TGACGTGCAAATCGGTCGTCCGACCTGGGT</td><td>170.0</td><td>-1.2</td><td>0.6</td><td>-2.1</td><td>3.6E-02</td><td>4.0E-02</td><td>BT20/MB468</td></tr>
<tr><td>>hg38_21_+_8433172_8446622_RNA45SN1_45S_50ntFlanks@10385.10405.21|CATGTGAACAGCAGTTGAACA</td><td>>hg38_21_+_8433172_8446622_RNA45SN1_45S_50ntFlanks</td><td>10385</td><td>10405</td><td>21</td><td>CATGTGAACAGCAGTTGAACA</td><td>151.9</td><td>-1.9</td><td>0.9</td><td>-2.1</td><td>3.6E-02</td><td>4.0E-02</td><td>BT20/MB468</td></tr>
<tr><td>>am_trna128_GlyGCC_6_-_27870686_27870756@1.32.32,trna133_GlyCCC_1_-_16872434_16872504@1.32.32,trna18_GlyGCC_16_+_70822597_70822667@1.32.32,trna19_GlyGCC_16_+_70823410_70823480@1.32.32,trna19_GlyGCC_2_-_157257659_157257729@1.32.32,trna24_GlyGCC_16_-_70812942_70813012@1.32.32,trna25_GlyGCC_16_-_70812114_70812184@1.32.32,trna4_GlyCCC_1_+_17188416_17188486@1.32.32,trna5_GlyGCC_17_+_8029064_8029134@1.32.32,trna68_GlyGCC_1_-_161493637_161493707@1.32.32|GCATTGGTGGTTCAGTGGTAGAATTCTCGCCT</td><td>>am_trna128_GlyGCC_6_-_27870686_27870756</td><td>1</td><td>32</td><td>32</td><td>GCATTGGTGGTTCAGTGGTAGAATTCTCGCCT</td><td>41952.4</td><td>-2.0</td><td>0.8</td><td>-2.5</td><td>1.1E-02</td><td>4.0E-02</td><td>BT20/MB231</td></tr>
<tr><td>>am_trnaMT_HisGTG_MT_+_12138_12206@19.38.20|AAACATCAGATTGTGAATCT</td><td>>am_trnaMT_HisGTG_MT_+_12138_12206</td><td>19</td><td>38</td><td>20</td><td>AAACATCAGATTGTGAATCT</td><td>350.8</td><td>2.0</td><td>0.8</td><td>2.5</td><td>1.1E-02</td><td>4.0E-02</td><td>BT20/MB231</td></tr>
<tr><td>>hg38_21_+_8433172_8446622_RNA45SN1_45S_50ntFlanks@12449.12486.38|GAATTCACCAAGCGTTGGATTGTTCACCCACTAATAGG</td><td>>hg38_21_+_8433172_8446622_RNA45SN1_45S_50ntFlanks</td><td>12449</td><td>12486</td><td>38</td><td>GAATTCACCAAGCGTTGGATTGTTCACCCACTAATAGG</td><td>200.2</td><td>-2.1</td><td>0.8</td><td>-2.5</td><td>1.1E-02</td><td>4.0E-02</td><td>BT20/MB231</td></tr>
<tr><td>>hg38_21_+_8433172_8446622_RNA45SN1_45S_50ntFlanks@11552.11590.39|CCTCGGCCGGCGCCTAGCAGCCGACTTAGAACTGGTGCG</td><td>>hg38_21_+_8433172_8446622_RNA45SN1_45S_50ntFlanks</td><td>11552</td><td>11590</td><td>39</td><td>CCTCGGCCGGCGCCTAGCAGCCGACTTAGAACTGGTGCG</td><td>198.2</td><td>-2.3</td><td>0.9</td><td>-2.5</td><td>1.1E-02</td><td>4.0E-02</td><td>BT20/MB231</td></tr>
<tr><td>>hg38_21_+_8433172_8446622_RNA45SN1_45S_50ntFlanks@6700.6719.20|CTAGCTGCGAGAATTAATGT</td><td>>hg38_21_+_8433172_8446622_RNA45SN1_45S_50ntFlanks</td><td>6700</td><td>6719</td><td>20</td><td>CTAGCTGCGAGAATTAATGT</td><td>434.2</td><td>-1.5</td><td>0.7</td><td>-2.1</td><td>3.4E-02</td><td>4.0E-02</td><td>MB231/MB468</td></tr>
<tr><td>>hg38_21_+_8433172_8446622_RNA45SN1_45S_50ntFlanks@6647.6676.30|CGTACGACTCTTAGCGGTGGATCACTCGGC</td><td>>hg38_21_+_8433172_8446622_RNA45SN1_45S_50ntFlanks</td><td>6647</td><td>6676</td><td>30</td><td>CGTACGACTCTTAGCGGTGGATCACTCGGC</td><td>416.4</td><td>2.0</td><td>0.8</td><td>2.5</td><td>1.1E-02</td><td>4.0E-02</td><td>BT20/MB231</td></tr>
<tr><td>>am_MI0016746|hsa-mir-548o-2&WithFlank&20|+|38516557|38516638@51.71.21(+1U);MI0006402|hsa-mir-548o&WithFlank&7|-|102405736|102405861@93.113.21(+1U)&[MIMAT0005919&hsa-miR-548o-3p&offsets|0|-1(+1U);m-431&20|+|38516607|38516628&offsets|0|-1(+1U)];[MIMAT0005919_1&hsa-miR-548o-3p&offsets|0|-1(+1U);m-430&7|-|102405748|102405769&offsets|0|-1(+1U)]|CCAAAACTGCAGTTACTTTTGT</td><td>>am_MI0016746|hsa-mir-548o-2&WithFlank&20|+|38516557|38516638</td><td>51</td><td>71</td><td>21(+1U)</td><td>CCAAAACTGCAGTTACTTTTGT</td><td>279.1</td><td>1.8</td><td>0.7</td><td>2.5</td><td>1.1E-02</td><td>4.0E-02</td><td>BT20/MB231</td></tr>
<tr><td>>MI0000077|hsa-mir-21&WithFlank&17|+|59841260|59841343@14.33.20(+1A)&[MIMAT0000076&hsa-miR-21-5p&offsets|0|-2(+1A);m-1&17|+|59841273|59841294&offsets|0|-2(+1A)]|TAGCTTATCAGACTGATGTTA</td><td>>MI0000077|hsa-mir-21&WithFlank&17|+|59841260|59841343</td><td>14</td><td>33</td><td>20(+1A)</td><td>TAGCTTATCAGACTGATGTTA</td><td>4744.2</td><td>1.9</td><td>0.8</td><td>2.5</td><td>1.1E-02</td><td>4.0E-02</td><td>BT20/MB231</td></tr>
<tr><td>>hg38_21_+_8433172_8446622_RNA45SN1_45S_50ntFlanks@6721.6760.40|AATTGCAGGACACATTGATCATCGACACTTCGAACGCACT</td><td>>hg38_21_+_8433172_8446622_RNA45SN1_45S_50ntFlanks</td><td>6721</td><td>6760</td><td>40</td><td>AATTGCAGGACACATTGATCATCGACACTTCGAACGCACT</td><td>372.9</td><td>1.1</td><td>0.5</td><td>2.1</td><td>3.5E-02</td><td>4.0E-02</td><td>MB231/MB468</td></tr>
<tr><td>>hg38_21_+_8433172_8446622_RNA45SN1_45S_50ntFlanks@10781.10799.19|AGGTGAACAGCCTCTGGCA</td><td>>hg38_21_+_8433172_8446622_RNA45SN1_45S_50ntFlanks</td><td>10781</td><td>10799</td><td>19</td><td>AGGTGAACAGCCTCTGGCA</td><td>162.0</td><td>-1.5</td><td>0.7</td><td>-2.1</td><td>3.6E-02</td><td>4.0E-02</td><td>BT20/MB468</td></tr>
<tr><td>>hg38_21_+_8433172_8446622_RNA45SN1_45S_50ntFlanks@9533.9569.37|GAGGAAACTCTGGTGGAGGTCCGTAGCGGTCCTGACG</td><td>>hg38_21_+_8433172_8446622_RNA45SN1_45S_50ntFlanks</td><td>9533</td><td>9569</td><td>37</td><td>GAGGAAACTCTGGTGGAGGTCCGTAGCGGTCCTGACG</td><td>198.4</td><td>-2.2</td><td>1.0</td><td>-2.1</td><td>3.6E-02</td><td>4.0E-02</td><td>BT20/MB468</td></tr>
<tr><td>>hg38_21_+_8433172_8446622_RNA45SN1_45S_50ntFlanks@8196.8224.29|CGTGGACGGTGTGAGGCCGGTAGCGGCCC</td><td>>hg38_21_+_8433172_8446622_RNA45SN1_45S_50ntFlanks</td><td>8196</td><td>8224</td><td>29</td><td>CGTGGACGGTGTGAGGCCGGTAGCGGCCC</td><td>489.8</td><td>-1.3</td><td>0.6</td><td>-2.1</td><td>3.5E-02</td><td>4.0E-02</td><td>MB231/MB468</td></tr>
<tr><td>>hg38_21_+_8433172_8446622_RNA45SN1_45S_50ntFlanks@11565.11588.24|CTAGCAGCCGACTTAGAACTGGTG</td><td>>hg38_21_+_8433172_8446622_RNA45SN1_45S_50ntFlanks</td><td>11565</td><td>11588</td><td>24</td><td>CTAGCAGCCGACTTAGAACTGGTG</td><td>148.7</td><td>-1.2</td><td>0.6</td><td>-2.1</td><td>3.5E-02</td><td>4.0E-02</td><td>MB231/MB468</td></tr>
</table>

| | | | | | | | | | | | | |
|---|---|---|---|---|---|---|---|---|---|---|---|---|
| >MI0000084\|hsa-mir-26b&WithFlank&2\|+\|218402640\|218402728@18.39.22(+1A)&[MIMAT0000083&hsa-miR-26b-5p&offsets\|0\|+1(+1A);m-53&2\|+\|218402657\|218402678&offsets\|0\|0(+1A)]\|TTCAAGTAATTCAGGATAGGTTA | >MI0000084\|hsa-mir-26b&WithFlank&2\|+\|218402640\|218402728 | 18 | 39 | 22(+1A) | TTCAAGTAATTCAGGATAGGTTA | 431.6 | 2.2 | 0.9 | 2.5 | 1.1E-02 | 4.0E-02 | BT20/MB231 |
| >MI0000108\|hsa-mir-103a-2&WithFlank&20\|+\|3917488\|3917577@54.76.23(+2U);MI0000109\|hsa-mir-103a-1&WithFlank&5\|-\|168560890\|168560979@54.76.23(+2U)&[MIMAT0000101&hsa-miR-103a-3p&offsets\|0\|0(+2U);m-12&20\|+\|3917541\|3917563&offsets\|0\|0(+2U)];[MIMAT0000101_1&hsa-miR-103a-3p&offsets\|0\|0(+2U);m-13&5\|-\|168560904\|168560926&offsets\|0\|0(+2U);m-44220&5\|-\|168560908\|168560926&offsets\|0\|+4(+2U)]\|AGCAGCATTGTACAGGGCTATGATT | >MI0000108\|hsa-mir-103a-2&WithFlank&20\|+\|3917488\|3917577 | 54 | 76 | 23(+2U) | AGCAGCATTGTACAGGGCTATGATT | 1608.3 | -1.8 | 0.7 | -2.5 | 1.1E-02 | 4.0E-02 | BT20/MB231 |
| >trnaMT_ValTAC_MT_+_1602_1670@54.72.19\|AACTTGACCGCTCTGACCA | >trnaMT_ValTAC_MT_+_1602_1670 | 54 | 72 | 19 | AACTTGACCGCTCTGACCA | 2400.8 | 1.8 | 0.7 | 2.5 | 1.1E-02 | 4.1E-02 | BT20/MB231 |
| >hg38_21_+_8433172_8446622_RNA45SN1_45S_50ntFlanks@12420.12456.37\|ATGTCGGCTCTTCCTATCATTGTGAAGCAGAATTCAC | >hg38_21_+_8433172_8446622_RNA45SN1_45S_50ntFlanks | 12420 | 12456 | 37 | ATGTCGGCTCTTCCTATCATTGTGAAGCAGAATTCAC | 990.3 | 1.3 | 0.6 | 2.1 | 3.5E-02 | 4.1E-02 | MB231/MB468 |
| >hg38_21_+_8433172_8446622_RNA45SN1_45S_50ntFlanks@9533.9562.30\|GAGGAAACTCTGGTGGAGGTCCGTAGCGGT | >hg38_21_+_8433172_8446622_RNA45SN1_45S_50ntFlanks | 9533 | 9562 | 30 | GAGGAAACTCTGGTGGAGGTCCGTAGCGGT | 151.1 | -1.0 | 0.5 | -2.1 | 3.5E-02 | 4.1E-02 | MB231/MB468 |
| >hg38_21_+_8433172_8446622_RNA45SN1_45S_50ntFlanks@12455.12476.22\|ACCAAGCGTTGGATTGTTCACC | >hg38_21_+_8433172_8446622_RNA45SN1_45S_50ntFlanks | 12455 | 12476 | 22 | ACCAAGCGTTGGATTGTTCACC | 819.0 | -1.6 | 0.8 | -2.1 | 3.7E-02 | 4.1E-02 | BT20/MB468 |
| >hg38_21_+_8433172_8446622_RNA45SN1_45S_50ntFlanks@10678.10705.28\|CGGTTCCGGCGGCGTCCGGTGAGCTCTC | >hg38_21_+_8433172_8446622_RNA45SN1_45S_50ntFlanks | 10678 | 10705 | 28 | CGGTTCCGGCGGCGTCCGGTGAGCTCTC | 600.8 | -1.9 | 0.9 | -2.1 | 3.7E-02 | 4.1E-02 | BT20/MB468 |
| >hg38_21_+_8433172_8446622_RNA45SN1_45S_50ntFlanks@9545.9569.25\|GTGGAGGTCCGTAGCGGTCCTGACG | >hg38_21_+_8433172_8446622_RNA45SN1_45S_50ntFlanks | 9545 | 9569 | 25 | GTGGAGGTCCGTAGCGGTCCTGACG | 404.0 | -1.4 | 0.6 | -2.1 | 3.5E-02 | 4.1E-02 | MB231/MB468 |
| >MI0000342\|hsa-mir-200b&WithFlank&1\|+\|1167098\|1167204@63.85.23(+1C)&[MIMAT0000318&hsa-miR-200b-3p&offsets\|0\|+1(+1C);m-93&1\|+\|1167160\|1167182&offsets\|0\|0(+1C)]\|TAATACTGCCTGGTAATGATGACC | >MI0000342\|hsa-mir-200b&WithFlank&1\|+\|1167098\|1167204 | 63 | 85 | 23(+1C) | TAATACTGCCTGGTAATGATGACC | 225.7 | -1.4 | 0.7 | -2.1 | 3.5E-02 | 4.1E-02 | MB231/MB468 |
| >MI0000650\|hsa-mir-200c&WithFlank&12\|+\|6963693\|6963772@50.72.23&[MIMAT0000617&hsa-miR-200c-3p&offsets\|0\|0;m-54&12\|+\|6963742\|6963764&offsets\|0\|0]\|TAATACTGCCGGGTAATGATGGA | >MI0000650\|hsa-mir-200c&WithFlank&12\|+\|6963693\|6963772 | 50 | 72 | 23 | TAATACTGCCGGGTAATGATGGA | 1583.6 | -1.5 | 0.7 | -2.1 | 3.5E-02 | 4.1E-02 | MB231/MB468 |
| >hg38_21_+_8433172_8446622_RNA45SN1_45S_50ntFlanks@12406.12432.27\|TTTTTGATCCTTCGATGTCGGCTCTTC | >hg38_21_+_8433172_8446622_RNA45SN1_45S_50ntFlanks | 12406 | 12432 | 27 | TTTTTGATCCTTCGATGTCGGCTCTTC | 155.6 | -1.1 | 0.5 | -2.1 | 3.5E-02 | 4.1E-02 | MB231/MB468 |
| >hg38_21_+_8433172_8446622_RNA45SN1_45S_50ntFlanks@3835.3868.34\|CTCCTCTCCTACTTGGATAACTGTGGTAATTCTA | >hg38_21_+_8433172_8446622_RNA45SN1_45S_50ntFlanks | 3835 | 3868 | 34 | CTCCTCTCCTACTTGGATAACTGTGGTAATTCTA | 101.5 | -1.4 | 0.7 | -2.1 | 3.7E-02 | 4.1E-02 | BT20/MB468 |
| >hg38_21_+_8433172_8446622_RNA45SN1_45S_50ntFlanks@8104.8138.35\|CCGCGGCGGGGCGCGGGACATGTGGCGTACGGAAG | >hg38_21_+_8433172_8446622_RNA45SN1_45S_50ntFlanks | 8104 | 8138 | 35 | CCGCGGCGGGGCGCGGGACATGTGGCGTACGGAAG | 216.7 | -1.5 | 0.7 | -2.1 | 3.5E-02 | 4.1E-02 | MB231/MB468 |
| >hg38_21_+_8433172_8446622_RNA45SN1_45S_50ntFlanks@3985.4024.40\|CGGCTTTGGTGACTCTAGATAACCTCGGGCCGATCGCACG | >hg38_21_+_8433172_8446622_RNA45SN1_45S_50ntFlanks | 3985 | 4024 | 40 | CGGCTTTGGTGACTCTAGATAACCTCGGGCCGATCGCACG | 286.0 | -4.4 | 2.1 | -2.1 | 3.5E-02 | 4.1E-02 | MB231/MB468 |
| >hg38_21_+_8433172_8446622_RNA45SN1_45S_50ntFlanks@6646.6663.18\|TCGTACGACTCTTAGCGG | >hg38_21_+_8433172_8446622_RNA45SN1_45S_50ntFlanks | 6646 | 6663 | 18 | TCGTACGACTCTTAGCGG | 6458.1 | 1.5 | 0.7 | 2.1 | 3.5E-02 | 4.1E-02 | MB231/MB468 |
| >hg38_21_+_8433172_8446622_RNA45SN1_45S_50ntFlanks@10767.10785.19\|GCAGCAGGTCTCCAAGGTG | >hg38_21_+_8433172_8446622_RNA45SN1_45S_50ntFlanks | 10767 | 10785 | 19 | GCAGCAGGTCTCCAAGGTG | 213.1 | -1.1 | 0.5 | -2.1 | 3.5E-02 | 4.1E-02 | MB231/MB468 |
| >hg38_21_+_8433172_8446622_RNA45SN1_45S_50ntFlanks@6651.6672.22\|CGACTCTTAGCGGTGGATCACT | >hg38_21_+_8433172_8446622_RNA45SN1_45S_50ntFlanks | 6651 | 6672 | 22 | CGACTCTTAGCGGTGGATCACT | 3389.7 | 1.9 | 0.8 | 2.5 | 1.2E-02 | 4.1E-02 | BT20/MB231 |
| >hg38_21_+_8433172_8446622_RNA45SN1_45S_50ntFlanks@10770.10796.27\|GCAGGTCTCCAAGGTGAACAGCCTCTG | >hg38_21_+_8433172_8446622_RNA45SN1_45S_50ntFlanks | 10770 | 10796 | 27 | GCAGGTCTCCAAGGTGAACAGCCTCTG | 290.4 | -2.3 | 0.9 | -2.5 | 1.2E-02 | 4.1E-02 | BT20/MB231 |
| >trnaMT_HisGTG_MT_+_12138_12206@20.55.36\|AACATCAGATTGTGAATCTGACAACAGAGGCTTACG | >trnaMT_HisGTG_MT_+_12138_12206 | 20 | 55 | 36 | AACATCAGATTGTGAATCTGACAACAGAGGCTTACG | 214.4 | 2.2 | 0.9 | 2.5 | 1.2E-02 | 4.1E-02 | BT20/MB231 |
| >hg38_21_+_8433172_8446622_RNA45SN1_45S_50ntFlanks@12995.13012.18\|CTCGCTGCGATCTATTGA | >hg38_21_+_8433172_8446622_RNA45SN1_45S_50ntFlanks | 12995 | 13012 | 18 | CTCGCTGCGATCTATTGA | 315.8 | 1.0 | 0.5 | 2.1 | 3.7E-02 | 4.1E-02 | BT20/MB468 |
| >MI0000270\|hsa-mir-181b-1&WithFlank&1\|-\|198858867\|198858988@42.61.20;MI0000683\|hsa-mir-181b-2&WithFlank&9\|+\|124693704\|124693804@22.41.20&[MIMAT0000257&hsa-miR-181b-5p&offsets\|0\|-3;m-73&1\|-\|198858925\|198858947&offsets\|0\|-3];[MIMAT0000257_1&hsa-miR-181b-5p&offsets\|0\|-3;m-72&9\|+\|124693725\|124693747&offsets\|0\|-3]\|AACATTCATTGCTGTCGGTG | >MI0000270\|hsa-mir-181b-1&WithFlank&1\|-\|198858867\|198858988 | 42 | 61 | 20 | AACATTCATTGCTGTCGGTG | 262.5 | -1.8 | 0.7 | -2.5 | 1.2E-02 | 4.1E-02 | BT20/MB231 |
| >hg38_21_+_8433172_8446622_RNA45SN1_45S_50ntFlanks@7980.8009.30\|CCTCAGATCAGACGTGGCGACCCGCTGAAT | >hg38_21_+_8433172_8446622_RNA45SN1_45S_50ntFlanks | 7980 | 8009 | 30 | CCTCAGATCAGACGTGGCGACCCGCTGAAT | 159.6 | -1.1 | 0.5 | -2.1 | 3.6E-02 | 4.1E-02 | MB231/MB468 |
| >hg38_21_+_8433172_8446622_RNA45SN1_45S_50ntFlanks@11920.11950.31\|ATGAGAGGTGTAGAATAAGTGGGAGGCCCCC | >hg38_21_+_8433172_8446622_RNA45SN1_45S_50ntFlanks | 11920 | 11950 | 31 | ATGAGAGGTGTAGAATAAGTGGGAGGCCCCC | 288.6 | 1.9 | 0.9 | 2.1 | 3.6E-02 | 4.1E-02 | MB231/MB468 |
| >hg38_21_+_8433172_8446622_RNA45SN1_45S_50ntFlanks@5467.5491.25\|GGCCCACGGCCCTGGCGGAGCGCTG | >hg38_21_+_8433172_8446622_RNA45SN1_45S_50ntFlanks | 5467 | 5491 | 25 | GGCCCACGGCCCTGGCGGAGCGCTG | 161.3 | -2.1 | 1.0 | -2.1 | 3.8E-02 | 4.1E-02 | BT20/MB468 |
| >hg38_21_+_8433172_8446622_RNA45SN1_45S_50ntFlanks@8105.8138.34\|CGCGGCGGGGCGCGGGACATGTGGCGTACGGAAG | >hg38_21_+_8433172_8446622_RNA45SN1_45S_50ntFlanks | 8105 | 8138 | 34 | CGCGGCGGGGCGCGGGACATGTGGCGTACGGAAG | 322.8 | -2.2 | 1.0 | -2.1 | 3.8E-02 | 4.2E-02 | BT20/MB468 |
| >hg38_21_+_8433172_8446622_RNA45SN1_45S_50ntFlanks@3854.3884.31\|ACTGTGGTAATTCTAGAGCTAATACATGCCG | >hg38_21_+_8433172_8446622_RNA45SN1_45S_50ntFlanks | 3854 | 3884 | 31 | ACTGTGGTAATTCTAGAGCTAATACATGCCG | 158.9 | -1.1 | 0.5 | -2.1 | 3.6E-02 | 4.2E-02 | MB231/MB468 |
| >hg38_21_+_8433172_8446622_RNA45SN1_45S_50ntFlanks@9534.9559.26\|AGGAAACTCTGGTGGAGGTCCGTAGC | >hg38_21_+_8433172_8446622_RNA45SN1_45S_50ntFlanks | 9534 | 9559 | 26 | AGGAAACTCTGGTGGAGGTCCGTAGC | 120.7 | -1.0 | 0.5 | -2.1 | 3.6E-02 | 4.2E-02 | MB231/MB468 |
| >hg38_21_+_8433172_8446622_RNA45SN1_45S_50ntFlanks@12990.13014.25\|GCTCCCTCGCTGCGATCTATTGAAA | >hg38_21_+_8433172_8446622_RNA45SN1_45S_50ntFlanks | 12990 | 13014 | 25 | GCTCCCTCGCTGCGATCTATTGAAA | 323.5 | -1.7 | 0.8 | -2.1 | 3.8E-02 | 4.2E-02 | BT20/MB468 |
| >hg38_21_+_8433172_8446622_RNA45SN1_45S_50ntFlanks@12476.12497.22\|CCACTAATAGGGAACGTGAGCT | >hg38_21_+_8433172_8446622_RNA45SN1_45S_50ntFlanks | 12476 | 12497 | 22 | CCACTAATAGGGAACGTGAGCT | 1628.7 | -1.2 | 0.6 | -2.1 | 3.6E-02 | 4.2E-02 | MB231/MB468 |
| >hg38_21_+_8433172_8446622_RNA45SN1_45S_50ntFlanks@6702.6720.19\|AGCTGCGAGAATTAATGTG | >hg38_21_+_8433172_8446622_RNA45SN1_45S_50ntFlanks | 6702 | 6720 | 19 | AGCTGCGAGAATTAATGTG | 982.4 | 1.2 | 0.6 | 2.1 | 3.6E-02 | 4.2E-02 | MB231/MB468 |
| >hg38_21_+_8433172_8446622_RNA45SN1_45S_50ntFlanks@12456.12491.36\|CCAAGCGTTGGATTGTTCACCCACTAATAGGGAACG | >hg38_21_+_8433172_8446622_RNA45SN1_45S_50ntFlanks | 12456 | 12491 | 36 | CCAAGCGTTGGATTGTTCACCCACTAATAGGGAACG | 100.3 | -1.5 | 0.7 | -2.1 | 3.6E-02 | 4.2E-02 | MB231/MB468 |
| >hg38_21_+_8433172_8446622_RNA45SN1_45S_50ntFlanks@482.507.26\|ATCGATGTGGTGACGTCGTGCTCTCC | >hg38_21_+_8433172_8446622_RNA45SN1_45S_50ntFlanks | 482 | 507 | 26 | ATCGATGTGGTGACGTCGTGCTCTCC | 2466.0 | 5.7 | 2.7 | 2.1 | 3.6E-02 | 4.2E-02 | MB231/MB468 |
| >hg38_21_+_8433172_8446622_RNA45SN1_45S_50ntFlanks@3854.3873.20\|ACTGTGGTAATTCTAGAGCT | >hg38_21_+_8433172_8446622_RNA45SN1_45S_50ntFlanks | 3854 | 3873 | 20 | ACTGTGGTAATTCTAGAGCT | 1199.9 | -1.9 | 0.9 | -2.1 | 3.8E-02 | 4.2E-02 | BT20/MB468 |
| >hg38_21_+_8433172_8446622_RNA45SN1_45S_50ntFlanks@12281.12321.41\|AATACAGACCGTGAAAGCGGGGCCTCACGATCCTTCTGACC | >hg38_21_+_8433172_8446622_RNA45SN1_45S_50ntFlanks | 12281 | 12321 | 41 | AATACAGACCGTGAAAGCGGGGCCTCACGATCCTTCTGACC | 151.3 | -2.4 | 1.0 | -2.5 | 1.2E-02 | 4.2E-02 | BT20/MB231 |
| >hg38_21_+_8433172_8446622_RNA45SN1_45S_50ntFlanks@12420.12443.24\|ATGTCGGCTCTTCCTATCATTGTG | >hg38_21_+_8433172_8446622_RNA45SN1_45S_50ntFlanks | 12420 | 12443 | 24 | ATGTCGGCTCTTCCTATCATTGTG | 2813.8 | -1.5 | 0.7 | -2.1 | 3.6E-02 | 4.2E-02 | MB231/MB468 |
| >MI0000299\|hsa-mir-222&WithFlank&X\|-\|45747009\|45747130@75.98.24(+1G)&[MIMAT0000279&hsa-miR-222-3p&offsets\|0\|+3(+1G);m-140&X\|-\|45747033\|45747056&offsets\|0\|0(+1G)]\|AGCTACATCTGGCTACTGGGTCTCG | >MI0000299\|hsa-mir-222&WithFlank&X\|-\|45747009\|45747130 | 75 | 98 | 24(+1G) | AGCTACATCTGGCTACTGGGTCTCG | 3467.6 | -1.8 | 0.7 | -2.5 | 1.2E-02 | 4.2E-02 | BT20/MB231 |
| >hg38_21_+_8433172_8446622_RNA45SN1_45S_50ntFlanks@9549.9571.23\|AGGTCCGTAGCGGTCCTGACGTG | >hg38_21_+_8433172_8446622_RNA45SN1_45S_50ntFlanks | 9549 | 9571 | 23 | AGGTCCGTAGCGGTCCTGACGTG | 257.1 | 1.0 | 0.5 | 2.1 | 3.7E-02 | 4.2E-02 | MB231/MB468 |
| >hg38_21_+_8433172_8446622_RNA45SN1_45S_50ntFlanks@12937.12960.24\|AACCATTCGTAGACGACCTGCTTC | >hg38_21_+_8433172_8446622_RNA45SN1_45S_50ntFlanks | 12937 | 12960 | 24 | AACCATTCGTAGACGACCTGCTTC | 352.5 | 1.3 | 0.6 | 2.1 | 3.7E-02 | 4.2E-02 | MB231/MB468 |
| >hg38_21_+_8433172_8446622_RNA45SN1_45S_50ntFlanks@4976.4993.18\|CCGGACACGGACAGGATT | >hg38_21_+_8433172_8446622_RNA45SN1_45S_50ntFlanks | 4976 | 4993 | 18 | CCGGACACGGACAGGATT | 242.5 | 1.8 | 0.7 | 2.5 | 1.2E-02 | 4.2E-02 | BT20/MB231 |
| >MI0000077\|hsa-mir-21&WithFlank&17\|+\|59841260\|59841343@14.34.21(+1U)&[MIMAT0000076&hsa-miR-21-5p&offsets\|0\|-1(+1U);m-1&17\|+\|59841273\|59841294&offsets\|0\|-1(+1U)]\|TAGCTTATCAGACTGATGTTGT | >MI0000077\|hsa-mir-21&WithFlank&17\|+\|59841260\|59841343 | 14 | 34 | 21(+1U) | TAGCTTATCAGACTGATGTTGT | 15588.4 | 2.3 | 0.9 | 2.5 | 1.2E-02 | 4.2E-02 | BT20/MB231 |
| >MI0000342\|hsa-mir-200b&WithFlank&1\|+\|1167098\|1167204@63.84.22(+1U)&[MIMAT0000318&hsa-miR-200b-3p&offsets\|0\|0(+1U);m-93&1\|+\|1167160\|1167182&offsets\|0\|-1(+1U)]\|TAATACTGCCTGGTAATGATGAT | >MI0000342\|hsa-mir-200b&WithFlank&1\|+\|1167098\|1167204 | 63 | 84 | 22(+1U) | TAATACTGCCTGGTAATGATGAT | 231.4 | -1.2 | 0.6 | -2.1 | 3.7E-02 | 4.2E-02 | MB231/MB468 |
| >hg38_21_+_8433172_8446622_RNA45SN1_45S_50ntFlanks@10680.10705.26\|GTTCCGGCGGCGTCCGGTGAGCTCTC | >hg38_21_+_8433172_8446622_RNA45SN1_45S_50ntFlanks | 10680 | 10705 | 26 | GTTCCGGCGGCGTCCGGTGAGCTCTC | 384.9 | -1.8 | 0.9 | -2.1 | 3.9E-02 | 4.3E-02 | BT20/MB468 |
| >hg38_21_+_8433172_8446622_RNA45SN1_45S_50ntFlanks@12913.12933.21\|ACGTTCGTGGGGAACCTGGCG | >hg38_21_+_8433172_8446622_RNA45SN1_45S_50ntFlanks | 12913 | 12933 | 21 | ACGTTCGTGGGGAACCTGGCG | 158.7 | -1.8 | 0.9 | -2.1 | 3.7E-02 | 4.3E-02 | MB231/MB468 |
| >hg38_21_+_8433172_8446622_RNA45SN1_45S_50ntFlanks@10348.10374.27\|AAACGAGAACTTTGAAGGCCGAAGTGG | >hg38_21_+_8433172_8446622_RNA45SN1_45S_50ntFlanks | 10348 | 10374 | 27 | AAACGAGAACTTTGAAGGCCGAAGTGG | 520.8 | -1.2 | 0.6 | -2.1 | 3.7E-02 | 4.3E-02 | MB231/MB468 |
| >hg38_21_+_8433172_8446622_RNA45SN1_45S_50ntFlanks@12396.12414.19\|GCGACGTCGCTTTTTGATC | >hg38_21_+_8433172_8446622_RNA45SN1_45S_50ntFlanks | 12396 | 12414 | 19 | GCGACGTCGCTTTTTGATC | 1323.3 | -1.0 | 0.5 | -2.1 | 3.9E-02 | 4.3E-02 | BT20/MB468 |

<table>
<tr><td>>hg38_21_+_8433172_8446622_RNA45SN1_45S_50ntFlanks@4498.4517.20|AAGCGTTTACTTTGAAAAAA</td><td>>hg38_21_+_8433172_8446622_RNA45SN1_45S_50ntFlanks</td><td>4498</td><td>4517</td><td>20</td><td>AAGCGTTTACTTTGAAAAAA</td><td>427.9</td><td>-1.4</td><td>0.7</td><td>-2.1</td><td>3.9E-02</td><td>4.3E-02</td><td>BT20/MB468</td></tr>
<tr><td>>hg38_21_+_8433172_8446622_RNA45SN1_45S_50ntFlanks@11672.11702.31|CAGTGCTCTGAATGTCAAAGTGAAGAAATTC</td><td>>hg38_21_+_8433172_8446622_RNA45SN1_45S_50ntFlanks</td><td>11672</td><td>11702</td><td>31</td><td>CAGTGCTCTGAATGTCAAAGTGAAGAAATTC</td><td>510.8</td><td>-2.1</td><td>0.8</td><td>-2.5</td><td>1.2E-02</td><td>4.3E-02</td><td>BT20/MB231</td></tr>
<tr><td>>hg38_21_+_8433172_8446622_RNA45SN1_45S_50ntFlanks@12399.12433.35|ACGTCGCTTTTTGATCCTTCGATGTCGGCTCTTCC</td><td>>hg38_21_+_8433172_8446622_RNA45SN1_45S_50ntFlanks</td><td>12399</td><td>12433</td><td>35</td><td>ACGTCGCTTTTTGATCCTTCGATGTCGGCTCTTCC</td><td>261.5</td><td>-2.7</td><td>1.1</td><td>-2.5</td><td>1.2E-02</td><td>4.3E-02</td><td>BT20/MB231</td></tr>
<tr><td>>MI0000433|hsa-let-7g&WithFlank&3|-|52268272|52268367@11.29.19&[MIMAT0000414&hsa-let-7g-5p&offsets|0|-3;m-58&3|-|52268336|52268357&offsets|0|-3]|TGAGGTAGTAGTTTGTACA</td><td>>MI0000433|hsa-let-7g&WithFlank&3|-|52268272|52268367</td><td>11</td><td>29</td><td>19</td><td>TGAGGTAGTAGTTTGTACA</td><td>308.9</td><td>1.7</td><td>0.7</td><td>2.5</td><td>1.2E-02</td><td>4.3E-02</td><td>BT20/MB231</td></tr>
<tr><td>>hg38_21_+_8433172_8446622_RNA45SN1_45S_50ntFlanks@4729.4767.39|TCAAGAACGAAAGTCGGAGGTTCGAAGACGATCAGATAC</td><td>>hg38_21_+_8433172_8446622_RNA45SN1_45S_50ntFlanks</td><td>4729</td><td>4767</td><td>39</td><td>TCAAGAACGAAAGTCGGAGGTTCGAAGACGATCAGATAC</td><td>159.4</td><td>-2.5</td><td>1.0</td><td>-2.5</td><td>1.2E-02</td><td>4.3E-02</td><td>BT20/MB231</td></tr>
<tr><td>>MI0000070|hsa-mir-16-1&WithFlank&13|-|50048967|50049067@20.39.20(+1U);MI0000115|hsa-mir-16-2&WithFlank&3|+|160404739|160404831@16.35.20(+1U)&[MIMAT0000069&hsa-miR-16-5p&offsets|0|-2(+1U);m-62&13|-|50049027|50049048&offsets|0|-2(+1U)];[MIMAT0000069_1&hsa-miR-16-5p&offsets|0|-2(+1U);m-61&3|+|160404754|160404775&offsets|0|-2(+1U)]|TAGCAGCACGTAAATATTGGT</td><td>>MI0000070|hsa-mir-16-1&WithFlank&13|-|50048967|50049067</td><td>20</td><td>39</td><td>20(+1U)</td><td>TAGCAGCACGTAAATATTGGT</td><td>1073.2</td><td>2.2</td><td>0.9</td><td>2.5</td><td>1.2E-02</td><td>4.3E-02</td><td>BT20/MB231</td></tr>
<tr><td>>MI0000469|hsa-mir-125a&WithFlank&19|+|51693248|51693345@21.41.21(+1G)&[MIMAT0000443&hsa-miR-125a-5p&offsets|0|-3(+1G);m-63&19|+|51693268|51693289&offsets|0|-1(+1G)]|TCCCTGAGACCCTTTAACCTGG</td><td>>MI0000469|hsa-mir-125a&WithFlank&19|+|51693248|51693345</td><td>21</td><td>41</td><td>21(+1G)</td><td>TCCCTGAGACCCTTTAACCTGG</td><td>300.4</td><td>2.0</td><td>0.8</td><td>2.5</td><td>1.2E-02</td><td>4.4E-02</td><td>BT20/MB231</td></tr>
<tr><td>>hg38_21_+_8433172_8446622_RNA45SN1_45S_50ntFlanks@11567.11588.22|AGCAGCCGACTTAGAACTGGTG</td><td>>hg38_21_+_8433172_8446622_RNA45SN1_45S_50ntFlanks</td><td>11567</td><td>11588</td><td>22</td><td>AGCAGCCGACTTAGAACTGGTG</td><td>305.9</td><td>-2.0</td><td>0.8</td><td>-2.5</td><td>1.3E-02</td><td>4.4E-02</td><td>BT20/MB231</td></tr>
<tr><td>>hg38_21_+_8433172_8446622_RNA45SN1_45S_50ntFlanks@9653.9677.25|AGGATAGCTGGCGCTCTCGCAGACC</td><td>>hg38_21_+_8433172_8446622_RNA45SN1_45S_50ntFlanks</td><td>9653</td><td>9677</td><td>25</td><td>AGGATAGCTGGCGCTCTCGCAGACC</td><td>231.4</td><td>1.7</td><td>0.8</td><td>2.1</td><td>3.8E-02</td><td>4.4E-02</td><td>MB231/MB468</td></tr>
<tr><td>>hg38_21_+_8433172_8446622_RNA45SN1_45S_50ntFlanks@4738.4758.21|AAAGTCGGAGGTTCGAAGACG</td><td>>hg38_21_+_8433172_8446622_RNA45SN1_45S_50ntFlanks</td><td>4738</td><td>4758</td><td>21</td><td>AAAGTCGGAGGTTCGAAGACG</td><td>3218.0</td><td>1.4</td><td>0.7</td><td>2.1</td><td>3.8E-02</td><td>4.4E-02</td><td>MB231/MB468</td></tr>
<tr><td>>hg38_21_+_8433172_8446622_RNA45SN1_45S_50ntFlanks@6652.6696.45|GACTCTTAGCGGTGGATCACTCGGCTCGTGCGTCGATGAAGAACG</td><td>>hg38_21_+_8433172_8446622_RNA45SN1_45S_50ntFlanks</td><td>6652</td><td>6696</td><td>45</td><td>GACTCTTAGCGGTGGATCACTCGGCTCGTGCGTCGATGAAGAACG</td><td>2071.1</td><td>-1.9</td><td>0.8</td><td>-2.5</td><td>1.3E-02</td><td>4.4E-02</td><td>BT20/MB231</td></tr>
<tr><td>>hg38_21_+_8433172_8446622_RNA45SN1_45S_50ntFlanks@4735.4758.24|ACGAAAGTCGGAGGTTCGAAGACG</td><td>>hg38_21_+_8433172_8446622_RNA45SN1_45S_50ntFlanks</td><td>4735</td><td>4758</td><td>24</td><td>ACGAAAGTCGGAGGTTCGAAGACG</td><td>584.9</td><td>-2.2</td><td>0.9</td><td>-2.5</td><td>1.3E-02</td><td>4.4E-02</td><td>BT20/MB231</td></tr>
<tr><td>>MI0000103|hsa-mir-101-1&WithFlank&1|-|65058428|65058514@53.74.22;MI0000739|hsa-mir-101-2&WithFlank&9|+|4850291|4850381@55.76.22&[MIMAT0000099&hsa-miR-101-3p&offsets|0|+1;m-17&1|-|65058443|65058462&offsets|0|+2];[MIMAT0000099_1&hsa-miR-101-3p&offsets|0|+1;m-16&9|+|4850345|4850364&offsets|0|+2]|TACAGTACTGTGATAACTGAAG</td><td>>MI0000103|hsa-mir-101-1&WithFlank&1|-|65058428|65058514</td><td>53</td><td>74</td><td>22</td><td>TACAGTACTGTGATAACTGAAG</td><td>13120.9</td><td>2.4</td><td>1.0</td><td>2.5</td><td>1.3E-02</td><td>4.4E-02</td><td>BT20/MB231</td></tr>
<tr><td>>MI0000298|hsa-mir-221&WithFlank&X|-|45746151|45746272@71.91.21(+1C)&[MIMAT0000278&hsa-miR-221-3p&offsets|0|-2(+1C);m-91&X|-|45746181|45746202&offsets|0|-1(+1C)]|AGCTACATTGTCTGCTGGGTTC</td><td>>MI0000298|hsa-mir-221&WithFlank&X|-|45746151|45746272</td><td>71</td><td>91</td><td>21(+1C)</td><td>AGCTACATTGTCTGCTGGGTTC</td><td>3378.0</td><td>-1.7</td><td>0.7</td><td>-2.5</td><td>1.3E-02</td><td>4.4E-02</td><td>BT20/MB231</td></tr>
<tr><td>>hg38_21_+_8433172_8446622_RNA45SN1_45S_50ntFlanks@10391.10429.39|AACAGCAGTTGAACATGGGTCAGTCGGTCCTGAGAGATG</td><td>>hg38_21_+_8433172_8446622_RNA45SN1_45S_50ntFlanks</td><td>10391</td><td>10429</td><td>39</td><td>AACAGCAGTTGAACATGGGTCAGTCGGTCCTGAGAGATG</td><td>858.0</td><td>-1.3</td><td>0.6</td><td>-2.1</td><td>3.9E-02</td><td>4.4E-02</td><td>MB231/MB468</td></tr>
<tr><td>>hg38_21_+_8433172_8446622_RNA45SN1_45S_50ntFlanks@10350.10371.22|ACGAGAACTTTGAAGGCCGAAG</td><td>>hg38_21_+_8433172_8446622_RNA45SN1_45S_50ntFlanks</td><td>10350</td><td>10371</td><td>22</td><td>ACGAGAACTTTGAAGGCCGAAG</td><td>332.9</td><td>-2.0</td><td>0.8</td><td>-2.5</td><td>1.3E-02</td><td>4.4E-02</td><td>BT20/MB231</td></tr>
<tr><td>>hg38_21_+_8433172_8446622_RNA45SN1_45S_50ntFlanks@12438.12473.36|ATTGTGAAGCAGAATTCACCAAGCGTTGGATTGTTC</td><td>>hg38_21_+_8433172_8446622_RNA45SN1_45S_50ntFlanks</td><td>12438</td><td>12473</td><td>36</td><td>ATTGTGAAGCAGAATTCACCAAGCGTTGGATTGTTC</td><td>447.5</td><td>-2.0</td><td>0.8</td><td>-2.5</td><td>1.3E-02</td><td>4.4E-02</td><td>BT20/MB231</td></tr>
<tr><td>>hg38_21_+_8433172_8446622_RNA45SN1_45S_50ntFlanks@6647.6669.23|CGTACGACTCTTAGCGGTGGATC</td><td>>hg38_21_+_8433172_8446622_RNA45SN1_45S_50ntFlanks</td><td>6647</td><td>6669</td><td>23</td><td>CGTACGACTCTTAGCGGTGGATC</td><td>3249.4</td><td>2.1</td><td>0.9</td><td>2.5</td><td>1.3E-02</td><td>4.4E-02</td><td>BT20/MB231</td></tr>
<tr><td>>hg38_21_+_8433172_8446622_RNA45SN1_45S_50ntFlanks@7988.8019.32|CAGACGTGGCGACCCGCTGAATTTAAGCATAT</td><td>>hg38_21_+_8433172_8446622_RNA45SN1_45S_50ntFlanks</td><td>7988</td><td>8019</td><td>32</td><td>CAGACGTGGCGACCCGCTGAATTTAAGCATAT</td><td>196.6</td><td>-2.4</td><td>1.0</td><td>-2.5</td><td>1.3E-02</td><td>4.4E-02</td><td>BT20/MB231</td></tr>
<tr><td>>MI0000289|hsa-mir-181a-1&WithFlank&1|-|198859038|198859159@30.53.24;MI0000269|hsa-mir-181a-2&WithFlank&9|+|124692436|124692557@45.68.24&[MIMAT0000256&hsa-miR-181a-5p&offsets|0|+1;m-32&1|-|198859108|198859130&offsets|0|+1];[MIMAT0000256_1&hsa-miR-181a-5p&offsets|0|+1;m-32&9|+|124692480|124692502&offsets|0|+1]|AACATTCAACGCTGTCGGTGAGTT</td><td>>MI0000289|hsa-mir-181a-1&WithFlank&1|-|198859038|198859159</td><td>30</td><td>53</td><td>24</td><td>AACATTCAACGCTGTCGGTGAGTT</td><td>1837.1</td><td>-2.0</td><td>0.8</td><td>-2.5</td><td>1.3E-02</td><td>4.4E-02</td><td>BT20/MB231</td></tr>
<tr><td>>hg38_1_-_228634819_228635039_RNA5S12_5S_50ntFlanks@51.87.37|GTCTACGGCCATACCACCCTGAACGCGCCCGATCTCG</td><td>>hg38_1_-_228634819_228635039_RNA5S12_5S_50ntFlanks</td><td>51</td><td>87</td><td>37</td><td>GTCTACGGCCATACCACCCTGAACGCGCCCGATCTCG</td><td>864.6</td><td>2.1</td><td>0.9</td><td>2.5</td><td>1.3E-02</td><td>4.4E-02</td><td>BT20/MB231</td></tr>
<tr><td>>hg38_21_+_8433172_8446622_RNA45SN1_45S_50ntFlanks@7974.8016.43|ACGCGACCTCAGATCAGACGTGGCGACCCGCTGAATTTAAGCA</td><td>>hg38_21_+_8433172_8446622_RNA45SN1_45S_50ntFlanks</td><td>7974</td><td>8016</td><td>43</td><td>ACGCGACCTCAGATCAGACGTGGCGACCCGCTGAATTTAAGCA</td><td>1338.9</td><td>-2.2</td><td>0.9</td><td>-2.5</td><td>1.3E-02</td><td>4.4E-02</td><td>BT20/MB231</td></tr>
<tr><td>>hg38_21_+_8433172_8446622_RNA45SN1_45S_50ntFlanks@11321.11340.20|CACGTCTCGTCGCGCGCGCG</td><td>>hg38_21_+_8433172_8446622_RNA45SN1_45S_50ntFlanks</td><td>11321</td><td>11340</td><td>20</td><td>CACGTCTCGTCGCGCGCGCG</td><td>151.0</td><td>-3.7</td><td>1.8</td><td>-2.1</td><td>3.9E-02</td><td>4.4E-02</td><td>MB231/MB468</td></tr>
<tr><td>>trna146_GlnCTG_6_-_27515531_27515602@1.30.30,trna1_GlnCTG_6_+_18836402_18836473@1.30.30,trna3_GlnCTG_17_+_8023070_8023141@1.30.30,trna42_GlnCTG_6_+_27263212_27263283@1.30.30,trna49_GlnCTG_6_+_27487308_27487379@1.30.30,trna7_GlnCTG_15_-_66161400_66161471@1.30.30,trna99_GlnCTG_6_-_28909378_28909449@1.30.30|GGTTCCATGGTGTAATGGTTAGCACTCTGG</td><td>>trna146_GlnCTG_6_-_27515531_27515602</td><td>1</td><td>30</td><td>30</td><td>GGTTCCATGGTGTAATGGTTAGCACTCTGG</td><td>504.4</td><td>2.1</td><td>1.0</td><td>2.1</td><td>3.9E-02</td><td>4.5E-02</td><td>MB231/MB468</td></tr>
<tr><td>>MI0000737|hsa-mir-200a&WithFlank&1|+|1167857|1167958@60.80.21&[MIMAT0000682&hsa-miR-200a-3p&offsets|0|-1;m-142&1|+|1167916|1167938&offsets|0|-2]|TAACACTGTCTGGTAACGATG</td><td>>MI0000737|hsa-mir-200a&WithFlank&1|+|1167857|1167958</td><td>60</td><td>80</td><td>21</td><td>TAACACTGTCTGGTAACGATG</td><td>115.9</td><td>-1.4</td><td>0.7</td><td>-2.0</td><td>4.1E-02</td><td>4.5E-02</td><td>BT20/MB468</td></tr>
<tr><td>>hg38_21_+_8433172_8446622_RNA45SN1_45S_50ntFlanks@7992.8017.26|CGTGGCGACCCGCTGAATTTAAGCAT</td><td>>hg38_21_+_8433172_8446622_RNA45SN1_45S_50ntFlanks</td><td>7992</td><td>8017</td><td>26</td><td>CGTGGCGACCCGCTGAATTTAAGCAT</td><td>147.9</td><td>-2.3</td><td>0.9</td><td>-2.5</td><td>1.3E-02</td><td>4.5E-02</td><td>BT20/MB231</td></tr>
<tr><td>>hg38_21_+_8433172_8446622_RNA45SN1_45S_50ntFlanks@11581.11604.24|AACTGGTGCGGACCAGGGGAATCC</td><td>>hg38_21_+_8433172_8446622_RNA45SN1_45S_50ntFlanks</td><td>11581</td><td>11604</td><td>24</td><td>AACTGGTGCGGACCAGGGGAATCC</td><td>209.9</td><td>-1.2</td><td>0.6</td><td>-2.0</td><td>4.1E-02</td><td>4.5E-02</td><td>BT20/MB468</td></tr>
<tr><td>>hg38_21_+_8433172_8446622_RNA45SN1_45S_50ntFlanks@10349.10383.35|AACGAGAACTTTGAAGGCCGAAGTGGAGAAGGGTT</td><td>>hg38_21_+_8433172_8446622_RNA45SN1_45S_50ntFlanks</td><td>10349</td><td>10383</td><td>35</td><td>AACGAGAACTTTGAAGGCCGAAGTGGAGAAGGGTT</td><td>2627.3</td><td>-1.4</td><td>0.7</td><td>-2.1</td><td>3.9E-02</td><td>4.5E-02</td><td>MB231/MB468</td></tr>
<tr><td>>MI0000542|hsa-mir-320a&WithFlank&8|-|22244960|22245043@50.70.21(+1U)&[MIMAT0000510&hsa-miR-320a-3p&offsets|+2|+1(+1U);m-60&8|-|22244975|22244996&offsets|+2|+1(+1U)]|AAGCTGGGTTGAGAGGGCGAAT</td><td>>MI0000542|hsa-mir-320a&WithFlank&8|-|22244960|22245043</td><td>50</td><td>70</td><td>21(+1U)</td><td>AAGCTGGGTTGAGAGGGCGAAT</td><td>224.9</td><td>1.8</td><td>0.7</td><td>2.5</td><td>1.3E-02</td><td>4.5E-02</td><td>BT20/MB231</td></tr>
<tr><td>>hg38_MT_+_1621_3279_RNR2_16S_50ntFlanks@1349.1388.40|ACAATAGGGTTTACGACCTCGATGTTGGATCAGGACATCC</td><td>>hg38_MT_+_1621_3279_RNR2_16S_50ntFlanks</td><td>1349</td><td>1388</td><td>40</td><td>ACAATAGGGTTTACGACCTCGATGTTGGATCAGGACATCC</td><td>1722.7</td><td>-2.4</td><td>1.0</td><td>-2.5</td><td>1.3E-02</td><td>4.5E-02</td><td>BT20/MB231</td></tr>
<tr><td>>hg38_21_+_8433172_8446622_RNA45SN1_45S_50ntFlanks@11209.11229.21|CGGCGACTCTGGACGCGAGCC</td><td>>hg38_21_+_8433172_8446622_RNA45SN1_45S_50ntFlanks</td><td>11209</td><td>11229</td><td>21</td><td>CGGCGACTCTGGACGCGAGCC</td><td>383.7</td><td>-1.3</td><td>0.6</td><td>-2.1</td><td>3.9E-02</td><td>4.5E-02</td><td>MB231/MB468</td></tr>
<tr><td>>hg38_21_+_8433172_8446622_RNA45SN1_45S_50ntFlanks@5486.5514.29|GCGCTGAGAAGACGGTCGAACTTGACTAT</td><td>>hg38_21_+_8433172_8446622_RNA45SN1_45S_50ntFlanks</td><td>5486</td><td>5514</td><td>29</td><td>GCGCTGAGAAGACGGTCGAACTTGACTAT</td><td>455.4</td><td>-1.6</td><td>0.8</td><td>-2.0</td><td>4.1E-02</td><td>4.5E-02</td><td>BT20/MB468</td></tr>
<tr><td>>trnaMT_SerGCT_MT_+_12207_12265@1.18.18|GAGAAAGCTCACAAGAAC</td><td>>trnaMT_SerGCT_MT_+_12207_12265</td><td>1</td><td>18</td><td>18</td><td>GAGAAAGCTCACAAGAAC</td><td>34717.1</td><td>2.7</td><td>1.1</td><td>2.5</td><td>1.3E-02</td><td>4.5E-02</td><td>BT20/MB231</td></tr>
<tr><td>>hg38_21_+_8433172_8446622_RNA45SN1_45S_50ntFlanks@10792.10816.25|CTCTGGCATGTTGGAACAATGTAGG</td><td>>hg38_21_+_8433172_8446622_RNA45SN1_45S_50ntFlanks</td><td>10792</td><td>10816</td><td>25</td><td>CTCTGGCATGTTGGAACAATGTAGG</td><td>226.0</td><td>-1.1</td><td>0.5</td><td>-2.1</td><td>4.0E-02</td><td>4.5E-02</td><td>MB231/MB468</td></tr>
<tr><td>>am_trnaMT_HisGTG_MT_+_12138_12206@18.38.21|AAAACATCAGATTGTGAATCT</td><td>>am_trnaMT_HisGTG_MT_+_12138_12206</td><td>18</td><td>38</td><td>21</td><td>AAAACATCAGATTGTGAATCT</td><td>423.2</td><td>2.0</td><td>0.8</td><td>2.5</td><td>1.3E-02</td><td>4.5E-02</td><td>BT20/MB231</td></tr>
<tr><td>>MI0000542|hsa-mir-320a&WithFlank&8|-|22244960|22245043@49.69.21(+2U)&[MIMAT0000510&hsa-miR-320a-3p&offsets|+1|0(+2U);m-60&8|-|22244975|22244996&offsets|+1|0(+2U)]|AAAGCTGGGTTGAGAGGGCGATT</td><td>>MI0000542|hsa-mir-320a&WithFlank&8|-|22244960|22245043</td><td>49</td><td>69</td><td>21(+2U)</td><td>AAAGCTGGGTTGAGAGGGCGATT</td><td>278.4</td><td>1.9</td><td>0.8</td><td>2.5</td><td>1.3E-02</td><td>4.6E-02</td><td>BT20/MB231</td></tr>
<tr><td>>hg38_21_+_8433172_8446622_RNA45SN1_45S_50ntFlanks@3999.4024.26|CTAGATAACCTCGGGCCGATCGCACG</td><td>>hg38_21_+_8433172_8446622_RNA45SN1_45S_50ntFlanks</td><td>3999</td><td>4024</td><td>26</td><td>CTAGATAACCTCGGGCCGATCGCACG</td><td>108.2</td><td>-1.1</td><td>0.5</td><td>-2.1</td><td>4.0E-02</td><td>4.6E-02</td><td>MB231/MB468</td></tr>
<tr><td>>hg38_21_+_8433172_8446622_RNA45SN1_45S_50ntFlanks@12416.12450.35|TTCGATGTCGGCTCTTCCTATCATTGTGAAGCAGA</td><td>>hg38_21_+_8433172_8446622_RNA45SN1_45S_50ntFlanks</td><td>12416</td><td>12450</td><td>35</td><td>TTCGATGTCGGCTCTTCCTATCATTGTGAAGCAGA</td><td>499.9</td><td>-2.1</td><td>0.9</td><td>-2.5</td><td>1.3E-02</td><td>4.6E-02</td><td>BT20/MB231</td></tr>
<tr><td>>hg38_21_+_8433172_8446622_RNA45SN1_45S_50ntFlanks@11727.11744.18|AGTAACTATGACTCTCTT</td><td>>hg38_21_+_8433172_8446622_RNA45SN1_45S_50ntFlanks</td><td>11727</td><td>11744</td><td>18</td><td>AGTAACTATGACTCTCTT</td><td>1018.6</td><td>1.4</td><td>0.7</td><td>2.1</td><td>4.0E-02</td><td>4.6E-02</td><td>MB231/MB468</td></tr>
<tr><td>>hg38_21_+_8433172_8446622_RNA45SN1_45S_50ntFlanks@12397.12418.22|CGACGTCGCTTTTTGATCCTTC</td><td>>hg38_21_+_8433172_8446622_RNA45SN1_45S_50ntFlanks</td><td>12397</td><td>12418</td><td>22</td><td>CGACGTCGCTTTTTGATCCTTC</td><td>572.0</td><td>-2.4</td><td>1.0</td><td>-2.5</td><td>1.3E-02</td><td>4.6E-02</td><td>BT20/MB231</td></tr>
</table>

<table>
<tr><td></td><td></td><td></td><td></td><td></td><td></td><td></td><td></td><td></td><td></td><td></td><td></td><td></td></tr>
<tr><td>>hg38_21_+_8433172_8446622_RNA45SN1_45S_50ntFlanks@9574.9601.28|AATCGGTCGTCCGACCTGGGTATAGGGG</td><td>>hg38_21_+_8433172_8446622_RNA45SN1_45S_50ntFlanks</td><td>9574</td><td>9601</td><td>28</td><td>AATCGGTCGTCCGACCTGGGTATAGGGG</td><td>460.1</td><td>1.0</td><td>0.5</td><td>2.1</td><td>4.0E-02</td><td>4.6E-02</td><td>MB231/MB468</td></tr>
<tr><td>>hg38_21_+_8433172_8446622_RNA45SN1_45S_50ntFlanks@6661.6689.29|CGGTGGATCACTCGGCTCGTGCGTCGATG</td><td>>hg38_21_+_8433172_8446622_RNA45SN1_45S_50ntFlanks</td><td>6661</td><td>6689</td><td>29</td><td>CGGTGGATCACTCGGCTCGTGCGTCGATG</td><td>1764.9</td><td>2.1</td><td>0.8</td><td>2.5</td><td>1.4E-02</td><td>4.6E-02</td><td>BT20/MB231</td></tr>
<tr><td>>hg38_21_+_8433172_8446622_RNA45SN1_45S_50ntFlanks@12419.12438.20|GATGTCGGCTCTTCCTATCA</td><td>>hg38_21_+_8433172_8446622_RNA45SN1_45S_50ntFlanks</td><td>12419</td><td>12438</td><td>20</td><td>GATGTCGGCTCTTCCTATCA</td><td>2131.7</td><td>-2.0</td><td>0.8</td><td>-2.5</td><td>1.4E-02</td><td>4.6E-02</td><td>BT20/MB231</td></tr>
<tr><td>>hg38_21_+_8433172_8446622_RNA45SN1_45S_50ntFlanks@11575.11594.20|ACTTAGAACTGGTGCGGACC</td><td>>hg38_21_+_8433172_8446622_RNA45SN1_45S_50ntFlanks</td><td>11575</td><td>11594</td><td>20</td><td>ACTTAGAACTGGTGCGGACC</td><td>1107.1</td><td>-1.1</td><td>0.5</td><td>-2.0</td><td>4.1E-02</td><td>4.6E-02</td><td>MB231/MB468</td></tr>
<tr><td>>hg38_21_+_8433172_8446622_RNA45SN1_45S_50ntFlanks@9534.9570.37|AGGAAACTCTGGTGGAGGTCCGTAGCGGTCCTGACGT</td><td>>hg38_21_+_8433172_8446622_RNA45SN1_45S_50ntFlanks</td><td>9534</td><td>9570</td><td>37</td><td>AGGAAACTCTGGTGGAGGTCCGTAGCGGTCCTGACGT</td><td>724.1</td><td>-1.0</td><td>0.5</td><td>-2.0</td><td>4.1E-02</td><td>4.7E-02</td><td>MB231/MB468</td></tr>
<tr><td>>hg38_21_+_8433172_8446622_RNA45SN1_45S_50ntFlanks@5495.5516.22|AGACGGTCGAACTTGACTATCT</td><td>>hg38_21_+_8433172_8446622_RNA45SN1_45S_50ntFlanks</td><td>5495</td><td>5516</td><td>22</td><td>AGACGGTCGAACTTGACTATCT</td><td>678.6</td><td>2.8</td><td>1.1</td><td>2.5</td><td>1.4E-02</td><td>4.7E-02</td><td>BT20/MB231</td></tr>
<tr><td>>hg38_21_+_8433172_8446622_RNA45SN1_45S_50ntFlanks@12999.13027.29|CTGCGATCTATTGAAAGTCAGCCCTCGAC</td><td>>hg38_21_+_8433172_8446622_RNA45SN1_45S_50ntFlanks</td><td>12999</td><td>13027</td><td>29</td><td>CTGCGATCTATTGAAAGTCAGCCCTCGAC</td><td>235.4</td><td>-2.5</td><td>1.0</td><td>-2.5</td><td>1.4E-02</td><td>4.7E-02</td><td>BT20/MB231</td></tr>
<tr><td>>hg38_21_+_8433172_8446622_RNA45SN1_45S_50ntFlanks@7975.8004.30|CGCGACCTCAGATCAGACGTGGCGACCCGC</td><td>>hg38_21_+_8433172_8446622_RNA45SN1_45S_50ntFlanks</td><td>7975</td><td>8004</td><td>30</td><td>CGCGACCTCAGATCAGACGTGGCGACCCGC</td><td>4704.8</td><td>1.8</td><td>0.7</td><td>2.5</td><td>1.4E-02</td><td>4.7E-02</td><td>BT20/MB231</td></tr>
<tr><td>>hg38_21_+_8433172_8446622_RNA45SN1_45S_50ntFlanks@12630.12667.38|GGGCGAAGCTACCATCTGTGGGATTATGACTGAACGCC</td><td>>hg38_21_+_8433172_8446622_RNA45SN1_45S_50ntFlanks</td><td>12630</td><td>12667</td><td>38</td><td>GGGCGAAGCTACCATCTGTGGGATTATGACTGAACGCC</td><td>155.7</td><td>-2.2</td><td>0.9</td><td>-2.5</td><td>1.4E-02</td><td>4.7E-02</td><td>BT20/MB231</td></tr>
<tr><td>>trna10_LysCTT_16_+_3241501_3241573@3.33.31,trna13_LysCTT_14_-_58706613_58706685@3.33.31,trna2_LysCTT_15_+_79152904_79152976@3.33.31|CCGGCTAGCTCAGTCGGTAGAGCATGGGACT</td><td>>trna10_LysCTT_16_+_3241501_3241573</td><td>3</td><td>33</td><td>31</td><td>CCGGCTAGCTCAGTCGGTAGAGCATGGGACT</td><td>605.9</td><td>-2.1</td><td>0.9</td><td>-2.5</td><td>1.4E-02</td><td>4.7E-02</td><td>BT20/MB231</td></tr>
<tr><td>>hg38_21_+_8433172_8446622_RNA45SN1_45S_50ntFlanks@5485.5510.26|AGCGCTGAGAAGACGGTCGAACTTGA</td><td>>hg38_21_+_8433172_8446622_RNA45SN1_45S_50ntFlanks</td><td>5485</td><td>5510</td><td>26</td><td>AGCGCTGAGAAGACGGTCGAACTTGA</td><td>134.0</td><td>-1.8</td><td>0.9</td><td>-2.0</td><td>4.3E-02</td><td>4.7E-02</td><td>BT20/MB468</td></tr>
<tr><td>>MI0005531|hsa-mir-450b&WithFlank&X|-|134540179|134540268@17.38.22&[MIMAT0004909&hsa-miR-450b-5p&offsets|0|0;m-266&X|-|134540232|134540252&offsets|0|+1]|TTTTGCAATATGTTCCTGAATA</td><td>>MI0005531|hsa-mir-450b&WithFlank&X|-|134540179|134540268</td><td>17</td><td>38</td><td>22</td><td>TTTTGCAATATGTTCCTGAATA</td><td>672.2</td><td>2.2</td><td>0.9</td><td>2.5</td><td>1.4E-02</td><td>4.7E-02</td><td>BT20/MB231</td></tr>
<tr><td>>hg38_21_+_8433172_8446622_RNA45SN1_45S_50ntFlanks@7988.8013.26|CAGACGTGGCGACCCGCTGAATTTAA</td><td>>hg38_21_+_8433172_8446622_RNA45SN1_45S_50ntFlanks</td><td>7988</td><td>8013</td><td>26</td><td>CAGACGTGGCGACCCGCTGAATTTAA</td><td>170.3</td><td>1.0</td><td>0.5</td><td>2.0</td><td>4.1E-02</td><td>4.7E-02</td><td>MB231/MB468</td></tr>
<tr><td>>hg38_21_+_8433172_8446622_RNA45SN1_45S_50ntFlanks@6645.6663.19|CTCGTACGACTCTTAGCGG</td><td>>hg38_21_+_8433172_8446622_RNA45SN1_45S_50ntFlanks</td><td>6645</td><td>6663</td><td>19</td><td>CTCGTACGACTCTTAGCGG</td><td>1707.3</td><td>2.1</td><td>0.9</td><td>2.5</td><td>1.4E-02</td><td>4.7E-02</td><td>BT20/MB231</td></tr>
<tr><td>>hg38_21_+_8433172_8446622_RNA45SN1_45S_50ntFlanks@6648.6681.34|GTACGACTCTTAGCGGTGGATCACTCGGCTCGTG</td><td>>hg38_21_+_8433172_8446622_RNA45SN1_45S_50ntFlanks</td><td>6648</td><td>6681</td><td>34</td><td>GTACGACTCTTAGCGGTGGATCACTCGGCTCGTG</td><td>1520.0</td><td>-2.0</td><td>0.8</td><td>-2.5</td><td>1.4E-02</td><td>4.7E-02</td><td>BT20/MB231</td></tr>
<tr><td>>hg38_21_+_8433172_8446622_RNA45SN1_45S_50ntFlanks@6653.6698.46|ACTCTTAGCGGTGGATCACTCGGCTCGTGCGTCGATGAAGAACGCA</td><td>>hg38_21_+_8433172_8446622_RNA45SN1_45S_50ntFlanks</td><td>6653</td><td>6698</td><td>46</td><td>ACTCTTAGCGGTGGATCACTCGGCTCGTGCGTCGATGAAGAACGCA</td><td>259.7</td><td>-1.9</td><td>0.8</td><td>-2.5</td><td>1.4E-02</td><td>4.7E-02</td><td>BT20/MB231</td></tr>
<tr><td>>hg38_21_+_8433172_8446622_RNA45SN1_45S_50ntFlanks@7989.8014.26|AGACGTGGCGACCCGCTGAATTTAAG</td><td>>hg38_21_+_8433172_8446622_RNA45SN1_45S_50ntFlanks</td><td>7989</td><td>8014</td><td>26</td><td>AGACGTGGCGACCCGCTGAATTTAAG</td><td>580.7</td><td>1.9</td><td>0.9</td><td>2.0</td><td>4.3E-02</td><td>4.7E-02</td><td>BT20/MB468</td></tr>
<tr><td>>hg38_21_+_8433172_8446622_RNA45SN1_45S_50ntFlanks@10830.10855.26|CAAGCCGGATCCGTAACTTCGGGATA</td><td>>hg38_21_+_8433172_8446622_RNA45SN1_45S_50ntFlanks</td><td>10830</td><td>10855</td><td>26</td><td>CAAGCCGGATCCGTAACTTCGGGATA</td><td>144.4</td><td>-3.8</td><td>1.9</td><td>-2.0</td><td>4.3E-02</td><td>4.7E-02</td><td>BT20/MB468</td></tr>
<tr><td>>hg38_21_+_8433172_8446622_RNA45SN1_45S_50ntFlanks@12455.12475.21|ACCAAGCGTTGGATTGTTCAC</td><td>>hg38_21_+_8433172_8446622_RNA45SN1_45S_50ntFlanks</td><td>12455</td><td>12475</td><td>21</td><td>ACCAAGCGTTGGATTGTTCAC</td><td>167.1</td><td>-1.3</td><td>0.6</td><td>-2.0</td><td>4.2E-02</td><td>4.8E-02</td><td>MB231/MB468</td></tr>
<tr><td>>hg38_21_+_8433172_8446622_RNA45SN1_45S_50ntFlanks@10798.10815.18|CATGTTGGAACAATGTAG</td><td>>hg38_21_+_8433172_8446622_RNA45SN1_45S_50ntFlanks</td><td>10798</td><td>10815</td><td>18</td><td>CATGTTGGAACAATGTAG</td><td>129.3</td><td>-3.8</td><td>1.9</td><td>-2.0</td><td>4.3E-02</td><td>4.8E-02</td><td>BT20/MB468</td></tr>
<tr><td>>hg38_21_+_8433172_8446622_RNA45SN1_45S_50ntFlanks@11877.11898.22|AGACCCTGTTGAGCTTGACTCT</td><td>>hg38_21_+_8433172_8446622_RNA45SN1_45S_50ntFlanks</td><td>11877</td><td>11898</td><td>22</td><td>AGACCCTGTTGAGCTTGACTCT</td><td>413.4</td><td>1.8</td><td>0.9</td><td>2.0</td><td>4.4E-02</td><td>4.8E-02</td><td>BT20/MB468</td></tr>
<tr><td>>hg38_21_+_8433172_8446622_RNA45SN1_45S_50ntFlanks@4735.4761.27|ACGAAAGTCGGAGGTTCGAAGACGATC</td><td>>hg38_21_+_8433172_8446622_RNA45SN1_45S_50ntFlanks</td><td>4735</td><td>4761</td><td>27</td><td>ACGAAAGTCGGAGGTTCGAAGACGATC</td><td>638.8</td><td>-1.7</td><td>0.7</td><td>-2.5</td><td>1.4E-02</td><td>4.8E-02</td><td>BT20/MB231</td></tr>
<tr><td>>hg38_21_+_8433172_8446622_RNA45SN1_45S_50ntFlanks@11208.11230.23|GCGGCGACTCTGGACGCGAGCCG</td><td>>hg38_21_+_8433172_8446622_RNA45SN1_45S_50ntFlanks</td><td>11208</td><td>11230</td><td>23</td><td>GCGGCGACTCTGGACGCGAGCCG</td><td>709.1</td><td>-2.2</td><td>1.1</td><td>-2.0</td><td>4.4E-02</td><td>4.8E-02</td><td>BT20/MB468</td></tr>
<tr><td>>hg38_21_+_8433172_8446622_RNA45SN1_45S_50ntFlanks@11671.11706.36|CCAGTGCTCTGAATGTCAAAGTGAAGAAATTCAATG</td><td>>hg38_21_+_8433172_8446622_RNA45SN1_45S_50ntFlanks</td><td>11671</td><td>11706</td><td>36</td><td>CCAGTGCTCTGAATGTCAAAGTGAAGAAATTCAATG</td><td>309.1</td><td>-1.8</td><td>0.9</td><td>-2.0</td><td>4.4E-02</td><td>4.8E-02</td><td>BT20/MB468</td></tr>
<tr><td>>hg38_21_+_8433172_8446622_RNA45SN1_45S_50ntFlanks@6732.6752.21|ACATTGATCATCGACACTTCG</td><td>>hg38_21_+_8433172_8446622_RNA45SN1_45S_50ntFlanks</td><td>6732</td><td>6752</td><td>21</td><td>ACATTGATCATCGACACTTCG</td><td>700.7</td><td>1.2</td><td>0.6</td><td>2.0</td><td>4.3E-02</td><td>4.9E-02</td><td>MB231/MB468</td></tr>
<tr><td>>hg38_21_+_8433172_8446622_RNA45SN1_45S_50ntFlanks@11200.11225.26|CGGCGGCGGCGGCGACTCTGGACGCG</td><td>>hg38_21_+_8433172_8446622_RNA45SN1_45S_50ntFlanks</td><td>11200</td><td>11225</td><td>26</td><td>CGGCGGCGGCGGCGACTCTGGACGCG</td><td>489.8</td><td>-4.0</td><td>2.0</td><td>-2.0</td><td>4.3E-02</td><td>4.9E-02</td><td>MB231/MB468</td></tr>
<tr><td>>hg38_21_+_8433172_8446622_RNA45SN1_45S_50ntFlanks@8105.8141.37|CGCGGCGGGGCGCGGGACATGTGGCGTACGGAAGACC</td><td>>hg38_21_+_8433172_8446622_RNA45SN1_45S_50ntFlanks</td><td>8105</td><td>8141</td><td>37</td><td>CGCGGCGGGGCGCGGGACATGTGGCGTACGGAAGACC</td><td>91.8</td><td>-1.8</td><td>0.9</td><td>-2.0</td><td>4.3E-02</td><td>4.9E-02</td><td>MB231/MB468</td></tr>
<tr><td>>hg38_21_+_8433172_8446622_RNA45SN1_45S_50ntFlanks@11579.11602.24|AGAACTGGTGCGGACCAGGGGAAT</td><td>>hg38_21_+_8433172_8446622_RNA45SN1_45S_50ntFlanks</td><td>11579</td><td>11602</td><td>24</td><td>AGAACTGGTGCGGACCAGGGGAAT</td><td>483.7</td><td>-1.7</td><td>0.8</td><td>-2.0</td><td>4.3E-02</td><td>4.9E-02</td><td>MB231/MB468</td></tr>
<tr><td>>trna111_HisGTG_1_-_147774845_147774916@3.31.29,trna118_HisGTG_1_-_145396881_145396952@3.31.29,trna16_HisGTG_1_+_146544773_146544844@3.31.29,trna1_HisGTG_15_+_45493349_45493420@3.31.29,trna21_HisGTG_1_+_147753471_147753542@3.31.29,trna33_HisGTG_6_+_27125906_27125977@3.31.29,trna7_HisGTG_9_-_14433938_14434009@3.31.29,trna8_HisGTG_15_-_45492611_45492682@3.31.29,trna9_HisGTG_15_-_45490804_45490875@3.31.29|CGTGATCGTATAGTGGTTAGTACTCTGCG</td><td>>trna111_HisGTG_1_-_147774845_147774916</td><td>3</td><td>31</td><td>29</td><td>CGTGATCGTATAGTGGTTAGTACTCTGCG</td><td>256.6</td><td>1.5</td><td>0.7</td><td>2.0</td><td>4.3E-02</td><td>4.9E-02</td><td>MB231/MB468</td></tr>
<tr><td>>trnaMT_GluTTC_MT_-_14674_14742@53.72.20,trnalookalike8_GluTTC_5_-_93905172_93905240@53.72.20|TAGTCCGTGCGAGAATACCA</td><td>>trnaMT_GluTTC_MT_-_14674_14742</td><td>53</td><td>72</td><td>20</td><td>TAGTCCGTGCGAGAATACCA</td><td>275.3</td><td>1.1</td><td>0.6</td><td>2.0</td><td>4.3E-02</td><td>4.9E-02</td><td>MB231/MB468</td></tr>
<tr><td>>hg38_21_+_8433172_8446622_RNA45SN1_45S_50ntFlanks@4297.4318.22|CATTGGAGGGCAAGTCTGGTGC</td><td>>hg38_21_+_8433172_8446622_RNA45SN1_45S_50ntFlanks</td><td>4297</td><td>4318</td><td>22</td><td>CATTGGAGGGCAAGTCTGGTGC</td><td>95.6</td><td>-1.7</td><td>0.9</td><td>-2.0</td><td>4.3E-02</td><td>4.9E-02</td><td>MB231/MB468</td></tr>
<tr><td>>hg38_1_-_228634819_228635039_RNA5S12_5S_50ntFlanks@144.171.28|CCTGGGAATACCGGGTGCTGTAGGCTTT</td><td>>hg38_1_-_228634819_228635039_RNA5S12_5S_50ntFlanks</td><td>144</td><td>171</td><td>28</td><td>CCTGGGAATACCGGGTGCTGTAGGCTTT</td><td>8567.6</td><td>-1.7</td><td>0.7</td><td>-2.4</td><td>1.5E-02</td><td>4.9E-02</td><td>BT20/MB231</td></tr>
<tr><td>>hg38_21_+_8433172_8446622_RNA45SN1_45S_50ntFlanks@8358.8377.20|AAAGAACTTTGAAGAGAGAG</td><td>>hg38_21_+_8433172_8446622_RNA45SN1_45S_50ntFlanks</td><td>8358</td><td>8377</td><td>20</td><td>AAAGAACTTTGAAGAGAGAG</td><td>165.4</td><td>-1.1</td><td>0.5</td><td>-2.0</td><td>4.5E-02</td><td>4.9E-02</td><td>BT20/MB468</td></tr>
<tr><td>>hg38_21_+_8433172_8446622_RNA45SN1_45S_50ntFlanks@12417.12448.32|TCGATGTCGGCTCTTCCTATCATTGTGAAGCA</td><td>>hg38_21_+_8433172_8446622_RNA45SN1_45S_50ntFlanks</td><td>12417</td><td>12448</td><td>32</td><td>TCGATGTCGGCTCTTCCTATCATTGTGAAGCA</td><td>869.4</td><td>-2.2</td><td>0.9</td><td>-2.4</td><td>1.5E-02</td><td>4.9E-02</td><td>BT20/MB231</td></tr>
<tr><td>>hg38_21_+_8433172_8446622_RNA45SN1_45S_50ntFlanks@7990.8018.29|GACGTGGCGACCCGCTGAATTTAAGCATA</td><td>>hg38_21_+_8433172_8446622_RNA45SN1_45S_50ntFlanks</td><td>7990</td><td>8018</td><td>29</td><td>GACGTGGCGACCCGCTGAATTTAAGCATA</td><td>204.4</td><td>-2.4</td><td>1.0</td><td>-2.4</td><td>1.5E-02</td><td>4.9E-02</td><td>BT20/MB231</td></tr>
<tr><td>>hg38_21_+_8433172_8446622_RNA45SN1_45S_50ntFlanks@12415.12447.33|CTTCGATGTCGGCTCTTCCTATCATTGTGAAGC</td><td>>hg38_21_+_8433172_8446622_RNA45SN1_45S_50ntFlanks</td><td>12415</td><td>12447</td><td>33</td><td>CTTCGATGTCGGCTCTTCCTATCATTGTGAAGC</td><td>514.2</td><td>-2.1</td><td>0.8</td><td>-2.4</td><td>1.5E-02</td><td>5.0E-02</td><td>BT20/MB231</td></tr>
<tr><td>>hg38_21_+_8433172_8446622_RNA45SN1_45S_50ntFlanks@4727.4752.26|AATCAAGAACGAAAGTCGGAGGTTCG</td><td>>hg38_21_+_8433172_8446622_RNA45SN1_45S_50ntFlanks</td><td>4727</td><td>4752</td><td>26</td><td>AATCAAGAACGAAAGTCGGAGGTTCG</td><td>141.8</td><td>-1.1</td><td>0.5</td><td>-2.0</td><td>4.5E-02</td><td>5.0E-02</td><td>BT20/MB468</td></tr>
<tr><td>>MI0000483|hsa-mir-186&WithFlank&1|-|71067625|71067722@21.41.21(+1G)&[MIMAT0000456&hsa-miR-186-5p&offsets|0|-1(+1G);m-84&1|-|71067681|71067702&offsets|0|-1(+1G)]|CAAAGAATTCTCCTTTTGGGCG</td><td>>MI0000483|hsa-mir-186&WithFlank&1|-|71067625|71067722</td><td>21</td><td>41</td><td>21(+1G)</td><td>CAAAGAATTCTCCTTTTGGGCG</td><td>349.7</td><td>2.4</td><td>1.0</td><td>2.4</td><td>1.5E-02</td><td>5.0E-02</td><td>BT20/MB231</td></tr>
<tr><td>>am_MI0000809|hsa-mir-151a&WithFlank&8|-|140732558|140732659@53.72.20&[MIMAT0000757&hsa-miR-151a-3p&offsets|0|-1;m-30&8|-|140732587|140732607&offsets|0|-1]|CTAGACTGAAGCTCCTTGAG</td><td>>am_MI0000809|hsa-mir-151a&WithFlank&8|-|140732558|140732659</td><td>53</td><td>72</td><td>20</td><td>CTAGACTGAAGCTCCTTGAG</td><td>5956.1</td><td>1.5</td><td>0.6</td><td>2.4</td><td>1.5E-02</td><td>5.0E-02</td><td>BT20/MB231</td></tr>
<tr><td>>hg38_21_+_8433172_8446622_RNA45SN1_45S_50ntFlanks@5468.5499.32|GCCCACGGCCCTGGCGGAGCGCTGAGAAGACG</td><td>>hg38_21_+_8433172_8446622_RNA45SN1_45S_50ntFlanks</td><td>5468</td><td>5499</td><td>32</td><td>GCCCACGGCCCTGGCGGAGCGCTGAGAAGACG</td><td>121.6</td><td>-1.7</td><td>0.8</td><td>-2.0</td><td>4.5E-02</td><td>5.0E-02</td><td>BT20/MB468</td></tr>
<tr><td>>hg38_21_+_8433172_8446622_RNA45SN1_45S_50ntFlanks@12008.12029.22|CCGCCGGTGAAATACCACTACT</td><td>>hg38_21_+_8433172_8446622_RNA45SN1_45S_50ntFlanks</td><td>12008</td><td>12029</td><td>22</td><td>CCGCCGGTGAAATACCACTACT</td><td>197.1</td><td>-2.1</td><td>1.1</td><td>-2.0</td><td>4.6E-02</td><td>5.0E-02</td><td>BT20/MB468</td></tr>
</table>
