## Supplementary material for "Single-nucleotide Differences and Cell Type Decide the Subcellular Localization of miRNA Isoforms (isomiRs), tRNA-derived Fragments (tRFs) and rRNA-derived Fragments (rRFs)": Supp. Table S5 (1 of 4)

### Nucleus

**Supplemental Table S5:** Data table of the top 10% most abundant short RNA from each cell line for each cell fraction based on average abundance of the cell line replicates. The annotation "am" is used to indicate abiguous short RNA, those that can be mapped to a noncoding RNA gene as well as somewhere else on the geneome. The four different cell fractions are separated by sheets (Nucleus, Cytoplasm, Mitochondria, Mitoplast) The annotation "multi" referrs to short RNA that map to locations in both the mitochondrial and nuclear genomes. The rRF are flanked by 50 nucleotdies upstream and downstream and their coordiantes reflect this. The coordiantes for the isomiR and rRF are from the hg38 human genome assembly while the coordintes for the tRF are from the hg19 human genome assembly. Columns H-J indicate whether or not a given short RNA is present in the top 10% of the short RNA in one of the three cell lines and is idicated by blue (TRUE) or white (FALSE) sorted from most abundant to least abundant.

### Nucleus

| | | | | | | | | | |
|---|---|---|---|---|---|---|---|---|---|
| >hg38_21_+_8433172_8446622_RNA45SN1_45S_50ntFlanks@3705.3746.42\|TACCTGGTTGATCCTGCCAGTAGCATATGCTTGTCTCAAAGA | >hg38_21_+_8433172_8446622_RNA45SN1_45S_50ntFlanks | 3705 | 3746 | 42 | TACCTGGTTGATCCTGCCAGTAGCATATGCTTGTCTCAAAGA | rRF | TRUE | FALSE | FALSE |
| >MI0000098\|hsa-mir-96&WithFlank&7\|-\|129774686\|129774775@15.37.23&[MIMAT0000095&hsa-miR-96-5p&offsets\|0\|0;m-248&7\|-\|129774739\|129774761&offsets\|0\|0]\|TTTGGCACTAGCACATTTTTGCT | >MI0000098\|hsa-mir-96&WithFlank&7\|-\|129774686\|129774775 | 15 | 37 | 23 | TTTGGCACTAGCACATTTTTGCT | isomiR | TRUE | FALSE | FALSE |
| >hg38_21_+_8433172_8446622_RNA45SN1_45S_50ntFlanks@7974.7994.21\|ACGCGACCTCAGATCAGACGT | >hg38_21_+_8433172_8446622_RNA45SN1_45S_50ntFlanks | 7974 | 7994 | 21 | ACGCGACCTCAGATCAGACGT | rRF | TRUE | FALSE | FALSE |
| >MI0000298\|hsa-mir-221&WithFlank&X\|-\|45746151\|45746272@71.93.23&[MIMAT0000278&hsa-miR-221-3p&offsets\|0\|0;m-91&X\|-\|45746181\|45746202&offsets\|0\|+1]\|AGCTACATTGTCTGCTGGGTTTC | >MI0000298\|hsa-mir-221&WithFlank&X\|-\|45746151\|45746272 | 71 | 93 | 23 | AGCTACATTGTCTGCTGGGTTTC | isomiR | TRUE | TRUE | FALSE |
| >hg38_1_-_228634819_228635039_RNA5S12_5S_50ntFlanks@51.84.34\|GTCTACGGCCATACCACCCTGAACGCGCCCGATC | >hg38_1_-_228634819_228635039_RNA5S12_5S_50ntFlanks | 51 | 84 | 34 | GTCTACGGCCATACCACCCTGAACGCGCCCGATC | rRF | TRUE | TRUE | FALSE |
| >am_MI0000809\|hsa-mir-151a&WithFlank&8\|-\|140732558\|140732659@53.73.21(+1U)&[MIMAT0000757&hsa-miR-151a-3p&offsets\|0\|0(+1U);m-30&8\|-\|140732587\|140732607&offsets\|0\|0(+1U)]\|CTAGACTGAAGCTCCTTGAGGT | >am_MI0000809\|hsa-mir-151a&WithFlank&8\|-\|140732558\|140732659 | 53 | 73 | 21(+1U) | CTAGACTGAAGCTCCTTGAGGT | isomiR | TRUE | TRUE | FALSE |
| >hg38_21_+_8433172_8446622_RNA45SN1_45S_50ntFlanks@7974.8007.34\|ACGCGACCTCAGATCAGACGTGGCGACCCGCTGA | >hg38_21_+_8433172_8446622_RNA45SN1_45S_50ntFlanks | 7974 | 8007 | 34 | ACGCGACCTCAGATCAGACGTGGCGACCCGCTGA | rRF | TRUE | FALSE | FALSE |
| >MI0000441\|hsa-mir-30b&WithFlank&8\|-\|134800514\|134800613@23.44.22&[MIMAT0000420&hsa-miR-30b-5p&offsets\|0\|0;m-76&8\|-\|134800570\|134800591&offsets\|0\|0]\|TGTAAACATCCTACACTCAGCT | >MI0000441\|hsa-mir-30b&WithFlank&8\|-\|134800514\|134800613 | 23 | 44 | 22 | TGTAAACATCCTACACTCAGCT | isomiR | TRUE | FALSE | FALSE |
| >MI0000542\|hsa-mir-320a&WithFlank&8\|-\|22244960\|22245043@48.70.23&[MIMAT0000510&hsa-miR-320a-3p&offsets\|0\|+1;m-60&8\|-\|22244975\|22244996&offsets\|0\|+1]\|AAAAGCTGGGTTGAGAGGGCGAA | >MI0000542\|hsa-mir-320a&WithFlank&8\|-\|22244960\|22245043 | 48 | 70 | 23 | AAAAGCTGGGTTGAGAGGGCGAA | isomiR | TRUE | FALSE | FALSE |
| >MI0000070\|hsa-mir-16-1&WithFlank&13\|-\|50048967\|50049067@20.41.22;MI0000115\|hsa-mir-16-2&WithFlank&3\|+\|160404739\|160404831@16.37.22&[MIMAT0000069&hsa-miR-16-5p&offsets\|0\|0;m-62&13\|-\|50049027\|50049048&offsets\|0\|0];[MIMAT0000069_1&hsa-miR-16-5p&offsets\|0\|0;m-61&3\|+\|160404754\|160404775&offsets\|0\|0]\|TAGCAGCACGTAAATATTGGCG | >MI0000070\|hsa-mir-16-1&WithFlank&13\|-\|50048967\|50049067 | 20 | 41 | 22 | TAGCAGCACGTAAATATTGGCG | isomiR | TRUE | FALSE | FALSE |
| >hg38_21_+_8433172_8446622_RNA45SN1_45S_50ntFlanks@6651.6684.34\|CGACTCTTAGCGGTGGATCACTCGGCTCGTGCGT | >hg38_21_+_8433172_8446622_RNA45SN1_45S_50ntFlanks | 6651 | 6684 | 34 | CGACTCTTAGCGGTGGATCACTCGGCTCGTGCGT | rRF | TRUE | TRUE | FALSE |
| >hg38_21_+_8433172_8446622_RNA45SN1_45S_50ntFlanks@475.496.22\|CTTCGTGATCGATGTGGTGACG | >hg38_21_+_8433172_8446622_RNA45SN1_45S_50ntFlanks | 475 | 496 | 22 | CTTCGTGATCGATGTGGTGACG | rRF | TRUE | TRUE | FALSE |
| >hg38_21_+_8433172_8446622_RNA45SN1_45S_50ntFlanks@3705.3748.44\|TACCTGGTTGATCCTGCCAGTAGCATATGCTTGTCTCAAAGATT | >hg38_21_+_8433172_8446622_RNA45SN1_45S_50ntFlanks | 3705 | 3748 | 44 | TACCTGGTTGATCCTGCCAGTAGCATATGCTTGTCTCAAAGATT | rRF | TRUE | FALSE | FALSE |
| >hg38_21_+_8433172_8446622_RNA45SN1_45S_50ntFlanks@7975.7996.22\|CGCGACCTCAGATCAGACGTGG | >hg38_21_+_8433172_8446622_RNA45SN1_45S_50ntFlanks | 7975 | 7996 | 22 | CGCGACCTCAGATCAGACGTGG | rRF | TRUE | FALSE | FALSE |
| >MI0000440\|hsa-mir-27b&WithFlank&9\|+\|95085439\|95085547@67.86.20&[MIMAT0000419&hsa-miR-27b-3p&offsets\|0\|-1;m-39&9\|+\|95085505\|95085525&offsets\|0\|-1]\|TTCACAGTGGCTAAGTTCTG | >MI0000440\|hsa-mir-27b&WithFlank&9\|+\|95085439\|95085547 | 67 | 86 | 20 | TTCACAGTGGCTAAGTTCTG | isomiR | TRUE | TRUE | FALSE |
| >am_MI0005763\|hsa-mir-941-1&WithFlank&20\|+\|63919443\|63919526@53.75.23;MI0005764\|hsa-mir-941-2&WithFlank&20\|+\|63919499\|63919582@53.75.23;MI0005765\|hsa-mir-941-3&WithFlank&20\|+\|63919555\|63919638@53.75.23;MI0005766\|hsa-mir-941-4&WithFlank&20\|+\|63919750\|63919833@53.75.23;MI0031520\|hsa-mir-941-5&WithFlank&20\|+\|63919862\|63919945@53.75.23&[MIMAT0004984&hsa-miR-941&offsets\|0\|0;m-263&20\|+\|63919495\|63919517&offsets\|0\|0];[MIMAT0004984_1&hsa-miR-941&offsets\|0\|0;m-263&20\|+\|63919551\|63919573&offsets\|0\|0];[MIMAT0004984_2&hsa-miR-941&offsets\|0\|0;m-263&20\|+\|63919607\|63919629&offsets\|0\|0];[MIMAT0004984_3&hsa-miR-941&offsets\|0\|0;m-263&20\|+\|63919802\|63919824&offsets\|0\|0];[MIMAT0004984_4&hsa-miR-941&offsets\|0\|0;m-263&20\|+\|63919914\|63919936&offsets\|0\|0]\|CACCCGGCTGTGTGCACATGTGC | >am_MI0005763\|hsa-mir-941-1&WithFlank&20\|+\|63919443\|63919526 | 53 | 75 | 23 | CACCCGGCTGTGTGCACATGTGC | isomiR | TRUE | TRUE | FALSE |
| >MI0000063\|hsa-let-7b&WithFlank&22\|+\|46113680\|46113774@12.34.23&[MIMAT0000063&hsa-let-7b-5p&offsets\|0\|+1;m-14&22\|+\|46113691\|46113712&offsets\|0\|+1]\|TGAGGTAGTAGGTTGTGTGGTTT | >MI0000063\|hsa-let-7b&WithFlank&22\|+\|46113680\|46113774 | 12 | 34 | 23 | TGAGGTAGTAGGTTGTGTGGTTT | isomiR | TRUE | FALSE | FALSE |
| >MI0000255\|hsa-mir-30d&WithFlank&8\|-\|134804870\|134804951@12.34.23(+1A)&[MIMAT0000245&hsa-miR-30d-5p&offsets\|0\|+1(+1A);m-21&8\|-\|134804919\|134804940&offsets\|0\|+1(+1A)]\|TGTAAACATCCCCGACTGGAAGCA | >MI0000255\|hsa-mir-30d&WithFlank&8\|-\|134804870\|134804951 | 12 | 34 | 23(+1A) | TGTAAACATCCCCGACTGGAAGCA | isomiR | TRUE | FALSE | FALSE |
| >hg38_21_+_8433172_8446622_RNA45SN1_45S_50ntFlanks@7975.8017.43\|CGCGACCTCAGATCAGACGTGGCGACCCGCTGAATTTAAGCAT | >hg38_21_+_8433172_8446622_RNA45SN1_45S_50ntFlanks | 7975 | 8017 | 43 | CGCGACCTCAGATCAGACGTGGCGACCCGCTGAATTTAAGCAT | rRF | TRUE | FALSE | FALSE |
| >hg38_21_+_8433172_8446622_RNA45SN1_45S_50ntFlanks@7974.8004.31\|ACGCGACCTCAGATCAGACGTGGCGACCCGC | >hg38_21_+_8433172_8446622_RNA45SN1_45S_50ntFlanks | 7974 | 8004 | 31 | ACGCGACCTCAGATCAGACGTGGCGACCCGC | rRF | TRUE | FALSE | FALSE |
| >hg38_21_+_8433172_8446622_RNA45SN1_45S_50ntFlanks@6651.6688.38\|CGACTCTTAGCGGTGGATCACTCGGCTCGTGCGTCGAT | >hg38_21_+_8433172_8446622_RNA45SN1_45S_50ntFlanks | 6651 | 6688 | 38 | CGACTCTTAGCGGTGGATCACTCGGCTCGTGCGTCGAT | rRF | TRUE | FALSE | FALSE |
| >hg38_21_+_8433172_8446622_RNA45SN1_45S_50ntFlanks@5485.5516.32\|AGCGCTGAGAAGACGGTCGAACTTGACTATCT | >hg38_21_+_8433172_8446622_RNA45SN1_45S_50ntFlanks | 5485 | 5516 | 32 | AGCGCTGAGAAGACGGTCGAACTTGACTATCT | rRF | TRUE | FALSE | FALSE |
| >MI0003137\|hsa-mir-193b&WithFlank&16\|+\|14303961\|14304055@57.78.22&[MIMAT0002819&hsa-miR-193b-3p&offsets\|0\|0;m-166&16\|+\|14304017\|14304038&offsets\|0\|0]\|AACTGGCCCTCAAAGTCCCGCT | >MI0003137\|hsa-mir-193b&WithFlank&16\|+\|14303961\|14304055 | 57 | 78 | 22 | AACTGGCCCTCAAAGTCCCGCT | isomiR | TRUE | FALSE | FALSE |
| >MI0000063\|hsa-let-7b&WithFlank&22\|+\|46113680\|46113774@12.32.21&[MIMAT0000063&hsa-let-7b-5p&offsets\|0\|-1;m-14&22\|+\|46113691\|46113712&offsets\|0\|-1]\|TGAGGTAGTAGGTTGTGTGGT | >MI0000063\|hsa-let-7b&WithFlank&22\|+\|46113680\|46113774 | 12 | 32 | 21 | TGAGGTAGTAGGTTGTGTGGT | isomiR | TRUE | FALSE | FALSE |
| >MI0000088\|hsa-mir-30a&WithFlank&6\|-\|71403545\|71403627@12.33.22(+1U)&[MIMAT0000087&hsa-miR-30a-5p&offsets\|0\|0(+1U);m-15&6\|-\|71403595\|71403616&offsets\|0\|0(+1U)]\|TGTAAACATCCTCGACTGGAAGT | >MI0000088\|hsa-mir-30a&WithFlank&6\|-\|71403545\|71403627 | 12 | 33 | 22(+1U) | TGTAAACATCCTCGACTGGAAGT | isomiR | TRUE | FALSE | FALSE |
| >MI0000079\|hsa-mir-23a&WithFlank&19\|-\|13836581\|13836665@54.72.19&[MIMAT0000078&hsa-miR-23a-3p&offsets\|+3\|+1;m-31&19\|-\|13836595\|13836615&offsets\|+3\|+1]\|ACATTGCCAGGGATTTCCA | >MI0000079\|hsa-mir-23a&WithFlank&19\|-\|13836581\|13836665 | 54 | 72 | 19 | ACATTGCCAGGGATTTCCA | isomiR | TRUE | FALSE | FALSE |

### Nucleus

| | | | | | | | | | |
|---|---|---|---|---|---|---|---|---|---|
| >hg38_21_+_8433172_8446622_RNA45SN1_45S_50ntFlanks@476.523.48\|TTCGTGATCGATGTGGTGACGTCGTGCTCTCCCGGGCCGGGTCCGAGC | >hg38_21_+_8433172_8446622_RNA45SN1_45S_50ntFlanks | 476 | 523 | 48 | TTCGTGATCGATGTGGTGACGTCGTGCTCTCCCGGGCCGGGTC | rRF | FALSE | TRUE | FALSE |
| >MI0000298\|hsa-mir-221&WithFlank&X\|-\|45746151\|45746272@71.93.23(+1U)&[MIMAT0000278&hsa-miR-221-3p&offsets\|0\|0(+1U);m-91&X\|-\|45746181\|45746202&offsets\|0\|+1(+1U)]\|AGCTACATTGTCTGCTGGGTTTCT | >MI0000298\|hsa-mir-221&WithFlank&X\|-\|45746151\|45746272 | 71 | 93 | 23(+1U) | AGCTACATTGTCTGCTGGGTTTCT | isomiR | FALSE | TRUE | FALSE |
| >hg38_21_+_8433172_8446622_RNA45SN1_45S_50ntFlanks@476.504.29\|TTCGTGATCGATGTGGTGACGTCGTGCTC | >hg38_21_+_8433172_8446622_RNA45SN1_45S_50ntFlanks | 476 | 504 | 29 | TTCGTGATCGATGTGGTGACGTCGTGCTC | rRF | FALSE | TRUE | FALSE |
| >hg38_21_+_8433172_8446622_RNA45SN1_45S_50ntFlanks@476.517.42\|TTCGTGATCGATGTGGTGACGTCGTGCTCTCCCGGGCCGGGT | >hg38_21_+_8433172_8446622_RNA45SN1_45S_50ntFlanks | 476 | 517 | 42 | TTCGTGATCGATGTGGTGACGTCGTGCTCTCCCGGGCCGGGT | rRF | FALSE | TRUE | FALSE |
| >MI0000446\|hsa-mir-125b-1&WithFlank&11\|-\|122099751\|122099850@21.41.21;MI0000470\|hsa-mir-125b-2&WithFlank&21\|+\|16590231\|16590331@23.43.21&[MIMAT0000423&hsa-miR-125b-5p&offsets\|0\|-1;m-67&11\|-\|122099809\|122099830&offsets\|0\|-1];[MIMAT0000423_1&hsa-miR-125b-5p&offsets\|0\|-1;m-68&21\|+\|16590253\|16590274&offsets\|0\|-1]\|TCCCTGAGACCCTAACTTGTG | >MI0000446\|hsa-mir-125b-1&WithFlank&11\|-\|122099751\|122099850 | 21 | 41 | 21 | TCCCTGAGACCCTAACTTGTG | isomiR | FALSE | TRUE | FALSE |
| >hg38_21_+_8433172_8446622_RNA45SN1_45S_50ntFlanks@6651.6675.25\|CGACTCTTAGCGGTGGATCACTCGG | >hg38_21_+_8433172_8446622_RNA45SN1_45S_50ntFlanks | 6651 | 6675 | 25 | CGACTCTTAGCGGTGGATCACTCGG | rRF | FALSE | TRUE | FALSE |
| >MI0000088\|hsa-mir-30a&WithFlank&6\|-\|71403545\|71403627@53.73.21(+1U)&[MIMAT0000088&hsa-miR-30a-3p&offsets\|0\|-1(+1U);m-38&6\|-\|71403554\|71403575&offsets\|0\|-1(+1U)]\|CTTTCAGTCGGATGTTTGCAGT | >MI0000088\|hsa-mir-30a&WithFlank&6\|-\|71403545\|71403627 | 53 | 73 | 21(+1U) | CTTTCAGTCGGATGTTTGCAGT | isomiR | FALSE | TRUE | FALSE |
| >trna111_HisGTG_1_-_147774845_147774916@-1G.34.35,trna118_HisGTG_1_-_145396881_145396952@-1G.34.35,trna16_HisGTG_1_+_146544773_146544844@-1G.34.35,trna1_HisGTG_15_+_45493349_45493420@-1G.34.35,trna21_HisGTG_1_+_147753471_147753542@-1G.34.35,trna33_HisGTG_6_+_27125906_27125977@-1G.34.35,trna7_HisGTG_9_-_14433938_14434009@-1G.34.35,trna8_HisGTG_15_-_45492611_45492682@-1G.34.35,trna9_HisGTG_15_-_45490804_45490875@-1G.34.35\|GGCCGTGATCGTATAGTGGTTAGTACTCTGCGTTG | >trna111_HisGTG_1_-_147774845_147774916 | -1G | 34 | 35 | GGCCGTGATCGTATAGTGGTTAGTACTCTGCGTTG | tRF | FALSE | TRUE | FALSE |
| >hg38_21_+_8433172_8446622_RNA45SN1_45S_50ntFlanks@476.498.23\|TTCGTGATCGATGTGGTGACGTC | >hg38_21_+_8433172_8446622_RNA45SN1_45S_50ntFlanks | 476 | 498 | 23 | TTCGTGATCGATGTGGTGACGTC | rRF | FALSE | TRUE | FALSE |
| >hg38_21_+_8433172_8446622_RNA45SN1_45S_50ntFlanks@476.524.49\|TTCGTGATCGATGTGGTGACGTCGTGCTCTCCCGGGCCGGGTCCGAGCC | >hg38_21_+_8433172_8446622_RNA45SN1_45S_50ntFlanks | 476 | 524 | 49 | TTCGTGATCGATGTGGTGACGTCGTGCTCTCCCGGGCCGGGTC | rRF | FALSE | TRUE | FALSE |
| >hg38_21_+_8433172_8446622_RNA45SN1_45S_50ntFlanks@6651.6698.48\|CGACTCTTAGCGGTGGATCACTCGGCTCGTGCGTCGATGAAGAACGCA | >hg38_21_+_8433172_8446622_RNA45SN1_45S_50ntFlanks | 6651 | 6698 | 48 | CGACTCTTAGCGGTGGATCACTCGGCTCGTGCGTCGATGAAGA | rRF | FALSE | TRUE | TRUE |
| >trnaMT_GluTTC_MT_-_14674_14742@37.72.36,trnalookalike8_GluTTC_5_-_93905172_93905240@37.72.36\|TCATTGGTCGTGGTTGTAGTCCGTGCGAGAATACCA | >trnaMT_GluTTC_MT_-_14674_14742 | 37 | 72 | 36 | TCATTGGTCGTGGTTGTAGTCCGTGCGAGAATACCA | tRF | FALSE | TRUE | FALSE |
| >trnaMT_GluTTC_MT_-_14674_14742@41.72.32,trnalookalike8_GluTTC_5_-_93905172_93905240@41.72.32\|TGGTCGTGGTTGTAGTCCGTGCGAGAATACCA | >trnaMT_GluTTC_MT_-_14674_14742 | 41 | 72 | 32 | TGGTCGTGGTTGTAGTCCGTGCGAGAATACCA | tRF | FALSE | TRUE | FALSE |
| >MI0000071\|hsa-mir-17&WithFlank&13\|+\|91350599\|91350694@20.42.23&[MIMAT0000070&hsa-miR-17-5p&offsets\|0\|0;m-88&13\|+\|91350618\|91350640&offsets\|0\|0]\|CAAAGTGCTTACAGTGCAGGTAG | >MI0000071\|hsa-mir-17&WithFlank&13\|+\|91350599\|91350694 | 20 | 42 | 23 | CAAAGTGCTTACAGTGCAGGTAG | isomiR | FALSE | TRUE | FALSE |
| >hg38_21_+_8433172_8446622_RNA45SN1_45S_50ntFlanks@12999.13039.41\|CTGCGATCTATTGAAAGTCAGCCCTCGACACAAGGGTTTGT | >hg38_21_+_8433172_8446622_RNA45SN1_45S_50ntFlanks | 12999 | 13039 | 41 | CTGCGATCTATTGAAAGTCAGCCCTCGACACAAGGGTTTGT | rRF | FALSE | TRUE | FALSE |
| >hg38_21_+_8433172_8446622_RNA45SN1_45S_50ntFlanks@6652.6681.30\|GACTCTTAGCGGTGGATCACTCGGCTCGTG | >hg38_21_+_8433172_8446622_RNA45SN1_45S_50ntFlanks | 6652 | 6681 | 30 | GACTCTTAGCGGTGGATCACTCGGCTCGTG | rRF | FALSE | TRUE | TRUE |
| >hg38_21_+_8433172_8446622_RNA45SN1_45S_50ntFlanks@6647.6681.35\|CGTACGACTCTTAGCGGTGGATCACTCGGCTCGTG | >hg38_21_+_8433172_8446622_RNA45SN1_45S_50ntFlanks | 6647 | 6681 | 35 | CGTACGACTCTTAGCGGTGGATCACTCGGCTCGTG | rRF | FALSE | TRUE | TRUE |
| >hg38_21_+_8433172_8446622_RNA45SN1_45S_50ntFlanks@475.509.35\|CTTCGTGATCGATGTGGTGACGTCGTGCTCTCCCG | >hg38_21_+_8433172_8446622_RNA45SN1_45S_50ntFlanks | 475 | 509 | 35 | CTTCGTGATCGATGTGGTGACGTCGTGCTCTCCCG | rRF | FALSE | TRUE | FALSE |
| >hg38_21_+_8433172_8446622_RNA45SN1_45S_50ntFlanks@485.520.36\|GATGTGGTGACGTCGTGCTCTCCCGGGCCGGGTCCG | >hg38_21_+_8433172_8446622_RNA45SN1_45S_50ntFlanks | 485 | 520 | 36 | GATGTGGTGACGTCGTGCTCTCCCGGGCCGGGTCCG | rRF | FALSE | TRUE | FALSE |
| >MI0000434\|hsa-let-7i&WithFlank&12\|+\|62603680\|62603775@12.32.21&[MIMAT0000415&hsa-let-7i-5p&offsets\|0\|-1;m-49&12\|+\|62603691\|62603712&offsets\|0\|-1]\|TGAGGTAGTAGTTTGTGCTGT | >MI0000434\|hsa-let-7i&WithFlank&12\|+\|62603680\|62603775 | 12 | 32 | 21 | TGAGGTAGTAGTTTGTGCTGT | isomiR | FALSE | TRUE | FALSE |
| >MI0000093\|hsa-mir-92a-1&WithFlank&13\|+\|91351308\|91351397@54.74.21(+1C);MI0000094\|hsa-mir-92a-2&WithFlank&X\|-\|134169532\|134169618@54.74.21(+1C)&[MIMAT0000092&hsa-miR-92a-3p&offsets\|0\|-1(+1C);m-9&13\|+\|91351361\|91351382&offsets\|0\|-1(+1C)];[MIMAT0000092_1&hsa-miR-92a-3p&offsets\|0\|-1(+1C);m-11&X\|-\|134169544\|134169565&offsets\|0\|-1(+1C)]\|TATTGCACTTGTCCCGGCCTGC | >MI0000093\|hsa-mir-92a-1&WithFlank&13\|+\|91351308\|91351397 | 54 | 74 | 21(+1C) | TATTGCACTTGTCCCGGCCTGC | isomiR | FALSE | TRUE | FALSE |
| >MI0000093\|hsa-mir-92a-1&WithFlank&13\|+\|91351308\|91351397@54.75.22(+1A);MI0000094\|hsa-mir-92a-2&WithFlank&X\|-\|134169532\|134169618@54.75.22(+1A)&[MIMAT0000092&hsa-miR-92a-3p&offsets\|0\|0(+1A);m-9&13\|+\|91351361\|91351382&offsets\|0\|0(+1A)];[MIMAT0000092_1&hsa-miR-92a-3p&offsets\|0\|0(+1A);m-11&X\|-\|134169544\|134169565&offsets\|0\|0(+1A)]\|TATTGCACTTGTCCCGGCCTGTA | >MI0000093\|hsa-mir-92a-1&WithFlank&13\|+\|91351308\|91351397 | 54 | 75 | 22(+1A) | TATTGCACTTGTCCCGGCCTGTA | isomiR | FALSE | TRUE | FALSE |
| >MI0000434\|hsa-let-7i&WithFlank&12\|+\|62603680\|62603775@12.33.22(+1U)&[MIMAT0000415&hsa-let-7i-5p&offsets\|0\|0(+1U);m-49&12\|+\|62603691\|62603712&offsets\|0\|0(+1U)]\|TGAGGTAGTAGTTTGTGCTGTTT | >MI0000434\|hsa-let-7i&WithFlank&12\|+\|62603680\|62603775 | 12 | 33 | 22(+1U) | TGAGGTAGTAGTTTGTGCTGTTT | isomiR | FALSE | TRUE | FALSE |
| >hg38_21_+_8433172_8446622_RNA45SN1_45S_50ntFlanks@475.524.50\|CTTCGTGATCGATGTGGTGACGTCGTGCTCTCCCGGGCCGGGTCCGAGCC | >hg38_21_+_8433172_8446622_RNA45SN1_45S_50ntFlanks | 475 | 524 | 50 | CTTCGTGATCGATGTGGTGACGTCGTGCTCTCCCGGGCCGGGT | rRF | FALSE | TRUE | FALSE |
| >hg38_21_+_8433172_8446622_RNA45SN1_45S_50ntFlanks@476.499.24\|TTCGTGATCGATGTGGTGACGTCG | >hg38_21_+_8433172_8446622_RNA45SN1_45S_50ntFlanks | 476 | 499 | 24 | TTCGTGATCGATGTGGTGACGTCG | rRF | FALSE | TRUE | FALSE |
| >MI0000102\|hsa-mir-100&WithFlank&11\|-\|122152223\|122152314@19.40.22(+1C)&[MIMAT0000098&hsa-miR-100-5p&offsets\|0\|0(+1C);m-27&11\|-\|122152276\|122152296&offsets\|0\|+1(+1C)]\|AACCCGTAGATCCGAACTTGTGC | >MI0000102\|hsa-mir-100&WithFlank&11\|-\|122152223\|122152314 | 19 | 40 | 22(+1C) | AACCCGTAGATCCGAACTTGTGC | isomiR | FALSE | TRUE | FALSE |
| >hg38_21_+_8433172_8446622_RNA45SN1_45S_50ntFlanks@6661.6683.23\|CGGTGGATCACTCGGCTCGTGCG | >hg38_21_+_8433172_8446622_RNA45SN1_45S_50ntFlanks | 6661 | 6683 | 23 | CGGTGGATCACTCGGCTCGTGCG | rRF | FALSE | TRUE | FALSE |

### Nucleus

<table>
<tr><td>>hg38_21_+_8433172_8446622_RNA45SN1_45S_50ntFlanks@485.521.37|GATGTGGTGACGTCGTGCTCTCCCGGGCCGGGTCCGA</td><td>>hg38_21_+_8433172_8446622_RNA45SN1_45S_50ntFlanks</td><td>485</td><td>521</td><td>37</td><td>GATGTGGTGACGTCGTGCTCTCCCGGGCCGGGTCCGA</td><td>rRF</td><td>FALSE</td><td>TRUE</td><td>FALSE</td></tr>
<tr><td>>hg38_21_+_8433172_8446622_RNA45SN1_45S_50ntFlanks@476.509.34|TTCGTGATCGATGTGGTGACGTCGTGCTCTCCCG</td><td>>hg38_21_+_8433172_8446622_RNA45SN1_45S_50ntFlanks</td><td>476</td><td>509</td><td>34</td><td>TTCGTGATCGATGTGGTGACGTCGTGCTCTCCCG</td><td>rRF</td><td>FALSE</td><td>TRUE</td><td>FALSE</td></tr>
<tr><td>>am_trna128_GlyGCC_6_-_27870686_27870756@1.32.32,trna133_GlyCCC_1_-_16872434_16872504@1.32.32,trna18_GlyGCC_16_+_70822597_70822667@1.32.32,trna19_GlyGCC_16_+_70823410_70823480@1.32.32,trna19_GlyGCC_2_-_157257659_157257729@1.32.32,trna24_GlyGCC_16_-_70812942_70813012@1.32.32,trna25_GlyGCC_16_-_70812114_70812184@1.32.32,trna4_GlyCCC_1_+_17188416_17188486@1.32.32,trna5_GlyGCC_17_+_8029064_8029134@1.32.32,trna68_GlyGCC_1_-_161493637_161493707@1.32.32|GCATTGGTGGTTCAGTGGTAGAATTCTCGCCT</td><td>>am_trna128_GlyGCC_6_-_27870686_27870756</td><td>1</td><td>32</td><td>32</td><td>GCATTGGTGGTTCAGTGGTAGAATTCTCGCCT</td><td>tRF</td><td>FALSE</td><td>TRUE</td><td>FALSE</td></tr>
<tr><td>>hg38_21_+_8433172_8446622_RNA45SN1_45S_50ntFlanks@6651.6692.42|CGACTCTTAGCGGTGGATCACTCGGCTCGTGCGTCGATGAAG</td><td>>hg38_21_+_8433172_8446622_RNA45SN1_45S_50ntFlanks</td><td>6651</td><td>6692</td><td>42</td><td>CGACTCTTAGCGGTGGATCACTCGGCTCGTGCGTCGATGAAG</td><td>rRF</td><td>FALSE</td><td>TRUE</td><td>FALSE</td></tr>
<tr><td>>hg38_21_+_8433172_8446622_RNA45SN1_45S_50ntFlanks@6648.6681.34|GTACGACTCTTAGCGGTGGATCACTCGGCTCGTG</td><td>>hg38_21_+_8433172_8446622_RNA45SN1_45S_50ntFlanks</td><td>6648</td><td>6681</td><td>34</td><td>GTACGACTCTTAGCGGTGGATCACTCGGCTCGTG</td><td>rRF</td><td>FALSE</td><td>TRUE</td><td>FALSE</td></tr>
<tr><td>>hg38_21_+_8433172_8446622_RNA45SN1_45S_50ntFlanks@6650.6683.34|ACGACTCTTAGCGGTGGATCACTCGGCTCGTGCG</td><td>>hg38_21_+_8433172_8446622_RNA45SN1_45S_50ntFlanks</td><td>6650</td><td>6683</td><td>34</td><td>ACGACTCTTAGCGGTGGATCACTCGGCTCGTGCG</td><td>rRF</td><td>FALSE</td><td>TRUE</td><td>FALSE</td></tr>
<tr><td>>trnaMT_GluTTC_MT_-_14674_14742@35.72.38,trnalookalike8_GluTTC_5_-_93905172_93905240@35.72.38|TATCATTGGTCGTGGTTGTAGTCCGTGCGAGAATACCA</td><td>>trnaMT_GluTTC_MT_-_14674_14742</td><td>35</td><td>72</td><td>38</td><td>TATCATTGGTCGTGGTTGTAGTCCGTGCGAGAATACCA</td><td>tRF</td><td>FALSE</td><td>TRUE</td><td>FALSE</td></tr>
<tr><td>>hg38_21_+_8433172_8446622_RNA45SN1_45S_50ntFlanks@6651.6679.29|CGACTCTTAGCGGTGGATCACTCGGCTCG</td><td>>hg38_21_+_8433172_8446622_RNA45SN1_45S_50ntFlanks</td><td>6651</td><td>6679</td><td>29</td><td>CGACTCTTAGCGGTGGATCACTCGGCTCG</td><td>rRF</td><td>FALSE</td><td>TRUE</td><td>TRUE</td></tr>
<tr><td>>MI0000088|hsa-mir-30a&WithFlank&6|-|71403545|71403627@12.34.23(+1A)&[MIMAT0000087&hsa-miR-30a-5p&offsets|0|+1(+1A);m-15&6|-|71403595|71403616&offsets|0|+1(+1A)]|TGTAAACATCCTCGACTGGAAGCA</td><td>>MI0000088|hsa-mir-30a&WithFlank&6|-|71403545|71403627</td><td>12</td><td>34</td><td>23(+1A)</td><td>TGTAAACATCCTCGACTGGAAGCA</td><td>isomiR</td><td>FALSE</td><td>TRUE</td><td>FALSE</td></tr>
<tr><td>>hg38_21_+_8433172_8446622_RNA45SN1_45S_50ntFlanks@482.505.24|ATCGATGTGGTGACGTCGTGCTCT</td><td>>hg38_21_+_8433172_8446622_RNA45SN1_45S_50ntFlanks</td><td>482</td><td>505</td><td>24</td><td>ATCGATGTGGTGACGTCGTGCTCT</td><td>rRF</td><td>FALSE</td><td>TRUE</td><td>FALSE</td></tr>
<tr><td>>MI0000299|hsa-mir-222&WithFlank&X|-|45747009|45747130@75.98.24(+1C)&[MIMAT0000279&hsa-miR-222-3p&offsets|0|+3(+1C);m-140&X|-|45747033|45747056&offsets|0|0(+1C)]|AGCTACATCTGGCTACTGGGTCTCC</td><td>>MI0000299|hsa-mir-222&WithFlank&X|-|45747009|45747130</td><td>75</td><td>98</td><td>24(+1C)</td><td>AGCTACATCTGGCTACTGGGTCTCC</td><td>isomiR</td><td>FALSE</td><td>TRUE</td><td>FALSE</td></tr>
<tr><td>>hg38_21_+_8433172_8446622_RNA45SN1_45S_50ntFlanks@485.525.41|GATGTGGTGACGTCGTGCTCTCCCGGGCCGGGTCCGAGCCG</td><td>>hg38_21_+_8433172_8446622_RNA45SN1_45S_50ntFlanks</td><td>485</td><td>525</td><td>41</td><td>GATGTGGTGACGTCGTGCTCTCCCGGGCCGGGTCCGAGCCG</td><td>rRF</td><td>FALSE</td><td>TRUE</td><td>FALSE</td></tr>
<tr><td>>MI0000102|hsa-mir-100&WithFlank&11|-|122152223|122152314@19.39.21(+1U)&[MIMAT0000098&hsa-miR-100-5p&offsets|0|-1(+1U);m-27&11|-|122152276|122152296&offsets|0|0(+1U)]|AACCCGTAGATCCGAACTTGTT</td><td>>MI0000102|hsa-mir-100&WithFlank&11|-|122152223|122152314</td><td>19</td><td>39</td><td>21(+1U)</td><td>AACCCGTAGATCCGAACTTGTT</td><td>isomiR</td><td>FALSE</td><td>TRUE</td><td>FALSE</td></tr>
<tr><td>>hg38_21_+_8433172_8446622_RNA45SN1_45S_50ntFlanks@475.495.21|CTTCGTGATCGATGTGGTGAC</td><td>>hg38_21_+_8433172_8446622_RNA45SN1_45S_50ntFlanks</td><td>475</td><td>495</td><td>21</td><td>CTTCGTGATCGATGTGGTGAC</td><td>rRF</td><td>FALSE</td><td>TRUE</td><td>FALSE</td></tr>
<tr><td>>MI0000734|hsa-mir-106b&WithFlank&7|-|100093987|100094080@58.79.22&[MIMAT0004672&hsa-miR-106b-3p&offsets|0|0;m-87&7|-|100094003|100094023&offsets|0|+1]|CCGCACTGTGGGTACTTGCTGC</td><td>>MI0000734|hsa-mir-106b&WithFlank&7|-|100093987|100094080</td><td>58</td><td>79</td><td>22</td><td>CCGCACTGTGGGTACTTGCTGC</td><td>isomiR</td><td>FALSE</td><td>TRUE</td><td>FALSE</td></tr>
<tr><td>>hg38_21_+_8433172_8446622_RNA45SN1_45S_50ntFlanks@6651.6682.32|CGACTCTTAGCGGTGGATCACTCGGCTCGTGC</td><td>>hg38_21_+_8433172_8446622_RNA45SN1_45S_50ntFlanks</td><td>6651</td><td>6682</td><td>32</td><td>CGACTCTTAGCGGTGGATCACTCGGCTCGTGC</td><td>rRF</td><td>FALSE</td><td>TRUE</td><td>FALSE</td></tr>
<tr><td>>hg38_21_+_8433172_8446622_RNA45SN1_45S_50ntFlanks@485.522.38|GATGTGGTGACGTCGTGCTCTCCCGGGCCGGGTCCGAG</td><td>>hg38_21_+_8433172_8446622_RNA45SN1_45S_50ntFlanks</td><td>485</td><td>522</td><td>38</td><td>GATGTGGTGACGTCGTGCTCTCCCGGGCCGGGTCCGAG</td><td>rRF</td><td>FALSE</td><td>TRUE</td><td>FALSE</td></tr>
<tr><td>>hg38_21_+_8433172_8446622_RNA45SN1_45S_50ntFlanks@476.518.43|TTCGTGATCGATGTGGTGACGTCGTGCTCTCCCGGGCCGGGTC</td><td>>hg38_21_+_8433172_8446622_RNA45SN1_45S_50ntFlanks</td><td>476</td><td>518</td><td>43</td><td>TTCGTGATCGATGTGGTGACGTCGTGCTCTCCCGGGCCGGGTC</td><td>rRF</td><td>FALSE</td><td>TRUE</td><td>FALSE</td></tr>
<tr><td>>MI0000736|hsa-mir-30c-1&WithFlank&1|+|40757278|40757378@23.44.22(+1U);MI0000254|hsa-mir-30c-2&WithFlank&6|-|71376954|71377037@13.34.22(+1U)&[MIMAT0000244&hsa-miR-30c-5p&offsets|0|-1(+1U);m-57&1|+|40757300|40757323&offsets|0|-2(+1U)];[MIMAT0000244_1&hsa-miR-30c-5p&offsets|0|-1(+1U);m-56&6|-|71377002|71377025&offsets|0|-2(+1U)]|TGTAAACATCCTACACTCTCAGT</td><td>>MI0000736|hsa-mir-30c-1&WithFlank&1|+|40757278|40757378</td><td>23</td><td>44</td><td>22(+1U)</td><td>TGTAAACATCCTACACTCTCAGT</td><td>isomiR</td><td>FALSE</td><td>TRUE</td><td>FALSE</td></tr>
<tr><td>>hg38_21_+_8433172_8446622_RNA45SN1_45S_50ntFlanks@6652.6698.47|GACTCTTAGCGGTGGATCACTCGGCTCGTGCGTCGATGAAGAACGCA</td><td>>hg38_21_+_8433172_8446622_RNA45SN1_45S_50ntFlanks</td><td>6652</td><td>6698</td><td>47</td><td>GACTCTTAGCGGTGGATCACTCGGCTCGTGCGTCGATGAAGAA</td><td>rRF</td><td>FALSE</td><td>TRUE</td><td>TRUE</td></tr>
<tr><td>>hg38_21_+_8433172_8446622_RNA45SN1_45S_50ntFlanks@476.495.20|TTCGTGATCGATGTGGTGAC</td><td>>hg38_21_+_8433172_8446622_RNA45SN1_45S_50ntFlanks</td><td>476</td><td>495</td><td>20</td><td>TTCGTGATCGATGTGGTGAC</td><td>rRF</td><td>FALSE</td><td>TRUE</td><td>FALSE</td></tr>
<tr><td>>hg38_21_+_8433172_8446622_RNA45SN1_45S_50ntFlanks@476.507.32|TTCGTGATCGATGTGGTGACGTCGTGCTCTCC</td><td>>hg38_21_+_8433172_8446622_RNA45SN1_45S_50ntFlanks</td><td>476</td><td>507</td><td>32</td><td>TTCGTGATCGATGTGGTGACGTCGTGCTCTCC</td><td>rRF</td><td>FALSE</td><td>TRUE</td><td>FALSE</td></tr>
<tr><td>>hg38_21_+_8433172_8446622_RNA45SN1_45S_50ntFlanks@6645.6662.18|CTCGTACGACTCTTAGCG</td><td>>hg38_21_+_8433172_8446622_RNA45SN1_45S_50ntFlanks</td><td>6645</td><td>6662</td><td>18</td><td>CTCGTACGACTCTTAGCG</td><td>rRF</td><td>FALSE</td><td>TRUE</td><td>FALSE</td></tr>
<tr><td>>hg38_21_+_8433172_8446622_RNA45SN1_45S_50ntFlanks@476.503.28|TTCGTGATCGATGTGGTGACGTCGTGCT</td><td>>hg38_21_+_8433172_8446622_RNA45SN1_45S_50ntFlanks</td><td>476</td><td>503</td><td>28</td><td>TTCGTGATCGATGTGGTGACGTCGTGCT</td><td>rRF</td><td>FALSE</td><td>TRUE</td><td>FALSE</td></tr>
<tr><td>>MI0000085|hsa-mir-27a&WithFlank&19|-|13836434|13836523@57.76.20(+1U)&[MIMAT0000084&hsa-miR-27a-3p&offsets|0|-1(+1U);m-43&19|-|13836447|13836467&offsets|0|-1(+1U)]|TTCACAGTGGCTAAGTTCCGT</td><td>>MI0000085|hsa-mir-27a&WithFlank&19|-|13836434|13836523</td><td>57</td><td>76</td><td>20(+1U)</td><td>TTCACAGTGGCTAAGTTCCGT</td><td>isomiR</td><td>FALSE</td><td>TRUE</td><td>FALSE</td></tr>
<tr><td>>trna111_HisGTG_1_-_147774845_147774916@-1T.23.24,trna118_HisGTG_1_-_145396881_145396952@-1T.23.24,trna16_HisGTG_1_+_146544773_146544844@-1T.23.24,trna1_HisGTG_15_+_45493349_45493420@-1T.23.24,trna21_HisGTG_1_+_147753471_147753542@-1T.23.24,trna33_HisGTG_6_+_27125906_27125977@-1T.23.24,trna7_HisGTG_9_-_14433938_14434009@-1T.23.24,trna8_HisGTG_15_-_45492611_45492682@-1T.23.24,trna9_HisGTG_15_-_45490804_45490875@-1T.23.24|TGCCGTGATCGTATAGTGGTTAGT</td><td>>trna111_HisGTG_1_-_147774845_147774916</td><td>-1T</td><td>23</td><td>24</td><td>TGCCGTGATCGTATAGTGGTTAGT</td><td>tRF</td><td>FALSE</td><td>TRUE</td><td>FALSE</td></tr>
<tr><td>>MI0000299|hsa-mir-222&WithFlank&X|-|45747009|45747130@75.97.23&[MIMAT0000279&hsa-miR-222-3p&offsets|0|+2;m-140&X|-|45747033|45747056&offsets|0|-1]|AGCTACATCTGGCTACTGGGTCT</td><td>>MI0000299|hsa-mir-222&WithFlank&X|-|45747009|45747130</td><td>75</td><td>97</td><td>23</td><td>AGCTACATCTGGCTACTGGGTCT</td><td>isomiR</td><td>FALSE</td><td>TRUE</td><td>FALSE</td></tr>
</table>

### Nucleus

| | | | | | | | | | |
|---|---|---|---|---|---|---|---|---|---|
| >trna116_GluCTC_1_-_145399233_145399304@37.75.39,trna134_GluTTC_1_-_16861774_16861845@37.75.39,trna71_GluCTC_1_-_161439189_161439260@37.75.39,trna74_GluCTC_1_-_161431809_161431880@37.75.39,trna77_GluCTC_1_-_161424398_161424469@37.75.39,trna77_GluCTC_6_+_28949976_28950047@37.75.39,trna80_GluCTC_1_-_161417018_161417089@37.75.39,trna84_GluTTC_1_-_161391883_161391954@37.75.39,trna87_GluCTC_6_-_126101393_126101464@37.75.39\|ACCGCCGCGGCCCGGGTTCGATTCCCGGTCAGGGAACCA | >trna116_GluCTC_1_-_145399233_145399304 | 37 | 75 | 39 | ACCGCCGCGGCCCGGGTTCGATTCCCGGTCAGGGAACCA | tRF | FALSE | TRUE | FALSE |
| >hg38_21_+_8433172_8446622_RNA45SN1_45S_50ntFlanks@485.516.32\|GATGTGGTGACGTCGTGCTCTCCCGGGCCGGG | >hg38_21_+_8433172_8446622_RNA45SN1_45S_50ntFlanks | 485 | 516 | 32 | GATGTGGTGACGTCGTGCTCTCCCGGGCCGGG | rRF | FALSE | TRUE | FALSE |
| >hg38_21_+_8433172_8446622_RNA45SN1_45S_50ntFlanks@12998.13039.42\|GCTGCGATCTATTGAAAGTCAGCCCTCGACACAAGGGTTTGT | >hg38_21_+_8433172_8446622_RNA45SN1_45S_50ntFlanks | 12998 | 13039 | 42 | GCTGCGATCTATTGAAAGTCAGCCCTCGACACAAGGGTTTGT | rRF | FALSE | TRUE | FALSE |
| >hg38_21_+_8433172_8446622_RNA45SN1_45S_50ntFlanks@476.510.35\|TTCGTGATCGATGTGGTGACGTCGTGCTCTCCCGG | >hg38_21_+_8433172_8446622_RNA45SN1_45S_50ntFlanks | 476 | 510 | 35 | TTCGTGATCGATGTGGTGACGTCGTGCTCTCCCGG | rRF | FALSE | TRUE | FALSE |
| >MI0000440\|hsa-mir-27b&WithFlank&9\|+\|95085439\|95085547@67.88.22&[MIMAT0000419&hsa-miR-27b-3p&offsets\|0\|+1;m-39&9\|+\|95085505\|95085525&offsets\|0\|+1]\|TTCACAGTGGCTAAGTTCTGCA | >MI0000440\|hsa-mir-27b&WithFlank&9\|+\|95085439\|95085547 | 67 | 88 | 22 | TTCACAGTGGCTAAGTTCTGCA | isomiR | FALSE | TRUE | FALSE |
| >hg38_21_+_8433172_8446622_RNA45SN1_45S_50ntFlanks@475.511.37\|CTTCGTGATCGATGTGGTGACGTCGTGCTCTCCCGGG | >hg38_21_+_8433172_8446622_RNA45SN1_45S_50ntFlanks | 475 | 511 | 37 | CTTCGTGATCGATGTGGTGACGTCGTGCTCTCCCGGG | rRF | FALSE | TRUE | FALSE |
| >hg38_21_+_8433172_8446622_RNA45SN1_45S_50ntFlanks@475.514.40\|CTTCGTGATCGATGTGGTGACGTCGTGCTCTCCCGGGCCG | >hg38_21_+_8433172_8446622_RNA45SN1_45S_50ntFlanks | 475 | 514 | 40 | CTTCGTGATCGATGTGGTGACGTCGTGCTCTCCCGGGCCG | rRF | FALSE | TRUE | FALSE |
| >MI0000082\|hsa-mir-25&WithFlank&7\|-\|100093554\|100093649@58.78.21&[MIMAT0000081&hsa-miR-25-3p&offsets\|0\|-1;m-23&7\|-\|100093571\|100093592&offsets\|0\|-1]\|CATTGCACTTGTCTCGGTCTG | >MI0000082\|hsa-mir-25&WithFlank&7\|-\|100093554\|100093649 | 58 | 78 | 21 | CATTGCACTTGTCTCGGTCTG | isomiR | FALSE | TRUE | FALSE |
| >hg38_21_+_8433172_8446622_RNA45SN1_45S_50ntFlanks@485.524.40\|GATGTGGTGACGTCGTGCTCTCCCGGGCCGGGTCCGAGCC | >hg38_21_+_8433172_8446622_RNA45SN1_45S_50ntFlanks | 485 | 524 | 40 | GATGTGGTGACGTCGTGCTCTCCCGGGCCGGGTCCGAGCC | rRF | FALSE | TRUE | FALSE |
| >hg38_21_+_8433172_8446622_RNA45SN1_45S_50ntFlanks@475.518.44\|CTTCGTGATCGATGTGGTGACGTCGTGCTCTCCCGGGCCGGGTC | >hg38_21_+_8433172_8446622_RNA45SN1_45S_50ntFlanks | 475 | 518 | 44 | CTTCGTGATCGATGTGGTGACGTCGTGCTCTCCCGGGCCGGGT | rRF | FALSE | TRUE | FALSE |
| >hg38_21_+_8433172_8446622_RNA45SN1_45S_50ntFlanks@475.507.33\|CTTCGTGATCGATGTGGTGACGTCGTGCTCTCC | >hg38_21_+_8433172_8446622_RNA45SN1_45S_50ntFlanks | 475 | 507 | 33 | CTTCGTGATCGATGTGGTGACGTCGTGCTCTCC | rRF | FALSE | TRUE | FALSE |
| >hg38_21_+_8433172_8446622_RNA45SN1_45S_50ntFlanks@482.531.50\|ATCGATGTGGTGACGTCGTGCTCTCCCGGGCCGGGTCCGAGCCGCGACGG | >hg38_21_+_8433172_8446622_RNA45SN1_45S_50ntFlanks | 482 | 531 | 50 | ATCGATGTGGTGACGTCGTGCTCTCCCGGGCCGGGTCCGAGC | rRF | FALSE | TRUE | FALSE |
| >MI0000093\|hsa-mir-92a-1&WithFlank&13\|+\|91351308\|91351397@54.76.23&[MIMAT0000092&hsa-miR-92a-3p&offsets\|0\|+1;m-9&13\|+\|91351361\|91351382&offsets\|0\|+1]\|TATTGCACTTGTCCCGGCCTGTT | >MI0000093\|hsa-mir-92a-1&WithFlank&13\|+\|91351308\|91351397 | 54 | 76 | 23 | TATTGCACTTGTCCCGGCCTGTT | isomiR | FALSE | TRUE | FALSE |
| >hg38_21_+_8433172_8446622_RNA45SN1_45S_50ntFlanks@6649.6683.35\|TACGACTCTTAGCGGTGGATCACTCGGCTCGTGCG | >hg38_21_+_8433172_8446622_RNA45SN1_45S_50ntFlanks | 6649 | 6683 | 35 | TACGACTCTTAGCGGTGGATCACTCGGCTCGTGCG | rRF | FALSE | TRUE | FALSE |
| >hg38_21_+_8433172_8446622_RNA45SN1_45S_50ntFlanks@6646.6675.30\|TCGTACGACTCTTAGCGGTGGATCACTCGG | >hg38_21_+_8433172_8446622_RNA45SN1_45S_50ntFlanks | 6646 | 6675 | 30 | TCGTACGACTCTTAGCGGTGGATCACTCGG | rRF | FALSE | TRUE | FALSE |
| >hg38_21_+_8433172_8446622_RNA45SN1_45S_50ntFlanks@6651.6700.50\|CGACTCTTAGCGGTGGATCACTCGGCTCGTGCGTCGATGAAGAACGCAGC | >hg38_21_+_8433172_8446622_RNA45SN1_45S_50ntFlanks | 6651 | 6700 | 50 | CGACTCTTAGCGGTGGATCACTCGGCTCGTGCGTCGATGAAGA | rRF | FALSE | TRUE | FALSE |
| >MI0000750\|hsa-mir-26a-2&WithFlank&12\|-\|57824603\|57824698@20.40.21(+1A);MI0000083\|hsa-mir-26a-1&WithFlank&3\|+\|37969398\|37969486@16.36.21(+1A)&[MIMAT0000082&hsa-miR-26a-5p&offsets\|0\|-1(+1A);m-33&12\|-\|57824658\|57824679&offsets\|0\|-1(+1A)];[MIMAT0000082_1&hsa-miR-26a-5p&offsets\|0\|-1(+1A);m-34&3\|+\|37969413\|37969434&offsets\|0\|-1(+1A)]\|TTCAAGTAATCCAGGATAGGCA | >MI0000750\|hsa-mir-26a-2&WithFlank&12\|-\|57824603\|57824698 | 20 | 40 | 21(+1A) | TTCAAGTAATCCAGGATAGGCA | isomiR | FALSE | TRUE | FALSE |
| >MI0001448\|hsa-mir-425&WithFlank&3\|-\|49020142\|49020240@20.44.25&[MIMAT0003393&hsa-miR-425-5p&offsets\|0\|+2;m-95&3\|-\|49020200\|49020221&offsets\|0\|+3]\|AATGACACGATCACTCCCGTTGAGT | >MI0001448\|hsa-mir-425&WithFlank&3\|-\|49020142\|49020240 | 20 | 44 | 25 | AATGACACGATCACTCCCGTTGAGT | isomiR | FALSE | TRUE | FALSE |
| >hg38_1_-_228634819_228635039_RNA5S12_5S_50ntFlanks@134.169.36\|TGGGAGACCGCCTGGGAATACCGGGTGCTGTAGGCT | >hg38_1_-_228634819_228635039_RNA5S12_5S_50ntFlanks | 134 | 169 | 36 | TGGGAGACCGCCTGGGAATACCGGGTGCTGTAGGCT | rRF | FALSE | TRUE | FALSE |
| >MI0000289\|hsa-mir-181a-1&WithFlank&1\|-\|198859038\|198859159@30.52.23;MI0000269\|hsa-mir-181a-2&WithFlank&9\|+\|124692436\|124692557@45.67.23&[MIMAT0000256&hsa-miR-181a-5p&offsets\|0\|0;m-32&1\|-\|198859108\|198859130&offsets\|0\|0];[MIMAT0000256_1&hsa-miR-181a-5p&offsets\|0\|0;m-32&9\|+\|124692480\|124692502&offsets\|0\|0]\|AACATTCAACGCTGTCGGTGAGT | >MI0000289\|hsa-mir-181a-1&WithFlank&1\|-\|198859038\|198859159 | 30 | 52 | 23 | AACATTCAACGCTGTCGGTGAGT | isomiR | FALSE | TRUE | FALSE |
| >trnaMT_ValTAC_MT_+_1602_1670@42.72.31\|AGATTTCAACTTAACTTGACCGCTCTGACCA | >trnaMT_ValTAC_MT_+_1602_1670 | 42 | 72 | 31 | AGATTTCAACTTAACTTGACCGCTCTGACCA | tRF | FALSE | TRUE | FALSE |
| >MI0000434\|hsa-let-7i&WithFlank&12\|+\|62603680\|62603775@12.33.22(+1A)&[MIMAT0000415&hsa-let-7i-5p&offsets\|0\|0(+1A);m-49&12\|+\|62603691\|62603712&offsets\|0\|0(+1A)]\|TGAGGTAGTAGTTTGTGCTGTTA | >MI0000434\|hsa-let-7i&WithFlank&12\|+\|62603680\|62603775 | 12 | 33 | 22(+1A) | TGAGGTAGTAGTTTGTGCTGTTA | isomiR | FALSE | TRUE | FALSE |
| >trnaMT_GluTTC_MT_-_14674_14742@38.72.35,trnalookalike8_GluTTC_5_-_93905172_93905240@38.72.35\|CATTGGTCGTGGTTGTAGTCCGTGCGAGAATACCA | >trnaMT_GluTTC_MT_-_14674_14742 | 38 | 72 | 35 | CATTGGTCGTGGTTGTAGTCCGTGCGAGAATACCA | tRF | FALSE | TRUE | FALSE |
| >hg38_21_+_8433172_8446622_RNA45SN1_45S_50ntFlanks@6652.6696.45\|GACTCTTAGCGGTGGATCACTCGGCTCGTGCGTCGATGAAGAACG | >hg38_21_+_8433172_8446622_RNA45SN1_45S_50ntFlanks | 6652 | 6696 | 45 | GACTCTTAGCGGTGGATCACTCGGCTCGTGCGTCGATGAAGAA | rRF | FALSE | TRUE | FALSE |
| >trna119_LysCTT_1_-_145395522_145395594@1.34.34,trna11_LysCTT_5_-_180648979_180649051@1.34.34,trna13_LysCTT_6_+_26556774_26556846@1.34.34,trna7_LysCTT_16_+_3225692_3225764@1.34.34,trna9_LysCTT_5_+_180634755_180634827@1.34.34\|GCCCGGCTAGCTCAGTCGGTAGAGCATGAGACTC | >trna119_LysCTT_1_-_145395522_145395594 | 1 | 34 | 34 | GCCCGGCTAGCTCAGTCGGTAGAGCATGAGACTC | tRF | FALSE | TRUE | FALSE |
| >hg38_1_-_228634819_228635039_RNA5S12_5S_50ntFlanks@144.171.28\|CCTGGGAATACCGGGTGCTGTAGGCTTT | >hg38_1_-_228634819_228635039_RNA5S12_5S_50ntFlanks | 144 | 171 | 28 | CCTGGGAATACCGGGTGCTGTAGGCTTT | rRF | FALSE | TRUE | FALSE |
| >hg38_21_+_8433172_8446622_RNA45SN1_45S_50ntFlanks@475.523.49\|CTTCGTGATCGATGTGGTGACGTCGTGCTCTCCCGGGCCGGGTCCGAGC | >hg38_21_+_8433172_8446622_RNA45SN1_45S_50ntFlanks | 475 | 523 | 49 | CTTCGTGATCGATGTGGTGACGTCGTGCTCTCCCGGGCCGGGT | rRF | FALSE | TRUE | FALSE |
| >MI0000434\|hsa-let-7i&WithFlank&12\|+\|62603680\|62603775@12.32.21(+1A)&[MIMAT0000415&hsa-let-7i-5p&offsets\|0\|-1(+1A);m-49&12\|+\|62603691\|62603712&offsets\|0\|-1(+1A)]\|TGAGGTAGTAGTTTGTGCTGTA | >MI0000434\|hsa-let-7i&WithFlank&12\|+\|62603680\|62603775 | 12 | 32 | 21(+1A) | TGAGGTAGTAGTTTGTGCTGTA | isomiR | FALSE | TRUE | FALSE |

### Nucleus

| | | | | | | | | | |
|---|---|---|---|---|---|---|---|---|---|
| >hg38_21_+_8433172_8446622_RNA45SN1_45S_50ntFlanks@11729.11749.21\|TAACTATGACTCTCTTAAGGT | >hg38_21_+_8433172_8446622_RNA45SN1_45S_50ntFlanks | 11729 | 11749 | 21 | TAACTATGACTCTCTTAAGGT | rRF | FALSE | FALSE | TRUE |
| >hg38_21_+_8433172_8446622_RNA45SN1_45S_50ntFlanks@5467.5499.33\|GGCCCACGGCCCTGGCGGAGCGCTGAGAAGACG | >hg38_21_+_8433172_8446622_RNA45SN1_45S_50ntFlanks | 5467 | 5499 | 33 | GGCCCACGGCCCTGGCGGAGCGCTGAGAAGACG | rRF | FALSE | FALSE | TRUE |
| >hg38_21_+_8433172_8446622_RNA45SN1_45S_50ntFlanks@6719.6744.26\|TGAATTGCAGGACACATTGATCATCG | >hg38_21_+_8433172_8446622_RNA45SN1_45S_50ntFlanks | 6719 | 6744 | 26 | TGAATTGCAGGACACATTGATCATCG | rRF | FALSE | FALSE | TRUE |
| >hg38_21_+_8433172_8446622_RNA45SN1_45S_50ntFlanks@11670.11702.33\|CCCAGTGCTCTGAATGTCAAAGTGAAGAAATTC | >hg38_21_+_8433172_8446622_RNA45SN1_45S_50ntFlanks | 11670 | 11702 | 33 | CCCAGTGCTCTGAATGTCAAAGTGAAGAAATTC | rRF | FALSE | FALSE | TRUE |
| >hg38_21_+_8433172_8446622_RNA45SN1_45S_50ntFlanks@12405.12429.25\|CTTTTTGATCCTTCGATGTCGGCTC | >hg38_21_+_8433172_8446622_RNA45SN1_45S_50ntFlanks | 12405 | 12429 | 25 | CTTTTTGATCCTTCGATGTCGGCTC | rRF | FALSE | FALSE | TRUE |
| >hg38_21_+_8433172_8446622_RNA45SN1_45S_50ntFlanks@12420.12441.22\|ATGTCGGCTCTTCCTATCATTG | >hg38_21_+_8433172_8446622_RNA45SN1_45S_50ntFlanks | 12420 | 12441 | 22 | ATGTCGGCTCTTCCTATCATTG | rRF | FALSE | FALSE | TRUE |
| >hg38_21_+_8433172_8446622_RNA45SN1_45S_50ntFlanks@11320.11344.25\|CCACGTCTCGTCGCGCGCGCGTCCG | >hg38_21_+_8433172_8446622_RNA45SN1_45S_50ntFlanks | 11320 | 11344 | 25 | CCACGTCTCGTCGCGCGCGCGTCCG | rRF | FALSE | FALSE | TRUE |
| >hg38_21_+_8433172_8446622_RNA45SN1_45S_50ntFlanks@10845.10865.21\|ACTTCGGGATAAGGATTGGCT | >hg38_21_+_8433172_8446622_RNA45SN1_45S_50ntFlanks | 10845 | 10865 | 21 | ACTTCGGGATAAGGATTGGCT | rRF | FALSE | FALSE | TRUE |
| >hg38_21_+_8433172_8446622_RNA45SN1_45S_50ntFlanks@11661.11685.25\|TGATTTCTGCCCAGTGCTCTGAATG | >hg38_21_+_8433172_8446622_RNA45SN1_45S_50ntFlanks | 11661 | 11685 | 25 | TGATTTCTGCCCAGTGCTCTGAATG | rRF | FALSE | FALSE | TRUE |
| >hg38_21_+_8433172_8446622_RNA45SN1_45S_50ntFlanks@12437.12455.19\|CATTGTGAAGCAGAATTCA | >hg38_21_+_8433172_8446622_RNA45SN1_45S_50ntFlanks | 12437 | 12455 | 19 | CATTGTGAAGCAGAATTCA | rRF | FALSE | FALSE | TRUE |
| >hg38_21_+_8433172_8446622_RNA45SN1_45S_50ntFlanks@12990.13013.24\|GCTCCCTCGCTGCGATCTATTGAA | >hg38_21_+_8433172_8446622_RNA45SN1_45S_50ntFlanks | 12990 | 13013 | 24 | GCTCCCTCGCTGCGATCTATTGAA | rRF | FALSE | FALSE | TRUE |
| >hg38_21_+_8433172_8446622_RNA45SN1_45S_50ntFlanks@11768.11794.27\|CTAATTAGTGACGCGCATGAATGGATG | >hg38_21_+_8433172_8446622_RNA45SN1_45S_50ntFlanks | 11768 | 11794 | 27 | CTAATTAGTGACGCGCATGAATGGATG | rRF | FALSE | FALSE | TRUE |
| >hg38_21_+_8433172_8446622_RNA45SN1_45S_50ntFlanks@11781.11804.24\|CGCATGAATGGATGAACGAGATTC | >hg38_21_+_8433172_8446622_RNA45SN1_45S_50ntFlanks | 11781 | 11804 | 24 | CGCATGAATGGATGAACGAGATTC | rRF | FALSE | FALSE | TRUE |
| >hg38_21_+_8433172_8446622_RNA45SN1_45S_50ntFlanks@10351.10384.34\|CGAGAACTTTGAAGGCCGAAGTGGAGAAGGGTTC | >hg38_21_+_8433172_8446622_RNA45SN1_45S_50ntFlanks | 10351 | 10384 | 34 | CGAGAACTTTGAAGGCCGAAGTGGAGAAGGGTTC | rRF | FALSE | FALSE | TRUE |
| >hg38_21_+_8433172_8446622_RNA45SN1_45S_50ntFlanks@6719.6740.22\|TGAATTGCAGGACACATTGATC | >hg38_21_+_8433172_8446622_RNA45SN1_45S_50ntFlanks | 6719 | 6740 | 22 | TGAATTGCAGGACACATTGATC | rRF | FALSE | FALSE | TRUE |
| >am_trna128_GlyGCC_6_-_27870686_27870756@1.30.30,trna133_GlyCCC_1_-_16872434_16872504@1.30.30,trna18_GlyGCC_16_+_70822597_70822667@1.30.30,trna19_GlyGCC_16_+_70823410_70823480@1.30.30,trna19_GlyGCC_2_-_157257659_157257729@1.30.30,trna24_GlyGCC_16_-_70812942_70813012@1.30.30,trna25_GlyGCC_16_-_70812114_70812184@1.30.30,trna4_GlyCCC_1_+_17188416_17188486@1.30.30,trna5_GlyGCC_17_+_8029064_8029134@1.30.30,trna68_GlyGCC_1_-_161493637_161493707@1.30.30\|GCATTGGTGGTTCAGTGGTAGAATTCTCGC | >am_trna128_GlyGCC_6_-_27870686_27870756 | 1 | 30 | 30 | GCATTGGTGGTTCAGTGGTAGAATTCTCGC | tRF | FALSE | FALSE | TRUE |
| >hg38_21_+_8433172_8446622_RNA45SN1_45S_50ntFlanks@11785.11803.19\|TGAATGGATGAACGAGATT | >hg38_21_+_8433172_8446622_RNA45SN1_45S_50ntFlanks | 11785 | 11803 | 19 | TGAATGGATGAACGAGATT | rRF | FALSE | FALSE | TRUE |
| >hg38_21_+_8433172_8446622_RNA45SN1_45S_50ntFlanks@3985.4002.18\|CGGCTTTGGTGACTCTAG | >hg38_21_+_8433172_8446622_RNA45SN1_45S_50ntFlanks | 3985 | 4002 | 18 | CGGCTTTGGTGACTCTAG | rRF | FALSE | FALSE | TRUE |
| >hg38_21_+_8433172_8446622_RNA45SN1_45S_50ntFlanks@10408.10429.22\|GGTCAGTCGGTCCTGAGAGATG | >hg38_21_+_8433172_8446622_RNA45SN1_45S_50ntFlanks | 10408 | 10429 | 22 | GGTCAGTCGGTCCTGAGAGATG | rRF | FALSE | FALSE | TRUE |
| >hg38_21_+_8433172_8446622_RNA45SN1_45S_50ntFlanks@10680.10712.33\|GTTCCGGCGGCGTCCGGTGAGCTCTCGCTGGCC | >hg38_21_+_8433172_8446622_RNA45SN1_45S_50ntFlanks | 10680 | 10712 | 33 | GTTCCGGCGGCGTCCGGTGAGCTCTCGCTGGCC | rRF | FALSE | FALSE | TRUE |
| >hg38_21_+_8433172_8446622_RNA45SN1_45S_50ntFlanks@11718.11740.23\|AACGGCGGGAGTAACTATGACTC | >hg38_21_+_8433172_8446622_RNA45SN1_45S_50ntFlanks | 11718 | 11740 | 23 | AACGGCGGGAGTAACTATGACTC | rRF | FALSE | FALSE | TRUE |
| >hg38_21_+_8433172_8446622_RNA45SN1_45S_50ntFlanks@11569.11600.32\|CAGCCGACTTAGAACTGGTGCGGACCAGGGGA | >hg38_21_+_8433172_8446622_RNA45SN1_45S_50ntFlanks | 11569 | 11600 | 32 | CAGCCGACTTAGAACTGGTGCGGACCAGGGGA | rRF | FALSE | FALSE | TRUE |
| >hg38_21_+_8433172_8446622_RNA45SN1_45S_50ntFlanks@11572.11600.29\|CCGACTTAGAACTGGTGCGGACCAGGGGA | >hg38_21_+_8433172_8446622_RNA45SN1_45S_50ntFlanks | 11572 | 11600 | 29 | CCGACTTAGAACTGGTGCGGACCAGGGGA | rRF | FALSE | FALSE | TRUE |
| >hg38_21_+_8433172_8446622_RNA45SN1_45S_50ntFlanks@12007.12032.26\|GCCGCCGGTGAAATACCACTACTCTG | >hg38_21_+_8433172_8446622_RNA45SN1_45S_50ntFlanks | 12007 | 12032 | 26 | GCCGCCGGTGAAATACCACTACTCTG | rRF | FALSE | FALSE | TRUE |
| >hg38_21_+_8433172_8446622_RNA45SN1_45S_50ntFlanks@10402.10429.28\|AACATGGGTCAGTCGGTCCTGAGAGATG | >hg38_21_+_8433172_8446622_RNA45SN1_45S_50ntFlanks | 10402 | 10429 | 28 | AACATGGGTCAGTCGGTCCTGAGAGATG | rRF | FALSE | FALSE | TRUE |
| >hg38_21_+_8433172_8446622_RNA45SN1_45S_50ntFlanks@12405.12444.40\|CTTTTTGATCCTTCGATGTCGGCTCTTCCTATCATTGTGA | >hg38_21_+_8433172_8446622_RNA45SN1_45S_50ntFlanks | 12405 | 12444 | 40 | CTTTTTGATCCTTCGATGTCGGCTCTTCCTATCATTGTGA | rRF | FALSE | FALSE | TRUE |
| >trna152_ValCAC_6_-_27248049_27248121@1.29.29\|GCTTCTGTAGTGTAGTGGTTATCACGTTC | >trna152_ValCAC_6_-_27248049_27248121 | 1 | 29 | 29 | GCTTCTGTAGTGTAGTGGTTATCACGTTC | tRF | FALSE | FALSE | TRUE |
| >trna111_HisGTG_1_-_147774845_147774916@-1T.29.30,trna118_HisGTG_1_-_145396881_145396952@-1T.29.30,trna16_HisGTG_1_+_146544773_146544844@-1T.29.30,trna1_HisGTG_15_+_45493349_45493420@-1T.29.30,trna21_HisGTG_1_+_147753471_147753542@-1T.29.30,trna33_HisGTG_6_+_27125906_27125977@-1T.29.30,trna7_HisGTG_9_-_14433938_14434009@-1T.29.30,trna8_HisGTG_15_-_45492611_45492682@-1T.29.30,trna9_HisGTG_15_-_45490804_45490875@-1T.29.30\|TGCCGTGATCGTATAGTGGTTAGTACTCTG | >trna111_HisGTG_1_-_147774845_147774916 | -1T | 29 | 30 | TGCCGTGATCGTATAGTGGTTAGTACTCTG | tRF | FALSE | FALSE | TRUE |
| >hg38_21_+_8433172_8446622_RNA45SN1_45S_50ntFlanks@7980.8006.27\|CCTCAGATCAGACGTGGCGACCCGCTG | >hg38_21_+_8433172_8446622_RNA45SN1_45S_50ntFlanks | 7980 | 8006 | 27 | CCTCAGATCAGACGTGGCGACCCGCTG | rRF | FALSE | FALSE | TRUE |
| >hg38_21_+_8433172_8446622_RNA45SN1_45S_50ntFlanks@8170.8212.43\|CAAGTCCTTCTGATCGAGGCCCAGCCCGTGGACGGTGTGAGGC | >hg38_21_+_8433172_8446622_RNA45SN1_45S_50ntFlanks | 8170 | 8212 | 43 | CAAGTCCTTCTGATCGAGGCCCAGCCCGTGGACGGTGTGAGG[illegible] | rRF | FALSE | FALSE | TRUE |
| >hg38_21_+_8433172_8446622_RNA45SN1_45S_50ntFlanks@12417.12435.19\|TCGATGTCGGCTCTTCCTA | >hg38_21_+_8433172_8446622_RNA45SN1_45S_50ntFlanks | 12417 | 12435 | 19 | TCGATGTCGGCTCTTCCTA | rRF | FALSE | FALSE | TRUE |
| >hg38_21_+_8433172_8446622_RNA45SN1_45S_50ntFlanks@8104.8143.40\|CCGCGGCGGGGCGCGGGACATGTGGCGTACGGAAGACCCG | >hg38_21_+_8433172_8446622_RNA45SN1_45S_50ntFlanks | 8104 | 8143 | 40 | CCGCGGCGGGGCGCGGGACATGTGGCGTACGGAAGACCCG | rRF | FALSE | FALSE | TRUE |
| >hg38_21_+_8433172_8446622_RNA45SN1_45S_50ntFlanks@10819.10840.22\|AGGGAAGTCGGCAAGCCGGATC | >hg38_21_+_8433172_8446622_RNA45SN1_45S_50ntFlanks | 10819 | 10840 | 22 | AGGGAAGTCGGCAAGCCGGATC | rRF | FALSE | FALSE | TRUE |
| >hg38_21_+_8433172_8446622_RNA45SN1_45S_50ntFlanks@11575.11603.29\|ACTTAGAACTGGTGCGGACCAGGGGAATC | >hg38_21_+_8433172_8446622_RNA45SN1_45S_50ntFlanks | 11575 | 11603 | 29 | ACTTAGAACTGGTGCGGACCAGGGGAATC | rRF | FALSE | FALSE | TRUE |

### Nucleus

| | | | | | | | | | |
|---|---|---|---|---|---|---|---|---|---|
| >hg38_21_+_8433172_8446622_RNA45SN1_45S_50ntFlanks@10690.10708.19\|CGTCCGGTGAGCTCTCGCT | >hg38_21_+_8433172_8446622_RNA45SN1_45S_50ntFlanks | 10690 | 10708 | 19 | CGTCCGGTGAGCTCTCGCT | rRF | FALSE | FALSE | TRUE |
| >hg38_21_+_8433172_8446622_RNA45SN1_45S_50ntFlanks@10684.10711.28\|CGGCGGCGTCCGGTGAGCTCTCGCTGGC | >hg38_21_+_8433172_8446622_RNA45SN1_45S_50ntFlanks | 10684 | 10711 | 28 | CGGCGGCGTCCGGTGAGCTCTCGCTGGC | rRF | FALSE | FALSE | TRUE |
| >hg38_21_+_8433172_8446622_RNA45SN1_45S_50ntFlanks@12605.12624.20\|TATGTGCTTGGCTGAGGAGC | >hg38_21_+_8433172_8446622_RNA45SN1_45S_50ntFlanks | 12605 | 12624 | 20 | TATGTGCTTGGCTGAGGAGC | rRF | FALSE | FALSE | TRUE |
| >hg38_21_+_8433172_8446622_RNA45SN1_45S_50ntFlanks@10350.10368.19\|ACGAGAACTTTGAAGGCCG | >hg38_21_+_8433172_8446622_RNA45SN1_45S_50ntFlanks | 10350 | 10368 | 19 | ACGAGAACTTTGAAGGCCG | rRF | FALSE | FALSE | TRUE |
| >hg38_21_+_8433172_8446622_RNA45SN1_45S_50ntFlanks@10411.10428.18\|CAGTCGGTCCTGAGAGAT | >hg38_21_+_8433172_8446622_RNA45SN1_45S_50ntFlanks | 10411 | 10428 | 18 | CAGTCGGTCCTGAGAGAT | rRF | FALSE | FALSE | TRUE |
| >hg38_21_+_8433172_8446622_RNA45SN1_45S_50ntFlanks@7983.8006.24\|CAGATCAGACGTGGCGACCCGCTG | >hg38_21_+_8433172_8446622_RNA45SN1_45S_50ntFlanks | 7983 | 8006 | 24 | CAGATCAGACGTGGCGACCCGCTG | rRF | FALSE | FALSE | TRUE |
| >hg38_21_+_8433172_8446622_RNA45SN1_45S_50ntFlanks@9544.9571.28\|GGTGGAGGTCCGTAGCGGTCCTGACGTG | >hg38_21_+_8433172_8446622_RNA45SN1_45S_50ntFlanks | 9544 | 9571 | 28 | GGTGGAGGTCCGTAGCGGTCCTGACGTG | rRF | FALSE | FALSE | TRUE |
| >hg38_21_+_8433172_8446622_RNA45SN1_45S_50ntFlanks@9534.9561.28\|AGGAAACTCTGGTGGAGGTCCGTAGCGG | >hg38_21_+_8433172_8446622_RNA45SN1_45S_50ntFlanks | 9534 | 9561 | 28 | AGGAAACTCTGGTGGAGGTCCGTAGCGG | rRF | FALSE | FALSE | TRUE |
| >trna134_GluTTC_1_-_16861774_16861845@1.31.31,trna5_GluTTC_1_+_17199078_17199149@1.31.31,trna84_GluTTC_1_-_161391883_161391954@1.31.31,trna94_GluTTC_1_-_149664355_149664427@1.31.31\|TCCCTGGTGGTCTAGTGGCTAGGATTCGGCG | >trna134_GluTTC_1_-_16861774_16861845 | 1 | 31 | 31 | TCCCTGGTGGTCTAGTGGCTAGGATTCGGCG | tRF | FALSE | FALSE | TRUE |
| >hg38_21_+_8433172_8446622_RNA45SN1_45S_50ntFlanks@10392.10410.19\|ACAGCAGTTGAACATGGGT | >hg38_21_+_8433172_8446622_RNA45SN1_45S_50ntFlanks | 10392 | 10410 | 19 | ACAGCAGTTGAACATGGGT | rRF | FALSE | FALSE | TRUE |
| >hg38_21_+_8433172_8446622_RNA45SN1_45S_50ntFlanks@12957.12982.26\|CTTCTGGGTCGGGGTTTCGTACGTAG | >hg38_21_+_8433172_8446622_RNA45SN1_45S_50ntFlanks | 12957 | 12982 | 26 | CTTCTGGGTCGGGGTTTCGTACGTAG | rRF | FALSE | FALSE | TRUE |
| >hg38_21_+_8433172_8446622_RNA45SN1_45S_50ntFlanks@12002.12032.31\|TGCGGGCCGCCGGTGAAATACCACTACTCTG | >hg38_21_+_8433172_8446622_RNA45SN1_45S_50ntFlanks | 12002 | 12032 | 31 | TGCGGGCCGCCGGTGAAATACCACTACTCTG | rRF | FALSE | FALSE | TRUE |
| >hg38_21_+_8433172_8446622_RNA45SN1_45S_50ntFlanks@10411.10431.21\|CAGTCGGTCCTGAGAGATGGG | >hg38_21_+_8433172_8446622_RNA45SN1_45S_50ntFlanks | 10411 | 10431 | 21 | CAGTCGGTCCTGAGAGATGGG | rRF | FALSE | FALSE | TRUE |
| >hg38_21_+_8433172_8446622_RNA45SN1_45S_50ntFlanks@12456.12485.30\|CCAAGCGTTGGATTGTTCACCCACTAATAG | >hg38_21_+_8433172_8446622_RNA45SN1_45S_50ntFlanks | 12456 | 12485 | 30 | CCAAGCGTTGGATTGTTCACCCACTAATAG | rRF | FALSE | FALSE | TRUE |
| >hg38_21_+_8433172_8446622_RNA45SN1_45S_50ntFlanks@5446.5466.21\|CGGATCGGCCCCGCCGGGGTC | >hg38_21_+_8433172_8446622_RNA45SN1_45S_50ntFlanks | 5446 | 5466 | 21 | CGGATCGGCCCCGCCGGGGTC | rRF | FALSE | FALSE | TRUE |
| >hg38_21_+_8433172_8446622_RNA45SN1_45S_50ntFlanks@12415.12432.18\|CTTCGATGTCGGCTCTTC | >hg38_21_+_8433172_8446622_RNA45SN1_45S_50ntFlanks | 12415 | 12432 | 18 | CTTCGATGTCGGCTCTTC | rRF | FALSE | FALSE | TRUE |
| >hg38_21_+_8433172_8446622_RNA45SN1_45S_50ntFlanks@12407.12436.30\|TTTTGATCCTTCGATGTCGGCTCTTCCTAT | >hg38_21_+_8433172_8446622_RNA45SN1_45S_50ntFlanks | 12407 | 12436 | 30 | TTTTGATCCTTCGATGTCGGCTCTTCCTAT | rRF | FALSE | FALSE | TRUE |
| >am_trna10_ValCAC_5_-_180649395_180649467@1.29.29,trna12_ValAAC_5_-_180645270_180645342@1.29.29,trna132_ValAAC_6_-_27721179_27721251@1.29.29,trna136_ValAAC_6_-_27648885_27648957@1.29.29,trna139_ValAAC_6_-_27618707_27618779@1.29.29,trna18_ValCAC_5_-_180529253_180529325@1.29.29,trna2_ValAAC_3_+_169490018_169490090@1.29.29,trna2_ValCAC_5_+_180524070_180524142@1.29.29,trna4_ValAAC_5_+_180591154_180591226@1.29.29,trna5_ValAAC_5_+_180596610_180596682@1.29.29,trna6_ValCAC_5_+_180600650_180600722@1.29.29,trna85_ValCAC_1_-_161369490_161369562@1.29.29,trna90_ValCAC_1_-_149684088_149684161@1.29.29,trna98_ValCAC_1_-_149298555_149298627@1.29.29,trna9_ValCAC_6_+_26538282_26538354@1.29.29\|GTTTCCGTAGTGTAGTGGTTATCACGTTC | >am_trna10_ValCAC_5_-_180649395_180649467 | 1 | 29 | 29 | GTTTCCGTAGTGTAGTGGTTATCACGTTC | tRF | FALSE | FALSE | TRUE |
| >trnaMT_ValTAC_MT_+_1602_1670@51.72.22\|CTTAACTTGACCGCTCTGACCA | >trnaMT_ValTAC_MT_+_1602_1670 | 51 | 72 | 22 | CTTAACTTGACCGCTCTGACCA | tRF | FALSE | FALSE | TRUE |
| >hg38_21_+_8433172_8446622_RNA45SN1_45S_50ntFlanks@10391.10410.20\|AACAGCAGTTGAACATGGGT | >hg38_21_+_8433172_8446622_RNA45SN1_45S_50ntFlanks | 10391 | 10410 | 20 | AACAGCAGTTGAACATGGGT | rRF | FALSE | FALSE | TRUE |
| >hg38_21_+_8433172_8446622_RNA45SN1_45S_50ntFlanks@12488.12509.22\|AACGTGAGCTGGGTTTAGACCG | >hg38_21_+_8433172_8446622_RNA45SN1_45S_50ntFlanks | 12488 | 12509 | 22 | AACGTGAGCTGGGTTTAGACCG | rRF | FALSE | FALSE | TRUE |
| >hg38_21_+_8433172_8446622_RNA45SN1_45S_50ntFlanks@9534.9570.37\|AGGAAACTCTGGTGGAGGTCCGTAGCGGTCCTGACGT | >hg38_21_+_8433172_8446622_RNA45SN1_45S_50ntFlanks | 9534 | 9570 | 37 | AGGAAACTCTGGTGGAGGTCCGTAGCGGTCCTGACGT | rRF | FALSE | FALSE | TRUE |
| >hg38_21_+_8433172_8446622_RNA45SN1_45S_50ntFlanks@4283.4317.35\|CTTTAACGAGGATCCATTGGAGGGCAAGTCTGGTG | >hg38_21_+_8433172_8446622_RNA45SN1_45S_50ntFlanks | 4283 | 4317 | 35 | CTTTAACGAGGATCCATTGGAGGGCAAGTCTGGTG | rRF | FALSE | FALSE | TRUE |
| >hg38_21_+_8433172_8446622_RNA45SN1_45S_50ntFlanks@12415.12435.21\|CTTCGATGTCGGCTCTTCCTA | >hg38_21_+_8433172_8446622_RNA45SN1_45S_50ntFlanks | 12415 | 12435 | 21 | CTTCGATGTCGGCTCTTCCTA | rRF | FALSE | FALSE | TRUE |
| >hg38_21_+_8433172_8446622_RNA45SN1_45S_50ntFlanks@10687.10706.20\|CGGCGTCCGGTGAGCTCTCG | >hg38_21_+_8433172_8446622_RNA45SN1_45S_50ntFlanks | 10687 | 10706 | 20 | CGGCGTCCGGTGAGCTCTCG | rRF | FALSE | FALSE | TRUE |
| >hg38_21_+_8433172_8446622_RNA45SN1_45S_50ntFlanks@11572.11605.34\|CCGACTTAGAACTGGTGCGGACCAGGGGAATCCG | >hg38_21_+_8433172_8446622_RNA45SN1_45S_50ntFlanks | 11572 | 11605 | 34 | CCGACTTAGAACTGGTGCGGACCAGGGGAATCCG | rRF | FALSE | FALSE | TRUE |
| >hg38_21_+_8433172_8446622_RNA45SN1_45S_50ntFlanks@9534.9564.31\|AGGAAACTCTGGTGGAGGTCCGTAGCGGTCC | >hg38_21_+_8433172_8446622_RNA45SN1_45S_50ntFlanks | 9534 | 9564 | 31 | AGGAAACTCTGGTGGAGGTCCGTAGCGGTCC | rRF | FALSE | FALSE | TRUE |
| >hg38_21_+_8433172_8446622_RNA45SN1_45S_50ntFlanks@6719.6736.18\|TGAATTGCAGGACACATT | >hg38_21_+_8433172_8446622_RNA45SN1_45S_50ntFlanks | 6719 | 6736 | 18 | TGAATTGCAGGACACATT | rRF | FALSE | FALSE | TRUE |
| >trna116_GluCTC_1_-_145399233_145399304@32.70.39,trna71_GluCTC_1_-_161439189_161439260@32.70.39,trna74_GluCTC_1_-_161431809_161431880@32.70.39,trna77_GluCTC_1_-_161424398_161424469@32.70.39,trna77_GluCTC_6_+_28949976_28950047@32.70.39,trna80_GluCTC_1_-_161417018_161417089@32.70.39,trna87_GluCTC_6_-_126101393_126101464@32.70.39\|CTCTCACCGCCGCGGCCCGGGTTCGATTCCCGGTCAGGG | >trna116_GluCTC_1_-_145399233_145399304 | 32 | 70 | 39 | CTCTCACCGCCGCGGCCCGGGTTCGATTCCCGGTCAGGG | tRF | FALSE | FALSE | TRUE |
| >hg38_21_+_8433172_8446622_RNA45SN1_45S_50ntFlanks@12654.12675.22\|TATGACTGAACGCCTCTAAGTC | >hg38_21_+_8433172_8446622_RNA45SN1_45S_50ntFlanks | 12654 | 12675 | 22 | TATGACTGAACGCCTCTAAGTC | rRF | FALSE | FALSE | TRUE |
| >hg38_21_+_8433172_8446622_RNA45SN1_45S_50ntFlanks@12991.13011.21\|CTCCCTCGCTGCGATCTATTG | >hg38_21_+_8433172_8446622_RNA45SN1_45S_50ntFlanks | 12991 | 13011 | 21 | CTCCCTCGCTGCGATCTATTG | rRF | FALSE | FALSE | TRUE |

### Nucleus

<table>
<tr><th></th><th></th><th></th><th></th><th></th><th></th><th></th><th></th><th></th><th></th></tr>
<tr><td>>hg38_21_+_8433172_8446622_RNA45SN1_45S_50ntFlanks@12415.12434.20|CTTCGATGTCGGCTCTTCCT</td><td>>hg38_21_+_8433172_8446622_RNA45SN1_45S_50ntFlanks</td><td>12415</td><td>12434</td><td>20</td><td>CTTCGATGTCGGCTCTTCCT</td><td>rRF</td><td>FALSE</td><td>FALSE</td><td>TRUE</td></tr>
<tr><td>>hg38_21_+_8433172_8446622_RNA45SN1_45S_50ntFlanks@10687.10711.25|CGGCGTCCGGTGAGCTCTCGCTGGC</td><td>>hg38_21_+_8433172_8446622_RNA45SN1_45S_50ntFlanks</td><td>10687</td><td>10711</td><td>25</td><td>CGGCGTCCGGTGAGCTCTCGCTGGC</td><td>rRF</td><td>FALSE</td><td>FALSE</td><td>TRUE</td></tr>
<tr><td>>hg38_21_+_8433172_8446622_RNA45SN1_45S_50ntFlanks@11319.11344.26|CCCACGTCTCGTCGCGCGCGCGTCCG</td><td>>hg38_21_+_8433172_8446622_RNA45SN1_45S_50ntFlanks</td><td>11319</td><td>11344</td><td>26</td><td>CCCACGTCTCGTCGCGCGCGCGTCCG</td><td>rRF</td><td>FALSE</td><td>FALSE</td><td>TRUE</td></tr>
<tr><td>>hg38_21_+_8433172_8446622_RNA45SN1_45S_50ntFlanks@10687.10710.24|CGGCGTCCGGTGAGCTCTCGCTGG</td><td>>hg38_21_+_8433172_8446622_RNA45SN1_45S_50ntFlanks</td><td>10687</td><td>10710</td><td>24</td><td>CGGCGTCCGGTGAGCTCTCGCTGG</td><td>rRF</td><td>FALSE</td><td>FALSE</td><td>TRUE</td></tr>
<tr><td>>hg38_21_+_8433172_8446622_RNA45SN1_45S_50ntFlanks@12407.12443.37|TTTTGATCCTTCGATGTCGGCTCTTCCTATCATTGTG</td><td>>hg38_21_+_8433172_8446622_RNA45SN1_45S_50ntFlanks</td><td>12407</td><td>12443</td><td>37</td><td>TTTTGATCCTTCGATGTCGGCTCTTCCTATCATTGTG</td><td>rRF</td><td>FALSE</td><td>FALSE</td><td>TRUE</td></tr>
<tr><td>>hg38_21_+_8433172_8446622_RNA45SN1_45S_50ntFlanks@8357.8377.21|AAAAGAACTTTGAAGAGAGAG</td><td>>hg38_21_+_8433172_8446622_RNA45SN1_45S_50ntFlanks</td><td>8357</td><td>8377</td><td>21</td><td>AAAAGAACTTTGAAGAGAGAG</td><td>rRF</td><td>FALSE</td><td>FALSE</td><td>TRUE</td></tr>
<tr><td>>hg38_21_+_8433172_8446622_RNA45SN1_45S_50ntFlanks@11784.11803.20|ATGAATGGATGAACGAGATT</td><td>>hg38_21_+_8433172_8446622_RNA45SN1_45S_50ntFlanks</td><td>11784</td><td>11803</td><td>20</td><td>ATGAATGGATGAACGAGATT</td><td>rRF</td><td>FALSE</td><td>FALSE</td><td>TRUE</td></tr>
<tr><td>>hg38_21_+_8433172_8446622_RNA45SN1_45S_50ntFlanks@12417.12434.18|TCGATGTCGGCTCTTCCT</td><td>>hg38_21_+_8433172_8446622_RNA45SN1_45S_50ntFlanks</td><td>12417</td><td>12434</td><td>18</td><td>TCGATGTCGGCTCTTCCT</td><td>rRF</td><td>FALSE</td><td>FALSE</td><td>TRUE</td></tr>
<tr><td>>hg38_21_+_8433172_8446622_RNA45SN1_45S_50ntFlanks@10677.10710.34|GCGGTTCCGGCGGCGTCCGGTGAGCTCTCGCTGG</td><td>>hg38_21_+_8433172_8446622_RNA45SN1_45S_50ntFlanks</td><td>10677</td><td>10710</td><td>34</td><td>GCGGTTCCGGCGGCGTCCGGTGAGCTCTCGCTGG</td><td>rRF</td><td>FALSE</td><td>FALSE</td><td>TRUE</td></tr>
<tr><td>>hg38_21_+_8433172_8446622_RNA45SN1_45S_50ntFlanks@7981.8006.26|CTCAGATCAGACGTGGCGACCCGCTG</td><td>>hg38_21_+_8433172_8446622_RNA45SN1_45S_50ntFlanks</td><td>7981</td><td>8006</td><td>26</td><td>CTCAGATCAGACGTGGCGACCCGCTG</td><td>rRF</td><td>FALSE</td><td>FALSE</td><td>TRUE</td></tr>
<tr><td>>hg38_21_+_8433172_8446622_RNA45SN1_45S_50ntFlanks@4726.4758.33|TAATCAAGAACGAAAGTCGGAGGTTCGAAGACG</td><td>>hg38_21_+_8433172_8446622_RNA45SN1_45S_50ntFlanks</td><td>4726</td><td>4758</td><td>33</td><td>TAATCAAGAACGAAAGTCGGAGGTTCGAAGACG</td><td>rRF</td><td>FALSE</td><td>FALSE</td><td>TRUE</td></tr>
<tr><td>>hg38_21_+_8433172_8446622_RNA45SN1_45S_50ntFlanks@12456.12477.22|CCAAGCGTTGGATTGTTCACCC</td><td>>hg38_21_+_8433172_8446622_RNA45SN1_45S_50ntFlanks</td><td>12456</td><td>12477</td><td>22</td><td>CCAAGCGTTGGATTGTTCACCC</td><td>rRF</td><td>FALSE</td><td>FALSE</td><td>TRUE</td></tr>
<tr><td>>hg38_21_+_8433172_8446622_RNA45SN1_45S_50ntFlanks@12625.12654.30|CAATGGGGCGAAGCTACCATCTGTGGGATT</td><td>>hg38_21_+_8433172_8446622_RNA45SN1_45S_50ntFlanks</td><td>12625</td><td>12654</td><td>30</td><td>CAATGGGGCGAAGCTACCATCTGTGGGATT</td><td>rRF</td><td>FALSE</td><td>FALSE</td><td>TRUE</td></tr>
<tr><td>>hg38_21_+_8433172_8446622_RNA45SN1_45S_50ntFlanks@10841.10864.24|CGTAACTTCGGGATAAGGATTGGC</td><td>>hg38_21_+_8433172_8446622_RNA45SN1_45S_50ntFlanks</td><td>10841</td><td>10864</td><td>24</td><td>CGTAACTTCGGGATAAGGATTGGC</td><td>rRF</td><td>FALSE</td><td>FALSE</td><td>TRUE</td></tr>
<tr><td>>hg38_21_+_8433172_8446622_RNA45SN1_45S_50ntFlanks@4722.4759.38|TCATTAATCAAGAACGAAAGTCGGAGGTTCGAAGACGA</td><td>>hg38_21_+_8433172_8446622_RNA45SN1_45S_50ntFlanks</td><td>4722</td><td>4759</td><td>38</td><td>TCATTAATCAAGAACGAAAGTCGGAGGTTCGAAGACGA</td><td>rRF</td><td>FALSE</td><td>FALSE</td><td>TRUE</td></tr>
<tr><td>>hg38_21_+_8433172_8446622_RNA45SN1_45S_50ntFlanks@12419.12436.18|GATGTCGGCTCTTCCTAT</td><td>>hg38_21_+_8433172_8446622_RNA45SN1_45S_50ntFlanks</td><td>12419</td><td>12436</td><td>18</td><td>GATGTCGGCTCTTCCTAT</td><td>rRF</td><td>FALSE</td><td>FALSE</td><td>TRUE</td></tr>
<tr><td>>hg38_21_+_8433172_8446622_RNA45SN1_45S_50ntFlanks@4724.4759.36|ATTAATCAAGAACGAAAGTCGGAGGTTCGAAGACGA</td><td>>hg38_21_+_8433172_8446622_RNA45SN1_45S_50ntFlanks</td><td>4724</td><td>4759</td><td>36</td><td>ATTAATCAAGAACGAAAGTCGGAGGTTCGAAGACGA</td><td>rRF</td><td>FALSE</td><td>FALSE</td><td>TRUE</td></tr>
<tr><td>>hg38_21_+_8433172_8446622_RNA45SN1_45S_50ntFlanks@9534.9551.18|AGGAAACTCTGGTGGAGG</td><td>>hg38_21_+_8433172_8446622_RNA45SN1_45S_50ntFlanks</td><td>9534</td><td>9551</td><td>18</td><td>AGGAAACTCTGGTGGAGG</td><td>rRF</td><td>FALSE</td><td>FALSE</td><td>TRUE</td></tr>
<tr><td>>hg38_21_+_8433172_8446622_RNA45SN1_45S_50ntFlanks@5484.5514.31|GAGCGCTGAGAAGACGGTCGAACTTGACTAT</td><td>>hg38_21_+_8433172_8446622_RNA45SN1_45S_50ntFlanks</td><td>5484</td><td>5514</td><td>31</td><td>GAGCGCTGAGAAGACGGTCGAACTTGACTAT</td><td>rRF</td><td>FALSE</td><td>FALSE</td><td>TRUE</td></tr>
<tr><td>>hg38_21_+_8433172_8446622_RNA45SN1_45S_50ntFlanks@12420.12455.36|ATGTCGGCTCTTCCTATCATTGTGAAGCAGAATTCA</td><td>>hg38_21_+_8433172_8446622_RNA45SN1_45S_50ntFlanks</td><td>12420</td><td>12455</td><td>36</td><td>ATGTCGGCTCTTCCTATCATTGTGAAGCAGAATTCA</td><td>rRF</td><td>FALSE</td><td>FALSE</td><td>TRUE</td></tr>
<tr><td>>hg38_21_+_8433172_8446622_RNA45SN1_45S_50ntFlanks@11316.11338.23|CACCCCACGTCTCGTCGCGCGCG</td><td>>hg38_21_+_8433172_8446622_RNA45SN1_45S_50ntFlanks</td><td>11316</td><td>11338</td><td>23</td><td>CACCCCACGTCTCGTCGCGCGCG</td><td>rRF</td><td>FALSE</td><td>FALSE</td><td>TRUE</td></tr>
<tr><td>>hg38_21_+_8433172_8446622_RNA45SN1_45S_50ntFlanks@9533.9563.31|GAGGAAACTCTGGTGGAGGTCCGTAGCGGTC</td><td>>hg38_21_+_8433172_8446622_RNA45SN1_45S_50ntFlanks</td><td>9533</td><td>9563</td><td>31</td><td>GAGGAAACTCTGGTGGAGGTCCGTAGCGGTC</td><td>rRF</td><td>FALSE</td><td>FALSE</td><td>TRUE</td></tr>
<tr><td>>hg38_21_+_8433172_8446622_RNA45SN1_45S_50ntFlanks@11564.11601.38|CCTAGCAGCCGACTTAGAACTGGTGCGGACCAGGGGAA</td><td>>hg38_21_+_8433172_8446622_RNA45SN1_45S_50ntFlanks</td><td>11564</td><td>11601</td><td>38</td><td>CCTAGCAGCCGACTTAGAACTGGTGCGGACCAGGGGAA</td><td>rRF</td><td>FALSE</td><td>FALSE</td><td>TRUE</td></tr>
<tr><td>>hg38_21_+_8433172_8446622_RNA45SN1_45S_50ntFlanks@11208.11233.26|GCGGCGACTCTGGACGCGAGCCGGGC</td><td>>hg38_21_+_8433172_8446622_RNA45SN1_45S_50ntFlanks</td><td>11208</td><td>11233</td><td>26</td><td>GCGGCGACTCTGGACGCGAGCCGGGC</td><td>rRF</td><td>FALSE</td><td>FALSE</td><td>TRUE</td></tr>
<tr><td>>hg38_21_+_8433172_8446622_RNA45SN1_45S_50ntFlanks@3984.4001.18|GCGGCTTTGGTGACTCTA</td><td>>hg38_21_+_8433172_8446622_RNA45SN1_45S_50ntFlanks</td><td>3984</td><td>4001</td><td>18</td><td>GCGGCTTTGGTGACTCTA</td><td>rRF</td><td>FALSE</td><td>FALSE</td><td>TRUE</td></tr>
<tr><td>>hg38_21_+_8433172_8446622_RNA45SN1_45S_50ntFlanks@12420.12444.25|ATGTCGGCTCTTCCTATCATTGTGA</td><td>>hg38_21_+_8433172_8446622_RNA45SN1_45S_50ntFlanks</td><td>12420</td><td>12444</td><td>25</td><td>ATGTCGGCTCTTCCTATCATTGTGA</td><td>rRF</td><td>FALSE</td><td>FALSE</td><td>TRUE</td></tr>
<tr><td>>MI0000342|hsa-mir-200b&WithFlank&1|+|1167098|1167204@63.85.23&[MIMAT0000318&hsa-miR-200b-3p&offsets|0|+1;m-93&1|+|1167160|1167182&offsets|0|0]|TAATACTGCCTGGTAATGATGAC</td><td>>MI0000342|hsa-mir-200b&WithFlank&1|+|1167098|1167204</td><td>63</td><td>85</td><td>23</td><td>TAATACTGCCTGGTAATGATGAC</td><td>isomiR</td><td>FALSE</td><td>FALSE</td><td>TRUE</td></tr>
<tr><td>>hg38_21_+_8433172_8446622_RNA45SN1_45S_50ntFlanks@12605.12626.22|TATGTGCTTGGCTGAGGAGCCA</td><td>>hg38_21_+_8433172_8446622_RNA45SN1_45S_50ntFlanks</td><td>12605</td><td>12626</td><td>22</td><td>TATGTGCTTGGCTGAGGAGCCA</td><td>rRF</td><td>FALSE</td><td>FALSE</td><td>TRUE</td></tr>
<tr><td>>hg38_21_+_8433172_8446622_RNA45SN1_45S_50ntFlanks@12001.12032.32|CTGCGGGCCGCCGGTGAAATACCACTACTCTG</td><td>>hg38_21_+_8433172_8446622_RNA45SN1_45S_50ntFlanks</td><td>12001</td><td>12032</td><td>32</td><td>CTGCGGGCCGCCGGTGAAATACCACTACTCTG</td><td>rRF</td><td>FALSE</td><td>FALSE</td><td>TRUE</td></tr>
<tr><td>>hg38_21_+_8433172_8446622_RNA45SN1_45S_50ntFlanks@10350.10384.35|ACGAGAACTTTGAAGGCCGAAGTGGAGAAGGGTTC</td><td>>hg38_21_+_8433172_8446622_RNA45SN1_45S_50ntFlanks</td><td>10350</td><td>10384</td><td>35</td><td>ACGAGAACTTTGAAGGCCGAAGTGGAGAAGGGTTC</td><td>rRF</td><td>FALSE</td><td>FALSE</td><td>TRUE</td></tr>
<tr><td>>hg38_21_+_8433172_8446622_RNA45SN1_45S_50ntFlanks@10458.10478.21|ATGGCCTCCGTTGCCCTCGGC</td><td>>hg38_21_+_8433172_8446622_RNA45SN1_45S_50ntFlanks</td><td>10458</td><td>10478</td><td>21</td><td>ATGGCCTCCGTTGCCCTCGGC</td><td>rRF</td><td>FALSE</td><td>FALSE</td><td>TRUE</td></tr>
<tr><td>>hg38_21_+_8433172_8446622_RNA45SN1_45S_50ntFlanks@11534.11551.18|GTTCCGCGCGGCGCCTCG</td><td>>hg38_21_+_8433172_8446622_RNA45SN1_45S_50ntFlanks</td><td>11534</td><td>11551</td><td>18</td><td>GTTCCGCGCGGCGCCTCG</td><td>rRF</td><td>FALSE</td><td>FALSE</td><td>TRUE</td></tr>
<tr><td>>hg38_21_+_8433172_8446622_RNA45SN1_45S_50ntFlanks@9534.9566.33|AGGAAACTCTGGTGGAGGTCCGTAGCGGTCCTG</td><td>>hg38_21_+_8433172_8446622_RNA45SN1_45S_50ntFlanks</td><td>9534</td><td>9566</td><td>33</td><td>AGGAAACTCTGGTGGAGGTCCGTAGCGGTCCTG</td><td>rRF</td><td>FALSE</td><td>FALSE</td><td>TRUE</td></tr>
<tr><td>>hg38_21_+_8433172_8446622_RNA45SN1_45S_50ntFlanks@12992.13013.22|TCCCTCGCTGCGATCTATTGAA</td><td>>hg38_21_+_8433172_8446622_RNA45SN1_45S_50ntFlanks</td><td>12992</td><td>13013</td><td>22</td><td>TCCCTCGCTGCGATCTATTGAA</td><td>rRF</td><td>FALSE</td><td>FALSE</td><td>TRUE</td></tr>
<tr><td>>hg38_21_+_8433172_8446622_RNA45SN1_45S_50ntFlanks@10410.10431.22|TCAGTCGGTCCTGAGAGATGGG</td><td>>hg38_21_+_8433172_8446622_RNA45SN1_45S_50ntFlanks</td><td>10410</td><td>10431</td><td>22</td><td>TCAGTCGGTCCTGAGAGATGGG</td><td>rRF</td><td>FALSE</td><td>FALSE</td><td>TRUE</td></tr>
<tr><td>>hg38_21_+_8433172_8446622_RNA45SN1_45S_50ntFlanks@11874.11893.20|AGAAGACCCTGTTGAGCTTG</td><td>>hg38_21_+_8433172_8446622_RNA45SN1_45S_50ntFlanks</td><td>11874</td><td>11893</td><td>20</td><td>AGAAGACCCTGTTGAGCTTG</td><td>rRF</td><td>FALSE</td><td>FALSE</td><td>TRUE</td></tr>
<tr><td>>hg38_21_+_8433172_8446622_RNA45SN1_45S_50ntFlanks@10362.10383.22|AAGGCCGAAGTGGAGAAGGGTT</td><td>>hg38_21_+_8433172_8446622_RNA45SN1_45S_50ntFlanks</td><td>10362</td><td>10383</td><td>22</td><td>AAGGCCGAAGTGGAGAAGGGTT</td><td>rRF</td><td>FALSE</td><td>FALSE</td><td>TRUE</td></tr>
<tr><td>>hg38_21_+_8433172_8446622_RNA45SN1_45S_50ntFlanks@3875.3898.24|ATACATGCCGACGGGCGCTGACCC</td><td>>hg38_21_+_8433172_8446622_RNA45SN1_45S_50ntFlanks</td><td>3875</td><td>3898</td><td>24</td><td>ATACATGCCGACGGGCGCTGACCC</td><td>rRF</td><td>FALSE</td><td>FALSE</td><td>TRUE</td></tr>
<tr><td>>hg38_21_+_8433172_8446622_RNA45SN1_45S_50ntFlanks@9554.9584.31|CGTAGCGGTCCTGACGTGCAAATCGGTCGTC</td><td>>hg38_21_+_8433172_8446622_RNA45SN1_45S_50ntFlanks</td><td>9554</td><td>9584</td><td>31</td><td>CGTAGCGGTCCTGACGTGCAAATCGGTCGTC</td><td>rRF</td><td>FALSE</td><td>FALSE</td><td>TRUE</td></tr>
</table>

### Nucleus

| | | | | | | | | | |
|---|---|---|---|---|---|---|---|---|---|
| >hg38_21_+_8433172_8446622_RNA45SN1_45S_50ntFlanks@13002.13028.27\|CGATCTATTGAAAGTCAGCCCTCGACA | >hg38_21_+_8433172_8446622_RNA45SN1_45S_50ntFlanks | 13002 | 13028 | 27 | CGATCTATTGAAAGTCAGCCCTCGACA | rRF | FALSE | FALSE | TRUE |
| >hg38_21_+_8433172_8446622_RNA45SN1_45S_50ntFlanks@11576.11603.28\|CTTAGAACTGGTGCGGACCAGGGGAATC | >hg38_21_+_8433172_8446622_RNA45SN1_45S_50ntFlanks | 11576 | 11603 | 28 | CTTAGAACTGGTGCGGACCAGGGGAATC | rRF | FALSE | FALSE | TRUE |
| >hg38_21_+_8433172_8446622_RNA45SN1_45S_50ntFlanks@4657.4689.33\|CTAGAGGTGAAATTCTTGGACCGGCGCAAGACG | >hg38_21_+_8433172_8446622_RNA45SN1_45S_50ntFlanks | 4657 | 4689 | 33 | CTAGAGGTGAAATTCTTGGACCGGCGCAAGACG | rRF | FALSE | FALSE | TRUE |
| >hg38_21_+_8433172_8446622_RNA45SN1_45S_50ntFlanks@10420.10456.37\|CTGAGAGATGGGCGAGCGCCGTTCCGAAGGGACGGGC | >hg38_21_+_8433172_8446622_RNA45SN1_45S_50ntFlanks | 10420 | 10456 | 37 | CTGAGAGATGGGCGAGCGCCGTTCCGAAGGGACGGGC | rRF | FALSE | FALSE | TRUE |
| >hg38_21_+_8433172_8446622_RNA45SN1_45S_50ntFlanks@12935.12956.22\|TAAACCATTCGTAGACGACCTG | >hg38_21_+_8433172_8446622_RNA45SN1_45S_50ntFlanks | 12935 | 12956 | 22 | TAAACCATTCGTAGACGACCTG | rRF | FALSE | FALSE | TRUE |
| >hg38_21_+_8433172_8446622_RNA45SN1_45S_50ntFlanks@3984.4002.19\|GCGGCTTTGGTGACTCTAG | >hg38_21_+_8433172_8446622_RNA45SN1_45S_50ntFlanks | 3984 | 4002 | 19 | GCGGCTTTGGTGACTCTAG | rRF | FALSE | FALSE | TRUE |
| >hg38_21_+_8433172_8446622_RNA45SN1_45S_50ntFlanks@11641.11670.30\|CGGCGGGTGTTGACGCGATGTGATTTCTGC | >hg38_21_+_8433172_8446622_RNA45SN1_45S_50ntFlanks | 11641 | 11670 | 30 | CGGCGGGTGTTGACGCGATGTGATTTCTGC | rRF | FALSE | FALSE | TRUE |
| >hg38_21_+_8433172_8446622_RNA45SN1_45S_50ntFlanks@10737.10755.19\|TAAATCTCGCGCCGGGCCG | >hg38_21_+_8433172_8446622_RNA45SN1_45S_50ntFlanks | 10737 | 10755 | 19 | TAAATCTCGCGCCGGGCCG | rRF | FALSE | FALSE | TRUE |
| >hg38_21_+_8433172_8446622_RNA45SN1_45S_50ntFlanks@10350.10390.41\|ACGAGAACTTTGAAGGCCGAAGTGGAGAAGGGTTCCATGTG | >hg38_21_+_8433172_8446622_RNA45SN1_45S_50ntFlanks | 10350 | 10390 | 41 | ACGAGAACTTTGAAGGCCGAAGTGGAGAAGGGTTCCATGTG | rRF | FALSE | FALSE | TRUE |
| >hg38_21_+_8433172_8446622_RNA45SN1_45S_50ntFlanks@10350.10377.28\|ACGAGAACTTTGAAGGCCGAAGTGGAGA | >hg38_21_+_8433172_8446622_RNA45SN1_45S_50ntFlanks | 10350 | 10377 | 28 | ACGAGAACTTTGAAGGCCGAAGTGGAGA | rRF | FALSE | FALSE | TRUE |
| >hg38_21_+_8433172_8446622_RNA45SN1_45S_50ntFlanks@12076.12095.20\|AGGGGCTCTCGCTTCTGGCG | >hg38_21_+_8433172_8446622_RNA45SN1_45S_50ntFlanks | 12076 | 12095 | 20 | AGGGGCTCTCGCTTCTGGCG | rRF | FALSE | FALSE | TRUE |
| >hg38_21_+_8433172_8446622_RNA45SN1_45S_50ntFlanks@3874.3898.25\|AATACATGCCGACGGGCGCTGACCC | >hg38_21_+_8433172_8446622_RNA45SN1_45S_50ntFlanks | 3874 | 3898 | 25 | AATACATGCCGACGGGCGCTGACCC | rRF | FALSE | FALSE | TRUE |
| >hg38_21_+_8433172_8446622_RNA45SN1_45S_50ntFlanks@12457.12476.20\|CAAGCGTTGGATTGTTCACC | >hg38_21_+_8433172_8446622_RNA45SN1_45S_50ntFlanks | 12457 | 12476 | 20 | CAAGCGTTGGATTGTTCACC | rRF | FALSE | FALSE | TRUE |
| >hg38_21_+_8433172_8446622_RNA45SN1_45S_50ntFlanks@12407.12434.28\|TTTTGATCCTTCGATGTCGGCTCTTCCT | >hg38_21_+_8433172_8446622_RNA45SN1_45S_50ntFlanks | 12407 | 12434 | 28 | TTTTGATCCTTCGATGTCGGCTCTTCCT | rRF | FALSE | FALSE | TRUE |
| >hg38_21_+_8433172_8446622_RNA45SN1_45S_50ntFlanks@11873.11893.21\|AAGAAGACCCTGTTGAGCTTG | >hg38_21_+_8433172_8446622_RNA45SN1_45S_50ntFlanks | 11873 | 11893 | 21 | AAGAAGACCCTGTTGAGCTTG | rRF | FALSE | FALSE | TRUE |
| >hg38_21_+_8433172_8446622_RNA45SN1_45S_50ntFlanks@4760.4780.21\|TCAGATACCGTCGTAGTTCCG | >hg38_21_+_8433172_8446622_RNA45SN1_45S_50ntFlanks | 4760 | 4780 | 21 | TCAGATACCGTCGTAGTTCCG | rRF | FALSE | FALSE | TRUE |
| >hg38_21_+_8433172_8446622_RNA45SN1_45S_50ntFlanks@10362.10388.27\|AAGGCCGAAGTGGAGAAGGGTTCCATG | >hg38_21_+_8433172_8446622_RNA45SN1_45S_50ntFlanks | 10362 | 10388 | 27 | AAGGCCGAAGTGGAGAAGGGTTCCATG | rRF | FALSE | FALSE | TRUE |
| >trna2_GlyGCC_21_-_18827107_18827177@1.30.30,trna35_GlyGCC_1_+_161413094_161413164@1.30.30,trna37_GlyGCC_1_+_161420467_161420537@1.30.30,trna39_GlyGCC_1_+_161427898_161427968@1.30.30,trna41_GlyGCC_1_+_161435258_161435328@1.30.30\|GCATGGGTGGTTCAGTGGTAGAATTCTCGC | >trna2_GlyGCC_21_-_18827107_18827177 | 1 | 30 | 30 | GCATGGGTGGTTCAGTGGTAGAATTCTCGC | tRF | FALSE | FALSE | TRUE |
| >hg38_21_+_8433172_8446622_RNA45SN1_45S_50ntFlanks@10681.10706.26\|TTCCGGCGGCGTCCGGTGAGCTCTCG | >hg38_21_+_8433172_8446622_RNA45SN1_45S_50ntFlanks | 10681 | 10706 | 26 | TTCCGGCGGCGTCCGGTGAGCTCTCG | rRF | FALSE | FALSE | TRUE |
| >hg38_21_+_8433172_8446622_RNA45SN1_45S_50ntFlanks@12418.12446.29\|CGATGTCGGCTCTTCCTATCATTGTGAAG | >hg38_21_+_8433172_8446622_RNA45SN1_45S_50ntFlanks | 12418 | 12446 | 29 | CGATGTCGGCTCTTCCTATCATTGTGAAG | rRF | FALSE | FALSE | TRUE |
| >hg38_21_+_8433172_8446622_RNA45SN1_45S_50ntFlanks@12418.12455.38\|CGATGTCGGCTCTTCCTATCATTGTGAAGCAGAATTCA | >hg38_21_+_8433172_8446622_RNA45SN1_45S_50ntFlanks | 12418 | 12455 | 38 | CGATGTCGGCTCTTCCTATCATTGTGAAGCAGAATTCA | rRF | FALSE | FALSE | TRUE |
| >hg38_21_+_8433172_8446622_RNA45SN1_45S_50ntFlanks@6721.6758.38\|AATTGCAGGACACATTGATCATCGACACTTCGAACGCA | >hg38_21_+_8433172_8446622_RNA45SN1_45S_50ntFlanks | 6721 | 6758 | 38 | AATTGCAGGACACATTGATCATCGACACTTCGAACGCA | rRF | FALSE | FALSE | TRUE |
| >hg38_21_+_8433172_8446622_RNA45SN1_45S_50ntFlanks@8176.8203.28\|CTTCTGATCGAGGCCCAGCCCGTGGACG | >hg38_21_+_8433172_8446622_RNA45SN1_45S_50ntFlanks | 8176 | 8203 | 28 | CTTCTGATCGAGGCCCAGCCCGTGGACG | rRF | FALSE | FALSE | TRUE |
| >hg38_21_+_8433172_8446622_RNA45SN1_45S_50ntFlanks@6661.6689.29\|CGGTGGATCACTCGGCTCGTGCGTCGATG | >hg38_21_+_8433172_8446622_RNA45SN1_45S_50ntFlanks | 6661 | 6689 | 29 | CGGTGGATCACTCGGCTCGTGCGTCGATG | rRF | FALSE | FALSE | TRUE |
| >hg38_21_+_8433172_8446622_RNA45SN1_45S_50ntFlanks@10389.10410.22\|TGAACAGCAGTTGAACATGGGT | >hg38_21_+_8433172_8446622_RNA45SN1_45S_50ntFlanks | 10389 | 10410 | 22 | TGAACAGCAGTTGAACATGGGT | rRF | FALSE | FALSE | TRUE |
| >hg38_21_+_8433172_8446622_RNA45SN1_45S_50ntFlanks@11703.11728.26\|AATGAAGCGCGGGTAAACGGCGGGAG | >hg38_21_+_8433172_8446622_RNA45SN1_45S_50ntFlanks | 11703 | 11728 | 26 | AATGAAGCGCGGGTAAACGGCGGGAG | rRF | FALSE | FALSE | TRUE |
| >hg38_21_+_8433172_8446622_RNA45SN1_45S_50ntFlanks@9534.9552.19\|AGGAAACTCTGGTGGAGGT | >hg38_21_+_8433172_8446622_RNA45SN1_45S_50ntFlanks | 9534 | 9552 | 19 | AGGAAACTCTGGTGGAGGT | rRF | FALSE | FALSE | TRUE |
| >hg38_21_+_8433172_8446622_RNA45SN1_45S_50ntFlanks@12539.12569.31\|ATGATGTGTTGTTGCCATGGTAATCCTGCTC | >hg38_21_+_8433172_8446622_RNA45SN1_45S_50ntFlanks | 12539 | 12569 | 31 | ATGATGTGTTGTTGCCATGGTAATCCTGCTC | rRF | FALSE | FALSE | TRUE |
| >hg38_21_+_8433172_8446622_RNA45SN1_45S_50ntFlanks@9565.9597.33\|TGACGTGCAAATCGGTCGTCCGACCTGGGTATA | >hg38_21_+_8433172_8446622_RNA45SN1_45S_50ntFlanks | 9565 | 9597 | 33 | TGACGTGCAAATCGGTCGTCCGACCTGGGTATA | rRF | FALSE | FALSE | TRUE |
| >hg38_21_+_8433172_8446622_RNA45SN1_45S_50ntFlanks@10841.10867.27\|CGTAACTTCGGGATAAGGATTGGCTCT | >hg38_21_+_8433172_8446622_RNA45SN1_45S_50ntFlanks | 10841 | 10867 | 27 | CGTAACTTCGGGATAAGGATTGGCTCT | rRF | FALSE | FALSE | TRUE |
| >hg38_21_+_8433172_8446622_RNA45SN1_45S_50ntFlanks@6719.6737.19\|TGAATTGCAGGACACATTG | >hg38_21_+_8433172_8446622_RNA45SN1_45S_50ntFlanks | 6719 | 6737 | 19 | TGAATTGCAGGACACATTG | rRF | FALSE | FALSE | TRUE |
| >hg38_21_+_8433172_8446622_RNA45SN1_45S_50ntFlanks@10392.10429.38\|ACAGCAGTTGAACATGGGTCAGTCGGTCCTGAGAGATG | >hg38_21_+_8433172_8446622_RNA45SN1_45S_50ntFlanks | 10392 | 10429 | 38 | ACAGCAGTTGAACATGGGTCAGTCGGTCCTGAGAGATG | rRF | FALSE | FALSE | TRUE |
| >hg38_21_+_8433172_8446622_RNA45SN1_45S_50ntFlanks@8170.8197.28\|CAAGTCCTTCTGATCGAGGCCCAGCCCG | >hg38_21_+_8433172_8446622_RNA45SN1_45S_50ntFlanks | 8170 | 8197 | 28 | CAAGTCCTTCTGATCGAGGCCCAGCCCG | rRF | FALSE | FALSE | TRUE |
| >hg38_21_+_8433172_8446622_RNA45SN1_45S_50ntFlanks@11317.11342.26\|ACCCCACGTCTCGTCGCGCGCGCGTC | >hg38_21_+_8433172_8446622_RNA45SN1_45S_50ntFlanks | 11317 | 11342 | 26 | ACCCCACGTCTCGTCGCGCGCGCGTC | rRF | FALSE | FALSE | TRUE |
| >hg38_21_+_8433172_8446622_RNA45SN1_45S_50ntFlanks@9567.9596.30\|ACGTGCAAATCGGTCGTCCGACCTGGGTAT | >hg38_21_+_8433172_8446622_RNA45SN1_45S_50ntFlanks | 9567 | 9596 | 30 | ACGTGCAAATCGGTCGTCCGACCTGGGTAT | rRF | FALSE | FALSE | TRUE |
| >hg38_21_+_8433172_8446622_RNA45SN1_45S_50ntFlanks@11768.11792.25\|CTAATTAGTGACGCGCATGAATGGA | >hg38_21_+_8433172_8446622_RNA45SN1_45S_50ntFlanks | 11768 | 11792 | 25 | CTAATTAGTGACGCGCATGAATGGA | rRF | FALSE | FALSE | TRUE |
| >hg38_21_+_8433172_8446622_RNA45SN1_45S_50ntFlanks@4728.4758.31\|ATCAAGAACGAAAGTCGGAGGTTCGAAGACG | >hg38_21_+_8433172_8446622_RNA45SN1_45S_50ntFlanks | 4728 | 4758 | 31 | ATCAAGAACGAAAGTCGGAGGTTCGAAGACG | rRF | FALSE | FALSE | TRUE |
| >hg38_21_+_8433172_8446622_RNA45SN1_45S_50ntFlanks@4047.4066.20\|ATTCGAACGTCTGCCCTATC | >hg38_21_+_8433172_8446622_RNA45SN1_45S_50ntFlanks | 4047 | 4066 | 20 | ATTCGAACGTCTGCCCTATC | rRF | FALSE | FALSE | TRUE |

### Nucleus

| | | | | | | | | | |
|---|---|---|---|---|---|---|---|---|---|
| >hg38_21_+_8433172_8446622_RNA45SN1_45S_50ntFlanks@5445.5466.22\|TCGGATCGGCCCCGCCGGGGTC | >hg38_21_+_8433172_8446622_RNA45SN1_45S_50ntFlanks | 5445 | 5466 | 22 | TCGGATCGGCCCCGCCGGGGTC | rRF | FALSE | FALSE | TRUE |
| >hg38_21_+_8433172_8446622_RNA45SN1_45S_50ntFlanks@3835.3860.26\|CTCCTCTCCTACTTGGATAACTGTGG | >hg38_21_+_8433172_8446622_RNA45SN1_45S_50ntFlanks | 3835 | 3860 | 26 | CTCCTCTCCTACTTGGATAACTGTGG | rRF | FALSE | FALSE | TRUE |
| >hg38_21_+_8433172_8446622_RNA45SN1_45S_50ntFlanks@9534.9562.29\|AGGAAACTCTGGTGGAGGTCCGTAGCGGT | >hg38_21_+_8433172_8446622_RNA45SN1_45S_50ntFlanks | 9534 | 9562 | 29 | AGGAAACTCTGGTGGAGGTCCGTAGCGGT | rRF | FALSE | FALSE | TRUE |
| >hg38_21_+_8433172_8446622_RNA45SN1_45S_50ntFlanks@11671.11702.32\|CCAGTGCTCTGAATGTCAAAGTGAAGAAATTC | >hg38_21_+_8433172_8446622_RNA45SN1_45S_50ntFlanks | 11671 | 11702 | 32 | CCAGTGCTCTGAATGTCAAAGTGAAGAAATTC | rRF | FALSE | FALSE | TRUE |
| >multi-am_trnaMT_ValTAC_MT_+_1602_1670@49.68.20\|AACTTAACTTGACCGCTCTG | >multi-am_trnaMT_ValTAC_MT_+_1602_1670 | 49 | 68 | 20 | AACTTAACTTGACCGCTCTG | tRF | FALSE | FALSE | TRUE |
| >hg38_21_+_8433172_8446622_RNA45SN1_45S_50ntFlanks@9539.9571.33\|ACTCTGGTGGAGGTCCGTAGCGGTCCTGACGTG | >hg38_21_+_8433172_8446622_RNA45SN1_45S_50ntFlanks | 9539 | 9571 | 33 | ACTCTGGTGGAGGTCCGTAGCGGTCCTGACGTG | rRF | FALSE | FALSE | TRUE |
| >hg38_21_+_8433172_8446622_RNA45SN1_45S_50ntFlanks@11561.11601.41\|GCGCCTAGCAGCCGACTTAGAACTGGTGCGGACCAGGGGAA | >hg38_21_+_8433172_8446622_RNA45SN1_45S_50ntFlanks | 11561 | 11601 | 41 | GCGCCTAGCAGCCGACTTAGAACTGGTGCGGACCAGGGGAA | rRF | FALSE | FALSE | TRUE |
| >hg38_21_+_8433172_8446622_RNA45SN1_45S_50ntFlanks@11316.11343.28\|CACCCCACGTCTCGTCGCGCGCGCGTCC | >hg38_21_+_8433172_8446622_RNA45SN1_45S_50ntFlanks | 11316 | 11343 | 28 | CACCCCACGTCTCGTCGCGCGCGCGTCC | rRF | FALSE | FALSE | TRUE |
| >hg38_21_+_8433172_8446622_RNA45SN1_45S_50ntFlanks@12418.12444.27\|CGATGTCGGCTCTTCCTATCATTGTGA | >hg38_21_+_8433172_8446622_RNA45SN1_45S_50ntFlanks | 12418 | 12444 | 27 | CGATGTCGGCTCTTCCTATCATTGTGA | rRF | FALSE | FALSE | TRUE |
| >hg38_21_+_8433172_8446622_RNA45SN1_45S_50ntFlanks@10402.10430.29\|AACATGGGTCAGTCGGTCCTGAGAGATGG | >hg38_21_+_8433172_8446622_RNA45SN1_45S_50ntFlanks | 10402 | 10430 | 29 | AACATGGGTCAGTCGGTCCTGAGAGATGG | rRF | FALSE | FALSE | TRUE |
| >hg38_21_+_8433172_8446622_RNA45SN1_45S_50ntFlanks@10735.10755.21\|TGTAAATCTCGCGCCGGGCCG | >hg38_21_+_8433172_8446622_RNA45SN1_45S_50ntFlanks | 10735 | 10755 | 21 | TGTAAATCTCGCGCCGGGCCG | rRF | FALSE | FALSE | TRUE |
| >hg38_21_+_8433172_8446622_RNA45SN1_45S_50ntFlanks@10686.10710.25\|GCGGCGTCCGGTGAGCTCTCGCTGG | >hg38_21_+_8433172_8446622_RNA45SN1_45S_50ntFlanks | 10686 | 10710 | 25 | GCGGCGTCCGGTGAGCTCTCGCTGG | rRF | FALSE | FALSE | TRUE |
| >hg38_21_+_8433172_8446622_RNA45SN1_45S_50ntFlanks@11641.11665.25\|CGGCGGGTGTTGACGCGATGTGATT | >hg38_21_+_8433172_8446622_RNA45SN1_45S_50ntFlanks | 11641 | 11665 | 25 | CGGCGGGTGTTGACGCGATGTGATT | rRF | FALSE | FALSE | TRUE |
| >hg38_21_+_8433172_8446622_RNA45SN1_45S_50ntFlanks@12417.12437.21\|TCGATGTCGGCTCTTCCTATC | >hg38_21_+_8433172_8446622_RNA45SN1_45S_50ntFlanks | 12417 | 12437 | 21 | TCGATGTCGGCTCTTCCTATC | rRF | FALSE | FALSE | TRUE |
| >hg38_21_+_8433172_8446622_RNA45SN1_45S_50ntFlanks@9566.9596.31\|GACGTGCAAATCGGTCGTCCGACCTGGGTAT | >hg38_21_+_8433172_8446622_RNA45SN1_45S_50ntFlanks | 9566 | 9596 | 31 | GACGTGCAAATCGGTCGTCCGACCTGGGTAT | rRF | FALSE | FALSE | TRUE |
| >hg38_21_+_8433172_8446622_RNA45SN1_45S_50ntFlanks@11999.12031.33\|CCCTGCGGGCCGCCGGTGAAATACCACTACTCT | >hg38_21_+_8433172_8446622_RNA45SN1_45S_50ntFlanks | 11999 | 12031 | 33 | CCCTGCGGGCCGCCGGTGAAATACCACTACTCT | rRF | FALSE | FALSE | TRUE |
| >hg38_21_+_8433172_8446622_RNA45SN1_45S_50ntFlanks@10408.10430.23\|GGTCAGTCGGTCCTGAGAGATGG | >hg38_21_+_8433172_8446622_RNA45SN1_45S_50ntFlanks | 10408 | 10430 | 23 | GGTCAGTCGGTCCTGAGAGATGG | rRF | FALSE | FALSE | TRUE |
| >hg38_21_+_8433172_8446622_RNA45SN1_45S_50ntFlanks@6646.6679.34\|TCGTACGACTCTTAGCGGTGGATCACTCGGCTCG | >hg38_21_+_8433172_8446622_RNA45SN1_45S_50ntFlanks | 6646 | 6679 | 34 | TCGTACGACTCTTAGCGGTGGATCACTCGGCTCG | rRF | FALSE | FALSE | TRUE |
| >hg38_21_+_8433172_8446622_RNA45SN1_45S_50ntFlanks@10679.10706.28\|GGTTCCGGCGGCGTCCGGTGAGCTCTCG | >hg38_21_+_8433172_8446622_RNA45SN1_45S_50ntFlanks | 10679 | 10706 | 28 | GGTTCCGGCGGCGTCCGGTGAGCTCTCG | rRF | FALSE | FALSE | TRUE |
| >hg38_21_+_8433172_8446622_RNA45SN1_45S_50ntFlanks@12628.12654.27\|TGGGGCGAAGCTACCATCTGTGGGATT | >hg38_21_+_8433172_8446622_RNA45SN1_45S_50ntFlanks | 12628 | 12654 | 27 | TGGGGCGAAGCTACCATCTGTGGGATT | rRF | FALSE | FALSE | TRUE |
| >hg38_21_+_8433172_8446622_RNA45SN1_45S_50ntFlanks@9534.9573.40\|AGGAAACTCTGGTGGAGGTCCGTAGCGGTCCTGACGTGCA | >hg38_21_+_8433172_8446622_RNA45SN1_45S_50ntFlanks | 9534 | 9573 | 40 | AGGAAACTCTGGTGGAGGTCCGTAGCGGTCCTGACGTGCA | rRF | FALSE | FALSE | TRUE |
| >hg38_21_+_8433172_8446622_RNA45SN1_45S_50ntFlanks@10393.10410.18\|CAGCAGTTGAACATGGGT | >hg38_21_+_8433172_8446622_RNA45SN1_45S_50ntFlanks | 10393 | 10410 | 18 | CAGCAGTTGAACATGGGT | rRF | FALSE | FALSE | TRUE |
| >hg38_21_+_8433172_8446622_RNA45SN1_45S_50ntFlanks@12415.12443.29\|CTTCGATGTCGGCTCTTCCTATCATTGTG | >hg38_21_+_8433172_8446622_RNA45SN1_45S_50ntFlanks | 12415 | 12443 | 29 | CTTCGATGTCGGCTCTTCCTATCATTGTG | rRF | FALSE | FALSE | TRUE |
| >hg38_21_+_8433172_8446622_RNA45SN1_45S_50ntFlanks@10869.10902.34\|AGGGCTGGGTCGGTCGGGCTGGGGCGCGAAGCGG | >hg38_21_+_8433172_8446622_RNA45SN1_45S_50ntFlanks | 10869 | 10902 | 34 | AGGGCTGGGTCGGTCGGGCTGGGGCGCGAAGCGG | rRF | FALSE | FALSE | TRUE |
| >hg38_21_+_8433172_8446622_RNA45SN1_45S_50ntFlanks@11572.11612.41\|CCGACTTAGAACTGGTGCGGACCAGGGGAATCCGACTGTTT | >hg38_21_+_8433172_8446622_RNA45SN1_45S_50ntFlanks | 11572 | 11612 | 41 | CCGACTTAGAACTGGTGCGGACCAGGGGAATCCGACTGTTT | rRF | FALSE | FALSE | TRUE |
| >hg38_21_+_8433172_8446622_RNA45SN1_45S_50ntFlanks@11561.11603.43\|GCGCCTAGCAGCCGACTTAGAACTGGTGCGGACCAGGGGAATC | >hg38_21_+_8433172_8446622_RNA45SN1_45S_50ntFlanks | 11561 | 11603 | 43 | GCGCCTAGCAGCCGACTTAGAACTGGTGCGGACCAGGGGAAT[illegible] | rRF | FALSE | FALSE | TRUE |
| >hg38_21_+_8433172_8446622_RNA45SN1_45S_50ntFlanks@4738.4758.21\|AAAGTCGGAGGTTCGAAGACG | >hg38_21_+_8433172_8446622_RNA45SN1_45S_50ntFlanks | 4738 | 4758 | 21 | AAAGTCGGAGGTTCGAAGACG | rRF | FALSE | FALSE | TRUE |
| >hg38_21_+_8433172_8446622_RNA45SN1_45S_50ntFlanks@9738.9765.28\|GAAACGATCTCAACCTATTCTCAAACTT | >hg38_21_+_8433172_8446622_RNA45SN1_45S_50ntFlanks | 9738 | 9765 | 28 | GAAACGATCTCAACCTATTCTCAAACTT | rRF | FALSE | FALSE | TRUE |
| >hg38_21_+_8433172_8446622_RNA45SN1_45S_50ntFlanks@12457.12478.22\|CAAGCGTTGGATTGTTCACCCA | >hg38_21_+_8433172_8446622_RNA45SN1_45S_50ntFlanks | 12457 | 12478 | 22 | CAAGCGTTGGATTGTTCACCCA | rRF | FALSE | FALSE | TRUE |
| >hg38_21_+_8433172_8446622_RNA45SN1_45S_50ntFlanks@10392.10424.33\|ACAGCAGTTGAACATGGGTCAGTCGGTCCTGAG | >hg38_21_+_8433172_8446622_RNA45SN1_45S_50ntFlanks | 10392 | 10424 | 33 | ACAGCAGTTGAACATGGGTCAGTCGGTCCTGAG | rRF | FALSE | FALSE | TRUE |
| >hg38_21_+_8433172_8446622_RNA45SN1_45S_50ntFlanks@4003.4024.22\|ATAACCTCGGGCCGATCGCACG | >hg38_21_+_8433172_8446622_RNA45SN1_45S_50ntFlanks | 4003 | 4024 | 22 | ATAACCTCGGGCCGATCGCACG | rRF | FALSE | FALSE | TRUE |
| >hg38_21_+_8433172_8446622_RNA45SN1_45S_50ntFlanks@11322.11343.22\|ACGTCTCGTCGCGCGCGCGTCC | >hg38_21_+_8433172_8446622_RNA45SN1_45S_50ntFlanks | 11322 | 11343 | 22 | ACGTCTCGTCGCGCGCGCGTCC | rRF | FALSE | FALSE | TRUE |
| >am_trna146_GlnCTG_6_-_27515531_27515602@1.29.29,trna1_GlnCTG_6_+_18836402_18836473@1.29.29,trna3_GlnCTG_17_+_8023070_8023141@1.29.29,trna42_GlnCTG_6_+_27263212_27263283@1.29.29,trna49_GlnCTG_6_+_27487308_27487379@1.29.29,trna7_GlnCTG_15_-_66161400_66161471@1.29.29,trna99_GlnCTG_6_-_28909378_28909449@1.29.29\|GGTTCCATGGTGTAATGGTTAGCACTCTG | >am_trna146_GlnCTG_6_-_27515531_27515602 | 1 | 29 | 29 | GGTTCCATGGTGTAATGGTTAGCACTCTG | tRF | FALSE | FALSE | TRUE |
| >hg38_21_+_8433172_8446622_RNA45SN1_45S_50ntFlanks@12417.12443.27\|TCGATGTCGGCTCTTCCTATCATTGTG | >hg38_21_+_8433172_8446622_RNA45SN1_45S_50ntFlanks | 12417 | 12443 | 27 | TCGATGTCGGCTCTTCCTATCATTGTG | rRF | FALSE | FALSE | TRUE |
| >hg38_21_+_8433172_8446622_RNA45SN1_45S_50ntFlanks@10846.10865.20\|CTTCGGGATAAGGATTGGCT | >hg38_21_+_8433172_8446622_RNA45SN1_45S_50ntFlanks | 10846 | 10865 | 20 | CTTCGGGATAAGGATTGGCT | rRF | FALSE | FALSE | TRUE |
| >hg38_21_+_8433172_8446622_RNA45SN1_45S_50ntFlanks@12933.12956.24\|GCTAAACCATTCGTAGACGACCTG | >hg38_21_+_8433172_8446622_RNA45SN1_45S_50ntFlanks | 12933 | 12956 | 24 | GCTAAACCATTCGTAGACGACCTG | rRF | FALSE | FALSE | TRUE |
| >hg38_21_+_8433172_8446622_RNA45SN1_45S_50ntFlanks@12415.12437.23\|CTTCGATGTCGGCTCTTCCTATC | >hg38_21_+_8433172_8446622_RNA45SN1_45S_50ntFlanks | 12415 | 12437 | 23 | CTTCGATGTCGGCTCTTCCTATC | rRF | FALSE | FALSE | TRUE |
| >hg38_21_+_8433172_8446622_RNA45SN1_45S_50ntFlanks@11586.11603.18\|GTGCGGACCAGGGGAATC | >hg38_21_+_8433172_8446622_RNA45SN1_45S_50ntFlanks | 11586 | 11603 | 18 | GTGCGGACCAGGGGAATC | rRF | FALSE | FALSE | TRUE |

### Nucleus

| | | | | | | | | | |
|---|---|---|---|---|---|---|---|---|---|
| >hg38_21_+_8433172_8446622_RNA45SN1_45S_50ntFlanks@6738.6771.34\|ATCATCGACACTTCGAACGCACTTGCGGCCCCGG | >hg38_21_+_8433172_8446622_RNA45SN1_45S_50ntFlanks | 6738 | 6771 | 34 | ATCATCGACACTTCGAACGCACTTGCGGCCCCGG | rRF | FALSE | FALSE | TRUE |
| >hg38_21_+_8433172_8446622_RNA45SN1_45S_50ntFlanks@12007.12043.37\|GCCGCCGGTGAAATACCACTACTCTGATCGTTTTTTC | >hg38_21_+_8433172_8446622_RNA45SN1_45S_50ntFlanks | 12007 | 12043 | 37 | GCCGCCGGTGAAATACCACTACTCTGATCGTTTTTTC | rRF | FALSE | FALSE | TRUE |
| >hg38_21_+_8433172_8446622_RNA45SN1_45S_50ntFlanks@11316.11336.21\|CACCCCACGTCTCGTCGCGCG | >hg38_21_+_8433172_8446622_RNA45SN1_45S_50ntFlanks | 11316 | 11336 | 21 | CACCCCACGTCTCGTCGCGCG | rRF | FALSE | FALSE | TRUE |
| >hg38_21_+_8433172_8446622_RNA45SN1_45S_50ntFlanks@11570.11601.32\|AGCCGACTTAGAACTGGTGCGGACCAGGGGAA | >hg38_21_+_8433172_8446622_RNA45SN1_45S_50ntFlanks | 11570 | 11601 | 32 | AGCCGACTTAGAACTGGTGCGGACCAGGGGAA | rRF | FALSE | FALSE | TRUE |
| >hg38_21_+_8433172_8446622_RNA45SN1_45S_50ntFlanks@10866.10902.37\|CTAAGGGCTGGGTCGGTCGGGCTGGGGCGCGAAGCGG | >hg38_21_+_8433172_8446622_RNA45SN1_45S_50ntFlanks | 10866 | 10902 | 37 | CTAAGGGCTGGGTCGGTCGGGCTGGGGCGCGAAGCGG | rRF | FALSE | FALSE | TRUE |
| >hg38_21_+_8433172_8446622_RNA45SN1_45S_50ntFlanks@10681.10712.32\|TTCCGGCGGCGTCCGGTGAGCTCTCGCTGGCC | >hg38_21_+_8433172_8446622_RNA45SN1_45S_50ntFlanks | 10681 | 10712 | 32 | TTCCGGCGGCGTCCGGTGAGCTCTCGCTGGCC | rRF | FALSE | FALSE | TRUE |
| >hg38_21_+_8433172_8446622_RNA45SN1_45S_50ntFlanks@9564.9595.32\|CTGACGTGCAAATCGGTCGTCCGACCTGGGTA | >hg38_21_+_8433172_8446622_RNA45SN1_45S_50ntFlanks | 9564 | 9595 | 32 | CTGACGTGCAAATCGGTCGTCCGACCTGGGTA | rRF | FALSE | FALSE | TRUE |
| >am_trna116_GluCTC_1_-_145399233_145399304@1.27.27,trna59_GluCTC_1_+_249168447_249168518@1.27.27,trna71_GluCTC_1_-_161439189_161439260@1.27.27,trna74_GluCTC_1_-_161431809_161431880@1.27.27,trna77_GluCTC_1_-_161424398_161424469@1.27.27,trna77_GluCTC_6_+_28949976_28950047@1.27.27,trna80_GluCTC_1_-_161417018_161417089@1.27.27,trna87_GluCTC_6_-_126101393_126101464@1.27.27\|TCCCTGGTGGTCTAGTGGTTAGGATTC | >am_trna116_GluCTC_1_-_145399233_145399304 | 1 | 27 | 27 | TCCCTGGTGGTCTAGTGGTTAGGATTC | tRF | FALSE | FALSE | TRUE |
| >hg38_21_+_8433172_8446622_RNA45SN1_45S_50ntFlanks@12938.12956.19\|ACCATTCGTAGACGACCTG | >hg38_21_+_8433172_8446622_RNA45SN1_45S_50ntFlanks | 12938 | 12956 | 19 | ACCATTCGTAGACGACCTG | rRF | FALSE | FALSE | TRUE |
| >hg38_21_+_8433172_8446622_RNA45SN1_45S_50ntFlanks@10350.10388.39\|ACGAGAACTTTGAAGGCCGAAGTGGAGAAGGGTTCCATG | >hg38_21_+_8433172_8446622_RNA45SN1_45S_50ntFlanks | 10350 | 10388 | 39 | ACGAGAACTTTGAAGGCCGAAGTGGAGAAGGGTTCCATG | rRF | FALSE | FALSE | TRUE |
| >hg38_21_+_8433172_8446622_RNA45SN1_45S_50ntFlanks@10513.10533.21\|CCGGAGTGGCGGAGATGGGCG | >hg38_21_+_8433172_8446622_RNA45SN1_45S_50ntFlanks | 10513 | 10533 | 21 | CCGGAGTGGCGGAGATGGGCG | rRF | FALSE | FALSE | TRUE |
| >hg38_21_+_8433172_8446622_RNA45SN1_45S_50ntFlanks@11781.11803.23\|CGCATGAATGGATGAACGAGATT | >hg38_21_+_8433172_8446622_RNA45SN1_45S_50ntFlanks | 11781 | 11803 | 23 | CGCATGAATGGATGAACGAGATT | rRF | FALSE | FALSE | TRUE |
| >hg38_21_+_8433172_8446622_RNA45SN1_45S_50ntFlanks@12457.12477.21\|CAAGCGTTGGATTGTTCACCC | >hg38_21_+_8433172_8446622_RNA45SN1_45S_50ntFlanks | 12457 | 12477 | 21 | CAAGCGTTGGATTGTTCACCC | rRF | FALSE | FALSE | TRUE |
| >hg38_21_+_8433172_8446622_RNA45SN1_45S_50ntFlanks@11027.11060.34\|CTCTCTCTCTCTCTCCCCCGCTCCCCGTCCTCCC | >hg38_21_+_8433172_8446622_RNA45SN1_45S_50ntFlanks | 11027 | 11060 | 34 | CTCTCTCTCTCTCTCCCCCGCTCCCCGTCCTCCC | rRF | FALSE | FALSE | TRUE |
| >hg38_21_+_8433172_8446622_RNA45SN1_45S_50ntFlanks@12457.12487.31\|CAAGCGTTGGATTGTTCACCCACTAATAGGG | >hg38_21_+_8433172_8446622_RNA45SN1_45S_50ntFlanks | 12457 | 12487 | 31 | CAAGCGTTGGATTGTTCACCCACTAATAGGG | rRF | FALSE | FALSE | TRUE |
| >hg38_21_+_8433172_8446622_RNA45SN1_45S_50ntFlanks@6699.6719.21\|GCTAGCTGCGAGAATTAATGT | >hg38_21_+_8433172_8446622_RNA45SN1_45S_50ntFlanks | 6699 | 6719 | 21 | GCTAGCTGCGAGAATTAATGT | rRF | FALSE | FALSE | TRUE |
| >hg38_21_+_8433172_8446622_RNA45SN1_45S_50ntFlanks@12405.12426.22\|CTTTTTGATCCTTCGATGTCGG | >hg38_21_+_8433172_8446622_RNA45SN1_45S_50ntFlanks | 12405 | 12426 | 22 | CTTTTTGATCCTTCGATGTCGG | rRF | FALSE | FALSE | TRUE |
| >hg38_21_+_8433172_8446622_RNA45SN1_45S_50ntFlanks@11564.11603.40\|CCTAGCAGCCGACTTAGAACTGGTGCGGACCAGGGGAATC | >hg38_21_+_8433172_8446622_RNA45SN1_45S_50ntFlanks | 11564 | 11603 | 40 | CCTAGCAGCCGACTTAGAACTGGTGCGGACCAGGGGAATC | rRF | FALSE | FALSE | TRUE |
| >hg38_21_+_8433172_8446622_RNA45SN1_45S_50ntFlanks@12626.12653.28\|AATGGGGCGAAGCTACCATCTGTGGGAT | >hg38_21_+_8433172_8446622_RNA45SN1_45S_50ntFlanks | 12626 | 12653 | 28 | AATGGGGCGAAGCTACCATCTGTGGGAT | rRF | FALSE | FALSE | TRUE |
| >hg38_21_+_8433172_8446622_RNA45SN1_45S_50ntFlanks@4003.4020.18\|ATAACCTCGGGCCGATCG | >hg38_21_+_8433172_8446622_RNA45SN1_45S_50ntFlanks | 4003 | 4020 | 18 | ATAACCTCGGGCCGATCG | rRF | FALSE | FALSE | TRUE |
| >hg38_21_+_8433172_8446622_RNA45SN1_45S_50ntFlanks@4730.4758.29\|CAAGAACGAAAGTCGGAGGTTCGAAGACG | >hg38_21_+_8433172_8446622_RNA45SN1_45S_50ntFlanks | 4730 | 4758 | 29 | CAAGAACGAAAGTCGGAGGTTCGAAGACG | rRF | FALSE | FALSE | TRUE |
| >hg38_21_+_8433172_8446622_RNA45SN1_45S_50ntFlanks@4046.4063.18\|CATTCGAACGTCTGCCCT | >hg38_21_+_8433172_8446622_RNA45SN1_45S_50ntFlanks | 4046 | 4063 | 18 | CATTCGAACGTCTGCCCT | rRF | FALSE | FALSE | TRUE |
| >hg38_21_+_8433172_8446622_RNA45SN1_45S_50ntFlanks@11668.11702.35\|TGCCCAGTGCTCTGAATGTCAAAGTGAAGAAATTC | >hg38_21_+_8433172_8446622_RNA45SN1_45S_50ntFlanks | 11668 | 11702 | 35 | TGCCCAGTGCTCTGAATGTCAAAGTGAAGAAATTC | rRF | FALSE | FALSE | TRUE |
| >hg38_21_+_8433172_8446622_RNA45SN1_45S_50ntFlanks@8170.8214.45\|CAAGTCCTTCTGATCGAGGCCCAGCCCGTGGACGGTGTGAGGCCG | >hg38_21_+_8433172_8446622_RNA45SN1_45S_50ntFlanks | 8170 | 8214 | 45 | CAAGTCCTTCTGATCGAGGCCCAGCCCGTGGACGGTGTGAGG | rRF | FALSE | FALSE | TRUE |
| >hg38_21_+_8433172_8446622_RNA45SN1_45S_50ntFlanks@12420.12445.26\|ATGTCGGCTCTTCCTATCATTGTGAA | >hg38_21_+_8433172_8446622_RNA45SN1_45S_50ntFlanks | 12420 | 12445 | 26 | ATGTCGGCTCTTCCTATCATTGTGAA | rRF | FALSE | FALSE | TRUE |
| >hg38_21_+_8433172_8446622_RNA45SN1_45S_50ntFlanks@10680.10709.30\|GTTCCGGCGGCGTCCGGTGAGCTCTCGCTG | >hg38_21_+_8433172_8446622_RNA45SN1_45S_50ntFlanks | 10680 | 10709 | 30 | GTTCCGGCGGCGTCCGGTGAGCTCTCGCTG | rRF | FALSE | FALSE | TRUE |
| >hg38_21_+_8433172_8446622_RNA45SN1_45S_50ntFlanks@10351.10374.24\|CGAGAACTTTGAAGGCCGAAGTGG | >hg38_21_+_8433172_8446622_RNA45SN1_45S_50ntFlanks | 10351 | 10374 | 24 | CGAGAACTTTGAAGGCCGAAGTGG | rRF | FALSE | FALSE | TRUE |
| >hg38_21_+_8433172_8446622_RNA45SN1_45S_50ntFlanks@11564.11582.19\|CCTAGCAGCCGACTTAGAA | >hg38_21_+_8433172_8446622_RNA45SN1_45S_50ntFlanks | 11564 | 11582 | 19 | CCTAGCAGCCGACTTAGAA | rRF | FALSE | FALSE | TRUE |
| >hg38_21_+_8433172_8446622_RNA45SN1_45S_50ntFlanks@11641.11660.20\|CGGCGGGTGTTGACGCGATG | >hg38_21_+_8433172_8446622_RNA45SN1_45S_50ntFlanks | 11641 | 11660 | 20 | CGGCGGGTGTTGACGCGATG | rRF | FALSE | FALSE | TRUE |
| >hg38_21_+_8433172_8446622_RNA45SN1_45S_50ntFlanks@9454.9503.50\|CCGCGCCGGGGAGGTGGAGCACGAGCGCACGTGTTAGGACCCGAAAGATG | >hg38_21_+_8433172_8446622_RNA45SN1_45S_50ntFlanks | 9454 | 9503 | 50 | CCGCGCCGGGGAGGTGGAGCACGAGCGCACGTGTTAGGACC | rRF | FALSE | FALSE | TRUE |
| >hg38_21_+_8433172_8446622_RNA45SN1_45S_50ntFlanks@11572.11590.19\|CCGACTTAGAACTGGTGCG | >hg38_21_+_8433172_8446622_RNA45SN1_45S_50ntFlanks | 11572 | 11590 | 19 | CCGACTTAGAACTGGTGCG | rRF | FALSE | FALSE | TRUE |
| >hg38_21_+_8433172_8446622_RNA45SN1_45S_50ntFlanks@11670.11701.32\|CCCAGTGCTCTGAATGTCAAAGTGAAGAAATT | >hg38_21_+_8433172_8446622_RNA45SN1_45S_50ntFlanks | 11670 | 11701 | 32 | CCCAGTGCTCTGAATGTCAAAGTGAAGAAATT | rRF | FALSE | FALSE | TRUE |
| >hg38_21_+_8433172_8446622_RNA45SN1_45S_50ntFlanks@8194.8215.22\|CCCGTGGACGGTGTGAGGCCGG | >hg38_21_+_8433172_8446622_RNA45SN1_45S_50ntFlanks | 8194 | 8215 | 22 | CCCGTGGACGGTGTGAGGCCGG | rRF | FALSE | FALSE | TRUE |
| >hg38_21_+_8433172_8446622_RNA45SN1_45S_50ntFlanks@3775.3792.18\|GCCGGTACAGTGAAACTG | >hg38_21_+_8433172_8446622_RNA45SN1_45S_50ntFlanks | 3775 | 3792 | 18 | GCCGGTACAGTGAAACTG | rRF | FALSE | FALSE | TRUE |
| >hg38_21_+_8433172_8446622_RNA45SN1_45S_50ntFlanks@10402.10424.23\|AACATGGGTCAGTCGGTCCTGAG | >hg38_21_+_8433172_8446622_RNA45SN1_45S_50ntFlanks | 10402 | 10424 | 23 | AACATGGGTCAGTCGGTCCTGAG | rRF | FALSE | FALSE | TRUE |
| >hg38_21_+_8433172_8446622_RNA45SN1_45S_50ntFlanks@10349.10368.20\|AACGAGAACTTTGAAGGCCG | >hg38_21_+_8433172_8446622_RNA45SN1_45S_50ntFlanks | 10349 | 10368 | 20 | AACGAGAACTTTGAAGGCCG | rRF | FALSE | FALSE | TRUE |
| >hg38_21_+_8433172_8446622_RNA45SN1_45S_50ntFlanks@9568.9596.29\|CGTGCAAATCGGTCGTCCGACCTGGGTAT | >hg38_21_+_8433172_8446622_RNA45SN1_45S_50ntFlanks | 9568 | 9596 | 29 | CGTGCAAATCGGTCGTCCGACCTGGGTAT | rRF | FALSE | FALSE | TRUE |

### Nucleus

| | | | | | | | | | |
|---|---|---|---|---|---|---|---|---|---|
| >hg38_21_+_8433172_8446622_RNA45SN1_45S_50ntFlanks@10512.10533.22\|TCCGGAGTGGCGGAGATGGGCG | >hg38_21_+_8433172_8446622_RNA45SN1_45S_50ntFlanks | 10512 | 10533 | 22 | TCCGGAGTGGCGGAGATGGGCG | rRF | FALSE | FALSE | TRUE |
| >hg38_21_+_8433172_8446622_RNA45SN1_45S_50ntFlanks@10682.10706.25\|TCCGGCGGCGTCCGGTGAGCTCTCG | >hg38_21_+_8433172_8446622_RNA45SN1_45S_50ntFlanks | 10682 | 10706 | 25 | TCCGGCGGCGTCCGGTGAGCTCTCG | rRF | FALSE | FALSE | TRUE |
| >hg38_21_+_8433172_8446622_RNA45SN1_45S_50ntFlanks@13002.13029.28\|CGATCTATTGAAAGTCAGCCCTCGACAC | >hg38_21_+_8433172_8446622_RNA45SN1_45S_50ntFlanks | 13002 | 13029 | 28 | CGATCTATTGAAAGTCAGCCCTCGACAC | rRF | FALSE | FALSE | TRUE |
| >am_trna11_ProAGG_16_+_3241989_3242060@1.29.29,trna12_ProAGG_6_+_26555498_26555569@1.29.29,trna12_ProTGG_11_-_75946869_75946940@1.29.29,trna14_ProTGG_5_-_180615854_180615925@1.29.29,trna22_ProAGG_14_-_21081560_21081631@1.29.29,trna23_ProAGG_14_-_21077495_21077566@1.29.29,trna28_ProTGG_16_-_3234133_3234204@1.29.29,trna29_ProAGG_16_-_3232635_3232706@1.29.29,trna2_ProAGG_7_+_128423504_128423575@1.29.29,trna30_ProCGG_6_+_27059521_27059592@1.29.29,trna37_ProCGG_17_-_8126151_8126222@1.29.29,trna3_ProTGG_16_+_3208923_3208994@1.29.29,trna52_ProCGG_1_+_167683962_167684033@1.29.29,trna65_ProAGG_1_-_167684725_167684796@1.29.29,trna6_ProCGG_16_+_3222049_3222120@1.29.29,trna6_ProTGG_14_+_21152175_21152246@1.29.29,trna8_ProTGG_16_+_3238094_3238165@1.29.29,trna9_ProAGG_11_+_75946557_75946628@1.29.29,trna9_ProAGG_16_+_3239634_3239705@1.29.29\|GGCTCGTTGGTCTAGGGGTATGATTCTCG | >am_trna11_ProAGG_16_+_3241989_3242060 | 1 | 29 | 29 | GGCTCGTTGGTCTAGGGGTATGATTCTCG | tRF | FALSE | FALSE | TRUE |
| >hg38_21_+_8433172_8446622_RNA45SN1_45S_50ntFlanks@12633.12654.22\|CGAAGCTACCATCTGTGGGATT | >hg38_21_+_8433172_8446622_RNA45SN1_45S_50ntFlanks | 12633 | 12654 | 22 | CGAAGCTACCATCTGTGGGATT | rRF | FALSE | FALSE | TRUE |
| >hg38_21_+_8433172_8446622_RNA45SN1_45S_50ntFlanks@11767.11794.28\|TCTAATTAGTGACGCGCATGAATGGATG | >hg38_21_+_8433172_8446622_RNA45SN1_45S_50ntFlanks | 11767 | 11794 | 28 | TCTAATTAGTGACGCGCATGAATGGATG | rRF | FALSE | FALSE | TRUE |
| >hg38_21_+_8433172_8446622_RNA45SN1_45S_50ntFlanks@11575.11594.20\|ACTTAGAACTGGTGCGGACC | >hg38_21_+_8433172_8446622_RNA45SN1_45S_50ntFlanks | 11575 | 11594 | 20 | ACTTAGAACTGGTGCGGACC | rRF | FALSE | FALSE | TRUE |
| >hg38_21_+_8433172_8446622_RNA45SN1_45S_50ntFlanks@4995.5024.30\|ACAGATTGATAGCTCTTTCTCGATTCCGTG | >hg38_21_+_8433172_8446622_RNA45SN1_45S_50ntFlanks | 4995 | 5024 | 30 | ACAGATTGATAGCTCTTTCTCGATTCCGTG | rRF | FALSE | FALSE | TRUE |
| >hg38_21_+_8433172_8446622_RNA45SN1_45S_50ntFlanks@12003.12032.30\|GCGGGCCGCCGGTGAAATACCACTACTCTG | >hg38_21_+_8433172_8446622_RNA45SN1_45S_50ntFlanks | 12003 | 12032 | 30 | GCGGGCCGCCGGTGAAATACCACTACTCTG | rRF | FALSE | FALSE | TRUE |
| >hg38_21_+_8433172_8446622_RNA45SN1_45S_50ntFlanks@11318.11342.25\|CCCCACGTCTCGTCGCGCGCGCGTC | >hg38_21_+_8433172_8446622_RNA45SN1_45S_50ntFlanks | 11318 | 11342 | 25 | CCCCACGTCTCGTCGCGCGCGCGTC | rRF | FALSE | FALSE | TRUE |
| >hg38_21_+_8433172_8446622_RNA45SN1_45S_50ntFlanks@11208.11234.27\|GCGGCGACTCTGGACGCGAGCCGGGCC | >hg38_21_+_8433172_8446622_RNA45SN1_45S_50ntFlanks | 11208 | 11234 | 27 | GCGGCGACTCTGGACGCGAGCCGGGCC | rRF | FALSE | FALSE | TRUE |
| >hg38_21_+_8433172_8446622_RNA45SN1_45S_50ntFlanks@6738.6770.33\|ATCATCGACACTTCGAACGCACTTGCGGCCCCG | >hg38_21_+_8433172_8446622_RNA45SN1_45S_50ntFlanks | 6738 | 6770 | 33 | ATCATCGACACTTCGAACGCACTTGCGGCCCCG | rRF | FALSE | FALSE | TRUE |
| >hg38_21_+_8433172_8446622_RNA45SN1_45S_50ntFlanks@9534.9565.32\|AGGAAACTCTGGTGGAGGTCCGTAGCGGTCCT | >hg38_21_+_8433172_8446622_RNA45SN1_45S_50ntFlanks | 9534 | 9565 | 32 | AGGAAACTCTGGTGGAGGTCCGTAGCGGTCCT | rRF | FALSE | FALSE | TRUE |
| >hg38_21_+_8433172_8446622_RNA45SN1_45S_50ntFlanks@12994.13013.20\|CCTCGCTGCGATCTATTGAA | >hg38_21_+_8433172_8446622_RNA45SN1_45S_50ntFlanks | 12994 | 13013 | 20 | CCTCGCTGCGATCTATTGAA | rRF | FALSE | FALSE | TRUE |
| >hg38_21_+_8433172_8446622_RNA45SN1_45S_50ntFlanks@12456.12483.28\|CCAAGCGTTGGATTGTTCACCCACTAAT | >hg38_21_+_8433172_8446622_RNA45SN1_45S_50ntFlanks | 12456 | 12483 | 28 | CCAAGCGTTGGATTGTTCACCCACTAAT | rRF | FALSE | FALSE | TRUE |
| >hg38_21_+_8433172_8446622_RNA45SN1_45S_50ntFlanks@6733.6771.39\|CATTGATCATCGACACTTCGAACGCACTTGCGGCCCCGG | >hg38_21_+_8433172_8446622_RNA45SN1_45S_50ntFlanks | 6733 | 6771 | 39 | CATTGATCATCGACACTTCGAACGCACTTGCGGCCCCGG | rRF | FALSE | FALSE | TRUE |
| >hg38_21_+_8433172_8446622_RNA45SN1_45S_50ntFlanks@12415.12441.27\|CTTCGATGTCGGCTCTTCCTATCATTG | >hg38_21_+_8433172_8446622_RNA45SN1_45S_50ntFlanks | 12415 | 12441 | 27 | CTTCGATGTCGGCTCTTCCTATCATTG | rRF | FALSE | FALSE | TRUE |
| >am_trna134_GluTTC_1_-_16861774_16861845@1.29.29,trna5_GluTTC_1_+_17199078_17199149@1.29.29,trna84_GluTTC_1_-_161391883_161391954@1.29.29,trna94_GluTTC_1_-_149664355_149664427@1.29.29\|TCCCTGGTGGTCTAGTGGCTAGGATTCGG | >am_trna134_GluTTC_1_-_16861774_16861845 | 1 | 29 | 29 | TCCCTGGTGGTCTAGTGGCTAGGATTCGG | tRF | FALSE | FALSE | TRUE |
| >hg38_21_+_8433172_8446622_RNA45SN1_45S_50ntFlanks@11199.11225.27\|CCGGCGGCGGCGGCGACTCTGGACGCG | >hg38_21_+_8433172_8446622_RNA45SN1_45S_50ntFlanks | 11199 | 11225 | 27 | CCGGCGGCGGCGGCGACTCTGGACGCG | rRF | FALSE | FALSE | TRUE |
| >hg38_21_+_8433172_8446622_RNA45SN1_45S_50ntFlanks@11581.11603.23\|AACTGGTGCGGACCAGGGGAATC | >hg38_21_+_8433172_8446622_RNA45SN1_45S_50ntFlanks | 11581 | 11603 | 23 | AACTGGTGCGGACCAGGGGAATC | rRF | FALSE | FALSE | TRUE |
| >hg38_21_+_8433172_8446622_RNA45SN1_45S_50ntFlanks@10391.10424.34\|AACAGCAGTTGAACATGGGTCAGTCGGTCCTGAG | >hg38_21_+_8433172_8446622_RNA45SN1_45S_50ntFlanks | 10391 | 10424 | 34 | AACAGCAGTTGAACATGGGTCAGTCGGTCCTGAG | rRF | FALSE | FALSE | TRUE |
| >hg38_21_+_8433172_8446622_RNA45SN1_45S_50ntFlanks@6733.6770.38\|CATTGATCATCGACACTTCGAACGCACTTGCGGCCCCG | >hg38_21_+_8433172_8446622_RNA45SN1_45S_50ntFlanks | 6733 | 6770 | 38 | CATTGATCATCGACACTTCGAACGCACTTGCGGCCCCG | rRF | FALSE | FALSE | TRUE |
| >hg38_21_+_8433172_8446622_RNA45SN1_45S_50ntFlanks@9567.9597.31\|ACGTGCAAATCGGTCGTCCGACCTGGGTATA | >hg38_21_+_8433172_8446622_RNA45SN1_45S_50ntFlanks | 9567 | 9597 | 31 | ACGTGCAAATCGGTCGTCCGACCTGGGTATA | rRF | FALSE | FALSE | TRUE |
| >hg38_21_+_8433172_8446622_RNA45SN1_45S_50ntFlanks@11575.11592.18\|ACTTAGAACTGGTGCGGA | >hg38_21_+_8433172_8446622_RNA45SN1_45S_50ntFlanks | 11575 | 11592 | 18 | ACTTAGAACTGGTGCGGA | rRF | FALSE | FALSE | TRUE |
| >hg38_21_+_8433172_8446622_RNA45SN1_45S_50ntFlanks@8171.8212.42\|AAGTCCTTCTGATCGAGGCCCAGCCCGTGGACGGTGTGAGGC | >hg38_21_+_8433172_8446622_RNA45SN1_45S_50ntFlanks | 8171 | 8212 | 42 | AAGTCCTTCTGATCGAGGCCCAGCCCGTGGACGGTGTGAGGC | rRF | FALSE | FALSE | TRUE |
| >hg38_21_+_8433172_8446622_RNA45SN1_45S_50ntFlanks@5471.5499.29\|CACGGCCCTGGCGGAGCGCTGAGAAGACG | >hg38_21_+_8433172_8446622_RNA45SN1_45S_50ntFlanks | 5471 | 5499 | 29 | CACGGCCCTGGCGGAGCGCTGAGAAGACG | rRF | FALSE | FALSE | TRUE |
| >hg38_21_+_8433172_8446622_RNA45SN1_45S_50ntFlanks@12437.12454.18\|CATTGTGAAGCAGAATTC | >hg38_21_+_8433172_8446622_RNA45SN1_45S_50ntFlanks | 12437 | 12454 | 18 | CATTGTGAAGCAGAATTC | rRF | FALSE | FALSE | TRUE |
| >hg38_21_+_8433172_8446622_RNA45SN1_45S_50ntFlanks@10689.10706.18\|GCGTCCGGTGAGCTCTCG | >hg38_21_+_8433172_8446622_RNA45SN1_45S_50ntFlanks | 10689 | 10706 | 18 | GCGTCCGGTGAGCTCTCG | rRF | FALSE | FALSE | TRUE |
| >hg38_21_+_8433172_8446622_RNA45SN1_45S_50ntFlanks@11769.11794.26\|TAATTAGTGACGCGCATGAATGGATG | >hg38_21_+_8433172_8446622_RNA45SN1_45S_50ntFlanks | 11769 | 11794 | 26 | TAATTAGTGACGCGCATGAATGGATG | rRF | FALSE | FALSE | TRUE |
| >hg38_21_+_8433172_8446622_RNA45SN1_45S_50ntFlanks@8356.8378.23\|GAAAAGAACTTTGAAGAGAGAGT | >hg38_21_+_8433172_8446622_RNA45SN1_45S_50ntFlanks | 8356 | 8378 | 23 | GAAAAGAACTTTGAAGAGAGAGT | rRF | FALSE | FALSE | TRUE |

### Nucleus

| | | | | | | | | | |
|---|---|---|---|---|---|---|---|---|---|
| >am_trna11_ProAGG_16_+_3241989_3242060@1.28.28,trna12_ProAGG_6_+_26555498_26555569@1.28.28,trna12_ProTGG_11_-_75946869_75946940@1.28.28,trna14_ProTGG_5_-_180615854_180615925@1.28.28,trna22_ProAGG_14_-_21081560_21081631@1.28.28,trna23_ProAGG_14_-_21077495_21077566@1.28.28,trna28_ProTGG_16_-_3234133_3234204@1.28.28,trna29_ProAGG_16_-_3232635_3232706@1.28.28,trna2_ProAGG_7_+_128423504_128423575@1.28.28,trna30_ProCGG_6_+_27059521_27059592@1.28.28,trna37_ProCGG_17_-_8126151_8126222@1.28.28,trna3_ProTGG_16_+_3208923_3208994@1.28.28,trna52_ProCGG_1_+_167683962_167684033@1.28.28,trna65_ProAGG_1_-_167684725_167684796@1.28.28,trna6_ProCGG_16_+_3222049_3222120@1.28.28,trna6_ProTGG_14_+_21152175_21152246@1.28.28,trna8_ProTGG_16_+_3238094_3238165@1.28.28,trna9_ProAGG_11_+_75946557_75946628@1.28.28,trna9_ProAGG_16_+_3239634_3239705@1.28.28\|GGCTCGTTGGTCTAGGGGTATGATTCTC | >am_trna11_ProAGG_16_+_3241989_3242060 | 1 | 28 | 28 | GGCTCGTTGGTCTAGGGGTATGATTCTC | tRF | FALSE | FALSE | TRUE |
| >hg38_21_+_8433172_8446622_RNA45SN1_45S_50ntFlanks@8357.8375.19\|AAAAGAACTTTGAAGAGAG | >hg38_21_+_8433172_8446622_RNA45SN1_45S_50ntFlanks | 8357 | 8375 | 19 | AAAAGAACTTTGAAGAGAG | rRF | FALSE | FALSE | TRUE |
| >hg38_21_+_8433172_8446622_RNA45SN1_45S_50ntFlanks@11564.11585.22\|CCTAGCAGCCGACTTAGAACTG | >hg38_21_+_8433172_8446622_RNA45SN1_45S_50ntFlanks | 11564 | 11585 | 22 | CCTAGCAGCCGACTTAGAACTG | rRF | FALSE | FALSE | TRUE |
| >hg38_21_+_8433172_8446622_RNA45SN1_45S_50ntFlanks@10362.10390.29\|AAGGCCGAAGTGGAGAAGGGTTCCATGTG | >hg38_21_+_8433172_8446622_RNA45SN1_45S_50ntFlanks | 10362 | 10390 | 29 | AAGGCCGAAGTGGAGAAGGGTTCCATGTG | rRF | FALSE | FALSE | TRUE |
| >hg38_21_+_8433172_8446622_RNA45SN1_45S_50ntFlanks@11580.11603.24\|GAACTGGTGCGGACCAGGGGAATC | >hg38_21_+_8433172_8446622_RNA45SN1_45S_50ntFlanks | 11580 | 11603 | 24 | GAACTGGTGCGGACCAGGGGAATC | rRF | FALSE | FALSE | TRUE |
| >hg38_21_+_8433172_8446622_RNA45SN1_45S_50ntFlanks@5485.5514.30\|AGCGCTGAGAAGACGGTCGAACTTGACTAT | >hg38_21_+_8433172_8446622_RNA45SN1_45S_50ntFlanks | 5485 | 5514 | 30 | AGCGCTGAGAAGACGGTCGAACTTGACTAT | rRF | FALSE | FALSE | TRUE |
| >hg38_21_+_8433172_8446622_RNA45SN1_45S_50ntFlanks@10350.10374.25\|ACGAGAACTTTGAAGGCCGAAGTGG | >hg38_21_+_8433172_8446622_RNA45SN1_45S_50ntFlanks | 10350 | 10374 | 25 | ACGAGAACTTTGAAGGCCGAAGTGG | rRF | FALSE | FALSE | TRUE |
| >hg38_21_+_8433172_8446622_RNA45SN1_45S_50ntFlanks@9568.9597.30\|CGTGCAAATCGGTCGTCCGACCTGGGTATA | >hg38_21_+_8433172_8446622_RNA45SN1_45S_50ntFlanks | 9568 | 9597 | 30 | CGTGCAAATCGGTCGTCCGACCTGGGTATA | rRF | FALSE | FALSE | TRUE |
| >hg38_21_+_8433172_8446622_RNA45SN1_45S_50ntFlanks@11322.11342.21\|ACGTCTCGTCGCGCGCGCGTC | >hg38_21_+_8433172_8446622_RNA45SN1_45S_50ntFlanks | 11322 | 11342 | 21 | ACGTCTCGTCGCGCGCGCGTC | rRF | FALSE | FALSE | TRUE |
| >hg38_21_+_8433172_8446622_RNA45SN1_45S_50ntFlanks@9504.9523.20\|GTGAACTATGCCTGGGCAGG | >hg38_21_+_8433172_8446622_RNA45SN1_45S_50ntFlanks | 9504 | 9523 | 20 | GTGAACTATGCCTGGGCAGG | rRF | FALSE | FALSE | TRUE |
| >hg38_21_+_8433172_8446622_RNA45SN1_45S_50ntFlanks@4002.4024.23\|GATAACCTCGGGCCGATCGCACG | >hg38_21_+_8433172_8446622_RNA45SN1_45S_50ntFlanks | 4002 | 4024 | 23 | GATAACCTCGGGCCGATCGCACG | rRF | FALSE | FALSE | TRUE |
| >hg38_21_+_8433172_8446622_RNA45SN1_45S_50ntFlanks@12992.13039.48\|TCCCTCGCTGCGATCTATTGAAAGTCAGCCCTCGACACAAGGGTTTGT | >hg38_21_+_8433172_8446622_RNA45SN1_45S_50ntFlanks | 12992 | 13039 | 48 | TCCCTCGCTGCGATCTATTGAAAGTCAGCCCTCGACACAAGGG | rRF | FALSE | FALSE | TRUE |
| >trnaMT_LeuTAA_MT_+_3230_3304@16.44.29\|CCCGGTAATCGCATAAAACTTAAAACTTT | >trnaMT_LeuTAA_MT_+_3230_3304 | 16 | 44 | 29 | CCCGGTAATCGCATAAAACTTAAAACTTT | tRF | FALSE | FALSE | TRUE |
| >hg38_21_+_8433172_8446622_RNA45SN1_45S_50ntFlanks@9554.9597.44\|CGTAGCGGTCCTGACGTGCAAATCGGTCGTCCGACCTGGGTATA | >hg38_21_+_8433172_8446622_RNA45SN1_45S_50ntFlanks | 9554 | 9597 | 44 | CGTAGCGGTCCTGACGTGCAAATCGGTCGTCCGACCTGGGTAT | rRF | FALSE | FALSE | TRUE |
| >hg38_21_+_8433172_8446622_RNA45SN1_45S_50ntFlanks@7986.8003.18\|ATCAGACGTGGCGACCCG | >hg38_21_+_8433172_8446622_RNA45SN1_45S_50ntFlanks | 7986 | 8003 | 18 | ATCAGACGTGGCGACCCG | rRF | FALSE | FALSE | TRUE |
| >multi-am_trnaMT_ArgTCG_MT_+_10405_10469@28.48.21\|ATTTCGACTCATTAAATTATG | >multi-am_trnaMT_ArgTCG_MT_+_10405_10469 | 28 | 48 | 21 | ATTTCGACTCATTAAATTATG | tRF | FALSE | FALSE | TRUE |
| >hg38_21_+_8433172_8446622_RNA45SN1_45S_50ntFlanks@5073.5098.26\|AATTCCGATAACGAACGAGACTCTGG | >hg38_21_+_8433172_8446622_RNA45SN1_45S_50ntFlanks | 5073 | 5098 | 26 | AATTCCGATAACGAACGAGACTCTGG | rRF | FALSE | FALSE | TRUE |
| >hg38_21_+_8433172_8446622_RNA45SN1_45S_50ntFlanks@10687.10708.22\|CGGCGTCCGGTGAGCTCTCGCT | >hg38_21_+_8433172_8446622_RNA45SN1_45S_50ntFlanks | 10687 | 10708 | 22 | CGGCGTCCGGTGAGCTCTCGCT | rRF | FALSE | FALSE | TRUE |
| >hg38_21_+_8433172_8446622_RNA45SN1_45S_50ntFlanks@11728.11751.24\|GTAACTATGACTCTCTTAAGGTAG | >hg38_21_+_8433172_8446622_RNA45SN1_45S_50ntFlanks | 11728 | 11751 | 24 | GTAACTATGACTCTCTTAAGGTAG | rRF | FALSE | FALSE | TRUE |
| >hg38_21_+_8433172_8446622_RNA45SN1_45S_50ntFlanks@10841.10863.23\|CGTAACTTCGGGATAAGGATTGG | >hg38_21_+_8433172_8446622_RNA45SN1_45S_50ntFlanks | 10841 | 10863 | 23 | CGTAACTTCGGGATAAGGATTGG | rRF | FALSE | FALSE | TRUE |
| >hg38_21_+_8433172_8446622_RNA45SN1_45S_50ntFlanks@6645.6681.37\|CTCGTACGACTCTTAGCGGTGGATCACTCGGCTCGTG | >hg38_21_+_8433172_8446622_RNA45SN1_45S_50ntFlanks | 6645 | 6681 | 37 | CTCGTACGACTCTTAGCGGTGGATCACTCGGCTCGTG | rRF | FALSE | FALSE | TRUE |
| >hg38_21_+_8433172_8446622_RNA45SN1_45S_50ntFlanks@10346.10368.23\|TCAAACGAGAACTTTGAAGGCCG | >hg38_21_+_8433172_8446622_RNA45SN1_45S_50ntFlanks | 10346 | 10368 | 23 | TCAAACGAGAACTTTGAAGGCCG | rRF | FALSE | FALSE | TRUE |
| >hg38_21_+_8433172_8446622_RNA45SN1_45S_50ntFlanks@6745.6770.26\|ACACTTCGAACGCACTTGCGGCCCCG | >hg38_21_+_8433172_8446622_RNA45SN1_45S_50ntFlanks | 6745 | 6770 | 26 | ACACTTCGAACGCACTTGCGGCCCCG | rRF | FALSE | FALSE | TRUE |
| >hg38_21_+_8433172_8446622_RNA45SN1_45S_50ntFlanks@10402.10431.30\|AACATGGGTCAGTCGGTCCTGAGAGATGGG | >hg38_21_+_8433172_8446622_RNA45SN1_45S_50ntFlanks | 10402 | 10431 | 30 | AACATGGGTCAGTCGGTCCTGAGAGATGGG | rRF | FALSE | FALSE | TRUE |
| >hg38_21_+_8433172_8446622_RNA45SN1_45S_50ntFlanks@11579.11603.25\|AGAACTGGTGCGGACCAGGGGAATC | >hg38_21_+_8433172_8446622_RNA45SN1_45S_50ntFlanks | 11579 | 11603 | 25 | AGAACTGGTGCGGACCAGGGGAATC | rRF | FALSE | FALSE | TRUE |
| >hg38_21_+_8433172_8446622_RNA45SN1_45S_50ntFlanks@4773.4793.21\|TAGTTCCGACCATAAACGATG | >hg38_21_+_8433172_8446622_RNA45SN1_45S_50ntFlanks | 4773 | 4793 | 21 | TAGTTCCGACCATAAACGATG | rRF | FALSE | FALSE | TRUE |
| >hg38_21_+_8433172_8446622_RNA45SN1_45S_50ntFlanks@12991.13016.26\|CTCCCTCGCTGCGATCTATTGAAAGT | >hg38_21_+_8433172_8446622_RNA45SN1_45S_50ntFlanks | 12991 | 13016 | 26 | CTCCCTCGCTGCGATCTATTGAAAGT | rRF | FALSE | FALSE | TRUE |
| >hg38_21_+_8433172_8446622_RNA45SN1_45S_50ntFlanks@10408.10435.28\|GGTCAGTCGGTCCTGAGAGATGGGCGAG | >hg38_21_+_8433172_8446622_RNA45SN1_45S_50ntFlanks | 10408 | 10435 | 28 | GGTCAGTCGGTCCTGAGAGATGGGCGAG | rRF | FALSE | FALSE | TRUE |
