## Supplementary material for "Single-nucleotide Differences and Cell Type Decide the Subcellular Localization of miRNA Isoforms (isomiRs), tRNA-derived Fragments (tRFs) and rRNA-derived Fragments (rRFs)": Supp. Table S5 (2 of 4)

### Cytoplasm

<table>
<tr><th></th><th></th><th></th><th></th><th></th><th></th><th></th><th></th><th></th><th></th></tr>
<tr><td>>trna11_GluTTC_15_-_26327381_26327452@1.33.33,trna3_GluTTC_13_-_45492062_45492133@1.33.33|TCCCACATGGTCTAGCGGTTAGGATTCCTGGTT</td><td>>trna11_GluTTC_15_-_26327381_26327452</td><td>1</td><td>33</td><td>33</td><td>TCCCACATGGTCTAGCGGTTAGGATTCCTGGTT</td><td>tRF</td><td>TRUE</td><td>TRUE</td><td>FALSE</td></tr>
<tr><td>>trna2_GlyGCC_21_-_18827107_18827177@1.28.28,trna35_GlyGCC_1_+_161413094_161413164@1.28.28,trna37_GlyGCC_1_+_161420467_161420537@1.28.28,trna39_GlyGCC_1_+_161427898_161427968@1.28.28,trna41_GlyGCC_1_+_161435258_161435328@1.28.28|GCATGGGTGGTTCAGTGGTAGAATTCTC</td><td>>trna2_GlyGCC_21_-_18827107_18827177</td><td>1</td><td>28</td><td>28</td><td>GCATGGGTGGTTCAGTGGTAGAATTCTC</td><td>tRF</td><td>TRUE</td><td>FALSE</td><td>TRUE</td></tr>
<tr><td>>hg38_21_+_8433172_8446622_RNA45SN1_45S_50ntFlanks@5473.5501.29|CGGCCCTGGCGGAGCGCTGAGAAGACGGT</td><td>>hg38_21_+_8433172_8446622_RNA45SN1_45S_50ntFlanks</td><td>5473</td><td>5501</td><td>29</td><td>CGGCCCTGGCGGAGCGCTGAGAAGACGGT</td><td>rRF</td><td>TRUE</td><td>FALSE</td><td>FALSE</td></tr>
<tr><td>>MI0000469|hsa-mir-125a&WithFlank&19|+|51693248|51693345@21.44.24&[MIMAT0000443&hsa-miR-125a-5p&offsets|0|0;m-63&19|+|51693268|51693289&offsets|0|+2]|TCCCTGAGACCCTTTAACCTGTGA</td><td>>MI0000469|hsa-mir-125a&WithFlank&19|+|51693248|51693345</td><td>21</td><td>44</td><td>24</td><td>TCCCTGAGACCCTTTAACCTGTGA</td><td>isomiR</td><td>TRUE</td><td>FALSE</td><td>FALSE</td></tr>
<tr><td>>MI0000093|hsa-mir-92a-1&WithFlank&13|+|91351308|91351397@54.74.21(+1C);MI0000094|hsa-mir-92a-2&WithFlank&X|-|134169532|134169618@54.74.21(+1C)&[MIMAT0000092&hsa-miR-92a-3p&offsets|0|-1(+1C);m-9&13|+|91351361|91351382&offsets|0|-1(+1C)];[MIMAT0000092_1&hsa-miR-92a-3p&offsets|0|-1(+1C);m-11&X|-|134169544|134169565&offsets|0|-1(+1C)]|TATTGCACTTGTCCCGGCCTGC</td><td>>MI0000093|hsa-mir-92a-1&WithFlank&13|+|91351308|91351397</td><td>54</td><td>74</td><td>21(+1C)</td><td>TATTGCACTTGTCCCGGCCTGC</td><td>isomiR</td><td>TRUE</td><td>TRUE</td><td>FALSE</td></tr>
<tr><td>>MI0000811|hsa-mir-148b&WithFlank&12|+|54337210|54337320@69.90.22&[MIMAT0000759&hsa-miR-148b-3p&offsets|0|0;m-77&12|+|54337278|54337299&offsets|0|0]|TCAGTGCATCACAGAACTTTGT</td><td>>MI0000811|hsa-mir-148b&WithFlank&12|+|54337210|54337320</td><td>69</td><td>90</td><td>22</td><td>TCAGTGCATCACAGAACTTTGT</td><td>isomiR</td><td>TRUE</td><td>TRUE</td><td>FALSE</td></tr>
<tr><td>>MI0000067|hsa-let-7f-1&WithFlank&9|+|94176341|94176439@13.30.18;MI0000068|hsa-let-7f-2&WithFlank&X|-|53557186|53557280@14.31.18&[MIMAT0000067&hsa-let-7f-5p&offsets|0|-4;m-20&9|+|94176353|94176374&offsets|0|-4];[MIMAT0000067_1&hsa-let-7f-5p&offsets|0|-4;m-19&X|-|53557246|53557267&offsets|0|-4]|TGAGGTAGTAGATTGTAT</td><td>>MI0000067|hsa-let-7f-1&WithFlank&9|+|94176341|94176439</td><td>13</td><td>30</td><td>18</td><td>TGAGGTAGTAGATTGTAT</td><td>isomiR</td><td>TRUE</td><td>FALSE</td><td>FALSE</td></tr>
<tr><td>>MI0000253|hsa-mir-148a&WithFlank&7|-|25949913|25949992@50.70.21(+1A)&[MIMAT0000243&hsa-miR-148a-3p&offsets|0|-1(+1A);m-7&7|-|25949922|25949943&offsets|0|-1(+1A)]|TCAGTGCACTACAGAACTTTGA</td><td>>MI0000253|hsa-mir-148a&WithFlank&7|-|25949913|25949992</td><td>50</td><td>70</td><td>21(+1A)</td><td>TCAGTGCACTACAGAACTTTGA</td><td>isomiR</td><td>TRUE</td><td>FALSE</td><td>FALSE</td></tr>
<tr><td>>trna116_GluCTC_1_-_145399233_145399304@1.36.36,trna59_GluCTC_1_+_249168447_249168518@1.36.36,trna71_GluCTC_1_-_161439189_161439260@1.36.36,trna74_GluCTC_1_-_161431809_161431880@1.36.36,trna77_GluCTC_1_-_161424398_161424469@1.36.36,trna77_GluCTC_6_+_28949976_28950047@1.36.36,trna80_GluCTC_1_-_161417018_161417089@1.36.36,trna87_GluCTC_6_-_126101393_126101464@1.36.36|TCCCTGGTGGTCTAGTGGTTAGGATTCGGCGCTCTC</td><td>>trna116_GluCTC_1_-_145399233_145399304</td><td>1</td><td>36</td><td>36</td><td>TCCCTGGTGGTCTAGTGGTTAGGATTCGGCGCTCTC</td><td>tRF</td><td>TRUE</td><td>TRUE</td><td>FALSE</td></tr>
<tr><td>>am_trna128_GlyGCC_6_-_27870686_27870756@1.32.32,trna133_GlyCCC_1_-_16872434_16872504@1.32.32,trna18_GlyGCC_16_+_70822597_70822667@1.32.32,trna19_GlyGCC_16_+_70823410_70823480@1.32.32,trna19_GlyGCC_2_-_157257659_157257729@1.32.32,trna24_GlyGCC_16_-_70812942_70813012@1.32.32,trna25_GlyGCC_16_-_70812114_70812184@1.32.32,trna4_GlyCCC_1_+_17188416_17188486@1.32.32,trna5_GlyGCC_17_+_8029064_8029134@1.32.32,trna68_GlyGCC_1_-_161493637_161493707@1.32.32|GCATTGGTGGTTCAGTGGTAGAATTCTCGCCT</td><td>>am_trna128_GlyGCC_6_-_27870686_27870756</td><td>1</td><td>32</td><td>32</td><td>GCATTGGTGGTTCAGTGGTAGAATTCTCGCCT</td><td>tRF</td><td>TRUE</td><td>TRUE</td><td>FALSE</td></tr>
<tr><td>>trna116_GluCTC_1_-_145399233_145399304@1.32.32,trna59_GluCTC_1_+_249168447_249168518@1.32.32,trna71_GluCTC_1_-_161439189_161439260@1.32.32,trna74_GluCTC_1_-_161431809_161431880@1.32.32,trna77_GluCTC_1_-_161424398_161424469@1.32.32,trna77_GluCTC_6_+_28949976_28950047@1.32.32,trna80_GluCTC_1_-_161417018_161417089@1.32.32,trna87_GluCTC_6_-_126101393_126101464@1.32.32|TCCCTGGTGGTCTAGTGGTTAGGATTCGGCGC</td><td>>trna116_GluCTC_1_-_145399233_145399304</td><td>1</td><td>32</td><td>32</td><td>TCCCTGGTGGTCTAGTGGTTAGGATTCGGCGC</td><td>tRF</td><td>TRUE</td><td>TRUE</td><td>FALSE</td></tr>
<tr><td>>hg38_21_+_8433172_8446622_RNA45SN1_45S_50ntFlanks@7974.8006.33|ACGCGACCTCAGATCAGACGTGGCGACCCGCTG</td><td>>hg38_21_+_8433172_8446622_RNA45SN1_45S_50ntFlanks</td><td>7974</td><td>8006</td><td>33</td><td>ACGCGACCTCAGATCAGACGTGGCGACCCGCTG</td><td>rRF</td><td>TRUE</td><td>FALSE</td><td>FALSE</td></tr>
<tr><td>>MI0000272|hsa-mir-182&WithFlank&7|-|129770377|129770498@29.50.22&[MIMAT0000259&hsa-miR-182-5p&offsets|0|-2;m-22&7|-|129770449|129770470&offsets|0|0]|TTTGGCAATGGTAGAACTCACA</td><td>>MI0000272|hsa-mir-182&WithFlank&7|-|129770377|129770498</td><td>29</td><td>50</td><td>22</td><td>TTTGGCAATGGTAGAACTCACA</td><td>isomiR</td><td>TRUE</td><td>FALSE</td><td>FALSE</td></tr>
<tr><td>>am_MI0000786|hsa-mir-378a&WithFlank&5|+|149732819|149732896@49.69.21(+1U)&[MIMAT0000732&hsa-miR-378a-3p&offsets|0|-1(+1U);m-44&5|+|149732867|149732888&offsets|0|-1(+1U)]|ACTGGACTTGGAGTCAGAAGGT</td><td>>am_MI0000786|hsa-mir-378a&WithFlank&5|+|149732819|149732896</td><td>49</td><td>69</td><td>21(+1U)</td><td>ACTGGACTTGGAGTCAGAAGGT</td><td>isomiR</td><td>TRUE</td><td>FALSE</td><td>FALSE</td></tr>
<tr><td>>MI0000264|hsa-mir-7-2&WithFlank&15|+|88611819|88611940@38.60.23(+1A);MI0000265|hsa-mir-7-3&WithFlank&19|+|4770664|4770785@37.59.23(+1A);MI0000263|hsa-mir-7-1&WithFlank&9|-|83969742|83969863@30.52.23(+1A)&[MIMAT0000252&hsa-miR-7-5p&offsets|0|-1(+1A);m-400&15|+|88611856|88611879&offsets|0|-1(+1A)];[MIMAT0000252_1&hsa-miR-7-5p&offsets|0|-1(+1A);m-402&19|+|4770700|4770723&offsets|0|-1(+1A)];[MIMAT0000252_2&hsa-miR-7-5p&offsets|0|-1(+1A);m-401&9|-|83969811|83969834&offsets|0|-1(+1A)]|TGGAAGACTAGTGATTTTGTTGTA</td><td>>MI0000264|hsa-mir-7-2&WithFlank&15|+|88611819|88611940</td><td>38</td><td>60</td><td>23(+1A)</td><td>TGGAAGACTAGTGATTTTGTTGTA</td><td>isomiR</td><td>TRUE</td><td>FALSE</td><td>FALSE</td></tr>
<tr><td>>hg38_21_+_8433172_8446622_RNA45SN1_45S_50ntFlanks@5488.5516.29|GCTGAGAAGACGGTCGAACTTGACTATCT</td><td>>hg38_21_+_8433172_8446622_RNA45SN1_45S_50ntFlanks</td><td>5488</td><td>5516</td><td>29</td><td>GCTGAGAAGACGGTCGAACTTGACTATCT</td><td>rRF</td><td>TRUE</td><td>FALSE</td><td>FALSE</td></tr>
<tr><td>>hg38_21_+_8433172_8446622_RNA45SN1_45S_50ntFlanks@5487.5516.30|CGCTGAGAAGACGGTCGAACTTGACTATCT</td><td>>hg38_21_+_8433172_8446622_RNA45SN1_45S_50ntFlanks</td><td>5487</td><td>5516</td><td>30</td><td>CGCTGAGAAGACGGTCGAACTTGACTATCT</td><td>rRF</td><td>TRUE</td><td>TRUE</td><td>FALSE</td></tr>
<tr><td>>MI0000279|hsa-mir-196a-2&WithFlank&12|+|53991732|53991853@31.52.22;MI0000238|hsa-mir-196a-1&WithFlank&17|-|48632484|48632565@13.34.22&[MIMAT0000226&hsa-miR-196a-5p&offsets|0|0;m-261&12|+|53991762|53991783&offsets|0|0];[MIMAT0000226_1&hsa-miR-196a-5p&offsets|0|0;m-268&17|-|48632532|48632553&offsets|0|0]|TAGGTAGTTTCATGTTGTTGGG</td><td>>MI0000279|hsa-mir-196a-2&WithFlank&12|+|53991732|53991853</td><td>31</td><td>52</td><td>22</td><td>TAGGTAGTTTCATGTTGTTGGG</td><td>isomiR</td><td>TRUE</td><td>FALSE</td><td>FALSE</td></tr>
<tr><td>>hg38_1_-_228634819_228635039_RNA5S12_5S_50ntFlanks@146.170.25|TGGGAATACCGGGTGCTGTAGGCTT</td><td>>hg38_1_-_228634819_228635039_RNA5S12_5S_50ntFlanks</td><td>146</td><td>170</td><td>25</td><td>TGGGAATACCGGGTGCTGTAGGCTT</td><td>rRF</td><td>TRUE</td><td>TRUE</td><td>FALSE</td></tr>
<tr><td>>hg38_1_-_228634819_228635039_RNA5S12_5S_50ntFlanks@51.94.44|GTCTACGGCCATACCACCCTGAACGCGCCCGATCTCGTCTGATC</td><td>>hg38_1_-_228634819_228635039_RNA5S12_5S_50ntFlanks</td><td>51</td><td>94</td><td>44</td><td>GTCTACGGCCATACCACCCTGAACGCGCCCGATCTCGTCTGATC</td><td>rRF</td><td>TRUE</td><td>FALSE</td><td>FALSE</td></tr>
<tr><td>>MI0000439|hsa-mir-23b&WithFlank&9|+|95085202|95085310@67.86.20&[MIMAT0000418&hsa-miR-23b-3p&offsets|+3|0;m-42&9|+|95085265|95085285&offsets|+3|+2]|ACATTGCCAGGGATTACCAC</td><td>>MI0000439|hsa-mir-23b&WithFlank&9|+|95085202|95085310</td><td>67</td><td>86</td><td>20</td><td>ACATTGCCAGGGATTACCAC</td><td>isomiR</td><td>TRUE</td><td>FALSE</td><td>FALSE</td></tr>
</table>

### Cytoplasm

| | | | | | | | | | |
|---|---|---|---|---|---|---|---|---|---|
| >hg38_21_+_8433172_8446622_RNA45SN1_45S_50ntFlanks@7975.8012.38\|CGCGACCTCAGATCAGACGTGGCGACCCGCTGAATTTA | >hg38_21_+_8433172_8446622_RNA45SN1_45S_50ntFlanks | 7975 | 8012 | 38 | CGCGACCTCAGATCAGACGTGGCGACCCGCTGAATTTA | rRF | TRUE | TRUE | FALSE |
| >MI0000298\|hsa-mir-221&WithFlank&X\|-\|45746151\|45746272@71.93.23(+1U)&[MIMAT0000278&hsa-miR-221-3p&offsets\|0\|0(+1U);m-91&X\|-\|45746181\|45746202&offsets\|0\|+1(+1U)]\|AGCTACATTGTCTGCTGGGTTTCT | >MI0000298\|hsa-mir-221&WithFlank&X\|-\|45746151\|45746272 | 71 | 93 | 23(+1U) | AGCTACATTGTCTGCTGGGTTTCT | isomiR | TRUE | TRUE | FALSE |
| >MI0000087\|hsa-mir-29a&WithFlank&7\|-\|130876741\|130876816@48.68.21&[MIMAT0000086&hsa-miR-29a-3p&offsets\|0\|-1;m-26&7\|-\|130876749\|130876769&offsets\|0\|0]\|TAGCACCATCTGAAATCGGTT | >MI0000087\|hsa-mir-29a&WithFlank&7\|-\|130876741\|130876816 | 48 | 68 | 21 | TAGCACCATCTGAAATCGGTT | isomiR | TRUE | FALSE | FALSE |
| >hg38_1_-_228634819_228635039_RNA5S12_5S_50ntFlanks@133.169.37\|ATGGGAGACCGCCTGGGAATACCGGGTGCTGTAGGCT | >hg38_1_-_228634819_228635039_RNA5S12_5S_50ntFlanks | 133 | 169 | 37 | ATGGGAGACCGCCTGGGAATACCGGGTGCTGTAGGCT | rRF | TRUE | FALSE | FALSE |
| >multi-am_trnaMT_ValTAC_MT_+_1602_1670@49.67.19\|AACTTAACTTGACCGCTCT | >multi-am_trnaMT_ValTAC_MT_+_1602_1670 | 49 | 67 | 19 | AACTTAACTTGACCGCTCT | tRF | TRUE | FALSE | FALSE |
| >am_MI0005763\|hsa-mir-941-1&WithFlank&20\|+\|63919443\|63919526@53.74.22(+1U);MI0005764\|hsa-mir-941-2&WithFlank&20\|+\|63919499\|63919582@53.74.22(+1U);MI0005765\|hsa-mir-941-3&WithFlank&20\|+\|63919555\|63919638@53.74.22(+1U);MI0005766\|hsa-mir-941-4&WithFlank&20\|+\|63919750\|63919833@53.74.22(+1U);MI0031520\|hsa-mir-941-5&WithFlank&20\|+\|63919862\|63919945@53.74.22(+1U)&[MIMAT0004984&hsa-miR-941&offsets\|0\|-1(+1U);m-263&20\|+\|63919495\|63919517&offsets\|0\|-1(+1U)];[MIMAT0004984_1&hsa-miR-941&offsets\|0\|-1(+1U);m-263&20\|+\|63919551\|63919573&offsets\|0\|-1(+1U)];[MIMAT0004984_2&hsa-miR-941&offsets\|0\|-1(+1U);m-263&20\|+\|63919607\|63919629&offsets\|0\|-1(+1U)];[MIMAT0004984_3&hsa-miR-941&offsets\|0\|-1(+1U);m-263&20\|+\|63919802\|63919824&offsets\|0\|-1(+1U)];[MIMAT0004984_4&hsa-miR-941&offsets\|0\|-1(+1U);m-263&20\|+\|63919914\|63919936&offsets\|0\|-1(+1U)]\|CACCCGGCTGTGTGCACATGTGT | >am_MI0005763\|hsa-mir-941-1&WithFlank&20\|+\|63919443\|63919526 | 53 | 74 | 22(+1U) | CACCCGGCTGTGTGCACATGTGT | isomiR | TRUE | FALSE | FALSE |
| >MI0000542\|hsa-mir-320a&WithFlank&8\|-\|22244960\|22245043@48.68.21(+3U)&[MIMAT0000510&hsa-miR-320a-3p&offsets\|0\|-1(+3U);m-60&8\|-\|22244975\|22244996&offsets\|0\|-1(+3U)]\|AAAAGCTGGGTTGAGAGGGCGTTT | >MI0000542\|hsa-mir-320a&WithFlank&8\|-\|22244960\|22245043 | 48 | 68 | 21(+3U) | AAAAGCTGGGTTGAGAGGGCGTTT | isomiR | TRUE | FALSE | FALSE |
| >am_MI0000809\|hsa-mir-151a&WithFlank&8\|-\|140732558\|140732659@53.72.20(+1U)&[MIMAT0000757&hsa-miR-151a-3p&offsets\|0\|-1(+1U);m-30&8\|-\|140732587\|140732607&offsets\|0\|-1(+1U)]\|CTAGACTGAAGCTCCTTGAGT | >am_MI0000809\|hsa-mir-151a&WithFlank&8\|-\|140732558\|140732659 | 53 | 72 | 20(+1U) | CTAGACTGAAGCTCCTTGAGT | isomiR | TRUE | FALSE | FALSE |
| >hg38_21_+_8433172_8446622_RNA45SN1_45S_50ntFlanks@5472.5502.31\|ACGGCCCTGGCGGAGCGCTGAGAAGACGGTC | >hg38_21_+_8433172_8446622_RNA45SN1_45S_50ntFlanks | 5472 | 5502 | 31 | ACGGCCCTGGCGGAGCGCTGAGAAGACGGTC | rRF | TRUE | FALSE | FALSE |
| >MI0000272\|hsa-mir-182&WithFlank&7\|-\|129770377\|129770498@29.54.26&[MIMAT0000259&hsa-miR-182-5p&offsets\|0\|+2;m-22&7\|-\|129770449\|129770470&offsets\|0\|+4]\|TTTGGCAATGGTAGAACTCACACTGG | >MI0000272\|hsa-mir-182&WithFlank&7\|-\|129770377\|129770498 | 29 | 54 | 26 | TTTGGCAATGGTAGAACTCACACTGG | isomiR | TRUE | FALSE | FALSE |
| >trna111_HisGTG_1_-_147774845_147774916@-1G.33.34,trna118_HisGTG_1_-_145396881_145396952@-1G.33.34,trna16_HisGTG_1_+_146544773_146544844@-1G.33.34,trna1_HisGTG_15_+_45493349_45493420@-1G.33.34,trna21_HisGTG_1_+_147753471_147753542@-1G.33.34,trna33_HisGTG_6_+_27125906_27125977@-1G.33.34,trna7_HisGTG_9_-_14433938_14434009@-1G.33.34,trna8_HisGTG_15_-_45492611_45492682@-1G.33.34,trna9_HisGTG_15_-_45490804_45490875@-1G.33.34\|GGCCGTGATCGTATAGTGGTTAGTACTCTGCGTT | >trna111_HisGTG_1_-_147774845_147774916 | -1G | 33 | 34 | GGCCGTGATCGTATAGTGGTTAGTACTCTGCGTT | tRF | TRUE | TRUE | FALSE |
| >hg38_21_+_8433172_8446622_RNA45SN1_45S_50ntFlanks@7975.7992.18\|CGCGACCTCAGATCAGAC | >hg38_21_+_8433172_8446622_RNA45SN1_45S_50ntFlanks | 7975 | 7992 | 18 | CGCGACCTCAGATCAGAC | rRF | TRUE | TRUE | FALSE |
| >hg38_21_+_8433172_8446622_RNA45SN1_45S_50ntFlanks@3705.3741.37\|TACCTGGTTGATCCTGCCAGTAGCATATGCTTGTCTC | >hg38_21_+_8433172_8446622_RNA45SN1_45S_50ntFlanks | 3705 | 3741 | 37 | TACCTGGTTGATCCTGCCAGTAGCATATGCTTGTCTC | rRF | TRUE | FALSE | FALSE |
| >hg38_1_-_228634819_228635039_RNA5S12_5S_50ntFlanks@147.170.24\|GGGAATACCGGGTGCTGTAGGCTT | >hg38_1_-_228634819_228635039_RNA5S12_5S_50ntFlanks | 147 | 170 | 24 | GGGAATACCGGGTGCTGTAGGCTT | rRF | TRUE | TRUE | FALSE |
| >hg38_1_-_228634819_228635039_RNA5S12_5S_50ntFlanks@140.171.32\|ACCGCCTGGGAATACCGGGTGCTGTAGGCTTT | >hg38_1_-_228634819_228635039_RNA5S12_5S_50ntFlanks | 140 | 171 | 32 | ACCGCCTGGGAATACCGGGTGCTGTAGGCTTT | rRF | TRUE | TRUE | FALSE |
| >MI0000255\|hsa-mir-30d&WithFlank&8\|-\|134804870\|134804951@12.33.22&[MIMAT0000245&hsa-miR-30d-5p&offsets\|0\|0;m-21&8\|-\|134804919\|134804940&offsets\|0\|0]\|TGTAAACATCCCCGACTGGAAG | >MI0000255\|hsa-mir-30d&WithFlank&8\|-\|134804870\|134804951 | 12 | 33 | 22 | TGTAAACATCCCCGACTGGAAG | isomiR | TRUE | FALSE | FALSE |
| >MI0000082\|hsa-mir-25&WithFlank&7\|-\|100093554\|100093649@58.78.21&[MIMAT0000081&hsa-miR-25-3p&offsets\|0\|-1;m-23&7\|-\|100093571\|100093592&offsets\|0\|-1]\|CATTGCACTTGTCTCGGTCTG | >MI0000082\|hsa-mir-25&WithFlank&7\|-\|100093554\|100093649 | 58 | 78 | 21 | CATTGCACTTGTCTCGGTCTG | isomiR | TRUE | FALSE | FALSE |
| >MI0000239\|hsa-mir-197&WithFlank&1\|+\|109598887\|109598973@54.74.21(+1U)&[MIMAT0000227&hsa-miR-197-3p&offsets\|0\|-1(+1U);m-99&1\|+\|109598940\|109598961&offsets\|0\|-1(+1U)]\|TTCACCACCTTCTCCACCCAGT | >MI0000239\|hsa-mir-197&WithFlank&1\|+\|109598887\|109598973 | 54 | 74 | 21(+1U) | TTCACCACCTTCTCCACCCAGT | isomiR | TRUE | FALSE | FALSE |
| >MI0000088\|hsa-mir-30a&WithFlank&6\|-\|71403545\|71403627@53.73.21(+1U)&[MIMAT0000088&hsa-miR-30a-3p&offsets\|0\|-1(+1U);m-38&6\|-\|71403554\|71403575&offsets\|0\|-1(+1U)]\|CTTTCAGTCGGATGTTTGCAGT | >MI0000088\|hsa-mir-30a&WithFlank&6\|-\|71403545\|71403627 | 53 | 73 | 21(+1U) | CTTTCAGTCGGATGTTTGCAGT | isomiR | TRUE | TRUE | FALSE |
| >MI0003776\|hsa-mir-320b-1&WithFlank&1\|+\|116671740\|116671823@48.68.21;MI0003839\|hsa-mir-320b-2&WithFlank&1\|-\|224257034\|224257116@47.67.21&[MIMAT0005792&hsa-miR-320b&offsets\|0\|-1;m-183&1\|+\|116671787\|116671806&offsets\|0\|+1];[MIMAT0005792_1&hsa-miR-320b&offsets\|0\|-1;m-183&1\|-\|224257051\|224257070&offsets\|0\|+1]\|AAAAGCTGGGTTGAGAGGGCA | >MI0003776\|hsa-mir-320b-1&WithFlank&1\|+\|116671740\|116671823 | 48 | 68 | 21 | AAAAGCTGGGTTGAGAGGGCA | isomiR | TRUE | FALSE | FALSE |
| >MI0000063\|hsa-let-7b&WithFlank&22\|+\|46113680\|46113774@12.32.21(+1A)&[MIMAT0000063&hsa-let-7b-5p&offsets\|0\|-1(+1A);m-14&22\|+\|46113691\|46113712&offsets\|0\|-1(+1A)]\|TGAGGTAGTAGGTTGTGTGGTA | >MI0000063\|hsa-let-7b&WithFlank&22\|+\|46113680\|46113774 | 12 | 32 | 21(+1A) | TGAGGTAGTAGGTTGTGTGGTA | isomiR | TRUE | FALSE | FALSE |
| >MI0000471\|hsa-mir-126&WithFlank&9\|+\|136670596\|136670692@58.79.22&[MIMAT0000445&hsa-miR-126-3p&offsets\|0\|0;m-40&9\|+\|136670653\|136670674&offsets\|0\|0]\|TCGTACCGTGAGTAATAATGCG | >MI0000471\|hsa-mir-126&WithFlank&9\|+\|136670596\|136670692 | 58 | 79 | 22 | TCGTACCGTGAGTAATAATGCG | isomiR | TRUE | FALSE | FALSE |
| >MI0006444\|hsa-mir-1307&WithFlank&10\|-\|103394247\|103394407@86.108.23&[MIMAT0005951&hsa-miR-1307-3p&offsets\|0\|+1;m-66&10\|-\|103394300\|103394322&offsets\|0\|0]\|ACTCGGCGTGGCGTCGGTCGTGG | >MI0006444\|hsa-mir-1307&WithFlank&10\|-\|103394247\|103394407 | 86 | 108 | 23 | ACTCGGCGTGGCGTCGGTCGTGG | isomiR | TRUE | FALSE | FALSE |
| >MI0000440\|hsa-mir-27b&WithFlank&9\|+\|95085439\|95085547@67.86.20(+1A)&[MIMAT0000419&hsa-miR-27b-3p&offsets\|0\|-1(+1A);m-39&9\|+\|95085505\|95085525&offsets\|0\|-1(+1A)]\|TTCACAGTGGCTAAGTTCTGA | >MI0000440\|hsa-mir-27b&WithFlank&9\|+\|95085439\|95085547 | 67 | 86 | 20(+1A) | TTCACAGTGGCTAAGTTCTGA | isomiR | TRUE | FALSE | FALSE |

### Cytoplasm

| | | | | | | | | | |
|---|---|---|---|---|---|---|---|---|---|
| >MI0000469\|hsa-mir-125a&WithFlank&19\|+\|51693248\|51693345@21.42.22(+1A)&[MIMAT0000443&hsa-miR-125a-5p&offsets\|0\|-2(+1A);m-63&19\|+\|51693268\|51693289&offsets\|0\|0(+1A)]\|TCCCTGAGACCCTTTAACCTGTA | >MI0000469\|hsa-mir-125a&WithFlank&19\|+\|51693248\|51693345 | 21 | 42 | 22(+1A) | TCCCTGAGACCCTTTAACCTGTA | isomiR | TRUE | FALSE | FALSE |
| >MI0000273\|hsa-mir-183&WithFlank&7\|-\|129774899\|129775020@34.54.21&[MIMAT0000261&hsa-miR-183-5p&offsets\|+1\|0;m-41&7\|-\|129774966\|129774987&offsets\|0\|-1]\|ATGGCACTGGTAGAATTCACT | >MI0000273\|hsa-mir-183&WithFlank&7\|-\|129774899\|129775020 | 34 | 54 | 21 | ATGGCACTGGTAGAATTCACT | isomiR | TRUE | FALSE | FALSE |
| >MI0000734\|hsa-mir-106b&WithFlank&7\|-\|100093987\|100094080@58.78.21&[MIMAT0004672&hsa-miR-106b-3p&offsets\|0\|-1;m-87&7\|-\|100094003\|100094023&offsets\|0\|0]\|CCGCACTGTGGGTACTTGCTG | >MI0000734\|hsa-mir-106b&WithFlank&7\|-\|100093987\|100094080 | 58 | 78 | 21 | CCGCACTGTGGGTACTTGCTG | isomiR | TRUE | FALSE | FALSE |
| >hg38_21_+_8433172_8446622_RNA45SN1_45S_50ntFlanks@475.506.32\|CTTCGTGATCGATGTGGTGACGTCGTGCTCTC | >hg38_21_+_8433172_8446622_RNA45SN1_45S_50ntFlanks | 475 | 506 | 32 | CTTCGTGATCGATGTGGTGACGTCGTGCTCTC | rRF | TRUE | FALSE | FALSE |
| >hg38_21_+_8433172_8446622_RNA45SN1_45S_50ntFlanks@6647.6665.19\|CGTACGACTCTTAGCGGTG | >hg38_21_+_8433172_8446622_RNA45SN1_45S_50ntFlanks | 6647 | 6665 | 19 | CGTACGACTCTTAGCGGTG | rRF | TRUE | FALSE | FALSE |
| >hg38_1_-_228634819_228635039_RNA5S12_5S_50ntFlanks@51.88.38\|GTCTACGGCCATACCACCCTGAACGCGCCCGATCTCGT | >hg38_1_-_228634819_228635039_RNA5S12_5S_50ntFlanks | 51 | 88 | 38 | GTCTACGGCCATACCACCCTGAACGCGCCCGATCTCGT | rRF | TRUE | TRUE | FALSE |
| >hg38_21_+_8433172_8446622_RNA45SN1_45S_50ntFlanks@5491.5512.22\|GAGAAGACGGTCGAACTTGACT | >hg38_21_+_8433172_8446622_RNA45SN1_45S_50ntFlanks | 5491 | 5512 | 22 | GAGAAGACGGTCGAACTTGACT | rRF | TRUE | FALSE | FALSE |
| >MI0003560\|hsa-mir-92b&WithFlank&1\|+\|155195171\|155195278@67.87.21(+1U)&[MIMAT0003218&hsa-miR-92b-3p&offsets\|0\|-1(+1U);m-115&1\|+\|155195237\|155195258&offsets\|0\|-1(+1U)]\|TATTGCACTCGTCCCGGCCTCT | >MI0003560\|hsa-mir-92b&WithFlank&1\|+\|155195171\|155195278 | 67 | 87 | 21(+1U) | TATTGCACTCGTCCCGGCCTCT | isomiR | TRUE | FALSE | FALSE |
| >hg38_21_+_8433172_8446622_RNA45SN1_45S_50ntFlanks@7974.7995.22\|ACGCGACCTCAGATCAGACGTG | >hg38_21_+_8433172_8446622_RNA45SN1_45S_50ntFlanks | 7974 | 7995 | 22 | ACGCGACCTCAGATCAGACGTG | rRF | TRUE | FALSE | FALSE |
| >hg38_1_-_228634819_228635039_RNA5S12_5S_50ntFlanks@133.171.39\|ATGGGAGACCGCCTGGGAATACCGGGTGCTGTAGGCTTT | >hg38_1_-_228634819_228635039_RNA5S12_5S_50ntFlanks | 133 | 171 | 39 | ATGGGAGACCGCCTGGGAATACCGGGTGCTGTAGGCTTT | rRF | TRUE | TRUE | FALSE |
| >hg38_21_+_8433172_8446622_RNA45SN1_45S_50ntFlanks@7974.8011.38\|ACGCGACCTCAGATCAGACGTGGCGACCCGCTGAATTT | >hg38_21_+_8433172_8446622_RNA45SN1_45S_50ntFlanks | 7974 | 8011 | 38 | ACGCGACCTCAGATCAGACGTGGCGACCCGCTGAATTT | rRF | TRUE | TRUE | FALSE |
| >MI0000650\|hsa-mir-200c&WithFlank&12\|+\|6963693\|6963772@50.70.21&[MIMAT0000617&hsa-miR-200c-3p&offsets\|0\|-2;m-54&12\|+\|6963742\|6963764&offsets\|0\|-2]\|TAATACTGCCGGGTAATGATG | >MI0000650\|hsa-mir-200c&WithFlank&12\|+\|6963693\|6963772 | 50 | 70 | 21 | TAATACTGCCGGGTAATGATG | isomiR | TRUE | FALSE | FALSE |
| >am_trna116_GluCTC_1_-_145399233_145399304@1.26.26,trna59_GluCTC_1_+_249168447_249168518@1.26.26,trna71_GluCTC_1_-_161439189_161439260@1.26.26,trna74_GluCTC_1_-_161431809_161431880@1.26.26,trna77_GluCTC_1_-_161424398_161424469@1.26.26,trna77_GluCTC_6_+_28949976_28950047@1.26.26,trna80_GluCTC_1_-_161417018_161417089@1.26.26,trna87_GluCTC_6_-_126101393_126101464@1.26.26\|TCCCTGGTGGTCTAGTGGTTAGGATT | >am_trna116_GluCTC_1_-_145399233_145399304 | 1 | 26 | 26 | TCCCTGGTGGTCTAGTGGTTAGGATT | tRF | TRUE | FALSE | FALSE |
| >MI0000067\|hsa-let-7f-1&WithFlank&9\|+\|94176341\|94176439@13.33.21(+1A);MI0000068\|hsa-let-7f-2&WithFlank&X\|-\|53557186\|53557280@14.34.21(+1A)&[MIMAT0000067&hsa-let-7f-5p&offsets\|0\|-1(+1A);m-20&9\|+\|94176353\|94176374&offsets\|0\|-1(+1A)];[MIMAT0000067_1&hsa-let-7f-5p&offsets\|0\|-1(+1A);m-19&X\|-\|53557246\|53557267&offsets\|0\|-1(+1A)]\|TGAGGTAGTAGATTGTATAGTA | >MI0000067\|hsa-let-7f-1&WithFlank&9\|+\|94176341\|94176439 | 13 | 33 | 21(+1A) | TGAGGTAGTAGATTGTATAGTA | isomiR | TRUE | FALSE | FALSE |
| >trna119_LysCTT_1_-_145395522_145395594@1.36.36,trna11_LysCTT_5_-_180648979_180649051@1.36.36,trna13_LysCTT_6_+_26556774_26556846@1.36.36,trna7_LysCTT_16_+_3225692_3225764@1.36.36,trna9_LysCTT_5_+_180634755_180634827@1.36.36\|GCCCGGCTAGCTCAGTCGGTAGAGCATGAGACTCTT | >trna119_LysCTT_1_-_145395522_145395594 | 1 | 36 | 36 | GCCCGGCTAGCTCAGTCGGTAGAGCATGAGACTCTT | tRF | TRUE | FALSE | FALSE |
| >MI0003776\|hsa-mir-320b-1&WithFlank&1\|+\|116671740\|116671823@48.67.20;MI0003839\|hsa-mir-320b-2&WithFlank&1\|-\|224257034\|224257116@47.66.20;MI0000542\|hsa-mir-320a&WithFlank&8\|-\|22244960\|22245043@48.67.20&[MIMAT0005792&hsa-miR-320b&offsets\|0\|-2;m-183&1\|+\|116671787\|116671806&offsets\|0\|0];[MIMAT0005792_1&hsa-miR-320b&offsets\|0\|-2;m-183&1\|-\|224257051\|224257070&offsets\|0\|0];[MIMAT0000510&hsa-miR-320a-3p&offsets\|0\|-2;m-60&8\|-\|22244975\|22244996&offsets\|0\|-2]\|AAAAGCTGGGTTGAGAGGGC | >MI0003776\|hsa-mir-320b-1&WithFlank&1\|+\|116671740\|116671823 | 48 | 67 | 20 | AAAAGCTGGGTTGAGAGGGC | isomiR | TRUE | FALSE | FALSE |
| >MI0000433\|hsa-let-7g&WithFlank&3\|-\|52268272\|52268367@11.31.21&[MIMAT0000414&hsa-let-7g-5p&offsets\|0\|-1;m-58&3\|-\|52268336\|52268357&offsets\|0\|-1]\|TGAGGTAGTAGTTTGTACAGT | >MI0000433\|hsa-let-7g&WithFlank&3\|-\|52268272\|52268367 | 11 | 31 | 21 | TGAGGTAGTAGTTTGTACAGT | isomiR | TRUE | FALSE | FALSE |
| >hg38_21_+_8433172_8446622_RNA45SN1_45S_50ntFlanks@5494.5516.23\|AAGACGGTCGAACTTGACTATCT | >hg38_21_+_8433172_8446622_RNA45SN1_45S_50ntFlanks | 5494 | 5516 | 23 | AAGACGGTCGAACTTGACTATCT | rRF | TRUE | TRUE | FALSE |
| >MI0000108\|hsa-mir-103a-2&WithFlank&20\|+\|3917488\|3917577@54.75.22;MI0000109\|hsa-mir-103a-1&WithFlank&5\|-\|168560890\|168560979@54.75.22&[MIMAT0000101&hsa-miR-103a-3p&offsets\|0\|-1;m-12&20\|+\|3917541\|3917563&offsets\|0\|-1];[MIMAT0000101_1&hsa-miR-103a-3p&offsets\|0\|-1;m-13&5\|-\|168560904\|168560926&offsets\|0\|-1;m-44220&5\|-\|168560908\|168560926&offsets\|0\|+3]\|AGCAGCATTGTACAGGGCTATG | >MI0000108\|hsa-mir-103a-2&WithFlank&20\|+\|3917488\|3917577 | 54 | 75 | 22 | AGCAGCATTGTACAGGGCTATG | isomiR | TRUE | FALSE | FALSE |
| >MI0000273\|hsa-mir-183&WithFlank&7\|-\|129774899\|129775020@33.55.23&[MIMAT0000261&hsa-miR-183-5p&offsets\|0\|+1;m-41&7\|-\|129774966\|129774987&offsets\|-1\|0]\|TATGGCACTGGTAGAATTCACTG | >MI0000273\|hsa-mir-183&WithFlank&7\|-\|129774899\|129775020 | 33 | 55 | 23 | TATGGCACTGGTAGAATTCACTG | isomiR | TRUE | FALSE | FALSE |
| >MI0000070\|hsa-mir-16-1&WithFlank&13\|-\|50048967\|50049067@20.41.22;MI0000115\|hsa-mir-16-2&WithFlank&3\|+\|160404739\|160404831@16.37.22&[MIMAT0000069&hsa-miR-16-5p&offsets\|0\|0;m-62&13\|-\|50049027\|50049048&offsets\|0\|0];[MIMAT0000069_1&hsa-miR-16-5p&offsets\|0\|0;m-61&3\|+\|160404754\|160404775&offsets\|0\|0]\|TAGCAGCACGTAAATATTGGCG | >MI0000070\|hsa-mir-16-1&WithFlank&13\|-\|50048967\|50049067 | 20 | 41 | 22 | TAGCAGCACGTAAATATTGGCG | isomiR | TRUE | FALSE | FALSE |
| >MI0000100\|hsa-mir-98&WithFlank&X\|-\|53556217\|53556347@28.49.22&[MIMAT0000096&hsa-miR-98-5p&offsets\|0\|0;m-143&X\|-\|53556299\|53556320&offsets\|0\|0]\|TGAGGTAGTAAGTTGTATTGTT | >MI0000100\|hsa-mir-98&WithFlank&X\|-\|53556217\|53556347 | 28 | 49 | 22 | TGAGGTAGTAAGTTGTATTGTT | isomiR | TRUE | FALSE | FALSE |
| >hg38_21_+_8433172_8446622_RNA45SN1_45S_50ntFlanks@6645.6669.25\|CTCGTACGACTCTTAGCGGTGGATC | >hg38_21_+_8433172_8446622_RNA45SN1_45S_50ntFlanks | 6645 | 6669 | 25 | CTCGTACGACTCTTAGCGGTGGATC | rRF | TRUE | FALSE | FALSE |
| >MI0003205\|hsa-mir-532&WithFlank&X\|+\|50003142\|50003244@26.47.22&[MIMAT0002888&hsa-miR-532-5p&offsets\|0\|0;m-65&X\|+\|50003167\|50003188&offsets\|0\|0]\|CATGCCTTGAGTGTAGGACCGT | >MI0003205\|hsa-mir-532&WithFlank&X\|+\|50003142\|50003244 | 26 | 47 | 22 | CATGCCTTGAGTGTAGGACCGT | isomiR | TRUE | FALSE | FALSE |

### Cytoplasm

<table>
<tr><td>>MI0000061|hsa-let-7a-2&WithFlank&11|-|122146516|122146599@11.28.18;MI0000064|hsa-let-7c&WithFlank&21|+|16539822|16539917@17.34.18;MI0000062|hsa-let-7a-3&WithFlank&22|+|46112743|46112828@10.27.18;MI0000060|hsa-let-7a-1&WithFlank&9|+|94175951|94176042@12.29.18&[MIMAT0000062&hsa-let-7a-5p&offsets|0|-4;m-6&11|-|122146568|122146589&offsets|0|-4];[MIMAT0000064&hsa-let-7c-5p&offsets|0|-4;m-37&21|+|16539838|16539859&offsets|0|-4];[MIMAT0000062_1&hsa-let-7a-5p&offsets|0|-4;m-5&22|+|46112752|46112773&offsets|0|-4];[MIMAT0000062_2&hsa-let-7a-5p&offsets|0|-4;m-4&9|+|94175962|94175983&offsets|0|-4]|TGAGGTAGTAGGTTGTAT</td><td>>MI0000061|hsa-let-7a-2&WithFlank&11|-|122146516|122146599</td><td>11</td><td>28</td><td>18</td><td>TGAGGTAGTAGGTTGTAT</td><td>isomiR</td><td>TRUE</td><td>FALSE</td><td>FALSE</td></tr>
<tr><td>>MI0000086|hsa-mir-28&WithFlank&3|+|188688775|188688872@60.81.22(+1A)&[MIMAT0004502&hsa-miR-28-3p&offsets|0|0(+1A);m-28&3|+|188688834|188688855&offsets|0|0(+1A)]|CACTAGATTGTGAGCTCCTGGAA</td><td>>MI0000086|hsa-mir-28&WithFlank&3|+|188688775|188688872</td><td>60</td><td>81</td><td>22(+1A)</td><td>CACTAGATTGTGAGCTCCTGGAA</td><td>isomiR</td><td>TRUE</td><td>FALSE</td><td>FALSE</td></tr>
<tr><td>>MI0000811|hsa-mir-148b&WithFlank&12|+|54337210|54337320@69.90.22(+1U)&[MIMAT0000759&hsa-miR-148b-3p&offsets|0|0(+1U);m-77&12|+|54337278|54337299&offsets|0|0(+1U)]|TCAGTGCATCACAGAACTTTGTT</td><td>>MI0000811|hsa-mir-148b&WithFlank&12|+|54337210|54337320</td><td>69</td><td>90</td><td>22(+1U)</td><td>TCAGTGCATCACAGAACTTTGTT</td><td>isomiR</td><td>TRUE</td><td>FALSE</td><td>FALSE</td></tr>
<tr><td>>MI0000066|hsa-let-7e&WithFlank&19|+|51692780|51692870@14.34.21&[MIMAT0000066&hsa-let-7e-5p&offsets|0|-1;m-50&19|+|51692793|51692814&offsets|0|-1]|TGAGGTAGGAGGTTGTATAGT</td><td>>MI0000066|hsa-let-7e&WithFlank&19|+|51692780|51692870</td><td>14</td><td>34</td><td>21</td><td>TGAGGTAGGAGGTTGTATAGT</td><td>isomiR</td><td>TRUE</td><td>FALSE</td><td>FALSE</td></tr>
<tr><td>>hg38_21_+_8433172_8446622_RNA45SN1_45S_50ntFlanks@12995.13036.42|CTCGCTGCGATCTATTGAAAGTCAGCCCTCGACACAAGGGTT</td><td>>hg38_21_+_8433172_8446622_RNA45SN1_45S_50ntFlanks</td><td>12995</td><td>13036</td><td>42</td><td>CTCGCTGCGATCTATTGAAAGTCAGCCCTCGACACAAGGGTT</td><td>rRF</td><td>TRUE</td><td>TRUE</td><td>FALSE</td></tr>
<tr><td>>MI0000650|hsa-mir-200c&WithFlank&12|+|6963693|6963772@50.70.21(+1A)&[MIMAT0000617&hsa-miR-200c-3p&offsets|0|-2(+1A);m-54&12|+|6963742|6963764&offsets|0|-2(+1A)]|TAATACTGCCGGGTAATGATGA</td><td>>MI0000650|hsa-mir-200c&WithFlank&12|+|6963693|6963772</td><td>50</td><td>70</td><td>21(+1A)</td><td>TAATACTGCCGGGTAATGATGA</td><td>isomiR</td><td>TRUE</td><td>FALSE</td><td>FALSE</td></tr>
<tr><td>>MI0000085|hsa-mir-27a&WithFlank&19|-|13836434|13836523@57.76.20&[MIMAT0000084&hsa-miR-27a-3p&offsets|0|-1;m-43&19|-|13836447|13836467&offsets|0|-1]|TTCACAGTGGCTAAGTTCCG</td><td>>MI0000085|hsa-mir-27a&WithFlank&19|-|13836434|13836523</td><td>57</td><td>76</td><td>20</td><td>TTCACAGTGGCTAAGTTCCG</td><td>isomiR</td><td>TRUE</td><td>FALSE</td><td>FALSE</td></tr>
<tr><td>>MI0000301|hsa-mir-224&WithFlank&X|-|151958572|151958664@14.37.24&[MIMAT0000281&hsa-miR-224-5p&offsets|+1|0;m-238&X|-|151958630|151958651&offsets|0|+2]|CAAGTCACTAGTGGTTCCGTTTAG</td><td>>MI0000301|hsa-mir-224&WithFlank&X|-|151958572|151958664</td><td>14</td><td>37</td><td>24</td><td>CAAGTCACTAGTGGTTCCGTTTAG</td><td>isomiR</td><td>TRUE</td><td>FALSE</td><td>FALSE</td></tr>
<tr><td>>MI0000746|hsa-mir-99b&WithFlank&19|+|51692606|51692687@13.34.22(+1U)&[MIMAT0000689&hsa-miR-99b-5p&offsets|0|0(+1U);m-10&19|+|51692618|51692639&offsets|0|0(+1U)]|CACCCGTAGAACCGACCTTGCGT</td><td>>MI0000746|hsa-mir-99b&WithFlank&19|+|51692606|51692687</td><td>13</td><td>34</td><td>22(+1U)</td><td>CACCCGTAGAACCGACCTTGCGT</td><td>isomiR</td><td>TRUE</td><td>FALSE</td><td>FALSE</td></tr>
<tr><td>>MI0000273|hsa-mir-183&WithFlank&7|-|129774899|129775020@34.55.22&[MIMAT0000261&hsa-miR-183-5p&offsets|+1|+1;m-41&7|-|129774966|129774987&offsets|0|0]|ATGGCACTGGTAGAATTCACTG</td><td>>MI0000273|hsa-mir-183&WithFlank&7|-|129774899|129775020</td><td>34</td><td>55</td><td>22</td><td>ATGGCACTGGTAGAATTCACTG</td><td>isomiR</td><td>TRUE</td><td>FALSE</td><td>FALSE</td></tr>
<tr><td>>hg38_21_+_8433172_8446622_RNA45SN1_45S_50ntFlanks@5472.5516.45|ACGGCCCTGGCGGAGCGCTGAGAAGACGGTCGAACTTGACTATCT</td><td>>hg38_21_+_8433172_8446622_RNA45SN1_45S_50ntFlanks</td><td>5472</td><td>5516</td><td>45</td><td>ACGGCCCTGGCGGAGCGCTGAGAAGACGGTCGAACTTGACTATCT</td><td>rRF</td><td>FALSE</td><td>TRUE</td><td>FALSE</td></tr>
<tr><td>>trna111_HisGTG_1_-_147774845_147774916@-1G.34.35,trna118_HisGTG_1_-_145396881_145396952@-1G.34.35,trna16_HisGTG_1_+_146544773_146544844@-1G.34.35,trna1_HisGTG_15_+_45493349_45493420@-1G.34.35,trna21_HisGTG_1_+_147753471_147753542@-1G.34.35,trna33_HisGTG_6_+_27125906_27125977@-1G.34.35,trna7_HisGTG_9_-_14433938_14434009@-1G.34.35,trna8_HisGTG_15_-_45492611_45492682@-1G.34.35,trna9_HisGTG_15_-_45490804_45490875@-1G.34.35|GGCCGTGATCGTATAGTGGTTAGTACTCTGCGTTG</td><td>>trna111_HisGTG_1_-_147774845_147774916</td><td>-1G</td><td>34</td><td>35</td><td>GGCCGTGATCGTATAGTGGTTAGTACTCTGCGTTG</td><td>tRF</td><td>FALSE</td><td>TRUE</td><td>FALSE</td></tr>
<tr><td>>MI0000102|hsa-mir-100&WithFlank&11|-|122152223|122152314@19.40.22(+1U)&[MIMAT0000098&hsa-miR-100-5p&offsets|0|0(+1U);m-27&11|-|122152276|122152296&offsets|0|+1(+1U)]|AACCCGTAGATCCGAACTTGTGT</td><td>>MI0000102|hsa-mir-100&WithFlank&11|-|122152223|122152314</td><td>19</td><td>40</td><td>22(+1U)</td><td>AACCCGTAGATCCGAACTTGTGT</td><td>isomiR</td><td>FALSE</td><td>TRUE</td><td>FALSE</td></tr>
<tr><td>>hg38_21_+_8433172_8446622_RNA45SN1_45S_50ntFlanks@12994.13039.46|CCTCGCTGCGATCTATTGAAAGTCAGCCCTCGACACAAGGGTTTGT</td><td>>hg38_21_+_8433172_8446622_RNA45SN1_45S_50ntFlanks</td><td>12994</td><td>13039</td><td>46</td><td>CCTCGCTGCGATCTATTGAAAGTCAGCCCTCGACACAAGGGTTTGT</td><td>rRF</td><td>FALSE</td><td>TRUE</td><td>FALSE</td></tr>
<tr><td>>hg38_21_+_8433172_8446622_RNA45SN1_45S_50ntFlanks@5473.5516.44|CGGCCCTGGCGGAGCGCTGAGAAGACGGTCGAACTTGACTATCT</td><td>>hg38_21_+_8433172_8446622_RNA45SN1_45S_50ntFlanks</td><td>5473</td><td>5516</td><td>44</td><td>CGGCCCTGGCGGAGCGCTGAGAAGACGGTCGAACTTGACTATCT</td><td>rRF</td><td>FALSE</td><td>TRUE</td><td>FALSE</td></tr>
<tr><td>>trna10_AspGTC_12_-_125424193_125424264@35.75.41,trna12_AspGTC_12_-_125411891_125411962@35.75.41,trna144_AspGTC_6_-_27551236_27551307@35.75.41,trna38_AspGTC_17_-_8125556_8125627@35.75.41,trna45_AspGTC_6_+_27447453_27447524@35.75.41,trna48_AspGTC_6_+_27471523_27471594@35.75.41,trna4_AspGTC_12_+_96429799_96429870@35.75.41,trna69_AspGTC_1_-_161440205_161440276@35.75.41,trna72_AspGTC_1_-_161432824_161432895@35.75.41,trna75_AspGTC_1_-_161425414_161425485@35.75.41,trna78_AspGTC_1_-_161418033_161418104@35.75.41,trna81_AspGTC_1_-_161410615_161410686@35.75.41|TCACGCGGGAGACCGGGGTTCGATTCCCCGACGGGGAGCCA</td><td>>trna10_AspGTC_12_-_125424193_125424264</td><td>35</td><td>75</td><td>41</td><td>TCACGCGGGAGACCGGGGTTCGATTCCCCGACGGGGAGCCA</td><td>tRF</td><td>FALSE</td><td>TRUE</td><td>FALSE</td></tr>
<tr><td>>trna119_LysCTT_1_-_145395522_145395594@1.33.33,trna11_LysCTT_5_-_180648979_180649051@1.33.33,trna13_LysCTT_6_+_26556774_26556846@1.33.33,trna7_LysCTT_16_+_3225692_3225764@1.33.33,trna9_LysCTT_5_+_180634755_180634827@1.33.33|GCCCGGCTAGCTCAGTCGGTAGAGCATGAGACT</td><td>>trna119_LysCTT_1_-_145395522_145395594</td><td>1</td><td>33</td><td>33</td><td>GCCCGGCTAGCTCAGTCGGTAGAGCATGAGACT</td><td>tRF</td><td>FALSE</td><td>TRUE</td><td>FALSE</td></tr>
<tr><td>>trna111_HisGTG_1_-_147774845_147774916@-1G.31.32,trna118_HisGTG_1_-_145396881_145396952@-1G.31.32,trna16_HisGTG_1_+_146544773_146544844@-1G.31.32,trna1_HisGTG_15_+_45493349_45493420@-1G.31.32,trna21_HisGTG_1_+_147753471_147753542@-1G.31.32,trna33_HisGTG_6_+_27125906_27125977@-1G.31.32,trna7_HisGTG_9_-_14433938_14434009@-1G.31.32,trna8_HisGTG_15_-_45492611_45492682@-1G.31.32,trna9_HisGTG_15_-_45490804_45490875@-1G.31.32|GGCCGTGATCGTATAGTGGTTAGTACTCTGCG</td><td>>trna111_HisGTG_1_-_147774845_147774916</td><td>-1G</td><td>31</td><td>32</td><td>GGCCGTGATCGTATAGTGGTTAGTACTCTGCG</td><td>tRF</td><td>FALSE</td><td>TRUE</td><td>TRUE</td></tr>
<tr><td>>MI0000102|hsa-mir-100&WithFlank&11|-|122152223|122152314@19.40.22(+1A)&[MIMAT0000098&hsa-miR-100-5p&offsets|0|0(+1A);m-27&11|-|122152276|122152296&offsets|0|+1(+1A)]|AACCCGTAGATCCGAACTTGTGA</td><td>>MI0000102|hsa-mir-100&WithFlank&11|-|122152223|122152314</td><td>19</td><td>40</td><td>22(+1A)</td><td>AACCCGTAGATCCGAACTTGTGA</td><td>isomiR</td><td>FALSE</td><td>TRUE</td><td>FALSE</td></tr>
</table>

### Cytoplasm

| | | | | | | | | | |
|---|---|---|---|---|---|---|---|---|---|
| >trna10_AspGTC_12_-_125424193_125424264@37.74.38,trna12_AspGTC_12_-_125411891_125411962@37.74.38,trna144_AspGTC_6_-_27551236_27551307@37.74.38,trna38_AspGTC_17_-_8125556_8125627@37.74.38,trna45_AspGTC_6_+_27447453_27447524@37.74.38,trna48_AspGTC_6_+_27471523_27471594@37.74.38,trna4_AspGTC_12_+_96429799_96429870@37.74.38,trna69_AspGTC_1_-_161440205_161440276@37.74.38,trna72_AspGTC_1_-_161432824_161432895@37.74.38,trna75_AspGTC_1_-_161425414_161425485@37.74.38,trna78_AspGTC_1_-_161418033_161418104@37.74.38,trna81_AspGTC_1_-_161410615_161410686@37.74.38\|ACGCGGGAGACCGGGGTTCGATTCCCCGACGGGGAGCC | >trna10_AspGTC_12_-_125424193_125424264 | 37 | 74 | 38 | ACGCGGGAGACCGGGGTTCGATTCCCCGACGGGGAGCC | tRF | FALSE | TRUE | FALSE |
| >MI0000102\|hsa-mir-100&WithFlank&11\|-\|122152223\|122152314@19.38.20&[MIMAT0000098&hsa-miR-100-5p&offsets\|0\|-2;m-27&11\|-\|122152276\|122152296&offsets\|0\|-1]\|AACCCGTAGATCCGAACTTG | >MI0000102\|hsa-mir-100&WithFlank&11\|-\|122152223\|122152314 | 19 | 38 | 20 | AACCCGTAGATCCGAACTTG | isomiR | FALSE | TRUE | FALSE |
| >trna10_AspGTC_12_-_125424193_125424264@37.73.37,trna12_AspGTC_12_-_125411891_125411962@37.73.37,trna144_AspGTC_6_-_27551236_27551307@37.73.37,trna38_AspGTC_17_-_8125556_8125627@37.73.37,trna45_AspGTC_6_+_27447453_27447524@37.73.37,trna48_AspGTC_6_+_27471523_27471594@37.73.37,trna4_AspGTC_12_+_96429799_96429870@37.73.37,trna69_AspGTC_1_-_161440205_161440276@37.73.37,trna72_AspGTC_1_-_161432824_161432895@37.73.37,trna75_AspGTC_1_-_161425414_161425485@37.73.37,trna78_AspGTC_1_-_161418033_161418104@37.73.37,trna81_AspGTC_1_-_161410615_161410686@37.73.37\|ACGCGGGAGACCGGGGTTCGATTCCCCGACGGGGAGC | >trna10_AspGTC_12_-_125424193_125424264 | 37 | 73 | 37 | ACGCGGGAGACCGGGGTTCGATTCCCCGACGGGGAGC | tRF | FALSE | TRUE | FALSE |
| >trna10_AspGTC_12_-_125424193_125424264@37.75.39,trna12_AspGTC_12_-_125411891_125411962@37.75.39,trna144_AspGTC_6_-_27551236_27551307@37.75.39,trna38_AspGTC_17_-_8125556_8125627@37.75.39,trna45_AspGTC_6_+_27447453_27447524@37.75.39,trna48_AspGTC_6_+_27471523_27471594@37.75.39,trna4_AspGTC_12_+_96429799_96429870@37.75.39,trna69_AspGTC_1_-_161440205_161440276@37.75.39,trna72_AspGTC_1_-_161432824_161432895@37.75.39,trna75_AspGTC_1_-_161425414_161425485@37.75.39,trna78_AspGTC_1_-_161418033_161418104@37.75.39,trna81_AspGTC_1_-_161410615_161410686@37.75.39\|ACGCGGGAGACCGGGGTTCGATTCCCCGACGGGGAGCCA | >trna10_AspGTC_12_-_125424193_125424264 | 37 | 75 | 39 | ACGCGGGAGACCGGGGTTCGATTCCCCGACGGGGAGCCA | tRF | FALSE | TRUE | FALSE |
| >trna10_AspGTC_12_-_125424193_125424264@36.75.40,trna12_AspGTC_12_-_125411891_125411962@36.75.40,trna144_AspGTC_6_-_27551236_27551307@36.75.40,trna38_AspGTC_17_-_8125556_8125627@36.75.40,trna45_AspGTC_6_+_27447453_27447524@36.75.40,trna48_AspGTC_6_+_27471523_27471594@36.75.40,trna4_AspGTC_12_+_96429799_96429870@36.75.40,trna69_AspGTC_1_-_161440205_161440276@36.75.40,trna72_AspGTC_1_-_161432824_161432895@36.75.40,trna75_AspGTC_1_-_161425414_161425485@36.75.40,trna78_AspGTC_1_-_161418033_161418104@36.75.40,trna81_AspGTC_1_-_161410615_161410686@36.75.40\|CACGCGGGAGACCGGGGTTCGATTCCCCGACGGGGAGCCA | >trna10_AspGTC_12_-_125424193_125424264 | 36 | 75 | 40 | CACGCGGGAGACCGGGGTTCGATTCCCCGACGGGGAGCCA | tRF | FALSE | TRUE | FALSE |
| >MI0000102\|hsa-mir-100&WithFlank&11\|-\|122152223\|122152314@19.38.20(+1C)&[MIMAT0000098&hsa-miR-100-5p&offsets\|0\|-2(+1C);m-27&11\|-\|122152276\|122152296&offsets\|0\|-1(+1C)]\|AACCCGTAGATCCGAACTTGC | >MI0000102\|hsa-mir-100&WithFlank&11\|-\|122152223\|122152314 | 19 | 38 | 20(+1C) | AACCCGTAGATCCGAACTTGC | isomiR | FALSE | TRUE | FALSE |
| >am_trna10_ValCAC_5_-_180649395_180649467@1.32.32,trna12_ValAAC_5_-_180645270_180645342@1.32.32,trna132_ValAAC_6_-_27721179_27721251@1.32.32,trna136_ValAAC_6_-_27648885_27648957@1.32.32,trna139_ValAAC_6_-_27618707_27618779@1.32.32,trna18_ValCAC_5_-_180529253_180529325@1.32.32,trna2_ValAAC_3_+_169490018_169490090@1.32.32,trna2_ValCAC_5_+_180524070_180524142@1.32.32,trna4_ValAAC_5_+_180591154_180591226@1.32.32,trna5_ValAAC_5_+_180596610_180596682@1.32.32,trna6_ValCAC_5_+_180600650_180600722@1.32.32,trna85_ValCAC_1_-_161369490_161369562@1.32.32,trna90_ValCAC_1_-_149684088_149684161@1.32.32,trna98_ValCAC_1_-_149298555_149298627@1.32.32,trna9_ValCAC_6_+_26538282_26538354@1.32.32\|GTTTCCGTAGTGTAGTGGTTATCACGTTCGCC | >am_trna10_ValCAC_5_-_180649395_180649467 | 1 | 32 | 32 | GTTTCCGTAGTGTAGTGGTTATCACGTTCGCC | tRF | FALSE | TRUE | FALSE |
| >trna116_GluCTC_1_-_145399233_145399304@37.75.39,trna134_GluTTC_1_-_16861774_16861845@37.75.39,trna71_GluCTC_1_-_161439189_161439260@37.75.39,trna74_GluCTC_1_-_161431809_161431880@37.75.39,trna77_GluCTC_1_-_161424398_161424469@37.75.39,trna77_GluCTC_6_+_28949976_28950047@37.75.39,trna80_GluCTC_1_-_161417018_161417089@37.75.39,trna84_GluTTC_1_-_161391883_161391954@37.75.39,trna87_GluCTC_6_-_126101393_126101464@37.75.39\|ACCGCCGCGGCCCGGGTTCGATTCCCGGTCAGGGAACCA | >trna116_GluCTC_1_-_145399233_145399304 | 37 | 75 | 39 | ACCGCCGCGGCCCGGGTTCGATTCCCGGTCAGGGAACCA | tRF | FALSE | TRUE | FALSE |
| >am_trna111_HisGTG_1_-_147774845_147774916@-1G.26.27,trna118_HisGTG_1_-_145396881_145396952@-1G.26.27,trna16_HisGTG_1_+_146544773_146544844@-1G.26.27,trna1_HisGTG_15_+_45493349_45493420@-1G.26.27,trna21_HisGTG_1_+_147753471_147753542@-1G.26.27,trna33_HisGTG_6_+_27125906_27125977@-1G.26.27,trna7_HisGTG_9_-_14433938_14434009@-1G.26.27,trna8_HisGTG_15_-_45492611_45492682@-1G.26.27,trna9_HisGTG_15_-_45490804_45490875@-1G.26.27\|GGCCGTGATCGTATAGTGGTTAGTACT | >am_trna111_HisGTG_1_-_147774845_147774916 | -1G | 26 | 27 | GGCCGTGATCGTATAGTGGTTAGTACT | tRF | FALSE | TRUE | FALSE |
| >trnaMT_GluTTC_MT_-_14674_14742@40.72.33,trnalookalike8_GluTTC_5_-_93905172_93905240@40.72.33\|TTGGTCGTGGTTGTAGTCCGTGCGAGAATACCA | >trnaMT_GluTTC_MT_-_14674_14742 | 40 | 72 | 33 | TTGGTCGTGGTTGTAGTCCGTGCGAGAATACCA | tRF | FALSE | TRUE | FALSE |
| >hg38_21_+_8433172_8446622_RNA45SN1_45S_50ntFlanks@7975.8013.39\|CGCGACCTCAGATCAGACGTGGCGACCCGCTGAATTTAA | >hg38_21_+_8433172_8446622_RNA45SN1_45S_50ntFlanks | 7975 | 8013 | 39 | CGCGACCTCAGATCAGACGTGGCGACCCGCTGAATTTAA | rRF | FALSE | TRUE | FALSE |

### Cytoplasm

| | | | | | | | | | |
|---|---|---|---|---|---|---|---|---|---|
| >trna116_GluCTC_1_-_145399233_145399304@37.74.38,trna134_GluTTC_1_-_16861774_16861845@37.74.38,trna71_GluCTC_1_-_161439189_161439260@37.74.38,trna74_GluCTC_1_-_161431809_161431880@37.74.38,trna77_GluCTC_1_-_161424398_161424469@37.74.38,trna77_GluCTC_6_+_28949976_28950047@37.74.38,trna80_GluCTC_1_-_161417018_161417089@37.74.38,trna84_GluTTC_1_-_161391883_161391954@37.74.38,trna87_GluCTC_6_-_126101393_126101464@37.74.38\|ACCGCCGCGGCCCGGGTTCGATTCCCGGTCAGGGAACC | >trna116_GluCTC_1_-_145399233_145399304 | 37 | 74 | 38 | ACCGCCGCGGCCCGGGTTCGATTCCCGGTCAGGGAACC | tRF | FALSE | TRUE | FALSE |
| >am_trna128_GlyGCC_6_-_27870686_27870756@1.30.30,trna133_GlyCCC_1_-_16872434_16872504@1.30.30,trna18_GlyGCC_16_+_70822597_70822667@1.30.30,trna19_GlyGCC_16_+_70823410_70823480@1.30.30,trna19_GlyGCC_2_-_157257659_157257729@1.30.30,trna24_GlyGCC_16_-_70812942_70813012@1.30.30,trna25_GlyGCC_16_-_70812114_70812184@1.30.30,trna4_GlyCCC_1_+_17188416_17188486@1.30.30,trna5_GlyGCC_17_+_8029064_8029134@1.30.30,trna68_GlyGCC_1_-_161493637_161493707@1.30.30\|GCATTGGTGGTTCAGTGGTAGAATTCTCGC | >am_trna128_GlyGCC_6_-_27870686_27870756 | 1 | 30 | 30 | GCATTGGTGGTTCAGTGGTAGAATTCTCGC | tRF | FALSE | TRUE | TRUE |
| >trna152_ValCAC_6_-_27248049_27248121@1.34.34\|GCTTCTGTAGTGTAGTGGTTATCACGTTCGCCTC | >trna152_ValCAC_6_-_27248049_27248121 | 1 | 34 | 34 | GCTTCTGTAGTGTAGTGGTTATCACGTTCGCCTC | tRF | FALSE | TRUE | FALSE |
| >hg38_21_+_8433172_8446622_RNA45SN1_45S_50ntFlanks@7975.8016.42\|CGCGACCTCAGATCAGACGTGGCGACCCGCTGAATTTAAGCA | >hg38_21_+_8433172_8446622_RNA45SN1_45S_50ntFlanks | 7975 | 8016 | 42 | CGCGACCTCAGATCAGACGTGGCGACCCGCTGAATTTAAGCA | rRF | FALSE | TRUE | FALSE |
| >trna119_LysCTT_1_-_145395522_145395594@1.34.34,trna11_LysCTT_5_-_180648979_180649051@1.34.34,trna13_LysCTT_6_+_26556774_26556846@1.34.34,trna7_LysCTT_16_+_3225692_3225764@1.34.34,trna9_LysCTT_5_+_180634755_180634827@1.34.34\|GCCCGGCTAGCTCAGTCGGTAGAGCATGAGACTC | >trna119_LysCTT_1_-_145395522_145395594 | 1 | 34 | 34 | GCCCGGCTAGCTCAGTCGGTAGAGCATGAGACTC | tRF | FALSE | TRUE | FALSE |
| >hg38_21_+_8433172_8446622_RNA45SN1_45S_50ntFlanks@12416.12433.18\|TTCGATGTCGGCTCTTCC | >hg38_21_+_8433172_8446622_RNA45SN1_45S_50ntFlanks | 12416 | 12433 | 18 | TTCGATGTCGGCTCTTCC | rRF | FALSE | TRUE | FALSE |
| >hg38_21_+_8433172_8446622_RNA45SN1_45S_50ntFlanks@9554.9571.18\|CGTAGCGGTCCTGACGTG | >hg38_21_+_8433172_8446622_RNA45SN1_45S_50ntFlanks | 9554 | 9571 | 18 | CGTAGCGGTCCTGACGTG | rRF | FALSE | TRUE | TRUE |
| >trna10_AspGTC_12_-_125424193_125424264@35.74.40,trna12_AspGTC_12_-_125411891_125411962@35.74.40,trna144_AspGTC_6_-_27551236_27551307@35.74.40,trna38_AspGTC_17_-_8125556_8125627@35.74.40,trna45_AspGTC_6_+_27447453_27447524@35.74.40,trna48_AspGTC_6_+_27471523_27471594@35.74.40,trna4_AspGTC_12_+_96429799_96429870@35.74.40,trna69_AspGTC_1_-_161440205_161440276@35.74.40,trna72_AspGTC_1_-_161432824_161432895@35.74.40,trna75_AspGTC_1_-_161425414_161425485@35.74.40,trna78_AspGTC_1_-_161418033_161418104@35.74.40,trna81_AspGTC_1_-_161410615_161410686@35.74.40\|TCACGCGGGAGACCGGGGTTCGATTCCCCGACGGGGAGCC | >trna10_AspGTC_12_-_125424193_125424264 | 35 | 74 | 40 | TCACGCGGGAGACCGGGGTTCGATTCCCCGACGGGGAGCC | tRF | FALSE | TRUE | FALSE |
| >trnaMT_GluTTC_MT_-_14674_14742@39.72.34,trnalookalike8_GluTTC_5_-_93905172_93905240@39.72.34\|ATTGGTCGTGGTTGTAGTCCGTGCGAGAATACCA | >trnaMT_GluTTC_MT_-_14674_14742 | 39 | 72 | 34 | ATTGGTCGTGGTTGTAGTCCGTGCGAGAATACCA | tRF | FALSE | TRUE | FALSE |
| >MI0000446\|hsa-mir-125b-1&WithFlank&11\|-\|122099751\|122099850@21.42.22;MI0000470\|hsa-mir-125b-2&WithFlank&21\|+\|16590231\|16590331@23.44.22&[MIMAT0000423&hsa-miR-125b-5p&offsets\|0\|0;m-67&11\|-\|122099809\|122099830&offsets\|0\|0];[MIMAT0000423_1&hsa-miR-125b-5p&offsets\|0\|0;m-68&21\|+\|16590253\|16590274&offsets\|0\|0]\|TCCCTGAGACCCTAACTTGTGA | >MI0000446\|hsa-mir-125b-1&WithFlank&11\|-\|122099751\|122099850 | 21 | 42 | 22 | TCCCTGAGACCCTAACTTGTGA | isomiR | FALSE | TRUE | FALSE |
| >am_trna128_GlyGCC_6_-_27870686_27870756@1.35.35,trna18_GlyGCC_16_+_70822597_70822667@1.35.35,trna19_GlyGCC_16_+_70823410_70823480@1.35.35,trna19_GlyGCC_2_-_157257659_157257729@1.35.35,trna24_GlyGCC_16_-_70812942_70813012@1.35.35,trna25_GlyGCC_16_-_70812114_70812184@1.35.35,trna5_GlyGCC_17_+_8029064_8029134@1.35.35,trna68_GlyGCC_1_-_161493637_161493707@1.35.35\|GCATTGGTGGTTCAGTGGTAGAATTCTCGCCTGCC | >am_trna128_GlyGCC_6_-_27870686_27870756 | 1 | 35 | 35 | GCATTGGTGGTTCAGTGGTAGAATTCTCGCCTGCC | tRF | FALSE | TRUE | FALSE |
| >trnaMT_GluTTC_MT_-_14674_14742@35.72.38,trnalookalike8_GluTTC_5_-_93905172_93905240@35.72.38\|TATCATTGGTCGTGGTTGTAGTCCGTGCGAGAATACCA | >trnaMT_GluTTC_MT_-_14674_14742 | 35 | 72 | 38 | TATCATTGGTCGTGGTTGTAGTCCGTGCGAGAATACCA | tRF | FALSE | TRUE | FALSE |
| >hg38_21_+_8433172_8446622_RNA45SN1_45S_50ntFlanks@12999.13039.41\|CTGCGATCTATTGAAAGTCAGCCCTCGACACAAGGGTTTGT | >hg38_21_+_8433172_8446622_RNA45SN1_45S_50ntFlanks | 12999 | 13039 | 41 | CTGCGATCTATTGAAAGTCAGCCCTCGACACAAGGGTTTGT | rRF | FALSE | TRUE | FALSE |
| >MI0000477\|hsa-mir-146a&WithFlank&5\|+\|160485346\|160485456@27.48.22&[MIMAT0000449&hsa-miR-146a-5p&offsets\|0\|0;m-129&5\|+\|160485372\|160485393&offsets\|0\|0]\|TGAGAACTGAATTCCATGGGTT | >MI0000477\|hsa-mir-146a&WithFlank&5\|+\|160485346\|160485456 | 27 | 48 | 22 | TGAGAACTGAATTCCATGGGTT | isomiR | FALSE | TRUE | FALSE |
| >trna10_AspGTC_12_-_125424193_125424264@36.74.39,trna12_AspGTC_12_-_125411891_125411962@36.74.39,trna144_AspGTC_6_-_27551236_27551307@36.74.39,trna38_AspGTC_17_-_8125556_8125627@36.74.39,trna45_AspGTC_6_+_27447453_27447524@36.74.39,trna48_AspGTC_6_+_27471523_27471594@36.74.39,trna4_AspGTC_12_+_96429799_96429870@36.74.39,trna69_AspGTC_1_-_161440205_161440276@36.74.39,trna72_AspGTC_1_-_161432824_161432895@36.74.39,trna75_AspGTC_1_-_161425414_161425485@36.74.39,trna78_AspGTC_1_-_161418033_161418104@36.74.39,trna81_AspGTC_1_-_161410615_161410686@36.74.39\|CACGCGGGAGACCGGGGTTCGATTCCCCGACGGGGAGCC | >trna10_AspGTC_12_-_125424193_125424264 | 36 | 74 | 39 | CACGCGGGAGACCGGGGTTCGATTCCCCGACGGGGAGCC | tRF | FALSE | TRUE | FALSE |
| >MI0000446\|hsa-mir-125b-1&WithFlank&11\|-\|122099751\|122099850@21.41.21;MI0000470\|hsa-mir-125b-2&WithFlank&21\|+\|16590231\|16590331@23.43.21&[MIMAT0000423&hsa-miR-125b-5p&offsets\|0\|-1;m-67&11\|-\|122099809\|122099830&offsets\|0\|-1];[MIMAT0000423_1&hsa-miR-125b-5p&offsets\|0\|-1;m-68&21\|+\|16590253\|16590274&offsets\|0\|-1]\|TCCCTGAGACCCTAACTTGTG | >MI0000446\|hsa-mir-125b-1&WithFlank&11\|-\|122099751\|122099850 | 21 | 41 | 21 | TCCCTGAGACCCTAACTTGTG | isomiR | FALSE | TRUE | FALSE |
| >hg38_21_+_8433172_8446622_RNA45SN1_45S_50ntFlanks@5471.5516.46\|CACGGCCCTGGCGGAGCGCTGAGAAGACGGTCGAACTTGACTATCT | >hg38_21_+_8433172_8446622_RNA45SN1_45S_50ntFlanks | 5471 | 5516 | 46 | CACGGCCCTGGCGGAGCGCTGAGAAGACGGTCGAACTTGACTATCT | rRF | FALSE | TRUE | FALSE |

### Cytoplasm

<table>
<tr><td>>trna111_HisGTG_1_-_147774845_147774916@1.34.34,trna118_HisGTG_1_-_145396881_145396952@1.34.34,trna16_HisGTG_1_+_146544773_146544844@1.34.34,trna1_HisGTG_15_+_45493349_45493420@1.34.34,trna21_HisGTG_1_+_147753471_147753542@1.34.34,trna33_HisGTG_6_+_27125906_27125977@1.34.34,trna7_HisGTG_9_-_14433938_14434009@1.34.34,trna8_HisGTG_15_-_45492611_45492682@1.34.34,trna9_HisGTG_15_-_45490804_45490875@1.34.34|GCCGTGATCGTATAGTGGTTAGTACTCTGCGTTG</td><td>>trna111_HisGTG_1_-_147774845_147774916</td><td>1</td><td>34</td><td>34</td><td>GCCGTGATCGTATAGTGGTTAGTACTCTGCGTTG</td><td>tRF</td><td>FALSE</td><td>TRUE</td><td>FALSE</td></tr>
<tr><td>>hg38_21_+_8433172_8446622_RNA45SN1_45S_50ntFlanks@8196.8213.18|CGTGGACGGTGTGAGGCC</td><td>>hg38_21_+_8433172_8446622_RNA45SN1_45S_50ntFlanks</td><td>8196</td><td>8213</td><td>18</td><td>CGTGGACGGTGTGAGGCC</td><td>rRF</td><td>FALSE</td><td>TRUE</td><td>FALSE</td></tr>
<tr><td>>hg38_21_+_8433172_8446622_RNA45SN1_45S_50ntFlanks@12419.12448.30|GATGTCGGCTCTTCCTATCATTGTGAAGCA</td><td>>hg38_21_+_8433172_8446622_RNA45SN1_45S_50ntFlanks</td><td>12419</td><td>12448</td><td>30</td><td>GATGTCGGCTCTTCCTATCATTGTGAAGCA</td><td>rRF</td><td>FALSE</td><td>TRUE</td><td>FALSE</td></tr>
<tr><td>>hg38_21_+_8433172_8446622_RNA45SN1_45S_50ntFlanks@12419.12437.19|GATGTCGGCTCTTCCTATC</td><td>>hg38_21_+_8433172_8446622_RNA45SN1_45S_50ntFlanks</td><td>12419</td><td>12437</td><td>19</td><td>GATGTCGGCTCTTCCTATC</td><td>rRF</td><td>FALSE</td><td>TRUE</td><td>FALSE</td></tr>
<tr><td>>trnaMT_GluTTC_MT_-_14674_14742@41.72.32,trnalookalike8_GluTTC_5_-_93905172_93905240@41.72.32|TGGTCGTGGTTGTAGTCCGTGCGAGAATACCA</td><td>>trnaMT_GluTTC_MT_-_14674_14742</td><td>41</td><td>72</td><td>32</td><td>TGGTCGTGGTTGTAGTCCGTGCGAGAATACCA</td><td>tRF</td><td>FALSE</td><td>TRUE</td><td>FALSE</td></tr>
<tr><td>>hg38_21_+_8433172_8446622_RNA45SN1_45S_50ntFlanks@5472.5512.41|ACGGCCCTGGCGGAGCGCTGAGAAGACGGTCGAACTTGACT</td><td>>hg38_21_+_8433172_8446622_RNA45SN1_45S_50ntFlanks</td><td>5472</td><td>5512</td><td>41</td><td>ACGGCCCTGGCGGAGCGCTGAGAAGACGGTCGAACTTGACT</td><td>rRF</td><td>FALSE</td><td>TRUE</td><td>FALSE</td></tr>
<tr><td>>trna111_HisGTG_1_-_147774845_147774916@1.31.31,trna118_HisGTG_1_-_145396881_145396952@1.31.31,trna16_HisGTG_1_+_146544773_146544844@1.31.31,trna1_HisGTG_15_+_45493349_45493420@1.31.31,trna21_HisGTG_1_+_147753471_147753542@1.31.31,trna33_HisGTG_6_+_27125906_27125977@1.31.31,trna7_HisGTG_9_-_14433938_14434009@1.31.31,trna8_HisGTG_15_-_45492611_45492682@1.31.31,trna9_HisGTG_15_-_45490804_45490875@1.31.31|GCCGTGATCGTATAGTGGTTAGTACTCTGCG</td><td>>trna111_HisGTG_1_-_147774845_147774916</td><td>1</td><td>31</td><td>31</td><td>GCCGTGATCGTATAGTGGTTAGTACTCTGCG</td><td>tRF</td><td>FALSE</td><td>TRUE</td><td>TRUE</td></tr>
<tr><td>>hg38_21_+_8433172_8446622_RNA45SN1_45S_50ntFlanks@12995.13038.44|CTCGCTGCGATCTATTGAAAGTCAGCCCTCGACACAAGGGTTTG</td><td>>hg38_21_+_8433172_8446622_RNA45SN1_45S_50ntFlanks</td><td>12995</td><td>13038</td><td>44</td><td>CTCGCTGCGATCTATTGAAAGTCAGCCCTCGACACAAGGGTTTG</td><td>rRF</td><td>FALSE</td><td>TRUE</td><td>FALSE</td></tr>
<tr><td>>hg38_21_+_8433172_8446622_RNA45SN1_45S_50ntFlanks@12416.12434.19|TTCGATGTCGGCTCTTCCT</td><td>>hg38_21_+_8433172_8446622_RNA45SN1_45S_50ntFlanks</td><td>12416</td><td>12434</td><td>19</td><td>TTCGATGTCGGCTCTTCCT</td><td>rRF</td><td>FALSE</td><td>TRUE</td><td>FALSE</td></tr>
<tr><td>>hg38_21_+_8433172_8446622_RNA45SN1_45S_50ntFlanks@11668.11702.35|TGCCCAGTGCTCTGAATGTCAAAGTGAAGAAATTC</td><td>>hg38_21_+_8433172_8446622_RNA45SN1_45S_50ntFlanks</td><td>11668</td><td>11702</td><td>35</td><td>TGCCCAGTGCTCTGAATGTCAAAGTGAAGAAATTC</td><td>rRF</td><td>FALSE</td><td>TRUE</td><td>FALSE</td></tr>
<tr><td>>MI0000299|hsa-mir-222&WithFlank&X|-|45747009|45747130@75.98.24(+1C)&[MIMAT0000279&hsa-miR-222-3p&offsets|0|+3(+1C);m-140&X|-|45747033|45747056&offsets|0|0(+1C)]|AGCTACATCTGGCTACTGGGTCTCC</td><td>>MI0000299|hsa-mir-222&WithFlank&X|-|45747009|45747130</td><td>75</td><td>98</td><td>24(+1C)</td><td>AGCTACATCTGGCTACTGGGTCTCC</td><td>isomiR</td><td>FALSE</td><td>TRUE</td><td>FALSE</td></tr>
<tr><td>>hg38_21_+_8433172_8446622_RNA45SN1_45S_50ntFlanks@12416.12437.22|TTCGATGTCGGCTCTTCCTATC</td><td>>hg38_21_+_8433172_8446622_RNA45SN1_45S_50ntFlanks</td><td>12416</td><td>12437</td><td>22</td><td>TTCGATGTCGGCTCTTCCTATC</td><td>rRF</td><td>FALSE</td><td>TRUE</td><td>FALSE</td></tr>
<tr><td>>hg38_21_+_8433172_8446622_RNA45SN1_45S_50ntFlanks@7975.8014.40|CGCGACCTCAGATCAGACGTGGCGACCCGCTGAATTTAAG</td><td>>hg38_21_+_8433172_8446622_RNA45SN1_45S_50ntFlanks</td><td>7975</td><td>8014</td><td>40</td><td>CGCGACCTCAGATCAGACGTGGCGACCCGCTGAATTTAAG</td><td>rRF</td><td>FALSE</td><td>TRUE</td><td>FALSE</td></tr>
<tr><td>>trna111_HisGTG_1_-_147774845_147774916@-1G.30.31,trna118_HisGTG_1_-_145396881_145396952@-1G.30.31,trna16_HisGTG_1_+_146544773_146544844@-1G.30.31,trna1_HisGTG_15_+_45493349_45493420@-1G.30.31,trna21_HisGTG_1_+_147753471_147753542@-1G.30.31,trna33_HisGTG_6_+_27125906_27125977@-1G.30.31,trna7_HisGTG_9_-_14433938_14434009@-1G.30.31,trna8_HisGTG_15_-_45492611_45492682@-1G.30.31,trna9_HisGTG_15_-_45490804_45490875@-1G.30.31|GGCCGTGATCGTATAGTGGTTAGTACTCTGC</td><td>>trna111_HisGTG_1_-_147774845_147774916</td><td>-1G</td><td>30</td><td>31</td><td>GGCCGTGATCGTATAGTGGTTAGTACTCTGC</td><td>tRF</td><td>FALSE</td><td>TRUE</td><td>FALSE</td></tr>
<tr><td>>am_trna10_ValCAC_5_-_180649395_180649467@1.33.33,trna12_ValAAC_5_-_180645270_180645342@1.33.33,trna132_ValAAC_6_-_27721179_27721251@1.33.33,trna136_ValAAC_6_-_27648885_27648957@1.33.33,trna139_ValAAC_6_-_27618707_27618779@1.33.33,trna18_ValCAC_5_-_180529253_180529325@1.33.33,trna2_ValAAC_3_+_169490018_169490090@1.33.33,trna2_ValCAC_5_+_180524070_180524142@1.33.33,trna4_ValAAC_5_+_180591154_180591226@1.33.33,trna5_ValAAC_5_+_180596610_180596682@1.33.33,trna6_ValCAC_5_+_180600650_180600722@1.33.33,trna85_ValCAC_1_-_161369490_161369562@1.33.33,trna90_ValCAC_1_-_149684088_149684161@1.33.33,trna98_ValCAC_1_-_149298555_149298627@1.33.33,trna9_ValCAC_6_+_26538282_26538354@1.33.33|GTTTCCGTAGTGTAGTGGTTATCACGTTCGCCT</td><td>>am_trna10_ValCAC_5_-_180649395_180649467</td><td>1</td><td>33</td><td>33</td><td>GTTTCCGTAGTGTAGTGGTTATCACGTTCGCCT</td><td>tRF</td><td>FALSE</td><td>TRUE</td><td>FALSE</td></tr>
<tr><td>>hg38_21_+_8433172_8446622_RNA45SN1_45S_50ntFlanks@6788.6806.19|CGCCTGTCTGAGCGTCGCT</td><td>>hg38_21_+_8433172_8446622_RNA45SN1_45S_50ntFlanks</td><td>6788</td><td>6806</td><td>19</td><td>CGCCTGTCTGAGCGTCGCT</td><td>rRF</td><td>FALSE</td><td>TRUE</td><td>FALSE</td></tr>
<tr><td>>hg38_21_+_8433172_8446622_RNA45SN1_45S_50ntFlanks@6651.6696.46|CGACTCTTAGCGGTGGATCACTCGGCTCGTGCGTCGATGAAGAACG</td><td>>hg38_21_+_8433172_8446622_RNA45SN1_45S_50ntFlanks</td><td>6651</td><td>6696</td><td>46</td><td>CGACTCTTAGCGGTGGATCACTCGGCTCGTGCGTCGATGAAGAACG</td><td>rRF</td><td>FALSE</td><td>TRUE</td><td>FALSE</td></tr>
<tr><td>>hg38_21_+_8433172_8446622_RNA45SN1_45S_50ntFlanks@12416.12448.33|TTCGATGTCGGCTCTTCCTATCATTGTGAAGCA</td><td>>hg38_21_+_8433172_8446622_RNA45SN1_45S_50ntFlanks</td><td>12416</td><td>12448</td><td>33</td><td>TTCGATGTCGGCTCTTCCTATCATTGTGAAGCA</td><td>rRF</td><td>FALSE</td><td>TRUE</td><td>FALSE</td></tr>
<tr><td>>hg38_1_-_228634819_228635039_RNA5S12_5S_50ntFlanks@130.170.41|TGGATGGGAGACCGCCTGGGAATACCGGGTGCTGTAGGCTT</td><td>>hg38_1_-_228634819_228635039_RNA5S12_5S_50ntFlanks</td><td>130</td><td>170</td><td>41</td><td>TGGATGGGAGACCGCCTGGGAATACCGGGTGCTGTAGGCTT</td><td>rRF</td><td>FALSE</td><td>TRUE</td><td>FALSE</td></tr>
<tr><td>>trna152_ValCAC_6_-_27248049_27248121@1.32.32|GCTTCTGTAGTGTAGTGGTTATCACGTTCGCC</td><td>>trna152_ValCAC_6_-_27248049_27248121</td><td>1</td><td>32</td><td>32</td><td>GCTTCTGTAGTGTAGTGGTTATCACGTTCGCC</td><td>tRF</td><td>FALSE</td><td>TRUE</td><td>FALSE</td></tr>
<tr><td>>hg38_21_+_8433172_8446622_RNA45SN1_45S_50ntFlanks@7975.8015.41|CGCGACCTCAGATCAGACGTGGCGACCCGCTGAATTTAAGC</td><td>>hg38_21_+_8433172_8446622_RNA45SN1_45S_50ntFlanks</td><td>7975</td><td>8015</td><td>41</td><td>CGCGACCTCAGATCAGACGTGGCGACCCGCTGAATTTAAGC</td><td>rRF</td><td>FALSE</td><td>TRUE</td><td>FALSE</td></tr>
<tr><td>>hg38_21_+_8433172_8446622_RNA45SN1_45S_50ntFlanks@7974.7991.18|ACGCGACCTCAGATCAGA</td><td>>hg38_21_+_8433172_8446622_RNA45SN1_45S_50ntFlanks</td><td>7974</td><td>7991</td><td>18</td><td>ACGCGACCTCAGATCAGA</td><td>rRF</td><td>FALSE</td><td>TRUE</td><td>FALSE</td></tr>
<tr><td>>hg38_21_+_8433172_8446622_RNA45SN1_45S_50ntFlanks@11669.11702.34|GCCCAGTGCTCTGAATGTCAAAGTGAAGAAATTC</td><td>>hg38_21_+_8433172_8446622_RNA45SN1_45S_50ntFlanks</td><td>11669</td><td>11702</td><td>34</td><td>GCCCAGTGCTCTGAATGTCAAAGTGAAGAAATTC</td><td>rRF</td><td>FALSE</td><td>TRUE</td><td>FALSE</td></tr>
<tr><td>>hg38_1_-_228634819_228635039_RNA5S12_5S_50ntFlanks@129.170.42|TTGGATGGGAGACCGCCTGGGAATACCGGGTGCTGTAGGCTT</td><td>>hg38_1_-_228634819_228635039_RNA5S12_5S_50ntFlanks</td><td>129</td><td>170</td><td>42</td><td>TTGGATGGGAGACCGCCTGGGAATACCGGGTGCTGTAGGCTT</td><td>rRF</td><td>FALSE</td><td>TRUE</td><td>FALSE</td></tr>
<tr><td>>trna5_AspGTC_12_+_98897281_98897352@35.75.41|TCACGCGGGAGACCGGGGTTCAATTCCCCGACGGGGAGCCA</td><td>>trna5_AspGTC_12_+_98897281_98897352</td><td>35</td><td>75</td><td>41</td><td>TCACGCGGGAGACCGGGGTTCAATTCCCCGACGGGGAGCCA</td><td>tRF</td><td>FALSE</td><td>TRUE</td><td>FALSE</td></tr>
<tr><td>>hg38_1_-_228634819_228635039_RNA5S12_5S_50ntFlanks@152.171.20|TACCGGGTGCTGTAGGCTTT</td><td>>hg38_1_-_228634819_228635039_RNA5S12_5S_50ntFlanks</td><td>152</td><td>171</td><td>20</td><td>TACCGGGTGCTGTAGGCTTT</td><td>rRF</td><td>FALSE</td><td>TRUE</td><td>FALSE</td></tr>
</table>

### Cytoplasm

<table>
<tr><td>>hg38_1_-_228634819_228635039_RNA5S12_5S_50ntFlanks@149.171.23|GAATACCGGGTGCTGTAGGCTTT</td><td>>hg38_1_-_228634819_228635039_RNA5S12_5S_50ntFlanks</td><td>149</td><td>171</td><td>23</td><td>GAATACCGGGTGCTGTAGGCTTT</td><td>rRF</td><td>FALSE</td><td>TRUE</td><td>FALSE</td></tr>
<tr><td>>trnaMT_GluTTC_MT_-_14674_14742@37.72.36,trnalookalike8_GluTTC_5_-_93905172_93905240@37.72.36|TCATTGGTCGTGGTTGTAGTCCGTGCGAGAATACCA</td><td>>trnaMT_GluTTC_MT_-_14674_14742</td><td>37</td><td>72</td><td>36</td><td>TCATTGGTCGTGGTTGTAGTCCGTGCGAGAATACCA</td><td>tRF</td><td>FALSE</td><td>TRUE</td><td>FALSE</td></tr>
<tr><td>>hg38_21_+_8433172_8446622_RNA45SN1_45S_50ntFlanks@12998.13039.42|GCTGCGATCTATTGAAAGTCAGCCCTCGACACAAGGGTTTGT</td><td>>hg38_21_+_8433172_8446622_RNA45SN1_45S_50ntFlanks</td><td>12998</td><td>13039</td><td>42</td><td>GCTGCGATCTATTGAAAGTCAGCCCTCGACACAAGGGTTTGT</td><td>rRF</td><td>FALSE</td><td>TRUE</td><td>FALSE</td></tr>
<tr><td>>hg38_1_-_228634819_228635039_RNA5S12_5S_50ntFlanks@144.171.28|CCTGGGAATACCGGGTGCTGTAGGCTTT</td><td>>hg38_1_-_228634819_228635039_RNA5S12_5S_50ntFlanks</td><td>144</td><td>171</td><td>28</td><td>CCTGGGAATACCGGGTGCTGTAGGCTTT</td><td>rRF</td><td>FALSE</td><td>TRUE</td><td>FALSE</td></tr>
<tr><td>>hg38_1_-_228634819_228635039_RNA5S12_5S_50ntFlanks@154.171.18|CCGGGTGCTGTAGGCTTT</td><td>>hg38_1_-_228634819_228635039_RNA5S12_5S_50ntFlanks</td><td>154</td><td>171</td><td>18</td><td>CCGGGTGCTGTAGGCTTT</td><td>rRF</td><td>FALSE</td><td>TRUE</td><td>FALSE</td></tr>
<tr><td>>hg38_1_-_228634819_228635039_RNA5S12_5S_50ntFlanks@150.171.22|AATACCGGGTGCTGTAGGCTTT</td><td>>hg38_1_-_228634819_228635039_RNA5S12_5S_50ntFlanks</td><td>150</td><td>171</td><td>22</td><td>AATACCGGGTGCTGTAGGCTTT</td><td>rRF</td><td>FALSE</td><td>TRUE</td><td>FALSE</td></tr>
<tr><td>>MI0000079|hsa-mir-23a&WithFlank&19|-|13836581|13836665@54.73.20(+1U)&[MIMAT0000078&hsa-miR-23a-3p&offsets|+3|+2(+1U);m-31&19|-|13836595|13836615&offsets|+3|+2(+1U)]|ACATTGCCAGGGATTTCCAAT</td><td>>MI0000079|hsa-mir-23a&WithFlank&19|-|13836581|13836665</td><td>54</td><td>73</td><td>20(+1U)</td><td>ACATTGCCAGGGATTTCCAAT</td><td>isomiR</td><td>FALSE</td><td>TRUE</td><td>FALSE</td></tr>
<tr><td>>hg38_1_-_228634819_228635039_RNA5S12_5S_50ntFlanks@148.171.24|GGAATACCGGGTGCTGTAGGCTTT</td><td>>hg38_1_-_228634819_228635039_RNA5S12_5S_50ntFlanks</td><td>148</td><td>171</td><td>24</td><td>GGAATACCGGGTGCTGTAGGCTTT</td><td>rRF</td><td>FALSE</td><td>TRUE</td><td>FALSE</td></tr>
<tr><td>>am_trna10_ValCAC_5_-_180649395_180649467@1.34.34,trna18_ValCAC_5_-_180529253_180529325@1.34.34,trna2_ValCAC_5_+_180524070_180524142@1.34.34,trna6_ValCAC_5_+_180600650_180600722@1.34.34,trna85_ValCAC_1_-_161369490_161369562@1.34.34,trna90_ValCAC_1_-_149684088_149684161@1.34.34,trna98_ValCAC_1_-_149298555_149298627@1.34.34,trna9_ValCAC_6_+_26538282_26538354@1.34.34|GTTTCCGTAGTGTAGTGGTTATCACGTTCGCCTC</td><td>>am_trna10_ValCAC_5_-_180649395_180649467</td><td>1</td><td>34</td><td>34</td><td>GTTTCCGTAGTGTAGTGGTTATCACGTTCGCCTC</td><td>tRF</td><td>FALSE</td><td>TRUE</td><td>FALSE</td></tr>
<tr><td>>trna152_ValCAC_6_-_27248049_27248121@1.33.33|GCTTCTGTAGTGTAGTGGTTATCACGTTCGCCT</td><td>>trna152_ValCAC_6_-_27248049_27248121</td><td>1</td><td>33</td><td>33</td><td>GCTTCTGTAGTGTAGTGGTTATCACGTTCGCCT</td><td>tRF</td><td>FALSE</td><td>TRUE</td><td>FALSE</td></tr>
<tr><td>>MI0000093|hsa-mir-92a-1&WithFlank&13|+|91351308|91351397@54.76.23&[MIMAT0000092&hsa-miR-92a-3p&offsets|0|+1;m-9&13|+|91351361|91351382&offsets|0|+1]|TATTGCACTTGTCCCGGCCTGTT</td><td>>MI0000093|hsa-mir-92a-1&WithFlank&13|+|91351308|91351397</td><td>54</td><td>76</td><td>23</td><td>TATTGCACTTGTCCCGGCCTGTT</td><td>isomiR</td><td>FALSE</td><td>TRUE</td><td>FALSE</td></tr>
<tr><td>>hg38_21_+_8433172_8446622_RNA45SN1_45S_50ntFlanks@4727.4767.41|AATCAAGAACGAAAGTCGGAGGTTCGAAGACGATCAGATAC</td><td>>hg38_21_+_8433172_8446622_RNA45SN1_45S_50ntFlanks</td><td>4727</td><td>4767</td><td>41</td><td>AATCAAGAACGAAAGTCGGAGGTTCGAAGACGATCAGATAC</td><td>rRF</td><td>FALSE</td><td>TRUE</td><td>FALSE</td></tr>
<tr><td>>hg38_1_-_228634819_228635039_RNA5S12_5S_50ntFlanks@147.171.25|GGGAATACCGGGTGCTGTAGGCTTT</td><td>>hg38_1_-_228634819_228635039_RNA5S12_5S_50ntFlanks</td><td>147</td><td>171</td><td>25</td><td>GGGAATACCGGGTGCTGTAGGCTTT</td><td>rRF</td><td>FALSE</td><td>TRUE</td><td>FALSE</td></tr>
<tr><td>>MI0000102|hsa-mir-100&WithFlank&11|-|122152223|122152314@19.40.22(+1C)&[MIMAT0000098&hsa-miR-100-5p&offsets|0|0(+1C);m-27&11|-|122152276|122152296&offsets|0|+1(+1C)]|AACCCGTAGATCCGAACTTGTGC</td><td>>MI0000102|hsa-mir-100&WithFlank&11|-|122152223|122152314</td><td>19</td><td>40</td><td>22(+1C)</td><td>AACCCGTAGATCCGAACTTGTGC</td><td>isomiR</td><td>FALSE</td><td>TRUE</td><td>FALSE</td></tr>
<tr><td>>hg38_21_+_8433172_8446622_RNA45SN1_45S_50ntFlanks@12997.13014.18|CGCTGCGATCTATTGAAA</td><td>>hg38_21_+_8433172_8446622_RNA45SN1_45S_50ntFlanks</td><td>12997</td><td>13014</td><td>18</td><td>CGCTGCGATCTATTGAAA</td><td>rRF</td><td>FALSE</td><td>TRUE</td><td>FALSE</td></tr>
<tr><td>>hg38_21_+_8433172_8446622_RNA45SN1_45S_50ntFlanks@11540.11584.45|CGCGGCGCCTCGCCTCGGCCGGCGCCTAGCAGCCGACTTAGAACT</td><td>>hg38_21_+_8433172_8446622_RNA45SN1_45S_50ntFlanks</td><td>11540</td><td>11584</td><td>45</td><td>CGCGGCGCCTCGCCTCGGCCGGCGCCTAGCAGCCGACTTAGAACT</td><td>rRF</td><td>FALSE</td><td>TRUE</td><td>FALSE</td></tr>
<tr><td>>hg38_21_+_8433172_8446622_RNA45SN1_45S_50ntFlanks@12997.13038.42|CGCTGCGATCTATTGAAAGTCAGCCCTCGACACAAGGGTTTG</td><td>>hg38_21_+_8433172_8446622_RNA45SN1_45S_50ntFlanks</td><td>12997</td><td>13038</td><td>42</td><td>CGCTGCGATCTATTGAAAGTCAGCCCTCGACACAAGGGTTTG</td><td>rRF</td><td>FALSE</td><td>TRUE</td><td>FALSE</td></tr>
<tr><td>>trna111_HisGTG_1_-_147774845_147774916@-1G.32.33,trna118_HisGTG_1_-_145396881_145396952@-1G.32.33,trna16_HisGTG_1_+_146544773_146544844@-1G.32.33,trna1_HisGTG_15_+_45493349_45493420@-1G.32.33,trna21_HisGTG_1_+_147753471_147753542@-1G.32.33,trna33_HisGTG_6_+_27125906_27125977@-1G.32.33,trna7_HisGTG_9_-_14433938_14434009@-1G.32.33,trna8_HisGTG_15_-_45492611_45492682@-1G.32.33,trna9_HisGTG_15_-_45490804_45490875@-1G.32.33|GGCCGTGATCGTATAGTGGTTAGTACTCTGCGT</td><td>>trna111_HisGTG_1_-_147774845_147774916</td><td>-1G</td><td>32</td><td>33</td><td>GGCCGTGATCGTATAGTGGTTAGTACTCTGCGT</td><td>tRF</td><td>FALSE</td><td>TRUE</td><td>FALSE</td></tr>
<tr><td>>hg38_21_+_8433172_8446622_RNA45SN1_45S_50ntFlanks@4731.4767.37|AAGAACGAAAGTCGGAGGTTCGAAGACGATCAGATAC</td><td>>hg38_21_+_8433172_8446622_RNA45SN1_45S_50ntFlanks</td><td>4731</td><td>4767</td><td>37</td><td>AAGAACGAAAGTCGGAGGTTCGAAGACGATCAGATAC</td><td>rRF</td><td>FALSE</td><td>TRUE</td><td>FALSE</td></tr>
<tr><td>>hg38_21_+_8433172_8446622_RNA45SN1_45S_50ntFlanks@8176.8195.20|CTTCTGATCGAGGCCCAGCC</td><td>>hg38_21_+_8433172_8446622_RNA45SN1_45S_50ntFlanks</td><td>8176</td><td>8195</td><td>20</td><td>CTTCTGATCGAGGCCCAGCC</td><td>rRF</td><td>FALSE</td><td>TRUE</td><td>FALSE</td></tr>
<tr><td>>hg38_1_-_228634819_228635039_RNA5S12_5S_50ntFlanks@152.170.19|TACCGGGTGCTGTAGGCTT</td><td>>hg38_1_-_228634819_228635039_RNA5S12_5S_50ntFlanks</td><td>152</td><td>170</td><td>19</td><td>TACCGGGTGCTGTAGGCTT</td><td>rRF</td><td>FALSE</td><td>TRUE</td><td>FALSE</td></tr>
<tr><td>>hg38_21_+_8433172_8446622_RNA45SN1_45S_50ntFlanks@12996.13013.18|TCGCTGCGATCTATTGAA</td><td>>hg38_21_+_8433172_8446622_RNA45SN1_45S_50ntFlanks</td><td>12996</td><td>13013</td><td>18</td><td>TCGCTGCGATCTATTGAA</td><td>rRF</td><td>FALSE</td><td>TRUE</td><td>FALSE</td></tr>
<tr><td>>hg38_21_+_8433172_8446622_RNA45SN1_45S_50ntFlanks@6645.6662.18|CTCGTACGACTCTTAGCG</td><td>>hg38_21_+_8433172_8446622_RNA45SN1_45S_50ntFlanks</td><td>6645</td><td>6662</td><td>18</td><td>CTCGTACGACTCTTAGCG</td><td>rRF</td><td>FALSE</td><td>TRUE</td><td>FALSE</td></tr>
<tr><td>>hg38_1_-_228634819_228635039_RNA5S12_5S_50ntFlanks@130.171.42|TGGATGGGAGACCGCCTGGGAATACCGGGTGCTGTAGGCTTT</td><td>>hg38_1_-_228634819_228635039_RNA5S12_5S_50ntFlanks</td><td>130</td><td>171</td><td>42</td><td>TGGATGGGAGACCGCCTGGGAATACCGGGTGCTGTAGGCTTT</td><td>rRF</td><td>FALSE</td><td>TRUE</td><td>FALSE</td></tr>
<tr><td>>hg38_1_-_228634819_228635039_RNA5S12_5S_50ntFlanks@141.171.31|CCGCCTGGGAATACCGGGTGCTGTAGGCTTT</td><td>>hg38_1_-_228634819_228635039_RNA5S12_5S_50ntFlanks</td><td>141</td><td>171</td><td>31</td><td>CCGCCTGGGAATACCGGGTGCTGTAGGCTTT</td><td>rRF</td><td>FALSE</td><td>TRUE</td><td>FALSE</td></tr>
<tr><td>>trna111_HisGTG_1_-_147774845_147774916@-1T.34.35,trna118_HisGTG_1_-_145396881_145396952@-1T.34.35,trna16_HisGTG_1_+_146544773_146544844@-1T.34.35,trna1_HisGTG_15_+_45493349_45493420@-1T.34.35,trna21_HisGTG_1_+_147753471_147753542@-1T.34.35,trna33_HisGTG_6_+_27125906_27125977@-1T.34.35,trna7_HisGTG_9_-_14433938_14434009@-1T.34.35,trna8_HisGTG_15_-_45492611_45492682@-1T.34.35,trna9_HisGTG_15_-_45490804_45490875@-1T.34.35|TGCCGTGATCGTATAGTGGTTAGTACTCTGCGTTG</td><td>>trna111_HisGTG_1_-_147774845_147774916</td><td>-1T</td><td>34</td><td>35</td><td>TGCCGTGATCGTATAGTGGTTAGTACTCTGCGTTG</td><td>tRF</td><td>FALSE</td><td>TRUE</td><td>FALSE</td></tr>
<tr><td>>hg38_1_-_228634819_228635039_RNA5S12_5S_50ntFlanks@151.171.21|ATACCGGGTGCTGTAGGCTTT</td><td>>hg38_1_-_228634819_228635039_RNA5S12_5S_50ntFlanks</td><td>151</td><td>171</td><td>21</td><td>ATACCGGGTGCTGTAGGCTTT</td><td>rRF</td><td>FALSE</td><td>TRUE</td><td>FALSE</td></tr>
</table>

### Cytoplasm

<table>
<tr><th></th><th></th><th></th><th></th><th></th><th></th><th></th><th></th><th></th><th></th></tr>
<tr><td>>trna10_AspGTC_12_-_125424193_125424264@36.73.38,trna12_AspGTC_12_-_125411891_125411962@36.73.38,trna144_AspGTC_6_-_27551236_27551307@36.73.38,trna38_AspGTC_17_-_8125556_8125627@36.73.38,trna45_AspGTC_6_+_27447453_27447524@36.73.38,trna48_AspGTC_6_+_27471523_27471594@36.73.38,trna4_AspGTC_12_+_96429799_96429870@36.73.38,trna69_AspGTC_1_-_161440205_161440276@36.73.38,trna72_AspGTC_1_-_161432824_161432895@36.73.38,trna75_AspGTC_1_-_161425414_161425485@36.73.38,trna78_AspGTC_1_-_161418033_161418104@36.73.38,trna81_AspGTC_1_-_161410615_161410686@36.73.38|CACGCGGGAGACCGGGGTTCGATTCCCCGACGGGGAGC</td><td>>trna10_AspGTC_12_-_125424193_125424264</td><td>36</td><td>73</td><td>38</td><td>CACGCGGGAGACCGGGGTTCGATTCCCCGACGGGGAGC</td><td>tRF</td><td>FALSE</td><td>TRUE</td><td>FALSE</td></tr>
<tr><td>>hg38_21_+_8433172_8446622_RNA45SN1_45S_50ntFlanks@12416.12438.23|TTCGATGTCGGCTCTTCCTATCA</td><td>>hg38_21_+_8433172_8446622_RNA45SN1_45S_50ntFlanks</td><td>12416</td><td>12438</td><td>23</td><td>TTCGATGTCGGCTCTTCCTATCA</td><td>rRF</td><td>FALSE</td><td>TRUE</td><td>FALSE</td></tr>
<tr><td>>hg38_21_+_8433172_8446622_RNA45SN1_45S_50ntFlanks@12416.12440.25|TTCGATGTCGGCTCTTCCTATCATT</td><td>>hg38_21_+_8433172_8446622_RNA45SN1_45S_50ntFlanks</td><td>12416</td><td>12440</td><td>25</td><td>TTCGATGTCGGCTCTTCCTATCATT</td><td>rRF</td><td>FALSE</td><td>TRUE</td><td>FALSE</td></tr>
<tr><td>>hg38_1_-_228634819_228635039_RNA5S12_5S_50ntFlanks@129.171.43|TTGGATGGGAGACCGCCTGGGAATACCGGGTGCTGTAGGCTTT</td><td>>hg38_1_-_228634819_228635039_RNA5S12_5S_50ntFlanks</td><td>129</td><td>171</td><td>43</td><td>TTGGATGGGAGACCGCCTGGGAATACCGGGTGCTGTAGGCTTT</td><td>rRF</td><td>FALSE</td><td>TRUE</td><td>FALSE</td></tr>
<tr><td>>hg38_21_+_8433172_8446622_RNA45SN1_45S_50ntFlanks@12419.12442.24|GATGTCGGCTCTTCCTATCATTGT</td><td>>hg38_21_+_8433172_8446622_RNA45SN1_45S_50ntFlanks</td><td>12419</td><td>12442</td><td>24</td><td>GATGTCGGCTCTTCCTATCATTGT</td><td>rRF</td><td>FALSE</td><td>TRUE</td><td>FALSE</td></tr>
<tr><td>>hg38_21_+_8433172_8446622_RNA45SN1_45S_50ntFlanks@7977.8011.35|CGACCTCAGATCAGACGTGGCGACCCGCTGAATTT</td><td>>hg38_21_+_8433172_8446622_RNA45SN1_45S_50ntFlanks</td><td>7977</td><td>8011</td><td>35</td><td>CGACCTCAGATCAGACGTGGCGACCCGCTGAATTT</td><td>rRF</td><td>FALSE</td><td>TRUE</td><td>FALSE</td></tr>
<tr><td>>MI0000299|hsa-mir-222&WithFlank&X|-|45747009|45747130@75.99.25(+1U)&[MIMAT0000279&hsa-miR-222-3p&offsets|0|+4(+1U);m-140&X|-|45747033|45747056&offsets|0|+1(+1U)]|AGCTACATCTGGCTACTGGGTCTCTT</td><td>>MI0000299|hsa-mir-222&WithFlank&X|-|45747009|45747130</td><td>75</td><td>99</td><td>25(+1U)</td><td>AGCTACATCTGGCTACTGGGTCTCTT</td><td>isomiR</td><td>FALSE</td><td>TRUE</td><td>FALSE</td></tr>
<tr><td>>hg38_1_-_228634819_228635039_RNA5S12_5S_50ntFlanks@126.170.45|TACTTGGATGGGAGACCGCCTGGGAATACCGGGTGCTGTAGGCTT</td><td>>hg38_1_-_228634819_228635039_RNA5S12_5S_50ntFlanks</td><td>126</td><td>170</td><td>45</td><td>TACTTGGATGGGAGACCGCCTGGGAATACCGGGTGCTGTAGGCTT</td><td>rRF</td><td>FALSE</td><td>TRUE</td><td>FALSE</td></tr>
<tr><td>>hg38_21_+_8433172_8446622_RNA45SN1_45S_50ntFlanks@11567.11584.18|AGCAGCCGACTTAGAACT</td><td>>hg38_21_+_8433172_8446622_RNA45SN1_45S_50ntFlanks</td><td>11567</td><td>11584</td><td>18</td><td>AGCAGCCGACTTAGAACT</td><td>rRF</td><td>FALSE</td><td>TRUE</td><td>FALSE</td></tr>
<tr><td>>MI0000102|hsa-mir-100&WithFlank&11|-|122152223|122152314@19.39.21(+1U)&[MIMAT0000098&hsa-miR-100-5p&offsets|0|-1(+1U);m-27&11|-|122152276|122152296&offsets|0|0(+1U)]|AACCCGTAGATCCGAACTTGTT</td><td>>MI0000102|hsa-mir-100&WithFlank&11|-|122152223|122152314</td><td>19</td><td>39</td><td>21(+1U)</td><td>AACCCGTAGATCCGAACTTGTT</td><td>isomiR</td><td>FALSE</td><td>TRUE</td><td>FALSE</td></tr>
<tr><td>>hg38_21_+_8433172_8446622_RNA45SN1_45S_50ntFlanks@12419.12438.20|GATGTCGGCTCTTCCTATCA</td><td>>hg38_21_+_8433172_8446622_RNA45SN1_45S_50ntFlanks</td><td>12419</td><td>12438</td><td>20</td><td>GATGTCGGCTCTTCCTATCA</td><td>rRF</td><td>FALSE</td><td>TRUE</td><td>FALSE</td></tr>
<tr><td>>MI0000102|hsa-mir-100&WithFlank&11|-|122152223|122152314@19.38.20(+1A)&[MIMAT0000098&hsa-miR-100-5p&offsets|0|-2(+1A);m-27&11|-|122152276|122152296&offsets|0|-1(+1A)]|AACCCGTAGATCCGAACTTGA</td><td>>MI0000102|hsa-mir-100&WithFlank&11|-|122152223|122152314</td><td>19</td><td>38</td><td>20(+1A)</td><td>AACCCGTAGATCCGAACTTGA</td><td>isomiR</td><td>FALSE</td><td>TRUE</td><td>FALSE</td></tr>
<tr><td>>hg38_21_+_8433172_8446622_RNA45SN1_45S_50ntFlanks@6651.6695.45|CGACTCTTAGCGGTGGATCACTCGGCTCGTGCGTCGATGAAGAAC</td><td>>hg38_21_+_8433172_8446622_RNA45SN1_45S_50ntFlanks</td><td>6651</td><td>6695</td><td>45</td><td>CGACTCTTAGCGGTGGATCACTCGGCTCGTGCGTCGATGAAGAAC</td><td>rRF</td><td>FALSE</td><td>TRUE</td><td>FALSE</td></tr>
<tr><td>>hg38_21_+_8433172_8446622_RNA45SN1_45S_50ntFlanks@5472.5514.43|ACGGCCCTGGCGGAGCGCTGAGAAGACGGTCGAACTTGACTAT</td><td>>hg38_21_+_8433172_8446622_RNA45SN1_45S_50ntFlanks</td><td>5472</td><td>5514</td><td>43</td><td>ACGGCCCTGGCGGAGCGCTGAGAAGACGGTCGAACTTGACTAT</td><td>rRF</td><td>FALSE</td><td>TRUE</td><td>FALSE</td></tr>
<tr><td>>MI0000299|hsa-mir-222&WithFlank&X|-|45747009|45747130@75.98.24(+1A)&[MIMAT0000279&hsa-miR-222-3p&offsets|0|+3(+1A);m-140&X|-|45747033|45747056&offsets|0|0(+1A)]|AGCTACATCTGGCTACTGGGTCTCA</td><td>>MI0000299|hsa-mir-222&WithFlank&X|-|45747009|45747130</td><td>75</td><td>98</td><td>24(+1A)</td><td>AGCTACATCTGGCTACTGGGTCTCA</td><td>isomiR</td><td>FALSE</td><td>TRUE</td><td>FALSE</td></tr>
<tr><td>>hg38_1_-_228634819_228635039_RNA5S12_5S_50ntFlanks@146.171.26|TGGGAATACCGGGTGCTGTAGGCTTT</td><td>>hg38_1_-_228634819_228635039_RNA5S12_5S_50ntFlanks</td><td>146</td><td>171</td><td>26</td><td>TGGGAATACCGGGTGCTGTAGGCTTT</td><td>rRF</td><td>FALSE</td><td>TRUE</td><td>FALSE</td></tr>
<tr><td>>trna10_AspGTC_12_-_125424193_125424264@35.73.39,trna12_AspGTC_12_-_125411891_125411962@35.73.39,trna144_AspGTC_6_-_27551236_27551307@35.73.39,trna38_AspGTC_17_-_8125556_8125627@35.73.39,trna45_AspGTC_6_+_27447453_27447524@35.73.39,trna48_AspGTC_6_+_27471523_27471594@35.73.39,trna4_AspGTC_12_+_96429799_96429870@35.73.39,trna69_AspGTC_1_-_161440205_161440276@35.73.39,trna72_AspGTC_1_-_161432824_161432895@35.73.39,trna75_AspGTC_1_-_161425414_161425485@35.73.39,trna78_AspGTC_1_-_161418033_161418104@35.73.39,trna81_AspGTC_1_-_161410615_161410686@35.73.39|TCACGCGGGAGACCGGGGTTCGATTCCCCGACGGGGAGC</td><td>>trna10_AspGTC_12_-_125424193_125424264</td><td>35</td><td>73</td><td>39</td><td>TCACGCGGGAGACCGGGGTTCGATTCCCCGACGGGGAGC</td><td>tRF</td><td>FALSE</td><td>TRUE</td><td>FALSE</td></tr>
<tr><td>>hg38_21_+_8433172_8446622_RNA45SN1_45S_50ntFlanks@12993.13039.47|CCCTCGCTGCGATCTATTGAAAGTCAGCCCTCGACACAAGGGTTTGT</td><td>>hg38_21_+_8433172_8446622_RNA45SN1_45S_50ntFlanks</td><td>12993</td><td>13039</td><td>47</td><td>CCCTCGCTGCGATCTATTGAAAGTCAGCCCTCGACACAAGGGTTTGT</td><td>rRF</td><td>FALSE</td><td>TRUE</td><td>FALSE</td></tr>
<tr><td>>hg38_1_-_228634819_228635039_RNA5S12_5S_50ntFlanks@145.171.27|CTGGGAATACCGGGTGCTGTAGGCTTT</td><td>>hg38_1_-_228634819_228635039_RNA5S12_5S_50ntFlanks</td><td>145</td><td>171</td><td>27</td><td>CTGGGAATACCGGGTGCTGTAGGCTTT</td><td>rRF</td><td>FALSE</td><td>TRUE</td><td>FALSE</td></tr>
<tr><td>>hg38_1_-_228634819_228635039_RNA5S12_5S_50ntFlanks@149.170.22|GAATACCGGGTGCTGTAGGCTT</td><td>>hg38_1_-_228634819_228635039_RNA5S12_5S_50ntFlanks</td><td>149</td><td>170</td><td>22</td><td>GAATACCGGGTGCTGTAGGCTT</td><td>rRF</td><td>FALSE</td><td>TRUE</td><td>FALSE</td></tr>
<tr><td>>trna32_LysCTT_16_-_3207406_3207478@1.33.33|GCCCGGCTAGCTCAGTCGGTAGAGCATGAGACC</td><td>>trna32_LysCTT_16_-_3207406_3207478</td><td>1</td><td>33</td><td>33</td><td>GCCCGGCTAGCTCAGTCGGTAGAGCATGAGACC</td><td>tRF</td><td>FALSE</td><td>TRUE</td><td>FALSE</td></tr>
<tr><td>>hg38_21_+_8433172_8446622_RNA45SN1_45S_50ntFlanks@8342.8365.24|CGTAAGGGAAAGTTGAAAAGAACT</td><td>>hg38_21_+_8433172_8446622_RNA45SN1_45S_50ntFlanks</td><td>8342</td><td>8365</td><td>24</td><td>CGTAAGGGAAAGTTGAAAAGAACT</td><td>rRF</td><td>FALSE</td><td>TRUE</td><td>FALSE</td></tr>
<tr><td>>hg38_1_-_228634819_228635039_RNA5S12_5S_50ntFlanks@150.170.21|AATACCGGGTGCTGTAGGCTT</td><td>>hg38_1_-_228634819_228635039_RNA5S12_5S_50ntFlanks</td><td>150</td><td>170</td><td>21</td><td>AATACCGGGTGCTGTAGGCTT</td><td>rRF</td><td>FALSE</td><td>TRUE</td><td>FALSE</td></tr>
<tr><td>>hg38_21_+_8433172_8446622_RNA45SN1_45S_50ntFlanks@6651.6681.31|CGACTCTTAGCGGTGGATCACTCGGCTCGTG</td><td>>hg38_21_+_8433172_8446622_RNA45SN1_45S_50ntFlanks</td><td>6651</td><td>6681</td><td>31</td><td>CGACTCTTAGCGGTGGATCACTCGGCTCGTG</td><td>rRF</td><td>FALSE</td><td>TRUE</td><td>TRUE</td></tr>
<tr><td>>trna111_HisGTG_1_-_147774845_147774916@-1G.35.36,trna118_HisGTG_1_-_145396881_145396952@-1G.35.36,trna16_HisGTG_1_+_146544773_146544844@-1G.35.36,trna1_HisGTG_15_+_45493349_45493420@-1G.35.36,trna21_HisGTG_1_+_147753471_147753542@-1G.35.36,trna33_HisGTG_6_+_27125906_27125977@-1G.35.36,trna7_HisGTG_9_-_14433938_14434009@-1G.35.36,trna8_HisGTG_15_-_45492611_45492682@-1G.35.36,trna9_HisGTG_15_-_45490804_45490875@-1G.35.36|GGCCGTGATCGTATAGTGGTTAGTACTCTGCGTTGT</td><td>>trna111_HisGTG_1_-_147774845_147774916</td><td>-1G</td><td>35</td><td>36</td><td>GGCCGTGATCGTATAGTGGTTAGTACTCTGCGTTGT</td><td>tRF</td><td>FALSE</td><td>TRUE</td><td>FALSE</td></tr>
</table>

### Cytoplasm

| | | | | | | | | | |
|---|---|---|---|---|---|---|---|---|---|
| >MI0000079\|hsa-mir-23a&WithFlank&19\|-\|13836581\|13836665@54.73.20&[MIMAT0000078&hsa-miR-23a-3p&offsets\|+3\|+2;m-31&19\|-\|13836595\|13836615&offsets\|+3\|+2]\|ACATTGCCAGGGATTTCCAA | >MI0000079\|hsa-mir-23a&WithFlank&19\|-\|13836581\|13836665 | 54 | 73 | 20 | ACATTGCCAGGGATTTCCAA | isomiR | FALSE | TRUE | FALSE |
| >trna119_LysCTT_1_-_145395522_145395594@1.32.32,trna11_LysCTT_5_-_180648979_180649051@1.32.32,trna13_LysCTT_6_+_26556774_26556846@1.32.32,trna32_LysCTT_16_-_3207406_3207478@1.32.32,trna7_LysCTT_16_+_3225692_3225764@1.32.32,trna9_LysCTT_5_+_180634755_180634827@1.32.32\|GCCCGGCTAGCTCAGTCGGTAGAGCATGAGAC | >trna119_LysCTT_1_-_145395522_145395594 | 1 | 32 | 32 | GCCCGGCTAGCTCAGTCGGTAGAGCATGAGAC | tRF | FALSE | TRUE | FALSE |
| >hg38_21_+_8433172_8446622_RNA45SN1_45S_50ntFlanks@12996.13038.43\|TCGCTGCGATCTATTGAAAGTCAGCCCTCGACACAAGGGTTTG | >hg38_21_+_8433172_8446622_RNA45SN1_45S_50ntFlanks | 12996 | 13038 | 43 | TCGCTGCGATCTATTGAAAGTCAGCCCTCGACACAAGGGTTTG | rRF | FALSE | TRUE | FALSE |
| >am_trna128_GlyGCC_6_-_27870686_27870756@1.33.33,trna18_GlyGCC_16_+_70822597_70822667@1.33.33,trna19_GlyGCC_16_+_70823410_70823480@1.33.33,trna19_GlyGCC_2_-_157257659_157257729@1.33.33,trna24_GlyGCC_16_-_70812942_70813012@1.33.33,trna25_GlyGCC_16_-_70812114_70812184@1.33.33,trna5_GlyGCC_17_+_8029064_8029134@1.33.33,trna68_GlyGCC_1_-_161493637_161493707@1.33.33\|GCATTGGTGGTTCAGTGGTAGAATTCTCGCCTG | >am_trna128_GlyGCC_6_-_27870686_27870756 | 1 | 33 | 33 | GCATTGGTGGTTCAGTGGTAGAATTCTCGCCTG | tRF | FALSE | TRUE | FALSE |
| >hg38_21_+_8433172_8446622_RNA45SN1_45S_50ntFlanks@4731.4768.38\|AAGAACGAAAGTCGGAGGTTCGAAGACGATCAGATACC | >hg38_21_+_8433172_8446622_RNA45SN1_45S_50ntFlanks | 4731 | 4768 | 38 | AAGAACGAAAGTCGGAGGTTCGAAGACGATCAGATACC | rRF | FALSE | TRUE | FALSE |
| >am_trna116_GluCTC_1_-_145399233_145399304@1.29.29,trna59_GluCTC_1_+_249168447_249168518@1.29.29,trna71_GluCTC_1_-_161439189_161439260@1.29.29,trna74_GluCTC_1_-_161431809_161431880@1.29.29,trna77_GluCTC_1_-_161424398_161424469@1.29.29,trna77_GluCTC_6_+_28949976_28950047@1.29.29,trna80_GluCTC_1_-_161417018_161417089@1.29.29,trna87_GluCTC_6_-_126101393_126101464@1.29.29\|TCCCTGGTGGTCTAGTGGTTAGGATTCGG | >am_trna116_GluCTC_1_-_145399233_145399304 | 1 | 29 | 29 | TCCCTGGTGGTCTAGTGGTTAGGATTCGG | tRF | FALSE | TRUE | TRUE |
| >trna10_AspGTC_12_-_125424193_125424264@39.75.37,trna12_AspGTC_12_-_125411891_125411962@39.75.37,trna144_AspGTC_6_-_27551236_27551307@39.75.37,trna38_AspGTC_17_-_8125556_8125627@39.75.37,trna45_AspGTC_6_+_27447453_27447524@39.75.37,trna48_AspGTC_6_+_27471523_27471594@39.75.37,trna4_AspGTC_12_+_96429799_96429870@39.75.37,trna69_AspGTC_1_-_161440205_161440276@39.75.37,trna72_AspGTC_1_-_161432824_161432895@39.75.37,trna75_AspGTC_1_-_161425414_161425485@39.75.37,trna78_AspGTC_1_-_161418033_161418104@39.75.37,trna81_AspGTC_1_-_161410615_161410686@39.75.37\|GCGGGAGACCGGGGTTCGATTCCCCGACGGGGAGCCA | >trna10_AspGTC_12_-_125424193_125424264 | 39 | 75 | 37 | GCGGGAGACCGGGGTTCGATTCCCCGACGGGGAGCCA | tRF | FALSE | TRUE | FALSE |
| >hg38_21_+_8433172_8446622_RNA45SN1_45S_50ntFlanks@10383.10424.42\|TCCATGTGAACAGCAGTTGAACATGGGTCAGTCGGTCCTGAG | >hg38_21_+_8433172_8446622_RNA45SN1_45S_50ntFlanks | 10383 | 10424 | 42 | TCCATGTGAACAGCAGTTGAACATGGGTCAGTCGGTCCTGAG | rRF | FALSE | TRUE | FALSE |
| >hg38_21_+_8433172_8446622_RNA45SN1_45S_50ntFlanks@9554.9573.20\|CGTAGCGGTCCTGACGTGCA | >hg38_21_+_8433172_8446622_RNA45SN1_45S_50ntFlanks | 9554 | 9573 | 20 | CGTAGCGGTCCTGACGTGCA | rRF | FALSE | TRUE | TRUE |
| >hg38_21_+_8433172_8446622_RNA45SN1_45S_50ntFlanks@12416.12444.29\|TTCGATGTCGGCTCTTCCTATCATTGTGA | >hg38_21_+_8433172_8446622_RNA45SN1_45S_50ntFlanks | 12416 | 12444 | 29 | TTCGATGTCGGCTCTTCCTATCATTGTGA | rRF | FALSE | TRUE | FALSE |
| >hg38_21_+_8433172_8446622_RNA45SN1_45S_50ntFlanks@10396.10424.29\|CAGTTGAACATGGGTCAGTCGGTCCTGAG | >hg38_21_+_8433172_8446622_RNA45SN1_45S_50ntFlanks | 10396 | 10424 | 29 | CAGTTGAACATGGGTCAGTCGGTCCTGAG | rRF | FALSE | TRUE | FALSE |
| >hg38_21_+_8433172_8446622_RNA45SN1_45S_50ntFlanks@6651.6678.28\|CGACTCTTAGCGGTGGATCACTCGGCTC | >hg38_21_+_8433172_8446622_RNA45SN1_45S_50ntFlanks | 6651 | 6678 | 28 | CGACTCTTAGCGGTGGATCACTCGGCTC | rRF | FALSE | TRUE | FALSE |
| >hg38_21_+_8433172_8446622_RNA45SN1_45S_50ntFlanks@12419.12447.29\|GATGTCGGCTCTTCCTATCATTGTGAAGC | >hg38_21_+_8433172_8446622_RNA45SN1_45S_50ntFlanks | 12419 | 12447 | 29 | GATGTCGGCTCTTCCTATCATTGTGAAGC | rRF | FALSE | TRUE | FALSE |
| >hg38_1_-_228634819_228635039_RNA5S12_5S_50ntFlanks@148.170.23\|GGAATACCGGGTGCTGTAGGCTT | >hg38_1_-_228634819_228635039_RNA5S12_5S_50ntFlanks | 148 | 170 | 23 | GGAATACCGGGTGCTGTAGGCTT | rRF | FALSE | TRUE | FALSE |
| >trna111_HisGTG_1_-_147774845_147774916@-1T.33.34,trna118_HisGTG_1_-_145396881_145396952@-1T.33.34,trna16_HisGTG_1_+_146544773_146544844@-1T.33.34,trna1_HisGTG_15_+_45493349_45493420@-1T.33.34,trna21_HisGTG_1_+_147753471_147753542@-1T.33.34,trna33_HisGTG_6_+_27125906_27125977@-1T.33.34,trna7_HisGTG_9_-_14433938_14434009@-1T.33.34,trna8_HisGTG_15_-_45492611_45492682@-1T.33.34,trna9_HisGTG_15_-_45490804_45490875@-1T.33.34\|TGCCGTGATCGTATAGTGGTTAGTACTCTGCGTT | >trna111_HisGTG_1_-_147774845_147774916 | -1T | 33 | 34 | TGCCGTGATCGTATAGTGGTTAGTACTCTGCGTT | tRF | FALSE | TRUE | FALSE |
| >MI0000446\|hsa-mir-125b-1&WithFlank&11\|-\|122099751\|122099850@21.42.22(+1A);MI0000470\|hsa-mir-125b-2&WithFlank&21\|+\|16590231\|16590331@23.44.22(+1A)&[MIMAT0000423&hsa-miR-125b-5p&offsets\|0\|0(+1A);m-67&11\|-\|122099809\|122099830&offsets\|0\|0(+1A)];[MIMAT0000423_1&hsa-miR-125b-5p&offsets\|0\|0(+1A);m-68&21\|+\|16590253\|16590274&offsets\|0\|0(+1A)]\|TCCCTGAGACCCTAACTTGTGAA | >MI0000446\|hsa-mir-125b-1&WithFlank&11\|-\|122099751\|122099850 | 21 | 42 | 22(+1A) | TCCCTGAGACCCTAACTTGTGAA | isomiR | FALSE | TRUE | FALSE |
| >hg38_21_+_8433172_8446622_RNA45SN1_45S_50ntFlanks@12419.12439.21\|GATGTCGGCTCTTCCTATCAT | >hg38_21_+_8433172_8446622_RNA45SN1_45S_50ntFlanks | 12419 | 12439 | 21 | GATGTCGGCTCTTCCTATCAT | rRF | FALSE | TRUE | FALSE |
| >hg38_21_+_8433172_8446622_RNA45SN1_45S_50ntFlanks@10841.10867.27\|CGTAACTTCGGGATAAGGATTGGCTCT | >hg38_21_+_8433172_8446622_RNA45SN1_45S_50ntFlanks | 10841 | 10867 | 27 | CGTAACTTCGGGATAAGGATTGGCTCT | rRF | FALSE | TRUE | TRUE |
| >hg38_1_-_228634819_228635039_RNA5S12_5S_50ntFlanks@127.170.44\|ACTTGGATGGGAGACCGCCTGGGAATACCGGGTGCTGTAGGCTT | >hg38_1_-_228634819_228635039_RNA5S12_5S_50ntFlanks | 127 | 170 | 44 | ACTTGGATGGGAGACCGCCTGGGAATACCGGGTGCTGTAGGCTT | rRF | FALSE | TRUE | FALSE |
| >hg38_21_+_8433172_8446622_RNA45SN1_45S_50ntFlanks@11668.11706.39\|TGCCCAGTGCTCTGAATGTCAAAGTGAAGAAATTCAATG | >hg38_21_+_8433172_8446622_RNA45SN1_45S_50ntFlanks | 11668 | 11706 | 39 | TGCCCAGTGCTCTGAATGTCAAAGTGAAGAAATTCAATG | rRF | FALSE | TRUE | FALSE |
| >hg38_21_+_8433172_8446622_RNA45SN1_45S_50ntFlanks@12996.13036.41\|TCGCTGCGATCTATTGAAAGTCAGCCCTCGACACAAGGGTT | >hg38_21_+_8433172_8446622_RNA45SN1_45S_50ntFlanks | 12996 | 13036 | 41 | TCGCTGCGATCTATTGAAAGTCAGCCCTCGACACAAGGGTT | rRF | FALSE | TRUE | FALSE |

### Cytoplasm

<table>
<tr><td>>am_trna128_GlyGCC_6_-_27870686_27870756@1.34.34,trna18_GlyGCC_16_+_70822597_70822667@1.34.34,trna19_GlyGCC_16_+_70823410_70823480@1.34.34,trna19_GlyGCC_2_-_157257659_157257729@1.34.34,trna24_GlyGCC_16_-_70812942_70813012@1.34.34,trna25_GlyGCC_16_-_70812114_70812184@1.34.34,trna5_GlyGCC_17_+_8029064_8029134@1.34.34,trna68_GlyGCC_1_-_161493637_161493707@1.34.34|GCATTGGTGGTTCAGTGGTAGAATTCTCGCCTGC</td><td>>am_trna128_GlyGCC_6_-_27870686_27870756</td><td>1</td><td>34</td><td>34</td><td>GCATTGGTGGTTCAGTGGTAGAATTCTCGCCTGC</td><td>tRF</td><td>FALSE</td><td>TRUE</td><td>FALSE</td></tr>
<tr><td>>hg38_1_-_228634819_228635039_RNA5S12_5S_50ntFlanks@151.170.20|ATACCGGGTGCTGTAGGCTT</td><td>>hg38_1_-_228634819_228635039_RNA5S12_5S_50ntFlanks</td><td>151</td><td>170</td><td>20</td><td>ATACCGGGTGCTGTAGGCTT</td><td>rRF</td><td>FALSE</td><td>TRUE</td><td>FALSE</td></tr>
<tr><td>>MI0003581|hsa-mir-574&WithFlank&4|+|38868026|38868133@67.88.22&[MIMAT0003239&hsa-miR-574-3p&offsets|0|0;m-149&4|+|38868092|38868113&offsets|0|0]|CACGCTCATGCACACACCCACA</td><td>>MI0003581|hsa-mir-574&WithFlank&4|+|38868026|38868133</td><td>67</td><td>88</td><td>22</td><td>CACGCTCATGCACACACCCACA</td><td>isomiR</td><td>FALSE</td><td>TRUE</td><td>FALSE</td></tr>
<tr><td>>hg38_21_+_8433172_8446622_RNA45SN1_45S_50ntFlanks@7989.8011.23|AGACGTGGCGACCCGCTGAATTT</td><td>>hg38_21_+_8433172_8446622_RNA45SN1_45S_50ntFlanks</td><td>7989</td><td>8011</td><td>23</td><td>AGACGTGGCGACCCGCTGAATTT</td><td>rRF</td><td>FALSE</td><td>TRUE</td><td>FALSE</td></tr>
<tr><td>>MI0000299|hsa-mir-222&WithFlank&X|-|45747009|45747130@75.97.23&[MIMAT0000279&hsa-miR-222-3p&offsets|0|+2;m-140&X|-|45747033|45747056&offsets|0|-1]|AGCTACATCTGGCTACTGGGTCT</td><td>>MI0000299|hsa-mir-222&WithFlank&X|-|45747009|45747130</td><td>75</td><td>97</td><td>23</td><td>AGCTACATCTGGCTACTGGGTCT</td><td>isomiR</td><td>FALSE</td><td>TRUE</td><td>FALSE</td></tr>
<tr><td>>hg38_21_+_8433172_8446622_RNA45SN1_45S_50ntFlanks@11669.11706.38|GCCCAGTGCTCTGAATGTCAAAGTGAAGAAATTCAATG</td><td>>hg38_21_+_8433172_8446622_RNA45SN1_45S_50ntFlanks</td><td>11669</td><td>11706</td><td>38</td><td>GCCCAGTGCTCTGAATGTCAAAGTGAAGAAATTCAATG</td><td>rRF</td><td>FALSE</td><td>TRUE</td><td>FALSE</td></tr>
<tr><td>>hg38_21_+_8433172_8446622_RNA45SN1_45S_50ntFlanks@4727.4768.42|AATCAAGAACGAAAGTCGGAGGTTCGAAGACGATCAGATACC</td><td>>hg38_21_+_8433172_8446622_RNA45SN1_45S_50ntFlanks</td><td>4727</td><td>4768</td><td>42</td><td>AATCAAGAACGAAAGTCGGAGGTTCGAAGACGATCAGATACC</td><td>rRF</td><td>FALSE</td><td>TRUE</td><td>FALSE</td></tr>
<tr><td>>hg38_21_+_8433172_8446622_RNA45SN1_45S_50ntFlanks@10678.10710.33|CGGTTCCGGCGGCGTCCGGTGAGCTCTCGCTGG</td><td>>hg38_21_+_8433172_8446622_RNA45SN1_45S_50ntFlanks</td><td>10678</td><td>10710</td><td>33</td><td>CGGTTCCGGCGGCGTCCGGTGAGCTCTCGCTGG</td><td>rRF</td><td>FALSE</td><td>FALSE</td><td>TRUE</td></tr>
<tr><td>>hg38_21_+_8433172_8446622_RNA45SN1_45S_50ntFlanks@9534.9553.20|AGGAAACTCTGGTGGAGGTC</td><td>>hg38_21_+_8433172_8446622_RNA45SN1_45S_50ntFlanks</td><td>9534</td><td>9553</td><td>20</td><td>AGGAAACTCTGGTGGAGGTC</td><td>rRF</td><td>FALSE</td><td>FALSE</td><td>TRUE</td></tr>
<tr><td>>hg38_21_+_8433172_8446622_RNA45SN1_45S_50ntFlanks@10841.10865.25|CGTAACTTCGGGATAAGGATTGGCT</td><td>>hg38_21_+_8433172_8446622_RNA45SN1_45S_50ntFlanks</td><td>10841</td><td>10865</td><td>25</td><td>CGTAACTTCGGGATAAGGATTGGCT</td><td>rRF</td><td>FALSE</td><td>FALSE</td><td>TRUE</td></tr>
<tr><td>>am_trna128_GlyGCC_6_-_27870686_27870756@1.28.28,trna133_GlyCCC_1_-_16872434_16872504@1.28.28,trna18_GlyGCC_16_+_70822597_70822667@1.28.28,trna19_GlyGCC_16_+_70823410_70823480@1.28.28,trna19_GlyGCC_2_-_157257659_157257729@1.28.28,trna24_GlyGCC_16_-_70812942_70813012@1.28.28,trna25_GlyGCC_16_-_70812114_70812184@1.28.28,trna4_GlyCCC_1_+_17188416_17188486@1.28.28,trna5_GlyGCC_17_+_8029064_8029134@1.28.28,trna68_GlyGCC_1_-_161493637_161493707@1.28.28|GCATTGGTGGTTCAGTGGTAGAATTCTC</td><td>>am_trna128_GlyGCC_6_-_27870686_27870756</td><td>1</td><td>28</td><td>28</td><td>GCATTGGTGGTTCAGTGGTAGAATTCTC</td><td>tRF</td><td>FALSE</td><td>FALSE</td><td>TRUE</td></tr>
<tr><td>>am_trna128_GlyGCC_6_-_27870686_27870756@1.29.29,trna133_GlyCCC_1_-_16872434_16872504@1.29.29,trna18_GlyGCC_16_+_70822597_70822667@1.29.29,trna19_GlyGCC_16_+_70823410_70823480@1.29.29,trna19_GlyGCC_2_-_157257659_157257729@1.29.29,trna24_GlyGCC_16_-_70812942_70813012@1.29.29,trna25_GlyGCC_16_-_70812114_70812184@1.29.29,trna4_GlyCCC_1_+_17188416_17188486@1.29.29,trna5_GlyGCC_17_+_8029064_8029134@1.29.29,trna68_GlyGCC_1_-_161493637_161493707@1.29.29|GCATTGGTGGTTCAGTGGTAGAATTCTCG</td><td>>am_trna128_GlyGCC_6_-_27870686_27870756</td><td>1</td><td>29</td><td>29</td><td>GCATTGGTGGTTCAGTGGTAGAATTCTCG</td><td>tRF</td><td>FALSE</td><td>FALSE</td><td>TRUE</td></tr>
<tr><td>>trna2_GlyGCC_21_-_18827107_18827177@1.29.29,trna35_GlyGCC_1_+_161413094_161413164@1.29.29,trna37_GlyGCC_1_+_161420467_161420537@1.29.29,trna39_GlyGCC_1_+_161427898_161427968@1.29.29,trna41_GlyGCC_1_+_161435258_161435328@1.29.29|GCATGGGTGGTTCAGTGGTAGAATTCTCG</td><td>>trna2_GlyGCC_21_-_18827107_18827177</td><td>1</td><td>29</td><td>29</td><td>GCATGGGTGGTTCAGTGGTAGAATTCTCG</td><td>tRF</td><td>FALSE</td><td>FALSE</td><td>TRUE</td></tr>
<tr><td>>hg38_21_+_8433172_8446622_RNA45SN1_45S_50ntFlanks@11729.11751.23|TAACTATGACTCTCTTAAGGTAG</td><td>>hg38_21_+_8433172_8446622_RNA45SN1_45S_50ntFlanks</td><td>11729</td><td>11751</td><td>23</td><td>TAACTATGACTCTCTTAAGGTAG</td><td>rRF</td><td>FALSE</td><td>FALSE</td><td>TRUE</td></tr>
<tr><td>>hg38_21_+_8433172_8446622_RNA45SN1_45S_50ntFlanks@12456.12487.32|CCAAGCGTTGGATTGTTCACCCACTAATAGGG</td><td>>hg38_21_+_8433172_8446622_RNA45SN1_45S_50ntFlanks</td><td>12456</td><td>12487</td><td>32</td><td>CCAAGCGTTGGATTGTTCACCCACTAATAGGG</td><td>rRF</td><td>FALSE</td><td>FALSE</td><td>TRUE</td></tr>
<tr><td>>hg38_21_+_8433172_8446622_RNA45SN1_45S_50ntFlanks@10678.10711.34|CGGTTCCGGCGGCGTCCGGTGAGCTCTCGCTGGC</td><td>>hg38_21_+_8433172_8446622_RNA45SN1_45S_50ntFlanks</td><td>10678</td><td>10711</td><td>34</td><td>CGGTTCCGGCGGCGTCCGGTGAGCTCTCGCTGGC</td><td>rRF</td><td>FALSE</td><td>FALSE</td><td>TRUE</td></tr>
<tr><td>>hg38_21_+_8433172_8446622_RNA45SN1_45S_50ntFlanks@6699.6720.22|GCTAGCTGCGAGAATTAATGTG</td><td>>hg38_21_+_8433172_8446622_RNA45SN1_45S_50ntFlanks</td><td>6699</td><td>6720</td><td>22</td><td>GCTAGCTGCGAGAATTAATGTG</td><td>rRF</td><td>FALSE</td><td>FALSE</td><td>TRUE</td></tr>
<tr><td>>trna119_LysCTT_1_-_145395522_145395594@1.30.30,trna11_LysCTT_5_-_180648979_180649051@1.30.30,trna13_LysCTT_6_+_26556774_26556846@1.30.30,trna32_LysCTT_16_-_3207406_3207478@1.30.30,trna7_LysCTT_16_+_3225692_3225764@1.30.30,trna9_LysCTT_5_+_180634755_180634827@1.30.30|GCCCGGCTAGCTCAGTCGGTAGAGCATGAG</td><td>>trna119_LysCTT_1_-_145395522_145395594</td><td>1</td><td>30</td><td>30</td><td>GCCCGGCTAGCTCAGTCGGTAGAGCATGAG</td><td>tRF</td><td>FALSE</td><td>FALSE</td><td>TRUE</td></tr>
<tr><td>>hg38_21_+_8433172_8446622_RNA45SN1_45S_50ntFlanks@12456.12476.21|CCAAGCGTTGGATTGTTCACC</td><td>>hg38_21_+_8433172_8446622_RNA45SN1_45S_50ntFlanks</td><td>12456</td><td>12476</td><td>21</td><td>CCAAGCGTTGGATTGTTCACC</td><td>rRF</td><td>FALSE</td><td>FALSE</td><td>TRUE</td></tr>
<tr><td>>hg38_21_+_8433172_8446622_RNA45SN1_45S_50ntFlanks@6721.6744.24|AATTGCAGGACACATTGATCATCG</td><td>>hg38_21_+_8433172_8446622_RNA45SN1_45S_50ntFlanks</td><td>6721</td><td>6744</td><td>24</td><td>AATTGCAGGACACATTGATCATCG</td><td>rRF</td><td>FALSE</td><td>FALSE</td><td>TRUE</td></tr>
<tr><td>>hg38_21_+_8433172_8446622_RNA45SN1_45S_50ntFlanks@9534.9571.38|AGGAAACTCTGGTGGAGGTCCGTAGCGGTCCTGACGTG</td><td>>hg38_21_+_8433172_8446622_RNA45SN1_45S_50ntFlanks</td><td>9534</td><td>9571</td><td>38</td><td>AGGAAACTCTGGTGGAGGTCCGTAGCGGTCCTGACGTG</td><td>rRF</td><td>FALSE</td><td>FALSE</td><td>TRUE</td></tr>
<tr><td>>hg38_21_+_8433172_8446622_RNA45SN1_45S_50ntFlanks@10679.10710.32|GGTTCCGGCGGCGTCCGGTGAGCTCTCGCTGG</td><td>>hg38_21_+_8433172_8446622_RNA45SN1_45S_50ntFlanks</td><td>10679</td><td>10710</td><td>32</td><td>GGTTCCGGCGGCGTCCGGTGAGCTCTCGCTGG</td><td>rRF</td><td>FALSE</td><td>FALSE</td><td>TRUE</td></tr>
<tr><td>>trna152_ValCAC_6_-_27248049_27248121@1.29.29|GCTTCTGTAGTGTAGTGGTTATCACGTTC</td><td>>trna152_ValCAC_6_-_27248049_27248121</td><td>1</td><td>29</td><td>29</td><td>GCTTCTGTAGTGTAGTGGTTATCACGTTC</td><td>tRF</td><td>FALSE</td><td>FALSE</td><td>TRUE</td></tr>
<tr><td>>hg38_21_+_8433172_8446622_RNA45SN1_45S_50ntFlanks@9545.9571.27|GTGGAGGTCCGTAGCGGTCCTGACGTG</td><td>>hg38_21_+_8433172_8446622_RNA45SN1_45S_50ntFlanks</td><td>9545</td><td>9571</td><td>27</td><td>GTGGAGGTCCGTAGCGGTCCTGACGTG</td><td>rRF</td><td>FALSE</td><td>FALSE</td><td>TRUE</td></tr>
<tr><td>>hg38_21_+_8433172_8446622_RNA45SN1_45S_50ntFlanks@8357.8379.23|AAAAGAACTTTGAAGAGAGAGTT</td><td>>hg38_21_+_8433172_8446622_RNA45SN1_45S_50ntFlanks</td><td>8357</td><td>8379</td><td>23</td><td>AAAAGAACTTTGAAGAGAGAGTT</td><td>rRF</td><td>FALSE</td><td>FALSE</td><td>TRUE</td></tr>
<tr><td>>hg38_21_+_8433172_8446622_RNA45SN1_45S_50ntFlanks@5444.5466.23|CTCGGATCGGCCCCGCCGGGGTC</td><td>>hg38_21_+_8433172_8446622_RNA45SN1_45S_50ntFlanks</td><td>5444</td><td>5466</td><td>23</td><td>CTCGGATCGGCCCCGCCGGGGTC</td><td>rRF</td><td>FALSE</td><td>FALSE</td><td>TRUE</td></tr>
</table>

### Cytoplasm

| | | | | | | | | | |
|---|---|---|---|---|---|---|---|---|---|
| >hg38_21_+_8433172_8446622_RNA45SN1_45S_50ntFlanks@11661.11685.25\|TGATTTCTGCCCAGTGCTCTGAATG | >hg38_21_+_8433172_8446622_RNA45SN1_45S_50ntFlanks | 11661 | 11685 | 25 | TGATTTCTGCCCAGTGCTCTGAATG | rRF | FALSE | FALSE | TRUE |
| >hg38_21_+_8433172_8446622_RNA45SN1_45S_50ntFlanks@11208.11233.26\|GCGGCGACTCTGGACGCGAGCCGGGC | >hg38_21_+_8433172_8446622_RNA45SN1_45S_50ntFlanks | 11208 | 11233 | 26 | GCGGCGACTCTGGACGCGAGCCGGGC | rRF | FALSE | FALSE | TRUE |
| >hg38_21_+_8433172_8446622_RNA45SN1_45S_50ntFlanks@10351.10388.38\|CGAGAACTTTGAAGGCCGAAGTGGAGAAGGGTTCCATG | >hg38_21_+_8433172_8446622_RNA45SN1_45S_50ntFlanks | 10351 | 10388 | 38 | CGAGAACTTTGAAGGCCGAAGTGGAGAAGGGTTCCATG | rRF | FALSE | FALSE | TRUE |
| >hg38_21_+_8433172_8446622_RNA45SN1_45S_50ntFlanks@12420.12437.18\|ATGTCGGCTCTTCCTATC | >hg38_21_+_8433172_8446622_RNA45SN1_45S_50ntFlanks | 12420 | 12437 | 18 | ATGTCGGCTCTTCCTATC | rRF | FALSE | FALSE | TRUE |
| >hg38_21_+_8433172_8446622_RNA45SN1_45S_50ntFlanks@10350.10383.34\|ACGAGAACTTTGAAGGCCGAAGTGGAGAAGGGTT | >hg38_21_+_8433172_8446622_RNA45SN1_45S_50ntFlanks | 10350 | 10383 | 34 | ACGAGAACTTTGAAGGCCGAAGTGGAGAAGGGTT | rRF | FALSE | FALSE | TRUE |
| >hg38_21_+_8433172_8446622_RNA45SN1_45S_50ntFlanks@10410.10430.21\|TCAGTCGGTCCTGAGAGATGG | >hg38_21_+_8433172_8446622_RNA45SN1_45S_50ntFlanks | 10410 | 10430 | 21 | TCAGTCGGTCCTGAGAGATGG | rRF | FALSE | FALSE | TRUE |
| >hg38_21_+_8433172_8446622_RNA45SN1_45S_50ntFlanks@11317.11344.28\|ACCCCACGTCTCGTCGCGCGCGCGTCCG | >hg38_21_+_8433172_8446622_RNA45SN1_45S_50ntFlanks | 11317 | 11344 | 28 | ACCCCACGTCTCGTCGCGCGCGCGTCCG | rRF | FALSE | FALSE | TRUE |
| >hg38_21_+_8433172_8446622_RNA45SN1_45S_50ntFlanks@11729.11747.19\|TAACTATGACTCTCTTAAG | >hg38_21_+_8433172_8446622_RNA45SN1_45S_50ntFlanks | 11729 | 11747 | 19 | TAACTATGACTCTCTTAAG | rRF | FALSE | FALSE | TRUE |
| >hg38_21_+_8433172_8446622_RNA45SN1_45S_50ntFlanks@12405.12426.22\|CTTTTTGATCCTTCGATGTCGG | >hg38_21_+_8433172_8446622_RNA45SN1_45S_50ntFlanks | 12405 | 12426 | 22 | CTTTTTGATCCTTCGATGTCGG | rRF | FALSE | FALSE | TRUE |
| >hg38_21_+_8433172_8446622_RNA45SN1_45S_50ntFlanks@12405.12429.25\|CTTTTTGATCCTTCGATGTCGGCTC | >hg38_21_+_8433172_8446622_RNA45SN1_45S_50ntFlanks | 12405 | 12429 | 25 | CTTTTTGATCCTTCGATGTCGGCTC | rRF | FALSE | FALSE | TRUE |
| >hg38_21_+_8433172_8446622_RNA45SN1_45S_50ntFlanks@12415.12432.18\|CTTCGATGTCGGCTCTTC | >hg38_21_+_8433172_8446622_RNA45SN1_45S_50ntFlanks | 12415 | 12432 | 18 | CTTCGATGTCGGCTCTTC | rRF | FALSE | FALSE | TRUE |
| >hg38_21_+_8433172_8446622_RNA45SN1_45S_50ntFlanks@12654.12675.22\|TATGACTGAACGCCTCTAAGTC | >hg38_21_+_8433172_8446622_RNA45SN1_45S_50ntFlanks | 12654 | 12675 | 22 | TATGACTGAACGCCTCTAAGTC | rRF | FALSE | FALSE | TRUE |
| >hg38_21_+_8433172_8446622_RNA45SN1_45S_50ntFlanks@11323.11343.21\|CGTCTCGTCGCGCGCGCGTCC | >hg38_21_+_8433172_8446622_RNA45SN1_45S_50ntFlanks | 11323 | 11343 | 21 | CGTCTCGTCGCGCGCGCGTCC | rRF | FALSE | FALSE | TRUE |
| >hg38_21_+_8433172_8446622_RNA45SN1_45S_50ntFlanks@10687.10711.25\|CGGCGTCCGGTGAGCTCTCGCTGGC | >hg38_21_+_8433172_8446622_RNA45SN1_45S_50ntFlanks | 10687 | 10711 | 25 | CGGCGTCCGGTGAGCTCTCGCTGGC | rRF | FALSE | FALSE | TRUE |
| >hg38_21_+_8433172_8446622_RNA45SN1_45S_50ntFlanks@4722.4758.37\|TCATTAATCAAGAACGAAAGTCGGAGGTTCGAAGACG | >hg38_21_+_8433172_8446622_RNA45SN1_45S_50ntFlanks | 4722 | 4758 | 37 | TCATTAATCAAGAACGAAAGTCGGAGGTTCGAAGACG | rRF | FALSE | FALSE | TRUE |
| >hg38_21_+_8433172_8446622_RNA45SN1_45S_50ntFlanks@10841.10864.24\|CGTAACTTCGGGATAAGGATTGGC | >hg38_21_+_8433172_8446622_RNA45SN1_45S_50ntFlanks | 10841 | 10864 | 24 | CGTAACTTCGGGATAAGGATTGGC | rRF | FALSE | FALSE | TRUE |
| >hg38_21_+_8433172_8446622_RNA45SN1_45S_50ntFlanks@12456.12488.33\|CCAAGCGTTGGATTGTTCACCCACTAATAGGGA | >hg38_21_+_8433172_8446622_RNA45SN1_45S_50ntFlanks | 12456 | 12488 | 33 | CCAAGCGTTGGATTGTTCACCCACTAATAGGGA | rRF | FALSE | FALSE | TRUE |
| >hg38_21_+_8433172_8446622_RNA45SN1_45S_50ntFlanks@12626.12654.29\|AATGGGGCGAAGCTACCATCTGTGGGATT | >hg38_21_+_8433172_8446622_RNA45SN1_45S_50ntFlanks | 12626 | 12654 | 29 | AATGGGGCGAAGCTACCATCTGTGGGATT | rRF | FALSE | FALSE | TRUE |
| >hg38_21_+_8433172_8446622_RNA45SN1_45S_50ntFlanks@12002.12032.31\|TGCGGGCCGCCGGTGAAATACCACTACTCTG | >hg38_21_+_8433172_8446622_RNA45SN1_45S_50ntFlanks | 12002 | 12032 | 31 | TGCGGGCCGCCGGTGAAATACCACTACTCTG | rRF | FALSE | FALSE | TRUE |
| >hg38_21_+_8433172_8446622_RNA45SN1_45S_50ntFlanks@12973.12991.19\|TCGTACGTAGCAGAGCAGC | >hg38_21_+_8433172_8446622_RNA45SN1_45S_50ntFlanks | 12973 | 12991 | 19 | TCGTACGTAGCAGAGCAGC | rRF | FALSE | FALSE | TRUE |
| >hg38_21_+_8433172_8446622_RNA45SN1_45S_50ntFlanks@4724.4758.35\|ATTAATCAAGAACGAAAGTCGGAGGTTCGAAGACG | >hg38_21_+_8433172_8446622_RNA45SN1_45S_50ntFlanks | 4724 | 4758 | 35 | ATTAATCAAGAACGAAAGTCGGAGGTTCGAAGACG | rRF | FALSE | FALSE | TRUE |
| >hg38_21_+_8433172_8446622_RNA45SN1_45S_50ntFlanks@6698.6718.21\|AGCTAGCTGCGAGAATTAATG | >hg38_21_+_8433172_8446622_RNA45SN1_45S_50ntFlanks | 6698 | 6718 | 21 | AGCTAGCTGCGAGAATTAATG | rRF | FALSE | FALSE | TRUE |
| >hg38_21_+_8433172_8446622_RNA45SN1_45S_50ntFlanks@12973.12990.18\|TCGTACGTAGCAGAGCAG | >hg38_21_+_8433172_8446622_RNA45SN1_45S_50ntFlanks | 12973 | 12990 | 18 | TCGTACGTAGCAGAGCAG | rRF | FALSE | FALSE | TRUE |
| >hg38_21_+_8433172_8446622_RNA45SN1_45S_50ntFlanks@9534.9551.18\|AGGAAACTCTGGTGGAGG | >hg38_21_+_8433172_8446622_RNA45SN1_45S_50ntFlanks | 9534 | 9551 | 18 | AGGAAACTCTGGTGGAGG | rRF | FALSE | FALSE | TRUE |
| >hg38_21_+_8433172_8446622_RNA45SN1_45S_50ntFlanks@11318.11344.27\|CCCCACGTCTCGTCGCGCGCGCGTCCG | >hg38_21_+_8433172_8446622_RNA45SN1_45S_50ntFlanks | 11318 | 11344 | 27 | CCCCACGTCTCGTCGCGCGCGCGTCCG | rRF | FALSE | FALSE | TRUE |
| >hg38_21_+_8433172_8446622_RNA45SN1_45S_50ntFlanks@12456.12486.31\|CCAAGCGTTGGATTGTTCACCCACTAATAGG | >hg38_21_+_8433172_8446622_RNA45SN1_45S_50ntFlanks | 12456 | 12486 | 31 | CCAAGCGTTGGATTGTTCACCCACTAATAGG | rRF | FALSE | FALSE | TRUE |
| >hg38_21_+_8433172_8446622_RNA45SN1_45S_50ntFlanks@10841.10862.22\|CGTAACTTCGGGATAAGGATTG | >hg38_21_+_8433172_8446622_RNA45SN1_45S_50ntFlanks | 10841 | 10862 | 22 | CGTAACTTCGGGATAAGGATTG | rRF | FALSE | FALSE | TRUE |
| >hg38_21_+_8433172_8446622_RNA45SN1_45S_50ntFlanks@10846.10865.20\|CTTCGGGATAAGGATTGGCT | >hg38_21_+_8433172_8446622_RNA45SN1_45S_50ntFlanks | 10846 | 10865 | 20 | CTTCGGGATAAGGATTGGCT | rRF | FALSE | FALSE | TRUE |
| >hg38_21_+_8433172_8446622_RNA45SN1_45S_50ntFlanks@12417.12435.19\|TCGATGTCGGCTCTTCCTA | >hg38_21_+_8433172_8446622_RNA45SN1_45S_50ntFlanks | 12417 | 12435 | 19 | TCGATGTCGGCTCTTCCTA | rRF | FALSE | FALSE | TRUE |
| >hg38_21_+_8433172_8446622_RNA45SN1_45S_50ntFlanks@10411.10431.21\|CAGTCGGTCCTGAGAGATGGG | >hg38_21_+_8433172_8446622_RNA45SN1_45S_50ntFlanks | 10411 | 10431 | 21 | CAGTCGGTCCTGAGAGATGGG | rRF | FALSE | FALSE | TRUE |
| >am_trna117_GlyTCC_1_-_145397864_145397935@1.30.30,trna45_GlyTCC_1_+_161500903_161500974@1.30.30,trna70_GlyTCC_1_-_161439547_161439618@1.30.30,trna73_GlyTCC_1_-_161432166_161432237@1.30.30,trna76_GlyTCC_1_-_161424756_161424827@1.30.30,trna79_GlyTCC_1_-_161417375_161417446@1.30.30\|GCGTTGGTGGTATAGTGGTGAGCATAGCTG | >am_trna117_GlyTCC_1_-_145397864_145397935 | 1 | 30 | 30 | GCGTTGGTGGTATAGTGGTGAGCATAGCTG | tRF | FALSE | FALSE | TRUE |
| >hg38_21_+_8433172_8446622_RNA45SN1_45S_50ntFlanks@6745.6770.26\|ACACTTCGAACGCACTTGCGGCCCCG | >hg38_21_+_8433172_8446622_RNA45SN1_45S_50ntFlanks | 6745 | 6770 | 26 | ACACTTCGAACGCACTTGCGGCCCCG | rRF | FALSE | FALSE | TRUE |
| >hg38_21_+_8433172_8446622_RNA45SN1_45S_50ntFlanks@6721.6743.23\|AATTGCAGGACACATTGATCATC | >hg38_21_+_8433172_8446622_RNA45SN1_45S_50ntFlanks | 6721 | 6743 | 23 | AATTGCAGGACACATTGATCATC | rRF | FALSE | FALSE | TRUE |
| >hg38_21_+_8433172_8446622_RNA45SN1_45S_50ntFlanks@5445.5466.22\|TCGGATCGGCCCCGCCGGGGTC | >hg38_21_+_8433172_8446622_RNA45SN1_45S_50ntFlanks | 5445 | 5466 | 22 | TCGGATCGGCCCCGCCGGGGTC | rRF | FALSE | FALSE | TRUE |
| >hg38_21_+_8433172_8446622_RNA45SN1_45S_50ntFlanks@12936.12956.21\|AAACCATTCGTAGACGACCTG | >hg38_21_+_8433172_8446622_RNA45SN1_45S_50ntFlanks | 12936 | 12956 | 21 | AAACCATTCGTAGACGACCTG | rRF | FALSE | FALSE | TRUE |
| >hg38_21_+_8433172_8446622_RNA45SN1_45S_50ntFlanks@12437.12473.37\|CATTGTGAAGCAGAATTCACCAAGCGTTGGATTGTTC | >hg38_21_+_8433172_8446622_RNA45SN1_45S_50ntFlanks | 12437 | 12473 | 37 | CATTGTGAAGCAGAATTCACCAAGCGTTGGATTGTTC | rRF | FALSE | FALSE | TRUE |
| >hg38_21_+_8433172_8446622_RNA45SN1_45S_50ntFlanks@11576.11603.28\|CTTAGAACTGGTGCGGACCAGGGGAATC | >hg38_21_+_8433172_8446622_RNA45SN1_45S_50ntFlanks | 11576 | 11603 | 28 | CTTAGAACTGGTGCGGACCAGGGGAATC | rRF | FALSE | FALSE | TRUE |
| >hg38_21_+_8433172_8446622_RNA45SN1_45S_50ntFlanks@9567.9596.30\|ACGTGCAAATCGGTCGTCCGACCTGGGTAT | >hg38_21_+_8433172_8446622_RNA45SN1_45S_50ntFlanks | 9567 | 9596 | 30 | ACGTGCAAATCGGTCGTCCGACCTGGGTAT | rRF | FALSE | FALSE | TRUE |
| >hg38_21_+_8433172_8446622_RNA45SN1_45S_50ntFlanks@11572.11602.31\|CCGACTTAGAACTGGTGCGGACCAGGGGAAT | >hg38_21_+_8433172_8446622_RNA45SN1_45S_50ntFlanks | 11572 | 11602 | 31 | CCGACTTAGAACTGGTGCGGACCAGGGGAAT | rRF | FALSE | FALSE | TRUE |
| >hg38_21_+_8433172_8446622_RNA45SN1_45S_50ntFlanks@10684.10706.23\|CGGCGGCGTCCGGTGAGCTCTCG | >hg38_21_+_8433172_8446622_RNA45SN1_45S_50ntFlanks | 10684 | 10706 | 23 | CGGCGGCGTCCGGTGAGCTCTCG | rRF | FALSE | FALSE | TRUE |

### Cytoplasm

| | | | | | | | | | |
|---|---|---|---|---|---|---|---|---|---|
| >hg38_21_+_8433172_8446622_RNA45SN1_45S_50ntFlanks@11575.11592.18\|ACTTAGAACTGGTGCGGA | >hg38_21_+_8433172_8446622_RNA45SN1_45S_50ntFlanks | 11575 | 11592 | 18 | ACTTAGAACTGGTGCGGA | rRF | FALSE | FALSE | TRUE |
| >hg38_21_+_8433172_8446622_RNA45SN1_45S_50ntFlanks@12418.12455.38\|CGATGTCGGCTCTTCCTATCATTGTGAAGCAGAATTCA | >hg38_21_+_8433172_8446622_RNA45SN1_45S_50ntFlanks | 12418 | 12455 | 38 | CGATGTCGGCTCTTCCTATCATTGTGAAGCAGAATTCA | rRF | FALSE | FALSE | TRUE |
| >hg38_21_+_8433172_8446622_RNA45SN1_45S_50ntFlanks@3875.3898.24\|ATACATGCCGACGGGCGCTGACCC | >hg38_21_+_8433172_8446622_RNA45SN1_45S_50ntFlanks | 3875 | 3898 | 24 | ATACATGCCGACGGGCGCTGACCC | rRF | FALSE | FALSE | TRUE |
| >hg38_21_+_8433172_8446622_RNA45SN1_45S_50ntFlanks@11718.11742.25\|AACGGCGGGAGTAACTATGACTCTC | >hg38_21_+_8433172_8446622_RNA45SN1_45S_50ntFlanks | 11718 | 11742 | 25 | AACGGCGGGAGTAACTATGACTCTC | rRF | FALSE | FALSE | TRUE |
| >hg38_21_+_8433172_8446622_RNA45SN1_45S_50ntFlanks@10392.10410.19\|ACAGCAGTTGAACATGGGT | >hg38_21_+_8433172_8446622_RNA45SN1_45S_50ntFlanks | 10392 | 10410 | 19 | ACAGCAGTTGAACATGGGT | rRF | FALSE | FALSE | TRUE |
| >hg38_21_+_8433172_8446622_RNA45SN1_45S_50ntFlanks@4046.4063.18\|CATTCGAACGTCTGCCCT | >hg38_21_+_8433172_8446622_RNA45SN1_45S_50ntFlanks | 4046 | 4063 | 18 | CATTCGAACGTCTGCCCT | rRF | FALSE | FALSE | TRUE |
| >hg38_21_+_8433172_8446622_RNA45SN1_45S_50ntFlanks@9567.9595.29\|ACGTGCAAATCGGTCGTCCGACCTGGGTA | >hg38_21_+_8433172_8446622_RNA45SN1_45S_50ntFlanks | 9567 | 9595 | 29 | ACGTGCAAATCGGTCGTCCGACCTGGGTA | rRF | FALSE | FALSE | TRUE |
| >hg38_21_+_8433172_8446622_RNA45SN1_45S_50ntFlanks@8198.8215.18\|TGGACGGTGTGAGGCCGG | >hg38_21_+_8433172_8446622_RNA45SN1_45S_50ntFlanks | 8198 | 8215 | 18 | TGGACGGTGTGAGGCCGG | rRF | FALSE | FALSE | TRUE |
| >hg38_21_+_8433172_8446622_RNA45SN1_45S_50ntFlanks@9738.9758.21\|GAAACGATCTCAACCTATTCT | >hg38_21_+_8433172_8446622_RNA45SN1_45S_50ntFlanks | 9738 | 9758 | 21 | GAAACGATCTCAACCTATTCT | rRF | FALSE | FALSE | TRUE |
| >hg38_21_+_8433172_8446622_RNA45SN1_45S_50ntFlanks@10798.10818.21\|CATGTTGGAACAATGTAGGTA | >hg38_21_+_8433172_8446622_RNA45SN1_45S_50ntFlanks | 10798 | 10818 | 21 | CATGTTGGAACAATGTAGGTA | rRF | FALSE | FALSE | TRUE |
| >hg38_21_+_8433172_8446622_RNA45SN1_45S_50ntFlanks@12415.12434.20\|CTTCGATGTCGGCTCTTCCT | >hg38_21_+_8433172_8446622_RNA45SN1_45S_50ntFlanks | 12415 | 12434 | 20 | CTTCGATGTCGGCTCTTCCT | rRF | FALSE | FALSE | TRUE |
| >hg38_21_+_8433172_8446622_RNA45SN1_45S_50ntFlanks@12488.12509.22\|AACGTGAGCTGGGTTTAGACCG | >hg38_21_+_8433172_8446622_RNA45SN1_45S_50ntFlanks | 12488 | 12509 | 22 | AACGTGAGCTGGGTTTAGACCG | rRF | FALSE | FALSE | TRUE |
| >am_trna117_GlyTCC_1_-_145397864_145397935@1.29.29,trna45_GlyTCC_1_+_161500903_161500974@1.29.29,trna70_GlyTCC_1_-_161439547_161439618@1.29.29,trna73_GlyTCC_1_-_161432166_161432237@1.29.29,trna76_GlyTCC_1_-_161424756_161424827@1.29.29,trna79_GlyTCC_1_-_161417375_161417446@1.29.29\|GCGTTGGTGGTATAGTGGTGAGCATAGCT | >am_trna117_GlyTCC_1_-_145397864_145397935 | 1 | 29 | 29 | GCGTTGGTGGTATAGTGGTGAGCATAGCT | tRF | FALSE | FALSE | TRUE |
| >hg38_21_+_8433172_8446622_RNA45SN1_45S_50ntFlanks@10411.10428.18\|CAGTCGGTCCTGAGAGAT | >hg38_21_+_8433172_8446622_RNA45SN1_45S_50ntFlanks | 10411 | 10428 | 18 | CAGTCGGTCCTGAGAGAT | rRF | FALSE | FALSE | TRUE |
| >hg38_21_+_8433172_8446622_RNA45SN1_45S_50ntFlanks@12458.12476.19\|AAGCGTTGGATTGTTCACC | >hg38_21_+_8433172_8446622_RNA45SN1_45S_50ntFlanks | 12458 | 12476 | 19 | AAGCGTTGGATTGTTCACC | rRF | FALSE | FALSE | TRUE |
| >hg38_21_+_8433172_8446622_RNA45SN1_45S_50ntFlanks@10840.10865.26\|CCGTAACTTCGGGATAAGGATTGGCT | >hg38_21_+_8433172_8446622_RNA45SN1_45S_50ntFlanks | 10840 | 10865 | 26 | CCGTAACTTCGGGATAAGGATTGGCT | rRF | FALSE | FALSE | TRUE |
| >hg38_21_+_8433172_8446622_RNA45SN1_45S_50ntFlanks@12615.12654.40\|GCTGAGGAGCCAATGGGGCGAAGCTACCATCTGTGGGATT | >hg38_21_+_8433172_8446622_RNA45SN1_45S_50ntFlanks | 12615 | 12654 | 40 | GCTGAGGAGCCAATGGGGCGAAGCTACCATCTGTGGGATT | rRF | FALSE | FALSE | TRUE |
| >hg38_21_+_8433172_8446622_RNA45SN1_45S_50ntFlanks@12418.12446.29\|CGATGTCGGCTCTTCCTATCATTGTGAAG | >hg38_21_+_8433172_8446622_RNA45SN1_45S_50ntFlanks | 12418 | 12446 | 29 | CGATGTCGGCTCTTCCTATCATTGTGAAG | rRF | FALSE | FALSE | TRUE |
| >hg38_21_+_8433172_8446622_RNA45SN1_45S_50ntFlanks@6651.6689.39\|CGACTCTTAGCGGTGGATCACTCGGCTCGTGCGTCGATG | >hg38_21_+_8433172_8446622_RNA45SN1_45S_50ntFlanks | 6651 | 6689 | 39 | CGACTCTTAGCGGTGGATCACTCGGCTCGTGCGTCGATG | rRF | FALSE | FALSE | TRUE |
| >hg38_21_+_8433172_8446622_RNA45SN1_45S_50ntFlanks@10402.10429.28\|AACATGGGTCAGTCGGTCCTGAGAGATG | >hg38_21_+_8433172_8446622_RNA45SN1_45S_50ntFlanks | 10402 | 10429 | 28 | AACATGGGTCAGTCGGTCCTGAGAGATG | rRF | FALSE | FALSE | TRUE |
| >hg38_21_+_8433172_8446622_RNA45SN1_45S_50ntFlanks@9574.9596.23\|AATCGGTCGTCCGACCTGGGTAT | >hg38_21_+_8433172_8446622_RNA45SN1_45S_50ntFlanks | 9574 | 9596 | 23 | AATCGGTCGTCCGACCTGGGTAT | rRF | FALSE | FALSE | TRUE |
| >hg38_21_+_8433172_8446622_RNA45SN1_45S_50ntFlanks@9574.9597.24\|AATCGGTCGTCCGACCTGGGTATA | >hg38_21_+_8433172_8446622_RNA45SN1_45S_50ntFlanks | 9574 | 9597 | 24 | AATCGGTCGTCCGACCTGGGTATA | rRF | FALSE | FALSE | TRUE |
| >hg38_21_+_8433172_8446622_RNA45SN1_45S_50ntFlanks@9568.9597.30\|CGTGCAAATCGGTCGTCCGACCTGGGTATA | >hg38_21_+_8433172_8446622_RNA45SN1_45S_50ntFlanks | 9568 | 9597 | 30 | CGTGCAAATCGGTCGTCCGACCTGGGTATA | rRF | FALSE | FALSE | TRUE |
| >hg38_21_+_8433172_8446622_RNA45SN1_45S_50ntFlanks@4722.4759.38\|TCATTAATCAAGAACGAAAGTCGGAGGTTCGAAGACGA | >hg38_21_+_8433172_8446622_RNA45SN1_45S_50ntFlanks | 4722 | 4759 | 38 | TCATTAATCAAGAACGAAAGTCGGAGGTTCGAAGACGA | rRF | FALSE | FALSE | TRUE |
| >hg38_21_+_8433172_8446622_RNA45SN1_45S_50ntFlanks@11212.11232.21\|CGACTCTGGACGCGAGCCGGG | >hg38_21_+_8433172_8446622_RNA45SN1_45S_50ntFlanks | 11212 | 11232 | 21 | CGACTCTGGACGCGAGCCGGG | rRF | FALSE | FALSE | TRUE |
| >hg38_21_+_8433172_8446622_RNA45SN1_45S_50ntFlanks@10680.10709.30\|GTTCCGGCGGCGTCCGGTGAGCTCTCGCTG | >hg38_21_+_8433172_8446622_RNA45SN1_45S_50ntFlanks | 10680 | 10709 | 30 | GTTCCGGCGGCGTCCGGTGAGCTCTCGCTG | rRF | FALSE | FALSE | TRUE |
| >hg38_21_+_8433172_8446622_RNA45SN1_45S_50ntFlanks@11787.11804.18\|AATGGATGAACGAGATTC | >hg38_21_+_8433172_8446622_RNA45SN1_45S_50ntFlanks | 11787 | 11804 | 18 | AATGGATGAACGAGATTC | rRF | FALSE | FALSE | TRUE |
| >hg38_21_+_8433172_8446622_RNA45SN1_45S_50ntFlanks@9568.9595.28\|CGTGCAAATCGGTCGTCCGACCTGGGTA | >hg38_21_+_8433172_8446622_RNA45SN1_45S_50ntFlanks | 9568 | 9595 | 28 | CGTGCAAATCGGTCGTCCGACCTGGGTA | rRF | FALSE | FALSE | TRUE |
| >trna111_HisGTG_1_-_147774845_147774916@-1A.31.32,trna118_HisGTG_1_-_145396881_145396952@-1A.31.32,trna16_HisGTG_1_+_146544773_146544844@-1A.31.32,trna1_HisGTG_15_+_45493349_45493420@-1A.31.32,trna21_HisGTG_1_+_147753471_147753542@-1A.31.32,trna33_HisGTG_6_+_27125906_27125977@-1A.31.32,trna7_HisGTG_9_-_14433938_14434009@-1A.31.32,trna8_HisGTG_15_-_45492611_45492682@-1A.31.32,trna9_HisGTG_15_-_45490804_45490875@-1A.31.32\|AGCCGTGATCGTATAGTGGTTAGTACTCTGCG | >trna111_HisGTG_1_-_147774845_147774916 | -1A | 31 | 32 | AGCCGTGATCGTATAGTGGTTAGTACTCTGCG | tRF | FALSE | FALSE | TRUE |
| >hg38_21_+_8433172_8446622_RNA45SN1_45S_50ntFlanks@11783.11803.21\|CATGAATGGATGAACGAGATT | >hg38_21_+_8433172_8446622_RNA45SN1_45S_50ntFlanks | 11783 | 11803 | 21 | CATGAATGGATGAACGAGATT | rRF | FALSE | FALSE | TRUE |
| >hg38_21_+_8433172_8446622_RNA45SN1_45S_50ntFlanks@10687.10709.23\|CGGCGTCCGGTGAGCTCTCGCTG | >hg38_21_+_8433172_8446622_RNA45SN1_45S_50ntFlanks | 10687 | 10709 | 23 | CGGCGTCCGGTGAGCTCTCGCTG | rRF | FALSE | FALSE | TRUE |
| >hg38_21_+_8433172_8446622_RNA45SN1_45S_50ntFlanks@11235.11254.20\|CTTCCCGTGGATCGCCCCAG | >hg38_21_+_8433172_8446622_RNA45SN1_45S_50ntFlanks | 11235 | 11254 | 20 | CTTCCCGTGGATCGCCCCAG | rRF | FALSE | FALSE | TRUE |
| >hg38_21_+_8433172_8446622_RNA45SN1_45S_50ntFlanks@12633.12654.22\|CGAAGCTACCATCTGTGGGATT | >hg38_21_+_8433172_8446622_RNA45SN1_45S_50ntFlanks | 12633 | 12654 | 22 | CGAAGCTACCATCTGTGGGATT | rRF | FALSE | FALSE | TRUE |
| >hg38_21_+_8433172_8446622_RNA45SN1_45S_50ntFlanks@11199.11225.27\|CCGGCGGCGGCGGCGACTCTGGACGCG | >hg38_21_+_8433172_8446622_RNA45SN1_45S_50ntFlanks | 11199 | 11225 | 27 | CCGGCGGCGGCGGCGACTCTGGACGCG | rRF | FALSE | FALSE | TRUE |
| >hg38_21_+_8433172_8446622_RNA45SN1_45S_50ntFlanks@11569.11601.33\|CAGCCGACTTAGAACTGGTGCGGACCAGGGGAA | >hg38_21_+_8433172_8446622_RNA45SN1_45S_50ntFlanks | 11569 | 11601 | 33 | CAGCCGACTTAGAACTGGTGCGGACCAGGGGAA | rRF | FALSE | FALSE | TRUE |
| >hg38_21_+_8433172_8446622_RNA45SN1_45S_50ntFlanks@12324.12344.21\|TTGGGTTTTAAGCAGGAGGTG | >hg38_21_+_8433172_8446622_RNA45SN1_45S_50ntFlanks | 12324 | 12344 | 21 | TTGGGTTTTAAGCAGGAGGTG | rRF | FALSE | FALSE | TRUE |
| >hg38_21_+_8433172_8446622_RNA45SN1_45S_50ntFlanks@12974.12991.18\|CGTACGTAGCAGAGCAGC | >hg38_21_+_8433172_8446622_RNA45SN1_45S_50ntFlanks | 12974 | 12991 | 18 | CGTACGTAGCAGAGCAGC | rRF | FALSE | FALSE | TRUE |

### Cytoplasm

| | | | | | | | | | |
|---|---|---|---|---|---|---|---|---|---|
| >hg38_21_+_8433172_8446622_RNA45SN1_45S_50ntFlanks@11316.11338.23\|CACCCCACGTCTCGTCGCGCGCG | >hg38_21_+_8433172_8446622_RNA45SN1_45S_50ntFlanks | 11316 | 11338 | 23 | CACCCCACGTCTCGTCGCGCGCG | rRF | FALSE | FALSE | TRUE |
| >hg38_21_+_8433172_8446622_RNA45SN1_45S_50ntFlanks@10346.10368.23\|TCAAACGAGAACTTTGAAGGCCG | >hg38_21_+_8433172_8446622_RNA45SN1_45S_50ntFlanks | 10346 | 10368 | 23 | TCAAACGAGAACTTTGAAGGCCG | rRF | FALSE | FALSE | TRUE |
| >hg38_21_+_8433172_8446622_RNA45SN1_45S_50ntFlanks@9504.9521.18\|GTGAACTATGCCTGGGCA | >hg38_21_+_8433172_8446622_RNA45SN1_45S_50ntFlanks | 9504 | 9521 | 18 | GTGAACTATGCCTGGGCA | rRF | FALSE | FALSE | TRUE |
| >hg38_21_+_8433172_8446622_RNA45SN1_45S_50ntFlanks@10350.10390.41\|ACGAGAACTTTGAAGGCCGAAGTGGAGAAGGGTTCCATGTG | >hg38_21_+_8433172_8446622_RNA45SN1_45S_50ntFlanks | 10350 | 10390 | 41 | ACGAGAACTTTGAAGGCCGAAGTGGAGAAGGGTTCCATGTG | rRF | FALSE | FALSE | TRUE |
| >hg38_21_+_8433172_8446622_RNA45SN1_45S_50ntFlanks@9534.9561.28\|AGGAAACTCTGGTGGAGGTCCGTAGCGG | >hg38_21_+_8433172_8446622_RNA45SN1_45S_50ntFlanks | 9534 | 9561 | 28 | AGGAAACTCTGGTGGAGGTCCGTAGCGG | rRF | FALSE | FALSE | TRUE |
| >hg38_21_+_8433172_8446622_RNA45SN1_45S_50ntFlanks@9566.9596.31\|GACGTGCAAATCGGTCGTCCGACCTGGGTAT | >hg38_21_+_8433172_8446622_RNA45SN1_45S_50ntFlanks | 9566 | 9596 | 31 | GACGTGCAAATCGGTCGTCCGACCTGGGTAT | rRF | FALSE | FALSE | TRUE |
| >hg38_21_+_8433172_8446622_RNA45SN1_45S_50ntFlanks@10681.10706.26\|TTCCGGCGGCGTCCGGTGAGCTCTCG | >hg38_21_+_8433172_8446622_RNA45SN1_45S_50ntFlanks | 10681 | 10706 | 26 | TTCCGGCGGCGTCCGGTGAGCTCTCG | rRF | FALSE | FALSE | TRUE |
| >hg38_21_+_8433172_8446622_RNA45SN1_45S_50ntFlanks@11768.11794.27\|CTAATTAGTGACGCGCATGAATGGATG | >hg38_21_+_8433172_8446622_RNA45SN1_45S_50ntFlanks | 11768 | 11794 | 27 | CTAATTAGTGACGCGCATGAATGGATG | rRF | FALSE | FALSE | TRUE |
| >hg38_21_+_8433172_8446622_RNA45SN1_45S_50ntFlanks@9554.9574.21\|CGTAGCGGTCCTGACGTGCAA | >hg38_21_+_8433172_8446622_RNA45SN1_45S_50ntFlanks | 9554 | 9574 | 21 | CGTAGCGGTCCTGACGTGCAA | rRF | FALSE | FALSE | TRUE |
| >hg38_21_+_8433172_8446622_RNA45SN1_45S_50ntFlanks@4002.4024.23\|GATAACCTCGGGCCGATCGCACG | >hg38_21_+_8433172_8446622_RNA45SN1_45S_50ntFlanks | 4002 | 4024 | 23 | GATAACCTCGGGCCGATCGCACG | rRF | FALSE | FALSE | TRUE |
| >hg38_21_+_8433172_8446622_RNA45SN1_45S_50ntFlanks@11208.11227.20\|GCGGCGACTCTGGACGCGAG | >hg38_21_+_8433172_8446622_RNA45SN1_45S_50ntFlanks | 11208 | 11227 | 20 | GCGGCGACTCTGGACGCGAG | rRF | FALSE | FALSE | TRUE |
| >hg38_21_+_8433172_8446622_RNA45SN1_45S_50ntFlanks@11209.11233.25\|CGGCGACTCTGGACGCGAGCCGGGC | >hg38_21_+_8433172_8446622_RNA45SN1_45S_50ntFlanks | 11209 | 11233 | 25 | CGGCGACTCTGGACGCGAGCCGGGC | rRF | FALSE | FALSE | TRUE |
| >hg38_21_+_8433172_8446622_RNA45SN1_45S_50ntFlanks@5471.5514.44\|CACGGCCCTGGCGGAGCGCTGAGAAGACGGTCGAACTTGACTAT | >hg38_21_+_8433172_8446622_RNA45SN1_45S_50ntFlanks | 5471 | 5514 | 44 | CACGGCCCTGGCGGAGCGCTGAGAAGACGGTCGAACTTGACTAT | rRF | FALSE | FALSE | TRUE |
| >hg38_21_+_8433172_8446622_RNA45SN1_45S_50ntFlanks@7983.8006.24\|CAGATCAGACGTGGCGACCCGCTG | >hg38_21_+_8433172_8446622_RNA45SN1_45S_50ntFlanks | 7983 | 8006 | 24 | CAGATCAGACGTGGCGACCCGCTG | rRF | FALSE | FALSE | TRUE |
| >hg38_21_+_8433172_8446622_RNA45SN1_45S_50ntFlanks@4003.4042.40\|ATAACCTCGGGCCGATCGCACGCCCCCCGTGGCGGCGACG | >hg38_21_+_8433172_8446622_RNA45SN1_45S_50ntFlanks | 4003 | 4042 | 40 | ATAACCTCGGGCCGATCGCACGCCCCCCGTGGCGGCGACG | rRF | FALSE | FALSE | TRUE |
| >hg38_21_+_8433172_8446622_RNA45SN1_45S_50ntFlanks@8176.8203.28\|CTTCTGATCGAGGCCCAGCCCGTGGACG | >hg38_21_+_8433172_8446622_RNA45SN1_45S_50ntFlanks | 8176 | 8203 | 28 | CTTCTGATCGAGGCCCAGCCCGTGGACG | rRF | FALSE | FALSE | TRUE |
| >hg38_21_+_8433172_8446622_RNA45SN1_45S_50ntFlanks@12072.12095.24\|CCCGAGGGGCTCTCGCTTCTGGCG | >hg38_21_+_8433172_8446622_RNA45SN1_45S_50ntFlanks | 12072 | 12095 | 24 | CCCGAGGGGCTCTCGCTTCTGGCG | rRF | FALSE | FALSE | TRUE |
| >hg38_21_+_8433172_8446622_RNA45SN1_45S_50ntFlanks@9570.9595.26\|TGCAAATCGGTCGTCCGACCTGGGTA | >hg38_21_+_8433172_8446622_RNA45SN1_45S_50ntFlanks | 9570 | 9595 | 26 | TGCAAATCGGTCGTCCGACCTGGGTA | rRF | FALSE | FALSE | TRUE |
| >am_trna10_ValCAC_5_-_180649395_180649467@1.23.23,trna12_ValAAC_5_-_180645270_180645342@1.23.23,trna132_ValAAC_6_-_27721179_27721251@1.23.23,trna136_ValAAC_6_-_27648885_27648957@1.23.23,trna139_ValAAC_6_-_27618707_27618779@1.23.23,trna18_ValCAC_5_-_180529253_180529325@1.23.23,trna2_ValAAC_3_+_169490018_169490090@1.23.23,trna2_ValCAC_5_+_180524070_180524142@1.23.23,trna37_ValAAC_6_+_27203288_27203360@1.23.23,trna4_ValAAC_5_+_180591154_180591226@1.23.23,trna5_ValAAC_5_+_180596610_180596682@1.23.23,trna6_ValCAC_5_+_180600650_180600722@1.23.23,trna85_ValCAC_1_-_161369490_161369562@1.23.23,trna90_ValCAC_1_-_149684088_149684161@1.23.23,trna98_ValCAC_1_-_149298555_149298627@1.23.23,trna9_ValCAC_6_+_26538282_26538354@1.23.23\|GTTTCCGTAGTGTAGTGGTTATC | >am_trna10_ValCAC_5_-_180649395_180649467 | 1 | 23 | 23 | GTTTCCGTAGTGTAGTGGTTATC | tRF | FALSE | FALSE | TRUE |
| >hg38_21_+_8433172_8446622_RNA45SN1_45S_50ntFlanks@12438.12455.18\|ATTGTGAAGCAGAATTCA | >hg38_21_+_8433172_8446622_RNA45SN1_45S_50ntFlanks | 12438 | 12455 | 18 | ATTGTGAAGCAGAATTCA | rRF | FALSE | FALSE | TRUE |
| >hg38_21_+_8433172_8446622_RNA45SN1_45S_50ntFlanks@13004.13027.24\|ATCTATTGAAAGTCAGCCCTCGAC | >hg38_21_+_8433172_8446622_RNA45SN1_45S_50ntFlanks | 13004 | 13027 | 24 | ATCTATTGAAAGTCAGCCCTCGAC | rRF | FALSE | FALSE | TRUE |
| >hg38_21_+_8433172_8446622_RNA45SN1_45S_50ntFlanks@9565.9597.33\|TGACGTGCAAATCGGTCGTCCGACCTGGGTATA | >hg38_21_+_8433172_8446622_RNA45SN1_45S_50ntFlanks | 9565 | 9597 | 33 | TGACGTGCAAATCGGTCGTCCGACCTGGGTATA | rRF | FALSE | FALSE | TRUE |
| >hg38_21_+_8433172_8446622_RNA45SN1_45S_50ntFlanks@11581.11605.25\|AACTGGTGCGGACCAGGGGAATCCG | >hg38_21_+_8433172_8446622_RNA45SN1_45S_50ntFlanks | 11581 | 11605 | 25 | AACTGGTGCGGACCAGGGGAATCCG | rRF | FALSE | FALSE | TRUE |
| >hg38_21_+_8433172_8446622_RNA45SN1_45S_50ntFlanks@10350.10388.39\|ACGAGAACTTTGAAGGCCGAAGTGGAGAAGGGTTCCATG | >hg38_21_+_8433172_8446622_RNA45SN1_45S_50ntFlanks | 10350 | 10388 | 39 | ACGAGAACTTTGAAGGCCGAAGTGGAGAAGGGTTCCATG | rRF | FALSE | FALSE | TRUE |
| >hg38_21_+_8433172_8446622_RNA45SN1_45S_50ntFlanks@9545.9572.28\|GTGGAGGTCCGTAGCGGTCCTGACGTGC | >hg38_21_+_8433172_8446622_RNA45SN1_45S_50ntFlanks | 9545 | 9572 | 28 | GTGGAGGTCCGTAGCGGTCCTGACGTGC | rRF | FALSE | FALSE | TRUE |
| >hg38_21_+_8433172_8446622_RNA45SN1_45S_50ntFlanks@10410.10431.22\|TCAGTCGGTCCTGAGAGATGGG | >hg38_21_+_8433172_8446622_RNA45SN1_45S_50ntFlanks | 10410 | 10431 | 22 | TCAGTCGGTCCTGAGAGATGGG | rRF | FALSE | FALSE | TRUE |
| >hg38_21_+_8433172_8446622_RNA45SN1_45S_50ntFlanks@6690.6720.31\|AAGAACGCAGCTAGCTGCGAGAATTAATGTG | >hg38_21_+_8433172_8446622_RNA45SN1_45S_50ntFlanks | 6690 | 6720 | 31 | AAGAACGCAGCTAGCTGCGAGAATTAATGTG | rRF | FALSE | FALSE | TRUE |
| >hg38_21_+_8433172_8446622_RNA45SN1_45S_50ntFlanks@10351.10384.34\|CGAGAACTTTGAAGGCCGAAGTGGAGAAGGGTTC | >hg38_21_+_8433172_8446622_RNA45SN1_45S_50ntFlanks | 10351 | 10384 | 34 | CGAGAACTTTGAAGGCCGAAGTGGAGAAGGGTTC | rRF | FALSE | FALSE | TRUE |
| >hg38_21_+_8433172_8446622_RNA45SN1_45S_50ntFlanks@9554.9584.31\|CGTAGCGGTCCTGACGTGCAAATCGGTCGTC | >hg38_21_+_8433172_8446622_RNA45SN1_45S_50ntFlanks | 9554 | 9584 | 31 | CGTAGCGGTCCTGACGTGCAAATCGGTCGTC | rRF | FALSE | FALSE | TRUE |
| >hg38_21_+_8433172_8446622_RNA45SN1_45S_50ntFlanks@11572.11605.34\|CCGACTTAGAACTGGTGCGGACCAGGGGAATCCG | >hg38_21_+_8433172_8446622_RNA45SN1_45S_50ntFlanks | 11572 | 11605 | 34 | CCGACTTAGAACTGGTGCGGACCAGGGGAATCCG | rRF | FALSE | FALSE | TRUE |
| >hg38_21_+_8433172_8446622_RNA45SN1_45S_50ntFlanks@13002.13039.38\|CGATCTATTGAAAGTCAGCCCTCGACACAAGGGTTTGT | >hg38_21_+_8433172_8446622_RNA45SN1_45S_50ntFlanks | 13002 | 13039 | 38 | CGATCTATTGAAAGTCAGCCCTCGACACAAGGGTTTGT | rRF | FALSE | FALSE | TRUE |
| >hg38_21_+_8433172_8446622_RNA45SN1_45S_50ntFlanks@6744.6770.27\|GACACTTCGAACGCACTTGCGGCCCCG | >hg38_21_+_8433172_8446622_RNA45SN1_45S_50ntFlanks | 6744 | 6770 | 27 | GACACTTCGAACGCACTTGCGGCCCCG | rRF | FALSE | FALSE | TRUE |
| >hg38_21_+_8433172_8446622_RNA45SN1_45S_50ntFlanks@8196.8214.19\|CGTGGACGGTGTGAGGCCG | >hg38_21_+_8433172_8446622_RNA45SN1_45S_50ntFlanks | 8196 | 8214 | 19 | CGTGGACGGTGTGAGGCCG | rRF | FALSE | FALSE | TRUE |
| >hg38_21_+_8433172_8446622_RNA45SN1_45S_50ntFlanks@9504.9523.20\|GTGAACTATGCCTGGGCAGG | >hg38_21_+_8433172_8446622_RNA45SN1_45S_50ntFlanks | 9504 | 9523 | 20 | GTGAACTATGCCTGGGCAGG | rRF | FALSE | FALSE | TRUE |
| >hg38_21_+_8433172_8446622_RNA45SN1_45S_50ntFlanks@10686.10710.25\|GCGGCGTCCGGTGAGCTCTCGCTGG | >hg38_21_+_8433172_8446622_RNA45SN1_45S_50ntFlanks | 10686 | 10710 | 25 | GCGGCGTCCGGTGAGCTCTCGCTGG | rRF | FALSE | FALSE | TRUE |
| >hg38_21_+_8433172_8446622_RNA45SN1_45S_50ntFlanks@11315.11344.30\|CCACCCCACGTCTCGTCGCGCGCGCGTCCG | >hg38_21_+_8433172_8446622_RNA45SN1_45S_50ntFlanks | 11315 | 11344 | 30 | CCACCCCACGTCTCGTCGCGCGCGCGTCCG | rRF | FALSE | FALSE | TRUE |
| >hg38_21_+_8433172_8446622_RNA45SN1_45S_50ntFlanks@10786.10816.31\|AACAGCCTCTGGCATGTTGGAACAATGTAGG | >hg38_21_+_8433172_8446622_RNA45SN1_45S_50ntFlanks | 10786 | 10816 | 31 | AACAGCCTCTGGCATGTTGGAACAATGTAGG | rRF | FALSE | FALSE | TRUE |

### Cytoplasm

| | | | | | | | | | |
|---|---|---|---|---|---|---|---|---|---|
| >hg38_21_+_8433172_8446622_RNA45SN1_45S_50ntFlanks@4588.4616.29\|CTATTTTGTTGGTTTTCGGAACTGAGGCC | >hg38_21_+_8433172_8446622_RNA45SN1_45S_50ntFlanks | 4588 | 4616 | 29 | CTATTTTGTTGGTTTTCGGAACTGAGGCC | rRF | FALSE | FALSE | TRUE |
| >hg38_21_+_8433172_8446622_RNA45SN1_45S_50ntFlanks@10389.10407.19\|TGAACAGCAGTTGAACATG | >hg38_21_+_8433172_8446622_RNA45SN1_45S_50ntFlanks | 10389 | 10407 | 19 | TGAACAGCAGTTGAACATG | rRF | FALSE | FALSE | TRUE |
| >hg38_21_+_8433172_8446622_RNA45SN1_45S_50ntFlanks@12008.12031.24\|CCGCCGGTGAAATACCACTACTCT | >hg38_21_+_8433172_8446622_RNA45SN1_45S_50ntFlanks | 12008 | 12031 | 24 | CCGCCGGTGAAATACCACTACTCT | rRF | FALSE | FALSE | TRUE |
| >hg38_21_+_8433172_8446622_RNA45SN1_45S_50ntFlanks@11161.11178.18\|CACGGTCCCCCGCGAGGG | >hg38_21_+_8433172_8446622_RNA45SN1_45S_50ntFlanks | 11161 | 11178 | 18 | CACGGTCCCCCGCGAGGG | rRF | FALSE | FALSE | TRUE |
| >hg38_21_+_8433172_8446622_RNA45SN1_45S_50ntFlanks@9545.9566.22\|GTGGAGGTCCGTAGCGGTCCTG | >hg38_21_+_8433172_8446622_RNA45SN1_45S_50ntFlanks | 9545 | 9566 | 22 | GTGGAGGTCCGTAGCGGTCCTG | rRF | FALSE | FALSE | TRUE |
| >hg38_21_+_8433172_8446622_RNA45SN1_45S_50ntFlanks@4724.4759.36\|ATTAATCAAGAACGAAAGTCGGAGGTTCGAAGACGA | >hg38_21_+_8433172_8446622_RNA45SN1_45S_50ntFlanks | 4724 | 4759 | 36 | ATTAATCAAGAACGAAAGTCGGAGGTTCGAAGACGA | rRF | FALSE | FALSE | TRUE |
| >hg38_21_+_8433172_8446622_RNA45SN1_45S_50ntFlanks@7979.8006.28\|ACCTCAGATCAGACGTGGCGACCCGCTG | >hg38_21_+_8433172_8446622_RNA45SN1_45S_50ntFlanks | 7979 | 8006 | 28 | ACCTCAGATCAGACGTGGCGACCCGCTG | rRF | FALSE | FALSE | TRUE |
| >trna119_LysCTT_1_-_145395522_145395594@1.29.29,trna11_LysCTT_5_-_180648979_180649051@1.29.29,trna13_LysCTT_6_+_26556774_26556846@1.29.29,trna32_LysCTT_16_-_3207406_3207478@1.29.29,trna7_LysCTT_16_+_3225692_3225764@1.29.29,trna9_LysCTT_5_+_180634755_180634827@1.29.29\|GCCCGGCTAGCTCAGTCGGTAGAGCATGA | >trna119_LysCTT_1_-_145395522_145395594 | 1 | 29 | 29 | GCCCGGCTAGCTCAGTCGGTAGAGCATGA | tRF | FALSE | FALSE | TRUE |
| >hg38_21_+_8433172_8446622_RNA45SN1_45S_50ntFlanks@12405.12424.20\|CTTTTTGATCCTTCGATGTC | >hg38_21_+_8433172_8446622_RNA45SN1_45S_50ntFlanks | 12405 | 12424 | 20 | CTTTTTGATCCTTCGATGTC | rRF | FALSE | FALSE | TRUE |
| >hg38_21_+_8433172_8446622_RNA45SN1_45S_50ntFlanks@12991.13015.25\|CTCCCTCGCTGCGATCTATTGAAAG | >hg38_21_+_8433172_8446622_RNA45SN1_45S_50ntFlanks | 12991 | 13015 | 25 | CTCCCTCGCTGCGATCTATTGAAAG | rRF | FALSE | FALSE | TRUE |
| >hg38_21_+_8433172_8446622_RNA45SN1_45S_50ntFlanks@12627.12653.27\|ATGGGGCGAAGCTACCATCTGTGGGAT | >hg38_21_+_8433172_8446622_RNA45SN1_45S_50ntFlanks | 12627 | 12653 | 27 | ATGGGGCGAAGCTACCATCTGTGGGAT | rRF | FALSE | FALSE | TRUE |
| >hg38_21_+_8433172_8446622_RNA45SN1_45S_50ntFlanks@6745.6771.27\|ACACTTCGAACGCACTTGCGGCCCCGG | >hg38_21_+_8433172_8446622_RNA45SN1_45S_50ntFlanks | 6745 | 6771 | 27 | ACACTTCGAACGCACTTGCGGCCCCGG | rRF | FALSE | FALSE | TRUE |
| >hg38_21_+_8433172_8446622_RNA45SN1_45S_50ntFlanks@11575.11605.31\|ACTTAGAACTGGTGCGGACCAGGGGAATCCG | >hg38_21_+_8433172_8446622_RNA45SN1_45S_50ntFlanks | 11575 | 11605 | 31 | ACTTAGAACTGGTGCGGACCAGGGGAATCCG | rRF | FALSE | FALSE | TRUE |
| >hg38_21_+_8433172_8446622_RNA45SN1_45S_50ntFlanks@12324.12346.23\|TTGGGTTTTAAGCAGGAGGTGTC | >hg38_21_+_8433172_8446622_RNA45SN1_45S_50ntFlanks | 12324 | 12346 | 23 | TTGGGTTTTAAGCAGGAGGTGTC | rRF | FALSE | FALSE | TRUE |
| >hg38_21_+_8433172_8446622_RNA45SN1_45S_50ntFlanks@10362.10390.29\|AAGGCCGAAGTGGAGAAGGGTTCCATGTG | >hg38_21_+_8433172_8446622_RNA45SN1_45S_50ntFlanks | 10362 | 10390 | 29 | AAGGCCGAAGTGGAGAAGGGTTCCATGTG | rRF | FALSE | FALSE | TRUE |
| >hg38_21_+_8433172_8446622_RNA45SN1_45S_50ntFlanks@11583.11603.21\|CTGGTGCGGACCAGGGGAATC | >hg38_21_+_8433172_8446622_RNA45SN1_45S_50ntFlanks | 11583 | 11603 | 21 | CTGGTGCGGACCAGGGGAATC | rRF | FALSE | FALSE | TRUE |
| >hg38_21_+_8433172_8446622_RNA45SN1_45S_50ntFlanks@12405.12441.37\|CTTTTTGATCCTTCGATGTCGGCTCTTCCTATCATTG | >hg38_21_+_8433172_8446622_RNA45SN1_45S_50ntFlanks | 12405 | 12441 | 37 | CTTTTTGATCCTTCGATGTCGGCTCTTCCTATCATTG | rRF | FALSE | FALSE | TRUE |
| >hg38_21_+_8433172_8446622_RNA45SN1_45S_50ntFlanks@8194.8215.22\|CCCGTGGACGGTGTGAGGCCGG | >hg38_21_+_8433172_8446622_RNA45SN1_45S_50ntFlanks | 8194 | 8215 | 22 | CCCGTGGACGGTGTGAGGCCGG | rRF | FALSE | FALSE | TRUE |
| >hg38_21_+_8433172_8446622_RNA45SN1_45S_50ntFlanks@6651.6699.49\|CGACTCTTAGCGGTGGATCACTCGGCTCGTGCGTCGATGAAGAACGCAG | >hg38_21_+_8433172_8446622_RNA45SN1_45S_50ntFlanks | 6651 | 6699 | 49 | CGACTCTTAGCGGTGGATCACTCGGCTCGTGCGTCGATGAAGAACGCAG | rRF | FALSE | FALSE | TRUE |
| >hg38_21_+_8433172_8446622_RNA45SN1_45S_50ntFlanks@9534.9566.33\|AGGAAACTCTGGTGGAGGTCCGTAGCGGTCCTG | >hg38_21_+_8433172_8446622_RNA45SN1_45S_50ntFlanks | 9534 | 9566 | 33 | AGGAAACTCTGGTGGAGGTCCGTAGCGGTCCTG | rRF | FALSE | FALSE | TRUE |
| >hg38_21_+_8433172_8446622_RNA45SN1_45S_50ntFlanks@11885.11904.20\|TTGAGCTTGACTCTAGTCTG | >hg38_21_+_8433172_8446622_RNA45SN1_45S_50ntFlanks | 11885 | 11904 | 20 | TTGAGCTTGACTCTAGTCTG | rRF | FALSE | FALSE | TRUE |
| >hg38_21_+_8433172_8446622_RNA45SN1_45S_50ntFlanks@8380.8400.21\|CAAGAGGGCGTGAAACCGTTA | >hg38_21_+_8433172_8446622_RNA45SN1_45S_50ntFlanks | 8380 | 8400 | 21 | CAAGAGGGCGTGAAACCGTTA | rRF | FALSE | FALSE | TRUE |
| >hg38_21_+_8433172_8446622_RNA45SN1_45S_50ntFlanks@10393.10410.18\|CAGCAGTTGAACATGGGT | >hg38_21_+_8433172_8446622_RNA45SN1_45S_50ntFlanks | 10393 | 10410 | 18 | CAGCAGTTGAACATGGGT | rRF | FALSE | FALSE | TRUE |
| >hg38_21_+_8433172_8446622_RNA45SN1_45S_50ntFlanks@3873.3898.26\|TAATACATGCCGACGGGCGCTGACCC | >hg38_21_+_8433172_8446622_RNA45SN1_45S_50ntFlanks | 3873 | 3898 | 26 | TAATACATGCCGACGGGCGCTGACCC | rRF | FALSE | FALSE | TRUE |
| >hg38_21_+_8433172_8446622_RNA45SN1_45S_50ntFlanks@10690.10709.20\|CGTCCGGTGAGCTCTCGCTG | >hg38_21_+_8433172_8446622_RNA45SN1_45S_50ntFlanks | 10690 | 10709 | 20 | CGTCCGGTGAGCTCTCGCTG | rRF | FALSE | FALSE | TRUE |
| >hg38_21_+_8433172_8446622_RNA45SN1_45S_50ntFlanks@11586.11611.26\|GTGCGGACCAGGGGAATCCGACTGTT | >hg38_21_+_8433172_8446622_RNA45SN1_45S_50ntFlanks | 11586 | 11611 | 26 | GTGCGGACCAGGGGAATCCGACTGTT | rRF | FALSE | FALSE | TRUE |
| >hg38_21_+_8433172_8446622_RNA45SN1_45S_50ntFlanks@12456.12480.25\|CCAAGCGTTGGATTGTTCACCCACT | >hg38_21_+_8433172_8446622_RNA45SN1_45S_50ntFlanks | 12456 | 12480 | 25 | CCAAGCGTTGGATTGTTCACCCACT | rRF | FALSE | FALSE | TRUE |
| >hg38_21_+_8433172_8446622_RNA45SN1_45S_50ntFlanks@3985.4002.18\|CGGCTTTGGTGACTCTAG | >hg38_21_+_8433172_8446622_RNA45SN1_45S_50ntFlanks | 3985 | 4002 | 18 | CGGCTTTGGTGACTCTAG | rRF | FALSE | FALSE | TRUE |
| >hg38_21_+_8433172_8446622_RNA45SN1_45S_50ntFlanks@10781.10818.38\|AGGTGAACAGCCTCTGGCATGTTGGAACAATGTAGGTA | >hg38_21_+_8433172_8446622_RNA45SN1_45S_50ntFlanks | 10781 | 10818 | 38 | AGGTGAACAGCCTCTGGCATGTTGGAACAATGTAGGTA | rRF | FALSE | FALSE | TRUE |
| >hg38_21_+_8433172_8446622_RNA45SN1_45S_50ntFlanks@10392.10429.38\|ACAGCAGTTGAACATGGGTCAGTCGGTCCTGAGAGATG | >hg38_21_+_8433172_8446622_RNA45SN1_45S_50ntFlanks | 10392 | 10429 | 38 | ACAGCAGTTGAACATGGGTCAGTCGGTCCTGAGAGATG | rRF | FALSE | FALSE | TRUE |
| >hg38_21_+_8433172_8446622_RNA45SN1_45S_50ntFlanks@12407.12431.25\|TTTTGATCCTTCGATGTCGGCTCTT | >hg38_21_+_8433172_8446622_RNA45SN1_45S_50ntFlanks | 12407 | 12431 | 25 | TTTTGATCCTTCGATGTCGGCTCTT | rRF | FALSE | FALSE | TRUE |
| >hg38_21_+_8433172_8446622_RNA45SN1_45S_50ntFlanks@6721.6745.25\|AATTGCAGGACACATTGATCATCGA | >hg38_21_+_8433172_8446622_RNA45SN1_45S_50ntFlanks | 6721 | 6745 | 25 | AATTGCAGGACACATTGATCATCGA | rRF | FALSE | FALSE | TRUE |
| >hg38_21_+_8433172_8446622_RNA45SN1_45S_50ntFlanks@10421.10456.36\|TGAGAGATGGGCGAGCGCCGTTCCGAAGGGACGGGC | >hg38_21_+_8433172_8446622_RNA45SN1_45S_50ntFlanks | 10421 | 10456 | 36 | TGAGAGATGGGCGAGCGCCGTTCCGAAGGGACGGGC | rRF | FALSE | FALSE | TRUE |
| >hg38_21_+_8433172_8446622_RNA45SN1_45S_50ntFlanks@11575.11593.19\|ACTTAGAACTGGTGCGGAC | >hg38_21_+_8433172_8446622_RNA45SN1_45S_50ntFlanks | 11575 | 11593 | 19 | ACTTAGAACTGGTGCGGAC | rRF | FALSE | FALSE | TRUE |
| >hg38_21_+_8433172_8446622_RNA45SN1_45S_50ntFlanks@10687.10708.22\|CGGCGTCCGGTGAGCTCTCGCT | >hg38_21_+_8433172_8446622_RNA45SN1_45S_50ntFlanks | 10687 | 10708 | 22 | CGGCGTCCGGTGAGCTCTCGCT | rRF | FALSE | FALSE | TRUE |
| >hg38_21_+_8433172_8446622_RNA45SN1_45S_50ntFlanks@9534.9570.37\|AGGAAACTCTGGTGGAGGTCCGTAGCGGTCCTGACGT | >hg38_21_+_8433172_8446622_RNA45SN1_45S_50ntFlanks | 9534 | 9570 | 37 | AGGAAACTCTGGTGGAGGTCCGTAGCGGTCCTGACGT | rRF | FALSE | FALSE | TRUE |
| >hg38_21_+_8433172_8446622_RNA45SN1_45S_50ntFlanks@12633.12653.21\|CGAAGCTACCATCTGTGGGAT | >hg38_21_+_8433172_8446622_RNA45SN1_45S_50ntFlanks | 12633 | 12653 | 21 | CGAAGCTACCATCTGTGGGAT | rRF | FALSE | FALSE | TRUE |
| >hg38_21_+_8433172_8446622_RNA45SN1_45S_50ntFlanks@12417.12446.30\|TCGATGTCGGCTCTTCCTATCATTGTGAAG | >hg38_21_+_8433172_8446622_RNA45SN1_45S_50ntFlanks | 12417 | 12446 | 30 | TCGATGTCGGCTCTTCCTATCATTGTGAAG | rRF | FALSE | FALSE | TRUE |
| >hg38_21_+_8433172_8446622_RNA45SN1_45S_50ntFlanks@6652.6681.30\|GACTCTTAGCGGTGGATCACTCGGCTCGTG | >hg38_21_+_8433172_8446622_RNA45SN1_45S_50ntFlanks | 6652 | 6681 | 30 | GACTCTTAGCGGTGGATCACTCGGCTCGTG | rRF | FALSE | FALSE | TRUE |
| >hg38_21_+_8433172_8446622_RNA45SN1_45S_50ntFlanks@11572.11590.19\|CCGACTTAGAACTGGTGCG | >hg38_21_+_8433172_8446622_RNA45SN1_45S_50ntFlanks | 11572 | 11590 | 19 | CCGACTTAGAACTGGTGCG | rRF | FALSE | FALSE | TRUE |
| >hg38_1_-_228634819_228635039_RNA5S12_5S_50ntFlanks@84.101.18\|CTCGTCTGATCTCGGAAG | >hg38_1_-_228634819_228635039_RNA5S12_5S_50ntFlanks | 84 | 101 | 18 | CTCGTCTGATCTCGGAAG | rRF | FALSE | FALSE | TRUE |

### Cytoplasm

| | | | | | | | | | |
|---|---|---|---|---|---|---|---|---|---|
| >hg38_21_+_8433172_8446622_RNA45SN1_45S_50ntFlanks@3775.3792.18\|GCCGGTACAGTGAAACTG | >hg38_21_+_8433172_8446622_RNA45SN1_45S_50ntFlanks | 3775 | 3792 | 18 | GCCGGTACAGTGAAACTG | rRF | FALSE | FALSE | TRUE |
| >hg38_21_+_8433172_8446622_RNA45SN1_45S_50ntFlanks@6743.6770.28\|CGACACTTCGAACGCACTTGCGGCCCCG | >hg38_21_+_8433172_8446622_RNA45SN1_45S_50ntFlanks | 6743 | 6770 | 28 | CGACACTTCGAACGCACTTGCGGCCCCG | rRF | FALSE | FALSE | TRUE |
| >hg38_21_+_8433172_8446622_RNA45SN1_45S_50ntFlanks@12415.12435.21\|CTTCGATGTCGGCTCTTCCTA | >hg38_21_+_8433172_8446622_RNA45SN1_45S_50ntFlanks | 12415 | 12435 | 21 | CTTCGATGTCGGCTCTTCCTA | rRF | FALSE | FALSE | TRUE |
| >hg38_21_+_8433172_8446622_RNA45SN1_45S_50ntFlanks@11208.11231.24\|GCGGCGACTCTGGACGCGAGCCGG | >hg38_21_+_8433172_8446622_RNA45SN1_45S_50ntFlanks | 11208 | 11231 | 24 | GCGGCGACTCTGGACGCGAGCCGG | rRF | FALSE | FALSE | TRUE |
| >hg38_21_+_8433172_8446622_RNA45SN1_45S_50ntFlanks@4045.4063.19\|CCATTCGAACGTCTGCCCT | >hg38_21_+_8433172_8446622_RNA45SN1_45S_50ntFlanks | 4045 | 4063 | 19 | CCATTCGAACGTCTGCCCT | rRF | FALSE | FALSE | TRUE |
| >hg38_21_+_8433172_8446622_RNA45SN1_45S_50ntFlanks@11768.11792.25\|CTAATTAGTGACGCGCATGAATGGA | >hg38_21_+_8433172_8446622_RNA45SN1_45S_50ntFlanks | 11768 | 11792 | 25 | CTAATTAGTGACGCGCATGAATGGA | rRF | FALSE | FALSE | TRUE |
| >trna11_GluTTC_15_-_26327381_26327452@1.31.31,trna3_GluTTC_13_-_45492062_45492133@1.31.31\|TCCCACATGGTCTAGCGGTTAGGATTCCTGG | >trna11_GluTTC_15_-_26327381_26327452 | 1 | 31 | 31 | TCCCACATGGTCTAGCGGTTAGGATTCCTGG | tRF | FALSE | FALSE | TRUE |
| >trna11_GluTTC_15_-_26327381_26327452@1.30.30,trna3_GluTTC_13_-_45492062_45492133@1.30.30\|TCCCACATGGTCTAGCGGTTAGGATTCCTG | >trna11_GluTTC_15_-_26327381_26327452 | 1 | 30 | 30 | TCCCACATGGTCTAGCGGTTAGGATTCCTG | tRF | FALSE | FALSE | TRUE |
| >hg38_21_+_8433172_8446622_RNA45SN1_45S_50ntFlanks@12938.12956.19\|ACCATTCGTAGACGACCTG | >hg38_21_+_8433172_8446622_RNA45SN1_45S_50ntFlanks | 12938 | 12956 | 19 | ACCATTCGTAGACGACCTG | rRF | FALSE | FALSE | TRUE |
| >hg38_21_+_8433172_8446622_RNA45SN1_45S_50ntFlanks@12476.12498.23\|CCACTAATAGGGAACGTGAGCTG | >hg38_21_+_8433172_8446622_RNA45SN1_45S_50ntFlanks | 12476 | 12498 | 23 | CCACTAATAGGGAACGTGAGCTG | rRF | FALSE | FALSE | TRUE |
| >hg38_21_+_8433172_8446622_RNA45SN1_45S_50ntFlanks@9544.9571.28\|GGTGGAGGTCCGTAGCGGTCCTGACGTG | >hg38_21_+_8433172_8446622_RNA45SN1_45S_50ntFlanks | 9544 | 9571 | 28 | GGTGGAGGTCCGTAGCGGTCCTGACGTG | rRF | FALSE | FALSE | TRUE |
| >hg38_21_+_8433172_8446622_RNA45SN1_45S_50ntFlanks@12073.12097.25\|CCGAGGGGCTCTCGCTTCTGGCGCC | >hg38_21_+_8433172_8446622_RNA45SN1_45S_50ntFlanks | 12073 | 12097 | 25 | CCGAGGGGCTCTCGCTTCTGGCGCC | rRF | FALSE | FALSE | TRUE |
| >hg38_21_+_8433172_8446622_RNA45SN1_45S_50ntFlanks@11316.11336.21\|CACCCCACGTCTCGTCGCGCG | >hg38_21_+_8433172_8446622_RNA45SN1_45S_50ntFlanks | 11316 | 11336 | 21 | CACCCCACGTCTCGTCGCGCG | rRF | FALSE | FALSE | TRUE |
| >hg38_21_+_8433172_8446622_RNA45SN1_45S_50ntFlanks@12991.13012.22\|CTCCCTCGCTGCGATCTATTGA | >hg38_21_+_8433172_8446622_RNA45SN1_45S_50ntFlanks | 12991 | 13012 | 22 | CTCCCTCGCTGCGATCTATTGA | rRF | FALSE | FALSE | TRUE |
| >hg38_21_+_8433172_8446622_RNA45SN1_45S_50ntFlanks@12366.12383.18\|TAACTGGCTTGTGGCGGC | >hg38_21_+_8433172_8446622_RNA45SN1_45S_50ntFlanks | 12366 | 12383 | 18 | TAACTGGCTTGTGGCGGC | rRF | FALSE | FALSE | TRUE |
| >hg38_21_+_8433172_8446622_RNA45SN1_45S_50ntFlanks@11321.11342.22\|CACGTCTCGTCGCGCGCGCGTC | >hg38_21_+_8433172_8446622_RNA45SN1_45S_50ntFlanks | 11321 | 11342 | 22 | CACGTCTCGTCGCGCGCGCGTC | rRF | FALSE | FALSE | TRUE |
| >hg38_21_+_8433172_8446622_RNA45SN1_45S_50ntFlanks@9739.9758.20\|AAACGATCTCAACCTATTCT | >hg38_21_+_8433172_8446622_RNA45SN1_45S_50ntFlanks | 9739 | 9758 | 20 | AAACGATCTCAACCTATTCT | rRF | FALSE | FALSE | TRUE |
| >hg38_21_+_8433172_8446622_RNA45SN1_45S_50ntFlanks@9533.9571.39\|GAGGAAACTCTGGTGGAGGTCCGTAGCGGTCCTGACGTG | >hg38_21_+_8433172_8446622_RNA45SN1_45S_50ntFlanks | 9533 | 9571 | 39 | GAGGAAACTCTGGTGGAGGTCCGTAGCGGTCCTGACGTG | rRF | FALSE | FALSE | TRUE |
| >hg38_21_+_8433172_8446622_RNA45SN1_45S_50ntFlanks@10351.10374.24\|CGAGAACTTTGAAGGCCGAAGTGG | >hg38_21_+_8433172_8446622_RNA45SN1_45S_50ntFlanks | 10351 | 10374 | 24 | CGAGAACTTTGAAGGCCGAAGTGG | rRF | FALSE | FALSE | TRUE |
| >hg38_21_+_8433172_8446622_RNA45SN1_45S_50ntFlanks@3835.3860.26\|CTCCTCTCCTACTTGGATAACTGTGG | >hg38_21_+_8433172_8446622_RNA45SN1_45S_50ntFlanks | 3835 | 3860 | 26 | CTCCTCTCCTACTTGGATAACTGTGG | rRF | FALSE | FALSE | TRUE |
| >hg38_21_+_8433172_8446622_RNA45SN1_45S_50ntFlanks@12456.12485.30\|CCAAGCGTTGGATTGTTCACCCACTAATAG | >hg38_21_+_8433172_8446622_RNA45SN1_45S_50ntFlanks | 12456 | 12485 | 30 | CCAAGCGTTGGATTGTTCACCCACTAATAG | rRF | FALSE | FALSE | TRUE |
| >hg38_21_+_8433172_8446622_RNA45SN1_45S_50ntFlanks@11891.11912.22\|TTGACTCTAGTCTGGCACGGTG | >hg38_21_+_8433172_8446622_RNA45SN1_45S_50ntFlanks | 11891 | 11912 | 22 | TTGACTCTAGTCTGGCACGGTG | rRF | FALSE | FALSE | TRUE |
| >hg38_21_+_8433172_8446622_RNA45SN1_45S_50ntFlanks@10681.10712.32\|TTCCGGCGGCGTCCGGTGAGCTCTCGCTGGCC | >hg38_21_+_8433172_8446622_RNA45SN1_45S_50ntFlanks | 10681 | 10712 | 32 | TTCCGGCGGCGTCCGGTGAGCTCTCGCTGGCC | rRF | FALSE | FALSE | TRUE |
| >hg38_21_+_8433172_8446622_RNA45SN1_45S_50ntFlanks@10683.10712.30\|CCGGCGGCGTCCGGTGAGCTCTCGCTGGCC | >hg38_21_+_8433172_8446622_RNA45SN1_45S_50ntFlanks | 10683 | 10712 | 30 | CCGGCGGCGTCCGGTGAGCTCTCGCTGGCC | rRF | FALSE | FALSE | TRUE |
| >hg38_21_+_8433172_8446622_RNA45SN1_45S_50ntFlanks@9570.9596.27\|TGCAAATCGGTCGTCCGACCTGGGTAT | >hg38_21_+_8433172_8446622_RNA45SN1_45S_50ntFlanks | 9570 | 9596 | 27 | TGCAAATCGGTCGTCCGACCTGGGTAT | rRF | FALSE | FALSE | TRUE |
| >trna111_HisGTG_1_-_147774845_147774916@1.30.30,trna118_HisGTG_1_-_145396881_145396952@1.30.30,trna16_HisGTG_1_+_146544773_146544844@1.30.30,trna1_HisGTG_15_+_45493349_45493420@1.30.30,trna21_HisGTG_1_+_147753471_147753542@1.30.30,trna33_HisGTG_6_+_27125906_27125977@1.30.30,trna7_HisGTG_9_-_14433938_14434009@1.30.30,trna8_HisGTG_15_-_45492611_45492682@1.30.30,trna9_HisGTG_15_-_45490804_45490875@1.30.30\|GCCGTGATCGTATAGTGGTTAGTACTCTGC | >trna111_HisGTG_1_-_147774845_147774916 | 1 | 30 | 30 | GCCGTGATCGTATAGTGGTTAGTACTCTGC | tRF | FALSE | FALSE | TRUE |
| >hg38_21_+_8433172_8446622_RNA45SN1_45S_50ntFlanks@10362.10383.22\|AAGGCCGAAGTGGAGAAGGGTT | >hg38_21_+_8433172_8446622_RNA45SN1_45S_50ntFlanks | 10362 | 10383 | 22 | AAGGCCGAAGTGGAGAAGGGTT | rRF | FALSE | FALSE | TRUE |
| >hg38_21_+_8433172_8446622_RNA45SN1_45S_50ntFlanks@12437.12454.18\|CATTGTGAAGCAGAATTC | >hg38_21_+_8433172_8446622_RNA45SN1_45S_50ntFlanks | 12437 | 12454 | 18 | CATTGTGAAGCAGAATTC | rRF | FALSE | FALSE | TRUE |
| >hg38_21_+_8433172_8446622_RNA45SN1_45S_50ntFlanks@6671.6699.29\|CTCGGCTCGTGCGTCGATGAAGAACGCAG | >hg38_21_+_8433172_8446622_RNA45SN1_45S_50ntFlanks | 6671 | 6699 | 29 | CTCGGCTCGTGCGTCGATGAAGAACGCAG | rRF | FALSE | FALSE | TRUE |
| >hg38_21_+_8433172_8446622_RNA45SN1_45S_50ntFlanks@11322.11342.21\|ACGTCTCGTCGCGCGCGCGTC | >hg38_21_+_8433172_8446622_RNA45SN1_45S_50ntFlanks | 11322 | 11342 | 21 | ACGTCTCGTCGCGCGCGCGTC | rRF | FALSE | FALSE | TRUE |
| >hg38_21_+_8433172_8446622_RNA45SN1_45S_50ntFlanks@12991.13039.49\|CTCCCTCGCTGCGATCTATTGAAAGTCAGCCCTCGACACAAGGGTTTGT | >hg38_21_+_8433172_8446622_RNA45SN1_45S_50ntFlanks | 12991 | 13039 | 49 | CTCCCTCGCTGCGATCTATTGAAAGTCAGCCCTCGACACAAGGGTTTGT | rRF | FALSE | FALSE | TRUE |
| >hg38_21_+_8433172_8446622_RNA45SN1_45S_50ntFlanks@9545.9570.26\|GTGGAGGTCCGTAGCGGTCCTGACGT | >hg38_21_+_8433172_8446622_RNA45SN1_45S_50ntFlanks | 9545 | 9570 | 26 | GTGGAGGTCCGTAGCGGTCCTGACGT | rRF | FALSE | FALSE | TRUE |
