## Supplementary material for "Single-nucleotide Differences and Cell Type Decide the Subcellular Localization of miRNA Isoforms (isomiRs), tRNA-derived Fragments (tRFs) and rRNA-derived Fragments (rRFs)": Supp. Table S5 (3 of 4)

### Mitochondria

<table>
<tr><td>>MI0000063|hsa-let-7b&WithFlank&22|+|46113680|46113774@12.32.21&[MIMAT0000063&hsa-let-7b-5p&offsets|0|-1;m-14&22|+|46113691|46113712&offsets|0|-1]|TGAGGTAGTAGGTTGTGTGGT</td><td>>MI0000063|hsa-let-7b&WithFlank&22|+|46113680|46113774</td><td>12</td><td>32</td><td>21</td><td>TGAGGTAGTAGGTTGTGTGGT</td><td>isomiR</td><td>TRUE</td><td>TRUE</td><td>FALSE</td></tr>
<tr><td>>multi-hg38_MT_+_1621_3279_RNR2_16S_50ntFlanks@51.84.34|GCTAAACCTAGCCCCAAACCCACTCCACCTTACT</td><td>>multi-hg38_MT_+_1621_3279_RNR2_16S_50ntFlanks</td><td>51</td><td>84</td><td>34</td><td>GCTAAACCTAGCCCCAAACCCACTCCACCTTACT</td><td>rRF</td><td>TRUE</td><td>TRUE</td><td>FALSE</td></tr>
<tr><td>>hg38_21_+_8433172_8446622_RNA45SN1_45S_50ntFlanks@6651.6676.26|CGACTCTTAGCGGTGGATCACTCGGC</td><td>>hg38_21_+_8433172_8446622_RNA45SN1_45S_50ntFlanks</td><td>6651</td><td>6676</td><td>26</td><td>CGACTCTTAGCGGTGGATCACTCGGC</td><td>rRF</td><td>TRUE</td><td>FALSE</td><td>FALSE</td></tr>
<tr><td>>trnaMT_SerGCT_MT_+_12207_12265@1.18.18|GAGAAAGCTCACAAGAAC</td><td>>trnaMT_SerGCT_MT_+_12207_12265</td><td>1</td><td>18</td><td>18</td><td>GAGAAAGCTCACAAGAAC</td><td>tRF</td><td>TRUE</td><td>TRUE</td><td>TRUE</td></tr>
<tr><td>>MI0000434|hsa-let-7i&WithFlank&12|+|62603680|62603775@12.33.22(+1A)&[MIMAT0000415&hsa-let-7i-5p&offsets|0|0(+1A);m-49&12|+|62603691|62603712&offsets|0|0(+1A)]|TGAGGTAGTAGTTTGTGCTGTTA</td><td>>MI0000434|hsa-let-7i&WithFlank&12|+|62603680|62603775</td><td>12</td><td>33</td><td>22(+1A)</td><td>TGAGGTAGTAGTTTGTGCTGTTA</td><td>isomiR</td><td>TRUE</td><td>TRUE</td><td>FALSE</td></tr>
<tr><td>>MI0000542|hsa-mir-320a&WithFlank&8|-|22244960|22245043@48.69.22(+1U)&[MIMAT0000510&hsa-miR-320a-3p&offsets|0|0(+1U);m-60&8|-|22244975|22244996&offsets|0|0(+1U)]|AAAAGCTGGGTTGAGAGGGCGAT</td><td>>MI0000542|hsa-mir-320a&WithFlank&8|-|22244960|22245043</td><td>48</td><td>69</td><td>22(+1U)</td><td>AAAAGCTGGGTTGAGAGGGCGAT</td><td>isomiR</td><td>TRUE</td><td>FALSE</td><td>FALSE</td></tr>
<tr><td>>trna111_HisGTG_1_-_147774845_147774916@-1G.33.34,trna118_HisGTG_1_-_145396881_145396952@-1G.33.34,trna16_HisGTG_1_+_146544773_146544844@-1G.33.34,trna1_HisGTG_15_+_45493349_45493420@-1G.33.34,trna21_HisGTG_1_+_147753471_147753542@-1G.33.34,trna33_HisGTG_6_+_27125906_27125977@-1G.33.34,trna7_HisGTG_9_-_14433938_14434009@-1G.33.34,trna8_HisGTG_15_-_45492611_45492682@-1G.33.34,trna9_HisGTG_15_-_45490804_45490875@-1G.33.34|GGCCGTGATCGTATAGTGGTTAGTACTCTGCGTT</td><td>>trna111_HisGTG_1_-_147774845_147774916</td><td>-1G</td><td>33</td><td>34</td><td>GGCCGTGATCGTATAGTGGTTAGTACTCTGCGTT</td><td>tRF</td><td>TRUE</td><td>TRUE</td><td>FALSE</td></tr>
<tr><td>>hg38_21_+_8433172_8446622_RNA45SN1_45S_50ntFlanks@7975.8004.30|CGCGACCTCAGATCAGACGTGGCGACCCGC</td><td>>hg38_21_+_8433172_8446622_RNA45SN1_45S_50ntFlanks</td><td>7975</td><td>8004</td><td>30</td><td>CGCGACCTCAGATCAGACGTGGCGACCCGC</td><td>rRF</td><td>TRUE</td><td>FALSE</td><td>FALSE</td></tr>
<tr><td>>MI0001445|hsa-mir-423&WithFlank&17|+|30117073|30117178@23.43.21&[MIMAT0004748&hsa-miR-423-5p&offsets|0|-2;m-141&17|+|30117095|30117117&offsets|0|-2]|TGAGGGGCAGAGAGCGAGACT</td><td>>MI0001445|hsa-mir-423&WithFlank&17|+|30117073|30117178</td><td>23</td><td>43</td><td>21</td><td>TGAGGGGCAGAGAGCGAGACT</td><td>isomiR</td><td>TRUE</td><td>TRUE</td><td>FALSE</td></tr>
<tr><td>>multi-hg38_MT_+_1621_3279_RNR2_16S_50ntFlanks@51.82.32|GCTAAACCTAGCCCCAAACCCACTCCACCTTA</td><td>>multi-hg38_MT_+_1621_3279_RNR2_16S_50ntFlanks</td><td>51</td><td>82</td><td>32</td><td>GCTAAACCTAGCCCCAAACCCACTCCACCTTA</td><td>rRF</td><td>TRUE</td><td>FALSE</td><td>FALSE</td></tr>
<tr><td>>hg38_21_+_8433172_8446622_RNA45SN1_45S_50ntFlanks@7974.8008.35|ACGCGACCTCAGATCAGACGTGGCGACCCGCTGAA</td><td>>hg38_21_+_8433172_8446622_RNA45SN1_45S_50ntFlanks</td><td>7974</td><td>8008</td><td>35</td><td>ACGCGACCTCAGATCAGACGTGGCGACCCGCTGAA</td><td>rRF</td><td>TRUE</td><td>FALSE</td><td>FALSE</td></tr>
<tr><td>>hg38_21_+_8433172_8446622_RNA45SN1_45S_50ntFlanks@7975.8002.28|CGCGACCTCAGATCAGACGTGGCGACCC</td><td>>hg38_21_+_8433172_8446622_RNA45SN1_45S_50ntFlanks</td><td>7975</td><td>8002</td><td>28</td><td>CGCGACCTCAGATCAGACGTGGCGACCC</td><td>rRF</td><td>TRUE</td><td>FALSE</td><td>FALSE</td></tr>
<tr><td>>MI0000070|hsa-mir-16-1&WithFlank&13|-|50048967|50049067@20.41.22;MI0000115|hsa-mir-16-2&WithFlank&3|+|160404739|160404831@16.37.22&[MIMAT0000069&hsa-miR-16-5p&offsets|0|0;m-62&13|-|50049027|50049048&offsets|0|0];[MIMAT0000069_1&hsa-miR-16-5p&offsets|0|0;m-61&3|+|160404754|160404775&offsets|0|0]|TAGCAGCACGTAAATATTGGCG</td><td>>MI0000070|hsa-mir-16-1&WithFlank&13|-|50048967|50049067</td><td>20</td><td>41</td><td>22</td><td>TAGCAGCACGTAAATATTGGCG</td><td>isomiR</td><td>TRUE</td><td>TRUE</td><td>FALSE</td></tr>
<tr><td>>trnaMT_HisGTG_MT_+_12138_12206@32.71.40|TGAATCTGACAACAGAGGCTTACGACCCCTTATTTACCCC</td><td>>trnaMT_HisGTG_MT_+_12138_12206</td><td>32</td><td>71</td><td>40</td><td>TGAATCTGACAACAGAGGCTTACGACCCCTTATTTACCCC</td><td>tRF</td><td>TRUE</td><td>TRUE</td><td>TRUE</td></tr>
<tr><td>>MI0000811|hsa-mir-148b&WithFlank&12|+|54337210|54337320@69.90.22&[MIMAT0000759&hsa-miR-148b-3p&offsets|0|0;m-77&12|+|54337278|54337299&offsets|0|0]|TCAGTGCATCACAGAACTTTGT</td><td>>MI0000811|hsa-mir-148b&WithFlank&12|+|54337210|54337320</td><td>69</td><td>90</td><td>22</td><td>TCAGTGCATCACAGAACTTTGT</td><td>isomiR</td><td>TRUE</td><td>TRUE</td><td>FALSE</td></tr>
<tr><td>>hg38_21_+_8433172_8446622_RNA45SN1_45S_50ntFlanks@6651.6681.31|CGACTCTTAGCGGTGGATCACTCGGCTCGTG</td><td>>hg38_21_+_8433172_8446622_RNA45SN1_45S_50ntFlanks</td><td>6651</td><td>6681</td><td>31</td><td>CGACTCTTAGCGGTGGATCACTCGGCTCGTG</td><td>rRF</td><td>TRUE</td><td>FALSE</td><td>TRUE</td></tr>
<tr><td>>hg38_21_+_8433172_8446622_RNA45SN1_45S_50ntFlanks@10396.10424.29|CAGTTGAACATGGGTCAGTCGGTCCTGAG</td><td>>hg38_21_+_8433172_8446622_RNA45SN1_45S_50ntFlanks</td><td>10396</td><td>10424</td><td>29</td><td>CAGTTGAACATGGGTCAGTCGGTCCTGAG</td><td>rRF</td><td>TRUE</td><td>FALSE</td><td>FALSE</td></tr>
<tr><td>>MI0001445|hsa-mir-423&WithFlank&17|+|30117073|30117178@23.44.22(+1C)&[MIMAT0004748&hsa-miR-423-5p&offsets|0|-1(+1C);m-141&17|+|30117095|30117117&offsets|0|-1(+1C)]|TGAGGGGCAGAGAGCGAGACTTC</td><td>>MI0001445|hsa-mir-423&WithFlank&17|+|30117073|30117178</td><td>23</td><td>44</td><td>22(+1C)</td><td>TGAGGGGCAGAGAGCGAGACTTC</td><td>isomiR</td><td>TRUE</td><td>FALSE</td><td>FALSE</td></tr>
<tr><td>>hg38_1_-_228634819_228635039_RNA5S12_5S_50ntFlanks@140.170.31|ACCGCCTGGGAATACCGGGTGCTGTAGGCTT</td><td>>hg38_1_-_228634819_228635039_RNA5S12_5S_50ntFlanks</td><td>140</td><td>170</td><td>31</td><td>ACCGCCTGGGAATACCGGGTGCTGTAGGCTT</td><td>rRF</td><td>TRUE</td><td>FALSE</td><td>TRUE</td></tr>
<tr><td>>hg38_21_+_8433172_8446622_RNA45SN1_45S_50ntFlanks@8357.8375.19|AAAAGAACTTTGAAGAGAG</td><td>>hg38_21_+_8433172_8446622_RNA45SN1_45S_50ntFlanks</td><td>8357</td><td>8375</td><td>19</td><td>AAAAGAACTTTGAAGAGAG</td><td>rRF</td><td>TRUE</td><td>FALSE</td><td>TRUE</td></tr>
<tr><td>>MI0000542|hsa-mir-320a&WithFlank&8|-|22244960|22245043@48.70.23(+1U)&[MIMAT0000510&hsa-miR-320a-3p&offsets|0|+1(+1U);m-60&8|-|22244975|22244996&offsets|0|+1(+1U)]|AAAAGCTGGGTTGAGAGGGCGAAT</td><td>>MI0000542|hsa-mir-320a&WithFlank&8|-|22244960|22245043</td><td>48</td><td>70</td><td>23(+1U)</td><td>AAAAGCTGGGTTGAGAGGGCGAAT</td><td>isomiR</td><td>TRUE</td><td>FALSE</td><td>FALSE</td></tr>
<tr><td>>hg38_21_+_8433172_8446622_RNA45SN1_45S_50ntFlanks@7975.7995.21|CGCGACCTCAGATCAGACGTG</td><td>>hg38_21_+_8433172_8446622_RNA45SN1_45S_50ntFlanks</td><td>7975</td><td>7995</td><td>21</td><td>CGCGACCTCAGATCAGACGTG</td><td>rRF</td><td>TRUE</td><td>FALSE</td><td>TRUE</td></tr>
<tr><td>>hg38_21_+_8433172_8446622_RNA45SN1_45S_50ntFlanks@6651.6669.19|CGACTCTTAGCGGTGGATC</td><td>>hg38_21_+_8433172_8446622_RNA45SN1_45S_50ntFlanks</td><td>6651</td><td>6669</td><td>19</td><td>CGACTCTTAGCGGTGGATC</td><td>rRF</td><td>TRUE</td><td>FALSE</td><td>TRUE</td></tr>
<tr><td>>multi-hg38_MT_+_1621_3279_RNR2_16S_50ntFlanks@51.79.29|GCTAAACCTAGCCCCAAACCCACTCCACC</td><td>>multi-hg38_MT_+_1621_3279_RNR2_16S_50ntFlanks</td><td>51</td><td>79</td><td>29</td><td>GCTAAACCTAGCCCCAAACCCACTCCACC</td><td>rRF</td><td>TRUE</td><td>FALSE</td><td>FALSE</td></tr>
<tr><td>>hg38_21_+_8433172_8446622_RNA45SN1_45S_50ntFlanks@6646.6667.22|TCGTACGACTCTTAGCGGTGGA</td><td>>hg38_21_+_8433172_8446622_RNA45SN1_45S_50ntFlanks</td><td>6646</td><td>6667</td><td>22</td><td>TCGTACGACTCTTAGCGGTGGA</td><td>rRF</td><td>TRUE</td><td>FALSE</td><td>FALSE</td></tr>
<tr><td>>hg38_21_+_8433172_8446622_RNA45SN1_45S_50ntFlanks@6661.6689.29|CGGTGGATCACTCGGCTCGTGCGTCGATG</td><td>>hg38_21_+_8433172_8446622_RNA45SN1_45S_50ntFlanks</td><td>6661</td><td>6689</td><td>29</td><td>CGGTGGATCACTCGGCTCGTGCGTCGATG</td><td>rRF</td><td>TRUE</td><td>FALSE</td><td>TRUE</td></tr>
<tr><td>>MI0000746|hsa-mir-99b&WithFlank&19|+|51692606|51692687@13.34.22&[MIMAT0000689&hsa-miR-99b-5p&offsets|0|0;m-10&19|+|51692618|51692639&offsets|0|0]|CACCCGTAGAACCGACCTTGCG</td><td>>MI0000746|hsa-mir-99b&WithFlank&19|+|51692606|51692687</td><td>13</td><td>34</td><td>22</td><td>CACCCGTAGAACCGACCTTGCG</td><td>isomiR</td><td>TRUE</td><td>TRUE</td><td>FALSE</td></tr>
<tr><td>>MI0000063|hsa-let-7b&WithFlank&22|+|46113680|46113774@12.32.21(+1C)&[MIMAT0000063&hsa-let-7b-5p&offsets|0|-1(+1C);m-14&22|+|46113691|46113712&offsets|0|-1(+1C)]|TGAGGTAGTAGGTTGTGTGGTC</td><td>>MI0000063|hsa-let-7b&WithFlank&22|+|46113680|46113774</td><td>12</td><td>32</td><td>21(+1C)</td><td>TGAGGTAGTAGGTTGTGTGGTC</td><td>isomiR</td><td>TRUE</td><td>TRUE</td><td>FALSE</td></tr>
</table>

### Mitochondria

<table>
<tr><th></th><th></th><th></th><th></th><th></th><th></th><th></th><th></th><th></th><th></th></tr>
<tr><td>>trna116_GluCTC_1_-_145399233_145399304@1.33.33,trna59_GluCTC_1_+_249168447_249168518@1.33.33,trna71_GluCTC_1_-_161439189_161439260@1.33.33,trna74_GluCTC_1_-_161431809_161431880@1.33.33,trna77_GluCTC_1_-_161424398_161424469@1.33.33,trna77_GluCTC_6_+_28949976_28950047@1.33.33,trna80_GluCTC_1_-_161417018_161417089@1.33.33,trna87_GluCTC_6_-_126101393_126101464@1.33.33|TCCCTGGTGGTCTAGTGGTTAGGATTCGGCGCT</td><td>>trna116_GluCTC_1_-_145399233_145399304</td><td>1</td><td>33</td><td>33</td><td>TCCCTGGTGGTCTAGTGGTTAGGATTCGGCGCT</td><td>tRF</td><td>TRUE</td><td>TRUE</td><td>FALSE</td></tr>
<tr><td>>MI0000272|hsa-mir-182&WithFlank&7|-|129770377|129770498@29.52.24&[MIMAT0000259&hsa-miR-182-5p&offsets|0|0;m-22&7|-|129770449|129770470&offsets|0|+2]|TTTGGCAATGGTAGAACTCACACT</td><td>>MI0000272|hsa-mir-182&WithFlank&7|-|129770377|129770498</td><td>29</td><td>52</td><td>24</td><td>TTTGGCAATGGTAGAACTCACACT</td><td>isomiR</td><td>TRUE</td><td>FALSE</td><td>FALSE</td></tr>
<tr><td>>hg38_21_+_8433172_8446622_RNA45SN1_45S_50ntFlanks@7974.7993.20|ACGCGACCTCAGATCAGACG</td><td>>hg38_21_+_8433172_8446622_RNA45SN1_45S_50ntFlanks</td><td>7974</td><td>7993</td><td>20</td><td>ACGCGACCTCAGATCAGACG</td><td>rRF</td><td>TRUE</td><td>FALSE</td><td>FALSE</td></tr>
<tr><td>>hg38_21_+_8433172_8446622_RNA45SN1_45S_50ntFlanks@10391.10424.34|AACAGCAGTTGAACATGGGTCAGTCGGTCCTGAG</td><td>>hg38_21_+_8433172_8446622_RNA45SN1_45S_50ntFlanks</td><td>10391</td><td>10424</td><td>34</td><td>AACAGCAGTTGAACATGGGTCAGTCGGTCCTGAG</td><td>rRF</td><td>TRUE</td><td>FALSE</td><td>FALSE</td></tr>
<tr><td>>hg38_21_+_8433172_8446622_RNA45SN1_45S_50ntFlanks@10401.10424.24|GAACATGGGTCAGTCGGTCCTGAG</td><td>>hg38_21_+_8433172_8446622_RNA45SN1_45S_50ntFlanks</td><td>10401</td><td>10424</td><td>24</td><td>GAACATGGGTCAGTCGGTCCTGAG</td><td>rRF</td><td>TRUE</td><td>TRUE</td><td>FALSE</td></tr>
<tr><td>>hg38_21_+_8433172_8446622_RNA45SN1_45S_50ntFlanks@10389.10424.36|TGAACAGCAGTTGAACATGGGTCAGTCGGTCCTGAG</td><td>>hg38_21_+_8433172_8446622_RNA45SN1_45S_50ntFlanks</td><td>10389</td><td>10424</td><td>36</td><td>TGAACAGCAGTTGAACATGGGTCAGTCGGTCCTGAG</td><td>rRF</td><td>TRUE</td><td>FALSE</td><td>FALSE</td></tr>
<tr><td>>trnaMT_GluTTC_MT_-_14674_14742@38.72.35,trnalookalike8_GluTTC_5_-_93905172_93905240@38.72.35|CATTGGTCGTGGTTGTAGTCCGTGCGAGAATACCA</td><td>>trnaMT_GluTTC_MT_-_14674_14742</td><td>38</td><td>72</td><td>35</td><td>CATTGGTCGTGGTTGTAGTCCGTGCGAGAATACCA</td><td>tRF</td><td>TRUE</td><td>TRUE</td><td>FALSE</td></tr>
<tr><td>>hg38_21_+_8433172_8446622_RNA45SN1_45S_50ntFlanks@6661.6685.25|CGGTGGATCACTCGGCTCGTGCGTC</td><td>>hg38_21_+_8433172_8446622_RNA45SN1_45S_50ntFlanks</td><td>6661</td><td>6685</td><td>25</td><td>CGGTGGATCACTCGGCTCGTGCGTC</td><td>rRF</td><td>TRUE</td><td>FALSE</td><td>FALSE</td></tr>
<tr><td>>MI0000088|hsa-mir-30a&WithFlank&6|-|71403545|71403627@12.33.22(+1U)&[MIMAT0000087&hsa-miR-30a-5p&offsets|0|0(+1U);m-15&6|-|71403595|71403616&offsets|0|0(+1U)]|TGTAAACATCCTCGACTGGAAGT</td><td>>MI0000088|hsa-mir-30a&WithFlank&6|-|71403545|71403627</td><td>12</td><td>33</td><td>22(+1U)</td><td>TGTAAACATCCTCGACTGGAAGT</td><td>isomiR</td><td>TRUE</td><td>TRUE</td><td>FALSE</td></tr>
<tr><td>>hg38_21_+_8433172_8446622_RNA45SN1_45S_50ntFlanks@10394.10424.31|AGCAGTTGAACATGGGTCAGTCGGTCCTGAG</td><td>>hg38_21_+_8433172_8446622_RNA45SN1_45S_50ntFlanks</td><td>10394</td><td>10424</td><td>31</td><td>AGCAGTTGAACATGGGTCAGTCGGTCCTGAG</td><td>rRF</td><td>TRUE</td><td>FALSE</td><td>FALSE</td></tr>
<tr><td>>hg38_21_+_8433172_8446622_RNA45SN1_45S_50ntFlanks@6651.6677.27|CGACTCTTAGCGGTGGATCACTCGGCT</td><td>>hg38_21_+_8433172_8446622_RNA45SN1_45S_50ntFlanks</td><td>6651</td><td>6677</td><td>27</td><td>CGACTCTTAGCGGTGGATCACTCGGCT</td><td>rRF</td><td>TRUE</td><td>FALSE</td><td>FALSE</td></tr>
<tr><td>>hg38_21_+_8433172_8446622_RNA45SN1_45S_50ntFlanks@10393.10424.32|CAGCAGTTGAACATGGGTCAGTCGGTCCTGAG</td><td>>hg38_21_+_8433172_8446622_RNA45SN1_45S_50ntFlanks</td><td>10393</td><td>10424</td><td>32</td><td>CAGCAGTTGAACATGGGTCAGTCGGTCCTGAG</td><td>rRF</td><td>TRUE</td><td>FALSE</td><td>FALSE</td></tr>
<tr><td>>hg38_21_+_8433172_8446622_RNA45SN1_45S_50ntFlanks@6651.6685.35|CGACTCTTAGCGGTGGATCACTCGGCTCGTGCGTC</td><td>>hg38_21_+_8433172_8446622_RNA45SN1_45S_50ntFlanks</td><td>6651</td><td>6685</td><td>35</td><td>CGACTCTTAGCGGTGGATCACTCGGCTCGTGCGTC</td><td>rRF</td><td>TRUE</td><td>FALSE</td><td>FALSE</td></tr>
<tr><td>>hg38_1_-_228634819_228635039_RNA5S12_5S_50ntFlanks@147.169.23|GGGAATACCGGGTGCTGTAGGCT</td><td>>hg38_1_-_228634819_228635039_RNA5S12_5S_50ntFlanks</td><td>147</td><td>169</td><td>23</td><td>GGGAATACCGGGTGCTGTAGGCT</td><td>rRF</td><td>TRUE</td><td>FALSE</td><td>FALSE</td></tr>
<tr><td>>MI0000746|hsa-mir-99b&WithFlank&19|+|51692606|51692687@13.32.20(+1U)&[MIMAT0000689&hsa-miR-99b-5p&offsets|0|-2(+1U);m-10&19|+|51692618|51692639&offsets|0|-2(+1U)]|CACCCGTAGAACCGACCTTGT</td><td>>MI0000746|hsa-mir-99b&WithFlank&19|+|51692606|51692687</td><td>13</td><td>32</td><td>20(+1U)</td><td>CACCCGTAGAACCGACCTTGT</td><td>isomiR</td><td>TRUE</td><td>FALSE</td><td>FALSE</td></tr>
<tr><td>>trnaMT_GluTTC_MT_-_14674_14742@41.72.32,trnalookalike8_GluTTC_5_-_93905172_93905240@41.72.32|TGGTCGTGGTTGTAGTCCGTGCGAGAATACCA</td><td>>trnaMT_GluTTC_MT_-_14674_14742</td><td>41</td><td>72</td><td>32</td><td>TGGTCGTGGTTGTAGTCCGTGCGAGAATACCA</td><td>tRF</td><td>TRUE</td><td>TRUE</td><td>FALSE</td></tr>
<tr><td>>hg38_21_+_8433172_8446622_RNA45SN1_45S_50ntFlanks@10403.10424.22|ACATGGGTCAGTCGGTCCTGAG</td><td>>hg38_21_+_8433172_8446622_RNA45SN1_45S_50ntFlanks</td><td>10403</td><td>10424</td><td>22</td><td>ACATGGGTCAGTCGGTCCTGAG</td><td>rRF</td><td>TRUE</td><td>FALSE</td><td>FALSE</td></tr>
<tr><td>>hg38_21_+_8433172_8446622_RNA45SN1_45S_50ntFlanks@6645.6662.18|CTCGTACGACTCTTAGCG</td><td>>hg38_21_+_8433172_8446622_RNA45SN1_45S_50ntFlanks</td><td>6645</td><td>6662</td><td>18</td><td>CTCGTACGACTCTTAGCG</td><td>rRF</td><td>TRUE</td><td>FALSE</td><td>FALSE</td></tr>
<tr><td>>hg38_21_+_8433172_8446622_RNA45SN1_45S_50ntFlanks@7975.8007.33|CGCGACCTCAGATCAGACGTGGCGACCCGCTGA</td><td>>hg38_21_+_8433172_8446622_RNA45SN1_45S_50ntFlanks</td><td>7975</td><td>8007</td><td>33</td><td>CGCGACCTCAGATCAGACGTGGCGACCCGCTGA</td><td>rRF</td><td>TRUE</td><td>FALSE</td><td>FALSE</td></tr>
<tr><td>>MI0000301|hsa-mir-224&WithFlank&X|-|151958572|151958664@14.35.22&[MIMAT0000281&hsa-miR-224-5p&offsets|+1|-2;m-238&X|-|151958630|151958651&offsets|0|0]|CAAGTCACTAGTGGTTCCGTTT</td><td>>MI0000301|hsa-mir-224&WithFlank&X|-|151958572|151958664</td><td>14</td><td>35</td><td>22</td><td>CAAGTCACTAGTGGTTCCGTTT</td><td>isomiR</td><td>TRUE</td><td>FALSE</td><td>FALSE</td></tr>
<tr><td>>hg38_21_+_8433172_8446622_RNA45SN1_45S_50ntFlanks@10404.10424.21|CATGGGTCAGTCGGTCCTGAG</td><td>>hg38_21_+_8433172_8446622_RNA45SN1_45S_50ntFlanks</td><td>10404</td><td>10424</td><td>21</td><td>CATGGGTCAGTCGGTCCTGAG</td><td>rRF</td><td>TRUE</td><td>TRUE</td><td>FALSE</td></tr>
<tr><td>>MI0000088|hsa-mir-30a&WithFlank&6|-|71403545|71403627@12.34.23(+1A)&[MIMAT0000087&hsa-miR-30a-5p&offsets|0|+1(+1A);m-15&6|-|71403595|71403616&offsets|0|+1(+1A)]|TGTAAACATCCTCGACTGGAAGCA</td><td>>MI0000088|hsa-mir-30a&WithFlank&6|-|71403545|71403627</td><td>12</td><td>34</td><td>23(+1A)</td><td>TGTAAACATCCTCGACTGGAAGCA</td><td>isomiR</td><td>TRUE</td><td>TRUE</td><td>FALSE</td></tr>
<tr><td>>hg38_21_+_8433172_8446622_RNA45SN1_45S_50ntFlanks@10388.10424.37|GTGAACAGCAGTTGAACATGGGTCAGTCGGTCCTGAG</td><td>>hg38_21_+_8433172_8446622_RNA45SN1_45S_50ntFlanks</td><td>10388</td><td>10424</td><td>37</td><td>GTGAACAGCAGTTGAACATGGGTCAGTCGGTCCTGAG</td><td>rRF</td><td>TRUE</td><td>FALSE</td><td>FALSE</td></tr>
<tr><td>>MI0000077|hsa-mir-21&WithFlank&17|+|59841260|59841343@14.34.21(+1G)&[MIMAT0000076&hsa-miR-21-5p&offsets|0|-1(+1G);m-1&17|+|59841273|59841294&offsets|0|-1(+1G)]|TAGCTTATCAGACTGATGTTGG</td><td>>MI0000077|hsa-mir-21&WithFlank&17|+|59841260|59841343</td><td>14</td><td>34</td><td>21(+1G)</td><td>TAGCTTATCAGACTGATGTTGG</td><td>isomiR</td><td>TRUE</td><td>TRUE</td><td>FALSE</td></tr>
<tr><td>>hg38_1_-_228634819_228635039_RNA5S12_5S_50ntFlanks@51.84.34|GTCTACGGCCATACCACCCTGAACGCGCCCGATC</td><td>>hg38_1_-_228634819_228635039_RNA5S12_5S_50ntFlanks</td><td>51</td><td>84</td><td>34</td><td>GTCTACGGCCATACCACCCTGAACGCGCCCGATC</td><td>rRF</td><td>TRUE</td><td>FALSE</td><td>FALSE</td></tr>
<tr><td>>hg38_21_+_8433172_8446622_RNA45SN1_45S_50ntFlanks@5484.5514.31|GAGCGCTGAGAAGACGGTCGAACTTGACTAT</td><td>>hg38_21_+_8433172_8446622_RNA45SN1_45S_50ntFlanks</td><td>5484</td><td>5514</td><td>31</td><td>GAGCGCTGAGAAGACGGTCGAACTTGACTAT</td><td>rRF</td><td>TRUE</td><td>FALSE</td><td>TRUE</td></tr>
<tr><td>>am_MI0000809|hsa-mir-151a&WithFlank&8|-|140732558|140732659@53.73.21(+1U)&[MIMAT0000757&hsa-miR-151a-3p&offsets|0|0(+1U);m-30&8|-|140732587|140732607&offsets|0|0(+1U)]|CTAGACTGAAGCTCCTTGAGGT</td><td>>am_MI0000809|hsa-mir-151a&WithFlank&8|-|140732558|140732659</td><td>53</td><td>73</td><td>21(+1U)</td><td>CTAGACTGAAGCTCCTTGAGGT</td><td>isomiR</td><td>TRUE</td><td>TRUE</td><td>FALSE</td></tr>
<tr><td>>MI0000434|hsa-let-7i&WithFlank&12|+|62603680|62603775@12.33.22(+1U)&[MIMAT0000415&hsa-let-7i-5p&offsets|0|0(+1U);m-49&12|+|62603691|62603712&offsets|0|0(+1U)]|TGAGGTAGTAGTTTGTGCTGTTT</td><td>>MI0000434|hsa-let-7i&WithFlank&12|+|62603680|62603775</td><td>12</td><td>33</td><td>22(+1U)</td><td>TGAGGTAGTAGTTTGTGCTGTTT</td><td>isomiR</td><td>TRUE</td><td>TRUE</td><td>FALSE</td></tr>
<tr><td>>hg38_21_+_8433172_8446622_RNA45SN1_45S_50ntFlanks@10402.10424.23|AACATGGGTCAGTCGGTCCTGAG</td><td>>hg38_21_+_8433172_8446622_RNA45SN1_45S_50ntFlanks</td><td>10402</td><td>10424</td><td>23</td><td>AACATGGGTCAGTCGGTCCTGAG</td><td>rRF</td><td>TRUE</td><td>FALSE</td><td>FALSE</td></tr>
<tr><td>>hg38_21_+_8433172_8446622_RNA45SN1_45S_50ntFlanks@6661.6700.40|CGGTGGATCACTCGGCTCGTGCGTCGATGAAGAACGCAGC</td><td>>hg38_21_+_8433172_8446622_RNA45SN1_45S_50ntFlanks</td><td>6661</td><td>6700</td><td>40</td><td>CGGTGGATCACTCGGCTCGTGCGTCGATGAAGAACGCAGC</td><td>rRF</td><td>TRUE</td><td>FALSE</td><td>FALSE</td></tr>
<tr><td>>trna10_LysCTT_16_+_3241501_3241573@1.36.36,trna13_LysCTT_14_-_58706613_58706685@1.36.36,trna2_LysCTT_15_+_79152904_79152976@1.36.36|GCCCGGCTAGCTCAGTCGGTAGAGCATGGGACTCTT</td><td>>trna10_LysCTT_16_+_3241501_3241573</td><td>1</td><td>36</td><td>36</td><td>GCCCGGCTAGCTCAGTCGGTAGAGCATGGGACTCTT</td><td>tRF</td><td>TRUE</td><td>TRUE</td><td>FALSE</td></tr>
</table>

### Mitochondria

<table>
<tr><th></th><th></th><th></th><th></th><th></th><th></th><th></th><th></th><th></th><th></th></tr>
<tr><td>>MI0000746|hsa-mir-99b&WithFlank&19|+|51692606|51692687@13.33.21&[MIMAT0000689&hsa-miR-99b-5p&offsets|0|-1;m-10&19|+|51692618|51692639&offsets|0|-1]|CACCCGTAGAACCGACCTTGC</td><td>>MI0000746|hsa-mir-99b&WithFlank&19|+|51692606|51692687</td><td>13</td><td>33</td><td>21</td><td>CACCCGTAGAACCGACCTTGC</td><td>isomiR</td><td>TRUE</td><td>TRUE</td><td>FALSE</td></tr>
<tr><td>>hg38_1_-_228634819_228635039_RNA5S12_5S_50ntFlanks@151.170.20|ATACCGGGTGCTGTAGGCTT</td><td>>hg38_1_-_228634819_228635039_RNA5S12_5S_50ntFlanks</td><td>151</td><td>170</td><td>20</td><td>ATACCGGGTGCTGTAGGCTT</td><td>rRF</td><td>TRUE</td><td>FALSE</td><td>TRUE</td></tr>
<tr><td>>MI0003591|hsa-mir-584&WithFlank&5|-|149062307|149062415@22.42.21&[MIMAT0003249&hsa-miR-584-5p&offsets|0|-1;m-167&5|-|149062374|149062394&offsets|0|0]|TTATGGTTTGCCTGGGACTGA</td><td>>MI0003591|hsa-mir-584&WithFlank&5|-|149062307|149062415</td><td>22</td><td>42</td><td>21</td><td>TTATGGTTTGCCTGGGACTGA</td><td>isomiR</td><td>TRUE</td><td>TRUE</td><td>FALSE</td></tr>
<tr><td>>MI0003137|hsa-mir-193b&WithFlank&16|+|14303961|14304055@57.78.22&[MIMAT0002819&hsa-miR-193b-3p&offsets|0|0;m-166&16|+|14304017|14304038&offsets|0|0]|AACTGGCCCTCAAAGTCCCGCT</td><td>>MI0003137|hsa-mir-193b&WithFlank&16|+|14303961|14304055</td><td>57</td><td>78</td><td>22</td><td>AACTGGCCCTCAAAGTCCCGCT</td><td>isomiR</td><td>TRUE</td><td>FALSE</td><td>FALSE</td></tr>
<tr><td>>MI0000086|hsa-mir-28&WithFlank&3|+|188688775|188688872@60.81.22&[MIMAT0004502&hsa-miR-28-3p&offsets|0|0;m-28&3|+|188688834|188688855&offsets|0|0]|CACTAGATTGTGAGCTCCTGGA</td><td>>MI0000086|hsa-mir-28&WithFlank&3|+|188688775|188688872</td><td>60</td><td>81</td><td>22</td><td>CACTAGATTGTGAGCTCCTGGA</td><td>isomiR</td><td>TRUE</td><td>TRUE</td><td>FALSE</td></tr>
<tr><td>>hg38_21_+_8433172_8446622_RNA45SN1_45S_50ntFlanks@475.498.24|CTTCGTGATCGATGTGGTGACGTC</td><td>>hg38_21_+_8433172_8446622_RNA45SN1_45S_50ntFlanks</td><td>475</td><td>498</td><td>24</td><td>CTTCGTGATCGATGTGGTGACGTC</td><td>rRF</td><td>TRUE</td><td>TRUE</td><td>FALSE</td></tr>
<tr><td>>MI0001445|hsa-mir-423&WithFlank&17|+|30117073|30117178@23.46.24&[MIMAT0004748&hsa-miR-423-5p&offsets|0|+1;m-141&17|+|30117095|30117117&offsets|0|+1]|TGAGGGGCAGAGAGCGAGACTTTT</td><td>>MI0001445|hsa-mir-423&WithFlank&17|+|30117073|30117178</td><td>23</td><td>46</td><td>24</td><td>TGAGGGGCAGAGAGCGAGACTTTT</td><td>isomiR</td><td>TRUE</td><td>FALSE</td><td>FALSE</td></tr>
<tr><td>>hg38_21_+_8433172_8446622_RNA45SN1_45S_50ntFlanks@10398.10424.27|GTTGAACATGGGTCAGTCGGTCCTGAG</td><td>>hg38_21_+_8433172_8446622_RNA45SN1_45S_50ntFlanks</td><td>10398</td><td>10424</td><td>27</td><td>GTTGAACATGGGTCAGTCGGTCCTGAG</td><td>rRF</td><td>TRUE</td><td>FALSE</td><td>FALSE</td></tr>
<tr><td>>hg38_21_+_8433172_8446622_RNA45SN1_45S_50ntFlanks@6652.6689.38|GACTCTTAGCGGTGGATCACTCGGCTCGTGCGTCGATG</td><td>>hg38_21_+_8433172_8446622_RNA45SN1_45S_50ntFlanks</td><td>6652</td><td>6689</td><td>38</td><td>GACTCTTAGCGGTGGATCACTCGGCTCGTGCGTCGATG</td><td>rRF</td><td>TRUE</td><td>FALSE</td><td>TRUE</td></tr>
<tr><td>>hg38_21_+_8433172_8446622_RNA45SN1_45S_50ntFlanks@7974.8005.32|ACGCGACCTCAGATCAGACGTGGCGACCCGCT</td><td>>hg38_21_+_8433172_8446622_RNA45SN1_45S_50ntFlanks</td><td>7974</td><td>8005</td><td>32</td><td>ACGCGACCTCAGATCAGACGTGGCGACCCGCT</td><td>rRF</td><td>TRUE</td><td>FALSE</td><td>FALSE</td></tr>
<tr><td>>am_MI0000786|hsa-mir-378a&WithFlank&5|+|149732819|149732896@49.70.22(+1U)&[MIMAT0000732&hsa-miR-378a-3p&offsets|0|0(+1U);m-44&5|+|149732867|149732888&offsets|0|0(+1U)]|ACTGGACTTGGAGTCAGAAGGCT</td><td>>am_MI0000786|hsa-mir-378a&WithFlank&5|+|149732819|149732896</td><td>49</td><td>70</td><td>22(+1U)</td><td>ACTGGACTTGGAGTCAGAAGGCT</td><td>isomiR</td><td>TRUE</td><td>TRUE</td><td>FALSE</td></tr>
<tr><td>>MI0000063|hsa-let-7b&WithFlank&22|+|46113680|46113774@12.33.22(+1A)&[MIMAT0000063&hsa-let-7b-5p&offsets|0|0(+1A);m-14&22|+|46113691|46113712&offsets|0|0(+1A)]|TGAGGTAGTAGGTTGTGTGGTTA</td><td>>MI0000063|hsa-let-7b&WithFlank&22|+|46113680|46113774</td><td>12</td><td>33</td><td>22(+1A)</td><td>TGAGGTAGTAGGTTGTGTGGTTA</td><td>isomiR</td><td>TRUE</td><td>FALSE</td><td>FALSE</td></tr>
<tr><td>>MI0000266|hsa-mir-10a&WithFlank&17|-|48579832|48579953@28.50.23&[MIMAT0000253&hsa-miR-10a-5p&offsets|0|0;m-18&17|-|48579905|48579926&offsets|0|+1]|TACCCTGTAGATCCGAATTTGTG</td><td>>MI0000266|hsa-mir-10a&WithFlank&17|-|48579832|48579953</td><td>28</td><td>50</td><td>23</td><td>TACCCTGTAGATCCGAATTTGTG</td><td>isomiR</td><td>TRUE</td><td>TRUE</td><td>FALSE</td></tr>
<tr><td>>hg38_21_+_8433172_8446622_RNA45SN1_45S_50ntFlanks@6647.6664.18|CGTACGACTCTTAGCGGT</td><td>>hg38_21_+_8433172_8446622_RNA45SN1_45S_50ntFlanks</td><td>6647</td><td>6664</td><td>18</td><td>CGTACGACTCTTAGCGGT</td><td>rRF</td><td>TRUE</td><td>FALSE</td><td>FALSE</td></tr>
<tr><td>>hg38_21_+_8433172_8446622_RNA45SN1_45S_50ntFlanks@10395.10424.30|GCAGTTGAACATGGGTCAGTCGGTCCTGAG</td><td>>hg38_21_+_8433172_8446622_RNA45SN1_45S_50ntFlanks</td><td>10395</td><td>10424</td><td>30</td><td>GCAGTTGAACATGGGTCAGTCGGTCCTGAG</td><td>rRF</td><td>TRUE</td><td>FALSE</td><td>FALSE</td></tr>
<tr><td>>hg38_21_+_8433172_8446622_RNA45SN1_45S_50ntFlanks@12997.13039.43|CGCTGCGATCTATTGAAAGTCAGCCCTCGACACAAGGGTTTGT</td><td>>hg38_21_+_8433172_8446622_RNA45SN1_45S_50ntFlanks</td><td>12997</td><td>13039</td><td>43</td><td>CGCTGCGATCTATTGAAAGTCAGCCCTCGACACAAGGGTTTGT</td><td>rRF</td><td>TRUE</td><td>TRUE</td><td>FALSE</td></tr>
<tr><td>>hg38_21_+_8433172_8446622_RNA45SN1_45S_50ntFlanks@8355.8375.21|TGAAAAGAACTTTGAAGAGAG</td><td>>hg38_21_+_8433172_8446622_RNA45SN1_45S_50ntFlanks</td><td>8355</td><td>8375</td><td>21</td><td>TGAAAAGAACTTTGAAGAGAG</td><td>rRF</td><td>TRUE</td><td>FALSE</td><td>FALSE</td></tr>
<tr><td>>hg38_1_-_228634819_228635039_RNA5S12_5S_50ntFlanks@133.171.39|ATGGGAGACCGCCTGGGAATACCGGGTGCTGTAGGCTTT</td><td>>hg38_1_-_228634819_228635039_RNA5S12_5S_50ntFlanks</td><td>133</td><td>171</td><td>39</td><td>ATGGGAGACCGCCTGGGAATACCGGGTGCTGTAGGCTTT</td><td>rRF</td><td>TRUE</td><td>FALSE</td><td>FALSE</td></tr>
<tr><td>>am_MI0000809|hsa-mir-151a&WithFlank&8|-|140732558|140732659@53.72.20&[MIMAT0000757&hsa-miR-151a-3p&offsets|0|-1;m-30&8|-|140732587|140732607&offsets|0|-1]|CTAGACTGAAGCTCCTTGAG</td><td>>am_MI0000809|hsa-mir-151a&WithFlank&8|-|140732558|140732659</td><td>53</td><td>72</td><td>20</td><td>CTAGACTGAAGCTCCTTGAG</td><td>isomiR</td><td>TRUE</td><td>FALSE</td><td>FALSE</td></tr>
<tr><td>>MI0003560|hsa-mir-92b&WithFlank&1|+|155195171|155195278@67.88.22&[MIMAT0003218&hsa-miR-92b-3p&offsets|0|0;m-115&1|+|155195237|155195258&offsets|0|0]|TATTGCACTCGTCCCGGCCTCC</td><td>>MI0003560|hsa-mir-92b&WithFlank&1|+|155195171|155195278</td><td>67</td><td>88</td><td>22</td><td>TATTGCACTCGTCCCGGCCTCC</td><td>isomiR</td><td>TRUE</td><td>TRUE</td><td>FALSE</td></tr>
<tr><td>>hg38_21_+_8433172_8446622_RNA45SN1_45S_50ntFlanks@7974.8010.37|ACGCGACCTCAGATCAGACGTGGCGACCCGCTGAATT</td><td>>hg38_21_+_8433172_8446622_RNA45SN1_45S_50ntFlanks</td><td>7974</td><td>8010</td><td>37</td><td>ACGCGACCTCAGATCAGACGTGGCGACCCGCTGAATT</td><td>rRF</td><td>TRUE</td><td>FALSE</td><td>FALSE</td></tr>
<tr><td>>trna116_GluCTC_1_-_145399233_145399304@1.31.31,trna59_GluCTC_1_+_249168447_249168518@1.31.31,trna71_GluCTC_1_-_161439189_161439260@1.31.31,trna74_GluCTC_1_-_161431809_161431880@1.31.31,trna77_GluCTC_1_-_161424398_161424469@1.31.31,trna77_GluCTC_6_+_28949976_28950047@1.31.31,trna80_GluCTC_1_-_161417018_161417089@1.31.31,trna87_GluCTC_6_-_126101393_126101464@1.31.31|TCCCTGGTGGTCTAGTGGTTAGGATTCGGCG</td><td>>trna116_GluCTC_1_-_145399233_145399304</td><td>1</td><td>31</td><td>31</td><td>TCCCTGGTGGTCTAGTGGTTAGGATTCGGCG</td><td>tRF</td><td>TRUE</td><td>FALSE</td><td>TRUE</td></tr>
<tr><td>>MI0001445|hsa-mir-423&WithFlank&17|+|30117073|30117178@23.44.22&[MIMAT0004748&hsa-miR-423-5p&offsets|0|-1;m-141&17|+|30117095|30117117&offsets|0|-1]|TGAGGGGCAGAGAGCGAGACTT</td><td>>MI0001445|hsa-mir-423&WithFlank&17|+|30117073|30117178</td><td>23</td><td>44</td><td>22</td><td>TGAGGGGCAGAGAGCGAGACTT</td><td>isomiR</td><td>TRUE</td><td>FALSE</td><td>FALSE</td></tr>
<tr><td>>MI0000264|hsa-mir-7-2&WithFlank&15|+|88611819|88611940@38.60.23;MI0000265|hsa-mir-7-3&WithFlank&19|+|4770664|4770785@37.59.23;MI0000263|hsa-mir-7-1&WithFlank&9|-|83969742|83969863@30.52.23&[MIMAT0000252&hsa-miR-7-5p&offsets|0|-1;m-400&15|+|88611856|88611879&offsets|0|-1];[MIMAT0000252_1&hsa-miR-7-5p&offsets|0|-1;m-402&19|+|4770700|4770723&offsets|0|-1];[MIMAT0000252_2&hsa-miR-7-5p&offsets|0|-1;m-401&9|-|83969811|83969834&offsets|0|-1]|TGGAAGACTAGTGATTTTGTTGT</td><td>>MI0000264|hsa-mir-7-2&WithFlank&15|+|88611819|88611940</td><td>38</td><td>60</td><td>23</td><td>TGGAAGACTAGTGATTTTGTTGT</td><td>isomiR</td><td>TRUE</td><td>TRUE</td><td>FALSE</td></tr>
<tr><td>>trna10_LysCTT_16_+_3241501_3241573@1.35.35,trna13_LysCTT_14_-_58706613_58706685@1.35.35,trna2_LysCTT_15_+_79152904_79152976@1.35.35|GCCCGGCTAGCTCAGTCGGTAGAGCATGGGACTCT</td><td>>trna10_LysCTT_16_+_3241501_3241573</td><td>1</td><td>35</td><td>35</td><td>GCCCGGCTAGCTCAGTCGGTAGAGCATGGGACTCT</td><td>tRF</td><td>TRUE</td><td>FALSE</td><td>FALSE</td></tr>
<tr><td>>MI0000734|hsa-mir-106b&WithFlank&7|-|100093987|100094080@58.79.22&[MIMAT0004672&hsa-miR-106b-3p&offsets|0|0;m-87&7|-|100094003|100094023&offsets|0|+1]|CCGCACTGTGGGTACTTGCTGC</td><td>>MI0000734|hsa-mir-106b&WithFlank&7|-|100093987|100094080</td><td>58</td><td>79</td><td>22</td><td>CCGCACTGTGGGTACTTGCTGC</td><td>isomiR</td><td>TRUE</td><td>FALSE</td><td>FALSE</td></tr>
<tr><td>>MI0000082|hsa-mir-25&WithFlank&7|-|100093554|100093649@58.78.21&[MIMAT0000081&hsa-miR-25-3p&offsets|0|-1;m-23&7|-|100093571|100093592&offsets|0|-1]|CATTGCACTTGTCTCGGTCTG</td><td>>MI0000082|hsa-mir-25&WithFlank&7|-|100093554|100093649</td><td>58</td><td>78</td><td>21</td><td>CATTGCACTTGTCTCGGTCTG</td><td>isomiR</td><td>TRUE</td><td>TRUE</td><td>FALSE</td></tr>
</table>

### Mitochondria

| | | | | | | | | | |
|---|---|---|---|---|---|---|---|---|---|
| >hg38_1_-_228634819_228635039_RNA5S12_5S_50ntFlanks@150.170.21\|AATACCGGGTGCTGTAGGCTT | >hg38_1_-_228634819_228635039_RNA5S12_5S_50ntFlanks | 150 | 170 | 21 | AATACCGGGTGCTGTAGGCTT | rRF | TRUE | FALSE | TRUE |
| >MI0000081\|hsa-mir-24-2&WithFlank&19\|-\|13836281\|13836365@56.76.21;MI0000080\|hsa-mir-24-1&WithFlank&9\|+\|95086015\|95086094@50.70.21&[MIMAT0000080&hsa-miR-24-3p&offsets\|0\|-1;m-36&19\|-\|13836288\|13836310&offsets\|0\|-2];[MIMAT0000080_1&hsa-miR-24-3p&offsets\|0\|-1;m-35&9\|+\|95086064\|95086086&offsets\|0\|-2]\|TGGCTCAGTTCAGCAGGAACA | >MI0000081\|hsa-mir-24-2&WithFlank&19\|-\|13836281\|13836365 | 56 | 76 | 21 | TGGCTCAGTTCAGCAGGAACA | isomiR | TRUE | TRUE | FALSE |
| >hg38_21_+_8433172_8446622_RNA45SN1_45S_50ntFlanks@6645.6664.20\|CTCGTACGACTCTTAGCGGT | >hg38_21_+_8433172_8446622_RNA45SN1_45S_50ntFlanks | 6645 | 6664 | 20 | CTCGTACGACTCTTAGCGGT | rRF | TRUE | FALSE | FALSE |
| >MI0005756\|hsa-mir-934&WithFlank&X\|+\|136550872\|136550966@21.41.21&[MIMAT0004977&hsa-miR-934&offsets\|0\|-1;m-943&X\|+\|136550892\|136550913&offsets\|0\|-1]\|TGTCTACTACTGGAGACACTG | >MI0005756\|hsa-mir-934&WithFlank&X\|+\|136550872\|136550966 | 21 | 41 | 21 | TGTCTACTACTGGAGACACTG | isomiR | TRUE | FALSE | FALSE |
| >hg38_21_+_8433172_8446622_RNA45SN1_45S_50ntFlanks@10400.10424.25\|TGAACATGGGTCAGTCGGTCCTGAG | >hg38_21_+_8433172_8446622_RNA45SN1_45S_50ntFlanks | 10400 | 10424 | 25 | TGAACATGGGTCAGTCGGTCCTGAG | rRF | TRUE | FALSE | FALSE |
| >MI0000079\|hsa-mir-23a&WithFlank&19\|-\|13836581\|13836665@54.71.18&[MIMAT0000078&hsa-miR-23a-3p&offsets\|+3\|0;m-31&19\|-\|13836595\|13836615&offsets\|+3\|0]\|ACATTGCCAGGGATTTCC | >MI0000079\|hsa-mir-23a&WithFlank&19\|-\|13836581\|13836665 | 54 | 71 | 18 | ACATTGCCAGGGATTTCC | isomiR | TRUE | TRUE | FALSE |
| >multi-hg38_MT_+_598_1651_RNR1_12S_50ntFlanks@52.74.23\|ATAGGTTTGGTCCTAGCCTTTCT | >multi-hg38_MT_+_598_1651_RNR1_12S_50ntFlanks | 52 | 74 | 23 | ATAGGTTTGGTCCTAGCCTTTCT | rRF | TRUE | FALSE | FALSE |
| >MI0000542\|hsa-mir-320a&WithFlank&8\|-\|22244960\|22245043@48.68.21(+2U)&[MIMAT0000510&hsa-miR-320a-3p&offsets\|0\|-1(+2U);m-60&8\|-\|22244975\|22244996&offsets\|0\|-1(+2U)]\|AAAAGCTGGGTTGAGAGGGCGTT | >MI0000542\|hsa-mir-320a&WithFlank&8\|-\|22244960\|22245043 | 48 | 68 | 21(+2U) | AAAAGCTGGGTTGAGAGGGCGTT | isomiR | TRUE | FALSE | FALSE |
| >MI0003643\|hsa-mir-629&WithFlank&15\|-\|70079366\|70079474@28.48.21&[MIMAT0004810&hsa-miR-629-5p&offsets\|0\|0;m-150&15\|-\|70079427\|70079447&offsets\|0\|0]\|TGGGTTTACGTTGGGAGAACT | >MI0003643\|hsa-mir-629&WithFlank&15\|-\|70079366\|70079474 | 28 | 48 | 21 | TGGGTTTACGTTGGGAGAACT | isomiR | TRUE | FALSE | FALSE |
| >am_MI0000786\|hsa-mir-378a&WithFlank&5\|+\|149732819\|149732896@49.70.22(+1A)&[MIMAT0000732&hsa-miR-378a-3p&offsets\|0\|0(+1A);m-44&5\|+\|149732867\|149732888&offsets\|0\|0(+1A)]\|ACTGGACTTGGAGTCAGAAGGCA | >am_MI0000786\|hsa-mir-378a&WithFlank&5\|+\|149732819\|149732896 | 49 | 70 | 22(+1A) | ACTGGACTTGGAGTCAGAAGGCA | isomiR | TRUE | TRUE | FALSE |
| >hg38_21_+_8433172_8446622_RNA45SN1_45S_50ntFlanks@6651.6668.18\|CGACTCTTAGCGGTGGAT | >hg38_21_+_8433172_8446622_RNA45SN1_45S_50ntFlanks | 6651 | 6668 | 18 | CGACTCTTAGCGGTGGAT | rRF | TRUE | FALSE | FALSE |
| >hg38_21_+_8433172_8446622_RNA45SN1_45S_50ntFlanks@8353.8375.23\|GTTGAAAAGAACTTTGAAGAGAG | >hg38_21_+_8433172_8446622_RNA45SN1_45S_50ntFlanks | 8353 | 8375 | 23 | GTTGAAAAGAACTTTGAAGAGAG | rRF | TRUE | FALSE | FALSE |
| >hg38_21_+_8433172_8446622_RNA45SN1_45S_50ntFlanks@6651.6689.39\|CGACTCTTAGCGGTGGATCACTCGGCTCGTGCGTCGATG | >hg38_21_+_8433172_8446622_RNA45SN1_45S_50ntFlanks | 6651 | 6689 | 39 | CGACTCTTAGCGGTGGATCACTCGGCTCGTGCGTCGATG | rRF | TRUE | FALSE | TRUE |
| >hg38_21_+_8433172_8446622_RNA45SN1_45S_50ntFlanks@8352.8375.24\|AGTTGAAAAGAACTTTGAAGAGAG | >hg38_21_+_8433172_8446622_RNA45SN1_45S_50ntFlanks | 8352 | 8375 | 24 | AGTTGAAAAGAACTTTGAAGAGAG | rRF | TRUE | FALSE | FALSE |
| >hg38_1_-_228634819_228635039_RNA5S12_5S_50ntFlanks@130.170.41\|TGGATGGGAGACCGCCTGGGAATACCGGGTGCTGTAGGCTT | >hg38_1_-_228634819_228635039_RNA5S12_5S_50ntFlanks | 130 | 170 | 41 | TGGATGGGAGACCGCCTGGGAATACCGGGTGCTGTAGGCTT | rRF | TRUE | FALSE | FALSE |
| >MI0003205\|hsa-mir-532&WithFlank&X\|+\|50003142\|50003244@26.47.22&[MIMAT0002888&hsa-miR-532-5p&offsets\|0\|0;m-65&X\|+\|50003167\|50003188&offsets\|0\|0]\|CATGCCTTGAGTGTAGGACCGT | >MI0003205\|hsa-mir-532&WithFlank&X\|+\|50003142\|50003244 | 26 | 47 | 22 | CATGCCTTGAGTGTAGGACCGT | isomiR | TRUE | TRUE | FALSE |
| >MI0000542\|hsa-mir-320a&WithFlank&8\|-\|22244960\|22245043@48.68.21(+1C)&[MIMAT0000510&hsa-miR-320a-3p&offsets\|0\|-1(+1C);m-60&8\|-\|22244975\|22244996&offsets\|0\|-1(+1C)]\|AAAAGCTGGGTTGAGAGGGCGC | >MI0000542\|hsa-mir-320a&WithFlank&8\|-\|22244960\|22245043 | 48 | 68 | 21(+1C) | AAAAGCTGGGTTGAGAGGGCGC | isomiR | TRUE | FALSE | FALSE |
| >am_trna116_GluCTC_1_-_145399233_145399304@1.30.30,trna59_GluCTC_1_+_249168447_249168518@1.30.30,trna71_GluCTC_1_-_161439189_161439260@1.30.30,trna74_GluCTC_1_-_161431809_161431880@1.30.30,trna77_GluCTC_1_-_161424398_161424469@1.30.30,trna77_GluCTC_6_+_28949976_28950047@1.30.30,trna80_GluCTC_1_-_161417018_161417089@1.30.30,trna87_GluCTC_6_-_126101393_126101464@1.30.30\|TCCCTGGTGGTCTAGTGGTTAGGATTCGGC | >am_trna116_GluCTC_1_-_145399233_145399304 | 1 | 30 | 30 | TCCCTGGTGGTCTAGTGGTTAGGATTCGGC | tRF | TRUE | FALSE | TRUE |
| >hg38_21_+_8433172_8446622_RNA45SN1_45S_50ntFlanks@10392.10424.33\|ACAGCAGTTGAACATGGGTCAGTCGGTCCTGAG | >hg38_21_+_8433172_8446622_RNA45SN1_45S_50ntFlanks | 10392 | 10424 | 33 | ACAGCAGTTGAACATGGGTCAGTCGGTCCTGAG | rRF | TRUE | FALSE | FALSE |
| >hg38_21_+_8433172_8446622_RNA45SN1_45S_50ntFlanks@5489.5516.28\|CTGAGAAGACGGTCGAACTTGACTATCT | >hg38_21_+_8433172_8446622_RNA45SN1_45S_50ntFlanks | 5489 | 5516 | 28 | CTGAGAAGACGGTCGAACTTGACTATCT | rRF | TRUE | FALSE | FALSE |
| >hg38_1_-_228634819_228635039_RNA5S12_5S_50ntFlanks@146.170.25\|TGGGAATACCGGGTGCTGTAGGCTT | >hg38_1_-_228634819_228635039_RNA5S12_5S_50ntFlanks | 146 | 170 | 25 | TGGGAATACCGGGTGCTGTAGGCTT | rRF | TRUE | FALSE | FALSE |
| >am_MI0005763\|hsa-mir-941-1&WithFlank&20\|+\|63919443\|63919526@53.75.23;MI0005764\|hsa-mir-941-2&WithFlank&20\|+\|63919499\|63919582@53.75.23;MI0005765\|hsa-mir-941-3&WithFlank&20\|+\|63919555\|63919638@53.75.23;MI0005766\|hsa-mir-941-4&WithFlank&20\|+\|63919750\|63919833@53.75.23;MI0031520\|hsa-mir-941-5&WithFlank&20\|+\|63919862\|63919945@53.75.23&[MIMAT0004984&hsa-miR-941&offsets\|0\|0;m-263&20\|+\|63919495\|63919517&offsets\|0\|0];[MIMAT0004984_1&hsa-miR-941&offsets\|0\|0;m-263&20\|+\|63919551\|63919573&offsets\|0\|0];[MIMAT0004984_2&hsa-miR-941&offsets\|0\|0;m-263&20\|+\|63919607\|63919629&offsets\|0\|0];[MIMAT0004984_3&hsa-miR-941&offsets\|0\|0;m-263&20\|+\|63919802\|63919824&offsets\|0\|0];[MIMAT0004984_4&hsa-miR-941&offsets\|0\|0;m-263&20\|+\|63919914\|63919936&offsets\|0\|0]\|CACCCGGCTGTGTGCACATGTGC | >am_MI0005763\|hsa-mir-941-1&WithFlank&20\|+\|63919443\|63919526 | 53 | 75 | 23 | CACCCGGCTGTGTGCACATGTGC | isomiR | TRUE | TRUE | FALSE |
| >MI0000085\|hsa-mir-27a&WithFlank&19\|-\|13836434\|13836523@57.76.20&[MIMAT0000084&hsa-miR-27a-3p&offsets\|0\|-1;m-43&19\|-\|13836447\|13836467&offsets\|0\|-1]\|TTCACAGTGGCTAAGTTCCG | >MI0000085\|hsa-mir-27a&WithFlank&19\|-\|13836434\|13836523 | 57 | 76 | 20 | TTCACAGTGGCTAAGTTCCG | isomiR | TRUE | TRUE | FALSE |
| >hg38_21_+_8433172_8446622_RNA45SN1_45S_50ntFlanks@5491.5516.26\|GAGAAGACGGTCGAACTTGACTATCT | >hg38_21_+_8433172_8446622_RNA45SN1_45S_50ntFlanks | 5491 | 5516 | 26 | GAGAAGACGGTCGAACTTGACTATCT | rRF | TRUE | FALSE | FALSE |
| >hg38_21_+_8433172_8446622_RNA45SN1_45S_50ntFlanks@12996.13039.44\|TCGCTGCGATCTATTGAAAGTCAGCCCTCGACACAAGGGTTTGT | >hg38_21_+_8433172_8446622_RNA45SN1_45S_50ntFlanks | 12996 | 13039 | 44 | TCGCTGCGATCTATTGAAAGTCAGCCCTCGACACAAGGGTTTG | rRF | TRUE | TRUE | FALSE |

### Mitochondria

<table>
<tr><th></th><th></th><th></th><th></th><th></th><th></th><th></th><th></th><th></th><th></th></tr>
<tr><td>>MI0000077|hsa-mir-21&WithFlank&17|+|59841260|59841343@14.34.21(+1U)&[MIMAT0000076&hsa-miR-21-5p&offsets|0|-1(+1U);m-1&17|+|59841273|59841294&offsets|0|-1(+1U)]|TAGCTTATCAGACTGATGTTGT</td><td>>MI0000077|hsa-mir-21&WithFlank&17|+|59841260|59841343</td><td>14</td><td>34</td><td>21(+1U)</td><td>TAGCTTATCAGACTGATGTTGT</td><td>isomiR</td><td>TRUE</td><td>TRUE</td><td>FALSE</td></tr>
<tr><td>>am_MI0000786|hsa-mir-378a&WithFlank&5|+|149732819|149732896@50.70.21&[MIMAT0000732&hsa-miR-378a-3p&offsets|+1|0;m-44&5|+|149732867|149732888&offsets|+1|0]|CTGGACTTGGAGTCAGAAGGC</td><td>>am_MI0000786|hsa-mir-378a&WithFlank&5|+|149732819|149732896</td><td>50</td><td>70</td><td>21</td><td>CTGGACTTGGAGTCAGAAGGC</td><td>isomiR</td><td>TRUE</td><td>TRUE</td><td>FALSE</td></tr>
<tr><td>>am_MI0000786|hsa-mir-378a&WithFlank&5|+|149732819|149732896@49.69.21&[MIMAT0000732&hsa-miR-378a-3p&offsets|0|-1;m-44&5|+|149732867|149732888&offsets|0|-1]|ACTGGACTTGGAGTCAGAAGG</td><td>>am_MI0000786|hsa-mir-378a&WithFlank&5|+|149732819|149732896</td><td>49</td><td>69</td><td>21</td><td>ACTGGACTTGGAGTCAGAAGG</td><td>isomiR</td><td>TRUE</td><td>TRUE</td><td>TRUE</td></tr>
<tr><td>>hg38_21_+_8433172_8446622_RNA45SN1_45S_50ntFlanks@485.505.21|GATGTGGTGACGTCGTGCTCT</td><td>>hg38_21_+_8433172_8446622_RNA45SN1_45S_50ntFlanks</td><td>485</td><td>505</td><td>21</td><td>GATGTGGTGACGTCGTGCTCT</td><td>rRF</td><td>TRUE</td><td>TRUE</td><td>FALSE</td></tr>
<tr><td>>MI0000440|hsa-mir-27b&WithFlank&9|+|95085439|95085547@67.86.20&[MIMAT0000419&hsa-miR-27b-3p&offsets|0|-1;m-39&9|+|95085505|95085525&offsets|0|-1]|TTCACAGTGGCTAAGTTCTG</td><td>>MI0000440|hsa-mir-27b&WithFlank&9|+|95085439|95085547</td><td>67</td><td>86</td><td>20</td><td>TTCACAGTGGCTAAGTTCTG</td><td>isomiR</td><td>TRUE</td><td>TRUE</td><td>FALSE</td></tr>
<tr><td>>multi-hg38_MT_+_1621_3279_RNR2_16S_50ntFlanks@51.74.24|GCTAAACCTAGCCCCAAACCCACT</td><td>>multi-hg38_MT_+_1621_3279_RNR2_16S_50ntFlanks</td><td>51</td><td>74</td><td>24</td><td>GCTAAACCTAGCCCCAAACCCACT</td><td>rRF</td><td>TRUE</td><td>TRUE</td><td>FALSE</td></tr>
<tr><td>>hg38_21_+_8433172_8446622_RNA45SN1_45S_50ntFlanks@8358.8375.18|AAAGAACTTTGAAGAGAG</td><td>>hg38_21_+_8433172_8446622_RNA45SN1_45S_50ntFlanks</td><td>8358</td><td>8375</td><td>18</td><td>AAAGAACTTTGAAGAGAG</td><td>rRF</td><td>TRUE</td><td>FALSE</td><td>FALSE</td></tr>
<tr><td>>MI0000108|hsa-mir-103a-2&WithFlank&20|+|3917488|3917577@54.76.23;MI0000109|hsa-mir-103a-1&WithFlank&5|-|168560890|168560979@54.76.23&[MIMAT0000101&hsa-miR-103a-3p&offsets|0|0;m-12&20|+|3917541|3917563&offsets|0|0];[MIMAT0000101_1&hsa-miR-103a-3p&offsets|0|0;m-13&5|-|168560904|168560926&offsets|0|0;m-44220&5|-|168560908|168560926&offsets|0|+4]|AGCAGCATTGTACAGGGCTATGA</td><td>>MI0000108|hsa-mir-103a-2&WithFlank&20|+|3917488|3917577</td><td>54</td><td>76</td><td>23</td><td>AGCAGCATTGTACAGGGCTATGA</td><td>isomiR</td><td>TRUE</td><td>TRUE</td><td>TRUE</td></tr>
<tr><td>>hg38_1_-_228634819_228635039_RNA5S12_5S_50ntFlanks@154.171.18|CCGGGTGCTGTAGGCTTT</td><td>>hg38_1_-_228634819_228635039_RNA5S12_5S_50ntFlanks</td><td>154</td><td>171</td><td>18</td><td>CCGGGTGCTGTAGGCTTT</td><td>rRF</td><td>TRUE</td><td>FALSE</td><td>TRUE</td></tr>
<tr><td>>MI0000285|hsa-mir-205&WithFlank&1|+|209432127|209432248@40.60.21&[MIMAT0000266&hsa-miR-205-5p&offsets|0|-1;m-351&1|+|209432166|209432187&offsets|0|-1]|TCCTTCATTCCACCGGAGTCT</td><td>>MI0000285|hsa-mir-205&WithFlank&1|+|209432127|209432248</td><td>40</td><td>60</td><td>21</td><td>TCCTTCATTCCACCGGAGTCT</td><td>isomiR</td><td>TRUE</td><td>FALSE</td><td>FALSE</td></tr>
<tr><td>>hg38_21_+_8433172_8446622_RNA45SN1_45S_50ntFlanks@6652.6678.27|GACTCTTAGCGGTGGATCACTCGGCTC</td><td>>hg38_21_+_8433172_8446622_RNA45SN1_45S_50ntFlanks</td><td>6652</td><td>6678</td><td>27</td><td>GACTCTTAGCGGTGGATCACTCGGCTC</td><td>rRF</td><td>TRUE</td><td>FALSE</td><td>FALSE</td></tr>
<tr><td>>MI0000447|hsa-mir-128-1&WithFlank&2|+|135665391|135665484@56.76.21;MI0000727|hsa-mir-128-2&WithFlank&3|+|35744470|35744565@58.78.21&[MIMAT0000424&hsa-miR-128-3p&offsets|0|0;m-108&2|+|135665446|135665466&offsets|0|0];[MIMAT0000424_1&hsa-miR-128-3p&offsets|0|0;m-117&3|+|35744527|35744547&offsets|0|0]|TCACAGTGAACCGGTCTCTTT</td><td>>MI0000447|hsa-mir-128-1&WithFlank&2|+|135665391|135665484</td><td>56</td><td>76</td><td>21</td><td>TCACAGTGAACCGGTCTCTTT</td><td>isomiR</td><td>TRUE</td><td>FALSE</td><td>FALSE</td></tr>
<tr><td>>am_MI0039740|hsa-mir-12136&WithFlank&1|-|632609|632691@7.32.26&[MIMAT0049032&hsa-miR-12136&offsets|0|+8]|GAAAAAGTCATGGAGGCCATGGGGTT</td><td>>am_MI0039740|hsa-mir-12136&WithFlank&1|-|632609|632691</td><td>7</td><td>32</td><td>26</td><td>GAAAAAGTCATGGAGGCCATGGGGTT</td><td>isomiR</td><td>TRUE</td><td>FALSE</td><td>FALSE</td></tr>
<tr><td>>multi-trnaMT_SerTGA_MT_-_7446_7514@1.26.26,trnalookalike6_SerTGA_1_-_567997_568065@1.26.26|GAAAAAGTCATGGAGGCCATGGGGTT</td><td>>multi-trnaMT_SerTGA_MT_-_7446_7514</td><td>1</td><td>26</td><td>26</td><td>GAAAAAGTCATGGAGGCCATGGGGTT</td><td>tRF</td><td>TRUE</td><td>FALSE</td><td>FALSE</td></tr>
<tr><td>>hg38_21_+_8433172_8446622_RNA45SN1_45S_50ntFlanks@12995.13039.45|CTCGCTGCGATCTATTGAAAGTCAGCCCTCGACACAAGGGTTTGT</td><td>>hg38_21_+_8433172_8446622_RNA45SN1_45S_50ntFlanks</td><td>12995</td><td>13039</td><td>45</td><td>CTCGCTGCGATCTATTGAAAGTCAGCCCTCGACACAAGGGTTT</td><td>rRF</td><td>TRUE</td><td>TRUE</td><td>FALSE</td></tr>
<tr><td>>trnaMT_LeuTAA_MT_+_3230_3304@1.30.30|GTTAAGATGGCAGAGCCCGGTAATCGCATA</td><td>>trnaMT_LeuTAA_MT_+_3230_3304</td><td>1</td><td>30</td><td>30</td><td>GTTAAGATGGCAGAGCCCGGTAATCGCATA</td><td>tRF</td><td>TRUE</td><td>FALSE</td><td>FALSE</td></tr>
<tr><td>>hg38_21_+_8433172_8446622_RNA45SN1_45S_50ntFlanks@6734.6772.39|ATTGATCATCGACACTTCGAACGCACTTGCGGCCCCGGG</td><td>>hg38_21_+_8433172_8446622_RNA45SN1_45S_50ntFlanks</td><td>6734</td><td>6772</td><td>39</td><td>ATTGATCATCGACACTTCGAACGCACTTGCGGCCCCGGG</td><td>rRF</td><td>TRUE</td><td>FALSE</td><td>FALSE</td></tr>
<tr><td>>hg38_21_+_8433172_8446622_RNA45SN1_45S_50ntFlanks@6646.6668.23|TCGTACGACTCTTAGCGGTGGAT</td><td>>hg38_21_+_8433172_8446622_RNA45SN1_45S_50ntFlanks</td><td>6646</td><td>6668</td><td>23</td><td>TCGTACGACTCTTAGCGGTGGAT</td><td>rRF</td><td>TRUE</td><td>FALSE</td><td>FALSE</td></tr>
<tr><td>>MI0000809|hsa-mir-151a&WithFlank&8|-|140732558|140732659@53.74.22(+1A)&[MIMAT0000757&hsa-miR-151a-3p&offsets|0|+1(+1A);m-30&8|-|140732587|140732607&offsets|0|+1(+1A)]|CTAGACTGAAGCTCCTTGAGGAA</td><td>>MI0000809|hsa-mir-151a&WithFlank&8|-|140732558|140732659</td><td>53</td><td>74</td><td>22(+1A)</td><td>CTAGACTGAAGCTCCTTGAGGAA</td><td>isomiR</td><td>TRUE</td><td>FALSE</td><td>FALSE</td></tr>
<tr><td>>hg38_21_+_8433172_8446622_RNA45SN1_45S_50ntFlanks@6699.6720.22|GCTAGCTGCGAGAATTAATGTG</td><td>>hg38_21_+_8433172_8446622_RNA45SN1_45S_50ntFlanks</td><td>6699</td><td>6720</td><td>22</td><td>GCTAGCTGCGAGAATTAATGTG</td><td>rRF</td><td>TRUE</td><td>FALSE</td><td>TRUE</td></tr>
<tr><td>>hg38_1_-_228634819_228635039_RNA5S12_5S_50ntFlanks@149.170.22|GAATACCGGGTGCTGTAGGCTT</td><td>>hg38_1_-_228634819_228635039_RNA5S12_5S_50ntFlanks</td><td>149</td><td>170</td><td>22</td><td>GAATACCGGGTGCTGTAGGCTT</td><td>rRF</td><td>TRUE</td><td>FALSE</td><td>TRUE</td></tr>
<tr><td>>hg38_21_+_8433172_8446622_RNA45SN1_45S_50ntFlanks@6651.6679.29|CGACTCTTAGCGGTGGATCACTCGGCTCG</td><td>>hg38_21_+_8433172_8446622_RNA45SN1_45S_50ntFlanks</td><td>6651</td><td>6679</td><td>29</td><td>CGACTCTTAGCGGTGGATCACTCGGCTCG</td><td>rRF</td><td>TRUE</td><td>FALSE</td><td>TRUE</td></tr>
<tr><td>>MI0000088|hsa-mir-30a&WithFlank&6|-|71403545|71403627@12.31.20&[MIMAT0000087&hsa-miR-30a-5p&offsets|0|-2;m-15&6|-|71403595|71403616&offsets|0|-2]|TGTAAACATCCTCGACTGGA</td><td>>MI0000088|hsa-mir-30a&WithFlank&6|-|71403545|71403627</td><td>12</td><td>31</td><td>20</td><td>TGTAAACATCCTCGACTGGA</td><td>isomiR</td><td>TRUE</td><td>TRUE</td><td>FALSE</td></tr>
<tr><td>>multi-hg38_MT_+_598_1651_RNR1_12S_50ntFlanks@52.83.32|ATAGGTTTGGTCCTAGCCTTTCTATTAGCTCT</td><td>>multi-hg38_MT_+_598_1651_RNR1_12S_50ntFlanks</td><td>52</td><td>83</td><td>32</td><td>ATAGGTTTGGTCCTAGCCTTTCTATTAGCTCT</td><td>rRF</td><td>TRUE</td><td>TRUE</td><td>FALSE</td></tr>
<tr><td>>trna10_LysCTT_16_+_3241501_3241573@1.34.34,trna13_LysCTT_14_-_58706613_58706685@1.34.34,trna2_LysCTT_15_+_79152904_79152976@1.34.34|GCCCGGCTAGCTCAGTCGGTAGAGCATGGGACTC</td><td>>trna10_LysCTT_16_+_3241501_3241573</td><td>1</td><td>34</td><td>34</td><td>GCCCGGCTAGCTCAGTCGGTAGAGCATGGGACTC</td><td>tRF</td><td>TRUE</td><td>TRUE</td><td>FALSE</td></tr>
<tr><td>>hg38_21_+_8433172_8446622_RNA45SN1_45S_50ntFlanks@8351.8375.25|AAGTTGAAAAGAACTTTGAAGAGAG</td><td>>hg38_21_+_8433172_8446622_RNA45SN1_45S_50ntFlanks</td><td>8351</td><td>8375</td><td>25</td><td>AAGTTGAAAAGAACTTTGAAGAGAG</td><td>rRF</td><td>TRUE</td><td>FALSE</td><td>FALSE</td></tr>
<tr><td>>am_MI0000809|hsa-mir-151a&WithFlank&8|-|140732558|140732659@53.72.20(+1U)&[MIMAT0000757&hsa-miR-151a-3p&offsets|0|-1(+1U);m-30&8|-|140732587|140732607&offsets|0|-1(+1U)]|CTAGACTGAAGCTCCTTGAGT</td><td>>am_MI0000809|hsa-mir-151a&WithFlank&8|-|140732558|140732659</td><td>53</td><td>72</td><td>20(+1U)</td><td>CTAGACTGAAGCTCCTTGAGT</td><td>isomiR</td><td>TRUE</td><td>TRUE</td><td>FALSE</td></tr>
<tr><td>>hg38_1_-_228634819_228635039_RNA5S12_5S_50ntFlanks@134.171.38|TGGGAGACCGCCTGGGAATACCGGGTGCTGTAGGCTTT</td><td>>hg38_1_-_228634819_228635039_RNA5S12_5S_50ntFlanks</td><td>134</td><td>171</td><td>38</td><td>TGGGAGACCGCCTGGGAATACCGGGTGCTGTAGGCTTT</td><td>rRF</td><td>TRUE</td><td>TRUE</td><td>FALSE</td></tr>
<tr><td>>trnaMT_PheGAA_MT_+_577_647@1.21.21|GTTTATGTAGCTTACCTCCTC</td><td>>trnaMT_PheGAA_MT_+_577_647</td><td>1</td><td>21</td><td>21</td><td>GTTTATGTAGCTTACCTCCTC</td><td>tRF</td><td>TRUE</td><td>FALSE</td><td>FALSE</td></tr>
<tr><td>>MI0000440|hsa-mir-27b&WithFlank&9|+|95085439|95085547@67.86.20(+1U)&[MIMAT0000419&hsa-miR-27b-3p&offsets|0|-1(+1U);m-39&9|+|95085505|95085525&offsets|0|-1(+1U)]|TTCACAGTGGCTAAGTTCTGT</td><td>>MI0000440|hsa-mir-27b&WithFlank&9|+|95085439|95085547</td><td>67</td><td>86</td><td>20(+1U)</td><td>TTCACAGTGGCTAAGTTCTGT</td><td>isomiR</td><td>TRUE</td><td>TRUE</td><td>FALSE</td></tr>
</table>

### Mitochondria

| | | | | | | | | | |
|---|---|---|---|---|---|---|---|---|---|
| >MI0000285\|hsa-mir-205&WithFlank&1\|+\|209432127\|209432248@40.60.21(+1A)&[MIMAT0000266&hsa-miR-205-5p&offsets\|0\|-1(+1A);m-351&1\|+\|209432166\|209432187&offsets\|0\|-1(+1A)]\|TCCTTCATTCCACCGGAGTCTA | >MI0000285\|hsa-mir-205&WithFlank&1\|+\|209432127\|209432248 | 40 | 60 | 21(+1A) | TCCTTCATTCCACCGGAGTCTA | isomiR | TRUE | FALSE | FALSE |
| >hg38_21_+_8433172_8446622_RNA45SN1_45S_50ntFlanks@7975.7996.22\|CGCGACCTCAGATCAGACGTGG | >hg38_21_+_8433172_8446622_RNA45SN1_45S_50ntFlanks | 7975 | 7996 | 22 | CGCGACCTCAGATCAGACGTGG | rRF | TRUE | FALSE | TRUE |
| >hg38_1_-_228634819_228635039_RNA5S12_5S_50ntFlanks@51.89.39\|GTCTACGGCCATACCACCCTGAACGCGCCCGATCTCGTC | >hg38_1_-_228634819_228635039_RNA5S12_5S_50ntFlanks | 51 | 89 | 39 | GTCTACGGCCATACCACCCTGAACGCGCCCGATCTCGTC | rRF | TRUE | TRUE | FALSE |
| >MI0000650\|hsa-mir-200c&WithFlank&12\|+\|6963693\|6963772@50.70.21&[MIMAT0000617&hsa-miR-200c-3p&offsets\|0\|-2;m-54&12\|+\|6963742\|6963764&offsets\|0\|-2]\|TAATACTGCCGGGTAATGATG | >MI0000650\|hsa-mir-200c&WithFlank&12\|+\|6963693\|6963772 | 50 | 70 | 21 | TAATACTGCCGGGTAATGATG | isomiR | TRUE | FALSE | FALSE |
| >MI0000102\|hsa-mir-100&WithFlank&11\|-\|122152223\|122152314@19.40.22&[MIMAT0000098&hsa-miR-100-5p&offsets\|0\|0;m-27&11\|-\|122152276\|122152296&offsets\|0\|+1]\|AACCCGTAGATCCGAACTTGTG | >MI0000102\|hsa-mir-100&WithFlank&11\|-\|122152223\|122152314 | 19 | 40 | 22 | AACCCGTAGATCCGAACTTGTG | isomiR | TRUE | TRUE | FALSE |
| >hg38_1_-_228634819_228635039_RNA5S12_5S_50ntFlanks@131.170.40\|GGATGGGAGACCGCCTGGGAATACCGGGTGCTGTAGGCTT | >hg38_1_-_228634819_228635039_RNA5S12_5S_50ntFlanks | 131 | 170 | 40 | GGATGGGAGACCGCCTGGGAATACCGGGTGCTGTAGGCTT | rRF | TRUE | FALSE | FALSE |
| >MI0000081\|hsa-mir-24-2&WithFlank&19\|-\|13836281\|13836365@56.76.21(+1A);MI0000080\|hsa-mir-24-1&WithFlank&9\|+\|95086015\|95086094@50.70.21(+1A)&[MIMAT0000080&hsa-miR-24-3p&offsets\|0\|-1(+1A);m-36&19\|-\|13836288\|13836310&offsets\|0\|-2(+1A)];[MIMAT0000080_1&hsa-miR-24-3p&offsets\|0\|-1(+1A);m-35&9\|+\|95086064\|95086086&offsets\|0\|-2(+1A)]\|TGGCTCAGTTCAGCAGGAACAA | >MI0000081\|hsa-mir-24-2&WithFlank&19\|-\|13836281\|13836365 | 56 | 76 | 21(+1A) | TGGCTCAGTTCAGCAGGAACAA | isomiR | TRUE | TRUE | FALSE |
| >MI0003776\|hsa-mir-320b-1&WithFlank&1\|+\|116671740\|116671823@48.68.21;MI0003839\|hsa-mir-320b-2&WithFlank&1\|-\|224257034\|224257116@47.67.21&[MIMAT0005792&hsa-miR-320b&offsets\|0\|-1;m-183&1\|+\|116671787\|116671806&offsets\|0\|+1];[MIMAT0005792_1&hsa-miR-320b&offsets\|0\|-1;m-183&1\|-\|224257051\|224257070&offsets\|0\|+1]\|AAAAGCTGGGTTGAGAGGGCA | >MI0003776\|hsa-mir-320b-1&WithFlank&1\|+\|116671740\|116671823 | 48 | 68 | 21 | AAAAGCTGGGTTGAGAGGGCA | isomiR | TRUE | FALSE | FALSE |
| >hg38_1_-_228634819_228635039_RNA5S12_5S_50ntFlanks@133.169.37\|ATGGGAGACCGCCTGGGAATACCGGGTGCTGTAGGCT | >hg38_1_-_228634819_228635039_RNA5S12_5S_50ntFlanks | 133 | 169 | 37 | ATGGGAGACCGCCTGGGAATACCGGGTGCTGTAGGCT | rRF | TRUE | FALSE | FALSE |
| >hg38_21_+_8433172_8446622_RNA45SN1_45S_50ntFlanks@8356.8375.20\|GAAAAGAACTTTGAAGAGAG | >hg38_21_+_8433172_8446622_RNA45SN1_45S_50ntFlanks | 8356 | 8375 | 20 | GAAAAGAACTTTGAAGAGAG | rRF | TRUE | FALSE | FALSE |
| >MI0000285\|hsa-mir-205&WithFlank&1\|+\|209432127\|209432248@40.62.23&[MIMAT0000266&hsa-miR-205-5p&offsets\|0\|+1;m-351&1\|+\|209432166\|209432187&offsets\|0\|+1]\|TCCTTCATTCCACCGGAGTCTGT | >MI0000285\|hsa-mir-205&WithFlank&1\|+\|209432127\|209432248 | 40 | 62 | 23 | TCCTTCATTCCACCGGAGTCTGT | isomiR | TRUE | FALSE | TRUE |
| >hg38_21_+_8433172_8446622_RNA45SN1_45S_50ntFlanks@10386.10424.39\|ATGTGAACAGCAGTTGAACATGGGTCAGTCGGTCCTGAG | >hg38_21_+_8433172_8446622_RNA45SN1_45S_50ntFlanks | 10386 | 10424 | 39 | ATGTGAACAGCAGTTGAACATGGGTCAGTCGGTCCTGAG | rRF | TRUE | FALSE | FALSE |
| >multi-hg38_MT_+_598_1651_RNR1_12S_50ntFlanks@52.75.24\|ATAGGTTTGGTCCTAGCCTTTCTA | >multi-hg38_MT_+_598_1651_RNR1_12S_50ntFlanks | 52 | 75 | 24 | ATAGGTTTGGTCCTAGCCTTTCTA | rRF | TRUE | FALSE | FALSE |
| >hg38_21_+_8433172_8446622_RNA45SN1_45S_50ntFlanks@6646.6689.44\|TCGTACGACTCTTAGCGGTGGATCACTCGGCTCGTGCGTCGATG | >hg38_21_+_8433172_8446622_RNA45SN1_45S_50ntFlanks | 6646 | 6689 | 44 | TCGTACGACTCTTAGCGGTGGATCACTCGGCTCGTGCGTCGAT | rRF | TRUE | FALSE | TRUE |
| >trna2_GlyGCC_21_-_18827107_18827177@1.28.28,trna35_GlyGCC_1_+_161413094_161413164@1.28.28,trna37_GlyGCC_1_+_161420467_161420537@1.28.28,trna39_GlyGCC_1_+_161427898_161427968@1.28.28,trna41_GlyGCC_1_+_161435258_161435328@1.28.28\|GCATGGGTGGTTCAGTGGTAGAATTCTC | >trna2_GlyGCC_21_-_18827107_18827177 | 1 | 28 | 28 | GCATGGGTGGTTCAGTGGTAGAATTCTC | tRF | TRUE | FALSE | TRUE |
| >hg38_21_+_8433172_8446622_RNA45SN1_45S_50ntFlanks@6646.6679.34\|TCGTACGACTCTTAGCGGTGGATCACTCGGCTCG | >hg38_21_+_8433172_8446622_RNA45SN1_45S_50ntFlanks | 6646 | 6679 | 34 | TCGTACGACTCTTAGCGGTGGATCACTCGGCTCG | rRF | TRUE | FALSE | FALSE |
| >am_trna10_ValCAC_5_-_180649395_180649467@1.23.23,trna12_ValAAC_5_-_180645270_180645342@1.23.23,trna132_ValAAC_6_-_27721179_27721251@1.23.23,trna136_ValAAC_6_-_27648885_27648957@1.23.23,trna139_ValAAC_6_-_27618707_27618779@1.23.23,trna18_ValCAC_5_-_180529253_180529325@1.23.23,trna2_ValAAC_3_+_169490018_169490090@1.23.23,trna2_ValCAC_5_+_180524070_180524142@1.23.23,trna37_ValAAC_6_+_27203288_27203360@1.23.23,trna4_ValAAC_5_+_180591154_180591226@1.23.23,trna5_ValAAC_5_+_180596610_180596682@1.23.23,trna6_ValCAC_5_+_180600650_180600722@1.23.23,trna85_ValCAC_1_-_161369490_161369562@1.23.23,trna90_ValCAC_1_-_149684088_149684161@1.23.23,trna98_ValCAC_1_-_149298555_149298627@1.23.23,trna9_ValCAC_6_+_26538282_26538354@1.23.23\|GTTTCCGTAGTGTAGTGGTTATC | >am_trna10_ValCAC_5_-_180649395_180649467 | 1 | 23 | 23 | GTTTCCGTAGTGTAGTGGTTATC | tRF | TRUE | FALSE | FALSE |
| >hg38_21_+_8433172_8446622_RNA45SN1_45S_50ntFlanks@6646.6681.36\|TCGTACGACTCTTAGCGGTGGATCACTCGGCTCGTG | >hg38_21_+_8433172_8446622_RNA45SN1_45S_50ntFlanks | 6646 | 6681 | 36 | TCGTACGACTCTTAGCGGTGGATCACTCGGCTCGTG | rRF | TRUE | FALSE | TRUE |
| >hg38_21_+_8433172_8446622_RNA45SN1_45S_50ntFlanks@10387.10424.38\|TGTGAACAGCAGTTGAACATGGGTCAGTCGGTCCTGAG | >hg38_21_+_8433172_8446622_RNA45SN1_45S_50ntFlanks | 10387 | 10424 | 38 | TGTGAACAGCAGTTGAACATGGGTCAGTCGGTCCTGAG | rRF | TRUE | FALSE | FALSE |
| >multi-hg38_MT_+_598_1651_RNR1_12S_50ntFlanks@52.73.22\|ATAGGTTTGGTCCTAGCCTTTC | >multi-hg38_MT_+_598_1651_RNR1_12S_50ntFlanks | 52 | 73 | 22 | ATAGGTTTGGTCCTAGCCTTTC | rRF | TRUE | FALSE | FALSE |
| >MI0000065\|hsa-let-7d&WithFlank&9\|+\|94178828\|94178926@68.89.22&[MIMAT0004484&hsa-let-7d-3p&offsets\|0\|0;m-98&9\|+\|94178895\|94178916&offsets\|0\|0]\|CTATACGACCTGCTGCCTTTCT | >MI0000065\|hsa-let-7d&WithFlank&9\|+\|94178828\|94178926 | 68 | 89 | 22 | CTATACGACCTGCTGCCTTTCT | isomiR | TRUE | TRUE | FALSE |
| >hg38_21_+_8433172_8446622_RNA45SN1_45S_50ntFlanks@4380.4404.25\|CGTAGTTGGATCTTGGGAGCGGGCG | >hg38_21_+_8433172_8446622_RNA45SN1_45S_50ntFlanks | 4380 | 4404 | 25 | CGTAGTTGGATCTTGGGAGCGGGCG | rRF | TRUE | FALSE | FALSE |
| >MI0000088\|hsa-mir-30a&WithFlank&6\|-\|71403545\|71403627@12.33.22&[MIMAT0000087&hsa-miR-30a-5p&offsets\|0\|0;m-15&6\|-\|71403595\|71403616&offsets\|0\|0]\|TGTAAACATCCTCGACTGGAAG | >MI0000088\|hsa-mir-30a&WithFlank&6\|-\|71403545\|71403627 | 12 | 33 | 22 | TGTAAACATCCTCGACTGGAAG | isomiR | TRUE | TRUE | FALSE |
| >hg38_21_+_8433172_8446622_RNA45SN1_45S_50ntFlanks@10240.10259.20\|CTACGCCGCGACGAGTAGGA | >hg38_21_+_8433172_8446622_RNA45SN1_45S_50ntFlanks | 10240 | 10259 | 20 | CTACGCCGCGACGAGTAGGA | rRF | TRUE | FALSE | FALSE |

### Mitochondria

<table>
<tr><td>>am_trna2_GlyGCC_21_-_18827107_18827177@1.25.25,trna35_GlyGCC_1_+_161413094_161413164@1.25.25,trna37_GlyGCC_1_+_161420467_161420537@1.25.25,trna39_GlyGCC_1_+_161427898_161427968@1.25.25,trna41_GlyGCC_1_+_161435258_161435328@1.25.25|GCATGGGTGGTTCAGTGGTAGAATT</td><td>>am_trna2_GlyGCC_21_-_18827107_18827177</td><td>1</td><td>25</td><td>25</td><td>GCATGGGTGGTTCAGTGGTAGAATT</td><td>tRF</td><td>TRUE</td><td>FALSE</td><td>FALSE</td></tr>
<tr><td>>trnaMT_PheGAA_MT_+_577_647@39.56.18|ATGTTTAGACGGGCTCAC</td><td>>trnaMT_PheGAA_MT_+_577_647</td><td>39</td><td>56</td><td>18</td><td>ATGTTTAGACGGGCTCAC</td><td>tRF</td><td>TRUE</td><td>FALSE</td><td>FALSE</td></tr>
<tr><td>>MI0003778|hsa-mir-320c-1&WithFlank&18|+|21683512|21683595@48.67.20;MI0008191|hsa-mir-320c-2&WithFlank&18|+|24321669|24321752@48.67.20&[MIMAT0005793&hsa-miR-320c&offsets|0|0;m-222&18|+|21683559|21683578&offsets|0|0];[MIMAT0005793_1&hsa-miR-320c&offsets|0|0;m-219&18|+|24321716|24321735&offsets|0|0]|AAAAGCTGGGTTGAGAGGGT</td><td>>MI0003778|hsa-mir-320c-1&WithFlank&18|+|21683512|21683595</td><td>48</td><td>67</td><td>20</td><td>AAAAGCTGGGTTGAGAGGGT</td><td>isomiR</td><td>TRUE</td><td>FALSE</td><td>FALSE</td></tr>
<tr><td>>hg38_21_+_8433172_8446622_RNA45SN1_45S_50ntFlanks@6646.6677.32|TCGTACGACTCTTAGCGGTGGATCACTCGGCT</td><td>>hg38_21_+_8433172_8446622_RNA45SN1_45S_50ntFlanks</td><td>6646</td><td>6677</td><td>32</td><td>TCGTACGACTCTTAGCGGTGGATCACTCGGCT</td><td>rRF</td><td>TRUE</td><td>FALSE</td><td>FALSE</td></tr>
<tr><td>>MI0000279|hsa-mir-196a-2&WithFlank&12|+|53991732|53991853@31.52.22;MI0000238|hsa-mir-196a-1&WithFlank&17|-|48632484|48632565@13.34.22&[MIMAT0000226&hsa-miR-196a-5p&offsets|0|0;m-261&12|+|53991762|53991783&offsets|0|0];[MIMAT0000226_1&hsa-miR-196a-5p&offsets|0|0;m-268&17|-|48632532|48632553&offsets|0|0]|TAGGTAGTTTCATGTTGTTGGG</td><td>>MI0000279|hsa-mir-196a-2&WithFlank&12|+|53991732|53991853</td><td>31</td><td>52</td><td>22</td><td>TAGGTAGTTTCATGTTGTTGGG</td><td>isomiR</td><td>TRUE</td><td>FALSE</td><td>FALSE</td></tr>
<tr><td>>hg38_1_-_228634819_228635039_RNA5S12_5S_50ntFlanks@135.171.37|GGGAGACCGCCTGGGAATACCGGGTGCTGTAGGCTTT</td><td>>hg38_1_-_228634819_228635039_RNA5S12_5S_50ntFlanks</td><td>135</td><td>171</td><td>37</td><td>GGGAGACCGCCTGGGAATACCGGGTGCTGTAGGCTTT</td><td>rRF</td><td>TRUE</td><td>TRUE</td><td>FALSE</td></tr>
<tr><td>>MI0000255|hsa-mir-30d&WithFlank&8|-|134804870|134804951@12.33.22(+1U)&[MIMAT0000245&hsa-miR-30d-5p&offsets|0|0(+1U);m-21&8|-|134804919|134804940&offsets|0|0(+1U)]|TGTAAACATCCCCGACTGGAAGT</td><td>>MI0000255|hsa-mir-30d&WithFlank&8|-|134804870|134804951</td><td>12</td><td>33</td><td>22(+1U)</td><td>TGTAAACATCCCCGACTGGAAGT</td><td>isomiR</td><td>TRUE</td><td>FALSE</td><td>FALSE</td></tr>
<tr><td>>multi-hg38_MT_+_1621_3279_RNR2_16S_50ntFlanks@51.88.38|GCTAAACCTAGCCCCAAACCCACTCCACCTTACTACCA</td><td>>multi-hg38_MT_+_1621_3279_RNR2_16S_50ntFlanks</td><td>51</td><td>88</td><td>38</td><td>GCTAAACCTAGCCCCAAACCCACTCCACCTTACTACCA</td><td>rRF</td><td>TRUE</td><td>TRUE</td><td>FALSE</td></tr>
<tr><td>>MI0000299|hsa-mir-222&WithFlank&X|-|45747009|45747130@75.95.21&[MIMAT0000279&hsa-miR-222-3p&offsets|0|0;m-140&X|-|45747033|45747056&offsets|0|-3]|AGCTACATCTGGCTACTGGGT</td><td>>MI0000299|hsa-mir-222&WithFlank&X|-|45747009|45747130</td><td>75</td><td>95</td><td>21</td><td>AGCTACATCTGGCTACTGGGT</td><td>isomiR</td><td>TRUE</td><td>TRUE</td><td>FALSE</td></tr>
<tr><td>>multi-hg38_MT_+_598_1651_RNR1_12S_50ntFlanks@52.81.30|ATAGGTTTGGTCCTAGCCTTTCTATTAGCT</td><td>>multi-hg38_MT_+_598_1651_RNR1_12S_50ntFlanks</td><td>52</td><td>81</td><td>30</td><td>ATAGGTTTGGTCCTAGCCTTTCTATTAGCT</td><td>rRF</td><td>TRUE</td><td>FALSE</td><td>FALSE</td></tr>
<tr><td>>MI0000434|hsa-let-7i&WithFlank&12|+|62603680|62603775@12.32.21&[MIMAT0000415&hsa-let-7i-5p&offsets|0|-1;m-49&12|+|62603691|62603712&offsets|0|-1]|TGAGGTAGTAGTTTGTGCTGT</td><td>>MI0000434|hsa-let-7i&WithFlank&12|+|62603680|62603775</td><td>12</td><td>32</td><td>21</td><td>TGAGGTAGTAGTTTGTGCTGT</td><td>isomiR</td><td>TRUE</td><td>TRUE</td><td>FALSE</td></tr>
<tr><td>>hg38_21_+_8433172_8446622_RNA45SN1_45S_50ntFlanks@6652.6677.26|GACTCTTAGCGGTGGATCACTCGGCT</td><td>>hg38_21_+_8433172_8446622_RNA45SN1_45S_50ntFlanks</td><td>6652</td><td>6677</td><td>26</td><td>GACTCTTAGCGGTGGATCACTCGGCT</td><td>rRF</td><td>TRUE</td><td>FALSE</td><td>FALSE</td></tr>
<tr><td>>MI0000465|hsa-mir-191&WithFlank&3|-|49020612|49020715@22.43.22&[MIMAT0000440&hsa-miR-191-5p&offsets|0|-1;m-48&3|-|49020672|49020694&offsets|0|-1]|CAACGGAATCCCAAAAGCAGCT</td><td>>MI0000465|hsa-mir-191&WithFlank&3|-|49020612|49020715</td><td>22</td><td>43</td><td>22</td><td>CAACGGAATCCCAAAAGCAGCT</td><td>isomiR</td><td>TRUE</td><td>TRUE</td><td>FALSE</td></tr>
<tr><td>>hg38_1_-_228634819_228635039_RNA5S12_5S_50ntFlanks@51.91.41|GTCTACGGCCATACCACCCTGAACGCGCCCGATCTCGTCTG</td><td>>hg38_1_-_228634819_228635039_RNA5S12_5S_50ntFlanks</td><td>51</td><td>91</td><td>41</td><td>GTCTACGGCCATACCACCCTGAACGCGCCCGATCTCGTCTG</td><td>rRF</td><td>TRUE</td><td>FALSE</td><td>FALSE</td></tr>
<tr><td>>MI0000542|hsa-mir-320a&WithFlank&8|-|22244960|22245043@49.69.21&[MIMAT0000510&hsa-miR-320a-3p&offsets|+1|0;m-60&8|-|22244975|22244996&offsets|+1|0]|AAAGCTGGGTTGAGAGGGCGA</td><td>>MI0000542|hsa-mir-320a&WithFlank&8|-|22244960|22245043</td><td>49</td><td>69</td><td>21</td><td>AAAGCTGGGTTGAGAGGGCGA</td><td>isomiR</td><td>TRUE</td><td>FALSE</td><td>FALSE</td></tr>
<tr><td>>hg38_21_+_8433172_8446622_RNA45SN1_45S_50ntFlanks@8349.8375.27|GAAAGTTGAAAAGAACTTTGAAGAGAG</td><td>>hg38_21_+_8433172_8446622_RNA45SN1_45S_50ntFlanks</td><td>8349</td><td>8375</td><td>27</td><td>GAAAGTTGAAAAGAACTTTGAAGAGAG</td><td>rRF</td><td>TRUE</td><td>FALSE</td><td>FALSE</td></tr>
<tr><td>>hg38_21_+_8433172_8446622_RNA45SN1_45S_50ntFlanks@5492.5516.25|AGAAGACGGTCGAACTTGACTATCT</td><td>>hg38_21_+_8433172_8446622_RNA45SN1_45S_50ntFlanks</td><td>5492</td><td>5516</td><td>25</td><td>AGAAGACGGTCGAACTTGACTATCT</td><td>rRF</td><td>TRUE</td><td>FALSE</td><td>FALSE</td></tr>
<tr><td>>hg38_21_+_8433172_8446622_RNA45SN1_45S_50ntFlanks@6652.6700.49|GACTCTTAGCGGTGGATCACTCGGCTCGTGCGTCGATGAAGAACGCAGC</td><td>>hg38_21_+_8433172_8446622_RNA45SN1_45S_50ntFlanks</td><td>6652</td><td>6700</td><td>49</td><td>GACTCTTAGCGGTGGATCACTCGGCTCGTGCGTCGATGAAGAA</td><td>rRF</td><td>TRUE</td><td>FALSE</td><td>FALSE</td></tr>
<tr><td>>am_MI0000786|hsa-mir-378a&WithFlank&5|+|149732819|149732896@49.69.21(+1A)&[MIMAT0000732&hsa-miR-378a-3p&offsets|0|-1(+1A);m-44&5|+|149732867|149732888&offsets|0|-1(+1A)]|ACTGGACTTGGAGTCAGAAGGA</td><td>>am_MI0000786|hsa-mir-378a&WithFlank&5|+|149732819|149732896</td><td>49</td><td>69</td><td>21(+1A)</td><td>ACTGGACTTGGAGTCAGAAGGA</td><td>isomiR</td><td>TRUE</td><td>TRUE</td><td>FALSE</td></tr>
<tr><td>>trna2_GlyGCC_21_-_18827107_18827177@1.30.30,trna35_GlyGCC_1_+_161413094_161413164@1.30.30,trna37_GlyGCC_1_+_161420467_161420537@1.30.30,trna39_GlyGCC_1_+_161427898_161427968@1.30.30,trna41_GlyGCC_1_+_161435258_161435328@1.30.30|GCATGGGTGGTTCAGTGGTAGAATTCTCGC</td><td>>trna2_GlyGCC_21_-_18827107_18827177</td><td>1</td><td>30</td><td>30</td><td>GCATGGGTGGTTCAGTGGTAGAATTCTCGC</td><td>tRF</td><td>TRUE</td><td>FALSE</td><td>TRUE</td></tr>
<tr><td>>trnaMT_HisGTG_MT_+_12138_12206@32.72.41|TGAATCTGACAACAGAGGCTTACGACCCCTTATTTACCCCA</td><td>>trnaMT_HisGTG_MT_+_12138_12206</td><td>32</td><td>72</td><td>41</td><td>TGAATCTGACAACAGAGGCTTACGACCCCTTATTTACCCCA</td><td>tRF</td><td>TRUE</td><td>FALSE</td><td>FALSE</td></tr>
<tr><td>>hg38_21_+_8433172_8446622_RNA45SN1_45S_50ntFlanks@6645.6663.19|CTCGTACGACTCTTAGCGG</td><td>>hg38_21_+_8433172_8446622_RNA45SN1_45S_50ntFlanks</td><td>6645</td><td>6663</td><td>19</td><td>CTCGTACGACTCTTAGCGG</td><td>rRF</td><td>TRUE</td><td>FALSE</td><td>FALSE</td></tr>
<tr><td>>hg38_21_+_8433172_8446622_RNA45SN1_45S_50ntFlanks@8348.8375.28|GGAAAGTTGAAAAGAACTTTGAAGAGAG</td><td>>hg38_21_+_8433172_8446622_RNA45SN1_45S_50ntFlanks</td><td>8348</td><td>8375</td><td>28</td><td>GGAAAGTTGAAAAGAACTTTGAAGAGAG</td><td>rRF</td><td>TRUE</td><td>FALSE</td><td>FALSE</td></tr>
<tr><td>>hg38_1_-_228634819_228635039_RNA5S12_5S_50ntFlanks@129.170.42|TTGGATGGGAGACCGCCTGGGAATACCGGGTGCTGTAGGCTT</td><td>>hg38_1_-_228634819_228635039_RNA5S12_5S_50ntFlanks</td><td>129</td><td>170</td><td>42</td><td>TTGGATGGGAGACCGCCTGGGAATACCGGGTGCTGTAGGCTT</td><td>rRF</td><td>TRUE</td><td>FALSE</td><td>FALSE</td></tr>
<tr><td>>hg38_21_+_8433172_8446622_RNA45SN1_45S_50ntFlanks@6646.6676.31|TCGTACGACTCTTAGCGGTGGATCACTCGGC</td><td>>hg38_21_+_8433172_8446622_RNA45SN1_45S_50ntFlanks</td><td>6646</td><td>6676</td><td>31</td><td>TCGTACGACTCTTAGCGGTGGATCACTCGGC</td><td>rRF</td><td>TRUE</td><td>FALSE</td><td>FALSE</td></tr>
<tr><td>>hg38_21_+_8433172_8446622_RNA45SN1_45S_50ntFlanks@7974.7992.19|ACGCGACCTCAGATCAGAC</td><td>>hg38_21_+_8433172_8446622_RNA45SN1_45S_50ntFlanks</td><td>7974</td><td>7992</td><td>19</td><td>ACGCGACCTCAGATCAGAC</td><td>rRF</td><td>TRUE</td><td>FALSE</td><td>FALSE</td></tr>
<tr><td>>trnaMT_GluTTC_MT_-_14674_14742@34.72.39,trnalookalike8_GluTTC_5_-_93905172_93905240@34.72.39|ATATCATTGGTCGTGGTTGTAGTCCGTGCGAGAATACCA</td><td>>trnaMT_GluTTC_MT_-_14674_14742</td><td>34</td><td>72</td><td>39</td><td>ATATCATTGGTCGTGGTTGTAGTCCGTGCGAGAATACCA</td><td>tRF</td><td>TRUE</td><td>TRUE</td><td>FALSE</td></tr>
<tr><td>>am_MI0000786|hsa-mir-378a&WithFlank&5|+|149732819|149732896@50.70.21(+1U)&[MIMAT0000732&hsa-miR-378a-3p&offsets|+1|0(+1U);m-44&5|+|149732867|149732888&offsets|+1|0(+1U)]|CTGGACTTGGAGTCAGAAGGCT</td><td>>am_MI0000786|hsa-mir-378a&WithFlank&5|+|149732819|149732896</td><td>50</td><td>70</td><td>21(+1U)</td><td>CTGGACTTGGAGTCAGAAGGCT</td><td>isomiR</td><td>TRUE</td><td>FALSE</td><td>FALSE</td></tr>
<tr><td>>hg38_21_+_8433172_8446622_RNA45SN1_45S_50ntFlanks@7976.8011.36|GCGACCTCAGATCAGACGTGGCGACCCGCTGAATTT</td><td>>hg38_21_+_8433172_8446622_RNA45SN1_45S_50ntFlanks</td><td>7976</td><td>8011</td><td>36</td><td>GCGACCTCAGATCAGACGTGGCGACCCGCTGAATTT</td><td>rRF</td><td>TRUE</td><td>FALSE</td><td>FALSE</td></tr>
</table>

### Mitochondria

| | | | | | | | | | |
|---|---|---|---|---|---|---|---|---|---|
| >hg38_21_+_8433172_8446622_RNA45SN1_45S_50ntFlanks@6661.6688.28\|CGGTGGATCACTCGGCTCGTGCGTCGAT | >hg38_21_+_8433172_8446622_RNA45SN1_45S_50ntFlanks | 6661 | 6688 | 28 | CGGTGGATCACTCGGCTCGTGCGTCGAT | rRF | TRUE | FALSE | FALSE |
| >MI0000650\|hsa-mir-200c&WithFlank&12\|+\|6963693\|6963772@50.72.23(+1A)&[MIMAT0000617&hsa-miR-200c-3p&offsets\|0\|0(+1A);m-54&12\|+\|6963742\|6963764&offsets\|0\|0(+1A)]\|TAATACTGCCGGGTAATGATGGAA | >MI0000650\|hsa-mir-200c&WithFlank&12\|+\|6963693\|6963772 | 50 | 72 | 23(+1A) | TAATACTGCCGGGTAATGATGGAA | isomiR | TRUE | FALSE | FALSE |
| >MI0000078\|hsa-mir-22&WithFlank&17\|-\|1713897\|1713993@59.79.21(+1C)&[MIMAT0000077&hsa-miR-22-3p&offsets\|0\|-1(+1C);m-2&17\|-\|1713914\|1713935&offsets\|0\|-1(+1C)]\|AAGCTGCCAGTTGAAGAACTGC | >MI0000078\|hsa-mir-22&WithFlank&17\|-\|1713897\|1713993 | 59 | 79 | 21(+1C) | AAGCTGCCAGTTGAAGAACTGC | isomiR | TRUE | TRUE | FALSE |
| >MI0000255\|hsa-mir-30d&WithFlank&8\|-\|134804870\|134804951@12.34.23(+1C)&[MIMAT0000245&hsa-miR-30d-5p&offsets\|0\|+1(+1C);m-21&8\|-\|134804919\|134804940&offsets\|0\|+1(+1C)]\|TGTAAACATCCCCGACTGGAAGCC | >MI0000255\|hsa-mir-30d&WithFlank&8\|-\|134804870\|134804951 | 12 | 34 | 23(+1C) | TGTAAACATCCCCGACTGGAAGCC | isomiR | TRUE | TRUE | FALSE |
| >hg38_21_+_8433172_8446622_RNA45SN1_45S_50ntFlanks@8196.8213.18\|CGTGGACGGTGTGAGGCC | >hg38_21_+_8433172_8446622_RNA45SN1_45S_50ntFlanks | 8196 | 8213 | 18 | CGTGGACGGTGTGAGGCC | rRF | TRUE | FALSE | FALSE |
| >hg38_21_+_8433172_8446622_RNA45SN1_45S_50ntFlanks@4297.4316.20\|CATTGGAGGGCAAGTCTGGT | >hg38_21_+_8433172_8446622_RNA45SN1_45S_50ntFlanks | 4297 | 4316 | 20 | CATTGGAGGGCAAGTCTGGT | rRF | TRUE | FALSE | FALSE |
| >hg38_21_+_8433172_8446622_RNA45SN1_45S_50ntFlanks@6646.6680.35\|TCGTACGACTCTTAGCGGTGGATCACTCGGCTCGT | >hg38_21_+_8433172_8446622_RNA45SN1_45S_50ntFlanks | 6646 | 6680 | 35 | TCGTACGACTCTTAGCGGTGGATCACTCGGCTCGT | rRF | TRUE | FALSE | FALSE |
| >hg38_21_+_8433172_8446622_RNA45SN1_45S_50ntFlanks@6645.6666.22\|CTCGTACGACTCTTAGCGGTGG | >hg38_21_+_8433172_8446622_RNA45SN1_45S_50ntFlanks | 6645 | 6666 | 22 | CTCGTACGACTCTTAGCGGTGG | rRF | TRUE | FALSE | FALSE |
| >MI0005559\|hsa-mir-744&WithFlank&17\|+\|12081893\|12082002@17.37.21&[MIMAT0004945&hsa-miR-744-5p&offsets\|0\|-1;m-172&17\|+\|12081909\|12081929&offsets\|0\|0]\|TGCGGGGCTAGGGCTAACAGC | >MI0005559\|hsa-mir-744&WithFlank&17\|+\|12081893\|12082002 | 17 | 37 | 21 | TGCGGGGCTAGGGCTAACAGC | isomiR | TRUE | FALSE | FALSE |
| >hg38_21_+_8433172_8446622_RNA45SN1_45S_50ntFlanks@5483.5514.32\|GGAGCGCTGAGAAGACGGTCGAACTTGACTAT | >hg38_21_+_8433172_8446622_RNA45SN1_45S_50ntFlanks | 5483 | 5514 | 32 | GGAGCGCTGAGAAGACGGTCGAACTTGACTAT | rRF | TRUE | FALSE | FALSE |
| >MI0000077\|hsa-mir-21&WithFlank&17\|+\|59841260\|59841343@14.36.23(+1C)&[MIMAT0000076&hsa-miR-21-5p&offsets\|0\|+1(+1C);m-1&17\|+\|59841273\|59841294&offsets\|0\|+1(+1C)]\|TAGCTTATCAGACTGATGTTGACC | >MI0000077\|hsa-mir-21&WithFlank&17\|+\|59841260\|59841343 | 14 | 36 | 23(+1C) | TAGCTTATCAGACTGATGTTGACC | isomiR | TRUE | TRUE | FALSE |
| >hg38_21_+_8433172_8446622_RNA45SN1_45S_50ntFlanks@8350.8375.26\|AAAGTTGAAAAGAACTTTGAAGAGAG | >hg38_21_+_8433172_8446622_RNA45SN1_45S_50ntFlanks | 8350 | 8375 | 26 | AAAGTTGAAAAGAACTTTGAAGAGAG | rRF | TRUE | FALSE | FALSE |
| >hg38_1_-_228634819_228635039_RNA5S12_5S_50ntFlanks@152.170.19\|TACCGGGTGCTGTAGGCTT | >hg38_1_-_228634819_228635039_RNA5S12_5S_50ntFlanks | 152 | 170 | 19 | TACCGGGTGCTGTAGGCTT | rRF | TRUE | FALSE | TRUE |
| >MI0000070\|hsa-mir-16-1&WithFlank&13\|-\|50048967\|50049067@20.42.23;MI0000115\|hsa-mir-16-2&WithFlank&3\|+\|160404739\|160404831@16.38.23&[MIMAT0000069&hsa-miR-16-5p&offsets\|0\|+1;m-62&13\|-\|50049027\|50049048&offsets\|0\|+1];[MIMAT0000069_1&hsa-miR-16-5p&offsets\|0\|+1;m-61&3\|+\|160404754\|160404775&offsets\|0\|+1]\|TAGCAGCACGTAAATATTGGCGT | >MI0000070\|hsa-mir-16-1&WithFlank&13\|-\|50048967\|50049067 | 20 | 42 | 23 | TAGCAGCACGTAAATATTGGCGT | isomiR | TRUE | TRUE | FALSE |
| >MI0000087\|hsa-mir-29a&WithFlank&7\|-\|130876741\|130876816@47.69.23&[MIMAT0000086&hsa-miR-29a-3p&offsets\|-1\|0;m-26&7\|-\|130876749\|130876769&offsets\|-1\|+1]\|CTAGCACCATCTGAAATCGGTTA | >MI0000087\|hsa-mir-29a&WithFlank&7\|-\|130876741\|130876816 | 47 | 69 | 23 | CTAGCACCATCTGAAATCGGTTA | isomiR | TRUE | TRUE | FALSE |
| >MI0000650\|hsa-mir-200c&WithFlank&12\|+\|6963693\|6963772@50.70.21(+1A)&[MIMAT0000617&hsa-miR-200c-3p&offsets\|0\|-2(+1A);m-54&12\|+\|6963742\|6963764&offsets\|0\|-2(+1A)]\|TAATACTGCCGGGTAATGATGA | >MI0000650\|hsa-mir-200c&WithFlank&12\|+\|6963693\|6963772 | 50 | 70 | 21(+1A) | TAATACTGCCGGGTAATGATGA | isomiR | TRUE | FALSE | FALSE |
| >MI0000471\|hsa-mir-126&WithFlank&9\|+\|136670596\|136670692@58.79.22&[MIMAT0000445&hsa-miR-126-3p&offsets\|0\|0;m-40&9\|+\|136670653\|136670674&offsets\|0\|0]\|TCGTACCGTGAGTAATAATGCG | >MI0000471\|hsa-mir-126&WithFlank&9\|+\|136670596\|136670692 | 58 | 79 | 22 | TCGTACCGTGAGTAATAATGCG | isomiR | TRUE | TRUE | FALSE |
| >hg38_21_+_8433172_8446622_RNA45SN1_45S_50ntFlanks@6652.6676.25\|GACTCTTAGCGGTGGATCACTCGGC | >hg38_21_+_8433172_8446622_RNA45SN1_45S_50ntFlanks | 6652 | 6676 | 25 | GACTCTTAGCGGTGGATCACTCGGC | rRF | TRUE | FALSE | FALSE |
| >MI0000065\|hsa-let-7d&WithFlank&9\|+\|94178828\|94178926@14.35.22&[MIMAT0000065&hsa-let-7d-5p&offsets\|0\|0;m-90&9\|+\|94178841\|94178862&offsets\|0\|0]\|AGAGGTAGTAGGTTGCATAGTT | >MI0000065\|hsa-let-7d&WithFlank&9\|+\|94178828\|94178926 | 14 | 35 | 22 | AGAGGTAGTAGGTTGCATAGTT | isomiR | TRUE | TRUE | FALSE |
| >MI0000255\|hsa-mir-30d&WithFlank&8\|-\|134804870\|134804951@12.31.20&[MIMAT0000245&hsa-miR-30d-5p&offsets\|0\|-2;m-21&8\|-\|134804919\|134804940&offsets\|0\|-2]\|TGTAAACATCCCCGACTGGA | >MI0000255\|hsa-mir-30d&WithFlank&8\|-\|134804870\|134804951 | 12 | 31 | 20 | TGTAAACATCCCCGACTGGA | isomiR | TRUE | FALSE | FALSE |
| >am_trna2_GlyGCC_21_-_18827107_18827177@1.27.27,trna35_GlyGCC_1_+_161413094_161413164@1.27.27,trna37_GlyGCC_1_+_161420467_161420537@1.27.27,trna39_GlyGCC_1_+_161427898_161427968@1.27.27,trna41_GlyGCC_1_+_161435258_161435328@1.27.27\|GCATGGGTGGTTCAGTGGTAGAATTCT | >am_trna2_GlyGCC_21_-_18827107_18827177 | 1 | 27 | 27 | GCATGGGTGGTTCAGTGGTAGAATTCT | tRF | TRUE | FALSE | FALSE |
| >hg38_21_+_8433172_8446622_RNA45SN1_45S_50ntFlanks@6646.6675.30\|TCGTACGACTCTTAGCGGTGGATCACTCGG | >hg38_21_+_8433172_8446622_RNA45SN1_45S_50ntFlanks | 6646 | 6675 | 30 | TCGTACGACTCTTAGCGGTGGATCACTCGG | rRF | TRUE | FALSE | FALSE |
| >hg38_21_+_8433172_8446622_RNA45SN1_45S_50ntFlanks@6651.6675.25\|CGACTCTTAGCGGTGGATCACTCGG | >hg38_21_+_8433172_8446622_RNA45SN1_45S_50ntFlanks | 6651 | 6675 | 25 | CGACTCTTAGCGGTGGATCACTCGG | rRF | TRUE | FALSE | FALSE |
| >hg38_21_+_8433172_8446622_RNA45SN1_45S_50ntFlanks@6646.6678.33\|TCGTACGACTCTTAGCGGTGGATCACTCGGCTC | >hg38_21_+_8433172_8446622_RNA45SN1_45S_50ntFlanks | 6646 | 6678 | 33 | TCGTACGACTCTTAGCGGTGGATCACTCGGCTC | rRF | TRUE | FALSE | FALSE |
| >hg38_21_+_8433172_8446622_RNA45SN1_45S_50ntFlanks@7974.7997.24\|ACGCGACCTCAGATCAGACGTGGC | >hg38_21_+_8433172_8446622_RNA45SN1_45S_50ntFlanks | 7974 | 7997 | 24 | ACGCGACCTCAGATCAGACGTGGC | rRF | TRUE | FALSE | FALSE |
| >hg38_21_+_8433172_8446622_RNA45SN1_45S_50ntFlanks@6652.6698.47\|GACTCTTAGCGGTGGATCACTCGGCTCGTGCGTCGATGAAGAACGCA | >hg38_21_+_8433172_8446622_RNA45SN1_45S_50ntFlanks | 6652 | 6698 | 47 | GACTCTTAGCGGTGGATCACTCGGCTCGTGCGTCGATGAAGAA | rRF | TRUE | FALSE | FALSE |
| >hg38_21_+_8433172_8446622_RNA45SN1_45S_50ntFlanks@6661.6699.39\|CGGTGGATCACTCGGCTCGTGCGTCGATGAAGAACGCAG | >hg38_21_+_8433172_8446622_RNA45SN1_45S_50ntFlanks | 6661 | 6699 | 39 | CGGTGGATCACTCGGCTCGTGCGTCGATGAAGAACGCAG | rRF | TRUE | FALSE | FALSE |
| >MI0000434\|hsa-let-7i&WithFlank&12\|+\|62603680\|62603775@12.32.21(+1A)&[MIMAT0000415&hsa-let-7i-5p&offsets\|0\|-1(+1A);m-49&12\|+\|62603691\|62603712&offsets\|0\|-1(+1A)]\|TGAGGTAGTAGTTTGTGCTGTA | >MI0000434\|hsa-let-7i&WithFlank&12\|+\|62603680\|62603775 | 12 | 32 | 21(+1A) | TGAGGTAGTAGTTTGTGCTGTA | isomiR | TRUE | TRUE | FALSE |
| >hg38_21_+_8433172_8446622_RNA45SN1_45S_50ntFlanks@6647.6666.20\|CGTACGACTCTTAGCGGTGG | >hg38_21_+_8433172_8446622_RNA45SN1_45S_50ntFlanks | 6647 | 6666 | 20 | CGTACGACTCTTAGCGGTGG | rRF | TRUE | FALSE | FALSE |

### Mitochondria

| | | | | | | | | | |
|---|---|---|---|---|---|---|---|---|---|
| >hg38_1_-_228634819_228635039_RNA5S12_5S_50ntFlanks@140.169.30\|ACCGCCTGGGAATACCGGGTGCTGTAGGCT | >hg38_1_-_228634819_228635039_RNA5S12_5S_50ntFlanks | 140 | 169 | 30 | ACCGCCTGGGAATACCGGGTGCTGTAGGCT | rRF | TRUE | FALSE | TRUE |
| >MI0000286\|hsa-mir-210&WithFlank&11\|-\|568083\|568204@72.93.22(+2A)&[MIMAT0000267&hsa-miR-210-3p&offsets\|0\|0(+2A);m-102&11\|-\|568112\|568133&offsets\|0\|0(+2A)]\|CTGTGCGTGTGACAGCGGCTGAAA | >MI0000286\|hsa-mir-210&WithFlank&11\|-\|568083\|568204 | 72 | 93 | 22(+2A) | CTGTGCGTGTGACAGCGGCTGAAA | isomiR | TRUE | FALSE | FALSE |
| >trnaMT_SerGCT_MT_+_12207_12265@1.20.20\|GAGAAAGCTCACAAGAACTG | >trnaMT_SerGCT_MT_+_12207_12265 | 1 | 20 | 20 | GAGAAAGCTCACAAGAACTG | tRF | TRUE | TRUE | FALSE |
| >hg38_21_+_8433172_8446622_RNA45SN1_45S_50ntFlanks@4243.4271.29\|CTTTCGAGGCCCTGTAATTGGAATGAGTC | >hg38_21_+_8433172_8446622_RNA45SN1_45S_50ntFlanks | 4243 | 4271 | 29 | CTTTCGAGGCCCTGTAATTGGAATGAGTC | rRF | TRUE | FALSE | FALSE |
| >hg38_21_+_8433172_8446622_RNA45SN1_45S_50ntFlanks@7974.8011.38\|ACGCGACCTCAGATCAGACGTGGCGACCCGCTGAATTT | >hg38_21_+_8433172_8446622_RNA45SN1_45S_50ntFlanks | 7974 | 8011 | 38 | ACGCGACCTCAGATCAGACGTGGCGACCCGCTGAATTT | rRF | TRUE | FALSE | FALSE |
| >am_MI0000809\|hsa-mir-151a&WithFlank&8\|-\|140732558\|140732659@53.70.18&[MIMAT0000757&hsa-miR-151a-3p&offsets\|0\|-3;m-30&8\|-\|140732587\|140732607&offsets\|0\|-3]\|CTAGACTGAAGCTCCTTG | >am_MI0000809\|hsa-mir-151a&WithFlank&8\|-\|140732558\|140732659 | 53 | 70 | 18 | CTAGACTGAAGCTCCTTG | isomiR | TRUE | FALSE | FALSE |
| >trnaMT_GluTTC_MT_-_14674_14742@35.72.38,trnalookalike8_GluTTC_5_-_93905172_93905240@35.72.38\|TATCATTGGTCGTGGTTGTAGTCCGTGCGAGAATACCA | >trnaMT_GluTTC_MT_-_14674_14742 | 35 | 72 | 38 | TATCATTGGTCGTGGTTGTAGTCCGTGCGAGAATACCA | tRF | TRUE | TRUE | FALSE |
| >MI0000469\|hsa-mir-125a&WithFlank&19\|+\|51693248\|51693345@21.43.23&[MIMAT0000443&hsa-miR-125a-5p&offsets\|0\|-1;m-63&19\|+\|51693268\|51693289&offsets\|0\|+1]\|TCCCTGAGACCCTTTAACCTGTG | >MI0000469\|hsa-mir-125a&WithFlank&19\|+\|51693248\|51693345 | 21 | 43 | 23 | TCCCTGAGACCCTTTAACCTGTG | isomiR | TRUE | FALSE | FALSE |
| >MI0000273\|hsa-mir-183&WithFlank&7\|-\|129774899\|129775020@33.54.22&[MIMAT0000261&hsa-miR-183-5p&offsets\|0\|0;m-41&7\|-\|129774966\|129774987&offsets\|-1\|-1]\|TATGGCACTGGTAGAATTCACT | >MI0000273\|hsa-mir-183&WithFlank&7\|-\|129774899\|129775020 | 33 | 54 | 22 | TATGGCACTGGTAGAATTCACT | isomiR | TRUE | FALSE | FALSE |
| >MI0000299\|hsa-mir-222&WithFlank&X\|-\|45747009\|45747130@75.98.24&[MIMAT0000279&hsa-miR-222-3p&offsets\|0\|+3;m-140&X\|-\|45747033\|45747056&offsets\|0\|0]\|AGCTACATCTGGCTACTGGGTCTC | >MI0000299\|hsa-mir-222&WithFlank&X\|-\|45747009\|45747130 | 75 | 98 | 24 | AGCTACATCTGGCTACTGGGTCTC | isomiR | TRUE | TRUE | FALSE |
| >hg38_21_+_8433172_8446622_RNA45SN1_45S_50ntFlanks@6661.6695.35\|CGGTGGATCACTCGGCTCGTGCGTCGATGAAGAAC | >hg38_21_+_8433172_8446622_RNA45SN1_45S_50ntFlanks | 6661 | 6695 | 35 | CGGTGGATCACTCGGCTCGTGCGTCGATGAAGAAC | rRF | TRUE | FALSE | FALSE |
| >multi-hg38_MT_+_1621_3279_RNR2_16S_50ntFlanks@51.81.31\|GCTAAACCTAGCCCCAAACCCACTCCACCTT | >multi-hg38_MT_+_1621_3279_RNR2_16S_50ntFlanks | 51 | 81 | 31 | GCTAAACCTAGCCCCAAACCCACTCCACCTT | rRF | TRUE | FALSE | FALSE |
| >MI0003560\|hsa-mir-92b&WithFlank&1\|+\|155195171\|155195278@67.87.21(+1U)&[MIMAT0003218&hsa-miR-92b-3p&offsets\|0\|-1(+1U);m-115&1\|+\|155195237\|155195258&offsets\|0\|-1(+1U)]\|TATTGCACTCGTCCCGGCCTCT | >MI0003560\|hsa-mir-92b&WithFlank&1\|+\|155195171\|155195278 | 67 | 87 | 21(+1U) | TATTGCACTCGTCCCGGCCTCT | isomiR | TRUE | FALSE | FALSE |
| >multi-hg38_MT_+_598_1651_RNR1_12S_50ntFlanks@52.93.42\|ATAGGTTTGGTCCTAGCCTTTCTATTAGCTCTTAGTAAGATT | >multi-hg38_MT_+_598_1651_RNR1_12S_50ntFlanks | 52 | 93 | 42 | ATAGGTTTGGTCCTAGCCTTTCTATTAGCTCTTAGTAAGATT | rRF | TRUE | FALSE | FALSE |
| >hg38_1_-_228634819_228635039_RNA5S12_5S_50ntFlanks@149.171.23\|GAATACCGGGTGCTGTAGGCTTT | >hg38_1_-_228634819_228635039_RNA5S12_5S_50ntFlanks | 149 | 171 | 23 | GAATACCGGGTGCTGTAGGCTTT | rRF | TRUE | FALSE | TRUE |
| >hg38_MT_+_1621_3279_RNR2_16S_50ntFlanks@51.90.40\|GCTAAACCTAGCCCCAAACCCACTCCACCTTACTACCAGA | >hg38_MT_+_1621_3279_RNR2_16S_50ntFlanks | 51 | 90 | 40 | GCTAAACCTAGCCCCAAACCCACTCCACCTTACTACCAGA | rRF | TRUE | FALSE | FALSE |
| >hg38_21_+_8433172_8446622_RNA45SN1_45S_50ntFlanks@6651.6700.50\|CGACTCTTAGCGGTGGATCACTCGGCTCGTGCGTCGATGAAGAACGCAGC | >hg38_21_+_8433172_8446622_RNA45SN1_45S_50ntFlanks | 6651 | 6700 | 50 | CGACTCTTAGCGGTGGATCACTCGGCTCGTGCGTCGATGAAG/ | rRF | TRUE | FALSE | FALSE |
| >MI0000286\|hsa-mir-210&WithFlank&11\|-\|568083\|568204@72.93.22&[MIMAT0000267&hsa-miR-210-3p&offsets\|0\|0;m-102&11\|-\|568112\|568133&offsets\|0\|0]\|CTGTGCGTGTGACAGCGGCTGA | >MI0000286\|hsa-mir-210&WithFlank&11\|-\|568083\|568204 | 72 | 93 | 22 | CTGTGCGTGTGACAGCGGCTGA | isomiR | TRUE | FALSE | FALSE |
| >MI0000285\|hsa-mir-205&WithFlank&1\|+\|209432127\|209432248@40.61.22(+1A)&[MIMAT0000266&hsa-miR-205-5p&offsets\|0\|0(+1A);m-351&1\|+\|209432166\|209432187&offsets\|0\|0(+1A)]\|TCCTTCATTCCACCGGAGTCTGA | >MI0000285\|hsa-mir-205&WithFlank&1\|+\|209432127\|209432248 | 40 | 61 | 22(+1A) | TCCTTCATTCCACCGGAGTCTGA | isomiR | TRUE | FALSE | TRUE |
| >trna14_LysTTT_11_-_59327808_59327880@1.33.33,trna2_LysTTT_17_+_8022473_8022545@1.33.33,trna54_LysTTT_1_+_204475655_204475727@1.33.33,trna5_LysTTT_11_+_59323902_59323974@1.33.33,trna62_LysTTT_1_-_204476158_204476230@1.33.33,trna76_LysTTT_6_+_28918806_28918878@1.33.33\|GCCCGGATAGCTCAGTCGGTAGAGCATCAGACT | >trna14_LysTTT_11_-_59327808_59327880 | 1 | 33 | 33 | GCCCGGATAGCTCAGTCGGTAGAGCATCAGACT | tRF | TRUE | TRUE | FALSE |
| >hg38_21_+_8433172_8446622_RNA45SN1_45S_50ntFlanks@7974.7994.21\|ACGCGACCTCAGATCAGACGT | >hg38_21_+_8433172_8446622_RNA45SN1_45S_50ntFlanks | 7974 | 7994 | 21 | ACGCGACCTCAGATCAGACGT | rRF | TRUE | FALSE | FALSE |
| >MI0000071\|hsa-mir-17&WithFlank&13\|+\|91350599\|91350694@20.42.23&[MIMAT0000070&hsa-miR-17-5p&offsets\|0\|0;m-88&13\|+\|91350618\|91350640&offsets\|0\|0]\|CAAAGTGCTTACAGTGCAGGTAG | >MI0000071\|hsa-mir-17&WithFlank&13\|+\|91350599\|91350694 | 20 | 42 | 23 | CAAAGTGCTTACAGTGCAGGTAG | isomiR | TRUE | TRUE | FALSE |
| >MI0000076\|hsa-mir-20a&WithFlank&13\|+\|91351059\|91351141@14.36.23&[MIMAT0000075&hsa-miR-20a-5p&offsets\|0\|0;m-96&13\|+\|91351072\|91351094&offsets\|0\|0]\|TAAAGTGCTTATAGTGCAGGTAG | >MI0000076\|hsa-mir-20a&WithFlank&13\|+\|91351059\|91351141 | 14 | 36 | 23 | TAAAGTGCTTATAGTGCAGGTAG | isomiR | TRUE | TRUE | TRUE |
| >trnaMT_SerGCT_MT_+_12207_12265@1.21.21\|GAGAAAGCTCACAAGAACTGC | >trnaMT_SerGCT_MT_+_12207_12265 | 1 | 21 | 21 | GAGAAAGCTCACAAGAACTGC | tRF | TRUE | TRUE | FALSE |
| >multi-hg38_MT_+_598_1651_RNR1_12S_50ntFlanks@52.80.29\|ATAGGTTTGGTCCTAGCCTTTCTATTAGC | >multi-hg38_MT_+_598_1651_RNR1_12S_50ntFlanks | 52 | 80 | 29 | ATAGGTTTGGTCCTAGCCTTTCTATTAGC | rRF | TRUE | FALSE | FALSE |
| >am_MI0039740\|hsa-mir-12136&WithFlank&1\|-\|632609\|632691@7.31.25&[MIMAT0049032&hsa-miR-12136&offsets\|0\|+7]\|GAAAAAGTCATGGAGGCCATGGGGT | >am_MI0039740\|hsa-mir-12136&WithFlank&1\|-\|632609\|632691 | 7 | 31 | 25 | GAAAAAGTCATGGAGGCCATGGGGT | isomiR | TRUE | FALSE | FALSE |
| >multi-trnaMT_SerTGA_MT_-_7446_7514@1.25.25,trnalookalike6_SerTGA_1_-_567997_568065@1.25.25\|GAAAAAGTCATGGAGGCCATGGGGT | >multi-trnaMT_SerTGA_MT_-_7446_7514 | 1 | 25 | 25 | GAAAAAGTCATGGAGGCCATGGGGT | tRF | TRUE | FALSE | FALSE |
| >hg38_21_+_8433172_8446622_RNA45SN1_45S_50ntFlanks@4733.4761.29\|GAACGAAAGTCGGAGGTTCGAAGACGATC | >hg38_21_+_8433172_8446622_RNA45SN1_45S_50ntFlanks | 4733 | 4761 | 29 | GAACGAAAGTCGGAGGTTCGAAGACGATC | rRF | TRUE | FALSE | FALSE |
| >hg38_1_-_228634819_228635039_RNA5S12_5S_50ntFlanks@134.169.36\|TGGGAGACCGCCTGGGAATACCGGGTGCTGTAGGCT | >hg38_1_-_228634819_228635039_RNA5S12_5S_50ntFlanks | 134 | 169 | 36 | TGGGAGACCGCCTGGGAATACCGGGTGCTGTAGGCT | rRF | TRUE | FALSE | FALSE |
| >MI0000088\|hsa-mir-30a&WithFlank&6\|-\|71403545\|71403627@12.35.24(+1U)&[MIMAT0000087&hsa-miR-30a-5p&offsets\|0\|+2(+1U);m-15&6\|-\|71403595\|71403616&offsets\|0\|+2(+1U)]\|TGTAAACATCCTCGACTGGAAGCTT | >MI0000088\|hsa-mir-30a&WithFlank&6\|-\|71403545\|71403627 | 12 | 35 | 24(+1U) | TGTAAACATCCTCGACTGGAAGCTT | isomiR | TRUE | TRUE | FALSE |

### Mitochondria

<table>
<tr><td>>multi-hg38_MT_+_1621_3279_RNR2_16S_50ntFlanks@51.71.21|GCTAAACCTAGCCCCAAACCC</td><td>>multi-hg38_MT_+_1621_3279_RNR2_16S_50ntFlanks</td><td>51</td><td>71</td><td>21</td><td>GCTAAACCTAGCCCCAAACCC</td><td>rRF</td><td>TRUE</td><td>TRUE</td><td>FALSE</td></tr>
<tr><td>>MI0000102|hsa-mir-100&WithFlank&11|-|122152223|122152314@19.39.21&[MIMAT0000098&hsa-miR-100-5p&offsets|0|-1;m-27&11|-|122152276|122152296&offsets|0|0]|AACCCGTAGATCCGAACTTGT</td><td>>MI0000102|hsa-mir-100&WithFlank&11|-|122152223|122152314</td><td>19</td><td>39</td><td>21</td><td>AACCCGTAGATCCGAACTTGT</td><td>isomiR</td><td>TRUE</td><td>TRUE</td><td>FALSE</td></tr>
<tr><td>>MI0000085|hsa-mir-27a&WithFlank&19|-|13836434|13836523@57.77.21&[MIMAT0000084&hsa-miR-27a-3p&offsets|0|0;m-43&19|-|13836447|13836467&offsets|0|0]|TTCACAGTGGCTAAGTTCCGC</td><td>>MI0000085|hsa-mir-27a&WithFlank&19|-|13836434|13836523</td><td>57</td><td>77</td><td>21</td><td>TTCACAGTGGCTAAGTTCCGC</td><td>isomiR</td><td>TRUE</td><td>TRUE</td><td>FALSE</td></tr>
<tr><td>>hg38_21_+_8433172_8446622_RNA45SN1_45S_50ntFlanks@4734.4758.25|AACGAAAGTCGGAGGTTCGAAGACG</td><td>>hg38_21_+_8433172_8446622_RNA45SN1_45S_50ntFlanks</td><td>4734</td><td>4758</td><td>25</td><td>AACGAAAGTCGGAGGTTCGAAGACG</td><td>rRF</td><td>TRUE</td><td>FALSE</td><td>TRUE</td></tr>
<tr><td>>MI0000093|hsa-mir-92a-1&WithFlank&13|+|91351308|91351397@54.74.21(+1C);MI0000094|hsa-mir-92a-2&WithFlank&X|-|134169532|134169618@54.74.21(+1C)&[MIMAT0000092&hsa-miR-92a-3p&offsets|0|-1(+1C);m-9&13|+|91351361|91351382&offsets|0|-1(+1C)];[MIMAT0000092_1&hsa-miR-92a-3p&offsets|0|-1(+1C);m-11&X|-|134169544|134169565&offsets|0|-1(+1C)]|TATTGCACTTGTCCCGGCCTGC</td><td>>MI0000093|hsa-mir-92a-1&WithFlank&13|+|91351308|91351397</td><td>54</td><td>74</td><td>21(+1C)</td><td>TATTGCACTTGTCCCGGCCTGC</td><td>isomiR</td><td>TRUE</td><td>TRUE</td><td>FALSE</td></tr>
<tr><td>>hg38_21_+_8433172_8446622_RNA45SN1_45S_50ntFlanks@11660.11685.26|GTGATTTCTGCCCAGTGCTCTGAATG</td><td>>hg38_21_+_8433172_8446622_RNA45SN1_45S_50ntFlanks</td><td>11660</td><td>11685</td><td>26</td><td>GTGATTTCTGCCCAGTGCTCTGAATG</td><td>rRF</td><td>TRUE</td><td>FALSE</td><td>FALSE</td></tr>
<tr><td>>hg38_21_+_8433172_8446622_RNA45SN1_45S_50ntFlanks@4731.4758.28|AAGAACGAAAGTCGGAGGTTCGAAGACG</td><td>>hg38_21_+_8433172_8446622_RNA45SN1_45S_50ntFlanks</td><td>4731</td><td>4758</td><td>28</td><td>AAGAACGAAAGTCGGAGGTTCGAAGACG</td><td>rRF</td><td>TRUE</td><td>FALSE</td><td>TRUE</td></tr>
<tr><td>>hg38_21_+_8433172_8446622_RNA45SN1_45S_50ntFlanks@8342.8375.34|CGTAAGGGAAAGTTGAAAAGAACTTTGAAGAGAG</td><td>>hg38_21_+_8433172_8446622_RNA45SN1_45S_50ntFlanks</td><td>8342</td><td>8375</td><td>34</td><td>CGTAAGGGAAAGTTGAAAAGAACTTTGAAGAGAG</td><td>rRF</td><td>TRUE</td><td>FALSE</td><td>FALSE</td></tr>
<tr><td>>MI0000542|hsa-mir-320a&WithFlank&8|-|22244960|22245043@48.71.24&[MIMAT0000510&hsa-miR-320a-3p&offsets|0|+2;m-60&8|-|22244975|22244996&offsets|0|+2]|AAAAGCTGGGTTGAGAGGGCGAAA</td><td>>MI0000542|hsa-mir-320a&WithFlank&8|-|22244960|22245043</td><td>48</td><td>71</td><td>24</td><td>AAAAGCTGGGTTGAGAGGGCGAAA</td><td>isomiR</td><td>TRUE</td><td>FALSE</td><td>FALSE</td></tr>
<tr><td>>hg38_21_+_8433172_8446622_RNA45SN1_45S_50ntFlanks@6652.6685.34|GACTCTTAGCGGTGGATCACTCGGCTCGTGCGTC</td><td>>hg38_21_+_8433172_8446622_RNA45SN1_45S_50ntFlanks</td><td>6652</td><td>6685</td><td>34</td><td>GACTCTTAGCGGTGGATCACTCGGCTCGTGCGTC</td><td>rRF</td><td>TRUE</td><td>FALSE</td><td>FALSE</td></tr>
<tr><td>>hg38_21_+_8433172_8446622_RNA45SN1_45S_50ntFlanks@7975.8015.41|CGCGACCTCAGATCAGACGTGGCGACCCGCTGAATTTAAGC</td><td>>hg38_21_+_8433172_8446622_RNA45SN1_45S_50ntFlanks</td><td>7975</td><td>8015</td><td>41</td><td>CGCGACCTCAGATCAGACGTGGCGACCCGCTGAATTTAAGC</td><td>rRF</td><td>TRUE</td><td>FALSE</td><td>FALSE</td></tr>
<tr><td>>trnaMT_PheGAA_MT_+_577_647@1.44.44|GTTTATGTAGCTTACCTCCTCAAAGCAATACACTGAAAATGTTT</td><td>>trnaMT_PheGAA_MT_+_577_647</td><td>1</td><td>44</td><td>44</td><td>GTTTATGTAGCTTACCTCCTCAAAGCAATACACTGAAAATGTTT</td><td>tRF</td><td>TRUE</td><td>FALSE</td><td>FALSE</td></tr>
<tr><td>>MI0000081|hsa-mir-24-2&WithFlank&19|-|13836281|13836365@56.74.19(+1U);MI0000080|hsa-mir-24-1&WithFlank&9|+|95086015|95086094@50.68.19(+1U)&[MIMAT0000080&hsa-miR-24-3p&offsets|0|-3(+1U);m-36&19|-|13836288|13836310&offsets|0|-4(+1U)];[MIMAT0000080_1&hsa-miR-24-3p&offsets|0|-3(+1U);m-35&9|+|95086064|95086086&offsets|0|-4(+1U)]|TGGCTCAGTTCAGCAGGAAT</td><td>>MI0000081|hsa-mir-24-2&WithFlank&19|-|13836281|13836365</td><td>56</td><td>74</td><td>19(+1U)</td><td>TGGCTCAGTTCAGCAGGAAT</td><td>isomiR</td><td>TRUE</td><td>TRUE</td><td>FALSE</td></tr>
<tr><td>>MI0000273|hsa-mir-183&WithFlank&7|-|129774899|129775020@33.55.23&[MIMAT0000261&hsa-miR-183-5p&offsets|0|+1;m-41&7|-|129774966|129774987&offsets|-1|0]|TATGGCACTGGTAGAATTCACTG</td><td>>MI0000273|hsa-mir-183&WithFlank&7|-|129774899|129775020</td><td>33</td><td>55</td><td>23</td><td>TATGGCACTGGTAGAATTCACTG</td><td>isomiR</td><td>TRUE</td><td>FALSE</td><td>FALSE</td></tr>
<tr><td>>hg38_21_+_8433172_8446622_RNA45SN1_45S_50ntFlanks@6652.6681.30|GACTCTTAGCGGTGGATCACTCGGCTCGTG</td><td>>hg38_21_+_8433172_8446622_RNA45SN1_45S_50ntFlanks</td><td>6652</td><td>6681</td><td>30</td><td>GACTCTTAGCGGTGGATCACTCGGCTCGTG</td><td>rRF</td><td>TRUE</td><td>FALSE</td><td>TRUE</td></tr>
<tr><td>>MI0000077|hsa-mir-21&WithFlank&17|+|59841260|59841343@14.32.19&[MIMAT0000076&hsa-miR-21-5p&offsets|0|-3;m-1&17|+|59841273|59841294&offsets|0|-3]|TAGCTTATCAGACTGATGT</td><td>>MI0000077|hsa-mir-21&WithFlank&17|+|59841260|59841343</td><td>14</td><td>32</td><td>19</td><td>TAGCTTATCAGACTGATGT</td><td>isomiR</td><td>TRUE</td><td>TRUE</td><td>FALSE</td></tr>
<tr><td>>MI0000063|hsa-let-7b&WithFlank&22|+|46113680|46113774@12.32.21(+1A)&[MIMAT0000063&hsa-let-7b-5p&offsets|0|-1(+1A);m-14&22|+|46113691|46113712&offsets|0|-1(+1A)]|TGAGGTAGTAGGTTGTGTGGTA</td><td>>MI0000063|hsa-let-7b&WithFlank&22|+|46113680|46113774</td><td>12</td><td>32</td><td>21(+1A)</td><td>TGAGGTAGTAGGTTGTGTGGTA</td><td>isomiR</td><td>TRUE</td><td>FALSE</td><td>FALSE</td></tr>
<tr><td>>hg38_21_+_8433172_8446622_RNA45SN1_45S_50ntFlanks@6651.6680.30|CGACTCTTAGCGGTGGATCACTCGGCTCGT</td><td>>hg38_21_+_8433172_8446622_RNA45SN1_45S_50ntFlanks</td><td>6651</td><td>6680</td><td>30</td><td>CGACTCTTAGCGGTGGATCACTCGGCTCGT</td><td>rRF</td><td>TRUE</td><td>FALSE</td><td>TRUE</td></tr>
<tr><td>>MI0000068|hsa-let-7f-2&WithFlank&X|-|53557186|53557280@14.36.23&[MIMAT0000067_1&hsa-let-7f-5p&offsets|0|+1;m-19&X|-|53557246|53557267&offsets|0|+1]|TGAGGTAGTAGATTGTATAGTTT</td><td>>MI0000068|hsa-let-7f-2&WithFlank&X|-|53557186|53557280</td><td>14</td><td>36</td><td>23</td><td>TGAGGTAGTAGATTGTATAGTTT</td><td>isomiR</td><td>TRUE</td><td>TRUE</td><td>FALSE</td></tr>
<tr><td>>MI0000082|hsa-mir-25&WithFlank&7|-|100093554|100093649@58.78.21(+1U)&[MIMAT0000081&hsa-miR-25-3p&offsets|0|-1(+1U);m-23&7|-|100093571|100093592&offsets|0|-1(+1U)]|CATTGCACTTGTCTCGGTCTGT</td><td>>MI0000082|hsa-mir-25&WithFlank&7|-|100093554|100093649</td><td>58</td><td>78</td><td>21(+1U)</td><td>CATTGCACTTGTCTCGGTCTGT</td><td>isomiR</td><td>TRUE</td><td>FALSE</td><td>FALSE</td></tr>
<tr><td>>MI0000087|hsa-mir-29a&WithFlank&7|-|130876741|130876816@47.68.22(+1U)&[MIMAT0000086&hsa-miR-29a-3p&offsets|-1|-1(+1U);m-26&7|-|130876749|130876769&offsets|-1|0(+1U)]|CTAGCACCATCTGAAATCGGTTT</td><td>>MI0000087|hsa-mir-29a&WithFlank&7|-|130876741|130876816</td><td>47</td><td>68</td><td>22(+1U)</td><td>CTAGCACCATCTGAAATCGGTTT</td><td>isomiR</td><td>TRUE</td><td>TRUE</td><td>FALSE</td></tr>
<tr><td>>MI0000077|hsa-mir-21&WithFlank&17|+|59841260|59841343@14.33.20(+1A)&[MIMAT0000076&hsa-miR-21-5p&offsets|0|-2(+1A);m-1&17|+|59841273|59841294&offsets|0|-2(+1A)]|TAGCTTATCAGACTGATGTTA</td><td>>MI0000077|hsa-mir-21&WithFlank&17|+|59841260|59841343</td><td>14</td><td>33</td><td>20(+1A)</td><td>TAGCTTATCAGACTGATGTTA</td><td>isomiR</td><td>TRUE</td><td>TRUE</td><td>FALSE</td></tr>
<tr><td>>MI0000299|hsa-mir-222&WithFlank&X|-|45747009|45747130@75.99.25&[MIMAT0000279&hsa-miR-222-3p&offsets|0|+4;m-140&X|-|45747033|45747056&offsets|0|+1]|AGCTACATCTGGCTACTGGGTCTCT</td><td>>MI0000299|hsa-mir-222&WithFlank&X|-|45747009|45747130</td><td>75</td><td>99</td><td>25</td><td>AGCTACATCTGGCTACTGGGTCTCT</td><td>isomiR</td><td>TRUE</td><td>TRUE</td><td>FALSE</td></tr>
<tr><td>>hg38_21_+_8433172_8446622_RNA45SN1_45S_50ntFlanks@6647.6669.23|CGTACGACTCTTAGCGGTGGATC</td><td>>hg38_21_+_8433172_8446622_RNA45SN1_45S_50ntFlanks</td><td>6647</td><td>6669</td><td>23</td><td>CGTACGACTCTTAGCGGTGGATC</td><td>rRF</td><td>TRUE</td><td>FALSE</td><td>FALSE</td></tr>
<tr><td>>hg38_1_-_228634819_228635039_RNA5S12_5S_50ntFlanks@99.135.37|AAGCTAAGCAGGGTCGGGCCTGGTTAGTACTTGGATG</td><td>>hg38_1_-_228634819_228635039_RNA5S12_5S_50ntFlanks</td><td>99</td><td>135</td><td>37</td><td>AAGCTAAGCAGGGTCGGGCCTGGTTAGTACTTGGATG</td><td>rRF</td><td>TRUE</td><td>FALSE</td><td>FALSE</td></tr>
<tr><td>>MI0001445|hsa-mir-423&WithFlank&17|+|30117073|30117178@59.78.20&[MIMAT0001340&hsa-miR-423-3p&offsets|0|-3;m-100&17|+|30117131|30117153&offsets|0|-3]|AGCTCGGTCTGAGGCCCCTC</td><td>>MI0001445|hsa-mir-423&WithFlank&17|+|30117073|30117178</td><td>59</td><td>78</td><td>20</td><td>AGCTCGGTCTGAGGCCCCTC</td><td>isomiR</td><td>TRUE</td><td>FALSE</td><td>FALSE</td></tr>
<tr><td>>MI0000809|hsa-mir-151a&WithFlank&8|-|140732558|140732659@51.73.23&[MIMAT0000757&hsa-miR-151a-3p&offsets|-2|0;m-30&8|-|140732587|140732607&offsets|-2|0]|TACTAGACTGAAGCTCCTTGAGG</td><td>>MI0000809|hsa-mir-151a&WithFlank&8|-|140732558|140732659</td><td>51</td><td>73</td><td>23</td><td>TACTAGACTGAAGCTCCTTGAGG</td><td>isomiR</td><td>TRUE</td><td>TRUE</td><td>FALSE</td></tr>
<tr><td>>MI0000077|hsa-mir-21&WithFlank&17|+|59841260|59841343@14.35.22(+1A)&[MIMAT0000076&hsa-miR-21-5p&offsets|0|0(+1A);m-1&17|+|59841273|59841294&offsets|0|0(+1A)]|TAGCTTATCAGACTGATGTTGAA</td><td>>MI0000077|hsa-mir-21&WithFlank&17|+|59841260|59841343</td><td>14</td><td>35</td><td>22(+1A)</td><td>TAGCTTATCAGACTGATGTTGAA</td><td>isomiR</td><td>TRUE</td><td>TRUE</td><td>FALSE</td></tr>
</table>

### Mitochondria

| | | | | | | | | | |
|---|---|---|---|---|---|---|---|---|---|
| >hg38_21_+_8433172_8446622_RNA45SN1_45S_50ntFlanks@10390.10424.35\|GAACAGCAGTTGAACATGGGTCAGTCGGTCCTGAG | >hg38_21_+_8433172_8446622_RNA45SN1_45S_50ntFlanks | 10390 | 10424 | 35 | GAACAGCAGTTGAACATGGGTCAGTCGGTCCTGAG | rRF | TRUE | FALSE | FALSE |
| >trnaMT_SerGCT_MT_+_12207_12265@1.23.23\|GAGAAAGCTCACAAGAACTGCTA | >trnaMT_SerGCT_MT_+_12207_12265 | 1 | 23 | 23 | GAGAAAGCTCACAAGAACTGCTA | tRF | TRUE | FALSE | FALSE |
| >MI0000063\|hsa-let-7b&WithFlank&22\|+\|46113680\|46113774@12.31.20&[MIMAT0000063&hsa-let-7b-5p&offsets\|0\|-2;m-14&22\|+\|46113691\|46113712&offsets\|0\|-2]\|TGAGGTAGTAGGTTGTGTGG | >MI0000063\|hsa-let-7b&WithFlank&22\|+\|46113680\|46113774 | 12 | 31 | 20 | TGAGGTAGTAGGTTGTGTGG | isomiR | TRUE | FALSE | FALSE |
| >hg38_21_+_8433172_8446622_RNA45SN1_45S_50ntFlanks@5485.5516.32\|AGCGCTGAGAAGACGGTCGAACTTGACTATCT | >hg38_21_+_8433172_8446622_RNA45SN1_45S_50ntFlanks | 5485 | 5516 | 32 | AGCGCTGAGAAGACGGTCGAACTTGACTATCT | rRF | TRUE | FALSE | FALSE |
| >MI0000273\|hsa-mir-183&WithFlank&7\|-\|129774899\|129775020@34.55.22&[MIMAT0000261&hsa-miR-183-5p&offsets\|+1\|+1;m-41&7\|-\|129774966\|129774987&offsets\|0\|0]\|ATGGCACTGGTAGAATTCACTG | >MI0000273\|hsa-mir-183&WithFlank&7\|-\|129774899\|129775020 | 34 | 55 | 22 | ATGGCACTGGTAGAATTCACTG | isomiR | TRUE | FALSE | FALSE |
| >trna116_GluCTC_1_-_145399233_145399304@1.34.34,trna59_GluCTC_1_+_249168447_249168518@1.34.34,trna71_GluCTC_1_-_161439189_161439260@1.34.34,trna74_GluCTC_1_-_161431809_161431880@1.34.34,trna77_GluCTC_1_-_161424398_161424469@1.34.34,trna77_GluCTC_6_+_28949976_28950047@1.34.34,trna80_GluCTC_1_-_161417018_161417089@1.34.34,trna87_GluCTC_6_-_126101393_126101464@1.34.34\|TCCCTGGTGGTCTAGTGGTTAGGATTCGGCGCTC | >trna116_GluCTC_1_-_145399233_145399304 | 1 | 34 | 34 | TCCCTGGTGGTCTAGTGGTTAGGATTCGGCGCTC | tRF | TRUE | FALSE | FALSE |
| >hg38_21_+_8433172_8446622_RNA45SN1_45S_50ntFlanks@8355.8373.19\|TGAAAAGAACTTTGAAGAG | >hg38_21_+_8433172_8446622_RNA45SN1_45S_50ntFlanks | 8355 | 8373 | 19 | TGAAAAGAACTTTGAAGAG | rRF | TRUE | FALSE | FALSE |
| >hg38_21_+_8433172_8446622_RNA45SN1_45S_50ntFlanks@6698.6720.23\|AGCTAGCTGCGAGAATTAATGTG | >hg38_21_+_8433172_8446622_RNA45SN1_45S_50ntFlanks | 6698 | 6720 | 23 | AGCTAGCTGCGAGAATTAATGTG | rRF | TRUE | FALSE | TRUE |
| >MI0000746\|hsa-mir-99b&WithFlank&19\|+\|51692606\|51692687@13.32.20&[MIMAT0000689&hsa-miR-99b-5p&offsets\|0\|-2;m-10&19\|+\|51692618\|51692639&offsets\|0\|-2]\|CACCCGTAGAACCGACCTTG | >MI0000746\|hsa-mir-99b&WithFlank&19\|+\|51692606\|51692687 | 13 | 32 | 20 | CACCCGTAGAACCGACCTTG | isomiR | TRUE | FALSE | FALSE |
| >hg38_21_+_8433172_8446622_RNA45SN1_45S_50ntFlanks@8347.8375.29\|GGGAAAGTTGAAAAGAACTTTGAAGAGAG | >hg38_21_+_8433172_8446622_RNA45SN1_45S_50ntFlanks | 8347 | 8375 | 29 | GGGAAAGTTGAAAAGAACTTTGAAGAGAG | rRF | TRUE | FALSE | FALSE |
| >hg38_21_+_8433172_8446622_RNA45SN1_45S_50ntFlanks@6647.6667.21\|CGTACGACTCTTAGCGGTGGA | >hg38_21_+_8433172_8446622_RNA45SN1_45S_50ntFlanks | 6647 | 6667 | 21 | CGTACGACTCTTAGCGGTGGA | rRF | TRUE | FALSE | FALSE |
| >hg38_21_+_8433172_8446622_RNA45SN1_45S_50ntFlanks@6645.6669.25\|CTCGTACGACTCTTAGCGGTGGATC | >hg38_21_+_8433172_8446622_RNA45SN1_45S_50ntFlanks | 6645 | 6669 | 25 | CTCGTACGACTCTTAGCGGTGGATC | rRF | TRUE | FALSE | FALSE |
| >am_MI0003776\|hsa-mir-320b-1&WithFlank&1\|+\|116671740\|116671823@48.65.18;MI0003839\|hsa-mir-320b-2&WithFlank&1\|-\|224257034\|224257116@47.64.18;MI0008190\|hsa-mir-320d-1&WithFlank&13\|-\|40727810\|40727893@48.65.18;MI0003778\|hsa-mir-320c-1&WithFlank&18\|+\|21683512\|21683595@48.65.18;MI0008191\|hsa-mir-320c-2&WithFlank&18\|+\|24321669\|24321752@48.65.18;MI0000542\|hsa-mir-320a&WithFlank&8\|-\|22244960\|22245043@48.65.18;MI0008192\|hsa-mir-320d-2&WithFlank&X\|-\|140926154\|140926237@48.65.18&[MIMAT0005792&hsa-miR-320b&offsets\|0\|-4;m-183&1\|+\|116671787\|116671806&offsets\|0\|-2];[MIMAT0005792_1&hsa-miR-320b&offsets\|0\|-4;m-183&1\|-\|224257051\|224257070&offsets\|0\|-2];[MIMAT0006764&hsa-miR-320d&offsets\|0\|-1;m-246&13\|-\|40727829\|40727846&offsets\|0\|0];[MIMAT0005793&hsa-miR-320c&offsets\|0\|-2;m-222&18\|+\|21683559\|21683578&offsets\|0\|-2];[MIMAT0005793_1&hsa-miR-320c&offsets\|0\|-2;m-219&18\|+\|24321716\|24321735&offsets\|0\|-2];[MIMAT0000510&hsa-miR-320a-3p&offsets\|0\|-4;m-60&8\|-\|22244975\|22244996&offsets\|0\|-4];[MIMAT0006764_1&hsa-miR-320d&offsets\|0\|-1;m-253&X\|-\|140926173\|140926190&offsets\|0\|0]\|AAAAGCTGGGTTGAGAGG | >am_MI0003776\|hsa-mir-320b-1&WithFlank&1\|+\|116671740\|116671823 | 48 | 65 | 18 | AAAAGCTGGGTTGAGAGG | isomiR | TRUE | FALSE | FALSE |
| >MI0000087\|hsa-mir-29a&WithFlank&7\|-\|130876741\|130876816@48.67.20(+1C)&[MIMAT0000086&hsa-miR-29a-3p&offsets\|0\|-2(+1C);m-26&7\|-\|130876749\|130876769&offsets\|0\|-1(+1C)]\|TAGCACCATCTGAAATCGGTC | >MI0000087\|hsa-mir-29a&WithFlank&7\|-\|130876741\|130876816 | 48 | 67 | 20(+1C) | TAGCACCATCTGAAATCGGTC | isomiR | TRUE | TRUE | FALSE |
| >MI0000465\|hsa-mir-191&WithFlank&3\|-\|49020612\|49020715@22.45.24&[MIMAT0000440&hsa-miR-191-5p&offsets\|0\|+1;m-48&3\|-\|49020672\|49020694&offsets\|0\|+1]\|CAACGGAATCCCAAAAGCAGCTGT | >MI0000465\|hsa-mir-191&WithFlank&3\|-\|49020612\|49020715 | 22 | 45 | 24 | CAACGGAATCCCAAAAGCAGCTGT | isomiR | TRUE | TRUE | FALSE |
| >trna111_HisGTG_1_-_147774845_147774916@1.33.33,trna118_HisGTG_1_-_145396881_145396952@1.33.33,trna16_HisGTG_1_+_146544773_146544844@1.33.33,trna1_HisGTG_15_+_45493349_45493420@1.33.33,trna21_HisGTG_1_+_147753471_147753542@1.33.33,trna33_HisGTG_6_+_27125906_27125977@1.33.33,trna7_HisGTG_9_-_14433938_14434009@1.33.33,trna8_HisGTG_15_-_45492611_45492682@1.33.33,trna9_HisGTG_15_-_45490804_45490875@1.33.33\|GCCGTGATCGTATAGTGGTTAGTACTCTGCGTT | >trna111_HisGTG_1_-_147774845_147774916 | 1 | 33 | 33 | GCCGTGATCGTATAGTGGTTAGTACTCTGCGTT | tRF | TRUE | FALSE | FALSE |
| >MI0003776\|hsa-mir-320b-1&WithFlank&1\|+\|116671740\|116671823@48.66.19;MI0003839\|hsa-mir-320b-2&WithFlank&1\|-\|224257034\|224257116@47.65.19;MI0003778\|hsa-mir-320c-1&WithFlank&18\|+\|21683512\|21683595@48.66.19;MI0008191\|hsa-mir-320c-2&WithFlank&18\|+\|24321669\|24321752@48.66.19;MI0000542\|hsa-mir-320a&WithFlank&8\|-\|22244960\|22245043@48.66.19&[MIMAT0005792&hsa-miR-320b&offsets\|0\|-3;m-183&1\|+\|116671787\|116671806&offsets\|0\|-1];[MIMAT0005792_1&hsa-miR-320b&offsets\|0\|-3;m-183&1\|-\|224257051\|224257070&offsets\|0\|-1];[MIMAT0005793&hsa-miR-320c&offsets\|0\|-1;m-222&18\|+\|21683559\|21683578&offsets\|0\|-1];[MIMAT0005793_1&hsa-miR-320c&offsets\|0\|-1;m-219&18\|+\|24321716\|24321735&offsets\|0\|-1];[MIMAT0000510&hsa-miR-320a-3p&offsets\|0\|-3;m-60&8\|-\|22244975\|22244996&offsets\|0\|-3]\|AAAAGCTGGGTTGAGAGGG | >MI0003776\|hsa-mir-320b-1&WithFlank&1\|+\|116671740\|116671823 | 48 | 66 | 19 | AAAAGCTGGGTTGAGAGGG | isomiR | TRUE | FALSE | FALSE |
| >hg38_21_+_8433172_8446622_RNA45SN1_45S_50ntFlanks@7975.8003.29\|CGCGACCTCAGATCAGACGTGGCGACCCG | >hg38_21_+_8433172_8446622_RNA45SN1_45S_50ntFlanks | 7975 | 8003 | 29 | CGCGACCTCAGATCAGACGTGGCGACCCG | rRF | TRUE | FALSE | FALSE |
| >MI0000085\|hsa-mir-27a&WithFlank&19\|-\|13836434\|13836523@57.76.20(+1U)&[MIMAT0000084&hsa-miR-27a-3p&offsets\|0\|-1(+1U);m-43&19\|-\|13836447\|13836467&offsets\|0\|-1(+1U)]\|TTCACAGTGGCTAAGTTCCGT | >MI0000085\|hsa-mir-27a&WithFlank&19\|-\|13836434\|13836523 | 57 | 76 | 20(+1U) | TTCACAGTGGCTAAGTTCCGT | isomiR | TRUE | TRUE | FALSE |
| >hg38_21_+_8433172_8446622_RNA45SN1_45S_50ntFlanks@6651.6674.24\|CGACTCTTAGCGGTGGATCACTCG | >hg38_21_+_8433172_8446622_RNA45SN1_45S_50ntFlanks | 6651 | 6674 | 24 | CGACTCTTAGCGGTGGATCACTCG | rRF | TRUE | FALSE | FALSE |
| >hg38_21_+_8433172_8446622_RNA45SN1_45S_50ntFlanks@8345.8375.31\|AAGGGAAAGTTGAAAAGAACTTTGAAGAGAG | >hg38_21_+_8433172_8446622_RNA45SN1_45S_50ntFlanks | 8345 | 8375 | 31 | AAGGGAAAGTTGAAAAGAACTTTGAAGAGAG | rRF | TRUE | FALSE | FALSE |

### Mitochondria

<table>
<tr><td>>MI0000061|hsa-let-7a-2&WithFlank&11|-|122146516|122146599@11.31.21;MI0000062|hsa-let-7a-3&WithFlank&22|+|46112743|46112828@10.30.21;MI0000060|hsa-let-7a-1&WithFlank&9|+|94175951|94176042@12.32.21&[MIMAT0000062&hsa-let-7a-5p&offsets|0|-1;m-6&11|-|122146568|122146589&offsets|0|-1];[MIMAT0000062_1&hsa-let-7a-5p&offsets|0|-1;m-5&22|+|46112752|46112773&offsets|0|-1];[MIMAT0000062_2&hsa-let-7a-5p&offsets|0|-1;m-4&9|+|94175962|94175983&offsets|0|-1]|TGAGGTAGTAGGTTGTATAGT</td><td>>MI0000061|hsa-let-7a-2&WithFlank&11|-|122146516|122146599</td><td>11</td><td>31</td><td>21</td><td>TGAGGTAGTAGGTTGTATAGT</td><td>isomiR</td><td>FALSE</td><td>TRUE</td><td>FALSE</td></tr>
<tr><td>>MI0000088|hsa-mir-30a&WithFlank&6|-|71403545|71403627@12.34.23(+1G)&[MIMAT0000087&hsa-miR-30a-5p&offsets|0|+1(+1G);m-15&6|-|71403595|71403616&offsets|0|+1(+1G)]|TGTAAACATCCTCGACTGGAAGCG</td><td>>MI0000088|hsa-mir-30a&WithFlank&6|-|71403545|71403627</td><td>12</td><td>34</td><td>23(+1G)</td><td>TGTAAACATCCTCGACTGGAAGCG</td><td>isomiR</td><td>FALSE</td><td>TRUE</td><td>FALSE</td></tr>
<tr><td>>MI0000088|hsa-mir-30a&WithFlank&6|-|71403545|71403627@53.73.21(+1U)&[MIMAT0000088&hsa-miR-30a-3p&offsets|0|-1(+1U);m-38&6|-|71403554|71403575&offsets|0|-1(+1U)]|CTTTCAGTCGGATGTTTGCAGT</td><td>>MI0000088|hsa-mir-30a&WithFlank&6|-|71403545|71403627</td><td>53</td><td>73</td><td>21(+1U)</td><td>CTTTCAGTCGGATGTTTGCAGT</td><td>isomiR</td><td>FALSE</td><td>TRUE</td><td>FALSE</td></tr>
<tr><td>>MI0000289|hsa-mir-181a-1&WithFlank&1|-|198859038|198859159@30.53.24;MI0000269|hsa-mir-181a-2&WithFlank&9|+|124692436|124692557@45.68.24&[MIMAT0000256&hsa-miR-181a-5p&offsets|0|+1;m-32&1|-|198859108|198859130&offsets|0|+1];[MIMAT0000256_1&hsa-miR-181a-5p&offsets|0|+1;m-32&9|+|124692480|124692502&offsets|0|+1]|AACATTCAACGCTGTCGGTGAGTT</td><td>>MI0000289|hsa-mir-181a-1&WithFlank&1|-|198859038|198859159</td><td>30</td><td>53</td><td>24</td><td>AACATTCAACGCTGTCGGTGAGTT</td><td>isomiR</td><td>FALSE</td><td>TRUE</td><td>FALSE</td></tr>
<tr><td>>MI0000298|hsa-mir-221&WithFlank&X|-|45746151|45746272@71.91.21&[MIMAT0000278&hsa-miR-221-3p&offsets|0|-2;m-91&X|-|45746181|45746202&offsets|0|-1]|AGCTACATTGTCTGCTGGGTT</td><td>>MI0000298|hsa-mir-221&WithFlank&X|-|45746151|45746272</td><td>71</td><td>91</td><td>21</td><td>AGCTACATTGTCTGCTGGGTT</td><td>isomiR</td><td>FALSE</td><td>TRUE</td><td>FALSE</td></tr>
<tr><td>>MI0000477|hsa-mir-146a&WithFlank&5|+|160485346|160485456@27.47.21(+1A)&[MIMAT0000449&hsa-miR-146a-5p&offsets|0|-1(+1A);m-129&5|+|160485372|160485393&offsets|0|-1(+1A)]|TGAGAACTGAATTCCATGGGTA</td><td>>MI0000477|hsa-mir-146a&WithFlank&5|+|160485346|160485456</td><td>27</td><td>47</td><td>21(+1A)</td><td>TGAGAACTGAATTCCATGGGTA</td><td>isomiR</td><td>FALSE</td><td>TRUE</td><td>FALSE</td></tr>
<tr><td>>MI0000264|hsa-mir-7-2&WithFlank&15|+|88611819|88611940@38.60.23(+1C);MI0000265|hsa-mir-7-3&WithFlank&19|+|4770664|4770785@37.59.23(+1C);MI0000263|hsa-mir-7-1&WithFlank&9|-|83969742|83969863@30.52.23(+1C)&[MIMAT0000252&hsa-miR-7-5p&offsets|0|-1(+1C);m-400&15|+|88611856|88611879&offsets|0|-1(+1C)];[MIMAT0000252_1&hsa-miR-7-5p&offsets|0|-1(+1C);m-402&19|+|4770700|4770723&offsets|0|-1(+1C)];[MIMAT0000252_2&hsa-miR-7-5p&offsets|0|-1(+1C);m-401&9|-|83969811|83969834&offsets|0|-1(+1C)]|TGGAAGACTAGTGATTTTGTTGTC</td><td>>MI0000264|hsa-mir-7-2&WithFlank&15|+|88611819|88611940</td><td>38</td><td>60</td><td>23(+1C)</td><td>TGGAAGACTAGTGATTTTGTTGTC</td><td>isomiR</td><td>FALSE</td><td>TRUE</td><td>FALSE</td></tr>
<tr><td>>MI0000446|hsa-mir-125b-1&WithFlank&11|-|122099751|122099850@21.41.21;MI0000470|hsa-mir-125b-2&WithFlank&21|+|16590231|16590331@23.43.21&[MIMAT0000423&hsa-miR-125b-5p&offsets|0|-1;m-67&11|-|122099809|122099830&offsets|0|-1];[MIMAT0000423_1&hsa-miR-125b-5p&offsets|0|-1;m-68&21|+|16590253|16590274&offsets|0|-1]|TCCCTGAGACCCTAACTTGTG</td><td>>MI0000446|hsa-mir-125b-1&WithFlank&11|-|122099751|122099850</td><td>21</td><td>41</td><td>21</td><td>TCCCTGAGACCCTAACTTGTG</td><td>isomiR</td><td>FALSE</td><td>TRUE</td><td>FALSE</td></tr>
<tr><td>>MI0000477|hsa-mir-146a&WithFlank&5|+|160485346|160485456@27.49.23&[MIMAT0000449&hsa-miR-146a-5p&offsets|0|+1;m-129&5|+|160485372|160485393&offsets|0|+1]|TGAGAACTGAATTCCATGGGTTG</td><td>>MI0000477|hsa-mir-146a&WithFlank&5|+|160485346|160485456</td><td>27</td><td>49</td><td>23</td><td>TGAGAACTGAATTCCATGGGTTG</td><td>isomiR</td><td>FALSE</td><td>TRUE</td><td>FALSE</td></tr>
<tr><td>>MI0000067|hsa-let-7f-1&WithFlank&9|+|94176341|94176439@13.33.21;MI0000068|hsa-let-7f-2&WithFlank&X|-|53557186|53557280@14.34.21&[MIMAT0000067&hsa-let-7f-5p&offsets|0|-1;m-20&9|+|94176353|94176374&offsets|0|-1];[MIMAT0000067_1&hsa-let-7f-5p&offsets|0|-1;m-19&X|-|53557246|53557267&offsets|0|-1]|TGAGGTAGTAGATTGTATAGT</td><td>>MI0000067|hsa-let-7f-1&WithFlank&9|+|94176341|94176439</td><td>13</td><td>33</td><td>21</td><td>TGAGGTAGTAGATTGTATAGT</td><td>isomiR</td><td>FALSE</td><td>TRUE</td><td>FALSE</td></tr>
<tr><td>>MI0000102|hsa-mir-100&WithFlank&11|-|122152223|122152314@19.37.19(+1A)&[MIMAT0000098&hsa-miR-100-5p&offsets|0|-3(+1A);m-27&11|-|122152276|122152296&offsets|0|-2(+1A)]|AACCCGTAGATCCGAACTTA</td><td>>MI0000102|hsa-mir-100&WithFlank&11|-|122152223|122152314</td><td>19</td><td>37</td><td>19(+1A)</td><td>AACCCGTAGATCCGAACTTA</td><td>isomiR</td><td>FALSE</td><td>TRUE</td><td>FALSE</td></tr>
<tr><td>>MI0000084|hsa-mir-26b&WithFlank&2|+|218402640|218402728@18.39.22&[MIMAT0000083&hsa-miR-26b-5p&offsets|0|+1;m-53&2|+|218402657|218402678&offsets|0|0]|TTCAAGTAATTCAGGATAGGTT</td><td>>MI0000084|hsa-mir-26b&WithFlank&2|+|218402640|218402728</td><td>18</td><td>39</td><td>22</td><td>TTCAAGTAATTCAGGATAGGTT</td><td>isomiR</td><td>FALSE</td><td>TRUE</td><td>FALSE</td></tr>
<tr><td>>MI0000736|hsa-mir-30c-1&WithFlank&1|+|40757278|40757378@23.45.23;MI0000254|hsa-mir-30c-2&WithFlank&6|-|71376954|71377037@13.35.23&[MIMAT0000244&hsa-miR-30c-5p&offsets|0|0;m-57&1|+|40757300|40757323&offsets|0|-1];[MIMAT0000244_1&hsa-miR-30c-5p&offsets|0|0;m-56&6|-|71377002|71377025&offsets|0|-1]|TGTAAACATCCTACACTCTCAGC</td><td>>MI0000736|hsa-mir-30c-1&WithFlank&1|+|40757278|40757378</td><td>23</td><td>45</td><td>23</td><td>TGTAAACATCCTACACTCTCAGC</td><td>isomiR</td><td>FALSE</td><td>TRUE</td><td>FALSE</td></tr>
<tr><td>>hg38_21_+_8433172_8446622_RNA45SN1_45S_50ntFlanks@475.497.23|CTTCGTGATCGATGTGGTGACGT</td><td>>hg38_21_+_8433172_8446622_RNA45SN1_45S_50ntFlanks</td><td>475</td><td>497</td><td>23</td><td>CTTCGTGATCGATGTGGTGACGT</td><td>rRF</td><td>FALSE</td><td>TRUE</td><td>FALSE</td></tr>
<tr><td>>MI0000750|hsa-mir-26a-2&WithFlank&12|-|57824603|57824698@20.41.22(+1U);MI0000083|hsa-mir-26a-1&WithFlank&3|+|37969398|37969486@16.37.22(+1U)&[MIMAT0000082&hsa-miR-26a-5p&offsets|0|0(+1U);m-33&12|-|57824658|57824679&offsets|0|0(+1U)];[MIMAT0000082_1&hsa-miR-26a-5p&offsets|0|0(+1U);m-34&3|+|37969413|37969434&offsets|0|0(+1U)]|TTCAAGTAATCCAGGATAGGCTT</td><td>>MI0000750|hsa-mir-26a-2&WithFlank&12|-|57824603|57824698</td><td>20</td><td>41</td><td>22(+1U)</td><td>TTCAAGTAATCCAGGATAGGCTT</td><td>isomiR</td><td>FALSE</td><td>TRUE</td><td>FALSE</td></tr>
<tr><td>>MI0000061|hsa-let-7a-2&WithFlank&11|-|122146516|122146599@11.33.23;MI0000062|hsa-let-7a-3&WithFlank&22|+|46112743|46112828@10.32.23;MI0000060|hsa-let-7a-1&WithFlank&9|+|94175951|94176042@12.34.23&[MIMAT0000062&hsa-let-7a-5p&offsets|0|+1;m-6&11|-|122146568|122146589&offsets|0|+1];[MIMAT0000062_1&hsa-let-7a-5p&offsets|0|+1;m-5&22|+|46112752|46112773&offsets|0|+1];[MIMAT0000062_2&hsa-let-7a-5p&offsets|0|+1;m-4&9|+|94175962|94175983&offsets|0|+1]|TGAGGTAGTAGGTTGTATAGTTT</td><td>>MI0000061|hsa-let-7a-2&WithFlank&11|-|122146516|122146599</td><td>11</td><td>33</td><td>23</td><td>TGAGGTAGTAGGTTGTATAGTTT</td><td>isomiR</td><td>FALSE</td><td>TRUE</td><td>FALSE</td></tr>
<tr><td>>MI0000289|hsa-mir-181a-1&WithFlank&1|-|198859038|198859159@30.54.25;MI0000269|hsa-mir-181a-2&WithFlank&9|+|124692436|124692557@45.69.25&[MIMAT0000256&hsa-miR-181a-5p&offsets|0|+2;m-32&1|-|198859108|198859130&offsets|0|+2];[MIMAT0000256_1&hsa-miR-181a-5p&offsets|0|+2;m-32&9|+|124692480|124692502&offsets|0|+2]|AACATTCAACGCTGTCGGTGAGTTT</td><td>>MI0000289|hsa-mir-181a-1&WithFlank&1|-|198859038|198859159</td><td>30</td><td>54</td><td>25</td><td>AACATTCAACGCTGTCGGTGAGTTT</td><td>isomiR</td><td>FALSE</td><td>TRUE</td><td>FALSE</td></tr>
<tr><td>>MI0000085|hsa-mir-27a&WithFlank&19|-|13836434|13836523@57.76.20(+1A)&[MIMAT0000084&hsa-miR-27a-3p&offsets|0|-1(+1A);m-43&19|-|13836447|13836467&offsets|0|-1(+1A)]|TTCACAGTGGCTAAGTTCCGA</td><td>>MI0000085|hsa-mir-27a&WithFlank&19|-|13836434|13836523</td><td>57</td><td>76</td><td>20(+1A)</td><td>TTCACAGTGGCTAAGTTCCGA</td><td>isomiR</td><td>FALSE</td><td>TRUE</td><td>FALSE</td></tr>
</table>

### Mitochondria

| | | | | | | | | | |
|---|---|---|---|---|---|---|---|---|---|
| >trna111_HisGTG_1_-_147774845_147774916@-1G.34.35,trna118_HisGTG_1_-_145396881_145396952@-1G.34.35,trna16_HisGTG_1_+_146544773_146544844@-1G.34.35,trna1_HisGTG_15_+_45493349_45493420@-1G.34.35,trna21_HisGTG_1_+_147753471_147753542@-1G.34.35,trna33_HisGTG_6_+_27125906_27125977@-1G.34.35,trna7_HisGTG_9_-_14433938_14434009@-1G.34.35,trna8_HisGTG_15_-_45492611_45492682@-1G.34.35,trna9_HisGTG_15_-_45490804_45490875@-1G.34.35\|GGCCGTGATCGTATAGTGGTTAGTACTCTGCGTTG | >trna111_HisGTG_1_-_147774845_147774916 | -1G | 34 | 35 | GGCCGTGATCGTATAGTGGTTAGTACTCTGCGTTG | tRF | FALSE | TRUE | FALSE |
| >trnaMT_ValTAC_MT_+_1602_1670@42.71.30\|AGATTTCAACTTAACTTGACCGCTCTGACC | >trnaMT_ValTAC_MT_+_1602_1670 | 42 | 71 | 30 | AGATTTCAACTTAACTTGACCGCTCTGACC | tRF | FALSE | TRUE | FALSE |
| >MI0000471\|hsa-mir-126&WithFlank&9\|+\|136670596\|136670692@58.78.21&[MIMAT0000445&hsa-miR-126-3p&offsets\|0\|-1;m-40&9\|+\|136670653\|136670674&offsets\|0\|-1]\|TCGTACCGTGAGTAATAATGC | >MI0000471\|hsa-mir-126&WithFlank&9\|+\|136670596\|136670692 | 58 | 78 | 21 | TCGTACCGTGAGTAATAATGC | isomiR | FALSE | TRUE | FALSE |
| >multi-hg38_MT_+_598_1651_RNR1_12S_50ntFlanks@905.929.25\|AGTATACTTCAAAGGACATTTAACT | >multi-hg38_MT_+_598_1651_RNR1_12S_50ntFlanks | 905 | 929 | 25 | AGTATACTTCAAAGGACATTTAACT | rRF | FALSE | TRUE | FALSE |
| >MI0000095\|hsa-mir-93&WithFlank&7\|-\|100093762\|100093853@17.37.21&[MIMAT0000093&hsa-miR-93-5p&offsets\|0\|-2;m-29&7\|-\|100093815\|100093837&offsets\|0\|-2]\|CAAAGTGCTGTTCGTGCAGGT | >MI0000095\|hsa-mir-93&WithFlank&7\|-\|100093762\|100093853 | 17 | 37 | 21 | CAAAGTGCTGTTCGTGCAGGT | isomiR | FALSE | TRUE | FALSE |
| >MI0000100\|hsa-mir-98&WithFlank&X\|-\|53556217\|53556347@28.49.22&[MIMAT0000096&hsa-miR-98-5p&offsets\|0\|0;m-143&X\|-\|53556299\|53556320&offsets\|0\|0]\|TGAGGTAGTAAGTTGTATTGTT | >MI0000100\|hsa-mir-98&WithFlank&X\|-\|53556217\|53556347 | 28 | 49 | 22 | TGAGGTAGTAAGTTGTATTGTT | isomiR | FALSE | TRUE | FALSE |
| >MI0000067\|hsa-let-7f-1&WithFlank&9\|+\|94176341\|94176439@13.30.18;MI0000068\|hsa-let-7f-2&WithFlank&X\|-\|53557186\|53557280@14.31.18&[MIMAT0000067&hsa-let-7f-5p&offsets\|0\|-4;m-20&9\|+\|94176353\|94176374&offsets\|0\|-4];[MIMAT0000067_1&hsa-let-7f-5p&offsets\|0\|-4;m-19&X\|-\|53557246\|53557267&offsets\|0\|-4]\|TGAGGTAGTAGATTGTAT | >MI0000067\|hsa-let-7f-1&WithFlank&9\|+\|94176341\|94176439 | 13 | 30 | 18 | TGAGGTAGTAGATTGTAT | isomiR | FALSE | TRUE | FALSE |
| >am_trna128_GlyGCC_6_-_27870686_27870756@1.33.33,trna18_GlyGCC_16_+_70822597_70822667@1.33.33,trna19_GlyGCC_16_+_70823410_70823480@1.33.33,trna19_GlyGCC_2_-_157257659_157257729@1.33.33,trna24_GlyGCC_16_-_70812942_70813012@1.33.33,trna25_GlyGCC_16_-_70812114_70812184@1.33.33,trna5_GlyGCC_17_+_8029064_8029134@1.33.33,trna68_GlyGCC_1_-_161493637_161493707@1.33.33\|GCATTGGTGGTTCAGTGGTAGAATTCTCGCCTG | >am_trna128_GlyGCC_6_-_27870686_27870756 | 1 | 33 | 33 | GCATTGGTGGTTCAGTGGTAGAATTCTCGCCTG | tRF | FALSE | TRUE | FALSE |
| >MI0000085\|hsa-mir-27a&WithFlank&19\|-\|13836434\|13836523@57.75.19(+1A)&[MIMAT0000084&hsa-miR-27a-3p&offsets\|0\|-2(+1A);m-43&19\|-\|13836447\|13836467&offsets\|0\|-2(+1A)]\|TTCACAGTGGCTAAGTTCCA | >MI0000085\|hsa-mir-27a&WithFlank&19\|-\|13836434\|13836523 | 57 | 75 | 19(+1A) | TTCACAGTGGCTAAGTTCCA | isomiR | FALSE | TRUE | FALSE |
| >MI0000085\|hsa-mir-27a&WithFlank&19\|-\|13836434\|13836523@57.76.20(+2U)&[MIMAT0000084&hsa-miR-27a-3p&offsets\|0\|-1(+2U);m-43&19\|-\|13836447\|13836467&offsets\|0\|-1(+2U)]\|TTCACAGTGGCTAAGTTCCGTT | >MI0000085\|hsa-mir-27a&WithFlank&19\|-\|13836434\|13836523 | 57 | 76 | 20(+2U) | TTCACAGTGGCTAAGTTCCGTT | isomiR | FALSE | TRUE | FALSE |
| >MI0000087\|hsa-mir-29a&WithFlank&7\|-\|130876741\|130876816@47.67.21(+1C)&[MIMAT0000086&hsa-miR-29a-3p&offsets\|-1\|-2(+1C);m-26&7\|-\|130876749\|130876769&offsets\|-1\|-1(+1C)]\|CTAGCACCATCTGAAATCGGTC | >MI0000087\|hsa-mir-29a&WithFlank&7\|-\|130876741\|130876816 | 47 | 67 | 21(+1C) | CTAGCACCATCTGAAATCGGTC | isomiR | FALSE | TRUE | FALSE |
| >multi-am_trnaMT_MetCAT_MT_+_4402_4469@1.21.21\|AGTAAGGTCAGCTAAATAAGC | >multi-am_trnaMT_MetCAT_MT_+_4402_4469 | 1 | 21 | 21 | AGTAAGGTCAGCTAAATAAGC | tRF | FALSE | TRUE | FALSE |
| >am_trna128_GlyGCC_6_-_27870686_27870756@1.35.35,trna18_GlyGCC_16_+_70822597_70822667@1.35.35,trna19_GlyGCC_16_+_70823410_70823480@1.35.35,trna19_GlyGCC_2_-_157257659_157257729@1.35.35,trna24_GlyGCC_16_-_70812942_70813012@1.35.35,trna25_GlyGCC_16_-_70812114_70812184@1.35.35,trna5_GlyGCC_17_+_8029064_8029134@1.35.35,trna68_GlyGCC_1_-_161493637_161493707@1.35.35\|GCATTGGTGGTTCAGTGGTAGAATTCTCGCCTGCC | >am_trna128_GlyGCC_6_-_27870686_27870756 | 1 | 35 | 35 | GCATTGGTGGTTCAGTGGTAGAATTCTCGCCTGCC | tRF | FALSE | TRUE | FALSE |
| >MI0000270\|hsa-mir-181b-1&WithFlank&1\|-\|198858867\|198858988@42.65.24;MI0000683\|hsa-mir-181b-2&WithFlank&9\|+\|124693704\|124693804@22.45.24&[MIMAT0000257&hsa-miR-181b-5p&offsets\|0\|+1;m-73&1\|-\|198858925\|198858947&offsets\|0\|+1];[MIMAT0000257_1&hsa-miR-181b-5p&offsets\|0\|+1;m-72&9\|+\|124693725\|124693747&offsets\|0\|+1]\|AACATTCATTGCTGTCGGTGGGTT | >MI0000270\|hsa-mir-181b-1&WithFlank&1\|-\|198858867\|198858988 | 42 | 65 | 24 | AACATTCATTGCTGTCGGTGGGTT | isomiR | FALSE | TRUE | FALSE |
| >MI0000088\|hsa-mir-30a&WithFlank&6\|-\|71403545\|71403627@12.33.22(+1A)&[MIMAT0000087&hsa-miR-30a-5p&offsets\|0\|0(+1A);m-15&6\|-\|71403595\|71403616&offsets\|0\|0(+1A)]\|TGTAAACATCCTCGACTGGAAGA | >MI0000088\|hsa-mir-30a&WithFlank&6\|-\|71403545\|71403627 | 12 | 33 | 22(+1A) | TGTAAACATCCTCGACTGGAAGA | isomiR | FALSE | TRUE | FALSE |
| >MI0000095\|hsa-mir-93&WithFlank&7\|-\|100093762\|100093853@17.38.22(+1A)&[MIMAT0000093&hsa-miR-93-5p&offsets\|0\|-1(+1A);m-29&7\|-\|100093815\|100093837&offsets\|0\|-1(+1A)]\|CAAAGTGCTGTTCGTGCAGGTAA | >MI0000095\|hsa-mir-93&WithFlank&7\|-\|100093762\|100093853 | 17 | 38 | 22(+1A) | CAAAGTGCTGTTCGTGCAGGTAA | isomiR | FALSE | TRUE | FALSE |
| >multi-am_trnaMT_ProTGG_MT_-_15956_16023@52.71.20\|AAAGACTTTTTCTCTGACCA | >multi-am_trnaMT_ProTGG_MT_-_15956_16023 | 52 | 71 | 20 | AAAGACTTTTTCTCTGACCA | tRF | FALSE | TRUE | FALSE |
| >MI0000088\|hsa-mir-30a&WithFlank&6\|-\|71403545\|71403627@12.35.24(+2U)&[MIMAT0000087&hsa-miR-30a-5p&offsets\|0\|+2(+2U);m-15&6\|-\|71403595\|71403616&offsets\|0\|+2(+2U)]\|TGTAAACATCCTCGACTGGAAGCTTT | >MI0000088\|hsa-mir-30a&WithFlank&6\|-\|71403545\|71403627 | 12 | 35 | 24(+2U) | TGTAAACATCCTCGACTGGAAGCTTT | isomiR | FALSE | TRUE | FALSE |
| >MI0000088\|hsa-mir-30a&WithFlank&6\|-\|71403545\|71403627@12.35.24(+1C)&[MIMAT0000087&hsa-miR-30a-5p&offsets\|0\|+2(+1C);m-15&6\|-\|71403595\|71403616&offsets\|0\|+2(+1C)]\|TGTAAACATCCTCGACTGGAAGCTC | >MI0000088\|hsa-mir-30a&WithFlank&6\|-\|71403545\|71403627 | 12 | 35 | 24(+1C) | TGTAAACATCCTCGACTGGAAGCTC | isomiR | FALSE | TRUE | FALSE |
| >trna111_HisGTG_1_-_147774845_147774916@-1G.32.33,trna118_HisGTG_1_-_145396881_145396952@-1G.32.33,trna16_HisGTG_1_+_146544773_146544844@-1G.32.33,trna1_HisGTG_15_+_45493349_45493420@-1G.32.33,trna21_HisGTG_1_+_147753471_147753542@-1G.32.33,trna33_HisGTG_6_+_27125906_27125977@-1G.32.33,trna7_HisGTG_9_-_14433938_14434009@-1G.32.33,trna8_HisGTG_15_-_45492611_45492682@-1G.32.33,trna9_HisGTG_15_-_45490804_45490875@-1G.32.33\|GGCCGTGATCGTATAGTGGTTAGTACTCTGCGT | >trna111_HisGTG_1_-_147774845_147774916 | -1G | 32 | 33 | GGCCGTGATCGTATAGTGGTTAGTACTCTGCGT | tRF | FALSE | TRUE | FALSE |
| >trnaMT_HisGTG_MT_+_12138_12206@32.70.39\|TGAATCTGACAACAGAGGCTTACGACCCCTTATTTACCC | >trnaMT_HisGTG_MT_+_12138_12206 | 32 | 70 | 39 | TGAATCTGACAACAGAGGCTTACGACCCCTTATTTACCC | tRF | FALSE | TRUE | FALSE |
| >trnaMT_ValTAC_MT_+_1602_1670@42.72.31\|AGATTTCAACTTAACTTGACCGCTCTGACCA | >trnaMT_ValTAC_MT_+_1602_1670 | 42 | 72 | 31 | AGATTTCAACTTAACTTGACCGCTCTGACCA | tRF | FALSE | TRUE | FALSE |
| >multi-hg38_MT_+_1621_3279_RNR2_16S_50ntFlanks@51.70.20\|GCTAAACCTAGCCCCAAACC | >multi-hg38_MT_+_1621_3279_RNR2_16S_50ntFlanks | 51 | 70 | 20 | GCTAAACCTAGCCCCAAACC | rRF | FALSE | TRUE | FALSE |

### Mitochondria

<table>
<tr><td>>MI0000749|hsa-mir-30e&WithFlank&1|+|40754349|40754452@23.45.23(+1C)&[MIMAT0000692&hsa-miR-30e-5p&offsets|0|+1(+1C);m-24&1|+|40754371|40754394&offsets|0|-1(+1C)]|TGTAAACATCCTTGACTGGAAGCC</td><td>>MI0000749|hsa-mir-30e&WithFlank&1|+|40754349|40754452</td><td>23</td><td>45</td><td>23(+1C)</td><td>TGTAAACATCCTTGACTGGAAGCC</td><td>isomiR</td><td>FALSE</td><td>TRUE</td><td>FALSE</td></tr>
<tr><td>>MI0000433|hsa-let-7g&WithFlank&3|-|52268272|52268367@11.31.21&[MIMAT0000414&hsa-let-7g-5p&offsets|0|-1;m-58&3|-|52268336|52268357&offsets|0|-1]|TGAGGTAGTAGTTTGTACAGT</td><td>>MI0000433|hsa-let-7g&WithFlank&3|-|52268272|52268367</td><td>11</td><td>31</td><td>21</td><td>TGAGGTAGTAGTTTGTACAGT</td><td>isomiR</td><td>FALSE</td><td>TRUE</td><td>FALSE</td></tr>
<tr><td>>MI0000078|hsa-mir-22&WithFlank&17|-|1713897|1713993@21.41.21&[MIMAT0004495&hsa-miR-22-5p&offsets|0|-1;m-217&17|-|1713953|1713973&offsets|0|0]|AGTTCTTCAGTGGCAAGCTTT</td><td>>MI0000078|hsa-mir-22&WithFlank&17|-|1713897|1713993</td><td>21</td><td>41</td><td>21</td><td>AGTTCTTCAGTGGCAAGCTTT</td><td>isomiR</td><td>FALSE</td><td>TRUE</td><td>FALSE</td></tr>
<tr><td>>MI0000088|hsa-mir-30a&WithFlank&6|-|71403545|71403627@54.74.21&[MIMAT0000088&hsa-miR-30a-3p&offsets|+1|0;m-38&6|-|71403554|71403575&offsets|+1|0]|TTTCAGTCGGATGTTTGCAGC</td><td>>MI0000088|hsa-mir-30a&WithFlank&6|-|71403545|71403627</td><td>54</td><td>74</td><td>21</td><td>TTTCAGTCGGATGTTTGCAGC</td><td>isomiR</td><td>FALSE</td><td>TRUE</td><td>FALSE</td></tr>
<tr><td>>MI0000093|hsa-mir-92a-1&WithFlank&13|+|91351308|91351397@54.75.22(+1A);MI0000094|hsa-mir-92a-2&WithFlank&X|-|134169532|134169618@54.75.22(+1A)&[MIMAT0000092&hsa-miR-92a-3p&offsets|0|0(+1A);m-9&13|+|91351361|91351382&offsets|0|0(+1A)];[MIMAT0000092_1&hsa-miR-92a-3p&offsets|0|0(+1A);m-11&X|-|134169544|134169565&offsets|0|0(+1A)]|TATTGCACTTGTCCCGGCCTGTA</td><td>>MI0000093|hsa-mir-92a-1&WithFlank&13|+|91351308|91351397</td><td>54</td><td>75</td><td>22(+1A)</td><td>TATTGCACTTGTCCCGGCCTGTA</td><td>isomiR</td><td>FALSE</td><td>TRUE</td><td>FALSE</td></tr>
<tr><td>>MI0000750|hsa-mir-26a-2&WithFlank&12|-|57824603|57824698@20.41.22(+2U);MI0000083|hsa-mir-26a-1&WithFlank&3|+|37969398|37969486@16.37.22(+2U)&[MIMAT0000082&hsa-miR-26a-5p&offsets|0|0(+2U);m-33&12|-|57824658|57824679&offsets|0|0(+2U)];[MIMAT0000082_1&hsa-miR-26a-5p&offsets|0|0(+2U);m-34&3|+|37969413|37969434&offsets|0|0(+2U)]|TTCAAGTAATCCAGGATAGGCTTT</td><td>>MI0000750|hsa-mir-26a-2&WithFlank&12|-|57824603|57824698</td><td>20</td><td>41</td><td>22(+2U)</td><td>TTCAAGTAATCCAGGATAGGCTTT</td><td>isomiR</td><td>FALSE</td><td>TRUE</td><td>FALSE</td></tr>
<tr><td>>MI0000076|hsa-mir-20a&WithFlank&13|+|91351059|91351141@14.35.22(+1A)&[MIMAT0000075&hsa-miR-20a-5p&offsets|0|-1(+1A);m-96&13|+|91351072|91351094&offsets|0|-1(+1A)]|TAAAGTGCTTATAGTGCAGGTAA</td><td>>MI0000076|hsa-mir-20a&WithFlank&13|+|91351059|91351141</td><td>14</td><td>35</td><td>22(+1A)</td><td>TAAAGTGCTTATAGTGCAGGTAA</td><td>isomiR</td><td>FALSE</td><td>TRUE</td><td>FALSE</td></tr>
<tr><td>>MI0000114|hsa-mir-107&WithFlank&10|-|89592741|89592833@56.76.21;MI0000108|hsa-mir-103a-2&WithFlank&20|+|3917488|3917577@54.74.21;MI0000109|hsa-mir-103a-1&WithFlank&5|-|168560890|168560979@54.74.21&[MIMAT0000104&hsa-miR-107&offsets|0|-2;m-78&10|-|89592758|89592778&offsets|0|0];[MIMAT0000101&hsa-miR-103a-3p&offsets|0|-2;m-12&20|+|3917541|3917563&offsets|0|-2];[MIMAT0000101_1&hsa-miR-103a-3p&offsets|0|-2;m-13&5|-|168560904|168560926&offsets|0|-2;m-44220&5|-|168560908|168560926&offsets|0|+2]|AGCAGCATTGTACAGGGCTAT</td><td>>MI0000114|hsa-mir-107&WithFlank&10|-|89592741|89592833</td><td>56</td><td>76</td><td>21</td><td>AGCAGCATTGTACAGGGCTAT</td><td>isomiR</td><td>FALSE</td><td>TRUE</td><td>FALSE</td></tr>
<tr><td>>hg38_21_+_8433172_8446622_RNA45SN1_45S_50ntFlanks@4738.4758.21|AAAGTCGGAGGTTCGAAGACG</td><td>>hg38_21_+_8433172_8446622_RNA45SN1_45S_50ntFlanks</td><td>4738</td><td>4758</td><td>21</td><td>AAAGTCGGAGGTTCGAAGACG</td><td>rRF</td><td>FALSE</td><td>TRUE</td><td>TRUE</td></tr>
<tr><td>>MI0000102|hsa-mir-100&WithFlank&11|-|122152223|122152314@19.38.20(+1G)&[MIMAT0000098&hsa-miR-100-5p&offsets|0|-2(+1G);m-27&11|-|122152276|122152296&offsets|0|-1(+1G)]|AACCCGTAGATCCGAACTTGG</td><td>>MI0000102|hsa-mir-100&WithFlank&11|-|122152223|122152314</td><td>19</td><td>38</td><td>20(+1G)</td><td>AACCCGTAGATCCGAACTTGG</td><td>isomiR</td><td>FALSE</td><td>TRUE</td><td>FALSE</td></tr>
<tr><td>>hg38_MT_+_1621_3279_RNR2_16S_50ntFlanks@51.95.45|GCTAAACCTAGCCCCAAACCCACTCCACCTTACTACCAGACAACC</td><td>>hg38_MT_+_1621_3279_RNR2_16S_50ntFlanks</td><td>51</td><td>95</td><td>45</td><td>GCTAAACCTAGCCCCAAACCCACTCCACCTTACTACCAGACA[illegible]</td><td>rRF</td><td>FALSE</td><td>TRUE</td><td>FALSE</td></tr>
<tr><td>>multi-am_trnaMT_ProTGG_MT_-_15956_16023@53.71.19|AAGACTTTTTCTCTGACCA</td><td>>multi-am_trnaMT_ProTGG_MT_-_15956_16023</td><td>53</td><td>71</td><td>19</td><td>AAGACTTTTTCTCTGACCA</td><td>tRF</td><td>FALSE</td><td>TRUE</td><td>FALSE</td></tr>
<tr><td>>MI0000088|hsa-mir-30a&WithFlank&6|-|71403545|71403627@54.73.20(+1U)&[MIMAT0000088&hsa-miR-30a-3p&offsets|+1|-1(+1U);m-38&6|-|71403554|71403575&offsets|+1|-1(+1U)]|TTTCAGTCGGATGTTTGCAGT</td><td>>MI0000088|hsa-mir-30a&WithFlank&6|-|71403545|71403627</td><td>54</td><td>73</td><td>20(+1U)</td><td>TTTCAGTCGGATGTTTGCAGT</td><td>isomiR</td><td>FALSE</td><td>TRUE</td><td>FALSE</td></tr>
<tr><td>>hg38_MT_+_1621_3279_RNR2_16S_50ntFlanks@51.92.42|GCTAAACCTAGCCCCAAACCCACTCCACCTTACTACCAGACA</td><td>>hg38_MT_+_1621_3279_RNR2_16S_50ntFlanks</td><td>51</td><td>92</td><td>42</td><td>GCTAAACCTAGCCCCAAACCCACTCCACCTTACTACCAGACA</td><td>rRF</td><td>FALSE</td><td>TRUE</td><td>FALSE</td></tr>
<tr><td>>trna119_LysCTT_1_-_145395522_145395594@1.33.33,trna11_LysCTT_5_-_180648979_180649051@1.33.33,trna13_LysCTT_6_+_26556774_26556846@1.33.33,trna7_LysCTT_16_+_3225692_3225764@1.33.33,trna9_LysCTT_5_+_180634755_180634827@1.33.33|GCCCGGCTAGCTCAGTCGGTAGAGCATGAGACT</td><td>>trna119_LysCTT_1_-_145395522_145395594</td><td>1</td><td>33</td><td>33</td><td>GCCCGGCTAGCTCAGTCGGTAGAGCATGAGACT</td><td>tRF</td><td>FALSE</td><td>TRUE</td><td>FALSE</td></tr>
<tr><td>>MI0000433|hsa-let-7g&WithFlank&3|-|52268272|52268367@11.28.18&[MIMAT0000414&hsa-let-7g-5p&offsets|0|-4;m-58&3|-|52268336|52268357&offsets|0|-4]|TGAGGTAGTAGTTTGTAC</td><td>>MI0000433|hsa-let-7g&WithFlank&3|-|52268272|52268367</td><td>11</td><td>28</td><td>18</td><td>TGAGGTAGTAGTTTGTAC</td><td>isomiR</td><td>FALSE</td><td>TRUE</td><td>FALSE</td></tr>
<tr><td>>MI0000811|hsa-mir-148b&WithFlank&12|+|54337210|54337320@69.90.22(+1U)&[MIMAT0000759&hsa-miR-148b-3p&offsets|0|0(+1U);m-77&12|+|54337278|54337299&offsets|0|0(+1U)]|TCAGTGCATCACAGAACTTTGTT</td><td>>MI0000811|hsa-mir-148b&WithFlank&12|+|54337210|54337320</td><td>69</td><td>90</td><td>22(+1U)</td><td>TCAGTGCATCACAGAACTTTGTT</td><td>isomiR</td><td>FALSE</td><td>TRUE</td><td>FALSE</td></tr>
<tr><td>>MI0000077|hsa-mir-21&WithFlank&17|+|59841260|59841343@13.35.23&[MIMAT0000076&hsa-miR-21-5p&offsets|-1|0;m-1&17|+|59841273|59841294&offsets|-1|0]|GTAGCTTATCAGACTGATGTTGA</td><td>>MI0000077|hsa-mir-21&WithFlank&17|+|59841260|59841343</td><td>13</td><td>35</td><td>23</td><td>GTAGCTTATCAGACTGATGTTGA</td><td>isomiR</td><td>FALSE</td><td>TRUE</td><td>FALSE</td></tr>
<tr><td>>MI0000093|hsa-mir-92a-1&WithFlank&13|+|91351308|91351397@54.76.23&[MIMAT0000092&hsa-miR-92a-3p&offsets|0|+1;m-9&13|+|91351361|91351382&offsets|0|+1]|TATTGCACTTGTCCCGGCCTGTT</td><td>>MI0000093|hsa-mir-92a-1&WithFlank&13|+|91351308|91351397</td><td>54</td><td>76</td><td>23</td><td>TATTGCACTTGTCCCGGCCTGTT</td><td>isomiR</td><td>FALSE</td><td>TRUE</td><td>FALSE</td></tr>
<tr><td>>MI0000299|hsa-mir-222&WithFlank&X|-|45747009|45747130@75.98.24(+1C)&[MIMAT0000279&hsa-miR-222-3p&offsets|0|+3(+1C);m-140&X|-|45747033|45747056&offsets|0|0(+1C)]|AGCTACATCTGGCTACTGGGTCTCC</td><td>>MI0000299|hsa-mir-222&WithFlank&X|-|45747009|45747130</td><td>75</td><td>98</td><td>24(+1C)</td><td>AGCTACATCTGGCTACTGGGTCTCC</td><td>isomiR</td><td>FALSE</td><td>TRUE</td><td>FALSE</td></tr>
<tr><td>>MI0000077|hsa-mir-21&WithFlank&17|+|59841260|59841343@14.33.20&[MIMAT0000076&hsa-miR-21-5p&offsets|0|-2;m-1&17|+|59841273|59841294&offsets|0|-2]|TAGCTTATCAGACTGATGTT</td><td>>MI0000077|hsa-mir-21&WithFlank&17|+|59841260|59841343</td><td>14</td><td>33</td><td>20</td><td>TAGCTTATCAGACTGATGTT</td><td>isomiR</td><td>FALSE</td><td>TRUE</td><td>FALSE</td></tr>
<tr><td>>MI0000477|hsa-mir-146a&WithFlank&5|+|160485346|160485456@27.48.22(+1A)&[MIMAT0000449&hsa-miR-146a-5p&offsets|0|0(+1A);m-129&5|+|160485372|160485393&offsets|0|0(+1A)]|TGAGAACTGAATTCCATGGGTTA</td><td>>MI0000477|hsa-mir-146a&WithFlank&5|+|160485346|160485456</td><td>27</td><td>48</td><td>22(+1A)</td><td>TGAGAACTGAATTCCATGGGTTA</td><td>isomiR</td><td>FALSE</td><td>TRUE</td><td>FALSE</td></tr>
<tr><td>>MI0000079|hsa-mir-23a&WithFlank&19|-|13836581|13836665@54.72.19&[MIMAT0000078&hsa-miR-23a-3p&offsets|+3|+1;m-31&19|-|13836595|13836615&offsets|+3|+1]|ACATTGCCAGGGATTTCCA</td><td>>MI0000079|hsa-mir-23a&WithFlank&19|-|13836581|13836665</td><td>54</td><td>72</td><td>19</td><td>ACATTGCCAGGGATTTCCA</td><td>isomiR</td><td>FALSE</td><td>TRUE</td><td>FALSE</td></tr>
<tr><td>>MI0000433|hsa-let-7g&WithFlank&3|-|52268272|52268367@11.33.23&[MIMAT0000414&hsa-let-7g-5p&offsets|0|+1;m-58&3|-|52268336|52268357&offsets|0|+1]|TGAGGTAGTAGTTTGTACAGTTT</td><td>>MI0000433|hsa-let-7g&WithFlank&3|-|52268272|52268367</td><td>11</td><td>33</td><td>23</td><td>TGAGGTAGTAGTTTGTACAGTTT</td><td>isomiR</td><td>FALSE</td><td>TRUE</td><td>FALSE</td></tr>
<tr><td>>hg38_21_+_8433172_8446622_RNA45SN1_45S_50ntFlanks@11641.11660.20|CGGCGGGTGTTGACGCGATG</td><td>>hg38_21_+_8433172_8446622_RNA45SN1_45S_50ntFlanks</td><td>11641</td><td>11660</td><td>20</td><td>CGGCGGGTGTTGACGCGATG</td><td>rRF</td><td>FALSE</td><td>FALSE</td><td>TRUE</td></tr>
</table>

### Mitochondria

| | | | | | | | | | |
|---|---|---|---|---|---|---|---|---|---|
| >hg38_21_+_8433172_8446622_RNA45SN1_45S_50ntFlanks@8114.8140.27\|GCGCGGGACATGTGGCGTACGGAAGAC | >hg38_21_+_8433172_8446622_RNA45SN1_45S_50ntFlanks | 8114 | 8140 | 27 | GCGCGGGACATGTGGCGTACGGAAGAC | rRF | FALSE | FALSE | TRUE |
| >hg38_21_+_8433172_8446622_RNA45SN1_45S_50ntFlanks@11579.11603.25\|AGAACTGGTGCGGACCAGGGGAATC | >hg38_21_+_8433172_8446622_RNA45SN1_45S_50ntFlanks | 11579 | 11603 | 25 | AGAACTGGTGCGGACCAGGGGAATC | rRF | FALSE | FALSE | TRUE |
| >hg38_21_+_8433172_8446622_RNA45SN1_45S_50ntFlanks@5487.5514.28\|CGCTGAGAAGACGGTCGAACTTGACTAT | >hg38_21_+_8433172_8446622_RNA45SN1_45S_50ntFlanks | 5487 | 5514 | 28 | CGCTGAGAAGACGGTCGAACTTGACTAT | rRF | FALSE | FALSE | TRUE |
| >hg38_21_+_8433172_8446622_RNA45SN1_45S_50ntFlanks@11572.11591.20\|CCGACTTAGAACTGGTGCGG | >hg38_21_+_8433172_8446622_RNA45SN1_45S_50ntFlanks | 11572 | 11591 | 20 | CCGACTTAGAACTGGTGCGG | rRF | FALSE | FALSE | TRUE |
| >hg38_21_+_8433172_8446622_RNA45SN1_45S_50ntFlanks@11787.11804.18\|AATGGATGAACGAGATTC | >hg38_21_+_8433172_8446622_RNA45SN1_45S_50ntFlanks | 11787 | 11804 | 18 | AATGGATGAACGAGATTC | rRF | FALSE | FALSE | TRUE |
| >hg38_21_+_8433172_8446622_RNA45SN1_45S_50ntFlanks@12456.12475.20\|CCAAGCGTTGGATTGTTCAC | >hg38_21_+_8433172_8446622_RNA45SN1_45S_50ntFlanks | 12456 | 12475 | 20 | CCAAGCGTTGGATTGTTCAC | rRF | FALSE | FALSE | TRUE |
| >hg38_21_+_8433172_8446622_RNA45SN1_45S_50ntFlanks@11641.11665.25\|CGGCGGGTGTTGACGCGATGTGATT | >hg38_21_+_8433172_8446622_RNA45SN1_45S_50ntFlanks | 11641 | 11665 | 25 | CGGCGGGTGTTGACGCGATGTGATT | rRF | FALSE | FALSE | TRUE |
| >hg38_21_+_8433172_8446622_RNA45SN1_45S_50ntFlanks@7984.8006.23\|AGATCAGACGTGGCGACCCGCTG | >hg38_21_+_8433172_8446622_RNA45SN1_45S_50ntFlanks | 7984 | 8006 | 23 | AGATCAGACGTGGCGACCCGCTG | rRF | FALSE | FALSE | TRUE |
| >hg38_21_+_8433172_8446622_RNA45SN1_45S_50ntFlanks@11572.11603.32\|CCGACTTAGAACTGGTGCGGACCAGGGGAATC | >hg38_21_+_8433172_8446622_RNA45SN1_45S_50ntFlanks | 11572 | 11603 | 32 | CCGACTTAGAACTGGTGCGGACCAGGGGAATC | rRF | FALSE | FALSE | TRUE |
| >hg38_21_+_8433172_8446622_RNA45SN1_45S_50ntFlanks@8357.8377.21\|AAAAGAACTTTGAAGAGAGAG | >hg38_21_+_8433172_8446622_RNA45SN1_45S_50ntFlanks | 8357 | 8377 | 21 | AAAAGAACTTTGAAGAGAGAG | rRF | FALSE | FALSE | TRUE |
| >hg38_21_+_8433172_8446622_RNA45SN1_45S_50ntFlanks@7991.8009.19\|ACGTGGCGACCCGCTGAAT | >hg38_21_+_8433172_8446622_RNA45SN1_45S_50ntFlanks | 7991 | 8009 | 19 | ACGTGGCGACCCGCTGAAT | rRF | FALSE | FALSE | TRUE |
| >hg38_21_+_8433172_8446622_RNA45SN1_45S_50ntFlanks@11641.11662.22\|CGGCGGGTGTTGACGCGATGTG | >hg38_21_+_8433172_8446622_RNA45SN1_45S_50ntFlanks | 11641 | 11662 | 22 | CGGCGGGTGTTGACGCGATGTG | rRF | FALSE | FALSE | TRUE |
| >hg38_21_+_8433172_8446622_RNA45SN1_45S_50ntFlanks@11208.11232.25\|GCGGCGACTCTGGACGCGAGCCGGG | >hg38_21_+_8433172_8446622_RNA45SN1_45S_50ntFlanks | 11208 | 11232 | 25 | GCGGCGACTCTGGACGCGAGCCGGG | rRF | FALSE | FALSE | TRUE |
| >hg38_21_+_8433172_8446622_RNA45SN1_45S_50ntFlanks@12163.12182.20\|ACTGGGGCGGTACACCTGTC | >hg38_21_+_8433172_8446622_RNA45SN1_45S_50ntFlanks | 12163 | 12182 | 20 | ACTGGGGCGGTACACCTGTC | rRF | FALSE | FALSE | TRUE |
| >hg38_1_-_228634819_228635039_RNA5S12_5S_50ntFlanks@151.171.21\|ATACCGGGTGCTGTAGGCTTT | >hg38_1_-_228634819_228635039_RNA5S12_5S_50ntFlanks | 151 | 171 | 21 | ATACCGGGTGCTGTAGGCTTT | rRF | FALSE | FALSE | TRUE |
| >hg38_21_+_8433172_8446622_RNA45SN1_45S_50ntFlanks@10510.10533.24\|AATCCGGAGTGGCGGAGATGGGCG | >hg38_21_+_8433172_8446622_RNA45SN1_45S_50ntFlanks | 10510 | 10533 | 24 | AATCCGGAGTGGCGGAGATGGGCG | rRF | FALSE | FALSE | TRUE |
| >hg38_21_+_8433172_8446622_RNA45SN1_45S_50ntFlanks@10679.10706.28\|GGTTCCGGCGGCGTCCGGTGAGCTCTCG | >hg38_21_+_8433172_8446622_RNA45SN1_45S_50ntFlanks | 10679 | 10706 | 28 | GGTTCCGGCGGCGTCCGGTGAGCTCTCG | rRF | FALSE | FALSE | TRUE |
| >hg38_21_+_8433172_8446622_RNA45SN1_45S_50ntFlanks@6745.6770.26\|ACACTTCGAACGCACTTGCGGCCCCG | >hg38_21_+_8433172_8446622_RNA45SN1_45S_50ntFlanks | 6745 | 6770 | 26 | ACACTTCGAACGCACTTGCGGCCCCG | rRF | FALSE | FALSE | TRUE |
| >hg38_21_+_8433172_8446622_RNA45SN1_45S_50ntFlanks@10350.10368.19\|ACGAGAACTTTGAAGGCCG | >hg38_21_+_8433172_8446622_RNA45SN1_45S_50ntFlanks | 10350 | 10368 | 19 | ACGAGAACTTTGAAGGCCG | rRF | FALSE | FALSE | TRUE |
| >hg38_21_+_8433172_8446622_RNA45SN1_45S_50ntFlanks@11323.11344.22\|CGTCTCGTCGCGCGCGCGTCCG | >hg38_21_+_8433172_8446622_RNA45SN1_45S_50ntFlanks | 11323 | 11344 | 22 | CGTCTCGTCGCGCGCGCGTCCG | rRF | FALSE | FALSE | TRUE |
| >hg38_21_+_8433172_8446622_RNA45SN1_45S_50ntFlanks@8105.8138.34\|CGCGGCGGGGCGCGGGACATGTGGCGTACGGAAG | >hg38_21_+_8433172_8446622_RNA45SN1_45S_50ntFlanks | 8105 | 8138 | 34 | CGCGGCGGGGCGCGGGACATGTGGCGTACGGAAG | rRF | FALSE | FALSE | TRUE |
| >hg38_21_+_8433172_8446622_RNA45SN1_45S_50ntFlanks@9572.9595.24\|CAAATCGGTCGTCCGACCTGGGTA | >hg38_21_+_8433172_8446622_RNA45SN1_45S_50ntFlanks | 9572 | 9595 | 24 | CAAATCGGTCGTCCGACCTGGGTA | rRF | FALSE | FALSE | TRUE |
| >hg38_21_+_8433172_8446622_RNA45SN1_45S_50ntFlanks@8104.8140.37\|CCGCGGCGGGGCGCGGGACATGTGGCGTACGGAAGAC | >hg38_21_+_8433172_8446622_RNA45SN1_45S_50ntFlanks | 8104 | 8140 | 37 | CCGCGGCGGGGCGCGGGACATGTGGCGTACGGAAGAC | rRF | FALSE | FALSE | TRUE |
| >hg38_21_+_8433172_8446622_RNA45SN1_45S_50ntFlanks@9574.9595.22\|AATCGGTCGTCCGACCTGGGTA | >hg38_21_+_8433172_8446622_RNA45SN1_45S_50ntFlanks | 9574 | 9595 | 22 | AATCGGTCGTCCGACCTGGGTA | rRF | FALSE | FALSE | TRUE |
| >hg38_21_+_8433172_8446622_RNA45SN1_45S_50ntFlanks@5492.5514.23\|AGAAGACGGTCGAACTTGACTAT | >hg38_21_+_8433172_8446622_RNA45SN1_45S_50ntFlanks | 5492 | 5514 | 23 | AGAAGACGGTCGAACTTGACTAT | rRF | FALSE | FALSE | TRUE |
| >hg38_21_+_8433172_8446622_RNA45SN1_45S_50ntFlanks@12626.12653.28\|AATGGGGCGAAGCTACCATCTGTGGGAT | >hg38_21_+_8433172_8446622_RNA45SN1_45S_50ntFlanks | 12626 | 12653 | 28 | AATGGGGCGAAGCTACCATCTGTGGGAT | rRF | FALSE | FALSE | TRUE |
| >hg38_21_+_8433172_8446622_RNA45SN1_45S_50ntFlanks@8176.8203.28\|CTTCTGATCGAGGCCCAGCCCGTGGACG | >hg38_21_+_8433172_8446622_RNA45SN1_45S_50ntFlanks | 8176 | 8203 | 28 | CTTCTGATCGAGGCCCAGCCCGTGGACG | rRF | FALSE | FALSE | TRUE |
| >hg38_21_+_8433172_8446622_RNA45SN1_45S_50ntFlanks@9575.9596.22\|ATCGGTCGTCCGACCTGGGTAT | >hg38_21_+_8433172_8446622_RNA45SN1_45S_50ntFlanks | 9575 | 9596 | 22 | ATCGGTCGTCCGACCTGGGTAT | rRF | FALSE | FALSE | TRUE |
| >hg38_21_+_8433172_8446622_RNA45SN1_45S_50ntFlanks@10351.10384.34\|CGAGAACTTTGAAGGCCGAAGTGGAGAAGGGTTC | >hg38_21_+_8433172_8446622_RNA45SN1_45S_50ntFlanks | 10351 | 10384 | 34 | CGAGAACTTTGAAGGCCGAAGTGGAGAAGGGTTC | rRF | FALSE | FALSE | TRUE |
| >hg38_21_+_8433172_8446622_RNA45SN1_45S_50ntFlanks@12456.12474.19\|CCAAGCGTTGGATTGTTCA | >hg38_21_+_8433172_8446622_RNA45SN1_45S_50ntFlanks | 12456 | 12474 | 19 | CCAAGCGTTGGATTGTTCA | rRF | FALSE | FALSE | TRUE |
| >hg38_21_+_8433172_8446622_RNA45SN1_45S_50ntFlanks@12627.12653.27\|ATGGGGCGAAGCTACCATCTGTGGGAT | >hg38_21_+_8433172_8446622_RNA45SN1_45S_50ntFlanks | 12627 | 12653 | 27 | ATGGGGCGAAGCTACCATCTGTGGGAT | rRF | FALSE | FALSE | TRUE |
| >hg38_21_+_8433172_8446622_RNA45SN1_45S_50ntFlanks@6700.6718.19\|CTAGCTGCGAGAATTAATG | >hg38_21_+_8433172_8446622_RNA45SN1_45S_50ntFlanks | 6700 | 6718 | 19 | CTAGCTGCGAGAATTAATG | rRF | FALSE | FALSE | TRUE |
| >hg38_21_+_8433172_8446622_RNA45SN1_45S_50ntFlanks@9573.9596.24\|AAATCGGTCGTCCGACCTGGGTAT | >hg38_21_+_8433172_8446622_RNA45SN1_45S_50ntFlanks | 9573 | 9596 | 24 | AAATCGGTCGTCCGACCTGGGTAT | rRF | FALSE | FALSE | TRUE |
| >hg38_21_+_8433172_8446622_RNA45SN1_45S_50ntFlanks@10351.10376.26\|CGAGAACTTTGAAGGCCGAAGTGGAG | >hg38_21_+_8433172_8446622_RNA45SN1_45S_50ntFlanks | 10351 | 10376 | 26 | CGAGAACTTTGAAGGCCGAAGTGGAG | rRF | FALSE | FALSE | TRUE |
| >hg38_21_+_8433172_8446622_RNA45SN1_45S_50ntFlanks@12635.12653.19\|AAGCTACCATCTGTGGGAT | >hg38_21_+_8433172_8446622_RNA45SN1_45S_50ntFlanks | 12635 | 12653 | 19 | AAGCTACCATCTGTGGGAT | rRF | FALSE | FALSE | TRUE |
| >hg38_21_+_8433172_8446622_RNA45SN1_45S_50ntFlanks@12163.12180.18\|ACTGGGGCGGTACACCTG | >hg38_21_+_8433172_8446622_RNA45SN1_45S_50ntFlanks | 12163 | 12180 | 18 | ACTGGGGCGGTACACCTG | rRF | FALSE | FALSE | TRUE |
| >hg38_21_+_8433172_8446622_RNA45SN1_45S_50ntFlanks@8356.8378.23\|GAAAAGAACTTTGAAGAGAGAGT | >hg38_21_+_8433172_8446622_RNA45SN1_45S_50ntFlanks | 8356 | 8378 | 23 | GAAAAGAACTTTGAAGAGAGAGT | rRF | FALSE | FALSE | TRUE |
| >hg38_21_+_8433172_8446622_RNA45SN1_45S_50ntFlanks@10819.10840.22\|AGGGAAGTCGGCAAGCCGGATC | >hg38_21_+_8433172_8446622_RNA45SN1_45S_50ntFlanks | 10819 | 10840 | 22 | AGGGAAGTCGGCAAGCCGGATC | rRF | FALSE | FALSE | TRUE |
| >hg38_21_+_8433172_8446622_RNA45SN1_45S_50ntFlanks@10351.10374.24\|CGAGAACTTTGAAGGCCGAAGTGG | >hg38_21_+_8433172_8446622_RNA45SN1_45S_50ntFlanks | 10351 | 10374 | 24 | CGAGAACTTTGAAGGCCGAAGTGG | rRF | FALSE | FALSE | TRUE |
| >hg38_21_+_8433172_8446622_RNA45SN1_45S_50ntFlanks@10678.10708.31\|CGGTTCCGGCGGCGTCCGGTGAGCTCTCGCT | >hg38_21_+_8433172_8446622_RNA45SN1_45S_50ntFlanks | 10678 | 10708 | 31 | CGGTTCCGGCGGCGTCCGGTGAGCTCTCGCT | rRF | FALSE | FALSE | TRUE |

### Mitochondria

| | | | | | | | | | |
|---|---|---|---|---|---|---|---|---|---|
| >am_trna146_GlnCTG_6_-_27515531_27515602@1.29.29,trna1_GlnCTG_6_+_18836402_18836473@1.29.29,trna3_GlnCTG_17_+_8023070_8023141@1.29.29,trna42_GlnCTG_6_+_27263212_27263283@1.29.29,trna49_GlnCTG_6_+_27487308_27487379@1.29.29,trna7_GlnCTG_15_-_66161400_66161471@1.29.29,trna99_GlnCTG_6_-_28909378_28909449@1.29.29\|GGTTCCATGGTGTAATGGTTAGCACTCTG | >am_trna146_GlnCTG_6_-_27515531_27515602 | 1 | 29 | 29 | GGTTCCATGGTGTAATGGTTAGCACTCTG | tRF | FALSE | FALSE | TRUE |
| >MI0000737\|hsa-mir-200a&WithFlank&1\|+\|1167857\|1167958@60.82.23&[MIMAT0000682&hsa-miR-200a-3p&offsets\|0\|+1;m-142&1\|+\|1167916\|1167938&offsets\|0\|0]\|TAACACTGTCTGGTAACGATGTT | >MI0000737\|hsa-mir-200a&WithFlank&1\|+\|1167857\|1167958 | 60 | 82 | 23 | TAACACTGTCTGGTAACGATGTT | isomiR | FALSE | FALSE | TRUE |
| >hg38_1_-_228634819_228635039_RNA5S12_5S_50ntFlanks@140.171.32\|ACCGCCTGGGAATACCGGGTGCTGTAGGCTTT | >hg38_1_-_228634819_228635039_RNA5S12_5S_50ntFlanks | 140 | 171 | 32 | ACCGCCTGGGAATACCGGGTGCTGTAGGCTTT | rRF | FALSE | FALSE | TRUE |
| >hg38_21_+_8433172_8446622_RNA45SN1_45S_50ntFlanks@10351.10388.38\|CGAGAACTTTGAAGGCCGAAGTGGAGAAGGGTTCCATG | >hg38_21_+_8433172_8446622_RNA45SN1_45S_50ntFlanks | 10351 | 10388 | 38 | CGAGAACTTTGAAGGCCGAAGTGGAGAAGGGTTCCATG | rRF | FALSE | FALSE | TRUE |
| >hg38_21_+_8433172_8446622_RNA45SN1_45S_50ntFlanks@7989.8007.19\|AGACGTGGCGACCCGCTGA | >hg38_21_+_8433172_8446622_RNA45SN1_45S_50ntFlanks | 7989 | 8007 | 19 | AGACGTGGCGACCCGCTGA | rRF | FALSE | FALSE | TRUE |
| >hg38_21_+_8433172_8446622_RNA45SN1_45S_50ntFlanks@11729.11751.23\|TAACTATGACTCTCTTAAGGTAG | >hg38_21_+_8433172_8446622_RNA45SN1_45S_50ntFlanks | 11729 | 11751 | 23 | TAACTATGACTCTCTTAAGGTAG | rRF | FALSE | FALSE | TRUE |
| >hg38_21_+_8433172_8446622_RNA45SN1_45S_50ntFlanks@11874.11893.20\|AGAAGACCCTGTTGAGCTTG | >hg38_21_+_8433172_8446622_RNA45SN1_45S_50ntFlanks | 11874 | 11893 | 20 | AGAAGACCCTGTTGAGCTTG | rRF | FALSE | FALSE | TRUE |
| >hg38_21_+_8433172_8446622_RNA45SN1_45S_50ntFlanks@9554.9572.19\|CGTAGCGGTCCTGACGTGC | >hg38_21_+_8433172_8446622_RNA45SN1_45S_50ntFlanks | 9554 | 9572 | 19 | CGTAGCGGTCCTGACGTGC | rRF | FALSE | FALSE | TRUE |
| >hg38_21_+_8433172_8446622_RNA45SN1_45S_50ntFlanks@6651.6699.49\|CGACTCTTAGCGGTGGATCACTCGGCTCGTGCGTCGATGAAGAACGCAG | >hg38_21_+_8433172_8446622_RNA45SN1_45S_50ntFlanks | 6651 | 6699 | 49 | CGACTCTTAGCGGTGGATCACTCGGCTCGTGCGTCGATGAAG[illegible] | rRF | FALSE | FALSE | TRUE |
| >hg38_21_+_8433172_8446622_RNA45SN1_45S_50ntFlanks@7981.8006.26\|CTCAGATCAGACGTGGCGACCCGCTG | >hg38_21_+_8433172_8446622_RNA45SN1_45S_50ntFlanks | 7981 | 8006 | 26 | CTCAGATCAGACGTGGCGACCCGCTG | rRF | FALSE | FALSE | TRUE |
| >hg38_21_+_8433172_8446622_RNA45SN1_45S_50ntFlanks@9564.9596.33\|CTGACGTGCAAATCGGTCGTCCGACCTGGGTAT | >hg38_21_+_8433172_8446622_RNA45SN1_45S_50ntFlanks | 9564 | 9596 | 33 | CTGACGTGCAAATCGGTCGTCCGACCTGGGTAT | rRF | FALSE | FALSE | TRUE |
| >hg38_21_+_8433172_8446622_RNA45SN1_45S_50ntFlanks@4759.4780.22\|ATCAGATACCGTCGTAGTTCCG | >hg38_21_+_8433172_8446622_RNA45SN1_45S_50ntFlanks | 4759 | 4780 | 22 | ATCAGATACCGTCGTAGTTCCG | rRF | FALSE | FALSE | TRUE |
| >hg38_21_+_8433172_8446622_RNA45SN1_45S_50ntFlanks@12456.12488.33\|CCAAGCGTTGGATTGTTCACCCACTAATAGGGA | >hg38_21_+_8433172_8446622_RNA45SN1_45S_50ntFlanks | 12456 | 12488 | 33 | CCAAGCGTTGGATTGTTCACCCACTAATAGGGA | rRF | FALSE | FALSE | TRUE |
| >hg38_21_+_8433172_8446622_RNA45SN1_45S_50ntFlanks@6663.6689.27\|GTGGATCACTCGGCTCGTGCGTCGATG | >hg38_21_+_8433172_8446622_RNA45SN1_45S_50ntFlanks | 6663 | 6689 | 27 | GTGGATCACTCGGCTCGTGCGTCGATG | rRF | FALSE | FALSE | TRUE |
| >hg38_21_+_8433172_8446622_RNA45SN1_45S_50ntFlanks@12438.12455.18\|ATTGTGAAGCAGAATTCA | >hg38_21_+_8433172_8446622_RNA45SN1_45S_50ntFlanks | 12438 | 12455 | 18 | ATTGTGAAGCAGAATTCA | rRF | FALSE | FALSE | TRUE |
| >hg38_21_+_8433172_8446622_RNA45SN1_45S_50ntFlanks@10786.10818.33\|AACAGCCTCTGGCATGTTGGAACAATGTAGGTA | >hg38_21_+_8433172_8446622_RNA45SN1_45S_50ntFlanks | 10786 | 10818 | 33 | AACAGCCTCTGGCATGTTGGAACAATGTAGGTA | rRF | FALSE | FALSE | TRUE |
| >hg38_21_+_8433172_8446622_RNA45SN1_45S_50ntFlanks@5489.5514.26\|CTGAGAAGACGGTCGAACTTGACTAT | >hg38_21_+_8433172_8446622_RNA45SN1_45S_50ntFlanks | 5489 | 5514 | 26 | CTGAGAAGACGGTCGAACTTGACTAT | rRF | FALSE | FALSE | TRUE |
| >hg38_21_+_8433172_8446622_RNA45SN1_45S_50ntFlanks@7979.8006.28\|ACCTCAGATCAGACGTGGCGACCCGCTG | >hg38_21_+_8433172_8446622_RNA45SN1_45S_50ntFlanks | 7979 | 8006 | 28 | ACCTCAGATCAGACGTGGCGACCCGCTG | rRF | FALSE | FALSE | TRUE |
| >hg38_21_+_8433172_8446622_RNA45SN1_45S_50ntFlanks@6671.6699.29\|CTCGGCTCGTGCGTCGATGAAGAACGCAG | >hg38_21_+_8433172_8446622_RNA45SN1_45S_50ntFlanks | 6671 | 6699 | 29 | CTCGGCTCGTGCGTCGATGAAGAACGCAG | rRF | FALSE | FALSE | TRUE |
| >hg38_21_+_8433172_8446622_RNA45SN1_45S_50ntFlanks@3985.4002.18\|CGGCTTTGGTGACTCTAG | >hg38_21_+_8433172_8446622_RNA45SN1_45S_50ntFlanks | 3985 | 4002 | 18 | CGGCTTTGGTGACTCTAG | rRF | FALSE | FALSE | TRUE |
| >hg38_21_+_8433172_8446622_RNA45SN1_45S_50ntFlanks@12159.12182.24\|TTTGACTGGGGCGGTACACCTGTC | >hg38_21_+_8433172_8446622_RNA45SN1_45S_50ntFlanks | 12159 | 12182 | 24 | TTTGACTGGGGCGGTACACCTGTC | rRF | FALSE | FALSE | TRUE |
| >hg38_21_+_8433172_8446622_RNA45SN1_45S_50ntFlanks@10690.10708.19\|CGTCCGGTGAGCTCTCGCT | >hg38_21_+_8433172_8446622_RNA45SN1_45S_50ntFlanks | 10690 | 10708 | 19 | CGTCCGGTGAGCTCTCGCT | rRF | FALSE | FALSE | TRUE |
| >hg38_21_+_8433172_8446622_RNA45SN1_45S_50ntFlanks@9572.9603.32\|CAAATCGGTCGTCCGACCTGGGTATAGGGGCG | >hg38_21_+_8433172_8446622_RNA45SN1_45S_50ntFlanks | 9572 | 9603 | 32 | CAAATCGGTCGTCCGACCTGGGTATAGGGGCG | rRF | FALSE | FALSE | TRUE |
| >hg38_21_+_8433172_8446622_RNA45SN1_45S_50ntFlanks@3899.3919.21\|CCTTCGCGGGGGGGATGCGTG | >hg38_21_+_8433172_8446622_RNA45SN1_45S_50ntFlanks | 3899 | 3919 | 21 | CCTTCGCGGGGGGGATGCGTG | rRF | FALSE | FALSE | TRUE |
| >hg38_21_+_8433172_8446622_RNA45SN1_45S_50ntFlanks@12456.12486.31\|CCAAGCGTTGGATTGTTCACCCACTAATAGG | >hg38_21_+_8433172_8446622_RNA45SN1_45S_50ntFlanks | 12456 | 12486 | 31 | CCAAGCGTTGGATTGTTCACCCACTAATAGG | rRF | FALSE | FALSE | TRUE |
| >hg38_21_+_8433172_8446622_RNA45SN1_45S_50ntFlanks@10350.10374.25\|ACGAGAACTTTGAAGGCCGAAGTGG | >hg38_21_+_8433172_8446622_RNA45SN1_45S_50ntFlanks | 10350 | 10374 | 25 | ACGAGAACTTTGAAGGCCGAAGTGG | rRF | FALSE | FALSE | TRUE |
| >hg38_1_-_228634819_228635039_RNA5S12_5S_50ntFlanks@152.171.20\|TACCGGGTGCTGTAGGCTTT | >hg38_1_-_228634819_228635039_RNA5S12_5S_50ntFlanks | 152 | 171 | 20 | TACCGGGTGCTGTAGGCTTT | rRF | FALSE | FALSE | TRUE |
| >hg38_21_+_8433172_8446622_RNA45SN1_45S_50ntFlanks@6721.6740.20\|AATTGCAGGACACATTGATC | >hg38_21_+_8433172_8446622_RNA45SN1_45S_50ntFlanks | 6721 | 6740 | 20 | AATTGCAGGACACATTGATC | rRF | FALSE | FALSE | TRUE |
| >hg38_21_+_8433172_8446622_RNA45SN1_45S_50ntFlanks@8114.8141.28\|GCGCGGGACATGTGGCGTACGGAAGACC | >hg38_21_+_8433172_8446622_RNA45SN1_45S_50ntFlanks | 8114 | 8141 | 28 | GCGCGGGACATGTGGCGTACGGAAGACC | rRF | FALSE | FALSE | TRUE |
| >hg38_21_+_8433172_8446622_RNA45SN1_45S_50ntFlanks@9573.9597.25\|AAATCGGTCGTCCGACCTGGGTATA | >hg38_21_+_8433172_8446622_RNA45SN1_45S_50ntFlanks | 9573 | 9597 | 25 | AAATCGGTCGTCCGACCTGGGTATA | rRF | FALSE | FALSE | TRUE |
| >hg38_21_+_8433172_8446622_RNA45SN1_45S_50ntFlanks@7989.8012.24\|AGACGTGGCGACCCGCTGAATTTA | >hg38_21_+_8433172_8446622_RNA45SN1_45S_50ntFlanks | 7989 | 8012 | 24 | AGACGTGGCGACCCGCTGAATTTA | rRF | FALSE | FALSE | TRUE |
| >hg38_21_+_8433172_8446622_RNA45SN1_45S_50ntFlanks@6647.6681.35\|CGTACGACTCTTAGCGGTGGATCACTCGGCTCGTG | >hg38_21_+_8433172_8446622_RNA45SN1_45S_50ntFlanks | 6647 | 6681 | 35 | CGTACGACTCTTAGCGGTGGATCACTCGGCTCGTG | rRF | FALSE | FALSE | TRUE |
| >hg38_21_+_8433172_8446622_RNA45SN1_45S_50ntFlanks@12008.12030.23\|CCGCCGGTGAAATACCACTACTC | >hg38_21_+_8433172_8446622_RNA45SN1_45S_50ntFlanks | 12008 | 12030 | 23 | CCGCCGGTGAAATACCACTACTC | rRF | FALSE | FALSE | TRUE |
| >hg38_21_+_8433172_8446622_RNA45SN1_45S_50ntFlanks@6745.6771.27\|ACACTTCGAACGCACTTGCGGCCCCGG | >hg38_21_+_8433172_8446622_RNA45SN1_45S_50ntFlanks | 6745 | 6771 | 27 | ACACTTCGAACGCACTTGCGGCCCCGG | rRF | FALSE | FALSE | TRUE |
| >hg38_21_+_8433172_8446622_RNA45SN1_45S_50ntFlanks@12003.12032.30\|GCGGGCCGCCGGTGAAATACCACTACTCTG | >hg38_21_+_8433172_8446622_RNA45SN1_45S_50ntFlanks | 12003 | 12032 | 30 | GCGGGCCGCCGGTGAAATACCACTACTCTG | rRF | FALSE | FALSE | TRUE |
| >hg38_21_+_8433172_8446622_RNA45SN1_45S_50ntFlanks@11781.11804.24\|CGCATGAATGGATGAACGAGATTC | >hg38_21_+_8433172_8446622_RNA45SN1_45S_50ntFlanks | 11781 | 11804 | 24 | CGCATGAATGGATGAACGAGATTC | rRF | FALSE | FALSE | TRUE |
| >hg38_21_+_8433172_8446622_RNA45SN1_45S_50ntFlanks@8115.8140.26\|CGCGGGACATGTGGCGTACGGAAGAC | >hg38_21_+_8433172_8446622_RNA45SN1_45S_50ntFlanks | 8115 | 8140 | 26 | CGCGGGACATGTGGCGTACGGAAGAC | rRF | FALSE | FALSE | TRUE |
| >hg38_21_+_8433172_8446622_RNA45SN1_45S_50ntFlanks@10349.10383.35\|AACGAGAACTTTGAAGGCCGAAGTGGAGAAGGGTT | >hg38_21_+_8433172_8446622_RNA45SN1_45S_50ntFlanks | 10349 | 10383 | 35 | AACGAGAACTTTGAAGGCCGAAGTGGAGAAGGGTT | rRF | FALSE | FALSE | TRUE |

### Mitochondria

| | | | | | | | | | |
|---|---|---|---|---|---|---|---|---|---|
| >hg38_21_+_8433172_8446622_RNA45SN1_45S_50ntFlanks@7988.8012.25\|CAGACGTGGCGACCCGCTGAATTTA | >hg38_21_+_8433172_8446622_RNA45SN1_45S_50ntFlanks | 7988 | 8012 | 25 | CAGACGTGGCGACCCGCTGAATTTA | rRF | FALSE | FALSE | TRUE |
| >hg38_21_+_8433172_8446622_RNA45SN1_45S_50ntFlanks@11583.11603.21\|CTGGTGCGGACCAGGGGAATC | >hg38_21_+_8433172_8446622_RNA45SN1_45S_50ntFlanks | 11583 | 11603 | 21 | CTGGTGCGGACCAGGGGAATC | rRF | FALSE | FALSE | TRUE |
| >hg38_21_+_8433172_8446622_RNA45SN1_45S_50ntFlanks@10411.10430.20\|CAGTCGGTCCTGAGAGATGG | >hg38_21_+_8433172_8446622_RNA45SN1_45S_50ntFlanks | 10411 | 10430 | 20 | CAGTCGGTCCTGAGAGATGG | rRF | FALSE | FALSE | TRUE |
| >hg38_21_+_8433172_8446622_RNA45SN1_45S_50ntFlanks@4728.4758.31\|ATCAAGAACGAAAGTCGGAGGTTCGAAGACG | >hg38_21_+_8433172_8446622_RNA45SN1_45S_50ntFlanks | 4728 | 4758 | 31 | ATCAAGAACGAAAGTCGGAGGTTCGAAGACG | rRF | FALSE | FALSE | TRUE |
| >hg38_21_+_8433172_8446622_RNA45SN1_45S_50ntFlanks@12636.12654.19\|AGCTACCATCTGTGGGATT | >hg38_21_+_8433172_8446622_RNA45SN1_45S_50ntFlanks | 12636 | 12654 | 19 | AGCTACCATCTGTGGGATT | rRF | FALSE | FALSE | TRUE |
| >hg38_21_+_8433172_8446622_RNA45SN1_45S_50ntFlanks@6663.6681.19\|GTGGATCACTCGGCTCGTG | >hg38_21_+_8433172_8446622_RNA45SN1_45S_50ntFlanks | 6663 | 6681 | 19 | GTGGATCACTCGGCTCGTG | rRF | FALSE | FALSE | TRUE |
| >hg38_21_+_8433172_8446622_RNA45SN1_45S_50ntFlanks@6664.6681.18\|TGGATCACTCGGCTCGTG | >hg38_21_+_8433172_8446622_RNA45SN1_45S_50ntFlanks | 6664 | 6681 | 18 | TGGATCACTCGGCTCGTG | rRF | FALSE | FALSE | TRUE |
| >hg38_21_+_8433172_8446622_RNA45SN1_45S_50ntFlanks@12635.12659.25\|AAGCTACCATCTGTGGGATTATGAC | >hg38_21_+_8433172_8446622_RNA45SN1_45S_50ntFlanks | 12635 | 12659 | 25 | AAGCTACCATCTGTGGGATTATGAC | rRF | FALSE | FALSE | TRUE |
| >hg38_21_+_8433172_8446622_RNA45SN1_45S_50ntFlanks@6721.6744.24\|AATTGCAGGACACATTGATCATCG | >hg38_21_+_8433172_8446622_RNA45SN1_45S_50ntFlanks | 6721 | 6744 | 24 | AATTGCAGGACACATTGATCATCG | rRF | FALSE | FALSE | TRUE |
| >hg38_21_+_8433172_8446622_RNA45SN1_45S_50ntFlanks@12437.12454.18\|CATTGTGAAGCAGAATTC | >hg38_21_+_8433172_8446622_RNA45SN1_45S_50ntFlanks | 12437 | 12454 | 18 | CATTGTGAAGCAGAATTC | rRF | FALSE | FALSE | TRUE |
| >hg38_21_+_8433172_8446622_RNA45SN1_45S_50ntFlanks@12405.12432.28\|CTTTTTGATCCTTCGATGTCGGCTCTTC | >hg38_21_+_8433172_8446622_RNA45SN1_45S_50ntFlanks | 12405 | 12432 | 28 | CTTTTTGATCCTTCGATGTCGGCTCTTC | rRF | FALSE | FALSE | TRUE |
| >hg38_21_+_8433172_8446622_RNA45SN1_45S_50ntFlanks@11661.11685.25\|TGATTTCTGCCCAGTGCTCTGAATG | >hg38_21_+_8433172_8446622_RNA45SN1_45S_50ntFlanks | 11661 | 11685 | 25 | TGATTTCTGCCCAGTGCTCTGAATG | rRF | FALSE | FALSE | TRUE |
| >hg38_21_+_8433172_8446622_RNA45SN1_45S_50ntFlanks@12163.12181.19\|ACTGGGGCGGTACACCTGT | >hg38_21_+_8433172_8446622_RNA45SN1_45S_50ntFlanks | 12163 | 12181 | 19 | ACTGGGGCGGTACACCTGT | rRF | FALSE | FALSE | TRUE |
| >hg38_21_+_8433172_8446622_RNA45SN1_45S_50ntFlanks@9575.9597.23\|ATCGGTCGTCCGACCTGGGTATA | >hg38_21_+_8433172_8446622_RNA45SN1_45S_50ntFlanks | 9575 | 9597 | 23 | ATCGGTCGTCCGACCTGGGTATA | rRF | FALSE | FALSE | TRUE |
| >hg38_21_+_8433172_8446622_RNA45SN1_45S_50ntFlanks@12638.12661.24\|CTACCATCTGTGGGATTATGACTG | >hg38_21_+_8433172_8446622_RNA45SN1_45S_50ntFlanks | 12638 | 12661 | 24 | CTACCATCTGTGGGATTATGACTG | rRF | FALSE | FALSE | TRUE |
| >hg38_21_+_8433172_8446622_RNA45SN1_45S_50ntFlanks@11569.11603.35\|CAGCCGACTTAGAACTGGTGCGGACCAGGGGAATC | >hg38_21_+_8433172_8446622_RNA45SN1_45S_50ntFlanks | 11569 | 11603 | 35 | CAGCCGACTTAGAACTGGTGCGGACCAGGGGAATC | rRF | FALSE | FALSE | TRUE |
| >hg38_21_+_8433172_8446622_RNA45SN1_45S_50ntFlanks@11643.11660.18\|GCGGGTGTTGACGCGATG | >hg38_21_+_8433172_8446622_RNA45SN1_45S_50ntFlanks | 11643 | 11660 | 18 | GCGGGTGTTGACGCGATG | rRF | FALSE | FALSE | TRUE |
| >hg38_21_+_8433172_8446622_RNA45SN1_45S_50ntFlanks@11575.11603.29\|ACTTAGAACTGGTGCGGACCAGGGGAATC | >hg38_21_+_8433172_8446622_RNA45SN1_45S_50ntFlanks | 11575 | 11603 | 29 | ACTTAGAACTGGTGCGGACCAGGGGAATC | rRF | FALSE | FALSE | TRUE |
| >hg38_21_+_8433172_8446622_RNA45SN1_45S_50ntFlanks@7991.8012.22\|ACGTGGCGACCCGCTGAATTTA | >hg38_21_+_8433172_8446622_RNA45SN1_45S_50ntFlanks | 7991 | 8012 | 22 | ACGTGGCGACCCGCTGAATTTA | rRF | FALSE | FALSE | TRUE |
| >hg38_21_+_8433172_8446622_RNA45SN1_45S_50ntFlanks@9574.9603.30\|AATCGGTCGTCCGACCTGGGTATAGGGGCG | >hg38_21_+_8433172_8446622_RNA45SN1_45S_50ntFlanks | 9574 | 9603 | 30 | AATCGGTCGTCCGACCTGGGTATAGGGGCG | rRF | FALSE | FALSE | TRUE |
| >hg38_21_+_8433172_8446622_RNA45SN1_45S_50ntFlanks@4730.4758.29\|CAAGAACGAAAGTCGGAGGTTCGAAGACG | >hg38_21_+_8433172_8446622_RNA45SN1_45S_50ntFlanks | 4730 | 4758 | 29 | CAAGAACGAAAGTCGGAGGTTCGAAGACG | rRF | FALSE | FALSE | TRUE |
| >hg38_21_+_8433172_8446622_RNA45SN1_45S_50ntFlanks@11572.11592.21\|CCGACTTAGAACTGGTGCGGA | >hg38_21_+_8433172_8446622_RNA45SN1_45S_50ntFlanks | 11572 | 11592 | 21 | CCGACTTAGAACTGGTGCGGA | rRF | FALSE | FALSE | TRUE |
| >hg38_21_+_8433172_8446622_RNA45SN1_45S_50ntFlanks@6650.6681.32\|ACGACTCTTAGCGGTGGATCACTCGGCTCGTG | >hg38_21_+_8433172_8446622_RNA45SN1_45S_50ntFlanks | 6650 | 6681 | 32 | ACGACTCTTAGCGGTGGATCACTCGGCTCGTG | rRF | FALSE | FALSE | TRUE |
| >am_trna116_GluCTC_1_-_145399233_145399304@1.28.28,trna59_GluCTC_1_+_249168447_249168518@1.28.28,trna71_GluCTC_1_-_161439189_161439260@1.28.28,trna74_GluCTC_1_-_161431809_161431880@1.28.28,trna77_GluCTC_1_-_161424398_161424469@1.28.28,trna77_GluCTC_6_+_28949976_28950047@1.28.28,trna80_GluCTC_1_-_161417018_161417089@1.28.28,trna87_GluCTC_6_-_126101393_126101464@1.28.28\|TCCCTGGTGGTCTAGTGGTTAGGATTCG | >am_trna116_GluCTC_1_-_145399233_145399304 | 1 | 28 | 28 | TCCCTGGTGGTCTAGTGGTTAGGATTCG | tRF | FALSE | FALSE | TRUE |
| >hg38_21_+_8433172_8446622_RNA45SN1_45S_50ntFlanks@8380.8405.26\|CAAGAGGGCGTGAAACCGTTAAGAGG | >hg38_21_+_8433172_8446622_RNA45SN1_45S_50ntFlanks | 8380 | 8405 | 26 | CAAGAGGGCGTGAAACCGTTAAGAGG | rRF | FALSE | FALSE | TRUE |
| >hg38_21_+_8433172_8446622_RNA45SN1_45S_50ntFlanks@9352.9370.19\|CCGTGGCGCAATGAAGGTG | >hg38_21_+_8433172_8446622_RNA45SN1_45S_50ntFlanks | 9352 | 9370 | 19 | CCGTGGCGCAATGAAGGTG | rRF | FALSE | FALSE | TRUE |
| >hg38_21_+_8433172_8446622_RNA45SN1_45S_50ntFlanks@9554.9573.20\|CGTAGCGGTCCTGACGTGCA | >hg38_21_+_8433172_8446622_RNA45SN1_45S_50ntFlanks | 9554 | 9573 | 20 | CGTAGCGGTCCTGACGTGCA | rRF | FALSE | FALSE | TRUE |
| >hg38_21_+_8433172_8446622_RNA45SN1_45S_50ntFlanks@7990.8008.19\|GACGTGGCGACCCGCTGAA | >hg38_21_+_8433172_8446622_RNA45SN1_45S_50ntFlanks | 7990 | 8008 | 19 | GACGTGGCGACCCGCTGAA | rRF | FALSE | FALSE | TRUE |
| >hg38_21_+_8433172_8446622_RNA45SN1_45S_50ntFlanks@10411.10443.33\|CAGTCGGTCCTGAGAGATGGGCGAGCGCCGTTC | >hg38_21_+_8433172_8446622_RNA45SN1_45S_50ntFlanks | 10411 | 10443 | 33 | CAGTCGGTCCTGAGAGATGGGCGAGCGCCGTTC | rRF | FALSE | FALSE | TRUE |
| >hg38_21_+_8433172_8446622_RNA45SN1_45S_50ntFlanks@8380.8400.21\|CAAGAGGGCGTGAAACCGTTA | >hg38_21_+_8433172_8446622_RNA45SN1_45S_50ntFlanks | 8380 | 8400 | 21 | CAAGAGGGCGTGAAACCGTTA | rRF | FALSE | FALSE | TRUE |
| >hg38_21_+_8433172_8446622_RNA45SN1_45S_50ntFlanks@4731.4759.29\|AAGAACGAAAGTCGGAGGTTCGAAGACGA | >hg38_21_+_8433172_8446622_RNA45SN1_45S_50ntFlanks | 4731 | 4759 | 29 | AAGAACGAAAGTCGGAGGTTCGAAGACGA | rRF | FALSE | FALSE | TRUE |
| >hg38_21_+_8433172_8446622_RNA45SN1_45S_50ntFlanks@9567.9596.30\|ACGTGCAAATCGGTCGTCCGACCTGGGTAT | >hg38_21_+_8433172_8446622_RNA45SN1_45S_50ntFlanks | 9567 | 9596 | 30 | ACGTGCAAATCGGTCGTCCGACCTGGGTAT | rRF | FALSE | FALSE | TRUE |
| >hg38_21_+_8433172_8446622_RNA45SN1_45S_50ntFlanks@7984.8008.25\|AGATCAGACGTGGCGACCCGCTGAA | >hg38_21_+_8433172_8446622_RNA45SN1_45S_50ntFlanks | 7984 | 8008 | 25 | AGATCAGACGTGGCGACCCGCTGAA | rRF | FALSE | FALSE | TRUE |
| >hg38_21_+_8433172_8446622_RNA45SN1_45S_50ntFlanks@12159.12181.23\|TTTGACTGGGGCGGTACACCTGT | >hg38_21_+_8433172_8446622_RNA45SN1_45S_50ntFlanks | 12159 | 12181 | 23 | TTTGACTGGGGCGGTACACCTGT | rRF | FALSE | FALSE | TRUE |
| >hg38_21_+_8433172_8446622_RNA45SN1_45S_50ntFlanks@6738.6770.33\|ATCATCGACACTTCGAACGCACTTGCGGCCCCG | >hg38_21_+_8433172_8446622_RNA45SN1_45S_50ntFlanks | 6738 | 6770 | 33 | ATCATCGACACTTCGAACGCACTTGCGGCCCCG | rRF | FALSE | FALSE | TRUE |
| >hg38_21_+_8433172_8446622_RNA45SN1_45S_50ntFlanks@7983.8008.26\|CAGATCAGACGTGGCGACCCGCTGAA | >hg38_21_+_8433172_8446622_RNA45SN1_45S_50ntFlanks | 7983 | 8008 | 26 | CAGATCAGACGTGGCGACCCGCTGAA | rRF | FALSE | FALSE | TRUE |
| >hg38_21_+_8433172_8446622_RNA45SN1_45S_50ntFlanks@12008.12043.36\|CCGCCGGTGAAATACCACTACTCTGATCGTTTTTTC | >hg38_21_+_8433172_8446622_RNA45SN1_45S_50ntFlanks | 12008 | 12043 | 36 | CCGCCGGTGAAATACCACTACTCTGATCGTTTTTTC | rRF | FALSE | FALSE | TRUE |
| >hg38_21_+_8433172_8446622_RNA45SN1_45S_50ntFlanks@10821.10840.20\|GGAAGTCGGCAAGCCGGATC | >hg38_21_+_8433172_8446622_RNA45SN1_45S_50ntFlanks | 10821 | 10840 | 20 | GGAAGTCGGCAAGCCGGATC | rRF | FALSE | FALSE | TRUE |

### Mitochondria

| | | | | | | | | | |
|---|---|---|---|---|---|---|---|---|---|
| >hg38_21_+_8433172_8446622_RNA45SN1_45S_50ntFlanks@10516.10533.18\|GAGTGGCGGAGATGGGCG | >hg38_21_+_8433172_8446622_RNA45SN1_45S_50ntFlanks | 10516 | 10533 | 18 | GAGTGGCGGAGATGGGCG | rRF | FALSE | FALSE | TRUE |
| >hg38_21_+_8433172_8446622_RNA45SN1_45S_50ntFlanks@12436.12455.20\|TCATTGTGAAGCAGAATTCA | >hg38_21_+_8433172_8446622_RNA45SN1_45S_50ntFlanks | 12436 | 12455 | 20 | TCATTGTGAAGCAGAATTCA | rRF | FALSE | FALSE | TRUE |
| >hg38_21_+_8433172_8446622_RNA45SN1_45S_50ntFlanks@12456.12484.29\|CCAAGCGTTGGATTGTTCACCCACTAATA | >hg38_21_+_8433172_8446622_RNA45SN1_45S_50ntFlanks | 12456 | 12484 | 29 | CCAAGCGTTGGATTGTTCACCCACTAATA | rRF | FALSE | FALSE | TRUE |
| >hg38_21_+_8433172_8446622_RNA45SN1_45S_50ntFlanks@6656.6681.26\|CTTAGCGGTGGATCACTCGGCTCGTG | >hg38_21_+_8433172_8446622_RNA45SN1_45S_50ntFlanks | 6656 | 6681 | 26 | CTTAGCGGTGGATCACTCGGCTCGTG | rRF | FALSE | FALSE | TRUE |
| >hg38_21_+_8433172_8446622_RNA45SN1_45S_50ntFlanks@9574.9594.21\|AATCGGTCGTCCGACCTGGGT | >hg38_21_+_8433172_8446622_RNA45SN1_45S_50ntFlanks | 9574 | 9594 | 21 | AATCGGTCGTCCGACCTGGGT | rRF | FALSE | FALSE | TRUE |
| >hg38_21_+_8433172_8446622_RNA45SN1_45S_50ntFlanks@10402.10429.28\|AACATGGGTCAGTCGGTCCTGAGAGATG | >hg38_21_+_8433172_8446622_RNA45SN1_45S_50ntFlanks | 10402 | 10429 | 28 | AACATGGGTCAGTCGGTCCTGAGAGATG | rRF | FALSE | FALSE | TRUE |
| >hg38_21_+_8433172_8446622_RNA45SN1_45S_50ntFlanks@7989.8009.21\|AGACGTGGCGACCCGCTGAAT | >hg38_21_+_8433172_8446622_RNA45SN1_45S_50ntFlanks | 7989 | 8009 | 21 | AGACGTGGCGACCCGCTGAAT | rRF | FALSE | FALSE | TRUE |
| >hg38_21_+_8433172_8446622_RNA45SN1_45S_50ntFlanks@9572.9594.23\|CAAATCGGTCGTCCGACCTGGGT | >hg38_21_+_8433172_8446622_RNA45SN1_45S_50ntFlanks | 9572 | 9594 | 23 | CAAATCGGTCGTCCGACCTGGGT | rRF | FALSE | FALSE | TRUE |
| >hg38_21_+_8433172_8446622_RNA45SN1_45S_50ntFlanks@8113.8140.28\|GGCGCGGGACATGTGGCGTACGGAAGAC | >hg38_21_+_8433172_8446622_RNA45SN1_45S_50ntFlanks | 8113 | 8140 | 28 | GGCGCGGGACATGTGGCGTACGGAAGAC | rRF | FALSE | FALSE | TRUE |
| >trna15_ValAAC_5_-_180615416_180615488@1.29.29\|GTTTCCGTAGTGTAGTGGTCATCACGTTC | >trna15_ValAAC_5_-_180615416_180615488 | 1 | 29 | 29 | GTTTCCGTAGTGTAGTGGTCATCACGTTC | tRF | FALSE | FALSE | TRUE |
| >hg38_21_+_8433172_8446622_RNA45SN1_45S_50ntFlanks@9533.9553.21\|GAGGAAACTCTGGTGGAGGTC | >hg38_21_+_8433172_8446622_RNA45SN1_45S_50ntFlanks | 9533 | 9553 | 21 | GAGGAAACTCTGGTGGAGGTC | rRF | FALSE | FALSE | TRUE |
| >am_trna111_HisGTG_1_-_147774845_147774916@-1G.29.30,trna118_HisGTG_1_-_145396881_145396952@-1G.29.30,trna16_HisGTG_1_+_146544773_146544844@-1G.29.30,trna1_HisGTG_15_+_45493349_45493420@-1G.29.30,trna21_HisGTG_1_+_147753471_147753542@-1G.29.30,trna33_HisGTG_6_+_27125906_27125977@-1G.29.30,trna7_HisGTG_9_-_14433938_14434009@-1G.29.30,trna8_HisGTG_15_-_45492611_45492682@-1G.29.30,trna9_HisGTG_15_-_45490804_45490875@-1G.29.30\|GGCCGTGATCGTATAGTGGTTAGTACTCTG | >am_trna111_HisGTG_1_-_147774845_147774916 | -1G | 29 | 30 | GGCCGTGATCGTATAGTGGTTAGTACTCTG | tRF | FALSE | FALSE | TRUE |
| >hg38_21_+_8433172_8446622_RNA45SN1_45S_50ntFlanks@9568.9596.29\|CGTGCAAATCGGTCGTCCGACCTGGGTAT | >hg38_21_+_8433172_8446622_RNA45SN1_45S_50ntFlanks | 9568 | 9596 | 29 | CGTGCAAATCGGTCGTCCGACCTGGGTAT | rRF | FALSE | FALSE | TRUE |
| >hg38_21_+_8433172_8446622_RNA45SN1_45S_50ntFlanks@12625.12654.30\|CAATGGGGCGAAGCTACCATCTGTGGGATT | >hg38_21_+_8433172_8446622_RNA45SN1_45S_50ntFlanks | 12625 | 12654 | 30 | CAATGGGGCGAAGCTACCATCTGTGGGATT | rRF | FALSE | FALSE | TRUE |
| >hg38_21_+_8433172_8446622_RNA45SN1_45S_50ntFlanks@11569.11592.24\|CAGCCGACTTAGAACTGGTGCGGA | >hg38_21_+_8433172_8446622_RNA45SN1_45S_50ntFlanks | 11569 | 11592 | 24 | CAGCCGACTTAGAACTGGTGCGGA | rRF | FALSE | FALSE | TRUE |
| >hg38_21_+_8433172_8446622_RNA45SN1_45S_50ntFlanks@10679.10709.31\|GGTTCCGGCGGCGTCCGGTGAGCTCTCGCTG | >hg38_21_+_8433172_8446622_RNA45SN1_45S_50ntFlanks | 10679 | 10709 | 31 | GGTTCCGGCGGCGTCCGGTGAGCTCTCGCTG | rRF | FALSE | FALSE | TRUE |
| >hg38_21_+_8433172_8446622_RNA45SN1_45S_50ntFlanks@4657.4689.33\|CTAGAGGTGAAATTCTTGGACCGGCGCAAGACG | >hg38_21_+_8433172_8446622_RNA45SN1_45S_50ntFlanks | 4657 | 4689 | 33 | CTAGAGGTGAAATTCTTGGACCGGCGCAAGACG | rRF | FALSE | FALSE | TRUE |
| >hg38_21_+_8433172_8446622_RNA45SN1_45S_50ntFlanks@6719.6743.25\|TGAATTGCAGGACACATTGATCATC | >hg38_21_+_8433172_8446622_RNA45SN1_45S_50ntFlanks | 6719 | 6743 | 25 | TGAATTGCAGGACACATTGATCATC | rRF | FALSE | FALSE | TRUE |
| >hg38_21_+_8433172_8446622_RNA45SN1_45S_50ntFlanks@12712.12742.31\|CGGAGCCTCGGTTGGCCTCGGATAGCCGGTC | >hg38_21_+_8433172_8446622_RNA45SN1_45S_50ntFlanks | 12712 | 12742 | 31 | CGGAGCCTCGGTTGGCCTCGGATAGCCGGTC | rRF | FALSE | FALSE | TRUE |
| >hg38_21_+_8433172_8446622_RNA45SN1_45S_50ntFlanks@6648.6681.34\|GTACGACTCTTAGCGGTGGATCACTCGGCTCGTG | >hg38_21_+_8433172_8446622_RNA45SN1_45S_50ntFlanks | 6648 | 6681 | 34 | GTACGACTCTTAGCGGTGGATCACTCGGCTCGTG | rRF | FALSE | FALSE | TRUE |
| >hg38_21_+_8433172_8446622_RNA45SN1_45S_50ntFlanks@10684.10706.23\|CGGCGGCGTCCGGTGAGCTCTCG | >hg38_21_+_8433172_8446622_RNA45SN1_45S_50ntFlanks | 10684 | 10706 | 23 | CGGCGGCGTCCGGTGAGCTCTCG | rRF | FALSE | FALSE | TRUE |
| >hg38_21_+_8433172_8446622_RNA45SN1_45S_50ntFlanks@7975.8012.38\|CGCGACCTCAGATCAGACGTGGCGACCCGCTGAATTTA | >hg38_21_+_8433172_8446622_RNA45SN1_45S_50ntFlanks | 7975 | 8012 | 38 | CGCGACCTCAGATCAGACGTGGCGACCCGCTGAATTTA | rRF | FALSE | FALSE | TRUE |
| >hg38_21_+_8433172_8446622_RNA45SN1_45S_50ntFlanks@9534.9551.18\|AGGAAACTCTGGTGGAGG | >hg38_21_+_8433172_8446622_RNA45SN1_45S_50ntFlanks | 9534 | 9551 | 18 | AGGAAACTCTGGTGGAGG | rRF | FALSE | FALSE | TRUE |
| >hg38_21_+_8433172_8446622_RNA45SN1_45S_50ntFlanks@11323.11342.20\|CGTCTCGTCGCGCGCGCGTC | >hg38_21_+_8433172_8446622_RNA45SN1_45S_50ntFlanks | 11323 | 11342 | 20 | CGTCTCGTCGCGCGCGCGTC | rRF | FALSE | FALSE | TRUE |
| >hg38_21_+_8433172_8446622_RNA45SN1_45S_50ntFlanks@3985.4024.40\|CGGCTTTGGTGACTCTAGATAACCTCGGGCCGATCGCACG | >hg38_21_+_8433172_8446622_RNA45SN1_45S_50ntFlanks | 3985 | 4024 | 40 | CGGCTTTGGTGACTCTAGATAACCTCGGGCCGATCGCACG | rRF | FALSE | FALSE | TRUE |
| >hg38_21_+_8433172_8446622_RNA45SN1_45S_50ntFlanks@8105.8141.37\|CGCGGCGGGGCGCGGGACATGTGGCGTACGGAAGACC | >hg38_21_+_8433172_8446622_RNA45SN1_45S_50ntFlanks | 8105 | 8141 | 37 | CGCGGCGGGGCGCGGGACATGTGGCGTACGGAAGACC | rRF | FALSE | FALSE | TRUE |
| >hg38_21_+_8433172_8446622_RNA45SN1_45S_50ntFlanks@7982.8006.25\|TCAGATCAGACGTGGCGACCCGCTG | >hg38_21_+_8433172_8446622_RNA45SN1_45S_50ntFlanks | 7982 | 8006 | 25 | TCAGATCAGACGTGGCGACCCGCTG | rRF | FALSE | FALSE | TRUE |
| >hg38_21_+_8433172_8446622_RNA45SN1_45S_50ntFlanks@10687.10710.24\|CGGCGTCCGGTGAGCTCTCGCTGG | >hg38_21_+_8433172_8446622_RNA45SN1_45S_50ntFlanks | 10687 | 10710 | 24 | CGGCGTCCGGTGAGCTCTCGCTGG | rRF | FALSE | FALSE | TRUE |
| >hg38_21_+_8433172_8446622_RNA45SN1_45S_50ntFlanks@9573.9595.23\|AAATCGGTCGTCCGACCTGGGTA | >hg38_21_+_8433172_8446622_RNA45SN1_45S_50ntFlanks | 9573 | 9595 | 23 | AAATCGGTCGTCCGACCTGGGTA | rRF | FALSE | FALSE | TRUE |
| >hg38_21_+_8433172_8446622_RNA45SN1_45S_50ntFlanks@4759.4782.24\|ATCAGATACCGTCGTAGTTCCGAC | >hg38_21_+_8433172_8446622_RNA45SN1_45S_50ntFlanks | 4759 | 4782 | 24 | ATCAGATACCGTCGTAGTTCCGAC | rRF | FALSE | FALSE | TRUE |
| >hg38_21_+_8433172_8446622_RNA45SN1_45S_50ntFlanks@11572.11590.19\|CCGACTTAGAACTGGTGCG | >hg38_21_+_8433172_8446622_RNA45SN1_45S_50ntFlanks | 11572 | 11590 | 19 | CCGACTTAGAACTGGTGCG | rRF | FALSE | FALSE | TRUE |
| >hg38_21_+_8433172_8446622_RNA45SN1_45S_50ntFlanks@9570.9596.27\|TGCAAATCGGTCGTCCGACCTGGGTAT | >hg38_21_+_8433172_8446622_RNA45SN1_45S_50ntFlanks | 9570 | 9596 | 27 | TGCAAATCGGTCGTCCGACCTGGGTAT | rRF | FALSE | FALSE | TRUE |
| >hg38_21_+_8433172_8446622_RNA45SN1_45S_50ntFlanks@3882.3899.18\|CCGACGGGCGCTGACCCC | >hg38_21_+_8433172_8446622_RNA45SN1_45S_50ntFlanks | 3882 | 3899 | 18 | CCGACGGGCGCTGACCCC | rRF | FALSE | FALSE | TRUE |
| >hg38_21_+_8433172_8446622_RNA45SN1_45S_50ntFlanks@12157.12182.26\|AGTTTGACTGGGGCGGTACACCTGTC | >hg38_21_+_8433172_8446622_RNA45SN1_45S_50ntFlanks | 12157 | 12182 | 26 | AGTTTGACTGGGGCGGTACACCTGTC | rRF | FALSE | FALSE | TRUE |
| >hg38_21_+_8433172_8446622_RNA45SN1_45S_50ntFlanks@12456.12480.25\|CCAAGCGTTGGATTGTTCACCCACT | >hg38_21_+_8433172_8446622_RNA45SN1_45S_50ntFlanks | 12456 | 12480 | 25 | CCAAGCGTTGGATTGTTCACCCACT | rRF | FALSE | FALSE | TRUE |
| >hg38_21_+_8433172_8446622_RNA45SN1_45S_50ntFlanks@6646.6670.25\|TCGTACGACTCTTAGCGGTGGATCA | >hg38_21_+_8433172_8446622_RNA45SN1_45S_50ntFlanks | 6646 | 6670 | 25 | TCGTACGACTCTTAGCGGTGGATCA | rRF | FALSE | FALSE | TRUE |
| >hg38_21_+_8433172_8446622_RNA45SN1_45S_50ntFlanks@8196.8215.20\|CGTGGACGGTGTGAGGCCGG | >hg38_21_+_8433172_8446622_RNA45SN1_45S_50ntFlanks | 8196 | 8215 | 20 | CGTGGACGGTGTGAGGCCGG | rRF | FALSE | FALSE | TRUE |
| >hg38_21_+_8433172_8446622_RNA45SN1_45S_50ntFlanks@9575.9595.21\|ATCGGTCGTCCGACCTGGGTA | >hg38_21_+_8433172_8446622_RNA45SN1_45S_50ntFlanks | 9575 | 9595 | 21 | ATCGGTCGTCCGACCTGGGTA | rRF | FALSE | FALSE | TRUE |

### Mitochondria

| | | | | | | | | | |
|---|---|---|---|---|---|---|---|---|---|
| >hg38_21_+_8433172_8446622_RNA45SN1_45S_50ntFlanks@9308.9338.31\|AGGAGTCTAACACGTGCGCGAGTCGGGGGCT | >hg38_21_+_8433172_8446622_RNA45SN1_45S_50ntFlanks | 9308 | 9338 | 31 | AGGAGTCTAACACGTGCGCGAGTCGGGGGCT | rRF | FALSE | FALSE | TRUE |
| >hg38_21_+_8433172_8446622_RNA45SN1_45S_50ntFlanks@11575.11592.18\|ACTTAGAACTGGTGCGGA | >hg38_21_+_8433172_8446622_RNA45SN1_45S_50ntFlanks | 11575 | 11592 | 18 | ACTTAGAACTGGTGCGGA | rRF | FALSE | FALSE | TRUE |
| >hg38_21_+_8433172_8446622_RNA45SN1_45S_50ntFlanks@5073.5098.26\|AATTCCGATAACGAACGAGACTCTGG | >hg38_21_+_8433172_8446622_RNA45SN1_45S_50ntFlanks | 5073 | 5098 | 26 | AATTCCGATAACGAACGAGACTCTGG | rRF | FALSE | FALSE | TRUE |
| >hg38_21_+_8433172_8446622_RNA45SN1_45S_50ntFlanks@8194.8211.18\|CCCGTGGACGGTGTGAGG | >hg38_21_+_8433172_8446622_RNA45SN1_45S_50ntFlanks | 8194 | 8211 | 18 | CCCGTGGACGGTGTGAGG | rRF | FALSE | FALSE | TRUE |
| >hg38_21_+_8433172_8446622_RNA45SN1_45S_50ntFlanks@6719.6744.26\|TGAATTGCAGGACACATTGATCATCG | >hg38_21_+_8433172_8446622_RNA45SN1_45S_50ntFlanks | 6719 | 6744 | 26 | TGAATTGCAGGACACATTGATCATCG | rRF | FALSE | FALSE | TRUE |
| >hg38_21_+_8433172_8446622_RNA45SN1_45S_50ntFlanks@10841.10868.28\|CGTAACTTCGGGATAAGGATTGGCTCTA | >hg38_21_+_8433172_8446622_RNA45SN1_45S_50ntFlanks | 10841 | 10868 | 28 | CGTAACTTCGGGATAAGGATTGGCTCTA | rRF | FALSE | FALSE | TRUE |
| >hg38_21_+_8433172_8446622_RNA45SN1_45S_50ntFlanks@6645.6681.37\|CTCGTACGACTCTTAGCGGTGGATCACTCGGCTCGTG | >hg38_21_+_8433172_8446622_RNA45SN1_45S_50ntFlanks | 6645 | 6681 | 37 | CTCGTACGACTCTTAGCGGTGGATCACTCGGCTCGTG | rRF | FALSE | FALSE | TRUE |
| >hg38_21_+_8433172_8446622_RNA45SN1_45S_50ntFlanks@12002.12032.31\|TGCGGGCCGCCGGTGAAATACCACTACTCTG | >hg38_21_+_8433172_8446622_RNA45SN1_45S_50ntFlanks | 12002 | 12032 | 31 | TGCGGGCCGCCGGTGAAATACCACTACTCTG | rRF | FALSE | FALSE | TRUE |
| >hg38_21_+_8433172_8446622_RNA45SN1_45S_50ntFlanks@12324.12346.23\|TTGGGTTTTAAGCAGGAGGTGTC | >hg38_21_+_8433172_8446622_RNA45SN1_45S_50ntFlanks | 12324 | 12346 | 23 | TTGGGTTTTAAGCAGGAGGTGTC | rRF | FALSE | FALSE | TRUE |
| >hg38_21_+_8433172_8446622_RNA45SN1_45S_50ntFlanks@12405.12431.27\|CTTTTTGATCCTTCGATGTCGGCTCTT | >hg38_21_+_8433172_8446622_RNA45SN1_45S_50ntFlanks | 12405 | 12431 | 27 | CTTTTTGATCCTTCGATGTCGGCTCTT | rRF | FALSE | FALSE | TRUE |
| >hg38_21_+_8433172_8446622_RNA45SN1_45S_50ntFlanks@7986.8008.23\|ATCAGACGTGGCGACCCGCTGAA | >hg38_21_+_8433172_8446622_RNA45SN1_45S_50ntFlanks | 7986 | 8008 | 23 | ATCAGACGTGGCGACCCGCTGAA | rRF | FALSE | FALSE | TRUE |
| >hg38_21_+_8433172_8446622_RNA45SN1_45S_50ntFlanks@10350.10376.27\|ACGAGAACTTTGAAGGCCGAAGTGGAG | >hg38_21_+_8433172_8446622_RNA45SN1_45S_50ntFlanks | 10350 | 10376 | 27 | ACGAGAACTTTGAAGGCCGAAGTGGAG | rRF | FALSE | FALSE | TRUE |
| >hg38_21_+_8433172_8446622_RNA45SN1_45S_50ntFlanks@10349.10368.20\|AACGAGAACTTTGAAGGCCG | >hg38_21_+_8433172_8446622_RNA45SN1_45S_50ntFlanks | 10349 | 10368 | 20 | AACGAGAACTTTGAAGGCCG | rRF | FALSE | FALSE | TRUE |
| >hg38_21_+_8433172_8446622_RNA45SN1_45S_50ntFlanks@12157.12180.24\|AGTTTGACTGGGGCGGTACACCTG | >hg38_21_+_8433172_8446622_RNA45SN1_45S_50ntFlanks | 12157 | 12180 | 24 | AGTTTGACTGGGGCGGTACACCTG | rRF | FALSE | FALSE | TRUE |
| >am_trna111_HisGTG_1_-_147774845_147774916@1.29.29,trna118_HisGTG_1_-_145396881_145396952@1.29.29,trna16_HisGTG_1_+_146544773_146544844@1.29.29,trna1_HisGTG_15_+_45493349_45493420@1.29.29,trna21_HisGTG_1_+_147753471_147753542@1.29.29,trna33_HisGTG_6_+_27125906_27125977@1.29.29,trna7_HisGTG_9_-_14433938_14434009@1.29.29,trna8_HisGTG_15_-_45492611_45492682@1.29.29,trna9_HisGTG_15_-_45490804_45490875@1.29.29\|GCCGTGATCGTATAGTGGTTAGTACTCTG | >am_trna111_HisGTG_1_-_147774845_147774916 | 1 | 29 | 29 | GCCGTGATCGTATAGTGGTTAGTACTCTG | tRF | FALSE | FALSE | TRUE |
| >hg38_21_+_8433172_8446622_RNA45SN1_45S_50ntFlanks@11569.11602.34\|CAGCCGACTTAGAACTGGTGCGGACCAGGGGAAT | >hg38_21_+_8433172_8446622_RNA45SN1_45S_50ntFlanks | 11569 | 11602 | 34 | CAGCCGACTTAGAACTGGTGCGGACCAGGGGAAT | rRF | FALSE | FALSE | TRUE |
| >hg38_21_+_8433172_8446622_RNA45SN1_45S_50ntFlanks@11564.11601.38\|CCTAGCAGCCGACTTAGAACTGGTGCGGACCAGGGGAA | >hg38_21_+_8433172_8446622_RNA45SN1_45S_50ntFlanks | 11564 | 11601 | 38 | CCTAGCAGCCGACTTAGAACTGGTGCGGACCAGGGGAA | rRF | FALSE | FALSE | TRUE |
| >hg38_21_+_8433172_8446622_RNA45SN1_45S_50ntFlanks@8113.8143.31\|GGCGCGGGACATGTGGCGTACGGAAGACCCG | >hg38_21_+_8433172_8446622_RNA45SN1_45S_50ntFlanks | 8113 | 8143 | 31 | GGCGCGGGACATGTGGCGTACGGAAGACCCG | rRF | FALSE | FALSE | TRUE |
| >hg38_21_+_8433172_8446622_RNA45SN1_45S_50ntFlanks@5444.5467.24\|CTCGGATCGGCCCCGCCGGGGTCG | >hg38_21_+_8433172_8446622_RNA45SN1_45S_50ntFlanks | 5444 | 5467 | 24 | CTCGGATCGGCCCCGCCGGGGTCG | rRF | FALSE | FALSE | TRUE |
| >hg38_21_+_8433172_8446622_RNA45SN1_45S_50ntFlanks@3984.4024.41\|GCGGCTTTGGTGACTCTAGATAACCTCGGGCCGATCGCACG | >hg38_21_+_8433172_8446622_RNA45SN1_45S_50ntFlanks | 3984 | 4024 | 41 | GCGGCTTTGGTGACTCTAGATAACCTCGGGCCGATCGCACG | rRF | FALSE | FALSE | TRUE |
| >hg38_21_+_8433172_8446622_RNA45SN1_45S_50ntFlanks@12632.12654.23\|GCGAAGCTACCATCTGTGGGATT | >hg38_21_+_8433172_8446622_RNA45SN1_45S_50ntFlanks | 12632 | 12654 | 23 | GCGAAGCTACCATCTGTGGGATT | rRF | FALSE | FALSE | TRUE |
| >hg38_21_+_8433172_8446622_RNA45SN1_45S_50ntFlanks@10841.10863.23\|CGTAACTTCGGGATAAGGATTGG | >hg38_21_+_8433172_8446622_RNA45SN1_45S_50ntFlanks | 10841 | 10863 | 23 | CGTAACTTCGGGATAAGGATTGG | rRF | FALSE | FALSE | TRUE |
| >hg38_21_+_8433172_8446622_RNA45SN1_45S_50ntFlanks@4283.4317.35\|CTTTAACGAGGATCCATTGGAGGGCAAGTCTGGTG | >hg38_21_+_8433172_8446622_RNA45SN1_45S_50ntFlanks | 4283 | 4317 | 35 | CTTTAACGAGGATCCATTGGAGGGCAAGTCTGGTG | rRF | FALSE | FALSE | TRUE |
| >hg38_21_+_8433172_8446622_RNA45SN1_45S_50ntFlanks@9568.9597.30\|CGTGCAAATCGGTCGTCCGACCTGGGTATA | >hg38_21_+_8433172_8446622_RNA45SN1_45S_50ntFlanks | 9568 | 9597 | 30 | CGTGCAAATCGGTCGTCCGACCTGGGTATA | rRF | FALSE | FALSE | TRUE |
| >hg38_21_+_8433172_8446622_RNA45SN1_45S_50ntFlanks@12934.12956.23\|CTAAACCATTCGTAGACGACCTG | >hg38_21_+_8433172_8446622_RNA45SN1_45S_50ntFlanks | 12934 | 12956 | 23 | CTAAACCATTCGTAGACGACCTG | rRF | FALSE | FALSE | TRUE |
| >hg38_21_+_8433172_8446622_RNA45SN1_45S_50ntFlanks@10346.10368.23\|TCAAACGAGAACTTTGAAGGCCG | >hg38_21_+_8433172_8446622_RNA45SN1_45S_50ntFlanks | 10346 | 10368 | 23 | TCAAACGAGAACTTTGAAGGCCG | rRF | FALSE | FALSE | TRUE |
| >hg38_21_+_8433172_8446622_RNA45SN1_45S_50ntFlanks@6721.6743.23\|AATTGCAGGACACATTGATCATC | >hg38_21_+_8433172_8446622_RNA45SN1_45S_50ntFlanks | 6721 | 6743 | 23 | AATTGCAGGACACATTGATCATC | rRF | FALSE | FALSE | TRUE |
| >hg38_21_+_8433172_8446622_RNA45SN1_45S_50ntFlanks@8104.8138.35\|CCGCGGCGGGGCGCGGGACATGTGGCGTACGGAAG | >hg38_21_+_8433172_8446622_RNA45SN1_45S_50ntFlanks | 8104 | 8138 | 35 | CCGCGGCGGGGCGCGGGACATGTGGCGTACGGAAG | rRF | FALSE | FALSE | TRUE |
| >hg38_1_-_228634819_228635039_RNA5S12_5S_50ntFlanks@150.169.20\|AATACCGGGTGCTGTAGGCT | >hg38_1_-_228634819_228635039_RNA5S12_5S_50ntFlanks | 150 | 169 | 20 | AATACCGGGTGCTGTAGGCT | rRF | FALSE | FALSE | TRUE |
| >hg38_21_+_8433172_8446622_RNA45SN1_45S_50ntFlanks@4732.4758.27\|AGAACGAAAGTCGGAGGTTCGAAGACG | >hg38_21_+_8433172_8446622_RNA45SN1_45S_50ntFlanks | 4732 | 4758 | 27 | AGAACGAAAGTCGGAGGTTCGAAGACG | rRF | FALSE | FALSE | TRUE |
| >MI0000101\|hsa-mir-99a&WithFlank&21\|+\|16539083\|16539175@19.39.21&[MIMAT0000097&hsa-miR-99a-5p&offsets\|0\|-1;m-64&21\|+\|16539101\|16539121&offsets\|0\|0]\|AACCCGTAGATCCGATCTTGT | >MI0000101\|hsa-mir-99a&WithFlank&21\|+\|16539083\|16539175 | 19 | 39 | 21 | AACCCGTAGATCCGATCTTGT | isomiR | FALSE | FALSE | TRUE |
| >hg38_21_+_8433172_8446622_RNA45SN1_45S_50ntFlanks@5486.5514.29\|GCGCTGAGAAGACGGTCGAACTTGACTAT | >hg38_21_+_8433172_8446622_RNA45SN1_45S_50ntFlanks | 5486 | 5514 | 29 | GCGCTGAGAAGACGGTCGAACTTGACTAT | rRF | FALSE | FALSE | TRUE |
| >hg38_21_+_8433172_8446622_RNA45SN1_45S_50ntFlanks@11876.11893.18\|AAGACCCTGTTGAGCTTG | >hg38_21_+_8433172_8446622_RNA45SN1_45S_50ntFlanks | 11876 | 11893 | 18 | AAGACCCTGTTGAGCTTG | rRF | FALSE | FALSE | TRUE |
| >hg38_21_+_8433172_8446622_RNA45SN1_45S_50ntFlanks@9567.9597.31\|ACGTGCAAATCGGTCGTCCGACCTGGGTATA | >hg38_21_+_8433172_8446622_RNA45SN1_45S_50ntFlanks | 9567 | 9597 | 31 | ACGTGCAAATCGGTCGTCCGACCTGGGTATA | rRF | FALSE | FALSE | TRUE |
| >hg38_21_+_8433172_8446622_RNA45SN1_45S_50ntFlanks@4729.4758.30\|TCAAGAACGAAAGTCGGAGGTTCGAAGACG | >hg38_21_+_8433172_8446622_RNA45SN1_45S_50ntFlanks | 4729 | 4758 | 30 | TCAAGAACGAAAGTCGGAGGTTCGAAGACG | rRF | FALSE | FALSE | TRUE |
| >hg38_21_+_8433172_8446622_RNA45SN1_45S_50ntFlanks@12636.12661.26\|AGCTACCATCTGTGGGATTATGACTG | >hg38_21_+_8433172_8446622_RNA45SN1_45S_50ntFlanks | 12636 | 12661 | 26 | AGCTACCATCTGTGGGATTATGACTG | rRF | FALSE | FALSE | TRUE |
| >hg38_21_+_8433172_8446622_RNA45SN1_45S_50ntFlanks@8196.8214.19\|CGTGGACGGTGTGAGGCCG | >hg38_21_+_8433172_8446622_RNA45SN1_45S_50ntFlanks | 8196 | 8214 | 19 | CGTGGACGGTGTGAGGCCG | rRF | FALSE | FALSE | TRUE |
| >hg38_21_+_8433172_8446622_RNA45SN1_45S_50ntFlanks@12936.12956.21\|AAACCATTCGTAGACGACCTG | >hg38_21_+_8433172_8446622_RNA45SN1_45S_50ntFlanks | 12936 | 12956 | 21 | AAACCATTCGTAGACGACCTG | rRF | FALSE | FALSE | TRUE |

### Mitochondria

| | | | | | | | | | |
|---|---|---|---|---|---|---|---|---|---|
| >trna119_LysCTT_1_-_145395522_145395594@1.30.30,trna11_LysCTT_5_-_180648979_180649051@1.30.30,trna13_LysCTT_6_+_26556774_26556846@1.30.30,trna32_LysCTT_16_-_3207406_3207478@1.30.30,trna7_LysCTT_16_+_3225692_3225764@1.30.30,trna9_LysCTT_5_+_180634755_180634827@1.30.30\|GCCCGGCTAGCTCAGTCGGTAGAGCATGAG | >trna119_LysCTT_1_-_145395522_145395594 | 1 | 30 | 30 | GCCCGGCTAGCTCAGTCGGTAGAGCATGAG | tRF | FALSE | FALSE | TRUE |
| >hg38_21_+_8433172_8446622_RNA45SN1_45S_50ntFlanks@10410.10429.20\|TCAGTCGGTCCTGAGAGATG | >hg38_21_+_8433172_8446622_RNA45SN1_45S_50ntFlanks | 10410 | 10429 | 20 | TCAGTCGGTCCTGAGAGATG | rRF | FALSE | FALSE | TRUE |
| >hg38_21_+_8433172_8446622_RNA45SN1_45S_50ntFlanks@11620.11640.21\|ACAAAGCATCGCGAAGGCCCG | >hg38_21_+_8433172_8446622_RNA45SN1_45S_50ntFlanks | 11620 | 11640 | 21 | ACAAAGCATCGCGAAGGCCCG | rRF | FALSE | FALSE | TRUE |
| >hg38_21_+_8433172_8446622_RNA45SN1_45S_50ntFlanks@4726.4758.33\|TAATCAAGAACGAAAGTCGGAGGTTCGAAGACG | >hg38_21_+_8433172_8446622_RNA45SN1_45S_50ntFlanks | 4726 | 4758 | 33 | TAATCAAGAACGAAAGTCGGAGGTTCGAAGACG | rRF | FALSE | FALSE | TRUE |
| >hg38_21_+_8433172_8446622_RNA45SN1_45S_50ntFlanks@10351.10382.32\|CGAGAACTTTGAAGGCCGAAGTGGAGAAGGGT | >hg38_21_+_8433172_8446622_RNA45SN1_45S_50ntFlanks | 10351 | 10382 | 32 | CGAGAACTTTGAAGGCCGAAGTGGAGAAGGGT | rRF | FALSE | FALSE | TRUE |
| >hg38_21_+_8433172_8446622_RNA45SN1_45S_50ntFlanks@10841.10864.24\|CGTAACTTCGGGATAAGGATTGGC | >hg38_21_+_8433172_8446622_RNA45SN1_45S_50ntFlanks | 10841 | 10864 | 24 | CGTAACTTCGGGATAAGGATTGGC | rRF | FALSE | FALSE | TRUE |
| >hg38_21_+_8433172_8446622_RNA45SN1_45S_50ntFlanks@11783.11806.24\|CATGAATGGATGAACGAGATTCCC | >hg38_21_+_8433172_8446622_RNA45SN1_45S_50ntFlanks | 11783 | 11806 | 24 | CATGAATGGATGAACGAGATTCCC | rRF | FALSE | FALSE | TRUE |
| >hg38_1_-_228634819_228635039_RNA5S12_5S_50ntFlanks@144.171.28\|CCTGGGAATACCGGGTGCTGTAGGCTTT | >hg38_1_-_228634819_228635039_RNA5S12_5S_50ntFlanks | 144 | 171 | 28 | CCTGGGAATACCGGGTGCTGTAGGCTTT | rRF | FALSE | FALSE | TRUE |
| >hg38_21_+_8433172_8446622_RNA45SN1_45S_50ntFlanks@8079.8104.26\|AAGAGCCCAGCGCCGAATCCCCGCCC | >hg38_21_+_8433172_8446622_RNA45SN1_45S_50ntFlanks | 8079 | 8104 | 26 | AAGAGCCCAGCGCCGAATCCCCGCCC | rRF | FALSE | FALSE | TRUE |
| >hg38_21_+_8433172_8446622_RNA45SN1_45S_50ntFlanks@8115.8141.27\|CGCGGGACATGTGGCGTACGGAAGACC | >hg38_21_+_8433172_8446622_RNA45SN1_45S_50ntFlanks | 8115 | 8141 | 27 | CGCGGGACATGTGGCGTACGGAAGACC | rRF | FALSE | FALSE | TRUE |
| >hg38_21_+_8433172_8446622_RNA45SN1_45S_50ntFlanks@10350.10384.35\|ACGAGAACTTTGAAGGCCGAAGTGGAGAAGGGTTC | >hg38_21_+_8433172_8446622_RNA45SN1_45S_50ntFlanks | 10350 | 10384 | 35 | ACGAGAACTTTGAAGGCCGAAGTGGAGAAGGGTTC | rRF | FALSE | FALSE | TRUE |
| >hg38_21_+_8433172_8446622_RNA45SN1_45S_50ntFlanks@11641.11664.24\|CGGCGGGTGTTGACGCGATGTGAT | >hg38_21_+_8433172_8446622_RNA45SN1_45S_50ntFlanks | 11641 | 11664 | 24 | CGGCGGGTGTTGACGCGATGTGAT | rRF | FALSE | FALSE | TRUE |
| >hg38_21_+_8433172_8446622_RNA45SN1_45S_50ntFlanks@6738.6771.34\|ATCATCGACACTTCGAACGCACTTGCGGCCCCGG | >hg38_21_+_8433172_8446622_RNA45SN1_45S_50ntFlanks | 6738 | 6771 | 34 | ATCATCGACACTTCGAACGCACTTGCGGCCCCGG | rRF | FALSE | FALSE | TRUE |
| >hg38_21_+_8433172_8446622_RNA45SN1_45S_50ntFlanks@10099.10117.19\|CGCGCGCGCGTGTGGTGTG | >hg38_21_+_8433172_8446622_RNA45SN1_45S_50ntFlanks | 10099 | 10117 | 19 | CGCGCGCGCGTGTGGTGTG | rRF | FALSE | FALSE | TRUE |
| >hg38_21_+_8433172_8446622_RNA45SN1_45S_50ntFlanks@11583.11611.29\|CTGGTGCGGACCAGGGGAATCCGACTGTT | >hg38_21_+_8433172_8446622_RNA45SN1_45S_50ntFlanks | 11583 | 11611 | 29 | CTGGTGCGGACCAGGGGAATCCGACTGTT | rRF | FALSE | FALSE | TRUE |
| >multi-am_trnaMT_ValTAC_MT_+_1602_1670@50.68.19\|ACTTAACTTGACCGCTCTG | >multi-am_trnaMT_ValTAC_MT_+_1602_1670 | 50 | 68 | 19 | ACTTAACTTGACCGCTCTG | tRF | FALSE | FALSE | TRUE |
| >hg38_21_+_8433172_8446622_RNA45SN1_45S_50ntFlanks@10362.10388.27\|AAGGCCGAAGTGGAGAAGGGTTCCATG | >hg38_21_+_8433172_8446622_RNA45SN1_45S_50ntFlanks | 10362 | 10388 | 27 | AAGGCCGAAGTGGAGAAGGGTTCCATG | rRF | FALSE | FALSE | TRUE |
| >hg38_21_+_8433172_8446622_RNA45SN1_45S_50ntFlanks@8114.8139.26\|GCGCGGGACATGTGGCGTACGGAAGA | >hg38_21_+_8433172_8446622_RNA45SN1_45S_50ntFlanks | 8114 | 8139 | 26 | GCGCGGGACATGTGGCGTACGGAAGA | rRF | FALSE | FALSE | TRUE |
| >hg38_21_+_8433172_8446622_RNA45SN1_45S_50ntFlanks@5471.5499.29\|CACGGCCCTGGCGGAGCGCTGAGAAGACG | >hg38_21_+_8433172_8446622_RNA45SN1_45S_50ntFlanks | 5471 | 5499 | 29 | CACGGCCCTGGCGGAGCGCTGAGAAGACG | rRF | FALSE | FALSE | TRUE |
| >hg38_21_+_8433172_8446622_RNA45SN1_45S_50ntFlanks@11569.11600.32\|CAGCCGACTTAGAACTGGTGCGGACCAGGGGA | >hg38_21_+_8433172_8446622_RNA45SN1_45S_50ntFlanks | 11569 | 11600 | 32 | CAGCCGACTTAGAACTGGTGCGGACCAGGGGA | rRF | FALSE | FALSE | TRUE |
| >hg38_21_+_8433172_8446622_RNA45SN1_45S_50ntFlanks@9504.9523.20\|GTGAACTATGCCTGGGCAGG | >hg38_21_+_8433172_8446622_RNA45SN1_45S_50ntFlanks | 9504 | 9523 | 20 | GTGAACTATGCCTGGGCAGG | rRF | FALSE | FALSE | TRUE |
| >hg38_21_+_8433172_8446622_RNA45SN1_45S_50ntFlanks@10687.10706.20\|CGGCGTCCGGTGAGCTCTCG | >hg38_21_+_8433172_8446622_RNA45SN1_45S_50ntFlanks | 10687 | 10706 | 20 | CGGCGTCCGGTGAGCTCTCG | rRF | FALSE | FALSE | TRUE |
| >hg38_21_+_8433172_8446622_RNA45SN1_45S_50ntFlanks@12637.12654.18\|GCTACCATCTGTGGGATT | >hg38_21_+_8433172_8446622_RNA45SN1_45S_50ntFlanks | 12637 | 12654 | 18 | GCTACCATCTGTGGGATT | rRF | FALSE | FALSE | TRUE |
| >hg38_21_+_8433172_8446622_RNA45SN1_45S_50ntFlanks@11562.11592.31\|CGCCTAGCAGCCGACTTAGAACTGGTGCGGA | >hg38_21_+_8433172_8446622_RNA45SN1_45S_50ntFlanks | 11562 | 11592 | 31 | CGCCTAGCAGCCGACTTAGAACTGGTGCGGA | rRF | FALSE | FALSE | TRUE |
| >hg38_21_+_8433172_8446622_RNA45SN1_45S_50ntFlanks@7974.8006.33\|ACGCGACCTCAGATCAGACGTGGCGACCCGCTG | >hg38_21_+_8433172_8446622_RNA45SN1_45S_50ntFlanks | 7974 | 8006 | 33 | ACGCGACCTCAGATCAGACGTGGCGACCCGCTG | rRF | FALSE | FALSE | TRUE |
| >hg38_21_+_8433172_8446622_RNA45SN1_45S_50ntFlanks@6661.6681.21\|CGGTGGATCACTCGGCTCGTG | >hg38_21_+_8433172_8446622_RNA45SN1_45S_50ntFlanks | 6661 | 6681 | 21 | CGGTGGATCACTCGGCTCGTG | rRF | FALSE | FALSE | TRUE |
| >hg38_21_+_8433172_8446622_RNA45SN1_45S_50ntFlanks@10392.10410.19\|ACAGCAGTTGAACATGGGT | >hg38_21_+_8433172_8446622_RNA45SN1_45S_50ntFlanks | 10392 | 10410 | 19 | ACAGCAGTTGAACATGGGT | rRF | FALSE | FALSE | TRUE |
| >hg38_21_+_8433172_8446622_RNA45SN1_45S_50ntFlanks@10684.10710.27\|CGGCGGCGTCCGGTGAGCTCTCGCTGG | >hg38_21_+_8433172_8446622_RNA45SN1_45S_50ntFlanks | 10684 | 10710 | 27 | CGGCGGCGTCCGGTGAGCTCTCGCTGG | rRF | FALSE | FALSE | TRUE |
| >hg38_21_+_8433172_8446622_RNA45SN1_45S_50ntFlanks@7988.8009.22\|CAGACGTGGCGACCCGCTGAAT | >hg38_21_+_8433172_8446622_RNA45SN1_45S_50ntFlanks | 7988 | 8009 | 22 | CAGACGTGGCGACCCGCTGAAT | rRF | FALSE | FALSE | TRUE |
| >hg38_21_+_8433172_8446622_RNA45SN1_45S_50ntFlanks@12626.12661.36\|AATGGGGCGAAGCTACCATCTGTGGGATTATGACTG | >hg38_21_+_8433172_8446622_RNA45SN1_45S_50ntFlanks | 12626 | 12661 | 36 | AATGGGGCGAAGCTACCATCTGTGGGATTATGACTG | rRF | FALSE | FALSE | TRUE |
| >hg38_21_+_8433172_8446622_RNA45SN1_45S_50ntFlanks@10679.10711.33\|GGTTCCGGCGGCGTCCGGTGAGCTCTCGCTGGC | >hg38_21_+_8433172_8446622_RNA45SN1_45S_50ntFlanks | 10679 | 10711 | 33 | GGTTCCGGCGGCGTCCGGTGAGCTCTCGCTGGC | rRF | FALSE | FALSE | TRUE |
| >hg38_21_+_8433172_8446622_RNA45SN1_45S_50ntFlanks@9504.9525.22\|GTGAACTATGCCTGGGCAGGGC | >hg38_21_+_8433172_8446622_RNA45SN1_45S_50ntFlanks | 9504 | 9525 | 22 | GTGAACTATGCCTGGGCAGGGC | rRF | FALSE | FALSE | TRUE |
| >MI0000342\|hsa-mir-200b&WithFlank&1\|+\|1167098\|1167204@63.84.22&[MIMAT0000318&hsa-miR-200b-3p&offsets\|0\|0;m-93&1\|+\|1167160\|1167182&offsets\|0\|-1]\|TAATACTGCCTGGTAATGATGA | >MI0000342\|hsa-mir-200b&WithFlank&1\|+\|1167098\|1167204 | 63 | 84 | 22 | TAATACTGCCTGGTAATGATGA | isomiR | FALSE | FALSE | TRUE |
| >hg38_21_+_8433172_8446622_RNA45SN1_45S_50ntFlanks@11703.11728.26\|AATGAAGCGCGGGTAAACGGCGGGAG | >hg38_21_+_8433172_8446622_RNA45SN1_45S_50ntFlanks | 11703 | 11728 | 26 | AATGAAGCGCGGGTAAACGGCGGGAG | rRF | FALSE | FALSE | TRUE |
| >hg38_21_+_8433172_8446622_RNA45SN1_45S_50ntFlanks@12628.12654.27\|TGGGGCGAAGCTACCATCTGTGGGATT | >hg38_21_+_8433172_8446622_RNA45SN1_45S_50ntFlanks | 12628 | 12654 | 27 | TGGGGCGAAGCTACCATCTGTGGGATT | rRF | FALSE | FALSE | TRUE |
| >hg38_21_+_8433172_8446622_RNA45SN1_45S_50ntFlanks@12973.12990.18\|TCGTACGTAGCAGAGCAG | >hg38_21_+_8433172_8446622_RNA45SN1_45S_50ntFlanks | 12973 | 12990 | 18 | TCGTACGTAGCAGAGCAG | rRF | FALSE | FALSE | TRUE |
| >hg38_21_+_8433172_8446622_RNA45SN1_45S_50ntFlanks@5485.5509.25\|AGCGCTGAGAAGACGGTCGAACTTG | >hg38_21_+_8433172_8446622_RNA45SN1_45S_50ntFlanks | 5485 | 5509 | 25 | AGCGCTGAGAAGACGGTCGAACTTG | rRF | FALSE | FALSE | TRUE |
| >hg38_21_+_8433172_8446622_RNA45SN1_45S_50ntFlanks@11576.11603.28\|CTTAGAACTGGTGCGGACCAGGGGAATC | >hg38_21_+_8433172_8446622_RNA45SN1_45S_50ntFlanks | 11576 | 11603 | 28 | CTTAGAACTGGTGCGGACCAGGGGAATC | rRF | FALSE | FALSE | TRUE |
| >hg38_21_+_8433172_8446622_RNA45SN1_45S_50ntFlanks@10797.10818.22\|GCATGTTGGAACAATGTAGGTA | >hg38_21_+_8433172_8446622_RNA45SN1_45S_50ntFlanks | 10797 | 10818 | 22 | GCATGTTGGAACAATGTAGGTA | rRF | FALSE | FALSE | TRUE |

### Mitochondria

| | | | | | | | | | |
|---|---|---|---|---|---|---|---|---|---|
| >hg38_21_+_8433172_8446622_RNA45SN1_45S_50ntFlanks@11641.11659.19\|CGGCGGGTGTTGACGCGAT | >hg38_21_+_8433172_8446622_RNA45SN1_45S_50ntFlanks | 11641 | 11659 | 19 | CGGCGGGTGTTGACGCGAT | rRF | FALSE | FALSE | TRUE |
| >hg38_21_+_8433172_8446622_RNA45SN1_45S_50ntFlanks@10362.10383.22\|AAGGCCGAAGTGGAGAAGGGTT | >hg38_21_+_8433172_8446622_RNA45SN1_45S_50ntFlanks | 10362 | 10383 | 22 | AAGGCCGAAGTGGAGAAGGGTT | rRF | FALSE | FALSE | TRUE |
| >hg38_21_+_8433172_8446622_RNA45SN1_45S_50ntFlanks@12457.12476.20\|CAAGCGTTGGATTGTTCACC | >hg38_21_+_8433172_8446622_RNA45SN1_45S_50ntFlanks | 12457 | 12476 | 20 | CAAGCGTTGGATTGTTCACC | rRF | FALSE | FALSE | TRUE |
| >hg38_21_+_8433172_8446622_RNA45SN1_45S_50ntFlanks@10841.10862.22\|CGTAACTTCGGGATAAGGATTG | >hg38_21_+_8433172_8446622_RNA45SN1_45S_50ntFlanks | 10841 | 10862 | 22 | CGTAACTTCGGGATAAGGATTG | rRF | FALSE | FALSE | TRUE |
| >hg38_21_+_8433172_8446622_RNA45SN1_45S_50ntFlanks@12073.12095.23\|CCGAGGGGCTCTCGCTTCTGGCG | >hg38_21_+_8433172_8446622_RNA45SN1_45S_50ntFlanks | 12073 | 12095 | 23 | CCGAGGGGCTCTCGCTTCTGGCG | rRF | FALSE | FALSE | TRUE |
| >hg38_21_+_8433172_8446622_RNA45SN1_45S_50ntFlanks@6651.6698.48\|CGACTCTTAGCGGTGGATCACTCGGCTCGTGCGTCGATGAAGAACGCA | >hg38_21_+_8433172_8446622_RNA45SN1_45S_50ntFlanks | 6651 | 6698 | 48 | CGACTCTTAGCGGTGGATCACTCGGCTCGTGCGTCGATGAAG[illegible] | rRF | FALSE | FALSE | TRUE |
| >hg38_21_+_8433172_8446622_RNA45SN1_45S_50ntFlanks@10680.10711.32\|GTTCCGGCGGCGTCCGGTGAGCTCTCGCTGGC | >hg38_21_+_8433172_8446622_RNA45SN1_45S_50ntFlanks | 10680 | 10711 | 32 | GTTCCGGCGGCGTCCGGTGAGCTCTCGCTGGC | rRF | FALSE | FALSE | TRUE |
| >hg38_21_+_8433172_8446622_RNA45SN1_45S_50ntFlanks@12605.12625.21\|TATGTGCTTGGCTGAGGAGCC | >hg38_21_+_8433172_8446622_RNA45SN1_45S_50ntFlanks | 12605 | 12625 | 21 | TATGTGCTTGGCTGAGGAGCC | rRF | FALSE | FALSE | TRUE |
| >hg38_21_+_8433172_8446622_RNA45SN1_45S_50ntFlanks@11208.11227.20\|GCGGCGACTCTGGACGCGAG | >hg38_21_+_8433172_8446622_RNA45SN1_45S_50ntFlanks | 11208 | 11227 | 20 | GCGGCGACTCTGGACGCGAG | rRF | FALSE | FALSE | TRUE |
| >hg38_21_+_8433172_8446622_RNA45SN1_45S_50ntFlanks@5449.5466.18\|ATCGGCCCCGCCGGGGTC | >hg38_21_+_8433172_8446622_RNA45SN1_45S_50ntFlanks | 5449 | 5466 | 18 | ATCGGCCCCGCCGGGGTC | rRF | FALSE | FALSE | TRUE |
| >hg38_21_+_8433172_8446622_RNA45SN1_45S_50ntFlanks@6671.6696.26\|CTCGGCTCGTGCGTCGATGAAGAACG | >hg38_21_+_8433172_8446622_RNA45SN1_45S_50ntFlanks | 6671 | 6696 | 26 | CTCGGCTCGTGCGTCGATGAAGAACG | rRF | FALSE | FALSE | TRUE |
| >hg38_21_+_8433172_8446622_RNA45SN1_45S_50ntFlanks@12405.12426.22\|CTTTTTGATCCTTCGATGTCGG | >hg38_21_+_8433172_8446622_RNA45SN1_45S_50ntFlanks | 12405 | 12426 | 22 | CTTTTTGATCCTTCGATGTCGG | rRF | FALSE | FALSE | TRUE |
| >hg38_21_+_8433172_8446622_RNA45SN1_45S_50ntFlanks@10411.10433.23\|CAGTCGGTCCTGAGAGATGGGCG | >hg38_21_+_8433172_8446622_RNA45SN1_45S_50ntFlanks | 10411 | 10433 | 23 | CAGTCGGTCCTGAGAGATGGGCG | rRF | FALSE | FALSE | TRUE |
| >hg38_21_+_8433172_8446622_RNA45SN1_45S_50ntFlanks@11999.12032.34\|CCCTGCGGGCCGCCGGTGAAATACCACTACTCTG | >hg38_21_+_8433172_8446622_RNA45SN1_45S_50ntFlanks | 11999 | 12032 | 34 | CCCTGCGGGCCGCCGGTGAAATACCACTACTCTG | rRF | FALSE | FALSE | TRUE |
| >hg38_21_+_8433172_8446622_RNA45SN1_45S_50ntFlanks@11621.11640.20\|CAAAGCATCGCGAAGGCCCG | >hg38_21_+_8433172_8446622_RNA45SN1_45S_50ntFlanks | 11621 | 11640 | 20 | CAAAGCATCGCGAAGGCCCG | rRF | FALSE | FALSE | TRUE |
| >hg38_1_-_228634819_228635039_RNA5S12_5S_50ntFlanks@144.170.27\|CCTGGGAATACCGGGTGCTGTAGGCTT | >hg38_1_-_228634819_228635039_RNA5S12_5S_50ntFlanks | 144 | 170 | 27 | CCTGGGAATACCGGGTGCTGTAGGCTT | rRF | FALSE | FALSE | TRUE |
| >hg38_21_+_8433172_8446622_RNA45SN1_45S_50ntFlanks@12676.12703.28\|AGAATCCCGCCCAGGCGGAACGATACGG | >hg38_21_+_8433172_8446622_RNA45SN1_45S_50ntFlanks | 12676 | 12703 | 28 | AGAATCCCGCCCAGGCGGAACGATACGG | rRF | FALSE | FALSE | TRUE |
| >hg38_21_+_8433172_8446622_RNA45SN1_45S_50ntFlanks@12640.12661.22\|ACCATCTGTGGGATTATGACTG | >hg38_21_+_8433172_8446622_RNA45SN1_45S_50ntFlanks | 12640 | 12661 | 22 | ACCATCTGTGGGATTATGACTG | rRF | FALSE | FALSE | TRUE |
| >hg38_21_+_8433172_8446622_RNA45SN1_45S_50ntFlanks@10845.10865.21\|ACTTCGGGATAAGGATTGGCT | >hg38_21_+_8433172_8446622_RNA45SN1_45S_50ntFlanks | 10845 | 10865 | 21 | ACTTCGGGATAAGGATTGGCT | rRF | FALSE | FALSE | TRUE |
| >hg38_21_+_8433172_8446622_RNA45SN1_45S_50ntFlanks@9534.9561.28\|AGGAAACTCTGGTGGAGGTCCGTAGCGG | >hg38_21_+_8433172_8446622_RNA45SN1_45S_50ntFlanks | 9534 | 9561 | 28 | AGGAAACTCTGGTGGAGGTCCGTAGCGG | rRF | FALSE | FALSE | TRUE |
| >hg38_21_+_8433172_8446622_RNA45SN1_45S_50ntFlanks@9604.9622.19\|AAAGACTAATCGAACCATC | >hg38_21_+_8433172_8446622_RNA45SN1_45S_50ntFlanks | 9604 | 9622 | 19 | AAAGACTAATCGAACCATC | rRF | FALSE | FALSE | TRUE |
| >hg38_21_+_8433172_8446622_RNA45SN1_45S_50ntFlanks@8114.8136.23\|GCGCGGGACATGTGGCGTACGGA | >hg38_21_+_8433172_8446622_RNA45SN1_45S_50ntFlanks | 8114 | 8136 | 23 | GCGCGGGACATGTGGCGTACGGA | rRF | FALSE | FALSE | TRUE |
| >hg38_21_+_8433172_8446622_RNA45SN1_45S_50ntFlanks@9305.9338.34\|CCAAGGAGTCTAACACGTGCGCGAGTCGGGGGCT | >hg38_21_+_8433172_8446622_RNA45SN1_45S_50ntFlanks | 9305 | 9338 | 34 | CCAAGGAGTCTAACACGTGCGCGAGTCGGGGGCT | rRF | FALSE | FALSE | TRUE |
| >hg38_21_+_8433172_8446622_RNA45SN1_45S_50ntFlanks@10788.10818.31\|CAGCCTCTGGCATGTTGGAACAATGTAGGTA | >hg38_21_+_8433172_8446622_RNA45SN1_45S_50ntFlanks | 10788 | 10818 | 31 | CAGCCTCTGGCATGTTGGAACAATGTAGGTA | rRF | FALSE | FALSE | TRUE |
| >hg38_21_+_8433172_8446622_RNA45SN1_45S_50ntFlanks@11666.11685.20\|TCTGCCCAGTGCTCTGAATG | >hg38_21_+_8433172_8446622_RNA45SN1_45S_50ntFlanks | 11666 | 11685 | 20 | TCTGCCCAGTGCTCTGAATG | rRF | FALSE | FALSE | TRUE |
| >hg38_21_+_8433172_8446622_RNA45SN1_45S_50ntFlanks@9570.9597.28\|TGCAAATCGGTCGTCCGACCTGGGTATA | >hg38_21_+_8433172_8446622_RNA45SN1_45S_50ntFlanks | 9570 | 9597 | 28 | TGCAAATCGGTCGTCCGACCTGGGTATA | rRF | FALSE | FALSE | TRUE |
| >hg38_21_+_8433172_8446622_RNA45SN1_45S_50ntFlanks@11581.11605.25\|AACTGGTGCGGACCAGGGGAATCCG | >hg38_21_+_8433172_8446622_RNA45SN1_45S_50ntFlanks | 11581 | 11605 | 25 | AACTGGTGCGGACCAGGGGAATCCG | rRF | FALSE | FALSE | TRUE |
| >hg38_21_+_8433172_8446622_RNA45SN1_45S_50ntFlanks@4724.4759.36\|ATTAATCAAGAACGAAAGTCGGAGGTTCGAAGACGA | >hg38_21_+_8433172_8446622_RNA45SN1_45S_50ntFlanks | 4724 | 4759 | 36 | ATTAATCAAGAACGAAAGTCGGAGGTTCGAAGACGA | rRF | FALSE | FALSE | TRUE |
| >hg38_21_+_8433172_8446622_RNA45SN1_45S_50ntFlanks@10351.10378.28\|CGAGAACTTTGAAGGCCGAAGTGGAGAA | >hg38_21_+_8433172_8446622_RNA45SN1_45S_50ntFlanks | 10351 | 10378 | 28 | CGAGAACTTTGAAGGCCGAAGTGGAGAA | rRF | FALSE | FALSE | TRUE |
| >hg38_21_+_8433172_8446622_RNA45SN1_45S_50ntFlanks@7986.8003.18\|ATCAGACGTGGCGACCCG | >hg38_21_+_8433172_8446622_RNA45SN1_45S_50ntFlanks | 7986 | 8003 | 18 | ATCAGACGTGGCGACCCG | rRF | FALSE | FALSE | TRUE |
| >hg38_21_+_8433172_8446622_RNA45SN1_45S_50ntFlanks@11593.11611.19\|CCAGGGGAATCCGACTGTT | >hg38_21_+_8433172_8446622_RNA45SN1_45S_50ntFlanks | 11593 | 11611 | 19 | CCAGGGGAATCCGACTGTT | rRF | FALSE | FALSE | TRUE |
| >hg38_21_+_8433172_8446622_RNA45SN1_45S_50ntFlanks@11572.11605.34\|CCGACTTAGAACTGGTGCGGACCAGGGGAATCCG | >hg38_21_+_8433172_8446622_RNA45SN1_45S_50ntFlanks | 11572 | 11605 | 34 | CCGACTTAGAACTGGTGCGGACCAGGGGAATCCG | rRF | FALSE | FALSE | TRUE |
| >hg38_21_+_8433172_8446622_RNA45SN1_45S_50ntFlanks@12633.12659.27\|CGAAGCTACCATCTGTGGGATTATGAC | >hg38_21_+_8433172_8446622_RNA45SN1_45S_50ntFlanks | 12633 | 12659 | 27 | CGAAGCTACCATCTGTGGGATTATGAC | rRF | FALSE | FALSE | TRUE |
| >hg38_21_+_8433172_8446622_RNA45SN1_45S_50ntFlanks@9564.9597.34\|CTGACGTGCAAATCGGTCGTCCGACCTGGGTATA | >hg38_21_+_8433172_8446622_RNA45SN1_45S_50ntFlanks | 9564 | 9597 | 34 | CTGACGTGCAAATCGGTCGTCCGACCTGGGTATA | rRF | FALSE | FALSE | TRUE |
| >hg38_21_+_8433172_8446622_RNA45SN1_45S_50ntFlanks@8176.8204.29\|CTTCTGATCGAGGCCCAGCCCGTGGACGG | >hg38_21_+_8433172_8446622_RNA45SN1_45S_50ntFlanks | 8176 | 8204 | 29 | CTTCTGATCGAGGCCCAGCCCGTGGACGG | rRF | FALSE | FALSE | TRUE |
| >hg38_21_+_8433172_8446622_RNA45SN1_45S_50ntFlanks@12126.12146.21\|CGACCCGCTCCGGGGACAGTG | >hg38_21_+_8433172_8446622_RNA45SN1_45S_50ntFlanks | 12126 | 12146 | 21 | CGACCCGCTCCGGGGACAGTG | rRF | FALSE | FALSE | TRUE |
| >hg38_21_+_8433172_8446622_RNA45SN1_45S_50ntFlanks@11322.11344.23\|ACGTCTCGTCGCGCGCGCGTCCG | >hg38_21_+_8433172_8446622_RNA45SN1_45S_50ntFlanks | 11322 | 11344 | 23 | ACGTCTCGTCGCGCGCGCGTCCG | rRF | FALSE | FALSE | TRUE |
| >hg38_21_+_8433172_8446622_RNA45SN1_45S_50ntFlanks@11613.11640.28\|AATTAAAACAAAGCATCGCGAAGGCCCG | >hg38_21_+_8433172_8446622_RNA45SN1_45S_50ntFlanks | 11613 | 11640 | 28 | AATTAAAACAAAGCATCGCGAAGGCCCG | rRF | FALSE | FALSE | TRUE |
| >hg38_21_+_8433172_8446622_RNA45SN1_45S_50ntFlanks@12627.12659.33\|ATGGGGCGAAGCTACCATCTGTGGGATTATGAC | >hg38_21_+_8433172_8446622_RNA45SN1_45S_50ntFlanks | 12627 | 12659 | 33 | ATGGGGCGAAGCTACCATCTGTGGGATTATGAC | rRF | FALSE | FALSE | TRUE |
| >hg38_21_+_8433172_8446622_RNA45SN1_45S_50ntFlanks@6721.6746.26\|AATTGCAGGACACATTGATCATCGAC | >hg38_21_+_8433172_8446622_RNA45SN1_45S_50ntFlanks | 6721 | 6746 | 26 | AATTGCAGGACACATTGATCATCGAC | rRF | FALSE | FALSE | TRUE |

### Mitochondria

| | | | | | | | | | |
|---|---|---|---|---|---|---|---|---|---|
| >hg38_21_+_8433172_8446622_RNA45SN1_45S_50ntFlanks@8198.8215.18\|TGGACGGTGTGAGGCCGG | >hg38_21_+_8433172_8446622_RNA45SN1_45S_50ntFlanks | 8198 | 8215 | 18 | TGGACGGTGTGAGGCCGG | rRF | FALSE | FALSE | TRUE |
| >hg38_21_+_8433172_8446622_RNA45SN1_45S_50ntFlanks@9570.9595.26\|TGCAAATCGGTCGTCCGACCTGGGTA | >hg38_21_+_8433172_8446622_RNA45SN1_45S_50ntFlanks | 9570 | 9595 | 26 | TGCAAATCGGTCGTCCGACCTGGGTA | rRF | FALSE | FALSE | TRUE |
| >hg38_21_+_8433172_8446622_RNA45SN1_45S_50ntFlanks@11580.11611.32\|GAACTGGTGCGGACCAGGGGAATCCGACTGTT | >hg38_21_+_8433172_8446622_RNA45SN1_45S_50ntFlanks | 11580 | 11611 | 32 | GAACTGGTGCGGACCAGGGGAATCCGACTGTT | rRF | FALSE | FALSE | TRUE |
| >hg38_21_+_8433172_8446622_RNA45SN1_45S_50ntFlanks@11845.11872.28\|GGAACGGGCTTGGCGGAATCAGCGGGGA | >hg38_21_+_8433172_8446622_RNA45SN1_45S_50ntFlanks | 11845 | 11872 | 28 | GGAACGGGCTTGGCGGAATCAGCGGGGA | rRF | FALSE | FALSE | TRUE |
| >hg38_21_+_8433172_8446622_RNA45SN1_45S_50ntFlanks@10348.10368.21\|AAACGAGAACTTTGAAGGCCG | >hg38_21_+_8433172_8446622_RNA45SN1_45S_50ntFlanks | 10348 | 10368 | 21 | AAACGAGAACTTTGAAGGCCG | rRF | FALSE | FALSE | TRUE |
| >hg38_21_+_8433172_8446622_RNA45SN1_45S_50ntFlanks@12627.12661.35\|ATGGGGCGAAGCTACCATCTGTGGGATTATGACTG | >hg38_21_+_8433172_8446622_RNA45SN1_45S_50ntFlanks | 12627 | 12661 | 35 | ATGGGGCGAAGCTACCATCTGTGGGATTATGACTG | rRF | FALSE | FALSE | TRUE |
| >hg38_21_+_8433172_8446622_RNA45SN1_45S_50ntFlanks@8104.8141.38\|CCGCGGCGGGGCGCGGGACATGTGGCGTACGGAAGACC | >hg38_21_+_8433172_8446622_RNA45SN1_45S_50ntFlanks | 8104 | 8141 | 38 | CCGCGGCGGGGCGCGGGACATGTGGCGTACGGAAGACC | rRF | FALSE | FALSE | TRUE |
| >hg38_21_+_8433172_8446622_RNA45SN1_45S_50ntFlanks@6743.6770.28\|CGACACTTCGAACGCACTTGCGGCCCCG | >hg38_21_+_8433172_8446622_RNA45SN1_45S_50ntFlanks | 6743 | 6770 | 28 | CGACACTTCGAACGCACTTGCGGCCCCG | rRF | FALSE | FALSE | TRUE |
| >hg38_21_+_8433172_8446622_RNA45SN1_45S_50ntFlanks@12937.12956.20\|AACCATTCGTAGACGACCTG | >hg38_21_+_8433172_8446622_RNA45SN1_45S_50ntFlanks | 12937 | 12956 | 20 | AACCATTCGTAGACGACCTG | rRF | FALSE | FALSE | TRUE |
| >hg38_21_+_8433172_8446622_RNA45SN1_45S_50ntFlanks@10349.10377.29\|AACGAGAACTTTGAAGGCCGAAGTGGAGA | >hg38_21_+_8433172_8446622_RNA45SN1_45S_50ntFlanks | 10349 | 10377 | 29 | AACGAGAACTTTGAAGGCCGAAGTGGAGA | rRF | FALSE | FALSE | TRUE |
| >trna111_HisGTG_1_-_147774845_147774916@-1T.29.30,trna118_HisGTG_1_-_145396881_145396952@-1T.29.30,trna16_HisGTG_1_+_146544773_146544844@-1T.29.30,trna1_HisGTG_15_+_45493349_45493420@-1T.29.30,trna21_HisGTG_1_+_147753471_147753542@-1T.29.30,trna33_HisGTG_6_+_27125906_27125977@-1T.29.30,trna7_HisGTG_9_-_14433938_14434009@-1T.29.30,trna8_HisGTG_15_-_45492611_45492682@-1T.29.30,trna9_HisGTG_15_-_45490804_45490875@-1T.29.30\|TGCCGTGATCGTATAGTGGTTAGTACTCTG | >trna111_HisGTG_1_-_147774845_147774916 | -1T | 29 | 30 | TGCCGTGATCGTATAGTGGTTAGTACTCTG | tRF | FALSE | FALSE | TRUE |
| >hg38_21_+_8433172_8446622_RNA45SN1_45S_50ntFlanks@11581.11609.29\|AACTGGTGCGGACCAGGGGAATCCGACTG | >hg38_21_+_8433172_8446622_RNA45SN1_45S_50ntFlanks | 11581 | 11609 | 29 | AACTGGTGCGGACCAGGGGAATCCGACTG | rRF | FALSE | FALSE | TRUE |
| >hg38_21_+_8433172_8446622_RNA45SN1_45S_50ntFlanks@11564.11591.28\|CCTAGCAGCCGACTTAGAACTGGTGCGG | >hg38_21_+_8433172_8446622_RNA45SN1_45S_50ntFlanks | 11564 | 11591 | 28 | CCTAGCAGCCGACTTAGAACTGGTGCGG | rRF | FALSE | FALSE | TRUE |
| >hg38_21_+_8433172_8446622_RNA45SN1_45S_50ntFlanks@9565.9596.32\|TGACGTGCAAATCGGTCGTCCGACCTGGGTAT | >hg38_21_+_8433172_8446622_RNA45SN1_45S_50ntFlanks | 9565 | 9596 | 32 | TGACGTGCAAATCGGTCGTCCGACCTGGGTAT | rRF | FALSE | FALSE | TRUE |
