## Supplementary material for "Single-nucleotide Differences and Cell Type Decide the Subcellular Localization of miRNA Isoforms (isomiRs), tRNA-derived Fragments (tRFs) and rRNA-derived Fragments (rRFs)": Supp. Table S5 (4 of 4)

### Mitoplast

| | | | | | | | | | |
|---|---|---|---|---|---|---|---|---|---|
| >hg38_21_+_8433172_8446622_RNA45SN1_45S_50ntFlanks@12002.12032.31\|TGCGGGCCGCCGGTGAAATACCACTACTCTG | >hg38_21_+_8433172_8446622_RNA45SN1_45S_50ntFlanks | 12002 | 12032 | 31 | TGCGGGCCGCCGGTGAAATACCACTACTCTG | rRF | FALSE | FALSE | TRUE |
| >hg38_21_+_8433172_8446622_RNA45SN1_45S_50ntFlanks@7991.8011.21\|ACGTGGCGACCCGCTGAATTT | >hg38_21_+_8433172_8446622_RNA45SN1_45S_50ntFlanks | 7991 | 8011 | 21 | ACGTGGCGACCCGCTGAATTT | rRF | FALSE | FALSE | TRUE |
| >hg38_21_+_8433172_8446622_RNA45SN1_45S_50ntFlanks@12306.12323.18\|CACGATCCTTCTGACCTT | >hg38_21_+_8433172_8446622_RNA45SN1_45S_50ntFlanks | 12306 | 12323 | 18 | CACGATCCTTCTGACCTT | rRF | FALSE | FALSE | TRUE |
| >hg38_21_+_8433172_8446622_RNA45SN1_45S_50ntFlanks@6671.6696.26\|CTCGGCTCGTGCGTCGATGAAGAACG | >hg38_21_+_8433172_8446622_RNA45SN1_45S_50ntFlanks | 6671 | 6696 | 26 | CTCGGCTCGTGCGTCGATGAAGAACG | rRF | FALSE | FALSE | TRUE |
| >hg38_21_+_8433172_8446622_RNA45SN1_45S_50ntFlanks@3985.4002.18\|CGGCTTTGGTGACTCTAG | >hg38_21_+_8433172_8446622_RNA45SN1_45S_50ntFlanks | 3985 | 4002 | 18 | CGGCTTTGGTGACTCTAG | rRF | FALSE | FALSE | TRUE |
| >hg38_21_+_8433172_8446622_RNA45SN1_45S_50ntFlanks@5487.5514.28\|CGCTGAGAAGACGGTCGAACTTGACTAT | >hg38_21_+_8433172_8446622_RNA45SN1_45S_50ntFlanks | 5487 | 5514 | 28 | CGCTGAGAAGACGGTCGAACTTGACTAT | rRF | FALSE | FALSE | TRUE |
| >hg38_21_+_8433172_8446622_RNA45SN1_45S_50ntFlanks@9563.9596.34\|CCTGACGTGCAAATCGGTCGTCCGACCTGGGTAT | >hg38_21_+_8433172_8446622_RNA45SN1_45S_50ntFlanks | 9563 | 9596 | 34 | CCTGACGTGCAAATCGGTCGTCCGACCTGGGTAT | rRF | FALSE | FALSE | TRUE |
| >hg38_21_+_8433172_8446622_RNA45SN1_45S_50ntFlanks@6698.6718.21\|AGCTAGCTGCGAGAATTAATG | >hg38_21_+_8433172_8446622_RNA45SN1_45S_50ntFlanks | 6698 | 6718 | 21 | AGCTAGCTGCGAGAATTAATG | rRF | FALSE | FALSE | TRUE |
| >hg38_21_+_8433172_8446622_RNA45SN1_45S_50ntFlanks@13002.13027.26\|CGATCTATTGAAAGTCAGCCCTCGAC | >hg38_21_+_8433172_8446622_RNA45SN1_45S_50ntFlanks | 13002 | 13027 | 26 | CGATCTATTGAAAGTCAGCCCTCGAC | rRF | FALSE | FALSE | TRUE |
| >hg38_21_+_8433172_8446622_RNA45SN1_45S_50ntFlanks@12641.12659.19\|CCATCTGTGGGATTATGAC | >hg38_21_+_8433172_8446622_RNA45SN1_45S_50ntFlanks | 12641 | 12659 | 19 | CCATCTGTGGGATTATGAC | rRF | FALSE | FALSE | TRUE |
| >hg38_21_+_8433172_8446622_RNA45SN1_45S_50ntFlanks@4047.4066.20\|ATTCGAACGTCTGCCCTATC | >hg38_21_+_8433172_8446622_RNA45SN1_45S_50ntFlanks | 4047 | 4066 | 20 | ATTCGAACGTCTGCCCTATC | rRF | FALSE | FALSE | TRUE |
| >hg38_21_+_8433172_8446622_RNA45SN1_45S_50ntFlanks@12636.12661.26\|AGCTACCATCTGTGGGATTATGACTG | >hg38_21_+_8433172_8446622_RNA45SN1_45S_50ntFlanks | 12636 | 12661 | 26 | AGCTACCATCTGTGGGATTATGACTG | rRF | FALSE | FALSE | TRUE |
| >hg38_21_+_8433172_8446622_RNA45SN1_45S_50ntFlanks@12405.12437.33\|CTTTTTGATCCTTCGATGTCGGCTCTTCCTATC | >hg38_21_+_8433172_8446622_RNA45SN1_45S_50ntFlanks | 12405 | 12437 | 33 | CTTTTTGATCCTTCGATGTCGGCTCTTCCTATC | rRF | FALSE | FALSE | TRUE |
| >hg38_21_+_8433172_8446622_RNA45SN1_45S_50ntFlanks@10690.10708.19\|CGTCCGGTGAGCTCTCGCT | >hg38_21_+_8433172_8446622_RNA45SN1_45S_50ntFlanks | 10690 | 10708 | 19 | CGTCCGGTGAGCTCTCGCT | rRF | FALSE | FALSE | TRUE |
| >hg38_21_+_8433172_8446622_RNA45SN1_45S_50ntFlanks@6740.6771.32\|CATCGACACTTCGAACGCACTTGCGGCCCCGG | >hg38_21_+_8433172_8446622_RNA45SN1_45S_50ntFlanks | 6740 | 6771 | 32 | CATCGACACTTCGAACGCACTTGCGGCCCCGG | rRF | FALSE | FALSE | TRUE |
| >hg38_21_+_8433172_8446622_RNA45SN1_45S_50ntFlanks@8114.8143.30\|GCGCGGGACATGTGGCGTACGGAAGACCCG | >hg38_21_+_8433172_8446622_RNA45SN1_45S_50ntFlanks | 8114 | 8143 | 30 | GCGCGGGACATGTGGCGTACGGAAGACCCG | rRF | FALSE | FALSE | TRUE |
| >hg38_21_+_8433172_8446622_RNA45SN1_45S_50ntFlanks@10684.10711.28\|CGGCGGCGTCCGGTGAGCTCTCGCTGGC | >hg38_21_+_8433172_8446622_RNA45SN1_45S_50ntFlanks | 10684 | 10711 | 28 | CGGCGGCGTCCGGTGAGCTCTCGCTGGC | rRF | FALSE | FALSE | TRUE |
| >hg38_21_+_8433172_8446622_RNA45SN1_45S_50ntFlanks@9554.9572.19\|CGTAGCGGTCCTGACGTGC | >hg38_21_+_8433172_8446622_RNA45SN1_45S_50ntFlanks | 9554 | 9572 | 19 | CGTAGCGGTCCTGACGTGC | rRF | FALSE | FALSE | TRUE |
| >hg38_21_+_8433172_8446622_RNA45SN1_45S_50ntFlanks@6663.6689.27\|GTGGATCACTCGGCTCGTGCGTCGATG | >hg38_21_+_8433172_8446622_RNA45SN1_45S_50ntFlanks | 6663 | 6689 | 27 | GTGGATCACTCGGCTCGTGCGTCGATG | rRF | FALSE | FALSE | TRUE |
| >hg38_21_+_8433172_8446622_RNA45SN1_45S_50ntFlanks@11564.11602.39\|CCTAGCAGCCGACTTAGAACTGGTGCGGACCAGGGGAAT | >hg38_21_+_8433172_8446622_RNA45SN1_45S_50ntFlanks | 11564 | 11602 | 39 | CCTAGCAGCCGACTTAGAACTGGTGCGGACCAGGGGAAT | rRF | FALSE | FALSE | TRUE |
| >hg38_21_+_8433172_8446622_RNA45SN1_45S_50ntFlanks@3899.3919.21\|CCTTCGCGGGGGGGATGCGTG | >hg38_21_+_8433172_8446622_RNA45SN1_45S_50ntFlanks | 3899 | 3919 | 21 | CCTTCGCGGGGGGGATGCGTG | rRF | FALSE | FALSE | TRUE |
| >hg38_21_+_8433172_8446622_RNA45SN1_45S_50ntFlanks@6652.6686.35\|GACTCTTAGCGGTGGATCACTCGGCTCGTGCGTCG | >hg38_21_+_8433172_8446622_RNA45SN1_45S_50ntFlanks | 6652 | 6686 | 35 | GACTCTTAGCGGTGGATCACTCGGCTCGTGCGTCG | rRF | FALSE | FALSE | TRUE |
| >hg38_21_+_8433172_8446622_RNA45SN1_45S_50ntFlanks@12628.12661.34\|TGGGGCGAAGCTACCATCTGTGGGATTATGACTG | >hg38_21_+_8433172_8446622_RNA45SN1_45S_50ntFlanks | 12628 | 12661 | 34 | TGGGGCGAAGCTACCATCTGTGGGATTATGACTG | rRF | FALSE | FALSE | TRUE |
| >hg38_21_+_8433172_8446622_RNA45SN1_45S_50ntFlanks@11315.11344.30\|CCACCCCACGTCTCGTCGCGCGCGCGTCCG | >hg38_21_+_8433172_8446622_RNA45SN1_45S_50ntFlanks | 11315 | 11344 | 30 | CCACCCCACGTCTCGTCGCGCGCGCGTCCG | rRF | FALSE | FALSE | TRUE |
| >hg38_21_+_8433172_8446622_RNA45SN1_45S_50ntFlanks@12007.12031.25\|GCCGCCGGTGAAATACCACTACTCT | >hg38_21_+_8433172_8446622_RNA45SN1_45S_50ntFlanks | 12007 | 12031 | 25 | GCCGCCGGTGAAATACCACTACTCT | rRF | FALSE | FALSE | TRUE |
| >hg38_21_+_8433172_8446622_RNA45SN1_45S_50ntFlanks@9504.9526.23\|GTGAACTATGCCTGGGCAGGGCG | >hg38_21_+_8433172_8446622_RNA45SN1_45S_50ntFlanks | 9504 | 9526 | 23 | GTGAACTATGCCTGGGCAGGGCG | rRF | FALSE | FALSE | TRUE |
| >hg38_21_+_8433172_8446622_RNA45SN1_45S_50ntFlanks@9004.9026.23\|TCTCCCCCGCGGGGGCGCGCCGG | >hg38_21_+_8433172_8446622_RNA45SN1_45S_50ntFlanks | 9004 | 9026 | 23 | TCTCCCCCGCGGGGGCGCGCCGG | rRF | FALSE | FALSE | TRUE |
| >hg38_21_+_8433172_8446622_RNA45SN1_45S_50ntFlanks@12893.12933.41\|CCTCGCCCGTCACGCACCGCACGTTCGTGGGGAACCTGGCG | >hg38_21_+_8433172_8446622_RNA45SN1_45S_50ntFlanks | 12893 | 12933 | 41 | CCTCGCCCGTCACGCACCGCACGTTCGTGGGGAACCTGGCG | rRF | FALSE | FALSE | TRUE |
| >hg38_21_+_8433172_8446622_RNA45SN1_45S_50ntFlanks@12456.12486.31\|CCAAGCGTTGGATTGTTCACCCACTAATAGG | >hg38_21_+_8433172_8446622_RNA45SN1_45S_50ntFlanks | 12456 | 12486 | 31 | CCAAGCGTTGGATTGTTCACCCACTAATAGG | rRF | FALSE | FALSE | TRUE |
| >hg38_21_+_8433172_8446622_RNA45SN1_45S_50ntFlanks@5445.5466.22\|TCGGATCGGCCCCGCCGGGGTC | >hg38_21_+_8433172_8446622_RNA45SN1_45S_50ntFlanks | 5445 | 5466 | 22 | TCGGATCGGCCCCGCCGGGGTC | rRF | FALSE | FALSE | TRUE |
| >hg38_21_+_8433172_8446622_RNA45SN1_45S_50ntFlanks@11569.11612.44\|CAGCCGACTTAGAACTGGTGCGGACCAGGGGAATCCGACTGTTT | >hg38_21_+_8433172_8446622_RNA45SN1_45S_50ntFlanks | 11569 | 11612 | 44 | CAGCCGACTTAGAACTGGTGCGGACCAGGGGAATCCGACTGTTT | rRF | FALSE | FALSE | TRUE |
| >hg38_21_+_8433172_8446622_RNA45SN1_45S_50ntFlanks@9575.9597.23\|ATCGGTCGTCCGACCTGGGTATA | >hg38_21_+_8433172_8446622_RNA45SN1_45S_50ntFlanks | 9575 | 9597 | 23 | ATCGGTCGTCCGACCTGGGTATA | rRF | FALSE | FALSE | TRUE |
| >hg38_21_+_8433172_8446622_RNA45SN1_45S_50ntFlanks@11641.11669.29\|CGGCGGGTGTTGACGCGATGTGATTTCTG | >hg38_21_+_8433172_8446622_RNA45SN1_45S_50ntFlanks | 11641 | 11669 | 29 | CGGCGGGTGTTGACGCGATGTGATTTCTG | rRF | FALSE | FALSE | TRUE |
| >hg38_21_+_8433172_8446622_RNA45SN1_45S_50ntFlanks@6650.6698.49\|ACGACTCTTAGCGGTGGATCACTCGGCTCGTGCGTCGATGAAGAACGCA | >hg38_21_+_8433172_8446622_RNA45SN1_45S_50ntFlanks | 6650 | 6698 | 49 | ACGACTCTTAGCGGTGGATCACTCGGCTCGTGCGTCGATGAAGAACGCA | rRF | FALSE | FALSE | TRUE |
| >trnaMT_ValTAC_MT_+_1602_1670@51.72.22\|CTTAACTTGACCGCTCTGACCA | >trnaMT_ValTAC_MT_+_1602_1670 | 51 | 72 | 22 | CTTAACTTGACCGCTCTGACCA | tRF | FALSE | FALSE | TRUE |
| >hg38_21_+_8433172_8446622_RNA45SN1_45S_50ntFlanks@9564.9595.32\|CTGACGTGCAAATCGGTCGTCCGACCTGGGTA | >hg38_21_+_8433172_8446622_RNA45SN1_45S_50ntFlanks | 9564 | 9595 | 32 | CTGACGTGCAAATCGGTCGTCCGACCTGGGTA | rRF | FALSE | FALSE | TRUE |
| >hg38_21_+_8433172_8446622_RNA45SN1_45S_50ntFlanks@11316.11338.23\|CACCCCACGTCTCGTCGCGCGCG | >hg38_21_+_8433172_8446622_RNA45SN1_45S_50ntFlanks | 11316 | 11338 | 23 | CACCCCACGTCTCGTCGCGCGCG | rRF | FALSE | FALSE | TRUE |
| >hg38_21_+_8433172_8446622_RNA45SN1_45S_50ntFlanks@12893.12932.40\|CCTCGCCCGTCACGCACCGCACGTTCGTGGGGAACCTGGC | >hg38_21_+_8433172_8446622_RNA45SN1_45S_50ntFlanks | 12893 | 12932 | 40 | CCTCGCCCGTCACGCACCGCACGTTCGTGGGGAACCTGGC | rRF | FALSE | FALSE | TRUE |
| >hg38_21_+_8433172_8446622_RNA45SN1_45S_50ntFlanks@12003.12032.30\|GCGGGCCGCCGGTGAAATACCACTACTCTG | >hg38_21_+_8433172_8446622_RNA45SN1_45S_50ntFlanks | 12003 | 12032 | 30 | GCGGGCCGCCGGTGAAATACCACTACTCTG | rRF | FALSE | FALSE | TRUE |
| >hg38_21_+_8433172_8446622_RNA45SN1_45S_50ntFlanks@11572.11602.31\|CCGACTTAGAACTGGTGCGGACCAGGGGAAT | >hg38_21_+_8433172_8446622_RNA45SN1_45S_50ntFlanks | 11572 | 11602 | 31 | CCGACTTAGAACTGGTGCGGACCAGGGGAAT | rRF | FALSE | FALSE | TRUE |
| >hg38_21_+_8433172_8446622_RNA45SN1_45S_50ntFlanks@6650.6679.30\|ACGACTCTTAGCGGTGGATCACTCGGCTCG | >hg38_21_+_8433172_8446622_RNA45SN1_45S_50ntFlanks | 6650 | 6679 | 30 | ACGACTCTTAGCGGTGGATCACTCGGCTCG | rRF | FALSE | FALSE | TRUE |
| >hg38_21_+_8433172_8446622_RNA45SN1_45S_50ntFlanks@6653.6689.37\|ACTCTTAGCGGTGGATCACTCGGCTCGTGCGTCGATG | >hg38_21_+_8433172_8446622_RNA45SN1_45S_50ntFlanks | 6653 | 6689 | 37 | ACTCTTAGCGGTGGATCACTCGGCTCGTGCGTCGATG | rRF | FALSE | FALSE | TRUE |
| >hg38_21_+_8433172_8446622_RNA45SN1_45S_50ntFlanks@12417.12443.27\|TCGATGTCGGCTCTTCCTATCATTGTG | >hg38_21_+_8433172_8446622_RNA45SN1_45S_50ntFlanks | 12417 | 12443 | 27 | TCGATGTCGGCTCTTCCTATCATTGTG | rRF | FALSE | FALSE | TRUE |

### Mitoplast

| | | | | | | | | | |
|---|---|---|---|---|---|---|---|---|---|
| >hg38_21_+_8433172_8446622_RNA45SN1_45S_50ntFlanks@9554.9597.44\|CGTAGCGGTCCTGACGTGCAAATCGGTCGTCCGACCTGGGTATA | >hg38_21_+_8433172_8446622_RNA45SN1_45S_50ntFlanks | 9554 | 9597 | 44 | CGTAGCGGTCCTGACGTGCAAATCGGTCGTCCGACCTGGGTATA | rRF | FALSE | FALSE | TRUE |
| >hg38_21_+_8433172_8446622_RNA45SN1_45S_50ntFlanks@6671.6697.27\|CTCGGCTCGTGCGTCGATGAAGAACGC | >hg38_21_+_8433172_8446622_RNA45SN1_45S_50ntFlanks | 6671 | 6697 | 27 | CTCGGCTCGTGCGTCGATGAAGAACGC | rRF | FALSE | FALSE | TRUE |
| >hg38_21_+_8433172_8446622_RNA45SN1_45S_50ntFlanks@11583.11611.29\|CTGGTGCGGACCAGGGGAATCCGACTGTT | >hg38_21_+_8433172_8446622_RNA45SN1_45S_50ntFlanks | 11583 | 11611 | 29 | CTGGTGCGGACCAGGGGAATCCGACTGTT | rRF | FALSE | FALSE | TRUE |
| >hg38_21_+_8433172_8446622_RNA45SN1_45S_50ntFlanks@8356.8379.24\|GAAAAGAACTTTGAAGAGAGAGTT | >hg38_21_+_8433172_8446622_RNA45SN1_45S_50ntFlanks | 8356 | 8379 | 24 | GAAAAGAACTTTGAAGAGAGAGTT | rRF | FALSE | FALSE | TRUE |
| >hg38_21_+_8433172_8446622_RNA45SN1_45S_50ntFlanks@6773.6803.31\|TTCCTCCCGGGGCTACGCCTGTCTGAGCGTC | >hg38_21_+_8433172_8446622_RNA45SN1_45S_50ntFlanks | 6773 | 6803 | 31 | TTCCTCCCGGGGCTACGCCTGTCTGAGCGTC | rRF | FALSE | FALSE | TRUE |
| >hg38_21_+_8433172_8446622_RNA45SN1_45S_50ntFlanks@10350.10390.41\|ACGAGAACTTTGAAGGCCGAAGTGGAGAAGGGTTCCATGTG | >hg38_21_+_8433172_8446622_RNA45SN1_45S_50ntFlanks | 10350 | 10390 | 41 | ACGAGAACTTTGAAGGCCGAAGTGGAGAAGGGTTCCATGTG | rRF | FALSE | FALSE | TRUE |
| >hg38_21_+_8433172_8446622_RNA45SN1_45S_50ntFlanks@6721.6744.24\|AATTGCAGGACACATTGATCATCG | >hg38_21_+_8433172_8446622_RNA45SN1_45S_50ntFlanks | 6721 | 6744 | 24 | AATTGCAGGACACATTGATCATCG | rRF | FALSE | FALSE | TRUE |
| >hg38_21_+_8433172_8446622_RNA45SN1_45S_50ntFlanks@11572.11611.40\|CCGACTTAGAACTGGTGCGGACCAGGGGAATCCGACTGTT | >hg38_21_+_8433172_8446622_RNA45SN1_45S_50ntFlanks | 11572 | 11611 | 40 | CCGACTTAGAACTGGTGCGGACCAGGGGAATCCGACTGTT | rRF | FALSE | FALSE | TRUE |
| >hg38_21_+_8433172_8446622_RNA45SN1_45S_50ntFlanks@8155.8181.27\|CGCTCGTGGGGGGCCCAAGTCCTTCTG | >hg38_21_+_8433172_8446622_RNA45SN1_45S_50ntFlanks | 8155 | 8181 | 27 | CGCTCGTGGGGGGCCCAAGTCCTTCTG | rRF | FALSE | FALSE | TRUE |
| >hg38_21_+_8433172_8446622_RNA45SN1_45S_50ntFlanks@8380.8402.23\|CAAGAGGGCGTGAAACCGTTAAG | >hg38_21_+_8433172_8446622_RNA45SN1_45S_50ntFlanks | 8380 | 8402 | 23 | CAAGAGGGCGTGAAACCGTTAAG | rRF | FALSE | FALSE | TRUE |
| >hg38_21_+_8433172_8446622_RNA45SN1_45S_50ntFlanks@10687.10711.25\|CGGCGTCCGGTGAGCTCTCGCTGGC | >hg38_21_+_8433172_8446622_RNA45SN1_45S_50ntFlanks | 10687 | 10711 | 25 | CGGCGTCCGGTGAGCTCTCGCTGGC | rRF | FALSE | FALSE | TRUE |
| >hg38_21_+_8433172_8446622_RNA45SN1_45S_50ntFlanks@12638.12659.22\|CTACCATCTGTGGGATTATGAC | >hg38_21_+_8433172_8446622_RNA45SN1_45S_50ntFlanks | 12638 | 12659 | 22 | CTACCATCTGTGGGATTATGAC | rRF | FALSE | FALSE | TRUE |
| >hg38_21_+_8433172_8446622_RNA45SN1_45S_50ntFlanks@11569.11602.34\|CAGCCGACTTAGAACTGGTGCGGACCAGGGGAAT | >hg38_21_+_8433172_8446622_RNA45SN1_45S_50ntFlanks | 11569 | 11602 | 34 | CAGCCGACTTAGAACTGGTGCGGACCAGGGGAAT | rRF | FALSE | FALSE | TRUE |
| >hg38_21_+_8433172_8446622_RNA45SN1_45S_50ntFlanks@12636.12659.24\|AGCTACCATCTGTGGGATTATGAC | >hg38_21_+_8433172_8446622_RNA45SN1_45S_50ntFlanks | 12636 | 12659 | 24 | AGCTACCATCTGTGGGATTATGAC | rRF | FALSE | FALSE | TRUE |
| >hg38_21_+_8433172_8446622_RNA45SN1_45S_50ntFlanks@11561.11601.41\|GCGCCTAGCAGCCGACTTAGAACTGGTGCGGACCAGGGGAA | >hg38_21_+_8433172_8446622_RNA45SN1_45S_50ntFlanks | 11561 | 11601 | 41 | GCGCCTAGCAGCCGACTTAGAACTGGTGCGGACCAGGGGAA | rRF | FALSE | FALSE | TRUE |
| >hg38_21_+_8433172_8446622_RNA45SN1_45S_50ntFlanks@6651.6683.33\|CGACTCTTAGCGGTGGATCACTCGGCTCGTGCG | >hg38_21_+_8433172_8446622_RNA45SN1_45S_50ntFlanks | 6651 | 6683 | 33 | CGACTCTTAGCGGTGGATCACTCGGCTCGTGCG | rRF | FALSE | FALSE | TRUE |
| >hg38_21_+_8433172_8446622_RNA45SN1_45S_50ntFlanks@7984.8008.25\|AGATCAGACGTGGCGACCCGCTGAA | >hg38_21_+_8433172_8446622_RNA45SN1_45S_50ntFlanks | 7984 | 8008 | 25 | AGATCAGACGTGGCGACCCGCTGAA | rRF | FALSE | FALSE | TRUE |
| >hg38_21_+_8433172_8446622_RNA45SN1_45S_50ntFlanks@6648.6679.32\|GTACGACTCTTAGCGGTGGATCACTCGGCTCG | >hg38_21_+_8433172_8446622_RNA45SN1_45S_50ntFlanks | 6648 | 6679 | 32 | GTACGACTCTTAGCGGTGGATCACTCGGCTCG | rRF | FALSE | FALSE | TRUE |
| >hg38_21_+_8433172_8446622_RNA45SN1_45S_50ntFlanks@6733.6770.38\|CATTGATCATCGACACTTCGAACGCACTTGCGGCCCCG | >hg38_21_+_8433172_8446622_RNA45SN1_45S_50ntFlanks | 6733 | 6770 | 38 | CATTGATCATCGACACTTCGAACGCACTTGCGGCCCCG | rRF | FALSE | FALSE | TRUE |
| >hg38_21_+_8433172_8446622_RNA45SN1_45S_50ntFlanks@12934.12956.23\|CTAAACCATTCGTAGACGACCTG | >hg38_21_+_8433172_8446622_RNA45SN1_45S_50ntFlanks | 12934 | 12956 | 23 | CTAAACCATTCGTAGACGACCTG | rRF | FALSE | FALSE | TRUE |
| >hg38_21_+_8433172_8446622_RNA45SN1_45S_50ntFlanks@6645.6679.35\|CTCGTACGACTCTTAGCGGTGGATCACTCGGCTCG | >hg38_21_+_8433172_8446622_RNA45SN1_45S_50ntFlanks | 6645 | 6679 | 35 | CTCGTACGACTCTTAGCGGTGGATCACTCGGCTCG | rRF | FALSE | FALSE | TRUE |
| >hg38_21_+_8433172_8446622_RNA45SN1_45S_50ntFlanks@12626.12659.34\|AATGGGGCGAAGCTACCATCTGTGGGATTATGAC | >hg38_21_+_8433172_8446622_RNA45SN1_45S_50ntFlanks | 12626 | 12659 | 34 | AATGGGGCGAAGCTACCATCTGTGGGATTATGAC | rRF | FALSE | FALSE | TRUE |
| >hg38_21_+_8433172_8446622_RNA45SN1_45S_50ntFlanks@9574.9603.30\|AATCGGTCGTCCGACCTGGGTATAGGGGCG | >hg38_21_+_8433172_8446622_RNA45SN1_45S_50ntFlanks | 9574 | 9603 | 30 | AATCGGTCGTCCGACCTGGGTATAGGGGCG | rRF | FALSE | FALSE | TRUE |
| >hg38_21_+_8433172_8446622_RNA45SN1_45S_50ntFlanks@9562.9596.35\|TCCTGACGTGCAAATCGGTCGTCCGACCTGGGTAT | >hg38_21_+_8433172_8446622_RNA45SN1_45S_50ntFlanks | 9562 | 9596 | 35 | TCCTGACGTGCAAATCGGTCGTCCGACCTGGGTAT | rRF | FALSE | FALSE | TRUE |
| >hg38_21_+_8433172_8446622_RNA45SN1_45S_50ntFlanks@8105.8140.36\|CGCGGCGGGGCGCGGGACATGTGGCGTACGGAAGAC | >hg38_21_+_8433172_8446622_RNA45SN1_45S_50ntFlanks | 8105 | 8140 | 36 | CGCGGCGGGGCGCGGGACATGTGGCGTACGGAAGAC | rRF | FALSE | FALSE | TRUE |
| >hg38_21_+_8433172_8446622_RNA45SN1_45S_50ntFlanks@12632.12654.23\|GCGAAGCTACCATCTGTGGGATT | >hg38_21_+_8433172_8446622_RNA45SN1_45S_50ntFlanks | 12632 | 12654 | 23 | GCGAAGCTACCATCTGTGGGATT | rRF | FALSE | FALSE | TRUE |
| >hg38_21_+_8433172_8446622_RNA45SN1_45S_50ntFlanks@3874.3898.25\|AATACATGCCGACGGGCGCTGACCC | >hg38_21_+_8433172_8446622_RNA45SN1_45S_50ntFlanks | 3874 | 3898 | 25 | AATACATGCCGACGGGCGCTGACCC | rRF | FALSE | FALSE | TRUE |
| >hg38_21_+_8433172_8446622_RNA45SN1_45S_50ntFlanks@10411.10430.20\|CAGTCGGTCCTGAGAGATGG | >hg38_21_+_8433172_8446622_RNA45SN1_45S_50ntFlanks | 10411 | 10430 | 20 | CAGTCGGTCCTGAGAGATGG | rRF | FALSE | FALSE | TRUE |
| >hg38_21_+_8433172_8446622_RNA45SN1_45S_50ntFlanks@9714.9737.24\|CGAATGATTAGAGGTCTTGGGGCC | >hg38_21_+_8433172_8446622_RNA45SN1_45S_50ntFlanks | 9714 | 9737 | 24 | CGAATGATTAGAGGTCTTGGGGCC | rRF | FALSE | FALSE | TRUE |
| >hg38_21_+_8433172_8446622_RNA45SN1_45S_50ntFlanks@12456.12488.33\|CCAAGCGTTGGATTGTTCACCCACTAATAGGGA | >hg38_21_+_8433172_8446622_RNA45SN1_45S_50ntFlanks | 12456 | 12488 | 33 | CCAAGCGTTGGATTGTTCACCCACTAATAGGGA | rRF | FALSE | FALSE | TRUE |
| >hg38_21_+_8433172_8446622_RNA45SN1_45S_50ntFlanks@9738.9759.22\|GAAACGATCTCAACCTATTCTC | >hg38_21_+_8433172_8446622_RNA45SN1_45S_50ntFlanks | 9738 | 9759 | 22 | GAAACGATCTCAACCTATTCTC | rRF | FALSE | FALSE | TRUE |
| >hg38_21_+_8433172_8446622_RNA45SN1_45S_50ntFlanks@12456.12477.22\|CCAAGCGTTGGATTGTTCACCC | >hg38_21_+_8433172_8446622_RNA45SN1_45S_50ntFlanks | 12456 | 12477 | 22 | CCAAGCGTTGGATTGTTCACCC | rRF | FALSE | FALSE | TRUE |
| >hg38_21_+_8433172_8446622_RNA45SN1_45S_50ntFlanks@3985.4026.42\|CGGCTTTGGTGACTCTAGATAACCTCGGGCCGATCGCACGCC | >hg38_21_+_8433172_8446622_RNA45SN1_45S_50ntFlanks | 3985 | 4026 | 42 | CGGCTTTGGTGACTCTAGATAACCTCGGGCCGATCGCACGCC | rRF | FALSE | FALSE | TRUE |
| >hg38_21_+_8433172_8446622_RNA45SN1_45S_50ntFlanks@10687.10706.20\|CGGCGTCCGGTGAGCTCTCG | >hg38_21_+_8433172_8446622_RNA45SN1_45S_50ntFlanks | 10687 | 10706 | 20 | CGGCGTCCGGTGAGCTCTCG | rRF | FALSE | FALSE | TRUE |
| >hg38_21_+_8433172_8446622_RNA45SN1_45S_50ntFlanks@6647.6689.43\|CGTACGACTCTTAGCGGTGGATCACTCGGCTCGTGCGTCGATG | >hg38_21_+_8433172_8446622_RNA45SN1_45S_50ntFlanks | 6647 | 6689 | 43 | CGTACGACTCTTAGCGGTGGATCACTCGGCTCGTGCGTCGATG | rRF | FALSE | FALSE | TRUE |
| >hg38_21_+_8433172_8446622_RNA45SN1_45S_50ntFlanks@10513.10533.21\|CCGGAGTGGCGGAGATGGGCG | >hg38_21_+_8433172_8446622_RNA45SN1_45S_50ntFlanks | 10513 | 10533 | 21 | CCGGAGTGGCGGAGATGGGCG | rRF | FALSE | FALSE | TRUE |
| >hg38_21_+_8433172_8446622_RNA45SN1_45S_50ntFlanks@6656.6681.26\|CTTAGCGGTGGATCACTCGGCTCGTG | >hg38_21_+_8433172_8446622_RNA45SN1_45S_50ntFlanks | 6656 | 6681 | 26 | CTTAGCGGTGGATCACTCGGCTCGTG | rRF | FALSE | FALSE | TRUE |
| >hg38_21_+_8433172_8446622_RNA45SN1_45S_50ntFlanks@10346.10368.23\|TCAAACGAGAACTTTGAAGGCCG | >hg38_21_+_8433172_8446622_RNA45SN1_45S_50ntFlanks | 10346 | 10368 | 23 | TCAAACGAGAACTTTGAAGGCCG | rRF | FALSE | FALSE | TRUE |
| >hg38_21_+_8433172_8446622_RNA45SN1_45S_50ntFlanks@6653.6681.29\|ACTCTTAGCGGTGGATCACTCGGCTCGTG | >hg38_21_+_8433172_8446622_RNA45SN1_45S_50ntFlanks | 6653 | 6681 | 29 | ACTCTTAGCGGTGGATCACTCGGCTCGTG | rRF | FALSE | FALSE | TRUE |
| >hg38_21_+_8433172_8446622_RNA45SN1_45S_50ntFlanks@9739.9759.21\|AAACGATCTCAACCTATTCTC | >hg38_21_+_8433172_8446622_RNA45SN1_45S_50ntFlanks | 9739 | 9759 | 21 | AAACGATCTCAACCTATTCTC | rRF | FALSE | FALSE | TRUE |
| >hg38_21_+_8433172_8446622_RNA45SN1_45S_50ntFlanks@10351.10377.27\|CGAGAACTTTGAAGGCCGAAGTGGAGA | >hg38_21_+_8433172_8446622_RNA45SN1_45S_50ntFlanks | 10351 | 10377 | 27 | CGAGAACTTTGAAGGCCGAAGTGGAGA | rRF | FALSE | FALSE | TRUE |
| >hg38_21_+_8433172_8446622_RNA45SN1_45S_50ntFlanks@9739.9757.19\|AAACGATCTCAACCTATTC | >hg38_21_+_8433172_8446622_RNA45SN1_45S_50ntFlanks | 9739 | 9757 | 19 | AAACGATCTCAACCTATTC | rRF | FALSE | FALSE | TRUE |

### Mitoplast

| | | | | | | | | | |
|---|---|---|---|---|---|---|---|---|---|
| >hg38_21_+_8433172_8446622_RNA45SN1_45S_50ntFlanks@9198.9222.25\|GTTCTCTCGGGGCCACGCGCGCGTC | >hg38_21_+_8433172_8446622_RNA45SN1_45S_50ntFlanks | 9198 | 9222 | 25 | GTTCTCTCGGGGCCACGCGCGCGTC | rRF | FALSE | FALSE | TRUE |
| >hg38_21_+_8433172_8446622_RNA45SN1_45S_50ntFlanks@4759.4782.24\|ATCAGATACCGTCGTAGTTCCGAC | >hg38_21_+_8433172_8446622_RNA45SN1_45S_50ntFlanks | 4759 | 4782 | 24 | ATCAGATACCGTCGTAGTTCCGAC | rRF | FALSE | FALSE | TRUE |
| >hg38_21_+_8433172_8446622_RNA45SN1_45S_50ntFlanks@9567.9595.29\|ACGTGCAAATCGGTCGTCCGACCTGGGTA | >hg38_21_+_8433172_8446622_RNA45SN1_45S_50ntFlanks | 9567 | 9595 | 29 | ACGTGCAAATCGGTCGTCCGACCTGGGTA | rRF | FALSE | FALSE | TRUE |
| >trnaMT_ValTAC_MT_+_1602_1670@55.72.18\|ACTTGACCGCTCTGACCA | >trnaMT_ValTAC_MT_+_1602_1670 | 55 | 72 | 18 | ACTTGACCGCTCTGACCA | tRF | FALSE | FALSE | TRUE |
| >hg38_21_+_8433172_8446622_RNA45SN1_45S_50ntFlanks@12636.12654.19\|AGCTACCATCTGTGGGATT | >hg38_21_+_8433172_8446622_RNA45SN1_45S_50ntFlanks | 12636 | 12654 | 19 | AGCTACCATCTGTGGGATT | rRF | FALSE | FALSE | TRUE |
| >hg38_21_+_8433172_8446622_RNA45SN1_45S_50ntFlanks@12405.12432.28\|CTTTTTGATCCTTCGATGTCGGCTCTTC | >hg38_21_+_8433172_8446622_RNA45SN1_45S_50ntFlanks | 12405 | 12432 | 28 | CTTTTTGATCCTTCGATGTCGGCTCTTC | rRF | FALSE | FALSE | TRUE |
| >hg38_21_+_8433172_8446622_RNA45SN1_45S_50ntFlanks@12001.12032.32\|CTGCGGGCCGCCGGTGAAATACCACTACTCTG | >hg38_21_+_8433172_8446622_RNA45SN1_45S_50ntFlanks | 12001 | 12032 | 32 | CTGCGGGCCGCCGGTGAAATACCACTACTCTG | rRF | FALSE | FALSE | TRUE |
| >hg38_21_+_8433172_8446622_RNA45SN1_45S_50ntFlanks@4722.4759.38\|TCATTAATCAAGAACGAAAGTCGGAGGTTCGAAGACGA | >hg38_21_+_8433172_8446622_RNA45SN1_45S_50ntFlanks | 4722 | 4759 | 38 | TCATTAATCAAGAACGAAAGTCGGAGGTTCGAAGACGA | rRF | FALSE | FALSE | TRUE |
| >hg38_21_+_8433172_8446622_RNA45SN1_45S_50ntFlanks@12615.12653.39\|GCTGAGGAGCCAATGGGGCGAAGCTACCATCTGTGGGAT | >hg38_21_+_8433172_8446622_RNA45SN1_45S_50ntFlanks | 12615 | 12653 | 39 | GCTGAGGAGCCAATGGGGCGAAGCTACCATCTGTGGGAT | rRF | FALSE | FALSE | TRUE |
| >hg38_21_+_8433172_8446622_RNA45SN1_45S_50ntFlanks@11316.11336.21\|CACCCCACGTCTCGTCGCGCG | >hg38_21_+_8433172_8446622_RNA45SN1_45S_50ntFlanks | 11316 | 11336 | 21 | CACCCCACGTCTCGTCGCGCG | rRF | FALSE | FALSE | TRUE |
| >hg38_21_+_8433172_8446622_RNA45SN1_45S_50ntFlanks@12289.12318.30\|CCGTGAAAGCGGGGCCTCACGATCCTTCTG | >hg38_21_+_8433172_8446622_RNA45SN1_45S_50ntFlanks | 12289 | 12318 | 30 | CCGTGAAAGCGGGGCCTCACGATCCTTCTG | rRF | FALSE | FALSE | TRUE |
| >hg38_21_+_8433172_8446622_RNA45SN1_45S_50ntFlanks@4726.4758.33\|TAATCAAGAACGAAAGTCGGAGGTTCGAAGACG | >hg38_21_+_8433172_8446622_RNA45SN1_45S_50ntFlanks | 4726 | 4758 | 33 | TAATCAAGAACGAAAGTCGGAGGTTCGAAGACG | rRF | FALSE | FALSE | TRUE |
| >hg38_21_+_8433172_8446622_RNA45SN1_45S_50ntFlanks@9575.9596.22\|ATCGGTCGTCCGACCTGGGTAT | >hg38_21_+_8433172_8446622_RNA45SN1_45S_50ntFlanks | 9575 | 9596 | 22 | ATCGGTCGTCCGACCTGGGTAT | rRF | FALSE | FALSE | TRUE |
| >hg38_21_+_8433172_8446622_RNA45SN1_45S_50ntFlanks@12936.12956.21\|AAACCATTCGTAGACGACCTG | >hg38_21_+_8433172_8446622_RNA45SN1_45S_50ntFlanks | 12936 | 12956 | 21 | AAACCATTCGTAGACGACCTG | rRF | FALSE | FALSE | TRUE |
| >hg38_21_+_8433172_8446622_RNA45SN1_45S_50ntFlanks@11581.11612.32\|AACTGGTGCGGACCAGGGGAATCCGACTGTTT | >hg38_21_+_8433172_8446622_RNA45SN1_45S_50ntFlanks | 11581 | 11612 | 32 | AACTGGTGCGGACCAGGGGAATCCGACTGTTT | rRF | FALSE | FALSE | TRUE |
| >hg38_21_+_8433172_8446622_RNA45SN1_45S_50ntFlanks@9739.9767.29\|AAACGATCTCAACCTATTCTCAAACTTTA | >hg38_21_+_8433172_8446622_RNA45SN1_45S_50ntFlanks | 9739 | 9767 | 29 | AAACGATCTCAACCTATTCTCAAACTTTA | rRF | FALSE | FALSE | TRUE |
| >hg38_21_+_8433172_8446622_RNA45SN1_45S_50ntFlanks@11580.11611.32\|GAACTGGTGCGGACCAGGGGAATCCGACTGTT | >hg38_21_+_8433172_8446622_RNA45SN1_45S_50ntFlanks | 11580 | 11611 | 32 | GAACTGGTGCGGACCAGGGGAATCCGACTGTT | rRF | FALSE | FALSE | TRUE |
| >hg38_21_+_8433172_8446622_RNA45SN1_45S_50ntFlanks@10350.10388.39\|ACGAGAACTTTGAAGGCCGAAGTGGAGAAGGGTTCCATG | >hg38_21_+_8433172_8446622_RNA45SN1_45S_50ntFlanks | 10350 | 10388 | 39 | ACGAGAACTTTGAAGGCCGAAGTGGAGAAGGGTTCCATG | rRF | FALSE | FALSE | TRUE |
| >hg38_21_+_8433172_8446622_RNA45SN1_45S_50ntFlanks@9352.9370.19\|CCGTGGCGCAATGAAGGTG | >hg38_21_+_8433172_8446622_RNA45SN1_45S_50ntFlanks | 9352 | 9370 | 19 | CCGTGGCGCAATGAAGGTG | rRF | FALSE | FALSE | TRUE |
| >hg38_21_+_8433172_8446622_RNA45SN1_45S_50ntFlanks@12405.12441.37\|CTTTTTGATCCTTCGATGTCGGCTCTTCCTATCATTG | >hg38_21_+_8433172_8446622_RNA45SN1_45S_50ntFlanks | 12405 | 12441 | 37 | CTTTTTGATCCTTCGATGTCGGCTCTTCCTATCATTG | rRF | FALSE | FALSE | TRUE |
| >hg38_21_+_8433172_8446622_RNA45SN1_45S_50ntFlanks@12624.12657.34\|CCAATGGGGCGAAGCTACCATCTGTGGGATTATG | >hg38_21_+_8433172_8446622_RNA45SN1_45S_50ntFlanks | 12624 | 12657 | 34 | CCAATGGGGCGAAGCTACCATCTGTGGGATTATG | rRF | FALSE | FALSE | TRUE |
| >hg38_21_+_8433172_8446622_RNA45SN1_45S_50ntFlanks@4730.4758.29\|CAAGAACGAAAGTCGGAGGTTCGAAGACG | >hg38_21_+_8433172_8446622_RNA45SN1_45S_50ntFlanks | 4730 | 4758 | 29 | CAAGAACGAAAGTCGGAGGTTCGAAGACG | rRF | FALSE | FALSE | TRUE |
| >hg38_21_+_8433172_8446622_RNA45SN1_45S_50ntFlanks@8357.8378.22\|AAAAGAACTTTGAAGAGAGAGT | >hg38_21_+_8433172_8446622_RNA45SN1_45S_50ntFlanks | 8357 | 8378 | 22 | AAAAGAACTTTGAAGAGAGAGT | rRF | FALSE | FALSE | TRUE |
| >hg38_21_+_8433172_8446622_RNA45SN1_45S_50ntFlanks@10808.10840.33\|CAATGTAGGTAAGGGAAGTCGGCAAGCCGGATC | >hg38_21_+_8433172_8446622_RNA45SN1_45S_50ntFlanks | 10808 | 10840 | 33 | CAATGTAGGTAAGGGAAGTCGGCAAGCCGGATC | rRF | FALSE | FALSE | TRUE |
| >trna2_GlyGCC_21_-_18827107_18827177@1.29.29,trna35_GlyGCC_1_+_161413094_161413164@1.29.29,trna37_GlyGCC_1_+_161420467_161420537@1.29.29,trna39_GlyGCC_1_+_161427898_161427968@1.29.29,trna41_GlyGCC_1_+_161435258_161435328@1.29.29\|GCATGGGTGGTTCAGTGGTAGAATTCTCG | >trna2_GlyGCC_21_-_18827107_18827177 | 1 | 29 | 29 | GCATGGGTGGTTCAGTGGTAGAATTCTCG | tRF | FALSE | FALSE | TRUE |
| >hg38_21_+_8433172_8446622_RNA45SN1_45S_50ntFlanks@9381.9409.29\|CGCTCGCCGGCCGAGGTGGGATCCCGAGG | >hg38_21_+_8433172_8446622_RNA45SN1_45S_50ntFlanks | 9381 | 9409 | 29 | CGCTCGCCGGCCGAGGTGGGATCCCGAGG | rRF | FALSE | FALSE | TRUE |
| >hg38_21_+_8433172_8446622_RNA45SN1_45S_50ntFlanks@6740.6770.31\|CATCGACACTTCGAACGCACTTGCGGCCCCG | >hg38_21_+_8433172_8446622_RNA45SN1_45S_50ntFlanks | 6740 | 6770 | 31 | CATCGACACTTCGAACGCACTTGCGGCCCCG | rRF | FALSE | FALSE | TRUE |
| >hg38_21_+_8433172_8446622_RNA45SN1_45S_50ntFlanks@10684.10709.26\|CGGCGGCGTCCGGTGAGCTCTCGCTG | >hg38_21_+_8433172_8446622_RNA45SN1_45S_50ntFlanks | 10684 | 10709 | 26 | CGGCGGCGTCCGGTGAGCTCTCGCTG | rRF | FALSE | FALSE | TRUE |
| >hg38_21_+_8433172_8446622_RNA45SN1_45S_50ntFlanks@10350.10368.19\|ACGAGAACTTTGAAGGCCG | >hg38_21_+_8433172_8446622_RNA45SN1_45S_50ntFlanks | 10350 | 10368 | 19 | ACGAGAACTTTGAAGGCCG | rRF | FALSE | FALSE | TRUE |
| >hg38_21_+_8433172_8446622_RNA45SN1_45S_50ntFlanks@8380.8405.26\|CAAGAGGGCGTGAAACCGTTAAGAGG | >hg38_21_+_8433172_8446622_RNA45SN1_45S_50ntFlanks | 8380 | 8405 | 26 | CAAGAGGGCGTGAAACCGTTAAGAGG | rRF | FALSE | FALSE | TRUE |
| >hg38_21_+_8433172_8446622_RNA45SN1_45S_50ntFlanks@12616.12654.39\|CTGAGGAGCCAATGGGGCGAAGCTACCATCTGTGGGATT | >hg38_21_+_8433172_8446622_RNA45SN1_45S_50ntFlanks | 12616 | 12654 | 39 | CTGAGGAGCCAATGGGGCGAAGCTACCATCTGTGGGATT | rRF | FALSE | FALSE | TRUE |
| >hg38_21_+_8433172_8446622_RNA45SN1_45S_50ntFlanks@9166.9197.32\|CCGGGCGGGTCGCGCCGTCGGGCCCGGGGGAG | >hg38_21_+_8433172_8446622_RNA45SN1_45S_50ntFlanks | 9166 | 9197 | 32 | CCGGGCGGGTCGCGCCGTCGGGCCCGGGGGAG | rRF | FALSE | FALSE | TRUE |
| >trna2_GlyGCC_21_-_18827107_18827177@1.28.28,trna35_GlyGCC_1_+_161413094_161413164@1.28.28,trna37_GlyGCC_1_+_161420467_161420537@1.28.28,trna39_GlyGCC_1_+_161427898_161427968@1.28.28,trna41_GlyGCC_1_+_161435258_161435328@1.28.28\|GCATGGGTGGTTCAGTGGTAGAATTCTC | >trna2_GlyGCC_21_-_18827107_18827177 | 1 | 28 | 28 | GCATGGGTGGTTCAGTGGTAGAATTCTC | tRF | FALSE | FALSE | TRUE |
| >hg38_21_+_8433172_8446622_RNA45SN1_45S_50ntFlanks@5084.5110.27\|CGAACGAGACTCTGGCATGCTAACTAG | >hg38_21_+_8433172_8446622_RNA45SN1_45S_50ntFlanks | 5084 | 5110 | 27 | CGAACGAGACTCTGGCATGCTAACTAG | rRF | FALSE | FALSE | TRUE |
| >hg38_21_+_8433172_8446622_RNA45SN1_45S_50ntFlanks@9736.9765.30\|CCGAAACGATCTCAACCTATTCTCAAACTT | >hg38_21_+_8433172_8446622_RNA45SN1_45S_50ntFlanks | 9736 | 9765 | 30 | CCGAAACGATCTCAACCTATTCTCAAACTT | rRF | FALSE | FALSE | TRUE |
| >hg38_21_+_8433172_8446622_RNA45SN1_45S_50ntFlanks@10687.10709.23\|CGGCGTCCGGTGAGCTCTCGCTG | >hg38_21_+_8433172_8446622_RNA45SN1_45S_50ntFlanks | 10687 | 10709 | 23 | CGGCGTCCGGTGAGCTCTCGCTG | rRF | FALSE | FALSE | TRUE |
| >hg38_21_+_8433172_8446622_RNA45SN1_45S_50ntFlanks@9568.9595.28\|CGTGCAAATCGGTCGTCCGACCTGGGTA | >hg38_21_+_8433172_8446622_RNA45SN1_45S_50ntFlanks | 9568 | 9595 | 28 | CGTGCAAATCGGTCGTCCGACCTGGGTA | rRF | FALSE | FALSE | TRUE |
| >hg38_21_+_8433172_8446622_RNA45SN1_45S_50ntFlanks@11575.11603.29\|ACTTAGAACTGGTGCGGACCAGGGGAATC | >hg38_21_+_8433172_8446622_RNA45SN1_45S_50ntFlanks | 11575 | 11603 | 29 | ACTTAGAACTGGTGCGGACCAGGGGAATC | rRF | FALSE | FALSE | TRUE |

### Mitoplast

| | | | | | | | | | |
|---|---|---|---|---|---|---|---|---|---|
| >am_trna116_GluCTC_1_-_145399233_145399304@1.28.28,trna59_GluCTC_1_+_249168447_249168518@1.28.28,trna71_GluCTC_1_-_161439189_161439260@1.28.28,trna74_GluCTC_1_-_161431809_161431880@1.28.28,trna77_GluCTC_1_-_161424398_161424469@1.28.28,trna77_GluCTC_6_+_28949976_28950047@1.28.28,trna80_GluCTC_1_-_161417018_161417089@1.28.28,trna87_GluCTC_6_-_126101393_126101464@1.28.28\|TCCTGGTGGTCTAGTGGTTAGGATTCG | >am_trna116_GluCTC_1_-_145399233_145399304 | 1 | 28 | 28 | TCCTGGTGGTCTAGTGGTTAGGATTCG | tRF | FALSE | FALSE | TRUE |
| >hg38_21_+_8433172_8446622_RNA45SN1_45S_50ntFlanks@10349.10383.35\|AACGAGAACTTTGAAGGCCGAAGTGGAGAAGGGTT | >hg38_21_+_8433172_8446622_RNA45SN1_45S_50ntFlanks | 10349 | 10383 | 35 | AACGAGAACTTTGAAGGCCGAAGTGGAGAAGGGTT | rRF | FALSE | FALSE | TRUE |
| >hg38_21_+_8433172_8446622_RNA45SN1_45S_50ntFlanks@12415.12437.23\|CTTCGATGTCGGCTCTTCCTATC | >hg38_21_+_8433172_8446622_RNA45SN1_45S_50ntFlanks | 12415 | 12437 | 23 | CTTCGATGTCGGCTCTTCCTATC | rRF | FALSE | FALSE | TRUE |
| >hg38_21_+_8433172_8446622_RNA45SN1_45S_50ntFlanks@11319.11344.26\|CCCACGTCTCGTCGCGCGCGCGTCCG | >hg38_21_+_8433172_8446622_RNA45SN1_45S_50ntFlanks | 11319 | 11344 | 26 | CCCACGTCTCGTCGCGCGCGCGTCCG | rRF | FALSE | FALSE | TRUE |
| >hg38_21_+_8433172_8446622_RNA45SN1_45S_50ntFlanks@10351.10384.34\|CGAGAACTTTGAAGGCCGAAGTGGAGAAGGGTTC | >hg38_21_+_8433172_8446622_RNA45SN1_45S_50ntFlanks | 10351 | 10384 | 34 | CGAGAACTTTGAAGGCCGAAGTGGAGAAGGGTTC | rRF | FALSE | FALSE | TRUE |
| >hg38_21_+_8433172_8446622_RNA45SN1_45S_50ntFlanks@6732.6771.40\|ACATTGATCATCGACACTTCGAACGCACTTGCGGCCCCGG | >hg38_21_+_8433172_8446622_RNA45SN1_45S_50ntFlanks | 6732 | 6771 | 40 | ACATTGATCATCGACACTTCGAACGCACTTGCGGCCCCGG | rRF | FALSE | FALSE | TRUE |
| >hg38_21_+_8433172_8446622_RNA45SN1_45S_50ntFlanks@11613.11640.28\|AATTAAAACAAAGCATCGCGAAGGCCCG | >hg38_21_+_8433172_8446622_RNA45SN1_45S_50ntFlanks | 11613 | 11640 | 28 | AATTAAAACAAAGCATCGCGAAGGCCCG | rRF | FALSE | FALSE | TRUE |
| >hg38_21_+_8433172_8446622_RNA45SN1_45S_50ntFlanks@3882.3899.18\|CCGACGGGCGCTGACCCC | >hg38_21_+_8433172_8446622_RNA45SN1_45S_50ntFlanks | 3882 | 3899 | 18 | CCGACGGGCGCTGACCCC | rRF | FALSE | FALSE | TRUE |
| >hg38_21_+_8433172_8446622_RNA45SN1_45S_50ntFlanks@6741.6771.31\|ATCGACACTTCGAACGCACTTGCGGCCCCGG | >hg38_21_+_8433172_8446622_RNA45SN1_45S_50ntFlanks | 6741 | 6771 | 31 | ATCGACACTTCGAACGCACTTGCGGCCCCGG | rRF | FALSE | FALSE | TRUE |
| >hg38_1_-_228634819_228635039_RNA5S12_5S_50ntFlanks@150.171.22\|AATACCGGGTGCTGTAGGCTTT | >hg38_1_-_228634819_228635039_RNA5S12_5S_50ntFlanks | 150 | 171 | 22 | AATACCGGGTGCTGTAGGCTTT | rRF | FALSE | FALSE | TRUE |
| >hg38_21_+_8433172_8446622_RNA45SN1_45S_50ntFlanks@3782.3801.20\|CAGTGAAACTGCGAATGGCT | >hg38_21_+_8433172_8446622_RNA45SN1_45S_50ntFlanks | 3782 | 3801 | 20 | CAGTGAAACTGCGAATGGCT | rRF | FALSE | FALSE | TRUE |
| >hg38_21_+_8433172_8446622_RNA45SN1_45S_50ntFlanks@12627.12657.31\|ATGGGGCGAAGCTACCATCTGTGGGATTATG | >hg38_21_+_8433172_8446622_RNA45SN1_45S_50ntFlanks | 12627 | 12657 | 31 | ATGGGGCGAAGCTACCATCTGTGGGATTATG | rRF | FALSE | FALSE | TRUE |
| >hg38_21_+_8433172_8446622_RNA45SN1_45S_50ntFlanks@12163.12180.18\|ACTGGGGCGGTACACCTG | >hg38_21_+_8433172_8446622_RNA45SN1_45S_50ntFlanks | 12163 | 12180 | 18 | ACTGGGGCGGTACACCTG | rRF | FALSE | FALSE | TRUE |
| >hg38_21_+_8433172_8446622_RNA45SN1_45S_50ntFlanks@6697.6718.22\|CAGCTAGCTGCGAGAATTAATG | >hg38_21_+_8433172_8446622_RNA45SN1_45S_50ntFlanks | 6697 | 6718 | 22 | CAGCTAGCTGCGAGAATTAATG | rRF | FALSE | FALSE | TRUE |
| >hg38_21_+_8433172_8446622_RNA45SN1_45S_50ntFlanks@6738.6772.35\|ATCATCGACACTTCGAACGCACTTGCGGCCCCGGG | >hg38_21_+_8433172_8446622_RNA45SN1_45S_50ntFlanks | 6738 | 6772 | 35 | ATCATCGACACTTCGAACGCACTTGCGGCCCCGGG | rRF | FALSE | FALSE | TRUE |
| >hg38_21_+_8433172_8446622_RNA45SN1_45S_50ntFlanks@4723.4759.37\|CATTAATCAAGAACGAAAGTCGGAGGTTCGAAGACGA | >hg38_21_+_8433172_8446622_RNA45SN1_45S_50ntFlanks | 4723 | 4759 | 37 | CATTAATCAAGAACGAAAGTCGGAGGTTCGAAGACGA | rRF | FALSE | FALSE | TRUE |
| >hg38_21_+_8433172_8446622_RNA45SN1_45S_50ntFlanks@12133.12150.18\|CTCCGGGGACAGTGCCAG | >hg38_21_+_8433172_8446622_RNA45SN1_45S_50ntFlanks | 12133 | 12150 | 18 | CTCCGGGGACAGTGCCAG | rRF | FALSE | FALSE | TRUE |
| >hg38_21_+_8433172_8446622_RNA45SN1_45S_50ntFlanks@10786.10826.41\|AACAGCCTCTGGCATGTTGGAACAATGTAGGTAAGGGAAGT | >hg38_21_+_8433172_8446622_RNA45SN1_45S_50ntFlanks | 10786 | 10826 | 41 | AACAGCCTCTGGCATGTTGGAACAATGTAGGTAAGGGAAGT | rRF | FALSE | FALSE | TRUE |
| >hg38_21_+_8433172_8446622_RNA45SN1_45S_50ntFlanks@9574.9595.22\|AATCGGTCGTCCGACCTGGGTA | >hg38_21_+_8433172_8446622_RNA45SN1_45S_50ntFlanks | 9574 | 9595 | 22 | AATCGGTCGTCCGACCTGGGTA | rRF | FALSE | FALSE | TRUE |
| >hg38_21_+_8433172_8446622_RNA45SN1_45S_50ntFlanks@13002.13028.27\|CGATCTATTGAAAGTCAGCCCTCGACA | >hg38_21_+_8433172_8446622_RNA45SN1_45S_50ntFlanks | 13002 | 13028 | 27 | CGATCTATTGAAAGTCAGCCCTCGACA | rRF | FALSE | FALSE | TRUE |
| >hg38_21_+_8433172_8446622_RNA45SN1_45S_50ntFlanks@12625.12653.29\|CAATGGGGCGAAGCTACCATCTGTGGGAT | >hg38_21_+_8433172_8446622_RNA45SN1_45S_50ntFlanks | 12625 | 12653 | 29 | CAATGGGGCGAAGCTACCATCTGTGGGAT | rRF | FALSE | FALSE | TRUE |
| >hg38_21_+_8433172_8446622_RNA45SN1_45S_50ntFlanks@11581.11603.23\|AACTGGTGCGGACCAGGGGAATC | >hg38_21_+_8433172_8446622_RNA45SN1_45S_50ntFlanks | 11581 | 11603 | 23 | AACTGGTGCGGACCAGGGGAATC | rRF | FALSE | FALSE | TRUE |
| >hg38_21_+_8433172_8446622_RNA45SN1_45S_50ntFlanks@3984.4002.19\|GCGGCTTTGGTGACTCTAG | >hg38_21_+_8433172_8446622_RNA45SN1_45S_50ntFlanks | 3984 | 4002 | 19 | GCGGCTTTGGTGACTCTAG | rRF | FALSE | FALSE | TRUE |
| >hg38_21_+_8433172_8446622_RNA45SN1_45S_50ntFlanks@7979.8008.30\|ACCTCAGATCAGACGTGGCGACCCGCTGAA | >hg38_21_+_8433172_8446622_RNA45SN1_45S_50ntFlanks | 7979 | 8008 | 30 | ACCTCAGATCAGACGTGGCGACCCGCTGAA | rRF | FALSE | FALSE | TRUE |
| >hg38_21_+_8433172_8446622_RNA45SN1_45S_50ntFlanks@10686.10710.25\|GCGGCGTCCGGTGAGCTCTCGCTGG | >hg38_21_+_8433172_8446622_RNA45SN1_45S_50ntFlanks | 10686 | 10710 | 25 | GCGGCGTCCGGTGAGCTCTCGCTGG | rRF | FALSE | FALSE | TRUE |
| >hg38_21_+_8433172_8446622_RNA45SN1_45S_50ntFlanks@12641.12661.21\|CCATCTGTGGGATTATGACTG | >hg38_21_+_8433172_8446622_RNA45SN1_45S_50ntFlanks | 12641 | 12661 | 21 | CCATCTGTGGGATTATGACTG | rRF | FALSE | FALSE | TRUE |
| >hg38_21_+_8433172_8446622_RNA45SN1_45S_50ntFlanks@6719.6744.26\|TGAATTGCAGGACACATTGATCATCG | >hg38_21_+_8433172_8446622_RNA45SN1_45S_50ntFlanks | 6719 | 6744 | 26 | TGAATTGCAGGACACATTGATCATCG | rRF | FALSE | FALSE | TRUE |
| >hg38_21_+_8433172_8446622_RNA45SN1_45S_50ntFlanks@10351.10374.24\|CGAGAACTTTGAAGGCCGAAGTGG | >hg38_21_+_8433172_8446622_RNA45SN1_45S_50ntFlanks | 10351 | 10374 | 24 | CGAGAACTTTGAAGGCCGAAGTGG | rRF | FALSE | FALSE | TRUE |
| >hg38_21_+_8433172_8446622_RNA45SN1_45S_50ntFlanks@7977.8006.30\|CGACCTCAGATCAGACGTGGCGACCCGCTG | >hg38_21_+_8433172_8446622_RNA45SN1_45S_50ntFlanks | 7977 | 8006 | 30 | CGACCTCAGATCAGACGTGGCGACCCGCTG | rRF | FALSE | FALSE | TRUE |
| >hg38_21_+_8433172_8446622_RNA45SN1_45S_50ntFlanks@12420.12437.18\|ATGTCGGCTCTTCCTATC | >hg38_21_+_8433172_8446622_RNA45SN1_45S_50ntFlanks | 12420 | 12437 | 18 | ATGTCGGCTCTTCCTATC | rRF | FALSE | FALSE | TRUE |
| >hg38_21_+_8433172_8446622_RNA45SN1_45S_50ntFlanks@12415.12441.27\|CTTCGATGTCGGCTCTTCCTATCATTG | >hg38_21_+_8433172_8446622_RNA45SN1_45S_50ntFlanks | 12415 | 12441 | 27 | CTTCGATGTCGGCTCTTCCTATCATTG | rRF | FALSE | FALSE | TRUE |
| >hg38_21_+_8433172_8446622_RNA45SN1_45S_50ntFlanks@12289.12323.35\|CCGTGAAAGCGGGGCCTCACGATCCTTCTGACCTT | >hg38_21_+_8433172_8446622_RNA45SN1_45S_50ntFlanks | 12289 | 12323 | 35 | CCGTGAAAGCGGGGCCTCACGATCCTTCTGACCTT | rRF | FALSE | FALSE | TRUE |
| >hg38_21_+_8433172_8446622_RNA45SN1_45S_50ntFlanks@9180.9197.18\|CCGTCGGGCCCGGGGGAG | >hg38_21_+_8433172_8446622_RNA45SN1_45S_50ntFlanks | 9180 | 9197 | 18 | CCGTCGGGCCCGGGGGAG | rRF | FALSE | FALSE | TRUE |
| >hg38_21_+_8433172_8446622_RNA45SN1_45S_50ntFlanks@12405.12431.27\|CTTTTTGATCCTTCGATGTCGGCTCTT | >hg38_21_+_8433172_8446622_RNA45SN1_45S_50ntFlanks | 12405 | 12431 | 27 | CTTTTTGATCCTTCGATGTCGGCTCTT | rRF | FALSE | FALSE | TRUE |
| >hg38_21_+_8433172_8446622_RNA45SN1_45S_50ntFlanks@12619.12654.36\|AGGAGCCAATGGGGCGAAGCTACCATCTGTGGGATT | >hg38_21_+_8433172_8446622_RNA45SN1_45S_50ntFlanks | 12619 | 12654 | 36 | AGGAGCCAATGGGGCGAAGCTACCATCTGTGGGATT | rRF | FALSE | FALSE | TRUE |
| >hg38_21_+_8433172_8446622_RNA45SN1_45S_50ntFlanks@6650.6699.50\|ACGACTCTTAGCGGTGGATCACTCGGCTCGTGCGTCGATGAAGAACGCAG | >hg38_21_+_8433172_8446622_RNA45SN1_45S_50ntFlanks | 6650 | 6699 | 50 | ACGACTCTTAGCGGTGGATCACTCGGCTCGTGCGTCGATGAAGAACGCAG | rRF | FALSE | FALSE | TRUE |
| >hg38_21_+_8433172_8446622_RNA45SN1_45S_50ntFlanks@8105.8138.34\|CGCGGCGGGGCGCGGGACATGTGGCGTACGGAAG | >hg38_21_+_8433172_8446622_RNA45SN1_45S_50ntFlanks | 8105 | 8138 | 34 | CGCGGCGGGGCGCGGGACATGTGGCGTACGGAAG | rRF | FALSE | FALSE | TRUE |
| >hg38_21_+_8433172_8446622_RNA45SN1_45S_50ntFlanks@11317.11342.26\|ACCCCACGTCTCGTCGCGCGCGCGTC | >hg38_21_+_8433172_8446622_RNA45SN1_45S_50ntFlanks | 11317 | 11342 | 26 | ACCCCACGTCTCGTCGCGCGCGCGTC | rRF | FALSE | FALSE | TRUE |
| >hg38_21_+_8433172_8446622_RNA45SN1_45S_50ntFlanks@12973.12990.18\|TCGTACGTAGCAGAGCAG | >hg38_21_+_8433172_8446622_RNA45SN1_45S_50ntFlanks | 12973 | 12990 | 18 | TCGTACGTAGCAGAGCAG | rRF | FALSE | FALSE | TRUE |

### Mitoplast

| | | | | | | | | | |
|---|---|---|---|---|---|---|---|---|---|
| >hg38_21_+_8433172_8446622_RNA45SN1_45S_50ntFlanks@12415.12435.21\|CTTCGATGTCGGCTCTTCCTA | >hg38_21_+_8433172_8446622_RNA45SN1_45S_50ntFlanks | 12415 | 12435 | 21 | CTTCGATGTCGGCTCTTCCTA | rRF | FALSE | FALSE | TRUE |
| >hg38_21_+_8433172_8446622_RNA45SN1_45S_50ntFlanks@9570.9595.26\|TGCAAATCGGTCGTCCGACCTGGGTA | >hg38_21_+_8433172_8446622_RNA45SN1_45S_50ntFlanks | 9570 | 9595 | 26 | TGCAAATCGGTCGTCCGACCTGGGTA | rRF | FALSE | FALSE | TRUE |
| >hg38_21_+_8433172_8446622_RNA45SN1_45S_50ntFlanks@12457.12476.20\|CAAGCGTTGGATTGTTCACC | >hg38_21_+_8433172_8446622_RNA45SN1_45S_50ntFlanks | 12457 | 12476 | 20 | CAAGCGTTGGATTGTTCACC | rRF | FALSE | FALSE | TRUE |
| >hg38_21_+_8433172_8446622_RNA45SN1_45S_50ntFlanks@6731.6771.41\|CACATTGATCATCGACACTTCGAACGCACTTGCGGCCCCGG | >hg38_21_+_8433172_8446622_RNA45SN1_45S_50ntFlanks | 6731 | 6771 | 41 | CACATTGATCATCGACACTTCGAACGCACTTGCGGCCCCGG | rRF | FALSE | FALSE | TRUE |
| >hg38_21_+_8433172_8446622_RNA45SN1_45S_50ntFlanks@6772.6807.36\|GTTCCTCCCGGGGCTACGCCTGTCTGAGCGTCGCTT | >hg38_21_+_8433172_8446622_RNA45SN1_45S_50ntFlanks | 6772 | 6807 | 36 | GTTCCTCCCGGGGCTACGCCTGTCTGAGCGTCGCTT | rRF | FALSE | FALSE | TRUE |
| >hg38_21_+_8433172_8446622_RNA45SN1_45S_50ntFlanks@6645.6670.26\|CTCGTACGACTCTTAGCGGTGGATCA | >hg38_21_+_8433172_8446622_RNA45SN1_45S_50ntFlanks | 6645 | 6670 | 26 | CTCGTACGACTCTTAGCGGTGGATCA | rRF | FALSE | FALSE | TRUE |
| >hg38_21_+_8433172_8446622_RNA45SN1_45S_50ntFlanks@6773.6806.34\|TTCCTCCCGGGGCTACGCCTGTCTGAGCGTCGCT | >hg38_21_+_8433172_8446622_RNA45SN1_45S_50ntFlanks | 6773 | 6806 | 34 | TTCCTCCCGGGGCTACGCCTGTCTGAGCGTCGCT | rRF | FALSE | FALSE | TRUE |
| >hg38_21_+_8433172_8446622_RNA45SN1_45S_50ntFlanks@4003.4024.22\|ATAACCTCGGGCCGATCGCACG | >hg38_21_+_8433172_8446622_RNA45SN1_45S_50ntFlanks | 4003 | 4024 | 22 | ATAACCTCGGGCCGATCGCACG | rRF | FALSE | FALSE | TRUE |
| >hg38_21_+_8433172_8446622_RNA45SN1_45S_50ntFlanks@11874.11906.33\|AGAAGACCCTGTTGAGCTTGACTCTAGTCTGGC | >hg38_21_+_8433172_8446622_RNA45SN1_45S_50ntFlanks | 11874 | 11906 | 33 | AGAAGACCCTGTTGAGCTTGACTCTAGTCTGGC | rRF | FALSE | FALSE | TRUE |
| >hg38_21_+_8433172_8446622_RNA45SN1_45S_50ntFlanks@11561.11585.25\|GCGCCTAGCAGCCGACTTAGAACTG | >hg38_21_+_8433172_8446622_RNA45SN1_45S_50ntFlanks | 11561 | 11585 | 25 | GCGCCTAGCAGCCGACTTAGAACTG | rRF | FALSE | FALSE | TRUE |
| >hg38_21_+_8433172_8446622_RNA45SN1_45S_50ntFlanks@8176.8203.28\|CTTCTGATCGAGGCCCAGCCCGTGGACG | >hg38_21_+_8433172_8446622_RNA45SN1_45S_50ntFlanks | 8176 | 8203 | 28 | CTTCTGATCGAGGCCCAGCCCGTGGACG | rRF | FALSE | FALSE | TRUE |
| >hg38_21_+_8433172_8446622_RNA45SN1_45S_50ntFlanks@4760.4780.21\|TCAGATACCGTCGTAGTTCCG | >hg38_21_+_8433172_8446622_RNA45SN1_45S_50ntFlanks | 4760 | 4780 | 21 | TCAGATACCGTCGTAGTTCCG | rRF | FALSE | FALSE | TRUE |
| >hg38_21_+_8433172_8446622_RNA45SN1_45S_50ntFlanks@9562.9597.36\|TCCTGACGTGCAAATCGGTCGTCCGACCTGGGTATA | >hg38_21_+_8433172_8446622_RNA45SN1_45S_50ntFlanks | 9562 | 9597 | 36 | TCCTGACGTGCAAATCGGTCGTCCGACCTGGGTATA | rRF | FALSE | FALSE | TRUE |
| >hg38_21_+_8433172_8446622_RNA45SN1_45S_50ntFlanks@10402.10429.28\|AACATGGGTCAGTCGGTCCTGAGAGATG | >hg38_21_+_8433172_8446622_RNA45SN1_45S_50ntFlanks | 10402 | 10429 | 28 | AACATGGGTCAGTCGGTCCTGAGAGATG | rRF | FALSE | FALSE | TRUE |
| >hg38_21_+_8433172_8446622_RNA45SN1_45S_50ntFlanks@9930.9959.30\|ATATAGACAGCAGGACGGTGGCCATGGAAG | >hg38_21_+_8433172_8446622_RNA45SN1_45S_50ntFlanks | 9930 | 9959 | 30 | ATATAGACAGCAGGACGGTGGCCATGGAAG | rRF | FALSE | FALSE | TRUE |
| >hg38_21_+_8433172_8446622_RNA45SN1_45S_50ntFlanks@11660.11685.26\|GTGATTTCTGCCCAGTGCTCTGAATG | >hg38_21_+_8433172_8446622_RNA45SN1_45S_50ntFlanks | 11660 | 11685 | 26 | GTGATTTCTGCCCAGTGCTCTGAATG | rRF | FALSE | FALSE | TRUE |
| >hg38_21_+_8433172_8446622_RNA45SN1_45S_50ntFlanks@12417.12455.39\|TCGATGTCGGCTCTTCCTATCATTGTGAAGCAGAATTCA | >hg38_21_+_8433172_8446622_RNA45SN1_45S_50ntFlanks | 12417 | 12455 | 39 | TCGATGTCGGCTCTTCCTATCATTGTGAAGCAGAATTCA | rRF | FALSE | FALSE | TRUE |
| >hg38_21_+_8433172_8446622_RNA45SN1_45S_50ntFlanks@11999.12043.45\|CCCTGCGGGCCGCCGGTGAAATACCACTACTCTGATCGTTTTTTC | >hg38_21_+_8433172_8446622_RNA45SN1_45S_50ntFlanks | 11999 | 12043 | 45 | CCCTGCGGGCCGCCGGTGAAATACCACTACTCTGATCGTTTTTTC | rRF | FALSE | FALSE | TRUE |
| >hg38_21_+_8433172_8446622_RNA45SN1_45S_50ntFlanks@3835.3860.26\|CTCCTCTCCTACTTGGATAACTGTGG | >hg38_21_+_8433172_8446622_RNA45SN1_45S_50ntFlanks | 3835 | 3860 | 26 | CTCCTCTCCTACTTGGATAACTGTGG | rRF | FALSE | FALSE | TRUE |
| >hg38_21_+_8433172_8446622_RNA45SN1_45S_50ntFlanks@3873.3898.26\|TAATACATGCCGACGGGCGCTGACCC | >hg38_21_+_8433172_8446622_RNA45SN1_45S_50ntFlanks | 3873 | 3898 | 26 | TAATACATGCCGACGGGCGCTGACCC | rRF | FALSE | FALSE | TRUE |
| >hg38_21_+_8433172_8446622_RNA45SN1_45S_50ntFlanks@12163.12181.19\|ACTGGGGCGGTACACCTGT | >hg38_21_+_8433172_8446622_RNA45SN1_45S_50ntFlanks | 12163 | 12181 | 19 | ACTGGGGCGGTACACCTGT | rRF | FALSE | FALSE | TRUE |
| >hg38_21_+_8433172_8446622_RNA45SN1_45S_50ntFlanks@6731.6770.40\|CACATTGATCATCGACACTTCGAACGCACTTGCGGCCCCG | >hg38_21_+_8433172_8446622_RNA45SN1_45S_50ntFlanks | 6731 | 6770 | 40 | CACATTGATCATCGACACTTCGAACGCACTTGCGGCCCCG | rRF | FALSE | FALSE | TRUE |
| >hg38_1_-_228634819_228635039_RNA5S12_5S_50ntFlanks@148.170.23\|GGAATACCGGGTGCTGTAGGCTT | >hg38_1_-_228634819_228635039_RNA5S12_5S_50ntFlanks | 148 | 170 | 23 | GGAATACCGGGTGCTGTAGGCTT | rRF | FALSE | FALSE | TRUE |
| >hg38_21_+_8433172_8446622_RNA45SN1_45S_50ntFlanks@12159.12179.21\|TTTGACTGGGGCGGTACACCT | >hg38_21_+_8433172_8446622_RNA45SN1_45S_50ntFlanks | 12159 | 12179 | 21 | TTTGACTGGGGCGGTACACCT | rRF | FALSE | FALSE | TRUE |
| >hg38_21_+_8433172_8446622_RNA45SN1_45S_50ntFlanks@6661.6681.21\|CGGTGGATCACTCGGCTCGTG | >hg38_21_+_8433172_8446622_RNA45SN1_45S_50ntFlanks | 6661 | 6681 | 21 | CGGTGGATCACTCGGCTCGTG | rRF | FALSE | FALSE | TRUE |
| >hg38_21_+_8433172_8446622_RNA45SN1_45S_50ntFlanks@11561.11602.42\|GCGCCTAGCAGCCGACTTAGAACTGGTGCGGACCAGGGGAAT | >hg38_21_+_8433172_8446622_RNA45SN1_45S_50ntFlanks | 11561 | 11602 | 42 | GCGCCTAGCAGCCGACTTAGAACTGGTGCGGACCAGGGGAAT | rRF | FALSE | FALSE | TRUE |
| >hg38_21_+_8433172_8446622_RNA45SN1_45S_50ntFlanks@6655.6681.27\|TCTTAGCGGTGGATCACTCGGCTCGTG | >hg38_21_+_8433172_8446622_RNA45SN1_45S_50ntFlanks | 6655 | 6681 | 27 | TCTTAGCGGTGGATCACTCGGCTCGTG | rRF | FALSE | FALSE | TRUE |
| >hg38_21_+_8433172_8446622_RNA45SN1_45S_50ntFlanks@12635.12653.19\|AAGCTACCATCTGTGGGAT | >hg38_21_+_8433172_8446622_RNA45SN1_45S_50ntFlanks | 12635 | 12653 | 19 | AAGCTACCATCTGTGGGAT | rRF | FALSE | FALSE | TRUE |
| >hg38_21_+_8433172_8446622_RNA45SN1_45S_50ntFlanks@7989.8007.19\|AGACGTGGCGACCCGCTGA | >hg38_21_+_8433172_8446622_RNA45SN1_45S_50ntFlanks | 7989 | 8007 | 19 | AGACGTGGCGACCCGCTGA | rRF | FALSE | FALSE | TRUE |
| >hg38_21_+_8433172_8446622_RNA45SN1_45S_50ntFlanks@9566.9597.32\|GACGTGCAAATCGGTCGTCCGACCTGGGTATA | >hg38_21_+_8433172_8446622_RNA45SN1_45S_50ntFlanks | 9566 | 9597 | 32 | GACGTGCAAATCGGTCGTCCGACCTGGGTATA | rRF | FALSE | FALSE | TRUE |
| >hg38_21_+_8433172_8446622_RNA45SN1_45S_50ntFlanks@6772.6799.28\|GTTCCTCCCGGGGCTACGCCTGTCTGAG | >hg38_21_+_8433172_8446622_RNA45SN1_45S_50ntFlanks | 6772 | 6799 | 28 | GTTCCTCCCGGGGCTACGCCTGTCTGAG | rRF | FALSE | FALSE | TRUE |
| >hg38_1_-_228634819_228635039_RNA5S12_5S_50ntFlanks@150.170.21\|AATACCGGGTGCTGTAGGCTT | >hg38_1_-_228634819_228635039_RNA5S12_5S_50ntFlanks | 150 | 170 | 21 | AATACCGGGTGCTGTAGGCTT | rRF | FALSE | FALSE | TRUE |
| >hg38_21_+_8433172_8446622_RNA45SN1_45S_50ntFlanks@11570.11603.34\|AGCCGACTTAGAACTGGTGCGGACCAGGGGAATC | >hg38_21_+_8433172_8446622_RNA45SN1_45S_50ntFlanks | 11570 | 11603 | 34 | AGCCGACTTAGAACTGGTGCGGACCAGGGGAATC | rRF | FALSE | FALSE | TRUE |
| >hg38_21_+_8433172_8446622_RNA45SN1_45S_50ntFlanks@8157.8181.25\|CTCGTGGGGGGCCCAAGTCCTTCTG | >hg38_21_+_8433172_8446622_RNA45SN1_45S_50ntFlanks | 8157 | 8181 | 25 | CTCGTGGGGGGCCCAAGTCCTTCTG | rRF | FALSE | FALSE | TRUE |
| >hg38_21_+_8433172_8446622_RNA45SN1_45S_50ntFlanks@8104.8140.37\|CCGCGGCGGGGCGCGGGACATGTGGCGTACGGAAGAC | >hg38_21_+_8433172_8446622_RNA45SN1_45S_50ntFlanks | 8104 | 8140 | 37 | CCGCGGCGGGGCGCGGGACATGTGGCGTACGGAAGAC | rRF | FALSE | FALSE | TRUE |
| >hg38_21_+_8433172_8446622_RNA45SN1_45S_50ntFlanks@11235.11254.20\|CTTCCCGTGGATCGCCCCAG | >hg38_21_+_8433172_8446622_RNA45SN1_45S_50ntFlanks | 11235 | 11254 | 20 | CTTCCCGTGGATCGCCCCAG | rRF | FALSE | FALSE | TRUE |
| >multi-am_trnaMT_ValTAC_MT_+_1602_1670@50.68.19\|ACTTAACTTGACCGCTCTG | >multi-am_trnaMT_ValTAC_MT_+_1602_1670 | 50 | 68 | 19 | ACTTAACTTGACCGCTCTG | tRF | FALSE | FALSE | TRUE |
| >hg38_21_+_8433172_8446622_RNA45SN1_45S_50ntFlanks@6649.6681.33\|TACGACTCTTAGCGGTGGATCACTCGGCTCGTG | >hg38_21_+_8433172_8446622_RNA45SN1_45S_50ntFlanks | 6649 | 6681 | 33 | TACGACTCTTAGCGGTGGATCACTCGGCTCGTG | rRF | FALSE | FALSE | TRUE |
| >hg38_21_+_8433172_8446622_RNA45SN1_45S_50ntFlanks@6743.6771.29\|CGACACTTCGAACGCACTTGCGGCCCCGG | >hg38_21_+_8433172_8446622_RNA45SN1_45S_50ntFlanks | 6743 | 6771 | 29 | CGACACTTCGAACGCACTTGCGGCCCCGG | rRF | FALSE | FALSE | TRUE |
| >hg38_21_+_8433172_8446622_RNA45SN1_45S_50ntFlanks@6776.6803.28\|CTCCCGGGGCTACGCCTGTCTGAGCGTC | >hg38_21_+_8433172_8446622_RNA45SN1_45S_50ntFlanks | 6776 | 6803 | 28 | CTCCCGGGGCTACGCCTGTCTGAGCGTC | rRF | FALSE | FALSE | TRUE |
| >hg38_21_+_8433172_8446622_RNA45SN1_45S_50ntFlanks@7983.8004.22\|CAGATCAGACGTGGCGACCCGC | >hg38_21_+_8433172_8446622_RNA45SN1_45S_50ntFlanks | 7983 | 8004 | 22 | CAGATCAGACGTGGCGACCCGC | rRF | FALSE | FALSE | TRUE |
| >hg38_21_+_8433172_8446622_RNA45SN1_45S_50ntFlanks@11579.11603.25\|AGAACTGGTGCGGACCAGGGGAATC | >hg38_21_+_8433172_8446622_RNA45SN1_45S_50ntFlanks | 11579 | 11603 | 25 | AGAACTGGTGCGGACCAGGGGAATC | rRF | FALSE | FALSE | TRUE |

### Mitoplast

<table>
<tr><th></th><th></th><th></th><th></th><th></th><th></th><th></th><th></th><th></th><th></th></tr>
<tr><td>>hg38_21_+_8433172_8446622_RNA45SN1_45S_50ntFlanks@10788.10818.31|CAGCCTCTGGCATGTTGGAACAATGTAGGTA</td><td>>hg38_21_+_8433172_8446622_RNA45SN1_45S_50ntFlanks</td><td>10788</td><td>10818</td><td>31</td><td>CAGCCTCTGGCATGTTGGAACAATGTAGGTA</td><td>rRF</td><td>FALSE</td><td>FALSE</td><td>TRUE</td></tr>
<tr><td>>am_trna111_HisGTG_1_-_147774845_147774916@-1G.29.30,trna118_HisGTG_1_-_145396881_145396952@-1G.29.30,trna16_HisGTG_1_+_146544773_146544844@-1G.29.30,trna1_HisGTG_15_+_45493349_45493420@-1G.29.30,trna21_HisGTG_1_+_147753471_147753542@-1G.29.30,trna33_HisGTG_6_+_27125906_27125977@-1G.29.30,trna7_HisGTG_9_-_14433938_14434009@-1G.29.30,trna8_HisGTG_15_-_45492611_45492682@-1G.29.30,trna9_HisGTG_15_-_45490804_45490875@-1G.29.30|GGCCGTGATCGTATAGTGGTTAGTACTCTG</td><td>>am_trna111_HisGTG_1_-_147774845_147774916</td><td>-1G</td><td>29</td><td>30</td><td>GGCCGTGATCGTATAGTGGTTAGTACTCTG</td><td>tRF</td><td>FALSE</td><td>FALSE</td><td>TRUE</td></tr>
<tr><td>>hg38_21_+_8433172_8446622_RNA45SN1_45S_50ntFlanks@11593.11611.19|CCAGGGGAATCCGACTGTT</td><td>>hg38_21_+_8433172_8446622_RNA45SN1_45S_50ntFlanks</td><td>11593</td><td>11611</td><td>19</td><td>CCAGGGGAATCCGACTGTT</td><td>rRF</td><td>FALSE</td><td>FALSE</td><td>TRUE</td></tr>
<tr><td>>hg38_21_+_8433172_8446622_RNA45SN1_45S_50ntFlanks@12893.12930.38|CCTCGCCCGTCACGCACCGCACGTTCGTGGGGAACCTG</td><td>>hg38_21_+_8433172_8446622_RNA45SN1_45S_50ntFlanks</td><td>12893</td><td>12930</td><td>38</td><td>CCTCGCCCGTCACGCACCGCACGTTCGTGGGGAACCTG</td><td>rRF</td><td>FALSE</td><td>FALSE</td><td>TRUE</td></tr>
</table>
