## Supplementary material for "Single-nucleotide Differences and Cell Type Decide the Subcellular Localization of miRNA Isoforms (isomiRs), tRNA-derived Fragments (tRFs) and rRNA-derived Fragments (rRFs)": Supp. Table S7

**Supplemental Table S7:** Data table of the differentially abundant (DA) short RNA for the cell line RNA between the three cell lines in Total RNA and Cytoplasmic RNA with FDR ≤ 0.05. The table is displayed as the output from DESeq2 where MedianFC (median fold change) is presented as raw reads . The table in decreasing order of FDR value. The annotation "am" is used to indicate abiguous short RNA, those that can be mapped to a noncoding RNA gene as well as somewhere else on the geneome. The annotation "multi" referrs to short RNA that map to locations in both the mitochondrial and nuclear genomes. The rRF are flanked by 50 nucleotdies upstream and downstream and their coordiantes reflect this. The coordiantes for the isomiR and rRF are from the hg38 human genome assembly while the coordintes for the tRF are from the hg19 human genome assembly.

| Sample | Combo | tRNA | Start | End | Length | Sequence | MedianFC | log2FC | lfcSE | stat | p-value | adjp-value | Numerator | Denominator |
|---|---|---|---|---|---|---|---|---|---|---|---|---|---|---|
| CYTO | BT20MB231 | chr1.trna119-LysCTT, chr16.trna1 | 1 | 27 | 27 | GCCCGGCTAGCTCAGTCGGTAGAGCAT | 136.9035 | -5.112242 | 1.322235 | -3.866364 | 0.00011 | 0.002676 | BT20 | MB231 |
| CYTO | BT20MB468 | chr1.trna119-LysCTT, chr16.trna1 | 1 | 27 | 27 | GCCCGGCTAGCTCAGTCGGTAGAGCAT | 221.7359 | -1.084059 | 1.004005 | -1.079735 | 0.28026 | 0.357457 | BT20 | MB468 |
| CYTO | MB231MB468 | chr1.trna119-LysCTT, chr16.trna1 | 1 | 27 | 27 | GCCCGGCTAGCTCAGTCGGTAGAGCAT | 97.47746 | 4.284318 | 0.989424 | 4.330115 | 1.49E-05 | 5.57E-05 | MB231 | MB468 |
| CYTO | BT20MB231 | chr1.trna119-LysCTT, chr16.trna1 | 1 | 28 | 28 | GCCCGGCTAGCTCAGTCGGTAGAGCATG | 345.1893 | 1.013837 | 1.039171 | 0.975621 | 0.329253 | 0.549992 | BT20 | MB231 |
| CYTO | BT20MB468 | chr1.trna119-LysCTT, chr16.trna1 | 1 | 28 | 28 | GCCCGGCTAGCTCAGTCGGTAGAGCATG | 6159.313 | 5.273845 | 0.887106 | 5.944998 | 2.76E-09 | 2.44E-08 | BT20 | MB468 |
| CYTO | MB231MB468 | chr1.trna119-LysCTT, chr16.trna1 | 1 | 28 | 28 | GCCCGGCTAGCTCAGTCGGTAGAGCATG | 6904.262 | 4.309082 | 0.912746 | 4.72101 | 2.35E-06 | 1.07E-05 | MB231 | MB468 |
| TOTAL | BT20MB231 | chr1.trna119-LysCTT, chr16.trna1 | 1 | 27 | 27 | GCCCGGCTAGCTCAGTCGGTAGAGCAT | 60.13546 | -2.238438 | 1.267317 | -1.766282 | 0.077349 | 0.293819 | BT20 | MB231 |
| TOTAL | BT20MB468 | chr1.trna119-LysCTT, chr16.trna1 | 1 | 27 | 27 | GCCCGGCTAGCTCAGTCGGTAGAGCAT | 143.2213 | -0.36165 | 1.078488 | -0.335331 | 0.737376 | 0.782309 | BT20 | MB468 |
| TOTAL | MB231MB468 | chr1.trna119-LysCTT, chr16.trna1 | 1 | 27 | 27 | GCCCGGCTAGCTCAGTCGGTAGAGCAT | 86.11266 | 1.353679 | 0.878771 | 1.540423 | 0.123457 | 0.158581 | MB231 | MB468 |
| TOTAL | BT20MB231 | chr1.trna119-LysCTT, chr16.trna1 | 1 | 28 | 28 | GCCCGGCTAGCTCAGTCGGTAGAGCATG | 155.2123 | 1.574368 | 0.984684 | 1.598856 | 0.109853 | 0.347699 | BT20 | MB231 |
| TOTAL | BT20MB468 | chr1.trna119-LysCTT, chr16.trna1 | 1 | 28 | 28 | GCCCGGCTAGCTCAGTCGGTAGAGCATG | 4133.494 | 5.722558 | 0.758645 | 7.543129 | 4.59E-14 | 6.09E-13 | BT20 | MB468 |
| TOTAL | MB231MB468 | chr1.trna119-LysCTT, chr16.trna1 | 1 | 28 | 28 | GCCCGGCTAGCTCAGTCGGTAGAGCATG | 3984.215 | 3.438232 | 0.825019 | 4.16746 | 3.08E-05 | 6.82E-05 | MB231 | MB468 |
| CYTO | BT20MB231 | chr1.trna119-LysCTT | 1 | 29 | 29 | GCCCGGCTAGCTCAGTCGGTAGAGCATGA | 167.7047 | 0.625458 | 1.161692 | 0.538403 | 0.590299 | 0.754816 | BT20 | MB231 |
| CYTO | BT20MB468 | chr1.trna119-LysCTT | 1 | 29 | 29 | GCCCGGCTAGCTCAGTCGGTAGAGCATGA | 2897.604 | 4.963657 | 0.978098 | 5.074805 | 3.88E-07 | 2.01E-06 | BT20 | MB468 |
| CYTO | MB231MB468 | chr1.trna119-LysCTT | 1 | 29 | 29 | GCCCGGCTAGCTCAGTCGGTAGAGCATGA | 3210.681 | 4.550426 | 0.870523 | 5.227232 | 1.72E-07 | 1.06E-06 | MB231 | MB468 |
| CYTO | BT20MB231 | chr1.trna119-LysCTT | 1 | 30 | 30 | GCCCGGCTAGCTCAGTCGGTAGAGCATGAG | 384.4622 | 0.410307 | 1.048506 | 0.391325 | 0.695557 | 0.827698 | BT20 | MB231 |
| CYTO | BT20MB468 | chr1.trna119-LysCTT | 1 | 30 | 30 | GCCCGGCTAGCTCAGTCGGTAGAGCATGAG | 39682.98 | 7.461396 | 0.970195 | 7.690618 | 1.46E-14 | 4.01E-13 | BT20 | MB468 |
| CYTO | MB231MB468 | chr1.trna119-LysCTT | 1 | 30 | 30 | GCCCGGCTAGCTCAGTCGGTAGAGCATGAG | 43710.31 | 7.130227 | 0.854432 | 8.344987 | 7.12E-17 | 7.77E-15 | MB231 | MB468 |
| CYTO | BT20MB231 | chr1.trna119-LysCTT | 1 | 31 | 31 | GCCCGGCTAGCTCAGTCGGTAGAGCATGAGA | 644.5306 | 0.89302 | 1.027222 | 0.869355 | 0.384653 | 0.595194 | BT20 | MB231 |
| CYTO | BT20MB468 | chr1.trna119-LysCTT | 1 | 31 | 31 | GCCCGGCTAGCTCAGTCGGTAGAGCATGAGA | 13747.07 | 5.428219 | 0.747338 | 7.263409 | 3.77E-13 | 7.69E-12 | BT20 | MB468 |
| CYTO | MB231MB468 | chr1.trna119-LysCTT | 1 | 31 | 31 | GCCCGGCTAGCTCAGTCGGTAGAGCATGAGA | 15279.51 | 4.540918 | 0.926 | 4.903799 | 9.40E-07 | 4.78E-06 | MB231 | MB468 |
| CYTO | BT20MB231 | chr1.trna119-LysCTT | 1 | 32 | 32 | GCCCGGCTAGCTCAGTCGGTAGAGCATGAGAC | 4628.617 | 2.149881 | 1.079989 | 1.990651 | 0.046519 | 0.186356 | BT20 | MB231 |
| CYTO | BT20MB468 | chr1.trna119-LysCTT | 1 | 32 | 32 | GCCCGGCTAGCTCAGTCGGTAGAGCATGAGAC | 2176.763 | 0.123623 | 0.74796 | 0.16528 | 0.868724 | 0.897914 | BT20 | MB468 |
| CYTO | MB231MB468 | chr1.trna119-LysCTT | 1 | 32 | 32 | GCCCGGCTAGCTCAGTCGGTAGAGCATGAGAC | 5751.361 | -2.037386 | 1.015927 | -2.005447 | 0.044915 | 0.072457 | MB231 | MB468 |
| CYTO | BT20MB231 | chr1.trna119-LysCTT | 1 | 33 | 33 | GCCCGGCTAGCTCAGTCGGTAGAGCATGAGACT | 34507.85 | 3.03805 | 1.029608 | 2.950686 | 0.003171 | 0.034405 | BT20 | MB231 |
| CYTO | BT20MB468 | chr1.trna119-LysCTT | 1 | 33 | 33 | GCCCGGCTAGCTCAGTCGGTAGAGCATGAGACT | 4069.59 | -4.513511 | 0.681818 | -6.619821 | 3.60E-11 | 4.86E-10 | BT20 | MB468 |
| CYTO | MB231MB468 | chr1.trna119-LysCTT | 1 | 33 | 33 | GCCCGGCTAGCTCAGTCGGTAGAGCATGAGACT | 35466.71 | -7.590695 | 1.040295 | -7.296673 | 2.95E-13 | 9.95E-12 | MB231 | MB468 |
| CYTO | BT20MB231 | chr1.trna119-LysCTT | 1 | 34 | 34 | GCCCGGCTAGCTCAGTCGGTAGAGCATGAGACTC | 13739.86 | 2.144061 | 1.107131 | 1.936592 | 0.052795 | 0.198828 | BT20 | MB231 |
| CYTO | BT20MB468 | chr1.trna119-LysCTT | 1 | 34 | 34 | GCCCGGCTAGCTCAGTCGGTAGAGCATGAGACTC | 2710.439 | -6.220041 | 0.787865 | -7.894804 | 2.91E-15 | 9.17E-14 | BT20 | MB468 |
| CYTO | MB231MB468 | chr1.trna119-LysCTT | 1 | 34 | 34 | GCCCGGCTAGCTCAGTCGGTAGAGCATGAGACTC | 13125.75 | -8.401187 | 1.138958 | -7.376205 | 1.63E-13 | 6.00E-12 | MB231 | MB468 |
| CYTO | BT20MB231 | chr1.trna119-LysCTT | 1 | 35 | 35 | GCCCGGCTAGCTCAGTCGGTAGAGCATGAGACTCT | 32676.48 | -2.720171 | 1.015168 | -2.679529 | 0.007373 | 0.059653 | BT20 | MB231 |
| CYTO | BT20MB468 | chr1.trna119-LysCTT | 1 | 35 | 35 | GCCCGGCTAGCTCAGTCGGTAGAGCATGAGACTCT | 30622.57 | -9.704852 | 0.868248 | -11.17751 | 5.25E-29 | 1.49E-26 | BT20 | MB468 |
| CYTO | MB231MB468 | chr1.trna119-LysCTT | 1 | 35 | 35 | GCCCGGCTAGCTCAGTCGGTAGAGCATGAGACTCT | 4672.388 | -6.855459 | 0.888419 | -7.716472 | 1.20E-14 | 6.18E-13 | MB231 | MB468 |
| CYTO | BT20MB231 | chr1.trna119-LysCTT | 1 | 36 | 36 | GCCCGGCTAGCTCAGTCGGTAGAGCATGAGACTCTT | 4722.106 | -2.682882 | 1.039908 | -2.579923 | 0.009882 | 0.071359 | BT20 | MB231 |
| CYTO | BT20MB468 | chr1.trna119-LysCTT | 1 | 36 | 36 | GCCCGGCTAGCTCAGTCGGTAGAGCATGAGACTCTT | 4262.027 | -7.519811 | 0.903566 | -8.322372 | 8.62E-17 | 3.75E-15 | BT20 | MB468 |
| CYTO | MB231MB468 | chr1.trna119-LysCTT | 1 | 36 | 36 | GCCCGGCTAGCTCAGTCGGTAGAGCATGAGACTCTT | 696.2377 | -4.729327 | 0.91869 | -5.147903 | 2.63E-07 | 1.56E-06 | MB231 | MB468 |
| CYTO | BT20MB231 | chr1.trna119-LysCTT | 1 | 37 | 37 | GCCCGGCTAGCTCAGTCGGTAGAGCATGAGACTCTTA | 77.10944 | -1.213926 | 1.429442 | -0.849231 | 0.395753 | 0.605917 | BT20 | MB231 |
| CYTO | BT20MB468 | chr1.trna119-LysCTT | 1 | 37 | 37 | GCCCGGCTAGCTCAGTCGGTAGAGCATGAGACTCTTA | 59.6051 | -6.100727 | 1.334778 | -4.570593 | 4.86E-06 | 1.94E-05 | BT20 | MB468 |
| CYTO | MB231MB468 | chr1.trna119-LysCTT | 1 | 37 | 37 | GCCCGGCTAGCTCAGTCGGTAGAGCATGAGACTCTTA | 24.74232 | -4.685547 | 1.739986 | -2.692864 | 0.007084 | 0.014031 | MB231 | MB468 |
| TOTAL | BT20MB231 | chr1.trna119-LysCTT | 1 | 29 | 29 | GCCCGGCTAGCTCAGTCGGTAGAGCATGA | 52.85889 | 1.866908 | 1.043865 | 1.788457 | 0.073702 | 0.287476 | BT20 | MB231 |
| TOTAL | BT20MB468 | chr1.trna119-LysCTT | 1 | 29 | 29 | GCCCGGCTAGCTCAGTCGGTAGAGCATGA | 2233.401 | 6.67234 | 0.668645 | 9.978895 | 1.89E-23 | 9.96E-22 | BT20 | MB468 |
| TOTAL | MB231MB468 | chr1.trna119-LysCTT | 1 | 29 | 29 | GCCCGGCTAGCTCAGTCGGTAGAGCATGA | 2101.573 | 4.235997 | 0.767842 | 5.516753 | 3.45E-08 | 1.19E-07 | MB231 | MB468 |
| TOTAL | BT20MB231 | chr1.trna119-LysCTT | 1 | 30 | 30 | GCCCGGCTAGCTCAGTCGGTAGAGCATGAG | 119.3296 | 2.034558 | 0.940239 | 2.163872 | 0.030474 | 0.182557 | BT20 | MB231 |
| TOTAL | BT20MB468 | chr1.trna119-LysCTT | 1 | 30 | 30 | GCCCGGCTAGCTCAGTCGGTAGAGCATGAG | 16978.19 | 8.560963 | 0.699187 | 12.24417 | 1.81E-34 | 2.66E-32 | BT20 | MB468 |
| TOTAL | MB231MB468 | chr1.trna119-LysCTT | 1 | 30 | 30 | GCCCGGCTAGCTCAGTCGGTAGAGCATGAG | 15852.55 | 5.828101 | 0.800096 | 7.284256 | 3.23E-13 | 3.11E-12 | MB231 | MB468 |
| TOTAL | BT20MB231 | chr1.trna119-LysCTT | 1 | 31 | 31 | GCCCGGCTAGCTCAGTCGGTAGAGCATGAGA | 436.1558 | 2.929107 | 0.966868 | 3.02948 | 0.00245 | 0.03815 | BT20 | MB231 |
| TOTAL | BT20MB468 | chr1.trna119-LysCTT | 1 | 31 | 31 | GCCCGGCTAGCTCAGTCGGTAGAGCATGAGA | 6092.82 | 5.886272 | 0.679173 | 8.66682 | 4.44E-18 | 1.16E-16 | BT20 | MB468 |
| TOTAL | MB231MB468 | chr1.trna119-LysCTT | 1 | 31 | 31 | GCCCGGCTAGCTCAGTCGGTAGAGCATGAGA | 6366.868 | 2.289027 | 0.859387 | 2.663559 | 0.007732 | 0.012407 | MB231 | MB468 |
| TOTAL | BT20MB231 | chr1.trna119-LysCTT | 1 | 32 | 32 | GCCCGGCTAGCTCAGTCGGTAGAGCATGAGAC | 2310.238 | 2.613408 | 0.969563 | 2.69545 | 0.007029 | 0.076874 | BT20 | MB231 |
| TOTAL | BT20MB468 | chr1.trna119-LysCTT | 1 | 32 | 32 | GCCCGGCTAGCTCAGTCGGTAGAGCATGAGAC | 955.067 | -0.438391 | 0.743105 | -0.589945 | 0.555227 | 0.616512 | BT20 | MB468 |
| TOTAL | MB231MB468 | chr1.trna119-LysCTT | 1 | 32 | 32 | GCCCGGCTAGCTCAGTCGGTAGAGCATGAGAC | 5101.984 | -3.844276 | 0.887086 | -4.333602 | 1.47E-05 | 3.39E-05 | MB231 | MB468 |
| TOTAL | BT20MB231 | chr1.trna119-LysCTT | 1 | 33 | 33 | GCCCGGCTAGCTCAGTCGGTAGAGCATGAGACT | 16796.04 | 3.648406 | 1.004325 | 3.632695 | 0.00028 | 0.007789 | BT20 | MB231 |
| TOTAL | BT20MB468 | chr1.trna119-LysCTT | 1 | 33 | 33 | GCCCGGCTAGCTCAGTCGGTAGAGCATGAGACT | 1802.272 | -4.312021 | 0.422579 | -10.20406 | 1.90E-24 | 1.12E-22 | BT20 | MB468 |
| TOTAL | MB231MB468 | chr1.trna119-LysCTT | 1 | 33 | 33 | GCCCGGCTAGCTCAGTCGGTAGAGCATGAGACT | 38959.38 | -8.969138 | 1.048718 | -8.552482 | 1.20E-17 | 2.81E-16 | MB231 | MB468 |
| TOTAL | BT20MB231 | chr1.trna119-LysCTT | 1 | 34 | 34 | GCCCGGCTAGCTCAGTCGGTAGAGCATGAGACTC | 6899.163 | 1.636718 | 1.035607 | 1.580443 | 0.114005 | 0.354522 | BT20 | MB231 |
| TOTAL | BT20MB468 | chr1.trna119-LysCTT | 1 | 34 | 34 | GCCCGGCTAGCTCAGTCGGTAGAGCATGAGACTC | 2664.719 | -6.983866 | 0.81833 | -8.534293 | 1.41E-17 | 3.41E-16 | BT20 | MB468 |
| TOTAL | MB231MB468 | chr1.trna119-LysCTT | 1 | 34 | 34 | GCCCGGCTAGCTCAGTCGGTAGAGCATGAGACTC | 12867.57 | -9.402845 | 1.077859 | -8.723634 | NA | NA | MB231 | MB468 |
| TOTAL | BT20MB231 | chr1.trna119-LysCTT | 1 | 35 | 35 | GCCCGGCTAGCTCAGTCGGTAGAGCATGAGACTCT | 18057.71 | -2.874459 | 0.920297 | -3.123402 | 0.001788 | 0.03062 | BT20 | MB231 |
| TOTAL | BT20MB468 | chr1.trna119-LysCTT | 1 | 35 | 35 | GCCCGGCTAGCTCAGTCGGTAGAGCATGAGACTCT | 26731.15 | -10.42058 | 0.852451 | -12.22426 | 2.31E-34 | 3.35E-32 | BT20 | MB468 |
| TOTAL | MB231MB468 | chr1.trna119-LysCTT | 1 | 35 | 35 | GCCCGGCTAGCTCAGTCGGTAGAGCATGAGACTCT | 4929.351 | -8.111503 | 0.795454 | -10.19733 | 2.04E-24 | 1.33E-22 | MB231 | MB468 |

| | | | | | | | | | | | | | | |
|---|---|---|---|---|---|---|---|---|---|---|---|---|---|---|
| TOTAL | BT20MB231 | chr1.trna119-LysCTT | 1 | 36 | 36 | GCCCGGCTAGCTCAGTCGGTAGAGCATGAGACTCTT | 2679.114 | -1.288839 | 0.950857 | -1.35545 | 0.175274 | 0.43282 | BT20 | MB231 |
| TOTAL | BT20MB468 | chr1.trna119-LysCTT | 1 | 36 | 36 | GCCCGGCTAGCTCAGTCGGTAGAGCATGAGACTCTT | 3062.754 | -7.928139 | 0.927197 | -8.550655 | 1.22E-17 | 2.98E-16 | BT20 | MB468 |
| TOTAL | MB231MB468 | chr1.trna119-LysCTT | 1 | 36 | 36 | GCCCGGCTAGCTCAGTCGGTAGAGCATGAGACTCTT | 1804.355 | -7.280072 | 0.855955 | -8.505203 | 1.81E-17 | 4.09E-16 | MB231 | MB468 |
| TOTAL | BT20MB231 | chr1.trna119-LysCTT | 1 | 37 | 37 | GCCCGGCTAGCTCAGTCGGTAGAGCATGAGACTCTTA | 26.00293 | -0.859364 | 1.024167 | -0.839086 | 0.401421 | NA | BT20 | MB231 |
| TOTAL | BT20MB468 | chr1.trna119-LysCTT | 1 | 37 | 37 | GCCCGGCTAGCTCAGTCGGTAGAGCATGAGACTCTTA | 24.21943 | -8.902296 | 1.448181 | -6.147224 | 7.89E-10 | 4.41E-09 | BT20 | MB468 |
| TOTAL | MB231MB468 | chr1.trna119-LysCTT | 1 | 37 | 37 | GCCCGGCTAGCTCAGTCGGTAGAGCATGAGACTCTTA | 19.91785 | -8.717078 | 1.482448 | -5.880191 | 4.10E-09 | 1.67E-08 | MB231 | MB468 |
| CYTO | BT20MB231 | chr16.trna10-LysCTT | 1 | 29 | 29 | GCCCGGCTAGCTCAGTCGGTAGAGCATGG | 27.72145 | 0.610129 | 1.644102 | 0.371102 | 0.710562 | NA | BT20 | MB231 |
| CYTO | BT20MB468 | chr16.trna10-LysCTT | 1 | 29 | 29 | GCCCGGCTAGCTCAGTCGGTAGAGCATGG | 1185.405 | 6.268885 | 1.15705 | 5.417989 | 6.03E-08 | 3.86E-07 | BT20 | MB468 |
| CYTO | MB231MB468 | chr16.trna10-LysCTT | 1 | 29 | 29 | GCCCGGCTAGCTCAGTCGGTAGAGCATGG | 1304.65 | 5.857657 | 0.849415 | 6.896111 | 5.34E-12 | 1.22E-10 | MB231 | MB468 |
| CYTO | BT20MB231 | chr16.trna10-LysCTT | 1 | 31 | 31 | GCCCGGCTAGCTCAGTCGGTAGAGCATGGGA | 3.168505 | -2.258015 | 3.826933 | -0.590032 | 0.555169 | NA | BT20 | MB231 |
| CYTO | BT20MB468 | chr16.trna10-LysCTT | 1 | 31 | 31 | GCCCGGCTAGCTCAGTCGGTAGAGCATGGGA | 112.9537 | 4.95412 | 1.328293 | 3.729688 | 0.000192 | 0.000534 | BT20 | MB468 |
| CYTO | MB231MB468 | chr16.trna10-LysCTT | 1 | 31 | 31 | GCCCGGCTAGCTCAGTCGGTAGAGCATGGGA | 121.2383 | 7.423438 | 1.200899 | 6.18157 | 6.35E-10 | 7.77E-09 | MB231 | MB468 |
| CYTO | BT20MB231 | chr16.trna10-LysCTT | 1 | 32 | 32 | GCCCGGCTAGCTCAGTCGGTAGAGCATGGGAC | 45.1024 | 0.74846 | 1.829972 | 0.409001 | 0.682539 | NA | BT20 | MB231 |
| CYTO | BT20MB468 | chr16.trna10-LysCTT | 1 | 32 | 32 | GCCCGGCTAGCTCAGTCGGTAGAGCATGGGAC | 34.06873 | -0.620788 | 1.349133 | -0.460138 | 0.645417 | 0.710052 | BT20 | MB468 |
| CYTO | MB231MB468 | chr16.trna10-LysCTT | 1 | 32 | 32 | GCCCGGCTAGCTCAGTCGGTAGAGCATGGGAC | 49.79864 | -1.315347 | 1.375356 | -0.956368 | 0.338886 | 0.4196 | MB231 | MB468 |
| CYTO | BT20MB231 | chr16.trna10-LysCTT | 1 | 33 | 33 | GCCCGGCTAGCTCAGTCGGTAGAGCATGGGACT | 1447.308 | 0.08225 | 1.144987 | 0.071835 | 0.942733 | 0.970213 | BT20 | MB231 |
| CYTO | BT20MB468 | chr16.trna10-LysCTT | 1 | 33 | 33 | GCCCGGCTAGCTCAGTCGGTAGAGCATGGGACT | 760.0952 | -6.495322 | 0.69734 | -9.31443 | 1.23E-20 | 1.04E-18 | BT20 | MB468 |
| CYTO | MB231MB468 | chr16.trna10-LysCTT | 1 | 33 | 33 | GCCCGGCTAGCTCAGTCGGTAGAGCATGGGACT | 883.0484 | -6.588163 | 1.144291 | -5.757417 | 8.54E-09 | 7.57E-08 | MB231 | MB468 |
| CYTO | BT20MB231 | chr16.trna10-LysCTT | 1 | 34 | 34 | GCCCGGCTAGCTCAGTCGGTAGAGCATGGGACTC | 4282.912 | 0.974044 | 1.203 | 0.809678 | 0.418125 | 0.626509 | BT20 | MB231 |
| CYTO | BT20MB468 | chr16.trna10-LysCTT | 1 | 34 | 34 | GCCCGGCTAGCTCAGTCGGTAGAGCATGGGACTC | 1559.817 | -6.621274 | 0.768498 | -8.615858 | 6.94E-18 | 3.75E-16 | BT20 | MB468 |
| CYTO | MB231MB468 | chr16.trna10-LysCTT | 1 | 34 | 34 | GCCCGGCTAGCTCAGTCGGTAGAGCATGGGACTC | 3391.053 | -7.631591 | 1.195121 | -6.385622 | 1.71E-10 | 2.43E-09 | MB231 | MB468 |
| CYTO | BT20MB231 | chr16.trna10-LysCTT | 1 | 35 | 35 | GCCCGGCTAGCTCAGTCGGTAGAGCATGGGACTCT | 208170.2 | -3.883649 | 1.171377 | -3.315455 | 0.000915 | 0.013962 | BT20 | MB231 |
| CYTO | BT20MB468 | chr16.trna10-LysCTT | 1 | 35 | 35 | GCCCGGCTAGCTCAGTCGGTAGAGCATGGGACTCT | 210782.8 | -11.69715 | 0.94748 | -12.34554 | 5.15E-35 | 3.51E-32 | BT20 | MB468 |
| CYTO | MB231MB468 | chr16.trna10-LysCTT | 1 | 35 | 35 | GCCCGGCTAGCTCAGTCGGTAGAGCATGGGACTCT | 14654.47 | -7.721846 | 1.046669 | -7.377545 | 1.61E-13 | 5.96E-12 | MB231 | MB468 |
| CYTO | BT20MB231 | chr16.trna10-LysCTT | 1 | 36 | 36 | GCCCGGCTAGCTCAGTCGGTAGAGCATGGGACTCTT | 7639.712 | -3.630328 | 0.992586 | -3.657446 | 0.000255 | 0.005177 | BT20 | MB231 |
| CYTO | BT20MB468 | chr16.trna10-LysCTT | 1 | 36 | 36 | GCCCGGCTAGCTCAGTCGGTAGAGCATGGGACTCTT | 7463.221 | -8.04506 | 0.820712 | -9.802536 | 1.10E-22 | 1.32E-20 | BT20 | MB468 |
| CYTO | MB231MB468 | chr16.trna10-LysCTT | 1 | 36 | 36 | GCCCGGCTAGCTCAGTCGGTAGAGCATGGGACTCTT | 636.0885 | -4.283704 | 0.86804 | -4.934916 | 8.02E-07 | 4.15E-06 | MB231 | MB468 |
| CYTO | BT20MB231 | chr16.trna10-LysCTT | 1 | 37 | 37 | GCCCGGCTAGCTCAGTCGGTAGAGCATGGGACTCTTA | 248.5715 | -2.704934 | 1.368026 | -1.977253 | 0.048013 | 0.188971 | BT20 | MB231 |
| CYTO | BT20MB468 | chr16.trna10-LysCTT | 1 | 37 | 37 | GCCCGGCTAGCTCAGTCGGTAGAGCATGGGACTCTTA | 235.8573 | -7.44173 | 1.753384 | -4.24421 | 2.19E-05 | 7.53E-05 | BT20 | MB468 |
| CYTO | MB231MB468 | chr16.trna10-LysCTT | 1 | 37 | 37 | GCCCGGCTAGCTCAGTCGGTAGAGCATGGGACTCTTA | 35.31841 | -4.560544 | 3.507203 | -1.300337 | 0.193486 | 0.259171 | MB231 | MB468 |
| TOTAL | BT20MB231 | chr16.trna10-LysCTT | 1 | 29 | 29 | GCCCGGCTAGCTCAGTCGGTAGAGCATGG | 14.01437 | 0.962359 | 1.437228 | 0.669594 | 0.503117 | NA | BT20 | MB231 |
| TOTAL | BT20MB468 | chr16.trna10-LysCTT | 1 | 29 | 29 | GCCCGGCTAGCTCAGTCGGTAGAGCATGG | 736.0133 | 6.438819 | 0.911622 | 7.063038 | 1.63E-12 | 1.58E-11 | BT20 | MB468 |
| TOTAL | MB231MB468 | chr16.trna10-LysCTT | 1 | 29 | 29 | GCCCGGCTAGCTCAGTCGGTAGAGCATGG | 683.6233 | 4.76339 | 0.73147 | 6.512074 | 7.41E-11 | 4.37E-10 | MB231 | MB468 |
| TOTAL | BT20MB231 | chr16.trna10-LysCTT | 1 | 32 | 32 | GCCCGGCTAGCTCAGTCGGTAGAGCATGGGAC | 27.8758 | 1.630184 | 1.350462 | 1.207131 | 0.227382 | NA | BT20 | MB231 |
| TOTAL | BT20MB468 | chr16.trna10-LysCTT | 1 | 32 | 32 | GCCCGGCTAGCTCAGTCGGTAGAGCATGGGAC | 18.84834 | -1.244904 | 1.800507 | -0.691418 | 0.489303 | 0.553504 | BT20 | MB468 |
| TOTAL | MB231MB468 | chr16.trna10-LysCTT | 1 | 32 | 32 | GCCCGGCTAGCTCAGTCGGTAGAGCATGGGAC | 54.52339 | -3.374919 | 1.345851 | -2.507646 | 0.012154 | 0.018918 | MB231 | MB468 |
| TOTAL | BT20MB231 | chr16.trna10-LysCTT | 1 | 33 | 33 | GCCCGGCTAGCTCAGTCGGTAGAGCATGGGACT | 609.0181 | 0.959744 | 1.061753 | 0.903924 | 0.366036 | 0.616327 | BT20 | MB231 |
| TOTAL | BT20MB468 | chr16.trna10-LysCTT | 1 | 33 | 33 | GCCCGGCTAGCTCAGTCGGTAGAGCATGGGACT | 345.014 | -6.171188 | 1.361572 | -4.532399 | 5.83E-06 | 1.50E-05 | BT20 | MB468 |
| TOTAL | MB231MB468 | chr16.trna10-LysCTT | 1 | 33 | 33 | GCCCGGCTAGCTCAGTCGGTAGAGCATGGGACT | 1014.853 | -7.900805 | 1.516745 | -5.209053 | 1.90E-07 | 5.75E-07 | MB231 | MB468 |
| TOTAL | BT20MB231 | chr16.trna10-LysCTT | 1 | 34 | 34 | GCCCGGCTAGCTCAGTCGGTAGAGCATGGGACTC | 4568.08 | 0.124852 | 1.134547 | 0.110045 | 0.912373 | 0.962345 | BT20 | MB231 |
| TOTAL | BT20MB468 | chr16.trna10-LysCTT | 1 | 34 | 34 | GCCCGGCTAGCTCAGTCGGTAGAGCATGGGACTC | 3850.159 | -8.929541 | 1.468128 | -6.082262 | 1.18E-09 | 6.39E-09 | BT20 | MB468 |
| TOTAL | MB231MB468 | chr16.trna10-LysCTT | 1 | 34 | 34 | GCCCGGCTAGCTCAGTCGGTAGAGCATGGGACTC | 5990.198 | -9.698439 | 1.524362 | -6.362294 | NA | NA | MB231 | MB468 |
| TOTAL | BT20MB231 | chr16.trna10-LysCTT | 1 | 35 | 35 | GCCCGGCTAGCTCAGTCGGTAGAGCATGGGACTCT | 185122.2 | -4.807024 | 1.132094 | -4.246135 | 2.17E-05 | 0.000984 | BT20 | MB231 |
| TOTAL | BT20MB468 | chr16.trna10-LysCTT | 1 | 35 | 35 | GCCCGGCTAGCTCAGTCGGTAGAGCATGGGACTCT | 323019.4 | -13.64385 | 1.191763 | -11.44846 | NA | NA | BT20 | MB468 |
| TOTAL | MB231MB468 | chr16.trna10-LysCTT | 1 | 35 | 35 | GCCCGGCTAGCTCAGTCGGTAGAGCATGGGACTCT | 15406.42 | -9.380204 | 1.007702 | -9.308505 | 1.30E-20 | 4.88E-19 | MB231 | MB468 |
| TOTAL | BT20MB231 | chr16.trna10-LysCTT | 1 | 36 | 36 | GCCCGGCTAGCTCAGTCGGTAGAGCATGGGACTCTT | 7925.733 | -2.758296 | 1.07009 | -2.577631 | 0.009948 | 0.094492 | BT20 | MB231 |
| TOTAL | BT20MB468 | chr16.trna10-LysCTT | 1 | 36 | 36 | GCCCGGCTAGCTCAGTCGGTAGAGCATGGGACTCTT | 12027.22 | -9.92965 | 1.114905 | -8.906272 | 5.28E-19 | 1.58E-17 | BT20 | MB468 |
| TOTAL | MB231MB468 | chr16.trna10-LysCTT | 1 | 36 | 36 | GCCCGGCTAGCTCAGTCGGTAGAGCATGGGACTCTT | 2416.073 | -7.736447 | 0.917294 | -8.433986 | 3.34E-17 | 7.15E-16 | MB231 | MB468 |
| TOTAL | BT20MB231 | chr16.trna10-LysCTT | 1 | 37 | 37 | GCCCGGCTAGCTCAGTCGGTAGAGCATGGGACTCTTA | 60.25761 | -1.896063 | 0.954456 | -1.986538 | 0.046974 | 0.22963 | BT20 | MB231 |
| TOTAL | BT20MB468 | chr16.trna10-LysCTT | 1 | 37 | 37 | GCCCGGCTAGCTCAGTCGGTAGAGCATGGGACTCTTA | 74.79299 | -10.53505 | 1.512609 | -6.964818 | 3.29E-12 | 3.01E-11 | BT20 | MB468 |
| TOTAL | MB231MB468 | chr16.trna10-LysCTT | 1 | 37 | 37 | GCCCGGCTAGCTCAGTCGGTAGAGCATGGGACTCTTA | 28.76255 | -9.257631 | 1.526796 | -6.063437 | 1.33E-09 | 6.01E-09 | MB231 | MB468 |
