## Supplementary material for "Single-nucleotide Differences and Cell Type Decide the Subcellular Localization of miRNA Isoforms (isomiRs), tRNA-derived Fragments (tRFs) and rRNA-derived Fragments (rRFs)": Supp. Table S8 (1 of 5)

### Supp Table S8.1.Columns_B_to_AL

**Supplemental Table S8:** The abundances (RPM) of each of the short RNA (Row 4) in publically available data (Column 1 contains SRA codes). The 150 short RNA are referenced in the paper and represent the top 17 most abundant short RNA from Figure 3, 20 LysCTT tRF from Figure 4, 111 short RNA from Figure 5, and 11 ETS1 rRF from Figure 6. Some RNA overlap across datasets from the figures. Conditional color formating ranges from 0 to 5000 RPM.

| | Top 17 short RNA (Figure 3) | | | | | | | | | | | | | | | | | 20 LysCTT tRF (Figure 4) | | | | | | | | | | | | | | | | | | | |
|---|---|---|---|---|---|---|---|---|---|---|---|---|---|---|---|---|---|---|---|---|---|---|---|---|---|---|---|---|---|---|---|---|---|---|---|---|---|
| | >hg38_21_+_8433172_8446622_RNA45SN1_45S_50ntFlanks@7975.8012.38\|CGCGACCTCAGATCAGACGTGGCGACCCGCTGAATTTA | >hg38_21_+_8433172_8446622_RNA45SN1_45S_50ntFlanks@7975.8008.34\|CGCGACCTCAGATCAGACGTGGCGACCCGCTGAA | >hg38_21_+_8433172_8446622_RNA45SN1_45S_50ntFlanks@7975.7994.20\|CGCGACCTCAGATCAGACGT | >hg38_1_-_228634819_228635039_RNA5S12_5S_50ntFlanks@132.170.39\|GATGGGAGACCGCCTGGGAATACCGGGTGCTGTAGGCTT | >MI0000077\|hsa-mir-21&WithFlank&17\|+\|59841260\|59841343@14.36.23(+1A)&[MIMAT0000076&hsa-miR-21-5p&offsets\|0\|+1(+1A);m-1&17\|+\|59841273\|59841294&offsets\|0\|+1(+1A)]\|TAGCTTATCAGACTGATGTTGACA | >MI0000077\|hsa-mir-21&WithFlank&17\|+\|59841260\|59841343@14.37.24&[MIMAT0000076&hsa-miR-21-5p&offsets\|0\|+2;m-1&17\|+\|59841273\|59841294&offsets\|0\|+2]\|TAGCTTATCAGACTGATGTTGACT | >MI0000077\|hsa-mir-21&WithFlank&17\|+\|59841260\|59841343@14.36.23&[MIMAT0000076&hsa-miR-21-5p&offsets\|0\|+1;m-1&17\|+\|59841273\|59841294&offsets\|0\|+1]\|TAGCTTATCAGACTGATGTTGAC | >MI0000077\|hsa-mir-21&WithFlank&17\|+\|59841260\|59841343@14.35.22&[MIMAT0000076&hsa-miR-21-5p&offsets\|0\|0;m-1&17\|+\|59841273\|59841294&offsets\|0\|0]\|TAGCTTATCAGACTGATGTTGA | >MI0000253\|hsa-mir-148a&WithFlank&7\|-\|25949913\|25949992@50.71.22&[MIMAT0000243&hsa-miR-148a-3p&offsets\|0\|0;m-7&7\|-\|25949922\|25949943&offsets\|0\|0]\|TCAGTGCACTACAGAACTTTGT | >hg38_21_+_8433172_8446622_RNA45SN1_45S_50ntFlanks@6646.6669.24\|TCGTACGACTCTTAGCGGTGGATC | >MI0000067\|hsa-let-7f-1&WithFlank&9\|+\|94176341\|94176439@13.34.22;MI0000068\|hsa-let-7f-2&WithFlank&X\|-\|53557186\|53557280@14.35.22&[MIMAT0000067&hsa-let-7f-5p&offsets\|0\|0;m-20&9\|+\|94176353\|94176374&offsets\|0\|0];[MIMAT0000067_1&hsa-let-7f-5p&offsets\|0\|0;m-19&X\|- | >MI0000061\|hsa-let-7a-2&WithFlank&11\|-\|122146516\|122146599@11.32.22;MI0000062\|hsa-let-7a-3&WithFlank&22\|+\|46112743\|46112828@10.31.22;MI0000060\|hsa-let-7a-1&WithFlank&9\|+\|94175951\|94176042@12.33.22&[MIMAT0000062&hsa-let-7a-5p&offsets\|0\|0;m-6&11\|-\|122146568\|122146589 | >MI0000433\|hsa-let-7g&WithFlank&3\|-\|52268272\|52268367@11.32.22&[MIMAT0000414&hsa-let-7g-5p&offsets\|0\|0;m-58&3\|-\|52268336\|52268357&offsets\|0\|0]\|TGAGGTAGTAGTTTGTACAGTT | >MI0000434\|hsa-let-7i&WithFlank&12\|+\|62603680\|62603775@12.33.22&[MIMAT0000415&hsa-let-7i-5p&offsets\|0\|0;m-49&12\|+\|62603691\|62603712&offsets\|0\|0]\|TGAGGTAGTAGTTTGTGCTGTT | >MI0000264\|hsa-mir-7-2&WithFlank&15\|+\|88611819\|88611940@38.61.24;MI0000265\|hsa-mir-7-3&WithFlank&19\|+\|4770664\|4770785@37.60.24;MI0000263\|hsa-mir-7-1&WithFlank&9\|-\|83969742\|83969863@30.53.24&[MIMAT0000252&hsa-miR-7-5p&offsets\|0\|0;m-400&15\|+\|88611856\|88 | >MI0000088\|hsa-mir-30a&WithFlank&6\|-\|71403545\|71403627@12.35.24&[MIMAT0000087&hsa-miR-30a-5p&offsets\|0\|+2;m-15&6\|-\|71403595\|71403616&offsets\|0\|+2]\|TGTAAACATCCTCGACTGGAAGCT | >MI0000750\|hsa-mir-26a-2&WithFlank&12\|-\|57824603\|57824698@20.41.22;MI0000083\|hsa-mir-26a-1&WithFlank&3\|+\|37969398\|37969486@16.37.22&[MIMAT0000082&hsa-miR-26a-5p&offsets\|0\|0;m-33&12\|-\|57824658\|57824679&offsets\|0\|0];[MIMAT0000082_1&hsa-miR-26a-5p&offsets\| | >trna119_LysCTT_1_-_145395522_145395594@1.37.37,trna11_LysCTT_5_-_180648979_180649051@1.37.37,trna13_LysCTT_6_+_26556774_26556846@1.37.37,trna7_LysCTT_16_+_3225692_3225764@1.37.37,trna9_LysCTT_5_+_180634755_180634827@1.37.37\|GCCCGGCTAGCTCAGTCGGTAGAGCAT | >trna119_LysCTT_1_-_145395522_145395594@1.36.36,trna11_LysCTT_5_-_180648979_180649051@1.36.36,trna13_LysCTT_6_+_26556774_26556846@1.36.36,trna7_LysCTT_16_+_3225692_3225764@1.36.36,trna9_LysCTT_5_+_180634755_180634827@1.36.36\|GCCCGGCTAGCTCAGTCGGTAGAGCAT | >trna119_LysCTT_1_-_145395522_145395594@1.35.35,trna11_LysCTT_5_-_180648979_180649051@1.35.35,trna13_LysCTT_6_+_26556774_26556846@1.35.35,trna7_LysCTT_16_+_3225692_3225764@1.35.35,trna9_LysCTT_5_+_180634755_180634827@1.35.35\|GCCCGGCTAGCTCAGTCGGTAGAGCAT | >trna119_LysCTT_1_-_145395522_145395594@1.34.34,trna11_LysCTT_5_-_180648979_180649051@1.34.34,trna13_LysCTT_6_+_26556774_26556846@1.34.34,trna7_LysCTT_16_+_3225692_3225764@1.34.34,trna9_LysCTT_5_+_180634755_180634827@1.34.34\|GCCCGGCTAGCTCAGTCGGTAGAGCAT | >trna119_LysCTT_1_-_145395522_145395594@1.33.33,trna11_LysCTT_5_-_180648979_180649051@1.33.33,trna13_LysCTT_6_+_26556774_26556846@1.33.33,trna7_LysCTT_16_+_3225692_3225764@1.33.33,trna9_LysCTT_5_+_180634755_180634827@1.33.33\|GCCCGGCTAGCTCAGTCGGTAGAGCAT | >trna119_LysCTT_1_-_145395522_145395594@1.32.32,trna11_LysCTT_5_-_180648979_180649051@1.32.32,trna13_LysCTT_6_+_26556774_26556846@1.32.32,trna32_LysCTT_16_-_3207406_3207478@1.32.32,trna7_LysCTT_16_+_3225692_3225764@1.32.32,trna9_LysCTT_5_+_180634755_180 | >trna119_LysCTT_1_-_145395522_145395594@1.31.31,trna11_LysCTT_5_-_180648979_180649051@1.31.31,trna13_LysCTT_6_+_26556774_26556846@1.31.31,trna32_LysCTT_16_-_3207406_3207478@1.31.31,trna7_LysCTT_16_+_3225692_3225764@1.31.31,trna9_LysCTT_5_+_180634755_180 | >trna119_LysCTT_1_-_145395522_145395594@1.30.30,trna11_LysCTT_5_-_180648979_180649051@1.30.30,trna13_LysCTT_6_+_26556774_26556846@1.30.30,trna32_LysCTT_16_-_3207406_3207478@1.30.30,trna7_LysCTT_16_+_3225692_3225764@1.30.30,trna9_LysCTT_5_+_180634755_180 | >trna119_LysCTT_1_-_145395522_145395594@1.29.29,trna11_LysCTT_5_-_180648979_180649051@1.29.29,trna13_LysCTT_6_+_26556774_26556846@1.29.29,trna32_LysCTT_16_-_3207406_3207478@1.29.29,trna7_LysCTT_16_+_3225692_3225764@1.29.29,trna9_LysCTT_5_+_180634755_180 | >trna10_LysCTT_16_+_3241501_3241573@1.37.37,trna13_LysCTT_14_-_58706613_58706685@1.37.37,trna2_LysCTT_15_+_79152904_79152976@1.37.37\|GCCCGGCTAGCTCAGTCGGTAGAGCATGGGACTCTTA | >trna10_LysCTT_16_+_3241501_3241573@1.36.36,trna13_LysCTT_14_-_58706613_58706685@1.36.36,trna2_LysCTT_15_+_79152904_79152976@1.36.36\|GCCCGGCTAGCTCAGTCGGTAGAGCATGGGACTCTT | >trna10_LysCTT_16_+_3241501_3241573@1.35.35,trna13_LysCTT_14_-_58706613_58706685@1.35.35,trna2_LysCTT_15_+_79152904_79152976@1.35.35\|GCCCGGCTAGCTCAGTCGGTAGAGCATGGGACTCT | >trna10_LysCTT_16_+_3241501_3241573@1.34.34,trna13_LysCTT_14_-_58706613_58706685@1.34.34,trna2_LysCTT_15_+_79152904_79152976@1.34.34\|GCCCGGCTAGCTCAGTCGGTAGAGCATGGGACTC | >trna10_LysCTT_16_+_3241501_3241573@1.33.33,trna13_LysCTT_14_-_58706613_58706685@1.33.33,trna2_LysCTT_15_+_79152904_79152976@1.33.33\|GCCCGGCTAGCTCAGTCGGTAGAGCATGGGACT | >trna10_LysCTT_16_+_3241501_3241573@1.32.32,trna13_LysCTT_14_-_58706613_58706685@1.32.32,trna2_LysCTT_15_+_79152904_79152976@1.32.32\|GCCCGGCTAGCTCAGTCGGTAGAGCATGGGAC | >trna10_LysCTT_16_+_3241501_3241573@1.31.31,trna13_LysCTT_14_-_58706613_58706685@1.31.31,trna2_LysCTT_15_+_79152904_79152976@1.31.31\|GCCCGGCTAGCTCAGTCGGTAGAGCATGGGA | >trna10_LysCTT_16_+_3241501_3241573@1.30.30,trna13_LysCTT_14_-_58706613_58706685@1.30.30,trna2_LysCTT_15_+_79152904_79152976@1.30.30\|GCCCGGCTAGCTCAGTCGGTAGAGCATGGG | >trna10_LysCTT_16_+_3241501_3241573@1.29.29,trna13_LysCTT_14_-_58706613_58706685@1.29.29,trna2_LysCTT_15_+_79152904_79152976@1.29.29\|GCCCGGCTAGCTCAGTCGGTAGAGCATGG | >trna10_LysCTT_16_+_3241501_3241573@1.28.28,trna119_LysCTT_1_-_145395522_145395594@1.28.28,trna11_LysCTT_5_-_180648979_180649051@1.28.28,trna13_LysCTT_14_-_58706613_58706685@1.28.28,trna13_LysCTT_6_+_26556774_26556846@1.28.28,trna2_LysCTT_15_+_79152904_79 | >trna10_LysCTT_16_+_3241501_3241573@1.27.27,trna119_LysCTT_1_-_145395522_145395594@1.27.27,trna11_LysCTT_5_-_180648979_180649051@1.27.27,trna13_LysCTT_14_-_58706613_58706685@1.27.27,trna13_LysCTT_6_+_26556774_26556846@1.27.27,trna2_LysCTT_15_+_79152904_79 |
| SRR10024150 | 2356 | 4983 | 128935 | 3409 | 32 | 39 | 170 | 166 | 4351 | 715 | 470 | 338 | 727 | 4717 | 348 | 49 | 3638 | 0 | 0 | 0 | 0 | 0 | 0 | 0 | 0 | 0 | 0 | 0 | 0 | 0 | 0 | 0 | 0 | 0 | 0 | 0 | 0 |
| SRR10024151 | 2446 | 7707 | 97782 | 3155 | 37 | 45 | 106 | 93 | 4106 | 340 | 2047 | 2035 | 2637 | 26866 | 1693 | 11 | 2082 | 0 | 0 | 0 | 0 | 0 | 0 | 0 | 0 | 0 | 0 | 0 | 0 | 0 | 0 | 0 | 0 | 0 | 0 | 0 | 0 |
| SRR10024152 | 1558 | 4180 | 92140 | 2797 | 30 | 41 | 125 | 132 | 6042 | 432 | 411 | 362 | 756 | 7199 | 406 | 11 | 2637 | 0 | 0 | 0 | 0 | 0 | 0 | 0 | 0 | 0 | 0 | 0 | 0 | 0 | 0 | 0 | 0 | 0 | 0 | 0 | 0 |
| SRR10024153 | 767 | 6452 | 133195 | 717 | 52 | 52 | 235 | 258 | 5890 | 831 | 792 | 628 | 1072 | 6477 | 637 | 22 | 4569 | 0 | 0 | 0 | 0 | 0 | 0 | 0 | 0 | 0 | 0 | 0 | 0 | 0 | 0 | 0 | 0 | 0 | 0 | 0 | 0 |
| SRR10024154 | 784 | 3567 | 97905 | 1148 | 38 | 36 | 171 | 255 | 3059 | 417 | 879 | 700 | 1502 | 11709 | 519 | 0 | 2314 | 0 | 0 | 0 | 0 | 0 | 0 | 0 | 0 | 0 | 0 | 0 | 0 | 0 | 0 | 0 | 0 | 0 | 0 | 0 | 0 |
| SRR10024155 | 7713 | 25963 | 40854 | 709 | 27 | 31 | 152 | 231 | 2122 | 1049 | 1626 | 1136 | 2300 | 13578 | 539 | 14 | 2652 | 0 | 0 | 0 | 0 | 0 | 0 | 0 | 0 | 0 | 0 | 0 | 16 | 0 | 0 | 0 | 0 | 0 | 0 | 0 | 0 |
| SRR10024156 | 866 | 5904 | 41148 | 4731 | 55 | 57 | 197 | 337 | 6592 | 323 | 2365 | 10 | 3199 | 21879 | 1607 | 11 | 2547 | 0 | 0 | 0 | 0 | 0 | 0 | 0 | 0 | 0 | 0 | 0 | 0 | 0 | 0 | 0 | 0 | 0 | 0 | 0 | 0 |
| SRR10024158 | 355 | 3299 | 112203 | 169 | 46 | 53 | 235 | 372 | 2990 | 437 | 1084 | 1117 | 2578 | 17203 | 631 | 36 | 3244 | 0 | 0 | 0 | 0 | 0 | 0 | 0 | 0 | 0 | 0 | 0 | 0 | 0 | 0 | 0 | 0 | 0 | 0 | 0 | 0 |
| SRR10024159 | 294 | 3171 | 69773 | 291 | 54 | 47 | 155 | 235 | 6936 | 568 | 1482 | 1346 | 1992 | 14708 | 1716 | 14 | 2168 | 0 | 0 | 0 | 0 | 0 | 0 | 0 | 0 | 0 | 0 | 0 | 0 | 0 | 0 | 0 | 0 | 0 | 0 | 0 | 0 |
| SRR10024161 | 2145 | 8676 | 123031 | 699 | 22 | 27 | 134 | 225 | 2239 | 1141 | 936 | 817 | 1651 | 9255 | 294 | 14 | 3368 | 0 | 0 | 0 | 0 | 0 | 0 | 0 | 0 | 0 | 0 | 0 | 0 | 0 | 0 | 0 | 0 | 0 | 0 | 0 | 0 |
| SRR10024162 | 1644 | 4649 | 117387 | 2043 | 38 | 37 | 143 | 183 | 3736 | 685 | 842 | 958 | 1587 | 12963 | 682 | 19 | 2319 | 0 | 0 | 0 | 0 | 0 | 0 | 0 | 0 | 0 | 0 | 0 | 0 | 0 | 0 | 0 | 0 | 0 | 0 | 0 | 0 |
| SRR10024163 | 1945 | 7767 | 112460 | 2906 | 25 | 27 | 85 | 124 | 3868 | 476 | 974 | 770 | 1168 | 13415 | 609 | 0 | 1857 | 0 | 0 | 0 | 0 | 0 | 0 | 0 | 0 | 0 | 0 | 0 | 0 | 0 | 0 | 0 | 0 | 0 | 0 | 0 | 0 |
| SRR10024164 | 156 | 2322 | 32983 | 468 | 56 | 63 | 203 | 160 | 7052 | 1187 | 476 | 313 | 790 | 6072 | 610 | 0 | 1100 | 0 | 0 | 0 | 0 | 0 | 0 | 0 | 0 | 0 | 0 | 0 | 0 | 0 | 0 | 0 | 0 | 0 | 0 | 0 | 0 |
| SRR10024166 | 277 | 2741 | 63240 | 615 | 17 | 22 | 69 | 42 | 8392 | 615 | 883 | 468 | 1095 | 8718 | 981 | 0 | 1053 | 0 | 0 | 0 | 0 | 0 | 0 | 0 | 0 | 0 | 0 | 0 | 0 | 0 | 0 | 0 | 0 | 0 | 0 | 0 | 0 |
| SRR10024167 | 102 | 1782 | 142926 | 106 | 20 | 20 | 106 | 124 | 3800 | 1069 | 765 | 940 | 1524 | 10285 | 319 | 36 | 1236 | 0 | 0 | 0 | 0 | 0 | 0 | 0 | 0 | 0 | 0 | 0 | 0 | 0 | 0 | 0 | 0 | 0 | 0 | 0 | 0 |
| SRR10024168 | 1995 | 9838 | 51977 | 1385 | 71 | 66 | 231 | 261 | 6482 | 1196 | 1315 | 824 | 1640 | 11810 | 901 | 24 | 2958 | 0 | 0 | 0 | 0 | 0 | 12 | 0 | 0 | 0 | 0 | 0 | 12 | 0 | 0 | 0 | 0 | 0 | 0 | 0 | 0 |
| SRR10024169 | 279 | 3248 | 139560 | 246 | 14 | 22 | 64 | 71 | 3728 | 521 | 1047 | 1536 | 1738 | 14898 | 884 | 13 | 1991 | 0 | 0 | 0 | 0 | 0 | 0 | 0 | 0 | 0 | 0 | 0 | 0 | 0 | 0 | 0 | 0 | 0 | 0 | 0 | 0 |
| SRR10024170 | 256 | 2811 | 55429 | 733 | 48 | 50 | 203 | 227 | 9782 | 408 | 830 | 588 | 1348 | 10414 | 998 | 0 | 1387 | 0 | 0 | 0 | 0 | 0 | 0 | 0 | 0 | 0 | 0 | 0 | 0 | 0 | 0 | 0 | 0 | 0 | 0 | 0 | 0 |
| SRR10024171 | 918 | 3167 | 75635 | 1258 | 25 | 24 | 116 | 186 | 3040 | 476 | 1143 | 1387 | 1974 | 11265 | 414 | 0 | 1738 | 0 | 0 | 0 | 0 | 0 | 0 | 0 | 0 | 0 | 0 | 0 | 0 | 0 | 0 | 0 | 0 | 0 | 0 | 0 | 0 |
| SRR10024173 | 509 | 3873 | 24042 | 995 | 76 | 77 | 299 | 450 | 5876 | 910 | 1038 | 708 | 1940 | 16712 | 1219 | 0 | 1579 | 0 | 0 | 0 | 0 | 0 | 0 | 0 | 0 | 0 | 0 | 0 | 13 | 0 | 0 | 0 | 0 | 0 | 0 | 0 | 0 |
| SRR10024176 | 233 | 2905 | 93065 | 128 | 15 | 14 | 75 | 88 | 1948 | 884 | 1260 | 1547 | 2088 | 8840 | 427 | 27 | 3922 | 0 | 0 | 0 | 0 | 0 | 0 | 0 | 0 | 0 | 0 | 0 | 0 | 0 | 0 | 0 | 0 | 0 | 0 | 0 | 0 |
| SRR10024178 | 1450 | 4417 | 72015 | 1386 | 11 | 15 | 41 | 78 | 1836 | 470 | 1778 | 1292 | 1885 | 16243 | 570 | 0 | 1749 | 0 | 0 | 0 | 0 | 0 | 0 | 0 | 0 | 0 | 0 | 0 | 0 | 0 | 0 | 0 | 0 | 0 | 0 | 0 | 0 |
| SRR10024179 | 1073 | 4612 | 56949 | 122 | 15 | 16 | 71 | 175 | 3178 | 551 | 1295 | 1685 | 1746 | 14650 | 293 | 0 | 1995 | 0 | 0 | 0 | 0 | 0 | 0 | 0 | 0 | 0 | 0 | 0 | 0 | 0 | 0 | 0 | 0 | 0 | 0 | 0 | 0 |
| SRR10024180 | 294 | 2838 | 151695 | 191 | 14 | 16 | 71 | 86 | 1967 | 858 | 650 | 633 | 900 | 5907 | 351 | 13 | 2446 | 0 | 0 | 0 | 0 | 0 | 0 | 0 | 0 | 0 | 0 | 0 | 0 | 0 | 0 | 0 | 0 | 0 | 0 | 0 | 0 |
| SRR10024181 | 1456 | 3161 | 119685 | 2213 | 39 | 39 | 180 | 262 | 4269 | 377 | 1389 | 1708 | 2321 | 14809 | 887 | 24 | 1886 | 0 | 0 | 0 | 0 | 0 | 0 | 0 | 0 | 0 | 0 | 0 | 0 | 0 | 0 | 0 | 0 | 0 | 0 | 0 | 0 |
| SRR10024182 | 1680 | 8357 | 70753 | 2491 | 24 | 24 | 77 | 132 | 3778 | 632 | 700 | 565 | 1009 | 11275 | 699 | 0 | 2537 | 0 | 0 | 0 | 0 | 0 | 0 | 0 | 0 | 0 | 0 | 0 | 0 | 0 | 0 | 0 | 0 | 0 | 0 | 0 | 0 |
| SRR10024183 | 678 | 3262 | 39042 | 1567 | 36 | 41 | 136 | 203 | 14017 | 608 | 1687 | 1268 | 2384 | 17305 | 1125 | 0 | 1291 | 0 | 0 | 0 | 0 | 0 | 0 | 0 | 0 | 0 | 0 | 0 | 0 | 0 | 0 | 0 | 0 | 0 | 0 | 0 | 0 |
| SRR10024185 | 328 | 2042 | 48758 | 4151 | 36 | 30 | 134 | 179 | 6992 | 392 | 1147 | 806 | 1872 | 11514 | 810 | 0 | 1769 | 0 | 0 | 0 | 0 | 0 | 0 | 0 | 0 | 0 | 0 | 0 | 0 | 0 | 0 | 0 | 0 | 0 | 0 | 0 | 0 |
| SRR10024186 | 523 | 2489 | 147411 | 341 | 24 | 23 | 106 | 137 | 4762 | 773 | 809 | 986 | 1270 | 9189 | 391 | 0 | 1672 | 0 | 0 | 0 | 0 | 0 | 0 | 0 | 0 | 0 | 0 | 0 | 0 | 0 | 0 | 0 | 0 | 0 | 0 | 0 | 0 |
| SRR10024187 | 164 | 1587 | 8936 | 1045 | 54 | 49 | 188 | 224 | 27430 | 567 | 609 | 409 | 620 | 5151 | 1137 | 10 | 1742 | 0 | 0 | 0 | 0 | 0 | 0 | 0 | 0 | 0 | 0 | 0 | 22 | 0 | 0 | 0 | 0 | 0 | 0 | 0 | 0 |
| SRR10024188 | 214 | 3403 | 58164 | 883 | 40 | 33 | 163 | 103 | 22434 | 460 | 983 | 246 | 800 | 10116 | 1039 | 21 | 1326 | 0 | 0 | 0 | 0 | 0 | 0 | 0 | 0 | 0 | 0 | 0 | 0 | 0 | 0 | 0 | 0 | 0 | 0 | 0 | 0 |
| SRR10024189 | 150 | 2160 | 51939 | 438 | 60 | 52 | 225 | 193 | 11802 | 759 | 1145 | 518 | 1535 | 12701 | 836 | 66 | 2024 | 0 | 0 | 0 | 0 | 0 | 0 | 0 | 0 | 0 | 0 | 0 | 0 | 0 | 0 | 0 | 0 | 0 | 0 | 0 | 0 |
| SRR10024190 | 362 | 10444 | 45681 | 667 | 31 | 27 | 130 | 37 | 27059 | 627 | 1139 | 173 | 562 | 11445 | 2018 | 0 | 454 | 0 | 0 | 0 | 0 | 0 | 0 | 0 | 0 | 0 | 0 | 0 | 0 | 0 | 0 | 0 | 0 | 0 | 0 | 0 | 0 |
| SRR10024191 | 531 | 2657 | 42362 | 651 | 40 | 47 | 146 | 324 | 2926 | 770 | 864 | 475 | 2004 | 13838 | 602 | 0 | 1436 | 0 | 0 | 0 | 0 | 0 | 0 | 0 | 0 | 0 | 0 | 0 | 0 | 0 | 0 | 0 | 0 | 0 | 0 | 0 | 0 |
| SRR10024192 | 320 | 4014 | 81378 | 811 | 37 | 42 | 172 | 184 | 5464 | 886 | 979 | 374 | 944 | 9712 | 702 | 21 | 1645 | 0 | 0 | 0 | 0 | 0 | 0 | 0 | 0 | 0 | 0 | 0 | 0 | 0 | 0 | 0 | 0 | 0 | 0 | 0 | 0 |
| SRR10024193 | 1276 | 6991 | 117650 | 678 | 33 | 37 | 182 | 280 | 2583 | 1192 | 1522 | 1286 | 2599 | 15229 | 616 | 15 | 3241 | 0 | 0 | 0 | 0 | 0 | 0 | 0 | 0 | 0 | 0 | 0 | 0 | 0 | 0 | 0 | 0 | 0 | 0 | 0 | 0 |
| SRR10024194 | 659 | 5909 | 97806 | 331 | 37 | 45 | 208 | 220 | 5215 | 829 | 792 | 746 | 1634 | 11301 | 483 | 31 | 2166 | 0 | 0 | 0 | 0 | 0 | 0 | 0 | 0 | 0 | 0 | 0 | 0 | 0 | 0 | 0 | 0 | 0 | 0 | 0 | 0 |
| SRR10024195 | 1100 | 4464 | 93349 | 2149 | 43 | 35 | 149 | 209 | 5341 | 444 | 1540 | 1299 | 1934 | 16197 | 595 | 11 | 2681 | 0 | 0 | 0 | 0 | 0 | 0 | 0 | 0 | 0 | 0 | 0 | 0 | 0 | 0 | 0 | 0 | 0 | 0 | 0 | 0 |
| SRR10024196 | 141 | 1463 | 34869 | 462 | 51 | 45 | 168 | 198 | 11869 | 712 | 1106 | 488 | 1405 | 9994 | 1142 | 0 | 1295 | 0 | 0 | 0 | 0 | 0 | 0 | 0 | 0 | 0 | 0 | 0 | 0 | 0 | 0 | 0 | 0 | 0 | 0 | 0 | 0 |
| SRR10024197 | 791 | 4242 | 158663 | 986 | 43 | 42 | 216 | 244 | 3646 | 774 | 841 | 699 | 1212 | 8473 | 233 | 0 | 2494 | 0 | 0 | 0 | 0 | 0 | 0 | 0 | 0 | 0 | 0 | 0 | 0 | 0 | 0 | 0 | 0 | 0 | 0 | 0 | 0 |
| SRR10024198 | 964 | 3894 | 57207 | 597 | 23 | 21 | 101 | 229 | 2872 | 406 | 1609 | 1652 | 2185 | 16118 | 417 | 0 | 2636 | 0 | 0 | 0 | 0 | 0 | 0 | 0 | 0 | 0 | 0 | 0 | 0 | 0 | 0 | 0 | 0 | 0 | 0 | 0 | 0 |
| SRR10024199 | 239 | 4107 | 16228 | 2060 | 88 | 68 | 218 | 276 | 15098 | 349 | 1373 | 578 | 1715 | 9100 | 2529 | 0 | 1437 | 0 | 0 | 0 | 0 | 0 | 0 | 0 | 0 | 0 | 0 | 0 | 0 | 0 | 0 | 0 | 0 | 0 | 0 | 0 | 0 |
| SRR10024201 | 442 | 3040 | 174917 | 356 | 20 | 15 | 75 | 109 | 2218 | 523 | 1201 | 1089 | 1398 | 10038 | 499 | 0 | 2011 | 0 | 0 | 0 | 0 | 0 | 0 | 0 | 0 | 0 | 0 | 0 | 0 | 0 | 0 | 0 | 0 | 0 | 0 | 0 | 0 |
| SRR10024202 | 311 | 6147 | 27849 | 2488 | 72 | 58 | 167 | 140 | 26169 | 694 | 813 | 377 | 1036 | 6027 | 1667 | 0 | 1698 | 0 | 0 | 0 | 0 | 0 | 0 | 0 | 0 | 0 | 0 | 0 | 0 | 0 | 0 | 0 | 0 | 0 | 0 | 0 | 0 |
| SRR10024203 | 2222 | 6081 | 160502 | 856 | 29 | 36 | 126 | 237 | 3927 | 501 | 1415 | 1559 | 2195 | 16535 | 1082 | 11 | 3399 | 0 | 0 | 0 | 0 | 0 | 0 | 0 | 0 | 0 | 0 | 0 | 0 | 0 | 0 | 0 | 0 | 0 | 0 | 0 | 0 |
| SRR10024204 | 150 | 2486 | 173621 | 169 | 22 | 33 | 144 | 171 | 5192 | 641 | 1127 | 907 | 1716 | 17944 | 494 | 11 | 3320 | 0 | 0 | 0 | 0 | 0 | 0 | 0 | 0 | 0 | 0 | 0 | 0 | 0 | 0 | 0 | 0 | 0 | 0 | 0 | 0 |
| SRR10024205 | 290 | 5634 | 90496 | 941 | 63 | 69 | 250 | 183 | 5751 | 767 | 1463 | 607 | 1954 | 13007 | 847 | 0 | 1748 | 0 | 0 | 0 | 0 | 0 | 0 | 0 | 0 | 0 | 0 | 0 | 0 | 0 | 0 | 0 | 0 | 0 | 0 | 0 | 0 |
| SRR10024206 | 506 | 6022 | 91948 | 515 | 40 | 42 | 162 | 225 | 3573 | 598 | 1241 | 972 | 2060 | 15942 | 587 | 0 | 3037 | 0 | 0 | 0 | 0 | 0 | 0 | 0 | 0 | 0 | 0 | 0 | 0 | 0 | 0 | 0 | 0 | 0 | 0 | 0 | 0 |
| SRR10024207 | 61 | 1050 | 30862 | 66 | 22 | 23 | 76 | 92 | 6616 | 785 | 710 | 555 | 1182 | 6817 | 693 | 65 | 1463 | 0 | 0 | 0 | 0 | 0 | 0 | 0 | 0 | 0 | 0 | 0 | 0 | 0 | 0 | 0 | 0 | 0 | 0 | 0 | 0 |
| SRR10024208 | 1919 | 9541 | 89714 | 1504 | 74 | 70 | 319 | 346 | 6377 | 874 | 687 | 410 | 955 | 6261 | 354 | 14 | 6014 | 0 | 0 | 0 | 0 | 0 | 0 | 0 | 0 | 0 | 0 | 0 | 0 | 0 | 0 | 0 | 0 | 0 | 0 | 0 | 0 |
| SRR1003025 | 104 | 63631 | 3759 | 824 | 886 | 9836 | 24698 | 18321 | 6126 | 58 | 10191 | 6800 | 2526 | 7076 | 0 | 3838 | 12042 | 0 | 0 | 10 | 2893 | 574 | 257 | 0 | 0 | 0 | 0 | 0 | 195 | 916 | 675 | 92 | 0 | 0 | 0 | 0 | 0 |
| SRR1005875 | 0 | 194 | 1784 | 233 | 64 | 1056 | 6353 | 5425 | 3516 | 0 | 3317 | 3232 | 1277 | 6647 | 0 | 689 | 70476 | 0 | 0 | 0 | 0 | 0 | 0 | 0 | 0 | 0 | 0 | 0 | 0 | 0 | 0 | 0 | 0 | 0 | 0 | 0 | 0 |
| SRR10072377 | 295 | 1732 | 24018 | 1785 | 733 | 688 | 8406 | 7874 | 10424 | 4134 | 17693 | 9388 | 10600 | 4525 | 194 | 3211 | 32626 | 0 | 100 | 68 | 281 | 442 | 178 | 23 | 0 | 0 | 0 | 108 | 155 | 179 | 125 | 0 | 0 | 0 | 0 | 0 | 0 |
| SRR10072378 | 237 | 1851 | 24052 | 1973 | 731 | 674 | 8410 | 7953 | 10566 | 4170 | 17755 | 9440 | 10902 | 4528 | 190 | 3248 | 32378 | 0 | 84 | 70 | 294 | 442 | 186 | 19 | 0 | 0 | 0 | 114 | 164 | 189 | 128 | 11 | 0 | 0 | 0 | 0 | 0 |
| SRR10072379 | 468 | 2925 | 13185 | 1353 | 583 | 463 | 5285 | 7393 | 11744 | 3739 | 12588 | 6635 | 7827 | 3293 | 108 | 3074 | 35349 | 0 | 216 | 64 | 50 | 98 | 28 | 11 | 0 | 0 | 0 | 140 | 137 | 24 | 48 | 0 | 0 | 0 | 0 | 0 | 0 |
| SRR10072380 | 511 | 2928 | 13217 | 1422 | 647 | 483 | 5375 | 7221 | 11893 | 3777 | 12622 | 6572 | 7931 | 3282 | 110 | 2994 | 35018 | 0 | 208 | 60 | 47 | 108 | 28 | 16 | 0 | 0 | 0 | 136 | 142 | 24 | 64 | 0 | 0 | 0 | 0 | 0 | 0 |
| SRR10072381 | 513 | 3024 | 22119 | 1245 | 516 | 490 | 5832 | 5024 | 11629 | 2953 | 14792 | 7133 | 7457 | 2726 | 86 | 3330 | 34690 | 0 | 252 | 100 | 364 | 718 | 167 | 21 | 0 | 0 | 0 | 181 | 224 | 159 | 196 | 0 | 0 | 0 | 0 | 17 | 0 |
| SRR10072382 | 540 | 2996 | 22240 | 1343 | 490 | 499 | 5856 | 5017 | 11673 | 3026 | 14800 | 7263 | 7484 | 2735 | 82 | 3244 | 34171 | 0 | 252 | 98 | 364 | 743 | 165 | 25 | 0 | 11 | 0 | 176 | 221 | 157 | 193 | 0 | 0 | 0 | 0 | 17 | 0 |
| SRR10072383 | 277 | 1403 | 13392 | 1084 | 753 | 557 | 5734 | 6786 | 10433 | 5934 | 12085 | 5783 | 6873 | 2741 | 89 | 3997 | 33485 | 0 | 141 | 33 | 117 | 635 | 117 | 15 | 0 | 0 | 0 | 72 | 66 | 52 | 138 | 0 | 0 | 0 | 0 | 0 | 0 |
| SRR10072384 | 291 | 1395 | 13242 | 1247 | 756 | 570 | 5716 | 6930 | 10573 | 5882 | 12237 | 5877 | 6844 | 2749 | 88 | 3887 | 33263 | 0 | 151 | 34 | 130 | 647 | 120 | 13 | 0 | 0 | 0 | 70 | 58 | 51 | 146 | 0 | 0 | 0 | 0 | 0 | 0 |
| SRR10072385 | 583 | 3322 | 37682 | 1769 | 563 | 497 | 5271 | 5523 | 13659 | 2751 | 15168 | 7764 | 9225 | 3709 | 112 | 3289 | 38237 | 0 | 125 | 111 | 696 | 1584 | 784 | 138 | 32 | 11 | 0 | 100 | 225 | 228 | 213 | 20 | 0 | 0 | 0 | 21 | 0 |
| SRR10072386 | 612 | 3325 | 37457 | 1730 | 554 | 505 | 5253 | 5535 | 13776 | 2838 | 15184 | 7785 | 9181 | 3653 | 106 | 3194 | 37801 | 0 | 124 | 109 | 720 | 1644 | 771 | 142 | 34 | 13 | 0 | 105 | 240 | 229 | 199 | 26 | 0 | 0 | 0 | 20 | 0 |
| SRR10072387 | 480 | 3026 | 27064 | 998 | 411 | 453 | 5170 | 4992 | 12572 | 3061 | 13190 | 7256 | 7455 | 2923 | 53 | 3073 | 39083 | 0 | 206 | 316 | 4888 | 3932 | 767 | 78 | 22 | 28 | 0 | 144 | 548 | 1646 | 523 | 24 | 0 | 0 | 0 | 47 | 0 |
| SRR10072388 | 499 | 3134 | 26963 | 1054 | 408 | 457 | 5023 | 5040 | 12725 | 2971 | 13227 | 7224 | 7368 | 2977 | 58 | 3059 | 38209 | 0 | 212 | 323 | 5097 | 4066 | 782 | 73 | 25 | 33 | 0 | 160 | 551 | 1704 | 543 | 24 | 0 | 0 | 0 | 44 | 0 |
| SRR10072389 | 130 | 1123 | 22388 | 1554 | 1274 | 1250 | 8196 | 9372 | 11065 | 8668 | 11168 | 5208 | 4813 | 2330 | 116 | 7099 | 21047 | 0 | 79 | 25 | 30 | 46 | 0 | 0 | 0 | 0 | 0 | 112 | 74 | 42 | 39 | 0 | 0 | 0 | 0 | 0 | 0 |
| SRR10072390 | 139 | 1154 | 22533 | 1723 | 1250 | 1275 | 8284 | 9491 | 11322 | 8693 | 11203 | 5213 | 4889 | 2342 | 113 | 7041 | 20890 | 0 | 76 | 26 | 27 | 43 | 0 | 0 | 0 | 0 | 0 | 110 | 78 | 36 | 41 | 0 | 0 | 0 | 0 | 0 | 0 |
| SRR10072391 | 304 | 1640 | 10591 | 1201 | 919 | 643 | 6145 | 8381 | 16162 | 5437 | 13195 | 6273 | 6640 | 2962 | 94 | 4734 | 37949 | 0 | 160 | 54 | 204 | 674 | 138 | 15 | 0 | 0 | 0 | 95 | 116 | 113 | 166 | 0 | 0 | 0 | 0 | 0 | 0 |
| SRR10072392 | 324 | 1626 | 10561 | 1186 | 920 | 630 | 6185 | 8630 | 16415 | 5381 | 13284 | 6285 | 6646 | 2943 | 88 | 4683 | 37757 | 0 | 172 | 55 | 207 | 704 | 153 | 20 | 0 | 0 | 0 | 100 | 122 | 123 | 167 | 0 | 0 | 0 | 0 | 11 | 0 |
| SRR10072393 | 309 | 1910 | 19966 | 1413 | 1062 | 773 | 10085 | 9030 | 11388 | 3744 | 13756 | 6392 | 8737 | 3893 | 88 | 4171 | 29411 | 0 | 216 | 106 | 157 | 1225 | 535 | 35 | 0 | 0 | 0 | 181 | 249 | 81 | 285 | 18 | 0 | 0 | 0 | 13 | 0 |
| SRR10072394 | 293 | 1994 | 19820 | 1548 | 1050 | 784 | 10126 | 9121 | 11519 | 3751 | 13821 | 6306 | 8709 | 3912 | 86 | 4081 | 29156 | 0 | 227 | 128 | 171 | 1280 | 556 | 39 | 0 | 0 | 0 | 189 | 270 | 80 | 291 | 17 | 0 | 0 | 0 | 12 | 0 |
| SRR10072395 | 238 | 1573 | 49872 | 436 | 2763 | 1636 | 14250 | 18747 | 6544 | 10751 | 17844 | 12 | 8808 | 5104 | 278 | 3614 | 34031 | 0 | 54 | 10 | 58 | 254 | 42 | 0 | 0 | 0 | 0 | 28 | 18 | 22 | 23 | 0 | 0 | 0 | 0 | 0 | 0 |
| SRR10072396 | 181 | 1636 | 51422 | 530 | 2765 | 1638 | 14318 | 18810 | 6695 | 11011 | 17865 | 7760 | 8894 | 5160 | 270 | 3577 | 33702 | 0 | 54 | 10 | 53 | 266 | 38 | 0 | 0 | 0 | 0 | 30 | 22 | 22 | 25 | 0 | 0 | 0 | 0 | 0 | 0 |
| SRR10072397 | 251 | 2108 | 59137 | 3200 | 960 | 816 | 7917 | 8788 | 21224 | 3186 | 13382 | 7127 | 8077 | 3458 | 105 | 5155 | 30439 | 0 | 206 | 113 | 212 | 1836 | 903 | 57 | 0 | 0 | 0 | 180 | 227 | 145 | 429 | 52 | 0 | 0 | 0 | 13 | 0 |
| SRR10072398 | 284 | 2108 | 58715 | 3239 | 985 | 833 | 7886 | 8826 | 21641 | 3154 | 13380 | 7175 | 8038 | 3466 | 116 | 4994 | 30094 | 0 | 206 | 123 | 224 | 1923 | 925 | 53 | 0 | 0 | 0 | 184 | 222 | 150 | 437 | 55 | 0 | 0 | 0 | 15 | 0 |
| SRR10072399 | 533 | 3058 | 16217 | 1363 | 567 | 432 | 5588 | 5728 | 14790 | 3533 | 16990 | 7408 | 10079 | 3138 | 80 | 2945 | 36285 | 0 | 247 | 187 | 403 | 1803 | 406 | 27 | 0 | 14 | 0 | 180 | 320 | 136 | 238 | 11 | 0 | 0 | 0 | 18 | 0 |
| SRR10072400 | 481 | 3091 | 15799 | 1460 | 584 | 437 | 5581 | 5682 | 15130 | 3537 | 17001 | 7440 | 10164 | 3177 | 68 | 2983 | 36181 | 0 | 259 | 184 | 432 | 1864 | 420 | 21 | 0 | 15 | 0 | 179 | 323 | 141 | 250 | 12 | 0 | 0 | 0 | 21 | 0 |
| SRR10077412 | 0 | 17241 | 1786 | 0 | 3135 | 26035 | 23775 | 14622 | 567 | 39 | 20753 | 51014 | 5280 | 8232 | 23 | 92381 | 24989 | 0 | 0 | 0 | 36 | 0 | 35 | 0 | 0 | 0 | 0 | 0 | 0 | 0 | 0 | 11 | 0 | 0 | 0 | 0 | 0 |
| SRR10077413 | 15 | 17410 | 1722 | 15 | 3187 | 26167 | 23845 | 14798 | 566 | 49 | 20644 | 50472 | 5222 | 8263 | 22 | 93898 | 23596 | 0 | 0 | 0 | 30 | 0 | 31 | 0 | 0 | 0 | 0 | 0 | 0 | 0 | 0 | 12 | 0 | 0 | 0 | 0 | 0 |
| SRR10077414 | 11 | 17711 | 1898 | 0 | 3089 | 25799 | 23610 | 14528 | 549 | 53 | 20899 | 96 | 5272 | 8423 | 25 | 92269 | 24838 | 0 | 0 | 0 | 36 | 0 | 29 | 0 | 0 | 0 | 0 | 0 | 0 | 0 | 0 | 15 | 0 | 0 | 0 | 0 | 0 |
| SRR10077415 | 0 | 18008 | 1711 | 0 | 3159 | 26390 | 23924 | 14792 | 567 | 55 | 20831 | 50643 | 5311 | 8365 | 26 | 94609 | 23458 | 0 | 0 | 0 | 34 | 0 | 30 | 0 | 0 | 0 | 0 | 0 | 0 | 0 | 0 | 12 | 0 | 0 | 0 | 0 | 0 |
| SRR10077416 | 0 | 4838 | 1290 | 0 | 3166 | 25940 | 26850 | 16369 | 674 | 72 | 18979 | 45258 | 5334 | 7271 | 33 | 100753 | 21909 | 0 | 0 | 0 | 11 | 0 | 0 | 0 | 0 | 0 | 0 | 0 | 0 | 0 | 0 | 0 | 0 | 0 | 0 | 0 | 0 |
| SRR10077417 | 0 | 4993 | 1389 | 0 | 3242 | 26169 | 27108 | 16535 | 686 | 64 | 18878 | 44949 | 5266 | 7213 | 35 | 101904 | 20509 | 0 | 0 | 0 | 0 | 0 | 0 | 0 | 0 | 0 | 0 | 0 | 0 | 0 | 0 | 0 | 0 | 0 | 0 | 0 | 0 |
| SRR10077418 | 0 | 4712 | 1310 | 0 | 3123 | 25590 | 26675 | 16280 | 698 | 58 | 19134 | 84 | 5246 | 7303 | 31 | 100494 | 21830 | 0 | 0 | 0 | 11 | 0 | 0 | 0 | 0 | 0 | 0 | 0 | 0 | 0 | 0 | 0 | 0 | 0 | 0 | 0 | 0 |
| SRR10077419 | 0 | 4852 | 1413 | 0 | 3172 | 26272 | 26932 | 16546 | 700 | 40 | 19076 | 45098 | 5240 | 7228 | 33 | 102690 | 20735 | 0 | 0 | 0 | 0 | 0 | 0 | 0 | 0 | 0 | 0 | 0 | 0 | 0 | 0 | 0 | 0 | 0 | 0 | 0 | 0 |
| SRR10077420 | 0 | 5007 | 1250 | 0 | 3796 | 28226 | 28238 | 17938 | 795 | 29 | 24625 | 43 | 6011 | 10664 | 35 | 103136 | 28736 | 0 | 0 | 0 | 16 | 0 | 0 | 0 | 0 | 0 | 0 | 0 | 0 | 0 | 0 | 0 | 0 | 0 | 0 | 0 | 0 |
| SRR10077421 | 0 | 4918 | 1254 | 0 | 3732 | 28500 | 28459 | 18008 | 806 | 51 | 24574 | 51194 | 6025 | 10718 | 34 | 104865 | 27150 | 0 | 0 | 0 | 16 | 0 | 10 | 0 | 0 | 0 | 0 | 0 | 0 | 0 | 0 | 0 | 0 | 0 | 0 | 0 | 0 |
| SRR10077422 | 0 | 5014 | 1409 | 0 | 3668 | 27910 | 27980 | 17689 | 807 | 0 | 24855 | 100 | 5934 | 10662 | 38 | 102906 | 28851 | 0 | 0 | 0 | 14 | 0 | 13 | 0 | 0 | 0 | 0 | 0 | 0 | 0 | 0 | 0 | 0 | 0 | 0 | 0 | 0 |
| SRR10077423 | 0 | 4889 | 1390 | 0 | 3737 | 28678 | 28516 | 18125 | 770 | 39 | 24760 | 51606 | 5966 | 10764 | 35 | 105432 | 27248 | 0 | 0 | 0 | 13 | 0 | 13 | 0 | 0 | 0 | 0 | 0 | 0 | 0 | 0 | 0 | 0 | 0 | 0 | 0 | 0 |
| SRR10077424 | 14 | 8502 | 6167 | 0 | 3886 | 32313 | 28631 | 19180 | 768 | 97 | 19536 | 45036 | 5268 | 7543 | 66 | 89772 | 21445 | 0 | 0 | 0 | 55 | 22 | 26 | 0 | 0 | 0 | 0 | 0 | 0 | 11 | 13 | 12 | 0 | 0 | 0 | 0 | 0 |
| SRR10077425 | 11 | 8697 | 5930 | 15 | 3890 | 32305 | 28587 | 19444 | 783 | 48 | 19403 | 92 | 5193 | 7596 | 65 | 90509 | 20432 | 0 | 0 | 0 | 56 | 22 | 26 | 0 | 0 | 0 | 0 | 0 | 0 | 0 | 13 | 13 | 0 | 0 | 0 | 0 | 0 |
| SRR10077426 | 12 | 8212 | 5835 | 12 | 3776 | 31812 | 28400 | 19115 | 801 | 67 | 19755 | 90 | 5270 | 7604 | 64 | 89084 | 21595 | 0 | 0 | 0 | 53 | 19 | 23 | 0 | 0 | 0 | 0 | 0 | 0 | 0 | 13 | 13 | 0 | 0 | 0 | 0 | 0 |
| SRR10077427 | 0 | 8575 | 5821 | 0 | 3871 | 32417 | 28666 | 19308 | 778 | 63 | 19487 | 44813 | 5289 | 7583 | 64 | 91564 | 20576 | 0 | 0 | 0 | 54 | 21 | 23 | 0 | 0 | 0 | 0 | 0 | 0 | 0 | 15 | 11 | 0 | 0 | 0 | 0 | 0 |

Supp Table S8.1.Columns_B_to_AL

| | | | | | | | | | | | | | | | | | | | | | | | | | | | | | | | | | | | | | |
|---|---|---|---|---|---|---|---|---|---|---|---|---|---|---|---|---|---|---|---|---|---|---|---|---|---|---|---|---|---|---|---|---|---|---|---|---|---|
| SRR10077428 | 0 | 63997 | 4426 | 21 | 4714 | 37476 | 33051 | 19972 | 732 | 28 | 19697 | 30 | 5274 | 7415 | 82 | 118949 | 21460 | 0 | 0 | 0 | 415 | 165 | 137 | 0 | 12 | 0 | 0 | 0 | 69 | 85 | 113 | 78 | 0 | 10 | 0 | 23 | 0 |
| SRR10077429 | 36 | 65177 | 4415 | 58 | 4800 | 37488 | 33305 | 20120 | 702 | 66 | 19796 | 43199 | 5242 | 7513 | 77 | 120543 | 20150 | 0 | 0 | 0 | 441 | 154 | 135 | 0 | 12 | 0 | 0 | 0 | 63 | 89 | 113 | 65 | 0 | 0 | 0 | 16 | 0 |
| SRR10077430 | 16 | 64857 | 4469 | 0 | 4755 | 36835 | 32799 | 20028 | 720 | 70 | 20068 | 78 | 5342 | 7510 | 78 | 118434 | 21439 | 0 | 0 | 0 | 393 | 146 | 149 | 0 | 13 | 0 | 0 | 0 | 61 | 80 | 106 | 67 | 0 | 13 | 0 | 14 | 0 |
| SRR10077431 | 29 | 64489 | 4460 | 0 | 4783 | 37622 | 33500 | 20175 | 722 | 73 | 19928 | 43610 | 5344 | 7470 | 93 | 121184 | 20098 | 0 | 0 | 0 | 407 | 163 | 156 | 12 | 14 | 0 | 0 | 0 | 59 | 90 | 105 | 67 | 0 | 11 | 0 | 17 | 0 |
| SRR10077432 | 0 | 15826 | 8074 | 0 | 4510 | 37933 | 32873 | 23556 | 658 | 56 | 21822 | 42 | 5861 | 8269 | 82 | 111404 | 26248 | 0 | 0 | 0 | 81 | 35 | 40 | 0 | 0 | 0 | 0 | 0 | 10 | 15 | 27 | 23 | 0 | 0 | 0 | 16 | 0 |
| SRR10077433 | 0 | 16567 | 7682 | 0 | 4485 | 38349 | 33253 | 23899 | 664 | 54 | 22125 | 51333 | 5866 | 8330 | 78 | 113195 | 24636 | 0 | 0 | 0 | 84 | 36 | 43 | 0 | 0 | 0 | 0 | 0 | 0 | 16 | 25 | 23 | 0 | 0 | 0 | 13 | 0 |
| SRR10077434 | 0 | 15848 | 7912 | 12 | 4394 | 37625 | 32595 | 23406 | 656 | 31 | 22089 | 98 | 5914 | 8271 | 76 | 110900 | 26220 | 0 | 0 | 0 | 70 | 32 | 36 | 0 | 0 | 0 | 0 | 0 | 10 | 12 | 24 | 22 | 0 | 0 | 0 | 11 | 0 |
| SRR10077435 | 0 | 16264 | 8077 | 0 | 4566 | 38476 | 33393 | 23925 | 658 | 33 | 22267 | 51725 | 5920 | 8322 | 74 | 113952 | 24839 | 0 | 0 | 0 | 79 | 34 | 42 | 0 | 0 | 0 | 0 | 0 | 10 | 17 | 21 | 21 | 0 | 0 | 0 | 14 | 0 |
| SRR10116271 | 0 | 1887 | 3277 | 13 | 5526 | 42807 | 59612 | 86704 | 19021 | 215 | 19232 | 24575 | 5035 | 11151 | 174 | 2245 | 56307 | 0 | 0 | 0 | 0 | 114 | 24 | 0 | 0 | 0 | 0 | 0 | 37 | 0 | 216 | 0 | 0 | 0 | 0 | 0 | 0 |
| SRR10116272 | 0 | 403 | 21335 | 13 | 301 | 1375 | 3121 | 4171 | 33331 | 152 | 10374 | 27342 | 4560 | 2554 | 53 | 9863 | 65170 | 0 | 0 | 0 | 0 | 69 | 26 | 0 | 0 | 0 | 0 | 10 | 52 | 10 | 135 | 0 | 0 | 0 | 0 | 0 | 0 |
| SRR10116273 | 0 | 1116 | 6375 | 10 | 4518 | 42119 | 53195 | 41870 | 14444 | 59 | 33985 | 21517 | 3475 | 4568 | 1461 | 1389 | 37731 | 0 | 0 | 0 | 0 | 0 | 0 | 0 | 0 | 0 | 0 | 0 | 0 | 0 | 12 | 0 | 0 | 0 | 0 | 0 | 0 |
| SRR10116274 | 0 | 706 | 16870 | 0 | 868 | 4814 | 9826 | 15871 | 43016 | 181 | 18765 | 24034 | 8287 | 3942 | 121 | 5576 | 65447 | 0 | 0 | 0 | 0 | 36 | 16 | 0 | 0 | 0 | 0 | 0 | 30 | 0 | 64 | 0 | 0 | 0 | 0 | 0 | 0 |
| SRR10116275 | 0 | 2691 | 17475 | 24 | 2355 | 11819 | 17218 | 17036 | 12803 | 220 | 24373 | 48418 | 9553 | 18348 | 201 | 4781 | 48458 | 0 | 0 | 0 | 34 | 273 | 144 | 32 | 0 | 0 | 0 | 0 | 70 | 36 | 219 | 0 | 0 | 0 | 0 | 0 | 0 |
| SRR10119011 | 134 | 2639 | 1234 | 625 | 51 | 383 | 350 | 465 | 1058 | 13 | 41373 | 84922 | 3266 | 8538 | 0 | 534 | 9192 | 0 | 0 | 0 | 577 | 288 | 66 | 0 | 0 | 0 | 0 | 0 | 11 | 219 | 199 | 39 | 0 | 0 | 0 | 0 | 0 |
| SRR10119012 | 82 | 2541 | 631 | 64 | 0 | 53 | 47 | 456 | 352 | 61 | 592 | 311 | 43 | 479 | 0 | 109 | 525 | 0 | 0 | 0 | 0 | 0 | 0 | 0 | 0 | 0 | 0 | 0 | 0 | 0 | 0 | 0 | 0 | 0 | 0 | 0 | 0 |
| SRR10119014 | 0 | 1209 | 256 | 22 | 0 | 0 | 0 | 0 | 407 | 0 | 4343 | 1448 | 193 | 2681 | 0 | 0 | 622 | 0 | 0 | 0 | 0 | 0 | 0 | 0 | 0 | 0 | 0 | 0 | 0 | 0 | 0 | 0 | 0 | 0 | 0 | 0 | 0 |
| SRR10119022 | 86 | 13832 | 794 | 322 | 19 | 152 | 148 | 222 | 468 | 63 | 43127 | 112557 | 6008 | 13592 | 0 | 588 | 6154 | 0 | 0 | 0 | 1020 | 271 | 35 | 0 | 0 | 0 | 0 | 0 | 46 | 598 | 224 | 31 | 0 | 0 | 0 | 0 | 0 |
| SRR10119033 | 67 | 4689 | 297 | 531 | 16 | 199 | 316 | 631 | 476 | 0 | 1703 | 706 | 139 | 1544 | 0 | 175 | 1569 | 0 | 0 | 0 | 0 | 0 | 0 | 0 | 0 | 0 | 0 | 0 | 0 | 0 | 0 | 0 | 0 | 0 | 0 | 0 | 0 |
| SRR10119034 | 45 | 1300 | 0 | 417 | 0 | 27 | 28 | 238 | 213 | 0 | 529 | 163 | 81 | 676 | 0 | 51 | 456 | 0 | 0 | 0 | 0 | 0 | 0 | 0 | 0 | 0 | 0 | 0 | 0 | 0 | 0 | 0 | 0 | 0 | 0 | 0 | 0 |
| SRR10119035 | 64 | 38434 | 887 | 209 | 0 | 40 | 67 | 69 | 406 | 29 | 18358 | 65872 | 3267 | 4827 | 0 | 498 | 4029 | 0 | 0 | 0 | 2176 | 684 | 170 | 0 | 0 | 0 | 0 | 0 | 48 | 466 | 282 | 63 | 0 | 0 | 0 | 0 | 0 |
| SRR10119037 | 0 | 1303 | 107 | 284 | 0 | 60 | 67 | 310 | 200 | 0 | 423 | 186 | 71 | 1129 | 0 | 195 | 653 | 0 | 0 | 0 | 0 | 0 | 0 | 0 | 0 | 0 | 0 | 0 | 0 | 0 | 0 | 0 | 0 | 0 | 0 | 0 | 0 |
| SRR10119040 | 495 | 9514 | 243 | 0 | 0 | 78 | 87 | 312 | 221 | 0 | 691 | 383 | 0 | 282 | 0 | 173 | 342 | 0 | 0 | 0 | 0 | 0 | 0 | 0 | 0 | 0 | 0 | 0 | 0 | 0 | 0 | 0 | 0 | 0 | 0 | 0 | 0 |
| SRR10119044 | 34 | 10310 | 1252 | 602 | 0 | 16 | 31 | 302 | 404 | 0 | 193 | 114 | 45 | 202 | 0 | 92 | 231 | 0 | 0 | 0 | 0 | 0 | 0 | 0 | 0 | 0 | 0 | 0 | 0 | 0 | 0 | 0 | 0 | 0 | 0 | 0 | 0 |
| SRR10119046 | 84 | 37187 | 932 | 295 | 0 | 42 | 40 | 71 | 388 | 28 | 18086 | 71356 | 3268 | 6102 | 0 | 443 | 4169 | 0 | 0 | 0 | 1476 | 499 | 121 | 0 | 0 | 0 | 0 | 0 | 35 | 354 | 204 | 42 | 0 | 0 | 0 | 0 | 0 |
| SRR10119048 | 75 | 9249 | 2536 | 199 | 0 | 307 | 284 | 442 | 608 | 0 | 850 | 745 | 102 | 1018 | 0 | 184 | 1512 | 0 | 0 | 0 | 0 | 0 | 0 | 0 | 0 | 0 | 0 | 0 | 0 | 0 | 0 | 0 | 0 | 0 | 0 | 0 | 0 |
| SRR10119051 | 0 | 1130 | 233 | 74 | 0 | 42 | 53 | 412 | 215 | 0 | 847 | 431 | 50 | 537 | 0 | 133 | 385 | 0 | 0 | 0 | 0 | 0 | 0 | 0 | 0 | 0 | 0 | 0 | 0 | 0 | 0 | 0 | 0 | 0 | 0 | 0 | 0 |
| SRR10119052 | 86 | 2523 | 534 | 17 | 0 | 86 | 83 | 333 | 204 | 0 | 900 | 462 | 40 | 531 | 0 | 69 | 382 | 0 | 0 | 0 | 0 | 0 | 0 | 0 | 0 | 0 | 0 | 0 | 0 | 0 | 0 | 0 | 0 | 0 | 0 | 0 | 0 |
| SRR10119056 | 0 | 2409 | 157 | 64 | 18 | 139 | 116 | 223 | 202 | 0 | 1014 | 558 | 76 | 906 | 0 | 26 | 387 | 0 | 0 | 0 | 0 | 0 | 0 | 0 | 0 | 0 | 0 | 0 | 0 | 0 | 0 | 0 | 0 | 0 | 0 | 0 | 0 |
| SRR10119057 | 67 | 3248 | 292 | 617 | 0 | 63 | 71 | 148 | 699 | 12 | 18744 | 68406 | 3213 | 5526 | 0 | 515 | 8048 | 0 | 0 | 0 | 54 | 37 | 0 | 0 | 0 | 0 | 0 | 0 | 0 | 32 | 30 | 0 | 0 | 0 | 0 | 0 | 0 |
| SRR10119058 | 65 | 3761 | 315 | 487 | 0 | 54 | 61 | 125 | 711 | 11 | 9043 | 55709 | 2229 | 3945 | 0 | 707 | 10886 | 0 | 0 | 0 | 34 | 29 | 0 | 0 | 0 | 0 | 0 | 0 | 0 | 20 | 19 | 0 | 0 | 0 | 0 | 0 | 0 |
| SRR10119059 | 91 | 10574 | 713 | 647 | 20 | 126 | 228 | 319 | 577 | 115 | 37175 | 139428 | 4752 | 21898 | 0 | 464 | 5737 | 0 | 0 | 0 | 792 | 304 | 25 | 0 | 0 | 0 | 0 | 0 | 35 | 665 | 259 | 49 | 0 | 0 | 0 | 0 | 0 |
| SRR10119063 | 0 | 9958 | 445 | 33 | 0 | 35 | 0 | 347 | 212 | 0 | 1418 | 352 | 189 | 914 | 0 | 0 | 909 | 0 | 0 | 0 | 0 | 0 | 0 | 0 | 0 | 0 | 0 | 0 | 0 | 11 | 34 | 0 | 0 | 0 | 0 | 0 | 0 |
| SRR10119066 | 45 | 12920 | 469 | 436 | 0 | 55 | 64 | 153 | 712 | 11 | 2040 | 32690 | 1600 | 2818 | 0 | 1640 | 11162 | 0 | 0 | 0 | 1787 | 733 | 149 | 0 | 0 | 0 | 0 | 0 | 40 | 697 | 448 | 83 | 0 | 0 | 0 | 0 | 0 |
| SRR10119067 | 46 | 14217 | 443 | 421 | 0 | 46 | 48 | 115 | 482 | 0 | 5592 | 49633 | 2990 | 3192 | 0 | 1603 | 7785 | 0 | 0 | 0 | 1480 | 549 | 91 | 0 | 0 | 0 | 0 | 0 | 30 | 626 | 357 | 59 | 0 | 0 | 0 | 0 | 0 |
| SRR10119068 | 140 | 16768 | 1265 | 392 | 0 | 30 | 48 | 71 | 316 | 24 | 7575 | 33643 | 1849 | 4267 | 0 | 658 | 3349 | 0 | 0 | 0 | 380 | 282 | 29 | 0 | 0 | 0 | 0 | 0 | 24 | 292 | 175 | 23 | 0 | 0 | 0 | 0 | 0 |
| SRR10119069 | 140 | 15375 | 1200 | 370 | 0 | 54 | 34 | 97 | 413 | 16 | 10487 | 36291 | 2126 | 4793 | 0 | 562 | 3685 | 0 | 0 | 0 | 307 | 259 | 23 | 0 | 0 | 0 | 0 | 0 | 28 | 253 | 164 | 18 | 0 | 0 | 0 | 0 | 0 |
| SRR10119070 | 51 | 2625 | 236 | 785 | 67 | 348 | 441 | 508 | 1280 | 21 | 22193 | 87786 | 3088 | 10396 | 0 | 628 | 6170 | 0 | 0 | 0 | 504 | 187 | 41 | 0 | 0 | 0 | 0 | 0 | 12 | 272 | 165 | 29 | 0 | 0 | 0 | 0 | 0 |
| SRR10119071 | 37 | 2989 | 157 | 547 | 0 | 127 | 87 | 259 | 486 | 0 | 21229 | 75423 | 3497 | 10514 | 0 | 932 | 5449 | 0 | 0 | 0 | 108 | 48 | 0 | 0 | 0 | 0 | 0 | 0 | 0 | 57 | 42 | 0 | 0 | 0 | 0 | 0 | 0 |
| SRR10119072 | 33 | 1381 | 139 | 1220 | 44 | 374 | 446 | 336 | 1079 | 0 | 37980 | 125799 | 4499 | 21554 | 0 | 903 | 9830 | 0 | 0 | 0 | 121 | 77 | 11 | 0 | 0 | 0 | 0 | 0 | 0 | 58 | 55 | 12 | 0 | 0 | 0 | 0 | 0 |
| SRR10119073 | 19 | 709 | 79 | 861 | 47 | 548 | 804 | 827 | 1571 | 0 | 59829 | 120741 | 3966 | 11830 | 0 | 855 | 11314 | 0 | 0 | 0 | 121 | 29 | 0 | 0 | 0 | 0 | 0 | 0 | 0 | 29 | 24 | 0 | 0 | 0 | 0 | 0 | 0 |
| SRR10119074 | 29 | 930 | 66 | 711 | 14 | 329 | 469 | 675 | 1478 | 0 | 57865 | 122306 | 4247 | 11197 | 0 | 456 | 8714 | 0 | 0 | 0 | 103 | 27 | 0 | 0 | 0 | 0 | 0 | 0 | 0 | 20 | 19 | 0 | 0 | 0 | 0 | 0 | 0 |
| SRR10119075 | 78 | 7231 | 969 | 517 | 0 | 111 | 167 | 168 | 564 | 32 | 24088 | 74659 | 3405 | 5956 | 0 | 392 | 7184 | 0 | 0 | 0 | 433 | 237 | 50 | 0 | 0 | 0 | 0 | 0 | 0 | 135 | 125 | 20 | 0 | 0 | 0 | 0 | 0 |
| SRR10119076 | 74 | 5018 | 759 | 690 | 23 | 291 | 372 | 495 | 899 | 41 | 38052 | 99626 | 4603 | 9647 | 0 | 416 | 9678 | 0 | 0 | 0 | 387 | 242 | 40 | 0 | 0 | 0 | 0 | 0 | 0 | 136 | 138 | 25 | 0 | 0 | 0 | 0 | 0 |
| SRR10119077 | 26 | 12966 | 667 | 249 | 0 | 19 | 18 | 61 | 299 | 0 | 901 | 14221 | 525 | 1236 | 0 | 591 | 5411 | 0 | 0 | 0 | 776 | 282 | 71 | 0 | 0 | 0 | 0 | 0 | 11 | 135 | 105 | 24 | 0 | 0 | 0 | 0 | 0 |
| SRR10119078 | 23 | 13465 | 562 | 263 | 0 | 17 | 26 | 41 | 278 | 22 | 627 | 11918 | 407 | 1420 | 0 | 715 | 3849 | 0 | 0 | 0 | 733 | 201 | 49 | 0 | 0 | 0 | 0 | 0 | 0 | 189 | 78 | 20 | 0 | 0 | 0 | 0 | 0 |
| SRR10119079 | 81 | 8108 | 1394 | 584 | 17 | 86 | 127 | 155 | 311 | 38 | 46302 | 97589 | 5200 | 13716 | 0 | 272 | 3056 | 0 | 0 | 0 | 1717 | 468 | 159 | 0 | 0 | 0 | 0 | 0 | 32 | 604 | 253 | 89 | 0 | 0 | 0 | 0 | 0 |
| SRR10119080 | 56 | 8751 | 1709 | 601 | 0 | 109 | 108 | 138 | 312 | 35 | 24350 | 86942 | 4131 | 10383 | 0 | 254 | 3549 | 0 | 0 | 0 | 1786 | 462 | 135 | 0 | 0 | 0 | 0 | 0 | 31 | 576 | 262 | 86 | 0 | 0 | 0 | 0 | 0 |
| SRR10119081 | 23 | 2258 | 146 | 570 | 0 | 84 | 117 | 207 | 397 | 0 | 9329 | 34183 | 1792 | 4285 | 0 | 679 | 3290 | 0 | 0 | 0 | 129 | 22 | 0 | 0 | 0 | 0 | 0 | 0 | 0 | 42 | 13 | 0 | 0 | 0 | 0 | 0 | 0 |
| SRR10119082 | 19 | 1692 | 162 | 770 | 0 | 207 | 211 | 283 | 554 | 10 | 30353 | 66680 | 2860 | 9696 | 0 | 542 | 4620 | 0 | 0 | 0 | 151 | 47 | 0 | 0 | 0 | 0 | 0 | 0 | 0 | 52 | 29 | 0 | 0 | 0 | 0 | 0 | 0 |
| SRR1014666 | 0 | 81 | 92571 | 0 | 227 | 2014 | 4039 | 6810 | 16247 | 81 | 21091 | 6072 | 1541 | 2723 | 77 | 0 | 8012 | 0 | 0 | 0 | 0 | 0 | 0 | 0 | 0 | 0 | 0 | 0 | 0 | 31 | 38 | 0 | 0 | 0 | 0 | 0 | 0 |
| SRR1014667 | 0 | 241 | 36610 | 0 | 396 | 2456 | 7030 | 11692 | 20140 | 96 | 13495 | 4418 | 1603 | 2490 | 78 | 0 | 9902 | 0 | 0 | 0 | 0 | 0 | 0 | 0 | 0 | 0 | 0 | 0 | 56 | 283 | 553 | 22 | 0 | 0 | 0 | 0 | 0 |
| SRR1014669 | 0 | 82 | 62039 | 0 | 1024 | 2388 | 15012 | 9256 | 14001 | 926 | 7097 | 4312 | 1157 | 1845 | 76 | 0 | 7012 | 0 | 0 | 0 | 0 | 0 | 0 | 0 | 0 | 0 | 0 | 0 | 0 | 47 | 767 | 27 | 0 | 0 | 0 | 0 | 0 |
| SRR10197184 | 32 | 898 | 46798 | 42 | 29517 | 14944 | 244638 | 44010 | 884 | 2408 | 31538 | 13957 | 19572 | 65227 | 2159 | 13500 | 8860 | 0 | 11 | 0 | 14 | 23 | 13 | 0 | 0 | 0 | 0 | 19 | 24 | 0 | 0 | 0 | 0 | 0 | 0 | 0 | 0 |
| SRR10197185 | 12 | 1306 | 104718 | 75 | 1911 | 3111 | 27875 | 894 | 59076 | 167 | 8497 | 2110 | 1067 | 44497 | 4154 | 4633 | 2706 | 0 | 0 | 0 | 0 | 0 | 0 | 0 | 0 | 0 | 0 | 0 | 0 | 0 | 0 | 0 | 0 | 0 | 0 | 0 | 0 |
| SRR10197186 | 29 | 530 | 96073 | 29 | 11946 | 10673 | 124441 | 6202 | 7368 | 6014 | 34349 | 11543 | 10951 | 90798 | 3329 | 10964 | 7277 | 0 | 0 | 0 | 0 | 14 | 0 | 0 | 0 | 0 | 0 | 0 | 0 | 0 | 0 | 0 | 0 | 0 | 0 | 0 | 0 |
| SRR10197187 | 27 | 550 | 38252 | 57 | 18966 | 12646 | 191606 | 16252 | 2958 | 3867 | 36254 | 12470 | 14730 | 86438 | 2551 | 13232 | 7136 | 0 | 0 | 0 | 0 | 0 | 0 | 0 | 0 | 0 | 0 | 0 | 0 | 0 | 0 | 0 | 0 | 0 | 0 | 20 | 0 |
| SRR10197188 | 20 | 463 | 33107 | 38 | 23689 | 13402 | 230554 | 28775 | 1264 | 2616 | 35504 | 13935 | 18047 | 70034 | 2157 | 14281 | 8653 | 0 | 0 | 0 | 0 | 11 | 0 | 0 | 0 | 0 | 0 | 0 | 12 | 0 | 0 | 0 | 0 | 0 | 0 | 40 | 0 |
| SRR10197189 | 13 | 284 | 95247 | 57 | 10910 | 11743 | 111902 | 5322 | 11857 | 1941 | 34528 | 11353 | 8224 | 86106 | 3005 | 9956 | 5786 | 0 | 0 | 0 | 0 | 0 | 0 | 0 | 0 | 0 | 0 | 0 | 0 | 0 | 0 | 0 | 0 | 0 | 0 | 0 | 0 |
| SRR10197190 | 0 | 86 | 16329 | 24 | 22436 | 10809 | 226846 | 35852 | 850 | 653 | 23347 | 10131 | 13564 | 55470 | 1653 | 11728 | 6459 | 0 | 0 | 0 | 0 | 11 | 0 | 0 | 0 | 0 | 0 | 0 | 0 | 0 | 0 | 0 | 0 | 0 | 0 | 10 | 0 |
| SRR10197191 | 18 | 160 | 17627 | 45 | 24219 | 11360 | 217250 | 30582 | 1908 | 1440 | 33868 | 13034 | 17615 | 75232 | 2279 | 12706 | 8900 | 0 | 0 | 0 | 0 | 14 | 10 | 0 | 0 | 0 | 0 | 0 | 0 | 0 | 0 | 0 | 0 | 0 | 0 | 0 | 0 |
| SRR10197192 | 10 | 117 | 21442 | 45 | 30811 | 11712 | 241707 | 46088 | 1056 | 276 | 36749 | 12 | 19709 | 83533 | 2153 | 12298 | 7725 | 0 | 0 | 0 | 0 | 19 | 11 | 0 | 0 | 0 | 0 | 0 | 0 | 0 | 0 | 0 | 0 | 0 | 0 | 0 | 0 |
| SRR10225092 | 0 | 59921 | 613 | 175 | 613 | 2058 | 4312 | 1423 | 700 | 175 | 591 | 30187 | 1467 | 1795 | 0 | 876 | 8625 | 0 | 0 | 0 | 101 | 25 | 13 | 0 | 0 | 0 | 0 | 0 | 44 | 76 | 19 | 0 | 0 | 0 | 0 | 0 | 0 |
| SRR10225093 | 583 | 20845 | 90712 | 469 | 1616 | 34495 | 20479 | 35542 | 3049 | 170 | 16018 | 11227 | 2845 | 9000 | 0 | 10243 | 10882 | 0 | 10 | 34 | 734 | 1006 | 105 | 0 | 0 | 0 | 0 | 52 | 523 | 301 | 611 | 72 | 0 | 0 | 0 | 0 | 0 |
| SRR10225094 | 981 | 98586 | 562 | 507 | 63 | 1112 | 633 | 760 | 118 | 44 | 1103 | 13509 | 868 | 1600 | 0 | 4774 | 4810 | 0 | 0 | 0 | 260 | 43 | 25 | 0 | 0 | 0 | 0 | 0 | 31 | 76 | 0 | 11 | 0 | 0 | 0 | 0 | 0 |
| SRR10225096 | 98 | 92960 | 1917 | 264 | 56 | 338 | 310 | 647 | 253 | 59 | 450 | 19335 | 366 | 2983 | 0 | 1914 | 3068 | 0 | 0 | 0 | 96 | 0 | 0 | 0 | 23 | 0 | 0 | 0 | 0 | 19 | 0 | 0 | 0 | 23 | 10 | 71 | 0 |
| SRR10282881 | 11 | 15424 | 115667 | 0 | 232 | 7814 | 5193 | 6282 | 9247 | 42 | 48942 | 6484 | 3438 | 7034 | 0 | 1884 | 68702 | 0 | 0 | 0 | 242 | 1534 | 23 | 0 | 0 | 0 | 0 | 36 | 628 | 232 | 1036 | 25 | 0 | 11 | 0 | 0 | 0 |
| SRR10282882 | 0 | 18344 | 133790 | 0 | 251 | 8540 | 5814 | 6779 | 9191 | 41 | 54388 | 7324 | 3771 | 7215 | 0 | 2014 | 56549 | 0 | 0 | 0 | 192 | 1222 | 18 | 0 | 0 | 0 | 0 | 26 | 518 | 189 | 850 | 17 | 0 | 0 | 0 | 0 | 0 |
| SRR10282883 | 82 | 15311 | 73154 | 16 | 398 | 18018 | 11318 | 9076 | 12926 | 20 | 77015 | 11346 | 4710 | 8139 | 0 | 2592 | 99417 | 0 | 0 | 15 | 337 | 897 | 16 | 0 | 0 | 0 | 0 | 196 | 985 | 241 | 911 | 22 | 0 | 0 | 0 | 0 | 0 |
| SRR10282884 | 106 | 17733 | 82422 | 18 | 413 | 19458 | 11765 | 9730 | 12688 | 20 | 85410 | 12709 | 5308 | 8276 | 0 | 2556 | 79825 | 0 | 0 | 10 | 275 | 707 | 12 | 0 | 0 | 0 | 0 | 162 | 812 | 193 | 734 | 20 | 0 | 0 | 0 | 0 | 0 |
| SRR10282885 | 23 | 18391 | 32398 | 0 | 468 | 21042 | 13158 | 12454 | 9824 | 37 | 19904 | 4494 | 3005 | 7834 | 0 | 2358 | 82272 | 0 | 0 | 21 | 345 | 829 | 20 | 0 | 0 | 0 | 0 | 86 | 582 | 280 | 1471 | 33 | 0 | 0 | 0 | 0 | 0 |
| SRR10282886 | 34 | 21143 | 37466 | 0 | 484 | 23074 | 13930 | 13096 | 9732 | 41 | 22827 | 4955 | 3511 | 8288 | 0 | 2291 | 64304 | 0 | 0 | 18 | 295 | 650 | 17 | 0 | 0 | 0 | 0 | 67 | 456 | 215 | 1135 | 24 | 0 | 0 | 0 | 0 | 0 |
| SRR10282890 | 0 | 15617 | 40538 | 0 | 616 | 20353 | 13551 | 9405 | 13603 | 0 | 89085 | 9006 | 5519 | 11487 | 0 | 2426 | 99717 | 0 | 0 | 10 | 398 | 1515 | 48 | 0 | 0 | 0 | 0 | 75 | 368 | 227 | 885 | 66 | 0 | 14 | 0 | 0 | 0 |
| SRR10282891 | 0 | 17971 | 45882 | 0 | 639 | 21755 | 14442 | 9885 | 13342 | 0 | 98958 | 10064 | 6120 | 11884 | 0 | 2503 | 79337 | 0 | 0 | 0 | 322 | 1212 | 35 | 0 | 0 | 0 | 0 | 63 | 306 | 184 | 712 | 50 | 0 | 10 | 0 | 0 | 0 |
| SRR10282892 | 0 | 11241 | 35368 | 0 | 602 | 23528 | 13778 | 10861 | 11390 | 0 | 29330 | 4643 | 2852 | 7853 | 0 | 1737 | 110089 | 0 | 0 | 0 | 74 | 284 | 19 | 0 | 0 | 0 | 0 | 19 | 223 | 156 | 1221 | 42 | 0 | 0 | 0 | 0 | 0 |
| SRR10282893 | 0 | 12944 | 40228 | 0 | 579 | 25356 | 14572 | 11434 | 11228 | 0 | 32498 | 5240 | 3318 | 8198 | 0 | 1740 | 88022 | 0 | 0 | 0 | 59 | 231 | 14 | 0 | 0 | 0 | 0 | 18 | 186 | 129 | 978 | 33 | 0 | 0 | 0 | 0 | 0 |
| SRR10282894 | 0 | 9519 | 4706 | 0 | 226 | 10302 | 5743 | 4088 | 6801 | 0 | 18439 | 3307 | 1558 | 3214 | 12 | 693 | 28423 | 0 | 0 | 0 | 49 | 168 | 0 | 0 | 0 | 0 | 0 | 13 | 59 | 64 | 405 | 14 | 0 | 0 | 0 | 0 | 0 |
| SRR10282895 | 0 | 11127 | 5318 | 0 | 248 | 11159 | 6116 | 4292 | 6760 | 0 | 20795 | 3682 | 1857 | 3363 | 11 | 699 | 21990 | 0 | 0 | 0 | 39 | 135 | 0 | 0 | 0 | 0 | 0 | 12 | 50 | 52 | 313 | 10 | 0 | 0 | 0 | 0 | 0 |
| SRR10323931 | 53 | 13976 | 69002 | 21 | 17870 | 11754 | 98795 | 25494 | 156429 | 4618 | 56 | 22 | 58 | 27 | 2273 | 8917 | 17657 | 0 | 0 | 0 | 11 | 18 | 77 | 48 | 13 | 0 | 0 | 0 | 11 | 0 | 0 | 0 | 0 | 0 | 0 | 0 | 0 |
| SRR10323932 | 58 | 9622 | 26418 | 45 | 12584 | 7912 | 71171 | 15686 | 79005 | 2687 | 35 | 31 | 35 | 20 | 1178 | 5002 | 14267 | 0 | 0 | 0 | 38 | 15 | 35 | 11 | 0 | 0 | 0 | 0 | 27 | 27 | 0 | 0 | 0 | 0 | 0 | 0 | 0 |
| SRR10323933 | 1623 | 110857 | 15203 | 117 | 8970 | 4328 | 40422 | 11228 | 54329 | 2828 | 31 | 26 | 22 | 16 | 2432 | 5200 | 12993 | 0 | 26 | 30 | 215 | 62 | 190 | 68 | 14 | 0 | 0 | 59 | 151 | 120 | 11 | 0 | 0 | 0 | 0 | 0 | 0 |
| SRR10323939 | 14 | 4168 | 70022 | 12 | 12919 | 7266 | 78960 | 17907 | 105223 | 7444 | 105 | 46 | 69 | 29 | 1754 | 13626 | 31182 | 0 | 0 | 0 | 0 | 0 | 11 | 0 | 0 | 0 | 0 | 0 | 0 | 0 | 0 | 0 | 0 | 0 | 0 | 0 | 0 |
| SRR10323940 | 202 | 12735 | 28834 | 55 | 11482 | 7954 | 58548 | 19847 | 87656 | 3516 | 102 | 76 | 65 | 41 | 2793 | 11140 | 29966 | 0 | 20 | 15 | 56 | 80 | 67 | 31 | 13 | 0 | 0 | 42 | 96 | 46 | 15 | 0 | 0 | 0 | 0 | 0 | 0 |
| SRR10323941 | 59 | 2343 | 21551 | 13 | 7785 | 6061 | 38521 | 13026 | 70057 | 2334 | 102 | 72 | 76 | 59 | 1711 | 6121 | 27148 | 0 | 14 | 11 | 33 | 59 | 44 | 15 | 16 | 0 | 0 | 35 | 70 | 36 | 0 | 0 | 0 | 0 | 0 | 0 | 0 |
| SRR10385125 | 20 | 192 | 231564 | 29 | 14230 | 23385 | 134439 | 21593 | 18058 | 4443 | 29700 | 15 | 33032 | 16440 | 6237 | 25 | 10441 | 0 | 30 | 37 | 37 | 104 | 35 | 13 | 0 | 0 | 0 | 20 | 24 | 0 | 11 | 0 | 0 | 0 | 0 | 0 | 0 |
| SRR10385126 | 81 | 781 | 139399 | 60 | 11940 | 15959 | 80268 | 12073 | 19708 | 4911 | 32786 | 16 | 37891 | 19276 | 13305 | 18 | 7741 | 0 | 0 | 16 | 24 | 55 | 62 | 31 | 14 | 0 | 0 | 0 | 12 | 0 | 0 | 0 | 0 | 0 | 0 | 13 | 0 |
| SRR10416439 | 1224 | 4525 | 270643 | 551 | 1010 | 297 | 10535 | 1423 | 472 | 92 | 23263 | 14996 | 115306 | 7316 | 2161 | 0 | 62097 | 0 | 0 | 0 | 0 | 16 | 172 | 27 | 63 | 0 | 0 | 0 | 0 | 0 | 0 | 20 | 0 | 0 | 0 | 28 | 12 |
| SRR10416440 | 1311 | 4690 | 248677 | 255 | 4881 | 2187 | 57544 | 7223 | 351 | 52 | 22261 | 15 | 69026 | 11431 | 2383 | 0 | 32383 | 0 | 0 | 0 | 0 | 146 | 196 | 33 | 63 | 0 | 0 | 19 | 0 | 11 | 20 | 38 | 0 | 0 | 17 | 67 | 22 |
| SRR10416441 | 1060 | 6698 | 242395 | 262 | 3924 | 1319 | 38942 | 7548 | 366 | 93 | 20718 | 12111 | 88048 | 14837 | 1814 | 0 | 37282 | 0 | 0 | 0 | 0 | 45 | 270 | 50 | 86 | 0 | 0 | 0 | 0 | 22 | 12 | 53 | 0 | 0 | 0 | 65 | 25 |
| SRR10416442 | 899 | 3566 | 136751 | 484 | 1232 | 401 | 7386 | 6589 | 1572 | 199 | 21535 | 14 | 100918 | 7404 | 1102 | 11 | 73893 | 0 | 0 | 0 | 14 | 22 | 142 | 31 | 33 | 0 | 0 | 0 | 0 | 15 | 0 | 20 | 0 | 0 | 0 | 25 | 0 |
| SRR10416496 | 4227 | 1262 | 780 | 1796 | 0 | 10 | 0 | 45 | 74 | 72 | 524 | 240 | 48 | 300 | 0 | 0 | 500 | 0 | 0 | 0 | 37 | 0 | 0 | 0 | 0 | 0 | 0 | 86 | 253 | 32 | 12 | 0 | 0 | 0 | 0 | 0 | 0 |
| SRR10416497 | 3812 | 1861 | 795 | 1861 | 0 | 0 | 0 | 31 | 46 | 18 | 143 | 110 | 0 | 129 | 0 | 0 | 354 | 0 | 0 | 0 | 15 | 0 | 0 | 0 | 0 | 0 | 0 | 33 | 86 | 13 | 0 | 0 | 0 | 0 | 0 | 0 | 0 |
| SRR10416498 | 6218 | 3281 | 834 | 925 | 0 | 0 | 0 | 39 | 84 | 0 | 456 | 376 | 57 | 373 | 0 | 0 | 876 | 0 | 0 | 0 | 18 | 0 | 0 | 0 | 0 | 0 | 0 | 51 | 135 | 11 | 0 | 0 | 0 | 0 | 0 | 0 | 0 |
| SRR10416499 | 5445 | 1695 | 587 | 2576 | 0 | 0 | 16 | 59 | 115 | 68 | 416 | 189 | 41 | 240 | 0 | 0 | 463 | 0 | 0 | 0 | 0 | 0 | 0 | 0 | 0 | 0 | 0 | 54 | 122 | 20 | 0 | 0 | 0 | 0 | 0 | 0 | 0 |
| SRR10416500 | 5294 | 3941 | 1011 | 1234 | 0 | 15 | 11 | 136 | 177 | 15 | 747 | 441 | 90 | 549 | 0 | 0 | 1006 | 0 | 0 | 0 | 14 | 0 | 0 | 0 | 0 | 0 | 0 | 36 | 73 | 10 | 0 | 0 | 0 | 0 | 0 | 0 | 0 |
| SRR10416501 | 3455 | 1772 | 298 | 1981 | 0 | 0 | 0 | 99 | 159 | 89 | 500 | 371 | 77 | 416 | 0 | 0 | 882 | 0 | 0 | 0 | 24 | 0 | 0 | 0 | 0 | 0 | 0 | 93 | 204 | 22 | 0 | 0 | 0 | 0 | 0 | 0 | 0 |
| SRR10416502 | 5839 | 2800 | 1053 | 1311 | 0 | 20 | 35 | 200 | 110 | 99 | 911 | 496 | 130 | 883 | 0 | 13 | 769 | 0 | 0 | 0 | 10 | 0 | 0 | 0 | 0 | 0 | 0 | 29 | 59 | 0 | 0 | 0 | 0 | 0 | 0 | 0 | 0 |
| SRR10416503 | 3202 | 2352 | 697 | 2766 | 0 | 25 | 19 | 191 | 203 | 152 | 703 | 350 | 114 | 1067 | 0 | 21 | 850 | 0 | 0 | 0 | 0 | 0 | 0 | 0 | 0 | 0 | 0 | 29 | 60 | 0 | 0 | 0 | 0 | 0 | 0 | 0 | 0 |
| SRR10416504 | 3701 | 2348 | 677 | 1438 | 0 | 13 | 15 | 123 | 167 | 42 | 476 | 301 | 78 | 455 | 0 | 0 | 553 | 0 | 0 | 0 | 0 | 0 | 0 | 0 | 0 | 0 | 0 | 18 | 23 | 0 | 0 | 0 | 0 | 0 | 0 | 0 | 0 |
| SRR10416505 | 3499 | 1892 | 357 | 1367 | 0 | 0 | 0 | 66 | 99 | 20 | 310 | 345 | 45 | 534 | 0 | 0 | 946 | 0 | 0 | 0 | 26 | 0 | 0 | 0 | 0 | 0 | 0 | 81 | 236 | 15 | 13 | 0 | 0 | 0 | 0 | 0 | 0 |
| SRR10416506 | 4537 | 2719 | 618 | 1818 | 0 | 24 | 38 | 225 | 318 | 18 | 896 | 501 | 160 | 1020 | 0 | 0 | 1234 | 0 | 0 | 0 | 19 | 0 | 0 | 0 | 0 | 0 | 0 | 42 | 82 | 13 | 0 | 0 | 0 | 0 | 0 | 0 | 0 |
| SRR10416507 | 5838 | 2323 | 625 | 1891 | 0 | 0 | 12 | 77 | 63 | 74 | 553 | 286 | 73 | 549 | 0 | 0 | 504 | 0 | 0 | 0 | 13 | 0 | 0 | 0 | 0 | 0 | 0 | 35 | 81 | 0 | 0 | 0 | 0 | 0 | 0 | 0 | 0 |
| SRR10416508 | 4773 | 2307 | 1991 | 860 | 0 | 11 | 17 | 107 | 235 | 68 | 414 | 345 | 70 | 414 | 0 | 0 | 586 | 0 | 0 | 0 | 0 | 0 | 0 | 0 | 0 | 0 | 0 | 19 | 42 | 0 | 0 | 0 | 0 | 0 | 0 | 0 | 0 |
| SRR10416509 | 4016 | 2482 | 815 | 1260 | 0 | 23 | 17 | 113 | 211 | 15 | 823 | 611 | 116 | 913 | 0 | 0 | 1383 | 0 | 0 | 0 | 25 | 0 | 0 | 0 | 0 | 0 | 0 | 64 | 155 | 0 | 0 | 0 | 0 | 0 | 0 | 0 | 0 |
| SRR10416510 | 567 | 850 | 409 | 3872 | 0 | 0 | 0 | 0 | 0 | 94 | 129 | 413 | 19 | 148 | 0 | 0 | 148 | 0 | 0 | 0 | 0 | 0 | 0 | 0 | 0 | 0 | 0 | 0 | 0 | 0 | 0 | 0 | 0 | 0 | 0 | 0 | 0 |
| SRR10416513 | 2897 | 1440 | 468 | 1200 | 0 | 16 | 0 | 116 | 163 | 29 | 1102 | 648 | 106 | 635 | 0 | 0 | 1796 | 0 | 0 | 10 | 96 | 15 | 0 | 0 | 0 | 0 | 0 | 165 | 689 | 41 | 20 | 0 | 0 | 0 | 0 | 0 | 0 |
| SRR10416514 | 6159 | 2974 | 562 | 2108 | 0 | 10 | 0 | 48 | 131 | 23 | 314 | 269 | 49 | 423 | 0 | 25 | 448 | 0 | 0 | 0 | 0 | 0 | 0 | 0 | 0 | 0 | 0 | 22 | 47 | 11 | 0 | 0 | 0 | 0 | 0 | 0 | 0 |
| SRR10416515 | 3514 | 2185 | 248 | 3761 | 0 | 19 | 29 | 119 | 215 | 68 | 868 | 458 | 152 | 732 | 0 | 0 | 1051 | 0 | 0 | 0 | 18 | 0 | 0 | 0 | 0 | 0 | 0 | 26 | 53 | 15 | 0 | 0 | 0 | 0 | 0 | 0 | 0 |
| SRR10416516 | 8901 | 1876 | 720 | 1876 | 0 | 0 | 0 | 47 | 128 | 131 | 584 | 308 | 54 | 463 | 0 | 0 | 569 | 0 | 0 | 0 | 27 | 0 | 0 | 0 | 0 | 0 | 0 | 77 | 137 | 16 | 0 | 0 | 0 | 0 | 0 | 0 | 0 |
| SRR10416517 | 4666 | 1940 | 246 | 3110 | 0 | 0 | 11 | 59 | 109 | 62 | 603 | 347 | 75 | 480 | 0 | 0 | 916 | 0 | 0 | 0 | 26 | 0 | 0 | 0 | 0 | 0 | 0 | 74 | 153 | 15 | 0 | 0 | 0 | 0 | 0 | 0 | 0 |
| SRR10416518 | 748 | 2217 | 914 | 4544 | 0 | 0 | 0 | 12 | 51 | 83 | 165 | 112 | 0 | 146 | 0 | 0 | 270 | 0 | 0 | 0 | 13 | 0 | 0 | 0 | 0 | 0 | 0 | 0 | 13 | 19 | 0 | 0 | 0 | 0 | 0 | 0 | 0 |
| SRR10416519 | 290 | 1370 | 258 | 6157 | 0 | 0 | 0 | 0 | 0 | 48 | 59 | 143 | 0 | 59 | 0 | 0 | 185 | 0 | 0 | 0 | 0 | 0 | 0 | 0 | 0 | 0 | 0 | 0 | 0 | 0 | 0 | 0 | 0 | 0 | 0 | 0 | 0 |
| SRR10416520 | 193 | 3135 | 232 | 2902 | 0 | 0 | 0 | 0 | 0 | 39 | 43 | 78 | 0 | 83 | 0 | 0 | 105 | 0 | 0 | 0 | 0 | 0 | 0 | 0 | 0 | 0 | 0 | 0 | 0 | 0 | 0 | 0 | 0 | 0 | 0 | 0 | 0 |
| SRR10416521 | 370 | 3919 | 444 | 5693 | 0 | 0 | 0 | 56 | 135 | 74 | 231 | 209 | 25 | 560 | 0 | 0 | 320 | 0 | 0 | 0 | 0 | 0 | 0 | 0 | 0 | 0 | 0 | 0 | 0 | 0 | 0 | 0 | 0 | 0 | 0 | 0 | 0 |
| SRR10416522 | 90 | 1219 | 361 | 2844 | 0 | 0 | 0 | 0 | 0 | 90 | 28 | 26 | 0 | 16 | 0 | 0 | 217 | 0 | 0 | 0 | 0 | 0 | 0 | 0 | 0 | 0 | 0 | 0 | 0 | 0 | 0 | 0 | 0 | 0 | 0 | 0 | 0 |
| SRR10416523 | 1488 | 1886 | 168 | 1907 | 0 | 0 | 0 | 0 | 11 | 42 | 13 | 24 | 0 | 14 | 0 | 0 | 97 | 0 | 0 | 0 | 0 | 0 | 0 | 0 | 0 | 0 | 0 | 0 | 0 | 0 | 0 | 0 | 0 | 0 | 0 | 0 | 0 |
| SRR10416524 | 148 | 1805 | 266 | 1183 | 0 | 0 | 0 | 13 | 10 | 0 | 72 | 108 | 22 | 71 | 0 | 0 | 174 | 0 | 0 | 0 | 0 | 0 | 0 | 0 | 0 | 0 | 0 | 0 | 0 | 0 | 0 | 0 | 0 | 0 | 0 | 0 | 0 |
| SRR10416525 | 345 | 5471 | 86 | 5126 | 0 | 0 | 0 | 0 | 0 | 0 | 0 | 17 | 0 | 0 | 0 | 0 | 38 | 0 | 0 | 0 | 0 | 0 | 0 | 0 | 0 | 0 | 0 | 0 | 0 | 0 | 0 | 0 | 0 | 0 | 0 | 0 | 0 |
| SRR10416526 | 309 | 1621 | 39 | 2432 | 0 | 0 | 0 | 14 | 0 | 0 | 356 | 314 | 36 | 46 | 0 | 0 | 882 | 0 | 0 | 0 | 0 | 0 | 0 | 0 | 0 | 0 | 0 | 0 | 0 | 0 | 0 | 0 | 0 | 0 | 0 | 0 | 0 |
| SRR10416527 | 475 | 4414 | 380 | 5173 | 0 | 0 | 0 | 0 | 12 | 0 | 56 | 128 | 22 | 48 | 0 | 0 | 206 | 0 | 0 | 0 | 0 | 0 | 0 | 0 | 0 | 0 | 0 | 0 | 0 | 0 | 0 | 0 | 0 | 0 | 0 | 0 | 0 |
| SRR10416528 | 163 | 3257 | 529 | 2484 | 0 | 0 | 0 | 0 | 0 | 0 | 52 | 83 | 0 | 42 | 0 | 0 | 171 | 0 | 0 | 0 | 0 | 0 | 0 | 0 | 0 | 0 | 0 | 0 | 0 | 0 | 0 | 0 | 0 | 0 | 0 | 0 | 0 |
| SRR10416529 | 382 | 2944 | 327 | 1418 | 0 | 0 | 0 | 0 | 0 | 55 | 81 | 162 | 17 | 108 | 0 | 0 | 421 | 0 | 0 | 0 | 0 | 0 | 0 | 0 | 0 | 0 | 0 | 0 | 0 | 0 | 0 | 0 | 0 | 0 | 0 | 0 | 0 |
| SRR10416530 | 417 | 6249 | 655 | 3511 | 0 | 0 | 0 | 0 | 0 | 0 | 0 | 26 | 0 | 0 | 0 | 0 | 24 | 0 | 0 | 0 | 0 | 0 | 0 | 0 | 0 | 0 | 0 | 0 | 0 | 0 | 0 | 0 | 0 | 0 | 0 | 0 | 0 |
| SRR10416531 | 298 | 4667 | 695 | 1986 | 0 | 0 | 0 | 0 | 28 | 0 | 44 | 85 | 11 | 147 | 0 | 0 | 402 | 0 | 0 | 0 | 0 | 0 | 0 | 0 | 0 | 0 | 0 | 0 | 0 | 0 | 0 | 0 | 0 | 0 | 0 | 0 | 0 |
| SRR10416532 | 276 | 2556 | 207 | 3869 | 0 | 0 | 0 | 0 | 17 | 69 | 15 | 40 | 0 | 19 | 0 | 0 | 110 | 0 | 0 | 0 | 0 | 0 | 0 | 0 | 0 | 0 | 0 | 0 | 0 | 0 | 0 | 0 | 0 | 0 | 0 | 0 | 0 |

### Supp Table S8.1.Columns_B_to_AL

| | | | | | | | | | | | | | | | | | | | | | | | | | | | | | | | | | | | | | |
|---|---|---|---|---|---|---|---|---|---|---|---|---|---|---|---|---|---|---|---|---|---|---|---|---|---|---|---|---|---|---|---|---|---|---|---|---|---|
| SRR10416533 | 569 | 1662 | 437 | 2187 | 0 | 0 | 0 | 24 | 21 | 131 | 84 | 77 | 10 | 73 | 0 | 0 | 627 | 0 | 0 | 0 | 0 | 0 | 0 | 0 | 0 | 0 | 0 | 0 | 0 | 0 | 0 | 0 | 0 | 0 | 0 | 0 | 0 |
| SRR10416534 | 426 | 6045 | 341 | 4172 | 0 | 0 | 0 | 0 | 0 | 0 | 0 | 0 | 0 | 14 | 0 | 0 | 135 | 0 | 0 | 0 | 0 | 0 | 0 | 0 | 0 | 0 | 0 | 0 | 0 | 0 | 0 | 0 | 0 | 0 | 0 | 0 | 0 |
| SRR10416535 | 548 | 8084 | 274 | 4567 | 0 | 0 | 0 | 0 | 0 | 0 | 0 | 22 | 0 | 0 | 0 | 0 | 11 | 0 | 0 | 0 | 0 | 0 | 0 | 0 | 0 | 0 | 0 | 0 | 0 | 0 | 0 | 0 | 0 | 0 | 0 | 0 | 0 |
| SRR10416536 | 797 | 6907 | 310 | 4383 | 0 | 0 | 0 | 0 | 0 | 44 | 0 | 0 | 0 | 0 | 0 | 0 | 0 | 0 | 0 | 0 | 0 | 0 | 0 | 0 | 0 | 0 | 0 | 0 | 0 | 0 | 0 | 0 | 0 | 0 | 0 | 0 | 0 |
| SRR10416537 | 269 | 6988 | 134 | 3315 | 0 | 0 | 0 | 0 | 0 | 45 | 0 | 25 | 0 | 0 | 0 | 0 | 33 | 0 | 0 | 0 | 11 | 0 | 0 | 0 | 0 | 0 | 0 | 0 | 0 | 0 | 0 | 0 | 0 | 0 | 0 | 0 | 0 |
| SRR10416538 | 205 | 3618 | 171 | 2458 | 0 | 0 | 0 | 0 | 0 | 34 | 24 | 61 | 0 | 32 | 0 | 0 | 57 | 0 | 0 | 0 | 0 | 0 | 0 | 0 | 0 | 0 | 0 | 0 | 0 | 0 | 0 | 0 | 0 | 0 | 0 | 0 | 0 |
| SRR10416539 | 134 | 2141 | 201 | 3278 | 0 | 17 | 0 | 82 | 61 | 0 | 345 | 456 | 135 | 510 | 0 | 0 | 1456 | 0 | 0 | 0 | 0 | 0 | 0 | 0 | 0 | 0 | 0 | 0 | 0 | 0 | 0 | 0 | 0 | 0 | 0 | 0 | 0 |
| SRR10416540 | 252 | 5504 | 252 | 2979 | 0 | 0 | 0 | 0 | 0 | 0 | 0 | 21 | 0 | 0 | 0 | 0 | 21 | 0 | 0 | 0 | 0 | 0 | 0 | 0 | 0 | 0 | 0 | 0 | 0 | 0 | 0 | 0 | 0 | 0 | 0 | 0 | 0 |
| SRR10416541 | 114 | 2623 | 0 | 2623 | 0 | 0 | 0 | 0 | 11 | 0 | 86 | 266 | 13 | 113 | 0 | 0 | 355 | 0 | 0 | 0 | 0 | 0 | 0 | 0 | 0 | 0 | 0 | 0 | 0 | 0 | 0 | 0 | 0 | 0 | 0 | 0 | 0 |
| SRR10416542 | 361 | 3973 | 632 | 3793 | 0 | 0 | 0 | 0 | 0 | 0 | 0 | 73 | 0 | 20 | 0 | 0 | 73 | 0 | 0 | 0 | 0 | 0 | 0 | 0 | 0 | 0 | 0 | 0 | 0 | 0 | 0 | 0 | 0 | 0 | 0 | 0 | 0 |
| SRR10416543 | 7489 | 1695 | 291 | 2247 | 0 | 23 | 33 | 103 | 237 | 31 | 114 | 73 | 14 | 123 | 0 | 0 | 1653 | 0 | 0 | 0 | 43 | 0 | 0 | 0 | 0 | 0 | 0 | 115 | 338 | 42 | 0 | 0 | 0 | 0 | 0 | 0 | 0 |
| SRR10416544 | 858 | 2106 | 624 | 5069 | 0 | 0 | 0 | 0 | 0 | 0 | 37 | 97 | 0 | 34 | 0 | 0 | 37 | 0 | 0 | 0 | 0 | 0 | 0 | 0 | 0 | 0 | 0 | 0 | 0 | 0 | 0 | 0 | 0 | 0 | 0 | 0 | 0 |
| SRR10416545 | 1297 | 1935 | 338 | 1954 | 0 | 0 | 0 | 0 | 15 | 56 | 32 | 25 | 0 | 0 | 0 | 0 | 332 | 0 | 0 | 0 | 0 | 0 | 0 | 0 | 0 | 0 | 0 | 0 | 0 | 0 | 0 | 0 | 0 | 0 | 0 | 0 | 0 |
| SRR10416546 | 1253 | 2770 | 198 | 1055 | 0 | 0 | 0 | 12 | 26 | 0 | 229 | 249 | 18 | 53 | 0 | 0 | 412 | 0 | 0 | 0 | 0 | 0 | 0 | 0 | 0 | 0 | 0 | 0 | 0 | 0 | 0 | 0 | 0 | 0 | 0 | 0 | 0 |
| SRR10416547 | 1354 | 2079 | 290 | 919 | 0 | 0 | 0 | 0 | 0 | 48 | 97 | 52 | 14 | 43 | 0 | 0 | 167 | 0 | 0 | 0 | 0 | 0 | 0 | 0 | 0 | 0 | 0 | 0 | 0 | 0 | 0 | 0 | 0 | 0 | 0 | 0 | 0 |
| SRR10416549 | 2226 | 4770 | 545 | 1840 | 0 | 0 | 0 | 0 | 0 | 23 | 0 | 24 | 0 | 0 | 0 | 0 | 49 | 0 | 0 | 0 | 0 | 0 | 0 | 0 | 0 | 0 | 0 | 0 | 0 | 0 | 0 | 0 | 0 | 0 | 0 | 0 | 0 |
| SRR10416550 | 1208 | 10892 | 479 | 1041 | 0 | 0 | 0 | 0 | 13 | 0 | 0 | 20 | 0 | 11 | 0 | 0 | 317 | 0 | 0 | 0 | 12 | 0 | 0 | 0 | 0 | 0 | 0 | 0 | 0 | 0 | 0 | 0 | 0 | 0 | 0 | 0 | 0 |
| SRR10416551 | 1617 | 3738 | 318 | 2837 | 0 | 0 | 0 | 0 | 14 | 0 | 26 | 26 | 0 | 20 | 0 | 0 | 130 | 0 | 0 | 0 | 14 | 0 | 0 | 0 | 0 | 0 | 0 | 0 | 0 | 0 | 0 | 0 | 0 | 0 | 0 | 0 | 0 |
| SRR10416553 | 130 | 4279 | 778 | 2658 | 0 | 0 | 0 | 12 | 0 | 65 | 33 | 72 | 0 | 74 | 0 | 0 | 269 | 0 | 0 | 0 | 0 | 0 | 0 | 0 | 0 | 0 | 0 | 0 | 0 | 0 | 0 | 0 | 0 | 0 | 0 | 0 | 0 |
| SRR10416554 | 708 | 2478 | 283 | 3540 | 0 | 11 | 14 | 74 | 34 | 142 | 786 | 460 | 78 | 411 | 0 | 0 | 804 | 0 | 0 | 0 | 0 | 0 | 0 | 0 | 0 | 0 | 0 | 0 | 0 | 0 | 0 | 0 | 0 | 0 | 0 | 0 | 0 |
| SRR10416555 | 343 | 5940 | 114 | 2856 | 0 | 0 | 0 | 0 | 14 | 114 | 11 | 39 | 0 | 14 | 0 | 0 | 151 | 0 | 0 | 0 | 0 | 0 | 0 | 0 | 0 | 0 | 0 | 0 | 0 | 0 | 0 | 0 | 0 | 0 | 0 | 0 | 0 |
| SRR10416557 | 692 | 6574 | 461 | 2076 | 0 | 0 | 0 | 0 | 0 | 115 | 0 | 0 | 0 | 0 | 0 | 0 | 37 | 0 | 0 | 0 | 0 | 0 | 0 | 0 | 0 | 0 | 0 | 0 | 0 | 0 | 0 | 0 | 0 | 0 | 0 | 0 | 0 |
| SRR10416558 | 355 | 1267 | 101 | 2737 | 0 | 0 | 0 | 0 | 0 | 0 | 20 | 36 | 0 | 34 | 0 | 0 | 73 | 0 | 0 | 0 | 0 | 0 | 0 | 0 | 0 | 0 | 0 | 0 | 0 | 0 | 0 | 0 | 0 | 0 | 0 | 0 | 0 |
| SRR10416559 | 156 | 2802 | 389 | 2452 | 0 | 0 | 0 | 0 | 13 | 117 | 25 | 120 | 0 | 53 | 0 | 0 | 330 | 0 | 0 | 0 | 0 | 0 | 0 | 0 | 0 | 0 | 0 | 0 | 0 | 0 | 0 | 0 | 0 | 0 | 0 | 0 | 0 |
| SRR10416560 | 276 | 1518 | 529 | 2254 | 0 | 0 | 0 | 0 | 0 | 23 | 42 | 124 | 0 | 21 | 0 | 0 | 67 | 0 | 0 | 0 | 0 | 0 | 0 | 0 | 0 | 0 | 0 | 0 | 0 | 0 | 0 | 0 | 0 | 0 | 0 | 0 | 0 |
| SRR10416562 | 485 | 1664 | 312 | 2288 | 0 | 0 | 0 | 0 | 0 | 35 | 74 | 126 | 0 | 41 | 0 | 0 | 42 | 0 | 0 | 0 | 0 | 0 | 0 | 0 | 0 | 0 | 0 | 0 | 0 | 0 | 0 | 0 | 0 | 0 | 0 | 0 | 0 |
| SRR10416563 | 345 | 1185 | 210 | 4665 | 0 | 0 | 0 | 0 | 0 | 30 | 0 | 45 | 0 | 0 | 0 | 0 | 28 | 0 | 0 | 0 | 0 | 0 | 0 | 0 | 0 | 0 | 0 | 0 | 0 | 0 | 0 | 0 | 0 | 0 | 0 | 0 | 0 |
| SRR10416564 | 1894 | 3315 | 474 | 1377 | 0 | 0 | 0 | 0 | 0 | 43 | 30 | 44 | 0 | 20 | 0 | 0 | 152 | 0 | 0 | 0 | 0 | 0 | 0 | 0 | 0 | 0 | 0 | 0 | 0 | 0 | 0 | 0 | 0 | 0 | 0 | 0 | 0 |
| SRR10416565 | 1495 | 3083 | 498 | 1557 | 0 | 0 | 0 | 0 | 0 | 31 | 10 | 0 | 0 | 12 | 0 | 0 | 83 | 0 | 0 | 0 | 0 | 0 | 0 | 0 | 0 | 0 | 0 | 0 | 0 | 0 | 0 | 0 | 0 | 0 | 0 | 0 | 0 |
| SRR10416566 | 360 | 5333 | 144 | 721 | 0 | 0 | 0 | 0 | 0 | 0 | 0 | 29 | 0 | 0 | 0 | 0 | 53 | 0 | 0 | 0 | 0 | 0 | 0 | 0 | 0 | 0 | 0 | 0 | 0 | 0 | 0 | 0 | 0 | 0 | 0 | 0 | 0 |
| SRR10416567 | 214 | 1711 | 1604 | 2353 | 0 | 0 | 0 | 0 | 23 | 0 | 84 | 76 | 10 | 102 | 0 | 0 | 90 | 0 | 0 | 0 | 0 | 0 | 0 | 0 | 0 | 0 | 0 | 0 | 0 | 0 | 0 | 0 | 0 | 0 | 0 | 0 | 0 |
| SRR10416568 | 552 | 6067 | 473 | 3467 | 0 | 0 | 0 | 0 | 0 | 0 | 19 | 39 | 0 | 39 | 0 | 0 | 81 | 0 | 0 | 0 | 0 | 0 | 0 | 0 | 0 | 0 | 0 | 0 | 0 | 0 | 0 | 0 | 0 | 0 | 0 | 0 | 0 |
| SRR10416569 | 620 | 2145 | 1001 | 3051 | 0 | 0 | 0 | 0 | 0 | 0 | 24 | 54 | 0 | 39 | 0 | 0 | 191 | 0 | 0 | 0 | 0 | 0 | 0 | 0 | 0 | 0 | 0 | 0 | 0 | 0 | 0 | 0 | 0 | 0 | 0 | 0 | 0 |
| SRR10416570 | 633 | 2955 | 950 | 4327 | 0 | 12 | 22 | 115 | 182 | 53 | 377 | 244 | 47 | 229 | 0 | 0 | 1323 | 0 | 0 | 0 | 0 | 0 | 0 | 0 | 0 | 0 | 0 | 0 | 0 | 0 | 0 | 0 | 0 | 0 | 0 | 0 | 0 |
| SRR10416571 | 795 | 6602 | 367 | 2140 | 0 | 0 | 0 | 0 | 0 | 122 | 0 | 30 | 0 | 13 | 0 | 0 | 64 | 0 | 0 | 0 | 0 | 0 | 0 | 0 | 0 | 0 | 0 | 0 | 0 | 0 | 0 | 0 | 0 | 0 | 0 | 0 | 0 |
| SRR10416572 | 262 | 1351 | 785 | 3269 | 0 | 0 | 0 | 45 | 51 | 131 | 217 | 162 | 17 | 293 | 0 | 0 | 326 | 0 | 0 | 0 | 0 | 0 | 0 | 0 | 0 | 0 | 0 | 0 | 0 | 0 | 0 | 0 | 0 | 0 | 0 | 0 | 0 |
| SRR10416573 | 716 | 6849 | 511 | 1738 | 0 | 0 | 0 | 0 | 0 | 0 | 0 | 14 | 0 | 0 | 0 | 0 | 98 | 0 | 0 | 0 | 0 | 0 | 0 | 0 | 0 | 0 | 0 | 0 | 0 | 0 | 0 | 0 | 0 | 0 | 0 | 0 | 0 |
| SRR10416574 | 383 | 5743 | 861 | 2872 | 0 | 0 | 0 | 0 | 0 | 0 | 0 | 14 | 0 | 0 | 0 | 0 | 65 | 0 | 0 | 0 | 0 | 0 | 0 | 0 | 0 | 0 | 0 | 0 | 0 | 0 | 0 | 0 | 0 | 0 | 0 | 0 | 0 |
| SRR10416575 | 340 | 935 | 552 | 3909 | 0 | 0 | 0 | 0 | 0 | 0 | 21 | 67 | 0 | 13 | 0 | 0 | 28 | 0 | 0 | 0 | 0 | 0 | 0 | 0 | 0 | 0 | 0 | 0 | 0 | 0 | 0 | 0 | 0 | 0 | 0 | 0 | 0 |
| SRR10416576 | 141 | 932 | 226 | 6185 | 0 | 0 | 0 | 0 | 0 | 28 | 31 | 163 | 0 | 62 | 0 | 0 | 84 | 0 | 0 | 0 | 0 | 0 | 0 | 0 | 0 | 0 | 0 | 0 | 0 | 0 | 0 | 0 | 0 | 0 | 0 | 0 | 0 |
| SRR10416577 | 855 | 4558 | 1496 | 2635 | 0 | 0 | 0 | 0 | 11 | 71 | 15 | 26 | 0 | 0 | 0 | 0 | 272 | 0 | 0 | 0 | 0 | 0 | 0 | 0 | 0 | 0 | 0 | 0 | 0 | 0 | 0 | 0 | 0 | 0 | 0 | 0 | 0 |
| SRR10416578 | 0 | 1169 | 730 | 1899 | 0 | 0 | 0 | 26 | 42 | 146 | 155 | 95 | 13 | 287 | 0 | 0 | 405 | 0 | 0 | 0 | 0 | 0 | 0 | 0 | 0 | 0 | 0 | 0 | 0 | 0 | 0 | 0 | 0 | 0 | 0 | 0 | 0 |
| SRR10416579 | 68 | 5942 | 68 | 4098 | 0 | 0 | 0 | 0 | 0 | 137 | 10 | 40 | 0 | 12 | 0 | 0 | 93 | 0 | 0 | 0 | 0 | 0 | 0 | 0 | 0 | 0 | 0 | 0 | 0 | 0 | 0 | 0 | 0 | 0 | 0 | 0 | 0 |
| SRR10416580 | 311 | 6124 | 415 | 1972 | 0 | 0 | 0 | 0 | 0 | 52 | 13 | 46 | 0 | 31 | 0 | 0 | 248 | 0 | 0 | 0 | 0 | 0 | 0 | 0 | 0 | 0 | 0 | 0 | 0 | 0 | 0 | 0 | 0 | 0 | 0 | 0 | 0 |
| SRR10416581 | 505 | 3857 | 404 | 4241 | 0 | 0 | 0 | 0 | 0 | 0 | 15 | 79 | 0 | 17 | 0 | 0 | 43 | 0 | 0 | 0 | 0 | 0 | 0 | 0 | 0 | 0 | 0 | 0 | 0 | 0 | 0 | 0 | 0 | 0 | 0 | 0 | 0 |
| SRR10416582 | 552 | 3863 | 736 | 2115 | 0 | 0 | 0 | 0 | 0 | 92 | 36 | 44 | 0 | 24 | 0 | 0 | 77 | 0 | 0 | 0 | 0 | 0 | 0 | 0 | 0 | 0 | 0 | 0 | 0 | 0 | 0 | 0 | 0 | 0 | 0 | 0 | 0 |
| SRR10416583 | 418 | 3013 | 670 | 2678 | 0 | 14 | 0 | 81 | 43 | 84 | 559 | 277 | 33 | 349 | 0 | 0 | 1404 | 0 | 0 | 0 | 0 | 0 | 0 | 0 | 0 | 0 | 0 | 0 | 0 | 0 | 0 | 0 | 0 | 0 | 0 | 0 | 0 |
| SRR10416584 | 445 | 3868 | 489 | 2490 | 0 | 0 | 0 | 0 | 12 | 0 | 70 | 79 | 12 | 41 | 0 | 0 | 187 | 0 | 0 | 0 | 0 | 0 | 0 | 0 | 0 | 0 | 0 | 0 | 0 | 0 | 0 | 0 | 0 | 0 | 0 | 0 | 0 |
| SRR10416585 | 659 | 3624 | 329 | 2142 | 0 | 0 | 0 | 0 | 0 | 0 | 0 | 31 | 0 | 14 | 0 | 0 | 111 | 0 | 0 | 0 | 0 | 0 | 0 | 0 | 0 | 0 | 0 | 0 | 0 | 0 | 0 | 0 | 0 | 0 | 0 | 0 | 0 |
| SRR10416586 | 94 | 6865 | 94 | 4326 | 0 | 0 | 0 | 0 | 0 | 94 | 0 | 16 | 0 | 0 | 0 | 0 | 74 | 0 | 0 | 0 | 0 | 0 | 0 | 0 | 0 | 0 | 0 | 0 | 0 | 0 | 0 | 0 | 0 | 0 | 0 | 0 | 0 |
| SRR10416587 | 645 | 2001 | 387 | 4002 | 0 | 0 | 0 | 12 | 16 | 65 | 52 | 52 | 0 | 49 | 0 | 0 | 371 | 0 | 0 | 0 | 0 | 0 | 0 | 0 | 0 | 0 | 0 | 0 | 0 | 0 | 0 | 0 | 0 | 0 | 0 | 0 | 0 |
| SRR10416588 | 932 | 4739 | 233 | 2486 | 0 | 0 | 0 | 0 | 0 | 0 | 0 | 24 | 0 | 37 | 0 | 0 | 117 | 0 | 0 | 0 | 0 | 0 | 0 | 0 | 0 | 0 | 0 | 0 | 11 | 0 | 0 | 0 | 0 | 0 | 0 | 0 | 0 |
| SRR10416589 | 295 | 2631 | 369 | 7376 | 0 | 0 | 0 | 0 | 15 | 25 | 13 | 37 | 0 | 43 | 0 | 0 | 206 | 0 | 0 | 0 | 0 | 0 | 0 | 0 | 0 | 0 | 0 | 0 | 0 | 0 | 0 | 0 | 0 | 0 | 0 | 0 | 0 |
| SRR10416590 | 202 | 5522 | 337 | 3569 | 0 | 0 | 0 | 0 | 0 | 0 | 0 | 22 | 0 | 40 | 0 | 0 | 97 | 0 | 0 | 0 | 14 | 0 | 0 | 0 | 0 | 0 | 0 | 0 | 14 | 0 | 0 | 0 | 0 | 0 | 0 | 0 | 0 |
| SRR10416591 | 1255 | 6635 | 90 | 2511 | 0 | 0 | 0 | 0 | 0 | 0 | 0 | 25 | 0 | 0 | 0 | 0 | 55 | 0 | 0 | 0 | 0 | 0 | 0 | 0 | 0 | 0 | 0 | 0 | 0 | 0 | 0 | 0 | 0 | 0 | 0 | 0 | 0 |
| SRR10416592 | 454 | 2495 | 907 | 4672 | 0 | 0 | 0 | 13 | 70 | 91 | 117 | 135 | 21 | 123 | 0 | 0 | 356 | 0 | 0 | 0 | 0 | 0 | 0 | 0 | 0 | 0 | 0 | 0 | 0 | 0 | 0 | 0 | 0 | 0 | 0 | 0 | 0 |
| SRR10416593 | 234 | 8136 | 140 | 5284 | 0 | 0 | 0 | 0 | 0 | 0 | 0 | 0 | 0 | 0 | 0 | 0 | 12 | 0 | 0 | 0 | 0 | 0 | 0 | 0 | 0 | 0 | 0 | 0 | 0 | 0 | 0 | 0 | 0 | 0 | 0 | 0 | 0 |
| SRR10416594 | 901 | 13883 | 750 | 2026 | 0 | 0 | 0 | 0 | 0 | 0 | 0 | 0 | 0 | 0 | 0 | 0 | 66 | 0 | 0 | 0 | 0 | 0 | 0 | 0 | 0 | 0 | 0 | 0 | 0 | 0 | 0 | 0 | 0 | 0 | 0 | 0 | 0 |
| SRR10416595 | 879 | 2554 | 251 | 3350 | 0 | 0 | 0 | 0 | 0 | 126 | 37 | 49 | 0 | 23 | 0 | 0 | 35 | 0 | 0 | 0 | 0 | 0 | 0 | 0 | 0 | 0 | 0 | 0 | 0 | 0 | 0 | 0 | 0 | 0 | 0 | 0 | 0 |
| SRR10416596 | 424 | 1330 | 154 | 3470 | 0 | 0 | 0 | 0 | 0 | 39 | 10 | 62 | 0 | 0 | 0 | 0 | 15 | 0 | 0 | 0 | 0 | 0 | 0 | 0 | 0 | 0 | 0 | 0 | 0 | 0 | 0 | 0 | 0 | 0 | 0 | 0 | 0 |
| SRR10416597 | 576 | 1563 | 219 | 6363 | 0 | 0 | 0 | 0 | 15 | 27 | 0 | 24 | 0 | 21 | 0 | 0 | 30 | 0 | 0 | 0 | 0 | 0 | 0 | 0 | 0 | 0 | 0 | 0 | 11 | 0 | 0 | 0 | 0 | 0 | 0 | 0 | 0 |
| SRR10416598 | 303 | 963 | 550 | 3743 | 0 | 0 | 0 | 0 | 0 | 55 | 28 | 60 | 0 | 89 | 0 | 0 | 48 | 0 | 0 | 0 | 0 | 0 | 0 | 0 | 0 | 0 | 0 | 0 | 0 | 0 | 0 | 0 | 0 | 0 | 0 | 0 | 0 |
| SRR10416599 | 252 | 1639 | 946 | 2648 | 0 | 0 | 0 | 25 | 46 | 0 | 101 | 68 | 0 | 140 | 0 | 0 | 300 | 0 | 0 | 0 | 0 | 0 | 0 | 0 | 0 | 0 | 0 | 0 | 0 | 0 | 0 | 0 | 0 | 0 | 0 | 0 | 0 |
| SRR10416600 | 100 | 1555 | 259 | 7378 | 0 | 0 | 0 | 0 | 0 | 100 | 35 | 111 | 14 | 91 | 0 | 0 | 79 | 0 | 0 | 0 | 0 | 0 | 0 | 0 | 0 | 0 | 0 | 0 | 0 | 0 | 0 | 0 | 0 | 0 | 0 | 0 | 0 |
| SRR10428553 | 76 | 6423 | 3443 | 1033 | 147 | 1343 | 1625 | 11256 | 16508 | 21 | 8612 | 25762 | 2301 | 10095 | 0 | 3847 | 36968 | 0 | 0 | 0 | 177 | 932 | 1343 | 74 | 80 | 0 | 0 | 0 | 21 | 86 | 679 | 1128 | 16 | 219 | 426 | 0 | 0 |
| SRR10428554 | 68 | 6738 | 3824 | 388 | 104 | 1349 | 1964 | 5149 | 9554 | 42 | 9416 | 17676 | 2684 | 15160 | 0 | 906 | 54126 | 0 | 0 | 0 | 176 | 734 | 245 | 0 | 0 | 0 | 0 | 0 | 31 | 98 | 780 | 254 | 0 | 10 | 10 | 0 | 0 |
| SRR10428555 | 90 | 4302 | 1759 | 793 | 75 | 669 | 1278 | 5399 | 19578 | 51 | 9171 | 17486 | 2467 | 14428 | 0 | 1097 | 91855 | 0 | 0 | 0 | 152 | 509 | 683 | 13 | 17 | 0 | 0 | 0 | 11 | 69 | 435 | 545 | 0 | 38 | 51 | 0 | 0 |
| SRR10428556 | 63 | 5204 | 3057 | 300 | 212 | 2057 | 2592 | 9042 | 15761 | 16 | 15078 | 22865 | 2898 | 13168 | 0 | 3072 | 49678 | 0 | 0 | 0 | 945 | 901 | 195 | 0 | 29 | 0 | 0 | 0 | 74 | 592 | 776 | 183 | 0 | 95 | 221 | 0 | 0 |
| SRR10428557 | 104 | 5684 | 1494 | 507 | 334 | 2208 | 4240 | 11291 | 3858 | 24 | 15564 | 34181 | 3886 | 7998 | 0 | 6435 | 83384 | 0 | 0 | 0 | 513 | 1678 | 434 | 0 | 0 | 0 | 0 | 0 | 34 | 168 | 1356 | 280 | 0 | 0 | 16 | 0 | 0 |
| SRR10428558 | 84 | 7820 | 4546 | 208 | 127 | 1244 | 1714 | 6536 | 4154 | 15 | 14376 | 33982 | 3144 | 8526 | 0 | 5048 | 85361 | 0 | 0 | 0 | 4816 | 2368 | 730 | 0 | 0 | 0 | 0 | 0 | 103 | 2003 | 2023 | 600 | 0 | 20 | 37 | 0 | 0 |
| SRR10428559 | 65 | 10219 | 3678 | 263 | 362 | 4337 | 5059 | 16122 | 14629 | 15 | 14014 | 27748 | 3142 | 9485 | 0 | 3925 | 75644 | 0 | 0 | 0 | 1340 | 2627 | 705 | 0 | 0 | 0 | 0 | 0 | 102 | 823 | 2677 | 839 | 0 | 34 | 48 | 0 | 0 |
| SRR10428560 | 32 | 5999 | 5919 | 101 | 171 | 1724 | 2278 | 6528 | 9466 | 0 | 14235 | 19535 | 3809 | 11365 | 0 | 3986 | 47201 | 0 | 0 | 0 | 1416 | 2402 | 602 | 16 | 15 | 0 | 0 | 0 | 114 | 1215 | 3143 | 845 | 0 | 62 | 102 | 0 | 0 |
| SRR10446207 | 199 | 361 | 16708 | 82 | 3446 | 1323 | 11946 | 26776 | 9847 | 3452 | 29612 | 34884 | 15205 | 2007 | 213 | 932 | 35241 | 0 | 0 | 0 | 37 | 46 | 0 | 0 | 0 | 0 | 0 | 0 | 0 | 27 | 25 | 0 | 0 | 0 | 0 | 0 | 0 |
| SRR10446208 | 224 | 726 | 29431 | 266 | 2408 | 1506 | 12186 | 21636 | 18110 | 3293 | 16 | 27 | 13184 | 3932 | 161 | 1424 | 44963 | 0 | 0 | 0 | 72 | 97 | 0 | 0 | 0 | 0 | 0 | 0 | 32 | 189 | 158 | 38 | 0 | 0 | 0 | 0 | 0 |
| SRR10446209 | 44 | 310 | 11429 | 388 | 1980 | 1376 | 9485 | 18792 | 29987 | 4172 | 19 | 17 | 12488 | 4021 | 144 | 945 | 43077 | 0 | 0 | 0 | 81 | 107 | 0 | 0 | 0 | 0 | 0 | 0 | 0 | 77 | 55 | 17 | 0 | 0 | 0 | 0 | 0 |
| SRR10446210 | 79 | 125 | 12189 | 243 | 14971 | 7115 | 58549 | 89424 | 20314 | 3989 | 27 | 23 | 12474 | 3028 | 307 | 1601 | 18386 | 0 | 0 | 0 | 0 | 16 | 0 | 0 | 0 | 0 | 0 | 0 | 0 | 14 | 47 | 24 | 0 | 0 | 0 | 0 | 0 |
| SRR10446211 | 141 | 745 | 6272 | 2322 | 5004 | 2684 | 34057 | 45038 | 7357 | 1423 | 12 | 34 | 12098 | 10695 | 138 | 983 | 60140 | 0 | 46 | 22 | 188 | 941 | 34 | 0 | 0 | 0 | 0 | 219 | 67 | 180 | 362 | 44 | 0 | 0 | 0 | 0 | 0 |
| SRR10446212 | 255 | 1168 | 12589 | 2043 | 3067 | 2424 | 17458 | 34648 | 20795 | 4038 | 22 | 38 | 12655 | 6829 | 230 | 487 | 25719 | 0 | 0 | 0 | 76 | 105 | 0 | 0 | 0 | 0 | 0 | 55 | 82 | 167 | 117 | 20 | 0 | 0 | 0 | 0 | 0 |
| SRR10531363 | 558 | 4864 | 3181 | 0 | 42 | 45 | 429 | 276 | 172962 | 0 | 1515 | 27 | 168 | 5474 | 51 | 60 | 237 | 0 | 0 | 0 | 0 | 0 | 0 | 0 | 0 | 0 | 0 | 0 | 0 | 0 | 0 | 0 | 0 | 0 | 0 | 0 | 0 |
| SRR10531364 | 1233 | 3375 | 2702 | 99 | 39 | 34 | 427 | 399 | 491479 | 0 | 3557 | 3385 | 192 | 2942 | 133 | 107 | 271 | 0 | 0 | 0 | 0 | 0 | 0 | 0 | 0 | 0 | 0 | 0 | 0 | 0 | 0 | 0 | 0 | 0 | 0 | 0 | 0 |
| SRR10531365 | 1377 | 3515 | 4237 | 128 | 0 | 0 | 193 | 109 | 682233 | 155 | 1407 | 1197 | 104 | 1302 | 71 | 35 | 108 | 0 | 0 | 0 | 0 | 0 | 0 | 0 | 0 | 0 | 0 | 0 | 0 | 0 | 0 | 0 | 0 | 0 | 0 | 0 | 0 |
| SRR10531367 | 1959 | 4922 | 6295 | 0 | 51 | 153 | 1041 | 181 | 220680 | 0 | 2619 | 3923 | 107 | 3974 | 549 | 36 | 423 | 0 | 0 | 0 | 0 | 0 | 0 | 0 | 0 | 0 | 0 | 0 | 0 | 0 | 0 | 0 | 0 | 0 | 0 | 0 | 0 |
| SRR10531368 | 2451 | 7222 | 6155 | 138 | 78 | 90 | 693 | 149 | 326886 | 0 | 1615 | 11 | 70 | 3752 | 201 | 74 | 438 | 0 | 0 | 0 | 0 | 0 | 0 | 0 | 0 | 0 | 0 | 0 | 0 | 0 | 0 | 0 | 0 | 0 | 0 | 0 | 0 |
| SRR10531369 | 1384 | 4844 | 10208 | 0 | 23 | 36 | 363 | 129 | 442945 | 0 | 2586 | 13 | 196 | 3515 | 183 | 72 | 203 | 0 | 0 | 0 | 0 | 0 | 0 | 0 | 0 | 0 | 0 | 0 | 0 | 0 | 0 | 0 | 0 | 0 | 0 | 0 | 0 |
| SRR10531370 | 1117 | 5297 | 8000 | 540 | 120 | 115 | 1868 | 358 | 210210 | 0 | 2120 | 16 | 134 | 10702 | 378 | 40 | 746 | 0 | 0 | 0 | 0 | 0 | 0 | 0 | 0 | 0 | 0 | 0 | 0 | 0 | 0 | 0 | 0 | 0 | 0 | 0 | 0 |
| SRR10544939 | 21 | 244 | 42 | 982 | 438 | 294 | 7050 | 1892 | 349 | 0 | 289 | 1090 | 62 | 27 | 276 | 0 | 1115 | 0 | 0 | 0 | 80 | 17 | 19 | 0 | 0 | 0 | 0 | 0 | 0 | 39 | 13 | 0 | 0 | 0 | 0 | 0 | 0 |
| SRR10544940 | 0 | 155 | 76 | 300 | 344 | 212 | 7336 | 1802 | 267 | 0 | 231 | 1003 | 77 | 28 | 153 | 0 | 966 | 0 | 0 | 0 | 131 | 41 | 32 | 0 | 0 | 0 | 0 | 0 | 0 | 62 | 30 | 0 | 0 | 0 | 0 | 0 | 0 |
| SRR10544941 | 23 | 336 | 127 | 404 | 727 | 381 | 14276 | 3575 | 263 | 21 | 412 | 1318 | 104 | 51 | 295 | 0 | 1616 | 0 | 0 | 0 | 188 | 59 | 60 | 13 | 15 | 0 | 0 | 0 | 0 | 107 | 46 | 15 | 0 | 0 | 0 | 0 | 0 |
| SRR10544942 | 0 | 156 | 65 | 574 | 151 | 75 | 2139 | 765 | 253 | 0 | 300 | 795 | 122 | 47 | 1115 | 0 | 1240 | 0 | 0 | 0 | 523 | 56 | 37 | 0 | 0 | 0 | 0 | 0 | 0 | 247 | 39 | 0 | 0 | 0 | 0 | 0 | 0 |
| SRR10544943 | 12 | 177 | 52 | 189 | 199 | 112 | 3812 | 1124 | 297 | 0 | 469 | 1903 | 91 | 46 | 358 | 0 | 1361 | 0 | 0 | 0 | 46 | 16 | 18 | 0 | 0 | 0 | 0 | 0 | 0 | 23 | 11 | 0 | 0 | 0 | 0 | 0 | 0 |
| SRR10544944 | 20 | 193 | 52 | 349 | 311 | 149 | 5371 | 2043 | 697 | 0 | 404 | 1369 | 115 | 34 | 657 | 0 | 2141 | 0 | 0 | 0 | 83 | 27 | 41 | 12 | 0 | 0 | 0 | 0 | 0 | 50 | 29 | 15 | 12 | 0 | 0 | 0 | 0 |
| SRR10544945 | 0 | 34 | 0 | 19 | 28 | 0 | 586 | 146 | 1151 | 0 | 844 | 2650 | 300 | 28 | 174 | 160 | 614 | 0 | 0 | 0 | 0 | 0 | 0 | 0 | 0 | 0 | 0 | 0 | 0 | 0 | 0 | 0 | 0 | 0 | 0 | 0 | 0 |
| SRR10544946 | 0 | 41 | 0 | 49 | 0 | 0 | 267 | 70 | 622 | 0 | 837 | 2616 | 197 | 67 | 434 | 106 | 334 | 0 | 0 | 0 | 11 | 11 | 11 | 0 | 0 | 0 | 0 | 0 | 0 | 0 | 0 | 0 | 0 | 0 | 0 | 0 | 0 |
| SRR10544947 | 0 | 59 | 0 | 42 | 32 | 0 | 352 | 127 | 1036 | 0 | 867 | 1783 | 271 | 67 | 152 | 148 | 497 | 0 | 0 | 0 | 11 | 0 | 0 | 0 | 0 | 0 | 0 | 0 | 0 | 0 | 0 | 0 | 0 | 0 | 0 | 0 | 0 |
| SRR10544948 | 0 | 46 | 0 | 59 | 0 | 0 | 224 | 58 | 605 | 0 | 828 | 1558 | 255 | 67 | 519 | 121 | 300 | 0 | 0 | 0 | 34 | 18 | 14 | 0 | 0 | 0 | 0 | 0 | 0 | 17 | 0 | 0 | 0 | 0 | 0 | 0 | 0 |
| SRR10544949 | 0 | 76 | 0 | 53 | 0 | 0 | 289 | 92 | 513 | 0 | 1184 | 4039 | 285 | 123 | 254 | 219 | 355 | 0 | 0 | 0 | 15 | 0 | 12 | 0 | 0 | 0 | 0 | 0 | 0 | 10 | 0 | 0 | 0 | 0 | 0 | 0 | 0 |
| SRR10544950 | 0 | 79 | 0 | 45 | 14 | 0 | 392 | 168 | 658 | 0 | 1261 | 3733 | 413 | 105 | 995 | 217 | 560 | 0 | 0 | 0 | 12 | 13 | 12 | 0 | 0 | 0 | 0 | 0 | 0 | 0 | 0 | 0 | 0 | 0 | 0 | 0 | 0 |
| SRR10756575 | 1246 | 40543 | 65991 | 966 | 6030 | 6655 | 32750 | 6239 | 13602 | 243 | 36025 | 17889 | 85338 | 48135 | 3120 | 43 | 104134 | 0 | 0 | 0 | 14 | 265 | 39 | 12 | 0 | 0 | 0 | 0 | 24 | 20 | 13 | 0 | 0 | 0 | 0 | 0 | 0 |
| SRR10756576 | 687 | 12925 | 63785 | 450 | 11434 | 11007 | 55617 | 10622 | 9483 | 251 | 42605 | 19902 | 79391 | 53087 | 3444 | 26 | 108759 | 0 | 0 | 24 | 25 | 404 | 52 | 17 | 0 | 0 | 0 | 13 | 79 | 33 | 16 | 0 | 0 | 0 | 0 | 0 | 0 |
| SRR10756577 | 196 | 9099 | 108790 | 161 | 8445 | 8931 | 47805 | 13166 | 12866 | 833 | 44070 | 18561 | 85466 | 49235 | 3429 | 41 | 99547 | 0 | 0 | 0 | 12 | 137 | 38 | 14 | 0 | 0 | 0 | 0 | 12 | 19 | 10 | 0 | 0 | 0 | 0 | 0 | 0 |
| SRR10756578 | 845 | 8603 | 98587 | 504 | 5935 | 6185 | 38799 | 11633 | 14332 | 867 | 51687 | 22083 | 92688 | 44064 | 2693 | 42 | 81730 | 0 | 0 | 0 | 21 | 72 | 38 | 0 | 0 | 0 | 0 | 15 | 100 | 49 | 18 | 0 | 0 | 0 | 0 | 0 | 0 |
| SRR10756579 | 700 | 13953 | 115923 | 596 | 10024 | 10222 | 52462 | 12475 | 17328 | 712 | 45169 | 17605 | 90439 | 49889 | 3624 | 39 | 85277 | 0 | 0 | 0 | 18 | 150 | 37 | 12 | 0 | 0 | 0 | 0 | 23 | 49 | 0 | 0 | 0 | 0 | 0 | 0 | 0 |
| SRR10756581 | 917 | 4189 | 59889 | 815 | 9974 | 11580 | 54017 | 9424 | 27130 | 1034 | 35963 | 19 | 66395 | 48691 | 3773 | 55 | 80811 | 0 | 0 | 0 | 29 | 95 | 42 | 0 | 0 | 0 | 0 | 24 | 79 | 159 | 23 | 0 | 0 | 0 | 0 | 0 | 0 |
| SRR10756582 | 1619 | 12961 | 39481 | 1427 | 9986 | 12476 | 48971 | 6605 | 36202 | 340 | 37362 | 18120 | 50075 | 61433 | 4363 | 53 | 92403 | 0 | 0 | 0 | 0 | 62 | 20 | 0 | 0 | 0 | 0 | 0 | 71 | 51 | 16 | 0 | 0 | 0 | 0 | 0 | 0 |
| SRR10756583 | 360 | 3192 | 81598 | 410 | 7225 | 9354 | 45524 | 8021 | 22467 | 469 | 31052 | 12765 | 29803 | 48465 | 2431 | 75 | 106782 | 0 | 0 | 0 | 0 | 45 | 12 | 0 | 0 | 0 | 0 | 0 | 29 | 19 | 13 | 0 | 0 | 0 | 0 | 0 | 0 |
| SRR10875785 | 231 | 5861 | 22281 | 526 | 3219 | 7260 | 16226 | 10020 | 476269 | 73 | 7321 | 2827 | 10064 | 3744 | 12350 | 17 | 16459 | 0 | 12 | 27 | 82 | 486 | 68 | 100 | 107 | 41 | 0 | 19 | 48 | 78 | 23 | 0 | 0 | 0 | 19 | 50 | 0 |
| SRR10875786 | 92 | 643 | 1671 | 496 | 7370 | 11850 | 42341 | 21488 | 422404 | 100 | 10707 | 3255 | 9955 | 3966 | 17878 | 68 | 14839 | 0 | 0 | 0 | 11 | 33 | 12 | 68 | 54 | 21 | 0 | 0 | 0 | 0 | 0 | 0 | 0 | 0 | 13 | 37 | 0 |
| SRR10953234 | 2175 | 7094 | 1794 | 4558 | 30836 | 26365 | 250620 | 94616 | 13630 | 1458 | 17595 | 3128 | 44452 | 19104 | 6645 | 32 | 13312 | 0 | 27 | 0 | 82 | 106 | 135 | 31 | 17 | 0 | 39 | 126 | 208 | 106 | 110 | 0 | 0 | 0 | 0 | 0 | 0 |
| SRR10953235 | 1881 | 5308 | 2064 | 4788 | 31663 | 30112 | 262109 | 76194 | 15243 | 1481 | 22642 | 4239 | 48322 | 20690 | 6379 | 33 | 14210 | 0 | 22 | 0 | 108 | 114 | 119 | 25 | 16 | 0 | 25 | 94 | 158 | 100 | 85 | 0 | 0 | 0 | 0 | 0 | 0 |
| SRR10953236 | 1738 | 5168 | 2297 | 4594 | 32253 | 29854 | 262974 | 82940 | 13158 | 1731 | 19297 | 3564 | 49143 | 18285 | 7300 | 37 | 15298 | 0 | 21 | 0 | 45 | 58 | 95 | 20 | 12 | 0 | 23 | 86 | 138 | 64 | 69 | 0 | 0 | 0 | 0 | 0 | 0 |
| SRR10953237 | 1660 | 4728 | 2472 | 6634 | 38394 | 34788 | 293659 | 81129 | 14486 | 1027 | 16758 | 2416 | 23527 | 11599 | 6850 | 34 | 13524 | 0 | 26 | 11 | 48 | 56 | 93 | 19 | 0 | 0 | 24 | 102 | 160 | 68 | 97 | 0 | 0 | 0 | 0 | 0 | 0 |
| SRR10953238 | 1675 | 5647 | 1796 | 5140 | 23454 | 21804 | 195664 | 62464 | 11365 | 1683 | 15385 | 3054 | 61856 | 24435 | 4986 | 26 | 21065 | 0 | 26 | 0 | 75 | 147 | 130 | 30 | 20 | 0 | 28 | 95 | 149 | 99 | 75 | 0 | 0 | 0 | 0 | 0 | 0 |
| SRR10953239 | 1606 | 6050 | 3233 | 6451 | 27576 | 25189 | 248925 | 51699 | 11061 | 1759 | 20383 | 3600 | 60792 | 22206 | 4930 | 26 | 18086 | 0 | 18 | 0 | 76 | 136 | 117 | 25 | 21 | 0 | 18 | 68 | 117 | 89 | 55 | 0 | 0 | 0 | 0 | 12 | 0 |
| SRR1107492 | 202 | 62015 | 5427 | 699 | 3441 | 49005 | 32231 | 45510 | 6455 | 0 | 27469 | 21481 | 22715 | 10321 | 12 | 61 | 48676 | 0 | 0 | 0 | 0 | 0 | 0 | 0 | 0 | 0 | 0 | 0 | 0 | 0 | 0 | 0 | 0 | 0 | 0 | 0 | 0 |
| SRR1107493 | 213 | 40260 | 4658 | 562 | 3049 | 44365 | 17915 | 39687 | 5632 | 0 | 26132 | 20350 | 20649 | 9153 | 14 | 69 | 31459 | 0 | 0 | 0 | 0 | 0 | 0 | 0 | 0 | 0 | 0 | 0 | 0 | 16 | 62 | 11 | 0 | 0 | 0 | 0 | 0 |
| SRR1107494 | 205 | 47649 | 3135 | 808 | 2087 | 26776 | 20396 | 32369 | 4065 | 18 | 18358 | 16958 | 16186 | 5893 | 13 | 48 | 55059 | 0 | 0 | 0 | 28 | 42 | 12 | 0 | 0 | 0 | 0 | 0 | 34 | 57 | 233 | 38 | 0 | 0 | 0 | 0 | 0 |
| SRR1107495 | 169 | 46696 | 3236 | 812 | 2098 | 28048 | 20778 | 32224 | 4102 | 19 | 19809 | 17982 | 16871 | 6547 | 11 | 48 | 57599 | 0 | 0 | 0 | 26 | 36 | 11 | 0 | 0 | 0 | 0 | 0 | 31 | 60 | 230 | 39 | 0 | 0 | 0 | 0 | 0 |
| SRR1107496 | 216 | 50581 | 2898 | 617 | 2818 | 32345 | 28115 | 35171 | 4468 | 0 | 17662 | 20918 | 15184 | 5380 | 0 | 44 | 57533 | 0 | 0 | 0 | 31 | 40 | 18 | 0 | 0 | 0 | 0 | 0 | 37 | 75 | 246 | 49 | 0 | 0 | 0 | 0 | 0 |
| SRR1107497 | 148 | 51815 | 3119 | 619 | 2825 | 32891 | 29140 | 36588 | 4447 | 13 | 19178 | 22430 | 15703 | 5600 | 0 | 46 | 60406 | 0 | 0 | 0 | 26 | 31 | 14 | 0 | 0 | 0 | 0 | 0 | 32 | 55 | 221 | 44 | 0 | 0 | 0 | 0 | 0 |
| SRR1159512 | 82 | 85419 | 3607 | 372 | 3860 | 35032 | 48315 | 38178 | 2859 | 48 | 24277 | 36988 | 4652 | 8902 | 54 | 49217 | 15547 | 0 | 0 | 0 | 287 | 137 | 63 | 0 | 0 | 0 | 0 | 0 | 27 | 78 | 102 | 29 | 0 | 0 | 0 | 0 | 0 |
| SRR1207050 | 66 | 3353 | 1171 | 0 | 402 | 121 | 1880 | 3362 | 19 | 0 | 60677 | 138 | 7255 | 7398 | 0 | 417 | 248 | 0 | 0 | 0 | 0 | 16 | 24 | 77 | 65 | 0 | 0 | 0 | 0 | 0 | 0 | 14 | 51 | 401 | 91 | 47 | 0 |
| SRR1207051 | 390 | 19243 | 36623 | 0 | 1529 | 611 | 7605 | 3820 | 12 | 31 | 78876 | 307356 | 8185 | 5280 | 22 | 370 | 226 | 0 | 0 | 0 | 0 | 18 | 31 | 20 | 15 | 0 | 0 | 0 | 0 | 0 | 0 | 10 | 23 | 86 | 62 | 25 | 0 |
| SRR1207053 | 31 | 14776 | 27972 | 0 | 1129 | 451 | 5924 | 4082 | 30 | 28 | 98208 | 222 | 8697 | 5924 | 21 | 115 | 227 | 0 | 0 | 0 | 0 | 11 | 44 | 130 | 58 | 16 | 0 | 0 | 0 | 0 | 0 | 0 | 11 | 78 | 133 | 130 | 0 |
| SRR1207054 | 66 | 1984 | 872 | 0 | 718 | 294 | 3974 | 3120 | 28 | 0 | 79364 | 216523 | 9542 | 7677 | 15 | 371 | 295 | 0 | 12 | 0 | 0 | 62 | 37 | 147 | 72 | 16 | 0 | 0 | 0 | 0 | 0 | 38 | 127 | 655 | 233 | 139 | 0 |

### Supp Table S8.1.Columns_B_to_AL

| | | | | | | | | | | | | | | | | | | | | | | | | | | | | | | | | | | | | | |
|---|---|---|---|---|---|---|---|---|---|---|---|---|---|---|---|---|---|---|---|---|---|---|---|---|---|---|---|---|---|---|---|---|---|---|---|---|---|
| SRR1207055 | 66 | 11301 | 31098 | 0 | 1051 | 462 | 5715 | 3737 | 18 | 122 | 91604 | 101 | 8541 | 5664 | 26 | 283 | 255 | 0 | 0 | 0 | 0 | 17 | 105 | 546 | 149 | 59 | 0 | 0 | 0 | 0 | 0 | 38 | 229 | 1186 | 390 | 666 | 0 |
| SRR1232072 | 112 | 15110 | 4294 | 3690 | 814 | 8922 | 6533 | 15012 | 14702 | 11 | 4575 | 4943 | 1722 | 1634 | 11 | 8233 | 13928 | 0 | 0 | 0 | 70 | 312 | 19 | 0 | 0 | 0 | 0 | 14 | 113 | 148 | 1393 | 119 | 0 | 36 | 0 | 0 | 0 |
| SRR1232073 | 186 | 20866 | 9897 | 2146 | 2721 | 43829 | 33409 | 36518 | 6371 | 13 | 15069 | 11624 | 3857 | 7741 | 0 | 2442 | 34179 | 0 | 0 | 0 | 90 | 181 | 16 | 0 | 0 | 0 | 0 | 12 | 99 | 222 | 1087 | 130 | 0 | 12 | 0 | 0 | 0 |
| SRR1240812 | 20 | 1571 | 258 | 118 | 5454 | 19474 | 37207 | 10026 | 2448 | 84 | 16886 | 62999 | 6282 | 15964 | 745 | 131928 | 10161 | 0 | 0 | 0 | 642 | 62 | 131 | 0 | 0 | 0 | 0 | 0 | 27 | 261 | 46 | 93 | 0 | 0 | 0 | 10 | 0 |
| SRR1240813 | 32 | 2107 | 194 | 105 | 3171 | 18280 | 26498 | 6164 | 1914 | 69 | 15532 | 69631 | 5587 | 14979 | 110 | 121796 | 9575 | 0 | 0 | 0 | 683 | 57 | 130 | 0 | 14 | 0 | 0 | 0 | 24 | 224 | 49 | 66 | 0 | 17 | 0 | 0 | 0 |
| SRR1240814 | 25 | 15208 | 305 | 119 | 7470 | 100573 | 72449 | 26931 | 63 | 36 | 9035 | 35877 | 2861 | 9863 | 41 | 69924 | 5930 | 0 | 0 | 0 | 315 | 16 | 53 | 0 | 0 | 0 | 0 | 0 | 20 | 94 | 25 | 29 | 0 | 0 | 0 | 0 | 0 |
| SRR1240815 | 33 | 14384 | 255 | 70 | 7185 | 96830 | 74234 | 24101 | 54 | 54 | 8451 | 37182 | 2726 | 9633 | 56 | 78801 | 5692 | 0 | 0 | 0 | 264 | 19 | 38 | 0 | 0 | 0 | 0 | 0 | 14 | 96 | 22 | 28 | 0 | 0 | 0 | 0 | 0 |
| SRR1240816 | 55 | 2828 | 217 | 113 | 115 | 608 | 926 | 262 | 13275 | 289 | 2379 | 7532 | 751 | 105 | 547 | 9223 | 4795 | 0 | 0 | 0 | 1296 | 93 | 368 | 14 | 33 | 0 | 0 | 0 | 25 | 352 | 44 | 209 | 0 | 36 | 0 | 0 | 0 |
| SRR1240817 | 62 | 2771 | 259 | 135 | 96 | 628 | 853 | 261 | 12466 | 298 | 2402 | 7508 | 742 | 99 | 566 | 8668 | 4679 | 0 | 0 | 0 | 1499 | 87 | 327 | 16 | 30 | 0 | 0 | 0 | 20 | 436 | 49 | 173 | 0 | 38 | 0 | 0 | 0 |
| SRR1240818 | 2245 | 27505 | 596 | 967 | 2020 | 23914 | 18133 | 10285 | 1769 | 16 | 15871 | 93636 | 3305 | 8639 | 11 | 142283 | 9660 | 0 | 17 | 33 | 2377 | 543 | 158 | 0 | 0 | 0 | 0 | 79 | 501 | 506 | 235 | 78 | 0 | 0 | 0 | 0 | 0 |
| SRR1241600 | 117 | 51227 | 1585 | 54 | 102 | 706 | 4468 | 1082 | 18912 | 47 | 2547 | 4825 | 512 | 827 | 177 | 455 | 3130 | 0 | 0 | 0 | 36 | 17 | 120 | 41 | 31 | 0 | 0 | 0 | 0 | 118 | 16 | 107 | 26 | 0 | 33 | 231 | 0 |
| SRR1241601 | 326 | 37748 | 936 | 67 | 577 | 6649 | 19241 | 4702 | 1486 | 80 | 1702 | 3245 | 115 | 2048 | 87 | 1370 | 3649 | 0 | 0 | 0 | 55 | 26 | 136 | 58 | 17 | 0 | 0 | 0 | 0 | 151 | 30 | 197 | 50 | 13 | 27 | 186 | 0 |
| SRR1241602 | 94 | 44598 | 1306 | 35 | 773 | 7901 | 23451 | 5556 | 2132 | 22 | 2292 | 1506 | 253 | 2811 | 40 | 2105 | 6782 | 0 | 0 | 0 | 18 | 17 | 43 | 0 | 0 | 0 | 0 | 0 | 0 | 74 | 46 | 138 | 19 | 0 | 20 | 227 | 0 |
| SRR1241603 | 0 | 6817 | 290 | 4631 | 67 | 15574 | 1271 | 376 | 7712 | 121 | 4563 | 2475 | 401 | 557 | 28 | 4296 | 8220 | 0 | 0 | 0 | 2043 | 397 | 610 | 0 | 45 | 0 | 0 | 10 | 111 | 385 | 254 | 197 | 0 | 33 | 20 | 52 | 0 |
| SRR1241604 | 0 | 17311 | 234 | 2631 | 110 | 2419 | 2577 | 665 | 26685 | 62 | 25822 | 4845 | 3439 | 6474 | 21 | 6875 | 20163 | 0 | 0 | 0 | 1665 | 459 | 527 | 0 | 32 | 0 | 0 | 0 | 89 | 315 | 280 | 180 | 0 | 27 | 17 | 29 | 0 |
| SRR1241605 | 0 | 10350 | 224 | 2893 | 86 | 1762 | 1876 | 468 | 9609 | 92 | 19285 | 7235 | 2444 | 5196 | 17 | 5659 | 26199 | 0 | 0 | 0 | 1893 | 534 | 764 | 10 | 37 | 0 | 0 | 0 | 61 | 300 | 238 | 124 | 0 | 24 | 11 | 21 | 0 |
| SRR1264353 | 0 | 147 | 198 | 73 | 9433 | 28580 | 94172 | 28841 | 78 | 34 | 23579 | 45286 | 16159 | 49449 | 1531 | 23618 | 19514 | 0 | 0 | 0 | 58 | 167 | 127 | 71 | 23 | 15 | 0 | 0 | 15 | 19 | 335 | 0 | 0 | 0 | 0 | 14 | 0 |
| SRR1264354 | 0 | 209 | 44 | 24 | 415 | 1116 | 3549 | 1151 | 18650 | 33 | 5904 | 6439 | 3456 | 519 | 6222 | 9554 | 27259 | 0 | 0 | 0 | 79 | 116 | 164 | 123 | 45 | 18 | 0 | 0 | 12 | 19 | 187 | 0 | 0 | 0 | 0 | 21 | 0 |
| SRR1270153 | 0 | 22 | 8530 | 22 | 114 | 34 | 1001 | 1829 | 268 | 170 | 55172 | 111737 | 14953 | 548 | 199 | 382 | 16417 | 0 | 0 | 0 | 0 | 91 | 59 | 0 | 0 | 11 | 0 | 16 | 16 | 0 | 0 | 0 | 0 | 0 | 94 | 0 | 0 |
| SRR1290784 | 17 | 3269 | 731 | 129 | 429 | 7567 | 7756 | 4646 | 3847 | 96 | 54575 | 25270 | 15013 | 10903 | 309 | 14 | 9327 | 0 | 0 | 0 | 482 | 33 | 53 | 0 | 0 | 0 | 0 | 0 | 27 | 151 | 51 | 38 | 0 | 19 | 0 | 0 | 0 |
| SRR1290785 | 53 | 1604 | 725 | 209 | 537 | 7864 | 9232 | 5003 | 2974 | 70 | 51460 | 27800 | 17506 | 11398 | 349 | 20 | 8282 | 0 | 0 | 0 | 487 | 29 | 41 | 0 | 0 | 0 | 0 | 0 | 27 | 168 | 51 | 35 | 0 | 17 | 0 | 0 | 0 |
| SRR1290786 | 45 | 1536 | 789 | 93 | 494 | 7759 | 8871 | 4572 | 2994 | 62 | 48864 | 23782 | 15396 | 10535 | 279 | 60 | 8721 | 0 | 0 | 0 | 437 | 29 | 55 | 0 | 0 | 0 | 0 | 0 | 22 | 141 | 48 | 44 | 0 | 21 | 0 | 0 | 0 |
| SRR1290787 | 16 | 3400 | 923 | 178 | 413 | 7876 | 8573 | 4845 | 3020 | 56 | 46471 | 23499 | 14803 | 11309 | 228 | 14 | 8449 | 0 | 0 | 0 | 840 | 75 | 91 | 0 | 11 | 0 | 0 | 0 | 150 | 337 | 142 | 75 | 0 | 24 | 0 | 0 | 0 |
| SRR1290788 | 11 | 2597 | 1193 | 169 | 440 | 8436 | 9415 | 4760 | 3334 | 73 | 52595 | 24613 | 15471 | 12357 | 305 | 13 | 8003 | 0 | 0 | 0 | 523 | 47 | 48 | 0 | 0 | 0 | 0 | 0 | 82 | 216 | 83 | 41 | 0 | 17 | 0 | 0 | 0 |
| SRR1290789 | 0 | 1849 | 1018 | 23 | 424 | 8136 | 8067 | 4538 | 2988 | 83 | 55697 | 26718 | 15923 | 11442 | 329 | 13 | 8501 | 0 | 0 | 0 | 426 | 35 | 42 | 0 | 0 | 0 | 0 | 0 | 27 | 152 | 52 | 34 | 0 | 23 | 11 | 0 | 0 |
| SRR1290790 | 11 | 1752 | 1134 | 104 | 338 | 6483 | 7431 | 4525 | 3163 | 66 | 53957 | 26131 | 15756 | 11603 | 250 | 13 | 8290 | 0 | 0 | 0 | 739 | 37 | 39 | 0 | 0 | 0 | 0 | 0 | 31 | 241 | 40 | 26 | 0 | 11 | 0 | 0 | 0 |
| SRR1290791 | 0 | 3667 | 823 | 147 | 566 | 7328 | 7325 | 6550 | 3709 | 113 | 57773 | 32336 | 17831 | 14100 | 389 | 11 | 7712 | 0 | 0 | 0 | 883 | 62 | 64 | 0 | 0 | 0 | 0 | 0 | 36 | 310 | 76 | 47 | 0 | 24 | 17 | 0 | 0 |
| SRR1290792 | 21 | 2563 | 947 | 87 | 447 | 7421 | 7967 | 5348 | 4078 | 56 | 44650 | 24202 | 14656 | 10796 | 241 | 14 | 9640 | 0 | 0 | 0 | 511 | 39 | 62 | 0 | 0 | 0 | 0 | 0 | 35 | 173 | 64 | 48 | 0 | 24 | 0 | 0 | 0 |
| SRR1290793 | 0 | 2137 | 1035 | 56 | 537 | 8300 | 8378 | 5440 | 3056 | 79 | 43577 | 25683 | 15450 | 9350 | 266 | 14 | 11056 | 0 | 0 | 0 | 439 | 25 | 59 | 0 | 0 | 0 | 0 | 0 | 15 | 140 | 43 | 49 | 0 | 25 | 11 | 0 | 0 |
| SRR1290794 | 0 | 1479 | 1054 | 70 | 718 | 11440 | 13731 | 3015 | 2006 | 71 | 65930 | 34677 | 20400 | 12226 | 325 | 12 | 7393 | 0 | 0 | 0 | 394 | 14 | 47 | 0 | 11 | 0 | 0 | 0 | 12 | 157 | 26 | 25 | 0 | 26 | 13 | 0 | 0 |
| SRR1290795 | 0 | 1650 | 881 | 173 | 638 | 10381 | 12292 | 3201 | 2164 | 68 | 55132 | 29162 | 16994 | 10390 | 246 | 12 | 6753 | 0 | 0 | 0 | 470 | 18 | 46 | 0 | 0 | 0 | 0 | 0 | 10 | 195 | 26 | 34 | 0 | 23 | 10 | 0 | 0 |
| SRR1290796 | 26 | 1556 | 1485 | 88 | 426 | 8346 | 9197 | 4444 | 2387 | 63 | 47053 | 24242 | 14904 | 10971 | 155 | 0 | 7117 | 0 | 0 | 0 | 325 | 25 | 37 | 0 | 0 | 0 | 0 | 0 | 23 | 140 | 41 | 36 | 0 | 14 | 0 | 0 | 0 |
| SRR1290797 | 33 | 2206 | 1274 | 294 | 472 | 8546 | 9132 | 5254 | 2887 | 70 | 42836 | 19633 | 12288 | 10652 | 161 | 11 | 5935 | 0 | 0 | 0 | 590 | 29 | 30 | 0 | 0 | 0 | 0 | 0 | 36 | 280 | 53 | 34 | 0 | 0 | 0 | 0 | 0 |
| SRR1290798 | 43 | 2017 | 580 | 241 | 575 | 9186 | 9554 | 5310 | 5124 | 52 | 37813 | 15687 | 9624 | 10274 | 191 | 0 | 4263 | 0 | 0 | 0 | 447 | 24 | 33 | 0 | 0 | 0 | 0 | 0 | 13 | 252 | 32 | 33 | 0 | 0 | 0 | 0 | 0 |
| SRR1290799 | 203 | 20423 | 845 | 318 | 340 | 6386 | 9702 | 3772 | 5599 | 27 | 30914 | 11000 | 6819 | 12602 | 119 | 0 | 3101 | 0 | 0 | 0 | 813 | 79 | 18 | 0 | 0 | 0 | 0 | 18 | 120 | 803 | 183 | 35 | 0 | 0 | 0 | 0 | 0 |
| SRR1290800 | 459 | 15558 | 647 | 358 | 361 | 6764 | 9200 | 4677 | 5803 | 20 | 33981 | 11998 | 8054 | 12810 | 129 | 0 | 3795 | 0 | 0 | 0 | 868 | 65 | 14 | 0 | 0 | 0 | 0 | 17 | 118 | 708 | 145 | 25 | 0 | 0 | 0 | 0 | 0 |
| SRR1290801 | 832 | 56860 | 1193 | 365 | 415 | 9416 | 8605 | 4825 | 2979 | 31 | 39639 | 21327 | 13540 | 9936 | 222 | 18 | 11950 | 0 | 0 | 0 | 3177 | 279 | 61 | 0 | 0 | 0 | 0 | 43 | 298 | 1324 | 452 | 56 | 0 | 14 | 11 | 0 | 0 |
| SRR1290802 | 84 | 3217 | 929 | 153 | 451 | 7780 | 8149 | 4954 | 2236 | 94 | 48659 | 27471 | 17340 | 8583 | 390 | 23 | 12216 | 0 | 0 | 0 | 419 | 29 | 58 | 0 | 0 | 0 | 0 | 0 | 27 | 96 | 28 | 31 | 0 | 12 | 0 | 0 | 0 |
| SRR1290803 | 86 | 2071 | 876 | 188 | 375 | 6892 | 7271 | 4343 | 2862 | 70 | 47670 | 27271 | 16545 | 9502 | 299 | 16 | 10164 | 0 | 0 | 0 | 617 | 33 | 58 | 0 | 0 | 0 | 0 | 0 | 35 | 175 | 39 | 39 | 0 | 16 | 0 | 0 | 0 |
| SRR1290804 | 67 | 1848 | 885 | 393 | 484 | 8056 | 9124 | 5119 | 3719 | 90 | 52995 | 33741 | 18107 | 11576 | 385 | 18 | 9224 | 0 | 0 | 0 | 577 | 40 | 65 | 0 | 0 | 0 | 0 | 0 | 84 | 222 | 66 | 49 | 0 | 17 | 0 | 0 | 0 |
| SRR1290805 | 48 | 2565 | 719 | 501 | 471 | 8494 | 9212 | 4810 | 2948 | 50 | 45136 | 30665 | 15985 | 9796 | 238 | 18 | 9851 | 0 | 0 | 0 | 1048 | 66 | 107 | 0 | 11 | 0 | 0 | 0 | 195 | 424 | 100 | 68 | 0 | 24 | 11 | 0 | 0 |
| SRR1290806 | 12 | 2708 | 994 | 138 | 493 | 8402 | 9031 | 4633 | 3529 | 70 | 47867 | 30362 | 16301 | 9851 | 384 | 25 | 9217 | 0 | 0 | 0 | 827 | 59 | 95 | 0 | 14 | 0 | 0 | 0 | 89 | 322 | 74 | 73 | 0 | 30 | 0 | 0 | 0 |
| SRR1290807 | 18 | 2855 | 989 | 327 | 501 | 7760 | 8002 | 4697 | 3164 | 76 | 51960 | 31553 | 18895 | 11358 | 428 | 16 | 8436 | 0 | 0 | 0 | 813 | 64 | 85 | 0 | 13 | 0 | 0 | 0 | 63 | 272 | 60 | 57 | 0 | 26 | 11 | 0 | 0 |
| SRR1290808 | 14 | 3888 | 789 | 231 | 521 | 8001 | 8253 | 4991 | 4211 | 114 | 53417 | 33853 | 17002 | 10972 | 412 | 15 | 11256 | 0 | 0 | 0 | 1266 | 74 | 88 | 0 | 0 | 0 | 0 | 0 | 73 | 453 | 97 | 60 | 0 | 29 | 13 | 0 | 0 |
| SRR1290809 | 52 | 4137 | 1026 | 528 | 364 | 6606 | 6162 | 4166 | 4018 | 99 | 52578 | 31315 | 16976 | 10461 | 347 | 38 | 10578 | 0 | 0 | 0 | 1019 | 59 | 56 | 0 | 0 | 0 | 0 | 0 | 54 | 330 | 57 | 37 | 0 | 20 | 0 | 0 | 0 |
| SRR1292461 | 15 | 1356 | 12088 | 0 | 90 | 24 | 546 | 289 | 286 | 1611 | 12 | 49 | 17 | 10021 | 643 | 0 | 953 | 0 | 0 | 0 | 0 | 15 | 20 | 0 | 0 | 0 | 0 | 0 | 0 | 89 | 38 | 76 | 88 | 199 | 27 | 15 | 18 |
| SRR1304304 | 0 | 942 | 1091 | 20 | 74 | 1042 | 2861 | 2996 | 27122 | 22 | 9828 | 3750 | 286 | 362 | 0 | 20218 | 14784 | 0 | 0 | 0 | 38 | 0 | 0 | 0 | 0 | 0 | 0 | 0 | 0 | 39 | 0 | 0 | 0 | 0 | 0 | 0 | 0 |
| SRR1304307 | 14 | 3174 | 230868 | 0 | 786 | 13846 | 20361 | 26087 | 10555 | 51 | 61898 | 5786 | 2518 | 29027 | 0 | 11844 | 8982 | 0 | 0 | 0 | 0 | 0 | 0 | 0 | 0 | 0 | 0 | 0 | 0 | 92 | 208 | 52 | 0 | 0 | 0 | 0 | 0 |
| SRR1304309 | 38 | 9508 | 9478 | 35 | 1724 | 27190 | 40196 | 19122 | 105 | 239 | 19317 | 13866 | 1273 | 42049 | 0 | 20524 | 6707 | 0 | 0 | 0 | 0 | 0 | 0 | 0 | 0 | 0 | 0 | 0 | 0 | 0 | 0 | 0 | 0 | 0 | 0 | 0 | 0 |
| SRR1304313 | 131 | 8625 | 2264 | 3424 | 1058 | 12498 | 25567 | 28447 | 3005 | 47 | 17145 | 9035 | 3886 | 14249 | 0 | 826 | 23372 | 0 | 0 | 0 | 493 | 151 | 138 | 0 | 0 | 0 | 0 | 0 | 25 | 151 | 237 | 76 | 0 | 0 | 0 | 0 | 0 |
| SRR1393733 | 0 | 21016 | 2047 | 0 | 344 | 4078 | 4225 | 2472 | 17011 | 311 | 10788 | 9451 | 1756 | 1726 | 3363 | 37463 | 58046 | 0 | 0 | 0 | 132 | 43 | 231 | 0 | 0 | 0 | 0 | 0 | 0 | 17 | 10 | 132 | 0 | 0 | 0 | 17 | 0 |
| SRR1510077 | 100 | 5937 | 348 | 1138 | 128 | 997 | 1186 | 1830 | 1888 | 25 | 18031 | 33078 | 5926 | 4129 | 1828 | 11147 | 148961 | 0 | 0 | 12 | 1396 | 2683 | 226 | 0 | 0 | 0 | 0 | 153 | 375 | 520 | 4157 | 206 | 0 | 0 | 16 | 0 | 0 |
| SRR1510078 | 111 | 2525 | 466 | 919 | 41 | 306 | 283 | 969 | 1536 | 15 | 21019 | 44599 | 7890 | 2817 | 4018 | 11027 | 66616 | 0 | 0 | 17 | 2764 | 875 | 340 | 0 | 0 | 0 | 0 | 72 | 706 | 1475 | 1151 | 253 | 0 | 0 | 88 | 0 | 0 |
| SRR1510079 | 105 | 4315 | 137 | 880 | 25 | 86 | 111 | 259 | 877 | 18 | 5763 | 29395 | 2434 | 1788 | 979 | 9113 | 320211 | 0 | 0 | 0 | 518 | 3178 | 319 | 0 | 0 | 0 | 0 | 45 | 177 | 671 | 5367 | 435 | 0 | 0 | 77 | 0 | 0 |
| SRR1510080 | 68 | 4775 | 143 | 604 | 17 | 144 | 171 | 499 | 534 | 22 | 24618 | 65785 | 7380 | 4038 | 1505 | 10322 | 75150 | 0 | 0 | 0 | 832 | 641 | 150 | 0 | 0 | 0 | 0 | 141 | 519 | 819 | 1177 | 195 | 0 | 11 | 283 | 13 | 0 |
| SRR1510081 | 59 | 1400 | 100 | 736 | 52 | 362 | 398 | 1357 | 3283 | 0 | 11096 | 30886 | 3398 | 1382 | 434 | 6815 | 104731 | 0 | 0 | 0 | 649 | 1291 | 375 | 0 | 0 | 0 | 0 | 88 | 338 | 499 | 2518 | 458 | 0 | 22 | 247 | 12 | 0 |
| SRR1510082 | 93 | 2931 | 347 | 1098 | 29 | 204 | 202 | 671 | 6416 | 12 | 10717 | 34299 | 4203 | 1694 | 749 | 8118 | 96441 | 0 | 18 | 31 | 1814 | 985 | 568 | 0 | 0 | 0 | 0 | 313 | 1034 | 916 | 1600 | 724 | 0 | 13 | 68 | 0 | 0 |
| SRR1510083 | 77 | 2357 | 188 | 608 | 21 | 226 | 244 | 412 | 1315 | 0 | 9956 | 30433 | 4352 | 1954 | 1245 | 9152 | 59585 | 0 | 14 | 32 | 1947 | 1048 | 125 | 0 | 0 | 0 | 0 | 376 | 1560 | 1333 | 2022 | 157 | 0 | 0 | 41 | 0 | 0 |
| SRR1510084 | 85 | 2084 | 107 | 1316 | 18 | 123 | 160 | 293 | 1554 | 11 | 17977 | 53156 | 5858 | 2097 | 1674 | 7007 | 156401 | 0 | 17 | 18 | 1215 | 2661 | 383 | 0 | 0 | 0 | 0 | 307 | 769 | 868 | 4330 | 368 | 0 | 0 | 172 | 0 | 0 |
| SRR1510085 | 37 | 2188 | 266 | 368 | 152 | 1370 | 1735 | 2594 | 1665 | 12 | 20496 | 12 | 6755 | 2800 | 1018 | 8781 | 98323 | 0 | 23 | 26 | 2300 | 1680 | 248 | 0 | 0 | 0 | 0 | 395 | 934 | 1016 | 2175 | 207 | 0 | 0 | 10 | 0 | 0 |
| SRR1510086 | 48 | 22887 | 314 | 2219 | 34 | 118 | 163 | 805 | 1328 | 10 | 11872 | 31420 | 3434 | 2781 | 810 | 11580 | 83585 | 0 | 18 | 47 | 1462 | 1019 | 293 | 13 | 0 | 0 | 18 | 417 | 2286 | 1680 | 3136 | 363 | 0 | 19 | 256 | 10 | 0 |
| SRR1510087 | 183 | 8475 | 126 | 947 | 11 | 77 | 119 | 156 | 630 | 0 | 12910 | 46951 | 5055 | 1309 | 1142 | 2460 | 95146 | 0 | 23 | 51 | 14097 | 1101 | 432 | 0 | 0 | 0 | 0 | 233 | 1304 | 2730 | 537 | 105 | 0 | 0 | 41 | 0 | 0 |
| SRR1510088 | 77 | 948 | 138 | 639 | 25 | 167 | 200 | 531 | 796 | 11 | 23847 | 44019 | 9255 | 2531 | 1535 | 13613 | 65833 | 0 | 15 | 29 | 1889 | 248 | 37 | 0 | 0 | 0 | 13 | 477 | 2142 | 1549 | 606 | 63 | 0 | 16 | 269 | 23 | 0 |
| SRR1510089 | 73 | 2694 | 204 | 1262 | 0 | 57 | 57 | 108 | 605 | 11 | 6982 | 20499 | 3197 | 3001 | 246 | 6424 | 179633 | 0 | 13 | 25 | 2580 | 3669 | 242 | 0 | 0 | 0 | 0 | 186 | 850 | 1979 | 3513 | 210 | 0 | 0 | 34 | 0 | 0 |
| SRR1510090 | 55 | 1464 | 151 | 668 | 54 | 273 | 430 | 1041 | 3392 | 60 | 19932 | 41507 | 7099 | 3178 | 769 | 9506 | 183055 | 0 | 29 | 155 | 17740 | 1262 | 546 | 0 | 0 | 0 | 0 | 773 | 7941 | 12810 | 3653 | 903 | 0 | 0 | 0 | 0 | 0 |
| SRR1510091 | 50 | 1129 | 65 | 1057 | 24 | 122 | 143 | 432 | 710 | 12 | 12616 | 31777 | 4797 | 1735 | 1530 | 5049 | 316946 | 0 | 0 | 10 | 1273 | 2313 | 174 | 0 | 0 | 0 | 0 | 201 | 698 | 1051 | 3780 | 161 | 0 | 13 | 273 | 14 | 0 |
| SRR1510092 | 76 | 5325 | 102 | 589 | 25 | 178 | 209 | 577 | 336 | 13 | 13480 | 42644 | 3698 | 964 | 264 | 1188 | 50695 | 0 | 14 | 57 | 12868 | 1503 | 192 | 0 | 0 | 0 | 0 | 146 | 1761 | 2470 | 860 | 51 | 0 | 0 | 0 | 0 | 0 |
| SRR1510093 | 83 | 2945 | 226 | 465 | 16 | 83 | 100 | 220 | 981 | 0 | 12807 | 39907 | 6082 | 2768 | 945 | 9722 | 78716 | 0 | 12 | 37 | 6000 | 1087 | 311 | 0 | 0 | 0 | 0 | 318 | 1923 | 3494 | 2584 | 375 | 0 | 0 | 24 | 0 | 0 |
| SRR1510094 | 68 | 4052 | 283 | 1767 | 16 | 158 | 217 | 596 | 433 | 27 | 12211 | 38721 | 4029 | 2846 | 281 | 6207 | 145948 | 0 | 0 | 0 | 833 | 971 | 144 | 0 | 0 | 0 | 0 | 98 | 374 | 817 | 1573 | 227 | 0 | 0 | 155 | 0 | 0 |
| SRR1510095 | 26 | 788 | 145 | 550 | 22 | 100 | 183 | 456 | 691 | 19 | 14701 | 29873 | 6015 | 2793 | 335 | 7945 | 249244 | 0 | 15 | 22 | 2173 | 4789 | 323 | 0 | 0 | 0 | 0 | 410 | 1367 | 1743 | 9909 | 426 | 0 | 0 | 63 | 0 | 0 |
| SRR1510096 | 57 | 1753 | 175 | 839 | 29 | 158 | 227 | 479 | 922 | 36 | 25492 | 41553 | 8570 | 3646 | 2247 | 7025 | 198245 | 0 | 19 | 35 | 5109 | 1291 | 140 | 0 | 0 | 0 | 0 | 262 | 1181 | 2118 | 1832 | 145 | 0 | 0 | 15 | 0 | 0 |
| SRR1510097 | 68 | 3263 | 190 | 680 | 56 | 970 | 484 | 787 | 10275 | 73 | 16249 | 32947 | 4672 | 5318 | 99 | 2561 | 58560 | 0 | 0 | 0 | 1803 | 694 | 160 | 0 | 0 | 0 | 0 | 0 | 143 | 770 | 739 | 96 | 0 | 0 | 0 | 0 | 0 |
| SRR1510098 | 198 | 8391 | 368 | 907 | 23 | 154 | 127 | 190 | 2760 | 63 | 16404 | 53565 | 5315 | 3200 | 425 | 3851 | 71477 | 0 | 0 | 0 | 1366 | 748 | 144 | 0 | 0 | 0 | 0 | 31 | 207 | 587 | 850 | 134 | 0 | 0 | 38 | 0 | 0 |
| SRR1510099 | 351 | 8568 | 159 | 1416 | 40 | 372 | 262 | 187 | 823 | 50 | 18437 | 46923 | 2106 | 1652 | 25 | 1224 | 36021 | 0 | 0 | 0 | 1535 | 491 | 173 | 0 | 0 | 0 | 0 | 31 | 216 | 354 | 232 | 66 | 0 | 0 | 0 | 0 | 0 |
| SRR1510100 | 330 | 6867 | 1255 | 284 | 0 | 56 | 46 | 96 | 1802 | 30 | 7061 | 30768 | 2218 | 2159 | 60 | 1421 | 32184 | 0 | 0 | 0 | 435 | 158 | 38 | 0 | 0 | 0 | 0 | 16 | 120 | 277 | 141 | 26 | 0 | 0 | 0 | 0 | 0 |
| SRR1510101 | 182 | 11181 | 231 | 258 | 27 | 147 | 145 | 290 | 2103 | 49 | 19930 | 58788 | 6718 | 3000 | 3337 | 3248 | 40008 | 0 | 26 | 133 | 10615 | 714 | 303 | 0 | 0 | 0 | 0 | 570 | 5596 | 4793 | 993 | 173 | 0 | 0 | 13 | 0 | 0 |
| SRR1510102 | 74 | 1863 | 348 | 1069 | 36 | 242 | 261 | 628 | 3978 | 31 | 20199 | 38256 | 8023 | 4458 | 2110 | 7643 | 74325 | 0 | 29 | 58 | 3802 | 653 | 172 | 0 | 0 | 0 | 0 | 770 | 3336 | 2561 | 1449 | 222 | 0 | 0 | 11 | 0 | 0 |
| SRR1510103 | 34 | 1218 | 266 | 542 | 29 | 128 | 182 | 352 | 1656 | 22 | 14810 | 34132 | 6071 | 3986 | 1776 | 8300 | 162707 | 0 | 13 | 16 | 2201 | 2530 | 188 | 0 | 0 | 0 | 0 | 256 | 751 | 1130 | 3392 | 167 | 0 | 13 | 152 | 19 | 0 |
| SRR1510104 | 93 | 1844 | 144 | 1171 | 11 | 51 | 67 | 126 | 825 | 11 | 6266 | 27850 | 2683 | 1166 | 167 | 2277 | 111080 | 0 | 0 | 12 | 1626 | 2375 | 606 | 0 | 0 | 0 | 0 | 164 | 481 | 517 | 1162 | 200 | 0 | 0 | 68 | 0 | 0 |
| SRR1510105 | 68 | 1084 | 245 | 345 | 43 | 235 | 354 | 890 | 2188 | 44 | 24231 | 40045 | 8920 | 3987 | 1569 | 8926 | 63544 | 0 | 0 | 0 | 1950 | 808 | 117 | 0 | 0 | 0 | 0 | 125 | 497 | 1080 | 1023 | 109 | 0 | 0 | 18 | 0 | 0 |
| SRR1510106 | 54 | 2145 | 106 | 1751 | 23 | 190 | 192 | 546 | 1031 | 0 | 13660 | 48812 | 4902 | 1659 | 2000 | 4326 | 68254 | 0 | 0 | 16 | 1693 | 1388 | 202 | 0 | 0 | 0 | 11 | 260 | 991 | 1459 | 2415 | 190 | 0 | 19 | 470 | 15 | 0 |
| SRR1510107 | 74 | 1933 | 158 | 451 | 33 | 209 | 321 | 419 | 1578 | 38 | 13584 | 35796 | 5556 | 3223 | 3223 | 9007 | 260749 | 0 | 16 | 36 | 6412 | 2563 | 437 | 0 | 0 | 0 | 0 | 278 | 1293 | 3670 | 3936 | 433 | 0 | 0 | 11 | 0 | 0 |
| SRR1510108 | 394 | 7460 | 388 | 1014 | 0 | 46 | 62 | 96 | 856 | 52 | 8169 | 21262 | 2883 | 1612 | 725 | 2750 | 204680 | 0 | 0 | 12 | 1381 | 2105 | 175 | 0 | 0 | 0 | 0 | 149 | 424 | 649 | 1304 | 88 | 0 | 0 | 79 | 0 | 0 |
| SRR1510109 | 126 | 4361 | 213 | 725 | 30 | 207 | 218 | 461 | 4275 | 61 | 13949 | 29438 | 6018 | 3977 | 662 | 12270 | 129359 | 0 | 17 | 34 | 2949 | 1592 | 253 | 0 | 0 | 0 | 0 | 292 | 1394 | 2233 | 2930 | 299 | 0 | 0 | 121 | 0 | 0 |
| SRR1510110 | 127 | 2375 | 127 | 826 | 17 | 56 | 74 | 153 | 695 | 0 | 11756 | 49687 | 4846 | 1212 | 281 | 2902 | 47361 | 0 | 17 | 25 | 1634 | 442 | 112 | 0 | 0 | 0 | 0 | 194 | 773 | 473 | 339 | 57 | 0 | 0 | 82 | 0 | 0 |
| SRR1510111 | 145 | 3303 | 222 | 780 | 38 | 127 | 216 | 476 | 1960 | 55 | 11584 | 23483 | 4912 | 3125 | 1495 | 5896 | 351336 | 0 | 19 | 26 | 1772 | 2858 | 253 | 0 | 0 | 0 | 0 | 331 | 1097 | 1396 | 4279 | 256 | 0 | 0 | 124 | 0 | 0 |
| SRR1510112 | 51 | 2550 | 30 | 1785 | 63 | 318 | 350 | 693 | 2624 | 13 | 16250 | 33728 | 4388 | 1784 | 3128 | 15277 | 115658 | 0 | 14 | 38 | 1566 | 1004 | 96 | 0 | 0 | 0 | 0 | 292 | 2131 | 1537 | 2176 | 94 | 0 | 11 | 221 | 0 | 0 |
| SRR1510113 | 35 | 1711 | 183 | 185 | 48 | 385 | 411 | 1236 | 668 | 0 | 20723 | 14 | 7884 | 2325 | 1165 | 10865 | 82487 | 0 | 18 | 36 | 7585 | 1011 | 247 | 0 | 0 | 0 | 0 | 370 | 1546 | 3913 | 1758 | 239 | 0 | 0 | 31 | 0 | 0 |
| SRR1510114 | 107 | 1940 | 196 | 1151 | 29 | 268 | 288 | 864 | 1370 | 24 | 18548 | 44433 | 6643 | 1978 | 2176 | 7653 | 82966 | 0 | 0 | 11 | 954 | 1026 | 249 | 0 | 0 | 0 | 0 | 144 | 610 | 697 | 2079 | 302 | 0 | 17 | 233 | 13 | 0 |
| SRR1510115 | 39 | 409 | 24 | 543 | 20 | 157 | 148 | 485 | 1121 | 0 | 22755 | 49560 | 8685 | 2777 | 3885 | 8620 | 164616 | 0 | 11 | 27 | 2272 | 649 | 102 | 0 | 0 | 0 | 0 | 296 | 1533 | 1797 | 1161 | 122 | 0 | 24 | 626 | 28 | 0 |
| SRR1510116 | 77 | 7979 | 374 | 685 | 0 | 49 | 53 | 114 | 924 | 109 | 8845 | 35041 | 3050 | 1510 | 304 | 3029 | 74078 | 0 | 12 | 12 | 1208 | 1125 | 198 | 0 | 0 | 0 | 0 | 136 | 287 | 512 | 756 | 113 | 0 | 0 | 87 | 0 | 0 |
| SRR1510117 | 221 | 6936 | 120 | 1360 | 21 | 89 | 144 | 189 | 1613 | 27 | 7422 | 46949 | 2498 | 783 | 294 | 2032 | 74059 | 0 | 22 | 43 | 3609 | 1045 | 152 | 0 | 0 | 0 | 0 | 221 | 961 | 1245 | 642 | 70 | 0 | 0 | 73 | 0 | 0 |
| SRR1510118 | 134 | 3569 | 206 | 1161 | 15 | 70 | 78 | 227 | 551 | 27 | 13237 | 40887 | 4927 | 1549 | 1164 | 3603 | 79491 | 0 | 0 | 22 | 1956 | 1381 | 236 | 0 | 0 | 0 | 0 | 162 | 713 | 818 | 1209 | 141 | 0 | 0 | 80 | 0 | 0 |
| SRR1510119 | 68 | 1844 | 185 | 549 | 24 | 141 | 219 | 561 | 654 | 12 | 25675 | 67267 | 9484 | 3139 | 835 | 10335 | 72916 | 0 | 22 | 40 | 3935 | 1116 | 328 | 0 | 0 | 0 | 0 | 586 | 2891 | 3276 | 2219 | 410 | 0 | 0 | 152 | 0 | 0 |
| SRR1510120 | 131 | 1747 | 269 | 636 | 26 | 181 | 214 | 646 | 1229 | 25 | 20704 | 49308 | 8744 | 2704 | 703 | 9831 | 75285 | 0 | 0 | 16 | 1851 | 1150 | 202 | 0 | 0 | 0 | 0 | 244 | 853 | 1226 | 2264 | 272 | 0 | 0 | 105 | 0 | 0 |
| SRR1510121 | 192 | 4134 | 239 | 642 | 61 | 592 | 816 | 852 | 1189 | 74 | 20967 | 40077 | 6143 | 3014 | 595 | 5756 | 263090 | 0 | 0 | 0 | 1469 | 517 | 115 | 0 | 0 | 0 | 0 | 50 | 338 | 1098 | 1118 | 171 | 0 | 0 | 67 | 0 | 0 |
| SRR1519906 | 0 | 325 | 1213 | 22 | 0 | 81 | 110 | 13013 | 37380 | 0 | 920 | 358 | 73 | 314 | 0 | 820 | 1556 | 0 | 0 | 0 | 0 | 0 | 0 | 0 | 0 | 0 | 0 | 0 | 0 | 0 | 29 | 55 | 0 | 0 | 0 | 0 | 0 |
| SRR1519919 | 43 | 18425 | 8593 | 53 | 53 | 1744 | 1508 | 7347 | 4587 | 43 | 7141 | 6200 | 770 | 2396 | 0 | 963 | 17559 | 0 | 0 | 0 | 15 | 310 | 26 | 0 | 0 | 0 | 0 | 0 | 32 | 43 | 430 | 55 | 0 | 0 | 0 | 0 | 0 |
| SRR1525198 | 328 | 17227 | 445 | 614 | 109 | 2617 | 1418 | 874 | 28166 | 49 | 14850 | 6801 | 365 | 358 | 0 | 10914 | 15381 | 0 | 0 | 0 | 241 | 126 | 200 | 23 | 22 | 42 | 0 | 59 | 92 | 95 | 105 | 93 | 0 | 12 | 0 | 11 | 0 |
| SRR1525199 | 142 | 28605 | 209 | 480 | 156 | 2889 | 1247 | 1277 | 34894 | 28 | 20375 | 8696 | 646 | 731 | 0 | 11840 | 16648 | 0 | 0 | 0 | 186 | 54 | 181 | 13 | 19 | 0 | 0 | 35 | 65 | 90 | 78 | 95 | 0 | 15 | 0 | 0 | 0 |
| SRR1525200 | 335 | 43243 | 396 | 656 | 304 | 6223 | 3299 | 2663 | 34887 | 48 | 15148 | 7807 | 447 | 501 | 0 | 16345 | 17682 | 0 | 0 | 12 | 295 | 103 | 204 | 19 | 20 | 19 | 0 | 62 | 105 | 128 | 175 | 114 | 0 | 19 | 14 | 11 | 0 |
| SRR1525201 | 353 | 45456 | 431 | 752 | 226 | 4728 | 1741 | 2476 | 30587 | 33 | 16262 | 10320 | 605 | 726 | 0 | 12384 | 16322 | 0 | 0 | 0 | 260 | 78 | 190 | 12 | 16 | 0 | 0 | 51 | 96 | 129 | 106 | 92 | 0 | 17 | 0 | 0 | 0 |
| SRR1525202 | 3854 | 10081 | 4198 | 304 | 123 | 3103 | 1654 | 1070 | 31640 | 40 | 15089 | 14286 | 496 | 320 | 75 | 15414 | 18560 | 0 | 0 | 0 | 221 | 146 | 153 | 21 | 14 | 66 | 0 | 51 | 93 | 85 | 108 | 65 | 0 | 12 | 14 | 0 | 0 |
| SRR1525203 | 268 | 52085 | 327 | 785 | 94 | 2005 | 835 | 886 | 24944 | 35 | 17157 | 7693 | 616 | 655 | 0 | 9822 | 15524 | 0 | 0 | 0 | 258 | 82 | 217 | 16 | 20 | 0 | 0 | 62 | 95 | 98 | 75 | 104 | 0 | 16 | 0 | 10 | 0 |
| SRR1563015 | 22 | 1369 | 5170 | 4178 | 803 | 20836 | 11012 | 11344 | 16615 | 23 | 2762 | 1470 | 254 | 53 | 0 | 0 | 23233 | 0 | 0 | 0 | 807 | 602 | 192 | 0 | 23 | 0 | 0 | 15 | 142 | 290 | 752 | 161 | 0 | 38 | 19 | 0 | 0 |
| SRR1563018 | 102 | 2097 | 953 | 2456 | 278 | 5959 | 7946 | 15432 | 0 | 13 | 11184 | 9183 | 4631 | 3472 | 0 | 22 | 33718 | 0 | 0 | 0 | 249 | 97 | 122 | 0 | 27 | 0 | 0 | 0 | 0 | 85 | 102 | 50 | 0 | 35 | 0 | 0 | 0 |
| SRR1563057 | 62 | 1244 | 1421 | 975 | 144 | 3225 | 3245 | 5624 | 14556 | 10 | 12520 | 6912 | 4330 | 3571 | 0 | 0 | 31797 | 0 | 0 | 0 | 355 | 243 | 236 | 0 | 35 | 0 | 0 | 0 | 11 | 107 | 167 | 77 | 0 | 50 | 18 | 0 | 0 |
| SRR1563058 | 948 | 22286 | 21096 | 888 | 445 | 6588 | 6883 | 9261 | 6235 | 35 | 15822 | 9846 | 2032 | 3233 | 0 | 709 | 28744 | 0 | 0 | 10 | 369 | 689 | 50 | 0 | 0 | 0 | 0 | 35 | 173 | 164 | 888 | 58 | 0 | 14 | 0 | 0 | 0 |
| SRR1563059 | 209 | 72456 | 5693 | 2632 | 1562 | 26794 | 20865 | 36243 | 19508 | 30 | 28297 | 8626 | 6914 | 14665 | 0 | 1016 | 8498 | 0 | 0 | 0 | 1140 | 580 | 242 | 0 | 32 | 0 | 0 | 0 | 24 | 297 | 341 | 132 | 0 | 63 | 15 | 0 | 0 |
| SRR1563060 | 208 | 28996 | 74124 | 473 | 1408 | 36027 | 29725 | 62621 | 14922 | 98 | 22733 | 6783 | 4464 | 10911 | 0 | 1719 | 5560 | 0 | 0 | 0 | 788 | 568 | 76 | 0 | 0 | 0 | 0 | 27 | 211 | 403 | 770 | 133 | 0 | 11 | 0 | 0 | 0 |
| SRR1563061 | 184 | 1833 | 274 | 7597 | 1204 | 38160 | 23371 | 37073 | 4261 | 29 | 13860 | 4573 | 429 | 432 | 0 | 542 | 5298 | 0 | 0 | 0 | 97 | 59 | 22 | 0 | 0 | 0 | 0 | 0 | 0 | 37 | 101 | 15 | 0 | 0 | 20 | 0 | 0 |
| SRR1563062 | 316 | 23244 | 41668 | 369 | 2599 | 43443 | 41564 | 78388 | 12827 | 86 | 3969 | 2669 | 680 | 1140 | 30 | 2304 | 3919 | 0 | 0 | 0 | 403 | 489 | 38 | 0 | 0 | 0 | 0 | 10 | 117 | 218 | 638 | 56 | 0 | 0 | 11 | 0 | 0 |
| SRR1563063 | 0 | 13848 | 8604 | 473 | 2978 | 57282 | 33437 | 35427 | 14366 | 58 | 24913 | 8445 | 4611 | 9561 | 26 | 24 | 8195 | 0 | 0 | 0 | 301 | 949 | 104 | 0 | 19 | 0 | 0 | 0 | 24 | 232 | 529 | 165 | 0 | 42 | 29 | 0 | 0 |
| SRR1563064 | 528 | 20823 | 51962 | 673 | 2306 | 42296 | 47086 | 67603 | 24645 | 75 | 24940 | 8313 | 3693 | 11254 | 0 | 431 | 6605 | 0 | 0 | 0 | 687 | 729 | 88 | 0 | 0 | 0 | 0 | 24 | 175 | 381 | 703 | 145 | 0 | 0 | 0 | 0 | 0 |
| SRR1575598 | 0 | 20 | 188 | 0 | 1059 | 10753 | 14422 | 9705 | 4417 | 12 | 39848 | 14101 | 5470 | 40762 | 144 | 67285 | 24517 | 0 | 0 | 0 | 0 | 0 | 0 | 0 | 32 | 0 | 0 | 0 | 0 | 0 | 0 | 0 | 0 | 16 | 15 | 17 | 0 |
| SRR1575599 | 0 | 0 | 85 | 0 | 856 | 11040 | 6781 | 7979 | 3708 | 0 | 43002 | 18542 | 4007 | 38220 | 19 | 62234 | 24746 | 0 | 0 | 0 | 0 | 0 | 39 | 59 | 2864 | 99 | 0 | 0 | 0 | 0 | 0 | 47 | 33 | 1206 | 1219 | 512 | 0 |
| SRR1575601 | 0 | 0 | 159 | 0 | 290 | 3035 | 9352 | 9864 | 3877 | 0 | 29100 | 12459 | 5325 | 22029 | 24 | 16177 | 27652 | 0 | 0 | 0 | 0 | 0 | 0 | 0 | 90 | 0 | 0 | 0 | 0 | 0 | 0 | 0 | 0 | 35 | 69 | 26 | 0 |
| SRR1575602 | 0 | 304 | 145 | 21 | 1205 | 15724 | 12052 | 12388 | 2873 | 0 | 26535 | 12674 | 4632 | 27654 | 46 | 67834 | 23730 | 0 | 0 | 0 | 69 | 18 | 82 | 54 | 2308 | 96 | 0 | 0 | 0 | 65 | 19 | 53 | 26 | 1265 | 1196 | 619 | 0 |
| SRR1592244 | 240 | 25281 | 136 | 1743 | 32 | 247 | 730 | 198 | 762 | 0 | 4311 | 18735 | 5742 | 36778 | 0 | 51 | 8406 | 0 | 0 | 0 | 0 | 0 | 0 | 0 | 0 | 0 | 0 | 0 | 0 | 0 | 0 | 0 | 0 | 0 | 0 | 0 | 0 |
| SRR1592245 | 166 | 42690 | 408 | 1340 | 48 | 418 | 952 | 314 | 1117 | 15 | 7920 | 37374 | 11505 | 49541 | 0 | 37 | 8875 | 0 | 0 | 0 | 99 | 0 | 0 | 0 | 0 | 0 | 0 | 0 | 29 | 78 | 0 | 0 | 0 | 0 | 0 | 0 | 0 |
| SRR1592246 | 163 | 42841 | 442 | 1230 | 21 | 193 | 458 | 110 | 760 | 11 | 3839 | 18639 | 7156 | 52706 | 0 | 31 | 6299 | 0 | 0 | 0 | 209 | 0 | 0 | 0 | 0 | 0 | 0 | 0 | 37 | 159 | 0 | 0 | 0 | 0 | 0 | 0 | 0 |
| SRR1592247 | 808 | 74038 | 620 | 558 | 34 | 376 | 489 | 240 | 809 | 0 | 5763 | 44930 | 11079 | 43412 | 0 | 42 | 16733 | 0 | 0 | 0 | 14 | 0 | 0 | 0 | 0 | 0 | 0 | 0 | 0 | 31 | 0 | 0 | 0 | 0 | 0 | 0 | 0 |
| SRR1592248 | 174 | 47821 | 752 | 768 | 59 | 781 | 1058 | 470 | 1505 | 0 | 9396 | 34310 | 8357 | 34783 | 0 | 72 | 13826 | 0 | 0 | 0 | 83 | 0 | 0 | 0 | 0 | 0 | 0 | 0 | 14 | 92 | 0 | 0 | 0 | 0 | 0 | 0 | 0 |
| SRR1596226 | 1075 | 14653 | 1383 | 1190 | 791 | 3691 | 6736 | 12105 | 14066 | 220 | 7949 | 19835 | 3162 | 5513 | 0 | 5946 | 53388 | 0 | 0 | 0 | 174 | 25 | 22 | 0 | 0 | 0 | 0 | 34 | 291 | 241 | 162 | 82 | 0 | 14 | 47 | 0 | 0 |

### Supp Table S8.1.Columns_B_to_AL

| | | | | | | | | | | | | | | | | | | | | | | | | | | | | | | | | | | | | |
|---|---|---|---|---|---|---|---|---|---|---|---|---|---|---|---|---|---|---|---|---|---|---|---|---|---|---|---|---|---|---|---|---|---|---|---|---|
| SRR1596227 | 1298 | 14101 | 1327 | 1560 | 743 | 2099 | 7630 | 12529 | 13015 | 219 | 7766 | 19032 | 3004 | 5454 | 0 | 3823 | 46556 | 0 | 0 | 0 | 126 | 23 | 12 | 0 | 0 | 0 | 0 | 17 | 232 | 223 | 135 | 65 | 0 | 0 | 26 | 23 | 0 |
| SRR1596228 | 204 | 2358 | 1499 | 2528 | 1540 | 6323 | 24227 | 22970 | 15993 | 291 | 14340 | 16021 | 4668 | 7159 | 0 | 3724 | 30603 | 0 | 0 | 11 | 195 | 81 | 86 | 0 | 0 | 0 | 0 | 29 | 475 | 623 | 631 | 892 | 12 | 91 | 99 | 21 | 0 |
| SRR1596229 | 215 | 2612 | 1462 | 2280 | 1716 | 5861 | 23016 | 24303 | 15144 | 277 | 15030 | 16539 | 4491 | 7577 | 0 | 3894 | 29228 | 0 | 0 | 11 | 202 | 72 | 60 | 0 | 0 | 0 | 0 | 25 | 442 | 574 | 628 | 876 | 14 | 90 | 103 | 16 | 0 |
| SRR1596230 | 294 | 4679 | 2309 | 470 | 1416 | 3643 | 11172 | 16302 | 21611 | 203 | 7398 | 11243 | 2579 | 3643 | 0 | 3234 | 51236 | 0 | 10 | 24 | 275 | 151 | 65 | 0 | 0 | 0 | 0 | 96 | 393 | 452 | 1560 | 642 | 0 | 55 | 126 | 32 | 0 |
| SRR1596231 | 403 | 5254 | 2770 | 546 | 1502 | 3639 | 11212 | 15971 | 21077 | 187 | 7300 | 10741 | 2638 | 3471 | 0 | 2903 | 53144 | 0 | 0 | 13 | 229 | 154 | 55 | 0 | 0 | 0 | 0 | 69 | 409 | 447 | 1406 | 546 | 0 | 35 | 112 | 19 | 0 |
| SRR1608128 | 0 | 89 | 3078 | 0 | 129 | 63 | 3987 | 40641 | 4213 | 480 | 94 | 499 | 38 | 267 | 0 | 29 | 155 | 0 | 0 | 0 | 0 | 0 | 0 | 0 | 0 | 0 | 0 | 0 | 0 | 0 | 0 | 0 | 0 | 0 | 0 | 0 | 0 |
| SRR1608138 | 0 | 52 | 5253 | 0 | 87 | 48 | 3478 | 39121 | 1009 | 510 | 335 | 1093 | 258 | 375 | 0 | 16 | 636 | 0 | 0 | 0 | 0 | 0 | 0 | 0 | 0 | 0 | 0 | 0 | 0 | 0 | 0 | 0 | 0 | 0 | 0 | 0 | 0 |
| SRR1608143 | 0 | 16 | 2120 | 0 | 0 | 0 | 347 | 3356 | 911 | 257 | 241 | 1080 | 63 | 183 | 0 | 0 | 161 | 0 | 0 | 0 | 0 | 0 | 0 | 0 | 0 | 0 | 0 | 0 | 0 | 0 | 0 | 0 | 0 | 0 | 0 | 0 | 0 |
| SRR1608150 | 0 | 62 | 4697 | 0 | 161 | 83 | 5896 | 67391 | 1520 | 705 | 214 | 654 | 58 | 330 | 0 | 38 | 321 | 0 | 0 | 0 | 0 | 0 | 0 | 0 | 0 | 0 | 0 | 0 | 0 | 0 | 0 | 0 | 0 | 0 | 0 | 0 | 0 |
| SRR1608158 | 0 | 20 | 2876 | 0 | 37 | 12 | 1092 | 13480 | 1777 | 213 | 203 | 870 | 113 | 325 | 0 | 102 | 226 | 0 | 0 | 0 | 0 | 0 | 0 | 0 | 0 | 0 | 0 | 0 | 0 | 0 | 0 | 0 | 0 | 0 | 0 | 0 | 0 |
| SRR1608160 | 0 | 36 | 2726 | 0 | 32 | 0 | 1160 | 16124 | 3464 | 466 | 113 | 657 | 59 | 394 | 0 | 39 | 176 | 0 | 0 | 0 | 0 | 0 | 0 | 0 | 0 | 0 | 0 | 0 | 0 | 0 | 0 | 0 | 0 | 0 | 0 | 0 | 0 |
| SRR1608163 | 0 | 52 | 5402 | 0 | 100 | 45 | 3327 | 41295 | 3323 | 1428 | 131 | 580 | 71 | 265 | 0 | 85 | 704 | 0 | 0 | 0 | 0 | 0 | 11 | 0 | 0 | 0 | 0 | 0 | 0 | 0 | 0 | 0 | 0 | 0 | 0 | 0 | 0 |
| SRR1608165 | 0 | 17 | 2314 | 0 | 91 | 42 | 3739 | 37552 | 7319 | 340 | 139 | 577 | 75 | 876 | 0 | 20 | 169 | 0 | 0 | 0 | 0 | 0 | 0 | 0 | 0 | 0 | 0 | 0 | 0 | 0 | 0 | 0 | 0 | 0 | 0 | 0 | 0 |
| SRR1608166 | 13 | 204 | 4153 | 0 | 69 | 35 | 2994 | 32805 | 2958 | 796 | 78 | 743 | 49 | 334 | 0 | 23 | 295 | 0 | 0 | 0 | 0 | 0 | 0 | 0 | 0 | 0 | 0 | 0 | 0 | 0 | 0 | 0 | 0 | 0 | 0 | 0 | 0 |
| SRR1608167 | 15 | 50 | 4375 | 0 | 112 | 48 | 4191 | 53416 | 2513 | 758 | 153 | 507 | 43 | 632 | 0 | 10 | 95 | 0 | 0 | 0 | 0 | 0 | 0 | 0 | 0 | 0 | 0 | 0 | 0 | 0 | 0 | 0 | 0 | 0 | 0 | 0 | 0 |
| SRR1608168 | 0 | 116 | 3161 | 0 | 74 | 22 | 2099 | 18854 | 4403 | 600 | 156 | 196 | 65 | 446 | 0 | 31 | 70 | 0 | 0 | 0 | 0 | 0 | 0 | 0 | 0 | 0 | 0 | 0 | 0 | 0 | 0 | 0 | 11 | 0 | 0 | 0 | 0 |
| SRR1608169 | 29 | 91 | 3903 | 0 | 105 | 39 | 3708 | 49124 | 2967 | 506 | 100 | 604 | 100 | 722 | 0 | 18 | 177 | 0 | 0 | 0 | 0 | 0 | 0 | 0 | 0 | 0 | 0 | 0 | 0 | 0 | 0 | 0 | 0 | 0 | 0 | 0 | 0 |
| SRR1608170 | 0 | 20 | 2321 | 0 | 70 | 49 | 3078 | 40603 | 3161 | 265 | 182 | 615 | 92 | 288 | 0 | 25 | 439 | 0 | 0 | 0 | 0 | 0 | 0 | 0 | 0 | 0 | 0 | 0 | 0 | 0 | 0 | 0 | 0 | 0 | 0 | 0 | 0 |
| SRR1608171 | 0 | 44 | 5136 | 0 | 54 | 23 | 2122 | 21837 | 1228 | 520 | 110 | 373 | 60 | 319 | 0 | 13 | 112 | 0 | 0 | 0 | 0 | 0 | 0 | 0 | 0 | 0 | 0 | 0 | 0 | 0 | 0 | 0 | 0 | 0 | 0 | 0 | 0 |
| SRR1608172 | 0 | 68 | 2056 | 0 | 219 | 107 | 8934 | 84185 | 1212 | 370 | 200 | 822 | 129 | 1431 | 0 | 15 | 324 | 0 | 0 | 0 | 0 | 0 | 0 | 0 | 0 | 0 | 0 | 0 | 0 | 0 | 0 | 0 | 0 | 0 | 0 | 0 | 0 |
| SRR1608174 | 0 | 40 | 2362 | 0 | 214 | 150 | 8538 | 96959 | 1080 | 628 | 156 | 937 | 43 | 642 | 0 | 44 | 92 | 0 | 0 | 0 | 0 | 0 | 0 | 0 | 0 | 0 | 0 | 0 | 0 | 0 | 0 | 0 | 0 | 0 | 0 | 0 | 0 |
| SRR1608175 | 19 | 100 | 5878 | 0 | 102 | 40 | 3923 | 40527 | 1720 | 2226 | 181 | 561 | 146 | 391 | 0 | 12 | 300 | 0 | 0 | 0 | 0 | 14 | 24 | 0 | 0 | 0 | 0 | 0 | 0 | 0 | 0 | 11 | 0 | 0 | 0 | 0 | 0 |
| SRR1608177 | 0 | 18 | 2420 | 0 | 114 | 75 | 5759 | 55174 | 4446 | 566 | 140 | 550 | 55 | 665 | 0 | 56 | 187 | 0 | 0 | 0 | 0 | 0 | 0 | 0 | 0 | 0 | 0 | 0 | 0 | 0 | 0 | 0 | 0 | 0 | 0 | 0 | 0 |
| SRR1608178 | 0 | 85 | 4254 | 0 | 200 | 103 | 8236 | 70505 | 3468 | 1931 | 218 | 736 | 89 | 528 | 0 | 16 | 186 | 0 | 0 | 0 | 0 | 0 | 0 | 0 | 0 | 0 | 0 | 0 | 0 | 0 | 0 | 0 | 0 | 0 | 0 | 0 | 0 |
| SRR1608179 | 50 | 134 | 5465 | 11 | 192 | 108 | 8391 | 81823 | 2375 | 1843 | 143 | 758 | 122 | 749 | 0 | 0 | 175 | 0 | 0 | 0 | 0 | 0 | 0 | 0 | 0 | 0 | 0 | 0 | 0 | 0 | 0 | 0 | 0 | 0 | 0 | 0 | 0 |
| SRR1608180 | 21 | 68 | 5408 | 0 | 119 | 95 | 5504 | 52723 | 3530 | 1190 | 169 | 853 | 85 | 881 | 0 | 24 | 130 | 0 | 0 | 0 | 0 | 0 | 0 | 0 | 0 | 0 | 0 | 0 | 0 | 0 | 0 | 0 | 0 | 0 | 0 | 0 | 0 |
| SRR1608181 | 0 | 35 | 2303 | 0 | 179 | 73 | 6734 | 66364 | 2128 | 380 | 308 | 980 | 156 | 539 | 0 | 39 | 279 | 0 | 0 | 0 | 0 | 0 | 0 | 0 | 0 | 0 | 0 | 0 | 0 | 0 | 0 | 0 | 0 | 0 | 0 | 0 | 0 |
| SRR1608182 | 0 | 53 | 3380 | 0 | 98 | 34 | 3366 | 41630 | 1017 | 441 | 102 | 377 | 64 | 420 | 0 | 0 | 130 | 0 | 0 | 0 | 0 | 0 | 0 | 0 | 0 | 0 | 0 | 0 | 0 | 0 | 0 | 0 | 0 | 0 | 0 | 0 | 0 |
| SRR1608184 | 0 | 21 | 4242 | 0 | 150 | 73 | 6103 | 57754 | 1187 | 462 | 227 | 971 | 155 | 705 | 0 | 17 | 105 | 0 | 0 | 0 | 0 | 0 | 0 | 0 | 0 | 0 | 0 | 0 | 0 | 0 | 0 | 0 | 0 | 0 | 0 | 0 | 0 |
| SRR1608185 | 0 | 90 | 4612 | 0 | 99 | 47 | 3349 | 39282 | 3691 | 433 | 254 | 697 | 103 | 355 | 0 | 25 | 482 | 0 | 0 | 0 | 0 | 0 | 0 | 0 | 0 | 0 | 0 | 0 | 0 | 0 | 0 | 0 | 0 | 0 | 0 | 0 | 0 |
| SRR1608186 | 0 | 44 | 3975 | 0 | 131 | 47 | 4670 | 49558 | 2445 | 915 | 102 | 512 | 70 | 401 | 0 | 11 | 90 | 0 | 0 | 0 | 0 | 0 | 0 | 0 | 0 | 0 | 0 | 0 | 0 | 0 | 0 | 0 | 0 | 0 | 0 | 0 | 0 |
| SRR1608189 | 0 | 21 | 3089 | 0 | 190 | 57 | 6159 | 71809 | 1443 | 382 | 223 | 574 | 92 | 351 | 0 | 10 | 192 | 0 | 0 | 0 | 0 | 0 | 0 | 0 | 0 | 0 | 0 | 0 | 0 | 0 | 0 | 0 | 0 | 0 | 0 | 0 | 0 |
| SRR1608190 | 0 | 41 | 4674 | 0 | 108 | 39 | 3107 | 39717 | 4733 | 1097 | 298 | 1393 | 197 | 494 | 0 | 60 | 374 | 0 | 0 | 0 | 0 | 0 | 0 | 0 | 0 | 0 | 0 | 0 | 0 | 0 | 0 | 0 | 0 | 0 | 0 | 0 | 0 |
| SRR1608191 | 0 | 15 | 5239 | 0 | 86 | 31 | 2678 | 26584 | 5120 | 510 | 402 | 1225 | 215 | 1270 | 0 | 31 | 749 | 0 | 0 | 0 | 0 | 0 | 0 | 0 | 0 | 0 | 0 | 0 | 0 | 0 | 0 | 0 | 0 | 0 | 0 | 0 | 0 |
| SRR1608192 | 0 | 43 | 3390 | 0 | 89 | 52 | 3714 | 48346 | 3862 | 894 | 238 | 820 | 79 | 919 | 0 | 24 | 205 | 0 | 0 | 0 | 0 | 0 | 0 | 0 | 0 | 0 | 0 | 0 | 0 | 0 | 0 | 0 | 0 | 0 | 0 | 0 | 0 |
| SRR1608193 | 0 | 68 | 5121 | 0 | 154 | 89 | 6242 | 60758 | 1627 | 1281 | 218 | 659 | 155 | 577 | 0 | 28 | 368 | 0 | 0 | 0 | 0 | 0 | 0 | 0 | 0 | 0 | 0 | 0 | 0 | 0 | 0 | 0 | 0 | 0 | 0 | 0 | 0 |
| SRR1608194 | 19 | 93 | 5882 | 0 | 134 | 62 | 5295 | 50562 | 1940 | 1461 | 194 | 624 | 161 | 507 | 0 | 21 | 293 | 0 | 0 | 0 | 0 | 0 | 0 | 0 | 0 | 0 | 0 | 0 | 0 | 0 | 0 | 0 | 0 | 0 | 0 | 0 | 0 |
| SRR1608196 | 0 | 62 | 4566 | 0 | 64 | 21 | 2330 | 27546 | 2546 | 485 | 206 | 739 | 66 | 334 | 0 | 27 | 337 | 0 | 0 | 0 | 0 | 0 | 0 | 0 | 0 | 0 | 0 | 0 | 0 | 0 | 0 | 0 | 0 | 0 | 0 | 0 | 0 |
| SRR1608201 | 0 | 21 | 6065 | 0 | 132 | 71 | 4912 | 57705 | 5590 | 1092 | 220 | 632 | 95 | 380 | 0 | 15 | 377 | 0 | 0 | 0 | 0 | 0 | 0 | 0 | 0 | 0 | 0 | 0 | 0 | 0 | 0 | 0 | 0 | 0 | 0 | 0 | 0 |
| SRR1608208 | 0 | 59 | 5058 | 0 | 134 | 59 | 4455 | 55566 | 2844 | 1287 | 347 | 1044 | 158 | 454 | 0 | 37 | 347 | 0 | 0 | 0 | 0 | 0 | 0 | 0 | 0 | 0 | 0 | 0 | 0 | 0 | 0 | 0 | 0 | 0 | 0 | 0 | 0 |
| SRR1608213 | 0 | 51 | 3480 | 0 | 165 | 96 | 6336 | 69455 | 2293 | 1198 | 237 | 878 | 121 | 605 | 0 | 30 | 224 | 0 | 0 | 0 | 0 | 0 | 0 | 0 | 0 | 0 | 0 | 0 | 0 | 0 | 0 | 0 | 0 | 0 | 0 | 0 | 0 |
| SRR1608214 | 0 | 24 | 5980 | 0 | 167 | 54 | 5422 | 54423 | 1212 | 954 | 155 | 627 | 66 | 633 | 0 | 11 | 192 | 0 | 0 | 0 | 0 | 0 | 0 | 0 | 0 | 0 | 0 | 0 | 0 | 0 | 0 | 0 | 0 | 0 | 0 | 0 | 0 |
| SRR1608222 | 0 | 11 | 11547 | 0 | 25 | 0 | 968 | 8446 | 779 | 37 | 918 | 1566 | 29 | 184 | 0 | 113 | 85 | 0 | 0 | 0 | 0 | 0 | 0 | 0 | 0 | 0 | 0 | 0 | 0 | 0 | 0 | 0 | 0 | 0 | 0 | 0 | 0 |
| SRR1608232 | 0 | 83 | 4593 | 0 | 69 | 28 | 2153 | 23279 | 1770 | 233 | 598 | 1426 | 48 | 357 | 0 | 133 | 117 | 0 | 0 | 0 | 0 | 0 | 0 | 0 | 0 | 0 | 0 | 0 | 0 | 0 | 0 | 0 | 0 | 0 | 0 | 0 | 0 |
| SRR1608237 | 0 | 38 | 8830 | 0 | 32 | 17 | 1079 | 11139 | 788 | 98 | 849 | 1425 | 66 | 311 | 0 | 148 | 164 | 0 | 0 | 0 | 0 | 0 | 0 | 0 | 0 | 0 | 0 | 0 | 0 | 0 | 0 | 0 | 0 | 0 | 0 | 0 | 0 |
| SRR1608254 | 0 | 71 | 2910 | 0 | 25 | 12 | 861 | 10154 | 1522 | 104 | 400 | 1181 | 60 | 300 | 0 | 117 | 162 | 0 | 0 | 0 | 0 | 0 | 0 | 0 | 0 | 0 | 0 | 0 | 0 | 0 | 0 | 0 | 0 | 0 | 0 | 0 | 0 |
| SRR1608255 | 0 | 23 | 2366 | 0 | 31 | 11 | 1015 | 10734 | 1173 | 83 | 518 | 1705 | 63 | 367 | 0 | 185 | 150 | 0 | 0 | 0 | 0 | 0 | 0 | 0 | 0 | 0 | 0 | 0 | 0 | 0 | 0 | 0 | 0 | 0 | 0 | 0 | 0 |
| SRR1608256 | 0 | 19 | 2871 | 0 | 34 | 14 | 1085 | 10934 | 1214 | 85 | 607 | 1517 | 65 | 307 | 0 | 175 | 193 | 0 | 0 | 0 | 0 | 0 | 0 | 0 | 0 | 0 | 0 | 0 | 0 | 0 | 0 | 0 | 0 | 0 | 0 | 0 | 0 |
| SRR1608257 | 0 | 97 | 4904 | 0 | 65 | 21 | 2014 | 20202 | 1331 | 282 | 270 | 823 | 81 | 424 | 0 | 77 | 267 | 0 | 0 | 0 | 0 | 0 | 0 | 0 | 0 | 0 | 0 | 0 | 0 | 0 | 0 | 0 | 0 | 0 | 0 | 0 | 0 |
| SRR1608260 | 0 | 73 | 2740 | 0 | 18 | 0 | 697 | 7705 | 1713 | 136 | 420 | 1409 | 80 | 380 | 0 | 184 | 281 | 0 | 0 | 0 | 0 | 0 | 0 | 0 | 0 | 0 | 0 | 0 | 0 | 0 | 0 | 0 | 0 | 0 | 0 | 0 | 0 |
| SRR1608263 | 0 | 36 | 9144 | 0 | 25 | 11 | 932 | 9654 | 833 | 83 | 385 | 1405 | 65 | 306 | 0 | 218 | 155 | 0 | 0 | 0 | 0 | 0 | 0 | 0 | 0 | 0 | 0 | 0 | 0 | 0 | 0 | 0 | 0 | 0 | 0 | 0 | 0 |
| SRR1608264 | 0 | 60 | 4581 | 0 | 18 | 0 | 710 | 7744 | 1069 | 101 | 534 | 1734 | 43 | 253 | 0 | 175 | 105 | 0 | 0 | 0 | 0 | 0 | 0 | 0 | 0 | 0 | 0 | 0 | 0 | 0 | 0 | 0 | 0 | 0 | 0 | 0 | 0 |
| SRR1608266 | 0 | 124 | 7031 | 0 | 40 | 15 | 1521 | 18228 | 1046 | 85 | 446 | 1299 | 39 | 267 | 0 | 84 | 77 | 0 | 0 | 0 | 0 | 0 | 0 | 0 | 0 | 0 | 0 | 0 | 0 | 0 | 0 | 0 | 0 | 0 | 0 | 0 | 0 |
| SRR1608267 | 0 | 20 | 7809 | 0 | 22 | 0 | 742 | 7817 | 803 | 59 | 527 | 1481 | 48 | 227 | 0 | 173 | 85 | 0 | 0 | 0 | 0 | 0 | 0 | 0 | 0 | 0 | 0 | 0 | 0 | 0 | 0 | 0 | 0 | 0 | 0 | 0 | 0 |
| SRR1608269 | 27 | 97 | 6904 | 0 | 22 | 0 | 721 | 7539 | 916 | 133 | 622 | 1496 | 64 | 286 | 0 | 160 | 142 | 0 | 0 | 0 | 0 | 0 | 0 | 0 | 0 | 0 | 0 | 0 | 0 | 0 | 0 | 0 | 0 | 0 | 0 | 0 | 0 |
| SRR1608270 | 0 | 79 | 6342 | 0 | 88 | 35 | 2670 | 26921 | 1401 | 249 | 409 | 941 | 54 | 401 | 0 | 70 | 92 | 0 | 0 | 0 | 0 | 0 | 0 | 0 | 0 | 0 | 0 | 0 | 0 | 0 | 0 | 0 | 0 | 0 | 0 | 0 | 0 |
| SRR1608271 | 0 | 75 | 4145 | 0 | 37 | 16 | 1265 | 15790 | 1929 | 220 | 386 | 1423 | 57 | 555 | 0 | 53 | 113 | 0 | 0 | 0 | 0 | 0 | 0 | 0 | 0 | 0 | 0 | 0 | 0 | 0 | 0 | 0 | 0 | 0 | 0 | 0 | 0 |
| SRR1608274 | 0 | 63 | 5345 | 0 | 54 | 24 | 1697 | 19334 | 1536 | 231 | 535 | 1273 | 77 | 370 | 0 | 106 | 174 | 0 | 0 | 0 | 0 | 0 | 0 | 0 | 0 | 0 | 0 | 0 | 0 | 0 | 0 | 0 | 0 | 0 | 0 | 0 | 0 |
| SRR1608276 | 38 | 198 | 5537 | 0 | 53 | 21 | 1798 | 21796 | 1197 | 292 | 480 | 1166 | 77 | 364 | 0 | 123 | 156 | 0 | 0 | 0 | 0 | 0 | 0 | 0 | 0 | 0 | 0 | 0 | 0 | 0 | 0 | 0 | 0 | 0 | 0 | 0 | 0 |
| SRR1608277 | 0 | 56 | 6883 | 0 | 61 | 22 | 2024 | 19690 | 1233 | 171 | 466 | 1345 | 73 | 372 | 0 | 157 | 156 | 0 | 0 | 0 | 0 | 0 | 0 | 0 | 0 | 0 | 0 | 0 | 0 | 0 | 0 | 0 | 0 | 0 | 0 | 0 | 0 |
| SRR1608278 | 0 | 10 | 8891 | 0 | 23 | 0 | 765 | 8269 | 913 | 44 | 670 | 1630 | 48 | 207 | 0 | 110 | 109 | 0 | 0 | 0 | 0 | 0 | 0 | 0 | 0 | 0 | 0 | 0 | 0 | 0 | 0 | 0 | 0 | 0 | 0 | 0 | 0 |
| SRR1608281 | 0 | 38 | 5153 | 0 | 30 | 12 | 1125 | 12681 | 1504 | 111 | 589 | 1827 | 66 | 345 | 0 | 197 | 126 | 0 | 0 | 0 | 0 | 0 | 0 | 0 | 0 | 0 | 0 | 0 | 0 | 0 | 0 | 0 | 0 | 0 | 0 | 0 | 0 |
| SRR1608282 | 0 | 65 | 5825 | 0 | 37 | 12 | 1090 | 12013 | 925 | 179 | 628 | 1585 | 59 | 267 | 0 | 180 | 155 | 0 | 0 | 0 | 0 | 0 | 0 | 0 | 0 | 0 | 0 | 0 | 0 | 0 | 0 | 0 | 0 | 0 | 0 | 0 | 0 |
| SRR1608283 | 0 | 32 | 11139 | 0 | 24 | 0 | 752 | 7936 | 812 | 108 | 683 | 1692 | 46 | 307 | 0 | 140 | 120 | 0 | 0 | 0 | 0 | 0 | 0 | 0 | 0 | 0 | 0 | 0 | 0 | 0 | 0 | 0 | 0 | 0 | 0 | 0 | 0 |
| SRR1608286 | 0 | 37 | 5272 | 0 | 38 | 15 | 1279 | 13481 | 986 | 52 | 680 | 1572 | 43 | 249 | 0 | 160 | 92 | 0 | 0 | 0 | 0 | 0 | 0 | 0 | 0 | 0 | 0 | 0 | 0 | 0 | 0 | 0 | 0 | 0 | 0 | 0 | 0 |
| SRR1608287 | 0 | 23 | 7767 | 0 | 29 | 14 | 1022 | 10504 | 987 | 147 | 691 | 1411 | 62 | 314 | 0 | 115 | 149 | 0 | 0 | 0 | 0 | 0 | 0 | 0 | 0 | 0 | 0 | 0 | 0 | 0 | 0 | 0 | 0 | 0 | 0 | 0 | 0 |
| SRR1608289 | 0 | 55 | 4333 | 0 | 33 | 11 | 1112 | 10430 | 1168 | 141 | 816 | 1742 | 57 | 292 | 0 | 134 | 128 | 0 | 0 | 0 | 0 | 0 | 0 | 0 | 0 | 0 | 0 | 0 | 0 | 0 | 0 | 0 | 0 | 0 | 0 | 0 | 0 |
| SRR1608290 | 0 | 41 | 5974 | 0 | 30 | 10 | 1116 | 11868 | 963 | 114 | 686 | 1700 | 46 | 329 | 0 | 158 | 110 | 0 | 0 | 0 | 0 | 0 | 0 | 0 | 0 | 0 | 0 | 0 | 0 | 0 | 0 | 0 | 0 | 0 | 0 | 0 | 0 |
| SRR1608302 | 0 | 21 | 8419 | 0 | 60 | 24 | 2148 | 23461 | 979 | 98 | 564 | 1165 | 46 | 240 | 0 | 94 | 73 | 0 | 0 | 0 | 0 | 0 | 0 | 0 | 0 | 0 | 0 | 0 | 0 | 0 | 0 | 0 | 0 | 0 | 0 | 0 | 0 |
| SRR1608307 | 0 | 13 | 7114 | 0 | 31 | 0 | 968 | 11994 | 772 | 95 | 631 | 1451 | 44 | 235 | 0 | 144 | 87 | 0 | 0 | 0 | 0 | 0 | 0 | 0 | 0 | 0 | 0 | 0 | 0 | 0 | 0 | 0 | 0 | 0 | 0 | 0 | 0 |
| SRR1608308 | 0 | 57 | 5637 | 0 | 31 | 13 | 1018 | 11060 | 991 | 191 | 515 | 1396 | 64 | 336 | 0 | 166 | 120 | 0 | 0 | 0 | 0 | 0 | 0 | 0 | 0 | 0 | 0 | 0 | 0 | 0 | 0 | 0 | 0 | 0 | 0 | 0 | 0 |
| SRR1634379 | 936 | 14427 | 1618 | 157 | 335 | 1570 | 4340 | 5756 | 7943 | 596 | 2333 | 4692 | 763 | 2084 | 0 | 3174 | 55514 | 0 | 0 | 21 | 640 | 248 | 153 | 0 | 0 | 0 | 0 | 41 | 355 | 598 | 423 | 397 | 0 | 0 | 35 | 19 | 0 |
| SRR1634381 | 1495 | 15377 | 2865 | 279 | 1172 | 3243 | 8943 | 7828 | 6918 | 796 | 3377 | 6483 | 1126 | 2303 | 0 | 6797 | 47332 | 0 | 14 | 36 | 868 | 221 | 81 | 0 | 0 | 0 | 0 | 101 | 604 | 540 | 314 | 150 | 0 | 15 | 27 | 44 | 0 |
| SRR1634384 | 2207 | 22424 | 881 | 391 | 263 | 395 | 1568 | 1740 | 1383 | 163 | 1818 | 4229 | 623 | 881 | 0 | 8128 | 39131 | 0 | 13 | 36 | 1371 | 242 | 93 | 0 | 0 | 0 | 0 | 78 | 498 | 447 | 219 | 107 | 0 | 0 | 14 | 22 | 0 |
| SRR1634388 | 1954 | 24822 | 1478 | 424 | 392 | 715 | 2710 | 2625 | 1110 | 316 | 1770 | 3373 | 520 | 922 | 0 | 3352 | 30113 | 0 | 16 | 46 | 5846 | 522 | 135 | 0 | 0 | 0 | 0 | 88 | 580 | 1748 | 458 | 141 | 0 | 0 | 18 | 31 | 0 |
| SRR1634390 | 665 | 9021 | 4036 | 331 | 1060 | 3038 | 9096 | 5962 | 6343 | 1820 | 6732 | 8136 | 1838 | 2864 | 0 | 7066 | 56479 | 0 | 18 | 41 | 1284 | 424 | 96 | 0 | 0 | 0 | 0 | 110 | 636 | 579 | 635 | 157 | 0 | 16 | 27 | 62 | 0 |
| SRR1634392 | 1196 | 18526 | 2211 | 231 | 489 | 927 | 2993 | 2566 | 1971 | 778 | 2712 | 4268 | 764 | 1275 | 0 | 4464 | 41677 | 0 | 17 | 41 | 1397 | 304 | 103 | 0 | 0 | 0 | 0 | 98 | 568 | 552 | 304 | 123 | 0 | 13 | 21 | 45 | 0 |
| SRR1634394 | 1192 | 18163 | 2290 | 182 | 440 | 1219 | 3884 | 2834 | 1818 | 787 | 2444 | 4177 | 762 | 960 | 0 | 2828 | 28903 | 0 | 15 | 45 | 2879 | 412 | 142 | 0 | 0 | 0 | 0 | 76 | 508 | 917 | 351 | 158 | 0 | 10 | 21 | 41 | 0 |
| SRR1634395 | 784 | 10508 | 4335 | 436 | 3247 | 15342 | 25088 | 30229 | 11896 | 1612 | 7272 | 8725 | 2852 | 5383 | 12 | 4120 | 51535 | 0 | 0 | 18 | 967 | 301 | 75 | 0 | 0 | 0 | 0 | 56 | 352 | 569 | 458 | 166 | 0 | 14 | 30 | 54 | 0 |
| SRR1634398 | 1185 | 11656 | 2444 | 414 | 5597 | 20611 | 58050 | 52524 | 5636 | 433 | 10623 | 10723 | 3977 | 9010 | 26 | 1698 | 38224 | 0 | 12 | 36 | 1872 | 505 | 120 | 0 | 10 | 0 | 0 | 98 | 535 | 1028 | 774 | 199 | 0 | 16 | 20 | 64 | 0 |
| SRR1634399 | 717 | 10038 | 3316 | 447 | 1751 | 4189 | 11153 | 10998 | 3829 | 1518 | 5538 | 10022 | 1904 | 3792 | 0 | 5179 | 62492 | 0 | 13 | 36 | 2332 | 582 | 178 | 0 | 0 | 0 | 0 | 80 | 509 | 1173 | 827 | 297 | 0 | 18 | 33 | 54 | 0 |
| SRR1634400 | 722 | 10189 | 3228 | 436 | 1698 | 4183 | 11109 | 10850 | 4121 | 1524 | 5492 | 10047 | 1879 | 3741 | 0 | 5297 | 62775 | 0 | 16 | 38 | 2307 | 552 | 181 | 0 | 0 | 0 | 0 | 82 | 512 | 1158 | 836 | 297 | 0 | 19 | 33 | 49 | 0 |
| SRR1634406 | 842 | 15118 | 2217 | 352 | 528 | 1157 | 3987 | 4179 | 2618 | 747 | 2954 | 5379 | 841 | 1431 | 0 | 4173 | 51182 | 0 | 0 | 26 | 658 | 177 | 63 | 0 | 0 | 0 | 0 | 68 | 453 | 398 | 245 | 122 | 0 | 13 | 19 | 30 | 0 |
| SRR1634409 | 615 | 9464 | 5619 | 303 | 5597 | 19378 | 53843 | 58480 | 8557 | 1380 | 9243 | 7877 | 3126 | 8774 | 20 | 3414 | 46110 | 0 | 0 | 21 | 578 | 154 | 56 | 0 | 0 | 0 | 0 | 60 | 347 | 396 | 280 | 112 | 0 | 11 | 22 | 53 | 0 |
| SRR1634411 | 406 | 6130 | 2489 | 980 | 1952 | 3834 | 14772 | 14166 | 4206 | 1057 | 3972 | 5385 | 1276 | 3554 | 0 | 8508 | 44225 | 0 | 0 | 29 | 3320 | 677 | 113 | 0 | 0 | 0 | 0 | 52 | 360 | 1791 | 1115 | 268 | 0 | 0 | 16 | 26 | 0 |
| SRR1634412 | 383 | 6015 | 2487 | 1022 | 1920 | 3993 | 14696 | 14165 | 4118 | 1077 | 3834 | 5372 | 1344 | 3406 | 0 | 8533 | 44670 | 0 | 0 | 25 | 3297 | 673 | 110 | 0 | 0 | 0 | 0 | 52 | 371 | 1758 | 1127 | 270 | 0 | 0 | 16 | 23 | 0 |
| SRR1634414 | 913 | 12569 | 4914 | 347 | 3783 | 14282 | 34785 | 26434 | 8171 | 948 | 8462 | 12085 | 2178 | 4583 | 0 | 4792 | 45873 | 0 | 12 | 25 | 611 | 159 | 55 | 0 | 0 | 0 | 0 | 67 | 354 | 303 | 215 | 96 | 0 | 13 | 16 | 47 | 0 |
| SRR1634415 | 671 | 11000 | 3540 | 335 | 3349 | 11451 | 25746 | 19333 | 4252 | 1208 | 4011 | 6098 | 1250 | 3195 | 0 | 3243 | 40172 | 0 | 0 | 19 | 694 | 222 | 68 | 0 | 0 | 0 | 0 | 76 | 472 | 544 | 428 | 185 | 0 | 13 | 24 | 35 | 0 |
| SRR1634417 | 655 | 10909 | 5730 | 398 | 3226 | 10824 | 28342 | 22083 | 78394 | 1551 | 7050 | 8423 | 2324 | 3160 | 0 | 15653 | 69652 | 0 | 0 | 12 | 319 | 160 | 40 | 0 | 0 | 0 | 0 | 74 | 414 | 431 | 424 | 162 | 0 | 20 | 32 | 46 | 0 |
| SRR1636959 | 1087 | 97066 | 1193 | 322 | 345 | 2834 | 3986 | 5929 | 5044 | 24 | 11628 | 21431 | 2263 | 2618 | 0 | 5859 | 29329 | 0 | 0 | 0 | 1820 | 31 | 61 | 0 | 0 | 0 | 0 | 0 | 27 | 669 | 28 | 16 | 0 | 0 | 13 | 0 | 0 |
| SRR1636960 | 436 | 103264 | 1015 | 111 | 638 | 5557 | 6295 | 8240 | 3513 | 19 | 20764 | 24135 | 4025 | 2417 | 0 | 6238 | 26729 | 0 | 0 | 0 | 979 | 31 | 51 | 0 | 0 | 0 | 0 | 0 | 33 | 397 | 35 | 22 | 0 | 11 | 13 | 0 | 0 |
| SRR1636961 | 332 | 104193 | 963 | 122 | 516 | 5199 | 5581 | 7593 | 3228 | 19 | 20659 | 24720 | 4193 | 2193 | 0 | 6583 | 30225 | 0 | 0 | 0 | 693 | 23 | 47 | 0 | 0 | 0 | 0 | 0 | 25 | 261 | 35 | 18 | 0 | 12 | 13 | 0 | 0 |
| SRR1636962 | 75 | 117377 | 193 | 410 | 225 | 2618 | 1890 | 2777 | 7634 | 0 | 1028 | 8304 | 432 | 63 | 0 | 8586 | 43693 | 0 | 0 | 0 | 119 | 0 | 28 | 0 | 0 | 0 | 0 | 0 | 35 | 60 | 13 | 13 | 0 | 10 | 11 | 0 | 0 |
| SRR1636963 | 57 | 97821 | 245 | 517 | 185 | 2082 | 1328 | 2252 | 7341 | 0 | 749 | 8117 | 287 | 64 | 0 | 6905 | 39237 | 0 | 0 | 0 | 127 | 0 | 38 | 0 | 0 | 0 | 0 | 0 | 31 | 68 | 15 | 25 | 0 | 16 | 20 | 0 | 0 |
| SRR1636964 | 66 | 105572 | 194 | 499 | 147 | 1592 | 1071 | 1764 | 5754 | 15 | 650 | 7751 | 269 | 65 | 0 | 5953 | 40619 | 0 | 0 | 0 | 157 | 10 | 38 | 0 | 10 | 0 | 0 | 0 | 29 | 74 | 14 | 20 | 0 | 17 | 33 | 0 | 0 |
| SRR1636965 | 1148 | 8852 | 562 | 2082 | 1007 | 7992 | 14365 | 8536 | 16155 | 15 | 1498 | 2258 | 286 | 150 | 0 | 7088 | 49152 | 0 | 0 | 0 | 1348 | 1104 | 96 | 0 | 0 | 0 | 0 | 0 | 150 | 874 | 1356 | 50 | 0 | 14 | 90 | 0 | 0 |
| SRR1636966 | 986 | 9646 | 153 | 511 | 371 | 2229 | 3419 | 4495 | 8880 | 15 | 66 | 834 | 11 | 0 | 0 | 2989 | 53360 | 0 | 0 | 0 | 387 | 420 | 39 | 0 | 0 | 0 | 0 | 0 | 97 | 290 | 882 | 34 | 0 | 20 | 423 | 0 | 0 |
| SRR1636967 | 1342 | 32648 | 157 | 235 | 209 | 1672 | 1555 | 2041 | 5404 | 0 | 63 | 903 | 11 | 0 | 14 | 3212 | 46906 | 0 | 0 | 0 | 570 | 307 | 49 | 0 | 0 | 0 | 0 | 0 | 114 | 348 | 602 | 43 | 0 | 19 | 381 | 0 | 0 |
| SRR1636968 | 209 | 70711 | 4389 | 340 | 1131 | 8141 | 11865 | 6605 | 37549 | 33 | 150 | 1692 | 31 | 0 | 14 | 12282 | 68747 | 0 | 0 | 0 | 6428 | 453 | 148 | 0 | 11 | 0 | 0 | 0 | 64 | 4304 | 1148 | 117 | 0 | 32 | 126 | 0 | 0 |
| SRR1636969 | 194 | 84010 | 3984 | 293 | 1205 | 12115 | 12578 | 12330 | 32212 | 32 | 132 | 1397 | 23 | 0 | 17 | 14339 | 46635 | 0 | 0 | 0 | 7885 | 295 | 109 | 0 | 0 | 0 | 0 | 0 | 52 | 4664 | 546 | 78 | 0 | 24 | 56 | 0 | 0 |
| SRR1636970 | 293 | 71361 | 3804 | 307 | 1246 | 10618 | 13316 | 12370 | 46016 | 25 | 111 | 820 | 18 | 0 | 16 | 13615 | 37479 | 0 | 0 | 0 | 12058 | 461 | 123 | 0 | 10 | 0 | 0 | 0 | 75 | 6092 | 645 | 69 | 0 | 18 | 62 | 0 | 0 |
| SRR1642941 | 499 | 1794 | 5273 | 1501 | 3734 | 1981 | 38472 | 23808 | 29964 | 1654 | 16147 | 13186 | 15148 | 4964 | 251 | 1981 | 28155 | 0 | 26 | 0 | 19 | 130 | 87 | 25 | 14 | 12 | 16 | 48 | 63 | 37 | 63 | 0 | 0 | 0 | 0 | 16 | 0 |
| SRR1642942 | 646 | 3674 | 10287 | 1752 | 2304 | 3187 | 30605 | 12939 | 33492 | 2644 | 2251 | 3336 | 1780 | 645 | 626 | 129 | 12641 | 0 | 51 | 11 | 20 | 79 | 55 | 35 | 16 | 26 | 0 | 45 | 33 | 17 | 21 | 0 | 0 | 0 | 0 | 37 | 0 |
| SRR1642943 | 332 | 1349 | 5902 | 1548 | 2025 | 1171 | 21067 | 17505 | 45843 | 2974 | 12316 | 6742 | 11919 | 3618 | 279 | 1506 | 8215 | 0 | 0 | 17 | 71 | 82 | 42 | 10 | 0 | 13 | 11 | 87 | 202 | 419 | 150 | 12 | 0 | 0 | 0 | 24 | 0 |
| SRR1642944 | 579 | 2561 | 8755 | 1689 | 8846 | 4198 | 102741 | 38985 | 13303 | 3605 | 10063 | 6124 | 10776 | 4729 | 120 | 371 | 16572 | 0 | 0 | 0 | 0 | 0 | 0 | 0 | 0 | 0 | 0 | 0 | 12 | 0 | 0 | 0 | 0 | 0 | 0 | 0 | 0 |
| SRR1642945 | 294 | 1222 | 4144 | 2013 | 2180 | 1434 | 19437 | 15383 | 61794 | 3592 | 10611 | 5623 | 10280 | 2337 | 133 | 943 | 14541 | 0 | 38 | 24 | 69 | 315 | 60 | 12 | 0 | 10 | 29 | 240 | 325 | 345 | 451 | 28 | 0 | 0 | 0 | 26 | 0 |
| SRR1642946 | 447 | 1666 | 6508 | 2609 | 4224 | 2815 | 50987 | 26059 | 52068 | 3972 | 3805 | 5114 | 2795 | 940 | 281 | 286 | 16865 | 0 | 0 | 0 | 54 | 203 | 233 | 73 | 26 | 26 | 15 | 32 | 52 | 132 | 75 | 26 | 0 | 0 | 16 | 48 | 0 |
| SRR1642947 | 450 | 1512 | 3163 | 1937 | 2216 | 1271 | 16549 | 12768 | 76115 | 2791 | 7994 | 4823 | 6669 | 1233 | 114 | 1407 | 10496 | 0 | 11 | 0 | 0 | 34 | 25 | 0 | 0 | 0 | 46 | 95 | 148 | 75 | 103 | 0 | 0 | 0 | 0 | 12 | 0 |
| SRR1642948 | 568 | 1941 | 6234 | 1866 | 7633 | 5096 | 81293 | 49344 | 57007 | 2128 | 11020 | 6138 | 9036 | 3004 | 214 | 1212 | 12542 | 0 | 24 | 0 | 0 | 78 | 29 | 23 | 0 | 17 | 13 | 99 | 85 | 39 | 74 | 0 | 0 | 0 | 0 | 24 | 0 |
| SRR1642949 | 469 | 1902 | 3859 | 1695 | 3182 | 1627 | 33559 | 22364 | 33367 | 2983 | 12046 | 9902 | 11913 | 3306 | 110 | 1086 | 21653 | 0 | 0 | 0 | 0 | 0 | 0 | 0 | 0 | 0 | 19 | 42 | 74 | 35 | 46 | 0 | 0 | 0 | 0 | 0 | 0 |
| SRR1642950 | 459 | 1606 | 3566 | 1809 | 4056 | 2928 | 38492 | 27568 | 60148 | 1624 | 9162 | 5904 | 8900 | 1783 | 159 | 967 | 8170 | 0 | 0 | 0 | 0 | 0 | 0 | 0 | 0 | 0 | 18 | 44 | 70 | 35 | 50 | 0 | 0 | 0 | 0 | 0 | 0 |
| SRR1642951 | 606 | 1897 | 4132 | 1069 | 2848 | 1379 | 29020 | 22980 | 27204 | 2415 | 11018 | 9056 | 11024 | 2642 | 214 | 1067 | 17629 | 0 | 0 | 0 | 0 | 44 | 13 | 0 | 0 | 12 | 18 | 44 | 62 | 37 | 49 | 0 | 0 | 0 | 0 | 12 | 0 |
| SRR1642952 | 418 | 2101 | 4235 | 2083 | 6118 | 4559 | 58803 | 37149 | 58381 | 4228 | 4568 | 2172 | 4209 | 1613 | 165 | 1134 | 3970 | 0 | 0 | 0 | 16 | 105 | 57 | 29 | 12 | 19 | 19 | 38 | 64 | 51 | 81 | 13 | 0 | 0 | 0 | 28 | 0 |
| SRR1642953 | 529 | 3242 | 10520 | 771 | 2067 | 1684 | 16991 | 13411 | 84244 | 3074 | 10406 | 6228 | 10425 | 2224 | 160 | 2085 | 15556 | 0 | 0 | 0 | 43 | 56 | 44 | 12 | 0 | 0 | 18 | 67 | 196 | 300 | 140 | 22 | 0 | 0 | 0 | 13 | 0 |
| SRR1642954 | 288 | 2272 | 11137 | 906 | 9990 | 6206 | 138221 | 67351 | 22672 | 3876 | 15301 | 7464 | 16044 | 5618 | 589 | 2286 | 13777 | 0 | 19 | 11 | 53 | 314 | 391 | 86 | 33 | 35 | 10 | 48 | 94 | 125 | 141 | 50 | 0 | 0 | 11 | 53 | 0 |
| SRR1642955 | 148 | 1171 | 7899 | 612 | 942 | 739 | 12659 | 8246 | 34571 | 5505 | 10289 | 5803 | 12370 | 4673 | 176 | 1451 | 7944 | 0 | 35 | 33 | 127 | 661 | 287 | 35 | 0 | 0 | 0 | 306 | 479 | 739 | 981 | 106 | 0 | 0 | 0 | 19 | 0 |
| SRR1642956 | 300 | 4417 | 6607 | 1078 | 5199 | 4055 | 72684 | 29980 | 29652 | 5632 | 3581 | 1436 | 3811 | 2550 | 4433 | 357 | 12808 | 0 | 0 | 0 | 34 | 579 | 553 | 116 | 42 | 82 | 24 | 41 | 122 | 97 | 136 | 63 | 0 | 0 | 16 | 92 | 0 |
| SRR1642957 | 449 | 3049 | 6022 | 909 | 8625 | 5213 | 76987 | 56711 | 52387 | 4213 | 5214 | 4534 | 5565 | 2638 | 367 | 1292 | 14733 | 0 | 34 | 29 | 33 | 192 | 51 | 37 | 12 | 26 | 21 | 210 | 360 | 167 | 216 | 15 | 0 | 0 | 0 | 41 | 0 |
| SRR1642958 | 425 | 3582 | 7016 | 1053 | 2817 | 1808 | 35772 | 21443 | 20985 | 3443 | 14687 | 10047 | 13969 | 3899 | 358 | 1289 | 21169 | 0 | 39 | 0 | 15 | 74 | 31 | 20 | 0 | 16 | 16 | 151 | 134 | 68 | 98 | 0 | 0 | 0 | 0 | 22 | 0 |
| SRR1642959 | 399 | 2571 | 7320 | 884 | 2969 | 1702 | 30446 | 22339 | 43330 | 2822 | 13655 | 8290 | 13776 | 3611 | 222 | 1017 | 18554 | 0 | 0 | 0 | 0 | 27 | 14 | 12 | 0 | 0 | 20 | 64 | 118 | 47 | 61 | 0 | 0 | 0 | 0 | 14 | 0 |
| SRR1642960 | 388 | 2519 | 6570 | 878 | 4320 | 2614 | 43301 | 27005 | 48186 | 2130 | 12652 | 6400 | 11317 | 3533 | 202 | 1099 | 9070 | 0 | 19 | 18 | 27 | 60 | 39 | 33 | 16 | 29 | 17 | 69 | 156 | 64 | 88 | 0 | 0 | 0 | 0 | 32 | 0 |
| SRR1642961 | 323 | 2072 | 8458 | 1046 | 1725 | 1500 | 17431 | 8753 | 92890 | 3785 | 10149 | 5230 | 9755 | 1585 | 154 | 1849 | 12696 | 0 | 32 | 22 | 74 | 265 | 198 | 25 | 0 | 0 | 19 | 209 | 329 | 319 | 361 | 78 | 0 | 0 | 0 | 23 | 0 |
| SRR1642962 | 477 | 2739 | 9568 | 572 | 3587 | 3893 | 52491 | 30383 | 74695 | 3797 | 14075 | 6696 | 12627 | 1540 | 205 | 1024 | 10459 | 0 | 0 | 0 | 0 | 61 | 34 | 14 | 0 | 0 | 16 | 68 | 121 | 65 | 107 | 12 | 0 | 0 | 0 | 16 | 0 |
| SRR1642963 | 795 | 4338 | 9742 | 769 | 2843 | 2081 | 19949 | 19599 | 85117 | 3031 | 11920 | 6463 | 11195 | 2118 | 178 | 4618 | 15548 | 0 | 0 | 0 | 12 | 37 | 27 | 15 | 0 | 10 | 38 | 138 | 254 | 134 | 207 | 0 | 0 | 0 | 0 | 17 | 0 |

### Supp Table S8.1.Columns_B_to_AL

| | | | | | | | | | | | | | | | | | | | | | | | | | | | | | | | | | | | | | |
|---|---|---|---|---|---|---|---|---|---|---|---|---|---|---|---|---|---|---|---|---|---|---|---|---|---|---|---|---|---|---|---|---|---|---|---|---|---|
| SRR1642964 | 392 | 3162 | 8621 | 1190 | 1248 | 1220 | 13988 | 9470 | 107500 | 6414 | 10159 | 7474 | 10468 | 854 | 116 | 492 | 7434 | 0 | 17 | 0 | 10 | 68 | 20 | 0 | 0 | 0 | 0 | 98 | 82 | 34 | 51 | 0 | 0 | 0 | 0 | 0 | 0 |
| SRR1642965 | 20 | 582 | 7035 | 46 | 3815 | 2665 | 41772 | 24013 | 31148 | 3691 | 10732 | 6807 | 9693 | 3872 | 170 | 1710 | 13554 | 0 | 0 | 0 | 0 | 22 | 13 | 12 | 0 | 14 | 0 | 0 | 22 | 18 | 25 | 0 | 0 | 0 | 0 | 21 | 0 |
| SRR1642966 | 21 | 596 | 5661 | 79 | 4695 | 2562 | 48827 | 23369 | 89923 | 2838 | 11833 | 4891 | 9463 | 3399 | 606 | 5765 | 7427 | 0 | 0 | 0 | 17 | 69 | 57 | 27 | 16 | 34 | 0 | 0 | 27 | 22 | 50 | 0 | 0 | 0 | 0 | 37 | 0 |
| SRR1642967 | 27 | 675 | 9213 | 55 | 3150 | 1602 | 28954 | 23832 | 26437 | 3893 | 12553 | 7782 | 11817 | 3543 | 246 | 1427 | 15647 | 0 | 0 | 0 | 21 | 259 | 67 | 17 | 0 | 20 | 0 | 26 | 61 | 42 | 131 | 11 | 0 | 0 | 0 | 26 | 0 |
| SRR1642968 | 19 | 703 | 8836 | 62 | 2827 | 1700 | 27397 | 24922 | 23408 | 4786 | 13782 | 9150 | 12666 | 4347 | 253 | 1430 | 18293 | 0 | 0 | 0 | 0 | 28 | 34 | 0 | 0 | 0 | 0 | 0 | 14 | 14 | 24 | 0 | 0 | 0 | 0 | 14 | 0 |
| SRR1642969 | 71 | 1142 | 15551 | 57 | 2211 | 1556 | 20787 | 17022 | 78775 | 5528 | 13165 | 6278 | 10984 | 1837 | 163 | 2665 | 15244 | 0 | 0 | 0 | 0 | 64 | 158 | 29 | 15 | 0 | 0 | 33 | 57 | 84 | 179 | 93 | 12 | 0 | 14 | 20 | 0 |
| SRR1642970 | 32 | 732 | 14765 | 66 | 6747 | 4963 | 70586 | 43832 | 73466 | 4049 | 5537 | 5882 | 4939 | 1551 | 203 | 1728 | 11607 | 0 | 14 | 0 | 12 | 190 | 98 | 20 | 0 | 20 | 0 | 88 | 50 | 43 | 186 | 30 | 0 | 0 | 0 | 28 | 0 |
| SRR1642971 | 36 | 709 | 11723 | 69 | 4149 | 2415 | 45020 | 28628 | 46570 | 5369 | 15173 | 8088 | 14378 | 3971 | 297 | 3396 | 15891 | 0 | 0 | 0 | 11 | 67 | 245 | 74 | 38 | 19 | 0 | 14 | 29 | 42 | 88 | 54 | 0 | 0 | 16 | 29 | 0 |
| SRR1642972 | 26 | 771 | 12260 | 64 | 6921 | 5407 | 85497 | 42632 | 56510 | 7046 | 18198 | 8453 | 12961 | 6978 | 301 | 1386 | 12699 | 0 | 0 | 0 | 0 | 22 | 18 | 14 | 0 | 13 | 0 | 20 | 30 | 20 | 44 | 0 | 0 | 0 | 0 | 25 | 0 |
| SRR1642973 | 24 | 656 | 11724 | 63 | 3390 | 1870 | 34417 | 27661 | 30915 | 3825 | 10603 | 6453 | 11635 | 3106 | 270 | 1959 | 15579 | 0 | 0 | 0 | 0 | 35 | 23 | 0 | 0 | 0 | 0 | 10 | 25 | 26 | 45 | 0 | 0 | 0 | 0 | 15 | 0 |
| SRR1642974 | 29 | 806 | 14342 | 53 | 3275 | 1984 | 41056 | 26335 | 14486 | 3663 | 15794 | 16439 | 14441 | 3164 | 155 | 577 | 22963 | 0 | 0 | 0 | 0 | 34 | 13 | 0 | 0 | 0 | 0 | 0 | 0 | 0 | 16 | 0 | 0 | 0 | 0 | 11 | 0 |
| SRR1642975 | 39 | 911 | 12560 | 69 | 4001 | 2045 | 28817 | 35471 | 41432 | 6446 | 11768 | 7955 | 11483 | 2576 | 182 | 1387 | 18608 | 0 | 0 | 0 | 0 | 21 | 24 | 12 | 0 | 0 | 0 | 0 | 17 | 16 | 49 | 0 | 0 | 0 | 0 | 12 | 0 |
| SRR1642976 | 22 | 542 | 10483 | 66 | 2642 | 1651 | 26901 | 14845 | 81647 | 5596 | 9892 | 6200 | 9387 | 2604 | 131 | 2566 | 10868 | 0 | 0 | 0 | 11 | 174 | 194 | 47 | 24 | 42 | 0 | 14 | 20 | 28 | 104 | 36 | 0 | 0 | 0 | 54 | 0 |
| SRR1642977 | 197 | 2668 | 11006 | 272 | 1664 | 996 | 18659 | 15509 | 24819 | 2877 | 23999 | 17706 | 23379 | 6995 | 308 | 1788 | 30100 | 0 | 137 | 60 | 83 | 732 | 146 | 26 | 11 | 20 | 0 | 316 | 345 | 88 | 252 | 0 | 0 | 0 | 0 | 25 | 0 |
| SRR1642978 | 210 | 1936 | 11009 | 364 | 3604 | 3466 | 35405 | 26151 | 93599 | 3703 | 15564 | 6705 | 12284 | 2355 | 356 | 511 | 9904 | 0 | 44 | 31 | 47 | 175 | 55 | 13 | 0 | 0 | 0 | 334 | 435 | 220 | 185 | 15 | 0 | 0 | 0 | 17 | 0 |
| SRR1642979 | 271 | 2863 | 8169 | 497 | 6569 | 4091 | 91445 | 63963 | 18118 | 3417 | 27817 | 27 | 21033 | 6295 | 407 | 1086 | 32505 | 0 | 0 | 0 | 0 | 24 | 18 | 12 | 0 | 0 | 0 | 26 | 70 | 37 | 85 | 0 | 0 | 0 | 0 | 19 | 0 |
| SRR1642980 | 170 | 2219 | 14283 | 575 | 7411 | 8900 | 124721 | 38296 | 60923 | 3893 | 9857 | 3702 | 9711 | 3867 | 496 | 553 | 5153 | 0 | 0 | 0 | 0 | 19 | 11 | 0 | 0 | 0 | 0 | 26 | 44 | 17 | 13 | 0 | 0 | 0 | 0 | 0 | 0 |
| SRR1642981 | 118 | 1535 | 7264 | 407 | 3522 | 1773 | 34229 | 25355 | 27766 | 3069 | 18940 | 12242 | 18040 | 5575 | 236 | 978 | 25600 | 0 | 52 | 41 | 43 | 335 | 79 | 25 | 13 | 14 | 0 | 196 | 258 | 59 | 114 | 0 | 0 | 0 | 0 | 26 | 0 |
| SRR1642982 | 165 | 1959 | 14107 | 355 | 4873 | 4007 | 56719 | 36627 | 54406 | 1841 | 15511 | 5991 | 15446 | 3045 | 245 | 1514 | 12699 | 0 | 84 | 29 | 37 | 742 | 124 | 16 | 0 | 11 | 0 | 947 | 643 | 159 | 741 | 49 | 0 | 0 | 0 | 23 | 0 |
| SRR1642983 | 227 | 3647 | 12038 | 269 | 2125 | 1576 | 18795 | 11108 | 57374 | 4664 | 13596 | 6647 | 11785 | 2078 | 132 | 911 | 11073 | 0 | 24 | 19 | 87 | 96 | 35 | 0 | 0 | 0 | 0 | 188 | 394 | 474 | 159 | 11 | 0 | 0 | 0 | 11 | 0 |
| SRR1642984 | 230 | 4419 | 12945 | 477 | 9830 | 7386 | 100248 | 68908 | 112685 | 7313 | 15204 | 4333 | 9959 | 1837 | 442 | 670 | 8128 | 0 | 16 | 0 | 15 | 121 | 34 | 0 | 0 | 0 | 0 | 118 | 93 | 60 | 134 | 0 | 0 | 0 | 0 | 13 | 0 |
| SRR1642985 | 166 | 2534 | 9574 | 336 | 1561 | 919 | 16799 | 12031 | 20762 | 2888 | 10456 | 8841 | 9932 | 2329 | 176 | 849 | 15396 | 0 | 17 | 13 | 91 | 279 | 115 | 13 | 0 | 0 | 0 | 62 | 109 | 133 | 112 | 14 | 0 | 0 | 0 | 13 | 0 |
| SRR1642986 | 298 | 3292 | 10819 | 338 | 1596 | 914 | 19377 | 12320 | 23227 | 2678 | 11856 | 9165 | 11081 | 2293 | 145 | 831 | 16751 | 0 | 0 | 0 | 0 | 73 | 18 | 0 | 0 | 0 | 0 | 36 | 60 | 23 | 41 | 0 | 0 | 0 | 0 | 0 | 0 |
| SRR1642987 | 148 | 1712 | 9119 | 389 | 1135 | 1109 | 14733 | 6122 | 71847 | 3193 | 11321 | 4944 | 9914 | 2150 | 138 | 890 | 13420 | 0 | 0 | 0 | 12 | 0 | 0 | 0 | 0 | 0 | 0 | 20 | 98 | 87 | 20 | 0 | 0 | 0 | 0 | 0 | 0 |
| SRR1642988 | 214 | 2173 | 14937 | 417 | 6030 | 2879 | 80912 | 46743 | 29212 | 6944 | 19354 | 9300 | 14298 | 13548 | 142 | 238 | 22185 | 0 | 0 | 0 | 14 | 66 | 123 | 25 | 12 | 10 | 0 | 33 | 41 | 42 | 32 | 19 | 0 | 0 | 0 | 20 | 0 |
| SRR1657194 | 20 | 12385 | 987 | 141 | 125 | 1342 | 1516 | 678 | 17085 | 145 | 4443 | 25708 | 736 | 169 | 88 | 20478 | 15381 | 0 | 0 | 0 | 3233 | 350 | 203 | 0 | 0 | 0 | 0 | 0 | 46 | 802 | 214 | 122 | 0 | 13 | 0 | 26 | 0 |
| SRR1657195 | 0 | 9383 | 1232 | 46 | 95 | 1609 | 1834 | 815 | 12558 | 346 | 2224 | 17117 | 274 | 45 | 82 | 25004 | 9208 | 0 | 0 | 0 | 2961 | 381 | 158 | 0 | 0 | 0 | 0 | 0 | 35 | 783 | 232 | 104 | 0 | 14 | 0 | 31 | 0 |
| SRR1657196 | 13 | 15869 | 838 | 45 | 122 | 1601 | 1489 | 783 | 15356 | 120 | 1782 | 22361 | 240 | 36 | 169 | 21311 | 14260 | 0 | 0 | 0 | 4338 | 471 | 242 | 0 | 15 | 0 | 0 | 0 | 52 | 1229 | 230 | 150 | 0 | 23 | 0 | 30 | 0 |
| SRR1657197 | 97 | 13721 | 1161 | 484 | 106 | 1452 | 1358 | 814 | 18632 | 152 | 5772 | 38520 | 757 | 119 | 12 | 25602 | 14664 | 0 | 0 | 0 | 3643 | 360 | 327 | 0 | 17 | 0 | 0 | 0 | 53 | 761 | 248 | 168 | 0 | 33 | 13 | 38 | 0 |
| SRR1657198 | 22 | 7564 | 3454 | 22 | 151 | 1566 | 1543 | 903 | 19991 | 156 | 7372 | 34650 | 926 | 141 | 12 | 28153 | 14082 | 0 | 0 | 0 | 353 | 233 | 80 | 0 | 0 | 0 | 0 | 0 | 15 | 68 | 135 | 52 | 0 | 0 | 0 | 27 | 0 |
| SRR1686806 | 956 | 61361 | 742 | 394 | 1357 | 48620 | 14420 | 14323 | 250 | 14 | 25683 | 129527 | 3704 | 18441 | 5097 | 32282 | 21787 | 0 | 0 | 0 | 507 | 227 | 22 | 0 | 0 | 0 | 0 | 0 | 44 | 135 | 167 | 0 | 0 | 0 | 0 | 0 | 0 |
| SRR1707574 | 0 | 1477 | 413 | 0 | 3570 | 48350 | 28982 | 30945 | 0 | 14 | 33 | 98 | 0 | 19 | 0 | 3437 | 15369 | 0 | 0 | 0 | 0 | 0 | 183 | 615 | 0 | 888 | 0 | 0 | 0 | 0 | 21 | 108 | 788 | 0 | 412 | 60 | 14 |
| SRR1707575 | 306 | 1677 | 140 | 387 | 3862 | 456 | 80867 | 23281 | 419 | 18 | 355 | 769 | 29 | 683 | 0 | 128 | 557 | 0 | 0 | 0 | 0 | 0 | 404 | 0 | 47 | 0 | 0 | 0 | 0 | 0 | 0 | 226 | 0 | 49 | 0 | 0 | 53 |
| SRR1708906 | 196 | 8879 | 635 | 322 | 6536 | 90379 | 58324 | 72034 | 1296 | 13 | 32494 | 30514 | 15234 | 4643 | 12 | 37 | 34724 | 0 | 0 | 0 | 96 | 16 | 30 | 0 | 0 | 0 | 0 | 0 | 18 | 83 | 63 | 38 | 0 | 17 | 0 | 0 | 0 |
| SRR1708907 | 36 | 5328 | 667 | 24 | 4825 | 64793 | 37772 | 38729 | 1067 | 46 | 32375 | 38999 | 15213 | 4998 | 11 | 119 | 35520 | 0 | 0 | 0 | 84 | 34 | 32 | 0 | 0 | 0 | 0 | 0 | 26 | 80 | 101 | 40 | 0 | 16 | 27 | 0 | 0 |
| SRR1708908 | 0 | 826 | 831 | 0 | 3974 | 58787 | 31614 | 31952 | 1227 | 41 | 38774 | 43974 | 19948 | 5477 | 13 | 35 | 32283 | 0 | 0 | 0 | 21 | 0 | 0 | 0 | 0 | 0 | 0 | 0 | 0 | 20 | 24 | 17 | 0 | 0 | 11 | 0 | 0 |
| SRR1708909 | 152 | 4232 | 554 | 430 | 2811 | 38890 | 29248 | 19843 | 402 | 61 | 33314 | 52142 | 17994 | 7147 | 14 | 131 | 31204 | 0 | 0 | 0 | 138 | 32 | 20 | 0 | 0 | 0 | 0 | 20 | 80 | 174 | 82 | 55 | 0 | 27 | 42 | 14 | 0 |
| SRR1708910 | 11 | 1628 | 474 | 17 | 3411 | 57180 | 33931 | 26166 | 1750 | 61 | 27579 | 41825 | 14889 | 4491 | 11 | 12 | 32269 | 0 | 0 | 0 | 33 | 13 | 20 | 0 | 0 | 0 | 0 | 0 | 0 | 27 | 29 | 25 | 0 | 21 | 42 | 0 | 0 |
| SRR1708911 | 24 | 1506 | 708 | 0 | 3610 | 56506 | 34076 | 29759 | 1602 | 37 | 31364 | 39096 | 18033 | 4532 | 12 | 15 | 43291 | 0 | 0 | 0 | 13 | 0 | 13 | 0 | 0 | 0 | 0 | 0 | 0 | 0 | 15 | 15 | 0 | 12 | 0 | 0 | 0 |
| SRR1708912 | 32 | 2052 | 600 | 12 | 6331 | 84059 | 56086 | 76569 | 1506 | 49 | 35462 | 27137 | 14772 | 4645 | 17 | 50 | 28616 | 0 | 0 | 0 | 66 | 12 | 34 | 0 | 0 | 0 | 0 | 0 | 0 | 41 | 37 | 32 | 0 | 15 | 0 | 0 | 0 |
| SRR1708913 | 0 | 681 | 595 | 0 | 5528 | 78826 | 46375 | 63914 | 1141 | 82 | 34937 | 34506 | 15901 | 5189 | 17 | 167 | 31109 | 0 | 0 | 0 | 28 | 11 | 16 | 0 | 0 | 0 | 0 | 0 | 0 | 19 | 13 | 16 | 0 | 13 | 18 | 0 | 0 |
| SRR1708914 | 50 | 1769 | 556 | 25 | 6070 | 89382 | 53861 | 87199 | 1094 | 124 | 31242 | 31063 | 15429 | 4011 | 16 | 71 | 26928 | 0 | 0 | 0 | 44 | 0 | 21 | 0 | 0 | 0 | 0 | 0 | 0 | 32 | 16 | 21 | 0 | 0 | 0 | 0 | 0 |
| SRR1708915 | 122 | 3662 | 942 | 139 | 6001 | 72930 | 51087 | 71534 | 3515 | 31 | 37579 | 29334 | 17166 | 6737 | 17 | 254 | 36259 | 0 | 0 | 0 | 58 | 11 | 19 | 0 | 0 | 0 | 0 | 0 | 15 | 79 | 45 | 32 | 0 | 12 | 0 | 0 | 0 |
| SRR1708916 | 0 | 1409 | 967 | 0 | 5820 | 85905 | 55769 | 62452 | 2894 | 27 | 32016 | 26279 | 16941 | 4873 | 20 | 21 | 32908 | 0 | 0 | 0 | 24 | 0 | 20 | 0 | 0 | 0 | 0 | 0 | 0 | 16 | 16 | 28 | 0 | 17 | 0 | 0 | 0 |
| SRR1708917 | 40 | 2303 | 500 | 49 | 5349 | 79466 | 52965 | 61258 | 1443 | 19 | 37560 | 34102 | 21027 | 4863 | 16 | 22 | 38912 | 0 | 0 | 0 | 73 | 0 | 23 | 0 | 0 | 0 | 0 | 0 | 0 | 54 | 16 | 32 | 0 | 19 | 0 | 0 | 0 |
| SRR1708918 | 16 | 712 | 1513 | 0 | 4416 | 68702 | 43365 | 82768 | 1005 | 49 | 31902 | 28513 | 15296 | 5047 | 0 | 111 | 29410 | 0 | 0 | 0 | 31 | 13 | 18 | 0 | 0 | 0 | 0 | 0 | 0 | 17 | 25 | 18 | 0 | 12 | 0 | 0 | 0 |
| SRR1708919 | 19 | 1245 | 918 | 30 | 7419 | 86722 | 61135 | 107126 | 1096 | 50 | 28531 | 25610 | 13441 | 4351 | 16 | 495 | 22715 | 0 | 0 | 0 | 73 | 30 | 21 | 0 | 0 | 0 | 0 | 0 | 17 | 75 | 64 | 35 | 0 | 0 | 0 | 0 | 0 |
| SRR1708920 | 30 | 1415 | 677 | 19 | 5101 | 72493 | 42367 | 76699 | 1495 | 37 | 35283 | 31221 | 18107 | 5532 | 13 | 163 | 24831 | 0 | 0 | 0 | 77 | 25 | 33 | 0 | 0 | 0 | 0 | 0 | 13 | 66 | 54 | 38 | 0 | 16 | 0 | 0 | 0 |
| SRR1708921 | 129 | 5036 | 995 | 49 | 5087 | 62323 | 42015 | 88095 | 2847 | 42 | 33150 | 27855 | 15291 | 6209 | 11 | 501 | 33688 | 0 | 0 | 0 | 47 | 14 | 17 | 0 | 0 | 0 | 0 | 0 | 15 | 72 | 61 | 30 | 0 | 0 | 0 | 0 | 0 |
| SRR1708922 | 67 | 1897 | 816 | 37 | 4925 | 78557 | 46342 | 76430 | 2642 | 23 | 31207 | 30988 | 16867 | 4814 | 18 | 47 | 31010 | 0 | 0 | 0 | 64 | 15 | 24 | 0 | 0 | 0 | 0 | 0 | 17 | 53 | 43 | 34 | 0 | 19 | 0 | 0 | 0 |
| SRR1708923 | 58 | 5802 | 1877 | 55 | 5280 | 69090 | 45880 | 73626 | 1360 | 43 | 41204 | 35670 | 21967 | 5392 | 12 | 44 | 35987 | 0 | 0 | 0 | 119 | 21 | 26 | 0 | 0 | 0 | 0 | 0 | 45 | 140 | 86 | 49 | 0 | 12 | 0 | 0 | 0 |
| SRR1708924 | 102 | 11193 | 995 | 79 | 6583 | 83001 | 57463 | 49091 | 733 | 58 | 37408 | 35487 | 15166 | 4903 | 10 | 35 | 32340 | 0 | 0 | 0 | 282 | 52 | 33 | 0 | 0 | 0 | 0 | 0 | 52 | 278 | 163 | 47 | 0 | 15 | 0 | 0 | 0 |
| SRR1708925 | 163 | 18844 | 656 | 140 | 6124 | 87591 | 52309 | 55407 | 1190 | 46 | 31589 | 37026 | 14530 | 5310 | 0 | 167 | 33520 | 0 | 0 | 0 | 201 | 47 | 28 | 0 | 0 | 0 | 0 | 0 | 41 | 332 | 231 | 53 | 0 | 13 | 0 | 0 | 0 |
| SRR1708926 | 232 | 8335 | 558 | 414 | 3273 | 51942 | 27211 | 30851 | 880 | 41 | 28244 | 42084 | 16281 | 5227 | 11 | 31 | 33170 | 0 | 0 | 0 | 212 | 72 | 35 | 0 | 0 | 0 | 0 | 0 | 90 | 219 | 170 | 54 | 0 | 23 | 28 | 0 | 0 |
| SRR1708927 | 150 | 9472 | 546 | 168 | 5571 | 72103 | 46950 | 56788 | 2295 | 32 | 44602 | 38665 | 18151 | 6799 | 17 | 166 | 37281 | 0 | 0 | 0 | 69 | 16 | 18 | 0 | 0 | 0 | 0 | 0 | 31 | 106 | 121 | 51 | 0 | 17 | 0 | 0 | 0 |
| SRR1708928 | 67 | 6171 | 1050 | 45 | 6210 | 106645 | 66065 | 64885 | 2816 | 60 | 35095 | 29387 | 15248 | 4219 | 22 | 13 | 31328 | 0 | 0 | 0 | 52 | 11 | 17 | 0 | 0 | 0 | 0 | 0 | 19 | 71 | 62 | 54 | 0 | 12 | 0 | 0 | 0 |
| SRR1708929 | 60 | 6347 | 889 | 30 | 5267 | 72975 | 55686 | 43924 | 1451 | 62 | 41329 | 37749 | 19395 | 5204 | 11 | 16 | 37623 | 0 | 0 | 0 | 50 | 11 | 20 | 0 | 0 | 0 | 0 | 0 | 13 | 56 | 53 | 42 | 0 | 15 | 0 | 0 | 0 |
| SRR1708930 | 227 | 7211 | 877 | 140 | 6327 | 71905 | 46113 | 80307 | 967 | 62 | 33681 | 32002 | 14550 | 5665 | 22 | 48 | 33461 | 0 | 0 | 0 | 126 | 51 | 61 | 0 | 12 | 0 | 0 | 0 | 14 | 109 | 95 | 61 | 0 | 32 | 12 | 0 | 0 |
| SRR1708931 | 197 | 5385 | 484 | 108 | 7138 | 79771 | 46594 | 69517 | 1593 | 31 | 31131 | 32261 | 14100 | 6148 | 18 | 184 | 37691 | 0 | 0 | 0 | 119 | 50 | 48 | 0 | 0 | 0 | 0 | 0 | 22 | 109 | 90 | 54 | 0 | 26 | 0 | 0 | 0 |
| SRR1708932 | 0 | 948 | 837 | 20 | 6463 | 65746 | 41817 | 96839 | 1559 | 62 | 33918 | 26904 | 15628 | 5539 | 17 | 44 | 31628 | 0 | 0 | 0 | 26 | 19 | 38 | 0 | 0 | 0 | 0 | 0 | 0 | 22 | 26 | 40 | 0 | 24 | 0 | 0 | 0 |
| SRR1708933 | 27 | 1855 | 1605 | 18 | 5872 | 56452 | 41878 | 71211 | 2535 | 94 | 30987 | 26559 | 14575 | 6423 | 40 | 188 | 34614 | 0 | 0 | 0 | 35 | 21 | 27 | 0 | 0 | 0 | 0 | 0 | 0 | 37 | 30 | 45 | 0 | 17 | 11 | 0 | 0 |
| SRR1708934 | 226 | 4790 | 1379 | 117 | 7547 | 86297 | 58549 | 67050 | 2510 | 56 | 30087 | 23920 | 13733 | 4912 | 34 | 18 | 29780 | 0 | 0 | 0 | 163 | 55 | 61 | 0 | 12 | 0 | 0 | 0 | 18 | 129 | 79 | 90 | 0 | 41 | 0 | 0 | 0 |
| SRR1708935 | 0 | 654 | 658 | 21 | 6760 | 77429 | 58296 | 80587 | 2021 | 31 | 33816 | 28303 | 18280 | 6208 | 23 | 26 | 30941 | 0 | 0 | 0 | 27 | 10 | 21 | 0 | 0 | 0 | 0 | 0 | 0 | 33 | 17 | 27 | 0 | 15 | 0 | 0 | 0 |
| SRR1708936 | 169 | 9014 | 988 | 118 | 3618 | 24393 | 22352 | 93800 | 761 | 39 | 32597 | 29424 | 16520 | 6733 | 30 | 35 | 35927 | 0 | 0 | 0 | 180 | 167 | 83 | 0 | 0 | 0 | 0 | 0 | 18 | 117 | 229 | 65 | 0 | 25 | 10 | 0 | 0 |
| SRR1708937 | 0 | 253 | 737 | 0 | 4579 | 28613 | 25729 | 109434 | 1106 | 30 | 34669 | 31118 | 16772 | 7018 | 23 | 131 | 38146 | 0 | 0 | 0 | 11 | 0 | 12 | 0 | 0 | 0 | 0 | 0 | 0 | 0 | 14 | 12 | 0 | 11 | 0 | 0 | 0 |
| SRR1708938 | 0 | 1003 | 1641 | 0 | 3415 | 28460 | 22745 | 90501 | 1376 | 87 | 33437 | 28821 | 17293 | 6731 | 24 | 60 | 34252 | 0 | 0 | 0 | 21 | 27 | 23 | 0 | 0 | 0 | 0 | 0 | 0 | 14 | 40 | 22 | 0 | 15 | 0 | 0 | 0 |
| SRR1708939 | 0 | 1958 | 2127 | 13 | 4589 | 29432 | 30244 | 86672 | 1269 | 91 | 28920 | 28753 | 14884 | 7370 | 34 | 293 | 35809 | 0 | 0 | 0 | 32 | 37 | 25 | 0 | 0 | 0 | 0 | 0 | 0 | 31 | 71 | 39 | 0 | 10 | 0 | 0 | 0 |
| SRR1708940 | 21 | 2896 | 1024 | 14 | 5492 | 47123 | 36996 | 94836 | 2653 | 29 | 31754 | 27402 | 16844 | 6783 | 37 | 31 | 28806 | 0 | 0 | 0 | 68 | 75 | 47 | 0 | 0 | 0 | 0 | 0 | 0 | 61 | 115 | 59 | 0 | 20 | 13 | 0 | 0 |
| SRR1708941 | 11 | 2221 | 1709 | 0 | 5135 | 39111 | 31248 | 70816 | 1373 | 49 | 31395 | 29350 | 19690 | 6835 | 27 | 26 | 38965 | 0 | 0 | 0 | 73 | 49 | 44 | 0 | 0 | 0 | 0 | 0 | 0 | 58 | 95 | 50 | 0 | 21 | 10 | 0 | 0 |
| SRR1708942 | 37 | 5457 | 1362 | 52 | 6357 | 73534 | 52912 | 52114 | 1148 | 63 | 34770 | 32204 | 14130 | 5014 | 11 | 30 | 32147 | 0 | 0 | 0 | 155 | 32 | 37 | 0 | 0 | 0 | 0 | 0 | 22 | 181 | 143 | 63 | 0 | 16 | 0 | 0 | 0 |
| SRR1708943 | 19 | 3116 | 959 | 0 | 2643 | 39233 | 23498 | 23479 | 592 | 38 | 29875 | 47160 | 14493 | 5756 | 0 | 77 | 33369 | 0 | 0 | 0 | 55 | 21 | 19 | 0 | 0 | 0 | 0 | 0 | 12 | 42 | 46 | 25 | 0 | 21 | 30 | 0 | 0 |
| SRR1708944 | 32 | 7984 | 1666 | 27 | 3422 | 48925 | 31080 | 27312 | 565 | 94 | 30312 | 49060 | 17601 | 5526 | 12 | 30 | 26220 | 0 | 0 | 0 | 135 | 45 | 32 | 0 | 0 | 0 | 0 | 0 | 34 | 167 | 128 | 61 | 0 | 23 | 42 | 12 | 0 |
| SRR1708945 | 374 | 20317 | 1248 | 326 | 3652 | 44781 | 33738 | 34035 | 1674 | 95 | 30834 | 39440 | 14554 | 7643 | 0 | 105 | 31683 | 0 | 0 | 0 | 258 | 62 | 30 | 0 | 0 | 0 | 0 | 16 | 111 | 408 | 256 | 91 | 0 | 18 | 42 | 26 | 0 |
| SRR1708946 | 66 | 7583 | 1981 | 23 | 5297 | 84960 | 51059 | 42360 | 2619 | 14 | 33681 | 32450 | 14784 | 4659 | 21 | 12 | 37857 | 0 | 0 | 0 | 110 | 30 | 27 | 0 | 0 | 0 | 0 | 0 | 26 | 191 | 152 | 81 | 0 | 14 | 11 | 0 | 0 |
| SRR1708947 | 418 | 22302 | 913 | 242 | 3142 | 45697 | 30096 | 24059 | 958 | 32 | 28534 | 42799 | 17494 | 4590 | 13 | 11 | 37636 | 0 | 0 | 0 | 333 | 80 | 42 | 0 | 0 | 0 | 0 | 0 | 104 | 348 | 181 | 59 | 0 | 39 | 58 | 12 | 0 |
| SRR1708948 | 30 | 4135 | 1205 | 0 | 6942 | 79777 | 49887 | 70576 | 840 | 34 | 32731 | 32671 | 15022 | 5318 | 19 | 49 | 32799 | 0 | 0 | 0 | 64 | 31 | 43 | 0 | 0 | 0 | 0 | 0 | 10 | 67 | 70 | 60 | 0 | 26 | 0 | 0 | 0 |
| SRR1708949 | 99 | 6743 | 831 | 43 | 4961 | 60223 | 31800 | 48315 | 746 | 79 | 32747 | 43658 | 14596 | 6043 | 24 | 130 | 34393 | 0 | 0 | 0 | 123 | 66 | 37 | 0 | 0 | 0 | 0 | 0 | 25 | 81 | 104 | 45 | 0 | 24 | 21 | 0 | 0 |
| SRR1708950 | 24 | 3096 | 807 | 0 | 6035 | 67040 | 41277 | 61389 | 1539 | 25 | 34682 | 33774 | 18279 | 6136 | 14 | 65 | 30211 | 0 | 0 | 0 | 95 | 47 | 45 | 0 | 11 | 0 | 0 | 0 | 17 | 114 | 108 | 58 | 0 | 26 | 16 | 0 | 0 |
| SRR1708951 | 217 | 10504 | 1390 | 113 | 5757 | 54228 | 38481 | 79878 | 2158 | 36 | 34620 | 28712 | 16481 | 7933 | 22 | 167 | 41171 | 0 | 0 | 0 | 60 | 42 | 30 | 0 | 0 | 0 | 0 | 0 | 23 | 90 | 104 | 46 | 0 | 20 | 0 | 0 | 0 |
| SRR1708952 | 64 | 9615 | 1420 | 17 | 6431 | 70252 | 45048 | 72600 | 2048 | 44 | 30768 | 26441 | 14983 | 5513 | 37 | 31 | 38811 | 0 | 0 | 0 | 99 | 57 | 46 | 0 | 0 | 0 | 0 | 0 | 14 | 96 | 113 | 55 | 0 | 36 | 11 | 0 | 0 |
| SRR1708953 | 55 | 5037 | 1088 | 24 | 6071 | 60761 | 44666 | 77436 | 1817 | 43 | 37256 | 29316 | 18971 | 6433 | 27 | 29 | 35236 | 0 | 0 | 0 | 85 | 41 | 49 | 0 | 0 | 0 | 0 | 0 | 16 | 90 | 84 | 63 | 0 | 20 | 0 | 0 | 0 |
| SRR1708954 | 0 | 4920 | 1604 | 13 | 4103 | 31352 | 28055 | 99088 | 718 | 24 | 34041 | 30886 | 15581 | 5934 | 32 | 41 | 33506 | 0 | 0 | 0 | 63 | 57 | 37 | 0 | 0 | 0 | 0 | 0 | 0 | 40 | 84 | 38 | 0 | 17 | 0 | 0 | 0 |
| SRR1708955 | 22 | 2637 | 841 | 14 | 5544 | 32178 | 31459 | 117022 | 1251 | 31 | 33920 | 25754 | 16624 | 7520 | 31 | 169 | 31953 | 0 | 0 | 0 | 66 | 49 | 49 | 0 | 12 | 0 | 0 | 0 | 0 | 65 | 78 | 39 | 0 | 19 | 10 | 0 | 0 |
| SRR1708956 | 69 | 6066 | 1705 | 59 | 3016 | 22672 | 20018 | 104010 | 1011 | 39 | 33019 | 27902 | 19976 | 7114 | 31 | 50 | 36794 | 0 | 0 | 0 | 70 | 43 | 40 | 0 | 0 | 0 | 0 | 0 | 0 | 69 | 84 | 41 | 0 | 20 | 0 | 0 | 0 |
| SRR1708957 | 113 | 6825 | 881 | 99 | 3196 | 15358 | 19417 | 98485 | 1485 | 60 | 30422 | 29079 | 17578 | 7732 | 52 | 116 | 37922 | 0 | 0 | 0 | 87 | 66 | 43 | 0 | 0 | 0 | 0 | 0 | 11 | 72 | 121 | 54 | 0 | 22 | 11 | 0 | 0 |
| SRR1708958 | 53 | 7464 | 1380 | 11 | 3194 | 17054 | 19421 | 102839 | 1307 | 20 | 34803 | 26937 | 18994 | 7737 | 54 | 16 | 34996 | 0 | 0 | 0 | 74 | 87 | 57 | 0 | 0 | 0 | 0 | 0 | 11 | 62 | 141 | 60 | 0 | 29 | 0 | 0 | 0 |
| SRR1708959 | 42 | 10864 | 1633 | 25 | 3155 | 19184 | 20597 | 97168 | 808 | 40 | 33258 | 27935 | 18938 | 7073 | 30 | 28 | 38833 | 0 | 0 | 0 | 112 | 66 | 54 | 0 | 0 | 0 | 0 | 0 | 0 | 84 | 100 | 57 | 0 | 35 | 0 | 0 | 0 |
| SRR1708960 | 220 | 13556 | 765 | 216 | 7673 | 94845 | 64362 | 89389 | 1265 | 29 | 35534 | 29758 | 13603 | 5178 | 10 | 34 | 27381 | 0 | 0 | 0 | 160 | 23 | 20 | 0 | 0 | 0 | 0 | 0 | 31 | 226 | 123 | 36 | 0 | 12 | 0 | 0 | 0 |
| SRR1708961 | 438 | 27921 | 884 | 235 | 8108 | 107713 | 74087 | 71130 | 1281 | 78 | 32576 | 31204 | 13405 | 4724 | 0 | 223 | 30669 | 0 | 0 | 0 | 231 | 52 | 28 | 0 | 0 | 0 | 0 | 0 | 83 | 298 | 269 | 65 | 0 | 16 | 0 | 0 | 0 |
| SRR1708962 | 35 | 10198 | 922 | 29 | 5428 | 77458 | 45769 | 56153 | 1533 | 51 | 45284 | 37304 | 18644 | 6369 | 0 | 36 | 31917 | 0 | 0 | 0 | 86 | 32 | 30 | 0 | 0 | 0 | 0 | 0 | 26 | 139 | 194 | 79 | 0 | 14 | 0 | 0 | 0 |
| SRR1708963 | 31 | 13219 | 869 | 43 | 3688 | 50099 | 30184 | 31378 | 1631 | 39 | 40626 | 45762 | 16316 | 7220 | 12 | 113 | 36498 | 0 | 0 | 0 | 70 | 32 | 16 | 0 | 0 | 0 | 0 | 0 | 29 | 99 | 140 | 42 | 0 | 17 | 16 | 0 | 0 |
| SRR1708964 | 139 | 17854 | 877 | 34 | 5345 | 84810 | 50086 | 51979 | 2282 | 19 | 23382 | 34288 | 11145 | 3527 | 20 | 0 | 31528 | 0 | 0 | 0 | 96 | 24 | 22 | 0 | 0 | 0 | 0 | 0 | 27 | 182 | 119 | 65 | 0 | 0 | 0 | 0 | 0 |
| SRR1708965 | 0 | 3345 | 994 | 0 | 5726 | 86783 | 63531 | 61269 | 2150 | 37 | 40292 | 33836 | 17796 | 6063 | 13 | 16 | 33557 | 0 | 0 | 0 | 40 | 0 | 15 | 0 | 0 | 0 | 0 | 0 | 0 | 77 | 57 | 46 | 0 | 0 | 0 | 0 | 0 |
| SRR1708966 | 450 | 18727 | 1015 | 238 | 6739 | 75456 | 48593 | 72876 | 1050 | 23 | 33430 | 30135 | 14462 | 5304 | 20 | 50 | 34938 | 0 | 0 | 0 | 175 | 73 | 57 | 0 | 0 | 0 | 0 | 0 | 27 | 146 | 151 | 73 | 0 | 30 | 0 | 0 | 0 |
| SRR1708967 | 13 | 2335 | 989 | 10 | 8050 | 83915 | 51552 | 82641 | 1626 | 42 | 33079 | 28401 | 13955 | 6277 | 16 | 157 | 34792 | 0 | 0 | 0 | 21 | 15 | 17 | 0 | 0 | 0 | 0 | 0 | 0 | 22 | 37 | 22 | 0 | 13 | 0 | 0 | 0 |
| SRR1708968 | 16 | 3284 | 1956 | 0 | 6585 | 67240 | 43578 | 84607 | 1676 | 66 | 33897 | 28079 | 17375 | 5555 | 20 | 62 | 32882 | 0 | 0 | 0 | 24 | 19 | 18 | 0 | 0 | 0 | 0 | 0 | 0 | 20 | 37 | 24 | 0 | 15 | 0 | 0 | 0 |
| SRR1708969 | 46 | 6137 | 1990 | 25 | 6785 | 56201 | 45650 | 87792 | 1877 | 68 | 30407 | 27271 | 14063 | 7067 | 33 | 176 | 34958 | 0 | 0 | 0 | 32 | 26 | 17 | 0 | 0 | 0 | 0 | 0 | 11 | 56 | 59 | 38 | 0 | 0 | 0 | 0 | 0 |
| SRR1708970 | 21 | 4062 | 1449 | 13 | 8126 | 75964 | 59607 | 79926 | 3167 | 29 | 35338 | 28214 | 17028 | 7324 | 35 | 37 | 30593 | 0 | 0 | 0 | 48 | 23 | 30 | 0 | 0 | 0 | 0 | 0 | 0 | 55 | 56 | 41 | 0 | 22 | 10 | 0 | 0 |
| SRR1708971 | 11 | 1744 | 2042 | 11 | 6523 | 59371 | 51633 | 92792 | 1843 | 64 | 25257 | 22294 | 15138 | 6184 | 19 | 40 | 31841 | 0 | 0 | 0 | 23 | 13 | 17 | 0 | 0 | 0 | 0 | 0 | 0 | 22 | 23 | 31 | 0 | 12 | 0 | 10 | 0 |
| SRR1708972 | 0 | 6690 | 3062 | 0 | 3626 | 21723 | 23626 | 97901 | 715 | 34 | 35370 | 27661 | 18083 | 7076 | 33 | 33 | 33526 | 0 | 0 | 0 | 52 | 82 | 60 | 0 | 11 | 0 | 0 | 0 | 0 | 36 | 100 | 52 | 0 | 22 | 0 | 0 | 0 |
| SRR1708973 | 0 | 1115 | 3566 | 0 | 4148 | 21745 | 25229 | 125909 | 788 | 44 | 33663 | 26494 | 17122 | 7022 | 28 | 98 | 39937 | 0 | 0 | 0 | 0 | 0 | 0 | 0 | 0 | 0 | 0 | 0 | 0 | 0 | 12 | 0 | 0 | 0 | 0 | 0 | 0 |
| SRR1708974 | 15 | 5678 | 3143 | 0 | 2532 | 14263 | 17393 | 108003 | 549 | 45 | 33431 | 26337 | 19629 | 7504 | 32 | 47 | 30419 | 0 | 0 | 0 | 47 | 57 | 37 | 0 | 0 | 0 | 0 | 0 | 0 | 32 | 84 | 41 | 0 | 23 | 0 | 0 | 0 |
| SRR1708975 | 21 | 4417 | 1874 | 19 | 2745 | 13145 | 18287 | 100826 | 1419 | 37 | 33748 | 27772 | 19022 | 9582 | 33 | 103 | 33596 | 0 | 0 | 0 | 36 | 40 | 35 | 0 | 0 | 0 | 0 | 0 | 0 | 41 | 65 | 32 | 0 | 18 | 11 | 0 | 0 |
| SRR1708976 | 44 | 16285 | 2392 | 14 | 3293 | 19462 | 19918 | 94692 | 1569 | 17 | 30632 | 25048 | 18049 | 7740 | 55 | 24 | 36871 | 0 | 0 | 0 | 79 | 97 | 55 | 0 | 10 | 0 | 0 | 0 | 0 | 64 | 174 | 63 | 0 | 33 | 0 | 0 | 0 |
| SRR1708977 | 203 | 24138 | 2343 | 195 | 3520 | 20733 | 22049 | 95872 | 947 | 21 | 32843 | 28226 | 19898 | 7140 | 40 | 22 | 37828 | 0 | 0 | 0 | 133 | 71 | 54 | 0 | 0 | 0 | 0 | 0 | 12 | 101 | 127 | 59 | 0 | 27 | 0 | 0 | 0 |
| SRR1708978 | 19 | 6887 | 1271 | 28 | 5109 | 70616 | 44737 | 40387 | 584 | 56 | 38046 | 43685 | 17522 | 5086 | 17 | 24 | 32910 | 0 | 0 | 0 | 86 | 28 | 38 | 0 | 0 | 0 | 0 | 0 | 13 | 49 | 67 | 48 | 0 | 16 | 17 | 0 | 0 |
| SRR1708979 | 112 | 14027 | 872 | 67 | 3019 | 52132 | 26608 | 25656 | 652 | 73 | 34894 | 53689 | 15234 | 5939 | 0 | 81 | 28830 | 0 | 0 | 0 | 132 | 45 | 23 | 0 | 0 | 0 | 0 | 0 | 47 | 92 | 98 | 35 | 0 | 20 | 26 | 14 | 0 |
| SRR1708980 | 23 | 3151 | 382 | 15 | 5898 | 88757 | 54354 | 73362 | 1861 | 33 | 37793 | 33860 | 19054 | 5332 | 11 | 51 | 29684 | 0 | 0 | 0 | 26 | 0 | 11 | 0 | 0 | 0 | 0 | 0 | 0 | 29 | 34 | 27 | 0 | 0 | 0 | 0 | 0 |
| SRR1708981 | 27 | 2337 | 1098 | 23 | 2921 | 35923 | 24411 | 25014 | 856 | 56 | 36916 | 46498 | 18338 | 7421 | 16 | 116 | 37578 | 0 | 0 | 0 | 55 | 26 | 19 | 0 | 0 | 0 | 0 | 0 | 27 | 44 | 49 | 33 | 0 | 36 | 39 | 0 | 0 |
| SRR1708982 | 64 | 7368 | 997 | 22 | 5335 | 91799 | 54365 | 53879 | 1990 | 51 | 30473 | 33513 | 14583 | 4096 | 22 | 12 | 36983 | 0 | 0 | 0 | 31 | 11 | 19 | 0 | 0 | 0 | 0 | 0 | 0 | 40 | 57 | 59 | 0 | 11 | 0 | 0 | 0 |
| SRR1708983 | 174 | 7828 | 619 | 139 | 2837 | 43300 | 28308 | 21882 | 872 | 50 | 35289 | 47593 | 20903 | 5631 | 11 | 0 | 33355 | 0 | 0 | 0 | 152 | 53 | 40 | 0 | 11 | 0 | 0 | 0 | 68 | 126 | 91 | 55 | 0 | 40 | 37 | 17 | 0 |
| SRR1708984 | 14 | 2471 | 711 | 33 | 6661 | 89794 | 53693 | 79039 | 1061 | 58 | 34117 | 33585 | 13064 | 4197 | 17 | 35 | 27106 | 0 | 0 | 0 | 76 | 18 | 33 | 0 | 0 | 0 | 0 | 0 | 0 | 46 | 26 | 35 | 0 | 15 | 0 | 0 | 0 |
| SRR1708985 | 269 | 2311 | 402 | 268 | 7778 | 80469 | 56413 | 80660 | 1950 | 42 | 33294 | 31815 | 15185 | 6069 | 21 | 225 | 28656 | 0 | 0 | 0 | 141 | 41 | 57 | 0 | 12 | 0 | 0 | 0 | 17 | 152 | 74 | 63 | 0 | 23 | 16 | 0 | 0 |
| SRR1708986 | 118 | 1631 | 555 | 94 | 6709 | 73011 | 54369 | 94639 | 1525 | 44 | 29285 | 27330 | 15775 | 5503 | 17 | 57 | 27102 | 0 | 0 | 0 | 111 | 31 | 41 | 0 | 0 | 0 | 0 | 0 | 0 | 104 | 40 | 59 | 0 | 18 | 0 | 0 | 0 |
| SRR1708987 | 135 | 4351 | 555 | 30 | 5215 | 49771 | 35781 | 76820 | 2539 | 41 | 30669 | 28545 | 14601 | 6599 | 27 | 179 | 43465 | 0 | 0 | 0 | 67 | 44 | 46 | 0 | 0 | 0 | 0 | 0 | 0 | 78 | 53 | 70 | 0 | 24 | 0 | 0 | 0 |
| SRR1708988 | 63 | 3974 | 1882 | 66 | 5129 | 46375 | 38273 | 74951 | 1270 | 39 | 27551 | 23566 | 14643 | 6019 | 55 | 15 | 39602 | 0 | 0 | 0 | 69 | 68 | 45 | 0 | 0 | 0 | 0 | 0 | 11 | 51 | 69 | 62 | 0 | 33 | 33 | 0 | 0 |
| SRR1708989 | 0 | 2346 | 1107 | 57 | 4339 | 38342 | 31560 | 84446 | 1353 | 46 | 33918 | 33192 | 17946 | 6743 | 28 | 16 | 40809 | 0 | 0 | 0 | 42 | 55 | 36 | 0 | 0 | 0 | 0 | 0 | 0 | 38 | 51 | 47 | 0 | 22 | 10 | 0 | 0 |
| SRR1708990 | 173 | 4723 | 1659 | 103 | 6572 | 66048 | 53192 | 83065 | 1553 | 33 | 35473 | 25847 | 15668 | 6443 | 0 | 93 | 23630 | 0 | 0 | 0 | 187 | 155 | 73 | 0 | 0 | 0 | 0 | 0 | 42 | 176 | 295 | 69 | 0 | 17 | 0 | 0 | 0 |
| SRR1708991 | 29 | 2074 | 704 | 29 | 5249 | 43281 | 35413 | 126182 | 1291 | 22 | 30352 | 24322 | 15119 | 5956 | 19 | 263 | 31807 | 0 | 0 | 0 | 86 | 112 | 41 | 0 | 0 | 0 | 0 | 0 | 0 | 84 | 145 | 45 | 0 | 12 | 0 | 0 | 0 |
| SRR1708992 | 359 | 7106 | 1874 | 178 | 4621 | 43176 | 32201 | 83874 | 1229 | 61 | 32511 | 25797 | 17450 | 6208 | 22 | 106 | 30948 | 0 | 0 | 0 | 154 | 128 | 62 | 0 | 0 | 0 | 0 | 0 | 23 | 98 | 206 | 62 | 0 | 24 | 0 | 0 | 0 |
| SRR1708993 | 72 | 3553 | 2402 | 0 | 3349 | 19031 | 23138 | 89497 | 1554 | 66 | 35079 | 29169 | 16925 | 8728 | 29 | 163 | 38723 | 0 | 0 | 0 | 44 | 85 | 40 | 0 | 0 | 0 | 0 | 0 | 0 | 39 | 130 | 57 | 0 | 14 | 0 | 0 | 0 |
| SRR1708994 | 25 | 3032 | 3623 | 156 | 2819 | 20438 | 21642 | 106406 | 1139 | 30 | 35993 | 24842 | 16416 | 7906 | 38 | 15 | 33718 | 0 | 0 | 0 | 35 | 39 | 26 | 0 | 0 | 0 | 0 | 0 | 0 | 32 | 46 | 29 | 0 | 14 | 34 | 0 | 0 |
| SRR1708995 | 96 | 14776 | 2590 | 186 | 1914 | 9796 | 15164 | 104754 | 499 | 39 | 30254 | 22983 | 17134 | 7730 | 35 | 10 | 30053 | 0 | 0 | 0 | 95 | 63 | 50 | 0 | 0 | 0 | 0 | 0 | 0 | 67 | 56 | 47 | 0 | 26 | 0 | 0 | 0 |
| SRR1708996 | 73 | 6971 | 627 | 64 | 6583 | 92378 | 60020 | 67748 | 1268 | 74 | 36746 | 34671 | 14844 | 4941 | 13 | 38 | 27989 | 0 | 0 | 0 | 111 | 17 | 23 | 0 | 0 | 0 | 0 | 0 | 11 | 114 | 63 | 47 | 0 | 15 | 10 | 0 | 0 |

### Supp Table S8.1.Columns_B_to_AL

| | | | | | | | | | | | | | | | | | | | | | | | | | | | | | | | | | | | | | |
|---|---|---|---|---|---|---|---|---|---|---|---|---|---|---|---|---|---|---|---|---|---|---|---|---|---|---|---|---|---|---|---|---|---|---|---|---|---|
| SRR1708997 | 604 | 45674 | 384 | 692 | 4788 | 77846 | 36842 | 40261 | 972 | 34 | 33243 | 45227 | 14118 | 4605 | 14 | 132 | 34293 | 0 | 0 | 0 | 167 | 42 | 22 | 0 | 0 | 0 | 0 | 0 | 27 | 167 | 143 | 46 | 0 | 14 | 11 | 0 | 0 |
| SRR1708998 | 62 | 9266 | 393 | 65 | 5113 | 75157 | 42320 | 53822 | 1680 | 65 | 34394 | 33012 | 17686 | 4708 | 11 | 45 | 38035 | 0 | 0 | 0 | 41 | 13 | 17 | 0 | 0 | 0 | 0 | 0 | 12 | 52 | 60 | 40 | 0 | 11 | 0 | 0 | 0 |
| SRR1708999 | 14 | 1094 | 672 | 0 | 2739 | 42756 | 23759 | 27675 | 1023 | 56 | 34505 | 53884 | 16267 | 6709 | 10 | 125 | 40513 | 0 | 0 | 0 | 15 | 0 | 0 | 0 | 0 | 0 | 0 | 0 | 0 | 13 | 19 | 11 | 0 | 12 | 13 | 0 | 0 |
| SRR1709000 | 137 | 4611 | 731 | 125 | 6810 | 82260 | 51194 | 59133 | 715 | 49 | 29473 | 33658 | 13925 | 4819 | 20 | 41 | 28065 | 0 | 0 | 0 | 123 | 43 | 56 | 0 | 0 | 0 | 0 | 0 | 15 | 93 | 51 | 52 | 0 | 26 | 12 | 0 | 0 |
| SRR1709001 | 0 | 970 | 508 | 15 | 6526 | 80968 | 47138 | 69132 | 2021 | 55 | 31010 | 28681 | 13573 | 5258 | 17 | 188 | 33772 | 0 | 0 | 0 | 39 | 18 | 28 | 0 | 0 | 0 | 0 | 0 | 0 | 33 | 31 | 31 | 0 | 19 | 0 | 0 | 0 |
| SRR1709002 | 234 | 1976 | 649 | 233 | 6602 | 71968 | 49336 | 73210 | 1391 | 51 | 34480 | 26233 | 15315 | 5116 | 22 | 58 | 27865 | 0 | 0 | 0 | 91 | 42 | 46 | 0 | 0 | 0 | 0 | 0 | 18 | 95 | 41 | 60 | 0 | 26 | 0 | 0 | 0 |
| SRR1709003 | 53 | 2358 | 1089 | 30 | 5688 | 53450 | 39088 | 86179 | 2108 | 50 | 37346 | 30301 | 16567 | 7858 | 26 | 140 | 40825 | 0 | 0 | 0 | 53 | 37 | 48 | 0 | 0 | 0 | 0 | 0 | 0 | 56 | 39 | 63 | 0 | 26 | 0 | 0 | 0 |
| SRR1709004 | 33 | 3564 | 761 | 16 | 3923 | 54047 | 32860 | 31737 | 569 | 59 | 32224 | 44946 | 14497 | 4778 | 10 | 24 | 31152 | 0 | 0 | 0 | 75 | 15 | 23 | 0 | 0 | 0 | 0 | 0 | 15 | 41 | 36 | 26 | 0 | 16 | 19 | 10 | 0 |
| SRR1709005 | 76 | 4691 | 411 | 117 | 5333 | 78290 | 45433 | 46010 | 1412 | 35 | 38144 | 43862 | 15923 | 6192 | 0 | 170 | 29733 | 0 | 0 | 0 | 117 | 22 | 28 | 0 | 0 | 0 | 0 | 0 | 23 | 116 | 83 | 45 | 0 | 13 | 20 | 0 | 0 |
| SRR1709006 | 190 | 9872 | 398 | 229 | 5033 | 72358 | 41587 | 51485 | 1076 | 74 | 34787 | 38861 | 17783 | 5131 | 12 | 41 | 32049 | 0 | 0 | 0 | 55 | 13 | 16 | 0 | 0 | 0 | 0 | 0 | 18 | 71 | 56 | 41 | 0 | 0 | 0 | 0 | 0 |
| SRR1709007 | 171 | 5521 | 430 | 235 | 4868 | 62949 | 39081 | 43769 | 2155 | 51 | 34935 | 35610 | 15835 | 6600 | 19 | 149 | 39775 | 0 | 0 | 0 | 42 | 0 | 13 | 0 | 0 | 0 | 0 | 0 | 24 | 56 | 63 | 24 | 0 | 0 | 0 | 0 | 0 |
| SRR1709008 | 39 | 2186 | 776 | 37 | 4621 | 74354 | 44346 | 36469 | 1659 | 77 | 43267 | 46862 | 18437 | 5375 | 21 | 0 | 30983 | 0 | 0 | 0 | 35 | 13 | 23 | 0 | 0 | 0 | 0 | 0 | 0 | 37 | 49 | 58 | 0 | 13 | 16 | 0 | 0 |
| SRR1709009 | 48 | 7420 | 752 | 72 | 4635 | 79974 | 46139 | 41750 | 1835 | 72 | 51650 | 45798 | 19954 | 5647 | 13 | 14 | 34934 | 0 | 0 | 0 | 36 | 11 | 16 | 0 | 0 | 0 | 0 | 0 | 0 | 35 | 48 | 29 | 0 | 0 | 0 | 0 | 0 |
| SRR1709010 | 18 | 1642 | 1060 | 28 | 6416 | 50904 | 48003 | 124596 | 1114 | 71 | 27034 | 21671 | 11169 | 6400 | 16 | 41 | 29433 | 0 | 0 | 0 | 68 | 24 | 37 | 0 | 0 | 0 | 0 | 0 | 0 | 62 | 39 | 69 | 0 | 21 | 14 | 13 | 0 |
| SRR1709011 | 100 | 1922 | 588 | 101 | 6508 | 48354 | 38777 | 85757 | 915 | 77 | 35089 | 31068 | 14336 | 8062 | 28 | 161 | 38267 | 0 | 0 | 0 | 119 | 66 | 67 | 0 | 14 | 0 | 0 | 0 | 0 | 79 | 54 | 72 | 0 | 40 | 0 | 0 | 0 |
| SRR1709012 | 160 | 1975 | 459 | 113 | 4607 | 41667 | 27982 | 79590 | 1211 | 64 | 35843 | 33894 | 18862 | 6504 | 32 | 48 | 38309 | 0 | 0 | 0 | 101 | 58 | 64 | 0 | 0 | 0 | 0 | 0 | 0 | 96 | 66 | 64 | 0 | 41 | 12 | 0 | 0 |
| SRR1709013 | 30 | 1108 | 882 | 27 | 5148 | 36404 | 30943 | 84716 | 1704 | 48 | 38694 | 29893 | 16843 | 8825 | 35 | 146 | 41758 | 0 | 0 | 0 | 32 | 32 | 41 | 0 | 0 | 0 | 0 | 0 | 0 | 45 | 35 | 61 | 0 | 19 | 0 | 0 | 0 |
| SRR1709014 | 59 | 1725 | 797 | 32 | 5805 | 60177 | 39765 | 80627 | 2038 | 74 | 31010 | 32433 | 15961 | 5789 | 37 | 17 | 37485 | 0 | 0 | 0 | 119 | 63 | 48 | 0 | 11 | 0 | 0 | 0 | 0 | 109 | 53 | 68 | 0 | 26 | 0 | 0 | 0 |
| SRR1709015 | 31 | 1027 | 667 | 20 | 5379 | 43781 | 38274 | 89426 | 1435 | 60 | 32678 | 25741 | 16474 | 6034 | 25 | 22 | 38191 | 0 | 0 | 0 | 53 | 31 | 42 | 0 | 0 | 0 | 0 | 0 | 0 | 48 | 25 | 53 | 0 | 31 | 0 | 0 | 0 |
| SRR1709016 | 120 | 4771 | 957 | 112 | 3228 | 14915 | 21280 | 121010 | 500 | 43 | 33636 | 26118 | 17978 | 7671 | 38 | 33 | 30887 | 0 | 0 | 0 | 129 | 135 | 74 | 0 | 13 | 0 | 0 | 0 | 17 | 89 | 143 | 68 | 0 | 25 | 12 | 0 | 0 |
| SRR1709017 | 40 | 1617 | 691 | 83 | 2955 | 9488 | 17057 | 171019 | 536 | 55 | 23075 | 17925 | 11952 | 7253 | 31 | 94 | 25252 | 0 | 0 | 0 | 86 | 34 | 34 | 0 | 0 | 0 | 0 | 0 | 0 | 66 | 58 | 42 | 0 | 13 | 0 | 13 | 0 |
| SRR1709018 | 22 | 2019 | 659 | 14 | 1408 | 5618 | 10568 | 106720 | 366 | 39 | 32575 | 24833 | 20102 | 8394 | 39 | 18 | 29712 | 0 | 0 | 0 | 68 | 69 | 88 | 16 | 18 | 0 | 0 | 0 | 0 | 37 | 67 | 62 | 0 | 31 | 15 | 0 | 0 |
| SRR1709019 | 0 | 1218 | 643 | 0 | 1573 | 6050 | 12819 | 114652 | 714 | 57 | 31024 | 21629 | 16076 | 8270 | 54 | 73 | 30377 | 0 | 0 | 0 | 34 | 26 | 27 | 0 | 0 | 0 | 0 | 0 | 0 | 28 | 26 | 26 | 0 | 13 | 0 | 0 | 0 |
| SRR1709020 | 115 | 2068 | 581 | 122 | 3055 | 17695 | 21549 | 104811 | 1123 | 30 | 31916 | 23327 | 16414 | 6612 | 46 | 17 | 32325 | 0 | 0 | 0 | 110 | 63 | 44 | 0 | 0 | 0 | 0 | 0 | 0 | 88 | 81 | 53 | 0 | 29 | 0 | 0 | 0 |
| SRR1709021 | 32 | 2122 | 750 | 28 | 2391 | 10210 | 17049 | 133199 | 793 | 45 | 25265 | 19672 | 15983 | 7629 | 30 | 13 | 27997 | 0 | 0 | 0 | 103 | 38 | 43 | 0 | 0 | 0 | 0 | 0 | 0 | 86 | 53 | 58 | 0 | 18 | 12 | 16 | 0 |
| SRR1732015 | 0 | 3963 | 644 | 190 | 6976 | 67520 | 83589 | 37105 | 3840 | 105 | 87708 | 210325 | 13805 | 29658 | 483 | 8887 | 5979 | 0 | 0 | 0 | 442 | 46 | 54 | 0 | 80 | 0 | 0 | 0 | 0 | 113 | 42 | 14 | 0 | 36 | 45 | 28 | 0 |
| SRR1732016 | 10 | 3167 | 411 | 811 | 11168 | 25282 | 102376 | 114782 | 4184 | 110 | 59163 | 122949 | 14276 | 10823 | 355 | 15687 | 6999 | 0 | 0 | 0 | 173 | 36 | 18 | 12 | 740 | 62 | 0 | 0 | 11 | 55 | 40 | 12 | 0 | 497 | 355 | 151 | 0 |
| SRR1748132 | 0 | 2608 | 2621 | 68 | 41 | 787 | 461 | 764 | 6004 | 309 | 4519 | 27343 | 1149 | 601 | 0 | 3842 | 72673 | 0 | 0 | 21 | 1405 | 7159 | 840 | 11 | 11 | 0 | 0 | 87 | 952 | 1166 | 6826 | 1599 | 0 | 55 | 57 | 0 | 0 |
| SRR1748133 | 0 | 5390 | 1372 | 242 | 4748 | 39018 | 35164 | 38289 | 8874 | 391 | 7708 | 14932 | 2145 | 4171 | 0 | 3357 | 111937 | 0 | 0 | 12 | 2426 | 2342 | 556 | 0 | 0 | 0 | 0 | 40 | 280 | 1111 | 2168 | 644 | 0 | 11 | 15 | 0 | 0 |
| SRR1759212 | 41 | 486 | 264 | 569 | 45 | 162 | 413 | 636 | 569 | 63 | 16303 | 27621 | 7744 | 2508 | 404 | 9377 | 148927 | 0 | 62 | 90 | 3851 | 4745 | 309 | 0 | 0 | 0 | 0 | 541 | 3306 | 2671 | 13650 | 620 | 0 | 0 | 0 | 0 | 0 |
| SRR1759213 | 314 | 2532 | 852 | 647 | 54 | 219 | 499 | 987 | 413 | 28 | 20263 | 31245 | 9511 | 3497 | 188 | 12853 | 77622 | 0 | 17 | 48 | 2015 | 288 | 105 | 0 | 0 | 0 | 0 | 123 | 1922 | 1064 | 1031 | 144 | 0 | 0 | 0 | 11 | 0 |
| SRR1759214 | 51 | 602 | 369 | 726 | 41 | 272 | 435 | 893 | 523 | 37 | 20547 | 35913 | 10333 | 2676 | 306 | 12088 | 82770 | 0 | 18 | 36 | 3231 | 647 | 217 | 0 | 0 | 0 | 0 | 131 | 1412 | 1474 | 1411 | 301 | 0 | 0 | 0 | 0 | 0 |
| SRR1759215 | 452 | 1480 | 912 | 989 | 51 | 310 | 901 | 956 | 428 | 60 | 21662 | 33690 | 10096 | 3111 | 282 | 12869 | 77366 | 0 | 34 | 39 | 2052 | 1320 | 292 | 0 | 0 | 0 | 0 | 336 | 1758 | 1069 | 3279 | 486 | 0 | 0 | 12 | 13 | 0 |
| SRR1759216 | 176 | 2472 | 594 | 224 | 33 | 183 | 325 | 742 | 532 | 39 | 21182 | 35644 | 9807 | 2442 | 313 | 12140 | 98455 | 0 | 35 | 42 | 1867 | 2302 | 236 | 0 | 0 | 0 | 0 | 352 | 1758 | 1159 | 5892 | 482 | 0 | 0 | 0 | 0 | 0 |
| SRR1759217 | 1602 | 546 | 357 | 2750 | 50 | 196 | 848 | 905 | 473 | 32 | 19217 | 21916 | 8281 | 6108 | 276 | 10366 | 70908 | 0 | 43 | 46 | 2312 | 1489 | 327 | 0 | 0 | 0 | 0 | 264 | 898 | 1092 | 3601 | 615 | 0 | 0 | 0 | 0 | 0 |
| SRR1759218 | 86 | 2087 | 1306 | 885 | 48 | 200 | 542 | 962 | 312 | 61 | 17648 | 26616 | 8111 | 3417 | 290 | 8758 | 73911 | 0 | 38 | 39 | 1714 | 1116 | 200 | 0 | 0 | 0 | 0 | 332 | 1311 | 1030 | 2650 | 342 | 0 | 0 | 0 | 0 | 0 |
| SRR1759219 | 304 | 2021 | 730 | 313 | 51 | 234 | 487 | 1136 | 415 | 41 | 20432 | 33627 | 9371 | 3051 | 315 | 10969 | 91263 | 0 | 56 | 99 | 5553 | 1210 | 330 | 0 | 0 | 0 | 0 | 503 | 3618 | 3065 | 3458 | 658 | 0 | 0 | 0 | 0 | 0 |
| SRR1759220 | 531 | 1102 | 932 | 632 | 42 | 207 | 489 | 925 | 585 | 30 | 17657 | 30951 | 8509 | 2595 | 368 | 10801 | 87734 | 0 | 31 | 30 | 735 | 2070 | 305 | 0 | 0 | 0 | 0 | 278 | 966 | 372 | 4875 | 575 | 0 | 0 | 0 | 0 | 0 |
| SRR1759221 | 21 | 429 | 173 | 789 | 27 | 140 | 205 | 681 | 883 | 11 | 27464 | 38786 | 9090 | 3850 | 283 | 12486 | 136216 | 0 | 19 | 19 | 3961 | 4101 | 473 | 0 | 0 | 0 | 0 | 248 | 552 | 1787 | 10222 | 580 | 0 | 0 | 22 | 0 | 0 |
| SRR1759222 | 0 | 1364 | 874 | 46 | 61 | 609 | 873 | 1893 | 820 | 28 | 24262 | 14 | 9166 | 2975 | 64 | 8919 | 62422 | 0 | 0 | 0 | 1437 | 221 | 88 | 0 | 0 | 0 | 0 | 44 | 237 | 696 | 545 | 66 | 0 | 0 | 12 | 0 | 0 |
| SRR1759223 | 42 | 761 | 91 | 708 | 52 | 350 | 487 | 1467 | 1187 | 0 | 26162 | 40094 | 9918 | 3360 | 307 | 16842 | 119621 | 0 | 40 | 64 | 6435 | 860 | 282 | 0 | 0 | 0 | 0 | 465 | 1841 | 2926 | 1817 | 452 | 0 | 0 | 22 | 0 | 0 |
| SRR1759224 | 0 | 528 | 241 | 322 | 29 | 181 | 267 | 716 | 1036 | 0 | 37284 | 53908 | 11591 | 3456 | 204 | 23660 | 92539 | 0 | 0 | 0 | 1130 | 1163 | 182 | 0 | 0 | 0 | 0 | 122 | 357 | 1533 | 3905 | 343 | 0 | 0 | 0 | 0 | 0 |
| SRR1759225 | 0 | 609 | 196 | 169 | 25 | 192 | 211 | 921 | 801 | 10 | 26008 | 39044 | 9918 | 2716 | 222 | 9795 | 80958 | 0 | 17 | 46 | 8448 | 717 | 379 | 0 | 0 | 0 | 0 | 326 | 1904 | 3962 | 1741 | 349 | 0 | 13 | 58 | 12 | 0 |
| SRR1759226 | 0 | 512 | 130 | 410 | 40 | 227 | 497 | 1086 | 1493 | 0 | 29483 | 25 | 7591 | 3584 | 103 | 21895 | 97708 | 0 | 0 | 0 | 3091 | 467 | 107 | 0 | 0 | 0 | 0 | 53 | 259 | 3820 | 1505 | 159 | 0 | 0 | 0 | 0 | 0 |
| SRR1759227 | 18 | 527 | 204 | 500 | 22 | 175 | 235 | 948 | 723 | 0 | 26550 | 41916 | 9407 | 3184 | 207 | 11340 | 108082 | 0 | 14 | 22 | 2790 | 5092 | 623 | 0 | 0 | 0 | 0 | 229 | 564 | 1294 | 13591 | 871 | 0 | 0 | 35 | 0 | 0 |
| SRR1759228 | 0 | 500 | 204 | 961 | 26 | 218 | 322 | 1377 | 595 | 0 | 18848 | 32526 | 5933 | 2504 | 64 | 7624 | 98765 | 0 | 12 | 19 | 1263 | 1917 | 408 | 0 | 0 | 0 | 0 | 276 | 1301 | 1285 | 5381 | 530 | 0 | 0 | 56 | 11 | 0 |
| SRR1759229 | 25 | 323 | 96 | 483 | 30 | 186 | 269 | 1096 | 1199 | 0 | 24010 | 35252 | 7981 | 3421 | 159 | 13070 | 106533 | 0 | 27 | 66 | 12667 | 1925 | 766 | 0 | 0 | 0 | 0 | 406 | 2011 | 5915 | 4618 | 1481 | 0 | 15 | 43 | 14 | 0 |
| SRR1759230 | 0 | 247 | 60 | 617 | 35 | 206 | 238 | 1003 | 1743 | 0 | 29681 | 43162 | 8681 | 2765 | 246 | 13282 | 121067 | 0 | 0 | 13 | 2124 | 691 | 136 | 0 | 0 | 0 | 0 | 131 | 633 | 1774 | 1979 | 222 | 0 | 14 | 81 | 12 | 0 |
| SRR1759231 | 12 | 1084 | 396 | 118 | 29 | 224 | 321 | 1261 | 374 | 24 | 22285 | 36062 | 8226 | 2778 | 194 | 8447 | 96486 | 0 | 19 | 63 | 12513 | 1555 | 407 | 0 | 0 | 0 | 0 | 426 | 3441 | 7107 | 6151 | 558 | 0 | 0 | 0 | 0 | 0 |
| SRR1759232 | 0 | 1044 | 213 | 428 | 43 | 327 | 321 | 1167 | 2518 | 0 | 28083 | 40326 | 9690 | 3015 | 196 | 12286 | 98199 | 0 | 0 | 19 | 2846 | 902 | 244 | 0 | 0 | 0 | 0 | 158 | 736 | 1646 | 2381 | 277 | 0 | 0 | 19 | 0 | 0 |
| SRR1759233 | 0 | 351 | 145 | 129 | 22 | 161 | 203 | 914 | 766 | 0 | 29160 | 20 | 9212 | 3204 | 233 | 12648 | 113559 | 0 | 0 | 0 | 1280 | 1348 | 298 | 0 | 0 | 0 | 0 | 63 | 232 | 901 | 3178 | 427 | 0 | 0 | 40 | 0 | 0 |
| SRR1759234 | 25 | 487 | 158 | 283 | 39 | 251 | 392 | 1348 | 629 | 15 | 24018 | 36056 | 9485 | 3234 | 269 | 9820 | 117196 | 0 | 58 | 148 | 20527 | 3446 | 777 | 0 | 0 | 0 | 0 | 717 | 4974 | 8533 | 6534 | 1395 | 0 | 13 | 27 | 0 | 0 |
| SRR1759235 | 72 | 818 | 714 | 862 | 29 | 216 | 442 | 1014 | 344 | 18 | 16300 | 29201 | 7455 | 2713 | 205 | 7837 | 75375 | 0 | 46 | 42 | 2733 | 1193 | 196 | 0 | 0 | 0 | 0 | 554 | 2202 | 2338 | 3052 | 489 | 0 | 11 | 50 | 38 | 0 |
| SRR1759236 | 47 | 644 | 236 | 205 | 46 | 319 | 399 | 1473 | 1049 | 17 | 28721 | 44094 | 10782 | 2976 | 330 | 14002 | 95245 | 0 | 16 | 26 | 4551 | 1067 | 200 | 0 | 0 | 0 | 0 | 242 | 924 | 2277 | 2290 | 324 | 0 | 0 | 12 | 0 | 0 |
| SRR1759237 | 38 | 663 | 247 | 439 | 36 | 258 | 423 | 1474 | 677 | 23 | 19000 | 33703 | 7793 | 2956 | 222 | 11044 | 133774 | 0 | 17 | 23 | 1238 | 948 | 98 | 0 | 0 | 0 | 0 | 291 | 962 | 1109 | 2242 | 198 | 0 | 0 | 28 | 0 | 0 |
| SRR1759238 | 17 | 563 | 130 | 127 | 42 | 184 | 307 | 916 | 701 | 21 | 21669 | 36614 | 8300 | 3071 | 276 | 13861 | 166523 | 0 | 26 | 35 | 5567 | 2154 | 285 | 0 | 0 | 0 | 0 | 369 | 1142 | 3050 | 4977 | 504 | 0 | 0 | 0 | 0 | 0 |
| SRR1759239 | 0 | 169 | 67 | 408 | 50 | 96 | 436 | 821 | 792 | 12 | 5745 | 11827 | 1919 | 3365 | 1553 | 11743 | 91328 | 0 | 35 | 65 | 1347 | 301 | 93 | 0 | 0 | 0 | 14 | 366 | 1324 | 1106 | 1099 | 227 | 0 | 15 | 486 | 49 | 0 |
| SRR1759240 | 47 | 1267 | 360 | 443 | 37 | 226 | 334 | 1096 | 628 | 19 | 27761 | 40232 | 10501 | 3900 | 399 | 12878 | 75131 | 0 | 28 | 37 | 3039 | 922 | 193 | 0 | 0 | 0 | 0 | 463 | 1623 | 1950 | 2270 | 398 | 0 | 0 | 41 | 0 | 0 |
| SRR1759241 | 48 | 1059 | 356 | 505 | 38 | 203 | 326 | 950 | 700 | 23 | 25379 | 39712 | 10296 | 3109 | 586 | 13338 | 71944 | 0 | 20 | 28 | 3137 | 737 | 135 | 0 | 0 | 0 | 0 | 284 | 928 | 1756 | 1493 | 212 | 0 | 11 | 72 | 18 | 0 |
| SRR1759242 | 32 | 1192 | 170 | 1117 | 21 | 175 | 311 | 1178 | 386 | 33 | 17724 | 37289 | 6539 | 2975 | 150 | 7407 | 109578 | 0 | 14 | 19 | 1131 | 2985 | 480 | 0 | 0 | 0 | 0 | 281 | 970 | 1260 | 8545 | 1155 | 0 | 0 | 47 | 12 | 0 |
| SRR1759243 | 15 | 943 | 459 | 216 | 37 | 206 | 397 | 1299 | 697 | 35 | 22592 | 35010 | 8587 | 3849 | 237 | 10352 | 84214 | 0 | 20 | 95 | 16794 | 1091 | 545 | 0 | 0 | 0 | 0 | 363 | 3881 | 10897 | 3113 | 1022 | 0 | 0 | 15 | 0 | 0 |
| SRR1759244 | 194 | 865 | 1085 | 1784 | 53 | 135 | 575 | 898 | 506 | 66 | 19194 | 23131 | 7932 | 5426 | 304 | 11445 | 71350 | 0 | 38 | 100 | 3415 | 1106 | 191 | 0 | 0 | 0 | 0 | 255 | 2254 | 1746 | 2713 | 307 | 0 | 0 | 0 | 0 | 0 |
| SRR1759245 | 53 | 530 | 492 | 453 | 40 | 231 | 484 | 735 | 374 | 43 | 22786 | 38264 | 11302 | 2873 | 504 | 12812 | 103197 | 0 | 58 | 39 | 3199 | 4721 | 668 | 0 | 0 | 0 | 0 | 512 | 1579 | 1818 | 11218 | 1344 | 0 | 0 | 16 | 10 | 0 |
| SRR1759246 | 28 | 932 | 280 | 79 | 116 | 528 | 1067 | 2843 | 1991 | 0 | 17287 | 26722 | 5587 | 2358 | 222 | 9908 | 87344 | 0 | 30 | 108 | 11223 | 1994 | 552 | 0 | 0 | 0 | 0 | 385 | 2494 | 3936 | 5996 | 643 | 0 | 0 | 0 | 0 | 0 |
| SRR1759247 | 11 | 456 | 223 | 507 | 34 | 227 | 252 | 1214 | 1935 | 11 | 23618 | 34447 | 8209 | 3048 | 220 | 11279 | 107132 | 0 | 0 | 13 | 1679 | 1780 | 576 | 0 | 0 | 0 | 0 | 187 | 614 | 1149 | 4866 | 822 | 0 | 0 | 32 | 0 | 0 |
| SRR1759248 | 36 | 915 | 222 | 570 | 58 | 313 | 480 | 1274 | 818 | 18 | 23110 | 37452 | 9448 | 3066 | 422 | 16554 | 112076 | 0 | 33 | 56 | 4560 | 1148 | 211 | 0 | 0 | 0 | 0 | 516 | 1926 | 2077 | 2645 | 280 | 0 | 0 | 32 | 0 | 0 |
| SRR1759249 | 31 | 675 | 220 | 88 | 52 | 301 | 379 | 1369 | 3158 | 15 | 26844 | 39556 | 9631 | 3998 | 226 | 19385 | 103192 | 0 | 0 | 96 | 5778 | 645 | 201 | 0 | 0 | 0 | 0 | 110 | 4079 | 3496 | 1897 | 369 | 0 | 0 | 0 | 0 | 0 |
| SRR1759250 | 39 | 2120 | 385 | 157 | 44 | 269 | 444 | 1484 | 687 | 44 | 18490 | 38593 | 7484 | 2895 | 232 | 13912 | 88216 | 0 | 0 | 64 | 6768 | 629 | 242 | 0 | 0 | 0 | 0 | 88 | 1671 | 2532 | 1164 | 260 | 0 | 0 | 0 | 0 | 0 |
| SRR1759251 | 0 | 907 | 145 | 260 | 49 | 304 | 470 | 1455 | 995 | 16 | 25394 | 40286 | 9523 | 3633 | 293 | 18256 | 105629 | 0 | 27 | 54 | 3249 | 3131 | 235 | 0 | 0 | 0 | 0 | 434 | 1842 | 2087 | 8074 | 508 | 0 | 0 | 0 | 0 | 0 |
| SRR1759252 | 12 | 1096 | 309 | 309 | 70 | 366 | 675 | 1724 | 1617 | 21 | 15979 | 26343 | 5809 | 2147 | 223 | 14165 | 148978 | 0 | 42 | 45 | 3059 | 4442 | 1137 | 0 | 0 | 0 | 0 | 429 | 1096 | 1105 | 9973 | 1821 | 0 | 19 | 19 | 0 | 0 |
| SRR1759253 | 38 | 742 | 275 | 187 | 55 | 506 | 861 | 1420 | 689 | 18 | 28935 | 37470 | 10070 | 3488 | 121 | 12288 | 83491 | 0 | 0 | 30 | 3070 | 727 | 200 | 0 | 0 | 0 | 0 | 111 | 706 | 985 | 1188 | 226 | 0 | 0 | 0 | 0 | 0 |
| SRR1759254 | 78 | 927 | 151 | 190 | 41 | 215 | 344 | 1097 | 1548 | 21 | 20043 | 34093 | 7574 | 3371 | 287 | 18690 | 121718 | 0 | 53 | 119 | 5933 | 1632 | 355 | 0 | 0 | 0 | 0 | 667 | 3673 | 3292 | 4101 | 628 | 0 | 17 | 14 | 0 | 0 |
| SRR1759255 | 42 | 629 | 329 | 197 | 32 | 221 | 302 | 1244 | 1029 | 0 | 32604 | 40854 | 9650 | 3662 | 176 | 15577 | 78914 | 0 | 0 | 46 | 2195 | 749 | 118 | 0 | 0 | 0 | 0 | 133 | 1802 | 1141 | 1745 | 191 | 0 | 0 | 0 | 0 | 0 |
| SRR1759256 | 68 | 864 | 398 | 162 | 33 | 221 | 302 | 1026 | 1724 | 36 | 27841 | 43694 | 11075 | 3008 | 405 | 14321 | 87140 | 0 | 0 | 11 | 2137 | 338 | 146 | 0 | 0 | 0 | 0 | 86 | 595 | 1349 | 921 | 304 | 0 | 0 | 19 | 0 | 0 |
| SRR1759257 | 17 | 917 | 385 | 62 | 34 | 187 | 342 | 1258 | 857 | 48 | 20181 | 29131 | 7903 | 3799 | 216 | 12571 | 80533 | 0 | 13 | 123 | 12632 | 989 | 234 | 0 | 0 | 0 | 0 | 234 | 5893 | 10745 | 3293 | 506 | 0 | 0 | 0 | 0 | 0 |
| SRR1759258 | 72 | 1512 | 191 | 918 | 73 | 221 | 406 | 1042 | 1459 | 13 | 16240 | 31172 | 5057 | 1771 | 217 | 8137 | 161725 | 0 | 19 | 40 | 981 | 1635 | 947 | 11 | 13 | 0 | 0 | 419 | 1604 | 600 | 4880 | 1343 | 0 | 72 | 205 | 15 | 0 |
| SRR1759259 | 35 | 2209 | 693 | 318 | 49 | 250 | 408 | 1088 | 866 | 28 | 21468 | 34449 | 8379 | 2855 | 267 | 12022 | 134374 | 0 | 0 | 19 | 577 | 853 | 173 | 0 | 0 | 0 | 0 | 169 | 711 | 388 | 1945 | 264 | 0 | 0 | 0 | 0 | 0 |
| SRR1759260 | 40 | 313 | 123 | 230 | 48 | 276 | 391 | 910 | 474 | 11 | 20100 | 31213 | 9896 | 2405 | 345 | 11131 | 83283 | 11 | 46 | 115 | 24456 | 1084 | 833 | 12 | 0 | 0 | 11 | 522 | 5656 | 13638 | 3020 | 1557 | 0 | 30 | 68 | 30 | 0 |
| SRR1759261 | 91 | 521 | 274 | 1018 | 31 | 165 | 263 | 895 | 1504 | 12 | 24349 | 37996 | 7708 | 3069 | 195 | 11042 | 151720 | 0 | 18 | 31 | 1482 | 2218 | 349 | 0 | 0 | 0 | 0 | 368 | 1214 | 1010 | 6048 | 403 | 0 | 0 | 52 | 0 | 0 |
| SRR1759262 | 31 | 454 | 101 | 544 | 41 | 190 | 388 | 957 | 814 | 19 | 16483 | 26256 | 8572 | 2659 | 263 | 10986 | 87158 | 0 | 95 | 100 | 10763 | 3957 | 962 | 11 | 0 | 0 | 0 | 853 | 3660 | 5545 | 10293 | 2171 | 0 | 14 | 48 | 24 | 0 |
| SRR1759263 | 0 | 1022 | 497 | 85 | 38 | 275 | 370 | 1083 | 1310 | 18 | 21086 | 10 | 8881 | 2757 | 225 | 9185 | 101862 | 0 | 0 | 34 | 5795 | 325 | 261 | 0 | 0 | 0 | 0 | 147 | 1482 | 2112 | 742 | 224 | 0 | 0 | 0 | 0 | 0 |
| SRR1759264 | 17 | 653 | 445 | 116 | 30 | 208 | 262 | 918 | 907 | 15 | 31331 | 43047 | 10161 | 3203 | 258 | 18375 | 93514 | 0 | 0 | 14 | 3178 | 1407 | 192 | 0 | 0 | 0 | 0 | 139 | 611 | 2234 | 3513 | 261 | 0 | 0 | 0 | 0 | 0 |
| SRR1759265 | 12 | 1610 | 1868 | 117 | 26 | 241 | 361 | 2081 | 943 | 16 | 22301 | 31791 | 5829 | 4027 | 67 | 9427 | 66655 | 0 | 10 | 88 | 9167 | 704 | 211 | 0 | 0 | 0 | 0 | 222 | 3532 | 7091 | 3133 | 318 | 0 | 0 | 0 | 0 | 0 |
| SRR1759266 | 41 | 734 | 601 | 142 | 58 | 224 | 506 | 998 | 686 | 52 | 19616 | 31586 | 9175 | 2797 | 417 | 14197 | 81573 | 0 | 36 | 267 | 30743 | 1130 | 1536 | 12 | 0 | 0 | 0 | 464 | 13433 | 22665 | 4883 | 3615 | 0 | 14 | 0 | 0 | 0 |
| SRR1759267 | 0 | 777 | 146 | 47 | 127 | 656 | 965 | 2087 | 4272 | 32 | 15775 | 28821 | 5539 | 2819 | 173 | 15375 | 155847 | 0 | 11 | 31 | 3612 | 1445 | 496 | 0 | 0 | 0 | 0 | 187 | 1414 | 2977 | 6109 | 940 | 0 | 11 | 25 | 0 | 0 |
| SRR1759268 | 12 | 1251 | 382 | 134 | 49 | 398 | 575 | 1654 | 5254 | 0 | 21548 | 21 | 5187 | 2848 | 74 | 15424 | 117913 | 0 | 12 | 30 | 2453 | 1873 | 563 | 0 | 0 | 0 | 0 | 179 | 743 | 1651 | 5344 | 789 | 0 | 0 | 0 | 0 | 0 |
| SRR1759269 | 22 | 883 | 447 | 165 | 31 | 332 | 548 | 1659 | 1099 | 27 | 25514 | 35682 | 7841 | 3674 | 117 | 11130 | 88173 | 0 | 11 | 55 | 3359 | 1435 | 132 | 0 | 0 | 0 | 0 | 134 | 1552 | 1680 | 3388 | 136 | 0 | 0 | 0 | 0 | 0 |
| SRR1759270 | 53 | 602 | 204 | 156 | 105 | 520 | 1062 | 1485 | 725 | 40 | 18825 | 32565 | 9319 | 2751 | 349 | 10312 | 109274 | 0 | 84 | 115 | 8353 | 4043 | 483 | 0 | 0 | 0 | 0 | 656 | 3912 | 3881 | 10928 | 885 | 0 | 0 | 0 | 0 | 0 |
| SRR1759271 | 0 | 842 | 446 | 200 | 27 | 238 | 258 | 1053 | 1045 | 0 | 27750 | 40903 | 10240 | 2630 | 172 | 12662 | 70506 | 0 | 0 | 18 | 2525 | 479 | 113 | 0 | 0 | 0 | 0 | 183 | 1212 | 1917 | 1507 | 149 | 0 | 0 | 26 | 0 | 0 |
| SRR1759272 | 0 | 759 | 278 | 66 | 39 | 292 | 475 | 1151 | 1081 | 21 | 30289 | 26 | 8564 | 3899 | 166 | 14694 | 131850 | 0 | 0 | 22 | 2588 | 1270 | 107 | 0 | 0 | 0 | 0 | 133 | 948 | 2557 | 3872 | 172 | 0 | 0 | 0 | 0 | 0 |
| SRR1759273 | 124 | 1128 | 297 | 483 | 64 | 430 | 789 | 1963 | 10624 | 60 | 16260 | 34931 | 5235 | 4714 | 139 | 19330 | 102505 | 0 | 18 | 54 | 3995 | 941 | 302 | 0 | 0 | 0 | 0 | 266 | 1632 | 4324 | 4749 | 725 | 0 | 0 | 10 | 0 | 0 |
| SRR1759274 | 181 | 1772 | 245 | 586 | 39 | 229 | 415 | 680 | 502 | 18 | 19931 | 30480 | 9403 | 2722 | 377 | 12810 | 119611 | 0 | 38 | 60 | 9097 | 1540 | 592 | 0 | 0 | 0 | 0 | 521 | 2565 | 4057 | 4169 | 953 | 0 | 14 | 42 | 21 | 0 |
| SRR1759275 | 98 | 3108 | 783 | 3015 | 50 | 266 | 705 | 986 | 437 | 31 | 19766 | 28258 | 9049 | 3830 | 371 | 7848 | 69577 | 0 | 77 | 94 | 13036 | 1868 | 721 | 0 | 0 | 0 | 0 | 645 | 2956 | 4310 | 3038 | 751 | 0 | 14 | 61 | 35 | 0 |
| SRR1777167 | 2848 | 17766 | 38316 | 3842 | 1634 | 621 | 23355 | 666 | 1000 | 411 | 4267 | 2451 | 4675 | 5921 | 25332 | 2006 | 7061 | 0 | 0 | 0 | 28 | 35 | 65 | 0 | 14 | 0 | 0 | 18 | 33 | 35 | 0 | 0 | 0 | 0 | 0 | 0 | 0 |
| SRR1820679 | 1081 | 6002 | 6188 | 962 | 976 | 706 | 13141 | 3177 | 1628 | 1868 | 9341 | 4555 | 4192 | 2523 | 42 | 3799 | 30218 | 0 | 0 | 0 | 264 | 94 | 21 | 0 | 0 | 0 | 0 | 0 | 11 | 50 | 0 | 0 | 0 | 0 | 0 | 0 | 0 |
| SRR1820680 | 24 | 1087 | 20897 | 0 | 74 | 53 | 1190 | 758 | 530 | 5052 | 7256 | 3193 | 3200 | 1481 | 30 | 1821 | 16689 | 0 | 0 | 0 | 11 | 0 | 0 | 0 | 0 | 0 | 0 | 0 | 0 | 0 | 0 | 0 | 0 | 0 | 0 | 0 | 0 |
| SRR1820681 | 1205 | 4095 | 3947 | 934 | 202 | 99 | 2282 | 628 | 686 | 3490 | 7111 | 2840 | 3288 | 1414 | 37 | 4026 | 20595 | 0 | 0 | 24 | 998 | 120 | 42 | 0 | 0 | 0 | 0 | 0 | 34 | 243 | 10 | 0 | 0 | 0 | 0 | 0 | 0 |
| SRR1820682 | 39 | 1243 | 22200 | 14 | 98 | 54 | 1226 | 803 | 595 | 7329 | 8560 | 3891 | 3920 | 1882 | 33 | 2333 | 20605 | 0 | 0 | 0 | 79 | 22 | 11 | 0 | 0 | 0 | 0 | 0 | 0 | 20 | 0 | 0 | 0 | 0 | 0 | 0 | 0 |
| SRR1820683 | 1184 | 4883 | 7453 | 848 | 221 | 119 | 2425 | 697 | 716 | 2594 | 8084 | 3436 | 3579 | 1894 | 55 | 3924 | 25922 | 0 | 0 | 15 | 812 | 166 | 42 | 0 | 0 | 0 | 0 | 0 | 25 | 170 | 13 | 0 | 0 | 0 | 0 | 0 | 0 |
| SRR1820684 | 26 | 1626 | 21969 | 0 | 239 | 130 | 3062 | 1481 | 1343 | 7156 | 12760 | 7005 | 8043 | 5683 | 63 | 1872 | 28571 | 0 | 0 | 0 | 21 | 19 | 0 | 0 | 0 | 0 | 0 | 0 | 0 | 0 | 0 | 0 | 0 | 0 | 0 | 0 | 0 |
| SRR1820685 | 1240 | 4461 | 4249 | 1214 | 730 | 439 | 8555 | 2535 | 1314 | 3343 | 8724 | 4006 | 4100 | 1637 | 30 | 3674 | 27799 | 0 | 0 | 0 | 67 | 69 | 13 | 0 | 0 | 0 | 0 | 0 | 0 | 17 | 0 | 0 | 0 | 0 | 0 | 0 | 0 |
| SRR1820686 | 38 | 1492 | 19568 | 10 | 158 | 90 | 2107 | 1273 | 748 | 7413 | 10566 | 4806 | 4891 | 1823 | 22 | 1936 | 23996 | 0 | 0 | 0 | 0 | 0 | 0 | 0 | 0 | 0 | 0 | 0 | 0 | 0 | 0 | 0 | 0 | 0 | 0 | 0 | 0 |
| SRR1820687 | 90 | 2912 | 14935 | 44 | 806 | 714 | 13265 | 2500 | 1962 | 6202 | 13247 | 5902 | 7345 | 3325 | 28 | 3932 | 26088 | 0 | 0 | 0 | 50 | 46 | 15 | 0 | 0 | 0 | 0 | 0 | 0 | 16 | 0 | 0 | 0 | 0 | 0 | 11 | 0 |
| SRR1820688 | 70 | 2485 | 22361 | 29 | 234 | 158 | 3205 | 712 | 1362 | 4711 | 11338 | 4283 | 5945 | 2491 | 33 | 3527 | 22838 | 0 | 0 | 0 | 306 | 63 | 18 | 0 | 0 | 0 | 0 | 0 | 11 | 101 | 0 | 0 | 0 | 0 | 0 | 16 | 0 |
| SRR1820689 | 13 | 555 | 24181 | 14 | 283 | 285 | 4696 | 889 | 1262 | 4482 | 16270 | 6834 | 8195 | 5531 | 92 | 5035 | 30991 | 0 | 0 | 0 | 209 | 64 | 31 | 0 | 0 | 0 | 0 | 0 | 0 | 54 | 0 | 0 | 0 | 0 | 0 | 20 | 0 |
| SRR1820690 | 19 | 583 | 16871 | 23 | 207 | 157 | 3407 | 837 | 1185 | 3539 | 11356 | 3450 | 4479 | 2386 | 64 | 3224 | 18439 | 0 | 0 | 0 | 84 | 44 | 26 | 0 | 0 | 0 | 0 | 0 | 0 | 35 | 0 | 0 | 0 | 0 | 0 | 13 | 0 |
| SRR1853804 | 139 | 35891 | 54 | 197 | 13 | 122 | 57 | 309 | 644 | 0 | 7136 | 24675 | 16207 | 1025 | 35 | 0 | 78108 | 0 | 0 | 0 | 48 | 73 | 26 | 0 | 0 | 0 | 0 | 0 | 0 | 22 | 23 | 18 | 0 | 39 | 26 | 0 | 0 |
| SRR1853805 | 162 | 29096 | 22 | 180 | 24 | 173 | 60 | 289 | 701 | 0 | 9808 | 24561 | 16255 | 832 | 16 | 0 | 78731 | 0 | 0 | 0 | 34 | 91 | 28 | 0 | 0 | 0 | 0 | 0 | 0 | 15 | 24 | 23 | 0 | 30 | 15 | 0 | 0 |
| SRR1853806 | 63 | 33575 | 41 | 185 | 194 | 1648 | 798 | 572 | 1104 | 0 | 10862 | 48193 | 18834 | 1572 | 13 | 0 | 67864 | 0 | 0 | 0 | 44 | 69 | 26 | 0 | 0 | 0 | 0 | 0 | 22 | 32 | 101 | 17 | 0 | 41 | 34 | 0 | 0 |
| SRR1853807 | 74 | 54678 | 61 | 214 | 23 | 149 | 70 | 177 | 870 | 0 | 15111 | 42446 | 13003 | 938 | 17 | 0 | 40094 | 0 | 0 | 0 | 10 | 88 | 0 | 0 | 0 | 0 | 0 | 0 | 13 | 0 | 15 | 0 | 0 | 13 | 56 | 0 | 0 |
| SRR1853808 | 128 | 135049 | 51 | 151 | 37 | 268 | 109 | 272 | 1004 | 0 | 8986 | 32710 | 11189 | 852 | 45 | 0 | 35962 | 0 | 0 | 0 | 34 | 79 | 0 | 0 | 0 | 0 | 0 | 0 | 41 | 14 | 82 | 0 | 0 | 23 | 93 | 0 | 0 |
| SRR1853809 | 287 | 70533 | 60 | 451 | 47 | 458 | 181 | 334 | 1543 | 0 | 12573 | 31671 | 9972 | 1112 | 65 | 0 | 43718 | 0 | 0 | 0 | 80 | 161 | 20 | 0 | 0 | 0 | 0 | 0 | 131 | 34 | 168 | 14 | 0 | 41 | 385 | 0 | 0 |
| SRR1917323 | 239 | 530 | 7350 | 6786 | 22862 | 9059 | 103722 | 13689 | 31450 | 1969 | 13943 | 14087 | 11720 | 20437 | 18335 | 59 | 12373 | 0 | 23 | 11 | 57 | 107 | 96 | 24 | 18 | 0 | 0 | 14 | 19 | 12 | 0 | 0 | 0 | 0 | 0 | 15 | 0 |
| SRR1917324 | 556 | 2063 | 13589 | 6428 | 21816 | 6209 | 145292 | 8328 | 23047 | 2202 | 13136 | 9574 | 8295 | 14694 | 20083 | 53 | 9693 | 0 | 16 | 0 | 51 | 76 | 86 | 20 | 19 | 0 | 0 | 0 | 14 | 0 | 0 | 0 | 0 | 0 | 0 | 13 | 0 |
| SRR1917325 | 199 | 601 | 7208 | 3627 | 19663 | 8845 | 89107 | 13438 | 26801 | 3896 | 18193 | 15805 | 14196 | 21037 | 20721 | 65 | 14296 | 0 | 32 | 15 | 65 | 108 | 97 | 25 | 16 | 0 | 0 | 20 | 27 | 12 | 0 | 0 | 0 | 0 | 0 | 11 | 0 |
| SRR1917326 | 0 | 836 | 33372 | 136 | 2762 | 1357 | 32284 | 7130 | 12334 | 675 | 11507 | 19948 | 7537 | 3964 | 928 | 181 | 8598 | 0 | 32 | 52 | 78 | 377 | 170 | 14 | 0 | 0 | 0 | 37 | 44 | 76 | 22 | 0 | 0 | 0 | 0 | 0 | 0 |
| SRR1917327 | 15 | 2001 | 36432 | 83 | 5545 | 2615 | 51980 | 5452 | 3956 | 628 | 13018 | 33174 | 4329 | 5037 | 1717 | 136 | 20028 | 0 | 59 | 89 | 199 | 777 | 360 | 36 | 14 | 0 | 0 | 56 | 61 | 65 | 29 | 0 | 0 | 0 | 0 | 12 | 0 |
| SRR1917328 | 58 | 2855 | 84007 | 29 | 1324 | 445 | 8746 | 755 | 4841 | 0 | 3705 | 8307 | 866 | 615 | 921 | 20 | 2926 | 0 | 14 | 0 | 10 | 186 | 21 | 0 | 0 | 0 | 0 | 25 | 11 | 0 | 0 | 0 | 0 | 0 | 0 | 0 | 0 |
| SRR1917329 | 76 | 781 | 4160 | 3306 | 21621 | 7175 | 99861 | 15091 | 33874 | 3771 | 14663 | 23 | 12834 | 13991 | 15225 | 84 | 13581 | 0 | 41 | 23 | 116 | 110 | 172 | 32 | 25 | 0 | 0 | 23 | 26 | 19 | 0 | 0 | 0 | 0 | 0 | 14 | 0 |
| SRR1917330 | 228 | 1398 | 4695 | 6308 | 22392 | 7376 | 116721 | 11240 | 30860 | 2768 | 12999 | 11889 | 10354 | 11949 | 16852 | 96 | 10390 | 0 | 36 | 24 | 100 | 114 | 157 | 26 | 24 | 0 | 0 | 21 | 25 | 20 | 0 | 0 | 0 | 0 | 0 | 14 | 0 |
| SRR1917331 | 184 | 747 | 6077 | 5168 | 17427 | 7554 | 76322 | 11145 | 30082 | 2008 | 12371 | 19 | 10842 | 10848 | 13201 | 82 | 14831 | 0 | 50 | 33 | 110 | 173 | 169 | 37 | 27 | 0 | 0 | 44 | 42 | 26 | 10 | 0 | 0 | 0 | 0 | 15 | 0 |
| SRR1917332 | 16 | 1667 | 100610 | 316 | 4193 | 1965 | 29412 | 3720 | 93022 | 455 | 20494 | 29590 | 5219 | 7206 | 991 | 21 | 6065 | 0 | 21 | 0 | 14 | 111 | 28 | 0 | 0 | 0 | 0 | 125 | 73 | 88 | 0 | 0 | 0 | 0 | 0 | 0 | 0 |
| SRR1917333 | 12 | 2676 | 79621 | 250 | 5599 | 1788 | 33478 | 2365 | 58924 | 203 | 18316 | 46407 | 3399 | 6553 | 1103 | 74 | 5896 | 0 | 17 | 12 | 18 | 124 | 21 | 0 | 0 | 0 | 0 | 79 | 61 | 41 | 0 | 0 | 0 | 0 | 0 | 0 | 0 |
| SRR1917334 | 46 | 2194 | 33260 | 458 | 7897 | 2902 | 50906 | 2527 | 53074 | 388 | 26361 | 64724 | 3661 | 11421 | 2517 | 66 | 9409 | 0 | 52 | 26 | 36 | 190 | 22 | 0 | 0 | 0 | 0 | 205 | 126 | 91 | 0 | 0 | 0 | 0 | 0 | 0 | 0 |

### Supp Table S8.1.Columns_B_to_AL

| | | | | | | | | | | | | | | | | | | | | | | | | | | | | | | | | | | | | | |
|---|---|---|---|---|---|---|---|---|---|---|---|---|---|---|---|---|---|---|---|---|---|---|---|---|---|---|---|---|---|---|---|---|---|---|---|---|---|
| SRR1917335 | 183 | 671 | 6813 | 5636 | 21070 | 9255 | 111604 | 15447 | 26789 | 3814 | 15757 | 23 | 12794 | 17044 | 17264 | 63 | 11437 | 0 | 43 | 19 | 83 | 101 | 112 | 32 | 24 | 0 | 0 | 34 | 35 | 21 | 0 | 0 | 0 | 0 | 0 | 17 | 0 |
| SRR1917336 | 181 | 601 | 10325 | 3992 | 17910 | 8246 | 87226 | 13113 | 30735 | 1389 | 15606 | 29 | 13263 | 19467 | 16450 | 58 | 15650 | 0 | 25 | 21 | 59 | 108 | 71 | 21 | 19 | 0 | 0 | 10 | 17 | 0 | 0 | 0 | 0 | 0 | 0 | 15 | 0 |
| SRR1917337 | 416 | 626 | 6909 | 3591 | 15015 | 7398 | 73489 | 15685 | 21035 | 1644 | 17039 | 17904 | 14017 | 15058 | 15481 | 54 | 17851 | 0 | 63 | 39 | 127 | 176 | 132 | 42 | 29 | 10 | 0 | 34 | 32 | 17 | 0 | 0 | 0 | 0 | 0 | 23 | 0 |
| SRR1917338 | 93 | 12344 | 143935 | 349 | 2719 | 1279 | 22749 | 1679 | 14659 | 121 | 5271 | 11756 | 1166 | 2616 | 664 | 52 | 5116 | 0 | 18 | 16 | 22 | 119 | 79 | 11 | 0 | 0 | 0 | 32 | 22 | 0 | 0 | 0 | 0 | 0 | 0 | 0 | 0 |
| SRR1917339 | 39 | 5744 | 95472 | 400 | 2210 | 1255 | 25196 | 1098 | 34446 | 206 | 9364 | 7633 | 2966 | 5946 | 1022 | 185 | 5433 | 0 | 0 | 0 | 0 | 30 | 21 | 0 | 0 | 0 | 0 | 0 | 0 | 0 | 0 | 0 | 0 | 0 | 0 | 0 | 0 |
| SRR1917340 | 50 | 3992 | 110038 | 210 | 4606 | 2043 | 38718 | 1410 | 27277 | 279 | 14771 | 12785 | 3058 | 6345 | 1313 | 155 | 5308 | 0 | 13 | 12 | 18 | 55 | 20 | 0 | 0 | 0 | 0 | 20 | 24 | 0 | 0 | 0 | 0 | 0 | 0 | 0 | 0 |
| SRR1922516 | 399 | 9137 | 997 | 14 | 124 | 1487 | 1203 | 2029 | 2826 | 21 | 14847 | 21630 | 2472 | 17818 | 0 | 3315 | 36704 | 0 | 0 | 0 | 58 | 16 | 0 | 0 | 0 | 0 | 0 | 0 | 0 | 0 | 0 | 0 | 0 | 0 | 0 | 0 | 0 |
| SRR1922517 | 289 | 8495 | 936 | 0 | 86 | 1305 | 1023 | 1848 | 3209 | 14 | 16139 | 22813 | 2549 | 19424 | 0 | 1970 | 40163 | 0 | 0 | 0 | 47 | 15 | 0 | 0 | 0 | 0 | 0 | 0 | 0 | 0 | 0 | 0 | 0 | 0 | 0 | 0 | 0 |
| SRR1922518 | 352 | 7588 | 1097 | 14 | 92 | 1574 | 1376 | 2118 | 3235 | 26 | 13202 | 18214 | 2241 | 16623 | 0 | 3387 | 39875 | 0 | 0 | 0 | 56 | 18 | 11 | 0 | 0 | 0 | 0 | 0 | 0 | 0 | 0 | 0 | 0 | 0 | 0 | 0 | 0 |
| SRR1922519 | 428 | 5935 | 1353 | 10 | 75 | 1401 | 1100 | 1938 | 2986 | 24 | 14499 | 22256 | 2877 | 18670 | 0 | 2177 | 41643 | 0 | 0 | 0 | 31 | 10 | 0 | 0 | 0 | 0 | 0 | 0 | 0 | 0 | 0 | 0 | 0 | 0 | 0 | 0 | 0 |
| SRR1922520 | 228 | 11548 | 568 | 0 | 94 | 1660 | 1210 | 1969 | 2346 | 14 | 12450 | 15119 | 1660 | 11746 | 0 | 2240 | 26027 | 0 | 0 | 0 | 21 | 0 | 0 | 0 | 0 | 0 | 0 | 0 | 0 | 0 | 0 | 0 | 0 | 0 | 0 | 0 | 0 |
| SRR1922521 | 401 | 9602 | 987 | 0 | 94 | 1481 | 1149 | 2107 | 3191 | 14 | 8253 | 11624 | 1380 | 11710 | 0 | 3572 | 40757 | 0 | 0 | 0 | 57 | 16 | 0 | 0 | 0 | 0 | 0 | 0 | 0 | 0 | 0 | 0 | 0 | 0 | 0 | 0 | 0 |
| SRR1922522 | 305 | 8944 | 1058 | 12 | 121 | 2077 | 1655 | 2911 | 2250 | 20 | 11572 | 16976 | 2220 | 15870 | 0 | 2836 | 30236 | 0 | 0 | 0 | 31 | 11 | 0 | 0 | 0 | 0 | 0 | 0 | 0 | 0 | 0 | 0 | 0 | 0 | 0 | 0 | 0 |
| SRR1922523 | 245 | 7360 | 1534 | 0 | 79 | 1585 | 1232 | 2297 | 3265 | 19 | 8197 | 12148 | 1526 | 13444 | 0 | 3716 | 48610 | 0 | 0 | 0 | 48 | 16 | 0 | 0 | 0 | 0 | 0 | 0 | 0 | 0 | 0 | 0 | 0 | 0 | 0 | 0 | 0 |
| SRR1922524 | 360 | 7063 | 1241 | 23 | 135 | 2165 | 1703 | 2664 | 2200 | 30 | 11314 | 16779 | 2168 | 16533 | 0 | 2673 | 30704 | 0 | 0 | 0 | 70 | 20 | 11 | 0 | 0 | 0 | 0 | 0 | 0 | 0 | 0 | 0 | 0 | 0 | 0 | 0 | 0 |
| SRR1922525 | 270 | 9135 | 1031 | 10 | 125 | 1861 | 1410 | 2552 | 2225 | 18 | 11026 | 16850 | 1992 | 14899 | 0 | 4094 | 31715 | 0 | 0 | 0 | 22 | 0 | 0 | 0 | 0 | 0 | 0 | 0 | 0 | 0 | 0 | 0 | 0 | 0 | 0 | 0 | 0 |
| SRR1922526 | 349 | 6252 | 1781 | 26 | 117 | 1974 | 1591 | 2992 | 1892 | 45 | 11411 | 18009 | 2153 | 17011 | 0 | 2804 | 29779 | 0 | 0 | 0 | 29 | 0 | 0 | 0 | 0 | 0 | 0 | 0 | 0 | 0 | 0 | 0 | 0 | 0 | 0 | 0 | 0 |
| SRR1922527 | 335 | 8589 | 1864 | 10 | 107 | 2175 | 1656 | 2881 | 1974 | 26 | 10296 | 18108 | 2261 | 17790 | 0 | 4939 | 30041 | 0 | 0 | 0 | 31 | 0 | 0 | 0 | 0 | 0 | 0 | 0 | 0 | 0 | 0 | 0 | 0 | 0 | 0 | 0 | 0 |
| SRR1973407 | 562 | 93400 | 16287 | 403 | 973 | 10771 | 15583 | 15672 | 8436 | 42 | 1280 | 4829 | 421 | 135 | 0 | 6151 | 30314 | 0 | 0 | 0 | 150 | 17 | 25 | 0 | 0 | 0 | 0 | 0 | 72 | 67 | 44 | 11 | 0 | 0 | 0 | 0 | 0 |
| SRR1973408 | 343 | 101201 | 23045 | 638 | 1364 | 14646 | 21507 | 21482 | 9521 | 12 | 1396 | 5112 | 429 | 153 | 0 | 6437 | 27241 | 0 | 0 | 0 | 126 | 14 | 18 | 0 | 0 | 0 | 0 | 0 | 56 | 93 | 43 | 11 | 0 | 0 | 0 | 0 | 0 |
| SRR1973409 | 325 | 85595 | 28125 | 423 | 1228 | 15913 | 19127 | 19828 | 9951 | 49 | 1030 | 5875 | 329 | 107 | 13 | 5384 | 60638 | 0 | 0 | 0 | 192 | 37 | 31 | 0 | 0 | 0 | 0 | 0 | 142 | 114 | 80 | 31 | 0 | 0 | 0 | 0 | 0 |
| SRR1973410 | 338 | 108628 | 33206 | 328 | 1449 | 15510 | 21572 | 22410 | 11020 | 79 | 864 | 5010 | 250 | 70 | 0 | 6037 | 51717 | 0 | 0 | 0 | 168 | 29 | 38 | 0 | 0 | 0 | 0 | 11 | 97 | 70 | 68 | 22 | 0 | 0 | 0 | 0 | 0 |
| SRR1973411 | 340 | 103829 | 29197 | 309 | 1250 | 16217 | 20103 | 24966 | 9542 | 124 | 839 | 4874 | 303 | 109 | 0 | 5555 | 50092 | 0 | 0 | 0 | 150 | 29 | 27 | 0 | 0 | 0 | 0 | 0 | 63 | 43 | 45 | 20 | 0 | 0 | 0 | 0 | 0 |
| SRR1973412 | 585 | 96757 | 37820 | 285 | 1422 | 19188 | 21600 | 26018 | 9234 | 90 | 985 | 6049 | 307 | 93 | 0 | 5149 | 53316 | 0 | 0 | 0 | 133 | 23 | 21 | 0 | 0 | 0 | 0 | 0 | 33 | 43 | 33 | 18 | 0 | 0 | 0 | 0 | 0 |
| SRR1973413 | 416 | 110220 | 34559 | 315 | 1840 | 21644 | 27636 | 29133 | 10237 | 22 | 1034 | 5678 | 346 | 90 | 0 | 6773 | 45219 | 0 | 0 | 0 | 86 | 13 | 24 | 0 | 0 | 0 | 0 | 0 | 25 | 24 | 28 | 0 | 0 | 0 | 0 | 0 | 0 |
| SRR1973414 | 352 | 106018 | 38114 | 191 | 1954 | 21970 | 32290 | 27576 | 13854 | 73 | 1031 | 6011 | 261 | 116 | 0 | 7928 | 36876 | 0 | 0 | 0 | 64 | 17 | 26 | 0 | 0 | 0 | 0 | 0 | 0 | 13 | 19 | 0 | 0 | 0 | 0 | 0 | 0 |
| SRR1973415 | 548 | 94644 | 40877 | 129 | 2418 | 24801 | 36745 | 38347 | 10304 | 113 | 1158 | 7051 | 358 | 119 | 0 | 7931 | 42487 | 0 | 0 | 0 | 15 | 0 | 0 | 0 | 0 | 0 | 0 | 0 | 0 | 0 | 0 | 0 | 0 | 0 | 0 | 0 | 0 |
| SRR1973952 | 77 | 181 | 464 | 1586 | 30523 | 34165 | 306391 | 110885 | 30372 | 3882 | 12246 | 2358 | 1493 | 9194 | 11 | 1201 | 3531 | 0 | 0 | 0 | 24 | 15 | 0 | 0 | 0 | 0 | 0 | 21 | 53 | 33 | 28 | 0 | 0 | 0 | 0 | 0 | 0 |
| SRR1973953 | 1436 | 5981 | 20307 | 5427 | 16397 | 27919 | 373346 | 132777 | 5418 | 2099 | 9799 | 2147 | 3522 | 19905 | 21295 | 272 | 6148 | 27 | 55 | 112 | 175 | 433 | 147 | 36 | 24 | 33 | 19 | 34 | 152 | 125 | 57 | 0 | 0 | 0 | 0 | 44 | 0 |
| SRR1973956 | 2963 | 5652 | 7897 | 1577 | 11392 | 7917 | 256034 | 41128 | 6951 | 651 | 38618 | 95385 | 4827 | 13323 | 772 | 579 | 19695 | 0 | 0 | 0 | 0 | 0 | 19 | 0 | 0 | 0 | 0 | 0 | 0 | 0 | 0 | 0 | 0 | 0 | 0 | 0 | 0 |
| SRR1973958 | 0 | 586 | 27175 | 634 | 3768 | 2979 | 236923 | 8149 | 43810 | 0 | 43810 | 9025 | 2278 | 7711 | 1840 | 1402 | 4819 | 0 | 0 | 16 | 129 | 60 | 690 | 76 | 0 | 0 | 0 | 0 | 166 | 47 | 0 | 28 | 0 | 0 | 0 | 0 | 0 |
| SRR1973964 | 0 | 0 | 1908 | 5752 | 2432 | 2270 | 418808 | 25537 | 36684 | 0 | 37292 | 5432 | 1338 | 22051 | 5391 | 284 | 4135 | 0 | 0 | 0 | 0 | 0 | 38 | 0 | 0 | 0 | 0 | 0 | 0 | 0 | 0 | 0 | 0 | 0 | 0 | 0 | 0 |
| SRR1973966 | 1579 | 6593 | 24559 | 1898 | 24287 | 23260 | 411877 | 63076 | 1371 | 3179 | 17552 | 10 | 6053 | 11646 | 1260 | 7381 | 11585 | 49 | 38 | 58 | 61 | 80 | 93 | 126 | 57 | 27 | 37 | 28 | 100 | 48 | 15 | 0 | 0 | 0 | 0 | 42 | 0 |
| SRR1973967 | 275 | 2747 | 13929 | 2970 | 7188 | 13566 | 113500 | 19524 | 1302 | 987 | 2045 | 907 | 1071 | 23133 | 3859 | 4364 | 1881 | 20 | 34 | 34 | 21 | 40 | 217 | 23 | 17 | 57 | 11 | 11 | 29 | 25 | 11 | 27 | 0 | 0 | 0 | 78 | 0 |
| SRR1979367 | 0 | 398 | 3048 | 0 | 423 | 447 | 28672 | 31234 | 1483 | 25 | 40859 | 18416 | 10 | 1778 | 1428 | 435 | 22477 | 0 | 0 | 0 | 0 | 0 | 0 | 0 | 0 | 0 | 0 | 0 | 0 | 0 | 0 | 45 | 0 | 0 | 0 | 0 | 0 |
| SRR1988280 | 30 | 143215 | 310 | 77 | 120 | 587 | 764 | 589 | 5552 | 0 | 53 | 135 | 11 | 0 | 12 | 3064 | 19203 | 0 | 0 | 0 | 316 | 18 | 31 | 0 | 0 | 0 | 0 | 0 | 44 | 80 | 15 | 21 | 0 | 0 | 0 | 0 | 0 |
| SRR1988281 | 29 | 137683 | 380 | 88 | 139 | 631 | 921 | 623 | 6733 | 16 | 50 | 187 | 0 | 0 | 12 | 3427 | 21619 | 0 | 0 | 0 | 326 | 20 | 33 | 0 | 0 | 0 | 0 | 0 | 48 | 86 | 15 | 23 | 0 | 0 | 0 | 0 | 0 |
| SRR1988282 | 26 | 149889 | 303 | 58 | 136 | 591 | 890 | 629 | 5569 | 11 | 43 | 110 | 12 | 0 | 12 | 2883 | 20756 | 0 | 0 | 0 | 502 | 31 | 54 | 0 | 0 | 0 | 0 | 0 | 59 | 108 | 24 | 26 | 0 | 0 | 0 | 0 | 0 |
| SRR1988283 | 25 | 41231 | 2350 | 67 | 2463 | 35298 | 31016 | 13219 | 771 | 13 | 19702 | 45593 | 4624 | 5222 | 100 | 10316 | 19710 | 0 | 0 | 24 | 721 | 58 | 70 | 0 | 0 | 0 | 0 | 41 | 419 | 154 | 28 | 31 | 0 | 0 | 0 | 0 | 0 |
| SRR1988284 | 33 | 42239 | 2953 | 77 | 2334 | 33307 | 30142 | 13520 | 816 | 0 | 20891 | 41227 | 4447 | 5620 | 57 | 9140 | 22158 | 0 | 0 | 17 | 784 | 61 | 68 | 0 | 0 | 0 | 0 | 34 | 354 | 147 | 21 | 27 | 0 | 0 | 0 | 0 | 0 |
| SRR1988285 | 23 | 37184 | 3401 | 64 | 2165 | 31622 | 28140 | 12904 | 734 | 0 | 21011 | 43021 | 4472 | 5318 | 78 | 8703 | 21620 | 0 | 0 | 15 | 614 | 57 | 65 | 0 | 0 | 0 | 0 | 34 | 304 | 121 | 21 | 23 | 0 | 0 | 0 | 0 | 0 |
| SRR1988286 | 16 | 35575 | 2022 | 98 | 2447 | 32796 | 30821 | 13437 | 842 | 13 | 19031 | 45736 | 4615 | 5382 | 110 | 9406 | 19476 | 0 | 0 | 45 | 1299 | 73 | 93 | 0 | 0 | 0 | 0 | 45 | 754 | 281 | 40 | 49 | 0 | 0 | 0 | 0 | 0 |
| SRR1988287 | 17 | 269031 | 5651 | 16 | 636 | 3108 | 7843 | 2660 | 20221 | 34 | 506 | 1053 | 94 | 107 | 24 | 2882 | 25409 | 0 | 0 | 0 | 7957 | 573 | 156 | 0 | 0 | 0 | 0 | 0 | 96 | 1657 | 320 | 54 | 0 | 0 | 0 | 0 | 0 |
| SRR1988288 | 0 | 278189 | 5809 | 24 | 702 | 3258 | 8585 | 3010 | 21985 | 24 | 461 | 1009 | 79 | 122 | 23 | 2849 | 25174 | 0 | 0 | 0 | 7899 | 603 | 159 | 0 | 0 | 0 | 0 | 0 | 88 | 1552 | 328 | 57 | 0 | 0 | 0 | 0 | 0 |
| SRR1988289 | 12 | 158512 | 3600 | 30 | 1080 | 8803 | 13260 | 5074 | 16686 | 12 | 3742 | 9126 | 856 | 1310 | 37 | 3980 | 23646 | 0 | 0 | 0 | 7439 | 628 | 145 | 0 | 0 | 0 | 0 | 0 | 138 | 1349 | 306 | 54 | 0 | 0 | 0 | 0 | 0 |
| SRR1988290 | 0 | 152269 | 3454 | 31 | 650 | 3089 | 8582 | 2725 | 22498 | 21 | 475 | 1025 | 88 | 101 | 23 | 2839 | 23133 | 0 | 0 | 11 | 7248 | 674 | 154 | 0 | 0 | 0 | 0 | 0 | 112 | 1472 | 346 | 57 | 0 | 0 | 0 | 0 | 0 |
| SRR1988291 | 13 | 88082 | 613 | 69 | 103 | 781 | 875 | 619 | 2535 | 10 | 31 | 48 | 12 | 0 | 0 | 2081 | 16511 | 0 | 0 | 0 | 147 | 16 | 19 | 0 | 0 | 0 | 0 | 0 | 25 | 43 | 15 | 15 | 0 | 0 | 0 | 0 | 0 |
| SRR1988292 | 16 | 76288 | 748 | 78 | 120 | 999 | 1108 | 766 | 2717 | 24 | 38 | 43 | 12 | 0 | 13 | 2559 | 18028 | 0 | 0 | 0 | 160 | 16 | 19 | 0 | 0 | 0 | 0 | 0 | 25 | 42 | 16 | 12 | 0 | 0 | 0 | 0 | 0 |
| SRR1988293 | 22 | 83275 | 714 | 72 | 96 | 721 | 861 | 574 | 2275 | 15 | 33 | 49 | 10 | 0 | 14 | 2097 | 16638 | 0 | 0 | 0 | 158 | 17 | 20 | 0 | 0 | 0 | 0 | 0 | 24 | 51 | 17 | 13 | 0 | 0 | 0 | 0 | 0 |
| SRR1988294 | 18 | 87682 | 745 | 84 | 82 | 730 | 782 | 567 | 2243 | 15 | 34 | 46 | 0 | 0 | 0 | 1979 | 15958 | 0 | 0 | 0 | 128 | 12 | 18 | 0 | 0 | 0 | 0 | 0 | 23 | 48 | 14 | 0 | 0 | 0 | 0 | 0 | 0 |
| SRR1988295 | 15 | 84357 | 6384 | 65 | 1825 | 16406 | 15753 | 19091 | 4282 | 11 | 9070 | 6965 | 1170 | 2896 | 13 | 7248 | 17983 | 0 | 0 | 0 | 5241 | 822 | 76 | 0 | 0 | 0 | 0 | 0 | 122 | 950 | 184 | 18 | 0 | 0 | 18 | 0 | 0 |
| SRR1988296 | 18 | 90147 | 5880 | 49 | 2183 | 21369 | 21320 | 20602 | 5081 | 0 | 7065 | 7618 | 1250 | 2258 | 0 | 9124 | 19285 | 0 | 0 | 0 | 6673 | 921 | 81 | 0 | 0 | 0 | 0 | 0 | 132 | 1340 | 245 | 27 | 0 | 0 | 24 | 0 | 0 |
| SRR1988297 | 14 | 103901 | 6571 | 60 | 2119 | 18808 | 19511 | 19384 | 5318 | 0 | 8366 | 7529 | 1341 | 2530 | 11 | 9151 | 21412 | 0 | 0 | 0 | 6451 | 929 | 78 | 0 | 0 | 0 | 0 | 0 | 112 | 1245 | 245 | 24 | 0 | 0 | 22 | 0 | 0 |
| SRR1988298 | 17 | 88317 | 7284 | 30 | 1799 | 17911 | 17816 | 17631 | 5401 | 12 | 6457 | 5770 | 1023 | 2137 | 0 | 8450 | 22114 | 0 | 0 | 11 | 3556 | 603 | 57 | 0 | 0 | 0 | 0 | 0 | 156 | 653 | 163 | 19 | 0 | 0 | 13 | 0 | 0 |
| SRR1988299 | 41 | 76397 | 915 | 63 | 120 | 1276 | 1236 | 1128 | 2575 | 10 | 70 | 115 | 0 | 13 | 20 | 10184 | 17889 | 0 | 0 | 0 | 69 | 13 | 10 | 0 | 0 | 0 | 0 | 0 | 36 | 39 | 14 | 12 | 0 | 0 | 0 | 0 | 0 |
| SRR1988300 | 26 | 62232 | 989 | 104 | 207 | 2067 | 1918 | 1666 | 3118 | 16 | 60 | 71 | 15 | 0 | 33 | 14154 | 21467 | 0 | 0 | 0 | 88 | 12 | 14 | 0 | 0 | 0 | 0 | 0 | 43 | 43 | 15 | 14 | 0 | 0 | 0 | 0 | 0 |
| SRR1988301 | 23 | 73048 | 724 | 111 | 127 | 1163 | 1184 | 1068 | 2011 | 11 | 67 | 94 | 0 | 0 | 29 | 9984 | 18295 | 0 | 0 | 0 | 72 | 11 | 12 | 0 | 0 | 0 | 0 | 0 | 40 | 39 | 12 | 14 | 0 | 0 | 0 | 0 | 0 |
| SRR1988302 | 31 | 67619 | 888 | 96 | 135 | 1354 | 1302 | 1013 | 1965 | 11 | 72 | 77 | 16 | 15 | 21 | 11506 | 19879 | 0 | 0 | 0 | 76 | 11 | 12 | 0 | 0 | 0 | 0 | 0 | 42 | 45 | 16 | 12 | 0 | 0 | 0 | 0 | 0 |
| SRR1988303 | 39 | 83702 | 761 | 81 | 118 | 1050 | 1035 | 1052 | 1611 | 14 | 73 | 103 | 15 | 0 | 40 | 5164 | 13443 | 0 | 0 | 0 | 67 | 11 | 0 | 0 | 0 | 0 | 0 | 0 | 36 | 29 | 14 | 0 | 0 | 0 | 0 | 0 | 0 |
| SRR1988304 | 40 | 65116 | 1450 | 97 | 127 | 1196 | 1166 | 1225 | 1972 | 27 | 81 | 90 | 11 | 0 | 26 | 5661 | 11340 | 0 | 0 | 0 | 52 | 0 | 0 | 0 | 0 | 0 | 0 | 0 | 27 | 31 | 0 | 0 | 0 | 0 | 0 | 0 | 0 |
| SRR1988305 | 23 | 76067 | 961 | 63 | 114 | 937 | 950 | 957 | 1519 | 0 | 75 | 107 | 11 | 0 | 30 | 4613 | 11925 | 0 | 0 | 0 | 61 | 11 | 0 | 0 | 0 | 0 | 0 | 0 | 30 | 33 | 11 | 0 | 0 | 0 | 0 | 0 | 0 |
| SRR1988306 | 42 | 73285 | 1474 | 109 | 135 | 1194 | 1158 | 1066 | 1700 | 15 | 65 | 84 | 0 | 0 | 27 | 6714 | 13148 | 0 | 0 | 0 | 59 | 0 | 0 | 0 | 0 | 0 | 0 | 0 | 48 | 29 | 12 | 0 | 0 | 0 | 0 | 0 | 0 |
| SRR1988307 | 52 | 66748 | 1168 | 105 | 2191 | 20453 | 25115 | 16219 | 4088 | 28 | 224 | 358 | 34 | 0 | 0 | 29789 | 35986 | 0 | 0 | 0 | 363 | 70 | 21 | 0 | 0 | 0 | 0 | 11 | 57 | 109 | 32 | 0 | 0 | 0 | 0 | 0 | 0 |
| SRR1988308 | 119 | 74035 | 1857 | 124 | 361 | 3923 | 4440 | 3230 | 5191 | 24 | 152 | 190 | 18 | 0 | 0 | 19201 | 49080 | 0 | 0 | 0 | 277 | 33 | 12 | 0 | 0 | 0 | 0 | 0 | 45 | 132 | 28 | 12 | 0 | 0 | 0 | 0 | 0 |
| SRR1988309 | 131 | 58612 | 2257 | 170 | 1518 | 14632 | 18749 | 10482 | 3724 | 28 | 174 | 240 | 14 | 0 | 0 | 21454 | 39180 | 0 | 0 | 0 | 205 | 38 | 15 | 0 | 0 | 0 | 0 | 0 | 34 | 73 | 17 | 11 | 0 | 0 | 0 | 0 | 0 |
| SRR1988310 | 120 | 66977 | 2307 | 93 | 502 | 3671 | 4716 | 3365 | 3005 | 22 | 166 | 388 | 25 | 0 | 20 | 9761 | 36616 | 0 | 0 | 0 | 200 | 34 | 32 | 0 | 0 | 0 | 0 | 0 | 51 | 70 | 22 | 22 | 0 | 0 | 0 | 0 | 0 |
| SRR1988311 | 98 | 66454 | 3449 | 121 | 95 | 638 | 1106 | 605 | 3275 | 20 | 114 | 274 | 15 | 0 | 11 | 4310 | 34804 | 0 | 0 | 0 | 372 | 22 | 18 | 0 | 0 | 0 | 0 | 0 | 19 | 104 | 17 | 0 | 0 | 0 | 0 | 0 | 0 |
| SRR1988312 | 90 | 55240 | 2958 | 134 | 93 | 695 | 1081 | 607 | 3982 | 38 | 116 | 283 | 16 | 0 | 11 | 4880 | 32609 | 0 | 0 | 0 | 322 | 15 | 14 | 0 | 0 | 0 | 0 | 0 | 17 | 130 | 16 | 0 | 0 | 0 | 0 | 0 | 0 |
| SRR2052873 | 941 | 31936 | 3653 | 357 | 526 | 8044 | 7936 | 5041 | 2935 | 83 | 11159 | 63908 | 5354 | 1916 | 216 | 7974 | 35730 | 0 | 0 | 0 | 318 | 214 | 57 | 0 | 0 | 0 | 0 | 0 | 66 | 66 | 79 | 22 | 0 | 0 | 11 | 10 | 0 |
| SRR2052874 | 878 | 34498 | 2949 | 415 | 548 | 8186 | 7638 | 5413 | 2571 | 64 | 9716 | 61135 | 4912 | 1726 | 356 | 7545 | 37578 | 0 | 0 | 0 | 258 | 198 | 50 | 0 | 0 | 0 | 0 | 0 | 48 | 53 | 75 | 21 | 0 | 0 | 0 | 0 | 0 |
| SRR2052875 | 1039 | 36539 | 1954 | 251 | 622 | 9170 | 8869 | 6791 | 2734 | 55 | 12584 | 58999 | 4875 | 3022 | 172 | 8939 | 38682 | 0 | 0 | 0 | 339 | 117 | 34 | 0 | 0 | 0 | 0 | 0 | 50 | 54 | 42 | 18 | 0 | 0 | 0 | 0 | 0 |
| SRR2052877 | 912 | 38754 | 4291 | 267 | 436 | 6822 | 5817 | 4870 | 1363 | 72 | 24470 | 17 | 11129 | 5698 | 53 | 6638 | 34615 | 0 | 0 | 0 | 347 | 121 | 51 | 0 | 0 | 0 | 0 | 0 | 42 | 45 | 49 | 19 | 0 | 0 | 18 | 0 | 0 |
| SRR2052878 | 546 | 46202 | 1659 | 270 | 804 | 11155 | 11452 | 7097 | 1981 | 33 | 10693 | 61149 | 4687 | 2969 | 73 | 10394 | 34911 | 0 | 0 | 0 | 330 | 158 | 45 | 0 | 0 | 0 | 0 | 0 | 48 | 72 | 53 | 14 | 0 | 0 | 0 | 0 | 0 |
| SRR2053027 | 648 | 15066 | 5215 | 514 | 254 | 3691 | 3413 | 2683 | 1439 | 113 | 11689 | 83004 | 6047 | 2777 | 170 | 8685 | 42132 | 0 | 0 | 0 | 471 | 124 | 175 | 0 | 0 | 0 | 0 | 0 | 64 | 63 | 46 | 66 | 0 | 14 | 11 | 11 | 0 |
| SRR2053028 | 704 | 16115 | 5880 | 511 | 329 | 4879 | 4450 | 3191 | 1574 | 115 | 12106 | 78940 | 6481 | 2736 | 115 | 11175 | 40366 | 0 | 0 | 0 | 523 | 141 | 172 | 0 | 12 | 0 | 0 | 0 | 68 | 98 | 57 | 65 | 0 | 19 | 17 | 13 | 0 |
| SRR2053029 | 747 | 13887 | 4057 | 563 | 279 | 4183 | 3752 | 2524 | 1709 | 144 | 13323 | 92280 | 6950 | 2551 | 209 | 9247 | 42513 | 0 | 0 | 0 | 404 | 89 | 93 | 0 | 0 | 0 | 0 | 0 | 71 | 70 | 32 | 33 | 0 | 0 | 0 | 0 | 0 |
| SRR2061931 | 248 | 29178 | 3303 | 1469 | 2686 | 27430 | 29907 | 80512 | 9346 | 21 | 28257 | 19807 | 6583 | 3350 | 0 | 3653 | 36910 | 0 | 0 | 12 | 10096 | 1497 | 87 | 0 | 0 | 0 | 0 | 0 | 211 | 2437 | 745 | 63 | 0 | 0 | 0 | 0 | 0 |
| SRR2061932 | 203 | 40897 | 2835 | 918 | 1099 | 16876 | 11393 | 42480 | 11942 | 41 | 17405 | 20721 | 4974 | 3021 | 0 | 1719 | 48055 | 0 | 0 | 14 | 5981 | 1692 | 59 | 0 | 0 | 0 | 0 | 0 | 195 | 2112 | 913 | 30 | 0 | 0 | 0 | 0 | 0 |
| SRR2061933 | 117 | 11879 | 1614 | 1135 | 2220 | 25603 | 20311 | 68451 | 11600 | 70 | 27748 | 27066 | 5977 | 2208 | 0 | 5392 | 42763 | 0 | 0 | 14 | 6555 | 1469 | 80 | 0 | 0 | 0 | 0 | 0 | 201 | 1982 | 738 | 52 | 0 | 0 | 0 | 0 | 0 |
| SRR2061934 | 207 | 43193 | 1982 | 853 | 1276 | 15581 | 10602 | 38300 | 5035 | 115 | 14501 | 19648 | 3127 | 1823 | 14 | 3268 | 59266 | 0 | 0 | 10 | 4567 | 2046 | 145 | 0 | 0 | 0 | 0 | 0 | 111 | 1031 | 1035 | 91 | 0 | 0 | 40 | 0 | 0 |
| SRR2061935 | 354 | 27119 | 3643 | 987 | 803 | 9830 | 7653 | 47121 | 3361 | 51 | 15052 | 17462 | 3826 | 2516 | 0 | 1670 | 45916 | 0 | 0 | 0 | 2121 | 960 | 57 | 0 | 11 | 0 | 0 | 0 | 85 | 664 | 539 | 44 | 0 | 0 | 19 | 0 | 0 |
| SRR2061936 | 171 | 8409 | 1694 | 2239 | 2640 | 18786 | 19064 | 73651 | 17558 | 37 | 32851 | 22564 | 3115 | 1616 | 0 | 7429 | 57609 | 0 | 0 | 0 | 2149 | 495 | 25 | 0 | 0 | 0 | 0 | 0 | 99 | 984 | 394 | 17 | 0 | 0 | 0 | 0 | 0 |
| SRR2061937 | 224 | 8756 | 887 | 1746 | 3118 | 27002 | 20411 | 61747 | 17753 | 14 | 41424 | 28624 | 3468 | 3008 | 0 | 11074 | 60443 | 0 | 0 | 10 | 2906 | 588 | 19 | 0 | 0 | 0 | 0 | 0 | 132 | 1438 | 451 | 13 | 0 | 0 | 0 | 0 | 0 |
| SRR2061938 | 147 | 8155 | 1273 | 1938 | 2192 | 25604 | 17624 | 67020 | 11519 | 44 | 27769 | 22411 | 3539 | 2238 | 0 | 8668 | 62673 | 0 | 0 | 0 | 3849 | 669 | 37 | 0 | 0 | 0 | 0 | 0 | 149 | 1214 | 448 | 16 | 0 | 0 | 0 | 0 | 0 |
| SRR2061940 | 142 | 19657 | 1298 | 1062 | 1088 | 12220 | 7292 | 40446 | 13319 | 47 | 15806 | 24519 | 2231 | 1854 | 11 | 2653 | 96221 | 0 | 0 | 0 | 1486 | 350 | 14 | 0 | 0 | 0 | 0 | 0 | 80 | 792 | 460 | 0 | 0 | 0 | 0 | 0 | 0 |
| SRR2061941 | 206 | 115746 | 2841 | 1761 | 3818 | 41388 | 54142 | 64769 | 7308 | 85 | 15447 | 24771 | 4310 | 9819 | 75 | 22435 | 18129 | 0 | 0 | 0 | 4850 | 499 | 531 | 0 | 0 | 0 | 0 | 0 | 48 | 1601 | 164 | 95 | 0 | 0 | 0 | 0 | 0 |
| SRR2061942 | 351 | 131306 | 1863 | 2110 | 2897 | 37586 | 44915 | 44640 | 6548 | 41 | 17319 | 36413 | 4816 | 11576 | 16 | 16010 | 26994 | 0 | 0 | 0 | 6183 | 509 | 1033 | 0 | 0 | 0 | 0 | 0 | 68 | 952 | 214 | 105 | 0 | 0 | 0 | 0 | 0 |
| SRR2061943 | 226 | 252020 | 1091 | 1163 | 2115 | 37323 | 36297 | 49460 | 5464 | 0 | 12428 | 26284 | 2445 | 5681 | 0 | 21322 | 40599 | 0 | 0 | 0 | 6040 | 807 | 1315 | 0 | 15 | 0 | 0 | 0 | 130 | 928 | 358 | 89 | 0 | 0 | 0 | 12 | 0 |
| SRR2061944 | 475 | 167604 | 1276 | 1800 | 3081 | 44119 | 48702 | 43825 | 7653 | 15 | 22243 | 45985 | 5482 | 14067 | 0 | 20064 | 25130 | 0 | 0 | 0 | 5415 | 310 | 822 | 0 | 0 | 0 | 0 | 0 | 64 | 916 | 205 | 69 | 0 | 0 | 0 | 0 | 0 |
| SRR2061945 | 450 | 147210 | 1819 | 1885 | 2761 | 38872 | 44215 | 43193 | 7230 | 24 | 17140 | 34473 | 4112 | 10482 | 0 | 17356 | 34307 | 0 | 0 | 12 | 6101 | 512 | 1059 | 0 | 0 | 0 | 0 | 0 | 85 | 938 | 252 | 100 | 0 | 0 | 0 | 0 | 0 |
| SRR2061946 | 474 | 156421 | 2341 | 1101 | 2045 | 32437 | 37220 | 57972 | 5089 | 22 | 11167 | 20429 | 2374 | 5603 | 0 | 20697 | 37242 | 0 | 0 | 16 | 3639 | 2292 | 913 | 0 | 11 | 0 | 0 | 0 | 129 | 712 | 1123 | 91 | 0 | 0 | 0 | 0 | 0 |
| SRR2061947 | 368 | 150700 | 2306 | 1533 | 3855 | 42595 | 59832 | 54007 | 8969 | 37 | 15884 | 25050 | 4069 | 9708 | 0 | 20355 | 22799 | 0 | 0 | 10 | 7687 | 541 | 1206 | 0 | 10 | 0 | 0 | 0 | 105 | 1652 | 267 | 180 | 0 | 0 | 0 | 0 | 0 |
| SRR2061948 | 478 | 166668 | 1541 | 1533 | 2490 | 39074 | 39326 | 40989 | 7109 | 0 | 16826 | 33298 | 3795 | 10034 | 0 | 18130 | 32258 | 0 | 0 | 12 | 5404 | 490 | 1075 | 0 | 12 | 0 | 0 | 0 | 70 | 798 | 276 | 89 | 0 | 13 | 0 | 0 | 0 |
| SRR2061949 | 329 | 244975 | 1655 | 1024 | 1958 | 36287 | 38718 | 50161 | 5723 | 11 | 12265 | 24752 | 2260 | 5219 | 0 | 19019 | 39677 | 0 | 0 | 0 | 3366 | 1161 | 1134 | 0 | 14 | 0 | 0 | 0 | 111 | 777 | 685 | 98 | 0 | 10 | 0 | 0 | 0 |
| SRR2061950 | 271 | 143625 | 3339 | 1550 | 2111 | 35496 | 27670 | 22363 | 1921 | 45 | 48072 | 85305 | 7642 | 51081 | 12 | 49895 | 16578 | 0 | 0 | 0 | 1675 | 162 | 894 | 0 | 0 | 0 | 0 | 0 | 38 | 314 | 122 | 95 | 0 | 0 | 0 | 0 | 0 |
| SRR2061951 | 190 | 101558 | 3442 | 1508 | 2546 | 33525 | 33286 | 25185 | 1969 | 10 | 37024 | 69241 | 7053 | 58836 | 23 | 52406 | 16316 | 0 | 0 | 0 | 1409 | 171 | 762 | 0 | 11 | 0 | 0 | 0 | 45 | 296 | 99 | 79 | 0 | 0 | 0 | 0 | 0 |
| SRR2061952 | 134 | 181936 | 1064 | 900 | 1997 | 36190 | 27442 | 21433 | 2034 | 13 | 28891 | 56929 | 4227 | 30503 | 25 | 58536 | 18863 | 0 | 0 | 0 | 1654 | 298 | 836 | 0 | 20 | 0 | 0 | 0 | 73 | 317 | 158 | 76 | 0 | 13 | 0 | 11 | 0 |
| SRR2061953 | 250 | 62279 | 4164 | 2124 | 3128 | 35205 | 36752 | 33886 | 2349 | 28 | 37060 | 49603 | 6764 | 51980 | 65 | 64410 | 16050 | 0 | 0 | 0 | 1542 | 225 | 731 | 0 | 12 | 0 | 0 | 0 | 45 | 372 | 81 | 135 | 0 | 0 | 0 | 0 | 0 |
| SRR2061954 | 396 | 87332 | 3413 | 1468 | 2252 | 38641 | 31665 | 27824 | 2077 | 0 | 31798 | 55466 | 5083 | 44155 | 16 | 56572 | 18866 | 0 | 0 | 0 | 1092 | 190 | 765 | 0 | 16 | 0 | 0 | 0 | 41 | 211 | 124 | 81 | 0 | 16 | 0 | 0 | 0 |
| SRR2061955 | 183 | 127628 | 1008 | 416 | 1877 | 37416 | 28211 | 23621 | 2117 | 11 | 24348 | 47293 | 3211 | 26923 | 12 | 54702 | 18140 | 0 | 0 | 0 | 966 | 239 | 494 | 0 | 11 | 0 | 0 | 0 | 54 | 227 | 175 | 48 | 0 | 0 | 0 | 0 | 0 |
| SRR2061956 | 255 | 59275 | 5039 | 3176 | 3066 | 38912 | 42688 | 36421 | 2313 | 13 | 36707 | 47689 | 6334 | 51867 | 27 | 56983 | 14755 | 0 | 0 | 0 | 1288 | 177 | 721 | 0 | 14 | 0 | 0 | 0 | 41 | 305 | 110 | 132 | 0 | 0 | 0 | 0 | 0 |
| SRR2061957 | 360 | 80013 | 3238 | 2159 | 2211 | 39413 | 32975 | 30637 | 2135 | 19 | 34221 | 51345 | 5071 | 49165 | 13 | 55895 | 17841 | 0 | 0 | 0 | 1011 | 160 | 642 | 0 | 12 | 0 | 0 | 0 | 35 | 200 | 101 | 80 | 0 | 0 | 0 | 0 | 0 |
| SRR2061958 | 185 | 138518 | 1477 | 940 | 1619 | 36929 | 26942 | 22013 | 1912 | 0 | 27748 | 47798 | 3765 | 29878 | 15 | 49945 | 18432 | 0 | 0 | 0 | 1125 | 271 | 701 | 0 | 12 | 0 | 0 | 0 | 31 | 202 | 138 | 56 | 0 | 0 | 0 | 0 | 0 |
| SRR2069629 | 0 | 20 | 198 | 1262 | 908 | 839 | 10910 | 1404 | 49218 | 3321 | 7420 | 31278 | 2836 | 956 | 32645 | 6165 | 14778 | 0 | 16 | 0 | 15 | 66 | 31 | 53 | 64 | 30 | 0 | 12 | 20 | 20 | 78 | 0 | 0 | 0 | 0 | 49 | 0 |
| SRR2069630 | 0 | 0 | 793 | 425 | 701 | 587 | 5804 | 1313 | 56083 | 1700 | 6739 | 14238 | 3116 | 647 | 5228 | 6393 | 5383 | 0 | 12 | 0 | 0 | 23 | 0 | 14 | 17 | 0 | 0 | 0 | 11 | 0 | 58 | 0 | 0 | 0 | 0 | 11 | 0 |
| SRR2084358 | 49 | 15354 | 2649 | 932 | 10866 | 55075 | 137274 | 103910 | 4003 | 49 | 80143 | 17400 | 9085 | 13890 | 2756 | 255 | 17326 | 0 | 0 | 11 | 267 | 55 | 422 | 213 | 281 | 41 | 0 | 0 | 267 | 111 | 51 | 0 | 0 | 0 | 0 | 26 | 0 |
| SRR2084359 | 106 | 18186 | 2044 | 2890 | 9760 | 58320 | 122870 | 83270 | 3078 | 106 | 90350 | 20685 | 8849 | 14505 | 994 | 241 | 22142 | 0 | 11 | 19 | 554 | 124 | 818 | 368 | 430 | 68 | 0 | 29 | 570 | 200 | 88 | 0 | 0 | 0 | 0 | 29 | 0 |
| SRR2085173 | 0 | 0 | 3289 | 0 | 5116 | 4888 | 80024 | 41296 | 74435 | 416 | 9698 | 57975 | 563 | 11 | 788 | 1442 | 1939 | 0 | 0 | 0 | 0 | 0 | 0 | 37 | 138 | 22 | 0 | 0 | 0 | 0 | 0 | 0 | 0 | 0 | 0 | 0 | 0 |
| SRR2103379 | 152 | 7660 | 9730 | 637 | 1338 | 19805 | 17194 | 14153 | 3498 | 11 | 814 | 2077 | 98 | 14 | 31 | 34 | 18895 | 0 | 0 | 0 | 674 | 329 | 336 | 83 | 51 | 10 | 0 | 0 | 49 | 95 | 126 | 44 | 0 | 0 | 0 | 0 | 0 |
| SRR2103380 | 166 | 9234 | 7944 | 674 | 1087 | 16912 | 14581 | 12843 | 3137 | 12 | 688 | 1765 | 89 | 14 | 26 | 30 | 17535 | 0 | 0 | 0 | 669 | 406 | 386 | 106 | 60 | 15 | 0 | 0 | 58 | 98 | 145 | 50 | 0 | 12 | 0 | 11 | 0 |
| SRR2103381 | 143 | 9464 | 8149 | 687 | 1247 | 19024 | 15985 | 13126 | 3241 | 12 | 688 | 1863 | 82 | 16 | 23 | 27 | 16067 | 0 | 0 | 0 | 683 | 360 | 347 | 85 | 48 | 12 | 0 | 0 | 60 | 111 | 147 | 53 | 0 | 13 | 0 | 10 | 0 |
| SRR2103382 | 338 | 18857 | 2974 | 189 | 2249 | 13098 | 16065 | 18485 | 11580 | 27 | 578 | 1494 | 62 | 11 | 0 | 20 | 32549 | 0 | 0 | 0 | 433 | 224 | 127 | 0 | 0 | 0 | 0 | 11 | 92 | 108 | 110 | 57 | 0 | 11 | 0 | 0 | 0 |
| SRR2103383 | 353 | 12156 | 3454 | 196 | 2219 | 16207 | 17130 | 16561 | 10513 | 24 | 540 | 1443 | 65 | 15 | 0 | 46 | 31901 | 0 | 0 | 0 | 534 | 191 | 114 | 0 | 0 | 0 | 0 | 13 | 115 | 127 | 118 | 53 | 0 | 0 | 0 | 0 | 0 |
| SRR2103384 | 430 | 18119 | 2592 | 353 | 1355 | 14145 | 12266 | 13046 | 8417 | 23 | 559 | 1587 | 74 | 11 | 13 | 29 | 23698 | 0 | 0 | 0 | 901 | 306 | 197 | 0 | 13 | 0 | 0 | 0 | 118 | 179 | 173 | 70 | 0 | 14 | 0 | 11 | 0 |
| SRR2105940 | 0 | 52 | 16 | 19 | 1046 | 19414 | 27067 | 13810 | 1468 | 27 | 94818 | 27941 | 1054 | 47483 | 0 | 287 | 4925 | 0 | 0 | 0 | 0 | 0 | 0 | 0 | 0 | 0 | 0 | 0 | 0 | 425 | 245 | 30 | 0 | 0 | 15 | 0 | 0 |
| SRR2105943 | 0 | 19 | 25 | 11 | 212 | 3097 | 5300 | 3956 | 286 | 12 | 76325 | 24669 | 951 | 18553 | 0 | 462 | 10729 | 0 | 0 | 0 | 0 | 0 | 0 | 0 | 0 | 0 | 0 | 0 | 0 | 48 | 22 | 0 | 0 | 0 | 0 | 0 | 0 |
| SRR2105944 | 0 | 18 | 37 | 0 | 619 | 7101 | 11782 | 6129 | 2879 | 15 | 120270 | 29661 | 617 | 7326 | 0 | 17868 | 5518 | 0 | 0 | 0 | 0 | 0 | 0 | 0 | 0 | 0 | 0 | 0 | 0 | 64 | 43 | 0 | 0 | 0 | 0 | 0 | 0 |
| SRR2105949 | 0 | 25 | 66 | 0 | 1171 | 10821 | 13923 | 9232 | 5209 | 16 | 88008 | 36160 | 1551 | 11962 | 0 | 4342 | 7218 | 0 | 0 | 0 | 0 | 0 | 0 | 0 | 0 | 0 | 0 | 0 | 0 | 0 | 44 | 0 | 0 | 0 | 0 | 0 | 0 |
| SRR2105951 | 0 | 1153 | 59 | 11 | 653 | 5070 | 7535 | 5645 | 1776 | 45 | 117704 | 57066 | 1742 | 14489 | 0 | 4092 | 5803 | 0 | 0 | 0 | 0 | 0 | 0 | 0 | 0 | 0 | 0 | 0 | 0 | 20 | 35 | 0 | 0 | 0 | 28 | 0 | 0 |
| SRR2106873 | 21 | 43046 | 600 | 53 | 2983 | 78913 | 48702 | 33551 | 5145 | 106 | 43945 | 14 | 5029 | 1943 | 11 | 11738 | 13303 | 0 | 0 | 0 | 439 | 89 | 169 | 0 | 17 | 0 | 0 | 0 | 16 | 117 | 69 | 142 | 0 | 27 | 0 | 32 | 0 |
| SRR2106874 | 0 | 67388 | 1105 | 51 | 3324 | 75503 | 56404 | 37679 | 6311 | 51 | 38483 | 17 | 5246 | 1904 | 0 | 12818 | 13956 | 0 | 0 | 0 | 185 | 13 | 85 | 0 | 0 | 0 | 0 | 0 | 0 | 35 | 18 | 71 | 0 | 0 | 0 | 0 | 0 |
| SRR2106875 | 23 | 146534 | 1032 | 197 | 3643 | 70386 | 58697 | 34698 | 5310 | 46 | 36339 | 10 | 5345 | 1685 | 12 | 12337 | 15355 | 0 | 0 | 0 | 478 | 23 | 137 | 0 | 0 | 0 | 0 | 0 | 29 | 86 | 33 | 100 | 0 | 11 | 0 | 10 | 0 |
| SRR2106876 | 81 | 237970 | 442 | 329 | 2208 | 47271 | 39043 | 23086 | 8463 | 46 | 31059 | 52703 | 6266 | 2601 | 0 | 16929 | 17527 | 0 | 0 | 0 | 847 | 43 | 155 | 0 | 0 | 0 | 0 | 0 | 26 | 157 | 36 | 86 | 0 | 0 | 0 | 23 | 0 |
| SRR2106877 | 98 | 127696 | 3140 | 189 | 2417 | 77482 | 44236 | 42413 | 6356 | 30 | 52732 | 83500 | 6883 | 1845 | 22 | 8860 | 14455 | 0 | 0 | 0 | 463 | 45 | 84 | 0 | 0 | 0 | 0 | 0 | 18 | 91 | 69 | 75 | 0 | 0 | 0 | 0 | 0 |

### Supp Table S8.1.Columns_B_to_AL

| | | | | | | | | | | | | | | | | | | | | | | | | | | | | | | | | | | | | | |
|---|---|---|---|---|---|---|---|---|---|---|---|---|---|---|---|---|---|---|---|---|---|---|---|---|---|---|---|---|---|---|---|---|---|---|---|---|---|
| SRR2106878 | 160 | 41009 | 909 | 1155 | 2012 | 82623 | 41678 | 38370 | 4256 | 43 | 24853 | 39810 | 4162 | 1930 | 15 | 10894 | 13303 | 0 | 0 | 0 | 461 | 52 | 128 | 0 | 0 | 0 | 0 | 0 | 41 | 93 | 53 | 81 | 0 | 0 | 0 | 12 | 0 |
| SRR2106879 | 12 | 101440 | 2576 | 139 | 1540 | 36776 | 22349 | 39959 | 4792 | 50 | 57811 | 15 | 6764 | 2147 | 0 | 8561 | 16746 | 0 | 0 | 0 | 229 | 27 | 45 | 0 | 0 | 0 | 0 | 0 | 14 | 41 | 29 | 24 | 0 | 0 | 0 | 12 | 0 |
| SRR2106880 | 0 | 66782 | 638 | 131 | 1771 | 44682 | 31739 | 26122 | 5887 | 66 | 30150 | 46405 | 4937 | 2164 | 0 | 10207 | 14648 | 0 | 0 | 0 | 460 | 50 | 166 | 0 | 11 | 0 | 0 | 0 | 21 | 113 | 44 | 134 | 0 | 12 | 0 | 16 | 0 |
| SRR2106881 | 0 | 126971 | 1136 | 118 | 1758 | 41400 | 25686 | 33156 | 6889 | 109 | 49914 | 79158 | 7464 | 1782 | 21 | 9914 | 17233 | 0 | 0 | 0 | 243 | 23 | 51 | 0 | 0 | 0 | 0 | 0 | 14 | 67 | 36 | 50 | 0 | 0 | 0 | 16 | 0 |
| SRR2106882 | 85 | 81487 | 634 | 320 | 2800 | 62970 | 51394 | 26802 | 5343 | 130 | 44846 | 52638 | 4527 | 2637 | 13 | 5057 | 14666 | 0 | 0 | 0 | 701 | 113 | 182 | 0 | 14 | 0 | 0 | 0 | 35 | 149 | 70 | 127 | 0 | 19 | 0 | 21 | 0 |
| SRR2106883 | 358 | 47814 | 1574 | 568 | 2311 | 44507 | 42820 | 19414 | 8270 | 222 | 44970 | 68881 | 5679 | 2658 | 32 | 7992 | 15642 | 0 | 0 | 0 | 527 | 40 | 92 | 0 | 0 | 0 | 0 | 0 | 26 | 82 | 40 | 60 | 0 | 0 | 0 | 0 | 0 |
| SRR2106884 | 169 | 74183 | 1022 | 419 | 2732 | 64104 | 50201 | 29813 | 6663 | 161 | 43681 | 13 | 4963 | 2603 | 21 | 6596 | 15331 | 0 | 0 | 0 | 565 | 73 | 126 | 0 | 13 | 0 | 0 | 0 | 25 | 108 | 52 | 86 | 0 | 14 | 0 | 13 | 0 |
| SRR2124760 | 667 | 21980 | 985 | 2267 | 185 | 1759 | 1340 | 4726 | 5270 | 0 | 8488 | 18825 | 2369 | 3272 | 0 | 4534 | 52127 | 0 | 0 | 0 | 5467 | 532 | 46 | 0 | 0 | 0 | 0 | 0 | 139 | 1749 | 286 | 26 | 0 | 0 | 0 | 0 | 0 |
| SRR2124761 | 498 | 10970 | 807 | 1391 | 164 | 1952 | 1232 | 3897 | 9726 | 13 | 11914 | 34008 | 3597 | 3232 | 0 | 4788 | 82601 | 0 | 0 | 11 | 9824 | 1629 | 211 | 0 | 0 | 0 | 0 | 0 | 465 | 5545 | 1843 | 176 | 0 | 0 | 23 | 0 | 0 |
| SRR2124762 | 501 | 16022 | 876 | 1243 | 206 | 2248 | 1601 | 5246 | 5666 | 0 | 12413 | 31826 | 11 | 2813 | 0 | 6683 | 80215 | 0 | 0 | 20 | 21206 | 2779 | 222 | 0 | 0 | 0 | 0 | 0 | 760 | 10884 | 2732 | 174 | 0 | 0 | 35 | 0 | 0 |
| SRR2124763 | 372 | 18998 | 488 | 1015 | 225 | 1952 | 1264 | 4239 | 7222 | 21 | 9416 | 31844 | 3163 | 2310 | 0 | 6658 | 125098 | 0 | 0 | 12 | 13644 | 2295 | 282 | 0 | 0 | 0 | 0 | 0 | 766 | 9427 | 3742 | 320 | 0 | 20 | 50 | 11 | 0 |
| SRR2124764 | 629 | 60575 | 880 | 1628 | 237 | 3137 | 1794 | 5798 | 4514 | 15 | 15574 | 30691 | 3928 | 2329 | 0 | 5293 | 68547 | 0 | 0 | 12 | 12776 | 1599 | 109 | 0 | 0 | 0 | 0 | 0 | 330 | 4200 | 987 | 66 | 0 | 0 | 0 | 0 | 0 |
| SRR2124765 | 522 | 19113 | 814 | 1664 | 238 | 2526 | 1169 | 4285 | 16842 | 0 | 14587 | 22119 | 21 | 1312 | 0 | 6418 | 70761 | 0 | 0 | 0 | 9377 | 638 | 37 | 0 | 0 | 0 | 0 | 0 | 374 | 4996 | 682 | 36 | 0 | 0 | 0 | 0 | 0 |
| SRR2124766 | 606 | 15663 | 892 | 2695 | 167 | 1712 | 729 | 2711 | 17196 | 13 | 9999 | 15467 | 2228 | 1603 | 0 | 4348 | 50492 | 0 | 0 | 11 | 9173 | 478 | 27 | 0 | 0 | 0 | 0 | 0 | 628 | 8046 | 800 | 39 | 0 | 0 | 0 | 0 | 0 |
| SRR2124767 | 653 | 14851 | 930 | 2217 | 221 | 3070 | 1202 | 4247 | 32673 | 0 | 17216 | 18853 | 3416 | 1753 | 0 | 6642 | 69908 | 0 | 0 | 16 | 12986 | 748 | 71 | 0 | 0 | 0 | 0 | 0 | 1212 | 14409 | 2093 | 147 | 0 | 0 | 13 | 0 | 0 |
| SRR2124768 | 571 | 14853 | 855 | 1089 | 224 | 2681 | 1595 | 6132 | 20606 | 15 | 18884 | 24085 | 4234 | 2806 | 0 | 6537 | 77854 | 0 | 0 | 13 | 14950 | 1441 | 138 | 0 | 0 | 0 | 0 | 0 | 919 | 12340 | 2545 | 167 | 0 | 0 | 22 | 0 | 0 |
| SRR2124769 | 790 | 12799 | 840 | 1078 | 276 | 3069 | 1476 | 5830 | 23859 | 17 | 18014 | 20869 | 25 | 1895 | 0 | 6594 | 78800 | 0 | 0 | 0 | 3723 | 348 | 26 | 0 | 0 | 0 | 0 | 0 | 255 | 2915 | 656 | 38 | 0 | 0 | 0 | 0 | 0 |
| SRR2124770 | 897 | 28859 | 1513 | 1326 | 343 | 4033 | 2406 | 7750 | 10426 | 0 | 20608 | 22967 | 3935 | 2398 | 0 | 9660 | 53025 | 0 | 0 | 13 | 20044 | 1716 | 145 | 0 | 0 | 0 | 0 | 0 | 717 | 8920 | 1626 | 123 | 0 | 0 | 15 | 0 | 0 |
| SRR2124771 | 900 | 12659 | 1009 | 1569 | 189 | 3317 | 1369 | 5145 | 25836 | 16 | 23661 | 30748 | 4591 | 4125 | 0 | 5929 | 65291 | 0 | 0 | 0 | 6442 | 627 | 75 | 0 | 0 | 0 | 0 | 0 | 384 | 5326 | 1307 | 147 | 0 | 0 | 0 | 0 | 0 |
| SRR2124772 | 706 | 21143 | 1123 | 1209 | 231 | 3101 | 1431 | 5205 | 26613 | 0 | 19278 | 21906 | 3847 | 3513 | 0 | 5795 | 51683 | 0 | 0 | 14 | 15686 | 676 | 50 | 0 | 0 | 0 | 0 | 0 | 679 | 10423 | 1112 | 48 | 0 | 0 | 0 | 0 | 0 |
| SRR2124773 | 705 | 10137 | 598 | 2398 | 212 | 2776 | 1378 | 4729 | 46485 | 25 | 17823 | 19054 | 28 | 1649 | 0 | 5859 | 56629 | 0 | 0 | 0 | 1697 | 77 | 0 | 0 | 0 | 0 | 0 | 0 | 157 | 1476 | 143 | 16 | 0 | 0 | 0 | 0 | 0 |
| SRR2124774 | 709 | 26498 | 843 | 1494 | 274 | 3606 | 1842 | 7080 | 23713 | 0 | 15861 | 18670 | 25 | 2131 | 0 | 5783 | 77690 | 0 | 0 | 17 | 15187 | 1228 | 105 | 0 | 0 | 0 | 0 | 0 | 1044 | 12475 | 2145 | 116 | 0 | 0 | 23 | 0 | 0 |
| SRR2124775 | 574 | 10137 | 637 | 1188 | 213 | 2757 | 1034 | 4154 | 27997 | 0 | 14733 | 21671 | 27 | 1769 | 0 | 6032 | 79944 | 0 | 0 | 0 | 8779 | 613 | 47 | 0 | 0 | 0 | 0 | 0 | 792 | 9299 | 1427 | 80 | 0 | 0 | 13 | 0 | 0 |
| SRR2124776 | 729 | 34377 | 1057 | 1495 | 162 | 1857 | 1053 | 3960 | 13556 | 0 | 15393 | 22136 | 16 | 5160 | 0 | 4585 | 44643 | 0 | 0 | 11 | 12026 | 661 | 44 | 0 | 0 | 0 | 0 | 0 | 501 | 7190 | 843 | 42 | 0 | 0 | 0 | 0 | 0 |
| SRR2124777 | 908 | 16645 | 1105 | 1039 | 169 | 2204 | 916 | 3524 | 26833 | 11 | 17587 | 22963 | 15 | 2923 | 0 | 5069 | 53299 | 0 | 0 | 0 | 5835 | 626 | 114 | 0 | 0 | 0 | 0 | 0 | 473 | 4812 | 1644 | 183 | 0 | 0 | 14 | 0 | 0 |
| SRR2124778 | 684 | 21139 | 953 | 1180 | 298 | 2863 | 1914 | 6212 | 13353 | 0 | 13488 | 22095 | 3054 | 1906 | 0 | 6950 | 46377 | 0 | 0 | 25 | 30050 | 875 | 45 | 0 | 0 | 0 | 0 | 0 | 689 | 11641 | 753 | 28 | 0 | 0 | 0 | 0 | 0 |
| SRR2124779 | 614 | 14373 | 938 | 1379 | 265 | 2491 | 1338 | 4827 | 10640 | 12 | 10926 | 23878 | 2838 | 2014 | 0 | 5381 | 66790 | 0 | 0 | 14 | 12201 | 2188 | 256 | 0 | 0 | 0 | 0 | 0 | 689 | 8842 | 3032 | 259 | 0 | 11 | 45 | 18 | 0 |
| SRR2126527 | 1911 | 71896 | 355 | 798 | 959 | 12184 | 7854 | 13071 | 54732 | 54 | 29524 | 134429 | 9450 | 1485 | 72 | 5498 | 17262 | 0 | 0 | 0 | 1472 | 183 | 242 | 0 | 32 | 0 | 0 | 0 | 86 | 374 | 165 | 101 | 0 | 38 | 20 | 0 | 0 |
| SRR2126528 | 1541 | 78468 | 345 | 509 | 999 | 12306 | 8037 | 13469 | 49919 | 65 | 29800 | 144923 | 9834 | 1489 | 80 | 5575 | 16294 | 0 | 0 | 0 | 2235 | 235 | 255 | 0 | 27 | 0 | 0 | 0 | 95 | 601 | 213 | 114 | 0 | 43 | 21 | 0 | 0 |
| SRR2126529 | 2341 | 75760 | 361 | 586 | 1024 | 12169 | 8076 | 12641 | 57711 | 70 | 28162 | 135802 | 9158 | 1433 | 81 | 5497 | 17101 | 0 | 0 | 0 | 1598 | 216 | 306 | 0 | 31 | 0 | 0 | 0 | 106 | 389 | 203 | 127 | 0 | 44 | 26 | 0 | 0 |
| SRR2126530 | 2926 | 52028 | 339 | 874 | 1192 | 15445 | 11683 | 17838 | 43709 | 77 | 26764 | 130221 | 8896 | 1433 | 65 | 4778 | 19410 | 0 | 0 | 0 | 1856 | 200 | 290 | 0 | 27 | 0 | 0 | 15 | 135 | 481 | 198 | 119 | 0 | 39 | 26 | 0 | 0 |
| SRR2126531 | 3805 | 56385 | 380 | 515 | 1178 | 14687 | 11516 | 16715 | 51875 | 66 | 29896 | 141529 | 9562 | 1647 | 51 | 4695 | 16906 | 0 | 0 | 0 | 2768 | 206 | 272 | 0 | 27 | 0 | 0 | 16 | 144 | 754 | 187 | 108 | 0 | 43 | 26 | 0 | 0 |
| SRR2126532 | 2534 | 63214 | 336 | 677 | 1116 | 13381 | 9728 | 14041 | 48449 | 78 | 27910 | 138122 | 9276 | 1385 | 73 | 5180 | 17198 | 0 | 0 | 0 | 1782 | 178 | 352 | 0 | 34 | 0 | 0 | 0 | 109 | 419 | 167 | 141 | 0 | 49 | 28 | 0 | 0 |
| SRR2135530 | 186 | 4072 | 31264 | 828 | 15019 | 16496 | 297989 | 9364 | 61963 | 1416 | 34343 | 7266 | 19100 | 25836 | 8896 | 51 | 12264 | 0 | 26 | 150 | 1129 | 457 | 468 | 47 | 27 | 13 | 0 | 28 | 322 | 297 | 25 | 21 | 0 | 0 | 12 | 54 | 0 |
| SRR2135531 | 584 | 4045 | 11837 | 1121 | 14750 | 10492 | 204209 | 21660 | 43853 | 2060 | 29604 | 8407 | 19266 | 22257 | 6005 | 47 | 12651 | 11 | 72 | 349 | 2558 | 513 | 642 | 111 | 49 | 24 | 23 | 82 | 565 | 666 | 65 | 31 | 0 | 0 | 17 | 80 | 0 |
| SRR2135532 | 164 | 2742 | 12262 | 218 | 11275 | 7288 | 143645 | 23502 | 32462 | 2832 | 34613 | 10581 | 22797 | 23810 | 5863 | 36 | 12759 | 0 | 73 | 356 | 3057 | 708 | 728 | 139 | 57 | 29 | 14 | 80 | 597 | 930 | 87 | 31 | 0 | 0 | 17 | 83 | 0 |
| SRR2229584 | 0 | 377 | 115 | 4412 | 121 | 2504 | 3575 | 441 | 273 | 11 | 5327 | 11158 | 354 | 421 | 1469 | 0 | 746 | 0 | 0 | 0 | 14 | 0 | 0 | 0 | 0 | 0 | 0 | 0 | 0 | 20 | 0 | 0 | 0 | 0 | 0 | 0 | 0 |
| SRR2229585 | 45 | 2322 | 253 | 2703 | 226 | 3307 | 4259 | 1407 | 354 | 14 | 2842 | 8015 | 462 | 274 | 1251 | 31 | 2609 | 0 | 0 | 0 | 61 | 12 | 14 | 0 | 0 | 0 | 0 | 0 | 0 | 31 | 0 | 12 | 0 | 0 | 0 | 0 | 0 |
| SRR2229586 | 75 | 3849 | 29 | 1923 | 0 | 0 | 0 | 0 | 0 | 0 | 0 | 0 | 0 | 0 | 0 | 0 | 0 | 0 | 0 | 0 | 0 | 0 | 0 | 0 | 0 | 0 | 0 | 0 | 0 | 0 | 0 | 0 | 0 | 0 | 0 | 0 | 0 |
| SRR2229587 | 22 | 1789 | 35 | 2724 | 0 | 0 | 0 | 23 | 0 | 0 | 0 | 0 | 0 | 0 | 0 | 0 | 0 | 0 | 0 | 0 | 0 | 0 | 0 | 0 | 0 | 0 | 0 | 0 | 0 | 0 | 0 | 0 | 0 | 0 | 0 | 0 | 0 |
| SRR2229588 | 348 | 12385 | 857 | 10769 | 429 | 1414 | 2666 | 1238 | 799 | 0 | 1001 | 3298 | 196 | 76 | 1452 | 0 | 4958 | 0 | 0 | 0 | 41 | 39 | 0 | 0 | 0 | 0 | 0 | 0 | 0 | 0 | 11 | 0 | 0 | 0 | 20 | 0 | 0 |
| SRR2229589 | 317 | 14068 | 1196 | 6353 | 705 | 1523 | 4790 | 1844 | 850 | 0 | 864 | 2949 | 250 | 57 | 1352 | 0 | 4347 | 0 | 0 | 0 | 27 | 35 | 0 | 0 | 0 | 0 | 0 | 0 | 0 | 0 | 15 | 0 | 0 | 0 | 24 | 0 | 0 |
| SRR2287868 | 35 | 5890 | 20 | 389 | 74 | 174 | 417 | 783 | 1681 | 0 | 13271 | 43565 | 4319 | 2603 | 1642 | 11560 | 139763 | 0 | 213 | 616 | 22113 | 5447 | 709 | 0 | 0 | 0 | 0 | 2288 | 11866 | 20327 | 19586 | 1344 | 0 | 0 | 149 | 0 | 0 |
| SRR2289624 | 581 | 21752 | 274 | 969 | 69 | 520 | 1182 | 820 | 1216 | 0 | 16037 | 58881 | 4437 | 2781 | 2404 | 10838 | 127070 | 0 | 14 | 57 | 4512 | 855 | 301 | 0 | 0 | 0 | 0 | 199 | 1267 | 3597 | 2389 | 580 | 0 | 10 | 142 | 0 | 0 |
| SRR2289625 | 12 | 1735 | 156 | 199 | 34 | 215 | 372 | 596 | 1614 | 0 | 18440 | 69438 | 7721 | 3006 | 1273 | 8860 | 97527 | 0 | 25 | 74 | 2411 | 2126 | 345 | 0 | 0 | 0 | 0 | 257 | 1362 | 1118 | 3227 | 364 | 0 | 13 | 313 | 0 | 0 |
| SRR2289626 | 0 | 1074 | 237 | 165 | 41 | 145 | 423 | 863 | 2501 | 0 | 10174 | 18580 | 3256 | 3523 | 649 | 7899 | 333327 | 0 | 0 | 32 | 1158 | 2421 | 93 | 0 | 0 | 0 | 0 | 125 | 1013 | 1463 | 8003 | 231 | 0 | 10 | 26 | 0 | 0 |
| SRR2294377 | 12 | 1666 | 125 | 77 | 41 | 146 | 404 | 1143 | 457 | 0 | 12444 | 42106 | 4172 | 2319 | 465 | 6696 | 194501 | 0 | 12 | 47 | 4192 | 898 | 146 | 0 | 0 | 0 | 0 | 129 | 1074 | 2884 | 3094 | 214 | 0 | 0 | 45 | 0 | 0 |
| SRR2294380 | 0 | 5035 | 848 | 570 | 46 | 195 | 692 | 1872 | 467 | 15 | 7512 | 30221 | 2496 | 3322 | 162 | 4672 | 168374 | 0 | 39 | 116 | 2140 | 5084 | 590 | 0 | 0 | 0 | 0 | 428 | 2259 | 1967 | 16597 | 1180 | 0 | 0 | 68 | 0 | 0 |
| SRR2294739 | 14 | 1086 | 60 | 716 | 31 | 109 | 408 | 981 | 341 | 0 | 7455 | 19090 | 2298 | 2997 | 29 | 4705 | 419827 | 0 | 116 | 318 | 8198 | 10420 | 534 | 0 | 0 | 0 | 0 | 1474 | 6634 | 11026 | 51792 | 1491 | 0 | 0 | 64 | 0 | 0 |
| SRR2294742 | 16 | 2930 | 379 | 333 | 42 | 182 | 594 | 1348 | 339 | 16 | 10479 | 31973 | 3066 | 2548 | 207 | 3196 | 253354 | 0 | 30 | 66 | 1710 | 3471 | 118 | 0 | 0 | 0 | 0 | 475 | 1764 | 1672 | 13013 | 307 | 0 | 0 | 36 | 0 | 0 |
| SRR2294746 | 19 | 1839 | 255 | 481 | 35 | 175 | 417 | 1148 | 362 | 15 | 10360 | 45220 | 3570 | 2982 | 33 | 3120 | 153095 | 0 | 12 | 27 | 982 | 3152 | 210 | 0 | 0 | 0 | 0 | 200 | 714 | 824 | 8403 | 373 | 0 | 0 | 166 | 0 | 0 |
| SRR2294747 | 26 | 1844 | 1496 | 220 | 32 | 179 | 457 | 1434 | 2415 | 0 | 10381 | 27103 | 2865 | 5256 | 301 | 6785 | 241768 | 0 | 0 | 31 | 792 | 1755 | 21 | 0 | 0 | 0 | 0 | 141 | 1062 | 1120 | 7278 | 33 | 0 | 0 | 0 | 0 | 0 |
| SRR2294748 | 14 | 2568 | 242 | 93 | 30 | 166 | 394 | 1276 | 313 | 0 | 10102 | 48259 | 3332 | 1774 | 39 | 2461 | 212848 | 0 | 0 | 19 | 614 | 1331 | 68 | 0 | 0 | 0 | 0 | 91 | 447 | 467 | 4387 | 120 | 0 | 0 | 41 | 0 | 0 |
| SRR2294820 | 0 | 2380 | 126 | 1200 | 65 | 444 | 1325 | 2183 | 290 | 15 | 7990 | 26268 | 2362 | 3569 | 61 | 4603 | 135862 | 0 | 15 | 41 | 1829 | 4145 | 395 | 0 | 0 | 0 | 0 | 140 | 680 | 1885 | 14853 | 1075 | 0 | 0 | 20 | 0 | 0 |
| SRR2296788 | 3408 | 14447 | 2218 | 1630 | 678 | 9132 | 10148 | 3269 | 4671 | 167 | 26376 | 20 | 5788 | 4624 | 24 | 1056 | 11209 | 0 | 0 | 0 | 485 | 94 | 92 | 0 | 0 | 0 | 0 | 0 | 38 | 163 | 42 | 48 | 0 | 11 | 0 | 0 | 0 |
| SRR2296789 | 1659 | 14883 | 2967 | 2115 | 547 | 6163 | 7694 | 1910 | 3546 | 152 | 24584 | 42582 | 4622 | 4927 | 15 | 1479 | 5848 | 0 | 0 | 0 | 669 | 73 | 120 | 0 | 15 | 0 | 0 | 0 | 26 | 172 | 22 | 58 | 0 | 16 | 11 | 0 | 0 |
| SRR2296790 | 4316 | 14042 | 2224 | 880 | 511 | 6444 | 7219 | 2340 | 2927 | 315 | 28766 | 66408 | 7087 | 5379 | 43 | 1045 | 12301 | 0 | 0 | 0 | 580 | 110 | 95 | 0 | 0 | 0 | 0 | 0 | 47 | 91 | 21 | 28 | 0 | 0 | 0 | 0 | 0 |
| SRR2296791 | 4083 | 10643 | 2028 | 1936 | 615 | 6836 | 8531 | 3089 | 3059 | 79 | 28158 | 57173 | 8266 | 4899 | 114 | 991 | 9183 | 0 | 0 | 0 | 504 | 80 | 62 | 0 | 0 | 0 | 0 | 0 | 47 | 158 | 42 | 51 | 0 | 0 | 0 | 0 | 0 |
| SRR2296792 | 2144 | 13221 | 1561 | 4200 | 972 | 9833 | 12627 | 2447 | 5437 | 260 | 13621 | 18443 | 4603 | 5345 | 20 | 2175 | 6681 | 0 | 0 | 0 | 690 | 82 | 131 | 0 | 19 | 0 | 0 | 0 | 29 | 271 | 21 | 50 | 0 | 12 | 0 | 0 | 0 |
| SRR2296793 | 4805 | 14216 | 2046 | 1133 | 682 | 8043 | 8239 | 3640 | 3583 | 199 | 27853 | 56276 | 6275 | 4116 | 38 | 967 | 13481 | 0 | 0 | 0 | 368 | 72 | 70 | 0 | 0 | 0 | 0 | 12 | 52 | 112 | 39 | 34 | 0 | 0 | 0 | 11 | 0 |
| SRR2296794 | 4333 | 13428 | 1689 | 1875 | 749 | 7756 | 9617 | 3493 | 3350 | 106 | 29786 | 61005 | 8569 | 5746 | 78 | 1371 | 8978 | 0 | 0 | 0 | 471 | 67 | 72 | 0 | 0 | 0 | 0 | 0 | 51 | 142 | 31 | 31 | 0 | 0 | 0 | 0 | 0 |
| SRR2296795 | 4964 | 15825 | 1809 | 2206 | 1027 | 13071 | 12902 | 7478 | 5442 | 217 | 27179 | 43815 | 6788 | 4793 | 16 | 1820 | 15704 | 0 | 0 | 0 | 422 | 75 | 115 | 0 | 0 | 0 | 0 | 0 | 56 | 114 | 23 | 69 | 0 | 0 | 0 | 15 | 0 |
| SRR2296797 | 3649 | 10415 | 2397 | 1074 | 774 | 7438 | 9227 | 4540 | 2443 | 122 | 25975 | 13 | 7432 | 7056 | 18 | 2442 | 11536 | 0 | 0 | 0 | 540 | 95 | 178 | 0 | 0 | 0 | 0 | 0 | 31 | 141 | 24 | 79 | 0 | 0 | 0 | 0 | 0 |
| SRR2296798 | 4742 | 16072 | 2441 | 974 | 561 | 7306 | 6821 | 2995 | 2828 | 152 | 22004 | 16 | 6593 | 4234 | 36 | 1066 | 12885 | 0 | 0 | 0 | 477 | 100 | 100 | 0 | 0 | 0 | 0 | 0 | 50 | 109 | 34 | 44 | 0 | 0 | 0 | 0 | 0 |
| SRR2296799 | 2239 | 13619 | 3634 | 1499 | 734 | 8824 | 8823 | 2941 | 3791 | 177 | 30759 | 55052 | 6028 | 5809 | 30 | 1196 | 9131 | 0 | 0 | 0 | 441 | 62 | 73 | 0 | 14 | 0 | 0 | 0 | 24 | 83 | 28 | 18 | 0 | 12 | 10 | 0 | 0 |
| SRR2296800 | 2094 | 15783 | 1630 | 3303 | 1058 | 11272 | 14378 | 3179 | 5575 | 169 | 25475 | 40466 | 5028 | 5797 | 11 | 2643 | 6433 | 0 | 0 | 0 | 584 | 76 | 113 | 0 | 11 | 0 | 0 | 0 | 38 | 235 | 31 | 53 | 0 | 14 | 11 | 0 | 0 |
| SRR2296801 | 3405 | 11281 | 1996 | 3945 | 1210 | 14707 | 17209 | 7164 | 5031 | 166 | 26538 | 10 | 4911 | 5447 | 14 | 2278 | 11363 | 0 | 0 | 0 | 403 | 67 | 83 | 0 | 13 | 0 | 0 | 11 | 38 | 133 | 25 | 51 | 0 | 16 | 11 | 0 | 0 |
| SRR2296802 | 1304 | 12218 | 4684 | 2584 | 1286 | 13697 | 20796 | 7362 | 5263 | 290 | 27410 | 42514 | 5400 | 6877 | 40 | 3228 | 6780 | 0 | 0 | 0 | 375 | 36 | 87 | 0 | 0 | 0 | 0 | 0 | 0 | 135 | 19 | 63 | 0 | 12 | 0 | 0 | 0 |
| SRR2296805 | 1333 | 14874 | 3323 | 1688 | 546 | 7309 | 7481 | 2504 | 3039 | 181 | 30164 | 21 | 6243 | 5961 | 20 | 971 | 9447 | 0 | 0 | 0 | 624 | 95 | 97 | 0 | 15 | 0 | 0 | 0 | 30 | 139 | 23 | 34 | 0 | 13 | 12 | 0 | 0 |
| SRR2296806 | 804 | 10384 | 2253 | 3295 | 1049 | 10884 | 15365 | 3292 | 6141 | 171 | 30111 | 57534 | 6583 | 6651 | 17 | 2083 | 7779 | 0 | 0 | 0 | 517 | 48 | 66 | 0 | 0 | 0 | 0 | 0 | 17 | 246 | 26 | 26 | 0 | 0 | 0 | 0 | 0 |
| SRR2296807 | 2656 | 10937 | 2431 | 2238 | 636 | 9045 | 9763 | 3224 | 4500 | 209 | 24926 | 65829 | 4840 | 5191 | 16 | 1180 | 10520 | 0 | 0 | 0 | 442 | 90 | 70 | 0 | 14 | 0 | 0 | 0 | 27 | 146 | 41 | 47 | 0 | 19 | 10 | 0 | 0 |
| SRR2296808 | 2101 | 12832 | 4688 | 3264 | 1236 | 11983 | 17291 | 3277 | 5953 | 295 | 32081 | 52488 | 5686 | 8210 | 0 | 2126 | 8064 | 0 | 0 | 0 | 357 | 52 | 50 | 0 | 11 | 0 | 0 | 0 | 32 | 191 | 18 | 27 | 0 | 0 | 0 | 0 | 0 |
| SRR2296809 | 4428 | 9653 | 2376 | 2619 | 1328 | 15206 | 19276 | 7361 | 4347 | 152 | 21707 | 49354 | 4449 | 5785 | 11 | 2844 | 12155 | 0 | 0 | 0 | 458 | 82 | 118 | 0 | 19 | 0 | 0 | 0 | 32 | 125 | 40 | 66 | 0 | 27 | 0 | 0 | 0 |
| SRR2296810 | 1531 | 14093 | 3976 | 2618 | 1358 | 12057 | 18309 | 6014 | 4184 | 252 | 31189 | 45268 | 5708 | 7389 | 20 | 3735 | 8055 | 0 | 0 | 0 | 487 | 54 | 107 | 0 | 12 | 0 | 0 | 0 | 30 | 171 | 19 | 52 | 0 | 17 | 0 | 0 | 0 |
| SRR2353417 | 54 | 799 | 444 | 336 | 35 | 128 | 299 | 599 | 662 | 38 | 22556 | 36714 | 10803 | 3027 | 369 | 12655 | 90430 | 0 | 88 | 156 | 2866 | 2443 | 301 | 0 | 0 | 0 | 0 | 781 | 5297 | 1697 | 7065 | 477 | 0 | 0 | 0 | 0 | 0 |
| SRR2353418 | 57 | 551 | 390 | 350 | 31 | 183 | 355 | 845 | 623 | 37 | 19066 | 29426 | 8944 | 2741 | 199 | 11842 | 108494 | 0 | 50 | 118 | 3265 | 2333 | 311 | 0 | 0 | 0 | 0 | 479 | 5589 | 2052 | 5254 | 471 | 0 | 0 | 0 | 0 | 0 |
| SRR2353419 | 63 | 1063 | 419 | 362 | 28 | 158 | 250 | 607 | 420 | 15 | 20620 | 29202 | 10031 | 2332 | 373 | 11326 | 101237 | 0 | 25 | 27 | 1623 | 1917 | 209 | 0 | 0 | 0 | 0 | 245 | 1030 | 723 | 4904 | 408 | 0 | 0 | 0 | 0 | 0 |
| SRR2353420 | 49 | 531 | 444 | 397 | 37 | 205 | 316 | 645 | 691 | 17 | 21906 | 34463 | 10355 | 2265 | 366 | 12934 | 81065 | 0 | 27 | 40 | 1913 | 362 | 154 | 0 | 0 | 0 | 0 | 248 | 1409 | 836 | 909 | 239 | 0 | 0 | 12 | 10 | 0 |
| SRR2353421 | 70 | 761 | 710 | 604 | 38 | 258 | 393 | 935 | 595 | 0 | 23139 | 36473 | 10805 | 2337 | 268 | 12083 | 86290 | 0 | 19 | 39 | 3655 | 1162 | 242 | 0 | 0 | 0 | 0 | 234 | 1964 | 2067 | 3245 | 483 | 0 | 0 | 0 | 0 | 0 |
| SRR2353422 | 57 | 619 | 772 | 352 | 32 | 188 | 356 | 732 | 519 | 41 | 21715 | 32138 | 10482 | 2497 | 338 | 13312 | 87940 | 0 | 29 | 101 | 7396 | 1027 | 602 | 0 | 0 | 0 | 0 | 264 | 4199 | 3542 | 2750 | 976 | 0 | 0 | 0 | 0 | 0 |
| SRR2353423 | 52 | 755 | 577 | 595 | 36 | 210 | 388 | 919 | 452 | 39 | 21965 | 35504 | 10686 | 2869 | 262 | 12793 | 82469 | 0 | 41 | 47 | 2435 | 1698 | 241 | 0 | 0 | 0 | 0 | 421 | 1823 | 1241 | 4702 | 470 | 0 | 0 | 0 | 0 | 0 |
| SRR2353424 | 51 | 982 | 467 | 622 | 48 | 236 | 491 | 1136 | 362 | 29 | 18289 | 30570 | 8167 | 2837 | 213 | 9769 | 96276 | 0 | 46 | 105 | 4573 | 2758 | 241 | 0 | 0 | 0 | 0 | 529 | 4308 | 2402 | 9284 | 441 | 0 | 0 | 0 | 0 | 0 |
| SRR2353425 | 66 | 16454 | 1126 | 874 | 34 | 277 | 500 | 656 | 564 | 0 | 24192 | 36580 | 11114 | 2918 | 376 | 13113 | 99574 | 0 | 24 | 55 | 19944 | 1863 | 1174 | 0 | 0 | 0 | 0 | 236 | 1946 | 6677 | 4146 | 1498 | 0 | 15 | 33 | 19 | 0 |
| SRR2353426 | 63 | 838 | 704 | 295 | 44 | 325 | 440 | 1048 | 612 | 18 | 24440 | 37564 | 11772 | 2669 | 271 | 12996 | 96723 | 0 | 30 | 48 | 5256 | 574 | 313 | 0 | 0 | 0 | 0 | 315 | 2121 | 2774 | 1616 | 569 | 0 | 0 | 0 | 0 | 0 |
| SRR2353427 | 23 | 1011 | 987 | 244 | 35 | 228 | 315 | 671 | 538 | 17 | 22531 | 32420 | 11135 | 2587 | 550 | 13123 | 89337 | 0 | 24 | 54 | 7527 | 891 | 523 | 0 | 0 | 0 | 0 | 238 | 2355 | 3639 | 2403 | 886 | 0 | 0 | 10 | 0 | 0 |
| SRR2353428 | 44 | 1179 | 552 | 655 | 50 | 290 | 579 | 972 | 410 | 41 | 19638 | 30408 | 9524 | 3093 | 317 | 11251 | 95269 | 0 | 56 | 120 | 18624 | 1920 | 1105 | 0 | 0 | 0 | 0 | 473 | 4080 | 8246 | 5344 | 1671 | 0 | 10 | 19 | 14 | 0 |
| SRR2353429 | 117 | 2143 | 634 | 446 | 54 | 327 | 547 | 962 | 761 | 23 | 24557 | 36813 | 10836 | 2738 | 282 | 12077 | 84819 | 11 | 70 | 79 | 9654 | 1537 | 491 | 0 | 0 | 0 | 0 | 759 | 3860 | 5094 | 4043 | 937 | 0 | 23 | 53 | 20 | 0 |
| SRR2353430 | 168 | 2513 | 727 | 354 | 37 | 196 | 327 | 766 | 529 | 40 | 16861 | 34288 | 7979 | 2523 | 209 | 10231 | 99722 | 0 | 90 | 215 | 30604 | 2779 | 1723 | 11 | 0 | 0 | 0 | 704 | 6961 | 13626 | 6497 | 2487 | 0 | 0 | 0 | 0 | 0 |
| SRR2353431 | 141 | 2304 | 623 | 241 | 33 | 176 | 305 | 620 | 543 | 26 | 20929 | 34811 | 10138 | 2508 | 402 | 11721 | 85467 | 0 | 25 | 30 | 2881 | 1189 | 390 | 0 | 0 | 0 | 0 | 276 | 1274 | 1391 | 3191 | 674 | 0 | 0 | 16 | 12 | 0 |
| SRR2353432 | 40 | 639 | 598 | 262 | 50 | 194 | 426 | 788 | 542 | 19 | 21215 | 31346 | 9970 | 2990 | 434 | 12299 | 84582 | 0 | 66 | 156 | 23067 | 1358 | 1166 | 0 | 0 | 0 | 0 | 668 | 6396 | 12023 | 3531 | 2113 | 0 | 0 | 0 | 0 | 0 |
| SRR2353433 | 45 | 818 | 323 | 216 | 59 | 314 | 680 | 939 | 556 | 52 | 20726 | 34926 | 10069 | 3162 | 271 | 10751 | 83455 | 0 | 70 | 151 | 31914 | 2042 | 2199 | 18 | 0 | 0 | 0 | 689 | 5129 | 15863 | 5498 | 4207 | 0 | 0 | 0 | 0 | 0 |
| SRR2353434 | 51 | 717 | 436 | 434 | 36 | 240 | 424 | 1076 | 756 | 21 | 16626 | 28315 | 8540 | 2405 | 266 | 11087 | 89029 | 0 | 63 | 81 | 6209 | 1609 | 356 | 0 | 0 | 0 | 0 | 526 | 3010 | 3067 | 4298 | 609 | 0 | 0 | 12 | 14 | 0 |
| SRR2353435 | 143 | 775 | 540 | 448 | 78 | 505 | 1168 | 1165 | 1180 | 40 | 18037 | 28983 | 8526 | 2307 | 250 | 10286 | 102497 | 0 | 26 | 32 | 2757 | 1291 | 242 | 0 | 0 | 0 | 0 | 241 | 1227 | 1490 | 2849 | 437 | 0 | 0 | 0 | 0 | 0 |
| SRR2353436 | 79 | 1218 | 604 | 652 | 76 | 348 | 861 | 1706 | 530 | 50 | 17400 | 27279 | 8387 | 3432 | 222 | 9347 | 95927 | 0 | 66 | 344 | 16107 | 3037 | 619 | 0 | 0 | 0 | 0 | 588 | 13217 | 10973 | 10222 | 1253 | 0 | 0 | 0 | 0 | 0 |
| SRR2353437 | 116 | 1287 | 811 | 811 | 33 | 177 | 336 | 744 | 580 | 16 | 22827 | 32700 | 9848 | 3021 | 248 | 12994 | 101680 | 0 | 29 | 71 | 3197 | 647 | 174 | 0 | 0 | 0 | 0 | 318 | 3272 | 1915 | 2105 | 311 | 0 | 0 | 0 | 0 | 0 |
| SRR2353438 | 61 | 1206 | 557 | 671 | 33 | 183 | 408 | 1031 | 329 | 52 | 15728 | 28934 | 7486 | 2287 | 228 | 8135 | 108631 | 0 | 25 | 82 | 2746 | 1383 | 117 | 0 | 0 | 0 | 0 | 332 | 4652 | 2279 | 5010 | 258 | 0 | 0 | 0 | 0 | 0 |
| SRR2353439 | 68 | 985 | 392 | 307 | 31 | 207 | 426 | 1225 | 479 | 28 | 15020 | 23992 | 7102 | 2290 | 176 | 8495 | 80071 | 0 | 43 | 271 | 19779 | 2046 | 917 | 0 | 0 | 0 | 0 | 360 | 11214 | 10295 | 5507 | 1547 | 0 | 0 | 0 | 0 | 0 |
| SRR2353440 | 129 | 868 | 313 | 347 | 82 | 446 | 818 | 1371 | 559 | 20 | 20022 | 31762 | 9321 | 2710 | 229 | 11751 | 86186 | 0 | 71 | 209 | 18539 | 1750 | 807 | 0 | 0 | 0 | 0 | 633 | 7803 | 8695 | 5100 | 1351 | 0 | 0 | 0 | 0 | 0 |
| SRR2353441 | 50 | 800 | 311 | 429 | 44 | 248 | 480 | 1239 | 360 | 45 | 16132 | 27061 | 7556 | 2437 | 194 | 9074 | 94671 | 0 | 81 | 142 | 19620 | 5182 | 856 | 0 | 0 | 0 | 0 | 825 | 5945 | 10407 | 17237 | 1716 | 0 | 0 | 0 | 0 | 0 |
| SRR2353442 | 84 | 1094 | 511 | 324 | 56 | 311 | 575 | 1267 | 708 | 37 | 17255 | 26534 | 8273 | 2411 | 336 | 10865 | 82746 | 0 | 66 | 228 | 14304 | 2027 | 453 | 0 | 0 | 0 | 0 | 709 | 9840 | 8129 | 7152 | 913 | 0 | 0 | 0 | 0 | 0 |
| SRR2353443 | 66 | 666 | 473 | 782 | 30 | 150 | 275 | 707 | 712 | 34 | 20002 | 30465 | 9854 | 2663 | 329 | 10337 | 110492 | 0 | 45 | 29 | 2489 | 6886 | 1138 | 0 | 0 | 0 | 0 | 432 | 1091 | 1275 | 15892 | 2642 | 0 | 0 | 0 | 0 | 0 |
| SRR2353444 | 55 | 981 | 554 | 597 | 44 | 219 | 469 | 974 | 397 | 35 | 18156 | 28942 | 8746 | 3212 | 248 | 11644 | 112758 | 0 | 83 | 157 | 14209 | 4052 | 976 | 0 | 0 | 0 | 0 | 463 | 4137 | 5354 | 7671 | 1483 | 0 | 0 | 0 | 0 | 0 |
| SRR2353445 | 55 | 691 | 441 | 560 | 95 | 500 | 1149 | 1220 | 638 | 35 | 19723 | 32335 | 9288 | 2743 | 263 | 11429 | 82789 | 0 | 46 | 228 | 18548 | 1179 | 584 | 0 | 0 | 0 | 0 | 370 | 7352 | 7995 | 3028 | 885 | 0 | 0 | 0 | 0 | 0 |
| SRR2422969 | 191 | 2101 | 5126 | 1210 | 3589 | 5694 | 31629 | 12932 | 31 | 32 | 9239 | 15256 | 4779 | 14998 | 958 | 4635 | 2508 | 0 | 34 | 24 | 170 | 277 | 157 | 0 | 0 | 0 | 0 | 27 | 25 | 45 | 194 | 44 | 0 | 0 | 0 | 0 | 0 |
| SRR2422970 | 333 | 2497 | 5694 | 1465 | 3505 | 5925 | 31560 | 13031 | 37 | 67 | 9228 | 15530 | 4791 | 15254 | 836 | 4855 | 2366 | 0 | 38 | 35 | 169 | 254 | 148 | 0 | 0 | 0 | 0 | 17 | 41 | 55 | 199 | 33 | 0 | 0 | 0 | 0 | 0 |
| SRR2422971 | 302 | 2662 | 5656 | 1543 | 3188 | 5594 | 31421 | 13016 | 50 | 30 | 9202 | 15129 | 4624 | 14549 | 945 | 4561 | 2348 | 0 | 41 | 17 | 179 | 253 | 137 | 0 | 0 | 0 | 0 | 29 | 36 | 62 | 223 | 38 | 0 | 0 | 0 | 0 | 0 |
| SRR2422972 | 217 | 2171 | 5396 | 1489 | 3424 | 5887 | 31012 | 12855 | 26 | 31 | 9246 | 15725 | 4917 | 15129 | 914 | 4737 | 2124 | 0 | 38 | 22 | 200 | 244 | 140 | 0 | 0 | 0 | 0 | 23 | 34 | 46 | 218 | 42 | 0 | 0 | 0 | 0 | 0 |
| SRR2454328 | 0 | 8265 | 4208 | 21 | 4218 | 61408 | 38940 | 26262 | 3750 | 49 | 28506 | 36839 | 5047 | 7313 | 67 | 43282 | 11292 | 0 | 0 | 0 | 304 | 646 | 258 | 0 | 0 | 0 | 0 | 0 | 29 | 97 | 791 | 210 | 0 | 22 | 23 | 0 | 0 |
| SRR2454329 | 0 | 7132 | 3956 | 15 | 4340 | 63263 | 41489 | 28166 | 4091 | 44 | 29833 | 38479 | 4997 | 7596 | 68 | 43382 | 11732 | 0 | 0 | 0 | 296 | 707 | 240 | 0 | 0 | 0 | 0 | 0 | 30 | 94 | 858 | 199 | 0 | 20 | 22 | 0 | 0 |
| SRR2454330 | 0 | 6430 | 4439 | 23 | 4957 | 52857 | 36111 | 26655 | 3587 | 48 | 30508 | 43954 | 4746 | 8096 | 77 | 48483 | 11122 | 0 | 0 | 0 | 174 | 418 | 209 | 0 | 0 | 0 | 0 | 0 | 18 | 56 | 507 | 180 | 0 | 16 | 16 | 0 | 0 |
| SRR2454331 | 0 | 7580 | 3796 | 25 | 5546 | 55688 | 39597 | 28384 | 3400 | 50 | 33327 | 48337 | 5123 | 8802 | 76 | 50983 | 11269 | 0 | 0 | 0 | 193 | 500 | 217 | 0 | 0 | 0 | 0 | 0 | 20 | 68 | 633 | 170 | 0 | 16 | 14 | 0 | 0 |
| SRR2454332 | 0 | 36995 | 2805 | 80 | 6644 | 86093 | 44940 | 22402 | 2976 | 62 | 39333 | 47676 | 6603 | 7244 | 153 | 41756 | 11707 | 0 | 0 | 0 | 534 | 801 | 753 | 0 | 21 | 0 | 0 | 0 | 51 | 116 | 1000 | 483 | 0 | 35 | 12 | 0 | 0 |
| SRR2454333 | 10 | 35174 | 2415 | 48 | 6778 | 85528 | 46309 | 25131 | 3046 | 62 | 35792 | 50119 | 6269 | 6840 | 180 | 41486 | 12995 | 0 | 0 | 0 | 506 | 959 | 758 | 0 | 15 | 0 | 0 | 0 | 42 | 100 | 1015 | 506 | 0 | 25 | 10 | 11 | 0 |
| SRR2454334 | 0 | 40131 | 2997 | 68 | 4804 | 81536 | 49438 | 25213 | 3993 | 60 | 32330 | 43315 | 5488 | 7545 | 240 | 46868 | 13486 | 0 | 0 | 0 | 286 | 494 | 683 | 0 | 0 | 0 | 0 | 0 | 25 | 60 | 558 | 488 | 0 | 16 | 0 | 0 | 0 |
| SRR2454335 | 19 | 8006 | 4382 | 0 | 4061 | 75806 | 47071 | 30244 | 3901 | 60 | 36811 | 54589 | 6683 | 8791 | 263 | 45574 | 16575 | 0 | 0 | 0 | 61 | 127 | 186 | 0 | 0 | 0 | 0 | 0 | 0 | 12 | 136 | 144 | 0 | 0 | 0 | 0 | 0 |
| SRR2454336 | 0 | 7402 | 4156 | 31 | 5031 | 65781 | 42983 | 27144 | 3481 | 101 | 37736 | 58314 | 6564 | 8677 | 186 | 43699 | 14850 | 0 | 0 | 0 | 59 | 131 | 173 | 0 | 0 | 0 | 0 | 0 | 0 | 12 | 137 | 135 | 0 | 11 | 0 | 0 | 0 |
| SRR2454337 | 0 | 5796 | 4424 | 28 | 5306 | 70588 | 48006 | 30395 | 3555 | 125 | 42722 | 64847 | 6924 | 9490 | 214 | 46588 | 15196 | 0 | 0 | 0 | 50 | 106 | 142 | 0 | 0 | 0 | 0 | 0 | 0 | 12 | 118 | 100 | 0 | 0 | 0 | 0 | 0 |
| SRR2487412 | 11 | 10504 | 4879 | 11 | 559 | 4301 | 3756 | 15598 | 4549 | 16 | 25703 | 21017 | 1956 | 3588 | 0 | 11556 | 49540 | 0 | 0 | 0 | 226 | 378 | 63 | 0 | 0 | 0 | 0 | 0 | 34 | 97 | 389 | 72 | 0 | 0 | 0 | 0 | 0 |
| SRR2487413 | 24 | 10662 | 4836 | 18 | 568 | 4207 | 3717 | 15801 | 4510 | 55 | 25655 | 13 | 1957 | 3765 | 0 | 11556 | 49646 | 0 | 0 | 0 | 226 | 388 | 63 | 0 | 0 | 0 | 0 | 0 | 40 | 89 | 395 | 72 | 0 | 0 | 0 | 0 | 0 |
| SRR2487414 | 26 | 10771 | 5026 | 26 | 575 | 4313 | 3732 | 15804 | 4493 | 31 | 25629 | 20677 | 1915 | 3652 | 0 | 11399 | 49702 | 0 | 0 | 0 | 236 | 386 | 57 | 0 | 0 | 0 | 0 | 0 | 32 | 79 | 404 | 62 | 0 | 0 | 0 | 0 | 0 |
| SRR2487415 | 69 | 8442 | 2107 | 28 | 152 | 1093 | 810 | 3285 | 8986 | 55 | 25348 | 18 | 4380 | 1424 | 0 | 39661 | 49952 | 0 | 0 | 0 | 70 | 102 | 18 | 0 | 0 | 0 | 0 | 0 | 21 | 44 | 194 | 25 | 0 | 0 | 0 | 0 | 0 |
| SRR2487416 | 48 | 8095 | 2200 | 48 | 135 | 1075 | 833 | 3288 | 8960 | 64 | 25367 | 14 | 4353 | 1435 | 0 | 40039 | 51068 | 0 | 0 | 0 | 67 | 103 | 14 | 0 | 0 | 0 | 0 | 0 | 16 | 43 | 190 | 30 | 0 | 0 | 0 | 0 | 0 |
| SRR2487417 | 96 | 8851 | 2339 | 164 | 150 | 1081 | 824 | 3271 | 9003 | 82 | 25089 | 11 | 4429 | 1415 | 0 | 39459 | 50022 | 0 | 0 | 0 | 73 | 99 | 23 | 0 | 0 | 0 | 0 | 0 | 24 | 44 | 200 | 32 | 0 | 0 | 0 | 0 | 0 |
| SRR2487418 | 201 | 11995 | 2552 | 221 | 725 | 5928 | 6051 | 25667 | 10063 | 181 | 28856 | 21793 | 1736 | 3662 | 0 | 13919 | 43574 | 0 | 0 | 0 | 245 | 779 | 98 | 0 | 0 | 0 | 0 | 0 | 25 | 75 | 577 | 58 | 0 | 0 | 0 | 0 | 0 |

### Supp Table S8.1.Columns_B_to_AL

| | | | | | | | | | | | | | | | | | | | | | | | | | | | | | | | | | | | | | |
|---|---|---|---|---|---|---|---|---|---|---|---|---|---|---|---|---|---|---|---|---|---|---|---|---|---|---|---|---|---|---|---|---|---|---|---|---|---|
| SRR2487419 | 184 | 12124 | 2278 | 253 | 701 | 5825 | 6027 | 25930 | 10033 | 138 | 29092 | 22094 | 1829 | 3877 | 0 | 14141 | 44226 | 0 | 0 | 0 | 267 | 733 | 87 | 0 | 0 | 0 | 0 | 0 | 32 | 63 | 536 | 52 | 0 | 0 | 0 | 0 | 0 |
| SRR2487420 | 99 | 12953 | 2467 | 179 | 670 | 5875 | 5868 | 25989 | 10109 | 159 | 28878 | 21906 | 1682 | 3682 | 0 | 13853 | 43349 | 0 | 0 | 0 | 248 | 768 | 98 | 0 | 0 | 0 | 0 | 0 | 32 | 59 | 653 | 52 | 0 | 0 | 0 | 0 | 0 |
| SRR2487422 | 245 | 17457 | 3972 | 49 | 1402 | 13572 | 10524 | 21277 | 6543 | 294 | 33072 | 30774 | 4141 | 3627 | 0 | 41268 | 48451 | 0 | 0 | 0 | 78 | 304 | 21 | 0 | 0 | 0 | 0 | 0 | 17 | 42 | 328 | 25 | 0 | 0 | 0 | 0 | 0 |
| SRR2487423 | 248 | 15948 | 4056 | 138 | 1385 | 13763 | 10363 | 21527 | 6225 | 359 | 33536 | 31102 | 4293 | 3839 | 0 | 41930 | 49319 | 0 | 0 | 0 | 81 | 295 | 29 | 0 | 0 | 0 | 0 | 0 | 22 | 44 | 317 | 25 | 0 | 0 | 0 | 0 | 0 |
| SRR2487424 | 215 | 17972 | 3250 | 191 | 1443 | 13890 | 10429 | 21327 | 6388 | 430 | 33019 | 30598 | 4260 | 3699 | 0 | 40988 | 48934 | 0 | 0 | 0 | 79 | 327 | 24 | 0 | 0 | 0 | 0 | 0 | 19 | 40 | 335 | 23 | 0 | 0 | 0 | 0 | 0 |
| SRR2487426 | 59 | 19995 | 2684 | 106 | 1320 | 17979 | 17290 | 65945 | 9116 | 24 | 28643 | 16 | 2980 | 7797 | 0 | 17718 | 47785 | 0 | 0 | 0 | 283 | 681 | 49 | 0 | 0 | 0 | 0 | 0 | 147 | 117 | 624 | 55 | 0 | 0 | 0 | 0 | 0 |
| SRR2487427 | 121 | 19875 | 2501 | 148 | 1350 | 17915 | 17292 | 66540 | 8900 | 67 | 29200 | 21052 | 3022 | 8103 | 0 | 17885 | 48880 | 0 | 0 | 0 | 272 | 670 | 53 | 0 | 0 | 0 | 0 | 0 | 129 | 115 | 593 | 53 | 0 | 0 | 0 | 0 | 0 |
| SRR2487428 | 105 | 20711 | 2374 | 199 | 1330 | 17981 | 17551 | 65572 | 8959 | 58 | 28685 | 21070 | 2878 | 7954 | 0 | 17674 | 47936 | 0 | 0 | 0 | 311 | 674 | 41 | 0 | 0 | 0 | 0 | 0 | 145 | 111 | 637 | 53 | 0 | 0 | 0 | 0 | 0 |
| SRR2487430 | 148 | 22448 | 3900 | 99 | 778 | 8293 | 6058 | 28206 | 4123 | 49 | 27474 | 23 | 3555 | 4608 | 0 | 28909 | 56409 | 0 | 0 | 0 | 252 | 997 | 189 | 0 | 0 | 0 | 0 | 0 | 44 | 165 | 1413 | 289 | 0 | 13 | 18 | 0 | 0 |
| SRR2487431 | 131 | 20797 | 3751 | 150 | 779 | 8376 | 6019 | 28385 | 4019 | 75 | 27786 | 16 | 3544 | 4702 | 0 | 29332 | 56673 | 0 | 0 | 0 | 235 | 998 | 174 | 0 | 0 | 0 | 0 | 0 | 52 | 162 | 1294 | 262 | 0 | 11 | 13 | 0 | 0 |
| SRR2487432 | 81 | 22324 | 3314 | 97 | 752 | 8337 | 6119 | 28066 | 4129 | 81 | 27486 | 32529 | 3520 | 4569 | 0 | 28980 | 56386 | 0 | 0 | 0 | 245 | 1004 | 194 | 0 | 0 | 0 | 0 | 0 | 49 | 168 | 1415 | 280 | 0 | 16 | 15 | 0 | 0 |
| SRR2487434 | 34 | 10094 | 3916 | 41 | 1179 | 10637 | 9285 | 56547 | 5082 | 223 | 23456 | 17336 | 2896 | 4003 | 0 | 9020 | 53947 | 0 | 0 | 0 | 303 | 2608 | 189 | 0 | 0 | 0 | 0 | 0 | 128 | 108 | 2825 | 165 | 0 | 21 | 0 | 0 | 0 |
| SRR2487435 | 39 | 9880 | 3511 | 16 | 1130 | 10761 | 9224 | 57392 | 4983 | 171 | 23805 | 17307 | 2918 | 4194 | 0 | 9268 | 54806 | 0 | 0 | 0 | 286 | 2529 | 192 | 0 | 0 | 0 | 0 | 0 | 129 | 111 | 2645 | 160 | 0 | 21 | 0 | 0 | 0 |
| SRR2487436 | 20 | 10057 | 3671 | 13 | 1140 | 10799 | 9314 | 56690 | 5021 | 152 | 23447 | 17205 | 2790 | 3969 | 0 | 9013 | 54004 | 0 | 0 | 0 | 286 | 2661 | 197 | 0 | 0 | 0 | 0 | 0 | 143 | 110 | 2911 | 164 | 0 | 20 | 0 | 0 | 0 |
| SRR2487438 | 78 | 11159 | 2779 | 89 | 418 | 3405 | 2682 | 11127 | 8405 | 145 | 21058 | 26005 | 4170 | 2069 | 0 | 38377 | 60116 | 0 | 0 | 0 | 395 | 1657 | 297 | 0 | 0 | 0 | 0 | 13 | 70 | 221 | 2472 | 387 | 0 | 47 | 62 | 12 | 0 |
| SRR2487439 | 64 | 10962 | 2833 | 76 | 403 | 3441 | 2616 | 11164 | 8214 | 254 | 21172 | 26236 | 4258 | 2135 | 0 | 39056 | 60198 | 0 | 0 | 0 | 389 | 1595 | 272 | 0 | 0 | 0 | 0 | 13 | 68 | 235 | 2300 | 382 | 0 | 50 | 63 | 0 | 0 |
| SRR2487441 | 719 | 34233 | 1490 | 810 | 703 | 4848 | 4553 | 15084 | 3160 | 78 | 37085 | 10 | 2594 | 5055 | 0 | 20984 | 58811 | 0 | 0 | 0 | 207 | 446 | 85 | 0 | 0 | 0 | 0 | 0 | 50 | 58 | 397 | 54 | 0 | 0 | 0 | 0 | 0 |
| SRR2487443 | 510 | 34580 | 1397 | 1128 | 686 | 5042 | 4576 | 15524 | 3110 | 67 | 37399 | 33952 | 2585 | 5158 | 0 | 21417 | 56559 | 0 | 0 | 0 | 229 | 483 | 84 | 0 | 0 | 0 | 0 | 0 | 60 | 63 | 394 | 58 | 0 | 0 | 0 | 0 | 0 |
| SRR2487445 | 741 | 35390 | 1583 | 691 | 701 | 4812 | 4575 | 15356 | 3100 | 168 | 38545 | 11 | 2680 | 5357 | 0 | 20409 | 57363 | 0 | 0 | 0 | 209 | 424 | 84 | 0 | 0 | 0 | 0 | 0 | 56 | 51 | 365 | 51 | 0 | 0 | 0 | 0 | 0 |
| SRR2487447 | 665 | 34615 | 1480 | 765 | 679 | 4733 | 4311 | 15788 | 2982 | 100 | 39357 | 35360 | 2762 | 5516 | 0 | 21091 | 57368 | 0 | 0 | 0 | 184 | 442 | 77 | 0 | 0 | 0 | 0 | 0 | 50 | 53 | 333 | 54 | 0 | 0 | 0 | 0 | 0 |
| SRR2487449 | 1087 | 10541 | 1367 | 482 | 174 | 1012 | 952 | 3473 | 4505 | 45 | 22259 | 28875 | 3787 | 1686 | 0 | 59461 | 57807 | 0 | 0 | 0 | 806 | 1150 | 151 | 0 | 0 | 0 | 0 | 82 | 344 | 535 | 2374 | 295 | 0 | 61 | 25 | 0 | 0 |
| SRR2487451 | 871 | 10684 | 1479 | 481 | 176 | 1053 | 956 | 3496 | 4481 | 23 | 22366 | 28989 | 3956 | 1747 | 0 | 60877 | 56694 | 0 | 0 | 0 | 801 | 1205 | 152 | 0 | 0 | 0 | 0 | 89 | 337 | 516 | 2421 | 292 | 0 | 65 | 21 | 0 | 0 |
| SRR2487453 | 790 | 10581 | 1536 | 388 | 156 | 1033 | 947 | 3550 | 4369 | 14 | 22998 | 30116 | 3931 | 1755 | 0 | 58991 | 57422 | 0 | 0 | 0 | 736 | 1106 | 147 | 0 | 0 | 0 | 0 | 90 | 295 | 485 | 2191 | 289 | 0 | 64 | 18 | 0 | 0 |
| SRR2487455 | 1120 | 10887 | 1503 | 383 | 162 | 1054 | 907 | 3646 | 4399 | 28 | 23336 | 30310 | 4046 | 1822 | 0 | 59936 | 57158 | 0 | 0 | 0 | 749 | 1095 | 144 | 0 | 0 | 0 | 0 | 80 | 304 | 464 | 2102 | 264 | 0 | 65 | 17 | 0 | 0 |
| SRR2487457 | 786 | 21233 | 2127 | 217 | 573 | 5405 | 5557 | 26717 | 14731 | 27 | 31444 | 24535 | 3028 | 4279 | 0 | 31866 | 57036 | 0 | 0 | 20 | 2513 | 1732 | 211 | 0 | 12 | 0 | 0 | 16 | 217 | 275 | 690 | 86 | 0 | 28 | 0 | 0 | 0 |
| SRR2487459 | 983 | 22219 | 1766 | 299 | 608 | 5612 | 5834 | 26643 | 14708 | 0 | 32170 | 24383 | 3247 | 4396 | 0 | 32566 | 54509 | 0 | 0 | 19 | 2648 | 1747 | 234 | 0 | 13 | 0 | 0 | 13 | 229 | 305 | 670 | 87 | 0 | 26 | 0 | 0 | 0 |
| SRR2487461 | 902 | 21892 | 2020 | 234 | 623 | 5490 | 5444 | 26562 | 14256 | 18 | 33319 | 12 | 3183 | 4537 | 0 | 31372 | 54583 | 0 | 0 | 12 | 2366 | 1563 | 222 | 0 | 0 | 0 | 0 | 14 | 201 | 251 | 609 | 89 | 0 | 29 | 0 | 0 | 0 |
| SRR2487463 | 919 | 22257 | 2245 | 194 | 568 | 5363 | 5483 | 27194 | 14426 | 0 | 33443 | 26058 | 3279 | 4675 | 0 | 31727 | 55084 | 0 | 0 | 16 | 2332 | 1490 | 196 | 0 | 11 | 0 | 0 | 0 | 221 | 246 | 641 | 83 | 0 | 26 | 0 | 0 | 0 |
| SRR2487465 | 531 | 8703 | 2441 | 316 | 115 | 728 | 649 | 2569 | 6609 | 50 | 25224 | 21829 | 3413 | 1753 | 0 | 58655 | 55007 | 0 | 0 | 0 | 474 | 600 | 219 | 0 | 57 | 0 | 0 | 54 | 183 | 319 | 1055 | 325 | 0 | 164 | 195 | 16 | 0 |
| SRR2487467 | 412 | 7839 | 1887 | 395 | 87 | 729 | 657 | 2641 | 6609 | 0 | 25399 | 21876 | 3482 | 1767 | 0 | 60008 | 52197 | 0 | 0 | 0 | 492 | 565 | 236 | 0 | 53 | 0 | 0 | 59 | 185 | 356 | 1042 | 335 | 0 | 172 | 209 | 11 | 0 |
| SRR2487469 | 709 | 9263 | 1934 | 365 | 103 | 772 | 619 | 2663 | 6279 | 43 | 26205 | 22962 | 3440 | 1837 | 0 | 57402 | 53780 | 0 | 0 | 0 | 466 | 556 | 211 | 0 | 61 | 0 | 0 | 57 | 168 | 314 | 966 | 311 | 0 | 156 | 194 | 11 | 0 |
| SRR2487471 | 492 | 8591 | 2180 | 299 | 97 | 687 | 645 | 2582 | 6258 | 85 | 26521 | 23011 | 3554 | 1824 | 0 | 58354 | 53417 | 0 | 0 | 0 | 442 | 537 | 192 | 0 | 57 | 0 | 0 | 53 | 166 | 316 | 939 | 277 | 0 | 169 | 182 | 10 | 0 |
| SRR2487473 | 576 | 34632 | 1797 | 475 | 471 | 4575 | 3876 | 18163 | 5824 | 34 | 27261 | 18012 | 1376 | 2674 | 0 | 39215 | 43981 | 0 | 0 | 0 | 198 | 64 | 18 | 0 | 0 | 0 | 0 | 0 | 42 | 30 | 46 | 14 | 0 | 0 | 0 | 0 | 0 |
| SRR2487475 | 470 | 33728 | 1689 | 766 | 495 | 4677 | 3924 | 18361 | 5690 | 70 | 27549 | 17827 | 1392 | 2749 | 0 | 40254 | 42736 | 0 | 0 | 0 | 198 | 64 | 19 | 0 | 0 | 0 | 0 | 0 | 40 | 31 | 47 | 15 | 0 | 0 | 0 | 0 | 0 |
| SRR2487477 | 525 | 34997 | 2211 | 569 | 488 | 4502 | 3822 | 18310 | 5334 | 66 | 28306 | 18658 | 1446 | 2718 | 0 | 38844 | 43026 | 0 | 0 | 0 | 185 | 61 | 16 | 0 | 0 | 0 | 0 | 0 | 40 | 28 | 48 | 10 | 0 | 0 | 0 | 0 | 0 |
| SRR2487479 | 630 | 35312 | 1717 | 608 | 445 | 4544 | 3835 | 18696 | 5346 | 109 | 28669 | 18837 | 1449 | 2837 | 0 | 39421 | 43329 | 0 | 0 | 0 | 177 | 60 | 20 | 0 | 0 | 0 | 0 | 0 | 39 | 26 | 45 | 0 | 0 | 0 | 0 | 0 | 0 |
| SRR2487481 | 364 | 5979 | 1092 | 322 | 222 | 1641 | 1299 | 5220 | 5162 | 56 | 25660 | 31977 | 4566 | 2166 | 0 | 67067 | 51141 | 0 | 0 | 0 | 534 | 304 | 44 | 0 | 0 | 0 | 0 | 50 | 186 | 296 | 529 | 129 | 0 | 0 | 0 | 0 | 0 |
| SRR2487483 | 589 | 6577 | 1321 | 273 | 216 | 1745 | 1420 | 5349 | 5265 | 29 | 25974 | 31762 | 4666 | 2080 | 0 | 69041 | 49349 | 0 | 0 | 0 | 570 | 307 | 64 | 0 | 0 | 0 | 0 | 48 | 202 | 318 | 526 | 112 | 0 | 0 | 0 | 0 | 0 |
| SRR2487485 | 541 | 6057 | 919 | 198 | 215 | 1645 | 1398 | 5311 | 4887 | 18 | 26638 | 33501 | 4704 | 2173 | 0 | 66970 | 50268 | 0 | 0 | 0 | 502 | 269 | 46 | 0 | 0 | 0 | 0 | 40 | 178 | 270 | 475 | 103 | 0 | 0 | 0 | 0 | 0 |
| SRR2487487 | 627 | 6613 | 1201 | 448 | 190 | 1688 | 1347 | 5495 | 5033 | 72 | 26820 | 11 | 4972 | 2134 | 0 | 67766 | 49448 | 0 | 0 | 0 | 505 | 279 | 46 | 0 | 0 | 0 | 0 | 44 | 146 | 292 | 466 | 99 | 0 | 0 | 0 | 0 | 0 |
| SRR2487489 | 1668 | 77769 | 12646 | 306 | 1091 | 15534 | 15731 | 49174 | 11374 | 22 | 19211 | 11434 | 2219 | 11525 | 0 | 25014 | 31171 | 0 | 10 | 54 | 2920 | 4815 | 201 | 0 | 0 | 0 | 0 | 70 | 998 | 738 | 3498 | 237 | 0 | 0 | 0 | 0 | 0 |
| SRR2487491 | 1818 | 77865 | 12766 | 316 | 1240 | 16343 | 16710 | 50101 | 11176 | 0 | 19262 | 11517 | 2253 | 11983 | 0 | 25850 | 30221 | 0 | 10 | 48 | 2932 | 4921 | 231 | 0 | 0 | 0 | 0 | 80 | 995 | 742 | 3511 | 238 | 0 | 0 | 0 | 0 | 0 |
| SRR2487493 | 1577 | 78181 | 13188 | 326 | 1120 | 16108 | 16376 | 49516 | 10361 | 16 | 19931 | 12463 | 2412 | 12058 | 0 | 24559 | 31254 | 0 | 0 | 45 | 2645 | 4484 | 203 | 0 | 0 | 0 | 0 | 81 | 884 | 673 | 3100 | 232 | 0 | 0 | 0 | 0 | 0 |
| SRR2487495 | 1744 | 79352 | 13369 | 308 | 1044 | 16208 | 15561 | 51532 | 10880 | 12 | 19936 | 12440 | 2277 | 12484 | 0 | 25658 | 31639 | 0 | 0 | 48 | 2641 | 4448 | 202 | 0 | 0 | 0 | 0 | 59 | 854 | 675 | 3111 | 196 | 0 | 0 | 0 | 0 | 0 |
| SRR2487497 | 4493 | 53445 | 2135 | 258 | 512 | 4434 | 4714 | 5089 | 3687 | 52 | 19423 | 21013 | 3836 | 2265 | 0 | 55348 | 51423 | 0 | 15 | 12 | 2488 | 2914 | 293 | 0 | 0 | 0 | 0 | 168 | 518 | 1113 | 2805 | 299 | 0 | 0 | 0 | 0 | 0 |
| SRR2487499 | 4419 | 53489 | 1908 | 231 | 461 | 4649 | 4903 | 5071 | 3710 | 0 | 19435 | 20526 | 3943 | 2275 | 0 | 57164 | 49776 | 0 | 16 | 13 | 2559 | 3053 | 301 | 0 | 0 | 0 | 0 | 172 | 524 | 1115 | 2725 | 287 | 0 | 0 | 10 | 0 | 0 |
| SRR2487501 | 3895 | 55128 | 2058 | 255 | 482 | 4302 | 4812 | 4982 | 3432 | 66 | 20143 | 21523 | 3992 | 2307 | 0 | 55042 | 50927 | 0 | 13 | 11 | 2304 | 2756 | 280 | 0 | 0 | 0 | 0 | 168 | 470 | 1005 | 2570 | 295 | 0 | 0 | 10 | 0 | 0 |
| SRR2487503 | 4216 | 54336 | 1937 | 242 | 437 | 4324 | 4633 | 5080 | 3517 | 33 | 20550 | 21960 | 3998 | 2394 | 0 | 55851 | 50600 | 0 | 10 | 0 | 2303 | 2727 | 281 | 0 | 0 | 0 | 0 | 157 | 486 | 1011 | 2496 | 272 | 0 | 0 | 0 | 0 | 0 |
| SRR2487504 | 540 | 25069 | 1259 | 660 | 823 | 5319 | 7145 | 26544 | 6564 | 120 | 28214 | 17405 | 2026 | 4474 | 0 | 17451 | 39507 | 0 | 0 | 15 | 926 | 558 | 372 | 0 | 168 | 0 | 0 | 0 | 119 | 141 | 335 | 231 | 0 | 277 | 40 | 0 | 0 |
| SRR2487505 | 826 | 46608 | 1901 | 826 | 666 | 4949 | 6507 | 30005 | 5753 | 83 | 35851 | 21406 | 2484 | 5392 | 0 | 20370 | 43339 | 0 | 0 | 12 | 957 | 609 | 435 | 0 | 153 | 0 | 0 | 0 | 134 | 151 | 295 | 215 | 0 | 255 | 58 | 0 | 0 |
| SRR2487507 | 703 | 43804 | 1612 | 1033 | 717 | 5133 | 6492 | 31419 | 6010 | 0 | 36242 | 21246 | 2623 | 5777 | 0 | 20451 | 43345 | 0 | 0 | 19 | 933 | 593 | 461 | 15 | 163 | 0 | 0 | 0 | 130 | 134 | 304 | 218 | 0 | 243 | 49 | 0 | 0 |
| SRR2487508 | 253 | 27184 | 1897 | 1138 | 858 | 5498 | 7281 | 27568 | 6518 | 0 | 28870 | 18189 | 2075 | 4604 | 0 | 17998 | 40867 | 0 | 0 | 15 | 851 | 564 | 433 | 13 | 160 | 0 | 0 | 0 | 122 | 130 | 304 | 197 | 0 | 328 | 54 | 0 | 0 |
| SRR2487509 | 66 | 28498 | 1649 | 1055 | 777 | 5362 | 6683 | 27508 | 6500 | 66 | 27900 | 16602 | 2163 | 4447 | 0 | 18132 | 40334 | 0 | 0 | 0 | 764 | 502 | 405 | 12 | 145 | 0 | 0 | 0 | 110 | 125 | 237 | 239 | 0 | 264 | 44 | 0 | 0 |
| SRR2487510 | 338 | 25738 | 1824 | 1216 | 705 | 5333 | 6863 | 28013 | 6540 | 68 | 28536 | 17195 | 1933 | 4621 | 0 | 18572 | 40545 | 0 | 0 | 11 | 732 | 504 | 376 | 0 | 132 | 0 | 0 | 0 | 118 | 120 | 244 | 159 | 0 | 249 | 48 | 0 | 0 |
| SRR2487512 | 639 | 43153 | 1502 | 991 | 808 | 5373 | 6593 | 29880 | 6167 | 32 | 33789 | 20077 | 2496 | 5300 | 0 | 20041 | 44317 | 0 | 0 | 15 | 1024 | 682 | 430 | 0 | 163 | 0 | 0 | 0 | 152 | 150 | 344 | 248 | 0 | 266 | 50 | 0 | 0 |
| SRR2487514 | 532 | 43341 | 832 | 1464 | 806 | 5534 | 6759 | 30809 | 6016 | 67 | 34118 | 20252 | 2490 | 5221 | 0 | 20783 | 43067 | 0 | 0 | 25 | 992 | 691 | 494 | 13 | 150 | 0 | 0 | 0 | 143 | 166 | 380 | 270 | 0 | 276 | 51 | 0 | 0 |
| SRR2487516 | 0 | 6835 | 1859 | 151 | 295 | 1939 | 1675 | 6501 | 5660 | 0 | 21583 | 14 | 4349 | 1668 | 0 | 46704 | 49410 | 0 | 0 | 0 | 270 | 418 | 31 | 0 | 0 | 0 | 0 | 22 | 63 | 111 | 733 | 88 | 0 | 0 | 0 | 0 | 0 |
| SRR2487517 | 163 | 11555 | 1893 | 359 | 294 | 1817 | 1503 | 7159 | 5032 | 0 | 26226 | 30319 | 5173 | 1927 | 0 | 54313 | 54140 | 0 | 0 | 0 | 256 | 411 | 36 | 0 | 0 | 0 | 0 | 16 | 61 | 133 | 746 | 92 | 0 | 0 | 0 | 0 | 0 |
| SRR2487519 | 291 | 10682 | 1651 | 356 | 222 | 1856 | 1489 | 7012 | 5196 | 65 | 26081 | 30651 | 5285 | 2088 | 0 | 54779 | 53568 | 0 | 0 | 0 | 252 | 422 | 44 | 0 | 0 | 0 | 0 | 13 | 81 | 153 | 753 | 87 | 0 | 0 | 0 | 0 | 0 |
| SRR2487520 | 0 | 5699 | 2105 | 257 | 255 | 2024 | 1727 | 6307 | 5612 | 103 | 21876 | 26687 | 4542 | 1724 | 0 | 48365 | 50853 | 0 | 0 | 0 | 254 | 349 | 34 | 0 | 0 | 0 | 0 | 0 | 69 | 131 | 704 | 94 | 0 | 0 | 0 | 0 | 0 |
| SRR2487521 | 279 | 6796 | 1838 | 223 | 266 | 1861 | 1607 | 6393 | 5755 | 167 | 20884 | 12 | 3919 | 1728 | 0 | 46673 | 49809 | 0 | 0 | 0 | 219 | 402 | 38 | 0 | 0 | 0 | 0 | 11 | 59 | 118 | 625 | 75 | 0 | 0 | 0 | 0 | 0 |
| SRR2487522 | 169 | 6014 | 2080 | 169 | 271 | 1788 | 1649 | 6367 | 5820 | 56 | 21576 | 25464 | 4493 | 1663 | 0 | 47678 | 49893 | 0 | 0 | 0 | 220 | 354 | 24 | 0 | 0 | 0 | 0 | 11 | 55 | 120 | 644 | 68 | 0 | 0 | 0 | 0 | 0 |
| SRR2487524 | 406 | 10201 | 1599 | 254 | 242 | 1825 | 1551 | 6774 | 5473 | 0 | 25227 | 29072 | 5056 | 1966 | 0 | 54452 | 54345 | 0 | 0 | 0 | 261 | 449 | 44 | 0 | 0 | 0 | 0 | 14 | 81 | 137 | 859 | 100 | 0 | 0 | 0 | 0 | 0 |
| SRR2487526 | 469 | 10810 | 1901 | 469 | 274 | 1855 | 1703 | 7008 | 5258 | 52 | 25604 | 29028 | 4997 | 2017 | 0 | 56150 | 52159 | 0 | 0 | 0 | 285 | 470 | 58 | 0 | 0 | 0 | 0 | 12 | 89 | 159 | 808 | 82 | 0 | 0 | 0 | 0 | 0 |
| SRR2487529 | 257 | 25666 | 1543 | 239 | 429 | 3591 | 3855 | 15596 | 7777 | 71 | 22202 | 24370 | 2730 | 2743 | 0 | 29598 | 48625 | 0 | 0 | 0 | 322 | 1144 | 183 | 0 | 0 | 0 | 0 | 0 | 71 | 73 | 893 | 120 | 0 | 13 | 0 | 0 | 0 |
| SRR2487531 | 348 | 25136 | 1401 | 211 | 447 | 3693 | 3928 | 15942 | 7694 | 82 | 22429 | 24017 | 2847 | 2726 | 0 | 30501 | 46531 | 0 | 0 | 0 | 326 | 1184 | 181 | 0 | 10 | 0 | 0 | 0 | 76 | 70 | 878 | 131 | 0 | 12 | 0 | 0 | 0 |
| SRR2487533 | 340 | 25480 | 1724 | 181 | 432 | 3604 | 3926 | 15687 | 7382 | 23 | 23230 | 25726 | 2874 | 2814 | 0 | 29753 | 47792 | 0 | 0 | 0 | 301 | 1049 | 168 | 0 | 0 | 0 | 0 | 0 | 68 | 73 | 823 | 113 | 0 | 11 | 0 | 0 | 0 |
| SRR2487535 | 347 | 26170 | 1837 | 258 | 422 | 3588 | 3789 | 15930 | 7563 | 45 | 23548 | 25531 | 3002 | 2892 | 0 | 29740 | 47489 | 0 | 0 | 0 | 290 | 1035 | 168 | 0 | 0 | 0 | 0 | 0 | 64 | 69 | 771 | 115 | 0 | 12 | 0 | 0 | 0 |
| SRR2487537 | 1169 | 13720 | 1487 | 304 | 188 | 1415 | 1348 | 6029 | 5277 | 15 | 21131 | 24977 | 4396 | 1828 | 0 | 51759 | 52597 | 0 | 0 | 0 | 1152 | 2118 | 303 | 0 | 0 | 0 | 0 | 54 | 285 | 744 | 3752 | 476 | 0 | 0 | 0 | 0 | 0 |
| SRR2487539 | 1450 | 14229 | 1198 | 299 | 214 | 1460 | 1351 | 6080 | 5279 | 16 | 21421 | 24833 | 4415 | 1923 | 0 | 53329 | 50822 | 0 | 0 | 0 | 1196 | 2274 | 298 | 0 | 0 | 0 | 0 | 55 | 293 | 795 | 3699 | 465 | 0 | 0 | 0 | 0 | 0 |
| SRR2487541 | 1251 | 14270 | 1662 | 293 | 193 | 1412 | 1328 | 5864 | 5057 | 39 | 21874 | 26059 | 4624 | 2026 | 0 | 51941 | 51513 | 0 | 0 | 0 | 1079 | 2014 | 277 | 0 | 0 | 0 | 0 | 51 | 263 | 675 | 3509 | 456 | 0 | 0 | 0 | 0 | 0 |
| SRR2487543 | 1278 | 14702 | 1646 | 291 | 202 | 1406 | 1295 | 6186 | 5044 | 0 | 22289 | 26334 | 4571 | 1972 | 0 | 52031 | 51729 | 0 | 0 | 0 | 1072 | 1997 | 288 | 0 | 0 | 0 | 0 | 54 | 270 | 708 | 3278 | 429 | 0 | 0 | 0 | 0 | 0 |
| SRR2487545 | 550 | 36845 | 2568 | 397 | 579 | 6212 | 5263 | 23199 | 6730 | 61 | 31846 | 24232 | 2226 | 4215 | 0 | 28657 | 53854 | 0 | 0 | 40 | 4773 | 5418 | 1744 | 11 | 164 | 0 | 0 | 0 | 527 | 1026 | 4357 | 1241 | 0 | 267 | 61 | 11 | 0 |
| SRR2487547 | 467 | 36825 | 2274 | 436 | 589 | 6484 | 5454 | 23785 | 6694 | 31 | 31769 | 24328 | 2330 | 4252 | 0 | 29317 | 51486 | 0 | 0 | 46 | 4936 | 5490 | 1749 | 12 | 178 | 0 | 0 | 0 | 552 | 1044 | 4412 | 1214 | 0 | 268 | 67 | 12 | 0 |
| SRR2487549 | 510 | 38102 | 2668 | 235 | 628 | 6142 | 5457 | 24202 | 6331 | 118 | 33173 | 25517 | 2331 | 4361 | 0 | 28936 | 53442 | 0 | 0 | 41 | 4470 | 5103 | 1687 | 10 | 158 | 0 | 0 | 0 | 489 | 956 | 3976 | 1175 | 0 | 255 | 63 | 0 | 0 |
| SRR2487551 | 506 | 38166 | 2765 | 273 | 605 | 6145 | 5134 | 23997 | 6259 | 39 | 33746 | 25591 | 2423 | 4631 | 0 | 28637 | 53190 | 0 | 0 | 33 | 4424 | 4953 | 1581 | 0 | 165 | 0 | 0 | 0 | 473 | 903 | 3823 | 1111 | 0 | 235 | 57 | 15 | 0 |
| SRR2487553 | 2063 | 15856 | 121 | 1315 | 793 | 4780 | 2809 | 7583 | 5228 | 0 | 18506 | 18001 | 3065 | 1165 | 0 | 99576 | 43181 | 0 | 0 | 18 | 6553 | 5870 | 1915 | 12 | 22 | 0 | 0 | 84 | 835 | 3577 | 10070 | 3004 | 10 | 93 | 82 | 24 | 0 |
| SRR2487555 | 1966 | 16512 | 103 | 766 | 741 | 5377 | 3108 | 7295 | 5215 | 0 | 19142 | 17666 | 3316 | 1222 | 0 | 102576 | 41841 | 0 | 10 | 18 | 6738 | 5862 | 1967 | 0 | 27 | 0 | 0 | 83 | 812 | 3711 | 9719 | 3035 | 0 | 86 | 79 | 19 | 0 |
| SRR2487560 | 1038 | 32718 | 3251 | 69 | 1410 | 10477 | 12320 | 42266 | 7343 | 69 | 29904 | 26285 | 2236 | 3320 | 0 | 29983 | 44210 | 0 | 0 | 15 | 10714 | 5115 | 1164 | 0 | 32 | 0 | 0 | 0 | 157 | 2007 | 3685 | 766 | 0 | 39 | 53 | 18 | 0 |
| SRR2487561 | 3991 | 49768 | 189 | 605 | 2352 | 16819 | 14026 | 38359 | 5289 | 19 | 32170 | 26184 | 2250 | 3439 | 0 | 52190 | 40915 | 0 | 0 | 25 | 24378 | 10052 | 2399 | 0 | 28 | 0 | 0 | 0 | 430 | 4486 | 7165 | 1440 | 0 | 60 | 72 | 26 | 0 |
| SRR2487563 | 4195 | 49977 | 155 | 602 | 2319 | 16599 | 13974 | 41436 | 5852 | 19 | 32926 | 26290 | 2171 | 3343 | 0 | 53132 | 43062 | 0 | 0 | 32 | 24172 | 9927 | 2267 | 0 | 24 | 0 | 0 | 0 | 419 | 4486 | 6723 | 1429 | 0 | 65 | 50 | 20 | 0 |
| SRR2487564 | 819 | 31406 | 3572 | 298 | 1531 | 10853 | 12133 | 42757 | 7183 | 74 | 31035 | 26942 | 2299 | 3529 | 0 | 30991 | 45436 | 0 | 0 | 15 | 10662 | 4777 | 1104 | 0 | 27 | 0 | 0 | 0 | 180 | 2064 | 3493 | 776 | 0 | 46 | 53 | 15 | 0 |
| SRR2487565 | 691 | 35451 | 2149 | 614 | 1426 | 10112 | 11927 | 41706 | 6921 | 0 | 29509 | 25538 | 2190 | 3248 | 0 | 30390 | 44098 | 0 | 0 | 15 | 9791 | 4725 | 1061 | 0 | 22 | 0 | 0 | 0 | 161 | 1889 | 3457 | 801 | 0 | 36 | 60 | 25 | 0 |
| SRR2487566 | 865 | 31596 | 2751 | 0 | 1429 | 10517 | 12225 | 43328 | 7282 | 0 | 29682 | 26364 | 2409 | 3428 | 0 | 30949 | 43375 | 0 | 0 | 11 | 9534 | 4506 | 1018 | 0 | 22 | 0 | 0 | 0 | 151 | 1801 | 3057 | 711 | 0 | 30 | 44 | 28 | 0 |
| SRR2487568 | 3543 | 46965 | 294 | 514 | 2400 | 16696 | 14001 | 40057 | 6260 | 0 | 30313 | 25123 | 2126 | 3202 | 0 | 53070 | 41654 | 0 | 0 | 33 | 26297 | 10876 | 2467 | 11 | 23 | 0 | 0 | 0 | 466 | 4837 | 7831 | 1555 | 0 | 61 | 60 | 23 | 0 |
| SRR2487570 | 4052 | 47011 | 320 | 609 | 2426 | 17282 | 14370 | 39690 | 6064 | 46 | 30721 | 25101 | 2383 | 3377 | 0 | 54447 | 40592 | 0 | 0 | 33 | 26593 | 11033 | 2538 | 12 | 31 | 0 | 0 | 0 | 460 | 4983 | 7577 | 1550 | 0 | 59 | 69 | 25 | 0 |
| SRR2487572 | 1266 | 14608 | 4058 | 325 | 242 | 1685 | 1369 | 4903 | 5609 | 130 | 19771 | 20047 | 3292 | 1386 | 0 | 57651 | 45010 | 0 | 0 | 0 | 3461 | 3035 | 1036 | 0 | 30 | 0 | 0 | 0 | 172 | 2045 | 6141 | 2066 | 0 | 128 | 184 | 49 | 0 |
| SRR2487573 | 6234 | 27411 | 358 | 954 | 368 | 2488 | 1432 | 4665 | 3939 | 0 | 20988 | 19715 | 2997 | 1263 | 0 | 93497 | 42192 | 0 | 0 | 17 | 8912 | 6661 | 2342 | 0 | 54 | 0 | 0 | 14 | 445 | 4967 | 12632 | 4162 | 0 | 200 | 237 | 42 | 0 |
| SRR2487575 | 5699 | 27115 | 420 | 690 | 420 | 2486 | 1472 | 4580 | 4168 | 0 | 21070 | 19223 | 3172 | 1408 | 0 | 96088 | 41170 | 0 | 0 | 0 | 8825 | 6632 | 2225 | 0 | 54 | 0 | 0 | 13 | 456 | 5049 | 12016 | 3945 | 11 | 167 | 246 | 41 | 0 |
| SRR2487576 | 1072 | 15176 | 4355 | 503 | 254 | 1640 | 1473 | 5234 | 5586 | 168 | 20408 | 20186 | 3368 | 1462 | 0 | 59074 | 46566 | 0 | 0 | 0 | 3494 | 3004 | 1025 | 0 | 32 | 0 | 0 | 0 | 152 | 2099 | 5830 | 2026 | 0 | 133 | 173 | 37 | 0 |
| SRR2487577 | 834 | 14750 | 3987 | 471 | 219 | 1464 | 1325 | 5118 | 5449 | 181 | 19403 | 12 | 3236 | 1406 | 0 | 56927 | 46071 | 0 | 0 | 0 | 3175 | 2753 | 969 | 0 | 31 | 0 | 0 | 0 | 135 | 1916 | 5349 | 1894 | 0 | 129 | 161 | 33 | 0 |
| SRR2487578 | 1167 | 12403 | 4487 | 474 | 219 | 1553 | 1324 | 5246 | 5434 | 109 | 19856 | 19927 | 3240 | 1468 | 0 | 59029 | 45697 | 0 | 0 | 0 | 3221 | 2646 | 957 | 0 | 31 | 0 | 0 | 0 | 156 | 1821 | 5111 | 1754 | 0 | 130 | 172 | 32 | 0 |
| SRR2487580 | 5419 | 26234 | 295 | 726 | 295 | 2406 | 1417 | 4894 | 4451 | 0 | 19995 | 18335 | 2997 | 1550 | 0 | 96943 | 43342 | 0 | 0 | 13 | 9534 | 7207 | 2467 | 0 | 55 | 0 | 0 | 10 | 462 | 5343 | 14003 | 4391 | 12 | 183 | 234 | 53 | 0 |
| SRR2487582 | 5261 | 24615 | 232 | 904 | 305 | 2521 | 1531 | 4654 | 4471 | 23 | 19958 | 18335 | 3146 | 1143 | 0 | 97662 | 41538 | 0 | 0 | 13 | 9770 | 7458 | 2491 | 0 | 48 | 0 | 0 | 14 | 525 | 5609 | 13744 | 4515 | 11 | 189 | 262 | 47 | 0 |
| SRR2487584 | 2620 | 9951 | 14483 | 221 | 1062 | 9923 | 10926 | 32788 | 6646 | 192 | 28512 | 28157 | 2977 | 7439 | 0 | 18401 | 46587 | 0 | 0 | 0 | 558 | 601 | 65 | 0 | 0 | 0 | 0 | 0 | 51 | 141 | 445 | 78 | 0 | 0 | 0 | 0 | 0 |
| SRR2487585 | 11752 | 15098 | 954 | 398 | 2081 | 15573 | 13179 | 30833 | 5862 | 50 | 34055 | 32108 | 3379 | 7451 | 0 | 34882 | 46875 | 0 | 0 | 0 | 1059 | 907 | 135 | 0 | 12 | 0 | 0 | 35 | 123 | 255 | 795 | 127 | 0 | 0 | 0 | 0 | 0 |
| SRR2487587 | 11424 | 15760 | 1013 | 426 | 2175 | 16226 | 12196 | 32218 | 5629 | 19 | 34499 | 30021 | 3049 | 7527 | 0 | 35224 | 46780 | 0 | 0 | 0 | 940 | 916 | 117 | 0 | 0 | 0 | 0 | 27 | 122 | 252 | 732 | 112 | 0 | 0 | 0 | 0 | 0 |
| SRR2487588 | 2517 | 9432 | 15446 | 190 | 1044 | 10045 | 11121 | 33085 | 6859 | 225 | 29368 | 17 | 3128 | 8184 | 0 | 18247 | 47385 | 0 | 0 | 0 | 563 | 575 | 68 | 0 | 0 | 0 | 0 | 12 | 54 | 158 | 468 | 66 | 0 | 0 | 0 | 0 | 0 |
| SRR2487589 | 2478 | 10366 | 15610 | 114 | 1013 | 9690 | 10451 | 33276 | 6713 | 203 | 28262 | 28063 | 2856 | 7411 | 0 | 17804 | 47038 | 0 | 0 | 0 | 475 | 528 | 76 | 0 | 0 | 0 | 0 | 11 | 58 | 135 | 392 | 78 | 0 | 0 | 0 | 0 | 0 |
| SRR2487590 | 2498 | 9886 | 15627 | 256 | 1099 | 9815 | 10590 | 33441 | 6991 | 248 | 29044 | 28378 | 2960 | 7719 | 0 | 18628 | 47172 | 0 | 0 | 0 | 500 | 534 | 64 | 0 | 0 | 0 | 0 | 10 | 40 | 140 | 392 | 74 | 0 | 0 | 0 | 0 | 0 |
| SRR2487592 | 11289 | 14936 | 908 | 408 | 1923 | 16438 | 12948 | 32200 | 6269 | 21 | 31719 | 29083 | 3722 | 7070 | 0 | 35388 | 49262 | 0 | 0 | 0 | 1104 | 989 | 170 | 0 | 0 | 0 | 0 | 29 | 105 | 289 | 876 | 140 | 0 | 0 | 0 | 0 | 0 |
| SRR2487594 | 10966 | 14894 | 1035 | 459 | 2182 | 16585 | 13590 | 33761 | 6120 | 26 | 31524 | 29897 | 3088 | 7581 | 0 | 35702 | 45169 | 0 | 0 | 0 | 1123 | 1044 | 162 | 0 | 0 | 0 | 0 | 31 | 112 | 339 | 817 | 127 | 0 | 0 | 0 | 0 | 0 |
| SRR2487597 | 1152 | 6806 | 5189 | 518 | 370 | 2289 | 2027 | 6280 | 3602 | 74 | 24773 | 26 | 5879 | 2182 | 0 | 51928 | 54115 | 0 | 0 | 0 | 411 | 308 | 55 | 0 | 0 | 0 | 0 | 26 | 113 | 185 | 389 | 79 | 0 | 11 | 0 | 0 | 0 |
| SRR2487598 | 1298 | 7463 | 4742 | 429 | 350 | 2111 | 1925 | 6185 | 3503 | 79 | 23500 | 30 | 5432 | 1900 | 0 | 50815 | 52910 | 0 | 0 | 0 | 375 | 273 | 61 | 0 | 0 | 0 | 0 | 22 | 113 | 160 | 373 | 70 | 0 | 0 | 0 | 0 | 0 |
| SRR2487599 | 1343 | 7296 | 4940 | 376 | 344 | 2213 | 1971 | 6480 | 3473 | 91 | 24267 | 26 | 5515 | 2038 | 0 | 51906 | 52725 | 0 | 0 | 0 | 385 | 272 | 60 | 0 | 0 | 0 | 0 | 21 | 106 | 165 | 362 | 71 | 0 | 0 | 0 | 0 | 0 |
| SRR2487600 | 6082 | 12431 | 425 | 1087 | 600 | 3175 | 2112 | 5691 | 2914 | 32 | 24198 | 34293 | 5250 | 2119 | 0 | 84344 | 49530 | 0 | 0 | 0 | 1601 | 974 | 165 | 0 | 0 | 0 | 0 | 109 | 496 | 642 | 1364 | 201 | 0 | 24 | 13 | 0 | 0 |
| SRR2487602 | 5482 | 12017 | 519 | 957 | 539 | 3578 | 2208 | 6446 | 2890 | 16 | 25026 | 33695 | 5218 | 2134 | 0 | 84219 | 48293 | 0 | 0 | 15 | 1725 | 1115 | 225 | 0 | 0 | 0 | 0 | 105 | 495 | 629 | 1293 | 259 | 0 | 29 | 13 | 0 | 0 |
| SRR2487604 | 5236 | 14077 | 367 | 672 | 566 | 3247 | 2171 | 5537 | 2673 | 0 | 24966 | 35201 | 5665 | 1998 | 0 | 81977 | 48674 | 0 | 0 | 0 | 1439 | 973 | 211 | 0 | 0 | 0 | 0 | 93 | 460 | 646 | 1226 | 247 | 0 | 38 | 15 | 11 | 0 |
| SRR2487606 | 5697 | 11767 | 435 | 994 | 523 | 3229 | 1889 | 5863 | 2980 | 21 | 26494 | 36677 | 5792 | 1907 | 0 | 82827 | 48057 | 0 | 0 | 13 | 1470 | 955 | 170 | 0 | 0 | 0 | 0 | 103 | 433 | 620 | 1125 | 227 | 0 | 34 | 13 | 11 | 0 |
| SRR2487607 | 1116 | 7694 | 4043 | 513 | 357 | 2218 | 2008 | 6074 | 3562 | 70 | 23882 | 32 | 5425 | 1900 | 0 | 51217 | 52332 | 0 | 0 | 0 | 418 | 298 | 52 | 0 | 0 | 0 | 0 | 22 | 125 | 171 | 433 | 73 | 0 | 14 | 0 | 0 | 0 |
| SRR2487609 | 495 | 32471 | 2293 | 3768 | 1133 | 11399 | 12311 | 49810 | 16694 | 35 | 15227 | 11111 | 1441 | 5306 | 0 | 19193 | 36423 | 0 | 0 | 0 | 2892 | 4340 | 592 | 0 | 10 | 0 | 0 | 0 | 196 | 487 | 2612 | 335 | 0 | 22 | 0 | 0 | 0 |
| SRR2487610 | 613 | 36499 | 2035 | 3007 | 1061 | 10437 | 11382 | 47987 | 16243 | 12 | 14041 | 10300 | 1317 | 4889 | 0 | 18950 | 35732 | 0 | 0 | 13 | 2736 | 4023 | 572 | 0 | 0 | 0 | 0 | 0 | 188 | 466 | 2464 | 308 | 0 | 14 | 0 | 0 | 0 |
| SRR2487611 | 516 | 34973 | 2206 | 2817 | 1043 | 10897 | 11829 | 49324 | 16552 | 12 | 14463 | 10507 | 1302 | 5073 | 0 | 19470 | 35509 | 0 | 0 | 0 | 2632 | 3883 | 557 | 0 | 0 | 0 | 0 | 0 | 184 | 431 | 2368 | 300 | 0 | 13 | 0 | 0 | 0 |
| SRR2487612 | 3055 | 64850 | 101 | 6445 | 2516 | 19241 | 13365 | 47976 | 14348 | 0 | 14662 | 10731 | 1612 | 5090 | 0 | 38679 | 37127 | 0 | 0 | 44 | 9142 | 12085 | 1545 | 0 | 14 | 0 | 0 | 16 | 720 | 1592 | 7072 | 881 | 0 | 33 | 0 | 0 | 0 |
| SRR2487614 | 3749 | 67374 | 210 | 7252 | 2195 | 20536 | 15838 | 48232 | 14874 | 35 | 14853 | 10894 | 1375 | 5047 | 0 | 37379 | 35122 | 0 | 0 | 46 | 9593 | 12159 | 1605 | 0 | 17 | 0 | 0 | 19 | 701 | 1605 | 7096 | 827 | 0 | 36 | 0 | 0 | 0 |
| SRR2487616 | 3040 | 67641 | 272 | 6896 | 1989 | 18519 | 15834 | 47427 | 14019 | 0 | 15933 | 11037 | 1317 | 4648 | 0 | 38007 | 36117 | 0 | 0 | 35 | 8430 | 11075 | 1652 | 0 | 16 | 0 | 0 | 14 | 667 | 1441 | 6697 | 729 | 0 | 19 | 14 | 0 | 0 |
| SRR2487618 | 2972 | 68904 | 137 | 7087 | 1670 | 18009 | 14282 | 47952 | 14959 | 0 | 16872 | 11909 | 1307 | 5204 | 0 | 37907 | 37423 | 0 | 0 | 54 | 8333 | 10584 | 1544 | 0 | 16 | 0 | 0 | 11 | 658 | 1299 | 6347 | 703 | 0 | 22 | 11 | 0 | 0 |
| SRR2487619 | 604 | 35257 | 1940 | 3032 | 1169 | 10830 | 12118 | 48216 | 16666 | 21 | 14297 | 10428 | 1333 | 4919 | 0 | 19318 | 35137 | 0 | 0 | 14 | 2941 | 4377 | 611 | 0 | 0 | 0 | 0 | 0 | 202 | 495 | 2713 | 318 | 0 | 15 | 0 | 0 | 0 |
| SRR2487621 | 1255 | 10526 | 3842 | 205 | 212 | 2440 | 2382 | 13399 | 5295 | 77 | 23276 | 19 | 3632 | 3158 | 0 | 42382 | 59687 | 0 | 0 | 0 | 518 | 840 | 330 | 0 | 0 | 0 | 0 | 0 | 48 | 274 | 1064 | 482 | 0 | 28 | 43 | 16 | 0 |
| SRR2487622 | 1281 | 10982 | 3733 | 382 | 216 | 2382 | 2189 | 13318 | 5074 | 27 | 22133 | 27759 | 3458 | 2868 | 0 | 41343 | 58000 | 0 | 0 | 0 | 434 | 755 | 329 | 0 | 0 | 0 | 0 | 0 | 50 | 268 | 1006 | 472 | 0 | 26 | 52 | 17 | 0 |
| SRR2487623 | 1300 | 10676 | 3955 | 360 | 218 | 2374 | 2300 | 13629 | 5151 | 28 | 22594 | 27924 | 3544 | 3076 | 0 | 42357 | 58149 | 0 | 0 | 0 | 450 | 756 | 308 | 0 | 13 | 0 | 0 | 0 | 45 | 261 | 912 | 423 | 0 | 26 | 47 | 17 | 0 |
| SRR2487624 | 5603 | 17559 | 214 | 643 | 268 | 3447 | 2377 | 11126 | 4307 | 0 | 21299 | 25111 | 3242 | 2485 | 0 | 67988 | 51947 | 0 | 0 | 0 | 2082 | 3042 | 1219 | 10 | 20 | 0 | 0 | 26 | 264 | 1123 | 4071 | 1675 | 0 | 68 | 94 | 23 | 0 |
| SRR2487626 | 5650 | 17391 | 315 | 693 | 315 | 3669 | 2472 | 11361 | 4352 | 16 | 21911 | 25120 | 3163 | 2614 | 0 | 69437 | 49852 | 0 | 0 | 0 | 2073 | 3014 | 1189 | 0 | 22 | 0 | 0 | 25 | 243 | 1183 | 3873 | 1621 | 0 | 74 | 91 | 25 | 0 |
| SRR2487628 | 6109 | 19062 | 377 | 516 | 271 | 3265 | 2373 | 11280 | 3947 | 20 | 22825 | 27024 | 3405 | 2844 | 0 | 67037 | 50816 | 0 | 0 | 0 | 1912 | 2819 | 1202 | 0 | 36 | 0 | 0 | 15 | 228 | 1059 | 3598 | 1520 | 0 | 76 | 69 | 29 | 0 |

### Supp Table S8.1.Columns_B_to_AL

| | | | | | | | | | | | | | | | | | | | | | | | | | | | | | | | | | | | | | |
|---|---|---|---|---|---|---|---|---|---|---|---|---|---|---|---|---|---|---|---|---|---|---|---|---|---|---|---|---|---|---|---|---|---|---|---|---|---|
| SRR2487630 | 5898 | 16782 | 357 | 616 | 228 | 3261 | 2402 | 11446 | 4074 | 0 | 23252 | 27281 | 3727 | 2864 | 0 | 68898 | 52136 | 0 | 0 | 0 | 1840 | 2906 | 1101 | 0 | 11 | 0 | 0 | 28 | 217 | 1084 | 3379 | 1440 | 0 | 63 | 99 | 28 | 0 |
| SRR2487631 | 1159 | 9685 | 3333 | 266 | 217 | 2426 | 2379 | 12996 | 5169 | 97 | 22622 | 28019 | 3479 | 2983 | 0 | 41996 | 57497 | 0 | 0 | 0 | 508 | 856 | 338 | 0 | 0 | 0 | 0 | 0 | 59 | 299 | 1092 | 544 | 0 | 32 | 40 | 20 | 0 |
| SRR2487633 | 941 | 116422 | 111 | 775 | 3173 | 31162 | 24569 | 62596 | 10870 | 0 | 23212 | 19111 | 2583 | 5308 | 0 | 33764 | 52200 | 0 | 0 | 50 | 10634 | 13662 | 2564 | 11 | 20 | 0 | 0 | 12 | 1071 | 2288 | 9968 | 1871 | 0 | 49 | 45 | 26 | 0 |
| SRR2487635 | 926 | 113281 | 309 | 637 | 3114 | 31538 | 25896 | 62570 | 10050 | 0 | 23689 | 19160 | 2734 | 5302 | 0 | 35800 | 49558 | 0 | 0 | 57 | 10990 | 14018 | 2591 | 14 | 18 | 0 | 0 | 11 | 1070 | 2354 | 9544 | 1928 | 0 | 38 | 39 | 34 | 0 |
| SRR2487637 | 952 | 117293 | 333 | 667 | 2907 | 30425 | 24555 | 62446 | 9656 | 0 | 24189 | 20313 | 2605 | 5268 | 0 | 34105 | 52944 | 0 | 0 | 47 | 10147 | 12715 | 2401 | 0 | 19 | 0 | 0 | 12 | 977 | 2144 | 9107 | 1842 | 0 | 40 | 46 | 39 | 0 |
| SRR2487639 | 1256 | 115645 | 266 | 725 | 3217 | 30111 | 23929 | 64020 | 10709 | 0 | 24415 | 20132 | 2707 | 6088 | 0 | 35383 | 51318 | 0 | 0 | 54 | 9765 | 12512 | 2263 | 0 | 15 | 0 | 0 | 14 | 976 | 2141 | 8495 | 1702 | 0 | 37 | 34 | 36 | 0 |
| SRR2487641 | 1669 | 21200 | 114 | 915 | 623 | 3204 | 2292 | 7585 | 4849 | 0 | 18772 | 17479 | 3134 | 1374 | 0 | 97341 | 44846 | 0 | 0 | 13 | 9046 | 7792 | 2573 | 0 | 43 | 0 | 0 | 30 | 488 | 4234 | 11987 | 3439 | 0 | 170 | 211 | 48 | 0 |
| SRR2487643 | 1995 | 20423 | 142 | 926 | 530 | 3559 | 2283 | 7884 | 4800 | 0 | 19827 | 17984 | 3523 | 1476 | 0 | 98569 | 43491 | 0 | 0 | 14 | 9221 | 7918 | 2593 | 0 | 49 | 0 | 0 | 32 | 486 | 4361 | 11657 | 3420 | 0 | 172 | 210 | 43 | 0 |
| SRR2487645 | 1998 | 21586 | 61 | 848 | 592 | 3531 | 2325 | 7238 | 4613 | 0 | 19578 | 18716 | 3670 | 1396 | 0 | 96144 | 43397 | 0 | 0 | 13 | 8436 | 7270 | 2364 | 0 | 39 | 0 | 0 | 30 | 440 | 3914 | 10880 | 3313 | 0 | 172 | 197 | 50 | 0 |
| SRR2487647 | 1597 | 21597 | 123 | 645 | 533 | 3356 | 2410 | 7394 | 5033 | 0 | 19707 | 18897 | 3626 | 1543 | 0 | 97718 | 44611 | 0 | 0 | 10 | 8264 | 7089 | 2401 | 11 | 35 | 0 | 0 | 30 | 427 | 3892 | 10294 | 3016 | 0 | 165 | 192 | 49 | 0 |
| SRR2487648 | 712 | 87709 | 2450 | 1445 | 1766 | 17953 | 24457 | 76111 | 4232 | 63 | 18456 | 13187 | 1955 | 4958 | 0 | 27129 | 36084 | 0 | 0 | 0 | 2500 | 2112 | 411 | 0 | 0 | 0 | 0 | 0 | 132 | 608 | 1510 | 339 | 0 | 12 | 16 | 0 | 0 |
| SRR2487649 | 1600 | 152009 | 167 | 2221 | 3218 | 29426 | 28064 | 74254 | 3448 | 0 | 20818 | 15224 | 2171 | 5194 | 0 | 53351 | 37004 | 0 | 0 | 24 | 8102 | 5668 | 1108 | 0 | 0 | 0 | 0 | 0 | 475 | 1840 | 4202 | 899 | 0 | 22 | 23 | 12 | 0 |
| SRR2487651 | 1497 | 153874 | 190 | 2732 | 2529 | 30476 | 28209 | 76763 | 3200 | 24 | 21638 | 14592 | 2152 | 5459 | 0 | 53382 | 36589 | 0 | 0 | 24 | 7969 | 5693 | 1107 | 0 | 0 | 0 | 0 | 0 | 478 | 1927 | 4026 | 841 | 0 | 26 | 26 | 0 | 0 |
| SRR2487652 | 400 | 85697 | 2592 | 1180 | 1724 | 18058 | 24156 | 76940 | 4199 | 63 | 18634 | 13534 | 2184 | 5151 | 0 | 27868 | 37307 | 0 | 0 | 0 | 2504 | 2069 | 377 | 0 | 0 | 0 | 0 | 0 | 126 | 634 | 1492 | 310 | 0 | 13 | 19 | 0 | 0 |
| SRR2487653 | 389 | 89146 | 2333 | 1098 | 1625 | 17371 | 23381 | 76582 | 4047 | 23 | 18070 | 12883 | 2041 | 5028 | 0 | 27189 | 36796 | 0 | 0 | 0 | 2332 | 1909 | 364 | 0 | 0 | 0 | 0 | 0 | 116 | 527 | 1408 | 298 | 0 | 0 | 22 | 11 | 0 |
| SRR2487654 | 229 | 86479 | 2776 | 1170 | 1623 | 17780 | 23329 | 78214 | 4095 | 0 | 18337 | 13008 | 2055 | 5204 | 0 | 27636 | 36358 | 0 | 0 | 0 | 2269 | 1839 | 353 | 0 | 0 | 0 | 0 | 0 | 103 | 553 | 1303 | 294 | 0 | 0 | 16 | 0 | 0 |
| SRR2487656 | 1870 | 147784 | 130 | 2758 | 2928 | 29058 | 28901 | 74649 | 3632 | 0 | 20836 | 13695 | 2162 | 4672 | 0 | 50988 | 36863 | 0 | 0 | 28 | 8633 | 6164 | 1166 | 0 | 0 | 0 | 0 | 0 | 496 | 1989 | 4546 | 926 | 0 | 19 | 24 | 17 | 0 |
| SRR2487658 | 1891 | 148423 | 172 | 2674 | 2813 | 30730 | 29422 | 76754 | 3647 | 0 | 20213 | 13450 | 1944 | 5004 | 0 | 53000 | 34391 | 0 | 0 | 28 | 9074 | 6378 | 1193 | 0 | 0 | 0 | 0 | 0 | 462 | 2102 | 4453 | 907 | 0 | 36 | 16 | 15 | 0 |
| SRR2487661 | 1413 | 15006 | 226 | 1413 | 254 | 1501 | 1556 | 4347 | 6346 | 28 | 23669 | 17386 | 2575 | 2533 | 0 | 82935 | 48886 | 0 | 0 | 0 | 3280 | 9419 | 1755 | 0 | 0 | 0 | 0 | 49 | 373 | 1726 | 12040 | 2519 | 0 | 33 | 54 | 11 | 0 |
| SRR2487663 | 1983 | 16094 | 437 | 1225 | 204 | 1663 | 1585 | 4712 | 6531 | 0 | 23767 | 17726 | 2705 | 2727 | 0 | 85922 | 46900 | 0 | 0 | 0 | 3361 | 9466 | 1790 | 0 | 0 | 0 | 0 | 44 | 391 | 1824 | 11762 | 2550 | 0 | 32 | 48 | 11 | 0 |
| SRR2487665 | 2076 | 16572 | 334 | 1223 | 222 | 1728 | 1469 | 4962 | 6335 | 0 | 24507 | 18171 | 2901 | 2964 | 0 | 83253 | 47661 | 0 | 0 | 0 | 3023 | 8697 | 1684 | 0 | 0 | 0 | 0 | 41 | 339 | 1582 | 10901 | 2345 | 0 | 39 | 44 | 0 | 0 |
| SRR2487667 | 2243 | 15515 | 262 | 1084 | 230 | 1849 | 1492 | 4567 | 6391 | 0 | 24870 | 18704 | 2871 | 2933 | 0 | 83848 | 47983 | 0 | 0 | 0 | 2953 | 8562 | 1606 | 0 | 0 | 0 | 0 | 41 | 332 | 1581 | 10388 | 2182 | 0 | 32 | 40 | 0 | 0 |
| SRR2487668 | 109 | 51987 | 4955 | 765 | 1283 | 10024 | 10213 | 31380 | 3832 | 73 | 18699 | 9367 | 1563 | 3783 | 0 | 14372 | 30123 | 0 | 0 | 0 | 6337 | 3863 | 692 | 0 | 0 | 0 | 0 | 0 | 193 | 1889 | 2983 | 602 | 0 | 20 | 20 | 0 | 0 |
| SRR2487669 | 530 | 106404 | 1191 | 1225 | 1663 | 12926 | 10653 | 32363 | 3374 | 33 | 20530 | 9877 | 1750 | 4174 | 0 | 22961 | 32141 | 0 | 0 | 27 | 17822 | 9411 | 1583 | 0 | 20 | 0 | 0 | 0 | 502 | 5200 | 6984 | 1326 | 0 | 22 | 22 | 0 | 0 |
| SRR2487671 | 467 | 104176 | 1134 | 1434 | 1611 | 13342 | 10027 | 33558 | 3530 | 0 | 20903 | 10382 | 1750 | 4355 | 0 | 23600 | 31339 | 0 | 0 | 21 | 17473 | 9420 | 1576 | 0 | 12 | 0 | 0 | 0 | 539 | 5135 | 6640 | 1349 | 0 | 30 | 22 | 0 | 0 |
| SRR2487672 | 225 | 53088 | 4277 | 488 | 1255 | 10220 | 10198 | 31984 | 3842 | 0 | 19604 | 9778 | 1688 | 3948 | 0 | 14881 | 30776 | 0 | 0 | 0 | 6267 | 3851 | 643 | 0 | 0 | 0 | 0 | 0 | 168 | 1969 | 2834 | 606 | 0 | 15 | 16 | 0 | 0 |
| SRR2487673 | 118 | 49489 | 4700 | 671 | 1216 | 9473 | 10043 | 31972 | 3800 | 79 | 18414 | 9130 | 1569 | 3796 | 0 | 14622 | 30145 | 0 | 0 | 0 | 5823 | 3603 | 582 | 0 | 11 | 0 | 0 | 0 | 145 | 1699 | 2720 | 578 | 0 | 17 | 16 | 0 | 0 |
| SRR2487674 | 80 | 52052 | 4830 | 599 | 1257 | 9950 | 10034 | 32627 | 3860 | 0 | 18926 | 9416 | 1721 | 3743 | 0 | 15042 | 30486 | 0 | 0 | 0 | 5692 | 3474 | 591 | 0 | 0 | 0 | 0 | 0 | 138 | 1700 | 2459 | 493 | 0 | 21 | 16 | 0 | 0 |
| SRR2487676 | 470 | 101983 | 913 | 1304 | 1513 | 13189 | 10521 | 32526 | 3364 | 26 | 19892 | 9506 | 1800 | 3645 | 0 | 24572 | 32015 | 0 | 0 | 25 | 18698 | 9982 | 1636 | 0 | 12 | 0 | 0 | 0 | 562 | 5349 | 7681 | 1322 | 0 | 43 | 26 | 0 | 0 |
| SRR2487678 | 484 | 101337 | 968 | 1856 | 1591 | 13357 | 10741 | 32703 | 3313 | 0 | 18919 | 9380 | 1847 | 4023 | 0 | 24204 | 31093 | 0 | 0 | 28 | 19321 | 10347 | 1729 | 0 | 19 | 0 | 0 | 0 | 617 | 5520 | 7532 | 1435 | 0 | 34 | 27 | 0 | 0 |
| SRR2487688 | 204 | 8473 | 2469 | 340 | 237 | 1387 | 1314 | 4328 | 4008 | 23 | 19336 | 22371 | 3241 | 1380 | 0 | 46242 | 49768 | 0 | 0 | 0 | 409 | 498 | 45 | 0 | 0 | 0 | 0 | 17 | 154 | 304 | 926 | 112 | 0 | 0 | 0 | 0 | 0 |
| SRR2487689 | 707 | 16134 | 353 | 648 | 336 | 1841 | 1506 | 4495 | 3271 | 0 | 19486 | 21338 | 3239 | 1538 | 11 | 68845 | 48622 | 0 | 0 | 0 | 1590 | 1622 | 146 | 0 | 0 | 0 | 0 | 73 | 566 | 1164 | 3223 | 342 | 0 | 13 | 0 | 0 | 0 |
| SRR2487691 | 717 | 17558 | 179 | 537 | 230 | 1923 | 1327 | 4598 | 3344 | 0 | 20377 | 21673 | 3563 | 1547 | 0 | 68415 | 49584 | 0 | 0 | 0 | 1628 | 1475 | 164 | 0 | 0 | 0 | 0 | 89 | 553 | 1239 | 2986 | 351 | 0 | 35 | 13 | 0 | 0 |
| SRR2487692 | 299 | 8593 | 2166 | 299 | 194 | 1433 | 1312 | 4515 | 4066 | 23 | 19948 | 22776 | 3253 | 1491 | 0 | 47655 | 51705 | 0 | 0 | 0 | 406 | 464 | 46 | 0 | 0 | 0 | 0 | 14 | 166 | 294 | 851 | 109 | 0 | 0 | 0 | 0 | 0 |
| SRR2487693 | 103 | 8860 | 1721 | 282 | 187 | 1397 | 1257 | 4407 | 4019 | 26 | 19092 | 13 | 3116 | 1331 | 0 | 46035 | 50027 | 0 | 0 | 0 | 362 | 427 | 51 | 0 | 0 | 0 | 0 | 19 | 146 | 259 | 824 | 87 | 0 | 0 | 0 | 0 | 0 |
| SRR2487694 | 102 | 7544 | 2600 | 306 | 211 | 1412 | 1272 | 4519 | 3970 | 0 | 19273 | 13 | 3136 | 1458 | 0 | 47473 | 50182 | 0 | 0 | 0 | 365 | 398 | 34 | 0 | 0 | 0 | 0 | 13 | 135 | 275 | 772 | 84 | 0 | 0 | 0 | 0 | 0 |
| SRR2487696 | 820 | 16582 | 410 | 501 | 255 | 1871 | 1293 | 4617 | 3580 | 46 | 19176 | 21650 | 3240 | 1310 | 0 | 70333 | 49185 | 0 | 0 | 14 | 1845 | 1759 | 194 | 0 | 0 | 0 | 0 | 113 | 743 | 1264 | 3624 | 435 | 0 | 23 | 0 | 0 | 0 |
| SRR2487698 | 783 | 16942 | 460 | 506 | 358 | 1984 | 1468 | 4850 | 3234 | 0 | 18737 | 20660 | 3417 | 1538 | 0 | 72160 | 49123 | 0 | 0 | 19 | 1806 | 1656 | 245 | 0 | 0 | 0 | 0 | 109 | 773 | 1338 | 3370 | 393 | 0 | 21 | 14 | 0 | 0 |
| SRR2487700 | 78 | 41428 | 3362 | 222 | 1523 | 12887 | 12804 | 44712 | 5538 | 26 | 20327 | 20941 | 2545 | 3466 | 0 | 17039 | 43430 | 0 | 0 | 0 | 634 | 1920 | 231 | 0 | 0 | 0 | 0 | 0 | 81 | 132 | 1318 | 147 | 0 | 24 | 0 | 0 | 0 |
| SRR2487701 | 414 | 72383 | 488 | 488 | 2441 | 21107 | 14476 | 42664 | 4504 | 0 | 22249 | 20719 | 2420 | 3561 | 10 | 33080 | 42276 | 0 | 0 | 0 | 2187 | 5956 | 697 | 0 | 0 | 0 | 0 | 11 | 322 | 528 | 4112 | 454 | 0 | 34 | 11 | 0 | 0 |
| SRR2487703 | 608 | 69826 | 292 | 438 | 2499 | 21153 | 14139 | 44252 | 4485 | 0 | 22111 | 20377 | 2630 | 3437 | 0 | 32572 | 41581 | 0 | 0 | 11 | 2183 | 5864 | 751 | 0 | 0 | 0 | 0 | 20 | 355 | 555 | 3989 | 418 | 0 | 35 | 0 | 0 | 0 |
| SRR2487704 | 105 | 39716 | 3487 | 144 | 1476 | 13174 | 12443 | 45523 | 5400 | 0 | 20688 | 20769 | 2639 | 3655 | 0 | 17578 | 45078 | 0 | 0 | 0 | 642 | 1861 | 219 | 0 | 0 | 0 | 0 | 0 | 81 | 138 | 1250 | 144 | 0 | 16 | 0 | 0 | 0 |
| SRR2487705 | 100 | 40488 | 3462 | 200 | 1424 | 12628 | 12282 | 45580 | 5528 | 29 | 20045 | 20474 | 2481 | 3523 | 0 | 16934 | 43643 | 0 | 0 | 0 | 588 | 1721 | 203 | 0 | 0 | 0 | 0 | 0 | 81 | 131 | 1209 | 157 | 0 | 20 | 0 | 0 | 0 |
| SRR2487706 | 57 | 39956 | 2981 | 258 | 1482 | 13036 | 12194 | 46225 | 5481 | 0 | 20471 | 20497 | 2529 | 3528 | 0 | 17388 | 43710 | 0 | 0 | 0 | 557 | 1683 | 187 | 0 | 0 | 0 | 0 | 0 | 75 | 129 | 1127 | 134 | 0 | 16 | 0 | 0 | 0 |
| SRR2487708 | 528 | 70209 | 302 | 264 | 2590 | 21574 | 14441 | 42788 | 4518 | 0 | 20728 | 19998 | 2372 | 3093 | 0 | 32881 | 42578 | 0 | 0 | 18 | 2493 | 6354 | 684 | 0 | 0 | 0 | 0 | 13 | 322 | 564 | 4441 | 472 | 0 | 48 | 0 | 0 | 0 |
| SRR2487710 | 329 | 69747 | 271 | 349 | 2471 | 21851 | 15409 | 42501 | 4542 | 19 | 20667 | 19235 | 2753 | 3383 | 0 | 34703 | 40788 | 0 | 0 | 10 | 2624 | 6539 | 813 | 0 | 10 | 0 | 0 | 18 | 394 | 538 | 4527 | 544 | 0 | 45 | 12 | 0 | 0 |
| SRR2529523 | 357 | 16521 | 922 | 511 | 3184 | 37942 | 31251 | 40660 | 3977 | 34 | 55026 | 39291 | 13377 | 9116 | 0 | 1042 | 49785 | 0 | 0 | 0 | 133 | 19 | 17 | 0 | 0 | 0 | 0 | 12 | 214 | 208 | 56 | 34 | 0 | 0 | 0 | 0 | 0 |
| SRR2529524 | 400 | 16408 | 961 | 514 | 3209 | 37660 | 30980 | 40802 | 3813 | 45 | 39 | 39578 | 13363 | 9204 | 0 | 1053 | 49317 | 0 | 0 | 0 | 141 | 17 | 20 | 0 | 0 | 0 | 0 | 15 | 201 | 186 | 58 | 32 | 0 | 0 | 0 | 0 | 0 |
| SRR2529525 | 614 | 9214 | 348 | 1118 | 4672 | 63316 | 49390 | 71716 | 4231 | 0 | 67722 | 30212 | 11715 | 8306 | 0 | 750 | 39205 | 0 | 0 | 0 | 159 | 17 | 30 | 0 | 0 | 0 | 0 | 20 | 140 | 109 | 51 | 35 | 0 | 12 | 0 | 0 | 0 |
| SRR2529526 | 648 | 9202 | 402 | 1224 | 4795 | 63190 | 49055 | 71418 | 4231 | 16 | 68155 | 29981 | 11527 | 8393 | 0 | 748 | 38918 | 0 | 0 | 0 | 167 | 23 | 25 | 0 | 0 | 0 | 0 | 23 | 137 | 109 | 44 | 40 | 0 | 0 | 0 | 0 | 0 |
| SRR2529527 | 458 | 44591 | 307 | 640 | 4606 | 62299 | 47129 | 61883 | 7062 | 33 | 51906 | 29490 | 11221 | 6477 | 12 | 1022 | 41466 | 0 | 0 | 12 | 579 | 132 | 111 | 0 | 0 | 0 | 0 | 30 | 987 | 683 | 471 | 189 | 0 | 14 | 0 | 0 | 0 |
| SRR2529528 | 492 | 44908 | 339 | 619 | 4483 | 61640 | 47028 | 61431 | 7177 | 14 | 52031 | 29461 | 11207 | 6445 | 14 | 1019 | 41218 | 0 | 0 | 12 | 576 | 132 | 100 | 0 | 0 | 0 | 0 | 24 | 943 | 663 | 456 | 182 | 0 | 14 | 0 | 0 | 0 |
| SRR2529529 | 562 | 16400 | 1285 | 336 | 2442 | 28626 | 22035 | 39878 | 3125 | 64 | 59072 | 33461 | 13057 | 9089 | 0 | 430 | 56847 | 0 | 0 | 0 | 131 | 14 | 15 | 0 | 0 | 0 | 0 | 18 | 219 | 245 | 62 | 26 | 0 | 0 | 0 | 0 | 0 |
| SRR2529530 | 674 | 16423 | 1216 | 324 | 2411 | 28591 | 21910 | 39658 | 3199 | 52 | 59045 | 33500 | 13018 | 9086 | 0 | 415 | 56380 | 0 | 0 | 0 | 137 | 13 | 12 | 0 | 0 | 0 | 0 | 20 | 216 | 228 | 62 | 26 | 0 | 0 | 0 | 0 | 0 |
| SRR2529531 | 548 | 11012 | 1350 | 335 | 2590 | 45109 | 36450 | 46436 | 5475 | 78 | 47685 | 31277 | 13083 | 7897 | 0 | 1292 | 41466 | 0 | 0 | 0 | 208 | 31 | 41 | 0 | 0 | 0 | 0 | 18 | 278 | 272 | 69 | 47 | 0 | 0 | 0 | 0 | 0 |
| SRR2529532 | 622 | 11026 | 1381 | 399 | 2509 | 44970 | 36338 | 46305 | 5406 | 72 | 47747 | 31128 | 13077 | 8027 | 0 | 1308 | 41059 | 0 | 0 | 0 | 206 | 25 | 40 | 0 | 0 | 0 | 0 | 19 | 274 | 276 | 67 | 46 | 0 | 0 | 0 | 0 | 0 |
| SRR2529533 | 521 | 10488 | 1259 | 333 | 3332 | 39521 | 33026 | 54945 | 1639 | 85 | 54806 | 36746 | 14007 | 8331 | 0 | 974 | 47762 | 0 | 0 | 0 | 104 | 0 | 18 | 0 | 0 | 0 | 0 | 13 | 165 | 164 | 43 | 26 | 0 | 0 | 0 | 0 | 0 |
| SRR2529534 | 548 | 10630 | 1342 | 311 | 3255 | 39201 | 32578 | 54685 | 1627 | 75 | 55011 | 36609 | 13982 | 8408 | 0 | 963 | 47518 | 0 | 0 | 0 | 117 | 0 | 16 | 0 | 0 | 0 | 0 | 14 | 167 | 160 | 49 | 27 | 0 | 0 | 0 | 0 | 0 |
| SRR2529535 | 234 | 71802 | 1163 | 626 | 5280 | 44142 | 40393 | 80632 | 4554 | 30 | 48912 | 26192 | 9933 | 11335 | 0 | 865 | 45615 | 0 | 0 | 0 | 374 | 438 | 97 | 0 | 0 | 0 | 0 | 90 | 415 | 701 | 2730 | 418 | 0 | 32 | 16 | 11 | 0 |
| SRR2529536 | 188 | 71298 | 1198 | 639 | 5171 | 43798 | 40443 | 80735 | 4509 | 36 | 20 | 25897 | 10021 | 11394 | 0 | 886 | 45475 | 0 | 0 | 0 | 372 | 415 | 94 | 0 | 0 | 0 | 0 | 87 | 411 | 658 | 2639 | 395 | 0 | 33 | 16 | 12 | 0 |
| SRR2529537 | 568 | 61985 | 556 | 487 | 4394 | 36076 | 31372 | 96161 | 3162 | 18 | 53842 | 29186 | 8553 | 9932 | 0 | 346 | 47223 | 0 | 0 | 0 | 349 | 1256 | 183 | 0 | 15 | 0 | 0 | 21 | 143 | 259 | 2742 | 230 | 0 | 30 | 16 | 0 | 0 |
| SRR2529538 | 561 | 62002 | 600 | 486 | 4305 | 36248 | 31106 | 96305 | 3153 | 18 | 53877 | 29118 | 8473 | 9969 | 0 | 344 | 46990 | 0 | 0 | 0 | 346 | 1201 | 179 | 0 | 13 | 0 | 0 | 23 | 137 | 259 | 2685 | 214 | 0 | 30 | 18 | 11 | 0 |
| SRR2529539 | 563 | 77818 | 391 | 523 | 3633 | 29115 | 27861 | 96529 | 3217 | 0 | 49960 | 22709 | 7843 | 14581 | 0 | 514 | 42074 | 0 | 0 | 0 | 334 | 693 | 158 | 0 | 18 | 0 | 0 | 23 | 196 | 235 | 1094 | 209 | 0 | 34 | 13 | 0 | 0 |
| SRR2529540 | 523 | 78222 | 423 | 490 | 3619 | 28861 | 27964 | 96387 | 3189 | 21 | 31 | 22632 | 7754 | 14614 | 0 | 521 | 41807 | 0 | 0 | 0 | 341 | 669 | 155 | 0 | 14 | 0 | 0 | 18 | 186 | 234 | 1046 | 206 | 0 | 35 | 11 | 12 | 0 |
| SRR2529541 | 306 | 113515 | 988 | 377 | 3096 | 29013 | 23736 | 42894 | 1987 | 26 | 60019 | 34050 | 11186 | 14447 | 0 | 303 | 63043 | 0 | 0 | 0 | 325 | 213 | 44 | 0 | 0 | 0 | 0 | 49 | 443 | 616 | 1150 | 148 | 0 | 12 | 0 | 0 | 0 |
| SRR2529542 | 320 | 113032 | 912 | 382 | 3084 | 29226 | 23524 | 42884 | 2027 | 39 | 59663 | 33898 | 11266 | 14328 | 0 | 293 | 62412 | 0 | 0 | 0 | 328 | 217 | 49 | 0 | 0 | 0 | 0 | 49 | 427 | 593 | 1147 | 131 | 0 | 11 | 0 | 0 | 0 |
| SRR2529543 | 453 | 75984 | 1757 | 411 | 3020 | 21406 | 22220 | 60011 | 2818 | 126 | 57948 | 35251 | 9866 | 14364 | 0 | 787 | 50118 | 0 | 0 | 0 | 290 | 541 | 127 | 0 | 0 | 0 | 0 | 24 | 93 | 235 | 1454 | 221 | 0 | 22 | 10 | 0 | 0 |
| SRR2529544 | 529 | 75810 | 1845 | 406 | 2867 | 21300 | 22148 | 59401 | 2826 | 127 | 57824 | 35237 | 10111 | 14544 | 0 | 730 | 49744 | 0 | 0 | 0 | 290 | 548 | 128 | 0 | 0 | 0 | 0 | 24 | 89 | 243 | 1412 | 215 | 0 | 18 | 0 | 0 | 0 |
| SRR2529545 | 502 | 121327 | 1241 | 343 | 3315 | 34939 | 30054 | 50876 | 1120 | 54 | 65881 | 42916 | 13764 | 13769 | 0 | 666 | 55938 | 0 | 0 | 0 | 269 | 72 | 19 | 0 | 0 | 0 | 0 | 35 | 433 | 477 | 371 | 43 | 0 | 0 | 0 | 0 | 0 |
| SRR2529546 | 535 | 121109 | 1237 | 326 | 3423 | 34881 | 29966 | 50808 | 1103 | 69 | 38 | 42912 | 13490 | 13891 | 0 | 667 | 55121 | 0 | 0 | 0 | 258 | 70 | 26 | 0 | 0 | 0 | 0 | 34 | 441 | 460 | 351 | 49 | 0 | 0 | 0 | 0 | 0 |
| SRR2529547 | 23 | 10773 | 555 | 448 | 4285 | 54827 | 51576 | 69182 | 6743 | 20 | 42427 | 24181 | 11889 | 7030 | 0 | 1248 | 41500 | 0 | 0 | 0 | 76 | 26 | 25 | 0 | 0 | 0 | 0 | 0 | 18 | 76 | 52 | 50 | 0 | 0 | 0 | 0 | 0 |
| SRR2529548 | 21 | 10486 | 530 | 345 | 4318 | 54388 | 51356 | 68916 | 6833 | 33 | 40 | 24271 | 11776 | 7079 | 0 | 1296 | 41274 | 0 | 0 | 0 | 82 | 26 | 23 | 0 | 0 | 0 | 0 | 0 | 18 | 64 | 51 | 49 | 0 | 0 | 0 | 0 | 0 |
| SRR2529549 | 655 | 16452 | 528 | 876 | 4720 | 62418 | 49454 | 75665 | 4319 | 20 | 74337 | 31652 | 11401 | 8937 | 0 | 741 | 38775 | 0 | 0 | 0 | 139 | 14 | 29 | 0 | 0 | 0 | 0 | 18 | 75 | 85 | 33 | 35 | 0 | 0 | 0 | 0 | 0 |
| SRR2529550 | 615 | 16740 | 484 | 919 | 4826 | 62118 | 49364 | 75252 | 4324 | 34 | 74507 | 31895 | 11369 | 8901 | 0 | 735 | 37972 | 0 | 0 | 0 | 148 | 17 | 26 | 0 | 0 | 0 | 0 | 20 | 81 | 86 | 34 | 38 | 0 | 11 | 0 | 0 | 0 |
| SRR2529551 | 629 | 23767 | 443 | 498 | 4325 | 57236 | 46053 | 64779 | 6948 | 17 | 57839 | 32474 | 12757 | 6583 | 14 | 1028 | 43943 | 0 | 0 | 0 | 157 | 32 | 48 | 0 | 0 | 0 | 0 | 21 | 111 | 103 | 95 | 52 | 0 | 16 | 0 | 0 | 0 |
| SRR2529552 | 627 | 23525 | 475 | 445 | 4315 | 57186 | 45633 | 64517 | 6876 | 30 | 46 | 32650 | 12825 | 6501 | 11 | 1035 | 43896 | 0 | 0 | 0 | 164 | 34 | 44 | 0 | 0 | 0 | 0 | 24 | 109 | 87 | 92 | 54 | 0 | 13 | 0 | 0 | 0 |
| SRR2529553 | 542 | 12145 | 1524 | 409 | 2233 | 24141 | 19914 | 39410 | 3137 | 75 | 64548 | 36067 | 13377 | 9917 | 0 | 423 | 60854 | 0 | 0 | 0 | 74 | 27 | 21 | 0 | 0 | 0 | 0 | 0 | 29 | 54 | 33 | 30 | 0 | 0 | 0 | 0 | 0 |
| SRR2529554 | 431 | 12018 | 1405 | 375 | 2238 | 24230 | 19753 | 39357 | 3086 | 96 | 45 | 35868 | 13198 | 10144 | 0 | 429 | 60340 | 0 | 0 | 0 | 73 | 17 | 19 | 0 | 0 | 0 | 0 | 0 | 25 | 55 | 39 | 28 | 0 | 0 | 0 | 0 | 0 |
| SRR2529555 | 597 | 12485 | 1368 | 453 | 2805 | 44260 | 34743 | 48300 | 5929 | 54 | 22776 | 13682 | 15316 | 8891 | 12 | 1635 | 48314 | 0 | 0 | 0 | 127 | 20 | 36 | 0 | 0 | 0 | 0 | 0 | 25 | 70 | 35 | 41 | 0 | 0 | 0 | 0 | 0 |
| SRR2529556 | 619 | 12538 | 1436 | 474 | 2814 | 43703 | 34610 | 48657 | 5851 | 70 | 23571 | 14094 | 15166 | 8791 | 0 | 1627 | 47733 | 0 | 0 | 0 | 123 | 20 | 38 | 0 | 0 | 0 | 0 | 0 | 24 | 65 | 33 | 45 | 0 | 0 | 0 | 0 | 0 |
| SRR2529557 | 634 | 7374 | 1261 | 374 | 3051 | 34099 | 28857 | 55250 | 1771 | 44 | 59979 | 39937 | 14571 | 9114 | 0 | 997 | 51522 | 0 | 0 | 0 | 81 | 0 | 22 | 0 | 0 | 0 | 0 | 0 | 23 | 37 | 18 | 38 | 0 | 0 | 0 | 0 | 0 |
| SRR2529558 | 638 | 7407 | 1273 | 296 | 3050 | 33978 | 28635 | 54917 | 1798 | 81 | 59860 | 39934 | 14690 | 8985 | 0 | 1022 | 51241 | 0 | 0 | 0 | 81 | 11 | 33 | 0 | 0 | 0 | 0 | 0 | 20 | 47 | 15 | 39 | 0 | 0 | 0 | 0 | 0 |
| SRR2529559 | 47 | 71483 | 569 | 448 | 5099 | 53768 | 47577 | 70172 | 5952 | 49 | 42600 | 25172 | 11958 | 7649 | 0 | 1233 | 44117 | 0 | 0 | 0 | 98 | 17 | 25 | 0 | 0 | 0 | 0 | 0 | 55 | 112 | 51 | 43 | 0 | 0 | 0 | 0 | 0 |
| SRR2529560 | 30 | 70788 | 620 | 444 | 5120 | 53562 | 47699 | 69923 | 5833 | 57 | 36 | 25305 | 11970 | 7643 | 0 | 1212 | 43830 | 0 | 0 | 0 | 91 | 14 | 22 | 0 | 0 | 0 | 0 | 0 | 53 | 111 | 48 | 41 | 0 | 0 | 0 | 0 | 0 |
| SRR2529561 | 672 | 11920 | 349 | 523 | 4224 | 58549 | 43645 | 75247 | 4843 | 18 | 62622 | 32087 | 12924 | 7645 | 12 | 813 | 43758 | 0 | 0 | 0 | 136 | 28 | 31 | 0 | 0 | 0 | 0 | 23 | 59 | 61 | 56 | 37 | 0 | 0 | 0 | 0 | 0 |
| SRR2529562 | 674 | 11658 | 357 | 599 | 4236 | 58039 | 43615 | 74526 | 4827 | 22 | 62653 | 31992 | 12808 | 7506 | 0 | 762 | 43170 | 0 | 0 | 0 | 128 | 27 | 37 | 0 | 0 | 0 | 0 | 20 | 56 | 56 | 52 | 30 | 0 | 14 | 0 | 0 | 0 |
| SRR2529563 | 680 | 33417 | 337 | 386 | 4225 | 56866 | 47891 | 69513 | 6912 | 14 | 52184 | 28438 | 11629 | 6556 | 15 | 1010 | 41485 | 0 | 0 | 0 | 165 | 29 | 48 | 0 | 0 | 0 | 0 | 40 | 99 | 93 | 52 | 48 | 0 | 14 | 0 | 0 | 0 |
| SRR2529564 | 669 | 33416 | 373 | 378 | 4240 | 56742 | 47690 | 69316 | 6869 | 16 | 45 | 28830 | 11732 | 6663 | 11 | 985 | 41239 | 0 | 0 | 0 | 175 | 21 | 44 | 0 | 0 | 0 | 0 | 35 | 103 | 70 | 52 | 46 | 0 | 13 | 0 | 0 | 0 |
| SRR2529565 | 356 | 35823 | 1043 | 526 | 2729 | 25006 | 20482 | 42242 | 2880 | 124 | 62228 | 36556 | 14152 | 10128 | 0 | 426 | 61608 | 0 | 0 | 0 | 57 | 0 | 11 | 0 | 0 | 0 | 0 | 0 | 29 | 55 | 21 | 21 | 0 | 0 | 0 | 0 | 0 |
| SRR2529566 | 393 | 35548 | 1070 | 591 | 2706 | 24904 | 20401 | 42082 | 2793 | 137 | 40 | 36513 | 13910 | 10223 | 0 | 449 | 61653 | 0 | 0 | 0 | 74 | 12 | 13 | 0 | 0 | 0 | 0 | 0 | 34 | 56 | 21 | 14 | 0 | 0 | 0 | 0 | 0 |
| SRR2529567 | 525 | 54087 | 2093 | 661 | 2350 | 28227 | 25302 | 40236 | 4484 | 74 | 56905 | 36598 | 13840 | 11365 | 0 | 1199 | 48295 | 0 | 0 | 0 | 145 | 48 | 46 | 0 | 0 | 0 | 0 | 0 | 42 | 110 | 79 | 67 | 0 | 0 | 0 | 0 | 0 |
| SRR2529568 | 461 | 54217 | 1987 | 624 | 2435 | 27823 | 24899 | 40066 | 4444 | 87 | 29 | 36337 | 13727 | 11249 | 0 | 1181 | 48400 | 0 | 0 | 0 | 144 | 47 | 44 | 0 | 0 | 0 | 0 | 0 | 34 | 110 | 74 | 60 | 0 | 10 | 0 | 0 | 0 |
| SRR2529569 | 698 | 48312 | 1503 | 421 | 3244 | 30520 | 26528 | 55163 | 1389 | 94 | 68246 | 43826 | 15985 | 10652 | 0 | 868 | 52235 | 0 | 0 | 0 | 56 | 0 | 11 | 0 | 0 | 0 | 0 | 0 | 34 | 44 | 0 | 12 | 0 | 0 | 0 | 0 | 0 |
| SRR2529570 | 624 | 47803 | 1630 | 391 | 3219 | 30311 | 26364 | 54950 | 1436 | 97 | 40 | 43779 | 16090 | 10906 | 0 | 904 | 52086 | 0 | 0 | 0 | 54 | 0 | 10 | 0 | 0 | 0 | 0 | 0 | 32 | 44 | 13 | 13 | 0 | 0 | 0 | 0 | 0 |
| SRR2529571 | 631 | 86584 | 1156 | 322 | 2646 | 33586 | 25617 | 39252 | 3789 | 53 | 54157 | 38825 | 13412 | 8333 | 0 | 972 | 49203 | 0 | 0 | 0 | 99 | 12 | 26 | 0 | 0 | 0 | 0 | 0 | 36 | 88 | 36 | 28 | 0 | 0 | 0 | 0 | 0 |
| SRR2529572 | 526 | 86468 | 1187 | 361 | 2667 | 33291 | 25313 | 39042 | 3778 | 65 | 39 | 38572 | 13428 | 8246 | 0 | 991 | 48850 | 0 | 0 | 0 | 93 | 16 | 23 | 0 | 0 | 0 | 0 | 0 | 45 | 78 | 35 | 29 | 0 | 0 | 0 | 0 | 0 |
| SRR2529573 | 682 | 25975 | 469 | 871 | 4923 | 61452 | 50739 | 76354 | 4714 | 34 | 69825 | 27327 | 10376 | 9186 | 0 | 730 | 37106 | 0 | 0 | 0 | 279 | 24 | 35 | 0 | 0 | 0 | 0 | 24 | 122 | 184 | 51 | 34 | 0 | 0 | 0 | 0 | 0 |
| SRR2529574 | 774 | 25614 | 404 | 841 | 4837 | 61086 | 50094 | 76167 | 4582 | 35 | 40 | 27300 | 10427 | 9053 | 0 | 728 | 36957 | 0 | 0 | 0 | 273 | 29 | 31 | 0 | 0 | 0 | 0 | 21 | 118 | 168 | 50 | 28 | 0 | 0 | 0 | 0 | 0 |
| SRR2529575 | 513 | 39813 | 338 | 425 | 4190 | 55066 | 42371 | 58781 | 7698 | 0 | 31564 | 19602 | 11242 | 6332 | 16 | 904 | 48681 | 0 | 0 | 0 | 172 | 54 | 45 | 0 | 0 | 0 | 0 | 23 | 141 | 90 | 131 | 42 | 0 | 13 | 0 | 0 | 0 |
| SRR2529576 | 539 | 39801 | 378 | 469 | 4240 | 54388 | 42146 | 58239 | 7676 | 14 | 32353 | 20407 | 11133 | 6344 | 15 | 863 | 48488 | 0 | 0 | 0 | 151 | 52 | 46 | 0 | 0 | 0 | 0 | 21 | 150 | 90 | 130 | 41 | 0 | 11 | 0 | 0 | 0 |
| SRR2529577 | 437 | 65335 | 1141 | 284 | 2161 | 26273 | 19404 | 36414 | 2830 | 66 | 58893 | 33120 | 12871 | 8795 | 0 | 428 | 59260 | 0 | 0 | 0 | 69 | 13 | 13 | 0 | 0 | 0 | 0 | 0 | 48 | 79 | 36 | 26 | 0 | 0 | 0 | 0 | 0 |
| SRR2529578 | 471 | 65710 | 1065 | 318 | 2235 | 26146 | 19196 | 36175 | 2847 | 57 | 31 | 32899 | 12820 | 8802 | 0 | 419 | 58624 | 0 | 0 | 0 | 75 | 13 | 11 | 0 | 0 | 0 | 0 | 0 | 48 | 81 | 39 | 22 | 0 | 0 | 0 | 0 | 0 |
| SRR2529579 | 384 | 15402 | 1310 | 333 | 2596 | 42229 | 33755 | 46240 | 5292 | 50 | 46068 | 29460 | 11729 | 7712 | 10 | 1330 | 43986 | 0 | 0 | 0 | 82 | 0 | 28 | 0 | 0 | 0 | 0 | 0 | 13 | 55 | 18 | 35 | 0 | 0 | 0 | 0 | 0 |
| SRR2529580 | 441 | 15190 | 1252 | 384 | 2627 | 42077 | 33614 | 46072 | 5241 | 64 | 46108 | 29497 | 11676 | 7842 | 0 | 1325 | 44293 | 0 | 0 | 0 | 85 | 0 | 28 | 0 | 0 | 0 | 0 | 0 | 11 | 51 | 19 | 36 | 0 | 0 | 0 | 0 | 0 |
| SRR2529581 | 642 | 48333 | 1278 | 344 | 3143 | 36726 | 30386 | 54017 | 1545 | 112 | 56292 | 37084 | 14501 | 8317 | 0 | 1004 | 48298 | 0 | 0 | 0 | 66 | 0 | 14 | 0 | 0 | 0 | 0 | 0 | 28 | 58 | 15 | 23 | 0 | 0 | 0 | 0 | 0 |
| SRR2529582 | 591 | 48600 | 1322 | 324 | 3063 | 36571 | 30344 | 53718 | 1520 | 112 | 56277 | 36926 | 14423 | 8420 | 0 | 957 | 48004 | 0 | 0 | 0 | 72 | 0 | 13 | 0 | 0 | 0 | 0 | 0 | 26 | 58 | 16 | 18 | 0 | 0 | 0 | 0 | 0 |
| SRR2529583 | 35 | 126392 | 444 | 349 | 3393 | 35794 | 38610 | 87975 | 4037 | 44 | 49875 | 28633 | 13053 | 10154 | 0 | 814 | 50874 | 0 | 0 | 0 | 134 | 31 | 34 | 0 | 0 | 0 | 0 | 0 | 31 | 150 | 90 | 71 | 0 | 10 | 0 | 0 | 0 |
| SRR2529584 | 47 | 125499 | 446 | 353 | 3394 | 35735 | 38306 | 87565 | 4143 | 33 | 49838 | 28392 | 12932 | 10166 | 0 | 830 | 50558 | 0 | 0 | 0 | 129 | 28 | 32 | 0 | 0 | 0 | 0 | 0 | 32 | 141 | 86 | 74 | 0 | 11 | 0 | 0 | 0 |
| SRR2529585 | 542 | 56729 | 369 | 688 | 1984 | 13946 | 17414 | 112322 | 1765 | 20 | 55201 | 25450 | 9408 | 11722 | 0 | 314 | 40712 | 0 | 0 | 0 | 200 | 83 | 94 | 0 | 13 | 0 | 0 | 19 | 43 | 123 | 73 | 81 | 0 | 22 | 0 | 0 | 0 |
| SRR2529586 | 576 | 56744 | 387 | 720 | 1995 | 13901 | 17251 | 111679 | 1835 | 25 | 54960 | 25414 | 9438 | 11710 | 0 | 340 | 40560 | 0 | 0 | 0 | 195 | 88 | 93 | 0 | 0 | 0 | 0 | 11 | 39 | 114 | 70 | 83 | 0 | 22 | 0 | 0 | 0 |
| SRR2529587 | 390 | 45878 | 356 | 563 | 1443 | 10121 | 13578 | 103338 | 1888 | 37 | 50139 | 23139 | 8397 | 11644 | 0 | 359 | 35751 | 0 | 0 | 0 | 176 | 96 | 116 | 0 | 14 | 0 | 0 | 17 | 70 | 116 | 76 | 86 | 0 | 31 | 0 | 0 | 0 |
| SRR2529588 | 417 | 45337 | 376 | 579 | 1422 | 10110 | 13439 | 103171 | 1911 | 21 | 49929 | 23298 | 8334 | 11705 | 0 | 362 | 35218 | 0 | 0 | 0 | 184 | 85 | 107 | 0 | 13 | 0 | 0 | 16 | 61 | 118 | 71 | 88 | 0 | 26 | 0 | 0 | 0 |
| SRR2529589 | 270 | 73589 | 871 | 383 | 1814 | 19112 | 18934 | 50628 | 1706 | 35 | 60732 | 32584 | 12897 | 13582 | 0 | 302 | 65771 | 0 | 0 | 0 | 88 | 18 | 28 | 0 | 0 | 0 | 0 | 0 | 19 | 87 | 57 | 37 | 0 | 0 | 0 | 0 | 0 |
| SRR2529590 | 261 | 73872 | 872 | 348 | 1812 | 18734 | 18712 | 50470 | 1728 | 41 | 60373 | 32301 | 12877 | 13510 | 0 | 303 | 65394 | 0 | 0 | 0 | 92 | 18 | 23 | 0 | 0 | 0 | 0 | 0 | 22 | 80 | 42 | 36 | 0 | 0 | 0 | 0 | 0 |
| SRR2529591 | 746 | 80776 | 1604 | 453 | 1460 | 18844 | 21078 | 58676 | 2231 | 53 | 64790 | 40196 | 12717 | 16761 | 0 | 674 | 51378 | 0 | 0 | 0 | 105 | 16 | 42 | 0 | 0 | 0 | 0 | 0 | 12 | 56 | 28 | 38 | 0 | 0 | 0 | 0 | 0 |
| SRR2529592 | 697 | 80707 | 1553 | 510 | 1420 | 18379 | 21329 | 58500 | 2187 | 74 | 40 | 40211 | 12768 | 16757 | 0 | 691 | 51154 | 0 | 0 | 0 | 98 | 21 | 42 | 0 | 0 | 0 | 0 | 0 | 11 | 55 | 19 | 33 | 0 | 0 | 0 | 0 | 0 |
| SRR2529593 | 547 | 70840 | 1515 | 322 | 2206 | 22769 | 23817 | 60095 | 938 | 38 | 64086 | 40442 | 15338 | 13905 | 0 | 553 | 60301 | 0 | 0 | 0 | 85 | 0 | 16 | 0 | 0 | 0 | 0 | 0 | 19 | 49 | 17 | 26 | 0 | 0 | 0 | 0 | 0 |
| SRR2529594 | 658 | 70574 | 1601 | 274 | 2120 | 22523 | 23972 | 60272 | 923 | 70 | 53 | 40497 | 15405 | 13871 | 0 | 578 | 60480 | 0 | 0 | 0 | 89 | 0 | 13 | 0 | 0 | 0 | 0 | 0 | 17 | 57 | 17 | 18 | 0 | 0 | 0 | 0 | 0 |
| SRR2648184 | 0 | 6077 | 5366 | 23 | 1867 | 22559 | 12924 | 16107 | 17624 | 35 | 718 | 1288 | 83 | 0 | 192 | 3361 | 33385 | 0 | 0 | 0 | 14 | 29 | 12 | 0 | 0 | 0 | 0 | 0 | 12 | 27 | 86 | 61 | 0 | 22 | 0 | 0 | 0 |
| SRR2648185 | 0 | 5259 | 3978 | 15 | 1682 | 21048 | 11724 | 15329 | 16404 | 32 | 654 | 1308 | 78 | 0 | 198 | 3339 | 34218 | 0 | 0 | 0 | 14 | 22 | 10 | 0 | 0 | 0 | 0 | 0 | 12 | 31 | 73 | 57 | 0 | 18 | 0 | 0 | 0 |
| SRR2648186 | 0 | 6650 | 3901 | 21 | 1420 | 17343 | 9669 | 11650 | 16198 | 39 | 366 | 946 | 50 | 0 | 113 | 3619 | 31058 | 0 | 0 | 0 | 17 | 30 | 16 | 0 | 0 | 0 | 0 | 0 | 15 | 39 | 89 | 82 | 0 | 24 | 0 | 0 | 0 |
| SRR2648187 | 0 | 73743 | 796 | 17 | 610 | 7690 | 4737 | 8336 | 24804 | 83 | 590 | 1915 | 83 | 0 | 109 | 2387 | 38188 | 0 | 0 | 0 | 30 | 0 | 16 | 0 | 0 | 0 | 0 | 0 | 17 | 94 | 48 | 88 | 0 | 18 | 0 | 0 | 0 |
| SRR2648188 | 0 | 84570 | 771 | 0 | 677 | 8141 | 5133 | 8845 | 24697 | 96 | 608 | 1795 | 84 | 0 | 116 | 2526 | 38462 | 0 | 0 | 0 | 33 | 0 | 18 | 0 | 0 | 0 | 0 | 0 | 18 | 103 | 52 | 110 | 0 | 21 | 0 | 0 | 0 |
| SRR2648189 | 0 | 65127 | 854 | 14 | 600 | 7164 | 4522 | 7567 | 23127 | 64 | 549 | 1477 | 68 | 0 | 109 | 2356 | 41004 | 0 | 0 | 0 | 32 | 0 | 18 | 0 | 0 | 0 | 0 | 0 | 17 | 87 | 50 | 95 | 0 | 23 | 0 | 0 | 0 |

### Supp Table S8.1.Columns_B_to_AL

| | | | | | | | | | | | | | | | | | | | | | | | | | | | | | | | | | | | | |
|---|---|---|---|---|---|---|---|---|---|---|---|---|---|---|---|---|---|---|---|---|---|---|---|---|---|---|---|---|---|---|---|---|---|---|---|---|---|
| SRR2728221 | 0 | 0 | 5325 | 0 | 0 | 0 | 0 | 0 | 0 | 155 | 1090 | 1918 | 337 | 2204 | 0 | 0 | 261 | 0 | 0 | 0 | 0 | 0 | 0 | 0 | 0 | 0 | 0 | 0 | 0 | 0 | 0 | 0 | 0 | 0 | 0 | 0 | 0 |
| SRR2731278 | 89 | 1358 | 166 | 7398 | 4222 | 5977 | 74188 | 64387 | 2618 | 1348 | 8992 | 4702 | 9061 | 14256 | 658 | 10199 | 26182 | 0 | 33 | 217 | 385 | 133 | 40 | 0 | 0 | 0 | 0 | 103 | 759 | 946 | 120 | 19 | 0 | 0 | 0 | 11 | 0 |
| SRR2731279 | 706 | 2181 | 547 | 4653 | 13795 | 10869 | 136096 | 48170 | 3858 | 1275 | 18888 | 13568 | 14530 | 30461 | 2000 | 6931 | 18656 | 80 | 148 | 250 | 2319 | 1669 | 1159 | 137 | 36 | 47 | 104 | 157 | 390 | 1679 | 873 | 161 | 0 | 0 | 13 | 95 | 0 |
| SRR2731280 | 3529 | 7154 | 28201 | 5108 | 8342 | 3305 | 66868 | 16084 | 5758 | 1997 | 19921 | 19 | 35 | 9997 | 293 | 8500 | 40221 | 48 | 81 | 40 | 241 | 276 | 135 | 44 | 33 | 28 | 51 | 135 | 160 | 265 | 74 | 22 | 0 | 0 | 14 | 69 | 11 |
| SRR2731281 | 931 | 3173 | 1331 | 3928 | 14374 | 6077 | 150204 | 101562 | 4823 | 3592 | 18267 | 16479 | 12270 | 17258 | 608 | 9888 | 30456 | 43 | 322 | 314 | 715 | 315 | 193 | 75 | 29 | 52 | 56 | 447 | 691 | 582 | 316 | 51 | 0 | 0 | 19 | 124 | 0 |
| SRR2731282 | 666 | 1691 | 602 | 8588 | 12051 | 11321 | 115838 | 40067 | 4888 | 1677 | 14945 | 12320 | 12785 | 20556 | 1655 | 10832 | 25570 | 63 | 340 | 1058 | 1326 | 752 | 325 | 90 | 34 | 56 | 71 | 486 | 1733 | 965 | 530 | 75 | 0 | 0 | 22 | 115 | 0 |
| SRR2731283 | 1118 | 2225 | 421 | 6079 | 24627 | 22414 | 263827 | 68174 | 5463 | 1807 | 12599 | 6906 | 7579 | 22590 | 1180 | 21686 | 10852 | 204 | 345 | 551 | 1016 | 1120 | 544 | 206 | 80 | 118 | 123 | 312 | 578 | 510 | 421 | 79 | 0 | 0 | 26 | 246 | 13 |
| SRR2731284 | 981 | 2751 | 427 | 2184 | 15680 | 10443 | 162188 | 79086 | 4267 | 1038 | 13456 | 8435 | 9072 | 17518 | 1003 | 4223 | 26712 | 89 | 202 | 228 | 415 | 476 | 137 | 91 | 32 | 47 | 99 | 258 | 369 | 317 | 386 | 36 | 0 | 0 | 21 | 120 | 0 |
| SRR2731285 | 680 | 2343 | 1077 | 3237 | 17484 | 11634 | 150004 | 93949 | 5552 | 2802 | 15626 | 8529 | 14600 | 16469 | 1098 | 12954 | 22160 | 117 | 420 | 331 | 495 | 939 | 217 | 103 | 43 | 62 | 114 | 468 | 570 | 337 | 529 | 42 | 0 | 0 | 27 | 154 | 12 |
| SRR2731286 | 399 | 2296 | 750 | 4869 | 5696 | 6127 | 89134 | 46760 | 1834 | 2120 | 13741 | 7095 | 8880 | 45450 | 4910 | 12683 | 12847 | 29 | 201 | 443 | 656 | 182 | 116 | 19 | 0 | 16 | 44 | 363 | 1051 | 1123 | 189 | 32 | 0 | 0 | 0 | 38 | 0 |
| SRR2731287 | 731 | 1180 | 1150 | 6056 | 16586 | 13564 | 183827 | 23487 | 4383 | 1408 | 15266 | 5738 | 9300 | 19907 | 4078 | 41087 | 31852 | 0 | 38 | 243 | 438 | 234 | 122 | 22 | 0 | 12 | 0 | 62 | 377 | 221 | 77 | 18 | 0 | 0 | 0 | 33 | 0 |
| SRR2731288 | 718 | 2542 | 1659 | 4816 | 6432 | 7662 | 80917 | 36534 | 5008 | 2721 | 16498 | 10482 | 10171 | 15581 | 908 | 28139 | 27429 | 42 | 1524 | 644 | 948 | 1834 | 440 | 89 | 32 | 67 | 70 | 3064 | 1450 | 983 | 1317 | 136 | 0 | 0 | 30 | 173 | 0 |
| SRR2731289 | 1706 | 2318 | 1840 | 10290 | 26922 | 25307 | 413340 | 18692 | 3013 | 2088 | 13720 | 4969 | 7241 | 23242 | 2156 | 12009 | 13349 | 12 | 148 | 192 | 451 | 273 | 143 | 29 | 17 | 0 | 12 | 108 | 371 | 282 | 58 | 20 | 0 | 0 | 0 | 43 | 0 |
| SRR2731290 | 2132 | 2936 | 1439 | 9041 | 20537 | 11115 | 157343 | 12999 | 9672 | 1207 | 22365 | 9698 | 10411 | 18158 | 1688 | 24542 | 28569 | 26 | 118 | 129 | 302 | 286 | 95 | 21 | 12 | 0 | 30 | 186 | 476 | 367 | 81 | 15 | 0 | 0 | 0 | 25 | 0 |
| SRR2731291 | 863 | 2073 | 3332 | 5690 | 17952 | 24722 | 288631 | 19327 | 9759 | 1946 | 13103 | 7421 | 7333 | 28167 | 1141 | 10850 | 25975 | 0 | 260 | 205 | 289 | 857 | 88 | 20 | 12 | 11 | 0 | 232 | 353 | 144 | 115 | 0 | 0 | 0 | 0 | 36 | 0 |
| SRR2731292 | 1477 | 2546 | 1870 | 15452 | 8495 | 8783 | 139742 | 4514 | 20951 | 1105 | 26274 | 8463 | 8607 | 29292 | 2039 | 24874 | 18918 | 0 | 51 | 66 | 41 | 318 | 46 | 0 | 0 | 0 | 0 | 60 | 128 | 34 | 54 | 0 | 0 | 0 | 0 | 0 | 0 |
| SRR2731293 | 940 | 2138 | 631 | 3603 | 11076 | 8967 | 120424 | 64172 | 6681 | 880 | 13923 | 8038 | 14955 | 26872 | 1696 | 11430 | 23362 | 54 | 186 | 634 | 1014 | 337 | 143 | 37 | 16 | 32 | 64 | 257 | 952 | 573 | 307 | 25 | 0 | 0 | 0 | 47 | 0 |
| SRR2731294 | 888 | 2497 | 1270 | 3701 | 11168 | 5419 | 95695 | 58021 | 4422 | 5091 | 21770 | 16191 | 14327 | 13668 | 905 | 20125 | 28608 | 162 | 196 | 265 | 352 | 476 | 261 | 115 | 51 | 68 | 134 | 179 | 346 | 318 | 450 | 44 | 0 | 0 | 21 | 113 | 0 |
| SRR2731295 | 706 | 2308 | 1558 | 4307 | 15295 | 14271 | 213505 | 82720 | 1854 | 7170 | 20036 | 11736 | 13160 | 12850 | 686 | 7898 | 20203 | 90 | 894 | 337 | 310 | 1201 | 191 | 96 | 41 | 57 | 94 | 1103 | 484 | 302 | 691 | 53 | 0 | 0 | 26 | 163 | 10 |
| SRR2731296 | 477 | 1849 | 506 | 4871 | 10661 | 7929 | 78535 | 46990 | 4337 | 1559 | 16665 | 14160 | 15319 | 17730 | 1190 | 18012 | 30277 | 46 | 238 | 348 | 455 | 503 | 139 | 79 | 27 | 56 | 43 | 320 | 683 | 403 | 370 | 34 | 0 | 0 | 19 | 96 | 0 |
| SRR2731297 | 519 | 1331 | 319 | 4067 | 8159 | 6374 | 68243 | 48912 | 3624 | 1229 | 16343 | 12602 | 13708 | 11117 | 923 | 22913 | 24648 | 47 | 157 | 416 | 561 | 331 | 111 | 42 | 16 | 27 | 71 | 272 | 750 | 460 | 285 | 31 | 0 | 0 | 10 | 47 | 0 |
| SRR2731298 | 848 | 2117 | 15672 | 4713 | 18990 | 14745 | 365874 | 45785 | 8800 | 641 | 16485 | 11887 | 15643 | 16586 | 1000 | 2501 | 13747 | 12 | 532 | 126 | 840 | 693 | 476 | 44 | 21 | 0 | 22 | 1286 | 390 | 787 | 479 | 179 | 0 | 0 | 22 | 65 | 0 |
| SRR2731299 | 659 | 2191 | 569 | 5523 | 15107 | 9873 | 94161 | 54082 | 11416 | 1808 | 17790 | 11907 | 11057 | 12521 | 623 | 13442 | 32442 | 70 | 1109 | 1729 | 2312 | 701 | 203 | 52 | 21 | 39 | 52 | 870 | 1871 | 1002 | 354 | 26 | 0 | 0 | 0 | 36 | 0 |
| SRR2731300 | 610 | 1951 | 903 | 2712 | 5763 | 3296 | 44549 | 28456 | 5873 | 2575 | 16334 | 16869 | 12784 | 14919 | 396 | 10778 | 34854 | 108 | 199 | 233 | 242 | 392 | 152 | 74 | 35 | 46 | 154 | 298 | 467 | 259 | 463 | 36 | 0 | 0 | 22 | 80 | 10 |
| SRR2731301 | 880 | 2262 | 884 | 3115 | 9242 | 3866 | 77105 | 48685 | 4675 | 2346 | 12365 | 14422 | 12256 | 14433 | 308 | 20017 | 28782 | 81 | 459 | 1634 | 3648 | 571 | 471 | 93 | 38 | 74 | 94 | 636 | 2580 | 2206 | 446 | 85 | 0 | 0 | 17 | 136 | 0 |
| SRR2731302 | 1133 | 2516 | 1077 | 4973 | 6251 | 4979 | 46738 | 18227 | 7501 | 2178 | 18366 | 16414 | 13162 | 13781 | 634 | 34152 | 45649 | 27 | 1175 | 2891 | 3571 | 2130 | 385 | 66 | 22 | 53 | 33 | 1121 | 5295 | 2072 | 891 | 61 | 0 | 0 | 13 | 66 | 0 |
| SRR2731303 | 2495 | 3009 | 15628 | 8454 | 11759 | 5917 | 133584 | 46168 | 18342 | 2499 | 20715 | 16 | 48 | 15288 | 320 | 5341 | 25720 | 32 | 985 | 229 | 1351 | 672 | 272 | 34 | 15 | 22 | 36 | 1551 | 777 | 1096 | 253 | 47 | 0 | 0 | 0 | 39 | 0 |
| SRR2731304 | 681 | 2781 | 1136 | 3564 | 35404 | 16786 | 247606 | 137427 | 5262 | 5906 | 13709 | 7907 | 8641 | 12581 | 854 | 11516 | 18955 | 169 | 433 | 945 | 1434 | 782 | 351 | 132 | 45 | 82 | 73 | 453 | 940 | 809 | 495 | 37 | 0 | 0 | 15 | 120 | 0 |
| SRR2731305 | 870 | 2187 | 575 | 4149 | 10110 | 6788 | 77219 | 41108 | 5346 | 1280 | 14341 | 11618 | 9521 | 12802 | 634 | 18433 | 35481 | 148 | 323 | 817 | 855 | 1624 | 387 | 97 | 30 | 42 | 109 | 235 | 723 | 354 | 722 | 46 | 0 | 0 | 12 | 70 | 0 |
| SRR2731306 | 868 | 1734 | 605 | 7271 | 16948 | 13484 | 117184 | 42679 | 6459 | 1136 | 18249 | 11979 | 15414 | 19574 | 1764 | 17206 | 29709 | 61 | 486 | 701 | 888 | 446 | 142 | 38 | 13 | 23 | 62 | 485 | 1317 | 565 | 289 | 24 | 0 | 0 | 0 | 33 | 0 |
| SRR2731307 | 506 | 1457 | 598 | 4321 | 7056 | 7668 | 46161 | 17697 | 21282 | 1889 | 32914 | 13794 | 13433 | 14794 | 680 | 25483 | 36789 | 30 | 89 | 121 | 135 | 242 | 103 | 27 | 10 | 12 | 54 | 144 | 373 | 235 | 357 | 37 | 0 | 0 | 10 | 37 | 0 |
| SRR2731308 | 1059 | 2743 | 604 | 4331 | 18070 | 13087 | 143271 | 57882 | 3361 | 970 | 14642 | 11285 | 16544 | 22535 | 1591 | 5397 | 28518 | 59 | 393 | 522 | 616 | 706 | 229 | 70 | 35 | 38 | 83 | 498 | 1207 | 503 | 419 | 65 | 0 | 0 | 35 | 121 | 0 |
| SRR2731309 | 1533 | 3068 | 663 | 3627 | 6070 | 4611 | 48749 | 28697 | 7571 | 1397 | 23191 | 16572 | 11786 | 10538 | 785 | 23485 | 43697 | 85 | 235 | 289 | 469 | 416 | 123 | 43 | 19 | 27 | 78 | 282 | 599 | 364 | 354 | 25 | 0 | 0 | 0 | 51 | 0 |
| SRR2731310 | 669 | 1903 | 662 | 5529 | 12348 | 9783 | 114654 | 55852 | 5222 | 2072 | 15194 | 15273 | 12557 | 18737 | 531 | 24027 | 33215 | 55 | 445 | 795 | 924 | 551 | 165 | 49 | 16 | 25 | 58 | 398 | 935 | 462 | 418 | 34 | 0 | 0 | 0 | 46 | 0 |
| SRR2731311 | 714 | 2200 | 661 | 3684 | 2231 | 3010 | 33854 | 22111 | 2300 | 1218 | 20533 | 11406 | 10080 | 15536 | 1375 | 23401 | 21774 | 31 | 166 | 700 | 1134 | 304 | 160 | 21 | 0 | 18 | 40 | 232 | 1210 | 1061 | 251 | 43 | 0 | 0 | 0 | 30 | 0 |
| SRR2731312 | 727 | 1861 | 1267 | 4674 | 6093 | 5468 | 66939 | 28232 | 5564 | 842 | 20147 | 16898 | 13766 | 15559 | 996 | 10970 | 30197 | 63 | 263 | 551 | 720 | 421 | 203 | 30 | 16 | 18 | 56 | 314 | 782 | 425 | 309 | 47 | 0 | 0 | 0 | 35 | 0 |
| SRR2731313 | 955 | 2404 | 1160 | 3806 | 10467 | 7104 | 95514 | 59801 | 3279 | 1808 | 21587 | 11656 | 12744 | 14607 | 677 | 7351 | 35005 | 53 | 1470 | 767 | 995 | 1078 | 293 | 41 | 16 | 25 | 78 | 2114 | 1415 | 724 | 914 | 101 | 0 | 0 | 11 | 56 | 0 |
| SRR2731314 | 635 | 2231 | 1542 | 2154 | 7798 | 4682 | 46100 | 49690 | 4761 | 3455 | 17316 | 12788 | 9189 | 7236 | 384 | 15749 | 32183 | 124 | 232 | 289 | 248 | 604 | 141 | 102 | 37 | 63 | 135 | 252 | 504 | 333 | 761 | 27 | 0 | 0 | 21 | 109 | 11 |
| SRR2731315 | 697 | 3968 | 837 | 2814 | 7904 | 4299 | 68580 | 49446 | 5146 | 1783 | 18831 | 15391 | 12778 | 14169 | 635 | 13436 | 32392 | 91 | 2988 | 1924 | 2701 | 1106 | 342 | 110 | 32 | 63 | 55 | 1804 | 1697 | 1229 | 675 | 58 | 0 | 0 | 0 | 79 | 0 |
| SRR2731316 | 675 | 2472 | 1061 | 2931 | 8247 | 5778 | 59231 | 46005 | 9724 | 1172 | 28868 | 18142 | 13738 | 18443 | 691 | 12472 | 33366 | 130 | 238 | 348 | 330 | 429 | 128 | 63 | 24 | 40 | 115 | 212 | 467 | 284 | 444 | 25 | 0 | 0 | 12 | 46 | 0 |
| SRR2731317 | 1011 | 2955 | 1558 | 3297 | 8916 | 5536 | 61814 | 52098 | 2420 | 1472 | 17255 | 14132 | 12430 | 16517 | 826 | 13024 | 37465 | 100 | 542 | 663 | 383 | 1548 | 255 | 92 | 27 | 52 | 95 | 465 | 787 | 264 | 797 | 54 | 0 | 0 | 15 | 77 | 0 |
| SRR2731318 | 182 | 797 | 509 | 3028 | 10613 | 9933 | 142043 | 69393 | 3263 | 1148 | 12273 | 7599 | 10039 | 28033 | 4692 | 14020 | 11212 | 87 | 255 | 437 | 595 | 281 | 178 | 59 | 26 | 37 | 73 | 381 | 587 | 550 | 219 | 30 | 0 | 0 | 0 | 78 | 0 |
| SRR2731319 | 1267 | 3639 | 1049 | 4039 | 11217 | 5395 | 87553 | 48660 | 3626 | 1632 | 16022 | 13734 | 13961 | 11156 | 1094 | 11184 | 32944 | 99 | 243 | 427 | 877 | 616 | 231 | 88 | 35 | 44 | 115 | 345 | 896 | 692 | 487 | 58 | 0 | 0 | 21 | 98 | 0 |
| SRR2731320 | 455 | 1461 | 2532 | 5505 | 8253 | 7936 | 80426 | 42616 | 4214 | 953 | 20934 | 19969 | 12513 | 12846 | 1087 | 23747 | 24522 | 49 | 163 | 354 | 350 | 398 | 249 | 62 | 28 | 23 | 78 | 272 | 613 | 364 | 452 | 82 | 0 | 0 | 14 | 51 | 0 |
| SRR2731321 | 346 | 958 | 1317 | 1419 | 6911 | 3428 | 65832 | 30071 | 10920 | 1027 | 15152 | 8077 | 6970 | 12014 | 604 | 17789 | 23520 | 139 | 297 | 344 | 331 | 471 | 247 | 83 | 38 | 41 | 84 | 213 | 291 | 233 | 280 | 49 | 0 | 0 | 11 | 95 | 10 |
| SRR2731322 | 3266 | 8214 | 3435 | 2169 | 7844 | 2786 | 61262 | 36016 | 4945 | 770 | 16413 | 11384 | 8245 | 6341 | 385 | 11516 | 18514 | 197 | 230 | 501 | 1080 | 618 | 502 | 212 | 102 | 114 | 169 | 286 | 654 | 808 | 450 | 107 | 10 | 0 | 39 | 188 | 10 |
| SRR2731323 | 587 | 2408 | 357 | 3740 | 5790 | 5039 | 76961 | 43415 | 1442 | 165 | 13076 | 3892 | 4487 | 32453 | 1868 | 27288 | 6889 | 104 | 443 | 425 | 252 | 361 | 71 | 37 | 13 | 27 | 109 | 422 | 544 | 467 | 303 | 15 | 0 | 0 | 0 | 56 | 0 |
| SRR2731324 | 433 | 1723 | 770 | 2582 | 10927 | 8382 | 108815 | 90695 | 9334 | 2818 | 11936 | 7143 | 8724 | 8549 | 674 | 7575 | 21412 | 87 | 304 | 620 | 591 | 1301 | 229 | 78 | 33 | 71 | 88 | 428 | 1149 | 661 | 1061 | 60 | 0 | 0 | 26 | 114 | 14 |
| SRR2731325 | 524 | 1977 | 819 | 3003 | 7058 | 4750 | 73222 | 77639 | 4592 | 3164 | 16534 | 17127 | 13398 | 15322 | 296 | 21452 | 33763 | 86 | 169 | 431 | 712 | 466 | 176 | 103 | 37 | 78 | 77 | 218 | 624 | 535 | 362 | 40 | 0 | 0 | 19 | 134 | 0 |
| SRR2731326 | 1088 | 3481 | 1433 | 3712 | 13388 | 9995 | 103932 | 51887 | 2653 | 1190 | 16464 | 13831 | 10793 | 13974 | 646 | 6387 | 26880 | 115 | 205 | 305 | 604 | 712 | 258 | 91 | 28 | 40 | 130 | 258 | 543 | 444 | 545 | 57 | 0 | 0 | 23 | 85 | 0 |
| SRR2731327 | 914 | 2253 | 1042 | 1884 | 6893 | 4024 | 50247 | 40257 | 3272 | 2086 | 21206 | 19144 | 12012 | 11900 | 2682 | 10228 | 32004 | 119 | 270 | 326 | 309 | 459 | 145 | 66 | 30 | 44 | 146 | 456 | 624 | 299 | 516 | 37 | 0 | 0 | 18 | 83 | 0 |
| SRR2731328 | 403 | 2066 | 650 | 4137 | 7834 | 7179 | 85777 | 58134 | 3084 | 2130 | 17918 | 13160 | 12333 | 25611 | 1220 | 4133 | 19561 | 39 | 359 | 571 | 778 | 830 | 194 | 49 | 16 | 29 | 53 | 575 | 887 | 740 | 669 | 43 | 0 | 0 | 0 | 52 | 0 |
| SRR2731329 | 1418 | 4558 | 1961 | 3397 | 29819 | 31031 | 263646 | 45502 | 1439 | 1413 | 13459 | 10299 | 11131 | 15585 | 2526 | 11709 | 18598 | 104 | 1878 | 645 | 649 | 2892 | 334 | 114 | 50 | 82 | 87 | 2061 | 906 | 375 | 898 | 55 | 0 | 0 | 47 | 130 | 0 |
| SRR2731330 | 265 | 1561 | 330 | 5759 | 2917 | 5882 | 63214 | 18210 | 865 | 719 | 7263 | 2878 | 3247 | 45345 | 6232 | 15060 | 3383 | 20 | 397 | 876 | 656 | 202 | 77 | 0 | 0 | 0 | 27 | 1382 | 1839 | 2594 | 198 | 21 | 0 | 0 | 0 | 20 | 0 |
| SRR2731331 | 921 | 1675 | 3672 | 4531 | 10543 | 6521 | 55565 | 45896 | 6971 | 4453 | 37700 | 24256 | 17740 | 25274 | 633 | 32350 | 37398 | 0 | 120 | 91 | 64 | 95 | 43 | 22 | 0 | 0 | 0 | 62 | 64 | 12 | 33 | 0 | 0 | 0 | 0 | 12 | 0 |
| SRR2731332 | 1450 | 3835 | 1922 | 6769 | 9410 | 7671 | 60469 | 41084 | 16461 | 1188 | 25035 | 19452 | 14699 | 19659 | 623 | 13286 | 49767 | 54 | 248 | 539 | 309 | 801 | 110 | 45 | 12 | 23 | 27 | 340 | 902 | 231 | 461 | 17 | 0 | 0 | 0 | 34 | 0 |
| SRR2731333 | 495 | 1799 | 1108 | 7831 | 17493 | 28526 | 186972 | 66668 | 11218 | 3636 | 14292 | 7969 | 7681 | 27420 | 2007 | 6919 | 14463 | 40 | 285 | 1956 | 1532 | 679 | 295 | 62 | 16 | 46 | 20 | 417 | 2526 | 1251 | 387 | 41 | 0 | 0 | 0 | 66 | 0 |
| SRR2731335 | 708 | 1930 | 393 | 4376 | 19686 | 13402 | 225786 | 118261 | 5591 | 1044 | 9770 | 5076 | 7918 | 9248 | 786 | 16784 | 11693 | 65 | 201 | 895 | 2063 | 326 | 247 | 72 | 29 | 52 | 55 | 204 | 1022 | 1057 | 211 | 50 | 0 | 0 | 11 | 85 | 0 |
| SRR2731336 | 1153 | 2900 | 560 | 7378 | 11069 | 11333 | 90893 | 47046 | 7003 | 1585 | 22319 | 6038 | 8842 | 72689 | 5840 | 22435 | 9056 | 0 | 283 | 276 | 235 | 144 | 65 | 14 | 0 | 0 | 0 | 215 | 228 | 77 | 48 | 0 | 0 | 0 | 0 | 11 | 0 |
| SRR2731337 | 796 | 3750 | 1891 | 2809 | 11267 | 6316 | 69118 | 44913 | 10076 | 9185 | 36733 | 20268 | 15129 | 22198 | 645 | 13180 | 46632 | 0 | 213 | 263 | 270 | 375 | 79 | 38 | 12 | 11 | 0 | 139 | 181 | 76 | 106 | 0 | 0 | 0 | 0 | 25 | 0 |
| SRR2731338 | 184 | 550 | 994 | 4022 | 12779 | 9241 | 66788 | 48819 | 11317 | 1283 | 26299 | 12621 | 12453 | 23194 | 1174 | 26245 | 21710 | 13 | 253 | 359 | 224 | 168 | 94 | 23 | 0 | 0 | 0 | 262 | 274 | 95 | 81 | 0 | 0 | 0 | 0 | 13 | 0 |
| SRR2731339 | 167 | 906 | 3339 | 518 | 5466 | 4274 | 63768 | 24023 | 3936 | 849 | 18264 | 17162 | 11404 | 20919 | 1215 | 9425 | 42849 | 11 | 38 | 56 | 66 | 55 | 50 | 12 | 11 | 0 | 13 | 65 | 96 | 46 | 55 | 11 | 0 | 0 | 0 | 19 | 0 |
| SRR2731340 | 450 | 2326 | 1384 | 5550 | 24649 | 18589 | 152011 | 69996 | 6903 | 3263 | 22857 | 12962 | 18406 | 38970 | 4574 | 12533 | 21311 | 32 | 210 | 316 | 525 | 3521 | 2191 | 323 | 79 | 56 | 17 | 321 | 535 | 420 | 960 | 178 | 0 | 0 | 30 | 97 | 0 |
| SRR2731341 | 563 | 2258 | 993 | 3117 | 7401 | 4047 | 44952 | 37285 | 4141 | 1291 | 18061 | 18244 | 12095 | 11114 | 459 | 17034 | 33942 | 130 | 259 | 554 | 420 | 422 | 132 | 67 | 25 | 29 | 112 | 280 | 800 | 356 | 451 | 36 | 0 | 0 | 13 | 52 | 0 |
| SRR2731342 | 597 | 1744 | 884 | 2561 | 6013 | 3345 | 40710 | 41072 | 6699 | 2174 | 24387 | 19096 | 13902 | 11858 | 499 | 14450 | 31933 | 136 | 260 | 418 | 460 | 408 | 157 | 80 | 45 | 55 | 83 | 259 | 512 | 296 | 333 | 23 | 0 | 0 | 19 | 79 | 0 |
| SRR2731343 | 376 | 1438 | 827 | 3042 | 10949 | 10358 | 113784 | 70409 | 4798 | 2452 | 26560 | 13652 | 15673 | 38109 | 1348 | 14929 | 17321 | 118 | 249 | 378 | 505 | 390 | 206 | 71 | 27 | 36 | 119 | 322 | 606 | 534 | 400 | 37 | 0 | 0 | 0 | 55 | 0 |
| SRR2731344 | 580 | 2136 | 2828 | 3612 | 8700 | 5611 | 86929 | 53441 | 4001 | 4149 | 15967 | 28369 | 11111 | 18992 | 249 | 17795 | 35984 | 152 | 277 | 459 | 388 | 1399 | 337 | 130 | 48 | 75 | 132 | 262 | 592 | 301 | 1069 | 55 | 0 | 0 | 21 | 144 | 14 |
| SRR2731345 | 1123 | 3064 | 28064 | 5610 | 2825 | 2160 | 33780 | 19387 | 18175 | 2644 | 35793 | 43430 | 35142 | 7808 | 399 | 12869 | 44399 | 25 | 70 | 96 | 148 | 152 | 80 | 16 | 11 | 0 | 20 | 114 | 236 | 107 | 79 | 10 | 0 | 0 | 0 | 19 | 0 |
| SRR2731346 | 214 | 1066 | 888 | 3795 | 7363 | 5892 | 66155 | 40664 | 3943 | 2135 | 15052 | 17114 | 13659 | 16568 | 485 | 6670 | 27122 | 23 | 249 | 1248 | 1449 | 479 | 138 | 41 | 18 | 37 | 41 | 554 | 2281 | 1224 | 502 | 34 | 0 | 0 | 15 | 70 | 0 |
| SRR2731347 | 441 | 1747 | 1225 | 5009 | 12123 | 7951 | 82047 | 54765 | 2224 | 5264 | 25736 | 17869 | 14762 | 19172 | 435 | 18515 | 27834 | 72 | 412 | 526 | 416 | 491 | 119 | 44 | 19 | 29 | 46 | 526 | 836 | 338 | 358 | 18 | 0 | 0 | 0 | 43 | 0 |
| SRR2731348 | 428 | 1876 | 884 | 2490 | 14289 | 6577 | 135703 | 106056 | 6806 | 3343 | 15314 | 14289 | 12678 | 14915 | 916 | 8743 | 23489 | 52 | 592 | 1056 | 2282 | 511 | 320 | 97 | 39 | 80 | 51 | 484 | 914 | 954 | 282 | 36 | 0 | 0 | 0 | 100 | 0 |
| SRR2731349 | 766 | 3156 | 13365 | 4289 | 8191 | 5142 | 104532 | 56558 | 6506 | 7455 | 29526 | 17 | 45 | 12792 | 728 | 11001 | 25804 | 20 | 309 | 641 | 2699 | 793 | 269 | 24 | 13 | 15 | 29 | 552 | 1864 | 2685 | 254 | 31 | 0 | 0 | 0 | 37 | 0 |
| SRR2731350 | 713 | 3111 | 1475 | 3347 | 10897 | 7964 | 125738 | 69444 | 1908 | 4122 | 13541 | 9930 | 14888 | 21483 | 688 | 19588 | 27731 | 50 | 866 | 1063 | 1917 | 613 | 227 | 89 | 36 | 77 | 39 | 592 | 1678 | 823 | 360 | 27 | 0 | 0 | 16 | 113 | 11 |
| SRR2910706 | 165 | 52292 | 2530 | 760 | 2650 | 33560 | 27032 | 36353 | 3883 | 15 | 21229 | 23365 | 18072 | 5681 | 17 | 48 | 68583 | 0 | 0 | 0 | 41 | 63 | 24 | 0 | 0 | 0 | 0 | 13 | 41 | 89 | 358 | 70 | 0 | 0 | 0 | 0 | 0 |
| SRR2910737 | 97 | 36270 | 3391 | 500 | 2261 | 30653 | 21656 | 35845 | 3558 | 25 | 24752 | 23655 | 21092 | 6603 | 12 | 46 | 62822 | 0 | 0 | 0 | 31 | 41 | 17 | 0 | 0 | 0 | 0 | 11 | 36 | 79 | 265 | 55 | 0 | 0 | 0 | 0 | 0 |
| SRR2910738 | 227 | 43755 | 3531 | 687 | 2593 | 34087 | 26908 | 37876 | 3590 | 11 | 21121 | 20332 | 17762 | 5552 | 11 | 57 | 57894 | 0 | 0 | 0 | 43 | 61 | 20 | 0 | 0 | 0 | 0 | 13 | 50 | 102 | 345 | 71 | 0 | 0 | 0 | 0 | 0 |
| SRR2912507 | 0 | 15 | 0 | 265 | 21 | 0 | 4093 | 26721 | 709 | 58 | 719 | 704 | 765 | 252 | 1268 | 10 | 1880 | 0 | 0 | 394 | 200 | 97 | 296 | 79 | 19 | 69 | 0 | 0 | 275 | 180 | 74 | 1197 | 0 | 15 | 58 | 35 | 25 |
| SRR2912508 | 13 | 134 | 12 | 2079 | 21 | 11 | 6239 | 34243 | 466 | 199 | 654 | 884 | 783 | 284 | 1501 | 27 | 1838 | 0 | 0 | 506 | 223 | 105 | 174 | 46 | 35 | 145 | 0 | 0 | 272 | 165 | 49 | 519 | 0 | 19 | 81 | 74 | 47 |
| SRR2912510 | 35 | 776 | 0 | 495 | 36 | 0 | 5404 | 34496 | 514 | 18 | 702 | 1450 | 707 | 321 | 1130 | 0 | 1155 | 0 | 0 | 574 | 69 | 130 | 2631 | 1957 | 51 | 444 | 0 | 0 | 128 | 63 | 54 | 2529 | 51 | 32 | 89 | 187 | 477 |
| SRR2912511 | 24 | 938 | 0 | 479 | 47 | 0 | 5005 | 38275 | 419 | 12 | 634 | 1610 | 605 | 250 | 831 | 12 | 1203 | 0 | 0 | 434 | 53 | 101 | 1908 | 2039 | 40 | 277 | 0 | 0 | 98 | 54 | 41 | 2017 | 103 | 23 | 68 | 143 | 376 |
| SRR2912513 | 0 | 216 | 0 | 279 | 59 | 12 | 10643 | 71313 | 44 | 0 | 1220 | 1729 | 1070 | 1673 | 549 | 883 | 1946 | 0 | 0 | 343 | 55 | 149 | 2477 | 2207 | 39 | 74 | 0 | 0 | 114 | 62 | 78 | 3589 | 124 | 47 | 38 | 51 | 51 |
| SRR2912514 | 0 | 54 | 0 | 519 | 32 | 0 | 6107 | 29340 | 75 | 81 | 1446 | 1073 | 1118 | 1273 | 971 | 438 | 1804 | 0 | 0 | 397 | 177 | 70 | 285 | 111 | 24 | 90 | 0 | 0 | 285 | 164 | 45 | 919 | 0 | 21 | 72 | 49 | 33 |
| SRR2912515 | 16 | 309 | 0 | 336 | 117 | 0 | 13324 | 80530 | 34 | 0 | 1134 | 1951 | 1041 | 1811 | 828 | 849 | 1295 | 0 | 0 | 462 | 58 | 175 | 1341 | 2156 | 49 | 100 | 0 | 0 | 135 | 63 | 75 | 1665 | 88 | 42 | 33 | 59 | 41 |
| SRR2913202 | 750 | 20599 | 716 | 2183 | 142 | 1562 | 962 | 4627 | 5285 | 14 | 25007 | 51021 | 7828 | 7473 | 0 | 4517 | 51847 | 0 | 0 | 12 | 5700 | 339 | 0 | 0 | 0 | 0 | 0 | 0 | 224 | 2343 | 219 | 0 | 0 | 0 | 10 | 0 | 0 |
| SRR2913203 | 482 | 27544 | 502 | 1876 | 79 | 559 | 355 | 1275 | 5552 | 13 | 20511 | 42456 | 21 | 11348 | 0 | 1658 | 23315 | 0 | 0 | 0 | 7541 | 913 | 69 | 0 | 0 | 0 | 0 | 0 | 300 | 4850 | 1071 | 70 | 0 | 0 | 0 | 0 | 0 |
| SRR2913204 | 605 | 17402 | 872 | 932 | 134 | 1167 | 1001 | 4069 | 5647 | 0 | 19208 | 44828 | 5781 | 6307 | 0 | 3553 | 68013 | 0 | 0 | 13 | 9969 | 1826 | 186 | 0 | 0 | 0 | 0 | 0 | 291 | 3142 | 985 | 91 | 0 | 0 | 0 | 0 | 0 |
| SRR2913205 | 856 | 18806 | 987 | 1283 | 93 | 1819 | 921 | 5163 | 10078 | 0 | 16766 | 27873 | 3917 | 2345 | 0 | 5016 | 57712 | 0 | 0 | 12 | 11785 | 1563 | 99 | 0 | 0 | 0 | 0 | 0 | 360 | 3911 | 939 | 53 | 0 | 0 | 0 | 0 | 0 |
| SRR2913206 | 1135 | 16357 | 1180 | 1422 | 157 | 1795 | 1113 | 4562 | 8764 | 0 | 26567 | 38209 | 5307 | 5213 | 0 | 4069 | 44935 | 0 | 0 | 0 | 6424 | 252 | 13 | 0 | 0 | 0 | 0 | 0 | 187 | 2243 | 203 | 11 | 0 | 0 | 0 | 0 | 0 |
| SRR2913207 | 704 | 16415 | 1162 | 709 | 133 | 1384 | 1164 | 4985 | 5465 | 0 | 12727 | 18430 | 2493 | 1971 | 0 | 4139 | 69556 | 0 | 0 | 42 | 34934 | 4910 | 441 | 0 | 0 | 0 | 0 | 0 | 1408 | 16651 | 4653 | 296 | 0 | 0 | 25 | 11 | 0 |
| SRR2913208 | 995 | 10950 | 1268 | 653 | 138 | 1420 | 844 | 4594 | 7906 | 0 | 16615 | 26816 | 16 | 2264 | 0 | 5077 | 33564 | 0 | 0 | 0 | 2532 | 442 | 40 | 0 | 0 | 0 | 0 | 0 | 82 | 710 | 261 | 21 | 0 | 0 | 0 | 0 | 0 |
| SRR2913209 | 665 | 11802 | 838 | 1478 | 77 | 897 | 670 | 3422 | 1961 | 11 | 12586 | 33278 | 2831 | 3570 | 0 | 3302 | 64720 | 0 | 0 | 0 | 1415 | 937 | 137 | 0 | 0 | 0 | 0 | 0 | 67 | 424 | 529 | 70 | 0 | 0 | 11 | 0 | 0 |
| SRR2913210 | 598 | 20132 | 612 | 1244 | 173 | 1694 | 1248 | 5679 | 3202 | 0 | 37774 | 75060 | 25 | 12010 | 0 | 4327 | 42374 | 0 | 0 | 0 | 3337 | 433 | 19 | 0 | 0 | 0 | 0 | 0 | 119 | 959 | 233 | 14 | 0 | 0 | 0 | 0 | 0 |
| SRR2913211 | 819 | 18839 | 865 | 826 | 266 | 1804 | 1515 | 6364 | 3716 | 0 | 22114 | 38167 | 5949 | 5394 | 0 | 4812 | 60631 | 0 | 0 | 0 | 1723 | 143 | 0 | 0 | 0 | 0 | 0 | 0 | 75 | 445 | 84 | 0 | 0 | 0 | 0 | 0 | 0 |
| SRR2913212 | 541 | 8793 | 527 | 2199 | 68 | 1417 | 952 | 3477 | 2691 | 19 | 27883 | 88984 | 8988 | 16649 | 0 | 2283 | 30963 | 0 | 0 | 0 | 2754 | 550 | 77 | 0 | 0 | 0 | 0 | 0 | 108 | 1437 | 567 | 89 | 0 | 0 | 0 | 0 | 0 |
| SRR2913213 | 443 | 18127 | 1365 | 2049 | 127 | 1490 | 817 | 3863 | 3539 | 13 | 21136 | 57360 | 5502 | 9086 | 0 | 3264 | 72637 | 0 | 0 | 0 | 1820 | 256 | 11 | 0 | 0 | 0 | 0 | 0 | 70 | 595 | 179 | 0 | 0 | 0 | 0 | 0 | 0 |
| SRR2913214 | 1120 | 12829 | 1222 | 1510 | 158 | 1802 | 805 | 2499 | 24907 | 0 | 10212 | 17734 | 3105 | 1237 | 0 | 4932 | 39735 | 0 | 0 | 0 | 1797 | 72 | 0 | 0 | 0 | 0 | 0 | 0 | 86 | 987 | 97 | 0 | 0 | 0 | 0 | 0 | 0 |
| SRR2913215 | 815 | 9314 | 1248 | 655 | 212 | 2731 | 1244 | 4796 | 21451 | 0 | 15827 | 27476 | 14 | 2623 | 0 | 5512 | 61384 | 0 | 0 | 0 | 6627 | 651 | 54 | 0 | 0 | 0 | 0 | 0 | 264 | 3390 | 544 | 43 | 0 | 0 | 0 | 0 | 0 |
| SRR2913216 | 623 | 18078 | 1135 | 1234 | 84 | 865 | 607 | 2128 | 2017 | 0 | 10760 | 33138 | 3698 | 4382 | 0 | 4110 | 64196 | 0 | 0 | 0 | 10824 | 1742 | 90 | 0 | 0 | 0 | 0 | 0 | 305 | 4245 | 965 | 50 | 0 | 0 | 0 | 0 | 0 |
| SRR2913217 | 899 | 13926 | 1258 | 727 | 83 | 1080 | 732 | 3380 | 6039 | 0 | 10006 | 22120 | 3339 | 3370 | 0 | 3131 | 38403 | 0 | 0 | 0 | 4077 | 453 | 34 | 0 | 0 | 0 | 0 | 0 | 158 | 1718 | 428 | 29 | 0 | 0 | 0 | 0 | 0 |
| SRR2913218 | 770 | 12133 | 1111 | 909 | 234 | 2811 | 1737 | 5109 | 16093 | 0 | 16429 | 24601 | 4194 | 2970 | 0 | 4380 | 50603 | 0 | 0 | 20 | 24610 | 1627 | 128 | 0 | 0 | 0 | 0 | 0 | 672 | 9883 | 1484 | 80 | 0 | 0 | 14 | 0 | 0 |
| SRR2913219 | 907 | 17037 | 1327 | 609 | 309 | 3194 | 1912 | 6217 | 12812 | 0 | 15676 | 23174 | 3348 | 2148 | 0 | 5688 | 57940 | 0 | 0 | 0 | 4871 | 170 | 10 | 0 | 0 | 0 | 0 | 0 | 156 | 1623 | 132 | 0 | 0 | 0 | 0 | 0 | 0 |
| SRR2913220 | 836 | 18169 | 1378 | 652 | 146 | 1469 | 1197 | 5181 | 6383 | 0 | 16128 | 32203 | 4296 | 4714 | 0 | 5794 | 52418 | 0 | 0 | 16 | 16147 | 1727 | 123 | 0 | 0 | 0 | 0 | 0 | 309 | 4689 | 1083 | 63 | 0 | 0 | 0 | 0 | 0 |
| SRR2913221 | 527 | 15719 | 1101 | 419 | 193 | 1538 | 1137 | 6421 | 7916 | 0 | 11959 | 20611 | 2855 | 2418 | 0 | 6212 | 63439 | 0 | 0 | 42 | 50097 | 4864 | 347 | 0 | 0 | 0 | 0 | 0 | 774 | 12368 | 2781 | 140 | 0 | 0 | 11 | 0 | 0 |
| SRR2913222 | 1082 | 19326 | 1790 | 669 | 171 | 2066 | 1200 | 4314 | 11987 | 13 | 19087 | 27265 | 3905 | 3887 | 0 | 5287 | 53260 | 0 | 0 | 0 | 9519 | 931 | 89 | 0 | 0 | 0 | 0 | 0 | 332 | 4076 | 855 | 61 | 0 | 0 | 0 | 0 | 0 |
| SRR2913223 | 832 | 21429 | 1200 | 1691 | 59 | 738 | 369 | 1143 | 3142 | 26 | 19055 | 36278 | 4903 | 9532 | 0 | 1320 | 14981 | 0 | 0 | 15 | 15198 | 1439 | 71 | 0 | 0 | 0 | 0 | 0 | 468 | 7651 | 1411 | 48 | 0 | 0 | 0 | 0 | 0 |
| SRR2913224 | 597 | 14725 | 906 | 634 | 135 | 1660 | 857 | 2966 | 14686 | 0 | 11385 | 24442 | 2994 | 2244 | 0 | 4365 | 60465 | 0 | 0 | 16 | 20230 | 1522 | 92 | 0 | 0 | 0 | 0 | 0 | 787 | 11894 | 2042 | 73 | 0 | 0 | 0 | 0 | 0 |
| SRR2913225 | 666 | 12489 | 1054 | 777 | 177 | 1961 | 932 | 3152 | 14044 | 0 | 17152 | 40458 | 12 | 4163 | 0 | 4830 | 59565 | 0 | 0 | 0 | 8363 | 839 | 119 | 0 | 0 | 0 | 0 | 0 | 666 | 5978 | 1469 | 183 | 0 | 10 | 12 | 0 | 0 |
| SRR2915342 | 899 | 4085 | 35728 | 1616 | 33535 | 18889 | 467169 | 16983 | 1940 | 3233 | 12778 | 12530 | 5485 | 14856 | 1246 | 13596 | 10501 | 0 | 12 | 13 | 164 | 405 | 457 | 86 | 80 | 29 | 20 | 14 | 49 | 52 | 37 | 11 | 0 | 0 | 15 | 94 | 0 |
| SRR2915343 | 1274 | 6852 | 38426 | 1107 | 30408 | 17977 | 436536 | 21286 | 1891 | 3551 | 13227 | 14891 | 6224 | 15186 | 1207 | 12416 | 12962 | 0 | 13 | 16 | 208 | 536 | 592 | 93 | 81 | 36 | 19 | 14 | 47 | 60 | 48 | 18 | 0 | 0 | 17 | 111 | 0 |
| SRR2915344 | 1016 | 6769 | 59362 | 787 | 32449 | 17582 | 397896 | 21807 | 1779 | 3175 | 14748 | 19826 | 7184 | 15867 | 1256 | 15137 | 14088 | 0 | 14 | 18 | 198 | 632 | 526 | 101 | 84 | 43 | 20 | 14 | 46 | 52 | 57 | 15 | 0 | 0 | 12 | 114 | 0 |
| SRR2915345 | 664 | 3248 | 36944 | 3299 | 31068 | 22448 | 397746 | 29980 | 186 | 3841 | 19503 | 26539 | 13171 | 31920 | 6250 | 8713 | 13973 | 31 | 44 | 31 | 257 | 237 | 712 | 196 | 140 | 68 | 66 | 45 | 122 | 140 | 102 | 22 | 0 | 0 | 20 | 195 | 16 |
| SRR2915346 | 981 | 4325 | 29960 | 3711 | 26277 | 18614 | 353638 | 31414 | 146 | 3855 | 23937 | 33813 | 14611 | 33503 | 6352 | 8708 | 16530 | 31 | 42 | 27 | 213 | 173 | 573 | 190 | 136 | 62 | 72 | 55 | 141 | 153 | 101 | 20 | 0 | 0 | 19 | 158 | 14 |
| SRR2915347 | 982 | 3631 | 28956 | 3459 | 26872 | 19189 | 329443 | 37272 | 173 | 3739 | 22410 | 30453 | 13704 | 32274 | 6981 | 8187 | 17403 | 44 | 52 | 40 | 282 | 497 | 787 | 242 | 149 | 81 | 83 | 58 | 152 | 177 | 146 | 26 | 0 | 0 | 20 | 204 | 15 |
| SRR2915348 | 1360 | 4046 | 44455 | 3707 | 61587 | 26163 | 549383 | 38963 | 1362 | 4088 | 11185 | 8986 | 4496 | 6740 | 656 | 10779 | 4090 | 0 | 13 | 10 | 52 | 27 | 90 | 45 | 34 | 24 | 37 | 26 | 82 | 65 | 53 | 0 | 0 | 0 | 16 | 83 | 0 |
| SRR2915349 | 511 | 2553 | 53534 | 1536 | 57636 | 23354 | 524242 | 39623 | 1277 | 5411 | 11877 | 11944 | 4942 | 7069 | 620 | 10811 | 5222 | 0 | 11 | 12 | 57 | 30 | 100 | 57 | 39 | 30 | 21 | 18 | 65 | 69 | 62 | 0 | 0 | 0 | 24 | 102 | 0 |
| SRR2915350 | 796 | 4385 | 41486 | 1858 | 60013 | 24610 | 502505 | 40722 | 1370 | 5260 | 13816 | 15063 | 5891 | 7909 | 633 | 10888 | 5589 | 0 | 12 | 13 | 68 | 40 | 100 | 53 | 40 | 19 | 20 | 15 | 55 | 56 | 40 | 0 | 0 | 0 | 16 | 76 | 0 |
| SRR2915905 | 21 | 1546 | 563 | 147 | 161 | 1494 | 1184 | 3018 | 8739 | 51 | 10658 | 17412 | 3370 | 1412 | 14 | 6306 | 80735 | 0 | 85 | 99 | 5643 | 3900 | 686 | 0 | 0 | 0 | 0 | 1200 | 4006 | 2123 | 4851 | 656 | 0 | 25 | 53 | 27 | 0 |
| SRR2915915 | 15 | 3361 | 788 | 59 | 87 | 717 | 930 | 3236 | 8526 | 60 | 8515 | 23636 | 2754 | 1570 | 0 | 1255 | 194294 | 0 | 16 | 11 | 1019 | 5134 | 1003 | 18 | 0 | 0 | 0 | 207 | 322 | 291 | 2316 | 540 | 0 | 14 | 37 | 0 | 0 |
| SRR2915918 | 370 | 29372 | 3583 | 432 | 2108 | 20559 | 24349 | 46468 | 3583 | 52 | 29568 | 32850 | 7873 | 4777 | 65 | 9034 | 54175 | 0 | 0 | 44 | 28481 | 12539 | 2503 | 38 | 27 | 0 | 0 | 16 | 1879 | 12634 | 21044 | 2597 | 0 | 71 | 133 | 31 | 0 |
| SRR2915921 | 481 | 4809 | 581 | 1432 | 2134 | 19944 | 30181 | 49548 | 6312 | 18 | 20698 | 22646 | 5719 | 6096 | 0 | 8957 | 72112 | 0 | 0 | 10 | 2725 | 2192 | 381 | 0 | 0 | 0 | 0 | 30 | 455 | 1259 | 3776 | 584 | 0 | 0 | 15 | 0 | 0 |
| SRR2915923 | 95 | 27763 | 1956 | 634 | 1013 | 9248 | 13140 | 26045 | 1636 | 23 | 14205 | 32474 | 4777 | 3072 | 0 | 11914 | 68558 | 0 | 0 | 186 | 17977 | 9221 | 1993 | 0 | 0 | 0 | 0 | 78 | 6377 | 9016 | 23276 | 2450 | 0 | 13 | 13 | 0 | 0 |
| SRR2915926 | 262 | 29803 | 9768 | 404 | 1048 | 13342 | 12597 | 18341 | 2388 | 31 | 22373 | 69410 | 7184 | 7795 | 15 | 14150 | 54922 | 0 | 0 | 85 | 14568 | 2608 | 1348 | 0 | 0 | 0 | 0 | 36 | 3262 | 11479 | 11051 | 3180 | 0 | 46 | 14 | 0 | 0 |
| SRR2915928 | 105 | 19299 | 8495 | 291 | 1701 | 11941 | 21955 | 50792 | 9355 | 88 | 17935 | 27760 | 4866 | 7669 | 0 | 11515 | 43707 | 0 | 0 | 26 | 1975 | 2655 | 316 | 0 | 0 | 0 | 0 | 71 | 1230 | 2048 | 8559 | 818 | 0 | 0 | 0 | 0 | 0 |
| SRR2960039 | 0 | 17 | 3116 | 0 | 968 | 938 | 75948 | 50114 | 2138 | 0 | 56289 | 21025 | 7278 | 3628 | 36 | 28 | 6107 | 0 | 0 | 0 | 0 | 0 | 0 | 0 | 0 | 0 | 0 | 0 | 0 | 0 | 0 | 11 | 0 | 0 | 0 | 0 | 0 |

### Supp Table S8.1.Columns_B_to_AL

| | | | | | | | | | | | | | | | | | | | | | | | | | | | | | | | | | | | | | |
|---|---|---|---|---|---|---|---|---|---|---|---|---|---|---|---|---|---|---|---|---|---|---|---|---|---|---|---|---|---|---|---|---|---|---|---|---|---|
| SRR2976280 | 0 | 0 | 243 | 0 | 68 | 447 | 1122 | 2166 | 29454 | 0 | 52123 | 13666 | 1329 | 8831 | 0 | 29325 | 9820 | 0 | 0 | 0 | 0 | 0 | 0 | 0 | 0 | 0 | 0 | 0 | 0 | 130 | 302 | 80 | 0 | 0 | 0 | 0 | 0 |
| SRR2976305 | 0 | 34 | 754 | 0 | 115 | 779 | 1915 | 1876 | 29343 | 21 | 121363 | 43321 | 2465 | 5625 | 0 | 22911 | 8166 | 0 | 0 | 0 | 0 | 0 | 0 | 0 | 0 | 0 | 0 | 0 | 0 | 0 | 0 | 21 | 0 | 0 | 0 | 0 | 0 |
| SRR3092114 | 0 | 17 | 252 | 0 | 809 | 936 | 6787 | 367 | 28362 | 49 | 8815 | 8842 | 3411 | 1633 | 9560 | 9663 | 2870 | 0 | 0 | 0 | 31 | 0 | 17 | 0 | 0 | 0 | 0 | 0 | 0 | 79 | 0 | 0 | 0 | 0 | 0 | 0 | 0 |
| SRR3092115 | 0 | 31 | 141 | 22 | 445 | 786 | 5988 | 259 | 28361 | 44 | 3499 | 3885 | 1014 | 491 | 5596 | 8757 | 3048 | 0 | 0 | 0 | 40 | 0 | 12 | 0 | 0 | 0 | 0 | 0 | 0 | 62 | 0 | 0 | 0 | 0 | 0 | 0 | 0 |
| SRR3103222 | 0 | 29505 | 6786 | 0 | 739 | 6998 | 6778 | 22793 | 15735 | 99 | 15088 | 23 | 5700 | 1849 | 0 | 4640 | 63124 | 0 | 0 | 0 | 234 | 305 | 166 | 33 | 27 | 0 | 0 | 0 | 28 | 178 | 474 | 252 | 0 | 96 | 218 | 0 | 0 |
| SRR3103223 | 28 | 20183 | 682 | 853 | 1381 | 12317 | 9830 | 33358 | 24935 | 0 | 8749 | 12471 | 3365 | 872 | 0 | 1392 | 62438 | 0 | 0 | 0 | 14 | 0 | 0 | 0 | 0 | 0 | 0 | 0 | 0 | 14 | 13 | 0 | 0 | 0 | 0 | 0 | 0 |
| SRR3103224 | 13 | 17335 | 810 | 161 | 710 | 11021 | 3850 | 16350 | 61380 | 13 | 7474 | 28 | 2996 | 497 | 0 | 692 | 52608 | 0 | 0 | 0 | 0 | 0 | 0 | 0 | 0 | 0 | 0 | 0 | 0 | 0 | 0 | 0 | 0 | 0 | 0 | 0 | 0 |
| SRR3103225 | 241 | 46884 | 1479 | 227 | 1513 | 23311 | 13661 | 34968 | 40999 | 19 | 16890 | 22195 | 5093 | 1391 | 0 | 6307 | 45095 | 0 | 0 | 15 | 4173 | 6635 | 604 | 27 | 11 | 0 | 0 | 75 | 992 | 4945 | 7750 | 852 | 0 | 43 | 92 | 13 | 0 |
| SRR3103226 | 85 | 38891 | 232 | 634 | 2395 | 24011 | 19176 | 51909 | 25509 | 21 | 8710 | 12055 | 3153 | 1246 | 0 | 1059 | 48132 | 0 | 0 | 0 | 0 | 0 | 0 | 0 | 0 | 0 | 0 | 0 | 0 | 0 | 0 | 0 | 0 | 0 | 0 | 0 | 0 |
| SRR3103227 | 61 | 32752 | 4494 | 394 | 1020 | 10534 | 6273 | 22981 | 51467 | 10 | 16514 | 27656 | 7712 | 1730 | 0 | 4194 | 35402 | 0 | 0 | 0 | 2192 | 1104 | 242 | 0 | 0 | 0 | 0 | 0 | 301 | 4427 | 4304 | 754 | 0 | 29 | 135 | 25 | 0 |
| SRR3103228 | 57 | 156865 | 6618 | 141 | 475 | 6568 | 4098 | 12102 | 45679 | 17 | 11854 | 24388 | 4602 | 1213 | 0 | 2125 | 26313 | 0 | 0 | 0 | 898 | 891 | 144 | 0 | 0 | 0 | 0 | 0 | 181 | 1057 | 1099 | 171 | 0 | 17 | 70 | 0 | 0 |
| SRR3103229 | 140 | 25264 | 1278 | 189 | 731 | 9148 | 6181 | 22095 | 18806 | 0 | 12357 | 19446 | 4607 | 2028 | 0 | 4019 | 50390 | 0 | 0 | 0 | 388 | 1289 | 118 | 0 | 0 | 0 | 0 | 26 | 165 | 483 | 2163 | 197 | 0 | 13 | 24 | 0 | 0 |
| SRR3103230 | 42 | 20611 | 5751 | 303 | 794 | 8352 | 5392 | 12979 | 79260 | 25 | 18391 | 27639 | 7807 | 1464 | 0 | 3306 | 29304 | 0 | 0 | 0 | 1360 | 1413 | 145 | 0 | 0 | 0 | 0 | 0 | 419 | 2670 | 5703 | 452 | 0 | 12 | 27 | 0 | 0 |
| SRR3103231 | 91 | 7973 | 2518 | 319 | 359 | 5282 | 2044 | 8455 | 50182 | 0 | 12031 | 21080 | 4707 | 890 | 0 | 4787 | 39185 | 0 | 0 | 0 | 1110 | 640 | 164 | 0 | 0 | 0 | 0 | 0 | 133 | 2036 | 2613 | 503 | 0 | 53 | 135 | 0 | 0 |
| SRR3103232 | 54 | 36490 | 4646 | 340 | 449 | 5275 | 3191 | 14494 | 48815 | 0 | 15641 | 27674 | 6074 | 1411 | 0 | 3029 | 23008 | 0 | 0 | 0 | 655 | 411 | 78 | 0 | 0 | 0 | 0 | 0 | 71 | 673 | 708 | 131 | 0 | 0 | 30 | 0 | 0 |
| SRR3103233 | 36 | 32083 | 3315 | 65 | 602 | 6190 | 3797 | 16943 | 41411 | 47 | 16599 | 31092 | 6096 | 1417 | 0 | 3283 | 28137 | 0 | 0 | 0 | 826 | 533 | 102 | 0 | 0 | 0 | 0 | 0 | 116 | 815 | 1202 | 154 | 0 | 19 | 63 | 0 | 0 |
| SRR3103234 | 115 | 23598 | 1099 | 264 | 571 | 6887 | 3185 | 12308 | 84745 | 15 | 12157 | 15253 | 3827 | 847 | 0 | 3402 | 38181 | 0 | 0 | 0 | 1128 | 2389 | 253 | 0 | 0 | 0 | 0 | 0 | 493 | 2997 | 6920 | 654 | 0 | 31 | 119 | 13 | 0 |
| SRR3103235 | 48 | 23315 | 1398 | 338 | 381 | 4328 | 2214 | 10078 | 75546 | 16 | 11922 | 18343 | 3885 | 1098 | 0 | 4562 | 55657 | 0 | 0 | 0 | 1976 | 1571 | 321 | 0 | 0 | 0 | 0 | 0 | 392 | 4371 | 4516 | 635 | 0 | 61 | 163 | 13 | 0 |
| SRR3103236 | 77 | 14217 | 739 | 121 | 441 | 4292 | 2488 | 11693 | 49667 | 44 | 13484 | 23011 | 5011 | 1190 | 0 | 4076 | 58528 | 0 | 0 | 0 | 142 | 322 | 181 | 0 | 0 | 0 | 0 | 0 | 67 | 396 | 1845 | 767 | 0 | 130 | 260 | 0 | 0 |
| SRR3103237 | 33 | 13284 | 523 | 233 | 512 | 6519 | 2576 | 8994 | 71252 | 29 | 11168 | 16959 | 3712 | 961 | 0 | 4781 | 69850 | 0 | 0 | 0 | 847 | 1440 | 119 | 0 | 0 | 0 | 0 | 34 | 605 | 2774 | 7349 | 427 | 0 | 19 | 47 | 0 | 0 |
| SRR3103238 | 26 | 29307 | 7302 | 461 | 444 | 3397 | 2877 | 9869 | 53598 | 53 | 10227 | 15426 | 3233 | 1883 | 0 | 4691 | 44270 | 0 | 0 | 0 | 1004 | 887 | 145 | 0 | 0 | 0 | 0 | 18 | 557 | 3741 | 6389 | 556 | 0 | 26 | 63 | 0 | 0 |
| SRR3103239 | 151 | 17556 | 1286 | 84 | 270 | 4059 | 1745 | 7562 | 42862 | 21 | 11467 | 23849 | 4629 | 967 | 0 | 4772 | 34854 | 0 | 0 | 0 | 95 | 86 | 14 | 0 | 0 | 0 | 0 | 0 | 16 | 129 | 166 | 33 | 0 | 18 | 34 | 0 | 0 |
| SRR3103240 | 171 | 18046 | 1238 | 162 | 958 | 41660 | 11797 | 36595 | 8686 | 42 | 8325 | 41010 | 3780 | 599 | 0 | 14634 | 55406 | 0 | 0 | 11 | 678 | 1574 | 108 | 0 | 0 | 0 | 0 | 16 | 436 | 668 | 2192 | 195 | 0 | 17 | 0 | 0 | 0 |
| SRR3103241 | 246 | 24020 | 977 | 214 | 1101 | 52711 | 13297 | 33898 | 8840 | 32 | 7854 | 45481 | 3860 | 646 | 0 | 18713 | 38650 | 0 | 0 | 0 | 186 | 628 | 39 | 0 | 0 | 0 | 0 | 0 | 78 | 212 | 804 | 83 | 0 | 13 | 0 | 0 | 0 |
| SRR3103242 | 241 | 90813 | 664 | 1415 | 1946 | 28262 | 18177 | 48262 | 15787 | 60 | 24069 | 22316 | 6676 | 1489 | 20 | 4437 | 22791 | 0 | 0 | 0 | 472 | 565 | 82 | 0 | 0 | 0 | 0 | 0 | 97 | 292 | 1105 | 83 | 0 | 0 | 15 | 0 | 0 |
| SRR3103243 | 0 | 8193 | 2143 | 314 | 2991 | 27864 | 30454 | 72230 | 25143 | 0 | 4626 | 17 | 1322 | 259 | 0 | 150 | 24275 | 0 | 0 | 0 | 0 | 0 | 0 | 0 | 0 | 0 | 0 | 0 | 0 | 0 | 0 | 0 | 0 | 0 | 0 | 0 | 0 |
| SRR3103244 | 65 | 57687 | 4595 | 891 | 2973 | 32425 | 28206 | 61513 | 22059 | 0 | 11237 | 14943 | 3812 | 1171 | 0 | 809 | 57857 | 0 | 0 | 0 | 16 | 51 | 0 | 0 | 0 | 0 | 0 | 0 | 0 | 16 | 24 | 0 | 0 | 0 | 0 | 0 | 0 |
| SRR3103245 | 21 | 166201 | 3858 | 268 | 4851 | 86421 | 50668 | 103481 | 11279 | 0 | 13679 | 15386 | 4120 | 1045 | 0 | 919 | 28228 | 0 | 0 | 0 | 0 | 26 | 0 | 0 | 0 | 0 | 0 | 0 | 0 | 0 | 11 | 0 | 0 | 0 | 0 | 0 | 0 |
| SRR3103246 | 204 | 72669 | 433 | 5648 | 656 | 13560 | 10901 | 16622 | 11902 | 0 | 1008 | 2286 | 363 | 168 | 42 | 324 | 34564 | 0 | 0 | 0 | 0 | 100 | 0 | 0 | 0 | 0 | 0 | 0 | 0 | 0 | 17 | 0 | 0 | 0 | 0 | 0 | 0 |
| SRR3103247 | 88 | 126757 | 2701 | 142 | 1509 | 24717 | 17720 | 46669 | 52390 | 108 | 19352 | 35915 | 8878 | 3353 | 0 | 3220 | 15666 | 0 | 0 | 0 | 224 | 690 | 51 | 0 | 0 | 0 | 0 | 16 | 96 | 184 | 776 | 58 | 0 | 0 | 0 | 0 | 0 |
| SRR3103248 | 69 | 211631 | 1908 | 826 | 1303 | 25590 | 18040 | 35192 | 26289 | 21 | 7059 | 12641 | 2423 | 1237 | 14 | 1754 | 23187 | 0 | 0 | 0 | 1107 | 4484 | 322 | 0 | 0 | 0 | 0 | 0 | 173 | 628 | 2555 | 302 | 0 | 0 | 0 | 0 | 0 |
| SRR3103249 | 186 | 15525 | 2024 | 2315 | 2577 | 24638 | 25802 | 39684 | 18740 | 20 | 6871 | 14354 | 2395 | 2331 | 37 | 3807 | 112992 | 0 | 0 | 13 | 695 | 3879 | 191 | 0 | 0 | 0 | 0 | 25 | 156 | 320 | 2216 | 149 | 0 | 0 | 0 | 0 | 0 |
| SRR3103250 | 47 | 216954 | 8889 | 304 | 1109 | 12174 | 10936 | 26742 | 40107 | 19 | 15076 | 30766 | 7103 | 2702 | 36 | 2319 | 21014 | 0 | 0 | 0 | 254 | 1247 | 58 | 0 | 0 | 0 | 0 | 0 | 111 | 369 | 1497 | 131 | 0 | 14 | 28 | 0 | 0 |
| SRR3103251 | 43 | 39147 | 6600 | 100 | 3291 | 35203 | 25636 | 45422 | 54279 | 22 | 32003 | 14 | 8984 | 1436 | 11 | 3722 | 15206 | 0 | 0 | 0 | 101 | 135 | 13 | 0 | 0 | 0 | 0 | 0 | 0 | 61 | 123 | 15 | 0 | 0 | 0 | 0 | 0 |
| SRR3103252 | 46 | 168155 | 3604 | 62 | 1727 | 13064 | 14573 | 40038 | 9586 | 46 | 26696 | 30456 | 10608 | 2045 | 12 | 538 | 12561 | 0 | 0 | 0 | 164 | 740 | 40 | 0 | 0 | 0 | 0 | 0 | 28 | 155 | 600 | 41 | 0 | 0 | 0 | 0 | 0 |
| SRR3103253 | 13 | 35772 | 3458 | 52 | 1763 | 12772 | 14679 | 48770 | 21450 | 43 | 21197 | 44010 | 6837 | 3129 | 0 | 3194 | 48866 | 0 | 0 | 0 | 1244 | 2230 | 452 | 0 | 0 | 0 | 0 | 0 | 54 | 672 | 2753 | 482 | 0 | 0 | 32 | 0 | 0 |
| SRR3103254 | 158 | 71047 | 523 | 300 | 1221 | 16619 | 14058 | 30478 | 65802 | 42 | 16561 | 28589 | 4966 | 2818 | 0 | 996 | 40515 | 0 | 0 | 0 | 1304 | 3937 | 300 | 0 | 0 | 0 | 0 | 0 | 124 | 1876 | 3667 | 396 | 0 | 20 | 78 | 0 | 0 |
| SRR3103255 | 29 | 204509 | 237 | 110 | 1278 | 13170 | 13379 | 25572 | 78559 | 33 | 6471 | 11496 | 2584 | 709 | 30 | 1284 | 25391 | 0 | 0 | 0 | 160 | 1044 | 61 | 0 | 0 | 0 | 0 | 0 | 23 | 249 | 504 | 63 | 0 | 0 | 14 | 0 | 0 |
| SRR3103256 | 74 | 154828 | 445 | 571 | 891 | 12457 | 6091 | 12957 | 45371 | 25 | 7619 | 14727 | 2794 | 771 | 0 | 9425 | 31590 | 0 | 0 | 0 | 1154 | 4537 | 406 | 0 | 0 | 0 | 0 | 17 | 280 | 768 | 5043 | 295 | 0 | 0 | 43 | 0 | 0 |
| SRR3103257 | 131 | 183617 | 3166 | 331 | 756 | 10298 | 9883 | 28242 | 19142 | 36 | 20702 | 32053 | 4495 | 2968 | 0 | 2444 | 27041 | 0 | 0 | 0 | 203 | 935 | 60 | 0 | 0 | 0 | 0 | 0 | 73 | 245 | 1046 | 61 | 0 | 0 | 0 | 0 | 0 |
| SRR3103258 | 201 | 95910 | 1168 | 342 | 872 | 11863 | 9674 | 22135 | 68657 | 101 | 34716 | 42418 | 6006 | 2051 | 27 | 4874 | 39818 | 0 | 0 | 0 | 578 | 4462 | 376 | 0 | 0 | 0 | 0 | 0 | 39 | 271 | 1235 | 155 | 0 | 0 | 0 | 0 | 0 |
| SRR3103259 | 126 | 8004 | 9923 | 265 | 475 | 6824 | 3057 | 9885 | 51829 | 0 | 12228 | 18050 | 4529 | 943 | 0 | 4021 | 36247 | 0 | 0 | 0 | 439 | 728 | 73 | 0 | 0 | 0 | 0 | 21 | 156 | 810 | 2228 | 255 | 0 | 32 | 34 | 0 | 0 |
| SRR3103260 | 378 | 55851 | 5152 | 1424 | 486 | 10852 | 6155 | 9250 | 31740 | 25 | 4382 | 9913 | 1457 | 675 | 30 | 886 | 55175 | 0 | 0 | 0 | 419 | 2251 | 85 | 0 | 0 | 0 | 0 | 0 | 64 | 251 | 941 | 58 | 0 | 0 | 0 | 0 | 0 |
| SRR3103261 | 315 | 36833 | 6604 | 1391 | 345 | 6541 | 4779 | 7663 | 34396 | 11 | 3783 | 6809 | 1179 | 834 | 12 | 635 | 55983 | 0 | 0 | 0 | 469 | 681 | 111 | 0 | 0 | 0 | 0 | 0 | 32 | 315 | 432 | 108 | 0 | 12 | 19 | 0 | 0 |
| SRR3103262 | 232 | 43538 | 2042 | 1390 | 1923 | 33679 | 19078 | 41993 | 26174 | 100 | 27354 | 24765 | 6948 | 1826 | 15 | 4251 | 15584 | 0 | 0 | 0 | 2159 | 1882 | 230 | 0 | 0 | 0 | 0 | 0 | 203 | 1202 | 2315 | 266 | 0 | 0 | 24 | 0 | 0 |
| SRR3103263 | 0 | 29147 | 4719 | 453 | 2144 | 46580 | 43717 | 129293 | 5870 | 0 | 4924 | 6380 | 1852 | 1331 | 0 | 409 | 31064 | 0 | 0 | 0 | 0 | 0 | 0 | 0 | 0 | 0 | 0 | 0 | 0 | 0 | 0 | 0 | 0 | 0 | 0 | 0 | 0 |
| SRR3103264 | 216 | 19374 | 1595 | 5176 | 1402 | 12439 | 11230 | 23369 | 14210 | 0 | 12033 | 12848 | 3194 | 743 | 0 | 846 | 51700 | 0 | 0 | 0 | 14 | 91 | 0 | 0 | 0 | 0 | 0 | 0 | 0 | 16 | 38 | 0 | 0 | 0 | 0 | 0 | 0 |
| SRR3103265 | 123 | 136178 | 8104 | 267 | 4072 | 72969 | 40198 | 91488 | 21548 | 20 | 21880 | 21813 | 5338 | 1040 | 0 | 2367 | 23075 | 0 | 0 | 12 | 2754 | 7879 | 366 | 16 | 0 | 0 | 0 | 19 | 225 | 1274 | 3585 | 219 | 0 | 14 | 22 | 0 | 0 |
| SRR3103266 | 163 | 124166 | 833 | 6040 | 1254 | 11190 | 18198 | 22467 | 10247 | 19 | 1858 | 5117 | 622 | 492 | 37 | 1032 | 39046 | 0 | 0 | 0 | 14 | 164 | 11 | 0 | 0 | 0 | 0 | 0 | 0 | 0 | 48 | 0 | 0 | 0 | 0 | 0 | 0 |
| SRR3103267 | 73 | 73036 | 34481 | 173 | 1634 | 43450 | 43197 | 98330 | 22132 | 42 | 18399 | 29033 | 5190 | 4111 | 0 | 2030 | 18838 | 0 | 0 | 0 | 243 | 3198 | 100 | 0 | 0 | 0 | 0 | 0 | 115 | 281 | 1919 | 118 | 0 | 0 | 0 | 0 | 0 |
| SRR3103268 | 52 | 127068 | 3201 | 949 | 1813 | 30560 | 24046 | 34590 | 19595 | 33 | 6229 | 12121 | 2147 | 1394 | 20 | 1382 | 37620 | 0 | 0 | 0 | 3126 | 10754 | 1136 | 28 | 0 | 0 | 0 | 0 | 217 | 1719 | 5785 | 939 | 0 | 12 | 49 | 16 | 0 |
| SRR3103269 | 228 | 24941 | 3026 | 758 | 820 | 13847 | 11676 | 34825 | 36777 | 60 | 5113 | 13385 | 2165 | 770 | 0 | 2604 | 25735 | 0 | 0 | 0 | 315 | 3449 | 149 | 0 | 0 | 0 | 0 | 0 | 159 | 325 | 4500 | 220 | 0 | 0 | 0 | 0 | 0 |
| SRR3103270 | 72 | 24401 | 2427 | 311 | 939 | 9746 | 7073 | 12442 | 66906 | 0 | 13138 | 22240 | 5634 | 1806 | 0 | 3218 | 38207 | 0 | 0 | 0 | 2027 | 2177 | 337 | 0 | 0 | 0 | 0 | 0 | 278 | 3107 | 5210 | 584 | 0 | 20 | 73 | 10 | 0 |
| SRR3103271 | 129 | 36923 | 4099 | 252 | 3283 | 38785 | 25781 | 37946 | 49678 | 19 | 31523 | 40544 | 8412 | 1764 | 14 | 5071 | 19147 | 0 | 0 | 15 | 5281 | 5227 | 877 | 29 | 11 | 0 | 0 | 0 | 78 | 1612 | 2314 | 396 | 0 | 14 | 27 | 0 | 0 |
| SRR3103272 | 38 | 144898 | 3586 | 134 | 1823 | 12936 | 14864 | 34174 | 14608 | 0 | 23135 | 26932 | 8363 | 2399 | 20 | 895 | 14900 | 0 | 0 | 0 | 1597 | 4865 | 465 | 18 | 0 | 0 | 0 | 0 | 249 | 1338 | 4343 | 484 | 0 | 0 | 28 | 0 | 0 |
| SRR3103273 | 199 | 100334 | 6404 | 216 | 3781 | 33674 | 28159 | 71326 | 36825 | 69 | 27115 | 35731 | 9879 | 2556 | 12 | 4056 | 12694 | 0 | 0 | 26 | 12310 | 7141 | 1210 | 14 | 0 | 0 | 0 | 0 | 292 | 2478 | 3409 | 573 | 0 | 0 | 10 | 0 | 0 |
| SRR3103274 | 92 | 116573 | 676 | 228 | 1292 | 20002 | 17774 | 37148 | 68211 | 54 | 17755 | 31527 | 4639 | 3250 | 0 | 813 | 55247 | 0 | 0 | 0 | 1744 | 4385 | 316 | 0 | 0 | 0 | 0 | 0 | 121 | 2397 | 2741 | 284 | 0 | 0 | 29 | 0 | 0 |
| SRR3103275 | 77 | 226929 | 1746 | 672 | 1225 | 22272 | 16655 | 26978 | 62803 | 47 | 3214 | 5447 | 1147 | 677 | 56 | 4308 | 48137 | 0 | 0 | 0 | 2159 | 6523 | 466 | 0 | 0 | 0 | 0 | 0 | 169 | 2880 | 5484 | 507 | 0 | 29 | 88 | 0 | 0 |
| SRR3103276 | 47 | 144996 | 1685 | 269 | 871 | 13834 | 6184 | 13419 | 34782 | 44 | 10099 | 18894 | 3664 | 1055 | 0 | 10189 | 27339 | 0 | 0 | 0 | 1680 | 3511 | 257 | 0 | 0 | 0 | 0 | 18 | 338 | 1733 | 5072 | 284 | 0 | 13 | 38 | 0 | 0 |
| SRR3103277 | 30 | 259401 | 3342 | 203 | 907 | 12992 | 11504 | 24373 | 13045 | 42 | 19052 | 25255 | 3538 | 4029 | 0 | 2741 | 48045 | 0 | 0 | 0 | 3680 | 4416 | 504 | 0 | 0 | 0 | 0 | 0 | 233 | 5409 | 6911 | 728 | 0 | 31 | 103 | 11 | 0 |
| SRR3103278 | 69 | 105601 | 1483 | 362 | 695 | 6548 | 7081 | 19301 | 34639 | 34 | 23980 | 28727 | 4354 | 2127 | 17 | 4196 | 87474 | 0 | 0 | 0 | 2359 | 6486 | 1403 | 33 | 15 | 0 | 0 | 0 | 57 | 1379 | 3419 | 853 | 0 | 41 | 166 | 25 | 0 |
| SRR3103279 | 49 | 14805 | 2259 | 79 | 327 | 4681 | 2123 | 8789 | 36046 | 22 | 10644 | 18683 | 4102 | 1262 | 0 | 3351 | 84219 | 0 | 0 | 0 | 1553 | 1092 | 206 | 0 | 0 | 0 | 0 | 0 | 325 | 5957 | 4535 | 660 | 0 | 54 | 162 | 14 | 0 |
| SRR3103280 | 108 | 103835 | 2521 | 72 | 1068 | 31806 | 13713 | 38710 | 20206 | 24 | 25228 | 27360 | 6954 | 1388 | 12 | 3665 | 31407 | 0 | 0 | 0 | 745 | 3109 | 163 | 0 | 0 | 0 | 0 | 0 | 60 | 433 | 1223 | 118 | 0 | 0 | 11 | 0 | 0 |
| SRR3103281 | 217 | 61993 | 2788 | 191 | 1908 | 36724 | 24237 | 61470 | 10346 | 34 | 21461 | 22553 | 5272 | 2484 | 11 | 2599 | 53201 | 0 | 0 | 0 | 812 | 3173 | 294 | 0 | 0 | 0 | 0 | 0 | 51 | 398 | 1271 | 186 | 0 | 11 | 12 | 0 | 0 |
| SRR3103386 | 0 | 18230 | 32 | 0 | 2565 | 9372 | 22607 | 21477 | 3615 | 0 | 20379 | 29315 | 9134 | 5269 | 0 | 132 | 53851 | 0 | 0 | 0 | 39 | 16 | 0 | 0 | 0 | 0 | 0 | 0 | 27 | 52 | 40 | 0 | 0 | 0 | 0 | 0 | 0 |
| SRR3103387 | 0 | 13629 | 189 | 0 | 3797 | 14310 | 29660 | 20794 | 3865 | 0 | 38427 | 72111 | 12734 | 4819 | 23 | 190 | 47001 | 0 | 0 | 0 | 34 | 0 | 0 | 0 | 0 | 0 | 0 | 0 | 0 | 28 | 12 | 0 | 0 | 0 | 0 | 0 | 0 |
| SRR3103388 | 0 | 20833 | 44 | 88 | 2002 | 7143 | 16454 | 13692 | 2768 | 0 | 35701 | 59021 | 11110 | 4033 | 0 | 78 | 43816 | 0 | 0 | 0 | 142 | 84 | 10 | 0 | 0 | 0 | 0 | 0 | 55 | 161 | 115 | 10 | 0 | 0 | 0 | 0 | 0 |
| SRR3103389 | 32 | 7736 | 380 | 95 | 1847 | 19555 | 22187 | 16739 | 3900 | 0 | 30928 | 36596 | 7696 | 7373 | 0 | 170 | 85232 | 0 | 0 | 0 | 131 | 74 | 0 | 0 | 0 | 0 | 0 | 0 | 92 | 238 | 162 | 11 | 0 | 0 | 0 | 0 | 0 |
| SRR3103390 | 28 | 22934 | 167 | 56 | 1351 | 14947 | 14845 | 11171 | 3109 | 0 | 50902 | 63251 | 12425 | 5382 | 0 | 260 | 87242 | 0 | 0 | 0 | 20 | 0 | 0 | 0 | 0 | 0 | 0 | 0 | 14 | 34 | 14 | 16 | 0 | 0 | 0 | 0 | 0 |
| SRR3103391 | 0 | 6248 | 62 | 42 | 2852 | 14138 | 30146 | 22661 | 5509 | 0 | 33264 | 54928 | 11590 | 5012 | 13 | 163 | 71781 | 0 | 0 | 0 | 48 | 15 | 0 | 0 | 0 | 0 | 0 | 0 | 24 | 87 | 27 | 14 | 0 | 0 | 0 | 0 | 0 |
| SRR3103392 | 58 | 5633 | 39 | 58 | 4851 | 16332 | 57077 | 22312 | 4151 | 19 | 25168 | 24445 | 7553 | 10012 | 0 | 255 | 73994 | 0 | 0 | 13 | 658 | 196 | 17 | 0 | 0 | 0 | 0 | 0 | 569 | 1593 | 1002 | 49 | 0 | 0 | 0 | 0 | 0 |
| SRR3103393 | 0 | 10119 | 0 | 0 | 2350 | 7250 | 19788 | 18750 | 4303 | 0 | 31523 | 39502 | 7671 | 4621 | 0 | 88 | 44373 | 0 | 0 | 0 | 0 | 0 | 0 | 0 | 0 | 0 | 0 | 0 | 0 | 17 | 0 | 0 | 0 | 0 | 0 | 0 | 0 |
| SRR3103394 | 17 | 14746 | 25 | 42 | 3122 | 12610 | 26123 | 24557 | 3481 | 0 | 23851 | 30036 | 17529 | 5839 | 24 | 171 | 60624 | 0 | 0 | 0 | 84 | 50 | 0 | 0 | 0 | 0 | 0 | 0 | 44 | 130 | 94 | 0 | 0 | 0 | 0 | 0 | 0 |
| SRR3103395 | 0 | 3487 | 42 | 0 | 7292 | 14682 | 74419 | 46486 | 7374 | 0 | 23758 | 40468 | 6291 | 5298 | 27 | 173 | 57139 | 0 | 0 | 0 | 48 | 0 | 0 | 0 | 0 | 0 | 0 | 0 | 61 | 132 | 43 | 15 | 0 | 0 | 0 | 0 | 0 |
| SRR3103396 | 24 | 22817 | 333 | 48 | 4057 | 14545 | 33922 | 26170 | 3542 | 0 | 29089 | 61028 | 8773 | 4331 | 17 | 82 | 48082 | 0 | 0 | 0 | 166 | 44 | 0 | 0 | 0 | 0 | 0 | 0 | 86 | 355 | 160 | 20 | 0 | 0 | 0 | 0 | 0 |
| SRR3103397 | 0 | 10648 | 70 | 47 | 2435 | 10555 | 22255 | 14977 | 3709 | 0 | 35962 | 56094 | 8293 | 4276 | 0 | 161 | 60407 | 0 | 0 | 0 | 64 | 19 | 0 | 0 | 0 | 0 | 0 | 0 | 64 | 93 | 57 | 11 | 0 | 0 | 0 | 0 | 0 |
| SRR3103400 | 83 | 23592 | 580 | 166 | 1376 | 9520 | 15478 | 12516 | 2915 | 0 | 23358 | 21423 | 15242 | 6824 | 0 | 131 | 105009 | 0 | 0 | 17 | 859 | 808 | 68 | 0 | 0 | 0 | 0 | 0 | 1079 | 767 | 1384 | 64 | 0 | 10 | 0 | 0 | 0 |
| SRR3103401 | 0 | 47786 | 437 | 219 | 2556 | 11540 | 23621 | 19102 | 2898 | 0 | 17781 | 17482 | 12588 | 5497 | 16 | 48 | 83927 | 0 | 0 | 0 | 346 | 143 | 38 | 0 | 0 | 0 | 0 | 0 | 530 | 410 | 456 | 51 | 0 | 14 | 0 | 0 | 0 |
| SRR3103402 | 244 | 46108 | 568 | 812 | 3696 | 20914 | 29759 | 17281 | 2453 | 0 | 20861 | 23761 | 7619 | 6769 | 0 | 67 | 91507 | 0 | 0 | 0 | 614 | 296 | 50 | 0 | 0 | 0 | 0 | 0 | 614 | 755 | 812 | 57 | 0 | 13 | 0 | 0 | 0 |
| SRR3103403 | 54 | 26686 | 539 | 216 | 3332 | 15479 | 31545 | 20179 | 2196 | 0 | 34384 | 26035 | 18867 | 10516 | 0 | 138 | 98808 | 0 | 0 | 0 | 690 | 357 | 62 | 0 | 0 | 0 | 0 | 0 | 636 | 714 | 728 | 65 | 0 | 18 | 0 | 0 | 0 |
| SRR3103404 | 57 | 16830 | 285 | 171 | 1984 | 12303 | 23219 | 12580 | 3755 | 57 | 21284 | 25698 | 7836 | 6073 | 0 | 206 | 133517 | 0 | 0 | 19 | 616 | 248 | 37 | 0 | 0 | 0 | 0 | 0 | 1209 | 686 | 876 | 50 | 0 | 0 | 0 | 0 | 0 |
| SRR3103405 | 0 | 13181 | 107 | 161 | 8090 | 15264 | 67777 | 37402 | 3897 | 54 | 30965 | 26533 | 22635 | 9783 | 0 | 103 | 79215 | 0 | 0 | 27 | 750 | 446 | 39 | 0 | 0 | 0 | 0 | 19 | 1541 | 968 | 1499 | 68 | 0 | 0 | 0 | 0 | 0 |
| SRR3103406 | 134 | 37419 | 7544 | 33 | 976 | 12266 | 13978 | 10081 | 2393 | 0 | 43790 | 29913 | 15710 | 10007 | 0 | 211 | 123817 | 0 | 0 | 0 | 213 | 177 | 28 | 0 | 0 | 0 | 0 | 0 | 159 | 142 | 212 | 31 | 0 | 0 | 0 | 0 | 0 |
| SRR3103407 | 117 | 49516 | 1166 | 292 | 1635 | 8523 | 15694 | 13293 | 1678 | 58 | 46135 | 37550 | 14691 | 11343 | 0 | 158 | 113558 | 0 | 0 | 19 | 897 | 940 | 83 | 13 | 0 | 0 | 0 | 0 | 726 | 647 | 1984 | 65 | 0 | 0 | 0 | 0 | 0 |
| SRR3103408 | 228 | 48767 | 914 | 228 | 2050 | 10846 | 18511 | 14307 | 1617 | 0 | 46222 | 38227 | 20182 | 9579 | 0 | 190 | 105225 | 0 | 0 | 0 | 230 | 136 | 22 | 0 | 0 | 0 | 0 | 0 | 184 | 156 | 211 | 27 | 0 | 11 | 0 | 0 | 0 |
| SRR3103409 | 62 | 40085 | 1492 | 62 | 1204 | 8397 | 12770 | 7482 | 1906 | 62 | 74814 | 59941 | 24830 | 12254 | 0 | 130 | 102959 | 0 | 0 | 0 | 100 | 68 | 17 | 0 | 0 | 0 | 0 | 0 | 68 | 51 | 67 | 11 | 0 | 0 | 0 | 0 | 0 |
| SRR3138807 | 40 | 3197 | 1723 | 9946 | 15 | 229 | 63091 | 266790 | 378 | 11 | 2195 | 1162 | 689 | 2519 | 142 | 170 | 727 | 0 | 0 | 117 | 68 | 201 | 1672 | 555 | 274 | 548 | 0 | 0 | 722 | 447 | 1129 | 548 | 205 | 17 | 23 | 181 | 541 |
| SRR3138808 | 14 | 2142 | 1227 | 10353 | 0 | 61 | 17259 | 314254 | 294 | 0 | 1487 | 992 | 705 | 2971 | 37 | 104 | 962 | 0 | 0 | 77 | 40 | 166 | 3039 | 887 | 359 | 876 | 0 | 0 | 394 | 168 | 615 | 455 | 219 | 15 | 25 | 240 | 892 |
| SRR3138809 | 14 | 2288 | 1320 | 9548 | 0 | 34 | 14727 | 306902 | 236 | 0 | 1277 | 923 | 642 | 2901 | 23 | 79 | 821 | 0 | 0 | 72 | 38 | 172 | 3526 | 1005 | 433 | 1046 | 0 | 0 | 333 | 129 | 631 | 476 | 225 | 10 | 21 | 266 | 1106 |
| SRR3138810 | 83 | 2772 | 2536 | 4078 | 31 | 311 | 71082 | 229558 | 390 | 13 | 2030 | 641 | 515 | 1970 | 194 | 341 | 474 | 0 | 0 | 75 | 32 | 107 | 680 | 273 | 149 | 317 | 0 | 0 | 369 | 254 | 413 | 180 | 63 | 0 | 0 | 93 | 259 |
| SRR3199254 | 14 | 42587 | 2267 | 1811 | 2992 | 4371 | 102448 | 23687 | 1393 | 530 | 6250 | 5291 | 6543 | 3541 | 1823 | 1819 | 8269 | 0 | 10 | 0 | 756 | 104 | 2534 | 688 | 149 | 78 | 0 | 25 | 64 | 1410 | 280 | 21 | 0 | 0 | 0 | 10 | 0 |
| SRR3199255 | 0 | 80413 | 11405 | 1796 | 2146 | 2470 | 56412 | 8200 | 4326 | 294 | 94 | 55 | 81 | 26 | 2574 | 11711 | 18511 | 0 | 0 | 0 | 328 | 108 | 3296 | 842 | 150 | 52 | 0 | 22 | 84 | 672 | 291 | 31 | 0 | 0 | 0 | 13 | 0 |
| SRR3199256 | 17 | 7725 | 17038 | 634 | 2121 | 2433 | 104419 | 18463 | 6489 | 1298 | 1028 | 1459 | 551 | 103 | 1716 | 5450 | 7240 | 0 | 11 | 0 | 284 | 130 | 716 | 120 | 173 | 32 | 0 | 28 | 59 | 571 | 442 | 21 | 0 | 0 | 0 | 0 | 0 |
| SRR3199257 | 0 | 9859 | 8725 | 331 | 216 | 815 | 8890 | 1613 | 251 | 181 | 1727 | 691 | 1285 | 1528 | 751 | 5418 | 40212 | 0 | 0 | 0 | 73 | 28 | 520 | 146 | 28 | 45 | 0 | 0 | 24 | 167 | 111 | 0 | 0 | 0 | 0 | 0 | 0 |
| SRR3199258 | 0 | 355 | 4198 | 363 | 312 | 793 | 29384 | 1992 | 274 | 198 | 1761 | 503 | 1395 | 1542 | 1284 | 4162 | 21107 | 0 | 0 | 0 | 723 | 115 | 1907 | 55 | 0 | 0 | 0 | 12 | 102 | 2336 | 857 | 134 | 0 | 0 | 0 | 0 | 0 |
| SRR3199259 | 0 | 559 | 1414 | 548 | 509 | 1206 | 8048 | 4718 | 259 | 96 | 3898 | 1299 | 5318 | 2724 | 1766 | 3380 | 17058 | 0 | 0 | 0 | 162 | 138 | 466 | 138 | 10 | 19 | 0 | 18 | 146 | 596 | 813 | 22 | 0 | 0 | 0 | 0 | 0 |
| SRR3199260 | 0 | 426 | 1553 | 795 | 566 | 1259 | 6841 | 6090 | 170 | 76 | 5981 | 1425 | 5372 | 3985 | 3383 | 4347 | 18549 | 0 | 0 | 0 | 161 | 143 | 452 | 188 | 27 | 32 | 0 | 31 | 137 | 614 | 1417 | 17 | 0 | 0 | 0 | 0 | 0 |
| SRR3199261 | 0 | 394 | 5942 | 531 | 182 | 511 | 5678 | 2501 | 325 | 76 | 8796 | 3600 | 6148 | 3703 | 957 | 2006 | 68079 | 0 | 0 | 0 | 510 | 190 | 968 | 74 | 0 | 33 | 0 | 17 | 115 | 958 | 613 | 43 | 0 | 0 | 0 | 0 | 0 |
| SRR3199262 | 0 | 495 | 2854 | 651 | 169 | 606 | 3689 | 2405 | 363 | 70 | 10927 | 4449 | 6334 | 4097 | 1766 | 2394 | 72328 | 0 | 0 | 0 | 161 | 112 | 471 | 122 | 24 | 21 | 0 | 14 | 76 | 360 | 377 | 15 | 0 | 0 | 0 | 0 | 0 |
| SRR3199263 | 0 | 18597 | 30253 | 756 | 9094 | 30784 | 97719 | 20992 | 163 | 133 | 10420 | 14574 | 6595 | 12184 | 919 | 2882 | 14439 | 0 | 0 | 0 | 59 | 23 | 552 | 185 | 21 | 112 | 0 | 14 | 17 | 518 | 211 | 17 | 0 | 0 | 0 | 23 | 0 |
| SRR3235777 | 166 | 4131 | 5529 | 544 | 22377 | 19097 | 305984 | 14677 | 577 | 100 | 25118 | 42979 | 13716 | 19095 | 6807 | 15783 | 21873 | 0 | 55 | 42 | 572 | 156 | 174 | 47 | 29 | 0 | 0 | 20 | 32 | 88 | 0 | 0 | 0 | 0 | 0 | 14 | 0 |
| SRR3235778 | 222 | 1483 | 6657 | 560 | 32015 | 30613 | 431528 | 19788 | 1219 | 61 | 11992 | 25 | 6181 | 14974 | 6130 | 9534 | 12869 | 0 | 17 | 14 | 269 | 112 | 82 | 21 | 12 | 0 | 0 | 0 | 11 | 40 | 0 | 0 | 0 | 0 | 0 | 0 | 0 |
| SRR3235779 | 53 | 189 | 2380 | 1957 | 23737 | 19728 | 164360 | 20148 | 6182 | 65 | 32261 | 48891 | 20559 | 48243 | 23429 | 23436 | 14674 | 0 | 48 | 37 | 78 | 43 | 89 | 32 | 18 | 0 | 0 | 14 | 32 | 17 | 0 | 0 | 0 | 0 | 0 | 11 | 0 |
| SRR3235780 | 96 | 845 | 1908 | 11712 | 26085 | 29370 | 243040 | 33540 | 161 | 69 | 16299 | 11319 | 15067 | 90728 | 9362 | 26279 | 21387 | 0 | 29 | 32 | 78 | 88 | 119 | 32 | 30 | 25 | 0 | 13 | 29 | 21 | 0 | 0 | 0 | 0 | 13 | 44 | 0 |
| SRR3235781 | 108 | 2744 | 3080 | 1662 | 16607 | 21080 | 158639 | 23696 | 5911 | 62 | 34900 | 24 | 30576 | 86129 | 12394 | 29473 | 18228 | 0 | 22 | 12 | 42 | 28 | 73 | 32 | 18 | 0 | 0 | 0 | 16 | 18 | 33 | 0 | 0 | 0 | 0 | 16 | 0 |
| SRR3279553 | 179 | 6128 | 770 | 197 | 1893 | 8994 | 33846 | 42367 | 6154 | 54 | 24852 | 16095 | 2367 | 7101 | 0 | 947 | 17515 | 0 | 0 | 0 | 256 | 137 | 0 | 0 | 0 | 0 | 0 | 0 | 27 | 247 | 146 | 0 | 0 | 0 | 46 | 0 | 0 |
| SRR3279554 | 0 | 59 | 1114 | 235 | 2498 | 12017 | 44074 | 47159 | 7493 | 469 | 16102 | 5759 | 1146 | 7404 | 0 | 411 | 19951 | 0 | 0 | 0 | 102 | 172 | 67 | 0 | 0 | 0 | 0 | 11 | 77 | 351 | 1003 | 414 | 0 | 21 | 112 | 0 | 0 |
| SRR3279555 | 0 | 609 | 47 | 84 | 2354 | 25307 | 35732 | 38843 | 4204 | 59 | 26652 | 17067 | 3447 | 4540 | 0 | 1345 | 14545 | 0 | 0 | 0 | 173 | 154 | 17 | 0 | 0 | 0 | 0 | 17 | 58 | 124 | 203 | 23 | 0 | 0 | 21 | 0 | 0 |
| SRR3279556 | 14 | 859 | 318 | 243 | 2850 | 19990 | 53789 | 56170 | 4334 | 429 | 22970 | 11440 | 2915 | 9618 | 104 | 677 | 23465 | 0 | 0 | 0 | 615 | 212 | 44 | 0 | 0 | 0 | 0 | 18 | 104 | 654 | 562 | 88 | 0 | 0 | 19 | 12 | 0 |
| SRR3279557 | 22 | 910 | 500 | 635 | 3526 | 19581 | 50014 | 35878 | 8737 | 675 | 9504 | 4436 | 1738 | 9536 | 28 | 1703 | 44363 | 0 | 0 | 10 | 327 | 364 | 157 | 0 | 0 | 0 | 0 | 19 | 89 | 487 | 1512 | 477 | 0 | 10 | 44 | 32 | 0 |
| SRR3279558 | 0 | 125 | 125 | 260 | 1883 | 13368 | 23647 | 27450 | 7456 | 77 | 13292 | 5460 | 2862 | 25078 | 75 | 1770 | 11861 | 0 | 0 | 11 | 131 | 116 | 26 | 0 | 0 | 0 | 0 | 0 | 64 | 685 | 1082 | 236 | 0 | 0 | 19 | 0 | 0 |
| SRR3279559 | 0 | 109 | 362 | 706 | 1412 | 18718 | 20624 | 24082 | 4232 | 193 | 12923 | 4828 | 964 | 3886 | 0 | 1122 | 17665 | 0 | 0 | 0 | 461 | 741 | 257 | 0 | 15 | 13 | 0 | 15 | 133 | 969 | 1916 | 462 | 0 | 43 | 499 | 56 | 0 |
| SRR3279560 | 0 | 270 | 344 | 209 | 1540 | 10477 | 21242 | 28842 | 5521 | 86 | 24638 | 6774 | 1912 | 4389 | 19 | 1179 | 22698 | 0 | 0 | 0 | 294 | 559 | 230 | 18 | 0 | 0 | 0 | 0 | 34 | 690 | 1373 | 463 | 0 | 19 | 390 | 70 | 0 |
| SRR3279561 | 0 | 193 | 241 | 333 | 2927 | 19128 | 34502 | 29033 | 2280 | 108 | 13466 | 3862 | 1221 | 6335 | 40 | 3387 | 21966 | 0 | 0 | 0 | 1660 | 1458 | 407 | 12 | 12 | 0 | 0 | 0 | 94 | 2555 | 4506 | 790 | 0 | 51 | 613 | 92 | 0 |
| SRR3279562 | 12 | 247 | 457 | 148 | 3593 | 43872 | 45250 | 32674 | 4052 | 173 | 32664 | 11208 | 1786 | 3144 | 92 | 347 | 13862 | 0 | 0 | 0 | 208 | 481 | 242 | 10 | 0 | 0 | 0 | 22 | 144 | 429 | 1373 | 488 | 0 | 53 | 464 | 41 | 0 |
| SRR3279563 | 0 | 141 | 164 | 59 | 2403 | 10041 | 34682 | 19836 | 5637 | 141 | 74817 | 19467 | 1478 | 5144 | 31 | 1571 | 14631 | 0 | 0 | 11 | 378 | 518 | 200 | 11 | 0 | 0 | 0 | 11 | 74 | 481 | 629 | 244 | 0 | 26 | 281 | 37 | 0 |
| SRR3279564 | 0 | 177 | 218 | 487 | 1554 | 18638 | 25684 | 28557 | 8584 | 194 | 7190 | 5333 | 1076 | 3371 | 0 | 1056 | 31529 | 0 | 0 | 0 | 559 | 987 | 172 | 0 | 0 | 0 | 0 | 0 | 40 | 596 | 2197 | 279 | 0 | 11 | 72 | 0 | 0 |
| SRR3279565 | 57 | 2612 | 928 | 556 | 1707 | 13238 | 24082 | 21810 | 7827 | 397 | 18696 | 13779 | 3090 | 9715 | 96 | 806 | 33905 | 0 | 0 | 0 | 563 | 506 | 117 | 0 | 0 | 0 | 0 | 20 | 154 | 1259 | 1714 | 480 | 0 | 38 | 623 | 75 | 0 |
| SRR3279566 | 35 | 539 | 283 | 92 | 1394 | 12712 | 17263 | 42851 | 3773 | 227 | 21651 | 8939 | 2214 | 5946 | 41 | 328 | 16935 | 0 | 0 | 0 | 134 | 256 | 72 | 0 | 0 | 0 | 0 | 0 | 25 | 136 | 484 | 88 | 0 | 0 | 62 | 0 | 0 |
| SRR3279567 | 0 | 48 | 176 | 346 | 1680 | 18555 | 32038 | 19363 | 1035 | 230 | 7694 | 1327 | 1953 | 18772 | 17 | 1690 | 8612 | 0 | 0 | 0 | 214 | 231 | 64 | 0 | 0 | 0 | 0 | 0 | 79 | 1552 | 3518 | 763 | 0 | 13 | 22 | 23 | 0 |
| SRR3279568 | 42 | 1559 | 1100 | 863 | 2218 | 19569 | 31274 | 35846 | 3771 | 97 | 13905 | 8172 | 1536 | 4897 | 17 | 154 | 23186 | 0 | 0 | 0 | 1070 | 673 | 165 | 0 | 11 | 0 | 0 | 0 | 79 | 1296 | 1192 | 297 | 0 | 47 | 547 | 55 | 0 |
| SRR3279569 | 0 | 177 | 321 | 353 | 1228 | 8694 | 22895 | 24258 | 6544 | 64 | 19652 | 6237 | 1267 | 5239 | 134 | 557 | 25870 | 0 | 0 | 0 | 367 | 625 | 188 | 0 | 0 | 0 | 0 | 0 | 33 | 704 | 1090 | 365 | 0 | 25 | 404 | 36 | 0 |
| SRR3279570 | 0 | 1600 | 16 | 398 | 788 | 4120 | 10718 | 13899 | 6889 | 0 | 12914 | 3491 | 1933 | 5256 | 56 | 338 | 14115 | 0 | 0 | 0 | 434 | 294 | 16 | 0 | 0 | 0 | 0 | 0 | 52 | 1266 | 313 | 27 | 0 | 0 | 0 | 0 | 0 |
| SRR3279571 | 0 | 354 | 422 | 191 | 2600 | 14777 | 33040 | 33800 | 7907 | 218 | 16562 | 5091 | 1797 | 7461 | 0 | 368 | 28788 | 0 | 0 | 0 | 115 | 221 | 125 | 0 | 0 | 0 | 0 | 0 | 37 | 393 | 1063 | 394 | 0 | 39 | 321 | 29 | 0 |

### Supp Table S8.1.Columns_B_to_AL

| | | | | | | | | | | | | | | | | | | | | | | | | | | | | | | | | | | | | | |
|---|---|---|---|---|---|---|---|---|---|---|---|---|---|---|---|---|---|---|---|---|---|---|---|---|---|---|---|---|---|---|---|---|---|---|---|---|---|
| SRR3279572 | 0 | 208 | 191 | 444 | 605 | 3766 | 6534 | 9604 | 7498 | 34 | 21974 | 4601 | 1504 | 4301 | 67 | 2132 | 24975 | 0 | 0 | 0 | 392 | 584 | 264 | 64 | 11 | 19 | 0 | 0 | 61 | 1007 | 2770 | 1013 | 0 | 52 | 778 | 277 | 0 |
| SRR3279573 | 0 | 428 | 187 | 141 | 975 | 7274 | 13613 | 21013 | 4467 | 203 | 23215 | 8360 | 1942 | 10444 | 134 | 495 | 12607 | 0 | 0 | 0 | 408 | 249 | 50 | 0 | 0 | 0 | 0 | 0 | 47 | 508 | 563 | 60 | 0 | 0 | 25 | 0 | 0 |
| SRR3279574 | 35 | 1128 | 282 | 203 | 1428 | 10369 | 20505 | 21652 | 4190 | 25 | 24812 | 11376 | 3979 | 4962 | 234 | 1358 | 18960 | 0 | 0 | 0 | 703 | 710 | 163 | 14 | 0 | 0 | 0 | 13 | 79 | 863 | 1250 | 278 | 0 | 0 | 114 | 11 | 0 |
| SRR3279575 | 32 | 1420 | 1014 | 375 | 1025 | 8878 | 16423 | 27423 | 2462 | 958 | 13807 | 6475 | 2653 | 11699 | 27 | 780 | 44726 | 0 | 0 | 0 | 465 | 420 | 161 | 0 | 0 | 0 | 0 | 25 | 111 | 871 | 2319 | 644 | 0 | 26 | 210 | 48 | 0 |
| SRR3279576 | 32 | 689 | 681 | 228 | 3534 | 24918 | 69974 | 70622 | 4032 | 806 | 18366 | 9428 | 2361 | 14741 | 25 | 624 | 22432 | 0 | 0 | 22 | 785 | 738 | 116 | 0 | 0 | 0 | 0 | 57 | 251 | 946 | 2207 | 281 | 0 | 11 | 54 | 13 | 0 |
| SRR3279577 | 39 | 1101 | 347 | 1291 | 1946 | 15999 | 27633 | 30547 | 5238 | 87 | 13496 | 5764 | 2188 | 6701 | 21 | 1336 | 38847 | 0 | 0 | 20 | 935 | 1413 | 390 | 0 | 20 | 0 | 0 | 35 | 157 | 576 | 1770 | 447 | 0 | 58 | 730 | 96 | 0 |
| SRR3279578 | 0 | 173 | 126 | 485 | 2175 | 25753 | 22023 | 20692 | 6931 | 42 | 20286 | 9347 | 1838 | 2927 | 27 | 492 | 29606 | 0 | 0 | 0 | 1091 | 656 | 214 | 0 | 0 | 0 | 0 | 19 | 180 | 1233 | 1150 | 280 | 0 | 32 | 488 | 101 | 0 |
| SRR3279579 | 23 | 1447 | 419 | 160 | 2614 | 30740 | 38582 | 26924 | 3450 | 69 | 37118 | 17095 | 2457 | 4391 | 209 | 366 | 15579 | 0 | 0 | 0 | 239 | 303 | 128 | 0 | 0 | 0 | 0 | 12 | 54 | 206 | 455 | 208 | 0 | 0 | 211 | 19 | 0 |
| SRR3279580 | 15 | 675 | 735 | 486 | 1386 | 11345 | 18293 | 30229 | 6840 | 426 | 7759 | 3345 | 1654 | 6916 | 0 | 935 | 43457 | 0 | 0 | 12 | 605 | 901 | 209 | 0 | 0 | 0 | 0 | 52 | 288 | 1166 | 5689 | 905 | 0 | 37 | 104 | 25 | 0 |
| SRR3279581 | 30 | 1292 | 575 | 1180 | 852 | 7267 | 11737 | 18959 | 4300 | 236 | 23324 | 10078 | 3136 | 10182 | 78 | 651 | 31405 | 0 | 0 | 0 | 745 | 1375 | 339 | 27 | 14 | 0 | 0 | 0 | 63 | 988 | 2921 | 870 | 0 | 74 | 594 | 79 | 0 |
| SRR3279582 | 28 | 1226 | 194 | 237 | 1418 | 11898 | 20903 | 29772 | 7992 | 42 | 16603 | 6742 | 2296 | 10007 | 45 | 2713 | 28962 | 0 | 0 | 0 | 219 | 911 | 448 | 38 | 13 | 0 | 0 | 0 | 23 | 232 | 1072 | 483 | 0 | 36 | 420 | 43 | 0 |
| SRR3279583 | 51 | 1251 | 391 | 368 | 1087 | 11013 | 18800 | 20676 | 9186 | 202 | 23085 | 7621 | 1481 | 4534 | 55 | 443 | 28990 | 0 | 0 | 0 | 675 | 697 | 143 | 0 | 0 | 0 | 0 | 0 | 123 | 1407 | 1681 | 369 | 0 | 39 | 472 | 86 | 0 |
| SRR3279584 | 206 | 2884 | 169 | 2151 | 2441 | 12296 | 31969 | 35631 | 5265 | 85 | 15766 | 8097 | 2012 | 8659 | 15 | 1691 | 64268 | 0 | 0 | 10 | 631 | 510 | 278 | 20 | 0 | 0 | 0 | 35 | 198 | 853 | 2067 | 923 | 0 | 29 | 130 | 22 | 0 |
| SRR3279585 | 30 | 1609 | 152 | 175 | 1055 | 8282 | 13526 | 19122 | 7355 | 23 | 28459 | 9721 | 2878 | 5756 | 96 | 416 | 22575 | 0 | 0 | 0 | 429 | 679 | 148 | 0 | 0 | 0 | 0 | 15 | 46 | 436 | 913 | 199 | 0 | 12 | 117 | 12 | 0 |
| SRR3279586 | 21 | 324 | 103 | 75 | 1473 | 13226 | 18702 | 22481 | 7109 | 25 | 30135 | 21264 | 3234 | 3779 | 96 | 1537 | 14379 | 0 | 0 | 0 | 439 | 342 | 91 | 0 | 0 | 0 | 0 | 0 | 42 | 429 | 624 | 159 | 0 | 0 | 124 | 22 | 0 |
| SRR3317221 | 0 | 251 | 87655 | 0 | 35053 | 42850 | 585923 | 51770 | 1943 | 22299 | 12245 | 3197 | 3067 | 5633 | 3538 | 111 | 4310 | 0 | 0 | 0 | 0 | 0 | 13 | 0 | 0 | 0 | 0 | 0 | 0 | 0 | 0 | 0 | 0 | 0 | 0 | 0 | 0 |
| SRR3477269 | 0 | 17 | 205 | 0 | 0 | 262 | 919 | 525 | 0 | 0 | 919 | 656 | 262 | 1050 | 0 | 1837 | 7611 | 0 | 0 | 0 | 37 | 136 | 17 | 0 | 0 | 0 | 0 | 0 | 0 | 0 | 27 | 0 | 0 | 0 | 0 | 0 | 0 |
| SRR3477271 | 0 | 12 | 191 | 0 | 0 | 140 | 560 | 140 | 140 | 0 | 560 | 140 | 140 | 140 | 0 | 1260 | 3360 | 0 | 0 | 0 | 35 | 111 | 13 | 0 | 0 | 0 | 0 | 0 | 0 | 0 | 11 | 0 | 0 | 0 | 0 | 0 | 0 |
| SRR3490374 | 0 | 148766 | 0 | 0 | 50 | 235 | 542 | 1327 | 7710 | 0 | 5841 | 12788 | 1548 | 1041 | 14 | 4515 | 102316 | 0 | 0 | 0 | 165 | 427 | 11 | 0 | 0 | 0 | 0 | 0 | 140 | 113 | 548 | 17 | 0 | 0 | 0 | 0 | 0 |
| SRR3490375 | 0 | 18556 | 290 | 0 | 43 | 187 | 498 | 771 | 18224 | 290 | 7339 | 7050 | 1179 | 911 | 0 | 7082 | 27406 | 0 | 0 | 0 | 220 | 79 | 0 | 0 | 0 | 0 | 0 | 0 | 133 | 214 | 137 | 0 | 0 | 0 | 0 | 0 | 0 |
| SRR3490376 | 133 | 261408 | 311 | 399 | 84 | 705 | 741 | 1801 | 8758 | 15 | 11407 | 27147 | 2654 | 1050 | 0 | 8316 | 38089 | 0 | 0 | 0 | 1297 | 1373 | 19 | 0 | 0 | 0 | 0 | 0 | 211 | 894 | 1858 | 39 | 0 | 0 | 0 | 0 | 0 |
| SRR3490377 | 126 | 201929 | 828 | 342 | 126 | 1160 | 1351 | 2882 | 15347 | 0 | 5638 | 9231 | 891 | 598 | 0 | 8460 | 39823 | 0 | 0 | 13 | 1280 | 1358 | 49 | 0 | 0 | 0 | 0 | 0 | 269 | 610 | 1188 | 36 | 0 | 0 | 0 | 0 | 0 |
| SRR3490378 | 809 | 82871 | 1857 | 476 | 37 | 508 | 610 | 1714 | 14178 | 0 | 6024 | 10487 | 1173 | 702 | 0 | 15199 | 41846 | 0 | 0 | 14 | 1090 | 107 | 35 | 0 | 0 | 0 | 0 | 0 | 252 | 792 | 170 | 37 | 0 | 0 | 0 | 0 | 0 |
| SRR3490379 | 275 | 43311 | 5494 | 183 | 68 | 345 | 476 | 1223 | 27568 | 0 | 4387 | 4379 | 856 | 594 | 0 | 19798 | 24084 | 0 | 0 | 0 | 124 | 38 | 15 | 0 | 0 | 0 | 0 | 0 | 101 | 297 | 98 | 22 | 0 | 0 | 0 | 0 | 0 |
| SRR3490380 | 685 | 43008 | 3552 | 171 | 111 | 1222 | 1520 | 2196 | 16549 | 43 | 3759 | 4591 | 848 | 660 | 0 | 12692 | 41518 | 0 | 0 | 0 | 438 | 49 | 13 | 0 | 0 | 0 | 0 | 0 | 125 | 401 | 127 | 13 | 0 | 0 | 0 | 0 | 0 |
| SRR3490382 | 202 | 24184 | 0 | 0 | 23 | 222 | 363 | 1169 | 18747 | 0 | 3427 | 5649 | 994 | 901 | 0 | 1123 | 47832 | 0 | 0 | 0 | 61 | 555 | 11 | 0 | 0 | 0 | 0 | 0 | 24 | 90 | 1031 | 16 | 0 | 0 | 0 | 0 | 0 |
| SRR3490383 | 906 | 41529 | 843 | 50 | 368 | 4733 | 3027 | 8640 | 21013 | 38 | 15657 | 28911 | 3482 | 1392 | 0 | 2317 | 70739 | 0 | 0 | 0 | 32 | 46 | 0 | 0 | 0 | 0 | 0 | 0 | 40 | 35 | 106 | 0 | 0 | 0 | 0 | 0 | 0 |
| SRR3490384 | 140 | 17777 | 698 | 233 | 475 | 6401 | 4461 | 19351 | 15333 | 140 | 14950 | 27975 | 3518 | 2418 | 0 | 2308 | 79195 | 0 | 0 | 0 | 239 | 1048 | 21 | 0 | 0 | 0 | 0 | 0 | 99 | 262 | 1530 | 58 | 0 | 0 | 0 | 0 | 0 |
| SRR3490385 | 51 | 9727 | 507 | 152 | 60 | 746 | 642 | 2020 | 17676 | 101 | 12046 | 21927 | 2694 | 813 | 0 | 3548 | 49496 | 0 | 0 | 0 | 11 | 18 | 0 | 0 | 0 | 0 | 0 | 0 | 13 | 19 | 53 | 0 | 0 | 0 | 0 | 0 | 0 |
| SRR3490386 | 50 | 14177 | 1498 | 300 | 140 | 1525 | 1001 | 4268 | 19458 | 0 | 15106 | 26243 | 3227 | 943 | 0 | 3017 | 68592 | 0 | 0 | 0 | 112 | 538 | 45 | 0 | 0 | 0 | 0 | 38 | 185 | 154 | 676 | 40 | 0 | 0 | 0 | 0 | 0 |
| SRR3490387 | 109 | 14853 | 581 | 690 | 248 | 2967 | 2141 | 6460 | 23125 | 36 | 13368 | 24376 | 2789 | 1289 | 0 | 5330 | 69003 | 0 | 0 | 0 | 68 | 1010 | 25 | 0 | 0 | 0 | 0 | 0 | 39 | 137 | 1563 | 60 | 0 | 0 | 0 | 0 | 0 |
| SRR3490388 | 200 | 9200 | 800 | 400 | 88 | 380 | 805 | 1829 | 24050 | 200 | 4769 | 16340 | 1097 | 1770 | 15 | 3248 | 87744 | 0 | 0 | 0 | 50 | 222 | 32 | 0 | 0 | 0 | 0 | 0 | 14 | 68 | 258 | 23 | 0 | 0 | 0 | 0 | 0 |
| SRR3490389 | 77 | 6665 | 460 | 230 | 147 | 786 | 1218 | 2998 | 19225 | 230 | 9485 | 10665 | 1452 | 1719 | 0 | 7820 | 26882 | 0 | 0 | 0 | 25 | 63 | 0 | 0 | 0 | 0 | 0 | 0 | 16 | 36 | 103 | 20 | 0 | 0 | 0 | 0 | 0 |
| SRR3490390 | 65 | 46344 | 648 | 130 | 44 | 175 | 524 | 1462 | 7486 | 65 | 2685 | 10585 | 786 | 895 | 0 | 1506 | 56594 | 0 | 0 | 0 | 482 | 964 | 53 | 0 | 0 | 0 | 0 | 0 | 147 | 488 | 982 | 18 | 0 | 0 | 0 | 0 | 0 |
| SRR3490391 | 0 | 33841 | 1209 | 0 | 71 | 583 | 559 | 2474 | 10705 | 0 | 6280 | 12430 | 1011 | 868 | 0 | 3414 | 45997 | 0 | 0 | 0 | 59 | 218 | 0 | 0 | 0 | 0 | 0 | 0 | 0 | 99 | 601 | 0 | 0 | 0 | 0 | 0 | 0 |
| SRR3490392 | 51 | 16730 | 1167 | 172 | 103 | 893 | 936 | 3878 | 11384 | 51 | 13288 | 33270 | 1956 | 2046 | 0 | 6311 | 36203 | 0 | 0 | 0 | 99 | 2140 | 20 | 0 | 0 | 0 | 0 | 0 | 26 | 59 | 1438 | 30 | 0 | 0 | 0 | 0 | 0 |
| SRR3490393 | 681 | 62052 | 869 | 329 | 342 | 4483 | 2971 | 10286 | 13811 | 0 | 16123 | 36465 | 3659 | 3125 | 0 | 5850 | 79178 | 0 | 0 | 13 | 373 | 1266 | 58 | 0 | 0 | 0 | 0 | 17 | 177 | 351 | 2546 | 58 | 0 | 0 | 0 | 0 | 0 |
| SRR3490394 | 66 | 49482 | 697 | 332 | 202 | 1322 | 1147 | 4868 | 13922 | 33 | 14382 | 23428 | 3513 | 2000 | 0 | 5641 | 84814 | 0 | 0 | 0 | 483 | 3613 | 83 | 0 | 0 | 0 | 0 | 0 | 166 | 478 | 6485 | 94 | 0 | 0 | 0 | 0 | 0 |
| SRR3493372 | 658 | 5433 | 49950 | 197 | 20503 | 36094 | 248977 | 32376 | 23511 | 44 | 22968 | 4404 | 4726 | 5407 | 12239 | 5540 | 8346 | 0 | 38 | 0 | 16 | 18 | 0 | 0 | 0 | 0 | 0 | 116 | 64 | 57 | 0 | 0 | 0 | 0 | 0 | 0 | 0 |
| SRR3493373 | 172 | 2191 | 66259 | 766 | 5855 | 19597 | 266313 | 44048 | 26203 | 58 | 13648 | 2262 | 1317 | 4824 | 3355 | 7793 | 8093 | 0 | 15 | 0 | 11 | 18 | 11 | 15 | 84 | 0 | 0 | 54 | 40 | 47 | 0 | 0 | 0 | 0 | 0 | 72 | 0 |
| SRR3534933 | 105 | 7127 | 1674 | 110 | 3734 | 56862 | 63378 | 48657 | 9596 | 64 | 30197 | 34215 | 7237 | 7480 | 300 | 3250 | 24536 | 0 | 0 | 0 | 305 | 104 | 160 | 93 | 959 | 45 | 0 | 0 | 34 | 64 | 26 | 39 | 0 | 65 | 30 | 18 | 0 |
| SRR3534934 | 105 | 8843 | 1636 | 59 | 3239 | 53632 | 56089 | 55647 | 9532 | 57 | 27193 | 33055 | 6995 | 6656 | 280 | 2630 | 24924 | 0 | 0 | 0 | 501 | 400 | 263 | 51 | 318 | 32 | 0 | 20 | 83 | 105 | 97 | 69 | 0 | 32 | 20 | 28 | 0 |
| SRR3534935 | 19 | 7525 | 3122 | 78 | 4640 | 61894 | 54244 | 40875 | 9108 | 32 | 32335 | 38611 | 6753 | 6871 | 134 | 8940 | 16470 | 0 | 0 | 0 | 732 | 178 | 267 | 31 | 90 | 0 | 0 | 15 | 71 | 137 | 55 | 85 | 0 | 22 | 0 | 14 | 0 |
| SRR3534936 | 70 | 8965 | 807 | 92 | 4213 | 58841 | 52343 | 37453 | 7862 | 42 | 28216 | 39203 | 6336 | 6350 | 169 | 9851 | 17197 | 0 | 0 | 0 | 533 | 148 | 246 | 29 | 84 | 0 | 0 | 13 | 65 | 99 | 49 | 77 | 0 | 18 | 0 | 14 | 0 |
| SRR3534937 | 354 | 71659 | 2220 | 279 | 6593 | 57581 | 48202 | 47726 | 12526 | 49 | 23072 | 31054 | 5514 | 4182 | 145 | 12166 | 18632 | 0 | 0 | 0 | 944 | 264 | 341 | 65 | 263 | 18 | 0 | 12 | 87 | 191 | 122 | 131 | 0 | 47 | 18 | 20 | 0 |
| SRR3534938 | 240 | 82856 | 1050 | 128 | 6242 | 57336 | 47293 | 44030 | 12178 | 94 | 24162 | 32849 | 5810 | 4309 | 154 | 12837 | 17863 | 0 | 0 | 0 | 770 | 195 | 301 | 31 | 108 | 0 | 0 | 13 | 81 | 160 | 100 | 120 | 0 | 35 | 14 | 18 | 0 |
| SRR3566024 | 0 | 1457 | 194 | 1053 | 137 | 811 | 1295 | 492 | 7525 | 25 | 6097 | 8194 | 2771 | 364 | 2700 | 3534 | 32191 | 0 | 0 | 0 | 0 | 14 | 14 | 11 | 0 | 0 | 0 | 0 | 0 | 0 | 0 | 0 | 0 | 0 | 0 | 0 | 0 |
| SRR3566026 | 19 | 3201 | 76 | 1401 | 199 | 704 | 1730 | 604 | 8229 | 38 | 5778 | 7504 | 2760 | 347 | 2575 | 3543 | 32348 | 0 | 0 | 0 | 0 | 11 | 24 | 0 | 0 | 0 | 0 | 0 | 0 | 0 | 0 | 0 | 0 | 0 | 0 | 0 | 0 |
| SRR3566150 | 47 | 3325 | 246 | 1986 | 62 | 765 | 817 | 427 | 8143 | 131 | 3694 | 4736 | 1404 | 253 | 1316 | 3510 | 37914 | 0 | 0 | 0 | 0 | 0 | 26 | 14 | 0 | 0 | 0 | 0 | 0 | 0 | 0 | 0 | 0 | 0 | 0 | 0 | 0 |
| SRR3566151 | 60 | 2961 | 115 | 1971 | 95 | 716 | 761 | 379 | 7901 | 22 | 3824 | 5739 | 1348 | 174 | 1208 | 3332 | 37655 | 0 | 0 | 0 | 0 | 13 | 17 | 15 | 0 | 0 | 0 | 0 | 0 | 0 | 13 | 0 | 0 | 0 | 0 | 0 | 0 |
| SRR3566152 | 109 | 2294 | 366 | 1921 | 54 | 629 | 870 | 355 | 8306 | 109 | 2644 | 4002 | 1011 | 228 | 790 | 3875 | 40778 | 0 | 0 | 0 | 0 | 0 | 22 | 0 | 0 | 0 | 0 | 0 | 0 | 0 | 16 | 0 | 0 | 0 | 0 | 0 | 0 |
| SRR3566153 | 13 | 2398 | 293 | 1825 | 134 | 773 | 343 | 273 | 7184 | 67 | 2354 | 3621 | 791 | 174 | 715 | 3057 | 42084 | 0 | 0 | 0 | 0 | 0 | 11 | 11 | 0 | 0 | 0 | 0 | 0 | 0 | 0 | 0 | 0 | 0 | 0 | 0 | 0 |
| SRR3566154 | 0 | 1745 | 67 | 939 | 0 | 1015 | 547 | 156 | 7418 | 403 | 1484 | 1015 | 547 | 0 | 312 | 5466 | 45210 | 0 | 0 | 0 | 0 | 0 | 0 | 15 | 0 | 0 | 0 | 0 | 0 | 0 | 0 | 0 | 0 | 0 | 0 | 0 | 0 |
| SRR3566155 | 0 | 1643 | 583 | 2015 | 67 | 737 | 1206 | 737 | 7236 | 159 | 1139 | 2345 | 268 | 603 | 469 | 5427 | 40399 | 0 | 0 | 0 | 0 | 0 | 35 | 12 | 0 | 0 | 0 | 0 | 0 | 0 | 0 | 0 | 0 | 0 | 0 | 0 | 0 |
| SRR3582392 | 53 | 1874 | 194 | 690 | 363 | 959 | 2831 | 925 | 9571 | 53 | 6008 | 7018 | 3302 | 573 | 3659 | 4119 | 28792 | 0 | 0 | 0 | 0 | 0 | 12 | 17 | 0 | 0 | 0 | 0 | 0 | 0 | 14 | 0 | 0 | 0 | 0 | 0 | 0 |
| SRR3582393 | 39 | 1969 | 212 | 734 | 325 | 1184 | 2426 | 801 | 9423 | 77 | 6007 | 6774 | 3528 | 417 | 3460 | 3974 | 26615 | 0 | 0 | 0 | 0 | 0 | 0 | 29 | 0 | 0 | 0 | 0 | 0 | 0 | 16 | 0 | 0 | 0 | 0 | 0 | 0 |
| SRR3623374 | 0 | 73 | 5120 | 21 | 57 | 98 | 2913 | 5720 | 382 | 238 | 2037 | 2388 | 1635 | 1644 | 0 | 19 | 3658 | 0 | 0 | 0 | 0 | 0 | 0 | 0 | 0 | 0 | 0 | 0 | 0 | 0 | 0 | 0 | 0 | 0 | 0 | 0 | 0 |
| SRR3624069 | 64 | 258 | 6375 | 2212 | 18726 | 6273 | 159633 | 45580 | 16402 | 4696 | 21763 | 8932 | 10562 | 6165 | 11088 | 986 | 11628 | 0 | 23 | 15 | 27 | 1099 | 285 | 153 | 77 | 190 | 17 | 54 | 70 | 40 | 113 | 37 | 0 | 0 | 63 | 402 | 15 |
| SRR3624070 | 123 | 793 | 14668 | 5519 | 27492 | 7986 | 204079 | 45205 | 19057 | 6843 | 19564 | 8328 | 10141 | 6182 | 11389 | 1002 | 11365 | 0 | 30 | 16 | 70 | 2731 | 1072 | 908 | 184 | 107 | 25 | 69 | 86 | 71 | 180 | 99 | 0 | 0 | 70 | 434 | 21 |
| SRR3624071 | 72 | 476 | 15692 | 2150 | 22649 | 6722 | 176655 | 37041 | 18364 | 6630 | 19351 | 9298 | 10594 | 6138 | 11212 | 936 | 13721 | 0 | 32 | 16 | 31 | 1646 | 356 | 239 | 97 | 53 | 23 | 53 | 72 | 56 | 146 | 77 | 0 | 0 | 91 | 471 | 17 |
| SRR3624072 | 214 | 960 | 36132 | 3920 | 31101 | 9515 | 249828 | 37749 | 17704 | 3102 | 15334 | 7158 | 8149 | 5376 | 7455 | 645 | 11612 | 0 | 32 | 18 | 34 | 3262 | 480 | 157 | 69 | 56 | 0 | 45 | 64 | 40 | 109 | 55 | 0 | 0 | 46 | 304 | 12 |
| SRR3624073 | 66 | 474 | 29401 | 1300 | 20207 | 7028 | 183222 | 53531 | 14713 | 6596 | 19872 | 9314 | 10378 | 5762 | 5007 | 592 | 13105 | 0 | 28 | 17 | 35 | 2677 | 394 | 155 | 59 | 40 | 18 | 57 | 80 | 54 | 170 | 63 | 0 | 0 | 30 | 235 | 0 |
| SRR3624074 | 70 | 546 | 49551 | 1122 | 25737 | 8233 | 190780 | 51508 | 14090 | 5885 | 20041 | 10025 | 11539 | 6533 | 6573 | 787 | 14224 | 0 | 31 | 18 | 31 | 1804 | 319 | 198 | 70 | 37 | 18 | 45 | 65 | 56 | 122 | 50 | 0 | 0 | 42 | 249 | 12 |
| SRR3675003 | 604 | 104838 | 55 | 576 | 0 | 3397 | 425 | 546 | 182 | 27 | 243 | 1031 | 0 | 61 | 0 | 667 | 1031 | 0 | 0 | 615 | 33030 | 26720 | 5819 | 208 | 37 | 0 | 0 | 13 | 4650 | 6507 | 23322 | 2189 | 12 | 19 | 15 | 35 | 0 |
| SRR3675004 | 106 | 108997 | 112 | 957 | 0 | 408 | 637 | 484 | 3312 | 16 | 8075 | 7107 | 3643 | 6649 | 25 | 866 | 4738 | 0 | 0 | 491 | 16518 | 26835 | 4277 | 202 | 29 | 0 | 0 | 10 | 2976 | 3029 | 20659 | 1646 | 12 | 15 | 11 | 34 | 0 |
| SRR3675005 | 44 | 95407 | 1316 | 783 | 0 | 17705 | 587 | 231 | 780 | 38 | 481 | 2821 | 164 | 260 | 0 | 3090 | 2561 | 0 | 0 | 0 | 1694 | 684 | 689 | 0 | 0 | 0 | 0 | 0 | 65 | 584 | 372 | 405 | 0 | 0 | 0 | 11 | 0 |
| SRR3675006 | 222 | 93620 | 46 | 1153 | 0 | 1418 | 253 | 152 | 304 | 27 | 203 | 912 | 101 | 51 | 0 | 456 | 152 | 0 | 0 | 403 | 20293 | 23126 | 5636 | 200 | 29 | 0 | 0 | 10 | 2344 | 4381 | 18951 | 2282 | 11 | 14 | 11 | 33 | 0 |
| SRR3675007 | 63 | 128225 | 58 | 331 | 0 | 670 | 765 | 96 | 96 | 47 | 287 | 574 | 287 | 0 | 0 | 478 | 478 | 0 | 0 | 470 | 10413 | 15589 | 2620 | 64 | 0 | 0 | 0 | 13 | 3136 | 3100 | 13545 | 1040 | 0 | 0 | 0 | 0 | 0 |
| SRR3675008 | 186 | 128622 | 174 | 1004 | 0 | 2138 | 231 | 173 | 0 | 14 | 289 | 1271 | 58 | 58 | 0 | 347 | 520 | 0 | 0 | 364 | 14767 | 20013 | 3794 | 160 | 20 | 0 | 0 | 11 | 2573 | 2813 | 15974 | 1592 | 10 | 11 | 0 | 25 | 0 |
| SRR3713941 | 115 | 864 | 4440 | 459 | 26524 | 17544 | 330603 | 25127 | 12378 | 2775 | 25783 | 21001 | 12497 | 16007 | 2391 | 4378 | 9153 | 0 | 0 | 0 | 19 | 13 | 115 | 769 | 1059 | 137 | 0 | 0 | 0 | 0 | 0 | 0 | 0 | 0 | 37 | 411 | 0 |
| SRR3713942 | 123 | 2496 | 8355 | 1966 | 411 | 324 | 15985 | 21911 | 6078 | 431 | 1492 | 1990 | 930 | 6749 | 130 | 1687 | 1795 | 0 | 12 | 20 | 20 | 164 | 16 | 0 | 25 | 0 | 0 | 0 | 0 | 0 | 0 | 0 | 0 | 0 | 0 | 0 | 0 |
| SRR3713943 | 83 | 552 | 5761 | 151 | 36380 | 18398 | 481525 | 64415 | 8207 | 680 | 26263 | 6640 | 5363 | 11695 | 19424 | 63 | 8008 | 0 | 0 | 0 | 22 | 0 | 93 | 401 | 911 | 90 | 0 | 0 | 0 | 17 | 0 | 0 | 0 | 0 | 28 | 325 | 0 |
| SRR3713945 | 733 | 3438 | 5128 | 3952 | 34059 | 15912 | 252120 | 18068 | 450 | 2863 | 25656 | 20768 | 11586 | 22491 | 2309 | 20921 | 5537 | 0 | 27 | 28 | 96 | 63 | 131 | 218 | 341 | 34 | 15 | 34 | 86 | 63 | 33 | 0 | 0 | 0 | 18 | 72 | 0 |
| SRR3713946 | 377 | 1480 | 7390 | 5087 | 51668 | 25913 | 271367 | 44239 | 536 | 905 | 36179 | 31134 | 17573 | 19805 | 1607 | 21930 | 6879 | 0 | 13 | 25 | 87 | 90 | 82 | 109 | 219 | 21 | 0 | 11 | 52 | 42 | 18 | 0 | 0 | 0 | 0 | 35 | 0 |
| SRR3879751 | 203 | 2227 | 176803 | 60 | 9059 | 6114 | 158056 | 16667 | 4049 | 428 | 26482 | 20225 | 4574 | 8038 | 2013 | 2103 | 16471 | 0 | 92 | 55 | 271 | 2441 | 487 | 89 | 37 | 0 | 0 | 108 | 99 | 158 | 157 | 27 | 0 | 0 | 0 | 86 | 0 |
| SRR3879752 | 170 | 3353 | 163307 | 47 | 7776 | 4575 | 141929 | 15483 | 3083 | 375 | 22525 | 19996 | 3835 | 8336 | 2078 | 2007 | 14209 | 0 | 84 | 73 | 224 | 2238 | 505 | 73 | 29 | 0 | 0 | 94 | 88 | 113 | 173 | 28 | 0 | 0 | 0 | 70 | 0 |
| SRR3879754 | 31 | 2752 | 113329 | 30 | 15039 | 8830 | 190156 | 25761 | 4073 | 206 | 17952 | 13228 | 8907 | 14627 | 3654 | 6419 | 11993 | 0 | 33 | 42 | 126 | 1704 | 286 | 88 | 39 | 0 | 0 | 36 | 52 | 96 | 154 | 15 | 0 | 0 | 0 | 37 | 0 |
| SRR3879755 | 27 | 2087 | 112116 | 22 | 15113 | 9577 | 185114 | 26326 | 4700 | 155 | 17629 | 10870 | 8880 | 14540 | 3934 | 7360 | 12512 | 0 | 28 | 25 | 99 | 1294 | 194 | 69 | 29 | 0 | 0 | 41 | 44 | 84 | 130 | 11 | 0 | 0 | 0 | 38 | 0 |
| SRR3879756 | 49 | 2552 | 115113 | 31 | 15270 | 8636 | 198534 | 18772 | 4162 | 351 | 17389 | 12060 | 9202 | 16650 | 3939 | 5544 | 11209 | 0 | 44 | 53 | 173 | 2323 | 331 | 100 | 58 | 0 | 0 | 57 | 74 | 138 | 147 | 15 | 0 | 0 | 0 | 79 | 0 |
| SRR3879757 | 39 | 4148 | 62145 | 0 | 12935 | 12544 | 229675 | 19292 | 3081 | 312 | 19170 | 17385 | 11932 | 11859 | 3252 | 1565 | 15405 | 0 | 17 | 16 | 43 | 413 | 104 | 20 | 12 | 0 | 0 | 13 | 14 | 27 | 42 | 0 | 0 | 0 | 0 | 17 | 0 |
| SRR3879758 | 22 | 3307 | 66542 | 16 | 13292 | 11998 | 214716 | 17051 | 2688 | 255 | 20113 | 20162 | 11873 | 11351 | 3136 | 1469 | 15508 | 0 | 24 | 13 | 38 | 517 | 119 | 23 | 17 | 0 | 0 | 15 | 21 | 26 | 30 | 0 | 0 | 0 | 0 | 15 | 0 |
| SRR3879759 | 99 | 7144 | 58633 | 32 | 15276 | 14065 | 180528 | 23708 | 4394 | 671 | 23689 | 31022 | 12751 | 13690 | 2131 | 1108 | 16422 | 0 | 40 | 42 | 138 | 1544 | 357 | 74 | 39 | 0 | 0 | 38 | 55 | 66 | 72 | 0 | 0 | 0 | 0 | 60 | 0 |
| SRR3879760 | 17 | 4272 | 77693 | 17 | 9681 | 8480 | 149924 | 12154 | 5295 | 221 | 23413 | 17979 | 9192 | 8871 | 2752 | 6706 | 15465 | 0 | 0 | 0 | 18 | 314 | 61 | 23 | 12 | 0 | 0 | 0 | 0 | 0 | 19 | 0 | 0 | 0 | 0 | 20 | 0 |
| SRR3879761 | 32 | 3916 | 64381 | 20 | 10728 | 9262 | 216553 | 6852 | 3540 | 282 | 29818 | 23183 | 11532 | 8676 | 3572 | 7666 | 15875 | 0 | 0 | 0 | 14 | 289 | 67 | 19 | 13 | 0 | 0 | 0 | 0 | 0 | 0 | 0 | 0 | 0 | 0 | 22 | 0 |
| SRR3879766 | 242 | 2530 | 38786 | 24 | 15321 | 7783 | 380016 | 10559 | 2544 | 3770 | 22876 | 8923 | 4135 | 7263 | 4648 | 2772 | 11277 | 0 | 0 | 0 | 15 | 71 | 154 | 41 | 53 | 0 | 0 | 0 | 0 | 0 | 0 | 0 | 0 | 0 | 0 | 48 | 12 |
| SRR3879767 | 48 | 626 | 65445 | 0 | 7926 | 5882 | 529591 | 691 | 3322 | 97 | 16016 | 10078 | 765 | 12359 | 29151 | 9847 | 922 | 0 | 0 | 0 | 0 | 0 | 0 | 0 | 0 | 0 | 0 | 0 | 0 | 0 | 0 | 0 | 0 | 0 | 0 | 0 | 0 |
| SRR3879768 | 195 | 1765 | 34466 | 14 | 25292 | 18522 | 630072 | 5891 | 1149 | 1426 | 12453 | 11852 | 3584 | 12751 | 10780 | 1071 | 3579 | 0 | 0 | 0 | 13 | 53 | 213 | 49 | 83 | 11 | 0 | 0 | 0 | 0 | 0 | 0 | 0 | 0 | 0 | 104 | 16 |
| SRR3879769 | 154 | 1170 | 25340 | 13 | 19109 | 12843 | 705059 | 1716 | 1790 | 635 | 20067 | 5665 | 1997 | 7122 | 15328 | 12222 | 1668 | 0 | 0 | 0 | 0 | 0 | 92 | 19 | 32 | 0 | 0 | 0 | 0 | 0 | 0 | 0 | 0 | 0 | 0 | 36 | 0 |
| SRR3879770 | 250 | 1666 | 59974 | 16 | 8189 | 6209 | 408146 | 444 | 5376 | 292 | 13429 | 8194 | 833 | 8623 | 5336 | 1860 | 665 | 0 | 0 | 0 | 0 | 32 | 35 | 0 | 0 | 0 | 0 | 0 | 0 | 0 | 0 | 0 | 0 | 0 | 0 | 11 | 0 |
| SRR3923813 | 0 | 657 | 26544 | 0 | 1292 | 14714 | 21306 | 6626 | 1663 | 45 | 28103 | 11694 | 320 | 2434 | 0 | 0 | 4123 | 0 | 0 | 0 | 0 | 0 | 0 | 0 | 0 | 0 | 0 | 0 | 0 | 32 | 587 | 21 | 0 | 0 | 0 | 0 | 0 |
| SRR3931357 | 26 | 2262 | 758 | 45 | 788 | 9195 | 13470 | 8750 | 21359 | 77 | 2279 | 62094 | 452 | 139 | 69 | 45834 | 39729 | 0 | 0 | 0 | 134 | 0 | 33 | 0 | 0 | 0 | 0 | 0 | 17 | 31 | 18 | 24 | 0 | 0 | 0 | 13 | 0 |
| SRR3931358 | 13 | 1937 | 689 | 20 | 723 | 7875 | 12148 | 8710 | 33458 | 60 | 1694 | 18 | 298 | 124 | 54 | 37079 | 30712 | 0 | 0 | 0 | 78 | 0 | 24 | 0 | 0 | 0 | 0 | 0 | 0 | 21 | 14 | 18 | 0 | 0 | 0 | 0 | 0 |
| SRR3931359 | 1000 | 26408 | 53 | 245 | 629 | 6278 | 9224 | 5897 | 35727 | 41 | 1384 | 35293 | 267 | 179 | 86 | 30445 | 29362 | 0 | 0 | 19 | 693 | 52 | 0 | 0 | 0 | 0 | 0 | 45 | 381 | 293 | 65 | 0 | 0 | 0 | 0 | 0 | 0 |
| SRR3931360 | 805 | 24277 | 59 | 370 | 673 | 6666 | 9931 | 5678 | 37923 | 38 | 1533 | 35886 | 242 | 198 | 104 | 37000 | 24292 | 0 | 0 | 13 | 634 | 41 | 0 | 0 | 0 | 0 | 0 | 34 | 332 | 303 | 50 | 0 | 0 | 0 | 0 | 0 | 0 |
| SRR3931984 | 0 | 24 | 5387 | 0 | 317 | 2221 | 4101 | 2121 | 4070 | 59 | 20467 | 20447 | 8358 | 9549 | 98 | 6713 | 47669 | 0 | 0 | 0 | 0 | 0 | 0 | 0 | 0 | 0 | 0 | 0 | 0 | 18 | 0 | 12 | 0 | 0 | 0 | 30 | 0 |
| SRR3931987 | 0 | 11 | 12929 | 0 | 194 | 1458 | 3231 | 2417 | 5899 | 30 | 21816 | 14834 | 6169 | 9597 | 32 | 7463 | 41702 | 0 | 0 | 0 | 0 | 0 | 0 | 0 | 0 | 0 | 0 | 0 | 0 | 25 | 0 | 0 | 0 | 0 | 0 | 35 | 0 |
| SRR3931988 | 0 | 39 | 11107 | 0 | 386 | 2764 | 5679 | 2328 | 3633 | 94 | 13324 | 13647 | 9359 | 10155 | 121 | 6246 | 33220 | 0 | 0 | 0 | 0 | 13 | 14 | 0 | 0 | 0 | 0 | 0 | 0 | 42 | 21 | 28 | 0 | 0 | 0 | 43 | 0 |
| SRR3992213 | 1129 | 57690 | 106378 | 74 | 6708 | 3777 | 58201 | 1919 | 23 | 572 | 27270 | 38 | 17603 | 47168 | 576 | 6216 | 20311 | 0 | 0 | 0 | 51 | 43 | 83 | 0 | 0 | 0 | 0 | 0 | 0 | 0 | 0 | 0 | 0 | 0 | 0 | 0 | 0 |
| SRR3992215 | 142 | 4703 | 71076 | 0 | 9976 | 6547 | 177773 | 8644 | 99 | 952 | 7242 | 32 | 9476 | 20556 | 78 | 3741 | 25416 | 0 | 0 | 0 | 16 | 38 | 37 | 0 | 0 | 0 | 0 | 0 | 0 | 0 | 0 | 0 | 0 | 0 | 0 | 0 | 0 |
| SRR3992474 | 156 | 15331 | 154 | 1081 | 1506 | 34704 | 23347 | 21625 | 33931 | 0 | 15120 | 5173 | 2099 | 6455 | 113 | 7543 | 9883 | 0 | 0 | 0 | 446 | 61 | 98 | 0 | 61 | 0 | 0 | 0 | 0 | 60 | 16 | 26 | 0 | 17 | 48 | 15 | 0 |
| SRR3992475 | 142 | 10045 | 130 | 1966 | 2047 | 34341 | 34614 | 27901 | 34186 | 0 | 13726 | 5613 | 2086 | 7189 | 112 | 6329 | 8998 | 0 | 0 | 0 | 279 | 32 | 58 | 0 | 34 | 0 | 0 | 0 | 0 | 40 | 10 | 17 | 0 | 14 | 35 | 15 | 0 |
| SRR3992476 | 168 | 1913 | 127 | 1016 | 17846 | 83000 | 108711 | 51907 | 618 | 19 | 20187 | 86876 | 5333 | 3049 | 77 | 13151 | 5894 | 0 | 0 | 0 | 113 | 29 | 11 | 0 | 56 | 12 | 0 | 0 | 11 | 41 | 36 | 0 | 0 | 72 | 274 | 70 | 0 |
| SRR3992477 | 27 | 3084 | 95 | 364 | 13842 | 102931 | 86839 | 45741 | 715 | 34 | 18817 | 77720 | 4745 | 2763 | 46 | 14244 | 5931 | 0 | 0 | 0 | 64 | 33 | 0 | 0 | 0 | 0 | 0 | 0 | 0 | 34 | 41 | 0 | 0 | 12 | 29 | 0 | 0 |
| SRR3996364 | 0 | 288335 | 483 | 2159 | 811 | 76611 | 13543 | 11870 | 1597 | 47 | 44463 | 36321 | 2462 | 41546 | 392 | 37864 | 85434 | 0 | 0 | 0 | 99 | 168 | 125 | 0 | 0 | 0 | 0 | 0 | 25 | 26 | 112 | 48 | 0 | 0 | 15 | 0 | 0 |
| SRR3996365 | 0 | 266560 | 472 | 1640 | 1416 | 98246 | 23677 | 15521 | 1985 | 54 | 49447 | 34861 | 2710 | 48261 | 385 | 56001 | 69726 | 0 | 0 | 0 | 129 | 168 | 118 | 0 | 0 | 0 | 0 | 0 | 32 | 41 | 113 | 59 | 0 | 0 | 21 | 0 | 0 |
| SRR3996366 | 17 | 240676 | 1172 | 2248 | 931 | 92236 | 15495 | 14509 | 1881 | 42 | 57154 | 46721 | 2905 | 55231 | 656 | 42521 | 57370 | 0 | 0 | 0 | 173 | 274 | 96 | 0 | 0 | 0 | 0 | 0 | 57 | 47 | 151 | 40 | 0 | 0 | 19 | 0 | 0 |
| SRR3996367 | 14 | 290303 | 1526 | 1561 | 1105 | 98118 | 19459 | 14842 | 1818 | 55 | 38439 | 37997 | 2445 | 52177 | 542 | 53061 | 63509 | 0 | 0 | 0 | 199 | 381 | 114 | 0 | 0 | 0 | 0 | 0 | 55 | 59 | 210 | 54 | 0 | 0 | 19 | 0 | 0 |
| SRR3996368 | 25 | 269046 | 300 | 641 | 1121 | 110414 | 21559 | 25982 | 2349 | 0 | 49493 | 30351 | 2624 | 40385 | 127 | 71255 | 58138 | 0 | 0 | 0 | 185 | 243 | 98 | 0 | 0 | 0 | 0 | 0 | 91 | 104 | 272 | 84 | 0 | 0 | 20 | 0 | 0 |
| SRR3996369 | 40 | 353488 | 352 | 1060 | 1288 | 132158 | 26176 | 27700 | 2444 | 0 | 49055 | 29529 | 2491 | 39296 | 160 | 88297 | 43429 | 0 | 0 | 0 | 238 | 224 | 70 | 0 | 0 | 0 | 0 | 0 | 88 | 127 | 235 | 67 | 0 | 0 | 18 | 0 | 0 |
| SRR4000625 | 83 | 1702 | 21219 | 32 | 13579 | 9979 | 241894 | 10231 | 10012 | 1417 | 12160 | 18191 | 4627 | 6525 | 294 | 1910 | 22270 | 0 | 0 | 0 | 92 | 29 | 170 | 17 | 22 | 0 | 0 | 0 | 0 | 17 | 0 | 0 | 0 | 0 | 0 | 10 | 0 |
| SRR4000626 | 79 | 1661 | 21249 | 28 | 13341 | 10157 | 242354 | 10475 | 9948 | 1392 | 12213 | 18294 | 4464 | 6859 | 290 | 1883 | 21900 | 0 | 0 | 0 | 104 | 31 | 168 | 20 | 26 | 0 | 0 | 0 | 0 | 18 | 0 | 0 | 0 | 0 | 0 | 13 | 0 |
| SRR4000627 | 17 | 177 | 32139 | 0 | 8518 | 6530 | 156867 | 3810 | 7411 | 1787 | 9357 | 17468 | 3259 | 4222 | 728 | 1403 | 21534 | 0 | 0 | 0 | 0 | 0 | 0 | 0 | 0 | 0 | 0 | 0 | 0 | 0 | 0 | 0 | 0 | 0 | 0 | 0 | 0 |
| SRR4000628 | 16 | 169 | 31902 | 0 | 8641 | 6272 | 154550 | 3882 | 7388 | 1781 | 9148 | 16651 | 3474 | 3894 | 766 | 1508 | 21785 | 0 | 0 | 0 | 0 | 0 | 0 | 0 | 0 | 0 | 0 | 0 | 0 | 0 | 0 | 0 | 0 | 0 | 0 | 0 | 0 |
| SRR4000629 | 80 | 1058 | 25851 | 19 | 9534 | 6814 | 172866 | 3331 | 10482 | 1451 | 13783 | 19843 | 4392 | 4434 | 1468 | 1772 | 15804 | 0 | 15 | 0 | 50 | 30 | 105 | 12 | 15 | 0 | 0 | 0 | 0 | 0 | 0 | 0 | 0 | 0 | 0 | 12 | 0 |
| SRR4000630 | 80 | 1080 | 25722 | 17 | 9646 | 7240 | 173364 | 3276 | 10645 | 1459 | 13793 | 19474 | 4269 | 4643 | 1376 | 1742 | 15659 | 0 | 13 | 0 | 50 | 31 | 98 | 17 | 16 | 0 | 0 | 0 | 0 | 0 | 0 | 0 | 0 | 0 | 0 | 12 | 0 |
| SRR4000631 | 76 | 2408 | 31621 | 0 | 8525 | 8897 | 151388 | 4478 | 8709 | 1339 | 10790 | 16865 | 3643 | 4271 | 1099 | 1691 | 17722 | 0 | 15 | 14 | 52 | 27 | 74 | 10 | 19 | 0 | 0 | 0 | 12 | 0 | 0 | 0 | 0 | 0 | 0 | 0 | 0 |
| SRR4000632 | 82 | 2346 | 31697 | 11 | 9062 | 8557 | 152065 | 4551 | 8390 | 1360 | 10781 | 17330 | 3410 | 4199 | 1037 | 1728 | 17853 | 0 | 14 | 14 | 56 | 25 | 79 | 0 | 17 | 0 | 0 | 0 | 13 | 0 | 0 | 0 | 0 | 0 | 0 | 0 | 0 |
| SRR4000633 | 96 | 1319 | 24648 | 23 | 16842 | 15309 | 311340 | 8561 | 12167 | 1613 | 30336 | 25135 | 7511 | 9835 | 465 | 1597 | 17238 | 0 | 0 | 0 | 46 | 19 | 112 | 12 | 21 | 0 | 0 | 0 | 0 | 0 | 0 | 0 | 0 | 0 | 0 | 11 | 0 |
| SRR4000634 | 92 | 1360 | 24494 | 22 | 17125 | 15510 | 312157 | 8349 | 12484 | 1578 | 29483 | 24902 | 7439 | 10136 | 472 | 1566 | 17384 | 0 | 0 | 0 | 44 | 24 | 92 | 13 | 18 | 0 | 0 | 0 | 0 | 0 | 0 | 0 | 0 | 0 | 0 | 11 | 0 |
| SRR4000635 | 28 | 464 | 17897 | 0 | 11756 | 12270 | 264864 | 11155 | 4309 | 1796 | 15568 | 25432 | 5553 | 7805 | 122 | 2262 | 18605 | 0 | 0 | 0 | 28 | 0 | 44 | 0 | 19 | 0 | 0 | 0 | 0 | 0 | 0 | 0 | 0 | 0 | 0 | 10 | 0 |
| SRR4000636 | 34 | 459 | 17878 | 11 | 11932 | 12272 | 265195 | 10899 | 4168 | 1790 | 15778 | 26024 | 5753 | 7729 | 125 | 2238 | 18348 | 0 | 0 | 0 | 32 | 0 | 50 | 12 | 19 | 0 | 0 | 0 | 0 | 0 | 0 | 0 | 0 | 0 | 0 | 11 | 0 |
| SRR4000638 | 44 | 328 | 35352 | 22 | 12555 | 11454 | 188533 | 12225 | 4699 | 2376 | 41008 | 34365 | 10412 | 10509 | 1564 | 2360 | 21439 | 0 | 0 | 0 | 15 | 0 | 11 | 0 | 0 | 0 | 0 | 0 | 0 | 0 | 0 | 0 | 0 | 0 | 0 | 0 | 0 |
| SRR4000639 | 45 | 199 | 37378 | 32 | 4104 | 3980 | 83999 | 4557 | 3488 | 1564 | 12185 | 22120 | 5021 | 4753 | 392 | 1658 | 23934 | 0 | 0 | 0 | 0 | 0 | 0 | 0 | 0 | 0 | 0 | 0 | 0 | 0 | 0 | 0 | 0 | 0 | 0 | 0 | 0 |
| SRR4000640 | 42 | 221 | 37293 | 27 | 4329 | 4114 | 82948 | 4585 | 3472 | 1563 | 12019 | 21583 | 5099 | 4808 | 367 | 1766 | 24166 | 0 | 0 | 0 | 0 | 0 | 0 | 0 | 0 | 0 | 0 | 0 | 0 | 0 | 0 | 0 | 0 | 0 | 0 | 0 | 0 |

### Supp Table S8.1.Columns_B_to_AL

| | | | | | | | | | | | | | | | | | | | | | | | | | | | | | | | | | | | | | |
|---|---|---|---|---|---|---|---|---|---|---|---|---|---|---|---|---|---|---|---|---|---|---|---|---|---|---|---|---|---|---|---|---|---|---|---|---|---|
| SRR4000641 | 112 | 355 | 39079 | 48 | 11076 | 11497 | 160488 | 6781 | 4742 | 1290 | 18565 | 23938 | 5818 | 4587 | 826 | 2111 | 19718 | 0 | 0 | 0 | 12 | 0 | 0 | 0 | 0 | 0 | 0 | 0 | 0 | 0 | 0 | 0 | 0 | 0 | 0 | 0 | 0 |
| SRR4000642 | 104 | 398 | 39213 | 49 | 11060 | 11415 | 157885 | 6677 | 4502 | 1304 | 18641 | 24016 | 5715 | 4491 | 885 | 2295 | 19894 | 0 | 0 | 0 | 13 | 0 | 0 | 0 | 0 | 0 | 0 | 0 | 0 | 0 | 0 | 0 | 0 | 0 | 0 | 0 | 0 |
| SRR4000643 | 167 | 559 | 28054 | 117 | 11918 | 10382 | 176932 | 10843 | 3194 | 1506 | 17779 | 19718 | 5825 | 4275 | 613 | 1262 | 20944 | 0 | 0 | 0 | 13 | 0 | 11 | 0 | 0 | 0 | 0 | 0 | 0 | 0 | 0 | 0 | 0 | 0 | 0 | 0 | 0 |
| SRR4000644 | 171 | 579 | 27995 | 115 | 11897 | 10572 | 175312 | 10263 | 3385 | 1458 | 16910 | 19229 | 6237 | 4299 | 569 | 1044 | 20712 | 0 | 0 | 0 | 13 | 0 | 0 | 0 | 0 | 0 | 0 | 0 | 0 | 0 | 0 | 0 | 0 | 0 | 0 | 0 | 0 |
| SRR4000645 | 15 | 152 | 37078 | 0 | 8851 | 10283 | 144664 | 5936 | 3960 | 1379 | 11938 | 17936 | 4289 | 5070 | 318 | 1647 | 17089 | 0 | 0 | 0 | 0 | 0 | 0 | 0 | 0 | 0 | 0 | 0 | 0 | 0 | 0 | 0 | 0 | 0 | 0 | 0 | 0 |
| SRR4000649 | 21 | 359 | 19982 | 14 | 18498 | 21775 | 317261 | 17845 | 8992 | 1608 | 21343 | 15488 | 6724 | 9943 | 1730 | 5293 | 24834 | 0 | 0 | 0 | 30 | 15 | 121 | 30 | 61 | 12 | 0 | 0 | 0 | 0 | 0 | 0 | 0 | 0 | 0 | 25 | 0 |
| SRR4000650 | 24 | 371 | 19948 | 17 | 18537 | 21749 | 316323 | 17843 | 8879 | 1568 | 21211 | 15558 | 6689 | 9986 | 1704 | 5296 | 24853 | 0 | 0 | 0 | 29 | 14 | 122 | 34 | 62 | 12 | 0 | 0 | 0 | 0 | 0 | 0 | 0 | 0 | 0 | 23 | 0 |
| SRR4011144 | 36 | 3931 | 893 | 0 | 845 | 3107 | 17977 | 8075 | 226 | 0 | 32 | 28 | 10542 | 3902 | 223 | 139 | 13828 | 0 | 0 | 0 | 0 | 0 | 0 | 0 | 0 | 0 | 0 | 0 | 0 | 0 | 0 | 0 | 0 | 0 | 0 | 0 | 0 |
| SRR4044175 | 91 | 7319 | 167 | 732 | 727 | 28317 | 20002 | 12454 | 2232 | 33 | 39068 | 32827 | 3074 | 8529 | 56 | 12154 | 28048 | 0 | 0 | 0 | 73 | 11 | 65 | 134 | 853 | 31 | 0 | 0 | 0 | 17 | 0 | 35 | 11 | 438 | 404 | 278 | 0 |
| SRR4044176 | 60 | 5789 | 160 | 1153 | 923 | 25238 | 17792 | 12529 | 1940 | 19 | 35720 | 30460 | 2832 | 6744 | 40 | 10766 | 19786 | 0 | 0 | 0 | 38 | 11 | 55 | 99 | 743 | 26 | 0 | 0 | 0 | 12 | 0 | 34 | 0 | 415 | 478 | 352 | 0 |
| SRR4044177 | 97 | 5366 | 106 | 4948 | 1259 | 29516 | 17176 | 15026 | 1757 | 13 | 37138 | 35758 | 2794 | 5800 | 26 | 8971 | 21858 | 0 | 17 | 0 | 317 | 160 | 51 | 27 | 230 | 0 | 0 | 131 | 110 | 96 | 208 | 38 | 0 | 254 | 368 | 145 | 0 |
| SRR4044178 | 40 | 5940 | 138 | 455 | 702 | 28129 | 20068 | 11251 | 2009 | 32 | 37208 | 30527 | 2841 | 8050 | 42 | 12902 | 27045 | 0 | 0 | 0 | 87 | 14 | 90 | 144 | 1267 | 48 | 0 | 0 | 0 | 20 | 0 | 52 | 14 | 615 | 631 | 478 | 0 |
| SRR4044179 | 298 | 6636 | 143 | 2272 | 1041 | 27391 | 18298 | 12009 | 1703 | 15 | 37695 | 29161 | 2742 | 6737 | 49 | 11794 | 19798 | 0 | 0 | 0 | 91 | 16 | 98 | 125 | 1379 | 45 | 0 | 0 | 0 | 20 | 10 | 55 | 16 | 741 | 937 | 578 | 0 |
| SRR4044180 | 0 | 3262 | 133 | 2592 | 1331 | 30934 | 18569 | 15449 | 1751 | 0 | 32671 | 36269 | 2826 | 6384 | 72 | 10040 | 21764 | 0 | 0 | 0 | 130 | 72 | 40 | 30 | 350 | 13 | 0 | 77 | 58 | 46 | 119 | 40 | 0 | 356 | 649 | 268 | 0 |
| SRR4044206 | 47 | 1006 | 62758 | 4378 | 3452 | 5637 | 18336 | 9761 | 155876 | 1150 | 2325 | 5053 | 8032 | 5643 | 613 | 5684 | 7810 | 0 | 108 | 160 | 102 | 510 | 75 | 0 | 0 | 19 | 55 | 1022 | 1236 | 1226 | 354 | 23 | 0 | 0 | 11 | 19 | 0 |
| SRR4044210 | 245 | 1690 | 10621 | 311 | 899 | 1947 | 69904 | 15620 | 24907 | 556 | 5617 | 5676 | 3329 | 23268 | 766 | 26630 | 6266 | 0 | 0 | 0 | 0 | 0 | 0 | 0 | 0 | 0 | 0 | 0 | 0 | 0 | 0 | 0 | 0 | 0 | 0 | 0 | 0 |
| SRR4044211 | 802 | 5166 | 33440 | 379 | 1942 | 2083 | 29666 | 10620 | 34224 | 244 | 20738 | 25437 | 8756 | 12452 | 486 | 19751 | 19594 | 0 | 0 | 0 | 13 | 46 | 0 | 0 | 0 | 0 | 0 | 0 | 23 | 0 | 0 | 0 | 0 | 0 | 0 | 0 | 0 |
| SRR4270461 | 2032 | 4845 | 10936 | 2443 | 28662 | 23594 | 353643 | 55082 | 7675 | 7273 | 21702 | 11317 | 8406 | 8176 | 2574 | 27748 | 5420 | 22 | 29 | 18 | 65 | 48 | 127 | 40 | 30 | 0 | 42 | 39 | 97 | 56 | 62 | 0 | 0 | 0 | 0 | 24 | 0 |
| SRR4270462 | 1142 | 4624 | 8986 | 1777 | 30042 | 25919 | 409901 | 44283 | 8430 | 7988 | 21822 | 9451 | 83 | 7898 | 2086 | 27402 | 5722 | 0 | 14 | 10 | 43 | 23 | 105 | 26 | 22 | 0 | 19 | 22 | 53 | 26 | 31 | 0 | 0 | 0 | 0 | 17 | 0 |
| SRR4292754 | 25 | 646 | 419 | 0 | 3969 | 59239 | 43228 | 9056 | 2205 | 133 | 15065 | 13653 | 3305 | 8205 | 107 | 222263 | 10499 | 0 | 0 | 0 | 23 | 16 | 14 | 14 | 32 | 11 | 0 | 0 | 0 | 0 | 0 | 0 | 0 | 19 | 0 | 95 | 0 |
| SRR4292755 | 20 | 311 | 485 | 11 | 4283 | 48745 | 50241 | 10207 | 2277 | 111 | 16319 | 12328 | 3641 | 9945 | 154 | 228369 | 10235 | 0 | 0 | 0 | 23 | 12 | 13 | 0 | 28 | 0 | 0 | 0 | 0 | 0 | 0 | 0 | 0 | 19 | 0 | 112 | 0 |
| SRR4292756 | 0 | 225 | 762 | 0 | 3986 | 76520 | 66607 | 16370 | 3239 | 332 | 15522 | 17970 | 3559 | 8902 | 112 | 178543 | 11409 | 0 | 0 | 0 | 40 | 23 | 39 | 0 | 0 | 0 | 0 | 0 | 0 | 0 | 17 | 19 | 0 | 0 | 0 | 47 | 0 |
| SRR4292757 | 0 | 276 | 528 | 0 | 3659 | 88342 | 55167 | 12112 | 3268 | 390 | 16790 | 20170 | 3667 | 9733 | 109 | 185700 | 9947 | 0 | 0 | 0 | 47 | 33 | 43 | 0 | 0 | 0 | 0 | 0 | 0 | 0 | 35 | 22 | 0 | 0 | 0 | 75 | 0 |
| SRR4296311 | 911 | 2535 | 35441 | 666 | 44779 | 63375 | 274645 | 75877 | 3132 | 2325 | 9716 | 8722 | 5321 | 57923 | 2083 | 11295 | 4436 | 0 | 18 | 12 | 14 | 42 | 15 | 0 | 12 | 0 | 0 | 26 | 77 | 21 | 13 | 0 | 0 | 0 | 0 | 0 | 0 |
| SRR4296312 | 845 | 2433 | 34886 | 710 | 43843 | 63393 | 271077 | 76569 | 3248 | 2399 | 9856 | 11 | 5469 | 57119 | 2301 | 11567 | 4508 | 0 | 11 | 16 | 13 | 59 | 12 | 0 | 11 | 0 | 0 | 26 | 87 | 21 | 14 | 0 | 0 | 0 | 0 | 0 | 0 |
| SRR4296313 | 744 | 2062 | 39974 | 647 | 42173 | 58849 | 235667 | 85992 | 1811 | 2033 | 8242 | 7764 | 5219 | 53587 | 2363 | 11218 | 3911 | 0 | 20 | 25 | 11 | 31 | 14 | 0 | 0 | 0 | 0 | 38 | 90 | 24 | 12 | 0 | 0 | 0 | 0 | 0 | 0 |
| SRR4296314 | 733 | 2286 | 40776 | 460 | 42636 | 59507 | 242556 | 86511 | 1686 | 2152 | 8538 | 15 | 5418 | 55064 | 2328 | 10236 | 3400 | 0 | 15 | 25 | 0 | 24 | 0 | 0 | 0 | 0 | 0 | 33 | 68 | 27 | 14 | 0 | 0 | 0 | 0 | 0 | 0 |
| SRR4296315 | 713 | 2850 | 69684 | 312 | 37623 | 54315 | 275345 | 92183 | 2817 | 3012 | 9170 | 17 | 5605 | 51237 | 659 | 9612 | 4735 | 0 | 16 | 18 | 0 | 18 | 0 | 0 | 0 | 0 | 0 | 33 | 82 | 24 | 0 | 0 | 13 | 0 | 0 | 0 | 0 |
| SRR4296316 | 565 | 3081 | 68564 | 377 | 38387 | 55360 | 273726 | 90712 | 2755 | 3181 | 9572 | 8911 | 5636 | 51610 | 701 | 9757 | 4979 | 0 | 17 | 18 | 0 | 15 | 0 | 0 | 0 | 0 | 0 | 25 | 81 | 34 | 0 | 0 | 12 | 0 | 0 | 0 | 0 |
| SRR4296317 | 701 | 3262 | 68400 | 386 | 40125 | 56122 | 273487 | 85909 | 2736 | 2368 | 9278 | 8022 | 5819 | 55171 | 635 | 10693 | 4529 | 0 | 18 | 12 | 11 | 25 | 0 | 0 | 0 | 0 | 0 | 24 | 76 | 27 | 12 | 0 | 0 | 0 | 0 | 0 | 0 |
| SRR4296318 | 626 | 3358 | 70503 | 459 | 41261 | 57264 | 274623 | 86359 | 2612 | 2352 | 9498 | 12 | 6010 | 56478 | 608 | 10040 | 4087 | 0 | 13 | 0 | 0 | 23 | 0 | 0 | 0 | 0 | 0 | 25 | 68 | 24 | 0 | 0 | 0 | 0 | 0 | 0 | 0 |
| SRR4296319 | 335 | 3002 | 87385 | 1090 | 42061 | 58549 | 312520 | 106072 | 2527 | 994 | 8586 | 6780 | 5090 | 44482 | 708 | 8223 | 3796 | 0 | 0 | 12 | 0 | 32 | 0 | 0 | 0 | 0 | 0 | 18 | 76 | 18 | 0 | 0 | 0 | 0 | 0 | 0 | 0 |
| SRR4296320 | 359 | 2709 | 87688 | 1088 | 41375 | 57945 | 310001 | 107599 | 2537 | 1102 | 8963 | 6951 | 5115 | 44331 | 697 | 8244 | 3941 | 0 | 0 | 12 | 0 | 30 | 0 | 0 | 0 | 0 | 0 | 25 | 70 | 19 | 13 | 0 | 0 | 0 | 0 | 0 | 0 |
| SRR4296321 | 493 | 3053 | 72809 | 1116 | 39954 | 53036 | 312641 | 115296 | 2741 | 1064 | 9428 | 14 | 5199 | 40701 | 646 | 8872 | 3893 | 0 | 12 | 18 | 0 | 27 | 0 | 0 | 0 | 0 | 0 | 27 | 92 | 21 | 0 | 0 | 0 | 0 | 0 | 0 | 0 |
| SRR4296322 | 394 | 3177 | 72118 | 1113 | 38961 | 52356 | 311596 | 117700 | 2738 | 967 | 9445 | 7756 | 5146 | 40685 | 646 | 8932 | 3986 | 0 | 12 | 15 | 10 | 29 | 0 | 0 | 0 | 0 | 0 | 25 | 100 | 20 | 0 | 0 | 0 | 0 | 0 | 0 | 0 |
| SRR4296979 | 260 | 17622 | 2965 | 1129 | 1309 | 24774 | 20649 | 8547 | 6986 | 0 | 94100 | 29711 | 6355 | 15515 | 18 | 11 | 14057 | 0 | 0 | 0 | 687 | 86 | 111 | 0 | 11 | 0 | 0 | 12 | 53 | 84 | 27 | 35 | 0 | 0 | 0 | 0 | 0 |
| SRR4296985 | 33 | 7753 | 5962 | 415 | 1433 | 9934 | 13745 | 17887 | 29891 | 0 | 14976 | 8831 | 7930 | 6130 | 15 | 953 | 14698 | 0 | 0 | 0 | 298 | 118 | 17 | 0 | 0 | 0 | 0 | 11 | 55 | 45 | 43 | 11 | 0 | 0 | 0 | 0 | 0 |
| SRR4296986 | 49 | 8062 | 13724 | 216 | 1097 | 11577 | 13093 | 15761 | 43371 | 0 | 17666 | 9647 | 6045 | 6412 | 26 | 980 | 14816 | 0 | 0 | 16 | 539 | 206 | 33 | 0 | 0 | 0 | 0 | 15 | 96 | 72 | 67 | 16 | 0 | 0 | 0 | 0 | 0 |
| SRR4296987 | 2051 | 22091 | 90178 | 363 | 2012 | 33943 | 35426 | 16000 | 9736 | 50 | 73852 | 19187 | 4092 | 9824 | 0 | 248 | 16963 | 0 | 28 | 74 | 5534 | 1851 | 342 | 0 | 0 | 0 | 0 | 163 | 1201 | 1153 | 691 | 177 | 0 | 14 | 0 | 0 | 0 |
| SRR4296988 | 1941 | 30247 | 91946 | 295 | 1886 | 32617 | 30931 | 17479 | 10567 | 37 | 77139 | 20124 | 4405 | 9987 | 0 | 349 | 18226 | 0 | 26 | 92 | 5686 | 1822 | 321 | 0 | 13 | 0 | 0 | 150 | 1291 | 979 | 574 | 157 | 0 | 19 | 0 | 0 | 0 |
| SRR4328127 | 52 | 29 | 113469 | 0 | 3705 | 5740 | 83459 | 3853 | 30981 | 11 | 19511 | 19116 | 6747 | 25301 | 484 | 267 | 43626 | 0 | 0 | 0 | 0 | 0 | 0 | 0 | 0 | 0 | 0 | 0 | 0 | 0 | 0 | 0 | 0 | 0 | 0 | 0 | 0 |
| SRR4328130 | 31 | 23 | 40104 | 136 | 1495 | 1913 | 31968 | 4050 | 9611 | 0 | 50082 | 26263 | 10415 | 10158 | 129 | 321 | 49536 | 0 | 0 | 0 | 0 | 0 | 0 | 0 | 0 | 0 | 0 | 0 | 0 | 0 | 0 | 0 | 0 | 0 | 0 | 0 | 0 |
| SRR4328135 | 30 | 42 | 144722 | 0 | 1950 | 2151 | 35388 | 3154 | 28390 | 0 | 12159 | 13421 | 3183 | 24232 | 344 | 0 | 16977 | 0 | 0 | 0 | 0 | 0 | 0 | 0 | 0 | 0 | 0 | 0 | 0 | 0 | 0 | 0 | 0 | 0 | 0 | 0 | 0 |
| SRR4328136 | 79 | 44 | 54780 | 13 | 3612 | 4319 | 73216 | 3574 | 20124 | 17 | 11983 | 9556 | 4243 | 20545 | 401 | 287 | 29642 | 0 | 0 | 0 | 0 | 20 | 0 | 0 | 0 | 0 | 0 | 0 | 0 | 0 | 0 | 0 | 0 | 0 | 0 | 0 | 0 |
| SRR4328137 | 74 | 34 | 36518 | 46 | 3555 | 7857 | 70716 | 2494 | 33861 | 24 | 9541 | 9167 | 7421 | 24632 | 748 | 624 | 23821 | 0 | 0 | 0 | 0 | 27 | 0 | 0 | 0 | 0 | 0 | 0 | 0 | 0 | 0 | 0 | 0 | 0 | 0 | 0 | 0 |
| SRR4409201 | 13 | 3977 | 137 | 322 | 137 | 331 | 1655 | 5135 | 11105 | 0 | 2647 | 16843 | 2142 | 4273 | 0 | 3486 | 195104 | 0 | 0 | 0 | 380 | 1273 | 137 | 0 | 0 | 0 | 0 | 0 | 94 | 412 | 2363 | 211 | 0 | 0 | 0 | 0 | 0 |
| SRR4409202 | 271 | 23640 | 0 | 324 | 60 | 806 | 439 | 645 | 3147 | 0 | 2751 | 48152 | 6705 | 3518 | 40 | 15 | 6583 | 0 | 0 | 0 | 91 | 22 | 16 | 0 | 0 | 0 | 0 | 0 | 0 | 16 | 0 | 0 | 0 | 0 | 0 | 0 | 0 |
| SRR4409203 | 0 | 5183 | 174 | 253 | 76 | 176 | 1009 | 1762 | 2940 | 0 | 2105 | 16650 | 1177 | 3169 | 0 | 2942 | 108318 | 0 | 0 | 0 | 502 | 1179 | 108 | 0 | 0 | 0 | 0 | 0 | 98 | 587 | 2489 | 225 | 0 | 0 | 0 | 11 | 0 |
| SRR4409204 | 1166 | 243959 | 0 | 229 | 87 | 1135 | 819 | 602 | 1452 | 0 | 3145 | 25884 | 3586 | 3121 | 74 | 43 | 10777 | 0 | 0 | 0 | 21 | 0 | 0 | 0 | 0 | 0 | 0 | 0 | 0 | 0 | 0 | 0 | 0 | 0 | 0 | 0 | 0 |
| SRR4417245 | 0 | 47 | 6225 | 0 | 1053 | 189 | 39924 | 46364 | 2671 | 582 | 40823 | 34253 | 30933 | 14907 | 184 | 85 | 36598 | 0 | 0 | 0 | 0 | 0 | 0 | 0 | 0 | 0 | 0 | 0 | 0 | 0 | 0 | 0 | 0 | 0 | 0 | 0 | 0 |
| SRR4417246 | 0 | 0 | 5203 | 0 | 605 | 76 | 26192 | 50212 | 2662 | 110 | 44306 | 33339 | 37904 | 13164 | 134 | 26 | 26291 | 0 | 0 | 0 | 0 | 0 | 0 | 0 | 0 | 0 | 0 | 0 | 0 | 0 | 0 | 0 | 0 | 0 | 0 | 0 | 0 |
| SRR4417247 | 0 | 52 | 1385 | 0 | 1129 | 109 | 20339 | 41511 | 1380 | 160 | 47540 | 30517 | 43539 | 11337 | 477 | 95 | 15105 | 0 | 0 | 0 | 0 | 0 | 0 | 0 | 0 | 19 | 0 | 0 | 0 | 0 | 0 | 0 | 0 | 0 | 0 | 0 | 0 |
| SRR4417248 | 0 | 22 | 2906 | 0 | 1789 | 180 | 29443 | 62447 | 2449 | 182 | 45854 | 35256 | 35239 | 10511 | 547 | 35 | 12574 | 0 | 0 | 0 | 0 | 0 | 0 | 23 | 0 | 15 | 0 | 0 | 0 | 0 | 0 | 0 | 0 | 0 | 0 | 0 | 0 |
| SRR4417249 | 0 | 0 | 5745 | 0 | 1061 | 89 | 23544 | 62920 | 1490 | 109 | 63386 | 47736 | 47990 | 12448 | 329 | 26 | 17471 | 0 | 0 | 0 | 0 | 0 | 0 | 0 | 0 | 0 | 0 | 0 | 0 | 0 | 0 | 0 | 0 | 0 | 0 | 0 | 0 |
| SRR4417250 | 25 | 49 | 1817 | 0 | 2840 | 230 | 32799 | 56012 | 1563 | 333 | 52607 | 37033 | 41344 | 11495 | 388 | 34 | 15744 | 0 | 0 | 0 | 0 | 0 | 0 | 14 | 0 | 24 | 0 | 0 | 0 | 0 | 0 | 0 | 0 | 0 | 0 | 0 | 0 |
| SRR4423247 | 698 | 56584 | 2590 | 283 | 847 | 26310 | 31791 | 34377 | 20062 | 16 | 19809 | 41640 | 6839 | 5917 | 143 | 21273 | 21952 | 0 | 0 | 0 | 4305 | 1196 | 267 | 0 | 38 | 0 | 0 | 0 | 154 | 2287 | 910 | 200 | 0 | 75 | 56 | 0 | 0 |
| SRR4423248 | 1809 | 49070 | 1551 | 423 | 2394 | 70228 | 66601 | 73356 | 170 | 45 | 24766 | 48408 | 5502 | 22047 | 91 | 19411 | 25172 | 0 | 0 | 0 | 4765 | 632 | 135 | 0 | 16 | 0 | 0 | 0 | 47 | 1541 | 535 | 63 | 0 | 16 | 18 | 0 | 0 |
| SRR4423250 | 1307 | 79919 | 693 | 979 | 2943 | 49438 | 56518 | 25021 | 2083 | 35 | 27213 | 14 | 6543 | 16761 | 110 | 63166 | 11538 | 0 | 0 | 10 | 1531 | 385 | 179 | 0 | 24 | 0 | 0 | 0 | 105 | 337 | 173 | 74 | 0 | 30 | 102 | 0 | 0 |
| SRR4423251 | 874 | 26885 | 722 | 606 | 1128 | 32643 | 35374 | 16891 | 200 | 25 | 9584 | 36813 | 3219 | 2011 | 129 | 14240 | 7744 | 0 | 0 | 0 | 970 | 258 | 76 | 0 | 10 | 0 | 0 | 15 | 80 | 217 | 86 | 22 | 0 | 0 | 12 | 0 | 0 |
| SRR4473670 | 440 | 39056 | 323 | 327 | 91 | 1604 | 1335 | 1230 | 1543 | 0 | 10890 | 61346 | 4160 | 3494 | 0 | 2707 | 26842 | 0 | 0 | 13 | 920 | 214 | 83 | 0 | 10 | 0 | 0 | 0 | 91 | 138 | 89 | 23 | 0 | 0 | 14 | 0 | 0 |
| SRR4734676 | 99 | 7033 | 629 | 22 | 1792 | 13641 | 17118 | 17937 | 1666 | 34 | 12418 | 10 | 9125 | 2517 | 19 | 108 | 63823 | 0 | 0 | 0 | 0 | 0 | 0 | 0 | 0 | 0 | 0 | 0 | 0 | 0 | 12 | 12 | 0 | 0 | 0 | 0 | 0 |
| SRR4734677 | 50 | 11064 | 855 | 17 | 1702 | 15093 | 17671 | 17772 | 4258 | 50 | 10983 | 11 | 8095 | 3285 | 0 | 159 | 53963 | 0 | 0 | 0 | 15 | 10 | 14 | 0 | 0 | 0 | 0 | 0 | 0 | 23 | 48 | 37 | 0 | 0 | 0 | 0 | 0 |
| SRR4734678 | 544 | 33556 | 610 | 13 | 290 | 2260 | 2642 | 2881 | 675 | 27 | 2667 | 14306 | 2125 | 560 | 0 | 15 | 30382 | 0 | 0 | 0 | 0 | 0 | 0 | 0 | 0 | 0 | 0 | 0 | 0 | 0 | 0 | 0 | 0 | 0 | 0 | 0 | 0 |
| SRR4734679 | 82 | 16868 | 602 | 27 | 230 | 2453 | 2816 | 2672 | 1105 | 14 | 9318 | 20780 | 5424 | 853 | 0 | 40 | 49704 | 0 | 0 | 0 | 0 | 0 | 0 | 0 | 0 | 0 | 0 | 0 | 0 | 0 | 0 | 0 | 0 | 0 | 0 | 0 | 0 |
| SRR4734680 | 169 | 15604 | 792 | 67 | 266 | 2730 | 3203 | 3277 | 1234 | 67 | 6016 | 19172 | 4520 | 826 | 0 | 31 | 49813 | 0 | 0 | 0 | 0 | 0 | 0 | 0 | 0 | 0 | 0 | 0 | 0 | 0 | 0 | 0 | 0 | 0 | 0 | 0 | 0 |
| SRR4734681 | 54 | 15748 | 675 | 31 | 989 | 9382 | 10662 | 11384 | 2764 | 15 | 16391 | 26419 | 10409 | 1657 | 19 | 100 | 76560 | 0 | 0 | 0 | 0 | 0 | 0 | 0 | 0 | 0 | 0 | 0 | 0 | 0 | 0 | 0 | 0 | 0 | 0 | 0 | 0 |
| SRR4734682 | 444 | 13233 | 808 | 40 | 417 | 3449 | 4954 | 5393 | 1402 | 48 | 9947 | 10 | 6986 | 1182 | 0 | 32 | 54036 | 0 | 0 | 0 | 0 | 0 | 0 | 0 | 0 | 0 | 0 | 0 | 0 | 0 | 0 | 0 | 0 | 0 | 0 | 0 | 0 |
| SRR4734683 | 359 | 19550 | 780 | 34 | 169 | 1891 | 2128 | 2291 | 586 | 24 | 3906 | 12217 | 3174 | 759 | 0 | 22 | 39851 | 0 | 0 | 0 | 0 | 0 | 0 | 0 | 0 | 0 | 0 | 0 | 0 | 0 | 0 | 0 | 0 | 0 | 0 | 0 | 0 |
| SRR4734684 | 124 | 16359 | 1258 | 49 | 676 | 5692 | 6793 | 7337 | 1752 | 31 | 12238 | 23549 | 8573 | 1570 | 0 | 36 | 66115 | 0 | 0 | 0 | 15 | 0 | 12 | 0 | 0 | 0 | 0 | 0 | 0 | 0 | 0 | 0 | 0 | 0 | 0 | 0 | 0 |
| SRR4734685 | 36 | 12490 | 472 | 126 | 205 | 1891 | 2157 | 3166 | 1326 | 29 | 5555 | 10996 | 4015 | 587 | 0 | 30 | 48908 | 0 | 0 | 0 | 0 | 0 | 0 | 0 | 0 | 0 | 0 | 0 | 0 | 0 | 0 | 0 | 0 | 0 | 0 | 0 | 0 |
| SRR4734686 | 0 | 13141 | 624 | 30 | 814 | 7522 | 8403 | 8542 | 2901 | 25 | 9827 | 18324 | 6240 | 2190 | 0 | 63 | 46893 | 0 | 0 | 0 | 0 | 0 | 0 | 0 | 0 | 0 | 0 | 0 | 0 | 0 | 0 | 0 | 0 | 0 | 0 | 0 | 0 |
| SRR4787084 | 829 | 691 | 9602 | 138 | 61 | 15 | 1049 | 1674 | 529 | 2003 | 30437 | 33 | 15342 | 10814 | 2622 | 5042 | 130928 | 0 | 212 | 131 | 199 | 545 | 53 | 0 | 11 | 13 | 0 | 424 | 372 | 155 | 510 | 33 | 0 | 0 | 13 | 442 | 0 |
| SRR4787085 | 1999 | 6497 | 2177 | 321 | 19 | 109 | 275 | 728 | 524 | 0 | 19727 | 16 | 7564 | 4178 | 109 | 9876 | 207051 | 0 | 0 | 10 | 578 | 270 | 24 | 0 | 0 | 0 | 0 | 126 | 606 | 266 | 503 | 24 | 0 | 29 | 43 | 43 | 0 |
| SRR4787086 | 902 | 580 | 11022 | 129 | 94 | 64 | 1249 | 1849 | 406 | 2256 | 29590 | 34557 | 15462 | 9374 | 2761 | 4977 | 119439 | 0 | 130 | 128 | 110 | 528 | 35 | 0 | 0 | 0 | 0 | 233 | 240 | 84 | 374 | 40 | 0 | 0 | 20 | 365 | 0 |
| SRR4787087 | 940 | 985 | 11104 | 45 | 102 | 68 | 1350 | 1899 | 552 | 2776 | 29799 | 31367 | 14385 | 9524 | 2618 | 5369 | 113587 | 0 | 120 | 114 | 91 | 341 | 49 | 0 | 17 | 0 | 0 | 263 | 244 | 147 | 331 | 43 | 0 | 0 | 12 | 323 | 0 |
| SRR4787088 | 1324 | 1523 | 7349 | 66 | 43 | 43 | 735 | 1495 | 294 | 1854 | 15 | 30 | 12146 | 6621 | 1954 | 4257 | 87649 | 0 | 111 | 123 | 106 | 689 | 65 | 0 | 18 | 0 | 0 | 254 | 266 | 171 | 1035 | 58 | 0 | 18 | 35 | 400 | 0 |
| SRR4787089 | 1783 | 1516 | 8648 | 0 | 72 | 48 | 781 | 1208 | 290 | 1159 | 22162 | 40 | 11560 | 6239 | 1618 | 4097 | 89645 | 0 | 113 | 80 | 86 | 607 | 73 | 0 | 13 | 0 | 0 | 166 | 179 | 116 | 918 | 43 | 0 | 0 | 30 | 345 | 0 |
| SRR4787090 | 1441 | 2829 | 13259 | 1137 | 251 | 144 | 2806 | 1688 | 2120 | 5425 | 18589 | 14 | 17501 | 11089 | 4922 | 6632 | 80729 | 0 | 214 | 94 | 185 | 160 | 35 | 13 | 16 | 11 | 0 | 592 | 527 | 122 | 36 | 0 | 0 | 0 | 0 | 43 | 0 |
| SRR4787091 | 1413 | 1413 | 8162 | 0 | 52 | 42 | 878 | 1515 | 303 | 942 | 14 | 41 | 11930 | 6184 | 1954 | 4137 | 96746 | 0 | 109 | 87 | 78 | 598 | 27 | 0 | 14 | 0 | 0 | 137 | 141 | 100 | 890 | 18 | 0 | 23 | 27 | 205 | 0 |
| SRR4787092 | 1534 | 3231 | 15194 | 1034 | 230 | 118 | 2648 | 1812 | 1872 | 5438 | 17885 | 17 | 17464 | 10605 | 4607 | 6328 | 80635 | 0 | 257 | 106 | 202 | 177 | 31 | 11 | 21 | 12 | 13 | 614 | 518 | 126 | 42 | 0 | 0 | 0 | 0 | 48 | 0 |
| SRR4787093 | 1742 | 3581 | 14744 | 1103 | 225 | 105 | 2793 | 1852 | 1989 | 5278 | 17766 | 14 | 17537 | 10640 | 4542 | 6602 | 75832 | 0 | 243 | 109 | 186 | 175 | 35 | 0 | 13 | 20 | 11 | 644 | 543 | 150 | 32 | 0 | 0 | 0 | 0 | 47 | 0 |
| SRR4787095 | 219 | 554 | 5756 | 3097 | 352 | 136 | 2614 | 4009 | 1089 | 3853 | 44002 | 50275 | 11609 | 17337 | 12994 | 7967 | 50859 | 0 | 279 | 216 | 1980 | 569 | 51 | 0 | 0 | 0 | 0 | 1537 | 1007 | 1970 | 863 | 118 | 0 | 0 | 0 | 13 | 0 |
| SRR4787096 | 184 | 343 | 3894 | 2777 | 398 | 178 | 2974 | 4631 | 1166 | 2905 | 44822 | 52231 | 12458 | 16631 | 15480 | 7474 | 50043 | 0 | 345 | 258 | 2277 | 654 | 53 | 0 | 0 | 0 | 13 | 2074 | 1394 | 2310 | 1039 | 146 | 0 | 0 | 0 | 12 | 0 |
| SRR4787097 | 200 | 3551 | 1799 | 2125 | 71 | 125 | 324 | 630 | 167 | 484 | 34178 | 147123 | 4956 | 11980 | 924 | 412 | 50843 | 0 | 0 | 19 | 92 | 378 | 22 | 0 | 0 | 0 | 0 | 45 | 52 | 73 | 302 | 0 | 0 | 0 | 0 | 0 | 0 |
| SRR4787098 | 237 | 3811 | 1887 | 2176 | 90 | 80 | 397 | 728 | 186 | 463 | 33790 | 23 | 4826 | 12170 | 916 | 345 | 47560 | 0 | 0 | 14 | 124 | 401 | 33 | 0 | 0 | 0 | 0 | 35 | 45 | 83 | 297 | 14 | 0 | 0 | 0 | 0 | 0 |
| SRR4787099 | 233 | 3887 | 1583 | 2477 | 60 | 79 | 313 | 704 | 202 | 331 | 34479 | 147545 | 4738 | 11930 | 895 | 456 | 47399 | 0 | 10 | 19 | 95 | 337 | 15 | 0 | 0 | 0 | 0 | 42 | 49 | 96 | 270 | 16 | 0 | 0 | 0 | 0 | 0 |
| SRR4787100 | 267 | 2022 | 6160 | 1782 | 268 | 204 | 1509 | 2633 | 361 | 501 | 37353 | 89334 | 10356 | 13432 | 2390 | 757 | 72579 | 0 | 87 | 112 | 358 | 1562 | 109 | 0 | 0 | 0 | 0 | 151 | 180 | 224 | 1095 | 45 | 0 | 0 | 0 | 14 | 0 |
| SRR4787101 | 253 | 2526 | 5912 | 1497 | 238 | 166 | 1269 | 2272 | 325 | 424 | 37612 | 95862 | 10019 | 13655 | 2168 | 687 | 74493 | 0 | 61 | 88 | 297 | 1597 | 94 | 0 | 0 | 0 | 0 | 133 | 158 | 242 | 1023 | 44 | 0 | 0 | 0 | 0 | 0 |
| SRR4787102 | 261 | 1978 | 7117 | 2039 | 261 | 188 | 1529 | 2762 | 390 | 975 | 37762 | 92428 | 10212 | 14096 | 2472 | 873 | 72816 | 0 | 71 | 97 | 317 | 1290 | 76 | 0 | 0 | 0 | 0 | 152 | 185 | 199 | 910 | 33 | 0 | 0 | 0 | 15 | 0 |
| SRR4787103 | 106 | 799 | 9802 | 475 | 602 | 392 | 4936 | 6771 | 638 | 1031 | 32309 | 58907 | 11651 | 10955 | 4903 | 1660 | 60320 | 0 | 163 | 190 | 366 | 1504 | 107 | 0 | 0 | 0 | 0 | 279 | 347 | 301 | 1560 | 86 | 0 | 0 | 0 | 24 | 0 |
| SRR4787104 | 145 | 1017 | 10727 | 554 | 557 | 341 | 4395 | 6457 | 646 | 1740 | 31811 | 15 | 12150 | 11716 | 4959 | 1720 | 63298 | 0 | 183 | 214 | 422 | 1473 | 99 | 0 | 0 | 0 | 0 | 321 | 412 | 343 | 1587 | 83 | 0 | 0 | 0 | 30 | 0 |
| SRR4787105 | 493 | 6909 | 8516 | 820 | 97 | 1166 | 1255 | 2450 | 540 | 29 | 39589 | 12 | 3188 | 9754 | 20 | 5549 | 20692 | 0 | 0 | 0 | 764 | 472 | 41 | 0 | 0 | 0 | 0 | 72 | 544 | 621 | 614 | 76 | 0 | 0 | 0 | 0 | 0 |
| SRR4787106 | 439 | 7020 | 8673 | 827 | 97 | 1229 | 1349 | 2085 | 424 | 51 | 38451 | 49048 | 3243 | 9099 | 29 | 5614 | 23812 | 0 | 0 | 0 | 751 | 480 | 47 | 0 | 0 | 0 | 0 | 79 | 555 | 616 | 646 | 87 | 0 | 0 | 0 | 0 | 0 |
| SRR4787107 | 555 | 7536 | 8315 | 697 | 99 | 1238 | 1444 | 1972 | 370 | 61 | 31992 | 14 | 5234 | 5743 | 73 | 6244 | 43384 | 0 | 13 | 15 | 1490 | 1106 | 107 | 0 | 0 | 0 | 0 | 150 | 1059 | 963 | 1287 | 160 | 0 | 0 | 11 | 0 | 0 |
| SRR4787108 | 426 | 5912 | 6781 | 1003 | 151 | 1338 | 2445 | 1770 | 352 | 53 | 42004 | 11 | 4880 | 7622 | 47 | 10172 | 22452 | 0 | 0 | 0 | 1246 | 509 | 57 | 0 | 0 | 0 | 0 | 126 | 671 | 1273 | 566 | 101 | 0 | 0 | 0 | 0 | 0 |
| SRR4787109 | 460 | 6937 | 9012 | 736 | 92 | 1059 | 1194 | 2102 | 349 | 57 | 38250 | 50408 | 3266 | 8767 | 24 | 5396 | 27001 | 0 | 0 | 0 | 882 | 566 | 55 | 0 | 0 | 0 | 0 | 84 | 625 | 637 | 742 | 77 | 0 | 0 | 0 | 0 | 0 |
| SRR4787110 | 412 | 7025 | 8547 | 698 | 92 | 1159 | 1367 | 2210 | 396 | 42 | 30241 | 51080 | 5431 | 5649 | 51 | 5681 | 38367 | 0 | 10 | 12 | 1290 | 914 | 95 | 0 | 0 | 0 | 0 | 119 | 928 | 865 | 1183 | 149 | 0 | 0 | 0 | 0 | 0 |
| SRR4787118 | 0 | 0 | 1635 | 12581 | 0 | 0 | 1812 | 0 | 0 | 1936 | 10585 | 21384 | 17 | 25899 | 3292 | 3144 | 36944 | 0 | 0 | 0 | 0 | 0 | 0 | 0 | 0 | 0 | 0 | 648 | 0 | 584 | 0 | 0 | 0 | 0 | 0 | 0 | 0 |
| SRR4787119 | 0 | 6616 | 0 | 10794 | 0 | 0 | 0 | 0 | 18 | 0 | 12003 | 18759 | 26 | 24624 | 7412 | 8114 | 31977 | 0 | 0 | 0 | 0 | 0 | 0 | 0 | 0 | 0 | 0 | 177 | 0 | 2136 | 0 | 0 | 0 | 0 | 0 | 0 | 0 |
| SRR4787120 | 221 | 437 | 9650 | 1215 | 418 | 160 | 3283 | 4891 | 1061 | 5743 | 43688 | 49793 | 13419 | 13310 | 13049 | 7588 | 53992 | 0 | 358 | 266 | 2026 | 697 | 73 | 0 | 0 | 0 | 14 | 1282 | 880 | 1737 | 924 | 213 | 0 | 0 | 0 | 19 | 0 |
| SRR4787121 | 245 | 1373 | 6090 | 10342 | 70 | 60 | 399 | 847 | 338 | 2177 | 49508 | 23 | 5337 | 15809 | 2853 | 2566 | 34102 | 0 | 19 | 53 | 847 | 187 | 17 | 0 | 0 | 0 | 0 | 250 | 128 | 1356 | 219 | 104 | 0 | 0 | 0 | 0 | 0 |
| SRR4787122 | 139 | 853 | 4201 | 8647 | 85 | 31 | 413 | 864 | 232 | 2400 | 54634 | 30 | 5660 | 14158 | 3007 | 2005 | 27591 | 0 | 30 | 47 | 872 | 159 | 22 | 0 | 0 | 0 | 0 | 282 | 169 | 1521 | 211 | 118 | 0 | 0 | 0 | 0 | 0 |
| SRR4787123 | 280 | 978 | 3490 | 7669 | 52 | 26 | 423 | 710 | 231 | 2027 | 50257 | 25 | 5193 | 14642 | 2698 | 2135 | 28190 | 0 | 17 | 32 | 814 | 158 | 15 | 0 | 0 | 0 | 0 | 283 | 164 | 1766 | 193 | 121 | 0 | 0 | 0 | 0 | 0 |
| SRR4787124 | 146 | 799 | 1651 | 12008 | 171 | 166 | 0 | 171 | 333 | 739 | 16885 | 53461 | 1412 | 25278 | 1972 | 4054 | 25881 | 0 | 0 | 20 | 288 | 101 | 0 | 0 | 0 | 0 | 0 | 123 | 107 | 1788 | 157 | 56 | 0 | 0 | 0 | 0 | 0 |
| SRR4787125 | 349 | 828 | 1841 | 10992 | 119 | 0 | 303 | 499 | 188 | 1050 | 18438 | 61435 | 1193 | 24831 | 986 | 6187 | 21930 | 0 | 0 | 20 | 331 | 32 | 12 | 0 | 0 | 0 | 0 | 300 | 125 | 2236 | 66 | 34 | 0 | 0 | 0 | 0 | 0 |
| SRR4787126 | 367 | 378 | 878 | 11577 | 0 | 0 | 130 | 424 | 268 | 663 | 18302 | 66549 | 600 | 19360 | 972 | 5099 | 19385 | 0 | 0 | 24 | 349 | 105 | 0 | 0 | 0 | 0 | 0 | 234 | 43 | 1623 | 24 | 24 | 0 | 0 | 0 | 0 | 0 |
| SRR4787127 | 0 | 0 | 3878 | 11234 | 0 | 0 | 0 | 0 | 13 | 0 | 10576 | 29773 | 25 | 34591 | 21 | 11518 | 20067 | 0 | 0 | 437 | 0 | 0 | 0 | 0 | 0 | 0 | 0 | 423 | 337 | 1288 | 0 | 414 | 0 | 0 | 0 | 0 | 0 |
| SRR4787132 | 2100 | 7195 | 2194 | 142 | 17 | 128 | 273 | 708 | 559 | 24 | 19207 | 35874 | 7287 | 4114 | 70 | 9678 | 199840 | 0 | 0 | 0 | 596 | 269 | 35 | 0 | 0 | 0 | 0 | 127 | 662 | 285 | 527 | 28 | 0 | 27 | 39 | 47 | 0 |
| SRR4787133 | 1698 | 7389 | 2463 | 233 | 17 | 166 | 316 | 620 | 582 | 0 | 18245 | 26 | 7355 | 4436 | 89 | 9803 | 215499 | 0 | 0 | 0 | 548 | 318 | 34 | 0 | 0 | 0 | 0 | 137 | 662 | 281 | 526 | 38 | 0 | 23 | 45 | 50 | 0 |
| SRR4787134 | 187 | 7414 | 1712 | 1452 | 23 | 97 | 232 | 390 | 254 | 377 | 26526 | 26 | 2177 | 13528 | 629 | 613 | 35675 | 0 | 0 | 12 | 100 | 246 | 0 | 0 | 0 | 0 | 0 | 37 | 34 | 101 | 214 | 0 | 0 | 0 | 0 | 0 | 0 |
| SRR4787135 | 148 | 698 | 10056 | 800 | 588 | 342 | 4594 | 7572 | 575 | 947 | 31615 | 21 | 11858 | 10710 | 4850 | 1562 | 59676 | 0 | 204 | 228 | 402 | 1937 | 142 | 0 | 0 | 0 | 0 | 349 | 418 | 335 | 1722 | 100 | 0 | 0 | 0 | 32 | 0 |
| SRR4787136 | 191 | 2414 | 7923 | 862 | 142 | 210 | 1076 | 1498 | 310 | 1806 | 38135 | 139549 | 5150 | 13250 | 2446 | 1409 | 56198 | 0 | 102 | 319 | 860 | 1214 | 117 | 0 | 0 | 0 | 0 | 597 | 582 | 1276 | 1713 | 382 | 13 | 14 | 0 | 43 | 0 |
| SRR4787137 | 275 | 5286 | 3694 | 1996 | 56 | 133 | 349 | 654 | 267 | 1234 | 26525 | 22 | 2808 | 16162 | 812 | 944 | 34101 | 0 | 0 | 0 | 91 | 187 | 0 | 0 | 0 | 0 | 0 | 49 | 43 | 105 | 181 | 0 | 0 | 0 | 0 | 0 | 0 |
| SRR4787138 | 248 | 4967 | 8315 | 982 | 103 | 189 | 449 | 978 | 360 | 1863 | 36172 | 44 | 6633 | 14952 | 1654 | 1027 | 66624 | 0 | 21 | 86 | 223 | 951 | 42 | 0 | 0 | 0 | 0 | 85 | 122 | 172 | 732 | 30 | 0 | 0 | 0 | 14 | 0 |
| SRR4787139 | 397 | 4226 | 10356 | 1051 | 159 | 249 | 762 | 1517 | 416 | 3527 | 36824 | 121781 | 7668 | 15613 | 2001 | 1122 | 71177 | 0 | 36 | 108 | 285 | 1023 | 51 | 0 | 0 | 0 | 0 | 122 | 179 | 246 | 851 | 41 | 0 | 0 | 0 | 16 | 0 |
| SRR4787140 | 295 | 2656 | 14185 | 575 | 358 | 358 | 2144 | 3168 | 583 | 5864 | 39070 | 22 | 11866 | 15918 | 4582 | 1690 | 69526 | 0 | 121 | 227 | 444 | 1654 | 95 | 0 | 0 | 0 | 0 | 299 | 396 | 276 | 1535 | 62 | 0 | 0 | 0 | 47 | 0 |
| SRR4787141 | 391 | 3891 | 11214 | 1448 | 187 | 245 | 849 | 1579 | 396 | 4564 | 37175 | 119088 | 7421 | 16262 | 2210 | 1039 | 66989 | 0 | 34 | 87 | 265 | 1034 | 51 | 0 | 0 | 0 | 0 | 119 | 173 | 241 | 793 | 36 | 0 | 0 | 0 | 17 | 0 |
| SRR4787142 | 353 | 2554 | 14272 | 711 | 394 | 372 | 2702 | 3824 | 623 | 5406 | 39261 | 26 | 12818 | 15244 | 4992 | 1658 | 65482 | 0 | 145 | 232 | 507 | 1631 | 87 | 0 | 0 | 0 | 0 | 323 | 428 | 305 | 1456 | 66 | 0 | 0 | 0 | 41 | 0 |
| SRR4787143 | 272 | 2900 | 14011 | 543 | 335 | 328 | 1869 | 2996 | 484 | 5018 | 42926 | 100446 | 12343 | 17092 | 4527 | 1502 | 69680 | 0 | 101 | 165 | 357 | 1508 | 71 | 0 | 0 | 0 | 0 | 203 | 262 | 183 | 1076 | 39 | 0 | 0 | 0 | 27 | 0 |
| SRR4836291 | 126 | 33867 | 651 | 51 | 1769 | 27810 | 20557 | 21821 | 51675 | 0 | 29036 | 34235 | 11564 | 10702 | 19 | 6731 | 28903 | 0 | 0 | 25 | 3220 | 103 | 234 | 0 | 0 | 0 | 0 | 21 | 567 | 1717 | 52 | 119 | 0 | 0 | 13 | 0 | 0 |
| SRR4836293 | 142 | 34456 | 678 | 57 | 1583 | 27695 | 20195 | 22087 | 52863 | 0 | 29436 | 35018 | 11398 | 10712 | 35 | 6252 | 28288 | 0 | 0 | 21 | 3345 | 116 | 224 | 0 | 0 | 0 | 0 | 16 | 530 | 1791 | 52 | 124 | 0 | 0 | 0 | 0 | 0 |
| SRR4836294 | 136 | 34423 | 655 | 58 | 1796 | 28743 | 20734 | 22094 | 54168 | 0 | 28618 | 34244 | 12184 | 10579 | 11 | 6554 | 27620 | 0 | 0 | 23 | 3098 | 82 | 217 | 0 | 0 | 0 | 0 | 17 | 526 | 1603 | 55 | 132 | 0 | 0 | 0 | 0 | 0 |
| SRR4836295 | 2086 | 35758 | 417 | 556 | 2777 | 9230 | 30516 | 21380 | 36924 | 36 | 4093 | 9756 | 1542 | 8318 | 19 | 4855 | 4962 | 0 | 0 | 29 | 2800 | 225 | 122 | 11 | 0 | 0 | 0 | 48 | 356 | 1045 | 170 | 58 | 0 | 0 | 0 | 0 | 0 |
| SRR4836297 | 2236 | 36115 | 479 | 523 | 2417 | 9390 | 30146 | 21259 | 37140 | 19 | 4115 | 9835 | 1486 | 8196 | 13 | 4751 | 5112 | 0 | 0 | 21 | 2726 | 238 | 108 | 12 | 0 | 0 | 0 | 29 | 357 | 1079 | 181 | 56 | 0 | 0 | 0 | 0 | 0 |

### Supp Table S8.1.Columns_B_to_AL

| | | | | | | | | | | | | | | | | | | | | | | | | | | | | | | | | | | | | | |
|---|---|---|---|---|---|---|---|---|---|---|---|---|---|---|---|---|---|---|---|---|---|---|---|---|---|---|---|---|---|---|---|---|---|---|---|---|---|
| SRR4836299 | 2176 | 36045 | 434 | 583 | 2908 | 9215 | 31377 | 21753 | 37081 | 22 | 4106 | 9537 | 1605 | 8370 | 15 | 4698 | 5099 | 0 | 0 | 30 | 2545 | 215 | 107 | 11 | 0 | 0 | 0 | 38 | 326 | 1045 | 157 | 55 | 0 | 0 | 0 | 0 | 0 |
| SRR4836301 | 1382 | 31845 | 605 | 169 | 2205 | 23140 | 31281 | 27047 | 71271 | 11 | 16657 | 18384 | 7290 | 5511 | 24 | 11787 | 20940 | 0 | 0 | 86 | 11601 | 743 | 559 | 0 | 0 | 0 | 0 | 12 | 1686 | 3688 | 446 | 217 | 0 | 0 | 15 | 0 | 0 |
| SRR4836302 | 1560 | 33329 | 615 | 200 | 2130 | 24075 | 31647 | 27651 | 71716 | 11 | 16885 | 18248 | 7654 | 5531 | 0 | 11498 | 20914 | 0 | 0 | 101 | 11599 | 711 | 559 | 13 | 0 | 0 | 0 | 13 | 1704 | 3687 | 475 | 206 | 0 | 0 | 20 | 0 | 0 |
| SRR4836303 | 1405 | 31912 | 640 | 173 | 2252 | 24265 | 31790 | 27462 | 73025 | 0 | 17096 | 17953 | 7516 | 5523 | 31 | 10995 | 20836 | 0 | 0 | 86 | 10985 | 684 | 508 | 0 | 0 | 0 | 0 | 10 | 1524 | 3472 | 435 | 199 | 0 | 0 | 16 | 13 | 0 |
| SRR4836305 | 2366 | 72267 | 1793 | 538 | 1133 | 18061 | 16738 | 24883 | 40804 | 0 | 32441 | 25676 | 13788 | 12500 | 0 | 7988 | 33721 | 0 | 0 | 159 | 21428 | 5274 | 2504 | 139 | 18 | 16 | 0 | 0 | 2868 | 7969 | 2041 | 723 | 0 | 0 | 37 | 11 | 0 |
| SRR4836307 | 2603 | 72511 | 1645 | 615 | 1137 | 18546 | 17085 | 24686 | 40766 | 18 | 32562 | 25983 | 14009 | 12355 | 0 | 7342 | 33485 | 0 | 0 | 142 | 21471 | 5245 | 2453 | 159 | 12 | 0 | 0 | 0 | 2847 | 8032 | 2002 | 726 | 0 | 0 | 21 | 11 | 0 |
| SRR4836308 | 2591 | 71814 | 1724 | 726 | 1294 | 18685 | 17313 | 25481 | 41336 | 15 | 32412 | 25848 | 13745 | 12478 | 14 | 7453 | 33668 | 0 | 0 | 132 | 20268 | 4933 | 2326 | 140 | 13 | 0 | 0 | 10 | 2647 | 7510 | 1983 | 688 | 0 | 0 | 28 | 11 | 0 |
| SRR4836310 | 997 | 60281 | 623 | 93 | 1941 | 12772 | 17905 | 14012 | 17033 | 0 | 8480 | 32318 | 3466 | 6494 | 51 | 8108 | 13476 | 0 | 0 | 156 | 10815 | 306 | 501 | 12 | 0 | 0 | 0 | 12 | 1855 | 2671 | 128 | 200 | 0 | 0 | 12 | 0 | 0 |
| SRR4836311 | 1014 | 60811 | 740 | 109 | 1695 | 12770 | 17497 | 14517 | 16710 | 0 | 8393 | 33106 | 3457 | 6214 | 51 | 7499 | 13230 | 0 | 0 | 135 | 10611 | 297 | 506 | 0 | 0 | 0 | 0 | 13 | 1966 | 2565 | 130 | 201 | 0 | 0 | 0 | 0 | 0 |
| SRR4836312 | 983 | 60117 | 673 | 80 | 1889 | 12741 | 17939 | 14011 | 17487 | 0 | 8416 | 32555 | 3428 | 6307 | 46 | 7591 | 13376 | 0 | 0 | 127 | 10158 | 263 | 467 | 15 | 0 | 0 | 0 | 15 | 1829 | 2503 | 130 | 182 | 0 | 0 | 14 | 0 | 0 |
| SRR4836313 | 196 | 36538 | 378 | 193 | 2135 | 34195 | 27672 | 28755 | 25798 | 10 | 26733 | 36121 | 9894 | 7369 | 68 | 6192 | 23180 | 0 | 0 | 0 | 446 | 166 | 149 | 19 | 0 | 0 | 0 | 18 | 93 | 127 | 45 | 39 | 0 | 0 | 0 | 0 | 0 |
| SRR4836314 | 211 | 35284 | 401 | 192 | 2219 | 34099 | 27219 | 28406 | 25417 | 0 | 26612 | 36198 | 9829 | 7397 | 64 | 6732 | 22503 | 0 | 0 | 0 | 415 | 155 | 148 | 22 | 13 | 11 | 0 | 14 | 96 | 137 | 58 | 34 | 0 | 0 | 11 | 0 | 0 |
| SRR4836315 | 213 | 36295 | 415 | 179 | 2155 | 35663 | 28489 | 29789 | 26121 | 0 | 27770 | 36603 | 10250 | 7536 | 62 | 5868 | 22307 | 0 | 0 | 0 | 384 | 141 | 126 | 23 | 16 | 0 | 0 | 13 | 97 | 121 | 47 | 33 | 0 | 0 | 12 | 0 | 0 |
| SRR4836316 | 80 | 38540 | 848 | 84 | 2666 | 21134 | 21021 | 20908 | 59819 | 0 | 24630 | 38470 | 7289 | 10316 | 55 | 15732 | 14913 | 0 | 0 | 51 | 10641 | 565 | 501 | 12 | 0 | 0 | 0 | 30 | 1084 | 3596 | 243 | 201 | 0 | 0 | 0 | 14 | 0 |
| SRR4836317 | 76 | 37704 | 840 | 101 | 2568 | 20721 | 20872 | 20636 | 59732 | 0 | 24207 | 38230 | 7219 | 10315 | 68 | 16765 | 14837 | 0 | 0 | 60 | 10717 | 535 | 471 | 15 | 0 | 0 | 0 | 23 | 1109 | 3616 | 222 | 201 | 0 | 0 | 0 | 10 | 0 |
| SRR4836318 | 93 | 38668 | 844 | 78 | 2492 | 21550 | 21554 | 21059 | 62493 | 13 | 24945 | 15 | 7492 | 10452 | 101 | 14825 | 15215 | 0 | 0 | 54 | 9525 | 470 | 417 | 20 | 0 | 0 | 0 | 21 | 963 | 3297 | 205 | 201 | 0 | 0 | 0 | 11 | 0 |
| SRR4836320 | 95 | 69985 | 534 | 128 | 1393 | 9629 | 15735 | 24281 | 36436 | 29 | 15168 | 16586 | 7986 | 28935 | 82 | 5466 | 10252 | 0 | 0 | 158 | 55255 | 6069 | 7032 | 220 | 16 | 13 | 0 | 0 | 1468 | 12141 | 2013 | 1772 | 0 | 0 | 21 | 23 | 0 |
| SRR4836321 | 92 | 67301 | 584 | 100 | 1401 | 9682 | 15640 | 24315 | 35996 | 12 | 14998 | 16648 | 7767 | 28787 | 74 | 5890 | 10074 | 0 | 0 | 163 | 54869 | 6164 | 7078 | 199 | 15 | 0 | 0 | 0 | 1429 | 11924 | 1991 | 1713 | 0 | 0 | 18 | 21 | 0 |
| SRR4836323 | 78 | 70457 | 437 | 125 | 1321 | 9932 | 16150 | 24687 | 36785 | 20 | 15227 | 16755 | 8110 | 29105 | 91 | 5328 | 10275 | 0 | 0 | 145 | 48259 | 5334 | 6217 | 210 | 19 | 0 | 0 | 0 | 1297 | 10653 | 1771 | 1571 | 0 | 0 | 11 | 16 | 0 |
| SRR4836325 | 259 | 26453 | 391 | 159 | 3656 | 47996 | 43258 | 26643 | 1071 | 11 | 5062 | 40159 | 2372 | 1778 | 87 | 10387 | 12678 | 0 | 0 | 0 | 425 | 61 | 50 | 0 | 0 | 0 | 0 | 27 | 160 | 124 | 28 | 20 | 0 | 0 | 0 | 0 | 0 |
| SRR4836326 | 272 | 26600 | 440 | 186 | 3701 | 47650 | 42953 | 26333 | 1073 | 0 | 5137 | 39742 | 2375 | 1753 | 87 | 11267 | 12389 | 0 | 0 | 0 | 425 | 59 | 54 | 0 | 0 | 0 | 0 | 26 | 157 | 124 | 31 | 24 | 0 | 0 | 10 | 0 | 0 |
| SRR4836327 | 311 | 28023 | 397 | 143 | 3522 | 48778 | 44433 | 26711 | 1091 | 0 | 5136 | 16 | 2364 | 1831 | 95 | 9500 | 12219 | 0 | 0 | 0 | 403 | 49 | 46 | 0 | 0 | 0 | 0 | 24 | 149 | 119 | 31 | 14 | 0 | 0 | 0 | 0 | 0 |
| SRR4836328 | 183 | 55087 | 1017 | 119 | 2429 | 21866 | 20646 | 22932 | 41230 | 10 | 19890 | 28453 | 4589 | 49003 | 20 | 3063 | 154185 | 0 | 0 | 11 | 1983 | 56 | 106 | 0 | 0 | 0 | 0 | 28 | 295 | 781 | 76 | 40 | 0 | 0 | 14 | 0 | 0 |
| SRR4836330 | 176 | 54342 | 982 | 149 | 2429 | 21330 | 20773 | 22571 | 41593 | 0 | 19733 | 27599 | 4616 | 49017 | 29 | 3349 | 151183 | 0 | 0 | 11 | 1951 | 63 | 112 | 0 | 0 | 0 | 0 | 29 | 288 | 749 | 64 | 53 | 0 | 0 | 14 | 0 | 0 |
| SRR4836332 | 194 | 56295 | 1024 | 142 | 2379 | 21912 | 21404 | 22940 | 42654 | 0 | 20239 | 28309 | 4607 | 49965 | 30 | 2971 | 150546 | 0 | 0 | 10 | 1744 | 52 | 92 | 0 | 0 | 0 | 0 | 26 | 266 | 654 | 57 | 42 | 0 | 0 | 12 | 0 | 0 |
| SRR4836334 | 164 | 28647 | 736 | 202 | 2569 | 44904 | 27487 | 33644 | 27423 | 0 | 19850 | 37708 | 6165 | 12101 | 76 | 11401 | 32577 | 0 | 0 | 0 | 345 | 109 | 49 | 0 | 0 | 0 | 0 | 22 | 197 | 201 | 103 | 15 | 0 | 0 | 27 | 0 | 0 |
| SRR4836335 | 143 | 28767 | 856 | 176 | 2559 | 43963 | 27413 | 32964 | 26932 | 12 | 19772 | 37767 | 5892 | 11825 | 73 | 12660 | 32640 | 0 | 0 | 0 | 357 | 117 | 41 | 0 | 0 | 0 | 0 | 23 | 187 | 183 | 95 | 21 | 0 | 0 | 27 | 0 | 0 |
| SRR4836336 | 147 | 29575 | 807 | 164 | 2446 | 45662 | 27684 | 34327 | 27592 | 10 | 20371 | 38018 | 5869 | 12018 | 90 | 10871 | 32308 | 0 | 0 | 0 | 295 | 83 | 32 | 0 | 0 | 0 | 0 | 23 | 178 | 165 | 80 | 17 | 0 | 0 | 31 | 0 | 0 |
| SRR4836337 | 3097 | 48993 | 331 | 348 | 4674 | 43369 | 48350 | 33624 | 8366 | 17 | 35193 | 49775 | 9365 | 7916 | 28 | 18195 | 14599 | 0 | 11 | 66 | 8278 | 471 | 455 | 15 | 0 | 0 | 0 | 72 | 975 | 2114 | 146 | 178 | 0 | 0 | 16 | 18 | 0 |
| SRR4836339 | 2912 | 50020 | 285 | 284 | 4590 | 46051 | 50568 | 33979 | 8506 | 14 | 36339 | 50729 | 9625 | 8040 | 33 | 14907 | 14625 | 0 | 0 | 55 | 7182 | 394 | 366 | 16 | 0 | 0 | 0 | 55 | 822 | 1849 | 123 | 151 | 0 | 0 | 23 | 11 | 0 |
| SRR4836341 | 3000 | 48370 | 294 | 314 | 4838 | 43242 | 48095 | 32860 | 8260 | 12 | 34941 | 48767 | 9173 | 7556 | 30 | 19498 | 14275 | 0 | 0 | 57 | 8306 | 462 | 470 | 16 | 0 | 0 | 0 | 73 | 932 | 2137 | 143 | 173 | 0 | 0 | 27 | 15 | 0 |
| SRR4836343 | 3137 | 48508 | 344 | 323 | 4725 | 45888 | 50746 | 34298 | 8497 | 0 | 35463 | 15 | 9487 | 7773 | 38 | 16971 | 14325 | 0 | 10 | 56 | 7231 | 382 | 407 | 13 | 0 | 0 | 0 | 59 | 817 | 1856 | 134 | 151 | 0 | 0 | 25 | 13 | 0 |
| SRR4836344 | 545 | 20380 | 642 | 316 | 2220 | 23121 | 22301 | 16592 | 23031 | 0 | 18289 | 44731 | 10650 | 5296 | 339 | 11502 | 35597 | 0 | 0 | 22 | 3476 | 272 | 294 | 0 | 0 | 0 | 0 | 38 | 335 | 924 | 59 | 83 | 0 | 0 | 0 | 0 | 0 |
| SRR4836345 | 600 | 20314 | 783 | 285 | 2119 | 23548 | 23315 | 16972 | 23774 | 15 | 18762 | 46488 | 10958 | 5613 | 354 | 9353 | 36176 | 0 | 0 | 16 | 2950 | 238 | 237 | 0 | 0 | 0 | 0 | 26 | 273 | 789 | 48 | 68 | 0 | 0 | 0 | 0 | 0 |
| SRR4836346 | 577 | 20055 | 604 | 332 | 2208 | 22543 | 22185 | 15947 | 22890 | 27 | 18284 | 44464 | 10435 | 5425 | 311 | 12499 | 34937 | 0 | 0 | 22 | 3556 | 281 | 271 | 12 | 0 | 0 | 0 | 45 | 323 | 923 | 71 | 71 | 0 | 0 | 0 | 0 | 0 |
| SRR4836347 | 665 | 20509 | 650 | 360 | 2189 | 23323 | 23218 | 16437 | 23146 | 15 | 18387 | 44826 | 10680 | 5528 | 382 | 10685 | 35227 | 0 | 0 | 18 | 2990 | 232 | 229 | 12 | 0 | 0 | 0 | 21 | 277 | 774 | 58 | 74 | 0 | 0 | 0 | 0 | 0 |
| SRR4836348 | 538 | 23290 | 791 | 469 | 4547 | 51391 | 51200 | 42789 | 7267 | 0 | 19189 | 43773 | 8154 | 18350 | 46 | 7306 | 20368 | 0 | 0 | 0 | 373 | 93 | 102 | 0 | 10 | 0 | 0 | 0 | 94 | 97 | 29 | 29 | 0 | 0 | 11 | 0 | 0 |
| SRR4836349 | 712 | 22731 | 802 | 450 | 4325 | 52621 | 52219 | 43862 | 7498 | 0 | 20280 | 45159 | 8382 | 18867 | 52 | 6039 | 20431 | 0 | 0 | 0 | 255 | 90 | 92 | 13 | 0 | 0 | 0 | 11 | 73 | 91 | 24 | 28 | 0 | 0 | 0 | 0 | 0 |
| SRR4836350 | 543 | 23252 | 829 | 521 | 4601 | 50950 | 50160 | 42802 | 7073 | 0 | 19452 | 44294 | 8159 | 18278 | 44 | 7835 | 19892 | 0 | 0 | 0 | 360 | 99 | 96 | 14 | 10 | 0 | 0 | 0 | 84 | 100 | 34 | 41 | 0 | 0 | 0 | 0 | 0 |
| SRR4836351 | 566 | 24595 | 725 | 543 | 4502 | 52775 | 52336 | 44153 | 7009 | 0 | 19769 | 44158 | 8193 | 18383 | 52 | 6942 | 19982 | 0 | 0 | 0 | 316 | 83 | 84 | 0 | 11 | 0 | 0 | 0 | 68 | 99 | 28 | 28 | 0 | 0 | 0 | 0 | 0 |
| SRR4836352 | 740 | 35383 | 371 | 441 | 909 | 15167 | 10357 | 14943 | 46424 | 0 | 48069 | 41150 | 9472 | 9236 | 58 | 5283 | 22071 | 0 | 0 | 32 | 5273 | 223 | 244 | 10 | 0 | 0 | 0 | 14 | 266 | 883 | 64 | 61 | 0 | 0 | 14 | 0 | 0 |
| SRR4836353 | 757 | 36075 | 396 | 427 | 899 | 15281 | 10630 | 15154 | 47412 | 0 | 50011 | 42519 | 9861 | 9645 | 55 | 4031 | 22755 | 0 | 0 | 21 | 4587 | 183 | 204 | 0 | 0 | 0 | 0 | 0 | 236 | 786 | 50 | 55 | 0 | 0 | 0 | 10 | 0 |
| SRR4836354 | 717 | 35475 | 388 | 467 | 871 | 14624 | 10143 | 14662 | 46121 | 0 | 47681 | 40514 | 9831 | 9340 | 53 | 5542 | 22187 | 0 | 0 | 28 | 5262 | 226 | 274 | 0 | 0 | 0 | 0 | 14 | 282 | 937 | 61 | 58 | 0 | 0 | 12 | 16 | 0 |
| SRR4836355 | 776 | 35972 | 383 | 480 | 805 | 15064 | 10281 | 14927 | 47603 | 0 | 49517 | 14 | 9947 | 9752 | 78 | 4680 | 22132 | 0 | 0 | 31 | 4544 | 189 | 213 | 0 | 0 | 0 | 0 | 0 | 251 | 783 | 48 | 45 | 0 | 0 | 11 | 12 | 0 |
| SRR4836356 | 557 | 8810 | 544 | 318 | 2353 | 24855 | 27333 | 43185 | 25242 | 0 | 13242 | 28501 | 7352 | 7551 | 0 | 1156 | 25063 | 0 | 0 | 18 | 2525 | 403 | 132 | 0 | 0 | 0 | 0 | 0 | 385 | 681 | 207 | 56 | 0 | 0 | 15 | 0 | 0 |
| SRR4836357 | 559 | 9087 | 629 | 272 | 2265 | 25517 | 28097 | 44658 | 25344 | 0 | 13618 | 29326 | 7586 | 7580 | 0 | 920 | 25766 | 0 | 0 | 16 | 2183 | 363 | 109 | 0 | 0 | 0 | 0 | 0 | 323 | 599 | 172 | 45 | 0 | 0 | 12 | 0 | 0 |
| SRR4836359 | 552 | 8786 | 576 | 278 | 2323 | 24508 | 27287 | 42631 | 24916 | 0 | 13028 | 28123 | 7359 | 7357 | 0 | 1184 | 24557 | 0 | 0 | 20 | 2585 | 419 | 135 | 0 | 0 | 0 | 0 | 11 | 368 | 725 | 199 | 51 | 0 | 0 | 16 | 0 | 0 |
| SRR4836361 | 551 | 8997 | 581 | 308 | 2186 | 25316 | 28166 | 44079 | 25381 | 0 | 13279 | 28751 | 7471 | 7398 | 0 | 1084 | 24754 | 0 | 0 | 18 | 2234 | 367 | 129 | 0 | 0 | 0 | 0 | 0 | 317 | 630 | 167 | 39 | 0 | 0 | 15 | 0 | 0 |
| SRR4836362 | 3865 | 20605 | 1231 | 968 | 4326 | 76014 | 40195 | 37040 | 4441 | 0 | 19053 | 45535 | 4564 | 8238 | 126 | 29885 | 9539 | 0 | 0 | 0 | 483 | 167 | 81 | 0 | 0 | 0 | 0 | 22 | 121 | 90 | 63 | 15 | 0 | 0 | 16 | 0 | 0 |
| SRR4836364 | 4029 | 21446 | 1329 | 875 | 4217 | 78519 | 41226 | 38324 | 4462 | 12 | 19323 | 47033 | 4788 | 8596 | 152 | 23983 | 9627 | 0 | 0 | 0 | 396 | 136 | 61 | 0 | 0 | 0 | 0 | 16 | 96 | 69 | 58 | 15 | 0 | 0 | 11 | 0 | 0 |
| SRR4836366 | 3737 | 20324 | 1231 | 1017 | 4400 | 74921 | 39297 | 36116 | 4336 | 16 | 18848 | 44933 | 4547 | 8240 | 109 | 32786 | 9541 | 0 | 0 | 0 | 481 | 173 | 88 | 0 | 0 | 0 | 0 | 22 | 114 | 92 | 65 | 21 | 0 | 0 | 13 | 0 | 0 |
| SRR4836368 | 3916 | 20985 | 1295 | 956 | 4226 | 77613 | 41293 | 37619 | 4445 | 13 | 19020 | 15 | 4497 | 8464 | 147 | 27993 | 9216 | 0 | 0 | 0 | 423 | 151 | 72 | 10 | 0 | 0 | 0 | 14 | 102 | 70 | 56 | 14 | 0 | 0 | 11 | 0 | 0 |
| SRR4836369 | 1010 | 13450 | 660 | 542 | 3183 | 36158 | 31097 | 33908 | 27587 | 0 | 32835 | 43196 | 9012 | 7791 | 0 | 7706 | 15465 | 0 | 0 | 0 | 469 | 176 | 101 | 12 | 0 | 0 | 0 | 35 | 163 | 169 | 84 | 33 | 0 | 0 | 14 | 0 | 0 |
| SRR4836370 | 1067 | 13410 | 691 | 567 | 2979 | 36588 | 31799 | 34574 | 28386 | 0 | 33882 | 44054 | 9288 | 8019 | 14 | 6388 | 15951 | 0 | 0 | 0 | 441 | 163 | 89 | 0 | 0 | 0 | 0 | 30 | 152 | 145 | 75 | 29 | 0 | 0 | 23 | 0 | 0 |
| SRR4836371 | 1100 | 13268 | 663 | 530 | 2779 | 36642 | 31230 | 33855 | 28034 | 0 | 34241 | 43806 | 9161 | 7795 | 13 | 7349 | 15338 | 0 | 0 | 0 | 473 | 178 | 110 | 0 | 0 | 0 | 0 | 32 | 145 | 157 | 72 | 33 | 0 | 0 | 22 | 0 | 0 |
| SRR4836373 | 1104 | 13553 | 674 | 539 | 3035 | 36553 | 30912 | 33830 | 27995 | 0 | 33161 | 42899 | 9065 | 7702 | 0 | 7453 | 15424 | 0 | 0 | 0 | 432 | 174 | 85 | 0 | 0 | 0 | 0 | 33 | 150 | 133 | 70 | 33 | 0 | 0 | 17 | 0 | 0 |
| SRR4836375 | 2341 | 27211 | 678 | 397 | 1263 | 11391 | 12960 | 25521 | 53869 | 17 | 30415 | 31456 | 15181 | 4806 | 49 | 4927 | 25258 | 0 | 0 | 14 | 2031 | 369 | 172 | 0 | 0 | 0 | 0 | 14 | 212 | 434 | 139 | 36 | 0 | 0 | 12 | 0 | 0 |
| SRR4836376 | 2383 | 27699 | 704 | 366 | 1307 | 11679 | 13153 | 26105 | 56073 | 10 | 31537 | 32622 | 15563 | 4868 | 48 | 4230 | 25970 | 0 | 0 | 13 | 1874 | 340 | 152 | 0 | 0 | 0 | 0 | 16 | 182 | 358 | 124 | 36 | 0 | 0 | 11 | 0 | 0 |
| SRR4836378 | 2371 | 28124 | 697 | 380 | 1101 | 11553 | 13247 | 25751 | 55235 | 0 | 30783 | 31899 | 15315 | 4955 | 48 | 4886 | 24774 | 0 | 0 | 0 | 2111 | 375 | 163 | 14 | 0 | 0 | 0 | 19 | 205 | 422 | 129 | 37 | 0 | 0 | 11 | 0 | 0 |
| SRR4836380 | 2463 | 27834 | 684 | 428 | 1252 | 11845 | 13258 | 26142 | 55615 | 11 | 30661 | 31313 | 15109 | 4880 | 60 | 4827 | 24701 | 0 | 0 | 14 | 1925 | 333 | 156 | 0 | 0 | 0 | 0 | 17 | 191 | 374 | 119 | 44 | 0 | 0 | 12 | 0 | 0 |
| SRR4836382 | 357 | 80276 | 130 | 56 | 401 | 14906 | 8140 | 13645 | 45779 | 0 | 20727 | 29222 | 6037 | 5630 | 46 | 7192 | 11179 | 0 | 0 | 31 | 3076 | 459 | 351 | 21 | 0 | 0 | 0 | 0 | 435 | 1020 | 150 | 76 | 0 | 0 | 64 | 0 | 0 |
| SRR4836383 | 381 | 82200 | 127 | 71 | 447 | 15311 | 8284 | 14187 | 47305 | 0 | 20889 | 30447 | 5924 | 5673 | 47 | 6120 | 10986 | 0 | 0 | 25 | 2908 | 434 | 313 | 19 | 10 | 0 | 0 | 0 | 443 | 939 | 120 | 67 | 0 | 0 | 52 | 10 | 0 |
| SRR4836384 | 399 | 81499 | 167 | 51 | 586 | 16829 | 10194 | 15423 | 45032 | 0 | 20569 | 31492 | 6250 | 5625 | 33 | 7005 | 11195 | 0 | 0 | 27 | 3066 | 430 | 368 | 15 | 0 | 0 | 0 | 0 | 447 | 1009 | 154 | 74 | 0 | 0 | 59 | 10 | 0 |
| SRR4836386 | 408 | 82800 | 162 | 73 | 475 | 15067 | 8210 | 13984 | 45827 | 0 | 21205 | 30085 | 5797 | 5618 | 42 | 6912 | 10938 | 0 | 0 | 27 | 2920 | 432 | 328 | 13 | 0 | 0 | 0 | 0 | 393 | 959 | 124 | 64 | 0 | 0 | 60 | 0 | 0 |
| SRR4836388 | 597 | 42337 | 245 | 143 | 2001 | 15575 | 18885 | 36885 | 36360 | 0 | 28502 | 38539 | 5564 | 3752 | 140 | 1141 | 16564 | 0 | 0 | 0 | 895 | 138 | 151 | 0 | 13 | 0 | 0 | 0 | 93 | 147 | 35 | 24 | 0 | 0 | 10 | 0 | 0 |
| SRR4836390 | 664 | 43228 | 283 | 125 | 1959 | 16010 | 19095 | 38514 | 37717 | 10 | 29401 | 38960 | 5877 | 3814 | 120 | 915 | 17031 | 0 | 0 | 0 | 768 | 116 | 161 | 0 | 11 | 0 | 0 | 0 | 76 | 143 | 45 | 21 | 0 | 0 | 19 | 0 | 0 |
| SRR4836391 | 632 | 43465 | 289 | 110 | 1786 | 15449 | 18662 | 37075 | 37219 | 17 | 28692 | 38583 | 5738 | 3864 | 121 | 1148 | 16887 | 0 | 0 | 0 | 875 | 134 | 150 | 13 | 10 | 0 | 0 | 0 | 89 | 153 | 39 | 22 | 0 | 0 | 11 | 0 | 0 |
| SRR4836392 | 698 | 43752 | 258 | 142 | 2051 | 15618 | 18880 | 37214 | 36983 | 13 | 28731 | 10 | 5654 | 3818 | 113 | 1078 | 16639 | 0 | 0 | 0 | 796 | 126 | 152 | 11 | 0 | 0 | 0 | 0 | 81 | 149 | 34 | 21 | 0 | 0 | 13 | 0 | 0 |
| SRR4836394 | 316 | 32570 | 4417 | 114 | 2334 | 33097 | 19446 | 33375 | 65911 | 0 | 28042 | 54064 | 9290 | 4194 | 10 | 6091 | 28643 | 0 | 0 | 0 | 1012 | 45 | 111 | 0 | 0 | 0 | 0 | 22 | 161 | 319 | 29 | 37 | 0 | 0 | 0 | 0 | 0 |
| SRR4836396 | 329 | 33404 | 4614 | 116 | 2301 | 33892 | 19802 | 33761 | 67335 | 0 | 28665 | 55493 | 9528 | 4348 | 0 | 5166 | 29339 | 0 | 0 | 0 | 928 | 40 | 86 | 0 | 0 | 0 | 0 | 17 | 139 | 317 | 25 | 33 | 0 | 0 | 10 | 0 | 0 |
| SRR4836398 | 366 | 33198 | 4445 | 97 | 2105 | 33458 | 19926 | 33819 | 66643 | 0 | 28403 | 54640 | 9369 | 4337 | 0 | 5688 | 28362 | 0 | 0 | 0 | 1036 | 45 | 110 | 0 | 0 | 0 | 0 | 19 | 159 | 341 | 26 | 32 | 0 | 0 | 0 | 0 | 0 |
| SRR4836399 | 373 | 33212 | 4515 | 136 | 2298 | 33240 | 19770 | 33793 | 66942 | 0 | 27990 | 53768 | 9354 | 4235 | 0 | 5994 | 28194 | 0 | 0 | 0 | 918 | 43 | 95 | 0 | 0 | 0 | 0 | 17 | 141 | 306 | 23 | 29 | 0 | 0 | 0 | 0 | 0 |
| SRR4836400 | 854 | 34588 | 254 | 84 | 3654 | 26155 | 25946 | 22306 | 47944 | 17 | 25643 | 37803 | 7081 | 21830 | 55 | 6470 | 16999 | 0 | 0 | 13 | 953 | 88 | 160 | 12 | 10 | 0 | 0 | 56 | 270 | 263 | 63 | 48 | 0 | 12 | 0 | 0 | 0 |
| SRR4836402 | 844 | 34813 | 259 | 108 | 3492 | 26513 | 26638 | 22996 | 49753 | 0 | 26356 | 38376 | 7127 | 22480 | 40 | 5597 | 16776 | 0 | 0 | 15 | 886 | 77 | 138 | 0 | 0 | 0 | 0 | 55 | 251 | 209 | 53 | 44 | 0 | 0 | 13 | 0 | 0 |
| SRR4836404 | 736 | 35727 | 242 | 100 | 3155 | 26151 | 25959 | 22503 | 48953 | 35 | 25988 | 38446 | 7112 | 22383 | 74 | 6240 | 16310 | 0 | 0 | 12 | 967 | 101 | 170 | 0 | 0 | 0 | 0 | 67 | 283 | 266 | 69 | 49 | 0 | 0 | 0 | 0 | 0 |
| SRR4836406 | 903 | 35380 | 262 | 81 | 3452 | 26001 | 25964 | 22713 | 48945 | 17 | 26077 | 15 | 7170 | 21931 | 80 | 6366 | 16510 | 0 | 0 | 14 | 858 | 76 | 136 | 0 | 0 | 0 | 0 | 66 | 260 | 237 | 55 | 41 | 0 | 0 | 0 | 0 | 0 |
| SRR4836407 | 423 | 21803 | 1653 | 232 | 3874 | 44394 | 47818 | 31956 | 64009 | 0 | 15647 | 37115 | 4389 | 14550 | 82 | 12848 | 15724 | 0 | 0 | 0 | 363 | 151 | 100 | 0 | 11 | 0 | 0 | 12 | 79 | 90 | 77 | 43 | 0 | 0 | 0 | 0 | 0 |
| SRR4836408 | 460 | 20764 | 1659 | 195 | 4286 | 42730 | 46415 | 30991 | 62793 | 0 | 15281 | 35844 | 4314 | 14107 | 86 | 15206 | 15486 | 0 | 0 | 0 | 342 | 159 | 123 | 11 | 11 | 0 | 0 | 18 | 81 | 81 | 67 | 35 | 0 | 0 | 0 | 0 | 0 |
| SRR4836410 | 391 | 21775 | 1724 | 227 | 4079 | 44189 | 47946 | 31690 | 64945 | 0 | 15432 | 36247 | 4414 | 14254 | 88 | 13261 | 15214 | 0 | 0 | 0 | 302 | 124 | 98 | 11 | 0 | 0 | 0 | 17 | 70 | 84 | 56 | 35 | 0 | 0 | 0 | 0 | 0 |
| SRR4836412 | 245 | 183747 | 3478 | 121 | 652 | 8408 | 8427 | 7591 | 46440 | 10 | 6916 | 15254 | 1591 | 1304 | 48 | 8768 | 15423 | 0 | 0 | 270 | 22343 | 444 | 1337 | 17 | 0 | 0 | 0 | 0 | 3694 | 6309 | 205 | 490 | 0 | 0 | 24 | 18 | 0 |
| SRR4836414 | 287 | 178306 | 3318 | 130 | 630 | 8155 | 8212 | 7403 | 45234 | 0 | 6936 | 14710 | 1513 | 1244 | 48 | 10032 | 15628 | 0 | 0 | 324 | 23144 | 472 | 1364 | 13 | 0 | 0 | 0 | 0 | 3793 | 6432 | 234 | 489 | 0 | 0 | 31 | 15 | 0 |
| SRR4836415 | 247 | 183341 | 3619 | 87 | 621 | 8880 | 8777 | 7504 | 47569 | 0 | 7234 | 14740 | 1582 | 1263 | 41 | 9288 | 15233 | 0 | 0 | 271 | 20265 | 421 | 1234 | 14 | 13 | 0 | 0 | 0 | 3302 | 5687 | 181 | 443 | 0 | 0 | 32 | 20 | 0 |
| SRR4836416 | 103 | 56748 | 4197 | 78 | 3504 | 38920 | 29932 | 22650 | 45870 | 0 | 25936 | 47341 | 8872 | 7899 | 56 | 11830 | 15379 | 0 | 0 | 0 | 562 | 72 | 86 | 0 | 0 | 0 | 0 | 0 | 41 | 122 | 26 | 24 | 0 | 0 | 0 | 0 | 0 |
| SRR4836417 | 117 | 55550 | 4068 | 96 | 3904 | 38276 | 29227 | 21932 | 44775 | 0 | 25114 | 45300 | 8524 | 7500 | 77 | 13703 | 14934 | 0 | 0 | 0 | 596 | 79 | 95 | 0 | 0 | 0 | 0 | 0 | 42 | 130 | 23 | 24 | 0 | 0 | 0 | 0 | 0 |
| SRR4836419 | 112 | 57902 | 4249 | 94 | 3843 | 39305 | 30191 | 22567 | 45998 | 0 | 25499 | 46294 | 8636 | 8009 | 79 | 12239 | 15035 | 0 | 0 | 0 | 507 | 67 | 82 | 0 | 0 | 0 | 0 | 0 | 41 | 119 | 20 | 22 | 0 | 0 | 0 | 0 | 0 |
| SRR4836421 | 2658 | 42253 | 384 | 1013 | 1895 | 17702 | 18060 | 24191 | 26678 | 23 | 35261 | 39325 | 9136 | 9226 | 29 | 5307 | 20797 | 0 | 0 | 15 | 1902 | 276 | 168 | 0 | 0 | 0 | 0 | 17 | 252 | 513 | 126 | 57 | 0 | 0 | 39 | 0 | 0 |
| SRR4836422 | 2556 | 40676 | 434 | 1106 | 2082 | 17118 | 17416 | 23470 | 26036 | 11 | 33897 | 38363 | 8536 | 8924 | 24 | 6526 | 20388 | 0 | 0 | 19 | 1988 | 288 | 184 | 14 | 0 | 0 | 0 | 13 | 237 | 540 | 131 | 51 | 0 | 0 | 39 | 14 | 0 |
| SRR4836423 | 2549 | 42155 | 444 | 1126 | 2104 | 17460 | 18083 | 24223 | 26660 | 0 | 35124 | 15 | 8731 | 9098 | 25 | 5678 | 20778 | 0 | 0 | 14 | 1722 | 272 | 152 | 0 | 12 | 0 | 0 | 19 | 255 | 481 | 123 | 45 | 0 | 0 | 34 | 12 | 0 |
| SRR4836425 | 3106 | 22747 | 419 | 841 | 2957 | 54394 | 28198 | 29058 | 1595 | 16 | 41334 | 65092 | 7430 | 7992 | 42 | 15557 | 15984 | 0 | 0 | 0 | 1032 | 96 | 66 | 0 | 0 | 0 | 0 | 41 | 241 | 301 | 42 | 27 | 0 | 0 | 16 | 0 | 0 |
| SRR4836427 | 2813 | 22218 | 430 | 823 | 3234 | 52443 | 27449 | 28196 | 1564 | 26 | 40283 | 63142 | 7237 | 7976 | 34 | 18339 | 15844 | 0 | 0 | 11 | 1047 | 117 | 70 | 0 | 0 | 0 | 0 | 39 | 250 | 309 | 56 | 34 | 0 | 0 | 14 | 11 | 0 |
| SRR4836428 | 3026 | 22381 | 391 | 744 | 3133 | 54458 | 28214 | 28990 | 1693 | 22 | 42533 | 31 | 7431 | 8157 | 46 | 15636 | 15598 | 0 | 0 | 0 | 901 | 93 | 64 | 0 | 0 | 0 | 0 | 38 | 217 | 292 | 53 | 28 | 0 | 0 | 14 | 0 | 0 |
| SRR4836429 | 360 | 51382 | 476 | 193 | 1305 | 18781 | 14499 | 31430 | 39645 | 20 | 29143 | 44901 | 10670 | 8202 | 16 | 20873 | 27080 | 0 | 0 | 24 | 3617 | 211 | 438 | 30 | 19 | 0 | 0 | 0 | 382 | 1209 | 63 | 108 | 0 | 0 | 0 | 15 | 0 |
| SRR4836431 | 373 | 48668 | 426 | 198 | 1482 | 18018 | 14228 | 31229 | 38618 | 14 | 28160 | 43531 | 10372 | 7897 | 0 | 24584 | 27322 | 0 | 0 | 19 | 3764 | 213 | 444 | 38 | 17 | 10 | 0 | 11 | 368 | 1248 | 67 | 112 | 0 | 0 | 0 | 14 | 0 |
| SRR4836433 | 336 | 51070 | 507 | 197 | 1465 | 18799 | 14539 | 31703 | 39982 | 17 | 29316 | 44343 | 10610 | 8129 | 0 | 21077 | 26756 | 0 | 0 | 22 | 3290 | 204 | 380 | 26 | 19 | 0 | 0 | 12 | 344 | 1104 | 56 | 94 | 0 | 0 | 0 | 12 | 0 |
| SRR4836434 | 42 | 53427 | 1319 | 149 | 2589 | 32423 | 29577 | 30656 | 332 | 0 | 54045 | 66089 | 9535 | 15104 | 11 | 9425 | 29676 | 0 | 0 | 0 | 1005 | 95 | 131 | 0 | 0 | 0 | 0 | 0 | 88 | 199 | 37 | 35 | 0 | 0 | 22 | 0 | 0 |
| SRR4836435 | 24 | 52096 | 1271 | 194 | 2607 | 32152 | 28807 | 30035 | 351 | 0 | 53421 | 65895 | 9501 | 14943 | 0 | 10161 | 29307 | 0 | 0 | 0 | 992 | 99 | 145 | 14 | 12 | 0 | 0 | 0 | 103 | 235 | 40 | 34 | 0 | 0 | 13 | 0 | 0 |
| SRR4836437 | 21 | 52414 | 1318 | 165 | 2684 | 32064 | 29076 | 29928 | 293 | 0 | 52139 | 65703 | 9444 | 14871 | 0 | 10064 | 29492 | 0 | 0 | 0 | 972 | 116 | 156 | 10 | 11 | 0 | 0 | 0 | 109 | 202 | 50 | 37 | 0 | 0 | 15 | 0 | 0 |
| SRR4836439 | 23 | 52790 | 1324 | 209 | 2683 | 32716 | 29499 | 30193 | 316 | 0 | 53263 | 65289 | 9417 | 14446 | 0 | 10181 | 27937 | 0 | 0 | 12 | 913 | 97 | 121 | 0 | 0 | 0 | 0 | 0 | 104 | 193 | 28 | 31 | 0 | 0 | 18 | 0 | 0 |
| SRR4836440 | 115 | 51083 | 787 | 470 | 3348 | 50765 | 38021 | 28199 | 5195 | 0 | 6336 | 54952 | 2549 | 408 | 84 | 9421 | 11613 | 0 | 0 | 0 | 291 | 189 | 134 | 22 | 11 | 0 | 0 | 43 | 139 | 111 | 50 | 31 | 0 | 12 | 0 | 0 | 0 |
| SRR4836441 | 66 | 49195 | 787 | 431 | 3317 | 49929 | 38243 | 28238 | 5096 | 28 | 6385 | 54721 | 2689 | 433 | 60 | 10272 | 11557 | 0 | 0 | 0 | 328 | 171 | 109 | 21 | 25 | 0 | 0 | 34 | 150 | 108 | 43 | 20 | 0 | 0 | 0 | 0 | 0 |
| SRR4836443 | 107 | 51653 | 920 | 629 | 3361 | 50190 | 38120 | 27973 | 5151 | 0 | 6106 | 54914 | 2660 | 424 | 68 | 10192 | 11156 | 0 | 0 | 0 | 343 | 194 | 115 | 22 | 16 | 0 | 0 | 38 | 152 | 93 | 48 | 29 | 0 | 0 | 0 | 0 | 0 |
| SRR4836445 | 122 | 50784 | 1116 | 451 | 3468 | 52887 | 39131 | 28703 | 5357 | 0 | 6613 | 54656 | 2642 | 392 | 70 | 10392 | 10750 | 0 | 0 | 0 | 316 | 178 | 110 | 13 | 17 | 0 | 0 | 36 | 161 | 116 | 40 | 36 | 0 | 0 | 0 | 0 | 0 |
| SRR4836446 | 170 | 49714 | 1339 | 224 | 809 | 5500 | 8548 | 19086 | 7661 | 11 | 17635 | 25665 | 11823 | 20197 | 57 | 12064 | 17560 | 0 | 0 | 23 | 4237 | 539 | 448 | 23 | 19 | 0 | 0 | 16 | 274 | 797 | 161 | 98 | 0 | 0 | 24 | 15 | 0 |
| SRR4836447 | 193 | 49234 | 1343 | 233 | 838 | 5601 | 8241 | 19198 | 7336 | 19 | 17634 | 25148 | 12110 | 19890 | 42 | 12874 | 17606 | 0 | 0 | 23 | 4197 | 584 | 431 | 24 | 12 | 12 | 0 | 15 | 252 | 820 | 183 | 104 | 0 | 0 | 26 | 0 | 0 |
| SRR4836448 | 152 | 49761 | 1390 | 246 | 797 | 5512 | 8817 | 19083 | 7385 | 13 | 17328 | 25140 | 11401 | 20288 | 39 | 13403 | 17500 | 0 | 0 | 25 | 4330 | 539 | 402 | 16 | 14 | 0 | 0 | 11 | 270 | 863 | 182 | 96 | 0 | 0 | 24 | 12 | 0 |
| SRR4836450 | 152 | 49645 | 1375 | 264 | 869 | 5553 | 8596 | 19392 | 7753 | 0 | 16650 | 24771 | 11744 | 19866 | 40 | 13413 | 16695 | 0 | 0 | 22 | 4278 | 514 | 413 | 15 | 18 | 0 | 0 | 11 | 269 | 842 | 157 | 88 | 0 | 0 | 19 | 10 | 0 |
| SRR4836452 | 84 | 46718 | 2650 | 128 | 4684 | 51105 | 36902 | 23239 | 35628 | 0 | 27501 | 51993 | 6460 | 12284 | 11 | 18213 | 14288 | 0 | 0 | 13 | 746 | 57 | 51 | 0 | 0 | 0 | 0 | 12 | 120 | 161 | 25 | 16 | 0 | 0 | 0 | 0 | 0 |
| SRR4836453 | 94 | 46073 | 2639 | 192 | 4838 | 50894 | 36443 | 23329 | 34749 | 0 | 27233 | 51459 | 6413 | 12113 | 19 | 19822 | 14017 | 0 | 0 | 0 | 741 | 45 | 70 | 0 | 0 | 0 | 0 | 15 | 115 | 159 | 30 | 15 | 0 | 0 | 0 | 0 | 0 |
| SRR4836454 | 67 | 45376 | 2820 | 143 | 4438 | 50383 | 36494 | 23565 | 35706 | 0 | 27243 | 51311 | 6522 | 11924 | 19 | 20167 | 14480 | 0 | 0 | 0 | 697 | 63 | 69 | 0 | 0 | 0 | 0 | 14 | 105 | 149 | 29 | 15 | 0 | 0 | 0 | 0 | 0 |
| SRR4836456 | 97 | 46091 | 2564 | 132 | 4536 | 50598 | 35749 | 24096 | 34749 | 0 | 27233 | 49841 | 6352 | 11586 | 11 | 20018 | 13559 | 0 | 0 | 10 | 755 | 65 | 55 | 0 | 0 | 0 | 0 | 16 | 127 | 171 | 30 | 11 | 0 | 0 | 0 | 0 | 0 |
| SRR4836458 | 23 | 31703 | 1181 | 132 | 3030 | 61456 | 32391 | 32679 | 3960 | 0 | 29756 | 49627 | 6319 | 16063 | 42 | 8886 | 23207 | 0 | 0 | 0 | 184 | 107 | 94 | 0 | 0 | 0 | 0 | 14 | 84 | 76 | 30 | 20 | 0 | 0 | 0 | 0 | 0 |
| SRR4836459 | 34 | 31134 | 1154 | 99 | 2871 | 61397 | 32111 | 32587 | 4105 | 0 | 29945 | 49871 | 6200 | 16000 | 28 | 9822 | 22640 | 0 | 0 | 0 | 199 | 132 | 106 | 13 | 10 | 0 | 0 | 10 | 71 | 71 | 39 | 38 | 0 | 0 | 0 | 0 | 0 |
| SRR4836461 | 14 | 31788 | 1333 | 119 | 2841 | 61141 | 32106 | 32037 | 3974 | 0 | 29239 | 49586 | 6293 | 16077 | 24 | 9489 | 22771 | 0 | 0 | 0 | 173 | 90 | 104 | 12 | 0 | 0 | 0 | 22 | 90 | 76 | 44 | 31 | 0 | 0 | 0 | 0 | 0 |
| SRR4836463 | 32 | 31456 | 1146 | 146 | 3094 | 62511 | 32667 | 32819 | 4158 | 0 | 30249 | 49566 | 6055 | 15989 | 36 | 9614 | 21755 | 0 | 0 | 0 | 189 | 96 | 105 | 0 | 11 | 0 | 0 | 18 | 81 | 84 | 33 | 30 | 0 | 0 | 0 | 0 | 0 |
| SRR4836464 | 24 | 89582 | 465 | 66 | 589 | 5383 | 5910 | 19449 | 436 | 0 | 23326 | 36158 | 7043 | 5834 | 227 | 9892 | 21375 | 0 | 0 | 38 | 2843 | 204 | 301 | 11 | 0 | 0 | 0 | 0 | 459 | 621 | 49 | 66 | 0 | 0 | 0 | 15 | 0 |
| SRR4836466 | 34 | 87821 | 458 | 61 | 609 | 5192 | 5877 | 19057 | 441 | 0 | 23224 | 35479 | 6828 | 5748 | 223 | 10515 | 20861 | 0 | 0 | 43 | 2841 | 204 | 269 | 11 | 0 | 0 | 0 | 0 | 442 | 632 | 52 | 79 | 0 | 0 | 0 | 0 | 0 |
| SRR4836468 | 14 | 87804 | 451 | 59 | 612 | 5140 | 5786 | 18916 | 406 | 0 | 22787 | 35296 | 6940 | 5763 | 205 | 10613 | 20840 | 0 | 0 | 39 | 2903 | 200 | 312 | 10 | 0 | 0 | 0 | 0 | 479 | 588 | 52 | 75 | 0 | 0 | 0 | 12 | 0 |
| SRR4836469 | 22 | 87508 | 469 | 54 | 585 | 5324 | 5929 | 19186 | 427 | 0 | 24268 | 35542 | 6969 | 5782 | 208 | 10881 | 20526 | 0 | 0 | 42 | 2796 | 190 | 293 | 13 | 0 | 0 | 0 | 0 | 453 | 574 | 45 | 71 | 0 | 0 | 0 | 13 | 0 |
| SRR4836471 | 54 | 32643 | 733 | 342 | 1446 | 24519 | 16973 | 30253 | 36964 | 11 | 14573 | 25510 | 6280 | 3325 | 0 | 8209 | 21043 | 0 | 0 | 0 | 836 | 265 | 190 | 17 | 14 | 0 | 0 | 0 | 158 | 346 | 121 | 51 | 0 | 0 | 25 | 0 | 0 |
| SRR4836473 | 48 | 31231 | 656 | 376 | 1524 | 23170 | 16069 | 28835 | 35378 | 0 | 13990 | 24100 | 6005 | 3263 | 0 | 10891 | 19988 | 0 | 0 | 0 | 984 | 309 | 208 | 24 | 23 | 0 | 0 | 0 | 182 | 366 | 132 | 61 | 0 | 13 | 29 | 15 | 0 |
| SRR4836474 | 47 | 32206 | 654 | 371 | 1484 | 23628 | 16507 | 28921 | 35196 | 0 | 13746 | 24065 | 5920 | 3207 | 0 | 11080 | 19128 | 0 | 0 | 0 | 947 | 291 | 208 | 25 | 24 | 0 | 0 | 0 | 185 | 406 | 142 | 65 | 0 | 11 | 32 | 0 | 0 |
| SRR4836475 | 104 | 29456 | 3428 | 71 | 2539 | 25037 | 25286 | 19996 | 34705 | 0 | 33264 | 47653 | 8671 | 15727 | 0 | 11031 | 19922 | 0 | 0 | 0 | 562 | 60 | 139 | 13 | 14 | 0 | 0 | 0 | 62 | 121 | 19 | 37 | 0 | 0 | 0 | 0 | 0 |
| SRR4836477 | 55 | 27173 | 3197 | 145 | 2613 | 23960 | 24445 | 19173 | 33121 | 12 | 31780 | 45580 | 8390 | 14826 | 0 | 15101 | 19009 | 0 | 0 | 0 | 668 | 69 | 167 | 16 | 22 | 0 | 0 | 13 | 95 | 135 | 29 | 36 | 0 | 0 | 0 | 0 | 0 |
| SRR4836479 | 66 | 27840 | 3138 | 122 | 2789 | 24000 | 24016 | 19371 | 34152 | 0 | 31531 | 45497 | 8529 | 15086 | 13 | 15493 | 18201 | 0 | 0 | 0 | 647 | 83 | 187 | 10 | 19 | 0 | 0 | 0 | 86 | 147 | 22 | 42 | 0 | 0 | 0 | 0 | 0 |

### Supp Table S8.1.Columns_B_to_AL

| | | | | | | | | | | | | | | | | | | | | | | | | | | | | | | | | | | | | | |
|---|---|---|---|---|---|---|---|---|---|---|---|---|---|---|---|---|---|---|---|---|---|---|---|---|---|---|---|---|---|---|---|---|---|---|---|---|---|
| SRR4836481 | 12 | 49323 | 223 | 25 | 597 | 7503 | 7193 | 7830 | 86554 | 0 | 15006 | 17107 | 3787 | 5961 | 138 | 3150 | 10294 | 0 | 0 | 0 | 1814 | 271 | 290 | 22 | 0 | 0 | 0 | 0 | 171 | 625 | 87 | 47 | 0 | 0 | 28 | 0 | 0 |
| SRR4836482 | 12 | 47192 | 260 | 21 | 674 | 7271 | 6703 | 7386 | 84465 | 0 | 14297 | 16200 | 3557 | 5739 | 115 | 4605 | 9791 | 0 | 0 | 17 | 2049 | 375 | 355 | 16 | 10 | 0 | 0 | 0 | 212 | 682 | 100 | 54 | 0 | 0 | 23 | 0 | 0 |
| SRR4836483 | 22 | 48411 | 227 | 25 | 727 | 7560 | 6972 | 7523 | 84931 | 0 | 14134 | 16052 | 3547 | 5595 | 86 | 4389 | 9413 | 0 | 0 | 17 | 2131 | 333 | 374 | 12 | 0 | 0 | 0 | 0 | 182 | 636 | 92 | 45 | 0 | 0 | 29 | 0 | 0 |
| SRR4836484 | 11 | 52232 | 2205 | 251 | 908 | 11511 | 8132 | 17755 | 21712 | 11 | 11883 | 26857 | 8879 | 17443 | 28 | 6844 | 39495 | 0 | 0 | 0 | 151 | 76 | 86 | 11 | 14 | 0 | 0 | 37 | 114 | 47 | 22 | 0 | 0 | 0 | 0 | 0 | 0 |
| SRR4836485 | 43 | 52688 | 2266 | 318 | 936 | 10935 | 7937 | 16994 | 20666 | 26 | 11279 | 25457 | 8604 | 16885 | 21 | 9187 | 38366 | 0 | 0 | 0 | 176 | 62 | 106 | 16 | 12 | 0 | 0 | 37 | 101 | 51 | 25 | 19 | 0 | 0 | 0 | 0 | 0 |
| SRR4836487 | 0 | 52383 | 2133 | 407 | 954 | 11094 | 8076 | 17269 | 21055 | 26 | 11963 | 25617 | 8719 | 17043 | 16 | 9429 | 36377 | 0 | 0 | 0 | 167 | 67 | 106 | 12 | 11 | 0 | 0 | 43 | 103 | 55 | 24 | 27 | 0 | 0 | 0 | 0 | 0 |
| SRR4836489 | 142 | 49206 | 882 | 44 | 4475 | 52639 | 44913 | 26327 | 7734 | 0 | 22755 | 47221 | 6676 | 6102 | 45 | 11345 | 15863 | 0 | 0 | 58 | 8358 | 414 | 248 | 0 | 0 | 0 | 0 | 0 | 990 | 2227 | 177 | 107 | 0 | 0 | 10 | 0 | 0 |
| SRR4836491 | 135 | 47459 | 796 | 47 | 4642 | 50484 | 43185 | 25281 | 7571 | 0 | 21976 | 45544 | 6457 | 5784 | 46 | 15221 | 15350 | 0 | 0 | 75 | 9655 | 481 | 308 | 0 | 0 | 0 | 0 | 0 | 1148 | 2604 | 209 | 122 | 0 | 0 | 11 | 0 | 0 |
| SRR4836492 | 156 | 48220 | 788 | 63 | 4716 | 51393 | 44114 | 25541 | 7713 | 0 | 21767 | 44916 | 6292 | 5817 | 41 | 15534 | 14846 | 0 | 0 | 80 | 9350 | 469 | 294 | 12 | 0 | 0 | 0 | 0 | 1126 | 2503 | 193 | 127 | 0 | 0 | 14 | 12 | 0 |
| SRR4836493 | 16 | 36082 | 936 | 68 | 5255 | 87203 | 61038 | 59247 | 7277 | 0 | 24497 | 44111 | 10361 | 7853 | 20 | 8664 | 19601 | 0 | 0 | 0 | 199 | 88 | 69 | 0 | 0 | 0 | 0 | 0 | 53 | 72 | 43 | 17 | 0 | 0 | 0 | 0 | 0 |
| SRR4836494 | 15 | 34694 | 897 | 69 | 5477 | 84269 | 58933 | 57072 | 7096 | 0 | 23655 | 42594 | 10176 | 7733 | 16 | 12346 | 19075 | 0 | 0 | 0 | 226 | 114 | 69 | 10 | 0 | 0 | 0 | 0 | 62 | 78 | 54 | 24 | 0 | 0 | 0 | 0 | 0 |
| SRR4836495 | 16 | 35284 | 928 | 79 | 5598 | 85236 | 59896 | 57706 | 7187 | 0 | 23505 | 42160 | 10027 | 7634 | 0 | 12603 | 18309 | 0 | 0 | 0 | 228 | 108 | 72 | 0 | 0 | 0 | 0 | 0 | 63 | 79 | 43 | 26 | 0 | 0 | 0 | 0 | 0 |
| SRR4836496 | 1335 | 23818 | 1783 | 389 | 6136 | 87330 | 56399 | 33874 | 11225 | 0 | 15698 | 50529 | 5213 | 6782 | 111 | 15036 | 14598 | 0 | 0 | 15 | 1462 | 270 | 98 | 0 | 0 | 0 | 0 | 18 | 200 | 415 | 105 | 36 | 0 | 0 | 0 | 0 | 0 |
| SRR4836498 | 1394 | 24470 | 1805 | 404 | 5916 | 88059 | 57004 | 33770 | 11323 | 0 | 15809 | 51226 | 5456 | 6847 | 85 | 12827 | 14556 | 0 | 0 | 10 | 1314 | 229 | 84 | 0 | 0 | 0 | 0 | 17 | 181 | 368 | 89 | 31 | 0 | 0 | 0 | 0 | 0 |
| SRR4836500 | 1369 | 24174 | 1750 | 389 | 5704 | 87467 | 56332 | 34047 | 11229 | 0 | 15563 | 50859 | 5425 | 6843 | 107 | 14547 | 14685 | 0 | 0 | 12 | 1433 | 281 | 102 | 0 | 0 | 0 | 0 | 20 | 195 | 399 | 99 | 31 | 0 | 0 | 0 | 0 | 0 |
| SRR4836502 | 1320 | 23316 | 1815 | 453 | 6268 | 87655 | 56786 | 33151 | 11491 | 0 | 15211 | 49704 | 5327 | 6776 | 84 | 17175 | 13639 | 0 | 0 | 12 | 1482 | 266 | 106 | 12 | 0 | 0 | 0 | 19 | 200 | 428 | 104 | 32 | 0 | 0 | 0 | 0 | 0 |
| SRR4836503 | 722 | 87977 | 850 | 224 | 2248 | 21214 | 22056 | 25689 | 7280 | 0 | 28883 | 36226 | 11631 | 11835 | 34 | 8718 | 32494 | 0 | 0 | 306 | 27647 | 1187 | 2222 | 26 | 0 | 17 | 0 | 21 | 4884 | 9713 | 423 | 627 | 0 | 0 | 37 | 21 | 0 |
| SRR4836504 | 690 | 89945 | 855 | 236 | 2344 | 22218 | 22773 | 26678 | 7939 | 0 | 30268 | 37051 | 12051 | 12232 | 24 | 7170 | 33311 | 0 | 0 | 292 | 24669 | 1066 | 1952 | 31 | 10 | 12 | 0 | 16 | 4366 | 8680 | 387 | 550 | 0 | 0 | 37 | 16 | 0 |
| SRR4836505 | 644 | 89522 | 929 | 283 | 2144 | 21623 | 22283 | 26738 | 7499 | 0 | 29330 | 36132 | 11731 | 12342 | 21 | 8069 | 32035 | 0 | 0 | 305 | 27326 | 1227 | 2216 | 35 | 11 | 10 | 0 | 28 | 4956 | 9747 | 416 | 619 | 0 | 0 | 38 | 16 | 0 |
| SRR4836507 | 656 | 86104 | 921 | 240 | 2355 | 21193 | 21886 | 25445 | 7549 | 0 | 28445 | 34834 | 11473 | 11634 | 18 | 9679 | 32238 | 0 | 0 | 330 | 28226 | 1246 | 2160 | 39 | 10 | 10 | 0 | 20 | 5022 | 9729 | 424 | 652 | 0 | 0 | 40 | 17 | 0 |
| SRR4836509 | 817 | 109491 | 131 | 142 | 1450 | 18610 | 26032 | 25642 | 49245 | 0 | 6304 | 23629 | 2501 | 1861 | 47 | 10591 | 15221 | 0 | 0 | 395 | 33323 | 1754 | 1938 | 51 | 0 | 0 | 0 | 0 | 5251 | 16072 | 681 | 609 | 0 | 0 | 23 | 21 | 0 |
| SRR4836511 | 861 | 110390 | 141 | 148 | 1422 | 19164 | 26637 | 26225 | 51057 | 14 | 6260 | 24311 | 2609 | 1958 | 60 | 9165 | 15719 | 0 | 0 | 358 | 29882 | 1552 | 1692 | 45 | 0 | 0 | 0 | 0 | 4742 | 14372 | 628 | 535 | 0 | 0 | 23 | 18 | 0 |
| SRR4836512 | 882 | 110746 | 132 | 159 | 1346 | 18835 | 27130 | 25731 | 49754 | 0 | 6646 | 24216 | 2561 | 1847 | 41 | 10274 | 15343 | 0 | 0 | 386 | 33566 | 1748 | 1929 | 47 | 0 | 12 | 0 | 0 | 5295 | 15905 | 672 | 611 | 0 | 0 | 24 | 18 | 0 |
| SRR4836513 | 795 | 107533 | 126 | 163 | 1458 | 18113 | 26747 | 24625 | 48392 | 18 | 6139 | 22833 | 2457 | 1871 | 32 | 12452 | 14569 | 0 | 0 | 393 | 33672 | 1737 | 1934 | 46 | 0 | 10 | 0 | 0 | 5368 | 16079 | 689 | 610 | 0 | 0 | 17 | 18 | 0 |
| SRR4836514 | 520 | 23932 | 576 | 275 | 386 | 5541 | 4961 | 15787 | 54912 | 0 | 23110 | 19722 | 10075 | 10889 | 57 | 7573 | 34256 | 0 | 0 | 0 | 412 | 234 | 188 | 22 | 16 | 0 | 0 | 0 | 48 | 124 | 83 | 50 | 0 | 0 | 0 | 0 | 0 |
| SRR4836515 | 516 | 24117 | 552 | 260 | 347 | 5640 | 5107 | 16320 | 56366 | 0 | 23896 | 20336 | 10495 | 10984 | 72 | 6293 | 35278 | 0 | 0 | 0 | 375 | 220 | 169 | 20 | 0 | 0 | 0 | 0 | 43 | 132 | 72 | 44 | 0 | 0 | 0 | 0 | 0 |
| SRR4836517 | 525 | 24418 | 611 | 249 | 329 | 5636 | 4943 | 16127 | 55089 | 0 | 23272 | 20096 | 10415 | 11081 | 54 | 7132 | 34133 | 0 | 0 | 0 | 421 | 247 | 163 | 23 | 17 | 0 | 0 | 10 | 43 | 147 | 80 | 45 | 0 | 0 | 0 | 0 | 0 |
| SRR4836519 | 487 | 23754 | 539 | 319 | 366 | 5568 | 4942 | 15804 | 55727 | 0 | 22620 | 19319 | 9711 | 10510 | 39 | 9113 | 32020 | 0 | 0 | 0 | 396 | 230 | 178 | 22 | 19 | 0 | 0 | 0 | 49 | 144 | 77 | 51 | 0 | 11 | 0 | 0 | 0 |
| SRR4836521 | 493 | 39657 | 1195 | 134 | 748 | 11123 | 14308 | 23071 | 24703 | 13 | 21277 | 69003 | 7704 | 14632 | 0 | 3033 | 27602 | 0 | 0 | 57 | 4680 | 314 | 285 | 12 | 0 | 0 | 0 | 36 | 1643 | 2465 | 261 | 203 | 0 | 0 | 0 | 0 | 0 |
| SRR4836522 | 473 | 40651 | 1077 | 122 | 739 | 11386 | 14655 | 23408 | 25253 | 0 | 21802 | 70235 | 7916 | 14871 | 0 | 2532 | 27809 | 0 | 0 | 54 | 4188 | 282 | 263 | 11 | 0 | 0 | 0 | 32 | 1440 | 2147 | 249 | 190 | 0 | 0 | 0 | 0 | 0 |
| SRR4836523 | 506 | 40409 | 974 | 151 | 699 | 11246 | 14541 | 23146 | 24893 | 13 | 21615 | 69316 | 7937 | 14546 | 0 | 2848 | 26906 | 0 | 0 | 48 | 4643 | 326 | 274 | 14 | 0 | 0 | 0 | 27 | 1623 | 2411 | 249 | 204 | 0 | 0 | 0 | 0 | 0 |
| SRR4836524 | 481 | 39142 | 1018 | 217 | 757 | 11012 | 14469 | 22646 | 25405 | 0 | 20804 | 67015 | 7704 | 14187 | 0 | 3603 | 25547 | 0 | 0 | 50 | 4794 | 309 | 291 | 11 | 0 | 0 | 0 | 39 | 1646 | 2495 | 285 | 214 | 0 | 0 | 0 | 0 | 0 |
| SRR4836525 | 409 | 27946 | 348 | 299 | 316 | 7140 | 6160 | 23760 | 48437 | 0 | 22601 | 19472 | 10205 | 11465 | 99 | 9591 | 25775 | 0 | 0 | 0 | 468 | 260 | 186 | 17 | 12 | 0 | 0 | 11 | 52 | 138 | 77 | 49 | 0 | 0 | 12 | 0 | 0 |
| SRR4836526 | 419 | 28862 | 344 | 311 | 353 | 7255 | 6344 | 24166 | 49702 | 0 | 23265 | 19573 | 10399 | 11872 | 99 | 8176 | 26545 | 0 | 0 | 0 | 421 | 264 | 164 | 11 | 13 | 0 | 0 | 14 | 50 | 142 | 76 | 54 | 0 | 0 | 0 | 0 | 0 |
| SRR4836528 | 428 | 28756 | 389 | 310 | 324 | 6990 | 6304 | 23703 | 48791 | 10 | 22715 | 19559 | 10580 | 11632 | 88 | 9222 | 25990 | 0 | 0 | 0 | 461 | 267 | 192 | 24 | 13 | 0 | 0 | 12 | 62 | 153 | 83 | 52 | 0 | 10 | 12 | 0 | 0 |
| SRR4836529 | 403 | 28166 | 338 | 349 | 363 | 6949 | 6162 | 23228 | 49054 | 0 | 21892 | 18522 | 9852 | 11169 | 76 | 10985 | 24658 | 0 | 0 | 0 | 492 | 292 | 175 | 17 | 13 | 0 | 0 | 16 | 59 | 156 | 92 | 53 | 0 | 0 | 0 | 0 | 0 |
| SRR4836531 | 706 | 100982 | 1718 | 272 | 1731 | 18344 | 17719 | 12568 | 26412 | 34 | 31680 | 37154 | 7647 | 9473 | 21 | 7089 | 15607 | 0 | 0 | 25 | 3637 | 808 | 224 | 12 | 0 | 0 | 0 | 24 | 273 | 904 | 309 | 60 | 0 | 0 | 0 | 0 | 0 |
| SRR4836533 | 779 | 98386 | 1610 | 244 | 1769 | 18180 | 17080 | 12140 | 25683 | 22 | 30742 | 35276 | 7026 | 9026 | 30 | 8160 | 15023 | 0 | 0 | 19 | 3747 | 863 | 222 | 0 | 0 | 0 | 0 | 27 | 295 | 955 | 332 | 75 | 0 | 0 | 0 | 0 | 0 |
| SRR4836534 | 774 | 100165 | 1661 | 227 | 1862 | 19085 | 18373 | 12717 | 27074 | 27 | 31265 | 36312 | 7232 | 9366 | 25 | 6914 | 15538 | 0 | 0 | 20 | 3331 | 773 | 202 | 10 | 0 | 0 | 0 | 19 | 261 | 842 | 294 | 57 | 0 | 0 | 0 | 0 | 0 |
| SRR4836535 | 53 | 89539 | 1906 | 100 | 2373 | 23990 | 25313 | 25452 | 231 | 12 | 31964 | 31970 | 8774 | 9099 | 30 | 17679 | 23353 | 0 | 0 | 130 | 23791 | 732 | 1349 | 16 | 0 | 0 | 0 | 0 | 1624 | 5384 | 187 | 275 | 0 | 0 | 0 | 0 | 0 |
| SRR4836536 | 56 | 86876 | 1826 | 95 | 2617 | 22695 | 24724 | 24474 | 124 | 12 | 30931 | 31183 | 8301 | 8616 | 31 | 20951 | 22835 | 0 | 0 | 124 | 24303 | 727 | 1391 | 15 | 0 | 0 | 0 | 0 | 1632 | 5593 | 188 | 286 | 0 | 0 | 0 | 0 | 0 |
| SRR4836537 | 68 | 89082 | 1818 | 106 | 2620 | 23282 | 25612 | 26060 | 375 | 0 | 32025 | 31461 | 8328 | 9232 | 18 | 18109 | 23182 | 0 | 0 | 113 | 21947 | 669 | 1256 | 12 | 0 | 0 | 0 | 0 | 1477 | 5011 | 166 | 263 | 0 | 0 | 11 | 0 | 0 |
| SRR4836539 | 121 | 104421 | 117 | 185 | 1523 | 13662 | 11257 | 11204 | 17409 | 23 | 29537 | 39628 | 9734 | 23308 | 49 | 17601 | 12945 | 0 | 0 | 235 | 13561 | 561 | 728 | 22 | 19 | 19 | 0 | 11 | 3330 | 4107 | 209 | 190 | 0 | 0 | 148 | 34 | 0 |
| SRR4836541 | 112 | 101308 | 157 | 183 | 1853 | 13170 | 10812 | 10781 | 17634 | 0 | 29429 | 38602 | 9251 | 22684 | 18 | 20361 | 12999 | 0 | 0 | 251 | 14172 | 613 | 727 | 21 | 18 | 19 | 0 | 0 | 3525 | 4265 | 219 | 174 | 0 | 0 | 147 | 26 | 0 |
| SRR4836543 | 121 | 104027 | 111 | 181 | 1718 | 13243 | 11227 | 11485 | 18228 | 14 | 30182 | 39213 | 9796 | 23033 | 42 | 17530 | 13380 | 0 | 0 | 235 | 12747 | 535 | 685 | 19 | 18 | 16 | 0 | 10 | 3120 | 3762 | 193 | 166 | 0 | 0 | 147 | 27 | 0 |
| SRR4836544 | 50 | 52692 | 1310 | 115 | 2869 | 60305 | 42613 | 63046 | 5304 | 0 | 35990 | 46015 | 6069 | 8714 | 0 | 5925 | 17275 | 0 | 0 | 0 | 273 | 19 | 23 | 0 | 0 | 0 | 0 | 0 | 0 | 13 | 0 | 0 | 0 | 0 | 0 | 0 | 0 |
| SRR4836545 | 29 | 51342 | 1240 | 127 | 3136 | 59126 | 41767 | 62284 | 5345 | 0 | 35263 | 45321 | 5741 | 8560 | 0 | 7398 | 17445 | 0 | 0 | 0 | 247 | 21 | 30 | 0 | 0 | 0 | 0 | 0 | 0 | 0 | 0 | 0 | 0 | 0 | 0 | 0 | 0 |
| SRR4836546 | 41 | 51901 | 1329 | 140 | 3217 | 60868 | 43038 | 62806 | 5350 | 0 | 36346 | 45999 | 5941 | 8811 | 0 | 6108 | 17707 | 0 | 0 | 0 | 221 | 19 | 24 | 0 | 0 | 0 | 0 | 0 | 0 | 0 | 0 | 0 | 0 | 0 | 0 | 0 | 0 |
| SRR4836548 | 125 | 60039 | 1106 | 138 | 2397 | 18227 | 21004 | 21199 | 61752 | 0 | 27633 | 64346 | 6115 | 5977 | 18 | 6274 | 15285 | 0 | 0 | 0 | 2492 | 190 | 242 | 15 | 0 | 0 | 0 | 15 | 148 | 514 | 55 | 58 | 0 | 0 | 0 | 13 | 0 |
| SRR4836550 | 128 | 58875 | 1166 | 110 | 2698 | 17593 | 20272 | 20203 | 60214 | 0 | 26520 | 62095 | 5835 | 5683 | 22 | 7530 | 15075 | 0 | 0 | 15 | 2595 | 187 | 244 | 15 | 0 | 0 | 0 | 18 | 142 | 516 | 55 | 67 | 0 | 0 | 0 | 10 | 0 |
| SRR4836552 | 79 | 61570 | 1163 | 145 | 2706 | 18262 | 21361 | 21161 | 62168 | 14 | 27032 | 63271 | 5951 | 5807 | 16 | 6538 | 15538 | 0 | 0 | 14 | 2351 | 168 | 233 | 12 | 11 | 0 | 0 | 17 | 140 | 461 | 44 | 61 | 0 | 0 | 0 | 12 | 0 |
| SRR4836553 | 355 | 92362 | 2735 | 346 | 1019 | 23141 | 15539 | 27150 | 44489 | 22 | 45506 | 35845 | 9925 | 17371 | 21 | 6311 | 38389 | 0 | 0 | 26 | 2975 | 171 | 216 | 0 | 0 | 0 | 0 | 0 | 426 | 1015 | 95 | 74 | 0 | 0 | 23 | 14 | 0 |
| SRR4836554 | 309 | 89895 | 2639 | 224 | 992 | 21914 | 15280 | 26459 | 43828 | 17 | 44434 | 34744 | 9635 | 16870 | 19 | 7095 | 39209 | 0 | 0 | 24 | 3030 | 186 | 218 | 0 | 11 | 0 | 0 | 0 | 472 | 1025 | 99 | 83 | 0 | 0 | 25 | 11 | 0 |
| SRR4836555 | 290 | 92823 | 2886 | 256 | 980 | 22393 | 15489 | 27407 | 44551 | 11 | 44941 | 35255 | 9844 | 16838 | 0 | 6032 | 39346 | 0 | 0 | 28 | 2738 | 167 | 206 | 0 | 0 | 0 | 0 | 0 | 465 | 906 | 101 | 62 | 0 | 0 | 23 | 12 | 0 |
| SRR4836556 | 0 | 6684 | 1034 | 121 | 1266 | 14881 | 16054 | 16487 | 7200 | 29 | 37953 | 35597 | 10299 | 6270 | 25 | 21128 | 24837 | 0 | 0 | 0 | 94 | 80 | 40 | 0 | 0 | 0 | 0 | 0 | 35 | 76 | 74 | 20 | 0 | 0 | 14 | 0 | 0 |
| SRR4836560 | 0 | 6438 | 992 | 147 | 1330 | 14523 | 15936 | 15890 | 6938 | 15 | 37870 | 35084 | 10436 | 6145 | 24 | 22617 | 23784 | 0 | 0 | 0 | 90 | 80 | 35 | 0 | 0 | 0 | 0 | 0 | 43 | 82 | 74 | 19 | 0 | 0 | 13 | 0 | 0 |
| SRR4836561 | 0 | 6563 | 1120 | 125 | 1222 | 14601 | 16272 | 16323 | 6942 | 16 | 37318 | 34634 | 10028 | 6087 | 33 | 22801 | 22881 | 0 | 0 | 0 | 82 | 80 | 33 | 0 | 0 | 0 | 0 | 0 | 39 | 78 | 83 | 24 | 0 | 0 | 13 | 0 | 0 |
| SRR4836562 | 52 | 81264 | 4010 | 44 | 143 | 1569 | 2172 | 10461 | 41482 | 0 | 16406 | 10111 | 10983 | 8868 | 12 | 11222 | 25542 | 0 | 0 | 20 | 4420 | 1260 | 602 | 36 | 36 | 0 | 0 | 0 | 388 | 1620 | 250 | 151 | 0 | 13 | 0 | 11 | 0 |
| SRR4836563 | 16 | 79466 | 4142 | 44 | 133 | 1522 | 2037 | 10164 | 41234 | 16 | 16307 | 10187 | 11015 | 8642 | 29 | 11862 | 25155 | 0 | 0 | 18 | 4469 | 1290 | 598 | 33 | 23 | 0 | 0 | 0 | 390 | 1612 | 255 | 157 | 0 | 13 | 0 | 0 | 0 |
| SRR4836565 | 35 | 79623 | 4000 | 39 | 120 | 1610 | 2193 | 10302 | 41207 | 23 | 16177 | 10170 | 10915 | 8682 | 13 | 11906 | 25289 | 0 | 0 | 24 | 4490 | 1266 | 595 | 42 | 30 | 0 | 0 | 0 | 372 | 1617 | 278 | 150 | 0 | 12 | 0 | 0 | 0 |
| SRR4836567 | 0 | 79973 | 4337 | 42 | 122 | 1606 | 2004 | 10421 | 41553 | 0 | 16070 | 9791 | 10778 | 8602 | 12 | 12355 | 24330 | 0 | 0 | 26 | 4394 | 1270 | 592 | 40 | 28 | 0 | 0 | 0 | 399 | 1533 | 258 | 149 | 0 | 15 | 0 | 0 | 0 |
| SRR4836569 | 36 | 17066 | 1867 | 33 | 3697 | 63511 | 56668 | 43866 | 6001 | 29 | 11499 | 58499 | 5599 | 5873 | 0 | 11757 | 12430 | 0 | 0 | 0 | 135 | 44 | 52 | 0 | 0 | 0 | 0 | 0 | 40 | 37 | 31 | 17 | 0 | 0 | 0 | 0 | 0 |
| SRR4836570 | 29 | 16927 | 1945 | 32 | 3685 | 63385 | 56449 | 43721 | 5947 | 11 | 11443 | 58007 | 5563 | 5828 | 0 | 12774 | 12225 | 0 | 0 | 0 | 140 | 39 | 50 | 0 | 0 | 0 | 0 | 0 | 41 | 41 | 34 | 16 | 0 | 0 | 0 | 0 | 0 |
| SRR4836571 | 43 | 16527 | 1843 | 29 | 3673 | 63383 | 56181 | 43502 | 6062 | 0 | 11361 | 57834 | 5596 | 5825 | 0 | 12798 | 12182 | 0 | 0 | 0 | 137 | 45 | 58 | 0 | 0 | 0 | 0 | 0 | 40 | 34 | 31 | 18 | 0 | 0 | 0 | 0 | 0 |
| SRR4836572 | 41 | 16809 | 1999 | 53 | 3731 | 64087 | 56462 | 43983 | 6068 | 18 | 11321 | 57183 | 5493 | 5787 | 0 | 13059 | 11799 | 0 | 0 | 0 | 137 | 43 | 47 | 0 | 0 | 0 | 0 | 0 | 40 | 36 | 31 | 19 | 0 | 0 | 0 | 0 | 0 |
| SRR4896694 | 160 | 2719 | 799 | 534 | 213 | 919 | 2358 | 3614 | 7436 | 244 | 9429 | 11295 | 2231 | 2576 | 86 | 2695 | 51336 | 0 | 0 | 29 | 1690 | 4130 | 387 | 0 | 0 | 0 | 0 | 51 | 577 | 6714 | 16770 | 1657 | 0 | 0 | 37 | 10 | 0 |
| SRR4896695 | 20 | 11386 | 486 | 157 | 427 | 1899 | 2461 | 4551 | 11531 | 269 | 8142 | 12738 | 2133 | 1235 | 29 | 2049 | 33364 | 0 | 0 | 0 | 88 | 170 | 34 | 0 | 0 | 0 | 0 | 0 | 30 | 322 | 563 | 93 | 0 | 0 | 0 | 0 | 0 |
| SRR4896698 | 16 | 6556 | 477 | 129 | 393 | 1980 | 3560 | 4429 | 8443 | 595 | 9989 | 12544 | 2501 | 1936 | 130 | 3500 | 27746 | 0 | 0 | 0 | 69 | 427 | 67 | 0 | 0 | 0 | 0 | 0 | 66 | 273 | 2369 | 351 | 0 | 0 | 0 | 0 | 0 |
| SRR4896701 | 65 | 17103 | 321 | 756 | 1668 | 5949 | 12612 | 20197 | 35550 | 594 | 7356 | 18 | 3554 | 2781 | 250 | 3123 | 25092 | 0 | 0 | 29 | 2072 | 1489 | 227 | 0 | 0 | 0 | 0 | 21 | 932 | 5342 | 3872 | 963 | 0 | 0 | 39 | 0 | 0 |
| SRR4896708 | 0 | 11540 | 384 | 153 | 3327 | 11734 | 37422 | 38665 | 11048 | 829 | 10110 | 7193 | 2565 | 4936 | 1048 | 2639 | 35136 | 0 | 0 | 23 | 434 | 2802 | 90 | 0 | 0 | 0 | 0 | 53 | 316 | 1526 | 8414 | 342 | 0 | 0 | 0 | 0 | 0 |
| SRR5022482 | 15 | 5113 | 12548 | 30 | 114 | 728 | 3879 | 1278 | 4329 | 170 | 2284 | 8731 | 1265 | 411 | 43 | 12 | 5200 | 0 | 0 | 0 | 384 | 132 | 61 | 0 | 0 | 0 | 0 | 17 | 111 | 504 | 375 | 131 | 0 | 0 | 0 | 0 | 0 |
| SRR5022483 | 258 | 29968 | 31398 | 156 | 78 | 935 | 1898 | 906 | 6027 | 53 | 2219 | 2984 | 711 | 510 | 14 | 65 | 8436 | 0 | 25 | 30 | 1728 | 1157 | 119 | 89 | 155 | 0 | 0 | 102 | 538 | 661 | 478 | 101 | 0 | 52 | 0 | 91 | 0 |
| SRR5022484 | 28 | 13348 | 41062 | 72 | 65 | 1323 | 2251 | 1845 | 6104 | 55 | 3710 | 6218 | 2201 | 590 | 44 | 18 | 9558 | 0 | 0 | 0 | 274 | 74 | 43 | 0 | 0 | 0 | 0 | 0 | 38 | 101 | 79 | 32 | 0 | 0 | 0 | 0 | 0 |
| SRR5022485 | 171 | 32420 | 21829 | 258 | 110 | 1328 | 2432 | 1454 | 4877 | 136 | 2218 | 2630 | 815 | 798 | 16 | 190 | 5907 | 0 | 0 | 12 | 1072 | 478 | 74 | 27 | 74 | 0 | 0 | 28 | 183 | 423 | 199 | 51 | 0 | 24 | 0 | 37 | 0 |
| SRR5029397 | 62 | 424 | 71663 | 30 | 3547 | 2294 | 38127 | 5216 | 9793 | 2550 | 17137 | 13 | 14919 | 19220 | 2901 | 3030 | 40262 | 0 | 0 | 0 | 14 | 107 | 49 | 16 | 0 | 18 | 0 | 0 | 0 | 60 | 19 | 0 | 0 | 0 | 0 | 37 | 0 |
| SRR5029398 | 49 | 332 | 55868 | 124 | 9254 | 10198 | 187178 | 13024 | 10540 | 1870 | 20607 | 8614 | 10677 | 10123 | 8380 | 1092 | 21987 | 0 | 0 | 0 | 0 | 0 | 0 | 0 | 0 | 0 | 0 | 0 | 0 | 18 | 0 | 0 | 0 | 0 | 0 | 19 | 0 |
| SRR5029399 | 528 | 3430 | 49655 | 218 | 3248 | 2011 | 29034 | 5575 | 4978 | 4383 | 9549 | 2740 | 4536 | 4302 | 654 | 1301 | 28305 | 0 | 0 | 0 | 48 | 502 | 348 | 24 | 11 | 15 | 0 | 27 | 102 | 196 | 182 | 41 | 0 | 0 | 0 | 31 | 0 |
| SRR5029400 | 38 | 234 | 49623 | 38 | 6733 | 4759 | 158016 | 18599 | 20130 | 3725 | 10120 | 1721 | 5311 | 20441 | 11568 | 1295 | 13221 | 0 | 0 | 0 | 53 | 123 | 88 | 11 | 0 | 15 | 0 | 22 | 47 | 297 | 37 | 0 | 0 | 0 | 13 | 55 | 0 |
| SRR5029401 | 69 | 422 | 48160 | 50 | 8949 | 7531 | 121825 | 13980 | 11859 | 4450 | 16170 | 4237 | 8571 | 11174 | 10813 | 1294 | 12482 | 0 | 0 | 0 | 0 | 0 | 0 | 0 | 0 | 15 | 0 | 0 | 0 | 53 | 0 | 0 | 0 | 0 | 0 | 12 | 0 |
| SRR5029402 | 15 | 186 | 8692 | 78 | 12193 | 6911 | 112809 | 11033 | 9996 | 1916 | 20926 | 4151 | 9343 | 9780 | 8646 | 1584 | 12171 | 0 | 0 | 0 | 0 | 18 | 14 | 0 | 0 | 0 | 0 | 0 | 0 | 116 | 0 | 0 | 0 | 0 | 0 | 0 | 0 |
| SRR5029403 | 15 | 236 | 49727 | 37 | 12539 | 10370 | 151469 | 29751 | 23316 | 5468 | 13079 | 4088 | 8162 | 20921 | 1152 | 1098 | 28236 | 0 | 0 | 0 | 0 | 36 | 26 | 0 | 0 | 0 | 0 | 0 | 0 | 22 | 12 | 0 | 0 | 0 | 0 | 25 | 0 |
| SRR5029404 | 12 | 146 | 35032 | 64 | 4578 | 3804 | 80033 | 12024 | 7706 | 1794 | 19007 | 5628 | 10027 | 11578 | 7445 | 1985 | 12076 | 0 | 0 | 0 | 0 | 0 | 0 | 0 | 0 | 0 | 0 | 0 | 0 | 17 | 0 | 0 | 0 | 0 | 0 | 0 | 0 |
| SRR5029405 | 43 | 348 | 18035 | 97 | 5456 | 4605 | 68631 | 6768 | 6269 | 1880 | 16455 | 4785 | 7236 | 6589 | 4469 | 1785 | 17014 | 0 | 0 | 0 | 0 | 0 | 0 | 0 | 0 | 0 | 0 | 0 | 0 | 93 | 0 | 0 | 0 | 0 | 0 | 0 | 0 |
| SRR5029406 | 75 | 1058 | 12332 | 150 | 12834 | 9946 | 145359 | 10823 | 11777 | 3111 | 22222 | 4116 | 9017 | 6694 | 5904 | 879 | 12982 | 0 | 0 | 0 | 37 | 15 | 0 | 0 | 0 | 0 | 0 | 16 | 21 | 574 | 13 | 0 | 0 | 0 | 0 | 17 | 0 |
| SRR5029407 | 12 | 133 | 66887 | 17 | 4260 | 3489 | 49213 | 14017 | 10104 | 4005 | 16001 | 6149 | 9337 | 16582 | 1218 | 2372 | 33490 | 0 | 0 | 0 | 0 | 0 | 16 | 0 | 0 | 0 | 0 | 0 | 0 | 0 | 0 | 0 | 0 | 0 | 0 | 13 | 0 |
| SRR5029408 | 27 | 149 | 77387 | 17 | 7452 | 6561 | 105991 | 12800 | 12000 | 2525 | 21104 | 8144 | 12434 | 9835 | 7371 | 1428 | 30225 | 0 | 0 | 0 | 0 | 0 | 0 | 0 | 0 | 0 | 0 | 0 | 0 | 0 | 0 | 0 | 0 | 0 | 0 | 23 | 0 |
| SRR5029409 | 17 | 156 | 24029 | 36 | 9246 | 6365 | 119888 | 20921 | 6240 | 4761 | 13715 | 4797 | 4986 | 22944 | 948 | 2018 | 14487 | 0 | 0 | 0 | 0 | 27 | 10 | 0 | 0 | 0 | 0 | 0 | 0 | 28 | 0 | 0 | 0 | 0 | 0 | 15 | 0 |
| SRR5029410 | 333 | 2452 | 28504 | 508 | 4758 | 2459 | 35313 | 8063 | 4707 | 3874 | 14111 | 11 | 6995 | 7621 | 1838 | 3643 | 23231 | 0 | 0 | 0 | 44 | 221 | 177 | 14 | 0 | 0 | 0 | 10 | 75 | 425 | 82 | 43 | 0 | 0 | 0 | 14 | 0 |
| SRR5029411 | 135 | 832 | 61075 | 99 | 16961 | 10193 | 178105 | 25252 | 39321 | 1135 | 21640 | 7762 | 10379 | 5055 | 1604 | 430 | 26996 | 0 | 0 | 0 | 0 | 0 | 0 | 0 | 0 | 0 | 0 | 0 | 0 | 34 | 0 | 0 | 0 | 0 | 0 | 0 | 0 |
| SRR5029412 | 68 | 941 | 45053 | 70 | 3234 | 1901 | 30428 | 8248 | 7569 | 3709 | 12538 | 4137 | 6429 | 9487 | 740 | 1491 | 42214 | 0 | 0 | 0 | 68 | 466 | 170 | 16 | 0 | 15 | 0 | 27 | 145 | 343 | 124 | 20 | 0 | 0 | 0 | 43 | 0 |
| SRR5029414 | 14 | 51 | 10645 | 147 | 7373 | 6587 | 126127 | 13137 | 4595 | 1596 | 15357 | 4936 | 6052 | 15571 | 2958 | 4956 | 9095 | 0 | 0 | 0 | 0 | 0 | 0 | 0 | 0 | 0 | 0 | 0 | 0 | 49 | 0 | 0 | 0 | 0 | 0 | 0 | 0 |
| SRR5029415 | 48 | 38 | 44890 | 287 | 6145 | 7591 | 124658 | 13576 | 7281 | 1839 | 15953 | 4989 | 7225 | 9161 | 9959 | 1475 | 19082 | 0 | 0 | 0 | 0 | 90 | 0 | 0 | 0 | 0 | 0 | 0 | 16 | 58 | 0 | 0 | 0 | 0 | 0 | 0 | 0 |
| SRR5029416 | 605 | 3686 | 46689 | 401 | 8631 | 6205 | 88315 | 12266 | 17482 | 4395 | 16212 | 6401 | 9292 | 7726 | 3914 | 1279 | 36528 | 0 | 0 | 0 | 42 | 787 | 531 | 62 | 33 | 55 | 0 | 12 | 60 | 133 | 210 | 45 | 0 | 0 | 30 | 105 | 0 |
| SRR5029417 | 18 | 92 | 71099 | 16 | 10531 | 9236 | 144834 | 24947 | 14179 | 5476 | 14920 | 4226 | 8352 | 11865 | 7114 | 1695 | 26184 | 0 | 0 | 0 | 0 | 12 | 0 | 0 | 0 | 11 | 0 | 0 | 0 | 11 | 0 | 0 | 0 | 0 | 0 | 22 | 0 |
| SRR5029418 | 93 | 1008 | 12794 | 308 | 4537 | 3942 | 60637 | 6703 | 6028 | 1059 | 13438 | 4236 | 7470 | 8904 | 6258 | 3647 | 18388 | 0 | 0 | 0 | 28 | 40 | 27 | 0 | 0 | 0 | 0 | 17 | 21 | 739 | 0 | 0 | 0 | 0 | 0 | 23 | 0 |
| SRR5029419 | 42 | 339 | 33804 | 134 | 8407 | 6572 | 113946 | 11361 | 15711 | 1418 | 20371 | 8705 | 11898 | 8831 | 4847 | 1012 | 24618 | 0 | 0 | 0 | 0 | 36 | 20 | 0 | 0 | 0 | 0 | 0 | 0 | 55 | 0 | 0 | 0 | 0 | 0 | 16 | 0 |
| SRR5029420 | 49 | 505 | 29552 | 97 | 26709 | 25332 | 348713 | 49743 | 6277 | 6846 | 22284 | 5591 | 9377 | 52996 | 1316 | 1211 | 19168 | 0 | 0 | 0 | 13 | 187 | 170 | 14 | 10 | 0 | 0 | 0 | 15 | 112 | 67 | 14 | 0 | 0 | 0 | 26 | 0 |
| SRR5029422 | 40 | 1707 | 57901 | 67 | 14124 | 12128 | 156902 | 30507 | 19922 | 6713 | 18885 | 7203 | 9621 | 6794 | 4319 | 1036 | 26405 | 0 | 0 | 0 | 0 | 0 | 0 | 0 | 0 | 0 | 0 | 0 | 0 | 13 | 0 | 0 | 0 | 0 | 0 | 15 | 0 |
| SRR5029424 | 27 | 144 | 30657 | 0 | 607 | 446 | 8419 | 2674 | 2094 | 419 | 29181 | 5934 | 17013 | 30539 | 1454 | 24 | 6850 | 0 | 0 | 0 | 0 | 0 | 0 | 0 | 0 | 0 | 0 | 0 | 0 | 0 | 0 | 0 | 0 | 0 | 0 | 0 | 0 |
| SRR5029425 | 25 | 189 | 68338 | 14 | 6076 | 6548 | 113098 | 13006 | 12679 | 3466 | 16493 | 4896 | 7955 | 10405 | 10358 | 2011 | 16166 | 0 | 0 | 0 | 0 | 11 | 0 | 0 | 0 | 0 | 0 | 0 | 0 | 23 | 0 | 0 | 0 | 0 | 0 | 31 | 0 |
| SRR5029426 | 40 | 243 | 52333 | 26 | 6105 | 3516 | 58425 | 14212 | 9855 | 2952 | 17441 | 14 | 11098 | 15432 | 945 | 2475 | 34649 | 0 | 0 | 0 | 0 | 17 | 27 | 0 | 0 | 0 | 0 | 0 | 0 | 0 | 0 | 0 | 0 | 0 | 0 | 33 | 0 |
| SRR5029427 | 130 | 1322 | 33125 | 123 | 6530 | 6454 | 91860 | 12645 | 9687 | 3629 | 13956 | 4507 | 8471 | 13241 | 5415 | 2828 | 18538 | 0 | 0 | 0 | 38 | 71 | 40 | 20 | 18 | 24 | 0 | 14 | 42 | 474 | 26 | 0 | 0 | 0 | 0 | 42 | 0 |
| SRR5029428 | 182 | 786 | 10597 | 0 | 656 | 356 | 5902 | 3517 | 1640 | 218 | 31683 | 7824 | 24068 | 30417 | 1244 | 11 | 6169 | 0 | 0 | 0 | 17 | 34 | 14 | 0 | 0 | 0 | 0 | 0 | 12 | 69 | 13 | 0 | 0 | 0 | 0 | 0 | 0 |
| SRR5029429 | 57 | 760 | 59653 | 57 | 3382 | 2085 | 33184 | 8893 | 5866 | 5984 | 8700 | 2409 | 4399 | 4028 | 535 | 1144 | 25802 | 0 | 0 | 0 | 12 | 94 | 71 | 0 | 0 | 0 | 0 | 0 | 17 | 60 | 36 | 0 | 0 | 0 | 0 | 22 | 0 |
| SRR5029430 | 148 | 1442 | 64526 | 90 | 13163 | 8359 | 176561 | 24296 | 7151 | 5032 | 12847 | 4771 | 7073 | 12903 | 3217 | 1228 | 29617 | 0 | 0 | 0 | 14 | 180 | 144 | 20 | 0 | 18 | 0 | 0 | 0 | 32 | 26 | 0 | 0 | 0 | 0 | 40 | 0 |
| SRR5029431 | 26 | 90 | 29363 | 104 | 3991 | 3794 | 66926 | 7144 | 6704 | 1147 | 17298 | 8142 | 9842 | 10592 | 3920 | 2482 | 18778 | 0 | 0 | 0 | 0 | 0 | 0 | 0 | 0 | 0 | 0 | 0 | 0 | 18 | 0 | 0 | 0 | 0 | 0 | 0 | 0 |
| SRR5029432 | 66 | 464 | 64128 | 70 | 4269 | 3669 | 47584 | 9791 | 14441 | 4865 | 14398 | 6632 | 10590 | 12171 | 1149 | 1643 | 54787 | 0 | 0 | 0 | 31 | 848 | 924 | 134 | 53 | 42 | 0 | 0 | 28 | 74 | 175 | 57 | 0 | 0 | 39 | 166 | 0 |
| SRR5029433 | 0 | 65 | 28907 | 19 | 6785 | 6151 | 87989 | 15455 | 23928 | 2616 | 22561 | 5607 | 10098 | 10531 | 1814 | 1355 | 18052 | 0 | 0 | 0 | 0 | 0 | 0 | 0 | 0 | 0 | 0 | 0 | 0 | 0 | 0 | 0 | 0 | 0 | 0 | 0 | 0 |
| SRR5029434 | 18 | 171 | 7235 | 58 | 5683 | 4395 | 71622 | 7516 | 17743 | 1691 | 16716 | 4305 | 8625 | 9307 | 3791 | 1311 | 7139 | 0 | 0 | 0 | 0 | 12 | 0 | 0 | 0 | 0 | 0 | 0 | 0 | 153 | 11 | 0 | 0 | 0 | 0 | 0 | 0 |
| SRR5029435 | 97 | 631 | 39679 | 75 | 5211 | 3150 | 56046 | 10594 | 7238 | 3914 | 14212 | 6686 | 7399 | 10938 | 1046 | 1761 | 30186 | 0 | 0 | 0 | 15 | 157 | 143 | 12 | 0 | 12 | 0 | 0 | 10 | 87 | 45 | 0 | 0 | 0 | 0 | 45 | 0 |
| SRR5029436 | 123 | 1187 | 37259 | 74 | 15681 | 13095 | 241386 | 33305 | 14372 | 4821 | 13722 | 5193 | 6978 | 9315 | 5150 | 1290 | 20109 | 0 | 0 | 0 | 64 | 336 | 283 | 42 | 27 | 27 | 0 | 16 | 13 | 273 | 73 | 16 | 0 | 0 | 11 | 102 | 0 |
| SRR5029437 | 0 | 91 | 87875 | 0 | 565 | 331 | 25864 | 5423 | 2502 | 36 | 1993 | 444 | 2489 | 50756 | 2266 | 88 | 2407 | 0 | 0 | 0 | 0 | 0 | 0 | 0 | 0 | 0 | 0 | 0 | 0 | 16 | 0 | 0 | 0 | 0 | 0 | 0 | 0 |
| SRR5029438 | 19 | 312 | 34828 | 36 | 11313 | 8622 | 129888 | 26833 | 25950 | 2984 | 19362 | 7306 | 11002 | 15568 | 2299 | 1197 | 29416 | 0 | 0 | 0 | 0 | 187 | 170 | 37 | 21 | 12 | 0 | 0 | 0 | 59 | 59 | 19 | 0 | 0 | 14 | 72 | 0 |
| SRR5029439 | 27 | 195 | 56435 | 27 | 15943 | 11413 | 196217 | 27990 | 17805 | 2639 | 17384 | 4977 | 7692 | 6523 | 2843 | 734 | 31561 | 0 | 0 | 0 | 65 | 152 | 124 | 15 | 0 | 0 | 0 | 12 | 28 | 104 | 23 | 0 | 0 | 0 | 0 | 32 | 0 |
| SRR5029440 | 0 | 135 | 20411 | 0 | 2445 | 1789 | 28590 | 3904 | 2221 | 68 | 17941 | 2344 | 14621 | 30200 | 1450 | 47 | 5348 | 0 | 0 | 0 | 0 | 0 | 0 | 0 | 0 | 0 | 0 | 0 | 14 | 16 | 0 | 0 | 0 | 0 | 0 | 0 | 0 |
| SRR5029441 | 36 | 257 | 49955 | 11 | 7740 | 5681 | 87728 | 13788 | 20680 | 6145 | 19014 | 5399 | 10005 | 15473 | 2877 | 2351 | 20146 | 0 | 0 | 0 | 0 | 0 | 0 | 0 | 0 | 0 | 0 | 0 | 0 | 32 | 0 | 0 | 0 | 0 | 0 | 24 | 0 |
| SRR5029442 | 23 | 244 | 36969 | 23 | 19768 | 17322 | 221178 | 37310 | 16581 | 4572 | 18385 | 4559 | 8497 | 6933 | 5271 | 755 | 19516 | 0 | 0 | 0 | 0 | 26 | 22 | 0 | 0 | 0 | 0 | 0 | 0 | 19 | 0 | 0 | 0 | 0 | 0 | 29 | 0 |
| SRR5029444 | 31 | 124 | 43978 | 93 | 5975 | 4659 | 190000 | 27082 | 14294 | 4035 | 8156 | 1821 | 5570 | 44729 | 2064 | 1271 | 20557 | 0 | 0 | 0 | 68 | 367 | 252 | 22 | 14 | 14 | 0 | 0 | 43 | 432 | 68 | 11 | 0 | 0 | 0 | 108 | 0 |
| SRR5029446 | 40 | 160 | 21413 | 0 | 838 | 645 | 11021 | 3006 | 1859 | 258 | 32622 | 8131 | 21611 | 27211 | 1486 | 28 | 8230 | 0 | 0 | 0 | 0 | 0 | 0 | 0 | 0 | 0 | 0 | 0 | 0 | 0 | 0 | 0 | 0 | 0 | 0 | 0 | 0 |
| SRR5029447 | 29 | 154 | 59725 | 16 | 7771 | 6800 | 100412 | 17617 | 11948 | 4756 | 16313 | 4775 | 8012 | 8187 | 4504 | 1849 | 29121 | 0 | 0 | 0 | 0 | 24 | 21 | 0 | 0 | 13 | 0 | 0 | 0 | 21 | 0 | 0 | 0 | 0 | 0 | 31 | 0 |
| SRR5029448 | 45 | 343 | 69821 | 24 | 5265 | 3372 | 57515 | 9484 | 6242 | 3125 | 11697 | 5259 | 6179 | 7228 | 1619 | 1601 | 29329 | 0 | 0 | 0 | 0 | 49 | 72 | 0 | 0 | 15 | 0 | 0 | 0 | 0 | 11 | 0 | 0 | 0 | 0 | 24 | 0 |
| SRR5029452 | 0 | 143 | 229892 | 0 | 312 | 127 | 20608 | 2938 | 2175 | 106 | 973 | 309 | 1722 | 42681 | 2263 | 39 | 2158 | 0 | 0 | 0 | 0 | 0 | 0 | 0 | 0 | 0 | 0 | 0 | 0 | 0 | 0 | 0 | 0 | 0 | 0 | 0 | 0 |
| SRR5029455 | 22 | 541 | 46688 | 41 | 18853 | 18470 | 201930 | 30839 | 18488 | 4011 | 15233 | 6337 | 8593 | 8897 | 5496 | 686 | 31808 | 0 | 0 | 0 | 15 | 324 | 198 | 42 | 18 | 16 | 0 | 0 | 0 | 11 | 20 | 0 | 0 | 0 | 0 | 49 | 0 |
| SRR5029456 | 26 | 149 | 44738 | 16 | 8982 | 6977 | 143338 | 25730 | 14735 | 5731 | 17401 | 4374 | 15309 | 25044 | 4074 | 2406 | 26587 | 0 | 0 | 0 | 0 | 24 | 45 | 0 | 0 | 0 | 0 | 0 | 0 | 49 | 0 | 0 | 0 | 0 | 0 | 33 | 0 |

### Supp Table S8.1.Columns_B_to_AL

| | | | | | | | | | | | | | | | | | | | | | | | | | | | | | | | | | | | | | |
|---|---|---|---|---|---|---|---|---|---|---|---|---|---|---|---|---|---|---|---|---|---|---|---|---|---|---|---|---|---|---|---|---|---|---|---|---|---|
| SRR5029457 | 37 | 288 | 77841 | 24 | 10971 | 5934 | 119010 | 35331 | 8054 | 5092 | 16853 | 10033 | 11362 | 26613 | 580 | 1582 | 42367 | 0 | 0 | 0 | 0 | 17 | 26 | 0 | 0 | 18 | 0 | 0 | 0 | 0 | 0 | 0 | 0 | 0 | 0 | 26 | 0 |
| SRR5029459 | 109 | 1266 | 38469 | 110 | 16116 | 14334 | 235200 | 35032 | 18800 | 8984 | 16476 | 4473 | 5047 | 13957 | 7994 | 1561 | 13819 | 0 | 0 | 0 | 20 | 319 | 291 | 37 | 13 | 22 | 0 | 0 | 19 | 90 | 65 | 16 | 0 | 0 | 0 | 80 | 0 |
| SRR5029460 | 13 | 129 | 39368 | 18 | 10621 | 7175 | 126540 | 24835 | 17417 | 5351 | 19287 | 13 | 9871 | 24088 | 775 | 2120 | 35786 | 0 | 0 | 0 | 24 | 30 | 19 | 0 | 0 | 16 | 0 | 0 | 20 | 100 | 0 | 0 | 0 | 0 | 0 | 23 | 0 |
| SRR5029462 | 579 | 3656 | 45270 | 282 | 7059 | 4488 | 78988 | 15392 | 7166 | 4474 | 16001 | 7382 | 8402 | 11171 | 1319 | 1825 | 30404 | 0 | 0 | 0 | 0 | 38 | 40 | 11 | 0 | 21 | 0 | 10 | 31 | 43 | 27 | 0 | 0 | 0 | 0 | 24 | 0 |
| SRR5029463 | 18 | 211 | 48183 | 35 | 9656 | 9196 | 169359 | 26296 | 15735 | 4437 | 13110 | 7436 | 6173 | 28520 | 351 | 1889 | 22473 | 0 | 0 | 0 | 0 | 58 | 42 | 0 | 0 | 13 | 0 | 0 | 0 | 25 | 15 | 0 | 0 | 0 | 0 | 31 | 0 |
| SRR5029465 | 31 | 1276 | 10544 | 302 | 371 | 169 | 4555 | 2688 | 1387 | 259 | 16 | 19 | 19413 | 29009 | 1083 | 21 | 9904 | 0 | 0 | 0 | 25 | 58 | 0 | 0 | 0 | 0 | 0 | 0 | 80 | 129 | 19 | 0 | 0 | 0 | 0 | 0 | 0 |
| SRR5029466 | 107 | 1125 | 37798 | 130 | 11201 | 8425 | 136254 | 21467 | 14624 | 6104 | 17849 | 4377 | 9009 | 11862 | 2610 | 1959 | 26728 | 0 | 0 | 0 | 16 | 55 | 19 | 0 | 0 | 0 | 0 | 16 | 65 | 145 | 24 | 0 | 0 | 0 | 0 | 26 | 0 |
| SRR5029467 | 12 | 132 | 48397 | 12 | 8069 | 7013 | 124455 | 15808 | 13186 | 2585 | 18444 | 4740 | 9009 | 10468 | 4538 | 1743 | 15306 | 0 | 0 | 0 | 0 | 0 | 0 | 0 | 0 | 0 | 0 | 0 | 0 | 18 | 0 | 0 | 0 | 0 | 0 | 15 | 0 |
| SRR5029469 | 17 | 105 | 31286 | 20 | 7531 | 7501 | 114469 | 20127 | 16546 | 5159 | 11 | 5752 | 9128 | 10058 | 8400 | 1189 | 17335 | 0 | 0 | 0 | 0 | 37 | 26 | 0 | 0 | 0 | 0 | 0 | 0 | 21 | 12 | 0 | 0 | 0 | 0 | 45 | 0 |
| SRR5029471 | 27 | 358 | 57556 | 55 | 11777 | 6447 | 172917 | 25973 | 9890 | 7453 | 16703 | 4927 | 9212 | 13061 | 4809 | 1388 | 26263 | 0 | 0 | 0 | 58 | 199 | 313 | 48 | 21 | 19 | 0 | 0 | 21 | 110 | 21 | 0 | 0 | 0 | 0 | 98 | 0 |
| SRR5029472 | 62 | 838 | 20983 | 101 | 5395 | 5779 | 80373 | 9024 | 6870 | 4374 | 17702 | 5981 | 7896 | 11196 | 7868 | 2724 | 14441 | 0 | 0 | 0 | 16 | 109 | 54 | 0 | 0 | 10 | 0 | 0 | 21 | 150 | 33 | 0 | 0 | 0 | 0 | 54 | 0 |
| SRR5029473 | 161 | 1433 | 46180 | 47 | 24546 | 17464 | 263058 | 26895 | 12721 | 6745 | 19271 | 4889 | 7891 | 10742 | 10284 | 720 | 15577 | 0 | 0 | 0 | 23 | 57 | 56 | 19 | 13 | 19 | 0 | 20 | 32 | 202 | 33 | 0 | 0 | 0 | 0 | 76 | 0 |
| SRR5029474 | 135 | 1292 | 54118 | 90 | 15202 | 13979 | 198256 | 32280 | 27101 | 5213 | 25747 | 8565 | 12760 | 18485 | 7441 | 1110 | 19914 | 0 | 0 | 0 | 26 | 152 | 174 | 18 | 0 | 0 | 0 | 0 | 21 | 99 | 53 | 20 | 0 | 0 | 0 | 38 | 0 |
| SRR5029476 | 152 | 1510 | 56368 | 58 | 11861 | 7923 | 151582 | 25781 | 10657 | 5620 | 13013 | 3620 | 5980 | 11448 | 2415 | 1401 | 23971 | 0 | 0 | 15 | 269 | 1112 | 1156 | 156 | 57 | 35 | 0 | 18 | 169 | 976 | 242 | 92 | 0 | 0 | 24 | 198 | 0 |
| SRR5029477 | 23 | 71 | 55970 | 20 | 5391 | 6288 | 127357 | 16377 | 14063 | 4796 | 17163 | 5560 | 9341 | 10685 | 4361 | 1115 | 26680 | 0 | 0 | 0 | 0 | 39 | 31 | 0 | 0 | 0 | 0 | 0 | 0 | 61 | 15 | 0 | 0 | 0 | 0 | 23 | 0 |
| SRR5029478 | 56 | 390 | 36248 | 55 | 12712 | 9774 | 193134 | 16137 | 9433 | 3274 | 19882 | 5744 | 9480 | 13087 | 9516 | 1115 | 16803 | 0 | 0 | 0 | 10 | 34 | 16 | 0 | 0 | 12 | 0 | 0 | 0 | 83 | 0 | 0 | 0 | 0 | 0 | 27 | 0 |
| SRR5029480 | 64 | 837 | 66306 | 48 | 5122 | 4169 | 56214 | 8542 | 10754 | 4908 | 11996 | 4105 | 7114 | 5016 | 1757 | 1094 | 29981 | 0 | 0 | 0 | 0 | 76 | 82 | 15 | 0 | 13 | 0 | 0 | 0 | 14 | 17 | 0 | 0 | 0 | 0 | 27 | 0 |
| SRR5029481 | 16 | 263 | 23053 | 36 | 9696 | 7997 | 110276 | 17796 | 19207 | 3545 | 25745 | 7336 | 9552 | 7229 | 3862 | 858 | 13755 | 0 | 0 | 0 | 0 | 0 | 0 | 0 | 0 | 0 | 0 | 0 | 0 | 18 | 0 | 0 | 0 | 0 | 0 | 14 | 0 |
| SRR5029482 | 0 | 73 | 30684 | 0 | 2997 | 2142 | 102549 | 21012 | 7781 | 5632 | 5249 | 980 | 2660 | 14295 | 7144 | 2105 | 8656 | 0 | 0 | 0 | 44 | 140 | 69 | 0 | 0 | 11 | 0 | 0 | 37 | 225 | 25 | 0 | 0 | 0 | 0 | 53 | 0 |
| SRR5029483 | 11 | 95 | 18636 | 25 | 4205 | 3510 | 57634 | 9350 | 7268 | 3159 | 20720 | 7325 | 10960 | 18033 | 2872 | 4365 | 19472 | 0 | 0 | 0 | 21 | 61 | 41 | 0 | 0 | 0 | 0 | 0 | 0 | 214 | 17 | 0 | 0 | 0 | 0 | 33 | 0 |
| SRR5029484 | 644 | 4066 | 63818 | 142 | 28652 | 17185 | 333670 | 51400 | 12530 | 10398 | 18061 | 5036 | 9564 | 11641 | 12307 | 550 | 21866 | 0 | 14 | 0 | 37 | 205 | 141 | 24 | 19 | 21 | 14 | 60 | 77 | 130 | 102 | 18 | 0 | 0 | 0 | 60 | 0 |
| SRR5029486 | 16 | 215 | 8954 | 92 | 26029 | 13660 | 282501 | 37915 | 4301 | 4216 | 17541 | 8004 | 8281 | 17716 | 12365 | 714 | 14734 | 0 | 0 | 0 | 51 | 322 | 324 | 58 | 30 | 30 | 0 | 12 | 16 | 102 | 52 | 17 | 0 | 0 | 14 | 113 | 0 |
| SRR5029487 | 22 | 115 | 60540 | 12 | 7914 | 5957 | 90745 | 16082 | 21465 | 4910 | 18541 | 5742 | 8999 | 8588 | 2374 | 1400 | 28989 | 0 | 0 | 0 | 0 | 0 | 0 | 0 | 0 | 0 | 0 | 0 | 0 | 19 | 0 | 0 | 0 | 0 | 0 | 0 | 0 |
| SRR5029488 | 147 | 338 | 63828 | 235 | 18990 | 18004 | 255471 | 26210 | 11978 | 5004 | 16602 | 6126 | 7494 | 16709 | 7332 | 1513 | 16698 | 0 | 0 | 0 | 17 | 34 | 40 | 14 | 0 | 0 | 0 | 0 | 0 | 79 | 26 | 0 | 0 | 0 | 0 | 40 | 0 |
| SRR5029489 | 122 | 1416 | 47250 | 70 | 20125 | 11505 | 162533 | 29704 | 31879 | 5782 | 16905 | 3743 | 7831 | 5694 | 2843 | 973 | 25487 | 0 | 0 | 0 | 0 | 46 | 16 | 0 | 0 | 0 | 0 | 14 | 27 | 58 | 30 | 0 | 0 | 0 | 0 | 17 | 0 |
| SRR5029490 | 96 | 916 | 42784 | 92 | 14288 | 11243 | 234141 | 29443 | 30056 | 3679 | 21207 | 5680 | 9789 | 9914 | 6459 | 928 | 19726 | 0 | 24 | 0 | 92 | 442 | 229 | 26 | 23 | 18 | 0 | 59 | 52 | 244 | 103 | 13 | 0 | 0 | 13 | 66 | 0 |
| SRR5029491 | 35 | 308 | 34704 | 77 | 13791 | 10762 | 201173 | 34199 | 20452 | 5886 | 20205 | 5263 | 9261 | 13311 | 5608 | 1460 | 20857 | 0 | 0 | 0 | 0 | 85 | 91 | 15 | 0 | 17 | 0 | 0 | 0 | 47 | 20 | 0 | 0 | 0 | 0 | 71 | 0 |
| SRR5029492 | 28 | 285 | 41448 | 39 | 12577 | 12676 | 244881 | 38655 | 10305 | 5263 | 14483 | 4479 | 7972 | 36761 | 1505 | 1423 | 23900 | 0 | 0 | 0 | 68 | 357 | 265 | 28 | 11 | 19 | 0 | 17 | 35 | 268 | 87 | 23 | 0 | 0 | 0 | 70 | 0 |
| SRR5029493 | 30 | 170 | 48366 | 26 | 15994 | 8912 | 156169 | 25121 | 23665 | 7191 | 24415 | 5635 | 12320 | 12296 | 6027 | 1236 | 24766 | 0 | 0 | 0 | 30 | 272 | 222 | 65 | 35 | 39 | 0 | 0 | 10 | 67 | 40 | 10 | 0 | 0 | 12 | 141 | 0 |
| SRR5029494 | 32 | 288 | 42670 | 32 | 15608 | 12816 | 224482 | 32343 | 14191 | 10594 | 17919 | 3854 | 7057 | 15907 | 8978 | 915 | 19014 | 0 | 0 | 0 | 164 | 270 | 232 | 37 | 12 | 12 | 0 | 18 | 60 | 466 | 69 | 27 | 0 | 0 | 0 | 59 | 0 |
| SRR5029495 | 95 | 1278 | 28360 | 114 | 10139 | 8845 | 139713 | 20050 | 18612 | 4934 | 16669 | 3804 | 7626 | 7239 | 4194 | 1254 | 16889 | 0 | 0 | 0 | 39 | 176 | 111 | 10 | 0 | 10 | 0 | 13 | 38 | 301 | 64 | 11 | 0 | 0 | 0 | 50 | 0 |
| SRR5029496 | 18 | 380 | 61157 | 46 | 12603 | 10625 | 144274 | 18596 | 15209 | 3710 | 18820 | 10 | 8668 | 5383 | 4272 | 853 | 28684 | 0 | 0 | 0 | 0 | 24 | 15 | 0 | 0 | 0 | 0 | 0 | 0 | 12 | 0 | 0 | 0 | 0 | 0 | 16 | 0 |
| SRR5029497 | 27 | 261 | 31146 | 36 | 8375 | 8015 | 135949 | 28390 | 12770 | 5698 | 13815 | 4092 | 6674 | 16001 | 4438 | 2119 | 17461 | 0 | 0 | 0 | 11 | 66 | 39 | 0 | 0 | 11 | 0 | 0 | 23 | 103 | 25 | 0 | 0 | 0 | 0 | 39 | 0 |
| SRR5029498 | 0 | 60 | 12985 | 38 | 8492 | 8397 | 152320 | 16954 | 10030 | 4050 | 26093 | 6427 | 11013 | 15786 | 6795 | 571 | 10161 | 0 | 0 | 0 | 0 | 12 | 13 | 0 | 0 | 0 | 0 | 0 | 0 | 15 | 0 | 0 | 0 | 0 | 0 | 26 | 0 |
| SRR5029499 | 27 | 209 | 38633 | 16 | 9270 | 7444 | 151067 | 23650 | 18663 | 6419 | 19163 | 4922 | 9117 | 17719 | 7943 | 1429 | 18614 | 0 | 0 | 0 | 80 | 400 | 455 | 110 | 46 | 23 | 0 | 0 | 11 | 272 | 92 | 39 | 0 | 0 | 17 | 121 | 0 |
| SRR5029500 | 32 | 148 | 30447 | 138 | 14139 | 14624 | 209690 | 18505 | 8469 | 3197 | 19823 | 5447 | 8605 | 23735 | 11516 | 1834 | 16549 | 0 | 0 | 0 | 16 | 182 | 115 | 27 | 34 | 72 | 0 | 0 | 0 | 38 | 0 | 0 | 0 | 0 | 153 | 636 | 29 |
| SRR5029501 | 20 | 129 | 73072 | 0 | 6884 | 6145 | 122074 | 18075 | 37965 | 5113 | 15819 | 3607 | 7314 | 7862 | 6217 | 1124 | 16490 | 0 | 0 | 0 | 26 | 48 | 37 | 0 | 0 | 0 | 0 | 0 | 19 | 61 | 13 | 0 | 0 | 0 | 0 | 26 | 0 |
| SRR5029502 | 67 | 634 | 38853 | 145 | 18504 | 27694 | 278905 | 31428 | 8462 | 5358 | 14622 | 3414 | 5879 | 17872 | 20948 | 1076 | 17534 | 0 | 0 | 0 | 94 | 2064 | 1542 | 223 | 131 | 104 | 0 | 14 | 42 | 244 | 109 | 58 | 0 | 0 | 33 | 409 | 14 |
| SRR5029503 | 11 | 116 | 39573 | 23 | 15101 | 9061 | 204536 | 37910 | 4047 | 8724 | 13008 | 3728 | 8156 | 12227 | 5768 | 383 | 23155 | 0 | 0 | 0 | 25 | 116 | 79 | 14 | 0 | 29 | 0 | 0 | 14 | 58 | 16 | 0 | 0 | 0 | 0 | 65 | 0 |
| SRR5029504 | 13 | 102 | 25957 | 20 | 10440 | 7298 | 128336 | 14418 | 28466 | 4137 | 18590 | 6053 | 8931 | 10707 | 6119 | 1734 | 13111 | 0 | 0 | 0 | 0 | 13 | 0 | 0 | 0 | 0 | 0 | 0 | 0 | 17 | 0 | 0 | 0 | 0 | 0 | 19 | 0 |
| SRR5029505 | 20 | 205 | 53970 | 21 | 10423 | 7963 | 110439 | 23870 | 19016 | 4919 | 19868 | 4511 | 8789 | 7801 | 4660 | 931 | 24796 | 0 | 0 | 0 | 0 | 13 | 11 | 0 | 0 | 0 | 0 | 0 | 0 | 17 | 0 | 0 | 0 | 0 | 0 | 23 | 0 |
| SRR5029506 | 507 | 3989 | 56285 | 198 | 22033 | 16184 | 292387 | 23876 | 58259 | 12092 | 21626 | 3762 | 8410 | 5469 | 23399 | 356 | 11196 | 0 | 12 | 0 | 48 | 243 | 239 | 39 | 30 | 26 | 0 | 38 | 64 | 110 | 79 | 24 | 0 | 0 | 12 | 85 | 0 |
| SRR5029507 | 74 | 776 | 34545 | 85 | 19304 | 16990 | 268369 | 27945 | 32156 | 3141 | 20855 | 6328 | 9501 | 6158 | 5078 | 391 | 14542 | 0 | 0 | 0 | 0 | 64 | 55 | 14 | 22 | 26 | 0 | 0 | 0 | 53 | 21 | 0 | 0 | 0 | 16 | 168 | 0 |
| SRR5029508 | 38 | 189 | 16835 | 27 | 9540 | 7319 | 146412 | 15901 | 13356 | 5230 | 19153 | 4032 | 7638 | 11064 | 6061 | 1352 | 7940 | 0 | 0 | 0 | 0 | 33 | 16 | 0 | 0 | 0 | 0 | 0 | 0 | 74 | 14 | 0 | 0 | 0 | 0 | 13 | 0 |
| SRR5029509 | 0 | 54 | 8674 | 44 | 17037 | 12032 | 249736 | 33259 | 23391 | 3448 | 17379 | 3944 | 10124 | 11912 | 11931 | 430 | 17560 | 0 | 0 | 0 | 34 | 396 | 238 | 26 | 21 | 30 | 0 | 26 | 17 | 73 | 26 | 0 | 0 | 0 | 0 | 119 | 0 |
| SRR5029510 | 32 | 451 | 51242 | 39 | 8585 | 8899 | 165306 | 28727 | 16584 | 5134 | 20081 | 4681 | 9685 | 10450 | 10844 | 896 | 15736 | 0 | 0 | 0 | 0 | 79 | 61 | 13 | 12 | 14 | 0 | 0 | 10 | 36 | 18 | 0 | 0 | 0 | 0 | 52 | 0 |
| SRR5029511 | 62 | 938 | 42155 | 65 | 17346 | 12253 | 137295 | 31590 | 11310 | 8574 | 17858 | 3862 | 8578 | 5706 | 1978 | 817 | 27279 | 0 | 0 | 0 | 10 | 27 | 17 | 0 | 0 | 0 | 0 | 11 | 31 | 62 | 21 | 0 | 0 | 0 | 0 | 30 | 0 |
| SRR5029512 | 127 | 1224 | 46190 | 104 | 9240 | 9834 | 142881 | 28025 | 12281 | 5866 | 16073 | 4404 | 7341 | 8064 | 9950 | 1464 | 18998 | 0 | 0 | 0 | 23 | 101 | 44 | 16 | 0 | 18 | 0 | 17 | 35 | 137 | 52 | 0 | 0 | 0 | 0 | 65 | 0 |
| SRR5029513 | 0 | 72 | 28961 | 12 | 22108 | 11217 | 277068 | 30756 | 14464 | 4755 | 17510 | 2607 | 6011 | 8774 | 6328 | 985 | 13351 | 0 | 0 | 0 | 49 | 86 | 150 | 28 | 26 | 60 | 0 | 0 | 13 | 165 | 20 | 12 | 0 | 0 | 22 | 202 | 0 |
| SRR5029514 | 11 | 78 | 32757 | 0 | 16116 | 12104 | 213764 | 40560 | 6404 | 5515 | 10429 | 3788 | 4510 | 12161 | 5788 | 970 | 16364 | 0 | 0 | 0 | 0 | 35 | 25 | 0 | 0 | 0 | 0 | 0 | 0 | 0 | 0 | 0 | 0 | 0 | 0 | 30 | 0 |
| SRR5029515 | 12 | 89 | 43578 | 14 | 19320 | 18277 | 331906 | 31355 | 10033 | 3662 | 18755 | 3895 | 7273 | 9020 | 11730 | 765 | 15189 | 0 | 0 | 0 | 0 | 71 | 51 | 0 | 11 | 15 | 0 | 0 | 0 | 39 | 0 | 0 | 0 | 0 | 0 | 50 | 0 |
| SRR5029516 | 356 | 3242 | 42616 | 254 | 13044 | 10371 | 125728 | 26583 | 57825 | 7467 | 19031 | 8077 | 10167 | 3092 | 1373 | 490 | 27908 | 0 | 0 | 0 | 21 | 74 | 42 | 0 | 0 | 0 | 0 | 26 | 52 | 58 | 73 | 0 | 0 | 0 | 0 | 17 | 0 |
| SRR5029870 | 874 | 76032 | 1040 | 638 | 195 | 4549 | 1713 | 4345 | 4762 | 83 | 39983 | 20160 | 4615 | 3080 | 10 | 0 | 39404 | 0 | 0 | 0 | 1036 | 447 | 82 | 0 | 0 | 0 | 0 | 0 | 92 | 1700 | 885 | 203 | 0 | 29 | 10 | 0 | 0 |
| SRR5029871 | 397 | 57637 | 2665 | 227 | 149 | 3971 | 1538 | 4384 | 5075 | 19 | 41063 | 21321 | 4718 | 3238 | 0 | 0 | 45172 | 0 | 0 | 0 | 531 | 242 | 52 | 0 | 0 | 0 | 0 | 0 | 43 | 893 | 545 | 110 | 0 | 17 | 0 | 0 | 0 |
| SRR5029872 | 1072 | 76900 | 934 | 613 | 201 | 4594 | 1726 | 4447 | 4961 | 46 | 40964 | 20172 | 4723 | 3045 | 0 | 0 | 37563 | 0 | 0 | 0 | 1082 | 447 | 76 | 0 | 0 | 0 | 0 | 0 | 78 | 1685 | 858 | 171 | 0 | 22 | 10 | 0 | 0 |
| SRR5029873 | 275 | 55890 | 2597 | 216 | 179 | 4068 | 1592 | 4624 | 5134 | 79 | 13 | 20973 | 4934 | 3304 | 0 | 0 | 39321 | 0 | 0 | 0 | 512 | 245 | 50 | 0 | 0 | 0 | 0 | 0 | 45 | 928 | 485 | 113 | 0 | 22 | 12 | 0 | 0 |
| SRR5029874 | 1024 | 78413 | 853 | 583 | 195 | 4493 | 1700 | 4236 | 4866 | 100 | 14 | 19922 | 4583 | 3087 | 11 | 0 | 39737 | 0 | 0 | 0 | 1039 | 457 | 83 | 0 | 0 | 0 | 0 | 0 | 101 | 1685 | 893 | 191 | 0 | 37 | 14 | 0 | 0 |
| SRR5029875 | 616 | 58142 | 2638 | 289 | 161 | 3907 | 1543 | 4359 | 5096 | 39 | 14 | 21025 | 4649 | 3134 | 0 | 0 | 45893 | 0 | 0 | 0 | 532 | 263 | 59 | 0 | 0 | 0 | 0 | 0 | 45 | 844 | 486 | 117 | 0 | 18 | 11 | 0 | 0 |
| SRR5029876 | 1049 | 74333 | 1259 | 584 | 221 | 4764 | 1694 | 4490 | 5023 | 120 | 41488 | 20230 | 4852 | 3241 | 0 | 0 | 37715 | 0 | 0 | 0 | 1004 | 463 | 81 | 0 | 0 | 0 | 0 | 0 | 110 | 1723 | 830 | 166 | 0 | 28 | 14 | 0 | 0 |
| SRR5029877 | 437 | 57403 | 2480 | 139 | 160 | 3972 | 1518 | 4699 | 5084 | 79 | 41988 | 21030 | 4970 | 3252 | 12 | 0 | 40926 | 0 | 0 | 0 | 539 | 244 | 49 | 0 | 0 | 0 | 0 | 0 | 55 | 914 | 496 | 92 | 0 | 15 | 0 | 0 | 0 |
| SRR5029878 | 111 | 278865 | 807 | 222 | 317 | 9727 | 4791 | 10331 | 145 | 0 | 55777 | 19348 | 5124 | 5149 | 0 | 1404 | 42109 | 0 | 0 | 0 | 460 | 1547 | 116 | 0 | 0 | 0 | 0 | 31 | 205 | 440 | 2118 | 183 | 0 | 15 | 23 | 0 | 0 |
| SRR5029879 | 50 | 238187 | 1716 | 151 | 274 | 8413 | 4361 | 10496 | 133 | 50 | 55743 | 20335 | 5307 | 5524 | 0 | 1082 | 46805 | 0 | 0 | 0 | 335 | 1177 | 101 | 0 | 0 | 0 | 0 | 27 | 160 | 318 | 1709 | 122 | 0 | 11 | 19 | 0 | 0 |
| SRR5029880 | 108 | 281722 | 961 | 162 | 321 | 9588 | 4775 | 10306 | 127 | 0 | 55366 | 19389 | 5167 | 5230 | 0 | 1298 | 40058 | 0 | 0 | 0 | 469 | 1519 | 111 | 0 | 0 | 0 | 0 | 40 | 232 | 453 | 2053 | 175 | 0 | 14 | 21 | 0 | 0 |
| SRR5029882 | 82 | 280227 | 946 | 288 | 356 | 9581 | 4830 | 10194 | 144 | 0 | 55769 | 19302 | 5275 | 5239 | 0 | 1315 | 42024 | 0 | 0 | 0 | 463 | 1548 | 108 | 0 | 0 | 0 | 0 | 28 | 228 | 444 | 2212 | 173 | 0 | 18 | 29 | 0 | 0 |
| SRR5029883 | 51 | 243752 | 2036 | 153 | 229 | 8039 | 4424 | 10576 | 135 | 0 | 56000 | 20015 | 5326 | 5288 | 0 | 1356 | 46455 | 0 | 0 | 0 | 325 | 1174 | 83 | 0 | 0 | 0 | 0 | 18 | 127 | 335 | 1681 | 122 | 0 | 0 | 15 | 0 | 0 |
| SRR5029884 | 63 | 279377 | 931 | 157 | 322 | 9780 | 4806 | 10255 | 131 | 10 | 55670 | 19318 | 5186 | 5282 | 0 | 1396 | 40588 | 0 | 0 | 0 | 455 | 1512 | 106 | 0 | 0 | 0 | 0 | 44 | 229 | 476 | 2117 | 162 | 0 | 15 | 27 | 0 | 0 |
| SRR5029885 | 105 | 248507 | 1991 | 157 | 255 | 8292 | 4458 | 10393 | 141 | 0 | 56188 | 20132 | 5548 | 5398 | 0 | 1105 | 43258 | 0 | 0 | 0 | 311 | 1166 | 75 | 0 | 0 | 0 | 0 | 17 | 135 | 348 | 1642 | 145 | 0 | 15 | 20 | 0 | 0 |
| SRR5029886 | 42 | 10572 | 83 | 583 | 514 | 5907 | 4135 | 5410 | 3270 | 21 | 70430 | 27551 | 5768 | 5444 | 16 | 20 | 47836 | 0 | 0 | 0 | 282 | 255 | 274 | 0 | 19 | 0 | 0 | 0 | 22 | 117 | 130 | 234 | 0 | 56 | 0 | 0 | 0 |
| SRR5029887 | 0 | 6582 | 203 | 249 | 423 | 5356 | 3950 | 5527 | 3231 | 0 | 70308 | 28454 | 5766 | 5630 | 19 | 19 | 53036 | 0 | 0 | 0 | 150 | 140 | 176 | 0 | 15 | 0 | 0 | 0 | 0 | 71 | 82 | 162 | 0 | 40 | 0 | 0 | 0 |
| SRR5029888 | 56 | 9615 | 111 | 611 | 495 | 6165 | 4242 | 5693 | 3239 | 11 | 70012 | 27639 | 5709 | 5406 | 26 | 21 | 44999 | 0 | 0 | 0 | 269 | 249 | 289 | 0 | 22 | 0 | 0 | 0 | 19 | 122 | 113 | 224 | 0 | 53 | 0 | 15 | 0 |
| SRR5029889 | 19 | 6133 | 143 | 182 | 419 | 5430 | 3935 | 5637 | 3269 | 0 | 13 | 28277 | 5835 | 5637 | 17 | 17 | 47299 | 0 | 0 | 0 | 157 | 147 | 174 | 0 | 17 | 0 | 0 | 0 | 11 | 64 | 71 | 152 | 0 | 46 | 0 | 12 | 0 |
| SRR5029890 | 85 | 10097 | 149 | 595 | 443 | 5865 | 4037 | 5394 | 3254 | 32 | 15 | 27501 | 5691 | 5595 | 29 | 21 | 47605 | 0 | 0 | 0 | 295 | 248 | 279 | 0 | 21 | 0 | 0 | 0 | 22 | 115 | 130 | 241 | 0 | 57 | 0 | 12 | 0 |
| SRR5029891 | 38 | 6234 | 274 | 170 | 390 | 5208 | 3947 | 5464 | 3242 | 0 | 18 | 28451 | 5670 | 5502 | 18 | 15 | 54181 | 0 | 0 | 0 | 149 | 140 | 186 | 0 | 16 | 0 | 0 | 0 | 0 | 66 | 64 | 156 | 0 | 38 | 0 | 12 | 0 |
| SRR5029892 | 54 | 10170 | 98 | 587 | 456 | 5929 | 4262 | 5664 | 3198 | 0 | 11 | 27700 | 5704 | 5508 | 33 | 19 | 45913 | 0 | 0 | 0 | 293 | 250 | 273 | 0 | 22 | 0 | 0 | 0 | 18 | 106 | 118 | 253 | 0 | 68 | 11 | 11 | 0 |
| SRR5029893 | 29 | 6263 | 163 | 183 | 422 | 5469 | 3925 | 5550 | 3221 | 0 | 11 | 28142 | 5916 | 5593 | 30 | 13 | 48522 | 0 | 0 | 0 | 170 | 136 | 190 | 0 | 16 | 0 | 0 | 0 | 12 | 60 | 70 | 163 | 0 | 44 | 0 | 11 | 0 |
| SRR5029894 | 1247 | 56384 | 424 | 485 | 150 | 3414 | 1469 | 3157 | 530 | 12 | 75359 | 21765 | 5968 | 2773 | 0 | 227 | 33524 | 0 | 0 | 0 | 1660 | 490 | 84 | 0 | 0 | 0 | 0 | 0 | 149 | 3077 | 1180 | 188 | 0 | 46 | 15 | 0 | 0 |
| SRR5029895 | 620 | 45012 | 604 | 168 | 130 | 3002 | 1281 | 3238 | 543 | 23 | 74524 | 22070 | 6027 | 2848 | 0 | 184 | 37349 | 0 | 0 | 0 | 1010 | 324 | 53 | 0 | 0 | 0 | 0 | 0 | 81 | 1888 | 759 | 131 | 0 | 35 | 17 | 0 | 0 |
| SRR5029896 | 1244 | 56328 | 450 | 490 | 166 | 3417 | 1450 | 3222 | 473 | 0 | 74932 | 21451 | 5928 | 2841 | 0 | 226 | 31208 | 0 | 0 | 0 | 1621 | 527 | 85 | 0 | 0 | 0 | 0 | 0 | 151 | 3060 | 1166 | 180 | 0 | 46 | 22 | 0 | 0 |
| SRR5029897 | 536 | 43918 | 855 | 272 | 139 | 3042 | 1344 | 3251 | 539 | 0 | 13 | 21816 | 6045 | 2862 | 0 | 208 | 32737 | 0 | 0 | 0 | 995 | 331 | 57 | 0 | 0 | 0 | 0 | 0 | 91 | 1830 | 743 | 124 | 0 | 36 | 20 | 0 | 0 |
| SRR5029898 | 1170 | 56607 | 365 | 453 | 149 | 3349 | 1482 | 3205 | 543 | 13 | 15 | 21534 | 6082 | 2753 | 0 | 216 | 33651 | 0 | 0 | 0 | 1652 | 503 | 84 | 0 | 0 | 0 | 0 | 0 | 162 | 3045 | 1158 | 201 | 0 | 44 | 22 | 0 | 0 |
| SRR5029899 | 476 | 44320 | 897 | 226 | 148 | 2954 | 1350 | 3192 | 520 | 31 | 15 | 21932 | 6084 | 2764 | 0 | 182 | 37758 | 0 | 0 | 0 | 944 | 341 | 50 | 0 | 0 | 0 | 0 | 0 | 76 | 1838 | 768 | 126 | 0 | 37 | 21 | 0 | 0 |
| SRR5029900 | 1248 | 56370 | 392 | 534 | 162 | 3474 | 1571 | 3302 | 510 | 0 | 74527 | 21624 | 5947 | 2779 | 0 | 196 | 31496 | 0 | 0 | 0 | 1680 | 552 | 85 | 0 | 0 | 0 | 0 | 0 | 169 | 3147 | 1121 | 188 | 0 | 40 | 17 | 0 | 0 |
| SRR5029901 | 550 | 46555 | 846 | 191 | 125 | 2982 | 1330 | 3284 | 535 | 24 | 74775 | 22120 | 6118 | 2934 | 0 | 211 | 33650 | 0 | 0 | 0 | 989 | 337 | 53 | 0 | 0 | 0 | 0 | 0 | 99 | 1876 | 779 | 129 | 0 | 44 | 13 | 0 | 0 |
| SRR5029902 | 109 | 49007 | 116 | 858 | 457 | 5645 | 3799 | 6489 | 313 | 0 | 32753 | 16568 | 5468 | 2576 | 0 | 0 | 45938 | 0 | 0 | 0 | 129 | 59 | 76 | 0 | 0 | 0 | 0 | 0 | 13 | 43 | 36 | 70 | 0 | 19 | 0 | 0 | 0 |
| SRR5029904 | 67 | 49120 | 127 | 1072 | 491 | 5604 | 3845 | 6618 | 319 | 22 | 32583 | 16555 | 5574 | 2511 | 0 | 0 | 43713 | 0 | 0 | 0 | 120 | 50 | 83 | 0 | 0 | 0 | 0 | 0 | 0 | 40 | 44 | 60 | 0 | 22 | 0 | 0 | 0 |
| SRR5029905 | 56 | 37642 | 390 | 502 | 434 | 5039 | 3530 | 6817 | 338 | 112 | 32843 | 17168 | 6023 | 2697 | 0 | 0 | 46173 | 0 | 0 | 0 | 110 | 30 | 62 | 0 | 0 | 0 | 0 | 0 | 27 | 33 | 21 | 33 | 0 | 0 | 0 | 0 | 0 |
| SRR5029906 | 70 | 48702 | 134 | 824 | 474 | 5646 | 3708 | 6350 | 290 | 0 | 32460 | 16636 | 5627 | 2501 | 0 | 0 | 46251 | 0 | 0 | 0 | 131 | 59 | 73 | 0 | 0 | 0 | 0 | 0 | 12 | 38 | 36 | 61 | 0 | 19 | 0 | 0 | 0 |
| SRR5029908 | 109 | 48495 | 130 | 890 | 482 | 5901 | 3831 | 6587 | 319 | 14 | 33236 | 16782 | 5710 | 2589 | 0 | 0 | 44493 | 0 | 0 | 0 | 127 | 51 | 72 | 0 | 0 | 0 | 0 | 0 | 15 | 35 | 38 | 65 | 0 | 15 | 0 | 0 | 0 |
| SRR5029910 | 61 | 30175 | 42343 | 307 | 114 | 5678 | 1401 | 5522 | 9518 | 20 | 54120 | 19372 | 3401 | 3217 | 11 | 0 | 52776 | 0 | 0 | 0 | 104 | 1745 | 488 | 0 | 0 | 0 | 0 | 0 | 17 | 158 | 3455 | 1581 | 0 | 64 | 38 | 0 | 0 |
| SRR5029911 | 64 | 24332 | 53116 | 127 | 92 | 5712 | 1326 | 5578 | 9206 | 0 | 13 | 18879 | 3467 | 3210 | 19 | 0 | 55825 | 0 | 0 | 0 | 83 | 1459 | 424 | 0 | 0 | 0 | 0 | 0 | 11 | 122 | 2845 | 1390 | 0 | 64 | 32 | 0 | 0 |
| SRR5029912 | 22 | 29044 | 40403 | 200 | 113 | 5724 | 1342 | 5528 | 9341 | 0 | 54188 | 19387 | 3538 | 3430 | 11 | 0 | 50677 | 0 | 0 | 0 | 100 | 1757 | 474 | 0 | 0 | 0 | 0 | 0 | 24 | 178 | 3305 | 1573 | 0 | 57 | 42 | 0 | 0 |
| SRR5029913 | 44 | 23176 | 53982 | 133 | 116 | 5722 | 1400 | 5514 | 9393 | 0 | 52217 | 18930 | 3519 | 3356 | 13 | 0 | 50497 | 0 | 0 | 0 | 81 | 1449 | 400 | 0 | 0 | 0 | 0 | 0 | 14 | 116 | 2882 | 1394 | 0 | 60 | 42 | 13 | 0 |
| SRR5029914 | 21 | 29168 | 43218 | 235 | 102 | 5765 | 1347 | 5616 | 9276 | 0 | 53905 | 19160 | 3364 | 3251 | 11 | 0 | 52433 | 0 | 0 | 0 | 88 | 1754 | 485 | 0 | 0 | 0 | 0 | 0 | 22 | 166 | 3587 | 1667 | 0 | 54 | 40 | 0 | 0 |
| SRR5029915 | 22 | 23097 | 54402 | 87 | 127 | 5699 | 1336 | 5453 | 9175 | 0 | 51871 | 18888 | 3428 | 3211 | 0 | 0 | 56579 | 0 | 0 | 0 | 68 | 1427 | 430 | 0 | 0 | 0 | 0 | 0 | 15 | 117 | 2790 | 1416 | 0 | 58 | 35 | 0 | 0 |
| SRR5029916 | 64 | 29479 | 42950 | 107 | 119 | 5989 | 1365 | 5619 | 9285 | 0 | 11 | 19343 | 3541 | 3235 | 22 | 0 | 51285 | 0 | 0 | 0 | 98 | 1779 | 450 | 0 | 0 | 0 | 0 | 0 | 21 | 153 | 3432 | 1591 | 0 | 54 | 42 | 0 | 0 |
| SRR5029917 | 44 | 24224 | 53149 | 131 | 118 | 5769 | 1406 | 5587 | 9204 | 22 | 52066 | 18335 | 3493 | 3381 | 14 | 15 | 51198 | 0 | 0 | 0 | 80 | 1424 | 420 | 0 | 0 | 0 | 0 | 0 | 15 | 135 | 2707 | 1456 | 0 | 62 | 40 | 0 | 0 |
| SRR5029918 | 66 | 77952 | 449 | 199 | 212 | 2870 | 1485 | 4460 | 67 | 0 | 11 | 27031 | 5053 | 6435 | 0 | 10 | 48177 | 0 | 0 | 0 | 239 | 77 | 90 | 0 | 12 | 0 | 0 | 0 | 30 | 72 | 62 | 58 | 0 | 15 | 0 | 0 | 0 |
| SRR5029919 | 81 | 58732 | 710 | 169 | 186 | 2528 | 1390 | 4409 | 71 | 0 | 13 | 28069 | 5151 | 6461 | 0 | 0 | 54562 | 0 | 0 | 0 | 134 | 48 | 58 | 0 | 0 | 0 | 0 | 0 | 12 | 33 | 36 | 38 | 0 | 14 | 0 | 0 | 0 |
| SRR5029920 | 89 | 79204 | 514 | 195 | 229 | 2921 | 1476 | 4413 | 78 | 18 | 42728 | 26862 | 4896 | 6433 | 0 | 0 | 45601 | 0 | 0 | 0 | 223 | 70 | 101 | 0 | 0 | 0 | 0 | 0 | 18 | 65 | 66 | 61 | 0 | 11 | 0 | 0 | 0 |
| SRR5029921 | 25 | 57288 | 915 | 76 | 190 | 2635 | 1381 | 4528 | 65 | 0 | 11 | 27891 | 5186 | 6798 | 0 | 0 | 47609 | 0 | 0 | 0 | 123 | 41 | 60 | 0 | 0 | 0 | 0 | 0 | 11 | 31 | 35 | 35 | 0 | 13 | 0 | 0 | 0 |
| SRR5029922 | 102 | 80351 | 373 | 305 | 219 | 2785 | 1538 | 4339 | 59 | 0 | 17 | 27308 | 4967 | 6213 | 0 | 0 | 48152 | 0 | 0 | 0 | 225 | 61 | 99 | 0 | 0 | 0 | 0 | 0 | 22 | 65 | 63 | 49 | 0 | 14 | 0 | 0 | 0 |
| SRR5029923 | 57 | 58420 | 752 | 106 | 197 | 2549 | 1327 | 4394 | 64 | 33 | 42285 | 27620 | 5033 | 6532 | 0 | 0 | 54872 | 0 | 0 | 0 | 109 | 45 | 56 | 0 | 0 | 0 | 0 | 0 | 0 | 36 | 42 | 41 | 0 | 10 | 0 | 0 | 0 |
| SRR5029924 | 139 | 79815 | 521 | 226 | 205 | 2841 | 1514 | 4374 | 85 | 0 | 42282 | 27504 | 4981 | 6544 | 0 | 11 | 46670 | 0 | 0 | 0 | 238 | 79 | 74 | 0 | 0 | 0 | 0 | 0 | 27 | 60 | 67 | 63 | 0 | 18 | 0 | 0 | 0 |
| SRR5029925 | 50 | 57443 | 778 | 151 | 191 | 2594 | 1371 | 4587 | 56 | 17 | 42869 | 28030 | 5157 | 6525 | 0 | 10 | 49557 | 0 | 0 | 0 | 107 | 51 | 63 | 0 | 0 | 0 | 0 | 0 | 0 | 40 | 34 | 40 | 0 | 0 | 0 | 0 | 0 |
| SRR5029926 | 706 | 124247 | 460 | 286 | 249 | 4311 | 2280 | 4411 | 9626 | 0 | 62803 | 23246 | 6353 | 4791 | 10 | 27 | 36386 | 0 | 0 | 0 | 205 | 552 | 195 | 0 | 0 | 0 | 0 | 0 | 65 | 238 | 405 | 306 | 0 | 46 | 13 | 0 | 0 |
| SRR5029927 | 239 | 84593 | 1677 | 78 | 234 | 3757 | 2127 | 4622 | 9970 | 33 | 23 | 24099 | 6402 | 4762 | 0 | 33 | 40645 | 0 | 0 | 0 | 74 | 280 | 129 | 0 | 11 | 0 | 0 | 0 | 23 | 94 | 203 | 178 | 0 | 44 | 0 | 0 | 0 |
| SRR5029928 | 706 | 124475 | 740 | 232 | 312 | 4436 | 2333 | 4720 | 9604 | 34 | 17 | 23337 | 6403 | 4876 | 12 | 22 | 34353 | 0 | 0 | 0 | 206 | 520 | 217 | 0 | 10 | 0 | 0 | 0 | 64 | 234 | 361 | 306 | 0 | 53 | 18 | 11 | 0 |
| SRR5029929 | 203 | 87204 | 1603 | 104 | 222 | 3886 | 2161 | 4511 | 9907 | 12 | 21 | 23761 | 6561 | 5053 | 0 | 29 | 35962 | 0 | 0 | 0 | 88 | 263 | 130 | 0 | 12 | 0 | 0 | 0 | 32 | 106 | 195 | 176 | 0 | 43 | 11 | 10 | 0 |
| SRR5029930 | 576 | 126066 | 633 | 235 | 262 | 4360 | 2392 | 4488 | 9535 | 0 | 15 | 23446 | 6264 | 4501 | 14 | 39 | 36392 | 0 | 0 | 0 | 205 | 518 | 210 | 0 | 0 | 0 | 0 | 0 | 80 | 215 | 409 | 304 | 0 | 63 | 0 | 0 | 0 |
| SRR5029931 | 129 | 87086 | 1809 | 62 | 253 | 3767 | 2037 | 4579 | 10018 | 17 | 62649 | 23660 | 6428 | 4892 | 0 | 30 | 41232 | 0 | 0 | 0 | 80 | 270 | 123 | 0 | 10 | 0 | 0 | 0 | 19 | 103 | 189 | 197 | 0 | 35 | 13 | 0 | 0 |
| SRR5029932 | 585 | 124467 | 643 | 267 | 270 | 4357 | 2381 | 4659 | 9825 | 33 | 63303 | 23151 | 6353 | 4779 | 0 | 40 | 35122 | 0 | 0 | 0 | 221 | 505 | 232 | 0 | 13 | 0 | 0 | 0 | 88 | 241 | 364 | 314 | 0 | 51 | 12 | 0 | 0 |
| SRR5029933 | 145 | 85719 | 1597 | 93 | 232 | 3826 | 2097 | 4656 | 9904 | 23 | 11 | 23842 | 6463 | 4783 | 0 | 28 | 37077 | 0 | 0 | 0 | 83 | 282 | 125 | 0 | 0 | 0 | 0 | 0 | 28 | 105 | 197 | 179 | 0 | 33 | 14 | 0 | 0 |
| SRR5029934 | 159 | 27103 | 1470 | 503 | 514 | 11188 | 5134 | 10143 | 2168 | 0 | 43991 | 22062 | 5355 | 5667 | 14 | 12 | 46873 | 0 | 0 | 0 | 1708 | 1578 | 171 | 0 | 0 | 0 | 0 | 0 | 148 | 1730 | 741 | 234 | 0 | 18 | 12 | 0 | 0 |
| SRR5029935 | 45 | 16908 | 3022 | 150 | 452 | 10047 | 4867 | 9967 | 2168 | 15 | 43053 | 22306 | 5460 | 5703 | 25 | 0 | 52090 | 0 | 0 | 0 | 1182 | 1166 | 148 | 0 | 0 | 0 | 0 | 0 | 100 | 1241 | 558 | 185 | 0 | 17 | 11 | 11 | 0 |
| SRR5029936 | 244 | 27124 | 1912 | 460 | 505 | 11145 | 4936 | 10279 | 2107 | 0 | 43637 | 21682 | 5347 | 5583 | 13 | 0 | 44669 | 0 | 0 | 0 | 1767 | 1549 | 156 | 0 | 0 | 0 | 0 | 0 | 156 | 1779 | 680 | 235 | 0 | 19 | 13 | 0 | 0 |
| SRR5029937 | 110 | 17116 | 2960 | 157 | 448 | 10080 | 4861 | 10083 | 2117 | 47 | 44092 | 22420 | 5432 | 5821 | 17 | 11 | 46093 | 0 | 0 | 0 | 1176 | 1177 | 148 | 0 | 0 | 0 | 0 | 0 | 121 | 1249 | 507 | 168 | 0 | 25 | 12 | 0 | 0 |
| SRR5029938 | 287 | 27394 | 1693 | 478 | 478 | 10892 | 5166 | 10032 | 2164 | 14 | 43742 | 21947 | 5365 | 5696 | 13 | 0 | 47353 | 0 | 0 | 0 | 1773 | 1580 | 159 | 0 | 0 | 0 | 0 | 11 | 178 | 1818 | 762 | 238 | 0 | 24 | 14 | 0 | 0 |
| SRR5029939 | 61 | 16915 | 2582 | 199 | 421 | 9813 | 4603 | 9952 | 2163 | 15 | 43129 | 21832 | 5327 | 5543 | 0 | 0 | 52855 | 0 | 0 | 0 | 1131 | 1154 | 158 | 0 | 0 | 0 | 0 | 0 | 102 | 1238 | 525 | 199 | 0 | 22 | 17 | 0 | 0 |
| SRR5029940 | 295 | 28330 | 1629 | 562 | 613 | 11250 | 5180 | 10151 | 2111 | 0 | 43774 | 21715 | 5334 | 5651 | 19 | 0 | 45168 | 0 | 0 | 0 | 1768 | 1535 | 181 | 0 | 0 | 0 | 0 | 0 | 139 | 1766 | 689 | 210 | 0 | 24 | 13 | 0 | 0 |
| SRR5029941 | 31 | 16752 | 2941 | 125 | 506 | 10186 | 4869 | 10021 | 2183 | 0 | 43533 | 22126 | 5584 | 5714 | 17 | 0 | 47504 | 0 | 0 | 0 | 1172 | 1150 | 122 | 0 | 0 | 0 | 0 | 0 | 121 | 1221 | 548 | 195 | 0 | 23 | 18 | 12 | 0 |
| SRR5029942 | 406 | 54819 | 1191 | 799 | 84 | 1834 | 659 | 1266 | 3866 | 56 | 44838 | 20227 | 4237 | 4244 | 36 | 29 | 30499 | 0 | 0 | 15 | 757 | 590 | 227 | 0 | 12 | 0 | 0 | 27 | 419 | 412 | 464 | 205 | 0 | 20 | 16 | 11 | 0 |
| SRR5029943 | 28 | 29295 | 2576 | 83 | 75 | 1634 | 753 | 1300 | 3933 | 0 | 46864 | 21763 | 4421 | 4574 | 29 | 13 | 35092 | 0 | 0 | 0 | 313 | 296 | 136 | 0 | 0 | 0 | 0 | 10 | 120 | 165 | 210 | 122 | 0 | 24 | 14 | 14 | 0 |
| SRR5029944 | 232 | 55267 | 1238 | 681 | 116 | 1986 | 834 | 1271 | 3787 | 0 | 10 | 20261 | 4198 | 4348 | 30 | 0 | 29084 | 0 | 0 | 12 | 809 | 615 | 247 | 0 | 0 | 0 | 0 | 18 | 375 | 412 | 409 | 204 | 0 | 29 | 12 | 0 | 0 |
| SRR5029945 | 98 | 30113 | 2512 | 161 | 78 | 1622 | 724 | 1296 | 4018 | 29 | 47084 | 21699 | 4492 | 4755 | 35 | 12 | 30922 | 0 | 0 | 0 | 305 | 285 | 137 | 0 | 0 | 0 | 0 | 0 | 130 | 158 | 194 | 115 | 0 | 23 | 11 | 11 | 0 |

### Supp Table S8.1.Columns_B_to_AL

| | | | | | | | | | | | | | | | | | | | | | | | | | | | | | | | | | | | | | |
|---|---|---|---|---|---|---|---|---|---|---|---|---|---|---|---|---|---|---|---|---|---|---|---|---|---|---|---|---|---|---|---|---|---|---|---|---|---|
| SRR5029946 | 276 | 56090 | 975 | 582 | 93 | 1896 | 840 | 1238 | 3577 | 15 | 45005 | 20187 | 4191 | 4270 | 42 | 0 | 30308 | 0 | 0 | 18 | 709 | 598 | 243 | 0 | 0 | 0 | 0 | 28 | 372 | 394 | 454 | 223 | 0 | 27 | 27 | 12 | 0 |
| SRR5029947 | 56 | 30394 | 2745 | 95 | 62 | 1619 | 698 | 1284 | 3929 | 11 | 11 | 21596 | 4409 | 4575 | 34 | 11 | 35328 | 0 | 0 | 0 | 302 | 311 | 142 | 0 | 0 | 0 | 0 | 0 | 152 | 143 | 214 | 117 | 0 | 19 | 14 | 13 | 0 |
| SRR5029948 | 409 | 54741 | 1274 | 667 | 71 | 1879 | 817 | 1356 | 3693 | 0 | 45171 | 20403 | 4332 | 4300 | 39 | 0 | 28838 | 0 | 0 | 12 | 718 | 638 | 218 | 0 | 0 | 0 | 0 | 20 | 420 | 363 | 434 | 206 | 0 | 16 | 19 | 12 | 0 |
| SRR5029949 | 115 | 29963 | 2665 | 167 | 70 | 1627 | 719 | 1349 | 4008 | 12 | 47468 | 21719 | 4533 | 4713 | 28 | 12 | 31991 | 0 | 0 | 0 | 302 | 291 | 151 | 0 | 0 | 0 | 0 | 0 | 136 | 173 | 227 | 101 | 0 | 20 | 13 | 0 | 0 |
| SRR5029950 | 308 | 28137 | 606 | 909 | 403 | 6537 | 3104 | 4684 | 7365 | 0 | 10 | 16223 | 8723 | 5905 | 58 | 0 | 49057 | 0 | 0 | 0 | 141 | 193 | 81 | 0 | 0 | 0 | 0 | 0 | 31 | 108 | 352 | 125 | 0 | 41 | 54 | 15 | 0 |
| SRR5029951 | 256 | 25893 | 470 | 513 | 356 | 6806 | 3009 | 4454 | 7393 | 0 | 41222 | 15379 | 8691 | 5822 | 28 | 0 | 52300 | 0 | 0 | 0 | 109 | 197 | 88 | 0 | 11 | 0 | 0 | 0 | 21 | 56 | 384 | 123 | 18 | 32 | 63 | 25 | 0 |
| SRR5029952 | 391 | 28460 | 581 | 992 | 411 | 6663 | 3091 | 4744 | 7174 | 15 | 42435 | 16310 | 8745 | 5819 | 59 | 0 | 47702 | 0 | 0 | 0 | 133 | 183 | 71 | 0 | 0 | 0 | 0 | 0 | 33 | 104 | 340 | 113 | 0 | 42 | 59 | 17 | 0 |
| SRR5029953 | 132 | 25507 | 573 | 705 | 489 | 6059 | 2979 | 4745 | 7680 | 0 | 42422 | 16506 | 8863 | 5526 | 73 | 0 | 47511 | 0 | 0 | 0 | 129 | 151 | 77 | 0 | 0 | 0 | 0 | 0 | 15 | 66 | 305 | 107 | 0 | 51 | 40 | 55 | 0 |
| SRR5029954 | 407 | 27734 | 474 | 862 | 415 | 6478 | 3102 | 4604 | 7224 | 0 | 42546 | 16254 | 8707 | 5878 | 56 | 0 | 49470 | 0 | 0 | 0 | 144 | 192 | 72 | 0 | 0 | 0 | 0 | 0 | 39 | 84 | 349 | 141 | 0 | 37 | 62 | 18 | 0 |
| SRR5029955 | 218 | 26509 | 523 | 698 | 376 | 6134 | 3138 | 4402 | 6646 | 0 | 41905 | 15791 | 8477 | 5290 | 36 | 14 | 52654 | 0 | 0 | 0 | 132 | 179 | 93 | 0 | 14 | 0 | 0 | 0 | 36 | 118 | 350 | 143 | 11 | 64 | 93 | 43 | 0 |
| SRR5029956 | 327 | 28438 | 566 | 903 | 410 | 6657 | 3043 | 4815 | 7327 | 0 | 42739 | 16230 | 8734 | 5766 | 59 | 0 | 47904 | 0 | 0 | 0 | 143 | 186 | 75 | 0 | 0 | 0 | 0 | 0 | 35 | 98 | 336 | 119 | 0 | 47 | 56 | 17 | 0 |
| SRR5029957 | 309 | 25703 | 397 | 574 | 404 | 6675 | 2995 | 4445 | 7729 | 44 | 41385 | 15775 | 8465 | 5643 | 36 | 0 | 48414 | 0 | 0 | 0 | 106 | 197 | 47 | 0 | 11 | 0 | 0 | 0 | 33 | 98 | 383 | 160 | 0 | 36 | 84 | 11 | 0 |
| SRR5029958 | 118 | 26420 | 1340 | 884 | 98 | 2422 | 957 | 1993 | 15582 | 0 | 71622 | 29734 | 4454 | 4777 | 0 | 0 | 46839 | 0 | 0 | 0 | 157 | 654 | 276 | 0 | 19 | 0 | 0 | 0 | 37 | 205 | 833 | 616 | 0 | 83 | 58 | 29 | 0 |
| SRR5029959 | 25 | 13220 | 2614 | 147 | 83 | 2035 | 835 | 1998 | 16306 | 0 | 72936 | 31063 | 4664 | 4870 | 16 | 0 | 53241 | 0 | 0 | 0 | 71 | 330 | 161 | 0 | 11 | 0 | 0 | 0 | 11 | 81 | 415 | 363 | 0 | 61 | 53 | 24 | 0 |
| SRR5029960 | 79 | 25369 | 1247 | 868 | 99 | 2427 | 916 | 1972 | 15116 | 16 | 13 | 28994 | 4527 | 4657 | 17 | 0 | 45307 | 0 | 0 | 0 | 174 | 612 | 306 | 0 | 16 | 0 | 0 | 0 | 21 | 181 | 798 | 598 | 0 | 78 | 62 | 21 | 0 |
| SRR5029961 | 34 | 12701 | 2694 | 119 | 77 | 2102 | 875 | 2058 | 16556 | 0 | 14 | 30714 | 4755 | 5094 | 0 | 0 | 48047 | 0 | 0 | 0 | 62 | 345 | 177 | 0 | 15 | 0 | 0 | 0 | 0 | 85 | 413 | 350 | 0 | 45 | 50 | 20 | 0 |
| SRR5029962 | 180 | 25263 | 930 | 465 | 95 | 2424 | 936 | 2001 | 15385 | 0 | 71346 | 29141 | 4497 | 4826 | 18 | 0 | 47047 | 0 | 0 | 0 | 169 | 626 | 317 | 0 | 14 | 0 | 0 | 0 | 32 | 187 | 818 | 589 | 0 | 80 | 54 | 29 | 0 |
| SRR5029963 | 17 | 13158 | 2582 | 183 | 96 | 2073 | 870 | 2027 | 16229 | 0 | 72728 | 30839 | 4540 | 4985 | 12 | 0 | 53696 | 0 | 0 | 0 | 71 | 326 | 183 | 0 | 14 | 0 | 0 | 0 | 0 | 73 | 395 | 361 | 0 | 59 | 54 | 23 | 0 |
| SRR5029964 | 139 | 25606 | 988 | 818 | 127 | 2344 | 1053 | 1915 | 15672 | 15 | 71516 | 29233 | 4461 | 4780 | 12 | 12 | 45963 | 0 | 0 | 0 | 170 | 641 | 296 | 0 | 18 | 0 | 0 | 0 | 24 | 185 | 777 | 588 | 0 | 60 | 53 | 17 | 0 |
| SRR5029965 | 42 | 12585 | 2992 | 178 | 87 | 2049 | 916 | 2079 | 16337 | 17 | 73359 | 30853 | 4756 | 5053 | 14 | 0 | 49191 | 0 | 0 | 0 | 79 | 324 | 191 | 0 | 14 | 0 | 0 | 0 | 0 | 77 | 394 | 370 | 0 | 62 | 49 | 25 | 0 |
| SRR5029966 | 149 | 47434 | 1029 | 721 | 204 | 4208 | 2395 | 2548 | 16926 | 21 | 46391 | 20708 | 4498 | 7921 | 15 | 1990 | 50309 | 0 | 0 | 0 | 281 | 289 | 184 | 0 | 10 | 0 | 0 | 12 | 91 | 372 | 602 | 541 | 0 | 74 | 29 | 10 | 0 |
| SRR5029967 | 33 | 24492 | 2851 | 160 | 195 | 3591 | 2096 | 2520 | 17503 | 0 | 47076 | 21680 | 4553 | 8048 | 15 | 1701 | 57388 | 0 | 0 | 0 | 107 | 125 | 105 | 0 | 11 | 0 | 0 | 0 | 27 | 144 | 294 | 300 | 0 | 59 | 17 | 12 | 0 |
| SRR5029968 | 105 | 46845 | 922 | 887 | 253 | 4360 | 2463 | 2415 | 16780 | 12 | 46927 | 20931 | 4652 | 8157 | 19 | 2114 | 48686 | 0 | 0 | 0 | 282 | 284 | 184 | 0 | 14 | 0 | 0 | 13 | 100 | 389 | 580 | 526 | 0 | 77 | 25 | 15 | 0 |
| SRR5029969 | 0 | 24438 | 2791 | 203 | 188 | 3748 | 2122 | 2528 | 17509 | 29 | 47660 | 21607 | 4729 | 8357 | 14 | 1745 | 51375 | 0 | 0 | 0 | 117 | 128 | 113 | 0 | 0 | 0 | 0 | 0 | 25 | 148 | 283 | 294 | 0 | 59 | 23 | 0 | 0 |
| SRR5029970 | 110 | 46919 | 904 | 816 | 235 | 4307 | 2342 | 2387 | 16964 | 0 | 46496 | 21027 | 4324 | 7925 | 0 | 1964 | 50794 | 0 | 0 | 0 | 255 | 260 | 177 | 0 | 11 | 0 | 0 | 0 | 113 | 369 | 610 | 544 | 0 | 86 | 18 | 13 | 0 |
| SRR5029971 | 40 | 24848 | 2757 | 175 | 190 | 3506 | 2076 | 2485 | 17305 | 28 | 46898 | 21763 | 4581 | 8125 | 12 | 1725 | 58185 | 0 | 0 | 0 | 110 | 127 | 103 | 0 | 10 | 0 | 0 | 0 | 33 | 140 | 291 | 283 | 0 | 61 | 21 | 18 | 0 |
| SRR5029972 | 181 | 44491 | 1055 | 703 | 237 | 4376 | 2414 | 2521 | 16985 | 11 | 47480 | 20726 | 4466 | 8088 | 19 | 2027 | 48969 | 0 | 0 | 0 | 281 | 250 | 179 | 0 | 0 | 0 | 0 | 11 | 112 | 395 | 589 | 498 | 0 | 71 | 29 | 20 | 0 |
| SRR5029973 | 23 | 25459 | 2691 | 204 | 190 | 3564 | 2153 | 2524 | 17794 | 23 | 11 | 21796 | 4545 | 8306 | 16 | 1781 | 53005 | 0 | 0 | 0 | 118 | 126 | 111 | 0 | 0 | 0 | 0 | 0 | 35 | 157 | 277 | 268 | 0 | 63 | 18 | 0 | 0 |
| SRR5029974 | 91 | 53558 | 304 | 2213 | 246 | 6220 | 2586 | 3962 | 5100 | 0 | 60425 | 23948 | 6103 | 5878 | 22 | 15 | 41613 | 0 | 0 | 0 | 195 | 125 | 152 | 0 | 14 | 0 | 0 | 0 | 45 | 142 | 155 | 231 | 0 | 51 | 0 | 0 | 0 |
| SRR5029975 | 12 | 25320 | 803 | 464 | 249 | 5559 | 2476 | 4149 | 5096 | 0 | 61520 | 24770 | 6166 | 6180 | 16 | 23 | 47093 | 0 | 0 | 0 | 69 | 72 | 83 | 0 | 0 | 0 | 0 | 0 | 12 | 57 | 74 | 142 | 0 | 41 | 0 | 0 | 0 |
| SRR5029976 | 99 | 53246 | 234 | 2018 | 238 | 6481 | 2648 | 4249 | 5083 | 0 | 62005 | 24549 | 6126 | 6194 | 19 | 22 | 40733 | 0 | 0 | 0 | 206 | 121 | 150 | 0 | 0 | 0 | 0 | 0 | 52 | 164 | 144 | 252 | 0 | 55 | 11 | 11 | 0 |
| SRR5029977 | 0 | 25078 | 706 | 452 | 259 | 5511 | 2481 | 4139 | 5287 | 0 | 20 | 24805 | 6244 | 6158 | 12 | 18 | 41932 | 0 | 0 | 0 | 82 | 62 | 77 | 0 | 0 | 0 | 0 | 0 | 16 | 64 | 68 | 137 | 0 | 40 | 0 | 0 | 0 |
| SRR5029978 | 78 | 53946 | 370 | 2308 | 288 | 6155 | 2581 | 4042 | 5147 | 0 | 23 | 24290 | 6169 | 5965 | 16 | 20 | 42655 | 0 | 0 | 0 | 199 | 119 | 170 | 0 | 0 | 0 | 0 | 0 | 45 | 169 | 158 | 245 | 0 | 60 | 0 | 0 | 0 |
| SRR5029979 | 31 | 26486 | 594 | 392 | 232 | 5495 | 2374 | 4038 | 5162 | 0 | 15 | 24753 | 6207 | 6157 | 18 | 20 | 47840 | 0 | 0 | 0 | 66 | 65 | 66 | 0 | 0 | 0 | 0 | 0 | 12 | 61 | 73 | 147 | 0 | 32 | 0 | 0 | 0 |
| SRR5029980 | 78 | 53549 | 340 | 2273 | 268 | 6307 | 2705 | 4122 | 5167 | 0 | 11 | 24014 | 6009 | 5839 | 16 | 15 | 40267 | 0 | 0 | 0 | 186 | 124 | 144 | 0 | 0 | 0 | 0 | 0 | 38 | 137 | 163 | 228 | 0 | 52 | 16 | 0 | 0 |
| SRR5029981 | 12 | 26623 | 615 | 522 | 245 | 5392 | 2385 | 4214 | 5274 | 0 | 61440 | 24688 | 6250 | 6168 | 13 | 19 | 43055 | 0 | 0 | 0 | 69 | 57 | 91 | 0 | 0 | 0 | 0 | 0 | 12 | 68 | 68 | 155 | 0 | 39 | 11 | 10 | 0 |
| SRR5029982 | 28 | 43579 | 1285 | 470 | 137 | 3255 | 1191 | 3209 | 3186 | 14 | 41515 | 15143 | 6194 | 4279 | 27 | 11 | 46016 | 0 | 0 | 0 | 629 | 280 | 31 | 0 | 0 | 0 | 0 | 0 | 139 | 798 | 265 | 71 | 0 | 25 | 13 | 0 | 0 |
| SRR5029983 | 0 | 41160 | 1029 | 655 | 150 | 3270 | 1221 | 3266 | 3173 | 0 | 40218 | 14763 | 6381 | 4123 | 29 | 0 | 49714 | 0 | 0 | 10 | 650 | 341 | 56 | 0 | 0 | 0 | 0 | 13 | 132 | 815 | 285 | 79 | 0 | 33 | 0 | 0 | 0 |
| SRR5029984 | 75 | 43440 | 1288 | 569 | 139 | 3165 | 1227 | 3301 | 3140 | 0 | 10 | 15233 | 6318 | 4265 | 26 | 0 | 44106 | 0 | 0 | 0 | 600 | 305 | 28 | 0 | 0 | 0 | 0 | 0 | 130 | 828 | 247 | 72 | 0 | 21 | 15 | 0 | 0 |
| SRR5029986 | 28 | 43838 | 1316 | 481 | 128 | 3177 | 1186 | 3256 | 3241 | 0 | 41255 | 15077 | 6258 | 4282 | 29 | 0 | 46012 | 0 | 0 | 0 | 663 | 283 | 36 | 0 | 0 | 0 | 0 | 0 | 136 | 788 | 265 | 67 | 0 | 23 | 15 | 0 | 0 |
| SRR5029988 | 73 | 42720 | 1125 | 555 | 132 | 3210 | 1260 | 3350 | 3304 | 15 | 42038 | 15110 | 6241 | 4332 | 26 | 11 | 44739 | 0 | 0 | 0 | 636 | 296 | 25 | 0 | 0 | 0 | 0 | 12 | 134 | 788 | 248 | 68 | 0 | 23 | 12 | 13 | 0 |
| SRR5029990 | 121 | 13212 | 1569 | 1075 | 229 | 3164 | 2169 | 3496 | 10847 | 0 | 34143 | 18264 | 3556 | 5468 | 36 | 26 | 43461 | 0 | 0 | 0 | 102 | 363 | 167 | 0 | 0 | 0 | 0 | 0 | 71 | 143 | 945 | 944 | 0 | 24 | 49 | 15 | 0 |
| SRR5029991 | 75 | 8577 | 2969 | 424 | 227 | 2907 | 1975 | 3687 | 11492 | 0 | 34885 | 18619 | 3547 | 5712 | 32 | 23 | 48430 | 0 | 0 | 0 | 57 | 253 | 117 | 0 | 0 | 0 | 0 | 0 | 38 | 94 | 637 | 732 | 0 | 20 | 54 | 13 | 0 |
| SRR5029992 | 164 | 13214 | 1531 | 1142 | 271 | 3546 | 2216 | 3606 | 10965 | 0 | 34565 | 18190 | 3510 | 5458 | 29 | 13 | 41668 | 0 | 0 | 0 | 111 | 355 | 155 | 0 | 0 | 0 | 0 | 11 | 58 | 140 | 874 | 927 | 0 | 24 | 41 | 17 | 0 |
| SRR5029993 | 42 | 8900 | 3297 | 404 | 201 | 2854 | 2008 | 3652 | 11813 | 0 | 35398 | 18795 | 3657 | 5783 | 43 | 24 | 43068 | 0 | 0 | 0 | 64 | 244 | 120 | 0 | 0 | 0 | 0 | 0 | 50 | 80 | 600 | 715 | 0 | 34 | 43 | 18 | 0 |
| SRR5029994 | 109 | 13580 | 1665 | 915 | 292 | 3473 | 2185 | 3434 | 11045 | 0 | 34091 | 18093 | 3362 | 5413 | 25 | 27 | 43527 | 0 | 0 | 0 | 122 | 364 | 167 | 0 | 0 | 0 | 0 | 0 | 60 | 137 | 933 | 955 | 0 | 26 | 34 | 15 | 0 |
| SRR5029995 | 51 | 8609 | 3393 | 432 | 196 | 2857 | 1914 | 3575 | 11381 | 0 | 34382 | 18596 | 3525 | 5650 | 25 | 14 | 49700 | 0 | 0 | 0 | 64 | 243 | 129 | 0 | 0 | 0 | 0 | 0 | 46 | 81 | 631 | 718 | 0 | 20 | 40 | 21 | 0 |
| SRR5029996 | 129 | 13504 | 1542 | 958 | 234 | 3475 | 2043 | 3635 | 11140 | 0 | 34932 | 17938 | 3582 | 5523 | 18 | 41 | 41861 | 0 | 0 | 0 | 103 | 361 | 167 | 0 | 0 | 0 | 0 | 0 | 64 | 134 | 878 | 938 | 0 | 26 | 49 | 17 | 0 |
| SRR5029997 | 58 | 8822 | 3187 | 513 | 194 | 2997 | 2034 | 3531 | 11583 | 16 | 34944 | 18749 | 3486 | 5781 | 31 | 28 | 45173 | 0 | 0 | 0 | 66 | 245 | 111 | 0 | 0 | 0 | 0 | 0 | 42 | 92 | 602 | 667 | 0 | 21 | 47 | 20 | 0 |
| SRR5029998 | 700 | 25856 | 2597 | 186 | 289 | 4369 | 2142 | 3912 | 7341 | 44 | 40170 | 8973 | 4355 | 6186 | 0 | 20 | 24792 | 0 | 0 | 0 | 1585 | 1295 | 128 | 0 | 13 | 0 | 0 | 0 | 167 | 2233 | 1075 | 125 | 0 | 21 | 30 | 0 | 0 |
| SRR5029999 | 235 | 18275 | 6088 | 112 | 229 | 3789 | 2083 | 3987 | 7729 | 48 | 41215 | 9329 | 4460 | 6704 | 0 | 14 | 27966 | 0 | 0 | 0 | 942 | 834 | 95 | 0 | 0 | 0 | 0 | 0 | 88 | 1349 | 662 | 90 | 0 | 18 | 29 | 0 | 0 |
| SRR5030000 | 783 | 26564 | 2452 | 189 | 280 | 4399 | 2192 | 3773 | 7239 | 38 | 39470 | 8886 | 4276 | 6260 | 0 | 0 | 23973 | 0 | 0 | 0 | 1602 | 1313 | 133 | 0 | 0 | 0 | 0 | 0 | 179 | 2205 | 1044 | 141 | 0 | 22 | 45 | 0 | 0 |
| SRR5030001 | 312 | 18340 | 6336 | 100 | 217 | 3729 | 2093 | 4034 | 7610 | 89 | 10 | 9471 | 4493 | 6818 | 0 | 11 | 25344 | 0 | 0 | 0 | 964 | 827 | 99 | 0 | 0 | 0 | 0 | 0 | 94 | 1317 | 676 | 107 | 0 | 18 | 29 | 10 | 0 |
| SRR5030002 | 623 | 25086 | 2673 | 190 | 285 | 4200 | 2193 | 3766 | 7344 | 45 | 39889 | 8751 | 4217 | 6287 | 0 | 10 | 25408 | 0 | 0 | 0 | 1635 | 1342 | 142 | 0 | 11 | 0 | 0 | 0 | 181 | 2166 | 1072 | 142 | 0 | 24 | 36 | 13 | 0 |
| SRR5030003 | 212 | 17926 | 6021 | 142 | 238 | 3645 | 2013 | 3925 | 7521 | 38 | 40755 | 9387 | 4428 | 6676 | 0 | 0 | 28450 | 0 | 0 | 0 | 945 | 802 | 107 | 0 | 0 | 0 | 0 | 0 | 92 | 1280 | 703 | 97 | 0 | 17 | 28 | 11 | 0 |
| SRR5030004 | 632 | 25858 | 2326 | 192 | 253 | 4382 | 2173 | 3976 | 7231 | 27 | 40366 | 8986 | 4148 | 6338 | 0 | 21 | 23783 | 0 | 0 | 0 | 1703 | 1296 | 141 | 0 | 0 | 0 | 0 | 0 | 168 | 2303 | 1038 | 127 | 0 | 19 | 32 | 12 | 0 |
| SRR5030005 | 239 | 17747 | 5995 | 105 | 227 | 3806 | 2106 | 3999 | 7714 | 44 | 41149 | 9323 | 4479 | 6815 | 0 | 14 | 26004 | 0 | 0 | 0 | 940 | 812 | 92 | 0 | 0 | 0 | 0 | 0 | 87 | 1297 | 671 | 94 | 0 | 14 | 31 | 0 | 0 |
| SRR5030006 | 159 | 31602 | 309 | 401 | 237 | 4681 | 2402 | 6317 | 269 | 0 | 17 | 14223 | 4280 | 3740 | 0 | 12 | 37687 | 0 | 0 | 0 | 555 | 455 | 331 | 0 | 15 | 0 | 0 | 0 | 39 | 162 | 390 | 224 | 0 | 52 | 0 | 10 | 0 |
| SRR5030008 | 54 | 32261 | 288 | 577 | 214 | 4789 | 2397 | 6313 | 245 | 0 | 16 | 14075 | 4298 | 3706 | 0 | 16 | 35118 | 0 | 0 | 0 | 563 | 428 | 333 | 0 | 16 | 0 | 0 | 0 | 35 | 162 | 387 | 228 | 0 | 45 | 13 | 0 | 0 |
| SRR5030009 | 0 | 26004 | 314 | 79 | 198 | 4271 | 2359 | 6385 | 262 | 0 | 34 | 14619 | 4279 | 3929 | 0 | 13 | 36755 | 0 | 0 | 0 | 399 | 319 | 299 | 0 | 26 | 0 | 0 | 0 | 29 | 125 | 305 | 200 | 0 | 37 | 17 | 11 | 0 |
| SRR5030010 | 95 | 30642 | 232 | 559 | 210 | 4618 | 2388 | 6326 | 238 | 17 | 55218 | 14366 | 4391 | 3723 | 0 | 18 | 37663 | 0 | 0 | 0 | 542 | 450 | 340 | 0 | 14 | 0 | 0 | 0 | 29 | 176 | 408 | 253 | 0 | 39 | 0 | 14 | 0 |
| SRR5030011 | 0 | 26501 | 465 | 232 | 153 | 4115 | 2310 | 6396 | 306 | 0 | 17 | 14432 | 4330 | 3665 | 0 | 21 | 42461 | 0 | 0 | 0 | 357 | 382 | 326 | 0 | 11 | 0 | 0 | 0 | 28 | 105 | 246 | 160 | 0 | 69 | 0 | 11 | 0 |
| SRR5030012 | 79 | 31188 | 254 | 473 | 226 | 4867 | 2414 | 6377 | 254 | 0 | 55487 | 14507 | 4299 | 3772 | 0 | 14 | 35743 | 0 | 0 | 0 | 582 | 438 | 329 | 0 | 20 | 0 | 0 | 0 | 38 | 172 | 393 | 220 | 0 | 48 | 11 | 0 | 0 |
| SRR5030014 | 55 | 37692 | 124 | 386 | 158 | 2302 | 1278 | 2203 | 452 | 28 | 12 | 24754 | 4675 | 8100 | 72 | 0 | 47543 | 0 | 0 | 0 | 260 | 5915 | 434 | 0 | 0 | 0 | 0 | 0 | 96 | 285 | 2611 | 441 | 0 | 45 | 29 | 21 | 0 |
| SRR5030015 | 0 | 20303 | 369 | 164 | 120 | 2079 | 1261 | 2273 | 468 | 14 | 55343 | 25292 | 4960 | 8516 | 52 | 12 | 54086 | 0 | 0 | 0 | 109 | 3063 | 261 | 0 | 0 | 0 | 0 | 0 | 49 | 126 | 1352 | 299 | 0 | 38 | 22 | 20 | 0 |
| SRR5030016 | 30 | 36214 | 122 | 456 | 150 | 2473 | 1352 | 2140 | 444 | 30 | 55366 | 24410 | 4773 | 8253 | 66 | 12 | 45573 | 0 | 0 | 0 | 271 | 5812 | 434 | 0 | 14 | 0 | 0 | 0 | 93 | 323 | 2496 | 434 | 0 | 66 | 27 | 26 | 0 |
| SRR5030017 | 0 | 19899 | 272 | 129 | 128 | 1915 | 1217 | 2372 | 459 | 29 | 56449 | 25824 | 5022 | 8575 | 49 | 0 | 48818 | 0 | 0 | 0 | 130 | 3046 | 276 | 0 | 12 | 0 | 0 | 0 | 27 | 130 | 1266 | 291 | 0 | 38 | 23 | 18 | 0 |
| SRR5030018 | 57 | 37217 | 115 | 401 | 168 | 2370 | 1318 | 2106 | 511 | 43 | 10 | 24223 | 4809 | 8090 | 62 | 0 | 48029 | 0 | 0 | 0 | 274 | 5981 | 450 | 0 | 0 | 0 | 0 | 0 | 95 | 294 | 2554 | 477 | 0 | 45 | 31 | 18 | 0 |
| SRR5030019 | 14 | 19282 | 320 | 167 | 119 | 1966 | 1173 | 2262 | 477 | 28 | 54747 | 25600 | 4840 | 8329 | 49 | 0 | 54791 | 0 | 0 | 0 | 113 | 3014 | 248 | 0 | 0 | 0 | 0 | 0 | 31 | 117 | 1328 | 290 | 0 | 44 | 28 | 22 | 0 |
| SRR5030020 | 88 | 36505 | 133 | 472 | 162 | 2537 | 1385 | 2432 | 463 | 29 | 55576 | 24612 | 4874 | 8312 | 61 | 0 | 46041 | 0 | 0 | 0 | 258 | 5901 | 468 | 0 | 0 | 0 | 0 | 0 | 94 | 296 | 2483 | 447 | 0 | 54 | 33 | 17 | 0 |
| SRR5030021 | 0 | 18750 | 474 | 101 | 128 | 1959 | 1182 | 2210 | 484 | 72 | 55948 | 25902 | 4986 | 8576 | 52 | 0 | 49736 | 0 | 0 | 0 | 130 | 2962 | 263 | 0 | 0 | 0 | 0 | 0 | 39 | 135 | 1328 | 305 | 0 | 37 | 21 | 18 | 0 |
| SRR5030022 | 109 | 89095 | 1101 | 278 | 205 | 4562 | 1522 | 3875 | 185 | 0 | 10 | 18247 | 2989 | 2101 | 0 | 78 | 39368 | 0 | 0 | 0 | 254 | 195 | 128 | 0 | 13 | 0 | 0 | 0 | 32 | 181 | 122 | 119 | 0 | 33 | 0 | 0 | 0 |
| SRR5030023 | 134 | 72365 | 1275 | 179 | 193 | 4525 | 1526 | 3866 | 190 | 0 | 19 | 17854 | 2961 | 2078 | 0 | 89 | 41785 | 0 | 0 | 0 | 205 | 154 | 119 | 0 | 17 | 0 | 0 | 0 | 18 | 106 | 83 | 92 | 0 | 33 | 10 | 0 | 0 |
| SRR5030024 | 65 | 88088 | 1049 | 207 | 206 | 4431 | 1578 | 3910 | 193 | 0 | 44996 | 17783 | 2961 | 2207 | 0 | 84 | 37522 | 0 | 0 | 0 | 266 | 203 | 151 | 0 | 17 | 0 | 0 | 0 | 33 | 161 | 119 | 116 | 0 | 35 | 13 | 0 | 0 |
| SRR5030025 | 70 | 74767 | 1279 | 70 | 209 | 4455 | 1626 | 3878 | 194 | 23 | 14 | 17520 | 2893 | 2099 | 0 | 84 | 37694 | 0 | 0 | 0 | 211 | 144 | 121 | 0 | 16 | 0 | 0 | 0 | 23 | 143 | 73 | 73 | 0 | 40 | 13 | 0 | 0 |
| SRR5030026 | 99 | 89272 | 1268 | 236 | 211 | 4395 | 1524 | 3946 | 194 | 12 | 15 | 18190 | 2880 | 2103 | 0 | 66 | 39140 | 0 | 0 | 0 | 271 | 205 | 141 | 0 | 14 | 0 | 0 | 0 | 34 | 168 | 116 | 125 | 0 | 39 | 11 | 0 | 0 |
| SRR5030027 | 23 | 73726 | 1373 | 135 | 196 | 4514 | 1592 | 3715 | 198 | 23 | 43981 | 17345 | 2896 | 2135 | 0 | 74 | 42483 | 0 | 0 | 0 | 183 | 156 | 110 | 0 | 13 | 0 | 0 | 0 | 14 | 121 | 82 | 81 | 0 | 32 | 0 | 0 | 0 |
| SRR5030028 | 76 | 87106 | 1148 | 315 | 220 | 4566 | 1555 | 3887 | 207 | 13 | 45601 | 18109 | 3035 | 2209 | 0 | 72 | 37783 | 0 | 0 | 0 | 264 | 204 | 151 | 0 | 14 | 0 | 0 | 0 | 28 | 150 | 111 | 114 | 0 | 40 | 0 | 0 | 0 |
| SRR5030030 | 180 | 108240 | 202 | 630 | 161 | 3240 | 1438 | 2139 | 3176 | 23 | 63202 | 21487 | 5674 | 6043 | 23 | 0 | 36815 | 0 | 0 | 0 | 173 | 171 | 117 | 0 | 11 | 0 | 0 | 0 | 113 | 154 | 168 | 174 | 0 | 88 | 25 | 0 | 0 |
| SRR5030031 | 23 | 53626 | 510 | 151 | 133 | 2713 | 1339 | 2229 | 3165 | 23 | 19 | 22757 | 5845 | 6325 | 21 | 0 | 41097 | 0 | 0 | 0 | 48 | 68 | 63 | 0 | 11 | 0 | 0 | 0 | 38 | 58 | 76 | 85 | 0 | 52 | 31 | 0 | 0 |
| SRR5030032 | 157 | 108678 | 339 | 533 | 193 | 3205 | 1528 | 2149 | 3113 | 0 | 62096 | 21204 | 5700 | 6077 | 26 | 0 | 34866 | 0 | 0 | 0 | 171 | 177 | 118 | 0 | 11 | 0 | 0 | 0 | 119 | 164 | 197 | 157 | 0 | 65 | 34 | 10 | 0 |
| SRR5030033 | 36 | 52341 | 783 | 120 | 148 | 2739 | 1359 | 2321 | 3245 | 48 | 20 | 22268 | 5974 | 6438 | 26 | 0 | 37050 | 0 | 0 | 0 | 61 | 79 | 65 | 0 | 0 | 0 | 0 | 0 | 39 | 65 | 70 | 84 | 0 | 45 | 33 | 0 | 0 |
| SRR5030034 | 207 | 107450 | 323 | 587 | 189 | 3190 | 1495 | 2165 | 3067 | 23 | 11 | 21752 | 5603 | 5921 | 31 | 0 | 36406 | 0 | 0 | 0 | 174 | 183 | 121 | 0 | 0 | 0 | 0 | 0 | 132 | 149 | 168 | 179 | 0 | 78 | 30 | 11 | 0 |
| SRR5030035 | 12 | 53324 | 626 | 130 | 137 | 2614 | 1378 | 2190 | 3257 | 24 | 63838 | 22664 | 5846 | 6229 | 24 | 0 | 41276 | 0 | 0 | 0 | 61 | 69 | 69 | 0 | 0 | 0 | 0 | 0 | 33 | 69 | 77 | 84 | 0 | 47 | 20 | 10 | 0 |
| SRR5030036 | 141 | 105742 | 259 | 482 | 150 | 3185 | 1505 | 2183 | 3044 | 24 | 19 | 21811 | 5766 | 5901 | 21 | 0 | 35192 | 0 | 0 | 0 | 161 | 163 | 117 | 0 | 0 | 0 | 0 | 0 | 132 | 164 | 168 | 169 | 0 | 62 | 33 | 10 | 0 |
| SRR5030038 | 102 | 33408 | 4647 | 190 | 237 | 5083 | 1899 | 4363 | 221 | 13 | 12 | 30829 | 5628 | 5328 | 0 | 0 | 59120 | 0 | 0 | 0 | 296 | 570 | 395 | 0 | 27 | 0 | 0 | 0 | 40 | 181 | 318 | 465 | 0 | 108 | 48 | 14 | 0 |
| SRR5030039 | 47 | 24882 | 5856 | 118 | 287 | 5011 | 1898 | 4517 | 173 | 47 | 11 | 29981 | 5573 | 5178 | 0 | 0 | 64094 | 0 | 0 | 0 | 207 | 441 | 363 | 0 | 26 | 0 | 0 | 0 | 23 | 101 | 252 | 385 | 0 | 91 | 60 | 15 | 0 |
| SRR5030040 | 121 | 32233 | 4338 | 403 | 266 | 5014 | 1858 | 4489 | 226 | 27 | 10 | 30453 | 5660 | 5269 | 0 | 0 | 56814 | 0 | 0 | 0 | 319 | 542 | 438 | 0 | 25 | 0 | 0 | 0 | 38 | 147 | 337 | 459 | 0 | 101 | 55 | 21 | 0 |
| SRR5030041 | 97 | 24833 | 5478 | 195 | 282 | 5073 | 1915 | 4441 | 223 | 73 | 51722 | 30066 | 5643 | 5301 | 0 | 0 | 56849 | 0 | 0 | 0 | 213 | 402 | 360 | 0 | 25 | 0 | 0 | 0 | 26 | 112 | 245 | 403 | 0 | 107 | 64 | 19 | 0 |
| SRR5030042 | 142 | 32597 | 4707 | 349 | 278 | 5030 | 2015 | 4503 | 181 | 26 | 13 | 30510 | 5634 | 5185 | 0 | 0 | 59913 | 0 | 0 | 0 | 310 | 545 | 425 | 0 | 26 | 0 | 0 | 0 | 34 | 159 | 340 | 467 | 0 | 116 | 50 | 16 | 0 |
| SRR5030043 | 24 | 25447 | 5504 | 95 | 222 | 4998 | 1922 | 4276 | 201 | 24 | 51382 | 29533 | 5481 | 5039 | 0 | 0 | 64283 | 0 | 0 | 0 | 227 | 431 | 334 | 0 | 23 | 0 | 0 | 0 | 17 | 111 | 271 | 400 | 0 | 93 | 52 | 19 | 0 |
| SRR5030044 | 145 | 32026 | 4590 | 303 | 260 | 5230 | 1977 | 4492 | 200 | 0 | 53158 | 30470 | 5616 | 5136 | 0 | 0 | 57403 | 0 | 0 | 0 | 324 | 556 | 404 | 0 | 24 | 0 | 0 | 0 | 29 | 163 | 326 | 469 | 0 | 102 | 50 | 16 | 0 |
| SRR5030045 | 24 | 24289 | 5633 | 219 | 278 | 5062 | 1921 | 4464 | 156 | 24 | 51924 | 29781 | 5412 | 5140 | 0 | 0 | 59098 | 0 | 0 | 0 | 211 | 421 | 352 | 0 | 23 | 0 | 0 | 0 | 22 | 98 | 254 | 384 | 0 | 101 | 43 | 18 | 0 |
| SRR5030046 | 750 | 36772 | 6307 | 988 | 322 | 8418 | 3009 | 5999 | 18170 | 0 | 36055 | 22740 | 3956 | 2788 | 15 | 0 | 40701 | 0 | 0 | 0 | 259 | 337 | 246 | 0 | 0 | 0 | 0 | 10 | 157 | 216 | 606 | 674 | 0 | 47 | 42 | 12 | 0 |
| SRR5030048 | 821 | 38374 | 6436 | 990 | 297 | 8498 | 3051 | 6106 | 17810 | 32 | 35745 | 22468 | 4031 | 2883 | 20 | 0 | 38216 | 0 | 0 | 0 | 301 | 345 | 241 | 0 | 0 | 0 | 0 | 0 | 159 | 215 | 608 | 664 | 0 | 57 | 56 | 0 | 0 |
| SRR5030050 | 709 | 39042 | 6142 | 937 | 304 | 8321 | 3040 | 6121 | 17937 | 15 | 35608 | 22775 | 3926 | 2759 | 20 | 0 | 40635 | 0 | 0 | 0 | 275 | 343 | 241 | 0 | 0 | 0 | 0 | 0 | 168 | 206 | 610 | 659 | 0 | 57 | 49 | 14 | 0 |
| SRR5030052 | 740 | 38414 | 6047 | 968 | 374 | 8572 | 3149 | 6149 | 18024 | 0 | 36013 | 22871 | 3969 | 2736 | 22 | 0 | 38433 | 0 | 0 | 0 | 275 | 359 | 251 | 0 | 0 | 0 | 0 | 10 | 139 | 228 | 604 | 668 | 0 | 59 | 45 | 12 | 0 |
| SRR5030053 | 411 | 30950 | 12462 | 593 | 248 | 7484 | 2955 | 6369 | 18546 | 0 | 36927 | 23578 | 4158 | 3009 | 11 | 11 | 41972 | 0 | 0 | 0 | 183 | 270 | 188 | 0 | 0 | 0 | 0 | 0 | 89 | 179 | 445 | 475 | 0 | 50 | 35 | 11 | 0 |
| SRR5030054 | 218 | 17359 | 641 | 728 | 127 | 3601 | 1231 | 2819 | 5866 | 15 | 58242 | 24647 | 5899 | 5171 | 14 | 0 | 36840 | 0 | 0 | 0 | 363 | 8305 | 745 | 0 | 21 | 0 | 0 | 0 | 84 | 257 | 5542 | 827 | 0 | 85 | 133 | 30 | 0 |
| SRR5030055 | 96 | 13508 | 1202 | 625 | 73 | 3431 | 1059 | 2741 | 5775 | 0 | 57707 | 25413 | 5817 | 5416 | 14 | 0 | 41794 | 0 | 0 | 0 | 260 | 6061 | 583 | 0 | 0 | 0 | 0 | 0 | 60 | 176 | 4125 | 680 | 0 | 82 | 115 | 24 | 0 |
| SRR5030056 | 235 | 17388 | 502 | 768 | 141 | 3697 | 1179 | 2798 | 5627 | 31 | 58287 | 24420 | 5969 | 5322 | 0 | 0 | 34811 | 0 | 0 | 0 | 361 | 8101 | 751 | 0 | 19 | 0 | 0 | 0 | 82 | 243 | 5367 | 844 | 0 | 86 | 140 | 28 | 0 |
| SRR5030057 | 102 | 12908 | 1122 | 153 | 86 | 3210 | 1077 | 2740 | 5814 | 0 | 57866 | 24686 | 6274 | 5303 | 17 | 0 | 36582 | 0 | 0 | 0 | 263 | 6067 | 576 | 0 | 19 | 0 | 0 | 0 | 60 | 197 | 3951 | 649 | 0 | 67 | 108 | 20 | 0 |
| SRR5030058 | 150 | 17793 | 600 | 495 | 117 | 3506 | 1174 | 2718 | 5745 | 0 | 58293 | 24604 | 5904 | 5343 | 13 | 0 | 37320 | 0 | 0 | 0 | 379 | 8435 | 813 | 0 | 22 | 0 | 0 | 0 | 84 | 253 | 5549 | 847 | 0 | 81 | 128 | 23 | 0 |
| SRR5030059 | 0 | 14566 | 1230 | 344 | 95 | 3134 | 1099 | 2755 | 5736 | 0 | 57279 | 24917 | 6042 | 5219 | 0 | 0 | 41897 | 0 | 0 | 0 | 236 | 5914 | 574 | 0 | 17 | 0 | 0 | 0 | 44 | 169 | 4074 | 723 | 0 | 61 | 143 | 22 | 0 |
| SRR5030060 | 277 | 17329 | 522 | 507 | 111 | 3680 | 1209 | 2781 | 5953 | 31 | 58388 | 24350 | 6006 | 5297 | 10 | 0 | 35000 | 0 | 0 | 0 | 389 | 8350 | 769 | 0 | 22 | 0 | 0 | 0 | 72 | 256 | 5376 | 853 | 0 | 77 | 128 | 35 | 0 |
| SRR5030061 | 0 | 12883 | 1459 | 352 | 110 | 3190 | 1028 | 2738 | 6031 | 0 | 57343 | 25238 | 6075 | 5355 | 0 | 0 | 37519 | 0 | 0 | 0 | 248 | 6121 | 655 | 0 | 15 | 0 | 0 | 0 | 61 | 175 | 3967 | 662 | 0 | 69 | 116 | 39 | 0 |
| SRR5030062 | 65 | 139696 | 3363 | 312 | 37 | 1747 | 614 | 1444 | 17811 | 26 | 54528 | 25067 | 4699 | 4084 | 0 | 0 | 48146 | 0 | 0 | 0 | 69 | 140 | 105 | 0 | 0 | 0 | 0 | 0 | 16 | 86 | 227 | 323 | 0 | 31 | 22 | 0 | 0 |
| SRR5030064 | 42 | 138675 | 3264 | 474 | 53 | 1758 | 583 | 1428 | 17566 | 14 | 13 | 24874 | 4773 | 4177 | 0 | 0 | 45911 | 0 | 0 | 0 | 63 | 141 | 105 | 0 | 0 | 0 | 0 | 0 | 14 | 85 | 228 | 312 | 0 | 31 | 23 | 0 | 0 |
| SRR5030066 | 120 | 140071 | 3703 | 428 | 33 | 1723 | 600 | 1406 | 17807 | 13 | 54318 | 24803 | 4647 | 4064 | 0 | 0 | 48125 | 0 | 0 | 0 | 56 | 130 | 117 | 0 | 0 | 0 | 0 | 0 | 22 | 82 | 218 | 345 | 0 | 29 | 28 | 0 | 0 |
| SRR5030068 | 81 | 139825 | 3255 | 511 | 38 | 1728 | 610 | 1499 | 17796 | 27 | 54388 | 25083 | 4712 | 4122 | 0 | 0 | 46199 | 0 | 0 | 0 | 58 | 134 | 114 | 0 | 0 | 0 | 0 | 0 | 17 | 90 | 210 | 315 | 0 | 38 | 25 | 0 | 0 |
| SRR5030069 | 63 | 119202 | 4813 | 250 | 43 | 1658 | 576 | 1322 | 17570 | 63 | 10 | 24798 | 4701 | 4007 | 0 | 0 | 46769 | 0 | 0 | 0 | 80 | 117 | 87 | 0 | 0 | 0 | 0 | 0 | 14 | 60 | 153 | 300 | 0 | 28 | 12 | 0 | 0 |
| SRR5030070 | 40 | 103268 | 278 | 418 | 444 | 9552 | 4118 | 11673 | 312 | 30 | 54250 | 27483 | 6317 | 5931 | 0 | 14 | 70504 | 0 | 0 | 0 | 297 | 1851 | 568 | 0 | 13 | 0 | 0 | 0 | 27 | 177 | 754 | 605 | 0 | 40 | 15 | 0 | 0 |
| SRR5030071 | 41 | 75434 | 637 | 205 | 365 | 8627 | 3730 | 11755 | 337 | 0 | 13 | 28051 | 6202 | 5961 | 0 | 0 | 78804 | 0 | 0 | 0 | 149 | 1105 | 370 | 0 | 0 | 0 | 0 | 0 | 14 | 97 | 497 | 386 | 0 | 34 | 10 | 0 | 0 |
| SRR5030072 | 97 | 103268 | 387 | 440 | 455 | 9955 | 4135 | 12101 | 322 | 11 | 54625 | 27284 | 6072 | 5833 | 0 | 0 | 67458 | 0 | 0 | 0 | 288 | 1759 | 603 | 0 | 15 | 0 | 0 | 0 | 39 | 155 | 754 | 581 | 0 | 40 | 0 | 0 | 0 |
| SRR5030073 | 0 | 76640 | 722 | 212 | 380 | 8764 | 3829 | 12141 | 324 | 0 | 55169 | 28153 | 6569 | 6304 | 0 | 0 | 70006 | 0 | 0 | 0 | 161 | 1085 | 397 | 0 | 0 | 0 | 0 | 0 | 18 | 107 | 437 | 385 | 0 | 35 | 13 | 0 | 0 |
| SRR5030074 | 92 | 103042 | 204 | 429 | 495 | 9640 | 4165 | 11518 | 325 | 20 | 53817 | 27475 | 6105 | 5960 | 0 | 0 | 70615 | 0 | 0 | 0 | 282 | 1807 | 588 | 0 | 0 | 0 | 0 | 0 | 31 | 173 | 731 | 602 | 0 | 41 | 17 | 0 | 0 |
| SRR5030076 | 104 | 102506 | 292 | 302 | 394 | 9614 | 4181 | 12034 | 294 | 10 | 54602 | 27426 | 6222 | 6071 | 0 | 12 | 67893 | 0 | 0 | 0 | 293 | 1732 | 600 | 0 | 15 | 0 | 0 | 0 | 35 | 184 | 714 | 594 | 0 | 47 | 10 | 0 | 0 |
| SRR5030077 | 85 | 77418 | 552 | 318 | 383 | 8681 | 3700 | 11958 | 314 | 0 | 54855 | 27814 | 6314 | 6082 | 0 | 11 | 71972 | 0 | 0 | 0 | 160 | 1088 | 415 | 0 | 21 | 0 | 0 | 0 | 17 | 98 | 471 | 416 | 0 | 32 | 11 | 0 | 0 |
| SRR5030078 | 561 | 9942 | 1809 | 1608 | 241 | 3986 | 2111 | 4603 | 32 | 11 | 35170 | 9326 | 3041 | 3213 | 0 | 0 | 35862 | 0 | 0 | 0 | 198 | 102 | 31 | 0 | 0 | 0 | 0 | 14 | 87 | 133 | 295 | 45 | 0 | 0 | 0 | 0 | 0 |
| SRR5030079 | 206 | 7022 | 3159 | 515 | 211 | 3788 | 2077 | 4628 | 47 | 34 | 35431 | 9810 | 3089 | 3254 | 0 | 0 | 40806 | 0 | 0 | 0 | 112 | 64 | 21 | 0 | 0 | 0 | 0 | 0 | 60 | 82 | 184 | 28 | 0 | 0 | 0 | 0 | 0 |
| SRR5030080 | 572 | 9946 | 1866 | 1568 | 249 | 4114 | 2044 | 4596 | 36 | 23 | 34959 | 9260 | 2951 | 3232 | 18 | 0 | 33792 | 0 | 0 | 0 | 200 | 118 | 32 | 0 | 0 | 0 | 0 | 17 | 84 | 141 | 320 | 41 | 0 | 0 | 0 | 0 | 0 |
| SRR5030081 | 107 | 6686 | 2878 | 626 | 201 | 3697 | 1972 | 4970 | 35 | 0 | 35560 | 9641 | 3022 | 3322 | 0 | 0 | 35138 | 0 | 0 | 0 | 106 | 81 | 19 | 0 | 0 | 0 | 0 | 10 | 36 | 75 | 191 | 29 | 0 | 0 | 0 | 0 | 0 |
| SRR5030082 | 491 | 10645 | 1550 | 1398 | 239 | 3998 | 2029 | 4699 | 30 | 22 | 35080 | 9317 | 3020 | 3122 | 0 | 0 | 36205 | 0 | 0 | 0 | 172 | 120 | 32 | 0 | 0 | 0 | 0 | 12 | 85 | 121 | 340 | 44 | 0 | 0 | 12 | 0 | 0 |
| SRR5030083 | 264 | 7242 | 3375 | 457 | 208 | 3668 | 1919 | 4773 | 34 | 35 | 35409 | 9557 | 2986 | 3125 | 0 | 0 | 41286 | 0 | 0 | 0 | 103 | 67 | 25 | 0 | 0 | 0 | 0 | 0 | 47 | 72 | 210 | 31 | 0 | 0 | 0 | 0 | 0 |

### Supp Table S8.1.Columns_B_to_AL

| | | | | | | | | | | | | | | | | | | | | | | | | | | | | | | | | | | | | | |
|---|---|---|---|---|---|---|---|---|---|---|---|---|---|---|---|---|---|---|---|---|---|---|---|---|---|---|---|---|---|---|---|---|---|---|---|---|---|
| SRR5030084 | 493 | 9812 | 1693 | 1379 | 253 | 4023 | 2131 | 4723 | 33 | 11 | 34982 | 9311 | 3064 | 3229 | 0 | 0 | 34401 | 0 | 0 | 0 | 186 | 125 | 37 | 0 | 0 | 0 | 0 | 13 | 82 | 138 | 318 | 43 | 0 | 0 | 0 | 0 | 0 |
| SRR5030085 | 231 | 6924 | 2883 | 481 | 230 | 3615 | 2127 | 4619 | 42 | 0 | 35701 | 9611 | 3166 | 3322 | 0 | 0 | 36773 | 0 | 0 | 0 | 99 | 68 | 19 | 0 | 0 | 0 | 0 | 0 | 50 | 74 | 185 | 33 | 0 | 0 | 0 | 0 | 0 |
| SRR5030086 | 1584 | 114286 | 525 | 218 | 113 | 2669 | 1295 | 2823 | 848 | 0 | 39361 | 18186 | 4096 | 2858 | 0 | 96 | 44328 | 0 | 0 | 0 | 419 | 250 | 71 | 0 | 0 | 0 | 0 | 0 | 100 | 333 | 278 | 67 | 0 | 16 | 0 | 0 | 0 |
| SRR5030087 | 767 | 102603 | 724 | 187 | 121 | 2719 | 1312 | 2743 | 798 | 0 | 37983 | 17957 | 4061 | 2707 | 0 | 84 | 47563 | 0 | 0 | 0 | 304 | 172 | 45 | 0 | 0 | 0 | 0 | 0 | 69 | 247 | 225 | 52 | 0 | 17 | 0 | 0 | 0 |
| SRR5030088 | 1679 | 115558 | 471 | 256 | 104 | 2701 | 1178 | 2937 | 812 | 10 | 38743 | 18031 | 4121 | 2862 | 0 | 82 | 42564 | 0 | 0 | 0 | 433 | 231 | 64 | 0 | 0 | 0 | 0 | 0 | 104 | 344 | 266 | 66 | 0 | 13 | 0 | 0 | 0 |
| SRR5030090 | 1564 | 114383 | 521 | 226 | 119 | 2615 | 1235 | 2925 | 878 | 40 | 38558 | 18097 | 4035 | 2811 | 0 | 89 | 44928 | 0 | 0 | 0 | 412 | 234 | 71 | 0 | 0 | 0 | 0 | 0 | 95 | 320 | 293 | 54 | 0 | 12 | 11 | 0 | 0 |
| SRR5030091 | 1025 | 101361 | 783 | 129 | 146 | 2613 | 1160 | 2828 | 791 | 0 | 37916 | 17483 | 3989 | 2716 | 0 | 89 | 47900 | 0 | 0 | 0 | 310 | 177 | 60 | 0 | 0 | 0 | 0 | 0 | 77 | 254 | 221 | 66 | 0 | 18 | 0 | 0 | 0 |
| SRR5030092 | 1628 | 114148 | 443 | 315 | 138 | 2665 | 1274 | 2911 | 798 | 15 | 38957 | 18185 | 4151 | 2909 | 0 | 89 | 43323 | 0 | 0 | 0 | 406 | 240 | 70 | 0 | 0 | 0 | 0 | 0 | 102 | 329 | 277 | 68 | 0 | 12 | 0 | 0 | 0 |
| SRR5030093 | 928 | 103298 | 813 | 88 | 140 | 2697 | 1180 | 2919 | 835 | 18 | 38181 | 17712 | 4043 | 2836 | 26 | 95 | 43638 | 0 | 0 | 0 | 302 | 173 | 53 | 0 | 0 | 0 | 0 | 0 | 74 | 248 | 228 | 53 | 0 | 14 | 0 | 0 | 0 |
| SRR5030094 | 2273 | 142314 | 108 | 401 | 420 | 5644 | 3325 | 7377 | 373 | 39 | 34423 | 10307 | 4426 | 4120 | 0 | 38 | 43800 | 0 | 0 | 0 | 2905 | 608 | 155 | 0 | 0 | 0 | 0 | 11 | 151 | 2453 | 931 | 153 | 0 | 24 | 17 | 0 | 0 |
| SRR5030095 | 1315 | 130629 | 226 | 206 | 377 | 4581 | 2759 | 7848 | 385 | 0 | 35483 | 11518 | 4712 | 4704 | 0 | 0 | 50522 | 0 | 0 | 0 | 2075 | 443 | 109 | 0 | 0 | 0 | 0 | 0 | 46 | 1715 | 803 | 167 | 0 | 38 | 0 | 0 | 0 |
| SRR5030096 | 2092 | 142955 | 110 | 443 | 401 | 5710 | 3385 | 7642 | 394 | 15 | 34231 | 10233 | 4396 | 4058 | 0 | 40 | 42043 | 0 | 0 | 0 | 2920 | 632 | 158 | 0 | 16 | 0 | 0 | 0 | 145 | 2532 | 883 | 160 | 0 | 28 | 14 | 0 | 0 |
| SRR5030097 | 1320 | 130479 | 319 | 362 | 276 | 4687 | 2904 | 7518 | 432 | 43 | 35779 | 10744 | 4843 | 4375 | 0 | 55 | 42791 | 0 | 0 | 0 | 1915 | 437 | 92 | 0 | 0 | 0 | 0 | 0 | 105 | 1748 | 590 | 166 | 0 | 22 | 26 | 0 | 0 |
| SRR5030098 | 2211 | 144134 | 131 | 394 | 408 | 5479 | 3305 | 7465 | 356 | 27 | 34819 | 10329 | 4367 | 4095 | 0 | 25 | 44064 | 0 | 0 | 0 | 2963 | 620 | 132 | 0 | 0 | 0 | 0 | 0 | 155 | 2545 | 919 | 146 | 0 | 32 | 15 | 0 | 0 |
| SRR5030100 | 2184 | 142594 | 122 | 425 | 417 | 5622 | 3299 | 7546 | 376 | 21 | 34354 | 10275 | 4566 | 4085 | 0 | 52 | 42422 | 0 | 0 | 0 | 2904 | 604 | 164 | 0 | 0 | 0 | 0 | 0 | 155 | 2560 | 876 | 168 | 0 | 26 | 19 | 0 | 0 |
| SRR5030101 | 1214 | 128443 | 405 | 213 | 372 | 4548 | 2724 | 7699 | 399 | 43 | 35878 | 10949 | 4812 | 4512 | 0 | 27 | 44285 | 0 | 0 | 0 | 1709 | 445 | 100 | 0 | 13 | 0 | 0 | 0 | 61 | 1713 | 636 | 113 | 0 | 22 | 0 | 0 | 0 |
| SRR5030102 | 470 | 27599 | 247 | 393 | 252 | 6183 | 2337 | 3687 | 12379 | 15 | 12993 | 25397 | 2362 | 1787 | 49 | 12 | 54757 | 0 | 0 | 0 | 285 | 286 | 223 | 0 | 20 | 0 | 0 | 0 | 47 | 153 | 163 | 223 | 0 | 71 | 14 | 13 | 0 |
| SRR5030103 | 225 | 19585 | 366 | 178 | 215 | 5487 | 2096 | 3715 | 12385 | 28 | 12970 | 25965 | 2426 | 1816 | 35 | 0 | 61293 | 0 | 0 | 0 | 118 | 167 | 121 | 0 | 28 | 0 | 0 | 0 | 20 | 74 | 104 | 160 | 0 | 62 | 0 | 0 | 0 |
| SRR5030104 | 352 | 27797 | 210 | 520 | 232 | 6378 | 2456 | 3701 | 12311 | 0 | 13140 | 25214 | 2309 | 1799 | 58 | 0 | 52273 | 0 | 0 | 0 | 260 | 280 | 219 | 0 | 23 | 0 | 0 | 0 | 53 | 158 | 154 | 235 | 0 | 66 | 10 | 0 | 0 |
| SRR5030105 | 185 | 19071 | 526 | 224 | 199 | 5575 | 2255 | 3885 | 12494 | 0 | 13294 | 25677 | 2430 | 1875 | 49 | 0 | 54329 | 0 | 0 | 0 | 131 | 185 | 158 | 0 | 15 | 0 | 0 | 0 | 25 | 68 | 97 | 141 | 0 | 55 | 0 | 13 | 0 |
| SRR5030106 | 531 | 28322 | 309 | 508 | 238 | 6123 | 2222 | 3676 | 11965 | 16 | 13089 | 25343 | 2389 | 1882 | 45 | 0 | 54826 | 0 | 0 | 0 | 249 | 293 | 217 | 0 | 21 | 0 | 0 | 0 | 60 | 146 | 165 | 248 | 0 | 80 | 0 | 0 | 0 |
| SRR5030107 | 304 | 19541 | 570 | 181 | 216 | 5430 | 2174 | 3692 | 12251 | 19 | 13017 | 25807 | 2299 | 1857 | 52 | 0 | 62130 | 0 | 0 | 0 | 137 | 164 | 144 | 0 | 17 | 0 | 0 | 0 | 24 | 74 | 94 | 159 | 0 | 58 | 12 | 0 | 0 |
| SRR5030108 | 516 | 28668 | 229 | 410 | 248 | 6237 | 2409 | 3644 | 12213 | 0 | 13455 | 25434 | 2286 | 1892 | 53 | 0 | 52567 | 0 | 0 | 0 | 267 | 290 | 223 | 0 | 18 | 0 | 0 | 0 | 39 | 168 | 149 | 238 | 0 | 83 | 11 | 10 | 0 |
| SRR5030109 | 147 | 19718 | 461 | 167 | 223 | 5655 | 2296 | 3830 | 12548 | 20 | 13353 | 25610 | 2298 | 1878 | 44 | 0 | 56062 | 0 | 0 | 0 | 146 | 156 | 148 | 0 | 17 | 0 | 0 | 0 | 25 | 93 | 97 | 142 | 0 | 60 | 12 | 0 | 0 |
| SRR5030110 | 129 | 24377 | 1373 | 945 | 123 | 2728 | 1291 | 2178 | 8077 | 0 | 57762 | 22216 | 4615 | 5572 | 18 | 13 | 36497 | 0 | 0 | 0 | 419 | 300 | 182 | 0 | 17 | 0 | 0 | 0 | 90 | 182 | 450 | 208 | 0 | 50 | 107 | 49 | 0 |
| SRR5030111 | 39 | 12696 | 3510 | 184 | 145 | 2422 | 1177 | 2262 | 8570 | 11 | 58586 | 22860 | 4901 | 5646 | 0 | 16 | 40135 | 0 | 0 | 0 | 165 | 156 | 104 | 0 | 14 | 0 | 0 | 0 | 36 | 76 | 245 | 137 | 0 | 36 | 80 | 30 | 0 |
| SRR5030112 | 160 | 24378 | 1394 | 926 | 143 | 2948 | 1324 | 2305 | 8229 | 0 | 57335 | 21474 | 4565 | 5187 | 0 | 11 | 34612 | 0 | 0 | 0 | 391 | 289 | 174 | 0 | 17 | 0 | 0 | 0 | 101 | 197 | 473 | 207 | 0 | 54 | 93 | 33 | 0 |
| SRR5030113 | 23 | 12466 | 3190 | 144 | 129 | 2395 | 1196 | 2386 | 8537 | 17 | 58341 | 22867 | 4925 | 5746 | 10 | 0 | 36353 | 0 | 0 | 0 | 156 | 147 | 105 | 0 | 15 | 0 | 0 | 0 | 29 | 80 | 224 | 120 | 0 | 39 | 89 | 34 | 0 |
| SRR5030114 | 81 | 26803 | 1365 | 855 | 151 | 2826 | 1249 | 2368 | 8077 | 0 | 11 | 21971 | 4627 | 5578 | 25 | 0 | 36054 | 0 | 0 | 0 | 385 | 280 | 167 | 0 | 10 | 0 | 0 | 0 | 101 | 200 | 481 | 217 | 0 | 67 | 99 | 50 | 0 |
| SRR5030115 | 34 | 12576 | 3236 | 181 | 116 | 2370 | 1099 | 2240 | 8476 | 0 | 15 | 22902 | 4816 | 5615 | 0 | 11 | 40354 | 0 | 0 | 0 | 152 | 155 | 101 | 0 | 14 | 0 | 0 | 0 | 37 | 68 | 246 | 130 | 0 | 48 | 78 | 40 | 0 |
| SRR5030116 | 114 | 25072 | 1297 | 1183 | 153 | 2777 | 1326 | 2260 | 8208 | 10 | 57535 | 22087 | 4543 | 5580 | 11 | 11 | 34709 | 0 | 0 | 0 | 380 | 306 | 173 | 0 | 15 | 0 | 0 | 0 | 96 | 189 | 473 | 214 | 0 | 41 | 89 | 58 | 0 |
| SRR5030117 | 23 | 12496 | 3161 | 172 | 130 | 2395 | 1168 | 2314 | 8570 | 11 | 58206 | 22866 | 4823 | 5735 | 0 | 12 | 36937 | 0 | 0 | 0 | 165 | 146 | 101 | 0 | 0 | 0 | 0 | 0 | 37 | 85 | 254 | 108 | 0 | 44 | 84 | 40 | 0 |
| SRR5030118 | 1340 | 132615 | 465 | 1576 | 174 | 3833 | 2488 | 5570 | 17063 | 21 | 50883 | 27239 | 5905 | 5962 | 11 | 11 | 42443 | 0 | 0 | 0 | 2563 | 1218 | 35 | 0 | 0 | 0 | 0 | 0 | 168 | 6842 | 1436 | 89 | 0 | 18 | 61 | 0 | 0 |
| SRR5030119 | 688 | 109389 | 552 | 786 | 154 | 3715 | 2583 | 5505 | 16989 | 25 | 48989 | 26996 | 5541 | 5683 | 21 | 0 | 45675 | 0 | 0 | 0 | 1969 | 1031 | 36 | 0 | 0 | 0 | 0 | 0 | 111 | 5160 | 1238 | 77 | 0 | 14 | 67 | 14 | 0 |
| SRR5030120 | 1355 | 133275 | 354 | 1608 | 191 | 3874 | 2364 | 5481 | 16801 | 34 | 50380 | 26647 | 5892 | 5984 | 17 | 0 | 41351 | 0 | 0 | 0 | 2633 | 1175 | 50 | 0 | 0 | 0 | 0 | 0 | 185 | 6822 | 1398 | 82 | 0 | 18 | 45 | 0 | 0 |
| SRR5030121 | 662 | 109058 | 559 | 797 | 174 | 3902 | 2559 | 5522 | 16909 | 33 | 48921 | 26520 | 5754 | 5901 | 0 | 0 | 41373 | 0 | 0 | 0 | 1909 | 984 | 38 | 0 | 0 | 0 | 0 | 0 | 104 | 5187 | 1167 | 81 | 0 | 20 | 56 | 0 | 0 |
| SRR5030122 | 1134 | 133587 | 392 | 1512 | 173 | 3862 | 2368 | 5642 | 16793 | 36 | 50087 | 27292 | 5939 | 5818 | 19 | 0 | 43215 | 0 | 0 | 0 | 2588 | 1241 | 48 | 0 | 0 | 0 | 0 | 12 | 139 | 6936 | 1406 | 71 | 0 | 16 | 65 | 12 | 0 |
| SRR5030123 | 708 | 110827 | 589 | 872 | 187 | 3861 | 2390 | 5418 | 17020 | 39 | 11 | 26413 | 5543 | 5530 | 20 | 0 | 45881 | 0 | 0 | 0 | 1919 | 942 | 48 | 0 | 0 | 0 | 0 | 0 | 126 | 5129 | 1135 | 67 | 0 | 14 | 70 | 11 | 0 |
| SRR5030124 | 1363 | 133427 | 413 | 1370 | 154 | 3736 | 2374 | 5674 | 17278 | 29 | 50058 | 26857 | 5841 | 6038 | 18 | 0 | 41647 | 0 | 0 | 0 | 2671 | 1212 | 34 | 0 | 0 | 0 | 0 | 0 | 176 | 6856 | 1387 | 75 | 0 | 17 | 56 | 0 | 0 |
| SRR5030125 | 767 | 109672 | 528 | 876 | 171 | 3779 | 2440 | 5452 | 16951 | 30 | 48918 | 26411 | 5608 | 5889 | 22 | 0 | 41968 | 0 | 0 | 0 | 1901 | 925 | 32 | 0 | 0 | 0 | 0 | 0 | 112 | 5140 | 1176 | 79 | 0 | 23 | 65 | 18 | 0 |
| SRR5030126 | 823 | 107116 | 1131 | 353 | 116 | 2832 | 990 | 2658 | 8773 | 22 | 31595 | 35570 | 4079 | 2208 | 20 | 67 | 45361 | 0 | 0 | 0 | 59 | 54 | 28 | 0 | 0 | 0 | 0 | 0 | 11 | 39 | 90 | 83 | 0 | 27 | 11 | 0 | 0 |
| SRR5030127 | 241 | 80995 | 1916 | 96 | 91 | 2878 | 934 | 2673 | 8707 | 34 | 30547 | 35098 | 3944 | 2202 | 0 | 64 | 48811 | 0 | 0 | 0 | 31 | 26 | 26 | 0 | 0 | 0 | 0 | 0 | 0 | 22 | 47 | 63 | 0 | 22 | 14 | 0 | 0 |
| SRR5030128 | 705 | 105253 | 1073 | 368 | 98 | 2947 | 988 | 2654 | 8555 | 31 | 31141 | 35769 | 4072 | 2269 | 0 | 65 | 43500 | 0 | 0 | 0 | 62 | 49 | 40 | 0 | 0 | 0 | 0 | 0 | 13 | 61 | 91 | 64 | 0 | 20 | 11 | 0 | 0 |
| SRR5030130 | 772 | 106522 | 1034 | 427 | 108 | 2831 | 1008 | 2635 | 8599 | 22 | 31281 | 35724 | 4246 | 2192 | 0 | 71 | 45752 | 0 | 0 | 0 | 62 | 42 | 42 | 0 | 0 | 0 | 0 | 0 | 15 | 52 | 83 | 79 | 0 | 21 | 12 | 0 | 0 |
| SRR5030131 | 203 | 80179 | 2027 | 119 | 97 | 2740 | 958 | 2604 | 8678 | 28 | 30354 | 35135 | 4067 | 2323 | 0 | 74 | 49493 | 0 | 0 | 0 | 36 | 27 | 32 | 0 | 0 | 0 | 0 | 0 | 0 | 25 | 49 | 50 | 0 | 25 | 13 | 0 | 0 |
| SRR5030132 | 829 | 105056 | 937 | 415 | 101 | 2980 | 1000 | 2584 | 8637 | 0 | 31225 | 35637 | 4089 | 2209 | 0 | 67 | 44180 | 0 | 0 | 0 | 64 | 46 | 37 | 0 | 0 | 0 | 0 | 0 | 14 | 58 | 83 | 75 | 0 | 19 | 0 | 0 | 0 |
| SRR5030133 | 309 | 80951 | 1984 | 187 | 99 | 2827 | 888 | 2688 | 8706 | 29 | 30647 | 34817 | 4098 | 2269 | 11 | 68 | 45060 | 0 | 0 | 0 | 27 | 27 | 32 | 0 | 0 | 0 | 0 | 0 | 0 | 32 | 53 | 59 | 0 | 23 | 0 | 0 | 0 |
| SRR5030134 | 3364 | 66641 | 63 | 639 | 519 | 14660 | 4771 | 9000 | 76 | 17 | 40796 | 17005 | 3887 | 4186 | 0 | 0 | 38897 | 0 | 0 | 0 | 471 | 361 | 83 | 0 | 0 | 0 | 0 | 0 | 128 | 199 | 128 | 43 | 0 | 12 | 30 | 11 | 0 |
| SRR5030135 | 1442 | 50904 | 133 | 321 | 458 | 12778 | 4737 | 9349 | 44 | 31 | 41828 | 17793 | 4086 | 4482 | 0 | 0 | 43846 | 0 | 0 | 0 | 310 | 250 | 56 | 0 | 0 | 0 | 0 | 0 | 71 | 142 | 106 | 30 | 0 | 17 | 32 | 12 | 0 |
| SRR5030136 | 3259 | 67002 | 52 | 622 | 540 | 14747 | 4872 | 8945 | 43 | 24 | 40264 | 16604 | 3997 | 4065 | 0 | 0 | 36630 | 0 | 0 | 0 | 475 | 368 | 77 | 0 | 0 | 0 | 0 | 0 | 130 | 210 | 119 | 42 | 0 | 12 | 34 | 12 | 0 |
| SRR5030137 | 1511 | 50979 | 138 | 237 | 441 | 12966 | 4512 | 9166 | 48 | 14 | 41218 | 17543 | 4014 | 4672 | 0 | 0 | 39920 | 0 | 0 | 0 | 328 | 233 | 53 | 0 | 11 | 0 | 0 | 0 | 76 | 150 | 91 | 38 | 0 | 14 | 24 | 0 | 0 |
| SRR5030138 | 3479 | 66788 | 54 | 627 | 532 | 14571 | 4657 | 8966 | 44 | 12 | 40705 | 16641 | 3874 | 4119 | 0 | 0 | 39396 | 0 | 0 | 0 | 473 | 357 | 80 | 0 | 0 | 0 | 0 | 0 | 145 | 212 | 140 | 40 | 0 | 12 | 37 | 10 | 0 |
| SRR5030139 | 1486 | 51239 | 118 | 305 | 421 | 12770 | 4360 | 8827 | 57 | 21 | 41317 | 17631 | 4165 | 4628 | 0 | 0 | 44320 | 0 | 0 | 0 | 323 | 253 | 42 | 0 | 0 | 0 | 0 | 0 | 85 | 131 | 97 | 28 | 0 | 16 | 34 | 0 | 0 |
| SRR5030140 | 3359 | 66568 | 38 | 645 | 603 | 14925 | 4856 | 8806 | 63 | 13 | 39889 | 16848 | 3963 | 4399 | 0 | 0 | 37208 | 0 | 0 | 0 | 501 | 348 | 82 | 0 | 0 | 0 | 0 | 0 | 132 | 201 | 129 | 48 | 0 | 11 | 26 | 0 | 0 |
| SRR5030141 | 1488 | 51669 | 103 | 266 | 449 | 13035 | 4428 | 9237 | 45 | 21 | 42264 | 17638 | 4247 | 4541 | 0 | 0 | 40817 | 0 | 0 | 0 | 298 | 251 | 55 | 0 | 0 | 0 | 0 | 0 | 70 | 136 | 77 | 30 | 0 | 0 | 38 | 0 | 0 |
| SRR5030142 | 3158 | 11649 | 223 | 1068 | 289 | 5319 | 2367 | 5245 | 123 | 28 | 5656 | 14359 | 1017 | 258 | 96 | 15 | 31180 | 0 | 0 | 0 | 2894 | 669 | 50 | 0 | 0 | 0 | 0 | 16 | 337 | 4029 | 763 | 59 | 0 | 13 | 22 | 0 | 0 |
| SRR5030143 | 1890 | 9688 | 210 | 690 | 213 | 4796 | 2288 | 5746 | 190 | 0 | 5754 | 15339 | 1011 | 296 | 46 | 0 | 36933 | 0 | 0 | 0 | 2230 | 540 | 45 | 0 | 0 | 0 | 0 | 21 | 194 | 3145 | 650 | 45 | 0 | 15 | 24 | 0 | 0 |
| SRR5030144 | 3176 | 11218 | 237 | 1043 | 281 | 5392 | 2457 | 5129 | 112 | 23 | 5793 | 14339 | 1040 | 287 | 92 | 13 | 29626 | 0 | 0 | 0 | 2844 | 700 | 60 | 0 | 0 | 0 | 0 | 22 | 297 | 3983 | 710 | 54 | 0 | 11 | 22 | 0 | 0 |
| SRR5030145 | 1627 | 9441 | 351 | 542 | 215 | 4649 | 2209 | 5857 | 191 | 0 | 5897 | 15514 | 1089 | 207 | 64 | 16 | 32784 | 0 | 0 | 0 | 2226 | 542 | 47 | 0 | 0 | 0 | 0 | 19 | 274 | 3097 | 632 | 59 | 0 | 0 | 16 | 0 | 0 |
| SRR5030146 | 3284 | 11371 | 247 | 1057 | 272 | 5234 | 2497 | 5099 | 140 | 14 | 5573 | 14369 | 1070 | 256 | 72 | 11 | 31200 | 0 | 0 | 0 | 2914 | 674 | 47 | 0 | 0 | 0 | 0 | 21 | 323 | 3988 | 800 | 67 | 0 | 12 | 25 | 0 | 0 |
| SRR5030147 | 2115 | 10234 | 249 | 591 | 177 | 4656 | 2182 | 5435 | 170 | 0 | 5643 | 15718 | 1079 | 254 | 123 | 0 | 37656 | 0 | 0 | 0 | 2271 | 495 | 48 | 0 | 0 | 0 | 0 | 27 | 202 | 3141 | 628 | 48 | 0 | 0 | 21 | 0 | 0 |
| SRR5030148 | 3064 | 11266 | 241 | 1065 | 267 | 5504 | 2415 | 5284 | 129 | 19 | 5647 | 14380 | 1056 | 305 | 77 | 13 | 30215 | 0 | 0 | 0 | 2816 | 652 | 59 | 0 | 0 | 0 | 0 | 22 | 311 | 4039 | 702 | 66 | 0 | 12 | 17 | 11 | 0 |
| SRR5030149 | 2231 | 10087 | 409 | 691 | 198 | 5049 | 2302 | 5866 | 183 | 31 | 5954 | 15456 | 921 | 318 | 64 | 0 | 34563 | 0 | 0 | 0 | 2133 | 587 | 50 | 0 | 0 | 0 | 0 | 12 | 245 | 2906 | 590 | 65 | 0 | 16 | 16 | 0 | 0 |
| SRR5030150 | 1079 | 19475 | 800 | 930 | 168 | 5394 | 2186 | 3959 | 4610 | 55 | 39208 | 19213 | 4222 | 4424 | 21 | 0 | 28954 | 0 | 0 | 0 | 1051 | 1834 | 79 | 0 | 0 | 0 | 0 | 12 | 132 | 3766 | 2173 | 172 | 0 | 30 | 177 | 27 | 0 |
| SRR5030151 | 709 | 17737 | 690 | 572 | 169 | 5327 | 2061 | 3831 | 4537 | 39 | 39965 | 19314 | 4396 | 4556 | 28 | 0 | 30647 | 0 | 0 | 0 | 915 | 1640 | 72 | 0 | 0 | 0 | 0 | 0 | 115 | 3297 | 2135 | 144 | 0 | 43 | 165 | 18 | 0 |
| SRR5030152 | 1066 | 19214 | 720 | 863 | 168 | 5351 | 2072 | 3984 | 4457 | 42 | 39461 | 18953 | 4428 | 4508 | 21 | 0 | 27697 | 0 | 0 | 0 | 1111 | 1772 | 79 | 0 | 0 | 0 | 0 | 12 | 137 | 3796 | 2141 | 169 | 0 | 22 | 173 | 33 | 0 |
| SRR5030153 | 758 | 17101 | 799 | 676 | 243 | 5536 | 2185 | 3613 | 4497 | 143 | 11 | 19115 | 4138 | 4478 | 19 | 0 | 28274 | 0 | 0 | 0 | 843 | 1720 | 67 | 0 | 0 | 0 | 0 | 11 | 93 | 3225 | 1783 | 130 | 0 | 15 | 134 | 22 | 0 |
| SRR5030154 | 1174 | 19727 | 677 | 899 | 162 | 5413 | 2053 | 3858 | 4540 | 56 | 39249 | 19235 | 4316 | 4486 | 14 | 0 | 29172 | 0 | 0 | 0 | 1069 | 1809 | 87 | 0 | 0 | 0 | 0 | 14 | 134 | 3867 | 2197 | 153 | 0 | 32 | 168 | 30 | 0 |
| SRR5030155 | 568 | 16801 | 1177 | 487 | 231 | 5236 | 2210 | 3603 | 4794 | 81 | 38372 | 19099 | 4467 | 4198 | 29 | 0 | 31483 | 0 | 0 | 0 | 902 | 1712 | 84 | 0 | 0 | 0 | 0 | 15 | 99 | 3293 | 2019 | 153 | 0 | 40 | 161 | 29 | 0 |
| SRR5030156 | 1059 | 19356 | 811 | 854 | 177 | 5429 | 2200 | 3964 | 4412 | 51 | 39924 | 19056 | 4428 | 4431 | 23 | 0 | 27640 | 0 | 0 | 0 | 1080 | 1737 | 84 | 0 | 0 | 0 | 0 | 15 | 144 | 3834 | 2116 | 148 | 0 | 25 | 172 | 35 | 0 |
| SRR5030157 | 808 | 17276 | 994 | 394 | 147 | 5336 | 2138 | 3619 | 4541 | 21 | 39763 | 18636 | 4394 | 4522 | 78 | 0 | 28140 | 0 | 0 | 0 | 918 | 1529 | 60 | 0 | 0 | 0 | 0 | 15 | 105 | 3399 | 1975 | 195 | 0 | 26 | 195 | 37 | 0 |
| SRR5030158 | 1631 | 14918 | 832 | 766 | 77 | 4167 | 1163 | 2649 | 11125 | 62 | 44701 | 19910 | 5575 | 4639 | 0 | 0 | 43495 | 0 | 0 | 0 | 5199 | 1996 | 76 | 0 | 0 | 0 | 0 | 13 | 229 | 8977 | 2528 | 148 | 0 | 20 | 71 | 16 | 0 |
| SRR5030159 | 637 | 9731 | 1910 | 409 | 91 | 3661 | 1086 | 2569 | 11213 | 64 | 44333 | 20983 | 5557 | 4734 | 11 | 0 | 48264 | 0 | 0 | 0 | 3282 | 1398 | 55 | 0 | 0 | 0 | 0 | 0 | 121 | 5526 | 1780 | 116 | 0 | 24 | 64 | 21 | 0 |
| SRR5030160 | 1746 | 15175 | 1096 | 760 | 100 | 4210 | 1131 | 2666 | 11049 | 53 | 44659 | 19964 | 5620 | 4477 | 0 | 0 | 40869 | 0 | 0 | 0 | 5292 | 1999 | 87 | 0 | 0 | 0 | 0 | 12 | 218 | 8918 | 2423 | 151 | 0 | 22 | 68 | 19 | 0 |
| SRR5030161 | 753 | 10128 | 1892 | 306 | 84 | 3762 | 1020 | 2694 | 11465 | 37 | 45106 | 20700 | 5861 | 4605 | 0 | 0 | 42124 | 0 | 0 | 0 | 3229 | 1372 | 58 | 0 | 0 | 0 | 0 | 0 | 147 | 5644 | 1696 | 116 | 0 | 23 | 58 | 14 | 0 |
| SRR5030162 | 1580 | 15136 | 950 | 736 | 103 | 4116 | 1128 | 2683 | 11268 | 55 | 44956 | 20087 | 5407 | 4506 | 0 | 0 | 43454 | 0 | 0 | 0 | 5307 | 1993 | 85 | 0 | 0 | 0 | 0 | 11 | 234 | 8914 | 2530 | 142 | 0 | 22 | 72 | 18 | 0 |
| SRR5030163 | 806 | 10550 | 1866 | 272 | 98 | 3618 | 1045 | 2552 | 11057 | 47 | 44408 | 20432 | 5719 | 4403 | 0 | 0 | 48769 | 0 | 0 | 0 | 3267 | 1361 | 64 | 0 | 0 | 0 | 0 | 0 | 138 | 5512 | 1726 | 120 | 0 | 22 | 66 | 13 | 0 |
| SRR5030164 | 1475 | 15067 | 903 | 770 | 84 | 4241 | 1069 | 2769 | 11007 | 52 | 44871 | 19954 | 5640 | 4638 | 0 | 0 | 41553 | 0 | 0 | 0 | 5259 | 1942 | 79 | 0 | 0 | 0 | 0 | 13 | 227 | 8907 | 2402 | 146 | 0 | 18 | 79 | 25 | 0 |
| SRR5030165 | 683 | 9947 | 1886 | 330 | 95 | 3877 | 1039 | 2767 | 11431 | 49 | 44910 | 20123 | 5669 | 4705 | 0 | 0 | 43505 | 0 | 0 | 0 | 3240 | 1309 | 56 | 0 | 0 | 0 | 0 | 0 | 108 | 5540 | 1599 | 118 | 0 | 19 | 70 | 18 | 0 |
| SRR5030166 | 234 | 17575 | 169 | 794 | 331 | 13698 | 3879 | 10711 | 14119 | 13 | 42823 | 10240 | 3691 | 5055 | 0 | 0 | 36575 | 0 | 0 | 0 | 311 | 405 | 239 | 0 | 20 | 0 | 0 | 13 | 78 | 149 | 498 | 338 | 0 | 71 | 38 | 18 | 0 |
| SRR5030167 | 38 | 12303 | 410 | 286 | 318 | 12281 | 3556 | 10502 | 14439 | 48 | 11 | 10291 | 3733 | 5009 | 0 | 0 | 41030 | 0 | 0 | 0 | 202 | 244 | 162 | 0 | 11 | 0 | 0 | 0 | 36 | 92 | 298 | 235 | 0 | 58 | 29 | 15 | 0 |
| SRR5030168 | 85 | 16833 | 156 | 778 | 354 | 14167 | 3978 | 10830 | 13807 | 14 | 43717 | 10086 | 3641 | 5044 | 0 | 0 | 35345 | 0 | 0 | 0 | 328 | 379 | 243 | 0 | 14 | 0 | 0 | 14 | 64 | 171 | 473 | 372 | 0 | 66 | 41 | 20 | 0 |
| SRR5030169 | 50 | 12004 | 321 | 421 | 289 | 12569 | 3619 | 10840 | 14301 | 0 | 42834 | 10378 | 3792 | 5122 | 0 | 0 | 36456 | 0 | 0 | 0 | 188 | 250 | 177 | 0 | 18 | 0 | 0 | 0 | 40 | 91 | 283 | 234 | 0 | 54 | 36 | 23 | 0 |
| SRR5030170 | 121 | 17749 | 283 | 727 | 365 | 13903 | 3723 | 10462 | 13972 | 13 | 43058 | 10047 | 3573 | 4899 | 0 | 0 | 36650 | 0 | 0 | 0 | 356 | 389 | 267 | 0 | 15 | 0 | 0 | 0 | 84 | 169 | 507 | 327 | 0 | 65 | 43 | 14 | 0 |
| SRR5030171 | 78 | 12499 | 370 | 341 | 310 | 12303 | 3430 | 10351 | 14173 | 39 | 12 | 10338 | 3630 | 4980 | 0 | 0 | 41746 | 0 | 0 | 0 | 190 | 241 | 160 | 0 | 17 | 0 | 0 | 0 | 45 | 90 | 296 | 255 | 0 | 56 | 32 | 15 | 0 |
| SRR5030172 | 250 | 17915 | 278 | 874 | 365 | 13855 | 3889 | 10812 | 14000 | 14 | 43670 | 10295 | 3710 | 4994 | 0 | 0 | 35709 | 0 | 0 | 0 | 337 | 396 | 244 | 0 | 17 | 0 | 0 | 14 | 68 | 163 | 436 | 345 | 0 | 71 | 34 | 19 | 0 |
| SRR5030173 | 59 | 12058 | 454 | 355 | 305 | 12471 | 3565 | 10648 | 14378 | 39 | 43344 | 10381 | 3711 | 5100 | 0 | 0 | 37630 | 0 | 0 | 0 | 203 | 246 | 172 | 0 | 15 | 0 | 0 | 0 | 34 | 101 | 280 | 241 | 0 | 53 | 32 | 22 | 0 |
| SRR5030174 | 1682 | 113007 | 400 | 1675 | 46 | 2132 | 670 | 1222 | 10003 | 15 | 36370 | 18747 | 4148 | 3184 | 0 | 0 | 39548 | 0 | 0 | 0 | 935 | 1990 | 80 | 0 | 0 | 0 | 0 | 18 | 171 | 2164 | 2056 | 125 | 0 | 21 | 202 | 38 | 0 |
| SRR5030175 | 781 | 86945 | 1046 | 709 | 41 | 1559 | 559 | 1263 | 10375 | 17 | 36914 | 19489 | 4324 | 3336 | 0 | 0 | 43636 | 0 | 0 | 0 | 603 | 1446 | 62 | 0 | 0 | 0 | 0 | 0 | 86 | 1434 | 1492 | 94 | 0 | 16 | 218 | 48 | 0 |
| SRR5030176 | 1576 | 112807 | 376 | 1606 | 41 | 2006 | 661 | 1156 | 9776 | 26 | 35567 | 17895 | 3947 | 3151 | 0 | 0 | 36638 | 0 | 0 | 0 | 919 | 1932 | 65 | 0 | 0 | 0 | 0 | 13 | 164 | 2148 | 2044 | 127 | 0 | 21 | 221 | 50 | 0 |
| SRR5030177 | 800 | 87489 | 1102 | 743 | 49 | 1641 | 596 | 1308 | 10109 | 25 | 35899 | 19138 | 4289 | 3408 | 0 | 0 | 39096 | 0 | 0 | 0 | 609 | 1393 | 52 | 0 | 0 | 0 | 0 | 0 | 80 | 1346 | 1421 | 103 | 0 | 16 | 218 | 42 | 0 |
| SRR5030178 | 1572 | 113034 | 383 | 1554 | 46 | 1822 | 570 | 1301 | 9781 | 0 | 36525 | 18623 | 4039 | 3204 | 0 | 0 | 38766 | 0 | 0 | 0 | 917 | 1941 | 71 | 0 | 0 | 0 | 0 | 14 | 177 | 2204 | 2081 | 134 | 0 | 25 | 229 | 52 | 0 |
| SRR5030179 | 748 | 87005 | 1110 | 657 | 46 | 1691 | 570 | 1272 | 10393 | 31 | 36348 | 19069 | 4366 | 3282 | 0 | 0 | 43508 | 0 | 0 | 0 | 617 | 1390 | 57 | 0 | 0 | 0 | 0 | 0 | 92 | 1391 | 1411 | 107 | 0 | 21 | 217 | 45 | 0 |
| SRR5030180 | 1607 | 112147 | 407 | 1625 | 74 | 1970 | 598 | 1310 | 9970 | 24 | 35872 | 18364 | 3932 | 3157 | 0 | 13 | 37424 | 0 | 0 | 0 | 957 | 1956 | 76 | 0 | 0 | 0 | 0 | 13 | 160 | 2158 | 2041 | 143 | 0 | 23 | 199 | 40 | 0 |
| SRR5030181 | 788 | 87018 | 1125 | 688 | 46 | 1609 | 559 | 1374 | 10412 | 37 | 36035 | 19072 | 4295 | 3255 | 0 | 0 | 40168 | 0 | 0 | 0 | 586 | 1429 | 53 | 0 | 0 | 0 | 0 | 0 | 115 | 1441 | 1423 | 95 | 0 | 23 | 197 | 49 | 0 |
| SRR5030182 | 247 | 52053 | 288 | 412 | 151 | 2126 | 1051 | 2226 | 9516 | 0 | 58885 | 24638 | 4567 | 4702 | 0 | 0 | 44509 | 0 | 0 | 0 | 99 | 248 | 84 | 0 | 0 | 0 | 0 | 0 | 41 | 93 | 319 | 270 | 0 | 51 | 15 | 0 | 0 |
| SRR5030183 | 46 | 36960 | 311 | 58 | 142 | 1960 | 954 | 2284 | 9717 | 12 | 20 | 25026 | 4455 | 4675 | 0 | 0 | 50068 | 0 | 0 | 0 | 44 | 140 | 60 | 0 | 0 | 0 | 0 | 0 | 16 | 54 | 199 | 180 | 0 | 44 | 12 | 0 | 0 |
| SRR5030184 | 119 | 51937 | 224 | 417 | 154 | 2158 | 1041 | 2232 | 9443 | 60 | 59343 | 24185 | 4401 | 4562 | 0 | 0 | 42337 | 0 | 0 | 0 | 102 | 237 | 106 | 0 | 0 | 0 | 0 | 0 | 32 | 102 | 289 | 249 | 0 | 45 | 14 | 0 | 0 |
| SRR5030185 | 60 | 36374 | 442 | 239 | 115 | 1975 | 1024 | 2227 | 9790 | 0 | 59319 | 24639 | 4581 | 4811 | 0 | 0 | 43427 | 0 | 0 | 0 | 44 | 145 | 55 | 0 | 0 | 0 | 0 | 0 | 15 | 64 | 203 | 161 | 0 | 40 | 10 | 0 | 0 |
| SRR5030186 | 241 | 53107 | 142 | 595 | 134 | 2118 | 1111 | 2166 | 9307 | 0 | 15 | 24692 | 4420 | 4565 | 0 | 0 | 45096 | 0 | 0 | 0 | 76 | 233 | 92 | 0 | 0 | 0 | 0 | 0 | 38 | 114 | 324 | 266 | 0 | 49 | 11 | 0 | 0 |
| SRR5030187 | 58 | 36839 | 489 | 221 | 134 | 1934 | 930 | 2209 | 9675 | 12 | 19 | 24862 | 4451 | 4655 | 0 | 0 | 50594 | 0 | 0 | 0 | 50 | 128 | 60 | 0 | 0 | 0 | 0 | 0 | 15 | 58 | 189 | 185 | 0 | 39 | 11 | 0 | 0 |
| SRR5030188 | 262 | 50843 | 145 | 466 | 157 | 2201 | 1100 | 2197 | 9674 | 15 | 13 | 24574 | 4460 | 4750 | 10 | 13 | 42867 | 0 | 0 | 0 | 95 | 224 | 93 | 0 | 0 | 0 | 0 | 0 | 36 | 118 | 327 | 245 | 0 | 49 | 22 | 0 | 0 |
| SRR5030189 | 60 | 37892 | 513 | 143 | 128 | 1959 | 981 | 2308 | 9840 | 0 | 58847 | 24720 | 4603 | 4737 | 0 | 0 | 45050 | 0 | 0 | 0 | 41 | 136 | 55 | 0 | 0 | 0 | 0 | 0 | 21 | 63 | 186 | 172 | 0 | 36 | 15 | 0 | 0 |
| SRR5030190 | 997 | 45901 | 332 | 129 | 165 | 5603 | 1700 | 5444 | 275 | 0 | 54089 | 42883 | 7276 | 9579 | 25 | 0 | 65016 | 0 | 0 | 0 | 1488 | 1038 | 73 | 0 | 0 | 0 | 0 | 22 | 376 | 1997 | 875 | 134 | 0 | 19 | 32 | 0 | 0 |
| SRR5030191 | 522 | 39709 | 416 | 80 | 134 | 5373 | 1745 | 5409 | 276 | 0 | 52938 | 42284 | 7038 | 9371 | 22 | 0 | 69343 | 0 | 0 | 0 | 1132 | 922 | 77 | 0 | 0 | 0 | 0 | 18 | 277 | 1522 | 776 | 119 | 0 | 18 | 34 | 0 | 0 |
| SRR5030192 | 990 | 45959 | 333 | 141 | 150 | 5454 | 1768 | 5577 | 260 | 0 | 54018 | 43017 | 7116 | 9467 | 15 | 0 | 62047 | 0 | 0 | 0 | 1462 | 1045 | 73 | 0 | 0 | 0 | 0 | 26 | 362 | 2008 | 834 | 131 | 0 | 24 | 28 | 0 | 0 |
| SRR5030193 | 459 | 41640 | 459 | 92 | 162 | 5580 | 1840 | 5940 | 287 | 0 | 52540 | 41694 | 7082 | 10281 | 23 | 14 | 61984 | 0 | 0 | 0 | 1172 | 892 | 78 | 0 | 12 | 0 | 0 | 15 | 227 | 1631 | 728 | 95 | 0 | 17 | 30 | 0 | 0 |
| SRR5030194 | 968 | 45979 | 336 | 112 | 161 | 5590 | 1716 | 5561 | 261 | 0 | 54044 | 42443 | 7327 | 9132 | 27 | 0 | 65343 | 0 | 0 | 0 | 1493 | 1006 | 71 | 0 | 0 | 0 | 0 | 18 | 376 | 2062 | 858 | 135 | 0 | 16 | 31 | 0 | 0 |
| SRR5030196 | 926 | 46286 | 299 | 156 | 145 | 5641 | 1738 | 5572 | 283 | 0 | 54259 | 42965 | 7141 | 9744 | 24 | 0 | 62831 | 0 | 0 | 0 | 1503 | 1021 | 64 | 0 | 0 | 0 | 0 | 16 | 387 | 1959 | 839 | 117 | 0 | 14 | 31 | 0 | 0 |
| SRR5030197 | 477 | 39977 | 468 | 73 | 149 | 5645 | 1781 | 5408 | 256 | 0 | 53347 | 42187 | 7333 | 9059 | 33 | 0 | 64801 | 0 | 0 | 0 | 1183 | 860 | 63 | 0 | 10 | 0 | 0 | 0 | 239 | 1557 | 741 | 125 | 0 | 25 | 39 | 0 | 0 |
| SRR5030198 | 2263 | 41416 | 100 | 419 | 934 | 23371 | 7235 | 10282 | 25611 | 46 | 68359 | 29673 | 5711 | 8220 | 12 | 20 | 43385 | 0 | 0 | 0 | 1013 | 511 | 81 | 0 | 0 | 0 | 0 | 20 | 347 | 691 | 321 | 87 | 0 | 11 | 34 | 0 | 0 |
| SRR5030199 | 945 | 32795 | 240 | 160 | 940 | 23355 | 7003 | 9988 | 25191 | 51 | 13 | 29045 | 5488 | 8161 | 15 | 15 | 46990 | 0 | 0 | 0 | 717 | 408 | 84 | 0 | 0 | 0 | 0 | 15 | 264 | 541 | 259 | 79 | 0 | 14 | 32 | 0 | 0 |
| SRR5030200 | 1996 | 42170 | 191 | 328 | 941 | 23525 | 7276 | 10230 | 25215 | 44 | 67815 | 29598 | 5654 | 8223 | 10 | 14 | 41719 | 0 | 0 | 0 | 987 | 542 | 79 | 0 | 0 | 0 | 0 | 38 | 337 | 663 | 303 | 79 | 0 | 16 | 29 | 0 | 0 |
| SRR5030201 | 797 | 33150 | 190 | 273 | 903 | 23238 | 7287 | 10088 | 25562 | 0 | 67232 | 28976 | 5642 | 8110 | 15 | 25 | 41668 | 0 | 0 | 0 | 757 | 403 | 82 | 0 | 0 | 0 | 0 | 0 | 234 | 514 | 256 | 87 | 0 | 0 | 38 | 0 | 0 |
| SRR5030202 | 2049 | 41692 | 182 | 359 | 901 | 23487 | 6932 | 10270 | 25375 | 37 | 11 | 29468 | 5707 | 8341 | 14 | 24 | 43334 | 0 | 0 | 0 | 969 | 536 | 87 | 0 | 0 | 0 | 0 | 26 | 355 | 716 | 329 | 81 | 0 | 13 | 35 | 0 | 0 |
| SRR5030203 | 1121 | 33586 | 260 | 252 | 874 | 23010 | 6789 | 9975 | 25042 | 37 | 66249 | 29093 | 5628 | 8098 | 11 | 21 | 46839 | 0 | 0 | 0 | 662 | 432 | 86 | 0 | 0 | 0 | 0 | 16 | 245 | 479 | 245 | 70 | 0 | 13 | 34 | 0 | 0 |
| SRR5030204 | 2158 | 41504 | 166 | 394 | 908 | 23468 | 7242 | 10394 | 25200 | 33 | 68202 | 29311 | 5866 | 8291 | 18 | 14 | 42099 | 0 | 0 | 0 | 984 | 526 | 86 | 0 | 0 | 0 | 0 | 21 | 357 | 728 | 316 | 91 | 0 | 12 | 33 | 0 | 0 |
| SRR5030205 | 1046 | 33648 | 258 | 258 | 902 | 23620 | 7087 | 10406 | 25547 | 23 | 66423 | 28950 | 5667 | 8263 | 25 | 17 | 43134 | 0 | 0 | 0 | 713 | 396 | 83 | 0 | 12 | 0 | 0 | 15 | 241 | 537 | 240 | 87 | 0 | 17 | 33 | 0 | 0 |
| SRR5030206 | 1466 | 28674 | 218 | 187 | 134 | 5953 | 1705 | 5774 | 160 | 12 | 11 | 24526 | 5855 | 2951 | 13 | 0 | 80292 | 0 | 0 | 0 | 794 | 348 | 79 | 0 | 0 | 0 | 0 | 0 | 58 | 576 | 241 | 89 | 0 | 25 | 13 | 0 | 0 |
| SRR5030207 | 752 | 24499 | 237 | 40 | 140 | 5681 | 1635 | 5380 | 173 | 0 | 53726 | 23971 | 5467 | 2945 | 0 | 0 | 85475 | 0 | 0 | 0 | 718 | 269 | 78 | 0 | 10 | 0 | 0 | 0 | 33 | 462 | 251 | 83 | 0 | 33 | 0 | 13 | 0 |
| SRR5030208 | 1455 | 29272 | 179 | 173 | 146 | 6041 | 1732 | 5915 | 163 | 27 | 56520 | 24498 | 5705 | 2933 | 17 | 0 | 76750 | 0 | 0 | 0 | 773 | 325 | 95 | 0 | 0 | 0 | 0 | 0 | 52 | 602 | 237 | 87 | 0 | 27 | 11 | 0 | 0 |
| SRR5030209 | 898 | 22080 | 245 | 41 | 172 | 5980 | 1664 | 5521 | 180 | 0 | 54601 | 24006 | 5851 | 2896 | 13 | 13 | 76938 | 0 | 0 | 0 | 637 | 298 | 73 | 0 | 10 | 0 | 0 | 0 | 31 | 504 | 185 | 89 | 0 | 0 | 18 | 0 | 0 |
| SRR5030210 | 1582 | 29269 | 179 | 192 | 132 | 5744 | 1713 | 5718 | 157 | 0 | 11 | 24560 | 5720 | 2915 | 15 | 0 | 80273 | 0 | 0 | 0 | 812 | 339 | 86 | 0 | 0 | 0 | 0 | 0 | 60 | 642 | 250 | 95 | 0 | 26 | 0 | 0 | 0 |
| SRR5030211 | 980 | 25238 | 123 | 163 | 126 | 5801 | 1656 | 5759 | 138 | 0 | 54274 | 23800 | 5361 | 2821 | 17 | 0 | 86998 | 0 | 0 | 0 | 729 | 316 | 71 | 0 | 13 | 0 | 0 | 0 | 51 | 484 | 194 | 74 | 0 | 36 | 10 | 0 | 0 |

### Supp Table S8.1.Columns_B_to_AL

| | | | | | | | | | | | | | | | | | | | | | | | | | | | | | | | | | | | | | |
|---|---|---|---|---|---|---|---|---|---|---|---|---|---|---|---|---|---|---|---|---|---|---|---|---|---|---|---|---|---|---|---|---|---|---|---|---|---|
| SRR5030212 | 1362 | 28888 | 187 | 187 | 142 | 5843 | 1823 | 5911 | 178 | 19 | 56718 | 24504 | 5864 | 2949 | 14 | 0 | 78036 | 0 | 0 | 0 | 795 | 357 | 88 | 0 | 0 | 0 | 0 | 0 | 59 | 647 | 231 | 98 | 0 | 27 | 10 | 0 | 0 |
| SRR5030213 | 831 | 23023 | 249 | 208 | 163 | 5922 | 1812 | 5600 | 154 | 83 | 55090 | 23969 | 5947 | 3081 | 0 | 0 | 79196 | 0 | 0 | 0 | 635 | 314 | 52 | 0 | 13 | 0 | 0 | 0 | 50 | 543 | 206 | 81 | 0 | 24 | 18 | 0 | 0 |
| SRR5030214 | 295 | 29599 | 234 | 2121 | 1206 | 16650 | 8352 | 21798 | 107 | 0 | 10 | 27148 | 7190 | 12121 | 17 | 0 | 49681 | 0 | 0 | 0 | 3072 | 2161 | 144 | 0 | 0 | 0 | 0 | 0 | 69 | 1956 | 755 | 104 | 0 | 14 | 0 | 0 | 0 |
| SRR5030216 | 304 | 29270 | 146 | 2108 | 1249 | 16773 | 8415 | 21694 | 123 | 15 | 62262 | 27403 | 7081 | 12153 | 11 | 0 | 47905 | 0 | 0 | 0 | 3083 | 2202 | 146 | 0 | 0 | 0 | 0 | 0 | 66 | 1936 | 746 | 106 | 0 | 16 | 0 | 0 | 0 |
| SRR5030217 | 221 | 27473 | 184 | 1475 | 1328 | 17513 | 8459 | 22082 | 100 | 37 | 32 | 26936 | 7487 | 12048 | 0 | 23 | 48038 | 0 | 0 | 0 | 2949 | 2301 | 140 | 0 | 0 | 0 | 0 | 12 | 56 | 1833 | 844 | 124 | 0 | 0 | 16 | 0 | 0 |
| SRR5030218 | 391 | 29850 | 199 | 2088 | 1159 | 16746 | 8480 | 21530 | 115 | 0 | 62732 | 27451 | 7118 | 11873 | 0 | 0 | 50252 | 0 | 0 | 0 | 3142 | 2221 | 138 | 0 | 0 | 0 | 0 | 0 | 72 | 1991 | 760 | 109 | 0 | 0 | 0 | 0 | 0 |
| SRR5030220 | 389 | 29666 | 198 | 2081 | 1195 | 16727 | 8470 | 22190 | 116 | 11 | 62659 | 27196 | 7010 | 12106 | 0 | 0 | 48201 | 0 | 0 | 0 | 3087 | 2177 | 148 | 0 | 0 | 0 | 0 | 0 | 89 | 1999 | 743 | 113 | 0 | 13 | 11 | 0 | 0 |
| SRR5030221 | 260 | 27348 | 223 | 967 | 1281 | 17005 | 8433 | 21946 | 130 | 0 | 60959 | 26865 | 7098 | 12262 | 0 | 0 | 48896 | 0 | 0 | 0 | 2973 | 2262 | 160 | 0 | 0 | 0 | 0 | 0 | 72 | 1962 | 791 | 128 | 0 | 16 | 12 | 0 | 0 |
| SRR5030222 | 147 | 13597 | 229 | 1458 | 125 | 2011 | 1149 | 3248 | 13593 | 33 | 15 | 20710 | 6170 | 10217 | 49 | 0 | 43956 | 0 | 0 | 0 | 2844 | 9043 | 334 | 0 | 0 | 0 | 0 | 0 | 379 | 2957 | 3622 | 296 | 0 | 38 | 35 | 0 | 0 |
| SRR5030223 | 42 | 8102 | 343 | 365 | 108 | 2008 | 1089 | 3164 | 13774 | 64 | 67187 | 20085 | 5955 | 10398 | 59 | 0 | 46857 | 0 | 0 | 0 | 1559 | 5943 | 303 | 0 | 0 | 0 | 0 | 0 | 203 | 1643 | 2406 | 243 | 0 | 36 | 40 | 13 | 0 |
| SRR5030224 | 158 | 13161 | 263 | 1523 | 107 | 2007 | 1121 | 3316 | 13723 | 53 | 14 | 20196 | 6078 | 10450 | 42 | 0 | 42443 | 0 | 0 | 0 | 2831 | 8950 | 377 | 0 | 16 | 0 | 0 | 0 | 393 | 2954 | 3624 | 279 | 0 | 42 | 30 | 11 | 0 |
| SRR5030225 | 0 | 7672 | 368 | 386 | 116 | 2027 | 1092 | 3170 | 13885 | 61 | 67505 | 20011 | 6065 | 10629 | 51 | 0 | 42588 | 0 | 0 | 0 | 1551 | 5932 | 279 | 0 | 12 | 0 | 0 | 0 | 193 | 1671 | 2403 | 237 | 0 | 41 | 42 | 15 | 0 |
| SRR5030226 | 100 | 13035 | 300 | 1434 | 131 | 2061 | 1073 | 3222 | 13843 | 100 | 21 | 20261 | 6043 | 10387 | 56 | 0 | 44920 | 0 | 0 | 0 | 2805 | 9043 | 422 | 0 | 14 | 0 | 0 | 0 | 386 | 2941 | 3668 | 268 | 0 | 50 | 33 | 14 | 0 |
| SRR5030227 | 47 | 7835 | 319 | 358 | 90 | 1954 | 1097 | 3113 | 13673 | 65 | 66674 | 19962 | 6023 | 10327 | 63 | 0 | 47348 | 0 | 0 | 0 | 1572 | 5947 | 283 | 0 | 12 | 0 | 0 | 0 | 196 | 1613 | 2417 | 232 | 0 | 40 | 41 | 14 | 0 |
| SRR5030229 | 0 | 7800 | 347 | 443 | 107 | 2027 | 1089 | 3232 | 13773 | 88 | 11 | 20102 | 5972 | 10464 | 51 | 0 | 43825 | 0 | 0 | 0 | 1592 | 5959 | 282 | 0 | 12 | 0 | 0 | 0 | 202 | 1662 | 2403 | 220 | 0 | 35 | 32 | 12 | 0 |
| SRR5030230 | 474 | 81637 | 373 | 677 | 179 | 3079 | 1567 | 3323 | 1731 | 0 | 13 | 13711 | 4974 | 6143 | 58 | 15 | 34896 | 0 | 0 | 0 | 683 | 937 | 106 | 0 | 0 | 0 | 0 | 11 | 75 | 957 | 917 | 100 | 0 | 18 | 20 | 0 | 0 |
| SRR5030231 | 131 | 45128 | 862 | 165 | 166 | 2696 | 1466 | 3221 | 1655 | 17 | 42641 | 14262 | 5214 | 6579 | 38 | 0 | 39075 | 0 | 0 | 0 | 287 | 455 | 56 | 0 | 0 | 0 | 0 | 0 | 31 | 367 | 418 | 55 | 0 | 15 | 0 | 0 | 0 |
| SRR5030232 | 435 | 82188 | 254 | 616 | 194 | 3348 | 1622 | 3178 | 1506 | 0 | 41970 | 13696 | 5168 | 6383 | 39 | 0 | 33345 | 0 | 0 | 0 | 685 | 1045 | 110 | 0 | 0 | 0 | 0 | 0 | 70 | 994 | 905 | 133 | 0 | 30 | 26 | 0 | 0 |
| SRR5030233 | 179 | 45483 | 863 | 157 | 168 | 2697 | 1511 | 3254 | 1641 | 17 | 42856 | 14230 | 5232 | 6694 | 35 | 0 | 35470 | 0 | 0 | 0 | 275 | 443 | 55 | 0 | 0 | 0 | 0 | 0 | 35 | 376 | 413 | 65 | 0 | 13 | 11 | 0 | 0 |
| SRR5030234 | 587 | 82422 | 380 | 794 | 242 | 3224 | 1647 | 3283 | 1558 | 35 | 11 | 13642 | 5046 | 6325 | 71 | 11 | 35111 | 0 | 0 | 0 | 706 | 973 | 93 | 0 | 0 | 0 | 0 | 0 | 79 | 939 | 889 | 131 | 0 | 25 | 18 | 0 | 0 |
| SRR5030235 | 120 | 45696 | 907 | 116 | 150 | 2648 | 1497 | 3185 | 1673 | 30 | 42504 | 14202 | 5185 | 6514 | 39 | 0 | 39484 | 0 | 0 | 0 | 266 | 452 | 46 | 0 | 0 | 0 | 0 | 0 | 29 | 374 | 427 | 66 | 0 | 0 | 0 | 0 | 0 |
| SRR5030236 | 633 | 80390 | 352 | 598 | 211 | 3275 | 1833 | 3098 | 1539 | 0 | 41619 | 13931 | 5194 | 6233 | 64 | 0 | 34230 | 0 | 0 | 0 | 672 | 960 | 101 | 0 | 0 | 0 | 0 | 0 | 108 | 898 | 924 | 110 | 0 | 11 | 0 | 0 | 0 |
| SRR5030237 | 157 | 45507 | 969 | 114 | 163 | 2717 | 1456 | 3266 | 1658 | 17 | 42488 | 14125 | 5256 | 6685 | 38 | 0 | 36402 | 0 | 0 | 0 | 272 | 446 | 56 | 0 | 0 | 0 | 0 | 0 | 34 | 369 | 418 | 55 | 0 | 15 | 10 | 0 | 0 |
| SRR5030238 | 1133 | 46428 | 1492 | 552 | 240 | 4295 | 1835 | 4414 | 83 | 14 | 17 | 30688 | 8060 | 6453 | 0 | 11 | 62155 | 0 | 0 | 0 | 473 | 139 | 118 | 0 | 13 | 0 | 0 | 20 | 108 | 325 | 116 | 136 | 0 | 29 | 17 | 0 | 0 |
| SRR5030239 | 620 | 37705 | 2468 | 322 | 226 | 4330 | 1767 | 4473 | 81 | 12 | 97656 | 29975 | 7764 | 6112 | 11 | 15 | 66792 | 0 | 0 | 0 | 344 | 131 | 92 | 0 | 0 | 0 | 0 | 11 | 54 | 224 | 99 | 110 | 0 | 30 | 22 | 0 | 0 |
| SRR5030240 | 1155 | 46251 | 1448 | 632 | 216 | 4362 | 1822 | 4562 | 66 | 31 | 24 | 30414 | 8275 | 6487 | 0 | 0 | 60136 | 0 | 0 | 0 | 452 | 139 | 102 | 0 | 0 | 0 | 0 | 23 | 103 | 316 | 140 | 120 | 0 | 27 | 22 | 0 | 0 |
| SRR5030241 | 618 | 37429 | 2137 | 258 | 242 | 4398 | 1857 | 4599 | 83 | 13 | 99595 | 29826 | 7887 | 6237 | 0 | 0 | 60398 | 0 | 0 | 0 | 302 | 94 | 75 | 0 | 0 | 0 | 0 | 0 | 69 | 250 | 96 | 98 | 0 | 30 | 18 | 11 | 0 |
| SRR5030242 | 1179 | 46993 | 1575 | 579 | 227 | 4247 | 1823 | 4582 | 75 | 44 | 100192 | 30369 | 8065 | 6381 | 0 | 0 | 63140 | 0 | 0 | 0 | 476 | 145 | 108 | 0 | 0 | 0 | 0 | 22 | 95 | 353 | 132 | 127 | 0 | 20 | 15 | 0 | 0 |
| SRR5030243 | 632 | 37491 | 2275 | 455 | 239 | 4139 | 1813 | 4455 | 66 | 13 | 20 | 30090 | 7962 | 6107 | 13 | 0 | 67405 | 0 | 0 | 0 | 318 | 118 | 91 | 0 | 0 | 0 | 0 | 0 | 61 | 257 | 98 | 87 | 0 | 22 | 19 | 0 | 0 |
| SRR5030244 | 1088 | 46361 | 1620 | 630 | 233 | 4189 | 1795 | 4570 | 80 | 30 | 102160 | 30260 | 8225 | 6454 | 0 | 0 | 60502 | 0 | 0 | 0 | 478 | 143 | 107 | 0 | 0 | 0 | 0 | 20 | 81 | 308 | 120 | 129 | 0 | 42 | 17 | 11 | 0 |
| SRR5030245 | 412 | 37341 | 2228 | 322 | 239 | 4407 | 1740 | 4552 | 66 | 77 | 99218 | 30063 | 7889 | 6347 | 0 | 0 | 61785 | 0 | 0 | 0 | 340 | 100 | 82 | 0 | 0 | 0 | 0 | 11 | 69 | 226 | 91 | 103 | 0 | 29 | 0 | 0 | 0 |
| SRR5030246 | 268 | 56392 | 3028 | 1894 | 949 | 18693 | 10761 | 21322 | 235 | 278 | 29854 | 7226 | 4092 | 4804 | 11 | 23 | 41868 | 0 | 0 | 0 | 3064 | 421 | 159 | 0 | 0 | 0 | 0 | 0 | 115 | 2140 | 725 | 175 | 0 | 30 | 11 | 0 | 0 |
| SRR5030247 | 76 | 31434 | 6470 | 435 | 844 | 16361 | 10469 | 21787 | 251 | 304 | 30186 | 7355 | 4276 | 4929 | 13 | 0 | 47251 | 0 | 0 | 0 | 1298 | 222 | 93 | 0 | 0 | 0 | 0 | 0 | 47 | 983 | 377 | 107 | 0 | 15 | 10 | 0 | 0 |
| SRR5030248 | 262 | 56160 | 3018 | 1822 | 889 | 18455 | 11093 | 21200 | 250 | 262 | 29544 | 7098 | 4198 | 4904 | 26 | 0 | 39947 | 0 | 0 | 0 | 3083 | 399 | 164 | 0 | 0 | 0 | 0 | 0 | 120 | 2281 | 693 | 139 | 0 | 14 | 13 | 0 | 0 |
| SRR5030249 | 93 | 32052 | 6464 | 546 | 811 | 16786 | 10474 | 22118 | 228 | 359 | 29958 | 7246 | 4171 | 5076 | 15 | 12 | 41853 | 0 | 0 | 0 | 1322 | 243 | 123 | 0 | 0 | 0 | 0 | 0 | 49 | 1008 | 366 | 89 | 0 | 23 | 0 | 0 | 0 |
| SRR5030250 | 281 | 58321 | 3136 | 1968 | 934 | 18146 | 10880 | 20786 | 216 | 227 | 12 | 7282 | 4085 | 4836 | 12 | 0 | 42592 | 0 | 0 | 0 | 3012 | 424 | 156 | 0 | 0 | 0 | 0 | 0 | 139 | 2263 | 816 | 176 | 0 | 27 | 15 | 0 | 0 |
| SRR5030251 | 77 | 32384 | 6443 | 442 | 812 | 16038 | 10163 | 21328 | 251 | 273 | 30014 | 7413 | 4157 | 4860 | 17 | 0 | 48139 | 0 | 0 | 0 | 1305 | 218 | 105 | 0 | 0 | 0 | 0 | 0 | 44 | 1010 | 372 | 104 | 0 | 26 | 11 | 0 | 0 |
| SRR5030252 | 345 | 56569 | 2959 | 1802 | 948 | 18455 | 10913 | 21600 | 220 | 267 | 29684 | 7094 | 4088 | 4802 | 16 | 0 | 40557 | 0 | 0 | 0 | 3005 | 410 | 158 | 0 | 0 | 0 | 0 | 0 | 103 | 2193 | 701 | 178 | 0 | 29 | 0 | 0 | 0 |
| SRR5030253 | 57 | 33274 | 6192 | 523 | 849 | 16362 | 10316 | 21926 | 230 | 265 | 29950 | 7317 | 4284 | 5009 | 14 | 0 | 43250 | 0 | 0 | 0 | 1334 | 227 | 103 | 0 | 0 | 0 | 0 | 0 | 53 | 976 | 365 | 89 | 0 | 16 | 12 | 0 | 0 |
| SRR5030254 | 24 | 1745 | 2773 | 41 | 42 | 1463 | 630 | 1550 | 7469 | 19 | 16707 | 11903 | 1670 | 1197 | 0 | 0 | 18075 | 0 | 0 | 0 | 24 | 0 | 12 | 0 | 0 | 0 | 0 | 0 | 0 | 0 | 0 | 0 | 0 | 0 | 0 | 0 | 0 |
| SRR5030255 | 26 | 1807 | 2760 | 47 | 36 | 1479 | 637 | 1530 | 7351 | 10 | 16462 | 11742 | 1652 | 1204 | 0 | 0 | 17374 | 0 | 0 | 0 | 20 | 0 | 0 | 0 | 0 | 0 | 0 | 0 | 0 | 0 | 0 | 0 | 0 | 0 | 0 | 0 | 0 |
| SRR5030256 | 31 | 1786 | 2698 | 41 | 40 | 1431 | 622 | 1501 | 7253 | 13 | 16474 | 11704 | 1611 | 1188 | 0 | 0 | 18236 | 0 | 0 | 0 | 22 | 0 | 12 | 0 | 0 | 0 | 0 | 0 | 0 | 0 | 10 | 0 | 0 | 0 | 0 | 0 | 0 |
| SRR5030257 | 24 | 1797 | 2773 | 46 | 41 | 1449 | 617 | 1541 | 7361 | 26 | 16611 | 11796 | 1658 | 1197 | 0 | 0 | 17250 | 0 | 0 | 0 | 25 | 0 | 0 | 0 | 0 | 0 | 0 | 0 | 0 | 0 | 0 | 0 | 0 | 0 | 0 | 0 | 0 |
| SRR5030258 | 17 | 17293 | 16512 | 31 | 282 | 5347 | 3772 | 5299 | 5045 | 113 | 8112 | 19210 | 1124 | 774 | 68 | 0 | 24544 | 0 | 0 | 0 | 42 | 0 | 16 | 0 | 0 | 0 | 0 | 0 | 0 | 41 | 23 | 27 | 0 | 0 | 0 | 0 | 0 |
| SRR5030259 | 32 | 17370 | 16700 | 43 | 260 | 5328 | 3856 | 5277 | 4942 | 108 | 8025 | 19210 | 1098 | 760 | 67 | 0 | 23541 | 0 | 0 | 0 | 37 | 0 | 18 | 0 | 0 | 0 | 0 | 0 | 13 | 38 | 28 | 23 | 0 | 0 | 0 | 0 | 0 |
| SRR5030260 | 17 | 17810 | 16365 | 28 | 287 | 5172 | 3701 | 5192 | 4991 | 80 | 7967 | 19112 | 1078 | 758 | 69 | 0 | 24398 | 0 | 0 | 0 | 40 | 0 | 16 | 0 | 0 | 0 | 0 | 0 | 13 | 35 | 33 | 24 | 0 | 0 | 0 | 0 | 0 |
| SRR5030261 | 31 | 17507 | 17068 | 28 | 279 | 5279 | 3846 | 5279 | 4993 | 104 | 7890 | 19068 | 1087 | 779 | 67 | 0 | 23424 | 0 | 0 | 0 | 36 | 0 | 17 | 0 | 0 | 0 | 0 | 0 | 0 | 35 | 24 | 24 | 0 | 0 | 0 | 0 | 0 |
| SRR5030262 | 25 | 6808 | 42806 | 56 | 121 | 2645 | 1189 | 1654 | 11421 | 61 | 5705 | 12259 | 289 | 98 | 0 | 0 | 16549 | 0 | 0 | 0 | 136 | 111 | 72 | 0 | 11 | 0 | 0 | 0 | 25 | 52 | 164 | 67 | 0 | 22 | 0 | 0 | 0 |
| SRR5030263 | 26 | 6890 | 42298 | 58 | 103 | 2672 | 1149 | 1694 | 11384 | 32 | 5638 | 12277 | 286 | 100 | 0 | 0 | 15847 | 0 | 0 | 0 | 129 | 114 | 79 | 0 | 12 | 0 | 0 | 0 | 31 | 52 | 136 | 54 | 0 | 21 | 11 | 0 | 0 |
| SRR5030265 | 10 | 6685 | 42069 | 83 | 107 | 2643 | 1158 | 1651 | 11349 | 46 | 5497 | 12099 | 290 | 105 | 0 | 0 | 15805 | 0 | 0 | 0 | 136 | 110 | 70 | 0 | 13 | 0 | 0 | 0 | 31 | 57 | 159 | 58 | 0 | 20 | 0 | 0 | 0 |
| SRR5030266 | 0 | 2375 | 9392 | 15 | 50 | 791 | 936 | 779 | 30320 | 61 | 55402 | 24189 | 6007 | 7402 | 57 | 0 | 11293 | 0 | 0 | 0 | 142 | 23 | 35 | 0 | 0 | 0 | 0 | 0 | 0 | 34 | 23 | 12 | 0 | 0 | 0 | 0 | 0 |
| SRR5030267 | 0 | 2700 | 9540 | 126 | 57 | 805 | 859 | 790 | 30551 | 63 | 54124 | 23864 | 5918 | 7475 | 34 | 0 | 11051 | 0 | 0 | 0 | 133 | 18 | 30 | 0 | 0 | 0 | 0 | 0 | 0 | 39 | 17 | 10 | 0 | 0 | 0 | 0 | 0 |
| SRR5030268 | 0 | 2777 | 9317 | 62 | 63 | 833 | 841 | 782 | 30344 | 69 | 54500 | 23843 | 5914 | 7333 | 42 | 0 | 11387 | 0 | 0 | 0 | 142 | 20 | 38 | 0 | 0 | 0 | 0 | 0 | 0 | 36 | 21 | 11 | 0 | 0 | 0 | 0 | 0 |
| SRR5030270 | 0 | 29860 | 11487 | 53 | 25 | 1968 | 1347 | 2122 | 13 | 105 | 21311 | 21901 | 1423 | 2473 | 46 | 0 | 33715 | 0 | 0 | 0 | 91 | 34 | 48 | 0 | 12 | 0 | 0 | 0 | 14 | 62 | 59 | 57 | 0 | 21 | 0 | 0 | 0 |
| SRR5030271 | 0 | 30281 | 11473 | 42 | 34 | 2024 | 1442 | 2165 | 10 | 125 | 21478 | 21781 | 1509 | 2475 | 43 | 0 | 32711 | 0 | 0 | 0 | 105 | 32 | 49 | 0 | 11 | 0 | 0 | 0 | 13 | 72 | 62 | 54 | 0 | 25 | 0 | 0 | 0 |
| SRR5030272 | 27 | 31229 | 11709 | 47 | 24 | 2031 | 1353 | 2103 | 14 | 147 | 21350 | 21845 | 1504 | 2367 | 37 | 0 | 34005 | 0 | 0 | 0 | 93 | 32 | 47 | 0 | 11 | 0 | 0 | 0 | 11 | 61 | 54 | 56 | 0 | 24 | 0 | 0 | 0 |
| SRR5030273 | 33 | 31379 | 11533 | 53 | 33 | 2074 | 1388 | 2138 | 12 | 120 | 21320 | 21694 | 1443 | 2493 | 43 | 0 | 32254 | 0 | 0 | 0 | 90 | 32 | 49 | 0 | 10 | 0 | 0 | 0 | 13 | 58 | 56 | 59 | 0 | 27 | 0 | 0 | 0 |
| SRR5030274 | 33 | 18916 | 27974 | 20 | 379 | 9488 | 3894 | 7641 | 3184 | 99 | 9375 | 14997 | 773 | 569 | 15 | 0 | 21561 | 0 | 0 | 0 | 96 | 16 | 17 | 0 | 0 | 0 | 0 | 0 | 0 | 39 | 12 | 14 | 0 | 0 | 0 | 0 | 0 |
| SRR5030275 | 35 | 18286 | 27374 | 42 | 401 | 9417 | 3930 | 7604 | 3145 | 56 | 9076 | 14679 | 810 | 613 | 13 | 0 | 20459 | 0 | 0 | 0 | 95 | 18 | 19 | 0 | 0 | 0 | 0 | 0 | 0 | 33 | 12 | 14 | 0 | 0 | 0 | 0 | 0 |
| SRR5030276 | 20 | 19025 | 29079 | 0 | 406 | 9279 | 3943 | 7402 | 3234 | 61 | 9344 | 14770 | 772 | 544 | 15 | 0 | 21391 | 0 | 0 | 0 | 92 | 18 | 22 | 0 | 0 | 0 | 0 | 0 | 0 | 37 | 13 | 15 | 0 | 0 | 0 | 0 | 0 |
| SRR5030277 | 14 | 17967 | 28419 | 20 | 406 | 9378 | 3828 | 7470 | 3158 | 47 | 9289 | 14744 | 775 | 570 | 11 | 0 | 20460 | 0 | 0 | 0 | 98 | 14 | 20 | 0 | 0 | 0 | 0 | 0 | 0 | 34 | 0 | 16 | 0 | 0 | 0 | 0 | 0 |
| SRR5030278 | 29 | 5862 | 29846 | 61 | 687 | 23072 | 10841 | 19912 | 0 | 46 | 10069 | 13498 | 1093 | 364 | 0 | 0 | 33743 | 0 | 0 | 0 | 15 | 14 | 0 | 0 | 0 | 0 | 0 | 0 | 49 | 50 | 106 | 39 | 0 | 15 | 0 | 0 | 0 |
| SRR5030279 | 44 | 5701 | 29556 | 82 | 682 | 23254 | 11005 | 20334 | 0 | 49 | 10038 | 13294 | 1019 | 373 | 0 | 0 | 32179 | 0 | 0 | 0 | 16 | 17 | 0 | 0 | 0 | 0 | 0 | 0 | 45 | 53 | 90 | 40 | 0 | 14 | 0 | 0 | 0 |
| SRR5030280 | 35 | 5735 | 30183 | 82 | 648 | 22974 | 10734 | 19650 | 0 | 60 | 9992 | 13373 | 993 | 362 | 0 | 0 | 33632 | 0 | 0 | 0 | 14 | 18 | 0 | 0 | 0 | 0 | 0 | 0 | 48 | 51 | 95 | 37 | 0 | 12 | 0 | 0 | 0 |
| SRR5030281 | 35 | 5855 | 29794 | 102 | 687 | 23162 | 10800 | 19911 | 0 | 55 | 10035 | 13313 | 1030 | 367 | 0 | 0 | 31992 | 0 | 0 | 0 | 15 | 16 | 0 | 0 | 0 | 0 | 0 | 0 | 45 | 49 | 91 | 38 | 0 | 10 | 0 | 0 | 0 |
| SRR5030282 | 0 | 7334 | 2161 | 62 | 14 | 335 | 147 | 623 | 6559 | 135 | 35228 | 21217 | 5324 | 3053 | 0 | 0 | 83721 | 0 | 0 | 0 | 13 | 15 | 14 | 0 | 0 | 0 | 0 | 0 | 0 | 0 | 12 | 24 | 0 | 15 | 0 | 0 | 0 |
| SRR5030283 | 33 | 7638 | 2155 | 54 | 17 | 348 | 164 | 641 | 6576 | 103 | 35090 | 21029 | 5294 | 3081 | 0 | 0 | 80842 | 0 | 0 | 0 | 15 | 13 | 15 | 0 | 0 | 0 | 0 | 0 | 0 | 0 | 13 | 21 | 0 | 14 | 0 | 0 | 0 |
| SRR5030284 | 63 | 7750 | 2095 | 48 | 13 | 326 | 162 | 618 | 6551 | 111 | 34814 | 20925 | 5259 | 3011 | 0 | 0 | 84060 | 0 | 0 | 0 | 15 | 15 | 16 | 0 | 0 | 0 | 0 | 0 | 0 | 10 | 11 | 24 | 0 | 15 | 0 | 0 | 0 |
| SRR5030285 | 47 | 7712 | 2121 | 53 | 14 | 342 | 158 | 627 | 6540 | 190 | 35034 | 21145 | 5372 | 3091 | 0 | 0 | 80386 | 0 | 0 | 0 | 13 | 13 | 17 | 0 | 0 | 0 | 0 | 0 | 0 | 0 | 12 | 20 | 0 | 13 | 0 | 0 | 0 |
| SRR5030287 | 0 | 5200 | 1745 | 26 | 26 | 750 | 320 | 1114 | 14514 | 61 | 35556 | 12451 | 4372 | 3398 | 0 | 0 | 83119 | 0 | 0 | 0 | 13 | 14 | 23 | 0 | 0 | 0 | 0 | 0 | 0 | 18 | 13 | 61 | 0 | 30 | 0 | 0 | 0 |
| SRR5030289 | 17 | 5231 | 1684 | 43 | 24 | 734 | 302 | 1112 | 14417 | 120 | 35705 | 12693 | 4301 | 3381 | 0 | 0 | 82652 | 0 | 0 | 0 | 17 | 11 | 23 | 0 | 0 | 0 | 0 | 0 | 0 | 17 | 13 | 65 | 0 | 32 | 0 | 0 | 0 |
| SRR5030290 | 16 | 29403 | 2181 | 31 | 90 | 1674 | 967 | 2316 | 4147 | 47 | 64170 | 38628 | 6377 | 4769 | 0 | 43 | 46061 | 0 | 0 | 0 | 68 | 105 | 94 | 0 | 14 | 0 | 0 | 0 | 0 | 27 | 62 | 73 | 0 | 36 | 14 | 0 | 0 |
| SRR5030291 | 22 | 29952 | 2269 | 22 | 88 | 1683 | 949 | 2346 | 4109 | 60 | 63749 | 38516 | 6320 | 4814 | 0 | 56 | 44142 | 0 | 0 | 0 | 67 | 97 | 104 | 0 | 14 | 0 | 0 | 0 | 0 | 27 | 53 | 74 | 0 | 33 | 11 | 0 | 0 |
| SRR5030292 | 21 | 29337 | 2029 | 48 | 91 | 1692 | 944 | 2318 | 4072 | 43 | 63465 | 38360 | 6371 | 4610 | 0 | 41 | 46063 | 0 | 0 | 0 | 73 | 91 | 101 | 0 | 17 | 0 | 0 | 0 | 0 | 22 | 55 | 61 | 0 | 29 | 13 | 11 | 0 |
| SRR5030293 | 32 | 29048 | 2185 | 27 | 79 | 1694 | 980 | 2247 | 4105 | 27 | 63264 | 38740 | 6382 | 4837 | 0 | 77 | 44157 | 0 | 0 | 0 | 73 | 102 | 109 | 0 | 13 | 0 | 0 | 0 | 0 | 22 | 63 | 70 | 0 | 37 | 12 | 0 | 0 |
| SRR5030294 | 40 | 37406 | 2110 | 46 | 72 | 1276 | 730 | 1699 | 9823 | 66 | 54378 | 22875 | 4497 | 4146 | 0 | 56 | 40903 | 0 | 0 | 0 | 38 | 79 | 54 | 0 | 0 | 0 | 0 | 0 | 0 | 14 | 54 | 47 | 0 | 18 | 0 | 0 | 0 |
| SRR5030295 | 28 | 36442 | 1809 | 28 | 72 | 1264 | 692 | 1703 | 9709 | 70 | 53937 | 22542 | 4365 | 4124 | 0 | 41 | 39193 | 0 | 0 | 0 | 40 | 74 | 51 | 0 | 0 | 0 | 0 | 0 | 0 | 16 | 56 | 34 | 0 | 19 | 0 | 0 | 0 |
| SRR5030296 | 0 | 37640 | 1914 | 27 | 70 | 1277 | 677 | 1644 | 9739 | 108 | 53465 | 22354 | 4462 | 4124 | 0 | 50 | 41183 | 0 | 0 | 0 | 44 | 79 | 48 | 0 | 0 | 0 | 0 | 0 | 0 | 18 | 47 | 41 | 0 | 18 | 0 | 0 | 0 |
| SRR5030297 | 47 | 36854 | 1972 | 27 | 74 | 1305 | 715 | 1677 | 9640 | 74 | 54064 | 22457 | 4475 | 4104 | 0 | 50 | 39195 | 0 | 0 | 0 | 47 | 85 | 44 | 0 | 0 | 0 | 0 | 0 | 0 | 19 | 42 | 39 | 0 | 22 | 0 | 0 | 0 |
| SRR5034616 | 99 | 4357 | 1234 | 420 | 56 | 1601 | 1603 | 2174 | 1882 | 25 | 3670 | 2149 | 404 | 1451 | 0 | 206 | 17923 | 0 | 0 | 0 | 0 | 10 | 0 | 0 | 0 | 0 | 0 | 0 | 0 | 11 | 20 | 0 | 0 | 0 | 0 | 0 | 0 |
| SRR5034617 | 164 | 19031 | 949 | 379 | 43 | 990 | 963 | 1218 | 1050 | 15 | 4896 | 4522 | 477 | 1906 | 0 | 195 | 22379 | 0 | 0 | 0 | 0 | 0 | 0 | 0 | 0 | 0 | 0 | 0 | 0 | 0 | 16 | 0 | 0 | 0 | 0 | 0 | 0 |
| SRR5034618 | 826 | 18724 | 1311 | 1030 | 180 | 3043 | 3762 | 2818 | 5719 | 13 | 22697 | 16623 | 1441 | 4708 | 0 | 339 | 44777 | 0 | 0 | 0 | 20 | 17 | 0 | 0 | 0 | 0 | 0 | 0 | 26 | 19 | 39 | 11 | 0 | 0 | 0 | 0 | 0 |
| SRR5034619 | 29 | 2250 | 772 | 124 | 97 | 2077 | 2194 | 2749 | 2456 | 0 | 8153 | 3760 | 623 | 1979 | 0 | 245 | 25598 | 0 | 0 | 0 | 0 | 0 | 0 | 0 | 0 | 0 | 0 | 0 | 0 | 0 | 18 | 0 | 0 | 0 | 0 | 0 | 0 |
| SRR5034620 | 209 | 6148 | 2107 | 283 | 30 | 553 | 536 | 1238 | 1975 | 0 | 2736 | 1092 | 182 | 644 | 0 | 116 | 6446 | 0 | 0 | 0 | 12 | 26 | 0 | 0 | 0 | 0 | 0 | 0 | 11 | 13 | 40 | 14 | 0 | 0 | 0 | 0 | 0 |
| SRR5034621 | 750 | 25441 | 1444 | 763 | 36 | 667 | 728 | 1090 | 1934 | 13 | 5905 | 2524 | 358 | 1080 | 0 | 58 | 8960 | 0 | 0 | 0 | 23 | 13 | 0 | 0 | 0 | 0 | 0 | 0 | 20 | 30 | 38 | 13 | 0 | 0 | 0 | 0 | 0 |
| SRR5034622 | 643 | 18152 | 1735 | 978 | 219 | 5093 | 5219 | 4050 | 4695 | 19 | 18298 | 21516 | 1980 | 6582 | 0 | 561 | 54321 | 0 | 0 | 0 | 14 | 13 | 0 | 0 | 0 | 0 | 0 | 0 | 15 | 22 | 37 | 0 | 0 | 0 | 0 | 0 | 0 |
| SRR5034623 | 0 | 1639 | 5418 | 168 | 0 | 150 | 215 | 1286 | 891 | 0 | 1155 | 403 | 127 | 434 | 0 | 60 | 2906 | 0 | 0 | 0 | 13 | 136 | 109 | 0 | 0 | 0 | 0 | 0 | 0 | 15 | 349 | 409 | 0 | 0 | 0 | 0 | 0 |
| SRR5034625 | 170 | 9717 | 9065 | 1043 | 67 | 1408 | 1641 | 3591 | 3805 | 0 | 5719 | 2387 | 459 | 1586 | 0 | 368 | 10630 | 0 | 0 | 0 | 16 | 248 | 117 | 0 | 0 | 0 | 0 | 0 | 31 | 15 | 685 | 647 | 0 | 25 | 0 | 0 | 0 |
| SRR5034626 | 60 | 16458 | 84339 | 822 | 22 | 356 | 554 | 3428 | 4543 | 0 | 2500 | 1049 | 564 | 763 | 0 | 174 | 7988 | 0 | 0 | 0 | 0 | 300 | 1044 | 0 | 0 | 0 | 0 | 0 | 18 | 19 | 954 | 3178 | 0 | 63 | 47 | 0 | 0 |
| SRR5034627 | 93 | 12112 | 5197 | 226 | 0 | 49 | 58 | 444 | 410 | 0 | 394 | 187 | 48 | 123 | 0 | 66 | 700 | 0 | 0 | 0 | 0 | 112 | 35 | 0 | 0 | 0 | 0 | 0 | 0 | 0 | 111 | 102 | 0 | 0 | 0 | 0 | 0 |
| SRR5034628 | 739 | 6988 | 3309 | 594 | 0 | 255 | 287 | 2951 | 2101 | 16 | 2020 | 771 | 124 | 435 | 0 | 216 | 2331 | 0 | 0 | 0 | 20 | 291 | 46 | 0 | 0 | 0 | 0 | 0 | 26 | 0 | 193 | 79 | 0 | 0 | 0 | 0 | 0 |
| SRR5034629 | 71 | 13228 | 10321 | 846 | 0 | 143 | 263 | 3963 | 1068 | 0 | 1033 | 567 | 140 | 1196 | 0 | 257 | 2179 | 0 | 0 | 0 | 14 | 418 | 88 | 0 | 0 | 0 | 0 | 0 | 10 | 11 | 1067 | 270 | 0 | 0 | 0 | 0 | 0 |
| SRR5034630 | 35 | 4756 | 6392 | 247 | 57 | 1508 | 1392 | 14338 | 3466 | 12 | 4367 | 2061 | 319 | 1652 | 0 | 711 | 15367 | 0 | 0 | 0 | 19 | 99 | 91 | 0 | 0 | 0 | 0 | 0 | 21 | 26 | 267 | 301 | 0 | 0 | 0 | 0 | 0 |
| SRR5034632 | 42 | 4721 | 5743 | 548 | 11 | 365 | 496 | 2280 | 1704 | 0 | 3125 | 1261 | 259 | 995 | 0 | 873 | 5046 | 0 | 0 | 0 | 0 | 71 | 117 | 0 | 0 | 0 | 0 | 0 | 14 | 12 | 115 | 483 | 0 | 62 | 0 | 0 | 0 |
| SRR5034634 | 56 | 6917 | 10459 | 298 | 21 | 1048 | 795 | 3046 | 2331 | 0 | 2964 | 1223 | 403 | 1094 | 0 | 739 | 12941 | 0 | 0 | 0 | 0 | 62 | 189 | 0 | 0 | 0 | 0 | 0 | 0 | 0 | 146 | 773 | 0 | 149 | 0 | 0 | 0 |
| SRR5034635 | 112 | 8734 | 10280 | 653 | 51 | 1397 | 1621 | 2388 | 1715 | 32 | 6379 | 2055 | 481 | 1624 | 0 | 522 | 9804 | 0 | 0 | 0 | 0 | 75 | 113 | 0 | 0 | 0 | 0 | 0 | 0 | 0 | 139 | 376 | 0 | 50 | 0 | 0 | 0 |
| SRR5034636 | 75 | 9592 | 6051 | 226 | 98 | 2131 | 1614 | 3122 | 2610 | 0 | 4969 | 1815 | 416 | 1773 | 0 | 480 | 22229 | 0 | 0 | 0 | 0 | 171 | 173 | 0 | 0 | 0 | 0 | 0 | 0 | 11 | 340 | 440 | 0 | 119 | 0 | 0 | 0 |
| SRR5034637 | 743 | 9608 | 2870 | 154 | 37 | 493 | 428 | 2127 | 2073 | 0 | 4900 | 1608 | 320 | 1364 | 0 | 491 | 6195 | 0 | 0 | 0 | 0 | 50 | 73 | 0 | 0 | 0 | 0 | 0 | 12 | 0 | 53 | 176 | 0 | 46 | 0 | 0 | 0 |
| SRR5034639 | 224 | 23729 | 6112 | 224 | 109 | 3411 | 2360 | 6063 | 3727 | 0 | 7731 | 2913 | 695 | 2221 | 0 | 1240 | 30671 | 0 | 0 | 0 | 14 | 107 | 94 | 0 | 0 | 0 | 0 | 0 | 31 | 20 | 174 | 305 | 0 | 63 | 0 | 0 | 0 |
| SRR5034641 | 42 | 1574 | 305 | 132 | 12 | 196 | 155 | 5077 | 1284 | 0 | 719 | 354 | 56 | 338 | 0 | 263 | 992 | 0 | 0 | 0 | 35 | 78 | 17 | 0 | 0 | 0 | 0 | 0 | 0 | 21 | 75 | 37 | 0 | 0 | 0 | 0 | 0 |
| SRR5034642 | 49 | 5361 | 2784 | 223 | 0 | 196 | 153 | 1345 | 681 | 0 | 722 | 327 | 46 | 150 | 0 | 81 | 1264 | 0 | 0 | 0 | 10 | 37 | 270 | 0 | 19 | 0 | 0 | 0 | 11 | 0 | 84 | 992 | 0 | 188 | 0 | 0 | 0 |
| SRR5034643 | 0 | 7373 | 7402 | 179 | 44 | 1220 | 1502 | 4478 | 3703 | 11 | 3671 | 1484 | 311 | 1215 | 0 | 197 | 11234 | 0 | 0 | 0 | 20 | 268 | 532 | 0 | 0 | 0 | 0 | 0 | 52 | 24 | 781 | 1498 | 0 | 19 | 0 | 0 | 0 |
| SRR5034644 | 52 | 13133 | 12457 | 606 | 95 | 1329 | 1449 | 4095 | 2786 | 35 | 7686 | 4728 | 986 | 1529 | 0 | 2123 | 14268 | 0 | 0 | 0 | 33 | 422 | 193 | 0 | 0 | 0 | 0 | 0 | 34 | 38 | 793 | 1011 | 0 | 15 | 23 | 0 | 0 |
| SRR5034645 | 190 | 10289 | 2983 | 361 | 0 | 84 | 83 | 709 | 525 | 0 | 570 | 264 | 46 | 156 | 0 | 81 | 1075 | 0 | 0 | 0 | 0 | 135 | 52 | 0 | 0 | 0 | 0 | 0 | 0 | 0 | 294 | 123 | 0 | 0 | 0 | 0 | 0 |
| SRR5034646 | 256 | 14029 | 5487 | 706 | 48 | 908 | 908 | 2582 | 2737 | 0 | 4842 | 1696 | 387 | 1674 | 0 | 333 | 9541 | 0 | 0 | 0 | 13 | 311 | 176 | 0 | 0 | 0 | 0 | 0 | 23 | 23 | 772 | 577 | 0 | 21 | 0 | 0 | 0 |
| SRR5034647 | 51 | 8951 | 15661 | 139 | 26 | 370 | 401 | 3554 | 16321 | 0 | 2561 | 1396 | 260 | 656 | 0 | 249 | 5542 | 0 | 0 | 0 | 0 | 216 | 509 | 0 | 0 | 0 | 0 | 0 | 15 | 12 | 2206 | 4074 | 0 | 41 | 28 | 0 | 0 |
| SRR5034648 | 576 | 7893 | 7999 | 2242 | 51 | 785 | 1102 | 3641 | 5322 | 15 | 9269 | 3903 | 647 | 1435 | 0 | 752 | 9809 | 0 | 0 | 0 | 18 | 205 | 86 | 0 | 0 | 0 | 0 | 0 | 34 | 17 | 321 | 254 | 0 | 27 | 0 | 0 | 0 |
| SRR5034649 | 47 | 13969 | 15433 | 610 | 53 | 1183 | 1280 | 3397 | 7097 | 0 | 4861 | 1629 | 302 | 1392 | 0 | 648 | 11382 | 0 | 0 | 0 | 10 | 140 | 219 | 0 | 13 | 0 | 0 | 0 | 23 | 11 | 473 | 2614 | 17 | 369 | 14 | 0 | 0 |
| SRR5034650 | 542 | 38767 | 5923 | 1452 | 11 | 261 | 269 | 1293 | 1518 | 25 | 2752 | 1257 | 379 | 1110 | 0 | 221 | 5767 | 0 | 0 | 0 | 20 | 774 | 134 | 0 | 0 | 0 | 0 | 0 | 25 | 34 | 971 | 374 | 0 | 12 | 15 | 0 | 0 |
| SRR5034651 | 216 | 11347 | 2957 | 870 | 25 | 372 | 528 | 1412 | 1124 | 0 | 4128 | 1636 | 374 | 1020 | 0 | 147 | 5406 | 0 | 0 | 0 | 16 | 172 | 170 | 0 | 0 | 0 | 0 | 0 | 18 | 0 | 167 | 255 | 0 | 0 | 0 | 0 | 0 |
| SRR5034652 | 123 | 16600 | 13334 | 778 | 14 | 275 | 290 | 1234 | 1176 | 0 | 1653 | 592 | 218 | 769 | 0 | 175 | 5126 | 0 | 0 | 0 | 14 | 271 | 374 | 0 | 0 | 0 | 0 | 0 | 16 | 14 | 526 | 690 | 0 | 40 | 25 | 0 | 0 |
| SRR5034653 | 111 | 21798 | 14075 | 586 | 22 | 345 | 420 | 1304 | 1460 | 0 | 2419 | 890 | 302 | 1085 | 0 | 188 | 7035 | 0 | 0 | 0 | 0 | 245 | 377 | 0 | 0 | 0 | 0 | 0 | 18 | 11 | 542 | 752 | 0 | 29 | 31 | 0 | 0 |
| SRR5034654 | 44 | 23243 | 19622 | 613 | 24 | 506 | 540 | 1502 | 1388 | 0 | 3227 | 1609 | 458 | 1292 | 0 | 315 | 12410 | 0 | 0 | 0 | 11 | 292 | 343 | 0 | 0 | 0 | 0 | 0 | 29 | 17 | 864 | 1109 | 0 | 100 | 47 | 0 | 0 |
| SRR5034655 | 65 | 11547 | 7943 | 855 | 12 | 428 | 394 | 1213 | 2001 | 0 | 2533 | 891 | 228 | 892 | 0 | 155 | 6210 | 0 | 0 | 0 | 0 | 283 | 388 | 0 | 0 | 0 | 0 | 0 | 28 | 0 | 579 | 1289 | 0 | 56 | 18 | 0 | 0 |
| SRR5034656 | 144 | 7601 | 6868 | 586 | 68 | 1073 | 1353 | 2854 | 2522 | 0 | 6401 | 2251 | 714 | 1991 | 0 | 330 | 7776 | 0 | 0 | 0 | 0 | 52 | 162 | 0 | 0 | 0 | 0 | 0 | 19 | 0 | 139 | 1079 | 0 | 47 | 11 | 0 | 0 |
| SRR5034657 | 39 | 23823 | 15185 | 357 | 15 | 293 | 327 | 2844 | 3247 | 0 | 2138 | 834 | 246 | 882 | 0 | 447 | 3680 | 0 | 0 | 0 | 0 | 157 | 212 | 0 | 0 | 0 | 0 | 0 | 12 | 12 | 476 | 1640 | 0 | 189 | 20 | 0 | 0 |
| SRR5034658 | 165 | 6661 | 56298 | 430 | 41 | 476 | 500 | 2425 | 3208 | 11 | 3857 | 1629 | 470 | 737 | 0 | 399 | 6080 | 0 | 0 | 0 | 10 | 297 | 926 | 0 | 0 | 0 | 0 | 0 | 28 | 11 | 364 | 1793 | 0 | 50 | 30 | 0 | 0 |
| SRR5034659 | 12 | 12432 | 61733 | 239 | 20 | 312 | 346 | 2176 | 2394 | 0 | 1564 | 776 | 285 | 395 | 0 | 469 | 5463 | 0 | 0 | 0 | 0 | 334 | 1446 | 0 | 0 | 0 | 0 | 0 | 11 | 0 | 477 | 3644 | 0 | 55 | 51 | 0 | 0 |
| SRR5034660 | 26 | 7516 | 61113 | 256 | 23 | 425 | 419 | 1995 | 2605 | 0 | 2524 | 1003 | 284 | 576 | 0 | 300 | 6130 | 0 | 0 | 0 | 0 | 219 | 1051 | 0 | 0 | 0 | 0 | 0 | 0 | 0 | 452 | 2850 | 0 | 60 | 35 | 0 | 0 |
| SRR5034661 | 11 | 4446 | 5854 | 812 | 30 | 697 | 389 | 400 | 5127 | 0 | 2588 | 1394 | 99 | 327 | 0 | 1401 | 1310 | 0 | 0 | 0 | 0 | 34 | 213 | 0 | 0 | 0 | 0 | 0 | 0 | 0 | 143 | 1417 | 0 | 44 | 0 | 0 | 0 |
| SRR5034662 | 105 | 14252 | 20744 | 402 | 70 | 874 | 1496 | 2125 | 3296 | 0 | 6454 | 2466 | 428 | 1202 | 0 | 187 | 9076 | 0 | 0 | 0 | 0 | 262 | 397 | 0 | 0 | 0 | 0 | 0 | 26 | 11 | 721 | 2049 | 0 | 27 | 14 | 0 | 0 |
| SRR5034663 | 99 | 25866 | 7749 | 669 | 31 | 487 | 381 | 2058 | 3481 | 0 | 2301 | 1262 | 273 | 650 | 0 | 944 | 8924 | 0 | 0 | 0 | 15 | 338 | 323 | 0 | 10 | 0 | 0 | 0 | 30 | 19 | 565 | 1664 | 0 | 174 | 16 | 0 | 0 |
| SRR5054304 | 11 | 119 | 194516 | 50 | 34134 | 44363 | 173310 | 23257 | 56678 | 1912 | 28213 | 9792 | 18794 | 26503 | 22821 | 13959 | 7203 | 0 | 0 | 0 | 0 | 15 | 21 | 0 | 0 | 0 | 0 | 0 | 0 | 0 | 0 | 0 | 0 | 0 | 0 | 0 | 0 |
| SRR5054305 | 39 | 651 | 151611 | 164 | 25811 | 42565 | 75339 | 11561 | 101782 | 528 | 24434 | 8380 | 17320 | 26121 | 45011 | 15042 | 8479 | 0 | 0 | 0 | 37 | 143 | 28 | 14 | 0 | 0 | 0 | 0 | 0 | 11 | 0 | 0 | 0 | 0 | 0 | 0 | 0 |

### Supp Table S8.1.Columns_B_to_AL

| | | | | | | | | | | | | | | | | | | | | | | | | | | | | | | | | | | | | | |
|---|---|---|---|---|---|---|---|---|---|---|---|---|---|---|---|---|---|---|---|---|---|---|---|---|---|---|---|---|---|---|---|---|---|---|---|---|---|
| SRR5087508 | 0 | 54290 | 54676 | 34 | 2669 | 33457 | 19918 | 39581 | 40123 | 48 | 5058 | 35711 | 1891 | 425 | 0 | 12018 | 30215 | 0 | 0 | 0 | 332 | 312 | 151 | 0 | 0 | 0 | 0 | 29 | 242 | 328 | 1454 | 539 | 0 | 42 | 13 | 0 | 0 |
| SRR5087510 | 0 | 54839 | 49239 | 43 | 2741 | 28320 | 21727 | 43153 | 46908 | 44 | 4247 | 31425 | 1676 | 409 | 0 | 11178 | 31082 | 0 | 0 | 0 | 400 | 363 | 180 | 0 | 0 | 0 | 0 | 34 | 287 | 385 | 1653 | 671 | 0 | 41 | 10 | 0 | 0 |
| SRR5087512 | 0 | 57893 | 52984 | 43 | 2538 | 28272 | 20221 | 37269 | 40304 | 53 | 4421 | 32101 | 1700 | 392 | 0 | 11109 | 33130 | 0 | 0 | 0 | 361 | 366 | 168 | 0 | 0 | 0 | 0 | 34 | 301 | 352 | 1645 | 574 | 0 | 43 | 13 | 0 | 0 |
| SRR5087516 | 0 | 12146 | 92878 | 57 | 2452 | 26995 | 21188 | 50667 | 41713 | 37 | 4994 | 32388 | 1854 | 602 | 0 | 14227 | 39180 | 0 | 0 | 0 | 329 | 469 | 150 | 0 | 0 | 0 | 0 | 69 | 438 | 445 | 2717 | 891 | 12 | 49 | 10 | 0 | 0 |
| SRR5087517 | 0 | 14725 | 99719 | 53 | 2195 | 26061 | 21546 | 50061 | 41796 | 60 | 4594 | 34782 | 1705 | 556 | 0 | 13299 | 36302 | 0 | 0 | 0 | 270 | 395 | 100 | 0 | 0 | 0 | 0 | 52 | 353 | 418 | 2539 | 642 | 0 | 43 | 0 | 0 | 0 |
| SRR5087519 | 0 | 15295 | 101696 | 45 | 2280 | 26032 | 22347 | 45355 | 42055 | 36 | 4632 | 34366 | 1688 | 519 | 0 | 12903 | 36851 | 0 | 0 | 0 | 304 | 424 | 109 | 0 | 0 | 0 | 0 | 61 | 400 | 437 | 2729 | 650 | 0 | 43 | 11 | 0 | 0 |
| SRR5087522 | 0 | 114004 | 6413 | 55 | 2546 | 30148 | 17173 | 36039 | 32069 | 63 | 5150 | 33686 | 2135 | 320 | 0 | 10946 | 32988 | 0 | 0 | 0 | 171 | 63 | 40 | 0 | 0 | 0 | 0 | 0 | 62 | 107 | 204 | 92 | 0 | 18 | 0 | 0 | 0 |
| SRR5087527 | 0 | 115582 | 8096 | 51 | 2734 | 30928 | 18956 | 38809 | 33601 | 93 | 5146 | 32134 | 2207 | 343 | 0 | 11390 | 30488 | 0 | 0 | 0 | 154 | 72 | 41 | 0 | 0 | 0 | 0 | 0 | 62 | 118 | 195 | 84 | 0 | 16 | 0 | 0 | 0 |
| SRR5087554 | 13 | 113874 | 7392 | 41 | 2505 | 29805 | 17580 | 37732 | 31923 | 256 | 5123 | 36141 | 2181 | 332 | 0 | 10718 | 30760 | 0 | 0 | 0 | 140 | 73 | 39 | 0 | 0 | 0 | 0 | 0 | 53 | 118 | 204 | 81 | 0 | 14 | 0 | 0 | 0 |
| SRR5090283 | 778 | 3874 | 15569 | 899 | 32267 | 23134 | 396630 | 32824 | 2471 | 1557 | 32235 | 11756 | 27883 | 19994 | 739 | 24 | 13530 | 0 | 0 | 0 | 13 | 10 | 16 | 0 | 0 | 0 | 13 | 29 | 50 | 25 | 0 | 0 | 0 | 0 | 0 | 0 | 0 |
| SRR5092303 | 59 | 197 | 169634 | 59 | 927 | 2569 | 6015 | 2581 | 45395 | 492 | 14 | 83 | 17 | 60862 | 288 | 88 | 19348 | 0 | 0 | 0 | 0 | 0 | 0 | 0 | 0 | 0 | 0 | 0 | 0 | 0 | 0 | 0 | 0 | 0 | 0 | 0 | 0 |
| SRR5092304 | 0 | 117 | 167273 | 117 | 947 | 2417 | 6615 | 2669 | 39645 | 630 | 14 | 69 | 16 | 75417 | 260 | 73 | 24553 | 0 | 0 | 0 | 0 | 0 | 0 | 0 | 0 | 0 | 0 | 0 | 0 | 0 | 0 | 0 | 0 | 0 | 0 | 0 | 0 |
| SRR5092305 | 0 | 176 | 102714 | 220 | 476 | 1304 | 3781 | 1954 | 38395 | 308 | 14430 | 12237 | 26538 | 53134 | 120 | 61 | 12721 | 0 | 0 | 0 | 0 | 0 | 0 | 0 | 0 | 0 | 0 | 0 | 0 | 0 | 0 | 0 | 0 | 0 | 0 | 0 | 0 |
| SRR5092836 | 114 | 26150 | 23341 | 190 | 7947 | 41314 | 51029 | 52886 | 1682 | 76 | 9505 | 5951 | 1612 | 17318 | 0 | 66754 | 10149 | 0 | 0 | 24 | 202 | 444 | 50 | 0 | 0 | 0 | 0 | 26 | 182 | 137 | 967 | 48 | 0 | 0 | 0 | 0 | 0 |
| SRR5100734 | 295 | 704 | 7597 | 123729 | 14824 | 19649 | 207608 | 29578 | 7482 | 25 | 30348 | 26991 | 11957 | 61604 | 22166 | 18111 | 4126 | 0 | 0 | 0 | 0 | 0 | 0 | 14 | 221 | 187 | 0 | 0 | 0 | 0 | 0 | 0 | 0 | 0 | 137 | 189 | 15 |
| SRR5100735 | 2466 | 8940 | 3780 | 87859 | 16596 | 29779 | 89684 | 6964 | 15990 | 240 | 46843 | 51018 | 6427 | 40797 | 6895 | 10481 | 4227 | 0 | 0 | 36 | 66 | 944 | 352 | 131 | 360 | 540 | 0 | 0 | 67 | 33 | 0 | 0 | 0 | 0 | 142 | 797 | 28 |
| SRR5104206 | 27 | 37898 | 1857 | 100 | 635 | 5804 | 5496 | 5441 | 220 | 17 | 22570 | 12 | 4591 | 3610 | 0 | 13569 | 10879 | 0 | 0 | 0 | 365 | 37 | 67 | 0 | 0 | 0 | 0 | 0 | 22 | 46 | 16 | 28 | 0 | 0 | 0 | 0 | 0 |
| SRR5104207 | 19 | 49798 | 6937 | 65 | 445 | 4155 | 3637 | 4224 | 214 | 27 | 38257 | 25 | 7260 | 3802 | 0 | 7806 | 10654 | 0 | 0 | 0 | 226 | 24 | 69 | 0 | 0 | 0 | 0 | 0 | 0 | 29 | 11 | 23 | 0 | 0 | 0 | 0 | 0 |
| SRR5104208 | 50 | 43142 | 7187 | 106 | 407 | 3551 | 3198 | 3628 | 203 | 43 | 34043 | 19 | 6673 | 4325 | 0 | 7260 | 11091 | 0 | 0 | 0 | 555 | 72 | 82 | 0 | 0 | 0 | 0 | 0 | 16 | 78 | 32 | 29 | 0 | 0 | 0 | 0 | 0 |
| SRR5104209 | 19 | 39742 | 6418 | 75 | 307 | 2998 | 2480 | 2296 | 152 | 33 | 34124 | 66916 | 5751 | 2908 | 0 | 6207 | 8897 | 0 | 0 | 0 | 386 | 42 | 58 | 0 | 0 | 0 | 0 | 0 | 11 | 58 | 18 | 26 | 0 | 0 | 0 | 0 | 0 |
| SRR5104210 | 27 | 42932 | 2508 | 201 | 52 | 957 | 672 | 2320 | 297 | 29 | 23863 | 15 | 4590 | 2009 | 0 | 1378 | 13410 | 0 | 0 | 0 | 247 | 49 | 42 | 0 | 0 | 0 | 0 | 0 | 14 | 48 | 30 | 20 | 0 | 0 | 0 | 0 | 0 |
| SRR5104211 | 14 | 39440 | 2495 | 121 | 31 | 569 | 465 | 1529 | 231 | 22 | 21461 | 15 | 3666 | 1429 | 0 | 906 | 10182 | 0 | 0 | 0 | 89 | 31 | 36 | 0 | 0 | 0 | 0 | 0 | 0 | 17 | 19 | 22 | 0 | 0 | 0 | 0 | 0 |
| SRR5104212 | 21 | 42077 | 7006 | 111 | 26 | 468 | 312 | 1331 | 250 | 24 | 20183 | 16 | 4323 | 1688 | 0 | 1231 | 8392 | 0 | 0 | 0 | 99 | 32 | 35 | 0 | 0 | 0 | 0 | 0 | 13 | 26 | 28 | 21 | 0 | 0 | 0 | 0 | 0 |
| SRR5104213 | 13 | 46968 | 6662 | 115 | 32 | 508 | 378 | 1458 | 284 | 21 | 22335 | 43723 | 4076 | 1744 | 0 | 1229 | 11008 | 0 | 0 | 0 | 206 | 46 | 39 | 0 | 0 | 0 | 0 | 0 | 12 | 64 | 38 | 24 | 0 | 0 | 0 | 0 | 0 |
| SRR5104214 | 16 | 22714 | 1219 | 168 | 0 | 141 | 139 | 428 | 195 | 36 | 10070 | 23573 | 1899 | 706 | 0 | 558 | 9962 | 0 | 0 | 0 | 246 | 296 | 39 | 0 | 0 | 0 | 0 | 0 | 16 | 48 | 206 | 24 | 0 | 0 | 0 | 0 | 0 |
| SRR5104215 | 25 | 22963 | 3337 | 108 | 0 | 177 | 165 | 492 | 330 | 29 | 12967 | 28853 | 2590 | 1207 | 0 | 527 | 6808 | 0 | 0 | 0 | 84 | 59 | 39 | 0 | 0 | 0 | 0 | 0 | 0 | 16 | 50 | 23 | 0 | 0 | 0 | 0 | 0 |
| SRR5104216 | 17 | 26512 | 4037 | 113 | 19 | 376 | 391 | 1334 | 762 | 41 | 18996 | 29648 | 3718 | 1687 | 0 | 731 | 11461 | 0 | 0 | 0 | 639 | 123 | 43 | 0 | 0 | 0 | 0 | 0 | 11 | 138 | 85 | 19 | 0 | 0 | 0 | 0 | 0 |
| SRR5104217 | 20 | 23044 | 4083 | 180 | 14 | 231 | 209 | 774 | 605 | 29 | 14547 | 27848 | 2888 | 1098 | 0 | 503 | 7565 | 0 | 0 | 0 | 3625 | 728 | 40 | 0 | 0 | 0 | 0 | 0 | 45 | 823 | 553 | 27 | 0 | 0 | 0 | 0 | 0 |
| SRR5104218 | 0 | 11104 | 1766 | 194 | 10 | 158 | 174 | 672 | 429 | 37 | 17361 | 11 | 3182 | 949 | 0 | 271 | 9663 | 0 | 0 | 0 | 121 | 125 | 57 | 0 | 0 | 0 | 0 | 0 | 0 | 23 | 82 | 29 | 0 | 0 | 0 | 0 | 0 |
| SRR5104219 | 19 | 13651 | 1634 | 160 | 10 | 159 | 180 | 671 | 517 | 32 | 16265 | 27711 | 3185 | 948 | 0 | 424 | 10541 | 0 | 0 | 0 | 96 | 94 | 46 | 0 | 0 | 0 | 0 | 0 | 0 | 18 | 62 | 25 | 0 | 0 | 0 | 0 | 0 |
| SRR5104220 | 18 | 19301 | 2884 | 93 | 0 | 217 | 231 | 855 | 786 | 32 | 21114 | 32835 | 3797 | 1397 | 0 | 567 | 11289 | 0 | 0 | 0 | 402 | 154 | 69 | 0 | 0 | 0 | 0 | 0 | 16 | 80 | 102 | 28 | 0 | 0 | 0 | 0 | 0 |
| SRR5104221 | 16 | 8533 | 949 | 182 | 0 | 184 | 183 | 946 | 629 | 27 | 13947 | 12 | 2740 | 846 | 0 | 446 | 9729 | 0 | 0 | 0 | 406 | 396 | 39 | 0 | 0 | 0 | 0 | 0 | 0 | 73 | 287 | 16 | 0 | 0 | 0 | 0 | 0 |
| SRR5104222 | 10 | 5872 | 3254 | 195 | 0 | 82 | 94 | 365 | 695 | 22 | 12022 | 24780 | 2721 | 798 | 0 | 349 | 7898 | 0 | 0 | 0 | 3449 | 1762 | 107 | 0 | 0 | 0 | 0 | 0 | 36 | 735 | 1258 | 57 | 0 | 0 | 0 | 0 | 0 |
| SRR5104223 | 19 | 6719 | 1536 | 205 | 0 | 95 | 109 | 322 | 392 | 30 | 11443 | 23715 | 2253 | 622 | 0 | 132 | 7551 | 0 | 0 | 0 | 445 | 369 | 57 | 0 | 0 | 0 | 0 | 0 | 0 | 57 | 198 | 24 | 0 | 0 | 0 | 0 | 0 |
| SRR5104224 | 0 | 3673 | 990 | 140 | 0 | 90 | 115 | 386 | 395 | 32 | 11426 | 25678 | 2045 | 713 | 0 | 372 | 8759 | 0 | 0 | 0 | 2024 | 2890 | 135 | 0 | 0 | 0 | 0 | 0 | 22 | 439 | 2246 | 87 | 0 | 0 | 0 | 0 | 0 |
| SRR5104225 | 18 | 6977 | 3668 | 216 | 0 | 114 | 145 | 474 | 893 | 40 | 12186 | 23930 | 2678 | 868 | 0 | 387 | 7917 | 0 | 0 | 0 | 301 | 858 | 63 | 0 | 0 | 0 | 0 | 0 | 0 | 66 | 581 | 34 | 0 | 0 | 0 | 0 | 0 |
| SRR5116489 | 0 | 3741 | 330 | 367 | 2384 | 33760 | 43504 | 19751 | 5825 | 367 | 27219 | 24386 | 2425 | 7769 | 0 | 14664 | 32076 | 0 | 0 | 11 | 972 | 186 | 61 | 0 | 0 | 0 | 0 | 28 | 191 | 139 | 112 | 29 | 0 | 0 | 0 | 15 | 0 |
| SRR5116490 | 0 | 41432 | 132 | 4491 | 2348 | 26935 | 24836 | 14638 | 4919 | 382 | 33069 | 25412 | 1968 | 7869 | 0 | 21974 | 33838 | 0 | 66 | 160 | 6540 | 810 | 215 | 0 | 0 | 0 | 0 | 212 | 1371 | 886 | 321 | 64 | 0 | 0 | 0 | 0 | 0 |
| SRR5116491 | 0 | 11701 | 107 | 1982 | 2558 | 32135 | 39310 | 18003 | 5821 | 171 | 24710 | 24354 | 2802 | 6868 | 0 | 9820 | 38090 | 0 | 172 | 287 | 14338 | 1896 | 285 | 0 | 11 | 0 | 0 | 573 | 3774 | 1829 | 1109 | 137 | 0 | 19 | 0 | 17 | 0 |
| SRR5116492 | 0 | 10279 | 297 | 2320 | 2939 | 37973 | 29130 | 11794 | 7265 | 348 | 34558 | 28035 | 2782 | 15426 | 0 | 16047 | 33149 | 0 | 0 | 12 | 1409 | 461 | 176 | 0 | 10 | 0 | 0 | 12 | 127 | 221 | 186 | 36 | 0 | 0 | 0 | 0 | 0 |
| SRR5116493 | 0 | 19457 | 126 | 1669 | 3372 | 33935 | 43877 | 24245 | 5681 | 173 | 37143 | 25495 | 2974 | 12762 | 0 | 18246 | 36508 | 0 | 16 | 32 | 3020 | 390 | 157 | 0 | 0 | 0 | 0 | 72 | 582 | 439 | 258 | 67 | 0 | 0 | 0 | 0 | 0 |
| SRR5116494 | 0 | 15128 | 584 | 628 | 2597 | 31802 | 24461 | 14468 | 5824 | 404 | 52321 | 33252 | 2604 | 15249 | 0 | 27882 | 33813 | 0 | 0 | 0 | 565 | 140 | 115 | 0 | 0 | 0 | 0 | 0 | 34 | 124 | 46 | 16 | 0 | 0 | 0 | 0 | 0 |
| SRR5116495 | 0 | 7004 | 181 | 945 | 2725 | 27680 | 29831 | 13272 | 5120 | 105 | 34620 | 29069 | 3003 | 12831 | 0 | 12399 | 37696 | 0 | 0 | 0 | 1899 | 320 | 222 | 0 | 12 | 0 | 0 | 11 | 164 | 281 | 179 | 50 | 0 | 0 | 0 | 0 | 0 |
| SRR5116496 | 0 | 7296 | 187 | 452 | 2626 | 40025 | 29287 | 13414 | 6371 | 177 | 33251 | 20210 | 2909 | 15149 | 0 | 17545 | 44794 | 0 | 0 | 0 | 562 | 159 | 149 | 0 | 14 | 0 | 0 | 0 | 32 | 82 | 69 | 28 | 0 | 0 | 0 | 0 | 0 |
| SRR5121477 | 1110 | 8766 | 10214 | 834 | 2930 | 19697 | 37870 | 33225 | 5687 | 38 | 55027 | 18582 | 11136 | 4254 | 317 | 59 | 1987 | 0 | 0 | 0 | 0 | 0 | 0 | 0 | 0 | 0 | 0 | 0 | 0 | 0 | 0 | 0 | 0 | 0 | 0 | 0 | 0 |
| SRR5121478 | 311 | 6341 | 3907 | 339 | 9204 | 85954 | 98143 | 65475 | 219 | 27 | 36360 | 21147 | 9162 | 8152 | 132 | 116 | 2138 | 0 | 0 | 0 | 0 | 0 | 0 | 0 | 0 | 0 | 0 | 0 | 0 | 0 | 0 | 0 | 0 | 0 | 0 | 0 | 0 |
| SRR5121479 | 0 | 7139 | 6408 | 95 | 98 | 3535 | 2317 | 1912 | 0 | 24 | 60601 | 68002 | 9140 | 5825 | 123 | 0 | 26613 | 0 | 0 | 0 | 395 | 70 | 83 | 0 | 21 | 0 | 0 | 0 | 38 | 112 | 111 | 40 | 0 | 37 | 55 | 0 | 0 |
| SRR5121480 | 83 | 118719 | 11349 | 180 | 6107 | 41433 | 37371 | 62420 | 22 | 19 | 25004 | 25276 | 6802 | 6748 | 167 | 0 | 5679 | 0 | 0 | 0 | 189 | 221 | 71 | 0 | 16 | 0 | 0 | 0 | 28 | 54 | 159 | 28 | 0 | 17 | 31 | 0 | 0 |
| SRR5121481 | 220 | 53233 | 2689 | 1152 | 28 | 1415 | 763 | 1453 | 33102 | 22 | 21730 | 27026 | 7281 | 4987 | 104 | 0 | 18749 | 0 | 0 | 0 | 266 | 171 | 62 | 0 | 14 | 0 | 0 | 0 | 29 | 145 | 275 | 56 | 0 | 32 | 50 | 0 | 0 |
| SRR5121482 | 50 | 10822 | 4169 | 1426 | 52 | 1063 | 1270 | 816 | 17596 | 12 | 41457 | 41813 | 3862 | 6755 | 65 | 12 | 8278 | 0 | 0 | 0 | 369 | 108 | 55 | 0 | 13 | 0 | 0 | 18 | 90 | 302 | 254 | 48 | 0 | 43 | 124 | 0 | 0 |
| SRR5121483 | 89 | 5467 | 2961 | 309 | 57 | 1096 | 842 | 858 | 6671 | 43 | 18545 | 9655 | 5313 | 2150 | 13 | 32 | 3294 | 0 | 0 | 0 | 42 | 14 | 0 | 0 | 0 | 0 | 0 | 0 | 17 | 14 | 0 | 0 | 0 | 0 | 0 | 0 | 0 |
| SRR5121484 | 29 | 5660 | 588 | 1253 | 385 | 7038 | 5921 | 4438 | 6254 | 40 | 29958 | 25257 | 3196 | 6450 | 1059 | 22 | 5322 | 0 | 0 | 10 | 389 | 48 | 37 | 0 | 0 | 0 | 0 | 34 | 100 | 67 | 15 | 0 | 0 | 0 | 11 | 0 | 0 |
| SRR5121485 | 33 | 5602 | 63 | 626 | 1128 | 12013 | 13934 | 16643 | 44900 | 16 | 21964 | 21997 | 3377 | 3323 | 130 | 3576 | 9606 | 0 | 0 | 0 | 51 | 50 | 19 | 0 | 0 | 0 | 0 | 0 | 16 | 64 | 50 | 29 | 0 | 0 | 26 | 0 | 0 |
| SRR5121486 | 699 | 61300 | 469 | 327 | 373 | 6452 | 4198 | 20688 | 10180 | 0 | 32835 | 25704 | 6529 | 4942 | 87 | 3243 | 26820 | 0 | 0 | 10 | 748 | 617 | 60 | 0 | 0 | 0 | 0 | 171 | 542 | 567 | 456 | 81 | 0 | 48 | 117 | 0 | 0 |
| SRR5121487 | 82 | 6435 | 102 | 1145 | 1116 | 11264 | 10597 | 22111 | 48598 | 20 | 28192 | 27963 | 4250 | 3317 | 136 | 2661 | 10387 | 0 | 0 | 0 | 48 | 153 | 22 | 0 | 0 | 0 | 0 | 0 | 18 | 62 | 191 | 43 | 0 | 0 | 61 | 0 | 0 |
| SRR5127200 | 393 | 31462 | 93 | 3231 | 428 | 4630 | 2705 | 17724 | 1649 | 100 | 41294 | 15481 | 14463 | 7384 | 0 | 32 | 72419 | 0 | 0 | 0 | 43 | 127 | 16 | 0 | 0 | 0 | 0 | 0 | 10 | 65 | 417 | 36 | 0 | 0 | 0 | 0 | 0 |
| SRR5127201 | 89 | 170765 | 805 | 212 | 4868 | 34977 | 53481 | 29975 | 28 | 31 | 15452 | 29068 | 3233 | 5760 | 11 | 38823 | 17873 | 0 | 0 | 0 | 2464 | 387 | 359 | 0 | 10 | 0 | 0 | 11 | 281 | 771 | 410 | 261 | 0 | 12 | 0 | 0 | 0 |
| SRR5127202 | 40 | 185729 | 1281 | 71 | 3622 | 46221 | 36215 | 15035 | 2559 | 0 | 24269 | 13 | 4340 | 6647 | 69 | 15837 | 23558 | 0 | 0 | 0 | 1299 | 424 | 445 | 0 | 10 | 0 | 0 | 0 | 96 | 577 | 285 | 228 | 0 | 11 | 0 | 11 | 0 |
| SRR5127203 | 153 | 131512 | 297 | 141 | 5416 | 61160 | 56703 | 22324 | 6434 | 61 | 21944 | 54889 | 4997 | 2316 | 0 | 19481 | 21541 | 0 | 0 | 0 | 3078 | 110 | 429 | 0 | 0 | 0 | 0 | 0 | 61 | 798 | 125 | 113 | 0 | 0 | 0 | 0 | 0 |
| SRR5127204 | 101 | 165493 | 434 | 110 | 3091 | 40913 | 38938 | 47138 | 1273 | 35 | 14470 | 22349 | 4120 | 2039 | 0 | 99107 | 19050 | 0 | 0 | 0 | 240 | 15 | 49 | 0 | 0 | 0 | 0 | 0 | 20 | 79 | 30 | 21 | 0 | 0 | 0 | 0 | 0 |
| SRR5127205 | 91 | 263816 | 421 | 82 | 2690 | 36047 | 23766 | 15950 | 3104 | 32 | 12310 | 18 | 2603 | 6193 | 0 | 13009 | 35264 | 0 | 0 | 0 | 1548 | 34 | 97 | 0 | 0 | 0 | 0 | 0 | 64 | 461 | 49 | 70 | 0 | 13 | 0 | 0 | 0 |
| SRR5127206 | 37 | 163225 | 3737 | 60 | 5489 | 87566 | 58778 | 34985 | 976 | 20 | 13595 | 39717 | 2469 | 7173 | 0 | 17401 | 29141 | 0 | 0 | 0 | 1105 | 189 | 198 | 0 | 0 | 0 | 0 | 0 | 64 | 430 | 316 | 103 | 0 | 10 | 0 | 0 | 0 |
| SRR5127207 | 16 | 666065 | 589 | 24 | 830 | 9537 | 3927 | 4970 | 4711 | 0 | 12984 | 14873 | 1244 | 6644 | 0 | 3679 | 18983 | 0 | 0 | 0 | 443 | 124 | 124 | 0 | 0 | 0 | 0 | 0 | 67 | 162 | 145 | 32 | 0 | 0 | 0 | 0 | 0 |
| SRR5127208 | 101 | 239519 | 222 | 88 | 6502 | 86369 | 64516 | 26778 | 6927 | 61 | 24034 | 15 | 5163 | 2239 | 0 | 11974 | 23273 | 0 | 0 | 0 | 3322 | 106 | 429 | 0 | 0 | 0 | 0 | 0 | 131 | 868 | 170 | 74 | 0 | 0 | 0 | 0 | 0 |
| SRR5127209 | 174 | 22550 | 1264 | 282 | 89 | 1436 | 769 | 1549 | 433 | 53 | 17418 | 22602 | 6209 | 4177 | 332 | 10706 | 47124 | 0 | 0 | 0 | 464 | 66 | 81 | 0 | 0 | 0 | 0 | 0 | 132 | 143 | 92 | 63 | 0 | 10 | 0 | 16 | 0 |
| SRR5127210 | 76 | 337850 | 182 | 83 | 7053 | 80057 | 71646 | 27812 | 1268 | 22 | 15835 | 49170 | 3464 | 8873 | 37 | 57270 | 13459 | 0 | 0 | 0 | 528 | 55 | 107 | 0 | 0 | 0 | 0 | 0 | 26 | 138 | 49 | 38 | 0 | 0 | 0 | 0 | 0 |
| SRR5127211 | 91 | 415494 | 8313 | 207 | 2083 | 31135 | 24401 | 35393 | 6665 | 23 | 33649 | 70875 | 4228 | 9222 | 0 | 13119 | 38190 | 0 | 0 | 0 | 1694 | 82 | 146 | 0 | 0 | 0 | 0 | 0 | 113 | 639 | 83 | 99 | 0 | 0 | 0 | 0 | 0 |
| SRR5127212 | 43 | 172027 | 431 | 132 | 5290 | 57073 | 48629 | 46678 | 4639 | 80 | 15459 | 32837 | 3417 | 1216 | 0 | 36449 | 19527 | 0 | 0 | 0 | 2355 | 69 | 544 | 0 | 0 | 0 | 0 | 0 | 120 | 935 | 130 | 131 | 0 | 0 | 0 | 0 | 0 |
| SRR5127213 | 45 | 335047 | 206 | 131 | 3783 | 38004 | 40911 | 20030 | 706 | 45 | 15972 | 19635 | 3394 | 6296 | 37 | 38597 | 17547 | 0 | 0 | 0 | 1039 | 120 | 204 | 0 | 12 | 0 | 0 | 0 | 83 | 275 | 159 | 119 | 0 | 14 | 0 | 0 | 0 |
| SRR5127214 | 44 | 330187 | 456 | 105 | 8084 | 84199 | 68108 | 44431 | 806 | 40 | 13340 | 13 | 2442 | 5036 | 0 | 53985 | 18653 | 0 | 0 | 0 | 536 | 73 | 112 | 0 | 0 | 0 | 0 | 0 | 79 | 271 | 80 | 82 | 0 | 10 | 0 | 0 | 0 |
| SRR5127215 | 82 | 238813 | 363 | 67 | 5401 | 63081 | 53377 | 42317 | 5092 | 22 | 17819 | 38550 | 3359 | 8341 | 0 | 8169 | 49706 | 0 | 0 | 0 | 1066 | 66 | 662 | 0 | 0 | 0 | 0 | 0 | 101 | 840 | 214 | 335 | 0 | 0 | 0 | 0 | 0 |
| SRR5127216 | 68 | 119816 | 377 | 97 | 4155 | 58223 | 50907 | 32718 | 6362 | 94 | 16716 | 27428 | 2835 | 1923 | 0 | 32819 | 17179 | 0 | 0 | 0 | 4393 | 89 | 631 | 11 | 0 | 0 | 0 | 0 | 139 | 847 | 75 | 102 | 0 | 0 | 0 | 14 | 0 |
| SRR5127217 | 30 | 385038 | 527 | 91 | 6519 | 70078 | 59639 | 39914 | 862 | 22 | 12499 | 13 | 2513 | 10068 | 0 | 15383 | 27934 | 0 | 0 | 0 | 1888 | 86 | 1632 | 0 | 0 | 0 | 0 | 0 | 85 | 1083 | 338 | 746 | 0 | 10 | 0 | 0 | 0 |
| SRR5127218 | 55 | 183040 | 1045 | 123 | 2858 | 26969 | 24847 | 16172 | 1055 | 25 | 18282 | 12 | 3124 | 4012 | 14 | 33949 | 27064 | 0 | 0 | 0 | 2628 | 162 | 221 | 0 | 10 | 0 | 0 | 0 | 111 | 603 | 166 | 126 | 0 | 16 | 0 | 11 | 0 |
| SRR5127219 | 294 | 10748 | 1086 | 1075 | 458 | 7862 | 5113 | 15357 | 14494 | 0 | 16900 | 7501 | 2520 | 1635 | 0 | 26 | 30068 | 0 | 0 | 0 | 535 | 816 | 146 | 0 | 0 | 0 | 0 | 65 | 382 | 1110 | 3947 | 554 | 0 | 50 | 119 | 30 | 0 |
| SRR5127220 | 90 | 176180 | 4295 | 117 | 3399 | 41946 | 43977 | 17036 | 743 | 31 | 17116 | 26536 | 3901 | 4696 | 91 | 12590 | 26514 | 0 | 0 | 0 | 1633 | 641 | 550 | 11 | 15 | 0 | 0 | 0 | 68 | 505 | 578 | 265 | 0 | 22 | 0 | 14 | 0 |
| SRR5127221 | 58 | 220456 | 4187 | 89 | 3742 | 37971 | 37973 | 14950 | 140 | 25 | 12915 | 35880 | 2316 | 3410 | 12 | 14592 | 19895 | 0 | 0 | 0 | 932 | 564 | 180 | 0 | 0 | 0 | 0 | 0 | 73 | 190 | 349 | 58 | 0 | 0 | 0 | 0 | 0 |
| SRR5127222 | 54 | 136165 | 2919 | 84 | 3475 | 42161 | 43163 | 35041 | 2179 | 28 | 17841 | 28895 | 2968 | 5313 | 21 | 8301 | 34495 | 0 | 0 | 0 | 583 | 124 | 399 | 0 | 0 | 0 | 0 | 0 | 50 | 304 | 172 | 185 | 0 | 0 | 0 | 0 | 0 |
| SRR5127223 | 60 | 135306 | 5615 | 168 | 5511 | 66399 | 67455 | 31413 | 236 | 39 | 13860 | 17232 | 2676 | 4293 | 22 | 12608 | 18232 | 0 | 0 | 0 | 798 | 684 | 325 | 0 | 0 | 0 | 0 | 0 | 62 | 296 | 548 | 116 | 0 | 0 | 0 | 0 | 0 |
| SRR5127224 | 57 | 102356 | 417 | 189 | 3381 | 31682 | 29276 | 26994 | 6565 | 40 | 17274 | 11 | 3721 | 2391 | 24 | 29341 | 22652 | 0 | 0 | 0 | 605 | 42 | 104 | 0 | 0 | 0 | 0 | 0 | 36 | 147 | 34 | 28 | 0 | 0 | 0 | 0 | 0 |
| SRR5127225 | 189 | 73538 | 1541 | 108 | 2365 | 46564 | 33295 | 16820 | 1818 | 22 | 15164 | 17 | 3087 | 5294 | 35 | 14415 | 36517 | 0 | 0 | 0 | 1112 | 132 | 152 | 0 | 0 | 0 | 0 | 0 | 47 | 289 | 118 | 96 | 0 | 18 | 0 | 0 | 0 |
| SRR5127226 | 44 | 127581 | 2769 | 83 | 2519 | 30743 | 27818 | 18053 | 1032 | 34 | 17884 | 59574 | 3426 | 3232 | 18 | 17511 | 32718 | 0 | 0 | 0 | 782 | 139 | 138 | 0 | 0 | 0 | 0 | 0 | 50 | 242 | 169 | 61 | 0 | 0 | 0 | 0 | 0 |
| SRR5127227 | 45 | 150632 | 1796 | 83 | 2683 | 40395 | 25654 | 14097 | 742 | 36 | 14681 | 74547 | 3493 | 2986 | 17 | 20858 | 29133 | 0 | 0 | 0 | 1609 | 100 | 235 | 0 | 13 | 0 | 0 | 0 | 75 | 361 | 123 | 123 | 0 | 17 | 0 | 11 | 0 |
| SRR5127228 | 116 | 203462 | 1848 | 890 | 2257 | 28006 | 24697 | 27949 | 973 | 68 | 31624 | 30742 | 2729 | 16869 | 0 | 24784 | 41361 | 0 | 0 | 0 | 373 | 335 | 91 | 0 | 0 | 0 | 0 | 0 | 40 | 67 | 136 | 33 | 0 | 0 | 0 | 0 | 0 |
| SRR5127229 | 50 | 85106 | 306 | 180 | 2557 | 33644 | 27046 | 15761 | 6006 | 16 | 14350 | 43814 | 2429 | 4510 | 20 | 4117 | 21781 | 0 | 0 | 0 | 1016 | 156 | 400 | 15 | 22 | 0 | 0 | 0 | 39 | 226 | 116 | 143 | 0 | 18 | 0 | 19 | 0 |
| SRR5127230 | 73 | 172578 | 896 | 63 | 3511 | 50856 | 52277 | 38822 | 428 | 29 | 12351 | 24229 | 3360 | 3542 | 0 | 141544 | 16824 | 0 | 0 | 0 | 663 | 51 | 147 | 0 | 0 | 0 | 0 | 0 | 53 | 196 | 55 | 57 | 0 | 0 | 0 | 0 | 0 |
| SRR5127231 | 55 | 382981 | 3542 | 746 | 1719 | 29395 | 15696 | 24069 | 4354 | 14 | 35733 | 68973 | 3033 | 16976 | 0 | 11629 | 29960 | 0 | 0 | 0 | 547 | 40 | 70 | 0 | 0 | 0 | 0 | 0 | 58 | 143 | 26 | 30 | 0 | 12 | 0 | 0 | 0 |
| SRR5127232 | 0 | 640 | 233 | 730 | 12 | 125 | 120 | 1982 | 538 | 11 | 12491 | 4581 | 1880 | 12738 | 0 | 0 | 11381 | 0 | 0 | 0 | 0 | 0 | 0 | 0 | 0 | 0 | 0 | 0 | 0 | 0 | 0 | 0 | 0 | 0 | 0 | 0 | 0 |
| SRR5127233 | 81 | 192364 | 9606 | 98 | 4119 | 53368 | 39472 | 23668 | 2918 | 24 | 11457 | 15 | 2034 | 6226 | 0 | 14680 | 27891 | 0 | 0 | 0 | 1525 | 1077 | 574 | 19 | 0 | 0 | 0 | 0 | 145 | 540 | 933 | 258 | 0 | 13 | 0 | 0 | 0 |
| SRR5127234 | 385 | 31498 | 7936 | 174 | 705 | 8430 | 8542 | 11413 | 7715 | 29 | 5692 | 7745 | 2979 | 1908 | 1106 | 21997 | 42859 | 0 | 0 | 10 | 596 | 328 | 95 | 0 | 0 | 0 | 0 | 25 | 444 | 261 | 678 | 131 | 0 | 0 | 0 | 0 | 0 |
| SRR5127235 | 117 | 121203 | 596 | 125 | 5954 | 81210 | 73487 | 35766 | 1506 | 30 | 11837 | 14623 | 2835 | 3617 | 18 | 86436 | 13184 | 0 | 0 | 0 | 456 | 78 | 71 | 0 | 0 | 0 | 0 | 0 | 23 | 94 | 54 | 33 | 0 | 0 | 0 | 0 | 0 |
| SRR5127236 | 0 | 9177 | 212 | 601 | 3229 | 36235 | 34128 | 23216 | 1269 | 0 | 10374 | 34975 | 1728 | 5305 | 10 | 25549 | 22516 | 0 | 0 | 0 | 570 | 39 | 45 | 0 | 0 | 0 | 0 | 0 | 30 | 124 | 42 | 16 | 0 | 0 | 0 | 0 | 0 |
| SRR5136285 | 188 | 3921 | 3444 | 43 | 391 | 8984 | 7265 | 18592 | 7499 | 0 | 45700 | 12265 | 4765 | 11796 | 0 | 6015 | 22811 | 0 | 0 | 0 | 0 | 0 | 0 | 0 | 0 | 0 | 0 | 0 | 13 | 0 | 15 | 0 | 0 | 0 | 0 | 0 | 0 |
| SRR5136286 | 157 | 4016 | 3512 | 63 | 570 | 9938 | 7494 | 18573 | 8716 | 0 | 50505 | 12463 | 3177 | 11975 | 0 | 4643 | 20202 | 0 | 0 | 0 | 0 | 0 | 0 | 0 | 0 | 0 | 0 | 0 | 0 | 0 | 14 | 0 | 0 | 0 | 0 | 0 | 0 |
| SRR5136287 | 181 | 3938 | 3581 | 51 | 152 | 7899 | 7292 | 17849 | 7292 | 0 | 51117 | 13444 | 4101 | 10102 | 0 | 6456 | 22178 | 0 | 0 | 0 | 0 | 0 | 0 | 0 | 0 | 0 | 0 | 0 | 0 | 0 | 15 | 0 | 0 | 0 | 0 | 0 | 0 |
| SRR5136288 | 149 | 4042 | 3511 | 50 | 403 | 11069 | 10062 | 25257 | 10968 | 0 | 60978 | 16402 | 3622 | 16603 | 0 | 9257 | 28477 | 0 | 0 | 0 | 0 | 0 | 0 | 0 | 0 | 0 | 0 | 0 | 0 | 0 | 16 | 0 | 0 | 0 | 0 | 0 | 0 |
| SRR5136289 | 151 | 3984 | 3574 | 59 | 329 | 7365 | 7628 | 16044 | 7299 | 0 | 35836 | 11113 | 2564 | 10981 | 0 | 5655 | 21962 | 0 | 0 | 0 | 0 | 11 | 0 | 0 | 0 | 0 | 0 | 0 | 12 | 0 | 0 | 0 | 0 | 0 | 0 | 0 | 0 |
| SRR5136290 | 161 | 3872 | 3704 | 51 | 655 | 7493 | 7565 | 17458 | 8293 | 0 | 42118 | 12221 | 3564 | 11130 | 0 | 5674 | 23496 | 0 | 0 | 0 | 12 | 10 | 0 | 0 | 0 | 0 | 0 | 0 | 0 | 0 | 15 | 0 | 0 | 0 | 0 | 0 | 0 |
| SRR5136291 | 185 | 4006 | 3579 | 55 | 742 | 6902 | 7050 | 16178 | 9796 | 0 | 46828 | 15213 | 3414 | 12245 | 0 | 6902 | 26642 | 0 | 0 | 0 | 0 | 10 | 0 | 0 | 0 | 0 | 0 | 0 | 11 | 0 | 18 | 0 | 0 | 0 | 0 | 0 | 0 |
| SRR5136292 | 162 | 3987 | 3517 | 64 | 185 | 9805 | 9157 | 23772 | 11747 | 0 | 51707 | 16927 | 3700 | 12950 | 0 | 7492 | 26362 | 0 | 0 | 0 | 0 | 12 | 0 | 0 | 0 | 0 | 0 | 0 | 0 | 0 | 17 | 0 | 0 | 0 | 0 | 0 | 0 |
| SRR5136293 | 389 | 5990 | 19032 | 139 | 51 | 2427 | 6183 | 24732 | 14205 | 0 | 51201 | 13337 | 4803 | 7103 | 0 | 4446 | 27133 | 0 | 0 | 0 | 0 | 27 | 0 | 0 | 0 | 0 | 0 | 0 | 0 | 0 | 44 | 12 | 0 | 0 | 0 | 0 | 0 |
| SRR5136294 | 364 | 5898 | 18938 | 152 | 249 | 2434 | 6445 | 27662 | 14052 | 0 | 57177 | 13416 | 4592 | 7828 | 0 | 4232 | 24757 | 0 | 0 | 0 | 0 | 25 | 0 | 0 | 0 | 0 | 0 | 0 | 0 | 0 | 39 | 0 | 0 | 0 | 0 | 0 | 0 |
| SRR5136295 | 316 | 6025 | 18841 | 160 | 282 | 2687 | 5911 | 26614 | 14331 | 0 | 53331 | 14766 | 5476 | 7984 | 0 | 4376 | 26333 | 0 | 0 | 0 | 0 | 28 | 0 | 0 | 0 | 0 | 0 | 0 | 0 | 0 | 41 | 0 | 0 | 0 | 0 | 0 | 0 |
| SRR5136296 | 382 | 6166 | 18790 | 189 | 110 | 2336 | 5937 | 26907 | 15886 | 0 | 56617 | 13632 | 5607 | 7943 | 0 | 4013 | 23114 | 0 | 0 | 0 | 0 | 23 | 0 | 0 | 0 | 0 | 0 | 0 | 10 | 0 | 58 | 0 | 0 | 0 | 0 | 0 | 0 |
| SRR5136297 | 383 | 5808 | 19352 | 171 | 184 | 2264 | 6108 | 29300 | 12505 | 0 | 50150 | 11873 | 5449 | 7345 | 0 | 4133 | 24799 | 0 | 0 | 0 | 0 | 19 | 0 | 0 | 0 | 0 | 0 | 0 | 0 | 0 | 31 | 0 | 0 | 0 | 0 | 0 | 0 |
| SRR5136298 | 387 | 5973 | 19768 | 149 | 216 | 2434 | 5815 | 27993 | 13469 | 16 | 51090 | 12063 | 5031 | 7952 | 0 | 4922 | 23124 | 0 | 0 | 0 | 0 | 23 | 0 | 0 | 0 | 0 | 0 | 0 | 0 | 0 | 36 | 12 | 0 | 0 | 0 | 0 | 0 |
| SRR5136299 | 365 | 6139 | 19314 | 178 | 180 | 2447 | 7056 | 29101 | 13624 | 0 | 51558 | 12645 | 5485 | 8241 | 0 | 4584 | 26320 | 0 | 0 | 0 | 0 | 21 | 0 | 0 | 0 | 0 | 0 | 0 | 0 | 0 | 39 | 0 | 0 | 0 | 0 | 0 | 0 |
| SRR5136300 | 391 | 5912 | 19486 | 151 | 111 | 2213 | 7191 | 31670 | 13830 | 0 | 53687 | 12143 | 5338 | 7717 | 0 | 4785 | 24479 | 0 | 0 | 0 | 0 | 21 | 0 | 0 | 0 | 0 | 0 | 0 | 0 | 0 | 34 | 0 | 0 | 0 | 0 | 0 | 0 |
| SRR5139121 | 173 | 288740 | 288 | 650 | 3640 | 31373 | 34481 | 32307 | 347 | 21 | 19543 | 29 | 4564 | 5239 | 20 | 41192 | 25662 | 0 | 0 | 0 | 428 | 69 | 159 | 0 | 16 | 0 | 0 | 0 | 27 | 57 | 36 | 63 | 0 | 16 | 0 | 0 | 0 |
| SRR5145238 | 831 | 5407 | 11792 | 5330 | 2725 | 2903 | 28444 | 3468 | 64480 | 8207 | 3939 | 4326 | 1469 | 348 | 5940 | 24240 | 18728 | 22 | 41 | 44 | 211 | 110 | 450 | 109 | 82 | 46 | 44 | 47 | 155 | 99 | 170 | 20 | 0 | 0 | 16 | 142 | 24 |
| SRR5145239 | 1276 | 5583 | 10819 | 5615 | 2713 | 2714 | 29245 | 2476 | 59608 | 8389 | 4766 | 4142 | 2618 | 519 | 5134 | 20179 | 16059 | 15 | 21 | 16 | 89 | 48 | 173 | 70 | 41 | 28 | 43 | 41 | 159 | 86 | 157 | 0 | 0 | 0 | 0 | 74 | 0 |
| SRR5145240 | 39 | 895 | 20429 | 1247 | 4646 | 6605 | 77382 | 7898 | 42356 | 5724 | 4347 | 4362 | 2192 | 779 | 19756 | 18653 | 12126 | 0 | 13 | 0 | 11 | 72 | 77 | 45 | 47 | 25 | 0 | 49 | 45 | 22 | 13 | 0 | 0 | 0 | 16 | 127 | 0 |
| SRR5145241 | 173 | 3239 | 27154 | 2565 | 4097 | 8197 | 58972 | 6341 | 32007 | 13044 | 2861 | 3120 | 2644 | 903 | 29070 | 19006 | 7693 | 0 | 96 | 73 | 72 | 612 | 405 | 137 | 83 | 43 | 25 | 207 | 392 | 131 | 89 | 37 | 0 | 0 | 17 | 122 | 0 |
| SRR5145242 | 39 | 687 | 7646 | 1474 | 2726 | 3184 | 31988 | 4824 | 41894 | 3249 | 3902 | 3748 | 2140 | 652 | 19171 | 13557 | 9888 | 0 | 14 | 0 | 0 | 15 | 0 | 0 | 0 | 0 | 0 | 33 | 45 | 16 | 0 | 0 | 0 | 0 | 0 | 11 | 0 |
| SRR5145243 | 54 | 1981 | 20019 | 510 | 4323 | 4350 | 40760 | 4707 | 42672 | 18312 | 2958 | 3890 | 2521 | 663 | 18417 | 12878 | 11429 | 0 | 0 | 12 | 37 | 47 | 192 | 103 | 78 | 47 | 0 | 0 | 47 | 32 | 52 | 12 | 0 | 0 | 19 | 137 | 12 |
| SRR5145244 | 150 | 1132 | 13929 | 3302 | 3142 | 2935 | 31431 | 4631 | 61668 | 3960 | 4535 | 4921 | 2283 | 571 | 21025 | 13903 | 17380 | 0 | 27 | 19 | 14 | 43 | 17 | 15 | 10 | 0 | 0 | 44 | 55 | 18 | 35 | 0 | 0 | 0 | 0 | 32 | 0 |
| SRR5145245 | 2053 | 12620 | 10410 | 6565 | 5431 | 3789 | 37208 | 3398 | 48392 | 6923 | 4510 | 4256 | 2796 | 614 | 24837 | 14193 | 14580 | 18 | 29 | 41 | 129 | 129 | 404 | 130 | 77 | 37 | 97 | 91 | 325 | 157 | 222 | 30 | 0 | 0 | 15 | 90 | 10 |
| SRR5163971 | 95 | 988 | 6017 | 2356 | 4375 | 4811 | 14948 | 13995 | 190410 | 1141 | 15972 | 11 | 19825 | 8383 | 429 | 3441 | 21189 | 0 | 17 | 27 | 70 | 441 | 101 | 22 | 0 | 0 | 0 | 179 | 1041 | 324 | 155 | 0 | 0 | 0 | 0 | 0 | 0 |
| SRR5163972 | 118 | 894 | 6324 | 2301 | 4493 | 4780 | 14796 | 13571 | 190896 | 1078 | 17812 | 6764 | 20363 | 8401 | 402 | 3476 | 21034 | 0 | 24 | 31 | 67 | 420 | 99 | 23 | 0 | 0 | 0 | 172 | 982 | 329 | 152 | 0 | 0 | 0 | 0 | 0 | 0 |
| SRR5163973 | 335 | 2055 | 55076 | 7920 | 22267 | 28730 | 96666 | 69995 | 50207 | 1233 | 18914 | 6445 | 15375 | 18119 | 515 | 4239 | 23102 | 0 | 25 | 37 | 91 | 2091 | 1313 | 161 | 22 | 37 | 0 | 91 | 349 | 155 | 199 | 34 | 0 | 0 | 14 | 34 | 0 |
| SRR5163974 | 281 | 2135 | 57835 | 8035 | 22234 | 28329 | 96004 | 69961 | 50605 | 1375 | 19313 | 11 | 15334 | 18108 | 552 | 4294 | 23084 | 0 | 22 | 31 | 92 | 2026 | 1307 | 158 | 21 | 35 | 0 | 84 | 342 | 160 | 197 | 30 | 0 | 0 | 11 | 33 | 0 |
| SRR5163975 | 209 | 2566 | 8009 | 4934 | 45603 | 46501 | 181702 | 63348 | 26445 | 1655 | 22577 | 5867 | 17309 | 23704 | 2511 | 6294 | 9606 | 0 | 14 | 15 | 73 | 1269 | 626 | 64 | 20 | 22 | 0 | 51 | 163 | 134 | 90 | 12 | 0 | 0 | 0 | 15 | 0 |
| SRR5163976 | 240 | 2287 | 8243 | 5237 | 45755 | 45947 | 181746 | 62875 | 26511 | 1577 | 11 | 15 | 17453 | 23700 | 2433 | 6135 | 9536 | 0 | 15 | 15 | 69 | 1257 | 598 | 69 | 15 | 21 | 0 | 48 | 158 | 120 | 81 | 0 | 0 | 0 | 0 | 14 | 0 |
| SRR5163977 | 247 | 988 | 7298 | 6750 | 18666 | 18621 | 107741 | 75052 | 129134 | 1068 | 7260 | 2442 | 6455 | 7300 | 4561 | 1302 | 9939 | 0 | 0 | 0 | 0 | 153 | 30 | 19 | 0 | 0 | 0 | 77 | 221 | 44 | 20 | 0 | 0 | 0 | 0 | 0 | 0 |

### Supp Table S8.1.Columns_B_to_AL

| | | | | | | | | | | | | | | | | | | | | | | | | | | | | | | | | | | | | |
|---|---|---|---|---|---|---|---|---|---|---|---|---|---|---|---|---|---|---|---|---|---|---|---|---|---|---|---|---|---|---|---|---|---|---|---|---|---|
| SRR5163978 | 230 | 976 | 7375 | 6659 | 18856 | 18286 | 107795 | 74778 | 129536 | 1084 | 7922 | 2737 | 6599 | 7272 | 4488 | 1302 | 9706 | 0 | 12 | 0 | 10 | 147 | 30 | 19 | 0 | 0 | 10 | 82 | 216 | 42 | 16 | 0 | 0 | 0 | 0 | 0 | 0 |
| SRR5163979 | 182 | 767 | 4998 | 2838 | 12735 | 13164 | 57177 | 44742 | 128543 | 987 | 21758 | 6356 | 16263 | 26437 | 893 | 1049 | 28821 | 0 | 0 | 0 | 37 | 600 | 202 | 18 | 0 | 0 | 0 | 53 | 155 | 87 | 76 | 0 | 0 | 0 | 0 | 0 | 0 |
| SRR5163980 | 127 | 714 | 5098 | 2829 | 12820 | 13049 | 57710 | 44618 | 129217 | 920 | 21429 | 13 | 16118 | 26271 | 880 | 1087 | 28651 | 0 | 10 | 10 | 26 | 583 | 215 | 20 | 0 | 0 | 0 | 61 | 160 | 75 | 79 | 0 | 0 | 0 | 0 | 0 | 0 |
| SRR5163981 | 122 | 876 | 17998 | 3963 | 2960 | 2647 | 10489 | 9011 | 177841 | 1046 | 16939 | 5299 | 13921 | 6392 | 376 | 3078 | 23106 | 0 | 0 | 20 | 68 | 142 | 99 | 19 | 0 | 0 | 12 | 228 | 2697 | 1668 | 286 | 20 | 0 | 0 | 15 | 0 | 0 |
| SRR5163982 | 301 | 1145 | 9578 | 6208 | 17858 | 21673 | 77222 | 53807 | 29337 | 828 | 25762 | 25 | 14026 | 17038 | 641 | 5078 | 21722 | 0 | 12 | 26 | 129 | 1235 | 371 | 48 | 0 | 0 | 0 | 42 | 378 | 338 | 163 | 15 | 0 | 0 | 0 | 0 | 0 |
| SRR5163983 | 112 | 453 | 4887 | 1366 | 4976 | 4363 | 14407 | 10905 | 211156 | 1270 | 19958 | 5880 | 16837 | 6494 | 425 | 3631 | 16251 | 0 | 0 | 15 | 74 | 304 | 78 | 11 | 0 | 0 | 0 | 128 | 756 | 498 | 162 | 0 | 0 | 0 | 0 | 0 | 0 |
| SRR5163984 | 75 | 373 | 5283 | 1086 | 4859 | 4369 | 14310 | 10798 | 213832 | 1233 | 19751 | 5769 | 16724 | 6560 | 458 | 3524 | 16137 | 0 | 0 | 13 | 88 | 308 | 74 | 11 | 0 | 0 | 0 | 109 | 705 | 489 | 171 | 0 | 0 | 0 | 0 | 0 | 0 |
| SRR5163985 | 111 | 2288 | 6038 | 5032 | 1535 | 1668 | 5083 | 1925 | 658321 | 550 | 3587 | 1168 | 2710 | 1840 | 478 | 670 | 3888 | 0 | 0 | 0 | 0 | 60 | 12 | 0 | 0 | 0 | 0 | 0 | 37 | 10 | 12 | 0 | 0 | 0 | 0 | 0 | 0 |
| SRR5163986 | 102 | 2291 | 5965 | 4624 | 1491 | 1631 | 5004 | 1862 | 658684 | 624 | 3604 | 1271 | 2659 | 1818 | 483 | 667 | 3833 | 0 | 0 | 0 | 0 | 55 | 12 | 0 | 0 | 0 | 0 | 0 | 36 | 11 | 0 | 0 | 0 | 0 | 0 | 0 | 0 |
| SRR5163987 | 162 | 914 | 9919 | 3641 | 5479 | 5467 | 18904 | 17263 | 194998 | 980 | 18763 | 5644 | 15082 | 10094 | 356 | 3472 | 19847 | 0 | 19 | 33 | 136 | 500 | 114 | 18 | 0 | 0 | 0 | 182 | 1244 | 933 | 170 | 0 | 0 | 0 | 0 | 0 | 0 |
| SRR5168444 | 43 | 799 | 14713 | 9904 | 991 | 1327 | 3322 | 3700 | 205885 | 16 | 40373 | 30 | 37685 | 49401 | 697 | 5594 | 52999 | 0 | 79 | 1043 | 2464 | 1661 | 497 | 216 | 24 | 13 | 0 | 203 | 3083 | 3214 | 397 | 20 | 0 | 0 | 19 | 26 | 0 |
| SRR5168445 | 1003 | 3057 | 2514 | 5137 | 2953 | 2259 | 29210 | 6507 | 40310 | 251 | 18567 | 19713 | 14572 | 17259 | 349 | 7566 | 86576 | 0 | 564 | 1705 | 9538 | 3573 | 6466 | 1490 | 380 | 35 | 0 | 768 | 5120 | 4708 | 912 | 576 | 23 | 0 | 70 | 189 | 0 |
| SRR5168447 | 310 | 1455 | 20043 | 3108 | 785 | 604 | 8701 | 2977 | 70079 | 1137 | 39863 | 30202 | 31235 | 22932 | 268 | 9071 | 78914 | 0 | 93 | 544 | 2150 | 1596 | 819 | 47 | 28 | 26 | 0 | 125 | 1531 | 1876 | 582 | 84 | 0 | 0 | 15 | 65 | 0 |
| SRR5168448 | 532 | 16148 | 48163 | 6998 | 1176 | 1533 | 6209 | 3607 | 90836 | 429 | 24076 | 10832 | 19327 | 39354 | 2053 | 11177 | 18019 | 0 | 322 | 495 | 737 | 1851 | 428 | 58 | 0 | 0 | 0 | 2235 | 2402 | 8493 | 1116 | 45 | 0 | 0 | 0 | 12 | 0 |
| SRR5168451 | 637 | 9043 | 9052 | 3734 | 2402 | 1354 | 29905 | 9019 | 35550 | 357 | 33577 | 13058 | 22885 | 13407 | 358 | 21475 | 41075 | 0 | 166 | 680 | 5866 | 8371 | 384 | 36 | 25 | 41 | 0 | 398 | 3752 | 12596 | 4425 | 71 | 0 | 0 | 29 | 126 | 0 |
| SRR5168454 | 35 | 127 | 443 | 363 | 639 | 522 | 12140 | 5214 | 48004 | 887 | 38848 | 23900 | 27885 | 15156 | 199 | 3185 | 46836 | 0 | 10 | 28 | 495 | 567 | 1133 | 93 | 140 | 52 | 0 | 18 | 111 | 484 | 333 | 195 | 0 | 0 | 130 | 62 | 0 |
| SRR5168458 | 80 | 364 | 954 | 5237 | 1294 | 1018 | 16255 | 5886 | 45278 | 1971 | 21564 | 17961 | 15103 | 10347 | 106 | 8303 | 46092 | 19 | 449 | 1204 | 25336 | 2620 | 4024 | 239 | 98 | 79 | 14 | 656 | 4353 | 17577 | 858 | 701 | 24 | 13 | 111 | 265 | 0 |
| SRR5168460 | 81 | 3465 | 18605 | 624 | 624 | 818 | 11562 | 3868 | 47160 | 193 | 25629 | 15110 | 17846 | 14366 | 351 | 7500 | 46373 | 0 | 17 | 122 | 1152 | 2480 | 5317 | 518 | 143 | 19 | 0 | 59 | 558 | 1493 | 1069 | 1280 | 78 | 32 | 52 | 86 | 0 |
| SRR5168465 | 274 | 3036 | 6640 | 1063 | 2198 | 2088 | 38945 | 18303 | 60562 | 6599 | 42133 | 23644 | 33625 | 14338 | 183 | 18356 | 64323 | 0 | 13 | 132 | 2626 | 1069 | 1339 | 80 | 28 | 54 | 0 | 26 | 581 | 2599 | 538 | 162 | 0 | 0 | 11 | 142 | 0 |
| SRR5168467 | 302 | 3210 | 86152 | 674 | 400 | 610 | 6803 | 4348 | 103201 | 609 | 11015 | 5195 | 9957 | 11659 | 142 | 14080 | 32047 | 0 | 0 | 183 | 488 | 6808 | 2668 | 94 | 30 | 12 | 0 | 24 | 923 | 752 | 2732 | 467 | 0 | 0 | 14 | 70 | 0 |
| SRR5168471 | 119 | 9588 | 43644 | 330 | 1926 | 1726 | 51085 | 19551 | 67367 | 1023 | 24886 | 8173 | 18767 | 13139 | 323 | 10053 | 61725 | 0 | 0 | 25 | 201 | 421 | 126 | 0 | 0 | 0 | 0 | 11 | 163 | 487 | 501 | 34 | 0 | 0 | 0 | 0 | 0 |
| SRR5168475 | 182 | 24673 | 45585 | 300 | 444 | 259 | 11176 | 3666 | 39520 | 377 | 11037 | 4212 | 8438 | 8925 | 296 | 9648 | 30720 | 0 | 0 | 32 | 510 | 253 | 564 | 50 | 22 | 0 | 0 | 10 | 142 | 1045 | 134 | 75 | 0 | 0 | 0 | 28 | 0 |
| SRR5171728 | 199 | 4642 | 264 | 5932 | 2317 | 9366 | 62197 | 10812 | 8178 | 130 | 7703 | 8762 | 7940 | 34405 | 1505 | 347 | 3346 | 0 | 0 | 0 | 76 | 25 | 30 | 0 | 0 | 0 | 0 | 0 | 16 | 86 | 34 | 53 | 0 | 0 | 0 | 0 | 0 |
| SRR5171729 | 503 | 11022 | 289 | 7055 | 2081 | 6563 | 49032 | 10106 | 15623 | 132 | 4994 | 4209 | 5814 | 44716 | 1700 | 321 | 2093 | 0 | 0 | 12 | 205 | 27 | 24 | 0 | 0 | 0 | 0 | 11 | 70 | 241 | 58 | 51 | 0 | 0 | 0 | 0 | 0 |
| SRR5171730 | 65 | 3852 | 491 | 3965 | 2941 | 10179 | 65775 | 12290 | 6101 | 146 | 21726 | 12203 | 15042 | 26196 | 1063 | 422 | 7528 | 0 | 0 | 0 | 112 | 43 | 48 | 0 | 0 | 0 | 0 | 0 | 21 | 89 | 21 | 39 | 0 | 0 | 0 | 0 | 0 |
| SRR5171731 | 453 | 6559 | 506 | 10056 | 2118 | 5783 | 51810 | 7574 | 15661 | 145 | 1268 | 1827 | 2676 | 33869 | 1513 | 582 | 2734 | 0 | 0 | 0 | 154 | 29 | 35 | 0 | 0 | 0 | 0 | 0 | 15 | 157 | 55 | 60 | 0 | 0 | 0 | 0 | 0 |
| SRR5171732 | 93 | 2177 | 173 | 7262 | 1804 | 10866 | 74619 | 7698 | 5694 | 66 | 4130 | 3609 | 6656 | 30714 | 682 | 882 | 3769 | 0 | 0 | 0 | 96 | 22 | 31 | 0 | 0 | 0 | 0 | 0 | 0 | 67 | 36 | 43 | 0 | 0 | 0 | 0 | 0 |
| SRR5171733 | 361 | 3265 | 131 | 16338 | 1992 | 9774 | 60984 | 6868 | 9513 | 121 | 2177 | 1524 | 4267 | 37518 | 1328 | 686 | 1720 | 0 | 0 | 0 | 133 | 47 | 44 | 0 | 0 | 0 | 0 | 0 | 33 | 94 | 42 | 63 | 0 | 0 | 0 | 0 | 0 |
| SRR5195030 | 14 | 43732 | 22549 | 192 | 2020 | 20443 | 17082 | 8383 | 968 | 0 | 16606 | 51085 | 2477 | 5905 | 14 | 6870 | 20510 | 0 | 0 | 0 | 525 | 546 | 80 | 0 | 0 | 0 | 0 | 0 | 137 | 128 | 307 | 39 | 0 | 0 | 0 | 0 | 0 |
| SRR5195032 | 0 | 65452 | 3350 | 44 | 815 | 7170 | 5457 | 2559 | 1164 | 10 | 8049 | 91641 | 1899 | 5082 | 0 | 5773 | 29532 | 0 | 0 | 0 | 319 | 101 | 70 | 0 | 0 | 0 | 0 | 0 | 39 | 62 | 81 | 22 | 0 | 0 | 0 | 0 | 0 |
| SRR5195033 | 18 | 122966 | 12290 | 148 | 1012 | 9426 | 6462 | 3328 | 59 | 35 | 13305 | 11 | 3175 | 2929 | 18 | 19822 | 20971 | 0 | 0 | 0 | 889 | 299 | 94 | 0 | 0 | 0 | 0 | 0 | 124 | 220 | 157 | 30 | 0 | 0 | 0 | 0 | 0 |
| SRR5195034 | 73 | 75730 | 2606 | 268 | 938 | 9775 | 7256 | 2530 | 1800 | 0 | 6960 | 57244 | 1547 | 3671 | 0 | 11664 | 26269 | 0 | 0 | 0 | 915 | 413 | 130 | 0 | 0 | 0 | 0 | 0 | 133 | 177 | 300 | 40 | 0 | 0 | 0 | 0 | 0 |
| SRR5224010 | 732 | 5254 | 18467 | 2882 | 2413 | 2205 | 40012 | 8993 | 33978 | 1755 | 28931 | 20 | 13786 | 14728 | 314 | 7091 | 45285 | 0 | 0 | 0 | 0 | 42 | 43 | 15 | 13 | 12 | 0 | 0 | 14 | 0 | 14 | 0 | 0 | 0 | 12 | 43 | 0 |
| SRR5224011 | 833 | 5713 | 11545 | 2656 | 2793 | 2237 | 40451 | 8966 | 31860 | 1752 | 39343 | 23616 | 16773 | 14382 | 271 | 7295 | 45016 | 0 | 0 | 0 | 0 | 55 | 59 | 18 | 14 | 11 | 0 | 0 | 13 | 0 | 12 | 0 | 0 | 0 | 14 | 46 | 0 |
| SRR5224012 | 1689 | 7958 | 13411 | 2094 | 3201 | 2426 | 38632 | 10105 | 27509 | 1132 | 51384 | 31250 | 20704 | 15650 | 294 | 7518 | 45353 | 0 | 0 | 0 | 15 | 143 | 113 | 19 | 12 | 0 | 0 | 0 | 15 | 13 | 25 | 0 | 0 | 0 | 0 | 22 | 0 |
| SRR5224013 | 941 | 16527 | 15252 | 2494 | 17343 | 10448 | 270970 | 9534 | 9271 | 4337 | 21599 | 15973 | 10423 | 32999 | 3250 | 12735 | 15150 | 0 | 0 | 0 | 62 | 46 | 185 | 49 | 32 | 29 | 13 | 12 | 37 | 26 | 13 | 0 | 0 | 0 | 18 | 89 | 0 |
| SRR5224014 | 635 | 4265 | 11357 | 4961 | 7218 | 7005 | 207520 | 6575 | 24518 | 2990 | 30731 | 19850 | 10900 | 23852 | 1247 | 7301 | 28900 | 0 | 0 | 0 | 0 | 30 | 38 | 52 | 50 | 45 | 0 | 0 | 12 | 0 | 10 | 0 | 0 | 0 | 35 | 172 | 0 |
| SRR5224015 | 1208 | 11615 | 10054 | 2907 | 24338 | 7585 | 296107 | 10372 | 4873 | 6159 | 23491 | 21387 | 12405 | 38163 | 3598 | 10716 | 10308 | 14 | 16 | 0 | 50 | 53 | 95 | 75 | 53 | 58 | 23 | 13 | 32 | 30 | 20 | 0 | 0 | 0 | 30 | 165 | 0 |
| SRR5224016 | 1071 | 6784 | 15092 | 3541 | 21784 | 7072 | 314951 | 10040 | 6815 | 7542 | 23288 | 12 | 9687 | 32775 | 2277 | 8547 | 12377 | 0 | 0 | 0 | 16 | 19 | 32 | 42 | 36 | 37 | 12 | 0 | 18 | 13 | 0 | 0 | 0 | 0 | 15 | 109 | 0 |
| SRR5224017 | 585 | 5916 | 16730 | 5937 | 12019 | 10457 | 304267 | 3081 | 5780 | 1835 | 15023 | 14408 | 4745 | 36289 | 5606 | 12436 | 7692 | 0 | 0 | 0 | 12 | 27 | 43 | 49 | 38 | 25 | 0 | 0 | 0 | 0 | 0 | 0 | 0 | 0 | 19 | 136 | 0 |
| SRR5224018 | 929 | 11744 | 12750 | 4026 | 17881 | 7580 | 291924 | 10657 | 6999 | 7796 | 24065 | 21101 | 10729 | 32558 | 2168 | 8232 | 11505 | 0 | 0 | 0 | 29 | 12 | 76 | 25 | 21 | 21 | 0 | 0 | 18 | 15 | 0 | 0 | 0 | 0 | 0 | 61 | 0 |
| SRR5224019 | 872 | 7103 | 19896 | 2211 | 15205 | 9820 | 227493 | 8152 | 4041 | 5174 | 24992 | 24898 | 12640 | 43045 | 4637 | 13942 | 11612 | 0 | 14 | 0 | 32 | 46 | 58 | 44 | 29 | 36 | 12 | 11 | 29 | 15 | 18 | 0 | 0 | 0 | 17 | 111 | 0 |
| SRR5224020 | 1138 | 9034 | 21117 | 2595 | 22891 | 8528 | 361748 | 8444 | 4073 | 5137 | 13557 | 14996 | 6518 | 27064 | 1872 | 11166 | 7138 | 0 | 0 | 0 | 30 | 27 | 66 | 50 | 37 | 40 | 0 | 0 | 17 | 14 | 0 | 0 | 0 | 0 | 19 | 122 | 0 |
| SRR5224021 | 922 | 11865 | 19661 | 2808 | 13250 | 8959 | 214513 | 7186 | 2544 | 5384 | 25386 | 12 | 14115 | 37122 | 3038 | 13491 | 11331 | 0 | 11 | 0 | 41 | 44 | 137 | 57 | 32 | 34 | 11 | 0 | 27 | 18 | 12 | 0 | 0 | 0 | 18 | 99 | 0 |
| SRR5224022 | 1056 | 10911 | 12215 | 2883 | 24060 | 8769 | 305682 | 9721 | 4744 | 5536 | 19172 | 21862 | 9584 | 33547 | 3589 | 12287 | 7785 | 12 | 14 | 0 | 43 | 43 | 100 | 71 | 50 | 53 | 20 | 13 | 32 | 25 | 13 | 0 | 0 | 0 | 25 | 155 | 0 |
| SRR5224023 | 679 | 5335 | 22469 | 4339 | 15842 | 12284 | 256636 | 7403 | 3044 | 4666 | 19802 | 13472 | 8113 | 39205 | 6123 | 16127 | 9428 | 0 | 15 | 0 | 33 | 54 | 49 | 51 | 34 | 42 | 17 | 19 | 43 | 21 | 21 | 0 | 0 | 0 | 19 | 127 | 10 |
| SRR5224024 | 1066 | 9164 | 14314 | 3431 | 21073 | 8724 | 291880 | 9893 | 1951 | 7968 | 17260 | 11 | 8512 | 30835 | 2737 | 10250 | 6688 | 0 | 0 | 0 | 46 | 37 | 138 | 72 | 60 | 58 | 15 | 14 | 35 | 26 | 13 | 0 | 0 | 0 | 21 | 164 | 0 |
| SRR5230550 | 104 | 1434 | 13443 | 6356 | 399 | 645 | 16276 | 5259 | 28236 | 114 | 3364 | 5687 | 3456 | 13174 | 267 | 147 | 17472 | 0 | 0 | 0 | 0 | 51 | 169 | 15 | 0 | 0 | 0 | 0 | 0 | 0 | 0 | 29 | 0 | 0 | 0 | 0 | 0 |
| SRR5230551 | 44 | 370 | 14172 | 1930 | 30 | 75 | 1689 | 2519 | 7511 | 159 | 517 | 500 | 1068 | 5030 | 87 | 282 | 2461 | 0 | 0 | 0 | 0 | 22 | 100 | 0 | 0 | 0 | 0 | 0 | 0 | 0 | 0 | 13 | 0 | 0 | 0 | 0 | 0 |
| SRR5230552 | 200 | 1312 | 27663 | 1680 | 582 | 1168 | 24805 | 3823 | 19001 | 791 | 6780 | 7648 | 7403 | 28448 | 538 | 180 | 33764 | 0 | 0 | 0 | 0 | 56 | 161 | 37 | 19 | 0 | 0 | 0 | 0 | 0 | 0 | 15 | 0 | 0 | 0 | 34 | 0 |
| SRR5230553 | 152 | 1753 | 23622 | 4799 | 761 | 1624 | 33544 | 14685 | 37112 | 113 | 6303 | 5704 | 3507 | 15438 | 206 | 314 | 28211 | 0 | 0 | 0 | 12 | 67 | 310 | 38 | 16 | 0 | 0 | 0 | 0 | 0 | 0 | 21 | 0 | 0 | 0 | 17 | 0 |
| SRR5230554 | 125 | 916 | 8674 | 5904 | 637 | 1081 | 23532 | 5395 | 22351 | 176 | 4242 | 5209 | 5134 | 20954 | 990 | 153 | 13658 | 0 | 0 | 0 | 0 | 199 | 317 | 0 | 0 | 0 | 0 | 0 | 20 | 12 | 52 | 32 | 0 | 0 | 0 | 0 | 0 |
| SRR5230555 | 174 | 1025 | 12827 | 5832 | 994 | 1454 | 32433 | 11960 | 34441 | 127 | 5689 | 5778 | 4131 | 13603 | 371 | 342 | 27726 | 0 | 0 | 0 | 0 | 62 | 162 | 15 | 0 | 0 | 0 | 0 | 0 | 0 | 11 | 36 | 0 | 0 | 0 | 0 | 0 |
| SRR5230556 | 473 | 3404 | 21279 | 7525 | 193 | 257 | 2879 | 1768 | 31140 | 116 | 753 | 1706 | 925 | 6766 | 215 | 320 | 8041 | 0 | 0 | 0 | 0 | 108 | 91 | 12 | 0 | 0 | 0 | 0 | 0 | 0 | 0 | 0 | 0 | 0 | 0 | 11 | 0 |
| SRR5230557 | 70 | 634 | 20756 | 2842 | 677 | 1270 | 23617 | 7490 | 27391 | 171 | 3736 | 4056 | 3385 | 19003 | 503 | 490 | 16189 | 0 | 0 | 0 | 0 | 71 | 103 | 10 | 0 | 0 | 0 | 0 | 0 | 0 | 0 | 0 | 0 | 0 | 0 | 0 | 0 |
| SRR5230558 | 84 | 1244 | 14804 | 1684 | 762 | 1067 | 18764 | 4852 | 32209 | 188 | 6493 | 9870 | 4781 | 19553 | 612 | 210 | 21868 | 0 | 0 | 0 | 0 | 73 | 146 | 0 | 0 | 0 | 0 | 0 | 10 | 0 | 17 | 16 | 0 | 0 | 0 | 0 | 0 |
| SRR5230559 | 215 | 1131 | 18735 | 6011 | 905 | 1871 | 29473 | 3624 | 21893 | 198 | 2223 | 3115 | 2512 | 16599 | 501 | 337 | 22876 | 0 | 0 | 0 | 0 | 257 | 343 | 13 | 0 | 0 | 0 | 0 | 10 | 0 | 24 | 27 | 0 | 0 | 0 | 14 | 0 |
| SRR5230560 | 216 | 1231 | 9553 | 7276 | 262 | 491 | 8122 | 2777 | 16138 | 107 | 1919 | 2773 | 2480 | 14447 | 334 | 179 | 9644 | 0 | 0 | 0 | 0 | 70 | 221 | 21 | 0 | 0 | 0 | 0 | 0 | 0 | 0 | 21 | 0 | 0 | 0 | 14 | 0 |
| SRR5230561 | 99 | 867 | 22836 | 2556 | 45 | 89 | 2276 | 3324 | 10243 | 133 | 241 | 328 | 539 | 3111 | 58 | 149 | 3570 | 0 | 0 | 0 | 0 | 19 | 101 | 12 | 0 | 0 | 0 | 0 | 0 | 0 | 0 | 11 | 0 | 0 | 0 | 10 | 0 |
| SRR5230562 | 199 | 1577 | 24817 | 8809 | 634 | 2186 | 35736 | 1832 | 38097 | 180 | 4983 | 6339 | 2933 | 23095 | 834 | 392 | 26908 | 0 | 0 | 0 | 0 | 147 | 135 | 12 | 0 | 0 | 0 | 0 | 0 | 0 | 0 | 0 | 0 | 0 | 0 | 11 | 0 |
| SRR5230563 | 420 | 2421 | 25778 | 4209 | 206 | 469 | 11107 | 1861 | 10681 | 161 | 1669 | 2324 | 2284 | 14564 | 454 | 381 | 14657 | 0 | 0 | 0 | 0 | 56 | 207 | 100 | 36 | 11 | 0 | 0 | 0 | 0 | 0 | 11 | 0 | 0 | 13 | 61 | 0 |
| SRR5230564 | 131 | 1351 | 14495 | 3339 | 582 | 1068 | 19479 | 3488 | 18230 | 88 | 4129 | 5980 | 2241 | 10849 | 322 | 398 | 20393 | 0 | 0 | 0 | 0 | 133 | 194 | 11 | 0 | 0 | 0 | 0 | 0 | 0 | 22 | 30 | 0 | 0 | 0 | 0 | 0 |
| SRR5230565 | 0 | 136 | 5912 | 2580 | 66 | 72 | 1405 | 1824 | 7808 | 64 | 8823 | 14 | 6640 | 37568 | 940 | 200 | 5046 | 0 | 0 | 0 | 0 | 26 | 24 | 0 | 0 | 0 | 0 | 0 | 0 | 0 | 0 | 0 | 0 | 0 | 0 | 0 | 0 |
| SRR5230566 | 193 | 1219 | 15348 | 4638 | 307 | 697 | 13054 | 2274 | 49432 | 92 | 2243 | 2985 | 2858 | 11578 | 386 | 276 | 12265 | 0 | 0 | 0 | 0 | 71 | 142 | 17 | 0 | 0 | 0 | 0 | 0 | 0 | 0 | 23 | 0 | 0 | 0 | 0 | 0 |
| SRR5230567 | 119 | 1161 | 22703 | 1630 | 127 | 225 | 5927 | 2136 | 15634 | 249 | 1940 | 2265 | 2221 | 10475 | 186 | 109 | 7300 | 0 | 0 | 0 | 16 | 58 | 144 | 25 | 13 | 0 | 0 | 0 | 0 | 0 | 0 | 10 | 0 | 0 | 0 | 25 | 0 |
| SRR5230568 | 521 | 2541 | 21948 | 5298 | 713 | 1369 | 37757 | 2244 | 48024 | 157 | 3810 | 3306 | 3196 | 30997 | 697 | 811 | 16169 | 0 | 0 | 0 | 0 | 66 | 84 | 0 | 0 | 0 | 0 | 0 | 0 | 0 | 0 | 0 | 0 | 0 | 0 | 0 | 0 |
| SRR5230569 | 294 | 4587 | 16681 | 4368 | 1140 | 2045 | 53987 | 9382 | 32318 | 90 | 6329 | 6036 | 3577 | 19271 | 488 | 430 | 33232 | 0 | 0 | 0 | 0 | 45 | 165 | 0 | 0 | 0 | 0 | 0 | 0 | 0 | 0 | 20 | 0 | 0 | 0 | 0 | 0 |
| SRR5230570 | 307 | 1139 | 19410 | 6449 | 143 | 238 | 6592 | 3183 | 7017 | 181 | 1254 | 1597 | 1410 | 6411 | 166 | 173 | 9988 | 0 | 0 | 0 | 0 | 96 | 224 | 16 | 0 | 0 | 0 | 0 | 0 | 0 | 0 | 25 | 0 | 0 | 0 | 15 | 0 |
| SRR5230571 | 89 | 1537 | 24242 | 1046 | 214 | 334 | 9236 | 4688 | 9481 | 590 | 2183 | 1887 | 5174 | 17495 | 250 | 83 | 15364 | 0 | 0 | 0 | 0 | 128 | 231 | 72 | 25 | 0 | 0 | 0 | 0 | 0 | 17 | 31 | 0 | 0 | 12 | 61 | 0 |
| SRR5230572 | 113 | 1676 | 7688 | 4670 | 664 | 1031 | 20423 | 5575 | 21482 | 187 | 9195 | 10372 | 7653 | 20813 | 399 | 131 | 22009 | 0 | 0 | 0 | 0 | 93 | 141 | 11 | 0 | 0 | 0 | 0 | 10 | 0 | 23 | 31 | 0 | 0 | 0 | 0 | 0 |
| SRR5230573 | 154 | 2445 | 12407 | 5481 | 246 | 352 | 7433 | 4972 | 14965 | 307 | 5445 | 6749 | 6756 | 14669 | 161 | 270 | 14758 | 0 | 0 | 0 | 0 | 98 | 184 | 28 | 0 | 0 | 0 | 0 | 0 | 0 | 15 | 35 | 0 | 0 | 0 | 16 | 0 |
| SRR5230574 | 150 | 2731 | 30681 | 1061 | 582 | 694 | 18835 | 4157 | 20130 | 181 | 4137 | 4690 | 2327 | 11973 | 214 | 434 | 22449 | 0 | 0 | 0 | 0 | 74 | 193 | 23 | 0 | 0 | 0 | 0 | 0 | 0 | 0 | 22 | 0 | 0 | 0 | 13 | 0 |
| SRR5230575 | 135 | 2834 | 30465 | 1055 | 606 | 748 | 19073 | 4308 | 20920 | 172 | 4106 | 4723 | 2266 | 12107 | 243 | 458 | 23677 | 0 | 0 | 0 | 0 | 64 | 184 | 22 | 0 | 0 | 0 | 0 | 0 | 0 | 0 | 22 | 0 | 0 | 0 | 13 | 0 |
| SRR5230576 | 90 | 1534 | 24610 | 1511 | 732 | 921 | 17936 | 12951 | 104825 | 283 | 3108 | 3685 | 3268 | 9178 | 383 | 253 | 19049 | 0 | 0 | 0 | 11 | 106 | 313 | 38 | 11 | 0 | 0 | 0 | 0 | 19 | 0 | 40 | 0 | 0 | 0 | 22 | 0 |
| SRR5230577 | 80 | 1434 | 24981 | 1343 | 736 | 997 | 18033 | 13260 | 107050 | 292 | 2956 | 3731 | 3234 | 8937 | 456 | 251 | 19704 | 0 | 0 | 0 | 12 | 98 | 295 | 42 | 0 | 0 | 0 | 0 | 12 | 15 | 0 | 46 | 0 | 0 | 0 | 22 | 0 |
| SRR5230578 | 72 | 420 | 7518 | 3781 | 421 | 743 | 10120 | 4209 | 9232 | 51 | 10994 | 12503 | 6416 | 28887 | 1103 | 60 | 13294 | 0 | 0 | 0 | 0 | 133 | 211 | 0 | 0 | 0 | 0 | 0 | 29 | 17 | 55 | 43 | 0 | 0 | 0 | 0 | 0 |
| SRR5230579 | 59 | 448 | 7347 | 3423 | 487 | 756 | 10348 | 4357 | 9958 | 43 | 11386 | 12774 | 6510 | 29356 | 1195 | 68 | 14064 | 0 | 0 | 0 | 0 | 116 | 196 | 0 | 0 | 0 | 0 | 0 | 30 | 11 | 47 | 41 | 0 | 0 | 0 | 0 | 0 |
| SRR5230580 | 242 | 1987 | 11128 | 3124 | 550 | 554 | 11044 | 10107 | 50467 | 71 | 2304 | 2715 | 1351 | 6746 | 134 | 397 | 10459 | 0 | 0 | 0 | 11 | 76 | 129 | 14 | 0 | 0 | 0 | 0 | 13 | 0 | 0 | 18 | 0 | 0 | 0 | 0 | 0 |
| SRR5230581 | 160 | 1610 | 11199 | 3003 | 508 | 578 | 11589 | 10512 | 53117 | 58 | 2336 | 2728 | 1391 | 7038 | 110 | 380 | 11378 | 0 | 0 | 0 | 16 | 81 | 102 | 0 | 0 | 0 | 0 | 0 | 22 | 0 | 0 | 0 | 0 | 0 | 0 | 11 | 0 |
| SRR5230582 | 355 | 2089 | 15002 | 5031 | 1367 | 2138 | 36815 | 8075 | 53326 | 121 | 12294 | 10359 | 7077 | 22148 | 585 | 301 | 23605 | 0 | 0 | 0 | 0 | 81 | 137 | 0 | 0 | 0 | 0 | 0 | 28 | 14 | 0 | 18 | 0 | 0 | 0 | 0 | 0 |
| SRR5230583 | 385 | 2091 | 14857 | 5086 | 1482 | 2032 | 37546 | 8148 | 54341 | 189 | 11746 | 10274 | 6738 | 22391 | 588 | 334 | 24253 | 0 | 0 | 0 | 0 | 62 | 116 | 0 | 0 | 0 | 0 | 0 | 29 | 13 | 0 | 16 | 0 | 0 | 0 | 0 | 0 |
| SRR5230584 | 118 | 857 | 23325 | 2148 | 1749 | 2988 | 45709 | 8834 | 85193 | 119 | 4118 | 3952 | 3222 | 28403 | 1207 | 547 | 17504 | 0 | 0 | 0 | 0 | 161 | 199 | 24 | 13 | 0 | 0 | 0 | 17 | 15 | 0 | 24 | 0 | 0 | 0 | 17 | 0 |
| SRR5230585 | 90 | 792 | 23031 | 2190 | 1771 | 2801 | 44657 | 8967 | 87261 | 181 | 3855 | 3891 | 3186 | 27977 | 1223 | 590 | 18170 | 0 | 0 | 0 | 0 | 152 | 190 | 32 | 11 | 0 | 0 | 0 | 20 | 16 | 0 | 23 | 0 | 0 | 0 | 13 | 0 |
| SRR5230586 | 289 | 3734 | 23541 | 4175 | 189 | 254 | 7805 | 2952 | 18857 | 164 | 3191 | 6081 | 3603 | 13556 | 333 | 386 | 23928 | 0 | 0 | 0 | 15 | 129 | 557 | 100 | 35 | 0 | 0 | 0 | 0 | 13 | 12 | 51 | 0 | 0 | 16 | 62 | 12 |
| SRR5230587 | 274 | 3847 | 23633 | 4284 | 186 | 295 | 7984 | 3106 | 19093 | 127 | 3267 | 6226 | 3713 | 14024 | 347 | 390 | 24482 | 0 | 0 | 0 | 11 | 127 | 556 | 97 | 38 | 0 | 0 | 0 | 0 | 13 | 0 | 48 | 0 | 0 | 0 | 59 | 0 |
| SRR5230588 | 170 | 1597 | 11304 | 7037 | 432 | 705 | 14072 | 6777 | 16433 | 114 | 2962 | 3342 | 1523 | 6460 | 146 | 338 | 18208 | 0 | 0 | 0 | 31 | 267 | 209 | 22 | 0 | 0 | 0 | 0 | 10 | 10 | 24 | 19 | 0 | 0 | 0 | 11 | 0 |
| SRR5230589 | 169 | 1669 | 11097 | 6491 | 405 | 751 | 14233 | 6610 | 17286 | 62 | 2856 | 3438 | 1556 | 6501 | 148 | 375 | 19094 | 0 | 0 | 0 | 29 | 235 | 186 | 28 | 0 | 0 | 0 | 0 | 0 | 0 | 21 | 12 | 0 | 0 | 0 | 0 | 0 |
| SRR5230590 | 14 | 551 | 26869 | 262 | 162 | 185 | 10120 | 2075 | 4399 | 578 | 2175 | 2011 | 2077 | 9024 | 131 | 113 | 15998 | 0 | 0 | 0 | 0 | 12 | 52 | 0 | 0 | 0 | 0 | 0 | 0 | 0 | 0 | 0 | 0 | 0 | 0 | 0 | 0 |
| SRR5230591 | 14 | 487 | 27470 | 88 | 133 | 197 | 10378 | 2259 | 4792 | 534 | 2213 | 2180 | 2069 | 9488 | 128 | 111 | 17047 | 0 | 0 | 0 | 0 | 0 | 35 | 0 | 0 | 0 | 0 | 0 | 0 | 0 | 0 | 0 | 0 | 0 | 0 | 0 | 0 |
| SRR5230592 | 0 | 436 | 36293 | 290 | 34 | 82 | 2087 | 2429 | 5133 | 532 | 7271 | 37 | 6792 | 21906 | 435 | 89 | 10217 | 0 | 0 | 0 | 0 | 15 | 100 | 22 | 0 | 0 | 0 | 0 | 0 | 0 | 0 | 0 | 0 | 0 | 0 | 20 | 0 |
| SRR5230593 | 0 | 307 | 36343 | 141 | 33 | 67 | 2194 | 2371 | 5763 | 871 | 7186 | 6631 | 6597 | 22539 | 497 | 95 | 11269 | 0 | 0 | 0 | 0 | 0 | 117 | 14 | 0 | 0 | 0 | 0 | 0 | 0 | 0 | 13 | 0 | 0 | 0 | 12 | 0 |
| SRR5230594 | 231 | 3263 | 26684 | 4020 | 262 | 377 | 8527 | 7281 | 13591 | 91 | 1931 | 1896 | 2358 | 9425 | 157 | 509 | 14944 | 0 | 0 | 0 | 11 | 107 | 314 | 66 | 18 | 0 | 0 | 0 | 0 | 0 | 11 | 36 | 0 | 0 | 0 | 36 | 0 |
| SRR5230595 | 261 | 3240 | 25595 | 3874 | 261 | 407 | 8552 | 7413 | 14215 | 113 | 2044 | 1772 | 2335 | 9260 | 153 | 617 | 15446 | 0 | 0 | 0 | 0 | 100 | 298 | 59 | 12 | 0 | 0 | 0 | 0 | 0 | 0 | 29 | 0 | 0 | 15 | 31 | 0 |
| SRR5230596 | 262 | 1572 | 33567 | 1203 | 927 | 1416 | 31172 | 3513 | 23792 | 454 | 8327 | 10466 | 4191 | 14228 | 296 | 295 | 40649 | 0 | 0 | 0 | 0 | 57 | 124 | 0 | 0 | 0 | 0 | 0 | 10 | 0 | 0 | 21 | 0 | 0 | 0 | 0 | 0 |
| SRR5230597 | 266 | 1522 | 33503 | 1059 | 899 | 1491 | 31509 | 3658 | 24539 | 474 | 8115 | 10162 | 4052 | 14312 | 336 | 317 | 42801 | 0 | 0 | 0 | 0 | 54 | 121 | 0 | 0 | 0 | 0 | 0 | 0 | 0 | 0 | 19 | 0 | 0 | 0 | 0 | 0 |
| SRR5230600 | 70 | 3125 | 61486 | 672 | 192 | 558 | 13608 | 4737 | 12923 | 148 | 2268 | 2608 | 1816 | 8325 | 197 | 326 | 16188 | 0 | 0 | 0 | 0 | 21 | 175 | 47 | 25 | 0 | 0 | 0 | 0 | 0 | 0 | 13 | 0 | 0 | 0 | 40 | 0 |
| SRR5230601 | 64 | 3216 | 60188 | 797 | 194 | 589 | 13763 | 4802 | 13227 | 148 | 2375 | 2477 | 1799 | 8103 | 228 | 340 | 16721 | 0 | 0 | 0 | 0 | 16 | 167 | 46 | 23 | 0 | 0 | 0 | 0 | 0 | 0 | 12 | 0 | 0 | 0 | 37 | 0 |
| SRR5230602 | 334 | 2491 | 26699 | 4060 | 384 | 804 | 17095 | 3561 | 8316 | 219 | 5891 | 5937 | 3567 | 19876 | 596 | 335 | 22271 | 0 | 0 | 0 | 14 | 266 | 289 | 32 | 16 | 0 | 0 | 0 | 0 | 10 | 20 | 22 | 0 | 0 | 0 | 18 | 0 |
| SRR5230603 | 324 | 2339 | 26004 | 4059 | 358 | 833 | 17387 | 3529 | 8601 | 213 | 5693 | 6127 | 3717 | 19622 | 570 | 311 | 23100 | 0 | 0 | 0 | 13 | 263 | 262 | 35 | 13 | 0 | 0 | 0 | 0 | 0 | 19 | 19 | 0 | 0 | 0 | 15 | 0 |
| SRR5230604 | 251 | 2141 | 39373 | 861 | 536 | 1196 | 24774 | 3899 | 22177 | 173 | 3957 | 4380 | 2702 | 10489 | 419 | 422 | 29988 | 0 | 0 | 0 | 0 | 71 | 206 | 22 | 12 | 0 | 0 | 0 | 0 | 0 | 0 | 18 | 0 | 0 | 0 | 18 | 0 |
| SRR5230605 | 243 | 2111 | 38989 | 857 | 526 | 1212 | 24880 | 3873 | 22484 | 211 | 4022 | 4121 | 2567 | 10392 | 454 | 405 | 30905 | 0 | 0 | 0 | 0 | 69 | 184 | 24 | 11 | 0 | 0 | 0 | 0 | 0 | 0 | 17 | 0 | 0 | 0 | 16 | 0 |
| SRR5230606 | 168 | 1527 | 28265 | 1026 | 1847 | 3794 | 64650 | 8121 | 25152 | 182 | 12547 | 10067 | 4142 | 21191 | 441 | 396 | 52668 | 0 | 0 | 0 | 0 | 127 | 267 | 11 | 0 | 0 | 0 | 0 | 0 | 0 | 16 | 23 | 0 | 0 | 0 | 0 | 0 |
| SRR5230607 | 215 | 1575 | 28273 | 1145 | 1841 | 3857 | 64316 | 7968 | 25881 | 169 | 11955 | 13 | 3886 | 20832 | 451 | 390 | 54992 | 0 | 0 | 0 | 0 | 119 | 215 | 11 | 0 | 0 | 0 | 0 | 0 | 0 | 12 | 21 | 0 | 0 | 0 | 0 | 0 |
| SRR5230608 | 360 | 1884 | 12452 | 7048 | 185 | 391 | 9381 | 3822 | 15848 | 126 | 6620 | 6214 | 4830 | 23022 | 912 | 207 | 16891 | 0 | 0 | 0 | 11 | 119 | 166 | 38 | 21 | 0 | 0 | 0 | 0 | 0 | 13 | 15 | 0 | 0 | 0 | 29 | 0 |
| SRR5230609 | 267 | 1784 | 12230 | 7233 | 195 | 429 | 9276 | 3987 | 16322 | 107 | 6704 | 6227 | 5201 | 22539 | 1019 | 221 | 17827 | 0 | 0 | 0 | 11 | 113 | 139 | 42 | 16 | 0 | 0 | 0 | 0 | 0 | 12 | 21 | 0 | 0 | 0 | 31 | 0 |
| SRR5230610 | 20 | 381 | 51502 | 107 | 476 | 989 | 22201 | 4334 | 25542 | 283 | 6298 | 14 | 3897 | 14879 | 289 | 373 | 25930 | 0 | 0 | 0 | 0 | 25 | 73 | 0 | 0 | 0 | 0 | 0 | 0 | 0 | 0 | 15 | 0 | 0 | 0 | 0 | 0 |
| SRR5230611 | 11 | 308 | 51400 | 123 | 454 | 1035 | 22545 | 4394 | 26687 | 306 | 6191 | 11 | 3716 | 14797 | 314 | 403 | 27208 | 0 | 0 | 0 | 0 | 38 | 79 | 0 | 0 | 0 | 0 | 0 | 0 | 0 | 0 | 10 | 0 | 0 | 0 | 14 | 0 |
| SRR5230613 | 14 | 622 | 61575 | 94 | 480 | 983 | 25315 | 8214 | 30859 | 525 | 3143 | 2737 | 2976 | 12064 | 198 | 980 | 28128 | 0 | 0 | 0 | 0 | 14 | 175 | 70 | 27 | 0 | 0 | 0 | 0 | 0 | 0 | 15 | 0 | 0 | 0 | 30 | 0 |
| SRR5230614 | 23 | 847 | 40303 | 113 | 94 | 232 | 5862 | 3834 | 24314 | 187 | 2265 | 2329 | 2694 | 8660 | 143 | 597 | 16753 | 0 | 0 | 12 | 110 | 55 | 121 | 22 | 17 | 0 | 0 | 0 | 11 | 0 | 0 | 0 | 0 | 0 | 0 | 27 | 0 |
| SRR5230615 | 33 | 768 | 40853 | 91 | 138 | 223 | 5920 | 3735 | 26249 | 180 | 2107 | 2277 | 2857 | 8660 | 160 | 636 | 17953 | 0 | 0 | 10 | 128 | 50 | 114 | 19 | 17 | 0 | 0 | 0 | 0 | 11 | 0 | 0 | 0 | 0 | 0 | 26 | 0 |
| SRR5230617 | 13 | 806 | 48331 | 128 | 1044 | 2079 | 53506 | 13381 | 33053 | 133 | 6180 | 10 | 2625 | 15288 | 193 | 597 | 32397 | 0 | 0 | 0 | 0 | 15 | 102 | 36 | 15 | 0 | 0 | 0 | 0 | 0 | 0 | 0 | 0 | 0 | 0 | 15 | 0 |
| SRR5230618 | 16 | 670 | 26548 | 313 | 717 | 1387 | 32122 | 6609 | 26664 | 835 | 9803 | 8605 | 7350 | 24261 | 587 | 195 | 28105 | 0 | 0 | 0 | 0 | 71 | 76 | 11 | 0 | 0 | 0 | 0 | 0 | 0 | 0 | 0 | 0 | 0 | 0 | 15 | 0 |
| SRR5230619 | 17 | 463 | 26356 | 196 | 816 | 1441 | 33441 | 6784 | 28215 | 1006 | 9960 | 22 | 7368 | 25091 | 644 | 201 | 29855 | 0 | 0 | 0 | 0 | 70 | 87 | 0 | 0 | 0 | 0 | 0 | 0 | 0 | 14 | 12 | 0 | 0 | 0 | 18 | 0 |
| SRR5230620 | 145 | 2811 | 48247 | 979 | 845 | 1528 | 31010 | 4236 | 17091 | 156 | 5438 | 5647 | 2681 | 10657 | 188 | 295 | 38006 | 0 | 0 | 0 | 0 | 106 | 226 | 20 | 0 | 0 | 0 | 0 | 0 | 0 | 0 | 16 | 0 | 0 | 0 | 14 | 0 |
| SRR5230621 | 145 | 2898 | 47378 | 920 | 860 | 1678 | 31544 | 4155 | 17482 | 172 | 5552 | 5537 | 2682 | 10243 | 220 | 296 | 39715 | 0 | 0 | 0 | 0 | 100 | 219 | 14 | 0 | 0 | 0 | 0 | 0 | 0 | 0 | 13 | 0 | 0 | 0 | 13 | 0 |
| SRR5230622 | 42 | 508 | 2907 | 5943 | 94 | 184 | 3357 | 3650 | 8616 | 33 | 346 | 218 | 917 | 5313 | 155 | 76 | 2125 | 0 | 0 | 0 | 0 | 66 | 67 | 0 | 0 | 0 | 0 | 0 | 0 | 11 | 0 | 10 | 0 | 0 | 0 | 0 | 0 |
| SRR5230623 | 201 | 1857 | 7403 | 4314 | 192 | 258 | 5157 | 2979 | 10100 | 148 | 2569 | 2992 | 6559 | 13404 | 216 | 617 | 7954 | 0 | 0 | 0 | 14 | 476 | 433 | 39 | 13 | 0 | 0 | 0 | 12 | 11 | 34 | 29 | 0 | 0 | 0 | 17 | 0 |
| SRR5230624 | 385 | 2096 | 7602 | 2946 | 849 | 1106 | 56769 | 10174 | 18265 | 162 | 5507 | 5209 | 4603 | 17410 | 205 | 345 | 43487 | 0 | 0 | 0 | 10 | 43 | 370 | 28 | 13 | 0 | 0 | 0 | 0 | 0 | 0 | 36 | 0 | 0 | 0 | 17 | 0 |
| SRR5230625 | 207 | 1796 | 8928 | 5784 | 2579 | 3627 | 85093 | 19468 | 20365 | 106 | 9865 | 7694 | 3285 | 15603 | 399 | 719 | 38754 | 0 | 11 | 0 | 29 | 216 | 393 | 41 | 23 | 0 | 0 | 21 | 20 | 11 | 20 | 32 | 0 | 0 | 0 | 32 | 0 |
| SRR5230626 | 386 | 2410 | 8696 | 3654 | 268 | 292 | 8353 | 4689 | 8258 | 332 | 3757 | 3920 | 3308 | 14285 | 351 | 143 | 11043 | 0 | 0 | 0 | 20 | 169 | 861 | 55 | 22 | 0 | 0 | 0 | 0 | 16 | 15 | 89 | 0 | 0 | 10 | 52 | 0 |
| SRR5230627 | 132 | 995 | 4074 | 4396 | 1879 | 2494 | 49952 | 9496 | 21964 | 88 | 10315 | 11272 | 5858 | 15770 | 580 | 531 | 29205 | 0 | 0 | 0 | 13 | 157 | 319 | 19 | 0 | 0 | 0 | 12 | 26 | 15 | 24 | 44 | 0 | 0 | 0 | 12 | 0 |
| SRR5230628 | 458 | 2432 | 8935 | 2588 | 636 | 635 | 22443 | 5836 | 10258 | 147 | 3581 | 2327 | 3029 | 16044 | 429 | 223 | 17431 | 0 | 0 | 0 | 12 | 46 | 550 | 67 | 24 | 0 | 0 | 0 | 0 | 0 | 0 | 46 | 0 | 0 | 0 | 22 | 0 |
| SRR5230629 | 371 | 2580 | 7467 | 2907 | 326 | 325 | 11149 | 5441 | 25264 | 363 | 8181 | 5938 | 8996 | 24153 | 865 | 78 | 19191 | 0 | 0 | 0 | 19 | 54 | 301 | 44 | 14 | 0 | 0 | 0 | 0 | 24 | 12 | 45 | 0 | 0 | 0 | 17 | 0 |
| SRR5230630 | 231 | 1618 | 8362 | 5224 | 1151 | 1220 | 26721 | 10279 | 76854 | 62 | 3189 | 2681 | 3681 | 12640 | 646 | 461 | 13287 | 0 | 0 | 0 | 0 | 64 | 214 | 13 | 0 | 0 | 0 | 0 | 0 | 19 | 16 | 46 | 0 | 0 | 0 | 0 | 0 |

### Supp Table S8.1.Columns_B_to_AL

| | | | | | | | | | | | | | | | | | | | | | | | | | | | | | | | | | | | | |
|---|---|---|---|---|---|---|---|---|---|---|---|---|---|---|---|---|---|---|---|---|---|---|---|---|---|---|---|---|---|---|---|---|---|---|---|---|---|
| SRR5230631 | 196 | 2096 | 8612 | 3891 | 665 | 1261 | 27273 | 7579 | 12737 | 55 | 3577 | 2537 | 1726 | 11171 | 161 | 290 | 19005 | 0 | 0 | 0 | 0 | 419 | 386 | 24 | 0 | 0 | 0 | 0 | 0 | 0 | 32 | 40 | 0 | 0 | 0 | 10 | 0 |
| SRR5230632 | 275 | 2466 | 9975 | 3590 | 560 | 680 | 21773 | 9167 | 19154 | 139 | 4489 | 4619 | 7317 | 20967 | 343 | 240 | 19152 | 0 | 0 | 0 | 16 | 102 | 420 | 76 | 23 | 0 | 0 | 0 | 15 | 19 | 0 | 44 | 0 | 0 | 13 | 40 | 0 |
| SRR5230633 | 177 | 931 | 3379 | 2766 | 732 | 935 | 27163 | 6855 | 12667 | 64 | 5127 | 5660 | 3967 | 12024 | 446 | 303 | 14614 | 0 | 0 | 0 | 20 | 79 | 287 | 31 | 12 | 0 | 0 | 0 | 0 | 12 | 12 | 27 | 0 | 0 | 0 | 13 | 0 |
| SRR5230634 | 28 | 197 | 25129 | 2939 | 665 | 1106 | 6816 | 7777 | 51693 | 105 | 12894 | 15597 | 9793 | 30063 | 881 | 211 | 10491 | 0 | 0 | 0 | 0 | 171 | 36 | 0 | 0 | 0 | 0 | 0 | 0 | 0 | 0 | 0 | 0 | 0 | 0 | 0 | 0 |
| SRR5230635 | 670 | 1164 | 66645 | 873 | 731 | 1133 | 9476 | 4756 | 145906 | 79 | 9659 | 7772 | 2579 | 13207 | 443 | 91 | 6119 | 0 | 0 | 0 | 0 | 67 | 22 | 0 | 0 | 0 | 0 | 0 | 0 | 0 | 0 | 0 | 0 | 0 | 0 | 0 | 0 |
| SRR5230636 | 596 | 3913 | 107986 | 3821 | 987 | 1437 | 10251 | 4115 | 64608 | 87 | 17012 | 14327 | 5388 | 21016 | 611 | 118 | 9137 | 0 | 0 | 0 | 0 | 424 | 79 | 0 | 0 | 0 | 0 | 0 | 13 | 0 | 0 | 0 | 0 | 0 | 0 | 0 | 0 |
| SRR5230637 | 467 | 2132 | 76187 | 467 | 1499 | 2573 | 18975 | 4638 | 152252 | 59 | 6143 | 8214 | 1385 | 23604 | 1025 | 73 | 11111 | 0 | 0 | 0 | 0 | 58 | 48 | 0 | 0 | 0 | 0 | 0 | 0 | 0 | 0 | 0 | 0 | 0 | 0 | 0 | 0 |
| SRR5230638 | 663 | 9194 | 69646 | 697 | 2963 | 5882 | 37480 | 6195 | 118610 | 77 | 22903 | 13 | 4689 | 40193 | 1292 | 165 | 31827 | 0 | 0 | 0 | 0 | 105 | 53 | 0 | 0 | 0 | 0 | 0 | 0 | 0 | 0 | 0 | 0 | 0 | 0 | 0 | 0 |
| SRR5230639 | 1054 | 4079 | 55348 | 875 | 3135 | 5786 | 35649 | 4943 | 94242 | 44 | 36156 | 38677 | 8107 | 43680 | 1188 | 225 | 34255 | 0 | 0 | 0 | 0 | 233 | 202 | 29 | 0 | 0 | 0 | 0 | 17 | 0 | 0 | 0 | 0 | 0 | 0 | 0 | 0 |
| SRR5230640 | 474 | 4963 | 33903 | 777 | 1511 | 2625 | 16958 | 4626 | 62104 | 32 | 9728 | 12364 | 1764 | 21549 | 506 | 80 | 15167 | 0 | 0 | 0 | 0 | 83 | 70 | 0 | 0 | 0 | 0 | 0 | 0 | 0 | 0 | 0 | 0 | 0 | 0 | 0 | 0 |
| SRR5230641 | 1011 | 15391 | 74438 | 371 | 1948 | 3578 | 19816 | 2861 | 32409 | 55 | 12337 | 22937 | 2494 | 23119 | 1998 | 213 | 19458 | 0 | 0 | 0 | 0 | 503 | 45 | 0 | 0 | 0 | 0 | 0 | 0 | 0 | 12 | 0 | 0 | 0 | 0 | 0 | 0 |
| SRR5230642 | 120 | 2371 | 103115 | 545 | 512 | 641 | 5077 | 8815 | 71136 | 75 | 20316 | 16 | 20725 | 33390 | 1204 | 69 | 5729 | 0 | 0 | 0 | 0 | 179 | 44 | 0 | 0 | 0 | 0 | 0 | 18 | 0 | 15 | 0 | 0 | 0 | 0 | 0 | 0 |
| SRR5230643 | 732 | 10234 | 74626 | 652 | 2221 | 4217 | 24005 | 5711 | 107783 | 99 | 28680 | 16 | 6116 | 32987 | 1065 | 215 | 27562 | 0 | 0 | 0 | 0 | 213 | 78 | 11 | 0 | 0 | 0 | 0 | 0 | 0 | 0 | 0 | 0 | 0 | 0 | 0 | 0 |
| SRR5230644 | 641 | 10841 | 88886 | 675 | 2568 | 4538 | 27669 | 6627 | 95606 | 118 | 20529 | 23342 | 4609 | 32451 | 911 | 185 | 26039 | 0 | 0 | 0 | 0 | 149 | 53 | 0 | 0 | 0 | 0 | 0 | 12 | 0 | 0 | 0 | 0 | 0 | 0 | 0 | 0 |
| SRR5230645 | 878 | 6978 | 82746 | 960 | 3643 | 6797 | 41421 | 6653 | 119073 | 106 | 48193 | 24 | 11826 | 54310 | 2123 | 276 | 39439 | 0 | 0 | 0 | 0 | 173 | 48 | 0 | 0 | 0 | 0 | 0 | 19 | 0 | 0 | 0 | 0 | 0 | 0 | 0 | 0 |
| SRR5230646 | 299 | 3763 | 61626 | 326 | 1805 | 3028 | 18637 | 5254 | 127721 | 99 | 26683 | 30275 | 5799 | 21643 | 886 | 186 | 19659 | 0 | 0 | 0 | 0 | 141 | 148 | 20 | 0 | 0 | 0 | 0 | 0 | 0 | 0 | 0 | 0 | 0 | 0 | 0 | 0 |
| SRR5230647 | 134 | 977 | 8566 | 128 | 262 | 494 | 3676 | 2529 | 38031 | 33 | 1030 | 1100 | 215 | 3887 | 110 | 29 | 2808 | 0 | 0 | 0 | 0 | 113 | 38 | 0 | 0 | 0 | 0 | 0 | 0 | 0 | 0 | 0 | 0 | 0 | 0 | 0 | 0 |
| SRR5230648 | 584 | 1192 | 53933 | 870 | 2236 | 3676 | 22894 | 4967 | 109246 | 59 | 28113 | 26706 | 6703 | 31564 | 952 | 191 | 21403 | 0 | 0 | 0 | 0 | 292 | 134 | 0 | 0 | 0 | 0 | 0 | 18 | 0 | 14 | 0 | 0 | 0 | 0 | 0 | 0 |
| SRR5230649 | 677 | 8258 | 87529 | 683 | 2971 | 4768 | 31301 | 8128 | 97776 | 53 | 22835 | 29101 | 3899 | 34138 | 1373 | 288 | 25790 | 0 | 0 | 0 | 0 | 228 | 48 | 0 | 0 | 0 | 0 | 0 | 21 | 0 | 0 | 0 | 0 | 0 | 0 | 0 | 0 |
| SRR5230650 | 431 | 2187 | 16735 | 620 | 56 | 100 | 479 | 1276 | 16138 | 33 | 577 | 1470 | 174 | 1831 | 85 | 22 | 1395 | 0 | 0 | 0 | 0 | 63 | 22 | 0 | 0 | 0 | 0 | 0 | 0 | 0 | 0 | 0 | 0 | 0 | 0 | 0 | 0 |
| SRR5230651 | 837 | 8735 | 91493 | 464 | 1309 | 2444 | 11640 | 6328 | 117590 | 87 | 23040 | 12 | 7532 | 24183 | 911 | 242 | 24485 | 0 | 0 | 0 | 0 | 209 | 72 | 0 | 0 | 0 | 0 | 0 | 11 | 0 | 11 | 0 | 0 | 0 | 0 | 0 | 0 |
| SRR5230652 | 254 | 1215 | 41753 | 629 | 1636 | 2260 | 13750 | 6161 | 174641 | 37 | 27881 | 29970 | 6834 | 18067 | 737 | 201 | 15127 | 0 | 0 | 0 | 0 | 275 | 54 | 0 | 0 | 0 | 0 | 0 | 13 | 0 | 0 | 0 | 0 | 0 | 0 | 0 | 0 |
| SRR5230653 | 412 | 3519 | 49486 | 270 | 1362 | 2610 | 13450 | 3719 | 52996 | 22 | 16382 | 24692 | 2985 | 33030 | 2464 | 231 | 18126 | 0 | 0 | 0 | 0 | 268 | 51 | 0 | 0 | 0 | 0 | 0 | 10 | 0 | 0 | 0 | 0 | 0 | 0 | 0 | 0 |
| SRR5230654 | 591 | 5830 | 109694 | 303 | 2325 | 3690 | 22344 | 3617 | 131458 | 92 | 15157 | 18595 | 2965 | 18068 | 1449 | 278 | 15824 | 0 | 0 | 0 | 0 | 360 | 52 | 0 | 0 | 0 | 0 | 0 | 0 | 0 | 11 | 0 | 0 | 0 | 0 | 0 | 0 |
| SRR5230655 | 615 | 5034 | 68865 | 709 | 1206 | 2364 | 12967 | 3183 | 60469 | 54 | 14008 | 23285 | 2493 | 21786 | 651 | 261 | 25000 | 0 | 0 | 0 | 0 | 158 | 64 | 11 | 0 | 0 | 0 | 0 | 0 | 0 | 0 | 0 | 0 | 0 | 0 | 0 | 0 |
| SRR5230656 | 309 | 983 | 64830 | 997 | 667 | 1081 | 6063 | 7568 | 97762 | 103 | 2338 | 3991 | 641 | 11077 | 275 | 203 | 9496 | 0 | 0 | 0 | 0 | 85 | 35 | 14 | 0 | 0 | 0 | 0 | 0 | 0 | 0 | 0 | 0 | 0 | 0 | 0 | 0 |
| SRR5230657 | 337 | 3370 | 62184 | 718 | 1214 | 2480 | 11655 | 4410 | 190989 | 39 | 6793 | 11602 | 1410 | 30553 | 947 | 282 | 22496 | 0 | 0 | 0 | 0 | 152 | 64 | 11 | 0 | 0 | 0 | 0 | 10 | 0 | 0 | 0 | 0 | 0 | 0 | 0 | 0 |
| SRR5230658 | 515 | 949 | 29651 | 1017 | 2122 | 3856 | 24202 | 6522 | 108055 | 73 | 21021 | 24643 | 3929 | 28047 | 564 | 240 | 33130 | 0 | 0 | 0 | 0 | 128 | 98 | 16 | 0 | 0 | 0 | 0 | 11 | 0 | 0 | 0 | 0 | 0 | 0 | 0 | 0 |
| SRR5230659 | 714 | 4267 | 41411 | 599 | 1026 | 1785 | 9993 | 5494 | 47537 | 40 | 11247 | 14491 | 3283 | 19556 | 1272 | 191 | 16177 | 0 | 0 | 0 | 0 | 382 | 245 | 27 | 0 | 0 | 0 | 0 | 15 | 0 | 12 | 0 | 0 | 0 | 0 | 0 | 0 |
| SRR5230660 | 468 | 1625 | 24734 | 1550 | 937 | 1528 | 10679 | 5660 | 59923 | 94 | 7534 | 8392 | 1703 | 19114 | 511 | 144 | 11825 | 0 | 0 | 0 | 0 | 112 | 107 | 15 | 0 | 0 | 0 | 0 | 13 | 0 | 0 | 0 | 0 | 0 | 0 | 0 | 0 |
| SRR5230661 | 760 | 1033 | 22123 | 2043 | 290 | 452 | 2373 | 2219 | 45462 | 0 | 8968 | 19 | 2520 | 22487 | 715 | 102 | 7022 | 0 | 0 | 0 | 0 | 111 | 71 | 17 | 0 | 0 | 0 | 0 | 0 | 0 | 0 | 0 | 0 | 0 | 0 | 0 | 0 |
| SRR5230662 | 111 | 393 | 5751 | 825 | 193 | 264 | 1789 | 2827 | 27312 | 35 | 2280 | 2991 | 664 | 8313 | 166 | 86 | 3009 | 0 | 0 | 0 | 0 | 44 | 32 | 0 | 0 | 0 | 0 | 0 | 0 | 0 | 0 | 0 | 0 | 0 | 0 | 0 | 0 |
| SRR5230663 | 969 | 3810 | 26777 | 453 | 2087 | 3099 | 19720 | 5994 | 123891 | 106 | 22890 | 15 | 4433 | 24206 | 1127 | 166 | 25134 | 0 | 0 | 0 | 0 | 101 | 30 | 0 | 0 | 0 | 0 | 0 | 27 | 11 | 12 | 0 | 0 | 0 | 0 | 0 | 0 |
| SRR5230664 | 153 | 724 | 4431 | 452 | 299 | 508 | 2656 | 2917 | 58216 | 13 | 4299 | 6719 | 1005 | 10635 | 209 | 109 | 5890 | 0 | 0 | 0 | 0 | 43 | 155 | 15 | 0 | 0 | 0 | 0 | 0 | 0 | 0 | 0 | 0 | 0 | 0 | 0 | 0 |
| SRR5230665 | 1910 | 3350 | 41770 | 848 | 3078 | 3966 | 28240 | 6346 | 84564 | 60 | 32718 | 37908 | 9237 | 36760 | 1600 | 438 | 27695 | 0 | 0 | 0 | 0 | 260 | 110 | 0 | 0 | 0 | 0 | 16 | 33 | 0 | 14 | 0 | 0 | 0 | 0 | 0 | 0 |
| SRR5230666 | 314 | 1034 | 32402 | 982 | 1661 | 2392 | 16013 | 8316 | 218401 | 102 | 14094 | 19620 | 2870 | 14780 | 624 | 229 | 14738 | 0 | 0 | 0 | 0 | 44 | 90 | 0 | 0 | 0 | 0 | 0 | 14 | 0 | 0 | 0 | 0 | 0 | 0 | 0 | 0 |
| SRR5230667 | 100 | 390 | 7512 | 549 | 45 | 58 | 355 | 1270 | 27610 | 0 | 857 | 1430 | 411 | 4671 | 77 | 47 | 1381 | 0 | 0 | 0 | 0 | 44 | 30 | 0 | 0 | 0 | 0 | 0 | 0 | 0 | 0 | 0 | 0 | 0 | 0 | 0 | 0 |
| SRR5230668 | 360 | 1564 | 15157 | 597 | 1156 | 1893 | 11608 | 3167 | 25504 | 42 | 15839 | 17061 | 4841 | 18554 | 632 | 181 | 14004 | 0 | 0 | 0 | 0 | 103 | 41 | 0 | 0 | 0 | 0 | 0 | 17 | 0 | 12 | 0 | 0 | 0 | 0 | 0 | 0 |
| SRR5230669 | 461 | 6655 | 71487 | 3359 | 880 | 1424 | 7032 | 3602 | 115081 | 174 | 7957 | 9736 | 2190 | 13353 | 446 | 144 | 21751 | 0 | 0 | 0 | 0 | 266 | 157 | 25 | 0 | 0 | 0 | 0 | 11 | 0 | 12 | 0 | 0 | 0 | 0 | 0 | 0 |
| SRR5230670 | 500 | 1240 | 18935 | 896 | 842 | 1400 | 9647 | 4469 | 57125 | 54 | 7559 | 7619 | 1497 | 12887 | 463 | 208 | 11024 | 0 | 0 | 0 | 0 | 93 | 584 | 93 | 17 | 0 | 0 | 0 | 0 | 0 | 0 | 17 | 0 | 0 | 0 | 12 | 0 |
| SRR5230671 | 235 | 737 | 13526 | 10783 | 202 | 271 | 2138 | 2495 | 22969 | 21 | 9964 | 10947 | 2466 | 30717 | 592 | 87 | 5619 | 0 | 0 | 0 | 0 | 46 | 46 | 0 | 0 | 0 | 0 | 0 | 0 | 0 | 0 | 0 | 0 | 0 | 0 | 0 | 0 |
| SRR5230673 | 447 | 3377 | 18305 | 1601 | 574 | 1037 | 6493 | 2060 | 27641 | 11 | 6996 | 11009 | 1439 | 11075 | 216 | 106 | 9231 | 0 | 0 | 0 | 0 | 106 | 95 | 21 | 0 | 0 | 0 | 0 | 0 | 0 | 0 | 0 | 0 | 0 | 0 | 0 | 0 |
| SRR5230674 | 810 | 1061 | 22235 | 1227 | 2439 | 3284 | 21188 | 5248 | 65948 | 26 | 15020 | 19049 | 2602 | 19483 | 566 | 341 | 21276 | 0 | 0 | 18 | 0 | 198 | 88 | 10 | 0 | 0 | 0 | 21 | 35 | 0 | 0 | 0 | 0 | 0 | 0 | 0 | 0 |
| SRR5230675 | 64 | 131 | 5985 | 1078 | 282 | 447 | 2907 | 4360 | 139376 | 40 | 9088 | 11541 | 2771 | 15164 | 414 | 43 | 6284 | 0 | 0 | 0 | 0 | 41 | 14 | 0 | 0 | 0 | 0 | 0 | 0 | 0 | 0 | 0 | 0 | 0 | 0 | 0 | 0 |
| SRR5230676 | 602 | 1766 | 26401 | 2219 | 1197 | 1656 | 10826 | 2558 | 137011 | 63 | 8063 | 10609 | 1613 | 14185 | 498 | 181 | 13622 | 0 | 0 | 0 | 0 | 100 | 45 | 0 | 0 | 0 | 0 | 0 | 11 | 0 | 0 | 0 | 0 | 0 | 0 | 0 | 0 |
| SRR5230677 | 333 | 1028 | 13859 | 1690 | 186 | 360 | 1993 | 1254 | 26209 | 17 | 1773 | 2871 | 525 | 5589 | 109 | 177 | 4576 | 0 | 0 | 0 | 0 | 50 | 185 | 33 | 0 | 0 | 0 | 0 | 0 | 0 | 0 | 0 | 0 | 0 | 0 | 0 | 0 |
| SRR5230678 | 806 | 2343 | 34074 | 1500 | 1772 | 2857 | 20759 | 3168 | 92003 | 47 | 16063 | 18253 | 2710 | 22254 | 596 | 189 | 20243 | 0 | 0 | 0 | 0 | 65 | 230 | 28 | 0 | 0 | 0 | 0 | 11 | 0 | 0 | 11 | 0 | 0 | 0 | 0 | 0 |
| SRR5230679 | 987 | 2728 | 31654 | 13023 | 750 | 1037 | 7677 | 3354 | 57266 | 38 | 7116 | 9825 | 1320 | 11259 | 262 | 486 | 11423 | 0 | 0 | 0 | 0 | 132 | 81 | 19 | 0 | 0 | 0 | 0 | 0 | 0 | 0 | 0 | 0 | 0 | 0 | 0 | 0 |
| SRR5230680 | 159 | 453 | 13880 | 762 | 141 | 202 | 1380 | 4139 | 36428 | 28 | 2444 | 2868 | 823 | 6828 | 153 | 62 | 2337 | 0 | 0 | 0 | 0 | 58 | 110 | 12 | 0 | 0 | 0 | 0 | 0 | 0 | 0 | 0 | 0 | 0 | 0 | 0 | 0 |
| SRR5230681 | 422 | 4199 | 35598 | 1470 | 891 | 1477 | 7611 | 2092 | 36320 | 56 | 5349 | 5228 | 1149 | 9147 | 1498 | 193 | 7488 | 0 | 0 | 0 | 0 | 349 | 84 | 0 | 0 | 0 | 0 | 0 | 16 | 0 | 18 | 0 | 0 | 0 | 0 | 0 | 0 |
| SRR5230682 | 1121 | 3503 | 29891 | 1318 | 913 | 1500 | 7199 | 851 | 18200 | 39 | 18492 | 41991 | 3540 | 17858 | 522 | 142 | 23825 | 0 | 0 | 0 | 0 | 222 | 114 | 0 | 0 | 0 | 0 | 0 | 0 | 0 | 0 | 0 | 0 | 0 | 0 | 0 | 0 |
| SRR5230683 | 679 | 1102 | 25028 | 2343 | 678 | 1318 | 8009 | 3301 | 48146 | 49 | 5235 | 10 | 1092 | 13533 | 249 | 106 | 8590 | 0 | 0 | 0 | 0 | 105 | 171 | 25 | 0 | 0 | 0 | 0 | 0 | 0 | 0 | 0 | 0 | 0 | 0 | 0 | 0 |
| SRR5230684 | 1887 | 6092 | 47068 | 2192 | 3898 | 4996 | 25194 | 4513 | 38359 | 118 | 13458 | 20277 | 2576 | 14797 | 708 | 959 | 27718 | 0 | 0 | 39 | 0 | 464 | 100 | 18 | 0 | 0 | 0 | 30 | 64 | 0 | 14 | 0 | 0 | 0 | 0 | 0 | 0 |
| SRR5230685 | 1065 | 1497 | 25184 | 2993 | 1237 | 1853 | 14054 | 3036 | 38520 | 89 | 11742 | 13 | 2348 | 15557 | 689 | 166 | 16975 | 0 | 0 | 0 | 0 | 92 | 84 | 13 | 0 | 0 | 0 | 0 | 0 | 0 | 0 | 0 | 0 | 0 | 0 | 0 | 0 |
| SRR5230686 | 1530 | 3512 | 38549 | 4502 | 1683 | 2751 | 15063 | 3161 | 73286 | 68 | 11694 | 16042 | 2197 | 15087 | 480 | 236 | 22983 | 0 | 0 | 0 | 0 | 171 | 157 | 18 | 0 | 0 | 0 | 0 | 0 | 0 | 0 | 0 | 0 | 0 | 0 | 0 | 0 |
| SRR5230687 | 1230 | 2041 | 26680 | 3295 | 2957 | 5014 | 29125 | 4650 | 35143 | 53 | 32320 | 21 | 5774 | 29395 | 661 | 305 | 53360 | 0 | 0 | 0 | 0 | 268 | 256 | 25 | 0 | 0 | 0 | 0 | 22 | 0 | 25 | 0 | 0 | 0 | 0 | 0 | 0 |
| SRR5230688 | 196 | 294 | 12842 | 742 | 211 | 332 | 2344 | 3656 | 22943 | 17 | 4818 | 16 | 1550 | 14095 | 375 | 47 | 4882 | 0 | 0 | 0 | 0 | 127 | 47 | 0 | 0 | 0 | 0 | 0 | 0 | 0 | 0 | 0 | 0 | 0 | 0 | 0 | 0 |
| SRR5230689 | 898 | 1689 | 48506 | 6389 | 135 | 208 | 1011 | 821 | 56596 | 96 | 897 | 1905 | 189 | 1555 | 64 | 251 | 2738 | 0 | 0 | 0 | 0 | 79 | 36 | 0 | 0 | 0 | 0 | 0 | 0 | 0 | 0 | 0 | 0 | 0 | 0 | 0 | 0 |
| SRR5230690 | 666 | 1589 | 27078 | 2492 | 2220 | 4419 | 21876 | 4672 | 69934 | 50 | 24362 | 17 | 4936 | 23515 | 921 | 320 | 36917 | 0 | 0 | 0 | 0 | 144 | 139 | 12 | 0 | 0 | 0 | 0 | 11 | 0 | 0 | 0 | 0 | 0 | 0 | 0 | 0 |
| SRR5230691 | 876 | 10618 | 17742 | 947 | 265 | 346 | 1855 | 848 | 8305 | 32 | 2596 | 5112 | 615 | 4036 | 113 | 62 | 4332 | 0 | 0 | 0 | 0 | 339 | 112 | 11 | 0 | 0 | 0 | 0 | 10 | 0 | 0 | 0 | 0 | 0 | 0 | 0 | 0 |
| SRR5230692 | 1119 | 5035 | 44437 | 1340 | 2043 | 3674 | 21437 | 13340 | 86362 | 87 | 18138 | 17 | 3960 | 22201 | 867 | 272 | 24798 | 0 | 0 | 0 | 0 | 271 | 119 | 0 | 0 | 0 | 0 | 12 | 27 | 0 | 20 | 0 | 0 | 0 | 0 | 0 | 0 |
| SRR5230693 | 1341 | 3062 | 30816 | 1927 | 1689 | 3038 | 15286 | 2968 | 51227 | 50 | 13404 | 19764 | 2279 | 13037 | 544 | 235 | 31345 | 0 | 0 | 0 | 0 | 148 | 234 | 42 | 0 | 0 | 0 | 0 | 0 | 0 | 0 | 0 | 0 | 0 | 0 | 0 | 0 |
| SRR5230694 | 743 | 2650 | 35701 | 12596 | 801 | 847 | 5178 | 4791 | 32855 | 49 | 11189 | 15188 | 4396 | 10853 | 273 | 32 | 6700 | 0 | 0 | 0 | 0 | 111 | 200 | 31 | 0 | 0 | 0 | 11 | 27 | 11 | 0 | 0 | 0 | 0 | 0 | 0 | 0 |
| SRR5230695 | 1016 | 1875 | 38186 | 2204 | 418 | 674 | 3605 | 1941 | 26516 | 91 | 3066 | 5880 | 733 | 6309 | 171 | 264 | 8538 | 0 | 0 | 0 | 0 | 126 | 68 | 15 | 0 | 0 | 0 | 0 | 13 | 0 | 0 | 0 | 0 | 0 | 0 | 0 | 0 |
| SRR5230696 | 634 | 3365 | 34571 | 4160 | 422 | 589 | 6147 | 4858 | 135726 | 83 | 3523 | 6374 | 700 | 8873 | 206 | 77 | 6209 | 0 | 0 | 0 | 0 | 103 | 36 | 0 | 0 | 0 | 0 | 0 | 10 | 0 | 0 | 0 | 0 | 0 | 0 | 0 | 0 |
| SRR5230697 | 930 | 2047 | 16215 | 1597 | 618 | 1026 | 5394 | 955 | 18097 | 61 | 5727 | 10306 | 957 | 6695 | 270 | 170 | 12608 | 0 | 0 | 0 | 0 | 130 | 142 | 21 | 0 | 0 | 0 | 0 | 13 | 0 | 0 | 0 | 0 | 0 | 0 | 0 | 0 |
| SRR5230698 | 1120 | 7361 | 58856 | 1363 | 888 | 1580 | 9451 | 1322 | 25412 | 30 | 10980 | 20415 | 1942 | 16571 | 337 | 188 | 19478 | 0 | 0 | 0 | 0 | 293 | 179 | 17 | 0 | 0 | 0 | 0 | 25 | 0 | 17 | 0 | 0 | 0 | 0 | 0 | 0 |
| SRR5230699 | 728 | 4093 | 56681 | 1466 | 1881 | 2770 | 14289 | 4557 | 151589 | 182 | 23437 | 35972 | 5074 | 21889 | 1434 | 117 | 22216 | 0 | 0 | 0 | 0 | 174 | 44 | 0 | 0 | 0 | 0 | 15 | 52 | 12 | 23 | 0 | 0 | 0 | 0 | 0 | 0 |
| SRR5230700 | 1337 | 2175 | 25292 | 2600 | 599 | 1033 | 6716 | 2308 | 29269 | 90 | 6758 | 10026 | 1518 | 10761 | 252 | 102 | 9260 | 0 | 0 | 0 | 0 | 144 | 52 | 0 | 0 | 0 | 0 | 0 | 0 | 0 | 0 | 0 | 0 | 0 | 0 | 0 | 0 |
| SRR5230701 | 944 | 9928 | 39780 | 1047 | 517 | 801 | 3897 | 2145 | 20536 | 37 | 9742 | 17211 | 2384 | 15713 | 746 | 116 | 8335 | 0 | 0 | 0 | 0 | 349 | 190 | 27 | 0 | 0 | 0 | 0 | 14 | 0 | 13 | 0 | 0 | 0 | 0 | 0 | 0 |
| SRR5230702 | 832 | 7257 | 59983 | 1202 | 2257 | 3868 | 24707 | 5706 | 81765 | 66 | 15445 | 16387 | 2455 | 22701 | 1015 | 242 | 24695 | 0 | 0 | 0 | 0 | 225 | 83 | 0 | 0 | 0 | 0 | 0 | 21 | 0 | 11 | 0 | 0 | 0 | 0 | 0 | 0 |
| SRR5230703 | 832 | 2528 | 26946 | 2230 | 784 | 1020 | 7537 | 2214 | 33825 | 27 | 17546 | 21311 | 5794 | 22256 | 866 | 72 | 9461 | 0 | 0 | 0 | 0 | 209 | 103 | 0 | 0 | 0 | 0 | 0 | 27 | 0 | 0 | 0 | 0 | 0 | 0 | 0 | 0 |
| SRR5230704 | 1045 | 19522 | 33075 | 1303 | 1087 | 1149 | 9173 | 1598 | 52608 | 42 | 8201 | 14183 | 1670 | 12486 | 796 | 185 | 7927 | 0 | 0 | 0 | 0 | 325 | 56 | 0 | 0 | 0 | 0 | 0 | 16 | 0 | 0 | 0 | 0 | 0 | 0 | 0 | 0 |
| SRR5230705 | 141 | 615 | 8052 | 390 | 76 | 88 | 714 | 1350 | 14348 | 11 | 983 | 1320 | 273 | 5161 | 124 | 22 | 1095 | 0 | 0 | 0 | 0 | 60 | 45 | 0 | 0 | 0 | 0 | 0 | 0 | 0 | 0 | 0 | 0 | 0 | 0 | 0 | 0 |
| SRR5230706 | 614 | 3515 | 23725 | 2312 | 618 | 938 | 5115 | 2741 | 17348 | 100 | 14818 | 22134 | 6065 | 20982 | 675 | 116 | 15869 | 0 | 0 | 0 | 0 | 183 | 57 | 0 | 0 | 0 | 0 | 0 | 36 | 0 | 21 | 0 | 0 | 0 | 0 | 0 | 0 |
| SRR5230707 | 579 | 680 | 13803 | 691 | 949 | 1776 | 10086 | 4924 | 44595 | 85 | 10123 | 12661 | 3561 | 15577 | 590 | 153 | 19602 | 0 | 0 | 0 | 0 | 224 | 137 | 0 | 0 | 0 | 0 | 0 | 19 | 0 | 21 | 0 | 0 | 0 | 0 | 0 | 0 |
| SRR5230708 | 1144 | 5474 | 65261 | 1561 | 3595 | 5431 | 32074 | 5526 | 101146 | 129 | 22945 | 20783 | 4662 | 25904 | 608 | 258 | 42088 | 0 | 0 | 0 | 0 | 127 | 85 | 0 | 0 | 0 | 0 | 0 | 20 | 0 | 0 | 0 | 0 | 0 | 0 | 0 | 0 |
| SRR5230709 | 2605 | 2282 | 28422 | 1794 | 1252 | 2162 | 11192 | 5709 | 122430 | 187 | 16289 | 19053 | 3458 | 14952 | 915 | 266 | 20478 | 0 | 0 | 0 | 0 | 133 | 89 | 15 | 0 | 0 | 0 | 0 | 17 | 0 | 0 | 0 | 0 | 0 | 0 | 0 | 0 |
| SRR5230710 | 845 | 10933 | 18890 | 323 | 1060 | 1812 | 9883 | 3475 | 24191 | 82 | 9943 | 11028 | 2595 | 12551 | 381 | 192 | 15432 | 0 | 0 | 0 | 0 | 207 | 40 | 0 | 0 | 0 | 0 | 0 | 18 | 0 | 11 | 0 | 0 | 0 | 0 | 0 | 0 |
| SRR5230711 | 1298 | 5446 | 37296 | 1710 | 1003 | 1166 | 8146 | 6237 | 35984 | 114 | 8435 | 9471 | 2039 | 9563 | 397 | 374 | 9193 | 0 | 0 | 0 | 0 | 184 | 84 | 16 | 0 | 0 | 0 | 0 | 11 | 0 | 0 | 0 | 0 | 0 | 0 | 0 | 0 |
| SRR5230712 | 428 | 630 | 13846 | 562 | 606 | 1046 | 5993 | 5835 | 33031 | 79 | 7479 | 14 | 1674 | 9491 | 152 | 114 | 13677 | 0 | 0 | 0 | 0 | 101 | 80 | 0 | 0 | 0 | 0 | 0 | 11 | 0 | 0 | 0 | 0 | 0 | 0 | 0 | 0 |
| SRR5230713 | 1001 | 10714 | 35545 | 404 | 1986 | 2273 | 12788 | 5107 | 34348 | 106 | 12102 | 14 | 3005 | 10978 | 558 | 394 | 20211 | 0 | 0 | 22 | 0 | 525 | 156 | 22 | 0 | 0 | 0 | 25 | 59 | 0 | 22 | 0 | 0 | 0 | 0 | 0 | 0 |
| SRR5230714 | 425 | 1627 | 14501 | 2220 | 425 | 633 | 2897 | 4459 | 40781 | 43 | 5787 | 7474 | 2131 | 6269 | 196 | 89 | 6974 | 0 | 0 | 0 | 0 | 172 | 87 | 0 | 0 | 0 | 0 | 0 | 30 | 0 | 19 | 0 | 0 | 0 | 0 | 0 | 0 |
| SRR5230716 | 699 | 1445 | 16059 | 8550 | 301 | 507 | 2789 | 3869 | 25928 | 137 | 2610 | 3228 | 923 | 4382 | 94 | 188 | 4898 | 0 | 0 | 0 | 0 | 56 | 67 | 0 | 0 | 0 | 0 | 0 | 15 | 0 | 0 | 0 | 0 | 0 | 0 | 0 | 0 |
| SRR5230717 | 576 | 1717 | 21382 | 736 | 547 | 748 | 4734 | 3319 | 53009 | 155 | 8404 | 11 | 2689 | 13575 | 346 | 259 | 13056 | 0 | 0 | 0 | 0 | 162 | 73 | 10 | 0 | 0 | 0 | 0 | 13 | 0 | 0 | 0 | 0 | 0 | 0 | 0 | 0 |
| SRR5230718 | 925 | 4140 | 35348 | 857 | 1133 | 2116 | 12830 | 4830 | 38361 | 204 | 9951 | 10337 | 2088 | 11752 | 408 | 396 | 20149 | 0 | 0 | 0 | 0 | 152 | 298 | 29 | 0 | 0 | 0 | 0 | 11 | 0 | 0 | 0 | 0 | 0 | 0 | 0 | 0 |
| SRR5230719 | 1138 | 8606 | 30247 | 710 | 441 | 774 | 3976 | 1892 | 21189 | 67 | 10559 | 16715 | 2942 | 10565 | 487 | 235 | 15394 | 0 | 0 | 0 | 0 | 178 | 90 | 14 | 0 | 0 | 0 | 0 | 16 | 0 | 10 | 0 | 0 | 0 | 0 | 0 | 0 |
| SRR5230720 | 1417 | 9523 | 25055 | 813 | 960 | 1583 | 7749 | 1917 | 29435 | 43 | 16569 | 26301 | 3401 | 16896 | 708 | 115 | 23791 | 0 | 0 | 0 | 0 | 341 | 123 | 0 | 0 | 0 | 0 | 0 | 25 | 0 | 15 | 0 | 0 | 0 | 0 | 0 | 0 |
| SRR5230721 | 937 | 3485 | 38745 | 1025 | 509 | 636 | 4123 | 3694 | 69800 | 74 | 7537 | 12006 | 1657 | 8356 | 237 | 236 | 9997 | 0 | 0 | 0 | 0 | 228 | 53 | 0 | 0 | 0 | 0 | 12 | 15 | 0 | 0 | 0 | 0 | 0 | 0 | 0 | 0 |
| SRR5230722 | 1320 | 15225 | 25673 | 499 | 828 | 1306 | 6845 | 3910 | 17324 | 39 | 9392 | 14718 | 1879 | 8034 | 273 | 357 | 18416 | 0 | 0 | 0 | 0 | 206 | 99 | 0 | 0 | 0 | 0 | 0 | 0 | 0 | 0 | 0 | 0 | 0 | 0 | 0 | 0 |
| SRR5230723 | 660 | 2667 | 31817 | 1124 | 1151 | 1838 | 10030 | 5433 | 111425 | 94 | 9818 | 11713 | 3137 | 13137 | 346 | 382 | 19527 | 0 | 0 | 0 | 0 | 87 | 378 | 58 | 11 | 0 | 0 | 0 | 11 | 0 | 0 | 19 | 0 | 0 | 0 | 0 | 0 |
| SRR5230724 | 503 | 1158 | 18266 | 576 | 201 | 256 | 1842 | 3469 | 23914 | 61 | 1827 | 2714 | 789 | 8323 | 85 | 120 | 3021 | 0 | 0 | 0 | 0 | 99 | 497 | 59 | 12 | 0 | 0 | 0 | 0 | 0 | 0 | 17 | 0 | 0 | 0 | 0 | 0 |
| SRR5230725 | 213 | 1122 | 28282 | 827 | 1127 | 1773 | 10689 | 8245 | 54038 | 51 | 9063 | 8998 | 2087 | 11088 | 198 | 180 | 14957 | 0 | 0 | 0 | 0 | 107 | 83 | 0 | 0 | 0 | 0 | 0 | 17 | 0 | 0 | 0 | 0 | 0 | 0 | 0 | 0 |
| SRR5230726 | 330 | 598 | 8703 | 794 | 509 | 652 | 4642 | 2916 | 37285 | 38 | 6083 | 8827 | 1684 | 8299 | 267 | 131 | 8618 | 0 | 0 | 0 | 0 | 32 | 77 | 12 | 0 | 0 | 0 | 0 | 0 | 0 | 0 | 0 | 0 | 0 | 0 | 0 | 0 |
| SRR5230727 | 514 | 1229 | 13053 | 825 | 111 | 169 | 1147 | 1618 | 18861 | 33 | 1664 | 2917 | 587 | 3002 | 76 | 73 | 2801 | 0 | 0 | 0 | 0 | 67 | 103 | 17 | 0 | 0 | 0 | 0 | 11 | 0 | 0 | 0 | 0 | 0 | 0 | 0 | 0 |
| SRR5230728 | 462 | 2414 | 34378 | 568 | 1108 | 1509 | 10254 | 7986 | 90314 | 69 | 12567 | 13568 | 3216 | 15011 | 663 | 260 | 16215 | 0 | 0 | 0 | 0 | 209 | 49 | 0 | 0 | 0 | 0 | 0 | 30 | 13 | 18 | 0 | 0 | 0 | 0 | 0 | 0 |
| SRR5230729 | 485 | 3245 | 38337 | 1618 | 1537 | 2621 | 15986 | 7053 | 69739 | 109 | 15461 | 16808 | 4298 | 20174 | 429 | 253 | 24543 | 0 | 0 | 0 | 0 | 195 | 93 | 0 | 0 | 0 | 0 | 10 | 30 | 0 | 11 | 0 | 0 | 0 | 0 | 0 | 0 |
| SRR5230732 | 1221 | 2184 | 42753 | 444 | 1571 | 2650 | 15886 | 7304 | 68181 | 119 | 17457 | 21485 | 5209 | 23886 | 964 | 228 | 26003 | 0 | 0 | 0 | 0 | 218 | 143 | 0 | 0 | 0 | 0 | 11 | 16 | 0 | 23 | 0 | 0 | 0 | 0 | 0 | 0 |
| SRR5230733 | 317 | 1386 | 22324 | 329 | 512 | 622 | 4191 | 3481 | 21355 | 46 | 4944 | 6157 | 1524 | 7387 | 172 | 257 | 6568 | 0 | 0 | 0 | 0 | 174 | 83 | 17 | 0 | 0 | 0 | 0 | 13 | 0 | 0 | 0 | 0 | 0 | 0 | 0 | 0 |
| SRR5230734 | 538 | 2118 | 50412 | 365 | 2841 | 4582 | 32267 | 7707 | 84113 | 123 | 2238 | 1181 | 935 | 11215 | 410 | 1141 | 16539 | 0 | 0 | 12 | 0 | 146 | 44 | 0 | 0 | 0 | 0 | 0 | 13 | 0 | 0 | 0 | 0 | 0 | 0 | 0 | 0 |
| SRR5230735 | 1524 | 5264 | 71225 | 1348 | 1439 | 2117 | 14638 | 6716 | 33712 | 163 | 17476 | 20837 | 4521 | 16738 | 550 | 542 | 26756 | 0 | 0 | 0 | 0 | 449 | 151 | 12 | 0 | 0 | 0 | 0 | 33 | 0 | 16 | 0 | 0 | 0 | 0 | 0 | 0 |
| SRR5230736 | 192 | 644 | 16082 | 429 | 180 | 315 | 1959 | 2393 | 11663 | 80 | 2816 | 3704 | 1098 | 7458 | 165 | 126 | 4833 | 0 | 0 | 0 | 0 | 90 | 113 | 12 | 0 | 0 | 0 | 0 | 0 | 0 | 0 | 0 | 0 | 0 | 0 | 0 | 0 |
| SRR5230737 | 525 | 2433 | 62929 | 338 | 2226 | 3823 | 24194 | 10081 | 138708 | 136 | 18228 | 11 | 3540 | 21751 | 623 | 394 | 30592 | 0 | 0 | 0 | 0 | 67 | 67 | 0 | 0 | 0 | 0 | 0 | 16 | 0 | 0 | 0 | 0 | 0 | 0 | 0 | 0 |
| SRR5230738 | 655 | 2877 | 39011 | 719 | 793 | 1325 | 8280 | 5269 | 25623 | 217 | 8283 | 8666 | 2427 | 13061 | 350 | 188 | 16472 | 0 | 0 | 0 | 0 | 133 | 103 | 14 | 0 | 0 | 0 | 0 | 0 | 0 | 0 | 0 | 0 | 0 | 0 | 0 | 0 |
| SRR5230739 | 256 | 5317 | 42197 | 204 | 1318 | 2011 | 12034 | 8756 | 89783 | 65 | 10626 | 11174 | 2403 | 13557 | 376 | 297 | 18361 | 0 | 0 | 0 | 0 | 49 | 154 | 39 | 13 | 0 | 0 | 0 | 0 | 0 | 0 | 0 | 0 | 0 | 0 | 0 | 0 |
| SRR5230740 | 261 | 2246 | 27677 | 305 | 866 | 1173 | 7822 | 4387 | 50759 | 67 | 18929 | 19779 | 5604 | 34391 | 1178 | 147 | 16905 | 0 | 0 | 0 | 0 | 121 | 34 | 0 | 0 | 0 | 0 | 0 | 14 | 0 | 0 | 0 | 0 | 0 | 0 | 0 | 0 |
| SRR5230741 | 1146 | 9620 | 35483 | 433 | 3560 | 5318 | 31331 | 5233 | 24305 | 59 | 19823 | 11 | 5879 | 27451 | 1153 | 1783 | 30351 | 0 | 0 | 0 | 0 | 163 | 35 | 0 | 0 | 0 | 0 | 0 | 21 | 0 | 0 | 0 | 0 | 0 | 0 | 0 | 0 |
| SRR5230742 | 301 | 980 | 67685 | 784 | 1142 | 1840 | 13573 | 12197 | 100306 | 168 | 10308 | 9365 | 3324 | 19270 | 328 | 496 | 21925 | 0 | 0 | 0 | 0 | 55 | 72 | 0 | 0 | 0 | 0 | 0 | 0 | 0 | 0 | 0 | 0 | 0 | 0 | 0 | 0 |
| SRR5230743 | 584 | 5155 | 50619 | 1257 | 1239 | 2151 | 11277 | 3760 | 70564 | 152 | 23400 | 21 | 8939 | 35585 | 1943 | 288 | 21644 | 0 | 0 | 0 | 0 | 186 | 81 | 0 | 0 | 0 | 0 | 0 | 31 | 0 | 14 | 0 | 0 | 0 | 0 | 0 | 0 |
| SRR5230744 | 1025 | 14875 | 37584 | 355 | 503 | 610 | 3808 | 2174 | 12381 | 94 | 8777 | 11 | 2121 | 6967 | 344 | 493 | 13632 | 0 | 0 | 0 | 0 | 317 | 76 | 0 | 0 | 0 | 0 | 0 | 23 | 0 | 12 | 0 | 0 | 0 | 0 | 0 | 0 |
| SRR5230745 | 970 | 3452 | 47719 | 636 | 2000 | 2657 | 16630 | 10835 | 103893 | 192 | 23855 | 23 | 5563 | 24339 | 1222 | 327 | 26825 | 0 | 0 | 0 | 0 | 102 | 59 | 0 | 0 | 0 | 0 | 13 | 17 | 0 | 0 | 0 | 0 | 0 | 0 | 0 | 0 |
| SRR5230746 | 1122 | 2330 | 50753 | 1432 | 1102 | 1516 | 9540 | 6533 | 59371 | 182 | 12974 | 13 | 3291 | 12758 | 426 | 394 | 20240 | 0 | 0 | 0 | 0 | 202 | 103 | 11 | 0 | 0 | 0 | 0 | 12 | 0 | 0 | 0 | 0 | 0 | 0 | 0 | 0 |
| SRR5230747 | 296 | 1326 | 29959 | 367 | 2270 | 3167 | 20367 | 5903 | 34830 | 67 | 13522 | 13578 | 2676 | 17636 | 407 | 356 | 27545 | 0 | 0 | 0 | 0 | 67 | 179 | 16 | 0 | 0 | 0 | 0 | 0 | 0 | 0 | 0 | 0 | 0 | 0 | 0 | 0 |
| SRR5230748 | 346 | 1178 | 41653 | 583 | 1970 | 3052 | 19967 | 6941 | 56563 | 146 | 17530 | 21244 | 4186 | 17957 | 488 | 272 | 31546 | 0 | 0 | 0 | 0 | 120 | 100 | 0 | 0 | 0 | 0 | 0 | 11 | 0 | 12 | 0 | 0 | 0 | 0 | 0 | 0 |
| SRR5230749 | 284 | 956 | 34197 | 423 | 1040 | 1703 | 9761 | 4323 | 29565 | 75 | 11918 | 12518 | 3117 | 16124 | 596 | 143 | 17132 | 0 | 0 | 0 | 0 | 143 | 125 | 12 | 0 | 0 | 0 | 0 | 10 | 0 | 0 | 0 | 0 | 0 | 0 | 0 | 0 |
| SRR5230750 | 233 | 920 | 20228 | 352 | 262 | 353 | 2690 | 3822 | 35292 | 82 | 2498 | 2770 | 858 | 5058 | 127 | 242 | 5674 | 0 | 0 | 0 | 0 | 42 | 79 | 12 | 0 | 0 | 0 | 0 | 0 | 0 | 0 | 0 | 0 | 0 | 0 | 0 | 0 |
| SRR5230751 | 112 | 529 | 20866 | 514 | 131 | 207 | 1330 | 2592 | 28366 | 62 | 1554 | 1992 | 832 | 5457 | 105 | 92 | 3036 | 0 | 0 | 0 | 0 | 38 | 445 | 68 | 18 | 0 | 0 | 0 | 0 | 0 | 0 | 0 | 0 | 0 | 0 | 12 | 0 |
| SRR5230752 | 956 | 1795 | 43998 | 1033 | 920 | 1411 | 6639 | 4674 | 72794 | 66 | 8033 | 11096 | 1869 | 7405 | 434 | 440 | 14342 | 0 | 0 | 0 | 0 | 130 | 77 | 0 | 0 | 0 | 0 | 14 | 21 | 0 | 0 | 0 | 0 | 0 | 0 | 0 | 0 |
| SRR5230753 | 1199 | 8716 | 59144 | 542 | 2508 | 4623 | 21608 | 7890 | 57226 | 177 | 22846 | 26622 | 4825 | 16155 | 724 | 519 | 46360 | 0 | 0 | 0 | 0 | 163 | 88 | 18 | 0 | 0 | 0 | 0 | 28 | 0 | 11 | 0 | 0 | 0 | 0 | 0 | 0 |
| SRR5231309 | 212 | 47535 | 20 | 3985 | 7377 | 64209 | 48783 | 25567 | 886 | 13 | 11990 | 6309 | 4132 | 13686 | 32 | 187588 | 11638 | 0 | 0 | 0 | 325 | 51 | 228 | 238 | 878 | 102 | 0 | 0 | 17 | 176 | 51 | 285 | 34 | 1093 | 1923 | 525 | 0 |
| SRR5231310 | 85 | 35250 | 46 | 3459 | 7543 | 59589 | 52376 | 37780 | 762 | 14 | 19139 | 9409 | 5297 | 13109 | 43 | 140974 | 14625 | 0 | 0 | 0 | 257 | 49 | 148 | 131 | 360 | 69 | 0 | 0 | 19 | 112 | 47 | 168 | 18 | 573 | 1552 | 385 | 0 |
| SRR5231311 | 131 | 50316 | 114 | 4053 | 7388 | 60271 | 58327 | 38901 | 947 | 15 | 16849 | 8043 | 4945 | 12331 | 16 | 148985 | 13681 | 0 | 0 | 0 | 184 | 34 | 103 | 154 | 462 | 52 | 0 | 0 | 24 | 118 | 55 | 229 | 17 | 593 | 737 | 460 | 0 |
| SRR5231312 | 57 | 13923 | 40 | 1616 | 6814 | 53031 | 55529 | 30819 | 1015 | 34 | 18510 | 8796 | 4784 | 17519 | 18 | 164556 | 10984 | 0 | 0 | 0 | 110 | 16 | 109 | 154 | 539 | 74 | 0 | 0 | 0 | 72 | 18 | 166 | 17 | 638 | 1068 | 338 | 0 |
| SRR5231313 | 79 | 17340 | 46 | 7482 | 8575 | 62948 | 62502 | 29650 | 820 | 23 | 18780 | 7020 | 3975 | 14962 | 22 | 193973 | 9944 | 0 | 0 | 0 | 193 | 28 | 91 | 120 | 335 | 53 | 0 | 0 | 18 | 79 | 26 | 159 | 18 | 575 | 1253 | 364 | 0 |

### Supp Table S8.1.Columns_B_to_AL

| | | | | | | | | | | | | | | | | | | | | | | | | | | | | | | | | | | | | | |
|---|---|---|---|---|---|---|---|---|---|---|---|---|---|---|---|---|---|---|---|---|---|---|---|---|---|---|---|---|---|---|---|---|---|---|---|---|---|
| SRR5231314 | 120 | 44796 | 90 | 4253 | 9605 | 63902 | 66767 | 33944 | 949 | 19 | 16334 | 7923 | 3768 | 11616 | 0 | 178040 | 10611 | 0 | 0 | 0 | 233 | 60 | 75 | 111 | 284 | 40 | 0 | 0 | 35 | 120 | 75 | 153 | 12 | 304 | 353 | 317 | 0 |
| SRR5239182 | 512 | 179734 | 39 | 384 | 508 | 5601 | 4477 | 1792 | 11524 | 0 | 80 | 148 | 0 | 34 | 14 | 4744 | 30959 | 0 | 0 | 0 | 776 | 69 | 42 | 0 | 0 | 0 | 0 | 0 | 49 | 134 | 19 | 17 | 0 | 0 | 49 | 0 | 0 |
| SRR5239183 | 790 | 164512 | 79 | 1007 | 97 | 2147 | 1358 | 499 | 6733 | 0 | 83 | 28 | 0 | 0 | 0 | 1482 | 7356 | 0 | 0 | 0 | 243 | 116 | 27 | 0 | 12 | 0 | 0 | 0 | 23 | 48 | 11 | 0 | 0 | 11 | 154 | 0 | 0 |
| SRR5239184 | 828 | 175016 | 31 | 960 | 82 | 1104 | 880 | 454 | 6978 | 0 | 12 | 63 | 0 | 0 | 22 | 837 | 13093 | 0 | 0 | 0 | 226 | 89 | 18 | 0 | 0 | 0 | 0 | 0 | 19 | 31 | 10 | 0 | 0 | 0 | 84 | 0 | 0 |
| SRR5239185 | 234 | 60117 | 19 | 400 | 548 | 8301 | 5617 | 2074 | 16871 | 0 | 353 | 293 | 14 | 136 | 0 | 5240 | 26474 | 0 | 0 | 0 | 198 | 57 | 10 | 0 | 0 | 0 | 0 | 0 | 19 | 58 | 12 | 0 | 0 | 0 | 109 | 0 | 0 |
| SRR5239186 | 420 | 162905 | 33 | 333 | 500 | 6177 | 5134 | 1790 | 13855 | 0 | 213 | 1065 | 18 | 129 | 26 | 1636 | 31001 | 0 | 0 | 0 | 1069 | 102 | 43 | 0 | 0 | 0 | 0 | 0 | 77 | 255 | 36 | 23 | 0 | 0 | 140 | 0 | 0 |
| SRR5239187 | 618 | 183859 | 40 | 2615 | 139 | 1936 | 1605 | 687 | 6471 | 0 | 300 | 1111 | 43 | 246 | 12 | 602 | 17158 | 0 | 0 | 0 | 388 | 129 | 36 | 0 | 12 | 0 | 0 | 0 | 36 | 60 | 15 | 0 | 0 | 0 | 183 | 0 | 0 |
| SRR5239188 | 828 | 150048 | 32 | 2017 | 898 | 11546 | 7735 | 2419 | 6316 | 0 | 356 | 1319 | 37 | 151 | 16 | 2653 | 24152 | 0 | 0 | 0 | 566 | 95 | 29 | 0 | 0 | 14 | 0 | 0 | 64 | 104 | 20 | 15 | 0 | 0 | 320 | 0 | 0 |
| SRR5275577 | 384 | 38748 | 319 | 230 | 116 | 1845 | 914 | 368 | 11810 | 125 | 2234 | 2821 | 960 | 133 | 340 | 13224 | 7718 | 0 | 0 | 0 | 114 | 29 | 81 | 82 | 938 | 152 | 0 | 0 | 13 | 30 | 21 | 59 | 0 | 455 | 716 | 269 | 0 |
| SRR5275578 | 333 | 30794 | 530 | 190 | 93 | 1690 | 840 | 405 | 12530 | 146 | 3094 | 3861 | 1036 | 149 | 324 | 12082 | 8466 | 0 | 0 | 0 | 82 | 21 | 56 | 59 | 668 | 98 | 0 | 0 | 12 | 23 | 16 | 44 | 0 | 349 | 539 | 178 | 0 |
| SRR5275579 | 399 | 33699 | 394 | 300 | 96 | 1806 | 841 | 368 | 12296 | 101 | 2815 | 3220 | 944 | 164 | 364 | 13152 | 8130 | 0 | 0 | 0 | 96 | 22 | 69 | 71 | 914 | 136 | 0 | 0 | 13 | 28 | 17 | 51 | 0 | 461 | 739 | 209 | 0 |
| SRR5275580 | 182 | 26649 | 602 | 141 | 59 | 1304 | 998 | 490 | 14267 | 195 | 4082 | 4188 | 1269 | 218 | 222 | 9187 | 9485 | 0 | 0 | 0 | 84 | 24 | 51 | 61 | 533 | 128 | 0 | 0 | 0 | 24 | 15 | 37 | 0 | 223 | 541 | 211 | 0 |
| SRR5275581 | 116 | 18995 | 759 | 101 | 51 | 1223 | 904 | 454 | 13117 | 152 | 4043 | 4201 | 1263 | 173 | 223 | 8460 | 9384 | 0 | 0 | 0 | 60 | 17 | 43 | 50 | 497 | 108 | 0 | 0 | 0 | 16 | 11 | 29 | 0 | 228 | 593 | 201 | 0 |
| SRR5275582 | 643 | 52101 | 311 | 673 | 57 | 1480 | 1016 | 468 | 12513 | 76 | 3844 | 3946 | 1281 | 178 | 235 | 9697 | 9013 | 0 | 0 | 0 | 184 | 45 | 96 | 107 | 850 | 158 | 0 | 0 | 20 | 48 | 27 | 63 | 0 | 386 | 767 | 206 | 0 |
| SRR5284195 | 71 | 3540 | 4590 | 356 | 22509 | 12467 | 149919 | 26542 | 10916 | 4152 | 18326 | 36 | 5620 | 28771 | 850 | 6747 | 12800 | 10 | 36 | 48 | 83 | 76 | 53 | 33 | 16 | 0 | 14 | 29 | 81 | 47 | 18 | 0 | 0 | 0 | 0 | 19 | 0 |
| SRR5284196 | 770 | 6921 | 3891 | 1810 | 21313 | 10388 | 159052 | 31089 | 10943 | 2731 | 23648 | 39 | 7388 | 32697 | 988 | 5887 | 15243 | 12 | 50 | 80 | 155 | 86 | 41 | 29 | 13 | 0 | 21 | 42 | 125 | 63 | 24 | 0 | 0 | 0 | 0 | 16 | 0 |
| SRR5284197 | 325 | 5080 | 6667 | 1604 | 15332 | 15111 | 96120 | 16999 | 8792 | 2799 | 20032 | 34873 | 6125 | 26410 | 983 | 12339 | 13451 | 10 | 46 | 64 | 73 | 91 | 42 | 27 | 12 | 0 | 11 | 36 | 115 | 31 | 16 | 0 | 0 | 0 | 0 | 12 | 0 |
| SRR5284198 | 340 | 4969 | 5494 | 1085 | 20119 | 10730 | 139660 | 25040 | 8182 | 4891 | 19026 | 29467 | 5275 | 20862 | 947 | 10477 | 10787 | 13 | 42 | 63 | 84 | 75 | 36 | 30 | 16 | 0 | 18 | 33 | 115 | 49 | 16 | 0 | 0 | 0 | 0 | 19 | 0 |
| SRR5284199 | 377 | 3029 | 2595 | 1790 | 29217 | 14256 | 209553 | 41894 | 5545 | 4213 | 16731 | 31 | 5289 | 36277 | 620 | 4186 | 11384 | 17 | 43 | 42 | 61 | 52 | 52 | 34 | 12 | 0 | 27 | 39 | 99 | 54 | 29 | 0 | 0 | 0 | 0 | 16 | 0 |
| SRR5284200 | 431 | 3063 | 1559 | 2160 | 25957 | 12416 | 194604 | 52103 | 6126 | 3464 | 15853 | 23552 | 5314 | 42773 | 353 | 3268 | 11628 | 19 | 53 | 42 | 64 | 47 | 52 | 25 | 10 | 0 | 21 | 37 | 97 | 47 | 22 | 0 | 0 | 0 | 0 | 12 | 0 |
| SRR5284201 | 356 | 4391 | 5048 | 1252 | 17179 | 17609 | 112227 | 19791 | 5515 | 4911 | 19391 | 27818 | 5599 | 39260 | 884 | 14207 | 12125 | 14 | 44 | 55 | 29 | 47 | 50 | 18 | 0 | 0 | 0 | 31 | 93 | 20 | 14 | 0 | 0 | 0 | 0 | 11 | 0 |
| SRR5284202 | 187 | 5886 | 8188 | 498 | 9337 | 10216 | 77277 | 14366 | 6805 | 4068 | 22380 | 41572 | 6518 | 39067 | 570 | 10821 | 13081 | 0 | 26 | 41 | 50 | 61 | 88 | 19 | 0 | 0 | 0 | 26 | 94 | 20 | 14 | 0 | 0 | 0 | 0 | 13 | 0 |
| SRR5284203 | 195 | 2549 | 2445 | 3176 | 20266 | 19762 | 125365 | 25421 | 8731 | 1662 | 21438 | 30869 | 5670 | 79290 | 582 | 3407 | 10483 | 0 | 36 | 32 | 32 | 66 | 19 | 0 | 0 | 0 | 14 | 39 | 114 | 24 | 18 | 0 | 0 | 0 | 0 | 0 | 0 |
| SRR5284204 | 184 | 4679 | 2911 | 886 | 30303 | 10455 | 190011 | 50346 | 7615 | 3985 | 18505 | 35 | 5261 | 52726 | 346 | 2072 | 10020 | 0 | 24 | 20 | 32 | 23 | 18 | 14 | 0 | 0 | 19 | 25 | 74 | 37 | 14 | 0 | 0 | 0 | 0 | 0 | 0 |
| SRR5284205 | 93 | 3351 | 8330 | 976 | 16903 | 14882 | 93549 | 18215 | 6913 | 4096 | 20607 | 23856 | 5262 | 73380 | 881 | 10248 | 10723 | 0 | 33 | 57 | 165 | 69 | 13 | 12 | 0 | 0 | 0 | 27 | 101 | 49 | 10 | 0 | 0 | 0 | 0 | 0 | 0 |
| SRR5284206 | 130 | 3913 | 6824 | 825 | 16096 | 11698 | 83644 | 20196 | 7452 | 3075 | 21201 | 30711 | 5897 | 57502 | 929 | 8908 | 12253 | 0 | 36 | 51 | 80 | 53 | 17 | 14 | 0 | 0 | 10 | 32 | 110 | 36 | 14 | 0 | 0 | 0 | 0 | 0 | 0 |
| SRR5317818 | 0 | 4565 | 504 | 660 | 1069 | 5269 | 6417 | 18368 | 7878 | 237 | 20463 | 19659 | 2744 | 6742 | 0 | 8536 | 52011 | 0 | 0 | 0 | 270 | 532 | 119 | 0 | 0 | 0 | 0 | 12 | 165 | 318 | 878 | 257 | 0 | 21 | 28 | 0 | 0 |
| SRR5317819 | 79 | 13429 | 857 | 2109 | 516 | 2877 | 3635 | 8584 | 8008 | 95 | 10556 | 26561 | 4297 | 6388 | 0 | 18843 | 42339 | 0 | 0 | 0 | 267 | 799 | 312 | 12 | 0 | 0 | 0 | 15 | 108 | 713 | 1779 | 1236 | 0 | 44 | 145 | 0 | 0 |
| SRR5317820 | 0 | 4498 | 797 | 274 | 391 | 2820 | 3322 | 11164 | 2266 | 146 | 25357 | 24025 | 3879 | 4495 | 0 | 8774 | 32011 | 0 | 0 | 0 | 62 | 316 | 68 | 0 | 0 | 0 | 0 | 0 | 31 | 58 | 369 | 70 | 0 | 0 | 14 | 0 | 0 |
| SRR5317821 | 0 | 40413 | 2768 | 89 | 70 | 360 | 754 | 3344 | 12153 | 25 | 13125 | 10029 | 2264 | 6189 | 23 | 11061 | 78389 | 0 | 0 | 0 | 11 | 111 | 157 | 13 | 0 | 0 | 0 | 0 | 26 | 110 | 1332 | 2252 | 16 | 66 | 39 | 0 | 0 |
| SRR5317822 | 13 | 7709 | 1685 | 594 | 975 | 6926 | 8480 | 22215 | 8624 | 93 | 23857 | 22603 | 3253 | 7410 | 0 | 9591 | 52403 | 0 | 0 | 0 | 838 | 1975 | 1085 | 21 | 25 | 0 | 0 | 23 | 230 | 920 | 3754 | 1897 | 0 | 140 | 355 | 23 | 0 |
| SRR5317823 | 45 | 15205 | 417 | 2277 | 151 | 546 | 990 | 2414 | 14491 | 51 | 15833 | 19135 | 2970 | 6699 | 37 | 23169 | 92047 | 0 | 0 | 0 | 130 | 747 | 691 | 19 | 11 | 0 | 0 | 32 | 128 | 279 | 3255 | 2241 | 10 | 56 | 220 | 0 | 0 |
| SRR5317824 | 12 | 2264 | 1432 | 1582 | 702 | 5326 | 7725 | 16912 | 3428 | 443 | 25180 | 23875 | 4646 | 13291 | 0 | 13318 | 34226 | 0 | 0 | 0 | 1689 | 1240 | 133 | 0 | 0 | 0 | 0 | 19 | 349 | 1564 | 1978 | 361 | 0 | 28 | 56 | 0 | 0 |
| SRR5317825 | 11 | 32052 | 154 | 2663 | 145 | 548 | 1242 | 2980 | 10340 | 43 | 5494 | 5015 | 1645 | 3339 | 148 | 28823 | 65068 | 0 | 0 | 0 | 667 | 4040 | 3376 | 21 | 0 | 0 | 0 | 11 | 112 | 1308 | 18129 | 19498 | 20 | 145 | 659 | 22 | 0 |
| SRR5317826 | 18 | 4923 | 981 | 1412 | 1501 | 8327 | 12884 | 34538 | 11577 | 169 | 31901 | 42482 | 5543 | 16284 | 0 | 12748 | 86261 | 0 | 0 | 27 | 2115 | 1589 | 293 | 0 | 0 | 0 | 0 | 212 | 1059 | 3331 | 4462 | 758 | 0 | 32 | 270 | 21 | 0 |
| SRR5317827 | 24 | 41420 | 942 | 977 | 122 | 441 | 756 | 1930 | 8445 | 119 | 14490 | 44127 | 3741 | 10682 | 14 | 18900 | 45438 | 0 | 0 | 15 | 675 | 427 | 141 | 0 | 0 | 0 | 0 | 127 | 460 | 1038 | 1155 | 403 | 0 | 12 | 98 | 0 | 0 |
| SRR5387779 | 426 | 3569 | 467 | 357 | 0 | 0 | 0 | 64 | 87 | 0 | 801 | 668 | 225 | 317 | 0 | 14 | 1357 | 0 | 0 | 0 | 0 | 0 | 0 | 0 | 0 | 0 | 0 | 0 | 0 | 0 | 0 | 0 | 0 | 0 | 0 | 0 | 0 |
| SRR5387780 | 324 | 3575 | 414 | 783 | 0 | 12 | 12 | 85 | 85 | 0 | 1268 | 1022 | 342 | 875 | 0 | 25 | 2149 | 0 | 0 | 0 | 0 | 0 | 0 | 0 | 0 | 0 | 0 | 0 | 0 | 0 | 0 | 0 | 0 | 0 | 0 | 0 | 0 |
| SRR5387781 | 727 | 4658 | 641 | 444 | 0 | 13 | 0 | 125 | 80 | 0 | 1018 | 1215 | 398 | 1346 | 0 | 0 | 1211 | 0 | 0 | 0 | 0 | 0 | 0 | 0 | 0 | 0 | 0 | 0 | 0 | 0 | 0 | 0 | 0 | 0 | 0 | 0 | 0 |
| SRR5387782 | 359 | 2496 | 273 | 323 | 0 | 33 | 25 | 276 | 146 | 0 | 4263 | 2298 | 1017 | 2289 | 0 | 0 | 2773 | 0 | 0 | 0 | 0 | 0 | 0 | 0 | 0 | 0 | 0 | 0 | 0 | 0 | 0 | 0 | 0 | 0 | 0 | 0 | 0 |
| SRR5387783 | 619 | 2191 | 594 | 427 | 0 | 16 | 18 | 101 | 168 | 0 | 728 | 579 | 165 | 272 | 0 | 0 | 1595 | 0 | 0 | 0 | 0 | 0 | 0 | 0 | 0 | 0 | 0 | 0 | 0 | 0 | 0 | 0 | 0 | 0 | 0 | 0 | 0 |
| SRR5387784 | 1136 | 2241 | 697 | 360 | 0 | 26 | 21 | 124 | 261 | 31 | 2585 | 2450 | 861 | 1035 | 0 | 0 | 2229 | 0 | 0 | 0 | 0 | 0 | 0 | 0 | 0 | 0 | 0 | 0 | 0 | 0 | 0 | 0 | 0 | 0 | 0 | 0 | 0 |
| SRR5387785 | 205 | 3048 | 317 | 594 | 0 | 18 | 16 | 150 | 236 | 13 | 1178 | 1161 | 398 | 2015 | 0 | 0 | 2841 | 0 | 0 | 0 | 0 | 0 | 0 | 0 | 0 | 0 | 0 | 0 | 0 | 0 | 0 | 0 | 0 | 0 | 0 | 0 | 0 |
| SRR5387786 | 502 | 4262 | 2204 | 705 | 0 | 0 | 0 | 50 | 86 | 41 | 252 | 432 | 138 | 557 | 0 | 0 | 1301 | 0 | 0 | 0 | 0 | 0 | 0 | 0 | 0 | 0 | 0 | 0 | 0 | 0 | 0 | 0 | 0 | 0 | 0 | 0 | 0 |
| SRR5387787 | 681 | 3558 | 905 | 457 | 0 | 0 | 0 | 40 | 70 | 18 | 505 | 686 | 210 | 415 | 0 | 0 | 1532 | 0 | 0 | 0 | 0 | 0 | 0 | 0 | 0 | 0 | 0 | 0 | 0 | 0 | 0 | 0 | 0 | 0 | 0 | 0 | 0 |
| SRR5387788 | 318 | 2625 | 490 | 262 | 0 | 16 | 20 | 138 | 144 | 0 | 3514 | 3484 | 1080 | 2143 | 0 | 0 | 2597 | 0 | 0 | 0 | 0 | 0 | 0 | 0 | 0 | 0 | 0 | 0 | 0 | 0 | 0 | 0 | 0 | 0 | 0 | 0 | 0 |
| SRR5387789 | 545 | 1799 | 535 | 545 | 0 | 0 | 10 | 73 | 188 | 19 | 805 | 800 | 274 | 648 | 0 | 0 | 2409 | 0 | 0 | 0 | 0 | 0 | 0 | 0 | 0 | 0 | 0 | 0 | 0 | 0 | 0 | 0 | 0 | 0 | 0 | 0 | 0 |
| SRR5387790 | 421 | 1513 | 343 | 336 | 0 | 12 | 11 | 66 | 162 | 14 | 2083 | 1075 | 362 | 690 | 0 | 0 | 2193 | 0 | 0 | 0 | 0 | 0 | 0 | 0 | 0 | 0 | 0 | 0 | 0 | 0 | 0 | 0 | 0 | 0 | 0 | 0 | 0 |
| SRR5387791 | 670 | 2255 | 1092 | 1040 | 0 | 0 | 0 | 27 | 65 | 0 | 160 | 215 | 53 | 367 | 0 | 0 | 988 | 0 | 0 | 0 | 0 | 0 | 0 | 0 | 0 | 0 | 0 | 0 | 0 | 0 | 0 | 0 | 0 | 0 | 0 | 0 | 0 |
| SRR5387792 | 369 | 1988 | 502 | 553 | 0 | 20 | 14 | 100 | 112 | 0 | 1191 | 906 | 310 | 773 | 0 | 0 | 2223 | 0 | 0 | 0 | 0 | 0 | 0 | 0 | 0 | 0 | 0 | 0 | 0 | 0 | 0 | 0 | 0 | 0 | 0 | 0 | 0 |
| SRR5387793 | 446 | 2602 | 705 | 503 | 0 | 0 | 0 | 60 | 79 | 0 | 411 | 487 | 152 | 678 | 0 | 0 | 977 | 0 | 0 | 0 | 0 | 0 | 0 | 0 | 0 | 0 | 0 | 0 | 0 | 0 | 0 | 0 | 0 | 0 | 0 | 0 | 0 |
| SRR5387794 | 150 | 3138 | 369 | 1157 | 0 | 17 | 15 | 101 | 140 | 0 | 2125 | 1893 | 588 | 2105 | 0 | 0 | 1700 | 0 | 0 | 0 | 0 | 0 | 0 | 0 | 0 | 0 | 0 | 0 | 0 | 0 | 0 | 0 | 0 | 0 | 0 | 0 | 0 |
| SRR5387795 | 506 | 3412 | 441 | 571 | 0 | 0 | 0 | 23 | 24 | 11 | 251 | 226 | 106 | 262 | 0 | 0 | 807 | 0 | 0 | 0 | 0 | 0 | 0 | 0 | 0 | 0 | 0 | 0 | 0 | 0 | 0 | 0 | 0 | 0 | 0 | 0 | 0 |
| SRR5387796 | 643 | 1715 | 549 | 429 | 0 | 0 | 0 | 38 | 36 | 0 | 339 | 419 | 162 | 362 | 0 | 0 | 1204 | 0 | 0 | 0 | 0 | 0 | 0 | 0 | 0 | 0 | 0 | 0 | 0 | 0 | 0 | 0 | 0 | 0 | 0 | 0 | 0 |
| SRR5387797 | 571 | 7051 | 408 | 686 | 0 | 37 | 27 | 138 | 210 | 0 | 1318 | 1509 | 513 | 1896 | 0 | 0 | 2432 | 0 | 0 | 0 | 0 | 0 | 0 | 0 | 0 | 0 | 0 | 0 | 0 | 0 | 0 | 0 | 0 | 0 | 0 | 0 | 0 |
| SRR5387798 | 1123 | 6048 | 766 | 576 | 0 | 46 | 42 | 163 | 147 | 0 | 2172 | 2069 | 744 | 2596 | 0 | 0 | 2252 | 0 | 0 | 0 | 0 | 0 | 0 | 0 | 0 | 0 | 0 | 0 | 0 | 0 | 0 | 0 | 0 | 0 | 0 | 0 | 0 |
| SRR5387799 | 584 | 3427 | 584 | 296 | 0 | 0 | 0 | 28 | 40 | 16 | 380 | 633 | 129 | 504 | 0 | 0 | 1070 | 0 | 0 | 0 | 0 | 0 | 0 | 0 | 0 | 0 | 0 | 0 | 0 | 0 | 0 | 0 | 0 | 0 | 0 | 0 | 0 |
| SRR5387800 | 564 | 1724 | 481 | 547 | 0 | 0 | 0 | 21 | 29 | 17 | 419 | 674 | 224 | 510 | 0 | 0 | 785 | 0 | 0 | 0 | 0 | 0 | 0 | 0 | 0 | 0 | 0 | 0 | 0 | 0 | 0 | 0 | 0 | 0 | 0 | 0 | 0 |
| SRR5387801 | 510 | 1746 | 481 | 623 | 0 | 0 | 0 | 32 | 65 | 0 | 1013 | 929 | 231 | 957 | 0 | 0 | 691 | 0 | 0 | 0 | 0 | 0 | 0 | 0 | 0 | 0 | 0 | 0 | 0 | 0 | 0 | 0 | 0 | 0 | 0 | 0 | 0 |
| SRR5387802 | 319 | 1851 | 421 | 523 | 0 | 0 | 0 | 85 | 74 | 0 | 1495 | 1600 | 401 | 2363 | 0 | 0 | 1287 | 0 | 0 | 0 | 0 | 0 | 0 | 0 | 0 | 0 | 0 | 0 | 0 | 0 | 0 | 0 | 0 | 0 | 0 | 0 | 0 |
| SRR5387803 | 719 | 2312 | 588 | 490 | 0 | 12 | 0 | 87 | 106 | 0 | 1195 | 994 | 294 | 1307 | 0 | 0 | 1836 | 0 | 0 | 0 | 0 | 0 | 0 | 0 | 0 | 0 | 0 | 0 | 0 | 0 | 0 | 0 | 0 | 0 | 0 | 0 | 0 |
| SRR5387804 | 495 | 2319 | 385 | 479 | 0 | 18 | 16 | 126 | 90 | 16 | 1878 | 1191 | 465 | 899 | 0 | 0 | 2809 | 0 | 0 | 0 | 0 | 0 | 0 | 0 | 0 | 0 | 0 | 0 | 0 | 0 | 0 | 0 | 0 | 0 | 0 | 0 | 0 |
| SRR5387805 | 195 | 2794 | 260 | 2395 | 0 | 0 | 0 | 65 | 55 | 0 | 1685 | 1137 | 423 | 2757 | 0 | 0 | 1149 | 0 | 0 | 0 | 0 | 0 | 0 | 0 | 0 | 0 | 0 | 0 | 0 | 0 | 0 | 0 | 0 | 0 | 0 | 0 | 0 |
| SRR5387806 | 598 | 3878 | 311 | 670 | 0 | 19 | 21 | 134 | 127 | 0 | 2340 | 1398 | 539 | 1331 | 0 | 0 | 1751 | 0 | 0 | 0 | 11 | 0 | 0 | 0 | 0 | 0 | 0 | 0 | 0 | 0 | 0 | 0 | 0 | 0 | 0 | 0 | 0 |
| SRR5387807 | 982 | 3929 | 580 | 639 | 0 | 19 | 15 | 93 | 92 | 12 | 958 | 1151 | 349 | 1745 | 0 | 20 | 2146 | 0 | 0 | 0 | 0 | 0 | 0 | 0 | 0 | 0 | 0 | 0 | 0 | 0 | 0 | 0 | 0 | 0 | 0 | 0 | 0 |
| SRR5387808 | 407 | 2492 | 536 | 437 | 0 | 0 | 14 | 61 | 118 | 0 | 723 | 941 | 240 | 861 | 0 | 12 | 1898 | 0 | 0 | 0 | 0 | 0 | 0 | 0 | 0 | 0 | 0 | 0 | 0 | 0 | 0 | 0 | 0 | 0 | 0 | 0 | 0 |
| SRR5387809 | 603 | 2697 | 763 | 568 | 0 | 0 | 0 | 28 | 20 | 0 | 468 | 359 | 123 | 295 | 0 | 0 | 1089 | 0 | 0 | 0 | 0 | 0 | 0 | 0 | 0 | 0 | 0 | 0 | 0 | 0 | 0 | 0 | 0 | 0 | 0 | 0 | 0 |
| SRR5387810 | 174 | 3013 | 300 | 412 | 0 | 31 | 34 | 274 | 135 | 0 | 3324 | 2388 | 1254 | 3030 | 0 | 0 | 3460 | 0 | 0 | 0 | 0 | 0 | 0 | 0 | 0 | 0 | 0 | 0 | 0 | 0 | 0 | 0 | 0 | 0 | 0 | 0 | 0 |
| SRR5387811 | 288 | 2540 | 725 | 623 | 0 | 19 | 16 | 94 | 93 | 0 | 2049 | 1352 | 460 | 1577 | 0 | 0 | 1483 | 0 | 0 | 0 | 0 | 0 | 0 | 0 | 0 | 0 | 0 | 0 | 0 | 0 | 0 | 0 | 0 | 0 | 0 | 0 | 0 |
| SRR5387812 | 251 | 2158 | 640 | 689 | 0 | 16 | 13 | 140 | 80 | 0 | 871 | 809 | 343 | 864 | 0 | 0 | 1757 | 0 | 0 | 0 | 0 | 0 | 0 | 0 | 0 | 0 | 0 | 0 | 0 | 0 | 0 | 0 | 0 | 0 | 0 | 0 | 0 |
| SRR5387813 | 424 | 4074 | 303 | 697 | 0 | 0 | 0 | 30 | 73 | 0 | 338 | 688 | 149 | 589 | 0 | 0 | 1073 | 0 | 0 | 0 | 0 | 0 | 0 | 0 | 0 | 0 | 0 | 0 | 0 | 0 | 0 | 0 | 0 | 0 | 0 | 0 | 0 |
| SRR5387814 | 999 | 3199 | 815 | 657 | 0 | 0 | 0 | 74 | 78 | 0 | 999 | 1085 | 345 | 1482 | 0 | 0 | 1492 | 0 | 0 | 0 | 0 | 0 | 0 | 0 | 0 | 0 | 0 | 0 | 0 | 0 | 0 | 0 | 0 | 0 | 0 | 0 | 0 |
| SRR5387815 | 411 | 2562 | 404 | 362 | 0 | 27 | 24 | 173 | 106 | 14 | 2830 | 2215 | 830 | 1267 | 0 | 0 | 2848 | 0 | 0 | 0 | 0 | 0 | 0 | 0 | 0 | 0 | 0 | 0 | 0 | 0 | 0 | 0 | 0 | 0 | 0 | 0 | 0 |
| SRR5387816 | 485 | 2564 | 811 | 476 | 0 | 0 | 0 | 56 | 60 | 19 | 791 | 805 | 272 | 519 | 0 | 0 | 1760 | 0 | 0 | 0 | 0 | 0 | 0 | 0 | 0 | 0 | 0 | 0 | 0 | 0 | 0 | 0 | 0 | 0 | 0 | 0 | 0 |
| SRR5387817 | 745 | 2092 | 510 | 850 | 0 | 13 | 0 | 63 | 98 | 0 | 455 | 469 | 170 | 935 | 0 | 0 | 2714 | 0 | 0 | 0 | 0 | 0 | 0 | 0 | 0 | 0 | 0 | 0 | 0 | 0 | 0 | 0 | 0 | 0 | 0 | 0 | 0 |
| SRR5387818 | 906 | 2683 | 488 | 2491 | 0 | 0 | 0 | 52 | 49 | 0 | 743 | 783 | 280 | 1321 | 0 | 13 | 2179 | 0 | 0 | 0 | 0 | 0 | 0 | 0 | 0 | 0 | 0 | 0 | 0 | 0 | 0 | 0 | 0 | 0 | 0 | 0 | 0 |
| SRR5387819 | 403 | 3921 | 445 | 445 | 0 | 11 | 0 | 84 | 99 | 0 | 919 | 770 | 334 | 571 | 0 | 19 | 2147 | 0 | 0 | 0 | 0 | 0 | 0 | 0 | 0 | 0 | 0 | 0 | 0 | 0 | 0 | 0 | 0 | 0 | 0 | 0 | 0 |
| SRR5387820 | 574 | 3173 | 487 | 526 | 0 | 11 | 0 | 90 | 64 | 19 | 747 | 837 | 335 | 694 | 0 | 21 | 1769 | 0 | 0 | 0 | 0 | 0 | 0 | 0 | 0 | 0 | 0 | 0 | 0 | 0 | 0 | 0 | 0 | 0 | 0 | 0 | 0 |
| SRR5387821 | 662 | 5548 | 801 | 549 | 0 | 18 | 20 | 138 | 127 | 0 | 1821 | 1652 | 690 | 2994 | 0 | 17 | 3176 | 0 | 0 | 0 | 0 | 0 | 0 | 0 | 0 | 0 | 0 | 0 | 0 | 0 | 0 | 0 | 0 | 0 | 0 | 0 | 0 |
| SRR5387822 | 390 | 6101 | 524 | 538 | 0 | 0 | 0 | 43 | 57 | 0 | 906 | 1893 | 576 | 3002 | 0 | 0 | 1526 | 0 | 0 | 0 | 0 | 0 | 0 | 0 | 0 | 0 | 0 | 0 | 0 | 0 | 0 | 0 | 0 | 0 | 0 | 0 | 0 |
| SRR5387823 | 252 | 2092 | 277 | 350 | 0 | 12 | 15 | 95 | 68 | 0 | 770 | 961 | 345 | 913 | 0 | 0 | 2002 | 0 | 0 | 0 | 0 | 0 | 0 | 0 | 0 | 0 | 0 | 0 | 0 | 0 | 0 | 0 | 0 | 0 | 0 | 0 | 0 |
| SRR5387824 | 491 | 2082 | 682 | 500 | 0 | 30 | 28 | 199 | 120 | 0 | 2629 | 1668 | 682 | 907 | 0 | 0 | 2408 | 0 | 0 | 0 | 0 | 0 | 0 | 0 | 0 | 0 | 0 | 0 | 0 | 0 | 0 | 0 | 0 | 0 | 0 | 0 | 0 |
| SRR5387825 | 946 | 2451 | 716 | 551 | 0 | 12 | 13 | 92 | 219 | 0 | 500 | 665 | 213 | 1229 | 0 | 0 | 1688 | 0 | 0 | 0 | 0 | 0 | 0 | 0 | 0 | 0 | 0 | 0 | 0 | 0 | 0 | 0 | 0 | 0 | 0 | 0 | 0 |
| SRR5387826 | 719 | 2059 | 682 | 487 | 0 | 17 | 14 | 113 | 132 | 24 | 1735 | 1163 | 395 | 951 | 0 | 22 | 1845 | 0 | 0 | 0 | 0 | 0 | 0 | 0 | 0 | 0 | 0 | 0 | 0 | 0 | 0 | 0 | 0 | 0 | 0 | 0 | 0 |
| SRR5387827 | 174 | 1563 | 237 | 537 | 0 | 10 | 0 | 71 | 128 | 16 | 1097 | 1296 | 342 | 1799 | 0 | 0 | 1485 | 0 | 0 | 0 | 0 | 0 | 0 | 0 | 0 | 0 | 0 | 0 | 0 | 0 | 0 | 0 | 0 | 0 | 0 | 0 | 0 |
| SRR5387828 | 861 | 2747 | 559 | 394 | 0 | 11 | 0 | 68 | 96 | 0 | 675 | 968 | 313 | 1159 | 0 | 0 | 1306 | 0 | 0 | 0 | 0 | 0 | 0 | 0 | 0 | 0 | 0 | 0 | 0 | 0 | 0 | 0 | 0 | 0 | 0 | 0 | 0 |
| SRR5387829 | 542 | 2301 | 849 | 588 | 0 | 0 | 0 | 22 | 20 | 0 | 316 | 588 | 150 | 599 | 0 | 0 | 646 | 0 | 0 | 0 | 0 | 0 | 0 | 0 | 0 | 0 | 0 | 0 | 0 | 0 | 0 | 0 | 0 | 0 | 0 | 0 | 0 |
| SRR5387830 | 388 | 1810 | 427 | 427 | 0 | 20 | 15 | 149 | 65 | 0 | 1552 | 1315 | 478 | 1629 | 0 | 0 | 2031 | 0 | 0 | 0 | 0 | 0 | 0 | 0 | 0 | 0 | 0 | 0 | 0 | 0 | 0 | 0 | 0 | 0 | 0 | 0 | 0 |
| SRR5387831 | 417 | 1749 | 379 | 565 | 0 | 0 | 0 | 20 | 34 | 19 | 167 | 303 | 80 | 574 | 0 | 11 | 639 | 0 | 0 | 0 | 0 | 0 | 0 | 0 | 0 | 0 | 0 | 0 | 0 | 0 | 0 | 0 | 0 | 0 | 0 | 0 | 0 |
| SRR5387832 | 445 | 1646 | 455 | 466 | 0 | 0 | 0 | 48 | 58 | 10 | 592 | 657 | 209 | 530 | 0 | 12 | 889 | 0 | 0 | 0 | 0 | 0 | 0 | 0 | 0 | 0 | 0 | 0 | 0 | 0 | 0 | 0 | 0 | 0 | 0 | 0 | 0 |
| SRR5387833 | 1073 | 3532 | 1016 | 554 | 0 | 0 | 0 | 42 | 89 | 0 | 219 | 338 | 107 | 414 | 0 | 0 | 1199 | 0 | 0 | 0 | 0 | 0 | 0 | 0 | 0 | 0 | 0 | 0 | 0 | 0 | 0 | 0 | 0 | 0 | 0 | 0 | 0 |
| SRR5387834 | 482 | 4342 | 573 | 346 | 0 | 0 | 0 | 61 | 63 | 0 | 898 | 921 | 295 | 481 | 0 | 0 | 1521 | 0 | 0 | 0 | 0 | 0 | 0 | 0 | 0 | 0 | 0 | 0 | 0 | 0 | 0 | 0 | 0 | 0 | 0 | 0 | 0 |
| SRR5387835 | 421 | 2117 | 400 | 407 | 0 | 20 | 20 | 166 | 192 | 0 | 1207 | 1495 | 566 | 2512 | 0 | 0 | 1505 | 0 | 0 | 0 | 0 | 0 | 0 | 0 | 0 | 0 | 0 | 0 | 0 | 0 | 0 | 0 | 0 | 0 | 0 | 0 | 0 |
| SRR5387836 | 525 | 2181 | 517 | 444 | 0 | 0 | 0 | 58 | 83 | 0 | 876 | 842 | 248 | 847 | 0 | 0 | 917 | 0 | 0 | 0 | 0 | 0 | 0 | 0 | 0 | 0 | 0 | 0 | 0 | 0 | 0 | 0 | 0 | 0 | 0 | 0 | 0 |
| SRR5387837 | 496 | 3044 | 628 | 496 | 0 | 28 | 20 | 158 | 158 | 22 | 1480 | 995 | 326 | 1091 | 0 | 0 | 1661 | 0 | 0 | 0 | 0 | 0 | 0 | 0 | 0 | 0 | 0 | 0 | 0 | 0 | 0 | 0 | 0 | 0 | 0 | 0 | 0 |
| SRR5387838 | 518 | 2980 | 480 | 391 | 0 | 16 | 14 | 96 | 114 | 19 | 1534 | 1022 | 329 | 1077 | 0 | 0 | 1508 | 0 | 0 | 0 | 0 | 0 | 0 | 0 | 0 | 0 | 0 | 0 | 0 | 0 | 0 | 0 | 0 | 0 | 0 | 0 | 0 |
| SRR5387839 | 600 | 2602 | 568 | 407 | 0 | 0 | 0 | 65 | 99 | 0 | 500 | 643 | 179 | 736 | 0 | 0 | 1314 | 0 | 0 | 0 | 0 | 0 | 0 | 0 | 0 | 0 | 0 | 0 | 0 | 0 | 0 | 0 | 0 | 0 | 0 | 0 | 0 |
| SRR5387840 | 501 | 2117 | 572 | 415 | 0 | 0 | 0 | 63 | 122 | 43 | 422 | 538 | 129 | 649 | 0 | 0 | 954 | 0 | 0 | 0 | 0 | 0 | 0 | 0 | 0 | 0 | 0 | 0 | 0 | 0 | 0 | 0 | 0 | 0 | 0 | 0 | 0 |
| SRR5387841 | 400 | 3089 | 490 | 613 | 0 | 0 | 0 | 23 | 23 | 0 | 375 | 472 | 164 | 268 | 0 | 0 | 1142 | 0 | 0 | 0 | 0 | 0 | 0 | 0 | 0 | 0 | 0 | 0 | 0 | 0 | 0 | 0 | 0 | 0 | 0 | 0 | 0 |
| SRR5387842 | 430 | 2007 | 508 | 745 | 0 | 0 | 0 | 76 | 50 | 0 | 975 | 660 | 233 | 952 | 0 | 0 | 1059 | 0 | 0 | 0 | 0 | 0 | 0 | 0 | 0 | 0 | 0 | 0 | 0 | 0 | 0 | 0 | 0 | 0 | 0 | 0 | 0 |
| SRR5387843 | 485 | 4373 | 629 | 728 | 0 | 23 | 21 | 107 | 217 | 20 | 3286 | 2077 | 713 | 1530 | 0 | 0 | 1600 | 0 | 0 | 0 | 0 | 0 | 0 | 0 | 0 | 0 | 0 | 0 | 0 | 0 | 0 | 0 | 0 | 0 | 0 | 0 | 0 |
| SRR5387844 | 238 | 3848 | 312 | 687 | 0 | 35 | 31 | 183 | 359 | 0 | 2574 | 1825 | 502 | 1599 | 0 | 0 | 2153 | 0 | 0 | 0 | 0 | 0 | 0 | 0 | 0 | 0 | 0 | 0 | 0 | 0 | 0 | 0 | 0 | 0 | 0 | 0 | 0 |
| SRR5387845 | 397 | 4388 | 367 | 509 | 0 | 25 | 21 | 202 | 163 | 0 | 1381 | 1006 | 382 | 944 | 0 | 12 | 2702 | 0 | 0 | 0 | 0 | 0 | 0 | 0 | 0 | 0 | 0 | 0 | 0 | 0 | 0 | 0 | 0 | 0 | 0 | 0 | 0 |
| SRR5387846 | 194 | 2653 | 439 | 701 | 0 | 13 | 17 | 106 | 68 | 0 | 882 | 951 | 344 | 485 | 0 | 13 | 1870 | 0 | 0 | 0 | 0 | 0 | 0 | 0 | 0 | 0 | 0 | 0 | 0 | 0 | 0 | 0 | 0 | 0 | 0 | 0 | 0 |
| SRR5387847 | 498 | 3052 | 509 | 619 | 0 | 13 | 11 | 72 | 102 | 11 | 861 | 786 | 252 | 525 | 0 | 0 | 2036 | 0 | 0 | 0 | 0 | 0 | 0 | 0 | 0 | 0 | 0 | 0 | 0 | 0 | 0 | 0 | 0 | 0 | 0 | 0 | 0 |
| SRR5387848 | 218 | 1925 | 347 | 993 | 0 | 31 | 26 | 120 | 178 | 0 | 1169 | 1758 | 579 | 1786 | 0 | 0 | 2964 | 0 | 0 | 0 | 17 | 0 | 0 | 0 | 0 | 0 | 0 | 0 | 0 | 0 | 0 | 0 | 0 | 0 | 0 | 0 | 0 |
| SRR5387849 | 239 | 2988 | 378 | 455 | 0 | 0 | 0 | 88 | 96 | 0 | 544 | 573 | 231 | 746 | 0 | 0 | 1213 | 0 | 0 | 0 | 0 | 0 | 0 | 0 | 0 | 0 | 0 | 0 | 0 | 0 | 0 | 0 | 0 | 0 | 0 | 0 | 0 |
| SRR5387850 | 256 | 3121 | 375 | 558 | 0 | 13 | 12 | 92 | 117 | 0 | 630 | 774 | 283 | 634 | 0 | 0 | 1250 | 0 | 0 | 0 | 0 | 0 | 0 | 0 | 0 | 0 | 0 | 0 | 0 | 0 | 0 | 0 | 0 | 0 | 0 | 0 | 0 |
| SRR5387851 | 427 | 1750 | 455 | 620 | 0 | 11 | 11 | 103 | 76 | 0 | 966 | 1172 | 320 | 1227 | 0 | 0 | 2553 | 0 | 0 | 0 | 0 | 0 | 0 | 0 | 0 | 0 | 0 | 0 | 0 | 0 | 0 | 0 | 0 | 0 | 0 | 0 | 0 |
| SRR5387852 | 384 | 2086 | 467 | 430 | 0 | 20 | 23 | 183 | 149 | 18 | 2203 | 2513 | 775 | 3498 | 0 | 0 | 3647 | 0 | 0 | 0 | 0 | 0 | 0 | 0 | 0 | 0 | 0 | 0 | 0 | 0 | 0 | 0 | 0 | 0 | 0 | 0 | 0 |
| SRR5387853 | 353 | 2017 | 457 | 468 | 0 | 15 | 10 | 117 | 142 | 0 | 804 | 892 | 292 | 1689 | 0 | 23 | 808 | 0 | 0 | 0 | 0 | 0 | 0 | 0 | 0 | 0 | 0 | 0 | 0 | 0 | 0 | 0 | 0 | 0 | 0 | 0 | 0 |
| SRR5387854 | 932 | 3720 | 657 | 579 | 0 | 0 | 0 | 71 | 80 | 0 | 778 | 1105 | 295 | 1708 | 0 | 21 | 915 | 0 | 0 | 0 | 0 | 0 | 0 | 0 | 0 | 0 | 0 | 0 | 0 | 0 | 0 | 0 | 0 | 0 | 0 | 0 | 0 |
| SRR5387855 | 827 | 2169 | 496 | 303 | 0 | 0 | 0 | 45 | 78 | 0 | 550 | 647 | 184 | 870 | 0 | 0 | 1178 | 0 | 0 | 0 | 0 | 0 | 0 | 0 | 0 | 0 | 0 | 0 | 0 | 0 | 0 | 0 | 0 | 0 | 0 | 0 | 0 |
| SRR5387856 | 567 | 1876 | 547 | 499 | 0 | 0 | 0 | 65 | 67 | 0 | 1153 | 652 | 298 | 1322 | 0 | 0 | 1194 | 0 | 0 | 0 | 0 | 0 | 0 | 0 | 0 | 0 | 0 | 0 | 0 | 0 | 0 | 0 | 0 | 0 | 0 | 0 | 0 |
| SRR5387857 | 411 | 2178 | 467 | 345 | 0 | 18 | 15 | 99 | 125 | 11 | 772 | 881 | 287 | 1005 | 0 | 0 | 2045 | 0 | 0 | 0 | 0 | 0 | 0 | 0 | 0 | 0 | 0 | 0 | 0 | 0 | 0 | 0 | 0 | 0 | 0 | 0 | 0 |
| SRR5387858 | 319 | 2337 | 400 | 662 | 0 | 14 | 11 | 74 | 114 | 0 | 1285 | 840 | 275 | 1099 | 0 | 0 | 1841 | 0 | 0 | 0 | 0 | 0 | 0 | 0 | 0 | 0 | 0 | 0 | 0 | 0 | 0 | 0 | 0 | 0 | 0 | 0 | 0 |
| SRR5387859 | 289 | 1587 | 131 | 446 | 0 | 0 | 0 | 65 | 45 | 0 | 674 | 535 | 193 | 308 | 0 | 0 | 1622 | 0 | 0 | 0 | 0 | 0 | 0 | 0 | 0 | 0 | 0 | 0 | 0 | 0 | 0 | 0 | 0 | 0 | 0 | 0 | 0 |
| SRR5387860 | 265 | 1845 | 562 | 668 | 0 | 0 | 12 | 89 | 71 | 21 | 1017 | 674 | 217 | 835 | 0 | 18 | 1468 | 0 | 0 | 0 | 0 | 0 | 0 | 0 | 0 | 0 | 0 | 0 | 0 | 0 | 0 | 0 | 0 | 0 | 0 | 0 | 0 |
| SRR5387861 | 885 | 3551 | 672 | 407 | 0 | 0 | 0 | 49 | 60 | 10 | 1021 | 896 | 256 | 643 | 0 | 0 | 1043 | 0 | 0 | 0 | 0 | 0 | 0 | 0 | 0 | 0 | 0 | 0 | 0 | 0 | 0 | 0 | 0 | 0 | 0 | 0 | 0 |
| SRR5387862 | 361 | 3589 | 479 | 580 | 0 | 16 | 14 | 126 | 100 | 0 | 1393 | 1531 | 501 | 2091 | 0 | 0 | 1389 | 0 | 0 | 0 | 0 | 0 | 0 | 0 | 0 | 0 | 0 | 0 | 0 | 0 | 0 | 0 | 0 | 0 | 0 | 0 | 0 |
| SRR5387863 | 305 | 7063 | 258 | 401 | 0 | 46 | 34 | 342 | 261 | 0 | 2283 | 2263 | 1038 | 3003 | 0 | 17 | 4333 | 0 | 0 | 0 | 0 | 0 | 0 | 0 | 0 | 0 | 0 | 0 | 0 | 0 | 0 | 0 | 0 | 0 | 0 | 0 | 0 |
| SRR5387864 | 441 | 5799 | 457 | 677 | 0 | 29 | 30 | 174 | 137 | 0 | 2268 | 2231 | 1046 | 2254 | 0 | 20 | 4437 | 0 | 0 | 0 | 0 | 0 | 0 | 0 | 0 | 0 | 0 | 0 | 0 | 0 | 0 | 0 | 0 | 0 | 0 | 0 | 0 |
| SRR5387865 | 276 | 1197 | 342 | 822 | 0 | 0 | 0 | 40 | 103 | 0 | 416 | 717 | 211 | 1495 | 0 | 0 | 1115 | 0 | 0 | 0 | 0 | 0 | 0 | 0 | 0 | 0 | 0 | 0 | 0 | 0 | 0 | 0 | 0 | 0 | 0 | 0 | 0 |
| SRR5387866 | 2197 | 7753 | 951 | 674 | 0 | 17 | 16 | 114 | 158 | 15 | 1248 | 898 | 350 | 1130 | 0 | 0 | 1544 | 0 | 0 | 0 | 0 | 0 | 0 | 0 | 0 | 0 | 0 | 0 | 0 | 0 | 0 | 0 | 0 | 0 | 0 | 0 | 0 |

### Supp Table S8.1.Columns_B_to_AL

| | | | | | | | | | | | | | | | | | | | | | | | | | | | | | | | | | | | | | |
|---|---|---|---|---|---|---|---|---|---|---|---|---|---|---|---|---|---|---|---|---|---|---|---|---|---|---|---|---|---|---|---|---|---|---|---|---|---|
| SRR5387867 | 269 | 3288 | 301 | 982 | 0 | 0 | 0 | 44 | 55 | 0 | 879 | 781 | 287 | 736 | 0 | 0 | 1244 | 0 | 0 | 0 | 0 | 0 | 0 | 0 | 0 | 0 | 0 | 0 | 0 | 0 | 0 | 0 | 0 | 0 | 0 | 0 | 0 |
| SRR5387868 | 605 | 2067 | 751 | 783 | 0 | 0 | 0 | 46 | 43 | 10 | 538 | 425 | 139 | 619 | 0 | 0 | 972 | 0 | 0 | 0 | 0 | 0 | 0 | 0 | 0 | 0 | 0 | 0 | 0 | 0 | 0 | 0 | 0 | 0 | 0 | 0 | 0 |
| SRR5387869 | 323 | 1916 | 280 | 344 | 0 | 16 | 13 | 85 | 90 | 11 | 1490 | 1023 | 404 | 395 | 0 | 0 | 1695 | 0 | 0 | 0 | 0 | 0 | 0 | 0 | 0 | 0 | 0 | 0 | 0 | 0 | 0 | 0 | 0 | 0 | 0 | 0 | 0 |
| SRR5387870 | 860 | 2404 | 415 | 538 | 0 | 12 | 0 | 81 | 93 | 23 | 2192 | 1698 | 515 | 1066 | 0 | 0 | 1520 | 0 | 0 | 0 | 0 | 0 | 0 | 0 | 0 | 0 | 0 | 0 | 0 | 0 | 0 | 0 | 0 | 0 | 0 | 0 | 0 |
| SRR5387871 | 569 | 2526 | 557 | 463 | 0 | 0 | 0 | 46 | 51 | 59 | 994 | 1009 | 294 | 600 | 0 | 0 | 1165 | 0 | 0 | 0 | 0 | 0 | 0 | 0 | 0 | 0 | 0 | 0 | 0 | 0 | 0 | 0 | 0 | 0 | 0 | 0 | 0 |
| SRR5387872 | 677 | 2359 | 587 | 488 | 0 | 19 | 15 | 137 | 94 | 0 | 2293 | 1895 | 676 | 1344 | 0 | 0 | 2111 | 0 | 0 | 0 | 0 | 0 | 0 | 0 | 0 | 0 | 0 | 0 | 0 | 0 | 0 | 0 | 0 | 0 | 0 | 0 | 0 |
| SRR5387873 | 581 | 4028 | 689 | 562 | 0 | 0 | 0 | 26 | 41 | 0 | 811 | 799 | 253 | 404 | 0 | 0 | 1683 | 0 | 0 | 0 | 0 | 0 | 0 | 0 | 0 | 0 | 0 | 0 | 0 | 0 | 0 | 0 | 0 | 0 | 0 | 0 | 0 |
| SRR5387874 | 305 | 2790 | 339 | 814 | 0 | 14 | 12 | 88 | 147 | 0 | 1105 | 1471 | 410 | 1190 | 0 | 0 | 2489 | 0 | 0 | 0 | 11 | 0 | 0 | 0 | 0 | 0 | 0 | 0 | 0 | 0 | 0 | 0 | 0 | 0 | 0 | 0 | 0 |
| SRR5387875 | 292 | 1330 | 300 | 316 | 0 | 19 | 14 | 132 | 150 | 0 | 1054 | 1783 | 377 | 1878 | 0 | 12 | 1500 | 0 | 0 | 0 | 0 | 0 | 0 | 0 | 0 | 0 | 0 | 0 | 0 | 0 | 0 | 0 | 0 | 0 | 0 | 0 | 0 |
| SRR5387876 | 725 | 2052 | 509 | 486 | 0 | 0 | 0 | 58 | 62 | 0 | 963 | 877 | 226 | 1194 | 0 | 0 | 1018 | 0 | 0 | 0 | 0 | 0 | 0 | 0 | 0 | 0 | 0 | 0 | 0 | 0 | 0 | 0 | 0 | 0 | 0 | 0 | 0 |
| SRR5387877 | 399 | 2634 | 650 | 513 | 0 | 15 | 14 | 104 | 200 | 11 | 721 | 709 | 191 | 671 | 0 | 0 | 1096 | 0 | 0 | 0 | 0 | 0 | 0 | 0 | 0 | 0 | 0 | 0 | 0 | 0 | 0 | 0 | 0 | 0 | 0 | 0 | 0 |
| SRR5387878 | 1848 | 6046 | 2304 | 958 | 0 | 13 | 12 | 87 | 86 | 0 | 770 | 723 | 251 | 1262 | 0 | 0 | 1269 | 0 | 0 | 0 | 0 | 0 | 0 | 0 | 0 | 0 | 0 | 0 | 0 | 0 | 0 | 0 | 0 | 0 | 0 | 0 | 0 |
| SRR5387879 | 788 | 2747 | 738 | 769 | 0 | 0 | 0 | 53 | 85 | 31 | 957 | 953 | 362 | 1732 | 0 | 0 | 1633 | 0 | 0 | 0 | 0 | 0 | 0 | 0 | 0 | 0 | 0 | 0 | 0 | 0 | 0 | 0 | 0 | 0 | 0 | 0 | 0 |
| SRR5387880 | 592 | 2614 | 485 | 436 | 0 | 0 | 0 | 53 | 74 | 0 | 888 | 827 | 310 | 669 | 0 | 0 | 1122 | 0 | 0 | 0 | 0 | 0 | 0 | 0 | 0 | 0 | 0 | 0 | 0 | 0 | 0 | 0 | 0 | 0 | 0 | 0 | 0 |
| SRR5387881 | 1171 | 3683 | 478 | 817 | 0 | 12 | 0 | 74 | 71 | 15 | 619 | 839 | 289 | 1343 | 0 | 0 | 2165 | 0 | 0 | 0 | 0 | 0 | 0 | 0 | 0 | 0 | 0 | 0 | 0 | 0 | 0 | 0 | 0 | 0 | 0 | 0 | 0 |
| SRR5387882 | 734 | 2069 | 613 | 637 | 0 | 12 | 12 | 91 | 127 | 24 | 623 | 835 | 354 | 1756 | 0 | 20 | 2252 | 0 | 0 | 0 | 0 | 0 | 0 | 0 | 0 | 0 | 0 | 0 | 0 | 0 | 0 | 0 | 0 | 0 | 0 | 0 | 0 |
| SRR5387883 | 105 | 1268 | 210 | 398 | 0 | 40 | 31 | 197 | 143 | 0 | 3144 | 2558 | 1012 | 2095 | 0 | 0 | 2453 | 0 | 0 | 0 | 0 | 0 | 0 | 0 | 0 | 0 | 0 | 0 | 0 | 0 | 0 | 0 | 0 | 0 | 0 | 0 | 0 |
| SRR5387884 | 548 | 2069 | 356 | 617 | 0 | 16 | 16 | 86 | 117 | 0 | 780 | 1245 | 438 | 883 | 0 | 0 | 1975 | 0 | 0 | 0 | 0 | 0 | 0 | 0 | 0 | 0 | 0 | 0 | 0 | 0 | 0 | 0 | 0 | 0 | 0 | 0 | 0 |
| SRR5387885 | 537 | 2884 | 796 | 517 | 0 | 18 | 18 | 144 | 260 | 0 | 814 | 1063 | 321 | 1479 | 0 | 0 | 1275 | 0 | 0 | 0 | 0 | 0 | 0 | 0 | 0 | 0 | 0 | 0 | 0 | 0 | 0 | 0 | 0 | 0 | 0 | 0 | 0 |
| SRR5387886 | 475 | 1874 | 549 | 585 | 0 | 0 | 0 | 18 | 82 | 0 | 454 | 484 | 84 | 557 | 0 | 0 | 591 | 0 | 0 | 0 | 0 | 0 | 0 | 0 | 0 | 0 | 0 | 0 | 0 | 0 | 0 | 0 | 0 | 0 | 0 | 0 | 0 |
| SRR5387887 | 524 | 2262 | 535 | 568 | 0 | 0 | 0 | 63 | 106 | 0 | 721 | 695 | 275 | 714 | 0 | 0 | 1905 | 0 | 0 | 0 | 0 | 0 | 0 | 0 | 0 | 0 | 0 | 0 | 0 | 0 | 0 | 0 | 0 | 0 | 0 | 0 | 0 |
| SRR5387888 | 596 | 2089 | 379 | 441 | 0 | 25 | 21 | 203 | 163 | 0 | 2669 | 1504 | 652 | 1339 | 0 | 0 | 2253 | 0 | 0 | 0 | 0 | 0 | 0 | 0 | 0 | 0 | 0 | 0 | 0 | 0 | 0 | 0 | 0 | 0 | 0 | 0 | 0 |
| SRR5387889 | 539 | 5374 | 607 | 645 | 0 | 0 | 0 | 21 | 24 | 0 | 328 | 581 | 140 | 450 | 0 | 0 | 1056 | 0 | 0 | 0 | 0 | 0 | 0 | 0 | 0 | 0 | 0 | 0 | 0 | 0 | 0 | 0 | 0 | 0 | 0 | 0 | 0 |
| SRR5387890 | 425 | 4166 | 453 | 501 | 0 | 22 | 14 | 118 | 72 | 0 | 690 | 760 | 282 | 1537 | 0 | 0 | 1893 | 0 | 0 | 0 | 0 | 0 | 0 | 0 | 0 | 0 | 0 | 0 | 0 | 0 | 0 | 0 | 0 | 0 | 0 | 0 | 0 |
| SRR5387891 | 726 | 3891 | 564 | 726 | 0 | 22 | 26 | 147 | 168 | 19 | 1161 | 1123 | 378 | 1293 | 0 | 0 | 3330 | 0 | 0 | 0 | 0 | 0 | 0 | 0 | 0 | 0 | 0 | 0 | 0 | 0 | 0 | 0 | 0 | 0 | 0 | 0 | 0 |
| SRR5387892 | 407 | 2382 | 475 | 638 | 0 | 25 | 23 | 199 | 139 | 0 | 1979 | 1867 | 633 | 2313 | 0 | 0 | 3477 | 0 | 0 | 0 | 0 | 0 | 0 | 0 | 0 | 0 | 0 | 0 | 0 | 0 | 0 | 0 | 0 | 0 | 0 | 0 | 0 |
| SRR5387893 | 492 | 2103 | 553 | 713 | 0 | 25 | 25 | 154 | 454 | 0 | 3089 | 2545 | 931 | 2557 | 0 | 0 | 2813 | 0 | 0 | 0 | 0 | 0 | 0 | 0 | 0 | 0 | 0 | 0 | 0 | 0 | 0 | 0 | 0 | 0 | 0 | 0 | 0 |
| SRR5387894 | 387 | 2582 | 310 | 445 | 0 | 22 | 20 | 104 | 104 | 0 | 2851 | 1744 | 811 | 2105 | 0 | 0 | 3443 | 0 | 0 | 0 | 0 | 0 | 0 | 0 | 0 | 0 | 0 | 0 | 0 | 0 | 0 | 0 | 0 | 0 | 0 | 0 | 0 |
| SRR5387895 | 407 | 2457 | 483 | 707 | 0 | 14 | 16 | 130 | 142 | 0 | 3577 | 2862 | 977 | 3931 | 0 | 0 | 1720 | 0 | 0 | 0 | 0 | 0 | 0 | 0 | 0 | 0 | 0 | 0 | 0 | 0 | 0 | 0 | 0 | 0 | 0 | 0 | 0 |
| SRR5387896 | 647 | 3394 | 528 | 462 | 0 | 14 | 14 | 106 | 130 | 20 | 1382 | 1561 | 471 | 1895 | 0 | 0 | 1410 | 0 | 0 | 0 | 0 | 0 | 0 | 0 | 0 | 0 | 0 | 0 | 0 | 0 | 0 | 0 | 0 | 0 | 0 | 0 | 0 |
| SRR5387897 | 1118 | 2636 | 738 | 380 | 0 | 11 | 11 | 58 | 94 | 0 | 1124 | 1020 | 333 | 442 | 0 | 0 | 1655 | 0 | 0 | 0 | 0 | 0 | 0 | 0 | 0 | 0 | 0 | 0 | 0 | 0 | 0 | 0 | 0 | 0 | 0 | 0 | 0 |
| SRR5387898 | 481 | 2673 | 498 | 412 | 0 | 23 | 19 | 113 | 125 | 0 | 1532 | 879 | 344 | 542 | 0 | 0 | 1754 | 0 | 0 | 0 | 0 | 0 | 0 | 0 | 0 | 0 | 0 | 0 | 0 | 0 | 0 | 0 | 0 | 0 | 0 | 0 | 0 |
| SRR5387899 | 964 | 2975 | 707 | 275 | 0 | 12 | 11 | 101 | 60 | 18 | 1682 | 1828 | 578 | 1853 | 0 | 0 | 1369 | 0 | 0 | 0 | 0 | 0 | 0 | 0 | 0 | 0 | 0 | 0 | 0 | 0 | 0 | 0 | 0 | 0 | 0 | 0 | 0 |
| SRR5387900 | 1061 | 4347 | 811 | 776 | 0 | 0 | 0 | 48 | 35 | 17 | 782 | 799 | 281 | 1789 | 0 | 0 | 868 | 0 | 0 | 0 | 0 | 0 | 0 | 0 | 0 | 0 | 0 | 0 | 0 | 0 | 0 | 0 | 0 | 0 | 0 | 0 | 0 |
| SRR5387901 | 266 | 1937 | 254 | 303 | 0 | 51 | 40 | 299 | 242 | 0 | 5784 | 4793 | 1914 | 6006 | 0 | 31 | 3095 | 0 | 0 | 0 | 0 | 0 | 0 | 0 | 0 | 0 | 0 | 0 | 0 | 0 | 0 | 0 | 0 | 0 | 0 | 0 | 0 |
| SRR5387902 | 239 | 2839 | 294 | 368 | 0 | 19 | 19 | 97 | 98 | 0 | 703 | 1992 | 772 | 2798 | 0 | 11 | 2143 | 0 | 0 | 0 | 10 | 0 | 0 | 0 | 0 | 0 | 0 | 0 | 0 | 0 | 0 | 0 | 0 | 0 | 0 | 0 | 0 |
| SRR5387903 | 897 | 2862 | 673 | 634 | 0 | 17 | 17 | 101 | 174 | 0 | 1859 | 2207 | 914 | 2208 | 0 | 0 | 3980 | 0 | 0 | 0 | 0 | 0 | 0 | 0 | 0 | 0 | 0 | 0 | 0 | 0 | 0 | 0 | 0 | 0 | 0 | 0 | 0 |
| SRR5387904 | 100 | 2755 | 536 | 1182 | 0 | 0 | 0 | 17 | 20 | 0 | 321 | 627 | 214 | 712 | 0 | 0 | 804 | 0 | 0 | 0 | 0 | 0 | 0 | 0 | 0 | 0 | 0 | 0 | 0 | 0 | 0 | 0 | 0 | 0 | 0 | 0 | 0 |
| SRR5387905 | 301 | 2077 | 419 | 696 | 0 | 0 | 0 | 52 | 122 | 0 | 1179 | 899 | 239 | 890 | 0 | 0 | 1309 | 0 | 0 | 0 | 0 | 0 | 0 | 0 | 0 | 0 | 0 | 0 | 0 | 0 | 0 | 0 | 0 | 0 | 0 | 0 | 0 |
| SRR5387906 | 744 | 2503 | 620 | 479 | 0 | 0 | 0 | 52 | 80 | 0 | 517 | 643 | 140 | 719 | 0 | 0 | 910 | 0 | 0 | 0 | 0 | 0 | 0 | 0 | 0 | 0 | 0 | 0 | 0 | 0 | 0 | 0 | 0 | 0 | 0 | 0 | 0 |
| SRR5387907 | 448 | 1199 | 786 | 170 | 0 | 0 | 0 | 53 | 75 | 0 | 1150 | 724 | 227 | 1128 | 0 | 0 | 1179 | 0 | 0 | 0 | 0 | 0 | 0 | 0 | 0 | 0 | 0 | 0 | 0 | 0 | 0 | 0 | 0 | 0 | 0 | 0 | 0 |
| SRR5387908 | 329 | 1493 | 322 | 410 | 0 | 19 | 14 | 106 | 180 | 0 | 1258 | 966 | 310 | 1101 | 0 | 0 | 1799 | 0 | 0 | 0 | 0 | 0 | 0 | 0 | 0 | 0 | 0 | 0 | 0 | 0 | 0 | 0 | 0 | 0 | 0 | 0 | 0 |
| SRR5387909 | 251 | 3075 | 448 | 287 | 0 | 0 | 0 | 30 | 38 | 27 | 413 | 367 | 103 | 210 | 0 | 0 | 1234 | 0 | 0 | 0 | 0 | 0 | 0 | 0 | 0 | 0 | 0 | 0 | 0 | 0 | 0 | 0 | 0 | 0 | 0 | 0 | 0 |
| SRR5387910 | 589 | 2467 | 580 | 424 | 0 | 0 | 0 | 53 | 74 | 0 | 1607 | 1306 | 361 | 734 | 0 | 0 | 1191 | 0 | 0 | 0 | 0 | 0 | 0 | 0 | 0 | 0 | 0 | 0 | 0 | 0 | 0 | 0 | 0 | 0 | 0 | 0 | 0 |
| SRR5387911 | 261 | 5384 | 404 | 639 | 0 | 12 | 0 | 75 | 59 | 0 | 901 | 780 | 289 | 1720 | 0 | 0 | 1310 | 0 | 0 | 0 | 0 | 0 | 0 | 0 | 0 | 0 | 0 | 0 | 0 | 0 | 0 | 0 | 0 | 0 | 0 | 0 | 0 |
| SRR5387912 | 601 | 3422 | 532 | 455 | 0 | 0 | 0 | 37 | 68 | 0 | 842 | 1146 | 296 | 1305 | 0 | 0 | 662 | 0 | 0 | 0 | 0 | 0 | 0 | 0 | 0 | 0 | 0 | 0 | 0 | 0 | 0 | 0 | 0 | 0 | 0 | 0 | 0 |
| SRR5387913 | 774 | 3283 | 601 | 324 | 0 | 12 | 11 | 72 | 75 | 0 | 1205 | 1551 | 508 | 1651 | 0 | 0 | 2028 | 0 | 0 | 0 | 0 | 0 | 0 | 0 | 0 | 0 | 0 | 0 | 0 | 0 | 0 | 0 | 0 | 0 | 0 | 0 | 0 |
| SRR5387914 | 635 | 2753 | 781 | 508 | 0 | 0 | 0 | 46 | 46 | 0 | 399 | 622 | 184 | 834 | 0 | 0 | 1142 | 0 | 0 | 0 | 0 | 0 | 0 | 0 | 0 | 0 | 0 | 0 | 0 | 0 | 0 | 0 | 0 | 0 | 0 | 0 | 0 |
| SRR5387915 | 269 | 3215 | 238 | 376 | 0 | 23 | 20 | 195 | 141 | 0 | 1356 | 1057 | 528 | 1525 | 0 | 0 | 2421 | 0 | 0 | 0 | 0 | 0 | 0 | 0 | 0 | 0 | 0 | 0 | 0 | 0 | 0 | 0 | 0 | 0 | 0 | 0 | 0 |
| SRR5387916 | 398 | 2607 | 407 | 725 | 0 | 0 | 0 | 87 | 81 | 0 | 1420 | 1214 | 371 | 1738 | 0 | 0 | 1236 | 0 | 0 | 0 | 0 | 0 | 0 | 0 | 0 | 0 | 0 | 0 | 0 | 0 | 0 | 0 | 0 | 0 | 0 | 0 | 0 |
| SRR5387917 | 501 | 2352 | 533 | 404 | 0 | 14 | 0 | 67 | 102 | 0 | 1361 | 1736 | 617 | 2057 | 0 | 0 | 1465 | 0 | 0 | 0 | 0 | 0 | 0 | 0 | 0 | 0 | 0 | 0 | 0 | 0 | 0 | 0 | 0 | 0 | 0 | 0 | 0 |
| SRR5387918 | 809 | 2934 | 760 | 825 | 0 | 0 | 0 | 17 | 26 | 16 | 239 | 434 | 110 | 455 | 0 | 0 | 621 | 0 | 0 | 0 | 0 | 0 | 0 | 0 | 0 | 0 | 0 | 0 | 0 | 0 | 0 | 0 | 0 | 0 | 0 | 0 | 0 |
| SRR5387919 | 379 | 3953 | 309 | 689 | 0 | 21 | 15 | 152 | 134 | 20 | 1082 | 1239 | 431 | 2065 | 0 | 0 | 2597 | 0 | 0 | 0 | 0 | 0 | 0 | 0 | 0 | 0 | 0 | 0 | 0 | 0 | 0 | 0 | 0 | 0 | 0 | 0 | 0 |
| SRR5387920 | 476 | 3259 | 549 | 903 | 0 | 0 | 0 | 53 | 92 | 12 | 538 | 849 | 247 | 1323 | 0 | 0 | 1877 | 0 | 0 | 0 | 0 | 0 | 0 | 0 | 0 | 0 | 0 | 0 | 0 | 0 | 0 | 0 | 0 | 0 | 0 | 0 | 0 |
| SRR5387921 | 461 | 2240 | 110 | 527 | 0 | 0 | 0 | 43 | 89 | 0 | 412 | 442 | 137 | 268 | 0 | 0 | 1523 | 0 | 0 | 0 | 0 | 0 | 0 | 0 | 0 | 0 | 0 | 0 | 0 | 0 | 0 | 0 | 0 | 0 | 0 | 0 | 0 |
| SRR5387922 | 725 | 2680 | 705 | 806 | 0 | 0 | 0 | 16 | 45 | 20 | 158 | 191 | 66 | 251 | 0 | 0 | 1106 | 0 | 0 | 0 | 0 | 0 | 0 | 0 | 0 | 0 | 0 | 0 | 0 | 0 | 0 | 0 | 0 | 0 | 0 | 0 | 0 |
| SRR5387923 | 350 | 3200 | 384 | 837 | 0 | 0 | 0 | 76 | 56 | 11 | 546 | 521 | 165 | 618 | 0 | 0 | 1244 | 0 | 0 | 0 | 0 | 0 | 0 | 0 | 0 | 0 | 0 | 0 | 0 | 0 | 0 | 0 | 0 | 0 | 0 | 0 | 0 |
| SRR5387924 | 460 | 2113 | 670 | 491 | 0 | 14 | 0 | 79 | 69 | 16 | 1132 | 965 | 331 | 1337 | 0 | 0 | 1595 | 0 | 0 | 0 | 0 | 0 | 0 | 0 | 0 | 0 | 0 | 0 | 0 | 0 | 0 | 0 | 0 | 0 | 0 | 0 | 0 |
| SRR5387925 | 308 | 1407 | 365 | 325 | 0 | 14 | 12 | 126 | 155 | 0 | 3277 | 2231 | 863 | 1895 | 0 | 23 | 2104 | 0 | 0 | 0 | 0 | 0 | 0 | 0 | 0 | 0 | 0 | 0 | 0 | 0 | 0 | 0 | 0 | 0 | 0 | 0 | 0 |
| SRR5387926 | 386 | 1791 | 292 | 465 | 0 | 17 | 17 | 127 | 114 | 0 | 1257 | 1572 | 543 | 1264 | 0 | 19 | 1943 | 0 | 0 | 0 | 12 | 0 | 0 | 0 | 0 | 0 | 0 | 0 | 0 | 0 | 0 | 0 | 0 | 0 | 0 | 0 | 0 |
| SRR5387927 | 427 | 2144 | 398 | 1337 | 0 | 0 | 0 | 87 | 101 | 0 | 590 | 810 | 326 | 1722 | 0 | 0 | 1785 | 0 | 0 | 0 | 0 | 0 | 0 | 0 | 0 | 0 | 0 | 0 | 0 | 0 | 0 | 0 | 0 | 0 | 0 | 0 | 0 |
| SRR5387928 | 431 | 1756 | 358 | 2016 | 0 | 0 | 10 | 90 | 112 | 0 | 987 | 1264 | 603 | 2844 | 0 | 11 | 1974 | 0 | 0 | 0 | 0 | 0 | 0 | 0 | 0 | 0 | 0 | 0 | 0 | 0 | 0 | 0 | 0 | 0 | 0 | 0 | 0 |
| SRR5387929 | 575 | 3551 | 637 | 913 | 0 | 12 | 10 | 78 | 131 | 15 | 931 | 878 | 258 | 1160 | 0 | 0 | 1311 | 0 | 0 | 0 | 0 | 0 | 0 | 0 | 0 | 0 | 0 | 0 | 0 | 0 | 0 | 0 | 0 | 0 | 0 | 0 | 0 |
| SRR5387930 | 344 | 2543 | 528 | 688 | 0 | 0 | 0 | 59 | 52 | 12 | 677 | 662 | 162 | 258 | 0 | 0 | 1966 | 0 | 0 | 0 | 0 | 0 | 0 | 0 | 0 | 0 | 0 | 0 | 0 | 0 | 0 | 0 | 0 | 0 | 0 | 0 | 0 |
| SRR5387931 | 707 | 2417 | 664 | 579 | 0 | 0 | 0 | 63 | 61 | 0 | 771 | 924 | 296 | 1513 | 0 | 0 | 1354 | 0 | 0 | 0 | 0 | 0 | 0 | 0 | 0 | 0 | 0 | 0 | 0 | 0 | 0 | 0 | 0 | 0 | 0 | 0 | 0 |
| SRR5387932 | 457 | 1529 | 615 | 591 | 0 | 12 | 10 | 79 | 99 | 16 | 1923 | 1068 | 423 | 1212 | 0 | 0 | 1295 | 0 | 0 | 0 | 0 | 0 | 0 | 0 | 0 | 0 | 0 | 0 | 0 | 0 | 0 | 0 | 0 | 0 | 0 | 0 | 0 |
| SRR5387933 | 561 | 2171 | 500 | 372 | 0 | 28 | 38 | 250 | 413 | 0 | 2451 | 1333 | 427 | 1971 | 0 | 0 | 1326 | 0 | 0 | 0 | 0 | 0 | 0 | 0 | 0 | 0 | 0 | 0 | 0 | 0 | 0 | 0 | 0 | 0 | 0 | 0 | 0 |
| SRR5387934 | 455 | 1901 | 368 | 249 | 0 | 29 | 28 | 225 | 216 | 0 | 1538 | 1625 | 721 | 1780 | 0 | 0 | 2491 | 0 | 0 | 0 | 0 | 0 | 0 | 0 | 0 | 0 | 0 | 0 | 0 | 0 | 0 | 0 | 0 | 0 | 0 | 0 | 0 |
| SRR5387935 | 627 | 2822 | 554 | 219 | 0 | 27 | 32 | 207 | 216 | 10 | 1644 | 1445 | 484 | 2125 | 0 | 0 | 1950 | 0 | 0 | 0 | 0 | 0 | 0 | 0 | 0 | 0 | 0 | 0 | 0 | 0 | 0 | 0 | 0 | 0 | 0 | 0 | 0 |
| SRR5387936 | 298 | 3448 | 406 | 804 | 0 | 15 | 11 | 63 | 67 | 17 | 692 | 588 | 236 | 668 | 0 | 0 | 1431 | 0 | 0 | 0 | 0 | 0 | 0 | 0 | 0 | 0 | 0 | 0 | 0 | 0 | 0 | 0 | 0 | 0 | 0 | 0 | 0 |
| SRR5387937 | 821 | 2176 | 746 | 499 | 0 | 0 | 0 | 49 | 42 | 27 | 1924 | 1501 | 549 | 1289 | 0 | 0 | 1835 | 0 | 0 | 0 | 0 | 0 | 0 | 0 | 0 | 0 | 0 | 0 | 0 | 0 | 0 | 0 | 0 | 0 | 0 | 0 | 0 |
| SRR5387938 | 568 | 2493 | 481 | 481 | 0 | 10 | 0 | 82 | 58 | 14 | 1190 | 1365 | 490 | 1796 | 0 | 0 | 1517 | 0 | 0 | 0 | 0 | 0 | 0 | 0 | 0 | 0 | 0 | 0 | 0 | 0 | 0 | 0 | 0 | 0 | 0 | 0 | 0 |
| SRR5387939 | 480 | 2953 | 581 | 379 | 0 | 13 | 13 | 96 | 223 | 0 | 1549 | 2160 | 642 | 1284 | 0 | 0 | 3420 | 0 | 0 | 0 | 0 | 0 | 0 | 0 | 0 | 0 | 0 | 0 | 0 | 0 | 0 | 0 | 0 | 0 | 0 | 0 | 0 |
| SRR5387940 | 1012 | 5773 | 1181 | 612 | 0 | 22 | 18 | 115 | 99 | 31 | 1332 | 1077 | 453 | 818 | 0 | 0 | 1746 | 0 | 0 | 0 | 0 | 0 | 0 | 0 | 0 | 0 | 0 | 0 | 0 | 0 | 0 | 0 | 0 | 0 | 0 | 0 | 0 |
| SRR5387941 | 403 | 3188 | 762 | 1282 | 0 | 0 | 0 | 12 | 24 | 0 | 176 | 324 | 85 | 280 | 0 | 0 | 680 | 0 | 0 | 0 | 0 | 0 | 0 | 0 | 0 | 0 | 0 | 0 | 0 | 0 | 0 | 0 | 0 | 0 | 0 | 0 | 0 |
| SRR5387942 | 435 | 1205 | 887 | 89 | 0 | 0 | 12 | 88 | 60 | 0 | 1476 | 954 | 335 | 437 | 0 | 0 | 2365 | 0 | 0 | 0 | 0 | 0 | 0 | 0 | 0 | 0 | 0 | 0 | 0 | 0 | 0 | 0 | 0 | 0 | 0 | 0 | 0 |
| SRR5387943 | 556 | 2033 | 397 | 635 | 0 | 0 | 0 | 42 | 132 | 0 | 548 | 606 | 224 | 924 | 0 | 0 | 1580 | 0 | 0 | 0 | 0 | 0 | 0 | 0 | 0 | 0 | 0 | 0 | 0 | 0 | 0 | 0 | 0 | 0 | 0 | 0 | 0 |
| SRR5387944 | 241 | 1929 | 209 | 531 | 0 | 11 | 0 | 68 | 168 | 0 | 539 | 655 | 207 | 628 | 0 | 0 | 1350 | 0 | 0 | 0 | 0 | 0 | 0 | 0 | 0 | 0 | 0 | 0 | 0 | 0 | 0 | 0 | 0 | 0 | 0 | 0 | 0 |
| SRR5387945 | 269 | 1529 | 529 | 510 | 0 | 12 | 11 | 113 | 190 | 0 | 936 | 831 | 387 | 1414 | 0 | 0 | 2221 | 0 | 0 | 0 | 0 | 0 | 0 | 0 | 0 | 0 | 0 | 0 | 0 | 0 | 0 | 0 | 0 | 0 | 0 | 0 | 0 |
| SRR5387946 | 438 | 1660 | 517 | 770 | 0 | 0 | 0 | 0 | 28 | 11 | 145 | 295 | 59 | 407 | 0 | 0 | 469 | 0 | 0 | 0 | 0 | 0 | 0 | 0 | 0 | 0 | 0 | 0 | 0 | 0 | 0 | 0 | 0 | 0 | 0 | 0 | 0 |
| SRR5387947 | 281 | 1594 | 393 | 516 | 0 | 0 | 0 | 42 | 52 | 11 | 1088 | 794 | 264 | 309 | 0 | 0 | 1710 | 0 | 0 | 0 | 0 | 0 | 0 | 0 | 0 | 0 | 0 | 0 | 0 | 0 | 0 | 0 | 0 | 0 | 0 | 0 | 0 |
| SRR5387948 | 716 | 3617 | 580 | 281 | 0 | 30 | 25 | 208 | 146 | 27 | 3461 | 2172 | 865 | 1374 | 0 | 0 | 2465 | 0 | 0 | 0 | 0 | 0 | 0 | 0 | 0 | 0 | 0 | 0 | 0 | 0 | 0 | 0 | 0 | 0 | 0 | 0 | 0 |
| SRR5387949 | 464 | 3044 | 658 | 542 | 0 | 20 | 20 | 123 | 151 | 0 | 999 | 993 | 330 | 972 | 0 | 0 | 2971 | 0 | 0 | 0 | 0 | 0 | 0 | 0 | 0 | 0 | 0 | 0 | 0 | 0 | 0 | 0 | 0 | 0 | 0 | 0 | 0 |
| SRR5387950 | 394 | 2645 | 634 | 541 | 0 | 0 | 0 | 35 | 52 | 0 | 1623 | 1565 | 380 | 957 | 0 | 0 | 1518 | 0 | 0 | 0 | 0 | 0 | 0 | 0 | 0 | 0 | 0 | 0 | 0 | 0 | 0 | 0 | 0 | 0 | 0 | 0 | 0 |
| SRR5387951 | 906 | 2706 | 707 | 1149 | 0 | 0 | 0 | 33 | 29 | 0 | 579 | 1079 | 285 | 1231 | 0 | 0 | 752 | 0 | 0 | 0 | 0 | 0 | 0 | 0 | 0 | 0 | 0 | 0 | 0 | 0 | 0 | 0 | 0 | 0 | 0 | 0 | 0 |
| SRR5387952 | 883 | 2426 | 741 | 569 | 0 | 16 | 15 | 78 | 96 | 10 | 1235 | 1131 | 406 | 1417 | 0 | 0 | 1434 | 0 | 0 | 0 | 0 | 0 | 0 | 0 | 0 | 0 | 0 | 0 | 0 | 0 | 0 | 0 | 0 | 0 | 0 | 0 | 0 |
| SRR5387953 | 221 | 1996 | 422 | 633 | 0 | 0 | 0 | 35 | 40 | 0 | 575 | 972 | 256 | 1072 | 0 | 0 | 855 | 0 | 0 | 0 | 0 | 0 | 0 | 0 | 0 | 0 | 0 | 0 | 0 | 0 | 0 | 0 | 0 | 0 | 0 | 0 | 0 |
| SRR5387954 | 415 | 2677 | 457 | 775 | 0 | 21 | 17 | 108 | 91 | 0 | 2119 | 1724 | 596 | 2721 | 0 | 0 | 1924 | 0 | 0 | 0 | 11 | 0 | 0 | 0 | 0 | 0 | 0 | 0 | 0 | 0 | 0 | 0 | 0 | 0 | 0 | 0 | 0 |
| SRR5387955 | 715 | 1739 | 528 | 682 | 0 | 0 | 0 | 59 | 92 | 0 | 422 | 617 | 152 | 637 | 0 | 0 | 1079 | 0 | 0 | 0 | 0 | 0 | 0 | 0 | 0 | 0 | 0 | 0 | 0 | 0 | 0 | 0 | 0 | 0 | 0 | 0 | 0 |
| SRR5387956 | 339 | 2496 | 618 | 890 | 0 | 16 | 15 | 100 | 91 | 0 | 1109 | 767 | 233 | 1514 | 0 | 0 | 1379 | 0 | 0 | 0 | 0 | 0 | 0 | 0 | 0 | 0 | 0 | 0 | 0 | 0 | 0 | 0 | 0 | 0 | 0 | 0 | 0 |
| SRR5387957 | 653 | 2388 | 669 | 871 | 0 | 16 | 17 | 135 | 260 | 16 | 847 | 1392 | 375 | 2076 | 0 | 0 | 1836 | 0 | 0 | 0 | 0 | 0 | 0 | 0 | 0 | 0 | 0 | 0 | 0 | 0 | 0 | 0 | 0 | 0 | 0 | 0 | 0 |
| SRR5387958 | 301 | 2079 | 353 | 3826 | 0 | 0 | 0 | 30 | 50 | 0 | 884 | 686 | 190 | 1281 | 0 | 0 | 603 | 0 | 0 | 0 | 0 | 0 | 0 | 0 | 0 | 0 | 0 | 0 | 0 | 0 | 0 | 0 | 0 | 0 | 0 | 0 | 0 |
| SRR5387959 | 335 | 2084 | 347 | 1495 | 0 | 0 | 0 | 72 | 94 | 0 | 997 | 806 | 363 | 1902 | 0 | 0 | 1486 | 0 | 0 | 0 | 0 | 0 | 0 | 0 | 0 | 0 | 0 | 0 | 0 | 0 | 0 | 0 | 0 | 0 | 0 | 0 | 0 |
| SRR5387960 | 493 | 1285 | 381 | 710 | 0 | 0 | 0 | 81 | 81 | 0 | 518 | 766 | 240 | 1413 | 0 | 0 | 1047 | 0 | 0 | 0 | 0 | 0 | 0 | 0 | 0 | 0 | 0 | 0 | 0 | 0 | 0 | 0 | 0 | 0 | 0 | 0 | 0 |
| SRR5387961 | 653 | 2958 | 617 | 245 | 0 | 15 | 14 | 103 | 139 | 0 | 906 | 1382 | 360 | 1157 | 0 | 0 | 1793 | 0 | 0 | 0 | 0 | 0 | 0 | 0 | 0 | 0 | 0 | 0 | 0 | 0 | 0 | 0 | 0 | 0 | 0 | 0 | 0 |
| SRR5387962 | 311 | 2734 | 528 | 423 | 0 | 43 | 37 | 390 | 357 | 0 | 2217 | 2052 | 880 | 2307 | 0 | 0 | 2928 | 0 | 0 | 0 | 0 | 0 | 0 | 0 | 0 | 0 | 0 | 0 | 0 | 0 | 0 | 0 | 0 | 0 | 0 | 0 | 0 |
| SRR5387963 | 443 | 1411 | 301 | 668 | 0 | 0 | 11 | 84 | 144 | 0 | 1160 | 1190 | 421 | 1466 | 0 | 0 | 1546 | 0 | 0 | 0 | 0 | 0 | 0 | 0 | 0 | 0 | 0 | 0 | 0 | 0 | 0 | 0 | 0 | 0 | 0 | 0 | 0 |
| SRR5387964 | 740 | 1683 | 592 | 435 | 0 | 0 | 0 | 72 | 144 | 0 | 1777 | 1705 | 634 | 1173 | 0 | 0 | 1113 | 0 | 0 | 0 | 0 | 0 | 0 | 0 | 0 | 0 | 0 | 0 | 0 | 0 | 0 | 0 | 0 | 0 | 0 | 0 | 0 |
| SRR5387965 | 657 | 1387 | 571 | 610 | 0 | 0 | 0 | 43 | 84 | 13 | 395 | 535 | 179 | 634 | 0 | 0 | 618 | 0 | 0 | 0 | 0 | 0 | 0 | 0 | 0 | 0 | 0 | 0 | 0 | 0 | 0 | 0 | 0 | 0 | 0 | 0 | 0 |
| SRR5387966 | 393 | 2328 | 464 | 503 | 0 | 11 | 0 | 85 | 110 | 16 | 647 | 668 | 266 | 1258 | 0 | 0 | 1628 | 0 | 0 | 0 | 0 | 0 | 0 | 0 | 0 | 0 | 0 | 0 | 0 | 0 | 0 | 0 | 0 | 0 | 0 | 0 | 0 |
| SRR5387967 | 475 | 3100 | 482 | 1050 | 0 | 0 | 0 | 53 | 201 | 0 | 1255 | 1139 | 290 | 1000 | 0 | 17 | 1124 | 0 | 0 | 0 | 11 | 0 | 0 | 0 | 0 | 0 | 0 | 0 | 0 | 0 | 0 | 0 | 0 | 0 | 0 | 0 | 0 |
| SRR5387968 | 663 | 3160 | 640 | 484 | 0 | 0 | 0 | 65 | 94 | 0 | 801 | 967 | 248 | 972 | 0 | 18 | 699 | 0 | 0 | 0 | 0 | 0 | 0 | 0 | 0 | 0 | 0 | 0 | 0 | 0 | 0 | 0 | 0 | 0 | 0 | 0 | 0 |
| SRR5387969 | 1240 | 3719 | 1384 | 824 | 0 | 12 | 12 | 62 | 96 | 16 | 1775 | 1602 | 527 | 1144 | 0 | 0 | 1624 | 0 | 0 | 0 | 10 | 0 | 0 | 0 | 0 | 0 | 0 | 0 | 0 | 0 | 0 | 0 | 0 | 0 | 0 | 0 | 0 |
| SRR5387970 | 368 | 3986 | 428 | 428 | 0 | 25 | 21 | 193 | 93 | 0 | 2508 | 1207 | 640 | 1267 | 0 | 0 | 2237 | 0 | 0 | 0 | 0 | 0 | 0 | 0 | 0 | 0 | 0 | 0 | 0 | 0 | 0 | 0 | 0 | 0 | 0 | 0 | 0 |
| SRR5387971 | 834 | 2943 | 947 | 792 | 0 | 20 | 21 | 93 | 118 | 31 | 787 | 847 | 289 | 1119 | 0 | 14 | 2726 | 0 | 0 | 0 | 0 | 0 | 0 | 0 | 0 | 0 | 0 | 0 | 0 | 0 | 0 | 0 | 0 | 0 | 0 | 0 | 0 |
| SRR5387972 | 515 | 1722 | 500 | 577 | 0 | 0 | 0 | 69 | 74 | 0 | 520 | 742 | 230 | 1236 | 0 | 13 | 1208 | 0 | 0 | 0 | 0 | 0 | 0 | 0 | 0 | 0 | 0 | 0 | 0 | 0 | 0 | 0 | 0 | 0 | 0 | 0 | 0 |
| SRR5387973 | 738 | 3167 | 769 | 461 | 0 | 15 | 19 | 112 | 126 | 10 | 895 | 895 | 255 | 759 | 0 | 0 | 1382 | 0 | 0 | 0 | 0 | 0 | 0 | 0 | 0 | 0 | 0 | 0 | 0 | 0 | 0 | 0 | 0 | 0 | 0 | 0 | 0 |
| SRR5387974 | 797 | 2191 | 737 | 388 | 0 | 23 | 27 | 149 | 95 | 20 | 1339 | 1189 | 559 | 1424 | 0 | 0 | 1872 | 0 | 0 | 0 | 0 | 0 | 0 | 0 | 0 | 0 | 0 | 0 | 0 | 0 | 0 | 0 | 0 | 0 | 0 | 0 | 0 |
| SRR5387975 | 310 | 3442 | 346 | 856 | 0 | 0 | 0 | 69 | 88 | 0 | 692 | 578 | 221 | 345 | 0 | 0 | 2072 | 0 | 0 | 0 | 0 | 0 | 0 | 0 | 0 | 0 | 0 | 0 | 0 | 0 | 0 | 0 | 0 | 0 | 0 | 0 | 0 |
| SRR5387976 | 662 | 2957 | 420 | 834 | 0 | 0 | 0 | 52 | 86 | 17 | 847 | 1244 | 377 | 1939 | 0 | 0 | 1428 | 0 | 0 | 0 | 0 | 0 | 0 | 0 | 0 | 0 | 0 | 0 | 0 | 0 | 0 | 0 | 0 | 0 | 0 | 0 | 0 |
| SRR5387977 | 452 | 3431 | 490 | 481 | 0 | 19 | 15 | 143 | 110 | 0 | 1402 | 1166 | 459 | 875 | 0 | 0 | 2941 | 0 | 0 | 0 | 0 | 0 | 0 | 0 | 0 | 0 | 0 | 0 | 0 | 0 | 0 | 0 | 0 | 0 | 0 | 0 | 0 |
| SRR5387978 | 444 | 2268 | 529 | 577 | 0 | 16 | 15 | 90 | 125 | 19 | 2972 | 1825 | 741 | 1100 | 0 | 0 | 3526 | 0 | 0 | 0 | 0 | 0 | 0 | 0 | 0 | 0 | 0 | 0 | 0 | 0 | 0 | 0 | 0 | 0 | 0 | 0 | 0 |
| SRR5387979 | 306 | 2508 | 379 | 685 | 0 | 0 | 0 | 47 | 79 | 0 | 488 | 303 | 109 | 127 | 0 | 0 | 1438 | 0 | 0 | 0 | 0 | 0 | 0 | 0 | 0 | 0 | 0 | 0 | 0 | 0 | 0 | 0 | 0 | 0 | 0 | 0 | 0 |
| SRR5387980 | 447 | 1735 | 429 | 984 | 0 | 0 | 0 | 28 | 94 | 0 | 1079 | 1025 | 267 | 604 | 0 | 0 | 1437 | 0 | 0 | 0 | 0 | 0 | 0 | 0 | 0 | 0 | 0 | 0 | 0 | 0 | 0 | 0 | 0 | 0 | 0 | 0 | 0 |
| SRR5387981 | 375 | 1790 | 458 | 583 | 0 | 0 | 0 | 43 | 61 | 0 | 396 | 432 | 158 | 289 | 0 | 0 | 1057 | 0 | 0 | 0 | 0 | 0 | 0 | 0 | 0 | 0 | 0 | 0 | 0 | 0 | 0 | 0 | 0 | 0 | 0 | 0 | 0 |
| SRR5387982 | 653 | 2048 | 563 | 450 | 0 | 12 | 10 | 87 | 92 | 34 | 612 | 747 | 238 | 880 | 0 | 0 | 1155 | 0 | 0 | 0 | 0 | 0 | 0 | 0 | 0 | 0 | 0 | 0 | 0 | 0 | 0 | 0 | 0 | 0 | 0 | 0 | 0 |
| SRR5387983 | 387 | 1894 | 600 | 489 | 0 | 10 | 15 | 92 | 154 | 0 | 710 | 732 | 240 | 591 | 0 | 0 | 1684 | 0 | 0 | 0 | 0 | 0 | 0 | 0 | 0 | 0 | 0 | 0 | 0 | 0 | 0 | 0 | 0 | 0 | 0 | 0 | 0 |
| SRR5387984 | 338 | 2500 | 610 | 730 | 0 | 0 | 0 | 63 | 79 | 16 | 411 | 492 | 155 | 988 | 0 | 0 | 1068 | 0 | 0 | 0 | 0 | 0 | 0 | 0 | 0 | 0 | 0 | 0 | 0 | 0 | 0 | 0 | 0 | 0 | 0 | 0 | 0 |
| SRR5387985 | 270 | 3482 | 565 | 571 | 0 | 35 | 40 | 319 | 317 | 0 | 2644 | 1422 | 781 | 3051 | 0 | 0 | 2704 | 0 | 0 | 0 | 13 | 0 | 0 | 0 | 0 | 0 | 0 | 0 | 0 | 0 | 0 | 0 | 0 | 0 | 0 | 0 | 0 |
| SRR5387986 | 336 | 3158 | 390 | 340 | 0 | 36 | 28 | 228 | 153 | 0 | 3832 | 3131 | 1157 | 3085 | 0 | 0 | 3489 | 0 | 0 | 0 | 13 | 0 | 0 | 0 | 0 | 0 | 0 | 0 | 0 | 0 | 0 | 0 | 0 | 0 | 0 | 0 | 0 |
| SRR5387987 | 342 | 2262 | 533 | 794 | 0 | 0 | 0 | 49 | 71 | 0 | 428 | 931 | 261 | 685 | 0 | 0 | 1387 | 0 | 0 | 0 | 0 | 0 | 0 | 0 | 0 | 0 | 0 | 0 | 0 | 0 | 0 | 0 | 0 | 0 | 0 | 0 | 0 |
| SRR5387988 | 1157 | 2365 | 835 | 435 | 0 | 0 | 0 | 12 | 26 | 17 | 482 | 1140 | 282 | 829 | 0 | 0 | 927 | 0 | 0 | 0 | 0 | 0 | 0 | 0 | 0 | 0 | 0 | 0 | 0 | 0 | 0 | 0 | 0 | 0 | 0 | 0 | 0 |
| SRR5387989 | 1312 | 3296 | 5198 | 2457 | 0 | 0 | 0 | 44 | 58 | 0 | 1188 | 1059 | 343 | 1318 | 0 | 0 | 820 | 0 | 0 | 0 | 0 | 0 | 0 | 0 | 0 | 0 | 0 | 0 | 0 | 0 | 0 | 0 | 0 | 0 | 0 | 0 | 0 |
| SRR5387990 | 1461 | 2839 | 1105 | 409 | 0 | 0 | 0 | 13 | 23 | 15 | 143 | 261 | 64 | 316 | 0 | 0 | 449 | 0 | 0 | 0 | 0 | 0 | 0 | 0 | 0 | 0 | 0 | 0 | 0 | 0 | 0 | 0 | 0 | 0 | 0 | 0 | 0 |

### Supp Table S8.1.Columns_B_to_AL

| | | | | | | | | | | | | | | | | | | | | | | | | | | | | | | | | | | | | | |
|---|---|---|---|---|---|---|---|---|---|---|---|---|---|---|---|---|---|---|---|---|---|---|---|---|---|---|---|---|---|---|---|---|---|---|---|---|---|
| SRR5387991 | 1042 | 7614 | 705 | 659 | 0 | 24 | 26 | 213 | 532 | 31 | 1165 | 1459 | 408 | 1492 | 0 | 15 | 1978 | 0 | 0 | 0 | 0 | 0 | 0 | 0 | 0 | 0 | 0 | 0 | 0 | 0 | 0 | 0 | 0 | 0 | 0 | 0 | 0 |
| SRR5387992 | 620 | 4311 | 401 | 367 | 0 | 11 | 12 | 107 | 145 | 0 | 1033 | 822 | 244 | 1093 | 0 | 21 | 985 | 0 | 0 | 0 | 0 | 0 | 0 | 0 | 0 | 0 | 0 | 0 | 0 | 0 | 0 | 0 | 0 | 0 | 0 | 0 | 0 |
| SRR5387993 | 791 | 2356 | 499 | 473 | 0 | 0 | 0 | 54 | 88 | 0 | 1135 | 1061 | 261 | 796 | 0 | 10 | 980 | 0 | 0 | 0 | 0 | 0 | 0 | 0 | 0 | 0 | 0 | 0 | 0 | 0 | 0 | 0 | 0 | 0 | 0 | 0 | 0 |
| SRR5387994 | 906 | 3934 | 359 | 379 | 0 | 15 | 14 | 105 | 109 | 20 | 1366 | 1300 | 409 | 1068 | 0 | 11 | 1642 | 0 | 0 | 0 | 0 | 0 | 0 | 0 | 0 | 0 | 0 | 0 | 0 | 0 | 0 | 0 | 0 | 0 | 0 | 0 | 0 |
| SRR5387995 | 974 | 2617 | 730 | 513 | 0 | 0 | 0 | 42 | 83 | 0 | 2143 | 1698 | 520 | 1198 | 0 | 0 | 1276 | 0 | 0 | 0 | 0 | 0 | 0 | 0 | 0 | 0 | 0 | 0 | 0 | 0 | 0 | 0 | 0 | 0 | 0 | 0 | 0 |
| SRR5387996 | 749 | 2863 | 665 | 728 | 0 | 14 | 15 | 98 | 90 | 0 | 1763 | 1895 | 609 | 2273 | 0 | 0 | 1681 | 0 | 0 | 0 | 0 | 0 | 0 | 0 | 0 | 0 | 0 | 0 | 0 | 0 | 0 | 0 | 0 | 0 | 0 | 0 | 0 |
| SRR5387997 | 238 | 2264 | 319 | 483 | 0 | 17 | 19 | 191 | 241 | 0 | 1463 | 1034 | 485 | 1674 | 0 | 0 | 2928 | 0 | 0 | 0 | 0 | 0 | 0 | 0 | 0 | 0 | 0 | 0 | 0 | 0 | 0 | 0 | 0 | 0 | 0 | 0 | 0 |
| SRR5387998 | 282 | 2586 | 304 | 682 | 0 | 18 | 17 | 113 | 87 | 0 | 1631 | 893 | 413 | 1441 | 0 | 0 | 2060 | 0 | 0 | 0 | 0 | 0 | 0 | 0 | 0 | 0 | 0 | 0 | 0 | 0 | 0 | 0 | 0 | 0 | 0 | 0 | 0 |
| SRR5387999 | 509 | 6207 | 538 | 737 | 0 | 10 | 0 | 61 | 36 | 0 | 370 | 542 | 228 | 657 | 0 | 0 | 1157 | 0 | 0 | 0 | 0 | 0 | 0 | 0 | 0 | 0 | 0 | 0 | 0 | 0 | 0 | 0 | 0 | 0 | 0 | 0 | 0 |
| SRR5388000 | 302 | 3043 | 721 | 478 | 0 | 15 | 12 | 72 | 63 | 0 | 367 | 565 | 192 | 680 | 0 | 0 | 1691 | 0 | 0 | 0 | 0 | 0 | 0 | 0 | 0 | 0 | 0 | 0 | 0 | 0 | 0 | 0 | 0 | 0 | 0 | 0 | 0 |
| SRR5388001 | 299 | 4870 | 359 | 635 | 0 | 22 | 23 | 125 | 291 | 15 | 1469 | 1346 | 466 | 1410 | 0 | 0 | 1832 | 0 | 0 | 0 | 0 | 0 | 0 | 0 | 0 | 0 | 0 | 0 | 0 | 0 | 0 | 0 | 0 | 0 | 0 | 0 | 0 |
| SRR5388002 | 362 | 10188 | 445 | 599 | 0 | 27 | 24 | 164 | 176 | 0 | 1228 | 1502 | 532 | 1722 | 0 | 0 | 2333 | 0 | 0 | 0 | 0 | 0 | 0 | 0 | 0 | 0 | 0 | 0 | 0 | 0 | 0 | 0 | 0 | 0 | 0 | 0 | 0 |
| SRR5388003 | 460 | 1261 | 749 | 342 | 0 | 0 | 0 | 23 | 45 | 0 | 363 | 637 | 146 | 287 | 0 | 0 | 1099 | 0 | 0 | 0 | 0 | 0 | 0 | 0 | 0 | 0 | 0 | 0 | 0 | 0 | 0 | 0 | 0 | 0 | 0 | 0 | 0 |
| SRR5388004 | 362 | 1338 | 336 | 459 | 0 | 15 | 12 | 80 | 92 | 0 | 484 | 606 | 214 | 910 | 0 | 0 | 1177 | 0 | 0 | 0 | 0 | 0 | 0 | 0 | 0 | 0 | 0 | 0 | 0 | 0 | 0 | 0 | 0 | 0 | 0 | 0 | 0 |
| SRR5388005 | 534 | 2102 | 430 | 791 | 0 | 13 | 0 | 103 | 109 | 28 | 794 | 1326 | 316 | 2346 | 0 | 0 | 2431 | 0 | 0 | 0 | 0 | 0 | 0 | 0 | 0 | 0 | 0 | 0 | 0 | 0 | 0 | 0 | 0 | 0 | 0 | 0 | 0 |
| SRR5388006 | 456 | 1440 | 439 | 431 | 0 | 27 | 21 | 223 | 152 | 0 | 2268 | 1690 | 495 | 3742 | 0 | 0 | 2714 | 0 | 0 | 0 | 0 | 0 | 0 | 0 | 0 | 0 | 0 | 0 | 0 | 0 | 0 | 0 | 0 | 0 | 0 | 0 | 0 |
| SRR5388007 | 311 | 1588 | 445 | 818 | 0 | 33 | 23 | 234 | 145 | 0 | 2584 | 2061 | 882 | 4232 | 0 | 0 | 5385 | 0 | 0 | 0 | 0 | 0 | 0 | 0 | 0 | 0 | 0 | 0 | 0 | 0 | 0 | 0 | 0 | 0 | 0 | 0 | 0 |
| SRR5388008 | 352 | 1258 | 579 | 1224 | 0 | 16 | 14 | 108 | 74 | 0 | 3359 | 2481 | 870 | 5157 | 0 | 0 | 2722 | 0 | 0 | 0 | 11 | 0 | 0 | 0 | 0 | 0 | 0 | 0 | 0 | 0 | 0 | 0 | 0 | 0 | 0 | 0 | 0 |
| SRR5388009 | 544 | 2423 | 513 | 288 | 0 | 0 | 12 | 66 | 141 | 0 | 1177 | 885 | 374 | 1139 | 0 | 0 | 1151 | 0 | 0 | 0 | 0 | 0 | 0 | 0 | 0 | 0 | 0 | 0 | 0 | 0 | 0 | 0 | 0 | 0 | 0 | 0 | 0 |
| SRR5388010 | 347 | 3224 | 550 | 482 | 0 | 19 | 17 | 85 | 80 | 23 | 2272 | 1080 | 587 | 1930 | 0 | 0 | 2244 | 0 | 0 | 0 | 0 | 0 | 0 | 0 | 0 | 0 | 0 | 0 | 0 | 0 | 0 | 0 | 0 | 0 | 0 | 0 | 0 |
| SRR5388011 | 575 | 11255 | 428 | 620 | 0 | 18 | 13 | 114 | 193 | 0 | 1050 | 1107 | 302 | 686 | 0 | 11 | 1696 | 0 | 0 | 0 | 0 | 0 | 0 | 0 | 0 | 0 | 0 | 0 | 0 | 0 | 0 | 0 | 0 | 0 | 0 | 0 | 0 |
| SRR5388012 | 450 | 12001 | 566 | 733 | 0 | 26 | 22 | 134 | 228 | 42 | 1485 | 1555 | 453 | 1137 | 0 | 15 | 1916 | 0 | 0 | 0 | 0 | 0 | 0 | 0 | 0 | 0 | 0 | 0 | 0 | 0 | 0 | 0 | 0 | 0 | 0 | 0 | 0 |
| SRR5388013 | 527 | 2498 | 353 | 481 | 0 | 31 | 30 | 231 | 202 | 0 | 1746 | 1675 | 651 | 2543 | 0 | 0 | 3306 | 0 | 0 | 0 | 22 | 0 | 0 | 0 | 0 | 0 | 0 | 0 | 0 | 18 | 0 | 0 | 0 | 0 | 0 | 0 | 0 |
| SRR5388014 | 1188 | 2954 | 725 | 538 | 0 | 33 | 26 | 181 | 193 | 12 | 3018 | 2167 | 779 | 2611 | 0 | 0 | 3323 | 0 | 0 | 0 | 0 | 0 | 0 | 0 | 0 | 0 | 0 | 0 | 0 | 0 | 0 | 0 | 0 | 0 | 0 | 0 | 0 |
| SRR5388015 | 670 | 2304 | 583 | 391 | 0 | 0 | 0 | 53 | 71 | 0 | 244 | 348 | 112 | 698 | 0 | 0 | 760 | 0 | 0 | 0 | 0 | 0 | 0 | 0 | 0 | 0 | 0 | 0 | 0 | 0 | 0 | 0 | 0 | 0 | 0 | 0 | 0 |
| SRR5388016 | 654 | 3347 | 502 | 467 | 0 | 60 | 51 | 329 | 243 | 0 | 4943 | 3316 | 1634 | 5865 | 0 | 0 | 4560 | 0 | 0 | 0 | 13 | 0 | 0 | 0 | 0 | 0 | 0 | 0 | 0 | 0 | 0 | 0 | 0 | 0 | 0 | 0 | 0 |
| SRR5388017 | 976 | 2401 | 775 | 564 | 0 | 0 | 0 | 22 | 49 | 0 | 379 | 394 | 129 | 314 | 0 | 0 | 1411 | 0 | 0 | 0 | 0 | 0 | 0 | 0 | 0 | 0 | 0 | 0 | 0 | 0 | 0 | 0 | 0 | 0 | 0 | 0 | 0 |
| SRR5388018 | 551 | 1894 | 492 | 418 | 0 | 0 | 0 | 43 | 103 | 15 | 391 | 438 | 164 | 360 | 0 | 0 | 1235 | 0 | 0 | 0 | 0 | 0 | 0 | 0 | 0 | 0 | 0 | 0 | 0 | 0 | 0 | 0 | 0 | 0 | 0 | 0 | 0 |
| SRR5388019 | 986 | 3777 | 639 | 493 | 0 | 13 | 0 | 63 | 154 | 0 | 1081 | 1012 | 356 | 647 | 0 | 0 | 2047 | 0 | 0 | 0 | 0 | 0 | 0 | 0 | 0 | 0 | 0 | 0 | 0 | 0 | 0 | 0 | 0 | 0 | 0 | 0 | 0 |
| SRR5388020 | 522 | 2499 | 578 | 457 | 0 | 0 | 0 | 66 | 119 | 0 | 1400 | 1032 | 278 | 466 | 0 | 0 | 1362 | 0 | 0 | 0 | 0 | 0 | 0 | 0 | 0 | 0 | 0 | 0 | 0 | 0 | 0 | 0 | 0 | 0 | 0 | 0 | 0 |
| SRR5388021 | 591 | 1899 | 475 | 916 | 0 | 11 | 11 | 98 | 179 | 17 | 1327 | 1492 | 493 | 2349 | 0 | 0 | 2068 | 0 | 0 | 0 | 0 | 0 | 0 | 0 | 0 | 0 | 0 | 0 | 0 | 0 | 0 | 0 | 0 | 0 | 0 | 0 | 0 |
| SRR5388022 | 353 | 2218 | 391 | 472 | 0 | 11 | 12 | 112 | 54 | 0 | 2551 | 2474 | 813 | 1406 | 0 | 0 | 1859 | 0 | 0 | 0 | 0 | 0 | 0 | 0 | 0 | 0 | 0 | 0 | 0 | 0 | 0 | 0 | 0 | 0 | 0 | 0 | 0 |
| SRR5388023 | 828 | 1918 | 650 | 293 | 0 | 12 | 15 | 109 | 89 | 10 | 2223 | 2266 | 895 | 2368 | 0 | 0 | 2578 | 0 | 0 | 0 | 0 | 0 | 0 | 0 | 0 | 0 | 0 | 0 | 0 | 0 | 0 | 0 | 0 | 0 | 0 | 0 | 0 |
| SRR5388024 | 344 | 1792 | 442 | 237 | 0 | 15 | 13 | 107 | 86 | 16 | 1593 | 1540 | 535 | 1147 | 0 | 0 | 1616 | 0 | 0 | 0 | 0 | 0 | 0 | 0 | 0 | 0 | 0 | 0 | 0 | 0 | 0 | 0 | 0 | 0 | 0 | 0 | 0 |
| SRR5388025 | 522 | 2890 | 522 | 294 | 0 | 0 | 0 | 56 | 105 | 16 | 725 | 1036 | 285 | 773 | 0 | 0 | 1901 | 0 | 0 | 0 | 0 | 0 | 0 | 0 | 0 | 0 | 0 | 0 | 0 | 0 | 0 | 0 | 0 | 0 | 0 | 0 | 0 |
| SRR5388026 | 628 | 2835 | 598 | 736 | 0 | 13 | 13 | 79 | 121 | 0 | 1099 | 981 | 298 | 1817 | 0 | 0 | 2010 | 0 | 0 | 0 | 0 | 0 | 0 | 0 | 0 | 0 | 0 | 0 | 0 | 0 | 0 | 0 | 0 | 0 | 0 | 0 | 0 |
| SRR5388027 | 466 | 4394 | 418 | 418 | 0 | 12 | 0 | 58 | 70 | 0 | 774 | 892 | 237 | 1119 | 0 | 0 | 1403 | 0 | 0 | 0 | 0 | 0 | 0 | 0 | 0 | 0 | 0 | 0 | 0 | 0 | 0 | 0 | 0 | 0 | 0 | 0 | 0 |
| SRR5388028 | 349 | 2897 | 547 | 585 | 0 | 24 | 24 | 159 | 108 | 0 | 1006 | 1142 | 439 | 2356 | 0 | 0 | 2188 | 0 | 0 | 0 | 0 | 0 | 0 | 0 | 0 | 0 | 0 | 0 | 0 | 0 | 0 | 0 | 0 | 0 | 0 | 0 | 0 |
| SRR5388029 | 971 | 2316 | 556 | 639 | 0 | 0 | 0 | 53 | 92 | 0 | 611 | 882 | 233 | 1419 | 0 | 0 | 920 | 0 | 0 | 0 | 0 | 0 | 0 | 0 | 0 | 0 | 0 | 0 | 0 | 0 | 0 | 0 | 0 | 0 | 0 | 0 | 0 |
| SRR5388030 | 755 | 2545 | 628 | 543 | 0 | 0 | 0 | 54 | 51 | 0 | 857 | 1009 | 287 | 1549 | 0 | 0 | 761 | 0 | 0 | 0 | 0 | 0 | 0 | 0 | 0 | 0 | 0 | 0 | 0 | 0 | 0 | 0 | 0 | 0 | 0 | 0 | 0 |
| SRR5388031 | 492 | 3410 | 380 | 552 | 0 | 10 | 0 | 63 | 141 | 0 | 1052 | 1864 | 418 | 2205 | 0 | 0 | 2825 | 0 | 0 | 0 | 0 | 0 | 0 | 0 | 0 | 0 | 0 | 0 | 0 | 0 | 0 | 0 | 0 | 0 | 0 | 0 | 0 |
| SRR5388032 | 652 | 2098 | 784 | 309 | 0 | 17 | 14 | 129 | 136 | 0 | 1792 | 1302 | 394 | 1508 | 0 | 0 | 1882 | 0 | 0 | 0 | 0 | 0 | 0 | 0 | 0 | 0 | 0 | 0 | 0 | 0 | 0 | 0 | 0 | 0 | 0 | 0 | 0 |
| SRR5388033 | 606 | 3100 | 509 | 1056 | 0 | 0 | 0 | 23 | 149 | 20 | 235 | 283 | 80 | 506 | 0 | 0 | 858 | 0 | 0 | 0 | 0 | 0 | 0 | 0 | 0 | 0 | 0 | 0 | 0 | 0 | 0 | 0 | 0 | 0 | 0 | 0 | 0 |
| SRR5388034 | 849 | 2240 | 608 | 558 | 0 | 22 | 17 | 117 | 166 | 0 | 1344 | 1409 | 428 | 1170 | 0 | 0 | 1829 | 0 | 0 | 0 | 0 | 0 | 0 | 0 | 0 | 0 | 0 | 0 | 0 | 0 | 0 | 0 | 0 | 0 | 0 | 0 | 0 |
| SRR5388035 | 768 | 5729 | 437 | 520 | 0 | 13 | 10 | 73 | 84 | 0 | 695 | 733 | 278 | 889 | 0 | 0 | 1386 | 0 | 0 | 0 | 0 | 0 | 0 | 0 | 0 | 0 | 0 | 0 | 0 | 0 | 0 | 0 | 0 | 0 | 0 | 0 | 0 |
| SRR5388036 | 589 | 3581 | 570 | 531 | 0 | 13 | 14 | 97 | 108 | 0 | 1379 | 960 | 315 | 1359 | 0 | 0 | 1216 | 0 | 0 | 0 | 0 | 0 | 0 | 0 | 0 | 0 | 0 | 0 | 0 | 0 | 0 | 0 | 0 | 0 | 0 | 0 | 0 |
| SRR5388037 | 849 | 2985 | 478 | 800 | 0 | 20 | 16 | 84 | 140 | 0 | 933 | 1237 | 365 | 650 | 0 | 0 | 2243 | 0 | 0 | 0 | 0 | 0 | 0 | 0 | 0 | 0 | 0 | 0 | 0 | 0 | 0 | 0 | 0 | 0 | 0 | 0 | 0 |
| SRR5388038 | 601 | 1793 | 547 | 973 | 0 | 28 | 22 | 166 | 218 | 22 | 1561 | 1065 | 270 | 1809 | 0 | 0 | 1929 | 0 | 0 | 0 | 0 | 0 | 0 | 0 | 0 | 0 | 0 | 0 | 0 | 0 | 0 | 0 | 0 | 0 | 0 | 0 | 0 |
| SRR5388039 | 470 | 6201 | 649 | 1602 | 0 | 0 | 0 | 13 | 21 | 0 | 65 | 94 | 17 | 155 | 0 | 0 | 358 | 0 | 0 | 0 | 0 | 0 | 0 | 0 | 0 | 0 | 0 | 0 | 0 | 0 | 0 | 0 | 0 | 0 | 0 | 0 | 0 |
| SRR5388040 | 440 | 3540 | 477 | 648 | 0 | 17 | 14 | 91 | 71 | 0 | 1526 | 1335 | 379 | 1056 | 0 | 0 | 1637 | 0 | 0 | 0 | 0 | 0 | 0 | 0 | 0 | 0 | 0 | 0 | 0 | 0 | 0 | 0 | 0 | 0 | 0 | 0 | 0 |
| SRR5388041 | 456 | 2880 | 538 | 683 | 0 | 0 | 10 | 63 | 225 | 18 | 494 | 536 | 181 | 956 | 0 | 0 | 1430 | 0 | 0 | 0 | 0 | 0 | 0 | 0 | 0 | 0 | 0 | 0 | 0 | 0 | 0 | 0 | 0 | 0 | 0 | 0 | 0 |
| SRR5388042 | 609 | 3001 | 652 | 669 | 0 | 0 | 0 | 27 | 38 | 0 | 450 | 515 | 158 | 818 | 0 | 0 | 1098 | 0 | 0 | 0 | 0 | 0 | 0 | 0 | 0 | 0 | 0 | 0 | 0 | 0 | 0 | 0 | 0 | 0 | 0 | 0 | 0 |
| SRR5388043 | 375 | 1120 | 641 | 109 | 0 | 21 | 19 | 151 | 195 | 0 | 2574 | 1454 | 493 | 1328 | 0 | 0 | 1672 | 0 | 0 | 0 | 0 | 0 | 0 | 0 | 0 | 0 | 0 | 0 | 0 | 0 | 0 | 0 | 0 | 0 | 0 | 0 | 0 |
| SRR5388044 | 626 | 2624 | 540 | 530 | 0 | 25 | 24 | 150 | 256 | 0 | 1638 | 1634 | 592 | 1318 | 0 | 0 | 2911 | 0 | 0 | 0 | 0 | 0 | 0 | 0 | 0 | 0 | 0 | 0 | 0 | 0 | 0 | 0 | 0 | 0 | 0 | 0 | 0 |
| SRR5388045 | 543 | 4567 | 1193 | 560 | 0 | 0 | 0 | 36 | 26 | 58 | 715 | 1449 | 502 | 1936 | 0 | 0 | 701 | 0 | 0 | 0 | 0 | 0 | 0 | 0 | 0 | 0 | 0 | 0 | 0 | 0 | 0 | 0 | 0 | 0 | 0 | 0 | 0 |
| SRR5388046 | 388 | 3959 | 557 | 675 | 0 | 11 | 15 | 102 | 101 | 0 | 1237 | 1035 | 387 | 1499 | 0 | 0 | 1315 | 0 | 0 | 0 | 0 | 0 | 0 | 0 | 0 | 0 | 0 | 0 | 0 | 0 | 0 | 0 | 0 | 0 | 0 | 0 | 0 |
| SRR5388047 | 568 | 2395 | 542 | 507 | 0 | 17 | 19 | 165 | 195 | 0 | 1317 | 1631 | 570 | 2320 | 0 | 0 | 1971 | 0 | 0 | 0 | 0 | 0 | 0 | 0 | 0 | 0 | 0 | 0 | 0 | 0 | 0 | 0 | 0 | 0 | 0 | 0 | 0 |
| SRR5388048 | 352 | 2544 | 303 | 360 | 0 | 18 | 15 | 143 | 145 | 0 | 1166 | 1185 | 476 | 1905 | 0 | 0 | 2022 | 0 | 0 | 0 | 0 | 0 | 0 | 0 | 0 | 0 | 0 | 0 | 0 | 0 | 0 | 0 | 0 | 0 | 0 | 0 | 0 |
| SRR5388049 | 298 | 3153 | 470 | 1051 | 0 | 16 | 22 | 131 | 283 | 0 | 2032 | 1728 | 622 | 1094 | 0 | 13 | 1609 | 0 | 0 | 0 | 0 | 0 | 0 | 0 | 0 | 0 | 0 | 0 | 0 | 0 | 0 | 0 | 0 | 0 | 0 | 0 | 0 |
| SRR5388050 | 3995 | 34564 | 1572 | 764 | 0 | 23 | 20 | 191 | 308 | 0 | 2585 | 2091 | 673 | 2193 | 0 | 12 | 1872 | 0 | 0 | 0 | 0 | 0 | 0 | 0 | 0 | 0 | 0 | 0 | 0 | 0 | 0 | 0 | 0 | 0 | 0 | 0 | 0 |
| SRR5388051 | 422 | 1784 | 340 | 422 | 0 | 19 | 13 | 147 | 85 | 0 | 2143 | 2172 | 673 | 2910 | 0 | 0 | 2230 | 0 | 0 | 0 | 0 | 0 | 0 | 0 | 0 | 0 | 0 | 0 | 0 | 0 | 0 | 0 | 0 | 0 | 0 | 0 | 0 |
| SRR5388052 | 263 | 1589 | 360 | 417 | 0 | 18 | 13 | 109 | 106 | 0 | 1835 | 1557 | 638 | 2973 | 0 | 0 | 2203 | 0 | 0 | 0 | 0 | 0 | 0 | 0 | 0 | 0 | 0 | 0 | 0 | 0 | 0 | 0 | 0 | 0 | 0 | 0 | 0 |
| SRR5388053 | 507 | 3150 | 550 | 556 | 0 | 13 | 11 | 85 | 107 | 0 | 1011 | 764 | 345 | 612 | 0 | 0 | 2326 | 0 | 0 | 0 | 0 | 0 | 0 | 0 | 0 | 0 | 0 | 0 | 0 | 0 | 0 | 0 | 0 | 0 | 0 | 0 | 0 |
| SRR5388054 | 702 | 2671 | 672 | 483 | 0 | 12 | 12 | 88 | 89 | 0 | 1450 | 1204 | 365 | 833 | 0 | 0 | 1370 | 0 | 0 | 0 | 0 | 0 | 0 | 0 | 0 | 0 | 0 | 0 | 0 | 0 | 0 | 0 | 0 | 0 | 0 | 0 | 0 |
| SRR5388055 | 797 | 3177 | 597 | 654 | 0 | 16 | 12 | 111 | 156 | 28 | 755 | 1100 | 290 | 975 | 0 | 0 | 1487 | 0 | 0 | 0 | 0 | 0 | 0 | 0 | 0 | 0 | 0 | 0 | 0 | 0 | 0 | 0 | 0 | 0 | 0 | 0 | 0 |
| SRR5388056 | 526 | 2405 | 441 | 363 | 0 | 24 | 22 | 154 | 180 | 0 | 1867 | 1409 | 440 | 1350 | 0 | 0 | 1787 | 0 | 0 | 0 | 0 | 0 | 0 | 0 | 0 | 0 | 0 | 0 | 0 | 0 | 0 | 0 | 0 | 0 | 0 | 0 | 0 |
| SRR5388057 | 264 | 1760 | 290 | 458 | 0 | 0 | 0 | 74 | 57 | 18 | 1158 | 1120 | 309 | 1066 | 0 | 0 | 1512 | 0 | 0 | 0 | 0 | 0 | 0 | 0 | 0 | 0 | 0 | 0 | 0 | 0 | 0 | 0 | 0 | 0 | 0 | 0 | 0 |
| SRR5388058 | 546 | 1615 | 431 | 334 | 0 | 19 | 16 | 162 | 101 | 18 | 3977 | 3393 | 1265 | 3777 | 0 | 0 | 2918 | 0 | 0 | 0 | 0 | 0 | 0 | 0 | 0 | 0 | 0 | 0 | 0 | 0 | 0 | 0 | 0 | 0 | 0 | 0 | 0 |
| SRR5388059 | 713 | 2031 | 807 | 418 | 0 | 10 | 14 | 116 | 160 | 36 | 1127 | 1202 | 373 | 1586 | 0 | 0 | 1477 | 0 | 0 | 0 | 0 | 0 | 0 | 0 | 0 | 0 | 0 | 0 | 0 | 0 | 0 | 0 | 0 | 0 | 0 | 0 | 0 |
| SRR5388060 | 565 | 1910 | 558 | 383 | 0 | 0 | 0 | 50 | 61 | 0 | 826 | 769 | 207 | 481 | 0 | 0 | 1174 | 0 | 0 | 0 | 0 | 0 | 0 | 0 | 0 | 0 | 0 | 0 | 0 | 0 | 0 | 0 | 0 | 0 | 0 | 0 | 0 |
| SRR5388061 | 592 | 2503 | 603 | 444 | 0 | 21 | 22 | 115 | 114 | 11 | 1678 | 1666 | 638 | 850 | 0 | 0 | 2577 | 0 | 0 | 0 | 0 | 0 | 0 | 0 | 0 | 0 | 0 | 0 | 0 | 0 | 0 | 0 | 0 | 0 | 0 | 0 | 0 |
| SRR5388062 | 399 | 2655 | 360 | 384 | 0 | 19 | 19 | 150 | 112 | 0 | 732 | 836 | 337 | 1434 | 0 | 0 | 1400 | 0 | 0 | 0 | 0 | 0 | 0 | 0 | 0 | 0 | 0 | 0 | 0 | 0 | 0 | 0 | 0 | 0 | 0 | 0 | 0 |
| SRR5388063 | 735 | 3102 | 725 | 927 | 0 | 0 | 0 | 51 | 93 | 11 | 361 | 356 | 138 | 614 | 0 | 0 | 1514 | 0 | 0 | 0 | 0 | 0 | 0 | 0 | 0 | 0 | 0 | 0 | 0 | 0 | 0 | 0 | 0 | 0 | 0 | 0 | 0 |
| SRR5388064 | 380 | 2024 | 574 | 521 | 0 | 13 | 13 | 73 | 111 | 18 | 1213 | 979 | 425 | 868 | 0 | 0 | 2057 | 0 | 0 | 0 | 10 | 0 | 0 | 0 | 0 | 0 | 0 | 0 | 0 | 0 | 0 | 0 | 0 | 0 | 0 | 0 | 0 |
| SRR5388065 | 625 | 3986 | 458 | 319 | 0 | 19 | 19 | 96 | 144 | 14 | 862 | 884 | 341 | 1153 | 0 | 0 | 1832 | 0 | 0 | 0 | 0 | 0 | 0 | 0 | 0 | 0 | 0 | 0 | 0 | 0 | 0 | 0 | 0 | 0 | 0 | 0 | 0 |
| SRR5388066 | 851 | 4055 | 485 | 412 | 0 | 15 | 11 | 75 | 90 | 0 | 915 | 1179 | 377 | 1241 | 0 | 0 | 1575 | 0 | 0 | 0 | 0 | 0 | 0 | 0 | 0 | 0 | 0 | 0 | 0 | 0 | 0 | 0 | 0 | 0 | 0 | 0 | 0 |
| SRR5388067 | 502 | 2441 | 397 | 472 | 0 | 30 | 28 | 186 | 128 | 0 | 3197 | 3063 | 1032 | 3584 | 0 | 0 | 3017 | 0 | 0 | 0 | 0 | 0 | 0 | 0 | 0 | 0 | 0 | 0 | 0 | 0 | 0 | 0 | 0 | 0 | 0 | 0 | 0 |
| SRR5388068 | 821 | 3155 | 522 | 693 | 0 | 33 | 33 | 180 | 192 | 0 | 3198 | 2409 | 878 | 2045 | 0 | 0 | 2864 | 0 | 0 | 0 | 13 | 0 | 0 | 0 | 0 | 0 | 0 | 0 | 0 | 0 | 0 | 0 | 0 | 0 | 0 | 0 | 0 |
| SRR5388069 | 562 | 2175 | 577 | 585 | 0 | 0 | 0 | 64 | 99 | 0 | 1479 | 1238 | 373 | 1438 | 0 | 0 | 1807 | 0 | 0 | 0 | 0 | 0 | 0 | 0 | 0 | 0 | 0 | 0 | 0 | 0 | 0 | 0 | 0 | 0 | 0 | 0 | 0 |
| SRR5388070 | 1239 | 2971 | 866 | 656 | 0 | 0 | 0 | 32 | 36 | 16 | 583 | 1035 | 240 | 937 | 0 | 0 | 1005 | 0 | 0 | 0 | 0 | 0 | 0 | 0 | 0 | 0 | 0 | 0 | 0 | 0 | 0 | 0 | 0 | 0 | 0 | 0 | 0 |
| SRR5388071 | 1370 | 4782 | 589 | 817 | 0 | 11 | 0 | 77 | 80 | 0 | 856 | 991 | 308 | 1606 | 0 | 0 | 2209 | 0 | 0 | 0 | 0 | 0 | 0 | 0 | 0 | 0 | 0 | 0 | 0 | 0 | 0 | 0 | 0 | 0 | 0 | 0 | 0 |
| SRR5388072 | 712 | 3215 | 643 | 643 | 0 | 28 | 27 | 204 | 245 | 0 | 1892 | 1217 | 442 | 1919 | 0 | 0 | 2382 | 0 | 0 | 0 | 0 | 0 | 0 | 0 | 0 | 0 | 0 | 0 | 0 | 0 | 0 | 0 | 0 | 0 | 0 | 0 | 0 |
| SRR5388073 | 917 | 4405 | 381 | 763 | 0 | 10 | 0 | 58 | 115 | 0 | 639 | 962 | 273 | 578 | 0 | 0 | 2033 | 0 | 0 | 0 | 0 | 0 | 0 | 0 | 0 | 0 | 0 | 0 | 0 | 0 | 0 | 0 | 0 | 0 | 0 | 0 | 0 |
| SRR5388074 | 510 | 2869 | 452 | 443 | 0 | 28 | 18 | 176 | 167 | 0 | 1572 | 1326 | 450 | 1666 | 0 | 0 | 1759 | 0 | 0 | 0 | 0 | 0 | 0 | 0 | 0 | 0 | 0 | 0 | 0 | 0 | 0 | 0 | 0 | 0 | 0 | 0 | 0 |
| SRR5388075 | 425 | 2484 | 491 | 498 | 0 | 38 | 35 | 101 | 183 | 0 | 1214 | 1368 | 617 | 1909 | 0 | 0 | 1556 | 0 | 0 | 0 | 0 | 0 | 0 | 0 | 0 | 0 | 0 | 0 | 0 | 0 | 0 | 0 | 0 | 0 | 0 | 0 | 0 |
| SRR5388076 | 542 | 3034 | 390 | 576 | 0 | 13 | 0 | 66 | 51 | 0 | 1488 | 1218 | 416 | 1243 | 0 | 0 | 1464 | 0 | 0 | 0 | 0 | 0 | 0 | 0 | 0 | 0 | 0 | 0 | 0 | 0 | 0 | 0 | 0 | 0 | 0 | 0 | 0 |
| SRR5388077 | 292 | 3854 | 470 | 366 | 0 | 18 | 18 | 109 | 111 | 0 | 1032 | 862 | 381 | 1149 | 0 | 0 | 1813 | 0 | 0 | 0 | 0 | 0 | 0 | 0 | 0 | 0 | 0 | 0 | 0 | 0 | 0 | 0 | 0 | 0 | 0 | 0 | 0 |
| SRR5388078 | 4155 | 12614 | 2372 | 1690 | 0 | 27 | 18 | 209 | 201 | 14 | 2423 | 2025 | 822 | 3638 | 0 | 0 | 3429 | 0 | 0 | 0 | 0 | 0 | 0 | 0 | 0 | 0 | 0 | 0 | 10 | 0 | 0 | 0 | 0 | 0 | 0 | 0 | 0 |
| SRR5388079 | 731 | 2510 | 635 | 519 | 0 | 0 | 0 | 57 | 131 | 29 | 443 | 483 | 131 | 959 | 0 | 0 | 881 | 0 | 0 | 0 | 0 | 0 | 0 | 0 | 0 | 0 | 0 | 0 | 0 | 0 | 0 | 0 | 0 | 0 | 0 | 0 | 0 |
| SRR5388080 | 351 | 2543 | 522 | 599 | 0 | 14 | 17 | 133 | 106 | 0 | 1969 | 1641 | 641 | 1177 | 0 | 0 | 1371 | 0 | 0 | 0 | 0 | 0 | 0 | 0 | 0 | 0 | 0 | 0 | 0 | 0 | 0 | 0 | 0 | 0 | 0 | 0 | 0 |
| SRR5388081 | 710 | 2670 | 650 | 505 | 0 | 17 | 19 | 129 | 170 | 43 | 1039 | 1045 | 316 | 1640 | 0 | 0 | 2177 | 0 | 0 | 0 | 0 | 0 | 0 | 0 | 0 | 0 | 0 | 0 | 0 | 0 | 0 | 0 | 0 | 0 | 0 | 0 | 0 |
| SRR5388082 | 352 | 2255 | 492 | 612 | 0 | 15 | 14 | 101 | 93 | 0 | 1112 | 1066 | 324 | 1318 | 0 | 0 | 1475 | 0 | 0 | 0 | 0 | 0 | 0 | 0 | 0 | 0 | 0 | 0 | 0 | 0 | 0 | 0 | 0 | 0 | 0 | 0 | 0 |
| SRR5388083 | 553 | 3752 | 599 | 499 | 0 | 20 | 17 | 125 | 188 | 0 | 2160 | 1317 | 509 | 1011 | 0 | 0 | 1783 | 0 | 0 | 0 | 13 | 0 | 0 | 0 | 0 | 0 | 0 | 0 | 0 | 0 | 0 | 0 | 0 | 0 | 0 | 0 | 0 |
| SRR5388084 | 586 | 3628 | 670 | 474 | 0 | 14 | 0 | 74 | 93 | 0 | 1269 | 986 | 313 | 733 | 0 | 0 | 1302 | 0 | 0 | 0 | 0 | 0 | 0 | 0 | 0 | 0 | 0 | 0 | 0 | 0 | 0 | 0 | 0 | 0 | 0 | 0 | 0 |
| SRR5388085 | 450 | 2942 | 440 | 1058 | 0 | 13 | 0 | 88 | 95 | 10 | 531 | 468 | 165 | 782 | 0 | 0 | 1935 | 0 | 0 | 0 | 0 | 0 | 0 | 0 | 0 | 0 | 0 | 0 | 0 | 0 | 0 | 0 | 0 | 0 | 0 | 0 | 0 |
| SRR5388086 | 549 | 1968 | 392 | 601 | 0 | 17 | 19 | 112 | 155 | 0 | 667 | 672 | 237 | 498 | 0 | 0 | 1796 | 0 | 0 | 0 | 12 | 0 | 0 | 0 | 0 | 0 | 0 | 0 | 0 | 0 | 0 | 0 | 0 | 0 | 0 | 0 | 0 |
| SRR5388087 | 541 | 5011 | 465 | 701 | 0 | 22 | 16 | 181 | 169 | 0 | 1211 | 930 | 455 | 1296 | 0 | 0 | 1921 | 0 | 0 | 0 | 0 | 0 | 0 | 0 | 0 | 0 | 0 | 0 | 0 | 0 | 0 | 0 | 0 | 0 | 0 | 0 | 0 |
| SRR5388088 | 930 | 4577 | 774 | 579 | 0 | 0 | 0 | 65 | 44 | 0 | 1162 | 1224 | 424 | 1373 | 0 | 0 | 867 | 0 | 0 | 0 | 0 | 0 | 0 | 0 | 0 | 0 | 0 | 0 | 0 | 0 | 0 | 0 | 0 | 0 | 0 | 0 | 0 |
| SRR5388089 | 700 | 4602 | 746 | 623 | 0 | 13 | 12 | 75 | 100 | 0 | 1141 | 1118 | 386 | 1128 | 0 | 0 | 2596 | 0 | 0 | 0 | 0 | 0 | 0 | 0 | 0 | 0 | 0 | 0 | 0 | 0 | 0 | 0 | 0 | 0 | 0 | 0 | 0 |
| SRR5388090 | 307 | 3625 | 584 | 602 | 0 | 16 | 14 | 133 | 189 | 0 | 1391 | 1490 | 546 | 1517 | 0 | 0 | 2925 | 0 | 0 | 0 | 0 | 0 | 0 | 0 | 0 | 0 | 0 | 0 | 0 | 0 | 0 | 0 | 0 | 0 | 0 | 0 | 0 |
| SRR5388091 | 348 | 1656 | 439 | 520 | 0 | 16 | 12 | 114 | 90 | 11 | 1626 | 938 | 310 | 1049 | 0 | 0 | 1692 | 0 | 0 | 0 | 0 | 0 | 0 | 0 | 0 | 0 | 0 | 0 | 0 | 0 | 0 | 0 | 0 | 0 | 0 | 0 | 0 |
| SRR5388092 | 289 | 2015 | 390 | 349 | 0 | 12 | 10 | 87 | 62 | 0 | 1171 | 1157 | 386 | 716 | 0 | 0 | 1775 | 0 | 0 | 0 | 0 | 0 | 0 | 0 | 0 | 0 | 0 | 0 | 0 | 0 | 0 | 0 | 0 | 0 | 0 | 0 | 0 |
| SRR5388093 | 2395 | 10567 | 4044 | 2719 | 0 | 0 | 0 | 22 | 22 | 0 | 172 | 732 | 164 | 1188 | 0 | 0 | 471 | 0 | 0 | 0 | 0 | 0 | 0 | 0 | 0 | 0 | 0 | 0 | 0 | 0 | 0 | 0 | 0 | 0 | 0 | 0 | 0 |
| SRR5388094 | 295 | 2565 | 491 | 322 | 0 | 24 | 24 | 208 | 142 | 25 | 1217 | 1139 | 563 | 2893 | 0 | 0 | 3105 | 0 | 0 | 0 | 0 | 0 | 0 | 0 | 0 | 0 | 0 | 0 | 0 | 0 | 0 | 0 | 0 | 0 | 0 | 0 | 0 |
| SRR5388095 | 248 | 2638 | 334 | 756 | 0 | 10 | 0 | 78 | 105 | 0 | 530 | 457 | 156 | 742 | 0 | 0 | 1045 | 0 | 0 | 0 | 0 | 0 | 0 | 0 | 0 | 0 | 0 | 0 | 0 | 0 | 0 | 0 | 0 | 0 | 0 | 0 | 0 |
| SRR5388096 | 360 | 1740 | 309 | 798 | 0 | 0 | 0 | 80 | 83 | 0 | 1284 | 1109 | 350 | 1698 | 0 | 0 | 1601 | 0 | 0 | 0 | 0 | 0 | 0 | 0 | 0 | 0 | 0 | 0 | 0 | 0 | 0 | 0 | 0 | 0 | 0 | 0 | 0 |
| SRR5388097 | 474 | 1971 | 502 | 512 | 0 | 14 | 0 | 94 | 90 | 0 | 904 | 1311 | 376 | 1409 | 0 | 0 | 1261 | 0 | 0 | 0 | 0 | 0 | 0 | 0 | 0 | 0 | 0 | 0 | 0 | 0 | 0 | 0 | 0 | 0 | 0 | 0 | 0 |
| SRR5388098 | 411 | 1924 | 528 | 540 | 0 | 0 | 0 | 60 | 86 | 0 | 1162 | 1153 | 312 | 1198 | 0 | 0 | 1396 | 0 | 0 | 0 | 0 | 0 | 0 | 0 | 0 | 0 | 0 | 0 | 0 | 0 | 0 | 0 | 0 | 0 | 0 | 0 | 0 |
| SRR5388099 | 678 | 3526 | 560 | 552 | 0 | 0 | 0 | 61 | 81 | 0 | 603 | 596 | 219 | 1101 | 0 | 0 | 1104 | 0 | 0 | 0 | 0 | 0 | 0 | 0 | 0 | 0 | 0 | 0 | 0 | 0 | 0 | 0 | 0 | 0 | 0 | 0 | 0 |
| SRR5388100 | 491 | 3635 | 327 | 426 | 0 | 12 | 13 | 91 | 103 | 0 | 1311 | 1771 | 606 | 2642 | 0 | 0 | 1970 | 0 | 0 | 0 | 0 | 0 | 0 | 0 | 0 | 0 | 0 | 0 | 0 | 0 | 0 | 0 | 0 | 0 | 0 | 0 | 0 |
| SRR5388101 | 468 | 1799 | 248 | 753 | 0 | 0 | 0 | 47 | 82 | 0 | 580 | 585 | 222 | 904 | 0 | 0 | 1493 | 0 | 0 | 0 | 0 | 0 | 0 | 0 | 0 | 0 | 0 | 0 | 0 | 0 | 0 | 0 | 0 | 0 | 0 | 0 | 0 |
| SRR5388102 | 380 | 1511 | 380 | 648 | 0 | 0 | 0 | 48 | 73 | 0 | 987 | 849 | 291 | 714 | 0 | 0 | 1540 | 0 | 0 | 0 | 0 | 0 | 0 | 0 | 0 | 0 | 0 | 0 | 0 | 0 | 0 | 0 | 0 | 0 | 0 | 0 | 0 |
| SRR5388103 | 245 | 4732 | 277 | 1481 | 0 | 11 | 0 | 66 | 148 | 32 | 418 | 311 | 147 | 449 | 0 | 0 | 1529 | 0 | 0 | 0 | 0 | 0 | 0 | 0 | 0 | 0 | 0 | 0 | 0 | 0 | 0 | 0 | 0 | 0 | 0 | 0 | 0 |
| SRR5388104 | 449 | 4701 | 441 | 530 | 0 | 0 | 0 | 53 | 68 | 15 | 564 | 622 | 217 | 375 | 0 | 0 | 1418 | 0 | 0 | 0 | 0 | 0 | 0 | 0 | 0 | 0 | 0 | 0 | 0 | 0 | 0 | 0 | 0 | 0 | 0 | 0 | 0 |
| SRR5388105 | 1021 | 4184 | 735 | 573 | 0 | 0 | 0 | 52 | 58 | 25 | 812 | 746 | 304 | 1089 | 0 | 0 | 1655 | 0 | 0 | 0 | 0 | 0 | 0 | 0 | 0 | 0 | 0 | 0 | 0 | 0 | 0 | 0 | 0 | 0 | 0 | 0 | 0 |
| SRR5388106 | 430 | 2764 | 389 | 480 | 0 | 0 | 0 | 58 | 96 | 0 | 1083 | 1306 | 472 | 1140 | 0 | 0 | 1408 | 0 | 0 | 0 | 0 | 0 | 0 | 0 | 0 | 0 | 0 | 0 | 0 | 0 | 0 | 0 | 0 | 0 | 0 | 0 | 0 |
| SRR5388107 | 397 | 2589 | 277 | 687 | 0 | 0 | 0 | 57 | 95 | 0 | 540 | 544 | 173 | 800 | 0 | 0 | 1058 | 0 | 0 | 0 | 0 | 0 | 0 | 0 | 0 | 0 | 0 | 0 | 0 | 0 | 0 | 0 | 0 | 0 | 0 | 0 | 0 |
| SRR5388108 | 578 | 2112 | 521 | 828 | 0 | 18 | 12 | 67 | 54 | 0 | 1007 | 589 | 204 | 943 | 0 | 0 | 1109 | 0 | 0 | 0 | 0 | 0 | 0 | 0 | 0 | 0 | 0 | 0 | 0 | 0 | 0 | 0 | 0 | 0 | 0 | 0 | 0 |
| SRR5388109 | 349 | 3819 | 389 | 1001 | 0 | 19 | 15 | 133 | 172 | 23 | 1169 | 1137 | 388 | 1902 | 0 | 11 | 1716 | 0 | 0 | 0 | 0 | 0 | 0 | 0 | 0 | 0 | 0 | 0 | 0 | 0 | 0 | 0 | 0 | 0 | 0 | 0 | 0 |
| SRR5388110 | 198 | 3029 | 371 | 931 | 0 | 25 | 17 | 113 | 139 | 0 | 1627 | 1867 | 883 | 2767 | 0 | 15 | 2278 | 0 | 0 | 0 | 16 | 0 | 0 | 0 | 0 | 0 | 0 | 0 | 0 | 0 | 0 | 0 | 0 | 0 | 0 | 0 | 0 |
| SRR5388111 | 341 | 2605 | 651 | 409 | 0 | 55 | 45 | 255 | 363 | 0 | 2709 | 2266 | 721 | 3342 | 0 | 0 | 2576 | 0 | 0 | 0 | 0 | 0 | 0 | 0 | 0 | 0 | 0 | 0 | 0 | 0 | 0 | 0 | 0 | 0 | 0 | 0 | 0 |
| SRR5388112 | 582 | 3583 | 763 | 944 | 0 | 29 | 25 | 153 | 267 | 11 | 1214 | 1280 | 299 | 1950 | 0 | 0 | 2162 | 0 | 0 | 0 | 0 | 0 | 0 | 0 | 0 | 0 | 0 | 0 | 0 | 0 | 0 | 0 | 0 | 0 | 0 | 0 | 0 |
| SRR5388113 | 441 | 1739 | 464 | 472 | 0 | 12 | 0 | 81 | 200 | 15 | 872 | 883 | 234 | 851 | 0 | 0 | 2241 | 0 | 0 | 0 | 0 | 0 | 0 | 0 | 0 | 0 | 0 | 0 | 0 | 0 | 0 | 0 | 0 | 0 | 0 | 0 | 0 |
| SRR5388114 | 622 | 2016 | 795 | 910 | 0 | 10 | 0 | 60 | 110 | 0 | 495 | 567 | 183 | 1409 | 0 | 0 | 1432 | 0 | 0 | 0 | 11 | 0 | 0 | 0 | 0 | 0 | 0 | 0 | 0 | 0 | 0 | 0 | 0 | 0 | 0 | 0 | 0 |

### Supp Table S8.1.Columns_B_to_AL

| | | | | | | | | | | | | | | | | | | | | | | | | | | | | | | | | | | | | | |
|---|---|---|---|---|---|---|---|---|---|---|---|---|---|---|---|---|---|---|---|---|---|---|---|---|---|---|---|---|---|---|---|---|---|---|---|---|---|
| SRR5388115 | 843 | 3808 | 598 | 500 | 0 | 46 | 36 | 233 | 213 | 0 | 1578 | 2010 | 665 | 2501 | 0 | 0 | 2707 | 0 | 0 | 0 | 12 | 0 | 0 | 0 | 0 | 0 | 0 | 0 | 0 | 0 | 0 | 0 | 0 | 0 | 0 | 0 | 0 |
| SRR5388116 | 337 | 3983 | 520 | 596 | 0 | 20 | 14 | 101 | 79 | 0 | 2036 | 1450 | 416 | 1193 | 0 | 0 | 1758 | 0 | 0 | 0 | 0 | 0 | 0 | 0 | 0 | 0 | 0 | 0 | 0 | 0 | 0 | 0 | 0 | 0 | 0 | 0 | 0 |
| SRR5388117 | 510 | 1415 | 358 | 613 | 0 | 11 | 0 | 87 | 137 | 0 | 2226 | 1381 | 541 | 2046 | 0 | 0 | 2613 | 0 | 0 | 0 | 11 | 0 | 0 | 0 | 0 | 0 | 0 | 0 | 0 | 0 | 0 | 0 | 0 | 0 | 0 | 0 | 0 |
| SRR5388118 | 446 | 1332 | 405 | 387 | 0 | 13 | 11 | 138 | 127 | 0 | 1592 | 1980 | 579 | 2691 | 0 | 0 | 1766 | 0 | 0 | 0 | 0 | 0 | 0 | 0 | 0 | 0 | 0 | 0 | 0 | 0 | 0 | 0 | 0 | 0 | 0 | 0 | 0 |
| SRR5388119 | 688 | 1884 | 446 | 1188 | 0 | 0 | 0 | 30 | 76 | 18 | 1371 | 1612 | 598 | 2220 | 0 | 0 | 1219 | 0 | 0 | 0 | 0 | 0 | 0 | 0 | 0 | 0 | 0 | 0 | 0 | 0 | 0 | 0 | 0 | 0 | 0 | 0 | 0 |
| SRR5388120 | 459 | 5133 | 1103 | 2189 | 0 | 0 | 0 | 31 | 63 | 0 | 510 | 1124 | 421 | 1240 | 0 | 0 | 774 | 0 | 0 | 0 | 0 | 0 | 0 | 0 | 0 | 0 | 0 | 0 | 0 | 0 | 0 | 0 | 0 | 0 | 0 | 0 | 0 |
| SRR5388121 | 798 | 4796 | 621 | 2512 | 0 | 0 | 0 | 0 | 16 | 15 | 117 | 286 | 66 | 187 | 0 | 0 | 475 | 0 | 0 | 0 | 0 | 0 | 0 | 0 | 0 | 0 | 0 | 0 | 0 | 0 | 0 | 0 | 0 | 0 | 0 | 0 | 0 |
| SRR5388122 | 809 | 3589 | 590 | 657 | 0 | 0 | 0 | 28 | 27 | 0 | 440 | 539 | 192 | 504 | 0 | 11 | 1257 | 0 | 0 | 0 | 0 | 0 | 0 | 0 | 0 | 0 | 0 | 0 | 0 | 0 | 0 | 0 | 0 | 0 | 0 | 0 | 0 |
| SRR5388123 | 331 | 2811 | 307 | 835 | 0 | 40 | 33 | 210 | 428 | 12 | 1185 | 681 | 364 | 1275 | 0 | 0 | 4296 | 0 | 0 | 0 | 0 | 0 | 0 | 0 | 0 | 0 | 0 | 0 | 0 | 0 | 0 | 0 | 0 | 0 | 0 | 0 | 0 |
| SRR5388124 | 370 | 2545 | 520 | 898 | 0 | 20 | 15 | 84 | 154 | 0 | 999 | 949 | 390 | 1187 | 0 | 0 | 2325 | 0 | 0 | 0 | 12 | 0 | 0 | 0 | 0 | 0 | 0 | 0 | 0 | 0 | 0 | 0 | 0 | 0 | 0 | 0 | 0 |
| SRR5388125 | 260 | 1751 | 369 | 1328 | 0 | 11 | 0 | 66 | 104 | 0 | 930 | 938 | 243 | 1116 | 0 | 0 | 1462 | 0 | 0 | 0 | 0 | 0 | 0 | 0 | 0 | 0 | 0 | 0 | 0 | 0 | 0 | 0 | 0 | 0 | 0 | 0 | 0 |
| SRR5388126 | 510 | 2611 | 385 | 311 | 0 | 12 | 0 | 79 | 65 | 0 | 1325 | 952 | 301 | 890 | 0 | 0 | 1896 | 0 | 0 | 0 | 0 | 0 | 0 | 0 | 0 | 0 | 0 | 0 | 0 | 0 | 0 | 0 | 0 | 0 | 0 | 0 | 0 |
| SRR5388127 | 721 | 1862 | 689 | 398 | 0 | 0 | 0 | 46 | 70 | 22 | 819 | 776 | 285 | 742 | 0 | 0 | 902 | 0 | 0 | 0 | 0 | 0 | 0 | 0 | 0 | 0 | 0 | 0 | 0 | 0 | 0 | 0 | 0 | 0 | 0 | 0 | 0 |
| SRR5388128 | 928 | 2378 | 795 | 476 | 0 | 13 | 12 | 88 | 83 | 0 | 1854 | 1610 | 687 | 975 | 0 | 0 | 2078 | 0 | 0 | 0 | 0 | 0 | 0 | 0 | 0 | 0 | 0 | 0 | 0 | 0 | 0 | 0 | 0 | 0 | 0 | 0 | 0 |
| SRR5388129 | 215 | 1658 | 563 | 450 | 0 | 0 | 0 | 68 | 60 | 0 | 539 | 835 | 229 | 1362 | 0 | 0 | 1167 | 0 | 0 | 0 | 0 | 0 | 0 | 0 | 0 | 0 | 0 | 0 | 0 | 0 | 0 | 0 | 0 | 0 | 0 | 0 | 0 |
| SRR5388130 | 827 | 1708 | 551 | 483 | 0 | 0 | 0 | 29 | 32 | 23 | 894 | 1002 | 382 | 891 | 0 | 0 | 1298 | 0 | 0 | 0 | 0 | 0 | 0 | 0 | 0 | 0 | 0 | 0 | 0 | 0 | 0 | 0 | 0 | 0 | 0 | 0 | 0 |
| SRR5388131 | 301 | 3853 | 458 | 688 | 0 | 22 | 19 | 171 | 188 | 14 | 1350 | 737 | 299 | 614 | 0 | 0 | 3819 | 0 | 0 | 0 | 0 | 0 | 0 | 0 | 0 | 0 | 0 | 0 | 0 | 0 | 0 | 0 | 0 | 0 | 0 | 0 | 0 |
| SRR5388132 | 650 | 3167 | 1908 | 950 | 0 | 36 | 30 | 201 | 258 | 21 | 3186 | 2278 | 842 | 1340 | 0 | 0 | 3484 | 0 | 0 | 0 | 0 | 0 | 0 | 0 | 0 | 0 | 0 | 0 | 0 | 0 | 0 | 0 | 0 | 0 | 0 | 0 | 0 |
| SRR5388133 | 455 | 4558 | 277 | 1384 | 0 | 21 | 26 | 150 | 294 | 20 | 1011 | 667 | 250 | 779 | 0 | 0 | 3681 | 0 | 0 | 0 | 0 | 0 | 0 | 0 | 0 | 0 | 0 | 0 | 0 | 0 | 0 | 0 | 0 | 0 | 0 | 0 | 0 |
| SRR5388134 | 477 | 3013 | 370 | 731 | 0 | 17 | 15 | 87 | 102 | 13 | 1412 | 1912 | 575 | 2587 | 0 | 0 | 1802 | 0 | 0 | 0 | 0 | 0 | 0 | 0 | 0 | 0 | 0 | 0 | 0 | 0 | 0 | 0 | 0 | 0 | 0 | 0 | 0 |
| SRR5388135 | 846 | 2351 | 349 | 341 | 0 | 10 | 10 | 59 | 79 | 0 | 1240 | 1443 | 532 | 1433 | 0 | 0 | 1960 | 0 | 0 | 0 | 0 | 0 | 0 | 0 | 0 | 0 | 0 | 0 | 0 | 0 | 0 | 0 | 0 | 0 | 0 | 0 | 0 |
| SRR5388136 | 618 | 1719 | 328 | 531 | 0 | 0 | 0 | 69 | 80 | 0 | 423 | 572 | 178 | 634 | 0 | 0 | 1062 | 0 | 0 | 0 | 0 | 0 | 0 | 0 | 0 | 0 | 0 | 0 | 0 | 0 | 0 | 0 | 0 | 0 | 0 | 0 | 0 |
| SRR5388137 | 808 | 3560 | 603 | 286 | 0 | 0 | 10 | 73 | 79 | 12 | 1861 | 1189 | 469 | 521 | 0 | 0 | 1743 | 0 | 0 | 0 | 0 | 0 | 0 | 0 | 0 | 0 | 0 | 0 | 0 | 0 | 0 | 0 | 0 | 0 | 0 | 0 | 0 |
| SRR5388138 | 551 | 2400 | 428 | 458 | 0 | 15 | 12 | 146 | 107 | 0 | 2171 | 1987 | 735 | 2386 | 0 | 0 | 1977 | 0 | 0 | 0 | 0 | 0 | 0 | 0 | 0 | 0 | 0 | 0 | 0 | 0 | 0 | 0 | 0 | 0 | 0 | 0 | 0 |
| SRR5388139 | 314 | 2281 | 453 | 490 | 0 | 0 | 0 | 64 | 106 | 18 | 1538 | 1067 | 331 | 551 | 0 | 0 | 1572 | 0 | 0 | 0 | 0 | 0 | 0 | 0 | 0 | 0 | 0 | 0 | 0 | 0 | 0 | 0 | 0 | 0 | 0 | 0 | 0 |
| SRR5388140 | 455 | 6117 | 527 | 741 | 0 | 32 | 26 | 148 | 175 | 20 | 1294 | 1362 | 580 | 1737 | 0 | 11 | 3292 | 0 | 0 | 0 | 11 | 0 | 0 | 0 | 0 | 0 | 0 | 0 | 0 | 0 | 0 | 0 | 0 | 0 | 0 | 0 | 0 |
| SRR5388141 | 257 | 2424 | 526 | 601 | 0 | 18 | 13 | 89 | 105 | 0 | 1484 | 1351 | 411 | 726 | 0 | 0 | 1496 | 0 | 0 | 0 | 0 | 0 | 0 | 0 | 0 | 0 | 0 | 0 | 0 | 0 | 0 | 0 | 0 | 0 | 0 | 0 | 0 |
| SRR5388142 | 569 | 2078 | 614 | 385 | 0 | 31 | 28 | 216 | 158 | 0 | 2259 | 2072 | 860 | 2498 | 0 | 0 | 2441 | 0 | 0 | 0 | 0 | 0 | 0 | 0 | 0 | 0 | 0 | 0 | 0 | 0 | 0 | 0 | 0 | 0 | 0 | 0 | 0 |
| SRR5388143 | 552 | 3085 | 439 | 633 | 0 | 0 | 0 | 43 | 64 | 10 | 530 | 850 | 223 | 491 | 0 | 0 | 1078 | 0 | 0 | 0 | 0 | 0 | 0 | 0 | 0 | 0 | 0 | 0 | 0 | 0 | 0 | 0 | 0 | 0 | 0 | 0 | 0 |
| SRR5388144 | 120 | 2913 | 312 | 731 | 0 | 0 | 0 | 60 | 94 | 0 | 1070 | 1030 | 289 | 881 | 0 | 0 | 1410 | 0 | 0 | 0 | 0 | 0 | 0 | 0 | 0 | 0 | 0 | 0 | 0 | 0 | 0 | 0 | 0 | 0 | 0 | 0 | 0 |
| SRR5388145 | 403 | 6254 | 370 | 501 | 0 | 12 | 0 | 78 | 112 | 11 | 879 | 1258 | 315 | 1219 | 0 | 0 | 1425 | 0 | 0 | 0 | 0 | 0 | 0 | 0 | 0 | 0 | 0 | 0 | 0 | 0 | 0 | 0 | 0 | 0 | 0 | 0 | 0 |
| SRR5388146 | 246 | 3520 | 524 | 738 | 0 | 28 | 27 | 183 | 139 | 21 | 1676 | 1040 | 447 | 949 | 0 | 0 | 1879 | 0 | 0 | 0 | 0 | 0 | 0 | 0 | 0 | 0 | 0 | 0 | 0 | 0 | 0 | 0 | 0 | 0 | 0 | 0 | 0 |
| SRR5388147 | 659 | 3093 | 609 | 609 | 0 | 26 | 24 | 164 | 138 | 0 | 2635 | 2397 | 921 | 4296 | 0 | 25 | 2160 | 0 | 0 | 0 | 0 | 0 | 0 | 0 | 0 | 0 | 0 | 0 | 0 | 0 | 0 | 0 | 0 | 0 | 0 | 0 | 0 |
| SRR5388148 | 502 | 3398 | 465 | 665 | 0 | 28 | 24 | 169 | 160 | 0 | 4278 | 2674 | 977 | 3243 | 0 | 27 | 2202 | 0 | 0 | 0 | 0 | 0 | 0 | 0 | 0 | 0 | 0 | 0 | 0 | 0 | 0 | 0 | 0 | 0 | 0 | 0 | 0 |
| SRR5388149 | 501 | 2056 | 431 | 422 | 0 | 26 | 18 | 161 | 209 | 0 | 1763 | 1508 | 519 | 1626 | 0 | 0 | 3573 | 0 | 0 | 0 | 0 | 0 | 0 | 0 | 0 | 0 | 0 | 0 | 0 | 0 | 0 | 0 | 0 | 0 | 0 | 0 | 0 |
| SRR5388150 | 358 | 2261 | 541 | 724 | 0 | 14 | 17 | 87 | 190 | 0 | 2405 | 1613 | 463 | 1267 | 0 | 0 | 1829 | 0 | 0 | 0 | 0 | 0 | 0 | 0 | 0 | 0 | 0 | 0 | 0 | 0 | 0 | 0 | 0 | 0 | 0 | 0 | 0 |
| SRR5388151 | 335 | 2878 | 463 | 483 | 0 | 10 | 0 | 82 | 56 | 0 | 644 | 763 | 227 | 920 | 0 | 0 | 1930 | 0 | 0 | 0 | 0 | 0 | 0 | 0 | 0 | 0 | 0 | 0 | 0 | 0 | 0 | 0 | 0 | 0 | 0 | 0 | 0 |
| SRR5388152 | 874 | 2162 | 503 | 697 | 0 | 0 | 0 | 39 | 47 | 0 | 1100 | 1292 | 383 | 1534 | 0 | 0 | 2060 | 0 | 0 | 0 | 0 | 0 | 0 | 0 | 0 | 0 | 0 | 0 | 0 | 0 | 0 | 0 | 0 | 0 | 0 | 0 | 0 |
| SRR5388153 | 317 | 2467 | 300 | 867 | 0 | 0 | 0 | 62 | 90 | 0 | 922 | 808 | 302 | 1255 | 0 | 0 | 1550 | 0 | 0 | 0 | 0 | 0 | 0 | 0 | 0 | 0 | 0 | 0 | 0 | 0 | 0 | 0 | 0 | 0 | 0 | 0 | 0 |
| SRR5388154 | 661 | 2234 | 623 | 1124 | 0 | 0 | 0 | 64 | 52 | 0 | 1372 | 1549 | 520 | 3021 | 0 | 20 | 1858 | 0 | 0 | 0 | 0 | 0 | 0 | 0 | 0 | 0 | 0 | 0 | 0 | 0 | 0 | 0 | 0 | 0 | 0 | 0 | 0 |
| SRR5388155 | 1505 | 6327 | 700 | 553 | 0 | 12 | 0 | 72 | 101 | 0 | 706 | 1204 | 327 | 1464 | 0 | 0 | 1829 | 0 | 0 | 0 | 0 | 0 | 0 | 0 | 0 | 0 | 0 | 0 | 0 | 0 | 0 | 0 | 0 | 0 | 0 | 0 | 0 |
| SRR5388156 | 476 | 2520 | 295 | 918 | 0 | 0 | 0 | 87 | 91 | 0 | 1438 | 1361 | 469 | 2563 | 0 | 0 | 1404 | 0 | 0 | 0 | 0 | 0 | 0 | 0 | 0 | 0 | 0 | 0 | 0 | 0 | 0 | 0 | 0 | 0 | 0 | 0 | 0 |
| SRR5388157 | 458 | 4941 | 505 | 599 | 0 | 0 | 0 | 27 | 48 | 0 | 377 | 711 | 171 | 565 | 0 | 0 | 661 | 0 | 0 | 0 | 0 | 0 | 0 | 0 | 0 | 0 | 0 | 0 | 0 | 0 | 0 | 0 | 0 | 0 | 0 | 0 | 0 |
| SRR5388158 | 712 | 3503 | 819 | 463 | 0 | 0 | 0 | 29 | 27 | 12 | 301 | 279 | 73 | 164 | 0 | 0 | 744 | 0 | 0 | 0 | 0 | 0 | 0 | 0 | 0 | 0 | 0 | 0 | 0 | 0 | 0 | 0 | 0 | 0 | 0 | 0 | 0 |
| SRR5388159 | 199 | 3076 | 502 | 513 | 0 | 0 | 0 | 53 | 54 | 0 | 380 | 373 | 107 | 360 | 0 | 0 | 1353 | 0 | 0 | 0 | 0 | 0 | 0 | 0 | 0 | 0 | 0 | 0 | 0 | 0 | 0 | 0 | 0 | 0 | 0 | 0 | 0 |
| SRR5388160 | 474 | 1572 | 332 | 656 | 0 | 0 | 0 | 30 | 54 | 0 | 1103 | 1805 | 508 | 1874 | 0 | 0 | 1566 | 0 | 0 | 0 | 0 | 0 | 0 | 0 | 0 | 0 | 0 | 0 | 0 | 0 | 0 | 0 | 0 | 0 | 0 | 0 | 0 |
| SRR5388161 | 225 | 3503 | 275 | 306 | 0 | 0 | 0 | 33 | 49 | 0 | 460 | 661 | 153 | 413 | 0 | 0 | 1458 | 0 | 0 | 0 | 0 | 0 | 0 | 0 | 0 | 0 | 0 | 0 | 0 | 0 | 0 | 0 | 0 | 0 | 0 | 0 | 0 |
| SRR5388162 | 594 | 2406 | 641 | 762 | 0 | 0 | 0 | 54 | 99 | 0 | 348 | 574 | 140 | 783 | 0 | 0 | 1628 | 0 | 0 | 0 | 0 | 0 | 0 | 0 | 0 | 0 | 0 | 0 | 0 | 0 | 0 | 0 | 0 | 0 | 0 | 0 | 0 |
| SRR5388163 | 565 | 5074 | 776 | 500 | 0 | 0 | 0 | 46 | 108 | 0 | 510 | 686 | 149 | 378 | 0 | 0 | 3105 | 0 | 0 | 0 | 0 | 0 | 0 | 0 | 0 | 0 | 0 | 0 | 0 | 0 | 0 | 0 | 0 | 0 | 0 | 0 | 0 |
| SRR5388164 | 462 | 3810 | 600 | 1424 | 0 | 0 | 0 | 25 | 60 | 0 | 261 | 342 | 59 | 374 | 0 | 0 | 770 | 0 | 0 | 0 | 0 | 0 | 0 | 0 | 0 | 0 | 0 | 0 | 0 | 0 | 0 | 0 | 0 | 0 | 0 | 0 | 0 |
| SRR5388165 | 149 | 2019 | 202 | 1977 | 0 | 0 | 0 | 50 | 73 | 0 | 249 | 237 | 122 | 875 | 0 | 0 | 882 | 0 | 0 | 0 | 0 | 0 | 0 | 0 | 0 | 0 | 0 | 0 | 0 | 0 | 0 | 0 | 0 | 0 | 0 | 0 | 0 |
| SRR5388166 | 483 | 3170 | 965 | 790 | 0 | 0 | 0 | 27 | 22 | 0 | 338 | 277 | 93 | 522 | 0 | 0 | 541 | 0 | 0 | 0 | 0 | 0 | 0 | 0 | 0 | 0 | 0 | 0 | 0 | 0 | 0 | 0 | 0 | 0 | 0 | 0 | 0 |
| SRR5388167 | 79 | 1093 | 171 | 3023 | 0 | 0 | 0 | 37 | 201 | 0 | 294 | 275 | 76 | 527 | 0 | 0 | 1348 | 0 | 0 | 0 | 0 | 0 | 0 | 0 | 0 | 0 | 0 | 0 | 0 | 0 | 0 | 0 | 0 | 0 | 0 | 0 | 0 |
| SRR5388168 | 331 | 3428 | 925 | 390 | 0 | 19 | 16 | 115 | 174 | 0 | 2174 | 1490 | 493 | 1353 | 0 | 0 | 3088 | 0 | 0 | 0 | 0 | 0 | 0 | 0 | 0 | 0 | 0 | 0 | 0 | 0 | 0 | 0 | 0 | 0 | 0 | 0 | 0 |
| SRR5388169 | 363 | 2028 | 428 | 2130 | 0 | 0 | 0 | 15 | 68 | 0 | 275 | 293 | 63 | 655 | 0 | 0 | 675 | 0 | 0 | 0 | 13 | 0 | 0 | 0 | 0 | 0 | 0 | 0 | 0 | 0 | 0 | 0 | 0 | 0 | 0 | 0 | 0 |
| SRR5388170 | 613 | 1810 | 620 | 1321 | 0 | 0 | 0 | 24 | 43 | 0 | 643 | 338 | 110 | 923 | 0 | 13 | 1523 | 0 | 0 | 0 | 0 | 0 | 0 | 0 | 0 | 0 | 0 | 0 | 0 | 0 | 0 | 0 | 0 | 0 | 0 | 0 | 0 |
| SRR5388171 | 673 | 2436 | 755 | 1656 | 0 | 0 | 0 | 0 | 11 | 12 | 140 | 122 | 18 | 183 | 0 | 0 | 206 | 0 | 0 | 0 | 0 | 0 | 0 | 0 | 0 | 0 | 0 | 0 | 0 | 0 | 0 | 0 | 0 | 0 | 0 | 0 | 0 |
| SRR5388172 | 476 | 3516 | 626 | 2273 | 0 | 0 | 0 | 0 | 0 | 0 | 96 | 133 | 19 | 180 | 0 | 0 | 196 | 0 | 0 | 0 | 0 | 0 | 0 | 0 | 0 | 0 | 0 | 0 | 0 | 0 | 0 | 0 | 0 | 0 | 0 | 0 | 0 |
| SRR5388173 | 444 | 3089 | 503 | 1007 | 0 | 0 | 0 | 18 | 40 | 0 | 174 | 325 | 75 | 499 | 0 | 0 | 495 | 0 | 0 | 0 | 0 | 0 | 0 | 0 | 0 | 0 | 0 | 0 | 0 | 0 | 0 | 0 | 0 | 0 | 0 | 0 | 0 |
| SRR5388174 | 367 | 3463 | 612 | 1187 | 0 | 0 | 0 | 38 | 52 | 0 | 400 | 432 | 122 | 656 | 0 | 0 | 1094 | 0 | 0 | 0 | 0 | 0 | 0 | 0 | 0 | 0 | 0 | 0 | 0 | 0 | 0 | 0 | 0 | 0 | 0 | 0 | 0 |
| SRR5388175 | 492 | 2285 | 430 | 872 | 0 | 0 | 0 | 18 | 37 | 0 | 177 | 366 | 90 | 553 | 0 | 0 | 545 | 0 | 0 | 0 | 0 | 0 | 0 | 0 | 0 | 0 | 0 | 0 | 0 | 0 | 0 | 0 | 0 | 0 | 0 | 0 | 0 |
| SRR5388176 | 404 | 3179 | 414 | 1388 | 0 | 0 | 0 | 33 | 42 | 21 | 386 | 374 | 114 | 672 | 0 | 0 | 934 | 0 | 0 | 0 | 0 | 0 | 0 | 0 | 0 | 0 | 0 | 0 | 0 | 0 | 0 | 0 | 0 | 0 | 0 | 0 | 0 |
| SRR5388177 | 475 | 2914 | 425 | 951 | 0 | 0 | 0 | 20 | 105 | 0 | 158 | 225 | 50 | 410 | 0 | 0 | 1089 | 0 | 0 | 0 | 0 | 0 | 0 | 0 | 0 | 0 | 0 | 0 | 0 | 0 | 0 | 0 | 0 | 0 | 0 | 0 | 0 |
| SRR5388178 | 698 | 2574 | 676 | 1102 | 0 | 0 | 0 | 0 | 35 | 22 | 88 | 167 | 33 | 184 | 0 | 0 | 280 | 0 | 0 | 0 | 0 | 0 | 0 | 0 | 0 | 0 | 0 | 0 | 0 | 0 | 0 | 0 | 0 | 0 | 0 | 0 | 0 |
| SRR5388179 | 321 | 2327 | 413 | 1186 | 0 | 0 | 0 | 38 | 99 | 15 | 929 | 701 | 153 | 1139 | 0 | 0 | 1212 | 0 | 0 | 0 | 0 | 0 | 0 | 0 | 0 | 0 | 0 | 0 | 0 | 0 | 0 | 0 | 0 | 0 | 0 | 0 | 0 |
| SRR5388180 | 300 | 1869 | 417 | 1179 | 0 | 0 | 0 | 31 | 81 | 0 | 745 | 707 | 145 | 1237 | 0 | 0 | 1713 | 0 | 0 | 0 | 0 | 0 | 0 | 0 | 0 | 0 | 0 | 0 | 0 | 0 | 0 | 0 | 0 | 0 | 0 | 0 | 0 |
| SRR5388181 | 523 | 1909 | 431 | 564 | 0 | 16 | 13 | 133 | 235 | 21 | 640 | 651 | 226 | 877 | 0 | 0 | 1096 | 0 | 0 | 0 | 0 | 0 | 0 | 0 | 0 | 0 | 0 | 0 | 0 | 0 | 0 | 0 | 0 | 0 | 0 | 0 | 0 |
| SRR5388182 | 693 | 2595 | 604 | 934 | 0 | 12 | 0 | 75 | 120 | 0 | 917 | 573 | 212 | 930 | 0 | 0 | 1519 | 0 | 0 | 0 | 0 | 0 | 0 | 0 | 0 | 0 | 0 | 0 | 0 | 0 | 0 | 0 | 0 | 0 | 0 | 0 | 0 |
| SRR5388183 | 347 | 3203 | 476 | 707 | 0 | 13 | 0 | 82 | 214 | 0 | 586 | 516 | 145 | 775 | 0 | 0 | 1831 | 0 | 0 | 0 | 0 | 0 | 0 | 0 | 0 | 0 | 0 | 0 | 0 | 0 | 0 | 0 | 0 | 0 | 0 | 0 | 0 |
| SRR5388184 | 306 | 2840 | 1016 | 1234 | 0 | 0 | 0 | 44 | 50 | 0 | 348 | 309 | 96 | 589 | 0 | 0 | 968 | 0 | 0 | 0 | 0 | 0 | 0 | 0 | 0 | 0 | 0 | 0 | 0 | 0 | 0 | 0 | 0 | 0 | 0 | 0 | 0 |
| SRR5388185 | 678 | 1947 | 553 | 709 | 0 | 0 | 14 | 64 | 127 | 39 | 799 | 761 | 237 | 546 | 0 | 0 | 3216 | 0 | 0 | 0 | 0 | 0 | 0 | 0 | 0 | 0 | 0 | 0 | 0 | 0 | 0 | 0 | 0 | 0 | 0 | 0 | 0 |
| SRR5388186 | 785 | 3079 | 674 | 1228 | 0 | 0 | 0 | 10 | 27 | 17 | 54 | 105 | 23 | 175 | 0 | 0 | 800 | 0 | 0 | 0 | 0 | 0 | 0 | 0 | 0 | 0 | 0 | 0 | 0 | 0 | 0 | 0 | 0 | 0 | 0 | 0 | 0 |
| SRR5388187 | 714 | 4211 | 811 | 1137 | 0 | 20 | 13 | 116 | 170 | 0 | 825 | 523 | 152 | 830 | 0 | 0 | 1623 | 0 | 0 | 0 | 0 | 0 | 0 | 0 | 0 | 0 | 0 | 0 | 0 | 0 | 0 | 0 | 0 | 0 | 0 | 0 | 0 |
| SRR5388188 | 644 | 3402 | 622 | 990 | 0 | 13 | 0 | 72 | 125 | 0 | 739 | 488 | 162 | 909 | 0 | 0 | 1970 | 0 | 0 | 0 | 0 | 0 | 0 | 0 | 0 | 0 | 0 | 0 | 0 | 0 | 0 | 0 | 0 | 0 | 0 | 0 | 0 |
| SRR5388189 | 348 | 3085 | 743 | 946 | 0 | 0 | 0 | 29 | 51 | 0 | 304 | 228 | 60 | 423 | 0 | 0 | 838 | 0 | 0 | 0 | 0 | 0 | 0 | 0 | 0 | 0 | 0 | 0 | 0 | 0 | 0 | 0 | 0 | 0 | 0 | 0 | 0 |
| SRR5388190 | 295 | 2619 | 426 | 1096 | 0 | 0 | 0 | 38 | 77 | 0 | 275 | 285 | 97 | 959 | 0 | 0 | 931 | 0 | 0 | 0 | 0 | 0 | 0 | 0 | 0 | 0 | 0 | 0 | 0 | 0 | 0 | 0 | 0 | 0 | 0 | 0 | 0 |
| SRR5388191 | 578 | 6213 | 543 | 648 | 0 | 11 | 0 | 60 | 105 | 0 | 562 | 701 | 191 | 689 | 0 | 0 | 1698 | 0 | 0 | 0 | 0 | 0 | 0 | 0 | 0 | 0 | 0 | 0 | 0 | 0 | 0 | 0 | 0 | 0 | 0 | 0 | 0 |
| SRR5388192 | 235 | 3459 | 458 | 830 | 0 | 27 | 25 | 151 | 276 | 0 | 997 | 675 | 217 | 1235 | 0 | 0 | 2346 | 0 | 0 | 0 | 0 | 0 | 0 | 0 | 0 | 0 | 0 | 0 | 0 | 0 | 0 | 0 | 0 | 0 | 0 | 0 | 0 |
| SRR5388193 | 792 | 3246 | 784 | 1229 | 0 | 10 | 0 | 45 | 91 | 21 | 619 | 643 | 154 | 942 | 0 | 13 | 1626 | 0 | 0 | 0 | 0 | 0 | 0 | 0 | 0 | 0 | 0 | 0 | 0 | 0 | 0 | 0 | 0 | 0 | 0 | 0 | 0 |
| SRR5388194 | 289 | 3428 | 568 | 2171 | 0 | 0 | 0 | 38 | 90 | 0 | 588 | 550 | 175 | 1208 | 0 | 12 | 1755 | 0 | 0 | 0 | 0 | 0 | 0 | 0 | 0 | 0 | 0 | 0 | 0 | 0 | 0 | 0 | 0 | 0 | 0 | 0 | 0 |
| SRR5388195 | 476 | 2500 | 546 | 1081 | 0 | 0 | 0 | 61 | 171 | 0 | 601 | 551 | 149 | 998 | 0 | 0 | 1206 | 0 | 0 | 0 | 0 | 0 | 0 | 0 | 0 | 0 | 0 | 0 | 0 | 0 | 0 | 0 | 0 | 0 | 0 | 0 | 0 |
| SRR5388196 | 348 | 2126 | 465 | 707 | 0 | 0 | 0 | 77 | 138 | 0 | 954 | 969 | 295 | 1423 | 0 | 0 | 1721 | 0 | 0 | 0 | 0 | 0 | 0 | 0 | 0 | 0 | 0 | 0 | 0 | 0 | 0 | 0 | 0 | 0 | 0 | 0 | 0 |
| SRR5388197 | 656 | 3266 | 920 | 1555 | 0 | 0 | 0 | 26 | 66 | 14 | 293 | 242 | 73 | 577 | 0 | 10 | 826 | 0 | 0 | 0 | 0 | 0 | 0 | 0 | 0 | 0 | 0 | 0 | 0 | 0 | 0 | 0 | 0 | 0 | 0 | 0 | 0 |
| SRR5388198 | 676 | 2657 | 935 | 1513 | 0 | 0 | 0 | 33 | 69 | 0 | 414 | 275 | 88 | 633 | 0 | 11 | 1115 | 0 | 0 | 0 | 0 | 0 | 0 | 0 | 0 | 0 | 0 | 0 | 0 | 0 | 0 | 0 | 0 | 0 | 0 | 0 | 0 |
| SRR5388199 | 61 | 531 | 58 | 4496 | 0 | 17 | 12 | 85 | 353 | 0 | 444 | 362 | 156 | 925 | 0 | 0 | 2178 | 0 | 0 | 0 | 0 | 0 | 0 | 0 | 0 | 0 | 0 | 0 | 0 | 0 | 0 | 0 | 0 | 0 | 0 | 0 | 0 |
| SRR5388200 | 367 | 2270 | 354 | 1008 | 0 | 0 | 0 | 39 | 101 | 0 | 422 | 437 | 123 | 741 | 0 | 0 | 1408 | 0 | 0 | 0 | 0 | 0 | 0 | 0 | 0 | 0 | 0 | 0 | 0 | 0 | 0 | 0 | 0 | 0 | 0 | 0 | 0 |
| SRR5388201 | 630 | 2802 | 735 | 1098 | 0 | 0 | 0 | 12 | 22 | 0 | 164 | 218 | 41 | 273 | 0 | 0 | 355 | 0 | 0 | 0 | 0 | 0 | 0 | 0 | 0 | 0 | 0 | 0 | 0 | 0 | 0 | 0 | 0 | 0 | 0 | 0 | 0 |
| SRR5388202 | 545 | 3029 | 479 | 883 | 0 | 15 | 13 | 81 | 147 | 0 | 911 | 735 | 224 | 732 | 0 | 0 | 1437 | 0 | 0 | 0 | 0 | 0 | 0 | 0 | 0 | 0 | 0 | 0 | 0 | 0 | 0 | 0 | 0 | 0 | 0 | 0 | 0 |
| SRR5388203 | 483 | 4150 | 762 | 644 | 0 | 0 | 0 | 46 | 77 | 0 | 380 | 374 | 114 | 522 | 0 | 0 | 1289 | 0 | 0 | 0 | 0 | 0 | 0 | 0 | 0 | 0 | 0 | 0 | 0 | 0 | 0 | 0 | 0 | 0 | 0 | 0 | 0 |
| SRR5388204 | 648 | 3253 | 1073 | 743 | 0 | 0 | 0 | 36 | 39 | 0 | 322 | 281 | 83 | 500 | 0 | 0 | 725 | 0 | 0 | 0 | 0 | 0 | 0 | 0 | 0 | 0 | 0 | 0 | 0 | 0 | 0 | 0 | 0 | 0 | 0 | 0 | 0 |
| SRR5388205 | 691 | 2630 | 629 | 1390 | 0 | 0 | 0 | 0 | 13 | 0 | 125 | 169 | 35 | 360 | 0 | 0 | 411 | 0 | 0 | 0 | 0 | 0 | 0 | 0 | 0 | 0 | 0 | 0 | 0 | 0 | 0 | 0 | 0 | 0 | 0 | 0 | 0 |
| SRR5388206 | 648 | 5328 | 1154 | 1707 | 0 | 0 | 0 | 0 | 20 | 0 | 75 | 117 | 24 | 155 | 0 | 13 | 749 | 0 | 0 | 0 | 0 | 0 | 0 | 0 | 0 | 0 | 0 | 0 | 0 | 0 | 0 | 0 | 0 | 0 | 0 | 0 | 0 |
| SRR5388207 | 259 | 2490 | 380 | 744 | 0 | 0 | 0 | 45 | 91 | 0 | 363 | 360 | 125 | 464 | 0 | 0 | 1429 | 0 | 0 | 0 | 0 | 0 | 0 | 0 | 0 | 0 | 0 | 0 | 0 | 0 | 0 | 0 | 0 | 0 | 0 | 0 | 0 |
| SRR5388208 | 492 | 2386 | 529 | 1392 | 0 | 0 | 0 | 12 | 41 | 0 | 151 | 244 | 52 | 414 | 0 | 0 | 542 | 0 | 0 | 0 | 0 | 0 | 0 | 0 | 0 | 0 | 0 | 0 | 0 | 0 | 0 | 0 | 0 | 0 | 0 | 0 | 0 |
| SRR5388209 | 555 | 4229 | 612 | 2079 | 0 | 0 | 0 | 13 | 15 | 0 | 197 | 142 | 39 | 261 | 0 | 0 | 343 | 0 | 0 | 0 | 0 | 0 | 0 | 0 | 0 | 0 | 0 | 0 | 0 | 0 | 0 | 0 | 0 | 0 | 0 | 0 | 0 |
| SRR5388210 | 388 | 5347 | 949 | 2436 | 0 | 0 | 0 | 14 | 16 | 0 | 134 | 138 | 39 | 230 | 0 | 0 | 403 | 0 | 0 | 0 | 0 | 0 | 0 | 0 | 0 | 0 | 0 | 0 | 0 | 0 | 0 | 0 | 0 | 0 | 0 | 0 | 0 |
| SRR5388211 | 427 | 1687 | 725 | 439 | 0 | 0 | 0 | 42 | 72 | 0 | 411 | 334 | 108 | 860 | 0 | 0 | 996 | 0 | 0 | 0 | 0 | 0 | 0 | 0 | 0 | 0 | 0 | 0 | 0 | 0 | 0 | 0 | 0 | 0 | 0 | 0 | 0 |
| SRR5388212 | 282 | 1465 | 646 | 883 | 0 | 0 | 0 | 46 | 60 | 0 | 401 | 424 | 157 | 726 | 0 | 0 | 1396 | 0 | 0 | 0 | 0 | 0 | 0 | 0 | 0 | 0 | 0 | 0 | 0 | 0 | 0 | 0 | 0 | 0 | 0 | 0 | 0 |
| SRR5388213 | 450 | 4846 | 994 | 1342 | 0 | 0 | 0 | 0 | 12 | 16 | 64 | 105 | 23 | 274 | 0 | 0 | 311 | 0 | 0 | 0 | 0 | 0 | 0 | 0 | 0 | 0 | 0 | 0 | 0 | 0 | 0 | 0 | 0 | 0 | 0 | 0 | 0 |
| SRR5388214 | 634 | 6464 | 898 | 1426 | 0 | 0 | 0 | 0 | 23 | 11 | 82 | 88 | 21 | 196 | 0 | 0 | 501 | 0 | 0 | 0 | 0 | 0 | 0 | 0 | 0 | 0 | 0 | 0 | 0 | 0 | 0 | 0 | 0 | 0 | 0 | 0 | 0 |
| SRR5388215 | 281 | 2677 | 365 | 938 | 0 | 0 | 0 | 33 | 73 | 10 | 429 | 649 | 135 | 505 | 0 | 0 | 1209 | 0 | 0 | 0 | 0 | 0 | 0 | 0 | 0 | 0 | 0 | 0 | 0 | 0 | 0 | 0 | 0 | 0 | 0 | 0 | 0 |
| SRR5388216 | 875 | 2331 | 586 | 453 | 0 | 17 | 18 | 93 | 204 | 31 | 658 | 772 | 232 | 840 | 0 | 0 | 1917 | 0 | 0 | 0 | 0 | 0 | 0 | 0 | 0 | 0 | 0 | 0 | 0 | 0 | 0 | 0 | 0 | 0 | 0 | 0 | 0 |
| SRR5388217 | 144 | 2286 | 270 | 3618 | 0 | 0 | 0 | 25 | 31 | 0 | 111 | 171 | 63 | 575 | 0 | 0 | 600 | 0 | 0 | 0 | 0 | 0 | 0 | 0 | 0 | 0 | 0 | 0 | 0 | 0 | 0 | 0 | 0 | 0 | 0 | 0 | 0 |
| SRR5388218 | 106 | 2220 | 550 | 1419 | 0 | 0 | 0 | 44 | 76 | 0 | 206 | 202 | 72 | 435 | 0 | 0 | 824 | 0 | 0 | 0 | 0 | 0 | 0 | 0 | 0 | 0 | 0 | 0 | 0 | 0 | 0 | 0 | 0 | 0 | 0 | 0 | 0 |
| SRR5388219 | 479 | 1969 | 727 | 497 | 0 | 14 | 0 | 72 | 100 | 0 | 1006 | 1102 | 319 | 1115 | 0 | 0 | 2357 | 0 | 0 | 0 | 0 | 0 | 0 | 0 | 0 | 0 | 0 | 0 | 0 | 0 | 0 | 0 | 0 | 0 | 0 | 0 | 0 |
| SRR5388220 | 755 | 2653 | 661 | 341 | 0 | 0 | 0 | 27 | 30 | 13 | 387 | 593 | 179 | 526 | 0 | 0 | 1807 | 0 | 0 | 0 | 0 | 0 | 0 | 0 | 0 | 0 | 0 | 0 | 0 | 0 | 0 | 0 | 0 | 0 | 0 | 0 | 0 |
| SRR5388221 | 401 | 2226 | 523 | 452 | 0 | 0 | 0 | 62 | 99 | 0 | 444 | 514 | 189 | 665 | 0 | 0 | 1354 | 0 | 0 | 0 | 0 | 0 | 0 | 0 | 0 | 0 | 0 | 0 | 0 | 0 | 0 | 0 | 0 | 0 | 0 | 0 | 0 |
| SRR5388222 | 620 | 1810 | 610 | 376 | 0 | 14 | 0 | 79 | 176 | 0 | 631 | 780 | 196 | 510 | 0 | 0 | 1836 | 0 | 0 | 0 | 0 | 0 | 0 | 0 | 0 | 0 | 0 | 0 | 0 | 0 | 0 | 0 | 0 | 0 | 0 | 0 | 0 |
| SRR5388223 | 328 | 4261 | 415 | 743 | 0 | 0 | 0 | 64 | 134 | 0 | 467 | 616 | 152 | 405 | 0 | 0 | 2022 | 0 | 0 | 0 | 0 | 0 | 0 | 0 | 0 | 0 | 0 | 0 | 0 | 0 | 0 | 0 | 0 | 0 | 0 | 0 | 0 |
| SRR5388224 | 450 | 2157 | 429 | 634 | 0 | 0 | 0 | 69 | 135 | 10 | 608 | 731 | 262 | 698 | 0 | 0 | 2688 | 0 | 0 | 0 | 0 | 0 | 0 | 0 | 0 | 0 | 0 | 0 | 0 | 0 | 0 | 0 | 0 | 0 | 0 | 0 | 0 |
| SRR5388225 | 972 | 2178 | 855 | 1017 | 0 | 0 | 0 | 23 | 24 | 54 | 369 | 469 | 97 | 636 | 0 | 0 | 442 | 0 | 0 | 0 | 0 | 0 | 0 | 0 | 0 | 0 | 0 | 0 | 0 | 0 | 0 | 0 | 0 | 0 | 0 | 0 | 0 |
| SRR5388226 | 1582 | 2766 | 919 | 1610 | 0 | 0 | 0 | 19 | 26 | 19 | 473 | 739 | 199 | 1048 | 0 | 15 | 636 | 0 | 0 | 0 | 0 | 0 | 0 | 0 | 0 | 0 | 0 | 0 | 0 | 0 | 0 | 0 | 0 | 0 | 0 | 0 | 0 |
| SRR5388227 | 252 | 2031 | 412 | 956 | 0 | 0 | 0 | 24 | 56 | 0 | 236 | 525 | 100 | 606 | 0 | 0 | 1253 | 0 | 0 | 0 | 0 | 0 | 0 | 0 | 0 | 0 | 0 | 0 | 0 | 0 | 0 | 0 | 0 | 0 | 0 | 0 | 0 |
| SRR5388228 | 290 | 1328 | 377 | 514 | 0 | 11 | 0 | 73 | 147 | 0 | 762 | 1404 | 266 | 1317 | 0 | 0 | 1855 | 0 | 0 | 0 | 0 | 0 | 0 | 0 | 0 | 0 | 0 | 0 | 0 | 0 | 0 | 0 | 0 | 0 | 0 | 0 | 0 |
| SRR5388229 | 284 | 2044 | 328 | 1432 | 0 | 0 | 0 | 14 | 33 | 22 | 129 | 216 | 48 | 327 | 0 | 0 | 748 | 0 | 0 | 0 | 0 | 0 | 0 | 0 | 0 | 0 | 0 | 0 | 0 | 0 | 0 | 0 | 0 | 0 | 0 | 0 | 0 |
| SRR5388230 | 401 | 1538 | 486 | 1109 | 0 | 0 | 0 | 24 | 54 | 0 | 152 | 279 | 53 | 316 | 0 | 0 | 848 | 0 | 0 | 0 | 0 | 0 | 0 | 0 | 0 | 0 | 0 | 0 | 0 | 0 | 0 | 0 | 0 | 0 | 0 | 0 | 0 |
| SRR5388231 | 544 | 5528 | 717 | 1129 | 0 | 0 | 0 | 45 | 94 | 16 | 271 | 325 | 82 | 486 | 0 | 0 | 1153 | 0 | 0 | 0 | 0 | 0 | 0 | 0 | 0 | 0 | 0 | 0 | 0 | 0 | 0 | 0 | 0 | 0 | 0 | 0 | 0 |
| SRR5388232 | 376 | 2818 | 519 | 743 | 0 | 0 | 0 | 53 | 68 | 0 | 567 | 879 | 226 | 844 | 0 | 0 | 1566 | 0 | 0 | 0 | 0 | 0 | 0 | 0 | 0 | 0 | 0 | 0 | 0 | 0 | 0 | 0 | 0 | 0 | 0 | 0 | 0 |
| SRR5388233 | 146 | 1430 | 333 | 455 | 0 | 21 | 14 | 128 | 181 | 0 | 1115 | 1087 | 320 | 1102 | 0 | 23 | 3510 | 0 | 0 | 0 | 0 | 0 | 0 | 0 | 0 | 0 | 0 | 0 | 0 | 0 | 0 | 0 | 0 | 0 | 0 | 0 | 0 |
| SRR5388234 | 912 | 2137 | 896 | 640 | 0 | 0 | 0 | 26 | 42 | 32 | 217 | 374 | 104 | 691 | 0 | 11 | 1061 | 0 | 0 | 0 | 0 | 0 | 0 | 0 | 0 | 0 | 0 | 0 | 0 | 0 | 0 | 0 | 0 | 0 | 0 | 0 | 0 |
| SRR5388235 | 591 | 2129 | 563 | 568 | 0 | 17 | 13 | 117 | 94 | 14 | 1074 | 1100 | 345 | 1389 | 0 | 0 | 1846 | 0 | 0 | 0 | 0 | 0 | 0 | 0 | 0 | 0 | 0 | 0 | 0 | 0 | 0 | 0 | 0 | 0 | 0 | 0 | 0 |
| SRR5388236 | 375 | 1658 | 375 | 568 | 0 | 14 | 0 | 64 | 85 | 0 | 451 | 499 | 149 | 661 | 0 | 0 | 1417 | 0 | 0 | 0 | 0 | 0 | 0 | 0 | 0 | 0 | 0 | 0 | 0 | 0 | 0 | 0 | 0 | 0 | 0 | 0 | 0 |
| SRR5388237 | 237 | 1429 | 397 | 1513 | 0 | 21 | 12 | 108 | 121 | 13 | 730 | 800 | 241 | 1725 | 0 | 0 | 2042 | 0 | 0 | 0 | 0 | 0 | 0 | 0 | 0 | 0 | 0 | 0 | 0 | 0 | 0 | 0 | 0 | 0 | 0 | 0 | 0 |
| SRR5388238 | 538 | 1893 | 799 | 673 | 0 | 15 | 12 | 80 | 73 | 0 | 594 | 475 | 207 | 566 | 0 | 0 | 2155 | 0 | 0 | 0 | 0 | 0 | 0 | 0 | 0 | 0 | 0 | 0 | 0 | 0 | 0 | 0 | 0 | 0 | 0 | 0 | 0 |

### Supp Table S8.1.Columns_B_to_AL

| | | | | | | | | | | | | | | | | | | | | | | | | | | | | | | | | | | | | | |
|---|---|---|---|---|---|---|---|---|---|---|---|---|---|---|---|---|---|---|---|---|---|---|---|---|---|---|---|---|---|---|---|---|---|---|---|---|---|
| SRR5388239 | 319 | 4383 | 625 | 938 | 0 | 13 | 0 | 82 | 82 | 0 | 660 | 822 | 256 | 1337 | 0 | 0 | 1834 | 0 | 0 | 0 | 0 | 0 | 0 | 0 | 0 | 0 | 0 | 0 | 0 | 0 | 0 | 0 | 0 | 0 | 0 | 0 | 0 |
| SRR5388240 | 316 | 3684 | 703 | 527 | 0 | 12 | 12 | 93 | 86 | 0 | 739 | 745 | 299 | 419 | 0 | 0 | 2180 | 0 | 0 | 0 | 0 | 0 | 0 | 0 | 0 | 0 | 0 | 0 | 0 | 0 | 0 | 0 | 0 | 0 | 0 | 0 | 0 |
| SRR5388241 | 157 | 1555 | 318 | 723 | 0 | 12 | 10 | 98 | 169 | 0 | 623 | 841 | 229 | 2190 | 0 | 0 | 2376 | 0 | 0 | 0 | 12 | 0 | 0 | 0 | 0 | 0 | 0 | 0 | 0 | 0 | 0 | 0 | 0 | 0 | 0 | 0 | 0 |
| SRR5388242 | 211 | 1772 | 324 | 530 | 0 | 24 | 21 | 282 | 316 | 0 | 1081 | 1301 | 473 | 2799 | 0 | 0 | 3543 | 0 | 0 | 0 | 12 | 0 | 0 | 0 | 0 | 0 | 0 | 0 | 0 | 0 | 0 | 0 | 0 | 0 | 0 | 0 | 0 |
| SRR5388243 | 203 | 2223 | 468 | 793 | 0 | 11 | 0 | 102 | 86 | 0 | 1418 | 987 | 372 | 1493 | 0 | 0 | 2122 | 0 | 0 | 0 | 0 | 0 | 0 | 0 | 0 | 0 | 0 | 0 | 0 | 0 | 0 | 0 | 0 | 0 | 0 | 0 | 0 |
| SRR5388244 | 827 | 2429 | 684 | 633 | 0 | 0 | 0 | 39 | 38 | 20 | 684 | 868 | 254 | 286 | 0 | 0 | 2461 | 0 | 0 | 0 | 0 | 0 | 0 | 0 | 0 | 0 | 0 | 0 | 0 | 0 | 0 | 0 | 0 | 0 | 0 | 0 | 0 |
| SRR5388245 | 213 | 2383 | 457 | 1211 | 0 | 0 | 0 | 13 | 52 | 23 | 113 | 126 | 24 | 204 | 0 | 0 | 213 | 0 | 0 | 0 | 0 | 0 | 0 | 0 | 0 | 0 | 0 | 0 | 0 | 0 | 0 | 0 | 0 | 0 | 0 | 0 | 0 |
| SRR5388246 | 539 | 1478 | 568 | 1177 | 0 | 0 | 0 | 21 | 33 | 0 | 327 | 464 | 130 | 419 | 0 | 0 | 716 | 0 | 0 | 0 | 0 | 0 | 0 | 0 | 0 | 0 | 0 | 0 | 0 | 0 | 0 | 0 | 0 | 0 | 0 | 0 | 0 |
| SRR5388247 | 711 | 2154 | 696 | 1264 | 0 | 0 | 0 | 15 | 77 | 22 | 144 | 226 | 40 | 259 | 0 | 11 | 744 | 0 | 0 | 0 | 0 | 0 | 0 | 0 | 0 | 0 | 0 | 0 | 0 | 0 | 0 | 0 | 0 | 0 | 0 | 0 | 0 |
| SRR5388248 | 868 | 1832 | 1125 | 604 | 0 | 0 | 0 | 24 | 32 | 19 | 273 | 451 | 96 | 505 | 0 | 0 | 1002 | 0 | 0 | 0 | 0 | 0 | 0 | 0 | 0 | 0 | 0 | 0 | 0 | 0 | 0 | 0 | 0 | 0 | 0 | 0 | 0 |
| SRR5388249 | 265 | 4423 | 246 | 1210 | 0 | 0 | 0 | 10 | 30 | 0 | 157 | 384 | 61 | 445 | 0 | 0 | 1070 | 0 | 0 | 0 | 0 | 0 | 0 | 0 | 0 | 0 | 0 | 0 | 0 | 0 | 0 | 0 | 0 | 0 | 0 | 0 | 0 |
| SRR5388250 | 505 | 2007 | 426 | 444 | 0 | 0 | 0 | 34 | 60 | 12 | 273 | 440 | 102 | 560 | 0 | 0 | 1082 | 0 | 0 | 0 | 0 | 0 | 0 | 0 | 0 | 0 | 0 | 0 | 0 | 0 | 0 | 0 | 0 | 0 | 0 | 0 | 0 |
| SRR5388251 | 446 | 2162 | 779 | 630 | 0 | 0 | 0 | 80 | 97 | 0 | 448 | 284 | 103 | 661 | 0 | 0 | 1184 | 0 | 0 | 0 | 0 | 0 | 0 | 0 | 0 | 0 | 0 | 0 | 0 | 0 | 0 | 0 | 0 | 0 | 0 | 0 | 0 |
| SRR5388252 | 547 | 1776 | 665 | 623 | 0 | 0 | 0 | 57 | 73 | 17 | 452 | 567 | 188 | 812 | 0 | 0 | 1580 | 0 | 0 | 0 | 0 | 0 | 0 | 0 | 0 | 0 | 0 | 0 | 0 | 0 | 0 | 0 | 0 | 0 | 0 | 0 | 0 |
| SRR5388253 | 267 | 1961 | 515 | 555 | 0 | 18 | 12 | 160 | 192 | 0 | 1474 | 851 | 233 | 1134 | 0 | 0 | 1041 | 0 | 0 | 0 | 0 | 0 | 0 | 0 | 0 | 0 | 0 | 0 | 0 | 0 | 0 | 0 | 0 | 0 | 0 | 0 | 0 |
| SRR5388254 | 195 | 1834 | 443 | 470 | 0 | 17 | 15 | 117 | 211 | 0 | 836 | 705 | 289 | 434 | 0 | 0 | 2534 | 0 | 0 | 0 | 0 | 0 | 0 | 0 | 0 | 0 | 0 | 0 | 0 | 0 | 0 | 0 | 0 | 0 | 0 | 0 | 0 |
| SRR5388255 | 606 | 3626 | 334 | 2236 | 0 | 0 | 0 | 0 | 18 | 21 | 59 | 144 | 20 | 155 | 0 | 0 | 205 | 0 | 0 | 0 | 0 | 0 | 0 | 0 | 0 | 0 | 0 | 0 | 0 | 0 | 0 | 0 | 0 | 0 | 0 | 0 | 0 |
| SRR5388256 | 442 | 2612 | 608 | 885 | 0 | 0 | 0 | 15 | 34 | 0 | 167 | 263 | 54 | 357 | 0 | 0 | 391 | 0 | 0 | 0 | 0 | 0 | 0 | 0 | 0 | 0 | 0 | 0 | 0 | 0 | 0 | 0 | 0 | 0 | 0 | 0 | 0 |
| SRR5388257 | 400 | 2262 | 260 | 909 | 0 | 0 | 0 | 0 | 11 | 0 | 82 | 195 | 28 | 207 | 0 | 0 | 308 | 0 | 0 | 0 | 0 | 0 | 0 | 0 | 0 | 0 | 0 | 0 | 0 | 0 | 0 | 0 | 0 | 0 | 0 | 0 | 0 |
| SRR5388258 | 1069 | 2730 | 1080 | 778 | 0 | 0 | 0 | 19 | 47 | 23 | 207 | 214 | 59 | 443 | 0 | 22 | 1216 | 0 | 0 | 0 | 0 | 0 | 0 | 0 | 0 | 0 | 0 | 0 | 0 | 0 | 0 | 0 | 0 | 0 | 0 | 0 | 0 |
| SRR5388259 | 703 | 2463 | 832 | 1051 | 0 | 0 | 0 | 37 | 69 | 0 | 327 | 496 | 103 | 626 | 0 | 0 | 983 | 0 | 0 | 0 | 0 | 0 | 0 | 0 | 0 | 0 | 0 | 0 | 0 | 0 | 0 | 0 | 0 | 0 | 0 | 0 | 0 |
| SRR5388260 | 784 | 2663 | 822 | 680 | 0 | 0 | 0 | 28 | 33 | 0 | 197 | 408 | 78 | 1182 | 0 | 0 | 926 | 0 | 0 | 0 | 0 | 0 | 0 | 0 | 0 | 0 | 0 | 0 | 0 | 0 | 0 | 0 | 0 | 0 | 0 | 0 | 0 |
| SRR5388261 | 358 | 3100 | 662 | 662 | 0 | 0 | 0 | 54 | 64 | 0 | 384 | 366 | 95 | 675 | 0 | 15 | 945 | 0 | 0 | 0 | 0 | 0 | 0 | 0 | 0 | 0 | 0 | 0 | 0 | 0 | 0 | 0 | 0 | 0 | 0 | 0 | 0 |
| SRR5388262 | 760 | 3591 | 709 | 684 | 0 | 15 | 13 | 75 | 69 | 0 | 573 | 712 | 280 | 1083 | 0 | 23 | 2095 | 0 | 0 | 0 | 0 | 0 | 0 | 0 | 0 | 0 | 0 | 0 | 0 | 0 | 0 | 0 | 0 | 0 | 0 | 0 | 0 |
| SRR5388263 | 440 | 3695 | 626 | 870 | 0 | 0 | 0 | 17 | 45 | 0 | 289 | 548 | 107 | 387 | 0 | 0 | 1064 | 0 | 0 | 0 | 0 | 0 | 0 | 0 | 0 | 0 | 0 | 0 | 0 | 0 | 0 | 0 | 0 | 0 | 0 | 0 | 0 |
| SRR5388264 | 494 | 2411 | 527 | 763 | 0 | 0 | 0 | 33 | 42 | 22 | 251 | 400 | 106 | 345 | 0 | 0 | 1347 | 0 | 0 | 0 | 0 | 0 | 0 | 0 | 0 | 0 | 0 | 0 | 0 | 0 | 0 | 0 | 0 | 0 | 0 | 0 | 0 |
| SRR5388265 | 365 | 2606 | 712 | 2097 | 0 | 0 | 0 | 0 | 12 | 0 | 46 | 110 | 16 | 107 | 0 | 16 | 159 | 0 | 0 | 0 | 12 | 0 | 0 | 0 | 0 | 0 | 0 | 0 | 0 | 0 | 0 | 0 | 0 | 0 | 0 | 0 | 0 |
| SRR5388266 | 1134 | 1868 | 563 | 849 | 0 | 0 | 0 | 0 | 19 | 15 | 58 | 190 | 30 | 100 | 0 | 0 | 268 | 0 | 0 | 0 | 0 | 0 | 0 | 0 | 0 | 0 | 0 | 0 | 0 | 0 | 0 | 0 | 0 | 0 | 0 | 0 | 0 |
| SRR5388267 | 552 | 1723 | 762 | 916 | 0 | 12 | 12 | 76 | 100 | 0 | 571 | 378 | 141 | 494 | 0 | 28 | 2027 | 0 | 0 | 0 | 0 | 0 | 0 | 0 | 0 | 0 | 0 | 0 | 0 | 0 | 0 | 0 | 0 | 0 | 0 | 0 | 0 |
| SRR5388268 | 297 | 2367 | 398 | 1029 | 0 | 0 | 0 | 41 | 87 | 0 | 336 | 468 | 113 | 649 | 0 | 0 | 943 | 0 | 0 | 0 | 0 | 0 | 0 | 0 | 0 | 0 | 0 | 0 | 0 | 0 | 0 | 0 | 0 | 0 | 0 | 0 | 0 |
| SRR5388269 | 636 | 1530 | 534 | 903 | 0 | 11 | 0 | 46 | 40 | 0 | 796 | 1108 | 364 | 2758 | 0 | 23 | 1817 | 0 | 0 | 0 | 0 | 0 | 0 | 0 | 0 | 0 | 0 | 0 | 0 | 0 | 0 | 0 | 0 | 0 | 0 | 0 | 0 |
| SRR5388270 | 408 | 2429 | 909 | 1539 | 0 | 0 | 0 | 0 | 23 | 0 | 67 | 94 | 20 | 116 | 0 | 19 | 423 | 0 | 0 | 0 | 0 | 0 | 0 | 0 | 0 | 0 | 0 | 0 | 0 | 0 | 0 | 0 | 0 | 0 | 0 | 0 | 0 |
| SRR5388271 | 712 | 4241 | 1157 | 801 | 0 | 0 | 0 | 38 | 84 | 30 | 578 | 866 | 199 | 599 | 0 | 0 | 1322 | 0 | 0 | 0 | 0 | 0 | 0 | 0 | 0 | 0 | 0 | 0 | 0 | 0 | 0 | 0 | 0 | 0 | 0 | 0 | 0 |
| SRR5388272 | 369 | 3415 | 437 | 1135 | 0 | 0 | 0 | 41 | 120 | 0 | 440 | 626 | 125 | 679 | 0 | 0 | 1098 | 0 | 0 | 0 | 0 | 0 | 0 | 0 | 0 | 0 | 0 | 0 | 0 | 0 | 0 | 0 | 0 | 0 | 0 | 0 | 0 |
| SRR5388273 | 498 | 2419 | 611 | 932 | 0 | 0 | 0 | 21 | 57 | 0 | 253 | 505 | 92 | 597 | 0 | 0 | 918 | 0 | 0 | 0 | 0 | 0 | 0 | 0 | 0 | 0 | 0 | 0 | 0 | 0 | 0 | 0 | 0 | 0 | 0 | 0 | 0 |
| SRR5388274 | 366 | 2084 | 337 | 590 | 0 | 0 | 0 | 67 | 146 | 0 | 433 | 1009 | 215 | 1108 | 0 | 0 | 1591 | 0 | 0 | 0 | 0 | 0 | 0 | 0 | 0 | 0 | 0 | 0 | 0 | 0 | 0 | 0 | 0 | 0 | 0 | 0 | 0 |
| SRR5388275 | 440 | 2688 | 328 | 1030 | 0 | 0 | 0 | 10 | 24 | 0 | 255 | 653 | 100 | 523 | 0 | 0 | 945 | 0 | 0 | 0 | 0 | 0 | 0 | 0 | 0 | 0 | 0 | 0 | 0 | 0 | 0 | 0 | 0 | 0 | 0 | 0 | 0 |
| SRR5388276 | 345 | 1359 | 385 | 793 | 0 | 11 | 0 | 83 | 100 | 11 | 462 | 468 | 149 | 955 | 0 | 0 | 1451 | 0 | 0 | 0 | 0 | 0 | 0 | 0 | 0 | 0 | 0 | 0 | 0 | 0 | 0 | 0 | 0 | 0 | 0 | 0 | 0 |
| SRR5388277 | 306 | 1530 | 377 | 398 | 0 | 0 | 0 | 33 | 58 | 0 | 289 | 559 | 150 | 960 | 0 | 0 | 759 | 0 | 0 | 0 | 0 | 0 | 0 | 0 | 0 | 0 | 0 | 0 | 0 | 0 | 0 | 0 | 0 | 0 | 0 | 0 | 0 |
| SRR5388278 | 342 | 2299 | 583 | 800 | 0 | 0 | 0 | 39 | 73 | 0 | 329 | 482 | 132 | 857 | 0 | 0 | 1169 | 0 | 0 | 0 | 0 | 0 | 0 | 0 | 0 | 0 | 0 | 0 | 0 | 0 | 0 | 0 | 0 | 0 | 0 | 0 | 0 |
| SRR5388279 | 409 | 2643 | 837 | 1744 | 0 | 0 | 0 | 0 | 28 | 18 | 74 | 134 | 19 | 133 | 0 | 0 | 455 | 0 | 0 | 0 | 0 | 0 | 0 | 0 | 0 | 0 | 0 | 0 | 0 | 0 | 0 | 0 | 0 | 0 | 0 | 0 | 0 |
| SRR5388280 | 308 | 1453 | 589 | 1065 | 0 | 0 | 0 | 12 | 24 | 0 | 91 | 185 | 32 | 168 | 0 | 0 | 425 | 0 | 0 | 0 | 0 | 0 | 0 | 0 | 0 | 0 | 0 | 0 | 0 | 0 | 0 | 0 | 0 | 0 | 0 | 0 | 0 |
| SRR5388281 | 1065 | 10456 | 920 | 1476 | 0 | 0 | 0 | 33 | 44 | 0 | 287 | 463 | 111 | 380 | 0 | 0 | 1311 | 0 | 0 | 0 | 0 | 0 | 0 | 0 | 0 | 0 | 0 | 0 | 0 | 0 | 0 | 0 | 0 | 0 | 0 | 0 | 0 |
| SRR5388282 | 1207 | 2826 | 994 | 497 | 0 | 0 | 0 | 11 | 20 | 14 | 183 | 390 | 101 | 223 | 0 | 0 | 1155 | 0 | 0 | 0 | 0 | 0 | 0 | 0 | 0 | 0 | 0 | 0 | 0 | 0 | 0 | 0 | 0 | 0 | 0 | 0 | 0 |
| SRR5388283 | 766 | 3367 | 649 | 1363 | 0 | 0 | 0 | 0 | 13 | 0 | 53 | 174 | 21 | 209 | 0 | 0 | 271 | 0 | 0 | 0 | 0 | 0 | 0 | 0 | 0 | 0 | 0 | 0 | 0 | 0 | 0 | 0 | 0 | 0 | 0 | 0 | 0 |
| SRR5388284 | 885 | 2004 | 876 | 666 | 0 | 0 | 0 | 26 | 35 | 19 | 391 | 592 | 142 | 844 | 0 | 0 | 920 | 0 | 0 | 0 | 0 | 0 | 0 | 0 | 0 | 0 | 0 | 0 | 0 | 0 | 0 | 0 | 0 | 0 | 0 | 0 | 0 |
| SRR5388285 | 420 | 3420 | 266 | 1205 | 0 | 0 | 0 | 46 | 150 | 0 | 271 | 389 | 112 | 804 | 0 | 0 | 1669 | 0 | 0 | 0 | 0 | 0 | 0 | 0 | 0 | 0 | 0 | 0 | 0 | 0 | 0 | 0 | 0 | 0 | 0 | 0 | 0 |
| SRR5388286 | 370 | 4082 | 547 | 1080 | 0 | 0 | 0 | 35 | 124 | 0 | 277 | 342 | 76 | 469 | 0 | 0 | 1072 | 0 | 0 | 0 | 0 | 0 | 0 | 0 | 0 | 0 | 0 | 0 | 0 | 0 | 0 | 0 | 0 | 0 | 0 | 0 | 0 |
| SRR5388287 | 791 | 5342 | 876 | 842 | 0 | 0 | 0 | 18 | 54 | 0 | 130 | 124 | 35 | 247 | 0 | 13 | 880 | 0 | 0 | 0 | 0 | 0 | 0 | 0 | 0 | 0 | 0 | 0 | 0 | 0 | 0 | 0 | 0 | 0 | 0 | 0 | 0 |
| SRR5388288 | 266 | 2016 | 393 | 1185 | 0 | 0 | 0 | 38 | 87 | 0 | 202 | 271 | 68 | 428 | 0 | 21 | 1221 | 0 | 0 | 0 | 0 | 0 | 0 | 0 | 0 | 0 | 0 | 0 | 0 | 0 | 0 | 0 | 0 | 0 | 0 | 0 | 0 |
| SRR5388289 | 605 | 3653 | 803 | 1861 | 0 | 12 | 0 | 57 | 111 | 0 | 705 | 491 | 182 | 1113 | 0 | 0 | 1269 | 0 | 0 | 0 | 14 | 0 | 0 | 0 | 0 | 0 | 0 | 0 | 0 | 0 | 0 | 0 | 0 | 0 | 0 | 0 | 0 |
| SRR5388290 | 722 | 2222 | 541 | 627 | 0 | 11 | 0 | 59 | 92 | 0 | 806 | 1028 | 296 | 817 | 0 | 0 | 1509 | 0 | 0 | 0 | 0 | 0 | 0 | 0 | 0 | 0 | 0 | 0 | 0 | 0 | 0 | 0 | 0 | 0 | 0 | 0 | 0 |
| SRR5388291 | 554 | 5398 | 585 | 1091 | 0 | 0 | 0 | 57 | 126 | 12 | 661 | 487 | 142 | 1072 | 0 | 0 | 1563 | 0 | 0 | 0 | 0 | 0 | 0 | 0 | 0 | 0 | 0 | 0 | 0 | 0 | 0 | 0 | 0 | 0 | 0 | 0 | 0 |
| SRR5388292 | 609 | 4514 | 399 | 760 | 0 | 11 | 11 | 79 | 172 | 0 | 623 | 917 | 313 | 852 | 0 | 0 | 1989 | 0 | 0 | 0 | 0 | 0 | 0 | 0 | 0 | 0 | 0 | 0 | 0 | 0 | 0 | 0 | 0 | 0 | 0 | 0 | 0 |
| SRR5388293 | 284 | 2627 | 429 | 745 | 0 | 0 | 0 | 62 | 95 | 0 | 559 | 479 | 127 | 1025 | 0 | 0 | 1216 | 0 | 0 | 0 | 0 | 0 | 0 | 0 | 0 | 0 | 0 | 0 | 0 | 0 | 0 | 0 | 0 | 0 | 0 | 0 | 0 |
| SRR5388294 | 368 | 2449 | 499 | 657 | 0 | 0 | 0 | 41 | 69 | 0 | 347 | 423 | 118 | 595 | 0 | 0 | 1319 | 0 | 0 | 0 | 0 | 0 | 0 | 0 | 0 | 0 | 0 | 0 | 0 | 0 | 0 | 0 | 0 | 0 | 0 | 0 | 0 |
| SRR5476966 | 0 | 2723 | 0 | 0 | 64 | 160 | 352 | 96 | 2111 | 0 | 1023 | 3934 | 320 | 64 | 96 | 1535 | 2495 | 0 | 0 | 0 | 43 | 0 | 0 | 0 | 0 | 0 | 0 | 0 | 0 | 0 | 0 | 0 | 0 | 0 | 0 | 0 | 0 |
| SRR5476967 | 0 | 5410 | 0 | 902 | 0 | 140 | 140 | 47 | 981 | 0 | 1355 | 2569 | 280 | 47 | 0 | 1075 | 2850 | 0 | 0 | 0 | 103 | 26 | 0 | 0 | 52 | 0 | 0 | 26 | 103 | 52 | 0 | 0 | 0 | 0 | 26 | 26 | 0 |
| SRR5476969 | 933 | 7463 | 933 | 0 | 0 | 0 | 0 | 0 | 0 | 0 | 0 | 27778 | 0 | 0 | 0 | 0 | 0 | 0 | 0 | 0 | 833 | 139 | 139 | 0 | 0 | 0 | 0 | 0 | 0 | 278 | 278 | 139 | 0 | 0 | 139 | 0 | 0 |
| SRR5517463 | 355 | 1187 | 27508 | 381 | 20033 | 24197 | 494051 | 17607 | 1047 | 201 | 11231 | 14 | 5225 | 28498 | 3603 | 45721 | 17230 | 0 | 0 | 0 | 0 | 0 | 0 | 0 | 0 | 0 | 0 | 0 | 0 | 0 | 0 | 0 | 0 | 0 | 0 | 0 | 0 |
| SRR5517464 | 1162 | 2001 | 16865 | 1794 | 21198 | 29588 | 405954 | 22863 | 1248 | 188 | 13401 | 15 | 7708 | 31097 | 3995 | 53915 | 17339 | 0 | 0 | 0 | 17 | 0 | 15 | 0 | 0 | 0 | 0 | 0 | 0 | 0 | 0 | 0 | 0 | 0 | 0 | 0 | 0 |
| SRR5517465 | 2568 | 3481 | 15350 | 5291 | 22470 | 27914 | 459852 | 19406 | 1138 | 131 | 12244 | 14 | 6102 | 30322 | 4050 | 48950 | 16667 | 0 | 0 | 0 | 25 | 13 | 25 | 0 | 0 | 0 | 0 | 0 | 14 | 0 | 0 | 0 | 0 | 0 | 0 | 0 | 0 |
| SRR5517466 | 632 | 1543 | 19387 | 769 | 21881 | 26603 | 471061 | 23805 | 1116 | 173 | 12397 | 17 | 5522 | 29030 | 3451 | 51345 | 17270 | 0 | 0 | 0 | 0 | 0 | 13 | 0 | 0 | 0 | 0 | 0 | 0 | 0 | 0 | 0 | 0 | 0 | 0 | 0 | 0 |
| SRR5517467 | 594 | 2825 | 25445 | 2113 | 17081 | 29973 | 425267 | 21706 | 1567 | 147 | 13169 | 11665 | 4699 | 23834 | 1804 | 46438 | 20328 | 0 | 0 | 13 | 41 | 21 | 73 | 10 | 11 | 0 | 0 | 0 | 19 | 13 | 0 | 0 | 0 | 0 | 0 | 0 | 0 |
| SRR5517468 | 645 | 3122 | 22685 | 2011 | 20559 | 33727 | 444570 | 27742 | 1776 | 136 | 12986 | 8756 | 6348 | 23651 | 1859 | 47574 | 17004 | 0 | 11 | 15 | 52 | 20 | 58 | 0 | 0 | 0 | 0 | 13 | 23 | 22 | 0 | 0 | 0 | 0 | 0 | 0 | 0 |
| SRR5517469 | 352 | 3074 | 35515 | 609 | 17967 | 32546 | 478587 | 22085 | 1760 | 84 | 12418 | 9618 | 4270 | 22363 | 1555 | 44506 | 16375 | 0 | 0 | 0 | 25 | 11 | 48 | 0 | 0 | 0 | 0 | 0 | 0 | 0 | 0 | 0 | 0 | 0 | 0 | 0 | 0 |
| SRR5517470 | 1008 | 4174 | 26410 | 3301 | 22073 | 38158 | 453348 | 25067 | 2183 | 80 | 12697 | 14 | 4872 | 25048 | 1831 | 47790 | 16181 | 0 | 0 | 11 | 40 | 20 | 51 | 0 | 0 | 0 | 0 | 11 | 22 | 17 | 0 | 0 | 0 | 0 | 0 | 0 | 0 |
| SRR5575011 | 35 | 999 | 95 | 1706 | 10515 | 27048 | 47043 | 19386 | 12 | 52 | 3328 | 852 | 1582 | 2055 | 75 | 16 | 4376 | 0 | 0 | 0 | 10 | 82 | 258 | 111 | 43 | 0 | 0 | 0 | 0 | 0 | 15 | 83 | 284 | 2642 | 61 | 61 | 38 |
| SRR5575012 | 47 | 845 | 73 | 1318 | 11350 | 33229 | 38861 | 9405 | 12 | 109 | 1582 | 650 | 1077 | 2052 | 69 | 23 | 3068 | 0 | 0 | 0 | 10 | 88 | 355 | 144 | 59 | 0 | 0 | 0 | 0 | 0 | 24 | 139 | 478 | 3900 | 81 | 61 | 51 |
| SRR5575013 | 211 | 439 | 131 | 1692 | 10012 | 24538 | 53562 | 20369 | 20 | 131 | 1952 | 722 | 1374 | 1851 | 49 | 14 | 2228 | 0 | 0 | 0 | 0 | 70 | 120 | 49 | 28 | 0 | 0 | 0 | 0 | 0 | 27 | 59 | 194 | 1393 | 42 | 31 | 26 |
| SRR5575014 | 59 | 1884 | 124 | 1460 | 9351 | 24456 | 58370 | 21648 | 12 | 100 | 2348 | 649 | 1687 | 1953 | 78 | 19 | 3253 | 0 | 0 | 0 | 13 | 69 | 309 | 112 | 51 | 0 | 0 | 0 | 0 | 13 | 32 | 150 | 382 | 2987 | 74 | 57 | 41 |
| SRR5575015 | 58 | 1091 | 144 | 1360 | 10927 | 24818 | 64956 | 27606 | 13 | 108 | 1878 | 657 | 1416 | 1697 | 54 | 15 | 2926 | 0 | 0 | 0 | 21 | 78 | 390 | 134 | 69 | 0 | 0 | 0 | 0 | 27 | 33 | 202 | 488 | 3394 | 69 | 57 | 51 |
| SRR5575016 | 220 | 874 | 139 | 1678 | 9902 | 24963 | 46393 | 17012 | 10 | 154 | 1486 | 542 | 1113 | 1765 | 35 | 18 | 2232 | 0 | 0 | 0 | 0 | 77 | 140 | 67 | 30 | 0 | 0 | 0 | 0 | 0 | 27 | 66 | 206 | 1542 | 54 | 26 | 25 |
| SRR5575017 | 42 | 811 | 126 | 1379 | 9692 | 24237 | 58914 | 24473 | 10 | 82 | 2102 | 659 | 1545 | 1798 | 70 | 25 | 3281 | 0 | 0 | 0 | 12 | 65 | 258 | 110 | 53 | 0 | 0 | 0 | 0 | 12 | 24 | 112 | 344 | 2642 | 62 | 49 | 38 |
| SRR5575018 | 90 | 1049 | 118 | 1400 | 11694 | 21405 | 80269 | 40956 | 16 | 84 | 2582 | 814 | 1932 | 1812 | 60 | 23 | 3739 | 0 | 0 | 0 | 34 | 89 | 426 | 146 | 78 | 0 | 0 | 0 | 0 | 34 | 50 | 267 | 465 | 3148 | 62 | 51 | 37 |
| SRR5575019 | 240 | 1118 | 243 | 2340 | 7984 | 23302 | 46547 | 17835 | 0 | 138 | 1435 | 701 | 1098 | 1856 | 42 | 16 | 1543 | 0 | 0 | 0 | 0 | 88 | 135 | 55 | 26 | 0 | 0 | 0 | 0 | 0 | 25 | 57 | 172 | 1701 | 51 | 31 | 29 |
| SRR5644067 | 138 | 49588 | 834 | 1176 | 1156 | 13893 | 8770 | 21962 | 49460 | 45 | 24453 | 25347 | 5206 | 719 | 15 | 3317 | 13219 | 0 | 0 | 31 | 4103 | 564 | 192 | 0 | 0 | 0 | 0 | 19 | 1756 | 3919 | 2427 | 414 | 0 | 14 | 68 | 13 | 0 |
| SRR5644068 | 91 | 9585 | 396 | 2602 | 1496 | 14597 | 15358 | 19656 | 17220 | 91 | 8367 | 10315 | 3228 | 2676 | 40 | 3344 | 105731 | 0 | 0 | 0 | 4824 | 11588 | 822 | 20 | 0 | 0 | 0 | 115 | 342 | 7348 | 9836 | 1733 | 11 | 27 | 42 | 30 | 0 |
| SRR5644069 | 86 | 14952 | 1038 | 173 | 1139 | 17551 | 8903 | 15845 | 121178 | 126 | 16406 | 22743 | 5933 | 865 | 0 | 5064 | 37985 | 0 | 0 | 0 | 2617 | 1203 | 194 | 0 | 0 | 0 | 0 | 70 | 728 | 7140 | 9943 | 1011 | 0 | 13 | 28 | 0 | 0 |
| SRR5644070 | 89 | 14042 | 337 | 3026 | 436 | 7430 | 3613 | 16630 | 18004 | 10 | 17980 | 27451 | 8986 | 818 | 0 | 5048 | 34278 | 0 | 0 | 0 | 362 | 356 | 82 | 0 | 0 | 0 | 0 | 21 | 231 | 827 | 1468 | 323 | 0 | 13 | 106 | 29 | 0 |
| SRR5644071 | 152 | 15916 | 2390 | 94 | 524 | 6795 | 4816 | 6913 | 32045 | 85 | 18919 | 25793 | 10671 | 7299 | 0 | 8906 | 54620 | 0 | 0 | 0 | 473 | 2898 | 181 | 0 | 0 | 0 | 0 | 0 | 214 | 413 | 2371 | 289 | 0 | 15 | 0 | 0 | 0 |
| SRR5644072 | 150 | 16898 | 561 | 2844 | 3803 | 18258 | 36620 | 71056 | 7425 | 41 | 11881 | 14236 | 6491 | 6631 | 11 | 3669 | 46222 | 0 | 0 | 0 | 898 | 2144 | 556 | 0 | 0 | 0 | 0 | 0 | 212 | 763 | 3803 | 925 | 0 | 14 | 35 | 27 | 0 |
| SRR5644073 | 153 | 5928 | 220 | 2149 | 520 | 4942 | 2521 | 5531 | 52582 | 32 | 10613 | 16461 | 4158 | 1003 | 0 | 6673 | 45109 | 0 | 0 | 27 | 3109 | 704 | 212 | 0 | 0 | 0 | 0 | 41 | 3213 | 7036 | 5911 | 807 | 0 | 24 | 181 | 26 | 0 |
| SRR5644074 | 98 | 6466 | 1647 | 250 | 496 | 7150 | 2720 | 8290 | 36606 | 56 | 9050 | 14877 | 4804 | 656 | 0 | 6771 | 45739 | 0 | 0 | 0 | 376 | 418 | 78 | 0 | 0 | 0 | 0 | 0 | 226 | 900 | 2258 | 445 | 0 | 37 | 39 | 0 | 0 |
| SRR5644075 | 78 | 5520 | 270 | 563 | 693 | 5063 | 4259 | 12241 | 26697 | 12 | 12804 | 18782 | 4803 | 2070 | 0 | 7125 | 48026 | 0 | 0 | 51 | 3489 | 822 | 173 | 0 | 0 | 0 | 0 | 111 | 4082 | 5546 | 4276 | 538 | 0 | 0 | 39 | 0 | 0 |
| SRR5644076 | 0 | 2376 | 936 | 258 | 553 | 5731 | 3104 | 8710 | 55488 | 85 | 9933 | 14431 | 4955 | 1161 | 0 | 4073 | 58958 | 0 | 0 | 0 | 767 | 388 | 85 | 0 | 0 | 0 | 0 | 88 | 990 | 2456 | 3128 | 586 | 0 | 17 | 57 | 21 | 0 |
| SRR5644077 | 35 | 6856 | 1896 | 330 | 1329 | 9484 | 9192 | 18844 | 29402 | 207 | 10545 | 15978 | 4613 | 2060 | 0 | 8051 | 43633 | 0 | 0 | 0 | 753 | 720 | 110 | 0 | 0 | 0 | 0 | 20 | 466 | 1109 | 2637 | 280 | 0 | 13 | 11 | 0 | 0 |
| SRR5644078 | 94 | 5300 | 767 | 149 | 565 | 7797 | 4195 | 10434 | 57667 | 61 | 8389 | 11043 | 4098 | 922 | 0 | 6012 | 51493 | 0 | 0 | 0 | 892 | 515 | 95 | 0 | 0 | 0 | 0 | 54 | 839 | 2243 | 2849 | 546 | 0 | 18 | 23 | 0 | 0 |
| SRR5644085 | 104 | 7605 | 871 | 205 | 2319 | 18744 | 25605 | 32927 | 22307 | 108 | 2020 | 7159 | 871 | 380 | 0 | 985 | 51647 | 0 | 0 | 0 | 154 | 781 | 33 | 0 | 0 | 0 | 0 | 0 | 148 | 229 | 1718 | 114 | 0 | 0 | 0 | 0 | 0 |
| SRR5644086 | 243 | 15228 | 845 | 1009 | 2982 | 21316 | 24555 | 37244 | 14807 | 124 | 18591 | 29732 | 5347 | 1439 | 0 | 5740 | 30445 | 0 | 0 | 16 | 6250 | 2561 | 774 | 25 | 0 | 0 | 0 | 0 | 304 | 2741 | 3273 | 616 | 0 | 20 | 110 | 51 | 0 |
| SRR5644087 | 110 | 9356 | 1242 | 1638 | 783 | 8788 | 4707 | 11240 | 47099 | 71 | 10485 | 15036 | 4176 | 1462 | 0 | 7309 | 61906 | 0 | 0 | 11 | 916 | 467 | 78 | 0 | 0 | 0 | 0 | 33 | 1919 | 3905 | 3985 | 399 | 0 | 0 | 41 | 0 | 0 |
| SRR5644088 | 109 | 9843 | 1896 | 1419 | 1136 | 7262 | 6802 | 11605 | 45949 | 48 | 9170 | 13664 | 3271 | 1428 | 0 | 4961 | 58621 | 0 | 0 | 0 | 900 | 568 | 100 | 0 | 0 | 0 | 0 | 220 | 1056 | 4650 | 8562 | 1006 | 0 | 24 | 63 | 0 | 0 |
| SRR5657537 | 47 | 7829 | 1019 | 808 | 179 | 1400 | 1260 | 5212 | 10069 | 111 | 27736 | 59618 | 6212 | 6985 | 40 | 19412 | 106429 | 0 | 0 | 0 | 877 | 1531 | 473 | 0 | 19 | 0 | 0 | 17 | 192 | 680 | 1363 | 449 | 0 | 72 | 374 | 19 | 0 |
| SRR5657538 | 0 | 2800 | 1058 | 310 | 1068 | 7208 | 9714 | 32149 | 5167 | 222 | 38287 | 38966 | 8246 | 13380 | 10 | 14175 | 66596 | 0 | 0 | 0 | 525 | 1863 | 394 | 0 | 0 | 0 | 0 | 16 | 190 | 490 | 4026 | 736 | 0 | 24 | 83 | 12 | 0 |
| SRR5657539 | 0 | 2384 | 1294 | 1008 | 198 | 1113 | 1444 | 5079 | 8229 | 153 | 16683 | 30869 | 3780 | 6663 | 112 | 21546 | 98659 | 0 | 0 | 0 | 132 | 297 | 238 | 14 | 14 | 0 | 0 | 0 | 60 | 193 | 996 | 945 | 0 | 133 | 512 | 0 | 0 |
| SRR5657540 | 0 | 2670 | 1589 | 148 | 398 | 2044 | 3824 | 11979 | 10298 | 456 | 18064 | 13600 | 1750 | 7310 | 0 | 30143 | 69008 | 0 | 0 | 0 | 341 | 472 | 84 | 0 | 0 | 0 | 0 | 0 | 14 | 868 | 913 | 288 | 0 | 0 | 28 | 0 | 0 |
| SRR5657541 | 0 | 6387 | 948 | 336 | 169 | 531 | 1040 | 4400 | 15118 | 154 | 13904 | 15172 | 3202 | 4122 | 55 | 12094 | 119364 | 0 | 0 | 0 | 153 | 1233 | 466 | 13 | 0 | 0 | 0 | 0 | 92 | 229 | 3176 | 960 | 0 | 94 | 265 | 0 | 0 |
| SRR5657542 | 0 | 1951 | 961 | 265 | 511 | 3001 | 4909 | 18974 | 47221 | 179 | 20445 | 16414 | 3357 | 3103 | 0 | 2206 | 93516 | 0 | 0 | 12 | 780 | 2305 | 322 | 0 | 0 | 0 | 0 | 37 | 512 | 1181 | 5302 | 710 | 0 | 0 | 11 | 0 | 0 |
| SRR5657543 | 0 | 924 | 948 | 85 | 207 | 1040 | 1479 | 7167 | 11349 | 231 | 16921 | 21407 | 4213 | 5123 | 141 | 16503 | 155920 | 0 | 0 | 0 | 148 | 1124 | 526 | 0 | 0 | 0 | 0 | 0 | 52 | 224 | 3460 | 1981 | 0 | 82 | 287 | 50 | 0 |
| SRR5657544 | 0 | 3217 | 437 | 421 | 708 | 1883 | 6217 | 13958 | 7495 | 117 | 6621 | 25874 | 1451 | 19444 | 15 | 8526 | 97676 | 0 | 0 | 0 | 50 | 374 | 179 | 11 | 0 | 0 | 0 | 0 | 22 | 109 | 1337 | 403 | 0 | 25 | 199 | 0 | 0 |
| SRR5657545 | 0 | 1806 | 6443 | 61 | 480 | 594 | 4363 | 9574 | 17740 | 1178 | 21493 | 11000 | 4109 | 12310 | 51 | 16807 | 46654 | 0 | 0 | 0 | 25 | 62 | 111 | 19 | 0 | 0 | 0 | 0 | 22 | 86 | 279 | 757 | 20 | 113 | 211 | 24 | 0 |
| SRR5657546 | 0 | 6497 | 8526 | 72 | 2234 | 6852 | 27124 | 48247 | 4086 | 2129 | 13107 | 8144 | 1848 | 21059 | 0 | 4945 | 23289 | 0 | 0 | 0 | 77 | 138 | 44 | 0 | 0 | 0 | 0 | 0 | 81 | 210 | 439 | 245 | 0 | 13 | 21 | 14 | 0 |
| SRR5658693 | 17 | 51803 | 497 | 182 | 237 | 2349 | 1800 | 6246 | 1304 | 0 | 26777 | 56252 | 15179 | 3777 | 22 | 20 | 40580 | 0 | 0 | 0 | 12 | 0 | 0 | 0 | 0 | 0 | 0 | 0 | 0 | 14 | 0 | 0 | 0 | 0 | 0 | 0 | 0 |
| SRR5658694 | 53 | 23211 | 337 | 416 | 438 | 4916 | 3710 | 8686 | 1230 | 22 | 30229 | 48602 | 17689 | 4715 | 37 | 21 | 46245 | 0 | 0 | 0 | 16 | 0 | 0 | 0 | 0 | 0 | 0 | 0 | 0 | 24 | 15 | 0 | 0 | 0 | 0 | 0 | 0 |
| SRR5658695 | 14 | 18916 | 880 | 134 | 421 | 4771 | 4337 | 6926 | 1213 | 18 | 22081 | 45811 | 13630 | 3569 | 18 | 26 | 42453 | 0 | 0 | 0 | 10 | 0 | 0 | 0 | 0 | 0 | 0 | 0 | 0 | 12 | 20 | 0 | 0 | 0 | 0 | 0 | 0 |
| SRR5658696 | 26 | 26752 | 365 | 232 | 458 | 5063 | 4177 | 9269 | 1705 | 14 | 27413 | 42675 | 17094 | 4281 | 32 | 30 | 51908 | 0 | 0 | 0 | 19 | 0 | 0 | 0 | 0 | 0 | 0 | 0 | 0 | 17 | 20 | 0 | 0 | 0 | 0 | 0 | 0 |
| SRR5658697 | 14 | 17153 | 1872 | 202 | 546 | 6870 | 5780 | 10754 | 1537 | 21 | 25067 | 34591 | 16856 | 5066 | 19 | 37 | 59030 | 0 | 0 | 0 | 17 | 0 | 0 | 0 | 0 | 0 | 0 | 0 | 0 | 21 | 29 | 12 | 0 | 0 | 0 | 0 | 0 |
| SRR5687455 | 12 | 6214 | 5049 | 207 | 3459 | 89002 | 62388 | 35565 | 178 | 13 | 29989 | 21744 | 2775 | 30521 | 0 | 51172 | 24360 | 0 | 0 | 0 | 114 | 45 | 18 | 0 | 0 | 0 | 0 | 0 | 31 | 59 | 81 | 16 | 0 | 0 | 0 | 0 | 0 |
| SRR5687456 | 46 | 9017 | 5327 | 409 | 3865 | 95609 | 66208 | 35441 | 201 | 0 | 32297 | 19795 | 2453 | 23414 | 0 | 57375 | 24616 | 0 | 0 | 0 | 117 | 47 | 18 | 0 | 0 | 0 | 0 | 0 | 43 | 63 | 88 | 18 | 0 | 0 | 0 | 0 | 0 |
| SRR5687457 | 31 | 11672 | 8039 | 363 | 3053 | 83292 | 55408 | 28243 | 134 | 20 | 32164 | 24181 | 2708 | 28863 | 0 | 50759 | 23793 | 0 | 0 | 0 | 101 | 50 | 17 | 0 | 0 | 0 | 0 | 0 | 30 | 49 | 84 | 15 | 0 | 0 | 0 | 0 | 0 |
| SRR5687458 | 13 | 6523 | 4350 | 225 | 3460 | 85773 | 64146 | 35163 | 195 | 12 | 32410 | 23896 | 2795 | 32504 | 0 | 47922 | 24597 | 0 | 0 | 0 | 104 | 48 | 18 | 0 | 0 | 0 | 0 | 0 | 27 | 53 | 72 | 18 | 0 | 0 | 0 | 0 | 0 |
| SRR5687459 | 0 | 5281 | 4333 | 145 | 3316 | 82773 | 60042 | 33447 | 190 | 13 | 32434 | 24462 | 2862 | 31850 | 0 | 49861 | 25531 | 0 | 0 | 0 | 105 | 45 | 18 | 0 | 0 | 0 | 0 | 0 | 22 | 49 | 66 | 15 | 0 | 0 | 0 | 0 | 0 |
| SRR5687461 | 0 | 2770 | 16760 | 42 | 2206 | 70653 | 75393 | 59310 | 316 | 0 | 36011 | 20061 | 3638 | 26707 | 0 | 55602 | 32851 | 0 | 0 | 0 | 63 | 61 | 17 | 0 | 0 | 0 | 0 | 0 | 26 | 66 | 199 | 34 | 0 | 0 | 0 | 0 | 0 |
| SRR5687462 | 0 | 2795 | 14789 | 47 | 1780 | 54620 | 56552 | 41201 | 239 | 12 | 34739 | 26502 | 3393 | 24770 | 0 | 45249 | 33721 | 0 | 0 | 0 | 83 | 77 | 23 | 0 | 0 | 0 | 0 | 0 | 29 | 62 | 152 | 27 | 0 | 0 | 0 | 0 | 0 |
| SRR5687463 | 0 | 2960 | 17256 | 41 | 1624 | 52115 | 54855 | 41085 | 233 | 0 | 34067 | 25911 | 3434 | 24682 | 0 | 43954 | 33898 | 0 | 0 | 0 | 82 | 76 | 22 | 0 | 0 | 0 | 0 | 0 | 25 | 56 | 148 | 27 | 0 | 0 | 0 | 0 | 0 |
| SRR5687464 | 0 | 1004 | 15583 | 24 | 1688 | 56057 | 62412 | 47205 | 226 | 0 | 38848 | 28937 | 3818 | 28502 | 0 | 46527 | 32390 | 0 | 0 | 0 | 21 | 18 | 0 | 0 | 0 | 0 | 0 | 0 | 0 | 18 | 40 | 0 | 0 | 0 | 0 | 0 | 0 |
| SRR5687465 | 15 | 9257 | 6055 | 277 | 2634 | 85688 | 58414 | 30608 | 64 | 13 | 32674 | 21874 | 2732 | 18780 | 0 | 49637 | 27149 | 0 | 0 | 0 | 398 | 179 | 115 | 0 | 0 | 0 | 0 | 0 | 75 | 167 | 230 | 120 | 0 | 0 | 0 | 0 | 0 |
| SRR5687466 | 0 | 5295 | 5364 | 237 | 2731 | 89060 | 65798 | 35791 | 56 | 12 | 33421 | 22211 | 2788 | 19638 | 0 | 56244 | 28083 | 0 | 0 | 0 | 215 | 119 | 67 | 0 | 0 | 0 | 0 | 0 | 49 | 108 | 173 | 82 | 0 | 0 | 0 | 0 | 0 |
| SRR5687467 | 0 | 7128 | 4647 | 202 | 2600 | 84422 | 59256 | 32081 | 57 | 14 | 33195 | 24566 | 2819 | 18825 | 0 | 51362 | 29272 | 0 | 0 | 0 | 279 | 141 | 82 | 0 | 0 | 0 | 0 | 0 | 54 | 128 | 187 | 91 | 0 | 0 | 0 | 0 | 0 |
| SRR5687468 | 0 | 4691 | 8445 | 149 | 3009 | 109502 | 70655 | 43649 | 52 | 12 | 37064 | 17769 | 2622 | 18489 | 0 | 74508 | 25700 | 0 | 0 | 0 | 96 | 76 | 41 | 0 | 0 | 0 | 0 | 0 | 26 | 81 | 168 | 61 | 0 | 0 | 0 | 0 | 0 |
| SRR5687469 | 0 | 4929 | 6683 | 152 | 2651 | 82535 | 63132 | 38983 | 57 | 13 | 25397 | 17467 | 3097 | 22737 | 0 | 53517 | 30364 | 0 | 0 | 0 | 193 | 110 | 65 | 0 | 0 | 0 | 0 | 0 | 47 | 100 | 176 | 71 | 0 | 0 | 0 | 0 | 0 |
| SRR5753916 | 7031 | 4539 | 58961 | 2143 | 12876 | 23615 | 117528 | 25136 | 7021 | 5067 | 45730 | 23080 | 19699 | 38953 | 24851 | 34699 | 10063 | 0 | 53 | 84 | 238 | 604 | 577 | 130 | 33 | 12 | 0 | 55 | 111 | 75 | 39 | 0 | 0 | 0 | 0 | 26 | 14 |
| SRR5755813 | 161 | 8681 | 176 | 212 | 465 | 3158 | 3000 | 22305 | 1603 | 45 | 17618 | 12525 | 24994 | 3746 | 14 | 0 | 118871 | 0 | 0 | 0 | 404 | 39 | 34 | 0 | 0 | 0 | 0 | 0 | 117 | 335 | 49 | 51 | 0 | 0 | 0 | 0 | 0 |
| SRR5755814 | 287 | 15585 | 652 | 78 | 365 | 2952 | 3587 | 13444 | 838 | 0 | 19042 | 14597 | 24582 | 5503 | 41 | 17 | 95807 | 0 | 0 | 0 | 27 | 0 | 0 | 0 | 0 | 0 | 0 | 0 | 19 | 26 | 22 | 13 | 0 | 0 | 0 | 0 | 0 |

### Supp Table S8.1.Columns_B_to_AL

| | | | | | | | | | | | | | | | | | | | | | | | | | | | | | | | | | | | | | |
|---|---|---|---|---|---|---|---|---|---|---|---|---|---|---|---|---|---|---|---|---|---|---|---|---|---|---|---|---|---|---|---|---|---|---|---|---|---|
| SRR5755815 | 523 | 22824 | 872 | 140 | 3766 | 33482 | 49189 | 29309 | 1480 | 70 | 42380 | 40922 | 10759 | 7838 | 31 | 39 | 77500 | 0 | 0 | 0 | 288 | 12 | 15 | 0 | 0 | 0 | 0 | 0 | 214 | 452 | 47 | 37 | 0 | 0 | 0 | 0 | 0 |
| SRR5755818 | 85 | 16229 | 116 | 205 | 156 | 1784 | 1522 | 6396 | 164 | 0 | 26715 | 15564 | 11258 | 3215 | 18 | 0 | 43450 | 0 | 0 | 0 | 437 | 28 | 26 | 0 | 0 | 0 | 0 | 0 | 188 | 570 | 62 | 69 | 0 | 0 | 0 | 0 | 0 |
| SRR5755819 | 101 | 11193 | 202 | 134 | 310 | 2618 | 2812 | 16272 | 1206 | 34 | 19130 | 14737 | 24025 | 5060 | 21 | 0 | 100169 | 0 | 0 | 0 | 50 | 0 | 0 | 0 | 0 | 0 | 0 | 0 | 26 | 32 | 13 | 12 | 0 | 0 | 0 | 0 | 0 |
| SRR5755820 | 177 | 23911 | 412 | 88 | 169 | 1232 | 1306 | 7279 | 664 | 0 | 18546 | 13418 | 17847 | 5301 | 17 | 0 | 88845 | 0 | 0 | 0 | 11 | 11 | 0 | 0 | 0 | 0 | 0 | 0 | 0 | 0 | 0 | 0 | 0 | 0 | 0 | 0 | 0 |
| SRR5755821 | 80 | 14299 | 607 | 119 | 8229 | 52504 | 95236 | 41757 | 5040 | 30 | 30641 | 27929 | 10119 | 12326 | 15 | 41 | 46670 | 0 | 0 | 0 | 1027 | 11 | 23 | 0 | 0 | 0 | 0 | 24 | 857 | 1845 | 90 | 76 | 0 | 0 | 0 | 0 | 0 |
| SRR5755824 | 85 | 25671 | 254 | 56 | 119 | 1497 | 1249 | 5359 | 158 | 0 | 21075 | 16639 | 10621 | 3324 | 38 | 11 | 42743 | 0 | 0 | 0 | 17 | 0 | 0 | 0 | 0 | 0 | 0 | 0 | 17 | 31 | 0 | 14 | 0 | 0 | 0 | 0 | 0 |
| SRR5755828 | 73 | 8345 | 193 | 105 | 6069 | 32720 | 62902 | 38616 | 2576 | 23 | 29871 | 26567 | 10297 | 9850 | 32 | 21 | 59900 | 0 | 0 | 0 | 151 | 70 | 16 | 0 | 0 | 0 | 0 | 237 | 429 | 445 | 1262 | 120 | 0 | 11 | 53 | 10 | 0 |
| SRR5755830 | 336 | 27061 | 2409 | 195 | 80 | 981 | 901 | 5542 | 449 | 33 | 19202 | 17865 | 12687 | 7349 | 0 | 156 | 101979 | 0 | 0 | 0 | 19 | 75 | 0 | 0 | 0 | 0 | 0 | 0 | 18 | 24 | 139 | 16 | 0 | 0 | 0 | 0 | 0 |
| SRR5755831 | 38 | 2864 | 382 | 272 | 259 | 1652 | 2322 | 7760 | 291 | 21 | 14415 | 8374 | 9748 | 4323 | 14 | 0 | 34732 | 0 | 0 | 0 | 137 | 45 | 0 | 0 | 0 | 0 | 0 | 173 | 293 | 275 | 299 | 53 | 0 | 0 | 0 | 0 | 0 |
| SRR5755833 | 118 | 8872 | 229 | 218 | 471 | 3473 | 3201 | 21854 | 2505 | 12 | 22256 | 12870 | 24271 | 4810 | 0 | 0 | 114062 | 0 | 0 | 0 | 218 | 34 | 17 | 0 | 0 | 0 | 0 | 19 | 137 | 185 | 49 | 23 | 0 | 0 | 0 | 0 | 0 |
| SRR5755835 | 469 | 26765 | 2314 | 525 | 4417 | 42895 | 58224 | 32644 | 1926 | 31 | 47266 | 41565 | 13569 | 8888 | 29 | 25 | 61721 | 0 | 0 | 0 | 824 | 85 | 31 | 0 | 0 | 0 | 0 | 101 | 616 | 1445 | 508 | 117 | 0 | 0 | 0 | 0 | 0 |
| SRR5755837 | 684 | 22976 | 314 | 332 | 142 | 1189 | 1005 | 6671 | 1043 | 19 | 24004 | 15869 | 18851 | 6848 | 10 | 140 | 87152 | 0 | 0 | 0 | 190 | 57 | 15 | 0 | 0 | 0 | 0 | 0 | 97 | 231 | 58 | 35 | 0 | 0 | 0 | 0 | 0 |
| SRR5755838 | 75 | 24243 | 226 | 335 | 278 | 4405 | 3114 | 10240 | 476 | 0 | 29841 | 25255 | 14040 | 3043 | 20 | 0 | 61606 | 0 | 0 | 0 | 678 | 173 | 27 | 0 | 0 | 0 | 0 | 57 | 355 | 762 | 364 | 61 | 0 | 0 | 12 | 0 | 0 |
| SRR5755842 | 871 | 53547 | 1531 | 225 | 7580 | 50933 | 65525 | 43698 | 2571 | 0 | 38582 | 40307 | 10953 | 5210 | 38 | 32 | 72916 | 0 | 0 | 0 | 474 | 77 | 31 | 0 | 0 | 0 | 0 | 26 | 249 | 625 | 306 | 78 | 0 | 0 | 12 | 0 | 0 |
| SRR5755844 | 666 | 46194 | 217 | 217 | 140 | 1255 | 1059 | 5349 | 1402 | 0 | 27783 | 25982 | 19420 | 6793 | 0 | 126 | 82166 | 0 | 0 | 0 | 102 | 127 | 18 | 0 | 0 | 0 | 0 | 0 | 26 | 104 | 72 | 19 | 0 | 0 | 0 | 0 | 0 |
| SRR5755847 | 1008 | 31897 | 374 | 112 | 516 | 4287 | 3945 | 19955 | 904 | 0 | 21763 | 17999 | 27906 | 5010 | 33 | 0 | 106632 | 0 | 0 | 0 | 110 | 40 | 0 | 0 | 0 | 0 | 0 | 0 | 26 | 67 | 53 | 12 | 0 | 0 | 0 | 0 | 0 |
| SRR5755849 | 544 | 26494 | 4162 | 264 | 11288 | 58350 | 83694 | 43463 | 2628 | 16 | 33280 | 33015 | 11060 | 7731 | 39 | 45 | 59810 | 0 | 0 | 0 | 507 | 53 | 16 | 0 | 0 | 0 | 0 | 63 | 406 | 757 | 274 | 55 | 0 | 0 | 0 | 0 | 0 |
| SRR5755851 | 522 | 39926 | 439 | 119 | 102 | 1058 | 1015 | 4846 | 689 | 0 | 30204 | 26074 | 20971 | 6925 | 27 | 32 | 93721 | 0 | 0 | 0 | 173 | 111 | 20 | 0 | 0 | 0 | 0 | 0 | 54 | 141 | 66 | 27 | 0 | 0 | 0 | 0 | 0 |
| SRR5755854 | 239 | 12938 | 137 | 222 | 622 | 4956 | 4743 | 22147 | 1369 | 0 | 20029 | 18189 | 23115 | 6156 | 18 | 0 | 113367 | 0 | 0 | 0 | 1213 | 281 | 74 | 0 | 0 | 0 | 0 | 0 | 287 | 840 | 220 | 90 | 0 | 0 | 0 | 0 | 0 |
| SRR5755855 | 162 | 25680 | 304 | 243 | 543 | 3426 | 3172 | 16853 | 1203 | 20 | 20550 | 16332 | 23315 | 8041 | 18 | 0 | 93048 | 0 | 0 | 0 | 622 | 132 | 29 | 0 | 0 | 0 | 0 | 0 | 188 | 500 | 135 | 42 | 0 | 0 | 0 | 0 | 0 |
| SRR5755856 | 801 | 33721 | 937 | 231 | 8170 | 36184 | 67485 | 30036 | 5046 | 41 | 33284 | 31874 | 10031 | 9840 | 11 | 28 | 60307 | 0 | 0 | 0 | 1623 | 100 | 148 | 0 | 0 | 0 | 0 | 35 | 696 | 2417 | 416 | 552 | 0 | 0 | 0 | 0 | 0 |
| SRR5755858 | 432 | 31053 | 277 | 122 | 129 | 895 | 813 | 5802 | 2052 | 0 | 24054 | 23840 | 20475 | 7605 | 19 | 123 | 75316 | 0 | 0 | 0 | 504 | 66 | 24 | 0 | 0 | 0 | 0 | 0 | 224 | 626 | 95 | 57 | 0 | 0 | 0 | 0 | 0 |
| SRR5755861 | 148 | 16577 | 132 | 275 | 470 | 2965 | 2743 | 11296 | 1040 | 16 | 9837 | 8623 | 16265 | 2941 | 24 | 0 | 129028 | 0 | 0 | 12 | 2476 | 211 | 163 | 0 | 0 | 0 | 0 | 23 | 747 | 1913 | 183 | 131 | 0 | 0 | 0 | 0 | 0 |
| SRR5755862 | 101 | 21094 | 101 | 395 | 248 | 1303 | 1496 | 6181 | 477 | 17 | 12917 | 10456 | 23529 | 3695 | 11 | 12 | 114183 | 0 | 0 | 14 | 2122 | 470 | 122 | 0 | 0 | 0 | 0 | 19 | 482 | 1241 | 319 | 95 | 0 | 0 | 0 | 0 | 0 |
| SRR5755863 | 454 | 40949 | 689 | 983 | 9312 | 43652 | 64334 | 23686 | 1584 | 25 | 25956 | 11 | 8668 | 6121 | 19 | 34 | 75883 | 0 | 0 | 20 | 6048 | 429 | 269 | 0 | 0 | 0 | 0 | 86 | 2965 | 9645 | 2223 | 677 | 0 | 0 | 32 | 0 | 0 |
| SRR5755865 | 245 | 47104 | 135 | 882 | 100 | 1298 | 1042 | 5639 | 2366 | 26 | 16863 | 15310 | 12210 | 6702 | 0 | 75 | 104807 | 0 | 0 | 0 | 1064 | 341 | 75 | 0 | 0 | 0 | 0 | 11 | 524 | 999 | 221 | 84 | 0 | 0 | 12 | 0 | 0 |
| SRR5755866 | 236 | 38180 | 94 | 495 | 245 | 1667 | 1121 | 3158 | 133 | 12 | 12297 | 11894 | 8080 | 1865 | 34 | 0 | 42611 | 0 | 0 | 14 | 2476 | 162 | 109 | 0 | 0 | 0 | 0 | 34 | 985 | 2329 | 216 | 153 | 0 | 0 | 29 | 0 | 0 |
| SRR5755868 | 353 | 12471 | 249 | 222 | 253 | 2864 | 2155 | 6704 | 386 | 0 | 5884 | 4522 | 7041 | 2035 | 0 | 0 | 64346 | 0 | 0 | 0 | 606 | 28 | 20 | 0 | 0 | 0 | 0 | 0 | 199 | 285 | 18 | 14 | 0 | 0 | 0 | 0 | 0 |
| SRR5755870 | 622 | 55568 | 510 | 1079 | 5857 | 41715 | 48238 | 27836 | 877 | 39 | 23453 | 28334 | 7483 | 5994 | 0 | 22 | 72210 | 0 | 0 | 32 | 8046 | 576 | 286 | 0 | 0 | 0 | 0 | 87 | 4297 | 11278 | 2064 | 697 | 0 | 0 | 21 | 0 | 0 |
| SRR5755875 | 60 | 7172 | 269 | 313 | 1126 | 6272 | 7678 | 27928 | 876 | 30 | 10758 | 10903 | 17757 | 6589 | 20 | 83 | 103237 | 0 | 0 | 19 | 3373 | 1040 | 89 | 0 | 0 | 0 | 0 | 32 | 1281 | 3375 | 1013 | 133 | 0 | 0 | 0 | 0 | 0 |
| SRR5755876 | 29 | 10418 | 133 | 258 | 467 | 3200 | 3494 | 12713 | 889 | 15 | 9823 | 8211 | 12350 | 5363 | 10 | 55 | 94615 | 0 | 0 | 14 | 2110 | 473 | 34 | 0 | 0 | 0 | 0 | 30 | 1294 | 2755 | 716 | 74 | 0 | 0 | 0 | 0 | 0 |
| SRR5755877 | 786 | 72753 | 1551 | 905 | 9016 | 48696 | 69987 | 32752 | 3606 | 27 | 26378 | 26035 | 8439 | 6771 | 36 | 3653 | 67736 | 0 | 0 | 42 | 8829 | 593 | 402 | 0 | 0 | 0 | 0 | 75 | 4659 | 11849 | 2752 | 1044 | 0 | 0 | 39 | 17 | 0 |
| SRR5755879 | 323 | 35446 | 313 | 231 | 102 | 1078 | 860 | 4693 | 1397 | 0 | 18135 | 11917 | 10438 | 5649 | 14 | 995 | 101267 | 0 | 0 | 0 | 915 | 256 | 88 | 0 | 0 | 0 | 0 | 10 | 423 | 936 | 194 | 149 | 0 | 0 | 11 | 0 | 0 |
| SRR5755880 | 85 | 24378 | 157 | 539 | 266 | 2380 | 1547 | 5523 | 172 | 17 | 10265 | 6794 | 7573 | 2958 | 16 | 1269 | 44005 | 0 | 0 | 24 | 3365 | 1132 | 134 | 0 | 0 | 0 | 0 | 113 | 2015 | 3782 | 2511 | 296 | 0 | 0 | 17 | 0 | 0 |
| SRR5755882 | 51 | 7865 | 242 | 268 | 529 | 4424 | 4808 | 19822 | 1295 | 13 | 17391 | 13189 | 18285 | 7207 | 16 | 0 | 87444 | 0 | 0 | 0 | 823 | 207 | 18 | 0 | 0 | 0 | 0 | 0 | 170 | 577 | 149 | 22 | 0 | 0 | 0 | 0 | 0 |
| SRR5755884 | 449 | 13819 | 772 | 336 | 6453 | 50494 | 61577 | 36380 | 2572 | 20 | 33107 | 34536 | 11090 | 5841 | 36 | 14 | 66991 | 0 | 0 | 0 | 636 | 61 | 47 | 0 | 0 | 0 | 0 | 49 | 295 | 710 | 281 | 127 | 0 | 19 | 11 | 0 | 0 |
| SRR5755887 | 65 | 25859 | 113 | 355 | 319 | 2886 | 2105 | 4970 | 97 | 16 | 13595 | 10506 | 7216 | 3104 | 18 | 0 | 35227 | 0 | 0 | 0 | 1050 | 187 | 18 | 0 | 0 | 0 | 0 | 20 | 302 | 837 | 262 | 29 | 0 | 0 | 0 | 0 | 0 |
| SRR5755889 | 219 | 21768 | 178 | 554 | 449 | 6045 | 4253 | 21569 | 1116 | 0 | 15142 | 10597 | 15182 | 7128 | 0 | 0 | 111141 | 0 | 0 | 0 | 248 | 180 | 18 | 0 | 0 | 0 | 0 | 12 | 135 | 230 | 145 | 26 | 0 | 0 | 0 | 0 | 0 |
| SRR5755890 | 147 | 28382 | 348 | 224 | 269 | 2749 | 1908 | 8433 | 963 | 0 | 15977 | 10747 | 13032 | 4923 | 17 | 0 | 79424 | 0 | 0 | 0 | 187 | 150 | 13 | 0 | 0 | 0 | 0 | 0 | 73 | 144 | 73 | 11 | 0 | 0 | 0 | 0 | 0 |
| SRR5755891 | 264 | 17933 | 1100 | 492 | 3020 | 19274 | 25389 | 9338 | 797 | 18 | 12365 | 18733 | 4824 | 4307 | 16 | 10 | 57834 | 0 | 0 | 0 | 868 | 146 | 30 | 0 | 0 | 0 | 0 | 66 | 783 | 874 | 381 | 52 | 0 | 0 | 0 | 0 | 0 |
| SRR5755894 | 111 | 25552 | 120 | 392 | 81 | 1377 | 599 | 1647 | 118 | 0 | 10683 | 8857 | 4701 | 1595 | 11 | 0 | 38898 | 0 | 0 | 0 | 214 | 50 | 10 | 0 | 0 | 0 | 0 | 0 | 118 | 134 | 35 | 0 | 0 | 0 | 0 | 0 | 0 |
| SRR5755896 | 64 | 14339 | 158 | 94 | 512 | 2898 | 3889 | 21499 | 1202 | 23 | 12482 | 9192 | 18426 | 4437 | 29 | 0 | 93335 | 0 | 0 | 0 | 1284 | 814 | 49 | 0 | 0 | 0 | 0 | 0 | 317 | 879 | 776 | 55 | 0 | 0 | 0 | 0 | 0 |
| SRR5755897 | 24 | 14202 | 136 | 128 | 274 | 2157 | 2555 | 11728 | 1042 | 0 | 11325 | 7841 | 12241 | 6383 | 22 | 12 | 78092 | 0 | 0 | 0 | 1620 | 697 | 39 | 0 | 0 | 0 | 0 | 0 | 637 | 1447 | 876 | 55 | 0 | 0 | 0 | 0 | 0 |
| SRR5755898 | 546 | 19478 | 718 | 230 | 5384 | 39360 | 47684 | 31799 | 3615 | 14 | 37960 | 17 | 11292 | 6818 | 27 | 20 | 75215 | 0 | 0 | 0 | 475 | 29 | 43 | 0 | 0 | 0 | 0 | 0 | 253 | 645 | 148 | 102 | 0 | 29 | 0 | 0 | 0 |
| SRR5755901 | 35 | 37454 | 84 | 190 | 218 | 1776 | 1354 | 5380 | 203 | 22 | 13642 | 9097 | 9286 | 2861 | 40 | 0 | 34074 | 0 | 0 | 11 | 1401 | 691 | 54 | 0 | 0 | 0 | 0 | 0 | 764 | 1608 | 1482 | 95 | 0 | 0 | 0 | 0 | 0 |
| SRR5755903 | 89 | 14180 | 169 | 115 | 535 | 3374 | 3595 | 23611 | 1427 | 21 | 14418 | 12647 | 19527 | 4115 | 18 | 0 | 101759 | 0 | 0 | 0 | 959 | 1182 | 66 | 0 | 0 | 0 | 0 | 13 | 241 | 734 | 1076 | 78 | 0 | 0 | 0 | 0 | 0 |
| SRR5755905 | 182 | 31442 | 999 | 303 | 7187 | 35439 | 57623 | 29645 | 2596 | 27 | 32016 | 30816 | 10615 | 7774 | 28 | 17 | 63332 | 0 | 0 | 12 | 1191 | 309 | 67 | 0 | 0 | 0 | 0 | 42 | 1396 | 1610 | 1663 | 192 | 0 | 10 | 11 | 0 | 0 |
| SRR5755908 | 56 | 24762 | 501 | 270 | 207 | 1610 | 1565 | 6271 | 274 | 0 | 11821 | 8913 | 6475 | 3752 | 21 | 0 | 44712 | 0 | 0 | 14 | 2166 | 496 | 30 | 0 | 0 | 0 | 0 | 0 | 925 | 2134 | 839 | 61 | 0 | 0 | 0 | 0 | 0 |
| SRR5755910 | 0 | 3493 | 162 | 86 | 617 | 2578 | 5538 | 28839 | 1078 | 32 | 7914 | 6210 | 14852 | 7284 | 18 | 0 | 82245 | 0 | 0 | 17 | 2825 | 2977 | 159 | 0 | 0 | 0 | 0 | 24 | 980 | 3849 | 5067 | 336 | 0 | 0 | 0 | 0 | 0 |
| SRR5755911 | 0 | 7685 | 517 | 188 | 146 | 550 | 2886 | 16278 | 1185 | 31 | 3970 | 2602 | 6072 | 10102 | 25 | 0 | 53066 | 0 | 0 | 13 | 2924 | 2131 | 102 | 0 | 0 | 0 | 0 | 0 | 843 | 3867 | 3510 | 225 | 0 | 0 | 0 | 0 | 0 |
| SRR5755912 | 214 | 52089 | 1141 | 225 | 6643 | 46898 | 62998 | 32008 | 2577 | 0 | 28888 | 32190 | 9380 | 7357 | 40 | 13 | 64473 | 0 | 0 | 0 | 1190 | 212 | 58 | 0 | 0 | 0 | 0 | 49 | 1036 | 1742 | 1261 | 177 | 0 | 12 | 20 | 0 | 0 |
| SRR5755914 | 695 | 44385 | 381 | 285 | 82 | 1001 | 768 | 6671 | 1239 | 13 | 22892 | 18107 | 16864 | 5678 | 19 | 44 | 96365 | 0 | 0 | 0 | 71 | 58 | 12 | 0 | 0 | 0 | 0 | 0 | 41 | 75 | 40 | 17 | 0 | 0 | 0 | 0 | 0 |
| SRR5755915 | 0 | 9561 | 819 | 347 | 171 | 691 | 3167 | 7944 | 420 | 40 | 5237 | 2731 | 2928 | 7329 | 13 | 0 | 22845 | 0 | 0 | 19 | 3280 | 1911 | 114 | 0 | 0 | 0 | 0 | 15 | 1830 | 6700 | 6178 | 407 | 0 | 0 | 0 | 0 | 0 |
| SRR5755917 | 229 | 15681 | 265 | 151 | 3641 | 26510 | 32127 | 32399 | 1800 | 0 | 22517 | 25888 | 18171 | 4099 | 33 | 0 | 84823 | 0 | 0 | 0 | 171 | 34 | 28 | 0 | 0 | 0 | 0 | 0 | 92 | 261 | 79 | 81 | 0 | 20 | 12 | 0 | 0 |
| SRR5755918 | 109 | 26477 | 176 | 319 | 202 | 2456 | 1822 | 7991 | 418 | 20 | 24056 | 17662 | 29486 | 4524 | 24 | 0 | 100782 | 0 | 0 | 0 | 111 | 69 | 22 | 0 | 0 | 0 | 0 | 0 | 45 | 96 | 81 | 34 | 0 | 0 | 0 | 0 | 0 |
| SRR5755919 | 525 | 17786 | 845 | 457 | 5782 | 52359 | 51226 | 30325 | 799 | 29 | 42356 | 34605 | 10843 | 5757 | 39 | 20 | 86418 | 0 | 0 | 0 | 237 | 20 | 26 | 0 | 0 | 0 | 0 | 11 | 144 | 287 | 86 | 90 | 0 | 16 | 11 | 0 | 0 |
| SRR5755922 | 137 | 32047 | 200 | 366 | 340 | 3976 | 2264 | 7702 | 242 | 29 | 30664 | 21133 | 12878 | 2272 | 41 | 0 | 55598 | 0 | 0 | 0 | 103 | 35 | 20 | 0 | 0 | 0 | 0 | 0 | 48 | 93 | 65 | 45 | 0 | 0 | 0 | 0 | 0 |
| SRR5755924 | 101 | 13781 | 205 | 341 | 348 | 3542 | 2619 | 16004 | 926 | 0 | 24057 | 21808 | 29245 | 3244 | 17 | 0 | 142673 | 0 | 0 | 0 | 62 | 28 | 32 | 0 | 0 | 0 | 0 | 0 | 28 | 34 | 26 | 36 | 0 | 12 | 0 | 0 | 0 |
| SRR5755925 | 176 | 30715 | 159 | 557 | 230 | 2527 | 1765 | 8457 | 585 | 23 | 25073 | 19461 | 26256 | 3500 | 16 | 0 | 115111 | 0 | 0 | 0 | 54 | 20 | 22 | 0 | 0 | 0 | 0 | 0 | 21 | 31 | 18 | 35 | 0 | 0 | 0 | 0 | 0 |
| SRR5755926 | 231 | 12230 | 704 | 856 | 5565 | 44546 | 46713 | 32383 | 1098 | 16 | 37899 | 36022 | 10623 | 5590 | 30 | 21 | 82070 | 0 | 0 | 0 | 368 | 27 | 24 | 0 | 0 | 0 | 0 | 11 | 218 | 504 | 104 | 85 | 0 | 14 | 11 | 0 | 0 |
| SRR5755928 | 414 | 25732 | 269 | 328 | 392 | 3354 | 2925 | 7477 | 779 | 0 | 31063 | 25488 | 20782 | 6393 | 18 | 85 | 104590 | 0 | 0 | 0 | 47 | 33 | 17 | 0 | 0 | 0 | 0 | 0 | 35 | 43 | 23 | 25 | 0 | 0 | 0 | 0 | 0 |
| SRR5755929 | 223 | 61504 | 111 | 384 | 268 | 3212 | 1884 | 6174 | 94 | 25 | 30492 | 21947 | 11586 | 2994 | 11 | 0 | 58120 | 0 | 0 | 0 | 44 | 0 | 0 | 0 | 0 | 0 | 0 | 0 | 12 | 29 | 12 | 23 | 0 | 0 | 0 | 0 | 0 |
| SRR5755931 | 278 | 30969 | 132 | 185 | 421 | 3429 | 3053 | 20583 | 901 | 0 | 17695 | 13512 | 23997 | 4356 | 15 | 0 | 126619 | 0 | 0 | 0 | 297 | 203 | 19 | 0 | 0 | 0 | 0 | 0 | 121 | 206 | 171 | 28 | 0 | 0 | 0 | 0 | 0 |
| SRR5755933 | 564 | 30311 | 726 | 413 | 5154 | 42370 | 50837 | 33024 | 1546 | 30 | 40548 | 36930 | 9723 | 7973 | 25 | 30 | 73612 | 0 | 0 | 0 | 364 | 63 | 18 | 0 | 0 | 0 | 0 | 18 | 312 | 428 | 213 | 58 | 0 | 0 | 11 | 0 | 0 |
| SRR5755936 | 240 | 33243 | 150 | 397 | 342 | 2441 | 1535 | 4718 | 283 | 26 | 19076 | 14928 | 10689 | 2768 | 24 | 0 | 52073 | 0 | 0 | 10 | 1744 | 581 | 41 | 0 | 0 | 0 | 0 | 0 | 896 | 1743 | 1068 | 74 | 0 | 0 | 0 | 0 | 0 |
| SRR5755938 | 119 | 15266 | 106 | 185 | 573 | 4711 | 4016 | 20005 | 805 | 0 | 22878 | 20271 | 27161 | 4027 | 18 | 0 | 128617 | 0 | 0 | 0 | 366 | 207 | 29 | 0 | 0 | 0 | 0 | 0 | 94 | 249 | 168 | 23 | 0 | 0 | 0 | 0 | 0 |
| SRR5755939 | 114 | 16819 | 89 | 326 | 300 | 2171 | 1857 | 8671 | 775 | 0 | 16977 | 15117 | 25876 | 4400 | 26 | 0 | 106225 | 0 | 0 | 0 | 1281 | 553 | 41 | 0 | 0 | 0 | 0 | 14 | 374 | 912 | 579 | 63 | 0 | 0 | 0 | 0 | 0 |
| SRR5755940 | 564 | 15470 | 1238 | 321 | 6311 | 39236 | 55298 | 30778 | 3282 | 0 | 37979 | 13 | 11316 | 6924 | 30 | 26 | 67034 | 0 | 0 | 0 | 440 | 50 | 31 | 0 | 0 | 0 | 0 | 29 | 352 | 793 | 270 | 131 | 0 | 18 | 22 | 0 | 0 |
| SRR5755943 | 192 | 24652 | 159 | 348 | 386 | 2960 | 1928 | 7383 | 343 | 0 | 24993 | 15493 | 12314 | 2694 | 28 | 0 | 42450 | 0 | 0 | 0 | 443 | 149 | 21 | 0 | 0 | 0 | 0 | 11 | 241 | 496 | 248 | 53 | 0 | 0 | 0 | 0 | 0 |
| SRR5755946 | 206 | 21359 | 190 | 228 | 132 | 1466 | 1037 | 6695 | 1115 | 0 | 24224 | 17466 | 26478 | 4633 | 18 | 0 | 102566 | 0 | 0 | 0 | 47 | 33 | 12 | 0 | 0 | 0 | 0 | 0 | 25 | 36 | 24 | 22 | 0 | 0 | 0 | 0 | 0 |
| SRR5755947 | 527 | 9530 | 642 | 313 | 4896 | 35862 | 47394 | 23352 | 3058 | 33 | 39177 | 41411 | 11550 | 7139 | 28 | 27 | 79562 | 0 | 0 | 0 | 245 | 22 | 36 | 0 | 0 | 0 | 0 | 0 | 170 | 332 | 97 | 91 | 0 | 13 | 12 | 0 | 0 |
| SRR5755950 | 196 | 35006 | 134 | 207 | 282 | 1935 | 1374 | 5416 | 205 | 10 | 23464 | 17912 | 12521 | 3207 | 64 | 17 | 54254 | 0 | 0 | 0 | 74 | 23 | 19 | 0 | 0 | 0 | 0 | 0 | 37 | 60 | 39 | 43 | 0 | 0 | 0 | 0 | 0 |
| SRR5755952 | 169 | 16840 | 151 | 113 | 484 | 3941 | 3790 | 25059 | 2006 | 0 | 21478 | 17150 | 30724 | 4882 | 17 | 0 | 103202 | 0 | 0 | 0 | 212 | 176 | 17 | 0 | 0 | 0 | 0 | 0 | 84 | 207 | 176 | 48 | 0 | 0 | 0 | 0 | 0 |
| SRR5755953 | 197 | 23986 | 166 | 228 | 367 | 3259 | 2582 | 13177 | 677 | 0 | 19171 | 14422 | 24251 | 5439 | 42 | 0 | 87896 | 0 | 0 | 0 | 634 | 517 | 30 | 0 | 0 | 0 | 0 | 0 | 270 | 645 | 455 | 37 | 0 | 0 | 0 | 0 | 0 |
| SRR5755954 | 848 | 30968 | 1180 | 199 | 5135 | 36952 | 49542 | 28214 | 3468 | 0 | 37407 | 38044 | 12541 | 7493 | 21 | 44 | 65011 | 0 | 0 | 0 | 755 | 97 | 31 | 0 | 0 | 0 | 0 | 32 | 625 | 1156 | 531 | 114 | 0 | 0 | 0 | 0 | 0 |
| SRR5755956 | 1854 | 27100 | 256 | 208 | 123 | 1376 | 1158 | 8033 | 1212 | 0 | 28030 | 21810 | 22701 | 6940 | 29 | 79 | 84019 | 0 | 0 | 0 | 100 | 68 | 13 | 0 | 0 | 0 | 0 | 0 | 58 | 112 | 40 | 22 | 0 | 0 | 0 | 0 | 0 |
| SRR5755957 | 146 | 34316 | 231 | 219 | 274 | 2354 | 1947 | 5996 | 151 | 12 | 18545 | 11685 | 9738 | 4590 | 47 | 0 | 39084 | 0 | 0 | 0 | 870 | 191 | 14 | 0 | 0 | 0 | 0 | 0 | 409 | 932 | 301 | 25 | 0 | 0 | 0 | 0 | 0 |
| SRR5755959 | 82 | 9990 | 104 | 99 | 397 | 2958 | 2966 | 17693 | 2242 | 0 | 17332 | 14396 | 25284 | 3540 | 11 | 0 | 113505 | 0 | 0 | 0 | 375 | 121 | 27 | 0 | 0 | 0 | 0 | 0 | 57 | 197 | 79 | 36 | 0 | 0 | 0 | 0 | 0 |
| SRR5755960 | 35 | 15475 | 132 | 247 | 208 | 1214 | 1426 | 5917 | 635 | 18 | 18181 | 10741 | 20755 | 5163 | 28 | 0 | 80474 | 0 | 0 | 0 | 860 | 224 | 22 | 0 | 0 | 0 | 0 | 0 | 115 | 424 | 116 | 26 | 0 | 0 | 0 | 0 | 0 |
| SRR5755961 | 782 | 8876 | 996 | 293 | 6535 | 40538 | 61144 | 25234 | 886 | 0 | 35143 | 36000 | 11069 | 5670 | 57 | 35 | 70621 | 0 | 0 | 0 | 567 | 46 | 42 | 0 | 0 | 0 | 0 | 19 | 195 | 453 | 132 | 81 | 0 | 0 | 0 | 0 | 0 |
| SRR5755963 | 738 | 21362 | 476 | 228 | 86 | 984 | 855 | 5742 | 1201 | 13 | 23134 | 18415 | 16965 | 6246 | 18 | 127 | 86639 | 0 | 0 | 0 | 50 | 81 | 21 | 0 | 0 | 0 | 0 | 0 | 13 | 21 | 21 | 18 | 0 | 0 | 0 | 0 | 0 |
| SRR5755964 | 152 | 23196 | 160 | 321 | 298 | 2418 | 2279 | 4856 | 205 | 18 | 17033 | 12115 | 8472 | 2982 | 33 | 30 | 51412 | 0 | 0 | 0 | 1172 | 204 | 27 | 0 | 0 | 0 | 0 | 0 | 284 | 845 | 204 | 41 | 0 | 0 | 0 | 0 | 0 |
| SRR5755966 | 241 | 12302 | 105 | 181 | 601 | 4869 | 4144 | 20812 | 1317 | 0 | 20982 | 18397 | 23988 | 4526 | 23 | 0 | 115373 | 0 | 0 | 0 | 252 | 177 | 24 | 0 | 0 | 0 | 0 | 15 | 131 | 259 | 163 | 36 | 0 | 0 | 0 | 0 | 0 |
| SRR5755967 | 286 | 23412 | 95 | 206 | 256 | 2323 | 1900 | 8174 | 929 | 0 | 20262 | 14929 | 20866 | 5316 | 53 | 0 | 103487 | 0 | 0 | 0 | 363 | 205 | 14 | 0 | 0 | 0 | 0 | 0 | 155 | 354 | 181 | 23 | 0 | 0 | 0 | 0 | 0 |
| SRR5755968 | 225 | 20240 | 2700 | 81 | 4460 | 34904 | 44380 | 30697 | 1545 | 12 | 42157 | 38890 | 11891 | 7606 | 0 | 16 | 76893 | 0 | 0 | 0 | 183 | 19 | 0 | 0 | 0 | 0 | 0 | 0 | 132 | 253 | 85 | 34 | 0 | 0 | 0 | 0 | 0 |
| SRR5755971 | 48 | 18032 | 88 | 42 | 185 | 1986 | 1303 | 5321 | 257 | 0 | 23410 | 16380 | 12787 | 2591 | 12 | 0 | 43639 | 0 | 0 | 0 | 208 | 83 | 12 | 0 | 0 | 0 | 0 | 0 | 97 | 253 | 136 | 29 | 0 | 0 | 0 | 0 | 0 |
| SRR5755973 | 68 | 5557 | 83 | 83 | 772 | 4917 | 5853 | 30873 | 2022 | 30 | 16311 | 12820 | 24876 | 7195 | 21 | 0 | 96632 | 0 | 0 | 0 | 1114 | 354 | 30 | 0 | 0 | 0 | 0 | 0 | 294 | 1030 | 350 | 53 | 0 | 0 | 0 | 0 | 0 |
| SRR5755974 | 16 | 5905 | 90 | 33 | 484 | 3987 | 4727 | 19615 | 1713 | 37 | 13926 | 10045 | 16272 | 9870 | 16 | 0 | 66444 | 0 | 0 | 10 | 1723 | 400 | 34 | 0 | 0 | 0 | 0 | 0 | 586 | 1859 | 518 | 61 | 0 | 0 | 0 | 0 | 0 |
| SRR5755975 | 41 | 4543 | 878 | 136 | 5119 | 35621 | 59276 | 28396 | 7932 | 21 | 32795 | 34739 | 11323 | 8593 | 28 | 19 | 61962 | 0 | 0 | 0 | 401 | 41 | 20 | 0 | 0 | 0 | 0 | 21 | 235 | 526 | 179 | 68 | 0 | 0 | 0 | 0 | 0 |
| SRR5755978 | 0 | 8101 | 184 | 126 | 488 | 3739 | 4837 | 13388 | 891 | 25 | 18052 | 12054 | 9988 | 5737 | 0 | 11 | 43500 | 0 | 0 | 0 | 1368 | 243 | 23 | 0 | 0 | 0 | 0 | 0 | 445 | 1405 | 308 | 42 | 0 | 0 | 0 | 0 | 0 |
| SRR5755980 | 24 | 7054 | 258 | 85 | 507 | 2863 | 3335 | 17325 | 1512 | 0 | 14597 | 11262 | 22248 | 5442 | 23 | 0 | 92763 | 0 | 0 | 0 | 288 | 15 | 23 | 0 | 0 | 0 | 0 | 0 | 147 | 228 | 19 | 31 | 0 | 0 | 0 | 0 | 0 |
| SRR5755981 | 30 | 10677 | 397 | 159 | 228 | 1362 | 1536 | 7488 | 763 | 20 | 16934 | 12874 | 25235 | 5434 | 29 | 0 | 84247 | 0 | 0 | 0 | 312 | 0 | 0 | 0 | 0 | 0 | 0 | 0 | 116 | 221 | 12 | 17 | 0 | 0 | 0 | 0 | 0 |
| SRR5755982 | 95 | 18427 | 392 | 128 | 5703 | 31393 | 48850 | 24550 | 4650 | 18 | 37035 | 32701 | 11850 | 8440 | 19 | 26 | 65258 | 0 | 0 | 0 | 884 | 0 | 18 | 0 | 0 | 0 | 0 | 0 | 761 | 1591 | 25 | 70 | 0 | 0 | 0 | 0 | 0 |
| SRR5755984 | 200 | 12557 | 283 | 132 | 78 | 1114 | 772 | 6709 | 1321 | 0 | 22673 | 20268 | 17265 | 6284 | 11 | 24 | 100654 | 0 | 0 | 0 | 248 | 97 | 38 | 0 | 0 | 0 | 0 | 0 | 193 | 222 | 46 | 47 | 0 | 0 | 0 | 0 | 0 |
| SRR5755985 | 0 | 13964 | 380 | 69 | 263 | 1650 | 1619 | 5079 | 272 | 24 | 18501 | 14365 | 11863 | 3864 | 12 | 0 | 43848 | 0 | 0 | 0 | 657 | 0 | 11 | 0 | 0 | 0 | 0 | 0 | 318 | 731 | 18 | 29 | 0 | 0 | 0 | 0 | 0 |
| SRR5755987 | 40 | 6802 | 190 | 53 | 819 | 5648 | 6471 | 31435 | 1082 | 18 | 15532 | 12576 | 19893 | 7883 | 26 | 12 | 82357 | 0 | 0 | 0 | 726 | 497 | 27 | 0 | 0 | 0 | 0 | 0 | 263 | 859 | 629 | 58 | 0 | 0 | 0 | 0 | 0 |
| SRR5755988 | 0 | 7945 | 273 | 49 | 402 | 3003 | 3553 | 16442 | 895 | 15 | 14270 | 10202 | 14761 | 8762 | 29 | 0 | 76052 | 0 | 0 | 0 | 1301 | 528 | 29 | 0 | 0 | 0 | 0 | 0 | 466 | 1695 | 768 | 70 | 0 | 0 | 0 | 0 | 0 |
| SRR5755989 | 443 | 13932 | 1136 | 268 | 8073 | 49308 | 70629 | 42865 | 2651 | 28 | 32944 | 15 | 10466 | 6924 | 28 | 23 | 64574 | 0 | 0 | 0 | 1351 | 121 | 58 | 0 | 0 | 0 | 0 | 33 | 644 | 2323 | 741 | 206 | 0 | 13 | 12 | 0 | 0 |
| SRR5755991 | 85 | 14916 | 291 | 108 | 50 | 671 | 523 | 3683 | 397 | 0 | 25279 | 21236 | 20785 | 6940 | 18 | 127 | 79234 | 0 | 0 | 0 | 127 | 40 | 20 | 0 | 0 | 0 | 0 | 0 | 75 | 174 | 44 | 38 | 0 | 0 | 0 | 0 | 0 |
| SRR5755992 | 0 | 14824 | 163 | 81 | 270 | 2371 | 1900 | 6547 | 174 | 0 | 15452 | 11702 | 8994 | 3272 | 15 | 0 | 42134 | 0 | 0 | 0 | 906 | 232 | 17 | 0 | 0 | 0 | 0 | 0 | 419 | 1136 | 451 | 49 | 0 | 0 | 0 | 0 | 0 |
| SRR5755994 | 162 | 14897 | 335 | 257 | 4660 | 29133 | 38303 | 38054 | 3199 | 45 | 21916 | 12 | 18682 | 7664 | 33 | 14 | 58899 | 0 | 0 | 0 | 290 | 55 | 37 | 0 | 0 | 0 | 0 | 0 | 96 | 387 | 82 | 87 | 0 | 18 | 0 | 0 | 0 |
| SRR5755995 | 130 | 30565 | 237 | 172 | 345 | 4390 | 3802 | 14270 | 991 | 14 | 21923 | 15028 | 23266 | 6383 | 38 | 0 | 83320 | 0 | 0 | 0 | 88 | 52 | 24 | 0 | 0 | 0 | 0 | 0 | 47 | 77 | 50 | 30 | 0 | 0 | 0 | 0 | 0 |
| SRR5755996 | 345 | 9749 | 563 | 587 | 5977 | 56050 | 69368 | 40674 | 3138 | 16 | 27970 | 34288 | 10890 | 6722 | 52 | 29 | 56577 | 0 | 0 | 0 | 697 | 19 | 46 | 0 | 0 | 0 | 0 | 13 | 219 | 1314 | 102 | 162 | 0 | 18 | 14 | 0 | 0 |
| SRR5755997 | 233 | 8565 | 438 | 223 | 430 | 4681 | 7493 | 5860 | 13412 | 19 | 5528 | 6614 | 2953 | 3662 | 44 | 37 | 88186 | 0 | 0 | 0 | 22 | 0 | 0 | 0 | 0 | 0 | 0 | 0 | 47 | 123 | 261 | 22 | 0 | 0 | 0 | 0 | 0 |
| SRR5755998 | 167 | 29120 | 249 | 152 | 68 | 1106 | 828 | 6985 | 1360 | 22 | 27171 | 23158 | 19320 | 7386 | 0 | 129 | 89990 | 0 | 0 | 0 | 53 | 128 | 21 | 0 | 0 | 0 | 0 | 0 | 29 | 34 | 58 | 27 | 0 | 0 | 0 | 0 | 0 |
| SRR5755999 | 94 | 41078 | 195 | 339 | 260 | 3639 | 2839 | 7144 | 215 | 11 | 24107 | 16892 | 12876 | 3013 | 23 | 0 | 43507 | 0 | 0 | 0 | 57 | 16 | 14 | 0 | 0 | 0 | 0 | 0 | 25 | 54 | 22 | 28 | 0 | 0 | 0 | 0 | 0 |
| SRR5756000 | 0 | 812 | 487 | 325 | 0 | 35 | 30 | 1094 | 438 | 0 | 8480 | 3068 | 1221 | 5047 | 0 | 0 | 9313 | 0 | 0 | 0 | 0 | 0 | 0 | 0 | 0 | 0 | 0 | 0 | 0 | 0 | 0 | 0 | 0 | 0 | 0 | 0 | 0 |
| SRR5756001 | 69 | 11555 | 291 | 161 | 720 | 5348 | 6724 | 26769 | 947 | 0 | 17807 | 14900 | 25709 | 5583 | 27 | 10 | 102917 | 0 | 0 | 0 | 436 | 69 | 30 | 0 | 0 | 0 | 0 | 12 | 150 | 416 | 96 | 55 | 0 | 0 | 0 | 0 | 0 |
| SRR5756002 | 97 | 18535 | 478 | 186 | 242 | 2185 | 2689 | 12782 | 646 | 32 | 19041 | 13940 | 26313 | 7181 | 56 | 0 | 87694 | 0 | 0 | 0 | 105 | 14 | 0 | 0 | 0 | 0 | 0 | 0 | 46 | 105 | 22 | 24 | 0 | 0 | 0 | 0 | 0 |
| SRR5756003 | 222 | 12905 | 1038 | 280 | 4473 | 38400 | 58216 | 27785 | 3116 | 34 | 34586 | 40265 | 11989 | 6621 | 34 | 33 | 71537 | 0 | 0 | 0 | 945 | 57 | 43 | 0 | 0 | 0 | 0 | 11 | 325 | 1465 | 283 | 147 | 0 | 0 | 0 | 0 | 0 |
| SRR5756005 | 94 | 18440 | 326 | 134 | 68 | 1135 | 812 | 5923 | 381 | 0 | 26855 | 26724 | 23734 | 6119 | 18 | 57 | 115961 | 0 | 0 | 0 | 41 | 22 | 12 | 0 | 0 | 0 | 0 | 0 | 21 | 42 | 13 | 22 | 0 | 0 | 0 | 0 | 0 |
| SRR5756006 | 53 | 38750 | 564 | 159 | 206 | 2080 | 2109 | 5633 | 182 | 33 | 21576 | 16366 | 13542 | 3229 | 20 | 12 | 45059 | 0 | 0 | 0 | 97 | 0 | 15 | 0 | 0 | 0 | 0 | 0 | 54 | 105 | 19 | 25 | 0 | 0 | 0 | 0 | 0 |
| SRR5756007 | 0 | 1042 | 0 | 0 | 0 | 13 | 13 | 1271 | 252 | 0 | 7236 | 2710 | 1238 | 5793 | 0 | 0 | 6767 | 0 | 0 | 0 | 0 | 0 | 0 | 0 | 0 | 0 | 0 | 0 | 0 | 0 | 0 | 0 | 0 | 0 | 0 | 0 | 0 |
| SRR5756008 | 12 | 17164 | 457 | 189 | 240 | 1784 | 3618 | 16973 | 2257 | 30 | 9566 | 6103 | 11139 | 12810 | 0 | 0 | 63661 | 0 | 0 | 0 | 973 | 119 | 16 | 0 | 0 | 0 | 0 | 0 | 210 | 1117 | 228 | 34 | 0 | 0 | 0 | 0 | 0 |
| SRR5756009 | 206 | 19637 | 1510 | 119 | 3852 | 32799 | 49308 | 21480 | 2827 | 11 | 39040 | 38163 | 11571 | 9682 | 29 | 28 | 68924 | 0 | 0 | 0 | 1146 | 36 | 64 | 0 | 0 | 0 | 0 | 0 | 404 | 1418 | 152 | 179 | 0 | 0 | 0 | 0 | 0 |
| SRR5756011 | 169 | 29936 | 345 | 134 | 75 | 808 | 803 | 4018 | 779 | 0 | 25704 | 23866 | 18850 | 7940 | 23 | 122 | 85151 | 0 | 0 | 0 | 290 | 37 | 25 | 0 | 0 | 0 | 0 | 0 | 113 | 296 | 41 | 41 | 0 | 0 | 0 | 0 | 0 |
| SRR5756012 | 0 | 18600 | 111 | 124 | 209 | 1758 | 2339 | 6404 | 516 | 37 | 14208 | 12437 | 8937 | 5098 | 11 | 0 | 39093 | 0 | 0 | 0 | 2589 | 331 | 28 | 0 | 0 | 0 | 0 | 26 | 818 | 3703 | 890 | 92 | 0 | 0 | 0 | 0 | 0 |
| SRR5756014 | 31 | 6530 | 323 | 202 | 627 | 4543 | 11825 | 43426 | 1608 | 81 | 7845 | 5307 | 10010 | 12124 | 16 | 20 | 56870 | 0 | 0 | 0 | 2348 | 1171 | 76 | 0 | 0 | 0 | 0 | 13 | 534 | 3363 | 2067 | 229 | 0 | 0 | 0 | 0 | 0 |
| SRR5756016 | 215 | 23209 | 619 | 272 | 5898 | 39462 | 65226 | 26705 | 3837 | 21 | 36177 | 38226 | 11794 | 8329 | 38 | 40 | 62983 | 0 | 0 | 0 | 1633 | 76 | 54 | 0 | 0 | 0 | 0 | 51 | 760 | 3276 | 537 | 241 | 0 | 0 | 10 | 0 | 0 |
| SRR5756017 | 81 | 6192 | 508 | 162 | 301 | 2581 | 9565 | 6075 | 19922 | 20 | 3882 | 4004 | 1950 | 4265 | 36 | 40 | 65082 | 0 | 0 | 0 | 61 | 11 | 0 | 0 | 0 | 0 | 0 | 0 | 38 | 248 | 143 | 11 | 0 | 0 | 0 | 0 | 0 |
| SRR5756018 | 139 | 30053 | 434 | 139 | 102 | 1406 | 1024 | 7774 | 1333 | 0 | 28764 | 22661 | 19633 | 7571 | 22 | 47 | 88457 | 0 | 0 | 0 | 103 | 61 | 16 | 0 | 0 | 0 | 0 | 0 | 46 | 113 | 37 | 21 | 0 | 0 | 0 | 0 | 0 |
| SRR5756019 | 65 | 45213 | 557 | 376 | 385 | 3185 | 6882 | 9598 | 997 | 13 | 9915 | 5665 | 5069 | 9607 | 0 | 45 | 28648 | 0 | 0 | 0 | 729 | 122 | 16 | 0 | 0 | 0 | 0 | 0 | 216 | 1341 | 487 | 69 | 0 | 0 | 0 | 0 | 0 |
| SRR5756021 | 37 | 8251 | 176 | 146 | 540 | 3566 | 4748 | 27065 | 983 | 11 | 12605 | 11372 | 18903 | 4936 | 23 | 0 | 83940 | 0 | 0 | 0 | 1160 | 474 | 47 | 0 | 0 | 0 | 0 | 11 | 345 | 875 | 498 | 68 | 0 | 0 | 0 | 0 | 0 |
| SRR5756022 | 31 | 12841 | 313 | 123 | 210 | 1915 | 2143 | 11468 | 644 | 43 | 13002 | 11451 | 16594 | 6517 | 44 | 0 | 75365 | 0 | 0 | 0 | 742 | 151 | 20 | 0 | 0 | 0 | 0 | 0 | 236 | 603 | 192 | 30 | 0 | 0 | 0 | 0 | 0 |

### Supp Table S8.1.Columns_B_to_AL

| | | | | | | | | | | | | | | | | | |
|---|---|---|---|---|---|---|---|---|---|---|---|---|---|---|---|---|---|
| SRR5756023 | 378 | 21264 | 1310 | 257 | 5401 | 45697 | 60533 | 39359 | 585 | 0 | 30545 | 36423 | 9889 | 6321 | 37 | 25 | 74099 |
| SRR5756025 | 188 | 35291 | 393 | 143 | 53 | 881 | 780 | 5094 | 788 | 22 | 26263 | 22139 | 15788 | 7308 | 25 | 106 | 86285 |
| SRR5756026 | 44 | 24999 | 383 | 94 | 185 | 1828 | 1528 | 6449 | 175 | 19 | 15780 | 12516 | 9001 | 3606 | 28 | 0 | 41205 |
| SRR5756027 | 47 | 4998 | 257 | 162 | 530 | 3821 | 5419 | 26144 | 1550 | 37 | 12464 | 10799 | 18456 | 6721 | 21 | 0 | 100736 |
| SRR5756028 | 54 | 13534 | 395 | 116 | 247 | 1918 | 2660 | 13527 | 2124 | 15 | 14612 | 8810 | 15430 | 9227 | 0 | 0 | 83373 |
| SRR5756029 | 291 | 21747 | 1681 | 196 | 6093 | 41549 | 68096 | 37172 | 2686 | 25 | 33592 | 34744 | 10263 | 8132 | 35 | 20 | 67471 |
| SRR5756031 | 160 | 20921 | 275 | 105 | 46 | 616 | 546 | 3541 | 769 | 11 | 23464 | 19809 | 18377 | 6059 | 17 | 77 | 84796 |
| SRR5756032 | 26 | 20265 | 370 | 218 | 189 | 1480 | 1614 | 5896 | 232 | 40 | 14435 | 11141 | 8992 | 4677 | 0 | 0 | 48877 |
| SRR5756033 | 47 | 10137 | 154 | 221 | 386 | 3208 | 2778 | 19894 | 2565 | 0 | 23080 | 17825 | 31419 | 4316 | 12 | 0 | 110510 |
| SRR5756034 | 59 | 24822 | 168 | 162 | 110 | 1158 | 925 | 7832 | 699 | 0 | 22371 | 18230 | 33324 | 3984 | 29 | 0 | 94212 |
| SRR5756035 | 349 | 27097 | 557 | 184 | 2702 | 25706 | 35079 | 19086 | 1884 | 12 | 43231 | 41564 | 13233 | 7779 | 32 | 21 | 73541 |
| SRR5756037 | 75 | 28138 | 259 | 90 | 55 | 788 | 657 | 5913 | 1742 | 0 | 28507 | 22286 | 23732 | 6695 | 21 | 53 | 82467 |
| SRR5756038 | 58 | 46792 | 111 | 48 | 122 | 1458 | 1076 | 6046 | 97 | 0 | 23671 | 18157 | 15397 | 2974 | 33 | 0 | 44807 |
| SRR5756040 | 64 | 13147 | 158 | 218 | 399 | 3870 | 3533 | 23156 | 754 | 0 | 19404 | 17838 | 26534 | 5228 | 18 | 0 | 115253 |
| SRR5756041 | 80 | 30905 | 477 | 102 | 377 | 4159 | 3179 | 15341 | 722 | 17 | 23918 | 16029 | 21820 | 7100 | 26 | 0 | 84303 |
| SRR5756042 | 422 | 9909 | 1609 | 258 | 4817 | 41171 | 62803 | 35216 | 1777 | 23 | 41445 | 36228 | 12651 | 7838 | 36 | 29 | 61608 |
| SRR5756044 | 189 | 15805 | 330 | 92 | 117 | 1608 | 1325 | 8235 | 712 | 12 | 25436 | 19858 | 25677 | 6138 | 26 | 59 | 77823 |
| SRR5756045 | 42 | 58985 | 399 | 112 | 177 | 2496 | 1678 | 6226 | 114 | 28 | 21427 | 17825 | 10194 | 4476 | 0 | 13 | 50750 |
| SRR5756047 | 73 | 13553 | 87 | 188 | 418 | 2868 | 2666 | 19817 | 2290 | 0 | 19174 | 14770 | 25486 | 3617 | 12 | 0 | 109791 |
| SRR5756048 | 74 | 17369 | 356 | 130 | 128 | 1081 | 1282 | 5353 | 793 | 0 | 14850 | 10599 | 19694 | 3851 | 53 | 0 | 68340 |
| SRR5756049 | 456 | 3942 | 954 | 317 | 5103 | 37133 | 60625 | 26610 | 1141 | 0 | 38047 | 37703 | 12743 | 6014 | 23 | 18 | 59844 |
| SRR5756051 | 357 | 22386 | 384 | 202 | 125 | 1238 | 1302 | 6639 | 4217 | 12 | 25098 | 18118 | 20113 | 6163 | 35 | 114 | 67211 |
| SRR5756052 | 82 | 23943 | 309 | 134 | 189 | 1791 | 1596 | 4101 | 162 | 41 | 19367 | 13929 | 10134 | 2679 | 0 | 0 | 45757 |
| SRR5756054 | 97 | 9018 | 187 | 155 | 467 | 3462 | 3547 | 23766 | 2078 | 0 | 17496 | 12875 | 23741 | 4858 | 0 | 0 | 100595 |
| SRR5756055 | 64 | 12551 | 480 | 339 | 188 | 1609 | 2007 | 8349 | 548 | 0 | 16005 | 11491 | 21894 | 4942 | 27 | 0 | 70894 |
| SRR5756056 | 393 | 5905 | 1216 | 365 | 6124 | 43503 | 75026 | 33800 | 2505 | 22 | 38340 | 49 | 10873 | 8850 | 26 | 12 | 54585 |
| SRR5756058 | 207 | 13208 | 380 | 167 | 80 | 861 | 920 | 5806 | 747 | 0 | 22014 | 17848 | 19920 | 5930 | 28 | 77 | 74424 |
| SRR5756059 | 93 | 25158 | 1091 | 252 | 261 | 2268 | 2031 | 5677 | 199 | 0 | 19225 | 16267 | 10613 | 3524 | 0 | 0 | 41023 |
| SRR5756061 | 106 | 25371 | 363 | 319 | 562 | 3474 | 4135 | 23482 | 2492 | 0 | 16661 | 11522 | 21101 | 5550 | 12 | 0 | 89867 |
| SRR5756062 | 68 | 12963 | 263 | 277 | 379 | 2368 | 3418 | 10479 | 1467 | 24 | 22635 | 14085 | 24947 | 8763 | 0 | 0 | 53648 |
| SRR5756063 | 515 | 10874 | 928 | 338 | 6597 | 40933 | 68963 | 34242 | 2866 | 23 | 35400 | 31597 | 11797 | 7076 | 23 | 13 | 59900 |
| SRR5756065 | 282 | 19603 | 331 | 290 | 119 | 1625 | 1624 | 9160 | 3751 | 12 | 26688 | 19145 | 20289 | 7474 | 17 | 81 | 62704 |
| SRR5756066 | 67 | 29695 | 124 | 545 | 213 | 1634 | 1804 | 5353 | 318 | 16 | 21207 | 14089 | 13161 | 4392 | 24 | 0 | 32107 |
| SRR5756068 | 58 | 6832 | 145 | 196 | 329 | 2726 | 3077 | 13069 | 1104 | 0 | 18198 | 15032 | 24221 | 4003 | 0 | 0 | 96839 |
| SRR5756069 | 61 | 12120 | 326 | 232 | 125 | 1175 | 1350 | 5944 | 721 | 16 | 18509 | 10 | 27442 | 4158 | 14 | 0 | 84052 |
| SRR5756070 | 376 | 12459 | 1127 | 220 | 5274 | 36975 | 50302 | 27238 | 2830 | 33 | 39066 | 21 | 12219 | 6218 | 17 | 27 | 71517 |
| SRR5756072 | 168 | 18705 | 572 | 162 | 93 | 1033 | 1073 | 6063 | 1203 | 0 | 23633 | 16973 | 20224 | 7576 | 21 | 57 | 70754 |
| SRR5756073 | 124 | 20705 | 1241 | 102 | 178 | 1462 | 1368 | 4242 | 173 | 15 | 24603 | 15729 | 12292 | 3290 | 26 | 0 | 52701 |
| SRR5756075 | 100 | 12034 | 150 | 125 | 426 | 3573 | 2980 | 20131 | 2742 | 0 | 20898 | 17474 | 30738 | 4298 | 12 | 0 | 120957 |
| SRR5756076 | 87 | 16615 | 225 | 223 | 140 | 1637 | 1499 | 6711 | 788 | 15 | 19914 | 14897 | 27570 | 4211 | 11 | 0 | 95389 |
| SRR5756077 | 207 | 11298 | 800 | 138 | 5447 | 38424 | 48552 | 30570 | 3738 | 28 | 34453 | 33127 | 10503 | 6395 | 22 | 19 | 71435 |
| SRR5756079 | 157 | 18816 | 516 | 153 | 90 | 994 | 951 | 5805 | 1061 | 18 | 27099 | 18993 | 20208 | 7838 | 31 | 27 | 84730 |
| SRR5756080 | 57 | 24653 | 205 | 188 | 179 | 1606 | 1336 | 5253 | 275 | 0 | 25258 | 17875 | 15360 | 3061 | 15 | 0 | 46743 |
| SRR5756082 | 97 | 8619 | 196 | 161 | 583 | 4951 | 5522 | 24732 | 766 | 15 | 20089 | 15533 | 31838 | 4420 | 16 | 0 | 94753 |
| SRR5756083 | 128 | 23154 | 258 | 133 | 494 | 5148 | 4445 | 18206 | 650 | 14 | 28718 | 18148 | 29120 | 5967 | 26 | 0 | 70716 |
| SRR5756084 | 394 | 14404 | 741 | 224 | 6705 | 48477 | 62767 | 41528 | 2698 | 15 | 38763 | 39106 | 13161 | 7087 | 33 | 37 | 63045 |
| SRR5756086 | 245 | 11227 | 455 | 127 | 169 | 1764 | 1859 | 9954 | 1150 | 17 | 22707 | 16067 | 20762 | 6254 | 17 | 71 | 73911 |
| SRR5756087 | 75 | 38993 | 236 | 179 | 352 | 4229 | 3636 | 10336 | 125 | 17 | 29031 | 23898 | 15790 | 3727 | 17 | 0 | 50435 |
| SRR5756089 | 35 | 14973 | 82 | 94 | 487 | 4269 | 3826 | 24476 | 2125 | 0 | 19465 | 15291 | 27833 | 4441 | 17 | 0 | 106267 |
| SRR5756090 | 66 | 22316 | 248 | 101 | 229 | 2000 | 2059 | 11500 | 816 | 11 | 19290 | 13880 | 30816 | 4722 | 54 | 0 | 72820 |
| SRR5756091 | 250 | 5293 | 1437 | 170 | 5192 | 39121 | 63598 | 30391 | 1741 | 30 | 46167 | 41891 | 16639 | 7581 | 46 | 19 | 54435 |
| SRR5756093 | 208 | 12711 | 419 | 84 | 117 | 1123 | 1050 | 6841 | 862 | 14 | 26781 | 19490 | 25147 | 7216 | 31 | 50 | 68246 |
| SRR5756094 | 72 | 35616 | 264 | 94 | 242 | 2191 | 1767 | 7152 | 273 | 14 | 25691 | 16519 | 16020 | 2836 | 28 | 0 | 39161 |
| SRR5756096 | 73 | 4177 | 407 | 174 | 821 | 5986 | 8051 | 31461 | 1141 | 69 | 15428 | 10712 | 22480 | 6236 | 22 | 0 | 75799 |
| SRR5756097 | 0 | 11828 | 1745 | 154 | 573 | 3900 | 6690 | 21761 | 825 | 81 | 13290 | 7905 | 14898 | 8380 | 40 | 0 | 71038 |
| SRR5756098 | 569 | 17274 | 575 | 260 | 5624 | 43855 | 61440 | 39258 | 3996 | 27 | 36110 | 37356 | 12323 | 7135 | 23 | 39 | 66828 |
| SRR5756100 | 176 | 23763 | 417 | 126 | 74 | 789 | 672 | 4492 | 409 | 17 | 12 | 20898 | 22242 | 9346 | 10 | 229 | 67344 |
| SRR5756101 | 167 | 34520 | 1844 | 131 | 440 | 3682 | 3622 | 8389 | 170 | 60 | 18949 | 12546 | 11358 | 3701 | 20 | 0 | 37441 |
| SRR5756103 | 111 | 14508 | 52 | 97 | 432 | 3784 | 3040 | 20219 | 1486 | 0 | 21219 | 17426 | 33457 | 4234 | 18 | 0 | 119700 |
| SRR5756104 | 108 | 30529 | 195 | 127 | 199 | 2185 | 1691 | 10871 | 404 | 12 | 12 | 19042 | 32544 | 4509 | 23 | 0 | 97902 |
| SRR5756105 | 870 | 8111 | 603 | 150 | 5269 | 44365 | 59049 | 35133 | 1126 | 80 | 42937 | 50342 | 16916 | 5575 | 22 | 38 | 64937 |
| SRR5756106 | 461 | 8952 | 515 | 118 | 743 | 8240 | 7767 | 5093 | 9586 | 27 | 10464 | 13412 | 4305 | 2636 | 12 | 86 | 142747 |
| SRR5756107 | 103 | 20970 | 228 | 64 | 99 | 1334 | 1059 | 9050 | 1400 | 12 | 27728 | 22914 | 28579 | 7431 | 12 | 60 | 82327 |
| SRR5756108 | 93 | 74291 | 176 | 119 | 173 | 1988 | 1676 | 5300 | 151 | 21 | 11 | 17844 | 16157 | 3547 | 18 | 16 | 43589 |
| SRR5756110 | 0 | 2088 | 672 | 106 | 0 | 22 | 35 | 211 | 388 | 35 | 448 | 481 | 170 | 348 | 0 | 44 | 2572 |
| SRR5756114 | 24 | 4102 | 996 | 213 | 0 | 37 | 60 | 295 | 529 | 0 | 817 | 805 | 272 | 544 | 0 | 0 | 2437 |
| SRR5756115 | 0 | 2982 | 877 | 351 | 0 | 22 | 33 | 228 | 420 | 0 | 519 | 583 | 192 | 293 | 0 | 29 | 2007 |
| SRR5756116 | 68 | 3995 | 677 | 68 | 0 | 26 | 53 | 240 | 243 | 0 | 778 | 726 | 202 | 205 | 0 | 21 | 1744 |
| SRR5756118 | 0 | 1965 | 513 | 171 | 0 | 15 | 27 | 208 | 238 | 0 | 586 | 510 | 171 | 286 | 0 | 0 | 1864 |
| SRR5756120 | 0 | 1871 | 922 | 163 | 0 | 35 | 68 | 287 | 340 | 27 | 721 | 926 | 272 | 376 | 0 | 0 | 2100 |
| SRR5756121 | 0 | 2593 | 791 | 352 | 0 | 23 | 43 | 201 | 181 | 0 | 324 | 434 | 137 | 303 | 0 | 0 | 1759 |
| SRR5756122 | 0 | 2101 | 718 | 133 | 0 | 51 | 71 | 507 | 328 | 0 | 937 | 966 | 356 | 1237 | 0 | 0 | 2190 |
| SRR5756123 | 0 | 2319 | 773 | 86 | 0 | 22 | 32 | 205 | 391 | 0 | 356 | 537 | 165 | 553 | 0 | 0 | 1248 |
| SRR5756125 | 16 | 1661 | 617 | 47 | 0 | 84 | 81 | 564 | 1279 | 16 | 1535 | 1618 | 398 | 1431 | 0 | 0 | 2835 |
| SRR5756128 | 0 | 2688 | 1761 | 278 | 0 | 34 | 62 | 294 | 201 | 0 | 403 | 308 | 101 | 83 | 0 | 0 | 2756 |
| SRR5756130 | 23 | 2345 | 722 | 113 | 0 | 43 | 61 | 442 | 527 | 0 | 988 | 983 | 349 | 1386 | 0 | 0 | 1960 |
| SRR5756131 | 0 | 2735 | 690 | 77 | 0 | 20 | 21 | 161 | 175 | 0 | 393 | 412 | 149 | 529 | 0 | 0 | 1297 |
| SRR5756132 | 0 | 2407 | 951 | 0 | 0 | 50 | 71 | 445 | 387 | 0 | 1240 | 1108 | 371 | 1030 | 0 | 0 | 1805 |
| SRR5756133 | 36 | 7831 | 1129 | 219 | 0 | 22 | 57 | 228 | 400 | 0 | 738 | 430 | 257 | 199 | 0 | 0 | 2726 |
| SRR5756135 | 0 | 3826 | 662 | 184 | 0 | 38 | 60 | 257 | 268 | 37 | 786 | 1015 | 349 | 366 | 0 | 0 | 2282 |
| SRR5756136 | 36 | 4085 | 904 | 578 | 0 | 61 | 122 | 440 | 358 | 0 | 596 | 443 | 206 | 286 | 0 | 59 | 4566 |
| SRR5756137 | 71 | 3957 | 570 | 499 | 0 | 55 | 88 | 371 | 743 | 18 | 940 | 973 | 495 | 733 | 0 | 0 | 1826 |
| SRR5756138 | 86 | 3834 | 486 | 286 | 0 | 23 | 38 | 168 | 230 | 0 | 502 | 624 | 203 | 285 | 0 | 0 | 2004 |
| SRR5756139 | 69 | 7214 | 794 | 690 | 0 | 29 | 41 | 287 | 441 | 0 | 811 | 861 | 321 | 308 | 0 | 0 | 2190 |
| SRR5756140 | 0 | 7669 | 492 | 463 | 0 | 37 | 51 | 267 | 344 | 29 | 712 | 732 | 307 | 833 | 0 | 38 | 1741 |
| SRR5756141 | 32 | 3093 | 335 | 335 | 0 | 53 | 81 | 419 | 511 | 0 | 1152 | 1150 | 474 | 573 | 0 | 56 | 2483 |
| SRR5756142 | 18 | 2182 | 273 | 509 | 0 | 39 | 60 | 445 | 467 | 0 | 1590 | 1249 | 535 | 826 | 0 | 54 | 2253 |
| SRR5756143 | 36 | 3224 | 522 | 504 | 0 | 51 | 77 | 509 | 498 | 0 | 1754 | 1718 | 592 | 850 | 0 | 0 | 2345 |
| SRR5756144 | 41 | 1816 | 530 | 490 | 0 | 29 | 46 | 305 | 455 | 0 | 866 | 861 | 304 | 500 | 0 | 0 | 2109 |
| SRR5756145 | 76 | 4204 | 764 | 268 | 0 | 33 | 51 | 236 | 188 | 0 | 774 | 632 | 241 | 323 | 0 | 0 | 1360 |
| SRR5756146 | 20 | 4112 | 731 | 237 | 0 | 39 | 66 | 300 | 419 | 0 | 510 | 558 | 202 | 140 | 0 | 0 | 3053 |
| SRR5756147 | 39 | 5185 | 507 | 429 | 0 | 25 | 45 | 200 | 295 | 0 | 538 | 389 | 159 | 119 | 0 | 0 | 2460 |
| SRR5756148 | 0 | 3127 | 674 | 270 | 0 | 31 | 44 | 230 | 242 | 27 | 601 | 590 | 238 | 394 | 0 | 0 | 1692 |
| SRR5756149 | 151 | 3588 | 680 | 302 | 0 | 38 | 60 | 308 | 419 | 0 | 625 | 634 | 266 | 411 | 0 | 0 | 3176 |
| SRR5756150 | 97 | 3647 | 582 | 737 | 0 | 45 | 55 | 315 | 494 | 0 | 826 | 744 | 336 | 590 | 0 | 0 | 1900 |
| SRR5756151 | 77 | 5930 | 689 | 612 | 0 | 31 | 47 | 388 | 230 | 0 | 1039 | 1198 | 412 | 356 | 0 | 0 | 2357 |
| SRR5756152 | 87 | 7879 | 503 | 417 | 0 | 38 | 49 | 252 | 623 | 0 | 868 | 876 | 250 | 222 | 0 | 0 | 1528 |
| SRR5756153 | 71 | 3026 | 544 | 544 | 0 | 28 | 42 | 216 | 268 | 24 | 661 | 677 | 240 | 276 | 0 | 0 | 1986 |
| SRR5756154 | 0 | 5245 | 497 | 588 | 0 | 29 | 40 | 249 | 271 | 0 | 526 | 447 | 174 | 114 | 0 | 0 | 2100 |
| SRR5756155 | 70 | 2873 | 363 | 321 | 0 | 23 | 33 | 209 | 182 | 28 | 449 | 420 | 177 | 188 | 0 | 0 | 1911 |
| SRR5756156 | 62 | 6008 | 165 | 391 | 0 | 44 | 63 | 428 | 851 | 21 | 1380 | 1572 | 503 | 527 | 0 | 0 | 2539 |
| SRR5756157 | 38 | 2869 | 551 | 437 | 0 | 31 | 48 | 375 | 422 | 19 | 917 | 962 | 375 | 776 | 0 | 0 | 1250 |
| SRR5756158 | 0 | 4272 | 520 | 294 | 0 | 43 | 58 | 347 | 438 | 0 | 1200 | 1395 | 463 | 792 | 0 | 0 | 2373 |
| SRR5756159 | 0 | 3793 | 337 | 506 | 0 | 61 | 93 | 464 | 645 | 0 | 1158 | 1026 | 570 | 1416 | 0 | 0 | 1552 |
| SRR5756160 | 0 | 2584 | 1213 | 53 | 0 | 42 | 89 | 471 | 518 | 0 | 1326 | 1121 | 478 | 521 | 0 | 0 | 2160 |
| SRR5756162 | 58 | 3370 | 1046 | 116 | 0 | 54 | 95 | 551 | 602 | 0 | 1792 | 1698 | 613 | 699 | 0 | 65 | 2217 |
| SRR5756164 | 0 | 1181 | 713 | 0 | 0 | 114 | 195 | 944 | 647 | 0 | 2418 | 1607 | 998 | 2546 | 0 | 0 | 3528 |
| SRR5756166 | 0 | 1955 | 551 | 25 | 0 | 41 | 65 | 425 | 443 | 25 | 1029 | 951 | 376 | 693 | 0 | 45 | 2307 |
| SRR5756167 | 0 | 1683 | 827 | 89 | 0 | 50 | 92 | 582 | 418 | 0 | 1835 | 1343 | 662 | 1945 | 0 | 0 | 2203 |
| SRR5756168 | 0 | 1421 | 639 | 36 | 0 | 56 | 97 | 504 | 371 | 36 | 1684 | 1894 | 651 | 561 | 0 | 0 | 2546 |
| SRR5756177 | 0 | 3180 | 658 | 110 | 0 | 25 | 52 | 259 | 218 | 0 | 495 | 391 | 143 | 194 | 0 | 0 | 2442 |
| SRR5756178 | 0 | 2320 | 892 | 268 | 0 | 42 | 79 | 307 | 201 | 0 | 393 | 439 | 138 | 80 | 0 | 0 | 2753 |
| SRR5756179 | 0 | 3467 | 743 | 83 | 0 | 35 | 56 | 361 | 310 | 0 | 1061 | 1163 | 344 | 500 | 0 | 0 | 2051 |
| SRR5756180 | 0 | 1726 | 1114 | 111 | 0 | 53 | 96 | 473 | 497 | 56 | 1429 | 1494 | 571 | 403 | 0 | 0 | 1637 |
| SRR5756182 | 0 | 1415 | 1082 | 0 | 0 | 25 | 50 | 216 | 167 | 0 | 660 | 544 | 189 | 266 | 0 | 0 | 2436 |
| SRR5756183 | 34 | 4184 | 898 | 102 | 0 | 35 | 63 | 235 | 240 | 11 | 811 | 629 | 227 | 152 | 0 | 0 | 2403 |
| SRR5756184 | 90 | 2268 | 1015 | 269 | 0 | 36 | 55 | 270 | 260 | 0 | 852 | 396 | 140 | 141 | 0 | 0 | 2570 |
| SRR5756185 | 35 | 1914 | 679 | 400 | 0 | 15 | 26 | 160 | 525 | 0 | 526 | 639 | 178 | 366 | 0 | 0 | 1150 |
| SRR5756186 | 33 | 2083 | 541 | 213 | 0 | 14 | 25 | 152 | 490 | 0 | 497 | 603 | 177 | 300 | 0 | 0 | 1396 |
| SRR5804909 | 1734 | 350 | 1802 | 132 | 6458 | 4347 | 47690 | 5837 | 1863 | 0 | 56508 | 81967 | 57005 | 153254 | 2360 | 5713 | 6955 |
| SRR5804912 | 605 | 193 | 734 | 144 | 12695 | 7579 | 156601 | 26148 | 332 | 0 | 36853 | 33063 | 33442 | 112690 | 4547 | 33348 | 6726 |
| SRR5804914 | 1323 | 669 | 3451 | 430 | 7045 | 9619 | 52026 | 9619 | 677 | 0 | 83864 | 86438 | 134941 | 126948 | 1897 | 10839 | 3252 |
| SRR5817623 | 213 | 321 | 128 | 2269 | 23511 | 8793 | 70271 | 76438 | 9681 | 398 | 19053 | 6604 | 11631 | 7132 | 44755 | 944 | 18303 |
| SRR5817624 | 145 | 690 | 435 | 2848 | 21807 | 7653 | 59260 | 85359 | 6927 | 471 | 19654 | 5646 | 11602 | 5511 | 30158 | 611 | 21751 |

| | | | | | | | | | | | | | | | | | | | |
|---|---|---|---|---|---|---|---|---|---|---|---|---|---|---|---|---|---|---|---|
| SRR5756023 | 0 | 0 | 0 | 1163 | 48 | 54 | 0 | 0 | 0 | 0 | 0 | 289 | 1011 | 182 | 134 | 0 | 0 | 0 | 0 | 0 |
| SRR5756025 | 0 | 0 | 0 | 117 | 115 | 23 | 0 | 0 | 0 | 0 | 0 | 48 | 89 | 47 | 29 | 0 | 0 | 0 | 0 | 0 |
| SRR5756026 | 0 | 0 | 0 | 845 | 116 | 20 | 0 | 0 | 0 | 0 | 14 | 460 | 964 | 308 | 45 | 0 | 0 | 0 | 0 | 0 |
| SRR5756027 | 0 | 0 | 13 | 2467 | 540 | 60 | 0 | 0 | 0 | 0 | 16 | 562 | 1910 | 616 | 94 | 0 | 0 | 0 | 0 | 0 |
| SRR5756028 | 0 | 0 | 0 | 125 | 13 | 0 | 0 | 0 | 0 | 0 | 0 | 73 | 94 | 16 | 11 | 0 | 0 | 0 | 0 | 0 |
| SRR5756029 | 0 | 0 | 0 | 2472 | 97 | 102 | 0 | 0 | 0 | 0 | 16 | 789 | 2891 | 433 | 294 | 0 | 0 | 15 | 0 | 0 |
| SRR5756031 | 0 | 0 | 0 | 233 | 43 | 24 | 0 | 0 | 0 | 0 | 0 | 92 | 244 | 45 | 39 | 0 | 0 | 0 | 0 | 0 |
| SRR5756032 | 0 | 0 | 0 | 1268 | 161 | 26 | 0 | 0 | 0 | 0 | 13 | 556 | 1312 | 345 | 50 | 0 | 0 | 0 | 0 | 0 |
| SRR5756033 | 0 | 0 | 0 | 297 | 28 | 25 | 0 | 0 | 0 | 0 | 13 | 111 | 253 | 31 | 36 | 0 | 0 | 0 | 0 | 0 |
| SRR5756034 | 0 | 0 | 0 | 223 | 37 | 24 | 0 | 0 | 0 | 0 | 0 | 72 | 134 | 28 | 25 | 0 | 0 | 0 | 0 | 0 |
| SRR5756035 | 0 | 0 | 0 | 1496 | 45 | 69 | 0 | 0 | 0 | 0 | 0 | 454 | 1877 | 171 | 195 | 0 | 0 | 0 | 0 | 0 |
| SRR5756037 | 0 | 0 | 0 | 93 | 105 | 18 | 0 | 0 | 0 | 0 | 0 | 27 | 59 | 49 | 26 | 0 | 0 | 0 | 0 | 0 |
| SRR5756038 | 0 | 0 | 0 | 146 | 12 | 19 | 0 | 0 | 0 | 0 | 0 | 29 | 100 | 13 | 29 | 0 | 0 | 0 | 0 | 0 |
| SRR5756040 | 0 | 0 | 0 | 457 | 79 | 30 | 0 | 0 | 0 | 0 | 0 | 127 | 316 | 73 | 36 | 0 | 0 | 0 | 0 | 0 |
| SRR5756041 | 0 | 0 | 0 | 146 | 27 | 12 | 0 | 0 | 0 | 0 | 0 | 61 | 142 | 30 | 26 | 0 | 0 | 0 | 0 | 0 |
| SRR5756042 | 0 | 0 | 0 | 758 | 26 | 37 | 0 | 0 | 0 | 0 | 0 | 367 | 1138 | 113 | 130 | 0 | 0 | 0 | 0 | 0 |
| SRR5756044 | 0 | 0 | 0 | 124 | 18 | 10 | 0 | 0 | 0 | 0 | 0 | 33 | 128 | 0 | 31 | 0 | 0 | 0 | 0 | 0 |
| SRR5756045 | 0 | 0 | 0 | 72 | 0 | 0 | 0 | 0 | 0 | 0 | 0 | 50 | 64 | 0 | 0 | 0 | 0 | 0 | 0 | 0 |
| SRR5756047 | 0 | 0 | 0 | 374 | 125 | 23 | 0 | 0 | 0 | 0 | 0 | 62 | 194 | 70 | 27 | 0 | 0 | 0 | 0 | 0 |
| SRR5756048 | 0 | 0 | 0 | 601 | 219 | 24 | 0 | 0 | 0 | 0 | 0 | 108 | 293 | 102 | 31 | 0 | 0 | 0 | 0 | 0 |
| SRR5756049 | 0 | 0 | 0 | 778 | 43 | 39 | 0 | 0 | 0 | 0 | 16 | 208 | 684 | 119 | 106 | 0 | 0 | 11 | 0 | 0 |
| SRR5756051 | 0 | 0 | 0 | 187 | 580 | 93 | 0 | 0 | 0 | 0 | 0 | 32 | 84 | 118 | 55 | 0 | 0 | 12 | 15 | 0 |
| SRR5756052 | 0 | 0 | 0 | 571 | 104 | 22 | 0 | 0 | 0 | 0 | 0 | 159 | 436 | 129 | 33 | 0 | 0 | 0 | 0 | 0 |
| SRR5756054 | 0 | 0 | 0 | 333 | 191 | 24 | 0 | 0 | 0 | 0 | 0 | 121 | 283 | 183 | 38 | 0 | 0 | 0 | 0 | 0 |
| SRR5756055 | 0 | 0 | 0 | 290 | 185 | 15 | 0 | 0 | 0 | 0 | 0 | 77 | 264 | 161 | 30 | 0 | 0 | 0 | 0 | 0 |
| SRR5756056 | 0 | 0 | 0 | 500 | 43 | 24 | 0 | 0 | 0 | 0 | 16 | 242 | 965 | 198 | 105 | 0 | 0 | 0 | 0 | 0 |
| SRR5756058 | 0 | 0 | 0 | 71 | 30 | 0 | 0 | 0 | 0 | 0 | 0 | 32 | 82 | 35 | 25 | 0 | 0 | 0 | 0 | 0 |
| SRR5756059 | 0 | 0 | 0 | 104 | 16 | 13 | 0 | 0 | 0 | 0 | 0 | 45 | 112 | 22 | 24 | 0 | 0 | 0 | 0 | 0 |
| SRR5756061 | 0 | 0 | 0 | 1042 | 285 | 47 | 0 | 0 | 0 | 0 | 12 | 315 | 1097 | 321 | 79 | 0 | 0 | 0 | 0 | 0 |
| SRR5756062 | 0 | 0 | 0 | 680 | 149 | 26 | 0 | 0 | 0 | 0 | 0 | 229 | 904 | 161 | 50 | 0 | 0 | 0 | 0 | 0 |
| SRR5756063 | 0 | 0 | 0 | 2363 | 88 | 96 | 0 | 0 | 0 | 0 | 20 | 769 | 4218 | 457 | 482 | 0 | 0 | 0 | 0 | 0 |
| SRR5756065 | 0 | 0 | 0 | 383 | 225 | 59 | 0 | 0 | 0 | 0 | 0 | 113 | 430 | 115 | 93 | 0 | 0 | 0 | 11 | 0 |
| SRR5756066 | 0 | 0 | 0 | 787 | 153 | 37 | 0 | 0 | 0 | 0 | 14 | 368 | 1080 | 290 | 108 | 0 | 0 | 15 | 0 | 0 |
| SRR5756068 | 0 | 0 | 0 | 684 | 116 | 38 | 0 | 0 | 0 | 0 | 12 | 202 | 623 | 100 | 66 | 0 | 0 | 0 | 0 | 0 |
| SRR5756069 | 0 | 0 | 0 | 345 | 71 | 18 | 0 | 0 | 0 | 0 | 0 | 79 | 271 | 64 | 27 | 0 | 0 | 0 | 0 | 0 |
| SRR5756070 | 0 | 0 | 0 | 1274 | 72 | 48 | 0 | 0 | 0 | 0 | 22 | 455 | 1794 | 318 | 168 | 0 | 11 | 12 | 0 | 0 |
| SRR5756072 | 0 | 0 | 0 | 204 | 47 | 19 | 0 | 0 | 0 | 0 | 0 | 75 | 231 | 33 | 48 | 0 | 0 | 0 | 0 | 0 |
| SRR5756073 | 0 | 0 | 0 | 69 | 11 | 0 | 0 | 0 | 0 | 0 | 0 | 31 | 68 | 24 | 24 | 0 | 0 | 0 | 0 | 0 |
| SRR5756075 | 0 | 0 | 0 | 383 | 65 | 28 | 0 | 0 | 0 | 0 | 12 | 127 | 322 | 58 | 40 | 0 | 0 | 0 | 0 | 0 |
| SRR5756076 | 0 | 0 | 0 | 303 | 64 | 28 | 0 | 0 | 0 | 0 | 0 | 86 | 239 | 65 | 39 | 0 | 0 | 0 | 0 | 0 |
| SRR5756077 | 0 | 0 | 0 | 1451 | 86 | 62 | 0 | 0 | 0 | 0 | 14 | 450 | 1829 | 448 | 198 | 0 | 11 | 0 | 0 | 0 |
| SRR5756079 | 0 | 0 | 0 | 123 | 43 | 14 | 0 | 0 | 0 | 0 | 0 | 38 | 145 | 31 | 30 | 0 | 0 | 0 | 0 | 0 |
| SRR5756080 | 0 | 0 | 0 | 196 | 66 | 16 | 0 | 0 | 0 | 0 | 0 | 79 | 212 | 89 | 45 | 0 | 0 | 0 | 0 | 0 |
| SRR5756082 | 0 | 0 | 0 | 242 | 28 | 22 | 0 | 0 | 0 | 0 | 0 | 87 | 218 | 30 | 33 | 0 | 0 | 0 | 0 | 0 |
| SRR5756083 | 0 | 0 | 0 | 150 | 19 | 11 | 0 | 0 | 0 | 0 | 0 | 54 | 155 | 17 | 36 | 0 | 0 | 0 | 0 | 0 |
| SRR5756084 | 0 | 0 | 0 | 1782 | 84 | 58 | 0 | 0 | 0 | 0 | 20 | 593 | 2516 | 465 | 232 | 0 | 16 | 17 | 0 | 0 |
| SRR5756086 | 0 | 0 | 0 | 154 | 13 | 15 | 0 | 0 | 0 | 0 | 0 | 50 | 187 | 12 | 29 | 0 | 0 | 0 | 0 | 0 |
| SRR5756087 | 0 | 0 | 0 | 66 | 0 | 0 | 0 | 0 | 0 | 0 | 0 | 20 | 75 | 14 | 18 | 0 | 0 | 0 | 0 | 0 |
| SRR5756089 | 0 | 0 | 0 | 119 | 32 | 17 | 0 | 0 | 0 | 0 | 0 | 51 | 88 | 44 | 24 | 0 | 0 | 0 | 0 | 0 |
| SRR5756090 | 0 | 0 | 0 | 132 | 39 | 19 | 0 | 0 | 0 | 0 | 0 | 54 | 105 | 37 | 35 | 0 | 0 | 0 | 0 | 0 |
| SRR5756091 | 0 | 0 | 0 | 253 | 11 | 19 | 0 | 0 | 0 | 0 | 0 | 82 | 382 | 45 | 57 | 0 | 0 | 0 | 0 | 0 |
| SRR5756093 | 0 | 0 | 0 | 72 | 32 | 12 | 0 | 0 | 0 | 0 | 0 | 25 | 67 | 23 | 26 | 0 | 0 | 0 | 0 | 0 |
| SRR5756094 | 0 | 0 | 0 | 72 | 11 | 14 | 0 | 0 | 0 | 0 | 0 | 35 | 72 | 26 | 44 | 0 | 0 | 0 | 0 | 0 |
| SRR5756096 | 0 | 0 | 0 | 1559 | 595 | 56 | 0 | 0 | 0 | 0 | 17 | 469 | 1493 | 488 | 100 | 0 | 0 | 0 | 0 | 0 |
| SRR5756097 | 0 | 0 | 0 | 228 | 29 | 0 | 0 | 0 | 0 | 0 | 0 | 148 | 225 | 25 | 25 | 0 | 0 | 0 | 0 | 0 |
| SRR5756098 | 0 | 0 | 0 | 963 | 48 | 38 | 0 | 0 | 0 | 0 | 0 | 373 | 1607 | 212 | 120 | 0 | 0 | 0 | 0 | 0 |
| SRR5756100 | 0 | 0 | 0 | 37 | 0 | 0 | 0 | 0 | 0 | 0 | 0 | 24 | 43 | 0 | 0 | 0 | 0 | 0 | 0 | 0 |
| SRR5756101 | 0 | 0 | 0 | 101 | 0 | 0 | 0 | 0 | 0 | 0 | 0 | 90 | 100 | 0 | 22 | 0 | 0 | 0 | 0 | 0 |
| SRR5756103 | 0 | 0 | 0 | 32 | 20 | 27 | 0 | 0 | 0 | 0 | 0 | 0 | 25 | 20 | 45 | 0 | 15 | 0 | 0 | 0 |
| SRR5756104 | 0 | 0 | 0 | 13 | 0 | 11 | 0 | 0 | 0 | 0 | 0 | 0 | 11 | 0 | 20 | 0 | 0 | 0 | 0 | 0 |
| SRR5756105 | 0 | 0 | 0 | 12 | 0 | 0 | 0 | 0 | 0 | 0 | 0 | 0 | 19 | 12 | 21 | 0 | 0 | 0 | 0 | 0 |
| SRR5756106 | 0 | 0 | 0 | 0 | 0 | 0 | 0 | 0 | 0 | 0 | 0 | 17 | 23 | 31 | 0 | 0 | 0 | 0 | 0 | 0 |
| SRR5756107 | 0 | 0 | 0 | 0 | 21 | 0 | 0 | 0 | 0 | 0 | 0 | 0 | 0 | 11 | 11 | 0 | 0 | 0 | 0 | 0 |
| SRR5756108 | 0 | 0 | 0 | 14 | 0 | 0 | 0 | 0 | 0 | 0 | 0 | 0 | 13 | 0 | 21 | 0 | 0 | 0 | 0 | 0 |
| SRR5756110 | 0 | 0 | 0 | 0 | 0 | 0 | 0 | 0 | 0 | 0 | 0 | 0 | 0 | 0 | 0 | 0 | 0 | 0 | 0 | 0 |
| SRR5756114 | 0 | 0 | 0 | 0 | 0 | 0 | 0 | 0 | 0 | 0 | 0 | 0 | 0 | 0 | 0 | 0 | 0 | 0 | 0 | 0 |
| SRR5756115 | 0 | 0 | 0 | 0 | 0 | 0 | 0 | 0 | 0 | 0 | 0 | 0 | 0 | 0 | 0 | 0 | 0 | 0 | 0 | 0 |
| SRR5756116 | 0 | 0 | 0 | 0 | 0 | 0 | 0 | 0 | 0 | 0 | 0 | 0 | 0 | 0 | 0 | 0 | 0 | 0 | 0 | 0 |
| SRR5756118 | 0 | 0 | 0 | 0 | 0 | 0 | 0 | 0 | 0 | 0 | 0 | 0 | 0 | 0 | 0 | 0 | 0 | 0 | 0 | 0 |
| SRR5756120 | 0 | 0 | 0 | 0 | 0 | 0 | 0 | 0 | 0 | 0 | 0 | 0 | 0 | 0 | 0 | 0 | 0 | 0 | 0 | 0 |
| SRR5756121 | 0 | 0 | 0 | 0 | 0 | 0 | 0 | 0 | 0 | 0 | 0 | 0 | 0 | 0 | 0 | 0 | 0 | 0 | 0 | 0 |
| SRR5756122 | 0 | 0 | 0 | 0 | 0 | 0 | 0 | 0 | 0 | 0 | 0 | 0 | 0 | 0 | 0 | 0 | 0 | 0 | 0 | 0 |
| SRR5756123 | 0 | 0 | 0 | 0 | 0 | 0 | 0 | 0 | 0 | 0 | 0 | 0 | 0 | 0 | 0 | 0 | 0 | 0 | 0 | 0 |
| SRR5756125 | 0 | 0 | 0 | 0 | 0 | 0 | 0 | 0 | 0 | 0 | 0 | 0 | 0 | 0 | 0 | 0 | 0 | 0 | 0 | 0 |
| SRR5756128 | 0 | 0 | 0 | 0 | 0 | 0 | 0 | 0 | 0 | 0 | 0 | 0 | 0 | 0 | 0 | 0 | 0 | 0 | 0 | 0 |
| SRR5756130 | 0 | 0 | 0 | 0 | 0 | 0 | 0 | 0 | 0 | 0 | 0 | 0 | 0 | 0 | 0 | 0 | 0 | 0 | 0 | 0 |
| SRR5756131 | 0 | 0 | 0 | 0 | 0 | 0 | 0 | 0 | 0 | 0 | 0 | 0 | 0 | 0 | 0 | 0 | 0 | 0 | 0 | 0 |
| SRR5756132 | 0 | 0 | 0 | 0 | 0 | 0 | 0 | 0 | 0 | 0 | 0 | 0 | 0 | 0 | 0 | 0 | 0 | 0 | 0 | 0 |
| SRR5756133 | 0 | 0 | 0 | 0 | 0 | 0 | 0 | 0 | 0 | 0 | 0 | 0 | 0 | 0 | 0 | 0 | 0 | 0 | 0 | 0 |
| SRR5756135 | 0 | 0 | 0 | 0 | 0 | 0 | 0 | 0 | 0 | 0 | 0 | 0 | 0 | 0 | 0 | 0 | 0 | 0 | 0 | 0 |
| SRR5756136 | 0 | 0 | 0 | 0 | 0 | 0 | 0 | 0 | 0 | 0 | 0 | 0 | 11 | 0 | 0 | 0 | 0 | 0 | 0 | 0 |
| SRR5756137 | 0 | 0 | 0 | 0 | 0 | 0 | 0 | 0 | 0 | 0 | 0 | 0 | 0 | 0 | 0 | 0 | 0 | 0 | 0 | 0 |
| SRR5756138 | 0 | 0 | 0 | 0 | 0 | 0 | 0 | 0 | 0 | 0 | 0 | 0 | 0 | 0 | 0 | 0 | 0 | 0 | 0 | 0 |
| SRR5756139 | 0 | 0 | 0 | 0 | 0 | 0 | 0 | 0 | 0 | 0 | 0 | 0 | 0 | 0 | 0 | 0 | 0 | 0 | 0 | 0 |
| SRR5756140 | 0 | 0 | 0 | 0 | 0 | 0 | 0 | 0 | 0 | 0 | 0 | 0 | 0 | 0 | 0 | 0 | 0 | 0 | 0 | 0 |
| SRR5756141 | 0 | 0 | 0 | 0 | 0 | 0 | 0 | 0 | 0 | 0 | 0 | 0 | 0 | 0 | 0 | 0 | 0 | 0 | 0 | 0 |
| SRR5756142 | 0 | 0 | 0 | 0 | 0 | 0 | 0 | 0 | 0 | 0 | 0 | 0 | 0 | 0 | 0 | 0 | 0 | 0 | 0 | 0 |
| SRR5756143 | 0 | 0 | 0 | 0 | 0 | 0 | 0 | 0 | 0 | 0 | 0 | 0 | 0 | 0 | 0 | 0 | 0 | 0 | 0 | 0 |
| SRR5756144 | 0 | 0 | 0 | 0 | 0 | 0 | 0 | 0 | 0 | 0 | 0 | 0 | 0 | 0 | 0 | 0 | 0 | 0 | 0 | 0 |
| SRR5756145 | 0 | 0 | 0 | 0 | 0 | 0 | 0 | 0 | 0 | 0 | 0 | 0 | 0 | 0 | 0 | 0 | 0 | 0 | 0 | 0 |
| SRR5756146 | 0 | 0 | 0 | 0 | 0 | 0 | 0 | 0 | 0 | 0 | 0 | 0 | 0 | 0 | 0 | 0 | 0 | 0 | 0 | 0 |
| SRR5756147 | 0 | 0 | 0 | 0 | 0 | 0 | 0 | 0 | 0 | 0 | 0 | 0 | 0 | 0 | 0 | 0 | 0 | 0 | 0 | 0 |
| SRR5756148 | 0 | 0 | 0 | 0 | 0 | 0 | 0 | 0 | 0 | 0 | 0 | 0 | 0 | 0 | 0 | 0 | 0 | 0 | 0 | 0 |
| SRR5756149 | 0 | 0 | 0 | 0 | 0 | 0 | 0 | 0 | 0 | 0 | 0 | 0 | 0 | 0 | 0 | 0 | 0 | 0 | 0 | 0 |
| SRR5756150 | 0 | 0 | 0 | 15 | 0 | 0 | 0 | 0 | 0 | 0 | 0 | 0 | 10 | 0 | 0 | 0 | 0 | 0 | 0 | 0 |
| SRR5756151 | 0 | 0 | 0 | 0 | 0 | 0 | 0 | 0 | 0 | 0 | 0 | 0 | 0 | 0 | 0 | 0 | 0 | 0 | 0 | 0 |
| SRR5756152 | 0 | 0 | 0 | 13 | 0 | 0 | 0 | 0 | 0 | 0 | 0 | 0 | 0 | 0 | 0 | 0 | 0 | 0 | 0 | 0 |
| SRR5756153 | 0 | 0 | 0 | 0 | 0 | 0 | 0 | 0 | 0 | 0 | 0 | 0 | 0 | 0 | 0 | 0 | 0 | 0 | 0 | 0 |
| SRR5756154 | 0 | 0 | 0 | 0 | 0 | 0 | 0 | 0 | 0 | 0 | 0 | 0 | 0 | 0 | 0 | 0 | 0 | 0 | 0 | 0 |
| SRR5756155 | 0 | 0 | 0 | 0 | 0 | 0 | 0 | 0 | 0 | 0 | 0 | 0 | 0 | 0 | 0 | 0 | 0 | 0 | 0 | 0 |
| SRR5756156 | 0 | 0 | 0 | 12 | 0 | 0 | 0 | 0 | 0 | 0 | 0 | 0 | 11 | 0 | 0 | 0 | 0 | 0 | 0 | 0 |
| SRR5756157 | 0 | 0 | 0 | 11 | 0 | 0 | 0 | 0 | 0 | 0 | 0 | 0 | 0 | 0 | 0 | 0 | 0 | 0 | 0 | 0 |
| SRR5756158 | 0 | 0 | 0 | 0 | 0 | 0 | 0 | 0 | 0 | 0 | 0 | 0 | 0 | 0 | 0 | 0 | 0 | 0 | 0 | 0 |
| SRR5756159 | 0 | 0 | 0 | 0 | 0 | 0 | 0 | 0 | 0 | 0 | 0 | 0 | 0 | 0 | 0 | 0 | 0 | 0 | 0 | 0 |
| SRR5756160 | 0 | 0 | 0 | 0 | 0 | 0 | 0 | 0 | 0 | 0 | 0 | 0 | 0 | 0 | 0 | 0 | 0 | 0 | 0 | 0 |
| SRR5756162 | 0 | 0 | 0 | 0 | 0 | 0 | 0 | 0 | 0 | 0 | 0 | 0 | 0 | 0 | 0 | 0 | 0 | 0 | 0 | 0 |
| SRR5756164 | 0 | 0 | 0 | 0 | 0 | 0 | 0 | 0 | 0 | 0 | 0 | 0 | 0 | 0 | 0 | 0 | 0 | 0 | 0 | 0 |
| SRR5756166 | 0 | 0 | 0 | 0 | 0 | 0 | 0 | 0 | 0 | 0 | 0 | 0 | 0 | 0 | 0 | 0 | 0 | 0 | 0 | 0 |
| SRR5756167 | 0 | 0 | 0 | 0 | 0 | 0 | 0 | 0 | 0 | 0 | 0 | 0 | 0 | 0 | 0 | 0 | 0 | 0 | 0 | 0 |
| SRR5756168 | 0 | 0 | 0 | 0 | 0 | 0 | 0 | 0 | 0 | 0 | 0 | 0 | 0 | 0 | 0 | 0 | 0 | 0 | 0 | 0 |
| SRR5756177 | 0 | 0 | 0 | 0 | 0 | 0 | 0 | 0 | 0 | 0 | 0 | 0 | 0 | 0 | 0 | 0 | 0 | 0 | 0 | 0 |
| SRR5756178 | 0 | 0 | 0 | 0 | 0 | 0 | 0 | 0 | 0 | 0 | 0 | 0 | 0 | 0 | 0 | 0 | 0 | 0 | 0 | 0 |
| SRR5756179 | 0 | 0 | 0 | 0 | 0 | 0 | 0 | 0 | 0 | 0 | 0 | 0 | 0 | 0 | 0 | 0 | 0 | 0 | 0 | 0 |
| SRR5756180 | 0 | 0 | 0 | 0 | 0 | 0 | 0 | 0 | 0 | 0 | 0 | 0 | 0 | 0 | 0 | 0 | 0 | 0 | 0 | 0 |
| SRR5756182 | 0 | 0 | 0 | 0 | 0 | 0 | 0 | 0 | 0 | 0 | 0 | 0 | 0 | 0 | 0 | 0 | 0 | 0 | 0 | 0 |
| SRR5756183 | 0 | 0 | 0 | 0 | 0 | 0 | 0 | 0 | 0 | 0 | 0 | 0 | 0 | 0 | 0 | 0 | 0 | 0 | 0 | 0 |
| SRR5756184 | 0 | 0 | 0 | 0 | 0 | 0 | 0 | 0 | 0 | 0 | 0 | 0 | 0 | 0 | 0 | 0 | 0 | 0 | 0 | 0 |
| SRR5756185 | 0 | 0 | 0 | 0 | 0 | 0 | 0 | 0 | 0 | 0 | 0 | 0 | 0 | 0 | 0 | 0 | 0 | 0 | 0 | 0 |
| SRR5756186 | 0 | 0 | 0 | 0 | 0 | 0 | 0 | 0 | 0 | 0 | 0 | 0 | 0 | 0 | 0 | 0 | 0 | 0 | 0 | 0 |
| SRR5804909 | 0 | 0 | 0 | 0 | 0 | 0 | 0 | 0 | 0 | 0 | 0 | 0 | 0 | 0 | 0 | 0 | 0 | 0 | 0 | 0 |
| SRR5804912 | 0 | 0 | 0 | 0 | 0 | 0 | 0 | 0 | 0 | 0 | 0 | 0 | 0 | 0 | 0 | 0 | 0 | 0 | 0 | 0 |
| SRR5804914 | 0 | 0 | 0 | 0 | 0 | 0 | 0 | 0 | 0 | 0 | 0 | 0 | 0 | 0 | 0 | 0 | 0 | 0 | 0 | 0 |
| SRR5817623 | 0 | 11 | 0 | 147 | 59 | 919 | 570 | 326 | 138 | 0 | 14 | 20 | 70 | 12 | 0 | 0 | 0 | 0 | 112 | 0 |
| SRR5817624 | 0 | 12 | 0 | 154 | 140 | 1426 | 770 | 429 | 124 | 0 | 16 | 24 | 72 | 24 | 0 | 0 | 0 | 0 | 101 | 0 |

### Supp Table S8.1.Columns_B_to_AL

| | | | | | | | | | | | | | | | | | | | | | | | | | | | | | | | | | | | | | |
|---|---|---|---|---|---|---|---|---|---|---|---|---|---|---|---|---|---|---|---|---|---|---|---|---|---|---|---|---|---|---|---|---|---|---|---|---|---|
| SRR5817625 | 129 | 190 | 77 | 1931 | 23359 | 8574 | 67920 | 73790 | 10016 | 322 | 20514 | 6969 | 12329 | 7020 | 42902 | 875 | 17897 | 0 | 0 | 0 | 95 | 60 | 1075 | 641 | 378 | 139 | 0 | 14 | 17 | 43 | 0 | 0 | 0 | 0 | 0 | 103 | 0 |
| SRR5817626 | 139 | 411 | 164 | 3310 | 23104 | 8683 | 63380 | 87284 | 8433 | 360 | 20872 | 5675 | 12335 | 5344 | 31016 | 616 | 21767 | 0 | 0 | 0 | 140 | 109 | 1397 | 764 | 451 | 108 | 0 | 13 | 20 | 63 | 15 | 0 | 0 | 0 | 0 | 93 | 0 |
| SRR5821803 | 140 | 2527 | 2022 | 3642 | 3275 | 47577 | 97668 | 40466 | 3498 | 63 | 12977 | 15407 | 4009 | 40097 | 27 | 43231 | 8857 | 0 | 0 | 0 | 41 | 42 | 0 | 0 | 0 | 0 | 0 | 0 | 0 | 29 | 49 | 0 | 0 | 0 | 0 | 0 | 0 |
| SRR5821804 | 14 | 29177 | 47845 | 0 | 2199 | 63056 | 35249 | 40715 | 3154 | 14 | 28952 | 48252 | 4541 | 9844 | 11 | 38856 | 22148 | 0 | 0 | 0 | 314 | 491 | 276 | 0 | 11 | 0 | 0 | 0 | 54 | 105 | 419 | 143 | 0 | 23 | 20 | 13 | 0 |
| SRR5821805 | 0 | 58142 | 1242 | 16 | 1729 | 33047 | 24722 | 17606 | 3184 | 0 | 26412 | 10 | 6921 | 7276 | 130 | 60250 | 15225 | 0 | 0 | 0 | 81 | 54 | 44 | 0 | 0 | 0 | 0 | 0 | 0 | 18 | 48 | 11 | 0 | 0 | 0 | 0 | 0 |
| SRR5832003 | 0 | 1036 | 333 | 333 | 19 | 249 | 364 | 1271 | 1118 | 0 | 40169 | 64504 | 18069 | 4516 | 1380 | 9082 | 62607 | 0 | 17 | 29 | 1727 | 455 | 89 | 0 | 0 | 0 | 0 | 394 | 1350 | 1298 | 780 | 121 | 0 | 14 | 132 | 23 | 0 |
| SRR5832004 | 187 | 1249 | 312 | 281 | 21 | 127 | 239 | 824 | 951 | 0 | 46907 | 69726 | 17631 | 4605 | 753 | 8393 | 66734 | 0 | 47 | 155 | 11807 | 1345 | 371 | 0 | 0 | 0 | 0 | 945 | 6370 | 6040 | 2813 | 447 | 0 | 13 | 67 | 11 | 0 |
| SRR5832005 | 182 | 630 | 98 | 462 | 18 | 219 | 265 | 933 | 1134 | 0 | 43915 | 65410 | 16055 | 4316 | 850 | 7607 | 71079 | 0 | 17 | 53 | 2835 | 535 | 99 | 0 | 0 | 0 | 0 | 394 | 1850 | 1931 | 1230 | 114 | 0 | 22 | 252 | 66 | 0 |
| SRR5832006 | 172 | 1290 | 602 | 258 | 43 | 130 | 246 | 810 | 1013 | 0 | 45587 | 70862 | 16131 | 4080 | 579 | 9722 | 73090 | 0 | 25 | 20 | 8938 | 1714 | 538 | 0 | 0 | 0 | 0 | 360 | 1556 | 2790 | 3131 | 454 | 0 | 15 | 40 | 15 | 0 |
| SRR5832007 | 126 | 1806 | 3192 | 420 | 26 | 132 | 132 | 713 | 1320 | 0 | 21546 | 37652 | 8951 | 3565 | 1083 | 8502 | 192301 | 0 | 0 | 0 | 1028 | 1663 | 664 | 21 | 26 | 0 | 0 | 40 | 293 | 544 | 5538 | 1582 | 10 | 207 | 295 | 21 | 0 |
| SRR5832008 | 319 | 1092 | 147 | 331 | 20 | 253 | 98 | 780 | 1131 | 0 | 45414 | 66532 | 16126 | 3978 | 858 | 9516 | 65225 | 0 | 18 | 80 | 5479 | 884 | 189 | 0 | 0 | 0 | 0 | 415 | 3986 | 4220 | 2587 | 354 | 0 | 31 | 191 | 29 | 0 |
| SRR5832009 | 107 | 2071 | 286 | 214 | 28 | 222 | 245 | 946 | 1498 | 0 | 47290 | 11 | 18063 | 5013 | 872 | 8448 | 64453 | 0 | 29 | 56 | 2922 | 2076 | 468 | 0 | 0 | 0 | 0 | 317 | 1265 | 1614 | 4093 | 583 | 0 | 13 | 112 | 20 | 0 |
| SRR5832010 | 0 | 1340 | 279 | 586 | 16 | 143 | 271 | 1062 | 999 | 0 | 38477 | 75223 | 16349 | 4047 | 765 | 8626 | 61407 | 0 | 14 | 37 | 1696 | 684 | 107 | 0 | 0 | 0 | 0 | 201 | 857 | 933 | 965 | 87 | 0 | 12 | 196 | 28 | 0 |
| SRR5832011 | 174 | 3372 | 756 | 58 | 20 | 229 | 424 | 881 | 1102 | 0 | 40858 | 61899 | 15966 | 5075 | 477 | 9539 | 70215 | 0 | 16 | 177 | 27335 | 1151 | 776 | 0 | 0 | 0 | 0 | 261 | 5180 | 12722 | 2605 | 864 | 0 | 0 | 18 | 0 | 0 |
| SRR5832012 | 130 | 2405 | 130 | 65 | 23 | 289 | 320 | 882 | 1529 | 0 | 41338 | 65499 | 14165 | 3735 | 784 | 10643 | 62061 | 0 | 20 | 140 | 27378 | 1158 | 896 | 0 | 0 | 0 | 0 | 417 | 4654 | 11075 | 2653 | 1051 | 0 | 0 | 12 | 0 | 0 |
| SRR5832013 | 209 | 2050 | 377 | 293 | 53 | 326 | 559 | 1701 | 1186 | 0 | 36865 | 61964 | 15291 | 4126 | 656 | 8753 | 75855 | 0 | 0 | 27 | 2528 | 603 | 105 | 0 | 0 | 0 | 0 | 134 | 752 | 1102 | 1411 | 112 | 0 | 0 | 56 | 0 | 0 |
| SRR5832014 | 183 | 2677 | 365 | 122 | 52 | 188 | 325 | 1110 | 890 | 0 | 38096 | 61875 | 15188 | 5935 | 526 | 8240 | 78108 | 0 | 46 | 177 | 10296 | 2516 | 366 | 0 | 0 | 0 | 0 | 609 | 4034 | 4587 | 4866 | 354 | 0 | 0 | 39 | 0 | 0 |
| SRR5832015 | 0 | 734 | 367 | 294 | 14 | 132 | 223 | 1003 | 1003 | 73 | 27364 | 41517 | 10942 | 3197 | 390 | 9145 | 59590 | 0 | 27 | 53 | 6844 | 748 | 191 | 0 | 0 | 0 | 0 | 293 | 2257 | 3365 | 1527 | 255 | 0 | 29 | 91 | 0 | 0 |
| SRR5832016 | 229 | 1487 | 286 | 515 | 25 | 167 | 125 | 702 | 852 | 0 | 23869 | 36598 | 9360 | 2382 | 610 | 9787 | 70897 | 0 | 29 | 156 | 10181 | 1511 | 311 | 0 | 0 | 0 | 0 | 610 | 6203 | 5883 | 3127 | 401 | 0 | 0 | 46 | 0 | 0 |
| SRR5832017 | 0 | 1861 | 604 | 151 | 52 | 197 | 343 | 1424 | 536 | 50 | 13512 | 23176 | 5332 | 2593 | 103 | 7616 | 53019 | 0 | 16 | 82 | 20840 | 1091 | 711 | 0 | 0 | 0 | 0 | 199 | 2858 | 12279 | 2523 | 1197 | 0 | 0 | 0 | 0 | 0 |
| SRR5832018 | 25 | 1398 | 399 | 50 | 52 | 152 | 258 | 922 | 985 | 25 | 25448 | 40410 | 10608 | 3281 | 315 | 13789 | 46469 | 0 | 30 | 126 | 13364 | 633 | 447 | 0 | 0 | 0 | 0 | 493 | 5201 | 8169 | 1435 | 710 | 0 | 0 | 13 | 0 | 0 |
| SRR5832019 | 73 | 1238 | 291 | 36 | 56 | 239 | 518 | 1080 | 779 | 0 | 20625 | 30189 | 7919 | 3647 | 317 | 14882 | 76023 | 0 | 0 | 155 | 15583 | 767 | 247 | 0 | 0 | 0 | 0 | 94 | 5012 | 10035 | 1564 | 333 | 0 | 0 | 0 | 0 | 0 |
| SRR5832020 | 94 | 1157 | 500 | 156 | 55 | 177 | 299 | 834 | 1612 | 0 | 24063 | 34951 | 9118 | 2894 | 377 | 12252 | 47993 | 0 | 32 | 155 | 37465 | 1094 | 833 | 0 | 0 | 0 | 0 | 306 | 4152 | 13762 | 1579 | 783 | 0 | 0 | 0 | 0 | 0 |
| SRR5832021 | 56 | 744 | 279 | 763 | 26 | 141 | 252 | 1007 | 901 | 19 | 21271 | 34871 | 8781 | 2267 | 192 | 10668 | 51566 | 0 | 10 | 31 | 1902 | 335 | 78 | 0 | 0 | 0 | 0 | 318 | 1333 | 1606 | 734 | 119 | 0 | 17 | 137 | 40 | 0 |
| SRR5832022 | 156 | 5781 | 21742 | 156 | 30 | 144 | 296 | 865 | 1304 | 52 | 13847 | 23054 | 5521 | 2912 | 129 | 12816 | 79687 | 0 | 16 | 72 | 5244 | 1314 | 201 | 0 | 0 | 0 | 0 | 286 | 2795 | 4318 | 3782 | 598 | 0 | 0 | 0 | 0 | 0 |
| SRR5832023 | 44 | 935 | 522 | 326 | 24 | 148 | 227 | 888 | 1008 | 0 | 11689 | 44481 | 10151 | 2684 | 310 | 11187 | 53338 | 0 | 20 | 45 | 4393 | 546 | 103 | 0 | 0 | 0 | 0 | 298 | 1659 | 3061 | 1227 | 238 | 0 | 0 | 91 | 26 | 0 |
| SRR5832024 | 56 | 1005 | 56 | 56 | 39 | 187 | 275 | 864 | 891 | 0 | 20530 | 30646 | 7844 | 3631 | 248 | 13395 | 113726 | 0 | 19 | 54 | 9662 | 2894 | 285 | 0 | 0 | 0 | 0 | 279 | 2342 | 4835 | 5215 | 421 | 0 | 0 | 0 | 0 | 0 |
| SRR5832025 | 125 | 317 | 68 | 532 | 44 | 150 | 273 | 679 | 891 | 0 | 23491 | 37406 | 9700 | 2478 | 300 | 14294 | 53129 | 0 | 0 | 24 | 2673 | 309 | 64 | 0 | 11 | 0 | 14 | 345 | 1965 | 2110 | 708 | 99 | 0 | 17 | 127 | 52 | 0 |
| SRR5832026 | 129 | 757 | 129 | 322 | 29 | 175 | 117 | 911 | 897 | 0 | 22589 | 33297 | 7909 | 2136 | 204 | 11583 | 52897 | 0 | 26 | 98 | 9753 | 787 | 279 | 0 | 0 | 0 | 11 | 590 | 5611 | 7048 | 1925 | 438 | 0 | 11 | 53 | 17 | 0 |
| SRR5832027 | 33 | 1305 | 294 | 228 | 31 | 180 | 227 | 792 | 829 | 33 | 24525 | 40025 | 10583 | 2636 | 435 | 12272 | 53846 | 0 | 0 | 25 | 4175 | 685 | 187 | 0 | 0 | 0 | 0 | 261 | 916 | 2211 | 1361 | 220 | 0 | 0 | 32 | 0 | 0 |
| SRR5832028 | 109 | 518 | 82 | 273 | 24 | 145 | 207 | 900 | 768 | 0 | 25244 | 37735 | 9494 | 2064 | 313 | 10409 | 48298 | 0 | 0 | 37 | 2575 | 372 | 85 | 0 | 0 | 0 | 0 | 214 | 1072 | 1521 | 834 | 170 | 0 | 0 | 51 | 19 | 0 |
| SRR5832029 | 49 | 1919 | 492 | 148 | 27 | 215 | 309 | 1019 | 555 | 0 | 23144 | 38857 | 8972 | 2556 | 434 | 11404 | 60081 | 0 | 22 | 50 | 10210 | 1801 | 240 | 0 | 0 | 0 | 0 | 286 | 2554 | 5440 | 4505 | 459 | 0 | 0 | 14 | 0 | 0 |
| SRR5868085 | 176 | 3508 | 55977 | 4894 | 67860 | 40152 | 288860 | 19098 | 11447 | 4958 | 15 | 22 | 99 | 31798 | 6006 | 82524 | 5557 | 0 | 0 | 0 | 0 | 11 | 20 | 0 | 0 | 0 | 0 | 0 | 28 | 0 | 0 | 0 | 0 | 0 | 0 | 0 | 0 |
| SRR5901584 | 100 | 19596 | 4913 | 685 | 2081 | 15437 | 56911 | 15515 | 1264 | 21 | 10716 | 5027 | 1812 | 9160 | 13 | 3510 | 18833 | 0 | 16 | 20 | 1839 | 342 | 412 | 48 | 39 | 0 | 0 | 17 | 153 | 611 | 234 | 171 | 0 | 21 | 0 | 17 | 0 |
| SRR5901585 | 115 | 20691 | 5570 | 534 | 2115 | 19392 | 54242 | 17646 | 1465 | 15 | 12292 | 4987 | 2015 | 11469 | 0 | 4129 | 18571 | 0 | 14 | 15 | 1732 | 264 | 280 | 28 | 34 | 0 | 0 | 13 | 147 | 573 | 196 | 124 | 0 | 18 | 0 | 15 | 0 |
| SRR5901586 | 163 | 19562 | 6868 | 686 | 2170 | 18236 | 57159 | 25538 | 3199 | 32 | 12325 | 5328 | 1894 | 12137 | 10 | 4105 | 15918 | 0 | 11 | 25 | 1976 | 208 | 227 | 30 | 22 | 0 | 0 | 15 | 163 | 619 | 136 | 120 | 0 | 11 | 0 | 10 | 0 |
| SRR5901587 | 37 | 11220 | 6151 | 361 | 1291 | 15205 | 45496 | 10832 | 2116 | 52 | 12857 | 5740 | 1838 | 9211 | 49 | 4837 | 19001 | 0 | 0 | 0 | 1060 | 221 | 335 | 35 | 26 | 0 | 0 | 10 | 88 | 339 | 165 | 184 | 0 | 18 | 0 | 17 | 0 |
| SRR5901588 | 46 | 9528 | 4531 | 343 | 1527 | 13601 | 43089 | 12210 | 1570 | 20 | 9461 | 4874 | 1738 | 8371 | 32 | 3538 | 21105 | 0 | 10 | 14 | 1290 | 324 | 536 | 51 | 53 | 0 | 0 | 11 | 113 | 407 | 275 | 225 | 0 | 31 | 0 | 24 | 0 |
| SRR5901589 | 54 | 12365 | 6543 | 391 | 1479 | 16281 | 50201 | 12274 | 2008 | 43 | 10879 | 4345 | 1572 | 8069 | 18 | 4679 | 19257 | 0 | 0 | 0 | 1111 | 219 | 322 | 32 | 27 | 0 | 0 | 0 | 84 | 390 | 173 | 187 | 0 | 17 | 0 | 15 | 0 |
| SRR5901590 | 16 | 30517 | 4780 | 489 | 160 | 1334 | 3383 | 754 | 18098 | 0 | 3990 | 1762 | 457 | 249 | 135 | 5708 | 26689 | 0 | 0 | 27 | 15312 | 7632 | 1422 | 153 | 105 | 24 | 0 | 0 | 340 | 7329 | 6445 | 1222 | 27 | 76 | 15 | 86 | 0 |
| SRR5901591 | 20 | 22081 | 3275 | 575 | 110 | 1277 | 2906 | 578 | 12411 | 25 | 2584 | 1379 | 464 | 169 | 209 | 5254 | 29311 | 0 | 0 | 29 | 23632 | 16787 | 2215 | 281 | 140 | 41 | 0 | 0 | 388 | 11248 | 14313 | 1569 | 31 | 106 | 26 | 102 | 0 |
| SRR5901592 | 11 | 9381 | 3804 | 240 | 138 | 1275 | 3062 | 635 | 14349 | 0 | 3682 | 1629 | 504 | 251 | 203 | 5689 | 27908 | 0 | 0 | 39 | 27009 | 19089 | 1882 | 222 | 134 | 44 | 0 | 0 | 498 | 13825 | 14317 | 1462 | 29 | 90 | 22 | 115 | 0 |
| SRR5901593 | 44 | 7217 | 4084 | 736 | 138 | 1067 | 3170 | 572 | 12635 | 33 | 5538 | 2496 | 1102 | 434 | 138 | 11146 | 23441 | 0 | 0 | 25 | 28747 | 4142 | 465 | 56 | 42 | 19 | 0 | 0 | 314 | 13631 | 3609 | 451 | 0 | 31 | 19 | 66 | 0 |
| SRR5901594 | 57 | 5975 | 3756 | 448 | 139 | 1178 | 3216 | 610 | 7513 | 40 | 4627 | 2001 | 1223 | 541 | 57 | 11264 | 22473 | 0 | 0 | 25 | 37544 | 7033 | 714 | 132 | 63 | 29 | 0 | 15 | 408 | 16352 | 5155 | 482 | 0 | 27 | 17 | 59 | 0 |
| SRR5901595 | 54 | 6412 | 2772 | 557 | 119 | 1020 | 2737 | 525 | 11030 | 40 | 4720 | 1785 | 929 | 324 | 131 | 13731 | 24067 | 0 | 0 | 30 | 30619 | 5107 | 575 | 81 | 46 | 21 | 0 | 0 | 446 | 12629 | 3577 | 412 | 0 | 22 | 17 | 71 | 0 |
| SRR5922753 | 81 | 21646 | 7159 | 246 | 5019 | 8463 | 114403 | 57953 | 264 | 3870 | 14030 | 23349 | 6550 | 5716 | 196 | 3790 | 38244 | 0 | 0 | 13 | 68 | 54 | 158 | 47 | 13 | 0 | 0 | 0 | 19 | 108 | 166 | 52 | 0 | 0 | 0 | 0 | 0 |
| SRR5922754 | 0 | 4862 | 15541 | 0 | 3251 | 4873 | 85541 | 67556 | 261 | 2852 | 15851 | 26541 | 8437 | 6098 | 128 | 3129 | 43622 | 0 | 0 | 0 | 0 | 0 | 30 | 0 | 0 | 0 | 0 | 0 | 0 | 16 | 29 | 0 | 0 | 0 | 0 | 0 | 0 |
| SRR5922756 | 0 | 154 | 6044 | 0 | 3304 | 3794 | 93011 | 69397 | 500 | 502 | 15650 | 26815 | 8358 | 8903 | 77 | 5026 | 30993 | 0 | 0 | 0 | 0 | 0 | 10 | 0 | 27 | 62 | 0 | 0 | 0 | 0 | 18 | 0 | 0 | 0 | 12 | 117 | 0 |
| SRR5925650 | 83 | 17064 | 158 | 542 | 2104 | 28238 | 26077 | 28365 | 5367 | 21 | 10129 | 142488 | 2384 | 1258 | 35 | 19187 | 10408 | 0 | 0 | 0 | 27 | 0 | 29 | 27 | 587 | 17 | 0 | 0 | 0 | 0 | 0 | 18 | 0 | 289 | 821 | 83 | 0 |
| SRR5925651 | 112 | 16064 | 425 | 325 | 5065 | 65483 | 64248 | 52704 | 8104 | 54 | 21480 | 57596 | 4050 | 5423 | 0 | 32773 | 18634 | 0 | 0 | 0 | 45 | 0 | 36 | 13 | 178 | 0 | 0 | 0 | 0 | 0 | 0 | 19 | 0 | 104 | 41 | 105 | 0 |
| SRR5925652 | 854 | 55131 | 2558 | 824 | 6487 | 35009 | 45928 | 39099 | 25251 | 109 | 38852 | 80191 | 8646 | 9601 | 24 | 27657 | 16907 | 0 | 0 | 0 | 503 | 109 | 103 | 0 | 15 | 0 | 0 | 0 | 95 | 113 | 122 | 55 | 0 | 16 | 27 | 51 | 0 |
| SRR5925665 | 79 | 115301 | 1347 | 173 | 2749 | 55789 | 20442 | 41295 | 18736 | 0 | 7780 | 31985 | 4239 | 739 | 996 | 128 | 75976 | 0 | 0 | 0 | 136 | 39 | 24 | 0 | 0 | 0 | 0 | 0 | 0 | 43 | 22 | 0 | 0 | 0 | 0 | 0 | 0 |
| SRR5925666 | 120 | 122092 | 1273 | 120 | 2602 | 55855 | 18436 | 37778 | 19527 | 12 | 7293 | 30358 | 3910 | 706 | 877 | 140 | 75171 | 0 | 0 | 0 | 152 | 42 | 35 | 0 | 0 | 0 | 0 | 0 | 0 | 37 | 24 | 10 | 0 | 0 | 12 | 0 | 0 |
| SRR5925667 | 101 | 128079 | 1053 | 174 | 2985 | 63786 | 22768 | 39233 | 18672 | 16 | 7500 | 30703 | 4204 | 568 | 1049 | 231 | 72935 | 0 | 0 | 0 | 137 | 38 | 24 | 0 | 0 | 0 | 0 | 0 | 13 | 28 | 20 | 0 | 0 | 0 | 0 | 0 | 0 |
| SRR5925668 | 145 | 124625 | 1398 | 155 | 2618 | 56678 | 20103 | 38306 | 20570 | 11 | 7154 | 30939 | 3888 | 595 | 1207 | 199 | 77322 | 0 | 0 | 0 | 183 | 44 | 31 | 0 | 0 | 0 | 0 | 0 | 10 | 36 | 25 | 0 | 0 | 0 | 0 | 0 | 0 |
| SRR5925669 | 122 | 129918 | 1032 | 196 | 2571 | 64009 | 21047 | 37585 | 19723 | 12 | 7028 | 28451 | 3888 | 588 | 1066 | 243 | 69845 | 0 | 0 | 0 | 131 | 33 | 19 | 0 | 0 | 0 | 0 | 0 | 0 | 22 | 20 | 0 | 0 | 0 | 0 | 0 | 0 |
| SRR5925670 | 145 | 123001 | 2033 | 153 | 3125 | 65068 | 23487 | 40914 | 20344 | 18 | 7230 | 30746 | 3877 | 579 | 1161 | 202 | 71268 | 0 | 0 | 0 | 137 | 25 | 29 | 0 | 0 | 0 | 0 | 0 | 0 | 27 | 19 | 0 | 0 | 0 | 0 | 0 | 0 |
| SRR5925671 | 191 | 117771 | 1085 | 187 | 2431 | 74862 | 20319 | 46215 | 18120 | 19 | 6132 | 26179 | 2877 | 397 | 2402 | 264 | 73413 | 0 | 0 | 0 | 189 | 46 | 30 | 0 | 0 | 0 | 0 | 0 | 13 | 41 | 21 | 0 | 0 | 0 | 0 | 0 | 0 |
| SRR5925672 | 82 | 156517 | 923 | 135 | 2378 | 74842 | 20071 | 46017 | 16757 | 0 | 5960 | 25459 | 2851 | 351 | 2271 | 315 | 76185 | 0 | 0 | 0 | 135 | 37 | 31 | 0 | 0 | 0 | 0 | 0 | 0 | 27 | 22 | 12 | 0 | 0 | 0 | 0 | 0 |
| SRR5925673 | 103 | 97996 | 832 | 186 | 2295 | 75316 | 14760 | 39841 | 24017 | 29 | 5953 | 25180 | 2859 | 353 | 446 | 117 | 74959 | 0 | 0 | 0 | 66 | 20 | 12 | 0 | 0 | 0 | 0 | 0 | 0 | 19 | 11 | 0 | 0 | 0 | 0 | 0 | 0 |
| SRR5925674 | 184 | 113760 | 664 | 177 | 2293 | 63274 | 14223 | 40789 | 23562 | 15 | 7156 | 27705 | 3301 | 443 | 580 | 102 | 79710 | 0 | 0 | 0 | 83 | 32 | 25 | 0 | 0 | 0 | 0 | 0 | 0 | 17 | 25 | 0 | 0 | 0 | 0 | 0 | 0 |
| SRR5925675 | 307 | 150006 | 1623 | 240 | 2350 | 64638 | 15900 | 32876 | 16197 | 0 | 4456 | 20322 | 2555 | 347 | 363 | 210 | 93898 | 0 | 0 | 0 | 192 | 69 | 32 | 0 | 0 | 0 | 0 | 0 | 17 | 58 | 51 | 10 | 0 | 0 | 11 | 0 | 0 |
| SRR5925676 | 187 | 178069 | 777 | 178 | 3274 | 82611 | 23137 | 48765 | 17051 | 0 | 6636 | 23404 | 3250 | 368 | 408 | 242 | 81866 | 0 | 0 | 0 | 145 | 39 | 30 | 0 | 0 | 0 | 0 | 0 | 13 | 30 | 22 | 10 | 0 | 0 | 0 | 0 | 0 |
| SRR5925677 | 125 | 167659 | 950 | 143 | 2964 | 79919 | 28413 | 49712 | 13865 | 18 | 6552 | 26171 | 3308 | 445 | 868 | 475 | 73006 | 0 | 0 | 0 | 121 | 30 | 33 | 0 | 0 | 0 | 0 | 0 | 0 | 19 | 21 | 13 | 0 | 0 | 0 | 0 | 0 |
| SRR5925678 | 219 | 164576 | 707 | 111 | 3071 | 81850 | 28093 | 45750 | 13738 | 24 | 6188 | 24923 | 3211 | 379 | 815 | 346 | 77841 | 0 | 0 | 0 | 139 | 41 | 43 | 0 | 0 | 0 | 0 | 0 | 0 | 20 | 17 | 15 | 0 | 0 | 0 | 0 | 0 |
| SRR5925679 | 176 | 158936 | 663 | 153 | 2780 | 75908 | 19600 | 42627 | 24532 | 42 | 11126 | 36853 | 6003 | 868 | 800 | 221 | 72118 | 0 | 0 | 0 | 94 | 36 | 25 | 0 | 0 | 0 | 0 | 0 | 13 | 22 | 22 | 10 | 0 | 0 | 0 | 0 | 0 |
| SRR5925680 | 145 | 179436 | 709 | 129 | 2316 | 68241 | 17253 | 38443 | 26579 | 26 | 10388 | 37123 | 5735 | 865 | 788 | 185 | 72823 | 0 | 0 | 0 | 98 | 36 | 29 | 0 | 0 | 0 | 0 | 0 | 12 | 20 | 23 | 0 | 0 | 0 | 0 | 0 | 0 |
| SRR5925681 | 121 | 123026 | 695 | 235 | 2245 | 54985 | 22739 | 35501 | 23088 | 17 | 7879 | 30512 | 3766 | 596 | 1355 | 64 | 85549 | 0 | 0 | 0 | 205 | 52 | 48 | 0 | 0 | 0 | 0 | 0 | 16 | 43 | 33 | 15 | 0 | 0 | 11 | 0 | 0 |
| SRR5925682 | 123 | 172060 | 900 | 127 | 2472 | 63615 | 23278 | 38232 | 20964 | 15 | 7672 | 26398 | 3739 | 576 | 1754 | 103 | 76644 | 0 | 0 | 0 | 219 | 49 | 47 | 0 | 0 | 0 | 0 | 0 | 17 | 38 | 30 | 12 | 0 | 0 | 11 | 0 | 0 |
| SRR5925683 | 121 | 113008 | 1149 | 116 | 3178 | 57298 | 20882 | 41423 | 20464 | 45 | 7225 | 31521 | 3993 | 542 | 1123 | 250 | 65601 | 0 | 0 | 0 | 168 | 37 | 30 | 0 | 0 | 0 | 0 | 0 | 10 | 31 | 26 | 12 | 0 | 0 | 0 | 0 | 0 |
| SRR5925684 | 134 | 94460 | 1483 | 160 | 3108 | 56122 | 20757 | 43137 | 21313 | 31 | 7430 | 31448 | 3866 | 554 | 1093 | 185 | 65739 | 0 | 0 | 0 | 217 | 48 | 25 | 0 | 0 | 0 | 0 | 0 | 10 | 63 | 28 | 0 | 0 | 0 | 0 | 0 | 0 |
| SRR5925685 | 124 | 20966 | 5663 | 593 | 172 | 5180 | 1809 | 1854 | 6327 | 14 | 1050 | 6642 | 127 | 22 | 573 | 0 | 17918 | 0 | 0 | 0 | 184 | 104 | 24 | 0 | 0 | 0 | 0 | 0 | 39 | 35 | 130 | 0 | 0 | 0 | 0 | 0 | 0 |
| SRR5925686 | 117 | 22770 | 5137 | 639 | 164 | 5119 | 1796 | 1740 | 6266 | 0 | 1050 | 6328 | 124 | 25 | 557 | 0 | 16898 | 0 | 0 | 0 | 181 | 98 | 22 | 0 | 0 | 0 | 0 | 0 | 41 | 32 | 110 | 0 | 0 | 0 | 0 | 0 | 0 |
| SRR5925687 | 87 | 34400 | 6683 | 336 | 134 | 4683 | 1371 | 1364 | 5680 | 17 | 880 | 5812 | 138 | 17 | 507 | 0 | 13947 | 0 | 0 | 0 | 365 | 104 | 25 | 0 | 0 | 0 | 0 | 0 | 42 | 73 | 109 | 0 | 0 | 0 | 0 | 0 | 0 |
| SRR5925688 | 73 | 29934 | 6306 | 297 | 148 | 4408 | 1356 | 1229 | 5619 | 0 | 800 | 5731 | 118 | 17 | 459 | 0 | 12388 | 0 | 0 | 0 | 279 | 78 | 20 | 0 | 0 | 0 | 0 | 0 | 28 | 67 | 86 | 0 | 0 | 0 | 0 | 0 | 0 |
| SRR5925689 | 115 | 31233 | 7721 | 284 | 168 | 4548 | 1518 | 1340 | 5249 | 22 | 989 | 6807 | 142 | 19 | 594 | 0 | 15213 | 0 | 0 | 0 | 424 | 125 | 22 | 0 | 0 | 0 | 0 | 0 | 47 | 77 | 128 | 0 | 0 | 0 | 0 | 0 | 0 |
| SRR5925690 | 131 | 30998 | 7401 | 302 | 132 | 4476 | 1400 | 1266 | 5666 | 15 | 842 | 6167 | 109 | 18 | 482 | 0 | 13962 | 0 | 0 | 0 | 319 | 92 | 20 | 0 | 0 | 0 | 0 | 0 | 40 | 70 | 106 | 0 | 0 | 0 | 0 | 0 | 0 |
| SRR5925691 | 140 | 25536 | 6402 | 406 | 239 | 20183 | 3362 | 2665 | 6214 | 0 | 681 | 5207 | 79 | 13 | 728 | 0 | 19733 | 0 | 0 | 0 | 305 | 131 | 28 | 0 | 0 | 0 | 0 | 11 | 39 | 51 | 135 | 0 | 0 | 0 | 11 | 0 | 0 |
| SRR5925692 | 167 | 22366 | 5020 | 327 | 279 | 19024 | 3643 | 2732 | 5793 | 0 | 660 | 5156 | 81 | 11 | 722 | 0 | 20587 | 0 | 0 | 0 | 282 | 121 | 22 | 0 | 0 | 0 | 0 | 0 | 38 | 53 | 142 | 0 | 0 | 0 | 0 | 0 | 0 |
| SRR5925693 | 120 | 19601 | 6030 | 465 | 179 | 10756 | 2001 | 1693 | 3992 | 29 | 606 | 4385 | 79 | 12 | 408 | 0 | 15182 | 0 | 0 | 0 | 198 | 116 | 23 | 0 | 0 | 0 | 0 | 0 | 45 | 46 | 136 | 0 | 0 | 0 | 0 | 0 | 0 |
| SRR5925694 | 181 | 20061 | 3794 | 590 | 164 | 9389 | 2249 | 1367 | 3593 | 0 | 643 | 5544 | 74 | 15 | 293 | 0 | 10770 | 0 | 0 | 0 | 218 | 126 | 25 | 0 | 0 | 0 | 0 | 0 | 62 | 50 | 173 | 12 | 0 | 0 | 12 | 0 | 0 |
| SRR5925695 | 132 | 41208 | 6598 | 524 | 138 | 6241 | 1495 | 1132 | 4745 | 24 | 717 | 5376 | 100 | 13 | 401 | 0 | 15026 | 0 | 0 | 0 | 645 | 146 | 31 | 0 | 0 | 0 | 0 | 0 | 26 | 143 | 87 | 0 | 0 | 0 | 0 | 0 | 0 |
| SRR5925696 | 174 | 45160 | 6480 | 1024 | 129 | 6523 | 1451 | 1180 | 4583 | 19 | 721 | 5015 | 94 | 13 | 437 | 0 | 14359 | 0 | 0 | 0 | 530 | 153 | 24 | 0 | 0 | 0 | 0 | 0 | 27 | 96 | 87 | 0 | 0 | 0 | 0 | 0 | 0 |
| SRR5925697 | 111 | 40320 | 5035 | 324 | 172 | 5115 | 2788 | 1092 | 4853 | 21 | 579 | 4641 | 81 | 21 | 509 | 0 | 8812 | 0 | 0 | 0 | 459 | 158 | 34 | 0 | 0 | 0 | 0 | 0 | 51 | 94 | 126 | 0 | 0 | 0 | 0 | 0 | 0 |
| SRR5925698 | 74 | 34797 | 5994 | 316 | 136 | 4626 | 1251 | 918 | 4259 | 0 | 655 | 4690 | 70 | 12 | 702 | 0 | 13605 | 0 | 0 | 0 | 222 | 93 | 26 | 0 | 0 | 0 | 0 | 0 | 27 | 47 | 89 | 0 | 0 | 0 | 0 | 0 | 0 |
| SRR5925699 | 140 | 52368 | 5633 | 341 | 187 | 6948 | 1922 | 1396 | 5101 | 16 | 1662 | 8800 | 357 | 54 | 379 | 0 | 15417 | 0 | 0 | 0 | 320 | 115 | 24 | 0 | 0 | 0 | 0 | 0 | 38 | 65 | 108 | 0 | 0 | 0 | 0 | 0 | 0 |
| SRR5925700 | 122 | 47490 | 5273 | 513 | 239 | 7005 | 2362 | 1517 | 5142 | 0 | 1633 | 8408 | 340 | 59 | 392 | 0 | 14936 | 0 | 0 | 0 | 268 | 116 | 26 | 0 | 0 | 0 | 0 | 0 | 49 | 61 | 118 | 0 | 0 | 0 | 0 | 0 | 0 |
| SRR5925701 | 128 | 21831 | 6678 | 1049 | 202 | 6673 | 2212 | 2056 | 7211 | 18 | 914 | 5845 | 109 | 21 | 408 | 0 | 20271 | 0 | 0 | 0 | 223 | 105 | 22 | 0 | 0 | 0 | 0 | 0 | 36 | 43 | 111 | 0 | 0 | 0 | 0 | 0 | 0 |
| SRR5925702 | 140 | 22999 | 6196 | 428 | 160 | 6419 | 1847 | 1711 | 6444 | 0 | 793 | 5553 | 86 | 19 | 336 | 0 | 18790 | 0 | 0 | 0 | 171 | 129 | 28 | 0 | 0 | 0 | 0 | 0 | 35 | 30 | 115 | 0 | 0 | 0 | 0 | 0 | 0 |
| SRR5925703 | 176 | 23998 | 6687 | 892 | 216 | 7157 | 2564 | 2046 | 5172 | 0 | 850 | 4584 | 83 | 14 | 468 | 0 | 14900 | 0 | 0 | 0 | 332 | 130 | 19 | 0 | 0 | 0 | 0 | 0 | 44 | 63 | 134 | 0 | 0 | 0 | 0 | 0 | 0 |
| SRR5925704 | 100 | 19074 | 5023 | 376 | 259 | 6597 | 2617 | 1968 | 4808 | 13 | 686 | 4915 | 88 | 14 | 356 | 0 | 14037 | 0 | 0 | 0 | 231 | 96 | 21 | 0 | 0 | 0 | 0 | 0 | 27 | 56 | 105 | 0 | 0 | 0 | 0 | 0 | 0 |
| SRR5925705 | 0 | 6750 | 54 | 3021 | 1649 | 50053 | 27023 | 21332 | 21538 | 0 | 5601 | 6484 | 1648 | 2405 | 817 | 167 | 70541 | 0 | 0 | 0 | 132 | 35 | 22 | 0 | 0 | 0 | 0 | 0 | 21 | 32 | 24 | 10 | 0 | 0 | 13 | 0 | 0 |
| SRR5925706 | 12 | 14226 | 337 | 3702 | 708 | 64783 | 11471 | 24686 | 27240 | 0 | 16568 | 11019 | 995 | 2064 | 688 | 197 | 62282 | 0 | 0 | 0 | 162 | 41 | 25 | 0 | 0 | 0 | 0 | 0 | 34 | 31 | 15 | 0 | 0 | 0 | 14 | 0 | 0 |
| SRR5925707 | 0 | 7870 | 182 | 2819 | 1455 | 52150 | 23155 | 25143 | 22199 | 0 | 5820 | 5704 | 1559 | 2599 | 797 | 151 | 76592 | 0 | 0 | 0 | 157 | 42 | 22 | 0 | 0 | 0 | 0 | 0 | 29 | 40 | 29 | 0 | 0 | 0 | 16 | 0 | 0 |
| SRR5925708 | 0 | 11153 | 292 | 3528 | 727 | 63829 | 11535 | 25392 | 25460 | 0 | 14247 | 9791 | 1008 | 2069 | 696 | 187 | 66686 | 0 | 0 | 0 | 207 | 52 | 26 | 0 | 0 | 0 | 0 | 0 | 43 | 32 | 17 | 0 | 0 | 0 | 20 | 0 | 0 |
| SRR5925709 | 16 | 24333 | 1233 | 872 | 1533 | 43006 | 22038 | 19477 | 36569 | 0 | 10398 | 6025 | 2475 | 3457 | 791 | 0 | 68850 | 0 | 0 | 0 | 74 | 23 | 19 | 0 | 0 | 0 | 0 | 0 | 0 | 38 | 21 | 12 | 0 | 0 | 10 | 0 | 0 |
| SRR5925710 | 0 | 17601 | 2779 | 2779 | 812 | 46324 | 15770 | 17335 | 33743 | 0 | 10842 | 5740 | 1218 | 2377 | 638 | 0 | 60703 | 0 | 0 | 0 | 0 | 27 | 0 | 0 | 0 | 0 | 0 | 27 | 0 | 0 | 53 | 27 | 0 | 0 | 0 | 0 | 0 |
| SRR5925711 | 0 | 14858 | 462 | 3958 | 543 | 72270 | 7218 | 21750 | 35395 | 0 | 12632 | 8470 | 967 | 1182 | 413 | 47 | 72473 | 0 | 0 | 0 | 106 | 43 | 18 | 0 | 0 | 0 | 0 | 10 | 36 | 23 | 21 | 0 | 0 | 0 | 13 | 0 | 0 |
| SRR5925712 | 0 | 2516 | 98 | 4451 | 1568 | 53191 | 50693 | 21479 | 22591 | 0 | 972 | 1755 | 424 | 1315 | 572 | 28 | 49653 | 0 | 0 | 0 | 83 | 39 | 18 | 0 | 0 | 0 | 0 | 0 | 34 | 39 | 35 | 0 | 0 | 0 | 217 | 0 | 0 |
| SRR5925713 | 0 | 13136 | 229 | 2972 | 1213 | 49373 | 13504 | 20574 | 22943 | 0 | 4769 | 4376 | 972 | 1193 | 237 | 76 | 90265 | 0 | 0 | 0 | 90 | 31 | 15 | 0 | 0 | 0 | 0 | 0 | 21 | 32 | 29 | 0 | 0 | 0 | 17 | 0 | 0 |
| SRR5925714 | 0 | 12515 | 122 | 3173 | 1377 | 49772 | 14675 | 21023 | 23778 | 0 | 4984 | 4984 | 1083 | 1269 | 280 | 80 | 90981 | 0 | 0 | 0 | 94 | 32 | 17 | 0 | 0 | 0 | 0 | 0 | 25 | 24 | 25 | 0 | 0 | 0 | 14 | 0 | 0 |
| SRR5925715 | 15 | 37147 | 901 | 3690 | 521 | 54082 | 9846 | 25601 | 31726 | 0 | 10069 | 6378 | 485 | 780 | 483 | 148 | 69970 | 0 | 0 | 0 | 148 | 39 | 22 | 0 | 0 | 0 | 0 | 0 | 23 | 31 | 21 | 0 | 0 | 0 | 17 | 0 | 0 |
| SRR5925716 | 0 | 29285 | 557 | 4185 | 498 | 55460 | 9703 | 25409 | 31944 | 0 | 10199 | 6229 | 492 | 745 | 501 | 157 | 71217 | 0 | 0 | 0 | 142 | 39 | 24 | 0 | 0 | 0 | 0 | 0 | 26 | 26 | 19 | 0 | 0 | 0 | 18 | 0 | 0 |
| SRR5925717 | 0 | 4237 | 701 | 4083 | 520 | 82346 | 7113 | 22488 | 31540 | 0 | 15649 | 11556 | 808 | 1171 | 261 | 77 | 64433 | 0 | 0 | 0 | 219 | 65 | 19 | 0 | 0 | 0 | 0 | 14 | 54 | 52 | 25 | 0 | 0 | 0 | 15 | 0 | 0 |
| SRR5925718 | 0 | 5004 | 888 | 4038 | 546 | 84719 | 7470 | 23272 | 31257 | 0 | 13959 | 10443 | 910 | 1161 | 251 | 70 | 68398 | 0 | 0 | 0 | 253 | 83 | 22 | 0 | 0 | 0 | 0 | 11 | 66 | 62 | 29 | 0 | 0 | 0 | 18 | 0 | 0 |
| SRR5925719 | 12 | 2354 | 7292 | 1716 | 67 | 6845 | 1315 | 1219 | 10125 | 10 | 2077 | 2700 | 41 | 50 | 201 | 0 | 14337 | 0 | 0 | 0 | 122 | 73 | 16 | 0 | 0 | 0 | 0 | 0 | 13 | 19 | 37 | 0 | 0 | 0 | 0 | 0 | 0 |
| SRR5925720 | 28 | 2713 | 7226 | 1674 | 70 | 6523 | 1342 | 1139 | 9697 | 0 | 1747 | 2375 | 41 | 45 | 193 | 0 | 14214 | 0 | 0 | 0 | 116 | 75 | 17 | 0 | 0 | 0 | 0 | 0 | 16 | 20 | 39 | 0 | 0 | 0 | 0 | 0 | 0 |
| SRR5925721 | 50 | 2522 | 3438 | 3286 | 71 | 6401 | 1350 | 1499 | 12137 | 0 | 2256 | 2716 | 44 | 44 | 172 | 0 | 17123 | 0 | 0 | 0 | 70 | 53 | 12 | 0 | 0 | 0 | 0 | 0 | 13 | 13 | 36 | 0 | 0 | 0 | 0 | 0 | 0 |
| SRR5925722 | 50 | 2583 | 3429 | 3133 | 81 | 6278 | 1437 | 1582 | 11792 | 0 | 1901 | 2514 | 46 | 49 | 183 | 0 | 17711 | 0 | 0 | 0 | 77 | 60 | 12 | 0 | 0 | 0 | 0 | 0 | 13 | 13 | 34 | 0 | 0 | 0 | 0 | 0 | 0 |
| SRR5925723 | 64 | 2991 | 1608 | 3189 | 122 | 16789 | 2068 | 2835 | 8917 | 0 | 1342 | 1894 | 28 | 36 | 217 | 0 | 12966 | 0 | 0 | 0 | 128 | 58 | 21 | 0 | 0 | 0 | 0 | 0 | 0 | 24 | 21 | 0 | 0 | 0 | 0 | 0 | 0 |
| SRR5925724 | 76 | 2870 | 1820 | 3366 | 137 | 16664 | 2061 | 3233 | 8416 | 0 | 1009 | 1588 | 31 | 30 | 225 | 0 | 14813 | 0 | 0 | 0 | 159 | 72 | 23 | 0 | 0 | 0 | 0 | 0 | 0 | 26 | 24 | 0 | 0 | 0 | 0 | 0 | 0 |
| SRR5925725 | 104 | 4451 | 1790 | 5312 | 85 | 11186 | 1261 | 2076 | 9504 | 0 | 1332 | 2071 | 27 | 30 | 152 | 0 | 18292 | 0 | 0 | 0 | 189 | 144 | 28 | 0 | 0 | 0 | 0 | 0 | 0 | 31 | 47 | 0 | 0 | 0 | 10 | 0 | 0 |
| SRR5925726 | 54 | 2937 | 2102 | 5700 | 77 | 11890 | 1342 | 2383 | 9618 | 0 | 1092 | 1817 | 26 | 32 | 165 | 0 | 20235 | 0 | 0 | 0 | 177 | 143 | 24 | 0 | 0 | 0 | 0 | 0 | 0 | 30 | 41 | 0 | 0 | 0 | 0 | 0 | 0 |
| SRR5925727 | 0 | 5484 | 1828 | 7313 | 447 | 31725 | 12064 | 11618 | 21448 | 0 | 4915 | 3128 | 0 | 1340 | 0 | 0 | 41555 | 0 | 0 | 0 | 208 | 0 | 0 | 0 | 0 | 0 | 0 | 0 | 0 | 0 | 0 | 0 | 0 | 0 | 0 | 0 | 0 |
| SRR5925728 | 24 | 2060 | 3982 | 2866 | 109 | 6421 | 949 | 1718 | 8025 | 0 | 1117 | 1437 | 20 | 23 | 271 | 0 | 19638 | 0 | 0 | 0 | 88 | 112 | 13 | 0 | 0 | 0 | 0 | 0 | 24 | 30 | 73 | 0 | 0 | 0 | 0 | 0 | 0 |
| SRR5925729 | 31 | 3280 | 4651 | 2819 | 94 | 14235 | 1807 | 2742 | 7696 | 0 | 1197 | 1717 | 25 | 27 | 154 | 0 | 14605 | 0 | 0 | 0 | 147 | 84 | 24 | 0 | 0 | 0 | 0 | 0 | 0 | 25 | 26 | 0 | 0 | 0 | 0 | 0 | 0 |
| SRR5925730 | 18 | 3563 | 5043 | 2756 | 85 | 13249 | 1747 | 2682 | 7250 | 0 | 965 | 1494 | 20 | 24 | 163 | 0 | 14834 | 0 | 0 | 0 | 152 | 79 | 24 | 0 | 0 | 0 | 0 | 0 | 11 | 25 | 28 | 0 | 0 | 0 | 0 | 0 | 0 |
| SRR5925731 | 32 | 2625 | 7976 | 2270 | 58 | 8882 | 857 | 1615 | 6638 | 0 | 1796 | 2631 | 32 | 33 | 168 | 0 | 11375 | 0 | 0 | 0 | 47 | 65 | 0 | 0 | 0 | 0 | 0 | 0 | 11 | 0 | 25 | 0 | 0 | 0 | 0 | 0 | 0 |
| SRR5925732 | 51 | 2092 | 10148 | 2132 | 80 | 10379 | 1049 | 2114 | 7494 | 10 | 1942 | 2728 | 32 | 43 | 197 | 0 | 13519 | 0 | 0 | 0 | 51 | 89 | 0 | 0 | 0 | 0 | 0 | 0 | 0 | 10 | 30 | 0 | 0 | 0 | 0 | 0 | 0 |
| SRR5930976 | 52 | 17479 | 12325 | 241 | 4765 | 36909 | 42512 | 40412 | 7745 | 39 | 34150 | 101549 | 9698 | 5775 | 34 | 49946 | 15232 | 0 | 0 | 0 | 296 | 83 | 91 | 0 | 0 | 0 | 0 | 0 | 23 | 86 | 63 | 46 | 0 | 12 | 0 | 0 | 0 |
| SRR5930977 | 87 | 10593 | 103 | 818 | 1193 | 11756 | 15774 | 11019 | 2099 | 11 | 10668 | 28617 | 1453 | 4423 | 0 | 7613 | 9781 | 0 | 0 | 0 | 35 | 0 | 0 | 0 | 0 | 0 | 0 | 0 | 0 | 0 | 0 | 0 | 0 | 0 | 86 | 0 | 0 |
| SRR5930978 | 36 | 7167 | 560 | 206 | 3429 | 42500 | 47326 | 46124 | 7499 | 66 | 12498 | 76581 | 4149 | 2222 | 0 | 27420 | 21005 | 0 | 0 | 0 | 27 | 0 | 72 | 31 | 1076 | 19 | 0 | 0 | 0 | 0 | 0 | 49 | 0 | 518 | 189 | 276 | 0 |
| SRR5936591 | 0 | 2427 | 446 | 91 | 3281 | 28235 | 30185 | 29545 | 9762 | 0 | 19270 | 118120 | 3886 | 1710 | 133 | 17150 | 5869 | 0 | 0 | 0 | 135 | 30 | 27 | 0 | 0 | 0 | 0 | 0 | 0 | 39 | 22 | 15 | 0 | 0 | 0 | 0 | 0 |
| SRR5936592 | 37 | 5888 | 110 | 80 | 3237 | 28501 | 34875 | 39850 | 0 | 68 | 7953 | 28358 | 1682 | 1709 | 386 | 21999 | 4205 | 0 | 0 | 0 | 0 | 0 | 12 | 10 | 104 | 0 | 0 | 0 | 0 | 0 | 0 | 14 | 0 | 160 | 899 | 60 | 0 |

### Supp Table S8.1.Columns_B_to_AL

| | | | | | | | | | | | | | | | | | | | | | | | | | | | | | | | | | | | | | |
|---|---|---|---|---|---|---|---|---|---|---|---|---|---|---|---|---|---|---|---|---|---|---|---|---|---|---|---|---|---|---|---|---|---|---|---|---|---|
| SRR5951878 | 0 | 33 | 139 | 2177 | 33621 | 22805 | 183386 | 60799 | 13788 | 174 | 11809 | 6949 | 12871 | 17422 | 31003 | 74 | 2701 | 0 | 31 | 13 | 76 | 30 | 80 | 36 | 29 | 14 | 0 | 0 | 14 | 16 | 15 | 0 | 0 | 0 | 0 | 45 | 0 |
| SRR5951879 | 0 | 162 | 73 | 3514 | 22036 | 11052 | 202441 | 79305 | 5694 | 28 | 7849 | 10196 | 240 | 8375 | 18220 | 55 | 4980 | 0 | 29 | 173 | 800 | 250 | 140 | 26 | 15 | 18 | 0 | 0 | 49 | 96 | 28 | 0 | 0 | 0 | 0 | 17 | 0 |
| SRR5951880 | 28542 | 1453 | 0 | 0 | 260648 | 164 | 66 | 18006 | 0 | 0 | 0 | 0 | 0 | 0 | 0 | 0 | 0 | 0 | 0 | 0 | 0 | 0 | 0 | 0 | 0 | 0 | 0 | 0 | 0 | 0 | 0 | 0 | 0 | 0 | 0 | 0 | 0 |
| SRR5951881 | 18 | 60 | 440 | 2564 | 31332 | 26878 | 224972 | 59357 | 9507 | 723 | 9423 | 7092 | 6807 | 32347 | 13015 | 304 | 4016 | 0 | 44 | 25 | 45 | 25 | 45 | 35 | 34 | 15 | 0 | 0 | 18 | 0 | 0 | 0 | 0 | 0 | 0 | 36 | 0 |
| SRR5951882 | 39 | 204 | 458 | 1934 | 28227 | 17390 | 217404 | 79290 | 4803 | 67 | 5846 | 8210 | 5202 | 23116 | 7815 | 288 | 7582 | 0 | 83 | 106 | 475 | 717 | 161 | 22 | 18 | 12 | 0 | 19 | 30 | 43 | 43 | 0 | 0 | 0 | 0 | 14 | 0 |
| SRR5951883 | 15418 | 525 | 0 | 0 | 257600 | 141 | 50 | 21361 | 0 | 0 | 0 | 0 | 0 | 0 | 0 | 0 | 0 | 56 | 0 | 0 | 0 | 0 | 0 | 15 | 0 | 11 | 29 | 0 | 0 | 0 | 0 | 0 | 0 | 0 | 0 | 0 | 0 |
| SRR5951884 | 69 | 838 | 719 | 2333 | 27979 | 16496 | 256035 | 19966 | 9198 | 30 | 11684 | 13962 | 11241 | 25713 | 23064 | 111 | 5618 | 0 | 0 | 0 | 0 | 0 | 30 | 0 | 0 | 0 | 0 | 0 | 0 | 0 | 0 | 0 | 0 | 0 | 0 | 0 | 0 |
| SRR5951885 | 83 | 996 | 591 | 2297 | 27315 | 16458 | 278188 | 19278 | 9162 | 24 | 11653 | 14 | 160 | 24108 | 23612 | 101 | 5502 | 0 | 0 | 0 | 0 | 0 | 27 | 0 | 0 | 0 | 0 | 0 | 0 | 0 | 0 | 0 | 0 | 0 | 0 | 0 | 0 |
| SRR5951886 | 135 | 1471 | 1020 | 2734 | 27890 | 16406 | 273812 | 16722 | 7750 | 34 | 12150 | 15 | 146 | 22639 | 24717 | 99 | 5698 | 0 | 0 | 0 | 12 | 24 | 52 | 0 | 0 | 0 | 0 | 0 | 0 | 0 | 0 | 0 | 0 | 0 | 0 | 0 | 0 |
| SRR5951887 | 68 | 765 | 493 | 1748 | 35362 | 17104 | 323079 | 28789 | 7739 | 16 | 9974 | 10496 | 106 | 17503 | 17587 | 12 | 5332 | 0 | 11 | 0 | 24 | 33 | 77 | 16 | 17 | 0 | 0 | 0 | 0 | 0 | 0 | 0 | 0 | 0 | 0 | 0 | 0 |
| SRR5951888 | 101 | 1094 | 775 | 2400 | 34316 | 17359 | 302766 | 23948 | 7386 | 18 | 11070 | 13 | 107 | 19222 | 18587 | 12 | 5226 | 0 | 0 | 0 | 22 | 35 | 69 | 12 | 16 | 0 | 0 | 0 | 0 | 0 | 0 | 0 | 0 | 0 | 0 | 0 | 0 |
| SRR5951889 | 58 | 889 | 710 | 2576 | 37864 | 18619 | 303642 | 27281 | 7787 | 24 | 10904 | 12113 | 107 | 20694 | 19071 | 11 | 5088 | 0 | 19 | 0 | 28 | 41 | 61 | 14 | 12 | 0 | 0 | 11 | 0 | 0 | 0 | 0 | 0 | 0 | 0 | 0 | 0 |
| SRR5951890 | 44 | 704 | 479 | 3204 | 30975 | 16512 | 257717 | 24109 | 11114 | 208 | 7593 | 10613 | 7523 | 19212 | 19238 | 162 | 2888 | 0 | 26 | 0 | 44 | 19 | 78 | 13 | 14 | 0 | 0 | 15 | 0 | 11 | 0 | 0 | 0 | 0 | 0 | 0 | 0 |
| SRR5951891 | 85 | 1113 | 880 | 1829 | 31525 | 16329 | 258794 | 20452 | 9962 | 70 | 8557 | 11 | 145 | 18567 | 18096 | 153 | 4761 | 0 | 20 | 0 | 44 | 29 | 78 | 14 | 14 | 0 | 0 | 13 | 0 | 12 | 0 | 0 | 0 | 0 | 0 | 0 | 0 |
| SRR5951892 | 61 | 721 | 941 | 1366 | 33896 | 16544 | 260988 | 23944 | 11497 | 37 | 9339 | 12 | 178 | 24118 | 23121 | 188 | 3989 | 0 | 16 | 0 | 34 | 38 | 73 | 14 | 13 | 0 | 0 | 0 | 0 | 0 | 0 | 0 | 0 | 0 | 0 | 0 | 0 |
| SRR5951893 | 426 | 11625 | 4985 | 2329 | 33033 | 19426 | 311670 | 23834 | 6623 | 71 | 9745 | 11966 | 9330 | 25570 | 19374 | 2012 | 7505 | 0 | 20 | 0 | 38 | 58 | 54 | 0 | 0 | 0 | 0 | 14 | 0 | 0 | 0 | 0 | 0 | 0 | 0 | 0 | 0 |
| SRR5951894 | 121 | 1973 | 3229 | 2525 | 30256 | 25315 | 323048 | 25690 | 6058 | 34 | 6165 | 8017 | 159 | 38571 | 18303 | 4680 | 7077 | 0 | 0 | 0 | 0 | 0 | 32 | 0 | 0 | 0 | 0 | 0 | 0 | 0 | 0 | 0 | 0 | 0 | 0 | 0 | 0 |
| SRR5951895 | 378 | 9499 | 3756 | 3211 | 31587 | 20491 | 311192 | 25645 | 7518 | 77 | 9722 | 12 | 215 | 27030 | 19558 | 2196 | 5978 | 0 | 20 | 0 | 53 | 40 | 44 | 0 | 0 | 0 | 0 | 14 | 13 | 16 | 0 | 0 | 0 | 0 | 0 | 0 | 0 |
| SRR5951896 | 51 | 2528 | 1901 | 1279 | 19712 | 17498 | 288389 | 14552 | 0 | 430 | 12982 | 17172 | 6356 | 61244 | 5018 | 24197 | 11523 | 0 | 0 | 10 | 42 | 36 | 92 | 40 | 43 | 0 | 0 | 0 | 13 | 0 | 0 | 0 | 0 | 0 | 0 | 13 | 0 |
| SRR5951897 | 44 | 3831 | 1022 | 2985 | 24446 | 15575 | 315237 | 29596 | 0 | 215 | 5171 | 8656 | 237 | 31312 | 1849 | 18314 | 7583 | 0 | 0 | 14 | 40 | 51 | 40 | 12 | 0 | 0 | 0 | 0 | 10 | 11 | 0 | 0 | 0 | 0 | 0 | 0 | 0 |
| SRR5951898 | 77909 | 16019 | 0 | 0 | 327609 | 191 | 47 | 3373 | 0 | 0 | 0 | 0 | 0 | 0 | 0 | 0 | 0 | 19 | 0 | 0 | 0 | 0 | 0 | 0 | 0 | 0 | 19 | 0 | 0 | 0 | 0 | 0 | 0 | 0 | 0 | 0 | 0 |
| SRR5951899 | 80 | 11140 | 1681 | 2227 | 26590 | 22187 | 256546 | 21437 | 0 | 1200 | 5675 | 6845 | 3282 | 62241 | 1820 | 13906 | 9568 | 0 | 0 | 0 | 39 | 19 | 38 | 19 | 17 | 0 | 0 | 0 | 15 | 12 | 0 | 0 | 0 | 0 | 0 | 0 | 0 |
| SRR5951900 | 27 | 6672 | 1828 | 4515 | 23293 | 17934 | 196250 | 32762 | 0 | 181 | 2210 | 6358 | 2126 | 36661 | 1557 | 18812 | 7562 | 0 | 0 | 20 | 44 | 50 | 46 | 13 | 11 | 0 | 0 | 0 | 14 | 12 | 10 | 0 | 0 | 0 | 0 | 0 | 0 |
| SRR5951901 | 55150 | 17065 | 0 | 13 | 225327 | 123 | 45 | 2727 | 0 | 0 | 0 | 0 | 0 | 0 | 0 | 0 | 0 | 15 | 0 | 0 | 0 | 0 | 0 | 0 | 0 | 0 | 17 | 0 | 0 | 0 | 0 | 0 | 0 | 0 | 0 | 0 | 0 |
| SRR5951902 | 60 | 1845 | 1414 | 1415 | 20115 | 17394 | 330535 | 16001 | 0 | 503 | 10458 | 14183 | 5513 | 63700 | 1289 | 20211 | 10863 | 0 | 0 | 12 | 48 | 21 | 39 | 22 | 20 | 0 | 0 | 0 | 20 | 11 | 0 | 0 | 0 | 0 | 0 | 0 | 0 |
| SRR5951903 | 78 | 1277 | 950 | 2436 | 24633 | 13636 | 317035 | 36089 | 0 | 158 | 8471 | 15 | 331 | 28963 | 503 | 25061 | 8884 | 0 | 0 | 0 | 34 | 15 | 37 | 14 | 11 | 0 | 0 | 0 | 0 | 0 | 0 | 0 | 0 | 0 | 0 | 0 | 0 |
| SRR5951904 | 98630 | 7861 | 0 | 0 | 258647 | 165 | 35 | 3431 | 0 | 0 | 0 | 0 | 0 | 0 | 0 | 0 | 0 | 20 | 0 | 0 | 0 | 0 | 0 | 0 | 0 | 0 | 24 | 0 | 0 | 0 | 0 | 0 | 0 | 0 | 0 | 0 | 0 |
| SRR5951905 | 115 | 4686 | 3564 | 1204 | 18352 | 15588 | 280414 | 16895 | 0 | 844 | 11402 | 19 | 5774 | 60319 | 1129 | 24836 | 14129 | 0 | 0 | 17 | 43 | 24 | 29 | 13 | 17 | 0 | 0 | 0 | 22 | 0 | 0 | 0 | 0 | 0 | 0 | 0 | 0 |
| SRR5951906 | 77 | 2972 | 1308 | 1660 | 21202 | 12599 | 299950 | 26977 | 0 | 312 | 7774 | 19 | 259 | 30637 | 414 | 28697 | 21578 | 0 | 14 | 30 | 63 | 18 | 42 | 15 | 17 | 0 | 0 | 0 | 24 | 0 | 0 | 0 | 0 | 0 | 0 | 0 | 0 |
| SRR5951907 | 113552 | 6147 | 0 | 0 | 293783 | 186 | 37 | 2953 | 0 | 0 | 0 | 0 | 0 | 0 | 0 | 0 | 0 | 25 | 0 | 0 | 0 | 0 | 0 | 0 | 0 | 0 | 32 | 0 | 0 | 0 | 0 | 0 | 0 | 0 | 0 | 0 | 0 |
| SRR5951908 | 736 | 45354 | 11711 | 3734 | 18075 | 11341 | 200438 | 6149 | 0 | 150 | 8438 | 21890 | 3219 | 55207 | 1973 | 14996 | 14506 | 0 | 0 | 0 | 19 | 12 | 46 | 11 | 0 | 0 | 0 | 0 | 0 | 0 | 0 | 0 | 0 | 0 | 0 | 0 | 0 |
| SRR5951909 | 1271 | 81844 | 17016 | 6051 | 17925 | 13591 | 248910 | 5311 | 0 | 276 | 6963 | 27 | 288 | 46046 | 2066 | 17259 | 10554 | 0 | 0 | 0 | 21 | 20 | 54 | 11 | 10 | 0 | 0 | 0 | 0 | 0 | 0 | 0 | 0 | 0 | 0 | 0 | 0 |
| SRR5951910 | 930 | 79909 | 14800 | 6646 | 19050 | 13404 | 229095 | 4693 | 0 | 202 | 7325 | 17493 | 3132 | 49863 | 2147 | 16834 | 9950 | 0 | 0 | 0 | 20 | 17 | 56 | 12 | 11 | 0 | 0 | 0 | 0 | 0 | 0 | 0 | 0 | 0 | 0 | 0 | 0 |
| SRR5951911 | 1084 | 64057 | 13820 | 4850 | 15536 | 14172 | 267477 | 5315 | 0 | 96 | 9571 | 26 | 244 | 34703 | 8380 | 17627 | 11663 | 0 | 17 | 0 | 17 | 15 | 34 | 0 | 0 | 0 | 0 | 16 | 10 | 0 | 0 | 0 | 0 | 0 | 0 | 0 | 0 |
| SRR5951912 | 1006 | 81598 | 13515 | 4067 | 16485 | 14936 | 229585 | 5088 | 0 | 137 | 9736 | 28 | 267 | 40545 | 7605 | 16662 | 12972 | 0 | 18 | 0 | 16 | 12 | 35 | 0 | 0 | 0 | 0 | 19 | 12 | 0 | 0 | 0 | 0 | 0 | 0 | 0 | 0 |
| SRR5951913 | 813 | 119496 | 17796 | 4591 | 18281 | 15474 | 245755 | 5294 | 0 | 314 | 9747 | 22 | 328 | 48879 | 6046 | 18022 | 11136 | 0 | 0 | 0 | 13 | 0 | 22 | 0 | 0 | 0 | 0 | 0 | 0 | 0 | 0 | 0 | 0 | 0 | 0 | 0 | 0 |
| SRR6020460 | 0 | 145 | 215 | 37 | 68 | 357 | 870 | 897 | 377 | 141 | 24636 | 14 | 2342 | 2393 | 0 | 14941 | 22057 | 0 | 0 | 0 | 365 | 85 | 0 | 0 | 0 | 0 | 0 | 0 | 0 | 415 | 264 | 30 | 0 | 0 | 0 | 0 | 0 |
| SRR6020461 | 0 | 314 | 111 | 65 | 154 | 285 | 1722 | 1370 | 508 | 104 | 8899 | 4521 | 804 | 3373 | 0 | 18044 | 18903 | 0 | 0 | 0 | 2327 | 351 | 39 | 0 | 0 | 0 | 0 | 0 | 61 | 2194 | 1075 | 110 | 0 | 0 | 0 | 0 | 0 |
| SRR6020462 | 0 | 1173 | 395 | 63 | 55 | 321 | 468 | 561 | 201 | 178 | 21669 | 58927 | 2621 | 1477 | 0 | 13158 | 23524 | 0 | 0 | 0 | 1102 | 338 | 32 | 0 | 0 | 0 | 0 | 0 | 22 | 677 | 569 | 50 | 0 | 0 | 0 | 0 | 0 |
| SRR6020463 | 0 | 1348 | 271 | 77 | 73 | 350 | 652 | 569 | 316 | 204 | 20147 | 47982 | 2362 | 1109 | 0 | 14958 | 21499 | 0 | 0 | 0 | 2593 | 374 | 51 | 0 | 0 | 0 | 0 | 0 | 57 | 2345 | 905 | 102 | 0 | 0 | 0 | 0 | 0 |
| SRR6020464 | 0 | 627 | 308 | 19 | 80 | 501 | 924 | 1137 | 597 | 269 | 37143 | 52 | 3514 | 1290 | 0 | 13719 | 35797 | 0 | 0 | 0 | 80 | 24 | 0 | 0 | 0 | 0 | 0 | 0 | 0 | 71 | 59 | 0 | 0 | 0 | 0 | 0 | 0 |
| SRR6020465 | 0 | 257 | 288 | 20 | 85 | 410 | 985 | 962 | 357 | 160 | 29939 | 51802 | 2815 | 1662 | 0 | 14016 | 34804 | 0 | 0 | 0 | 301 | 39 | 0 | 0 | 0 | 0 | 0 | 0 | 0 | 246 | 102 | 10 | 0 | 0 | 0 | 0 | 0 |
| SRR6020466 | 0 | 845 | 196 | 135 | 149 | 558 | 879 | 916 | 516 | 183 | 26649 | 49022 | 2275 | 1248 | 0 | 22613 | 31697 | 0 | 0 | 0 | 924 | 123 | 12 | 0 | 0 | 0 | 0 | 0 | 32 | 1180 | 357 | 23 | 0 | 0 | 0 | 0 | 0 |
| SRR6020467 | 0 | 378 | 330 | 30 | 56 | 381 | 666 | 883 | 430 | 227 | 29571 | 13 | 3658 | 1388 | 0 | 15187 | 32374 | 0 | 0 | 0 | 501 | 42 | 0 | 0 | 0 | 0 | 0 | 0 | 11 | 311 | 98 | 0 | 0 | 0 | 0 | 0 | 0 |
| SRR6020468 | 0 | 449 | 473 | 33 | 42 | 296 | 540 | 792 | 402 | 187 | 18348 | 51888 | 2729 | 1307 | 0 | 11280 | 33065 | 0 | 0 | 0 | 1621 | 149 | 13 | 0 | 0 | 0 | 0 | 0 | 20 | 958 | 314 | 30 | 0 | 0 | 0 | 0 | 0 |
| SRR6020469 | 0 | 979 | 345 | 127 | 92 | 491 | 709 | 901 | 398 | 179 | 18816 | 42562 | 3457 | 1526 | 0 | 21408 | 26633 | 0 | 0 | 0 | 2615 | 424 | 44 | 0 | 0 | 0 | 0 | 0 | 43 | 1625 | 997 | 95 | 0 | 0 | 0 | 0 | 0 |
| SRR6020470 | 0 | 253 | 349 | 0 | 42 | 263 | 513 | 564 | 296 | 130 | 17757 | 18 | 2628 | 1112 | 0 | 12370 | 28573 | 0 | 0 | 0 | 99 | 15 | 0 | 0 | 0 | 0 | 0 | 0 | 0 | 92 | 37 | 0 | 0 | 0 | 0 | 0 | 0 |
| SRR6020471 | 0 | 252 | 130 | 39 | 125 | 345 | 1140 | 879 | 309 | 103 | 18695 | 49587 | 1676 | 2157 | 0 | 12052 | 22110 | 0 | 0 | 0 | 764 | 185 | 23 | 0 | 0 | 0 | 0 | 0 | 14 | 633 | 454 | 40 | 0 | 0 | 0 | 0 | 0 |
| SRR6020472 | 0 | 84 | 480 | 0 | 96 | 394 | 847 | 1780 | 701 | 89 | 29138 | 13 | 3242 | 2951 | 0 | 10177 | 79611 | 0 | 0 | 0 | 118 | 50 | 11 | 0 | 0 | 0 | 0 | 0 | 0 | 95 | 115 | 28 | 0 | 0 | 0 | 0 | 0 |
| SRR6020473 | 0 | 449 | 479 | 10 | 76 | 419 | 768 | 880 | 391 | 195 | 19091 | 50567 | 3204 | 1485 | 0 | 14149 | 21063 | 0 | 0 | 0 | 427 | 91 | 0 | 0 | 0 | 0 | 0 | 0 | 10 | 269 | 190 | 16 | 0 | 0 | 0 | 0 | 0 |
| SRR6020474 | 0 | 358 | 289 | 41 | 56 | 242 | 504 | 456 | 174 | 142 | 15094 | 45 | 1764 | 1108 | 0 | 12485 | 17093 | 0 | 0 | 0 | 2675 | 321 | 32 | 0 | 0 | 0 | 0 | 0 | 28 | 1828 | 755 | 59 | 0 | 0 | 0 | 0 | 0 |
| SRR6020475 | 0 | 220 | 348 | 27 | 65 | 277 | 627 | 700 | 251 | 114 | 20999 | 61289 | 1747 | 1080 | 0 | 8982 | 26647 | 0 | 0 | 0 | 279 | 78 | 0 | 0 | 0 | 0 | 0 | 0 | 0 | 227 | 181 | 14 | 0 | 0 | 0 | 0 | 0 |
| SRR6020476 | 0 | 807 | 157 | 197 | 90 | 270 | 665 | 458 | 179 | 198 | 21562 | 65226 | 1699 | 996 | 0 | 10813 | 15447 | 0 | 0 | 0 | 1626 | 126 | 14 | 0 | 0 | 0 | 0 | 0 | 28 | 1061 | 245 | 16 | 0 | 0 | 0 | 0 | 0 |
| SRR6020477 | 0 | 613 | 280 | 39 | 87 | 416 | 802 | 852 | 410 | 214 | 25171 | 23 | 3336 | 1468 | 0 | 13847 | 28296 | 0 | 0 | 0 | 224 | 61 | 0 | 0 | 0 | 0 | 0 | 0 | 0 | 180 | 146 | 12 | 0 | 0 | 0 | 0 | 0 |
| SRR6020478 | 0 | 192 | 411 | 28 | 100 | 690 | 1100 | 1695 | 474 | 216 | 34060 | 12 | 3746 | 1838 | 0 | 16977 | 36392 | 0 | 0 | 0 | 669 | 71 | 0 | 0 | 0 | 0 | 0 | 0 | 13 | 561 | 202 | 22 | 0 | 0 | 0 | 0 | 0 |
| SRR6020479 | 0 | 335 | 323 | 62 | 157 | 787 | 1525 | 2301 | 1054 | 301 | 32151 | 65446 | 3634 | 2268 | 0 | 17655 | 58120 | 0 | 0 | 0 | 278 | 78 | 0 | 0 | 0 | 0 | 0 | 0 | 0 | 190 | 145 | 11 | 0 | 0 | 0 | 0 | 0 |
| SRR6020480 | 0 | 537 | 124 | 95 | 220 | 616 | 1907 | 1527 | 519 | 224 | 26562 | 27414 | 2188 | 3037 | 0 | 23902 | 22814 | 0 | 0 | 0 | 1731 | 147 | 14 | 0 | 0 | 0 | 0 | 0 | 39 | 1592 | 452 | 33 | 0 | 0 | 10 | 0 | 0 |
| SRR6020481 | 0 | 404 | 446 | 20 | 53 | 329 | 561 | 736 | 409 | 114 | 23874 | 55929 | 2204 | 1234 | 0 | 12129 | 30475 | 0 | 0 | 0 | 1094 | 116 | 10 | 0 | 0 | 0 | 0 | 0 | 15 | 807 | 248 | 17 | 0 | 0 | 0 | 0 | 0 |
| SRR6020482 | 0 | 1335 | 344 | 86 | 102 | 621 | 914 | 971 | 411 | 247 | 25268 | 11 | 3441 | 1177 | 0 | 21423 | 38509 | 0 | 0 | 0 | 3500 | 450 | 45 | 0 | 0 | 0 | 0 | 0 | 71 | 2614 | 1114 | 91 | 0 | 0 | 0 | 0 | 0 |
| SRR6020483 | 0 | 272 | 575 | 0 | 64 | 408 | 613 | 780 | 336 | 109 | 25866 | 44 | 3639 | 1705 | 0 | 12469 | 28425 | 0 | 0 | 0 | 206 | 41 | 0 | 0 | 0 | 0 | 0 | 0 | 0 | 150 | 80 | 0 | 0 | 0 | 0 | 0 | 0 |
| SRR6020484 | 0 | 224 | 281 | 37 | 126 | 546 | 1162 | 1293 | 472 | 209 | 27994 | 43 | 2386 | 2052 | 0 | 18269 | 52882 | 0 | 0 | 0 | 838 | 121 | 15 | 0 | 0 | 0 | 0 | 0 | 14 | 764 | 315 | 40 | 0 | 0 | 0 | 0 | 0 |
| SRR6020485 | 0 | 589 | 345 | 69 | 116 | 671 | 922 | 966 | 413 | 140 | 26602 | 60868 | 3416 | 1377 | 0 | 19809 | 46066 | 0 | 0 | 0 | 1120 | 114 | 15 | 0 | 0 | 0 | 0 | 0 | 28 | 1127 | 306 | 30 | 0 | 0 | 0 | 0 | 0 |
| SRR6020486 | 0 | 189 | 402 | 39 | 67 | 353 | 947 | 863 | 340 | 116 | 27388 | 48297 | 2237 | 2126 | 0 | 12358 | 28860 | 0 | 0 | 0 | 676 | 84 | 10 | 0 | 0 | 0 | 0 | 0 | 14 | 636 | 251 | 29 | 0 | 0 | 0 | 0 | 0 |
| SRR6020487 | 0 | 271 | 308 | 22 | 67 | 406 | 730 | 840 | 521 | 132 | 26885 | 68212 | 2854 | 1775 | 0 | 12451 | 36007 | 0 | 0 | 0 | 637 | 95 | 11 | 0 | 0 | 0 | 0 | 0 | 12 | 483 | 223 | 21 | 0 | 0 | 0 | 0 | 0 |
| SRR6020488 | 0 | 448 | 307 | 52 | 61 | 378 | 541 | 676 | 362 | 166 | 14761 | 46214 | 2201 | 1030 | 0 | 14430 | 22102 | 0 | 0 | 0 | 4066 | 403 | 29 | 0 | 0 | 0 | 0 | 0 | 52 | 2693 | 827 | 55 | 0 | 0 | 0 | 0 | 0 |
| SRR6020489 | 0 | 52 | 164 | 11 | 71 | 241 | 1177 | 1203 | 535 | 116 | 13888 | 4355 | 795 | 4820 | 0 | 10200 | 17858 | 0 | 0 | 0 | 164 | 13 | 0 | 0 | 0 | 0 | 0 | 0 | 0 | 175 | 50 | 0 | 0 | 0 | 0 | 0 | 0 |
| SRR6020490 | 0 | 129 | 311 | 33 | 86 | 365 | 900 | 1244 | 392 | 143 | 28408 | 42130 | 3072 | 3327 | 0 | 13915 | 19874 | 0 | 0 | 0 | 1557 | 79 | 0 | 0 | 0 | 0 | 0 | 0 | 24 | 1266 | 174 | 14 | 0 | 0 | 0 | 0 | 0 |
| SRR6020491 | 0 | 184 | 382 | 16 | 31 | 252 | 435 | 513 | 221 | 144 | 20602 | 11 | 2831 | 1300 | 0 | 12333 | 19386 | 0 | 0 | 0 | 417 | 68 | 0 | 0 | 0 | 0 | 0 | 0 | 0 | 288 | 180 | 0 | 0 | 0 | 0 | 0 | 0 |
| SRR6020492 | 0 | 986 | 98 | 254 | 122 | 389 | 850 | 1014 | 363 | 166 | 16442 | 47674 | 1828 | 1241 | 0 | 17469 | 21698 | 0 | 0 | 0 | 3672 | 272 | 27 | 0 | 0 | 0 | 0 | 0 | 97 | 2751 | 633 | 54 | 0 | 0 | 0 | 0 | 0 |
| SRR6020493 | 0 | 220 | 293 | 27 | 75 | 425 | 710 | 767 | 457 | 194 | 25894 | 40 | 2370 | 1341 | 0 | 14433 | 29242 | 0 | 0 | 0 | 155 | 18 | 0 | 0 | 0 | 0 | 0 | 0 | 0 | 154 | 60 | 0 | 0 | 0 | 0 | 0 | 0 |
| SRR6020494 | 0 | 107 | 226 | 12 | 78 | 333 | 1137 | 1130 | 433 | 128 | 27540 | 47795 | 2654 | 2354 | 0 | 13072 | 23816 | 0 | 0 | 0 | 922 | 57 | 0 | 0 | 0 | 0 | 0 | 0 | 16 | 770 | 163 | 14 | 0 | 0 | 0 | 0 | 0 |
| SRR6020495 | 0 | 146 | 125 | 65 | 63 | 428 | 852 | 531 | 208 | 72 | 6685 | 12965 | 530 | 1022 | 0 | 10731 | 5805 | 0 | 0 | 0 | 107 | 13 | 0 | 0 | 0 | 0 | 0 | 0 | 0 | 114 | 36 | 0 | 0 | 0 | 0 | 0 | 0 |
| SRR6020496 | 0 | 120 | 291 | 38 | 74 | 460 | 843 | 996 | 548 | 156 | 24651 | 43267 | 1811 | 1625 | 0 | 13781 | 19486 | 0 | 0 | 0 | 76 | 12 | 0 | 0 | 0 | 0 | 0 | 0 | 0 | 85 | 55 | 0 | 0 | 0 | 0 | 0 | 0 |
| SRR6020497 | 0 | 115 | 627 | 0 | 78 | 451 | 818 | 1301 | 549 | 121 | 27249 | 41377 | 4172 | 1995 | 0 | 17753 | 28105 | 0 | 0 | 0 | 652 | 50 | 0 | 0 | 0 | 0 | 0 | 0 | 14 | 684 | 133 | 19 | 0 | 0 | 15 | 0 | 0 |
| SRR6020498 | 0 | 1788 | 77 | 335 | 173 | 360 | 1481 | 952 | 474 | 109 | 13097 | 8186 | 790 | 2968 | 0 | 15590 | 18236 | 0 | 0 | 0 | 2021 | 376 | 63 | 0 | 0 | 0 | 0 | 0 | 57 | 1931 | 1129 | 149 | 0 | 0 | 0 | 0 | 0 |
| SRR6020499 | 0 | 398 | 311 | 22 | 83 | 406 | 733 | 949 | 435 | 127 | 23190 | 51259 | 2860 | 1557 | 0 | 13018 | 41672 | 0 | 0 | 0 | 761 | 125 | 19 | 0 | 0 | 0 | 0 | 0 | 15 | 525 | 283 | 42 | 0 | 0 | 0 | 0 | 0 |
| SRR6020500 | 13 | 2338 | 155 | 447 | 106 | 610 | 900 | 734 | 269 | 105 | 20887 | 37073 | 2210 | 1103 | 0 | 15234 | 21870 | 0 | 0 | 0 | 1657 | 197 | 27 | 0 | 0 | 0 | 0 | 0 | 61 | 1251 | 491 | 54 | 0 | 15 | 0 | 0 | 0 |
| SRR6020501 | 0 | 283 | 467 | 20 | 72 | 348 | 673 | 662 | 283 | 153 | 20170 | 33 | 1871 | 1208 | 0 | 10168 | 20529 | 0 | 0 | 0 | 194 | 55 | 0 | 0 | 0 | 0 | 0 | 0 | 0 | 160 | 135 | 12 | 0 | 0 | 0 | 0 | 0 |
| SRR6020502 | 0 | 240 | 167 | 84 | 102 | 302 | 1277 | 755 | 197 | 107 | 8008 | 8612 | 586 | 1903 | 0 | 14341 | 13815 | 0 | 0 | 0 | 1640 | 134 | 11 | 0 | 0 | 0 | 0 | 0 | 38 | 1371 | 403 | 27 | 0 | 0 | 0 | 0 | 0 |
| SRR6020503 | 0 | 797 | 741 | 58 | 74 | 602 | 834 | 1423 | 437 | 206 | 25177 | 46182 | 3233 | 1544 | 0 | 14523 | 31901 | 0 | 0 | 0 | 659 | 162 | 12 | 0 | 0 | 0 | 0 | 0 | 13 | 360 | 257 | 16 | 0 | 0 | 0 | 0 | 0 |
| SRR6020504 | 0 | 343 | 280 | 18 | 281 | 1347 | 2694 | 3292 | 466 | 165 | 26733 | 18 | 3463 | 2039 | 0 | 15834 | 24409 | 0 | 0 | 0 | 169 | 30 | 0 | 0 | 0 | 0 | 0 | 0 | 0 | 124 | 83 | 11 | 0 | 0 | 0 | 0 | 0 |
| SRR6020505 | 0 | 504 | 329 | 38 | 54 | 315 | 584 | 674 | 293 | 145 | 13145 | 11 | 2367 | 1059 | 0 | 12840 | 18926 | 0 | 0 | 0 | 1337 | 172 | 14 | 0 | 0 | 0 | 0 | 0 | 22 | 909 | 365 | 22 | 0 | 0 | 0 | 0 | 0 |
| SRR6020506 | 0 | 376 | 303 | 39 | 71 | 384 | 769 | 881 | 370 | 201 | 22702 | 59661 | 2947 | 1384 | 0 | 13340 | 30963 | 0 | 0 | 0 | 177 | 35 | 0 | 0 | 0 | 0 | 0 | 0 | 0 | 151 | 84 | 0 | 0 | 0 | 0 | 0 | 0 |
| SRR6020507 | 0 | 298 | 233 | 0 | 126 | 275 | 1416 | 1242 | 397 | 97 | 11488 | 12941 | 966 | 2340 | 0 | 11817 | 8464 | 0 | 0 | 0 | 537 | 64 | 0 | 0 | 0 | 0 | 0 | 0 | 0 | 419 | 152 | 12 | 0 | 0 | 0 | 0 | 0 |
| SRR6020508 | 0 | 682 | 205 | 56 | 125 | 539 | 1012 | 1070 | 459 | 162 | 25661 | 48630 | 2730 | 1391 | 0 | 18743 | 41915 | 0 | 0 | 0 | 1514 | 267 | 31 | 0 | 0 | 0 | 0 | 0 | 35 | 1308 | 590 | 68 | 0 | 0 | 0 | 0 | 0 |
| SRR6020509 | 0 | 171 | 286 | 16 | 93 | 486 | 963 | 1357 | 432 | 123 | 28668 | 67433 | 2650 | 1871 | 0 | 9941 | 36919 | 0 | 0 | 0 | 275 | 55 | 0 | 0 | 0 | 0 | 0 | 0 | 0 | 291 | 181 | 20 | 0 | 0 | 0 | 0 | 0 |
| SRR6020510 | 0 | 133 | 451 | 0 | 64 | 408 | 806 | 1170 | 536 | 165 | 30101 | 65406 | 3602 | 2017 | 0 | 11496 | 38026 | 0 | 0 | 0 | 436 | 69 | 10 | 0 | 0 | 0 | 0 | 0 | 0 | 375 | 172 | 29 | 0 | 0 | 0 | 0 | 0 |
| SRR6020511 | 0 | 204 | 254 | 28 | 97 | 383 | 909 | 1049 | 556 | 147 | 19729 | 33 | 1072 | 1276 | 0 | 16519 | 27482 | 0 | 0 | 0 | 587 | 49 | 0 | 0 | 0 | 0 | 0 | 0 | 0 | 464 | 84 | 18 | 0 | 0 | 0 | 0 | 0 |
| SRR6020512 | 0 | 174 | 495 | 14 | 44 | 333 | 519 | 832 | 425 | 174 | 28236 | 61340 | 3978 | 1603 | 0 | 12328 | 33748 | 0 | 0 | 0 | 1250 | 162 | 18 | 0 | 0 | 0 | 0 | 0 | 20 | 795 | 376 | 34 | 0 | 0 | 0 | 0 | 0 |
| SRR6020513 | 0 | 213 | 391 | 16 | 58 | 391 | 711 | 852 | 340 | 149 | 26483 | 54290 | 2688 | 2030 | 0 | 11020 | 24016 | 0 | 0 | 0 | 91 | 18 | 0 | 0 | 0 | 0 | 0 | 0 | 0 | 91 | 45 | 0 | 0 | 0 | 0 | 0 | 0 |
| SRR6020514 | 0 | 116 | 212 | 12 | 82 | 275 | 873 | 611 | 207 | 130 | 24836 | 71956 | 1774 | 1616 | 0 | 10820 | 16813 | 0 | 0 | 0 | 338 | 28 | 0 | 0 | 0 | 0 | 0 | 0 | 0 | 253 | 77 | 10 | 0 | 0 | 0 | 0 | 0 |
| SRR6020515 | 0 | 284 | 736 | 16 | 62 | 433 | 638 | 882 | 375 | 153 | 24541 | 50253 | 4088 | 1201 | 0 | 15989 | 25229 | 0 | 0 | 0 | 195 | 30 | 0 | 0 | 0 | 0 | 0 | 0 | 0 | 161 | 66 | 0 | 0 | 0 | 0 | 0 | 0 |
| SRR6020516 | 0 | 1279 | 241 | 113 | 82 | 367 | 787 | 678 | 258 | 186 | 29340 | 49174 | 2339 | 1349 | 0 | 12326 | 36142 | 0 | 0 | 0 | 590 | 190 | 30 | 0 | 0 | 0 | 0 | 0 | 18 | 529 | 492 | 57 | 0 | 0 | 0 | 0 | 0 |
| SRR6020517 | 0 | 918 | 165 | 214 | 144 | 560 | 1116 | 1162 | 612 | 180 | 25551 | 42440 | 3137 | 1935 | 0 | 21374 | 33350 | 0 | 0 | 0 | 3371 | 294 | 38 | 0 | 0 | 0 | 0 | 0 | 79 | 2408 | 703 | 68 | 0 | 0 | 0 | 0 | 0 |
| SRR6020518 | 0 | 319 | 446 | 22 | 19 | 123 | 224 | 187 | 103 | 88 | 8456 | 48070 | 824 | 776 | 0 | 4988 | 7423 | 0 | 0 | 0 | 132 | 29 | 0 | 0 | 0 | 0 | 0 | 0 | 0 | 189 | 100 | 0 | 0 | 0 | 0 | 0 | 0 |
| SRR6020519 | 0 | 662 | 272 | 35 | 69 | 409 | 635 | 649 | 317 | 177 | 20877 | 46 | 3226 | 1209 | 0 | 15244 | 34153 | 0 | 0 | 0 | 173 | 44 | 0 | 0 | 0 | 0 | 0 | 0 | 0 | 131 | 95 | 13 | 0 | 0 | 0 | 0 | 0 |
| SRR6020520 | 0 | 780 | 508 | 31 | 69 | 343 | 560 | 837 | 425 | 187 | 38795 | 70 | 4216 | 1392 | 0 | 15141 | 36451 | 0 | 0 | 0 | 2192 | 147 | 17 | 0 | 0 | 0 | 0 | 0 | 44 | 1213 | 290 | 25 | 0 | 0 | 0 | 0 | 0 |
| SRR6020521 | 0 | 558 | 406 | 58 | 56 | 371 | 569 | 774 | 356 | 161 | 23015 | 57291 | 3408 | 1880 | 0 | 13718 | 37360 | 0 | 0 | 0 | 551 | 170 | 16 | 0 | 0 | 0 | 0 | 0 | 12 | 376 | 359 | 21 | 0 | 0 | 0 | 0 | 0 |
| SRR6020522 | 0 | 320 | 288 | 28 | 75 | 490 | 803 | 1207 | 390 | 178 | 27921 | 15 | 3816 | 1719 | 0 | 13688 | 35009 | 0 | 0 | 0 | 483 | 59 | 0 | 0 | 0 | 0 | 0 | 0 | 12 | 464 | 162 | 18 | 0 | 0 | 0 | 0 | 0 |
| SRR6020523 | 0 | 400 | 282 | 93 | 100 | 484 | 878 | 1460 | 434 | 206 | 27543 | 57052 | 4041 | 2560 | 0 | 15398 | 37457 | 0 | 0 | 0 | 279 | 66 | 11 | 0 | 0 | 0 | 0 | 0 | 12 | 259 | 184 | 29 | 0 | 0 | 0 | 0 | 0 |
| SRR6020524 | 0 | 126 | 472 | 11 | 56 | 402 | 627 | 787 | 456 | 154 | 25943 | 14 | 3767 | 1675 | 0 | 13370 | 41444 | 0 | 0 | 0 | 464 | 28 | 0 | 0 | 0 | 0 | 0 | 0 | 0 | 373 | 60 | 11 | 0 | 0 | 0 | 0 | 0 |
| SRR6020525 | 0 | 143 | 171 | 14 | 142 | 306 | 1562 | 1476 | 710 | 181 | 21262 | 25390 | 1409 | 3654 | 0 | 15115 | 29610 | 0 | 0 | 0 | 432 | 60 | 0 | 0 | 0 | 0 | 0 | 0 | 11 | 383 | 113 | 22 | 0 | 0 | 0 | 0 | 0 |
| SRR6020526 | 0 | 417 | 432 | 74 | 76 | 393 | 688 | 963 | 468 | 206 | 44404 | 14 | 3495 | 2009 | 0 | 16611 | 31227 | 0 | 0 | 0 | 801 | 161 | 23 | 0 | 0 | 0 | 0 | 0 | 18 | 579 | 369 | 29 | 0 | 0 | 0 | 0 | 0 |
| SRR6020527 | 0 | 107 | 101 | 54 | 174 | 449 | 1750 | 1061 | 335 | 191 | 14865 | 5226 | 943 | 4215 | 0 | 24006 | 22949 | 0 | 0 | 0 | 852 | 56 | 0 | 0 | 0 | 0 | 0 | 0 | 19 | 795 | 196 | 13 | 0 | 0 | 0 | 0 | 0 |
| SRR6020528 | 15 | 1257 | 72 | 476 | 227 | 471 | 1891 | 982 | 248 | 65 | 11444 | 13 | 824 | 1445 | 0 | 13799 | 13134 | 0 | 0 | 0 | 1608 | 207 | 21 | 0 | 0 | 0 | 0 | 0 | 40 | 1434 | 592 | 38 | 0 | 0 | 0 | 0 | 0 |
| SRR6020529 | 0 | 279 | 441 | 10 | 69 | 326 | 549 | 565 | 297 | 128 | 22224 | 55444 | 2368 | 1171 | 0 | 11726 | 22756 | 0 | 0 | 0 | 556 | 75 | 0 | 0 | 0 | 0 | 0 | 0 | 14 | 469 | 180 | 16 | 0 | 0 | 0 | 0 | 0 |
| SRR6020530 | 0 | 150 | 449 | 0 | 58 | 371 | 551 | 786 | 456 | 73 | 23619 | 42 | 3768 | 1568 | 0 | 13008 | 37546 | 0 | 0 | 0 | 133 | 44 | 0 | 0 | 0 | 0 | 0 | 0 | 0 | 91 | 96 | 14 | 0 | 0 | 0 | 0 | 0 |
| SRR6020531 | 0 | 137 | 339 | 14 | 85 | 434 | 902 | 935 | 386 | 173 | 31317 | 67303 | 3073 | 1940 | 0 | 12856 | 22637 | 0 | 0 | 0 | 66 | 0 | 0 | 0 | 0 | 0 | 0 | 0 | 0 | 51 | 17 | 0 | 0 | 0 | 0 | 0 | 0 |
| SRR6020532 | 0 | 152 | 211 | 25 | 91 | 359 | 795 | 913 | 443 | 163 | 16343 | 18 | 1506 | 949 | 0 | 13142 | 27855 | 0 | 0 | 0 | 534 | 58 | 0 | 0 | 0 | 0 | 0 | 0 | 0 | 467 | 136 | 13 | 0 | 0 | 0 | 0 | 0 |
| SRR6025030 | 2050 | 1369 | 20226 | 2867 | 6046 | 7490 | 68525 | 16981 | 45009 | 1540 | 10138 | 5596 | 16 | 3953 | 378 | 1595 | 23285 | 0 | 523 | 80 | 343 | 4706 | 703 | 61 | 16 | 21 | 16 | 2151 | 582 | 589 | 941 | 87 | 0 | 0 | 0 | 39 | 0 |
| SRR6025031 | 0 | 706 | 26 | 5188 | 3819 | 3432 | 77260 | 25228 | 2412 | 0 | 41584 | 41430 | 23692 | 12587 | 344 | 648 | 40969 | 0 | 0 | 0 | 0 | 0 | 0 | 0 | 0 | 0 | 0 | 35 | 27 | 17 | 0 | 0 | 0 | 0 | 0 | 0 | 0 |
| SRR6025033 | 1842 | 348 | 198 | 10213 | 4942 | 2789 | 75469 | 29914 | 3006 | 14 | 30349 | 24751 | 22572 | 11371 | 325 | 677 | 46584 | 0 | 0 | 0 | 0 | 0 | 0 | 0 | 0 | 0 | 0 | 0 | 0 | 0 | 0 | 0 | 0 | 0 | 0 | 0 | 0 |
| SRR6025035 | 577 | 2569 | 1795 | 1243 | 637 | 13616 | 14733 | 10801 | 26463 | 131 | 16763 | 17938 | 5116 | 2480 | 0 | 7295 | 92540 | 0 | 14 | 12 | 1510 | 4006 | 394 | 0 | 0 | 0 | 0 | 461 | 877 | 1678 | 6559 | 1178 | 0 | 29 | 14 | 0 | 0 |
| SRR6025036 | 105 | 2920 | 552 | 1012 | 517 | 4544 | 7038 | 7405 | 2704 | 137 | 2621 | 10576 | 895 | 574 | 0 | 14277 | 65349 | 0 | 18 | 38 | 3424 | 652 | 217 | 0 | 0 | 0 | 0 | 151 | 539 | 901 | 395 | 279 | 0 | 35 | 22 | 11 | 0 |
| SRR6025037 | 807 | 2176 | 633 | 1221 | 804 | 16606 | 17895 | 11347 | 28908 | 103 | 15401 | 16522 | 4611 | 2323 | 0 | 9130 | 102868 | 0 | 13 | 13 | 1485 | 3886 | 386 | 0 | 0 | 0 | 0 | 465 | 876 | 1873 | 6602 | 1196 | 0 | 28 | 14 | 0 | 0 |
| SRR6025038 | 722 | 0 | 2771 | 8368 | 4022 | 3431 | 70838 | 33300 | 2544 | 0 | 29004 | 30663 | 17242 | 9727 | 209 | 1220 | 44734 | 0 | 0 | 0 | 0 | 36 | 48 | 68 | 0 | 0 | 0 | 72 | 0 | 0 | 10 | 0 | 0 | 0 | 0 | 0 | 0 |
| SRR6025041 | 134 | 1342 | 23550 | 74 | 229 | 135 | 4053 | 301 | 1754 | 521 | 13396 | 27 | 5775 | 20395 | 4501 | 3038 | 56846 | 0 | 22 | 12 | 44 | 189 | 36 | 0 | 0 | 0 | 0 | 50 | 80 | 16 | 0 | 0 | 0 | 0 | 0 | 0 | 0 |
| SRR6025042 | 1797 | 13 | 2183 | 2953 | 4458 | 3836 | 78804 | 31238 | 2639 | 213 | 33384 | 22703 | 24457 | 10819 | 165 | 447 | 46035 | 0 | 0 | 0 | 0 | 0 | 0 | 0 | 0 | 0 | 0 | 0 | 0 | 0 | 0 | 0 | 0 | 0 | 0 | 0 | 0 |
| SRR6025043 | 410 | 3448 | 1573 | 1079 | 757 | 15131 | 16844 | 11280 | 29430 | 170 | 14886 | 17036 | 4898 | 2517 | 0 | 7916 | 96570 | 0 | 15 | 16 | 1771 | 4831 | 528 | 0 | 0 | 0 | 0 | 469 | 1003 | 2000 | 8171 | 1565 | 0 | 31 | 16 | 0 | 0 |
| SRR6025044 | 136 | 6099 | 1087 | 301 | 36 | 323 | 540 | 755 | 723 | 131 | 20532 | 42608 | 9773 | 3852 | 152 | 19595 | 121193 | 0 | 38 | 51 | 4444 | 1876 | 255 | 0 | 0 | 0 | 0 | 1029 | 2970 | 2566 | 2160 | 445 | 0 | 15 | 67 | 0 | 0 |
| SRR6025047 | 926 | 2447 | 45744 | 1024 | 4374 | 2658 | 51315 | 5247 | 5178 | 1887 | 2954 | 16382 | 1332 | 3042 | 362 | 3080 | 46558 | 0 | 173 | 64 | 264 | 485 | 365 | 39 | 23 | 15 | 0 | 85 | 116 | 38 | 10 | 0 | 0 | 0 | 0 | 28 | 0 |
| SRR6025048 | 3669 | 5664 | 14917 | 3152 | 5840 | 3470 | 81029 | 4881 | 12319 | 930 | 2613 | 14174 | 1344 | 4387 | 554 | 4801 | 28694 | 0 | 113 | 45 | 192 | 316 | 157 | 21 | 12 | 0 | 0 | 143 | 196 | 31 | 0 | 0 | 0 | 0 | 0 | 17 | 0 |
| SRR6025050 | 3251 | 1595 | 47812 | 5295 | 5471 | 6615 | 117892 | 4295 | 35751 | 2393 | 9472 | 3901 | 15 | 3514 | 575 | 1653 | 16330 | 0 | 195 | 15 | 99 | 2585 | 467 | 42 | 0 | 0 | 0 | 952 | 213 | 300 | 313 | 60 | 0 | 0 | 0 | 19 | 0 |
| SRR6025051 | 48 | 3765 | 870 | 533 | 489 | 3857 | 6085 | 7648 | 2864 | 251 | 2353 | 9733 | 778 | 554 | 0 | 12594 | 65337 | 0 | 13 | 33 | 3005 | 438 | 216 | 0 | 0 | 0 | 0 | 104 | 463 | 850 | 370 | 260 | 0 | 31 | 25 | 0 | 0 |

### Supp Table S8.1.Columns_B_to_AL

| | | | | | | | | | | | | | | | | | | | | | | | | | | | | | | | | | | | | | |
|---|---|---|---|---|---|---|---|---|---|---|---|---|---|---|---|---|---|---|---|---|---|---|---|---|---|---|---|---|---|---|---|---|---|---|---|---|---|
| SRR6025053 | 348 | 4205 | 570 | 314 | 36 | 337 | 555 | 698 | 652 | 69 | 20205 | 46177 | 9264 | 3813 | 97 | 18298 | 139893 | 0 | 42 | 58 | 4167 | 1644 | 230 | 0 | 0 | 0 | 0 | 910 | 2811 | 2324 | 2101 | 342 | 0 | 16 | 45 | 0 | 0 |
| SRR6025055 | 839 | 1754 | 14406 | 1313 | 377 | 149 | 3533 | 2211 | 1600 | 1616 | 14775 | 18 | 38 | 8578 | 5588 | 3818 | 54039 | 66 | 1383 | 249 | 1076 | 1621 | 335 | 24 | 0 | 0 | 181 | 2946 | 1276 | 672 | 215 | 21 | 0 | 0 | 0 | 23 | 0 |
| SRR6025056 | 576 | 1662 | 19270 | 1463 | 3308 | 2051 | 21075 | 14886 | 4015 | 1539 | 1614 | 3762 | 2147 | 912 | 300 | 2909 | 30481 | 0 | 416 | 156 | 510 | 391 | 227 | 41 | 19 | 17 | 0 | 357 | 213 | 146 | 57 | 12 | 0 | 0 | 0 | 20 | 0 |
| SRR6025058 | 431 | 6503 | 510 | 510 | 36 | 337 | 537 | 654 | 652 | 66 | 20639 | 44720 | 9053 | 3921 | 98 | 21081 | 138505 | 0 | 43 | 71 | 5187 | 2116 | 279 | 0 | 0 | 0 | 0 | 1204 | 3790 | 2894 | 2500 | 395 | 0 | 16 | 45 | 0 | 0 |
| SRR6025061 | 592 | 4808 | 1156 | 398 | 33 | 205 | 368 | 1121 | 801 | 122 | 19848 | 44767 | 8422 | 4142 | 55 | 14341 | 117805 | 0 | 0 | 18 | 1310 | 400 | 57 | 0 | 0 | 0 | 0 | 207 | 745 | 917 | 489 | 90 | 0 | 0 | 0 | 0 | 0 |
| SRR6025062 | 470 | 10752 | 883 | 608 | 38 | 306 | 431 | 935 | 838 | 67 | 19418 | 10 | 94 | 3391 | 80 | 16065 | 144932 | 0 | 16 | 40 | 2724 | 1148 | 161 | 0 | 0 | 0 | 0 | 454 | 1687 | 2098 | 1553 | 214 | 0 | 0 | 18 | 10 | 0 |
| SRR6025065 | 1710 | 3333 | 1584 | 827 | 714 | 10883 | 11227 | 16709 | 32590 | 141 | 14490 | 15775 | 3872 | 2333 | 0 | 5914 | 101003 | 0 | 0 | 0 | 650 | 1347 | 140 | 0 | 0 | 0 | 0 | 129 | 301 | 966 | 2267 | 444 | 0 | 0 | 0 | 0 | 0 |
| SRR6025066 | 815 | 247 | 524 | 448 | 152 | 35 | 931 | 2119 | 59 | 318 | 42086 | 13 | 22939 | 10649 | 3583 | 1410 | 22051 | 0 | 0 | 0 | 0 | 0 | 0 | 0 | 0 | 0 | 0 | 39 | 22 | 0 | 0 | 0 | 0 | 0 | 0 | 0 | 0 |
| SRR6025067 | 1710 | 4347 | 587 | 1209 | 1058 | 16463 | 13951 | 14527 | 28797 | 49 | 14818 | 15375 | 3920 | 2028 | 0 | 7837 | 120354 | 0 | 0 | 0 | 1096 | 2696 | 260 | 0 | 0 | 0 | 0 | 252 | 588 | 1715 | 5218 | 784 | 0 | 14 | 0 | 15 | 10 |
| SRR6025068 | 818 | 3607 | 706 | 1147 | 711 | 4894 | 6718 | 11007 | 4676 | 61 | 2502 | 9338 | 12 | 831 | 0 | 14314 | 59207 | 0 | 29 | 68 | 3573 | 789 | 285 | 0 | 0 | 0 | 0 | 282 | 1020 | 1570 | 699 | 211 | 0 | 10 | 0 | 0 | 0 |
| SRR6025069 | 1081 | 3343 | 918 | 785 | 594 | 3685 | 5288 | 12025 | 5017 | 146 | 2334 | 9350 | 701 | 918 | 0 | 10605 | 54283 | 0 | 12 | 34 | 1511 | 379 | 128 | 0 | 0 | 0 | 0 | 103 | 379 | 625 | 277 | 93 | 0 | 0 | 0 | 0 | 0 |
| SRR6025070 | 2297 | 3407 | 52498 | 1369 | 2644 | 3392 | 52514 | 3170 | 15091 | 1932 | 6445 | 10874 | 2648 | 3978 | 190 | 742 | 22348 | 0 | 49 | 13 | 75 | 1473 | 534 | 38 | 12 | 0 | 0 | 101 | 83 | 43 | 88 | 21 | 0 | 0 | 0 | 21 | 0 |
| SRR6025071 | 5805 | 5203 | 27962 | 2410 | 2834 | 3046 | 66347 | 2412 | 25806 | 1218 | 5287 | 8270 | 2065 | 4225 | 170 | 688 | 9056 | 0 | 21 | 0 | 40 | 617 | 244 | 15 | 0 | 0 | 0 | 93 | 81 | 31 | 18 | 0 | 0 | 0 | 0 | 11 | 0 |
| SRR6025072 | 722 | 4774 | 1942 | 1511 | 16 | 200 | 236 | 703 | 437 | 66 | 49945 | 62051 | 5970 | 11158 | 12 | 9397 | 97554 | 0 | 13 | 36 | 2283 | 1022 | 145 | 0 | 0 | 0 | 0 | 391 | 1414 | 886 | 1065 | 173 | 0 | 0 | 0 | 10 | 0 |
| SRR6025073 | 2178 | 5637 | 1218 | 3601 | 489 | 5467 | 4966 | 10176 | 3473 | 24 | 7127 | 15627 | 778 | 1511 | 0 | 12303 | 47238 | 0 | 47 | 93 | 6380 | 1636 | 595 | 0 | 13 | 0 | 0 | 414 | 1482 | 1187 | 781 | 308 | 0 | 16 | 0 | 14 | 0 |
| SRR6025075 | 2190 | 5556 | 2723 | 1475 | 426 | 4504 | 4236 | 11522 | 4602 | 41 | 8455 | 15580 | 674 | 1802 | 0 | 10781 | 35191 | 0 | 18 | 46 | 2928 | 736 | 247 | 0 | 0 | 0 | 0 | 165 | 571 | 738 | 353 | 129 | 0 | 0 | 0 | 0 | 0 |
| SRR6025079 | 9968 | 8756 | 2716 | 5862 | 4409 | 5345 | 71645 | 1908 | 19230 | 354 | 5989 | 5662 | 2301 | 5737 | 631 | 1370 | 8608 | 0 | 38 | 0 | 73 | 1099 | 398 | 19 | 0 | 0 | 0 | 198 | 164 | 54 | 39 | 11 | 0 | 0 | 0 | 0 | 0 |
| SRR6025080 | 1563 | 1581 | 17914 | 4929 | 7317 | 4640 | 62492 | 17680 | 10893 | 4242 | 2042 | 3037 | 36 | 1559 | 492 | 5489 | 28660 | 0 | 1139 | 380 | 820 | 785 | 418 | 46 | 18 | 28 | 0 | 1086 | 884 | 317 | 55 | 14 | 0 | 0 | 0 | 34 | 12 |
| SRR6025082 | 808 | 3992 | 8281 | 825 | 368 | 216 | 5816 | 1412 | 1937 | 1569 | 24815 | 14 | 20441 | 15623 | 7797 | 8141 | 68777 | 16 | 512 | 117 | 370 | 361 | 143 | 0 | 0 | 0 | 71 | 1745 | 1571 | 423 | 73 | 22 | 0 | 11 | 0 | 34 | 0 |
| SRR6025083 | 1812 | 1521 | 21474 | 4749 | 8173 | 10481 | 112794 | 17957 | 53035 | 3791 | 9559 | 4550 | 9627 | 3990 | 421 | 1785 | 23681 | 0 | 291 | 59 | 282 | 2755 | 681 | 48 | 11 | 20 | 0 | 1098 | 534 | 443 | 359 | 74 | 0 | 0 | 0 | 32 | 0 |
| SRR6025084 | 558 | 2445 | 22463 | 1586 | 478 | 273 | 5832 | 1606 | 2334 | 2952 | 14327 | 13465 | 16352 | 9559 | 6904 | 6641 | 71030 | 14 | 714 | 254 | 633 | 1079 | 259 | 15 | 0 | 0 | 38 | 1375 | 1248 | 342 | 101 | 16 | 0 | 0 | 0 | 15 | 0 |
| SRR6025085 | 1024 | 3136 | 12814 | 1368 | 6665 | 8910 | 139087 | 18897 | 52720 | 3491 | 16570 | 6628 | 14495 | 6267 | 431 | 2768 | 31483 | 0 | 166 | 32 | 162 | 957 | 526 | 0 | 21 | 28 | 0 | 929 | 569 | 382 | 189 | 58 | 10 | 0 | 0 | 96 | 0 |
| SRR6025086 | 1708 | 341 | 1181 | 875 | 4342 | 1560 | 29254 | 57524 | 1075 | 568 | 8080 | 45088 | 4640 | 2410 | 298 | 3807 | 24702 | 0 | 42 | 32 | 21 | 72 | 20 | 0 | 0 | 0 | 0 | 110 | 65 | 13 | 14 | 0 | 0 | 0 | 0 | 0 | 0 |
| SRR6025087 | 3773 | 325 | 1831 | 723 | 4311 | 3230 | 48130 | 55686 | 4249 | 572 | 36382 | 55846 | 21025 | 5709 | 219 | 1112 | 22237 | 0 | 12 | 0 | 0 | 175 | 23 | 0 | 0 | 0 | 0 | 103 | 38 | 16 | 91 | 0 | 0 | 0 | 0 | 0 | 0 |
| SRR6025088 | 857 | 2718 | 6964 | 1700 | 5552 | 3947 | 67150 | 13873 | 10283 | 2690 | 2748 | 3056 | 2491 | 2268 | 507 | 7767 | 32370 | 0 | 730 | 164 | 460 | 412 | 242 | 0 | 20 | 23 | 0 | 1268 | 1225 | 350 | 110 | 22 | 0 | 0 | 0 | 92 | 11 |
| SRR6025091 | 555 | 2676 | 34717 | 687 | 200 | 92 | 2765 | 292 | 497 | 638 | 16408 | 53 | 110 | 14924 | 4696 | 2328 | 76276 | 0 | 106 | 28 | 113 | 353 | 124 | 0 | 0 | 0 | 0 | 137 | 145 | 38 | 21 | 0 | 0 | 0 | 0 | 0 | 0 |
| SRR6025092 | 4512 | 5632 | 2220 | 2512 | 679 | 16689 | 12298 | 15824 | 24015 | 31 | 47452 | 28945 | 4584 | 5790 | 0 | 6765 | 56221 | 0 | 0 | 0 | 1394 | 3665 | 418 | 0 | 0 | 0 | 0 | 278 | 632 | 1078 | 4524 | 936 | 0 | 19 | 0 | 0 | 0 |
| SRR6025093 | 2563 | 4813 | 32207 | 2730 | 2347 | 3867 | 65775 | 5621 | 22043 | 1314 | 8390 | 13629 | 3661 | 5655 | 242 | 1274 | 21725 | 0 | 46 | 17 | 63 | 561 | 341 | 0 | 15 | 0 | 0 | 157 | 128 | 47 | 65 | 23 | 0 | 0 | 0 | 65 | 0 |
| SRR6025094 | 2679 | 6609 | 2576 | 1808 | 753 | 15517 | 10846 | 16226 | 32733 | 53 | 51061 | 24916 | 3316 | 5038 | 0 | 7061 | 46774 | 0 | 0 | 0 | 817 | 2063 | 224 | 0 | 0 | 0 | 0 | 138 | 344 | 899 | 2598 | 465 | 0 | 13 | 0 | 0 | 0 |
| SRR6025095 | 4520 | 4470 | 60696 | 914 | 3454 | 3237 | 56604 | 2765 | 20316 | 963 | 5073 | 7983 | 2074 | 3568 | 150 | 532 | 11095 | 0 | 22 | 0 | 56 | 1318 | 369 | 28 | 0 | 0 | 0 | 89 | 67 | 40 | 40 | 13 | 0 | 0 | 0 | 19 | 0 |
| SRR6025096 | 874 | 4128 | 32333 | 1516 | 186 | 104 | 2806 | 307 | 577 | 928 | 16908 | 45162 | 7873 | 17081 | 5171 | 2678 | 79080 | 0 | 142 | 48 | 149 | 430 | 144 | 0 | 0 | 0 | 0 | 187 | 226 | 52 | 25 | 0 | 0 | 0 | 0 | 0 | 0 |
| SRR6025097 | 1700 | 7929 | 26719 | 1658 | 183 | 100 | 3956 | 434 | 1008 | 659 | 21204 | 33 | 8678 | 25564 | 6276 | 4629 | 56911 | 0 | 96 | 44 | 108 | 112 | 57 | 0 | 0 | 0 | 0 | 297 | 327 | 62 | 15 | 0 | 0 | 0 | 0 | 12 | 0 |
| SRR6025099 | 267 | 725 | 270 | 2055 | 3556 | 2432 | 50997 | 25138 | 1671 | 0 | 36384 | 56737 | 27197 | 9410 | 995 | 928 | 42041 | 0 | 26 | 0 | 0 | 0 | 0 | 15 | 0 | 0 | 0 | 64 | 0 | 0 | 0 | 0 | 0 | 0 | 0 | 0 | 0 |
| SRR6025101 | 4219 | 12810 | 2691 | 2301 | 428 | 248 | 6747 | 311 | 1585 | 189 | 21768 | 48261 | 9082 | 39076 | 25757 | 7439 | 50115 | 0 | 96 | 26 | 121 | 268 | 58 | 0 | 0 | 0 | 0 | 310 | 359 | 49 | 0 | 0 | 0 | 0 | 0 | 0 | 0 |
| SRR6025103 | 3016 | 5956 | 27467 | 2475 | 5695 | 2839 | 81155 | 5165 | 12195 | 1392 | 3025 | 14919 | 1260 | 3997 | 445 | 3842 | 26301 | 0 | 114 | 40 | 197 | 285 | 149 | 17 | 13 | 0 | 0 | 132 | 186 | 33 | 0 | 0 | 0 | 0 | 0 | 17 | 0 |
| SRR6025104 | 183 | 1595 | 25563 | 212 | 3186 | 1567 | 55112 | 3248 | 16122 | 1603 | 4047 | 16742 | 978 | 3111 | 253 | 3177 | 15203 | 0 | 11 | 0 | 34 | 58 | 38 | 0 | 0 | 0 | 0 | 21 | 34 | 0 | 0 | 0 | 0 | 0 | 0 | 0 | 0 |
| SRR6025105 | 777 | 4815 | 1744 | 869 | 11 | 175 | 225 | 996 | 632 | 85 | 37373 | 52436 | 4711 | 11327 | 11 | 9897 | 61282 | 0 | 0 | 11 | 702 | 261 | 38 | 0 | 0 | 0 | 0 | 97 | 358 | 402 | 262 | 43 | 0 | 0 | 0 | 0 | 0 |
| SRR6025106 | 1549 | 4962 | 21417 | 1127 | 234 | 106 | 4406 | 307 | 1595 | 649 | 16676 | 45422 | 6609 | 25108 | 5842 | 3494 | 48847 | 0 | 48 | 21 | 78 | 178 | 42 | 0 | 0 | 0 | 0 | 147 | 208 | 26 | 0 | 0 | 0 | 0 | 0 | 0 | 0 |
| SRR6025108 | 2544 | 334 | 1831 | 6281 | 4660 | 4152 | 75173 | 31745 | 3761 | 0 | 32988 | 23417 | 21255 | 10759 | 363 | 301 | 45234 | 0 | 0 | 0 | 0 | 0 | 0 | 0 | 0 | 0 | 0 | 23 | 26 | 0 | 21 | 0 | 0 | 0 | 0 | 0 | 0 |
| SRR6025111 | 731 | 1760 | 19809 | 1449 | 296 | 177 | 2658 | 1488 | 1601 | 1188 | 17185 | 17852 | 19371 | 10470 | 6167 | 4273 | 61226 | 43 | 1322 | 272 | 771 | 1709 | 238 | 11 | 0 | 0 | 125 | 2880 | 1295 | 422 | 211 | 12 | 0 | 0 | 0 | 16 | 0 |
| SRR6025113 | 877 | 2025 | 21038 | 1736 | 4558 | 2797 | 47285 | 7587 | 4947 | 1100 | 1982 | 3229 | 2034 | 812 | 662 | 3893 | 28984 | 0 | 342 | 83 | 302 | 347 | 144 | 26 | 13 | 0 | 0 | 289 | 143 | 108 | 24 | 0 | 0 | 0 | 0 | 12 | 0 |
| SRR6025114 | 1039 | 2510 | 18130 | 1701 | 6261 | 2708 | 63707 | 9474 | 4065 | 1543 | 1787 | 2949 | 1865 | 725 | 532 | 3590 | 27301 | 0 | 316 | 63 | 406 | 268 | 159 | 30 | 10 | 0 | 0 | 271 | 149 | 156 | 21 | 13 | 0 | 0 | 0 | 15 | 0 |
| SRR6025116 | 4968 | 2679 | 63203 | 5962 | 5103 | 3955 | 158154 | 1222 | 41110 | 1906 | 10090 | 2880 | 4116 | 3630 | 878 | 1417 | 8592 | 0 | 58 | 0 | 25 | 984 | 309 | 21 | 0 | 0 | 0 | 318 | 70 | 113 | 73 | 23 | 0 | 0 | 0 | 12 | 0 |
| SRR6025119 | 5423 | 5050 | 32166 | 2112 | 2660 | 3000 | 64071 | 2408 | 25648 | 1110 | 4724 | 8063 | 1912 | 4142 | 171 | 705 | 9213 | 0 | 21 | 0 | 42 | 671 | 263 | 15 | 0 | 0 | 0 | 99 | 90 | 29 | 19 | 0 | 0 | 0 | 0 | 11 | 0 |
| SRR6025120 | 810 | 7469 | 2250 | 1214 | 21 | 200 | 249 | 755 | 576 | 71 | 59440 | 10 | 108 | 10339 | 17 | 10658 | 58659 | 0 | 0 | 19 | 1482 | 597 | 93 | 0 | 0 | 0 | 0 | 254 | 875 | 893 | 617 | 80 | 0 | 0 | 0 | 0 | 0 |
| SRR6025121 | 1968 | 4426 | 2027 | 933 | 360 | 3788 | 3776 | 13429 | 4999 | 67 | 4964 | 14464 | 581 | 1865 | 0 | 8943 | 33630 | 0 | 0 | 18 | 893 | 239 | 75 | 0 | 0 | 0 | 0 | 41 | 160 | 237 | 120 | 36 | 0 | 0 | 0 | 0 | 0 |
| SRR6025123 | 3023 | 3090 | 35283 | 2230 | 4201 | 2601 | 53206 | 5199 | 5553 | 792 | 2833 | 16 | 10 | 2854 | 414 | 3214 | 46264 | 0 | 322 | 95 | 328 | 567 | 411 | 39 | 24 | 13 | 0 | 167 | 172 | 52 | 18 | 0 | 0 | 0 | 0 | 22 | 0 |
| SRR6025125 | 1782 | 5294 | 17387 | 1507 | 269 | 131 | 4955 | 349 | 1666 | 601 | 18305 | 28 | 7689 | 26394 | 6399 | 3789 | 52712 | 0 | 56 | 22 | 92 | 176 | 44 | 0 | 0 | 0 | 0 | 173 | 221 | 31 | 0 | 0 | 0 | 0 | 0 | 0 | 0 |
| SRR6025126 | 3471 | 5238 | 20974 | 3229 | 6428 | 2987 | 78953 | 4922 | 11502 | 1139 | 2827 | 15004 | 1281 | 4032 | 467 | 4034 | 26605 | 0 | 101 | 31 | 163 | 239 | 124 | 15 | 10 | 0 | 0 | 117 | 166 | 27 | 0 | 0 | 0 | 0 | 0 | 12 | 0 |
| SRR6025128 | 4769 | 3378 | 49753 | 2559 | 2487 | 3264 | 54273 | 3465 | 15694 | 1238 | 6922 | 11177 | 2747 | 3848 | 188 | 684 | 21684 | 0 | 64 | 15 | 78 | 1294 | 518 | 35 | 12 | 0 | 0 | 129 | 81 | 49 | 88 | 24 | 0 | 0 | 0 | 20 | 0 |
| SRR6040044 | 188 | 3516 | 36118 | 107 | 42268 | 17570 | 82570 | 16112 | 78887 | 9334 | 9909 | 7669 | 4687 | 1920 | 32191 | 14550 | 8425 | 0 | 134 | 75 | 138 | 136 | 363 | 111 | 20 | 0 | 0 | 192 | 263 | 89 | 15 | 0 | 0 | 0 | 0 | 16 | 0 |
| SRR6040045 | 88 | 3444 | 51353 | 50 | 44059 | 16620 | 83118 | 16440 | 83243 | 12567 | 9783 | 7684 | 4565 | 1838 | 31674 | 13524 | 7970 | 0 | 89 | 55 | 127 | 129 | 370 | 113 | 19 | 0 | 0 | 125 | 184 | 76 | 14 | 0 | 0 | 0 | 0 | 15 | 0 |
| SRR6076852 | 297 | 2025 | 39805 | 39 | 1487 | 1267 | 5953 | 3376 | 36020 | 63 | 11698 | 19114 | 2275 | 25062 | 1002 | 224 | 18503 | 0 | 0 | 0 | 0 | 0 | 0 | 0 | 0 | 0 | 0 | 0 | 0 | 0 | 0 | 0 | 0 | 0 | 0 | 0 | 0 |
| SRR6076853 | 790 | 2965 | 54265 | 49 | 1183 | 1047 | 6029 | 3405 | 62805 | 83 | 11513 | 17750 | 2168 | 27171 | 1108 | 175 | 12982 | 0 | 0 | 0 | 0 | 15 | 0 | 0 | 0 | 0 | 0 | 0 | 0 | 0 | 0 | 0 | 0 | 0 | 0 | 0 | 0 |
| SRR6076854 | 3929 | 10095 | 42765 | 266 | 1105 | 853 | 2615 | 902 | 46008 | 170 | 3727 | 7238 | 711 | 3974 | 325 | 133 | 5224 | 0 | 0 | 0 | 0 | 15 | 0 | 0 | 0 | 0 | 0 | 0 | 0 | 0 | 0 | 0 | 0 | 0 | 0 | 0 | 0 |
| SRR6076855 | 192 | 1132 | 101089 | 22 | 1726 | 1749 | 7648 | 2111 | 41499 | 133 | 12293 | 19557 | 2837 | 21273 | 600 | 185 | 20839 | 0 | 0 | 0 | 0 | 12 | 0 | 0 | 0 | 0 | 0 | 0 | 0 | 0 | 0 | 0 | 0 | 0 | 0 | 0 | 0 |
| SRR6076856 | 956 | 5802 | 60692 | 39 | 1346 | 1469 | 7358 | 1605 | 86253 | 307 | 14906 | 26298 | 2837 | 19825 | 494 | 172 | 15222 | 0 | 0 | 0 | 0 | 16 | 0 | 0 | 0 | 0 | 0 | 0 | 0 | 0 | 0 | 0 | 0 | 0 | 0 | 0 | 0 |
| SRR6076858 | 79 | 384 | 82657 | 22 | 1024 | 896 | 5078 | 2834 | 22666 | 175 | 9281 | 22 | 2055 | 27077 | 912 | 255 | 12966 | 0 | 0 | 0 | 0 | 0 | 0 | 0 | 0 | 0 | 0 | 0 | 0 | 0 | 0 | 0 | 0 | 0 | 0 | 0 | 0 |
| SRR6076861 | 1967 | 11019 | 81862 | 103 | 1731 | 1936 | 10388 | 3011 | 41565 | 195 | 13601 | 20 | 3099 | 29268 | 1517 | 238 | 18486 | 0 | 0 | 0 | 0 | 91 | 0 | 0 | 0 | 0 | 0 | 0 | 0 | 0 | 0 | 0 | 0 | 0 | 0 | 0 | 0 |
| SRR6153965 | 18 | 86358 | 1014 | 798 | 289 | 6052 | 11176 | 15083 | 11408 | 526 | 2835 | 2557 | 478 | 4180 | 0 | 206 | 27136 | 0 | 0 | 0 | 560 | 59 | 28 | 0 | 0 | 0 | 0 | 17 | 136 | 187 | 63 | 27 | 0 | 0 | 0 | 0 | 0 |
| SRR6153968 | 95 | 180500 | 813 | 2096 | 125 | 1291 | 1771 | 1232 | 1876 | 241 | 618 | 1717 | 136 | 1453 | 18 | 233 | 5269 | 0 | 34 | 64 | 1585 | 294 | 81 | 0 | 0 | 0 | 0 | 136 | 773 | 435 | 555 | 80 | 0 | 0 | 12 | 0 | 0 |
| SRR6153976 | 0 | 86291 | 1201 | 176 | 37 | 554 | 1156 | 2015 | 2246 | 33 | 456 | 962 | 49 | 773 | 0 | 116 | 2270 | 0 | 12 | 13 | 446 | 111 | 13 | 0 | 0 | 0 | 0 | 97 | 288 | 258 | 134 | 19 | 0 | 0 | 0 | 0 | 0 |
| SRR6153984 | 15 | 217160 | 534 | 390 | 36 | 333 | 684 | 414 | 747 | 80 | 279 | 2366 | 0 | 234 | 0 | 45 | 1331 | 0 | 0 | 0 | 296 | 32 | 13 | 0 | 0 | 0 | 0 | 14 | 64 | 75 | 46 | 11 | 0 | 0 | 0 | 0 | 0 |
| SRR6153998 | 48 | 42378 | 603 | 3417 | 505 | 7303 | 9076 | 5873 | 10973 | 664 | 3438 | 6259 | 330 | 3277 | 0 | 1247 | 5552 | 0 | 17 | 20 | 590 | 114 | 19 | 0 | 0 | 0 | 0 | 91 | 328 | 259 | 135 | 18 | 0 | 0 | 0 | 0 | 0 |
| SRR6154002 | 0 | 284022 | 2211 | 419 | 1726 | 31322 | 39929 | 37490 | 18796 | 440 | 12825 | 17126 | 2442 | 4715 | 16 | 285 | 19485 | 0 | 0 | 20 | 1003 | 27 | 12 | 0 | 0 | 0 | 0 | 16 | 599 | 1447 | 85 | 49 | 0 | 0 | 0 | 0 | 0 |
| SRR6154003 | 424 | 16394 | 1543 | 452 | 582 | 9208 | 18132 | 38773 | 4686 | 69 | 4963 | 16499 | 2389 | 2612 | 0 | 85 | 11845 | 0 | 0 | 0 | 701 | 779 | 99 | 0 | 0 | 0 | 0 | 33 | 367 | 548 | 1275 | 280 | 0 | 0 | 0 | 0 | 0 |
| SRR6154004 | 33 | 81656 | 1078 | 2672 | 535 | 2495 | 9958 | 5968 | 5860 | 423 | 8611 | 8472 | 2193 | 3252 | 0 | 136 | 12556 | 0 | 39 | 48 | 1157 | 136 | 33 | 0 | 0 | 0 | 0 | 213 | 533 | 506 | 286 | 74 | 0 | 0 | 0 | 0 | 0 |
| SRR6154005 | 0 | 70400 | 1101 | 884 | 161 | 4154 | 8491 | 7511 | 5764 | 526 | 1357 | 2338 | 278 | 1523 | 0 | 155 | 10043 | 0 | 41 | 58 | 2387 | 177 | 69 | 0 | 0 | 0 | 0 | 180 | 844 | 985 | 394 | 92 | 0 | 0 | 0 | 0 | 0 |
| SRR6154006 | 465 | 34817 | 632 | 227 | 86 | 1579 | 3258 | 10228 | 1849 | 49 | 864 | 2982 | 485 | 2877 | 0 | 69 | 4014 | 0 | 0 | 13 | 748 | 825 | 212 | 0 | 0 | 0 | 0 | 37 | 402 | 685 | 2976 | 552 | 0 | 28 | 27 | 0 | 0 |
| SRR6203722 | 1649 | 1411 | 3310 | 649 | 43 | 202 | 340 | 1452 | 1191 | 25 | 23208 | 44810 | 6154 | 3907 | 77 | 11217 | 107275 | 0 | 0 | 13 | 1087 | 447 | 52 | 0 | 0 | 0 | 0 | 122 | 527 | 1089 | 949 | 66 | 0 | 0 | 0 | 0 | 0 |
| SRR6203757 | 1039 | 3624 | 2262 | 1247 | 27 | 232 | 602 | 4412 | 2558 | 46 | 18314 | 35398 | 4711 | 3655 | 98 | 14989 | 98799 | 0 | 0 | 16 | 1564 | 652 | 93 | 0 | 0 | 0 | 0 | 177 | 719 | 1654 | 1435 | 137 | 0 | 0 | 0 | 0 | 0 |
| SRR6203760 | 239 | 5381 | 2033 | 1435 | 0 | 0 | 95 | 1436 | 1793 | 0 | 1009 | 799 | 61 | 3932 | 0 | 2128 | 712 | 0 | 0 | 0 | 0 | 13 | 17 | 0 | 0 | 0 | 0 | 0 | 0 | 0 | 15 | 33 | 0 | 0 | 0 | 0 | 0 |
| SRR6203761 | 176 | 4234 | 1588 | 1588 | 0 | 17 | 71 | 1325 | 1589 | 0 | 1032 | 952 | 101 | 4102 | 0 | 1942 | 633 | 0 | 0 | 0 | 11 | 15 | 18 | 0 | 0 | 0 | 0 | 0 | 0 | 0 | 14 | 39 | 0 | 0 | 0 | 0 | 0 |
| SRR6203772 | 843 | 3903 | 2596 | 881 | 61 | 272 | 776 | 4517 | 1109 | 47 | 20191 | 39324 | 5136 | 4422 | 117 | 9419 | 101231 | 0 | 0 | 17 | 1158 | 519 | 61 | 0 | 0 | 0 | 0 | 152 | 650 | 1481 | 1116 | 84 | 0 | 0 | 0 | 0 | 0 |
| SRR6203807 | 974 | 1194 | 1674 | 247 | 64 | 1069 | 3529 | 26947 | 691 | 27 | 24291 | 22392 | 9733 | 13828 | 42 | 3941 | 60768 | 0 | 0 | 0 | 497 | 371 | 48 | 0 | 0 | 0 | 0 | 48 | 244 | 622 | 620 | 87 | 0 | 0 | 0 | 0 | 0 |
| SRR6224558 | 0 | 11 | 6284 | 0 | 1291 | 1180 | 32088 | 80084 | 71784 | 129 | 7421 | 8070 | 8801 | 9421 | 43 | 47 | 30679 | 0 | 0 | 0 | 0 | 0 | 0 | 0 | 0 | 0 | 0 | 0 | 0 | 0 | 0 | 0 | 0 | 0 | 0 | 0 | 0 |
| SRR6226650 | 0 | 27 | 173825 | 15 | 2961 | 1629 | 9477 | 6021 | 115504 | 279 | 14191 | 13 | 9442 | 38876 | 602 | 2965 | 45617 | 0 | 0 | 0 | 0 | 47 | 45 | 15 | 0 | 0 | 0 | 0 | 0 | 50 | 0 | 0 | 0 | 0 | 0 | 0 | 0 |
| SRR6226651 | 0 | 35 | 116734 | 18 | 9151 | 8861 | 48098 | 10732 | 142067 | 589 | 14043 | 5728 | 11928 | 44519 | 1757 | 2895 | 30539 | 0 | 0 | 0 | 30 | 179 | 172 | 41 | 11 | 0 | 0 | 0 | 0 | 208 | 16 | 0 | 0 | 0 | 0 | 13 | 0 |
| SRR6226652 | 0 | 239 | 189692 | 57 | 1115 | 613 | 3409 | 3041 | 107234 | 380 | 12206 | 13 | 12244 | 34097 | 447 | 2088 | 49035 | 0 | 0 | 0 | 14 | 53 | 46 | 23 | 0 | 0 | 0 | 0 | 0 | 138 | 0 | 0 | 0 | 0 | 0 | 0 | 0 |
| SRR6226653 | 0 | 166 | 164040 | 36 | 7899 | 7481 | 42529 | 26069 | 119204 | 604 | 18724 | 12698 | 12128 | 48279 | 406 | 1685 | 34592 | 0 | 0 | 0 | 61 | 154 | 541 | 206 | 47 | 0 | 0 | 0 | 14 | 126 | 0 | 11 | 0 | 0 | 0 | 44 | 0 |
| SRR6226654 | 0 | 318 | 49128 | 29 | 1707 | 974 | 6417 | 10275 | 59967 | 898 | 12886 | 19 | 12795 | 24713 | 313 | 2110 | 48025 | 0 | 0 | 0 | 60 | 136 | 379 | 163 | 34 | 0 | 0 | 0 | 44 | 120 | 25 | 13 | 0 | 0 | 15 | 14 | 0 |
| SRR6226655 | 0 | 462 | 71032 | 19 | 3579 | 2161 | 11948 | 10708 | 125518 | 915 | 16342 | 9468 | 14520 | 38574 | 510 | 2027 | 43257 | 0 | 0 | 0 | 117 | 352 | 523 | 206 | 37 | 0 | 0 | 0 | 28 | 112 | 15 | 10 | 0 | 0 | 11 | 18 | 0 |
| SRR6226656 | 0 | 293 | 96740 | 0 | 2156 | 1805 | 7893 | 4875 | 234198 | 338 | 6868 | 3624 | 6733 | 40019 | 753 | 2320 | 31986 | 0 | 0 | 0 | 35 | 108 | 262 | 236 | 38 | 0 | 0 | 0 | 12 | 108 | 0 | 0 | 0 | 0 | 23 | 18 | 0 |
| SRR6226657 | 0 | 1739 | 120094 | 100 | 4055 | 2601 | 11568 | 6700 | 184604 | 268 | 9013 | 7019 | 9465 | 41375 | 416 | 2054 | 33981 | 0 | 0 | 0 | 125 | 347 | 531 | 296 | 44 | 0 | 0 | 0 | 50 | 401 | 17 | 14 | 0 | 0 | 16 | 19 | 0 |
| SRR6230411 | 48 | 3545 | 31154 | 1055 | 38986 | 102679 | 251409 | 26766 | 113589 | 2119 | 20591 | 8100 | 6732 | 25423 | 18035 | 5782 | 3275 | 0 | 0 | 35 | 17 | 89 | 32 | 11 | 0 | 0 | 0 | 26 | 72 | 13 | 0 | 0 | 0 | 0 | 0 | 0 | 0 |
| SRR6230412 | 80 | 626 | 7788 | 1260 | 36529 | 80369 | 273579 | 42055 | 46734 | 2208 | 27881 | 11150 | 8869 | 33187 | 11662 | 5930 | 2405 | 0 | 43 | 46 | 30 | 117 | 43 | 13 | 0 | 0 | 0 | 88 | 241 | 24 | 0 | 0 | 0 | 0 | 0 | 0 | 0 |
| SRR6230413 | 47 | 1128 | 31260 | 1040 | 38654 | 85723 | 302996 | 48889 | 32230 | 4688 | 23677 | 12723 | 7864 | 27262 | 10956 | 5463 | 3178 | 0 | 14 | 13 | 42 | 45 | 102 | 31 | 18 | 0 | 0 | 36 | 92 | 35 | 0 | 0 | 0 | 0 | 0 | 0 | 0 |
| SRR6246305 | 313 | 5053 | 426 | 2132 | 188 | 3354 | 6066 | 10916 | 3893 | 51 | 12370 | 24168 | 2571 | 6285 | 0 | 6184 | 22867 | 0 | 0 | 231 | 747 | 12506 | 184 | 0 | 0 | 0 | 0 | 28 | 3236 | 701 | 16357 | 186 | 0 | 0 | 0 | 0 | 0 |
| SRR6246306 | 314 | 4987 | 399 | 2107 | 169 | 3548 | 6299 | 11079 | 4021 | 51 | 12491 | 24367 | 2449 | 6229 | 0 | 6130 | 22665 | 0 | 0 | 230 | 753 | 12432 | 178 | 0 | 0 | 0 | 0 | 26 | 3261 | 723 | 16202 | 186 | 0 | 0 | 0 | 0 | 0 |
| SRR6246307 | 317 | 5088 | 382 | 2106 | 168 | 3584 | 5967 | 11055 | 3842 | 50 | 12316 | 24215 | 2562 | 6183 | 0 | 6099 | 23199 | 0 | 0 | 234 | 752 | 12528 | 180 | 0 | 0 | 0 | 0 | 28 | 3190 | 723 | 16584 | 198 | 0 | 0 | 0 | 0 | 0 |
| SRR6246308 | 328 | 4909 | 422 | 1945 | 198 | 3473 | 6256 | 11323 | 3925 | 48 | 12441 | 24411 | 2672 | 6247 | 0 | 5962 | 22491 | 0 | 0 | 219 | 761 | 12519 | 169 | 0 | 0 | 0 | 0 | 26 | 3158 | 712 | 16187 | 175 | 0 | 0 | 0 | 0 | 0 |
| SRR6246309 | 91 | 5554 | 14628 | 1525 | 462 | 13829 | 14129 | 35110 | 10115 | 129 | 20976 | 14780 | 2956 | 4993 | 0 | 12521 | 31022 | 0 | 19 | 48 | 936 | 803 | 65 | 0 | 0 | 0 | 0 | 254 | 1296 | 1233 | 2180 | 207 | 0 | 0 | 0 | 0 | 0 |
| SRR6246310 | 94 | 5523 | 14858 | 1537 | 441 | 14099 | 14263 | 36163 | 10164 | 129 | 21071 | 14847 | 2867 | 4995 | 0 | 12194 | 29059 | 0 | 22 | 45 | 929 | 792 | 62 | 0 | 0 | 0 | 0 | 258 | 1295 | 1223 | 2107 | 203 | 0 | 0 | 0 | 0 | 0 |
| SRR6246311 | 95 | 5540 | 14721 | 1463 | 469 | 13737 | 14005 | 35632 | 10109 | 127 | 21078 | 14788 | 2819 | 4888 | 0 | 12449 | 31080 | 0 | 18 | 47 | 925 | 800 | 65 | 0 | 0 | 0 | 0 | 263 | 1287 | 1233 | 2163 | 199 | 0 | 0 | 0 | 0 | 0 |
| SRR6246312 | 91 | 5575 | 14926 | 1607 | 429 | 13736 | 14379 | 36224 | 10288 | 119 | 21330 | 14851 | 2850 | 5121 | 0 | 12544 | 29830 | 0 | 23 | 48 | 962 | 801 | 67 | 0 | 0 | 0 | 0 | 256 | 1295 | 1256 | 2159 | 209 | 0 | 0 | 0 | 0 | 0 |
| SRR6246313 | 52 | 6450 | 19611 | 559 | 612 | 13991 | 14326 | 21506 | 5866 | 90 | 16249 | 10333 | 1923 | 4642 | 0 | 14332 | 26959 | 0 | 17 | 70 | 823 | 1092 | 46 | 0 | 0 | 0 | 0 | 161 | 1165 | 1234 | 2253 | 118 | 0 | 0 | 0 | 0 | 0 |
| SRR6246314 | 53 | 6412 | 19219 | 615 | 630 | 13942 | 14132 | 21744 | 6022 | 107 | 16697 | 10168 | 2104 | 4790 | 0 | 14673 | 24424 | 0 | 14 | 63 | 832 | 1057 | 43 | 0 | 0 | 0 | 0 | 167 | 1154 | 1230 | 2189 | 126 | 0 | 0 | 0 | 0 | 0 |
| SRR6246315 | 55 | 6497 | 19317 | 555 | 657 | 14234 | 13778 | 21001 | 5867 | 115 | 17209 | 10383 | 2058 | 4715 | 0 | 14530 | 26910 | 0 | 18 | 78 | 813 | 1083 | 40 | 0 | 0 | 0 | 0 | 177 | 1140 | 1266 | 2338 | 131 | 0 | 0 | 0 | 0 | 0 |
| SRR6246316 | 56 | 6689 | 19104 | 543 | 593 | 14333 | 14060 | 21251 | 5834 | 112 | 16944 | 10415 | 2140 | 4803 | 0 | 14722 | 24572 | 0 | 13 | 69 | 807 | 1070 | 46 | 0 | 0 | 0 | 0 | 171 | 1147 | 1241 | 2254 | 124 | 0 | 0 | 0 | 0 | 0 |
| SRR6246317 | 278 | 20484 | 2860 | 349 | 693 | 25114 | 12572 | 26423 | 6526 | 0 | 39649 | 32257 | 3032 | 4274 | 0 | 20320 | 33469 | 0 | 0 | 12 | 256 | 287 | 26 | 0 | 0 | 0 | 0 | 66 | 221 | 139 | 274 | 40 | 0 | 21 | 0 | 0 | 0 |
| SRR6246318 | 335 | 21357 | 2653 | 393 | 672 | 25043 | 14226 | 26742 | 6303 | 0 | 38606 | 31128 | 3132 | 4060 | 0 | 19876 | 31306 | 0 | 0 | 20 | 291 | 320 | 20 | 0 | 0 | 0 | 0 | 55 | 216 | 131 | 235 | 31 | 0 | 12 | 0 | 0 | 0 |
| SRR6246319 | 280 | 20267 | 2764 | 341 | 688 | 25898 | 13588 | 26140 | 6315 | 0 | 38498 | 32086 | 2983 | 4378 | 0 | 19845 | 35127 | 0 | 0 | 12 | 256 | 332 | 30 | 0 | 0 | 0 | 0 | 49 | 251 | 118 | 278 | 35 | 0 | 18 | 0 | 0 | 0 |
| SRR6246320 | 254 | 20604 | 2760 | 418 | 756 | 24816 | 14117 | 26476 | 6317 | 0 | 39316 | 31329 | 3222 | 4411 | 0 | 20640 | 32144 | 0 | 0 | 13 | 246 | 295 | 12 | 10 | 0 | 0 | 0 | 57 | 224 | 164 | 260 | 30 | 0 | 13 | 0 | 0 | 0 |
| SRR6246321 | 2918 | 14876 | 5771 | 1941 | 179 | 5788 | 3114 | 6998 | 6400 | 0 | 39851 | 39313 | 2323 | 2898 | 0 | 4511 | 31038 | 0 | 0 | 28 | 499 | 1284 | 99 | 0 | 0 | 0 | 0 | 21 | 594 | 161 | 1312 | 68 | 0 | 0 | 0 | 0 | 0 |
| SRR6246322 | 2844 | 14845 | 5729 | 1958 | 295 | 5983 | 3286 | 7231 | 6797 | 12 | 39480 | 39286 | 2364 | 2767 | 0 | 4588 | 29397 | 0 | 0 | 17 | 489 | 1325 | 111 | 0 | 0 | 0 | 0 | 10 | 639 | 165 | 1235 | 63 | 0 | 0 | 0 | 0 | 0 |
| SRR6246323 | 2945 | 14887 | 5695 | 1852 | 258 | 5756 | 3117 | 7259 | 6714 | 12 | 39414 | 40564 | 2469 | 2896 | 0 | 4297 | 32059 | 0 | 0 | 22 | 492 | 1302 | 98 | 0 | 0 | 0 | 0 | 22 | 617 | 142 | 1340 | 86 | 0 | 0 | 0 | 0 | 0 |
| SRR6246324 | 2876 | 14822 | 5845 | 1912 | 214 | 6036 | 3183 | 7406 | 6473 | 16 | 39470 | 41291 | 2563 | 2800 | 0 | 4284 | 30412 | 0 | 0 | 21 | 509 | 1321 | 111 | 0 | 0 | 0 | 0 | 20 | 613 | 126 | 1282 | 65 | 0 | 0 | 0 | 0 | 0 |
| SRR6246325 | 138 | 9297 | 18032 | 1003 | 702 | 11698 | 12027 | 27378 | 12930 | 141 | 30957 | 19444 | 2990 | 4497 | 0 | 9379 | 20621 | 0 | 13 | 39 | 544 | 2695 | 366 | 0 | 0 | 0 | 0 | 137 | 970 | 415 | 6764 | 630 | 0 | 0 | 0 | 0 | 0 |
| SRR6246326 | 97 | 9640 | 17926 | 1050 | 501 | 11549 | 11985 | 27472 | 13056 | 169 | 30214 | 19319 | 3194 | 4769 | 0 | 9688 | 19650 | 0 | 0 | 49 | 532 | 2685 | 347 | 0 | 0 | 0 | 0 | 144 | 945 | 433 | 6518 | 652 | 0 | 0 | 0 | 0 | 0 |
| SRR6246327 | 123 | 9269 | 18386 | 976 | 532 | 11657 | 12394 | 27170 | 13081 | 131 | 31004 | 19215 | 2883 | 4311 | 0 | 9227 | 20589 | 0 | 12 | 46 | 549 | 2714 | 351 | 0 | 0 | 0 | 0 | 136 | 960 | 452 | 6820 | 635 | 0 | 0 | 0 | 0 | 0 |
| SRR6246328 | 142 | 9450 | 18412 | 1025 | 536 | 12151 | 12323 | 28134 | 13039 | 148 | 31171 | 19609 | 3265 | 4333 | 0 | 9386 | 19949 | 0 | 12 | 40 | 541 | 2664 | 351 | 0 | 0 | 0 | 0 | 129 | 953 | 423 | 6423 | 598 | 0 | 0 | 0 | 0 | 0 |
| SRR6246329 | 476 | 29768 | 25342 | 442 | 592 | 14881 | 9268 | 22535 | 8141 | 100 | 47845 | 16009 | 2174 | 3908 | 0 | 10009 | 30011 | 0 | 29 | 87 | 902 | 3006 | 241 | 0 | 0 | 0 | 0 | 153 | 1298 | 337 | 4839 | 284 | 0 | 11 | 0 | 0 | 0 |
| SRR6246330 | 435 | 29546 | 25082 | 385 | 602 | 15692 | 9645 | 23590 | 8764 | 127 | 48285 | 16239 | 2090 | 4141 | 0 | 10441 | 28213 | 0 | 26 | 86 | 916 | 3020 | 241 | 0 | 0 | 0 | 0 | 161 | 1313 | 349 | 4570 | 267 | 0 | 13 | 0 | 0 | 0 |
| SRR6246331 | 528 | 30239 | 25490 | 390 | 511 | 15397 | 8957 | 23692 | 8328 | 100 | 48514 | 15899 | 2350 | 4275 | 0 | 10030 | 29253 | 0 | 21 | 82 | 933 | 2992 | 257 | 0 | 0 | 0 | 0 | 157 | 1289 | 375 | 4836 | 284 | 0 | 15 | 0 | 0 | 0 |
| SRR6246332 | 453 | 29521 | 25562 | 400 | 544 | 15189 | 9536 | 23342 | 8987 | 102 | 49209 | 16463 | 2133 | 4099 | 0 | 10683 | 27852 | 0 | 24 | 85 | 920 | 3009 | 260 | 0 | 0 | 0 | 0 | 163 | 1310 | 362 | 4538 | 264 | 0 | 11 | 0 | 0 | 0 |
| SRR6246333 | 646 | 11470 | 6853 | 337 | 604 | 23376 | 10605 | 23494 | 8632 | 0 | 39505 | 30815 | 2092 | 3093 | 0 | 15186 | 29842 | 0 | 0 | 12 | 247 | 664 | 239 | 0 | 0 | 0 | 0 | 49 | 256 | 74 | 1310 | 301 | 0 | 29 | 0 | 0 | 0 |
| SRR6246334 | 577 | 11796 | 7118 | 319 | 646 | 24761 | 10612 | 24238 | 8982 | 20 | 39418 | 30052 | 2507 | 2891 | 0 | 15287 | 27929 | 0 | 0 | 16 | 251 | 704 | 229 | 0 | 0 | 0 | 0 | 62 | 261 | 84 | 1255 | 285 | 0 | 22 | 0 | 0 | 0 |
| SRR6246335 | 465 | 11197 | 6984 | 199 | 753 | 24108 | 10533 | 23036 | 8505 | 0 | 38872 | 27991 | 2434 | 3318 | 0 | 14329 | 30976 | 0 | 0 | 23 | 187 | 671 | 250 | 0 | 0 | 0 | 0 | 76 | 311 | 74 | 1370 | 355 | 0 | 34 | 0 | 0 | 0 |
| SRR6246336 | 432 | 11447 | 7023 | 196 | 664 | 23701 | 10651 | 23701 | 8343 | 0 | 39330 | 31546 | 2776 | 3470 | 15 | 14438 | 28076 | 0 | 12 | 0 | 225 | 690 | 253 | 0 | 12 | 0 | 0 | 49 | 273 | 73 | 1277 | 304 | 0 | 28 | 0 | 0 | 0 |
| SRR6246337 | 1430 | 17767 | 1183 | 497 | 1259 | 15141 | 25315 | 34224 | 11039 | 13 | 17298 | 38528 | 5082 | 4377 | 0 | 7211 | 34971 | 0 | 0 | 0 | 230 | 680 | 71 | 0 | 0 | 0 | 0 | 0 | 116 | 40 | 441 | 34 | 0 | 0 | 0 | 0 | 0 |
| SRR6246338 | 1446 | 17729 | 1217 | 477 | 1477 | 15738 | 23741 | 34445 | 11015 | 11 | 16847 | 38667 | 4786 | 4752 | 0 | 7028 | 34522 | 0 | 0 | 0 | 257 | 708 | 77 | 0 | 0 | 0 | 0 | 0 | 113 | 49 | 403 | 36 | 0 | 0 | 0 | 0 | 0 |
| SRR6246339 | 1457 | 17850 | 1108 | 475 | 1152 | 15833 | 24749 | 32850 | 11261 | 11 | 17588 | 39421 | 5180 | 4780 | 0 | 7021 | 35505 | 0 | 0 | 0 | 237 | 668 | 75 | 0 | 0 | 0 | 0 | 0 | 101 | 45 | 410 | 33 | 0 | 0 | 0 | 0 | 0 |
| SRR6246340 | 1442 | 17622 | 1150 | 478 | 1264 | 15239 | 24606 | 34833 | 11185 | 11 | 17020 | 39859 | 4956 | 4857 | 0 | 7465 | 33884 | 0 | 0 | 0 | 231 | 659 | 83 | 0 | 0 | 0 | 0 | 0 | 122 | 46 | 409 | 31 | 0 | 0 | 0 | 0 | 0 |
| SRR6246341 | 494 | 43949 | 37817 | 1174 | 1598 | 26152 | 28035 | 62961 | 12498 | 38 | 20702 | 11371 | 2197 | 13782 | 0 | 6178 | 21914 | 0 | 18 | 55 | 1458 | 2992 | 166 | 0 | 0 | 0 | 0 | 70 | 621 | 419 | 2720 | 128 | 0 | 0 | 0 | 0 | 0 |

### Supp Table S8.1.Columns_B_to_AL

| | | | | | | | | | | | | | | | | | | | | | | | | | | | | | | | | | | | | | |
|---|---|---|---|---|---|---|---|---|---|---|---|---|---|---|---|---|---|---|---|---|---|---|---|---|---|---|---|---|---|---|---|---|---|---|---|---|---|
| SRR6246342 | 538 | 43827 | 38389 | 1295 | 1930 | 25522 | 25537 | 65735 | 12555 | 43 | 20099 | 11567 | 1886 | 13881 | 0 | 6454 | 20984 | 0 | 19 | 52 | 1433 | 2937 | 148 | 0 | 0 | 0 | 0 | 83 | 622 | 420 | 2597 | 115 | 0 | 0 | 0 | 0 | 0 |
| SRR6246343 | 453 | 43967 | 37779 | 1190 | 1818 | 26328 | 28216 | 64627 | 12908 | 31 | 20491 | 12056 | 2002 | 13988 | 0 | 6972 | 23118 | 0 | 19 | 47 | 1483 | 3056 | 163 | 0 | 0 | 0 | 0 | 67 | 626 | 413 | 2724 | 126 | 0 | 0 | 0 | 0 | 0 |
| SRR6246344 | 520 | 44021 | 37922 | 1242 | 1989 | 27152 | 26673 | 64177 | 13286 | 39 | 20778 | 11354 | 1539 | 14491 | 0 | 6665 | 19950 | 0 | 17 | 55 | 1429 | 3041 | 158 | 0 | 0 | 0 | 0 | 74 | 604 | 420 | 2584 | 107 | 0 | 0 | 0 | 0 | 0 |
| SRR6246345 | 907 | 39299 | 51042 | 995 | 2261 | 33916 | 25073 | 58788 | 7336 | 22 | 32057 | 12160 | 1357 | 8492 | 0 | 8090 | 29143 | 0 | 27 | 36 | 1853 | 2078 | 198 | 0 | 0 | 0 | 0 | 31 | 522 | 292 | 1241 | 99 | 0 | 27 | 0 | 18 | 0 |
| SRR6246346 | 975 | 40886 | 48846 | 1021 | 2511 | 32170 | 24376 | 61568 | 7689 | 0 | 29398 | 12764 | 889 | 8736 | 0 | 8579 | 28247 | 0 | 0 | 70 | 1579 | 2095 | 183 | 0 | 0 | 0 | 0 | 66 | 492 | 323 | 1284 | 84 | 0 | 0 | 0 | 0 | 0 |
| SRR6246347 | 658 | 40655 | 49295 | 789 | 1947 | 34343 | 23411 | 60550 | 9385 | 44 | 29252 | 12979 | 1248 | 8486 | 0 | 8686 | 27854 | 0 | 18 | 35 | 1664 | 2076 | 182 | 0 | 0 | 0 | 0 | 40 | 484 | 293 | 1269 | 67 | 0 | 0 | 13 | 0 | 0 |
| SRR6246348 | 779 | 41540 | 50407 | 939 | 2141 | 33648 | 23197 | 61178 | 9279 | 46 | 29773 | 13051 | 1173 | 7800 | 0 | 6882 | 25389 | 0 | 46 | 55 | 1656 | 2238 | 170 | 0 | 0 | 0 | 0 | 64 | 436 | 239 | 1220 | 83 | 0 | 14 | 0 | 18 | 0 |
| SRR6246349 | 912 | 29448 | 15595 | 539 | 2208 | 30545 | 29931 | 72089 | 11490 | 41 | 19545 | 13207 | 1268 | 6379 | 0 | 6501 | 28705 | 0 | 0 | 0 | 592 | 488 | 113 | 0 | 0 | 0 | 0 | 0 | 225 | 141 | 329 | 85 | 0 | 0 | 19 | 0 | 0 |
| SRR6246350 | 390 | 27884 | 16626 | 779 | 2297 | 30293 | 28974 | 70328 | 10424 | 0 | 21401 | 11785 | 1489 | 6510 | 0 | 7531 | 28165 | 0 | 0 | 0 | 684 | 410 | 59 | 0 | 0 | 0 | 0 | 0 | 176 | 68 | 391 | 68 | 0 | 0 | 0 | 0 | 0 |
| SRR6246351 | 856 | 27590 | 16301 | 367 | 2306 | 30589 | 27311 | 74246 | 10803 | 0 | 21485 | 11896 | 1295 | 6069 | 0 | 7688 | 27797 | 0 | 0 | 18 | 671 | 561 | 55 | 0 | 0 | 0 | 0 | 37 | 212 | 37 | 405 | 55 | 0 | 0 | 18 | 0 | 0 |
| SRR6246352 | 802 | 27242 | 15247 | 591 | 2163 | 30819 | 29862 | 73698 | 10689 | 0 | 23083 | 11937 | 1165 | 6904 | 0 | 6488 | 27533 | 0 | 0 | 0 | 488 | 603 | 38 | 19 | 0 | 0 | 0 | 0 | 86 | 153 | 345 | 86 | 0 | 0 | 0 | 0 | 0 |
| SRR6246353 | 428 | 21539 | 1205 | 534 | 1315 | 7704 | 18769 | 8942 | 6809 | 117 | 5274 | 10061 | 2707 | 20553 | 0 | 3754 | 12667 | 0 | 30 | 411 | 1430 | 7281 | 98 | 0 | 0 | 0 | 0 | 91 | 2693 | 799 | 9207 | 96 | 0 | 0 | 0 | 11 | 0 |
| SRR6246354 | 417 | 21592 | 1196 | 546 | 1206 | 8002 | 18888 | 9194 | 6844 | 123 | 5112 | 10030 | 2807 | 20683 | 0 | 4023 | 11821 | 0 | 28 | 420 | 1440 | 7272 | 90 | 0 | 0 | 0 | 0 | 85 | 2701 | 784 | 9159 | 111 | 0 | 0 | 0 | 0 | 0 |
| SRR6246355 | 436 | 21471 | 1171 | 534 | 1193 | 7560 | 18818 | 9083 | 6817 | 123 | 5011 | 10583 | 2702 | 20097 | 0 | 3722 | 12700 | 0 | 32 | 422 | 1431 | 7276 | 99 | 0 | 0 | 0 | 0 | 91 | 2710 | 786 | 9244 | 109 | 0 | 0 | 0 | 12 | 0 |
| SRR6246356 | 401 | 21508 | 1250 | 552 | 1220 | 7791 | 18731 | 9286 | 6980 | 132 | 5441 | 10617 | 3164 | 20699 | 0 | 3845 | 12397 | 0 | 35 | 421 | 1466 | 7262 | 95 | 0 | 0 | 0 | 0 | 95 | 2741 | 798 | 9199 | 97 | 0 | 0 | 0 | 11 | 0 |
| SRR6246358 | 197 | 53923 | 19179 | 142 | 1769 | 13711 | 11057 | 24547 | 14153 | 66 | 7519 | 6192 | 3096 | 10173 | 0 | 5307 | 19903 | 0 | 0 | 16 | 279 | 1468 | 93 | 0 | 0 | 0 | 0 | 0 | 25 | 136 | 440 | 54 | 0 | 0 | 0 | 0 | 0 |
| SRR6246359 | 149 | 53992 | 20453 | 117 | 1902 | 13945 | 15424 | 20917 | 9930 | 42 | 6972 | 6550 | 1479 | 8874 | 0 | 2958 | 22396 | 0 | 0 | 0 | 281 | 1417 | 113 | 0 | 0 | 0 | 0 | 0 | 35 | 146 | 458 | 58 | 0 | 0 | 0 | 0 | 0 |
| SRR6246360 | 152 | 54355 | 20367 | 141 | 1530 | 8745 | 6559 | 21207 | 11150 | 33 | 6996 | 5903 | 1968 | 9620 | 0 | 3717 | 21863 | 0 | 0 | 0 | 362 | 1490 | 130 | 0 | 0 | 0 | 0 | 13 | 20 | 115 | 393 | 54 | 0 | 0 | 0 | 0 | 0 |
| SRR6246361 | 314 | 66053 | 13384 | 434 | 1761 | 35940 | 20197 | 32160 | 10823 | 48 | 16934 | 8234 | 1346 | 6836 | 0 | 7872 | 33454 | 0 | 18 | 39 | 1586 | 2606 | 328 | 0 | 11 | 0 | 0 | 61 | 431 | 613 | 1347 | 103 | 0 | 0 | 11 | 0 | 0 |
| SRR6246362 | 332 | 66590 | 13360 | 590 | 2019 | 33427 | 21683 | 34065 | 9353 | 0 | 15305 | 7706 | 1594 | 7865 | 0 | 7068 | 28538 | 0 | 0 | 59 | 1465 | 2381 | 278 | 0 | 15 | 0 | 0 | 44 | 363 | 667 | 1223 | 103 | 0 | 11 | 15 | 0 | 0 |
| SRR6246363 | 335 | 67108 | 13405 | 419 | 1549 | 32984 | 21525 | 32674 | 9291 | 72 | 13679 | 8052 | 1910 | 7020 | 0 | 8517 | 30506 | 0 | 0 | 36 | 1778 | 2654 | 268 | 0 | 0 | 0 | 0 | 57 | 426 | 673 | 1370 | 114 | 0 | 0 | 0 | 0 | 0 |
| SRR6246364 | 492 | 66405 | 14235 | 381 | 2530 | 36844 | 21453 | 34788 | 10173 | 12 | 17921 | 8223 | 1370 | 7959 | 0 | 7748 | 27883 | 0 | 0 | 58 | 1501 | 2722 | 258 | 15 | 0 | 0 | 0 | 73 | 422 | 778 | 1257 | 87 | 0 | 0 | 0 | 0 | 0 |
| SRR6246365 | 59 | 24661 | 999 | 353 | 1313 | 42000 | 25934 | 40462 | 10862 | 29 | 26748 | 17425 | 1675 | 6336 | 0 | 11179 | 32451 | 0 | 0 | 0 | 571 | 444 | 205 | 0 | 0 | 0 | 0 | 21 | 169 | 127 | 261 | 92 | 0 | 0 | 0 | 0 | 0 |
| SRR6246366 | 150 | 24684 | 1203 | 391 | 2111 | 44187 | 26154 | 42259 | 10737 | 0 | 24135 | 18262 | 1560 | 5598 | 0 | 11196 | 31568 | 0 | 0 | 22 | 457 | 435 | 189 | 0 | 0 | 0 | 0 | 36 | 145 | 167 | 290 | 102 | 0 | 15 | 0 | 0 | 0 |
| SRR6246367 | 58 | 24368 | 1485 | 378 | 2036 | 41670 | 25654 | 41490 | 10044 | 0 | 24975 | 19093 | 2036 | 6470 | 0 | 11402 | 34341 | 0 | 0 | 0 | 502 | 346 | 240 | 0 | 0 | 0 | 0 | 0 | 155 | 191 | 283 | 113 | 0 | 0 | 0 | 0 | 0 |
| SRR6246368 | 300 | 22565 | 1049 | 390 | 1552 | 42684 | 27619 | 43278 | 9222 | 0 | 25748 | 15978 | 1872 | 6117 | 0 | 11824 | 31408 | 0 | 0 | 0 | 463 | 470 | 202 | 0 | 0 | 0 | 0 | 14 | 130 | 195 | 340 | 87 | 0 | 0 | 0 | 0 | 0 |
| SRR6246369 | 0 | 121225 | 16723 | 471 | 770 | 10495 | 17082 | 11436 | 8128 | 154 | 3622 | 4278 | 3108 | 27520 | 0 | 11977 | 8327 | 0 | 0 | 146 | 235 | 8136 | 155 | 0 | 0 | 0 | 0 | 28 | 2157 | 282 | 19826 | 348 | 0 | 0 | 0 | 0 | 0 |
| SRR6246370 | 0 | 122276 | 16676 | 453 | 910 | 10214 | 17815 | 11241 | 8335 | 161 | 3874 | 4314 | 3082 | 27530 | 0 | 12151 | 8130 | 0 | 0 | 146 | 246 | 8184 | 163 | 0 | 0 | 0 | 0 | 28 | 2168 | 296 | 19531 | 339 | 0 | 0 | 0 | 10 | 0 |
| SRR6246371 | 0 | 121593 | 16835 | 428 | 899 | 9280 | 17458 | 12267 | 8584 | 190 | 4263 | 4495 | 2929 | 27347 | 0 | 10759 | 8613 | 0 | 0 | 147 | 251 | 8124 | 153 | 0 | 0 | 0 | 0 | 27 | 2191 | 284 | 19684 | 347 | 0 | 0 | 0 | 0 | 0 |
| SRR6246372 | 0 | 121970 | 16717 | 465 | 1079 | 9621 | 16123 | 10846 | 7755 | 162 | 3440 | 3965 | 3528 | 28631 | 0 | 10962 | 7639 | 0 | 0 | 133 | 254 | 8183 | 145 | 0 | 0 | 0 | 0 | 25 | 2211 | 281 | 19458 | 331 | 0 | 0 | 0 | 0 | 0 |
| SRR6246373 | 23 | 106892 | 28574 | 288 | 955 | 13641 | 12686 | 27827 | 14732 | 83 | 17187 | 10094 | 1637 | 9821 | 0 | 8184 | 18688 | 0 | 0 | 29 | 1228 | 2547 | 439 | 0 | 0 | 0 | 0 | 51 | 288 | 662 | 2145 | 425 | 0 | 0 | 0 | 0 | 0 |
| SRR6246374 | 31 | 109083 | 28611 | 273 | 983 | 17689 | 13758 | 31447 | 16285 | 70 | 16005 | 10529 | 1544 | 12916 | 0 | 6879 | 18110 | 0 | 12 | 19 | 1237 | 2518 | 421 | 0 | 0 | 0 | 0 | 48 | 311 | 679 | 2113 | 368 | 0 | 0 | 26 | 0 | 0 |
| SRR6246375 | 15 | 107518 | 28787 | 258 | 811 | 14732 | 11758 | 29599 | 15137 | 99 | 16489 | 8380 | 2838 | 10137 | 0 | 8785 | 20408 | 0 | 0 | 29 | 1168 | 2588 | 435 | 10 | 0 | 0 | 0 | 31 | 344 | 632 | 2234 | 418 | 0 | 12 | 12 | 0 | 0 |
| SRR6246376 | 31 | 107700 | 28228 | 384 | 1140 | 16814 | 12682 | 31063 | 12112 | 108 | 15247 | 8692 | 2137 | 9832 | 0 | 10687 | 19379 | 0 | 0 | 14 | 1233 | 2321 | 415 | 0 | 0 | 0 | 0 | 49 | 363 | 647 | 2172 | 385 | 0 | 0 | 14 | 10 | 0 |
| SRR6246379 | 0 | 214677 | 7403 | 322 | 0 | 33069 | 17196 | 22487 | 5291 | 0 | 21164 | 22487 | 0 | 1323 | 0 | 17196 | 25132 | 0 | 0 | 58 | 526 | 760 | 0 | 0 | 0 | 0 | 0 | 0 | 58 | 292 | 468 | 0 | 0 | 0 | 0 | 0 | 0 |
| SRR6246385 | 417 | 19281 | 2524 | 486 | 1786 | 16802 | 31555 | 19037 | 9240 | 38 | 9407 | 25214 | 3700 | 20265 | 0 | 3870 | 26066 | 0 | 29 | 357 | 1609 | 8009 | 302 | 0 | 0 | 0 | 0 | 94 | 2511 | 702 | 7790 | 255 | 0 | 0 | 0 | 0 | 0 |
| SRR6246386 | 402 | 19323 | 2457 | 500 | 1806 | 16782 | 32253 | 19666 | 9155 | 45 | 9236 | 25335 | 3792 | 20085 | 0 | 3905 | 25377 | 0 | 31 | 354 | 1616 | 7973 | 317 | 0 | 0 | 0 | 0 | 90 | 2507 | 699 | 7583 | 251 | 0 | 0 | 0 | 0 | 0 |
| SRR6246387 | 409 | 19357 | 2478 | 504 | 1778 | 16624 | 31946 | 19163 | 9299 | 42 | 9185 | 25333 | 3740 | 20235 | 10 | 4050 | 26509 | 0 | 29 | 374 | 1606 | 7988 | 292 | 0 | 0 | 0 | 0 | 100 | 2482 | 690 | 7665 | 270 | 0 | 0 | 0 | 0 | 0 |
| SRR6246388 | 426 | 19233 | 2494 | 477 | 1822 | 17153 | 32055 | 18932 | 9097 | 36 | 9457 | 25481 | 4007 | 20198 | 0 | 3936 | 25380 | 0 | 33 | 356 | 1620 | 7935 | 296 | 0 | 0 | 0 | 0 | 95 | 2497 | 686 | 7657 | 253 | 0 | 0 | 0 | 0 | 0 |
| SRR6246389 | 77 | 48138 | 49770 | 455 | 1203 | 22338 | 19485 | 54160 | 9416 | 112 | 10447 | 6529 | 2062 | 9244 | 0 | 3815 | 20551 | 0 | 0 | 13 | 695 | 2546 | 181 | 0 | 0 | 0 | 0 | 41 | 191 | 604 | 1450 | 124 | 0 | 0 | 0 | 0 | 0 |
| SRR6246390 | 62 | 46694 | 49596 | 512 | 1920 | 23004 | 19057 | 56745 | 9209 | 97 | 10986 | 7786 | 1849 | 8817 | 0 | 3804 | 17279 | 0 | 0 | 12 | 632 | 2612 | 165 | 0 | 0 | 0 | 0 | 25 | 176 | 588 | 1357 | 118 | 0 | 0 | 0 | 0 | 0 |
| SRR6246391 | 43 | 47673 | 50323 | 443 | 1830 | 24485 | 19615 | 55151 | 10533 | 85 | 10015 | 6907 | 2072 | 9911 | 0 | 4386 | 18510 | 0 | 0 | 17 | 630 | 2651 | 168 | 0 | 0 | 0 | 0 | 25 | 171 | 587 | 1479 | 127 | 0 | 0 | 0 | 0 | 0 |
| SRR6246392 | 70 | 47081 | 49934 | 436 | 1675 | 22158 | 20448 | 57366 | 9666 | 105 | 10713 | 6560 | 2163 | 9038 | 0 | 4222 | 17238 | 0 | 0 | 21 | 648 | 2582 | 179 | 11 | 0 | 0 | 0 | 27 | 207 | 558 | 1405 | 131 | 0 | 0 | 0 | 0 | 0 |
| SRR6246393 | 87 | 46628 | 11251 | 869 | 1641 | 30259 | 24061 | 38644 | 7352 | 29 | 18289 | 7231 | 2309 | 7777 | 0 | 6319 | 32568 | 0 | 0 | 0 | 368 | 1437 | 117 | 0 | 0 | 0 | 0 | 22 | 103 | 157 | 435 | 81 | 0 | 0 | 0 | 0 | 0 |
| SRR6246394 | 75 | 47277 | 10160 | 806 | 2572 | 28227 | 24276 | 42404 | 8155 | 0 | 19320 | 8092 | 1882 | 7151 | 0 | 5959 | 27914 | 0 | 0 | 0 | 556 | 1127 | 204 | 0 | 0 | 0 | 0 | 28 | 65 | 162 | 399 | 46 | 0 | 0 | 0 | 0 | 0 |
| SRR6246395 | 44 | 46809 | 11082 | 890 | 2928 | 30232 | 24437 | 41764 | 9440 | 15 | 19179 | 7349 | 2151 | 6871 | 0 | 6214 | 31846 | 0 | 0 | 23 | 541 | 1127 | 158 | 0 | 0 | 0 | 0 | 14 | 68 | 221 | 563 | 63 | 0 | 0 | 0 | 0 | 0 |
| SRR6246396 | 45 | 47861 | 10949 | 1202 | 2375 | 30191 | 22690 | 44443 | 6938 | 45 | 19315 | 7563 | 2313 | 7876 | 0 | 6438 | 32004 | 0 | 0 | 0 | 505 | 1266 | 156 | 0 | 0 | 0 | 0 | 0 | 115 | 206 | 500 | 50 | 0 | 0 | 0 | 0 | 0 |
| SRR6246397 | 363 | 9764 | 1308 | 239 | 2080 | 39885 | 25588 | 64395 | 8567 | 0 | 21938 | 12387 | 1362 | 6260 | 0 | 11366 | 39337 | 0 | 0 | 0 | 198 | 178 | 32 | 0 | 0 | 0 | 0 | 0 | 46 | 65 | 144 | 16 | 0 | 0 | 0 | 0 | 0 |
| SRR6246398 | 487 | 9618 | 1465 | 224 | 2148 | 41901 | 26843 | 68823 | 8632 | 23 | 22310 | 12377 | 1537 | 6346 | 0 | 11372 | 35397 | 0 | 0 | 0 | 209 | 226 | 29 | 0 | 0 | 0 | 0 | 18 | 46 | 93 | 131 | 20 | 0 | 0 | 0 | 0 | 0 |
| SRR6246399 | 369 | 9626 | 1332 | 191 | 1922 | 41352 | 26338 | 68266 | 8516 | 0 | 21397 | 11958 | 1596 | 6094 | 0 | 11169 | 41410 | 0 | 0 | 0 | 239 | 209 | 22 | 0 | 0 | 0 | 0 | 15 | 50 | 84 | 167 | 10 | 0 | 0 | 0 | 0 | 0 |
| SRR6246400 | 382 | 9888 | 1450 | 225 | 1992 | 42540 | 26491 | 66943 | 8083 | 0 | 20903 | 11873 | 1450 | 5782 | 0 | 11853 | 38711 | 0 | 0 | 0 | 224 | 230 | 30 | 0 | 0 | 0 | 0 | 0 | 50 | 62 | 150 | 0 | 0 | 0 | 0 | 0 | 0 |
| SRR6265480 | 10 | 67832 | 1328 | 234 | 3029 | 53629 | 32139 | 87945 | 9218 | 20 | 20716 | 16562 | 8669 | 4642 | 37 | 0 | 26190 | 0 | 0 | 0 | 728 | 171 | 52 | 0 | 0 | 0 | 0 | 0 | 136 | 263 | 280 | 30 | 0 | 25 | 27 | 0 | 0 |
| SRR6265481 | 96 | 111032 | 1365 | 416 | 3724 | 48031 | 36409 | 95540 | 5633 | 18 | 17110 | 14362 | 8261 | 4465 | 48 | 0 | 24651 | 0 | 0 | 0 | 311 | 248 | 34 | 0 | 0 | 0 | 0 | 0 | 73 | 177 | 604 | 27 | 0 | 15 | 13 | 0 | 0 |
| SRR6265482 | 0 | 19874 | 1240 | 109 | 4756 | 93263 | 51202 | 93452 | 8299 | 32 | 21335 | 15717 | 7193 | 3156 | 53 | 0 | 18970 | 0 | 0 | 0 | 171 | 88 | 26 | 0 | 0 | 0 | 0 | 0 | 23 | 93 | 131 | 14 | 0 | 15 | 15 | 0 | 0 |
| SRR6265483 | 0 | 32928 | 1241 | 190 | 3778 | 58152 | 32627 | 77975 | 12313 | 15 | 20845 | 18696 | 5687 | 2448 | 32 | 0 | 18392 | 0 | 0 | 0 | 305 | 69 | 40 | 0 | 0 | 0 | 0 | 0 | 45 | 129 | 148 | 25 | 0 | 18 | 29 | 0 | 0 |
| SRR6265484 | 27 | 49674 | 1155 | 423 | 3530 | 81139 | 39624 | 73323 | 5350 | 14 | 36179 | 32002 | 6211 | 1974 | 36 | 0 | 18173 | 0 | 0 | 0 | 427 | 103 | 65 | 0 | 10 | 0 | 0 | 0 | 45 | 140 | 151 | 36 | 0 | 28 | 15 | 0 | 0 |
| SRR6265485 | 37 | 27599 | 866 | 616 | 3607 | 66894 | 39222 | 82734 | 13235 | 18 | 26252 | 21941 | 7153 | 2773 | 15 | 0 | 21418 | 0 | 0 | 0 | 570 | 73 | 46 | 0 | 0 | 0 | 0 | 0 | 120 | 221 | 113 | 26 | 0 | 29 | 15 | 0 | 0 |
| SRR6265486 | 0 | 19520 | 711 | 102 | 2598 | 56007 | 29143 | 60403 | 13808 | 10 | 24562 | 20057 | 6176 | 2559 | 70 | 0 | 16784 | 0 | 0 | 0 | 432 | 79 | 66 | 0 | 13 | 0 | 0 | 0 | 27 | 124 | 45 | 22 | 0 | 33 | 21 | 0 | 0 |
| SRR6265487 | 13 | 186712 | 364 | 232 | 1026 | 14326 | 11825 | 65793 | 15864 | 13 | 23117 | 26967 | 9049 | 4158 | 19 | 0 | 28070 | 0 | 0 | 0 | 197 | 44 | 29 | 0 | 0 | 0 | 0 | 0 | 44 | 225 | 171 | 31 | 0 | 19 | 21 | 0 | 0 |
| SRR6265488 | 0 | 77470 | 1116 | 323 | 3150 | 50318 | 40097 | 78079 | 11772 | 26 | 23524 | 19694 | 7109 | 2651 | 47 | 0 | 23533 | 0 | 0 | 0 | 388 | 154 | 61 | 0 | 0 | 0 | 0 | 0 | 37 | 155 | 161 | 34 | 0 | 26 | 17 | 0 | 0 |
| SRR6265489 | 0 | 8609 | 0 | 23 | 2022 | 26684 | 21702 | 74605 | 9102 | 0 | 17355 | 16526 | 5602 | 2220 | 17 | 0 | 18832 | 0 | 0 | 0 | 76 | 10 | 0 | 0 | 0 | 0 | 0 | 0 | 17 | 25 | 0 | 0 | 0 | 0 | 11 | 0 | 0 |
| SRR6265490 | 61 | 74331 | 228 | 683 | 2675 | 53838 | 26941 | 64826 | 11424 | 14 | 31016 | 27316 | 7113 | 1977 | 30 | 0 | 20878 | 0 | 0 | 0 | 444 | 77 | 69 | 0 | 0 | 0 | 0 | 0 | 56 | 217 | 88 | 37 | 0 | 25 | 22 | 0 | 0 |
| SRR6265491 | 0 | 53203 | 402 | 332 | 3231 | 48142 | 31736 | 84704 | 9997 | 17 | 27695 | 24942 | 9003 | 2802 | 24 | 0 | 23488 | 0 | 0 | 0 | 208 | 38 | 36 | 0 | 0 | 0 | 0 | 0 | 20 | 123 | 72 | 22 | 0 | 20 | 24 | 0 | 0 |
| SRR6306014 | 0 | 32390 | 3908 | 620 | 3339 | 67381 | 45938 | 26020 | 490 | 30 | 51276 | 31887 | 1910 | 10055 | 0 | 27295 | 15428 | 0 | 0 | 0 | 1626 | 492 | 201 | 0 | 0 | 0 | 0 | 0 | 40 | 402 | 134 | 76 | 0 | 0 | 0 | 0 | 0 |
| SRR6306015 | 0 | 22539 | 2648 | 658 | 3695 | 69131 | 48156 | 31270 | 585 | 33 | 51989 | 31338 | 2016 | 12059 | 0 | 29651 | 13983 | 0 | 0 | 0 | 4085 | 1265 | 309 | 0 | 0 | 0 | 0 | 0 | 61 | 1131 | 319 | 129 | 0 | 0 | 0 | 0 | 0 |
| SRR6313948 | 180 | 2603 | 16929 | 325 | 81862 | 40208 | 285052 | 28508 | 11095 | 2692 | 38212 | 7421 | 21804 | 69516 | 8245 | 45234 | 5976 | 0 | 0 | 0 | 11 | 46 | 34 | 16 | 14 | 165 | 0 | 12 | 28 | 0 | 0 | 0 | 0 | 0 | 0 | 61 | 0 |
| SRR6313949 | 21 | 802 | 8958 | 18 | 68030 | 26140 | 310039 | 30470 | 5060 | 4580 | 46883 | 12689 | 28486 | 56466 | 7083 | 14031 | 8465 | 0 | 0 | 0 | 26 | 24 | 66 | 20 | 12 | 17 | 0 | 13 | 56 | 20 | 0 | 0 | 0 | 0 | 0 | 0 | 0 |
| SRR6317622 | 789 | 74322 | 2464 | 634 | 3535 | 48059 | 34014 | 30346 | 39 | 70 | 9391 | 50946 | 2203 | 7325 | 15 | 7765 | 15548 | 0 | 0 | 0 | 30 | 0 | 0 | 0 | 0 | 0 | 0 | 0 | 0 | 0 | 0 | 0 | 0 | 0 | 0 | 0 | 0 |
| SRR6317623 | 416 | 75550 | 2554 | 293 | 3331 | 44163 | 34100 | 29223 | 40 | 38 | 9184 | 52765 | 2177 | 6499 | 30 | 9848 | 13899 | 0 | 0 | 0 | 0 | 0 | 0 | 0 | 0 | 0 | 0 | 0 | 0 | 0 | 0 | 0 | 0 | 0 | 0 | 0 | 0 |
| SRR6317624 | 373 | 56496 | 2718 | 877 | 4018 | 53294 | 48467 | 35072 | 46 | 67 | 8961 | 44151 | 2064 | 9194 | 19 | 9032 | 15663 | 0 | 0 | 0 | 24 | 0 | 0 | 0 | 0 | 0 | 0 | 0 | 0 | 0 | 0 | 0 | 0 | 0 | 0 | 0 | 0 |
| SRR6317625 | 249 | 52374 | 2018 | 1052 | 4024 | 51881 | 51955 | 34354 | 37 | 43 | 8469 | 44953 | 1953 | 10861 | 11 | 8303 | 15964 | 0 | 0 | 0 | 25 | 0 | 0 | 0 | 0 | 0 | 0 | 0 | 0 | 0 | 0 | 0 | 0 | 0 | 0 | 0 | 0 |
| SRR6317626 | 151 | 239249 | 2553 | 298 | 3900 | 55903 | 49304 | 44346 | 20 | 27 | 4924 | 24190 | 1060 | 2724 | 0 | 5548 | 12185 | 0 | 0 | 0 | 21 | 0 | 0 | 0 | 0 | 0 | 0 | 0 | 0 | 0 | 0 | 0 | 0 | 0 | 0 | 0 | 0 |
| SRR6317627 | 180 | 109251 | 4352 | 2221 | 4488 | 59719 | 54401 | 55903 | 12 | 35 | 5145 | 21050 | 978 | 4015 | 0 | 6399 | 10737 | 0 | 0 | 0 | 37 | 0 | 0 | 0 | 0 | 0 | 0 | 0 | 14 | 23 | 17 | 0 | 0 | 0 | 0 | 0 | 0 |
| SRR6317628 | 326 | 86195 | 1034 | 209 | 3495 | 58561 | 46010 | 38749 | 34 | 33 | 9547 | 49735 | 2242 | 5116 | 0 | 5406 | 16720 | 0 | 0 | 0 | 10 | 0 | 0 | 0 | 0 | 0 | 0 | 0 | 0 | 0 | 0 | 0 | 0 | 0 | 0 | 0 | 0 |
| SRR6317629 | 225 | 229650 | 2569 | 278 | 3997 | 63575 | 48084 | 49217 | 23 | 50 | 7638 | 40896 | 1566 | 3603 | 0 | 6795 | 14769 | 0 | 0 | 0 | 13 | 0 | 0 | 0 | 0 | 0 | 0 | 0 | 0 | 0 | 0 | 0 | 0 | 0 | 0 | 0 | 0 |
| SRR6317630 | 806 | 97993 | 2405 | 420 | 3882 | 53724 | 38525 | 33378 | 54 | 31 | 9192 | 50204 | 2335 | 6721 | 25 | 8732 | 18782 | 0 | 0 | 0 | 27 | 0 | 0 | 0 | 0 | 0 | 0 | 0 | 0 | 0 | 0 | 0 | 0 | 0 | 0 | 0 | 0 |
| SRR6317631 | 557 | 69217 | 1948 | 1024 | 4035 | 47413 | 41798 | 34433 | 51 | 56 | 9479 | 49258 | 2253 | 8193 | 17 | 9187 | 18155 | 0 | 0 | 0 | 17 | 0 | 0 | 0 | 0 | 0 | 0 | 0 | 0 | 0 | 0 | 0 | 0 | 0 | 0 | 0 | 0 |
| SRR6317632 | 288 | 66909 | 2150 | 814 | 4096 | 53859 | 49061 | 34705 | 43 | 41 | 8733 | 44756 | 2203 | 8787 | 22 | 8880 | 16768 | 0 | 0 | 0 | 21 | 0 | 0 | 0 | 0 | 0 | 0 | 0 | 0 | 0 | 0 | 0 | 0 | 0 | 0 | 0 | 0 |
| SRR6317633 | 195 | 32826 | 2396 | 1159 | 3812 | 49499 | 51314 | 33723 | 48 | 42 | 8658 | 41965 | 2066 | 10892 | 16 | 9007 | 15810 | 0 | 0 | 0 | 27 | 0 | 0 | 0 | 0 | 0 | 0 | 0 | 0 | 0 | 0 | 0 | 0 | 0 | 0 | 0 | 0 |
| SRR6317634 | 171 | 220438 | 3487 | 311 | 4145 | 59651 | 53716 | 43736 | 23 | 44 | 5415 | 24846 | 1176 | 3058 | 0 | 6383 | 13106 | 0 | 0 | 0 | 27 | 0 | 0 | 0 | 0 | 0 | 0 | 0 | 0 | 10 | 13 | 0 | 0 | 0 | 0 | 0 | 0 |
| SRR6317635 | 258 | 235527 | 4590 | 1575 | 5098 | 68048 | 56429 | 60356 | 21 | 29 | 5460 | 25594 | 1200 | 3700 | 0 | 8092 | 12281 | 0 | 0 | 0 | 60 | 0 | 0 | 0 | 0 | 0 | 0 | 0 | 19 | 37 | 22 | 0 | 0 | 0 | 0 | 0 | 0 |
| SRR6317636 | 452 | 113391 | 1356 | 229 | 3315 | 56015 | 43060 | 37294 | 32 | 76 | 9090 | 49223 | 2064 | 5207 | 0 | 8528 | 19090 | 0 | 0 | 0 | 17 | 0 | 0 | 0 | 0 | 0 | 0 | 0 | 0 | 0 | 0 | 0 | 0 | 0 | 0 | 0 | 0 |
| SRR6317637 | 175 | 136115 | 2374 | 295 | 4244 | 67887 | 54388 | 46805 | 22 | 27 | 7357 | 37171 | 1525 | 3519 | 0 | 7926 | 14461 | 0 | 0 | 0 | 14 | 0 | 0 | 0 | 0 | 0 | 0 | 0 | 0 | 0 | 0 | 0 | 0 | 0 | 0 | 0 | 0 |
| SRR6317638 | 625 | 93738 | 2865 | 365 | 3400 | 48635 | 36080 | 30373 | 43 | 43 | 10280 | 49889 | 2300 | 6723 | 20 | 8750 | 15496 | 0 | 0 | 0 | 19 | 0 | 0 | 0 | 0 | 0 | 0 | 0 | 0 | 0 | 0 | 0 | 0 | 0 | 0 | 0 | 0 |
| SRR6317639 | 463 | 82727 | 2532 | 154 | 3709 | 48523 | 36362 | 32602 | 40 | 60 | 9974 | 51493 | 2354 | 6277 | 31 | 10522 | 16306 | 0 | 0 | 0 | 16 | 0 | 0 | 0 | 0 | 0 | 0 | 0 | 0 | 0 | 0 | 0 | 0 | 0 | 0 | 0 | 0 |
| SRR6317640 | 494 | 58558 | 2291 | 829 | 3977 | 52753 | 46911 | 32572 | 41 | 44 | 9810 | 45606 | 2279 | 8106 | 22 | 8864 | 15617 | 0 | 0 | 0 | 23 | 0 | 0 | 0 | 0 | 0 | 0 | 0 | 0 | 0 | 0 | 0 | 0 | 0 | 0 | 0 | 0 |
| SRR6317641 | 156 | 24877 | 2797 | 990 | 4379 | 48953 | 57801 | 33515 | 45 | 69 | 8508 | 39964 | 1955 | 12638 | 12 | 7667 | 15453 | 0 | 0 | 0 | 23 | 0 | 0 | 0 | 0 | 0 | 0 | 0 | 0 | 0 | 0 | 0 | 0 | 0 | 0 | 0 | 0 |
| SRR6317642 | 213 | 229871 | 4049 | 462 | 4005 | 60642 | 50804 | 45101 | 19 | 40 | 5621 | 25724 | 1114 | 2736 | 0 | 5980 | 13103 | 0 | 0 | 0 | 27 | 0 | 0 | 0 | 0 | 0 | 0 | 0 | 0 | 11 | 12 | 0 | 0 | 0 | 0 | 0 | 0 |
| SRR6317643 | 169 | 105948 | 13895 | 1027 | 4736 | 63341 | 53439 | 65685 | 20 | 53 | 5881 | 25985 | 1265 | 3649 | 0 | 5914 | 12242 | 0 | 0 | 0 | 38 | 0 | 0 | 0 | 0 | 0 | 0 | 0 | 0 | 18 | 13 | 0 | 0 | 0 | 0 | 0 | 0 |
| SRR6317644 | 500 | 140116 | 964 | 215 | 3024 | 54823 | 40346 | 34948 | 26 | 30 | 9409 | 51394 | 2153 | 4932 | 10 | 9130 | 17929 | 0 | 0 | 0 | 18 | 0 | 0 | 0 | 0 | 0 | 0 | 0 | 0 | 0 | 0 | 0 | 0 | 0 | 0 | 0 | 0 |
| SRR6317645 | 302 | 226110 | 2463 | 367 | 4189 | 65261 | 55192 | 45036 | 23 | 50 | 7505 | 37985 | 1548 | 3454 | 0 | 7807 | 14907 | 0 | 0 | 0 | 25 | 0 | 0 | 0 | 0 | 0 | 0 | 0 | 0 | 0 | 0 | 0 | 0 | 0 | 0 | 0 | 0 |
| SRR6324194 | 0 | 40671 | 8143 | 0 | 1821 | 22968 | 21235 | 14212 | 1781 | 13 | 8420 | 75934 | 1551 | 3955 | 16 | 6295 | 6755 | 0 | 0 | 0 | 58 | 95 | 27 | 0 | 0 | 0 | 0 | 0 | 0 | 32 | 51 | 0 | 0 | 0 | 0 | 0 | 0 |
| SRR6324195 | 19 | 208336 | 14323 | 37 | 2243 | 28128 | 22229 | 15246 | 5892 | 17 | 10148 | 98020 | 1616 | 4620 | 11 | 6190 | 7883 | 0 | 0 | 0 | 229 | 321 | 104 | 13 | 14 | 0 | 0 | 0 | 18 | 157 | 165 | 33 | 0 | 12 | 11 | 14 | 10 |
| SRR6363052 | 43 | 34903 | 1485 | 165 | 3939 | 36506 | 44365 | 20621 | 105 | 31 | 18864 | 65553 | 3434 | 5174 | 71 | 37216 | 4651 | 0 | 0 | 0 | 284 | 31 | 67 | 13 | 18 | 0 | 0 | 0 | 0 | 49 | 12 | 17 | 0 | 0 | 0 | 12 | 0 |
| SRR6363053 | 22 | 28427 | 1891 | 143 | 4161 | 39848 | 43916 | 21323 | 94 | 31 | 22106 | 70056 | 3730 | 5311 | 70 | 35098 | 5022 | 0 | 0 | 0 | 314 | 40 | 65 | 15 | 22 | 0 | 0 | 0 | 15 | 49 | 16 | 11 | 0 | 0 | 0 | 12 | 0 |
| SRR6363054 | 19 | 11805 | 3511 | 89 | 5365 | 56724 | 50397 | 27471 | 93 | 19 | 20045 | 50324 | 3142 | 4512 | 81 | 35173 | 4266 | 0 | 0 | 0 | 204 | 28 | 47 | 11 | 16 | 0 | 0 | 0 | 0 | 46 | 18 | 13 | 0 | 0 | 0 | 11 | 0 |
| SRR6370846 | 0 | 70 | 444 | 101 | 120 | 347 | 174 | 595 | 1157 | 15 | 4643 | 2445 | 5784 | 14739 | 730 | 0 | 945 | 0 | 0 | 0 | 0 | 0 | 0 | 0 | 0 | 0 | 0 | 0 | 0 | 0 | 0 | 0 | 0 | 0 | 0 | 0 | 0 |
| SRR6370847 | 269 | 1437 | 20767 | 34 | 197 | 205 | 573 | 1768 | 9329 | 378 | 14110 | 23 | 19368 | 49721 | 2392 | 0 | 2801 | 0 | 0 | 0 | 0 | 0 | 0 | 0 | 0 | 0 | 0 | 0 | 0 | 0 | 0 | 0 | 0 | 0 | 0 | 0 | 0 |
| SRR6370848 | 43 | 383 | 5831 | 12 | 294 | 455 | 686 | 2291 | 12451 | 128 | 9036 | 4526 | 17880 | 76135 | 3629 | 275 | 2999 | 0 | 0 | 0 | 0 | 0 | 0 | 0 | 0 | 0 | 0 | 0 | 0 | 0 | 0 | 0 | 0 | 0 | 0 | 0 | 0 |
| SRR6370849 | 138 | 2730 | 12141 | 50 | 293 | 715 | 810 | 1635 | 2096 | 228 | 3499 | 28 | 9720 | 57719 | 1525 | 70 | 1672 | 0 | 0 | 0 | 0 | 0 | 0 | 0 | 0 | 0 | 0 | 0 | 0 | 0 | 0 | 0 | 0 | 0 | 0 | 0 | 0 |
| SRR6370852 | 173 | 2393 | 9727 | 91 | 427 | 862 | 1162 | 2132 | 2301 | 473 | 5003 | 42 | 14317 | 56157 | 2664 | 104 | 2324 | 0 | 0 | 0 | 0 | 0 | 0 | 0 | 0 | 0 | 0 | 0 | 0 | 0 | 0 | 0 | 0 | 0 | 0 | 0 | 0 |
| SRR6370853 | 0 | 113 | 22832 | 0 | 252 | 472 | 828 | 1831 | 9500 | 199 | 13031 | 21 | 16048 | 57289 | 2675 | 0 | 2906 | 0 | 0 | 0 | 0 | 0 | 0 | 0 | 0 | 0 | 0 | 0 | 0 | 0 | 0 | 0 | 0 | 0 | 0 | 0 | 0 |
| SRR6370856 | 199 | 1106 | 15736 | 139 | 380 | 731 | 1250 | 1487 | 6660 | 360 | 7988 | 22 | 14921 | 66118 | 2016 | 94 | 2271 | 0 | 0 | 0 | 0 | 0 | 0 | 0 | 0 | 0 | 0 | 0 | 0 | 0 | 0 | 0 | 0 | 0 | 0 | 0 | 0 |
| SRR6372849 | 125 | 12482 | 22504 | 755 | 4154 | 2383 | 59811 | 2134 | 35698 | 2081 | 266 | 230 | 129 | 327 | 1634 | 1019 | 12611 | 0 | 29 | 19 | 79 | 59 | 94 | 63 | 32 | 15 | 0 | 42 | 57 | 41 | 32 | 0 | 0 | 0 | 0 | 26 | 0 |
| SRR6372850 | 101 | 7639 | 38608 | 109 | 2853 | 2003 | 50466 | 2004 | 22963 | 1137 | 222 | 216 | 90 | 252 | 2201 | 918 | 14182 | 0 | 0 | 12 | 122 | 60 | 120 | 34 | 38 | 17 | 0 | 0 | 13 | 18 | 0 | 0 | 0 | 0 | 0 | 41 | 0 |
| SRR6372851 | 83 | 16404 | 28051 | 156 | 5673 | 1617 | 76859 | 2413 | 29114 | 561 | 137 | 162 | 51 | 129 | 882 | 803 | 11964 | 0 | 0 | 16 | 467 | 60 | 149 | 34 | 39 | 11 | 0 | 0 | 22 | 127 | 0 | 0 | 0 | 0 | 0 | 32 | 0 |
| SRR6372852 | 519 | 17066 | 11467 | 1478 | 1372 | 904 | 15375 | 632 | 16628 | 2536 | 57 | 462 | 27 | 49 | 2879 | 780 | 17345 | 0 | 40 | 47 | 200 | 130 | 161 | 60 | 70 | 22 | 0 | 12 | 44 | 27 | 21 | 0 | 0 | 0 | 15 | 78 | 14 |
| SRR6372853 | 969 | 16991 | 8967 | 2345 | 2274 | 915 | 36211 | 2415 | 30132 | 399 | 75 | 275 | 31 | 23 | 1277 | 1396 | 20619 | 0 | 0 | 16 | 234 | 30 | 201 | 38 | 32 | 0 | 0 | 0 | 24 | 25 | 0 | 0 | 0 | 0 | 0 | 12 | 0 |
| SRR6372854 | 109 | 13425 | 55070 | 212 | 1884 | 1213 | 29868 | 998 | 33919 | 383 | 55 | 305 | 16 | 52 | 1670 | 2331 | 33906 | 0 | 0 | 0 | 83 | 26 | 107 | 12 | 15 | 0 | 0 | 0 | 10 | 11 | 0 | 0 | 0 | 0 | 0 | 17 | 0 |
| SRR6372855 | 23 | 484 | 22943 | 0 | 7185 | 5133 | 127637 | 11798 | 24019 | 344 | 405 | 593 | 152 | 71 | 369 | 1625 | 27425 | 0 | 0 | 0 | 0 | 0 | 0 | 0 | 0 | 0 | 0 | 0 | 0 | 0 | 0 | 0 | 0 | 0 | 0 | 0 | 0 |
| SRR6372856 | 315 | 6784 | 23324 | 106 | 13723 | 6172 | 186190 | 14682 | 26512 | 684 | 892 | 1624 | 267 | 452 | 414 | 2895 | 35169 | 0 | 13 | 21 | 49 | 93 | 61 | 49 | 50 | 14 | 0 | 30 | 96 | 59 | 53 | 0 | 0 | 0 | 12 | 63 | 0 |
| SRR6372857 | 56 | 3771 | 26157 | 99 | 11083 | 7365 | 138033 | 14894 | 21601 | 1421 | 729 | 2845 | 626 | 265 | 628 | 4383 | 54586 | 0 | 13 | 34 | 53 | 133 | 105 | 128 | 98 | 33 | 0 | 17 | 57 | 44 | 54 | 0 | 0 | 0 | 19 | 73 | 12 |
| SRR6389795 | 2051 | 4147 | 32 | 6663 | 3147 | 43996 | 27313 | 17895 | 5912 | 30 | 63414 | 35773 | 2815 | 8805 | 0 | 27869 | 19335 | 0 | 0 | 11 | 3766 | 491 | 144 | 0 | 19 | 0 | 0 | 78 | 530 | 1452 | 383 | 48 | 0 | 12 | 24 | 0 | 0 |
| SRR6389796 | 864 | 14514 | 2907 | 2883 | 2295 | 32650 | 17896 | 6691 | 38 | 27 | 51576 | 33468 | 2017 | 24671 | 0 | 99236 | 11914 | 0 | 0 | 84 | 48688 | 4202 | 336 | 0 | 20 | 0 | 0 | 84 | 1959 | 14246 | 2316 | 169 | 0 | 11 | 11 | 13 | 0 |
| SRR6389797 | 455 | 47532 | 295 | 2117 | 2527 | 40283 | 23989 | 10292 | 30 | 18 | 29565 | 24919 | 2065 | 57360 | 12 | 70562 | 78823 | 0 | 0 | 49 | 7829 | 1773 | 302 | 0 | 17 | 0 | 0 | 34 | 889 | 1670 | 887 | 131 | 0 | 14 | 14 | 12 | 0 |
| SRR6389798 | 241 | 164903 | 4742 | 1542 | 226 | 4792 | 3305 | 5280 | 6220 | 99 | 21884 | 18192 | 2015 | 4518 | 0 | 5921 | 13475 | 0 | 0 | 113 | 21457 | 4671 | 1409 | 12 | 18 | 0 | 0 | 14 | 1437 | 3588 | 1797 | 479 | 0 | 15 | 15 | 13 | 0 |
| SRR6389799 | 419 | 28266 | 534 | 2560 | 1635 | 13963 | 10094 | 4966 | 100 | 17 | 27888 | 21138 | 3270 | 9043 | 0 | 163283 | 17607 | 0 | 13 | 94 | 46529 | 4467 | 415 | 0 | 20 | 0 | 0 | 109 | 1942 | 10913 | 1753 | 180 | 0 | 18 | 27 | 18 | 0 |
| SRR6389800 | 238 | 55058 | 231 | 2074 | 5188 | 69305 | 47041 | 33144 | 82 | 45 | 35991 | 19094 | 1255 | 2797 | 0 | 97927 | 34956 | 0 | 0 | 73 | 8815 | 4602 | 769 | 0 | 29 | 0 | 0 | 60 | 1320 | 1776 | 2339 | 352 | 0 | 29 | 16 | 14 | 0 |
| SRR6389801 | 21 | 6781 | 2720 | 275 | 83 | 769 | 742 | 372 | 17003 | 26 | 66619 | 21268 | 2703 | 12034 | 0 | 97 | 6649 | 0 | 0 | 0 | 5322 | 1288 | 196 | 0 | 0 | 0 | 0 | 0 | 37 | 1793 | 547 | 93 | 0 | 0 | 0 | 0 | 0 |
| SRR6389802 | 12 | 19453 | 2104 | 834 | 7458 | 124050 | 78569 | 42534 | 683 | 71 | 79809 | 40294 | 1688 | 21624 | 0 | 65536 | 9939 | 0 | 0 | 0 | 546 | 628 | 74 | 0 | 0 | 0 | 0 | 14 | 114 | 212 | 238 | 38 | 0 | 0 | 0 | 0 | 0 |
| SRR6389803 | 37 | 11096 | 1131 | 1051 | 480 | 7477 | 3767 | 2627 | 12620 | 43 | 88005 | 48191 | 2988 | 15703 | 0 | 1133 | 7901 | 0 | 0 | 80 | 7786 | 1254 | 458 | 0 | 22 | 0 | 0 | 13 | 749 | 799 | 354 | 107 | 0 | 11 | 13 | 11 | 0 |
| SRR6389804 | 186 | 8117 | 3026 | 612 | 2876 | 38562 | 28847 | 22821 | 3618 | 61 | 78771 | 42874 | 2995 | 37645 | 0 | 85448 | 7957 | 0 | 0 | 0 | 605 | 428 | 194 | 0 | 18 | 0 | 0 | 32 | 172 | 187 | 358 | 104 | 0 | 24 | 0 | 11 | 0 |
| SRR6389805 | 73 | 17458 | 9311 | 3186 | 399 | 4795 | 3355 | 2810 | 5109 | 22 | 109944 | 57451 | 3592 | 14205 | 0 | 496 | 19822 | 0 | 0 | 16 | 533 | 415 | 151 | 0 | 11 | 0 | 0 | 25 | 438 | 177 | 381 | 119 | 0 | 13 | 32 | 16 | 0 |
| SRR6389806 | 164 | 7420 | 1074 | 1616 | 1311 | 9088 | 10414 | 5211 | 16864 | 27 | 36628 | 40551 | 1231 | 5452 | 0 | 28558 | 21653 | 0 | 0 | 50 | 2893 | 1359 | 208 | 0 | 0 | 0 | 0 | 52 | 740 | 445 | 431 | 74 | 0 | 0 | 0 | 0 | 0 |

### Supp Table S8.1.Columns_B_to_AL

| | | | | | | | | | | | | | | | | | | | | | | | | | | | | | | | | | | | | | |
|---|---|---|---|---|---|---|---|---|---|---|---|---|---|---|---|---|---|---|---|---|---|---|---|---|---|---|---|---|---|---|---|---|---|---|---|---|---|
| SRR6389807 | 208 | 3154 | 198 | 4207 | 7517 | 37944 | 31181 | 16625 | 9624 | 19 | 69292 | 30656 | 2988 | 8552 | 75 | 63820 | 11206 | 0 | 21 | 95 | 91281 | 18801 | 2706 | 29 | 54 | 16 | 0 | 125 | 1228 | 17829 | 11286 | 971 | 0 | 37 | 129 | 59 | 0 |
| SRR6389808 | 429 | 19478 | 198 | 1681 | 6256 | 135410 | 32658 | 9248 | 1673 | 16 | 57054 | 21707 | 1900 | 7547 | 16 | 85909 | 7350 | 0 | 18 | 72 | 22688 | 1983 | 215 | 0 | 12 | 0 | 0 | 123 | 1368 | 5423 | 1187 | 107 | 0 | 0 | 11 | 14 | 0 |
| SRR6389809 | 1643 | 12031 | 1104 | 2375 | 5148 | 40259 | 30616 | 25381 | 2972 | 28 | 65342 | 38874 | 2138 | 8383 | 14 | 23474 | 15529 | 0 | 0 | 117 | 11199 | 2912 | 513 | 0 | 28 | 0 | 0 | 25 | 2179 | 2904 | 1883 | 268 | 0 | 29 | 30 | 19 | 0 |
| SRR6389810 | 327 | 61191 | 521 | 501 | 3204 | 39795 | 30562 | 22909 | 5485 | 51 | 19798 | 22484 | 1064 | 2844 | 0 | 47853 | 13668 | 0 | 0 | 28 | 2630 | 845 | 222 | 0 | 10 | 0 | 0 | 15 | 377 | 417 | 311 | 75 | 0 | 0 | 11 | 10 | 0 |
| SRR6389811 | 260 | 42895 | 4030 | 3380 | 5616 | 51631 | 33999 | 15423 | 142 | 24 | 39632 | 37700 | 1923 | 29577 | 0 | 19263 | 11297 | 0 | 0 | 29 | 3945 | 729 | 169 | 0 | 0 | 0 | 0 | 27 | 559 | 749 | 232 | 60 | 0 | 0 | 0 | 0 | 0 |
| SRR6389812 | 20 | 10398 | 1122 | 374 | 3538 | 94868 | 39776 | 23512 | 3178 | 0 | 95707 | 26635 | 879 | 14981 | 0 | 24923 | 18626 | 0 | 0 | 42 | 2028 | 2364 | 471 | 0 | 14 | 0 | 0 | 11 | 674 | 307 | 729 | 165 | 0 | 0 | 14 | 11 | 0 |
| SRR6389813 | 137 | 33871 | 1068 | 288 | 2126 | 42876 | 25261 | 11934 | 3702 | 36 | 73689 | 40447 | 3845 | 20601 | 0 | 95063 | 16072 | 0 | 0 | 14 | 2663 | 1629 | 174 | 0 | 0 | 0 | 0 | 24 | 376 | 1160 | 534 | 102 | 0 | 0 | 12 | 0 | 0 |
| SRR6389814 | 27 | 45508 | 531 | 361 | 2563 | 20357 | 16386 | 11871 | 2244 | 0 | 131628 | 38328 | 2154 | 8015 | 0 | 22249 | 13080 | 0 | 0 | 16 | 1465 | 5949 | 690 | 0 | 19 | 0 | 0 | 0 | 120 | 126 | 1103 | 136 | 0 | 0 | 11 | 11 | 0 |
| SRR6389815 | 76 | 3437 | 225 | 584 | 494 | 6042 | 4696 | 6446 | 11943 | 21 | 49154 | 25 | 3271 | 10708 | 21 | 6540 | 35575 | 0 | 0 | 16 | 1629 | 239 | 97 | 0 | 0 | 0 | 0 | 0 | 543 | 550 | 174 | 63 | 0 | 0 | 10 | 0 | 0 |
| SRR6389816 | 17 | 2891 | 70 | 17216 | 3475 | 37824 | 23328 | 11453 | 15272 | 0 | 54800 | 20775 | 2724 | 17041 | 96 | 71681 | 21892 | 0 | 0 | 74 | 2478 | 10222 | 1012 | 17 | 27 | 0 | 0 | 29 | 1171 | 836 | 3226 | 517 | 0 | 20 | 43 | 22 | 0 |
| SRR6389817 | 71 | 12484 | 498 | 1437 | 608 | 7233 | 6434 | 9469 | 7366 | 19 | 115983 | 24634 | 2298 | 20039 | 0 | 25925 | 22365 | 0 | 0 | 0 | 773 | 754 | 184 | 0 | 0 | 0 | 0 | 0 | 159 | 198 | 552 | 88 | 0 | 0 | 12 | 0 | 0 |
| SRR6389818 | 385 | 7822 | 948 | 3269 | 1264 | 19589 | 12777 | 6462 | 1008 | 33 | 98165 | 52881 | 3470 | 13924 | 0 | 18125 | 25234 | 0 | 0 | 34 | 3754 | 10830 | 985 | 0 | 24 | 0 | 0 | 46 | 512 | 743 | 3218 | 334 | 0 | 19 | 19 | 16 | 0 |
| SRR6389819 | 51 | 9195 | 833 | 747 | 464 | 7694 | 6337 | 10480 | 179 | 34 | 33554 | 12640 | 907 | 2978 | 0 | 10667 | 50313 | 0 | 0 | 12 | 4059 | 3207 | 1011 | 17 | 74 | 0 | 0 | 11 | 234 | 814 | 2228 | 430 | 0 | 61 | 28 | 26 | 0 |
| SRR6389820 | 60 | 19227 | 390 | 583 | 2800 | 37912 | 23917 | 12758 | 7541 | 28 | 62524 | 21715 | 2245 | 7596 | 76 | 969 | 15278 | 0 | 0 | 56 | 3612 | 723 | 241 | 0 | 17 | 0 | 0 | 17 | 1222 | 908 | 408 | 124 | 0 | 15 | 13 | 11 | 0 |
| SRR6409751 | 0 | 8373 | 0 | 63 | 0 | 0 | 135 | 278 | 6146 | 0 | 5314 | 3629 | 0 | 801 | 0 | 7129 | 363 | 0 | 0 | 0 | 17 | 26 | 0 | 0 | 16 | 0 | 0 | 0 | 0 | 0 | 25 | 0 | 0 | 0 | 0 | 0 | 0 |
| SRR6409760 | 70 | 25206 | 0 | 98 | 0 | 0 | 199 | 486 | 3488 | 0 | 9444 | 4342 | 69 | 656 | 0 | 5433 | 518 | 0 | 0 | 0 | 0 | 0 | 64 | 29 | 0 | 0 | 0 | 0 | 14 | 0 | 0 | 28 | 0 | 0 | 40 | 0 | 0 |
| SRR6409798 | 0 | 17366 | 206 | 167 | 73 | 255 | 998 | 1049 | 1262 | 0 | 12954 | 34659 | 2748 | 2490 | 0 | 22018 | 108289 | 0 | 0 | 0 | 1351 | 609 | 96 | 0 | 0 | 0 | 0 | 46 | 140 | 1504 | 509 | 37 | 0 | 0 | 67 | 0 | 0 |
| SRR6409800 | 0 | 15594 | 0 | 0 | 0 | 253 | 0 | 291 | 3170 | 0 | 5164 | 2966 | 0 | 0 | 0 | 1210 | 522 | 0 | 0 | 0 | 0 | 0 | 0 | 0 | 0 | 0 | 0 | 0 | 0 | 0 | 0 | 0 | 0 | 0 | 0 | 0 | 0 |
| SRR6409827 | 0 | 18695 | 0 | 0 | 209 | 0 | 323 | 246 | 500 | 0 | 10692 | 32352 | 3584 | 2292 | 315 | 24016 | 104723 | 0 | 0 | 0 | 711 | 130 | 0 | 0 | 0 | 0 | 0 | 26 | 106 | 1169 | 235 | 0 | 0 | 0 | 0 | 0 | 0 |
| SRR6409832 | 0 | 20750 | 0 | 438 | 0 | 0 | 97 | 0 | 296 | 0 | 1176 | 617 | 0 | 653 | 0 | 358 | 223 | 0 | 0 | 0 | 0 | 44 | 0 | 0 | 0 | 0 | 0 | 0 | 0 | 0 | 0 | 0 | 0 | 0 | 0 | 0 | 0 |
| SRR6409833 | 0 | 2068 | 0 | 0 | 135 | 119 | 390 | 1203 | 1103 | 0 | 13540 | 31404 | 3648 | 954 | 0 | 28447 | 107690 | 0 | 16 | 0 | 1107 | 216 | 46 | 0 | 0 | 0 | 0 | 17 | 244 | 1273 | 270 | 35 | 0 | 0 | 35 | 0 | 0 |
| SRR6409834 | 0 | 2403 | 0 | 0 | 0 | 419 | 516 | 532 | 1182 | 0 | 12899 | 37 | 2797 | 2032 | 76 | 16735 | 100374 | 0 | 0 | 0 | 963 | 248 | 86 | 53 | 0 | 0 | 0 | 0 | 82 | 710 | 411 | 42 | 0 | 0 | 126 | 0 | 0 |
| SRR6409845 | 0 | 1387 | 0 | 0 | 0 | 0 | 62 | 109 | 151 | 0 | 162 | 152 | 43 | 439 | 0 | 157 | 127 | 0 | 0 | 0 | 0 | 0 | 0 | 0 | 0 | 0 | 0 | 0 | 0 | 0 | 0 | 0 | 0 | 0 | 35 | 0 | 0 |
| SRR6409851 | 0 | 413 | 0 | 114 | 59 | 225 | 830 | 677 | 759 | 0 | 15845 | 12 | 3757 | 2442 | 39 | 23721 | 132924 | 0 | 0 | 0 | 1852 | 681 | 150 | 0 | 0 | 0 | 0 | 49 | 290 | 2074 | 848 | 72 | 0 | 0 | 24 | 0 | 0 |
| SRR6409853 | 0 | 854 | 0 | 0 | 0 | 300 | 705 | 428 | 979 | 0 | 13547 | 40143 | 3349 | 2672 | 69 | 23111 | 115933 | 0 | 0 | 0 | 1450 | 400 | 72 | 0 | 0 | 0 | 0 | 26 | 153 | 1647 | 561 | 0 | 0 | 0 | 0 | 0 | 0 |
| SRR6409859 | 0 | 0 | 0 | 0 | 0 | 301 | 658 | 683 | 1379 | 0 | 11013 | 34340 | 3688 | 2185 | 35 | 23013 | 120294 | 0 | 0 | 0 | 1348 | 540 | 0 | 0 | 0 | 0 | 0 | 58 | 322 | 1589 | 662 | 0 | 0 | 0 | 0 | 0 | 0 |
| SRR6409860 | 0 | 1190 | 0 | 0 | 139 | 555 | 749 | 647 | 938 | 0 | 14808 | 40975 | 3378 | 2575 | 76 | 22226 | 112680 | 0 | 0 | 18 | 1220 | 298 | 0 | 0 | 0 | 0 | 0 | 54 | 192 | 1683 | 229 | 0 | 0 | 0 | 33 | 0 | 0 |
| SRR6409863 | 0 | 2931 | 0 | 0 | 95 | 133 | 591 | 421 | 1473 | 0 | 10666 | 29095 | 2362 | 1582 | 0 | 14792 | 81091 | 0 | 0 | 0 | 433 | 129 | 88 | 0 | 0 | 0 | 0 | 0 | 49 | 533 | 143 | 0 | 0 | 0 | 0 | 0 | 0 |
| SRR6409866 | 0 | 1724 | 0 | 149 | 121 | 0 | 377 | 248 | 616 | 0 | 7390 | 23305 | 2199 | 1266 | 41 | 13911 | 65244 | 0 | 0 | 0 | 1202 | 346 | 75 | 0 | 0 | 0 | 0 | 22 | 128 | 1206 | 444 | 0 | 0 | 0 | 0 | 0 | 0 |
| SRR6409867 | 0 | 6018 | 0 | 0 | 58 | 290 | 783 | 617 | 816 | 0 | 14720 | 38293 | 3809 | 2471 | 64 | 23042 | 111912 | 0 | 0 | 0 | 1667 | 583 | 0 | 0 | 0 | 0 | 0 | 98 | 259 | 2328 | 695 | 135 | 0 | 0 | 27 | 0 | 0 |
| SRR6409868 | 0 | 2868 | 0 | 0 | 73 | 0 | 410 | 158 | 838 | 0 | 10774 | 29329 | 2503 | 2045 | 0 | 19799 | 87094 | 0 | 0 | 0 | 605 | 147 | 97 | 0 | 0 | 0 | 0 | 0 | 66 | 400 | 413 | 0 | 0 | 0 | 0 | 0 | 0 |
| SRR6450851 | 892 | 4348 | 31665 | 4125 | 27133 | 15395 | 195318 | 34830 | 770 | 3791 | 51443 | 38 | 36177 | 39897 | 10263 | 35407 | 9108 | 19 | 0 | 0 | 57 | 19 | 153 | 114 | 114 | 0 | 0 | 95 | 76 | 134 | 19 | 0 | 0 | 0 | 0 | 0 | 0 |
| SRR6450852 | 317 | 3480 | 17212 | 3122 | 26730 | 12297 | 184059 | 28360 | 67 | 1464 | 48510 | 18430 | 33525 | 40814 | 9004 | 33185 | 7797 | 0 | 12 | 0 | 17 | 13 | 93 | 116 | 136 | 19 | 0 | 36 | 51 | 19 | 0 | 0 | 0 | 0 | 0 | 36 | 0 |
| SRR6450853 | 281 | 2776 | 18819 | 2809 | 25338 | 12663 | 175416 | 27697 | 193 | 2495 | 45705 | 16719 | 32861 | 39036 | 8907 | 32476 | 8257 | 0 | 14 | 0 | 21 | 10 | 83 | 93 | 97 | 14 | 14 | 42 | 97 | 14 | 0 | 0 | 10 | 0 | 0 | 24 | 0 |
| SRR6450854 | 151 | 1755 | 20408 | 1790 | 24703 | 11457 | 176140 | 27150 | 60 | 2348 | 44271 | 16071 | 31138 | 38048 | 8164 | 33237 | 7590 | 0 | 15 | 0 | 18 | 0 | 90 | 136 | 209 | 20 | 0 | 42 | 83 | 35 | 11 | 0 | 0 | 0 | 0 | 33 | 0 |
| SRR6450855 | 199 | 3445 | 32267 | 2584 | 22692 | 11282 | 167398 | 28907 | 213 | 3379 | 48661 | 17796 | 31334 | 36996 | 10388 | 31760 | 7408 | 0 | 13 | 0 | 0 | 13 | 76 | 64 | 64 | 0 | 0 | 13 | 76 | 13 | 0 | 0 | 0 | 0 | 13 | 13 | 0 |
| SRR6450856 | 660 | 4692 | 21888 | 4140 | 23882 | 11176 | 168969 | 30774 | 77 | 2964 | 40289 | 17145 | 29628 | 32459 | 8253 | 29774 | 8007 | 0 | 33 | 13 | 26 | 28 | 164 | 122 | 175 | 24 | 22 | 72 | 129 | 35 | 13 | 0 | 0 | 0 | 0 | 28 | 0 |
| SRR6450857 | 638 | 3918 | 20866 | 4647 | 28244 | 13433 | 178760 | 29803 | 120 | 2369 | 43596 | 16371 | 31063 | 35020 | 9715 | 36220 | 8995 | 0 | 17 | 0 | 35 | 0 | 104 | 70 | 139 | 17 | 0 | 87 | 157 | 52 | 17 | 0 | 0 | 0 | 0 | 17 | 0 |
| SRR6450858 | 325 | 3949 | 23169 | 4494 | 23747 | 11682 | 170384 | 23632 | 61 | 2381 | 44372 | 18 | 30765 | 37085 | 9373 | 29400 | 7547 | 0 | 14 | 11 | 0 | 0 | 99 | 80 | 108 | 0 | 0 | 39 | 69 | 28 | 0 | 0 | 0 | 0 | 0 | 18 | 0 |
| SRR6450859 | 332 | 4316 | 19088 | 4482 | 23768 | 12551 | 167163 | 31565 | 137 | 2545 | 20 | 16005 | 30745 | 31429 | 10260 | 32489 | 7660 | 0 | 40 | 50 | 20 | 0 | 150 | 100 | 100 | 30 | 0 | 100 | 130 | 60 | 30 | 0 | 0 | 0 | 0 | 60 | 0 |
| SRR6450860 | 278 | 3756 | 22166 | 3478 | 23595 | 10371 | 166155 | 32183 | 27 | 3710 | 18 | 27 | 30098 | 35091 | 7902 | 28342 | 9136 | 0 | 0 | 18 | 18 | 0 | 142 | 80 | 62 | 0 | 0 | 80 | 115 | 44 | 18 | 0 | 0 | 0 | 0 | 27 | 0 |
| SRR6450861 | 249 | 1931 | 23955 | 1371 | 23881 | 10889 | 171489 | 27438 | 138 | 2928 | 43418 | 11 | 31212 | 36204 | 7764 | 29600 | 7724 | 0 | 0 | 0 | 22 | 0 | 77 | 176 | 209 | 22 | 22 | 17 | 61 | 11 | 0 | 0 | 11 | 11 | 0 | 44 | 0 |
| SRR6450862 | 175 | 2405 | 20534 | 1632 | 24537 | 12019 | 173685 | 31444 | 72 | 2816 | 43864 | 17088 | 31734 | 36827 | 8280 | 31703 | 8578 | 0 | 12 | 0 | 20 | 0 | 111 | 332 | 647 | 52 | 0 | 29 | 58 | 21 | 0 | 0 | 0 | 0 | 12 | 86 | 0 |
| SRR6450863 | 188 | 2405 | 15712 | 2773 | 26926 | 13689 | 189743 | 28908 | 113 | 1159 | 54719 | 12 | 38438 | 48383 | 10005 | 34950 | 8189 | 0 | 12 | 0 | 12 | 14 | 93 | 95 | 100 | 18 | 0 | 18 | 35 | 18 | 0 | 0 | 0 | 0 | 0 | 39 | 0 |
| SRR6450867 | 28 | 7352 | 1714 | 617 | 30 | 20 | 316 | 110 | 326 | 0 | 8633 | 3860 | 5224 | 276 | 52347 | 1003 | 3159 | 0 | 0 | 0 | 0 | 0 | 55 | 225 | 265 | 76 | 0 | 0 | 0 | 0 | 0 | 0 | 109 | 116 | 115 | 468 | 0 |
| SRR6450868 | 0 | 289 | 4752 | 28 | 369 | 275 | 5149 | 7080 | 0 | 0 | 72479 | 72445 | 31657 | 39441 | 12088 | 3929 | 2809 | 0 | 0 | 0 | 0 | 0 | 0 | 0 | 0 | 0 | 0 | 0 | 0 | 0 | 0 | 0 | 0 | 0 | 16 | 25 | 0 |
| SRR6450869 | 0 | 289 | 4811 | 30 | 451 | 279 | 5388 | 6397 | 21 | 0 | 82818 | 72450 | 33166 | 40615 | 12215 | 4143 | 2619 | 0 | 0 | 0 | 0 | 0 | 0 | 0 | 18 | 0 | 0 | 0 | 0 | 0 | 0 | 0 | 0 | 21 | 16 | 26 | 0 |
| SRR6453341 | 588 | 2693 | 17564 | 464 | 139 | 96 | 1715 | 293 | 4586 | 318 | 5649 | 2699 | 4168 | 14100 | 457 | 46 | 4809 | 0 | 0 | 0 | 0 | 0 | 0 | 0 | 0 | 0 | 0 | 0 | 0 | 0 | 0 | 0 | 0 | 0 | 0 | 0 | 0 |
| SRR6453342 | 571 | 3647 | 13065 | 365 | 113 | 92 | 1766 | 423 | 9940 | 882 | 7405 | 3364 | 6633 | 11300 | 277 | 0 | 3744 | 0 | 0 | 0 | 0 | 0 | 0 | 0 | 0 | 0 | 0 | 0 | 0 | 0 | 0 | 0 | 0 | 0 | 0 | 0 | 0 |
| SRR6453343 | 394 | 2813 | 15492 | 385 | 124 | 95 | 1550 | 581 | 5255 | 948 | 5149 | 2980 | 6152 | 12135 | 432 | 0 | 4200 | 0 | 0 | 0 | 0 | 0 | 0 | 0 | 0 | 0 | 0 | 0 | 0 | 0 | 0 | 0 | 0 | 0 | 0 | 0 | 0 |
| SRR6453344 | 544 | 3650 | 16106 | 445 | 120 | 89 | 1534 | 429 | 4618 | 1383 | 3044 | 2098 | 4574 | 4626 | 298 | 0 | 4202 | 0 | 0 | 0 | 0 | 0 | 0 | 0 | 0 | 0 | 0 | 0 | 0 | 0 | 0 | 0 | 0 | 0 | 0 | 0 | 0 |
| SRR6453345 | 545 | 2970 | 18511 | 404 | 222 | 141 | 2261 | 692 | 8715 | 1009 | 5463 | 2861 | 5768 | 5775 | 524 | 13 | 7036 | 0 | 0 | 0 | 0 | 0 | 0 | 0 | 0 | 0 | 0 | 0 | 0 | 0 | 0 | 0 | 0 | 0 | 0 | 0 | 0 |
| SRR6453346 | 446 | 2439 | 14266 | 603 | 202 | 141 | 2125 | 704 | 14785 | 672 | 5757 | 2487 | 5773 | 10668 | 569 | 0 | 6329 | 0 | 0 | 0 | 0 | 0 | 0 | 0 | 0 | 0 | 0 | 0 | 0 | 0 | 0 | 0 | 0 | 0 | 0 | 0 | 0 |
| SRR6453347 | 431 | 3315 | 12988 | 297 | 82 | 71 | 1302 | 331 | 5383 | 1202 | 3847 | 2067 | 5128 | 3923 | 268 | 17 | 3706 | 0 | 0 | 0 | 0 | 0 | 0 | 0 | 0 | 0 | 0 | 0 | 0 | 0 | 0 | 0 | 0 | 0 | 0 | 0 | 0 |
| SRR6453348 | 543 | 2997 | 20241 | 547 | 243 | 151 | 3009 | 672 | 16662 | 815 | 6840 | 2659 | 7650 | 9256 | 372 | 0 | 4117 | 0 | 0 | 0 | 0 | 0 | 0 | 0 | 0 | 0 | 0 | 0 | 0 | 0 | 0 | 0 | 0 | 0 | 0 | 0 | 0 |
| SRR6453349 | 660 | 2995 | 18087 | 569 | 137 | 81 | 1436 | 570 | 8435 | 720 | 5534 | 2924 | 5924 | 7901 | 371 | 0 | 4085 | 0 | 0 | 0 | 0 | 0 | 0 | 0 | 0 | 0 | 0 | 0 | 0 | 0 | 0 | 0 | 0 | 0 | 0 | 0 | 0 |
| SRR6453350 | 518 | 3066 | 18905 | 399 | 103 | 66 | 1053 | 365 | 7043 | 1155 | 3133 | 1738 | 3880 | 5180 | 185 | 11 | 6955 | 0 | 0 | 0 | 0 | 0 | 0 | 0 | 0 | 0 | 0 | 0 | 0 | 0 | 0 | 0 | 0 | 0 | 0 | 0 | 0 |
| SRR6453351 | 464 | 2992 | 18614 | 505 | 169 | 125 | 2247 | 902 | 12752 | 898 | 6202 | 3187 | 7679 | 18250 | 572 | 0 | 5860 | 0 | 0 | 0 | 0 | 0 | 0 | 0 | 0 | 0 | 0 | 0 | 0 | 0 | 0 | 0 | 0 | 0 | 0 | 0 | 0 |
| SRR6453352 | 442 | 3394 | 14840 | 276 | 100 | 73 | 1265 | 409 | 6455 | 890 | 4281 | 3194 | 4587 | 9193 | 265 | 11 | 4612 | 0 | 0 | 0 | 0 | 0 | 0 | 0 | 0 | 0 | 0 | 0 | 0 | 0 | 0 | 0 | 0 | 0 | 0 | 0 | 0 |
| SRR6453353 | 398 | 2700 | 12695 | 340 | 191 | 119 | 2827 | 545 | 14117 | 707 | 6481 | 2821 | 6021 | 9738 | 605 | 0 | 3897 | 0 | 0 | 0 | 0 | 0 | 0 | 0 | 0 | 0 | 0 | 0 | 0 | 0 | 0 | 0 | 0 | 0 | 0 | 0 | 0 |
| SRR6453354 | 474 | 3340 | 12016 | 241 | 105 | 65 | 1060 | 532 | 6086 | 1009 | 4185 | 2674 | 4439 | 8688 | 305 | 0 | 4295 | 0 | 0 | 0 | 0 | 0 | 0 | 0 | 0 | 0 | 0 | 0 | 0 | 0 | 0 | 0 | 0 | 0 | 0 | 0 | 0 |
| SRR6453355 | 668 | 3798 | 12123 | 352 | 125 | 74 | 1323 | 686 | 7034 | 1364 | 3084 | 2439 | 3284 | 5330 | 305 | 43 | 3741 | 0 | 0 | 0 | 0 | 0 | 12 | 0 | 0 | 0 | 0 | 0 | 0 | 0 | 0 | 0 | 0 | 0 | 0 | 0 | 0 |
| SRR6453356 | 517 | 2878 | 15218 | 365 | 120 | 64 | 1411 | 456 | 5029 | 1066 | 2498 | 1253 | 2591 | 4624 | 222 | 12 | 4766 | 0 | 0 | 0 | 0 | 0 | 15 | 0 | 0 | 0 | 0 | 0 | 0 | 0 | 0 | 0 | 0 | 0 | 0 | 0 | 0 |
| SRR6453357 | 720 | 4743 | 14522 | 469 | 145 | 95 | 1916 | 449 | 8295 | 907 | 6729 | 3113 | 7738 | 8505 | 459 | 36 | 6631 | 0 | 0 | 0 | 0 | 0 | 0 | 0 | 0 | 0 | 0 | 0 | 0 | 0 | 0 | 0 | 0 | 0 | 0 | 0 | 0 |
| SRR6453358 | 828 | 4332 | 17132 | 365 | 108 | 58 | 1145 | 364 | 6180 | 1001 | 2913 | 1970 | 3604 | 5954 | 322 | 0 | 4290 | 0 | 0 | 0 | 0 | 0 | 0 | 0 | 0 | 0 | 0 | 0 | 0 | 0 | 0 | 0 | 0 | 0 | 0 | 0 | 0 |
| SRR6453359 | 402 | 2829 | 14658 | 221 | 143 | 94 | 1632 | 721 | 6246 | 1527 | 2982 | 1565 | 3489 | 7150 | 307 | 48 | 4478 | 0 | 0 | 0 | 0 | 0 | 12 | 0 | 0 | 0 | 0 | 0 | 0 | 0 | 0 | 0 | 0 | 0 | 0 | 0 | 0 |
| SRR6453360 | 416 | 3056 | 14631 | 452 | 103 | 80 | 1565 | 330 | 10233 | 647 | 2706 | 1985 | 2682 | 5367 | 224 | 0 | 5684 | 0 | 0 | 0 | 0 | 0 | 0 | 0 | 0 | 0 | 0 | 0 | 0 | 0 | 0 | 0 | 0 | 0 | 0 | 0 | 0 |
| SRR6453361 | 536 | 2901 | 13613 | 459 | 217 | 114 | 2263 | 769 | 9088 | 1253 | 5732 | 2889 | 5896 | 15823 | 402 | 12 | 5168 | 0 | 0 | 0 | 0 | 0 | 13 | 0 | 0 | 0 | 0 | 0 | 0 | 0 | 0 | 0 | 0 | 0 | 0 | 0 | 0 |
| SRR6453362 | 578 | 3778 | 13766 | 388 | 89 | 59 | 1212 | 452 | 6475 | 1836 | 2571 | 1680 | 3421 | 5681 | 221 | 32 | 3799 | 0 | 0 | 0 | 0 | 0 | 14 | 0 | 0 | 0 | 0 | 0 | 0 | 0 | 0 | 0 | 0 | 0 | 0 | 0 | 0 |
| SRR6453363 | 485 | 3196 | 16388 | 454 | 301 | 204 | 4248 | 818 | 23224 | 1195 | 4999 | 2518 | 5508 | 10434 | 857 | 16 | 5056 | 0 | 0 | 0 | 0 | 0 | 0 | 0 | 0 | 0 | 0 | 0 | 0 | 0 | 0 | 0 | 0 | 0 | 0 | 0 | 0 |
| SRR6453364 | 493 | 3274 | 14395 | 464 | 78 | 49 | 962 | 231 | 6369 | 673 | 3187 | 3337 | 3237 | 9074 | 285 | 0 | 5520 | 0 | 0 | 0 | 0 | 0 | 0 | 0 | 0 | 0 | 0 | 0 | 0 | 0 | 0 | 0 | 0 | 0 | 0 | 0 | 0 |
| SRR6453365 | 402 | 2413 | 13565 | 263 | 190 | 114 | 2401 | 454 | 14721 | 736 | 4872 | 2214 | 4594 | 8343 | 513 | 10 | 5669 | 0 | 0 | 0 | 0 | 0 | 11 | 0 | 0 | 0 | 0 | 0 | 0 | 0 | 0 | 0 | 0 | 0 | 0 | 0 | 0 |
| SRR6453366 | 487 | 3217 | 12443 | 304 | 91 | 60 | 1188 | 463 | 8128 | 1620 | 4209 | 2903 | 4960 | 11078 | 324 | 0 | 3940 | 0 | 0 | 0 | 0 | 0 | 0 | 0 | 0 | 0 | 0 | 0 | 0 | 0 | 0 | 0 | 0 | 0 | 0 | 0 | 0 |
| SRR6453367 | 371 | 1887 | 11561 | 998 | 78 | 53 | 745 | 276 | 5811 | 2974 | 3413 | 2647 | 3985 | 6829 | 276 | 15 | 4029 | 0 | 0 | 0 | 0 | 0 | 0 | 0 | 0 | 0 | 0 | 0 | 0 | 0 | 0 | 0 | 0 | 0 | 0 | 0 | 0 |
| SRR6453368 | 409 | 1779 | 13139 | 903 | 71 | 54 | 755 | 322 | 7425 | 2486 | 3724 | 2751 | 3527 | 7984 | 485 | 56 | 3406 | 0 | 0 | 0 | 0 | 0 | 0 | 0 | 0 | 0 | 0 | 0 | 0 | 0 | 0 | 0 | 0 | 0 | 0 | 0 | 0 |
| SRR6453369 | 516 | 2973 | 15356 | 405 | 97 | 67 | 1240 | 349 | 7692 | 640 | 5072 | 3279 | 5843 | 10478 | 405 | 0 | 4183 | 0 | 0 | 0 | 0 | 0 | 0 | 0 | 0 | 0 | 0 | 0 | 0 | 0 | 0 | 0 | 0 | 0 | 0 | 0 | 0 |
| SRR6453370 | 691 | 4464 | 11072 | 498 | 104 | 63 | 1316 | 302 | 7028 | 949 | 4719 | 2756 | 4710 | 5389 | 226 | 10 | 3709 | 0 | 0 | 0 | 0 | 0 | 0 | 0 | 0 | 0 | 0 | 0 | 0 | 0 | 0 | 0 | 0 | 0 | 0 | 0 | 0 |
| SRR6453371 | 441 | 2594 | 17201 | 389 | 255 | 217 | 4894 | 479 | 17775 | 672 | 6387 | 3573 | 6893 | 12173 | 822 | 0 | 5736 | 0 | 0 | 0 | 0 | 0 | 0 | 0 | 0 | 0 | 0 | 0 | 0 | 0 | 0 | 0 | 0 | 0 | 0 | 0 | 0 |
| SRR6453372 | 543 | 3026 | 14565 | 408 | 88 | 69 | 1488 | 212 | 7232 | 454 | 3004 | 1822 | 3751 | 9806 | 429 | 63 | 3137 | 0 | 0 | 0 | 0 | 0 | 0 | 0 | 0 | 0 | 0 | 0 | 0 | 0 | 0 | 0 | 0 | 0 | 0 | 0 | 0 |
| SRR6453373 | 491 | 2798 | 18690 | 367 | 116 | 117 | 2093 | 249 | 7695 | 681 | 2505 | 1392 | 3012 | 9950 | 461 | 26 | 5004 | 0 | 0 | 0 | 0 | 0 | 13 | 0 | 0 | 0 | 0 | 0 | 0 | 0 | 0 | 0 | 0 | 0 | 0 | 0 | 0 |
| SRR6453374 | 423 | 2215 | 18684 | 385 | 172 | 155 | 2932 | 310 | 6456 | 427 | 4108 | 2443 | 4975 | 15364 | 685 | 27 | 5429 | 0 | 0 | 0 | 0 | 0 | 0 | 0 | 0 | 0 | 0 | 0 | 0 | 0 | 0 | 0 | 0 | 0 | 0 | 0 | 0 |
| SRR6453375 | 450 | 2159 | 15879 | 274 | 151 | 142 | 3119 | 432 | 8090 | 595 | 7127 | 3133 | 7110 | 8884 | 780 | 137 | 4969 | 0 | 0 | 0 | 0 | 0 | 0 | 0 | 0 | 0 | 0 | 0 | 0 | 0 | 0 | 0 | 0 | 0 | 0 | 0 | 0 |
| SRR6453376 | 274 | 1039 | 18368 | 330 | 87 | 86 | 990 | 353 | 3740 | 718 | 1097 | 808 | 1284 | 1902 | 152 | 0 | 3666 | 0 | 0 | 0 | 0 | 0 | 0 | 0 | 0 | 0 | 0 | 0 | 0 | 0 | 0 | 0 | 0 | 0 | 0 | 0 | 0 |
| SRR6453377 | 418 | 2544 | 15145 | 406 | 171 | 126 | 2312 | 358 | 7137 | 602 | 1597 | 718 | 1233 | 2439 | 184 | 11 | 5550 | 0 | 0 | 0 | 0 | 0 | 0 | 0 | 0 | 0 | 0 | 0 | 0 | 0 | 0 | 0 | 0 | 0 | 0 | 0 | 0 |
| SRR6453378 | 352 | 2277 | 14768 | 277 | 127 | 96 | 2091 | 270 | 5675 | 508 | 3196 | 1599 | 3086 | 4870 | 266 | 43 | 4245 | 0 | 0 | 0 | 0 | 0 | 0 | 0 | 0 | 0 | 0 | 0 | 0 | 0 | 0 | 0 | 0 | 0 | 0 | 0 | 0 |
| SRR6453379 | 517 | 2990 | 16954 | 473 | 329 | 202 | 4723 | 281 | 13918 | 819 | 2367 | 1228 | 2322 | 4420 | 512 | 15 | 3107 | 0 | 0 | 0 | 0 | 0 | 13 | 0 | 0 | 0 | 0 | 0 | 0 | 0 | 0 | 0 | 0 | 0 | 0 | 0 | 0 |
| SRR6453380 | 310 | 1905 | 15358 | 203 | 245 | 192 | 3549 | 405 | 11732 | 649 | 2733 | 1389 | 2303 | 7557 | 530 | 16 | 6074 | 0 | 0 | 0 | 0 | 0 | 0 | 0 | 0 | 0 | 0 | 0 | 0 | 0 | 0 | 0 | 0 | 0 | 0 | 0 | 0 |
| SRR6453381 | 388 | 2267 | 13689 | 393 | 101 | 73 | 1433 | 283 | 7970 | 619 | 4683 | 2285 | 4706 | 7794 | 460 | 11 | 4135 | 0 | 0 | 0 | 0 | 0 | 0 | 0 | 0 | 0 | 0 | 0 | 0 | 0 | 0 | 0 | 0 | 0 | 0 | 0 | 0 |
| SRR6453382 | 495 | 3553 | 13593 | 330 | 130 | 101 | 1751 | 472 | 6911 | 1244 | 4980 | 1887 | 7100 | 5221 | 301 | 0 | 5369 | 0 | 0 | 0 | 0 | 0 | 12 | 0 | 0 | 0 | 0 | 0 | 0 | 0 | 0 | 0 | 0 | 0 | 0 | 0 | 0 |
| SRR6453383 | 410 | 2330 | 17688 | 538 | 181 | 115 | 2497 | 367 | 10838 | 683 | 2095 | 1013 | 2168 | 4835 | 428 | 12 | 3819 | 0 | 0 | 0 | 0 | 0 | 0 | 0 | 0 | 0 | 0 | 0 | 0 | 0 | 0 | 0 | 0 | 0 | 0 | 0 | 0 |
| SRR6453384 | 470 | 2565 | 15813 | 376 | 175 | 119 | 2808 | 379 | 12781 | 704 | 4029 | 1689 | 4231 | 6020 | 652 | 10 | 4476 | 0 | 0 | 0 | 0 | 0 | 0 | 0 | 0 | 0 | 0 | 0 | 0 | 0 | 0 | 0 | 0 | 0 | 0 | 0 | 0 |
| SRR6453385 | 509 | 2340 | 21411 | 586 | 258 | 177 | 3104 | 459 | 3531 | 546 | 7195 | 5179 | 8477 | 20981 | 895 | 0 | 6704 | 0 | 0 | 0 | 0 | 0 | 0 | 0 | 0 | 0 | 0 | 0 | 0 | 0 | 0 | 0 | 0 | 0 | 0 | 0 | 0 |
| SRR6453386 | 1537 | 3556 | 20891 | 156 | 2705 | 2557 | 61651 | 929 | 5262 | 210 | 8915 | 30593 | 16815 | 117793 | 145 | 73 | 19716 | 0 | 0 | 0 | 0 | 0 | 0 | 0 | 0 | 0 | 0 | 0 | 0 | 0 | 0 | 0 | 0 | 0 | 0 | 0 | 0 |
| SRR6453387 | 1234 | 5891 | 17023 | 71 | 2217 | 1934 | 42566 | 1094 | 3993 | 406 | 12418 | 32145 | 38239 | 114794 | 110 | 70 | 24400 | 0 | 0 | 0 | 0 | 0 | 0 | 0 | 0 | 0 | 0 | 0 | 0 | 0 | 0 | 0 | 0 | 0 | 0 | 0 | 0 |
| SRR6453388 | 778 | 3835 | 10871 | 912 | 6541 | 4687 | 94015 | 3445 | 10907 | 1597 | 20469 | 27304 | 38818 | 59073 | 361 | 154 | 25817 | 0 | 0 | 0 | 0 | 0 | 0 | 0 | 0 | 0 | 0 | 0 | 0 | 0 | 0 | 0 | 0 | 0 | 0 | 0 | 0 |
| SRR6453389 | 847 | 3590 | 20819 | 923 | 6946 | 7385 | 135965 | 4086 | 11121 | 1743 | 14082 | 21862 | 27631 | 75108 | 312 | 153 | 32850 | 0 | 0 | 0 | 0 | 0 | 0 | 0 | 0 | 0 | 0 | 0 | 0 | 0 | 0 | 0 | 0 | 0 | 0 | 0 | 0 |
| SRR6453390 | 657 | 3345 | 17872 | 611 | 4908 | 3441 | 53966 | 3531 | 5196 | 2892 | 12056 | 31104 | 33735 | 60202 | 220 | 139 | 27939 | 0 | 0 | 0 | 0 | 0 | 0 | 0 | 0 | 0 | 0 | 0 | 0 | 0 | 0 | 0 | 0 | 0 | 0 | 0 | 0 |
| SRR6453391 | 735 | 3715 | 16610 | 543 | 4077 | 3422 | 68388 | 1973 | 11185 | 1255 | 11809 | 18644 | 31660 | 71100 | 220 | 152 | 29561 | 0 | 0 | 0 | 0 | 0 | 0 | 0 | 0 | 0 | 0 | 0 | 0 | 0 | 0 | 0 | 0 | 0 | 0 | 0 | 0 |
| SRR6453392 | 804 | 4331 | 14584 | 571 | 2541 | 2299 | 41134 | 2393 | 8493 | 2660 | 16674 | 27904 | 28632 | 34015 | 214 | 99 | 27634 | 0 | 0 | 0 | 0 | 0 | 0 | 0 | 0 | 0 | 0 | 0 | 0 | 0 | 0 | 0 | 0 | 0 | 0 | 0 | 0 |
| SRR6453393 | 563 | 2674 | 15364 | 840 | 4630 | 3751 | 68400 | 2955 | 9101 | 4616 | 11984 | 23273 | 33130 | 69494 | 202 | 139 | 26972 | 0 | 0 | 0 | 0 | 0 | 0 | 0 | 0 | 0 | 0 | 0 | 0 | 0 | 0 | 0 | 0 | 0 | 0 | 0 | 0 |
| SRR6453394 | 908 | 4001 | 18376 | 608 | 11424 | 8789 | 156924 | 5748 | 8630 | 1562 | 16245 | 20411 | 21944 | 37360 | 383 | 45 | 30895 | 0 | 0 | 0 | 0 | 0 | 0 | 0 | 0 | 0 | 0 | 0 | 0 | 0 | 0 | 0 | 0 | 0 | 0 | 0 | 0 |
| SRR6453395 | 857 | 4413 | 14791 | 518 | 3989 | 3142 | 56874 | 2607 | 7924 | 2547 | 13647 | 20837 | 37761 | 64852 | 185 | 84 | 25298 | 0 | 0 | 0 | 0 | 0 | 0 | 0 | 0 | 0 | 0 | 0 | 0 | 0 | 0 | 0 | 0 | 0 | 0 | 0 | 0 |
| SRR6453396 | 891 | 3889 | 20571 | 504 | 4503 | 3496 | 72120 | 2614 | 7174 | 2625 | 11759 | 18657 | 28812 | 59257 | 254 | 77 | 22297 | 0 | 0 | 0 | 0 | 0 | 0 | 0 | 0 | 0 | 0 | 0 | 0 | 0 | 0 | 0 | 0 | 0 | 0 | 0 | 0 |
| SRR6453397 | 740 | 4019 | 12572 | 368 | 5435 | 4221 | 84717 | 2562 | 8700 | 823 | 12003 | 21422 | 26955 | 68489 | 252 | 66 | 26753 | 0 | 0 | 0 | 0 | 0 | 0 | 0 | 0 | 0 | 0 | 0 | 0 | 0 | 0 | 0 | 0 | 0 | 0 | 0 | 0 |
| SRR6453398 | 1304 | 5480 | 21771 | 342 | 8793 | 7422 | 154628 | 5133 | 10063 | 752 | 16157 | 14678 | 18497 | 39046 | 402 | 136 | 28537 | 0 | 0 | 0 | 0 | 0 | 0 | 0 | 0 | 0 | 0 | 0 | 0 | 0 | 0 | 0 | 0 | 0 | 0 | 0 | 0 |
| SRR6453399 | 786 | 3324 | 9878 | 1954 | 8704 | 8441 | 175705 | 4990 | 13293 | 874 | 27002 | 24045 | 23767 | 48742 | 415 | 189 | 36582 | 0 | 0 | 0 | 0 | 0 | 0 | 0 | 0 | 0 | 0 | 0 | 0 | 0 | 0 | 0 | 0 | 0 | 0 | 0 | 0 |
| SRR6453400 | 690 | 3593 | 14397 | 348 | 4201 | 4170 | 78166 | 2899 | 6583 | 1437 | 14716 | 19175 | 28810 | 59663 | 234 | 57 | 28763 | 0 | 0 | 0 | 0 | 0 | 0 | 0 | 0 | 0 | 0 | 0 | 0 | 0 | 0 | 0 | 0 | 0 | 0 | 0 | 0 |
| SRR6453401 | 785 | 4637 | 12946 | 630 | 4079 | 3210 | 59979 | 2776 | 6515 | 1956 | 13282 | 23368 | 23226 | 42784 | 212 | 106 | 25180 | 0 | 0 | 0 | 0 | 0 | 0 | 0 | 0 | 0 | 0 | 0 | 0 | 0 | 0 | 0 | 0 | 0 | 0 | 0 | 0 |
| SRR6453402 | 970 | 5484 | 9107 | 496 | 3270 | 2187 | 51700 | 1755 | 5963 | 1843 | 25770 | 45134 | 40437 | 76778 | 221 | 111 | 15980 | 0 | 0 | 0 | 0 | 0 | 0 | 0 | 0 | 0 | 0 | 0 | 0 | 0 | 0 | 0 | 0 | 0 | 0 | 0 | 0 |
| SRR6453403 | 820 | 4328 | 11489 | 812 | 2040 | 1857 | 42766 | 1057 | 6864 | 1899 | 13957 | 27431 | 32321 | 97092 | 214 | 159 | 16314 | 0 | 0 | 0 | 0 | 0 | 0 | 0 | 0 | 0 | 0 | 0 | 0 | 0 | 0 | 0 | 0 | 0 | 0 | 0 | 0 |
| SRR6453404 | 1452 | 5155 | 12917 | 1504 | 6112 | 6516 | 143024 | 3107 | 11994 | 877 | 32673 | 11 | 23234 | 61854 | 362 | 73 | 21733 | 0 | 0 | 0 | 0 | 0 | 0 | 0 | 0 | 0 | 0 | 0 | 0 | 0 | 0 | 0 | 0 | 0 | 0 | 0 | 0 |
| SRR6453405 | 1060 | 3910 | 20525 | 715 | 3862 | 3009 | 61628 | 1588 | 7863 | 1038 | 15025 | 26819 | 30806 | 80379 | 150 | 53 | 27163 | 0 | 0 | 0 | 0 | 0 | 0 | 0 | 0 | 0 | 0 | 0 | 0 | 0 | 0 | 0 | 0 | 0 | 0 | 0 | 0 |
| SRR6453406 | 1069 | 5146 | 13505 | 742 | 3640 | 2779 | 62148 | 2043 | 5239 | 1351 | 23265 | 10 | 45274 | 88776 | 205 | 97 | 15783 | 0 | 0 | 0 | 0 | 0 | 0 | 0 | 0 | 0 | 0 | 0 | 0 | 0 | 0 | 0 | 0 | 0 | 0 | 0 | 0 |
| SRR6453407 | 1035 | 5598 | 9250 | 999 | 3977 | 4256 | 99886 | 2085 | 9011 | 1330 | 19156 | 19735 | 12204 | 30800 | 241 | 117 | 19373 | 0 | 0 | 0 | 0 | 0 | 0 | 0 | 0 | 0 | 0 | 0 | 0 | 0 | 0 | 0 | 0 | 0 | 0 | 0 | 0 |
| SRR6453408 | 1326 | 4234 | 21069 | 1155 | 6413 | 4909 | 109096 | 1889 | 11777 | 771 | 20435 | 20342 | 16399 | 55108 | 352 | 198 | 24550 | 0 | 0 | 0 | 0 | 0 | 0 | 0 | 0 | 0 | 0 | 0 | 0 | 0 | 0 | 0 | 0 | 0 | 0 | 0 | 0 |
| SRR6453409 | 1948 | 6332 | 14242 | 1214 | 5552 | 4421 | 92148 | 2082 | 7941 | 690 | 17955 | 26869 | 23022 | 59457 | 211 | 67 | 17549 | 0 | 0 | 0 | 0 | 0 | 0 | 0 | 0 | 0 | 0 | 0 | 0 | 0 | 0 | 0 | 0 | 0 | 0 | 0 | 0 |
| SRR6453410 | 992 | 5365 | 10846 | 1219 | 6961 | 5187 | 124413 | 2520 | 11391 | 1040 | 26389 | 35170 | 21814 | 46359 | 374 | 164 | 21260 | 0 | 0 | 0 | 0 | 0 | 0 | 0 | 0 | 0 | 0 | 0 | 0 | 0 | 0 | 0 | 0 | 0 | 0 | 0 | 0 |
| SRR6453411 | 1236 | 5796 | 19410 | 670 | 4885 | 3531 | 75577 | 2243 | 6996 | 1587 | 18887 | 31851 | 33416 | 74569 | 259 | 122 | 24571 | 0 | 0 | 0 | 0 | 0 | 0 | 0 | 0 | 0 | 0 | 0 | 0 | 0 | 0 | 0 | 0 | 0 | 0 | 0 | 0 |
| SRR6453412 | 1315 | 3485 | 18745 | 185 | 3114 | 2707 | 58460 | 1021 | 7169 | 314 | 11904 | 31545 | 18151 | 142699 | 196 | 31 | 21231 | 0 | 0 | 0 | 0 | 0 | 0 | 0 | 0 | 0 | 0 | 0 | 0 | 0 | 0 | 0 | 0 | 0 | 0 | 0 | 0 |
| SRR6453413 | 1074 | 5072 | 18555 | 109 | 4667 | 3180 | 69326 | 1746 | 5416 | 407 | 13273 | 51242 | 37379 | 111472 | 178 | 50 | 21809 | 0 | 0 | 0 | 0 | 0 | 0 | 0 | 0 | 0 | 0 | 0 | 0 | 0 | 0 | 0 | 0 | 0 | 0 | 0 | 0 |
| SRR6453414 | 488 | 2201 | 16540 | 607 | 3225 | 2697 | 48746 | 3320 | 6255 | 2835 | 23646 | 11 | 34215 | 50040 | 219 | 186 | 19022 | 0 | 0 | 0 | 0 | 0 | 0 | 0 | 0 | 0 | 0 | 0 | 0 | 0 | 0 | 0 | 0 | 0 | 0 | 0 | 0 |
| SRR6453415 | 384 | 1649 | 13056 | 1834 | 7021 | 5805 | 95070 | 5400 | 14433 | 974 | 28413 | 11 | 27931 | 42396 | 504 | 202 | 26751 | 0 | 0 | 0 | 0 | 0 | 0 | 0 | 0 | 0 | 0 | 0 | 0 | 0 | 0 | 0 | 0 | 0 | 0 | 0 | 0 |
| SRR6453416 | 211 | 1296 | 11992 | 456 | 4616 | 4600 | 62145 | 3413 | 12882 | 2868 | 16743 | 17535 | 19182 | 46493 | 464 | 397 | 25982 | 0 | 0 | 0 | 0 | 0 | 0 | 0 | 0 | 0 | 0 | 0 | 57 | 0 | 0 | 0 | 0 | 0 | 0 | 0 | 0 |
| SRR6453417 | 790 | 2079 | 11640 | 896 | 5244 | 3855 | 68841 | 3752 | 7044 | 2258 | 22339 | 35173 | 31001 | 47675 | 291 | 150 | 22043 | 0 | 0 | 0 | 0 | 0 | 0 | 0 | 0 | 0 | 0 | 0 | 0 | 0 | 0 | 0 | 0 | 0 | 0 | 0 | 0 |

### Supp Table S8.1.Columns_B_to_AL

| | | | | | | | | | | | | | | | | | | | | | | | | | | | | | | | | | | | | | |
|---|---|---|---|---|---|---|---|---|---|---|---|---|---|---|---|---|---|---|---|---|---|---|---|---|---|---|---|---|---|---|---|---|---|---|---|---|---|
| SRR6453418 | 1150 | 4617 | 20069 | 728 | 6091 | 5662 | 91223 | 4630 | 10262 | 1483 | 15627 | 17278 | 24970 | 56718 | 299 | 139 | 29997 | 0 | 0 | 0 | 0 | 0 | 0 | 0 | 0 | 0 | 0 | 0 | 0 | 0 | 0 | 0 | 0 | 0 | 0 | 0 | 0 |
| SRR6453419 | 981 | 3272 | 17927 | 878 | 9943 | 6031 | 134500 | 4879 | 10135 | 1270 | 27397 | 29866 | 30029 | 52040 | 391 | 100 | 22339 | 0 | 0 | 0 | 0 | 0 | 0 | 0 | 0 | 0 | 0 | 0 | 0 | 0 | 0 | 0 | 0 | 0 | 0 | 0 | 0 |
| SRR6453420 | 1030 | 4341 | 16801 | 715 | 6228 | 4239 | 96054 | 2866 | 6123 | 1011 | 26998 | 34714 | 31662 | 54284 | 370 | 95 | 20486 | 0 | 0 | 0 | 0 | 0 | 0 | 0 | 0 | 0 | 0 | 0 | 0 | 0 | 0 | 0 | 0 | 0 | 0 | 0 | 0 |
| SRR6453421 | 1198 | 3784 | 27960 | 833 | 6478 | 4252 | 91865 | 1857 | 19311 | 1000 | 16035 | 21448 | 44105 | 89026 | 207 | 356 | 79653 | 0 | 0 | 0 | 0 | 0 | 0 | 0 | 0 | 0 | 0 | 0 | 12 | 0 | 0 | 0 | 0 | 0 | 0 | 0 | 0 |
| SRR6453422 | 1200 | 4711 | 15698 | 99 | 3156 | 2723 | 65630 | 1071 | 7915 | 527 | 13399 | 22892 | 35264 | 111470 | 303 | 157 | 14610 | 0 | 0 | 0 | 0 | 0 | 0 | 0 | 0 | 0 | 0 | 0 | 0 | 0 | 0 | 0 | 0 | 0 | 0 | 0 | 0 |
| SRR6453423 | 1262 | 4263 | 25020 | 75 | 2814 | 2589 | 51089 | 1508 | 5807 | 425 | 10772 | 20432 | 39208 | 115524 | 136 | 65 | 24842 | 0 | 0 | 0 | 0 | 0 | 0 | 0 | 0 | 0 | 0 | 0 | 0 | 0 | 0 | 0 | 0 | 0 | 0 | 0 | 0 |
| SRR6453424 | 786 | 2113 | 15717 | 1190 | 3491 | 3224 | 55622 | 2721 | 6985 | 2228 | 22117 | 34785 | 25432 | 31681 | 312 | 147 | 21112 | 0 | 0 | 0 | 0 | 0 | 0 | 0 | 0 | 0 | 0 | 0 | 0 | 0 | 0 | 0 | 0 | 0 | 0 | 0 | 0 |
| SRR6453425 | 421 | 1480 | 13141 | 2024 | 3198 | 5783 | 69631 | 4134 | 11342 | 1587 | 36087 | 40544 | 32807 | 40561 | 497 | 205 | 30254 | 0 | 0 | 0 | 0 | 0 | 0 | 0 | 0 | 0 | 0 | 0 | 0 | 0 | 0 | 0 | 0 | 0 | 0 | 0 | 0 |
| SRR6453426 | 557 | 1961 | 20124 | 1084 | 5221 | 4277 | 72477 | 2598 | 11841 | 1147 | 13913 | 21193 | 22293 | 54250 | 372 | 143 | 21693 | 0 | 0 | 0 | 0 | 0 | 0 | 0 | 0 | 0 | 0 | 0 | 0 | 0 | 0 | 0 | 0 | 0 | 0 | 0 | 0 |
| SRR6453427 | 356 | 1643 | 15305 | 838 | 4737 | 4262 | 66179 | 3505 | 8881 | 1087 | 15491 | 17728 | 22242 | 45899 | 353 | 172 | 22054 | 0 | 0 | 0 | 0 | 0 | 0 | 0 | 0 | 0 | 0 | 0 | 0 | 0 | 0 | 0 | 0 | 0 | 0 | 0 | 0 |
| SRR6453428 | 459 | 2188 | 18665 | 615 | 6199 | 4788 | 75756 | 4799 | 4087 | 1266 | 15796 | 18993 | 24054 | 45629 | 287 | 165 | 21598 | 0 | 0 | 0 | 0 | 0 | 0 | 0 | 0 | 0 | 0 | 0 | 0 | 0 | 0 | 0 | 0 | 0 | 0 | 0 | 0 |
| SRR6454160 | 0 | 15077 | 239 | 213 | 2404 | 13899 | 18078 | 21141 | 19216 | 53 | 15685 | 39350 | 7150 | 2544 | 16 | 8952 | 40818 | 0 | 0 | 20 | 6559 | 2524 | 369 | 16 | 0 | 0 | 0 | 0 | 413 | 7662 | 9349 | 821 | 0 | 12 | 22 | 0 | 0 |
| SRR6454161 | 17 | 23452 | 848 | 1285 | 1426 | 18235 | 16951 | 39504 | 12912 | 18 | 24802 | 18122 | 58 | 16757 | 0 | 10920 | 45984 | 0 | 0 | 14 | 4763 | 7551 | 2183 | 88 | 27 | 0 | 0 | 0 | 564 | 4970 | 14855 | 4142 | 16 | 68 | 61 | 44 | 0 |
| SRR6454162 | 123 | 7937 | 766 | 0 | 0 | 0 | 0 | 56 | 80 | 0 | 627 | 1107 | 278 | 1028 | 0 | 0 | 883 | 0 | 0 | 0 | 0 | 0 | 0 | 0 | 0 | 0 | 0 | 0 | 0 | 0 | 0 | 0 | 0 | 0 | 0 | 0 | 0 |
| SRR6454163 | 0 | 7021 | 888 | 929 | 627 | 9987 | 7005 | 30034 | 13406 | 62 | 46754 | 11 | 3658 | 10373 | 0 | 8558 | 41622 | 0 | 0 | 12 | 7370 | 4716 | 1511 | 121 | 31 | 31 | 0 | 0 | 505 | 6021 | 7281 | 1573 | 0 | 28 | 64 | 50 | 0 |
| SRR6454164 | 153 | 9651 | 674 | 92 | 0 | 0 | 0 | 56 | 68 | 31 | 749 | 1468 | 345 | 1210 | 0 | 0 | 996 | 0 | 0 | 0 | 0 | 0 | 0 | 0 | 0 | 0 | 0 | 0 | 0 | 0 | 0 | 0 | 0 | 0 | 0 | 0 | 0 |
| SRR6454165 | 0 | 9711 | 936 | 407 | 1024 | 16812 | 16185 | 48413 | 10839 | 46 | 45458 | 26897 | 5027 | 11333 | 0 | 13180 | 44492 | 0 | 0 | 0 | 4306 | 2512 | 1374 | 154 | 39 | 24 | 0 | 0 | 286 | 3636 | 3817 | 1700 | 0 | 36 | 46 | 44 | 0 |
| SRR6454166 | 127 | 5851 | 506 | 253 | 0 | 10 | 11 | 63 | 106 | 63 | 689 | 1341 | 328 | 769 | 0 | 0 | 673 | 0 | 0 | 0 | 0 | 0 | 0 | 0 | 0 | 0 | 0 | 0 | 0 | 0 | 0 | 0 | 0 | 0 | 0 | 0 | 0 |
| SRR6454167 | 0 | 8403 | 1160 | 944 | 859 | 13052 | 10322 | 27735 | 18282 | 72 | 64401 | 27582 | 5066 | 11016 | 0 | 10666 | 38117 | 0 | 0 | 0 | 5145 | 3804 | 930 | 54 | 16 | 12 | 0 | 0 | 355 | 4876 | 7427 | 1405 | 0 | 19 | 36 | 21 | 0 |
| SRR6454169 | 0 | 6144 | 985 | 400 | 1322 | 16561 | 15781 | 43949 | 12038 | 72 | 44591 | 20251 | 4646 | 14287 | 0 | 12903 | 54486 | 0 | 0 | 0 | 2446 | 2890 | 867 | 99 | 20 | 17 | 0 | 0 | 193 | 2340 | 4514 | 1217 | 0 | 17 | 34 | 30 | 0 |
| SRR6454170 | 265 | 8859 | 572 | 633 | 0 | 0 | 11 | 70 | 62 | 0 | 696 | 942 | 290 | 603 | 0 | 0 | 1295 | 0 | 0 | 0 | 0 | 0 | 0 | 0 | 0 | 0 | 0 | 0 | 0 | 0 | 0 | 0 | 0 | 0 | 0 | 0 | 0 |
| SRR6454171 | 35 | 7319 | 1650 | 435 | 846 | 13927 | 10419 | 37350 | 13949 | 106 | 39465 | 19709 | 5338 | 11147 | 0 | 11522 | 33303 | 0 | 0 | 16 | 10800 | 8169 | 1684 | 74 | 19 | 23 | 0 | 0 | 642 | 9339 | 13361 | 2094 | 0 | 24 | 43 | 43 | 0 |
| SRR6454172 | 558 | 10299 | 804 | 607 | 0 | 0 | 0 | 57 | 65 | 33 | 465 | 607 | 204 | 341 | 0 | 0 | 1284 | 0 | 0 | 0 | 0 | 0 | 0 | 0 | 0 | 0 | 0 | 0 | 0 | 0 | 0 | 0 | 0 | 0 | 0 | 0 | 0 |
| SRR6454173 | 16 | 8318 | 592 | 382 | 1166 | 21313 | 15646 | 53265 | 13508 | 23 | 35998 | 27843 | 12 | 7373 | 0 | 12496 | 43891 | 0 | 0 | 0 | 4350 | 3655 | 1429 | 239 | 42 | 17 | 0 | 0 | 268 | 3273 | 5600 | 1866 | 11 | 30 | 27 | 24 | 0 |
| SRR6454174 | 397 | 4924 | 1048 | 540 | 0 | 0 | 0 | 31 | 65 | 16 | 226 | 318 | 67 | 113 | 0 | 0 | 881 | 0 | 0 | 0 | 11 | 0 | 0 | 0 | 0 | 0 | 0 | 0 | 0 | 0 | 0 | 0 | 0 | 0 | 0 | 0 | 0 |
| SRR6454175 | 0 | 4058 | 102 | 1022 | 898 | 10350 | 10956 | 25009 | 13170 | 0 | 20123 | 18722 | 167 | 7095 | 0 | 8321 | 52508 | 0 | 0 | 14 | 10642 | 6159 | 1507 | 48 | 34 | 0 | 0 | 0 | 498 | 8035 | 8559 | 2147 | 15 | 72 | 65 | 27 | 0 |
| SRR6454176 | 250 | 4415 | 736 | 342 | 0 | 11 | 10 | 70 | 73 | 79 | 638 | 573 | 208 | 608 | 0 | 0 | 1090 | 0 | 0 | 0 | 0 | 0 | 0 | 0 | 0 | 0 | 0 | 0 | 0 | 0 | 0 | 0 | 0 | 0 | 0 | 0 | 0 |
| SRR6454177 | 39 | 29991 | 925 | 495 | 818 | 13672 | 10537 | 42769 | 10526 | 0 | 55877 | 11 | 4022 | 9611 | 0 | 12244 | 36490 | 0 | 0 | 0 | 3453 | 2602 | 1040 | 60 | 15 | 11 | 0 | 0 | 308 | 3025 | 3906 | 1216 | 0 | 23 | 43 | 38 | 0 |
| SRR6454178 | 242 | 6524 | 450 | 381 | 0 | 0 | 0 | 63 | 70 | 0 | 461 | 559 | 172 | 447 | 0 | 0 | 812 | 0 | 0 | 0 | 0 | 0 | 0 | 0 | 0 | 0 | 0 | 0 | 0 | 0 | 0 | 0 | 0 | 0 | 0 | 0 | 0 |
| SRR6454179 | 25 | 5971 | 792 | 277 | 1372 | 21111 | 21339 | 57125 | 13077 | 13 | 51111 | 26524 | 4582 | 9870 | 0 | 13155 | 50867 | 0 | 0 | 0 | 3300 | 4353 | 1830 | 207 | 42 | 44 | 0 | 0 | 297 | 3539 | 7051 | 2488 | 0 | 38 | 75 | 78 | 0 |
| SRR6454180 | 280 | 5345 | 391 | 256 | 0 | 0 | 0 | 105 | 224 | 24 | 552 | 481 | 203 | 659 | 0 | 0 | 908 | 0 | 0 | 0 | 0 | 0 | 0 | 0 | 0 | 0 | 0 | 0 | 0 | 0 | 0 | 0 | 0 | 0 | 0 | 0 | 0 |
| SRR6454181 | 30 | 11521 | 364 | 2221 | 659 | 12970 | 9892 | 23965 | 13641 | 10 | 52166 | 24810 | 3700 | 9909 | 0 | 10821 | 38705 | 0 | 0 | 15 | 10353 | 6470 | 1463 | 55 | 21 | 14 | 0 | 0 | 715 | 9731 | 11087 | 2194 | 0 | 38 | 76 | 55 | 0 |
| SRR6454182 | 175 | 7201 | 639 | 161 | 0 | 0 | 0 | 41 | 105 | 27 | 324 | 367 | 147 | 503 | 0 | 0 | 913 | 0 | 0 | 0 | 11 | 0 | 0 | 0 | 0 | 0 | 0 | 0 | 0 | 0 | 0 | 0 | 0 | 0 | 0 | 0 | 0 |
| SRR6454183 | 21 | 5373 | 628 | 374 | 1102 | 20055 | 17060 | 41271 | 12568 | 14 | 42485 | 24099 | 5127 | 9676 | 0 | 11175 | 41946 | 0 | 0 | 0 | 4832 | 6515 | 2153 | 148 | 67 | 22 | 0 | 0 | 346 | 5158 | 10593 | 3770 | 14 | 114 | 135 | 104 | 0 |
| SRR6454184 | 97 | 5357 | 516 | 387 | 0 | 0 | 0 | 42 | 62 | 16 | 279 | 425 | 111 | 300 | 0 | 0 | 506 | 0 | 0 | 0 | 0 | 0 | 0 | 0 | 0 | 0 | 0 | 0 | 0 | 0 | 0 | 0 | 0 | 0 | 0 | 0 | 0 |
| SRR6454185 | 95 | 9363 | 115 | 664 | 1122 | 17961 | 12637 | 23748 | 18521 | 0 | 49153 | 48536 | 8743 | 5845 | 0 | 13507 | 34564 | 0 | 0 | 13 | 7168 | 3908 | 1168 | 25 | 15 | 0 | 0 | 0 | 494 | 4841 | 7418 | 1849 | 0 | 30 | 56 | 34 | 0 |
| SRR6454186 | 466 | 5876 | 420 | 839 | 0 | 0 | 0 | 40 | 64 | 16 | 364 | 526 | 133 | 562 | 0 | 0 | 501 | 0 | 0 | 0 | 0 | 0 | 0 | 0 | 0 | 0 | 0 | 0 | 0 | 0 | 0 | 0 | 0 | 0 | 0 | 0 | 0 |
| SRR6454187 | 123 | 4149 | 2196 | 1737 | 1635 | 30234 | 24225 | 48013 | 12838 | 18 | 62767 | 31434 | 4643 | 13667 | 0 | 12582 | 37877 | 0 | 0 | 15 | 9195 | 8171 | 1662 | 111 | 26 | 20 | 0 | 0 | 820 | 8656 | 11582 | 2305 | 0 | 35 | 53 | 52 | 0 |
| SRR6454188 | 83 | 12954 | 372 | 83 | 0 | 19 | 17 | 126 | 185 | 0 | 732 | 884 | 317 | 1114 | 0 | 0 | 1135 | 0 | 0 | 0 | 0 | 0 | 0 | 0 | 0 | 0 | 0 | 0 | 0 | 0 | 0 | 0 | 0 | 0 | 0 | 0 | 0 |
| SRR6454189 | 45 | 24954 | 491 | 268 | 826 | 18914 | 12537 | 70153 | 12148 | 0 | 65617 | 16889 | 3808 | 10354 | 0 | 13899 | 33041 | 0 | 0 | 0 | 2118 | 1792 | 447 | 23 | 0 | 0 | 0 | 0 | 274 | 4076 | 6232 | 1241 | 0 | 18 | 31 | 24 | 0 |
| SRR6454190 | 125 | 6233 | 499 | 299 | 0 | 16 | 14 | 122 | 133 | 0 | 740 | 860 | 274 | 945 | 0 | 0 | 1010 | 0 | 0 | 0 | 0 | 0 | 0 | 0 | 0 | 0 | 0 | 0 | 0 | 0 | 0 | 0 | 0 | 0 | 0 | 0 | 0 |
| SRR6454191 | 23 | 9997 | 503 | 175 | 1106 | 18551 | 16307 | 53811 | 13177 | 0 | 44796 | 29971 | 6756 | 10294 | 0 | 13127 | 42845 | 0 | 0 | 0 | 2148 | 3690 | 1311 | 148 | 34 | 16 | 0 | 0 | 202 | 2358 | 5646 | 2137 | 0 | 37 | 42 | 38 | 0 |
| SRR6454193 | 0 | 8994 | 70 | 545 | 1254 | 17772 | 15231 | 50999 | 18976 | 0 | 69667 | 38896 | 6726 | 10427 | 0 | 14734 | 30613 | 0 | 0 | 0 | 3276 | 2456 | 602 | 24 | 0 | 0 | 0 | 0 | 306 | 3982 | 6842 | 1440 | 0 | 12 | 28 | 24 | 0 |
| SRR6454194 | 123 | 11463 | 666 | 666 | 0 | 15 | 14 | 79 | 118 | 0 | 438 | 543 | 184 | 956 | 0 | 0 | 723 | 0 | 0 | 0 | 0 | 0 | 0 | 0 | 0 | 0 | 0 | 0 | 0 | 0 | 0 | 0 | 0 | 0 | 0 | 0 | 0 |
| SRR6454195 | 18 | 10554 | 1368 | 160 | 1215 | 20640 | 19716 | 63462 | 12017 | 0 | 51441 | 25349 | 5443 | 15033 | 0 | 16290 | 43912 | 0 | 0 | 0 | 2233 | 2347 | 1114 | 74 | 18 | 0 | 0 | 0 | 205 | 2979 | 5039 | 2256 | 0 | 31 | 41 | 40 | 0 |
| SRR6454196 | 248 | 6393 | 173 | 124 | 0 | 0 | 0 | 15 | 23 | 0 | 46 | 75 | 13 | 53 | 0 | 0 | 201 | 0 | 0 | 0 | 0 | 0 | 0 | 0 | 0 | 0 | 0 | 0 | 0 | 0 | 0 | 0 | 0 | 0 | 0 | 0 | 0 |
| SRR6454197 | 35 | 6397 | 49 | 1243 | 1029 | 8021 | 9677 | 18588 | 17947 | 14 | 22550 | 25224 | 6841 | 6133 | 0 | 7725 | 48254 | 0 | 0 | 28 | 10222 | 7429 | 2014 | 32 | 18 | 0 | 0 | 0 | 1021 | 8028 | 18964 | 2891 | 0 | 56 | 131 | 58 | 0 |
| SRR6454198 | 863 | 8608 | 757 | 505 | 0 | 0 | 0 | 42 | 117 | 27 | 262 | 275 | 91 | 274 | 0 | 0 | 624 | 0 | 0 | 0 | 0 | 0 | 0 | 0 | 0 | 0 | 0 | 0 | 0 | 0 | 0 | 0 | 0 | 0 | 0 | 0 | 0 |
| SRR6454199 | 0 | 3528 | 55 | 868 | 1104 | 8109 | 12543 | 29335 | 13795 | 0 | 20783 | 19978 | 148 | 9784 | 0 | 10514 | 49729 | 0 | 0 | 16 | 5761 | 7055 | 1947 | 49 | 15 | 0 | 0 | 0 | 568 | 5342 | 14832 | 2835 | 11 | 40 | 76 | 32 | 0 |
| SRR6454201 | 66 | 12321 | 1518 | 358 | 357 | 4701 | 4087 | 30987 | 10498 | 0 | 23614 | 27706 | 3507 | 5632 | 0 | 5643 | 70795 | 0 | 0 | 0 | 3602 | 5690 | 1762 | 35 | 11 | 0 | 0 | 0 | 234 | 2805 | 9245 | 2691 | 0 | 28 | 40 | 15 | 0 |
| SRR6454203 | 68 | 5623 | 508 | 423 | 1056 | 17313 | 14507 | 59552 | 11664 | 0 | 52883 | 30536 | 5378 | 12155 | 0 | 13662 | 53120 | 0 | 0 | 0 | 2230 | 3207 | 1351 | 124 | 29 | 22 | 0 | 0 | 299 | 3265 | 7503 | 2814 | 0 | 59 | 125 | 86 | 0 |
| SRR6454205 | 37 | 6885 | 320 | 1721 | 689 | 10842 | 8318 | 46355 | 15574 | 74 | 60330 | 26212 | 4469 | 10966 | 0 | 12557 | 38148 | 0 | 0 | 0 | 3784 | 3346 | 999 | 82 | 16 | 16 | 0 | 0 | 308 | 3923 | 6222 | 1421 | 0 | 22 | 47 | 30 | 0 |
| SRR6454206 | 304 | 8299 | 1134 | 332 | 0 | 0 | 0 | 16 | 24 | 28 | 148 | 245 | 82 | 201 | 0 | 0 | 823 | 0 | 0 | 0 | 0 | 0 | 0 | 0 | 0 | 0 | 0 | 0 | 0 | 0 | 0 | 0 | 0 | 0 | 0 | 0 | 0 |
| SRR6454207 | 23 | 6297 | 683 | 1036 | 1120 | 16302 | 17924 | 58359 | 16064 | 34 | 62417 | 27969 | 4890 | 13696 | 0 | 17279 | 51764 | 0 | 0 | 0 | 1975 | 1742 | 979 | 139 | 34 | 19 | 0 | 0 | 205 | 2206 | 2165 | 1224 | 0 | 24 | 28 | 25 | 0 |
| SRR6454208 | 317 | 10395 | 1152 | 202 | 0 | 0 | 0 | 31 | 68 | 29 | 148 | 212 | 73 | 209 | 0 | 0 | 1145 | 0 | 0 | 0 | 0 | 0 | 0 | 0 | 0 | 0 | 0 | 0 | 0 | 0 | 0 | 0 | 0 | 0 | 0 | 0 | 0 |
| SRR6454209 | 25 | 8211 | 370 | 2463 | 695 | 10815 | 9523 | 40915 | 11336 | 59 | 57861 | 26352 | 4491 | 13056 | 0 | 12515 | 41541 | 0 | 0 | 0 | 2912 | 1938 | 560 | 15 | 0 | 0 | 0 | 0 | 235 | 3023 | 4373 | 1143 | 0 | 20 | 32 | 19 | 0 |
| SRR6454211 | 22 | 3537 | 597 | 1296 | 1160 | 16601 | 17269 | 43543 | 9692 | 0 | 39724 | 18662 | 4320 | 14532 | 0 | 13799 | 48072 | 0 | 0 | 0 | 4528 | 5951 | 2003 | 55 | 30 | 13 | 0 | 0 | 404 | 5451 | 10748 | 4132 | 0 | 80 | 160 | 96 | 0 |
| SRR6454213 | 45 | 9031 | 700 | 1355 | 1012 | 18112 | 13277 | 48199 | 14413 | 45 | 59634 | 23202 | 4726 | 11622 | 0 | 14065 | 35484 | 0 | 0 | 0 | 4085 | 2480 | 614 | 19 | 10 | 0 | 0 | 0 | 420 | 5395 | 6239 | 1588 | 0 | 25 | 38 | 28 | 0 |
| SRR6454214 | 242 | 5341 | 1020 | 161 | 0 | 0 | 0 | 28 | 39 | 0 | 161 | 270 | 85 | 329 | 0 | 0 | 698 | 0 | 0 | 0 | 0 | 0 | 0 | 0 | 0 | 0 | 0 | 0 | 0 | 0 | 0 | 0 | 0 | 0 | 0 | 0 | 0 |
| SRR6454215 | 65 | 6344 | 1113 | 678 | 1413 | 22136 | 18737 | 63146 | 14045 | 60 | 67519 | 22839 | 5485 | 14008 | 0 | 14802 | 40627 | 0 | 0 | 0 | 4551 | 3774 | 1091 | 42 | 18 | 0 | 0 | 0 | 439 | 5926 | 9123 | 2914 | 0 | 46 | 67 | 55 | 0 |
| SRR6454216 | 98 | 6908 | 686 | 49 | 0 | 10 | 0 | 43 | 83 | 0 | 286 | 530 | 98 | 206 | 0 | 0 | 612 | 0 | 0 | 0 | 0 | 0 | 0 | 0 | 0 | 0 | 0 | 0 | 0 | 0 | 10 | 0 | 0 | 0 | 0 | 0 | 0 |
| SRR6454217 | 25 | 12656 | 58 | 1080 | 1681 | 14583 | 14968 | 25906 | 27874 | 33 | 20061 | 23823 | 7016 | 3025 | 0 | 9890 | 39887 | 0 | 0 | 11 | 4470 | 2983 | 1771 | 67 | 41 | 0 | 0 | 0 | 482 | 3919 | 7122 | 2586 | 22 | 98 | 69 | 26 | 0 |
| SRR6454218 | 108 | 8661 | 1218 | 81 | 0 | 12 | 12 | 54 | 149 | 27 | 370 | 356 | 115 | 263 | 0 | 0 | 1150 | 0 | 0 | 0 | 0 | 0 | 0 | 0 | 0 | 0 | 0 | 0 | 0 | 0 | 11 | 0 | 0 | 0 | 0 | 0 | 0 |
| SRR6454219 | 35 | 9385 | 174 | 1304 | 961 | 9934 | 10992 | 27065 | 9422 | 0 | 18022 | 15512 | 4953 | 12909 | 0 | 8527 | 67189 | 0 | 0 | 18 | 8411 | 6182 | 2138 | 39 | 28 | 0 | 0 | 0 | 567 | 7311 | 13283 | 3702 | 13 | 87 | 116 | 56 | 0 |
| SRR6454220 | 22 | 5671 | 960 | 112 | 0 | 0 | 0 | 34 | 67 | 0 | 210 | 251 | 64 | 238 | 0 | 0 | 684 | 0 | 0 | 0 | 0 | 0 | 0 | 0 | 0 | 0 | 0 | 0 | 0 | 0 | 0 | 0 | 0 | 0 | 0 | 0 | 0 |
| SRR6454221 | 23 | 6622 | 79 | 1309 | 619 | 5777 | 5938 | 23232 | 17148 | 0 | 30566 | 27805 | 6464 | 7641 | 0 | 5698 | 48099 | 0 | 0 | 27 | 12714 | 3276 | 776 | 0 | 0 | 0 | 0 | 0 | 989 | 10729 | 7623 | 1053 | 0 | 21 | 29 | 10 | 0 |
| SRR6454222 | 0 | 4807 | 917 | 25 | 0 | 12 | 0 | 50 | 82 | 0 | 304 | 387 | 99 | 486 | 0 | 0 | 552 | 0 | 0 | 0 | 10 | 0 | 0 | 0 | 0 | 0 | 0 | 0 | 0 | 0 | 13 | 0 | 0 | 0 | 0 | 0 | 0 |
| SRR6454223 | 0 | 10918 | 72 | 941 | 1583 | 15527 | 16291 | 35454 | 12670 | 0 | 25505 | 21001 | 7140 | 10471 | 0 | 6143 | 54236 | 0 | 0 | 19 | 6800 | 6084 | 1740 | 46 | 23 | 0 | 0 | 0 | 692 | 6638 | 11869 | 2668 | 15 | 63 | 95 | 52 | 0 |
| SRR6454224 | 78 | 3607 | 234 | 26 | 0 | 0 | 0 | 11 | 22 | 0 | 135 | 182 | 42 | 101 | 0 | 0 | 286 | 0 | 0 | 0 | 0 | 0 | 0 | 0 | 0 | 0 | 0 | 0 | 0 | 0 | 0 | 0 | 0 | 0 | 0 | 0 | 0 |
| SRR6454225 | 0 | 6535 | 0 | 1564 | 1030 | 10374 | 9710 | 18275 | 21949 | 0 | 32612 | 21775 | 6970 | 6955 | 0 | 8197 | 39797 | 0 | 0 | 31 | 14259 | 5160 | 966 | 11 | 0 | 0 | 0 | 0 | 919 | 8724 | 10325 | 1119 | 0 | 18 | 45 | 19 | 0 |
| SRR6454226 | 0 | 3224 | 735 | 24 | 0 | 10 | 0 | 34 | 51 | 0 | 283 | 231 | 79 | 202 | 0 | 0 | 283 | 0 | 0 | 0 | 13 | 0 | 0 | 0 | 0 | 0 | 0 | 0 | 0 | 0 | 0 | 0 | 0 | 0 | 0 | 0 | 0 |
| SRR6454227 | 0 | 5920 | 146 | 1273 | 1161 | 9344 | 11418 | 26946 | 9408 | 0 | 17391 | 17448 | 5939 | 10540 | 0 | 8529 | 72855 | 0 | 0 | 18 | 8618 | 4469 | 939 | 19 | 16 | 0 | 0 | 0 | 505 | 5902 | 8720 | 1448 | 0 | 36 | 54 | 20 | 0 |
| SRR6454228 | 52 | 9017 | 538 | 52 | 0 | 0 | 0 | 19 | 33 | 0 | 190 | 256 | 69 | 212 | 0 | 0 | 451 | 0 | 0 | 0 | 0 | 0 | 0 | 0 | 0 | 0 | 0 | 0 | 0 | 0 | 0 | 0 | 0 | 0 | 0 | 0 | 0 |
| SRR6454229 | 0 | 7149 | 33 | 1750 | 907 | 9950 | 8399 | 18488 | 15515 | 0 | 34406 | 25664 | 140 | 7234 | 0 | 10923 | 36347 | 0 | 0 | 24 | 16074 | 3791 | 673 | 0 | 0 | 0 | 0 | 0 | 990 | 10892 | 7291 | 925 | 0 | 19 | 63 | 20 | 0 |
| SRR6454230 | 57 | 5906 | 705 | 38 | 0 | 0 | 0 | 28 | 48 | 0 | 188 | 255 | 80 | 261 | 0 | 0 | 503 | 0 | 0 | 0 | 0 | 0 | 0 | 0 | 0 | 0 | 0 | 0 | 0 | 0 | 0 | 0 | 0 | 0 | 0 | 0 | 0 |
| SRR6454231 | 13 | 7343 | 52 | 1373 | 1468 | 17279 | 16658 | 32024 | 12591 | 0 | 26337 | 23965 | 126 | 8329 | 0 | 7170 | 49683 | 0 | 0 | 18 | 8572 | 4211 | 1227 | 19 | 16 | 0 | 0 | 0 | 576 | 7128 | 7606 | 1893 | 0 | 44 | 67 | 36 | 0 |
| SRR6454232 | 16 | 2542 | 923 | 130 | 0 | 0 | 0 | 25 | 62 | 32 | 212 | 208 | 49 | 129 | 0 | 0 | 708 | 0 | 0 | 0 | 0 | 0 | 0 | 0 | 0 | 0 | 0 | 0 | 0 | 0 | 0 | 0 | 0 | 0 | 0 | 0 | 0 |
| SRR6454233 | 16 | 3110 | 259 | 550 | 1775 | 18866 | 20325 | 36614 | 23502 | 24 | 28571 | 13765 | 69 | 6351 | 0 | 7817 | 41098 | 0 | 0 | 13 | 5132 | 5895 | 1449 | 41 | 23 | 0 | 0 | 0 | 467 | 5616 | 10995 | 2531 | 15 | 56 | 53 | 25 | 0 |
| SRR6454234 | 0 | 4549 | 968 | 508 | 883 | 12370 | 10239 | 33516 | 17456 | 28 | 22713 | 20167 | 123 | 4617 | 0 | 10841 | 48081 | 0 | 0 | 17 | 8297 | 5762 | 1770 | 68 | 39 | 0 | 0 | 0 | 738 | 7650 | 12076 | 3326 | 13 | 99 | 100 | 34 | 0 |
| SRR6454235 | 16 | 5060 | 1188 | 98 | 0 | 0 | 0 | 43 | 123 | 33 | 262 | 406 | 109 | 416 | 0 | 0 | 1264 | 0 | 0 | 0 | 0 | 0 | 0 | 0 | 0 | 0 | 0 | 0 | 0 | 0 | 0 | 0 | 0 | 0 | 0 | 0 | 0 |
| SRR6454236 | 24 | 3441 | 405 | 319 | 1161 | 10289 | 12985 | 27504 | 12660 | 19 | 17633 | 14291 | 58 | 8759 | 0 | 7370 | 45554 | 0 | 0 | 12 | 4047 | 3201 | 1684 | 46 | 33 | 0 | 0 | 0 | 395 | 4206 | 9244 | 3528 | 15 | 132 | 229 | 70 | 0 |
| SRR6454237 | 58 | 2707 | 1500 | 39 | 0 | 0 | 0 | 44 | 66 | 19 | 377 | 489 | 152 | 500 | 0 | 0 | 867 | 0 | 0 | 0 | 0 | 0 | 0 | 0 | 0 | 0 | 0 | 0 | 0 | 0 | 0 | 0 | 0 | 0 | 0 | 0 | 0 |
| SRR6454238 | 0 | 2785 | 482 | 225 | 1313 | 15111 | 15519 | 40120 | 17757 | 72 | 18506 | 6199 | 4046 | 5759 | 0 | 7173 | 47703 | 0 | 0 | 21 | 8160 | 6283 | 1519 | 32 | 29 | 0 | 0 | 0 | 611 | 7314 | 13466 | 2962 | 14 | 72 | 95 | 58 | 0 |
| SRR6454239 | 10 | 1841 | 617 | 62 | 0 | 24 | 19 | 166 | 241 | 0 | 1255 | 1189 | 561 | 1775 | 0 | 58 | 1907 | 0 | 0 | 0 | 0 | 0 | 0 | 0 | 0 | 0 | 0 | 0 | 0 | 0 | 0 | 0 | 0 | 0 | 0 | 0 | 0 |
| SRR6454240 | 16 | 6071 | 451 | 231 | 1128 | 11192 | 9832 | 22948 | 10363 | 59 | 13682 | 19794 | 5832 | 3253 | 0 | 8006 | 46547 | 0 | 0 | 10 | 5366 | 2732 | 716 | 24 | 12 | 0 | 0 | 0 | 422 | 4651 | 7070 | 1483 | 11 | 54 | 77 | 21 | 0 |
| SRR6454241 | 0 | 3697 | 649 | 35 | 0 | 15 | 0 | 74 | 135 | 0 | 411 | 411 | 167 | 558 | 0 | 0 | 959 | 0 | 0 | 0 | 0 | 0 | 0 | 0 | 0 | 0 | 0 | 0 | 0 | 0 | 0 | 0 | 0 | 0 | 0 | 0 | 0 |
| SRR6454242 | 20 | 5234 | 170 | 291 | 1384 | 12808 | 13775 | 29339 | 19241 | 20 | 20515 | 17938 | 59 | 6147 | 0 | 7390 | 54637 | 0 | 0 | 14 | 3818 | 3473 | 2451 | 64 | 40 | 0 | 0 | 0 | 359 | 3796 | 6978 | 4891 | 25 | 169 | 146 | 57 | 0 |
| SRR6454244 | 0 | 12875 | 41 | 936 | 1097 | 10899 | 13215 | 30715 | 16680 | 14 | 22822 | 17504 | 89 | 14570 | 0 | 8264 | 50300 | 0 | 0 | 15 | 5434 | 5628 | 1141 | 28 | 19 | 0 | 0 | 0 | 483 | 5902 | 10891 | 2189 | 11 | 58 | 89 | 40 | 0 |
| SRR6454245 | 0 | 2497 | 933 | 110 | 0 | 0 | 11 | 42 | 117 | 0 | 183 | 236 | 70 | 216 | 0 | 0 | 723 | 0 | 0 | 0 | 0 | 0 | 0 | 0 | 0 | 0 | 0 | 0 | 0 | 0 | 0 | 0 | 0 | 0 | 0 | 0 | 0 |
| SRR6454246 | 0 | 5495 | 25 | 938 | 1079 | 15484 | 15028 | 35317 | 14952 | 0 | 32032 | 24254 | 119 | 11100 | 0 | 10813 | 46092 | 0 | 0 | 18 | 16352 | 13989 | 2001 | 54 | 29 | 0 | 0 | 0 | 740 | 13698 | 14882 | 3002 | 18 | 60 | 41 | 28 | 0 |
| SRR6454247 | 0 | 7655 | 34 | 411 | 995 | 8423 | 10507 | 25428 | 11810 | 0 | 15387 | 17774 | 146 | 6474 | 0 | 5169 | 49698 | 0 | 0 | 17 | 6022 | 2845 | 866 | 36 | 30 | 0 | 0 | 0 | 447 | 6108 | 6453 | 1419 | 11 | 73 | 78 | 26 | 0 |
| SRR6454249 | 0 | 5697 | 158 | 2048 | 716 | 7673 | 7543 | 19249 | 17067 | 0 | 32306 | 24906 | 131 | 7218 | 0 | 7694 | 38371 | 0 | 0 | 41 | 22866 | 10418 | 1612 | 21 | 15 | 0 | 0 | 0 | 1239 | 14896 | 17085 | 1943 | 0 | 28 | 80 | 39 | 0 |
| SRR6454251 | 0 | 4546 | 92 | 1044 | 1408 | 21348 | 17300 | 35459 | 9915 | 0 | 24503 | 20554 | 9027 | 5383 | 0 | 14812 | 36272 | 0 | 0 | 13 | 7416 | 5301 | 1305 | 38 | 27 | 0 | 0 | 0 | 611 | 7219 | 10426 | 2280 | 14 | 68 | 88 | 25 | 0 |
| SRR6454252 | 46 | 879 | 121 | 67 | 0 | 0 | 0 | 14 | 19 | 0 | 35 | 52 | 11 | 65 | 0 | 0 | 146 | 0 | 0 | 0 | 0 | 0 | 0 | 0 | 0 | 0 | 0 | 0 | 0 | 0 | 0 | 0 | 0 | 0 | 0 | 0 | 0 |
| SRR6454253 | 14 | 8583 | 259 | 734 | 707 | 10414 | 9207 | 24306 | 9286 | 0 | 23073 | 17663 | 127 | 8978 | 0 | 6371 | 58209 | 0 | 0 | 0 | 3969 | 3671 | 1647 | 51 | 45 | 0 | 0 | 0 | 372 | 3488 | 7342 | 3146 | 20 | 139 | 114 | 30 | 0 |
| SRR6454254 | 364 | 5603 | 812 | 238 | 0 | 10 | 12 | 61 | 107 | 42 | 373 | 459 | 135 | 335 | 0 | 0 | 721 | 0 | 0 | 0 | 11 | 0 | 0 | 0 | 0 | 0 | 0 | 0 | 0 | 0 | 0 | 0 | 0 | 0 | 0 | 0 | 0 |
| SRR6454255 | 0 | 6508 | 195 | 671 | 880 | 11784 | 9581 | 25543 | 9043 | 0 | 19268 | 20788 | 6727 | 4826 | 0 | 8013 | 48258 | 0 | 0 | 0 | 6098 | 5651 | 1744 | 46 | 47 | 0 | 0 | 0 | 389 | 5363 | 8741 | 3040 | 20 | 140 | 221 | 47 | 0 |
| SRR6454256 | 316 | 5253 | 1392 | 285 | 0 | 0 | 0 | 16 | 18 | 0 | 46 | 83 | 14 | 85 | 0 | 10 | 219 | 0 | 0 | 0 | 0 | 0 | 0 | 0 | 0 | 0 | 0 | 0 | 0 | 0 | 0 | 0 | 0 | 0 | 0 | 0 | 0 |
| SRR6454257 | 57 | 10914 | 194 | 638 | 895 | 13540 | 11168 | 31713 | 14660 | 0 | 15536 | 22416 | 109 | 4200 | 0 | 9349 | 46871 | 0 | 0 | 13 | 8587 | 5494 | 1961 | 66 | 36 | 0 | 0 | 0 | 500 | 6775 | 10245 | 2905 | 22 | 82 | 82 | 22 | 0 |
| SRR6454258 | 0 | 6501 | 49 | 509 | 696 | 7885 | 7539 | 21580 | 19635 | 0 | 28973 | 29014 | 283 | 9142 | 0 | 7274 | 34361 | 0 | 0 | 19 | 9726 | 5073 | 1207 | 42 | 22 | 0 | 0 | 0 | 586 | 9092 | 11722 | 1648 | 0 | 33 | 53 | 24 | 0 |
| SRR6454259 | 0 | 8697 | 166 | 351 | 1103 | 13345 | 13248 | 32509 | 17377 | 0 | 26025 | 21913 | 6827 | 8272 | 0 | 11685 | 42496 | 0 | 0 | 13 | 4969 | 4874 | 1621 | 183 | 71 | 11 | 0 | 0 | 429 | 4359 | 7447 | 2030 | 17 | 94 | 68 | 33 | 0 |
| SRR6454260 | 0 | 5703 | 125 | 316 | 687 | 6887 | 6217 | 13634 | 15759 | 19 | 28830 | 26171 | 369 | 5738 | 0 | 10111 | 31212 | 0 | 0 | 12 | 7144 | 4939 | 1112 | 47 | 17 | 11 | 0 | 0 | 487 | 6884 | 12235 | 1666 | 0 | 47 | 110 | 35 | 0 |
| SRR6454261 | 0 | 3167 | 753 | 275 | 1393 | 13261 | 16337 | 32805 | 11825 | 57 | 23429 | 17088 | 6193 | 11504 | 0 | 14960 | 45449 | 0 | 0 | 28 | 13956 | 9879 | 3371 | 148 | 87 | 31 | 0 | 0 | 983 | 12712 | 19468 | 5190 | 25 | 221 | 359 | 141 | 0 |
| SRR6454262 | 0 | 4849 | 63 | 203 | 982 | 12688 | 11963 | 26532 | 15622 | 0 | 27584 | 18641 | 7389 | 6778 | 0 | 8480 | 39278 | 0 | 0 | 18 | 9066 | 8196 | 2150 | 198 | 84 | 19 | 0 | 0 | 680 | 8713 | 16845 | 3229 | 30 | 155 | 234 | 62 | 0 |
| SRR6454263 | 32 | 28295 | 158 | 95 | 1295 | 15131 | 13248 | 26717 | 25172 | 13 | 16692 | 32695 | 8646 | 2371 | 0 | 10391 | 35226 | 0 | 0 | 10 | 4319 | 3172 | 1212 | 110 | 45 | 0 | 0 | 0 | 405 | 3082 | 6227 | 1709 | 16 | 78 | 54 | 18 | 0 |
| SRR6454264 | 0 | 4959 | 0 | 161 | 877 | 10708 | 10572 | 27443 | 13657 | 0 | 19914 | 21914 | 7718 | 6087 | 0 | 7909 | 40831 | 0 | 0 | 11 | 4538 | 3590 | 1410 | 93 | 57 | 11 | 0 | 0 | 352 | 4054 | 8265 | 2327 | 18 | 117 | 145 | 38 | 0 |
| SRR6454265 | 0 | 4084 | 255 | 135 | 847 | 8677 | 10053 | 25525 | 11849 | 0 | 10 | 18467 | 195 | 10947 | 0 | 5875 | 37175 | 0 | 0 | 11 | 6519 | 3617 | 942 | 57 | 26 | 0 | 0 | 0 | 343 | 6041 | 7716 | 1631 | 0 | 62 | 107 | 28 | 0 |
| SRR6454266 | 0 | 4615 | 130 | 125 | 757 | 8262 | 8263 | 29491 | 22469 | 0 | 19623 | 24298 | 8007 | 3735 | 0 | 8862 | 39884 | 0 | 0 | 0 | 2047 | 2653 | 1209 | 168 | 82 | 0 | 0 | 0 | 256 | 2440 | 7890 | 2443 | 27 | 155 | 107 | 32 | 0 |
| SRR6454267 | 11 | 12138 | 126 | 163 | 1593 | 16093 | 12870 | 29353 | 24157 | 11 | 18407 | 12 | 9492 | 4013 | 0 | 10073 | 33890 | 0 | 0 | 0 | 3944 | 3374 | 1207 | 117 | 63 | 13 | 0 | 0 | 263 | 3415 | 7016 | 2083 | 22 | 125 | 162 | 55 | 0 |
| SRR6454268 | 0 | 11108 | 27 | 589 | 585 | 7788 | 6129 | 17834 | 19041 | 0 | 12 | 25628 | 63 | 9874 | 0 | 6375 | 31502 | 0 | 0 | 17 | 5917 | 2303 | 456 | 18 | 0 | 0 | 0 | 0 | 493 | 6949 | 6376 | 871 | 0 | 20 | 39 | 15 | 0 |
| SRR6454269 | 0 | 10843 | 212 | 238 | 1078 | 11360 | 11307 | 26667 | 14404 | 26 | 21096 | 25438 | 67 | 7835 | 0 | 5784 | 43971 | 0 | 0 | 20 | 8182 | 7998 | 2009 | 120 | 53 | 14 | 0 | 0 | 800 | 8275 | 18612 | 3576 | 17 | 107 | 162 | 47 | 0 |
| SRR6454270 | 14 | 5895 | 195 | 84 | 1372 | 16578 | 15861 | 40447 | 17018 | 14 | 25199 | 18784 | 10600 | 8565 | 0 | 7149 | 32673 | 0 | 0 | 10 | 4119 | 4142 | 1197 | 122 | 82 | 14 | 0 | 0 | 351 | 5081 | 10918 | 2701 | 29 | 178 | 105 | 42 | 0 |
| SRR6454271 | 0 | 5238 | 0 | 2562 | 586 | 4533 | 5163 | 11843 | 8005 | 25 | 15491 | 12 | 4937 | 5136 | 0 | 6458 | 36274 | 0 | 0 | 0 | 3526 | 2217 | 179 | 0 | 0 | 0 | 0 | 0 | 179 | 3391 | 3893 | 347 | 0 | 0 | 0 | 0 | 0 |
| SRR6454272 | 16 | 1769 | 198 | 2245 | 2511 | 12608 | 21497 | 31252 | 5697 | 0 | 11900 | 13862 | 7188 | 6537 | 0 | 8820 | 31524 | 0 | 0 | 15 | 7093 | 3988 | 207 | 0 | 0 | 0 | 0 | 0 | 321 | 7268 | 6742 | 509 | 0 | 0 | 0 | 0 | 0 |
| SRR6454273 | 222 | 31471 | 540 | 2481 | 1786 | 20541 | 20002 | 13042 | 3318 | 62 | 26499 | 15736 | 4013 | 6795 | 0 | 515 | 98136 | 0 | 0 | 0 | 1217 | 661 | 44 | 0 | 0 | 0 | 0 | 35 | 362 | 1641 | 1227 | 112 | 0 | 0 | 0 | 0 | 0 |
| SRR6454274 | 200 | 23890 | 1645 | 1845 | 1371 | 16245 | 16900 | 9891 | 2839 | 86 | 25270 | 14162 | 4148 | 7327 | 0 | 484 | 98542 | 0 | 0 | 0 | 692 | 455 | 41 | 0 | 0 | 0 | 0 | 27 | 238 | 869 | 751 | 103 | 0 | 0 | 0 | 0 | 0 |
| SRR6454275 | 75 | 15078 | 330 | 842 | 655 | 6535 | 6795 | 29880 | 7339 | 11 | 19195 | 18691 | 3149 | 5275 | 0 | 9002 | 40333 | 0 | 0 | 11 | 8343 | 8029 | 1251 | 0 | 0 | 0 | 0 | 0 | 323 | 6081 | 12348 | 1915 | 0 | 13 | 54 | 22 | 0 |
| SRR6454276 | 80 | 10581 | 221 | 1489 | 1151 | 13831 | 11938 | 24909 | 9002 | 10 | 16000 | 20975 | 3651 | 3123 | 0 | 12217 | 37164 | 0 | 0 | 0 | 4389 | 1941 | 353 | 0 | 0 | 0 | 0 | 0 | 192 | 2381 | 1881 | 353 | 0 | 12 | 0 | 0 | 0 |
| SRR6454277 | 79 | 19720 | 163 | 1339 | 869 | 12044 | 9907 | 18897 | 9946 | 15 | 18939 | 13891 | 2808 | 4264 | 0 | 12919 | 32789 | 0 | 0 | 0 | 5046 | 2958 | 216 | 0 | 0 | 0 | 0 | 0 | 252 | 3537 | 3181 | 279 | 0 | 0 | 0 | 0 | 0 |
| SRR6454278 | 25 | 9003 | 221 | 1392 | 1549 | 18610 | 15764 | 26890 | 7003 | 10 | 17111 | 21840 | 4830 | 5805 | 0 | 7807 | 30489 | 0 | 0 | 0 | 8558 | 3608 | 276 | 0 | 0 | 0 | 0 | 0 | 341 | 8507 | 4944 | 573 | 0 | 0 | 24 | 0 | 0 |
| SRR6454279 | 0 | 10616 | 232 | 473 | 625 | 8321 | 7530 | 20230 | 12213 | 0 | 24421 | 20419 | 4510 | 8488 | 0 | 9973 | 34275 | 0 | 0 | 13 | 7014 | 6329 | 1099 | 13 | 0 | 0 | 0 | 0 | 331 | 5709 | 10131 | 1854 | 0 | 11 | 38 | 26 | 0 |
| SRR6454280 | 0 | 9087 | 1399 | 252 | 1243 | 15043 | 15366 | 30099 | 10866 | 0 | 16944 | 16881 | 3638 | 5964 | 0 | 7136 | 56821 | 0 | 0 | 0 | 7100 | 4680 | 704 | 0 | 0 | 0 | 0 | 0 | 222 | 4938 | 6293 | 1075 | 0 | 38 | 50 | 0 | 0 |
| SRR6454281 | 0 | 3630 | 2311 | 322 | 704 | 7082 | 8288 | 30677 | 8938 | 0 | 19425 | 25053 | 5751 | 6830 | 0 | 5111 | 41781 | 0 | 0 | 16 | 12742 | 11803 | 1363 | 15 | 0 | 0 | 0 | 0 | 525 | 12242 | 19933 | 2272 | 0 | 16 | 52 | 22 | 0 |
| SRR6454282 | 0 | 3856 | 98 | 459 | 817 | 7211 | 8071 | 24360 | 10183 | 0 | 20036 | 31335 | 7000 | 5139 | 11 | 5607 | 36483 | 0 | 0 | 0 | 6497 | 4615 | 500 | 10 | 0 | 0 | 0 | 0 | 230 | 5651 | 7515 | 1002 | 0 | 0 | 32 | 15 | 0 |

### Supp Table S8.1.Columns_B_to_AL

| | | | | | | | | | | | | | | | | | | | | | | | | | | | | | | | | | | | | | |
|---|---|---|---|---|---|---|---|---|---|---|---|---|---|---|---|---|---|---|---|---|---|---|---|---|---|---|---|---|---|---|---|---|---|---|---|---|---|
| SRR6454283 | 0 | 3070 | 303 | 485 | 386 | 4265 | 4401 | 24498 | 10683 | 0 | 20064 | 23560 | 4771 | 7043 | 0 | 4865 | 44450 | 0 | 0 | 13 | 9022 | 10584 | 2107 | 26 | 0 | 0 | 0 | 0 | 421 | 7359 | 20349 | 3651 | 0 | 30 | 132 | 70 | 0 |
| SRR6454284 | 16 | 7030 | 303 | 205 | 1176 | 11593 | 12511 | 27346 | 14281 | 22 | 14266 | 35012 | 5829 | 1773 | 0 | 7653 | 44083 | 0 | 0 | 0 | 5125 | 3207 | 435 | 19 | 0 | 0 | 0 | 0 | 167 | 4328 | 5909 | 744 | 0 | 20 | 11 | 0 | 0 |
| SRR6454285 | 66 | 6819 | 546 | 521 | 238 | 4653 | 2906 | 12297 | 5500 | 30 | 16863 | 18430 | 2253 | 5789 | 0 | 3876 | 79509 | 0 | 0 | 0 | 8321 | 14417 | 1424 | 0 | 0 | 0 | 0 | 0 | 222 | 5188 | 13943 | 1911 | 0 | 0 | 13 | 0 | 0 |
| SRR6454286 | 0 | 4735 | 105 | 531 | 1207 | 10621 | 12936 | 23454 | 11985 | 18 | 10187 | 23825 | 3121 | 2009 | 0 | 9983 | 57974 | 0 | 0 | 0 | 6538 | 1937 | 230 | 12 | 0 | 0 | 0 | 0 | 210 | 9401 | 3869 | 466 | 0 | 18 | 16 | 0 | 0 |
| SRR6454287 | 243 | 32742 | 135 | 1431 | 1445 | 18638 | 17206 | 12895 | 3095 | 54 | 43542 | 26532 | 6742 | 4607 | 0 | 353 | 81900 | 0 | 0 | 0 | 25 | 0 | 0 | 0 | 0 | 0 | 0 | 0 | 0 | 22 | 12 | 15 | 0 | 0 | 0 | 0 | 0 |
| SRR6454288 | 206 | 15683 | 537 | 1073 | 1151 | 18864 | 17581 | 13235 | 3584 | 41 | 48986 | 32662 | 8258 | 4120 | 0 | 405 | 102931 | 0 | 0 | 0 | 17 | 0 | 0 | 0 | 0 | 0 | 0 | 0 | 0 | 0 | 0 | 0 | 0 | 0 | 0 | 0 | 0 |
| SRR6454289 | 975 | 34327 | 387 | 790 | 1034 | 15174 | 16036 | 10160 | 2834 | 0 | 40694 | 34385 | 7822 | 3428 | 0 | 492 | 102205 | 0 | 0 | 0 | 22 | 0 | 0 | 0 | 0 | 0 | 0 | 0 | 0 | 25 | 17 | 0 | 0 | 0 | 0 | 0 | 0 |
| SRR6454290 | 547 | 24685 | 257 | 1396 | 2570 | 20318 | 25192 | 14266 | 2256 | 22 | 33424 | 14215 | 5586 | 6337 | 0 | 287 | 73746 | 0 | 0 | 0 | 100 | 19 | 0 | 0 | 0 | 0 | 0 | 0 | 42 | 120 | 48 | 16 | 0 | 0 | 0 | 0 | 0 |
| SRR6454291 | 20 | 9684 | 484 | 121 | 1548 | 15691 | 22696 | 15178 | 6918 | 101 | 32012 | 32004 | 7657 | 5270 | 12 | 375 | 91621 | 0 | 0 | 0 | 111 | 35 | 0 | 0 | 0 | 0 | 0 | 0 | 25 | 108 | 64 | 20 | 0 | 0 | 0 | 0 | 0 |
| SRR6454292 | 20 | 4629 | 363 | 128 | 1518 | 15682 | 24980 | 15008 | 6766 | 49 | 26854 | 26402 | 6203 | 6555 | 0 | 421 | 92329 | 0 | 0 | 0 | 136 | 31 | 12 | 0 | 0 | 0 | 0 | 0 | 34 | 182 | 58 | 32 | 0 | 0 | 0 | 0 | 0 |
| SRR6454293 | 0 | 24613 | 280 | 280 | 814 | 11534 | 13345 | 13492 | 3473 | 35 | 37039 | 29088 | 9647 | 4320 | 0 | 240 | 94296 | 0 | 0 | 0 | 17 | 14 | 0 | 0 | 0 | 0 | 0 | 0 | 0 | 14 | 17 | 0 | 0 | 0 | 0 | 0 | 0 |
| SRR6454294 | 24 | 14376 | 659 | 537 | 1066 | 15170 | 17477 | 13824 | 4114 | 0 | 41836 | 27873 | 5342 | 8095 | 0 | 349 | 101054 | 0 | 0 | 0 | 31 | 13 | 0 | 0 | 0 | 0 | 0 | 0 | 0 | 22 | 20 | 11 | 0 | 0 | 0 | 0 | 0 |
| SRR6454295 | 26 | 18206 | 972 | 276 | 1777 | 17237 | 23185 | 18589 | 3220 | 66 | 36270 | 27644 | 10025 | 5219 | 0 | 206 | 88268 | 0 | 0 | 0 | 107 | 37 | 14 | 0 | 0 | 0 | 0 | 0 | 26 | 128 | 69 | 26 | 0 | 0 | 0 | 0 | 0 |
| SRR6454296 | 53 | 14405 | 677 | 267 | 1444 | 15653 | 21530 | 16560 | 4497 | 53 | 32171 | 27731 | 7930 | 5282 | 0 | 212 | 89127 | 0 | 0 | 0 | 33 | 0 | 10 | 0 | 0 | 0 | 0 | 0 | 0 | 41 | 24 | 25 | 0 | 0 | 0 | 0 | 0 |
| SRR6454298 | 0 | 256 | 93 | 0 | 1147 | 10436 | 15190 | 23127 | 18394 | 16 | 8879 | 8395 | 2117 | 4461 | 0 | 9777 | 68059 | 0 | 0 | 0 | 122 | 368 | 254 | 21 | 10 | 0 | 0 | 0 | 0 | 344 | 950 | 449 | 0 | 32 | 115 | 39 | 0 |
| SRR6454299 | 0 | 1950 | 1357 | 36 | 1514 | 32525 | 23973 | 14339 | 3015 | 107 | 47721 | 30325 | 9980 | 7397 | 22 | 1109 | 101315 | 0 | 0 | 0 | 23 | 0 | 0 | 0 | 0 | 0 | 0 | 0 | 0 | 48 | 13 | 0 | 0 | 0 | 0 | 0 | 0 |
| SRR6454300 | 0 | 1228 | 283 | 22 | 2406 | 31008 | 32517 | 19584 | 4344 | 73 | 28059 | 20287 | 9243 | 6386 | 20 | 836 | 94722 | 0 | 0 | 0 | 62 | 18 | 0 | 0 | 0 | 0 | 0 | 0 | 21 | 157 | 31 | 12 | 0 | 0 | 0 | 0 | 0 |
| SRR6454301 | 13 | 6390 | 320 | 875 | 519 | 5932 | 5807 | 20108 | 9491 | 0 | 20518 | 20129 | 3336 | 4749 | 0 | 8460 | 43331 | 0 | 0 | 0 | 4945 | 2301 | 291 | 0 | 0 | 0 | 0 | 0 | 189 | 4170 | 3763 | 475 | 0 | 0 | 13 | 0 | 0 |
| SRR6454302 | 16 | 7069 | 206 | 303 | 1030 | 13357 | 13068 | 29499 | 12230 | 23 | 12253 | 33745 | 5380 | 1313 | 0 | 12274 | 53557 | 0 | 0 | 0 | 2228 | 1213 | 262 | 12 | 13 | 0 | 0 | 0 | 73 | 1503 | 1243 | 344 | 0 | 34 | 17 | 0 | 0 |
| SRR6454303 | 22 | 6252 | 283 | 244 | 813 | 8883 | 7736 | 20292 | 10524 | 0 | 11545 | 25086 | 3058 | 1503 | 0 | 7386 | 47868 | 0 | 0 | 17 | 11318 | 8125 | 1146 | 17 | 0 | 0 | 0 | 0 | 500 | 6713 | 11785 | 1428 | 0 | 0 | 56 | 19 | 0 |
| SRR6454304 | 86 | 16771 | 301 | 403 | 859 | 12118 | 12413 | 23079 | 5720 | 0 | 14216 | 19306 | 4452 | 5110 | 0 | 12429 | 41693 | 0 | 0 | 14 | 5736 | 10221 | 2092 | 27 | 12 | 0 | 0 | 0 | 429 | 3266 | 8540 | 2518 | 0 | 20 | 28 | 15 | 0 |
| SRR6454305 | 11 | 19203 | 705 | 371 | 486 | 8726 | 6354 | 17486 | 10171 | 32 | 19736 | 11 | 2311 | 6009 | 0 | 6257 | 59471 | 0 | 0 | 12 | 8856 | 8603 | 1650 | 0 | 0 | 0 | 0 | 0 | 289 | 4926 | 8696 | 2053 | 0 | 0 | 10 | 0 | 0 |
| SRR6454306 | 0 | 5948 | 423 | 104 | 1042 | 11401 | 12948 | 28870 | 7518 | 0 | 13124 | 16584 | 3911 | 5232 | 0 | 10831 | 77005 | 0 | 0 | 0 | 4333 | 6744 | 1716 | 26 | 0 | 0 | 0 | 0 | 191 | 2932 | 8430 | 2268 | 0 | 17 | 15 | 0 | 0 |
| SRR6454307 | 0 | 3371 | 196 | 387 | 517 | 6439 | 6252 | 24212 | 10818 | 0 | 22218 | 23538 | 4122 | 6232 | 0 | 5736 | 42561 | 0 | 0 | 0 | 6192 | 4566 | 523 | 0 | 0 | 0 | 0 | 0 | 241 | 5536 | 6561 | 829 | 0 | 11 | 17 | 0 | 0 |
| SRR6454308 | 17 | 13996 | 309 | 144 | 1550 | 17568 | 15702 | 24273 | 12237 | 0 | 17137 | 20 | 7825 | 1682 | 17 | 9289 | 32823 | 0 | 0 | 0 | 7783 | 5371 | 559 | 17 | 0 | 0 | 0 | 0 | 261 | 4896 | 6404 | 818 | 0 | 14 | 20 | 0 | 0 |
| SRR6454309 | 51 | 20985 | 228 | 948 | 698 | 6208 | 6828 | 19632 | 7841 | 67 | 15281 | 17071 | 3685 | 4375 | 0 | 7588 | 40106 | 0 | 0 | 0 | 1953 | 1240 | 180 | 0 | 0 | 0 | 0 | 0 | 119 | 1245 | 1852 | 275 | 0 | 0 | 0 | 0 | 0 |
| SRR6454310 | 47 | 6657 | 197 | 158 | 1575 | 16011 | 17577 | 29581 | 9699 | 20 | 9637 | 27639 | 4620 | 954 | 13 | 12340 | 53125 | 0 | 0 | 0 | 5303 | 1826 | 281 | 23 | 14 | 0 | 0 | 0 | 178 | 3308 | 2689 | 415 | 0 | 43 | 33 | 13 | 0 |
| SRR6454311 | 119 | 39126 | 314 | 714 | 1326 | 18489 | 17069 | 15775 | 2740 | 11 | 35673 | 23828 | 7798 | 4289 | 0 | 229 | 78695 | 0 | 0 | 0 | 21 | 10 | 0 | 0 | 0 | 0 | 0 | 0 | 0 | 25 | 22 | 16 | 0 | 0 | 0 | 0 | 0 |
| SRR6454312 | 623 | 33043 | 660 | 363 | 1547 | 18564 | 19754 | 16107 | 4307 | 22 | 28204 | 24534 | 6642 | 3715 | 12 | 297 | 100752 | 0 | 0 | 0 | 60 | 37 | 14 | 0 | 0 | 0 | 0 | 0 | 39 | 96 | 152 | 49 | 0 | 0 | 12 | 0 | 0 |
| SRR6454313 | 35 | 17169 | 940 | 371 | 1056 | 17081 | 18097 | 14027 | 3525 | 70 | 15 | 23642 | 9020 | 3019 | 0 | 267 | 96703 | 0 | 0 | 0 | 17 | 0 | 0 | 0 | 0 | 0 | 0 | 0 | 0 | 17 | 13 | 0 | 0 | 0 | 0 | 0 | 0 |
| SRR6454314 | 1027 | 50685 | 1474 | 409 | 880 | 15489 | 14976 | 13199 | 5216 | 0 | 37570 | 25490 | 6455 | 4423 | 0 | 291 | 94445 | 0 | 0 | 0 | 70 | 25 | 14 | 0 | 0 | 0 | 0 | 0 | 28 | 117 | 115 | 60 | 0 | 0 | 0 | 0 | 0 |
| SRR6454315 | 173 | 17265 | 743 | 314 | 1266 | 17765 | 18434 | 16333 | 8441 | 26 | 18 | 13 | 9545 | 3984 | 0 | 458 | 90117 | 0 | 0 | 0 | 34 | 16 | 13 | 0 | 0 | 0 | 0 | 0 | 0 | 34 | 22 | 34 | 0 | 0 | 0 | 0 | 0 |
| SRR6454316 | 171 | 15190 | 665 | 273 | 1051 | 15225 | 15798 | 14068 | 6631 | 26 | 46848 | 35969 | 10288 | 4133 | 0 | 380 | 97859 | 0 | 0 | 0 | 40 | 15 | 18 | 0 | 0 | 0 | 0 | 0 | 0 | 35 | 24 | 41 | 0 | 0 | 0 | 0 | 0 |
| SRR6454317 | 20 | 22723 | 5207 | 112 | 983 | 16064 | 15321 | 11646 | 5230 | 0 | 48216 | 44646 | 11453 | 4059 | 0 | 199 | 111827 | 0 | 0 | 0 | 19 | 0 | 0 | 0 | 0 | 0 | 0 | 0 | 0 | 14 | 0 | 17 | 0 | 0 | 0 | 0 | 0 |
| SRR6454318 | 43 | 18444 | 9206 | 150 | 912 | 16020 | 15552 | 12268 | 5048 | 21 | 47177 | 40204 | 11898 | 4171 | 0 | 236 | 108701 | 0 | 0 | 0 | 17 | 0 | 0 | 0 | 0 | 0 | 0 | 0 | 0 | 14 | 0 | 0 | 0 | 0 | 0 | 0 | 0 |
| SRR6454319 | 37 | 18083 | 1110 | 209 | 1472 | 20378 | 22408 | 16345 | 6825 | 30 | 51276 | 49566 | 12978 | 4199 | 0 | 519 | 89607 | 0 | 0 | 0 | 33 | 16 | 12 | 0 | 0 | 0 | 0 | 0 | 0 | 30 | 33 | 26 | 0 | 0 | 0 | 0 | 0 |
| SRR6454320 | 68 | 16757 | 1153 | 536 | 1736 | 20797 | 21104 | 19225 | 4806 | 46 | 42875 | 29018 | 7035 | 5621 | 0 | 299 | 94546 | 0 | 0 | 0 | 31 | 11 | 11 | 0 | 0 | 0 | 0 | 0 | 0 | 25 | 12 | 35 | 0 | 11 | 0 | 0 | 0 |
| SRR6454321 | 30 | 28777 | 378 | 438 | 789 | 15568 | 14114 | 12438 | 4052 | 30 | 41092 | 26931 | 9191 | 3080 | 0 | 395 | 110561 | 0 | 0 | 0 | 14 | 0 | 0 | 0 | 0 | 0 | 0 | 0 | 0 | 11 | 12 | 0 | 0 | 0 | 0 | 0 | 0 |
| SRR6454322 | 93 | 23969 | 691 | 572 | 894 | 14593 | 14094 | 12411 | 3642 | 13 | 33335 | 24993 | 7610 | 2891 | 0 | 388 | 110367 | 0 | 0 | 0 | 16 | 14 | 0 | 0 | 0 | 0 | 0 | 0 | 0 | 15 | 18 | 0 | 0 | 0 | 0 | 0 | 0 |
| SRR6454323 | 2236 | 106221 | 1141 | 274 | 1565 | 20151 | 20027 | 17665 | 3919 | 14 | 29934 | 23421 | 5610 | 3430 | 0 | 437 | 101297 | 0 | 0 | 0 | 33 | 17 | 0 | 0 | 0 | 0 | 0 | 0 | 23 | 60 | 48 | 22 | 0 | 0 | 0 | 0 | 0 |
| SRR6454324 | 72 | 21003 | 1938 | 404 | 1304 | 20864 | 20284 | 17866 | 2583 | 42 | 31140 | 27094 | 8771 | 2936 | 0 | 517 | 101193 | 0 | 0 | 0 | 20 | 0 | 0 | 0 | 0 | 0 | 0 | 0 | 0 | 13 | 0 | 12 | 0 | 0 | 0 | 0 | 0 |
| SRR6454325 | 36 | 13687 | 176 | 641 | 447 | 5454 | 5456 | 20256 | 7244 | 16 | 15824 | 11 | 3951 | 4707 | 0 | 6388 | 69726 | 0 | 0 | 14 | 7748 | 4531 | 700 | 10 | 0 | 0 | 0 | 0 | 286 | 4182 | 6669 | 845 | 0 | 0 | 36 | 12 | 0 |
| SRR6454326 | 127 | 15931 | 251 | 149 | 1892 | 13511 | 16333 | 20534 | 9149 | 13 | 10735 | 20 | 4757 | 1309 | 24 | 10330 | 36965 | 0 | 0 | 16 | 3033 | 738 | 151 | 0 | 0 | 0 | 0 | 0 | 409 | 1357 | 1127 | 200 | 0 | 0 | 0 | 0 | 0 |
| SRR6454331 | 107 | 31383 | 620 | 417 | 1527 | 19790 | 21526 | 16223 | 2503 | 32 | 34303 | 18 | 8686 | 3203 | 0 | 280 | 106627 | 0 | 0 | 0 | 27 | 0 | 0 | 0 | 0 | 0 | 0 | 0 | 0 | 18 | 14 | 12 | 0 | 0 | 0 | 0 | 0 |
| SRR6454332 | 564 | 16731 | 711 | 361 | 1367 | 18529 | 22059 | 15800 | 3176 | 45 | 33557 | 28020 | 6878 | 3589 | 0 | 305 | 107392 | 0 | 0 | 0 | 39 | 0 | 0 | 0 | 0 | 0 | 0 | 0 | 12 | 33 | 19 | 20 | 0 | 0 | 0 | 0 | 0 |
| SRR6454335 | 34 | 40894 | 1058 | 0 | 1017 | 17791 | 12281 | 9600 | 2816 | 102 | 40463 | 37737 | 9442 | 4698 | 18 | 824 | 123888 | 0 | 0 | 0 | 705 | 224 | 26 | 0 | 0 | 0 | 0 | 0 | 0 | 480 | 245 | 44 | 0 | 20 | 0 | 0 | 0 |
| SRR6454337 | 0 | 641 | 437 | 50 | 1704 | 15050 | 20917 | 49833 | 19666 | 27 | 9110 | 9872 | 2370 | 3832 | 0 | 12745 | 67770 | 0 | 0 | 0 | 190 | 601 | 255 | 12 | 0 | 0 | 0 | 0 | 17 | 516 | 1597 | 570 | 0 | 15 | 29 | 15 | 0 |
| SRR6454338 | 0 | 124 | 82 | 26 | 1514 | 10263 | 16799 | 30231 | 9866 | 31 | 7773 | 9501 | 2852 | 3633 | 0 | 8074 | 53253 | 0 | 0 | 0 | 69 | 296 | 181 | 15 | 0 | 0 | 0 | 0 | 0 | 44 | 469 | 191 | 0 | 0 | 32 | 15 | 0 |
| SRR6454339 | 0 | 89 | 120 | 28 | 2043 | 12697 | 15642 | 33358 | 14512 | 0 | 10868 | 12857 | 11 | 4606 | 0 | 7860 | 45998 | 0 | 0 | 0 | 296 | 327 | 209 | 13 | 0 | 0 | 0 | 0 | 40 | 471 | 1011 | 377 | 0 | 0 | 23 | 22 | 0 |
| SRR6454340 | 0 | 398 | 314 | 0 | 1779 | 18140 | 20978 | 26237 | 21579 | 90 | 4876 | 9578 | 1992 | 678 | 0 | 12822 | 55865 | 0 | 0 | 0 | 522 | 215 | 39 | 0 | 0 | 0 | 0 | 0 | 17 | 1081 | 519 | 76 | 0 | 0 | 0 | 0 | 0 |
| SRR6454341 | 0 | 103 | 52 | 11 | 1053 | 6856 | 10232 | 14238 | 13515 | 15 | 4207 | 6797 | 1171 | 1470 | 0 | 7989 | 68712 | 0 | 0 | 0 | 73 | 150 | 101 | 10 | 0 | 0 | 0 | 0 | 13 | 173 | 388 | 183 | 0 | 0 | 18 | 12 | 0 |
| SRR6454342 | 0 | 123 | 59 | 19 | 1651 | 11894 | 16361 | 17940 | 23586 | 31 | 4784 | 8776 | 1612 | 1144 | 0 | 8202 | 60332 | 0 | 0 | 0 | 116 | 313 | 167 | 20 | 0 | 0 | 0 | 0 | 17 | 332 | 1041 | 296 | 0 | 0 | 27 | 21 | 0 |
| SRR6454343 | 0 | 794 | 106 | 44 | 1693 | 10851 | 14825 | 16809 | 38044 | 20 | 8068 | 14150 | 2525 | 2049 | 0 | 10759 | 32087 | 0 | 0 | 0 | 322 | 897 | 392 | 29 | 0 | 0 | 0 | 0 | 55 | 942 | 3103 | 657 | 0 | 0 | 21 | 15 | 0 |
| SRR6454345 | 41 | 4404 | 678 | 50 | 1674 | 27909 | 24893 | 11401 | 2319 | 25 | 56851 | 26039 | 9770 | 8140 | 0 | 673 | 90271 | 0 | 0 | 0 | 0 | 0 | 0 | 0 | 0 | 0 | 0 | 0 | 0 | 21 | 0 | 11 | 0 | 0 | 0 | 0 | 0 |
| SRR6454347 | 0 | 7834 | 394 | 0 | 1385 | 20400 | 13722 | 9014 | 2412 | 17 | 41748 | 14177 | 4834 | 8583 | 0 | 338 | 139691 | 0 | 0 | 0 | 50 | 21 | 20 | 0 | 0 | 0 | 0 | 0 | 0 | 58 | 46 | 72 | 0 | 31 | 23 | 0 | 0 |
| SRR6454348 | 17 | 26126 | 660 | 0 | 1081 | 18215 | 12240 | 10452 | 2996 | 23 | 41163 | 27708 | 8990 | 4794 | 20 | 330 | 128086 | 0 | 0 | 0 | 41 | 26 | 16 | 0 | 0 | 0 | 0 | 0 | 0 | 23 | 22 | 32 | 0 | 0 | 13 | 0 | 0 |
| SRR6454349 | 14 | 24085 | 563 | 0 | 535 | 11226 | 7023 | 4820 | 3201 | 19 | 54442 | 44805 | 11984 | 4145 | 12 | 227 | 135164 | 0 | 0 | 0 | 39 | 13 | 18 | 0 | 0 | 0 | 0 | 0 | 0 | 25 | 18 | 28 | 0 | 11 | 0 | 0 | 0 |
| SRR6454350 | 19 | 15043 | 626 | 0 | 631 | 16220 | 10640 | 6616 | 3007 | 0 | 50505 | 24176 | 4284 | 6746 | 0 | 425 | 137948 | 0 | 0 | 0 | 23 | 0 | 0 | 0 | 0 | 0 | 0 | 0 | 0 | 21 | 15 | 22 | 0 | 11 | 0 | 0 | 0 |
| SRR6454351 | 0 | 4827 | 195 | 0 | 1929 | 17354 | 15081 | 25137 | 9124 | 15 | 14655 | 33205 | 6125 | 1889 | 27 | 13444 | 38173 | 0 | 0 | 0 | 2929 | 3122 | 687 | 31 | 20 | 18 | 0 | 0 | 0 | 3013 | 8787 | 1472 | 23 | 55 | 180 | 74 | 28 |
| SRR6454352 | 0 | 2510 | 118 | 0 | 1516 | 20809 | 13662 | 37590 | 9083 | 0 | 17108 | 21834 | 5429 | 3576 | 17 | 18931 | 49291 | 0 | 0 | 0 | 1240 | 4938 | 1956 | 74 | 52 | 54 | 0 | 0 | 0 | 2079 | 10757 | 4875 | 65 | 209 | 1138 | 545 | 42 |
| SRR6454354 | 0 | 2719 | 77 | 0 | 1149 | 17189 | 9963 | 26345 | 7618 | 0 | 17649 | 19261 | 5053 | 3890 | 0 | 14075 | 36719 | 0 | 0 | 0 | 1890 | 3076 | 1165 | 44 | 59 | 39 | 0 | 0 | 0 | 2540 | 7302 | 2223 | 27 | 223 | 801 | 333 | 27 |
| SRR6496456 | 24 | 57 | 468 | 385 | 26681 | 19139 | 213179 | 82167 | 511 | 65 | 8459 | 5170 | 6503 | 10321 | 5261 | 28098 | 13151 | 11 | 117 | 20 | 102 | 153 | 115 | 821 | 781 | 2102 | 0 | 285 | 72 | 25 | 55 | 0 | 0 | 0 | 315 | 544 | 0 |
| SRR6496457 | 22 | 62 | 325 | 644 | 21715 | 19346 | 167809 | 67491 | 492 | 38 | 10815 | 4785 | 7320 | 12730 | 5753 | 27192 | 20652 | 0 | 75 | 10 | 70 | 136 | 147 | 859 | 1329 | 2598 | 0 | 192 | 52 | 11 | 42 | 0 | 0 | 0 | 380 | 577 | 0 |
| SRR6496458 | 25 | 68 | 422 | 554 | 24041 | 18976 | 179933 | 70568 | 637 | 73 | 8167 | 3194 | 6726 | 10917 | 5942 | 25974 | 16831 | 17 | 119 | 13 | 97 | 168 | 152 | 962 | 680 | 2534 | 0 | 319 | 77 | 23 | 56 | 0 | 0 | 0 | 227 | 358 | 0 |
| SRR6496459 | 0 | 44 | 247 | 213 | 22210 | 19391 | 185386 | 72576 | 601 | 40 | 9476 | 4565 | 7522 | 14090 | 7171 | 28964 | 15116 | 0 | 35 | 0 | 45 | 68 | 62 | 511 | 680 | 1768 | 0 | 99 | 30 | 0 | 25 | 0 | 0 | 0 | 229 | 416 | 0 |
| SRR6496460 | 0 | 61 | 323 | 201 | 24310 | 21645 | 219783 | 71937 | 564 | 53 | 6258 | 4113 | 6111 | 12160 | 6449 | 31489 | 13822 | 0 | 62 | 0 | 65 | 104 | 120 | 588 | 457 | 1543 | 0 | 159 | 52 | 17 | 36 | 0 | 0 | 0 | 139 | 286 | 0 |
| SRR6496461 | 281 | 33257 | 485 | 1308 | 12520 | 85673 | 85659 | 41023 | 506 | 22 | 29373 | 28631 | 6015 | 7049 | 68 | 106731 | 12886 | 0 | 0 | 0 | 688 | 142 | 55 | 0 | 19 | 56 | 0 | 0 | 79 | 193 | 109 | 33 | 0 | 49 | 259 | 133 | 0 |
| SRR6496462 | 281 | 33291 | 463 | 1291 | 12607 | 86182 | 85869 | 41237 | 611 | 18 | 28644 | 28252 | 6014 | 6888 | 66 | 104967 | 12868 | 0 | 0 | 0 | 620 | 138 | 51 | 0 | 17 | 50 | 0 | 0 | 79 | 185 | 105 | 31 | 0 | 40 | 253 | 127 | 0 |
| SRR6496463 | 285 | 32827 | 509 | 1233 | 12458 | 86542 | 86062 | 41220 | 590 | 21 | 27585 | 26984 | 5858 | 6823 | 52 | 107176 | 13802 | 0 | 0 | 0 | 661 | 146 | 55 | 0 | 20 | 49 | 0 | 0 | 69 | 186 | 106 | 35 | 0 | 41 | 233 | 124 | 0 |
| SRR6496464 | 303 | 33231 | 452 | 1264 | 12648 | 85661 | 85480 | 41123 | 557 | 23 | 28692 | 27847 | 5957 | 6952 | 50 | 105781 | 12925 | 0 | 0 | 0 | 658 | 127 | 54 | 0 | 16 | 52 | 0 | 0 | 79 | 202 | 116 | 35 | 0 | 44 | 254 | 134 | 0 |
| SRR6496465 | 359 | 32514 | 373 | 1315 | 12871 | 86244 | 85951 | 41521 | 524 | 14 | 27919 | 26704 | 5752 | 7013 | 45 | 107170 | 13405 | 0 | 0 | 0 | 629 | 121 | 39 | 0 | 11 | 55 | 0 | 0 | 63 | 215 | 106 | 27 | 0 | 42 | 241 | 100 | 0 |
| SRR6498854 | 112 | 231123 | 205 | 497 | 2613 | 30909 | 19255 | 13670 | 44 | 20 | 22860 | 56017 | 3707 | 5706 | 0 | 32247 | 35990 | 0 | 0 | 0 | 698 | 223 | 561 | 16 | 10 | 0 | 0 | 0 | 41 | 226 | 138 | 131 | 0 | 11 | 0 | 0 | 0 |
| SRR6498855 | 98 | 229248 | 358 | 499 | 1893 | 29767 | 28701 | 10548 | 65 | 17 | 17163 | 49384 | 3088 | 5334 | 0 | 36063 | 28047 | 0 | 0 | 0 | 526 | 168 | 361 | 14 | 0 | 0 | 0 | 0 | 41 | 159 | 93 | 76 | 0 | 0 | 0 | 0 | 0 |
| SRR6498856 | 153 | 204323 | 312 | 507 | 2456 | 32866 | 19344 | 16260 | 70 | 13 | 25254 | 43099 | 3039 | 8037 | 0 | 33614 | 37266 | 0 | 0 | 0 | 656 | 281 | 520 | 18 | 11 | 0 | 0 | 0 | 38 | 228 | 144 | 121 | 0 | 12 | 0 | 0 | 0 |
| SRR6498857 | 66 | 168998 | 257 | 1240 | 1691 | 26557 | 26146 | 11133 | 98 | 19 | 13092 | 48669 | 3029 | 5668 | 11 | 31332 | 31149 | 0 | 0 | 0 | 671 | 318 | 537 | 25 | 16 | 0 | 0 | 0 | 74 | 211 | 190 | 163 | 0 | 17 | 0 | 0 | 0 |
| SRR6498858 | 154 | 128127 | 186 | 959 | 2322 | 24832 | 16044 | 11185 | 44 | 24 | 22638 | 59842 | 3945 | 5374 | 14 | 27108 | 45326 | 0 | 0 | 0 | 703 | 469 | 537 | 20 | 11 | 0 | 0 | 0 | 38 | 223 | 203 | 130 | 0 | 0 | 0 | 0 | 0 |
| SRR6498860 | 73 | 189307 | 317 | 381 | 1971 | 23935 | 16507 | 9483 | 56 | 17 | 20254 | 44852 | 3060 | 5286 | 0 | 29679 | 37206 | 0 | 0 | 0 | 555 | 190 | 402 | 14 | 0 | 0 | 0 | 0 | 53 | 184 | 113 | 97 | 0 | 0 | 0 | 0 | 0 |
| SRR6498861 | 49 | 164057 | 327 | 733 | 2682 | 49889 | 61193 | 15514 | 63 | 33 | 26543 | 69924 | 5615 | 7618 | 16 | 73345 | 24532 | 0 | 0 | 0 | 361 | 118 | 147 | 16 | 0 | 0 | 0 | 0 | 31 | 170 | 72 | 59 | 0 | 0 | 0 | 0 | 0 |
| SRR6498862 | 118 | 110437 | 165 | 880 | 2168 | 24801 | 20273 | 13924 | 101 | 17 | 17656 | 42669 | 3264 | 6243 | 0 | 23903 | 41707 | 0 | 0 | 0 | 571 | 347 | 346 | 15 | 0 | 0 | 0 | 0 | 52 | 163 | 207 | 119 | 0 | 11 | 0 | 0 | 0 |
| SRR6498863 | 115 | 93583 | 149 | 575 | 2097 | 31638 | 38345 | 11296 | 56 | 0 | 18821 | 63816 | 3646 | 6025 | 25 | 39674 | 41954 | 0 | 0 | 0 | 497 | 308 | 351 | 19 | 14 | 0 | 0 | 0 | 54 | 158 | 169 | 100 | 0 | 18 | 15 | 0 | 0 |
| SRR6498864 | 251 | 64483 | 91 | 683 | 2166 | 21799 | 17644 | 9214 | 40 | 24 | 18857 | 66139 | 4291 | 5454 | 15 | 24125 | 44343 | 0 | 0 | 0 | 616 | 615 | 397 | 13 | 0 | 0 | 0 | 0 | 33 | 204 | 319 | 113 | 0 | 0 | 11 | 0 | 0 |
| SRR6498865 | 150 | 54135 | 141 | 713 | 2238 | 25458 | 30445 | 10072 | 85 | 0 | 20798 | 76132 | 4863 | 6630 | 26 | 33370 | 29562 | 0 | 0 | 0 | 395 | 405 | 365 | 25 | 0 | 0 | 0 | 0 | 25 | 147 | 238 | 119 | 0 | 0 | 30 | 18 | 0 |
| SRR6498866 | 102 | 178792 | 311 | 428 | 1840 | 22137 | 12656 | 9212 | 20 | 10 | 25504 | 66472 | 3981 | 5675 | 21 | 25341 | 27312 | 0 | 0 | 0 | 663 | 297 | 694 | 21 | 0 | 0 | 0 | 0 | 41 | 131 | 92 | 114 | 0 | 0 | 0 | 0 | 0 |
| SRR6498867 | 70 | 116616 | 531 | 947 | 1188 | 18326 | 20234 | 7437 | 22 | 11 | 18649 | 90448 | 3448 | 4903 | 0 | 28159 | 21556 | 0 | 0 | 0 | 159 | 84 | 156 | 0 | 0 | 0 | 0 | 0 | 12 | 33 | 20 | 19 | 0 | 0 | 0 | 0 | 0 |
| SRR6498868 | 150 | 104891 | 273 | 1195 | 1347 | 14457 | 9169 | 6416 | 25 | 11 | 21716 | 88097 | 3475 | 4243 | 15 | 18887 | 29971 | 0 | 0 | 0 | 328 | 252 | 457 | 12 | 0 | 0 | 0 | 0 | 26 | 45 | 52 | 52 | 0 | 0 | 0 | 0 | 0 |
| SRR6498869 | 115 | 121824 | 433 | 589 | 1731 | 22748 | 24006 | 10133 | 45 | 22 | 15382 | 72104 | 3252 | 4799 | 22 | 28741 | 26818 | 0 | 0 | 0 | 490 | 374 | 540 | 12 | 12 | 0 | 0 | 0 | 28 | 111 | 133 | 95 | 0 | 0 | 0 | 0 | 0 |
| SRR6498870 | 188 | 98092 | 287 | 1054 | 1765 | 22858 | 12858 | 7250 | 14 | 17 | 32753 | 85796 | 5179 | 6621 | 22 | 33151 | 29153 | 0 | 0 | 0 | 449 | 412 | 344 | 0 | 0 | 0 | 0 | 0 | 28 | 92 | 108 | 61 | 0 | 0 | 0 | 0 | 0 |
| SRR6498871 | 129 | 129716 | 334 | 623 | 1089 | 16989 | 18854 | 7517 | 28 | 22 | 10630 | 67499 | 2006 | 3943 | 12 | 17443 | 22679 | 0 | 0 | 0 | 394 | 453 | 676 | 13 | 11 | 0 | 0 | 0 | 25 | 80 | 130 | 100 | 0 | 0 | 14 | 0 | 0 |
| SRR6705804 | 0 | 90 | 924 | 0 | 620 | 8821 | 9277 | 7666 | 9177 | 181 | 7297 | 8840 | 4093 | 5802 | 302 | 1650 | 39187 | 0 | 0 | 0 | 32 | 480 | 39 | 0 | 0 | 0 | 0 | 13 | 609 | 183 | 1272 | 69 | 0 | 0 | 0 | 19 | 10 |
| SRR6705805 | 0 | 87 | 282 | 0 | 205 | 2794 | 3514 | 2629 | 3469 | 97 | 2313 | 2430 | 1751 | 3173 | 145 | 1605 | 18361 | 0 | 0 | 0 | 36 | 479 | 37 | 0 | 0 | 0 | 0 | 16 | 1119 | 396 | 5565 | 184 | 0 | 0 | 0 | 0 | 0 |
| SRR6705806 | 0 | 0 | 0 | 0 | 1225 | 18382 | 18382 | 12255 | 7353 | 0 | 7353 | 4902 | 3676 | 4902 | 0 | 0 | 20833 | 0 | 0 | 0 | 0 | 2370 | 0 | 0 | 0 | 0 | 0 | 0 | 395 | 0 | 3752 | 0 | 0 | 0 | 0 | 0 | 0 |
| SRR6705808 | 0 | 3143 | 1403 | 0 | 329 | 3375 | 5841 | 6175 | 4956 | 214 | 24067 | 25275 | 12024 | 3654 | 463 | 1355 | 74611 | 0 | 0 | 13 | 211 | 9242 | 981 | 16 | 0 | 0 | 0 | 31 | 550 | 153 | 3462 | 222 | 0 | 0 | 43 | 92 | 0 |
| SRR6705809 | 0 | 213 | 1172 | 0 | 1326 | 18260 | 13646 | 11588 | 4298 | 80 | 9972 | 10742 | 6053 | 6358 | 1562 | 1005 | 51058 | 0 | 0 | 0 | 15 | 789 | 101 | 0 | 0 | 0 | 0 | 0 | 69 | 46 | 913 | 36 | 0 | 0 | 26 | 23 | 0 |
| SRR6705810 | 0 | 6405 | 1032 | 0 | 744 | 10227 | 11311 | 9292 | 2312 | 110 | 7711 | 6940 | 7936 | 5679 | 511 | 2636 | 31171 | 0 | 19 | 110 | 1326 | 12102 | 209 | 0 | 0 | 0 | 0 | 81 | 1826 | 483 | 4167 | 73 | 0 | 0 | 11 | 29 | 10 |
| SRR6705811 | 0 | 2737 | 249 | 0 | 386 | 2963 | 6741 | 7181 | 492 | 55 | 3171 | 3705 | 1943 | 4200 | 64 | 829 | 13116 | 0 | 19 | 95 | 2150 | 12142 | 481 | 0 | 0 | 0 | 0 | 135 | 2502 | 1250 | 3839 | 96 | 0 | 0 | 0 | 22 | 0 |
| SRR6705812 | 0 | 103 | 334 | 11 | 764 | 6250 | 14602 | 7790 | 4787 | 448 | 7638 | 3340 | 4164 | 4742 | 1536 | 2267 | 23287 | 0 | 0 | 12 | 59 | 331 | 15 | 0 | 0 | 0 | 0 | 24 | 249 | 55 | 1157 | 58 | 0 | 0 | 0 | 14 | 0 |
| SRR6705813 | 0 | 121 | 1383 | 0 | 444 | 3790 | 9469 | 7013 | 4414 | 685 | 9588 | 5706 | 5678 | 4234 | 426 | 1771 | 35983 | 0 | 0 | 10 | 52 | 332 | 21 | 0 | 0 | 0 | 0 | 27 | 182 | 63 | 460 | 39 | 0 | 0 | 0 | 24 | 0 |
| SRR6705814 | 23 | 334 | 1166 | 394 | 264 | 964 | 29681 | 23183 | 5152 | 100 | 3294 | 1458 | 2266 | 9504 | 1329 | 2763 | 13818 | 0 | 0 | 29 | 699 | 18170 | 1280 | 0 | 0 | 0 | 0 | 30 | 228 | 552 | 9968 | 1513 | 0 | 0 | 32 | 43 | 83 |
| SRR6705815 | 22 | 516 | 777 | 955 | 2765 | 6913 | 50587 | 33693 | 4486 | 386 | 27020 | 5983 | 12291 | 4705 | 1061 | 1014 | 45049 | 0 | 41 | 54 | 1510 | 7452 | 522 | 0 | 0 | 0 | 0 | 253 | 346 | 1056 | 5552 | 864 | 0 | 0 | 17 | 54 | 49 |
| SRR6705816 | 0 | 241 | 653 | 30 | 1505 | 4514 | 30287 | 13687 | 4347 | 310 | 15237 | 5894 | 9243 | 6141 | 1118 | 1771 | 42228 | 0 | 19 | 64 | 1856 | 8442 | 294 | 0 | 0 | 0 | 0 | 124 | 591 | 1283 | 3267 | 272 | 0 | 0 | 0 | 46 | 11 |
| SRR6705817 | 0 | 1762 | 1122 | 0 | 5931 | 29306 | 106204 | 42025 | 4698 | 163 | 28892 | 15891 | 13979 | 15268 | 11677 | 612 | 29619 | 0 | 14 | 30 | 346 | 5178 | 112 | 0 | 0 | 0 | 0 | 78 | 182 | 213 | 1940 | 97 | 0 | 0 | 0 | 28 | 0 |
| SRR6705818 | 52 | 834 | 1128 | 330 | 2045 | 7695 | 33993 | 19242 | 4852 | 285 | 18517 | 9789 | 10961 | 7267 | 1011 | 2337 | 33807 | 0 | 0 | 54 | 1002 | 6168 | 171 | 0 | 0 | 0 | 0 | 25 | 329 | 422 | 1903 | 158 | 0 | 0 | 0 | 27 | 0 |
| SRR6705819 | 62 | 1980 | 1312 | 1191 | 4801 | 21966 | 75065 | 51594 | 3439 | 175 | 22317 | 16576 | 10692 | 12340 | 2130 | 948 | 28870 | 0 | 32 | 117 | 1800 | 7180 | 157 | 0 | 0 | 0 | 0 | 81 | 508 | 666 | 1471 | 77 | 0 | 0 | 0 | 16 | 0 |
| SRR6705820 | 0 | 247 | 150 | 21 | 1806 | 4243 | 33387 | 19764 | 7254 | 101 | 19565 | 7222 | 8264 | 4312 | 1430 | 1706 | 45160 | 0 | 0 | 22 | 858 | 4066 | 303 | 0 | 0 | 0 | 0 | 50 | 173 | 588 | 1985 | 400 | 0 | 0 | 0 | 14 | 25 |
| SRR6705821 | 32 | 676 | 351 | 127 | 4735 | 8496 | 77996 | 51080 | 6931 | 265 | 25494 | 11063 | 10401 | 10798 | 1898 | 2511 | 33491 | 0 | 27 | 80 | 2046 | 7129 | 402 | 0 | 0 | 0 | 0 | 150 | 330 | 1768 | 3579 | 732 | 0 | 0 | 0 | 68 | 86 |
| SRR6705822 | 0 | 162 | 233 | 52 | 2738 | 5629 | 34234 | 11943 | 3072 | 152 | 13283 | 4898 | 3780 | 1917 | 3960 | 975 | 37836 | 0 | 0 | 13 | 179 | 686 | 53 | 0 | 0 | 0 | 0 | 33 | 51 | 600 | 1214 | 134 | 0 | 0 | 0 | 0 | 45 |
| SRR6705823 | 16 | 3566 | 310 | 77 | 13131 | 24786 | 95119 | 35550 | 7969 | 191 | 31558 | 15260 | 13069 | 5721 | 10884 | 1023 | 28430 | 0 | 21 | 89 | 922 | 3601 | 212 | 0 | 0 | 0 | 0 | 62 | 101 | 216 | 570 | 69 | 0 | 0 | 0 | 33 | 12 |
| SRR6705824 | 0 | 466 | 797 | 24 | 134 | 2127 | 3632 | 3329 | 4121 | 0 | 7323 | 9891 | 3119 | 3457 | 154 | 1483 | 30913 | 0 | 0 | 0 | 60 | 1911 | 139 | 0 | 0 | 0 | 0 | 0 | 139 | 459 | 5354 | 287 | 0 | 0 | 0 | 17 | 15 |
| SRR6705825 | 0 | 277 | 1402 | 14 | 1055 | 5477 | 18693 | 10183 | 1878 | 127 | 9702 | 11949 | 5760 | 5317 | 462 | 1087 | 23599 | 0 | 0 | 0 | 91 | 1470 | 263 | 0 | 0 | 0 | 0 | 0 | 131 | 777 | 9404 | 224 | 0 | 0 | 0 | 23 | 15 |
| SRR6705826 | 0 | 20 | 139 | 35 | 249 | 2037 | 7045 | 4744 | 3175 | 50 | 6973 | 5884 | 4762 | 6801 | 723 | 2382 | 59604 | 0 | 0 | 0 | 112 | 3866 | 161 | 0 | 0 | 0 | 0 | 11 | 128 | 185 | 7494 | 121 | 0 | 13 | 0 | 21 | 103 |
| SRR6705827 | 0 | 361 | 1352 | 67 | 717 | 6183 | 12998 | 11381 | 10781 | 25 | 11330 | 11560 | 9720 | 5676 | 288 | 1255 | 33001 | 0 | 0 | 12 | 613 | 6244 | 436 | 10 | 0 | 0 | 0 | 118 | 794 | 887 | 13233 | 239 | 0 | 0 | 10 | 19 | 28 |
| SRR6705828 | 0 | 557 | 2267 | 28 | 613 | 5191 | 7237 | 6197 | 5400 | 62 | 8734 | 6831 | 7357 | 4725 | 412 | 1266 | 57616 | 0 | 0 | 0 | 36 | 281 | 18 | 0 | 0 | 0 | 0 | 0 | 29 | 31 | 2262 | 56 | 0 | 0 | 0 | 27 | 0 |
| SRR6705829 | 0 | 1580 | 499 | 26 | 1153 | 8226 | 22731 | 14182 | 3704 | 38 | 17686 | 16310 | 13657 | 9240 | 2178 | 1044 | 17322 | 0 | 0 | 11 | 838 | 3297 | 177 | 0 | 0 | 0 | 0 | 77 | 817 | 633 | 3624 | 39 | 0 | 0 | 0 | 0 | 57 |
| SRR6705830 | 24 | 1193 | 570 | 0 | 1136 | 8116 | 15920 | 26382 | 10923 | 286 | 7773 | 5961 | 4638 | 8237 | 500 | 4911 | 16662 | 0 | 0 | 20 | 135 | 6678 | 915 | 37 | 0 | 0 | 0 | 24 | 334 | 82 | 4958 | 469 | 0 | 0 | 14 | 15 | 0 |
| SRR6705831 | 51 | 1146 | 814 | 25 | 1761 | 12629 | 33407 | 29076 | 4914 | 357 | 12341 | 9837 | 6904 | 14600 | 872 | 1937 | 28619 | 0 | 15 | 28 | 223 | 741 | 60 | 0 | 0 | 0 | 0 | 51 | 388 | 83 | 575 | 41 | 0 | 0 | 0 | 0 | 0 |
| SRR6705832 | 0 | 1128 | 566 | 0 | 1321 | 11288 | 17314 | 13652 | 4866 | 63 | 12859 | 14266 | 7148 | 8067 | 465 | 1704 | 31553 | 0 | 0 | 14 | 53 | 794 | 61 | 0 | 0 | 0 | 0 | 48 | 461 | 68 | 601 | 46 | 0 | 0 | 0 | 0 | 0 |
| SRR6705833 | 21 | 656 | 683 | 12 | 2348 | 15180 | 30679 | 29778 | 7219 | 328 | 14938 | 10908 | 7827 | 7447 | 770 | 3172 | 24589 | 0 | 0 | 18 | 84 | 2369 | 244 | 11 | 0 | 0 | 0 | 52 | 464 | 76 | 2530 | 196 | 0 | 0 | 0 | 0 | 13 |
| SRR6705834 | 11 | 2383 | 799 | 31 | 7574 | 15652 | 115149 | 50869 | 4871 | 180 | 39115 | 14846 | 16600 | 9427 | 4709 | 658 | 33480 | 0 | 87 | 175 | 5392 | 11019 | 456 | 0 | 0 | 0 | 0 | 259 | 586 | 1356 | 2070 | 152 | 0 | 0 | 0 | 23 | 0 |
| SRR6705835 | 0 | 612 | 549 | 97 | 1398 | 3305 | 25150 | 13270 | 4092 | 586 | 17767 | 6912 | 9246 | 4114 | 1118 | 1813 | 52830 | 0 | 26 | 140 | 3033 | 13071 | 630 | 0 | 0 | 0 | 0 | 134 | 848 | 1642 | 6456 | 661 | 0 | 0 | 0 | 41 | 13 |
| SRR6705837 | 0 | 130 | 262 | 24 | 263 | 725 | 10611 | 4266 | 1555 | 195 | 8193 | 3399 | 3448 | 4166 | 480 | 2151 | 17629 | 0 | 0 | 0 | 555 | 5791 | 282 | 0 | 0 | 0 | 0 | 11 | 78 | 510 | 4450 | 593 | 0 | 0 | 0 | 16 | 36 |
| SRR6705838 | 0 | 1094 | 530 | 19 | 3840 | 11629 | 59672 | 34816 | 3028 | 244 | 15567 | 7562 | 7035 | 8693 | 2264 | 814 | 18267 | 0 | 73 | 117 | 4442 | 18472 | 360 | 0 | 0 | 0 | 0 | 373 | 505 | 2018 | 2966 | 163 | 0 | 0 | 0 | 39 | 22 |
| SRR6705839 | 0 | 96 | 170 | 0 | 1218 | 4307 | 24251 | 16145 | 3977 | 176 | 21126 | 6191 | 11544 | 4869 | 1533 | 1795 | 60512 | 0 | 0 | 13 | 317 | 8987 | 307 | 0 | 0 | 0 | 0 | 22 | 64 | 191 | 3314 | 326 | 0 | 0 | 12 | 38 | 17 |
| SRR6705840 | 0 | 4681 | 1751 | 51 | 1981 | 11400 | 38772 | 31205 | 7848 | 130 | 23048 | 19705 | 12577 | 14840 | 2815 | 1073 | 30187 | 0 | 32 | 57 | 1730 | 8543 | 186 | 0 | 0 | 36 | 0 | 106 | 313 | 764 | 1507 | 81 | 0 | 0 | 158 | 989 | 0 |
| SRR6705841 | 0 | 594 | 196 | 16 | 1599 | 5107 | 49660 | 20394 | 4652 | 176 | 17088 | 5168 | 9457 | 10861 | 8593 | 2497 | 19293 | 0 | 0 | 60 | 2746 | 11758 | 240 | 0 | 0 | 0 | 0 | 37 | 364 | 2039 | 4113 | 244 | 0 | 0 | 39 | 166 | 42 |

### Supp Table S8.1.Columns_B_to_AL

| | | | | | | | | | | | | | | | | | | | | | | | | | | | | | | | | | | | | | |
|---|---|---|---|---|---|---|---|---|---|---|---|---|---|---|---|---|---|---|---|---|---|---|---|---|---|---|---|---|---|---|---|---|---|---|---|---|---|
| SRR6705842 | 0 | 536 | 293 | 12 | 2761 | 8985 | 59203 | 28309 | 2208 | 219 | 12015 | 6096 | 4911 | 7686 | 570 | 1114 | 20333 | 0 | 220 | 220 | 6023 | 13401 | 438 | 0 | 0 | 0 | 0 | 1165 | 1239 | 3682 | 3587 | 346 | 0 | 0 | 14 | 40 | 42 |
| SRR6705843 | 13 | 486 | 690 | 51 | 1612 | 5523 | 34699 | 19334 | 7252 | 305 | 15791 | 8712 | 8852 | 8900 | 1310 | 3723 | 45571 | 0 | 25 | 20 | 892 | 22980 | 709 | 0 | 0 | 0 | 0 | 127 | 172 | 531 | 7321 | 481 | 0 | 0 | 24 | 50 | 0 |
| SRR6705844 | 69 | 1545 | 669 | 169 | 781 | 2531 | 16585 | 13463 | 4336 | 19 | 8694 | 10721 | 6440 | 5050 | 607 | 1040 | 41065 | 0 | 0 | 16 | 232 | 11730 | 1044 | 10 | 0 | 0 | 0 | 19 | 104 | 191 | 44379 | 273 | 0 | 0 | 90 | 69 | 44 |
| SRR6705845 | 84 | 3617 | 992 | 118 | 2024 | 4715 | 25642 | 26530 | 4696 | 131 | 22174 | 25951 | 12931 | 5941 | 2794 | 393 | 48549 | 0 | 25 | 44 | 413 | 3766 | 314 | 0 | 0 | 0 | 0 | 39 | 209 | 230 | 6694 | 64 | 0 | 0 | 35 | 62 | 11 |
| SRR6705846 | 23 | 3019 | 621 | 114 | 612 | 3179 | 10648 | 11403 | 5985 | 68 | 15991 | 23742 | 9978 | 4487 | 786 | 667 | 64109 | 0 | 24 | 41 | 361 | 4186 | 479 | 0 | 0 | 0 | 0 | 66 | 229 | 190 | 13950 | 174 | 0 | 0 | 30 | 32 | 16 |
| SRR6705847 | 118 | 4798 | 3553 | 196 | 985 | 2400 | 12127 | 23047 | 4272 | 104 | 19423 | 27588 | 8104 | 4533 | 347 | 497 | 60200 | 0 | 13 | 25 | 295 | 6034 | 609 | 0 | 0 | 0 | 0 | 42 | 164 | 163 | 13373 | 139 | 0 | 0 | 14 | 29 | 0 |
| SRR6705848 | 11 | 933 | 216 | 62 | 734 | 3026 | 9735 | 11453 | 5454 | 22 | 11120 | 19035 | 8659 | 4257 | 401 | 594 | 78197 | 0 | 0 | 12 | 129 | 1626 | 145 | 0 | 0 | 0 | 0 | 16 | 67 | 78 | 5573 | 49 | 0 | 0 | 16 | 12 | 10 |
| SRR6705849 | 0 | 800 | 477 | 106 | 991 | 6475 | 18979 | 16614 | 27379 | 15 | 16013 | 22878 | 8717 | 6142 | 1332 | 585 | 39444 | 0 | 24 | 53 | 414 | 7829 | 1080 | 59 | 33 | 23 | 0 | 64 | 354 | 295 | 27202 | 208 | 26 | 39 | 396 | 171 | 0 |
| SRR6705850 | 49 | 1883 | 840 | 51 | 825 | 3277 | 10837 | 12947 | 4371 | 57 | 14965 | 24067 | 11155 | 4170 | 667 | 677 | 63540 | 0 | 0 | 15 | 151 | 1380 | 78 | 0 | 0 | 0 | 0 | 12 | 67 | 96 | 2216 | 38 | 0 | 0 | 14 | 24 | 0 |
| SRR6705851 | 26 | 2924 | 235 | 74 | 1626 | 6157 | 23324 | 33609 | 6157 | 31 | 13463 | 20987 | 10830 | 8209 | 1083 | 692 | 53339 | 0 | 0 | 24 | 278 | 1607 | 112 | 0 | 0 | 0 | 0 | 26 | 166 | 165 | 3758 | 55 | 0 | 0 | 22 | 38 | 0 |
| SRR6705852 | 32 | 1594 | 431 | 47 | 515 | 2255 | 7602 | 9667 | 4309 | 42 | 12142 | 17894 | 8596 | 3903 | 380 | 585 | 63624 | 0 | 11 | 39 | 341 | 5931 | 357 | 0 | 0 | 0 | 0 | 24 | 344 | 235 | 17220 | 70 | 0 | 0 | 55 | 161 | 14 |
| SRR6705853 | 37 | 3117 | 602 | 90 | 2134 | 7840 | 39209 | 31797 | 6353 | 51 | 14260 | 17801 | 6444 | 7067 | 2815 | 647 | 40055 | 0 | 40 | 79 | 570 | 6352 | 756 | 28 | 10 | 15 | 0 | 52 | 278 | 237 | 14630 | 113 | 0 | 0 | 67 | 171 | 0 |
| SRR6705854 | 90 | 5877 | 2698 | 188 | 1185 | 4924 | 19164 | 25656 | 6185 | 51 | 14243 | 23707 | 9140 | 6771 | 597 | 495 | 72045 | 0 | 63 | 127 | 510 | 9647 | 402 | 0 | 0 | 0 | 0 | 116 | 503 | 223 | 20065 | 101 | 0 | 0 | 0 | 26 | 0 |
| SRR6705855 | 0 | 1763 | 622 | 59 | 898 | 2877 | 14763 | 18798 | 5392 | 73 | 17779 | 22284 | 11465 | 4832 | 848 | 641 | 50147 | 0 | 74 | 126 | 595 | 8327 | 370 | 0 | 0 | 0 | 0 | 116 | 486 | 337 | 15395 | 131 | 0 | 0 | 37 | 65 | 17 |
| SRR6705856 | 33 | 3749 | 1590 | 140 | 3291 | 13184 | 48328 | 24701 | 1993 | 313 | 20658 | 14399 | 9058 | 9679 | 697 | 1244 | 20272 | 0 | 23 | 106 | 2160 | 14558 | 675 | 0 | 0 | 0 | 0 | 103 | 591 | 1100 | 5615 | 492 | 0 | 0 | 22 | 36 | 0 |
| SRR6705857 | 35 | 371 | 496 | 271 | 2467 | 5882 | 35106 | 22240 | 3198 | 293 | 19664 | 6624 | 9694 | 5664 | 1676 | 1847 | 33382 | 0 | 24 | 94 | 1972 | 8403 | 489 | 0 | 0 | 0 | 0 | 75 | 488 | 1077 | 6902 | 689 | 0 | 0 | 15 | 32 | 31 |
| SRR6705859 | 15 | 484 | 795 | 1176 | 711 | 3211 | 21969 | 15115 | 4985 | 138 | 22922 | 9412 | 12510 | 9255 | 1092 | 2187 | 48701 | 0 | 0 | 11 | 231 | 13743 | 1126 | 0 | 0 | 0 | 0 | 14 | 76 | 228 | 8806 | 1526 | 0 | 0 | 41 | 60 | 43 |
| SRR6705860 | 0 | 2695 | 1017 | 45 | 2101 | 19838 | 34180 | 25622 | 12147 | 72 | 22318 | 25181 | 13475 | 6646 | 1198 | 1095 | 29098 | 0 | 16 | 209 | 4030 | 7664 | 204 | 0 | 0 | 0 | 0 | 68 | 1316 | 967 | 1126 | 69 | 0 | 0 | 67 | 146 | 24 |
| SRR6705861 | 0 | 130 | 67 | 45 | 864 | 2869 | 26309 | 12978 | 5415 | 60 | 4041 | 1757 | 2884 | 5761 | 789 | 2741 | 22997 | 0 | 0 | 18 | 1079 | 9493 | 367 | 0 | 0 | 0 | 0 | 28 | 163 | 455 | 4613 | 370 | 0 | 0 | 17 | 18 | 18 |
| SRR6705862 | 0 | 160 | 206 | 33 | 1975 | 4617 | 56757 | 32687 | 3043 | 132 | 9594 | 3938 | 3796 | 10126 | 1379 | 1369 | 16255 | 0 | 112 | 179 | 7185 | 9324 | 231 | 0 | 0 | 0 | 0 | 507 | 1056 | 2550 | 2767 | 153 | 0 | 0 | 0 | 71 | 48 |
| SRR6705863 | 11 | 162 | 206 | 168 | 1562 | 3114 | 53423 | 24236 | 8484 | 580 | 8769 | 1267 | 3275 | 4633 | 3004 | 2826 | 29676 | 0 | 21 | 21 | 2025 | 9478 | 419 | 0 | 0 | 0 | 0 | 169 | 190 | 1575 | 5173 | 601 | 0 | 0 | 25 | 57 | 37 |
| SRR6705866 | 74 | 2906 | 631 | 62 | 937 | 5249 | 22916 | 19542 | 3655 | 74 | 11997 | 19776 | 8904 | 6514 | 562 | 890 | 42973 | 0 | 0 | 0 | 159 | 799 | 36 | 0 | 0 | 0 | 0 | 0 | 50 | 102 | 796 | 29 | 0 | 0 | 0 | 17 | 0 |
| SRR6705867 | 210 | 6808 | 1188 | 248 | 2786 | 7124 | 34669 | 37952 | 4639 | 171 | 11672 | 18600 | 7922 | 7470 | 602 | 798 | 36537 | 0 | 0 | 24 | 378 | 2194 | 207 | 0 | 0 | 0 | 0 | 31 | 119 | 185 | 2050 | 99 | 0 | 0 | 0 | 28 | 0 |
| SRR6705868 | 54 | 4111 | 750 | 142 | 1670 | 5180 | 26994 | 20029 | 4973 | 187 | 12145 | 15600 | 8130 | 7440 | 1448 | 996 | 30327 | 0 | 11 | 29 | 323 | 1638 | 84 | 0 | 0 | 0 | 0 | 33 | 117 | 212 | 1469 | 74 | 0 | 0 | 12 | 34 | 0 |
| SRR6705869 | 232 | 13298 | 864 | 269 | 3669 | 8895 | 38825 | 26699 | 4943 | 144 | 13864 | 17623 | 7930 | 8329 | 1120 | 708 | 29003 | 0 | 28 | 59 | 986 | 4565 | 249 | 0 | 0 | 0 | 0 | 80 | 311 | 548 | 3434 | 168 | 0 | 0 | 44 | 58 | 0 |
| SRR6705870 | 63 | 2990 | 441 | 142 | 901 | 3261 | 16636 | 14314 | 4002 | 92 | 12362 | 16475 | 8332 | 4737 | 456 | 971 | 49172 | 0 | 15 | 46 | 397 | 3238 | 415 | 20 | 0 | 0 | 0 | 31 | 195 | 246 | 6625 | 135 | 26 | 15 | 33 | 54 | 30 |
| SRR6705871 | 111 | 5486 | 823 | 214 | 1663 | 5854 | 24431 | 29387 | 7992 | 113 | 18051 | 26711 | 9603 | 5858 | 1531 | 474 | 50299 | 0 | 13 | 56 | 417 | 2760 | 357 | 12 | 18 | 0 | 0 | 44 | 222 | 213 | 3694 | 79 | 0 | 0 | 138 | 83 | 0 |
| SRR6705874 | 0 | 54 | 13 | 0 | 107 | 549 | 1654 | 1251 | 661 | 0 | 758 | 421 | 614 | 1133 | 177 | 569 | 3776 | 0 | 0 | 0 | 25 | 78 | 0 | 0 | 0 | 0 | 0 | 0 | 19 | 14 | 136 | 0 | 0 | 0 | 0 | 0 | 13 |
| SRR6705875 | 0 | 361 | 58 | 12 | 2365 | 12533 | 31512 | 16497 | 2284 | 14 | 4129 | 4942 | 3277 | 24383 | 286 | 2493 | 13312 | 0 | 0 | 0 | 105 | 1143 | 45 | 0 | 0 | 0 | 0 | 0 | 111 | 59 | 975 | 48 | 0 | 0 | 0 | 0 | 30 |
| SRR6705876 | 0 | 110 | 100 | 34 | 1215 | 9859 | 12463 | 22287 | 10137 | 0 | 3576 | 7637 | 2430 | 12602 | 347 | 5589 | 5520 | 0 | 0 | 0 | 0 | 196 | 105 | 0 | 0 | 0 | 0 | 0 | 14 | 0 | 205 | 162 | 0 | 0 | 0 | 0 | 0 |
| SRR6705877 | 0 | 44 | 408 | 0 | 165 | 741 | 3706 | 81302 | 14246 | 0 | 1866 | 1235 | 1125 | 11199 | 82 | 4447 | 2498 | 0 | 0 | 0 | 0 | 49 | 66 | 0 | 0 | 0 | 0 | 0 | 0 | 0 | 464 | 224 | 0 | 0 | 0 | 0 | 0 |
| SRR6705878 | 0 | 418 | 1154 | 15 | 186 | 4343 | 3991 | 5446 | 6845 | 0 | 3288 | 5379 | 3873 | 9682 | 601 | 2940 | 9662 | 0 | 0 | 0 | 53 | 1560 | 177 | 0 | 0 | 0 | 0 | 0 | 58 | 20 | 1456 | 189 | 0 | 0 | 0 | 0 | 86 |
| SRR6705879 | 0 | 787 | 181 | 29 | 835 | 6828 | 11667 | 8727 | 9186 | 12 | 14006 | 18941 | 10394 | 8770 | 314 | 955 | 20241 | 0 | 0 | 0 | 52 | 445 | 13 | 0 | 0 | 0 | 0 | 0 | 165 | 29 | 314 | 0 | 0 | 0 | 0 | 13 | 62 |
| SRR6705880 | 0 | 92 | 409 | 0 | 1256 | 3341 | 12699 | 13473 | 4400 | 30 | 5789 | 1871 | 4381 | 2726 | 518 | 1525 | 17597 | 0 | 0 | 0 | 46 | 687 | 53 | 0 | 0 | 0 | 0 | 0 | 61 | 123 | 1680 | 72 | 0 | 0 | 0 | 0 | 0 |
| SRR6705881 | 0 | 54 | 1527 | 0 | 4619 | 12635 | 47140 | 32147 | 7608 | 79 | 11858 | 5385 | 6503 | 3651 | 1382 | 809 | 26753 | 0 | 0 | 0 | 45 | 302 | 28 | 0 | 0 | 0 | 0 | 0 | 34 | 66 | 739 | 47 | 0 | 0 | 0 | 0 | 0 |
| SRR6705882 | 0 | 15 | 196 | 0 | 1565 | 3240 | 22822 | 15918 | 6522 | 77 | 3808 | 571 | 2358 | 2963 | 892 | 2519 | 16307 | 0 | 0 | 0 | 70 | 283 | 34 | 0 | 0 | 0 | 0 | 0 | 50 | 100 | 725 | 36 | 0 | 0 | 0 | 0 | 0 |
| SRR6705883 | 0 | 190 | 592 | 13 | 2497 | 9152 | 26405 | 15661 | 8751 | 49 | 8548 | 6333 | 5428 | 3231 | 743 | 612 | 22463 | 0 | 0 | 0 | 161 | 1678 | 130 | 0 | 0 | 0 | 0 | 32 | 187 | 317 | 2307 | 91 | 0 | 0 | 0 | 21 | 0 |
| SRR6705884 | 18 | 1566 | 224 | 51 | 1563 | 7798 | 21256 | 24054 | 6266 | 15 | 18723 | 28522 | 8867 | 8494 | 985 | 584 | 46009 | 0 | 14 | 41 | 347 | 4370 | 565 | 23 | 12 | 19 | 0 | 37 | 273 | 229 | 9954 | 76 | 0 | 0 | 202 | 265 | 0 |
| SRR6705885 | 153 | 2076 | 1187 | 119 | 942 | 3123 | 14664 | 13317 | 4774 | 53 | 16001 | 24831 | 11110 | 5052 | 835 | 709 | 49312 | 0 | 0 | 25 | 292 | 4734 | 576 | 16 | 23 | 15 | 0 | 24 | 195 | 197 | 15851 | 138 | 12 | 20 | 410 | 412 | 22 |
| SRR6705886 | 142 | 4492 | 741 | 88 | 1747 | 6705 | 24865 | 29874 | 4621 | 45 | 13889 | 21979 | 7585 | 8258 | 453 | 621 | 47957 | 0 | 13 | 19 | 267 | 1678 | 132 | 0 | 0 | 0 | 0 | 36 | 170 | 179 | 2086 | 65 | 0 | 0 | 48 | 51 | 0 |
| SRR6705887 | 164 | 4592 | 1667 | 78 | 1326 | 4726 | 24741 | 23306 | 3978 | 125 | 17022 | 23529 | 9946 | 6274 | 510 | 792 | 38744 | 0 | 0 | 17 | 336 | 1603 | 125 | 0 | 0 | 0 | 0 | 37 | 136 | 191 | 1358 | 80 | 0 | 0 | 13 | 37 | 0 |
| SRR6705888 | 11 | 634 | 252 | 26 | 1727 | 4600 | 26264 | 27249 | 5713 | 35 | 21113 | 20250 | 13641 | 5489 | 697 | 601 | 39860 | 0 | 0 | 13 | 160 | 1106 | 52 | 0 | 0 | 0 | 0 | 28 | 76 | 86 | 1717 | 33 | 0 | 0 | 11 | 16 | 0 |
| SRR6705889 | 56 | 1452 | 872 | 96 | 686 | 2221 | 11579 | 14689 | 2053 | 62 | 11870 | 18420 | 18657 | 8707 | 192 | 548 | 70181 | 0 | 11 | 25 | 230 | 1520 | 129 | 0 | 0 | 0 | 0 | 66 | 175 | 204 | 3256 | 60 | 0 | 0 | 32 | 32 | 0 |
| SRR6705890 | 137 | 11709 | 776 | 180 | 1025 | 6045 | 14071 | 14663 | 5667 | 76 | 17225 | 23701 | 9267 | 3693 | 839 | 442 | 57012 | 0 | 11 | 28 | 363 | 4301 | 508 | 29 | 0 | 0 | 0 | 25 | 117 | 145 | 3892 | 67 | 0 | 0 | 30 | 66 | 0 |
| SRR6705891 | 12 | 2211 | 609 | 88 | 823 | 2607 | 10028 | 11638 | 3916 | 61 | 18436 | 28095 | 13160 | 3704 | 405 | 541 | 60038 | 0 | 0 | 40 | 243 | 2545 | 337 | 0 | 0 | 0 | 0 | 24 | 198 | 151 | 5385 | 92 | 0 | 0 | 20 | 31 | 28 |
| SRR6705892 | 0 | 2415 | 637 | 69 | 1385 | 4231 | 22444 | 26458 | 4311 | 53 | 16376 | 27050 | 10110 | 6211 | 1365 | 272 | 53296 | 0 | 50 | 70 | 629 | 14385 | 1979 | 97 | 27 | 22 | 0 | 87 | 265 | 296 | 38944 | 253 | 44 | 33 | 221 | 301 | 0 |
| SRR6705893 | 77 | 3410 | 619 | 86 | 919 | 3249 | 13353 | 13541 | 3384 | 114 | 13838 | 22733 | 10099 | 4285 | 538 | 797 | 64890 | 0 | 11 | 30 | 341 | 2537 | 233 | 0 | 0 | 0 | 0 | 38 | 136 | 200 | 4563 | 91 | 0 | 0 | 65 | 74 | 13 |
| SRR6705894 | 0 | 950 | 694 | 0 | 1196 | 18579 | 23339 | 48235 | 11609 | 82 | 10730 | 12267 | 7930 | 11940 | 518 | 1110 | 58417 | 0 | 0 | 0 | 420 | 3314 | 122 | 0 | 0 | 0 | 0 | 39 | 531 | 310 | 4661 | 97 | 0 | 0 | 11 | 32 | 22 |
| SRR6705895 | 0 | 221 | 534 | 0 | 471 | 5785 | 10021 | 8581 | 4865 | 39 | 6214 | 3389 | 7192 | 2785 | 1466 | 1590 | 50561 | 0 | 0 | 0 | 154 | 2016 | 137 | 0 | 0 | 0 | 0 | 0 | 186 | 152 | 5302 | 98 | 0 | 0 | 0 | 11 | 58 |
| SRR6705897 | 0 | 637 | 592 | 19 | 129 | 1532 | 2260 | 3427 | 1125 | 140 | 3783 | 2564 | 3653 | 1545 | 673 | 1206 | 27768 | 0 | 0 | 0 | 112 | 1117 | 80 | 0 | 0 | 0 | 0 | 0 | 231 | 199 | 4950 | 89 | 0 | 0 | 0 | 0 | 62 |
| SRR6705898 | 0 | 16 | 12 | 0 | 178 | 1247 | 4394 | 2790 | 459 | 26 | 922 | 1366 | 673 | 3230 | 72 | 890 | 1963 | 0 | 0 | 10 | 338 | 1162 | 21 | 0 | 0 | 0 | 0 | 25 | 451 | 373 | 1727 | 43 | 0 | 0 | 0 | 0 | 46 |
| SRR6705899 | 0 | 364 | 48 | 0 | 30 | 202 | 377 | 401 | 148 | 14 | 543 | 818 | 369 | 901 | 0 | 482 | 2713 | 0 | 0 | 0 | 36 | 297 | 11 | 0 | 0 | 0 | 0 | 0 | 66 | 0 | 187 | 0 | 0 | 0 | 0 | 0 | 0 |
| SRR6705900 | 0 | 29 | 0 | 0 | 127 | 769 | 2894 | 1648 | 1036 | 0 | 637 | 633 | 580 | 2043 | 197 | 748 | 2104 | 0 | 0 | 0 | 111 | 244 | 0 | 0 | 0 | 0 | 0 | 0 | 105 | 73 | 358 | 13 | 0 | 0 | 0 | 0 | 0 |
| SRR6705901 | 0 | 23 | 13 | 0 | 155 | 994 | 3638 | 1982 | 191 | 40 | 622 | 417 | 478 | 2553 | 124 | 548 | 2423 | 0 | 0 | 22 | 990 | 2432 | 60 | 0 | 0 | 0 | 0 | 30 | 785 | 903 | 5467 | 92 | 0 | 0 | 0 | 0 | 62 |
| SRR6705902 | 0 | 688 | 212 | 173 | 1243 | 3666 | 37854 | 25683 | 5992 | 205 | 17524 | 4082 | 8431 | 6201 | 1729 | 1836 | 30170 | 0 | 24 | 18 | 1138 | 19056 | 1328 | 0 | 0 | 0 | 0 | 171 | 143 | 788 | 9268 | 1548 | 0 | 0 | 32 | 44 | 12 |
| SRR6705903 | 11 | 4334 | 816 | 159 | 2468 | 10057 | 35304 | 36244 | 3374 | 92 | 30706 | 27221 | 18835 | 9750 | 2221 | 653 | 35128 | 0 | 230 | 83 | 2254 | 4553 | 194 | 0 | 0 | 0 | 0 | 868 | 452 | 754 | 850 | 79 | 0 | 0 | 11 | 25 | 12 |
| SRR6726498 | 0 | 49433 | 511 | 20 | 495 | 5409 | 4608 | 3081 | 4430 | 0 | 7194 | 53081 | 1106 | 2827 | 16 | 607 | 2941 | 0 | 0 | 0 | 41 | 20 | 16 | 0 | 0 | 0 | 0 | 0 | 0 | 19 | 13 | 0 | 0 | 0 | 0 | 0 | 0 |
| SRR6726499 | 70 | 218710 | 13206 | 90 | 892 | 8700 | 7683 | 6559 | 4292 | 0 | 7015 | 64208 | 1234 | 2744 | 18 | 1261 | 3577 | 0 | 0 | 0 | 33 | 25 | 13 | 0 | 0 | 0 | 0 | 0 | 0 | 28 | 11 | 0 | 0 | 0 | 0 | 0 | 0 |
| SRR6757230 | 926 | 152096 | 10164 | 272 | 190 | 3236 | 3046 | 3210 | 6810 | 31 | 5113 | 8600 | 1030 | 309 | 13 | 33 | 14480 | 0 | 0 | 0 | 322 | 166 | 82 | 0 | 0 | 0 | 0 | 0 | 56 | 74 | 93 | 34 | 0 | 11 | 0 | 0 | 0 |
| SRR6757231 | 953 | 157521 | 10457 | 291 | 197 | 2992 | 2876 | 2960 | 6738 | 30 | 5081 | 8586 | 1040 | 325 | 0 | 31 | 12330 | 0 | 0 | 0 | 339 | 182 | 85 | 0 | 0 | 0 | 0 | 11 | 59 | 84 | 94 | 38 | 0 | 10 | 0 | 0 | 0 |
| SRR6757232 | 469 | 142502 | 13356 | 271 | 219 | 3392 | 3377 | 4420 | 5526 | 35 | 5274 | 7940 | 1080 | 380 | 10 | 37 | 18531 | 0 | 0 | 0 | 346 | 173 | 68 | 0 | 0 | 0 | 0 | 14 | 76 | 87 | 108 | 35 | 0 | 0 | 0 | 0 | 0 |
| SRR6757233 | 501 | 148437 | 13836 | 277 | 209 | 3226 | 3088 | 4183 | 5410 | 23 | 5256 | 8247 | 1058 | 347 | 15 | 34 | 16201 | 0 | 0 | 0 | 350 | 185 | 72 | 0 | 0 | 0 | 0 | 17 | 82 | 102 | 105 | 39 | 0 | 0 | 0 | 0 | 0 |
| SRR6757234 | 440 | 125746 | 9393 | 193 | 218 | 3274 | 3266 | 3700 | 7387 | 13 | 6215 | 9755 | 1200 | 481 | 0 | 45 | 21434 | 0 | 0 | 0 | 493 | 221 | 73 | 0 | 0 | 0 | 0 | 29 | 83 | 115 | 113 | 32 | 0 | 10 | 0 | 0 | 0 |
| SRR6757235 | 453 | 132576 | 9644 | 192 | 226 | 3043 | 3094 | 3443 | 7296 | 15 | 6193 | 9723 | 1268 | 455 | 14 | 33 | 18302 | 0 | 0 | 0 | 515 | 230 | 74 | 0 | 0 | 0 | 0 | 33 | 93 | 116 | 130 | 32 | 0 | 0 | 0 | 0 | 0 |
| SRR6757236 | 502 | 137720 | 7869 | 179 | 270 | 3986 | 4269 | 4776 | 5443 | 28 | 5488 | 9130 | 1255 | 446 | 13 | 33 | 18863 | 0 | 0 | 0 | 527 | 172 | 93 | 0 | 0 | 0 | 0 | 0 | 78 | 93 | 77 | 36 | 0 | 0 | 0 | 0 | 0 |
| SRR6757237 | 496 | 145205 | 8119 | 160 | 195 | 3716 | 3888 | 4363 | 5393 | 23 | 5698 | 9335 | 1348 | 520 | 0 | 23 | 16466 | 0 | 0 | 0 | 552 | 189 | 88 | 0 | 0 | 0 | 0 | 0 | 82 | 96 | 98 | 33 | 0 | 11 | 0 | 0 | 0 |
| SRR6757238 | 892 | 233373 | 772 | 442 | 349 | 5653 | 3620 | 2826 | 3583 | 12 | 2494 | 4589 | 500 | 159 | 15 | 69 | 13966 | 0 | 0 | 0 | 822 | 388 | 175 | 0 | 13 | 0 | 0 | 29 | 150 | 201 | 227 | 65 | 0 | 18 | 0 | 0 | 0 |
| SRR6757239 | 949 | 238223 | 910 | 401 | 316 | 5059 | 3394 | 2790 | 3658 | 10 | 4076 | 5758 | 746 | 302 | 18 | 62 | 11410 | 0 | 0 | 0 | 868 | 396 | 180 | 0 | 13 | 0 | 0 | 30 | 173 | 195 | 229 | 79 | 0 | 22 | 0 | 0 | 0 |
| SRR6757240 | 877 | 131678 | 416 | 416 | 289 | 4301 | 3022 | 2558 | 5741 | 15 | 2698 | 6120 | 769 | 207 | 20 | 60 | 12742 | 0 | 0 | 12 | 983 | 460 | 148 | 0 | 0 | 0 | 0 | 45 | 215 | 244 | 307 | 79 | 0 | 19 | 0 | 0 | 0 |
| SRR6757241 | 918 | 135928 | 498 | 430 | 268 | 4024 | 2805 | 2435 | 5738 | 11 | 2720 | 6014 | 709 | 203 | 25 | 38 | 11005 | 0 | 0 | 13 | 1035 | 483 | 156 | 0 | 0 | 0 | 0 | 54 | 246 | 252 | 306 | 85 | 0 | 16 | 0 | 0 | 0 |
| SRR6757242 | 622 | 231926 | 2300 | 234 | 254 | 4314 | 3865 | 3896 | 5778 | 23 | 4464 | 8302 | 1002 | 295 | 11 | 23 | 15371 | 0 | 0 | 0 | 315 | 213 | 73 | 0 | 0 | 0 | 0 | 13 | 79 | 85 | 127 | 33 | 0 | 10 | 0 | 0 | 0 |
| SRR6757243 | 606 | 229792 | 2380 | 247 | 255 | 4302 | 3740 | 3918 | 6193 | 26 | 4707 | 8987 | 1112 | 333 | 11 | 27 | 15935 | 0 | 0 | 0 | 330 | 237 | 82 | 0 | 0 | 0 | 0 | 16 | 85 | 90 | 138 | 40 | 0 | 12 | 0 | 0 | 0 |
| SRR6757244 | 1310 | 150295 | 3445 | 448 | 311 | 4560 | 3554 | 2541 | 4364 | 23 | 4102 | 7738 | 949 | 261 | 15 | 26 | 14259 | 0 | 0 | 0 | 504 | 231 | 123 | 0 | 0 | 0 | 0 | 29 | 97 | 121 | 169 | 54 | 0 | 17 | 0 | 0 | 0 |
| SRR6757245 | 1307 | 148588 | 3402 | 473 | 322 | 4456 | 3539 | 2589 | 4654 | 22 | 4397 | 8392 | 982 | 280 | 19 | 20 | 14828 | 0 | 0 | 0 | 529 | 241 | 127 | 0 | 11 | 0 | 0 | 27 | 104 | 137 | 179 | 61 | 0 | 16 | 0 | 0 | 0 |
| SRR6757246 | 700 | 117253 | 6229 | 266 | 171 | 2600 | 2816 | 3595 | 7281 | 17 | 4672 | 8036 | 956 | 310 | 0 | 10 | 14962 | 0 | 0 | 0 | 311 | 92 | 58 | 0 | 0 | 0 | 0 | 0 | 50 | 68 | 59 | 23 | 0 | 0 | 0 | 0 | 0 |
| SRR6757247 | 674 | 115472 | 6389 | 282 | 184 | 2575 | 2755 | 3485 | 7688 | 18 | 4842 | 8604 | 1040 | 344 | 0 | 14 | 15376 | 0 | 0 | 0 | 334 | 95 | 58 | 0 | 0 | 0 | 0 | 0 | 54 | 75 | 62 | 22 | 0 | 0 | 0 | 0 | 0 |
| SRR6757248 | 646 | 185278 | 5246 | 283 | 199 | 3590 | 3261 | 4406 | 5286 | 18 | 5038 | 9602 | 1049 | 266 | 0 | 22 | 15604 | 0 | 0 | 0 | 315 | 122 | 57 | 0 | 0 | 0 | 0 | 11 | 62 | 72 | 86 | 28 | 0 | 0 | 0 | 0 | 0 |
| SRR6757249 | 636 | 183594 | 5277 | 316 | 213 | 3564 | 3312 | 4387 | 5657 | 16 | 5299 | 10371 | 1110 | 299 | 0 | 20 | 15951 | 0 | 0 | 0 | 340 | 118 | 61 | 0 | 0 | 0 | 0 | 10 | 81 | 77 | 85 | 29 | 0 | 0 | 0 | 0 | 0 |
| SRR6757250 | 669 | 143191 | 8282 | 290 | 184 | 2990 | 2991 | 3418 | 6413 | 24 | 5039 | 9389 | 1177 | 373 | 10 | 18 | 14602 | 0 | 0 | 0 | 259 | 120 | 46 | 0 | 0 | 0 | 0 | 0 | 48 | 73 | 81 | 24 | 0 | 0 | 0 | 0 | 0 |
| SRR6757251 | 682 | 141552 | 8179 | 324 | 179 | 2989 | 2952 | 3415 | 6854 | 19 | 5484 | 9981 | 1268 | 416 | 12 | 16 | 14850 | 0 | 0 | 0 | 283 | 128 | 53 | 0 | 0 | 0 | 0 | 0 | 53 | 82 | 84 | 25 | 0 | 0 | 0 | 0 | 0 |
| SRR6757252 | 921 | 181309 | 5132 | 368 | 286 | 5239 | 4587 | 4471 | 4929 | 39 | 3763 | 7161 | 875 | 281 | 0 | 35 | 12767 | 0 | 0 | 0 | 270 | 97 | 58 | 0 | 0 | 0 | 0 | 0 | 40 | 73 | 60 | 26 | 0 | 0 | 0 | 0 | 0 |
| SRR6757253 | 971 | 180173 | 5051 | 448 | 286 | 5185 | 4490 | 4451 | 5409 | 40 | 4091 | 7670 | 918 | 309 | 0 | 40 | 13069 | 0 | 0 | 0 | 287 | 99 | 69 | 0 | 0 | 0 | 0 | 0 | 47 | 80 | 59 | 31 | 0 | 11 | 0 | 0 | 0 |
| SRR6757254 | 853 | 173091 | 19358 | 406 | 245 | 3501 | 2838 | 4112 | 6716 | 38 | 35898 | 25417 | 5139 | 4428 | 0 | 54 | 16359 | 0 | 0 | 0 | 383 | 242 | 104 | 0 | 0 | 0 | 0 | 20 | 97 | 112 | 189 | 62 | 0 | 14 | 0 | 0 | 0 |
| SRR6757255 | 880 | 180511 | 19816 | 394 | 227 | 3344 | 2602 | 3868 | 6688 | 34 | 35624 | 25047 | 5035 | 4383 | 0 | 55 | 14086 | 0 | 0 | 0 | 422 | 252 | 102 | 0 | 0 | 0 | 0 | 22 | 98 | 111 | 202 | 64 | 0 | 14 | 0 | 0 | 0 |
| SRR6757256 | 515 | 233106 | 11675 | 234 | 261 | 3669 | 3492 | 5792 | 4592 | 22 | 26221 | 21565 | 4067 | 3729 | 0 | 37 | 15075 | 0 | 0 | 0 | 322 | 198 | 67 | 0 | 0 | 0 | 0 | 26 | 75 | 103 | 159 | 39 | 0 | 0 | 0 | 0 | 0 |
| SRR6757257 | 506 | 242974 | 11979 | 213 | 238 | 3450 | 3320 | 5399 | 4573 | 19 | 26047 | 21401 | 3959 | 3773 | 0 | 45 | 12954 | 0 | 0 | 0 | 348 | 212 | 68 | 0 | 0 | 0 | 0 | 26 | 84 | 99 | 167 | 46 | 0 | 14 | 0 | 0 | 0 |
| SRR6757258 | 828 | 150775 | 17402 | 434 | 129 | 1791 | 1545 | 2975 | 4360 | 22 | 24079 | 22541 | 4628 | 3633 | 0 | 48 | 14554 | 0 | 0 | 0 | 343 | 246 | 56 | 0 | 0 | 0 | 0 | 22 | 101 | 132 | 174 | 37 | 0 | 11 | 0 | 0 | 0 |
| SRR6757259 | 800 | 154646 | 17931 | 425 | 127 | 1715 | 1446 | 2808 | 4321 | 20 | 23376 | 21736 | 4517 | 3558 | 0 | 39 | 12220 | 0 | 0 | 0 | 357 | 251 | 61 | 0 | 0 | 0 | 0 | 19 | 103 | 139 | 174 | 39 | 0 | 11 | 0 | 0 | 0 |
| SRR6757260 | 742 | 97896 | 13543 | 493 | 280 | 3822 | 3728 | 4868 | 4485 | 23 | 31871 | 25997 | 4636 | 3907 | 0 | 0 | 13119 | 0 | 0 | 0 | 378 | 159 | 85 | 0 | 0 | 0 | 0 | 23 | 66 | 99 | 124 | 47 | 0 | 12 | 0 | 0 | 0 |
| SRR6757261 | 783 | 104127 | 13943 | 439 | 261 | 3628 | 3487 | 4641 | 4499 | 31 | 31836 | 26056 | 4576 | 3875 | 0 | 0 | 11016 | 0 | 0 | 0 | 402 | 166 | 90 | 0 | 0 | 0 | 0 | 28 | 80 | 103 | 136 | 45 | 0 | 0 | 0 | 0 | 0 |
| SRR6757262 | 1247 | 125235 | 9175 | 497 | 289 | 4234 | 3114 | 3208 | 4047 | 32 | 26735 | 22586 | 3934 | 3128 | 12 | 23 | 10711 | 0 | 0 | 0 | 557 | 281 | 135 | 0 | 0 | 0 | 0 | 28 | 98 | 130 | 182 | 73 | 0 | 16 | 0 | 0 | 0 |
| SRR6757263 | 1258 | 130175 | 9550 | 455 | 234 | 3875 | 2841 | 2969 | 3954 | 41 | 26257 | 22356 | 3868 | 3130 | 0 | 19 | 8901 | 0 | 0 | 0 | 567 | 300 | 143 | 0 | 11 | 0 | 0 | 26 | 103 | 137 | 202 | 76 | 0 | 19 | 0 | 0 | 0 |
| SRR6757264 | 1165 | 122829 | 21751 | 314 | 199 | 2751 | 2937 | 6377 | 6767 | 20 | 29241 | 27532 | 3694 | 5640 | 0 | 35 | 12928 | 0 | 0 | 0 | 273 | 178 | 57 | 0 | 0 | 0 | 0 | 0 | 57 | 93 | 135 | 35 | 0 | 0 | 0 | 0 | 0 |
| SRR6757265 | 1144 | 121936 | 21830 | 348 | 196 | 2773 | 2980 | 6306 | 7214 | 23 | 31666 | 29465 | 3801 | 6007 | 0 | 38 | 12932 | 0 | 0 | 0 | 301 | 191 | 62 | 0 | 0 | 0 | 0 | 0 | 60 | 108 | 146 | 36 | 0 | 0 | 0 | 0 | 0 |
| SRR6757266 | 1563 | 148771 | 1149 | 515 | 230 | 3034 | 2050 | 2185 | 3424 | 16 | 17166 | 17298 | 2901 | 2342 | 0 | 0 | 11405 | 0 | 0 | 0 | 606 | 238 | 156 | 0 | 0 | 0 | 0 | 34 | 111 | 162 | 202 | 68 | 0 | 17 | 11 | 0 | 0 |
| SRR6757267 | 1668 | 156209 | 1230 | 506 | 224 | 2788 | 1915 | 1917 | 3532 | 12 | 17204 | 16988 | 2836 | 2328 | 0 | 0 | 9355 | 0 | 0 | 0 | 625 | 242 | 143 | 0 | 10 | 0 | 0 | 30 | 116 | 155 | 205 | 73 | 0 | 18 | 11 | 0 | 0 |
| SRR6757268 | 1549 | 142560 | 9196 | 493 | 216 | 2923 | 2234 | 2476 | 2841 | 38 | 21944 | 23486 | 4347 | 3137 | 0 | 50 | 11233 | 0 | 0 | 0 | 367 | 158 | 94 | 0 | 0 | 0 | 0 | 12 | 61 | 89 | 137 | 46 | 0 | 13 | 0 | 0 | 0 |
| SRR6757269 | 1522 | 141239 | 9291 | 503 | 190 | 2937 | 2176 | 2429 | 3047 | 48 | 23729 | 25607 | 4521 | 3335 | 0 | 56 | 11444 | 0 | 0 | 0 | 388 | 181 | 98 | 0 | 0 | 0 | 0 | 13 | 73 | 102 | 143 | 50 | 0 | 14 | 0 | 0 | 0 |
| SRR6757270 | 782 | 166092 | 19352 | 381 | 282 | 4321 | 3580 | 5284 | 3735 | 21 | 30424 | 28235 | 5189 | 4170 | 0 | 15 | 12636 | 0 | 0 | 0 | 300 | 119 | 56 | 0 | 0 | 0 | 0 | 11 | 63 | 96 | 115 | 33 | 0 | 0 | 0 | 0 | 0 |
| SRR6757271 | 798 | 164533 | 19478 | 405 | 296 | 4332 | 3559 | 5298 | 3945 | 24 | 32789 | 30433 | 5576 | 4567 | 0 | 16 | 12763 | 0 | 0 | 0 | 306 | 134 | 60 | 0 | 0 | 0 | 0 | 12 | 71 | 105 | 125 | 35 | 0 | 0 | 0 | 0 | 0 |
| SRR6757272 | 936 | 104360 | 18467 | 408 | 183 | 2435 | 2208 | 3391 | 4241 | 21 | 27938 | 27265 | 4314 | 3803 | 0 | 54 | 13492 | 0 | 0 | 0 | 332 | 144 | 66 | 0 | 0 | 0 | 0 | 12 | 64 | 97 | 118 | 38 | 0 | 0 | 0 | 0 | 0 |
| SRR6757273 | 922 | 103409 | 18515 | 446 | 175 | 2416 | 2227 | 3360 | 4609 | 35 | 29790 | 29183 | 4591 | 4122 | 0 | 56 | 13457 | 0 | 0 | 0 | 361 | 163 | 77 | 0 | 0 | 0 | 0 | 14 | 67 | 110 | 127 | 38 | 0 | 10 | 0 | 0 | 0 |
| SRR6757274 | 865 | 80807 | 6762 | 386 | 110 | 1729 | 1486 | 2166 | 3492 | 23 | 20415 | 21491 | 3529 | 3005 | 0 | 15 | 11647 | 0 | 0 | 0 | 325 | 183 | 60 | 0 | 0 | 0 | 0 | 16 | 66 | 90 | 128 | 31 | 0 | 0 | 0 | 0 | 0 |
| SRR6757275 | 886 | 79851 | 6780 | 401 | 111 | 1750 | 1571 | 2145 | 3747 | 18 | 22046 | 22917 | 3806 | 3271 | 0 | 11 | 11812 | 0 | 0 | 0 | 364 | 204 | 60 | 0 | 0 | 0 | 0 | 13 | 70 | 91 | 139 | 32 | 0 | 0 | 0 | 0 | 0 |
| SRR6757276 | 1339 | 93887 | 16037 | 623 | 188 | 2382 | 2061 | 2782 | 3418 | 57 | 16759 | 18066 | 3202 | 2591 | 0 | 57 | 12745 | 0 | 0 | 0 | 232 | 116 | 51 | 0 | 0 | 0 | 0 | 0 | 51 | 84 | 88 | 30 | 0 | 10 | 0 | 0 | 0 |
| SRR6757277 | 1381 | 93237 | 16159 | 643 | 188 | 2364 | 2040 | 2747 | 3713 | 38 | 18086 | 19549 | 3476 | 2702 | 0 | 56 | 12954 | 0 | 0 | 0 | 253 | 124 | 60 | 0 | 0 | 0 | 0 | 0 | 50 | 83 | 94 | 38 | 0 | 10 | 0 | 0 | 0 |
| SRR6757373 | 0 | 843 | 2292 | 0 | 492 | 4063 | 4104 | 17458 | 9789 | 23 | 15086 | 25013 | 4189 | 2656 | 0 | 11537 | 106503 | 0 | 0 | 0 | 136 | 412 | 141 | 0 | 0 | 0 | 0 | 0 | 0 | 94 | 750 | 227 | 0 | 37 | 242 | 33 | 0 |
| SRR6757374 | 0 | 724 | 1861 | 0 | 1195 | 10446 | 15077 | 50722 | 12779 | 44 | 14994 | 21126 | 4375 | 2823 | 0 | 3520 | 193184 | 0 | 0 | 0 | 51 | 252 | 137 | 0 | 0 | 0 | 0 | 0 | 0 | 16 | 173 | 104 | 0 | 15 | 53 | 14 | 0 |
| SRR6757375 | 0 | 147 | 4420 | 0 | 134 | 1173 | 1349 | 8479 | 6844 | 27 | 14208 | 15 | 4177 | 2131 | 0 | 12365 | 74047 | 0 | 0 | 0 | 0 | 0 | 0 | 0 | 0 | 0 | 0 | 0 | 0 | 0 | 10 | 0 | 0 | 0 | 0 | 0 | 0 |
| SRR6757376 | 0 | 592 | 3699 | 0 | 1117 | 9182 | 13335 | 47191 | 7935 | 118 | 17838 | 17648 | 5068 | 4395 | 0 | 847 | 88568 | 0 | 0 | 0 | 23 | 190 | 46 | 0 | 0 | 0 | 0 | 0 | 0 | 14 | 212 | 74 | 0 | 0 | 0 | 0 | 0 |
| SRR6757377 | 0 | 126 | 2383 | 0 | 245 | 2018 | 2335 | 10987 | 9529 | 33 | 13129 | 24446 | 3273 | 2361 | 0 | 16719 | 67462 | 0 | 0 | 0 | 14 | 11 | 0 | 0 | 0 | 0 | 0 | 0 | 0 | 19 | 21 | 0 | 0 | 0 | 0 | 0 | 0 |
| SRR6757378 | 32 | 3378 | 1439 | 19 | 2493 | 18491 | 25361 | 99257 | 9510 | 235 | 11 | 21024 | 4644 | 4110 | 0 | 2579 | 59383 | 0 | 0 | 0 | 399 | 1800 | 442 | 0 | 10 | 0 | 0 | 0 | 39 | 284 | 2122 | 698 | 0 | 42 | 137 | 41 | 0 |
| SRR6757379 | 0 | 1412 | 1040 | 0 | 388 | 3312 | 2923 | 11831 | 11291 | 12 | 17040 | 11 | 3861 | 2572 | 0 | 38047 | 87295 | 0 | 0 | 0 | 130 | 184 | 66 | 0 | 0 | 0 | 0 | 0 | 0 | 150 | 314 | 136 | 0 | 11 | 38 | 0 | 0 |
| SRR6757380 | 0 | 3397 | 248 | 13 | 1853 | 16998 | 18761 | 56029 | 14978 | 13 | 19168 | 24459 | 5076 | 4521 | 13 | 11980 | 117805 | 0 | 0 | 0 | 188 | 735 | 239 | 0 | 0 | 0 | 0 | 0 | 0 | 202 | 1049 | 462 | 0 | 31 | 91 | 28 | 0 |
| SRR6757381 | 0 | 6074 | 1029 | 0 | 674 | 6021 | 4826 | 17601 | 13194 | 76 | 13398 | 29369 | 3392 | 2630 | 0 | 19429 | 60606 | 0 | 0 | 0 | 108 | 173 | 99 | 0 | 0 | 0 | 0 | 0 | 0 | 120 | 307 | 207 | 0 | 18 | 27 | 0 | 0 |
| SRR6757382 | 0 | 1183 | 1731 | 0 | 1530 | 13704 | 19003 | 64980 | 37959 | 134 | 15968 | 40255 | 4044 | 2240 | 0 | 3648 | 37602 | 0 | 0 | 0 | 77 | 195 | 80 | 0 | 0 | 0 | 0 | 0 | 0 | 31 | 124 | 65 | 0 | 0 | 0 | 0 | 0 |
| SRR6757383 | 0 | 736 | 3084 | 0 | 296 | 2223 | 2265 | 10293 | 12978 | 145 | 14185 | 21 | 4146 | 2107 | 0 | 11661 | 110015 | 0 | 0 | 0 | 38 | 38 | 16 | 0 | 0 | 0 | 0 | 0 | 0 | 25 | 77 | 28 | 0 | 0 | 13 | 0 | 0 |
| SRR6757384 | 0 | 1149 | 1244 | 12 | 606 | 5513 | 7359 | 28230 | 107720 | 124 | 39278 | 27 | 7410 | 2053 | 12 | 1092 | 68724 | 0 | 0 | 0 | 0 | 17 | 0 | 0 | 0 | 0 | 0 | 0 | 0 | 0 | 28 | 0 | 0 | 0 | 0 | 0 | 0 |
| SRR6757385 | 0 | 283 | 11552 | 0 | 247 | 1421 | 2287 | 11599 | 7461 | 14 | 11075 | 22473 | 2841 | 2315 | 0 | 10422 | 93373 | 0 | 0 | 0 | 44 | 120 | 50 | 0 | 0 | 0 | 0 | 0 | 0 | 38 | 270 | 109 | 0 | 0 | 25 | 0 | 0 |
| SRR6757386 | 0 | 229 | 1257 | 0 | 572 | 4464 | 7557 | 32762 | 32976 | 47 | 13873 | 27053 | 3430 | 2898 | 0 | 5309 | 120101 | 0 | 0 | 0 | 220 | 864 | 333 | 0 | 10 | 0 | 0 | 0 | 17 | 111 | 1228 | 411 | 0 | 42 | 176 | 97 | 0 |
| SRR6757387 | 0 | 424 | 3554 | 0 | 1336 | 11541 | 14506 | 46779 | 12704 | 104 | 12620 | 19407 | 6429 | 5550 | 0 | 5364 | 61623 | 0 | 0 | 0 | 15 | 58 | 17 | 0 | 0 | 0 | 0 | 0 | 0 | 12 | 73 | 24 | 0 | 0 | 15 | 0 | 0 |
| SRR6757388 | 0 | 654 | 3085 | 0 | 1533 | 20863 | 26582 | 71729 | 24214 | 150 | 21935 | 40838 | 8405 | 3470 | 0 | 2288 | 66376 | 0 | 0 | 0 | 279 | 1176 | 268 | 0 | 11 | 0 | 0 | 0 | 14 | 77 | 832 | 199 | 0 | 19 | 60 | 36 | 0 |
| SRR6757389 | 0 | 2263 | 1575 | 0 | 282 | 2254 | 2138 | 8426 | 7686 | 45 | 14499 | 11 | 3196 | 2106 | 0 | 19134 | 89585 | 0 | 0 | 0 | 55 | 75 | 19 | 0 | 0 | 0 | 0 | 0 | 0 | 91 | 154 | 34 | 0 | 0 | 35 | 0 | 0 |
| SRR6757390 | 0 | 4338 | 423 | 157 | 2174 | 12841 | 23470 | 59729 | 21522 | 257 | 9729 | 18695 | 2098 | 2113 | 0 | 1775 | 96443 | 0 | 0 | 0 | 90 | 2358 | 598 | 0 | 13 | 0 | 0 | 0 | 14 | 70 | 1847 | 545 | 0 | 39 | 147 | 0 | 0 |

### Supp Table S8.1.Columns_B_to_AL

| | | | | | | | | | | | | | | | | | | | | | | | | | | | | | | | | | | | | | |
|---|---|---|---|---|---|---|---|---|---|---|---|---|---|---|---|---|---|---|---|---|---|---|---|---|---|---|---|---|---|---|---|---|---|---|---|---|---|
| SRR6757391 | 0 | 614 | 2170 | 0 | 292 | 2255 | 2422 | 11042 | 10670 | 14 | 16322 | 23621 | 4943 | 2369 | 0 | 14124 | 102375 | 0 | 0 | 0 | 22 | 82 | 35 | 0 | 0 | 0 | 0 | 0 | 0 | 16 | 152 | 63 | 0 | 0 | 0 | 0 | 0 |
| SRR6757392 | 0 | 766 | 1246 | 0 | 2149 | 24066 | 29541 | 83661 | 29358 | 46 | 19266 | 28537 | 5559 | 3037 | 0 | 3637 | 83052 | 0 | 0 | 0 | 366 | 1685 | 333 | 0 | 0 | 0 | 0 | 0 | 39 | 175 | 1471 | 339 | 0 | 26 | 76 | 45 | 0 |
| SRR6757393 | 0 | 400 | 4969 | 0 | 193 | 1666 | 1858 | 10579 | 7863 | 0 | 14456 | 25233 | 3875 | 2097 | 0 | 12326 | 104885 | 0 | 0 | 0 | 26 | 54 | 23 | 0 | 0 | 0 | 0 | 0 | 0 | 18 | 104 | 36 | 0 | 0 | 42 | 0 | 0 |
| SRR6757394 | 0 | 456 | 4947 | 0 | 402 | 5257 | 10165 | 57310 | 4200 | 16 | 13520 | 15 | 5585 | 4426 | 0 | 3037 | 41649 | 0 | 0 | 0 | 35 | 374 | 16 | 0 | 0 | 0 | 0 | 0 | 21 | 27 | 468 | 29 | 0 | 0 | 0 | 0 | 0 |
| SRR6757395 | 0 | 7811 | 900 | 0 | 306 | 2346 | 2028 | 7683 | 9109 | 0 | 14067 | 28528 | 3578 | 1872 | 0 | 18425 | 103274 | 0 | 0 | 0 | 1060 | 661 | 216 | 0 | 0 | 0 | 0 | 0 | 39 | 541 | 986 | 324 | 0 | 26 | 80 | 11 | 0 |
| SRR6757396 | 0 | 2767 | 803 | 0 | 1042 | 7328 | 9532 | 37628 | 36867 | 15 | 21627 | 40952 | 5606 | 2704 | 0 | 8188 | 88867 | 0 | 0 | 0 | 596 | 1367 | 301 | 0 | 0 | 0 | 0 | 0 | 29 | 396 | 2290 | 491 | 0 | 28 | 75 | 18 | 0 |
| SRR6757397 | 0 | 696 | 2718 | 0 | 310 | 2789 | 2145 | 10260 | 12506 | 0 | 20358 | 35975 | 5656 | 2481 | 0 | 13164 | 84931 | 0 | 0 | 0 | 31 | 69 | 24 | 0 | 0 | 0 | 0 | 0 | 0 | 24 | 92 | 34 | 0 | 0 | 34 | 0 | 0 |
| SRR6757398 | 0 | 2660 | 2024 | 0 | 1469 | 14730 | 18336 | 77902 | 20524 | 47 | 16247 | 31129 | 3602 | 3908 | 0 | 817 | 79211 | 0 | 0 | 0 | 19 | 210 | 42 | 0 | 0 | 0 | 0 | 0 | 0 | 0 | 76 | 27 | 0 | 0 | 0 | 0 | 0 |
| SRR6757399 | 0 | 606 | 5851 | 0 | 262 | 2560 | 2171 | 10600 | 10168 | 10 | 15856 | 12 | 3294 | 2521 | 0 | 19934 | 71672 | 0 | 0 | 0 | 0 | 23 | 0 | 0 | 0 | 0 | 0 | 0 | 0 | 0 | 45 | 18 | 0 | 0 | 0 | 0 | 0 |
| SRR6757400 | 0 | 1074 | 1393 | 0 | 2206 | 15268 | 22984 | 68123 | 38564 | 42 | 8612 | 14192 | 2847 | 3424 | 0 | 5217 | 94592 | 0 | 0 | 0 | 58 | 406 | 152 | 0 | 0 | 0 | 0 | 0 | 0 | 51 | 657 | 228 | 0 | 13 | 41 | 0 | 0 |
| SRR6757401 | 0 | 1353 | 2648 | 0 | 250 | 2126 | 1886 | 8332 | 9434 | 10 | 16623 | 28608 | 4396 | 2013 | 0 | 17002 | 102422 | 0 | 0 | 0 | 52 | 144 | 95 | 0 | 0 | 0 | 0 | 0 | 0 | 40 | 309 | 158 | 0 | 31 | 148 | 16 | 0 |
| SRR6757402 | 0 | 977 | 1413 | 0 | 856 | 6886 | 8336 | 37572 | 10749 | 15 | 17042 | 33467 | 4019 | 2230 | 0 | 5640 | 104673 | 0 | 0 | 0 | 185 | 530 | 184 | 0 | 0 | 0 | 0 | 0 | 0 | 115 | 699 | 245 | 0 | 30 | 201 | 64 | 0 |
| SRR6757403 | 0 | 809 | 1684 | 0 | 272 | 1898 | 1755 | 8461 | 9525 | 91 | 14235 | 12 | 3816 | 2081 | 0 | 11959 | 104209 | 0 | 0 | 0 | 73 | 92 | 39 | 0 | 0 | 0 | 0 | 0 | 0 | 56 | 210 | 70 | 0 | 15 | 76 | 0 | 0 |
| SRR6757404 | 0 | 2242 | 829 | 0 | 1288 | 11511 | 13137 | 56143 | 28655 | 59 | 14694 | 26894 | 4615 | 3096 | 0 | 3171 | 132109 | 0 | 0 | 0 | 259 | 3176 | 666 | 12 | 15 | 0 | 0 | 0 | 24 | 63 | 1731 | 375 | 0 | 23 | 61 | 29 | 0 |
| SRR6757405 | 0 | 565 | 1086 | 0 | 314 | 2890 | 2312 | 11572 | 10411 | 0 | 17362 | 14 | 3636 | 2347 | 0 | 20199 | 103822 | 0 | 0 | 0 | 82 | 96 | 33 | 0 | 0 | 0 | 0 | 0 | 0 | 83 | 146 | 56 | 0 | 0 | 18 | 0 | 0 |
| SRR6757406 | 0 | 664 | 819 | 0 | 782 | 6522 | 6638 | 30224 | 23671 | 0 | 16662 | 24157 | 3611 | 2298 | 0 | 12755 | 108156 | 0 | 0 | 0 | 58 | 162 | 60 | 0 | 0 | 0 | 0 | 0 | 0 | 81 | 316 | 111 | 0 | 0 | 66 | 12 | 0 |
| SRR6757407 | 0 | 630 | 2333 | 0 | 281 | 2707 | 2176 | 9223 | 9148 | 0 | 5467 | 12194 | 4355 | 2239 | 0 | 13681 | 97738 | 0 | 0 | 0 | 25 | 24 | 11 | 0 | 0 | 0 | 0 | 0 | 0 | 17 | 45 | 16 | 0 | 0 | 13 | 0 | 0 |
| SRR6757408 | 0 | 269 | 2939 | 0 | 1004 | 10093 | 12249 | 54000 | 13663 | 0 | 12468 | 24909 | 4795 | 12436 | 0 | 3167 | 218388 | 0 | 0 | 0 | 125 | 341 | 121 | 0 | 0 | 0 | 0 | 0 | 0 | 51 | 360 | 100 | 0 | 17 | 121 | 48 | 0 |
| SRR6757409 | 0 | 327 | 3621 | 0 | 255 | 1919 | 2400 | 11484 | 7630 | 19 | 7233 | 12972 | 4530 | 2380 | 0 | 12513 | 129042 | 0 | 0 | 0 | 115 | 249 | 101 | 0 | 0 | 0 | 0 | 0 | 0 | 110 | 656 | 209 | 0 | 24 | 71 | 14 | 0 |
| SRR6757410 | 0 | 1125 | 1234 | 12 | 796 | 4633 | 6220 | 28198 | 19971 | 30 | 18798 | 34623 | 5538 | 2547 | 0 | 7805 | 125753 | 0 | 0 | 0 | 95 | 227 | 97 | 0 | 0 | 0 | 0 | 0 | 0 | 55 | 453 | 151 | 0 | 22 | 108 | 40 | 0 |
| SRR6757411 | 0 | 1362 | 1421 | 0 | 323 | 2462 | 2138 | 8649 | 11280 | 38 | 13384 | 14 | 3203 | 1946 | 0 | 20383 | 98423 | 0 | 0 | 0 | 72 | 117 | 48 | 0 | 0 | 0 | 0 | 0 | 0 | 77 | 246 | 88 | 0 | 12 | 62 | 0 | 0 |
| SRR6757412 | 0 | 8526 | 561 | 10 | 1685 | 16157 | 24420 | 64977 | 73268 | 26 | 13567 | 15320 | 3305 | 4624 | 0 | 2908 | 81427 | 0 | 0 | 0 | 106 | 581 | 222 | 0 | 0 | 0 | 0 | 0 | 0 | 80 | 516 | 188 | 0 | 16 | 51 | 12 | 0 |
| SRR6757413 | 0 | 5491 | 7234 | 18 | 254 | 1915 | 1897 | 6630 | 7752 | 30 | 15040 | 29555 | 3932 | 2057 | 0 | 23818 | 114084 | 0 | 0 | 0 | 170 | 564 | 166 | 0 | 0 | 0 | 0 | 0 | 0 | 139 | 1156 | 345 | 0 | 22 | 84 | 15 | 0 |
| SRR6757414 | 0 | 775 | 1507 | 0 | 866 | 10117 | 11078 | 55220 | 14109 | 15 | 18502 | 27039 | 5330 | 3139 | 0 | 3567 | 108478 | 0 | 0 | 0 | 136 | 478 | 127 | 0 | 0 | 0 | 0 | 0 | 11 | 78 | 612 | 162 | 0 | 15 | 64 | 29 | 0 |
| SRR6757415 | 0 | 728 | 3660 | 0 | 250 | 2325 | 1785 | 11595 | 12520 | 54 | 18772 | 35484 | 5291 | 2069 | 0 | 11199 | 86215 | 0 | 0 | 0 | 15 | 15 | 0 | 0 | 0 | 0 | 0 | 0 | 0 | 0 | 24 | 13 | 0 | 0 | 0 | 0 | 0 |
| SRR6757416 | 0 | 385 | 2802 | 0 | 449 | 3985 | 5006 | 25556 | 17174 | 42 | 26277 | 36939 | 7556 | 2672 | 0 | 5277 | 102901 | 0 | 0 | 0 | 56 | 173 | 54 | 0 | 0 | 0 | 0 | 0 | 0 | 30 | 233 | 73 | 0 | 0 | 62 | 27 | 0 |
| SRR6757417 | 0 | 1198 | 1703 | 0 | 305 | 2358 | 2338 | 9697 | 12992 | 13 | 14154 | 19 | 3339 | 1920 | 0 | 20133 | 98099 | 0 | 0 | 0 | 209 | 355 | 181 | 0 | 0 | 0 | 0 | 0 | 0 | 196 | 698 | 265 | 0 | 43 | 144 | 14 | 0 |
| SRR6757418 | 0 | 3103 | 722 | 19 | 1497 | 9887 | 13009 | 40529 | 22814 | 48 | 15354 | 13 | 3451 | 2075 | 0 | 18488 | 110529 | 0 | 0 | 0 | 1208 | 3383 | 1774 | 20 | 64 | 30 | 0 | 0 | 56 | 1153 | 5351 | 2703 | 24 | 366 | 963 | 233 | 0 |
| SRR6757419 | 0 | 2983 | 3563 | 0 | 553 | 4894 | 4937 | 15470 | 10899 | 0 | 16023 | 20777 | 3655 | 2631 | 0 | 15714 | 79487 | 0 | 0 | 0 | 804 | 924 | 330 | 0 | 0 | 0 | 0 | 0 | 32 | 693 | 1444 | 494 | 0 | 46 | 75 | 0 | 0 |
| SRR6757420 | 0 | 7537 | 801 | 0 | 1711 | 13042 | 18237 | 63731 | 21083 | 17 | 18979 | 16 | 4979 | 3215 | 0 | 6058 | 86362 | 0 | 0 | 0 | 1341 | 2716 | 1107 | 0 | 38 | 15 | 0 | 0 | 60 | 788 | 2956 | 1190 | 0 | 156 | 348 | 81 | 0 |
| SRR6757421 | 0 | 396 | 3237 | 0 | 878 | 7295 | 7433 | 26338 | 16353 | 14 | 16764 | 20 | 3764 | 2686 | 0 | 20268 | 91089 | 0 | 0 | 0 | 32 | 59 | 28 | 0 | 0 | 0 | 0 | 0 | 0 | 37 | 155 | 58 | 0 | 16 | 48 | 0 | 0 |
| SRR6757422 | 0 | 597 | 2705 | 0 | 1084 | 7598 | 10948 | 47772 | 33926 | 15 | 13132 | 26620 | 3366 | 2585 | 0 | 10775 | 84364 | 0 | 0 | 0 | 71 | 379 | 131 | 0 | 0 | 0 | 0 | 0 | 0 | 54 | 468 | 167 | 0 | 25 | 82 | 0 | 0 |
| SRR6757423 | 0 | 2944 | 956 | 231 | 286 | 1936 | 2205 | 10992 | 7100 | 0 | 13678 | 31350 | 3188 | 1902 | 0 | 15037 | 123837 | 0 | 0 | 0 | 604 | 1219 | 319 | 0 | 0 | 0 | 0 | 0 | 29 | 517 | 2184 | 544 | 0 | 32 | 122 | 12 | 0 |
| SRR6757424 | 17 | 23974 | 1606 | 73 | 1667 | 19894 | 15637 | 55385 | 17618 | 13 | 14960 | 26670 | 4390 | 3120 | 0 | 9232 | 95979 | 0 | 0 | 0 | 734 | 2156 | 578 | 0 | 0 | 0 | 0 | 0 | 62 | 508 | 3006 | 887 | 0 | 30 | 94 | 14 | 0 |
| SRR6757425 | 0 | 1812 | 3955 | 11 | 446 | 4410 | 3950 | 12598 | 11728 | 0 | 15285 | 25394 | 3804 | 2159 | 0 | 19356 | 97164 | 0 | 0 | 0 | 62 | 58 | 14 | 0 | 0 | 0 | 0 | 0 | 0 | 61 | 99 | 31 | 0 | 0 | 0 | 0 | 0 |
| SRR6757426 | 0 | 1387 | 4635 | 0 | 1675 | 21791 | 21918 | 69523 | 38743 | 21 | 22427 | 33119 | 5010 | 4092 | 0 | 4381 | 67055 | 0 | 0 | 0 | 482 | 1436 | 237 | 0 | 0 | 0 | 0 | 0 | 35 | 265 | 1462 | 303 | 0 | 11 | 17 | 11 | 0 |
| SRR6757427 | 10 | 3979 | 2549 | 23 | 278 | 2280 | 2257 | 11197 | 7578 | 14 | 11706 | 24947 | 3296 | 2170 | 0 | 18541 | 136522 | 0 | 0 | 0 | 825 | 998 | 288 | 0 | 0 | 0 | 0 | 0 | 48 | 757 | 2271 | 607 | 0 | 17 | 47 | 0 | 0 |
| SRR6757428 | 0 | 2785 | 1190 | 0 | 2022 | 24133 | 25742 | 68084 | 30143 | 68 | 11950 | 25519 | 3353 | 3633 | 0 | 4170 | 100355 | 0 | 0 | 0 | 208 | 3175 | 1082 | 12 | 14 | 0 | 0 | 0 | 23 | 88 | 1436 | 635 | 0 | 23 | 12 | 0 | 0 |
| SRR6757429 | 0 | 279 | 1274 | 0 | 193 | 1798 | 1580 | 7447 | 7974 | 0 | 15169 | 27588 | 3942 | 1870 | 0 | 20791 | 93230 | 0 | 0 | 0 | 0 | 18 | 0 | 0 | 0 | 0 | 0 | 0 | 0 | 0 | 31 | 15 | 0 | 0 | 0 | 0 | 0 |
| SRR6757430 | 0 | 462 | 530 | 19 | 1172 | 12307 | 12023 | 50878 | 15470 | 0 | 14662 | 12 | 4322 | 3263 | 0 | 7285 | 91275 | 0 | 0 | 0 | 54 | 353 | 59 | 0 | 0 | 0 | 0 | 0 | 0 | 32 | 190 | 64 | 0 | 0 | 20 | 0 | 0 |
| SRR6757431 | 0 | 546 | 3946 | 0 | 176 | 1619 | 1598 | 9926 | 9393 | 0 | 12250 | 24609 | 3543 | 1658 | 0 | 9373 | 92374 | 0 | 0 | 0 | 46 | 35 | 0 | 0 | 0 | 0 | 0 | 0 | 0 | 50 | 63 | 11 | 0 | 0 | 0 | 0 | 0 |
| SRR6757432 | 0 | 2319 | 1080 | 0 | 1074 | 8950 | 11643 | 35011 | 12085 | 47 | 14731 | 20074 | 3771 | 3289 | 0 | 12564 | 144903 | 0 | 0 | 0 | 772 | 2760 | 760 | 0 | 11 | 0 | 0 | 0 | 33 | 419 | 2356 | 1042 | 0 | 49 | 156 | 110 | 0 |
| SRR6757433 | 0 | 978 | 3335 | 0 | 241 | 2468 | 1855 | 10226 | 8662 | 0 | 15938 | 22 | 4708 | 1666 | 0 | 10802 | 78260 | 0 | 0 | 0 | 44 | 46 | 0 | 0 | 0 | 0 | 0 | 0 | 0 | 33 | 63 | 13 | 0 | 0 | 0 | 0 | 0 |
| SRR6757434 | 11 | 2075 | 2576 | 29 | 1386 | 15243 | 17442 | 41705 | 11232 | 51 | 13643 | 17589 | 5931 | 4079 | 56 | 3668 | 86504 | 0 | 0 | 0 | 71 | 1869 | 280 | 10 | 0 | 0 | 0 | 0 | 0 | 33 | 781 | 229 | 0 | 0 | 28 | 31 | 0 |
| SRR6757435 | 0 | 1385 | 2256 | 0 | 348 | 3034 | 2504 | 11148 | 9455 | 14 | 13553 | 10 | 3518 | 2165 | 0 | 24710 | 103646 | 0 | 0 | 0 | 99 | 338 | 173 | 0 | 0 | 0 | 0 | 0 | 0 | 77 | 668 | 320 | 0 | 33 | 163 | 35 | 0 |
| SRR6757436 | 0 | 2480 | 1991 | 19 | 1889 | 21016 | 24862 | 67852 | 15133 | 106 | 23181 | 30622 | 4741 | 3971 | 0 | 4672 | 88498 | 0 | 0 | 0 | 281 | 1547 | 740 | 11 | 17 | 0 | 0 | 0 | 13 | 163 | 1510 | 888 | 0 | 62 | 224 | 108 | 0 |
| SRR6757437 | 0 | 3509 | 1810 | 0 | 406 | 3196 | 2869 | 12612 | 7751 | 0 | 17938 | 10 | 3272 | 2231 | 0 | 22071 | 108405 | 0 | 0 | 0 | 2029 | 1611 | 536 | 0 | 0 | 0 | 0 | 0 | 69 | 1319 | 2905 | 972 | 0 | 44 | 191 | 68 | 0 |
| SRR6757438 | 0 | 3311 | 417 | 23 | 2542 | 13955 | 21254 | 61556 | 28207 | 76 | 13779 | 18297 | 2508 | 2984 | 0 | 13267 | 119207 | 0 | 0 | 0 | 337 | 1569 | 678 | 0 | 0 | 0 | 0 | 0 | 20 | 184 | 1491 | 691 | 0 | 38 | 125 | 51 | 0 |
| SRR6757439 | 0 | 10339 | 1363 | 40 | 625 | 4854 | 4272 | 12646 | 9323 | 19 | 16980 | 27377 | 3420 | 2129 | 0 | 31378 | 100719 | 0 | 0 | 0 | 549 | 690 | 317 | 0 | 0 | 0 | 0 | 0 | 33 | 432 | 1275 | 709 | 0 | 26 | 24 | 0 | 0 |
| SRR6757440 | 23 | 18514 | 338 | 92 | 2119 | 22434 | 24279 | 59847 | 39328 | 37 | 15509 | 19360 | 3843 | 2707 | 0 | 2706 | 57684 | 0 | 0 | 0 | 1422 | 4326 | 1315 | 0 | 10 | 0 | 0 | 0 | 63 | 362 | 3150 | 989 | 0 | 31 | 64 | 21 | 0 |
| SRR6757441 | 0 | 7186 | 458 | 44 | 748 | 4987 | 4948 | 9992 | 12152 | 23 | 12669 | 22917 | 2627 | 2748 | 0 | 65501 | 53965 | 0 | 0 | 0 | 394 | 657 | 328 | 0 | 0 | 0 | 0 | 0 | 20 | 628 | 1303 | 811 | 0 | 54 | 61 | 0 | 0 |
| SRR6757442 | 0 | 1404 | 498 | 40 | 2914 | 20763 | 39059 | 87687 | 27520 | 69 | 13522 | 16589 | 2528 | 10137 | 0 | 4292 | 65639 | 0 | 0 | 0 | 70 | 1057 | 240 | 0 | 0 | 0 | 0 | 0 | 20 | 57 | 1397 | 376 | 0 | 0 | 0 | 0 | 0 |
| SRR6757443 | 0 | 1752 | 1897 | 0 | 572 | 5192 | 4056 | 16208 | 13714 | 45 | 16085 | 14 | 3702 | 2345 | 0 | 36841 | 76845 | 0 | 0 | 0 | 32 | 99 | 39 | 0 | 0 | 0 | 0 | 0 | 0 | 43 | 185 | 88 | 0 | 10 | 26 | 0 | 0 |
| SRR6757444 | 18 | 4487 | 1135 | 20 | 3541 | 37685 | 39716 | 91577 | 44684 | 134 | 21171 | 21963 | 3878 | 3971 | 0 | 4917 | 72639 | 0 | 0 | 0 | 217 | 1010 | 215 | 0 | 0 | 0 | 0 | 0 | 35 | 122 | 848 | 192 | 0 | 0 | 13 | 0 | 0 |
| SRR6757445 | 0 | 1188 | 2662 | 0 | 494 | 3930 | 3650 | 12533 | 15609 | 0 | 16751 | 18 | 3620 | 2292 | 0 | 25835 | 78950 | 0 | 0 | 0 | 165 | 190 | 96 | 0 | 0 | 0 | 0 | 0 | 0 | 184 | 334 | 186 | 0 | 36 | 108 | 0 | 0 |
| SRR6757446 | 0 | 798 | 3591 | 0 | 1452 | 15194 | 19188 | 51854 | 32112 | 45 | 15366 | 28196 | 3895 | 2784 | 0 | 10159 | 55177 | 0 | 0 | 0 | 64 | 72 | 30 | 0 | 0 | 0 | 0 | 0 | 0 | 44 | 50 | 25 | 0 | 0 | 13 | 0 | 0 |
| SRR6757447 | 0 | 938 | 2896 | 0 | 441 | 3553 | 3198 | 10802 | 13388 | 13 | 20495 | 17 | 4286 | 2451 | 0 | 24597 | 86123 | 0 | 0 | 0 | 111 | 83 | 31 | 0 | 0 | 0 | 0 | 0 | 0 | 119 | 148 | 60 | 0 | 0 | 24 | 0 | 0 |
| SRR6757448 | 13 | 2130 | 1426 | 27 | 3320 | 37995 | 45829 | 96238 | 13537 | 54 | 21019 | 12 | 5056 | 4590 | 0 | 2627 | 73349 | 0 | 0 | 0 | 855 | 3733 | 937 | 16 | 50 | 14 | 0 | 0 | 36 | 385 | 1629 | 637 | 0 | 81 | 237 | 45 | 0 |
| SRR6757449 | 0 | 673 | 2255 | 0 | 428 | 4365 | 4495 | 20788 | 9603 | 12 | 14455 | 31618 | 3739 | 3001 | 0 | 5797 | 66134 | 0 | 0 | 0 | 67 | 34 | 0 | 0 | 0 | 0 | 0 | 0 | 0 | 59 | 62 | 19 | 0 | 0 | 0 | 0 | 0 |
| SRR6757450 | 0 | 3577 | 1109 | 24 | 1175 | 12553 | 11471 | 38949 | 12618 | 91 | 21853 | 41345 | 7390 | 2615 | 0 | 6956 | 70875 | 0 | 0 | 0 | 87 | 98 | 18 | 0 | 0 | 0 | 0 | 0 | 0 | 84 | 153 | 32 | 0 | 0 | 0 | 0 | 0 |
| SRR6757451 | 0 | 2233 | 1423 | 15 | 636 | 4731 | 4673 | 15080 | 9903 | 130 | 10967 | 21772 | 2913 | 2297 | 0 | 18338 | 153968 | 0 | 0 | 0 | 259 | 1155 | 601 | 0 | 12 | 0 | 0 | 0 | 18 | 316 | 3602 | 1125 | 0 | 105 | 480 | 86 | 0 |
| SRR6757452 | 32 | 22653 | 6752 | 26 | 1647 | 26615 | 26283 | 41805 | 28724 | 306 | 12636 | 20435 | 3057 | 2731 | 0 | 9790 | 66531 | 0 | 0 | 0 | 211 | 1758 | 392 | 0 | 0 | 0 | 0 | 0 | 26 | 121 | 1516 | 442 | 0 | 17 | 34 | 35 | 0 |
| SRR6757453 | 0 | 2353 | 1822 | 0 | 266 | 1721 | 1740 | 6557 | 7920 | 0 | 10197 | 20858 | 2866 | 1801 | 0 | 20493 | 170007 | 0 | 0 | 0 | 772 | 1686 | 687 | 0 | 11 | 0 | 0 | 0 | 36 | 917 | 5175 | 1282 | 0 | 96 | 582 | 126 | 0 |
| SRR6757454 | 0 | 10552 | 1338 | 33 | 1720 | 15997 | 16672 | 56871 | 31330 | 0 | 16654 | 32086 | 4841 | 4106 | 0 | 8047 | 100843 | 0 | 0 | 0 | 1050 | 1230 | 330 | 0 | 0 | 0 | 0 | 0 | 43 | 705 | 1930 | 518 | 0 | 31 | 137 | 28 | 0 |
| SRR6757455 | 0 | 399 | 4032 | 0 | 215 | 1870 | 1836 | 8535 | 6665 | 71 | 12954 | 28304 | 3586 | 2025 | 0 | 14722 | 93428 | 0 | 0 | 0 | 14 | 36 | 13 | 0 | 0 | 0 | 0 | 0 | 0 | 12 | 92 | 32 | 0 | 0 | 0 | 0 | 0 |
| SRR6757456 | 0 | 854 | 1500 | 14 | 1811 | 28909 | 30027 | 91126 | 16331 | 180 | 15509 | 10 | 4892 | 2528 | 0 | 1996 | 98693 | 0 | 0 | 0 | 43 | 739 | 287 | 0 | 11 | 0 | 0 | 0 | 0 | 27 | 1083 | 488 | 0 | 49 | 130 | 39 | 0 |
| SRR6757457 | 0 | 287 | 2707 | 0 | 185 | 1502 | 1589 | 6919 | 5991 | 52 | 13806 | 26299 | 4228 | 2204 | 0 | 14082 | 72758 | 0 | 0 | 0 | 44 | 76 | 21 | 0 | 0 | 0 | 0 | 0 | 0 | 38 | 153 | 38 | 0 | 0 | 17 | 0 | 0 |
| SRR6757458 | 0 | 135 | 1656 | 12 | 586 | 5230 | 7337 | 41257 | 19754 | 96 | 21019 | 31545 | 5962 | 2341 | 0 | 3196 | 99155 | 0 | 0 | 0 | 26 | 146 | 43 | 0 | 0 | 0 | 0 | 0 | 0 | 17 | 174 | 61 | 0 | 0 | 27 | 0 | 0 |
| SRR6757459 | 0 | 827 | 1757 | 0 | 479 | 4508 | 3412 | 12359 | 7820 | 43 | 15576 | 33345 | 4272 | 2539 | 0 | 20718 | 120023 | 0 | 0 | 0 | 366 | 787 | 265 | 0 | 0 | 0 | 0 | 0 | 12 | 291 | 1637 | 465 | 0 | 78 | 343 | 30 | 0 |
| SRR6757460 | 0 | 1206 | 2128 | 16 | 1141 | 12260 | 16287 | 49547 | 53318 | 274 | 12200 | 17845 | 3843 | 4005 | 0 | 2052 | 108809 | 0 | 0 | 0 | 50 | 580 | 252 | 0 | 11 | 0 | 0 | 0 | 0 | 34 | 1003 | 392 | 0 | 46 | 120 | 33 | 0 |
| SRR6757461 | 0 | 186 | 4905 | 0 | 228 | 1396 | 1655 | 8502 | 8608 | 63 | 10581 | 26093 | 3066 | 1988 | 0 | 16605 | 105822 | 0 | 0 | 0 | 15 | 64 | 53 | 0 | 0 | 0 | 0 | 0 | 0 | 11 | 161 | 81 | 0 | 18 | 125 | 0 | 0 |
| SRR6757462 | 0 | 420 | 8472 | 0 | 1067 | 8254 | 12596 | 56845 | 20806 | 174 | 13221 | 30 | 5256 | 4504 | 0 | 3242 | 76763 | 0 | 0 | 0 | 207 | 1328 | 400 | 0 | 15 | 0 | 0 | 0 | 0 | 72 | 978 | 331 | 0 | 34 | 51 | 0 | 0 |
| SRR6757463 | 0 | 1524 | 8200 | 0 | 508 | 4485 | 3762 | 15334 | 12603 | 58 | 16929 | 35856 | 4918 | 2072 | 0 | 19307 | 80461 | 0 | 0 | 0 | 32 | 209 | 160 | 0 | 0 | 0 | 0 | 0 | 0 | 35 | 453 | 326 | 0 | 44 | 117 | 0 | 0 |
| SRR6757464 | 0 | 4461 | 2470 | 60 | 3165 | 46220 | 45716 | 64454 | 13899 | 257 | 18795 | 25302 | 6259 | 4244 | 48 | 3779 | 57080 | 0 | 0 | 0 | 1804 | 8443 | 2173 | 74 | 181 | 49 | 0 | 0 | 24 | 220 | 1749 | 609 | 0 | 133 | 448 | 494 | 0 |
| SRR6757465 | 0 | 1255 | 1496 | 0 | 364 | 3067 | 2617 | 11881 | 16484 | 111 | 7017 | 13 | 4057 | 1835 | 0 | 18975 | 97334 | 0 | 0 | 0 | 100 | 314 | 161 | 0 | 0 | 0 | 0 | 0 | 0 | 76 | 599 | 229 | 0 | 56 | 269 | 29 | 0 |
| SRR6757466 | 0 | 6801 | 1355 | 19 | 1008 | 14858 | 18744 | 51216 | 52837 | 243 | 14515 | 32301 | 7474 | 2858 | 18 | 2490 | 99288 | 0 | 0 | 0 | 132 | 4509 | 667 | 20 | 27 | 0 | 0 | 0 | 15 | 52 | 1329 | 385 | 0 | 52 | 113 | 58 | 0 |
| SRR6757467 | 0 | 345 | 11459 | 15 | 192 | 1597 | 2130 | 12056 | 13200 | 65 | 7182 | 11 | 3782 | 2382 | 0 | 8313 | 84859 | 0 | 0 | 0 | 25 | 67 | 42 | 0 | 0 | 0 | 0 | 0 | 0 | 26 | 186 | 89 | 0 | 13 | 61 | 0 | 0 |
| SRR6757468 | 0 | 516 | 3326 | 0 | 532 | 4567 | 7594 | 30284 | 17884 | 103 | 17143 | 34592 | 4713 | 2264 | 0 | 6256 | 108375 | 0 | 0 | 0 | 46 | 171 | 53 | 0 | 0 | 0 | 0 | 0 | 0 | 31 | 187 | 68 | 0 | 0 | 32 | 0 | 0 |
| SRR6761159 | 85 | 4686 | 1704 | 3749 | 0 | 137 | 73 | 487 | 2449 | 0 | 1642 | 2663 | 378 | 1299 | 0 | 634 | 10959 | 0 | 0 | 0 | 0 | 11 | 0 | 0 | 0 | 0 | 0 | 0 | 0 | 0 | 0 | 0 | 0 | 0 | 0 | 0 | 0 |
| SRR6761162 | 302 | 4832 | 1208 | 302 | 0 | 0 | 0 | 200 | 543 | 0 | 194 | 261 | 49 | 331 | 0 | 48 | 807 | 0 | 0 | 0 | 0 | 0 | 0 | 0 | 0 | 0 | 0 | 0 | 0 | 0 | 0 | 0 | 0 | 0 | 0 | 0 | 0 |
| SRR6761163 | 216 | 4752 | 972 | 540 | 0 | 30 | 39 | 309 | 847 | 0 | 1251 | 1636 | 316 | 810 | 0 | 216 | 6145 | 0 | 0 | 0 | 0 | 0 | 0 | 0 | 0 | 0 | 0 | 0 | 0 | 0 | 0 | 0 | 0 | 0 | 0 | 0 | 0 |
| SRR6761165 | 0 | 5553 | 2508 | 1881 | 0 | 98 | 0 | 430 | 1550 | 0 | 1897 | 2256 | 323 | 534 | 0 | 302 | 7653 | 0 | 0 | 0 | 0 | 0 | 0 | 0 | 0 | 0 | 0 | 0 | 0 | 0 | 0 | 0 | 0 | 0 | 0 | 0 | 0 |
| SRR6761167 | 0 | 14214 | 4610 | 1153 | 0 | 88 | 23 | 389 | 1538 | 0 | 792 | 820 | 88 | 709 | 0 | 264 | 4082 | 0 | 0 | 0 | 0 | 0 | 0 | 0 | 0 | 0 | 0 | 0 | 0 | 0 | 0 | 0 | 0 | 0 | 0 | 0 | 0 |
| SRR6761175 | 172 | 6108 | 2151 | 3699 | 0 | 54 | 18 | 215 | 476 | 0 | 734 | 1283 | 96 | 334 | 0 | 211 | 4304 | 0 | 0 | 0 | 0 | 0 | 0 | 0 | 0 | 0 | 0 | 0 | 0 | 0 | 0 | 0 | 0 | 0 | 0 | 0 | 0 |
| SRR6761179 | 0 | 6171 | 1543 | 5999 | 0 | 32 | 13 | 194 | 605 | 0 | 637 | 853 | 219 | 386 | 0 | 140 | 5279 | 0 | 0 | 0 | 0 | 0 | 0 | 0 | 0 | 0 | 0 | 0 | 0 | 0 | 0 | 0 | 0 | 0 | 0 | 0 | 0 |
| SRR6761181 | 0 | 2173 | 1114 | 2618 | 0 | 100 | 73 | 517 | 750 | 0 | 1020 | 1053 | 563 | 1405 | 0 | 212 | 7499 | 0 | 0 | 0 | 0 | 0 | 0 | 0 | 0 | 0 | 0 | 0 | 0 | 0 | 0 | 0 | 0 | 0 | 0 | 0 | 0 |
| SRR6761182 | 42 | 1761 | 1300 | 3229 | 0 | 66 | 32 | 428 | 1055 | 0 | 826 | 883 | 263 | 541 | 0 | 128 | 5174 | 0 | 0 | 0 | 0 | 0 | 0 | 0 | 0 | 0 | 0 | 0 | 0 | 0 | 0 | 0 | 0 | 0 | 0 | 0 | 0 |
| SRR6761183 | 0 | 4257 | 3275 | 6277 | 0 | 103 | 19 | 347 | 1515 | 0 | 2089 | 1644 | 428 | 880 | 0 | 268 | 3639 | 0 | 0 | 0 | 0 | 0 | 0 | 0 | 0 | 0 | 0 | 0 | 0 | 0 | 0 | 0 | 0 | 0 | 0 | 0 | 0 |
| SRR6761184 | 32 | 2218 | 1800 | 6942 | 0 | 70 | 19 | 533 | 1504 | 0 | 1407 | 1067 | 379 | 1156 | 0 | 150 | 2035 | 0 | 0 | 0 | 0 | 0 | 0 | 0 | 0 | 0 | 0 | 0 | 0 | 0 | 0 | 0 | 0 | 0 | 0 | 0 | 0 |
| SRR6761188 | 0 | 2477 | 976 | 8031 | 0 | 223 | 65 | 779 | 2810 | 0 | 4531 | 3333 | 455 | 997 | 0 | 377 | 6806 | 0 | 0 | 0 | 0 | 0 | 0 | 0 | 0 | 0 | 0 | 0 | 0 | 0 | 0 | 0 | 0 | 0 | 0 | 0 | 0 |
| SRR6761192 | 340 | 3063 | 3404 | 3063 | 0 | 154 | 120 | 1081 | 6639 | 0 | 5782 | 5953 | 721 | 1973 | 0 | 429 | 5644 | 0 | 0 | 0 | 70 | 23 | 0 | 0 | 0 | 0 | 0 | 0 | 0 | 13 | 17 | 0 | 0 | 0 | 0 | 0 | 0 |
| SRR6761198 | 18 | 1323 | 917 | 7338 | 13 | 249 | 88 | 948 | 3822 | 0 | 7204 | 7287 | 769 | 1443 | 12 | 691 | 9352 | 0 | 0 | 0 | 0 | 0 | 0 | 0 | 0 | 0 | 0 | 0 | 0 | 0 | 0 | 0 | 0 | 0 | 0 | 0 | 0 |
| SRR6767691 | 50 | 871 | 261 | 1668 | 0 | 235 | 356 | 882 | 765 | 0 | 1618 | 1116 | 286 | 858 | 0 | 483 | 6818 | 0 | 0 | 0 | 0 | 0 | 0 | 0 | 0 | 0 | 0 | 0 | 0 | 0 | 0 | 0 | 0 | 0 | 0 | 0 | 0 |
| SRR6767701 | 0 | 86 | 62 | 1483 | 0 | 30 | 25 | 308 | 46 | 0 | 146 | 64 | 105 | 550 | 0 | 20 | 367 | 0 | 0 | 0 | 0 | 0 | 0 | 0 | 0 | 0 | 0 | 0 | 0 | 0 | 0 | 0 | 0 | 0 | 0 | 0 | 0 |
| SRR6767712 | 0 | 481 | 67 | 2484 | 0 | 42 | 66 | 514 | 280 | 0 | 422 | 411 | 178 | 528 | 0 | 359 | 1531 | 0 | 0 | 0 | 0 | 0 | 0 | 0 | 0 | 0 | 0 | 0 | 0 | 0 | 0 | 0 | 0 | 0 | 0 | 0 | 0 |
| SRR6767713 | 0 | 782 | 219 | 1097 | 0 | 57 | 132 | 836 | 338 | 0 | 599 | 438 | 267 | 642 | 0 | 407 | 5100 | 0 | 0 | 0 | 0 | 0 | 0 | 0 | 0 | 0 | 0 | 0 | 0 | 0 | 0 | 0 | 0 | 0 | 0 | 0 | 0 |
| SRR6821085 | 197 | 11095 | 4560 | 171 | 3378 | 11548 | 47237 | 61464 | 3324 | 92 | 9924 | 18776 | 2993 | 3437 | 0 | 3412 | 25012 | 0 | 0 | 0 | 1844 | 1793 | 990 | 12 | 0 | 0 | 0 | 0 | 138 | 886 | 2587 | 1261 | 0 | 18 | 18 | 13 | 0 |
| SRR6821086 | 81 | 21932 | 6244 | 314 | 5834 | 4927 | 58842 | 123327 | 6807 | 156 | 9544 | 12907 | 2998 | 5142 | 0 | 2068 | 29018 | 0 | 0 | 50 | 2204 | 1431 | 1029 | 13 | 0 | 0 | 0 | 0 | 541 | 1519 | 6078 | 2203 | 0 | 45 | 63 | 14 | 0 |
| SRR6821087 | 22 | 3321 | 8042 | 212 | 4569 | 4617 | 58825 | 116054 | 3047 | 268 | 6290 | 13301 | 2050 | 3525 | 0 | 1205 | 13942 | 0 | 0 | 0 | 177 | 276 | 194 | 0 | 0 | 0 | 0 | 0 | 52 | 220 | 1333 | 887 | 0 | 0 | 0 | 0 | 0 |
| SRR6821088 | 26 | 1648 | 1210 | 75 | 29 | 148 | 246 | 698 | 356 | 29 | 21440 | 47189 | 9342 | 4594 | 69 | 11865 | 172250 | 0 | 0 | 13 | 752 | 868 | 206 | 0 | 0 | 0 | 0 | 121 | 381 | 220 | 823 | 193 | 0 | 0 | 0 | 0 | 0 |
| SRR6821089 | 18 | 459 | 872 | 45 | 38 | 177 | 370 | 1033 | 863 | 22 | 25085 | 11 | 21 | 1090 | 147 | 438 | 132806 | 0 | 12 | 0 | 309 | 1090 | 112 | 0 | 0 | 0 | 0 | 60 | 152 | 20 | 138 | 20 | 0 | 0 | 0 | 0 | 0 |
| SRR6821090 | 1312 | 2316 | 3262 | 67 | 32 | 164 | 275 | 824 | 327 | 43 | 14238 | 46021 | 9447 | 4224 | 140 | 8681 | 166793 | 0 | 23 | 50 | 2208 | 790 | 193 | 0 | 0 | 0 | 0 | 313 | 1552 | 509 | 568 | 96 | 0 | 0 | 0 | 0 | 0 |
| SRR6823984 | 819 | 3138 | 32230 | 368 | 17867 | 15040 | 176991 | 43680 | 8998 | 5393 | 30867 | 18182 | 13711 | 49326 | 10860 | 39308 | 4886 | 0 | 79 | 50 | 77 | 46 | 44 | 11 | 11 | 0 | 14 | 112 | 221 | 61 | 35 | 0 | 0 | 0 | 0 | 0 | 0 |
| SRR6823985 | 158 | 2574 | 65840 | 838 | 19183 | 12744 | 172374 | 35853 | 16062 | 12357 | 38178 | 33 | 17656 | 65801 | 12647 | 34639 | 5114 | 0 | 120 | 67 | 79 | 29 | 43 | 0 | 0 | 0 | 0 | 111 | 167 | 38 | 15 | 0 | 0 | 0 | 0 | 0 | 0 |
| SRR6823986 | 435 | 1181 | 29131 | 661 | 13864 | 20798 | 128213 | 67611 | 102247 | 7431 | 28245 | 32 | 9334 | 16481 | 9789 | 17613 | 7245 | 0 | 31 | 14 | 51 | 97 | 80 | 24 | 15 | 23 | 18 | 106 | 205 | 104 | 79 | 0 | 0 | 0 | 0 | 17 | 0 |
| SRR6823987 | 571 | 2019 | 59643 | 778 | 12584 | 18016 | 114971 | 57398 | 97312 | 11014 | 33047 | 26 | 11171 | 20800 | 8918 | 16456 | 6626 | 0 | 34 | 18 | 46 | 56 | 61 | 44 | 77 | 0 | 18 | 79 | 156 | 69 | 41 | 0 | 0 | 0 | 0 | 16 | 0 |
| SRR6823990 | 747 | 3569 | 74854 | 810 | 52099 | 43876 | 267009 | 50505 | 2957 | 2197 | 23450 | 6667 | 14921 | 37450 | 25173 | 845 | 4172 | 0 | 18 | 19 | 16 | 56 | 17 | 0 | 0 | 0 | 0 | 28 | 53 | 20 | 13 | 0 | 0 | 0 | 0 | 0 | 0 |
| SRR6823991 | 783 | 2789 | 87944 | 1601 | 47997 | 37192 | 227090 | 58903 | 4616 | 3751 | 26158 | 7417 | 16907 | 37385 | 24490 | 889 | 4463 | 0 | 20 | 30 | 0 | 15 | 0 | 28 | 37 | 0 | 0 | 34 | 77 | 27 | 13 | 0 | 17 | 12 | 0 | 0 | 0 |
| SRR6829958 | 65 | 3743 | 13054 | 864 | 3966 | 2623 | 17560 | 3452 | 170792 | 4048 | 17 | 27 | 26497 | 60506 | 6070 | 1529 | 25998 | 0 | 411 | 371 | 472 | 2271 | 1157 | 171 | 56 | 30 | 0 | 894 | 2131 | 496 | 59 | 12 | 0 | 0 | 12 | 57 | 0 |
| SRR6829959 | 222 | 5587 | 7701 | 1081 | 3520 | 2739 | 16729 | 3858 | 151435 | 5139 | 18 | 41 | 31635 | 28441 | 817 | 2598 | 36117 | 0 | 500 | 237 | 556 | 635 | 248 | 25 | 11 | 0 | 13 | 1733 | 3896 | 759 | 60 | 0 | 0 | 0 | 0 | 10 | 0 |
| SRR6829960 | 91 | 3406 | 19309 | 279 | 3613 | 2586 | 17838 | 4251 | 159178 | 4750 | 39620 | 13182 | 21373 | 50684 | 5481 | 1307 | 26548 | 0 | 446 | 295 | 386 | 1911 | 1030 | 155 | 46 | 23 | 0 | 862 | 1729 | 316 | 66 | 12 | 0 | 0 | 0 | 50 | 0 |
| SRR6829961 | 144 | 4395 | 21180 | 317 | 2883 | 2122 | 16062 | 4181 | 182052 | 12380 | 46983 | 19529 | 25462 | 20734 | 459 | 2416 | 39118 | 0 | 521 | 246 | 377 | 440 | 206 | 19 | 0 | 0 | 0 | 1748 | 4173 | 467 | 50 | 0 | 0 | 0 | 0 | 10 | 0 |
| SRR6829962 | 76 | 4086 | 26804 | 220 | 3235 | 2477 | 17524 | 3283 | 136870 | 5239 | 17 | 34 | 26909 | 59407 | 5403 | 1055 | 21228 | 0 | 217 | 182 | 173 | 1169 | 718 | 113 | 46 | 20 | 0 | 411 | 844 | 102 | 23 | 0 | 0 | 0 | 0 | 43 | 0 |
| SRR6829963 | 31 | 3636 | 35157 | 31 | 2071 | 1377 | 10119 | 2234 | 331532 | 8482 | 11 | 41 | 18963 | 19437 | 321 | 1654 | 31262 | 0 | 61 | 29 | 57 | 111 | 41 | 0 | 0 | 0 | 0 | 232 | 390 | 56 | 0 | 0 | 0 | 0 | 0 | 0 | 0 |
| SRR6831609 | 65 | 908 | 65844 | 22 | 7947 | 16110 | 83715 | 11357 | 23964 | 2032 | 40558 | 20 | 33200 | 46212 | 1070 | 259 | 127127 | 0 | 0 | 0 | 0 | 0 | 0 | 0 | 0 | 0 | 0 | 0 | 0 | 0 | 0 | 0 | 0 | 0 | 0 | 0 | 0 |
| SRR6831611 | 39 | 1141 | 127940 | 334 | 7553 | 15185 | 60728 | 6198 | 28502 | 1160 | 24214 | 13576 | 17931 | 59135 | 1433 | 335 | 112358 | 0 | 0 | 0 | 0 | 0 | 0 | 0 | 0 | 0 | 0 | 0 | 0 | 0 | 0 | 0 | 0 | 0 | 0 | 0 | 0 |
| SRR6831613 | 29 | 860 | 71938 | 115 | 9165 | 19128 | 76945 | 7672 | 15057 | 1118 | 24674 | 19111 | 14504 | 61883 | 1944 | 225 | 104509 | 0 | 0 | 0 | 0 | 0 | 0 | 0 | 0 | 0 | 0 | 0 | 0 | 0 | 0 | 0 | 0 | 0 | 0 | 0 | 0 |
| SRR6837863 | 0 | 54 | 2241 | 0 | 963 | 6705 | 13879 | 5401 | 1110 | 82 | 25964 | 16641 | 3780 | 29769 | 0 | 30334 | 29854 | 0 | 0 | 0 | 0 | 0 | 0 | 0 | 0 | 0 | 0 | 0 | 0 | 0 | 0 | 11 | 0 | 0 | 0 | 94 | 0 |
| SRR6837867 | 0 | 97 | 1901 | 0 | 934 | 6202 | 14305 | 5975 | 1124 | 65 | 27989 | 20890 | 3671 | 31992 | 0 | 27913 | 30980 | 0 | 0 | 0 | 0 | 0 | 0 | 0 | 0 | 0 | 0 | 0 | 0 | 0 | 0 | 0 | 0 | 0 | 0 | 52 | 0 |
| SRR6837868 | 0 | 275 | 1854 | 0 | 927 | 5486 | 12650 | 5741 | 1202 | 89 | 30239 | 21664 | 3849 | 31844 | 0 | 29679 | 30581 | 0 | 0 | 0 | 0 | 0 | 13 | 0 | 0 | 0 | 0 | 0 | 0 | 0 | 0 | 0 | 0 | 0 | 0 | 111 | 0 |
| SRR6880326 | 44 | 4775 | 249 | 3459 | 242 | 14775 | 8058 | 1504 | 2380 | 0 | 59909 | 17576 | 13928 | 17668 | 83 | 968 | 90142 | 0 | 0 | 0 | 0 | 0 | 0 | 0 | 0 | 0 | 0 | 0 | 0 | 0 | 0 | 0 | 0 | 0 | 0 | 0 | 0 |
| SRR6880328 | 46 | 833 | 105 | 16894 | 13 | 182 | 195 | 436 | 353 | 0 | 42343 | 9253 | 10877 | 17478 | 302 | 14 | 8734 | 0 | 0 | 0 | 0 | 0 | 0 | 0 | 0 | 0 | 0 | 0 | 0 | 0 | 0 | 0 | 0 | 0 | 10 | 0 | 0 |
| SRR6880329 | 63 | 981 | 164 | 14651 | 13 | 169 | 181 | 473 | 349 | 0 | 41161 | 9161 | 12076 | 16382 | 386 | 14 | 7908 | 0 | 0 | 0 | 0 | 0 | 0 | 0 | 0 | 0 | 0 | 0 | 0 | 0 | 0 | 0 | 0 | 0 | 0 | 0 | 0 |
| SRR6880330 | 93 | 1121 | 101 | 7706 | 12 | 201 | 189 | 384 | 325 | 0 | 48805 | 12268 | 11449 | 17120 | 342 | 17 | 7221 | 0 | 0 | 0 | 0 | 0 | 0 | 0 | 0 | 0 | 0 | 0 | 0 | 0 | 0 | 0 | 0 | 0 | 0 | 0 | 0 |

### Supp Table S8.1.Columns_B_to_AL

| | | | | | | | | | | | | | | | | | |
|---|---|---|---|---|---|---|---|---|---|---|---|---|---|---|---|---|---|
| SRR6880331 | 100 | 931 | 102 | 7208 | 0 | 130 | 106 | 272 | 312 | 0 | 50388 | 16752 | 11697 | 15095 | 361 | 12 | 6408 |
| SRR6880333 | 54 | 1733 | 257 | 1851 | 1221 | 13971 | 15643 | 5703 | 4412 | 12 | 86717 | 12928 | 52001 | 13875 | 387 | 214 | 75869 |
| SRR6880334 | 71 | 1992 | 223 | 2393 | 668 | 8690 | 9697 | 4624 | 2987 | 0 | 62531 | 6966 | 40142 | 10348 | 508 | 247 | 77160 |
| SRR6880335 | 80 | 2175 | 273 | 2930 | 1058 | 10373 | 13523 | 5385 | 6215 | 17 | 64100 | 9226 | 51132 | 11603 | 579 | 164 | 71403 |
| SRR6880336 | 0 | 229 | 673 | 16271 | 0 | 1842 | 3157 | 1710 | 6885 | 0 | 16970 | 5701 | 11664 | 15479 | 329 | 1359 | 18396 |
| SRR6880337 | 191 | 1971 | 393 | 4843 | 377 | 3208 | 6047 | 2898 | 17110 | 38 | 37075 | 24356 | 22697 | 7600 | 1038 | 912 | 106396 |
| SRR6880338 | 35 | 2019 | 268 | 4079 | 215 | 12818 | 6723 | 1390 | 1797 | 0 | 62697 | 19536 | 11187 | 13892 | 87 | 926 | 85669 |
| SRR6880339 | 24 | 2394 | 299 | 5795 | 208 | 13006 | 7482 | 1349 | 2363 | 16 | 51570 | 15420 | 10445 | 16783 | 87 | 878 | 75132 |
| SRR6880341 | 49 | 2825 | 363 | 8383 | 65 | 4743 | 2366 | 920 | 1734 | 0 | 31129 | 19503 | 2297 | 4185 | 50 | 358 | 27964 |
| SRR6880342 | 25 | 1779 | 624 | 8457 | 57 | 2685 | 1476 | 900 | 2226 | 0 | 23839 | 15643 | 3374 | 4980 | 31 | 634 | 22229 |
| SRR6880343 | 28 | 2203 | 987 | 7868 | 55 | 2975 | 1945 | 1025 | 2143 | 0 | 23385 | 12980 | 2706 | 4738 | 35 | 391 | 25301 |
| SRR6880345 | 51 | 647 | 433 | 11366 | 1138 | 5572 | 22585 | 5778 | 15269 | 28 | 57156 | 26418 | 37382 | 7817 | 922 | 536 | 65013 |
| SRR6880351 | 20 | 1626 | 765 | 4136 | 81 | 2307 | 2863 | 1993 | 6158 | 20 | 16667 | 11466 | 2405 | 1681 | 19 | 39258 | 20222 |
| SRR6880356 | 66 | 1916 | 529 | 2630 | 165 | 2316 | 3539 | 2668 | 31613 | 15 | 24247 | 18325 | 14501 | 4673 | 485 | 1999 | 87010 |
| SRR6880357 | 63 | 1648 | 483 | 2902 | 76 | 1501 | 4066 | 3740 | 23341 | 34 | 17571 | 14131 | 11074 | 5141 | 219 | 1046 | 81654 |
| SRR6898464 | 263 | 19114 | 14758 | 369 | 678 | 7102 | 7355 | 8632 | 3732 | 37 | 13652 | 69930 | 8590 | 1090 | 16 | 17 | 21861 |
| SRR6914752 | 0 | 2834 | 147 | 0 | 7606 | 4970 | 119975 | 36977 | 19 | 84 | 12085 | 5578 | 1672 | 1675 | 27578 | 130 | 3276 |
| SRR6914753 | 38 | 916 | 356 | 51 | 7663 | 3446 | 224279 | 45522 | 0 | 7577 | 5925 | 2873 | 2062 | 6772 | 28330 | 153 | 4039 |
| SRR6914754 | 67 | 10797 | 17909 | 679 | 4866 | 4691 | 147890 | 45044 | 53 | 226 | 12055 | 7345 | 1536 | 953 | 22503 | 327 | 4578 |
| SRR6914755 | 83 | 4644 | 18167 | 53 | 5428 | 2889 | 179438 | 52121 | 30 | 4294 | 8461 | 4570 | 2815 | 5162 | 20833 | 248 | 5555 |
| SRR6928219 | 2589 | 4951 | 9172 | 385 | 48352 | 49386 | 369281 | 30826 | 3931 | 16362 | 5664 | 17 | 6048 | 45182 | 4998 | 8950 | 2310 |
| SRR6928220 | 4241 | 6427 | 6592 | 436 | 42273 | 39110 | 250072 | 25429 | 100 | 14653 | 3922 | 4833 | 5197 | 44921 | 10480 | 13688 | 1372 |
| SRR6928221 | 631 | 9666 | 9924 | 538 | 109754 | 58220 | 126241 | 22700 | 4483 | 9114 | 3647 | 2640 | 3182 | 19210 | 6076 | 9709 | 1874 |
| SRR6928222 | 1646 | 3765 | 12221 | 417 | 96774 | 28198 | 162934 | 27362 | 1643 | 15471 | 3557 | 3351 | 4157 | 20193 | 4881 | 6141 | 1428 |
| SRR6928223 | 1316 | 2390 | 7743 | 399 | 37643 | 34378 | 259731 | 21828 | 1925 | 13708 | 8259 | 18 | 7684 | 50340 | 13152 | 9014 | 2579 |
| SRR6928224 | 3512 | 7062 | 5924 | 422 | 34004 | 52227 | 187001 | 21147 | 78 | 14151 | 3348 | 19 | 4895 | 54827 | 10781 | 15962 | 1284 |
| SRR6928225 | 1015 | 5661 | 10205 | 557 | 107076 | 54636 | 156706 | 18795 | 6154 | 11173 | 3401 | 2972 | 3243 | 21588 | 12811 | 9379 | 1927 |
| SRR6928226 | 1989 | 3901 | 11870 | 456 | 104258 | 35227 | 165275 | 30499 | 2207 | 16822 | 4025 | 3773 | 4669 | 21221 | 5051 | 7154 | 1554 |
| SRR6930936 | 350 | 920 | 1699 | 646 | 40272 | 27150 | 450773 | 72307 | 2292 | 2528 | 9099 | 8864 | 6394 | 12973 | 1122 | 4069 | 5433 |
| SRR6930937 | 272 | 865 | 4021 | 429 | 28305 | 34327 | 246226 | 49212 | 3430 | 1347 | 14100 | 16422 | 11329 | 21948 | 1763 | 5678 | 12724 |
| SRR6930939 | 73 | 238 | 1456 | 1202 | 19213 | 34173 | 257349 | 54695 | 47 | 2690 | 10375 | 15 | 10001 | 30728 | 688 | 5864 | 6313 |
| SRR6930941 | 241 | 745 | 2513 | 1230 | 27284 | 34081 | 327768 | 64352 | 31 | 3916 | 11346 | 21 | 7994 | 31727 | 796 | 7875 | 5414 |
| SRR6932941 | 0 | 29 | 288 | 0 | 5398 | 28746 | 29852 | 28140 | 1883 | 22 | 15831 | 34237 | 5613 | 10865 | 925 | 265 | 18670 |
| SRR6932942 | 0 | 70 | 368 | 40 | 2282 | 13129 | 12498 | 16320 | 787 | 20 | 7589 | 17882 | 2604 | 6426 | 355 | 620 | 13340 |
| SRR6932943 | 0 | 0 | 2655 | 0 | 9036 | 34983 | 131285 | 37098 | 576 | 404 | 10332 | 20093 | 6823 | 65043 | 514 | 2774 | 21709 |
| SRR6932946 | 0 | 28 | 3154 | 0 | 4908 | 24228 | 34072 | 27841 | 550 | 134 | 9404 | 20941 | 6382 | 24531 | 343 | 690 | 8176 |
| SRR6932949 | 0 | 0 | 2547 | 18 | 9142 | 10116 | 101422 | 14841 | 929 | 148 | 26391 | 49452 | 10341 | 16698 | 183 | 425 | 3701 |
| SRR6932950 | 0 | 0 | 2693 | 15 | 9820 | 9972 | 115643 | 18399 | 1107 | 159 | 30388 | 52107 | 11654 | 18839 | 245 | 357 | 3255 |
| SRR6932951 | 0 | 0 | 2730 | 28 | 8934 | 8708 | 126699 | 18964 | 1103 | 118 | 29073 | 11 | 12069 | 18391 | 185 | 326 | 3973 |
| SRR6932952 | 0 | 1323 | 1648 | 0 | 8100 | 10860 | 114239 | 11297 | 1397 | 114 | 23291 | 38854 | 8275 | 22759 | 50 | 1364 | 8728 |
| SRR6932953 | 0 | 1391 | 3207 | 62 | 9541 | 14614 | 150192 | 15547 | 1414 | 92 | 21504 | 34976 | 8944 | 32666 | 45 | 2485 | 8502 |
| SRR6932954 | 0 | 639 | 3990 | 0 | 9113 | 22482 | 135000 | 15676 | 1408 | 119 | 36953 | 49637 | 11248 | 45182 | 72 | 3169 | 6259 |
| SRR6932955 | 0 | 0 | 2785 | 10 | 6420 | 28992 | 64194 | 26240 | 594 | 237 | 12641 | 21468 | 5711 | 13183 | 212 | 439 | 11983 |
| SRR7062776 | 1021 | 3660 | 499 | 586 | 2669 | 4035 | 44297 | 21295 | 64 | 566 | 1455 | 1745 | 1288 | 1705 | 1001 | 41928 | 29629 |
| SRR7062777 | 836 | 3508 | 544 | 591 | 2711 | 4150 | 47017 | 21980 | 66 | 639 | 1501 | 1721 | 1391 | 1850 | 1196 | 39621 | 28995 |
| SRR7062778 | 632 | 2401 | 184 | 742 | 2557 | 3787 | 40831 | 20501 | 69 | 577 | 1391 | 1526 | 1193 | 1430 | 1884 | 41710 | 29694 |
| SRR7062779 | 599 | 2482 | 182 | 689 | 2669 | 4234 | 44580 | 20799 | 84 | 535 | 1458 | 1635 | 1270 | 1580 | 1555 | 43594 | 30036 |
| SRR7062780 | 462 | 1951 | 234 | 1206 | 3207 | 4381 | 47962 | 23305 | 45 | 416 | 1369 | 1515 | 1170 | 1375 | 1542 | 44783 | 30599 |
| SRR7062781 | 410 | 2097 | 313 | 1049 | 3192 | 4337 | 45699 | 23125 | 61 | 525 | 1275 | 1577 | 1120 | 1357 | 1639 | 46013 | 31294 |
| SRR7081766 | 165 | 5104 | 1123 | 269 | 0 | 83 | 364 | 1008 | 761 | 0 | 1505 | 658 | 359 | 1681 | 0 | 193 | 4067 |
| SRR7081767 | 40 | 2952 | 1152 | 425 | 0 | 100 | 549 | 1648 | 1045 | 0 | 2850 | 1193 | 693 | 2165 | 0 | 641 | 5942 |
| SRR7081768 | 12 | 2987 | 19917 | 238 | 28 | 368 | 2244 | 4894 | 3200 | 0 | 7400 | 3336 | 2033 | 5078 | 0 | 893 | 14871 |
| SRR7081769 | 43 | 11655 | 20366 | 181 | 15 | 185 | 1616 | 3615 | 19504 | 60 | 3358 | 1904 | 823 | 2103 | 0 | 397 | 14083 |
| SRR7081770 | 78 | 4545 | 1190 | 332 | 0 | 97 | 266 | 790 | 919 | 39 | 1160 | 559 | 332 | 779 | 0 | 311 | 2638 |
| SRR7081771 | 31 | 1957 | 1998 | 312 | 0 | 59 | 363 | 965 | 1028 | 31 | 2127 | 815 | 738 | 1756 | 0 | 241 | 6285 |
| SRR7081772 | 25 | 1991 | 3335 | 100 | 0 | 101 | 674 | 925 | 752 | 25 | 2049 | 853 | 504 | 1635 | 0 | 498 | 3550 |
| SRR7081773 | 107 | 1876 | 3532 | 180 | 0 | 92 | 166 | 1704 | 1817 | 0 | 705 | 376 | 89 | 360 | 0 | 261 | 774 |
| SRR7081774 | 95 | 1359 | 7903 | 353 | 0 | 96 | 669 | 1762 | 1196 | 0 | 1163 | 573 | 283 | 1143 | 0 | 399 | 2460 |
| SRR7081775 | 139 | 1658 | 2908 | 382 | 0 | 57 | 598 | 1397 | 1289 | 26 | 1451 | 771 | 225 | 1018 | 0 | 112 | 3118 |
| SRR7081776 | 46 | 579 | 305 | 411 | 0 | 54 | 152 | 623 | 593 | 0 | 830 | 448 | 232 | 1197 | 0 | 124 | 1301 |
| SRR7081777 | 73 | 2367 | 679 | 1156 | 0 | 146 | 628 | 1276 | 1726 | 0 | 1916 | 899 | 399 | 2011 | 0 | 332 | 5135 |
| SRR7107416 | 0 | 7799 | 0 | 836 | 0 | 0 | 0 | 0 | 0 | 100 | 584 | 313 | 30 | 497 | 0 | 664 | 795 |
| SRR7107417 | 0 | 7147 | 0 | 718 | 0 | 0 | 0 | 0 | 0 | 100 | 591 | 274 | 25 | 479 | 0 | 729 | 788 |
| SRR7107418 | 717 | 7197 | 1019 | 0 | 5245 | 83347 | 51167 | 53232 | 166 | 78 | 1740 | 2344 | 1354 | 21004 | 0 | 28508 | 0 |
| SRR7107419 | 711 | 8310 | 1459 | 0 | 5011 | 76876 | 49289 | 52130 | 163 | 36 | 8223 | 10474 | 1342 | 19660 | 0 | 28847 | 0 |
| SRR7107420 | 113 | 9178 | 0 | 0 | 3590 | 0 | 104166 | 94365 | 0 | 0 | 0 | 0 | 0 | 0 | 0 | 0 | 41 |
| SRR7107421 | 116 | 10159 | 0 | 0 | 3483 | 0 | 100445 | 93237 | 0 | 21 | 0 | 0 | 0 | 0 | 0 | 0 | 38 |
| SRR7126083 | 468 | 1067 | 52975 | 1813 | 368 | 812 | 2303 | 1465 | 17843 | 15 | 645 | 1725 | 586 | 24943 | 218 | 109 | 5526 |
| SRR7126084 | 771 | 3021 | 85645 | 1611 | 17992 | 18847 | 88726 | 13269 | 217 | 1283 | 44839 | 22 | 26 | 183788 | 6079 | 13391 | 14292 |
| SRR7126085 | 803 | 3914 | 98260 | 1572 | 19221 | 27311 | 97455 | 14252 | 242 | 1444 | 35054 | 17126 | 18291 | 189487 | 7043 | 16173 | 13012 |
| SRR7126086 | 453 | 3438 | 65645 | 2098 | 18118 | 34284 | 80416 | 13650 | 381 | 588 | 23103 | 15395 | 17 | 197718 | 6873 | 14055 | 12166 |
| SRR7126087 | 413 | 2106 | 92276 | 1307 | 18436 | 28402 | 101243 | 19488 | 351 | 558 | 34553 | 19090 | 19234 | 165905 | 5801 | 14312 | 13384 |
| SRR7126088 | 369 | 2993 | 80421 | 2204 | 16601 | 27997 | 81139 | 11052 | 357 | 346 | 32070 | 17985 | 20483 | 201880 | 6889 | 12592 | 13364 |
| SRR7126089 | 674 | 4488 | 79534 | 2740 | 14309 | 20470 | 78655 | 9240 | 373 | 780 | 41574 | 18 | 23 | 209556 | 5913 | 10250 | 15050 |
| SRR7126090 | 723 | 3853 | 111879 | 1097 | 19178 | 26854 | 87793 | 11680 | 706 | 1097 | 46840 | 21895 | 21279 | 196684 | 7182 | 10391 | 13713 |
| SRR7126091 | 722 | 2580 | 79359 | 1253 | 20136 | 23624 | 103909 | 16891 | 349 | 912 | 44335 | 22324 | 23771 | 180208 | 6249 | 14839 | 12730 |
| SRR7126092 | 470 | 3129 | 69949 | 2596 | 15229 | 21537 | 79101 | 11431 | 278 | 504 | 39090 | 18 | 26 | 195088 | 6057 | 11882 | 14897 |
| SRR7126093 | 356 | 2997 | 64285 | 2334 | 23324 | 36218 | 98827 | 14039 | 329 | 568 | 25747 | 15 | 22 | 183613 | 6351 | 14135 | 11835 |
| SRR7153253 | 102 | 27211 | 298 | 260 | 421 | 3794 | 3878 | 8290 | 3945 | 15 | 1683 | 9692 | 607 | 304 | 0 | 9376 | 63032 |
| SRR7153254 | 77 | 28267 | 338 | 425 | 324 | 3652 | 2952 | 7913 | 3795 | 0 | 2797 | 11486 | 991 | 478 | 0 | 7563 | 48736 |
| SRR7153255 | 101 | 55449 | 236 | 189 | 319 | 3752 | 2869 | 7942 | 4326 | 0 | 1696 | 12630 | 594 | 255 | 0 | 9559 | 46688 |
| SRR7153256 | 109 | 23934 | 383 | 233 | 427 | 4357 | 4073 | 8130 | 5745 | 21 | 2530 | 16011 | 914 | 567 | 0 | 11491 | 53969 |
| SRR7153257 | 192 | 21652 | 183 | 334 | 323 | 3242 | 2500 | 7473 | 4786 | 13 | 1659 | 7791 | 612 | 309 | 0 | 7897 | 46704 |
| SRR7153258 | 111 | 18537 | 213 | 250 | 293 | 2747 | 2692 | 7802 | 4282 | 12 | 1842 | 11742 | 734 | 333 | 0 | 8078 | 47696 |
| SRR7153259 | 167 | 24883 | 405 | 304 | 377 | 3648 | 3175 | 7515 | 3825 | 16 | 2154 | 10908 | 853 | 482 | 0 | 9233 | 58676 |
| SRR7153260 | 78 | 18933 | 320 | 389 | 499 | 4262 | 4396 | 11294 | 4269 | 11 | 1184 | 13393 | 414 | 280 | 0 | 10715 | 74683 |
| SRR7153261 | 144 | 15320 | 67 | 389 | 538 | 4631 | 4572 | 6842 | 5419 | 14 | 1221 | 9490 | 444 | 256 | 0 | 14620 | 67895 |
| SRR7153262 | 323 | 18987 | 188 | 316 | 317 | 3909 | 3120 | 7169 | 4089 | 0 | 1370 | 10732 | 568 | 214 | 0 | 10803 | 47200 |
| SRR7153263 | 51 | 32784 | 188 | 162 | 361 | 2989 | 3089 | 7018 | 4232 | 0 | 1645 | 13116 | 672 | 370 | 0 | 11493 | 57890 |
| SRR7153264 | 110 | 21958 | 325 | 416 | 562 | 4430 | 4793 | 9063 | 4168 | 41 | 1810 | 12214 | 506 | 435 | 0 | 9908 | 75220 |
| SRR7153265 | 109 | 14841 | 185 | 221 | 442 | 4381 | 3909 | 9613 | 6085 | 0 | 1739 | 13982 | 647 | 296 | 0 | 11379 | 48411 |
| SRR7153266 | 183 | 11920 | 130 | 296 | 337 | 3179 | 2649 | 6794 | 6205 | 13 | 1477 | 11319 | 428 | 215 | 0 | 11812 | 65172 |
| SRR7153267 | 122 | 12894 | 261 | 255 | 314 | 3110 | 2715 | 7828 | 6577 | 0 | 1402 | 12226 | 464 | 253 | 0 | 8348 | 55254 |
| SRR7153268 | 134 | 19899 | 15 | 273 | 549 | 3792 | 3230 | 5587 | 4422 | 0 | 1088 | 7384 | 380 | 279 | 0 | 14643 | 93573 |
| SRR7153269 | 173 | 11537 | 386 | 192 | 463 | 4489 | 3809 | 11089 | 6051 | 11 | 2006 | 9527 | 600 | 387 | 0 | 8487 | 59958 |
| SRR7153270 | 132 | 27552 | 153 | 333 | 361 | 3720 | 3028 | 8352 | 3780 | 0 | 1348 | 12402 | 460 | 231 | 0 | 10437 | 54340 |
| SRR7153271 | 115 | 26213 | 103 | 305 | 305 | 3339 | 2847 | 8158 | 2947 | 0 | 1591 | 9871 | 616 | 292 | 0 | 8789 | 49867 |
| SRR7153272 | 135 | 19579 | 320 | 237 | 247 | 2629 | 2330 | 6969 | 3081 | 23 | 1639 | 8783 | 559 | 278 | 0 | 8249 | 47230 |
| SRR7153273 | 35 | 2352 | 245 | 281 | 367 | 2833 | 3606 | 6703 | 4991 | 81 | 1997 | 8585 | 598 | 559 | 0 | 15420 | 50938 |
| SRR7153274 | 138 | 23230 | 253 | 332 | 279 | 3261 | 2310 | 6743 | 4918 | 0 | 2052 | 9140 | 675 | 472 | 0 | 8351 | 34745 |
| SRR7153275 | 187 | 50070 | 189 | 587 | 331 | 3853 | 2865 | 7757 | 5371 | 0 | 1382 | 8811 | 526 | 267 | 0 | 9887 | 51340 |
| SRR7153276 | 139 | 24408 | 171 | 194 | 300 | 3134 | 2675 | 7811 | 6038 | 0 | 1599 | 10581 | 621 | 329 | 0 | 9831 | 45472 |
| SRR7153277 | 87 | 24221 | 300 | 180 | 189 | 2016 | 1692 | 4825 | 3883 | 0 | 1183 | 6386 | 430 | 240 | 0 | 6390 | 48154 |
| SRR7153278 | 1265 | 37451 | 1415 | 655 | 400 | 4980 | 3778 | 7934 | 6655 | 14 | 2518 | 13370 | 935 | 404 | 0 | 8788 | 46121 |
| SRR7153279 | 44 | 5303 | 492 | 241 | 336 | 2914 | 3333 | 7070 | 4839 | 33 | 1522 | 8490 | 534 | 288 | 0 | 15294 | 42074 |
| SRR7153280 | 176 | 38737 | 301 | 253 | 258 | 2858 | 2048 | 6054 | 5793 | 0 | 1655 | 8344 | 582 | 291 | 0 | 8393 | 43746 |
| SRR7153281 | 133 | 55386 | 314 | 476 | 376 | 4036 | 3133 | 8748 | 5629 | 23 | 1589 | 8228 | 554 | 295 | 0 | 8183 | 76014 |
| SRR7153282 | 113 | 11623 | 345 | 244 | 235 | 2994 | 2127 | 7115 | 2389 | 20 | 1213 | 7977 | 412 | 216 | 0 | 8354 | 42428 |
| SRR7153283 | 186 | 46200 | 680 | 182 | 232 | 2738 | 2093 | 5965 | 7081 | 13 | 1535 | 7808 | 568 | 300 | 0 | 6530 | 50429 |
| SRR7153284 | 117 | 15945 | 106 | 694 | 313 | 2215 | 2271 | 5355 | 4126 | 16 | 868 | 7069 | 269 | 172 | 0 | 5722 | 92939 |
| SRR7153285 | 91 | 17374 | 237 | 214 | 224 | 2010 | 1788 | 5631 | 4080 | 18 | 1926 | 8022 | 695 | 370 | 0 | 6837 | 52627 |
| SRR7153286 | 230 | 32462 | 853 | 197 | 262 | 2991 | 2367 | 7104 | 5646 | 11 | 1673 | 8426 | 576 | 409 | 0 | 6633 | 41403 |
| SRR7153287 | 201 | 15184 | 71 | 535 | 572 | 3997 | 4035 | 10194 | 7261 | 0 | 2026 | 9727 | 626 | 564 | 0 | 8041 | 68369 |
| SRR7153288 | 218 | 50736 | 407 | 302 | 382 | 4318 | 2986 | 8300 | 4089 | 0 | 1632 | 8399 | 561 | 329 | 0 | 5908 | 58273 |
| SRR7153289 | 148 | 28614 | 398 | 252 | 282 | 2910 | 2472 | 6629 | 7613 | 26 | 1342 | 7116 | 492 | 286 | 0 | 7466 | 56542 |
| SRR7153290 | 94 | 33909 | 217 | 149 | 310 | 3402 | 2534 | 7043 | 6083 | 11 | 1921 | 9132 | 678 | 441 | 0 | 8142 | 46899 |
| SRR7153291 | 54 | 4912 | 523 | 289 | 353 | 3308 | 3815 | 8522 | 5293 | 29 | 1599 | 7895 | 457 | 331 | 0 | 13012 | 43331 |
| SRR7153292 | 58 | 3269 | 401 | 391 | 328 | 2937 | 3689 | 7124 | 3450 | 65 | 1479 | 6784 | 438 | 358 | 0 | 13421 | 51742 |
| SRR7153293 | 117 | 8838 | 182 | 599 | 570 | 5936 | 5820 | 9661 | 5009 | 11 | 1494 | 6938 | 489 | 324 | 0 | 16890 | 31395 |
| SRR7153294 | 112 | 14313 | 143 | 352 | 236 | 2210 | 1976 | 5688 | 3683 | 0 | 1136 | 5995 | 474 | 264 | 0 | 5549 | 45597 |
| SRR7153295 | 90 | 13294 | 140 | 730 | 453 | 3525 | 4048 | 8041 | 4994 | 21 | 1472 | 8109 | 438 | 320 | 0 | 8600 | 90942 |
| SRR7153296 | 132 | 11931 | 241 | 146 | 161 | 1553 | 1309 | 4365 | 4679 | 0 | 1539 | 6052 | 642 | 260 | 0 | 5313 | 41330 |
| SRR7153297 | 1887 | 10197 | 136 | 1444 | 306 | 3551 | 3188 | 7452 | 5411 | 17 | 1471 | 7052 | 494 | 313 | 0 | 5699 | 47430 |
| SRR7153298 | 166 | 18724 | 266 | 159 | 413 | 3753 | 2717 | 8669 | 4672 | 11 | 1302 | 4892 | 411 | 278 | 0 | 7810 | 50676 |

| | | | | | | | | | | | | | | | | | | | |
|---|---|---|---|---|---|---|---|---|---|---|---|---|---|---|---|---|---|---|---|
| SRR6880331 | 0 | 0 | 0 | 0 | 0 | 0 | 0 | 0 | 0 | 0 | 0 | 0 | 0 | 0 | 0 | 0 | 0 | 0 | 0 | 0 |
| SRR6880333 | 0 | 0 | 0 | 0 | 0 | 16 | 12 | 0 | 18 | 0 | 0 | 86 | 25 | 13 | 0 | 0 | 0 | 19 | 18 | 0 |
| SRR6880334 | 0 | 0 | 0 | 0 | 0 | 21 | 17 | 0 | 33 | 0 | 0 | 42 | 13 | 0 | 0 | 0 | 0 | 28 | 26 | 0 |
| SRR6880335 | 0 | 0 | 0 | 0 | 0 | 22 | 14 | 0 | 26 | 0 | 0 | 149 | 31 | 13 | 0 | 0 | 0 | 29 | 27 | 0 |
| SRR6880336 | 0 | 0 | 0 | 0 | 0 | 0 | 0 | 0 | 0 | 0 | 0 | 0 | 0 | 0 | 0 | 0 | 0 | 0 | 0 | 0 |
| SRR6880337 | 0 | 0 | 0 | 0 | 0 | 0 | 0 | 0 | 0 | 0 | 12 | 46 | 0 | 0 | 0 | 0 | 0 | 11 | 0 | 0 |
| SRR6880338 | 0 | 0 | 0 | 0 | 0 | 0 | 0 | 0 | 0 | 0 | 0 | 0 | 0 | 0 | 0 | 0 | 0 | 0 | 0 | 0 |
| SRR6880339 | 0 | 0 | 0 | 0 | 0 | 0 | 0 | 0 | 0 | 0 | 0 | 0 | 0 | 0 | 0 | 0 | 0 | 0 | 0 | 0 |
| SRR6880341 | 0 | 0 | 0 | 0 | 0 | 40 | 0 | 0 | 0 | 0 | 0 | 14 | 0 | 24 | 0 | 0 | 0 | 0 | 0 | 0 |
| SRR6880342 | 0 | 0 | 0 | 0 | 12 | 119 | 24 | 0 | 0 | 0 | 0 | 0 | 0 | 21 | 0 | 0 | 0 | 0 | 10 | 0 |
| SRR6880343 | 0 | 0 | 0 | 0 | 27 | 136 | 29 | 0 | 0 | 0 | 0 | 13 | 0 | 45 | 0 | 0 | 0 | 0 | 0 | 0 |
| SRR6880345 | 0 | 13 | 0 | 17 | 18 | 0 | 0 | 0 | 0 | 0 | 67 | 169 | 12 | 19 | 0 | 0 | 0 | 0 | 0 | 0 |
| SRR6880351 | 0 | 0 | 0 | 0 | 15 | 12 | 0 | 0 | 0 | 0 | 24 | 134 | 12 | 62 | 0 | 0 | 0 | 0 | 0 | 0 |
| SRR6880356 | 0 | 0 | 0 | 0 | 0 | 0 | 0 | 0 | 0 | 0 | 0 | 34 | 0 | 0 | 0 | 0 | 0 | 0 | 0 | 0 |
| SRR6880357 | 0 | 0 | 0 | 0 | 0 | 10 | 0 | 0 | 0 | 0 | 0 | 16 | 0 | 0 | 0 | 0 | 0 | 11 | 0 | 0 |
| SRR6898464 | 0 | 0 | 0 | 250 | 192 | 36 | 0 | 0 | 0 | 0 | 0 | 45 | 70 | 222 | 26 | 0 | 0 | 0 | 0 | 0 |
| SRR6914752 | 0 | 0 | 0 | 34 | 27 | 63 | 58 | 0 | 10 | 0 | 0 | 10 | 52 | 76 | 0 | 0 | 0 | 0 | 0 | 0 |
| SRR6914753 | 0 | 0 | 0 | 367 | 268 | 598 | 154 | 12 | 43 | 0 | 12 | 30 | 522 | 704 | 17 | 0 | 0 | 0 | 0 | 0 |
| SRR6914754 | 0 | 61 | 18 | 394 | 198 | 170 | 42 | 0 | 14 | 0 | 243 | 335 | 627 | 915 | 20 | 0 | 0 | 0 | 0 | 0 |
| SRR6914755 | 0 | 67 | 24 | 649 | 376 | 958 | 164 | 15 | 36 | 0 | 196 | 174 | 1156 | 1448 | 47 | 0 | 0 | 0 | 0 | 0 |
| SRR6928219 | 0 | 101 | 34 | 30 | 61 | 42 | 15 | 0 | 0 | 31 | 261 | 389 | 102 | 22 | 0 | 0 | 0 | 0 | 0 | 0 |
| SRR6928220 | 0 | 94 | 54 | 35 | 112 | 59 | 18 | 0 | 0 | 34 | 219 | 513 | 88 | 25 | 0 | 0 | 0 | 0 | 0 | 0 |
| SRR6928221 | 18 | 386 | 200 | 134 | 297 | 139 | 44 | 22 | 0 | 112 | 1282 | 2259 | 330 | 136 | 0 | 0 | 0 | 0 | 16 | 0 |
| SRR6928222 | 23 | 321 | 150 | 190 | 445 | 386 | 92 | 34 | 0 | 133 | 1218 | 2001 | 447 | 104 | 16 | 0 | 0 | 0 | 20 | 0 |
| SRR6928223 | 0 | 64 | 24 | 28 | 35 | 30 | 0 | 0 | 0 | 26 | 247 | 357 | 86 | 20 | 0 | 0 | 0 | 0 | 0 | 0 |
| SRR6928224 | 11 | 100 | 78 | 36 | 117 | 70 | 22 | 0 | 0 | 28 | 157 | 500 | 59 | 20 | 0 | 0 | 0 | 0 | 11 | 0 |
| SRR6928225 | 18 | 358 | 127 | 141 | 136 | 163 | 46 | 14 | 0 | 123 | 1726 | 2278 | 409 | 117 | 0 | 0 | 0 | 0 | 0 | 0 |
| SRR6928226 | 24 | 305 | 133 | 139 | 333 | 333 | 95 | 32 | 12 | 141 | 1396 | 1985 | 418 | 131 | 14 | 0 | 0 | 0 | 21 | 0 |
| SRR6930936 | 0 | 24 | 10 | 18 | 12 | 24 | 12 | 0 | 0 | 16 | 43 | 68 | 33 | 17 | 0 | 0 | 0 | 0 | 0 | 0 |
| SRR6930937 | 0 | 25 | 12 | 14 | 14 | 23 | 0 | 0 | 0 | 0 | 33 | 53 | 0 | 16 | 0 | 0 | 0 | 0 | 0 | 0 |
| SRR6930939 | 0 | 22 | 0 | 10 | 21 | 0 | 0 | 0 | 0 | 13 | 39 | 71 | 16 | 17 | 0 | 0 | 0 | 0 | 0 | 0 |
| SRR6930941 | 0 | 30 | 14 | 15 | 29 | 17 | 0 | 0 | 0 | 19 | 63 | 128 | 33 | 25 | 0 | 0 | 0 | 0 | 0 | 0 |
| SRR6932941 | 0 | 0 | 0 | 0 | 0 | 0 | 0 | 14 | 0 | 0 | 0 | 0 | 0 | 0 | 0 | 0 | 11 | 0 | 0 | 0 |
| SRR6932942 | 0 | 0 | 0 | 18 | 0 | 0 | 0 | 23 | 0 | 0 | 0 | 0 | 0 | 0 | 0 | 0 | 48 | 0 | 11 | 0 |
| SRR6932943 | 0 | 0 | 0 | 0 | 0 | 19 | 0 | 0 | 0 | 0 | 0 | 0 | 0 | 0 | 38 | 0 | 22 | 0 | 10 | 0 |
| SRR6932946 | 0 | 0 | 0 | 0 | 23 | 27 | 0 | 0 | 0 | 0 | 0 | 0 | 0 | 60 | 55 | 0 | 0 | 0 | 0 | 0 |
| SRR6932949 | 0 | 0 | 0 | 0 | 0 | 0 | 0 | 0 | 0 | 0 | 0 | 0 | 0 | 0 | 0 | 0 | 0 | 0 | 0 | 0 |
| SRR6932950 | 0 | 0 | 0 | 0 | 0 | 0 | 0 | 0 | 0 | 0 | 0 | 0 | 0 | 0 | 0 | 0 | 0 | 0 | 0 | 0 |
| SRR6932951 | 0 | 0 | 0 | 0 | 0 | 0 | 0 | 0 | 0 | 0 | 0 | 0 | 0 | 0 | 0 | 0 | 0 | 0 | 0 | 0 |
| SRR6932952 | 0 | 23 | 26 | 2016 | 402 | 296 | 26 | 47 | 0 | 0 | 218 | 1187 | 3409 | 401 | 734 | 13 | 39 | 13 | 0 | 0 |
| SRR6932953 | 0 | 0 | 19 | 1380 | 371 | 211 | 12 | 35 | 0 | 0 | 112 | 732 | 2493 | 538 | 852 | 13 | 41 | 0 | 0 | 0 |
| SRR6932954 | 0 | 0 | 0 | 414 | 100 | 33 | 0 | 0 | 0 | 0 | 23 | 163 | 1114 | 121 | 292 | 0 | 15 | 0 | 0 | 0 |
| SRR6932955 | 0 | 0 | 0 | 0 | 0 | 27 | 10 | 11 | 0 | 0 | 0 | 0 | 0 | 0 | 58 | 0 | 33 | 20 | 21 | 0 |
| SRR7062776 | 0 | 37 | 148 | 1342 | 728 | 181 | 0 | 0 | 0 | 0 | 84 | 1030 | 1734 | 770 | 391 | 0 | 0 | 0 | 52 | 0 |
| SRR7062777 | 0 | 33 | 144 | 1482 | 808 | 187 | 0 | 0 | 0 | 0 | 74 | 1002 | 2043 | 936 | 443 | 0 | 0 | 0 | 70 | 0 |
| SRR7062778 | 0 | 36 | 278 | 2103 | 705 | 357 | 0 | 0 | 0 | 0 | 82 | 1645 | 2772 | 978 | 888 | 0 | 0 | 0 | 85 | 0 |
| SRR7062779 | 0 | 33 | 236 | 2183 | 771 | 342 | 0 | 0 | 0 | 0 | 80 | 1618 | 2698 | 1035 | 863 | 0 | 0 | 0 | 82 | 0 |
| SRR7062780 | 0 | 41 | 477 | 1773 | 612 | 328 | 0 | 0 | 0 | 0 | 76 | 2725 | 2232 | 799 | 755 | 0 | 0 | 0 | 255 | 0 |
| SRR7062781 | 0 | 35 | 474 | 1875 | 615 | 325 | 0 | 0 | 0 | 0 | 64 | 2789 | 2265 | 756 | 792 | 0 | 0 | 0 | 196 | 0 |
| SRR7081766 | 0 | 0 | 0 | 15 | 0 | 0 | 0 | 0 | 0 | 0 | 0 | 0 | 18 | 0 | 13 | 0 | 0 | 0 | 0 | 0 |
| SRR7081767 | 0 | 0 | 0 | 10 | 0 | 0 | 0 | 0 | 0 | 0 | 0 | 0 | 10 | 0 | 0 | 0 | 0 | 0 | 0 | 0 |
| SRR7081768 | 0 | 0 | 0 | 14 | 0 | 0 | 0 | 0 | 0 | 0 | 0 | 0 | 18 | 0 | 16 | 0 | 0 | 0 | 0 | 0 |
| SRR7081769 | 0 | 0 | 0 | 0 | 0 | 0 | 0 | 0 | 0 | 0 | 0 | 0 | 0 | 0 | 0 | 0 | 0 | 0 | 0 | 0 |
| SRR7081770 | 0 | 0 | 0 | 28 | 0 | 0 | 0 | 0 | 0 | 0 | 0 | 0 | 27 | 0 | 20 | 0 | 0 | 0 | 0 | 0 |
| SRR7081771 | 0 | 0 | 0 | 62 | 0 | 16 | 0 | 0 | 0 | 0 | 0 | 0 | 64 | 0 | 31 | 0 | 12 | 0 | 0 | 0 |
| SRR7081772 | 0 | 0 | 0 | 19 | 0 | 0 | 0 | 0 | 0 | 0 | 0 | 0 | 12 | 0 | 0 | 0 | 0 | 0 | 0 | 0 |
| SRR7081773 | 0 | 0 | 0 | 38 | 0 | 0 | 0 | 0 | 0 | 0 | 0 | 0 | 23 | 0 | 11 | 0 | 0 | 0 | 0 | 0 |
| SRR7081774 | 0 | 0 | 0 | 16 | 0 | 0 | 0 | 0 | 0 | 0 | 0 | 0 | 11 | 0 | 0 | 0 | 0 | 0 | 0 | 0 |
| SRR7081775 | 0 | 0 | 0 | 0 | 0 | 0 | 0 | 0 | 0 | 0 | 0 | 0 | 0 | 0 | 0 | 0 | 0 | 0 | 0 | 0 |
| SRR7081776 | 0 | 0 | 0 | 23 | 0 | 0 | 0 | 0 | 0 | 0 | 0 | 0 | 0 | 0 | 0 | 0 | 0 | 0 | 0 | 0 |
| SRR7081777 | 0 | 0 | 0 | 16 | 0 | 0 | 0 | 0 | 0 | 0 | 0 | 0 | 0 | 0 | 0 | 0 | 0 | 0 | 0 | 0 |
| SRR7107416 | 0 | 0 | 0 | 446 | 0 | 232 | 0 | 14 | 0 | 0 | 0 | 0 | 137 | 0 | 126 | 0 | 0 | 0 | 10 | 0 |
| SRR7107417 | 0 | 0 | 0 | 305 | 0 | 180 | 0 | 0 | 0 | 0 | 0 | 0 | 110 | 0 | 120 | 0 | 0 | 0 | 0 | 0 |
| SRR7107418 | 0 | 35 | 115 | 5295 | 1140 | 133 | 0 | 0 | 12 | 0 | 228 | 2675 | 2535 | 1903 | 78 | 0 | 0 | 0 | 0 | 0 |
| SRR7107419 | 0 | 36 | 123 | 5801 | 1225 | 128 | 0 | 0 | 0 | 0 | 254 | 2953 | 2582 | 1909 | 69 | 0 | 0 | 0 | 0 | 0 |
| SRR7107420 | 17 | 0 | 0 | 378 | 0 | 264 | 0 | 19 | 0 | 12 | 0 | 0 | 121 | 0 | 154 | 0 | 12 | 0 | 0 | 0 |
| SRR7107421 | 13 | 0 | 0 | 372 | 0 | 249 | 0 | 18 | 0 | 0 | 0 | 0 | 111 | 0 | 146 | 0 | 13 | 0 | 0 | 0 |
| SRR7126083 | 0 | 0 | 0 | 0 | 91 | 82 | 0 | 0 | 0 | 0 | 0 | 0 | 0 | 0 | 0 | 0 | 0 | 0 | 0 | 0 |
| SRR7126084 | 0 | 84 | 220 | 1218 | 3164 | 560 | 87 | 32 | 22 | 0 | 74 | 219 | 288 | 46 | 0 | 0 | 0 | 0 | 36 | 0 |
| SRR7126085 | 0 | 102 | 231 | 784 | 2142 | 397 | 72 | 22 | 16 | 0 | 72 | 175 | 186 | 31 | 0 | 0 | 0 | 0 | 28 | 0 |
| SRR7126086 | 0 | 85 | 277 | 658 | 1750 | 638 | 80 | 28 | 10 | 0 | 54 | 138 | 103 | 25 | 0 | 0 | 0 | 0 | 25 | 0 |
| SRR7126087 | 0 | 72 | 273 | 1624 | 3076 | 1225 | 96 | 32 | 16 | 0 | 55 | 215 | 317 | 60 | 0 | 0 | 0 | 0 | 34 | 0 |
| SRR7126088 | 0 | 68 | 250 | 936 | 1796 | 451 | 61 | 22 | 0 | 0 | 48 | 113 | 138 | 21 | 0 | 0 | 0 | 0 | 20 | 0 |
| SRR7126089 | 0 | 56 | 200 | 1351 | 2592 | 637 | 82 | 23 | 0 | 0 | 49 | 125 | 227 | 29 | 0 | 0 | 0 | 0 | 25 | 0 |
| SRR7126090 | 0 | 69 | 212 | 1261 | 2603 | 380 | 35 | 14 | 0 | 0 | 58 | 145 | 299 | 32 | 0 | 0 | 0 | 0 | 19 | 0 |
| SRR7126091 | 0 | 118 | 251 | 1819 | 3074 | 570 | 62 | 24 | 12 | 0 | 88 | 271 | 393 | 54 | 0 | 0 | 0 | 0 | 28 | 0 |
| SRR7126092 | 0 | 73 | 248 | 1381 | 3619 | 813 | 96 | 28 | 11 | 0 | 62 | 155 | 254 | 51 | 0 | 0 | 0 | 0 | 37 | 0 |
| SRR7126093 | 0 | 99 | 352 | 994 | 2707 | 607 | 75 | 22 | 12 | 0 | 67 | 150 | 150 | 32 | 0 | 0 | 0 | 0 | 24 | 0 |
| SRR7153253 | 0 | 153 | 364 | 15474 | 1435 | 2257 | 0 | 0 | 0 | 0 | 994 | 1810 | 3359 | 4702 | 1024 | 0 | 12 | 0 | 0 | 0 |
| SRR7153254 | 0 | 105 | 267 | 12014 | 1307 | 1430 | 0 | 0 | 0 | 0 | 518 | 974 | 1770 | 2679 | 446 | 0 | 0 | 0 | 0 | 0 |
| SRR7153255 | 0 | 63 | 179 | 6547 | 754 | 697 | 0 | 0 | 0 | 0 | 368 | 826 | 1323 | 2063 | 322 | 0 | 0 | 0 | 0 | 0 |
| SRR7153256 | 0 | 61 | 177 | 7341 | 748 | 817 | 0 | 0 | 0 | 0 | 485 | 1024 | 1744 | 2447 | 362 | 0 | 0 | 0 | 0 | 0 |
| SRR7153257 | 0 | 230 | 557 | 19867 | 2010 | 2141 | 0 | 0 | 0 | 0 | 1276 | 2224 | 2956 | 4596 | 676 | 0 | 0 | 0 | 0 | 0 |
| SRR7153258 | 0 | 105 | 262 | 9772 | 1005 | 1432 | 0 | 0 | 0 | 0 | 845 | 1422 | 2096 | 3332 | 629 | 0 | 0 | 0 | 0 | 0 |
| SRR7153259 | 0 | 202 | 559 | 21987 | 2173 | 2937 | 0 | 0 | 0 | 0 | 1085 | 2227 | 3488 | 5701 | 1009 | 0 | 14 | 0 | 0 | 0 |
| SRR7153260 | 0 | 180 | 532 | 14593 | 2200 | 2614 | 0 | 0 | 0 | 0 | 987 | 2119 | 3445 | 7788 | 1190 | 0 | 17 | 0 | 0 | 0 |
| SRR7153261 | 0 | 414 | 1074 | 27506 | 3478 | 3596 | 0 | 0 | 0 | 0 | 2662 | 4649 | 6005 | 11368 | 1650 | 0 | 20 | 0 | 0 | 0 |
| SRR7153262 | 0 | 61 | 159 | 8861 | 807 | 1247 | 0 | 0 | 0 | 0 | 471 | 961 | 1950 | 2505 | 550 | 0 | 0 | 0 | 0 | 0 |
| SRR7153263 | 0 | 71 | 218 | 9988 | 856 | 1301 | 0 | 0 | 0 | 0 | 371 | 870 | 1990 | 2813 | 587 | 0 | 0 | 0 | 0 | 0 |
| SRR7153264 | 0 | 263 | 595 | 18422 | 2483 | 2728 | 0 | 0 | 0 | 0 | 1606 | 2677 | 3462 | 7099 | 1039 | 0 | 20 | 15 | 0 | 0 |
| SRR7153265 | 0 | 71 | 266 | 12088 | 1100 | 1881 | 0 | 0 | 0 | 0 | 414 | 1002 | 2180 | 3016 | 694 | 0 | 0 | 0 | 0 | 0 |
| SRR7153266 | 0 | 167 | 441 | 18743 | 1657 | 2111 | 0 | 0 | 0 | 0 | 1112 | 2173 | 3716 | 4916 | 916 | 0 | 13 | 0 | 0 | 0 |
| SRR7153267 | 0 | 107 | 327 | 14338 | 1252 | 2184 | 0 | 0 | 0 | 0 | 760 | 1547 | 2768 | 3704 | 858 | 0 | 13 | 0 | 0 | 0 |
| SRR7153268 | 0 | 382 | 835 | 34872 | 3411 | 5826 | 21 | 0 | 0 | 0 | 2251 | 3692 | 7574 | 11447 | 2543 | 0 | 49 | 23 | 0 | 0 |
| SRR7153269 | 0 | 97 | 333 | 20593 | 1587 | 2940 | 0 | 0 | 0 | 0 | 532 | 1419 | 3866 | 3797 | 1024 | 0 | 16 | 12 | 0 | 0 |
| SRR7153270 | 0 | 401 | 906 | 24502 | 3073 | 2496 | 0 | 0 | 0 | 0 | 1776 | 3029 | 3589 | 6245 | 770 | 0 | 11 | 0 | 0 | 0 |
| SRR7153271 | 0 | 106 | 292 | 10889 | 1164 | 1260 | 0 | 0 | 0 | 0 | 379 | 737 | 1234 | 1840 | 314 | 0 | 0 | 0 | 0 | 0 |
| SRR7153272 | 0 | 128 | 326 | 13486 | 1400 | 1780 | 0 | 0 | 0 | 0 | 683 | 1302 | 2169 | 3358 | 633 | 0 | 0 | 0 | 0 | 0 |
| SRR7153273 | 0 | 31 | 47 | 2873 | 750 | 346 | 0 | 0 | 0 | 0 | 290 | 1514 | 1704 | 1675 | 678 | 0 | 12 | 0 | 0 | 0 |
| SRR7153274 | 0 | 37 | 133 | 6572 | 535 | 920 | 0 | 0 | 0 | 0 | 260 | 615 | 1190 | 1369 | 326 | 0 | 0 | 0 | 0 | 0 |
| SRR7153275 | 0 | 157 | 428 | 18776 | 1633 | 3331 | 11 | 0 | 0 | 0 | 873 | 1838 | 3103 | 4023 | 1150 | 0 | 26 | 20 | 0 | 0 |
| SRR7153276 | 0 | 0 | 30 | 1166 | 259 | 348 | 0 | 0 | 0 | 0 | 44 | 105 | 213 | 531 | 135 | 0 | 0 | 0 | 0 | 0 |
| SRR7153277 | 0 | 71 | 228 | 9728 | 970 | 1498 | 0 | 0 | 0 | 0 | 496 | 1007 | 1909 | 2810 | 615 | 0 | 0 | 0 | 0 | 0 |
| SRR7153278 | 0 | 41 | 149 | 6622 | 683 | 974 | 0 | 0 | 0 | 0 | 224 | 578 | 1117 | 1666 | 367 | 0 | 0 | 0 | 0 | 0 |
| SRR7153279 | 0 | 15 | 26 | 2262 | 693 | 327 | 0 | 0 | 0 | 0 | 133 | 792 | 1127 | 1139 | 498 | 0 | 0 | 0 | 0 | 0 |
| SRR7153280 | 0 | 45 | 156 | 6975 | 603 | 952 | 0 | 0 | 0 | 0 | 387 | 1016 | 1688 | 2075 | 433 | 0 | 0 | 0 | 0 | 0 |
| SRR7153281 | 0 | 135 | 367 | 18448 | 1477 | 3990 | 20 | 11 | 0 | 0 | 678 | 1523 | 3480 | 3830 | 1588 | 0 | 54 | 51 | 14 | 0 |
| SRR7153282 | 0 | 53 | 143 | 6536 | 803 | 919 | 0 | 0 | 0 | 0 | 291 | 540 | 926 | 1479 | 274 | 0 | 0 | 0 | 0 | 0 |
| SRR7153283 | 0 | 79 | 196 | 12945 | 1224 | 2645 | 0 | 0 | 0 | 0 | 334 | 676 | 1600 | 2171 | 714 | 0 | 15 | 14 | 0 | 0 |
| SRR7153284 | 0 | 144 | 363 | 17915 | 2168 | 4129 | 17 | 0 | 0 | 0 | 629 | 1316 | 3001 | 5562 | 1455 | 0 | 28 | 17 | 0 | 0 |
| SRR7153285 | 0 | 67 | 224 | 12496 | 895 | 1782 | 0 | 0 | 0 | 0 | 407 | 1021 | 2169 | 2323 | 621 | 0 | 11 | 0 | 0 | 0 |
| SRR7153286 | 0 | 33 | 94 | 3798 | 578 | 986 | 0 | 0 | 0 | 0 | 151 | 358 | 726 | 1282 | 402 | 0 | 0 | 10 | 0 | 0 |
| SRR7153287 | 0 | 394 | 844 | 26573 | 2447 | 3030 | 0 | 0 | 0 | 0 | 2822 | 4561 | 6502 | 8985 | 1554 | 0 | 25 | 13 | 0 | 0 |
| SRR7153288 | 0 | 149 | 429 | 20037 | 1596 | 3195 | 0 | 0 | 0 | 0 | 786 | 1827 | 3448 | 4112 | 1270 | 0 | 25 | 17 | 0 | 0 |
| SRR7153289 | 0 | 103 | 302 | 14794 | 1184 | 2454 | 0 | 0 | 0 | 0 | 550 | 1298 | 2446 | 2969 | 853 | 0 | 17 | 12 | 0 | 0 |
| SRR7153290 | 0 | 50 | 195 | 9328 | 839 | 1879 | 0 | 0 | 0 | 0 | 414 | 1084 | 1885 | 2574 | 728 | 0 | 15 | 0 | 0 | 0 |
| SRR7153291 | 0 | 15 | 26 | 1726 | 502 | 240 | 0 | 0 | 0 | 0 | 127 | 727 | 857 | 940 | 402 | 0 | 0 | 0 | 0 | 0 |
| SRR7153292 | 0 | 36 | 58 | 3873 | 1087 | 700 | 18 | 0 | 0 | 0 | 330 | 1677 | 2075 | 1983 | 1172 | 0 | 39 | 31 | 25 | 0 |
| SRR7153293 | 0 | 24 | 36 | 2542 | 829 | 433 | 0 | 0 | 0 | 0 | 174 | 918 | 1132 | 1211 | 657 | 0 | 13 | 0 | 0 | 0 |
| SRR7153294 | 0 | 238 | 480 | 16139 | 1703 | 3829 | 11 | 0 | 0 | 0 | 1270 | 1930 | 2793 | 4952 | 1386 | 0 | 24 | 20 | 10 | 0 |
| SRR7153295 | 0 | 375 | 949 | 31866 | 3015 | 5702 | 27 | 11 | 0 | 0 | 1715 | 3430 | 5910 | 8328 | 2197 | 0 | 56 | 54 | 29 | 0 |
| SRR7153296 | 0 | 31 | 77 | 3849 | 367 | 604 | 0 | 0 | 0 | 0 | 218 | 409 | 672 | 932 | 208 | 0 | 0 | 0 | 0 | 0 |
| SRR7153297 | 0 | 24 | 83 | 3819 | 368 | 650 | 0 | 0 | 0 | 0 | 149 | 357 | 735 | 993 | 250 | 0 | 0 | 0 | 0 | 0 |
| SRR7153298 | 0 | 85 | 271 | 13601 | 1049 | 1661 | 0 | 0 | 0 | 0 | 471 | 1170 | 2346 | 2678 | 597 | 0 | 10 | 0 | 0 | 0 |

### Supp Table S8.1.Columns_B_to_AL

| | | | | | | | | | | | | | | | | | | | | | | | | | | | | | | | | | | | | | |
|---|---|---|---|---|---|---|---|---|---|---|---|---|---|---|---|---|---|---|---|---|---|---|---|---|---|---|---|---|---|---|---|---|---|---|---|---|---|
| SRR7153299 | 183 | 53255 | 449 | 224 | 292 | 4176 | 2724 | 8093 | 5628 | 0 | 1416 | 10696 | 468 | 206 | 0 | 9553 | 50003 | 0 | 142 | 456 | 18101 | 1804 | 2132 | 0 | 0 | 0 | 0 | 819 | 2011 | 3154 | 4384 | 773 | 0 | 14 | 0 | 0 | 0 |
| SRR7153300 | 268 | 35682 | 850 | 427 | 274 | 3705 | 2488 | 7238 | 4228 | 11 | 1289 | 8681 | 362 | 203 | 0 | 6299 | 44942 | 0 | 82 | 279 | 10627 | 1186 | 1537 | 0 | 0 | 0 | 0 | 435 | 944 | 1528 | 2221 | 463 | 0 | 0 | 0 | 0 | 0 |
| SRR7153301 | 1142 | 43670 | 2579 | 210 | 497 | 5074 | 4353 | 11360 | 6708 | 31 | 2520 | 13083 | 957 | 599 | 0 | 7165 | 63659 | 0 | 44 | 144 | 9318 | 767 | 1034 | 0 | 0 | 0 | 0 | 342 | 714 | 1924 | 2158 | 416 | 0 | 12 | 0 | 0 | 0 |
| SRR7153302 | 113 | 12713 | 318 | 180 | 268 | 3246 | 2361 | 7451 | 4803 | 11 | 1020 | 7451 | 340 | 176 | 0 | 6594 | 47696 | 0 | 107 | 300 | 12153 | 1143 | 1743 | 0 | 0 | 0 | 0 | 527 | 1074 | 1774 | 2651 | 577 | 0 | 0 | 0 | 0 | 0 |
| SRR7153303 | 55 | 11123 | 967 | 207 | 495 | 6335 | 5519 | 11428 | 5256 | 23 | 1443 | 6764 | 404 | 296 | 0 | 10244 | 35947 | 0 | 0 | 0 | 662 | 250 | 80 | 0 | 0 | 0 | 0 | 27 | 151 | 299 | 341 | 105 | 0 | 0 | 0 | 0 | 0 |
| SRR7153304 | 313 | 53226 | 307 | 275 | 426 | 4300 | 2972 | 8219 | 10698 | 0 | 1168 | 6126 | 392 | 194 | 0 | 9027 | 50723 | 0 | 189 | 548 | 23917 | 2182 | 4510 | 16 | 12 | 0 | 0 | 1091 | 2529 | 4254 | 5544 | 1693 | 0 | 45 | 47 | 19 | 0 |
| SRR7153305 | 141 | 33629 | 362 | 167 | 282 | 3302 | 2311 | 7428 | 3553 | 0 | 1537 | 8987 | 501 | 255 | 0 | 7968 | 43781 | 0 | 50 | 166 | 8560 | 764 | 1324 | 0 | 0 | 0 | 0 | 269 | 593 | 1136 | 1408 | 363 | 0 | 0 | 0 | 0 | 0 |
| SRR7153306 | 214 | 40173 | 285 | 393 | 442 | 4258 | 3397 | 7850 | 6510 | 20 | 1162 | 8698 | 395 | 247 | 0 | 7608 | 61803 | 0 | 179 | 452 | 16293 | 1738 | 2889 | 0 | 0 | 0 | 0 | 893 | 1695 | 2359 | 3940 | 896 | 0 | 19 | 15 | 0 | 0 |
| SRR7153307 | 134 | 40964 | 332 | 274 | 339 | 3366 | 2918 | 7704 | 5099 | 15 | 1387 | 8628 | 469 | 248 | 0 | 8286 | 44831 | 0 | 117 | 377 | 19430 | 1638 | 2642 | 0 | 0 | 0 | 0 | 711 | 1743 | 3579 | 4651 | 1040 | 0 | 24 | 12 | 0 | 0 |
| SRR7153308 | 178 | 29707 | 538 | 147 | 259 | 3034 | 2130 | 6706 | 6625 | 0 | 1384 | 8360 | 428 | 208 | 0 | 7471 | 39423 | 0 | 67 | 183 | 7822 | 751 | 1386 | 0 | 0 | 0 | 0 | 423 | 759 | 1234 | 1698 | 440 | 0 | 0 | 0 | 0 | 0 |
| SRR7153309 | 121 | 16618 | 275 | 218 | 303 | 3151 | 2628 | 7793 | 4432 | 0 | 1186 | 10103 | 455 | 242 | 0 | 8795 | 45166 | 0 | 114 | 348 | 13180 | 1431 | 2478 | 0 | 0 | 0 | 0 | 518 | 1091 | 2070 | 3333 | 788 | 0 | 13 | 12 | 0 | 0 |
| SRR7153310 | 297 | 66605 | 1827 | 124 | 403 | 5005 | 3535 | 10321 | 4115 | 15 | 1790 | 8822 | 650 | 357 | 0 | 6940 | 50805 | 0 | 64 | 173 | 8283 | 1076 | 1306 | 0 | 0 | 0 | 0 | 468 | 995 | 1570 | 2875 | 559 | 0 | 0 | 0 | 0 | 0 |
| SRR7153311 | 177 | 28340 | 343 | 108 | 281 | 3611 | 2516 | 7821 | 4246 | 0 | 1400 | 8572 | 467 | 192 | 0 | 7671 | 43288 | 0 | 37 | 93 | 5943 | 548 | 979 | 0 | 0 | 0 | 0 | 157 | 331 | 773 | 989 | 292 | 0 | 0 | 0 | 0 | 0 |
| SRR7153312 | 58 | 14973 | 512 | 343 | 347 | 4206 | 3671 | 7375 | 4336 | 27 | 1259 | 6583 | 399 | 233 | 0 | 13340 | 34897 | 0 | 12 | 20 | 1897 | 521 | 274 | 0 | 0 | 0 | 0 | 105 | 570 | 959 | 874 | 460 | 0 | 0 | 0 | 0 | 0 |
| SRR7153313 | 84 | 10038 | 1291 | 390 | 595 | 5497 | 6659 | 11662 | 3977 | 28 | 1331 | 6423 | 409 | 459 | 0 | 8884 | 62758 | 0 | 26 | 42 | 4775 | 1681 | 1300 | 30 | 14 | 0 | 0 | 193 | 1085 | 2355 | 2938 | 2097 | 0 | 94 | 91 | 25 | 0 |
| SRR7153314 | 174 | 30810 | 710 | 258 | 335 | 3277 | 2713 | 6800 | 5579 | 15 | 1511 | 6750 | 501 | 341 | 0 | 6995 | 54844 | 0 | 184 | 484 | 24591 | 1701 | 5046 | 22 | 12 | 0 | 0 | 855 | 1828 | 3894 | 3931 | 1602 | 0 | 41 | 38 | 15 | 0 |
| SRR7153315 | 372 | 17157 | 212 | 211 | 201 | 1991 | 1715 | 5200 | 4167 | 26 | 1384 | 7726 | 514 | 223 | 0 | 6839 | 47425 | 0 | 74 | 209 | 10729 | 811 | 1823 | 0 | 0 | 0 | 0 | 339 | 750 | 1591 | 1793 | 490 | 0 | 0 | 0 | 0 | 0 |
| SRR7153316 | 199 | 16957 | 69 | 298 | 292 | 3127 | 2079 | 6578 | 4279 | 0 | 1909 | 6199 | 669 | 399 | 0 | 8122 | 38384 | 0 | 64 | 213 | 10103 | 689 | 1208 | 0 | 0 | 0 | 0 | 509 | 1078 | 1942 | 1688 | 506 | 0 | 0 | 0 | 0 | 0 |
| SRR7153317 | 60 | 7809 | 867 | 264 | 397 | 3647 | 4400 | 8617 | 4076 | 38 | 1839 | 10469 | 658 | 392 | 0 | 18612 | 36600 | 0 | 33 | 51 | 1857 | 744 | 154 | 0 | 0 | 0 | 0 | 363 | 1616 | 1142 | 1504 | 310 | 0 | 0 | 0 | 0 | 0 |
| SRR7153318 | 1208 | 9473 | 72 | 2085 | 635 | 4723 | 5870 | 12893 | 4030 | 24 | 2009 | 14159 | 690 | 561 | 0 | 10932 | 72300 | 0 | 57 | 167 | 6248 | 770 | 997 | 0 | 0 | 0 | 0 | 493 | 983 | 1746 | 3245 | 582 | 0 | 11 | 12 | 0 | 0 |
| SRR7153319 | 50 | 4719 | 402 | 188 | 438 | 4334 | 4502 | 9948 | 4655 | 17 | 1106 | 6509 | 299 | 168 | 0 | 13255 | 33701 | 0 | 0 | 14 | 1878 | 400 | 257 | 0 | 0 | 0 | 0 | 78 | 413 | 892 | 554 | 324 | 0 | 13 | 10 | 0 | 0 |
| SRR7153320 | 49 | 8723 | 495 | 263 | 340 | 3232 | 3595 | 7006 | 4667 | 24 | 1177 | 7104 | 342 | 212 | 0 | 13725 | 33927 | 0 | 11 | 17 | 1219 | 350 | 140 | 0 | 0 | 0 | 0 | 77 | 455 | 539 | 503 | 182 | 0 | 0 | 0 | 0 | 0 |
| SRR7153321 | 156 | 13746 | 332 | 633 | 675 | 7519 | 6647 | 13632 | 8202 | 12 | 1829 | 7700 | 480 | 321 | 0 | 12982 | 35888 | 0 | 20 | 32 | 6022 | 1230 | 555 | 0 | 0 | 0 | 0 | 197 | 923 | 2860 | 1820 | 768 | 0 | 14 | 0 | 0 | 0 |
| SRR7153322 | 40 | 9195 | 1535 | 101 | 601 | 6601 | 6684 | 13570 | 9506 | 52 | 1112 | 6312 | 278 | 228 | 0 | 11738 | 55289 | 0 | 23 | 41 | 6079 | 1314 | 1028 | 21 | 0 | 0 | 0 | 207 | 1153 | 3054 | 2319 | 1592 | 0 | 45 | 38 | 10 | 0 |
| SRR7153323 | 68 | 6982 | 815 | 339 | 429 | 4253 | 4524 | 9058 | 4388 | 32 | 1980 | 8618 | 552 | 522 | 0 | 13166 | 43435 | 0 | 18 | 22 | 1540 | 448 | 240 | 0 | 0 | 0 | 0 | 137 | 690 | 736 | 754 | 353 | 0 | 0 | 0 | 0 | 0 |
| SRR7153324 | 168 | 35560 | 607 | 331 | 381 | 3149 | 3547 | 9032 | 9121 | 24 | 1919 | 9325 | 590 | 530 | 0 | 7744 | 89700 | 0 | 113 | 406 | 20557 | 1673 | 6774 | 60 | 29 | 11 | 0 | 711 | 1823 | 3863 | 5055 | 2457 | 11 | 129 | 212 | 112 | 0 |
| SRR7153325 | 44 | 4390 | 655 | 311 | 488 | 4477 | 5197 | 9819 | 4802 | 66 | 1799 | 8565 | 599 | 428 | 0 | 17488 | 38731 | 0 | 14 | 28 | 2195 | 570 | 228 | 0 | 0 | 0 | 0 | 163 | 944 | 1314 | 1128 | 414 | 0 | 0 | 0 | 0 | 0 |
| SRR7153326 | 98 | 11363 | 792 | 359 | 681 | 7427 | 6516 | 15252 | 7289 | 25 | 2309 | 6129 | 634 | 546 | 0 | 8136 | 37586 | 0 | 16 | 24 | 2087 | 821 | 330 | 0 | 0 | 0 | 0 | 150 | 724 | 945 | 1449 | 503 | 0 | 0 | 0 | 0 | 0 |
| SRR7153327 | 62 | 5332 | 1132 | 315 | 358 | 3870 | 4321 | 9161 | 5755 | 27 | 1325 | 7258 | 423 | 256 | 0 | 14544 | 38278 | 0 | 0 | 12 | 1197 | 374 | 120 | 0 | 0 | 0 | 0 | 64 | 411 | 660 | 563 | 174 | 0 | 0 | 0 | 0 | 0 |
| SRR7153328 | 43 | 4017 | 533 | 412 | 415 | 3871 | 4336 | 8328 | 5532 | 65 | 1500 | 8814 | 413 | 439 | 0 | 12619 | 56561 | 0 | 46 | 62 | 3274 | 1100 | 449 | 0 | 0 | 0 | 0 | 313 | 1266 | 1354 | 1852 | 659 | 0 | 13 | 0 | 0 | 0 |
| SRR7153329 | 70 | 9719 | 606 | 287 | 449 | 4066 | 4501 | 10051 | 2751 | 20 | 1619 | 7936 | 498 | 330 | 0 | 14128 | 39616 | 0 | 10 | 20 | 1603 | 446 | 241 | 0 | 0 | 0 | 0 | 85 | 431 | 605 | 636 | 303 | 0 | 0 | 0 | 0 | 0 |
| SRR7153330 | 120 | 6469 | 329 | 575 | 441 | 4633 | 4719 | 9349 | 4561 | 21 | 1463 | 6994 | 419 | 313 | 0 | 14534 | 36938 | 0 | 16 | 25 | 1798 | 532 | 251 | 0 | 0 | 0 | 0 | 153 | 808 | 1022 | 1098 | 439 | 0 | 0 | 0 | 0 | 0 |
| SRR7153331 | 265 | 6239 | 610 | 482 | 509 | 5582 | 5616 | 10775 | 4404 | 27 | 1789 | 6221 | 550 | 343 | 0 | 14177 | 31086 | 0 | 0 | 13 | 847 | 335 | 100 | 0 | 0 | 0 | 0 | 90 | 394 | 489 | 541 | 169 | 0 | 0 | 0 | 0 | 0 |
| SRR7162919 | 0 | 144 | 352 | 0 | 16519 | 6300 | 163451 | 101042 | 2214 | 65 | 8960 | 13083 | 4416 | 4847 | 10100 | 85 | 3371 | 0 | 0 | 0 | 0 | 0 | 0 | 0 | 0 | 0 | 0 | 0 | 0 | 0 | 0 | 0 | 0 | 0 | 0 | 0 | 0 |
| SRR7162920 | 0 | 72 | 106 | 0 | 8351 | 2221 | 54261 | 66782 | 1332 | 0 | 3762 | 5045 | 2688 | 3721 | 8060 | 46 | 2967 | 0 | 0 | 0 | 0 | 0 | 0 | 0 | 0 | 0 | 0 | 0 | 0 | 0 | 0 | 0 | 0 | 0 | 0 | 0 | 0 |
| SRR7162921 | 0 | 125 | 171 | 0 | 11838 | 4116 | 113975 | 82932 | 668 | 40 | 9853 | 15123 | 4014 | 6204 | 8887 | 112 | 4336 | 0 | 0 | 0 | 0 | 0 | 0 | 0 | 0 | 0 | 0 | 0 | 0 | 0 | 0 | 0 | 0 | 0 | 0 | 0 | 0 |
| SRR7178585 | 184 | 4764 | 251 | 554 | 437 | 4250 | 5605 | 8074 | 7247 | 32 | 11814 | 21690 | 3767 | 1392 | 17 | 1424 | 67108 | 0 | 0 | 0 | 1530 | 6451 | 957 | 14 | 0 | 0 | 0 | 0 | 47 | 1118 | 6256 | 1093 | 0 | 24 | 59 | 41 | 0 |
| SRR7178586 | 175 | 4695 | 284 | 610 | 423 | 4075 | 5441 | 7601 | 6987 | 59 | 11138 | 20831 | 3624 | 1350 | 11 | 1534 | 73047 | 0 | 0 | 0 | 1634 | 6837 | 1017 | 13 | 0 | 0 | 0 | 0 | 50 | 1187 | 6682 | 1152 | 0 | 23 | 66 | 46 | 0 |
| SRR7178587 | 222 | 4799 | 254 | 651 | 469 | 5068 | 6443 | 9424 | 7633 | 79 | 12990 | 23580 | 4040 | 1422 | 17 | 881 | 55777 | 0 | 0 | 0 | 1150 | 4685 | 683 | 0 | 0 | 0 | 0 | 0 | 31 | 808 | 4454 | 756 | 0 | 17 | 41 | 33 | 0 |
| SRR7178588 | 124 | 3981 | 195 | 548 | 485 | 4838 | 6238 | 9034 | 8222 | 71 | 14360 | 25785 | 4343 | 1587 | 0 | 622 | 42261 | 0 | 0 | 0 | 1093 | 4525 | 655 | 0 | 0 | 0 | 0 | 0 | 38 | 768 | 4334 | 739 | 0 | 15 | 33 | 26 | 0 |
| SRR7178589 | 70 | 25126 | 413 | 687 | 1490 | 17176 | 24156 | 35018 | 17675 | 45 | 13431 | 18189 | 4228 | 3708 | 37 | 2062 | 21935 | 0 | 0 | 0 | 6089 | 966 | 27 | 0 | 0 | 0 | 0 | 0 | 143 | 5043 | 601 | 30 | 0 | 0 | 21 | 25 | 0 |
| SRR7178590 | 63 | 24503 | 506 | 649 | 1462 | 16670 | 23260 | 33606 | 17350 | 51 | 12926 | 17626 | 4056 | 3637 | 40 | 2248 | 24529 | 0 | 0 | 0 | 6430 | 1015 | 26 | 0 | 0 | 0 | 0 | 0 | 158 | 5369 | 643 | 31 | 0 | 0 | 20 | 25 | 0 |
| SRR7178591 | 28 | 23916 | 431 | 675 | 1891 | 21134 | 28069 | 43105 | 18273 | 141 | 13289 | 18435 | 4190 | 3641 | 27 | 1166 | 18665 | 0 | 0 | 0 | 4481 | 794 | 21 | 0 | 0 | 0 | 0 | 0 | 101 | 3759 | 447 | 22 | 0 | 0 | 15 | 13 | 0 |
| SRR7178592 | 55 | 22373 | 415 | 720 | 1870 | 20850 | 27583 | 42613 | 19596 | 33 | 14489 | 19856 | 4462 | 3874 | 17 | 817 | 13377 | 0 | 0 | 0 | 4311 | 758 | 20 | 0 | 0 | 0 | 0 | 0 | 114 | 3700 | 449 | 22 | 0 | 0 | 16 | 17 | 0 |
| SRR7178593 | 415 | 14305 | 345 | 482 | 864 | 9105 | 13867 | 28645 | 29606 | 44 | 9440 | 13037 | 3001 | 3398 | 22 | 1425 | 39172 | 0 | 0 | 16 | 9597 | 5773 | 297 | 0 | 0 | 0 | 0 | 0 | 341 | 6463 | 6310 | 270 | 0 | 0 | 0 | 0 | 0 |
| SRR7178594 | 326 | 15084 | 390 | 420 | 851 | 9019 | 13874 | 28399 | 29302 | 30 | 9276 | 12671 | 2929 | 3360 | 20 | 1356 | 39775 | 0 | 0 | 19 | 9916 | 5959 | 314 | 0 | 0 | 0 | 0 | 0 | 362 | 6784 | 6612 | 286 | 0 | 0 | 0 | 0 | 0 |
| SRR7178595 | 380 | 13279 | 294 | 415 | 947 | 10215 | 15233 | 32396 | 30168 | 35 | 9096 | 12781 | 2924 | 3284 | 24 | 1096 | 42186 | 0 | 0 | 16 | 6459 | 4265 | 225 | 0 | 0 | 0 | 0 | 0 | 236 | 4582 | 4457 | 185 | 0 | 0 | 0 | 0 | 0 |
| SRR7178596 | 386 | 15358 | 313 | 699 | 835 | 9696 | 14009 | 30388 | 34090 | 24 | 10772 | 14877 | 3405 | 3789 | 14 | 1085 | 38851 | 0 | 0 | 15 | 6184 | 4015 | 205 | 0 | 0 | 0 | 0 | 0 | 252 | 4420 | 4265 | 188 | 0 | 0 | 0 | 0 | 0 |
| SRR7178597 | 114 | 14860 | 570 | 386 | 1027 | 10747 | 17609 | 36625 | 25988 | 48 | 10126 | 13284 | 3019 | 6401 | 20 | 1778 | 33952 | 0 | 0 | 12 | 9275 | 5304 | 202 | 0 | 0 | 0 | 0 | 0 | 308 | 6471 | 3522 | 178 | 0 | 0 | 0 | 0 | 0 |
| SRR7178598 | 114 | 16909 | 341 | 573 | 1440 | 17303 | 21008 | 42482 | 15716 | 16 | 13068 | 17730 | 4585 | 5562 | 46 | 1700 | 28858 | 0 | 0 | 0 | 12598 | 2455 | 62 | 0 | 0 | 0 | 0 | 0 | 396 | 8446 | 1925 | 67 | 0 | 0 | 0 | 0 | 0 |
| SRR7178599 | 63 | 16856 | 371 | 544 | 1534 | 18234 | 22195 | 44802 | 15835 | 29 | 13420 | 18162 | 4747 | 5702 | 43 | 1530 | 27844 | 0 | 0 | 0 | 12527 | 2443 | 60 | 0 | 0 | 0 | 0 | 0 | 390 | 8437 | 1906 | 67 | 0 | 0 | 0 | 0 | 0 |
| SRR7178600 | 12 | 3857 | 702 | 1187 | 2023 | 24622 | 60495 | 16869 | 9792 | 140 | 18852 | 21108 | 5194 | 5956 | 42 | 3805 | 24548 | 0 | 0 | 0 | 4035 | 921 | 40 | 0 | 0 | 0 | 0 | 0 | 113 | 7708 | 982 | 63 | 0 | 0 | 13 | 20 | 0 |
| SRR7178601 | 18 | 3735 | 667 | 1130 | 1934 | 23777 | 57707 | 16179 | 9664 | 120 | 18157 | 20330 | 5040 | 5811 | 57 | 4149 | 27690 | 0 | 0 | 0 | 4228 | 952 | 41 | 0 | 0 | 0 | 0 | 0 | 115 | 8177 | 1033 | 74 | 0 | 0 | 17 | 22 | 0 |
| SRR7178602 | 16 | 4078 | 629 | 1039 | 2373 | 27725 | 64234 | 20513 | 9262 | 213 | 20165 | 22876 | 5399 | 6105 | 31 | 2139 | 20134 | 0 | 0 | 0 | 3278 | 731 | 27 | 0 | 0 | 0 | 0 | 0 | 87 | 6349 | 769 | 52 | 0 | 0 | 13 | 13 | 0 |
| SRR7178603 | 13 | 3647 | 717 | 1719 | 2445 | 27729 | 63464 | 20210 | 10185 | 152 | 21675 | 24298 | 5855 | 6264 | 17 | 1573 | 14025 | 0 | 0 | 0 | 3077 | 713 | 28 | 0 | 0 | 0 | 0 | 0 | 95 | 6160 | 774 | 43 | 0 | 0 | 12 | 18 | 0 |
| SRR7178604 | 80 | 3301 | 135 | 891 | 433 | 4104 | 6492 | 7411 | 7171 | 22 | 8375 | 13896 | 3004 | 1184 | 17 | 1349 | 72903 | 0 | 0 | 0 | 1047 | 3852 | 1078 | 26 | 18 | 0 | 0 | 0 | 56 | 1136 | 5986 | 1754 | 0 | 106 | 296 | 240 | 0 |
| SRR7178605 | 101 | 3247 | 110 | 858 | 462 | 4175 | 6438 | 7295 | 7157 | 28 | 8283 | 13477 | 2906 | 1154 | 11 | 1252 | 73362 | 0 | 0 | 0 | 1119 | 4058 | 1165 | 21 | 16 | 0 | 0 | 0 | 58 | 1179 | 6387 | 1868 | 0 | 100 | 322 | 242 | 0 |
| SRR7178606 | 139 | 2844 | 111 | 943 | 423 | 4565 | 6631 | 7916 | 7018 | 42 | 8829 | 13640 | 3084 | 1166 | 18 | 1045 | 74128 | 0 | 0 | 0 | 743 | 2614 | 737 | 19 | 11 | 0 | 0 | 0 | 36 | 755 | 3912 | 1110 | 0 | 59 | 189 | 147 | 0 |
| SRR7178607 | 136 | 3498 | 214 | 816 | 456 | 4378 | 6325 | 7933 | 8312 | 58 | 10616 | 17006 | 3850 | 1425 | 0 | 1195 | 70540 | 0 | 0 | 0 | 721 | 2531 | 679 | 14 | 13 | 0 | 0 | 0 | 35 | 751 | 3843 | 1084 | 0 | 60 | 162 | 133 | 0 |
| SRR7178608 | 268 | 13626 | 676 | 557 | 1344 | 16410 | 19212 | 41824 | 18701 | 44 | 9572 | 16431 | 3434 | 3117 | 29 | 1199 | 22468 | 0 | 0 | 0 | 11524 | 3843 | 85 | 0 | 0 | 0 | 0 | 0 | 249 | 9580 | 2625 | 92 | 0 | 0 | 13 | 0 | 0 |
| SRR7178609 | 268 | 13314 | 638 | 527 | 1227 | 15613 | 18336 | 39901 | 18254 | 17 | 9193 | 15726 | 3257 | 3011 | 26 | 1267 | 24900 | 0 | 0 | 12 | 12198 | 4036 | 93 | 0 | 0 | 0 | 0 | 0 | 269 | 10157 | 2821 | 99 | 0 | 0 | 16 | 0 | 0 |
| SRR7178610 | 228 | 13398 | 607 | 522 | 1684 | 20410 | 23739 | 53202 | 19231 | 47 | 10424 | 17705 | 3495 | 3213 | 24 | 706 | 18864 | 0 | 0 | 0 | 8936 | 2932 | 61 | 0 | 0 | 0 | 0 | 0 | 177 | 7267 | 1991 | 75 | 0 | 0 | 0 | 0 | 0 |
| SRR7178611 | 119 | 11574 | 412 | 488 | 1668 | 20811 | 23960 | 53707 | 20789 | 87 | 11423 | 19392 | 3963 | 3588 | 18 | 513 | 13918 | 0 | 0 | 0 | 8598 | 2876 | 57 | 0 | 0 | 0 | 0 | 0 | 192 | 7166 | 1958 | 63 | 0 | 0 | 10 | 0 | 0 |
| SRR7178612 | 246 | 13951 | 341 | 280 | 1423 | 16430 | 18089 | 35831 | 23805 | 28 | 9685 | 16713 | 3596 | 3869 | 29 | 1496 | 24096 | 0 | 0 | 0 | 4250 | 562 | 18 | 0 | 0 | 0 | 0 | 0 | 117 | 3304 | 480 | 20 | 0 | 0 | 46 | 21 | 0 |
| SRR7178613 | 209 | 14285 | 342 | 292 | 1422 | 16248 | 18059 | 35683 | 23709 | 22 | 9599 | 16596 | 3524 | 3845 | 29 | 1425 | 24498 | 0 | 0 | 0 | 4396 | 596 | 23 | 0 | 0 | 0 | 0 | 0 | 118 | 3430 | 490 | 23 | 0 | 0 | 49 | 21 | 0 |
| SRR7178614 | 239 | 13198 | 258 | 182 | 1638 | 18591 | 20363 | 40791 | 23076 | 0 | 9926 | 16348 | 3448 | 3665 | 28 | 1131 | 23641 | 0 | 0 | 0 | 3005 | 387 | 14 | 0 | 0 | 0 | 0 | 0 | 80 | 2282 | 328 | 14 | 0 | 0 | 33 | 0 | 0 |
| SRR7178615 | 239 | 14765 | 345 | 305 | 1537 | 17093 | 18869 | 37862 | 25514 | 13 | 11502 | 19425 | 4143 | 4345 | 17 | 1231 | 22046 | 0 | 0 | 0 | 2969 | 390 | 15 | 0 | 0 | 0 | 0 | 0 | 82 | 2277 | 332 | 16 | 0 | 0 | 27 | 16 | 0 |
| SRR7178616 | 110 | 5394 | 240 | 533 | 1603 | 14440 | 19797 | 31197 | 16127 | 25 | 13580 | 18935 | 5337 | 3803 | 57 | 1842 | 29573 | 0 | 0 | 0 | 5379 | 2095 | 69 | 0 | 0 | 0 | 0 | 0 | 255 | 5730 | 1864 | 90 | 0 | 17 | 85 | 132 | 0 |
| SRR7178617 | 111 | 5085 | 241 | 518 | 1527 | 14017 | 19278 | 30302 | 15680 | 32 | 13115 | 18316 | 5224 | 3693 | 41 | 1998 | 32749 | 0 | 0 | 0 | 5706 | 2188 | 60 | 0 | 0 | 0 | 0 | 0 | 272 | 6068 | 1983 | 100 | 0 | 17 | 102 | 138 | 0 |
| SRR7178618 | 71 | 5218 | 294 | 548 | 1960 | 18050 | 24391 | 38299 | 16256 | 47 | 14494 | 20181 | 5598 | 3889 | 43 | 1060 | 25040 | 0 | 0 | 0 | 4342 | 1627 | 57 | 0 | 0 | 0 | 0 | 0 | 196 | 4424 | 1496 | 65 | 0 | 12 | 68 | 83 | 0 |
| SRR7178619 | 115 | 5299 | 224 | 666 | 1969 | 17844 | 24555 | 39638 | 17771 | 20 | 15979 | 21789 | 6091 | 4244 | 26 | 734 | 18252 | 0 | 0 | 0 | 4061 | 1599 | 49 | 0 | 0 | 0 | 0 | 0 | 197 | 4336 | 1363 | 75 | 0 | 0 | 64 | 95 | 0 |
| SRR7178620 | 81 | 8392 | 178 | 190 | 1587 | 17556 | 25092 | 34159 | 17143 | 50 | 13246 | 19994 | 4304 | 4624 | 34 | 2693 | 36351 | 0 | 0 | 10 | 18693 | 5592 | 206 | 0 | 0 | 0 | 0 | 0 | 517 | 15601 | 4650 | 242 | 0 | 0 | 0 | 0 | 0 |
| SRR7178621 | 92 | 8608 | 200 | 142 | 1618 | 17430 | 24918 | 34176 | 16977 | 44 | 13206 | 19669 | 4385 | 4628 | 41 | 2548 | 36815 | 0 | 0 | 14 | 19364 | 5877 | 205 | 0 | 0 | 0 | 0 | 0 | 538 | 16114 | 4784 | 265 | 0 | 0 | 0 | 0 | 0 |
| SRR7178622 | 55 | 7614 | 198 | 198 | 1734 | 18329 | 24551 | 38655 | 16502 | 22 | 13926 | 20586 | 4510 | 4587 | 42 | 2119 | 36386 | 0 | 0 | 0 | 13396 | 3937 | 139 | 0 | 0 | 0 | 0 | 0 | 372 | 11497 | 3366 | 167 | 0 | 0 | 0 | 0 | 0 |
| SRR7178623 | 152 | 8346 | 182 | 137 | 1652 | 17250 | 23114 | 36615 | 17928 | 46 | 16355 | 24879 | 5210 | 5487 | 27 | 2242 | 33447 | 0 | 0 | 0 | 12939 | 3871 | 136 | 0 | 0 | 0 | 0 | 0 | 353 | 11289 | 3199 | 191 | 0 | 0 | 0 | 0 | 0 |
| SRR7178624 | 273 | 25073 | 560 | 457 | 1581 | 21722 | 27021 | 39042 | 17295 | 23 | 14129 | 18895 | 4889 | 4851 | 81 | 1915 | 28075 | 0 | 0 | 12 | 18357 | 3801 | 87 | 0 | 0 | 0 | 0 | 0 | 385 | 8328 | 1653 | 54 | 0 | 0 | 34 | 57 | 0 |
| SRR7178625 | 241 | 24447 | 489 | 429 | 1529 | 20643 | 25807 | 37560 | 16807 | 40 | 13652 | 18212 | 4578 | 4713 | 85 | 2057 | 31252 | 0 | 0 | 15 | 19141 | 3977 | 96 | 0 | 0 | 0 | 0 | 0 | 388 | 8785 | 1755 | 68 | 0 | 0 | 32 | 71 | 0 |
| SRR7178626 | 240 | 25323 | 519 | 392 | 2086 | 27549 | 33493 | 51172 | 19381 | 25 | 15442 | 20266 | 5263 | 5029 | 69 | 1145 | 23540 | 0 | 0 | 11 | 13835 | 3008 | 67 | 0 | 0 | 0 | 0 | 0 | 307 | 6455 | 1285 | 49 | 0 | 0 | 22 | 41 | 0 |
| SRR7178627 | 236 | 22927 | 333 | 458 | 2075 | 28231 | 34262 | 51733 | 21255 | 42 | 17276 | 22245 | 5801 | 5586 | 46 | 834 | 18091 | 0 | 0 | 11 | 13276 | 2931 | 62 | 0 | 0 | 0 | 0 | 0 | 288 | 6364 | 1278 | 45 | 0 | 0 | 19 | 35 | 0 |
| SRR7178628 | 129 | 6233 | 404 | 289 | 1420 | 13323 | 20658 | 29731 | 26595 | 47 | 11966 | 16790 | 4508 | 4327 | 20 | 2065 | 37409 | 0 | 0 | 16 | 21274 | 3492 | 120 | 0 | 0 | 0 | 0 | 0 | 775 | 20852 | 4487 | 175 | 0 | 0 | 0 | 0 | 0 |
| SRR7178629 | 93 | 6504 | 416 | 308 | 1565 | 13992 | 21266 | 32680 | 25594 | 30 | 11803 | 16425 | 4388 | 4008 | 15 | 2129 | 33231 | 0 | 0 | 13 | 18531 | 3111 | 99 | 0 | 0 | 0 | 0 | 0 | 669 | 18420 | 3929 | 156 | 0 | 0 | 0 | 0 | 0 |
| SRR7178630 | 98 | 6550 | 417 | 249 | 1631 | 14798 | 21918 | 33932 | 26048 | 43 | 12013 | 16872 | 4498 | 4182 | 17 | 1990 | 31947 | 0 | 0 | 13 | 18535 | 3038 | 105 | 0 | 0 | 0 | 0 | 0 | 671 | 18398 | 3915 | 155 | 0 | 0 | 0 | 0 | 0 |
| SRR7178631 | 129 | 5902 | 302 | 588 | 1323 | 13239 | 20052 | 33402 | 15433 | 42 | 11793 | 18501 | 4956 | 5281 | 30 | 2096 | 32927 | 0 | 0 | 0 | 10884 | 3336 | 112 | 0 | 0 | 0 | 0 | 0 | 498 | 11529 | 3581 | 164 | 0 | 0 | 11 | 0 | 0 |
| SRR7178632 | 157 | 5838 | 294 | 603 | 1269 | 13007 | 19148 | 32176 | 15108 | 32 | 11381 | 17963 | 4858 | 5048 | 18 | 2298 | 36476 | 0 | 0 | 0 | 11317 | 3522 | 116 | 0 | 0 | 0 | 0 | 0 | 491 | 12072 | 3758 | 180 | 0 | 0 | 12 | 12 | 0 |
| SRR7178633 | 131 | 6122 | 372 | 513 | 1577 | 16416 | 24098 | 41352 | 15829 | 50 | 14204 | 19332 | 5312 | 5664 | 18 | 1148 | 27598 | 0 | 0 | 0 | 8398 | 2566 | 85 | 0 | 0 | 0 | 0 | 0 | 406 | 8921 | 2787 | 119 | 0 | 0 | 13 | 0 | 0 |
| SRR7178634 | 171 | 5638 | 274 | 879 | 1665 | 16507 | 23932 | 40760 | 17266 | 57 | 15728 | 21353 | 5732 | 5986 | 11 | 847 | 20457 | 0 | 0 | 0 | 8124 | 2473 | 75 | 0 | 0 | 0 | 0 | 0 | 378 | 8604 | 2686 | 115 | 0 | 0 | 0 | 10 | 0 |
| SRR7178635 | 224 | 25933 | 412 | 292 | 1075 | 13357 | 12731 | 26264 | 19927 | 11 | 15200 | 21897 | 5323 | 3510 | 65 | 1748 | 28391 | 0 | 0 | 0 | 9061 | 3613 | 96 | 0 | 0 | 0 | 0 | 0 | 282 | 5278 | 2136 | 79 | 0 | 0 | 13 | 28 | 0 |
| SRR7178636 | 266 | 26658 | 434 | 214 | 1087 | 13364 | 12654 | 26289 | 19650 | 18 | 15026 | 21627 | 5206 | 3543 | 66 | 1650 | 28928 | 0 | 0 | 0 | 9362 | 3754 | 104 | 0 | 0 | 0 | 0 | 0 | 296 | 5439 | 2190 | 84 | 0 | 0 | 11 | 26 | 0 |
| SRR7178637 | 255 | 24359 | 395 | 308 | 1226 | 15192 | 14203 | 29953 | 19545 | 40 | 15743 | 21710 | 5392 | 3488 | 64 | 1387 | 28435 | 0 | 0 | 0 | 6374 | 2590 | 65 | 0 | 0 | 0 | 0 | 0 | 202 | 3738 | 1490 | 59 | 0 | 0 | 0 | 14 | 0 |
| SRR7178638 | 305 | 27554 | 545 | 259 | 1188 | 14552 | 13496 | 28198 | 22003 | 46 | 18530 | 26063 | 6568 | 4149 | 63 | 1478 | 26574 | 0 | 0 | 0 | 6276 | 2542 | 72 | 0 | 0 | 0 | 0 | 0 | 197 | 3659 | 1474 | 56 | 0 | 0 | 0 | 18 | 0 |
| SRR7178639 | 65 | 21921 | 223 | 316 | 1079 | 14059 | 14179 | 32400 | 11415 | 34 | 12152 | 19123 | 3024 | 6723 | 23 | 1825 | 31598 | 0 | 0 | 0 | 7090 | 1484 | 36 | 0 | 0 | 0 | 0 | 0 | 159 | 4196 | 832 | 22 | 0 | 0 | 0 | 0 | 0 |
| SRR7178640 | 87 | 21377 | 239 | 329 | 1188 | 15087 | 15229 | 35621 | 11195 | 19 | 11789 | 18443 | 2894 | 6366 | 23 | 1876 | 27569 | 0 | 0 | 0 | 6209 | 1264 | 28 | 0 | 0 | 0 | 0 | 0 | 131 | 3636 | 716 | 23 | 0 | 0 | 0 | 0 | 0 |
| SRR7178641 | 57 | 21477 | 217 | 355 | 1205 | 15794 | 15682 | 36942 | 11238 | 30 | 12130 | 18847 | 2927 | 6385 | 17 | 1760 | 26340 | 0 | 0 | 0 | 6093 | 1280 | 28 | 0 | 0 | 0 | 0 | 0 | 129 | 3619 | 707 | 20 | 0 | 0 | 0 | 0 | 0 |
| SRR7178642 | 142 | 8260 | 204 | 131 | 1138 | 10393 | 13553 | 30174 | 19295 | 13 | 9217 | 16990 | 3899 | 4088 | 26 | 1428 | 46469 | 0 | 0 | 14 | 10636 | 3142 | 77 | 0 | 0 | 0 | 0 | 0 | 480 | 9122 | 4093 | 97 | 0 | 0 | 0 | 0 | 0 |
| SRR7178643 | 143 | 8171 | 157 | 89 | 1206 | 11172 | 14499 | 32804 | 19073 | 19 | 9100 | 16439 | 3878 | 3924 | 32 | 1481 | 41438 | 0 | 0 | 12 | 9145 | 2760 | 73 | 0 | 0 | 0 | 0 | 0 | 405 | 7828 | 3462 | 89 | 0 | 0 | 0 | 0 | 0 |
| SRR7178644 | 149 | 8054 | 244 | 117 | 1293 | 11843 | 15497 | 34566 | 19248 | 0 | 9469 | 16922 | 3984 | 3960 | 28 | 1402 | 39848 | 0 | 0 | 12 | 9120 | 2742 | 71 | 0 | 0 | 0 | 0 | 0 | 405 | 7853 | 3506 | 82 | 0 | 0 | 0 | 0 | 0 |
| SRR7178645 | 63 | 8464 | 225 | 251 | 1089 | 10787 | 14176 | 36147 | 17160 | 53 | 9604 | 18706 | 3571 | 4262 | 21 | 1133 | 46102 | 0 | 0 | 12 | 6643 | 4512 | 242 | 0 | 0 | 0 | 0 | 0 | 302 | 4734 | 5967 | 301 | 0 | 0 | 11 | 0 | 0 |
| SRR7178646 | 65 | 8483 | 201 | 259 | 1027 | 10167 | 13451 | 34457 | 16548 | 25 | 9200 | 17885 | 3515 | 4123 | 19 | 1243 | 50495 | 0 | 0 | 0 | 7014 | 4763 | 254 | 0 | 0 | 0 | 0 | 0 | 305 | 5062 | 6345 | 313 | 0 | 0 | 12 | 0 | 0 |
| SRR7178647 | 27 | 8210 | 242 | 215 | 1353 | 13835 | 17851 | 46788 | 18053 | 0 | 10750 | 20471 | 3714 | 4389 | 11 | 671 | 39342 | 0 | 0 | 0 | 4939 | 3345 | 187 | 0 | 0 | 0 | 0 | 0 | 203 | 3471 | 4298 | 197 | 0 | 0 | 0 | 0 | 0 |
| SRR7178648 | 61 | 7850 | 151 | 333 | 1414 | 14637 | 18446 | 48879 | 19775 | 45 | 12002 | 22676 | 4384 | 4846 | 16 | 508 | 29741 | 0 | 0 | 0 | 4749 | 3175 | 164 | 0 | 0 | 0 | 0 | 0 | 207 | 3323 | 4091 | 205 | 0 | 0 | 0 | 0 | 0 |
| SRR7178649 | 148 | 4774 | 379 | 917 | 421 | 4448 | 6267 | 7088 | 5824 | 40 | 9571 | 16923 | 3978 | 1145 | 19 | 1623 | 58992 | 0 | 0 | 0 | 1812 | 5720 | 1237 | 19 | 0 | 0 | 0 | 0 | 72 | 1198 | 6458 | 1482 | 0 | 46 | 145 | 144 | 0 |
| SRR7178650 | 187 | 4415 | 282 | 871 | 459 | 4656 | 6691 | 7443 | 5871 | 53 | 9620 | 16451 | 3856 | 1085 | 22 | 1795 | 53884 | 0 | 0 | 0 | 1572 | 4769 | 1020 | 18 | 10 | 0 | 0 | 0 | 56 | 988 | 5301 | 1222 | 0 | 39 | 118 | 115 | 0 |
| SRR7178651 | 191 | 4768 | 337 | 932 | 473 | 4763 | 6884 | 7761 | 6023 | 70 | 9881 | 17011 | 3924 | 1106 | 17 | 1567 | 52109 | 0 | 0 | 0 | 1555 | 4792 | 1023 | 21 | 0 | 0 | 0 | 0 | 53 | 983 | 5264 | 1185 | 0 | 34 | 120 | 110 | 0 |
| SRR7178652 | 54 | 9084 | 170 | 410 | 1397 | 12863 | 19003 | 34255 | 15097 | 34 | 10743 | 19443 | 4185 | 6180 | 23 | 2456 | 44015 | 0 | 0 | 0 | 11949 | 4274 | 162 | 0 | 0 | 0 | 0 | 0 | 433 | 11450 | 4128 | 223 | 0 | 0 | 14 | 14 | 0 |
| SRR7178653 | 74 | 9179 | 212 | 339 | 1365 | 12696 | 18894 | 34099 | 14921 | 48 | 10521 | 19243 | 4130 | 6093 | 25 | 2318 | 44519 | 0 | 0 | 11 | 12323 | 4428 | 166 | 0 | 0 | 0 | 0 | 0 | 472 | 11803 | 4280 | 243 | 0 | 0 | 11 | 16 | 0 |
| SRR7178654 | 104 | 8794 | 177 | 446 | 1531 | 14286 | 21011 | 38799 | 14922 | 31 | 12624 | 19509 | 4241 | 6042 | 27 | 1965 | 45326 | 0 | 0 | 0 | 8473 | 3075 | 110 | 0 | 0 | 0 | 0 | 0 | 332 | 8005 | 2824 | 148 | 0 | 0 | 0 | 14 | 0 |
| SRR7178655 | 116 | 9960 | 102 | 363 | 1454 | 13591 | 19775 | 36626 | 16470 | 102 | 14850 | 23364 | 4865 | 7207 | 31 | 2215 | 41740 | 0 | 0 | 0 | 8278 | 2967 | 112 | 0 | 0 | 0 | 0 | 0 | 324 | 7928 | 2792 | 142 | 0 | 0 | 0 | 0 | 0 |
| SRR7178656 | 107 | 12823 | 366 | 394 | 1316 | 11612 | 15677 | 37877 | 8981 | 38 | 10153 | 17228 | 3993 | 5973 | 17 | 2245 | 40655 | 0 | 0 | 12 | 15279 | 3117 | 65 | 0 | 0 | 0 | 0 | 0 | 454 | 10284 | 2338 | 54 | 0 | 0 | 0 | 0 | 0 |
| SRR7178657 | 129 | 13124 | 358 | 329 | 1292 | 11677 | 15688 | 37882 | 8894 | 39 | 9949 | 16966 | 3904 | 5948 | 16 | 2147 | 41556 | 0 | 0 | 15 | 15750 | 3194 | 68 | 0 | 0 | 0 | 0 | 0 | 478 | 10610 | 2400 | 62 | 0 | 0 | 0 | 0 | 0 |
| SRR7178658 | 98 | 11881 | 449 | 449 | 1397 | 12848 | 17432 | 42542 | 8915 | 55 | 10008 | 16206 | 3797 | 5537 | 17 | 1744 | 39795 | 0 | 0 | 0 | 10443 | 2228 | 46 | 0 | 0 | 0 | 0 | 0 | 309 | 7284 | 1633 | 36 | 0 | 0 | 0 | 0 | 0 |
| SRR7178659 | 105 | 13262 | 467 | 347 | 1306 | 11778 | 15746 | 38172 | 9662 | 45 | 11283 | 18841 | 4430 | 6411 | 19 | 1738 | 35582 | 0 | 0 | 0 | 10104 | 2218 | 51 | 0 | 0 | 0 | 0 | 0 | 318 | 7031 | 1607 | 41 | 0 | 0 | 0 | 0 | 0 |
| SRR7178660 | 129 | 3636 | 179 | 519 | 1322 | 13400 | 20361 | 25187 | 17993 | 74 | 9110 | 16598 | 3536 | 3252 | 40 | 1774 | 54775 | 0 | 0 | 10 | 12019 | 4702 | 189 | 0 | 0 | 0 | 0 | 0 | 515 | 11726 | 5414 | 297 | 0 | 0 | 0 | 11 | 0 |
| SRR7178661 | 111 | 3466 | 210 | 529 | 1459 | 14535 | 22198 | 27401 | 18135 | 71 | 9364 | 16030 | 3412 | 3117 | 36 | 1843 | 50336 | 0 | 0 | 0 | 10508 | 4080 | 157 | 0 | 0 | 0 | 0 | 0 | 435 | 10128 | 4563 | 264 | 0 | 0 | 0 | 0 | 0 |
| SRR7178662 | 142 | 3397 | 213 | 580 | 1529 | 15527 | 23320 | 29038 | 18260 | 69 | 9783 | 16712 | 3549 | 3220 | 35 | 1705 | 48619 | 0 | 0 | 0 | 10486 | 4077 | 161 | 0 | 0 | 0 | 0 | 0 | 436 | 10180 | 4574 | 255 | 0 | 0 | 0 | 0 | 0 |
| SRR7178663 | 20 | 24376 | 330 | 285 | 964 | 13035 | 14258 | 20190 | 14561 | 12 | 13600 | 21111 | 4550 | 2440 | 36 | 1188 | 44585 | 0 | 0 | 0 | 10192 | 6680 | 343 | 0 | 0 | 0 | 0 | 0 | 333 | 8341 | 5130 | 391 | 0 | 0 | 19 | 12 | 0 |
| SRR7178664 | 42 | 23358 | 378 | 280 | 893 | 12208 | 13444 | 18905 | 13947 | 23 | 12939 | 20014 | 4287 | 2307 | 34 | 1306 | 48120 | 0 | 0 | 0 | 10850 | 7100 | 357 | 0 | 0 | 0 | 0 | 0 | 344 | 8838 | 5478 | 416 | 0 | 10 | 19 | 13 | 0 |
| SRR7178665 | 0 | 24541 | 243 | 299 | 1259 | 16811 | 18841 | 26923 | 15338 | 93 | 14951 | 23349 | 4937 | 2584 | 22 | 684 | 39560 | 0 | 0 | 0 | 7877 | 5139 | 265 | 0 | 0 | 0 | 0 | 0 | 270 | 6291 | 3852 | 290 | 0 | 0 | 13 | 10 | 0 |
| SRR7178666 | 44 | 22466 | 283 | 457 | 1281 | 17300 | 19096 | 27690 | 17160 | 22 | 17538 | 26065 | 5477 | 2808 | 16 | 570 | 30048 | 0 | 0 | 0 | 7494 | 4905 | 226 | 0 | 0 | 0 | 0 | 0 | 239 | 6194 | 3708 | 292 | 0 | 0 | 14 | 10 | 0 |
| SRR7178667 | 36 | 2902 | 131 | 521 | 556 | 5093 | 7750 | 6072 | 4971 | 60 | 14392 | 26480 | 3707 | 1504 | 21 | 1683 | 38039 | 0 | 0 | 0 | 6396 | 1559 | 58 | 0 | 0 | 0 | 0 | 0 | 180 | 7171 | 1378 | 70 | 0 | 0 | 0 | 0 | 0 |
| SRR7178668 | 38 | 2807 | 135 | 614 | 548 | 4787 | 7358 | 5752 | 4828 | 55 | 13688 | 25486 | 3655 | 1416 | 27 | 1764 | 42071 | 0 | 0 | 0 | 6809 | 1662 | 65 | 0 | 0 | 0 | 0 | 0 | 197 | 7602 | 1465 | 81 | 0 | 0 | 0 | 0 | 0 |
| SRR7178669 | 53 | 2939 | 148 | 539 | 701 | 6163 | 9625 | 7460 | 5079 | 53 | 15034 | 27853 | 3894 | 1523 | 17 | 1000 | 32145 | 0 | 0 | 0 | 4769 | 1200 | 41 | 0 | 0 | 0 | 0 | 0 | 140 | 5447 | 1057 | 59 | 0 | 0 | 0 | 0 | 0 |
| SRR7178670 | 37 | 2789 | 158 | 755 | 663 | 6084 | 9518 | 7530 | 5430 | 73 | 16307 | 29552 | 4227 | 1566 | 16 | 694 | 23605 | 0 | 0 | 0 | 4625 | 1172 | 50 | 0 | 0 | 0 | 0 | 0 | 155 | 5255 | 1011 | 55 | 0 | 0 | 0 | 0 | 0 |
| SRR7178671 | 201 | 12303 | 207 | 329 | 1449 | 12540 | 18509 | 42265 | 16146 | 46 | 9178 | 16057 | 3864 | 6145 | 26 | 1893 | 42834 | 0 | 0 | 0 | 7927 | 3466 | 96 | 0 | 0 | 0 | 0 | 0 | 388 | 7140 | 3674 | 150 | 0 | 0 | 0 | 0 | 0 |
| SRR7178672 | 205 | 12548 | 276 | 289 | 1408 | 12457 | 18368 | 41670 | 15982 | 26 | 9047 | 16008 | 3795 | 6045 | 21 | 1756 | 43646 | 0 | 0 | 11 | 8215 | 3620 | 98 | 0 | 0 | 0 | 0 | 0 | 395 | 7500 | 3849 | 158 | 0 | 0 | 0 | 0 | 0 |

### Supp Table S8.1.Columns_B_to_AL

| | | | | | | | | | | | | | | | | | | | | | | | | | | | | | | | | | | | | | |
|---|---|---|---|---|---|---|---|---|---|---|---|---|---|---|---|---|---|---|---|---|---|---|---|---|---|---|---|---|---|---|---|---|---|---|---|---|---|
| SRR7178673 | 152 | 11749 | 163 | 413 | 1695 | 14725 | 20927 | 49945 | 16092 | 0 | 9490 | 16222 | 3896 | 5777 | 17 | 1532 | 44569 | 0 | 0 | 0 | 5385 | 2348 | 67 | 0 | 0 | 0 | 0 | 0 | 241 | 4765 | 2482 | 99 | 0 | 0 | 0 | 0 | 0 |
| SRR7178674 | 235 | 11408 | 279 | 440 | 1536 | 13763 | 19529 | 46714 | 18208 | 29 | 11244 | 19425 | 4379 | 6903 | 28 | 1597 | 40739 | 0 | 0 | 0 | 5098 | 2275 | 68 | 0 | 0 | 0 | 0 | 0 | 260 | 4649 | 2338 | 86 | 0 | 0 | 0 | 0 | 0 |
| SRR7178675 | 161 | 9534 | 234 | 661 | 1308 | 13165 | 18286 | 43250 | 13309 | 14 | 7812 | 14012 | 2439 | 5489 | 0 | 2453 | 31286 | 0 | 0 | 0 | 5215 | 2090 | 75 | 0 | 0 | 0 | 0 | 0 | 137 | 4651 | 2135 | 97 | 0 | 19 | 304 | 86 | 0 |
| SRR7178676 | 171 | 9180 | 282 | 631 | 1201 | 12524 | 17370 | 41184 | 13093 | 13 | 7381 | 13289 | 2351 | 5186 | 13 | 2624 | 34622 | 0 | 0 | 0 | 5466 | 2207 | 83 | 0 | 0 | 0 | 0 | 0 | 136 | 4894 | 2310 | 92 | 0 | 23 | 313 | 94 | 0 |
| SRR7178677 | 194 | 9406 | 272 | 428 | 1484 | 16606 | 22247 | 55150 | 13473 | 19 | 8445 | 14512 | 2463 | 5428 | 0 | 1444 | 26551 | 0 | 0 | 0 | 3962 | 1618 | 62 | 0 | 0 | 0 | 0 | 0 | 93 | 3408 | 1609 | 78 | 0 | 17 | 216 | 66 | 0 |
| SRR7178678 | 155 | 8409 | 288 | 843 | 1626 | 16882 | 22218 | 55026 | 14429 | 0 | 9459 | 15526 | 2745 | 6098 | 0 | 904 | 20129 | 0 | 0 | 0 | 3880 | 1531 | 48 | 0 | 0 | 0 | 0 | 0 | 105 | 3325 | 1580 | 70 | 0 | 20 | 224 | 56 | 0 |
| SRR7178679 | 208 | 8959 | 186 | 422 | 1402 | 14259 | 19090 | 33498 | 18951 | 35 | 9397 | 16207 | 3407 | 3859 | 25 | 1265 | 35356 | 0 | 0 | 17 | 17583 | 6658 | 194 | 0 | 0 | 0 | 0 | 0 | 623 | 13322 | 6508 | 215 | 0 | 0 | 0 | 0 | 0 |
| SRR7178680 | 228 | 8597 | 290 | 526 | 1542 | 15893 | 20958 | 36957 | 19108 | 27 | 9095 | 15981 | 3366 | 3651 | 26 | 1328 | 31318 | 0 | 0 | 17 | 15190 | 5709 | 160 | 0 | 0 | 0 | 0 | 0 | 526 | 11273 | 5513 | 170 | 0 | 0 | 0 | 0 | 0 |
| SRR7178681 | 216 | 9333 | 244 | 545 | 1621 | 16664 | 21942 | 38891 | 19445 | 36 | 9476 | 16371 | 3421 | 3772 | 30 | 1182 | 30322 | 0 | 0 | 18 | 15225 | 5697 | 159 | 0 | 0 | 0 | 0 | 0 | 523 | 11234 | 5449 | 179 | 0 | 0 | 0 | 0 | 0 |
| SRR7178682 | 329 | 36753 | 443 | 440 | 823 | 11871 | 11378 | 23835 | 19927 | 26 | 11009 | 18849 | 3934 | 2355 | 19 | 988 | 34239 | 0 | 0 | 0 | 7176 | 3386 | 94 | 0 | 0 | 0 | 0 | 0 | 232 | 5692 | 2752 | 117 | 0 | 0 | 0 | 0 | 0 |
| SRR7178683 | 369 | 35823 | 442 | 405 | 799 | 11293 | 10754 | 22522 | 19205 | 27 | 10410 | 17880 | 3711 | 2252 | 22 | 1069 | 37829 | 0 | 0 | 0 | 7559 | 3625 | 104 | 0 | 0 | 0 | 0 | 0 | 241 | 6023 | 2907 | 122 | 0 | 0 | 0 | 0 | 0 |
| SRR7178684 | 415 | 35619 | 422 | 451 | 1081 | 14500 | 14602 | 32337 | 19110 | 87 | 11182 | 19412 | 4027 | 2482 | 23 | 634 | 31449 | 0 | 0 | 0 | 5335 | 2564 | 67 | 0 | 0 | 0 | 0 | 0 | 181 | 4323 | 2085 | 90 | 0 | 0 | 0 | 0 | 0 |
| SRR7178685 | 357 | 33379 | 433 | 662 | 1151 | 15137 | 14863 | 33704 | 21319 | 51 | 13010 | 21941 | 4498 | 2867 | 13 | 441 | 23709 | 0 | 0 | 0 | 5109 | 2462 | 67 | 0 | 0 | 0 | 0 | 0 | 183 | 4167 | 2078 | 80 | 0 | 0 | 0 | 0 | 0 |
| SRR7178686 | 59 | 6077 | 296 | 666 | 433 | 4658 | 6473 | 7139 | 5995 | 41 | 10372 | 17892 | 3442 | 1056 | 12 | 2082 | 39186 | 0 | 0 | 0 | 644 | 628 | 63 | 0 | 0 | 0 | 0 | 0 | 35 | 641 | 848 | 94 | 0 | 0 | 0 | 0 | 0 |
| SRR7178687 | 55 | 6168 | 206 | 610 | 439 | 4605 | 6453 | 7031 | 5923 | 40 | 10233 | 17505 | 3384 | 1057 | 10 | 1968 | 39383 | 0 | 0 | 0 | 652 | 658 | 63 | 0 | 0 | 0 | 0 | 0 | 36 | 659 | 869 | 96 | 0 | 0 | 0 | 0 | 0 |
| SRR7178688 | 68 | 5994 | 330 | 686 | 497 | 5206 | 7310 | 8254 | 5727 | 68 | 10397 | 17002 | 3296 | 974 | 13 | 1599 | 40118 | 0 | 0 | 0 | 432 | 450 | 45 | 0 | 0 | 0 | 0 | 0 | 27 | 432 | 584 | 60 | 0 | 0 | 0 | 0 | 0 |
| SRR7178689 | 94 | 6583 | 211 | 751 | 473 | 4926 | 7015 | 7962 | 6619 | 35 | 12509 | 20914 | 4055 | 1197 | 0 | 1874 | 37724 | 0 | 0 | 0 | 445 | 456 | 40 | 0 | 0 | 0 | 0 | 0 | 23 | 439 | 586 | 69 | 0 | 0 | 0 | 0 | 0 |
| SRR7178690 | 138 | 6952 | 174 | 395 | 1171 | 11123 | 16651 | 33229 | 16359 | 36 | 9646 | 16705 | 3498 | 4877 | 27 | 2012 | 65126 | 0 | 0 | 0 | 4948 | 3052 | 117 | 0 | 0 | 0 | 0 | 0 | 301 | 4045 | 3654 | 189 | 0 | 0 | 56 | 32 | 0 |
| SRR7178691 | 195 | 14362 | 514 | 409 | 1192 | 11824 | 18730 | 40370 | 25914 | 29 | 9706 | 12802 | 2896 | 6086 | 18 | 1916 | 30668 | 0 | 0 | 10 | 7895 | 4697 | 185 | 0 | 0 | 0 | 0 | 0 | 268 | 5530 | 3024 | 148 | 0 | 0 | 0 | 0 | 0 |
| SRR7178692 | 123 | 14773 | 448 | 409 | 1209 | 12341 | 19510 | 42105 | 26374 | 48 | 9901 | 13039 | 2961 | 6247 | 15 | 1714 | 29555 | 0 | 0 | 11 | 7848 | 4656 | 183 | 0 | 0 | 0 | 0 | 0 | 267 | 5539 | 2992 | 147 | 0 | 0 | 0 | 0 | 0 |
| SRR7178693 | 133 | 14007 | 437 | 271 | 1153 | 16916 | 16011 | 34432 | 22829 | 23 | 11284 | 19877 | 4347 | 3593 | 32 | 1335 | 35639 | 0 | 0 | 0 | 8422 | 2017 | 57 | 0 | 0 | 0 | 0 | 0 | 254 | 6768 | 1733 | 74 | 0 | 0 | 27 | 18 | 0 |
| SRR7178694 | 150 | 14397 | 460 | 265 | 1167 | 16792 | 15852 | 34300 | 22433 | 36 | 11036 | 19534 | 4140 | 3523 | 28 | 1214 | 36130 | 0 | 0 | 0 | 8791 | 2069 | 61 | 0 | 0 | 0 | 0 | 0 | 251 | 7049 | 1797 | 82 | 0 | 0 | 33 | 22 | 0 |
| SRR7178695 | 119 | 13986 | 451 | 307 | 1439 | 19083 | 18266 | 42605 | 20924 | 17 | 11205 | 19565 | 4290 | 3794 | 33 | 1046 | 38794 | 0 | 0 | 0 | 5887 | 1424 | 45 | 0 | 0 | 0 | 0 | 0 | 188 | 4829 | 1225 | 52 | 0 | 0 | 16 | 12 | 0 |
| SRR7178696 | 94 | 14776 | 578 | 283 | 1380 | 18352 | 17835 | 41114 | 23769 | 12 | 13308 | 23186 | 5177 | 4487 | 37 | 1172 | 37089 | 0 | 0 | 0 | 5791 | 1370 | 44 | 0 | 0 | 0 | 0 | 0 | 186 | 4722 | 1220 | 53 | 0 | 0 | 22 | 13 | 0 |
| SRR7178697 | 93 | 16996 | 319 | 536 | 1271 | 15648 | 19136 | 38502 | 15782 | 26 | 13303 | 18244 | 4823 | 5871 | 42 | 1599 | 32099 | 0 | 0 | 12 | 14580 | 2832 | 67 | 0 | 0 | 0 | 0 | 0 | 466 | 9748 | 2282 | 76 | 0 | 0 | 0 | 0 | 0 |
| SRR7178698 | 140 | 6852 | 154 | 409 | 1308 | 12293 | 18096 | 36372 | 16572 | 39 | 9438 | 16244 | 3505 | 4608 | 31 | 2113 | 58717 | 0 | 0 | 0 | 4178 | 2717 | 104 | 0 | 0 | 0 | 0 | 0 | 256 | 3488 | 3121 | 164 | 0 | 0 | 45 | 23 | 0 |
| SRR7178699 | 153 | 7033 | 155 | 433 | 1323 | 12833 | 19003 | 38233 | 16912 | 32 | 9823 | 16791 | 3547 | 4775 | 23 | 1984 | 56750 | 0 | 0 | 0 | 4246 | 2713 | 99 | 0 | 0 | 0 | 0 | 0 | 248 | 3503 | 3131 | 162 | 0 | 0 | 46 | 26 | 0 |
| SRR7178700 | 74 | 5405 | 235 | 321 | 1206 | 13510 | 17977 | 24177 | 17929 | 53 | 12481 | 19898 | 3442 | 3477 | 36 | 1621 | 44552 | 0 | 0 | 0 | 6189 | 2761 | 151 | 0 | 0 | 0 | 0 | 0 | 243 | 6124 | 3080 | 243 | 0 | 0 | 18 | 24 | 0 |
| SRR7178701 | 51 | 5690 | 196 | 391 | 1135 | 13213 | 17906 | 24018 | 17770 | 60 | 12238 | 19403 | 3377 | 3403 | 25 | 1464 | 44619 | 0 | 0 | 0 | 6430 | 2890 | 165 | 0 | 0 | 0 | 0 | 0 | 251 | 6312 | 3185 | 274 | 0 | 0 | 18 | 23 | 0 |
| SRR7178702 | 53 | 4760 | 158 | 338 | 1418 | 15725 | 20475 | 28769 | 18158 | 95 | 13191 | 19888 | 3552 | 3501 | 24 | 1257 | 46450 | 0 | 0 | 0 | 4283 | 1904 | 113 | 0 | 0 | 0 | 0 | 0 | 172 | 4331 | 2131 | 176 | 0 | 0 | 12 | 13 | 0 |
| SRR7178703 | 147 | 5409 | 324 | 486 | 1270 | 14833 | 20012 | 27775 | 20487 | 133 | 15747 | 24490 | 4278 | 4095 | 23 | 1332 | 44297 | 0 | 0 | 0 | 4210 | 1893 | 105 | 0 | 0 | 0 | 0 | 0 | 175 | 4274 | 2109 | 173 | 0 | 0 | 13 | 12 | 0 |
| SRR7178704 | 244 | 6908 | 216 | 371 | 314 | 3243 | 3965 | 4321 | 2858 | 30 | 12707 | 26045 | 3453 | 1034 | 13 | 1763 | 60497 | 0 | 0 | 0 | 2451 | 2197 | 184 | 0 | 0 | 0 | 0 | 0 | 108 | 2060 | 1785 | 200 | 0 | 0 | 0 | 0 | 0 |
| SRR7178705 | 246 | 6854 | 281 | 460 | 340 | 3523 | 4372 | 4714 | 2874 | 35 | 12286 | 25231 | 3392 | 1016 | 0 | 1878 | 55691 | 0 | 0 | 0 | 2135 | 1928 | 155 | 0 | 0 | 0 | 0 | 0 | 95 | 1780 | 1481 | 166 | 0 | 0 | 0 | 0 | 0 |
| SRR7178706 | 267 | 6981 | 269 | 453 | 366 | 3734 | 4532 | 5028 | 2922 | 18 | 12703 | 25999 | 3483 | 1046 | 0 | 1704 | 53822 | 0 | 0 | 0 | 2080 | 1924 | 148 | 0 | 0 | 0 | 0 | 0 | 92 | 1751 | 1524 | 177 | 0 | 0 | 0 | 0 | 0 |
| SRR7178707 | 13 | 3760 | 96 | 108 | 907 | 7498 | 14290 | 33713 | 14736 | 83 | 6036 | 14288 | 1965 | 3999 | 0 | 2104 | 75394 | 0 | 0 | 0 | 2917 | 1867 | 106 | 0 | 0 | 0 | 0 | 0 | 135 | 3015 | 3034 | 218 | 0 | 0 | 36 | 12 | 0 |
| SRR7178708 | 31 | 4069 | 123 | 141 | 881 | 7516 | 14180 | 33228 | 14925 | 55 | 5995 | 14192 | 1996 | 3940 | 0 | 1961 | 75885 | 0 | 0 | 0 | 3033 | 2000 | 117 | 0 | 0 | 0 | 0 | 0 | 137 | 3151 | 3220 | 220 | 0 | 0 | 36 | 11 | 0 |
| SRR7178709 | 50 | 3270 | 75 | 176 | 1037 | 8865 | 16110 | 39648 | 14389 | 75 | 6312 | 14017 | 1915 | 3826 | 14 | 1584 | 78160 | 0 | 0 | 0 | 1950 | 1231 | 71 | 0 | 0 | 0 | 0 | 0 | 84 | 1954 | 1996 | 143 | 0 | 0 | 22 | 0 | 0 |
| SRR7178710 | 0 | 4052 | 68 | 204 | 1009 | 8305 | 14685 | 37135 | 16264 | 136 | 7339 | 16902 | 2490 | 4653 | 0 | 1905 | 72374 | 0 | 0 | 0 | 1931 | 1221 | 68 | 0 | 0 | 0 | 0 | 0 | 81 | 1900 | 1923 | 136 | 0 | 0 | 20 | 0 | 0 |
| SRR7178711 | 205 | 23107 | 353 | 191 | 806 | 8505 | 9293 | 22562 | 15000 | 29 | 8117 | 16070 | 2696 | 2305 | 14 | 1092 | 48472 | 0 | 0 | 0 | 2234 | 901 | 30 | 0 | 0 | 0 | 0 | 0 | 94 | 2520 | 1232 | 45 | 0 | 0 | 0 | 0 | 0 |
| SRR7178712 | 168 | 22287 | 239 | 252 | 885 | 9403 | 10186 | 25367 | 14887 | 35 | 8022 | 15629 | 2605 | 2277 | 13 | 1146 | 45239 | 0 | 0 | 0 | 1982 | 801 | 22 | 0 | 0 | 0 | 0 | 0 | 82 | 2166 | 1033 | 41 | 0 | 0 | 0 | 0 | 0 |
| SRR7178713 | 169 | 22698 | 311 | 178 | 964 | 10034 | 10854 | 26882 | 15069 | 32 | 8340 | 16155 | 2713 | 2321 | 17 | 1064 | 43906 | 0 | 0 | 0 | 2000 | 785 | 23 | 0 | 0 | 0 | 0 | 0 | 87 | 2148 | 1037 | 39 | 0 | 0 | 0 | 0 | 0 |
| SRR7178714 | 175 | 22903 | 258 | 235 | 1519 | 17212 | 22196 | 45336 | 22157 | 34 | 8864 | 14189 | 3109 | 5978 | 25 | 1401 | 19704 | 0 | 0 | 11 | 7826 | 2893 | 101 | 0 | 0 | 0 | 0 | 0 | 252 | 7489 | 2550 | 114 | 0 | 0 | 0 | 0 | 0 |
| SRR7178715 | 113 | 22280 | 271 | 222 | 1421 | 16396 | 21136 | 43703 | 21478 | 28 | 8483 | 13605 | 2910 | 5761 | 19 | 1528 | 21748 | 0 | 0 | 0 | 8279 | 3097 | 101 | 0 | 0 | 0 | 0 | 0 | 260 | 7972 | 2770 | 114 | 0 | 0 | 0 | 0 | 0 |
| SRR7178716 | 171 | 21288 | 295 | 311 | 1898 | 21456 | 27457 | 57477 | 22680 | 0 | 9404 | 14534 | 3066 | 6129 | 20 | 841 | 16542 | 0 | 0 | 0 | 5997 | 2215 | 66 | 0 | 0 | 0 | 0 | 0 | 195 | 5727 | 1962 | 62 | 0 | 0 | 0 | 0 | 0 |
| SRR7178717 | 193 | 20934 | 245 | 210 | 1938 | 21690 | 27670 | 57822 | 24608 | 35 | 10107 | 15631 | 3411 | 6615 | 10 | 566 | 12194 | 0 | 0 | 0 | 5823 | 2186 | 73 | 0 | 0 | 0 | 0 | 0 | 191 | 5590 | 1918 | 81 | 0 | 0 | 0 | 0 | 0 |
| SRR7178718 | 201 | 16047 | 318 | 135 | 1268 | 12880 | 17052 | 39440 | 24885 | 25 | 8566 | 15684 | 3131 | 4268 | 22 | 1332 | 35160 | 0 | 0 | 0 | 8708 | 4230 | 142 | 0 | 0 | 0 | 0 | 0 | 289 | 9755 | 4597 | 192 | 0 | 0 | 0 | 0 | 0 |
| SRR7178719 | 194 | 16088 | 282 | 167 | 1437 | 13976 | 18674 | 45102 | 23662 | 42 | 8165 | 14942 | 3021 | 4089 | 21 | 1414 | 33034 | 0 | 0 | 0 | 7260 | 3637 | 116 | 0 | 0 | 0 | 0 | 0 | 264 | 8326 | 3946 | 162 | 0 | 0 | 0 | 0 | 0 |
| SRR7178720 | 188 | 15720 | 332 | 156 | 1523 | 14807 | 19514 | 47566 | 24036 | 26 | 8449 | 15453 | 3101 | 4244 | 21 | 1282 | 32197 | 0 | 0 | 0 | 7264 | 3589 | 109 | 0 | 0 | 0 | 0 | 0 | 263 | 8236 | 3907 | 166 | 0 | 0 | 0 | 0 | 0 |
| SRR7178721 | 92 | 5632 | 184 | 352 | 842 | 8958 | 12296 | 14435 | 20755 | 32 | 5800 | 10996 | 1857 | 2086 | 28 | 2073 | 50934 | 0 | 0 | 0 | 2512 | 2005 | 115 | 0 | 0 | 0 | 0 | 0 | 116 | 2379 | 2437 | 171 | 0 | 0 | 44 | 20 | 0 |
| SRR7178722 | 74 | 5663 | 140 | 357 | 826 | 8822 | 12100 | 14368 | 20431 | 35 | 5736 | 10825 | 1823 | 2037 | 27 | 1954 | 51306 | 0 | 0 | 0 | 2642 | 2044 | 110 | 0 | 0 | 0 | 0 | 0 | 110 | 2476 | 2552 | 156 | 0 | 0 | 46 | 25 | 0 |
| SRR7178723 | 47 | 4805 | 124 | 420 | 947 | 10492 | 14192 | 16839 | 20665 | 78 | 5972 | 10478 | 1773 | 2025 | 32 | 1539 | 51977 | 0 | 0 | 0 | 1827 | 1428 | 76 | 0 | 0 | 0 | 0 | 0 | 90 | 1664 | 1687 | 106 | 0 | 0 | 36 | 14 | 0 |
| SRR7178724 | 22 | 5424 | 151 | 452 | 921 | 9951 | 13610 | 15882 | 23742 | 108 | 7231 | 13014 | 2252 | 2539 | 22 | 1649 | 49478 | 0 | 0 | 0 | 1782 | 1404 | 81 | 0 | 0 | 0 | 0 | 0 | 73 | 1643 | 1654 | 111 | 0 | 0 | 26 | 10 | 0 |
| SRR7213733 | 146 | 753 | 9947 | 589 | 550 | 668 | 38161 | 9996 | 15960 | 491 | 3031 | 2377 | 3737 | 19354 | 111 | 720 | 46594 | 0 | 0 | 0 | 0 | 0 | 21 | 0 | 0 | 0 | 0 | 0 | 0 | 0 | 0 | 0 | 0 | 0 | 0 | 0 | 0 |
| SRR7213734 | 167 | 987 | 11779 | 1147 | 320 | 456 | 41457 | 10332 | 22754 | 397 | 2762 | 1883 | 4083 | 19071 | 111 | 724 | 41168 | 0 | 0 | 0 | 0 | 0 | 14 | 0 | 0 | 0 | 0 | 0 | 0 | 0 | 0 | 0 | 0 | 0 | 0 | 0 | 0 |
| SRR7213735 | 241 | 891 | 12076 | 402 | 499 | 760 | 49703 | 11808 | 24642 | 365 | 4158 | 2972 | 4593 | 21712 | 106 | 772 | 45852 | 0 | 0 | 0 | 0 | 0 | 16 | 0 | 0 | 0 | 0 | 0 | 0 | 0 | 0 | 0 | 0 | 0 | 0 | 0 | 0 |
| SRR7226512 | 388 | 1660 | 17211 | 5942 | 1477 | 2168 | 17609 | 5865 | 36471 | 8059 | 14928 | 20273 | 8831 | 2983 | 199 | 1172 | 13890 | 0 | 13 | 14 | 171 | 22 | 0 | 0 | 0 | 0 | 15 | 601 | 329 | 1081 | 130 | 77 | 0 | 0 | 0 | 17 | 0 |
| SRR7226513 | 1711 | 6193 | 47665 | 1392 | 2086 | 1429 | 10047 | 11456 | 21491 | 3983 | 16128 | 32889 | 13810 | 7926 | 72 | 1734 | 16092 | 0 | 0 | 0 | 126 | 21 | 0 | 0 | 0 | 0 | 0 | 65 | 38 | 160 | 28 | 16 | 0 | 0 | 0 | 0 | 0 |
| SRR7226514 | 275 | 4005 | 17181 | 3940 | 6781 | 5458 | 73860 | 73925 | 1949 | 13635 | 45858 | 13 | 16088 | 11346 | 443 | 12252 | 22238 | 0 | 0 | 0 | 94 | 19 | 0 | 0 | 0 | 0 | 0 | 37 | 40 | 119 | 45 | 59 | 0 | 0 | 0 | 0 | 0 |
| SRR7226515 | 75 | 3436 | 34298 | 1480 | 7435 | 8384 | 53160 | 44332 | 7678 | 18498 | 49154 | 86082 | 12146 | 19792 | 446 | 8240 | 25620 | 0 | 0 | 0 | 13 | 13 | 0 | 0 | 0 | 0 | 0 | 0 | 0 | 20 | 11 | 0 | 0 | 0 | 0 | 0 | 0 |
| SRR7226516 | 212 | 4293 | 31488 | 2267 | 7488 | 4576 | 82051 | 76205 | 10819 | 14927 | 34787 | 47592 | 14371 | 17430 | 1817 | 3707 | 20537 | 0 | 0 | 13 | 105 | 70 | 0 | 0 | 0 | 0 | 0 | 27 | 47 | 147 | 63 | 33 | 0 | 0 | 0 | 0 | 0 |
| SRR7226517 | 104 | 2652 | 41752 | 3293 | 7394 | 5037 | 60420 | 77718 | 18813 | 3731 | 49925 | 41401 | 15977 | 19328 | 392 | 6060 | 21276 | 0 | 78 | 111 | 6136 | 266 | 35 | 0 | 0 | 0 | 0 | 253 | 284 | 4465 | 328 | 238 | 0 | 0 | 0 | 47 | 0 |
| SRR7226518 | 299 | 1985 | 36387 | 4362 | 6529 | 4617 | 53351 | 67015 | 18218 | 1745 | 55857 | 44391 | 16352 | 25655 | 434 | 5859 | 24181 | 0 | 54 | 62 | 1197 | 206 | 22 | 0 | 0 | 0 | 0 | 139 | 127 | 650 | 161 | 83 | 0 | 0 | 0 | 18 | 0 |
| SRR7226519 | 382 | 2322 | 28001 | 4753 | 7096 | 5041 | 62501 | 74736 | 18895 | 2319 | 45296 | 34059 | 13014 | 33252 | 286 | 5700 | 22433 | 0 | 35 | 39 | 687 | 112 | 11 | 0 | 0 | 0 | 0 | 126 | 102 | 333 | 81 | 41 | 0 | 0 | 0 | 0 | 0 |
| SRR7226520 | 583 | 3310 | 34665 | 2097 | 559 | 428 | 3358 | 3192 | 14666 | 8935 | 21331 | 19663 | 8111 | 8880 | 66 | 464 | 16400 | 0 | 0 | 17 | 144 | 23 | 0 | 0 | 0 | 0 | 0 | 82 | 133 | 244 | 98 | 107 | 0 | 0 | 0 | 0 | 0 |
| SRR7226521 | 200 | 3609 | 37897 | 1471 | 779 | 592 | 4998 | 6325 | 19288 | 8331 | 31260 | 24559 | 10975 | 11318 | 259 | 1118 | 25294 | 0 | 12 | 15 | 90 | 64 | 11 | 0 | 0 | 0 | 0 | 94 | 125 | 280 | 174 | 176 | 0 | 0 | 0 | 18 | 0 |
| SRR7226522 | 363 | 2909 | 26362 | 3076 | 724 | 475 | 5071 | 4130 | 20277 | 15252 | 24442 | 19940 | 7229 | 12257 | 84 | 536 | 14057 | 0 | 0 | 0 | 48 | 13 | 0 | 0 | 0 | 0 | 0 | 85 | 88 | 205 | 59 | 71 | 0 | 0 | 0 | 0 | 0 |
| SRR7226523 | 276 | 4382 | 15446 | 3505 | 8752 | 8590 | 117410 | 95255 | 3101 | 10387 | 47152 | 40329 | 19444 | 15816 | 455 | 7795 | 32661 | 0 | 22 | 29 | 321 | 56 | 14 | 0 | 0 | 0 | 0 | 88 | 127 | 290 | 134 | 173 | 0 | 0 | 0 | 24 | 0 |
| SRR7226524 | 483 | 1284 | 1067 | 4422 | 17239 | 12942 | 179495 | 88912 | 4378 | 937 | 44926 | 26533 | 14854 | 18765 | 521 | 10294 | 19125 | 0 | 0 | 21 | 275 | 20 | 0 | 0 | 0 | 0 | 0 | 46 | 55 | 168 | 12 | 0 | 0 | 0 | 0 | 0 | 0 |
| SRR7226525 | 195 | 4297 | 25666 | 3137 | 11356 | 10350 | 137906 | 105942 | 3304 | 13856 | 46976 | 29753 | 16916 | 15888 | 349 | 8606 | 30514 | 0 | 0 | 14 | 284 | 48 | 0 | 0 | 0 | 0 | 0 | 64 | 88 | 418 | 94 | 108 | 0 | 0 | 0 | 0 | 0 |
| SRR7226526 | 216 | 4519 | 25232 | 2643 | 17251 | 16008 | 196691 | 76712 | 3759 | 5788 | 44613 | 34330 | 16239 | 17139 | 946 | 9313 | 27112 | 0 | 33 | 45 | 426 | 95 | 25 | 0 | 0 | 0 | 0 | 135 | 172 | 464 | 197 | 234 | 0 | 0 | 0 | 25 | 0 |
| SRR7233553 | 57 | 2785 | 30956 | 44 | 0 | 0 | 47 | 27 | 1519 | 2067 | 961 | 987 | 1051 | 3885 | 187 | 0 | 1091 | 0 | 0 | 0 | 0 | 0 | 0 | 0 | 0 | 0 | 0 | 0 | 0 | 0 | 0 | 0 | 0 | 0 | 0 | 0 | 0 |
| SRR7233554 | 63 | 3106 | 24637 | 61 | 10 | 0 | 92 | 60 | 1415 | 1493 | 1113 | 1117 | 1178 | 4705 | 206 | 0 | 2049 | 0 | 0 | 0 | 0 | 0 | 0 | 0 | 0 | 0 | 0 | 0 | 0 | 0 | 0 | 0 | 0 | 0 | 0 | 0 | 0 |
| SRR7233556 | 51 | 2255 | 20620 | 63 | 0 | 0 | 51 | 37 | 848 | 1431 | 1086 | 1146 | 1036 | 3975 | 221 | 0 | 1341 | 0 | 0 | 0 | 0 | 0 | 0 | 0 | 0 | 0 | 0 | 0 | 0 | 0 | 0 | 0 | 0 | 0 | 0 | 0 | 0 |
| SRR7233557 | 39 | 1144 | 23607 | 28 | 0 | 0 | 13 | 0 | 402 | 985 | 682 | 727 | 630 | 2090 | 105 | 0 | 704 | 0 | 0 | 0 | 0 | 0 | 0 | 0 | 0 | 0 | 0 | 0 | 0 | 0 | 0 | 0 | 0 | 0 | 0 | 0 | 0 |
| SRR7233558 | 91 | 3909 | 22385 | 67 | 0 | 0 | 33 | 12 | 525 | 1929 | 539 | 678 | 495 | 2254 | 80 | 0 | 1336 | 0 | 0 | 0 | 0 | 0 | 0 | 0 | 0 | 0 | 0 | 0 | 0 | 0 | 0 | 0 | 0 | 0 | 0 | 0 | 0 |
| SRR7233559 | 24 | 1047 | 30850 | 21 | 0 | 0 | 40 | 15 | 934 | 1616 | 1051 | 1092 | 1033 | 3894 | 144 | 0 | 1094 | 0 | 0 | 0 | 0 | 0 | 0 | 0 | 0 | 0 | 0 | 0 | 0 | 0 | 0 | 0 | 0 | 0 | 0 | 0 | 0 |
| SRR7233560 | 24 | 1002 | 24991 | 17 | 0 | 0 | 54 | 28 | 993 | 1233 | 1069 | 1008 | 887 | 3796 | 227 | 0 | 1325 | 0 | 0 | 0 | 0 | 0 | 0 | 0 | 0 | 0 | 0 | 0 | 0 | 0 | 0 | 0 | 0 | 0 | 0 | 0 | 0 |
| SRR7233561 | 129 | 3268 | 30870 | 119 | 0 | 0 | 40 | 19 | 878 | 3174 | 766 | 876 | 822 | 2772 | 170 | 25 | 1428 | 0 | 0 | 0 | 0 | 0 | 0 | 0 | 0 | 0 | 0 | 0 | 0 | 0 | 0 | 0 | 0 | 0 | 0 | 0 | 0 |
| SRR7233562 | 54 | 1755 | 25835 | 49 | 0 | 0 | 24 | 15 | 908 | 1220 | 859 | 748 | 735 | 2817 | 160 | 23 | 910 | 0 | 0 | 0 | 0 | 0 | 0 | 0 | 0 | 0 | 0 | 0 | 0 | 0 | 0 | 0 | 0 | 0 | 0 | 0 | 0 |
| SRR7233563 | 182 | 5693 | 23887 | 123 | 12 | 0 | 130 | 66 | 2011 | 2177 | 1005 | 659 | 861 | 3148 | 372 | 0 | 1472 | 0 | 0 | 0 | 0 | 0 | 0 | 0 | 0 | 0 | 0 | 0 | 0 | 0 | 0 | 0 | 0 | 0 | 0 | 0 | 0 |
| SRR7233564 | 122 | 5007 | 27645 | 123 | 12 | 0 | 105 | 48 | 1406 | 2877 | 972 | 625 | 838 | 2904 | 358 | 0 | 1570 | 0 | 0 | 0 | 0 | 0 | 0 | 0 | 0 | 0 | 0 | 0 | 0 | 0 | 0 | 0 | 0 | 0 | 0 | 0 | 0 |
| SRR7233565 | 177 | 6043 | 21881 | 109 | 0 | 0 | 14 | 12 | 204 | 2622 | 660 | 543 | 583 | 1560 | 173 | 0 | 752 | 0 | 0 | 0 | 0 | 0 | 0 | 0 | 0 | 0 | 0 | 0 | 0 | 0 | 0 | 0 | 0 | 0 | 0 | 0 | 0 |
| SRR7233566 | 210 | 6315 | 22888 | 158 | 0 | 0 | 22 | 13 | 287 | 1813 | 373 | 372 | 372 | 1559 | 115 | 0 | 970 | 0 | 0 | 0 | 0 | 0 | 0 | 0 | 0 | 0 | 0 | 0 | 0 | 0 | 0 | 0 | 0 | 0 | 0 | 0 | 0 |
| SRR7233568 | 59 | 2065 | 22437 | 67 | 11 | 0 | 72 | 48 | 1588 | 1771 | 1171 | 918 | 961 | 2859 | 347 | 11 | 1875 | 0 | 0 | 0 | 0 | 0 | 0 | 0 | 0 | 0 | 0 | 0 | 0 | 0 | 0 | 0 | 0 | 0 | 0 | 0 | 0 |
| SRR7233570 | 49 | 1883 | 24620 | 35 | 15 | 0 | 109 | 50 | 1433 | 1556 | 1244 | 745 | 1053 | 2931 | 456 | 45 | 1340 | 0 | 0 | 0 | 0 | 0 | 0 | 0 | 0 | 0 | 0 | 0 | 0 | 0 | 0 | 0 | 0 | 0 | 0 | 0 | 0 |
| SRR7233571 | 95 | 4004 | 35026 | 94 | 0 | 0 | 31 | 13 | 918 | 2634 | 874 | 737 | 847 | 2060 | 164 | 10 | 1187 | 0 | 0 | 0 | 0 | 0 | 0 | 0 | 0 | 0 | 0 | 0 | 0 | 0 | 0 | 0 | 0 | 0 | 0 | 0 | 0 |
| SRR7233572 | 38 | 1361 | 23631 | 49 | 0 | 0 | 39 | 21 | 764 | 1596 | 666 | 530 | 661 | 1844 | 155 | 0 | 1343 | 0 | 0 | 0 | 0 | 0 | 0 | 0 | 0 | 0 | 0 | 0 | 0 | 0 | 0 | 0 | 0 | 0 | 0 | 0 | 0 |
| SRR7262826 | 25 | 676 | 25931 | 435 | 1771 | 2088 | 6404 | 5645 | 66248 | 1551 | 10675 | 32 | 9949 | 21850 | 220 | 568 | 48204 | 0 | 0 | 22 | 141 | 1084 | 478 | 84 | 13 | 0 | 0 | 13 | 83 | 307 | 130 | 16 | 0 | 0 | 0 | 14 | 0 |
| SRR7262827 | 61 | 1220 | 30316 | 1182 | 2163 | 3018 | 9953 | 6543 | 46174 | 1506 | 8847 | 17672 | 12035 | 27842 | 341 | 765 | 49458 | 0 | 18 | 49 | 232 | 1230 | 438 | 75 | 0 | 11 | 0 | 83 | 265 | 745 | 201 | 19 | 0 | 0 | 0 | 23 | 0 |
| SRR7262828 | 47 | 1442 | 17110 | 1194 | 3443 | 3870 | 18001 | 8287 | 44956 | 1539 | 8901 | 14469 | 11907 | 29637 | 514 | 969 | 41984 | 0 | 23 | 67 | 354 | 1255 | 703 | 97 | 16 | 10 | 0 | 126 | 417 | 1471 | 225 | 36 | 0 | 0 | 0 | 33 | 0 |
| SRR7262829 | 190 | 1271 | 20544 | 2650 | 2294 | 2091 | 7813 | 5458 | 93253 | 906 | 10522 | 26 | 9018 | 21697 | 232 | 413 | 41765 | 0 | 0 | 38 | 239 | 1614 | 565 | 106 | 18 | 0 | 0 | 19 | 105 | 507 | 113 | 19 | 0 | 0 | 0 | 22 | 0 |
| SRR7262830 | 22 | 1014 | 25393 | 456 | 2096 | 2837 | 9391 | 5476 | 45245 | 1856 | 8155 | 34 | 11487 | 27476 | 387 | 887 | 50495 | 0 | 0 | 38 | 207 | 1178 | 482 | 82 | 13 | 14 | 0 | 50 | 198 | 671 | 186 | 22 | 0 | 0 | 0 | 29 | 0 |
| SRR7288164 | 31 | 276 | 15 | 0 | 0 | 13 | 55 | 1325 | 1068 | 0 | 193 | 116 | 113 | 388 | 0 | 97 | 1883 | 0 | 0 | 0 | 0 | 0 | 0 | 0 | 0 | 0 | 0 | 0 | 0 | 0 | 0 | 0 | 0 | 0 | 0 | 0 | 0 |
| SRR7288166 | 46 | 275 | 15 | 31 | 0 | 23 | 62 | 1320 | 1108 | 0 | 222 | 89 | 126 | 363 | 0 | 90 | 1801 | 0 | 0 | 0 | 0 | 0 | 0 | 0 | 0 | 0 | 0 | 0 | 0 | 0 | 0 | 0 | 0 | 0 | 0 | 0 | 0 |
| SRR7288167 | 16 | 331 | 16 | 31 | 0 | 20 | 50 | 1131 | 1076 | 0 | 215 | 115 | 113 | 391 | 0 | 74 | 1593 | 0 | 0 | 0 | 0 | 0 | 0 | 0 | 0 | 0 | 0 | 0 | 0 | 0 | 0 | 0 | 0 | 0 | 0 | 0 | 0 |
| SRR7288236 | 16 | 1179 | 0 | 0 | 0 | 0 | 26 | 277 | 466 | 0 | 231 | 57 | 20 | 294 | 0 | 0 | 506 | 0 | 0 | 0 | 0 | 0 | 0 | 0 | 0 | 0 | 0 | 0 | 0 | 0 | 0 | 0 | 0 | 0 | 0 | 0 | 0 |
| SRR7288237 | 16 | 1605 | 0 | 0 | 0 | 0 | 26 | 213 | 515 | 0 | 215 | 31 | 44 | 241 | 0 | 0 | 478 | 0 | 0 | 0 | 0 | 0 | 0 | 0 | 0 | 0 | 0 | 0 | 0 | 0 | 0 | 0 | 0 | 0 | 0 | 0 | 0 |
| SRR7288238 | 15 | 1853 | 0 | 0 | 0 | 0 | 30 | 184 | 491 | 0 | 235 | 47 | 26 | 248 | 0 | 0 | 547 | 0 | 0 | 0 | 0 | 0 | 0 | 0 | 0 | 0 | 0 | 0 | 0 | 0 | 0 | 0 | 0 | 0 | 0 | 0 | 0 |
| SRR7288239 | 0 | 1900 | 15 | 0 | 0 | 0 | 23 | 267 | 452 | 0 | 215 | 42 | 38 | 250 | 0 | 0 | 507 | 0 | 0 | 0 | 0 | 0 | 0 | 0 | 0 | 0 | 0 | 0 | 0 | 0 | 0 | 0 | 0 | 0 | 0 | 0 | 0 |
| SRR7288244 | 0 | 2558 | 853 | 0 | 0 | 55 | 0 | 992 | 1322 | 0 | 606 | 331 | 55 | 276 | 0 | 276 | 1818 | 0 | 0 | 0 | 12 | 0 | 0 | 0 | 0 | 0 | 0 | 0 | 0 | 12 | 0 | 0 | 0 | 12 | 0 | 0 | 0 |
| SRR7288264 | 0 | 270 | 472 | 22 | 0 | 59 | 273 | 3087 | 1962 | 0 | 1003 | 309 | 496 | 1508 | 0 | 151 | 10198 | 0 | 0 | 0 | 0 | 0 | 0 | 0 | 0 | 0 | 0 | 0 | 0 | 0 | 0 | 0 | 0 | 0 | 0 | 0 | 0 |
| SRR7288265 | 23 | 274 | 457 | 137 | 0 | 54 | 269 | 2968 | 1948 | 0 | 1069 | 290 | 562 | 1492 | 0 | 175 | 8881 | 0 | 0 | 0 | 0 | 0 | 0 | 0 | 0 | 0 | 0 | 0 | 0 | 0 | 0 | 0 | 0 | 0 | 0 | 0 | 0 |
| SRR7288266 | 0 | 309 | 508 | 66 | 0 | 56 | 263 | 3190 | 2014 | 0 | 1069 | 263 | 552 | 1554 | 0 | 151 | 10279 | 0 | 0 | 0 | 0 | 0 | 0 | 0 | 0 | 0 | 0 | 0 | 0 | 0 | 0 | 0 | 0 | 0 | 0 | 0 | 0 |
| SRR7288267 | 0 | 182 | 410 | 0 | 0 | 33 | 261 | 3332 | 1905 | 0 | 1062 | 249 | 495 | 1560 | 0 | 180 | 9193 | 0 | 0 | 0 | 0 | 0 | 0 | 0 | 0 | 0 | 0 | 0 | 0 | 0 | 0 | 0 | 0 | 0 | 0 | 0 | 0 |
| SRR7288277 | 0 | 1400 | 910 | 140 | 11 | 115 | 115 | 2881 | 1228 | 0 | 666 | 356 | 34 | 195 | 0 | 264 | 1354 | 0 | 0 | 0 | 0 | 12 | 0 | 0 | 0 | 0 | 0 | 0 | 0 | 0 | 0 | 0 | 0 | 0 | 0 | 0 | 0 |
| SRR7288300 | 0 | 1234 | 76 | 25 | 0 | 48 | 270 | 2841 | 3123 | 0 | 1146 | 333 | 422 | 1352 | 0 | 92 | 10545 | 0 | 0 | 0 | 33 | 22 | 0 | 0 | 0 | 0 | 0 | 0 | 0 | 11 | 25 | 0 | 0 | 0 | 0 | 0 | 0 |
| SRR7288301 | 26 | 1108 | 26 | 52 | 0 | 49 | 286 | 2875 | 3060 | 0 | 954 | 325 | 519 | 1438 | 0 | 94 | 9366 | 0 | 0 | 0 | 30 | 18 | 0 | 0 | 0 | 0 | 0 | 0 | 0 | 0 | 27 | 0 | 0 | 0 | 0 | 0 | 0 |
| SRR7288302 | 0 | 928 | 25 | 0 | 0 | 51 | 291 | 2879 | 3196 | 0 | 909 | 374 | 478 | 1289 | 0 | 79 | 10214 | 0 | 0 | 0 | 31 | 29 | 0 | 0 | 0 | 0 | 0 | 0 | 0 | 13 | 31 | 0 | 0 | 0 | 0 | 0 | 0 |
| SRR7288303 | 26 | 907 | 52 | 26 | 0 | 33 | 287 | 2909 | 3046 | 0 | 829 | 362 | 424 | 1329 | 0 | 75 | 9348 | 0 | 0 | 0 | 34 | 22 | 0 | 0 | 0 | 0 | 0 | 0 | 0 | 0 | 21 | 10 | 0 | 0 | 0 | 0 | 0 |
| SRR7288308 | 322 | 752 | 72 | 0 | 0 | 108 | 405 | 5872 | 6441 | 0 | 2375 | 904 | 288 | 1837 | 0 | 389 | 7336 | 0 | 0 | 0 | 0 | 0 | 0 | 0 | 0 | 0 | 0 | 0 | 0 | 0 | 0 | 0 | 0 | 0 | 0 | 0 | 0 |
| SRR7288309 | 187 | 859 | 299 | 37 | 23 | 131 | 446 | 6038 | 6278 | 0 | 2274 | 925 | 308 | 1782 | 0 | 473 | 6321 | 0 | 0 | 0 | 0 | 0 | 0 | 0 | 0 | 0 | 0 | 0 | 0 | 0 | 0 | 0 | 0 | 0 | 0 | 0 | 0 |
| SRR7288310 | 106 | 568 | 106 | 0 | 19 | 146 | 414 | 5860 | 6137 | 0 | 2440 | 865 | 286 | 1777 | 0 | 392 | 7400 | 0 | 0 | 0 | 0 | 0 | 0 | 0 | 0 | 0 | 0 | 0 | 0 | 0 | 0 | 0 | 0 | 0 | 0 | 0 | 0 |
| SRR7288311 | 111 | 297 | 111 | 37 | 16 | 126 | 368 | 6180 | 6352 | 0 | 2381 | 992 | 349 | 1577 | 0 | 407 | 6510 | 0 | 0 | 0 | 0 | 0 | 0 | 0 | 0 | 0 | 0 | 0 | 0 | 0 | 0 | 0 | 0 | 0 | 0 | 0 | 0 |
| SRR7288320 | 74 | 271 | 172 | 0 | 0 | 42 | 519 | 3759 | 5628 | 0 | 1186 | 517 | 472 | 3340 | 0 | 266 | 8804 | 0 | 0 | 0 | 0 | 0 | 0 | 0 | 0 | 0 | 0 | 0 | 0 | 0 | 0 | 0 | 0 | 0 | 0 | 0 | 0 |
| SRR7288321 | 130 | 391 | 52 | 0 | 0 | 75 | 633 | 3835 | 5646 | 0 | 1275 | 395 | 417 | 3406 | 0 | 296 | 7985 | 0 | 0 | 0 | 0 | 0 | 0 | 0 | 0 | 0 | 0 | 0 | 0 | 0 | 0 | 0 | 0 | 0 | 0 | 0 | 0 |
| SRR7288322 | 24 | 462 | 243 | 0 | 0 | 49 | 573 | 3805 | 5819 | 0 | 1332 | 427 | 539 | 3297 | 0 | 359 | 9152 | 0 | 0 | 0 | 12 | 0 | 0 | 0 | 0 | 0 | 0 | 0 | 0 | 0 | 0 | 0 | 0 | 0 | 0 | 0 | 0 |
| SRR7288323 | 77 | 154 | 179 | 0 | 0 | 68 | 616 | 3877 | 5753 | 0 | 1309 | 415 | 570 | 3524 | 0 | 288 | 7934 | 0 | 0 | 0 | 0 | 0 | 0 | 0 | 0 | 0 | 0 | 0 | 0 | 0 | 0 | 0 | 0 | 0 | 0 | 0 | 0 |
| SRR7288376 | 237 | 355 | 118 | 0 | 0 | 22 | 78 | 2414 | 2256 | 59 | 336 | 151 | 159 | 587 | 0 | 96 | 1757 | 0 | 0 | 0 | 0 | 0 | 0 | 0 | 0 | 0 | 0 | 0 | 0 | 0 | 0 | 0 | 0 | 0 | 0 | 0 | 0 |
| SRR7288377 | 61 | 550 | 61 | 0 | 0 | 19 | 66 | 2564 | 2298 | 61 | 289 | 77 | 116 | 528 | 0 | 154 | 1361 | 0 | 0 | 0 | 0 | 0 | 0 | 0 | 0 | 0 | 0 | 0 | 0 | 0 | 0 | 0 | 0 | 0 | 0 | 0 | 0 |
| SRR7288378 | 59 | 411 | 0 | 0 | 0 | 22 | 102 | 2428 | 2260 | 0 | 318 | 110 | 150 | 573 | 0 | 172 | 1592 | 0 | 0 | 0 | 0 | 0 | 0 | 0 | 0 | 0 | 0 | 0 | 0 | 0 | 0 | 0 | 0 | 0 | 0 | 0 | 0 |
| SRR7288379 | 123 | 368 | 123 | 61 | 0 | 38 | 80 | 2622 | 2220 | 0 | 395 | 110 | 133 | 516 | 0 | 144 | 1484 | 0 | 0 | 0 | 0 | 0 | 0 | 0 | 0 | 0 | 0 | 0 | 0 | 0 | 0 | 0 | 0 | 0 | 0 | 0 | 0 |
| SRR7288380 | 22 | 1039 | 22 | 22 | 0 | 46 | 129 | 1062 | 1464 | 0 | 1524 | 293 | 241 | 640 | 0 | 57 | 2877 | 0 | 0 | 0 | 0 | 0 | 0 | 0 | 0 | 0 | 0 | 0 | 0 | 0 | 0 | 0 | 0 | 0 | 0 | 0 | 0 |
| SRR7288381 | 23 | 1175 | 0 | 0 | 0 | 32 | 124 | 1039 | 1732 | 0 | 1403 | 315 | 283 | 589 | 0 | 72 | 2445 | 0 | 0 | 0 | 0 | 0 | 0 | 0 | 0 | 0 | 0 | 0 | 0 | 0 | 0 | 0 | 0 | 0 | 0 | 0 | 0 |
| SRR7288382 | 43 | 1233 | 0 | 0 | 0 | 34 | 118 | 1066 | 1564 | 0 | 1429 | 267 | 200 | 740 | 0 | 96 | 2767 | 0 | 0 | 0 | 0 | 0 | 0 | 0 | 0 | 0 | 0 | 0 | 0 | 0 | 0 | 0 | 0 | 0 | 0 | 0 | 0 |

### Supp Table S8.1.Columns_B_to_AL

| | | | | | | | | | | | | | | | | | | | | | | | | | | | | | | | | | | | | | |
|---|---|---|---|---|---|---|---|---|---|---|---|---|---|---|---|---|---|---|---|---|---|---|---|---|---|---|---|---|---|---|---|---|---|---|---|---|---|
| SRR7288383 | 43 | 1010 | 21 | 0 | 0 | 25 | 153 | 1058 | 1628 | 0 | 1303 | 333 | 236 | 661 | 0 | 94 | 2456 | 0 | 0 | 0 | 0 | 0 | 0 | 0 | 0 | 0 | 0 | 0 | 0 | 0 | 0 | 0 | 0 | 0 | 0 | 0 | 0 |
| SRR7288384 | 120 | 499 | 0 | 17 | 51 | 590 | 1089 | 8289 | 4001 | 0 | 9109 | 2022 | 714 | 2824 | 0 | 1138 | 9954 | 0 | 0 | 0 | 0 | 0 | 0 | 0 | 0 | 0 | 0 | 0 | 0 | 0 | 0 | 0 | 0 | 0 | 0 | 0 | 0 |
| SRR7288385 | 52 | 595 | 17 | 0 | 37 | 589 | 1170 | 8597 | 4261 | 0 | 9349 | 2101 | 721 | 2758 | 0 | 1301 | 8475 | 0 | 0 | 0 | 0 | 0 | 0 | 0 | 0 | 0 | 0 | 0 | 0 | 0 | 0 | 0 | 0 | 0 | 0 | 0 | 0 |
| SRR7288386 | 17 | 470 | 17 | 34 | 33 | 622 | 1163 | 8194 | 3913 | 0 | 9280 | 2182 | 719 | 2809 | 0 | 1193 | 9473 | 0 | 0 | 0 | 0 | 0 | 0 | 0 | 0 | 0 | 0 | 0 | 0 | 0 | 0 | 0 | 0 | 0 | 0 | 0 | 0 |
| SRR7288387 | 134 | 401 | 0 | 33 | 29 | 451 | 1208 | 8719 | 4017 | 0 | 9398 | 2139 | 635 | 2707 | 0 | 1279 | 8488 | 0 | 0 | 0 | 0 | 0 | 0 | 0 | 0 | 0 | 0 | 0 | 0 | 0 | 0 | 0 | 0 | 0 | 0 | 0 | 0 |
| SRR7288400 | 0 | 971 | 971 | 971 | 0 | 92 | 69 | 3298 | 8396 | 0 | 231 | 185 | 46 | 323 | 0 | 277 | 2076 | 0 | 0 | 0 | 0 | 0 | 0 | 0 | 0 | 0 | 0 | 0 | 0 | 0 | 0 | 0 | 0 | 0 | 0 | 0 | 0 |
| SRR7288402 | 0 | 1951 | 0 | 0 | 0 | 22 | 22 | 3166 | 8028 | 0 | 602 | 178 | 156 | 468 | 0 | 201 | 1717 | 0 | 0 | 0 | 0 | 0 | 0 | 0 | 0 | 0 | 0 | 0 | 0 | 0 | 0 | 0 | 0 | 0 | 0 | 0 | 0 |
| SRR7288472 | 0 | 398 | 0 | 0 | 0 | 25 | 82 | 943 | 2131 | 0 | 361 | 148 | 131 | 443 | 0 | 41 | 2680 | 0 | 0 | 0 | 0 | 0 | 0 | 0 | 0 | 0 | 0 | 0 | 0 | 0 | 0 | 0 | 0 | 0 | 0 | 0 | 0 |
| SRR7288474 | 0 | 388 | 0 | 0 | 0 | 24 | 32 | 988 | 1945 | 0 | 397 | 130 | 178 | 494 | 0 | 122 | 2771 | 0 | 0 | 0 | 0 | 0 | 0 | 0 | 0 | 0 | 0 | 0 | 0 | 0 | 0 | 0 | 0 | 0 | 0 | 0 | 0 |
| SRR7288475 | 0 | 515 | 0 | 0 | 0 | 51 | 85 | 1012 | 1760 | 0 | 442 | 179 | 145 | 400 | 0 | 60 | 2244 | 0 | 0 | 0 | 0 | 0 | 0 | 0 | 0 | 0 | 0 | 0 | 0 | 0 | 0 | 0 | 0 | 0 | 0 | 0 | 0 |
| SRR7288512 | 25 | 39090 | 50 | 0 | 0 | 16 | 162 | 890 | 1088 | 0 | 1487 | 246 | 242 | 550 | 0 | 63 | 3614 | 0 | 0 | 0 | 0 | 0 | 0 | 0 | 0 | 0 | 0 | 0 | 0 | 0 | 0 | 0 | 0 | 0 | 0 | 0 | 0 |
| SRR7288513 | 25 | 37178 | 0 | 0 | 0 | 23 | 172 | 920 | 1131 | 0 | 1443 | 251 | 272 | 522 | 0 | 79 | 3098 | 0 | 0 | 0 | 0 | 0 | 0 | 0 | 0 | 0 | 0 | 0 | 0 | 0 | 0 | 0 | 0 | 0 | 0 | 0 | 0 |
| SRR7288514 | 0 | 36977 | 24 | 0 | 0 | 18 | 157 | 806 | 1116 | 0 | 1420 | 252 | 292 | 555 | 0 | 69 | 3436 | 0 | 0 | 0 | 0 | 0 | 0 | 0 | 0 | 0 | 0 | 0 | 0 | 0 | 0 | 0 | 0 | 0 | 0 | 0 | 0 |
| SRR7288515 | 48 | 35976 | 0 | 24 | 0 | 14 | 171 | 902 | 1136 | 0 | 1497 | 256 | 288 | 619 | 0 | 69 | 3249 | 0 | 0 | 0 | 0 | 0 | 0 | 0 | 0 | 0 | 0 | 0 | 0 | 0 | 0 | 0 | 0 | 0 | 0 | 0 | 0 |
| SRR7293222 | 29 | 40876 | 8510 | 447 | 115 | 761 | 757 | 1196 | 240 | 25 | 14173 | 6948 | 885 | 8904 | 0 | 25 | 27071 | 0 | 0 | 0 | 433 | 277 | 84 | 0 | 0 | 0 | 0 | 10 | 99 | 136 | 178 | 55 | 0 | 0 | 0 | 0 | 0 |
| SRR7293223 | 31 | 29720 | 6762 | 448 | 103 | 744 | 704 | 1013 | 157 | 27 | 13359 | 6703 | 798 | 7840 | 0 | 25 | 25720 | 0 | 0 | 0 | 277 | 146 | 62 | 0 | 0 | 0 | 0 | 0 | 72 | 90 | 128 | 39 | 0 | 0 | 0 | 0 | 0 |
| SRR7293224 | 54 | 15911 | 1615 | 419 | 31 | 205 | 250 | 339 | 879 | 0 | 6862 | 5472 | 409 | 2397 | 0 | 0 | 25557 | 0 | 0 | 0 | 1027 | 186 | 95 | 0 | 0 | 0 | 0 | 15 | 164 | 525 | 107 | 78 | 0 | 17 | 0 | 0 | 0 |
| SRR7293225 | 76 | 19678 | 42549 | 396 | 56 | 371 | 454 | 984 | 715 | 15 | 8625 | 4935 | 491 | 3825 | 0 | 14 | 31194 | 0 | 0 | 0 | 75 | 117 | 13 | 0 | 0 | 0 | 0 | 0 | 104 | 53 | 156 | 24 | 0 | 0 | 0 | 0 | 0 |
| SRR7293226 | 60 | 13149 | 31889 | 555 | 85 | 556 | 626 | 1130 | 710 | 15 | 8849 | 5153 | 636 | 5319 | 0 | 20 | 29433 | 0 | 0 | 0 | 304 | 390 | 85 | 0 | 0 | 0 | 0 | 13 | 152 | 159 | 332 | 82 | 0 | 13 | 0 | 0 | 0 |
| SRR7293227 | 91 | 12351 | 18662 | 367 | 72 | 495 | 586 | 1041 | 387 | 12 | 8355 | 5094 | 636 | 4736 | 0 | 18 | 27642 | 0 | 0 | 0 | 244 | 341 | 64 | 0 | 0 | 0 | 0 | 13 | 159 | 126 | 349 | 118 | 0 | 23 | 0 | 0 | 0 |
| SRR7293228 | 0 | 6627 | 2243 | 136 | 75 | 503 | 504 | 591 | 223 | 0 | 15451 | 10207 | 838 | 6724 | 0 | 10 | 31187 | 0 | 0 | 0 | 1356 | 244 | 101 | 0 | 10 | 0 | 0 | 0 | 42 | 584 | 130 | 59 | 0 | 16 | 0 | 0 | 0 |
| SRR7293229 | 0 | 20362 | 29950 | 75 | 78 | 545 | 617 | 1273 | 291 | 13 | 7447 | 3338 | 677 | 6121 | 0 | 17 | 32484 | 0 | 0 | 0 | 43 | 48 | 11 | 0 | 0 | 0 | 0 | 0 | 35 | 26 | 61 | 16 | 0 | 0 | 0 | 0 | 0 |
| SRR7293230 | 81 | 9215 | 50422 | 729 | 68 | 786 | 691 | 2053 | 70 | 45 | 8646 | 4730 | 914 | 9445 | 0 | 27 | 59261 | 0 | 0 | 0 | 255 | 220 | 69 | 0 | 12 | 0 | 0 | 0 | 93 | 97 | 173 | 122 | 0 | 23 | 0 | 0 | 0 |
| SRR7293231 | 21 | 9065 | 49457 | 590 | 64 | 789 | 634 | 1869 | 42 | 57 | 15961 | 6175 | 900 | 8824 | 0 | 22 | 44420 | 0 | 0 | 0 | 199 | 164 | 53 | 0 | 0 | 0 | 0 | 0 | 76 | 90 | 132 | 137 | 0 | 24 | 0 | 0 | 0 |
| SRR7293232 | 0 | 5679 | 2373 | 491 | 49 | 658 | 506 | 1261 | 41 | 0 | 18115 | 8165 | 905 | 10282 | 0 | 13 | 47631 | 0 | 0 | 0 | 1463 | 108 | 34 | 0 | 0 | 0 | 0 | 0 | 81 | 677 | 60 | 30 | 0 | 0 | 0 | 0 | 0 |
| SRR7293233 | 121 | 14710 | 63851 | 1487 | 53 | 686 | 596 | 2135 | 62 | 34 | 17234 | 5908 | 980 | 7235 | 0 | 39 | 45128 | 0 | 0 | 0 | 198 | 113 | 23 | 0 | 0 | 0 | 0 | 13 | 126 | 109 | 141 | 32 | 0 | 0 | 0 | 0 | 0 |
| SRR7343626 | 62 | 1506 | 1258 | 227 | 0 | 0 | 0 | 0 | 171 | 41 | 206 | 301 | 65 | 315 | 0 | 19 | 1416 | 0 | 0 | 0 | 0 | 0 | 0 | 0 | 0 | 0 | 0 | 0 | 0 | 0 | 0 | 0 | 0 | 0 | 0 | 0 | 0 |
| SRR7343627 | 149 | 5880 | 3936 | 149 | 0 | 0 | 0 | 0 | 99 | 0 | 61 | 180 | 20 | 94 | 0 | 0 | 1181 | 0 | 0 | 0 | 0 | 0 | 0 | 0 | 0 | 0 | 0 | 0 | 0 | 0 | 0 | 0 | 0 | 0 | 0 | 0 | 0 |
| SRR7343629 | 100 | 4826 | 398 | 149 | 0 | 0 | 0 | 0 | 96 | 0 | 48 | 114 | 15 | 82 | 0 | 32 | 1362 | 0 | 0 | 0 | 0 | 0 | 0 | 0 | 0 | 0 | 0 | 0 | 0 | 0 | 0 | 0 | 0 | 0 | 0 | 0 | 0 |
| SRR7343630 | 0 | 754 | 935 | 73 | 0 | 13 | 0 | 33 | 551 | 45 | 276 | 342 | 86 | 681 | 12 | 0 | 3039 | 0 | 0 | 0 | 0 | 0 | 0 | 0 | 0 | 0 | 0 | 0 | 0 | 0 | 0 | 0 | 0 | 0 | 0 | 0 | 0 |
| SRR7343631 | 21 | 11012 | 406 | 64 | 0 | 0 | 0 | 0 | 98 | 21 | 72 | 118 | 17 | 144 | 0 | 0 | 1134 | 0 | 0 | 0 | 0 | 0 | 0 | 0 | 0 | 0 | 0 | 0 | 0 | 0 | 0 | 0 | 0 | 0 | 0 | 0 | 0 |
| SRR7343632 | 83 | 4391 | 1077 | 166 | 0 | 0 | 0 | 13 | 130 | 0 | 90 | 288 | 27 | 106 | 0 | 0 | 1243 | 0 | 0 | 0 | 0 | 0 | 0 | 0 | 0 | 0 | 0 | 0 | 0 | 0 | 0 | 0 | 0 | 0 | 0 | 0 | 0 |
| SRR7343633 | 175 | 6985 | 918 | 279 | 0 | 18 | 0 | 27 | 353 | 0 | 256 | 541 | 104 | 365 | 0 | 30 | 3300 | 0 | 0 | 0 | 0 | 0 | 0 | 0 | 0 | 0 | 0 | 0 | 0 | 0 | 0 | 0 | 0 | 0 | 0 | 0 | 0 |
| SRR7343635 | 0 | 891 | 697 | 155 | 0 | 22 | 10 | 48 | 360 | 0 | 429 | 652 | 155 | 609 | 16 | 0 | 2932 | 0 | 0 | 0 | 0 | 0 | 0 | 0 | 0 | 0 | 0 | 0 | 0 | 0 | 0 | 0 | 0 | 0 | 0 | 0 | 0 |
| SRR7343636 | 12 | 2119 | 512 | 195 | 0 | 13 | 0 | 23 | 299 | 12 | 229 | 505 | 93 | 346 | 0 | 41 | 3039 | 0 | 0 | 0 | 0 | 0 | 0 | 0 | 0 | 0 | 0 | 0 | 0 | 0 | 0 | 0 | 0 | 0 | 0 | 0 | 0 |
| SRR7343637 | 0 | 9036 | 635 | 254 | 0 | 20 | 0 | 40 | 401 | 0 | 363 | 697 | 156 | 802 | 24 | 0 | 3030 | 0 | 0 | 0 | 0 | 0 | 0 | 0 | 0 | 0 | 0 | 0 | 0 | 0 | 0 | 0 | 0 | 0 | 0 | 0 | 0 |
| SRR7343638 | 78 | 7119 | 2504 | 156 | 0 | 15 | 0 | 22 | 131 | 0 | 250 | 346 | 66 | 258 | 11 | 0 | 1208 | 0 | 0 | 0 | 0 | 0 | 0 | 0 | 0 | 0 | 0 | 0 | 0 | 0 | 0 | 0 | 0 | 0 | 0 | 0 | 0 |
| SRR7343639 | 44 | 2300 | 3184 | 88 | 0 | 0 | 0 | 0 | 49 | 44 | 64 | 162 | 24 | 83 | 0 | 18 | 488 | 0 | 0 | 0 | 0 | 0 | 0 | 0 | 0 | 0 | 0 | 0 | 0 | 0 | 0 | 0 | 0 | 0 | 0 | 0 | 0 |
| SRR7343640 | 19 | 2563 | 829 | 636 | 0 | 0 | 0 | 0 | 188 | 0 | 124 | 220 | 30 | 102 | 0 | 0 | 2146 | 0 | 0 | 0 | 0 | 0 | 0 | 0 | 0 | 0 | 0 | 0 | 0 | 0 | 0 | 0 | 0 | 0 | 0 | 0 | 0 |
| SRR7412243 | 483 | 5372 | 298 | 192 | 1533 | 1742 | 19505 | 29575 | 5403 | 0 | 25651 | 31907 | 15722 | 6438 | 21 | 11 | 44676 | 0 | 0 | 0 | 31 | 0 | 14 | 0 | 0 | 0 | 0 | 0 | 0 | 42 | 27 | 38 | 0 | 0 | 0 | 0 | 0 |
| SRR7412244 | 1152 | 14390 | 608 | 221 | 1624 | 1752 | 17372 | 31467 | 4538 | 0 | 26963 | 40233 | 17421 | 6229 | 42 | 17 | 39185 | 0 | 0 | 0 | 23 | 0 | 0 | 0 | 0 | 0 | 0 | 0 | 0 | 36 | 15 | 34 | 0 | 0 | 0 | 0 | 0 |
| SRR7412245 | 584 | 9226 | 257 | 273 | 1610 | 2091 | 19472 | 36125 | 6222 | 0 | 30137 | 34988 | 16898 | 5195 | 31 | 23 | 39381 | 0 | 0 | 0 | 31 | 0 | 18 | 0 | 0 | 0 | 0 | 0 | 0 | 39 | 21 | 25 | 0 | 0 | 0 | 0 | 0 |
| SRR7412246 | 346 | 7444 | 269 | 199 | 3671 | 2921 | 36907 | 46932 | 4616 | 0 | 25323 | 33433 | 13697 | 5683 | 26 | 12 | 38075 | 0 | 0 | 0 | 41 | 0 | 27 | 0 | 0 | 0 | 0 | 0 | 0 | 61 | 18 | 41 | 0 | 0 | 0 | 0 | 0 |
| SRR7412247 | 615 | 9290 | 453 | 131 | 1079 | 1508 | 13370 | 25603 | 6269 | 0 | 29023 | 32113 | 19511 | 6354 | 15 | 10 | 39808 | 0 | 0 | 0 | 34 | 0 | 15 | 0 | 0 | 0 | 0 | 0 | 24 | 74 | 38 | 27 | 0 | 0 | 0 | 0 | 0 |
| SRR7412248 | 730 | 4985 | 365 | 136 | 2713 | 3013 | 27702 | 41705 | 3053 | 0 | 33389 | 39190 | 14105 | 6992 | 17 | 0 | 30569 | 0 | 0 | 0 | 28 | 11 | 19 | 0 | 0 | 0 | 0 | 0 | 0 | 46 | 51 | 38 | 0 | 0 | 0 | 0 | 0 |
| SRR7412249 | 744 | 8142 | 416 | 136 | 1067 | 1291 | 11433 | 29873 | 4704 | 0 | 26045 | 33847 | 18828 | 5355 | 21 | 0 | 31948 | 0 | 0 | 0 | 28 | 0 | 22 | 0 | 0 | 0 | 0 | 0 | 0 | 31 | 10 | 30 | 0 | 0 | 0 | 0 | 0 |
| SRR7412250 | 604 | 6063 | 363 | 82 | 1646 | 1995 | 18719 | 31075 | 2967 | 14 | 24928 | 42830 | 13853 | 4962 | 30 | 22 | 38333 | 0 | 0 | 0 | 0 | 0 | 0 | 0 | 0 | 0 | 0 | 0 | 0 | 0 | 0 | 0 | 0 | 0 | 0 | 0 | 0 |
| SRR7412251 | 512 | 16685 | 264 | 66 | 742 | 2375 | 9999 | 16226 | 8245 | 0 | 25874 | 28206 | 19873 | 4070 | 37 | 74 | 62922 | 0 | 0 | 0 | 0 | 11 | 0 | 0 | 0 | 0 | 0 | 0 | 0 | 22 | 0 | 10 | 0 | 0 | 0 | 0 | 0 |
| SRR7412252 | 827 | 3489 | 356 | 111 | 1530 | 1528 | 15480 | 31172 | 4677 | 0 | 25545 | 29269 | 17877 | 5823 | 28 | 28 | 29389 | 0 | 0 | 0 | 21 | 0 | 0 | 0 | 0 | 0 | 0 | 0 | 0 | 20 | 47 | 30 | 0 | 0 | 0 | 0 | 0 |
| SRR7412253 | 795 | 12930 | 646 | 441 | 1881 | 7258 | 22456 | 39461 | 7011 | 0 | 23313 | 26033 | 14371 | 8291 | 18 | 58 | 34797 | 0 | 0 | 0 | 0 | 12 | 0 | 0 | 0 | 0 | 0 | 0 | 0 | 23 | 20 | 15 | 0 | 0 | 0 | 0 | 0 |
| SRR7412254 | 303 | 10674 | 581 | 449 | 1652 | 5750 | 25054 | 25672 | 4955 | 13 | 38066 | 42180 | 15600 | 7754 | 12 | 234 | 37585 | 0 | 0 | 0 | 0 | 0 | 0 | 0 | 0 | 0 | 0 | 0 | 0 | 15 | 0 | 13 | 0 | 0 | 0 | 0 | 0 |
| SRR7412255 | 481 | 5779 | 522 | 93 | 1327 | 1895 | 16218 | 29109 | 5046 | 0 | 27736 | 37987 | 14535 | 5209 | 14 | 15 | 37002 | 0 | 0 | 0 | 0 | 0 | 0 | 0 | 0 | 0 | 0 | 0 | 0 | 19 | 51 | 12 | 0 | 0 | 0 | 0 | 0 |
| SRR7412256 | 239 | 18964 | 509 | 301 | 1845 | 2129 | 21997 | 29316 | 4600 | 0 | 25234 | 30710 | 12870 | 5274 | 34 | 0 | 43203 | 0 | 0 | 0 | 36 | 0 | 22 | 0 | 0 | 0 | 0 | 0 | 0 | 29 | 12 | 32 | 0 | 0 | 0 | 0 | 0 |
| SRR7412257 | 277 | 13654 | 405 | 378 | 2328 | 3855 | 22829 | 29138 | 6719 | 20 | 22141 | 20288 | 15055 | 5516 | 25 | 24 | 44537 | 0 | 0 | 0 | 33 | 0 | 35 | 0 | 0 | 0 | 0 | 0 | 0 | 59 | 24 | 56 | 0 | 0 | 0 | 0 | 0 |
| SRR7412258 | 144 | 7176 | 541 | 72 | 1986 | 1824 | 23119 | 20882 | 3740 | 0 | 30225 | 69100 | 6991 | 5662 | 15 | 81 | 27610 | 0 | 0 | 0 | 18 | 0 | 11 | 0 | 0 | 0 | 0 | 0 | 0 | 20 | 14 | 23 | 0 | 0 | 0 | 0 | 0 |
| SRR7412259 | 404 | 28659 | 231 | 180 | 900 | 2735 | 9848 | 20305 | 7777 | 19 | 26326 | 30806 | 17472 | 4171 | 35 | 23 | 44090 | 0 | 0 | 0 | 17 | 0 | 14 | 0 | 0 | 0 | 0 | 0 | 0 | 38 | 28 | 23 | 0 | 0 | 0 | 0 | 0 |
| SRR7412260 | 108 | 6998 | 342 | 126 | 793 | 1051 | 10909 | 10120 | 4063 | 18 | 32076 | 81317 | 10193 | 5545 | 0 | 36 | 25355 | 0 | 0 | 0 | 0 | 0 | 0 | 0 | 0 | 0 | 0 | 0 | 0 | 15 | 21 | 14 | 0 | 0 | 0 | 0 | 0 |
| SRR7412261 | 208 | 10568 | 402 | 83 | 892 | 1159 | 10190 | 32801 | 3055 | 0 | 27073 | 39506 | 20437 | 5669 | 32 | 0 | 32620 | 0 | 0 | 0 | 32 | 0 | 18 | 0 | 0 | 0 | 0 | 0 | 0 | 38 | 54 | 29 | 0 | 0 | 0 | 0 | 0 |
| SRR7412262 | 323 | 9258 | 494 | 145 | 1452 | 1682 | 15957 | 33360 | 3457 | 0 | 27938 | 31791 | 19500 | 6142 | 16 | 15 | 40142 | 0 | 0 | 0 | 31 | 11 | 16 | 0 | 0 | 0 | 0 | 0 | 0 | 24 | 28 | 28 | 0 | 0 | 0 | 0 | 0 |
| SRR7412263 | 189 | 14992 | 402 | 686 | 2901 | 7014 | 36190 | 35404 | 7374 | 0 | 23775 | 29668 | 8429 | 8447 | 0 | 46 | 19152 | 0 | 0 | 0 | 29 | 12 | 0 | 0 | 0 | 0 | 0 | 0 | 0 | 36 | 29 | 33 | 0 | 0 | 0 | 0 | 0 |
| SRR7412264 | 145 | 11614 | 263 | 289 | 2247 | 4020 | 28266 | 27976 | 10158 | 0 | 20299 | 23155 | 13618 | 7475 | 15 | 35 | 28583 | 0 | 0 | 0 | 30 | 0 | 24 | 0 | 0 | 0 | 0 | 0 | 0 | 35 | 25 | 27 | 0 | 0 | 0 | 0 | 0 |
| SRR7412265 | 194 | 7685 | 480 | 404 | 889 | 1749 | 11470 | 24657 | 2827 | 0 | 36145 | 32850 | 26598 | 7471 | 15 | 36 | 39541 | 0 | 0 | 0 | 40 | 0 | 16 | 0 | 0 | 0 | 0 | 0 | 0 | 46 | 0 | 38 | 0 | 0 | 0 | 0 | 0 |
| SRR7412266 | 332 | 28117 | 945 | 701 | 1740 | 6078 | 22382 | 37946 | 5812 | 30 | 47869 | 44688 | 25842 | 10090 | 40 | 108 | 30194 | 0 | 0 | 0 | 48 | 41 | 23 | 0 | 0 | 0 | 0 | 0 | 0 | 51 | 35 | 41 | 0 | 0 | 0 | 0 | 0 |
| SRR7412267 | 334 | 10748 | 634 | 367 | 937 | 3024 | 14938 | 26397 | 5150 | 0 | 32810 | 35857 | 19561 | 5816 | 23 | 66 | 40056 | 0 | 0 | 0 | 25 | 0 | 0 | 0 | 0 | 0 | 0 | 0 | 0 | 29 | 0 | 24 | 0 | 0 | 0 | 0 | 0 |
| SRR7412268 | 307 | 13264 | 564 | 451 | 1040 | 4168 | 13973 | 28656 | 2989 | 13 | 40137 | 45780 | 23181 | 5850 | 38 | 70 | 41558 | 0 | 0 | 0 | 0 | 0 | 0 | 0 | 0 | 0 | 0 | 0 | 0 | 15 | 22 | 12 | 0 | 0 | 0 | 0 | 0 |
| SRR7412269 | 242 | 10164 | 532 | 447 | 1468 | 5739 | 21413 | 30731 | 5758 | 12 | 21867 | 15346 | 14630 | 8992 | 17 | 75 | 30746 | 0 | 0 | 0 | 24 | 11 | 0 | 0 | 0 | 0 | 0 | 0 | 0 | 49 | 72 | 42 | 0 | 0 | 0 | 0 | 0 |
| SRR7412270 | 223 | 19629 | 446 | 506 | 1221 | 4915 | 17750 | 32199 | 4636 | 0 | 32971 | 31167 | 22144 | 8074 | 23 | 23 | 33766 | 0 | 0 | 0 | 18 | 0 | 0 | 0 | 0 | 0 | 0 | 0 | 0 | 31 | 0 | 15 | 0 | 0 | 0 | 0 | 0 |
| SRR7412271 | 411 | 11931 | 343 | 525 | 1595 | 3987 | 20792 | 21534 | 10494 | 0 | 36023 | 35638 | 17710 | 5549 | 11 | 55 | 58585 | 0 | 0 | 0 | 24 | 0 | 0 | 0 | 0 | 0 | 0 | 0 | 0 | 26 | 12 | 13 | 0 | 0 | 0 | 0 | 0 |
| SRR7412272 | 463 | 15803 | 357 | 174 | 1006 | 3173 | 10244 | 20990 | 4299 | 19 | 29750 | 44603 | 16948 | 4798 | 14 | 41 | 58142 | 0 | 0 | 0 | 19 | 0 | 17 | 0 | 0 | 0 | 0 | 0 | 0 | 26 | 0 | 29 | 0 | 0 | 0 | 0 | 0 |
| SRR7412273 | 284 | 12212 | 288 | 369 | 3116 | 6816 | 24229 | 34258 | 6669 | 0 | 20402 | 22472 | 13747 | 6117 | 11 | 14 | 58820 | 0 | 0 | 0 | 58 | 25 | 22 | 0 | 0 | 0 | 0 | 0 | 10 | 87 | 70 | 48 | 0 | 0 | 0 | 0 | 0 |
| SRR7412274 | 391 | 22370 | 615 | 557 | 1641 | 4519 | 15432 | 25114 | 7793 | 0 | 20946 | 22490 | 15569 | 5560 | 12 | 17 | 50878 | 0 | 0 | 0 | 39 | 35 | 18 | 0 | 0 | 0 | 0 | 0 | 21 | 69 | 26 | 39 | 0 | 0 | 0 | 0 | 0 |
| SRR7412275 | 375 | 18222 | 336 | 439 | 999 | 3028 | 12571 | 20014 | 4436 | 0 | 37884 | 48089 | 16462 | 5672 | 0 | 41 | 49482 | 0 | 0 | 0 | 18 | 15 | 0 | 0 | 0 | 0 | 0 | 0 | 0 | 40 | 31 | 21 | 0 | 0 | 0 | 0 | 0 |
| SRR7412276 | 223 | 16985 | 205 | 205 | 1426 | 2816 | 14539 | 30413 | 4066 | 0 | 38976 | 37943 | 24657 | 5784 | 21 | 15 | 43062 | 0 | 0 | 0 | 25 | 0 | 10 | 0 | 0 | 0 | 0 | 0 | 0 | 23 | 0 | 30 | 0 | 0 | 0 | 0 | 0 |
| SRR7412277 | 410 | 15024 | 579 | 515 | 967 | 4955 | 12653 | 11349 | 7264 | 24 | 25880 | 38203 | 12397 | 4599 | 0 | 168 | 45197 | 0 | 0 | 0 | 29 | 0 | 0 | 0 | 0 | 0 | 0 | 0 | 0 | 33 | 18 | 29 | 0 | 0 | 0 | 0 | 0 |
| SRR7412278 | 226 | 17902 | 113 | 113 | 1158 | 1531 | 11906 | 16357 | 5903 | 0 | 31672 | 45050 | 18185 | 4650 | 15 | 21 | 37077 | 0 | 0 | 0 | 23 | 0 | 19 | 0 | 0 | 0 | 0 | 0 | 0 | 34 | 19 | 20 | 0 | 0 | 0 | 0 | 0 |
| SRR7412279 | 278 | 11327 | 229 | 351 | 1297 | 2880 | 17159 | 18651 | 9449 | 0 | 32069 | 30309 | 16979 | 6057 | 27 | 18 | 47512 | 0 | 0 | 0 | 19 | 0 | 11 | 0 | 0 | 0 | 0 | 0 | 0 | 18 | 19 | 17 | 0 | 0 | 0 | 0 | 0 |
| SRR7412280 | 291 | 4943 | 373 | 136 | 1060 | 2033 | 14510 | 22567 | 5516 | 18 | 19028 | 32486 | 8777 | 4359 | 15 | 40 | 26022 | 0 | 0 | 0 | 14 | 0 | 0 | 0 | 0 | 0 | 0 | 0 | 0 | 23 | 16 | 21 | 0 | 0 | 0 | 0 | 0 |
| SRR7412281 | 364 | 5695 | 472 | 334 | 1324 | 2087 | 18088 | 25916 | 3748 | 0 | 34745 | 50567 | 15071 | 5734 | 11 | 29 | 48293 | 0 | 0 | 0 | 33 | 0 | 24 | 0 | 0 | 0 | 0 | 0 | 0 | 26 | 25 | 35 | 0 | 0 | 0 | 0 | 0 |
| SRR7412282 | 262 | 5440 | 365 | 148 | 1026 | 1637 | 13081 | 21497 | 4913 | 0 | 25564 | 38166 | 12579 | 5618 | 16 | 23 | 41218 | 0 | 0 | 0 | 17 | 0 | 0 | 0 | 0 | 0 | 0 | 0 | 0 | 15 | 0 | 14 | 0 | 0 | 0 | 0 | 0 |
| SRR7412283 | 271 | 9211 | 394 | 114 | 880 | 1506 | 12049 | 16955 | 3982 | 26 | 28489 | 46677 | 12855 | 5241 | 23 | 35 | 54573 | 0 | 0 | 0 | 23 | 0 | 0 | 0 | 0 | 0 | 0 | 0 | 0 | 23 | 11 | 14 | 0 | 0 | 0 | 0 | 0 |
| SRR7412284 | 573 | 4473 | 483 | 98 | 1421 | 2013 | 16796 | 22771 | 5649 | 18 | 30041 | 38203 | 10435 | 6682 | 11 | 17 | 31149 | 0 | 0 | 0 | 13 | 0 | 12 | 0 | 0 | 0 | 0 | 0 | 0 | 19 | 10 | 14 | 0 | 0 | 0 | 0 | 0 |
| SRR7412285 | 240 | 6473 | 351 | 181 | 3060 | 2546 | 28720 | 31261 | 7648 | 18 | 15785 | 24316 | 7079 | 8721 | 12 | 11 | 33267 | 0 | 0 | 0 | 19 | 11 | 10 | 0 | 0 | 0 | 0 | 0 | 0 | 60 | 99 | 43 | 0 | 0 | 0 | 0 | 0 |
| SRR7412286 | 289 | 5314 | 217 | 105 | 1710 | 2547 | 20825 | 28422 | 5479 | 0 | 25125 | 41558 | 11678 | 5554 | 30 | 28 | 47452 | 0 | 0 | 0 | 12 | 0 | 0 | 0 | 0 | 0 | 0 | 0 | 15 | 15 | 0 | 0 | 0 | 0 | 0 | 0 | 0 |
| SRR7412287 | 343 | 15610 | 155 | 444 | 1676 | 3020 | 15781 | 23244 | 5654 | 0 | 33843 | 30666 | 22427 | 4487 | 19 | 0 | 54753 | 0 | 0 | 0 | 20 | 0 | 19 | 0 | 0 | 0 | 0 | 0 | 11 | 33 | 10 | 15 | 0 | 0 | 0 | 0 | 0 |
| SRR7412288 | 288 | 6759 | 337 | 134 | 1838 | 3589 | 29586 | 24514 | 8713 | 0 | 22543 | 26096 | 8607 | 7156 | 13 | 52 | 26957 | 0 | 0 | 0 | 0 | 0 | 0 | 0 | 0 | 0 | 0 | 0 | 0 | 0 | 0 | 0 | 0 | 0 | 0 | 0 | 0 |
| SRR7412289 | 276 | 10290 | 440 | 153 | 1148 | 1799 | 14940 | 19697 | 7165 | 0 | 27072 | 33437 | 14015 | 6256 | 29 | 70 | 43279 | 0 | 0 | 0 | 0 | 0 | 0 | 0 | 0 | 0 | 0 | 0 | 0 | 27 | 0 | 0 | 0 | 0 | 0 | 0 | 0 |
| SRR7412290 | 139 | 6565 | 222 | 153 | 1728 | 1345 | 19094 | 16543 | 3978 | 0 | 27184 | 33009 | 11050 | 3011 | 19 | 0 | 35067 | 0 | 0 | 0 | 12 | 0 | 13 | 0 | 0 | 0 | 0 | 0 | 0 | 26 | 0 | 20 | 0 | 0 | 0 | 0 | 0 |
| SRR7412291 | 359 | 5260 | 215 | 104 | 874 | 1495 | 10793 | 21475 | 3545 | 0 | 26427 | 41740 | 16063 | 4510 | 18 | 0 | 54470 | 0 | 0 | 0 | 29 | 0 | 13 | 0 | 0 | 0 | 0 | 0 | 0 | 45 | 51 | 39 | 0 | 0 | 0 | 0 | 0 |
| SRR7412292 | 457 | 2736 | 348 | 119 | 2134 | 2342 | 22993 | 31817 | 2706 | 0 | 29590 | 50992 | 12880 | 5719 | 18 | 0 | 40853 | 0 | 0 | 0 | 32 | 0 | 28 | 0 | 0 | 0 | 0 | 0 | 0 | 32 | 12 | 51 | 0 | 0 | 0 | 0 | 0 |
| SRR7412293 | 368 | 10833 | 515 | 423 | 924 | 2512 | 14448 | 21520 | 7031 | 0 | 25430 | 35698 | 10820 | 6009 | 14 | 59 | 44738 | 0 | 0 | 0 | 0 | 0 | 0 | 0 | 0 | 0 | 0 | 0 | 0 | 0 | 0 | 0 | 0 | 0 | 0 | 0 | 0 |
| SRR7412294 | 235 | 12334 | 1731 | 71 | 3695 | 3550 | 31646 | 31248 | 5950 | 192 | 15168 | 19489 | 8629 | 7266 | 14 | 44 | 44117 | 0 | 0 | 0 | 22 | 0 | 11 | 0 | 0 | 0 | 0 | 0 | 0 | 40 | 17 | 35 | 0 | 0 | 0 | 0 | 0 |
| SRR7412295 | 254 | 7130 | 831 | 385 | 2377 | 2304 | 26600 | 36078 | 4546 | 18 | 18024 | 26341 | 9367 | 8453 | 16 | 29 | 29050 | 0 | 0 | 0 | 17 | 0 | 0 | 0 | 0 | 0 | 0 | 0 | 0 | 31 | 16 | 12 | 0 | 0 | 0 | 0 | 0 |
| SRR7412296 | 548 | 20280 | 280 | 61 | 965 | 2727 | 9395 | 15818 | 6219 | 0 | 28643 | 26234 | 19910 | 3722 | 41 | 0 | 53102 | 0 | 0 | 0 | 18 | 17 | 26 | 0 | 0 | 0 | 0 | 0 | 0 | 26 | 26 | 29 | 0 | 0 | 0 | 0 | 0 |
| SRR7412297 | 180 | 13284 | 131 | 305 | 784 | 4184 | 8889 | 16026 | 6128 | 0 | 31781 | 36638 | 16689 | 4188 | 13 | 69 | 55219 | 0 | 0 | 0 | 35 | 14 | 0 | 0 | 0 | 0 | 0 | 0 | 0 | 53 | 61 | 28 | 0 | 0 | 0 | 0 | 0 |
| SRR7412298 | 166 | 4538 | 316 | 191 | 1707 | 1974 | 20432 | 19087 | 6374 | 0 | 15425 | 24546 | 5344 | 6770 | 0 | 44 | 22888 | 0 | 0 | 0 | 0 | 0 | 0 | 0 | 0 | 0 | 0 | 0 | 0 | 12 | 0 | 12 | 0 | 0 | 0 | 0 | 0 |
| SRR7412299 | 125 | 9678 | 267 | 53 | 883 | 1931 | 17277 | 12181 | 8659 | 0 | 39450 | 57592 | 7663 | 5360 | 0 | 61 | 39428 | 0 | 0 | 0 | 0 | 0 | 0 | 0 | 0 | 0 | 0 | 0 | 0 | 18 | 0 | 0 | 0 | 0 | 0 | 0 | 0 |
| SRR7412300 | 368 | 14172 | 490 | 479 | 928 | 2131 | 13395 | 13958 | 5477 | 11 | 34412 | 58110 | 7841 | 4504 | 11 | 45 | 43671 | 0 | 0 | 0 | 15 | 0 | 0 | 0 | 0 | 0 | 0 | 0 | 0 | 19 | 15 | 0 | 0 | 0 | 0 | 0 | 0 |
| SRR7412301 | 204 | 11358 | 113 | 170 | 2615 | 4506 | 23716 | 25867 | 13497 | 0 | 27157 | 31023 | 12258 | 8362 | 0 | 40 | 53714 | 0 | 0 | 0 | 10 | 0 | 0 | 0 | 0 | 0 | 0 | 0 | 14 | 51 | 11 | 12 | 0 | 0 | 0 | 0 | 0 |
| SRR7412302 | 483 | 8677 | 338 | 227 | 905 | 1842 | 14078 | 22212 | 10343 | 0 | 22007 | 21595 | 10125 | 6489 | 10 | 15 | 20945 | 0 | 0 | 0 | 11 | 0 | 0 | 0 | 0 | 0 | 0 | 0 | 0 | 35 | 20 | 21 | 0 | 0 | 0 | 0 | 0 |
| SRR7412303 | 196 | 19161 | 153 | 477 | 780 | 2933 | 8336 | 15885 | 6904 | 12 | 34345 | 32214 | 18685 | 7471 | 0 | 0 | 49862 | 0 | 0 | 0 | 22 | 0 | 11 | 0 | 0 | 0 | 0 | 0 | 0 | 52 | 16 | 33 | 0 | 0 | 0 | 0 | 0 |
| SRR7412304 | 1313 | 9181 | 744 | 326 | 896 | 1983 | 12714 | 20820 | 6410 | 0 | 34328 | 28507 | 13058 | 6328 | 0 | 20 | 19312 | 0 | 0 | 0 | 23 | 0 | 13 | 0 | 0 | 0 | 0 | 0 | 0 | 35 | 12 | 29 | 0 | 0 | 0 | 0 | 0 |
| SRR7412305 | 275 | 11637 | 654 | 183 | 1677 | 2460 | 23152 | 23969 | 6600 | 0 | 25017 | 41213 | 8155 | 5985 | 0 | 85 | 40097 | 0 | 0 | 0 | 10 | 0 | 0 | 0 | 0 | 0 | 0 | 0 | 0 | 29 | 0 | 17 | 0 | 0 | 0 | 0 | 0 |
| SRR7412306 | 367 | 23226 | 580 | 244 | 916 | 1600 | 11494 | 26808 | 3921 | 10 | 23376 | 39338 | 17461 | 4608 | 23 | 19 | 45952 | 0 | 0 | 0 | 29 | 0 | 12 | 0 | 0 | 0 | 0 | 0 | 0 | 31 | 11 | 33 | 0 | 0 | 0 | 0 | 0 |
| SRR7412307 | 230 | 4722 | 262 | 164 | 1860 | 1782 | 21778 | 23996 | 3054 | 0 | 28836 | 52789 | 10540 | 5792 | 11 | 27 | 36437 | 0 | 0 | 0 | 0 | 0 | 0 | 0 | 0 | 0 | 0 | 0 | 0 | 23 | 37 | 19 | 0 | 0 | 0 | 0 | 0 |
| SRR7412308 | 535 | 6798 | 220 | 124 | 1335 | 1522 | 15735 | 23713 | 4712 | 0 | 26963 | 28802 | 15017 | 5540 | 0 | 18 | 40470 | 0 | 0 | 0 | 24 | 0 | 12 | 0 | 0 | 0 | 0 | 0 | 11 | 31 | 24 | 18 | 0 | 0 | 0 | 0 | 0 |
| SRR7412309 | 518 | 8949 | 326 | 217 | 1827 | 2107 | 20274 | 35332 | 4155 | 0 | 21821 | 27755 | 12867 | 4780 | 21 | 11 | 33176 | 0 | 0 | 0 | 50 | 0 | 40 | 0 | 0 | 0 | 0 | 0 | 0 | 45 | 40 | 34 | 0 | 0 | 0 | 0 | 0 |
| SRR7412310 | 310 | 3303 | 180 | 46 | 2791 | 2015 | 24833 | 33585 | 3719 | 0 | 20408 | 33255 | 11321 | 5692 | 25 | 13 | 28686 | 0 | 0 | 0 | 32 | 0 | 27 | 0 | 0 | 0 | 0 | 0 | 0 | 42 | 15 | 56 | 0 | 15 | 0 | 0 | 0 |
| SRR7412311 | 321 | 4849 | 279 | 90 | 1789 | 1882 | 19236 | 35210 | 2819 | 0 | 27082 | 37575 | 16445 | 5080 | 17 | 10 | 35164 | 0 | 0 | 0 | 29 | 0 | 30 | 0 | 0 | 0 | 0 | 0 | 0 | 28 | 0 | 48 | 0 | 0 | 0 | 0 | 0 |
| SRR7412312 | 704 | 6671 | 416 | 42 | 999 | 1508 | 13450 | 25495 | 4112 | 0 | 33205 | 48330 | 16950 | 5674 | 0 | 18 | 36894 | 0 | 0 | 0 | 0 | 0 | 0 | 0 | 0 | 0 | 0 | 0 | 0 | 0 | 15 | 0 | 0 | 0 | 0 | 0 | 0 |
| SRR7412313 | 439 | 6456 | 497 | 334 | 553 | 1837 | 9666 | 19195 | 7045 | 0 | 43962 | 31513 | 24848 | 5414 | 15 | 66 | 41619 | 0 | 0 | 0 | 11 | 0 | 0 | 0 | 0 | 0 | 0 | 0 | 0 | 28 | 29 | 16 | 0 | 0 | 0 | 0 | 0 |
| SRR7412314 | 304 | 16763 | 238 | 653 | 890 | 1733 | 11323 | 10818 | 6549 | 15 | 36733 | 42175 | 11879 | 3764 | 19 | 33 | 53522 | 0 | 0 | 0 | 0 | 0 | 0 | 0 | 0 | 0 | 0 | 0 | 0 | 18 | 11 | 0 | 0 | 0 | 14 | 0 | 0 |
| SRR7412315 | 588 | 12119 | 576 | 141 | 700 | 1884 | 10320 | 16538 | 5679 | 24 | 36170 | 34468 | 25623 | 4655 | 31 | 16 | 61474 | 0 | 0 | 0 | 15 | 14 | 15 | 0 | 0 | 0 | 0 | 0 | 0 | 31 | 37 | 22 | 0 | 0 | 0 | 0 | 0 |
| SRR7412316 | 228 | 11282 | 519 | 483 | 1444 | 4386 | 20902 | 23420 | 5886 | 0 | 33461 | 26868 | 20679 | 7160 | 14 | 94 | 40224 | 0 | 0 | 0 | 0 | 0 | 0 | 0 | 0 | 0 | 0 | 0 | 0 | 26 | 14 | 23 | 0 | 0 | 0 | 0 | 0 |
| SRR7412317 | 265 | 15887 | 305 | 81 | 1634 | 1869 | 16001 | 21009 | 5509 | 17 | 21778 | 39411 | 9646 | 4137 | 21 | 17 | 43149 | 0 | 0 | 0 | 14 | 0 | 13 | 0 | 0 | 0 | 0 | 0 | 0 | 59 | 88 | 22 | 0 | 0 | 0 | 0 | 0 |
| SRR7412318 | 178 | 9635 | 331 | 127 | 1334 | 1759 | 15132 | 13882 | 5545 | 0 | 28834 | 46451 | 8810 | 4904 | 12 | 48 | 36943 | 0 | 0 | 0 | 0 | 0 | 0 | 0 | 0 | 0 | 0 | 0 | 0 | 17 | 0 | 0 | 0 | 0 | 0 | 0 | 0 |
| SRR7412319 | 433 | 16715 | 405 | 121 | 2950 | 3296 | 28579 | 29003 | 9959 | 0 | 27545 | 43197 | 10989 | 6309 | 24 | 26 | 36185 | 0 | 0 | 0 | 22 | 0 | 15 | 0 | 0 | 0 | 0 | 0 | 0 | 58 | 0 | 37 | 0 | 0 | 0 | 0 | 0 |
| SRR7412320 | 211 | 3508 | 638 | 146 | 2205 | 2690 | 27201 | 31154 | 8695 | 10 | 22290 | 33557 | 9767 | 6786 | 16 | 31 | 25519 | 0 | 0 | 0 | 0 | 0 | 0 | 0 | 0 | 0 | 0 | 0 | 0 | 0 | 0 | 10 | 0 | 0 | 0 | 0 | 0 |
| SRR7412321 | 502 | 6437 | 521 | 228 | 831 | 1991 | 11709 | 19539 | 6243 | 0 | 33883 | 33386 | 26173 | 5458 | 16 | 80 | 30943 | 0 | 0 | 0 | 14 | 0 | 0 | 0 | 0 | 0 | 0 | 0 | 0 | 18 | 0 | 18 | 0 | 0 | 0 | 0 | 0 |
| SRR7412322 | 292 | 9543 | 573 | 435 | 1424 | 2682 | 16716 | 25976 | 4907 | 0 | 37881 | 44369 | 18544 | 7387 | 14 | 0 | 35123 | 0 | 0 | 0 | 51 | 11 | 20 | 0 | 0 | 0 | 0 | 0 | 0 | 45 | 26 | 39 | 0 | 10 | 0 | 0 | 0 |
| SRR7412323 | 66 | 2574 | 648 | 475 | 2492 | 7103 | 41790 | 20458 | 7888 | 0 | 37887 | 27744 | 9789 | 13750 | 24 | 92 | 13987 | 0 | 0 | 0 | 14 | 0 | 0 | 0 | 0 | 0 | 0 | 0 | 0 | 29 | 14 | 19 | 0 | 0 | 0 | 0 | 0 |
| SRR7412324 | 64 | 2475 | 637 | 446 | 2347 | 4188 | 29563 | 24370 | 8941 | 18 | 27111 | 31755 | 11192 | 8648 | 13 | 35 | 15719 | 0 | 0 | 0 | 24 | 0 | 0 | 0 | 0 | 0 | 0 | 0 | 0 | 30 | 0 | 26 | 0 | 0 | 0 | 0 | 0 |
| SRR7412325 | 260 | 13811 | 715 | 163 | 2200 | 2015 | 20339 | 31556 | 2966 | 0 | 29045 | 59987 | 13573 | 6256 | 25 | 36 | 36701 | 0 | 0 | 0 | 11 | 0 | 0 | 0 | 0 | 0 | 0 | 0 | 0 | 16 | 0 | 0 | 0 | 0 | 0 | 0 | 0 |
| SRR7412326 | 536 | 9608 | 467 | 133 | 1336 | 1782 | 16497 | 28443 | 5770 | 0 | 27662 | 30959 | 15831 | 5796 | 16 | 24 | 40121 | 0 | 0 | 0 | 20 | 13 | 10 | 0 | 0 | 0 | 0 | 0 | 0 | 44 | 40 | 36 | 0 | 0 | 0 | 0 | 0 |
| SRR7412327 | 360 | 9655 | 432 | 190 | 1219 | 1193 | 12713 | 22597 | 3947 | 0 | 26604 | 30741 | 15446 | 5132 | 17 | 16 | 33986 | 0 | 0 | 0 | 19 | 0 | 22 | 0 | 0 | 0 | 0 | 0 | 0 | 23 | 24 | 36 | 0 | 0 | 0 | 0 | 0 |

### Supp Table S8.1.Columns_B_to_AL

| | | | | | | | | | | | | | | | | | | | | | | | | | | | | | | | | | | | | | |
|---|---|---|---|---|---|---|---|---|---|---|---|---|---|---|---|---|---|---|---|---|---|---|---|---|---|---|---|---|---|---|---|---|---|---|---|---|---|
| SRR7412328 | 317 | 6521 | 485 | 710 | 961 | 2595 | 13854 | 18096 | 1950 | 12 | 23333 | 20314 | 12936 | 4650 | 14 | 51 | 21417 | 0 | 0 | 0 | 29 | 11 | 0 | 0 | 0 | 0 | 0 | 0 | 0 | 71 | 54 | 41 | 0 | 0 | 0 | 0 | 0 |
| SRR7412329 | 224 | 5114 | 926 | 496 | 2832 | 8558 | 35401 | 33989 | 5924 | 112 | 19558 | 13447 | 10327 | 14065 | 0 | 140 | 27761 | 0 | 0 | 0 | 25 | 12 | 13 | 0 | 0 | 0 | 0 | 0 | 16 | 84 | 77 | 36 | 0 | 15 | 0 | 0 | 0 |
| SRR7412330 | 343 | 11438 | 3557 | 630 | 2987 | 6172 | 24469 | 24786 | 2232 | 702 | 20575 | 17913 | 13336 | 5560 | 0 | 0 | 35657 | 0 | 0 | 0 | 33 | 0 | 0 | 0 | 0 | 0 | 0 | 0 | 11 | 37 | 0 | 0 | 0 | 0 | 0 | 0 | 0 |
| SRR7412331 | 669 | 14226 | 438 | 877 | 2255 | 6167 | 30575 | 30827 | 11849 | 32 | 25687 | 27114 | 15458 | 7903 | 0 | 92 | 31176 | 0 | 0 | 0 | 14 | 0 | 0 | 0 | 0 | 0 | 0 | 0 | 0 | 70 | 30 | 32 | 0 | 0 | 0 | 0 | 0 |
| SRR7412332 | 312 | 28860 | 265 | 227 | 1146 | 3360 | 11638 | 22170 | 5786 | 0 | 29891 | 36304 | 19791 | 4703 | 20 | 17 | 43246 | 0 | 0 | 0 | 35 | 0 | 12 | 0 | 0 | 0 | 0 | 0 | 0 | 37 | 19 | 22 | 0 | 0 | 0 | 0 | 0 |
| SRR7412333 | 326 | 14793 | 331 | 297 | 1019 | 2942 | 11530 | 16866 | 4624 | 0 | 31534 | 38212 | 17398 | 4982 | 17 | 40 | 50823 | 0 | 0 | 0 | 27 | 0 | 17 | 0 | 0 | 0 | 0 | 0 | 0 | 55 | 19 | 29 | 0 | 0 | 0 | 0 | 0 |
| SRR7412334 | 496 | 24216 | 398 | 328 | 584 | 2331 | 6695 | 13537 | 4164 | 0 | 35784 | 43311 | 18576 | 4260 | 16 | 24 | 53794 | 0 | 0 | 0 | 36 | 81 | 22 | 0 | 0 | 0 | 0 | 0 | 19 | 35 | 50 | 19 | 0 | 0 | 0 | 0 | 0 |
| SRR7412335 | 977 | 23643 | 240 | 376 | 1295 | 2965 | 14251 | 15717 | 5433 | 30 | 30160 | 41879 | 13183 | 4689 | 16 | 66 | 45916 | 0 | 0 | 0 | 17 | 18 | 19 | 0 | 0 | 0 | 0 | 0 | 0 | 53 | 55 | 17 | 0 | 0 | 0 | 0 | 0 |
| SRR7412336 | 525 | 9763 | 947 | 469 | 1180 | 4970 | 15686 | 17964 | 4611 | 0 | 46985 | 29666 | 26816 | 8033 | 13 | 88 | 40791 | 0 | 0 | 0 | 0 | 0 | 0 | 0 | 0 | 0 | 0 | 0 | 0 | 22 | 0 | 0 | 0 | 0 | 0 | 0 | 0 |
| SRR7412337 | 158 | 15171 | 346 | 444 | 2591 | 5841 | 27229 | 23321 | 5889 | 30 | 29311 | 48567 | 10466 | 4872 | 30 | 14 | 43540 | 0 | 0 | 0 | 12 | 0 | 0 | 0 | 0 | 0 | 0 | 0 | 0 | 17 | 0 | 10 | 0 | 0 | 0 | 0 | 0 |
| SRR7412338 | 211 | 11766 | 271 | 135 | 2534 | 4674 | 34536 | 20360 | 13485 | 0 | 23886 | 23168 | 13493 | 7584 | 19 | 35 | 25045 | 0 | 0 | 0 | 0 | 0 | 0 | 0 | 0 | 0 | 0 | 0 | 0 | 34 | 0 | 0 | 0 | 0 | 0 | 0 | 0 |
| SRR7412339 | 432 | 8053 | 567 | 1295 | 1639 | 3427 | 24708 | 23202 | 6706 | 13 | 45104 | 36466 | 18114 | 8268 | 26 | 117 | 33537 | 0 | 0 | 0 | 10 | 0 | 0 | 0 | 0 | 0 | 0 | 0 | 0 | 22 | 0 | 0 | 0 | 0 | 0 | 0 | 0 |
| SRR7412340 | 96 | 6664 | 613 | 1155 | 1610 | 3248 | 21978 | 13414 | 3768 | 48 | 66081 | 41366 | 18318 | 8146 | 16 | 25 | 19898 | 0 | 0 | 0 | 30 | 0 | 0 | 0 | 0 | 0 | 0 | 0 | 0 | 15 | 0 | 0 | 0 | 0 | 0 | 0 | 0 |
| SRR7412341 | 169 | 6351 | 1349 | 1602 | 695 | 2647 | 13857 | 8933 | 3541 | 47 | 68663 | 35405 | 21500 | 8379 | 14 | 0 | 17195 | 0 | 0 | 0 | 19 | 0 | 0 | 0 | 0 | 0 | 0 | 0 | 0 | 15 | 0 | 0 | 0 | 0 | 0 | 0 | 0 |
| SRR7412342 | 414 | 9977 | 354 | 60 | 1331 | 2394 | 13113 | 17820 | 7003 | 0 | 21530 | 23341 | 15416 | 4334 | 59 | 26 | 51615 | 0 | 0 | 0 | 12 | 0 | 17 | 0 | 0 | 0 | 0 | 0 | 0 | 55 | 36 | 22 | 0 | 0 | 0 | 0 | 0 |
| SRR7412343 | 818 | 21530 | 576 | 442 | 1298 | 2373 | 15330 | 24754 | 14912 | 33 | 22139 | 27062 | 9329 | 4761 | 15 | 22 | 53594 | 0 | 0 | 0 | 0 | 0 | 0 | 0 | 0 | 0 | 0 | 0 | 0 | 22 | 0 | 0 | 0 | 0 | 0 | 0 | 0 |
| SRR7412344 | 90 | 5711 | 711 | 487 | 839 | 4300 | 14588 | 16468 | 4522 | 0 | 53110 | 22002 | 30190 | 10313 | 0 | 25 | 20540 | 0 | 0 | 0 | 15 | 0 | 0 | 0 | 0 | 0 | 0 | 0 | 0 | 21 | 0 | 24 | 0 | 0 | 0 | 0 | 0 |
| SRR7412345 | 820 | 10510 | 606 | 215 | 1303 | 2336 | 14870 | 24395 | 10158 | 39 | 35861 | 32660 | 23106 | 5029 | 21 | 12 | 42246 | 0 | 0 | 0 | 32 | 0 | 14 | 0 | 0 | 0 | 0 | 0 | 0 | 49 | 16 | 32 | 0 | 0 | 0 | 0 | 0 |
| SRR7466734 | 1580 | 27097 | 2194 | 199 | 1444 | 13973 | 17991 | 14293 | 17658 | 51 | 17308 | 38982 | 5555 | 15159 | 223 | 7630 | 15649 | 0 | 0 | 0 | 833 | 142 | 182 | 0 | 15 | 0 | 0 | 0 | 15 | 102 | 37 | 43 | 0 | 0 | 0 | 0 | 0 |
| SRR7466735 | 1488 | 27108 | 2176 | 236 | 1526 | 14247 | 18051 | 14603 | 17925 | 48 | 17153 | 39146 | 5692 | 15083 | 237 | 7732 | 14637 | 0 | 0 | 0 | 860 | 145 | 173 | 0 | 18 | 0 | 0 | 0 | 16 | 105 | 33 | 31 | 0 | 0 | 0 | 0 | 0 |
| SRR7466736 | 1345 | 19660 | 7295 | 749 | 1288 | 14326 | 19295 | 16700 | 18005 | 99 | 19597 | 37407 | 5293 | 24100 | 148 | 10093 | 13349 | 0 | 0 | 0 | 992 | 171 | 135 | 0 | 0 | 75 | 0 | 0 | 24 | 113 | 78 | 42 | 0 | 0 | 11 | 12 | 0 |
| SRR7466737 | 1340 | 19820 | 7105 | 684 | 1366 | 14243 | 19417 | 16969 | 17745 | 74 | 19647 | 36709 | 5249 | 24394 | 149 | 10307 | 12764 | 0 | 0 | 0 | 990 | 160 | 135 | 0 | 0 | 74 | 0 | 0 | 30 | 116 | 75 | 44 | 0 | 0 | 0 | 0 | 0 |
| SRR7466738 | 1367 | 20057 | 6962 | 641 | 1244 | 14263 | 19002 | 16756 | 17582 | 77 | 19564 | 36648 | 5178 | 23906 | 144 | 9967 | 13131 | 0 | 0 | 0 | 985 | 151 | 122 | 0 | 13 | 81 | 0 | 0 | 32 | 106 | 80 | 43 | 0 | 0 | 10 | 0 | 0 |
| SRR7466739 | 1271 | 20103 | 7105 | 658 | 1337 | 14218 | 19380 | 17015 | 18246 | 79 | 19595 | 37275 | 5228 | 24486 | 154 | 10217 | 13079 | 0 | 0 | 0 | 969 | 171 | 118 | 0 | 0 | 79 | 0 | 0 | 28 | 109 | 72 | 41 | 0 | 0 | 0 | 10 | 0 |
| SRR7466740 | 1523 | 25611 | 7970 | 379 | 1216 | 11242 | 15736 | 17524 | 15042 | 55 | 19898 | 41539 | 5871 | 18041 | 137 | 8160 | 16314 | 0 | 0 | 0 | 1047 | 168 | 171 | 0 | 12 | 82 | 0 | 0 | 29 | 133 | 54 | 45 | 0 | 0 | 14 | 11 | 0 |
| SRR7466741 | 1581 | 25806 | 8268 | 339 | 1298 | 11358 | 15424 | 17394 | 14950 | 81 | 19945 | 41369 | 6010 | 18549 | 141 | 8087 | 15485 | 0 | 0 | 0 | 1058 | 177 | 180 | 0 | 19 | 79 | 0 | 0 | 28 | 138 | 63 | 60 | 0 | 0 | 17 | 11 | 0 |
| SRR7466742 | 1580 | 25791 | 8240 | 332 | 1221 | 11467 | 15800 | 17592 | 15079 | 61 | 20057 | 41234 | 5799 | 18354 | 122 | 8024 | 16455 | 0 | 0 | 0 | 1111 | 174 | 177 | 0 | 15 | 73 | 0 | 0 | 32 | 136 | 60 | 55 | 0 | 0 | 12 | 0 | 0 |
| SRR7466743 | 1534 | 26025 | 8261 | 337 | 1228 | 11356 | 15186 | 17756 | 15420 | 74 | 20022 | 41105 | 5807 | 18435 | 132 | 8198 | 15621 | 0 | 0 | 0 | 1008 | 180 | 166 | 0 | 14 | 84 | 0 | 0 | 24 | 139 | 57 | 61 | 0 | 0 | 16 | 11 | 0 |
| SRR7466744 | 883 | 33381 | 4695 | 520 | 1260 | 9004 | 13961 | 14467 | 12940 | 87 | 24087 | 46278 | 5545 | 13901 | 134 | 4292 | 14795 | 0 | 0 | 0 | 1225 | 188 | 185 | 0 | 14 | 55 | 0 | 0 | 25 | 132 | 46 | 31 | 0 | 0 | 0 | 15 | 0 |
| SRR7466745 | 758 | 32803 | 5007 | 604 | 1252 | 8993 | 13784 | 14051 | 13224 | 84 | 23823 | 46200 | 5332 | 13847 | 151 | 4214 | 15071 | 0 | 0 | 0 | 1213 | 160 | 179 | 0 | 14 | 59 | 0 | 0 | 27 | 116 | 41 | 26 | 0 | 0 | 0 | 12 | 0 |
| SRR7466746 | 739 | 23209 | 4187 | 405 | 1606 | 12849 | 16767 | 14918 | 14106 | 48 | 22414 | 42619 | 6232 | 14242 | 189 | 6127 | 14193 | 0 | 0 | 0 | 879 | 159 | 173 | 0 | 15 | 21 | 0 | 0 | 21 | 102 | 30 | 37 | 0 | 0 | 0 | 0 | 0 |
| SRR7466747 | 896 | 23756 | 4331 | 360 | 1573 | 12644 | 16909 | 14578 | 14129 | 65 | 22184 | 42526 | 6202 | 13893 | 191 | 6085 | 14657 | 0 | 0 | 0 | 863 | 149 | 174 | 0 | 0 | 16 | 0 | 0 | 22 | 89 | 30 | 29 | 0 | 0 | 0 | 0 | 0 |
| SRR7466748 | 717 | 24026 | 4214 | 294 | 1575 | 12909 | 16985 | 14811 | 14044 | 59 | 22258 | 42429 | 6191 | 13814 | 192 | 6365 | 14384 | 0 | 0 | 0 | 884 | 151 | 154 | 0 | 19 | 23 | 0 | 0 | 23 | 92 | 30 | 38 | 0 | 0 | 0 | 0 | 0 |
| SRR7466749 | 764 | 23604 | 4163 | 338 | 1602 | 12645 | 16817 | 14722 | 13770 | 60 | 22368 | 42447 | 6039 | 13801 | 180 | 6063 | 15075 | 0 | 0 | 0 | 860 | 157 | 159 | 0 | 13 | 19 | 0 | 0 | 15 | 97 | 39 | 40 | 0 | 0 | 0 | 0 | 0 |
| SRR7466750 | 1540 | 17745 | 7902 | 555 | 1138 | 11786 | 15555 | 14282 | 13147 | 93 | 21772 | 46046 | 5732 | 17665 | 145 | 6705 | 14502 | 0 | 0 | 0 | 853 | 140 | 140 | 0 | 12 | 93 | 0 | 0 | 25 | 100 | 29 | 31 | 0 | 0 | 18 | 0 | 0 |
| SRR7466751 | 1463 | 17399 | 8177 | 497 | 1228 | 11547 | 15357 | 13876 | 13113 | 87 | 21570 | 45664 | 5807 | 17412 | 159 | 6521 | 15397 | 0 | 0 | 0 | 804 | 133 | 150 | 0 | 0 | 123 | 0 | 0 | 24 | 119 | 31 | 37 | 0 | 0 | 15 | 0 | 0 |
| SRR7466752 | 1626 | 17914 | 8337 | 461 | 1131 | 11520 | 15335 | 14268 | 13345 | 89 | 21576 | 45537 | 5666 | 17207 | 145 | 6679 | 14832 | 0 | 0 | 0 | 776 | 124 | 143 | 0 | 0 | 95 | 0 | 0 | 25 | 112 | 32 | 32 | 0 | 0 | 12 | 0 | 0 |
| SRR7466753 | 1588 | 17365 | 8270 | 518 | 1168 | 11727 | 15008 | 14105 | 13074 | 69 | 21719 | 45799 | 5701 | 17106 | 120 | 6515 | 15545 | 0 | 0 | 0 | 769 | 122 | 142 | 0 | 0 | 100 | 0 | 0 | 28 | 104 | 35 | 34 | 0 | 0 | 18 | 12 | 0 |
| SRR7466754 | 683 | 32171 | 3445 | 383 | 1218 | 14540 | 17409 | 14937 | 16279 | 99 | 18642 | 40475 | 5571 | 19261 | 191 | 7414 | 15037 | 0 | 0 | 0 | 1074 | 190 | 155 | 0 | 0 | 35 | 0 | 0 | 38 | 126 | 57 | 46 | 0 | 0 | 0 | 10 | 0 |
| SRR7466755 | 643 | 33277 | 3645 | 447 | 1244 | 14685 | 17745 | 15499 | 16256 | 87 | 18854 | 40427 | 5673 | 19765 | 217 | 7677 | 14334 | 0 | 0 | 0 | 1104 | 174 | 159 | 0 | 10 | 44 | 0 | 0 | 32 | 116 | 67 | 48 | 0 | 0 | 0 | 11 | 0 |
| SRR7466756 | 632 | 32760 | 3352 | 368 | 1157 | 14370 | 17319 | 15245 | 16354 | 74 | 18584 | 40290 | 5551 | 18786 | 211 | 7825 | 15104 | 0 | 0 | 0 | 1114 | 184 | 142 | 0 | 0 | 30 | 0 | 0 | 43 | 133 | 55 | 38 | 0 | 0 | 0 | 0 | 0 |
| SRR7466757 | 692 | 32630 | 3475 | 360 | 1159 | 14493 | 17661 | 15367 | 16424 | 49 | 18779 | 40270 | 5740 | 19877 | 217 | 7758 | 14177 | 0 | 0 | 0 | 1122 | 188 | 133 | 0 | 13 | 31 | 0 | 0 | 33 | 127 | 61 | 45 | 0 | 0 | 0 | 0 | 0 |
| SRR7466758 | 242 | 41103 | 1545 | 195 | 1180 | 12477 | 14554 | 14552 | 15661 | 47 | 17583 | 35460 | 4264 | 18536 | 108 | 6355 | 14423 | 0 | 0 | 0 | 1047 | 171 | 188 | 0 | 22 | 0 | 0 | 0 | 27 | 103 | 44 | 29 | 0 | 12 | 0 | 0 | 0 |
| SRR7466759 | 256 | 41289 | 1667 | 344 | 1109 | 13065 | 14998 | 15108 | 15638 | 47 | 17575 | 35611 | 4440 | 19485 | 123 | 6354 | 13889 | 0 | 0 | 0 | 1104 | 180 | 172 | 0 | 24 | 0 | 0 | 0 | 25 | 130 | 35 | 36 | 0 | 11 | 0 | 0 | 0 |
| SRR7466760 | 296 | 40462 | 1478 | 334 | 1033 | 13034 | 14781 | 14467 | 15520 | 45 | 17565 | 35158 | 4175 | 18756 | 132 | 6431 | 15027 | 0 | 0 | 0 | 1072 | 177 | 202 | 0 | 13 | 0 | 0 | 0 | 30 | 137 | 37 | 44 | 0 | 0 | 0 | 0 | 0 |
| SRR7466761 | 197 | 40198 | 1275 | 362 | 1006 | 12847 | 14743 | 14832 | 15792 | 59 | 17540 | 35459 | 4414 | 18908 | 109 | 6259 | 14179 | 0 | 0 | 0 | 1021 | 193 | 214 | 0 | 21 | 0 | 0 | 0 | 35 | 116 | 32 | 36 | 0 | 0 | 0 | 0 | 0 |
| SRR7466762 | 1566 | 26828 | 2248 | 270 | 1444 | 14197 | 18072 | 14226 | 17958 | 41 | 17384 | 39068 | 5597 | 14994 | 223 | 7688 | 15550 | 0 | 0 | 0 | 870 | 137 | 186 | 0 | 16 | 13 | 0 | 0 | 17 | 106 | 37 | 37 | 0 | 0 | 0 | 0 | 0 |
| SRR7466763 | 1574 | 27456 | 2218 | 235 | 1501 | 14047 | 17956 | 14596 | 17672 | 45 | 17530 | 39228 | 5663 | 15268 | 252 | 7673 | 14681 | 0 | 0 | 0 | 869 | 147 | 204 | 0 | 18 | 0 | 0 | 0 | 13 | 93 | 28 | 36 | 0 | 0 | 0 | 11 | 0 |
| SRR7466764 | 830 | 33177 | 4718 | 574 | 1283 | 8835 | 13976 | 14090 | 13153 | 90 | 23789 | 46299 | 5501 | 13769 | 168 | 4346 | 15269 | 0 | 0 | 0 | 1196 | 177 | 172 | 0 | 13 | 55 | 0 | 0 | 28 | 121 | 48 | 34 | 0 | 0 | 0 | 0 | 0 |
| SRR7466765 | 782 | 33261 | 4890 | 522 | 1301 | 8942 | 13801 | 14434 | 13140 | 72 | 23880 | 46483 | 5672 | 13726 | 165 | 4312 | 14898 | 0 | 0 | 0 | 1147 | 189 | 172 | 0 | 11 | 51 | 0 | 0 | 24 | 128 | 46 | 30 | 0 | 0 | 0 | 0 | 0 |
| SRR7541164 | 31 | 19164 | 502 | 19 | 50 | 594 | 419 | 1751 | 191285 | 52 | 38240 | 41677 | 23065 | 5541 | 40 | 80 | 51195 | 0 | 0 | 0 | 1033 | 3804 | 95 | 0 | 0 | 0 | 0 | 0 | 14 | 367 | 2893 | 72 | 0 | 0 | 0 | 0 | 0 |
| SRR7541165 | 0 | 3021 | 123 | 93 | 11 | 307 | 212 | 1199 | 38121 | 22 | 10002 | 64657 | 28659 | 14248 | 13 | 88 | 19333 | 0 | 0 | 0 | 48 | 2651 | 89 | 0 | 0 | 0 | 0 | 0 | 0 | 35 | 1147 | 70 | 0 | 0 | 12 | 0 | 0 |
| SRR7541166 | 41 | 9045 | 389 | 67 | 50 | 778 | 657 | 2516 | 133743 | 13 | 18167 | 16075 | 16679 | 8005 | 36 | 578 | 24785 | 0 | 0 | 0 | 218 | 678 | 13 | 0 | 0 | 0 | 0 | 0 | 37 | 662 | 1665 | 59 | 0 | 0 | 0 | 0 | 0 |
| SRR7541167 | 14 | 2568 | 258 | 97 | 114 | 1222 | 934 | 3464 | 68660 | 25 | 14418 | 14930 | 17358 | 13590 | 29 | 537 | 32603 | 0 | 0 | 0 | 93 | 1271 | 45 | 0 | 0 | 0 | 0 | 0 | 16 | 325 | 4857 | 242 | 0 | 16 | 41 | 13 | 0 |
| SRR7541168 | 47 | 15918 | 4736 | 114 | 48 | 656 | 699 | 1555 | 328024 | 250 | 9562 | 24559 | 5524 | 2127 | 48 | 868 | 9043 | 0 | 0 | 0 | 871 | 2115 | 95 | 0 | 0 | 0 | 0 | 0 | 42 | 1061 | 1352 | 115 | 0 | 12 | 41 | 35 | 0 |
| SRR7541169 | 398 | 73131 | 328 | 116 | 37 | 577 | 604 | 3279 | 83372 | 28 | 16530 | 11657 | 27591 | 9428 | 50 | 541 | 17638 | 0 | 0 | 0 | 16 | 122 | 0 | 0 | 0 | 0 | 0 | 0 | 10 | 52 | 541 | 49 | 0 | 0 | 29 | 0 | 0 |
| SRR7541170 | 12 | 3774 | 496 | 105 | 44 | 993 | 708 | 3690 | 86911 | 30 | 19415 | 23616 | 13071 | 6354 | 35 | 1629 | 36370 | 0 | 0 | 0 | 99 | 1205 | 67 | 0 | 0 | 0 | 0 | 0 | 11 | 103 | 2446 | 158 | 0 | 13 | 32 | 0 | 0 |
| SRR7541171 | 19 | 35421 | 2445 | 32 | 117 | 4720 | 1503 | 3338 | 120198 | 13 | 19484 | 23061 | 15823 | 4344 | 43 | 4717 | 53165 | 0 | 0 | 0 | 243 | 111 | 12 | 0 | 0 | 0 | 0 | 0 | 28 | 262 | 186 | 25 | 0 | 0 | 0 | 0 | 0 |
| SRR7541172 | 0 | 7114 | 664 | 42 | 71 | 1432 | 862 | 3055 | 68801 | 15 | 34505 | 32437 | 29150 | 7648 | 49 | 1938 | 29873 | 0 | 0 | 0 | 26 | 60 | 11 | 0 | 0 | 0 | 0 | 0 | 0 | 52 | 195 | 42 | 0 | 11 | 0 | 0 | 0 |
| SRR7541173 | 0 | 4474 | 489 | 28 | 214 | 5790 | 2960 | 18633 | 119076 | 20 | 27503 | 30832 | 21157 | 4886 | 76 | 858 | 51063 | 0 | 0 | 0 | 58 | 793 | 45 | 0 | 0 | 0 | 0 | 0 | 0 | 39 | 592 | 46 | 0 | 0 | 27 | 0 | 0 |
| SRR7541174 | 21 | 22865 | 314 | 107 | 78 | 1049 | 566 | 2278 | 118235 | 47 | 35068 | 44658 | 27656 | 3345 | 28 | 2496 | 21039 | 0 | 0 | 0 | 112 | 1316 | 51 | 0 | 0 | 0 | 0 | 0 | 0 | 101 | 1340 | 79 | 0 | 13 | 21 | 0 | 0 |
| SRR7541175 | 40 | 23491 | 481 | 31 | 104 | 2070 | 963 | 3429 | 178166 | 29 | 20366 | 18933 | 12232 | 6358 | 20 | 5631 | 48949 | 0 | 0 | 0 | 112 | 225 | 27 | 0 | 0 | 0 | 0 | 0 | 13 | 81 | 391 | 51 | 0 | 0 | 0 | 0 | 0 |
| SRR7541176 | 13 | 10231 | 476 | 38 | 51 | 531 | 375 | 1949 | 195176 | 0 | 40639 | 79070 | 21834 | 7274 | 18 | 3056 | 37052 | 0 | 0 | 0 | 77 | 380 | 32 | 0 | 0 | 0 | 0 | 0 | 0 | 55 | 704 | 38 | 0 | 0 | 39 | 0 | 0 |
| SRR7541177 | 24 | 20418 | 1005 | 0 | 92 | 3123 | 791 | 3487 | 103167 | 58 | 20885 | 29883 | 22518 | 7718 | 39 | 4798 | 56397 | 0 | 0 | 0 | 143 | 1652 | 109 | 0 | 0 | 0 | 0 | 0 | 0 | 62 | 1121 | 74 | 0 | 19 | 56 | 0 | 0 |
| SRR7541178 | 249 | 19397 | 1630 | 49 | 466 | 10677 | 4318 | 15848 | 144873 | 102 | 46441 | 33548 | 21943 | 5389 | 67 | 379 | 36962 | 0 | 0 | 0 | 113 | 324 | 28 | 0 | 0 | 0 | 0 | 0 | 21 | 108 | 237 | 48 | 0 | 10 | 0 | 0 | 0 |
| SRR7541179 | 0 | 10671 | 1012 | 16 | 40 | 795 | 799 | 7467 | 86868 | 24 | 22171 | 17234 | 28190 | 2933 | 31 | 916 | 33368 | 0 | 0 | 0 | 94 | 281 | 26 | 0 | 0 | 0 | 0 | 0 | 29 | 210 | 1128 | 113 | 0 | 11 | 14 | 0 | 0 |
| SRR7541180 | 15 | 5887 | 674 | 43 | 27 | 415 | 332 | 1342 | 171524 | 50 | 27135 | 53573 | 19908 | 9492 | 37 | 2996 | 24185 | 0 | 0 | 0 | 1701 | 6421 | 367 | 0 | 0 | 0 | 0 | 0 | 78 | 980 | 3293 | 299 | 0 | 19 | 83 | 13 | 0 |
| SRR7541181 | 0 | 16758 | 3427 | 10 | 27 | 385 | 314 | 2896 | 98283 | 70 | 20648 | 31574 | 13473 | 8996 | 36 | 1511 | 19211 | 0 | 0 | 0 | 31 | 225 | 15 | 0 | 0 | 0 | 0 | 0 | 0 | 32 | 164 | 25 | 0 | 0 | 13 | 0 | 0 |
| SRR7541182 | 16 | 21841 | 827 | 41 | 61 | 669 | 513 | 2286 | 148798 | 19 | 18967 | 18471 | 15518 | 6432 | 15 | 2644 | 46571 | 0 | 0 | 0 | 394 | 1739 | 124 | 0 | 0 | 0 | 0 | 0 | 17 | 183 | 1248 | 110 | 0 | 12 | 19 | 0 | 0 |
| SRR7541183 | 0 | 27747 | 412 | 26 | 57 | 879 | 719 | 2647 | 177953 | 20 | 18813 | 20401 | 24408 | 9527 | 35 | 330 | 17215 | 0 | 0 | 0 | 67 | 581 | 35 | 0 | 0 | 0 | 0 | 0 | 12 | 131 | 1275 | 119 | 0 | 12 | 15 | 0 | 0 |
| SRR7541184 | 0 | 7975 | 603 | 25 | 109 | 2425 | 1760 | 11957 | 85208 | 18 | 28919 | 22127 | 34727 | 8176 | 104 | 1072 | 28069 | 0 | 0 | 0 | 45 | 4571 | 132 | 0 | 0 | 0 | 0 | 0 | 0 | 78 | 4758 | 279 | 0 | 26 | 85 | 13 | 0 |
| SRR7541185 | 12 | 8179 | 733 | 35 | 66 | 873 | 882 | 4288 | 45568 | 31 | 21365 | 16664 | 27026 | 7416 | 763 | 78 | 30010 | 0 | 0 | 0 | 79 | 1179 | 96 | 0 | 0 | 0 | 0 | 0 | 0 | 65 | 912 | 132 | 0 | 19 | 0 | 0 | 0 |
| SRR7541186 | 0 | 18488 | 385 | 18 | 30 | 964 | 524 | 3415 | 140655 | 22 | 33917 | 25061 | 27095 | 7727 | 16 | 164 | 24701 | 0 | 0 | 0 | 68 | 513 | 19 | 0 | 0 | 0 | 0 | 0 | 0 | 196 | 1323 | 64 | 0 | 14 | 22 | 0 | 0 |
| SRR7541187 | 205 | 171954 | 870 | 22 | 109 | 2535 | 1495 | 3581 | 125731 | 14 | 22082 | 16108 | 16747 | 8404 | 27 | 326 | 37227 | 0 | 0 | 0 | 34 | 56 | 0 | 0 | 0 | 0 | 0 | 0 | 11 | 65 | 149 | 27 | 0 | 0 | 15 | 0 | 0 |
| SRR7541188 | 32 | 17177 | 818 | 11 | 97 | 2473 | 1139 | 5978 | 143820 | 0 | 21128 | 24897 | 18655 | 5594 | 0 | 11580 | 50508 | 0 | 0 | 0 | 78 | 200 | 22 | 0 | 0 | 0 | 0 | 0 | 0 | 65 | 305 | 40 | 0 | 0 | 0 | 0 | 0 |
| SRR7541189 | 0 | 2771 | 405 | 0 | 121 | 3091 | 1767 | 17322 | 101702 | 63 | 23887 | 15901 | 23963 | 11525 | 120 | 33 | 19198 | 0 | 0 | 0 | 150 | 353 | 35 | 0 | 0 | 0 | 0 | 0 | 0 | 40 | 154 | 25 | 0 | 0 | 0 | 0 | 0 |
| SRR7541190 | 54 | 15172 | 1411 | 18 | 139 | 5005 | 1615 | 8071 | 37748 | 41 | 31817 | 27530 | 19592 | 6354 | 33 | 0 | 27583 | 0 | 0 | 0 | 29 | 1033 | 37 | 0 | 0 | 0 | 0 | 0 | 0 | 58 | 507 | 66 | 0 | 18 | 40 | 0 | 0 |
| SRR7541191 | 0 | 10492 | 702 | 28 | 62 | 1168 | 624 | 3309 | 113024 | 25 | 21918 | 35425 | 19596 | 3037 | 64 | 824 | 38747 | 0 | 0 | 0 | 48 | 1039 | 51 | 0 | 0 | 0 | 0 | 0 | 0 | 49 | 1862 | 114 | 0 | 0 | 36 | 0 | 0 |
| SRR7541192 | 31 | 21680 | 1303 | 22 | 219 | 1909 | 1192 | 5232 | 169407 | 27 | 24540 | 23946 | 13040 | 6437 | 0 | 14239 | 64513 | 0 | 0 | 0 | 94 | 75 | 23 | 0 | 0 | 0 | 0 | 0 | 25 | 135 | 260 | 51 | 0 | 12 | 11 | 0 | 0 |
| SRR7541193 | 74 | 24637 | 1554 | 29 | 162 | 1750 | 1181 | 7540 | 138681 | 25 | 39087 | 28537 | 28688 | 7870 | 81 | 795 | 66284 | 0 | 0 | 0 | 25 | 90 | 0 | 0 | 0 | 0 | 0 | 0 | 0 | 24 | 164 | 20 | 0 | 0 | 0 | 0 | 0 |
| SRR7547853 | 643 | 2793 | 4132 | 3119 | 16577 | 19521 | 177508 | 79914 | 25135 | 3862 | 11175 | 10831 | 19975 | 10398 | 2357 | 7089 | 28920 | 0 | 50 | 71 | 232 | 6499 | 1349 | 256 | 103 | 163 | 26 | 77 | 232 | 170 | 525 | 73 | 0 | 0 | 39 | 322 | 0 |
| SRR7547854 | 1520 | 5159 | 7336 | 4851 | 10325 | 4635 | 99424 | 47804 | 96017 | 2692 | 20531 | 24183 | 17661 | 6702 | 3434 | 10548 | 31705 | 0 | 61 | 23 | 76 | 918 | 356 | 47 | 26 | 21 | 17 | 86 | 102 | 75 | 204 | 29 | 0 | 0 | 0 | 64 | 0 |
| SRR7547855 | 1109 | 4859 | 4761 | 1875 | 5938 | 3090 | 60625 | 38235 | 19217 | 2042 | 15411 | 13381 | 13014 | 15358 | 223 | 3889 | 44228 | 0 | 22 | 68 | 1326 | 2588 | 2492 | 223 | 94 | 71 | 11 | 28 | 167 | 632 | 339 | 144 | 0 | 0 | 28 | 182 | 0 |
| SRR7547856 | 761 | 3354 | 6284 | 4604 | 10260 | 11156 | 128692 | 50184 | 15722 | 4725 | 16661 | 17165 | 15142 | 33746 | 3369 | 10514 | 17666 | 0 | 67 | 62 | 176 | 2150 | 1055 | 157 | 50 | 48 | 25 | 59 | 155 | 94 | 304 | 38 | 0 | 0 | 0 | 103 | 0 |
| SRR7547857 | 800 | 3870 | 9941 | 3752 | 18998 | 18873 | 232830 | 81133 | 24568 | 4361 | 12244 | 8073 | 9506 | 8686 | 8271 | 2194 | 19258 | 0 | 78 | 223 | 445 | 28888 | 6430 | 752 | 183 | 87 | 0 | 49 | 293 | 141 | 1106 | 270 | 0 | 0 | 24 | 349 | 0 |
| SRR7547858 | 857 | 3604 | 6951 | 3522 | 12575 | 13151 | 130899 | 60520 | 56221 | 4597 | 19105 | 16 | 15724 | 10444 | 607 | 8717 | 41663 | 0 | 16 | 38 | 109 | 4655 | 2968 | 348 | 120 | 73 | 0 | 27 | 100 | 75 | 329 | 185 | 0 | 0 | 37 | 200 | 0 |
| SRR7547859 | 459 | 2563 | 6985 | 2311 | 19103 | 15321 | 280304 | 100862 | 19255 | 3787 | 10739 | 6275 | 7967 | 8849 | 630 | 2842 | 24008 | 0 | 26 | 109 | 1671 | 3255 | 7631 | 965 | 236 | 145 | 16 | 38 | 228 | 1262 | 427 | 536 | 25 | 0 | 52 | 291 | 0 |
| SRR7547860 | 1296 | 5620 | 7514 | 2467 | 43335 | 13757 | 220784 | 90458 | 37389 | 4298 | 11476 | 11620 | 13155 | 8145 | 2587 | 5783 | 24917 | 0 | 39 | 42 | 138 | 2162 | 907 | 160 | 68 | 40 | 19 | 42 | 102 | 76 | 185 | 43 | 0 | 0 | 0 | 107 | 0 |
| SRR7547861 | 1400 | 7599 | 7884 | 3766 | 17185 | 16244 | 267534 | 71276 | 25486 | 4735 | 15630 | 12027 | 11001 | 9655 | 2721 | 13694 | 22586 | 0 | 39 | 31 | 140 | 744 | 600 | 119 | 57 | 20 | 24 | 53 | 109 | 96 | 144 | 53 | 0 | 0 | 12 | 93 | 0 |
| SRR7547862 | 369 | 1914 | 2177 | 2063 | 19612 | 17956 | 158875 | 101728 | 33865 | 4004 | 15655 | 7262 | 16394 | 13540 | 3184 | 1131 | 23724 | 0 | 79 | 157 | 835 | 5144 | 1971 | 136 | 38 | 31 | 17 | 131 | 630 | 616 | 847 | 153 | 0 | 0 | 16 | 84 | 0 |
| SRR7547863 | 649 | 3833 | 4716 | 2320 | 15799 | 11305 | 169957 | 80577 | 27349 | 5454 | 5718 | 8363 | 6385 | 8253 | 1272 | 10334 | 18450 | 0 | 659 | 358 | 1061 | 10224 | 3675 | 660 | 105 | 139 | 29 | 503 | 680 | 565 | 880 | 183 | 11 | 0 | 21 | 331 | 0 |
| SRR7547864 | 414 | 2370 | 21444 | 910 | 2196 | 1707 | 23146 | 20188 | 37911 | 4531 | 16554 | 23 | 17196 | 9728 | 307 | 4758 | 66284 | 0 | 10 | 21 | 144 | 955 | 769 | 59 | 25 | 23 | 0 | 10 | 40 | 50 | 141 | 49 | 0 | 0 | 0 | 60 | 0 |
| SRR7547865 | 368 | 1698 | 12506 | 2532 | 6790 | 6688 | 57115 | 31767 | 19855 | 2214 | 23604 | 10656 | 55688 | 21739 | 2067 | 2706 | 50495 | 0 | 19 | 36 | 204 | 3108 | 3536 | 546 | 139 | 78 | 10 | 39 | 125 | 163 | 437 | 339 | 0 | 0 | 116 | 211 | 14 |
| SRR7547866 | 372 | 2344 | 2685 | 1629 | 18724 | 14782 | 120648 | 100506 | 15028 | 6135 | 14924 | 8319 | 15567 | 18718 | 2031 | 7653 | 35025 | 0 | 17 | 16 | 96 | 1712 | 1219 | 241 | 63 | 83 | 15 | 39 | 124 | 155 | 296 | 97 | 0 | 0 | 37 | 164 | 0 |
| SRR7547867 | 871 | 3001 | 4645 | 1415 | 16035 | 21476 | 182596 | 59516 | 49326 | 2417 | 28069 | 13443 | 18528 | 11039 | 1159 | 11266 | 20856 | 0 | 84 | 69 | 377 | 2933 | 1828 | 161 | 69 | 49 | 0 | 73 | 82 | 58 | 133 | 46 | 0 | 0 | 14 | 112 | 0 |
| SRR7547868 | 1119 | 5203 | 7029 | 1592 | 14330 | 10102 | 114525 | 77837 | 16299 | 1791 | 10600 | 17991 | 10114 | 7650 | 416 | 8965 | 33370 | 0 | 22 | 57 | 471 | 3795 | 3612 | 983 | 398 | 123 | 24 | 36 | 152 | 201 | 234 | 119 | 0 | 0 | 105 | 249 | 0 |
| SRR7547869 | 1035 | 5654 | 6508 | 4114 | 17699 | 17013 | 202192 | 69846 | 32281 | 1912 | 14727 | 9072 | 11153 | 9280 | 3087 | 25799 | 28713 | 0 | 21 | 39 | 374 | 2429 | 1411 | 215 | 94 | 38 | 25 | 56 | 222 | 283 | 487 | 142 | 0 | 0 | 44 | 124 | 0 |
| SRR7547870 | 1218 | 5252 | 7089 | 3502 | 12955 | 13128 | 186428 | 79130 | 53564 | 2117 | 13816 | 10154 | 12412 | 13117 | 1880 | 4905 | 27607 | 0 | 66 | 70 | 247 | 4928 | 1994 | 362 | 131 | 122 | 23 | 78 | 186 | 138 | 318 | 119 | 0 | 0 | 39 | 253 | 0 |
| SRR7547871 | 581 | 2540 | 8815 | 4133 | 19092 | 13466 | 179979 | 89410 | 34307 | 1226 | 13309 | 8476 | 12478 | 5204 | 534 | 11575 | 36386 | 0 | 27 | 81 | 684 | 6943 | 3134 | 196 | 69 | 80 | 13 | 42 | 248 | 420 | 860 | 304 | 0 | 0 | 28 | 182 | 0 |
| SRR7547872 | 746 | 3582 | 10125 | 2055 | 19605 | 21189 | 236201 | 87161 | 15362 | 8662 | 17272 | 11854 | 12303 | 9740 | 773 | 6019 | 27553 | 0 | 32 | 71 | 122 | 5492 | 1697 | 186 | 54 | 60 | 0 | 49 | 253 | 95 | 597 | 116 | 0 | 0 | 14 | 134 | 0 |
| SRR7547873 | 499 | 2191 | 3520 | 5079 | 6576 | 5586 | 60256 | 35556 | 120205 | 7025 | 17238 | 9666 | 19159 | 11459 | 3634 | 19319 | 12086 | 13 | 76 | 93 | 631 | 9945 | 8716 | 1861 | 350 | 382 | 20 | 64 | 176 | 267 | 380 | 337 | 21 | 0 | 106 | 874 | 37 |
| SRR7547874 | 1004 | 4635 | 7466 | 2606 | 11542 | 14413 | 141492 | 55526 | 41189 | 2410 | 23709 | 14560 | 12871 | 11133 | 649 | 16342 | 41851 | 0 | 21 | 100 | 278 | 5704 | 1475 | 133 | 56 | 49 | 17 | 39 | 228 | 110 | 475 | 59 | 0 | 0 | 0 | 127 | 0 |
| SRR7547875 | 673 | 2775 | 7936 | 3682 | 17045 | 17625 | 194140 | 77693 | 29478 | 1661 | 15738 | 12427 | 14280 | 18328 | 906 | 1835 | 32735 | 0 | 55 | 131 | 291 | 3838 | 1336 | 120 | 34 | 49 | 0 | 50 | 230 | 119 | 379 | 74 | 0 | 0 | 0 | 94 | 0 |
| SRR7547876 | 398 | 2068 | 6259 | 862 | 9302 | 9304 | 100585 | 65319 | 26026 | 6061 | 16372 | 11 | 16268 | 12915 | 407 | 1586 | 29971 | 0 | 12 | 20 | 108 | 583 | 340 | 81 | 46 | 45 | 12 | 25 | 80 | 83 | 160 | 21 | 0 | 0 | 14 | 87 | 0 |
| SRR7547877 | 763 | 2934 | 6789 | 2044 | 15588 | 17051 | 148394 | 69144 | 42095 | 2574 | 22479 | 13928 | 17277 | 10257 | 1299 | 5052 | 31288 | 0 | 71 | 39 | 57 | 1543 | 364 | 62 | 28 | 17 | 22 | 96 | 136 | 52 | 275 | 25 | 0 | 0 | 0 | 57 | 0 |
| SRR7547878 | 555 | 2280 | 14011 | 1895 | 10949 | 8977 | 188811 | 76710 | 21113 | 16403 | 13848 | 6731 | 10008 | 10457 | 2148 | 3317 | 20328 | 0 | 31 | 36 | 151 | 2377 | 1644 | 226 | 69 | 94 | 10 | 27 | 67 | 70 | 255 | 108 | 0 | 0 | 19 | 215 | 0 |
| SRR7547879 | 477 | 4988 | 22514 | 1499 | 5904 | 6563 | 106300 | 57133 | 56422 | 9038 | 22699 | 14333 | 10793 | 10851 | 1668 | 3943 | 20585 | 0 | 25 | 16 | 65 | 1136 | 697 | 76 | 30 | 31 | 0 | 45 | 78 | 93 | 183 | 70 | 0 | 0 | 15 | 77 | 0 |
| SRR7547880 | 437 | 2953 | 27286 | 561 | 4688 | 3297 | 79005 | 52091 | 13097 | 6761 | 19866 | 17 | 15419 | 14332 | 330 | 4427 | 37654 | 0 | 0 | 25 | 836 | 745 | 895 | 182 | 73 | 137 | 0 | 16 | 64 | 318 | 109 | 68 | 0 | 0 | 32 | 256 | 0 |
| SRR7547881 | 511 | 2727 | 17772 | 1578 | 10620 | 8430 | 167579 | 85563 | 25595 | 6725 | 13669 | 12766 | 12401 | 16141 | 668 | 3650 | 25554 | 0 | 11 | 21 | 106 | 1868 | 1064 | 173 | 80 | 143 | 0 | 14 | 40 | 59 | 141 | 63 | 0 | 0 | 20 | 321 | 0 |
| SRR7547882 | 1551 | 3538 | 4411 | 2254 | 488 | 318 | 4317 | 1959 | 5283 | 2914 | 4463 | 18148 | 7697 | 4803 | 62 | 1635 | 57072 | 0 | 0 | 0 | 57 | 441 | 213 | 19 | 0 | 0 | 0 | 0 | 19 | 17 | 24 | 0 | 0 | 0 | 0 | 19 | 0 |
| SRR7547883 | 1746 | 7489 | 5862 | 827 | 809 | 855 | 8888 | 4113 | 6193 | 1035 | 10470 | 32962 | 15614 | 19240 | 240 | 1535 | 35340 | 0 | 0 | 13 | 177 | 395 | 304 | 30 | 10 | 11 | 0 | 0 | 10 | 19 | 22 | 0 | 0 | 0 | 0 | 20 | 0 |
| SRR7547884 | 1057 | 2984 | 6196 | 3047 | 380 | 384 | 4243 | 1970 | 6153 | 2995 | 3477 | 17555 | 6234 | 5529 | 48 | 2654 | 42812 | 0 | 15 | 26 | 360 | 1287 | 1292 | 44 | 16 | 0 | 0 | 15 | 54 | 120 | 86 | 43 | 0 | 0 | 0 | 30 | 0 |
| SRR7547885 | 534 | 2056 | 14439 | 1707 | 12629 | 10261 | 156499 | 73664 | 26516 | 14452 | 24161 | 10 | 45 | 7234 | 4796 | 6705 | 37106 | 0 | 166 | 90 | 336 | 7092 | 2997 | 344 | 112 | 123 | 0 | 93 | 126 | 120 | 568 | 131 | 0 | 0 | 22 | 248 | 0 |
| SRR7547886 | 433 | 2305 | 18996 | 2854 | 4436 | 7789 | 130086 | 64597 | 8051 | 10757 | 17035 | 9413 | 77 | 6919 | 1593 | 3330 | 14506 | 0 | 10 | 15 | 129 | 2058 | 944 | 132 | 58 | 54 | 0 | 12 | 42 | 74 | 138 | 75 | 0 | 0 | 22 | 171 | 0 |
| SRR7547887 | 519 | 2752 | 19202 | 1328 | 9346 | 10571 | 157195 | 64316 | 37325 | 7885 | 9195 | 11701 | 37 | 5880 | 1498 | 4002 | 36582 | 0 | 27 | 45 | 195 | 2849 | 1750 | 233 | 71 | 75 | 0 | 30 | 95 | 89 | 265 | 103 | 0 | 0 | 20 | 202 | 0 |
| SRR7547888 | 396 | 2083 | 19235 | 2272 | 7906 | 7643 | 146798 | 59240 | 24860 | 6009 | 19343 | 8048 | 54 | 8874 | 2384 | 472 | 17613 | 0 | 24 | 31 | 161 | 3527 | 2034 | 583 | 130 | 189 | 0 | 24 | 55 | 65 | 195 | 118 | 0 | 0 | 36 | 446 | 15 |
| SRR7547889 | 755 | 2902 | 13246 | 2694 | 9614 | 9112 | 140388 | 47137 | 47983 | 7600 | 25888 | 14088 | 54 | 13193 | 3850 | 7550 | 30700 | 0 | 14 | 17 | 41 | 395 | 247 | 50 | 26 | 26 | 12 | 21 | 54 | 41 | 82 | 15 | 0 | 0 | 0 | 64 | 0 |
| SRR7547890 | 439 | 2154 | 21173 | 1974 | 5747 | 5881 | 111505 | 49596 | 76189 | 11298 | 17486 | 9699 | 43 | 14512 | 2084 | 774 | 24182 | 0 | 0 | 11 | 96 | 1033 | 491 | 75 | 28 | 25 | 11 | 16 | 64 | 114 | 183 | 62 | 0 | 0 | 18 | 97 | 0 |
| SRR7547891 | 570 | 2881 | 13409 | 952 | 8084 | 10341 | 128506 | 66823 | 25070 | 8270 | 17255 | 17421 | 12486 | 9637 | 1138 | 1722 | 33538 | 0 | 0 | 15 | 196 | 765 | 540 | 77 | 32 | 41 | 0 | 0 | 37 | 95 | 104 | 26 | 0 | 0 | 0 | 94 | 0 |
| SRR7547892 | 405 | 2268 | 15484 | 2221 | 5557 | 4070 | 80639 | 31329 | 111401 | 9435 | 14615 | 15087 | 10376 | 7254 | 218 | 5691 | 17195 | 0 | 35 | 22 | 103 | 756 | 1002 | 131 | 68 | 62 | 0 | 32 | 53 | 63 | 112 | 48 | 0 | 0 | 42 | 110 | 0 |
| SRR7547893 | 698 | 2815 | 16625 | 2293 | 15644 | 16346 | 292138 | 123180 | 9715 | 8504 | 10370 | 8248 | 43 | 3574 | 727 | 4490 | 13134 | 0 | 26 | 16 | 99 | 1153 | 507 | 85 | 39 | 31 | 0 | 28 | 41 | 57 | 105 | 33 | 0 | 0 | 0 | 100 | 0 |
| SRR7547894 | 363 | 2008 | 11435 | 2478 | 6882 | 7312 | 102141 | 43041 | 67493 | 7070 | 11178 | 13783 | 39 | 9685 | 534 | 1036 | 27554 | 0 | 40 | 120 | 717 | 19297 | 9958 | 1415 | 264 | 300 | 0 | 48 | 267 | 610 | 960 | 752 | 14 | 0 | 92 | 823 | 22 |
| SRR7547895 | 614 | 2470 | 20389 | 947 | 10686 | 17206 | 220577 | 64653 | 15536 | 6526 | 19857 | 9990 | 71 | 7449 | 2159 | 1036 | 24776 | 0 | 11 | 13 | 103 | 2288 | 717 | 125 | 73 | 59 | 12 | 22 | 51 | 76 | 185 | 65 | 0 | 0 | 17 | 177 | 0 |
| SRR7547896 | 394 | 2186 | 5048 | 1024 | 6826 | 5383 | 79814 | 61037 | 16937 | 6928 | 18263 | 9829 | 22654 | 14417 | 370 | 1391 | 33971 | 0 | 17 | 18 | 107 | 226 | 309 | 69 | 29 | 27 | 19 | 30 | 76 | 95 | 116 | 22 | 0 | 0 | 16 | 74 | 0 |

Supp Table S8.1.Columns_B_to_AL

| | | | | | | | | | | | | | | | | | | | | | | | | | | | | | | | | | | | | | |
|---|---|---|---|---|---|---|---|---|---|---|---|---|---|---|---|---|---|---|---|---|---|---|---|---|---|---|---|---|---|---|---|---|---|---|---|---|---|
| SRR7547897 | 705 | 2985 | 16180 | 1525 | 4428 | 2887 | 61792 | 58168 | 6889 | 5279 | 12908 | 16371 | 19938 | 7332 | 2790 | 3482 | 66665 | 0 | 0 | 0 | 67 | 246 | 193 | 29 | 21 | 11 | 0 | 11 | 33 | 65 | 75 | 24 | 0 | 0 | 12 | 51 | 0 |
| SRR7547898 | 640 | 3321 | 7805 | 1404 | 11939 | 10713 | 126656 | 78209 | 24827 | 7883 | 18021 | 12794 | 18999 | 12653 | 3817 | 1552 | 29235 | 0 | 0 | 0 | 75 | 296 | 189 | 35 | 15 | 11 | 25 | 32 | 102 | 123 | 236 | 26 | 0 | 0 | 0 | 38 | 0 |
| SRR7547899 | 652 | 3414 | 21433 | 3042 | 5402 | 5803 | 80138 | 47032 | 39737 | 5292 | 18427 | 24 | 217 | 8486 | 2090 | 13377 | 54560 | 0 | 0 | 10 | 107 | 1580 | 865 | 206 | 126 | 62 | 0 | 0 | 33 | 67 | 155 | 72 | 0 | 0 | 40 | 205 | 0 |
| SRR7547900 | 708 | 2954 | 21243 | 1307 | 10065 | 10436 | 158308 | 86865 | 17033 | 4520 | 17146 | 12422 | 10337 | 9785 | 1252 | 1585 | 33469 | 0 | 42 | 33 | 109 | 811 | 235 | 67 | 31 | 53 | 0 | 38 | 70 | 54 | 85 | 12 | 0 | 0 | 0 | 117 | 0 |
| SRR7547901 | 476 | 2716 | 8190 | 1204 | 14049 | 8411 | 155451 | 114569 | 8773 | 8305 | 7082 | 9478 | 85 | 7840 | 2825 | 1592 | 26888 | 0 | 0 | 20 | 307 | 1184 | 2815 | 555 | 211 | 136 | 18 | 21 | 73 | 179 | 221 | 174 | 0 | 0 | 104 | 326 | 0 |
| SRR7547902 | 362 | 2623 | 8287 | 1287 | 26191 | 14409 | 173871 | 97003 | 18596 | 3843 | 11031 | 12963 | 10110 | 4439 | 2167 | 5349 | 27453 | 0 | 555 | 472 | 2631 | 5361 | 3951 | 520 | 107 | 122 | 14 | 652 | 1025 | 1383 | 683 | 265 | 0 | 0 | 79 | 279 | 0 |
| SRR7547903 | 480 | 1845 | 9506 | 1718 | 7525 | 6955 | 80160 | 60585 | 119133 | 6786 | 17117 | 12997 | 22659 | 10498 | 3219 | 3715 | 28931 | 0 | 72 | 29 | 111 | 981 | 199 | 50 | 25 | 34 | 16 | 110 | 144 | 118 | 200 | 21 | 0 | 0 | 12 | 101 | 0 |
| SRR7547904 | 592 | 2154 | 11327 | 1134 | 15983 | 11181 | 151827 | 100285 | 37280 | 4384 | 17846 | 16872 | 7478 | 7215 | 585 | 8331 | 31160 | 0 | 12 | 22 | 241 | 1205 | 769 | 129 | 80 | 41 | 14 | 26 | 97 | 153 | 203 | 30 | 0 | 0 | 12 | 120 | 0 |
| SRR7547905 | 603 | 2608 | 17953 | 1199 | 15624 | 10407 | 204114 | 92006 | 21311 | 4427 | 12012 | 14545 | 9820 | 6286 | 2455 | 5836 | 27134 | 0 | 27 | 27 | 80 | 1052 | 379 | 60 | 32 | 28 | 0 | 29 | 66 | 51 | 146 | 18 | 0 | 0 | 0 | 80 | 0 |
| SRR7547906 | 851 | 3144 | 14037 | 725 | 376 | 341 | 5796 | 2619 | 6470 | 2847 | 3350 | 7162 | 5693 | 10257 | 117 | 1789 | 30387 | 0 | 0 | 0 | 101 | 746 | 500 | 15 | 10 | 13 | 0 | 0 | 13 | 21 | 44 | 14 | 0 | 0 | 0 | 29 | 0 |
| SRR7547907 | 897 | 3752 | 16692 | 652 | 382 | 198 | 5753 | 2169 | 2652 | 4899 | 5333 | 26626 | 17 | 8923 | 86 | 842 | 27394 | 0 | 0 | 13 | 389 | 1044 | 1590 | 59 | 26 | 27 | 0 | 0 | 13 | 52 | 46 | 38 | 0 | 0 | 0 | 83 | 0 |
| SRR7547908 | 1168 | 3516 | 9847 | 1036 | 407 | 247 | 3532 | 2959 | 7266 | 5772 | 3281 | 14501 | 4727 | 2629 | 34 | 2503 | 33184 | 0 | 0 | 0 | 46 | 183 | 360 | 21 | 0 | 0 | 0 | 18 | 25 | 39 | 31 | 17 | 0 | 0 | 0 | 14 | 0 |
| SRR7547909 | 469 | 2461 | 17365 | 2398 | 7938 | 6875 | 121179 | 54832 | 28665 | 8216 | 18686 | 11390 | 14820 | 17614 | 643 | 10136 | 39244 | 0 | 90 | 51 | 407 | 5034 | 3858 | 905 | 201 | 269 | 0 | 108 | 112 | 264 | 847 | 595 | 12 | 0 | 192 | 867 | 40 |
| SRR7547910 | 429 | 2024 | 17178 | 1798 | 8176 | 6904 | 104106 | 57515 | 20561 | 3943 | 21670 | 8638 | 20350 | 10815 | 1721 | 1340 | 46397 | 0 | 32 | 46 | 793 | 9169 | 6795 | 669 | 236 | 262 | 0 | 33 | 93 | 450 | 865 | 584 | 0 | 0 | 158 | 616 | 23 |
| SRR7547911 | 541 | 2376 | 16681 | 919 | 5442 | 6399 | 82530 | 57464 | 19940 | 3718 | 20904 | 31 | 16870 | 6495 | 1087 | 15408 | 58465 | 0 | 0 | 14 | 149 | 1665 | 1145 | 149 | 62 | 109 | 0 | 17 | 81 | 194 | 348 | 226 | 0 | 0 | 85 | 254 | 13 |
| SRR7547912 | 707 | 3764 | 14954 | 1594 | 10112 | 12918 | 195451 | 65223 | 32050 | 4727 | 13847 | 12 | 11240 | 10849 | 462 | 2033 | 39084 | 0 | 0 | 12 | 175 | 3158 | 3099 | 384 | 163 | 150 | 0 | 0 | 26 | 75 | 172 | 172 | 0 | 0 | 68 | 407 | 17 |
| SRR7547913 | 749 | 3318 | 14652 | 1187 | 10873 | 11632 | 165694 | 87222 | 19816 | 3729 | 16661 | 12 | 9203 | 6779 | 512 | 3270 | 23697 | 0 | 0 | 0 | 95 | 568 | 404 | 45 | 30 | 22 | 13 | 17 | 52 | 109 | 174 | 55 | 0 | 0 | 19 | 77 | 0 |
| SRR7547914 | 459 | 2378 | 12537 | 796 | 6249 | 5159 | 71379 | 50161 | 41421 | 4399 | 21588 | 20170 | 14229 | 9090 | 443 | 3335 | 45698 | 0 | 0 | 11 | 76 | 1815 | 1377 | 211 | 77 | 147 | 0 | 0 | 31 | 48 | 138 | 97 | 0 | 0 | 28 | 288 | 0 |
| SRR7547915 | 492 | 2238 | 10495 | 1063 | 8774 | 9795 | 111809 | 74285 | 29811 | 4485 | 16430 | 6859 | 14321 | 9535 | 1114 | 11872 | 30254 | 0 | 83 | 195 | 411 | 13026 | 4582 | 1039 | 130 | 190 | 14 | 83 | 428 | 237 | 1062 | 249 | 0 | 0 | 28 | 429 | 0 |
| SRR7547916 | 417 | 2320 | 8336 | 933 | 8055 | 8710 | 99966 | 57618 | 51004 | 4210 | 15440 | 12 | 12680 | 7292 | 827 | 4746 | 35525 | 0 | 11 | 33 | 116 | 1178 | 489 | 99 | 47 | 54 | 12 | 16 | 75 | 70 | 344 | 30 | 0 | 0 | 10 | 107 | 0 |
| SRR7547917 | 863 | 2886 | 14336 | 608 | 225 | 148 | 2464 | 1692 | 6453 | 6149 | 3050 | 11 | 5250 | 3491 | 11 | 1752 | 50980 | 0 | 24 | 38 | 105 | 1029 | 584 | 39 | 18 | 23 | 0 | 33 | 120 | 57 | 94 | 23 | 0 | 0 | 0 | 36 | 0 |
| SRR7547918 | 454 | 2294 | 15381 | 222 | 259 | 255 | 4293 | 2199 | 3658 | 2446 | 2371 | 17782 | 4878 | 4693 | 13 | 1231 | 37563 | 0 | 0 | 0 | 76 | 533 | 585 | 27 | 12 | 22 | 0 | 0 | 0 | 19 | 52 | 17 | 0 | 0 | 0 | 48 | 0 |
| SRR7547919 | 872 | 3313 | 19377 | 244 | 309 | 253 | 4485 | 2062 | 3849 | 2066 | 3096 | 18657 | 5841 | 8392 | 33 | 1162 | 24940 | 0 | 0 | 0 | 183 | 300 | 189 | 20 | 10 | 18 | 0 | 0 | 0 | 25 | 19 | 0 | 0 | 0 | 0 | 37 | 0 |
| SRR7547920 | 800 | 2238 | 5504 | 1341 | 567 | 451 | 5981 | 3231 | 12378 | 5051 | 4653 | 15452 | 5780 | 4267 | 41 | 2599 | 36695 | 0 | 15 | 0 | 23 | 169 | 131 | 19 | 0 | 13 | 0 | 21 | 24 | 13 | 17 | 0 | 0 | 0 | 0 | 21 | 0 |
| SRR7585373 | 98 | 2677 | 3289 | 250 | 165 | 2793 | 1899 | 2750 | 6710 | 0 | 8081 | 14364 | 31 | 1914 | 1985 | 302 | 28066 | 0 | 0 | 17 | 1780 | 1008 | 268 | 0 | 0 | 0 | 0 | 35 | 434 | 771 | 1698 | 243 | 0 | 13 | 0 | 0 | 0 |
| SRR7586110 | 1399 | 6177 | 28855 | 1020 | 4357 | 4686 | 52855 | 15486 | 40187 | 1903 | 42938 | 10134 | 21258 | 22807 | 3224 | 0 | 7487 | 0 | 0 | 0 | 122 | 159 | 136 | 17 | 21 | 0 | 0 | 0 | 0 | 19 | 0 | 0 | 0 | 0 | 0 | 0 | 0 |
| SRR7586111 | 1488 | 6137 | 28983 | 1016 | 4333 | 4814 | 53590 | 15483 | 39116 | 1882 | 42541 | 9778 | 21325 | 22594 | 3159 | 11 | 8110 | 0 | 0 | 0 | 123 | 158 | 135 | 15 | 23 | 0 | 0 | 0 | 0 | 18 | 0 | 0 | 0 | 0 | 0 | 0 | 0 |
| SRR7586113 | 1478 | 5468 | 16550 | 2476 | 10064 | 7586 | 98806 | 19778 | 34725 | 3442 | 42110 | 12799 | 23289 | 26252 | 4019 | 13 | 16620 | 0 | 0 | 0 | 78 | 171 | 218 | 33 | 35 | 0 | 0 | 0 | 11 | 18 | 10 | 0 | 0 | 0 | 0 | 20 | 0 |
| SRR7586114 | 1576 | 5804 | 17180 | 2426 | 9941 | 7615 | 98747 | 20115 | 33777 | 3438 | 41977 | 12717 | 23301 | 25991 | 3915 | 14 | 17661 | 0 | 0 | 0 | 82 | 179 | 226 | 34 | 35 | 0 | 0 | 0 | 12 | 19 | 12 | 0 | 0 | 0 | 0 | 19 | 0 |
| SRR7586116 | 1962 | 5598 | 7214 | 1661 | 20581 | 12772 | 231346 | 42044 | 16702 | 7705 | 39249 | 9231 | 23240 | 20613 | 4483 | 37 | 9196 | 0 | 20 | 16 | 176 | 80 | 200 | 36 | 31 | 0 | 0 | 18 | 32 | 39 | 0 | 0 | 0 | 0 | 0 | 16 | 0 |
| SRR7586117 | 2006 | 5854 | 7340 | 1539 | 20765 | 13038 | 232353 | 42010 | 16374 | 7552 | 39076 | 9100 | 23348 | 20289 | 4480 | 30 | 9852 | 0 | 20 | 17 | 192 | 84 | 206 | 40 | 36 | 0 | 0 | 18 | 35 | 42 | 0 | 0 | 0 | 0 | 0 | 16 | 0 |
| SRR7586118 | 1971 | 5735 | 7147 | 1577 | 20447 | 12846 | 230484 | 41558 | 16372 | 7672 | 39204 | 9121 | 23371 | 20556 | 4483 | 40 | 9916 | 0 | 21 | 18 | 185 | 85 | 210 | 38 | 35 | 0 | 0 | 20 | 33 | 43 | 0 | 0 | 0 | 0 | 0 | 16 | 0 |
| SRR7586119 | 1960 | 7329 | 10191 | 2141 | 13796 | 8782 | 100286 | 30803 | 46388 | 7151 | 43122 | 9815 | 28963 | 23322 | 3980 | 15 | 13469 | 10 | 25 | 16 | 336 | 301 | 297 | 56 | 49 | 0 | 22 | 37 | 60 | 143 | 28 | 0 | 0 | 0 | 0 | 24 | 0 |
| SRR7586120 | 1977 | 7497 | 10158 | 2107 | 13908 | 8904 | 100195 | 30304 | 46132 | 7058 | 43269 | 9569 | 28987 | 23005 | 3811 | 16 | 14216 | 10 | 25 | 16 | 345 | 323 | 297 | 56 | 49 | 0 | 22 | 36 | 62 | 144 | 27 | 0 | 0 | 0 | 0 | 25 | 0 |
| SRR7586122 | 2582 | 7639 | 8702 | 2739 | 18278 | 12913 | 220083 | 30442 | 61564 | 3569 | 34622 | 9802 | 17239 | 19984 | 6117 | 136 | 7686 | 0 | 13 | 0 | 88 | 87 | 242 | 47 | 45 | 0 | 0 | 10 | 19 | 16 | 0 | 0 | 0 | 0 | 0 | 24 | 0 |
| SRR7586123 | 2685 | 7971 | 8749 | 2672 | 18054 | 13108 | 219992 | 30260 | 61389 | 3498 | 34373 | 9814 | 17388 | 20013 | 5934 | 121 | 8244 | 0 | 13 | 11 | 89 | 87 | 257 | 46 | 46 | 0 | 0 | 0 | 16 | 18 | 0 | 0 | 0 | 0 | 0 | 23 | 0 |
| SRR7586125 | 2210 | 8488 | 13695 | 3176 | 14513 | 10147 | 181191 | 32547 | 37595 | 4379 | 32821 | 8741 | 18570 | 23487 | 4252 | 47 | 7430 | 0 | 13 | 14 | 145 | 142 | 376 | 63 | 63 | 0 | 0 | 17 | 31 | 34 | 14 | 11 | 0 | 0 | 0 | 26 | 0 |
| SRR7586126 | 2288 | 8738 | 13822 | 3033 | 14658 | 10342 | 183934 | 32985 | 36631 | 4401 | 32224 | 8664 | 18458 | 22868 | 4122 | 48 | 7863 | 0 | 11 | 14 | 158 | 153 | 392 | 59 | 65 | 0 | 0 | 13 | 31 | 35 | 14 | 13 | 0 | 0 | 0 | 28 | 0 |
| SRR7586128 | 3125 | 10904 | 13368 | 2649 | 19616 | 12022 | 196087 | 26723 | 40233 | 4029 | 37942 | 10065 | 18977 | 22293 | 5361 | 14 | 8832 | 0 | 35 | 34 | 379 | 549 | 442 | 76 | 74 | 13 | 0 | 29 | 53 | 73 | 21 | 12 | 0 | 0 | 0 | 42 | 0 |
| SRR7586129 | 3366 | 11832 | 13840 | 2562 | 19820 | 11826 | 196337 | 26481 | 38919 | 3888 | 37954 | 10283 | 18919 | 22149 | 5584 | 16 | 9429 | 0 | 37 | 34 | 393 | 560 | 458 | 73 | 69 | 13 | 0 | 30 | 56 | 77 | 22 | 13 | 0 | 0 | 0 | 49 | 0 |
| SRR7586131 | 2611 | 7501 | 13369 | 3514 | 12248 | 12415 | 184899 | 26486 | 41483 | 2191 | 35706 | 9204 | 18334 | 26684 | 5864 | 98 | 9826 | 0 | 0 | 0 | 79 | 139 | 224 | 38 | 39 | 0 | 0 | 0 | 12 | 13 | 0 | 0 | 0 | 0 | 0 | 23 | 0 |
| SRR7586132 | 2729 | 7914 | 13587 | 3455 | 12268 | 12717 | 183512 | 26618 | 40813 | 2185 | 35253 | 9380 | 18004 | 26605 | 5664 | 76 | 10558 | 0 | 11 | 0 | 79 | 138 | 234 | 38 | 37 | 0 | 0 | 0 | 13 | 17 | 0 | 0 | 0 | 0 | 0 | 20 | 0 |
| SRR7586134 | 3298 | 13380 | 10839 | 2464 | 14607 | 10356 | 168757 | 29380 | 50686 | 4953 | 36141 | 10761 | 19542 | 25516 | 5293 | 17 | 7991 | 0 | 15 | 12 | 241 | 211 | 370 | 63 | 70 | 11 | 0 | 14 | 25 | 49 | 13 | 11 | 0 | 0 | 0 | 34 | 0 |
| SRR7586135 | 3386 | 13441 | 10802 | 2369 | 14631 | 10598 | 170803 | 29603 | 50736 | 4920 | 36176 | 10907 | 19602 | 25533 | 5169 | 14 | 8790 | 0 | 15 | 12 | 269 | 227 | 375 | 68 | 69 | 0 | 0 | 15 | 25 | 51 | 12 | 12 | 0 | 0 | 0 | 35 | 0 |
| SRR7586137 | 1723 | 5869 | 11026 | 3002 | 8807 | 10293 | 123300 | 22724 | 54846 | 2313 | 37225 | 10415 | 22074 | 30232 | 5816 | 22 | 8514 | 0 | 0 | 0 | 96 | 159 | 99 | 17 | 18 | 0 | 0 | 0 | 0 | 19 | 0 | 0 | 0 | 0 | 0 | 0 | 0 |
| SRR7586138 | 1775 | 6017 | 10967 | 2972 | 8677 | 10373 | 122113 | 22622 | 54658 | 2241 | 37269 | 10470 | 22099 | 29774 | 5663 | 24 | 9370 | 0 | 0 | 0 | 92 | 164 | 102 | 16 | 20 | 0 | 0 | 0 | 0 | 20 | 0 | 0 | 0 | 0 | 0 | 12 | 0 |
| SRR7586140 | 2974 | 14947 | 12564 | 3550 | 14708 | 10037 | 160399 | 31004 | 72959 | 3810 | 29179 | 7653 | 16998 | 22468 | 4214 | 34 | 8763 | 0 | 15 | 13 | 283 | 237 | 283 | 55 | 65 | 11 | 0 | 14 | 26 | 59 | 15 | 0 | 0 | 0 | 10 | 39 | 0 |
| SRR7586141 | 2973 | 15109 | 12379 | 3450 | 14836 | 10124 | 159984 | 31168 | 71503 | 3726 | 29320 | 7705 | 17169 | 22253 | 4215 | 25 | 9394 | 0 | 14 | 11 | 287 | 239 | 299 | 57 | 61 | 11 | 0 | 16 | 26 | 64 | 16 | 11 | 0 | 0 | 11 | 42 | 0 |
| SRR7586143 | 2787 | 8353 | 13566 | 4638 | 24905 | 12657 | 262289 | 22637 | 56010 | 3273 | 31854 | 8992 | 18558 | 27344 | 7876 | 45 | 10385 | 0 | 0 | 0 | 160 | 147 | 345 | 64 | 73 | 0 | 0 | 0 | 13 | 30 | 0 | 0 | 0 | 0 | 0 | 34 | 0 |
| SRR7586144 | 2840 | 8688 | 13921 | 4495 | 24877 | 12808 | 264110 | 22536 | 55045 | 3228 | 31530 | 8931 | 18309 | 26631 | 7768 | 36 | 11264 | 0 | 0 | 0 | 162 | 151 | 377 | 63 | 73 | 0 | 0 | 0 | 15 | 32 | 0 | 11 | 0 | 0 | 0 | 36 | 0 |
| SRR7586146 | 897 | 7400 | 42654 | 1135 | 7872 | 9649 | 121193 | 18289 | 47778 | 5888 | 51457 | 15065 | 29404 | 29537 | 3914 | 22 | 16818 | 0 | 15 | 16 | 132 | 172 | 253 | 37 | 35 | 0 | 0 | 11 | 24 | 25 | 0 | 0 | 0 | 0 | 0 | 21 | 0 |
| SRR7586147 | 877 | 7803 | 42975 | 1081 | 7895 | 9671 | 119762 | 18091 | 46835 | 5845 | 50967 | 11 | 29091 | 29147 | 3835 | 18 | 17399 | 0 | 14 | 15 | 133 | 181 | 249 | 39 | 43 | 0 | 0 | 11 | 28 | 26 | 11 | 0 | 0 | 0 | 0 | 17 | 0 |
| SRR7586149 | 1383 | 7361 | 27171 | 3049 | 7401 | 5784 | 91374 | 18307 | 140540 | 5104 | 31781 | 8763 | 22749 | 31018 | 7530 | 26 | 9777 | 0 | 0 | 0 | 74 | 180 | 277 | 45 | 44 | 0 | 0 | 10 | 23 | 22 | 12 | 0 | 0 | 0 | 0 | 29 | 0 |
| SRR7586150 | 1454 | 7444 | 27429 | 2992 | 7495 | 5923 | 90887 | 18297 | 139747 | 5069 | 31738 | 9037 | 22816 | 30750 | 7496 | 28 | 10235 | 0 | 0 | 0 | 79 | 195 | 275 | 48 | 46 | 0 | 0 | 11 | 22 | 18 | 13 | 0 | 0 | 0 | 0 | 30 | 0 |
| SRR7586152 | 1430 | 7185 | 27571 | 3341 | 14620 | 14989 | 163420 | 15696 | 27833 | 4309 | 45212 | 11982 | 25075 | 24991 | 2590 | 12 | 14846 | 0 | 12 | 11 | 103 | 82 | 171 | 31 | 29 | 0 | 0 | 11 | 29 | 26 | 11 | 0 | 0 | 0 | 0 | 24 | 0 |
| SRR7586153 | 1528 | 7327 | 27587 | 3285 | 14538 | 15095 | 163267 | 15343 | 27132 | 4307 | 44699 | 12064 | 24916 | 24667 | 2552 | 17 | 15780 | 0 | 13 | 12 | 108 | 87 | 178 | 30 | 31 | 0 | 0 | 10 | 30 | 26 | 12 | 0 | 0 | 0 | 0 | 21 | 0 |
| SRR7586155 | 887 | 5751 | 42474 | 1398 | 12664 | 10254 | 118914 | 14465 | 60418 | 5404 | 37301 | 11766 | 18247 | 19473 | 3690 | 0 | 11762 | 0 | 14 | 16 | 168 | 230 | 259 | 39 | 40 | 0 | 0 | 11 | 25 | 31 | 13 | 0 | 0 | 0 | 0 | 27 | 0 |
| SRR7586156 | 915 | 5945 | 43219 | 1436 | 12595 | 10283 | 118842 | 14451 | 59140 | 5404 | 37483 | 11807 | 18105 | 19343 | 3727 | 15 | 12509 | 0 | 17 | 19 | 171 | 244 | 285 | 51 | 43 | 11 | 0 | 13 | 24 | 30 | 13 | 0 | 0 | 0 | 0 | 25 | 0 |
| SRR7586158 | 2535 | 10348 | 18829 | 4001 | 16827 | 14322 | 140305 | 14600 | 113941 | 4568 | 40649 | 11600 | 23340 | 24801 | 6974 | 15 | 9538 | 0 | 19 | 20 | 254 | 302 | 360 | 56 | 55 | 0 | 0 | 21 | 42 | 65 | 18 | 10 | 0 | 0 | 0 | 26 | 0 |
| SRR7586159 | 2591 | 10607 | 18806 | 3964 | 16701 | 14410 | 138682 | 14246 | 111780 | 4521 | 40105 | 11560 | 23027 | 24454 | 6751 | 17 | 10113 | 0 | 21 | 21 | 260 | 319 | 363 | 63 | 62 | 0 | 0 | 17 | 46 | 65 | 20 | 0 | 0 | 0 | 0 | 26 | 0 |
| SRR7609258 | 16 | 190 | 444458 | 74 | 11792 | 22010 | 106182 | 32091 | 24295 | 3157 | 17511 | 6472 | 44377 | 32377 | 2362 | 255 | 39509 | 0 | 0 | 0 | 16 | 101 | 23 | 0 | 0 | 0 | 0 | 0 | 11 | 30 | 0 | 0 | 0 | 0 | 0 | 10 | 0 |
| SRR7609259 | 95 | 4463 | 188107 | 116 | 14973 | 18754 | 111032 | 41442 | 53504 | 2249 | 16878 | 18671 | 8832 | 31775 | 5489 | 1501 | 63871 | 0 | 0 | 28 | 624 | 3194 | 1186 | 154 | 61 | 37 | 0 | 0 | 26 | 192 | 76 | 11 | 0 | 0 | 19 | 147 | 0 |
| SRR7609260 | 141 | 4190 | 161192 | 101 | 13942 | 19574 | 85530 | 27880 | 113148 | 1492 | 15041 | 28 | 8162 | 17523 | 1395 | 3942 | 48600 | 0 | 0 | 56 | 825 | 4863 | 662 | 107 | 38 | 20 | 0 | 0 | 69 | 325 | 99 | 0 | 0 | 0 | 0 | 117 | 0 |
| SRR7609261 | 54 | 645 | 73790 | 163 | 5416 | 9738 | 47950 | 25976 | 109039 | 1176 | 31051 | 25 | 8921 | 9603 | 2985 | 3095 | 54710 | 0 | 0 | 17 | 165 | 2283 | 929 | 201 | 55 | 29 | 0 | 0 | 61 | 137 | 77 | 12 | 0 | 0 | 24 | 215 | 15 |
| SRR7609262 | 20 | 428 | 121189 | 32 | 2133 | 3545 | 17000 | 7322 | 15797 | 1150 | 14279 | 31 | 9543 | 37303 | 1289 | 242 | 26664 | 0 | 0 | 0 | 63 | 562 | 321 | 63 | 29 | 15 | 0 | 0 | 0 | 22 | 14 | 0 | 0 | 0 | 0 | 72 | 0 |
| SRR7609263 | 28 | 326 | 95956 | 61 | 4775 | 7520 | 48632 | 28478 | 34188 | 1306 | 27022 | 26 | 10299 | 24203 | 2848 | 1845 | 67751 | 0 | 0 | 28 | 446 | 1934 | 354 | 46 | 23 | 21 | 0 | 0 | 28 | 144 | 63 | 0 | 0 | 0 | 12 | 126 | 12 |
| SRR7609264 | 55 | 413 | 86838 | 58 | 11808 | 20312 | 126941 | 52406 | 79366 | 1678 | 16416 | 24 | 7729 | 18658 | 4067 | 674 | 53032 | 0 | 0 | 31 | 850 | 3281 | 667 | 72 | 36 | 21 | 0 | 0 | 42 | 267 | 52 | 0 | 0 | 0 | 0 | 158 | 0 |
| SRR7609265 | 21 | 256 | 105045 | 20 | 4171 | 7330 | 39576 | 13043 | 82916 | 3078 | 22696 | 11354 | 16900 | 13992 | 2014 | 2539 | 45353 | 0 | 0 | 24 | 385 | 1399 | 312 | 44 | 23 | 21 | 0 | 0 | 23 | 112 | 20 | 0 | 0 | 0 | 0 | 128 | 0 |
| SRR7609266 | 13 | 242 | 39115 | 18 | 3489 | 6112 | 28086 | 15480 | 138339 | 680 | 18624 | 20887 | 10105 | 18496 | 838 | 1353 | 62587 | 0 | 0 | 12 | 221 | 1695 | 428 | 82 | 40 | 62 | 0 | 0 | 16 | 80 | 41 | 0 | 0 | 0 | 24 | 214 | 24 |
| SRR7609267 | 32 | 236 | 131541 | 32 | 6803 | 11535 | 71633 | 48433 | 21600 | 1814 | 22717 | 23567 | 8533 | 18403 | 3801 | 871 | 52942 | 0 | 0 | 25 | 473 | 3178 | 1969 | 386 | 157 | 138 | 0 | 0 | 26 | 130 | 114 | 20 | 0 | 0 | 66 | 558 | 61 |
| SRR7609268 | 141 | 556 | 80507 | 42 | 1601 | 2672 | 12172 | 6362 | 157419 | 1936 | 15347 | 21010 | 7976 | 8293 | 567 | 981 | 39264 | 0 | 0 | 38 | 598 | 2123 | 1013 | 184 | 91 | 102 | 0 | 0 | 109 | 325 | 83 | 0 | 0 | 0 | 49 | 420 | 33 |
| SRR7609269 | 12 | 171 | 75881 | 10 | 6317 | 11048 | 70000 | 27247 | 105755 | 4464 | 21696 | 12 | 13669 | 38695 | 2772 | 2555 | 51478 | 0 | 0 | 0 | 81 | 802 | 848 | 225 | 91 | 142 | 0 | 0 | 15 | 75 | 45 | 12 | 0 | 0 | 126 | 771 | 67 |
| SRR7609270 | 14 | 701 | 78659 | 85 | 18399 | 22729 | 124587 | 42716 | 183757 | 1399 | 19730 | 9359 | 11202 | 49785 | 6979 | 135 | 34088 | 0 | 0 | 0 | 55 | 2975 | 1613 | 467 | 241 | 376 | 0 | 0 | 0 | 36 | 41 | 14 | 0 | 0 | 181 | 1591 | 131 |
| SRR7609271 | 20 | 154 | 99221 | 27 | 3563 | 4745 | 31590 | 20758 | 31760 | 759 | 20343 | 22770 | 12856 | 30189 | 5171 | 1185 | 81843 | 0 | 0 | 11 | 181 | 881 | 230 | 37 | 19 | 12 | 0 | 0 | 17 | 72 | 28 | 0 | 0 | 0 | 0 | 83 | 0 |
| SRR7609272 | 15 | 260 | 82899 | 0 | 2060 | 3282 | 14738 | 6243 | 195106 | 5398 | 12363 | 26 | 7949 | 8041 | 911 | 2515 | 26087 | 0 | 0 | 17 | 275 | 1504 | 1141 | 286 | 91 | 98 | 0 | 0 | 27 | 106 | 28 | 0 | 0 | 0 | 24 | 339 | 24 |
| SRR7609273 | 0 | 246 | 256692 | 20 | 8829 | 13009 | 73488 | 36724 | 151336 | 3957 | 26634 | 44008 | 18669 | 13626 | 2384 | 1898 | 42181 | 0 | 0 | 0 | 0 | 189 | 68 | 14 | 0 | 0 | 0 | 0 | 0 | 0 | 0 | 0 | 0 | 0 | 0 | 22 | 0 |
| SRR7609274 | 24 | 296 | 184496 | 62 | 6535 | 10320 | 60972 | 32028 | 68530 | 3438 | 18439 | 21711 | 9567 | 19687 | 1165 | 460 | 45688 | 0 | 0 | 0 | 24 | 764 | 320 | 73 | 34 | 21 | 0 | 0 | 0 | 12 | 21 | 0 | 0 | 0 | 0 | 121 | 0 |
| SRR7609275 | 14 | 662 | 142141 | 43 | 6385 | 7192 | 45632 | 17152 | 382809 | 3070 | 12255 | 30 | 4928 | 7524 | 2104 | 226 | 36452 | 0 | 0 | 20 | 270 | 1710 | 1139 | 306 | 90 | 64 | 0 | 0 | 23 | 89 | 46 | 0 | 0 | 0 | 27 | 311 | 25 |
| SRR7609276 | 39 | 168 | 62890 | 21 | 1237 | 1600 | 6865 | 3981 | 105696 | 5891 | 8204 | 13 | 5687 | 4565 | 131 | 406 | 17975 | 0 | 0 | 0 | 17 | 30 | 0 | 0 | 0 | 0 | 0 | 0 | 0 | 22 | 0 | 0 | 0 | 0 | 0 | 0 | 0 |
| SRR7609277 | 18 | 610 | 160182 | 33 | 8131 | 14783 | 105368 | 29679 | 77087 | 1288 | 15336 | 17732 | 9966 | 19510 | 1982 | 2508 | 40110 | 0 | 0 | 51 | 1016 | 5091 | 1877 | 382 | 150 | 129 | 0 | 0 | 36 | 236 | 94 | 13 | 0 | 0 | 51 | 538 | 33 |
| SRR7609278 | 48 | 879 | 112958 | 47 | 11575 | 12800 | 86357 | 38279 | 96483 | 778 | 13806 | 19801 | 10525 | 26861 | 1449 | 1819 | 55807 | 0 | 0 | 16 | 288 | 335 | 29 | 0 | 0 | 0 | 0 | 0 | 26 | 118 | 12 | 0 | 0 | 0 | 0 | 13 | 0 |
| SRR7609279 | 0 | 275 | 90239 | 0 | 4645 | 4739 | 31036 | 19601 | 250689 | 1392 | 15224 | 13348 | 11753 | 19171 | 2101 | 860 | 60996 | 0 | 0 | 0 | 42 | 977 | 413 | 83 | 32 | 21 | 0 | 0 | 0 | 18 | 29 | 0 | 0 | 0 | 11 | 118 | 0 |
| SRR7609280 | 14 | 224 | 74229 | 0 | 2393 | 4098 | 23738 | 14493 | 174280 | 1361 | 20483 | 23171 | 9733 | 12255 | 728 | 1489 | 65082 | 0 | 0 | 0 | 155 | 1956 | 1382 | 316 | 119 | 71 | 0 | 0 | 11 | 46 | 33 | 10 | 0 | 0 | 30 | 444 | 21 |
| SRR7609281 | 20 | 449 | 92498 | 17 | 10564 | 22349 | 112094 | 39754 | 106974 | 1452 | 17586 | 17645 | 8725 | 27209 | 1630 | 961 | 53293 | 0 | 0 | 0 | 111 | 1403 | 245 | 21 | 13 | 0 | 0 | 0 | 12 | 49 | 43 | 0 | 0 | 0 | 0 | 54 | 0 |
| SRR7609282 | 20 | 142 | 96772 | 18 | 12184 | 15658 | 82939 | 55242 | 62218 | 1471 | 21711 | 14442 | 5521 | 19148 | 1670 | 1027 | 76985 | 0 | 0 | 0 | 242 | 2614 | 1039 | 205 | 100 | 67 | 0 | 0 | 0 | 58 | 37 | 0 | 0 | 0 | 20 | 351 | 19 |
| SRR7609283 | 13 | 964 | 100448 | 25 | 2979 | 3990 | 22410 | 9531 | 132462 | 2386 | 16280 | 60 | 7780 | 24672 | 1019 | 824 | 70505 | 0 | 0 | 0 | 32 | 1418 | 470 | 43 | 21 | 11 | 0 | 0 | 0 | 0 | 21 | 0 | 0 | 0 | 0 | 70 | 0 |
| SRR7609284 | 45 | 356 | 82861 | 39 | 2934 | 5431 | 35244 | 16054 | 125786 | 4702 | 26702 | 35 | 12488 | 11544 | 3898 | 329 | 38434 | 0 | 0 | 19 | 353 | 1817 | 474 | 61 | 27 | 14 | 0 | 0 | 24 | 102 | 26 | 0 | 0 | 0 | 0 | 91 | 0 |
| SRR7609285 | 44 | 1033 | 99765 | 28 | 5570 | 8046 | 41956 | 14502 | 104861 | 2020 | 21447 | 32646 | 8174 | 20001 | 1644 | 833 | 71092 | 0 | 0 | 0 | 58 | 2083 | 424 | 51 | 25 | 12 | 0 | 0 | 0 | 0 | 25 | 0 | 0 | 0 | 0 | 94 | 0 |
| SRR7609286 | 11 | 439 | 133892 | 47 | 2605 | 3569 | 19680 | 5499 | 193173 | 5284 | 12887 | 27319 | 9141 | 23304 | 624 | 2032 | 61406 | 0 | 0 | 0 | 33 | 1127 | 404 | 44 | 16 | 11 | 0 | 0 | 0 | 12 | 25 | 0 | 0 | 0 | 0 | 63 | 0 |
| SRR7609287 | 23 | 1464 | 211117 | 125 | 12512 | 21862 | 136754 | 66216 | 93773 | 3680 | 11852 | 11922 | 5805 | 27595 | 3638 | 1294 | 44697 | 0 | 0 | 0 | 116 | 2763 | 1620 | 291 | 101 | 45 | 0 | 0 | 0 | 55 | 91 | 18 | 0 | 0 | 23 | 203 | 0 |
| SRR7609288 | 35 | 608 | 225420 | 36 | 6312 | 9839 | 54797 | 26235 | 71995 | 12119 | 11871 | 11 | 13128 | 18568 | 4133 | 5938 | 53601 | 0 | 0 | 17 | 505 | 4757 | 1254 | 193 | 74 | 88 | 0 | 0 | 28 | 273 | 237 | 14 | 0 | 0 | 24 | 219 | 13 |
| SRR7609289 | 46 | 751 | 61044 | 110 | 9737 | 30289 | 161797 | 53078 | 88523 | 1018 | 17256 | 21 | 11579 | 36630 | 3789 | 1084 | 39919 | 0 | 0 | 12 | 196 | 1394 | 577 | 170 | 55 | 16 | 0 | 0 | 13 | 86 | 28 | 0 | 0 | 0 | 0 | 138 | 0 |
| SRR7609290 | 40 | 1049 | 138655 | 83 | 5632 | 7802 | 42644 | 27882 | 179708 | 3226 | 12468 | 14 | 9874 | 18841 | 13071 | 1557 | 38602 | 0 | 0 | 15 | 471 | 6199 | 3095 | 721 | 289 | 301 | 0 | 0 | 20 | 181 | 121 | 28 | 0 | 0 | 127 | 1087 | 92 |
| SRR7609291 | 0 | 513 | 225462 | 18 | 1555 | 2882 | 21496 | 8401 | 140162 | 310 | 1805 | 1906 | 2552 | 33807 | 1147 | 2076 | 28214 | 0 | 0 | 0 | 175 | 1525 | 1299 | 321 | 65 | 12 | 0 | 0 | 15 | 363 | 85 | 28 | 0 | 0 | 11 | 152 | 0 |
| SRR7609292 | 35 | 529 | 106515 | 13 | 2203 | 4440 | 20106 | 11412 | 199018 | 1921 | 19906 | 45 | 9876 | 12368 | 558 | 1509 | 47780 | 0 | 0 | 29 | 644 | 3235 | 1549 | 270 | 138 | 153 | 0 | 0 | 66 | 358 | 87 | 15 | 0 | 0 | 104 | 746 | 60 |
| SRR7609293 | 22 | 491 | 101390 | 53 | 11387 | 15081 | 106589 | 46320 | 96823 | 2264 | 15225 | 25 | 8197 | 25605 | 15252 | 217 | 74348 | 0 | 0 | 0 | 99 | 1656 | 639 | 96 | 43 | 21 | 0 | 0 | 0 | 29 | 44 | 0 | 0 | 0 | 0 | 137 | 0 |
| SRR7609294 | 28 | 994 | 227385 | 86 | 3060 | 4000 | 23070 | 12382 | 249128 | 650 | 13600 | 17746 | 8242 | 17143 | 572 | 663 | 71900 | 0 | 0 | 15 | 175 | 4888 | 1720 | 219 | 70 | 31 | 0 | 0 | 26 | 139 | 69 | 20 | 0 | 0 | 14 | 193 | 0 |
| SRR7609295 | 54 | 495 | 75827 | 170 | 10530 | 16023 | 114348 | 41866 | 67110 | 497 | 8317 | 11769 | 5704 | 19074 | 16570 | 1880 | 38981 | 0 | 0 | 197 | 3994 | 9530 | 1725 | 226 | 80 | 68 | 0 | 0 | 163 | 1354 | 190 | 14 | 0 | 0 | 38 | 268 | 11 |
| SRR7609296 | 0 | 114 | 77402 | 51 | 3468 | 4553 | 40064 | 25765 | 242458 | 1355 | 10610 | 19707 | 7735 | 18498 | 218 | 2722 | 48870 | 0 | 0 | 0 | 12 | 307 | 129 | 0 | 0 | 0 | 0 | 0 | 0 | 0 | 0 | 0 | 0 | 0 | 0 | 21 | 0 |
| SRR7609297 | 53 | 627 | 96586 | 167 | 1675 | 2346 | 16865 | 10931 | 57120 | 1120 | 19761 | 40175 | 12248 | 19326 | 924 | 1577 | 75483 | 0 | 0 | 0 | 72 | 925 | 283 | 42 | 13 | 0 | 0 | 0 | 16 | 45 | 42 | 0 | 0 | 0 | 16 | 71 | 0 |
| SRR7609298 | 99 | 457 | 62238 | 51 | 2193 | 2819 | 17934 | 8425 | 66700 | 648 | 15435 | 24 | 12534 | 10007 | 534 | 3458 | 53659 | 0 | 0 | 11 | 111 | 692 | 142 | 25 | 0 | 0 | 0 | 0 | 16 | 43 | 15 | 0 | 0 | 0 | 0 | 30 | 0 |
| SRR7609299 | 44 | 449 | 66096 | 74 | 3742 | 5809 | 51296 | 25157 | 263096 | 1153 | 15931 | 19 | 9262 | 14978 | 7691 | 949 | 36749 | 0 | 0 | 22 | 742 | 4835 | 1272 | 168 | 65 | 63 | 0 | 0 | 21 | 195 | 119 | 0 | 0 | 0 | 36 | 227 | 14 |
| SRR7609300 | 99 | 895 | 95969 | 386 | 3409 | 3708 | 36437 | 38242 | 94571 | 1415 | 8945 | 14660 | 6316 | 26483 | 919 | 258 | 37644 | 0 | 0 | 0 | 77 | 551 | 178 | 28 | 10 | 0 | 0 | 0 | 0 | 38 | 17 | 0 | 0 | 0 | 0 | 30 | 0 |
| SRR7609301 | 125 | 1284 | 170756 | 461 | 6485 | 9643 | 75890 | 32967 | 43836 | 892 | 25630 | 15 | 14506 | 23392 | 3821 | 563 | 55696 | 0 | 0 | 68 | 1375 | 10730 | 4246 | 703 | 291 | 209 | 0 | 0 | 60 | 384 | 223 | 34 | 0 | 0 | 100 | 616 | 31 |
| SRR7609302 | 193 | 806 | 72856 | 227 | 1744 | 1865 | 11326 | 7244 | 75891 | 254 | 12445 | 31 | 7127 | 10844 | 551 | 721 | 73012 | 0 | 0 | 58 | 758 | 1595 | 188 | 27 | 0 | 0 | 0 | 0 | 63 | 258 | 54 | 0 | 0 | 0 | 0 | 27 | 0 |
| SRR7644105 | 240 | 690 | 11048 | 210 | 0 | 0 | 68 | 96 | 118 | 961 | 207 | 700 | 46 | 557 | 0 | 43 | 450 | 0 | 0 | 0 | 0 | 0 | 0 | 0 | 0 | 0 | 0 | 0 | 0 | 0 | 0 | 0 | 0 | 0 | 0 | 0 | 0 |
| SRR7644106 | 228 | 943 | 10304 | 683 | 0 | 0 | 56 | 165 | 109 | 1040 | 165 | 622 | 64 | 701 | 0 | 15 | 435 | 0 | 0 | 0 | 0 | 0 | 0 | 0 | 0 | 0 | 0 | 0 | 0 | 0 | 0 | 0 | 0 | 0 | 0 | 0 | 0 |
| SRR7644107 | 304 | 1095 | 11134 | 456 | 0 | 14 | 61 | 155 | 169 | 700 | 191 | 651 | 72 | 720 | 0 | 32 | 385 | 0 | 0 | 0 | 0 | 0 | 0 | 0 | 0 | 0 | 0 | 0 | 0 | 0 | 0 | 0 | 0 | 0 | 0 | 0 | 0 |
| SRR7644108 | 289 | 931 | 10302 | 642 | 0 | 0 | 61 | 183 | 103 | 899 | 198 | 650 | 91 | 590 | 0 | 15 | 456 | 0 | 0 | 0 | 0 | 0 | 0 | 0 | 0 | 0 | 0 | 0 | 0 | 0 | 0 | 0 | 0 | 0 | 0 | 0 | 0 |
| SRR7644109 | 53 | 801 | 7472 | 747 | 0 | 54 | 89 | 214 | 228 | 640 | 278 | 710 | 89 | 1149 | 0 | 107 | 667 | 0 | 0 | 0 | 0 | 0 | 0 | 0 | 0 | 0 | 0 | 0 | 0 | 0 | 0 | 0 | 0 | 0 | 0 | 0 | 0 |
| SRR7644110 | 87 | 954 | 7578 | 781 | 0 | 42 | 112 | 239 | 262 | 810 | 197 | 999 | 81 | 991 | 0 | 89 | 617 | 0 | 0 | 0 | 0 | 0 | 0 | 0 | 0 | 0 | 0 | 0 | 0 | 0 | 0 | 0 | 0 | 0 | 0 | 0 | 0 |
| SRR7644111 | 190 | 925 | 7100 | 653 | 0 | 19 | 104 | 189 | 208 | 925 | 315 | 853 | 74 | 1160 | 0 | 119 | 601 | 0 | 0 | 0 | 0 | 0 | 0 | 0 | 0 | 0 | 0 | 0 | 0 | 0 | 0 | 0 | 0 | 0 | 0 | 0 | 0 |
| SRR7644112 | 171 | 940 | 7352 | 1026 | 0 | 27 | 86 | 176 | 282 | 570 | 258 | 876 | 78 | 1084 | 0 | 121 | 747 | 0 | 0 | 0 | 0 | 0 | 0 | 0 | 0 | 0 | 0 | 0 | 0 | 0 | 0 | 0 | 0 | 0 | 0 | 0 | 0 |
| SRR7644113 | 266 | 1242 | 9555 | 562 | 0 | 42 | 131 | 232 | 264 | 296 | 618 | 1719 | 124 | 1075 | 0 | 86 | 909 | 0 | 0 | 0 | 0 | 0 | 0 | 0 | 0 | 0 | 0 | 0 | 0 | 0 | 0 | 0 | 0 | 0 | 0 | 0 | 0 |
| SRR7644114 | 283 | 1258 | 8932 | 723 | 13 | 49 | 175 | 245 | 198 | 283 | 669 | 1821 | 117 | 1078 | 0 | 130 | 969 | 0 | 0 | 0 | 0 | 0 | 0 | 0 | 0 | 0 | 0 | 0 | 0 | 0 | 0 | 0 | 0 | 0 | 0 | 0 | 0 |
| SRR7644115 | 387 | 1309 | 10172 | 773 | 0 | 55 | 117 | 186 | 251 | 446 | 603 | 1715 | 159 | 1072 | 0 | 89 | 1087 | 0 | 0 | 0 | 0 | 0 | 0 | 0 | 0 | 0 | 0 | 0 | 0 | 0 | 0 | 0 | 0 | 0 | 0 | 0 | 0 |
| SRR7644116 | 312 | 1092 | 9704 | 905 | 0 | 47 | 147 | 265 | 236 | 686 | 601 | 1726 | 144 | 1065 | 0 | 94 | 1094 | 0 | 0 | 0 | 0 | 0 | 0 | 0 | 0 | 0 | 0 | 0 | 0 | 0 | 0 | 0 | 0 | 0 | 0 | 0 | 0 |
| SRR7644117 | 239 | 1484 | 5593 | 848 | 0 | 47 | 128 | 204 | 262 | 424 | 536 | 1504 | 122 | 1102 | 0 | 93 | 886 | 0 | 0 | 0 | 0 | 0 | 0 | 0 | 0 | 0 | 0 | 0 | 0 | 0 | 0 | 0 | 0 | 0 | 0 | 0 | 0 |
| SRR7644118 | 142 | 1557 | 6142 | 679 | 0 | 25 | 181 | 168 | 380 | 594 | 554 | 1793 | 75 | 1164 | 0 | 50 | 934 | 0 | 0 | 0 | 0 | 0 | 0 | 0 | 0 | 0 | 0 | 0 | 0 | 0 | 0 | 0 | 0 | 0 | 0 | 0 | 0 |
| SRR7644119 | 186 | 1221 | 6214 | 823 | 0 | 24 | 172 | 195 | 302 | 292 | 604 | 1663 | 112 | 1373 | 0 | 95 | 799 | 0 | 0 | 0 | 0 | 0 | 0 | 0 | 0 | 0 | 0 | 0 | 0 | 0 | 0 | 0 | 0 | 0 | 0 | 0 | 0 |
| SRR7644120 | 112 | 1179 | 6315 | 758 | 13 | 94 | 176 | 213 | 276 | 477 | 565 | 1826 | 132 | 1412 | 0 | 94 | 979 | 0 | 0 | 0 | 0 | 0 | 0 | 0 | 0 | 0 | 0 | 0 | 0 | 0 | 0 | 0 | 0 | 0 | 0 | 0 | 0 |
| SRR7644121 | 236 | 735 | 15084 | 761 | 0 | 46 | 228 | 311 | 161 | 603 | 774 | 2348 | 93 | 1835 | 0 | 118 | 1346 | 0 | 0 | 0 | 0 | 0 | 0 | 0 | 0 | 0 | 0 | 0 | 0 | 0 | 0 | 0 | 0 | 0 | 0 | 0 | 0 |
| SRR7644122 | 144 | 719 | 15061 | 632 | 19 | 50 | 213 | 348 | 197 | 920 | 847 | 2515 | 93 | 1842 | 0 | 93 | 1358 | 0 | 0 | 0 | 0 | 0 | 0 | 0 | 0 | 0 | 0 | 0 | 0 | 0 | 0 | 0 | 0 | 0 | 0 | 0 | 0 |
| SRR7644123 | 295 | 643 | 14987 | 456 | 18 | 33 | 267 | 354 | 208 | 858 | 774 | 2224 | 95 | 1932 | 0 | 121 | 1443 | 0 | 0 | 0 | 0 | 0 | 0 | 0 | 0 | 0 | 0 | 0 | 0 | 0 | 0 | 0 | 0 | 0 | 0 | 0 | 0 |

### Supp Table S8.1.Columns_B_to_AL

| | | | | | | | | | | | | | | | | | | | | | | | | | | | | | | | | | | | | |
|---|---|---|---|---|---|---|---|---|---|---|---|---|---|---|---|---|---|---|---|---|---|---|---|---|---|---|---|---|---|---|---|---|---|---|---|---|
| SRR7644124 | 226 | 792 | 14990 | 368 | 0 | 66 | 252 | 372 | 268 | 566 | 745 | 2436 | 97 | 1966 | 0 | 109 | 1435 | 0 | 0 | 0 | 0 | 0 | 0 | 0 | 0 | 0 | 0 | 0 | 0 | 0 | 0 | 0 | 0 | 0 | 0 | 0 | 0 |
| SRR7644125 | 92 | 916 | 8699 | 1168 | 25 | 37 | 286 | 197 | 151 | 572 | 758 | 2753 | 80 | 2497 | 0 | 59 | 890 | 0 | 0 | 0 | 0 | 0 | 0 | 0 | 0 | 0 | 0 | 0 | 0 | 0 | 0 | 0 | 0 | 0 | 0 | 0 | 0 |
| SRR7644126 | 25 | 792 | 7850 | 1808 | 0 | 50 | 222 | 212 | 173 | 693 | 766 | 2824 | 93 | 2731 | 0 | 76 | 922 | 0 | 0 | 0 | 0 | 0 | 0 | 0 | 0 | 0 | 0 | 0 | 0 | 0 | 0 | 0 | 0 | 0 | 0 | 0 | 0 |
| SRR7644128 | 69 | 805 | 8050 | 1633 | 0 | 35 | 192 | 182 | 160 | 736 | 823 | 2876 | 119 | 2503 | 0 | 66 | 961 | 0 | 0 | 0 | 0 | 0 | 0 | 0 | 0 | 0 | 0 | 0 | 0 | 0 | 0 | 0 | 0 | 0 | 0 | 0 | 0 |
| SRR7644129 | 50 | 743 | 8448 | 1759 | 0 | 61 | 212 | 195 | 131 | 941 | 761 | 2766 | 101 | 2817 | 0 | 54 | 858 | 0 | 0 | 0 | 0 | 0 | 0 | 0 | 0 | 0 | 0 | 0 | 0 | 0 | 0 | 0 | 0 | 0 | 0 | 0 | 0 |
| SRR7663662 | 51 | 1080 | 189 | 191 | 677 | 10176 | 12152 | 3619 | 0 | 0 | 79488 | 12954 | 1818 | 44982 | 0 | 66 | 6245 | 0 | 0 | 0 | 0 | 0 | 0 | 0 | 12 | 0 | 0 | 0 | 0 | 0 | 0 | 0 | 0 | 0 | 0 | 0 | 0 |
| SRR7663663 | 187 | 2229 | 158 | 912 | 1027 | 16240 | 22298 | 7502 | 0 | 0 | 11671 | 3586 | 322 | 10470 | 0 | 17 | 9060 | 0 | 0 | 0 | 0 | 0 | 0 | 0 | 0 | 0 | 0 | 0 | 0 | 0 | 0 | 0 | 0 | 0 | 0 | 0 | 0 |
| SRR7664346 | 0 | 315732 | 4728 | 0 | 7016 | 72177 | 74715 | 43355 | 509 | 61 | 13362 | 32 | 3555 | 10320 | 28 | 21335 | 23337 | 0 | 0 | 0 | 473 | 173 | 297 | 0 | 0 | 0 | 0 | 0 | 44 | 456 | 484 | 481 | 0 | 0 | 0 | 0 | 0 |
| SRR7664347 | 0 | 176812 | 13842 | 0 | 4944 | 58361 | 64173 | 44069 | 523 | 75 | 13781 | 58460 | 3749 | 11532 | 14 | 15976 | 25123 | 0 | 0 | 0 | 155 | 72 | 99 | 0 | 0 | 0 | 0 | 0 | 0 | 149 | 165 | 120 | 0 | 0 | 0 | 0 | 0 |
| SRR7690697 | 231 | 15280 | 1120 | 941 | 647 | 7603 | 9288 | 10773 | 4581 | 71 | 7273 | 24753 | 3926 | 1135 | 15 | 1749 | 41518 | 0 | 0 | 0 | 4857 | 2029 | 180 | 0 | 0 | 0 | 0 | 0 | 212 | 3461 | 1826 | 450 | 0 | 0 | 0 | 0 | 0 |
| SRR7690698 | 416 | 12341 | 1007 | 352 | 430 | 5353 | 6777 | 8342 | 5923 | 51 | 6467 | 28878 | 3807 | 971 | 10 | 1604 | 21842 | 0 | 0 | 0 | 4531 | 1621 | 122 | 0 | 0 | 0 | 0 | 0 | 250 | 4686 | 1927 | 665 | 0 | 0 | 0 | 0 | 0 |
| SRR7690699 | 346 | 36460 | 781 | 499 | 445 | 5043 | 5841 | 7500 | 4633 | 90 | 8081 | 24221 | 4929 | 1010 | 17 | 938 | 48114 | 0 | 0 | 0 | 5413 | 1572 | 193 | 0 | 0 | 0 | 0 | 0 | 285 | 3935 | 1466 | 342 | 0 | 16 | 0 | 0 | 0 |
| SRR7690700 | 226 | 2985 | 13553 | 394 | 6775 | 7250 | 91908 | 49255 | 29711 | 2891 | 12666 | 8514 | 12529 | 5434 | 3417 | 508 | 26315 | 0 | 12 | 21 | 538 | 921 | 343 | 120 | 46 | 81 | 18 | 23 | 96 | 494 | 150 | 41 | 0 | 0 | 25 | 127 | 0 |
| SRR7690701 | 81 | 1741 | 9981 | 65 | 7933 | 5203 | 96226 | 49737 | 29135 | 2791 | 14558 | 17 | 18 | 5900 | 3256 | 610 | 22938 | 0 | 0 | 13 | 377 | 260 | 151 | 90 | 40 | 79 | 0 | 10 | 51 | 358 | 67 | 12 | 0 | 0 | 14 | 96 | 0 |
| SRR7690702 | 303 | 2996 | 11186 | 366 | 7403 | 5700 | 87405 | 49514 | 28463 | 3304 | 10152 | 6976 | 11966 | 4671 | 1757 | 673 | 25833 | 0 | 11 | 14 | 218 | 237 | 152 | 94 | 33 | 54 | 20 | 20 | 60 | 303 | 94 | 22 | 0 | 0 | 22 | 98 | 0 |
| SRR7690703 | 102 | 2336 | 10314 | 93 | 5886 | 5131 | 91832 | 48821 | 22061 | 3073 | 13585 | 8823 | 13657 | 5940 | 1800 | 693 | 25119 | 0 | 13 | 23 | 65 | 198 | 67 | 65 | 29 | 51 | 11 | 40 | 138 | 142 | 79 | 13 | 0 | 0 | 29 | 89 | 0 |
| SRR7690704 | 55 | 1940 | 17317 | 78 | 7335 | 7355 | 83729 | 41099 | 33374 | 2190 | 11 | 12 | 13260 | 5806 | 2878 | 781 | 32952 | 0 | 0 | 10 | 256 | 359 | 138 | 69 | 29 | 43 | 0 | 0 | 41 | 255 | 74 | 18 | 0 | 0 | 15 | 91 | 0 |
| SRR7690705 | 62 | 1727 | 21067 | 35 | 5592 | 5417 | 88833 | 38830 | 27636 | 2649 | 14041 | 9141 | 16122 | 6961 | 1936 | 715 | 35586 | 0 | 0 | 0 | 56 | 95 | 69 | 49 | 28 | 37 | 0 | 0 | 12 | 52 | 20 | 0 | 0 | 0 | 0 | 58 | 0 |
| SRR7690706 | 86 | 1602 | 11248 | 264 | 2913 | 3986 | 33851 | 15175 | 37876 | 2849 | 6403 | 6887 | 10498 | 4430 | 830 | 1132 | 40732 | 0 | 0 | 0 | 214 | 384 | 115 | 71 | 35 | 47 | 0 | 0 | 35 | 212 | 59 | 11 | 0 | 0 | 11 | 82 | 0 |
| SRR7690707 | 1380 | 16539 | 650 | 378 | 388 | 5729 | 5641 | 6156 | 6094 | 95 | 8195 | 26962 | 5591 | 1265 | 12 | 1185 | 41480 | 0 | 0 | 0 | 2363 | 727 | 68 | 0 | 0 | 0 | 0 | 0 | 111 | 1427 | 450 | 106 | 0 | 0 | 13 | 0 | 0 |
| SRR7690708 | 123 | 2341 | 11761 | 582 | 7500 | 7151 | 77936 | 29959 | 34453 | 3185 | 7189 | 5984 | 9168 | 4602 | 1830 | 1131 | 30536 | 0 | 0 | 0 | 150 | 183 | 78 | 42 | 23 | 23 | 0 | 0 | 28 | 113 | 23 | 0 | 0 | 0 | 0 | 37 | 0 |
| SRR7690713 | 108 | 1789 | 9535 | 622 | 4869 | 5403 | 45882 | 25573 | 32538 | 2960 | 9558 | 13 | 14 | 3546 | 1034 | 982 | 42188 | 0 | 0 | 13 | 225 | 618 | 199 | 49 | 17 | 29 | 0 | 0 | 56 | 246 | 132 | 28 | 0 | 0 | 0 | 52 | 0 |
| SRR7690714 | 275 | 2604 | 18970 | 469 | 3618 | 3501 | 38206 | 15144 | 19476 | 1187 | 6575 | 12 | 10391 | 4093 | 615 | 1095 | 45503 | 0 | 0 | 0 | 76 | 190 | 73 | 37 | 19 | 18 | 0 | 0 | 21 | 59 | 31 | 0 | 0 | 0 | 0 | 41 | 0 |
| SRR7690715 | 1191 | 8226 | 1311 | 451 | 524 | 9080 | 10527 | 11693 | 9182 | 89 | 8553 | 15273 | 3888 | 1164 | 24 | 1508 | 36306 | 0 | 0 | 0 | 1076 | 1154 | 58 | 0 | 0 | 0 | 0 | 0 | 52 | 786 | 485 | 141 | 0 | 0 | 0 | 0 | 0 |
| SRR7690716 | 1072 | 13736 | 622 | 616 | 514 | 8145 | 7262 | 11280 | 7859 | 55 | 8033 | 17829 | 3968 | 835 | 20 | 1263 | 42720 | 0 | 0 | 0 | 4143 | 1845 | 155 | 0 | 0 | 0 | 0 | 0 | 159 | 2113 | 1297 | 192 | 0 | 0 | 0 | 0 | 0 |
| SRR7690717 | 97 | 3589 | 13153 | 135 | 7721 | 10151 | 97408 | 41272 | 31964 | 3275 | 10 | 8138 | 13198 | 4352 | 1947 | 409 | 31086 | 0 | 0 | 12 | 256 | 413 | 245 | 173 | 72 | 108 | 0 | 13 | 59 | 229 | 93 | 16 | 0 | 0 | 22 | 149 | 0 |
| SRR7690718 | 282 | 3144 | 7901 | 270 | 3902 | 4782 | 54067 | 30394 | 22588 | 1367 | 18140 | 11156 | 18016 | 3708 | 1373 | 215 | 43819 | 0 | 11 | 12 | 253 | 418 | 204 | 139 | 57 | 93 | 15 | 30 | 84 | 323 | 117 | 20 | 0 | 0 | 23 | 110 | 0 |
| SRR7690719 | 107 | 3415 | 10121 | 211 | 6121 | 6770 | 84433 | 41243 | 24577 | 3405 | 10851 | 7795 | 11297 | 4259 | 1994 | 1089 | 32228 | 0 | 0 | 0 | 114 | 363 | 232 | 146 | 62 | 68 | 0 | 12 | 39 | 161 | 84 | 30 | 0 | 0 | 29 | 128 | 0 |
| SRR7690720 | 99 | 1770 | 5793 | 400 | 6256 | 7044 | 63711 | 31521 | 27193 | 5206 | 7973 | 5064 | 8116 | 3534 | 1179 | 875 | 28369 | 0 | 0 | 15 | 134 | 201 | 86 | 38 | 20 | 38 | 0 | 11 | 56 | 154 | 56 | 18 | 0 | 0 | 0 | 61 | 0 |
| SRR7690721 | 77 | 1844 | 5530 | 782 | 5583 | 11509 | 53920 | 26607 | 21068 | 3241 | 5234 | 4895 | 6755 | 4584 | 1840 | 1085 | 19846 | 0 | 0 | 32 | 208 | 383 | 151 | 58 | 28 | 41 | 0 | 0 | 74 | 184 | 73 | 21 | 0 | 0 | 16 | 67 | 0 |
| SRR7690722 | 298 | 7015 | 39359 | 1216 | 9025 | 9196 | 44045 | 22229 | 113777 | 4443 | 18537 | 19 | 19311 | 9898 | 2606 | 585 | 37568 | 0 | 0 | 32 | 221 | 1030 | 132 | 21 | 0 | 0 | 0 | 11 | 108 | 236 | 35 | 0 | 0 | 0 | 0 | 16 | 0 |
| SRR7690723 | 102 | 1910 | 56382 | 398 | 6177 | 7709 | 33235 | 12144 | 106279 | 2696 | 17706 | 8912 | 16992 | 7766 | 2853 | 378 | 38499 | 0 | 0 | 0 | 25 | 163 | 25 | 0 | 0 | 0 | 0 | 0 | 15 | 22 | 0 | 0 | 0 | 0 | 0 | 0 | 0 |
| SRR7690724 | 102 | 6190 | 35305 | 767 | 10569 | 13330 | 51668 | 21758 | 202118 | 2491 | 11349 | 5681 | 12779 | 8544 | 7066 | 526 | 24886 | 0 | 0 | 38 | 198 | 3461 | 462 | 57 | 16 | 11 | 0 | 0 | 97 | 224 | 151 | 0 | 0 | 0 | 0 | 46 | 0 |
| SRR7690726 | 162 | 4509 | 30782 | 726 | 8318 | 8536 | 44691 | 22227 | 96586 | 6142 | 19601 | 19 | 18473 | 10267 | 1968 | 583 | 41804 | 0 | 0 | 46 | 291 | 1317 | 241 | 43 | 10 | 16 | 0 | 12 | 187 | 330 | 72 | 0 | 0 | 0 | 0 | 34 | 0 |
| SRR7690727 | 192 | 3753 | 12765 | 112 | 5468 | 4889 | 56534 | 30518 | 33982 | 2260 | 12 | 12227 | 19838 | 4909 | 1318 | 329 | 38420 | 0 | 0 | 15 | 334 | 468 | 222 | 104 | 39 | 65 | 10 | 24 | 81 | 347 | 117 | 16 | 0 | 0 | 18 | 79 | 0 |
| SRR7690728 | 81 | 1910 | 9962 | 47 | 5639 | 2749 | 56043 | 24145 | 25617 | 3175 | 23980 | 16 | 14 | 3088 | 1198 | 145 | 31401 | 0 | 0 | 11 | 415 | 252 | 167 | 95 | 42 | 69 | 0 | 13 | 53 | 316 | 54 | 12 | 0 | 0 | 20 | 95 | 0 |
| SRR7690729 | 98 | 2571 | 19408 | 47 | 10673 | 9036 | 105881 | 53340 | 92834 | 2571 | 14859 | 9662 | 14301 | 5822 | 2530 | 579 | 27719 | 0 | 0 | 0 | 265 | 203 | 65 | 61 | 25 | 48 | 0 | 14 | 47 | 258 | 49 | 0 | 0 | 0 | 12 | 66 | 0 |
| SRR7690730 | 244 | 3552 | 30186 | 40 | 4349 | 3892 | 53099 | 19546 | 26357 | 3460 | 7708 | 5707 | 9470 | 3316 | 616 | 500 | 33152 | 0 | 0 | 0 | 92 | 72 | 26 | 20 | 0 | 11 | 0 | 0 | 19 | 101 | 22 | 0 | 0 | 0 | 0 | 25 | 0 |
| SRR7690731 | 59 | 1584 | 10579 | 59 | 5684 | 6422 | 56488 | 26132 | 28103 | 3589 | 12508 | 8745 | 14008 | 5311 | 1493 | 864 | 32854 | 0 | 0 | 12 | 362 | 316 | 121 | 91 | 45 | 84 | 0 | 0 | 58 | 412 | 87 | 19 | 0 | 0 | 28 | 108 | 0 |
| SRR7690732 | 112 | 2420 | 9736 | 666 | 3776 | 4896 | 43846 | 23234 | 23158 | 3652 | 5986 | 11 | 8354 | 3396 | 576 | 969 | 28407 | 0 | 0 | 0 | 125 | 362 | 170 | 49 | 18 | 20 | 0 | 0 | 40 | 128 | 85 | 28 | 0 | 0 | 15 | 41 | 0 |
| SRR7690733 | 40 | 1812 | 8528 | 376 | 10100 | 11229 | 108788 | 38095 | 31587 | 5033 | 7530 | 4906 | 8482 | 4184 | 2412 | 864 | 23732 | 0 | 0 | 0 | 40 | 72 | 44 | 42 | 19 | 20 | 0 | 0 | 26 | 40 | 21 | 0 | 0 | 0 | 14 | 41 | 0 |
| SRR7690734 | 253 | 4037 | 66917 | 1677 | 17412 | 16626 | 95922 | 23461 | 172827 | 1288 | 16545 | 11 | 12881 | 10404 | 4465 | 492 | 15427 | 0 | 0 | 32 | 22 | 120 | 13 | 0 | 0 | 0 | 0 | 26 | 105 | 55 | 0 | 0 | 0 | 0 | 0 | 11 | 0 |
| SRR7690736 | 246 | 7925 | 51259 | 1273 | 19540 | 11906 | 89719 | 23929 | 84953 | 3389 | 15192 | 15 | 15273 | 9113 | 3474 | 1209 | 25685 | 0 | 0 | 27 | 189 | 461 | 143 | 36 | 0 | 0 | 0 | 0 | 113 | 195 | 20 | 0 | 0 | 0 | 0 | 17 | 0 |
| SRR7690737 | 51 | 4433 | 86299 | 105 | 17270 | 16432 | 89456 | 25079 | 156287 | 8159 | 14472 | 6217 | 15745 | 8727 | 4177 | 332 | 26930 | 0 | 0 | 0 | 103 | 569 | 101 | 26 | 0 | 0 | 0 | 0 | 37 | 100 | 20 | 0 | 0 | 0 | 0 | 18 | 0 |
| SRR7690739 | 301 | 17376 | 85770 | 452 | 11677 | 8492 | 46884 | 18313 | 87834 | 6083 | 15967 | 7242 | 18341 | 8701 | 2803 | 582 | 29498 | 0 | 0 | 14 | 65 | 473 | 113 | 29 | 0 | 0 | 0 | 0 | 43 | 60 | 26 | 0 | 0 | 0 | 0 | 0 | 0 |
| SRR7690740 | 411 | 11712 | 104668 | 580 | 12202 | 8833 | 48072 | 16774 | 106168 | 6113 | 17500 | 16 | 17561 | 9039 | 3723 | 574 | 29470 | 0 | 0 | 12 | 51 | 511 | 115 | 34 | 0 | 0 | 0 | 0 | 45 | 45 | 24 | 0 | 0 | 0 | 0 | 0 | 0 |
| SRR7690741 | 228 | 13313 | 772 | 368 | 662 | 6688 | 7215 | 9296 | 5557 | 59 | 9314 | 31294 | 6259 | 1203 | 17 | 1518 | 40342 | 0 | 0 | 0 | 3272 | 827 | 60 | 0 | 0 | 0 | 0 | 0 | 127 | 2596 | 817 | 147 | 0 | 0 | 0 | 0 | 0 |
| SRR7690742 | 166 | 11268 | 675 | 207 | 568 | 4882 | 6278 | 6640 | 6433 | 86 | 7760 | 30167 | 4985 | 883 | 12 | 1677 | 42223 | 0 | 0 | 0 | 5501 | 1389 | 110 | 0 | 0 | 0 | 0 | 0 | 220 | 3099 | 1192 | 157 | 0 | 0 | 0 | 0 | 0 |
| SRR7690743 | 587 | 10289 | 953 | 151 | 517 | 5456 | 5852 | 9368 | 4640 | 112 | 8146 | 21265 | 4667 | 748 | 0 | 1328 | 47110 | 0 | 0 | 0 | 2226 | 442 | 34 | 0 | 0 | 0 | 0 | 0 | 110 | 1298 | 355 | 54 | 0 | 0 | 0 | 0 | 0 |
| SRR7690744 | 473 | 4102 | 16863 | 635 | 8270 | 5335 | 75657 | 49643 | 22727 | 3727 | 9568 | 13 | 16 | 3694 | 1054 | 526 | 31623 | 0 | 0 | 39 | 1479 | 563 | 217 | 90 | 33 | 69 | 18 | 24 | 136 | 1304 | 158 | 31 | 0 | 0 | 24 | 121 | 0 |
| SRR7690745 | 72 | 1781 | 6369 | 203 | 9871 | 6166 | 69447 | 39517 | 38134 | 4067 | 11071 | 7508 | 12584 | 4598 | 1928 | 655 | 25226 | 0 | 0 | 23 | 612 | 361 | 148 | 82 | 36 | 59 | 0 | 14 | 67 | 409 | 128 | 19 | 0 | 0 | 27 | 115 | 0 |
| SRR7690746 | 385 | 2991 | 14218 | 306 | 8795 | 6077 | 95169 | 26780 | 29728 | 1136 | 11117 | 6600 | 11680 | 3819 | 1201 | 849 | 36475 | 0 | 0 | 11 | 269 | 155 | 78 | 39 | 23 | 31 | 0 | 11 | 41 | 207 | 35 | 0 | 0 | 0 | 14 | 71 | 0 |
| SRR7690747 | 77 | 1845 | 26232 | 64 | 5422 | 5323 | 54650 | 29118 | 26193 | 3308 | 11 | 8676 | 14583 | 5407 | 1415 | 704 | 44162 | 0 | 0 | 13 | 299 | 333 | 122 | 53 | 25 | 77 | 0 | 0 | 44 | 212 | 71 | 12 | 0 | 0 | 16 | 85 | 0 |
| SRR7690748 | 262 | 3893 | 10511 | 518 | 6530 | 7310 | 78232 | 30936 | 30369 | 3370 | 8311 | 6871 | 9809 | 4166 | 1411 | 1028 | 35451 | 0 | 0 | 0 | 93 | 182 | 115 | 101 | 42 | 39 | 0 | 11 | 38 | 98 | 31 | 13 | 0 | 0 | 19 | 84 | 0 |
| SRR7690749 | 588 | 3746 | 11407 | 420 | 2187 | 1783 | 28800 | 6480 | 19026 | 2690 | 4861 | 18873 | 13702 | 6312 | 247 | 1090 | 41310 | 0 | 0 | 0 | 118 | 314 | 143 | 90 | 47 | 55 | 0 | 0 | 32 | 124 | 54 | 16 | 0 | 0 | 15 | 102 | 0 |
| SRR7690750 | 220 | 3017 | 8563 | 210 | 6855 | 4776 | 64451 | 34560 | 31380 | 2568 | 16346 | 11007 | 17141 | 5092 | 1379 | 435 | 34193 | 0 | 15 | 25 | 837 | 879 | 388 | 158 | 59 | 136 | 22 | 32 | 123 | 827 | 194 | 28 | 0 | 0 | 28 | 164 | 0 |
| SRR7690751 | 356 | 2927 | 8047 | 320 | 7071 | 5363 | 90097 | 33540 | 38372 | 2949 | 14231 | 12 | 15 | 5652 | 1459 | 557 | 26392 | 0 | 16 | 23 | 577 | 508 | 229 | 142 | 53 | 132 | 30 | 32 | 123 | 678 | 135 | 22 | 0 | 0 | 29 | 169 | 0 |
| SRR7690752 | 40 | 1158 | 13700 | 61 | 6926 | 5848 | 69061 | 32294 | 46753 | 3540 | 12715 | 15 | 21 | 5448 | 1228 | 636 | 35542 | 0 | 0 | 0 | 227 | 244 | 85 | 55 | 29 | 69 | 0 | 0 | 39 | 250 | 59 | 11 | 0 | 0 | 14 | 95 | 0 |
| SRR7690753 | 283 | 2914 | 4717 | 1206 | 6546 | 5632 | 64583 | 30720 | 22069 | 3952 | 5946 | 4069 | 6204 | 3016 | 1023 | 1277 | 27582 | 0 | 11 | 22 | 280 | 248 | 101 | 49 | 19 | 44 | 14 | 18 | 75 | 286 | 65 | 10 | 0 | 0 | 0 | 84 | 0 |
| SRR7690754 | 185 | 3489 | 6255 | 609 | 8638 | 6085 | 64353 | 24218 | 30073 | 4930 | 10623 | 7730 | 11205 | 4135 | 1142 | 1086 | 38482 | 0 | 0 | 10 | 61 | 95 | 46 | 30 | 13 | 21 | 0 | 16 | 39 | 68 | 38 | 0 | 0 | 0 | 0 | 45 | 0 |
| SRR7690755 | 124 | 2465 | 3975 | 1504 | 5234 | 10660 | 43217 | 27656 | 19266 | 3617 | 5074 | 4344 | 6487 | 4241 | 1172 | 881 | 17391 | 0 | 0 | 16 | 95 | 96 | 51 | 29 | 12 | 25 | 0 | 11 | 37 | 104 | 29 | 11 | 0 | 0 | 0 | 42 | 0 |
| SRR7690756 | 23 | 1029 | 6912 | 124 | 7812 | 8792 | 75074 | 34118 | 30515 | 4930 | 7104 | 11 | 7659 | 3849 | 1977 | 1418 | 28460 | 0 | 0 | 12 | 183 | 349 | 105 | 42 | 18 | 38 | 0 | 0 | 37 | 215 | 108 | 27 | 0 | 0 | 11 | 83 | 0 |
| SRR7690757 | 181 | 1700 | 5085 | 1097 | 11643 | 10705 | 117478 | 49464 | 27847 | 4882 | 7208 | 4290 | 6673 | 3658 | 3941 | 863 | 21585 | 0 | 12 | 20 | 254 | 343 | 155 | 162 | 41 | 48 | 12 | 22 | 85 | 287 | 75 | 40 | 0 | 0 | 21 | 96 | 0 |
| SRR7690758 | 223 | 11278 | 36732 | 195 | 9921 | 6811 | 49826 | 22579 | 68355 | 8416 | 14951 | 5105 | 13705 | 8649 | 2388 | 717 | 25168 | 0 | 0 | 31 | 163 | 4861 | 1482 | 87 | 14 | 0 | 0 | 0 | 112 | 210 | 275 | 20 | 0 | 0 | 0 | 53 | 0 |
| SRR7690759 | 207 | 9944 | 32826 | 1388 | 13393 | 8455 | 58070 | 14657 | 65395 | 3443 | 20043 | 17 | 19091 | 13307 | 4347 | 1115 | 39893 | 0 | 0 | 15 | 192 | 315 | 57 | 18 | 0 | 0 | 0 | 0 | 61 | 162 | 11 | 0 | 0 | 0 | 0 | 11 | 0 |
| SRR7690762 | 293 | 6818 | 70626 | 436 | 9745 | 8963 | 46998 | 15425 | 86117 | 5232 | 18482 | 19 | 19091 | 10069 | 4104 | 660 | 32656 | 0 | 0 | 12 | 52 | 308 | 56 | 15 | 0 | 0 | 0 | 0 | 46 | 40 | 16 | 0 | 0 | 0 | 0 | 0 | 0 |
| SRR7690764 | 1448 | 18331 | 651 | 312 | 534 | 6497 | 6170 | 6566 | 7265 | 126 | 9117 | 28893 | 5547 | 1330 | 24 | 1458 | 48578 | 0 | 0 | 0 | 6079 | 1338 | 67 | 0 | 0 | 0 | 0 | 0 | 216 | 3559 | 901 | 110 | 0 | 0 | 0 | 0 | 0 |
| SRR7690765 | 980 | 7709 | 728 | 240 | 535 | 7297 | 7968 | 9926 | 6429 | 61 | 8824 | 19299 | 4396 | 850 | 15 | 1550 | 44816 | 0 | 0 | 0 | 4931 | 957 | 65 | 0 | 0 | 0 | 0 | 0 | 171 | 2860 | 699 | 94 | 0 | 0 | 0 | 0 | 0 |
| SRR7690766 | 318 | 2537 | 10572 | 463 | 4914 | 6762 | 61567 | 33728 | 36077 | 2323 | 11049 | 6598 | 11962 | 4533 | 1117 | 686 | 32055 | 0 | 12 | 20 | 214 | 605 | 168 | 83 | 31 | 89 | 10 | 23 | 85 | 275 | 133 | 11 | 0 | 0 | 14 | 96 | 0 |
| SRR7690767 | 173 | 2270 | 10002 | 198 | 5008 | 4826 | 60318 | 32179 | 26607 | 2231 | 15834 | 9637 | 15011 | 4669 | 1526 | 492 | 34001 | 0 | 0 | 17 | 521 | 536 | 232 | 149 | 52 | 128 | 12 | 17 | 82 | 560 | 128 | 15 | 0 | 0 | 23 | 159 | 0 |
| SRR7690768 | 76 | 1917 | 7170 | 384 | 5412 | 6483 | 53960 | 34121 | 26396 | 4428 | 7914 | 13 | 9579 | 3794 | 856 | 996 | 34122 | 0 | 0 | 0 | 109 | 348 | 101 | 36 | 18 | 27 | 0 | 0 | 36 | 114 | 71 | 17 | 0 | 0 | 12 | 56 | 0 |
| SRR7690769 | 239 | 7980 | 34744 | 1154 | 8251 | 9067 | 39152 | 15539 | 87750 | 2563 | 18029 | 8283 | 16818 | 7921 | 2455 | 614 | 35602 | 0 | 0 | 0 | 32 | 187 | 17 | 0 | 0 | 0 | 0 | 0 | 33 | 29 | 11 | 0 | 0 | 0 | 0 | 0 | 0 |
| SRR7690770 | 197 | 6620 | 30666 | 1127 | 7447 | 8798 | 35491 | 18886 | 100956 | 3343 | 19350 | 8694 | 20520 | 9340 | 2157 | 688 | 41528 | 0 | 0 | 41 | 422 | 1216 | 118 | 18 | 0 | 0 | 0 | 0 | 156 | 472 | 52 | 0 | 0 | 0 | 0 | 18 | 0 |
| SRR7690773 | 61 | 3477 | 23492 | 57 | 9647 | 8793 | 56621 | 28714 | 97340 | 13805 | 13465 | 14 | 12594 | 8836 | 4956 | 507 | 21370 | 0 | 0 | 13 | 42 | 407 | 37 | 0 | 0 | 0 | 0 | 0 | 47 | 50 | 26 | 0 | 0 | 0 | 0 | 16 | 0 |
| SRR7690775 | 110 | 2835 | 9227 | 614 | 4742 | 7379 | 40940 | 29343 | 18951 | 3575 | 6410 | 5089 | 8065 | 3169 | 1066 | 757 | 21600 | 0 | 0 | 16 | 169 | 163 | 90 | 57 | 27 | 40 | 0 | 0 | 39 | 182 | 57 | 14 | 0 | 0 | 17 | 73 | 0 |
| SRR7690777 | 109 | 2073 | 13697 | 251 | 6409 | 5778 | 58901 | 31988 | 36713 | 2618 | 10462 | 6237 | 11684 | 5682 | 2497 | 978 | 28833 | 0 | 11 | 32 | 233 | 645 | 300 | 74 | 29 | 65 | 0 | 18 | 80 | 182 | 214 | 39 | 0 | 0 | 21 | 92 | 0 |
| SRR7690778 | 475 | 3844 | 5856 | 835 | 2050 | 1943 | 22281 | 9672 | 12680 | 3174 | 2763 | 11003 | 9169 | 5323 | 192 | 1479 | 21373 | 0 | 0 | 0 | 226 | 143 | 111 | 97 | 54 | 53 | 0 | 0 | 27 | 239 | 34 | 14 | 0 | 0 | 14 | 80 | 0 |
| SRR7690779 | 70 | 1847 | 4689 | 2390 | 4989 | 13943 | 58671 | 14670 | 15112 | 2798 | 3622 | 3348 | 4001 | 4491 | 1314 | 1040 | 8423 | 0 | 0 | 12 | 55 | 74 | 41 | 17 | 10 | 17 | 0 | 0 | 16 | 40 | 12 | 0 | 0 | 0 | 0 | 38 | 0 |
| SRR7690780 | 121 | 1442 | 6598 | 449 | 6921 | 6654 | 68866 | 24378 | 17028 | 4169 | 10468 | 13 | 10976 | 5137 | 1637 | 885 | 27174 | 0 | 0 | 17 | 232 | 243 | 99 | 68 | 30 | 49 | 16 | 18 | 94 | 330 | 103 | 21 | 0 | 0 | 20 | 101 | 0 |
| SRR7690781 | 187 | 2644 | 6720 | 686 | 7173 | 5539 | 58713 | 26842 | 22026 | 3988 | 9657 | 6303 | 14 | 4641 | 1505 | 1545 | 41043 | 0 | 15 | 43 | 1283 | 1359 | 326 | 74 | 31 | 61 | 0 | 17 | 176 | 1368 | 276 | 47 | 0 | 0 | 13 | 117 | 0 |
| SRR7690783 | 388 | 21269 | 37578 | 1290 | 12929 | 9269 | 57212 | 18310 | 93639 | 6938 | 13896 | 11 | 15003 | 7575 | 2614 | 751 | 29067 | 0 | 0 | 11 | 108 | 378 | 69 | 15 | 0 | 0 | 0 | 0 | 50 | 133 | 20 | 0 | 0 | 0 | 0 | 0 | 0 |
| SRR7690788 | 416 | 4885 | 39024 | 1453 | 6354 | 6013 | 31045 | 10966 | 77342 | 5083 | 12633 | 23 | 15932 | 9136 | 1089 | 1438 | 42918 | 0 | 0 | 53 | 233 | 862 | 107 | 29 | 0 | 18 | 0 | 20 | 294 | 396 | 86 | 0 | 0 | 0 | 0 | 33 | 0 |
| SRR7690789 | 396 | 9876 | 18059 | 2904 | 12988 | 11382 | 47592 | 18675 | 96970 | 3247 | 17223 | 7488 | 18624 | 9366 | 3683 | 917 | 36438 | 0 | 0 | 43 | 201 | 4371 | 2311 | 223 | 45 | 27 | 0 | 22 | 155 | 221 | 285 | 42 | 0 | 0 | 17 | 57 | 0 |
| SRR7690790 | 297 | 6080 | 4571 | 2298 | 6072 | 8567 | 51247 | 34250 | 22164 | 3120 | 6848 | 5015 | 8458 | 3113 | 1227 | 816 | 25006 | 0 | 14 | 68 | 1003 | 1150 | 420 | 155 | 53 | 76 | 32 | 37 | 218 | 944 | 350 | 77 | 0 | 0 | 30 | 125 | 0 |
| SRR7690791 | 278 | 2811 | 9721 | 635 | 5324 | 3507 | 50472 | 25969 | 24548 | 3204 | 13887 | 11074 | 13935 | 3295 | 1445 | 211 | 39790 | 0 | 11 | 28 | 397 | 722 | 584 | 111 | 42 | 75 | 13 | 23 | 77 | 192 | 133 | 28 | 0 | 0 | 22 | 108 | 0 |
| SRR7690792 | 96 | 1355 | 8276 | 286 | 7792 | 6799 | 78205 | 40442 | 38378 | 2311 | 10675 | 5540 | 10938 | 5229 | 2337 | 613 | 23097 | 0 | 10 | 65 | 620 | 2859 | 658 | 86 | 33 | 79 | 0 | 17 | 157 | 483 | 578 | 82 | 0 | 0 | 19 | 129 | 0 |
| SRR7690793 | 300 | 2647 | 13294 | 425 | 5773 | 5077 | 71296 | 14035 | 31149 | 1581 | 3536 | 7626 | 8206 | 3846 | 710 | 1050 | 31383 | 0 | 0 | 11 | 355 | 324 | 157 | 123 | 71 | 83 | 0 | 12 | 40 | 316 | 50 | 13 | 0 | 0 | 14 | 130 | 0 |
| SRR7690794 | 148 | 1971 | 13914 | 130 | 1778 | 1221 | 21460 | 4289 | 16232 | 1993 | 3720 | 19657 | 8407 | 5105 | 234 | 622 | 50298 | 0 | 0 | 0 | 174 | 313 | 152 | 65 | 41 | 46 | 0 | 0 | 20 | 149 | 48 | 13 | 0 | 0 | 0 | 64 | 0 |
| SRR7690795 | 908 | 3097 | 5606 | 1962 | 2058 | 1546 | 25455 | 7103 | 12980 | 2099 | 2982 | 12812 | 6053 | 3684 | 219 | 595 | 39887 | 0 | 12 | 11 | 189 | 224 | 102 | 84 | 48 | 41 | 0 | 19 | 40 | 154 | 39 | 12 | 0 | 0 | 0 | 56 | 0 |
| SRR7690796 | 169 | 2126 | 9092 | 718 | 5955 | 5419 | 56409 | 25778 | 25310 | 3495 | 9745 | 6385 | 10416 | 3628 | 1024 | 1037 | 40932 | 0 | 11 | 18 | 454 | 460 | 109 | 46 | 20 | 37 | 0 | 20 | 87 | 553 | 112 | 16 | 0 | 0 | 11 | 70 | 0 |
| SRR7690797 | 782 | 21823 | 29018 | 413 | 9237 | 5990 | 40513 | 16813 | 63433 | 5726 | 18352 | 7304 | 14953 | 6670 | 2306 | 589 | 26520 | 0 | 0 | 0 | 40 | 113 | 25 | 0 | 0 | 0 | 0 | 0 | 32 | 31 | 0 | 0 | 0 | 0 | 0 | 0 | 0 |
| SRR7690798 | 357 | 17478 | 59601 | 579 | 8344 | 8767 | 34975 | 14928 | 93760 | 4766 | 15306 | 17 | 16798 | 7413 | 2067 | 698 | 34346 | 0 | 0 | 0 | 41 | 531 | 64 | 0 | 0 | 0 | 0 | 0 | 43 | 51 | 29 | 0 | 0 | 0 | 0 | 0 | 0 |
| SRR7690800 | 170 | 9180 | 40892 | 1374 | 5975 | 6789 | 31526 | 10241 | 87877 | 2594 | 19107 | 23 | 16320 | 8737 | 3209 | 671 | 45788 | 0 | 0 | 14 | 28 | 246 | 25 | 0 | 0 | 0 | 0 | 10 | 34 | 23 | 0 | 0 | 0 | 0 | 0 | 11 | 0 |
| SRR7690801 | 106 | 3421 | 61179 | 190 | 8535 | 9623 | 48782 | 19328 | 99367 | 11468 | 17229 | 15 | 17968 | 10288 | 4085 | 695 | 32670 | 0 | 0 | 27 | 133 | 1024 | 155 | 24 | 0 | 13 | 0 | 0 | 101 | 185 | 48 | 0 | 0 | 0 | 0 | 27 | 0 |
| SRR7690802 | 57 | 1545 | 50593 | 139 | 8614 | 8240 | 58402 | 19184 | 72628 | 4177 | 24199 | 11609 | 21124 | 8495 | 3460 | 511 | 38043 | 0 | 0 | 0 | 35 | 115 | 33 | 12 | 0 | 0 | 0 | 0 | 0 | 28 | 0 | 0 | 0 | 0 | 0 | 10 | 0 |
| SRR7690803 | 280 | 6080 | 38724 | 851 | 7899 | 6564 | 40094 | 25391 | 70953 | 4979 | 16006 | 16 | 17207 | 6985 | 1484 | 535 | 42877 | 0 | 0 | 11 | 92 | 293 | 48 | 14 | 0 | 0 | 0 | 11 | 58 | 136 | 24 | 0 | 0 | 0 | 0 | 12 | 0 |
| SRR7819267 | 0 | 10 | 88463 | 0 | 173 | 236 | 1714 | 7615 | 42336 | 185 | 921 | 1441 | 658 | 5761 | 27 | 54 | 16488 | 0 | 0 | 0 | 0 | 0 | 0 | 0 | 0 | 0 | 0 | 0 | 0 | 0 | 0 | 0 | 0 | 0 | 0 | 0 | 0 |
| SRR7819269 | 0 | 57 | 106703 | 0 | 442 | 924 | 5440 | 11987 | 79539 | 314 | 1327 | 1271 | 1099 | 17337 | 39 | 110 | 50540 | 0 | 0 | 0 | 0 | 0 | 0 | 0 | 0 | 0 | 0 | 0 | 0 | 0 | 0 | 0 | 0 | 0 | 0 | 0 | 0 |
| SRR7819272 | 0 | 105 | 124245 | 72 | 315 | 446 | 3838 | 13638 | 84919 | 138 | 1161 | 1531 | 940 | 13666 | 59 | 74 | 23698 | 0 | 0 | 0 | 0 | 0 | 0 | 0 | 0 | 0 | 0 | 0 | 0 | 0 | 0 | 0 | 0 | 0 | 0 | 0 | 0 |
| SRR7851651 | 0 | 72 | 5284 | 0 | 10544 | 9916 | 228070 | 68083 | 3165 | 312 | 9971 | 33225 | 4599 | 18344 | 170 | 2486 | 14937 | 0 | 0 | 0 | 0 | 0 | 19 | 12 | 87 | 88 | 0 | 0 | 0 | 0 | 0 | 0 | 0 | 0 | 18 | 145 | 0 |
| SRR7851652 | 0 | 36 | 5429 | 0 | 10419 | 11132 | 211634 | 65249 | 3640 | 198 | 9324 | 26042 | 4486 | 16812 | 213 | 2522 | 14118 | 0 | 0 | 0 | 0 | 0 | 14 | 17 | 204 | 188 | 0 | 0 | 0 | 0 | 0 | 0 | 0 | 0 | 41 | 264 | 0 |
| SRR7851653 | 0 | 18 | 4688 | 0 | 6685 | 7977 | 197484 | 66569 | 3399 | 127 | 7958 | 25492 | 3893 | 16748 | 101 | 1830 | 15529 | 0 | 0 | 0 | 0 | 0 | 0 | 0 | 40 | 32 | 0 | 0 | 0 | 0 | 0 | 0 | 0 | 0 | 0 | 45 | 0 |
| SRR7851655 | 0 | 41 | 5651 | 0 | 10666 | 10938 | 214866 | 65814 | 3454 | 225 | 10023 | 28866 | 4786 | 17901 | 213 | 2494 | 14106 | 0 | 0 | 0 | 0 | 0 | 14 | 0 | 99 | 106 | 0 | 0 | 0 | 0 | 0 | 0 | 0 | 0 | 25 | 188 | 0 |
| SRR7851656 | 0 | 81 | 7382 | 0 | 8679 | 9008 | 200274 | 64313 | 3382 | 285 | 10906 | 33392 | 4945 | 18089 | 191 | 2172 | 14578 | 0 | 0 | 0 | 0 | 0 | 19 | 0 | 51 | 42 | 0 | 0 | 0 | 0 | 0 | 0 | 0 | 0 | 0 | 67 | 0 |
| SRR7901057 | 772 | 3563 | 8405 | 363 | 4534 | 3564 | 44331 | 997 | 91 | 108 | 20487 | 31851 | 7408 | 60015 | 7908 | 1012 | 7446 | 208 | 253 | 84 | 273 | 348 | 360 | 187 | 236 | 60 | 385 | 280 | 445 | 196 | 67 | 18 | 0 | 0 | 30 | 308 | 58 |
| SRR7901058 | 428 | 2956 | 8816 | 192 | 4700 | 3410 | 46589 | 1210 | 125 | 96 | 21723 | 29166 | 7912 | 60175 | 7526 | 819 | 4973 | 156 | 146 | 53 | 190 | 206 | 237 | 132 | 136 | 47 | 232 | 194 | 212 | 97 | 40 | 0 | 0 | 0 | 16 | 182 | 39 |
| SRR7901059 | 619 | 4007 | 12965 | 155 | 4619 | 3646 | 44956 | 1104 | 118 | 66 | 17883 | 29706 | 6131 | 60288 | 6744 | 1185 | 8353 | 372 | 442 | 153 | 427 | 390 | 513 | 282 | 305 | 92 | 517 | 489 | 463 | 203 | 37 | 13 | 0 | 0 | 24 | 357 | 64 |
| SRR7903701 | 10511 | 21257 | 96198 | 618 | 28902 | 23953 | 85697 | 15362 | 29056 | 12323 | 3943 | 1857 | 4876 | 3178 | 3254 | 206 | 12221 | 0 | 90 | 133 | 96 | 344 | 133 | 42 | 14 | 0 | 0 | 87 | 123 | 37 | 29 | 0 | 0 | 0 | 0 | 12 | 0 |
| SRR7903702 | 8337 | 13818 | 102244 | 555 | 28702 | 25189 | 85412 | 14343 | 32292 | 13900 | 3853 | 1878 | 6241 | 3967 | 3248 | 200 | 13125 | 0 | 97 | 120 | 84 | 378 | 127 | 34 | 11 | 0 | 0 | 89 | 116 | 37 | 33 | 0 | 0 | 0 | 0 | 11 | 0 |
| SRR7903703 | 8363 | 15835 | 93765 | 500 | 29630 | 25010 | 83596 | 15847 | 28770 | 14984 | 4253 | 2002 | 5478 | 3396 | 3224 | 217 | 12546 | 0 | 109 | 133 | 102 | 366 | 136 | 40 | 13 | 0 | 10 | 100 | 148 | 42 | 33 | 0 | 0 | 0 | 0 | 12 | 0 |
| SRR7903704 | 7815 | 7589 | 13819 | 789 | 3813 | 3256 | 9213 | 1456 | 3649 | 641 | 15496 | 15396 | 12353 | 8911 | 6710 | 59 | 34511 | 624 | 1587 | 7501 | 2227 | 4954 | 1739 | 2427 | 895 | 935 | 736 | 2081 | 10800 | 1117 | 2567 | 85 | 123 | 99 | 202 | 907 | 349 |
| SRR7903705 | 5721 | 5522 | 24659 | 799 | 2994 | 2317 | 7077 | 1239 | 2614 | 441 | 16040 | 10 | 12468 | 7920 | 5605 | 50 | 41861 | 344 | 887 | 4156 | 1484 | 2778 | 1071 | 1152 | 755 | 701 | 247 | 872 | 4373 | 363 | 240 | 12 | 0 | 0 | 89 | 606 | 140 |
| SRR7903706 | 4624 | 4961 | 25582 | 731 | 3167 | 3174 | 7723 | 1278 | 3301 | 449 | 13305 | 14431 | 10313 | 8209 | 5745 | 44 | 26885 | 819 | 1763 | 8058 | 2881 | 4779 | 1823 | 1983 | 1040 | 1007 | 565 | 1905 | 9237 | 739 | 436 | 28 | 19 | 0 | 148 | 950 | 261 |
| SRR8063607 | 270 | 3557 | 132090 | 128 | 1614 | 952 | 13682 | 1194 | 68583 | 9480 | 9147 | 4395 | 10093 | 1941 | 88730 | 8513 | 7676 | 0 | 40 | 30 | 67 | 17 | 40 | 15 | 13 | 0 | 14 | 110 | 346 | 99 | 17 | 0 | 0 | 0 | 0 | 0 | 0 |
| SRR8063608 | 214 | 3601 | 135114 | 127 | 1617 | 969 | 13319 | 1204 | 70538 | 9109 | 8971 | 4343 | 10289 | 2038 | 30 | 8528 | 7703 | 0 | 37 | 23 | 79 | 20 | 52 | 15 | 16 | 0 | 11 | 97 | 391 | 99 | 18 | 0 | 0 | 0 | 0 | 0 | 0 |
| SRR8063609 | 595 | 4797 | 121949 | 345 | 1748 | 916 | 14407 | 1297 | 67641 | 10233 | 9100 | 4267 | 9941 | 1992 | 23 | 8968 | 7526 | 0 | 51 | 42 | 85 | 19 | 52 | 12 | 12 | 0 | 30 | 170 | 533 | 130 | 25 | 0 | 0 | 0 | 0 | 0 | 0 |
| SRR8063610 | 61 | 5854 | 38311 | 604 | 2204 | 1073 | 15518 | 1472 | 58927 | 2031 | 8641 | 4019 | 8222 | 1671 | 49034 | 8167 | 7257 | 0 | 353 | 965 | 3669 | 163 | 91 | 13 | 0 | 0 | 10 | 350 | 2782 | 2553 | 53 | 0 | 0 | 0 | 0 | 11 | 0 |
| SRR8063611 | 48 | 7471 | 37692 | 764 | 2136 | 1066 | 14930 | 1456 | 64272 | 2051 | 8226 | 3957 | 8014 | 1425 | 49137 | 7832 | 7262 | 0 | 451 | 1141 | 3856 | 171 | 92 | 0 | 0 | 0 | 15 | 485 | 3357 | 2747 | 54 | 0 | 0 | 0 | 0 | 0 | 0 |
| SRR8063612 | 130 | 7388 | 34713 | 1564 | 2005 | 1040 | 15232 | 1481 | 62355 | 1688 | 8586 | 4041 | 8093 | 1566 | 50964 | 7819 | 6928 | 12 | 544 | 1258 | 3938 | 167 | 89 | 12 | 0 | 0 | 18 | 654 | 3895 | 2918 | 57 | 0 | 0 | 0 | 0 | 0 | 0 |
| SRR8063613 | 1777 | 35709 | 17201 | 838 | 1161 | 670 | 5993 | 525 | 57109 | 7343 | 17331 | 14 | 12289 | 2045 | 16 | 1689 | 10030 | 0 | 0 | 25 | 78 | 0 | 0 | 0 | 0 | 0 | 0 | 0 | 29 | 0 | 0 | 0 | 0 | 0 | 0 | 0 | 0 |
| SRR8063614 | 5848 | 18705 | 18301 | 1069 | 1170 | 556 | 12861 | 1199 | 36933 | 3127 | 8772 | 10353 | 8200 | 1066 | 10 | 5038 | 7437 | 0 | 0 | 38 | 161 | 0 | 0 | 0 | 0 | 0 | 0 | 0 | 88 | 39 | 0 | 0 | 0 | 0 | 0 | 0 | 0 |
| SRR8063615 | 1089 | 22551 | 23567 | 180 | 1117 | 577 | 12830 | 1253 | 35469 | 4586 | 8790 | 10388 | 8283 | 1025 | 44029 | 4814 | 7578 | 0 | 0 | 29 | 175 | 0 | 16 | 0 | 0 | 0 | 0 | 0 | 72 | 42 | 0 | 0 | 0 | 0 | 0 | 0 | 0 |
| SRR8074870 | 0 | 16 | 1762 | 0 | 7419 | 4545 | 115358 | 32114 | 2517 | 180 | 41600 | 48429 | 25997 | 39270 | 9514 | 45067 | 6560 | 0 | 0 | 10 | 15 | 0 | 0 | 28 | 177 | 56 | 0 | 0 | 55 | 26 | 0 | 0 | 0 | 0 | 0 | 0 | 0 |
| SRR8074871 | 0 | 0 | 1695 | 0 | 7560 | 4613 | 115833 | 32681 | 2530 | 201 | 41423 | 16 | 25818 | 39138 | 9435 | 44447 | 6431 | 0 | 0 | 10 | 15 | 0 | 0 | 29 | 177 | 54 | 0 | 0 | 56 | 26 | 0 | 0 | 0 | 0 | 0 | 11 | 0 |

### Supp Table S8.1.Columns_B_to_AL

| | | | | | | | | | | | | | | | | | | | | | | | | | | | | | | | | | | | | | |
|---|---|---|---|---|---|---|---|---|---|---|---|---|---|---|---|---|---|---|---|---|---|---|---|---|---|---|---|---|---|---|---|---|---|---|---|---|---|
| SRR8074873 | 0 | 52 | 1080 | 0 | 28582 | 7144 | 169082 | 31068 | 2335 | 141 | 32564 | 38775 | 19483 | 30533 | 6325 | 60083 | 6318 | 0 | 0 | 17 | 42 | 0 | 47 | 251 | 989 | 121 | 0 | 0 | 112 | 100 | 0 | 0 | 0 | 0 | 0 | 12 | 0 |
| SRR8074874 | 0 | 131 | 912 | 23 | 38860 | 7852 | 185710 | 28180 | 2987 | 142 | 27931 | 31116 | 17618 | 18423 | 9678 | 52780 | 5149 | 0 | 58 | 115 | 162 | 15 | 77 | 252 | 946 | 113 | 0 | 80 | 554 | 296 | 0 | 0 | 0 | 0 | 0 | 15 | 0 |
| SRR8074875 | 0 | 118 | 848 | 26 | 38076 | 7919 | 191441 | 29461 | 3117 | 128 | 28025 | 10 | 17643 | 18380 | 9727 | 51592 | 5081 | 0 | 47 | 96 | 144 | 14 | 66 | 225 | 884 | 104 | 0 | 66 | 499 | 259 | 0 | 0 | 0 | 0 | 0 | 15 | 0 |
| SRR8094756 | 456 | 89570 | 24444 | 419 | 19815 | 17921 | 82770 | 18859 | 7066 | 638 | 5601 | 38 | 5936 | 16617 | 735 | 2216 | 15333 | 0 | 51 | 51 | 313 | 729 | 218 | 106 | 132 | 22 | 0 | 53 | 139 | 79 | 25 | 0 | 0 | 0 | 0 | 29 | 0 |
| SRR8094757 | 2100 | 13908 | 5224 | 815 | 27372 | 21656 | 98507 | 21756 | 16036 | 73 | 12563 | 10462 | 10837 | 25501 | 3476 | 2194 | 16167 | 15 | 194 | 204 | 329 | 1450 | 260 | 490 | 3790 | 898 | 0 | 185 | 577 | 81 | 40 | 0 | 0 | 29 | 254 | 818 | 24 |
| SRR8094758 | 290 | 35460 | 8803 | 596 | 46775 | 23441 | 149266 | 26535 | 48826 | 97 | 21932 | 18 | 13359 | 90988 | 1818 | 1439 | 13375 | 0 | 14 | 33 | 199 | 702 | 78 | 85 | 75 | 0 | 0 | 16 | 94 | 51 | 15 | 0 | 0 | 0 | 0 | 16 | 0 |
| SRR8094759 | 114 | 387 | 3526 | 2688 | 4132 | 4334 | 8715 | 10834 | 192665 | 594 | 14007 | 7582 | 12527 | 4403 | 572 | 3262 | 30839 | 0 | 65 | 163 | 2031 | 808 | 189 | 29 | 0 | 0 | 0 | 671 | 5261 | 13354 | 1672 | 35 | 0 | 0 | 27 | 27 | 0 |
| SRR8094760 | 57 | 1122 | 7269 | 611 | 4276 | 4504 | 8106 | 9057 | 334363 | 456 | 10838 | 4455 | 7966 | 2877 | 271 | 1906 | 24891 | 0 | 0 | 0 | 31 | 51 | 38 | 18 | 0 | 0 | 0 | 0 | 55 | 87 | 17 | 0 | 0 | 0 | 0 | 0 | 0 |
| SRR8094761 | 45 | 306 | 1782 | 585 | 4341 | 3553 | 9059 | 10987 | 297164 | 858 | 8461 | 4911 | 9519 | 1392 | 309 | 2062 | 17692 | 0 | 0 | 0 | 28 | 23 | 16 | 0 | 0 | 0 | 0 | 85 | 307 | 123 | 30 | 0 | 0 | 0 | 0 | 0 | 0 |
| SRR8094762 | 679 | 9862 | 5094 | 1965 | 7110 | 4914 | 14523 | 14347 | 297157 | 136 | 11552 | 5479 | 14171 | 4962 | 930 | 2051 | 20912 | 0 | 31 | 42 | 132 | 158 | 55 | 13 | 0 | 0 | 15 | 285 | 1424 | 395 | 104 | 0 | 0 | 0 | 0 | 0 | 0 |
| SRR8094763 | 343 | 16832 | 135021 | 5224 | 15272 | 16234 | 56327 | 55073 | 78383 | 528 | 2318 | 5843 | 1943 | 252 | 915 | 34 | 17100 | 0 | 42 | 60 | 313 | 507 | 51 | 14 | 15 | 128 | 0 | 70 | 328 | 421 | 127 | 0 | 0 | 0 | 71 | 12 | 0 |
| SRR8094764 | 469 | 22464 | 46046 | 11433 | 15949 | 14398 | 68505 | 62020 | 28791 | 1243 | 14972 | 13639 | 16584 | 2669 | 1114 | 53 | 27060 | 0 | 12 | 39 | 138 | 64 | 25 | 15 | 24 | 244 | 0 | 24 | 254 | 130 | 20 | 0 | 0 | 0 | 104 | 23 | 0 |
| SRR8094765 | 871 | 11895 | 69826 | 41697 | 17348 | 18799 | 76342 | 88838 | 32833 | 373 | 13945 | 12844 | 15714 | 3205 | 836 | 40 | 19479 | 0 | 16 | 62 | 106 | 57 | 0 | 0 | 12 | 195 | 0 | 29 | 318 | 104 | 13 | 0 | 0 | 0 | 56 | 22 | 0 |
| SRR8094766 | 250 | 18121 | 66404 | 3205 | 11108 | 11064 | 36023 | 43910 | 35542 | 664 | 11462 | 14 | 12330 | 2215 | 1139 | 127 | 28835 | 0 | 26 | 27 | 121 | 230 | 26 | 0 | 0 | 69 | 0 | 35 | 141 | 90 | 35 | 0 | 0 | 0 | 37 | 0 | 0 |
| SRR8094767 | 81 | 6724 | 88069 | 599 | 14143 | 15143 | 65181 | 69417 | 93123 | 1570 | 17248 | 20 | 14379 | 7190 | 509 | 31 | 29782 | 0 | 0 | 0 | 0 | 20 | 0 | 56 | 55 | 0 | 0 | 0 | 24 | 12 | 0 | 0 | 0 | 0 | 0 | 16 | 0 |
| SRR8094768 | 626 | 29919 | 22487 | 6558 | 13843 | 13568 | 49592 | 79090 | 39030 | 1522 | 29205 | 27 | 25721 | 6205 | 1125 | 77 | 24220 | 0 | 14 | 38 | 158 | 47 | 40 | 25 | 22 | 229 | 0 | 34 | 224 | 162 | 20 | 0 | 0 | 0 | 110 | 18 | 0 |
| SRR8094769 | 189 | 20499 | 187943 | 1422 | 11368 | 16322 | 49633 | 61222 | 131310 | 558 | 24611 | 21215 | 22066 | 5742 | 819 | 85 | 25750 | 0 | 11 | 15 | 42 | 55 | 18 | 30 | 17 | 104 | 0 | 16 | 61 | 37 | 12 | 0 | 0 | 0 | 48 | 12 | 0 |
| SRR8094770 | 554 | 10285 | 18295 | 6043 | 46094 | 58666 | 280500 | 61308 | 80172 | 740 | 2172 | 1631 | 3339 | 1097 | 7795 | 34 | 5734 | 0 | 0 | 10 | 17 | 35 | 21 | 10 | 0 | 1451 | 0 | 0 | 12 | 0 | 0 | 0 | 0 | 0 | 115 | 67 | 0 |
| SRR8137033 | 527 | 2189 | 10963 | 689 | 1149 | 1077 | 8447 | 862 | 204976 | 2796 | 9632 | 13 | 8359 | 4812 | 22 | 5151 | 6033 | 0 | 0 | 0 | 133 | 85 | 211 | 62 | 35 | 0 | 0 | 18 | 51 | 36 | 13 | 0 | 0 | 0 | 0 | 0 | 0 |
| SRR8137034 | 238 | 852 | 2797 | 203 | 1094 | 1229 | 5968 | 550 | 177398 | 797 | 6552 | 7532 | 6404 | 6011 | 114655 | 1946 | 18279 | 0 | 0 | 35 | 69 | 189 | 214 | 74 | 26 | 0 | 0 | 11 | 27 | 13 | 0 | 0 | 0 | 0 | 0 | 23 | 0 |
| SRR8137035 | 781 | 3405 | 11012 | 1499 | 941 | 1116 | 11351 | 131 | 347228 | 1308 | 7018 | 1990 | 1562 | 4084 | 38 | 1119 | 2586 | 0 | 0 | 0 | 27 | 115 | 117 | 37 | 20 | 0 | 0 | 0 | 15 | 0 | 0 | 0 | 0 | 0 | 0 | 14 | 0 |
| SRR8137036 | 1277 | 3179 | 15330 | 775 | 1574 | 2332 | 20871 | 468 | 188320 | 5774 | 8804 | 4783 | 6433 | 7002 | 41 | 2655 | 5381 | 0 | 0 | 0 | 58 | 80 | 232 | 77 | 35 | 0 | 0 | 14 | 35 | 16 | 0 | 0 | 0 | 0 | 0 | 22 | 0 |
| SRR8137037 | 255 | 18412 | 13174 | 380 | 36033 | 40702 | 118731 | 27436 | 17009 | 72 | 15350 | 7041 | 9851 | 53295 | 6451 | 7925 | 28547 | 0 | 24 | 99 | 55 | 1488 | 122 | 39 | 16 | 0 | 0 | 14 | 34 | 14 | 18 | 0 | 0 | 0 | 0 | 14 | 0 |
| SRR8203922 | 0 | 152782 | 14066 | 14 | 35 | 153 | 703 | 681 | 20013 | 0 | 393 | 265 | 131 | 118 | 16 | 10425 | 1067 | 0 | 52 | 55 | 307 | 997 | 123 | 1774 | 7570 | 221 | 0 | 0 | 10 | 35 | 138 | 25 | 52 | 294 | 172 | 139 | 0 |
| SRR8203923 | 0 | 152614 | 13904 | 0 | 33 | 197 | 706 | 807 | 20219 | 0 | 351 | 268 | 120 | 114 | 10 | 10516 | 917 | 0 | 48 | 54 | 310 | 1008 | 127 | 1755 | 7606 | 231 | 0 | 0 | 13 | 37 | 137 | 23 | 47 | 328 | 181 | 135 | 0 |
| SRR8203924 | 0 | 152892 | 13920 | 0 | 30 | 148 | 774 | 817 | 20170 | 0 | 344 | 207 | 190 | 112 | 20 | 10607 | 984 | 0 | 52 | 62 | 278 | 1015 | 140 | 1794 | 7617 | 235 | 0 | 0 | 12 | 35 | 136 | 24 | 49 | 316 | 168 | 138 | 0 |
| SRR8203925 | 0 | 152988 | 14006 | 0 | 40 | 144 | 683 | 820 | 20389 | 0 | 358 | 274 | 144 | 107 | 17 | 10159 | 994 | 0 | 48 | 52 | 297 | 1054 | 132 | 1854 | 7686 | 236 | 0 | 0 | 0 | 33 | 134 | 22 | 49 | 306 | 173 | 141 | 0 |
| SRR8203926 | 18 | 109464 | 5949 | 15 | 56 | 200 | 896 | 963 | 20817 | 15 | 539 | 390 | 165 | 163 | 43 | 9941 | 766 | 28 | 68 | 74 | 461 | 1065 | 143 | 2082 | 7034 | 234 | 0 | 14 | 59 | 148 | 310 | 94 | 109 | 735 | 148 | 223 | 0 |
| SRR8203927 | 11 | 109418 | 5756 | 22 | 56 | 160 | 876 | 1080 | 20050 | 14 | 565 | 397 | 151 | 174 | 34 | 9819 | 784 | 28 | 69 | 78 | 455 | 1065 | 150 | 2105 | 7036 | 231 | 0 | 0 | 67 | 143 | 286 | 90 | 103 | 763 | 137 | 232 | 0 |
| SRR8203928 | 11 | 109096 | 5850 | 26 | 47 | 170 | 895 | 1073 | 20476 | 21 | 619 | 342 | 161 | 183 | 33 | 9914 | 824 | 29 | 73 | 79 | 478 | 1077 | 144 | 2114 | 6932 | 239 | 0 | 14 | 60 | 151 | 328 | 83 | 119 | 729 | 133 | 211 | 0 |
| SRR8203929 | 19 | 109671 | 6050 | 19 | 61 | 179 | 899 | 1108 | 20754 | 0 | 542 | 360 | 167 | 162 | 31 | 9647 | 703 | 32 | 74 | 76 | 462 | 1082 | 143 | 2105 | 6983 | 234 | 0 | 14 | 54 | 158 | 302 | 89 | 111 | 748 | 144 | 224 | 0 |
| SRR8203930 | 0 | 134179 | 6248 | 34 | 32 | 109 | 717 | 1159 | 29875 | 0 | 526 | 292 | 288 | 135 | 0 | 15217 | 968 | 0 | 24 | 35 | 429 | 666 | 70 | 1222 | 3546 | 122 | 0 | 15 | 54 | 181 | 292 | 58 | 50 | 405 | 80 | 134 | 0 |
| SRR8203931 | 12 | 132698 | 6246 | 26 | 41 | 130 | 699 | 1142 | 29837 | 0 | 475 | 311 | 308 | 149 | 11 | 14902 | 891 | 0 | 27 | 35 | 425 | 668 | 87 | 1291 | 3536 | 120 | 0 | 13 | 56 | 173 | 279 | 62 | 55 | 412 | 90 | 130 | 0 |
| SRR8203932 | 0 | 133738 | 6188 | 37 | 34 | 102 | 699 | 1177 | 29372 | 18 | 470 | 285 | 293 | 130 | 10 | 14939 | 955 | 0 | 28 | 30 | 427 | 668 | 84 | 1236 | 3614 | 130 | 0 | 12 | 64 | 166 | 268 | 68 | 51 | 409 | 89 | 138 | 0 |
| SRR8203933 | 0 | 134380 | 6184 | 26 | 31 | 121 | 716 | 1273 | 29735 | 24 | 496 | 323 | 338 | 127 | 11 | 14816 | 857 | 0 | 27 | 32 | 416 | 632 | 74 | 1234 | 3573 | 118 | 0 | 0 | 56 | 171 | 270 | 57 | 47 | 423 | 88 | 143 | 0 |
| SRR8203934 | 15 | 190309 | 950 | 38 | 57 | 212 | 829 | 1205 | 21324 | 20 | 740 | 625 | 212 | 129 | 33 | 14297 | 1623 | 60 | 86 | 95 | 1485 | 1467 | 243 | 1515 | 13548 | 322 | 0 | 36 | 184 | 368 | 501 | 164 | 104 | 1316 | 238 | 625 | 0 |
| SRR8203935 | 0 | 191074 | 975 | 44 | 49 | 217 | 841 | 1112 | 21390 | 0 | 757 | 552 | 251 | 156 | 36 | 14503 | 1541 | 62 | 85 | 98 | 1517 | 1465 | 240 | 1560 | 13636 | 325 | 0 | 36 | 177 | 369 | 493 | 164 | 97 | 1353 | 239 | 625 | 0 |
| SRR8203936 | 12 | 190304 | 896 | 31 | 68 | 221 | 822 | 1181 | 21512 | 18 | 740 | 641 | 217 | 124 | 35 | 14274 | 1686 | 56 | 87 | 97 | 1505 | 1464 | 242 | 1577 | 13607 | 310 | 0 | 35 | 192 | 373 | 501 | 173 | 111 | 1334 | 241 | 617 | 0 |
| SRR8203937 | 13 | 190238 | 892 | 57 | 41 | 223 | 827 | 1137 | 21281 | 15 | 716 | 568 | 255 | 152 | 25 | 14371 | 1617 | 63 | 81 | 98 | 1527 | 1460 | 253 | 1544 | 13681 | 317 | 0 | 38 | 187 | 370 | 481 | 168 | 105 | 1339 | 235 | 626 | 0 |
| SRR8203938 | 0 | 107861 | 2280 | 43 | 63 | 233 | 1084 | 1365 | 25977 | 0 | 687 | 623 | 252 | 166 | 0 | 15584 | 2288 | 20 | 37 | 46 | 1246 | 1042 | 183 | 1028 | 11016 | 244 | 0 | 28 | 151 | 349 | 452 | 147 | 51 | 932 | 259 | 500 | 0 |
| SRR8203939 | 14 | 107999 | 2416 | 47 | 73 | 272 | 1068 | 1397 | 25927 | 12 | 709 | 630 | 246 | 196 | 22 | 15851 | 2360 | 16 | 35 | 59 | 1278 | 1014 | 179 | 1020 | 11029 | 240 | 0 | 26 | 143 | 347 | 453 | 154 | 51 | 931 | 263 | 483 | 0 |
| SRR8203940 | 13 | 108348 | 2368 | 38 | 71 | 254 | 1069 | 1408 | 25898 | 11 | 779 | 599 | 282 | 189 | 23 | 15628 | 2263 | 18 | 41 | 47 | 1264 | 1043 | 176 | 1035 | 11069 | 245 | 0 | 27 | 142 | 352 | 443 | 152 | 57 | 933 | 255 | 496 | 0 |
| SRR8203941 | 14 | 108432 | 2383 | 50 | 68 | 298 | 1055 | 1276 | 26123 | 21 | 731 | 605 | 272 | 212 | 0 | 15780 | 2190 | 17 | 31 | 52 | 1270 | 1040 | 190 | 1051 | 11105 | 248 | 0 | 25 | 143 | 355 | 419 | 139 | 53 | 959 | 263 | 489 | 0 |
| SRR8203942 | 0 | 35845 | 960 | 199 | 41 | 87 | 775 | 581 | 14016 | 51 | 806 | 301 | 148 | 219 | 31 | 25406 | 765 | 32 | 76 | 49 | 817 | 1148 | 128 | 942 | 4949 | 92 | 0 | 15 | 55 | 384 | 167 | 48 | 49 | 864 | 153 | 234 | 0 |
| SRR8203943 | 0 | 36867 | 794 | 129 | 27 | 133 | 662 | 630 | 13689 | 38 | 891 | 341 | 171 | 208 | 32 | 25885 | 694 | 45 | 65 | 57 | 862 | 1137 | 124 | 949 | 4945 | 99 | 0 | 16 | 51 | 378 | 164 | 50 | 42 | 826 | 149 | 245 | 0 |
| SRR8203944 | 0 | 37016 | 879 | 105 | 10 | 152 | 806 | 628 | 14165 | 26 | 932 | 272 | 236 | 199 | 31 | 25299 | 728 | 42 | 66 | 46 | 820 | 1089 | 125 | 915 | 4997 | 97 | 0 | 0 | 60 | 359 | 151 | 44 | 47 | 820 | 148 | 234 | 0 |
| SRR8203945 | 0 | 36453 | 872 | 155 | 16 | 149 | 690 | 546 | 13987 | 32 | 875 | 350 | 212 | 239 | 32 | 26329 | 610 | 44 | 79 | 55 | 859 | 1154 | 114 | 948 | 4946 | 92 | 0 | 0 | 56 | 395 | 142 | 46 | 52 | 860 | 152 | 232 | 0 |
| SRR8203946 | 0 | 50715 | 1965 | 186 | 32 | 200 | 947 | 770 | 18044 | 44 | 915 | 524 | 385 | 276 | 11 | 27072 | 1483 | 15 | 21 | 15 | 952 | 568 | 83 | 583 | 3708 | 60 | 0 | 23 | 77 | 341 | 210 | 74 | 32 | 570 | 127 | 237 | 0 |
| SRR8203947 | 0 | 50801 | 2074 | 193 | 51 | 181 | 983 | 839 | 18122 | 48 | 994 | 455 | 355 | 261 | 0 | 27515 | 1402 | 0 | 24 | 15 | 964 | 570 | 73 | 581 | 3682 | 53 | 0 | 20 | 84 | 345 | 213 | 78 | 26 | 607 | 134 | 244 | 0 |
| SRR8203948 | 13 | 50955 | 1997 | 171 | 37 | 198 | 931 | 807 | 18093 | 30 | 983 | 467 | 381 | 267 | 0 | 27194 | 1473 | 0 | 20 | 15 | 909 | 572 | 76 | 553 | 3714 | 55 | 0 | 21 | 79 | 362 | 200 | 79 | 25 | 595 | 123 | 236 | 0 |
| SRR8203949 | 0 | 50812 | 2073 | 178 | 40 | 148 | 892 | 826 | 18399 | 33 | 901 | 518 | 361 | 297 | 0 | 27709 | 1438 | 0 | 23 | 13 | 937 | 587 | 80 | 558 | 3619 | 43 | 0 | 20 | 79 | 370 | 189 | 80 | 26 | 599 | 128 | 255 | 0 |
| SRR8248787 | 1422 | 12471 | 53965 | 1429 | 13111 | 17326 | 37620 | 4373 | 8185 | 1168 | 16570 | 28520 | 14411 | 42480 | 17275 | 737 | 19560 | 0 | 21 | 52 | 24 | 147 | 45 | 16 | 10 | 0 | 0 | 30 | 75 | 12 | 0 | 0 | 0 | 0 | 0 | 10 | 0 |
| SRR8248788 | 881 | 26344 | 118248 | 1186 | 13996 | 17805 | 42080 | 4749 | 10845 | 1966 | 20213 | 31794 | 15233 | 41321 | 20188 | 595 | 19292 | 0 | 18 | 62 | 25 | 169 | 46 | 17 | 11 | 0 | 0 | 28 | 80 | 12 | 0 | 0 | 0 | 0 | 0 | 0 | 0 |
| SRR8248789 | 1075 | 26092 | 98776 | 1385 | 14352 | 18140 | 41122 | 4144 | 12350 | 2136 | 17628 | 51 | 12976 | 44411 | 22294 | 571 | 19139 | 0 | 17 | 53 | 18 | 153 | 39 | 16 | 11 | 0 | 0 | 25 | 78 | 14 | 0 | 0 | 0 | 0 | 0 | 12 | 0 |
| SRR8248790 | 521 | 7768 | 21314 | 742 | 11905 | 15912 | 38902 | 3142 | 11412 | 532 | 15794 | 28 | 11004 | 40583 | 19204 | 381 | 16257 | 0 | 0 | 26 | 10 | 94 | 19 | 0 | 0 | 0 | 0 | 0 | 24 | 0 | 0 | 0 | 0 | 0 | 0 | 0 | 0 |
| SRR8248791 | 230 | 15537 | 161091 | 328 | 15922 | 21740 | 48730 | 4183 | 13861 | 3002 | 15636 | 25311 | 13192 | 46557 | 26168 | 589 | 17178 | 0 | 0 | 29 | 0 | 136 | 35 | 16 | 10 | 0 | 0 | 0 | 30 | 0 | 0 | 0 | 0 | 0 | 0 | 14 | 0 |
| SRR8248792 | 245 | 21537 | 105593 | 468 | 13136 | 17220 | 47314 | 2296 | 16483 | 2724 | 15928 | 21015 | 9681 | 50158 | 26507 | 649 | 9816 | 0 | 0 | 21 | 0 | 82 | 26 | 0 | 0 | 0 | 0 | 0 | 23 | 0 | 0 | 0 | 0 | 0 | 0 | 0 | 0 |
| SRR8272058 | 0 | 0 | 1057 | 0 | 2441 | 27620 | 54036 | 17908 | 2368 | 68 | 101904 | 57518 | 1681 | 32775 | 0 | 87545 | 3803 | 0 | 0 | 0 | 0 | 0 | 0 | 0 | 0 | 0 | 0 | 0 | 0 | 0 | 0 | 0 | 0 | 0 | 0 | 0 | 0 |
| SRR8280405 | 131 | 36585 | 1748 | 874 | 0 | 91 | 16 | 84 | 513 | 0 | 513 | 609 | 39 | 1102 | 0 | 33 | 1159 | 0 | 0 | 0 | 0 | 56 | 0 | 0 | 0 | 0 | 0 | 0 | 0 | 0 | 81 | 14 | 0 | 0 | 0 | 0 | 0 |
| SRR8280408 | 0 | 34963 | 829 | 552 | 0 | 304 | 48 | 167 | 1221 | 0 | 1859 | 2604 | 264 | 2943 | 0 | 44 | 4010 | 0 | 0 | 0 | 0 | 79 | 0 | 0 | 0 | 0 | 0 | 0 | 0 | 0 | 96 | 18 | 0 | 0 | 0 | 0 | 0 |
| SRR8280414 | 224 | 6063 | 425 | 2528 | 0 | 90 | 25 | 184 | 635 | 0 | 585 | 655 | 125 | 2219 | 0 | 140 | 2423 | 0 | 0 | 0 | 0 | 26 | 0 | 0 | 0 | 0 | 0 | 0 | 0 | 0 | 48 | 11 | 0 | 0 | 0 | 0 | 0 |
| SRR8280415 | 108 | 12186 | 216 | 683 | 0 | 261 | 75 | 179 | 364 | 0 | 930 | 1283 | 108 | 1812 | 0 | 57 | 3419 | 0 | 0 | 0 | 0 | 16 | 0 | 0 | 0 | 0 | 0 | 0 | 0 | 0 | 43 | 13 | 0 | 0 | 0 | 0 | 0 |
| SRR8280416 | 0 | 18208 | 583 | 4758 | 0 | 120 | 46 | 131 | 310 | 0 | 310 | 523 | 90 | 1631 | 11 | 30 | 1925 | 0 | 0 | 0 | 0 | 16 | 0 | 0 | 0 | 0 | 0 | 0 | 0 | 0 | 35 | 0 | 0 | 0 | 11 | 0 | 0 |
| SRR8280417 | 103 | 12077 | 531 | 976 | 0 | 135 | 62 | 332 | 1372 | 0 | 617 | 793 | 133 | 2095 | 0 | 300 | 3836 | 0 | 0 | 0 | 0 | 0 | 0 | 0 | 0 | 0 | 0 | 0 | 0 | 0 | 15 | 18 | 0 | 0 | 0 | 0 | 0 |
| SRR8280423 | 0 | 18780 | 258 | 2452 | 0 | 44 | 11 | 20 | 235 | 0 | 181 | 347 | 20 | 880 | 0 | 0 | 531 | 0 | 0 | 0 | 0 | 19 | 0 | 0 | 0 | 0 | 0 | 0 | 0 | 0 | 24 | 0 | 0 | 0 | 0 | 0 | 0 |
| SRR8280425 | 0 | 10070 | 879 | 743 | 0 | 97 | 0 | 26 | 254 | 0 | 404 | 503 | 28 | 860 | 0 | 26 | 953 | 0 | 0 | 0 | 0 | 24 | 0 | 0 | 0 | 0 | 0 | 0 | 0 | 0 | 39 | 0 | 0 | 0 | 0 | 0 | 0 |
| SRR8280432 | 220 | 14292 | 550 | 3848 | 0 | 94 | 24 | 29 | 343 | 0 | 428 | 623 | 51 | 1316 | 0 | 22 | 1110 | 0 | 0 | 0 | 0 | 0 | 0 | 0 | 0 | 0 | 0 | 0 | 0 | 0 | 37 | 0 | 0 | 0 | 0 | 0 | 0 |
| SRR8280440 | 28 | 3955 | 1244 | 9586 | 0 | 122 | 33 | 173 | 3708 | 0 | 126 | 114 | 28 | 1922 | 0 | 44 | 1316 | 0 | 0 | 0 | 0 | 59 | 0 | 0 | 0 | 0 | 0 | 0 | 0 | 0 | 51 | 12 | 0 | 0 | 0 | 0 | 0 |
| SRR8280442 | 18 | 8081 | 886 | 3260 | 0 | 126 | 21 | 89 | 700 | 0 | 191 | 232 | 29 | 506 | 0 | 50 | 1394 | 0 | 0 | 0 | 0 | 61 | 0 | 0 | 0 | 0 | 0 | 0 | 0 | 0 | 60 | 0 | 0 | 0 | 0 | 0 | 0 |
| SRR8280452 | 0 | 6184 | 358 | 1434 | 0 | 31 | 0 | 0 | 116 | 0 | 81 | 170 | 46 | 336 | 0 | 0 | 294 | 0 | 0 | 0 | 0 | 0 | 11 | 0 | 0 | 0 | 0 | 0 | 0 | 0 | 12 | 0 | 0 | 0 | 0 | 0 | 0 |
| SRR8280460 | 14 | 14387 | 472 | 1988 | 0 | 101 | 11 | 115 | 837 | 0 | 247 | 417 | 46 | 1044 | 0 | 60 | 1316 | 0 | 0 | 0 | 0 | 65 | 0 | 0 | 0 | 0 | 0 | 0 | 0 | 0 | 104 | 14 | 0 | 0 | 0 | 0 | 0 |
| SRR8280461 | 0 | 10447 | 540 | 660 | 0 | 35 | 0 | 46 | 272 | 0 | 217 | 305 | 11 | 608 | 0 | 15 | 592 | 0 | 0 | 0 | 0 | 17 | 0 | 0 | 0 | 0 | 0 | 0 | 0 | 0 | 34 | 0 | 0 | 0 | 0 | 0 | 0 |
| SRR8280463 | 32 | 7831 | 554 | 981 | 0 | 16 | 0 | 33 | 107 | 0 | 21 | 40 | 0 | 199 | 0 | 0 | 196 | 0 | 0 | 0 | 0 | 12 | 0 | 0 | 0 | 0 | 0 | 0 | 0 | 0 | 32 | 0 | 0 | 0 | 0 | 0 | 0 |
| SRR8280464 | 46 | 12280 | 1385 | 277 | 0 | 426 | 84 | 248 | 1382 | 0 | 1890 | 2685 | 220 | 2394 | 0 | 56 | 5218 | 0 | 0 | 0 | 0 | 33 | 0 | 0 | 0 | 0 | 0 | 0 | 0 | 0 | 90 | 16 | 0 | 0 | 0 | 0 | 0 |
| SRR8280467 | 25 | 12270 | 499 | 1298 | 0 | 191 | 37 | 126 | 490 | 0 | 790 | 1627 | 279 | 2099 | 0 | 33 | 2647 | 0 | 0 | 0 | 0 | 218 | 17 | 0 | 0 | 0 | 0 | 0 | 0 | 0 | 218 | 17 | 0 | 0 | 0 | 0 | 0 |
| SRR8280468 | 0 | 8259 | 948 | 135 | 0 | 92 | 31 | 18 | 460 | 0 | 1236 | 2099 | 197 | 2174 | 0 | 26 | 1420 | 0 | 0 | 0 | 0 | 36 | 0 | 0 | 0 | 0 | 0 | 0 | 0 | 0 | 36 | 0 | 0 | 0 | 0 | 0 | 0 |
| SRR8280472 | 0 | 8066 | 0 | 684 | 0 | 158 | 0 | 34 | 293 | 0 | 675 | 1238 | 68 | 2318 | 23 | 0 | 1092 | 0 | 0 | 0 | 0 | 51 | 0 | 0 | 0 | 0 | 0 | 0 | 0 | 0 | 33 | 0 | 0 | 0 | 0 | 0 | 0 |
| SRR8280473 | 96 | 3305 | 527 | 96 | 0 | 223 | 40 | 83 | 478 | 0 | 2501 | 3635 | 286 | 2885 | 0 | 0 | 2584 | 0 | 0 | 0 | 0 | 34 | 0 | 0 | 0 | 0 | 0 | 0 | 0 | 0 | 31 | 0 | 0 | 0 | 0 | 0 | 0 |
| SRR8280476 | 0 | 3973 | 1398 | 221 | 0 | 95 | 14 | 90 | 367 | 0 | 703 | 1286 | 65 | 979 | 0 | 19 | 1025 | 0 | 0 | 0 | 0 | 27 | 0 | 0 | 0 | 0 | 0 | 0 | 0 | 0 | 23 | 0 | 0 | 0 | 0 | 0 | 0 |
| SRR8280479 | 0 | 3643 | 442 | 1104 | 0 | 114 | 33 | 83 | 245 | 0 | 298 | 364 | 37 | 1017 | 0 | 33 | 1683 | 0 | 0 | 0 | 0 | 13 | 0 | 0 | 0 | 0 | 0 | 0 | 0 | 0 | 27 | 15 | 0 | 0 | 0 | 0 | 0 |
| SRR8280481 | 0 | 4124 | 773 | 773 | 0 | 135 | 17 | 153 | 310 | 0 | 310 | 877 | 105 | 1584 | 0 | 39 | 3098 | 0 | 0 | 0 | 0 | 22 | 0 | 0 | 0 | 0 | 0 | 0 | 0 | 0 | 49 | 0 | 0 | 0 | 0 | 0 | 0 |
| SRR8280483 | 0 | 6871 | 204 | 476 | 0 | 63 | 30 | 120 | 455 | 0 | 273 | 315 | 93 | 1642 | 0 | 57 | 1998 | 0 | 0 | 0 | 0 | 13 | 0 | 0 | 0 | 0 | 0 | 0 | 0 | 0 | 0 | 12 | 0 | 0 | 0 | 0 | 0 |
| SRR8280485 | 109 | 10127 | 176 | 2525 | 0 | 31 | 0 | 47 | 176 | 0 | 134 | 144 | 41 | 445 | 0 | 16 | 466 | 0 | 0 | 0 | 0 | 14 | 0 | 0 | 0 | 0 | 0 | 0 | 0 | 0 | 0 | 13 | 0 | 0 | 0 | 0 | 0 |
| SRR8280487 | 0 | 14996 | 2499 | 1428 | 0 | 180 | 23 | 75 | 406 | 0 | 353 | 626 | 93 | 1981 | 12 | 12 | 1489 | 0 | 0 | 0 | 0 | 62 | 0 | 0 | 0 | 0 | 0 | 0 | 0 | 0 | 51 | 0 | 0 | 0 | 0 | 0 | 0 |
| SRR8280490 | 0 | 6827 | 1280 | 427 | 0 | 386 | 80 | 106 | 799 | 0 | 2423 | 4725 | 293 | 6256 | 13 | 40 | 4779 | 0 | 0 | 0 | 0 | 22 | 0 | 0 | 0 | 0 | 0 | 0 | 0 | 0 | 29 | 0 | 0 | 0 | 0 | 0 | 0 |
| SRR8280494 | 0 | 13282 | 1932 | 3019 | 0 | 110 | 0 | 58 | 291 | 0 | 265 | 375 | 58 | 1635 | 0 | 19 | 1053 | 0 | 0 | 0 | 0 | 11 | 0 | 0 | 0 | 0 | 0 | 0 | 0 | 0 | 16 | 0 | 0 | 0 | 0 | 0 | 0 |
| SRR8280497 | 151 | 4301 | 129 | 981 | 0 | 179 | 96 | 145 | 1036 | 0 | 434 | 662 | 83 | 1619 | 0 | 99 | 1655 | 0 | 0 | 0 | 0 | 13 | 0 | 0 | 0 | 0 | 0 | 0 | 0 | 16 | 36 | 13 | 0 | 0 | 0 | 0 | 0 |
| SRR8280498 | 170 | 8826 | 311 | 1980 | 0 | 274 | 62 | 154 | 638 | 0 | 566 | 617 | 117 | 2097 | 0 | 76 | 3461 | 0 | 0 | 0 | 0 | 0 | 0 | 0 | 0 | 0 | 0 | 0 | 0 | 0 | 31 | 0 | 0 | 0 | 0 | 0 | 0 |
| SRR8280500 | 76 | 22031 | 719 | 4189 | 0 | 105 | 35 | 308 | 3150 | 0 | 461 | 546 | 207 | 1882 | 0 | 171 | 4163 | 0 | 0 | 0 | 0 | 0 | 0 | 0 | 0 | 0 | 0 | 0 | 0 | 0 | 0 | 48 | 0 | 0 | 0 | 0 | 0 |
| SRR8280501 | 25 | 2935 | 29 | 157 | 372 | 8494 | 5476 | 4691 | 37447 | 0 | 23542 | 72221 | 6612 | 4352 | 46 | 170 | 55415 | 0 | 0 | 0 | 92 | 229 | 12 | 0 | 0 | 0 | 0 | 86 | 177 | 215 | 739 | 12 | 0 | 0 | 0 | 0 | 0 |
| SRR8280502 | 414 | 8908 | 0 | 1036 | 0 | 248 | 23 | 137 | 463 | 0 | 398 | 574 | 127 | 2178 | 0 | 62 | 3386 | 0 | 0 | 0 | 0 | 11 | 0 | 0 | 0 | 0 | 0 | 0 | 0 | 0 | 18 | 0 | 0 | 0 | 0 | 0 | 0 |
| SRR8280503 | 48 | 14114 | 224 | 214 | 254 | 6462 | 5364 | 5476 | 33777 | 33 | 19130 | 42155 | 5193 | 6713 | 27 | 178 | 36574 | 0 | 0 | 0 | 103 | 208 | 13 | 0 | 0 | 0 | 0 | 61 | 160 | 185 | 682 | 14 | 0 | 0 | 0 | 0 | 0 |
| SRR8280504 | 0 | 3335 | 834 | 538 | 0 | 451 | 117 | 309 | 1016 | 0 | 543 | 875 | 92 | 1638 | 0 | 130 | 5877 | 0 | 0 | 0 | 0 | 0 | 0 | 0 | 0 | 0 | 0 | 0 | 0 | 0 | 29 | 16 | 0 | 0 | 0 | 0 | 0 |
| SRR8280505 | 14 | 4554 | 176 | 854 | 799 | 12043 | 13031 | 5977 | 35300 | 97 | 19717 | 43853 | 4964 | 4514 | 45 | 78 | 52628 | 0 | 11 | 14 | 162 | 226 | 0 | 0 | 0 | 0 | 0 | 152 | 242 | 331 | 993 | 0 | 0 | 0 | 0 | 0 | 0 |
| SRR8280506 | 128 | 10344 | 513 | 2137 | 0 | 119 | 20 | 93 | 406 | 0 | 192 | 317 | 49 | 773 | 0 | 59 | 1490 | 0 | 0 | 0 | 0 | 13 | 0 | 0 | 0 | 0 | 0 | 0 | 0 | 0 | 22 | 13 | 0 | 0 | 0 | 0 | 0 |
| SRR8280507 | 60 | 7472 | 52 | 144 | 583 | 11840 | 7209 | 4603 | 35622 | 19 | 24571 | 63239 | 6620 | 4987 | 27 | 165 | 43593 | 0 | 0 | 13 | 231 | 363 | 19 | 0 | 0 | 0 | 0 | 168 | 348 | 398 | 911 | 16 | 0 | 0 | 0 | 0 | 0 |
| SRR8280509 | 32 | 2957 | 85 | 150 | 788 | 11226 | 10142 | 6005 | 33774 | 22 | 18670 | 41816 | 5444 | 5634 | 69 | 79 | 49143 | 0 | 0 | 0 | 101 | 133 | 0 | 0 | 0 | 0 | 0 | 114 | 168 | 170 | 539 | 0 | 0 | 0 | 0 | 0 | 0 |
| SRR8280510 | 59 | 7629 | 591 | 2484 | 0 | 163 | 38 | 313 | 803 | 0 | 327 | 438 | 167 | 2155 | 10 | 111 | 3743 | 0 | 0 | 0 | 0 | 11 | 0 | 0 | 0 | 0 | 0 | 0 | 0 | 0 | 20 | 12 | 0 | 0 | 0 | 0 | 0 |
| SRR8280511 | 38 | 3104 | 55 | 130 | 522 | 10258 | 6707 | 5778 | 32858 | 64 | 18674 | 67314 | 5972 | 5099 | 76 | 132 | 59210 | 0 | 11 | 14 | 81 | 299 | 10 | 0 | 0 | 0 | 0 | 183 | 309 | 184 | 1074 | 0 | 0 | 0 | 0 | 0 | 0 |
| SRR8280512 | 0 | 9264 | 463 | 602 | 0 | 280 | 47 | 140 | 718 | 0 | 539 | 1227 | 57 | 1690 | 0 | 89 | 4203 | 0 | 0 | 0 | 0 | 12 | 0 | 0 | 0 | 0 | 0 | 0 | 0 | 0 | 55 | 0 | 0 | 0 | 0 | 0 | 0 |
| SRR8280513 | 0 | 5321 | 100 | 210 | 109 | 2138 | 4876 | 5592 | 31417 | 0 | 17132 | 76377 | 7077 | 6529 | 17 | 188 | 27702 | 0 | 0 | 0 | 60 | 87 | 0 | 0 | 0 | 0 | 0 | 78 | 101 | 78 | 275 | 0 | 0 | 0 | 0 | 0 | 0 |
| SRR8280515 | 245 | 10535 | 245 | 1531 | 0 | 127 | 14 | 213 | 325 | 0 | 357 | 543 | 108 | 1545 | 0 | 50 | 2274 | 0 | 0 | 0 | 0 | 26 | 0 | 0 | 0 | 0 | 0 | 0 | 0 | 0 | 50 | 0 | 0 | 0 | 0 | 0 | 0 |
| SRR8280519 | 37 | 4715 | 3582 | 621 | 0 | 376 | 131 | 615 | 2137 | 0 | 645 | 444 | 87 | 1290 | 0 | 48 | 3876 | 0 | 0 | 0 | 0 | 32 | 0 | 0 | 0 | 0 | 0 | 0 | 0 | 0 | 58 | 0 | 0 | 0 | 0 | 0 | 0 |
| SRR8280521 | 0 | 2824 | 1213 | 2815 | 0 | 1037 | 178 | 296 | 2549 | 0 | 916 | 1406 | 217 | 4929 | 0 | 157 | 9174 | 0 | 0 | 0 | 0 | 78 | 10 | 0 | 0 | 0 | 0 | 0 | 0 | 0 | 140 | 15 | 0 | 0 | 0 | 0 | 0 |
| SRR8280525 | 376 | 14094 | 0 | 2349 | 18 | 510 | 108 | 204 | 749 | 0 | 2422 | 2908 | 306 | 4142 | 0 | 54 | 4250 | 0 | 0 | 0 | 0 | 114 | 0 | 0 | 0 | 0 | 0 | 0 | 0 | 0 | 106 | 10 | 0 | 0 | 0 | 0 | 0 |
| SRR8280531 | 217 | 12953 | 163 | 1409 | 0 | 209 | 63 | 125 | 395 | 0 | 1088 | 1304 | 85 | 1563 | 0 | 35 | 1638 | 0 | 0 | 0 | 0 | 64 | 0 | 0 | 0 | 0 | 0 | 0 | 0 | 0 | 74 | 25 | 0 | 0 | 0 | 0 | 0 |
| SRR8280533 | 301 | 10289 | 481 | 1685 | 0 | 159 | 32 | 114 | 452 | 0 | 723 | 779 | 58 | 960 | 0 | 40 | 1289 | 0 | 0 | 0 | 0 | 30 | 0 | 0 | 0 | 0 | 0 | 0 | 0 | 0 | 28 | 11 | 0 | 0 | 0 | 0 | 0 |
| SRR8280536 | 16 | 16518 | 855 | 3048 | 0 | 72 | 14 | 49 | 341 | 0 | 291 | 376 | 54 | 867 | 0 | 22 | 687 | 0 | 0 | 0 | 0 | 69 | 0 | 0 | 0 | 0 | 0 | 0 | 0 | 0 | 73 | 0 | 0 | 0 | 0 | 0 | 0 |
| SRR8280539 | 0 | 6942 | 449 | 404 | 0 | 435 | 71 | 142 | 777 | 0 | 1266 | 1614 | 139 | 2914 | 0 | 41 | 3302 | 0 | 0 | 0 | 0 | 45 | 0 | 0 | 0 | 0 | 0 | 0 | 0 | 0 | 75 | 15 | 0 | 0 | 0 | 0 | 0 |
| SRR8280541 | 37 | 11105 | 840 | 1388 | 0 | 476 | 71 | 155 | 735 | 0 | 2179 | 3017 | 186 | 4609 | 0 | 20 | 3736 | 0 | 0 | 0 | 0 | 42 | 0 | 0 | 0 | 0 | 0 | 0 | 0 | 0 | 73 | 12 | 0 | 0 | 0 | 0 | 0 |
| SRR8280543 | 0 | 3322 | 894 | 767 | 0 | 77 | 0 | 130 | 314 | 0 | 468 | 644 | 77 | 1380 | 0 | 69 | 1119 | 0 | 0 | 0 | 0 | 36 | 0 | 0 | 0 | 0 | 0 | 0 | 0 | 0 | 0 | 0 | 0 | 0 | 0 | 0 | 0 |
| SRR8280544 | 0 | 14234 | 1379 | 314 | 0 | 115 | 14 | 28 | 177 | 0 | 606 | 894 | 126 | 1290 | 0 | 26 | 1040 | 0 | 0 | 0 | 0 | 14 | 0 | 0 | 0 | 0 | 0 | 0 | 0 | 0 | 17 | 0 | 0 | 0 | 0 | 0 | 0 |
| SRR8280546 | 0 | 4822 | 1999 | 0 | 0 | 41 | 11 | 67 | 253 | 0 | 354 | 422 | 17 | 786 | 0 | 33 | 832 | 0 | 0 | 0 | 0 | 22 | 0 | 0 | 0 | 0 | 0 | 0 | 0 | 0 | 13 | 0 | 0 | 0 | 0 | 0 | 0 |
| SRR8280548 | 193 | 10246 | 1933 | 773 | 0 | 92 | 12 | 69 | 190 | 0 | 639 | 1002 | 81 | 1520 | 12 | 29 | 1059 | 0 | 0 | 0 | 0 | 17 | 0 | 0 | 0 | 0 | 0 | 0 | 0 | 0 | 0 | 0 | 0 | 0 | 0 | 0 | 0 |
| SRR8280550 | 0 | 16590 | 1326 | 1943 | 0 | 97 | 14 | 51 | 295 | 0 | 173 | 270 | 35 | 933 | 0 | 17 | 880 | 0 | 0 | 0 | 0 | 35 | 0 | 0 | 0 | 0 | 0 | 0 | 0 | 0 | 55 | 0 | 0 | 0 | 0 | 0 | 0 |
| SRR8280551 | 0 | 5616 | 105 | 632 | 0 | 15 | 0 | 43 | 195 | 0 | 197 | 180 | 12 | 538 | 0 | 11 | 440 | 0 | 0 | 0 | 0 | 18 | 0 | 0 | 0 | 0 | 0 | 0 | 0 | 0 | 16 | 0 | 0 | 0 | 0 | 0 | 0 |
| SRR8280552 | 0 | 9133 | 496 | 1390 | 0 | 101 | 31 | 21 | 182 | 0 | 273 | 371 | 66 | 1616 | 0 | 0 | 822 | 0 | 0 | 0 | 0 | 29 | 0 | 0 | 0 | 0 | 0 | 0 | 0 | 0 | 26 | 12 | 0 | 0 | 0 | 0 | 0 |
| SRR8280554 | 0 | 5355 | 487 | 765 | 0 | 548 | 42 | 73 | 898 | 0 | 2281 | 2881 | 245 | 4885 | 0 | 21 | 4243 | 0 | 0 | 0 | 0 | 50 | 0 | 0 | 0 | 0 | 0 | 0 | 0 | 0 | 54 | 0 | 0 | 0 | 0 | 0 | 0 |
| SRR8280558 | 0 | 9214 | 110 | 1536 | 0 | 115 | 12 | 15 | 280 | 0 | 403 | 676 | 119 | 2055 | 0 | 12 | 980 | 0 | 0 | 0 | 0 | 12 | 0 | 0 | 0 | 0 | 0 | 0 | 0 | 0 | 24 | 0 | 0 | 0 | 0 | 0 | 0 |
| SRR8280560 | 73 | 12402 | 294 | 1651 | 0 | 28 | 0 | 23 | 110 | 0 | 68 | 125 | 12 | 475 | 0 | 17 | 335 | 0 | 0 | 0 | 0 | 49 | 0 | 0 | 0 | 0 | 0 | 0 | 0 | 0 | 56 | 0 | 0 | 0 | 0 | 0 | 0 |
| SRR8280561 | 156 | 9370 | 311 | 1339 | 0 | 50 | 0 | 47 | 125 | 0 | 144 | 200 | 35 | 738 | 0 | 11 | 478 | 0 | 0 | 0 | 0 | 27 | 0 | 0 | 0 | 0 | 0 | 0 | 0 | 0 | 25 | 0 | 0 | 0 | 0 | 0 | 0 |
| SRR8280562 | 426 | 28323 | 994 | 426 | 0 | 189 | 31 | 118 | 989 | 0 | 968 | 1423 | 191 | 2053 | 0 | 92 | 2378 | 0 | 0 | 0 | 0 | 119 | 0 | 0 | 0 | 0 | 0 | 0 | 0 | 10 | 212 | 23 | 0 | 0 | 0 | 0 | 0 |
| SRR8280564 | 0 | 29269 | 461 | 1238 | 0 | 108 | 11 | 27 | 454 | 0 | 170 | 311 | 66 | 977 | 0 | 27 | 991 | 0 | 0 | 0 | 0 | 52 | 12 | 0 | 0 | 0 | 0 | 0 | 0 | 0 | 109 | 12 | 0 | 0 | 0 | 0 | 0 |
| SRR8280566 | 208 | 12801 | 675 | 1740 | 0 | 194 | 36 | 150 | 1289 | 0 | 785 | 1082 | 115 | 1668 | 0 | 35 | 1954 | 0 | 0 | 0 | 0 | 49 | 15 | 0 | 0 | 0 | 0 | 0 | 0 | 0 | 118 | 33 | 0 | 0 | 0 | 0 | 0 |

### Supp Table S8.1.Columns_B_to_AL

| | | | | | | | | | | | | | | | | | |
|---|---|---|---|---|---|---|---|---|---|---|---|---|---|---|---|---|---|
| SRR8280568 | 383 | 20362 | 639 | 1235 | 0 | 156 | 14 | 64 | 507 | 0 | 817 | 1118 | 113 | 1669 | 0 | 27 | 1148 |
| SRR8280572 | 29 | 19189 | 685 | 1626 | 0 | 87 | 22 | 158 | 513 | 0 | 263 | 459 | 92 | 1184 | 0 | 88 | 1559 |
| SRR8280573 | 115 | 15740 | 614 | 1651 | 0 | 509 | 78 | 201 | 947 | 0 | 1098 | 1592 | 187 | 2643 | 0 | 109 | 5436 |
| SRR8280574 | 76 | 11589 | 76 | 3050 | 0 | 199 | 40 | 81 | 530 | 0 | 664 | 664 | 197 | 1742 | 0 | 53 | 2511 |
| SRR8280576 | 0 | 5587 | 848 | 4889 | 0 | 50 | 18 | 91 | 590 | 0 | 33 | 63 | 20 | 648 | 0 | 86 | 901 |
| SRR8280580 | 40 | 5602 | 202 | 1491 | 0 | 120 | 27 | 156 | 601 | 0 | 334 | 538 | 79 | 1696 | 0 | 44 | 2852 |
| SRR8280582 | 64 | 16036 | 1018 | 3882 | 0 | 408 | 59 | 157 | 730 | 0 | 599 | 1139 | 128 | 2293 | 0 | 126 | 5117 |
| SRR8280583 | 30 | 4832 | 59 | 1340 | 122 | 2536 | 4713 | 4847 | 25260 | 12 | 20064 | 41720 | 4691 | 9253 | 22 | 153 | 46970 |
| SRR8280584 | 49 | 13620 | 98 | 2156 | 0 | 261 | 77 | 204 | 536 | 0 | 510 | 1145 | 131 | 2122 | 0 | 85 | 3832 |
| SRR8280589 | 0 | 3640 | 331 | 1390 | 0 | 67 | 0 | 50 | 326 | 0 | 134 | 130 | 20 | 733 | 0 | 20 | 640 |
| SRR8280590 | 0 | 9961 | 294 | 1360 | 0 | 306 | 73 | 191 | 886 | 37 | 342 | 309 | 113 | 1609 | 0 | 11 | 2091 |
| SRR8280592 | 116 | 5081 | 116 | 1241 | 0 | 230 | 44 | 85 | 496 | 0 | 762 | 798 | 125 | 1871 | 16 | 85 | 2556 |
| SRR8280593 | 0 | 5767 | 166 | 1027 | 0 | 75 | 16 | 38 | 284 | 0 | 312 | 516 | 49 | 1017 | 0 | 18 | 974 |
| SRR8311061 | 31 | 298 | 9547 | 31 | 0 | 0 | 100 | 855 | 169 | 130 | 536 | 446 | 404 | 5121 | 0 | 32 | 3624 |
| SRR8311062 | 162 | 659 | 4945 | 952 | 0 | 35 | 138 | 1372 | 572 | 162 | 481 | 771 | 375 | 4653 | 15 | 253 | 4411 |
| SRR8311063 | 113 | 816 | 7160 | 568 | 16 | 29 | 289 | 3695 | 1843 | 309 | 1290 | 1655 | 629 | 8654 | 22 | 715 | 6166 |
| SRR8311064 | 0 | 259 | 10853 | 0 | 62 | 157 | 917 | 2554 | 670 | 394 | 2362 | 2593 | 648 | 4951 | 0 | 140 | 11920 |
| SRR8311070 | 30 | 2509 | 6025 | 125 | 97 | 167 | 963 | 3289 | 789 | 180 | 3581 | 3295 | 1735 | 4571 | 82 | 175 | 6917 |
| SRR8311071 | 0 | 961 | 16054 | 599 | 41 | 31 | 282 | 3638 | 486 | 295 | 2146 | 3025 | 650 | 5783 | 36 | 314 | 5685 |
| SRR8311072 | 0 | 697 | 8968 | 634 | 0 | 15 | 238 | 1464 | 518 | 523 | 1290 | 1657 | 584 | 6102 | 15 | 92 | 4949 |
| SRR8311073 | 122 | 588 | 5340 | 614 | 0 | 41 | 742 | 1489 | 1059 | 0 | 1764 | 1868 | 530 | 7846 | 17 | 423 | 6485 |
| SRR8311074 | 275 | 310 | 9304 | 512 | 73 | 150 | 1518 | 5542 | 1048 | 310 | 5036 | 7317 | 399 | 5088 | 78 | 641 | 11842 |
| SRR8311076 | 0 | 548 | 2687 | 329 | 14 | 0 | 277 | 1128 | 525 | 126 | 791 | 1028 | 524 | 6778 | 12 | 70 | 4069 |
| SRR8311078 | 76 | 607 | 4529 | 451 | 31 | 48 | 468 | 1993 | 564 | 442 | 1227 | 2099 | 523 | 6671 | 0 | 211 | 6229 |
| SRR8311080 | 0 | 639 | 2575 | 474 | 12 | 13 | 73 | 742 | 340 | 0 | 735 | 967 | 285 | 5940 | 0 | 58 | 3291 |
| SRR8311081 | 0 | 0 | 3983 | 0 | 0 | 19 | 124 | 780 | 137 | 0 | 1083 | 1271 | 340 | 5055 | 0 | 59 | 3705 |
| SRR8311082 | 0 | 285 | 3921 | 171 | 0 | 11 | 117 | 2045 | 527 | 0 | 1210 | 1514 | 471 | 5782 | 16 | 309 | 2562 |
| SRR8311085 | 167 | 412 | 19247 | 39 | 67 | 16 | 848 | 6533 | 162 | 930 | 2556 | 4493 | 897 | 2882 | 55 | 262 | 6165 |
| SRR8311087 | 0 | 983 | 16399 | 392 | 30 | 60 | 660 | 4614 | 328 | 772 | 1780 | 3385 | 760 | 4623 | 26 | 385 | 9813 |
| SRR8311089 | 112 | 248 | 10546 | 0 | 11 | 14 | 356 | 1912 | 140 | 248 | 2311 | 4122 | 821 | 2523 | 28 | 111 | 4725 |
| SRR8311092 | 0 | 274 | 9678 | 173 | 46 | 110 | 1308 | 6127 | 1890 | 208 | 2533 | 3000 | 492 | 10610 | 0 | 602 | 13805 |
| SRR8311093 | 30 | 947 | 6104 | 243 | 0 | 25 | 137 | 1040 | 480 | 0 | 615 | 767 | 253 | 5378 | 0 | 146 | 3432 |
| SRR8311095 | 136 | 528 | 7577 | 82 | 18 | 22 | 387 | 2579 | 467 | 615 | 1986 | 2782 | 520 | 5252 | 14 | 182 | 3648 |
| SRR8311098 | 113 | 482 | 5785 | 116 | 19 | 105 | 583 | 3405 | 2442 | 531 | 2346 | 4267 | 558 | 10666 | 0 | 725 | 10229 |
| SRR8311105 | 0 | 177 | 6588 | 0 | 32 | 34 | 407 | 3069 | 485 | 29 | 1950 | 2077 | 999 | 7041 | 28 | 181 | 7081 |
| SRR8311106 | 0 | 511 | 10509 | 0 | 35 | 26 | 665 | 5592 | 687 | 632 | 1514 | 2780 | 380 | 5407 | 0 | 775 | 7824 |
| SRR835324 | 1217 | 4496 | 20583 | 0 | 21 | 347 | 313 | 427 | 211 | 4471 | 177 | 97 | 249 | 545 | 127 | 308 | 1166 |
| SRR835325 | 1264 | 2956 | 13867 | 0 | 46 | 366 | 738 | 705 | 548 | 3311 | 346 | 300 | 353 | 627 | 124 | 424 | 2291 |
| SRR8354354 | 984 | 2769 | 1998 | 426 | 4191 | 5759 | 60268 | 60632 | 64608 | 0 | 35797 | 9408 | 18749 | 20317 | 1605 | 5715 | 15918 |
| SRR837775 | 14 | 9002 | 779 | 438 | 913 | 8734 | 5608 | 6751 | 18117 | 78 | 48934 | 13 | 6707 | 3054 | 730 | 4808 | 34888 |
| SRR837776 | 0 | 6528 | 780 | 97 | 306 | 5691 | 3629 | 8082 | 6805 | 106 | 45202 | 16 | 7499 | 2278 | 149 | 3271 | 21800 |
| SRR837777 | 0 | 529 | 8158 | 0 | 2350 | 31478 | 41950 | 19086 | 30 | 41 | 16462 | 51714 | 5972 | 20894 | 28 | 14482 | 8418 |
| SRR837778 | 0 | 542 | 6695 | 0 | 1525 | 18597 | 25698 | 12533 | 1098 | 56 | 11153 | 43760 | 2418 | 7324 | 0 | 16910 | 14851 |
| SRR837779 | 0 | 6528 | 780 | 97 | 306 | 5691 | 3629 | 8082 | 6805 | 106 | 45202 | 16 | 7499 | 2278 | 149 | 3271 | 21800 |
| SRR837780 | 14 | 9002 | 779 | 438 | 913 | 8734 | 5608 | 6751 | 18117 | 78 | 48934 | 13 | 6707 | 3054 | 730 | 4808 | 34888 |
| SRR8380550 | 57 | 1765 | 22234 | 28 | 779 | 746 | 18222 | 23865 | 54020 | 10112 | 14077 | 1713 | 14 | 5895 | 237 | 1831 | 3270 |
| SRR8380552 | 1382 | 6861 | 143639 | 122 | 10876 | 12088 | 75759 | 16813 | 353627 | 2903 | 22721 | 6855 | 17588 | 15976 | 357 | 1108 | 14441 |
| SRR8380553 | 1052 | 9834 | 198364 | 338 | 8162 | 7389 | 49538 | 7801 | 254680 | 2578 | 17532 | 8023 | 15653 | 17307 | 993 | 1495 | 39785 |
| SRR8380554 | 177 | 5138 | 32015 | 172 | 774 | 454 | 3473 | 1189 | 233718 | 3979 | 17158 | 6850 | 11124 | 4694 | 76 | 1101 | 9497 |
| SRR8380555 | 281 | 3945 | 31461 | 10 | 3772 | 3280 | 36816 | 47613 | 34334 | 8528 | 44378 | 15 | 79 | 21270 | 684 | 391 | 24327 |
| SRR8380722 | 241 | 693 | 47897 | 2154 | 825 | 572 | 2704 | 3428 | 12801 | 1432 | 26414 | 21 | 18880 | 24379 | 20 | 6817 | 35475 |
| SRR8380723 | 25 | 421 | 85219 | 66 | 485 | 400 | 2309 | 3410 | 6993 | 5911 | 36795 | 26351 | 25778 | 17791 | 28940 | 8846 | 49324 |
| SRR8380724 | 717 | 1130 | 9884 | 387 | 523 | 302 | 2305 | 2570 | 6046 | 2198 | 30210 | 21 | 23643 | 18193 | 15 | 11346 | 56953 |
| SRR8380725 | 33 | 380 | 31347 | 48 | 552 | 418 | 2321 | 3781 | 11771 | 2382 | 12986 | 34 | 12868 | 8980 | 12 | 4573 | 40640 |
| SRR8380726 | 49 | 352 | 54185 | 318 | 651 | 479 | 2036 | 2928 | 23122 | 2009 | 16204 | 29 | 14795 | 18831 | 29 | 3369 | 31428 |
| SRR8380727 | 20 | 280 | 37753 | 71 | 648 | 548 | 3234 | 4515 | 6995 | 3517 | 27674 | 20803 | 26827 | 17171 | 18424 | 4182 | 58221 |
| SRR8400006 | 0 | 91 | 1753 | 26 | 6291 | 1190 | 189279 | 36377 | 841 | 484 | 8879 | 6404 | 7052 | 12550 | 2091 | 105 | 15513 |
| SRR8400007 | 0 | 0 | 581 | 0 | 3740 | 855 | 143898 | 29686 | 1802 | 155 | 8639 | 4950 | 9164 | 11117 | 1787 | 269 | 14660 |
| SRR8400008 | 0 | 12 | 269 | 34 | 9155 | 1269 | 275569 | 47238 | 851 | 534 | 5984 | 3941 | 6771 | 7609 | 1179 | 161 | 11986 |
| SRR8400009 | 0 | 17 | 506 | 15 | 1395 | 130 | 36496 | 13641 | 11449 | 45 | 8178 | 6147 | 13758 | 7363 | 4201 | 436 | 18085 |
| SRR8400010 | 0 | 20 | 1140 | 0 | 1554 | 199 | 41985 | 7147 | 1083 | 208 | 9408 | 8370 | 9210 | 5322 | 320 | 317 | 27129 |
| SRR8417876 | 21 | 45205 | 2900 | 319 | 2464 | 29531 | 23838 | 11621 | 626 | 12 | 13209 | 49006 | 3197 | 5015 | 49 | 34723 | 15165 |
| SRR8417877 | 102 | 72732 | 4206 | 307 | 2210 | 26156 | 18808 | 11893 | 864 | 47 | 12237 | 52634 | 3130 | 3917 | 29 | 34318 | 19726 |
| SRR8417878 | 711 | 78806 | 3781 | 170 | 2277 | 25981 | 18185 | 12077 | 1029 | 374 | 12427 | 53520 | 2977 | 4493 | 33 | 38388 | 20884 |
| SRR8435171 | 0 | 69 | 65948 | 29 | 17097 | 36028 | 197484 | 31333 | 14374 | 1222 | 29963 | 25 | 21619 | 12331 | 1743 | 3269 | 24559 |
| SRR8435172 | 0 | 29 | 23957 | 19 | 17666 | 34399 | 167016 | 26011 | 24475 | 296 | 41950 | 15616 | 25529 | 14545 | 1850 | 3360 | 30444 |
| SRR8481810 | 0 | 1131 | 1586 | 299 | 3832 | 63166 | 67675 | 91785 | 811 | 52 | 54486 | 26446 | 17020 | 5499 | 12 | 227 | 20510 |
| SRR8481811 | 13 | 1150 | 1209 | 227 | 3621 | 61100 | 61678 | 108250 | 954 | 31 | 72803 | 27250 | 16110 | 7519 | 0 | 294 | 16694 |
| SRR8481812 | 0 | 1147 | 3449 | 319 | 3575 | 58361 | 59264 | 110225 | 816 | 73 | 62418 | 27384 | 17489 | 5634 | 0 | 236 | 21606 |
| SRR8481813 | 21 | 1937 | 579 | 183 | 2881 | 55835 | 51873 | 117358 | 784 | 25 | 56903 | 23319 | 10983 | 10098 | 0 | 437 | 20619 |
| SRR8481814 | 18 | 1694 | 7610 | 246 | 4706 | 84264 | 71111 | 133844 | 1053 | 31 | 56179 | 25121 | 14820 | 4649 | 0 | 228 | 17834 |
| SRR8481815 | 0 | 2530 | 5090 | 108 | 3950 | 79278 | 58235 | 134200 | 956 | 49 | 45273 | 25146 | 14055 | 4606 | 0 | 267 | 22955 |
| SRR8481816 | 53 | 1916 | 6037 | 687 | 3430 | 80283 | 61698 | 136813 | 613 | 36 | 51945 | 33273 | 18781 | 4546 | 0 | 305 | 20303 |
| SRR8481817 | 55 | 2350 | 934 | 144 | 2975 | 69929 | 52642 | 142288 | 1194 | 18 | 33765 | 19831 | 9166 | 6297 | 0 | 683 | 28715 |
| SRR8481818 | 45 | 1609 | 1604 | 2540 | 3946 | 68175 | 56507 | 141147 | 938 | 11 | 48026 | 29166 | 13645 | 4718 | 0 | 193 | 26331 |
| SRR8481819 | 127 | 2866 | 2081 | 652 | 3752 | 66521 | 50963 | 132356 | 1176 | 47 | 35504 | 27138 | 11695 | 5220 | 11 | 264 | 29416 |
| SRR8481820 | 98 | 2679 | 947 | 746 | 3278 | 57612 | 46705 | 134783 | 1330 | 13 | 39496 | 32187 | 11592 | 4322 | 0 | 287 | 30016 |
| SRR8481821 | 81 | 2909 | 282 | 179 | 2811 | 56248 | 41576 | 138957 | 1276 | 16 | 35216 | 31321 | 8948 | 7746 | 0 | 457 | 24987 |
| SRR8485055 | 491 | 1661 | 12374 | 1063 | 4595 | 2929 | 41914 | 64558 | 8618 | 5033 | 36193 | 50695 | 17348 | 15809 | 196 | 1695 | 50341 |
| SRR8485056 | 363 | 2206 | 15098 | 859 | 1530 | 696 | 11899 | 25464 | 2893 | 4880 | 33871 | 58638 | 28676 | 28547 | 369 | 1862 | 70718 |
| SRR8485057 | 350 | 2561 | 30334 | 761 | 3942 | 2186 | 36584 | 40916 | 7598 | 8484 | 29834 | 49868 | 21149 | 31234 | 146 | 3627 | 29731 |
| SRR8485058 | 552 | 2502 | 13860 | 932 | 3260 | 1790 | 21855 | 33039 | 6375 | 12038 | 30104 | 62181 | 17244 | 18036 | 108 | 1582 | 40224 |
| SRR8485059 | 443 | 7833 | 19809 | 644 | 1181 | 527 | 9950 | 15996 | 4587 | 8991 | 14186 | 32308 | 15483 | 25300 | 129 | 1595 | 17044 |
| SRR8485060 | 715 | 3946 | 13148 | 607 | 2370 | 1206 | 15010 | 21087 | 5159 | 4804 | 19026 | 53065 | 17683 | 18976 | 164 | 1417 | 38887 |
| SRR8485061 | 345 | 2118 | 20150 | 1478 | 4339 | 2230 | 31067 | 43821 | 5106 | 3190 | 41385 | 43369 | 27633 | 29749 | 400 | 2401 | 73923 |
| SRR8485062 | 551 | 2339 | 10736 | 927 | 1311 | 1044 | 8067 | 11185 | 24476 | 6052 | 25980 | 39944 | 14656 | 6977 | 135 | 2064 | 55828 |
| SRR8485063 | 358 | 2623 | 13090 | 2224 | 5469 | 4221 | 52207 | 60808 | 4905 | 8067 | 34034 | 29498 | 12764 | 20105 | 385 | 2340 | 36079 |
| SRR8485064 | 368 | 1616 | 11357 | 1125 | 7076 | 3926 | 56773 | 75580 | 5780 | 8193 | 36528 | 43886 | 15841 | 15387 | 228 | 2439 | 43977 |
| SRR8485065 | 494 | 7141 | 27411 | 981 | 3452 | 2024 | 31913 | 46830 | 6039 | 4695 | 39451 | 56612 | 17148 | 22362 | 219 | 2657 | 31607 |
| SRR8485066 | 921 | 3136 | 18853 | 1465 | 1797 | 1485 | 14464 | 14882 | 18364 | 4968 | 39462 | 55320 | 18441 | 9404 | 226 | 1664 | 51889 |
| SRR8485067 | 265 | 1594 | 8978 | 4811 | 4496 | 3282 | 48514 | 68720 | 8855 | 1577 | 42638 | 39485 | 16218 | 15679 | 242 | 2927 | 60738 |
| SRR8487217 | 698 | 3735 | 25084 | 899 | 9744 | 5637 | 73187 | 87801 | 9683 | 4083 | 27634 | 27742 | 9856 | 19421 | 192 | 3406 | 46300 |
| SRR8487218 | 135 | 747 | 33038 | 543 | 6553 | 3856 | 68789 | 92626 | 4818 | 7673 | 25877 | 40515 | 13980 | 46122 | 530 | 929 | 54541 |
| SRR8487219 | 416 | 2306 | 10528 | 369 | 7942 | 6219 | 84229 | 85730 | 10987 | 9358 | 32727 | 37971 | 11204 | 23900 | 219 | 2124 | 48458 |
| SRR8487220 | 190 | 885 | 6209 | 580 | 5222 | 3773 | 53972 | 76733 | 9342 | 15078 | 20303 | 23545 | 10503 | 22902 | 189 | 2888 | 72821 |
| SRR8487221 | 256 | 2094 | 9617 | 487 | 1314 | 902 | 16345 | 28750 | 3913 | 7782 | 13518 | 12163 | 11202 | 37999 | 133 | 4326 | 69939 |
| SRR8487222 | 595 | 2888 | 9304 | 345 | 451 | 612 | 5851 | 6863 | 23724 | 6747 | 7776 | 5017 | 3997 | 7022 | 264 | 3415 | 35482 |
| SRR8487223 | 203 | 3396 | 14626 | 1123 | 1907 | 2537 | 42958 | 49528 | 4137 | 4659 | 6602 | 4359 | 3061 | 29521 | 1029 | 3209 | 10015 |
| SRR8487224 | 163 | 4990 | 19345 | 4518 | 2988 | 2092 | 51997 | 95472 | 5498 | 2762 | 27770 | 27371 | 12519 | 20789 | 200 | 4364 | 48067 |
| SRR8586304 | 0 | 1101 | 2642 | 93 | 156 | 4293 | 2807 | 17652 | 3302 | 0 | 6209 | 10558 | 2261 | 1503 | 95 | 2027 | 41513 |
| SRR871609 | 0 | 0 | 27913 | 0 | 4161 | 2332 | 37879 | 16915 | 24408 | 477 | 19682 | 20661 | 1938 | 169 | 551 | 820 | 2156 |
| SRR871671 | 0 | 15 | 3264 | 0 | 103 | 70 | 690 | 823 | 10997 | 76 | 71658 | 35 | 2087 | 73 | 12374 | 27 | 526 |
| SRR873381 | 20 | 38921 | 52763 | 609 | 814 | 7478 | 11371 | 11141 | 15959 | 22 | 44133 | 11436 | 3888 | 8634 | 123 | 1824 | 26646 |
| SRR873401 | 113 | 70986 | 1639 | 100 | 144 | 1415 | 2126 | 1453 | 10594 | 0 | 20579 | 44 | 2139 | 3782 | 59 | 401 | 28607 |
| SRR873410 | 223 | 33588 | 2115 | 648 | 94 | 950 | 1639 | 1626 | 33108 | 20 | 20201 | 8449 | 3582 | 5443 | 104 | 572 | 38358 |
| SRR8834226 | 0 | 0 | 0 | 102 | 0 | 180 | 6085 | 0 | 109 | 0 | 27905 | 19430 | 2375 | 818 | 284 | 0 | 77 |
| SRR8834227 | 0 | 0 | 0 | 0 | 0 | 74 | 2405 | 0 | 11 | 0 | 62463 | 26122 | 17194 | 8959 | 1134 | 0 | 105 |
| SRR8839811 | 349 | 4066 | 3701 | 10205 | 3240 | 3811 | 25713 | 37512 | 35466 | 19 | 3679 | 4999 | 6917 | 48272 | 796 | 2478 | 4738 |
| SRR8839812 | 30 | 9978 | 253290 | 1725 | 2488 | 3448 | 44535 | 49867 | 120171 | 148 | 4976 | 3341 | 9526 | 94331 | 853 | 10308 | 6647 |
| SRR8850891 | 4470 | 23554 | 36247 | 1884 | 16015 | 16381 | 89149 | 21449 | 12646 | 3710 | 12556 | 16231 | 8463 | 47416 | 4311 | 1563 | 21108 |
| SRR8850892 | 5088 | 30306 | 54674 | 2435 | 18003 | 16923 | 92728 | 22564 | 12693 | 5411 | 14248 | 36 | 9045 | 48424 | 4298 | 2168 | 21204 |
| SRR8850893 | 7178 | 29320 | 49511 | 816 | 19151 | 18285 | 102920 | 26237 | 11878 | 8612 | 13593 | 18586 | 9498 | 48869 | 3926 | 2096 | 18857 |
| SRR8850894 | 4857 | 16184 | 48243 | 963 | 24436 | 23950 | 131401 | 35745 | 9844 | 6941 | 13213 | 17718 | 8442 | 45503 | 3218 | 2293 | 19423 |
| SRR8850895 | 3503 | 7926 | 39781 | 991 | 16757 | 17389 | 83031 | 27326 | 5088 | 2809 | 12311 | 17509 | 8289 | 37292 | 4988 | 1210 | 17499 |
| SRR8850896 | 2432 | 9095 | 29144 | 692 | 18272 | 20204 | 84066 | 27876 | 6636 | 1156 | 14877 | 24 | 7156 | 38907 | 6281 | 1428 | 16429 |
| SRR8850897 | 3209 | 10028 | 13656 | 2292 | 18349 | 18123 | 92762 | 24411 | 9383 | 2558 | 12163 | 17603 | 7888 | 42204 | 5682 | 1688 | 19232 |
| SRR8850898 | 5412 | 9065 | 42760 | 686 | 18664 | 18078 | 101936 | 33013 | 5901 | 7523 | 10033 | 10960 | 7121 | 33009 | 4357 | 1938 | 18093 |
| SRR8850899 | 4753 | 16751 | 81087 | 707 | 23874 | 24147 | 113542 | 26515 | 3133 | 10346 | 17801 | 27224 | 12663 | 42229 | 6622 | 1347 | 19160 |
| SRR8850900 | 4265 | 11930 | 97370 | 603 | 18988 | 24749 | 78297 | 22062 | 3124 | 9172 | 16437 | 57 | 12286 | 37891 | 6040 | 1393 | 20869 |
| SRR8850901 | 4989 | 16245 | 65961 | 506 | 24467 | 25775 | 129720 | 32249 | 3893 | 11296 | 15355 | 44 | 11005 | 41376 | 5545 | 2497 | 18623 |
| SRR8850902 | 2830 | 13297 | 67177 | 351 | 22659 | 25962 | 117513 | 29570 | 3471 | 12681 | 15538 | 24802 | 11537 | 42554 | 5317 | 2471 | 20524 |
| SRR8858246 | 0 | 116 | 1665 | 0 | 1271 | 21636 | 29582 | 8654 | 13724 | 811 | 145563 | 76792 | 1873 | 11844 | 0 | 317 | 18739 |
| SRR8858247 | 0 | 143 | 1225 | 0 | 1176 | 28276 | 24423 | 9979 | 18505 | 332 | 140041 | 75398 | 1881 | 8434 | 0 | 1791 | 25375 |
| SRR8858248 | 0 | 109 | 1358 | 0 | 1399 | 17544 | 28836 | 9139 | 17951 | 591 | 92900 | 57552 | 2395 | 11062 | 0 | 1177 | 24434 |

| | | | | | | | | | | | | | | | | | | | | |
|---|---|---|---|---|---|---|---|---|---|---|---|---|---|---|---|---|---|---|---|---|
| SRR8280568 | 0 | 0 | 0 | 0 | 50 | 0 | 0 | 0 | 0 | 0 | 0 | 0 | 0 | 67 | 12 | 0 | 0 | 0 | 0 | 0 |
| SRR8280572 | 0 | 0 | 0 | 0 | 13 | 0 | 0 | 0 | 0 | 0 | 0 | 0 | 0 | 47 | 0 | 0 | 0 | 0 | 0 | 0 |
| SRR8280573 | 0 | 0 | 0 | 0 | 21 | 0 | 0 | 0 | 0 | 0 | 0 | 0 | 0 | 57 | 16 | 0 | 0 | 0 | 0 | 0 |
| SRR8280574 | 0 | 0 | 0 | 0 | 0 | 0 | 0 | 0 | 0 | 0 | 0 | 0 | 0 | 0 | 0 | 0 | 0 | 0 | 0 | 0 |
| SRR8280576 | 0 | 0 | 0 | 0 | 0 | 0 | 0 | 0 | 0 | 0 | 0 | 0 | 0 | 0 | 0 | 0 | 0 | 0 | 0 | 0 |
| SRR8280580 | 0 | 0 | 0 | 0 | 11 | 0 | 0 | 0 | 0 | 0 | 0 | 0 | 0 | 24 | 17 | 0 | 0 | 0 | 0 | 0 |
| SRR8280582 | 0 | 0 | 0 | 0 | 18 | 0 | 0 | 0 | 0 | 0 | 0 | 0 | 0 | 64 | 40 | 0 | 0 | 10 | 0 | 0 |
| SRR8280583 | 0 | 0 | 0 | 34 | 138 | 0 | 0 | 0 | 0 | 0 | 12 | 41 | 47 | 316 | 0 | 0 | 0 | 0 | 0 | 0 |
| SRR8280584 | 0 | 0 | 0 | 0 | 14 | 0 | 0 | 0 | 0 | 0 | 0 | 0 | 0 | 55 | 15 | 0 | 0 | 0 | 0 | 0 |
| SRR8280589 | 0 | 0 | 0 | 0 | 32 | 0 | 0 | 0 | 0 | 0 | 0 | 0 | 0 | 17 | 0 | 0 | 0 | 0 | 0 | 0 |
| SRR8280590 | 0 | 0 | 0 | 0 | 51 | 0 | 0 | 0 | 0 | 0 | 0 | 0 | 0 | 72 | 35 | 0 | 0 | 0 | 0 | 0 |
| SRR8280592 | 0 | 0 | 0 | 0 | 44 | 0 | 0 | 0 | 0 | 0 | 0 | 0 | 0 | 28 | 0 | 0 | 0 | 0 | 0 | 0 |
| SRR8280593 | 0 | 0 | 0 | 0 | 35 | 0 | 0 | 0 | 0 | 0 | 0 | 0 | 0 | 38 | 0 | 0 | 0 | 0 | 0 | 0 |
| SRR8311061 | 0 | 0 | 0 | 0 | 0 | 0 | 0 | 0 | 0 | 0 | 0 | 0 | 0 | 0 | 0 | 0 | 0 | 0 | 0 | 0 |
| SRR8311062 | 0 | 0 | 0 | 0 | 0 | 0 | 0 | 0 | 0 | 0 | 0 | 0 | 0 | 0 | 0 | 0 | 0 | 0 | 0 | 0 |
| SRR8311063 | 0 | 0 | 0 | 0 | 0 | 0 | 0 | 0 | 0 | 0 | 0 | 0 | 0 | 0 | 0 | 0 | 0 | 0 | 0 | 0 |
| SRR8311064 | 0 | 0 | 0 | 0 | 0 | 0 | 0 | 0 | 0 | 0 | 0 | 0 | 0 | 0 | 0 | 0 | 0 | 0 | 0 | 0 |
| SRR8311070 | 0 | 0 | 0 | 0 | 0 | 0 | 0 | 0 | 0 | 0 | 0 | 0 | 0 | 0 | 0 | 0 | 0 | 0 | 0 | 0 |
| SRR8311071 | 0 | 0 | 0 | 0 | 0 | 0 | 0 | 0 | 0 | 0 | 0 | 0 | 0 | 0 | 0 | 0 | 0 | 0 | 0 | 0 |
| SRR8311072 | 0 | 0 | 0 | 0 | 0 | 0 | 0 | 0 | 0 | 0 | 0 | 0 | 0 | 0 | 0 | 0 | 0 | 0 | 0 | 0 |
| SRR8311073 | 0 | 0 | 0 | 0 | 0 | 0 | 0 | 0 | 0 | 0 | 0 | 0 | 0 | 0 | 0 | 0 | 0 | 0 | 0 | 0 |
| SRR8311074 | 0 | 0 | 0 | 0 | 0 | 0 | 0 | 0 | 0 | 0 | 0 | 0 | 0 | 0 | 0 | 0 | 0 | 0 | 0 | 0 |
| SRR8311076 | 0 | 0 | 0 | 0 | 0 | 0 | 0 | 0 | 0 | 0 | 0 | 0 | 0 | 0 | 0 | 0 | 0 | 0 | 0 | 0 |
| SRR8311078 | 0 | 0 | 0 | 0 | 0 | 0 | 0 | 0 | 0 | 0 | 0 | 0 | 0 | 0 | 0 | 0 | 0 | 0 | 0 | 0 |
| SRR8311080 | 0 | 0 | 0 | 0 | 0 | 0 | 0 | 0 | 0 | 0 | 0 | 0 | 0 | 0 | 0 | 0 | 0 | 0 | 0 | 0 |
| SRR8311081 | 0 | 0 | 0 | 0 | 0 | 0 | 0 | 0 | 0 | 0 | 0 | 0 | 0 | 0 | 0 | 0 | 0 | 0 | 0 | 0 |
| SRR8311082 | 0 | 0 | 0 | 0 | 0 | 0 | 0 | 0 | 0 | 0 | 0 | 0 | 0 | 0 | 0 | 0 | 0 | 0 | 0 | 0 |
| SRR8311085 | 0 | 0 | 0 | 0 | 0 | 0 | 0 | 0 | 0 | 0 | 0 | 0 | 0 | 0 | 0 | 0 | 0 | 0 | 0 | 0 |
| SRR8311087 | 0 | 0 | 0 | 0 | 0 | 0 | 0 | 0 | 0 | 0 | 0 | 0 | 0 | 0 | 0 | 0 | 0 | 0 | 0 | 0 |
| SRR8311089 | 0 | 0 | 0 | 0 | 0 | 0 | 0 | 0 | 0 | 0 | 0 | 0 | 0 | 0 | 0 | 0 | 0 | 0 | 0 | 0 |
| SRR8311092 | 0 | 0 | 0 | 0 | 0 | 0 | 0 | 0 | 0 | 0 | 0 | 0 | 0 | 0 | 0 | 0 | 0 | 0 | 0 | 0 |
| SRR8311093 | 0 | 0 | 0 | 0 | 0 | 0 | 0 | 0 | 0 | 0 | 0 | 0 | 0 | 0 | 0 | 0 | 0 | 0 | 0 | 0 |
| SRR8311095 | 0 | 0 | 0 | 0 | 0 | 0 | 0 | 0 | 0 | 0 | 0 | 0 | 0 | 0 | 0 | 0 | 0 | 0 | 0 | 0 |
| SRR8311098 | 0 | 0 | 0 | 0 | 0 | 0 | 0 | 0 | 0 | 0 | 0 | 0 | 0 | 0 | 0 | 0 | 0 | 0 | 0 | 0 |
| SRR8311105 | 0 | 0 | 0 | 0 | 0 | 0 | 0 | 0 | 0 | 0 | 0 | 0 | 0 | 0 | 0 | 0 | 0 | 0 | 0 | 0 |
| SRR8311106 | 0 | 0 | 0 | 0 | 0 | 0 | 0 | 0 | 0 | 0 | 0 | 0 | 0 | 0 | 0 | 0 | 0 | 0 | 0 | 0 |
| SRR835324 | 0 | 0 | 0 | 0 | 21 | 0 | 10 | 0 | 0 | 0 | 13 | 20 | 14 | 75 | 0 | 0 | 0 | 0 | 18 | 0 |
| SRR835325 | 0 | 0 | 0 | 0 | 0 | 0 | 0 | 0 | 0 | 0 | 0 | 12 | 0 | 18 | 0 | 0 | 0 | 0 | 19 | 0 |
| SRR8354354 | 0 | 0 | 0 | 52 | 18 | 35 | 10 | 0 | 0 | 0 | 30 | 23 | 0 | 17 | 0 | 0 | 0 | 0 | 0 | 0 |
| SRR837775 | 0 | 0 | 0 | 209 | 43 | 90 | 0 | 32 | 0 | 0 | 0 | 0 | 43 | 13 | 17 | 0 | 27 | 39 | 0 | 0 |
| SRR837776 | 0 | 0 | 0 | 44 | 12 | 24 | 0 | 0 | 0 | 0 | 0 | 0 | 0 | 0 | 0 | 0 | 13 | 15 | 0 | 0 |
| SRR837777 | 0 | 0 | 0 | 10 | 0 | 0 | 0 | 0 | 0 | 0 | 0 | 0 | 0 | 0 | 0 | 0 | 0 | 0 | 0 | 0 |
| SRR837778 | 0 | 0 | 0 | 28 | 0 | 0 | 0 | 0 | 0 | 0 | 0 | 0 | 0 | 0 | 0 | 0 | 0 | 0 | 0 | 0 |
| SRR837779 | 0 | 0 | 0 | 44 | 12 | 24 | 0 | 0 | 0 | 0 | 0 | 0 | 0 | 0 | 0 | 0 | 13 | 15 | 0 | 0 |
| SRR837780 | 0 | 0 | 0 | 209 | 43 | 90 | 0 | 32 | 0 | 0 | 0 | 0 | 43 | 13 | 17 | 0 | 27 | 39 | 0 | 0 |
| SRR8380550 | 0 | 0 | 67 | 182 | 82 | 39 | 0 | 0 | 0 | 0 | 182 | 3291 | 3928 | 286 | 23 | 0 | 0 | 0 | 0 | 0 |
| SRR8380552 | 0 | 0 | 67 | 104 | 712 | 251 | 30 | 11 | 0 | 0 | 0 | 241 | 171 | 10 | 0 | 0 | 0 | 0 | 25 | 0 |
| SRR8380553 | 0 | 0 | 50 | 248 | 668 | 488 | 91 | 27 | 10 | 0 | 36 | 304 | 625 | 10 | 0 | 0 | 0 | 0 | 38 | 0 |
| SRR8380554 | 0 | 0 | 23 | 123 | 48 | 57 | 0 | 0 | 0 | 0 | 67 | 623 | 741 | 0 | 0 | 0 | 0 | 0 | 0 | 0 |
| SRR8380555 | 0 | 41 | 205 | 700 | 545 | 460 | 32 | 18 | 14 | 0 | 153 | 1690 | 1637 | 112 | 13 | 0 | 0 | 0 | 35 | 0 |
| SRR8380722 | 0 | 72 | 113 | 210 | 3135 | 251 | 0 | 0 | 0 | 0 | 296 | 689 | 416 | 795 | 11 | 0 | 0 | 0 | 14 | 0 |
| SRR8380723 | 0 | 26 | 92 | 55 | 2417 | 144 | 11 | 0 | 0 | 0 | 77 | 734 | 111 | 1022 | 0 | 0 | 0 | 0 | 24 | 0 |
| SRR8380724 | 0 | 282 | 934 | 389 | 13475 | 651 | 25 | 13 | 0 | 34 | 677 | 10384 | 708 | 5360 | 22 | 0 | 0 | 0 | 60 | 0 |
| SRR8380725 | 0 | 23 | 93 | 136 | 1978 | 503 | 19 | 10 | 0 | 0 | 37 | 400 | 125 | 948 | 16 | 0 | 0 | 0 | 25 | 0 |
| SRR8380726 | 0 | 51 | 116 | 367 | 3486 | 423 | 23 | 11 | 0 | 0 | 54 | 299 | 195 | 431 | 0 | 0 | 0 | 0 | 32 | 0 |
| SRR8380727 | 0 | 60 | 175 | 69 | 3494 | 105 | 0 | 0 | 0 | 0 | 113 | 1205 | 96 | 1484 | 0 | 0 | 0 | 0 | 28 | 0 |
| SRR8400006 | 0 | 60 | 169 | 1904 | 12190 | 1494 | 540 | 117 | 49 | 0 | 11 | 88 | 303 | 588 | 29 | 31 | 13 | 128 | 131 | 12 |
| SRR8400007 | 0 | 0 | 0 | 20 | 121 | 45 | 17 | 11 | 13 | 0 | 0 | 0 | 0 | 0 | 0 | 0 | 0 | 30 | 54 | 0 |
| SRR8400008 | 0 | 11 | 20 | 139 | 1230 | 359 | 128 | 53 | 52 | 0 | 0 | 0 | 20 | 56 | 0 | 0 | 0 | 96 | 140 | 13 |
| SRR8400009 | 0 | 0 | 14 | 171 | 2533 | 307 | 48 | 13 | 0 | 0 | 0 | 12 | 41 | 145 | 0 | 0 | 0 | 20 | 33 | 0 |
| SRR8400010 | 0 | 0 | 0 | 85 | 259 | 90 | 28 | 15 | 24 | 0 | 0 | 0 | 0 | 11 | 0 | 0 | 0 | 25 | 62 | 42 |
| SRR8417876 | 0 | 0 | 0 | 314 | 147 | 96 | 10 | 0 | 13 | 0 | 0 | 29 | 188 | 106 | 49 | 0 | 0 | 41 | 0 | 0 |
| SRR8417877 | 0 | 0 | 0 | 599 | 187 | 142 | 0 | 0 | 13 | 0 | 0 | 41 | 366 | 145 | 70 | 0 | 0 | 71 | 11 | 0 |
| SRR8417878 | 0 | 0 | 0 | 792 | 213 | 156 | 0 | 0 | 0 | 0 | 0 | 45 | 515 | 167 | 78 | 0 | 12 | 82 | 15 | 0 |
| SRR8435171 | 0 | 0 | 0 | 45 | 212 | 76 | 19 | 22 | 17 | 0 | 0 | 98 | 119 | 38 | 0 | 0 | 0 | 11 | 74 | 0 |
| SRR8435172 | 0 | 0 | 0 | 16 | 82 | 23 | 0 | 0 | 0 | 0 | 0 | 21 | 37 | 0 | 0 | 0 | 0 | 0 | 37 | 0 |
| SRR8481810 | 0 | 0 | 0 | 136 | 0 | 37 | 0 | 0 | 0 | 0 | 0 | 0 | 63 | 17 | 45 | 0 | 0 | 0 | 0 | 0 |
| SRR8481811 | 0 | 0 | 0 | 123 | 0 | 25 | 0 | 0 | 0 | 0 | 0 | 18 | 57 | 25 | 47 | 0 | 0 | 0 | 0 | 0 |
| SRR8481812 | 0 | 0 | 0 | 143 | 0 | 36 | 0 | 0 | 0 | 0 | 0 | 0 | 48 | 16 | 43 | 0 | 0 | 0 | 0 | 0 |
| SRR8481813 | 0 | 0 | 0 | 117 | 31 | 23 | 13 | 0 | 0 | 0 | 0 | 27 | 99 | 89 | 66 | 0 | 19 | 0 | 0 | 0 |
| SRR8481814 | 0 | 0 | 0 | 136 | 16 | 25 | 0 | 0 | 0 | 0 | 0 | 12 | 56 | 27 | 24 | 0 | 0 | 0 | 0 | 0 |
| SRR8481815 | 0 | 0 | 0 | 140 | 19 | 36 | 0 | 0 | 0 | 0 | 0 | 19 | 55 | 41 | 34 | 0 | 12 | 0 | 0 | 0 |
| SRR8481816 | 0 | 0 | 0 | 314 | 32 | 86 | 0 | 0 | 0 | 0 | 0 | 16 | 79 | 44 | 80 | 0 | 16 | 0 | 0 | 0 |
| SRR8481817 | 0 | 0 | 0 | 181 | 44 | 42 | 12 | 0 | 0 | 0 | 0 | 29 | 104 | 105 | 75 | 0 | 17 | 0 | 0 | 0 |
| SRR8481818 | 0 | 0 | 0 | 178 | 20 | 39 | 0 | 0 | 0 | 0 | 0 | 16 | 74 | 54 | 39 | 0 | 0 | 0 | 0 | 0 |
| SRR8481819 | 0 | 0 | 0 | 181 | 30 | 57 | 0 | 0 | 0 | 0 | 0 | 23 | 83 | 68 | 50 | 0 | 0 | 0 | 0 | 0 |
| SRR8481820 | 0 | 0 | 0 | 170 | 24 | 76 | 0 | 0 | 0 | 0 | 0 | 13 | 82 | 49 | 40 | 0 | 0 | 0 | 0 | 0 |
| SRR8481821 | 0 | 0 | 0 | 72 | 25 | 32 | 0 | 0 | 0 | 0 | 0 | 16 | 76 | 90 | 47 | 0 | 17 | 16 | 0 | 0 |
| SRR8485055 | 0 | 76 | 410 | 2057 | 2188 | 650 | 13 | 0 | 0 | 0 | 114 | 575 | 524 | 922 | 678 | 0 | 0 | 0 | 42 | 0 |
| SRR8485056 | 0 | 0 | 46 | 1481 | 1425 | 363 | 0 | 0 | 0 | 0 | 24 | 87 | 518 | 549 | 462 | 0 | 0 | 18 | 52 | 0 |
| SRR8485057 | 0 | 10 | 55 | 807 | 883 | 122 | 0 | 0 | 0 | 0 | 42 | 137 | 323 | 400 | 254 | 0 | 0 | 13 | 31 | 0 |
| SRR8485058 | 0 | 139 | 124 | 824 | 1544 | 104 | 0 | 0 | 0 | 0 | 281 | 194 | 191 | 902 | 114 | 0 | 0 | 0 | 23 | 0 |
| SRR8485059 | 0 | 0 | 11 | 636 | 2109 | 410 | 21 | 0 | 0 | 0 | 13 | 21 | 195 | 219 | 337 | 0 | 0 | 0 | 49 | 0 |
| SRR8485060 | 0 | 41 | 31 | 349 | 86 | 11 | 0 | 0 | 0 | 0 | 73 | 51 | 78 | 42 | 18 | 0 | 0 | 0 | 14 | 0 |
| SRR8485061 | 0 | 23 | 99 | 1857 | 1073 | 302 | 13 | 0 | 0 | 0 | 32 | 110 | 356 | 342 | 308 | 0 | 0 | 12 | 35 | 0 |
| SRR8485062 | 0 | 14 | 34 | 487 | 185 | 32 | 0 | 0 | 0 | 0 | 94 | 208 | 476 | 334 | 116 | 0 | 0 | 0 | 15 | 0 |
| SRR8485063 | 0 | 33 | 56 | 821 | 1433 | 191 | 0 | 0 | 0 | 0 | 40 | 63 | 164 | 279 | 140 | 0 | 0 | 0 | 51 | 0 |
| SRR8485064 | 0 | 51 | 67 | 608 | 1006 | 38 | 0 | 0 | 0 | 0 | 59 | 72 | 128 | 196 | 64 | 0 | 0 | 0 | 44 | 0 |
| SRR8485065 | 0 | 36 | 220 | 3034 | 5146 | 245 | 11 | 0 | 0 | 0 | 82 | 379 | 1157 | 1385 | 438 | 0 | 0 | 0 | 79 | 0 |
| SRR8485066 | 0 | 13 | 19 | 152 | 174 | 194 | 11 | 12 | 0 | 0 | 63 | 95 | 141 | 230 | 925 | 13 | 20 | 96 | 39 | 0 |
| SRR8485067 | 0 | 204 | 198 | 711 | 916 | 518 | 21 | 0 | 0 | 0 | 185 | 180 | 166 | 382 | 709 | 0 | 0 | 0 | 12 | 0 |
| SRR8487217 | 0 | 57 | 82 | 912 | 802 | 171 | 0 | 0 | 0 | 0 | 149 | 210 | 480 | 387 | 391 | 0 | 0 | 0 | 44 | 0 |
| SRR8487218 | 0 | 0 | 0 | 398 | 1026 | 471 | 13 | 0 | 0 | 0 | 13 | 16 | 111 | 274 | 568 | 0 | 0 | 25 | 30 | 0 |
| SRR8487219 | 0 | 26 | 46 | 1931 | 1026 | 297 | 0 | 0 | 0 | 0 | 89 | 126 | 916 | 552 | 570 | 0 | 0 | 46 | 174 | 0 |
| SRR8487220 | 0 | 111 | 124 | 563 | 989 | 674 | 52 | 17 | 0 | 0 | 174 | 184 | 149 | 249 | 1100 | 15 | 0 | 12 | 53 | 0 |
| SRR8487221 | 0 | 55 | 42 | 198 | 1034 | 52 | 0 | 0 | 0 | 0 | 571 | 406 | 260 | 1179 | 255 | 0 | 0 | 0 | 23 | 0 |
| SRR8487222 | 0 | 0 | 0 | 50 | 71 | 18 | 0 | 0 | 0 | 0 | 84 | 138 | 227 | 151 | 217 | 0 | 0 | 0 | 0 | 0 |
| SRR8487223 | 0 | 17 | 25 | 569 | 1895 | 562 | 11 | 0 | 0 | 0 | 61 | 81 | 344 | 1264 | 1203 | 0 | 0 | 33 | 73 | 0 |
| SRR8487224 | 0 | 0 | 17 | 425 | 504 | 185 | 0 | 0 | 0 | 0 | 19 | 24 | 97 | 136 | 193 | 0 | 0 | 0 | 34 | 0 |
| SRR8586304 | 0 | 0 | 0 | 19 | 1345 | 124 | 54 | 0 | 0 | 0 | 13 | 125 | 28 | 3590 | 33 | 0 | 0 | 0 | 0 | 0 |
| SRR871609 | 0 | 0 | 0 | 0 | 0 | 0 | 0 | 0 | 0 | 0 | 0 | 0 | 0 | 0 | 0 | 0 | 0 | 0 | 0 | 0 |
| SRR871671 | 0 | 0 | 0 | 0 | 0 | 0 | 0 | 0 | 0 | 0 | 0 | 0 | 0 | 0 | 0 | 0 | 0 | 0 | 0 | 0 |
| SRR873381 | 0 | 0 | 0 | 96 | 13 | 20 | 0 | 0 | 0 | 0 | 0 | 11 | 74 | 21 | 41 | 0 | 0 | 0 | 0 | 0 |
| SRR873401 | 0 | 0 | 0 | 77 | 45 | 11 | 0 | 0 | 0 | 0 | 0 | 15 | 44 | 28 | 0 | 0 | 0 | 0 | 0 | 0 |
| SRR873410 | 0 | 0 | 0 | 145 | 106 | 14 | 0 | 0 | 0 | 0 | 0 | 36 | 114 | 176 | 20 | 0 | 0 | 23 | 0 | 0 |
| SRR8834226 | 0 | 0 | 0 | 0 | 0 | 0 | 0 | 0 | 0 | 0 | 0 | 0 | 0 | 0 | 0 | 0 | 0 | 0 | 0 | 0 |
| SRR8834227 | 0 | 0 | 0 | 0 | 0 | 0 | 0 | 0 | 0 | 0 | 0 | 0 | 0 | 0 | 0 | 0 | 0 | 0 | 0 | 0 |
| SRR8839811 | 0 | 0 | 17 | 0 | 84 | 0 | 0 | 0 | 0 | 0 | 11 | 16 | 0 | 0 | 0 | 0 | 0 | 0 | 0 | 0 |
| SRR8839812 | 0 | 0 | 0 | 0 | 356 | 424 | 73 | 0 | 0 | 0 | 0 | 0 | 0 | 0 | 0 | 0 | 0 | 0 | 0 | 0 |
| SRR8850891 | 0 | 24 | 23 | 33 | 76 | 83 | 41 | 20 | 0 | 0 | 23 | 54 | 13 | 15 | 0 | 0 | 0 | 0 | 15 | 0 |
| SRR8850892 | 0 | 16 | 11 | 37 | 80 | 91 | 36 | 21 | 0 | 0 | 12 | 33 | 12 | 0 | 0 | 0 | 0 | 0 | 14 | 0 |
| SRR8850893 | 0 | 85 | 113 | 91 | 197 | 206 | 127 | 47 | 11 | 16 | 89 | 224 | 43 | 31 | 0 | 0 | 0 | 0 | 29 | 0 |
| SRR8850894 | 0 | 69 | 105 | 71 | 108 | 140 | 91 | 31 | 0 | 0 | 56 | 204 | 20 | 12 | 0 | 0 | 0 | 0 | 22 | 0 |
| SRR8850895 | 0 | 13 | 12 | 31 | 79 | 62 | 29 | 14 | 0 | 0 | 23 | 39 | 13 | 0 | 0 | 0 | 0 | 0 | 12 | 0 |
| SRR8850896 | 0 | 11 | 14 | 28 | 87 | 60 | 31 | 14 | 0 | 0 | 14 | 26 | 0 | 0 | 0 | 0 | 0 | 0 | 10 | 0 |
| SRR8850897 | 0 | 52 | 65 | 107 | 228 | 220 | 176 | 70 | 11 | 0 | 50 | 103 | 49 | 16 | 0 | 0 | 0 | 0 | 33 | 0 |
| SRR8850898 | 13 | 219 | 192 | 346 | 476 | 328 | 161 | 65 | 15 | 19 | 174 | 357 | 114 | 56 | 0 | 0 | 0 | 0 | 46 | 0 |
| SRR8850899 | 0 | 57 | 22 | 40 | 164 | 98 | 47 | 31 | 20 | 24 | 97 | 116 | 47 | 45 | 0 | 0 | 0 | 0 | 32 | 0 |
| SRR8850900 | 0 | 51 | 22 | 29 | 135 | 74 | 36 | 16 | 0 | 18 | 90 | 130 | 31 | 48 | 0 | 0 | 0 | 0 | 12 | 0 |
| SRR8850901 | 0 | 92 | 70 | 87 | 319 | 363 | 188 | 66 | 16 | 30 | 126 | 217 | 62 | 45 | 0 | 0 | 0 | 0 | 40 | 0 |
| SRR8850902 | 0 | 114 | 97 | 79 | 235 | 245 | 157 | 59 | 14 | 24 | 123 | 227 | 57 | 42 | 0 | 0 | 0 | 0 | 36 | 0 |
| SRR8858246 | 0 | 0 | 0 | 0 | 0 | 0 | 0 | 0 | 0 | 0 | 0 | 0 | 0 | 75 | 0 | 0 | 0 | 0 | 0 | 0 |
| SRR8858247 | 0 | 0 | 0 | 0 | 0 | 0 | 0 | 0 | 0 | 0 | 0 | 0 | 46 | 55 | 0 | 0 | 0 | 0 | 0 | 0 |
| SRR8858248 | 0 | 0 | 0 | 0 | 0 | 0 | 0 | 0 | 0 | 0 | 0 | 0 | 0 | 23 | 0 | 0 | 0 | 0 | 0 | 0 |

### Supp Table S8.1.Columns_B_to_AL

| | | | | | | | | | | | | | | | | | | | | | | | | | | | | | | | | | | | | | |
|---|---|---|---|---|---|---|---|---|---|---|---|---|---|---|---|---|---|---|---|---|---|---|---|---|---|---|---|---|---|---|---|---|---|---|---|---|---|
| SRR8858254 | 0 | 676 | 189 | 177 | 463 | 5101 | 4997 | 2561 | 11944 | 101 | 42135 | 44600 | 1401 | 8706 | 0 | 545 | 38717 | 0 | 0 | 0 | 0 | 0 | 0 | 0 | 0 | 0 | 0 | 0 | 0 | 0 | 26 | 20 | 0 | 0 | 0 | 0 | 0 |
| SRR8858258 | 0 | 77 | 372 | 0 | 248 | 1881 | 3267 | 2150 | 12777 | 52 | 93408 | 15 | 1518 | 5374 | 0 | 1628 | 26636 | 0 | 0 | 0 | 0 | 0 | 0 | 0 | 0 | 0 | 0 | 0 | 0 | 0 | 0 | 0 | 0 | 0 | 0 | 0 | 0 |
| SRR8907824 | 134 | 2885 | 152 | 1450 | 6744 | 12863 | 63825 | 2303 | 4118 | 96 | 5451 | 4694 | 3480 | 13594 | 394 | 3175 | 3441 | 0 | 0 | 0 | 11 | 272 | 131 | 33 | 121 | 62 | 0 | 0 | 0 | 0 | 0 | 0 | 0 | 0 | 61 | 260 | 25 |
| SRR8932122 | 0 | 860 | 747 | 322 | 2152 | 23503 | 27958 | 62850 | 13652 | 349 | 5516 | 10811 | 1420 | 8387 | 0 | 2365 | 112362 | 0 | 0 | 0 | 734 | 3577 | 438 | 0 | 0 | 0 | 0 | 50 | 160 | 935 | 7735 | 1100 | 0 | 43 | 205 | 22 | 0 |
| SRR8932123 | 0 | 402 | 628 | 179 | 244 | 1443 | 3445 | 5333 | 1080 | 196 | 3171 | 4317 | 832 | 985 | 0 | 1127 | 33702 | 0 | 0 | 14 | 741 | 1084 | 121 | 0 | 12 | 0 | 0 | 15 | 232 | 505 | 1451 | 165 | 0 | 32 | 295 | 101 | 0 |
| SRR8932124 | 0 | 3835 | 637 | 195 | 1380 | 10133 | 14189 | 28306 | 12035 | 358 | 6188 | 15132 | 2026 | 2132 | 52 | 1403 | 78429 | 0 | 0 | 0 | 746 | 2664 | 206 | 0 | 0 | 0 | 0 | 10 | 85 | 892 | 3877 | 429 | 0 | 18 | 85 | 19 | 0 |
| SRR8932125 | 14 | 1344 | 1408 | 380 | 648 | 8662 | 8210 | 16351 | 10817 | 277 | 9236 | 13241 | 3335 | 1674 | 16 | 2448 | 42168 | 0 | 0 | 0 | 105 | 526 | 93 | 0 | 0 | 0 | 0 | 0 | 23 | 130 | 772 | 237 | 0 | 0 | 43 | 12 | 0 |
| SRR8944248 | 0 | 9631 | 4606 | 0 | 0 | 79 | 103 | 84 | 75 | 0 | 965 | 3780 | 84 | 217 | 0 | 16 | 1473 | 0 | 0 | 0 | 0 | 0 | 0 | 0 | 0 | 0 | 0 | 0 | 0 | 0 | 0 | 0 | 0 | 0 | 0 | 0 | 0 |
| SRR8944250 | 0 | 27384 | 1494 | 0 | 0 | 334 | 198 | 144 | 84 | 0 | 2808 | 23109 | 319 | 532 | 0 | 19 | 2263 | 0 | 0 | 0 | 0 | 0 | 19 | 0 | 0 | 0 | 0 | 0 | 0 | 0 | 0 | 0 | 0 | 0 | 0 | 0 | 0 |
| SRR8944251 | 0 | 4610 | 3353 | 0 | 0 | 248 | 160 | 196 | 125 | 0 | 2611 | 14127 | 190 | 440 | 0 | 26 | 2892 | 0 | 0 | 0 | 0 | 0 | 0 | 0 | 0 | 0 | 0 | 0 | 0 | 0 | 0 | 0 | 0 | 0 | 0 | 0 | 0 |
| SRR8944252 | 0 | 6609 | 590 | 0 | 23 | 559 | 579 | 360 | 371 | 0 | 6219 | 34065 | 444 | 1069 | 0 | 53 | 4919 | 0 | 0 | 0 | 0 | 0 | 0 | 0 | 0 | 0 | 0 | 0 | 0 | 0 | 0 | 0 | 0 | 0 | 0 | 0 | 0 |
| SRR8944257 | 0 | 8186 | 1320 | 0 | 15 | 219 | 133 | 150 | 90 | 0 | 2417 | 10725 | 177 | 407 | 22 | 58 | 2072 | 0 | 0 | 0 | 0 | 0 | 0 | 0 | 0 | 0 | 0 | 0 | 0 | 0 | 0 | 0 | 0 | 0 | 0 | 0 | 0 |
| SRR8944259 | 0 | 8533 | 125 | 0 | 18 | 736 | 424 | 212 | 205 | 0 | 9307 | 56167 | 1017 | 1177 | 0 | 69 | 6146 | 0 | 0 | 0 | 0 | 0 | 0 | 0 | 0 | 0 | 0 | 0 | 0 | 0 | 0 | 0 | 0 | 0 | 0 | 0 | 0 |
| SRR8956086 | 316 | 6999 | 500739 | 211 | 40413 | 54534 | 213252 | 12115 | 115 | 134 | 2917 | 28785 | 2232 | 28104 | 17487 | 11267 | 14335 | 0 | 16 | 108 | 76 | 670 | 46 | 19 | 0 | 0 | 0 | 33 | 82 | 29 | 11 | 0 | 0 | 0 | 0 | 0 | 0 |
| SRR8956087 | 253 | 5899 | 495106 | 72 | 37455 | 48745 | 185669 | 8093 | 56 | 101 | 2323 | 29629 | 1806 | 18739 | 31443 | 10736 | 11264 | 0 | 15 | 86 | 63 | 631 | 42 | 17 | 0 | 0 | 0 | 29 | 58 | 20 | 0 | 0 | 0 | 0 | 0 | 0 | 0 |
| SRR8956089 | 512 | 10313 | 530725 | 336 | 31914 | 61382 | 203968 | 15339 | 66 | 264 | 3409 | 18085 | 2401 | 18995 | 28402 | 21156 | 21044 | 0 | 22 | 150 | 22 | 879 | 62 | 27 | 29 | 0 | 0 | 61 | 169 | 23 | 18 | 0 | 0 | 0 | 0 | 14 | 0 |
| SRR8956090 | 577 | 12773 | 518770 | 695 | 32895 | 55222 | 238151 | 12668 | 64 | 448 | 3428 | 9628 | 2077 | 18143 | 37453 | 28651 | 12339 | 0 | 16 | 125 | 20 | 601 | 49 | 18 | 33 | 0 | 0 | 76 | 223 | 34 | 14 | 0 | 0 | 0 | 0 | 11 | 0 |
| SRR8956091 | 527 | 11712 | 482836 | 526 | 29733 | 57540 | 231340 | 12368 | 50 | 439 | 3571 | 13721 | 2282 | 20396 | 31953 | 26502 | 15282 | 0 | 19 | 114 | 18 | 665 | 58 | 18 | 27 | 0 | 0 | 79 | 217 | 22 | 14 | 0 | 0 | 0 | 0 | 11 | 0 |
| SRR8992368 | 219 | 1009 | 39419 | 16750 | 26 | 36 | 280 | 2217 | 20906 | 51 | 554 | 1241 | 470 | 28326 | 223 | 2347 | 6970 | 0 | 0 | 0 | 10 | 27 | 13 | 24 | 0 | 0 | 0 | 17 | 15 | 0 | 0 | 0 | 0 | 0 | 0 | 0 | 0 |
| SRR8992369 | 131 | 2033 | 30940 | 36169 | 56 | 83 | 977 | 5419 | 26178 | 58 | 756 | 1038 | 1004 | 90329 | 326 | 4653 | 7717 | 0 | 30 | 49 | 70 | 510 | 392 | 371 | 45 | 0 | 0 | 99 | 52 | 51 | 36 | 0 | 0 | 0 | 0 | 26 | 0 |
| SRR8992370 | 217 | 3099 | 21571 | 36475 | 37 | 52 | 864 | 3940 | 32762 | 36 | 792 | 1154 | 1167 | 73008 | 335 | 3141 | 6745 | 0 | 27 | 41 | 72 | 478 | 279 | 333 | 50 | 0 | 0 | 104 | 54 | 55 | 27 | 0 | 0 | 0 | 0 | 27 | 0 |
| SRR8992371 | 140 | 1958 | 24560 | 29877 | 52 | 85 | 1147 | 4443 | 25225 | 42 | 837 | 1051 | 1112 | 67377 | 342 | 4232 | 9346 | 0 | 40 | 52 | 96 | 573 | 364 | 355 | 58 | 0 | 0 | 119 | 72 | 68 | 51 | 0 | 0 | 0 | 11 | 31 | 0 |
| SRR8992378 | 272 | 3303 | 25485 | 48815 | 93 | 159 | 1085 | 3373 | 28665 | 43 | 854 | 1276 | 1039 | 82399 | 347 | 3815 | 9145 | 0 | 25 | 49 | 64 | 479 | 336 | 337 | 49 | 0 | 0 | 99 | 67 | 49 | 29 | 0 | 0 | 0 | 0 | 30 | 0 |
| SRR8992379 | 397 | 1421 | 26297 | 23935 | 27 | 32 | 475 | 3633 | 29337 | 83 | 475 | 901 | 414 | 40507 | 191 | 1750 | 5993 | 0 | 0 | 0 | 0 | 19 | 10 | 20 | 0 | 0 | 0 | 11 | 0 | 0 | 0 | 0 | 0 | 0 | 0 | 0 | 0 |
| SRR8992380 | 128 | 2137 | 26906 | 46594 | 55 | 108 | 1433 | 4187 | 25869 | 38 | 723 | 941 | 1556 | 114377 | 476 | 5140 | 10839 | 0 | 35 | 48 | 75 | 471 | 312 | 282 | 45 | 0 | 0 | 99 | 68 | 70 | 38 | 0 | 0 | 0 | 0 | 30 | 0 |
| SRR8992381 | 461 | 1535 | 49661 | 34951 | 39 | 33 | 387 | 2297 | 24069 | 77 | 475 | 1069 | 587 | 34021 | 214 | 2148 | 7673 | 0 | 0 | 12 | 54 | 25 | 0 | 17 | 0 | 0 | 0 | 10 | 0 | 0 | 0 | 0 | 0 | 0 | 0 | 0 | 0 |
| SRR8992382 | 1002 | 2634 | 27300 | 38503 | 37 | 46 | 321 | 2554 | 23398 | 61 | 508 | 1249 | 315 | 28008 | 222 | 1817 | 6136 | 0 | 0 | 16 | 18 | 34 | 18 | 38 | 0 | 0 | 0 | 19 | 17 | 0 | 0 | 0 | 0 | 0 | 0 | 0 | 0 |
| SRR8992383 | 614 | 1423 | 40329 | 42885 | 29 | 39 | 368 | 2225 | 20349 | 79 | 451 | 949 | 501 | 36974 | 202 | 2160 | 6835 | 0 | 0 | 16 | 13 | 33 | 15 | 26 | 0 | 0 | 0 | 22 | 20 | 0 | 0 | 0 | 0 | 0 | 0 | 0 | 0 |
| SRR8992384 | 296 | 3217 | 21183 | 56998 | 198 | 345 | 1857 | 3971 | 29532 | 66 | 1064 | 1425 | 1106 | 80437 | 475 | 4247 | 10750 | 0 | 27 | 38 | 61 | 339 | 275 | 292 | 38 | 0 | 0 | 93 | 56 | 44 | 22 | 0 | 0 | 0 | 0 | 22 | 0 |
| SRR8992385 | 238 | 2667 | 25433 | 43708 | 64 | 93 | 1167 | 4689 | 32073 | 51 | 714 | 1085 | 955 | 97422 | 385 | 4183 | 7455 | 0 | 19 | 33 | 52 | 283 | 238 | 239 | 35 | 0 | 0 | 66 | 46 | 35 | 20 | 0 | 0 | 0 | 0 | 20 | 0 |
| SRR8992387 | 214 | 3015 | 30003 | 37685 | 81 | 127 | 1371 | 4478 | 31751 | 49 | 708 | 1049 | 1087 | 91927 | 374 | 3443 | 10441 | 0 | 21 | 46 | 65 | 373 | 282 | 286 | 40 | 0 | 0 | 76 | 52 | 42 | 23 | 0 | 0 | 0 | 0 | 21 | 0 |
| SRR8992388 | 110 | 2017 | 22126 | 51192 | 59 | 152 | 1371 | 4554 | 26360 | 33 | 351 | 498 | 694 | 120538 | 495 | 5043 | 5463 | 0 | 19 | 36 | 46 | 242 | 190 | 192 | 28 | 0 | 0 | 68 | 49 | 36 | 16 | 0 | 0 | 0 | 0 | 17 | 0 |
| SRR8992389 | 187 | 2032 | 37097 | 29877 | 43 | 77 | 1227 | 8084 | 31950 | 39 | 974 | 1296 | 1346 | 75213 | 286 | 4333 | 9206 | 0 | 21 | 28 | 50 | 195 | 217 | 178 | 25 | 0 | 0 | 91 | 56 | 52 | 22 | 0 | 0 | 0 | 0 | 14 | 0 |
| SRR8992390 | 294 | 3353 | 25601 | 53388 | 178 | 314 | 1893 | 5595 | 29192 | 45 | 1061 | 1200 | 923 | 88884 | 381 | 3579 | 10386 | 0 | 29 | 55 | 72 | 413 | 293 | 302 | 39 | 0 | 0 | 92 | 54 | 43 | 26 | 0 | 0 | 0 | 0 | 24 | 0 |
| SRR8992391 | 187 | 1740 | 20491 | 40004 | 258 | 612 | 1985 | 4855 | 29659 | 40 | 1603 | 2252 | 1009 | 26182 | 345 | 1559 | 8487 | 0 | 16 | 31 | 39 | 271 | 286 | 264 | 36 | 0 | 0 | 52 | 35 | 32 | 23 | 0 | 0 | 0 | 0 | 20 | 0 |
| SRR9003330 | 1715 | 6024 | 196736 | 1305 | 33370 | 36734 | 206248 | 13722 | 85591 | 7314 | 3919 | 2591 | 1293 | 585 | 27153 | 15 | 11639 | 0 | 16 | 38 | 21 | 37 | 30 | 0 | 0 | 0 | 0 | 115 | 155 | 37 | 0 | 0 | 0 | 0 | 0 | 0 | 0 |
| SRR9003332 | 0 | 0 | 112801 | 0 | 18706 | 24818 | 182753 | 49737 | 56863 | 752 | 3267 | 1354 | 1278 | 470 | 18604 | 28 | 21390 | 0 | 0 | 0 | 0 | 0 | 0 | 0 | 0 | 0 | 0 | 0 | 0 | 0 | 0 | 0 | 0 | 0 | 0 | 0 | 0 |
| SRR9003333 | 986 | 5158 | 221531 | 324 | 38739 | 48178 | 263414 | 24255 | 97001 | 8429 | 3596 | 1734 | 1554 | 606 | 34818 | 19 | 8103 | 0 | 13 | 25 | 16 | 31 | 20 | 0 | 0 | 0 | 0 | 116 | 161 | 35 | 0 | 0 | 0 | 0 | 0 | 0 | 0 |
| SRR9003334 | 0 | 10 | 288485 | 0 | 22741 | 27378 | 200000 | 36662 | 75301 | 2393 | 3340 | 1503 | 1078 | 564 | 24 | 23 | 15172 | 0 | 0 | 0 | 0 | 0 | 0 | 0 | 0 | 0 | 0 | 0 | 0 | 0 | 0 | 0 | 0 | 0 | 0 | 0 | 0 |
| SRR9003336 | 832 | 5249 | 202942 | 85 | 39221 | 45300 | 256570 | 19921 | 95350 | 9747 | 3636 | 2220 | 1422 | 621 | 38396 | 21 | 9332 | 0 | 14 | 28 | 23 | 36 | 28 | 11 | 0 | 0 | 0 | 118 | 157 | 44 | 0 | 0 | 0 | 0 | 0 | 0 | 0 |
| SRR9003338 | 0 | 0 | 288148 | 0 | 31191 | 31689 | 217324 | 34655 | 81478 | 1847 | 2981 | 1820 | 773 | 556 | 22 | 27 | 18044 | 0 | 0 | 0 | 0 | 0 | 0 | 0 | 0 | 0 | 0 | 0 | 0 | 0 | 0 | 0 | 0 | 0 | 0 | 0 | 0 |
| SRR9003339 | 1090 | 6014 | 182639 | 230 | 37212 | 42661 | 239820 | 17207 | 90034 | 10023 | 4002 | 2230 | 1695 | 598 | 41841 | 17 | 7472 | 0 | 22 | 42 | 25 | 40 | 28 | 0 | 0 | 0 | 0 | 160 | 212 | 48 | 0 | 0 | 0 | 0 | 0 | 0 | 0 |
| SRR9003341 | 0 | 29 | 140229 | 0 | 27330 | 24632 | 192486 | 31234 | 73961 | 6071 | 3086 | 2008 | 971 | 503 | 24 | 33 | 15676 | 0 | 0 | 0 | 0 | 0 | 0 | 0 | 0 | 0 | 0 | 0 | 0 | 0 | 0 | 0 | 0 | 0 | 0 | 0 | 0 |
| SRR9003342 | 932 | 5945 | 196448 | 79 | 33957 | 35106 | 207670 | 15466 | 95541 | 13543 | 3509 | 1890 | 1427 | 546 | 35505 | 17 | 9194 | 0 | 15 | 33 | 31 | 36 | 36 | 12 | 0 | 0 | 0 | 107 | 154 | 49 | 0 | 0 | 0 | 0 | 0 | 0 | 0 |
| SRR9018258 | 907 | 4176 | 80254 | 625 | 1337 | 1096 | 12364 | 1162 | 55270 | 9251 | 21563 | 8477 | 44226 | 26192 | 63811 | 6330 | 6977 | 0 | 31 | 0 | 26 | 10 | 52 | 161 | 151 | 45 | 13 | 55 | 90 | 46 | 0 | 0 | 0 | 0 | 15 | 70 | 0 |
| SRR9018259 | 1173 | 6066 | 91148 | 240 | 982 | 456 | 9225 | 672 | 51456 | 16097 | 17246 | 6474 | 35733 | 35158 | 64671 | 6893 | 8027 | 0 | 52 | 20 | 37 | 12 | 36 | 20 | 22 | 0 | 22 | 91 | 148 | 70 | 0 | 0 | 0 | 0 | 0 | 0 | 0 |
| SRR9018260 | 2216 | 5301 | 32783 | 343 | 622 | 312 | 3692 | 242 | 53537 | 16829 | 24542 | 19428 | 31937 | 27796 | 57782 | 983 | 10467 | 0 | 0 | 0 | 0 | 0 | 0 | 0 | 0 | 0 | 0 | 0 | 0 | 0 | 0 | 0 | 0 | 0 | 0 | 0 | 0 |
| SRR9018261 | 405 | 1256 | 24619 | 7071 | 1255 | 618 | 6970 | 449 | 128676 | 3025 | 21571 | 6314 | 40391 | 50592 | 17 | 4087 | 8951 | 0 | 0 | 0 | 14 | 0 | 29 | 71 | 92 | 15 | 0 | 20 | 40 | 0 | 0 | 0 | 0 | 0 | 0 | 24 | 0 |
| SRR952248 | 301 | 79287 | 4047 | 740 | 1799 | 31700 | 28793 | 24342 | 12632 | 16 | 11081 | 69431 | 3679 | 2273 | 37 | 3914 | 29358 | 0 | 0 | 0 | 412 | 119 | 100 | 0 | 12 | 0 | 0 | 0 | 55 | 76 | 88 | 35 | 0 | 20 | 21 | 0 | 0 |
| SRR952249 | 174 | 80079 | 3698 | 275 | 1841 | 33416 | 29725 | 26941 | 11650 | 20 | 10926 | 16 | 4219 | 2165 | 33 | 5309 | 24352 | 0 | 0 | 12 | 594 | 150 | 133 | 0 | 22 | 0 | 0 | 13 | 122 | 123 | 93 | 49 | 0 | 27 | 41 | 0 | 0 |
| SRR952288 | 182 | 79668 | 3723 | 279 | 1838 | 33529 | 29835 | 26883 | 11618 | 23 | 10900 | 81434 | 4230 | 2167 | 36 | 5326 | 24341 | 0 | 0 | 12 | 604 | 150 | 132 | 0 | 22 | 0 | 0 | 12 | 124 | 121 | 92 | 49 | 0 | 28 | 42 | 0 | 0 |
| SRR952289 | 72 | 83923 | 6046 | 274 | 1997 | 36498 | 30462 | 31125 | 12338 | 25 | 11090 | 16 | 3759 | 2069 | 29 | 5131 | 25706 | 0 | 0 | 0 | 486 | 142 | 118 | 0 | 18 | 0 | 0 | 0 | 88 | 96 | 99 | 45 | 0 | 28 | 41 | 0 | 0 |
| SRR952290 | 74 | 83577 | 6078 | 270 | 2004 | 36536 | 30535 | 31150 | 12318 | 22 | 11006 | 76823 | 3753 | 2062 | 29 | 5193 | 25682 | 0 | 0 | 0 | 484 | 140 | 120 | 0 | 18 | 0 | 0 | 0 | 90 | 96 | 100 | 46 | 0 | 28 | 42 | 0 | 0 |
| SRR952309 | 28 | 12921 | 3871 | 81 | 2316 | 46711 | 51302 | 19416 | 10865 | 73 | 14192 | 49671 | 4040 | 2696 | 26 | 4765 | 20360 | 0 | 0 | 0 | 156 | 28 | 47 | 0 | 0 | 0 | 0 | 0 | 19 | 41 | 29 | 26 | 0 | 12 | 0 | 0 | 0 |
| SRR952310 | 44 | 16125 | 3455 | 143 | 2976 | 48293 | 61386 | 19920 | 11506 | 56 | 13557 | 52058 | 4133 | 2283 | 32 | 4633 | 20887 | 0 | 0 | 0 | 405 | 56 | 99 | 0 | 13 | 0 | 0 | 0 | 46 | 102 | 45 | 55 | 0 | 18 | 0 | 0 | 0 |
| SRR952311 | 50 | 29890 | 9665 | 216 | 2545 | 51440 | 54714 | 21560 | 14218 | 69 | 12144 | 50010 | 3899 | 2108 | 27 | 5741 | 22979 | 0 | 0 | 0 | 475 | 152 | 68 | 0 | 0 | 0 | 0 | 19 | 145 | 134 | 177 | 44 | 0 | 14 | 0 | 0 | 0 |
| SRR952312 | 42 | 67928 | 5483 | 203 | 1222 | 17199 | 15937 | 14732 | 1888 | 18 | 3891 | 41704 | 1725 | 1096 | 32 | 1252 | 9856 | 0 | 0 | 0 | 146 | 56 | 43 | 0 | 0 | 0 | 0 | 0 | 13 | 49 | 46 | 13 | 0 | 12 | 23 | 0 | 0 |
| SRR952313 | 66 | 74940 | 4558 | 323 | 1330 | 20075 | 18494 | 15449 | 2414 | 39 | 5199 | 51814 | 1937 | 1289 | 35 | 1514 | 8773 | 0 | 0 | 0 | 135 | 50 | 33 | 0 | 0 | 0 | 0 | 0 | 19 | 62 | 42 | 13 | 0 | 0 | 17 | 0 | 0 |
| SRR952314 | 75 | 38428 | 1133 | 433 | 330 | 4509 | 5155 | 2816 | 3177 | 39 | 6495 | 63182 | 2163 | 1301 | 57 | 3974 | 10677 | 0 | 0 | 0 | 268 | 32 | 74 | 0 | 16 | 0 | 0 | 0 | 0 | 77 | 16 | 31 | 0 | 21 | 31 | 0 | 0 |
| SRR952315 | 121 | 33606 | 845 | 321 | 304 | 4137 | 4445 | 2723 | 2469 | 26 | 5584 | 62309 | 2116 | 1175 | 71 | 3075 | 12091 | 0 | 0 | 0 | 286 | 35 | 84 | 0 | 21 | 0 | 0 | 0 | 11 | 63 | 22 | 26 | 0 | 29 | 35 | 0 | 0 |
| SRR952316 | 0 | 17410 | 521 | 2325 | 0 | 2297 | 2256 | 1417 | 4163 | 0 | 3620 | 14965 | 6264 | 3237 | 225 | 393 | 1306 | 0 | 0 | 20 | 3084 | 1108 | 1031 | 0 | 257 | 0 | 0 | 41 | 420 | 1084 | 725 | 298 | 17 | 264 | 633 | 40 | 0 |
| SRR952317 | 22 | 19780 | 164 | 3803 | 233 | 5141 | 3671 | 3294 | 4087 | 0 | 3848 | 17582 | 3545 | 2163 | 102 | 665 | 2643 | 0 | 0 | 21 | 2666 | 1000 | 662 | 0 | 124 | 0 | 0 | 36 | 368 | 1419 | 893 | 280 | 0 | 199 | 413 | 0 | 0 |
| SRR952318 | 60 | 16882 | 13 | 4735 | 120 | 1165 | 1717 | 488 | 4059 | 0 | 4740 | 19266 | 3154 | 1632 | 42 | 1032 | 2666 | 0 | 0 | 14 | 4080 | 484 | 1202 | 18 | 273 | 0 | 0 | 0 | 131 | 1375 | 243 | 458 | 30 | 421 | 615 | 25 | 0 |
| SRR952319 | 0 | 14097 | 55 | 3647 | 0 | 269 | 241 | 571 | 2659 | 0 | 4952 | 16242 | 5775 | 988 | 0 | 263 | 803 | 0 | 0 | 25 | 4198 | 651 | 1738 | 20 | 567 | 15 | 0 | 16 | 128 | 1210 | 344 | 721 | 0 | 692 | 662 | 24 | 0 |
| SRR9588764 | 77 | 471 | 13549 | 1382 | 3328 | 7475 | 47396 | 13524 | 1187 | 1959 | 33288 | 21795 | 23554 | 17295 | 2206 | 10534 | 48088 | 0 | 52 | 13 | 33 | 1669 | 94 | 0 | 0 | 0 | 0 | 148 | 651 | 253 | 863 | 18 | 0 | 0 | 0 | 24 | 0 |
| SRR9588765 | 203 | 734 | 24489 | 743 | 468 | 959 | 8655 | 1974 | 162 | 1403 | 23245 | 20 | 20482 | 8680 | 21854 | 8105 | 57353 | 0 | 160 | 0 | 26 | 812 | 258 | 11 | 0 | 10 | 0 | 205 | 202 | 77 | 225 | 20 | 0 | 0 | 0 | 17 | 0 |
| SRR9588766 | 117 | 389 | 13945 | 1783 | 11367 | 28563 | 199587 | 43189 | 4708 | 2245 | 22499 | 12188 | 10150 | 17393 | 573 | 8081 | 10077 | 0 | 407 | 33 | 10 | 155 | 18 | 0 | 0 | 28 | 0 | 2713 | 1050 | 118 | 137 | 0 | 0 | 0 | 0 | 56 | 0 |
| SRR9588767 | 151 | 635 | 15588 | 1098 | 251 | 454 | 4239 | 1905 | 3210 | 2181 | 24140 | 23 | 14236 | 9097 | 9404 | 6948 | 35444 | 0 | 33 | 21 | 115 | 2454 | 1012 | 32 | 16 | 50 | 0 | 66 | 210 | 156 | 858 | 92 | 0 | 0 | 0 | 57 | 0 |
| SRR9618218 | 222 | 6364 | 683 | 295 | 3617 | 30568 | 35429 | 25697 | 4433 | 11 | 36053 | 41871 | 13776 | 6221 | 13 | 507 | 55873 | 0 | 0 | 0 | 83 | 0 | 21 | 0 | 0 | 0 | 0 | 0 | 33 | 154 | 31 | 90 | 0 | 20 | 11 | 0 | 0 |
| SRR9618219 | 0 | 7473 | 1269 | 23 | 8101 | 30818 | 60373 | 34622 | 3215 | 110 | 61738 | 11 | 29627 | 14571 | 728 | 63 | 52831 | 0 | 0 | 0 | 47 | 0 | 24 | 0 | 0 | 0 | 0 | 0 | 11 | 54 | 49 | 89 | 0 | 24 | 15 | 0 | 0 |
| SRR9618220 | 291 | 8731 | 451 | 561 | 4785 | 47243 | 48533 | 33037 | 4850 | 10 | 36381 | 36995 | 12240 | 6267 | 16 | 405 | 50231 | 0 | 0 | 0 | 94 | 0 | 31 | 0 | 0 | 0 | 0 | 0 | 40 | 186 | 30 | 130 | 0 | 20 | 0 | 0 | 0 |
| SRR9618221 | 115 | 4289 | 488 | 766 | 4401 | 41863 | 42543 | 31490 | 4389 | 26 | 32221 | 39727 | 12152 | 5823 | 19 | 488 | 51874 | 0 | 0 | 0 | 67 | 0 | 21 | 0 | 0 | 0 | 0 | 0 | 35 | 162 | 28 | 96 | 0 | 21 | 0 | 0 | 0 |
| SRR9618222 | 17 | 14458 | 1383 | 227 | 8012 | 27277 | 65957 | 45413 | 6893 | 73 | 48626 | 71883 | 16844 | 15040 | 448 | 38 | 33374 | 0 | 0 | 0 | 62 | 11 | 33 | 0 | 0 | 0 | 0 | 0 | 17 | 109 | 34 | 81 | 0 | 13 | 0 | 0 | 0 |
| SRR9627609 | 584 | 7646 | 998 | 914 | 228 | 2541 | 2169 | 6152 | 277 | 49 | 57261 | 14 | 40893 | 10704 | 593 | 17 | 52755 | 0 | 0 | 0 | 0 | 0 | 0 | 0 | 0 | 0 | 0 | 0 | 18 | 0 | 0 | 0 | 0 | 0 | 0 | 0 | 0 |
| SRR9627610 | 201 | 4401 | 1227 | 259 | 226 | 2514 | 2169 | 6500 | 317 | 63 | 58026 | 82025 | 41673 | 10777 | 567 | 19 | 54201 | 0 | 0 | 0 | 0 | 0 | 0 | 0 | 0 | 0 | 0 | 0 | 0 | 0 | 0 | 0 | 0 | 0 | 0 | 0 | 0 |
| SRR9627611 | 383 | 5352 | 977 | 453 | 222 | 2668 | 2387 | 6828 | 316 | 40 | 55136 | 11 | 41369 | 10728 | 552 | 25 | 54119 | 0 | 0 | 0 | 0 | 0 | 0 | 0 | 0 | 0 | 0 | 0 | 0 | 0 | 0 | 0 | 0 | 0 | 0 | 0 | 0 |
| SRR9627612 | 640 | 4877 | 897 | 510 | 140 | 1736 | 1301 | 4747 | 329 | 30 | 53429 | 12 | 44402 | 9967 | 474 | 23 | 69040 | 0 | 0 | 0 | 12 | 11 | 0 | 0 | 0 | 0 | 0 | 0 | 0 | 0 | 0 | 0 | 0 | 0 | 0 | 0 | 0 |
| SRR9627613 | 542 | 4337 | 779 | 450 | 147 | 1694 | 1280 | 4917 | 320 | 26 | 54400 | 76843 | 45959 | 10316 | 476 | 23 | 65345 | 0 | 0 | 0 | 0 | 0 | 0 | 0 | 0 | 0 | 0 | 0 | 0 | 0 | 0 | 0 | 0 | 0 | 0 | 0 | 0 |
| SRR9627614 | 185 | 3040 | 1084 | 159 | 186 | 1698 | 1545 | 5579 | 304 | 62 | 66779 | 19 | 46835 | 12088 | 562 | 24 | 55007 | 0 | 0 | 0 | 0 | 0 | 0 | 0 | 0 | 0 | 0 | 0 | 0 | 0 | 0 | 0 | 0 | 0 | 0 | 0 | 0 |
| SRR9627651 | 314 | 4235 | 5179 | 534 | 892 | 556 | 15779 | 14116 | 170 | 827 | 77647 | 73386 | 127802 | 16247 | 1617 | 0 | 14963 | 0 | 0 | 0 | 0 | 0 | 0 | 0 | 0 | 0 | 0 | 125 | 23 | 23 | 0 | 0 | 0 | 0 | 0 | 0 | 0 |
| SRR9627653 | 44 | 1176 | 8596 | 113 | 810 | 439 | 15561 | 13745 | 156 | 1531 | 83691 | 14 | 12 | 18387 | 1761 | 0 | 10674 | 0 | 0 | 0 | 0 | 0 | 0 | 0 | 0 | 0 | 0 | 19 | 0 | 0 | 0 | 0 | 0 | 0 | 0 | 0 | 0 |
| SRR9627654 | 150 | 1779 | 4447 | 183 | 650 | 328 | 9992 | 11723 | 165 | 841 | 78317 | 14 | 13 | 16517 | 1574 | 0 | 14332 | 0 | 0 | 0 | 0 | 0 | 0 | 0 | 0 | 0 | 0 | 48 | 0 | 18 | 0 | 0 | 0 | 0 | 0 | 0 | 0 |
| SRR9627655 | 73 | 956 | 5959 | 102 | 503 | 247 | 8647 | 10447 | 140 | 1081 | 81475 | 78403 | 152528 | 17194 | 1353 | 0 | 13216 | 0 | 0 | 0 | 0 | 0 | 0 | 0 | 0 | 0 | 0 | 20 | 0 | 0 | 0 | 0 | 0 | 0 | 0 | 0 | 0 |
| SRR9627656 | 317 | 2796 | 3770 | 462 | 673 | 332 | 10605 | 11482 | 154 | 927 | 80713 | 75590 | 148786 | 17458 | 1726 | 0 | 13537 | 0 | 0 | 0 | 0 | 0 | 0 | 0 | 0 | 0 | 0 | 93 | 16 | 22 | 0 | 0 | 0 | 0 | 0 | 0 | 0 |
| SRR9627693 | 1534 | 4423 | 6166 | 1329 | 696 | 870 | 4183 | 9445 | 1308 | 1307 | 44046 | 43277 | 45354 | 22306 | 616 | 0 | 105094 | 0 | 0 | 0 | 0 | 0 | 0 | 0 | 0 | 0 | 0 | 34 | 13 | 0 | 0 | 0 | 0 | 0 | 0 | 0 | 0 |
| SRR9627694 | 1619 | 4529 | 6058 | 1351 | 787 | 878 | 4850 | 10245 | 1240 | 1747 | 44821 | 42714 | 47443 | 23083 | 725 | 0 | 97730 | 0 | 0 | 0 | 0 | 0 | 0 | 0 | 0 | 0 | 0 | 40 | 15 | 0 | 12 | 0 | 0 | 0 | 0 | 0 | 0 |
| SRR9627695 | 1624 | 4553 | 6164 | 1300 | 704 | 796 | 4328 | 9316 | 1260 | 1821 | 44955 | 42001 | 46531 | 22765 | 705 | 0 | 107844 | 0 | 0 | 0 | 0 | 0 | 0 | 0 | 0 | 0 | 0 | 39 | 14 | 0 | 13 | 0 | 0 | 0 | 0 | 0 | 0 |
| SRR9627696 | 1932 | 4209 | 2310 | 1034 | 445 | 494 | 2501 | 6182 | 1092 | 1153 | 55203 | 58335 | 55395 | 22976 | 654 | 0 | 115316 | 0 | 0 | 0 | 0 | 0 | 0 | 0 | 0 | 0 | 0 | 13 | 0 | 0 | 0 | 0 | 0 | 0 | 0 | 0 | 0 |
| SRR9627697 | 2053 | 3844 | 2654 | 1257 | 417 | 586 | 2374 | 6392 | 1317 | 596 | 46553 | 49241 | 48792 | 23531 | 578 | 0 | 120128 | 0 | 0 | 0 | 0 | 0 | 0 | 0 | 0 | 0 | 0 | 20 | 0 | 0 | 0 | 0 | 0 | 0 | 0 | 0 | 0 |
| SRR9627698 | 2052 | 3111 | 4088 | 744 | 466 | 594 | 2570 | 6815 | 936 | 723 | 54207 | 57299 | 51852 | 21471 | 621 | 0 | 130431 | 0 | 0 | 0 | 0 | 0 | 0 | 0 | 0 | 0 | 0 | 0 | 0 | 0 | 0 | 0 | 0 | 0 | 0 | 0 | 0 |
| SRR9648387 | 0 | 651 | 246459 | 361 | 2211 | 2233 | 17992 | 18619 | 221546 | 67 | 29771 | 22435 | 12559 | 7764 | 913 | 363 | 8479 | 0 | 0 | 0 | 0 | 887 | 910 | 47 | 14 | 0 | 0 | 0 | 0 | 0 | 0 | 0 | 0 | 0 | 0 | 25 | 0 |
| SRR9648388 | 20 | 3629 | 6129 | 167403 | 834 | 668 | 1502 | 501 | 52403 | 40 | 4005 | 9513 | 1001 | 1836 | 167 | 167 | 1001 | 0 | 0 | 0 | 0 | 13 | 41 | 250 | 410 | 15 | 0 | 0 | 0 | 0 | 0 | 0 | 0 | 0 | 18 | 50 | 0 |
| SRR9648389 | 0 | 1315 | 26852 | 50230 | 498 | 747 | 8087 | 13064 | 102893 | 23 | 6843 | 5910 | 4230 | 16610 | 560 | 1555 | 4977 | 0 | 0 | 0 | 0 | 11 | 45 | 183 | 201 | 12 | 0 | 0 | 0 | 0 | 0 | 0 | 0 | 0 | 0 | 36 | 0 |
| SRR965653 | 1697 | 18477 | 732 | 17444 | 2144 | 30109 | 36242 | 35902 | 1366 | 101 | 17805 | 10181 | 2804 | 11859 | 0 | 290 | 8502 | 0 | 0 | 0 | 24 | 0 | 0 | 0 | 0 | 0 | 0 | 0 | 0 | 0 | 0 | 0 | 0 | 0 | 0 | 0 | 0 |
| SRR965655 | 135 | 8616 | 2267 | 3400 | 1018 | 12540 | 25617 | 28516 | 2976 | 37 | 17479 | 9200 | 3954 | 14510 | 0 | 810 | 23418 | 0 | 0 | 0 | 495 | 151 | 133 | 0 | 0 | 0 | 0 | 0 | 26 | 155 | 233 | 75 | 0 | 0 | 0 | 0 | 0 |
| SRR9833150 | 12 | 21152 | 689 | 0 | 285 | 1898 | 3029 | 1734 | 25062 | 17 | 87 | 999 | 21 | 0 | 0 | 6010 | 53548 | 0 | 0 | 0 | 1307 | 149 | 43 | 0 | 0 | 0 | 0 | 0 | 45 | 495 | 155 | 28 | 0 | 0 | 23 | 0 | 0 |
| SRR9833151 | 0 | 12491 | 729 | 0 | 250 | 3426 | 3859 | 2064 | 26950 | 15 | 94 | 1138 | 16 | 11 | 0 | 5924 | 47072 | 0 | 0 | 0 | 224 | 35 | 19 | 0 | 0 | 0 | 0 | 0 | 11 | 105 | 32 | 11 | 0 | 0 | 0 | 0 | 0 |
| SRR9833152 | 15 | 18412 | 720 | 0 | 377 | 2753 | 3967 | 2515 | 29484 | 14 | 116 | 1058 | 21 | 0 | 0 | 6198 | 54465 | 0 | 0 | 0 | 607 | 69 | 34 | 0 | 0 | 0 | 0 | 0 | 26 | 216 | 77 | 23 | 0 | 0 | 17 | 0 | 0 |
| SRR9833153 | 0 | 11709 | 1169 | 0 | 164 | 1669 | 1863 | 1534 | 26387 | 42 | 117 | 1450 | 12 | 12 | 0 | 3898 | 43362 | 0 | 0 | 0 | 147 | 22 | 17 | 0 | 0 | 0 | 0 | 0 | 12 | 76 | 32 | 13 | 0 | 0 | 0 | 0 | 0 |
| SRR9833154 | 0 | 12883 | 449 | 0 | 211 | 2604 | 3036 | 1826 | 25673 | 14 | 73 | 1428 | 0 | 0 | 15 | 4913 | 46702 | 0 | 0 | 0 | 147 | 18 | 22 | 0 | 0 | 0 | 0 | 0 | 10 | 60 | 22 | 13 | 0 | 0 | 0 | 0 | 0 |
| SRR9833155 | 14 | 23225 | 685 | 0 | 221 | 1667 | 2429 | 1951 | 26291 | 16 | 100 | 1348 | 17 | 0 | 0 | 4146 | 47145 | 0 | 0 | 0 | 272 | 38 | 29 | 0 | 0 | 0 | 0 | 0 | 34 | 140 | 66 | 26 | 0 | 0 | 0 | 0 | 0 |
| SRR9833156 | 15 | 55429 | 788 | 23 | 230 | 2166 | 2750 | 1937 | 26825 | 10 | 119 | 1232 | 20 | 0 | 0 | 4689 | 53176 | 0 | 0 | 0 | 562 | 55 | 40 | 0 | 0 | 0 | 0 | 16 | 85 | 193 | 64 | 25 | 0 | 0 | 0 | 0 | 0 |
| SRR9833157 | 29 | 54355 | 591 | 45 | 307 | 3326 | 3786 | 2308 | 24482 | 12 | 91 | 1024 | 19 | 0 | 0 | 7598 | 61824 | 0 | 0 | 0 | 1041 | 71 | 46 | 0 | 0 | 0 | 0 | 12 | 82 | 439 | 84 | 29 | 0 | 0 | 0 | 0 | 0 |
| SRR9833158 | 994 | 39418 | 770 | 250 | 419 | 3075 | 3994 | 2656 | 27788 | 10 | 117 | 806 | 21 | 0 | 0 | 6925 | 54424 | 0 | 0 | 0 | 1507 | 87 | 39 | 0 | 0 | 0 | 0 | 23 | 128 | 537 | 97 | 25 | 0 | 0 | 12 | 0 | 0 |
| SRR9833159 | 5574 | 19358 | 519 | 540 | 611 | 4423 | 4664 | 4694 | 38958 | 31 | 109 | 543 | 25 | 0 | 0 | 10040 | 62522 | 0 | 0 | 0 | 272 | 20 | 11 | 0 | 0 | 0 | 0 | 15 | 105 | 148 | 35 | 0 | 0 | 0 | 19 | 0 | 0 |
| SRR9833160 | 4645 | 29292 | 1584 | 986 | 615 | 5345 | 5191 | 4020 | 40155 | 19 | 173 | 699 | 31 | 0 | 0 | 13154 | 58091 | 0 | 0 | 0 | 575 | 31 | 19 | 0 | 0 | 0 | 0 | 21 | 147 | 312 | 41 | 13 | 0 | 0 | 61 | 0 | 0 |
| SRR9833161 | 3496 | 30766 | 865 | 732 | 705 | 5413 | 5346 | 4662 | 60445 | 39 | 305 | 837 | 56 | 0 | 45 | 13331 | 26586 | 0 | 0 | 0 | 157 | 0 | 0 | 0 | 0 | 0 | 0 | 13 | 61 | 125 | 0 | 0 | 0 | 0 | 29 | 0 | 0 |
| SRR9833162 | 3467 | 32988 | 1889 | 423 | 529 | 5221 | 4737 | 4491 | 23316 | 29 | 220 | 808 | 39 | 0 | 0 | 10801 | 65798 | 0 | 0 | 0 | 381 | 17 | 15 | 0 | 0 | 0 | 0 | 11 | 75 | 229 | 26 | 10 | 0 | 0 | 26 | 0 | 0 |
| SRR9833163 | 3578 | 30714 | 1246 | 417 | 575 | 3954 | 4445 | 4192 | 25159 | 26 | 173 | 844 | 33 | 0 | 10 | 9280 | 64234 | 0 | 0 | 0 | 416 | 24 | 15 | 0 | 0 | 0 | 0 | 15 | 94 | 226 | 35 | 11 | 0 | 0 | 27 | 0 | 0 |
| SRR9833164 | 1764 | 23262 | 1395 | 919 | 513 | 2883 | 4954 | 4477 | 30134 | 21 | 224 | 747 | 34 | 0 | 0 | 11303 | 71061 | 0 | 0 | 0 | 428 | 24 | 16 | 0 | 0 | 0 | 0 | 0 | 73 | 168 | 24 | 0 | 0 | 0 | 28 | 0 | 0 |
| SRR9856157 | 195 | 2076 | 12292 | 320 | 1700 | 743 | 14563 | 7837 | 3843 | 2552 | 7002 | 2681 | 2194 | 1129 | 76 | 3907 | 10280 | 0 | 439 | 46 | 39 | 80 | 45 | 0 | 0 | 0 | 0 | 580 | 204 | 32 | 54 | 0 | 0 | 0 | 0 | 0 | 0 |
| SRR9856158 | 34 | 1121 | 26220 | 242 | 1065 | 825 | 5453 | 7388 | 3022 | 2966 | 8552 | 3893 | 3996 | 2283 | 143 | 4040 | 19101 | 0 | 362 | 108 | 129 | 854 | 152 | 0 | 0 | 0 | 0 | 541 | 471 | 150 | 354 | 0 | 0 | 0 | 0 | 0 | 0 |
| SRR9856159 | 106 | 1283 | 10029 | 958 | 1794 | 671 | 12196 | 12797 | 5021 | 4671 | 8197 | 3830 | 3759 | 2206 | 74 | 6016 | 19753 | 0 | 319 | 68 | 123 | 475 | 164 | 11 | 0 | 0 | 0 | 625 | 387 | 133 | 130 | 0 | 0 | 0 | 0 | 13 | 0 |
| SRR9856160 | 109 | 4508 | 31341 | 917 | 618 | 504 | 8199 | 371 | 16072 | 936 | 2212 | 2023 | 380 | 226 | 92 | 6689 | 6441 | 0 | 81 | 17 | 69 | 299 | 37 | 0 | 0 | 0 | 0 | 172 | 131 | 74 | 16 | 0 | 0 | 0 | 0 | 0 | 0 |
| SRR9856161 | 18 | 580 | 9174 | 383 | 556 | 444 | 3742 | 4474 | 3460 | 2424 | 7430 | 2850 | 2335 | 1276 | 84 | 4569 | 15726 | 0 | 398 | 102 | 152 | 614 | 61 | 0 | 0 | 0 | 0 | 634 | 513 | 172 | 204 | 0 | 0 | 0 | 0 | 0 | 0 |
| SRR9856162 | 23 | 843 | 7776 | 332 | 441 | 348 | 2992 | 4121 | 1972 | 6512 | 6653 | 2847 | 2083 | 959 | 73 | 3888 | 13025 | 0 | 683 | 137 | 94 | 223 | 24 | 0 | 0 | 0 | 0 | 957 | 531 | 83 | 121 | 0 | 0 | 0 | 0 | 0 | 0 |
| SRR9856163 | 101 | 816 | 6129 | 1816 | 468 | 356 | 2882 | 5155 | 1631 | 2439 | 7442 | 3319 | 2451 | 1055 | 63 | 2568 | 16991 | 0 | 321 | 103 | 93 | 520 | 80 | 0 | 0 | 0 | 0 | 530 | 473 | 108 | 263 | 0 | 0 | 0 | 0 | 0 | 0 |
| SRR9856164 | 83 | 975 | 15508 | 523 | 1109 | 825 | 6786 | 11083 | 3070 | 1824 | 8019 | 3201 | 3457 | 1775 | 90 | 3968 | 15557 | 0 | 530 | 118 | 96 | 698 | 113 | 0 | 0 | 0 | 0 | 821 | 551 | 99 | 203 | 0 | 0 | 0 | 0 | 0 | 0 |
| SRR9856165 | 88 | 1618 | 14310 | 516 | 318 | 304 | 2870 | 2218 | 2301 | 2473 | 6392 | 2438 | 2117 | 984 | 90 | 2979 | 9281 | 0 | 168 | 24 | 27 | 73 | 17 | 0 | 0 | 0 | 0 | 257 | 140 | 23 | 47 | 0 | 0 | 0 | 0 | 0 | 0 |
| SRR9856166 | 22 | 863 | 9355 | 380 | 306 | 230 | 2172 | 2364 | 1187 | 3947 | 5742 | 2162 | 1609 | 751 | 51 | 2156 | 7716 | 0 | 257 | 43 | 42 | 143 | 30 | 0 | 0 | 0 | 0 | 406 | 233 | 44 | 123 | 0 | 0 | 0 | 0 | 0 | 0 |
| SRR9856167 | 171 | 2199 | 23589 | 843 | 565 | 723 | 8310 | 2172 | 3056 | 1124 | 5041 | 2996 | 966 | 590 | 49 | 5635 | 8473 | 0 | 125 | 18 | 107 | 103 | 16 | 0 | 0 | 0 | 0 | 249 | 124 | 83 | 24 | 0 | 0 | 0 | 0 | 0 | 0 |
| SRR9856168 | 118 | 1439 | 16691 | 1514 | 903 | 688 | 5695 | 5647 | 4930 | 4108 | 11770 | 3999 | 3072 | 2113 | 83 | 4194 | 13034 | 0 | 201 | 35 | 160 | 449 | 139 | 0 | 0 | 0 | 0 | 293 | 155 | 130 | 96 | 0 | 0 | 0 | 0 | 0 | 0 |
| SRR9856169 | 97 | 1551 | 12524 | 1211 | 798 | 773 | 7896 | 7344 | 3513 | 3096 | 8063 | 3231 | 3028 | 1791 | 83 | 5246 | 12655 | 0 | 618 | 85 | 337 | 662 | 178 | 0 | 0 | 0 | 0 | 1142 | 561 | 367 | 159 | 0 | 0 | 0 | 0 | 0 | 0 |
| SRR9856170 | 0 | 634 | 34977 | 56 | 765 | 631 | 3668 | 5988 | 5673 | 1934 | 10498 | 5107 | 5080 | 2454 | 76 | 4357 | 20876 | 0 | 145 | 32 | 25 | 191 | 27 | 0 | 0 | 0 | 0 | 197 | 126 | 17 | 45 | 0 | 0 | 0 | 0 | 0 | 0 |

### Supp Table S8.1.Columns_B_to_AL

| | | | | | | | | | | | | | | | | | | |
|---|---|---|---|---|---|---|---|---|---|---|---|---|---|---|---|---|---|---|
| SRR9856171 | 14 | 198 | 25101 | 49 | 471 | 372 | 3268 | 4826 | 2504 | 1164 | 8635 | 3859 | 2964 | 1409 | 52 | 5888 | 16234 | 0 |
| SRR9856172 | 29 | 484 | 7212 | 158 | 730 | 639 | 5524 | 7192 | 3170 | 4302 | 7222 | 3426 | 1940 | 1199 | 46 | 4653 | 13016 | 0 |
| SRR9856173 | 33 | 694 | 13326 | 738 | 2019 | 1623 | 13064 | 11190 | 6237 | 2436 | 9747 | 3435 | 3755 | 2404 | 122 | 7025 | 16727 | 0 |
| SRR9856174 | 51 | 1286 | 25450 | 229 | 2790 | 2811 | 21450 | 5620 | 13024 | 2124 | 8341 | 2965 | 2975 | 2230 | 104 | 6006 | 15409 | 0 |
| SRR9856175 | 179 | 1165 | 15200 | 1394 | 1195 | 917 | 7883 | 6909 | 5600 | 2750 | 10825 | 4000 | 4117 | 2184 | 126 | 5768 | 16906 | 0 |
| SRR9856176 | 102 | 1197 | 13480 | 508 | 2068 | 795 | 13936 | 11538 | 8385 | 4452 | 10907 | 4081 | 4362 | 2874 | 97 | 5637 | 16591 | 0 |
| SRR9856177 | 60 | 514 | 8144 | 788 | 4815 | 2459 | 40558 | 37835 | 3818 | 2330 | 9222 | 3469 | 5447 | 3538 | 141 | 4517 | 15048 | 0 |
| SRR9856178 | 408 | 3715 | 27489 | 1693 | 290 | 298 | 3953 | 268 | 10052 | 1167 | 2453 | 2423 | 454 | 200 | 69 | 7300 | 5922 | 0 |
| SRR9856179 | 43 | 368 | 6936 | 639 | 196 | 157 | 1293 | 1893 | 1888 | 1199 | 4342 | 2141 | 1198 | 714 | 64 | 2466 | 13456 | 0 |
| SRR9856180 | 100 | 724 | 11210 | 598 | 267 | 216 | 1891 | 2158 | 2477 | 1306 | 4946 | 2220 | 1348 | 771 | 70 | 2497 | 11482 | 0 |
| SRR9856181 | 84 | 1327 | 9184 | 607 | 337 | 264 | 2484 | 2459 | 2913 | 2696 | 5691 | 2153 | 1544 | 776 | 94 | 2879 | 11980 | 0 |
| SRR9856182 | 139 | 1292 | 8164 | 857 | 216 | 178 | 1773 | 1726 | 5750 | 1789 | 5257 | 2174 | 1099 | 668 | 63 | 3404 | 11263 | 0 |
| SRR9856183 | 27 | 438 | 14027 | 290 | 284 | 309 | 2394 | 1851 | 7034 | 3028 | 5389 | 2903 | 1500 | 693 | 68 | 5524 | 13172 | 0 |
| SRR9856184 | 145 | 1221 | 10099 | 1003 | 381 | 336 | 2979 | 1899 | 13274 | 1808 | 6863 | 3777 | 1912 | 966 | 90 | 4129 | 14724 | 0 |
| SRR9856185 | 29 | 1647 | 20938 | 348 | 125 | 115 | 955 | 406 | 3609 | 2126 | 3271 | 1519 | 349 | 263 | 29 | 2499 | 3930 | 0 |
| SRR9856186 | 80 | 1232 | 30173 | 264 | 440 | 353 | 3193 | 1728 | 12228 | 2053 | 6366 | 3483 | 26 | 1202 | 85 | 4892 | 12840 | 0 |
| SRR990360 | 0 | 11 | 1124 | 0 | 276 | 1783 | 5313 | 9951 | 4777 | 128 | 4001 | 4519 | 2644 | 2859 | 13 | 2643 | 40627 | 0 |
| SRR9924500 | 307 | 92667 | 522 | 726 | 2410 | 32895 | 27935 | 19984 | 17580 | 34 | 950 | 1106 | 73 | 0 | 176 | 153 | 36089 | 0 |
| SRR9924502 | 144 | 43988 | 927 | 335 | 2133 | 30783 | 27111 | 19343 | 15733 | 50 | 818 | 890 | 70 | 0 | 149 | 185 | 36232 | 0 |
| SRR9924511 | 347 | 56047 | 4441 | 632 | 2867 | 45943 | 46418 | 22676 | 13126 | 41 | 663 | 720 | 64 | 0 | 93 | 111 | 14487 | 0 |
| SRR9924512 | 292 | 72355 | 5365 | 472 | 2936 | 56167 | 47336 | 23557 | 5749 | 30 | 783 | 937 | 70 | 10 | 184 | 123 | 13169 | 0 |
| SRR9924514 | 493 | 29147 | 984 | 559 | 2827 | 48276 | 47457 | 23046 | 10197 | 47 | 733 | 797 | 60 | 13 | 104 | 118 | 16383 | 0 |
| SRR9924515 | 569 | 35708 | 2405 | 296 | 3112 | 49434 | 49751 | 23563 | 14251 | 42 | 622 | 660 | 49 | 13 | 100 | 119 | 12651 | 0 |
| SRR9924516 | 56 | 16643 | 1097 | 231 | 2746 | 46661 | 47666 | 23471 | 11574 | 45 | 569 | 643 | 51 | 0 | 80 | 118 | 14253 | 0 |
| SRR9924517 | 284 | 26699 | 1117 | 760 | 2970 | 51261 | 50638 | 25452 | 10630 | 63 | 648 | 758 | 52 | 11 | 92 | 119 | 14559 | 0 |
| SRR9924518 | 238 | 68508 | 1416 | 504 | 2409 | 40331 | 36646 | 20142 | 13748 | 38 | 998 | 1104 | 82 | 0 | 133 | 134 | 33054 | 0 |
| SRR9924519 | 567 | 60491 | 960 | 1139 | 2365 | 43601 | 36827 | 18073 | 13208 | 47 | 871 | 992 | 63 | 0 | 100 | 153 | 34402 | 0 |
| SRR9924520 | 480 | 37931 | 632 | 1095 | 2308 | 35255 | 33510 | 17946 | 11033 | 46 | 1012 | 1049 | 73 | 0 | 132 | 162 | 36496 | 0 |
| SRR9924521 | 387 | 85406 | 1293 | 696 | 2207 | 39503 | 34508 | 19264 | 15459 | 29 | 1034 | 1071 | 77 | 0 | 121 | 137 | 33488 | 0 |
| SRR9924522 | 131 | 22212 | 996 | 442 | 2083 | 33345 | 32312 | 18320 | 11980 | 65 | 903 | 986 | 64 | 0 | 114 | 158 | 36555 | 0 |
| SRR9924523 | 249 | 22417 | 545 | 867 | 2310 | 35595 | 35498 | 19542 | 10574 | 26 | 910 | 977 | 62 | 0 | 120 | 172 | 35132 | 0 |
| SRR9924524 | 347 | 180981 | 1200 | 481 | 2550 | 48373 | 35397 | 22703 | 16718 | 35 | 871 | 917 | 55 | 0 | 112 | 188 | 31532 | 0 |
| SRR9924525 | 362 | 82496 | 478 | 678 | 1997 | 27393 | 24539 | 18301 | 15308 | 21 | 973 | 1007 | 70 | 0 | 151 | 154 | 33061 | 0 |
| SRR9924526 | 277 | 251526 | 1979 | 595 | 2337 | 43743 | 32508 | 21720 | 17048 | 17 | 925 | 950 | 64 | 0 | 109 | 183 | 32187 | 0 |
| SRR9924527 | 263 | 184086 | 1674 | 769 | 2340 | 44885 | 33986 | 22633 | 17014 | 28 | 964 | 1008 | 63 | 0 | 103 | 170 | 34251 | 0 |

| | | | | | | | | | | | | | | | | | | | |
|---|---|---|---|---|---|---|---|---|---|---|---|---|---|---|---|---|---|---|---|
| SRR9856171 | 128 | 26 | 13 | 27 | 0 | 0 | 0 | 0 | 0 | 204 | 115 | 14 | 12 | 0 | 0 | 0 | 0 | 0 | 0 |
| SRR9856172 | 55 | 13 | 41 | 109 | 25 | 0 | 0 | 0 | 0 | 84 | 64 | 53 | 72 | 0 | 0 | 0 | 0 | 0 | 0 |
| SRR9856173 | 270 | 81 | 54 | 444 | 97 | 11 | 0 | 0 | 0 | 530 | 437 | 70 | 168 | 0 | 0 | 0 | 0 | 0 | 0 |
| SRR9856174 | 81 | 20 | 38 | 386 | 74 | 0 | 0 | 0 | 0 | 149 | 108 | 37 | 45 | 0 | 0 | 0 | 0 | 0 | 0 |
| SRR9856175 | 720 | 146 | 554 | 1456 | 392 | 12 | 0 | 0 | 0 | 1096 | 729 | 437 | 237 | 18 | 0 | 0 | 0 | 0 | 0 |
| SRR9856176 | 137 | 34 | 208 | 415 | 206 | 13 | 0 | 0 | 0 | 248 | 227 | 207 | 106 | 0 | 0 | 0 | 0 | 12 | 0 |
| SRR9856177 | 257 | 85 | 638 | 1835 | 1334 | 59 | 11 | 33 | 12 | 727 | 726 | 1263 | 393 | 88 | 0 | 0 | 0 | 39 | 0 |
| SRR9856178 | 22 | 0 | 0 | 33 | 0 | 0 | 0 | 0 | 0 | 38 | 22 | 0 | 12 | 0 | 0 | 0 | 0 | 0 | 0 |
| SRR9856179 | 65 | 16 | 10 | 62 | 0 | 0 | 0 | 0 | 0 | 110 | 80 | 16 | 110 | 0 | 0 | 0 | 0 | 0 | 0 |
| SRR9856180 | 358 | 70 | 53 | 121 | 29 | 0 | 0 | 0 | 0 | 434 | 243 | 53 | 91 | 0 | 0 | 0 | 0 | 0 | 0 |
| SRR9856181 | 377 | 77 | 54 | 189 | 63 | 0 | 0 | 0 | 0 | 456 | 289 | 51 | 88 | 0 | 0 | 0 | 0 | 0 | 0 |
| SRR9856182 | 687 | 118 | 38 | 73 | 21 | 0 | 0 | 0 | 0 | 718 | 374 | 32 | 63 | 0 | 0 | 0 | 0 | 0 | 0 |
| SRR9856183 | 230 | 62 | 41 | 401 | 40 | 0 | 0 | 0 | 0 | 398 | 341 | 50 | 109 | 0 | 0 | 0 | 0 | 0 | 0 |
| SRR9856184 | 424 | 66 | 17 | 36 | 0 | 0 | 0 | 0 | 0 | 583 | 267 | 11 | 16 | 0 | 0 | 0 | 0 | 0 | 0 |
| SRR9856185 | 0 | 0 | 0 | 17 | 0 | 0 | 0 | 0 | 0 | 0 | 0 | 0 | 0 | 0 | 0 | 0 | 0 | 0 | 0 |
| SRR9856186 | 116 | 45 | 120 | 560 | 151 | 0 | 0 | 0 | 0 | 152 | 144 | 52 | 38 | 0 | 0 | 0 | 0 | 0 | 0 |
| SRR990360 | 0 | 0 | 0 | 0 | 0 | 0 | 0 | 0 | 0 | 0 | 0 | 0 | 0 | 0 | 0 | 0 | 0 | 0 | 0 |
| SRR9924500 | 0 | 0 | 276 | 44 | 73 | 23 | 156 | 0 | 0 | 0 | 24 | 70 | 36 | 62 | 0 | 165 | 119 | 194 | 0 |
| SRR9924502 | 0 | 0 | 211 | 33 | 51 | 0 | 94 | 0 | 0 | 0 | 20 | 56 | 30 | 35 | 0 | 93 | 53 | 96 | 0 |
| SRR9924511 | 0 | 0 | 153 | 56 | 38 | 0 | 48 | 0 | 0 | 0 | 21 | 56 | 71 | 31 | 0 | 36 | 24 | 66 | 0 |
| SRR9924512 | 0 | 0 | 142 | 50 | 30 | 10 | 52 | 0 | 0 | 0 | 16 | 49 | 65 | 24 | 0 | 56 | 53 | 101 | 0 |
| SRR9924514 | 0 | 0 | 167 | 39 | 44 | 0 | 13 | 0 | 0 | 0 | 16 | 51 | 26 | 26 | 0 | 13 | 0 | 11 | 0 |
| SRR9924515 | 0 | 0 | 130 | 43 | 34 | 0 | 34 | 0 | 0 | 0 | 19 | 52 | 54 | 24 | 0 | 26 | 18 | 38 | 0 |
| SRR9924516 | 0 | 0 | 91 | 24 | 30 | 0 | 11 | 0 | 0 | 0 | 0 | 25 | 16 | 19 | 0 | 0 | 0 | 13 | 0 |
| SRR9924517 | 0 | 0 | 145 | 38 | 41 | 0 | 0 | 0 | 0 | 0 | 12 | 51 | 23 | 24 | 0 | 0 | 0 | 0 | 0 |
| SRR9924518 | 0 | 0 | 254 | 41 | 89 | 54 | 300 | 16 | 0 | 0 | 25 | 79 | 39 | 93 | 12 | 293 | 214 | 564 | 0 |
| SRR9924519 | 0 | 0 | 314 | 43 | 84 | 48 | 260 | 18 | 0 | 0 | 33 | 99 | 40 | 87 | 12 | 252 | 182 | 454 | 0 |
| SRR9924520 | 0 | 0 | 249 | 40 | 50 | 0 | 36 | 0 | 0 | 0 | 30 | 64 | 32 | 34 | 0 | 27 | 16 | 21 | 0 |
| SRR9924521 | 0 | 0 | 279 | 37 | 87 | 57 | 272 | 21 | 0 | 0 | 29 | 85 | 36 | 85 | 12 | 270 | 210 | 500 | 0 |
| SRR9924522 | 0 | 0 | 134 | 26 | 35 | 0 | 31 | 0 | 0 | 0 | 13 | 39 | 19 | 23 | 0 | 29 | 16 | 34 | 0 |
| SRR9924523 | 0 | 0 | 175 | 25 | 41 | 0 | 40 | 0 | 0 | 0 | 18 | 43 | 20 | 26 | 0 | 37 | 20 | 34 | 0 |
| SRR9924524 | 0 | 0 | 413 | 100 | 76 | 19 | 82 | 0 | 0 | 0 | 68 | 141 | 100 | 66 | 0 | 104 | 80 | 143 | 0 |
| SRR9924525 | 0 | 0 | 253 | 32 | 71 | 77 | 178 | 23 | 0 | 0 | 26 | 73 | 33 | 49 | 0 | 134 | 90 | 189 | 0 |
| SRR9924526 | 0 | 0 | 439 | 96 | 83 | 25 | 144 | 0 | 0 | 0 | 64 | 149 | 120 | 74 | 0 | 162 | 117 | 198 | 0 |
| SRR9924527 | 0 | 0 | 387 | 74 | 77 | 12 | 130 | 0 | 0 | 0 | 54 | 127 | 77 | 77 | 0 | 150 | 104 | 188 | 0 |
